## Supplemental Table 5 for "Differential CpG methylation at *Nnat* in the early establishment of beta cell heterogeneity"

| baseMean | log2FoldChange | lfcSE | stat | pvalue | padj | Symbol | GeneName | non β | non β | non β | NNAT$^{-}$ β | NNAT$^{-}$ β | NNAT$^{-}$ β | NNAT$^{-}$ β | NNAT$^{+}$ β | NNAT$^{+}$ β | NNAT$^{+}$ β | NNAT$^{+}$ β |
|---|---|---|---|---|---|---|---|---|---|---|---|---|---|---|---|---|---|---|
| 24738 | -1.45 | 0.07 | -19.91 | 0.00 | 8.45E-84 | Gpx3 | glutathione peroxidase 3 | 74603 | 80954 | 81455 | 7340 | 7042 | 4594 | 6617 | 3030 | 2797 | 2181 | 1509 |
| 7268 | -1.38 | 0.10 | -13.91 | 0.00 | 7.35E-40 | Rbp4 | retinol binding protein 4, plasma | 16770 | 25466 | 23187 | 2675 | 2811 | 2534 | 2465 | 1071 | 963 | 984 | 1021 |
| 24420 | -1.31 | 0.12 | -11.24 | 0.00 | 2.33E-25 | Pyy | peptide YY | 51527 | 86483 | 86871 | 7801 | 8777 | 6781 | 7612 | 4117 | 3327 | 2307 | 3019 |
| 1894 | -1.71 | 0.15 | -11.06 | 0.00 | 1.26E-24 | Eef1a2 | eukaryotic translation elongation factor 1 | 5439 | 6265 | 7712 | 260 | 334 | 191 | 301 | 67 | 112 | 91 | 61 |
| 13546 | -1.49 | 0.14 | -10.38 | 0.00 | 1.67E-21 | Ppy | pancreatic polypeptide | 22180 | 44993 | 55737 | 4562 | 6418 | 4307 | 3883 | 1937 | 1837 | 1054 | 2104 |
| 4497 | -1.11 | 0.11 | -10.17 | 0.00 | 1.24E-20 | Anpep | alanyl (membrane) aminopeptidase | 4223 | 4754 | 3336 | 7103 | 4983 | 5379 | 7701 | 3661 | 2147 | 4069 | 2108 |
| 2895 | -1.36 | 0.15 | -9.17 | 0.00 | 1.79E-16 | Cd24a | CD24a antigen | 7573 | 9490 | 9073 | 984 | 1180 | 780 | 1166 | 447 | 441 | 308 | 403 |
| 2 | -29.98 | 3.58 | -8.39 | 0.00 | 1.68E-13 | H2ac10 | H2A clustered histone 10 | 0 | 0 | 5 | 14 | 2 | 3 | 0 | 0 | 0 | 0 | 0 |
| 2 | -29.51 | 3.58 | -8.24 | 0.00 | 4.84E-13 | Gm43598 | | 0 | 10 | 0 | 0 | 6 | 9 | 0 | 0 | 0 | 0 | 0 |
| 2 | 29.50 | 3.58 | 8.23 | 0.00 | 4.84E-13 | Gm44064 | | 0 | 11 | 0 | 0 | 0 | 0 | 0 | 0 | 5 | 0 | 8 |
| 1 | -29.09 | 3.60 | -8.08 | 0.00 | 1.60E-12 | Gm38699 | | 0 | 11 | 0 | 0 | 4 | 1 | 0 | 0 | 0 | 0 | 0 |
| 3464 | -1.02 | 0.13 | -8.02 | 0.00 | 2.32E-12 | Cd81 | CD81 antigen | 10555 | 9751 | 10093 | 1497 | 1126 | 1178 | 1330 | 643 | 629 | 837 | 460 |
| 2 | -28.39 | 3.61 | -7.86 | 0.00 | 7.89E-12 | 9630002D21Rik | | 0 | 14 | 0 | 0 | 2 | 1 | 0 | 0 | 0 | 0 | 0 |
| 7 | 28.55 | 3.64 | 7.84 | 0.00 | 8.26E-12 | Gm33466 | | 0 | 0 | 0 | 0 | 0 | 0 | 0 | 0 | 0 | 73 | 0 |
| 1 | -28.27 | 3.63 | -7.79 | 0.00 | 1.14E-11 | Gm43608 | | 0 | 8 | 0 | 0 | 1 | 2 | 0 | 0 | 0 | 0 | 0 |
| 4 | 28.05 | 3.64 | 7.71 | 0.00 | 2.16E-11 | Gm48075 | | 0 | 0 | 0 | 0 | 0 | 0 | 0 | 0 | 0 | 44 | 0 |
| 2 | 27.38 | 3.64 | 7.52 | 0.00 | 8.01E-11 | Gm44626 | | 0 | 0 | 0 | 0 | 0 | 0 | 0 | 0 | 0 | 22 | 0 |
| 2 | 27.38 | 3.64 | 7.52 | 0.00 | 8.01E-11 | 1810046K07Ril | RIKEN cDNA 1810046K07 gene | 0 | 0 | 0 | 0 | 0 | 0 | 0 | 0 | 0 | 22 | 0 |
| 925 | -0.85 | 0.11 | -7.50 | 0.00 | 8.69E-11 | Scn3a | sodium channel, voltage-gated, type III, al | 1440 | 1796 | 1627 | 838 | 699 | 1198 | 689 | 447 | 417 | 389 | 632 |
| 4225 | -1.12 | 0.15 | -7.46 | 0.00 | 1.19E-10 | Mafb | v-maf musculoaponeurotic fibrosarcoma ( | 13704 | 14863 | 13185 | 814 | 692 | 964 | 742 | 389 | 368 | 210 | 549 |
| 2 | 26.76 | 3.64 | 7.35 | 0.00 | 2.39E-10 | Mir6399 | | 0 | 0 | 0 | 0 | 0 | 0 | 0 | 0 | 0 | 17 | 0 |
| 2 | 26.76 | 3.64 | 7.35 | 0.00 | 2.39E-10 | Gm4768 | | 0 | 0 | 0 | 0 | 0 | 0 | 0 | 0 | 0 | 17 | 0 |
| 1 | -26.38 | 3.60 | -7.32 | 0.00 | 2.88E-10 | Gm30524 | | 0 | 0 | 3 | 4 | 0 | 1 | 0 | 0 | 0 | 0 | 0 |
| 5 | 26.47 | 3.63 | 7.30 | 0.00 | 3.22E-10 | Gm10805 | | 0 | 13 | 18 | 0 | 0 | 0 | 0 | 24 | 0 | 0 | 0 |
| 2 | 26.42 | 3.64 | 7.26 | 0.00 | 3.80E-10 | Gm5842 | | 0 | 0 | 0 | 0 | 0 | 0 | 0 | 0 | 0 | 18 | 0 |
| 2 | 26.42 | 3.64 | 7.26 | 0.00 | 3.80E-10 | Gm15857 | | 0 | 0 | 0 | 0 | 0 | 0 | 0 | 0 | 0 | 18 | 0 |
| 2 | 26.42 | 3.64 | 7.26 | 0.00 | 3.80E-10 | Gm47099 | | 0 | 0 | 0 | 0 | 0 | 0 | 0 | 0 | 0 | 18 | 0 |
| 2 | 26.42 | 3.64 | 7.26 | 0.00 | 3.80E-10 | Gm12243 | | 0 | 0 | 0 | 0 | 0 | 0 | 0 | 0 | 0 | 18 | 0 |
| 1 | -26.11 | 3.64 | -7.17 | 0.00 | 6.94E-10 | Gm13564 | | 0 | 0 | 0 | 0 | 0 | 0 | 10 | 0 | 0 | 0 | 0 |
| 3 | -25.51 | 3.58 | -7.12 | 0.00 | 9.83E-10 | Tcerg1l | transcription elongation regulator 1-like | 0 | 0 | 21 | 7 | 0 | 2 | 0 | 0 | 0 | 0 | 0 |
| 145348 | -1.33 | 0.19 | -7.07 | 0.00 | 1.35E-09 | Sst | somatostatin | 254964 | 487895 | 470603 | 72280 | 86743 | 48926 | 66376 | 38660 | 28332 | 19486 | 24568 |
| 14 | -22.01 | 3.18 | -6.92 | 0.00 | 3.67E-09 | Vip | vasoactive intestinal polypeptide | 50 | 0 | 18 | 83 | 0 | 0 | 0 | 0 | 0 | 0 | 0 |
| 3 | -24.73 | 3.58 | -6.91 | 0.00 | 4.02E-09 | 4921501I09Rik | | 22 | 0 | 0 | 4 | 4 | 0 | 0 | 0 | 0 | 0 | 0 |
| 1 | -24.70 | 3.59 | -6.88 | 0.00 | 4.66E-09 | 4930456G14Rik | | 0 | 5 | 0 | 0 | 4 | 3 | 0 | 0 | 0 | 0 | 0 |
| 5 | 24.72 | 3.61 | 6.85 | 0.00 | 5.54E-09 | 1700016L21Rik | | 0 | 43 | 0 | 0 | 0 | 0 | 0 | 0 | 9 | 0 | 0 |
| 3 | 24.72 | 3.61 | 6.85 | 0.00 | 5.54E-09 | Gm38058 | | 14 | 0 | 0 | 0 | 0 | 0 | 0 | 20 | 0 | 0 | 0 |
| 13961 | -0.54 | 0.08 | -6.76 | 0.00 | 1.00E-08 | Txnip | thioredoxin interacting protein | 24456 | 20560 | 22853 | 8357 | 8122 | 22329 | 11207 | 5291 | 5500 | 7615 | 17277 |
| 621 | -1.83 | 0.27 | -6.70 | 0.00 | 1.49E-08 | Sgce | sarcoglycan, epsilon | 2008 | 2113 | 2181 | 84 | 126 | 108 | 96 | 27 | 39 | 13 | 39 |
| 3 | 23.63 | 3.54 | 6.67 | 0.00 | 1.75E-08 | Gm49312 | | 0 | 8 | 0 | 0 | 0 | 0 | 0 | 0 | 11 | 8 | 11 |
| 3 | -23.39 | 3.55 | -6.60 | 0.00 | 2.78E-08 | Gm6410 | | 0 | 0 | 9 | 11 | 0 | 3 | 5 | 0 | 0 | 0 | 0 |
| 2 | -23.57 | 3.61 | -6.53 | 0.00 | 4.34E-08 | Gm20478 | | 5 | 0 | 0 | 12 | 0 | 0 | 0 | 0 | 0 | 0 | 0 |
| 3 | -23.12 | 3.55 | -6.52 | 0.00 | 4.44E-08 | Gm50304 | | 0 | 7 | 0 | 0 | 9 | 3 | 9 | 0 | 0 | 0 | 0 |
| 1 | -23.65 | 3.64 | -6.50 | 0.00 | 5.12E-08 | Gm17748 | | 0 | 0 | 0 | 0 | 0 | 0 | 11 | 0 | 0 | 0 | 0 |
| 1472 | -1.59 | 0.25 | -6.45 | 0.00 | 6.58E-08 | Sorcs2 | sortilin-related VPS10 domain containing | 4247 | 4927 | 4646 | 386 | 473 | 389 | 519 | 70 | 229 | 155 | 154 |
| 4 | -23.23 | 3.61 | -6.43 | 0.00 | 7.63E-08 | Btnl5-ps | | 0 | 45 | 0 | 0 | 5 | 0 | 0 | 0 | 0 | 0 | 0 |
| 6 | -19.36 | 3.02 | -6.42 | 0.00 | 7.89E-08 | Phox2b | paired-like homeobox 2b | 27 | 10 | 23 | 11 | 0 | 0 | 0 | 0 | 0 | 0 | 0 |
| 2 | 23.28 | 3.64 | 6.40 | 0.00 | 8.88E-08 | 4930512P04Rik | | 0 | 0 | 0 | 0 | 0 | 0 | 0 | 21 | 0 | 0 | 0 |
| 2 | 23.28 | 3.64 | 6.40 | 0.00 | 8.88E-08 | Gm47066 | | 0 | 0 | 0 | 0 | 0 | 0 | 0 | 21 | 0 | 0 | 0 |
| 1799 | -1.24 | 0.19 | -6.38 | 0.00 | 9.33E-08 | Arg1 | arginase, liver | 4270 | 5818 | 5162 | 843 | 768 | 654 | 900 | 451 | 363 | 200 | 356 |
| 3 | 23.22 | 3.64 | 6.38 | 0.00 | 9.33E-08 | Slc16a8 | solute carrier family 16 (monocarboxylic a | 0 | 0 | 0 | 0 | 0 | 0 | 0 | 28 | 0 | 0 | 0 |
| 1 | -23.25 | 3.64 | -6.38 | 0.00 | 9.33E-08 | Gm37981 | | 0 | 0 | 0 | 12 | 0 | 0 | 0 | 0 | 0 | 0 | 0 |
| 526 | -1.25 | 0.20 | -6.36 | 0.00 | 1.02E-07 | Basp1 | brain abundant, membrane attached sign | 1570 | 1492 | 1909 | 123 | 164 | 147 | 140 | 65 | 63 | 52 | 62 |
| 1 | 23.01 | 3.62 | 6.35 | 0.00 | 1.05E-07 | Gm44175 | | 0 | 2 | 0 | 0 | 0 | 0 | 0 | 0 | 6 | 0 | 0 |
| 1 | 23.01 | 3.62 | 6.35 | 0.00 | 1.05E-07 | Gm7691 | | 0 | 2 | 0 | 0 | 0 | 0 | 0 | 0 | 6 | 0 | 0 |
| 904 | -1.42 | 0.22 | -6.35 | 0.00 | 1.07E-07 | Ehf | ets homologous factor | 2285 | 2717 | 2856 | 422 | 408 | 320 | 352 | 141 | 184 | 73 | 181 |
| 2 | 23.04 | 3.63 | 6.34 | 0.00 | 1.09E-07 | Mcf2 | mcf.2 transforming sequence | 0 | 16 | 4 | 0 | 0 | 0 | 0 | 0 | 7 | 0 | 0 |
| 3 | 22.97 | 3.64 | 6.31 | 0.00 | 1.29E-07 | AI314278 | | 0 | 0 | 0 | 0 | 0 | 0 | 0 | 36 | 0 | 0 | 0 |
| 1 | 22.96 | 3.64 | 6.31 | 0.00 | 1.31E-07 | 4930553I04Rik | | 0 | 0 | 0 | 0 | 0 | 0 | 0 | 16 | 0 | 0 | 0 |
| 5 | 20.48 | 3.25 | 6.29 | 0.00 | 1.41E-07 | Gm48684 | | 6 | 2 | 18 | 0 | 0 | 0 | 0 | 35 | 0 | 0 | 0 |

| | | | | | | | | | | | | | | | | | | |
|---|---|---|---|---|---|---|---|---|---|---|---|---|---|---|---|---|---|---|
| 6 | -19.34 | 3.08 | -6.28 | 0.00 | 1.49E-07 | Gm37805 | | 7 | 10 | 37 | 12 | 0 | 0 | 0 | 0 | 0 | 0 | 0 |
| 3 | 22.85 | 3.64 | 6.28 | 0.00 | 1.49E-07 | Hecw1 | HECT, C2 and WW domain containing E3 | 0 | 0 | 0 | 0 | 0 | 0 | 0 | 0 | 0 | 37 | 0 |
| 2 | 22.62 | 3.61 | 6.28 | 0.00 | 1.49E-07 | Rps19-ps9 | | 0 | 15 | 0 | 0 | 0 | 0 | 0 | 0 | 3 | 1 | 0 |
| 2 | -22.37 | 3.58 | -6.24 | 0.00 | 1.82E-07 | Gm18337 | | 0 | 11 | 0 | 0 | 5 | 0 | 3 | 0 | 0 | 0 | 0 |
| 3 | 22.72 | 3.64 | 6.24 | 0.00 | 1.82E-07 | Rlbp1 | retinaldehyde binding protein 1 | 0 | 0 | 0 | 0 | 0 | 0 | 0 | 0 | 0 | 30 | 0 |
| 1 | -22.39 | 3.61 | -6.20 | 0.00 | 2.36E-07 | Chil5 | chitinase-like 5 | 3 | 0 | 0 | 11 | 0 | 0 | 0 | 0 | 0 | 0 | 0 |
| 1 | -22.27 | 3.61 | -6.16 | 0.00 | 2.85E-07 | Chrna3 | cholinergic receptor, nicotinic, alpha poly | 2 | 0 | 0 | 12 | 0 | 0 | 0 | 0 | 0 | 0 | 0 |
| 1 | -22.28 | 3.61 | -6.16 | 0.00 | 2.85E-07 | Gm37446 | | 3 | 0 | 0 | 0 | 9 | 0 | 0 | 0 | 0 | 0 | 0 |
| 2 | 22.32 | 3.64 | 6.14 | 0.00 | 3.22E-07 | Gm15384 | | 0 | 13 | 2 | 0 | 0 | 0 | 0 | 0 | 5 | 0 | 0 |
| 2 | 22.13 | 3.62 | 6.12 | 0.00 | 3.49E-07 | Ighv5-9 | | 11 | 0 | 0 | 0 | 0 | 0 | 0 | 6 | 0 | 0 | 0 |
| 2 | -22.11 | 3.61 | -6.12 | 0.00 | 3.49E-07 | Gm45869 | | 0 | 7 | 0 | 0 | 9 | 0 | 0 | 0 | 0 | 0 | 0 |
| 1 | -22.15 | 3.62 | -6.11 | 0.00 | 3.49E-07 | Gm44081 | | 0 | 13 | 0 | 0 | 3 | 0 | 0 | 0 | 0 | 0 | 0 |
| 1 | 22.12 | 3.62 | 6.12 | 0.00 | 3.49E-07 | Gm45620 | | 8 | 0 | 0 | 0 | 0 | 0 | 0 | 0 | 6 | 0 | 0 |
| 1 | 22.12 | 3.62 | 6.12 | 0.00 | 3.49E-07 | 2810433D01Rik | | 8 | 0 | 0 | 0 | 0 | 0 | 0 | 0 | 6 | 0 | 0 |
| 1 | -22.26 | 3.64 | -6.12 | 0.00 | 3.49E-07 | Gm8000 | | 0 | 1 | 3 | 7 | 0 | 0 | 0 | 0 | 0 | 0 | 0 |
| 3 | 22.25 | 3.64 | 6.11 | 0.00 | 3.51E-07 | Dsc1 | desmocollin 1 | 0 | 0 | 0 | 0 | 0 | 0 | 0 | 32 | 0 | 0 | 0 |
| 1 | -22.24 | 3.64 | -6.11 | 0.00 | 3.56E-07 | Gm16325 | | 0 | 0 | 0 | 0 | 11 | 0 | 0 | 0 | 0 | 0 | 0 |
| 1017 | -1.06 | 0.17 | -6.10 | 0.00 | 3.65E-07 | Tspan8 | tetraspanin 8 | 1995 | 2958 | 2573 | 649 | 599 | 615 | 595 | 263 | 264 | 255 | 417 |
| 2 | 22.18 | 3.64 | 6.09 | 0.00 | 3.79E-07 | 4930479H17Rik | | 0 | 0 | 0 | 0 | 0 | 0 | 0 | 0 | 0 | 19 | 0 |
| 2 | 22.18 | 3.64 | 6.09 | 0.00 | 3.79E-07 | H2bc12 | H2B clustered histone 12 | 0 | 0 | 0 | 0 | 0 | 0 | 0 | 0 | 0 | 19 | 0 |
| 1 | -22.16 | 3.64 | -6.09 | 0.00 | 3.83E-07 | Gm41562 | | 0 | 0 | 0 | 0 | 0 | 0 | 13 | 0 | 0 | 0 | 0 |
| 1 | 21.97 | 3.62 | 6.07 | 0.00 | 4.12E-07 | Alpk2 | alpha-kinase 2 | 9 | 0 | 0 | 0 | 0 | 0 | 0 | 5 | 0 | 0 | 0 |
| 1 | -21.95 | 3.62 | -6.07 | 0.00 | 4.12E-07 | Gata1 | GATA binding protein 1 | 6 | 0 | 0 | 0 | 6 | 0 | 0 | 0 | 0 | 0 | 0 |
| 1 | -22.00 | 3.62 | -6.07 | 0.00 | 4.12E-07 | Bcas3os2 | | 0 | 0 | 8 | 3 | 0 | 0 | 0 | 0 | 0 | 0 | 0 |
| 1 | -21.90 | 3.61 | -6.06 | 0.00 | 4.34E-07 | Gm15731 | | 0 | 0 | 5 | 7 | 0 | 0 | 0 | 0 | 0 | 0 | 0 |
| 2 | -21.89 | 3.62 | -6.05 | 0.00 | 4.51E-07 | Pou2f3 | POU domain, class 2, transcription factor | 0 | 22 | 0 | 0 | 4 | 0 | 0 | 0 | 0 | 0 | 0 |
| 1 | 21.81 | 3.62 | 6.03 | 0.00 | 5.01E-07 | Gm49801 | | 5 | 0 | 0 | 0 | 0 | 0 | 0 | 0 | 7 | 0 | 0 |
| 1 | -21.77 | 3.61 | -6.02 | 0.00 | 5.26E-07 | Gm6871 | zinc finger protein 431-like | 0 | 6 | 0 | 0 | 7 | 0 | 0 | 0 | 0 | 0 | 0 |
| 2 | -21.90 | 3.64 | -6.02 | 0.00 | 5.39E-07 | Gm18990 | | 0 | 0 | 0 | 0 | 0 | 0 | 19 | 0 | 0 | 0 | 0 |
| 2976 | -1.04 | 0.17 | -6.00 | 0.00 | 5.91E-07 | Adgrf5 | adhesion G protein-coupled receptor F5 | 9098 | 10460 | 11113 | 343 | 242 | 539 | 252 | 200 | 127 | 77 | 285 |
| 459 | 21.84 | 3.64 | 6.00 | 0.00 | 5.91E-07 | Igkv9-120 | | 5051 | 0 | 0 | 0 | 0 | 0 | 0 | 0 | 1 | 0 | 0 |
| 1 | -21.62 | 3.61 | -5.98 | 0.00 | 6.45E-07 | Gm15486 | | 5 | 0 | 0 | 8 | 0 | 0 | 0 | 0 | 0 | 0 | 0 |
| 1 | -21.62 | 3.62 | -5.98 | 0.00 | 6.48E-07 | Gm9936 | | 2 | 0 | 0 | 10 | 0 | 0 | 0 | 0 | 0 | 0 | 0 |
| 1 | 21.56 | 3.62 | 5.96 | 0.00 | 7.15E-07 | Gm31282 | | 6 | 0 | 0 | 0 | 0 | 0 | 0 | 0 | 7 | 0 | 0 |
| 1 | -21.50 | 3.62 | -5.94 | 0.00 | 7.95E-07 | Gm2990 | | 3 | 0 | 0 | 0 | 7 | 0 | 0 | 0 | 0 | 0 | 0 |
| 2832 | -1.00 | 0.17 | -5.91 | 0.00 | 9.85E-07 | Ace2 | angiotensin I converting enzyme (peptidy | 8386 | 10082 | 9572 | 470 | 535 | 364 | 708 | 338 | 261 | 227 | 207 |
| 312 | -1.15 | 0.20 | -5.88 | 0.00 | 1.11E-06 | Agap2 | ArfGAP with GTPase domain, ankyrin rep | 483 | 416 | 695 | 248 | 393 | 292 | 327 | 116 | 210 | 129 | 120 |
| 460 | -1.56 | 0.27 | -5.80 | 0.00 | 1.86E-06 | Gata6 | GATA binding protein 6 | 1286 | 1137 | 1256 | 217 | 222 | 326 | 252 | 36 | 102 | 84 | 146 |
| 743 | -0.92 | 0.16 | -5.77 | 0.00 | 2.19E-06 | Hdac7 | histone deacetylase 7 | 1980 | 1683 | 1968 | 399 | 422 | 456 | 381 | 166 | 244 | 193 | 285 |
| 1063 | -1.49 | 0.26 | -5.76 | 0.00 | 2.26E-06 | Serpine2 | serine (or cysteine) peptidase inhibitor, cl | 3801 | 3108 | 4098 | 98 | 150 | 97 | 163 | 39 | 80 | 24 | 39 |
| 2 | -20.85 | 3.62 | -5.76 | 0.00 | 2.27E-06 | Gm10425 | | 0 | 0 | 24 | 3 | 0 | 0 | 0 | 0 | 0 | 0 | 0 |
| 1 | -20.79 | 3.62 | -5.74 | 0.00 | 2.48E-06 | Gm46555 | | 2 | 0 | 0 | 6 | 0 | 0 | 0 | 0 | 0 | 0 | 0 |
| 1 | -20.69 | 3.62 | -5.71 | 0.00 | 2.96E-06 | Rxfp3 | relaxin family peptide receptor 3 | 3 | 0 | 0 | 0 | 4 | 0 | 0 | 0 | 0 | 0 | 0 |
| 1 | -20.69 | 3.63 | -5.70 | 0.00 | 3.15E-06 | AU023070 | | 0 | 7 | 0 | 0 | 2 | 0 | 0 | 0 | 0 | 0 | 0 |
| 1 | -20.69 | 3.63 | -5.70 | 0.00 | 3.15E-06 | Gm44287 | | 0 | 7 | 0 | 0 | 2 | 0 | 0 | 0 | 0 | 0 | 0 |
| 1101 | -1.25 | 0.22 | -5.69 | 0.00 | 3.18E-06 | Tspan12 | tetraspanin 12 | 3783 | 4189 | 3124 | 137 | 178 | 172 | 233 | 63 | 81 | 63 | 92 |
| 1 | 20.57 | 3.61 | 5.69 | 0.00 | 3.18E-06 | Gm4940 | | 6 | 0 | 0 | 0 | 0 | 0 | 0 | 0 | 8 | 0 | 0 |
| 1 | -20.59 | 3.62 | -5.69 | 0.00 | 3.18E-06 | Gm13179 | | 3 | 0 | 0 | 0 | 6 | 0 | 0 | 0 | 0 | 0 | 0 |
| 782 | -1.48 | 0.26 | -5.69 | 0.00 | 3.19E-06 | Efnb3 | ephrin B3 | 1639 | 2491 | 2210 | 402 | 471 | 370 | 398 | 211 | 157 | 64 | 194 |
| 3 | 20.39 | 3.59 | 5.68 | 0.00 | 3.31E-06 | Gm42697 | | 0 | 0 | 5 | 0 | 0 | 0 | 0 | 0 | 0 | 26 | 1 |
| 1 | -20.55 | 3.62 | -5.68 | 0.00 | 3.31E-06 | Gm10403 | | 8 | 0 | 0 | 0 | 4 | 0 | 0 | 0 | 0 | 0 | 0 |
| 3168 | -0.80 | 0.14 | -5.66 | 0.00 | 3.53E-06 | Vwde | von Willebrand factor D and EGF domains | 8892 | 11262 | 12172 | 410 | 441 | 306 | 420 | 263 | 330 | 227 | 122 |
| 970 | -1.18 | 0.21 | -5.66 | 0.00 | 3.53E-06 | Spred1 | sprouty protein with EVH-1 domain 1, rela | 3608 | 2826 | 2857 | 306 | 197 | 223 | 225 | 80 | 103 | 139 | 101 |
| 1 | -20.52 | 3.62 | -5.66 | 0.00 | 3.53E-06 | Slc5a7 | solute carrier family 5 (choline transporter | 4 | 0 | 0 | 4 | 0 | 0 | 0 | 0 | 0 | 0 | 0 |
| 1 | 20.49 | 3.62 | 5.66 | 0.00 | 3.57E-06 | Gm43311 | | 0 | 8 | 0 | 0 | 0 | 0 | 0 | 0 | 4 | 0 | 0 |
| 1 | 20.46 | 3.62 | 5.65 | 0.00 | 3.70E-06 | Gm49593 | | 3 | 0 | 0 | 0 | 0 | 0 | 0 | 7 | 0 | 0 | 0 |
| 1 | -20.40 | 3.62 | -5.63 | 0.00 | 4.19E-06 | Ell3 | elongation factor RNA polymerase II-like | 0 | 0 | 3 | 4 | 0 | 0 | 0 | 0 | 0 | 0 | 0 |
| 264 | -2.43 | 0.43 | -5.59 | 0.00 | 5.20E-06 | Cacna1h | calcium channel, voltage-dependent, T ty | 888 | 787 | 428 | 157 | 157 | 216 | 127 | 34 | 26 | 8 | 73 |
| 1 | 20.23 | 3.62 | 5.58 | 0.00 | 5.28E-06 | Fgf23 | fibroblast growth factor 23 | 3 | 0 | 0 | 0 | 0 | 0 | 0 | 0 | 5 | 0 | 0 |
| 1 | -20.18 | 3.62 | -5.57 | 0.00 | 5.59E-06 | Gm3734 | | 4 | 0 | 0 | 4 | 0 | 0 | 0 | 0 | 0 | 0 | 0 |

| | | | | | | | | | | | | | | | | | | |
|---|---|---|---|---|---|---|---|---|---|---|---|---|---|---|---|---|---|---|
| 2 | -19.87 | 3.58 | -5.55 | 0.00 | 6.38E-06 | Gm4829 | | 0 | 10 | 0 | 0 | 0 | 6 | 6 | 0 | 0 | 0 | 0 |
| 1 | -20.02 | 3.61 | -5.55 | 0.00 | 6.44E-06 | Gm12643 | | 0 | 5 | 0 | 0 | 1 | 0 | 2 | 0 | 0 | 0 | 0 |
| 1 | -20.00 | 3.62 | -5.52 | 0.00 | 7.31E-06 | Gm50275 | | 0 | 4 | 0 | 0 | 4 | 0 | 0 | 0 | 0 | 0 | 0 |
| 1183 | -1.05 | 0.19 | -5.52 | 0.00 | 7.42E-06 | Ndst4 | N-deacetylase/N-sulfotransferase (hepari | 3209 | 5825 | 2850 | 195 | 125 | 233 | 211 | 104 | 62 | 89 | 115 |
| 36 | 20.05 | 3.64 | 5.50 | 0.00 | 7.94E-06 | Gabrp | gamma-aminobutyric acid (GABA) A rece | 146 | 78 | 155 | 0 | 0 | 0 | 0 | 0 | 21 | 0 | 0 |
| 1 | -19.86 | 3.61 | -5.50 | 0.00 | 8.06E-06 | Sypl2 | synaptophysin-like 2 | 3 | 0 | 0 | 6 | 0 | 0 | 1 | 0 | 0 | 0 | 0 |
| 1 | -19.89 | 3.62 | -5.49 | 0.00 | 8.28E-06 | Rspo2 | R-spondin 2 | 12 | 0 | 0 | 4 | 0 | 0 | 0 | 0 | 0 | 0 | 0 |
| 1 | 19.90 | 3.63 | 5.48 | 0.00 | 8.73E-06 | Gm27167 | | 0 | 4 | 0 | 0 | 0 | 0 | 0 | 0 | 3 | 0 | 0 |
| 8513 | -0.70 | 0.13 | -5.47 | 0.00 | 9.38E-06 | Itpr1 | inositol 1,4,5-trisphosphate receptor 1 | 18402 | 27333 | 18805 | 4105 | 3704 | 5846 | 4395 | 2599 | 2954 | 2134 | 3369 |
| 3 | -19.53 | 3.58 | -5.45 | 0.00 | 1.03E-05 | Gm48153 | | 0 | 5 | 15 | 3 | 0 | 0 | 11 | 0 | 0 | 0 | 0 |
| 11 | -17.48 | 3.21 | -5.44 | 0.00 | 1.06E-05 | Gimap7 | GTPase, IMAP family member 7 | 97 | 19 | 0 | 0 | 3 | 0 | 0 | 0 | 0 | 0 | 0 |
| 1 | -19.60 | 3.62 | -5.42 | 0.00 | 1.24E-05 | Gm26777 | | 0 | 5 | 0 | 0 | 5 | 0 | 0 | 0 | 0 | 0 | 0 |
| 2 | 19.52 | 3.61 | 5.41 | 0.00 | 1.29E-05 | Gm11536 | | 0 | 3 | 0 | 0 | 0 | 0 | 0 | 0 | 0 | 22 | 0 |
| 2 | 19.39 | 3.61 | 5.37 | 0.00 | 1.58E-05 | Tctex1d4 | Tctex1 domain containing 4 | 0 | 3 | 0 | 0 | 0 | 0 | 0 | 0 | 0 | 19 | 0 |
| 1 | -19.48 | 3.63 | -5.36 | 0.00 | 1.65E-05 | Gm35558 | | 9 | 0 | 0 | 2 | 0 | 0 | 0 | 0 | 0 | 0 | 0 |
| 7 | 19.48 | 3.64 | 5.35 | 0.00 | 1.75E-05 | Slc13a2 | solute carrier family 13 (sodium-depende | 0 | 15 | 39 | 0 | 0 | 0 | 0 | 0 | 0 | 23 | 0 |
| 1 | -19.32 | 3.62 | -5.33 | 0.00 | 1.91E-05 | Gm20703 | | 0 | 8 | 0 | 0 | 3 | 0 | 0 | 0 | 0 | 0 | 0 |
| 3855 | -0.53 | 0.10 | -5.33 | 0.00 | 1.94E-05 | 1700086L19Rik | | 10547 | 12325 | 12367 | 978 | 950 | 1343 | 966 | 618 | 659 | 751 | 903 |
| 55 | -19.39 | 3.64 | -5.32 | 0.00 | 1.94E-05 | Stmn2 | stathmin-like 2 | 233 | 152 | 181 | 35 | 0 | 0 | 0 | 0 | 0 | 0 | 0 |
| 32 | -19.39 | 3.64 | -5.32 | 0.00 | 1.94E-05 | Dnah1 | dynein, axonemal, heavy chain 1 | 92 | 129 | 115 | 19 | 0 | 0 | 0 | 0 | 0 | 0 | 0 |
| 2 | 19.23 | 3.62 | 5.31 | 0.00 | 2.09E-05 | Mir5136 | | 19 | 0 | 0 | 0 | 0 | 0 | 0 | 0 | 3 | 0 | 0 |
| 0 | -19.24 | 3.63 | -5.30 | 0.00 | 2.15E-05 | Cyp11a1 | cytochrome P450, family 11, subfamily a, | 2 | 0 | 0 | 4 | 0 | 0 | 0 | 0 | 0 | 0 | 0 |
| 2 | -18.97 | 3.58 | -5.30 | 0.00 | 2.17E-05 | Csf3 | colony stimulating factor 3 (granulocyte) | 0 | 0 | 11 | 0 | 0 | 5 | 9 | 0 | 0 | 0 | 0 |
| 1 | -19.13 | 3.62 | -5.29 | 0.00 | 2.26E-05 | Gm7019 | | 4 | 0 | 0 | 0 | 8 | 0 | 0 | 0 | 0 | 0 | 0 |
| 546 | -2.13 | 0.40 | -5.28 | 0.00 | 2.42E-05 | Ntn4 | netrin 4 | 2240 | 1759 | 1676 | 72 | 60 | 86 | 48 | 31 | 5 | 8 | 21 |
| 822 | -2.09 | 0.40 | -5.27 | 0.00 | 2.46E-05 | Pros1 | protein S (alpha) | 4142 | 2144 | 2492 | 36 | 97 | 30 | 57 | 10 | 20 | 5 | 12 |
| 3 | 18.86 | 3.58 | 5.27 | 0.00 | 2.52E-05 | Gm47621 | | 0 | 0 | 3 | 0 | 0 | 0 | 0 | 0 | 0 | 24 | 3 |
| 4 | -18.14 | 3.45 | -5.27 | 0.00 | 2.54E-05 | Ovol1 | ovo like zinc finger 1 | 20 | 1 | 22 | 6 | 0 | 0 | 0 | 0 | 0 | 0 | 0 |
| 1 | 19.00 | 3.62 | 5.25 | 0.00 | 2.69E-05 | Gm48505 | | 9 | 0 | 0 | 0 | 0 | 0 | 0 | 0 | 5 | 0 | 0 |
| 1 | -19.07 | 3.63 | -5.25 | 0.00 | 2.69E-05 | Tmem114 | transmembrane protein 114 | 6 | 1 | 0 | 6 | 0 | 0 | 0 | 0 | 0 | 0 | 0 |
| 1 | -18.95 | 3.62 | -5.24 | 0.00 | 2.84E-05 | Gm9850 | | 7 | 0 | 0 | 0 | 6 | 0 | 0 | 0 | 0 | 0 | 0 |
| 52 | 19.07 | 3.64 | 5.23 | 0.00 | 2.91E-05 | Ptpn22 | protein tyrosine phosphatase, non-recept | 415 | 111 | 35 | 0 | 0 | 0 | 0 | 0 | 9 | 0 | 0 |
| 0 | 19.05 | 3.64 | 5.23 | 0.00 | 2.91E-05 | Gm6899 | | 0 | 3 | 0 | 0 | 0 | 0 | 0 | 0 | 2 | 0 | 0 |
| 2 | -18.70 | 3.58 | -5.22 | 0.00 | 3.07E-05 | 2310081O03Rik | | 0 | 0 | 15 | 0 | 0 | 5 | 5 | 0 | 0 | 0 | 0 |
| 1 | -18.86 | 3.61 | -5.22 | 0.00 | 3.07E-05 | Gm8625 | | 1 | 0 | 9 | 1 | 0 | 0 | 3 | 0 | 0 | 0 | 0 |
| 1 | -18.92 | 3.62 | -5.22 | 0.00 | 3.07E-05 | Gm24853 | | 0 | 3 | 0 | 0 | 4 | 0 | 0 | 0 | 0 | 0 | 0 |
| 1 | 18.94 | 3.64 | 5.21 | 0.00 | 3.28E-05 | 2700069I18Rik | | 0 | 4 | 0 | 0 | 0 | 0 | 0 | 0 | 2 | 0 | 0 |
| 2 | -18.82 | 3.62 | -5.20 | 0.00 | 3.36E-05 | Gimd1 | GIMAP family P-loop NTPase domain cor | 0 | 15 | 0 | 0 | 4 | 0 | 0 | 0 | 0 | 0 | 0 |
| 78 | -18.91 | 3.64 | -5.19 | 0.00 | 3.49E-05 | Sncg | synuclein, gamma | 367 | 259 | 216 | 16 | 0 | 0 | 0 | 0 | 0 | 0 | 0 |
| 1 | 18.92 | 3.64 | 5.19 | 0.00 | 3.49E-05 | Serpinb1b | serine (or cysteine) peptidase inhibitor, cl | 1 | 2 | 0 | 0 | 0 | 0 | 0 | 0 | 8 | 0 | 0 |
| 2 | 18.60 | 3.58 | 5.19 | 0.00 | 3.52E-05 | Mogat1 | monoacylglycerol O-acyltransferase 1 | 10 | 0 | 0 | 0 | 0 | 0 | 0 | 0 | 5 | 0 | 8 |
| 88 | -4.82 | 0.93 | -5.19 | 0.00 | 3.53E-05 | Nlgn1 | neuroligin 1 | 183 | 299 | 189 | 105 | 41 | 58 | 61 | 0 | 2 | 0 | 25 |
| 641 | -0.96 | 0.18 | -5.18 | 0.00 | 3.61E-05 | Uchl1 | ubiquitin carboxy-terminal hydrolase L1 | 1341 | 1387 | 1729 | 342 | 467 | 359 | 545 | 166 | 253 | 252 | 209 |
| 4 | -18.69 | 3.61 | -5.17 | 0.00 | 3.80E-05 | Slc10a5 | solute carrier family 10 (sodium/bile acid ( | 3 | 24 | 0 | 0 | 0 | 0 | 18 | 0 | 0 | 0 | 0 |
| 0 | 18.74 | 3.64 | 5.16 | 0.00 | 4.11E-05 | Gm15880 | | 2 | 0 | 0 | 0 | 0 | 0 | 0 | 0 | 3 | 0 | 0 |
| 1 | -18.64 | 3.62 | -5.15 | 0.00 | 4.25E-05 | Gm14942 | | 0 | 0 | 12 | 0 | 0 | 0 | 3 | 0 | 0 | 0 | 0 |
| 2 | -18.46 | 3.59 | -5.14 | 0.00 | 4.29E-05 | Gm26839 | | 0 | 14 | 0 | 0 | 0 | 3 | 3 | 0 | 0 | 0 | 0 |
| 1 | 18.60 | 3.61 | 5.14 | 0.00 | 4.29E-05 | Gm43837 | | 0 | 7 | 0 | 0 | 0 | 0 | 0 | 0 | 0 | 8 | 0 |
| 1 | 18.75 | 3.65 | 5.14 | 0.00 | 4.29E-05 | Gm46203 | | 1 | 0 | 1 | 0 | 0 | 0 | 0 | 8 | 0 | 0 | 0 |
| 8709 | -0.29 | 0.06 | -5.13 | 0.00 | 4.52E-05 | Map3k15 | mitogen-activated protein kinase kinase k | 13032 | 14466 | 13930 | 6403 | 6192 | 11587 | 5735 | 5042 | 5472 | 4434 | 9504 |
| 2 | -18.40 | 3.59 | -5.13 | 0.00 | 4.56E-05 | Gm11465 | | 0 | 0 | 10 | 0 | 0 | 2 | 7 | 0 | 0 | 0 | 0 |
| 2 | 18.57 | 3.62 | 5.12 | 0.00 | 4.68E-05 | Gm7065 | | 1 | 3 | 0 | 0 | 0 | 0 | 0 | 0 | 0 | 18 | 0 |
| 532 | -1.35 | 0.26 | -5.11 | 0.00 | 4.98E-05 | Cldn6 | claudin 6 | 1191 | 1383 | 1573 | 439 | 286 | 247 | 237 | 134 | 210 | 82 | 72 |
| 65 | -18.58 | 3.64 | -5.10 | 0.00 | 5.22E-05 | Tmem255b | transmembrane protein 255B | 257 | 172 | 275 | 12 | 0 | 0 | 0 | 0 | 0 | 0 | 0 |
| 1 | -18.22 | 3.58 | -5.08 | 0.00 | 5.67E-05 | Mir344c | | 0 | 0 | 5 | 0 | 0 | 3 | 7 | 0 | 0 | 0 | 0 |
| 2 | 18.23 | 3.59 | 5.08 | 0.00 | 5.68E-05 | Cdh26 | cadherin-like 26 | 12 | 0 | 0 | 0 | 0 | 0 | 0 | 0 | 8 | 0 | 2 |
| 392 | -1.44 | 0.28 | -5.08 | 0.00 | 5.75E-05 | Ache | acetylcholinesterase | 1192 | 1059 | 1002 | 147 | 215 | 166 | 240 | 76 | 46 | 66 | 97 |
| 23248 | -0.24 | 0.05 | -5.08 | 0.00 | 5.78E-05 | Cltrn | collectrin, amino acid transport regulator | 38742 | 45841 | 43946 | 20072 | 18338 | 13686 | 16918 | 16709 | 15946 | 13223 | 12302 |
| 2 | -18.19 | 3.59 | -5.07 | 0.00 | 5.90E-05 | Gm28731 | | 0 | 9 | 0 | 0 | 0 | 2 | 7 | 0 | 0 | 0 | 0 |
| 1 | -18.24 | 3.62 | -5.04 | 0.00 | 6.88E-05 | Ostn | osteocrin | 0 | 10 | 0 | 0 | 0 | 0 | 4 | 0 | 0 | 0 | 0 |

| | | | | | | | | | | | | | | | | | | |
|---|---|---|---|---|---|---|---|---|---|---|---|---|---|---|---|---|---|---|
| 1 | -18.29 | 3.63 | -5.04 | 0.00 | 6.88E-05 | Gm19287 | | 3 | 0 | 0 | 3 | 0 | 0 | 0 | 0 | 0 | 0 | 0 |
| 131 | -4.99 | 0.99 | -5.03 | 0.00 | 7.16E-05 | Nrxn2 | neurexin II | 360 | 522 | 390 | 31 | 54 | 18 | 64 | 0 | 1 | 0 | 6 |
| 6 | -18.28 | 3.64 | -5.02 | 0.00 | 7.74E-05 | Nefl | neurofilament, light polypeptide | 45 | 0 | 8 | 7 | 0 | 0 | 0 | 0 | 0 | 0 | 0 |
| 626 | -1.86 | 0.37 | -5.00 | 0.00 | 8.40E-05 | AW551984 | expressed sequence AW551984 | 1583 | 2154 | 1754 | 268 | 277 | 249 | 270 | 35 | 129 | 35 | 134 |
| 1 | -18.08 | 3.62 | -5.00 | 0.00 | 8.40E-05 | Cckbr | cholecystokinin B receptor | 0 | 0 | 7 | 0 | 0 | 0 | 5 | 0 | 0 | 0 | 0 |
| 1 | -17.96 | 3.60 | -4.99 | 0.00 | 8.73E-05 | Gm16867 | predicted gene, 16867 | 0 | 9 | 0 | 0 | 0 | 3 | 2 | 0 | 0 | 0 | 0 |
| 1 | 17.95 | 3.60 | 4.99 | 0.00 | 8.73E-05 | Gm15750 | | 7 | 0 | 0 | 0 | 0 | 0 | 0 | 0 | 6 | 0 | 1 |
| 1 | -17.98 | 3.61 | -4.98 | 0.00 | 9.23E-05 | Gm4152 | | 0 | 2 | 0 | 0 | 0 | 0 | 12 | 0 | 0 | 0 | 0 |
| 3 | -18.13 | 3.65 | -4.97 | 0.00 | 9.50E-05 | Gm36723 | | 23 | 5 | 0 | 0 | 5 | 0 | 0 | 0 | 0 | 0 | 0 |
| 1 | -18.07 | 3.64 | -4.97 | 0.00 | 9.50E-05 | Efna3 | ephrin A3 | 0 | 0 | 4 | 2 | 0 | 0 | 0 | 0 | 0 | 0 | 0 |
| 2 | 18.13 | 3.65 | 4.97 | 0.00 | 9.59E-05 | Mpl | myeloproliferative leukemia virus oncoge | 15 | 7 | 0 | 0 | 0 | 0 | 0 | 0 | 4 | 0 | 0 |
| 175 | -18.07 | 3.64 | -4.96 | 0.00 | 9.64E-05 | Ms4a6c | membrane-spanning 4-domains, subfami | 934 | 374 | 586 | 0 | 0 | 0 | 31 | 0 | 0 | 0 | 0 |
| 1 | -17.79 | 3.59 | -4.96 | 0.00 | 9.81E-05 | Gm42577 | | 6 | 0 | 0 | 0 | 5 | 3 | 0 | 0 | 0 | 0 | 0 |
| 0 | 18.03 | 3.64 | 4.95 | 0.00 | 0.0001 | Gm48899 | | 3 | 0 | 0 | 0 | 0 | 0 | 0 | 2 | 0 | 0 | 0 |
| 1 | -17.90 | 3.62 | -4.95 | 0.00 | 0.0001 | Gm26153 | | 0 | 8 | 0 | 0 | 0 | 0 | 5 | 0 | 0 | 0 | 0 |
| 2 | -17.73 | 3.59 | -4.94 | 0.00 | 0.0001 | Gm45007 | | 8 | 0 | 0 | 9 | 0 | 2 | 0 | 0 | 0 | 0 | 0 |
| 0 | 17.99 | 3.64 | 4.94 | 0.00 | 0.0001 | 9330154J02Rik | | 3 | 0 | 0 | 0 | 0 | 0 | 0 | 0 | 2 | 0 | 0 |
| 1 | -17.76 | 3.60 | -4.94 | 0.00 | 0.00011 | Dppa3-ps | | 0 | 6 | 0 | 0 | 0 | 1 | 6 | 0 | 0 | 0 | 0 |
| 1 | -17.81 | 3.61 | -4.93 | 0.00 | 0.00011 | Gm34299 | | 0 | 3 | 0 | 0 | 0 | 0 | 9 | 0 | 0 | 0 | 0 |
| 3 | -17.86 | 3.63 | -4.93 | 0.00 | 0.00011 | Gm16344 | | 0 | 7 | 16 | 0 | 0 | 1 | 9 | 0 | 0 | 0 | 0 |
| 3 | 17.95 | 3.65 | 4.92 | 0.00 | 0.00011 | Kcnj16 | potassium inwardly-rectifying channel, su | 6 | 22 | 1 | 0 | 0 | 0 | 0 | 0 | 5 | 0 | 0 |
| 2 | 17.67 | 3.59 | 4.92 | 0.00 | 0.00012 | Scn5a | sodium channel, voltage-gated, type V, al | 12 | 0 | 0 | 0 | 0 | 0 | 0 | 0 | 3 | 0 | 4 |
| 124 | -5.72 | 1.16 | -4.91 | 0.00 | 0.00012 | Slc25a33 | solute carrier family 25, member 33 | 445 | 387 | 487 | 4 | 14 | 8 | 24 | 0 | 0 | 0 | 0 |
| 2 | -17.79 | 3.64 | -4.88 | 0.00 | 0.00014 | Ccdc187 | coiled-coil domain containing 187 | 0 | 5 | 10 | 0 | 0 | 0 | 10 | 0 | 0 | 0 | 0 |
| 1 | -17.79 | 3.64 | -4.88 | 0.00 | 0.00014 | Fabp1 | fatty acid binding protein 1, liver | 0 | 1 | 5 | 0 | 0 | 0 | 10 | 0 | 0 | 0 | 0 |
| 272 | -1.81 | 0.37 | -4.88 | 0.00 | 0.00014 | Shisal1 | shisa like 1 | 582 | 891 | 583 | 199 | 98 | 224 | 174 | 54 | 33 | 18 | 130 |
| 2 | -17.61 | 3.61 | -4.87 | 0.00 | 0.00014 | D1Pas1 | DNA segment, Chr 1, Pasteur Institute 1 | 0 | 0 | 14 | 0 | 0 | 0 | 7 | 0 | 0 | 0 | 0 |
| 1 | 17.61 | 3.61 | 4.87 | 0.00 | 0.00014 | Gm9959 | | 0 | 5 | 0 | 0 | 0 | 0 | 0 | 0 | 0 | 10 | 0 |
| 1 | -17.69 | 3.63 | -4.87 | 0.00 | 0.00014 | Gm7074 | | 0 | 0 | 10 | 0 | 0 | 0 | 2 | 0 | 0 | 0 | 0 |
| 2 | -17.70 | 3.65 | -4.85 | 0.00 | 0.00015 | Ugt1a1 | UDP glucuronosyltransferase 1 family, pc | 8 | 0 | 5 | 5 | 0 | 0 | 0 | 0 | 0 | 0 | 0 |
| 2 | -17.38 | 3.59 | -4.84 | 0.00 | 0.00016 | Gpr52 | G protein-coupled receptor 52 | 15 | 0 | 0 | 4 | 0 | 3 | 0 | 0 | 0 | 0 | 0 |
| 2 | -17.68 | 3.65 | -4.84 | 0.00 | 0.00016 | Gm2822 | | 4 | 2 | 8 | 4 | 0 | 0 | 0 | 0 | 0 | 0 | 0 |
| 2 | -17.53 | 3.63 | -4.83 | 0.00 | 0.00017 | Nat8f2 | N-acetyltransferase 8 (GCN5-related) fan | 3 | 0 | 5 | 0 | 0 | 0 | 14 | 0 | 0 | 0 | 0 |
| 1 | -17.43 | 3.61 | -4.83 | 0.00 | 0.00017 | Saa2 | serum amyloid A 2 | 0 | 0 | 7 | 0 | 0 | 0 | 8 | 0 | 0 | 0 | 0 |
| 2 | -17.58 | 3.65 | -4.81 | 0.00 | 0.00018 | Zfp36l1-ps | | 8 | 5 | 1 | 0 | 3 | 0 | 0 | 0 | 0 | 0 | 0 |
| 2 | -17.54 | 3.65 | -4.81 | 0.00 | 0.00019 | 1700006J14Rik | | 7 | 15 | 0 | 0 | 5 | 0 | 0 | 0 | 0 | 0 | 0 |
| 1 | -17.33 | 3.61 | -4.81 | 0.00 | 0.00019 | Gm12275 | | 0 | 8 | 0 | 0 | 0 | 1 | 3 | 0 | 0 | 0 | 0 |
| 953 | -1.12 | 0.23 | -4.79 | 0.00 | 0.0002 | Irx2 | Iroquois homeobox 2 | 2949 | 3569 | 3092 | 112 | 163 | 209 | 112 | 35 | 84 | 55 | 106 |
| 1 | -17.18 | 3.60 | -4.78 | 0.00 | 0.00021 | Slco1c1 | solute carrier organic anion transporter fa | 7 | 0 | 0 | 0 | 3 | 3 | 0 | 0 | 0 | 0 | 0 |
| 1 | -17.29 | 3.62 | -4.78 | 0.00 | 0.00021 | Gm15917 | | 0 | 3 | 0 | 0 | 6 | 0 | 0 | 0 | 0 | 0 | 0 |
| 1 | -17.25 | 3.61 | -4.77 | 0.00 | 0.00022 | Cfc1 | cripto, FRL-1, cryptic family 1 | 0 | 0 | 4 | 0 | 0 | 0 | 8 | 0 | 0 | 0 | 0 |
| 1 | -17.26 | 3.62 | -4.77 | 0.00 | 0.00022 | Lrp1b | low density lipoprotein-related protein 1B | 0 | 0 | 3 | 0 | 0 | 0 | 6 | 0 | 0 | 0 | 0 |
| 185 | -17.38 | 3.64 | -4.77 | 0.00 | 0.00022 | Ms4a4d | membrane-spanning 4-domains, subfami | 844 | 563 | 608 | 16 | 0 | 0 | 0 | 0 | 0 | 0 | 0 |
| 3 | -17.38 | 3.65 | -4.76 | 0.00 | 0.00023 | G430095P16Rik | | 5 | 1 | 21 | 4 | 0 | 0 | 0 | 0 | 0 | 0 | 0 |
| 2 | -17.16 | 3.60 | -4.76 | 0.00 | 0.00023 | Gm17017 | | 22 | 0 | 0 | 0 | 3 | 1 | 0 | 0 | 0 | 0 | 0 |
| 286 | -1.32 | 0.28 | -4.76 | 0.00 | 0.00023 | Ppp2r2c | protein phosphatase 2, regulatory subuni | 208 | 248 | 381 | 434 | 297 | 548 | 374 | 151 | 156 | 161 | 187 |
| 1 | -17.18 | 3.62 | -4.75 | 0.00 | 0.00024 | Atp6v1c2 | ATPase, H+ transporting, lysosomal V1 su | 0 | 0 | 6 | 0 | 0 | 1 | 2 | 0 | 0 | 0 | 0 |
| 719 | -1.02 | 0.22 | -4.73 | 0.00 | 0.00026 | Tenm4 | teneurin transmembrane protein 4 | 1604 | 1914 | 1560 | 469 | 401 | 418 | 580 | 331 | 205 | 301 | 127 |
| 1 | -17.09 | 3.61 | -4.73 | 0.00 | 0.00026 | Gm19554 | | 0 | 0 | 8 | 0 | 0 | 0 | 7 | 0 | 0 | 0 | 0 |
| 311 | -5.35 | 1.14 | -4.71 | 0.00 | 0.00029 | Thy1 | thymus cell antigen 1, theta | 1714 | 612 | 1052 | 17 | 5 | 3 | 16 | 0 | 0 | 0 | 0 |
| 1 | -17.19 | 3.65 | -4.71 | 0.00 | 0.00029 | 1500035N22Rik | | 3 | 4 | 0 | 0 | 3 | 0 | 0 | 0 | 0 | 0 | 0 |
| 1 | -17.02 | 3.63 | -4.69 | 0.00 | 0.00032 | 4933412O06Rik | | 0 | 0 | 7 | 0 | 0 | 0 | 2 | 0 | 0 | 0 | 0 |
| 346 | -17.00 | 3.64 | -4.67 | 0.00 | 0.00035 | Inpp5d | inositol polyphosphate-5-phosphatase D | 1818 | 918 | 1045 | 0 | 0 | 0 | 24 | 0 | 0 | 0 | 0 |
| 1 | -16.90 | 3.63 | -4.66 | 0.00 | 0.00036 | Gm13243 | | 0 | 4 | 0 | 0 | 0 | 0 | 3 | 0 | 0 | 0 | 0 |
| 1 | 16.87 | 3.63 | 4.65 | 0.00 | 0.00038 | Lrrc71 | leucine rich repeat containing 71 | 10 | 0 | 0 | 0 | 0 | 0 | 0 | 0 | 1 | 0 | 2 |
| 1 | -16.61 | 3.61 | -4.61 | 0.00 | 0.00046 | Atp13a5 | ATPase type 13A5 | 9 | 0 | 0 | 0 | 2 | 2 | 0 | 0 | 0 | 0 | 0 |
| 1803 | -0.74 | 0.16 | -4.61 | 0.00 | 0.00046 | Bcam | basal cell adhesion molecule | 6376 | 4811 | 5734 | 557 | 481 | 361 | 424 | 315 | 261 | 249 | 259 |
| 2994 | -1.31 | 0.29 | -4.59 | 0.00 | 0.0005 | Plpp3 | phospholipid phosphatase 3 | 11136 | 9676 | 11871 | 46 | 25 | 38 | 68 | 21 | 24 | 18 | 8 |
| 1 | -16.58 | 3.62 | -4.58 | 0.00 | 0.00052 | Dnm3os | | 0 | 5 | 0 | 0 | 0 | 0 | 4 | 0 | 0 | 0 | 0 |
| 2808 | -0.79 | 0.17 | -4.55 | 0.00 | 0.00059 | Serpinh1 | serine (or cysteine) peptidase inhibitor, cl | 11704 | 8175 | 9202 | 270 | 192 | 415 | 247 | 102 | 148 | 126 | 304 |

| | | | | | | | | | | | | | | | | | | |
|---|---|---|---|---|---|---|---|---|---|---|---|---|---|---|---|---|---|---|
| 7517 | -0.68 | 0.15 | -4.55 | 0.00 | 0.0006 | Dbpht2 | DNA binding protein with his-thr domain | 19816 | 23722 | 26186 | 1742 | 1794 | 2315 | 2058 | 1402 | 1423 | 775 | 1454 |
| 380 | -1.30 | 0.29 | -4.55 | 0.00 | 0.0006 | Grem2 | gremlin 2, DAN family BMP antagonist | 1097 | 1093 | 1042 | 153 | 211 | 106 | 206 | 46 | 107 | 55 | 63 |
| 494 | -0.96 | 0.21 | -4.55 | 0.00 | 0.0006 | Mest | mesoderm specific transcript | 1319 | 1715 | 1402 | 211 | 112 | 212 | 130 | 86 | 67 | 85 | 97 |
| 1 | -16.50 | 3.64 | -4.54 | 0.00 | 0.00063 | Kcnh4 | potassium voltage-gated channel, subfam | 1 | 5 | 0 | 0 | 0 | 0 | 4 | 0 | 0 | 0 | 0 |
| 360 | -2.46 | 0.54 | -4.51 | 0.00 | 0.00069 | Pcp4l1 | Purkinje cell protein 4-like 1 | 1162 | 1425 | 924 | 104 | 43 | 135 | 81 | 12 | 21 | 0 | 48 |
| 6856 | -0.78 | 0.17 | -4.51 | 0.00 | 0.0007 | Lrp1 | low density lipoprotein receptor-related pr | 31699 | 19833 | 20136 | 496 | 480 | 661 | 752 | 366 | 368 | 256 | 367 |
| 478 | -5.07 | 1.13 | -4.49 | 0.00 | 0.00076 | Vtn | vitronectin | 1867 | 1336 | 2016 | 16 | 5 | 1 | 16 | 0 | 0 | 0 | 0 |
| 484 | -1.77 | 0.39 | -4.49 | 0.00 | 0.00077 | Tfpi | tissue factor pathway inhibitor | 1809 | 1693 | 1590 | 34 | 77 | 37 | 30 | 21 | 17 | 9 | 6 |
| 86 | -3.87 | 0.87 | -4.47 | 0.00 | 0.00082 | Mob3b | MOB kinase activator 3B | 306 | 167 | 282 | 65 | 38 | 28 | 44 | 0 | 9 | 0 | 5 |
| 197 | -4.99 | 1.12 | -4.44 | 0.00 | 0.00094 | Fcgr2b | Fc receptor, IgG, low affinity IIb | 1066 | 559 | 509 | 4 | 4 | 4 | 19 | 0 | 0 | 0 | 0 |
| 329 | -16.21 | 3.65 | -4.45 | 0.00 | 0.00094 | Igkv15-103 | | 3613 | 0 | 0 | 0 | 0 | 0 | 3 | 0 | 0 | 0 | 0 |
| 35 | -4.90 | 1.11 | -4.43 | 0.00 | 0.00102 | Opcml | opioid binding protein/cell adhesion mole | 118 | 53 | 98 | 12 | 26 | 41 | 25 | 0 | 0 | 0 | 7 |
| 1 | -15.94 | 3.62 | -4.40 | 0.00 | 0.00114 | Gm48591 | | 0 | 0 | 5 | 0 | 0 | 0 | 4 | 0 | 0 | 0 | 0 |
| 1431 | -0.52 | 0.12 | -4.37 | 0.00 | 0.00133 | Enpep | glutamyl aminopeptidase | 1665 | 1773 | 2038 | 1764 | 1820 | 1208 | 1235 | 1276 | 1291 | 944 | 726 |
| 618 | -1.92 | 0.44 | -4.36 | 0.00 | 0.00135 | Hhex | hematopoietically expressed homeobox | 1512 | 1911 | 1786 | 344 | 228 | 325 | 314 | 79 | 127 | 13 | 159 |
| 245 | -4.80 | 1.11 | -4.34 | 0.00 | 0.00152 | Gm8995 | | 1177 | 745 | 748 | 5 | 7 | 4 | 7 | 0 | 0 | 0 | 0 |
| 271 | -2.57 | 0.60 | -4.30 | 0.00 | 0.00176 | Bmp6 | bone morphogenetic protein 6 | 1096 | 896 | 742 | 49 | 45 | 40 | 74 | 28 | 7 | 0 | 6 |
| 290 | -1.27 | 0.30 | -4.26 | 0.00 | 0.00214 | Neurog3 | neurogenin 3 | 625 | 819 | 762 | 154 | 219 | 134 | 174 | 96 | 66 | 100 | 35 |
| 769 | -0.99 | 0.23 | -4.23 | 0.00 | 0.00241 | Ambp | alpha 1 microglobulin/bikunin precursor | 1127 | 1961 | 2243 | 527 | 450 | 661 | 418 | 159 | 298 | 214 | 399 |
| 2351 | -0.48 | 0.11 | -4.22 | 0.00 | 0.00253 | Man1c1 | mannosidase, alpha, class 1C, member 1 | 5407 | 6134 | 5638 | 1293 | 1132 | 1047 | 1588 | 992 | 846 | 1001 | 781 |
| 171 | -4.66 | 1.11 | -4.19 | 0.00 | 0.0028 | Masp1 | mannan-binding lectin serine peptidase 1 | 998 | 413 | 372 | 38 | 16 | 17 | 18 | 0 | 0 | 0 | 10 |
| 657 | -0.79 | 0.19 | -4.19 | 0.00 | 0.00286 | Slc2a13 | solute carrier family 2 (facilitated glucose | 315 | 224 | 379 | 1123 | 1161 | 712 | 957 | 597 | 878 | 517 | 370 |
| 207 | -2.50 | 0.60 | -4.18 | 0.00 | 0.00297 | Pik3cd | phosphatidylinositol-4,5-bisphosphate 3-k | 1042 | 528 | 504 | 46 | 52 | 25 | 49 | 0 | 5 | 21 | 11 |
| 5937 | -0.53 | 0.13 | -4.13 | 0.00 | 0.00367 | Fosb | FBJ osteosarcoma oncogene B | 17298 | 8863 | 8108 | 568 | 645 | 15878 | 1260 | 346 | 577 | 735 | 11031 |
| 13837 | -0.51 | 0.12 | -4.12 | 0.00 | 0.00389 | Jun | jun proto-oncogene | 35519 | 21754 | 27360 | 2096 | 2416 | 28866 | 4145 | 1058 | 1741 | 3170 | 24084 |
| 376 | -1.17 | 0.28 | -4.11 | 0.00 | 0.00403 | Gucy1a1 | guanylate cyclase 1, soluble, alpha 1 | 1726 | 970 | 1024 | 67 | 58 | 96 | 68 | 17 | 26 | 50 | 40 |
| 1694 | 0.50 | 0.12 | 4.10 | 0.00 | 0.00411 | Gucy2c | guanylate cyclase 2c | 234 | 379 | 482 | 1889 | 2020 | 2093 | 1259 | 2982 | 2498 | 1811 | 2989 |
| 27 | -5.48 | 1.34 | -4.09 | 0.00 | 0.00421 | Fam166c | family with sequence similarity 166, meml | 52 | 119 | 89 | 20 | 5 | 8 | 8 | 0 | 0 | 0 | 0 |
| 2 | -14.88 | 3.64 | -4.09 | 0.00 | 0.00436 | Tnnt2 | troponin T2, cardiac | 10 | 0 | 0 | 0 | 0 | 8 | 0 | 0 | 0 | 0 | 0 |
| 422 | -0.85 | 0.21 | -4.08 | 0.00 | 0.00437 | Cobl | cordon-bleu WH2 repeat | 1102 | 1265 | 1012 | 266 | 150 | 210 | 187 | 111 | 111 | 95 | 126 |
| 982 | -0.79 | 0.19 | -4.08 | 0.00 | 0.00439 | Kcnk3 | potassium channel, subfamily K, member | 2893 | 3793 | 2781 | 246 | 232 | 207 | 164 | 114 | 125 | 147 | 103 |
| 173 | -3.88 | 0.95 | -4.08 | 0.00 | 0.00441 | Scin | scinderin | 689 | 584 | 561 | 8 | 31 | 10 | 14 | 0 | 3 | 0 | 1 |
| 4126 | 0.28 | 0.07 | 4.07 | 0.00 | 0.00455 | Lrpprc | leucine-rich PPR-motif containing | 3468 | 3725 | 3516 | 4093 | 3893 | 3394 | 4257 | 5478 | 4602 | 4859 | 4105 |
| 1 | 14.86 | 3.65 | 4.07 | 0.00 | 0.00455 | Gm48998 | | 0 | 9 | 0 | 0 | 0 | 0 | 0 | 0 | 0 | 0 | 2 |
| 1461 | -1.28 | 0.32 | -4.07 | 0.00 | 0.00459 | Prss23 | protease, serine 23 | 5893 | 3101 | 4210 | 677 | 602 | 417 | 322 | 292 | 301 | 103 | 151 |
| 1298 | -14.79 | 3.64 | -4.06 | 0.00 | 0.0047 | Emcn | endomucin | 4977 | 4388 | 4891 | 0 | 0 | 0 | 25 | 0 | 0 | 0 | 0 |
| 335 | -1.79 | 0.44 | -4.06 | 0.00 | 0.00478 | Fbp1 | fructose bisphosphatase 1 | 707 | 1368 | 1104 | 187 | 83 | 56 | 71 | 12 | 35 | 40 | 18 |
| 1 | 14.78 | 3.65 | 4.05 | 0.00 | 0.00483 | Bmp8a | bone morphogenetic protein 8a | 9 | 0 | 0 | 0 | 0 | 0 | 0 | 0 | 0 | 0 | 3 |
| 1 | 14.78 | 3.65 | 4.05 | 0.00 | 0.00483 | Mir301 | | 9 | 0 | 0 | 0 | 0 | 0 | 0 | 0 | 0 | 0 | 3 |
| 325 | -1.13 | 0.28 | -4.03 | 0.00 | 0.00523 | Ffar4 | free fatty acid receptor 4 | 847 | 974 | 941 | 165 | 96 | 206 | 94 | 44 | 72 | 48 | 89 |
| 396 | -3.62 | 0.90 | -4.03 | 0.00 | 0.00523 | Spry1 | sprouty RTK signaling antagonist 1 | 1657 | 1164 | 1478 | 10 | 15 | 27 | 3 | 0 | 0 | 0 | 6 |
| 41 | -5.35 | 1.33 | -4.03 | 0.00 | 0.00523 | Galnt3 | polypeptide N-acetylgalactosaminyltransf | 115 | 115 | 180 | 4 | 26 | 14 | 2 | 0 | 0 | 0 | 0 |
| 133 | 3.09 | 0.77 | 4.03 | 0.00 | 0.00533 | Gbp5 | guanylate binding protein 5 | 670 | 312 | 368 | 0 | 5 | 8 | 1 | 37 | 11 | 31 | 22 |
| 27 | -5.35 | 1.33 | -4.02 | 0.00 | 0.00554 | Frat1 | frequently rearranged in advanced T cell | 127 | 38 | 90 | 4 | 17 | 6 | 10 | 0 | 0 | 0 | 0 |
| 1 | 14.65 | 3.65 | 4.01 | 0.00 | 0.00563 | Jakmip3 | janus kinase and microtubule interacting | 8 | 0 | 0 | 0 | 0 | 0 | 0 | 0 | 0 | 0 | 2 |
| 1 | -14.61 | 3.64 | -4.01 | 0.00 | 0.00569 | Gm25363 | | 9 | 0 | 0 | 0 | 0 | 4 | 0 | 0 | 0 | 0 | 0 |
| 553 | -6.05 | 1.51 | -4.00 | 0.00 | 0.00588 | Nrp2 | neuropilin 2 | 2693 | 1559 | 1639 | 18 | 65 | 36 | 45 | 0 | 0 | 0 | 24 |
| 68 | -3.45 | 0.86 | -4.00 | 0.00 | 0.00588 | Sftpd | surfactant associated protein D | 132 | 198 | 273 | 30 | 50 | 19 | 33 | 0 | 9 | 0 | 6 |
| 372 | -3.55 | 0.89 | -3.99 | 0.00 | 0.00599 | Id3 | inhibitor of DNA binding 3 | 1721 | 1078 | 1247 | 6 | 6 | 22 | 9 | 0 | 0 | 0 | 4 |
| 1 | -14.55 | 3.65 | -3.99 | 0.00 | 0.00605 | 2900022M07Rik | | 9 | 0 | 0 | 0 | 0 | 3 | 0 | 0 | 0 | 0 | 0 |
| 26 | -4.62 | 1.16 | -3.98 | 0.00 | 0.00643 | C78197 | | 100 | 20 | 30 | 32 | 22 | 41 | 27 | 0 | 1 | 0 | 10 |
| 1 | 14.51 | 3.65 | 3.97 | 0.00 | 0.00643 | Gm39384 | | 9 | 0 | 0 | 0 | 0 | 0 | 0 | 0 | 0 | 0 | 2 |
| 1 | 14.51 | 3.65 | 3.97 | 0.00 | 0.00643 | Serpina3e-ps | | 9 | 0 | 0 | 0 | 0 | 0 | 0 | 0 | 0 | 0 | 2 |
| 137 | -4.76 | 1.20 | -3.97 | 0.00 | 0.00658 | Ifi27l2a | interferon, alpha-inducible protein 27 like | 668 | 318 | 489 | 12 | 6 | 0 | 18 | 0 | 0 | 0 | 0 |
| 4030 | -0.59 | 0.15 | -3.96 | 0.00 | 0.00665 | Ocrl | OCRL, inositol polyphosphate-5-phospha | 6663 | 8639 | 7475 | 2577 | 3186 | 4472 | 2610 | 1532 | 2726 | 1456 | 2997 |
| 3724 | -0.62 | 0.16 | -3.96 | 0.00 | 0.00667 | Zfp36 | zinc finger protein 36 | 18250 | 8313 | 8689 | 441 | 484 | 1874 | 654 | 251 | 386 | 376 | 1243 |
| 22 | -5.59 | 1.41 | -3.96 | 0.00 | 0.00672 | Anks1b | ankyrin repeat and sterile alpha motif dom | 72 | 101 | 28 | 7 | 10 | 15 | 7 | 0 | 0 | 0 | 0 |
| 281 | -2.21 | 0.57 | -3.91 | 0.00 | 0.00812 | Clmp | CXADR-like membrane protein | 1036 | 940 | 820 | 34 | 63 | 59 | 74 | 0 | 17 | 10 | 36 |
| 4520 | 0.48 | 0.12 | 3.89 | 0.00 | 0.00891 | Cntn1 | contactin 1 | 481 | 636 | 757 | 6364 | 4833 | 3324 | 5526 | 8449 | 7569 | 6815 | 4967 |
| 103 | -4.25 | 1.10 | -3.88 | 0.00 | 0.00939 | Cfi | complement component factor i | 248 | 514 | 293 | 22 | 9 | 20 | 20 | 0 | 0 | 0 | 9 |

| | | | | | | | | | | | | | | | | | |
|---|---|---|---|---|---|---|---|---|---|---|---|---|---|---|---|---|---|
| 50 | -4.93 | 1.27 | -3.87 | 0.00 | 0.00966 | Fabp7 | fatty acid binding protein 7, brain | 220 | 84 | 213 | 5 | 23 | 1 | 9 | 0 | 0 | 0 | 0 |
| 1 | -14.14 | 3.66 | -3.86 | 0.00 | 0.00989 | Cabp4 | calcium binding protein 4 | 8 | 0 | 0 | 0 | 0 | 1 | 0 | 0 | 0 | 0 | 0 |
| 3221 | -0.32 | 0.08 | -3.86 | 0.00 | 0.00998 | Prom1 | prominin 1 | 1717 | 2348 | 2004 | 4615 | 3421 | 4688 | 3577 | 3872 | 2931 | 2840 | 3419 |
| 1378 | -0.73 | 0.19 | -3.85 | 0.00 | 0.01016 | Ptprg | protein tyrosine phosphatase, receptor typ | 4600 | 4701 | 4358 | 265 | 163 | 296 | 216 | 107 | 120 | 183 | 153 |
| 142 | -4.46 | 1.16 | -3.85 | 0.00 | 0.01033 | Pdlim1 | PDZ and LIM domain 1 (elfin) | 549 | 405 | 558 | 16 | 5 | 8 | 22 | 0 | 2 | 0 | 0 |
| 525 | -0.72 | 0.19 | -3.83 | 0.00 | 0.0114 | Ston2 | stonin 2 | 1011 | 1283 | 1055 | 330 | 341 | 506 | 322 | 229 | 148 | 215 | 339 |
| 64890 | 0.27 | 0.07 | 3.82 | 0.00 | 0.01159 | Ccnd2 | cyclin D2 | 30966 | 35458 | 38091 | 70960 | 73039 | 58062 | 73836 | 98771 | 87016 | 79042 | 68546 |
| 1720 | -0.49 | 0.13 | -3.82 | 0.00 | 0.01169 | Gnai1 | guanine nucleotide binding protein (G pro | 3572 | 4543 | 3977 | 995 | 972 | 923 | 1077 | 797 | 809 | 721 | 534 |
| 58 | -4.61 | 1.21 | -3.81 | 0.00 | 0.01206 | Anxa13 | annexin A13 | 227 | 95 | 274 | 4 | 13 | 7 | 13 | 0 | 0 | 0 | 1 |
| 591 | -1.20 | 0.32 | -3.81 | 0.00 | 0.01212 | Myrf | myelin regulatory factor | 1904 | 1917 | 1634 | 210 | 201 | 148 | 154 | 72 | 155 | 39 | 63 |
| 510 | -0.82 | 0.21 | -3.80 | 0.00 | 0.01243 | Unc5c | unc-5 netrin receptor C | 813 | 1171 | 822 | 427 | 298 | 637 | 418 | 368 | 155 | 202 | 302 |
| 2201 | -0.44 | 0.12 | -3.80 | 0.00 | 0.01252 | Dapk1 | death associated protein kinase 1 | 3495 | 3503 | 3105 | 2206 | 1699 | 2085 | 2132 | 1298 | 1536 | 1504 | 1645 |
| 139 | -4.59 | 1.21 | -3.79 | 0.00 | 0.01298 | Cmah | cytidine monophospho-N-acetylneuramin | 848 | 283 | 375 | 8 | 9 | 2 | 3 | 0 | 0 | 0 | 0 |
| 295 | -1.54 | 0.41 | -3.78 | 0.00 | 0.01308 | Gap43 | growth associated protein 43 | 683 | 939 | 789 | 183 | 157 | 128 | 149 | 19 | 54 | 91 | 56 |
| 118 | -1.43 | 0.38 | -3.78 | 0.00 | 0.01329 | Bik | BCL2-interacting killer | 204 | 177 | 220 | 188 | 89 | 90 | 141 | 63 | 49 | 36 | 38 |
| 576 | -0.50 | 0.13 | -3.77 | 0.00 | 0.01356 | 2010300C02Ril | RIKEN cDNA 2010300C02 gene | 653 | 753 | 647 | 663 | 634 | 570 | 642 | 496 | 433 | 459 | 388 |
| 1582 | -0.53 | 0.14 | -3.74 | 0.00 | 0.01567 | Cadm1 | cell adhesion molecule 1 | 3860 | 4283 | 4454 | 708 | 655 | 606 | 878 | 448 | 573 | 491 | 451 |
| 688 | -0.84 | 0.23 | -3.74 | 0.00 | 0.0157 | Crip2 | cysteine rich protein 2 | 2572 | 1964 | 2476 | 101 | 126 | 54 | 79 | 56 | 53 | 62 | 30 |
| 152 | -1.37 | 0.37 | -3.73 | 0.00 | 0.01596 | Als2cl | ALS2 C-terminal like | 434 | 499 | 416 | 52 | 34 | 114 | 39 | 23 | 19 | 17 | 25 |
| 239 | -1.70 | 0.46 | -3.73 | 0.00 | 0.01601 | Kbtbd11 | kelch repeat and BTB (POZ) domain conta | 756 | 744 | 648 | 73 | 79 | 73 | 129 | 8 | 35 | 71 | 13 |
| 596 | -0.77 | 0.21 | -3.73 | 0.00 | 0.01609 | Dock6 | dedicator of cytokinesis 6 | 1682 | 1622 | 1602 | 155 | 187 | 554 | 145 | 87 | 137 | 71 | 312 |
| 200 | -4.44 | 1.19 | -3.73 | 0.00 | 0.01609 | Cd72 | CD72 antigen | 746 | 368 | 1063 | 7 | 5 | 5 | 1 | 0 | 0 | 0 | 0 |
| 32 | -4.69 | 1.26 | -3.72 | 0.00 | 0.01636 | 1700016P03Rik | | 56 | 21 | 54 | 17 | 21 | 95 | 18 | 0 | 0 | 0 | 67 |
| 238 | -2.77 | 0.74 | -3.72 | 0.00 | 0.01659 | Adamts18 | a disintegrin-like and metallopeptidase (r | 799 | 942 | 650 | 11 | 26 | 77 | 71 | 0 | 7 | 1 | 36 |
| 20 | -5.40 | 1.45 | -3.72 | 0.00 | 0.01659 | Myo16 | myosin XVI | 30 | 80 | 76 | 7 | 7 | 6 | 16 | 0 | 0 | 0 | 0 |
| 1913 | -0.37 | 0.10 | -3.70 | 0.00 | 0.01791 | Celsr3 | cadherin, EGF LAG seven-pass G-type re | 1935 | 1725 | 2010 | 1332 | 1542 | 4214 | 1629 | 1036 | 1403 | 1096 | 3126 |
| 2296 | 0.65 | 0.18 | 3.69 | 0.00 | 0.01859 | Nnat | neuronatin | 511 | 1142 | 673 | 1694 | 1662 | 3740 | 1819 | 3075 | 2424 | 2822 | 5695 |
| 302 | -1.36 | 0.37 | -3.67 | 0.00 | 0.01966 | Gpr153 | G protein-coupled receptor 153 | 1180 | 843 | 1049 | 39 | 30 | 41 | 71 | 16 | 13 | 37 | 8 |
| 186 | -1.21 | 0.33 | -3.66 | 0.00 | 0.02023 | Rtkn | rhotekin | 478 | 350 | 455 | 156 | 113 | 127 | 127 | 37 | 54 | 57 | 87 |
| 2520 | -5.73 | 1.56 | -3.66 | 0.00 | 0.02026 | Lyz2 | lysozyme 2 | 16193 | 6142 | 5226 | 8 | 36 | 18 | 83 | 0 | 0 | 0 | 18 |
| 295 | -2.60 | 0.71 | -3.66 | 0.00 | 0.02029 | Sned1 | sushi, nidogen and EGF-like domains 1 | 1370 | 931 | 764 | 10 | 20 | 105 | 16 | 0 | 7 | 0 | 21 |
| 297 | 1.45 | 0.39 | 3.66 | 0.00 | 0.02029 | Rai14 | retinoic acid induced 14 | 1085 | 941 | 976 | 5 | 18 | 41 | 14 | 43 | 50 | 39 | 55 |
| 2871 | 0.24 | 0.07 | 3.66 | 0.00 | 0.02051 | Tacc2 | transforming, acidic coiled-coil containing | 1339 | 2028 | 1742 | 3873 | 3306 | 2454 | 2464 | 4672 | 4032 | 2865 | 2809 |
| 985 | -1.19 | 0.32 | -3.65 | 0.00 | 0.02086 | Fmo2 | flavin containing monooxygenase 2 | 4790 | 2704 | 2767 | 94 | 146 | 68 | 84 | 34 | 94 | 36 | 21 |
| 31 | -3.49 | 0.96 | -3.62 | 0.00 | 0.02373 | Cfap44 | cilia and flagella associated protein 44 | 36 | 48 | 36 | 34 | 12 | 124 | 16 | 0 | 3 | 0 | 29 |
| 37 | -5.03 | 1.39 | -3.62 | 0.00 | 0.02381 | Gm38345 | | 126 | 81 | 90 | 22 | 11 | 9 | 40 | 0 | 0 | 0 | 21 |
| 68 | -4.78 | 1.33 | -3.61 | 0.00 | 0.02436 | Tbxas1 | thromboxane A synthase 1, platelet | 365 | 214 | 134 | 18 | 0 | 5 | 8 | 0 | 0 | 0 | 0 |
| 362 | -1.00 | 0.28 | -3.61 | 0.00 | 0.02461 | Nxph1 | neurexophilin 1 | 933 | 1311 | 746 | 204 | 131 | 127 | 206 | 87 | 105 | 64 | 70 |
| 26 | -4.16 | 1.16 | -3.59 | 0.00 | 0.02554 | Pcdhb6 | protocadherin beta 6 | 48 | 42 | 47 | 39 | 35 | 14 | 46 | 0 | 3 | 0 | 12 |
| 172 | -2.06 | 0.57 | -3.59 | 0.00 | 0.02626 | Proser2 | proline and serine rich 2 | 485 | 568 | 518 | 108 | 60 | 38 | 44 | 37 | 18 | 0 | 12 |
| 196 | -1.35 | 0.38 | -3.58 | 0.00 | 0.0267 | Slc2a3 | solute carrier family 2 (facilitated glucose | 579 | 754 | 633 | 52 | 32 | 25 | 27 | 16 | 11 | 9 | 16 |
| 1287 | -0.43 | 0.12 | -3.58 | 0.00 | 0.02721 | Klhdc8b | kelch domain containing 8B | 2294 | 3383 | 2914 | 1026 | 663 | 824 | 688 | 731 | 503 | 513 | 616 |
| 116 | -3.22 | 0.90 | -3.57 | 0.00 | 0.02769 | Tifab | TRAF-interacting protein with forkhead-as | 595 | 301 | 317 | 18 | 11 | 9 | 19 | 0 | 0 | 3 | 5 |
| 13867 | 0.25 | 0.07 | 3.57 | 0.00 | 0.02788 | Atp6ap2 | ATPase, H+ transporting, lysosomal acce | 6838 | 6828 | 7030 | 16499 | 15534 | 11984 | 16072 | 21834 | 16814 | 18410 | 14698 |
| 344 | -4.45 | 1.25 | -3.56 | 0.00 | 0.02916 | Cd83 | CD83 antigen | 1995 | 822 | 786 | 25 | 45 | 30 | 51 | 0 | 2 | 0 | 31 |
| 328 | -3.03 | 0.85 | -3.55 | 0.00 | 0.02992 | Podn | podocan | 2160 | 658 | 736 | 15 | 6 | 12 | 14 | 0 | 0 | 1 | 6 |
| 360 | -1.45 | 0.41 | -3.53 | 0.00 | 0.03169 | F5 | coagulation factor V | 789 | 1171 | 977 | 243 | 199 | 207 | 94 | 21 | 97 | 58 | 100 |
| 28 | -4.94 | 1.40 | -3.53 | 0.00 | 0.03185 | Oas3 | 2'-5' oligoadenylate synthetase 3 | 72 | 69 | 109 | 39 | 8 | 1 | 13 | 0 | 0 | 0 | 1 |
| 128 | -4.51 | 1.28 | -3.53 | 0.00 | 0.03189 | Epha3 | Eph receptor A3 | 866 | 271 | 243 | 15 | 0 | 2 | 11 | 0 | 0 | 0 | 0 |
| 254 | -0.77 | 0.22 | -3.52 | 0.00 | 0.0325 | Wfdc16 | WAP four-disulfide core domain 16 | 488 | 636 | 699 | 179 | 195 | 98 | 137 | 117 | 112 | 72 | 58 |
| 3328 | -0.37 | 0.11 | -3.52 | 0.00 | 0.03302 | Jup | junction plakoglobin | 7311 | 7071 | 6572 | 2538 | 2453 | 1905 | 1918 | 1934 | 1918 | 1659 | 1332 |
| 160 | 1.58 | 0.45 | 3.51 | 0.00 | 0.03393 | Mfng | MFNG O-fucosylpeptide 3-beta-N-acetylg | 507 | 325 | 351 | 14 | 33 | 83 | 28 | 126 | 84 | 57 | 146 |
| 923 | 0.45 | 0.13 | 3.51 | 0.00 | 0.03402 | Adgrl3 | adhesion G protein-coupled receptor L3 | 147 | 113 | 148 | 899 | 911 | 1207 | 1115 | 1331 | 1236 | 1511 | 1534 |
| 1063 | -0.64 | 0.18 | -3.51 | 0.00 | 0.03403 | Wwc1 | WW, C2 and coiled-coil domain containin | 2621 | 2496 | 2814 | 710 | 642 | 212 | 648 | 509 | 566 | 386 | 87 |
| 1968 | -0.56 | 0.16 | -3.50 | 0.00 | 0.03438 | Dpp6 | dipeptidylpeptidase 6 | 3543 | 5665 | 5162 | 889 | 1275 | 1006 | 1157 | 818 | 818 | 631 | 681 |
| 67 | -3.06 | 0.87 | -3.50 | 0.00 | 0.03438 | Tox2 | TOX high mobility group box family memb | 148 | 211 | 242 | 41 | 24 | 36 | 12 | 0 | 6 | 0 | 14 |
| 81 | -4.60 | 1.31 | -3.50 | 0.00 | 0.03438 | Ier5l | immediate early response 5-like | 377 | 223 | 234 | 31 | 16 | 5 | 3 | 0 | 0 | 0 | 2 |
| 857 | -0.60 | 0.17 | -3.50 | 0.00 | 0.03483 | Pon2 | paraoxonase 2 | 1458 | 1252 | 1278 | 724 | 795 | 565 | 1154 | 407 | 597 | 867 | 329 |
| 864 | -0.75 | 0.22 | -3.50 | 0.00 | 0.03493 | s | single-minded family bHLH transcription f | 1531 | 2010 | 1781 | 664 | 442 | 880 | 637 | 497 | 402 | 238 | 425 |
| 608 | -4.22 | 1.21 | -3.50 | 0.00 | 0.03493 | Plbd1 | phospholipase B domain containing 1 | 3514 | 1586 | 1539 | 4 | 17 | 0 | 29 | 0 | 1 | 0 | 0 |

| | | | | | | | | | | | | | | | | | | |
|---|---|---|---|---|---|---|---|---|---|---|---|---|---|---|---|---|---|---|
| 161 | -1.67 | 0.48 | -3.50 | 0.00 | 0.03502 | Cntnap5b | contactin associated protein-like 5B | 409 | 800 | 247 | 39 | 62 | 46 | 83 | 29 | 12 | 30 | 10 |
| 225 | -1.46 | 0.42 | -3.49 | 0.00 | 0.03518 | Asic2 | acid-sensing (proton-gated) ion channel 2 | 402 | 638 | 667 | 179 | 82 | 139 | 151 | 27 | 48 | 43 | 101 |
| 763 | -0.51 | 0.15 | -3.49 | 0.00 | 0.03562 | Plaat3 | phospholipase A and acyltransferase 3 | 1429 | 1504 | 1598 | 656 | 598 | 571 | 443 | 397 | 458 | 349 | 388 |
| 63 | -3.64 | 1.04 | -3.49 | 0.00 | 0.03562 | Oit1 | oncoprotein induced transcript 1 | 86 | 233 | 282 | 12 | 6 | 30 | 35 | 0 | 3 | 0 | 5 |
| 4324 | -0.71 | 0.20 | -3.49 | 0.00 | 0.03569 | Serping1 | serine (or cysteine) peptidase inhibitor, cl | 20978 | 10380 | 10484 | 988 | 944 | 844 | 766 | 545 | 613 | 504 | 516 |
| 193 | -1.42 | 0.41 | -3.48 | 0.00 | 0.03699 | Aldh3b1 | aldehyde dehydrogenase 3 family, memb | 571 | 415 | 350 | 216 | 92 | 148 | 118 | 39 | 78 | 48 | 46 |
| 137 | 3.89 | 1.12 | 3.48 | 0.00 | 0.03699 | Tmem88 | transmembrane protein 88 | 456 | 405 | 600 | 0 | 1 | 2 | 0 | 19 | 1 | 15 | 7 |
| 8 | -6.04 | 1.74 | -3.46 | 0.00 | 0.03859 | 4933427E11Rik | | 5 | 13 | 12 | 7 | 11 | 15 | 21 | 0 | 0 | 0 | 0 |
| 155 | -1.38 | 0.40 | -3.45 | 0.00 | 0.04084 | Tmie | transmembrane inner ear | 408 | 519 | 435 | 59 | 49 | 53 | 82 | 43 | 13 | 30 | 14 |
| 37 | -5.23 | 1.52 | -3.45 | 0.00 | 0.04087 | Grik2 | glutamate receptor, ionotropic, kainate 2 ( | 116 | 146 | 107 | 0 | 12 | 10 | 13 | 0 | 0 | 0 | 0 |
| 55 | -4.29 | 1.25 | -3.43 | 0.00 | 0.04261 | Ccl22 | chemokine (C-C motif) ligand 22 | 263 | 215 | 91 | 15 | 4 | 3 | 8 | 0 | 0 | 0 | 1 |
| 1234 | -0.59 | 0.17 | -3.43 | 0.00 | 0.04353 | Fgfr1 | fibroblast growth factor receptor 1 | 4919 | 2759 | 3399 | 417 | 315 | 340 | 418 | 271 | 212 | 319 | 200 |
| 321 | -3.35 | 0.98 | -3.43 | 0.00 | 0.04353 | Sh3pxd2b | SH3 and PX domains 2B | 1509 | 895 | 1070 | 22 | 3 | 6 | 16 | 1 | 0 | 0 | 3 |
| 1547 | -0.63 | 0.18 | -3.42 | 0.00 | 0.04449 | Nfix | nuclear factor I/X | 3143 | 1780 | 1979 | 1702 | 1552 | 1220 | 1611 | 1075 | 1259 | 1077 | 618 |
| 2006 | -0.46 | 0.13 | -3.41 | 0.00 | 0.04597 | Calb1 | calbindin 1 | 5584 | 5905 | 5704 | 768 | 655 | 538 | 865 | 641 | 558 | 450 | 403 |
| 400 | -1.85 | 0.54 | -3.41 | 0.00 | 0.04598 | Thsd7a | thrombospondin, type I, domain containin | 1887 | 1113 | 1223 | 28 | 7 | 47 | 52 | 0 | 5 | 22 | 13 |
| 233 | -1.43 | 0.42 | -3.41 | 0.00 | 0.04643 | Abcc3 | ATP-binding cassette, sub-family C (CFTF | 857 | 565 | 934 | 41 | 45 | 49 | 16 | 13 | 17 | 8 | 18 |
| 2248 | -0.58 | 0.17 | -3.39 | 0.00 | 0.04941 | Wnk3 | WNK lysine deficient protein kinase 3 | 5329 | 7846 | 6402 | 742 | 628 | 878 | 837 | 416 | 455 | 631 | 567 |
| 171 | -1.33 | 0.39 | -3.39 | 0.00 | 0.04941 | Kcnq1 | potassium voltage-gated channel, subfam | 518 | 464 | 507 | 64 | 55 | 96 | 61 | 15 | 48 | 19 | 30 |
| 2615 | -0.53 | 0.16 | -3.39 | 0.00 | 0.04977 | Smarca1 | SWI/SNF related, matrix associated, actin | 5948 | 8936 | 7035 | 928 | 985 | 1062 | 1074 | 579 | 801 | 741 | 680 |
| 262 | 0.94 | 0.28 | 3.38 | 0.00 | 0.05076 | Npy | neuropeptide Y | 45 | 75 | 44 | 244 | 268 | 196 | 184 | 718 | 506 | 249 | 348 |
| 386 | -0.98 | 0.29 | -3.37 | 0.00 | 0.05215 | Sema3g | sema domain, immunoglobulin domain (Ig | 1223 | 1081 | 1482 | 88 | 60 | 41 | 113 | 36 | 51 | 45 | 21 |
| 11849 | 0.16 | 0.05 | 3.37 | 0.00 | 0.05286 | Rnf130 | ring finger protein 130 | 11854 | 13211 | 11738 | 11770 | 11118 | 9599 | 11716 | 13121 | 12472 | 12881 | 10863 |
| 69 | -4.30 | 1.28 | -3.36 | 0.00 | 0.05407 | Impa2 | inositol (myo)-1(or 4)-monophosphatase 2 | 384 | 199 | 153 | 5 | 10 | 0 | 8 | 0 | 0 | 0 | 0 |
| 1231 | 0.37 | 0.11 | 3.34 | 0.00 | 0.05671 | Mis12 | MIS12 kinetochore complex component | 997 | 1030 | 1018 | 1197 | 1086 | 1276 | 1018 | 1607 | 1277 | 1492 | 1547 |
| 1087 | -0.49 | 0.15 | -3.35 | 0.00 | 0.05671 | Fam149a | family with sequence similarity 149, meml | 1789 | 1801 | 1853 | 1009 | 997 | 800 | 969 | 857 | 642 | 802 | 442 |
| 297 | -0.85 | 0.25 | -3.34 | 0.00 | 0.05671 | Spock3 | sparc/osteonectin, cwcv and kazal-like do | 800 | 1010 | 745 | 110 | 111 | 132 | 103 | 50 | 51 | 73 | 82 |
| 308 | -3.86 | 1.15 | -3.34 | 0.00 | 0.05671 | S100a16 | S100 calcium binding protein A16 | 1125 | 1068 | 1019 | 42 | 28 | 27 | 61 | 3 | 19 | 0 | 0 |
| 65 | -3.41 | 1.02 | -3.34 | 0.00 | 0.05671 | F730043M19Rik | | 232 | 189 | 113 | 5 | 54 | 57 | 29 | 0 | 4 | 0 | 29 |
| 1320 | -0.53 | 0.16 | -3.34 | 0.00 | 0.05797 | Nbl1 | NBL1, DAN family BMP antagonist | 4262 | 2125 | 2489 | 839 | 920 | 455 | 1059 | 649 | 575 | 882 | 262 |
| 456 | -3.13 | 0.94 | -3.33 | 0.00 | 0.05913 | Vcan | versican | 1761 | 1464 | 1243 | 62 | 104 | 128 | 150 | 9 | 11 | 0 | 88 |
| 272 | -1.16 | 0.35 | -3.33 | 0.00 | 0.05913 | Cacng4 | calcium channel, voltage-dependent, gan | 852 | 783 | 1011 | 39 | 58 | 56 | 83 | 10 | 34 | 28 | 35 |
| 204 | -1.19 | 0.36 | -3.33 | 0.00 | 0.05916 | Fxyd3 | FXYD domain-containing ion transport re | 326 | 394 | 478 | 237 | 180 | 196 | 103 | 114 | 104 | 23 | 92 |
| 7459 | 0.18 | 0.05 | 3.33 | 0.00 | 0.05968 | Ddx50 | DEAD (Asp-Glu-Ala-Asp) box polypeptide | 4440 | 4167 | 4737 | 7030 | 8059 | 9378 | 7748 | 7982 | 8564 | 8952 | 10991 |
| 491 | -0.83 | 0.25 | -3.32 | 0.00 | 0.0602 | Tiam1 | T cell lymphoma invasion and metastasis | 1482 | 1067 | 1358 | 304 | 276 | 199 | 181 | 151 | 153 | 97 | 135 |
| 1513 | -0.50 | 0.15 | -3.32 | 0.00 | 0.06091 | Nynrin | NYN domain and retroviral integrase cont | 3479 | 3286 | 3267 | 983 | 910 | 701 | 1265 | 534 | 674 | 1029 | 519 |
| 9909 | -0.29 | 0.09 | -3.31 | 0.00 | 0.06237 | Ucp2 | uncoupling protein 2 (mitochondrial, protc | 14599 | 15332 | 15653 | 8789 | 9727 | 7591 | 8620 | 6161 | 8605 | 8009 | 5907 |
| 146 | -3.78 | 1.15 | -3.28 | 0.00 | 0.06919 | Tlr4 | toll-like receptor 4 | 631 | 428 | 514 | 5 | 23 | 0 | 4 | 0 | 1 | 0 | 0 |
| 267 | -0.88 | 0.27 | -3.27 | 0.00 | 0.07213 | Creg2 | cellular repressor of E1A-stimulated gene | 480 | 658 | 692 | 141 | 207 | 214 | 157 | 76 | 82 | 112 | 118 |
| 1621 | -0.47 | 0.15 | -3.26 | 0.00 | 0.07301 | Arhgap29 | Rho GTPase activating protein 29 | 5705 | 5351 | 4290 | 397 | 299 | 453 | 296 | 320 | 206 | 210 | 305 |
| 332 | -0.70 | 0.21 | -3.26 | 0.00 | 0.07301 | Scn3b | sodium channel, voltage-gated, type III, b | 908 | 917 | 717 | 162 | 164 | 191 | 169 | 129 | 81 | 113 | 105 |
| 6 | 10.70 | 3.28 | 3.26 | 0.00 | 0.07301 | Nox1 | NADPH oxidase 1 | 0 | 31 | 0 | 1 | 0 | 0 | 0 | 0 | 3 | 30 | 0 |
| 133 | -4.03 | 1.24 | -3.26 | 0.00 | 0.07424 | Il16 | interleukin 16 | 807 | 250 | 382 | 0 | 2 | 2 | 24 | 0 | 0 | 0 | 0 |
| 1748 | -0.89 | 0.27 | -3.26 | 0.00 | 0.07446 | Pdk4 | pyruvate dehydrogenase kinase, isoenzy | 4453 | 6657 | 7260 | 107 | 95 | 168 | 179 | 60 | 98 | 43 | 106 |
| 130 | -2.02 | 0.62 | -3.26 | 0.00 | 0.07446 | Hgf | hepatocyte growth factor | 540 | 273 | 386 | 45 | 73 | 10 | 54 | 0 | 23 | 21 | 5 |
| 3105 | -0.86 | 0.26 | -3.26 | 0.00 | 0.07446 | Epas1 | endothelial PAS domain protein 1 | 11762 | 11367 | 10816 | 22 | 20 | 48 | 45 | 14 | 17 | 8 | 35 |
| 969 | -2.08 | 0.64 | -3.25 | 0.00 | 0.07529 | Mef2c | myocyte enhancer factor 2C | 3626 | 2952 | 2816 | 210 | 284 | 220 | 276 | 49 | 101 | 9 | 114 |
| 16 | -4.70 | 1.45 | -3.25 | 0.00 | 0.07529 | Gm36172 | | 67 | 23 | 43 | 13 | 5 | 4 | 16 | 0 | 1 | 0 | 0 |
| 88 | -2.80 | 0.87 | -3.23 | 0.00 | 0.08059 | Ephb3 | Eph receptor B3 | 338 | 163 | 329 | 37 | 18 | 17 | 36 | 0 | 11 | 15 | 0 |
| 3911 | -0.33 | 0.10 | -3.23 | 0.00 | 0.08178 | Fat1 | FAT atypical cadherin 1 | 5667 | 6933 | 5700 | 3510 | 3838 | 3136 | 3278 | 2629 | 3313 | 2584 | 2427 |
| 14 | -4.95 | 1.53 | -3.22 | 0.00 | 0.08182 | Iqcd | IQ motif containing D | 15 | 26 | 25 | 23 | 15 | 16 | 16 | 0 | 0 | 0 | 21 |
| 1385 | -0.45 | 0.14 | -3.22 | 0.00 | 0.08202 | Rgs16 | regulator of G-protein signaling 16 | 728 | 737 | 1009 | 1168 | 2021 | 2302 | 1843 | 898 | 1688 | 1115 | 1724 |
| 78 | -4.00 | 1.24 | -3.22 | 0.00 | 0.08227 | Pde1a | phosphodiesterase 1A, calmodulin-deper | 343 | 225 | 218 | 16 | 12 | 39 | 0 | 0 | 0 | 0 | 9 |
| 57 | -3.97 | 1.24 | -3.21 | 0.00 | 0.08607 | Cecr2 | CECR2, histone acetyl-lysine reader | 196 | 148 | 233 | 0 | 3 | 33 | 18 | 0 | 1 | 0 | 1 |
| 664 | -0.58 | 0.18 | -3.21 | 0.00 | 0.08664 | Cldn3 | claudin 3 | 741 | 996 | 1204 | 828 | 723 | 423 | 656 | 506 | 427 | 486 | 310 |
| 199 | -2.57 | 0.80 | -3.20 | 0.00 | 0.08769 | Amy1 | amylase 1, salivary | 238 | 1083 | 737 | 18 | 12 | 47 | 11 | 3 | 0 | 0 | 35 |
| 3858 | -0.28 | 0.09 | -3.20 | 0.00 | 0.08775 | Sfrp5 | secreted frizzled-related sequence protei | 8295 | 8851 | 9728 | 2126 | 2866 | 1318 | 2241 | 1981 | 2011 | 1953 | 1068 |
| 135 | -2.42 | 0.76 | -3.19 | 0.00 | 0.09036 | Fam13c | family with sequence similarity 13, memb | 398 | 554 | 382 | 20 | 17 | 24 | 57 | 7 | 2 | 0 | 22 |
| 4132 | -0.36 | 0.11 | -3.19 | 0.00 | 0.09129 | Man2a1 | mannosidase 2, alpha 1 | 9803 | 7744 | 7020 | 3622 | 2367 | 2558 | 3217 | 2785 | 2047 | 2267 | 2021 |
| 131 | -1.96 | 0.61 | -3.19 | 0.00 | 0.09129 | Zfhx4 | zinc finger homeodomain 4 | 502 | 427 | 265 | 37 | 54 | 52 | 47 | 8 | 26 | 1 | 17 |

| | | | | | | | | | | | | | | | | | | |
|---|---|---|---|---|---|---|---|---|---|---|---|---|---|---|---|---|---|---|
| 174 | -0.98 | 0.31 | -3.18 | 0.00 | 0.09467 | Gm42372 | predicted gene, 42372 | 314 | 496 | 317 | 159 | 143 | 125 | 99 | 58 | 78 | 71 | 56 |
| 344 | -1.38 | 0.44 | -3.17 | 0.00 | 0.09474 | Ptprz1 | protein tyrosine phosphatase, receptor typ | 691 | 1058 | 974 | 130 | 177 | 228 | 200 | 148 | 26 | 40 | 115 |
| 68 | -1.76 | 0.55 | -3.17 | 0.00 | 0.09474 | Pde8b | phosphodiesterase 8B | 181 | 236 | 136 | 31 | 26 | 10 | 81 | 12 | 7 | 21 | 3 |
| 254 | -1.18 | 0.37 | -3.17 | 0.00 | 0.09587 | Cyth4 | cytohesin 4 | 1276 | 558 | 660 | 62 | 37 | 69 | 39 | 19 | 18 | 31 | 26 |
| 667 | -1.36 | 0.43 | -3.16 | 0.00 | 0.09863 | Irx1 | Iroquois homeobox 1 | 2188 | 2606 | 2251 | 30 | 60 | 61 | 59 | 35 | 23 | 3 | 26 |
| 4233 | -0.26 | 0.08 | -3.15 | 0.00 | 0.10146 | Tbc1d16 | TBC1 domain family, member 16 | 3731 | 3581 | 3485 | 4996 | 4764 | 4116 | 5579 | 3918 | 4667 | 4607 | 3117 |
| 368 | -1.82 | 0.58 | -3.15 | 0.00 | 0.103 | Lrrk1 | leucine-rich repeat kinase 1 | 1724 | 985 | 1048 | 81 | 60 | 22 | 46 | 1 | 13 | 54 | 9 |
| 3045 | 0.21 | 0.07 | 3.15 | 0.00 | 0.10311 | Tomm70a | translocase of outer mitochondrial membr | 2833 | 2497 | 2911 | 2675 | 3002 | 2945 | 3099 | 3213 | 3375 | 3583 | 3366 |
| 16 | -4.73 | 1.51 | -3.14 | 0.00 | 0.10377 | Pgr | progesterone receptor | 24 | 75 | 37 | 4 | 25 | 7 | 5 | 0 | 0 | 0 | 1 |
| 602 | -4.69 | 1.49 | -3.14 | 0.00 | 0.10464 | Loxl1 | lysyl oxidase-like 1 | 3016 | 1436 | 2115 | 21 | 16 | 6 | 9 | 0 | 0 | 0 | 7 |
| 4612 | 0.33 | 0.11 | 3.14 | 0.00 | 0.10587 | P2ry1 | purinergic receptor P2Y, G-protein couple | 539 | 1076 | 602 | 6466 | 2780 | 8798 | 3565 | 6793 | 3799 | 4747 | 11567 |
| 657 | -0.74 | 0.23 | -3.14 | 0.00 | 0.10599 | Arid5b | AT rich interactive domain 5B (MRF1-like) | 1789 | 1367 | 1581 | 319 | 398 | 407 | 426 | 251 | 324 | 157 | 205 |
| 135 | -2.94 | 0.94 | -3.13 | 0.00 | 0.10787 | Ptger3 | prostaglandin E receptor 3 (subtype EP3) | 330 | 516 | 394 | 70 | 9 | 89 | 37 | 0 | 18 | 0 | 19 |
| 153 | 1.73 | 0.56 | 3.11 | 0.00 | 0.11294 | Gm15564 | | 33 | 21 | 155 | 36 | 76 | 63 | 153 | 144 | 241 | 635 | 128 |
| 26 | -4.59 | 1.47 | -3.11 | 0.00 | 0.11294 | Serpina1d | serine (or cysteine) peptidase inhibitor, cl | 66 | 102 | 82 | 8 | 7 | 0 | 18 | 0 | 0 | 0 | 0 |
| 34 | -4.58 | 1.47 | -3.11 | 0.00 | 0.1133 | Tlcd5 | TLC domain containing 5 | 96 | 116 | 121 | 15 | 0 | 9 | 14 | 0 | 0 | 0 | 1 |
| 58 | 2.96 | 0.95 | 3.11 | 0.00 | 0.11347 | Tead3 | TEA domain family member 3 | 272 | 157 | 125 | 2 | 0 | 12 | 2 | 25 | 22 | 15 | 7 |
| 252 | -1.69 | 0.54 | -3.11 | 0.00 | 0.11437 | Ghsr | growth hormone secretagogue receptor | 473 | 1055 | 678 | 94 | 104 | 127 | 88 | 3 | 39 | 28 | 83 |
| 37083 | -0.12 | 0.04 | -3.11 | 0.00 | 0.11493 | Pam | peptidylglycine alpha-amidating monooxy | 56137 | 57170 | 52916 | 33608 | 32126 | 25303 | 34710 | 31860 | 29694 | 30588 | 23797 |
| 213 | -1.76 | 0.57 | -3.11 | 0.00 | 0.11527 | Zbtb46 | zinc finger and BTB domain containing 46 | 896 | 748 | 587 | 17 | 26 | 16 | 29 | 0 | 6 | 15 | 7 |
| 925 | -1.15 | 0.37 | -3.10 | 0.00 | 0.11575 | Klhl13 | kelch-like 13 | 3341 | 2875 | 3226 | 138 | 99 | 103 | 154 | 29 | 108 | 27 | 78 |
| 4840 | 0.23 | 0.07 | 3.10 | 0.00 | 0.11772 | Lifr | LIF receptor alpha | 2608 | 2519 | 2855 | 5548 | 4995 | 5447 | 4844 | 6325 | 6180 | 5562 | 6358 |
| 16 | -4.66 | 1.52 | -3.07 | 0.00 | 0.1279 | Tent5b | terminal nucleotidyltransferase 5B | 88 | 25 | 9 | 24 | 7 | 6 | 13 | 0 | 0 | 0 | 3 |
| 168 | -1.36 | 0.44 | -3.07 | 0.00 | 0.12976 | D16Ertd472e | DNA segment, Chr 16, ERATO Doi 472, e | 459 | 456 | 359 | 96 | 84 | 99 | 100 | 27 | 26 | 21 | 120 |
| 3824 | 0.21 | 0.07 | 3.06 | 0.00 | 0.13343 | Yipf4 | Yip1 domain family, member 4 | 3305 | 3608 | 3564 | 3875 | 3977 | 2931 | 3810 | 4952 | 4353 | 4444 | 3241 |
| 220 | -1.92 | 0.63 | -3.06 | 0.00 | 0.13451 | Sh3bgrl2 | SH3 domain binding glutamic acid-rich pr | 444 | 856 | 784 | 27 | 78 | 90 | 62 | 0 | 39 | 12 | 31 |
| 6765 | 0.40 | 0.13 | 3.05 | 0.00 | 0.136 | Spc25 | SPC25, NDC80 kinetochore complex con | 1208 | 1047 | 2207 | 6672 | 8883 | 6826 | 7881 | 9775 | 11039 | 10092 | 8790 |
| 994 | 0.33 | 0.11 | 3.05 | 0.00 | 0.13651 | Clcc1 | chloride channel CLIC-like 1 | 971 | 956 | 869 | 988 | 778 | 974 | 865 | 1250 | 1001 | 1135 | 1152 |
| 69 | -3.01 | 0.99 | -3.04 | 0.00 | 0.13935 | Myo5c | myosin VC | 173 | 343 | 191 | 25 | 3 | 16 | 3 | 0 | 1 | 0 | 5 |
| 434 | -0.95 | 0.31 | -3.04 | 0.00 | 0.13963 | Drc1 | dynein regulatory complex subunit 1 | 1139 | 1834 | 1271 | 65 | 81 | 105 | 97 | 32 | 65 | 35 | 49 |
| 100 | 1.60 | 0.53 | 3.04 | 0.00 | 0.13963 | Cxcl10 | chemokine (C-X-C motif) ligand 10 | 327 | 141 | 228 | 14 | 29 | 4 | 55 | 22 | 166 | 85 | 24 |
| 14 | -4.49 | 1.48 | -3.04 | 0.00 | 0.13969 | Rad51b | RAD51 paralog B | 32 | 21 | 48 | 11 | 11 | 8 | 16 | 0 | 0 | 0 | 7 |
| 116 | -3.91 | 1.29 | -3.04 | 0.00 | 0.13969 | Adap2 | ArfGAP with dual PH domains 2 | 619 | 298 | 338 | 8 | 0 | 1 | 8 | 0 | 0 | 0 | 0 |
| 2733 | 0.26 | 0.09 | 3.04 | 0.00 | 0.14063 | Rer1 | retention in endoplasmic reticulum sorting | 2679 | 2644 | 2333 | 2818 | 2490 | 2308 | 2546 | 3645 | 2654 | 3268 | 2681 |
| 35 | -3.56 | 1.17 | -3.04 | 0.00 | 0.14063 | Frmpd4 | FERM and PDZ domain containing 4 | 33 | 105 | 77 | 37 | 20 | 48 | 34 | 0 | 27 | 0 | 3 |
| 139 | -3.76 | 1.24 | -3.04 | 0.00 | 0.14063 | Ptafr | platelet-activating factor receptor | 817 | 291 | 405 | 0 | 3 | 1 | 16 | 0 | 0 | 0 | 0 |
| 507 | -0.48 | 0.16 | -3.03 | 0.00 | 0.14461 | Zfp618 | zinc finger protein 618 | 806 | 940 | 723 | 475 | 358 | 555 | 427 | 357 | 294 | 250 | 397 |
| 456 | -0.84 | 0.28 | -3.02 | 0.00 | 0.14649 | Celsr1 | cadherin, EGF LAG seven-pass G-type re | 566 | 487 | 608 | 242 | 941 | 385 | 601 | 225 | 494 | 298 | 172 |
| 134 | -1.99 | 0.66 | -3.01 | 0.00 | 0.15285 | Dscam | DS cell adhesion molecule | 295 | 360 | 281 | 81 | 108 | 130 | 92 | 42 | 32 | 0 | 51 |
| 2109 | -2.83 | 0.94 | -3.01 | 0.00 | 0.15285 | Tns1 | tensin 1 | 9879 | 6334 | 6675 | 95 | 87 | 52 | 30 | 0 | 35 | 8 | 7 |
| 4516 | -0.31 | 0.10 | -3.01 | 0.00 | 0.15342 | Lamc1 | laminin, gamma 1 | 15135 | 11596 | 12488 | 1648 | 1417 | 1205 | 1498 | 1196 | 1306 | 1265 | 919 |
| 4384 | 0.18 | 0.06 | 2.99 | 0.00 | 0.15913 | Ascc3 | activating signal cointegrator 1 complex s | 2493 | 2561 | 2726 | 4580 | 5071 | 4839 | 4492 | 5088 | 5824 | 4879 | 5674 |
| 9 | -5.06 | 1.70 | -2.98 | 0.00 | 0.16549 | Prdm6 | PR domain containing 6 | 39 | 9 | 23 | 12 | 10 | 2 | 7 | 0 | 0 | 0 | 0 |
| 2276 | -0.51 | 0.17 | -2.98 | 0.00 | 0.1667 | Myh14 | myosin, heavy polypeptide 14 | 2211 | 2142 | 2761 | 2566 | 2849 | 2204 | 2772 | 2745 | 1882 | 1774 | 1136 |
| 15763 | -0.27 | 0.09 | -2.98 | 0.00 | 0.16671 | 5330417C22Ril | RIKEN cDNA 5330417C22 gene | 20978 | 30374 | 24701 | 14772 | 13315 | 11238 | 13779 | 11414 | 12558 | 11294 | 8977 |
| 275 | -1.34 | 0.45 | -2.97 | 0.00 | 0.17239 | Lrrk2 | leucine-rich repeat kinase 2 | 1111 | 748 | 733 | 35 | 94 | 101 | 85 | 36 | 38 | 9 | 38 |
| 61 | 3.17 | 1.07 | 2.96 | 0.00 | 0.17296 | Gm37206 | | 11 | 0 | 9 | 0 | 5 | 244 | 2 | 28 | 9 | 48 | 319 |
| 6749 | 0.30 | 0.10 | 2.96 | 0.00 | 0.17802 | Syt4 | synaptotagmin IV | 2069 | 2839 | 2610 | 7643 | 7329 | 6630 | 8332 | 10322 | 8598 | 9304 | 8562 |
| 61 | -3.05 | 1.03 | -2.95 | 0.00 | 0.17802 | Chrnb1 | cholinergic receptor, nicotinic, beta polyp | 259 | 144 | 176 | 9 | 15 | 14 | 31 | 0 | 2 | 0 | 19 |
| 20 | -4.59 | 1.55 | -2.95 | 0.00 | 0.17802 | Sprn | shadow of prion protein | 104 | 22 | 74 | 11 | 6 | 3 | 2 | 0 | 0 | 0 | 0 |
| 16 | -3.68 | 1.25 | -2.95 | 0.00 | 0.18259 | Nrk | Nik related kinase | 39 | 23 | 30 | 8 | 30 | 12 | 26 | 0 | 7 | 0 | 1 |
| 51 | -3.06 | 1.04 | -2.94 | 0.00 | 0.18259 | Slc28a3 | solute carrier family 28 (sodium-coupled r | 105 | 110 | 281 | 33 | 13 | 7 | 5 | 0 | 4 | 0 | 3 |
| 27 | -3.20 | 1.09 | -2.94 | 0.00 | 0.18663 | Sstr1 | somatostatin receptor 1 | 27 | 89 | 75 | 33 | 23 | 14 | 18 | 0 | 5 | 0 | 11 |
| 76 | -4.01 | 1.37 | -2.94 | 0.00 | 0.18663 | Efs | embryonal Fyn-associated substrate | 478 | 149 | 184 | 0 | 13 | 1 | 7 | 0 | 0 | 0 | 0 |
| 110 | -2.60 | 0.89 | -2.93 | 0.00 | 0.19043 | Lrrn3 | leucine rich repeat protein 3, neuronal | 234 | 479 | 426 | 6 | 6 | 29 | 15 | 0 | 2 | 0 | 12 |
| 223 | -1.74 | 0.60 | -2.92 | 0.00 | 0.19365 | Ncam2 | neural cell adhesion molecule 2 | 471 | 824 | 597 | 103 | 130 | 104 | 64 | 34 | 50 | 0 | 76 |
| 137 | -2.92 | 1.00 | -2.92 | 0.00 | 0.19371 | Trip6 | thyroid hormone receptor interactor 6 | 457 | 355 | 420 | 8 | 10 | 148 | 3 | 0 | 0 | 0 | 107 |
| 203 | -3.56 | 1.22 | -2.92 | 0.00 | 0.19461 | Il33 | interleukin 33 | 1416 | 558 | 237 | 11 | 0 | 5 | 4 | 0 | 0 | 0 | 1 |
| 321 | -1.89 | 0.65 | -2.92 | 0.00 | 0.19507 | Lhfpl2 | lipoma HMGIC fusion partner-like 2 | 1265 | 1034 | 1065 | 7 | 21 | 36 | 61 | 0 | 3 | 13 | 28 |
| 89 | -2.89 | 0.99 | -2.92 | 0.00 | 0.19507 | Apobr | apolipoprotein B receptor | 532 | 188 | 211 | 13 | 4 | 6 | 19 | 5 | 1 | 0 | 0 |

| | | | | | | | | | | | | | | | | | | |
|---|---|---|---|---|---|---|---|---|---|---|---|---|---|---|---|---|---|---|
| 5 | 7.41 | 2.54 | 2.92 | 0.00 | 0.19642 | Gm13241 | | 0 | 11 | 0 | 0 | 0 | 8 | 0 | 9 | 7 | 19 | 3 |
| 1265 | -0.49 | 0.17 | -2.91 | 0.00 | 0.19768 | Slc29a4 | solute carrier family 29 (nucleoside transp | 2115 | 2960 | 2115 | 958 | 857 | 1104 | 1011 | 554 | 682 | 790 | 766 |
| 408 | -0.97 | 0.33 | -2.91 | 0.00 | 0.20143 | Ptk7 | PTK7 protein tyrosine kinase 7 | 1533 | 991 | 974 | 178 | 168 | 122 | 171 | 137 | 115 | 58 | 38 |
| 530 | -3.15 | 1.08 | -2.90 | 0.00 | 0.20262 | Higd1a | HIG1 domain family, member 1A | 1790 | 1821 | 2030 | 25 | 33 | 56 | 48 | 0 | 18 | 0 | 12 |
| 128 | -1.32 | 0.46 | -2.90 | 0.00 | 0.2046 | Pak7 | p21 (RAC1) activated kinase 7 | 332 | 366 | 325 | 54 | 51 | 71 | 97 | 48 | 24 | 13 | 27 |
| 586 | -4.78 | 1.65 | -2.90 | 0.00 | 0.2046 | Ccn2 | cellular communication network factor 2 | 3630 | 1243 | 1545 | 10 | 0 | 8 | 11 | 0 | 0 | 0 | 0 |
| 16128 | 0.16 | 0.06 | 2.90 | 0.00 | 0.20564 | Rtn4 | reticulon 4 | 8679 | 7492 | 8509 | 19491 | 19263 | 12289 | 21082 | 21942 | 20764 | 24074 | 13821 |
| 260 | -4.41 | 1.53 | -2.88 | 0.00 | 0.2141 | Slit2 | slit guidance ligand 2 | 1445 | 636 | 733 | 14 | 22 | 8 | 2 | 0 | 0 | 0 | 5 |
| 110 | -1.24 | 0.43 | -2.88 | 0.00 | 0.21481 | Slc2a10 | solute carrier family 2 (facilitated glucose | 315 | 407 | 230 | 58 | 36 | 41 | 44 | 28 | 16 | 31 | 7 |
| 341 | -0.66 | 0.23 | -2.88 | 0.00 | 0.21595 | Caskin1 | CASK interacting protein 1 | 243 | 302 | 282 | 441 | 399 | 426 | 521 | 217 | 332 | 287 | 299 |
| 439 | -0.55 | 0.19 | -2.87 | 0.00 | 0.21856 | Ddo | D-aspartate oxidase | 323 | 563 | 556 | 593 | 503 | 566 | 364 | 355 | 339 | 317 | 346 |
| 224 | -1.76 | 0.61 | -2.88 | 0.00 | 0.21856 | Ccn3 | cellular communication network factor 3 | 984 | 565 | 679 | 53 | 58 | 21 | 40 | 9 | 46 | 1 | 7 |
| 16 | -5.24 | 1.82 | -2.88 | 0.00 | 0.21856 | A830011K09Rik | | 65 | 50 | 21 | 0 | 16 | 7 | 14 | 0 | 0 | 0 | 0 |
| 1050 | -0.35 | 0.12 | -2.87 | 0.00 | 0.22032 | Ttc30b | tetratricopeptide repeat domain 30B | 549 | 761 | 729 | 1443 | 1210 | 1582 | 1044 | 1066 | 961 | 729 | 1472 |
| 98 | 2.47 | 0.86 | 2.87 | 0.00 | 0.22057 | Gbp4 | guanylate binding protein 4 | 474 | 295 | 232 | 4 | 0 | 4 | 2 | 35 | 2 | 22 | 7 |
| 141 | -2.16 | 0.75 | -2.87 | 0.00 | 0.22184 | Acot7 | acyl-CoA thioesterase 7 | 479 | 534 | 337 | 50 | 52 | 32 | 26 | 0 | 4 | 30 | 13 |
| 1659 | -0.46 | 0.16 | -2.86 | 0.00 | 0.2247 | Jam2 | junction adhesion molecule 2 | 2816 | 3456 | 3263 | 1545 | 1060 | 725 | 1651 | 1097 | 991 | 1226 | 413 |
| 3901 | -3.45 | 1.21 | -2.86 | 0.00 | 0.22788 | Vim | vimentin | 23894 | 9465 | 9306 | 25 | 60 | 21 | 96 | 1 | 39 | 0 | 1 |
| 136 | -1.80 | 0.63 | -2.86 | 0.00 | 0.22788 | Ttbk1 | tau tubulin kinase 1 | 311 | 388 | 403 | 83 | 56 | 91 | 58 | 22 | 40 | 0 | 39 |
| 5059 | 0.22 | 0.08 | 2.85 | 0.00 | 0.2302 | Uba5 | ubiquitin-like modifier activating enzyme 5 | 2231 | 1810 | 2833 | 3500 | 5609 | 8540 | 5022 | 4740 | 6277 | 5401 | 9683 |
| 613 | -3.38 | 1.19 | -2.85 | 0.00 | 0.23108 | Antxr2 | anthrax toxin receptor 2 | 2666 | 1867 | 2115 | 38 | 22 | 1 | 31 | 0 | 7 | 0 | 1 |
| 47 | -3.40 | 1.20 | -2.84 | 0.00 | 0.23691 | Tmeff2 | transmembrane protein with EGF-like anc | 240 | 155 | 65 | 20 | 9 | 3 | 15 | 0 | 5 | 0 | 0 |
| 91 | -2.05 | 0.72 | -2.84 | 0.00 | 0.23767 | Slc38a11 | solute carrier family 38, member 11 | 304 | 344 | 237 | 26 | 29 | 25 | 11 | 16 | 3 | 0 | 7 |
| 147 | -0.99 | 0.35 | -2.84 | 0.00 | 0.23871 | Gk | glycerol kinase | 369 | 371 | 337 | 108 | 69 | 134 | 54 | 34 | 49 | 39 | 56 |
| 343 | -1.83 | 0.64 | -2.84 | 0.00 | 0.23969 | Ntn1 | netrin 1 | 2220 | 626 | 756 | 31 | 9 | 47 | 32 | 0 | 4 | 45 | 7 |
| 756 | 0.39 | 0.14 | 2.83 | 0.00 | 0.24879 | Mpp3 | membrane protein, palmitoylated 3 (MAG | 214 | 270 | 261 | 887 | 760 | 798 | 830 | 1101 | 1020 | 1161 | 1010 |
| 668 | -0.45 | 0.16 | -2.82 | 0.00 | 0.24879 | Samd14 | sterile alpha motif domain containing 14 | 1482 | 1783 | 1368 | 402 | 357 | 341 | 468 | 313 | 266 | 278 | 287 |
| 57 | -2.86 | 1.02 | -2.82 | 0.00 | 0.25273 | Fat3 | FAT atypical cadherin 3 | 57 | 206 | 200 | 16 | 36 | 46 | 30 | 0 | 29 | 0 | 7 |
| 1619 | -0.32 | 0.11 | -2.82 | 0.00 | 0.2538 | Rcn1 | reticulocalbin 1 | 3503 | 3407 | 3842 | 1000 | 1189 | 769 | 967 | 865 | 893 | 674 | 698 |
| 153 | -0.86 | 0.30 | -2.82 | 0.00 | 0.25412 | Clcn2 | chloride channel, voltage-sensitive 2 | 124 | 105 | 179 | 111 | 127 | 410 | 147 | 67 | 71 | 57 | 283 |
| 3259 | -0.31 | 0.11 | -2.81 | 0.00 | 0.25423 | Serinc5 | serine incorporator 5 | 6059 | 5307 | 5683 | 3151 | 2711 | 1844 | 2667 | 2007 | 2582 | 2387 | 1448 |
| 20 | 3.48 | 1.24 | 2.82 | 0.00 | 0.25423 | Gm13562 | | 6 | 24 | 29 | 0 | 6 | 18 | 1 | 51 | 10 | 54 | 22 |
| 404 | -4.05 | 1.44 | -2.81 | 0.00 | 0.25481 | Fscn1 | fascin actin-bundling protein 1 | 1986 | 1049 | 1387 | 5 | 2 | 0 | 19 | 0 | 0 | 0 | 0 |
| 799 | 0.36 | 0.13 | 2.81 | 0.00 | 0.25643 | Surf6 | surfeit gene 6 | 697 | 796 | 752 | 744 | 663 | 834 | 621 | 1040 | 771 | 878 | 991 |
| 20 | -3.91 | 1.39 | -2.81 | 0.00 | 0.25643 | Gm43672 | | 93 | 42 | 47 | 7 | 12 | 9 | 4 | 0 | 0 | 0 | 4 |
| 41 | -4.16 | 1.48 | -2.81 | 0.00 | 0.25665 | Chst7 | carbohydrate (N-acetylglucosamino) sulfo | 171 | 112 | 149 | 11 | 6 | 2 | 0 | 0 | 0 | 0 | 0 |
| 15690 | 0.16 | 0.06 | 2.81 | 0.01 | 0.25695 | Selenot | selenoprotein T | 8618 | 9375 | 9365 | 19074 | 17168 | 13832 | 18697 | 21350 | 19039 | 20085 | 15985 |
| 418 | -0.66 | 0.24 | -2.81 | 0.01 | 0.25695 | Ube2l6 | ubiquitin-conjugating enzyme E2L 6 | 1146 | 893 | 1188 | 235 | 244 | 156 | 202 | 115 | 195 | 100 | 123 |
| 41 | -2.97 | 1.06 | -2.81 | 0.00 | 0.25695 | Plekhg6 | pleckstrin homology domain containing, fa | 112 | 111 | 91 | 42 | 20 | 19 | 29 | 0 | 17 | 0 | 4 |
| 7 | -4.68 | 1.67 | -2.81 | 0.01 | 0.25695 | Chrna7 | cholinergic receptor, nicotinic, alpha poly | 13 | 9 | 12 | 3 | 15 | 9 | 10 | 0 | 0 | 0 | 1 |
| 277 | -0.63 | 0.23 | -2.80 | 0.01 | 0.25826 | Smoc1 | SPARC related modular calcium binding | 471 | 518 | 457 | 279 | 177 | 263 | 253 | 166 | 114 | 187 | 160 |
| 546 | -3.39 | 1.21 | -2.80 | 0.01 | 0.25985 | Efnb2 | ephrin B2 | 1757 | 2275 | 1722 | 67 | 27 | 56 | 39 | 0 | 8 | 0 | 50 |
| 176 | -2.67 | 0.95 | -2.80 | 0.01 | 0.25985 | Prrg3 | proline rich Gla (G-carboxyglutamic acid) | 801 | 468 | 584 | 23 | 8 | 9 | 24 | 0 | 16 | 0 | 1 |
| 10 | -5.15 | 1.84 | -2.80 | 0.01 | 0.26095 | Gm35721 | | 15 | 0 | 9 | 27 | 23 | 10 | 14 | 0 | 0 | 10 | 0 |
| 433 | -0.64 | 0.23 | -2.79 | 0.01 | 0.26414 | Znrf3 | zinc and ring finger 3 | 420 | 413 | 422 | 537 | 436 | 646 | 497 | 314 | 394 | 206 | 478 |
| 33 | 2.64 | 0.94 | 2.79 | 0.01 | 0.26414 | Xlr4a | X-linked lymphocyte-regulated 4A | 10 | 16 | 2 | 9 | 0 | 67 | 6 | 147 | 14 | 14 | 76 |
| 161 | -1.73 | 0.62 | -2.79 | 0.01 | 0.26492 | Olfm3 | olfactomedin 3 | 263 | 877 | 299 | 69 | 51 | 63 | 44 | 48 | 10 | 0 | 48 |
| 3146 | 0.27 | 0.10 | 2.78 | 0.01 | 0.27004 | Rnf128 | ring finger protein 128 | 1890 | 1869 | 2056 | 3320 | 3121 | 2785 | 3828 | 4464 | 3906 | 3957 | 3405 |
| 1348 | -0.39 | 0.14 | -2.78 | 0.01 | 0.27004 | Lrp8 | low density lipoprotein receptor-related pr | 677 | 915 | 814 | 1321 | 1442 | 2714 | 1563 | 989 | 1249 | 1063 | 2085 |
| 137 | -1.17 | 0.42 | -2.79 | 0.01 | 0.27004 | Erich5 | glutamate rich 5 | 198 | 455 | 329 | 140 | 61 | 85 | 79 | 37 | 23 | 55 | 42 |
| 15 | -3.72 | 1.34 | -2.78 | 0.01 | 0.27004 | 2900060N12Rik | | 5 | 12 | 36 | 10 | 29 | 19 | 28 | 0 | 3 | 0 | 26 |
| 241 | -3.06 | 1.10 | -2.78 | 0.01 | 0.27317 | Plekhg1 | pleckstrin homology domain containing, fa | 742 | 925 | 604 | 120 | 70 | 83 | 43 | 1 | 25 | 0 | 44 |
| 121 | -3.43 | 1.23 | -2.78 | 0.01 | 0.27461 | Mamdc2 | MAM domain containing 2 | 630 | 387 | 300 | 3 | 0 | 2 | 6 | 0 | 0 | 0 | 0 |
| 342 | -0.95 | 0.34 | -2.77 | 0.01 | 0.27786 | Sema3e | sema domain, immunoglobulin domain (Ig | 844 | 1171 | 1122 | 86 | 95 | 101 | 126 | 29 | 84 | 71 | 37 |
| 33 | -4.23 | 1.52 | -2.77 | 0.01 | 0.27786 | Rgs6 | regulator of G-protein signaling 6 | 168 | 49 | 133 | 5 | 6 | 5 | 0 | 0 | 0 | 0 | 0 |
| 6371 | 0.20 | 0.07 | 2.77 | 0.01 | 0.28156 | Ttc28 | tetratricopeptide repeat domain 28 | 2265 | 2050 | 1857 | 8138 | 7488 | 5195 | 8759 | 9552 | 9485 | 10059 | 5237 |
| 1750 | 0.30 | 0.11 | 2.77 | 0.01 | 0.28156 | Lysmd2 | LysM, putative peptidoglycan-binding, do | 1271 | 1236 | 1249 | 1694 | 1744 | 1731 | 1767 | 2519 | 1990 | 2133 | 1918 |
| 149 | -1.53 | 0.55 | -2.76 | 0.01 | 0.28156 | Epsti1 | epithelial stromal interaction 1 (breast) | 719 | 347 | 378 | 18 | 34 | 47 | 40 | 0 | 12 | 27 | 17 |
| 18 | -4.52 | 1.63 | -2.76 | 0.01 | 0.28156 | Kif26b | kinesin family member 26B | 88 | 42 | 46 | 14 | 9 | 1 | 1 | 0 | 0 | 0 | 0 |
| 2781 | -0.30 | 0.11 | -2.76 | 0.01 | 0.28476 | Furin | furin (paired basic amino acid cleaving er | 3103 | 3628 | 3260 | 2986 | 3153 | 2496 | 2709 | 2165 | 2784 | 2517 | 1793 |

| | | | | | | | | | | | | | | | | | |
|---|---|---|---|---|---|---|---|---|---|---|---|---|---|---|---|---|---|
| 119 | -1.45 | 0.53 | -2.76 | 0.01 | 0.28476 | Zfp521 | zinc finger protein 521 | 565 | 311 | 344 | 7 | 9 | 24 | 27 | 3 | 4 | 5 | 11 |
| 20 | -4.70 | 1.70 | -2.76 | 0.01 | 0.2852 | Gabrg3 | gamma-aminobutyric acid (GABA) A rece | 52 | 71 | 64 | 0 | 4 | 8 | 16 | 0 | 0 | 0 | 0 |
| 1527 | -0.38 | 0.14 | -2.76 | 0.01 | 0.28537 | Klf11 | Kruppel-like factor 11 | 2190 | 1946 | 2290 | 1048 | 1039 | 2281 | 1416 | 741 | 975 | 830 | 2039 |
| 649 | 0.58 | 0.21 | 2.76 | 0.01 | 0.28537 | Rras2 | related RAS viral (r-ras) oncogene 2 | 1021 | 797 | 664 | 528 | 426 | 509 | 388 | 977 | 565 | 663 | 605 |
| 1776 | -0.34 | 0.12 | -2.75 | 0.01 | 0.28725 | Myo1d | myosin ID | 2820 | 2355 | 2300 | 1753 | 1866 | 1155 | 1953 | 1277 | 1742 | 1394 | 923 |
| 6 | -5.02 | 1.83 | -2.75 | 0.01 | 0.2918 | Gm46633 | | 10 | 9 | 17 | 10 | 15 | 3 | 3 | 0 | 0 | 0 | 0 |
| 170 | -2.27 | 0.83 | -2.75 | 0.01 | 0.29484 | BC067074 | cDNA sequence BC067074 | 444 | 712 | 484 | 26 | 32 | 41 | 73 | 0 | 48 | 4 | 7 |
| 33 | -3.15 | 1.15 | -2.74 | 0.01 | 0.29711 | Adcy8 | adenylate cyclase 8 | 80 | 57 | 171 | 10 | 10 | 7 | 23 | 0 | 2 | 0 | 8 |
| 1855 | -0.26 | 0.09 | -2.74 | 0.01 | 0.29934 | Dennd4a | DENN/MADD domain containing 4A | 1682 | 1624 | 1536 | 2261 | 2002 | 2243 | 1963 | 1709 | 1745 | 1635 | 2011 |
| 182 | -2.16 | 0.79 | -2.74 | 0.01 | 0.2994 | Ugt8a | UDP galactosyltransferase 8A | 523 | 740 | 669 | 9 | 13 | 12 | 17 | 0 | 9 | 0 | 4 |
| 229 | -1.51 | 0.55 | -2.73 | 0.01 | 0.30239 | Cd79a | CD79A antigen (immunoglobulin-associa | 579 | 135 | 77 | 279 | 365 | 386 | 258 | 107 | 136 | 58 | 144 |
| 121 | -1.59 | 0.58 | -2.73 | 0.01 | 0.30465 | Medag | mesenteric estrogen dependent adipoger | 582 | 213 | 320 | 56 | 42 | 18 | 45 | 6 | 8 | 21 | 23 |
| 954 | -0.36 | 0.13 | -2.72 | 0.01 | 0.31076 | Rtn1 | reticulon 1 | 1930 | 1927 | 1543 | 843 | 677 | 536 | 820 | 655 | 588 | 529 | 449 |
| 168 | -1.85 | 0.68 | -2.72 | 0.01 | 0.31076 | Cacna2d2 | calcium channel, voltage-dependent, alpl | 358 | 462 | 457 | 62 | 166 | 120 | 83 | 58 | 56 | 0 | 30 |
| 1608 | -0.37 | 0.13 | -2.72 | 0.01 | 0.31076 | Gpld1 | glycosylphosphatidylinositol specific phos | 2646 | 3457 | 3078 | 1433 | 1086 | 837 | 1443 | 1215 | 894 | 878 | 723 |
| 404 | -0.73 | 0.27 | -2.72 | 0.01 | 0.3123 | Aldh1a3 | aldehyde dehydrogenase family 1, subfar | 448 | 157 | 213 | 536 | 524 | 178 | 911 | 285 | 505 | 620 | 63 |
| 757 | 0.43 | 0.16 | 2.72 | 0.01 | 0.31631 | E2f1 | E2F transcription factor 1 | 301 | 349 | 304 | 952 | 926 | 537 | 680 | 1591 | 1162 | 863 | 662 |
| 32536 | -0.52 | 0.19 | -2.71 | 0.01 | 0.317 | Pappa2 | pappalysin 2 | 80888 | 97942 | 82765 | 13146 | 12342 | 8739 | 22407 | 9852 | 10549 | 13875 | 5388 |
| 868 | 0.47 | 0.17 | 2.71 | 0.01 | 0.31711 | Myo15b | | 188 | 426 | 300 | 882 | 808 | 1372 | 609 | 1205 | 1087 | 979 | 1691 |
| 50 | -2.73 | 1.01 | -2.71 | 0.01 | 0.31711 | Fam83f | family with sequence similarity 83, memb | 165 | 167 | 117 | 15 | 15 | 10 | 44 | 0 | 6 | 0 | 8 |
| 8 | 4.27 | 1.57 | 2.71 | 0.01 | 0.31739 | Gm6286 | | 21 | 5 | 2 | 1 | 0 | 3 | 0 | 28 | 7 | 12 | 5 |
| 4087 | 0.19 | 0.07 | 2.71 | 0.01 | 0.3177 | Rabep1 | rabaptin, RAB GTPase binding effector pr | 3046 | 3114 | 3265 | 4362 | 4292 | 3552 | 4368 | 5290 | 4565 | 4964 | 4140 |
| 898 | -0.32 | 0.12 | -2.71 | 0.01 | 0.3177 | Tlnrd1 | talin rod domain containing 1 | 1525 | 1548 | 1381 | 843 | 782 | 605 | 772 | 722 | 622 | 596 | 477 |
| 124 | -2.97 | 1.09 | -2.71 | 0.01 | 0.3177 | Pygm | muscle glycogen phosphorylase | 650 | 233 | 458 | 0 | 6 | 7 | 6 | 0 | 0 | 0 | 2 |
| 710 | -4.69 | 1.73 | -2.71 | 0.01 | 0.31809 | Srgn | serglycin | 3284 | 2121 | 2353 | 0 | 14 | 4 | 32 | 0 | 0 | 0 | 1 |
| 1942 | 0.24 | 0.09 | 2.71 | 0.01 | 0.31888 | Fam114a2 | family with sequence similarity 114, meml | 1644 | 1756 | 1792 | 1794 | 1754 | 1916 | 1953 | 2168 | 2099 | 2121 | 2368 |
| 8725 | 0.19 | 0.07 | 2.70 | 0.01 | 0.32074 | Eprs | glutamyl-prolyl-tRNA synthetase | 8266 | 6997 | 7338 | 7580 | 9646 | 8187 | 8913 | 9311 | 10021 | 9865 | 9852 |
| 207 | -0.80 | 0.30 | -2.71 | 0.01 | 0.32074 | Oplah | 5-oxoprolinase (ATP-hydrolysing) | 547 | 522 | 528 | 111 | 79 | 95 | 147 | 44 | 64 | 71 | 67 |
| 59 | -1.98 | 0.73 | -2.70 | 0.01 | 0.32418 | Slc44a4 | solute carrier family 44, member 4 | 122 | 188 | 181 | 20 | 38 | 31 | 30 | 21 | 9 | 0 | 7 |
| 4924 | 0.29 | 0.11 | 2.70 | 0.01 | 0.32702 | Eif5b | eukaryotic translation initiation factor 5B | 4409 | 4226 | 4185 | 4213 | 4680 | 4921 | 4735 | 6786 | 5327 | 4862 | 5817 |
| 52 | -2.12 | 0.79 | -2.70 | 0.01 | 0.32785 | Nphs1 | nephrosis 1, nephrin | 127 | 136 | 186 | 25 | 13 | 27 | 29 | 0 | 5 | 5 | 20 |
| 18306 | 0.36 | 0.13 | 2.69 | 0.01 | 0.32838 | Tceal9 | transcription elongation factor A like 9 | 10504 | 11531 | 11780 | 19521 | 18554 | 13786 | 20492 | 36271 | 18764 | 23281 | 16887 |
| 400 | -1.08 | 0.40 | -2.69 | 0.01 | 0.32838 | Fmnl1 | formin-like 1 | 1765 | 1134 | 795 | 131 | 104 | 82 | 169 | 38 | 80 | 31 | 74 |
| 1470 | 0.48 | 0.18 | 2.69 | 0.01 | 0.32856 | Th | tyrosine hydroxylase | 150 | 166 | 240 | 1653 | 1212 | 2117 | 1537 | 2471 | 1785 | 1950 | 2885 |
| 461 | -6.01 | 2.23 | -2.69 | 0.01 | 0.32979 | Retnla | resistin like alpha | 3346 | 909 | 709 | 0 | 70 | 2 | 39 | 0 | 0 | 0 | 0 |
| 688 | 0.70 | 0.26 | 2.69 | 0.01 | 0.32981 | Reln | reelin | 1866 | 253 | 527 | 190 | 258 | 714 | 798 | 461 | 331 | 1154 | 1011 |
| 12365 | -0.25 | 0.09 | -2.69 | 0.01 | 0.33034 | Lpl | lipoprotein lipase | 21660 | 33270 | 25766 | 9235 | 6023 | 7260 | 7374 | 8435 | 4952 | 5724 | 6313 |
| 649 | -0.49 | 0.18 | -2.68 | 0.01 | 0.33399 | Tln2 | talin 2 | 617 | 448 | 379 | 1018 | 702 | 772 | 842 | 553 | 597 | 639 | 568 |
| 1032 | 1.09 | 0.41 | 2.68 | 0.01 | 0.33399 | Penk | preproenkephalin | 5835 | 2430 | 2949 | 14 | 4 | 7 | 24 | 28 | 26 | 15 | 21 |
| 6 | -7.74 | 2.88 | -2.68 | 0.01 | 0.33399 | Gm16306 | | 0 | 0 | 19 | 33 | 0 | 10 | 3 | 0 | 0 | 0 | 2 |
| 700 | -5.24 | 1.95 | -2.68 | 0.01 | 0.33418 | Col6a5 | collagen, type VI, alpha 5 | 4468 | 2657 | 534 | 0 | 7 | 16 | 16 | 0 | 0 | 0 | 0 |
| 393 | -0.53 | 0.20 | -2.68 | 0.01 | 0.33664 | Arl4c | ADP-ribosylation factor-like 4C | 1107 | 1009 | 952 | 186 | 151 | 190 | 214 | 133 | 137 | 129 | 112 |
| 1859 | 0.26 | 0.10 | 2.68 | 0.01 | 0.33686 | Naa30 | N(alpha)-acetyltransferase 30, NatC catal | 1193 | 1949 | 1475 | 2118 | 1653 | 1864 | 1560 | 2689 | 1887 | 1949 | 2117 |
| 908 | -2.79 | 1.04 | -2.68 | 0.01 | 0.33758 | Cd36 | CD36 molecule | 3608 | 3004 | 3241 | 28 | 35 | 23 | 21 | 0 | 12 | 0 | 12 |
| 139 | 0.88 | 0.33 | 2.68 | 0.01 | 0.33955 | Atp23 | ATP23 metallopeptidase and ATP syntha | 134 | 125 | 101 | 123 | 59 | 115 | 109 | 185 | 122 | 300 | 154 |
| 8 | -3.89 | 1.45 | -2.68 | 0.01 | 0.33955 | E330011M16Rik | | 2 | 17 | 8 | 11 | 11 | 19 | 9 | 0 | 2 | 0 | 2 |
| 1603 | 0.38 | 0.14 | 2.67 | 0.01 | 0.34036 | Tsc22d3 | TSC22 domain family, member 3 | 3485 | 2678 | 2863 | 854 | 854 | 1127 | 891 | 1364 | 877 | 1220 | 1421 |
| 7697 | -0.16 | 0.06 | -2.67 | 0.01 | 0.34246 | Scd2 | stearoyl-Coenzyme A desaturase 2 | 5642 | 5291 | 5487 | 9535 | 8433 | 8380 | 9628 | 8006 | 7928 | 8588 | 7745 |
| 2164 | 0.28 | 0.11 | 2.67 | 0.01 | 0.34246 | Pabpc4 | poly(A) binding protein, cytoplasmic 4 | 1820 | 1833 | 2111 | 2117 | 2188 | 1997 | 1823 | 2967 | 2494 | 2243 | 2211 |
| 182 | -1.61 | 0.61 | -2.66 | 0.01 | 0.34948 | Muc1 | mucin 1, transmembrane | 398 | 691 | 812 | 21 | 22 | 17 | 13 | 12 | 3 | 0 | 13 |
| 177 | -0.65 | 0.24 | -2.66 | 0.01 | 0.35114 | Kctd8 | potassium channel tetramerisation domai | 148 | 157 | 136 | 253 | 203 | 298 | 166 | 154 | 117 | 122 | 192 |
| 66 | -2.62 | 0.98 | -2.66 | 0.01 | 0.35114 | Pacsin3 | protein kinase C and casein kinase subst | 253 | 175 | 179 | 36 | 37 | 13 | 11 | 0 | 10 | 0 | 12 |
| 1949 | -0.22 | 0.08 | -2.66 | 0.01 | 0.35133 | Aip | aryl-hydrocarbon receptor-interacting pro | 2626 | 2564 | 2460 | 1964 | 1963 | 1569 | 1908 | 1642 | 1732 | 1735 | 1276 |
| 730 | -1.82 | 0.68 | -2.66 | 0.01 | 0.35133 | Hk1 | hexokinase 1 | 3080 | 2473 | 2243 | 18 | 32 | 57 | 78 | 15 | 26 | 8 | 4 |
| 104 | -3.35 | 1.26 | -2.66 | 0.01 | 0.35133 | Angptl4 | angiopoietin-like 4 | 499 | 199 | 423 | 8 | 9 | 4 | 0 | 0 | 2 | 0 | 0 |
| 533 | -0.46 | 0.17 | -2.66 | 0.01 | 0.35358 | Shisal2b | shisa like 2B | 722 | 796 | 947 | 375 | 627 | 493 | 479 | 327 | 385 | 296 | 414 |
| 54 | -2.01 | 0.76 | -2.65 | 0.01 | 0.35543 | Npl | N-acetylneuraminate pyruvate lyase | 219 | 92 | 147 | 26 | 28 | 20 | 25 | 0 | 14 | 17 | 3 |
| 487 | -0.84 | 0.32 | -2.65 | 0.01 | 0.3595 | Aqp4 | aquaporin 4 | 94 | 199 | 246 | 623 | 680 | 878 | 882 | 242 | 487 | 461 | 570 |
| 741 | -0.45 | 0.17 | -2.65 | 0.01 | 0.3595 | Celsr2 | cadherin, EGF LAG seven-pass G-type re | 577 | 524 | 814 | 930 | 892 | 952 | 809 | 514 | 796 | 689 | 651 |
| 400 | 0.52 | 0.20 | 2.65 | 0.01 | 0.3595 | Nat14 | N-acetyltransferase 14 | 406 | 344 | 415 | 309 | 308 | 381 | 328 | 520 | 399 | 541 | 445 |

| | | | | | | | | | | | | | | | | | | |
|---|---|---|---|---|---|---|---|---|---|---|---|---|---|---|---|---|---|---|
| 278 | -0.70 | 0.26 | -2.65 | 0.01 | 0.3595 | Stimate | STIM activating enhancer | 558 | 530 | 507 | 213 | 196 | 245 | 247 | 114 | 183 | 103 | 161 |
| 9 | 5.78 | 2.18 | 2.65 | 0.01 | 0.3595 | Gm40773 | | 0 | 0 | 0 | 0 | 0 | 14 | 0 | 23 | 36 | 9 | 18 |
| 745 | -2.28 | 0.86 | -2.65 | 0.01 | 0.36006 | Ecm1 | extracellular matrix protein 1 | 4371 | 1769 | 1953 | 5 | 4 | 43 | 26 | 1 | 1 | 0 | 27 |
| 777 | -1.05 | 0.40 | -2.64 | 0.01 | 0.36149 | Adgre5 | adhesion G protein-coupled receptor E5 | 3473 | 2397 | 2032 | 110 | 92 | 91 | 139 | 107 | 58 | 15 | 39 |
| 75 | -2.06 | 0.78 | -2.64 | 0.01 | 0.36149 | Rab29 | RAB29, member RAS oncogene family | 275 | 262 | 144 | 31 | 13 | 28 | 31 | 0 | 3 | 27 | 5 |
| 173 | -3.36 | 1.27 | -2.64 | 0.01 | 0.36505 | Plac8 | placenta-specific 8 | 942 | 419 | 529 | 10 | 0 | 3 | 4 | 0 | 0 | 0 | 1 |
| 2668 | -0.35 | 0.13 | -2.64 | 0.01 | 0.36624 | Tshz2 | teashirt zinc finger family member 2 | 5208 | 4866 | 4201 | 2505 | 2325 | 1547 | 2022 | 1970 | 2113 | 1512 | 1084 |
| 204 | -1.71 | 0.65 | -2.64 | 0.01 | 0.36707 | Sertm1 | serine rich and transmembrane domain c | 558 | 928 | 633 | 47 | 21 | 12 | 16 | 0 | 5 | 12 | 10 |
| 358 | -1.14 | 0.43 | -2.63 | 0.01 | 0.36977 | Spns2 | spinster homolog 2 | 1308 | 951 | 1195 | 131 | 105 | 46 | 48 | 20 | 75 | 22 | 36 |
| 3320 | -0.28 | 0.11 | -2.63 | 0.01 | 0.37228 | Fryl | FRY like transcription coactivator | 3532 | 3750 | 3371 | 3270 | 2522 | 5581 | 2835 | 2271 | 2654 | 2147 | 4592 |
| 108 | -3.34 | 1.27 | -2.63 | 0.01 | 0.3759 | Chdh | choline dehydrogenase | 324 | 324 | 523 | 4 | 16 | 1 | 0 | 0 | 0 | 0 | 1 |
| 96 | -2.74 | 1.04 | -2.63 | 0.01 | 0.37599 | Ptpro | protein tyrosine phosphatase, receptor ty | 462 | 329 | 197 | 21 | 0 | 15 | 15 | 0 | 1 | 0 | 11 |
| 7 | -6.03 | 2.30 | -2.63 | 0.01 | 0.37599 | Gm43307 | | 0 | 0 | 11 | 8 | 3 | 28 | 7 | 0 | 0 | 0 | 21 |
| 11 | -4.46 | 1.70 | -2.62 | 0.01 | 0.38044 | Gm39929 | | 0 | 0 | 27 | 20 | 22 | 24 | 4 | 0 | 20 | 0 | 1 |
| 2442 | -0.28 | 0.11 | -2.62 | 0.01 | 0.38405 | Cald1 | caldesmon 1 | 9276 | 5532 | 5102 | 845 | 628 | 850 | 1540 | 636 | 620 | 1095 | 744 |
| 13503 | -0.18 | 0.07 | -2.61 | 0.01 | 0.38819 | Insrr | insulin receptor-related receptor | 1836 | 2037 | 2697 | 15089 | 18514 | 27552 | 14690 | 12483 | 16811 | 14262 | 22563 |
| 2026 | 0.29 | 0.11 | 2.61 | 0.01 | 0.39005 | Rbm17 | RNA binding motif protein 17 | 1736 | 1325 | 1616 | 1782 | 1948 | 2347 | 1878 | 2464 | 2233 | 2281 | 2681 |
| 8 | -4.62 | 1.77 | -2.61 | 0.01 | 0.39005 | Gm8170 | | 1 | 12 | 4 | 4 | 22 | 25 | 10 | 0 | 0 | 0 | 14 |
| 4750 | -0.35 | 0.13 | -2.61 | 0.01 | 0.39156 | Alcam | activated leukocyte cell adhesion molecu | 8178 | 10141 | 8893 | 3158 | 3486 | 3363 | 3994 | 2110 | 3084 | 3136 | 2705 |
| 844 | -0.64 | 0.25 | -2.60 | 0.01 | 0.3943 | Thbs3 | thrombospondin 3 | 2170 | 1498 | 1850 | 515 | 484 | 930 | 317 | 171 | 343 | 308 | 701 |
| 558 | -2.99 | 1.15 | -2.61 | 0.01 | 0.3943 | Gja1 | gap junction protein, alpha 1 | 2349 | 1971 | 1804 | 6 | 5 | 0 | 4 | 0 | 1 | 0 | 0 |
| 29 | -3.86 | 1.48 | -2.60 | 0.01 | 0.3943 | Ccdc125 | coiled-coil domain containing 125 | 82 | 73 | 114 | 33 | 4 | 2 | 4 | 0 | 0 | 0 | 4 |
| 1531 | 0.26 | 0.10 | 2.60 | 0.01 | 0.39661 | Mfap3 | microfibrillar-associated protein 3 | 1573 | 1558 | 1462 | 1297 | 1331 | 1475 | 1479 | 1795 | 1603 | 1620 | 1653 |
| 370 | -2.95 | 1.14 | -2.60 | 0.01 | 0.39727 | Cavin3 | caveolae associated 3 | 1749 | 985 | 1274 | 32 | 6 | 6 | 16 | 0 | 4 | 0 | 3 |
| 78 | -3.08 | 1.19 | -2.60 | 0.01 | 0.39847 | Ripor2 | RHO family interacting cell polarization re | 370 | 217 | 190 | 18 | 9 | 10 | 18 | 0 | 0 | 30 | 0 |
| 112 | -2.23 | 0.86 | -2.60 | 0.01 | 0.40111 | Scn2b | sodium channel, voltage-gated, type II, be | 470 | 373 | 220 | 23 | 15 | 54 | 32 | 0 | 11 | 0 | 32 |
| 137 | -2.25 | 0.87 | -2.59 | 0.01 | 0.40622 | Otof | otoferlin | 349 | 577 | 424 | 19 | 15 | 37 | 42 | 0 | 13 | 0 | 25 |
| 3398 | 0.21 | 0.08 | 2.59 | 0.01 | 0.40637 | Nmt2 | N-myristoyltransferase 2 | 1585 | 1692 | 1586 | 3409 | 3277 | 4473 | 3910 | 3834 | 3925 | 4593 | 5094 |
| 2004 | 0.28 | 0.11 | 2.59 | 0.01 | 0.40637 | Birc2 | baculoviral IAP repeat-containing 2 | 1712 | 1458 | 1352 | 1489 | 1654 | 3295 | 1505 | 1748 | 1833 | 2213 | 3783 |
| 218 | -0.88 | 0.34 | -2.58 | 0.01 | 0.41381 | Ly6h | lymphocyte antigen 6 complex, locus H | 457 | 709 | 715 | 51 | 123 | 81 | 82 | 28 | 58 | 41 | 55 |
| 676 | -0.92 | 0.36 | -2.58 | 0.01 | 0.41393 | Sulf2 | sulfatase 2 | 3849 | 1377 | 1520 | 144 | 86 | 112 | 101 | 104 | 43 | 72 | 31 |
| 1394 | -0.28 | 0.11 | -2.58 | 0.01 | 0.41592 | Dab2ip | disabled 2 interacting protein | 3059 | 2466 | 2561 | 976 | 999 | 939 | 1061 | 804 | 906 | 823 | 734 |
| 688 | -0.60 | 0.23 | -2.58 | 0.01 | 0.4208 | Arrdc4 | arrestin domain containing 4 | 1527 | 1034 | 1411 | 526 | 351 | 798 | 506 | 365 | 341 | 237 | 468 |
| 45 | -2.57 | 1.00 | -2.58 | 0.01 | 0.4208 | Gbp8 | guanylate-binding protein 8 | 182 | 108 | 61 | 27 | 23 | 24 | 19 | 1 | 4 | 0 | 44 |
| 1194 | -0.27 | 0.10 | -2.57 | 0.01 | 0.4215 | G6pc3 | glucose 6 phosphatase, catalytic, 3 | 958 | 1065 | 1077 | 1364 | 1451 | 1238 | 1426 | 1185 | 1122 | 1215 | 1030 |
| 642 | -3.92 | 1.53 | -2.57 | 0.01 | 0.42583 | Dab2 | disabled 2, mitogen-responsive phospho | 3000 | 1799 | 2225 | 0 | 8 | 8 | 21 | 0 | 0 | 0 | 2 |
| 352 | -0.46 | 0.18 | -2.57 | 0.01 | 0.4266 | Adam23 | a disintegrin and metallopeptidase domai | 823 | 568 | 663 | 238 | 241 | 233 | 342 | 153 | 183 | 241 | 189 |
| 2119 | -0.32 | 0.12 | -2.56 | 0.01 | 0.42974 | Ppp1r9a | protein phosphatase 1, regulatory subuni | 2301 | 2963 | 2622 | 2067 | 2030 | 2342 | 2064 | 1694 | 1864 | 1240 | 2120 |
| 512 | -0.49 | 0.19 | -2.56 | 0.01 | 0.42974 | Spire2 | spire type actin nucleation factor 2 | 450 | 637 | 629 | 738 | 515 | 489 | 557 | 409 | 396 | 399 | 418 |
| 65 | -1.66 | 0.65 | -2.56 | 0.01 | 0.42974 | Ninl | ninein-like | 238 | 206 | 125 | 17 | 12 | 24 | 52 | 7 | 2 | 6 | 21 |
| 238 | -4.46 | 1.74 | -2.57 | 0.01 | 0.42974 | Ccl9 | chemokine (C-C motif) ligand 9 | 1488 | 530 | 568 | 0 | 7 | 2 | 26 | 0 | 0 | 0 | 0 |
| 9 | -4.53 | 1.77 | -2.56 | 0.01 | 0.42974 | Pcdha3 | protocadherin alpha 3 | 2 | 15 | 3 | 20 | 4 | 10 | 20 | 0 | 0 | 0 | 22 |
| 150 | -3.00 | 1.17 | -2.56 | 0.01 | 0.42982 | Folr2 | folate receptor 2 (fetal) | 955 | 263 | 406 | 0 | 10 | 5 | 4 | 0 | 0 | 0 | 2 |
| 22 | -2.91 | 1.13 | -2.56 | 0.01 | 0.4301 | Fam217a | family with sequence similarity 217, meml | 47 | 64 | 82 | 4 | 7 | 17 | 14 | 0 | 3 | 0 | 3 |
| 124 | -0.81 | 0.32 | -2.56 | 0.01 | 0.43135 | C230013L11Rik | | 120 | 159 | 179 | 141 | 184 | 144 | 107 | 59 | 133 | 70 | 73 |
| 176 | -0.87 | 0.34 | -2.56 | 0.01 | 0.43144 | Me3 | malic enzyme 3, NADP(+)-dependent, mit | 137 | 175 | 221 | 366 | 190 | 192 | 152 | 200 | 145 | 55 | 108 |
| 2623 | -0.61 | 0.24 | -2.56 | 0.01 | 0.43208 | Ace | angiotensin I converting enzyme (peptidy | 11009 | 8114 | 9380 | 40 | 55 | 14 | 104 | 31 | 29 | 57 | 15 |
| 32 | -3.78 | 1.48 | -2.56 | 0.01 | 0.43208 | Rerg | RAS-like, estrogen-regulated, growth-inhi | 193 | 46 | 79 | 11 | 8 | 2 | 6 | 0 | 0 | 0 | 6 |
| 3 | -6.72 | 2.63 | -2.56 | 0.01 | 0.43208 | 1700010B13Rik | | 4 | 0 | 0 | 12 | 9 | 8 | 2 | 0 | 0 | 0 | 2 |
| 91 | -1.30 | 0.51 | -2.56 | 0.01 | 0.43402 | Mospd1 | motile sperm domain containing 1 | 294 | 280 | 244 | 41 | 20 | 17 | 52 | 12 | 13 | 9 | 14 |
| 28010 | -0.10 | 0.04 | -2.56 | 0.01 | 0.4343 | Atp1a1 | ATPase, Na+/K+ transporting, alpha 1 pol | 22975 | 25133 | 25614 | 32559 | 31911 | 25971 | 30928 | 29615 | 30223 | 29198 | 23987 |
| 1994 | 0.29 | 0.11 | 2.55 | 0.01 | 0.43528 | Mtf1 | metal response element binding transcrip | 1287 | 1697 | 1438 | 2029 | 1906 | 1964 | 1924 | 3018 | 2289 | 2392 | 1984 |
| 70 | -2.86 | 1.12 | -2.55 | 0.01 | 0.43728 | Mmp17 | matrix metallopeptidase 17 | 320 | 267 | 130 | 25 | 12 | 1 | 9 | 0 | 2 | 0 | 3 |
| 39 | -3.11 | 1.22 | -2.55 | 0.01 | 0.43728 | Gm38357 | | 16 | 26 | 14 | 15 | 13 | 194 | 3 | 0 | 1 | 0 | 141 |
| 1205 | -0.30 | 0.12 | -2.55 | 0.01 | 0.43746 | Rtn4rl1 | reticulon 4 receptor-like 1 | 966 | 1186 | 958 | 1577 | 1427 | 1053 | 1532 | 1194 | 1141 | 1358 | 859 |
| 10 | -4.25 | 1.67 | -2.55 | 0.01 | 0.43746 | Gm49805 | | 0 | 0 | 9 | 25 | 26 | 14 | 19 | 0 | 14 | 0 | 6 |
| 36 | -3.60 | 1.41 | -2.55 | 0.01 | 0.43876 | Lrrtm3 | leucine rich repeat transmembrane neurc | 175 | 96 | 114 | 3 | 0 | 5 | 4 | 0 | 0 | 0 | 0 |
| 1017 | 0.43 | 0.17 | 2.55 | 0.01 | 0.43918 | Hmgn5 | high-mobility group nucleosome binding | 1510 | 1376 | 1299 | 657 | 715 | 866 | 735 | 1287 | 817 | 875 | 1045 |
| 1466 | -2.18 | 0.86 | -2.55 | 0.01 | 0.44101 | Dlc1 | deleted in liver cancer 1 | 5860 | 4853 | 5213 | 27 | 54 | 37 | 23 | 5 | 9 | 0 | 51 |
| 178 | -1.40 | 0.55 | -2.54 | 0.01 | 0.44538 | Rab15 | RAB15, member RAS oncogene family | 450 | 469 | 512 | 112 | 95 | 50 | 115 | 5 | 66 | 53 | 29 |

| | | | | | | | | | | | | | | | | | | |
|---|---|---|---|---|---|---|---|---|---|---|---|---|---|---|---|---|---|---|
| 25 | 1.93 | 0.76 | 2.54 | 0.01 | 0.44657 | Tex11 | testis expressed gene 11 | 16 | 5 | 14 | 12 | 7 | 32 | 5 | 38 | 55 | 26 | 65 |
| 117 | -1.43 | 0.56 | -2.54 | 0.01 | 0.44684 | Adamtsl4 | ADAMTS-like 4 | 522 | 245 | 339 | 49 | 29 | 21 | 28 | 3 | 21 | 22 | 7 |
| 3999 | 0.20 | 0.08 | 2.54 | 0.01 | 0.44696 | Cmpk1 | cytidine monophosphate (UMP-CMP) kin | 3321 | 2911 | 2986 | 4064 | 4395 | 3246 | 4490 | 5032 | 4366 | 5127 | 4050 |
| 1242 | -0.30 | 0.12 | -2.54 | 0.01 | 0.44696 | Fnbp1l | formin binding protein 1-like | 1639 | 1784 | 1610 | 1116 | 1266 | 1247 | 1126 | 931 | 918 | 837 | 1192 |
| 950 | -0.36 | 0.14 | -2.54 | 0.01 | 0.44696 | Agt | angiotensinogen (serpin peptidase inhibi | 1343 | 1473 | 1965 | 735 | 1132 | 605 | 717 | 736 | 784 | 492 | 469 |
| 24 | -3.83 | 1.51 | -2.54 | 0.01 | 0.44696 | Tmem200a | transmembrane protein 200A | 76 | 81 | 63 | 0 | 7 | 10 | 19 | 0 | 0 | 0 | 3 |
| 145 | -2.33 | 0.92 | -2.53 | 0.01 | 0.45163 | Evc2 | EvC ciliary complex subunit 2 | 658 | 487 | 314 | 17 | 36 | 8 | 44 | 0 | 26 | 0 | 3 |
| 338 | -0.87 | 0.34 | -2.53 | 0.01 | 0.4556 | Bmp3 | bone morphogenetic protein 3 | 714 | 1268 | 816 | 130 | 107 | 129 | 222 | 122 | 67 | 50 | 91 |
| 10 | -4.40 | 1.74 | -2.53 | 0.01 | 0.45609 | 4933429H19Rik | | 1 | 8 | 11 | 22 | 19 | 11 | 4 | 0 | 0 | 0 | 30 |
| 114 | -3.44 | 1.36 | -2.53 | 0.01 | 0.45702 | Lrp4 | low density lipoprotein receptor-related p | 546 | 404 | 272 | 9 | 9 | 0 | 8 | 0 | 0 | 0 | 2 |
| 3449 | 0.19 | 0.07 | 2.53 | 0.01 | 0.45756 | Eif3e | eukaryotic translation initiation factor 3, su | 5387 | 4579 | 5041 | 2356 | 2865 | 2494 | 3019 | 2954 | 2991 | 3510 | 2742 |
| 1056 | -0.32 | 0.13 | -2.52 | 0.01 | 0.4598 | Caskin2 | CASK-interacting protein 2 | 2330 | 2165 | 2385 | 620 | 644 | 753 | 612 | 492 | 606 | 432 | 578 |
| 55 | -2.81 | 1.11 | -2.52 | 0.01 | 0.4598 | Galm | galactose mutarotase | 182 | 106 | 266 | 18 | 2 | 5 | 16 | 0 | 3 | 0 | 3 |
| 87 | -2.36 | 0.94 | -2.52 | 0.01 | 0.46147 | Col8a2 | collagen, type VIII, alpha 2 | 411 | 235 | 243 | 23 | 17 | 13 | 5 | 0 | 7 | 0 | 6 |
| 81 | -3.32 | 1.32 | -2.52 | 0.01 | 0.46401 | Slc38a3 | solute carrier family 38, member 3 | 81 | 484 | 264 | 47 | 0 | 4 | 3 | 0 | 0 | 0 | 4 |
| 2868 | -0.31 | 0.12 | -2.51 | 0.01 | 0.46954 | Aatk | apoptosis-associated tyrosine kinase | 4282 | 4404 | 4407 | 2606 | 2183 | 2092 | 3301 | 2486 | 1848 | 2442 | 1496 |
| 2197 | 0.24 | 0.10 | 2.51 | 0.01 | 0.46954 | Notch1 | notch 1 | 4770 | 5235 | 4411 | 1065 | 963 | 1360 | 1092 | 1351 | 1182 | 1363 | 1372 |
| 439 | -0.55 | 0.22 | -2.51 | 0.01 | 0.46954 | Ror1 | receptor tyrosine kinase-like orphan rece | 770 | 491 | 445 | 444 | 433 | 405 | 570 | 204 | 363 | 393 | 314 |
| 259 | -0.69 | 0.28 | -2.51 | 0.01 | 0.46954 | Large1 | LARGE xylosyl- and glucuronyltransferas | 807 | 734 | 689 | 120 | 86 | 69 | 105 | 52 | 46 | 94 | 48 |
| 517 | -0.72 | 0.29 | -2.51 | 0.01 | 0.46954 | Galnt13 | polypeptide N-acetylgalactosaminyltransf | 1973 | 1718 | 1701 | 28 | 40 | 50 | 70 | 30 | 20 | 37 | 24 |
| 178 | -1.15 | 0.46 | -2.51 | 0.01 | 0.46954 | Kcna5 | potassium voltage-gated channel, shaker | 573 | 669 | 346 | 75 | 77 | 46 | 52 | 20 | 23 | 21 | 59 |
| 104 | -3.23 | 1.28 | -2.51 | 0.01 | 0.46954 | Acp5 | acid phosphatase 5, tartrate resistant | 583 | 303 | 248 | 4 | 0 | 1 | 4 | 0 | 0 | 0 | 0 |
| 2636 | -0.53 | 0.21 | -2.51 | 0.01 | 0.46963 | Gas6 | growth arrest specific 6 | 13977 | 5479 | 7660 | 327 | 301 | 169 | 311 | 191 | 253 | 215 | 114 |
| 63 | 1.55 | 0.62 | 2.51 | 0.01 | 0.47077 | Gm4332 | | 90 | 72 | 155 | 16 | 24 | 20 | 31 | 144 | 59 | 32 | 52 |
| 3087 | -0.25 | 0.10 | -2.51 | 0.01 | 0.47199 | Tspan2 | tetraspanin 2 | 2573 | 3070 | 2377 | 4234 | 3442 | 3038 | 3445 | 2916 | 3107 | 2908 | 2851 |
| 113 | 2.28 | 0.91 | 2.51 | 0.01 | 0.4723 | Mme | membrane metallo endopeptidase | 404 | 243 | 547 | 2 | 4 | 1 | 0 | 0 | 27 | 14 | 1 |
| 560 | 0.48 | 0.19 | 2.50 | 0.01 | 0.47406 | Igsf11 | immunoglobulin superfamily, member 11 | 63 | 81 | 123 | 587 | 735 | 423 | 725 | 862 | 968 | 974 | 624 |
| 320 | -0.59 | 0.24 | -2.50 | 0.01 | 0.47406 | Dtx1 | deltex 1, E3 ubiquitin ligase | 456 | 545 | 399 | 424 | 299 | 248 | 289 | 283 | 260 | 216 | 108 |
| 216 | -2.26 | 0.90 | -2.50 | 0.01 | 0.47406 | Sema3b | sema domain, immunoglobulin domain (I | 1042 | 540 | 760 | 8 | 3 | 8 | 9 | 0 | 5 | 0 | 1 |
| 1703 | -0.34 | 0.14 | -2.50 | 0.01 | 0.47803 | Septin8 | septin 8 | 3954 | 2998 | 3061 | 1336 | 1202 | 1018 | 1327 | 1043 | 1070 | 877 | 851 |
| 69450 | -0.14 | 0.05 | -2.50 | 0.01 | 0.48116 | Gnas | GNAS (guanine nucleotide binding protei | 123400 | 151911 | 150946 | 47955 | 48480 | 35154 | 45253 | 43461 | 43135 | 42269 | 31982 |
| 854 | 0.34 | 0.14 | 2.50 | 0.01 | 0.48116 | Sord | sorbitol dehydrogenase | 1108 | 1204 | 832 | 854 | 522 | 532 | 832 | 1182 | 761 | 1010 | 553 |
| 618 | -4.88 | 1.95 | -2.50 | 0.01 | 0.48116 | Spink1 | serine peptidase inhibitor, Kazal type 1 | 843 | 2843 | 2941 | 120 | 24 | 0 | 9 | 0 | 1 | 0 | 18 |
| 2922 | 0.23 | 0.09 | 2.50 | 0.01 | 0.48227 | Snw1 | SNW domain containing 1 | 3217 | 2920 | 3037 | 2800 | 2621 | 2438 | 2693 | 3543 | 2709 | 3250 | 2911 |
| 2570 | -0.20 | 0.08 | -2.49 | 0.01 | 0.48433 | Arhgef2 | rho/rac guanine nucleotide exchange fac | 4128 | 3799 | 3373 | 2453 | 1986 | 2013 | 2616 | 2153 | 1926 | 2205 | 1621 |
| 51 | -2.54 | 1.02 | -2.49 | 0.01 | 0.48707 | Calhm5 | calcium homeostasis modulator family me | 266 | 157 | 95 | 9 | 13 | 5 | 12 | 0 | 3 | 0 | 4 |
| 29 | -4.13 | 1.66 | -2.49 | 0.01 | 0.48707 | Cd2 | CD2 antigen | 244 | 54 | 5 | 0 | 4 | 6 | 8 | 0 | 0 | 0 | 0 |
| 61 | -2.10 | 0.84 | -2.49 | 0.01 | 0.4881 | 4930452B06Ril | RIKEN cDNA 4930452B06 gene | 133 | 158 | 156 | 38 | 69 | 54 | 7 | 0 | 22 | 3 | 26 |
| 8 | -4.06 | 1.63 | -2.48 | 0.01 | 0.49436 | Marchf11 | membrane associated ring-CH-type finge | 9 | 15 | 36 | 7 | 2 | 13 | 7 | 0 | 0 | 0 | 2 |
| 488 | -0.83 | 0.33 | -2.48 | 0.01 | 0.49516 | F2r | coagulation factor II (thrombin) receptor | 2023 | 1368 | 1725 | 47 | 31 | 29 | 56 | 22 | 23 | 14 | 29 |
| 1719 | 0.20 | 0.08 | 2.48 | 0.01 | 0.50088 | Abhd17c | abhydrolase domain containing 17C | 957 | 1228 | 1047 | 2028 | 1867 | 1668 | 1717 | 2417 | 2030 | 2007 | 1941 |
| 72 | -2.65 | 1.07 | -2.48 | 0.01 | 0.50088 | Nox4 | NADPH oxidase 4 | 467 | 116 | 169 | 8 | 12 | 4 | 5 | 0 | 2 | 0 | 3 |
| 174 | -1.27 | 0.51 | -2.48 | 0.01 | 0.50257 | Fgf14 | fibroblast growth factor 14 | 400 | 624 | 491 | 61 | 86 | 65 | 61 | 7 | 31 | 58 | 25 |
| 214 | -2.41 | 0.97 | -2.47 | 0.01 | 0.50528 | Cbr2 | carbonyl reductase 2 | 1160 | 586 | 583 | 0 | 2 | 8 | 12 | 0 | 0 | 0 | 4 |
| 1810 | -0.37 | 0.15 | -2.47 | 0.01 | 0.50565 | Fgf12 | fibroblast growth factor 12 | 1998 | 2160 | 2600 | 1953 | 1986 | 1625 | 1812 | 1036 | 1753 | 1529 | 1462 |
| 36 | -2.72 | 1.10 | -2.47 | 0.01 | 0.50871 | Rin1 | Ras and Rab interactor 1 | 120 | 103 | 113 | 3 | 3 | 28 | 7 | 0 | 0 | 0 | 14 |
| 89 | 2.39 | 0.97 | 2.47 | 0.01 | 0.51105 | Hey1 | hairy/enhancer-of-split related with YRPW | 320 | 256 | 319 | 0 | 7 | 3 | 1 | 42 | 5 | 19 | 2 |
| 506 | -0.52 | 0.21 | -2.47 | 0.01 | 0.51255 | Litaf | LPS-induced TN factor | 1630 | 1314 | 1303 | 277 | 203 | 94 | 206 | 195 | 152 | 131 | 65 |
| 205 | -2.14 | 0.87 | -2.46 | 0.01 | 0.515 | Dram1 | DNA-damage regulated autophagy modu | 927 | 718 | 548 | 16 | 3 | 13 | 12 | 13 | 1 | 0 | 0 |
| 168 | -0.91 | 0.37 | -2.46 | 0.01 | 0.51632 | Mapk4 | mitogen-activated protein kinase 4 | 321 | 776 | 306 | 83 | 65 | 64 | 83 | 37 | 40 | 40 | 38 |
| 78 | 1.29 | 0.52 | 2.46 | 0.01 | 0.52495 | Gpx7 | glutathione peroxidase 7 | 229 | 144 | 193 | 13 | 50 | 15 | 13 | 41 | 50 | 76 | 40 |
| 7 | -3.92 | 1.60 | -2.46 | 0.01 | 0.52495 | Gm10778 | predicted gene 10778 | 19 | 13 | 17 | 6 | 4 | 8 | 7 | 0 | 0 | 0 | 2 |
| 3888 | -0.17 | 0.07 | -2.46 | 0.01 | 0.52631 | Pip5k1c | phosphatidylinositol-4-phosphate 5-kinas | 4879 | 4372 | 4581 | 3832 | 3992 | 3292 | 4198 | 3113 | 3701 | 3701 | 3109 |
| 1704 | -1.51 | 0.62 | -2.45 | 0.01 | 0.52667 | Cilp | cartilage intermediate layer protein, nucle | 9640 | 2403 | 5154 | 222 | 455 | 323 | 143 | 90 | 176 | 32 | 106 |
| 1009 | -0.32 | 0.13 | -2.45 | 0.01 | 0.53107 | Elmsan1 | ELM2 and Myb/SANT-like domain contair | 1544 | 1273 | 1321 | 852 | 1018 | 971 | 1012 | 652 | 928 | 814 | 715 |
| 52 | -3.51 | 1.43 | -2.45 | 0.01 | 0.53107 | Gpr132 | G protein-coupled receptor 132 | 411 | 90 | 53 | 8 | 0 | 2 | 3 | 0 | 0 | 0 | 0 |
| 15 | -2.63 | 1.07 | -2.45 | 0.01 | 0.53138 | Nfyc-ps | | 11 | 7 | 35 | 21 | 27 | 20 | 23 | 0 | 3 | 5 | 16 |
| 321 | -0.65 | 0.27 | -2.45 | 0.01 | 0.53157 | Ankrd6 | ankyrin repeat domain 6 | 667 | 1002 | 900 | 102 | 131 | 193 | 163 | 81 | 97 | 79 | 116 |
| 20 | -3.66 | 1.49 | -2.45 | 0.01 | 0.53376 | Muc20 | mucin 20 | 75 | 11 | 104 | 9 | 10 | 8 | 2 | 0 | 0 | 0 | 6 |
| 210 | -2.23 | 0.91 | -2.45 | 0.01 | 0.53555 | Il1b | interleukin 1 beta | 1372 | 377 | 522 | 4 | 8 | 3 | 16 | 5 | 1 | 0 | 0 |

| | | | | | | | | | | | | | | | | | | |
|---|---|---|---|---|---|---|---|---|---|---|---|---|---|---|---|---|---|---|
| 5648 | -0.14 | 0.06 | -2.44 | 0.01 | 0.5356 | Rpl32 | ribosomal protein L32 | 9981 | 9060 | 9834 | 4435 | 4993 | 3214 | 4822 | 4157 | 4156 | 4519 | 2955 |
| 14658 | 0.16 | 0.06 | 2.44 | 0.01 | 0.53653 | Kif5b | kinesin family member 5B | 9412 | 9066 | 9058 | 16675 | 15942 | 14349 | 16232 | 19814 | 18181 | 16186 | 16323 |
| 101 | -2.21 | 0.90 | -2.44 | 0.01 | 0.53705 | Kcnk2 | potassium channel, subfamily K, member | 318 | 282 | 349 | 26 | 33 | 23 | 25 | 48 | 2 | 0 | 3 |
| 4 | 6.96 | 2.85 | 2.44 | 0.01 | 0.53786 | Gm49539 | | 0 | 2 | 12 | 0 | 0 | 3 | 0 | 15 | 0 | 13 | 1 |
| 3789 | -0.23 | 0.09 | -2.44 | 0.01 | 0.53872 | Thrap3 | thyroid hormone receptor associated prot | 3445 | 3190 | 3569 | 3841 | 4187 | 4892 | 3956 | 2733 | 4027 | 3296 | 4544 |
| 37 | -2.34 | 0.96 | -2.44 | 0.01 | 0.5419 | Vwa3b | von Willebrand factor A domain containin | 51 | 120 | 16 | 58 | 53 | 19 | 46 | 13 | 9 | 0 | 21 |
| 476 | 4.51 | 1.85 | 2.44 | 0.01 | 0.5419 | Cnn2 | calponin 2 | 2605 | 1214 | 1374 | 0 | 0 | 0 | 2 | 35 | 9 | 0 | 2 |
| 4 | 8.56 | 3.51 | 2.44 | 0.01 | 0.5419 | Ct55 | cancer/testis antigen 55 | 5 | 0 | 0 | 0 | 0 | 3 | 0 | 29 | 9 | 0 | 0 |
| 735 | -0.38 | 0.16 | -2.43 | 0.01 | 0.54719 | Slc20a2 | solute carrier family 20, member 2 | 634 | 532 | 580 | 868 | 813 | 815 | 1083 | 575 | 842 | 781 | 569 |
| 111 | -0.81 | 0.33 | -2.43 | 0.01 | 0.54719 | Arhgef4 | Rho guanine nucleotide exchange factor | 129 | 108 | 160 | 127 | 132 | 103 | 162 | 83 | 72 | 67 | 73 |
| 16 | 4.24 | 1.74 | 2.43 | 0.02 | 0.54849 | Ankrd55 | ankyrin repeat domain 55 | 37 | 77 | 42 | 0 | 0 | 1 | 0 | 0 | 13 | 9 | 2 |
| 9613 | 0.13 | 0.05 | 2.43 | 0.02 | 0.5492 | Eif1 | eukaryotic translation initiation factor 1 | 8648 | 8739 | 8924 | 9779 | 10001 | 8253 | 9919 | 10369 | 10471 | 11499 | 9144 |
| 990 | -0.49 | 0.20 | -2.43 | 0.02 | 0.5492 | Sult1d1 | sulfotransferase family 1D, member 1 | 847 | 1634 | 1101 | 1072 | 1035 | 1406 | 649 | 767 | 891 | 303 | 1184 |
| 1354 | -1.31 | 0.54 | -2.43 | 0.02 | 0.5492 | Rcn3 | reticulocalbin 3, EF-hand calcium binding | 6044 | 3722 | 4150 | 152 | 168 | 120 | 238 | 155 | 77 | 27 | 38 |
| 199 | -1.55 | 0.64 | -2.43 | 0.02 | 0.5492 | Wnt5a | wingless-type MMTV integration site famil | 1080 | 443 | 518 | 19 | 27 | 41 | 14 | 0 | 10 | 27 | 8 |
| 624 | -0.49 | 0.20 | -2.43 | 0.02 | 0.55096 | Dock1 | dedicator of cytokinesis 1 | 1854 | 1839 | 1736 | 232 | 219 | 183 | 201 | 165 | 198 | 104 | 132 |
| 1870 | -0.39 | 0.16 | -2.43 | 0.02 | 0.55186 | Selenon | selenoprotein N | 3655 | 3464 | 3239 | 1421 | 1382 | 1184 | 1801 | 755 | 1287 | 1285 | 1094 |
| 5 | -6.56 | 2.70 | -2.42 | 0.02 | 0.55266 | Gm50022 | | 0 | 3 | 6 | 30 | 0 | 2 | 9 | 0 | 0 | 0 | 2 |
| 1103 | -0.44 | 0.18 | -2.42 | 0.02 | 0.55311 | Ptprk | protein tyrosine phosphatase, receptor ty | 3651 | 3786 | 3350 | 191 | 175 | 277 | 135 | 129 | 158 | 89 | 196 |
| 8745 | 0.17 | 0.07 | 2.42 | 0.02 | 0.55322 | Sars | seryl-aminoacyl-tRNA synthetase | 5216 | 4870 | 5770 | 9297 | 10785 | 7732 | 9909 | 12030 | 11276 | 11103 | 8210 |
| 364 | -0.49 | 0.20 | -2.42 | 0.02 | 0.55551 | Dnase2a | deoxyribonuclease II alpha | 651 | 542 | 621 | 365 | 322 | 304 | 285 | 262 | 200 | 214 | 234 |
| 67 | -2.13 | 0.88 | -2.42 | 0.02 | 0.55551 | Lck | lymphocyte protein tyrosine kinase | 252 | 139 | 248 | 39 | 6 | 15 | 16 | 19 | 1 | 3 | 1 |
| 179 | -0.64 | 0.26 | -2.42 | 0.02 | 0.55583 | Hykk | hydroxylysine kinase 1 | 246 | 255 | 236 | 196 | 166 | 171 | 219 | 98 | 128 | 119 | 136 |
| 27 | -2.34 | 0.97 | -2.42 | 0.02 | 0.55583 | Gm16897 | | 37 | 6 | 23 | 32 | 46 | 36 | 60 | 0 | 22 | 5 | 28 |
| 7278 | 0.21 | 0.09 | 2.42 | 0.02 | 0.55613 | Atp6v1e1 | ATPase, H+ transporting, lysosomal V1 su | 3518 | 4234 | 4438 | 8579 | 8146 | 6103 | 8576 | 11254 | 9168 | 8685 | 7357 |
| 89 | -2.28 | 0.94 | -2.42 | 0.02 | 0.559 | Bmp7 | bone morphogenetic protein 7 | 117 | 389 | 378 | 13 | 17 | 14 | 20 | 0 | 7 | 0 | 19 |
| 139 | -2.19 | 0.91 | -2.42 | 0.02 | 0.559 | Mertk | MER proto-oncogene tyrosine kinase | 746 | 410 | 336 | 8 | 9 | 4 | 11 | 0 | 2 | 0 | 6 |
| 562 | -0.65 | 0.27 | -2.42 | 0.02 | 0.5595 | Klb | klotho beta | 1950 | 1603 | 1810 | 142 | 105 | 87 | 152 | 60 | 58 | 158 | 53 |
| 46 | -2.40 | 1.00 | -2.41 | 0.02 | 0.56155 | Slc1a1 | solute carrier family 1 (neuronal/epithelial | 97 | 191 | 153 | 5 | 22 | 12 | 16 | 0 | 8 | 0 | 5 |
| 5582 | -0.36 | 0.15 | -2.41 | 0.02 | 0.5635 | Dusp1 | dual specificity phosphatase 1 | 19196 | 10572 | 12120 | 634 | 629 | 7793 | 1540 | 346 | 661 | 1106 | 6809 |
| 18184 | 0.24 | 0.10 | 2.41 | 0.02 | 0.56484 | Camk2n1 | calcium/calmodulin-dependent protein kir | 7090 | 9975 | 9297 | 19746 | 22084 | 15278 | 22455 | 27339 | 23968 | 25102 | 17686 |
| 13813 | 0.29 | 0.12 | 2.41 | 0.02 | 0.56484 | Ftl1 | ferritin light polypeptide 1 | 8545 | 6984 | 8270 | 15258 | 15920 | 9929 | 16139 | 24812 | 16777 | 18328 | 10978 |
| 287 | -0.61 | 0.25 | -2.41 | 0.02 | 0.56484 | Ldlrad3 | low density lipoprotein receptor class A d | 664 | 521 | 579 | 201 | 177 | 218 | 232 | 126 | 81 | 152 | 202 |
| 6 | -6.73 | 2.80 | -2.41 | 0.02 | 0.56484 | Stkld1 | serine/threonine kinase-like domain conta | 0 | 0 | 4 | 13 | 0 | 19 | 9 | 0 | 0 | 0 | 19 |
| 237 | -2.26 | 0.94 | -2.41 | 0.02 | 0.56544 | Arhgap30 | Rho GTPase activating protein 30 | 1499 | 551 | 514 | 3 | 10 | 6 | 15 | 0 | 1 | 0 | 12 |
| 29 | -3.57 | 1.48 | -2.41 | 0.02 | 0.56544 | Gbgt1 | globoside alpha-1,3-N-acetylgalactosami | 117 | 47 | 116 | 10 | 0 | 12 | 10 | 0 | 0 | 0 | 5 |
| 132 | 3.01 | 1.25 | 2.41 | 0.02 | 0.56544 | Ndufa4l2 | Ndufa4, mitochondrial complex associate | 638 | 325 | 477 | 0 | 0 | 1 | 0 | 0 | 6 | 0 | 5 |
| 58 | 2.57 | 1.07 | 2.41 | 0.02 | 0.56544 | Tnfsf13b | tumor necrosis factor (ligand) superfamily | 274 | 112 | 213 | 0 | 1 | 6 | 0 | 0 | 12 | 9 | 8 |
| 7 | -4.14 | 1.72 | -2.41 | 0.02 | 0.56566 | 4932435O22Rik | | 33 | 8 | 8 | 7 | 11 | 4 | 4 | 0 | 0 | 0 | 2 |
| 266 | -5.32 | 2.21 | -2.40 | 0.02 | 0.56777 | Try10 | trypsin 10 | 212 | 1409 | 1243 | 25 | 9 | 0 | 22 | 0 | 0 | 0 | 9 |
| 3786 | 0.19 | 0.08 | 2.39 | 0.02 | 0.58119 | Atp11b | ATPase, class VI, type 11B | 3285 | 3769 | 2962 | 4001 | 3569 | 4089 | 3127 | 4665 | 3705 | 3939 | 4534 |
| 1397 | -0.61 | 0.26 | -2.40 | 0.02 | 0.58119 | Cd93 | CD93 antigen | 4990 | 4738 | 4201 | 281 | 244 | 117 | 227 | 231 | 146 | 72 | 117 |
| 16 | -3.58 | 1.50 | -2.39 | 0.02 | 0.58119 | Adgrb2 | adhesion G protein-coupled receptor B2 | 107 | 24 | 17 | 11 | 1 | 4 | 10 | 0 | 0 | 0 | 2 |
| 180 | -2.22 | 0.93 | -2.39 | 0.02 | 0.58119 | Pgm5 | phosphoglucomutase 5 | 847 | 590 | 500 | 18 | 3 | 11 | 4 | 0 | 5 | 0 | 2 |
| 68 | -1.80 | 0.75 | -2.39 | 0.02 | 0.58139 | Gm16011 | | 164 | 235 | 165 | 23 | 23 | 71 | 15 | 0 | 16 | 3 | 28 |
| 2279 | -0.37 | 0.16 | -2.39 | 0.02 | 0.58234 | Tmem132b | transmembrane protein 132B | 1441 | 2023 | 1875 | 2901 | 2378 | 2359 | 3490 | 2237 | 1913 | 2679 | 1768 |
| 486 | -0.48 | 0.20 | -2.39 | 0.02 | 0.58234 | Usb1 | U6 snRNA biogenesis 1 | 552 | 519 | 566 | 602 | 513 | 491 | 545 | 280 | 449 | 441 | 385 |
| 227 | -0.63 | 0.26 | -2.39 | 0.02 | 0.58234 | Sowahb | sosondowah ankyrin repeat domain famil | 165 | 237 | 186 | 423 | 285 | 176 | 291 | 178 | 227 | 176 | 149 |
| 151 | -2.04 | 0.85 | -2.39 | 0.02 | 0.58234 | Minar1 | membrane integral NOTCH2 associated r | 412 | 617 | 524 | 6 | 12 | 30 | 32 | 0 | 9 | 0 | 19 |
| 8 | -3.88 | 1.62 | -2.39 | 0.02 | 0.58234 | Klhl31 | kelch-like 31 | 2 | 0 | 4 | 18 | 20 | 19 | 13 | 0 | 6 | 8 | 0 |
| 46 | -3.00 | 1.26 | -2.39 | 0.02 | 0.58234 | Dennd2c | DENN/MADD domain containing 2C | 166 | 130 | 167 | 0 | 9 | 14 | 9 | 0 | 0 | 0 | 9 |
| 303 | -2.88 | 1.21 | -2.39 | 0.02 | 0.58234 | Srpx | sushi-repeat-containing protein | 1546 | 820 | 956 | 0 | 1 | 2 | 4 | 0 | 0 | 0 | 0 |
| 1387 | 3.37 | 1.41 | 2.39 | 0.02 | 0.58234 | Ccdc80 | coiled-coil domain containing 80 | 9279 | 2924 | 3032 | 0 | 0 | 2 | 0 | 0 | 8 | 5 | 5 |
| 245 | 0.54 | 0.23 | 2.39 | 0.02 | 0.58385 | Cdkn2b | cyclin dependent kinase inhibitor 2B | 197 | 135 | 130 | 242 | 253 | 168 | 250 | 372 | 330 | 362 | 260 |
| 360 | -0.48 | 0.20 | -2.38 | 0.02 | 0.58613 | Kif5a | kinesin family member 5A | 572 | 606 | 612 | 276 | 298 | 370 | 319 | 184 | 268 | 236 | 216 |
| 336 | -0.55 | 0.23 | -2.39 | 0.02 | 0.58613 | Gfod1 | glucose-fructose oxidoreductase domain | 806 | 743 | 920 | 183 | 192 | 149 | 208 | 156 | 121 | 102 | 118 |
| 233 | -0.69 | 0.29 | -2.38 | 0.02 | 0.58613 | Cpne8 | copine VIII | 581 | 645 | 615 | 151 | 76 | 119 | 101 | 101 | 55 | 71 | 53 |
| 366 | -3.64 | 1.53 | -2.38 | 0.02 | 0.58613 | Ggta1 | glycoprotein galactosyltransferase alpha | 1741 | 834 | 1424 | 11 | 6 | 1 | 5 | 0 | 0 | 0 | 2 |
| 25 | -2.56 | 1.07 | -2.38 | 0.02 | 0.58613 | Tstd1 | thiosulfate sulfurtransferase (rhodanese)- | 66 | 63 | 74 | 11 | 9 | 28 | 4 | 0 | 2 | 0 | 14 |
| 7971 | 0.16 | 0.07 | 2.38 | 0.02 | 0.58681 | Ufm1 | ubiquitin-fold modifier 1 | 2889 | 3171 | 3434 | 8596 | 9916 | 8778 | 9582 | 10813 | 10626 | 9898 | 9976 |

| | | | | | | | | | | | | | | | | | |
|---|---|---|---|---|---|---|---|---|---|---|---|---|---|---|---|---|---|
| 27 | -3.83 | 1.61 | -2.38 | 0.02 | 0.58681 | Lrrc75b | leucine rich repeat containing 75B | 122 | 100 | 62 | 11 | 5 | 1 | 0 | 0 | 0 | 0 | 0 |
| 4 | 7.58 | 3.18 | 2.38 | 0.02 | 0.58762 | Mir132 | | 4 | 0 | 0 | 0 | 0 | 8 | 0 | 16 | 8 | 0 | 4 |
| 545 | -0.75 | 0.32 | -2.38 | 0.02 | 0.59557 | Camk1d | calcium/calmodulin-dependent protein kir | 1260 | 1071 | 1028 | 327 | 631 | 267 | 389 | 161 | 536 | 148 | 178 |
| 911 | -0.34 | 0.14 | -2.37 | 0.02 | 0.59768 | Lonrf2 | LON peptidase N-terminal domain and rir | 1047 | 1527 | 1431 | 943 | 834 | 704 | 852 | 812 | 769 | 634 | 466 |
| 654 | -0.40 | 0.17 | -2.37 | 0.02 | 0.59768 | Zc3hav1l | zinc finger CCCH-type, antiviral 1-like | 697 | 609 | 581 | 896 | 738 | 656 | 736 | 629 | 652 | 479 | 520 |
| 1795 | 0.24 | 0.10 | 2.37 | 0.02 | 0.60549 | Ubr7 | ubiquitin protein ligase E3 component n-r | 1591 | 1536 | 1488 | 1754 | 1900 | 1571 | 1702 | 2211 | 1920 | 2294 | 1781 |
| 139 | -0.84 | 0.36 | -2.37 | 0.02 | 0.60549 | Trnp1 | TMF1-regulated nuclear protein 1 | 376 | 56 | 49 | 58 | 30 | 117 | 483 | 43 | 14 | 250 | 59 |
| 1334 | -2.70 | 1.14 | -2.37 | 0.02 | 0.60549 | Mgll | monoglyceride lipase | 3933 | 5532 | 5138 | 23 | 3 | 17 | 19 | 0 | 4 | 0 | 9 |
| 6 | -6.14 | 2.59 | -2.37 | 0.02 | 0.60549 | Gm43081 | | 0 | 0 | 25 | 7 | 0 | 10 | 15 | 0 | 1 | 0 | 5 |
| 634 | -0.41 | 0.17 | -2.37 | 0.02 | 0.60727 | Tubb4a | tubulin, beta 4A class IVA | 1181 | 1194 | 1235 | 560 | 487 | 369 | 477 | 506 | 374 | 376 | 213 |
| 331 | -0.81 | 0.34 | -2.36 | 0.02 | 0.60962 | Nyap1 | neuronal tyrosine-phosphorylated phospl | 907 | 1123 | 932 | 64 | 139 | 124 | 101 | 92 | 45 | 48 | 65 |
| 6 | 5.64 | 2.39 | 2.36 | 0.02 | 0.61092 | Pou4f1 | POU domain, class 4, transcription factor | 0 | 0 | 0 | 0 | 0 | 1 | 0 | 30 | 3 | 19 | 8 |
| 381 | -0.65 | 0.27 | -2.36 | 0.02 | 0.61356 | Cdyl | chromodomain protein, Y chromosome-lil | 447 | 360 | 497 | 497 | 375 | 379 | 490 | 221 | 415 | 210 | 297 |
| 3675 | 0.17 | 0.07 | 2.36 | 0.02 | 0.61398 | Trim37 | tripartite motif-containing 37 | 2555 | 2567 | 2568 | 3935 | 3955 | 3265 | 4286 | 4735 | 4181 | 4591 | 3789 |
| 2386 | 0.31 | 0.13 | 2.36 | 0.02 | 0.61398 | Srp9 | signal recognition particle 9 | 1484 | 1833 | 1861 | 2998 | 2305 | 2024 | 1999 | 4377 | 2515 | 2518 | 2329 |
| 140 | -0.93 | 0.39 | -2.36 | 0.02 | 0.61398 | Zfp101 | zinc finger protein 101 | 168 | 175 | 155 | 144 | 109 | 277 | 102 | 102 | 38 | 31 | 233 |
| 70 | -1.05 | 0.45 | -2.36 | 0.02 | 0.61789 | Rnf43 | ring finger protein 43 | 86 | 97 | 137 | 105 | 63 | 43 | 91 | 57 | 36 | 28 | 24 |
| 12 | -3.62 | 1.54 | -2.35 | 0.02 | 0.6208 | Cbln3 | cerebellin 3 precursor protein | 2 | 5 | 19 | 4 | 9 | 18 | 51 | 0 | 2 | 0 | 21 |
| 239 | -4.41 | 1.87 | -2.35 | 0.02 | 0.6208 | Zfp366 | zinc finger protein 366 | 919 | 962 | 722 | 0 | 7 | 2 | 18 | 0 | 0 | 0 | 0 |
| 5140 | 0.18 | 0.08 | 2.35 | 0.02 | 0.6243 | Hspd1 | heat shock protein 1 (chaperonin) | 4980 | 4624 | 5120 | 4653 | 4325 | 5735 | 4916 | 5590 | 5140 | 5061 | 6394 |
| 3967 | -0.26 | 0.11 | -2.35 | 0.02 | 0.6243 | Prox1 | prospero homeobox 1 | 5469 | 7636 | 7011 | 3554 | 2951 | 2903 | 3382 | 2777 | 3084 | 2443 | 2427 |
| 274 | -0.94 | 0.40 | -2.35 | 0.02 | 0.6243 | Acer2 | alkaline ceramidase 2 | 760 | 857 | 695 | 132 | 104 | 91 | 128 | 25 | 84 | 45 | 95 |
| 606 | -0.71 | 0.30 | -2.35 | 0.02 | 0.6243 | Ctsh | cathepsin H | 2633 | 1248 | 2125 | 123 | 108 | 61 | 112 | 29 | 84 | 89 | 56 |
| 34 | 1.44 | 0.62 | 2.35 | 0.02 | 0.6243 | Glra1 | glycine receptor, alpha 1 subunit | 2 | 6 | 0 | 32 | 11 | 28 | 27 | 89 | 28 | 68 | 86 |
| 9956 | -2.92 | 1.24 | -2.35 | 0.02 | 0.6243 | Col1a1 | collagen, type I, alpha 1 | 56890 | 20714 | 31880 | 4 | 9 | 1 | 19 | 0 | 1 | 0 | 2 |
| 153 | -2.35 | 1.00 | -2.35 | 0.02 | 0.6243 | Dkk3 | dickkopf WNT signaling pathway inhibitor | 1132 | 392 | 114 | 18 | 5 | 4 | 7 | 0 | 2 | 0 | 4 |
| 67 | 3.27 | 1.39 | 2.35 | 0.02 | 0.6243 | Klra2 | killer cell lectin-like receptor, subfamily A, | 280 | 75 | 369 | 0 | 0 | 0 | 0 | 0 | 6 | 0 | 3 |
| 69 | -2.21 | 0.94 | -2.35 | 0.02 | 0.62674 | Gprin2 | G protein regulated inducer of neurite out | 139 | 297 | 226 | 35 | 15 | 9 | 14 | 0 | 11 | 0 | 8 |
| 199 | 0.69 | 0.30 | 2.34 | 0.02 | 0.62674 | Pop1 | processing of precursor 1, ribonuclease F | 167 | 236 | 244 | 137 | 138 | 201 | 119 | 266 | 174 | 268 | 241 |
| 515 | -0.47 | 0.20 | -2.34 | 0.02 | 0.62742 | Vasn | vasorin | 1032 | 527 | 829 | 384 | 717 | 317 | 494 | 306 | 463 | 366 | 227 |
| 12 | -3.37 | 1.44 | -2.34 | 0.02 | 0.62786 | Ttc36 | tetratricopeptide repeat domain 36 | 3 | 23 | 14 | 23 | 27 | 8 | 16 | 0 | 4 | 0 | 19 |
| 2489 | 0.23 | 0.10 | 2.34 | 0.02 | 0.62797 | Tars | threonyl-tRNA synthetase | 1899 | 1899 | 2158 | 2065 | 2784 | 2825 | 2232 | 2789 | 2973 | 2874 | 2884 |
| 2450 | 0.21 | 0.09 | 2.34 | 0.02 | 0.63061 | Lrrc47 | leucine rich repeat containing 47 | 2203 | 2559 | 2338 | 2193 | 2328 | 2319 | 2334 | 2987 | 2663 | 2556 | 2474 |
| 55 | -2.77 | 1.18 | -2.34 | 0.02 | 0.63371 | Zfp423 | zinc finger protein 423 | 243 | 168 | 167 | 0 | 5 | 14 | 6 | 0 | 0 | 0 | 5 |
| 3319 | 0.20 | 0.08 | 2.34 | 0.02 | 0.6341 | Ppp1r2 | protein phosphatase 1, regulatory inhibito | 3369 | 2890 | 3124 | 3188 | 3312 | 2580 | 3560 | 3859 | 3296 | 4287 | 3046 |
| 1665 | 0.25 | 0.11 | 2.34 | 0.02 | 0.6341 | Smim1 | small integral membrane protein 1 | 888 | 920 | 968 | 1651 | 1850 | 1694 | 1883 | 2310 | 1998 | 2307 | 1845 |
| 780 | -0.49 | 0.21 | -2.34 | 0.02 | 0.6341 | Cadps2 | Ca2+-dependent activator protein for secr | 736 | 972 | 742 | 1081 | 954 | 684 | 878 | 481 | 774 | 650 | 627 |
| 26 | 3.90 | 1.67 | 2.34 | 0.02 | 0.6341 | Sh3bp1 | SH3-domain binding protein 1 | 154 | 43 | 56 | 0 | 0 | 2 | 0 | 21 | 0 | 10 | 1 |
| 2097 | 0.26 | 0.11 | 2.33 | 0.02 | 0.63574 | Fpgt | fucose-1-phosphate guanylyltransferase | 1048 | 1258 | 1060 | 2068 | 1878 | 2781 | 2183 | 3097 | 2106 | 2221 | 3363 |
| 3154 | 0.21 | 0.09 | 2.33 | 0.02 | 0.63833 | Yipf5 | Yip1 domain family, member 5 | 2038 | 2107 | 2219 | 2909 | 3734 | 3227 | 3258 | 4048 | 3791 | 3810 | 3551 |
| 43 | 2.48 | 1.06 | 2.33 | 0.02 | 0.63833 | Crp | C-reactive protein, pentraxin-related | 93 | 108 | 219 | 5 | 0 | 0 | 4 | 13 | 4 | 18 | 7 |
| 19 | -3.53 | 1.52 | -2.33 | 0.02 | 0.64448 | Kctd16 | potassium channel tetramerisation domai | 26 | 75 | 34 | 11 | 0 | 19 | 35 | 0 | 2 | 0 | 3 |
| 9088 | 0.17 | 0.07 | 2.32 | 0.02 | 0.6532 | Gars | glycyl-tRNA synthetase | 7056 | 7677 | 7778 | 8518 | 10558 | 7577 | 9856 | 11050 | 11155 | 10288 | 8453 |
| 1770 | 0.20 | 0.09 | 2.32 | 0.02 | 0.6532 | Rsl1d1 | ribosomal L1 domain containing 1 | 1898 | 1765 | 1733 | 1384 | 1598 | 1926 | 1631 | 1730 | 1764 | 1775 | 2264 |
| 5058 | -0.22 | 0.10 | -2.32 | 0.02 | 0.66103 | Slc6a17 | solute carrier family 6 (neurotransmitter tra | 4417 | 4989 | 4597 | 5681 | 6532 | 3970 | 6067 | 4798 | 6169 | 5431 | 2993 |
| 925 | -0.37 | 0.16 | -2.32 | 0.02 | 0.66161 | Sema6d | sema domain, transmembrane domain (T | 2282 | 2073 | 2059 | 410 | 407 | 839 | 486 | 356 | 407 | 294 | 562 |
| 13 | -4.06 | 1.75 | -2.32 | 0.02 | 0.66296 | Gm7224 | | 15 | 41 | 6 | 26 | 0 | 13 | 26 | 0 | 0 | 0 | 15 |
| 3591 | 0.22 | 0.09 | 2.31 | 0.02 | 0.66664 | Ctdspl | CTD (carboxy-terminal domain, RNA poly | 2916 | 2821 | 3251 | 3148 | 4203 | 3347 | 3394 | 4455 | 4659 | 3954 | 3352 |
| 3056 | 0.19 | 0.08 | 2.31 | 0.02 | 0.66664 | Vamp4 | vesicle-associated membrane protein 4 | 2051 | 2219 | 2337 | 3056 | 3051 | 3129 | 3389 | 3649 | 3629 | 3543 | 3566 |
| 66 | 0.96 | 0.41 | 2.31 | 0.02 | 0.66664 | Ube2cbp | ubiquitin-conjugating enzyme E2C bindin | 63 | 64 | 39 | 44 | 53 | 47 | 43 | 108 | 95 | 98 | 68 |
| 13 | 3.37 | 1.46 | 2.31 | 0.02 | 0.66664 | Gm47246 | | 5 | 0 | 11 | 0 | 5 | 30 | 0 | 14 | 26 | 5 | 47 |
| 41 | -2.68 | 1.16 | -2.31 | 0.02 | 0.66695 | Khdrbs2 | KH domain containing, RNA binding, sigr | 109 | 159 | 63 | 27 | 8 | 30 | 35 | 0 | 18 | 0 | 5 |
| 12443 | -2.08 | 0.90 | -2.31 | 0.02 | 0.66797 | Cpa2 | carboxypeptidase A2, pancreatic | 17169 | 62871 | 52961 | 2390 | 260 | 202 | 214 | 80 | 108 | 35 | 580 |
| 5337 | -0.16 | 0.07 | -2.31 | 0.02 | 0.67044 | Stim1 | stromal interaction molecule 1 | 4000 | 3673 | 3938 | 7085 | 6815 | 4579 | 6380 | 6064 | 6437 | 5634 | 4106 |
| 784 | 0.28 | 0.12 | 2.31 | 0.02 | 0.67044 | Apobec1 | apolipoprotein B mRNA editing enzyme, c | 1041 | 655 | 898 | 486 | 585 | 927 | 713 | 589 | 727 | 791 | 1211 |
| 89 | 0.90 | 0.39 | 2.31 | 0.02 | 0.67044 | Asphd2 | aspartate beta-hydroxylase domain conta | 106 | 111 | 72 | 64 | 42 | 36 | 87 | 181 | 97 | 144 | 38 |
| 46 | -2.76 | 1.20 | -2.31 | 0.02 | 0.67044 | Gstt3 | glutathione S-transferase, theta 3 | 208 | 129 | 135 | 5 | 5 | 3 | 14 | 0 | 5 | 0 | 0 |
| 2910 | -0.26 | 0.11 | -2.31 | 0.02 | 0.67204 | Llgl2 | LLGL2 scribble cell polarity complex com | 2068 | 2780 | 2171 | 3248 | 3438 | 3830 | 3095 | 2507 | 2917 | 2736 | 3221 |
| 9944 | -0.13 | 0.05 | -2.30 | 0.02 | 0.67214 | Camk2b | calcium/calmodulin-dependent protein kir | 8870 | 11032 | 12087 | 10846 | 10780 | 9601 | 9128 | 9651 | 10407 | 8375 | 8604 |
| 767 | 0.33 | 0.14 | 2.30 | 0.02 | 0.6747 | Ttc38 | tetratricopeptide repeat domain 38 | 934 | 1017 | 748 | 770 | 641 | 525 | 596 | 904 | 753 | 947 | 603 |

| | | | | | | | | | | | | | | | | | | |
|---|---|---|---|---|---|---|---|---|---|---|---|---|---|---|---|---|---|---|
| 2227 | -0.21 | 0.09 | -2.30 | 0.02 | 0.67785 | Galnt10 | polypeptide N-acetylgalactosaminyltransf | 2099 | 1700 | 2117 | 2634 | 2602 | 2156 | 2562 | 2163 | 2327 | 2339 | 1796 |
| 75 | 2.95 | 1.28 | 2.30 | 0.02 | 0.67785 | Bin2 | bridging integrator 2 | 384 | 203 | 203 | 0 | 0 | 1 | 1 | 0 | 3 | 27 | 0 |
| 8 | -3.63 | 1.58 | -2.30 | 0.02 | 0.68092 | Hsh2d | hematopoietic SH2 domain containing | 34 | 3 | 10 | 15 | 6 | 6 | 10 | 0 | 1 | 0 | 3 |
| 7 | 3.66 | 1.59 | 2.30 | 0.02 | 0.68092 | Gm5491 | | 8 | 11 | 12 | 1 | 0 | 3 | 0 | 23 | 9 | 1 | 5 |
| 1072 | -0.28 | 0.12 | -2.30 | 0.02 | 0.68119 | Car8 | carbonic anhydrase 8 | 1256 | 1894 | 2643 | 917 | 1111 | 631 | 613 | 642 | 1048 | 501 | 538 |
| 96 | -2.40 | 1.04 | -2.30 | 0.02 | 0.68119 | Gabra3 | gamma-aminobutyric acid (GABA) A rece | 875 | 33 | 101 | 8 | 7 | 3 | 10 | 0 | 1 | 19 | 0 |
| 3931 | -2.84 | 1.24 | -2.30 | 0.02 | 0.68119 | Col6a2 | collagen, type VI, alpha 2 | 18484 | 10859 | 13858 | 9 | 12 | 4 | 7 | 1 | 0 | 0 | 9 |
| 23 | -3.51 | 1.53 | -2.30 | 0.02 | 0.68119 | Acy3 | aspartoacylase (aminoacylase) 3 | 85 | 68 | 81 | 11 | 0 | 5 | 4 | 1 | 0 | 0 | 0 |
| 14 | -3.01 | 1.31 | -2.30 | 0.02 | 0.68156 | Sbk2 | SH3-binding domain kinase family, memb | 8 | 44 | 10 | 24 | 12 | 21 | 7 | 0 | 2 | 0 | 22 |
| 3982 | 0.22 | 0.09 | 2.29 | 0.02 | 0.68277 | Naca | nascent polypeptide-associated complex | 5928 | 5480 | 6250 | 2939 | 3213 | 2592 | 3351 | 4108 | 3442 | 3543 | 2956 |
| 108 | -2.91 | 1.27 | -2.29 | 0.02 | 0.68361 | Pdlim2 | PDZ and LIM domain 2 | 674 | 194 | 316 | 0 | 2 | 3 | 2 | 0 | 0 | 0 | 0 |
| 20 | 3.32 | 1.45 | 2.29 | 0.02 | 0.68611 | Gm9926 | | 35 | 65 | 75 | 0 | 0 | 2 | 2 | 9 | 10 | 23 | 0 |
| 3 | -7.60 | 3.32 | -2.29 | 0.02 | 0.69012 | Gm42588 | | 0 | 0 | 12 | 8 | 0 | 4 | 10 | 0 | 0 | 0 | 2 |
| 1185 | -0.38 | 0.17 | -2.29 | 0.02 | 0.69472 | Zfp236 | zinc finger protein 236 | 1373 | 1504 | 1132 | 1079 | 899 | 1888 | 1195 | 709 | 903 | 705 | 1651 |
| 7 | -4.55 | 1.99 | -2.28 | 0.02 | 0.69472 | Gm973 | predicted gene 973 | 0 | 3 | 7 | 7 | 4 | 25 | 19 | 0 | 0 | 0 | 9 |
| 5 | -3.64 | 1.59 | -2.28 | 0.02 | 0.69472 | Npm3-ps1 | | 3 | 6 | 6 | 18 | 8 | 3 | 10 | 0 | 2 | 1 | 0 |
| 12 | 2.48 | 1.09 | 2.28 | 0.02 | 0.70114 | Styxl1 | serine/threonine/tyrosine interacting-like ' | 9 | 16 | 8 | 2 | 6 | 1 | 5 | 58 | 6 | 13 | 12 |
| 200 | -0.76 | 0.33 | -2.28 | 0.02 | 0.70249 | Peli2 | pellino 2 | 679 | 549 | 581 | 79 | 43 | 41 | 86 | 28 | 30 | 53 | 35 |
| 2416 | -4.67 | 2.05 | -2.28 | 0.02 | 0.70249 | Esm1 | endothelial cell-specific molecule 1 | 8123 | 9084 | 9330 | 0 | 2 | 5 | 35 | 0 | 0 | 0 | 0 |
| 69 | 3.14 | 1.38 | 2.28 | 0.02 | 0.70249 | Tlr13 | toll-like receptor 13 | 405 | 197 | 138 | 0 | 1 | 0 | 0 | 0 | 6 | 14 | 0 |
| 15 | -3.39 | 1.49 | -2.28 | 0.02 | 0.70536 | H1f11-ps | | 62 | 27 | 45 | 1 | 9 | 5 | 12 | 0 | 0 | 0 | 5 |
| 2440 | -0.26 | 0.11 | -2.28 | 0.02 | 0.70558 | Bace1 | beta-site APP cleaving enzyme 1 | 2926 | 3399 | 2563 | 2607 | 2329 | 2277 | 2573 | 1863 | 2170 | 2132 | 1997 |
| 13 | -3.01 | 1.32 | -2.28 | 0.02 | 0.70558 | Spats1 | spermatogenesis associated, serine-rich | 9 | 42 | 21 | 14 | 13 | 19 | 12 | 0 | 11 | 0 | 2 |
| 204 | -1.87 | 0.82 | -2.27 | 0.02 | 0.70626 | Ramp3 | receptor (calcitonin) activity modifying pro | 722 | 559 | 909 | 20 | 7 | 6 | 6 | 0 | 7 | 0 | 4 |
| 5783 | -0.15 | 0.07 | -2.27 | 0.02 | 0.70998 | Lpgat1 | lysophosphatidylglycerol acyltransferase | 3976 | 3700 | 3909 | 6611 | 6561 | 6996 | 7206 | 5487 | 6636 | 6497 | 6033 |
| 1809 | 0.23 | 0.10 | 2.27 | 0.02 | 0.70998 | Usp14 | ubiquitin specific peptidase 14 | 1892 | 1558 | 1912 | 1525 | 1523 | 1923 | 1720 | 1942 | 1772 | 1999 | 2128 |
| 4964 | 0.17 | 0.07 | 2.27 | 0.02 | 0.71109 | Ube3a | ubiquitin protein ligase E3A | 4036 | 4523 | 3859 | 5166 | 4804 | 5105 | 4758 | 6341 | 5519 | 4753 | 5745 |
| 67 | -2.01 | 0.88 | -2.27 | 0.02 | 0.71192 | St8sia2 | ST8 alpha-N-acetyl-neuraminide alpha-2 | 220 | 138 | 184 | 37 | 50 | 14 | 28 | 0 | 9 | 57 | 2 |
| 398 | 0.52 | 0.23 | 2.27 | 0.02 | 0.71247 | Fastkd1 | FAST kinase domains 1 | 323 | 204 | 222 | 340 | 295 | 500 | 373 | 560 | 502 | 589 | 471 |
| 1166 | -0.25 | 0.11 | -2.27 | 0.02 | 0.71446 | Csad | cysteine sulfinic acid decarboxylase | 1220 | 1182 | 1345 | 1191 | 1345 | 1106 | 1274 | 1143 | 1118 | 1063 | 840 |
| 21 | -2.88 | 1.27 | -2.27 | 0.02 | 0.71446 | Apob | apolipoprotein B | 5 | 31 | 19 | 24 | 59 | 20 | 32 | 0 | 40 | 0 | 5 |
| 41 | -3.25 | 1.43 | -2.27 | 0.02 | 0.71613 | Cadm2 | cell adhesion molecule 2 | 144 | 114 | 160 | 12 | 0 | 2 | 14 | 0 | 3 | 0 | 0 |
| 108 | -1.50 | 0.66 | -2.27 | 0.02 | 0.71674 | Slc24a3 | solute carrier family 24 (sodium/potassiur | 365 | 362 | 336 | 31 | 15 | 25 | 23 | 0 | 6 | 19 | 12 |
| 12567 | -0.24 | 0.11 | -2.26 | 0.02 | 0.71802 | Sez6l | seizure related 6 homolog like | 5137 | 8350 | 6857 | 16480 | 18303 | 13126 | 15845 | 13737 | 16265 | 13721 | 10418 |
| 1543 | 0.24 | 0.10 | 2.26 | 0.02 | 0.71802 | Slc35a1 | solute carrier family 35 (CMP-sialic acid tr | 1307 | 1364 | 1299 | 1580 | 1593 | 1136 | 1678 | 2040 | 1694 | 1772 | 1513 |
| 923 | -0.30 | 0.13 | -2.26 | 0.02 | 0.71802 | Gga3 | golgi associated, gamma adaptin ear con | 700 | 725 | 758 | 1046 | 894 | 1727 | 738 | 783 | 816 | 578 | 1390 |
| 865 | 0.36 | 0.16 | 2.26 | 0.02 | 0.71802 | Thoc2l | THO complex subunit 2-like | 611 | 796 | 754 | 403 | 404 | 2044 | 454 | 630 | 623 | 451 | 2350 |
| 420 | -0.41 | 0.18 | -2.26 | 0.02 | 0.71802 | Dclk2 | doublecortin-like kinase 2 | 668 | 865 | 720 | 251 | 221 | 657 | 228 | 217 | 190 | 135 | 472 |
| 1119 | -1.21 | 0.53 | -2.26 | 0.02 | 0.71802 | Ptgfrn | prostaglandin F2 receptor negative regula | 5336 | 3257 | 3198 | 126 | 74 | 76 | 76 | 19 | 72 | 22 | 49 |
| 224 | -3.90 | 1.73 | -2.26 | 0.02 | 0.71802 | Pdzd2 | PDZ domain containing 2 | 886 | 744 | 803 | 0 | 17 | 3 | 13 | 0 | 0 | 0 | 2 |
| 40 | 3.77 | 1.66 | 2.26 | 0.02 | 0.71802 | Pax5 | paired box 5 | 255 | 153 | 3 | 0 | 0 | 3 | 0 | 12 | 4 | 5 | 0 |
| 4466 | 0.18 | 0.08 | 2.26 | 0.02 | 0.71828 | Larp4b | La ribonucleoprotein domain family, mem | 2907 | 3618 | 3569 | 4451 | 4589 | 4835 | 4375 | 5824 | 4979 | 4562 | 5417 |
| 637 | -0.41 | 0.18 | -2.26 | 0.02 | 0.72102 | Hnf1b | HNF1 homeobox B | 651 | 695 | 807 | 544 | 596 | 878 | 743 | 361 | 594 | 497 | 641 |
| 34 | -2.81 | 1.25 | -2.26 | 0.02 | 0.72102 | Gm43681 | | 24 | 34 | 27 | 37 | 1 | 110 | 23 | 0 | 3 | 0 | 114 |
| 4 | -4.39 | 1.94 | -2.26 | 0.02 | 0.72102 | Gm4540 | | 1 | 3 | 3 | 16 | 2 | 8 | 9 | 0 | 0 | 0 | 2 |
| 19 | -1.96 | 0.87 | -2.26 | 0.02 | 0.72141 | Gm10167 | | 18 | 14 | 26 | 35 | 48 | 15 | 19 | 1 | 22 | 5 | 7 |
| 7 | -4.79 | 2.12 | -2.26 | 0.02 | 0.72192 | Gm31645 | | 0 | 0 | 17 | 18 | 13 | 5 | 8 | 0 | 14 | 0 | 0 |
| 3774 | 0.17 | 0.08 | 2.25 | 0.02 | 0.72372 | Cpeb1 | cytoplasmic polyadenylation element binc | 1059 | 1267 | 1096 | 4897 | 3625 | 5374 | 4030 | 5828 | 4093 | 4463 | 5780 |
| 2970 | 0.31 | 0.14 | 2.25 | 0.02 | 0.72372 | Rpl5 | ribosomal protein L5 | 4596 | 3788 | 5192 | 2042 | 2236 | 1929 | 2321 | 3194 | 2372 | 2451 | 2554 |
| 35 | -3.51 | 1.56 | -2.25 | 0.02 | 0.72398 | Lypd1 | Ly6/Plaur domain containing 1 | 182 | 117 | 30 | 0 | 11 | 7 | 29 | 0 | 4 | 0 | 0 |
| 96326 | 0.13 | 0.06 | 2.25 | 0.02 | 0.7246 | Atp2a2 | ATPase, Ca++ transporting, cardiac musc | 22687 | 21167 | 25951 | 126158 | 124165 | 90701 | 131708 | 133288 | 143475 | 141227 | 99065 |
| 713 | 0.42 | 0.19 | 2.25 | 0.02 | 0.7246 | Cybc1 | cytochrome b 245 chaperone 1 | 1070 | 934 | 895 | 510 | 490 | 600 | 496 | 922 | 545 | 754 | 621 |
| 50 | -2.78 | 1.23 | -2.25 | 0.02 | 0.7246 | Cldn9 | claudin 9 | 106 | 166 | 260 | 0 | 9 | 7 | 3 | 0 | 0 | 0 | 3 |
| 3135 | 0.16 | 0.07 | 2.25 | 0.02 | 0.72761 | Otud4 | OTU domain containing 4 | 2281 | 2493 | 2117 | 3318 | 2845 | 4232 | 2650 | 3689 | 3060 | 3133 | 4667 |
| 409 | 0.55 | 0.25 | 2.25 | 0.02 | 0.72828 | Psmb9 | proteasome (prosome, macropain) subun | 1330 | 614 | 868 | 85 | 225 | 164 | 217 | 202 | 275 | 344 | 173 |
| 4 | -7.14 | 3.17 | -2.25 | 0.02 | 0.72828 | Gm33228 | | 0 | 13 | 0 | 0 | 8 | 11 | 6 | 0 | 0 | 0 | 4 |
| 231 | -0.74 | 0.33 | -2.25 | 0.02 | 0.72915 | Prkg1 | protein kinase, cGMP-dependent, type I | 761 | 611 | 377 | 179 | 82 | 71 | 169 | 82 | 97 | 76 | 35 |
| 244 | -0.65 | 0.29 | -2.25 | 0.02 | 0.72943 | Vangl1 | VANGL planar cell polarity 1 | 583 | 644 | 660 | 134 | 80 | 148 | 122 | 86 | 84 | 49 | 90 |
| 1037 | 0.29 | 0.13 | 2.25 | 0.02 | 0.72992 | Rhou | ras homolog family member U | 3045 | 2652 | 3260 | 274 | 331 | 195 | 300 | 361 | 434 | 344 | 213 |
| 503 | -3.69 | 1.64 | -2.25 | 0.02 | 0.72992 | Lsp1 | lymphocyte specific 1 | 2938 | 1331 | 1214 | 16 | 0 | 4 | 31 | 3 | 0 | 0 | 0 |

| | | | | | | | | | | | | | | | | | | |
|---|---|---|---|---|---|---|---|---|---|---|---|---|---|---|---|---|---|---|
| 11 | -4.53 | 2.02 | -2.25 | 0.02 | 0.73074 | Gm44292 | | 9 | 1 | 26 | 7 | 0 | 24 | 9 | 0 | 0 | 0 | 47 |
| 89 | -0.80 | 0.36 | -2.24 | 0.02 | 0.73325 | Nyap2 | neuronal tyrosine-phophorylated phosph | 69 | 95 | 56 | 159 | 80 | 150 | 85 | 81 | 47 | 40 | 115 |
| 212 | 0.78 | 0.35 | 2.24 | 0.02 | 0.73409 | Sergef | secretion regulating guanine nucleotide e | 344 | 234 | 226 | 153 | 181 | 149 | 81 | 308 | 240 | 219 | 199 |
| 1391 | 0.28 | 0.12 | 2.24 | 0.02 | 0.73445 | Klhl12 | kelch-like 12 | 1042 | 1147 | 1109 | 1351 | 1557 | 1269 | 1242 | 1990 | 1532 | 1606 | 1458 |
| 80 | -2.84 | 1.27 | -2.24 | 0.02 | 0.73461 | Chst1 | carbohydrate sulfotransferase 1 | 539 | 170 | 133 | 11 | 1 | 3 | 10 | 0 | 0 | 0 | 9 |
| 12 | 3.95 | 1.76 | 2.24 | 0.02 | 0.73463 | Gm15494 | | 3 | 6 | 11 | 0 | 0 | 36 | 0 | 0 | 17 | 10 | 47 |
| 25766 | -0.19 | 0.09 | -2.24 | 0.03 | 0.74328 | Ece1 | endothelin converting enzyme 1 | 18754 | 20755 | 20085 | 31630 | 30391 | 22513 | 34756 | 23640 | 28728 | 32131 | 20037 |
| 784 | 0.32 | 0.14 | 2.24 | 0.03 | 0.74334 | Rab4a | RAB4A, member RAS oncogene family | 419 | 472 | 496 | 731 | 903 | 626 | 961 | 1064 | 1112 | 1096 | 742 |
| 5048 | -0.32 | 0.14 | -2.23 | 0.03 | 0.74422 | Timp2 | tissue inhibitor of metalloproteinase 2 | 25392 | 11615 | 11856 | 1065 | 849 | 593 | 1172 | 704 | 609 | 1117 | 560 |
| 7 | -5.38 | 2.41 | -2.24 | 0.03 | 0.74422 | A530041M06Rik | | 0 | 20 | 0 | 2 | 1 | 17 | 12 | 0 | 0 | 0 | 24 |
| 590 | -4.31 | 1.93 | -2.23 | 0.03 | 0.74504 | Cd248 | CD248 antigen, endosialin | 3125 | 1353 | 1980 | 0 | 17 | 9 | 1 | 0 | 0 | 0 | 0 |
| 78 | -0.86 | 0.39 | -2.23 | 0.03 | 0.74626 | 2310040G24Rik | | 55 | 48 | 64 | 122 | 124 | 99 | 100 | 71 | 80 | 36 | 62 |
| 7448 | 0.21 | 0.09 | 2.23 | 0.03 | 0.75015 | Sv2a | synaptic vesicle glycoprotein 2 a | 2173 | 3003 | 2610 | 9819 | 8814 | 6313 | 9203 | 13652 | 9086 | 10153 | 7106 |
| 7669 | 0.13 | 0.06 | 2.23 | 0.03 | 0.75109 | Dnajc10 | DnaJ heat shock protein family (Hsp40) m | 5931 | 5599 | 4925 | 8516 | 7985 | 7624 | 8315 | 8900 | 8574 | 9618 | 8369 |
| 335 | -0.43 | 0.19 | -2.23 | 0.03 | 0.75109 | Gm17690 | | 306 | 286 | 260 | 403 | 392 | 404 | 422 | 265 | 292 | 274 | 378 |
| 192 | -0.64 | 0.29 | -2.23 | 0.03 | 0.75109 | Kirrel2 | kirre like nephrin family adhesion molecu | 450 | 438 | 514 | 116 | 100 | 116 | 97 | 70 | 73 | 43 | 97 |
| 6517 | -0.19 | 0.09 | -2.23 | 0.03 | 0.7529 | Polr2a | polymerase (RNA) II (DNA directed) polyp | 5164 | 4807 | 4722 | 8355 | 8171 | 6308 | 7507 | 7136 | 7987 | 6161 | 5368 |
| 25 | 2.46 | 1.10 | 2.23 | 0.03 | 0.75346 | Gm43071 | | 28 | 3 | 12 | 0 | 3 | 59 | 12 | 50 | 14 | 21 | 74 |
| 736 | -0.30 | 0.13 | -2.23 | 0.03 | 0.75377 | Abtb1 | ankyrin repeat and BTB (POZ) domain co | 649 | 665 | 660 | 820 | 793 | 1019 | 739 | 696 | 686 | 515 | 855 |
| 453 | 0.37 | 0.17 | 2.23 | 0.03 | 0.75377 | Dars2 | aspartyl-tRNA synthetase 2 (mitochondria | 474 | 401 | 475 | 382 | 357 | 441 | 399 | 431 | 452 | 595 | 576 |
| 62 | -1.79 | 0.80 | -2.23 | 0.03 | 0.75377 | Shmt1 | serine hydroxymethyltransferase 1 (solub | 262 | 121 | 98 | 59 | 33 | 23 | 42 | 0 | 18 | 15 | 15 |
| 3965 | 0.21 | 0.09 | 2.22 | 0.03 | 0.75837 | Atp6v1d | ATPase, H+ transporting, lysosomal V1 su | 2407 | 2488 | 2592 | 4367 | 3915 | 3517 | 4985 | 5429 | 4912 | 4801 | 4204 |
| 5620 | -0.21 | 0.09 | -2.22 | 0.03 | 0.75837 | Gls | glutaminase | 9977 | 11917 | 9631 | 4101 | 3720 | 4410 | 3945 | 3188 | 3459 | 3048 | 4426 |
| 1815 | -0.30 | 0.13 | -2.22 | 0.03 | 0.75837 | Slc35g1 | solute carrier family 35, member G1 | 1325 | 1820 | 1559 | 2291 | 1959 | 1713 | 2475 | 1672 | 1832 | 1824 | 1500 |
| 885 | -0.31 | 0.14 | -2.22 | 0.03 | 0.75837 | Tulp3 | tubby-like protein 3 | 1123 | 1035 | 1072 | 959 | 948 | 849 | 837 | 616 | 842 | 742 | 711 |
| 826 | -0.57 | 0.25 | -2.22 | 0.03 | 0.75837 | Mxra8 | matrix-remodelling associated 8 | 3037 | 1416 | 1771 | 371 | 584 | 254 | 486 | 182 | 501 | 277 | 206 |
| 706 | -0.73 | 0.33 | -2.22 | 0.03 | 0.75837 | Gfra1 | glial cell line derived neurotrophic factor f | 2433 | 2479 | 1823 | 79 | 175 | 129 | 244 | 120 | 64 | 167 | 50 |
| 307 | 0.59 | 0.26 | 2.22 | 0.03 | 0.75837 | Zfand1 | zinc finger, AN1-type domain 1 | 219 | 252 | 340 | 187 | 152 | 552 | 188 | 327 | 253 | 309 | 595 |
| 166 | -0.67 | 0.30 | -2.22 | 0.03 | 0.75837 | Csrnp2 | cysteine-serine-rich nuclear protein 2 | 149 | 191 | 165 | 144 | 265 | 229 | 175 | 136 | 160 | 89 | 126 |
| 73 | -1.37 | 0.62 | -2.22 | 0.03 | 0.75837 | Cacna1i | calcium channel, voltage-dependent, alpl | 184 | 309 | 99 | 28 | 16 | 51 | 47 | 29 | 4 | 4 | 31 |
| 102 | -1.59 | 0.72 | -2.22 | 0.03 | 0.75837 | Tubb1 | tubulin, beta 1 class VI | 257 | 447 | 347 | 11 | 9 | 9 | 24 | 0 | 7 | 6 | 4 |
| 62 | -2.07 | 0.93 | -2.22 | 0.03 | 0.75837 | Map3k6 | mitogen-activated protein kinase kinase k | 269 | 166 | 159 | 11 | 27 | 13 | 13 | 0 | 12 | 0 | 8 |
| 61 | -1.97 | 0.89 | -2.22 | 0.03 | 0.75837 | Egr4 | early growth response 4 | 224 | 145 | 99 | 7 | 15 | 107 | 12 | 0 | 7 | 0 | 58 |
| 78 | -2.57 | 1.16 | -2.22 | 0.03 | 0.75837 | Homer2 | homer scaffolding protein 2 | 130 | 373 | 292 | 46 | 4 | 0 | 4 | 0 | 5 | 0 | 0 |
| 75 | -2.05 | 0.92 | -2.22 | 0.03 | 0.75837 | Fermt1 | fermitin family member 1 | 186 | 288 | 243 | 20 | 37 | 19 | 4 | 0 | 17 | 0 | 9 |
| 368 | -1.04 | 0.47 | -2.22 | 0.03 | 0.75851 | Bcl2 | B cell leukemia/lymphoma 2 | 1370 | 1121 | 1066 | 52 | 83 | 84 | 93 | 9 | 86 | 19 | 64 |
| 47 | -2.74 | 1.24 | -2.21 | 0.03 | 0.76594 | Fam184b | family with sequence similarity 184, meml | 154 | 168 | 161 | 7 | 0 | 8 | 13 | 0 | 3 | 0 | 1 |
| 5 | 5.69 | 2.57 | 2.21 | 0.03 | 0.76681 | Gm15577 | | 0 | 15 | 0 | 3 | 0 | 5 | 0 | 0 | 5 | 23 | 7 |
| 11691 | -0.60 | 0.27 | -2.21 | 0.03 | 0.76724 | Egr1 | early growth response 1 | 35511 | 20055 | 23122 | 1444 | 2187 | 20889 | 4054 | 564 | 1660 | 2983 | 16134 |
| 3386 | 0.19 | 0.09 | 2.21 | 0.03 | 0.76724 | Ddx3y | DEAD (Asp-Glu-Ala-Asp) box polypeptide | 3076 | 2706 | 2714 | 2836 | 3350 | 4435 | 2795 | 3303 | 3497 | 3438 | 5092 |
| 325 | -0.57 | 0.26 | -2.21 | 0.03 | 0.76724 | Ephb2 | Eph receptor B2 | 803 | 787 | 739 | 203 | 134 | 183 | 227 | 130 | 158 | 116 | 98 |
| 43 | 1.94 | 0.88 | 2.21 | 0.03 | 0.76724 | Castor1 | cytosolic arginine sensor for mTORC1 sul | 135 | 101 | 140 | 4 | 0 | 19 | 4 | 31 | 5 | 15 | 21 |
| 15 | -3.73 | 1.69 | -2.21 | 0.03 | 0.77004 | Mstn | myostatin | 28 | 40 | 80 | 5 | 4 | 4 | 0 | 0 | 0 | 0 | 0 |
| 6 | -4.55 | 2.06 | -2.21 | 0.03 | 0.77282 | 1700093J21Rik | | 15 | 0 | 0 | 17 | 9 | 5 | 13 | 0 | 0 | 1 | 4 |
| 1002 | -0.33 | 0.15 | -2.20 | 0.03 | 0.77325 | Rbms1 | RNA binding motif, single stranded intera | 3034 | 2367 | 2349 | 392 | 424 | 527 | 480 | 306 | 415 | 380 | 350 |
| 350 | 0.41 | 0.19 | 2.21 | 0.03 | 0.77325 | Gm6548 | | 232 | 442 | 531 | 370 | 357 | 218 | 198 | 467 | 449 | 282 | 302 |
| 5009 | 0.19 | 0.09 | 2.20 | 0.03 | 0.78051 | Pacc1 | proton activated chloride channel 1 | 1461 | 1510 | 1787 | 6203 | 6090 | 4799 | 6435 | 7133 | 6637 | 7500 | 5546 |
| 32 | -2.50 | 1.14 | -2.20 | 0.03 | 0.78051 | Rtl4 | retrotransposon Gag like 4 | 44 | 77 | 88 | 25 | 14 | 26 | 39 | 0 | 13 | 0 | 25 |
| 152 | 1.30 | 0.59 | 2.20 | 0.03 | 0.78051 | Rasa4 | RAS p21 protein activator 4 | 650 | 428 | 399 | 9 | 10 | 23 | 11 | 78 | 27 | 13 | 21 |
| 3208 | 0.22 | 0.10 | 2.20 | 0.03 | 0.78158 | Zdhhc3 | zinc finger, DHHC domain containing 3 | 2308 | 1967 | 2244 | 3289 | 3759 | 2808 | 3415 | 4469 | 3680 | 4149 | 3197 |
| 1189 | 0.24 | 0.11 | 2.20 | 0.03 | 0.78158 | Fbxw7 | F-box and WD-40 domain protein 7 | 836 | 676 | 720 | 1090 | 1230 | 1507 | 1146 | 1344 | 1336 | 1539 | 1658 |
| 17 | -2.69 | 1.22 | -2.20 | 0.03 | 0.78158 | Gm39157 | | 11 | 16 | 26 | 11 | 32 | 22 | 36 | 0 | 16 | 0 | 12 |
| 183 | 0.74 | 0.34 | 2.20 | 0.03 | 0.78213 | Paqr4 | progestin and adipoQ receptor family mer | 412 | 416 | 361 | 55 | 79 | 62 | 99 | 165 | 103 | 194 | 63 |
| 78 | -2.13 | 0.97 | -2.20 | 0.03 | 0.78213 | Rasgrp1 | RAS guanyl releasing protein 1 | 333 | 261 | 135 | 13 | 25 | 41 | 19 | 0 | 22 | 0 | 11 |
| 2064 | -0.29 | 0.13 | -2.20 | 0.03 | 0.78235 | Gna12 | guanine nucleotide binding protein, alpha | 3618 | 2226 | 2970 | 2170 | 2012 | 1395 | 2066 | 1638 | 1705 | 1754 | 1152 |
| 5 | -3.93 | 1.79 | -2.20 | 0.03 | 0.78299 | Gm11574 | | 10 | 4 | 4 | 12 | 8 | 2 | 12 | 0 | 2 | 0 | 0 |
| 1544 | -0.27 | 0.12 | -2.19 | 0.03 | 0.7839 | Snx18 | sorting nexin 18 | 3116 | 2996 | 3206 | 1168 | 909 | 1119 | 995 | 981 | 876 | 745 | 875 |
| 227 | -0.48 | 0.22 | -2.19 | 0.03 | 0.7839 | Cspg5 | chondroitin sulfate proteoglycan 5 | 323 | 227 | 299 | 233 | 286 | 156 | 289 | 147 | 217 | 188 | 132 |
| 53 | -3.09 | 1.41 | -2.19 | 0.03 | 0.7839 | Alox5 | arachidonate 5-lipoxygenase | 294 | 164 | 116 | 0 | 0 | 7 | 5 | 0 | 0 | 0 | 0 |
| 465 | -0.39 | 0.18 | -2.19 | 0.03 | 0.78481 | Slc25a29 | solute carrier family 25 (mitochondrial car | 167 | 201 | 190 | 702 | 725 | 527 | 632 | 513 | 444 | 599 | 410 |

| | | | | | | | | | | | | | | | | | | |
|---|---|---|---|---|---|---|---|---|---|---|---|---|---|---|---|---|---|---|
| 458 | -0.40 | 0.18 | -2.19 | 0.03 | 0.78481 | Snhg18 | | 619 | 575 | 533 | 599 | 464 | 295 | 503 | 524 | 355 | 390 | 185 |
| 73 | -1.23 | 0.56 | -2.19 | 0.03 | 0.78681 | Zfp85 | zinc finger protein 85 | 74 | 48 | 136 | 109 | 75 | 104 | 68 | 14 | 72 | 24 | 76 |
| 4551 | -0.23 | 0.11 | -2.19 | 0.03 | 0.78917 | Rps27 | ribosomal protein S27 | 7808 | 7120 | 7968 | 3587 | 3925 | 2779 | 4355 | 2405 | 3814 | 3688 | 2611 |
| 957 | 0.40 | 0.18 | 2.19 | 0.03 | 0.78917 | Khk | ketohexokinase | 595 | 714 | 730 | 748 | 903 | 975 | 1032 | 1416 | 1036 | 1325 | 1055 |
| 62 | -1.91 | 0.87 | -2.19 | 0.03 | 0.78917 | Tuba8 | tubulin, alpha 8 | 13 | 35 | 60 | 88 | 129 | 65 | 126 | 83 | 57 | 0 | 23 |
| 2374 | -3.86 | 1.76 | -2.19 | 0.03 | 0.78917 | Eng | endoglin | 9303 | 7963 | 8806 | 0 | 6 | 5 | 23 | 2 | 0 | 0 | 0 |
| 174 | -1.65 | 0.75 | -2.19 | 0.03 | 0.78917 | Itgb3 | integrin beta 3 | 693 | 654 | 455 | 42 | 4 | 17 | 18 | 6 | 0 | 23 | 3 |
| 969 | 0.31 | 0.14 | 2.19 | 0.03 | 0.7914 | Dennd1b | DENN/MADD domain containing 1B | 1126 | 765 | 851 | 803 | 790 | 1072 | 878 | 1079 | 1050 | 1010 | 1233 |
| 494 | -0.77 | 0.35 | -2.19 | 0.03 | 0.79232 | Peg10 | paternally expressed 10 | 953 | 1794 | 2042 | 122 | 110 | 164 | 27 | 54 | 41 | 54 | 68 |
| 267 | -0.77 | 0.35 | -2.18 | 0.03 | 0.79498 | Lpar6 | lysophosphatidic acid receptor 6 | 511 | 435 | 498 | 244 | 181 | 276 | 190 | 115 | 134 | 53 | 299 |
| 585 | -0.62 | 0.28 | -2.18 | 0.03 | 0.79498 | Zyx | zyxin | 2328 | 1584 | 1484 | 188 | 163 | 167 | 112 | 80 | 143 | 84 | 107 |
| 708 | -1.79 | 0.82 | -2.18 | 0.03 | 0.79498 | Ldha | lactate dehydrogenase A | 3632 | 1803 | 2254 | 23 | 16 | 13 | 17 | 0 | 2 | 13 | 12 |
| 7 | -3.19 | 1.46 | -2.18 | 0.03 | 0.79498 | Gm17771 | | 7 | 11 | 12 | 4 | 11 | 12 | 17 | 0 | 3 | 0 | 3 |
| 39 | -2.44 | 1.12 | -2.18 | 0.03 | 0.79498 | Fjx1 | four jointed box 1 | 156 | 78 | 162 | 0 | 8 | 8 | 14 | 0 | 2 | 0 | 3 |
| 269 | -2.69 | 1.23 | -2.18 | 0.03 | 0.79498 | Ppic | peptidylprolyl isomerase C | 1261 | 767 | 919 | 9 | 3 | 0 | 1 | 0 | 1 | 0 | 0 |
| 1195 | -0.39 | 0.18 | -2.18 | 0.03 | 0.79667 | Mpp2 | membrane protein, palmitoylated 2 (MAG | 2191 | 3455 | 3066 | 652 | 591 | 625 | 642 | 387 | 587 | 462 | 483 |
| 1240 | -0.35 | 0.16 | -2.18 | 0.03 | 0.79747 | Prdx4 | peroxiredoxin 4 | 2724 | 3086 | 2748 | 859 | 755 | 479 | 761 | 468 | 650 | 689 | 420 |
| 887 | -2.90 | 1.33 | -2.18 | 0.03 | 0.80048 | Ltbp1 | latent transforming growth factor beta binc | 4004 | 2237 | 3476 | 12 | 9 | 3 | 8 | 0 | 10 | 0 | 0 |
| 746 | -0.45 | 0.21 | -2.18 | 0.03 | 0.80119 | Folr1 | folate receptor 1 (adult) | 913 | 1599 | 1559 | 705 | 663 | 426 | 528 | 810 | 409 | 339 | 258 |
| 653 | -0.39 | 0.18 | -2.18 | 0.03 | 0.80159 | 2310022A10Ril | RIKEN cDNA 2310022A10 gene | 750 | 840 | 758 | 667 | 776 | 665 | 626 | 415 | 604 | 612 | 467 |
| 2315 | -0.22 | 0.10 | -2.18 | 0.03 | 0.8032 | Kifc3 | kinesin family member C3 | 3653 | 3582 | 4277 | 1826 | 1881 | 2301 | 1511 | 1542 | 1498 | 1526 | 1871 |
| 7110 | 0.12 | 0.06 | 2.17 | 0.03 | 0.80748 | Rab14 | RAB14, member RAS oncogene family | 7052 | 7103 | 6756 | 6989 | 6559 | 6501 | 7397 | 7892 | 7062 | 7457 | 7446 |
| 69 | -0.87 | 0.40 | -2.17 | 0.03 | 0.8079 | Rgn | regucalcin | 71 | 114 | 86 | 90 | 95 | 63 | 70 | 46 | 50 | 28 | 50 |
| 19 | 2.17 | 1.00 | 2.17 | 0.03 | 0.8079 | Gm8355 | | 15 | 10 | 10 | 3 | 14 | 14 | 2 | 76 | 12 | 23 | 30 |
| 34 | -3.62 | 1.67 | -2.17 | 0.03 | 0.8111 | Lsamp | limbic system-associated membrane prot | 135 | 123 | 92 | 21 | 4 | 0 | 0 | 0 | 0 | 0 | 0 |
| 718 | 0.41 | 0.19 | 2.17 | 0.03 | 0.81672 | Optn | optineurin | 568 | 538 | 424 | 535 | 594 | 744 | 885 | 959 | 940 | 853 | 862 |
| 10430 | 0.30 | 0.14 | 2.16 | 0.03 | 0.8229 | Rps3a1 | ribosomal protein S3A1 | 18387 | 15449 | 19623 | 6690 | 7488 | 4849 | 8406 | 11813 | 7535 | 8289 | 6202 |
| 1466 | -0.31 | 0.14 | -2.16 | 0.03 | 0.8229 | Klf3 | Kruppel-like factor 3 (basic) | 2508 | 2160 | 2462 | 1308 | 1173 | 1122 | 1375 | 1020 | 1227 | 844 | 934 |
| 118 | -1.26 | 0.58 | -2.16 | 0.03 | 0.8229 | Col20a1 | collagen, type XX, alpha 1 | 217 | 273 | 310 | 25 | 19 | 241 | 26 | 10 | 17 | 0 | 159 |
| 229 | 0.64 | 0.30 | 2.16 | 0.03 | 0.82378 | Zfp189 | zinc finger protein 189 | 248 | 213 | 165 | 202 | 174 | 224 | 132 | 438 | 219 | 240 | 268 |
| 260 | 0.65 | 0.30 | 2.16 | 0.03 | 0.82378 | Msantd2 | Myb/SANT-like DNA-binding domain cont | 182 | 195 | 234 | 89 | 121 | 739 | 48 | 159 | 138 | 146 | 814 |
| 1181 | -0.28 | 0.13 | -2.16 | 0.03 | 0.82432 | Pygb | brain glycogen phosphorylase | 3531 | 2269 | 3037 | 543 | 708 | 382 | 642 | 382 | 625 | 549 | 326 |
| 184 | 0.68 | 0.31 | 2.16 | 0.03 | 0.82491 | Gm20604 | predicted gene 20604 | 124 | 138 | 206 | 126 | 147 | 134 | 197 | 266 | 196 | 265 | 229 |
| 23 | 1.48 | 0.69 | 2.16 | 0.03 | 0.82491 | 2810039B14Rik | | 38 | 49 | 32 | 17 | 4 | 5 | 10 | 43 | 21 | 24 | 10 |
| 4892 | 0.13 | 0.06 | 2.16 | 0.03 | 0.82641 | Smim10l1 | small integral membrane protein 10 like 1 | 3470 | 3753 | 3884 | 5547 | 5424 | 3946 | 5481 | 6094 | 5870 | 5965 | 4384 |
| 1677 | -0.30 | 0.14 | -2.16 | 0.03 | 0.82763 | Cd151 | CD151 antigen | 4328 | 3601 | 3557 | 1119 | 1088 | 656 | 975 | 692 | 970 | 891 | 569 |
| 372 | -0.48 | 0.22 | -2.16 | 0.03 | 0.82763 | Dclre1a | DNA cross-link repair 1A | 351 | 467 | 468 | 413 | 333 | 545 | 328 | 228 | 348 | 182 | 434 |
| 585 | 0.43 | 0.20 | 2.16 | 0.03 | 0.82817 | Btbd8 | RIKEN cDNA A830010M20 gene | 409 | 505 | 284 | 417 | 437 | 764 | 629 | 663 | 628 | 754 | 948 |
| 986 | 0.27 | 0.12 | 2.16 | 0.03 | 0.82846 | Smim20 | small integral membrane protein 20 | 592 | 646 | 655 | 1111 | 1083 | 822 | 1040 | 1457 | 1108 | 1328 | 1003 |
| 63 | -2.23 | 1.03 | -2.16 | 0.03 | 0.82846 | Pdlim3 | PDZ and LIM domain 3 | 440 | 74 | 80 | 21 | 14 | 21 | 18 | 0 | 9 | 0 | 14 |
| 16 | -2.42 | 1.12 | -2.16 | 0.03 | 0.82846 | Lor | loricrin | 29 | 21 | 42 | 14 | 12 | 19 | 15 | 1 | 17 | 1 | 1 |
| 235 | -1.86 | 0.87 | -2.15 | 0.03 | 0.83237 | Adamtsl2 | ADAMTS-like 2 | 1262 | 474 | 814 | 0 | 3 | 9 | 17 | 0 | 2 | 0 | 5 |
| 19 | -3.11 | 1.45 | -2.15 | 0.03 | 0.83386 | Krt90 | keratin 90 | 39 | 77 | 44 | 5 | 30 | 9 | 0 | 0 | 1 | 0 | 4 |
| 6651 | 0.13 | 0.06 | 2.15 | 0.03 | 0.83417 | Ywhab | tyrosine 3-monooxygenase/tryptophan 5- | 8707 | 7579 | 8711 | 5664 | 6233 | 5034 | 6098 | 6505 | 6428 | 6768 | 5436 |
| 16 | -3.25 | 1.51 | -2.15 | 0.03 | 0.83451 | Gm44829 | | 15 | 0 | 19 | 11 | 9 | 40 | 26 | 0 | 4 | 0 | 52 |
| 9155 | 0.22 | 0.10 | 2.15 | 0.03 | 0.8358 | Sf3b2 | splicing factor 3b, subunit 2 | 9573 | 8970 | 9437 | 7860 | 8924 | 7903 | 8844 | 12143 | 9232 | 9194 | 8627 |
| 1620 | -0.25 | 0.11 | -2.15 | 0.03 | 0.8358 | Acadl | acyl-Coenzyme A dehydrogenase, long-c | 2178 | 2673 | 2544 | 1946 | 1366 | 1034 | 1289 | 1686 | 1227 | 1073 | 806 |
| 3401 | 0.26 | 0.12 | 2.15 | 0.03 | 0.83731 | Rps6 | ribosomal protein S6 | 4531 | 4491 | 5191 | 2620 | 2905 | 2048 | 2909 | 4073 | 2986 | 3501 | 2160 |
| 17 | -2.36 | 1.10 | -2.15 | 0.03 | 0.84112 | Adm2 | adrenomedullin 2 | 18 | 48 | 19 | 11 | 33 | 7 | 33 | 5 | 12 | 0 | 2 |
| 177 | -0.84 | 0.39 | -2.14 | 0.03 | 0.84421 | Nsun5 | NOL1/NOP2/Sun domain family, member | 223 | 184 | 135 | 155 | 162 | 407 | 117 | 44 | 109 | 59 | 355 |
| 2901 | 0.15 | 0.07 | 2.14 | 0.03 | 0.84448 | Afg3l2 | AFG3-like AAA ATPase 2 | 2246 | 2232 | 2134 | 3068 | 2821 | 3130 | 2986 | 3242 | 3306 | 3189 | 3556 |
| 249 | -2.94 | 1.37 | -2.14 | 0.03 | 0.84448 | Cmtm3 | CKLF-like MARVEL transmembrane dom | 1263 | 767 | 684 | 0 | 12 | 4 | 11 | 0 | 1 | 0 | 2 |
| 2629 | 0.16 | 0.07 | 2.14 | 0.03 | 0.84676 | Inpp5f | inositol polyphosphate-5-phosphatase F | 1645 | 1770 | 1459 | 2998 | 2746 | 2713 | 2907 | 3259 | 2929 | 3282 | 3211 |
| 1992 | 0.22 | 0.11 | 2.14 | 0.03 | 0.8538 | Rala | v-ral simian leukemia viral oncogene A (ra | 2348 | 2088 | 2457 | 1745 | 1856 | 1506 | 1819 | 2336 | 1998 | 1985 | 1770 |
| 973 | 0.31 | 0.14 | 2.14 | 0.03 | 0.85616 | 4930453N24Ril | RIKEN cDNA 4930453N24 gene | 720 | 612 | 727 | 1069 | 1016 | 829 | 948 | 1389 | 1275 | 1015 | 1099 |
| 118 | -1.10 | 0.52 | -2.14 | 0.03 | 0.85616 | Cpne4 | copine IV | 115 | 201 | 120 | 162 | 235 | 74 | 115 | 32 | 124 | 36 | 84 |
| 3636 | 0.19 | 0.09 | 2.13 | 0.03 | 0.85771 | Svip | small VCP/p97-interacting protein | 1162 | 1496 | 1435 | 4346 | 4414 | 3478 | 4530 | 5293 | 4822 | 4963 | 4060 |
| 1963 | 0.20 | 0.09 | 2.13 | 0.03 | 0.85771 | Rhot1 | ras homolog family member T1 | 1789 | 1393 | 1445 | 1792 | 1926 | 2199 | 2000 | 2146 | 2232 | 2357 | 2317 |
| 1927 | -0.19 | 0.09 | -2.13 | 0.03 | 0.85771 | Sipa1l1 | signal-induced proliferation-associated 1 | 2648 | 2295 | 2588 | 1771 | 1993 | 1526 | 1981 | 1387 | 1901 | 1767 | 1339 |
| 604 | 0.33 | 0.15 | 2.13 | 0.03 | 0.85771 | Wdyhv1 | WDYHV motif containing 1 | 438 | 497 | 482 | 514 | 580 | 576 | 645 | 789 | 687 | 755 | 681 |

| | | | | | | | | | | | | | | | | | | |
|---|---|---|---|---|---|---|---|---|---|---|---|---|---|---|---|---|---|---|
| 666 | 0.49 | 0.23 | 2.13 | 0.03 | 0.85771 | Tbcc | tubulin-specific chaperone C | 656 | 645 | 870 | 636 | 519 | 442 | 526 | 1090 | 554 | 835 | 552 |
| 6 | -3.62 | 1.70 | -2.13 | 0.03 | 0.86501 | Gm12830 | | 7 | 12 | 15 | 9 | 9 | 5 | 7 | 0 | 5 | 0 | 0 |
| 458 | 0.64 | 0.30 | 2.13 | 0.03 | 0.86506 | Hpgds | hematopoietic prostaglandin D synthase | 331 | 132 | 435 | 331 | 432 | 574 | 306 | 542 | 547 | 725 | 682 |
| 5 | -5.54 | 2.60 | -2.13 | 0.03 | 0.86545 | Actc1 | actin, alpha, cardiac muscle 1 | 2 | 34 | 0 | 0 | 2 | 7 | 9 | 0 | 0 | 0 | 4 |
| 15 | -5.12 | 2.41 | -2.13 | 0.03 | 0.86572 | Cdh23 | cadherin 23 (otocadherin) | 52 | 11 | 0 | 16 | 0 | 17 | 43 | 0 | 24 | 0 | 0 |
| 915 | -0.36 | 0.17 | -2.13 | 0.03 | 0.86615 | Ubald1 | UBA-like domain containing 1 | 758 | 911 | 794 | 852 | 965 | 1468 | 967 | 526 | 847 | 757 | 1221 |
| 228 | -3.67 | 1.73 | -2.13 | 0.03 | 0.86615 | Tspan18 | tetraspanin 18 | 922 | 742 | 804 | 0 | 9 | 3 | 19 | 0 | 0 | 0 | 3 |
| 1553 | 0.24 | 0.11 | 2.13 | 0.03 | 0.86665 | Ppp4r2 | protein phosphatase 4, regulatory subuni | 1449 | 1945 | 1445 | 1364 | 1269 | 1546 | 1413 | 1920 | 1502 | 1471 | 1757 |
| 836 | 0.39 | 0.18 | 2.13 | 0.03 | 0.86665 | Cryzl2 | crystallin zeta like 2 | 583 | 495 | 672 | 646 | 936 | 759 | 902 | 1261 | 994 | 1024 | 921 |
| 1136 | -0.29 | 0.14 | -2.13 | 0.03 | 0.86665 | Ciz1 | CDKN1A interacting zinc finger protein 1 | 957 | 1378 | 1172 | 1289 | 1212 | 1676 | 809 | 880 | 994 | 840 | 1290 |
| 7 | -3.58 | 1.68 | -2.13 | 0.03 | 0.86665 | Gm45619 | | 0 | 5 | 6 | 11 | 15 | 13 | 19 | 0 | 7 | 0 | 3 |
| 3301 | 0.15 | 0.07 | 2.12 | 0.03 | 0.86801 | Gmps | guanine monophosphate synthetase | 2918 | 2697 | 3103 | 2973 | 3568 | 3163 | 3375 | 3544 | 3695 | 3536 | 3739 |
| 2091 | -0.20 | 0.09 | -2.12 | 0.03 | 0.86801 | Itga3 | integrin alpha 3 | 2503 | 2473 | 2600 | 2138 | 2222 | 1443 | 2449 | 2119 | 1815 | 1964 | 1272 |
| 2120 | 0.25 | 0.12 | 2.12 | 0.03 | 0.86801 | Txndc17 | thioredoxin domain containing 17 | 1953 | 1853 | 2144 | 1879 | 2283 | 1528 | 2246 | 2803 | 2239 | 2577 | 1815 |
| 20 | -2.60 | 1.23 | -2.12 | 0.03 | 0.86801 | Gm14964 | | 45 | 74 | 27 | 25 | 7 | 16 | 13 | 0 | 9 | 0 | 3 |
| 3 | -7.35 | 3.46 | -2.12 | 0.03 | 0.86801 | Gm39383 | | 10 | 0 | 0 | 8 | 6 | 4 | 0 | 0 | 0 | 0 | 1 |
| 721 | -2.30 | 1.09 | -2.12 | 0.03 | 0.86836 | Itpripl2 | inositol 1,4,5-triphosphate receptor intera | 2955 | 2143 | 2770 | 13 | 3 | 4 | 32 | 0 | 7 | 6 | 0 |
| 13175 | 0.12 | 0.06 | 2.12 | 0.03 | 0.87121 | Sar1a | secretion associated Ras related GTPase | 9566 | 9104 | 10389 | 13163 | 15118 | 11754 | 15558 | 15484 | 15975 | 16413 | 12398 |
| 40 | -1.99 | 0.94 | -2.12 | 0.03 | 0.87121 | Gm648 | predicted gene 648 | 75 | 154 | 127 | 31 | 4 | 13 | 19 | 13 | 1 | 0 | 7 |
| 15 | 3.49 | 1.65 | 2.12 | 0.03 | 0.87121 | Gm31045 | | 27 | 60 | 37 | 0 | 0 | 4 | 0 | 27 | 4 | 0 | 5 |
| 3223 | -0.21 | 0.10 | -2.12 | 0.03 | 0.87449 | Zfp704 | zinc finger protein 704 | 3883 | 5071 | 3349 | 3475 | 2814 | 2999 | 3056 | 3502 | 2548 | 2273 | 2482 |
| 2226 | -0.24 | 0.11 | -2.12 | 0.03 | 0.87449 | Slc37a1 | solute carrier family 37 (glycerol-3-phospl | 1501 | 2140 | 1956 | 2782 | 2492 | 2296 | 2668 | 2214 | 2284 | 2272 | 1884 |
| 586 | 0.33 | 0.16 | 2.12 | 0.03 | 0.87449 | Mrps35 | mitochondrial ribosomal protein S35 | 613 | 633 | 635 | 550 | 426 | 525 | 513 | 766 | 524 | 649 | 608 |
| 236 | -0.83 | 0.39 | -2.12 | 0.03 | 0.87449 | Mvb12b | multivesicular body subunit 12B | 756 | 630 | 899 | 79 | 43 | 44 | 34 | 16 | 32 | 30 | 34 |
| 20 | 1.48 | 0.70 | 2.12 | 0.03 | 0.87449 | Gm15492 | | 15 | 7 | 6 | 17 | 6 | 17 | 11 | 35 | 21 | 44 | 38 |
| 4427 | 0.13 | 0.06 | 2.11 | 0.03 | 0.87756 | Selenos | selenoprotein S | 2489 | 2025 | 2761 | 4066 | 5584 | 5296 | 4885 | 4716 | 5905 | 5433 | 5535 |
| 167 | -1.73 | 0.82 | -2.11 | 0.03 | 0.87756 | Galnt16 | polypeptide N-acetylgalactosaminyltransf | 746 | 455 | 545 | 15 | 14 | 11 | 27 | 0 | 13 | 0 | 13 |
| 126 | -0.68 | 0.32 | -2.11 | 0.03 | 0.87759 | Sass6 | SAS-6 centriolar assembly protein | 221 | 216 | 149 | 119 | 86 | 159 | 129 | 61 | 78 | 62 | 102 |
| 1684 | -0.27 | 0.13 | -2.11 | 0.03 | 0.87855 | Galnt4 | polypeptide N-acetylgalactosaminyltransf | 1793 | 2262 | 1961 | 1810 | 1907 | 1314 | 1730 | 1636 | 1634 | 1641 | 835 |
| 19 | -2.37 | 1.12 | -2.11 | 0.03 | 0.87855 | Gm36026 | | 8 | 18 | 52 | 72 | 16 | 16 | 5 | 14 | 1 | 0 | 11 |
| 1635 | 0.21 | 0.10 | 2.11 | 0.03 | 0.87921 | Gtpbp4 | GTP binding protein 4 | 1641 | 1960 | 1902 | 1511 | 1433 | 1568 | 1273 | 1819 | 1451 | 1489 | 1939 |
| 33 | -1.90 | 0.90 | -2.11 | 0.03 | 0.87921 | Gnal | guanine nucleotide binding protein, alpha | 57 | 4 | 42 | 50 | 40 | 49 | 60 | 8 | 23 | 3 | 30 |
| 101 | -0.85 | 0.40 | -2.11 | 0.03 | 0.88184 | Pcdhga12 | protocadherin gamma subfamily A, 12 | 117 | 149 | 160 | 130 | 86 | 79 | 148 | 46 | 60 | 62 | 74 |
| 2603 | 0.18 | 0.09 | 2.11 | 0.04 | 0.88428 | Psmd13 | proteasome (prosome, macropain) 26S su | 2933 | 2728 | 2895 | 2196 | 2584 | 2105 | 2513 | 2840 | 2564 | 2924 | 2349 |
| 195 | -0.91 | 0.43 | -2.11 | 0.04 | 0.88428 | Shroom4 | shroom family member 4 | 770 | 478 | 421 | 106 | 102 | 57 | 45 | 56 | 61 | 27 | 24 |
| 275 | -0.59 | 0.28 | -2.11 | 0.04 | 0.88454 | Card10 | caspase recruitment domain family, meml | 926 | 761 | 950 | 75 | 60 | 46 | 54 | 36 | 41 | 45 | 34 |
| 480 | -2.63 | 1.25 | -2.11 | 0.04 | 0.88476 | Ildr2 | immunoglobulin-like domain containing re | 3094 | 1329 | 802 | 19 | 9 | 8 | 7 | 0 | 2 | 0 | 13 |
| 622 | 0.44 | 0.21 | 2.11 | 0.04 | 0.88565 | Mir670hg | | 155 | 137 | 229 | 676 | 745 | 728 | 537 | 945 | 932 | 1030 | 726 |
| 17 | -3.59 | 1.71 | -2.11 | 0.04 | 0.88565 | Insc | INSC spindle orientation adaptor protein | 73 | 47 | 58 | 0 | 6 | 1 | 8 | 0 | 0 | 0 | 0 |
| 148 | -1.88 | 0.89 | -2.11 | 0.04 | 0.88649 | Tm6sf1 | transmembrane 6 superfamily member 1 | 705 | 373 | 503 | 12 | 4 | 19 | 2 | 0 | 3 | 0 | 9 |
| 4954 | 0.20 | 0.09 | 2.10 | 0.04 | 0.88925 | Riok3 | RIO kinase 3 | 3831 | 3469 | 4060 | 4574 | 5128 | 5549 | 4856 | 6381 | 5555 | 5118 | 5970 |
| 2184 | -0.17 | 0.08 | -2.10 | 0.04 | 0.88925 | Pomgnt1 | protein O-linked mannose beta 1,2-N-ace | 1838 | 1775 | 2035 | 2389 | 2658 | 2089 | 2604 | 2289 | 2430 | 2073 | 1843 |
| 855 | 0.24 | 0.12 | 2.10 | 0.04 | 0.88925 | Tyw1 | tRNA-yW synthesizing protein 1 homolog | 864 | 876 | 999 | 740 | 842 | 709 | 762 | 963 | 929 | 911 | 812 |
| 180 | -0.92 | 0.44 | -2.10 | 0.04 | 0.88925 | Amot | angiomotin | 492 | 501 | 434 | 116 | 86 | 76 | 77 | 24 | 28 | 97 | 54 |
| 731 | -0.48 | 0.23 | -2.10 | 0.04 | 0.88925 | Lama5 | laminin, alpha 5 | 2817 | 2337 | 2453 | 56 | 84 | 35 | 75 | 49 | 38 | 68 | 25 |
| 101 | 1.12 | 0.53 | 2.10 | 0.04 | 0.88925 | Rpl19-ps11 | | 136 | 195 | 102 | 46 | 86 | 40 | 29 | 231 | 55 | 119 | 76 |
| 85 | 1.24 | 0.59 | 2.10 | 0.04 | 0.88925 | Hs3st3b1 | heparan sulfate (glucosamine) 3-O-sulfot | 172 | 210 | 94 | 32 | 63 | 14 | 27 | 37 | 80 | 162 | 46 |
| 10 | -3.83 | 1.82 | -2.10 | 0.04 | 0.88925 | Gm5668 | | 0 | 0 | 7 | 22 | 3 | 34 | 14 | 0 | 12 | 0 | 23 |
| 14 | -2.91 | 1.39 | -2.10 | 0.04 | 0.89302 | Cacng2 | calcium channel, voltage-dependent, gan | 19 | 28 | 34 | 4 | 24 | 7 | 23 | 0 | 14 | 0 | 1 |
| 4505 | 0.15 | 0.07 | 2.09 | 0.04 | 0.89671 | Ccng1 | cyclin G1 | 3956 | 3586 | 3393 | 4792 | 4595 | 3821 | 5121 | 5604 | 5173 | 5101 | 4418 |
| 3393 | 0.17 | 0.08 | 2.09 | 0.04 | 0.89671 | Rheb | Ras homolog enriched in brain | 3349 | 3161 | 3303 | 3457 | 3310 | 2735 | 3450 | 4180 | 3492 | 3635 | 3249 |
| 2381 | 0.18 | 0.08 | 2.10 | 0.04 | 0.89671 | Creb1 | cAMP responsive element binding proteir | 2784 | 2834 | 2375 | 2266 | 1823 | 2492 | 1992 | 2445 | 2235 | 2378 | 2572 |
| 253 | -0.46 | 0.22 | -2.10 | 0.04 | 0.89671 | Traf3ip1 | TRAF3 interacting protein 1 | 268 | 301 | 292 | 221 | 202 | 500 | 196 | 136 | 186 | 146 | 334 |
| 73 | -1.07 | 0.51 | -2.09 | 0.04 | 0.89671 | Calr3 | calreticulin 3 | 149 | 123 | 76 | 46 | 89 | 49 | 137 | 27 | 43 | 30 | 41 |
| 13 | -3.27 | 1.56 | -2.10 | 0.04 | 0.89671 | Gm26704 | | 57 | 15 | 37 | 6 | 1 | 9 | 11 | 0 | 0 | 0 | 10 |
| 3 | -6.86 | 3.28 | -2.09 | 0.04 | 0.89671 | Gm37472 | | 0 | 0 | 5 | 8 | 0 | 3 | 7 | 0 | 0 | 0 | 14 |
| 945 | 0.30 | 0.14 | 2.09 | 0.04 | 0.89795 | Usp18 | ubiquitin specific peptidase 18 | 668 | 710 | 939 | 929 | 1171 | 653 | 880 | 1266 | 1191 | 1240 | 749 |
| 67 | -2.04 | 0.97 | -2.09 | 0.04 | 0.89795 | Prr16 | proline rich 16 | 290 | 204 | 124 | 25 | 12 | 25 | 25 | 0 | 10 | 0 | 26 |
| 69 | -2.89 | 1.38 | -2.09 | 0.04 | 0.89795 | Ly9 | lymphocyte antigen 9 | 430 | 194 | 125 | 0 | 0 | 3 | 7 | 0 | 0 | 0 | 0 |
| 81 | -0.75 | 0.36 | -2.09 | 0.04 | 0.89988 | Mrpl48-ps | | 61 | 78 | 128 | 131 | 139 | 48 | 76 | 73 | 88 | 39 | 34 |

| | | | | | | | | | | | | | | | | | |
|---|---|---|---|---|---|---|---|---|---|---|---|---|---|---|---|---|---|
| 10 | -2.79 | 1.33 | -2.09 | 0.04 | 0.89988 | 2010109A12Ril | RIKEN cDNA 2010109A12 gene | 6 | 3 | 12 | 9 | 33 | 15 | 8 | 0 | 6 | 0 | 14 |
| 53 | -2.99 | 1.43 | -2.09 | 0.04 | 0.89988 | Il1rl2 | interleukin 1 receptor-like 2 | 282 | 129 | 166 | 0 | 0 | 5 | 5 | 0 | 0 | 0 | 0 |
| 2992 | -2.97 | 1.42 | -2.09 | 0.04 | 0.90421 | Col14a1 | collagen, type XIV, alpha 1 | 19055 | 5653 | 8136 | 3 | 31 | 2 | 14 | 0 | 2 | 0 | 11 |
| 19471 | 0.12 | 0.06 | 2.09 | 0.04 | 0.90468 | Eif4a2 | eukaryotic translation initiation factor 4A2 | 14174 | 16303 | 15087 | 20027 | 18891 | 21583 | 20317 | 23371 | 19813 | 20743 | 23876 |
| 3621 | 0.16 | 0.07 | 2.09 | 0.04 | 0.90468 | Rragd | Ras-related GTP binding D | 1422 | 1601 | 1439 | 4989 | 3846 | 3193 | 4729 | 5430 | 4536 | 5039 | 3603 |
| 2271 | 0.18 | 0.09 | 2.09 | 0.04 | 0.90468 | Sptssa | serine palmitoyltransferase, small subunit | 2590 | 2340 | 2316 | 2360 | 2040 | 1517 | 2381 | 2719 | 2146 | 2817 | 1750 |
| 516 | -0.81 | 0.39 | -2.09 | 0.04 | 0.90468 | Pik3ap1 | phosphoinositide-3-kinase adaptor protei | 2028 | 1648 | 1767 | 39 | 27 | 47 | 31 | 32 | 10 | 36 | 10 |
| 1951 | 0.28 | 0.13 | 2.09 | 0.04 | 0.90479 | Rpl22l1 | ribosomal protein L22 like 1 | 2822 | 2950 | 2729 | 1365 | 1213 | 1620 | 1636 | 2209 | 1386 | 1645 | 1885 |
| 8 | 4.17 | 2.00 | 2.09 | 0.04 | 0.90479 | 1700010K24Rik | | 9 | 9 | 19 | 0 | 0 | 3 | 0 | 0 | 7 | 41 | 3 |
| 41 | -2.16 | 1.04 | -2.08 | 0.04 | 0.90738 | Plcd3 | phospholipase C, delta 3 | 169 | 82 | 125 | 4 | 24 | 8 | 18 | 0 | 0 | 8 | 9 |
| 10 | -3.82 | 1.83 | -2.08 | 0.04 | 0.90864 | 8430406P12Rik | | 7 | 18 | 0 | 4 | 3 | 30 | 16 | 0 | 0 | 0 | 29 |
| 5445 | 0.19 | 0.09 | 2.08 | 0.04 | 0.91171 | Selenok | selenoprotein K | 3415 | 3470 | 3654 | 5810 | 6102 | 5203 | 5921 | 7364 | 5793 | 7338 | 5826 |
| 22903 | -0.69 | 0.33 | -2.08 | 0.04 | 0.91261 | Fos | FBJ osteosarcoma oncogene | 69329 | 42818 | 48820 | 3303 | 4071 | 35728 | 8320 | 1002 | 2778 | 6233 | 29529 |
| 7293 | 0.21 | 0.10 | 2.08 | 0.04 | 0.91261 | Cct5 | chaperonin containing Tcp1, subunit 5 (e | 8615 | 7343 | 8232 | 6210 | 6821 | 6384 | 6556 | 8871 | 6991 | 6873 | 7330 |
| 1813 | -0.30 | 0.15 | -2.08 | 0.04 | 0.91261 | Jmy | junction-mediating and regulatory protein | 1720 | 1512 | 2208 | 1806 | 1998 | 1979 | 2137 | 1211 | 2057 | 1328 | 1986 |
| 1659 | 0.23 | 0.11 | 2.08 | 0.04 | 0.91261 | Rhobtb3 | Rho-related BTB domain containing 3 | 1378 | 1272 | 1433 | 1371 | 1674 | 1614 | 1872 | 1920 | 2075 | 1892 | 1744 |
| 993 | -0.27 | 0.13 | -2.08 | 0.04 | 0.91261 | Hspa2 | heat shock protein 2 | 1415 | 1358 | 1322 | 859 | 995 | 878 | 994 | 791 | 680 | 940 | 692 |
| 153 | -1.72 | 0.83 | -2.08 | 0.04 | 0.91261 | Hr | lysine demethylase and nuclear receptor | 660 | 355 | 499 | 35 | 36 | 22 | 17 | 50 | 8 | 0 | 2 |
| 2574 | 0.16 | 0.08 | 2.08 | 0.04 | 0.91312 | Bet1 | Bet1 golgi vesicular membrane trafficking | 1827 | 1771 | 1896 | 2268 | 2877 | 2929 | 2707 | 2601 | 3137 | 2973 | 3334 |
| 1145 | -0.26 | 0.12 | -2.08 | 0.04 | 0.91467 | Nsmf | NMDA receptor synaptonuclear signaling | 1322 | 1023 | 1312 | 1140 | 1300 | 1193 | 1236 | 1044 | 1137 | 921 | 966 |
| 2789 | -0.33 | 0.16 | -2.07 | 0.04 | 0.91686 | Etv1 | ets variant 1 | 5779 | 8707 | 6735 | 1539 | 961 | 1105 | 1707 | 1087 | 1037 | 1130 | 895 |
| 717 | -0.52 | 0.25 | -2.07 | 0.04 | 0.91686 | Arc | activity regulated cytoskeletal-associated | 1226 | 739 | 958 | 94 | 87 | 2478 | 218 | 36 | 88 | 161 | 1805 |
| 435 | -0.86 | 0.42 | -2.07 | 0.04 | 0.91686 | Pde3a | phosphodiesterase 3A, cGMP inhibited | 1755 | 850 | 1035 | 134 | 134 | 274 | 185 | 82 | 125 | 26 | 184 |
| 107 | 0.68 | 0.33 | 2.07 | 0.04 | 0.91686 | Gm13331 | | 160 | 40 | 171 | 41 | 110 | 41 | 115 | 63 | 160 | 211 | 66 |
| 455 | -2.11 | 1.02 | -2.07 | 0.04 | 0.91686 | Fev | FEV transcription factor, ETS family memb | 1114 | 1616 | 1638 | 160 | 148 | 105 | 87 | 0 | 43 | 62 | 26 |
| 238 | -1.18 | 0.57 | -2.07 | 0.04 | 0.91686 | Itga4 | integrin alpha 4 | 1552 | 392 | 402 | 19 | 53 | 36 | 69 | 0 | 18 | 43 | 40 |
| 363 | -3.43 | 1.65 | -2.07 | 0.04 | 0.91686 | Ccl6 | chemokine (C-C motif) ligand 6 | 2194 | 889 | 848 | 0 | 5 | 13 | 32 | 0 | 0 | 0 | 10 |
| 462 | -2.08 | 1.00 | -2.07 | 0.04 | 0.91686 | Kirrel | kirre like nephrin family adhesion molecu | 2351 | 1097 | 1547 | 5 | 12 | 23 | 25 | 0 | 7 | 0 | 16 |
| 280 | 1.54 | 0.74 | 2.07 | 0.04 | 0.91686 | Mgst1 | microsomal glutathione S-transferase 1 | 1418 | 695 | 897 | 2 | 6 | 1 | 5 | 0 | 15 | 36 | 1 |
| 1658 | -2.90 | 1.40 | -2.07 | 0.04 | 0.91702 | Abca8a | ATP-binding cassette, sub-family A (ABC | 7704 | 4562 | 5915 | 10 | 20 | 4 | 9 | 0 | 18 | 0 | 0 |
| 659 | -0.50 | 0.24 | -2.07 | 0.04 | 0.9193 | P3h3 | prolyl 3-hydroxylase 3 | 2125 | 1220 | 1163 | 435 | 450 | 369 | 341 | 221 | 346 | 314 | 262 |
| 5268 | 0.16 | 0.08 | 2.07 | 0.04 | 0.92228 | Ddx1 | DEAD (Asp-Glu-Ala-Asp) box polypeptide | 4359 | 3923 | 4821 | 4975 | 5796 | 5042 | 5424 | 6115 | 5838 | 6139 | 5517 |
| 74 | -2.49 | 1.20 | -2.07 | 0.04 | 0.92228 | Gdpd5 | glycerophosphodiester phosphodiesteras | 314 | 215 | 229 | 25 | 16 | 4 | 0 | 0 | 4 | 0 | 3 |
| 1315 | 0.23 | 0.11 | 2.07 | 0.04 | 0.92325 | Bfar | bifunctional apoptosis regulator | 1473 | 1425 | 1308 | 1158 | 1129 | 1399 | 1049 | 1453 | 1260 | 1360 | 1447 |
| 11 | -3.89 | 1.88 | -2.07 | 0.04 | 0.92341 | Kirrel3 | kirre like nephrin family adhesion molecu | 53 | 17 | 25 | 5 | 15 | 3 | 0 | 0 | 0 | 0 | 1 |
| 236 | -0.62 | 0.30 | -2.07 | 0.04 | 0.92599 | Pla2g2d | phospholipase A2, group IID | 896 | 496 | 342 | 170 | 92 | 96 | 168 | 109 | 60 | 82 | 81 |
| 218 | -0.75 | 0.36 | -2.07 | 0.04 | 0.92599 | Il17rd | interleukin 17 receptor D | 623 | 739 | 652 | 100 | 33 | 45 | 65 | 35 | 25 | 54 | 26 |
| 1029 | 0.27 | 0.13 | 2.07 | 0.04 | 0.92602 | Slco3a1 | solute carrier organic anion transporter fa | 1742 | 1305 | 1326 | 861 | 642 | 710 | 941 | 1030 | 904 | 1019 | 834 |
| 85 | -0.82 | 0.40 | -2.06 | 0.04 | 0.92648 | 1700003E16Ril | RIKEN cDNA 1700003E16 gene | 88 | 175 | 114 | 139 | 77 | 62 | 74 | 83 | 39 | 64 | 22 |
| 451 | -0.39 | 0.19 | -2.06 | 0.04 | 0.92788 | Zbtb45 | zinc finger and BTB domain containing 45 | 539 | 475 | 494 | 529 | 565 | 425 | 437 | 422 | 404 | 301 | 367 |
| 2204 | -0.18 | 0.09 | -2.06 | 0.04 | 0.92802 | Tspan15 | tetraspanin 15 | 1906 | 2501 | 2375 | 2511 | 2457 | 2136 | 2172 | 2062 | 2322 | 1941 | 1862 |
| 346 | -0.76 | 0.37 | -2.06 | 0.04 | 0.92802 | Lats2 | large tumor suppressor 2 | 1383 | 1058 | 959 | 63 | 47 | 57 | 80 | 76 | 34 | 32 | 15 |
| 13 | -2.78 | 1.35 | -2.06 | 0.04 | 0.929 | | | 9 | 24 | 14 | 5 | 12 | 27 | 34 | 0 | 7 | 0 | 9 |
| 19100 | 0.15 | 0.08 | 2.06 | 0.04 | 0.93216 | Glul | glutamate-ammonia ligase (glutamine syr | 11124 | 11913 | 12474 | 23910 | 19950 | 16944 | 21600 | 28945 | 22376 | 22608 | 18256 |
| 6183 | -0.12 | 0.06 | -2.06 | 0.04 | 0.93216 | Zc3h7b | zinc finger CCCH type containing 7B | 6873 | 6940 | 7078 | 6240 | 6658 | 4942 | 6703 | 5541 | 6125 | 6127 | 4780 |
| 1167 | 0.25 | 0.12 | 2.06 | 0.04 | 0.93216 | Fbxo7 | F-box protein 7 | 791 | 801 | 799 | 1062 | 1395 | 1027 | 1295 | 1527 | 1565 | 1361 | 1219 |
| 717 | 0.27 | 0.13 | 2.06 | 0.04 | 0.93216 | Fam199x | family with sequence similarity 199, X-link | 715 | 791 | 494 | 690 | 511 | 778 | 689 | 852 | 604 | 912 | 850 |
| 496 | -0.43 | 0.21 | -2.06 | 0.04 | 0.93216 | Nhsl1 | NHS-like 1 | 1273 | 1461 | 1202 | 285 | 193 | 215 | 172 | 232 | 170 | 137 | 112 |
| 6064 | -0.17 | 0.08 | -2.06 | 0.04 | 0.93275 | Actg1 | actin, gamma, cytoplasmic 1 | 10724 | 10172 | 8875 | 4917 | 4785 | 4909 | 4907 | 4072 | 4398 | 5029 | 3921 |
| 4346 | 0.15 | 0.07 | 2.06 | 0.04 | 0.93275 | Becn1 | beclin 1, autophagy related | 3876 | 3555 | 4025 | 4123 | 4373 | 4358 | 4405 | 4793 | 4572 | 5156 | 4572 |
| 217 | -1.55 | 0.75 | -2.06 | 0.04 | 0.93436 | Fkbp10 | FK506 binding protein 10 | 1027 | 626 | 616 | 31 | 28 | 13 | 8 | 22 | 2 | 0 | 9 |
| 2488 | -0.22 | 0.11 | -2.06 | 0.04 | 0.93534 | Ehd4 | EH-domain containing 4 | 8235 | 6721 | 7672 | 682 | 692 | 510 | 677 | 551 | 620 | 520 | 491 |
| 10 | -3.23 | 1.57 | -2.05 | 0.04 | 0.93664 | AA467197 | expressed sequence AA467197 | 14 | 16 | 39 | 9 | 9 | 5 | 8 | 0 | 13 | 0 | 0 |
| 171 | 1.82 | 0.89 | 2.05 | 0.04 | 0.93664 | Cryab | crystallin, alpha B | 814 | 508 | 469 | 3 | 11 | 1 | 3 | 46 | 12 | 0 | 8 |
| 3042 | 0.20 | 0.10 | 2.05 | 0.04 | 0.93747 | Elp2 | elongator acetyltransferase complex subu | 2571 | 2595 | 2809 | 2666 | 2980 | 3196 | 3029 | 3757 | 3199 | 3174 | 3487 |
| 486 | -0.51 | 0.25 | -2.05 | 0.04 | 0.9387 | Hirip3 | HIRA interacting protein 3 | 663 | 397 | 492 | 459 | 479 | 702 | 546 | 285 | 406 | 241 | 682 |
| 75156 | 0.10 | 0.05 | 2.05 | 0.04 | 0.94162 | Hspa5 | heat shock protein 5 | 46739 | 40727 | 51433 | 72100 | 91071 | 87944 | 81665 | 82329 | 91992 | 83631 | 97087 |
| 4 | -4.45 | 2.17 | -2.05 | 0.04 | 0.94162 | Gm15603 | | 0 | 0 | 3 | 18 | 5 | 1 | 19 | 0 | 2 | 0 | 1 |
| 205 | -1.80 | 0.88 | -2.05 | 0.04 | 0.94184 | Adamts15 | a disintegrin-like and metallopeptidase (r | 882 | 595 | 733 | 18 | 7 | 5 | 4 | 0 | 3 | 0 | 9 |
| 25 | -2.52 | 1.23 | -2.05 | 0.04 | 0.94201 | Gm10416 | | 58 | 61 | 31 | 42 | 9 | 28 | 17 | 0 | 16 | 0 | 7 |

| | | | | | | | | | | | | | | | | | | |
|---|---|---|---|---|---|---|---|---|---|---|---|---|---|---|---|---|---|---|
| 334 | -0.67 | 0.33 | -2.05 | 0.04 | 0.94488 | Ramp1 | receptor (calcitonin) activity modifying pro | 506 | 728 | 766 | 307 | 316 | 227 | 184 | 100 | 156 | 224 | 163 |
| 6 | -4.90 | 2.39 | -2.05 | 0.04 | 0.94644 | 4930470G03Rik | | 5 | 15 | 0 | 0 | 2 | 14 | 14 | 0 | 0 | 0 | 20 |
| 217 | 0.54 | 0.27 | 2.05 | 0.04 | 0.9473 | Zfp747 | zinc finger protein 747 | 297 | 229 | 217 | 185 | 125 | 204 | 156 | 256 | 229 | 286 | 201 |
| 133 | -0.78 | 0.38 | -2.05 | 0.04 | 0.94755 | Mex3a | mex3 RNA binding family member A | 257 | 225 | 316 | 118 | 69 | 98 | 129 | 49 | 51 | 102 | 48 |
| 3782 | 0.24 | 0.12 | 2.04 | 0.04 | 0.94793 | Prkcb | protein kinase C, beta | 2160 | 1990 | 1764 | 4235 | 4142 | 3544 | 4281 | 6635 | 4462 | 4620 | 3767 |
| 1783 | 0.22 | 0.11 | 2.04 | 0.04 | 0.94793 | Pex7 | peroxisomal biogenesis factor 7 | 1232 | 1274 | 1112 | 1850 | 1726 | 1824 | 1957 | 2505 | 1824 | 2451 | 1859 |
| 1862 | -0.36 | 0.17 | -2.04 | 0.04 | 0.94793 | Mbd6 | methyl-CpG binding domain protein 6 | 1626 | 1812 | 1665 | 1845 | 1739 | 3161 | 1809 | 914 | 1701 | 1579 | 2628 |
| 1148 | -0.32 | 0.15 | -2.04 | 0.04 | 0.94793 | Vsig10 | V-set and immunoglobulin domain contai | 1462 | 1623 | 1942 | 1174 | 1066 | 980 | 976 | 1121 | 816 | 632 | 841 |
| 984 | -0.31 | 0.15 | -2.04 | 0.04 | 0.94793 | Leo1 | Leo1, Paf1/RNA polymerase II complex co | 1071 | 1090 | 1506 | 941 | 1144 | 946 | 884 | 614 | 1073 | 607 | 951 |
| 680 | -0.35 | 0.17 | -2.04 | 0.04 | 0.94793 | Olfm1 | olfactomedin 1 | 1602 | 1360 | 1324 | 404 | 470 | 382 | 537 | 393 | 384 | 313 | 308 |
| 411 | -0.37 | 0.18 | -2.04 | 0.04 | 0.94793 | Tmem63c | transmembrane protein 63c | 224 | 301 | 274 | 513 | 520 | 521 | 536 | 455 | 438 | 322 | 416 |
| 447 | -0.55 | 0.27 | -2.05 | 0.04 | 0.94793 | Dclk1 | doublecortin-like kinase 1 | 1487 | 1192 | 747 | 298 | 147 | 284 | 168 | 131 | 137 | 120 | 210 |
| 80 | -1.73 | 0.85 | -2.04 | 0.04 | 0.94793 | Ccdc39 | coiled-coil domain containing 39 | 86 | 83 | 61 | 59 | 35 | 276 | 40 | 5 | 38 | 0 | 196 |
| 41 | -2.62 | 1.28 | -2.04 | 0.04 | 0.94793 | Adgrg3 | adhesion G protein-coupled receptor G3 | 102 | 22 | 72 | 3 | 0 | 138 | 13 | 0 | 0 | 0 | 100 |
| 65 | -1.41 | 0.69 | -2.04 | 0.04 | 0.94859 | Prox1os | | 178 | 229 | 143 | 32 | 33 | 31 | 22 | 21 | 17 | 0 | 12 |
| 96 | -2.14 | 1.05 | -2.04 | 0.04 | 0.9507 | Fbln7 | fibulin 7 | 358 | 243 | 317 | 66 | 18 | 25 | 3 | 0 | 13 | 0 | 18 |
| 17858 | -0.11 | 0.06 | -2.04 | 0.04 | 0.9512 | App | amyloid beta (A4) precursor protein | 42491 | 38711 | 39071 | 10109 | 10392 | 7200 | 11715 | 9008 | 9876 | 11638 | 6229 |
| 2885 | 0.14 | 0.07 | 2.04 | 0.04 | 0.9512 | Csnk2a1 | casein kinase 2, alpha 1 polypeptide | 2400 | 2580 | 2649 | 2696 | 2924 | 3126 | 2722 | 3156 | 3241 | 2988 | 3247 |
| 9490 | -0.12 | 0.06 | -2.04 | 0.04 | 0.95641 | Ank | progressive ankylosis | 8375 | 9299 | 9264 | 10388 | 10566 | 8929 | 10470 | 9001 | 10003 | 9561 | 8537 |
| 12363 | 0.13 | 0.07 | 2.03 | 0.04 | 0.95874 | Rab2a | RAB2A, member RAS oncogene family | 9446 | 9965 | 10054 | 12851 | 12568 | 11110 | 14254 | 15500 | 13391 | 14478 | 12373 |
| 258 | -0.58 | 0.28 | -2.04 | 0.04 | 0.95874 | Lrp8os3 | | 134 | 220 | 163 | 486 | 332 | 301 | 263 | 224 | 292 | 126 | 295 |
| 484 | -3.25 | 1.60 | -2.03 | 0.04 | 0.95874 | Adgre1 | adhesion G protein-coupled receptor E1 | 2415 | 1111 | 1696 | 0 | 20 | 14 | 57 | 0 | 9 | 0 | 1 |
| 5 | -3.76 | 1.85 | -2.04 | 0.04 | 0.95874 | Gm14487 | | 0 | 2 | 2 | 20 | 9 | 4 | 10 | 0 | 3 | 0 | 1 |
| 9844 | 0.17 | 0.08 | 2.03 | 0.04 | 0.95978 | Hspa9 | heat shock protein 9 | 7813 | 6784 | 7710 | 9531 | 10330 | 9340 | 11337 | 12794 | 11009 | 11376 | 10256 |
| 4647 | 0.14 | 0.07 | 2.03 | 0.04 | 0.95978 | Rnf141 | ring finger protein 141 | 3416 | 4077 | 2965 | 5323 | 4440 | 4298 | 5211 | 6051 | 4683 | 6050 | 4603 |
| 3254 | -0.19 | 0.09 | -2.03 | 0.04 | 0.95978 | Rcc2 | regulator of chromosome condensation 2 | 4676 | 3823 | 4495 | 3211 | 3411 | 2718 | 2807 | 2546 | 2975 | 2705 | 2421 |
| 1659 | -0.24 | 0.12 | -2.03 | 0.04 | 0.95978 | Pik3r2 | phosphoinositide-3-kinase regulatory sub | 1953 | 1809 | 1572 | 1761 | 1903 | 1545 | 1782 | 1468 | 1665 | 1412 | 1374 |
| 1039 | 0.48 | 0.24 | 2.03 | 0.04 | 0.95978 | Nme1 | NME/NM23 nucleoside diphosphate kina | 810 | 518 | 894 | 884 | 945 | 867 | 1141 | 1934 | 1148 | 1230 | 1059 |
| 1377 | -2.05 | 1.01 | -2.03 | 0.04 | 0.95978 | Cav1 | caveolin 1, caveolae protein | 6692 | 4570 | 3799 | 10 | 24 | 3 | 20 | 0 | 4 | 22 | 0 |
| 194 | -1.43 | 0.71 | -2.03 | 0.04 | 0.95978 | Itih2 | inter-alpha trypsin inhibitor, heavy chain 2 | 719 | 766 | 565 | 9 | 31 | 8 | 13 | 0 | 13 | 13 | 1 |
| 187 | -2.18 | 1.07 | -2.03 | 0.04 | 0.95978 | Fyb | FYN binding protein | 998 | 504 | 539 | 5 | 5 | 3 | 3 | 0 | 4 | 0 | 0 |
| 12 | -3.64 | 1.79 | -2.03 | 0.04 | 0.95978 | Trpc5 | transient receptor potential cation channe | 33 | 55 | 8 | 9 | 10 | 8 | 0 | 0 | 0 | 0 | 4 |
| 5 | 6.38 | 3.14 | 2.03 | 0.04 | 0.95978 | Vamp9 | vesicle-associated membrane protein 9 | 7 | 0 | 6 | 0 | 0 | 3 | 0 | 36 | 4 | 0 | 0 |
| 5084 | -0.27 | 0.14 | -2.03 | 0.04 | 0.95993 | Rps11 | ribosomal protein S11 | 9884 | 8468 | 7994 | 3936 | 4340 | 2902 | 4841 | 2339 | 3916 | 4892 | 2408 |
| 3455 | -0.21 | 0.10 | -2.03 | 0.04 | 0.95993 | Kat6a | K(lysine) acetyltransferase 6A | 3801 | 4094 | 3383 | 3848 | 3395 | 3612 | 3467 | 3397 | 3366 | 2499 | 3144 |
| 60 | -1.60 | 0.79 | -2.03 | 0.04 | 0.95993 | Cer1 | cerberus 1, DAN family BMP antagonist | 122 | 189 | 141 | 52 | 26 | 36 | 26 | 37 | 4 | 0 | 28 |
| 530 | -0.34 | 0.17 | -2.03 | 0.04 | 0.96045 | Fbxo30 | F-box protein 30 | 752 | 901 | 715 | 507 | 335 | 636 | 448 | 368 | 320 | 287 | 563 |
| 8799 | 0.15 | 0.07 | 2.03 | 0.04 | 0.9606 | Npc2 | NPC intracellular cholesterol transporter 2 | 13793 | 12101 | 13309 | 7588 | 7248 | 4734 | 7683 | 9131 | 7526 | 8526 | 5149 |
| 3057 | -0.16 | 0.08 | -2.03 | 0.04 | 0.9606 | Eloa | elongin A | 3598 | 3136 | 3302 | 2870 | 3436 | 2967 | 3154 | 2478 | 2921 | 2996 | 2772 |
| 1238 | -0.39 | 0.19 | -2.03 | 0.04 | 0.9606 | Khnyn | KH and NYN domain containing | 1597 | 1268 | 1502 | 1152 | 1271 | 1755 | 993 | 558 | 1314 | 802 | 1401 |
| 2424 | -0.23 | 0.11 | -2.03 | 0.04 | 0.9606 | Klf6 | Kruppel-like factor 6 | 8468 | 4860 | 4838 | 543 | 584 | 2640 | 800 | 511 | 478 | 609 | 2333 |
| 18 | -2.46 | 1.21 | -2.03 | 0.04 | 0.9606 | Mfsd4b3-ps | | 27 | 15 | 42 | 11 | 30 | 12 | 24 | 0 | 11 | 0 | 20 |
| 6 | -5.53 | 2.73 | -2.03 | 0.04 | 0.9606 | Gm38157 | | 3 | 0 | 0 | 28 | 0 | 12 | 3 | 0 | 0 | 0 | 16 |
| 39 | -2.78 | 1.37 | -2.03 | 0.04 | 0.96103 | Gm9899 | | 78 | 190 | 122 | 0 | 13 | 8 | 11 | 0 | 1 | 0 | 10 |
| 687 | 0.46 | 0.23 | 2.03 | 0.04 | 0.96112 | Nckipsd | NCK interacting protein with SH3 domain | 651 | 563 | 833 | 445 | 549 | 755 | 592 | 936 | 721 | 845 | 669 |
| 305 | -1.29 | 0.64 | -2.03 | 0.04 | 0.96112 | Msrb3 | methionine sulfoxide reductase B3 | 1562 | 708 | 879 | 39 | 12 | 32 | 57 | 15 | 11 | 0 | 36 |
| 5036 | -0.17 | 0.08 | -2.02 | 0.04 | 0.96439 | Hipk1 | homeodomain interacting protein kinase 1 | 6622 | 6474 | 6686 | 4709 | 4974 | 4208 | 4909 | 3583 | 4714 | 4722 | 3794 |
| 2579 | 0.26 | 0.13 | 2.02 | 0.04 | 0.96439 | Ift20 | intraflagellar transport 20 | 1476 | 1298 | 1800 | 2276 | 2851 | 3087 | 2640 | 3157 | 2655 | 3479 | 3651 |
| 1714 | -0.21 | 0.10 | -2.02 | 0.04 | 0.96439 | Kdm3b | KDM3B lysine (K)-specific demethylase 3 | 1998 | 2004 | 1845 | 1637 | 1706 | 2016 | 1582 | 1299 | 1461 | 1319 | 1983 |
| 85 | -2.63 | 1.30 | -2.02 | 0.04 | 0.96439 | Fcgr1 | Fc receptor, IgG, high affinity I | 315 | 155 | 443 | 0 | 3 | 0 | 17 | 0 | 1 | 0 | 0 |
| 13 | -2.59 | 1.28 | -2.02 | 0.04 | 0.96439 | 4921504A21Rik | | 26 | 35 | 31 | 22 | 9 | 4 | 9 | 0 | 6 | 0 | 2 |
| 10 | -3.67 | 1.82 | -2.02 | 0.04 | 0.96439 | Gm11734 | | 4 | 45 | 47 | 2 | 6 | 12 | 0 | 0 | 0 | 0 | 0 |
| 3303 | 0.28 | 0.14 | 2.02 | 0.04 | 0.96534 | Eif3m | eukaryotic translation initiation factor 3, su | 2973 | 2633 | 3103 | 3054 | 3364 | 2964 | 3077 | 4872 | 3313 | 3250 | 3734 |
| 7 | 3.58 | 1.77 | 2.02 | 0.04 | 0.96739 | Gm44672 | | 0 | 0 | 10 | 0 | 5 | 7 | 1 | 14 | 9 | 12 | 22 |
| 6136 | -0.16 | 0.08 | -2.02 | 0.04 | 0.96739 | Tjp1 | tight junction protein 1 | 4562 | 4859 | 4977 | 7323 | 6771 | 7742 | 6182 | 5552 | 6856 | 5893 | 6779 |
| 853 | -0.48 | 0.24 | -2.02 | 0.04 | 0.96739 | Plekhb1 | pleckstrin homology domain containing, fa | 1506 | 2249 | 2091 | 522 | 637 | 416 | 487 | 366 | 279 | 535 | 301 |
| 313 | -2.40 | 1.19 | -2.02 | 0.04 | 0.96739 | Rasip1 | Ras interacting protein 1 | 1259 | 958 | 1111 | 36 | 19 | 11 | 22 | 0 | 15 | 0 | 9 |
| 432 | -0.79 | 0.39 | -2.02 | 0.04 | 0.96863 | Syt1 | synaptotagmin I | 847 | 1172 | 1262 | 229 | 257 | 165 | 263 | 41 | 156 | 189 | 169 |
| 8580 | 0.20 | 0.10 | 2.02 | 0.04 | 0.96924 | Maob | monoamine oxidase B | 2984 | 3467 | 4034 | 10060 | 9666 | 8601 | 10747 | 11954 | 10612 | 11760 | 10496 |
| 348 | -0.36 | 0.18 | -2.02 | 0.04 | 0.96924 | Nr6a1 | nuclear receptor subfamily 6, group A, me | 310 | 268 | 255 | 319 | 306 | 729 | 309 | 251 | 214 | 246 | 614 |

| | | | | | | | | | | | | | | | | | | |
|---|---|---|---|---|---|---|---|---|---|---|---|---|---|---|---|---|---|---|
| 701 | -0.49 | 0.24 | -2.02 | 0.04 | 0.96924 | Npas4 | neuronal PAS domain protein 4 | 949 | 651 | 813 | 99 | 171 | 2387 | 285 | 52 | 118 | 220 | 1960 |
| 5 | -4.29 | 2.13 | -2.02 | 0.04 | 0.96924 | Vsig1 | V-set and immunoglobulin domain contai | 8 | 0 | 6 | 9 | 6 | 18 | 6 | 0 | 8 | 0 | 0 |
| 5 | -4.33 | 2.15 | -2.02 | 0.04 | 0.96924 | | | 13 | 1 | 19 | 5 | 12 | 2 | 1 | 0 | 0 | 0 | 0 |
| 3525 | 0.22 | 0.11 | 2.01 | 0.04 | 0.96977 | Cdh8 | cadherin 8 | 1425 | 1967 | 1673 | 4129 | 3534 | 3648 | 4299 | 4610 | 4466 | 4387 | 4642 |
| 449 | -0.55 | 0.27 | -2.01 | 0.04 | 0.96977 | Msi1 | musashi RNA-binding protein 1 | 450 | 768 | 564 | 437 | 410 | 471 | 554 | 231 | 326 | 434 | 293 |
| 220 | -0.64 | 0.32 | -2.01 | 0.04 | 0.96977 | Tspyl3 | TSPY-like 3 | 384 | 354 | 477 | 205 | 201 | 101 | 229 | 190 | 106 | 89 | 82 |
| 711 | -0.38 | 0.19 | -2.01 | 0.04 | 0.96977 | Vwa1 | von Willebrand factor A domain containin | 2107 | 1973 | 2413 | 190 | 213 | 152 | 192 | 161 | 119 | 176 | 120 |
| 402 | -0.61 | 0.30 | -2.01 | 0.04 | 0.96977 | Plekho2 | pleckstrin homology domain containing, f | 2093 | 1130 | 888 | 49 | 50 | 33 | 53 | 22 | 31 | 45 | 25 |
| 6 | -4.32 | 2.15 | -2.01 | 0.04 | 0.96977 | BC023105 | | 5 | 2 | 0 | 12 | 9 | 10 | 19 | 0 | 13 | 0 | 0 |
| 115 | -2.00 | 0.99 | -2.01 | 0.04 | 0.96977 | Trabd2b | TraB domain containing 2B | 540 | 321 | 342 | 39 | 0 | 3 | 11 | 6 | 3 | 0 | 1 |
| 4799 | -0.14 | 0.07 | -2.01 | 0.04 | 0.96988 | Rhob | ras homolog family member B | 12048 | 10429 | 11633 | 1728 | 1905 | 3995 | 2121 | 1420 | 1738 | 1993 | 3775 |
| 454 | -0.52 | 0.26 | -2.01 | 0.04 | 0.96988 | Nrsn1 | neurensin 1 | 546 | 799 | 692 | 498 | 528 | 232 | 495 | 213 | 418 | 359 | 210 |
| 213 | -0.67 | 0.33 | -2.01 | 0.04 | 0.96988 | 2900076A07Rik | | 57 | 141 | 135 | 184 | 278 | 527 | 137 | 102 | 203 | 48 | 532 |
| 819 | -3.29 | 1.64 | -2.01 | 0.04 | 0.96988 | Tgfbi | transforming growth factor, beta induced | 4452 | 1786 | 2738 | 0 | 8 | 5 | 18 | 0 | 0 | 0 | 6 |
| 114 | 2.75 | 1.37 | 2.01 | 0.04 | 0.96988 | Tnfrsf19 | tumor necrosis factor receptor superfamily | 564 | 327 | 354 | 0 | 0 | 0 | 0 | 0 | 2 | 0 | 3 |
| 63 | -2.72 | 1.35 | -2.01 | 0.04 | 0.96988 | Gsta3 | glutathione S-transferase, alpha 3 | 315 | 146 | 224 | 0 | 2 | 8 | 0 | 0 | 0 | 0 | 0 |
| 445 | -0.43 | 0.21 | -2.01 | 0.04 | 0.96993 | N4bp2os | | 375 | 600 | 541 | 437 | 529 | 421 | 541 | 415 | 420 | 285 | 331 |
| 10856 | -0.10 | 0.05 | -2.01 | 0.04 | 0.97064 | Slc6a6 | solute carrier family 6 (neurotransmitter tra | 8578 | 7986 | 9519 | 12215 | 13006 | 10883 | 12142 | 10829 | 12931 | 11361 | 9963 |
| 2290 | 0.17 | 0.09 | 2.01 | 0.04 | 0.97346 | Mfsd14b | major facilitator superfamily domain conta | 2318 | 2576 | 2417 | 2048 | 2162 | 1984 | 2217 | 2280 | 2517 | 2455 | 2218 |
| 11816 | 0.15 | 0.08 | 2.01 | 0.04 | 0.97353 | Scgn | secretagogin, EF-hand calcium binding p | 11597 | 13220 | 12674 | 11001 | 11133 | 10107 | 11501 | 13911 | 10893 | 12646 | 11297 |
| 1133 | 0.33 | 0.17 | 2.01 | 0.04 | 0.9744 | Ptprd | protein tyrosine phosphatase, receptor typ | 2277 | 3187 | 2329 | 466 | 405 | 562 | 632 | 719 | 515 | 670 | 698 |
| 341 | -0.39 | 0.19 | -2.01 | 0.04 | 0.9744 | Fmo5 | flavin containing monooxygenase 5 | 727 | 547 | 420 | 277 | 248 | 197 | 456 | 239 | 178 | 285 | 177 |
| 8 | -2.84 | 1.42 | -2.01 | 0.04 | 0.97447 | Gm19510 | | 6 | 13 | 12 | 25 | 4 | 17 | 6 | 0 | 4 | 0 | 5 |
| 2143 | -0.26 | 0.13 | -2.01 | 0.04 | 0.97574 | Bhlhb9 | basic helix-loop-helix domain containing, | 2451 | 3052 | 2813 | 2082 | 1889 | 2120 | 2197 | 1448 | 1738 | 1640 | 2146 |
| 636 | 0.35 | 0.18 | 2.01 | 0.04 | 0.97574 | Mrpl16 | mitochondrial ribosomal protein L16 | 573 | 678 | 671 | 611 | 639 | 490 | 478 | 897 | 637 | 792 | 533 |
| 939 | -0.30 | 0.15 | -2.00 | 0.04 | 0.97633 | Klf10 | Kruppel-like factor 10 | 1381 | 1283 | 1235 | 704 | 808 | 959 | 1055 | 520 | 617 | 799 | 963 |
| 9 | -2.89 | 1.44 | -2.00 | 0.04 | 0.97633 | Gm34006 | | 6 | 12 | 8 | 4 | 9 | 21 | 17 | 0 | 2 | 0 | 16 |
| 4601 | 0.18 | 0.09 | 2.00 | 0.05 | 0.97641 | Psip1 | PC4 and SFRS1 interacting protein 1 | 4850 | 4612 | 4552 | 4098 | 4073 | 4499 | 4481 | 5487 | 4347 | 4409 | 5200 |
| 7853 | 0.15 | 0.07 | 2.00 | 0.05 | 0.97725 | Ubxn4 | UBX domain protein 4 | 4393 | 4558 | 5410 | 6911 | 9447 | 9858 | 8066 | 8974 | 9577 | 8643 | 10548 |
| 7 | -3.48 | 1.74 | -2.00 | 0.05 | 0.97725 | F420014N23Rik | | 18 | 0 | 3 | 11 | 7 | 11 | 20 | 0 | 3 | 0 | 2 |
| 8 | -4.19 | 2.09 | -2.00 | 0.05 | 0.97725 | Ascl1 | achaete-scute family bHLH transcription f | 21 | 22 | 19 | 13 | 1 | 10 | 0 | 0 | 0 | 0 | 0 |
| 1543 | -0.21 | 0.11 | -2.00 | 0.05 | 0.97791 | Npc1 | NPC intracellular cholesterol transporter | 1635 | 1555 | 1506 | 1938 | 1714 | 1342 | 1573 | 1679 | 1455 | 1508 | 1066 |
| 386 | -3.23 | 1.61 | -2.00 | 0.05 | 0.97791 | Cd163 | CD163 antigen | 2504 | 679 | 1007 | 0 | 21 | 3 | 29 | 0 | 5 | 0 | 0 |
| 2659 | -0.19 | 0.10 | -2.00 | 0.05 | 0.98092 | Selenoi | selenoprotein I | 2798 | 4433 | 3813 | 2486 | 2628 | 2497 | 2081 | 2178 | 2346 | 1716 | 2277 |
| 1192 | 0.28 | 0.14 | 2.00 | 0.05 | 0.98092 | Cox5a | cytochrome c oxidase subunit 5A | 1164 | 1151 | 1183 | 1066 | 1093 | 951 | 1210 | 1683 | 1206 | 1403 | 1002 |
| 405 | -0.45 | 0.23 | -2.00 | 0.05 | 0.98092 | Zfp954 | zinc finger protein 954 | 331 | 517 | 476 | 409 | 418 | 469 | 510 | 313 | 285 | 359 | 367 |
| 529 | -0.40 | 0.20 | -2.00 | 0.05 | 0.98092 | Slc41a2 | solute carrier family 41, member 2 | 887 | 1150 | 1162 | 312 | 443 | 287 | 454 | 279 | 306 | 282 | 254 |
| 33 | 1.61 | 0.80 | 2.00 | 0.05 | 0.98092 | AI593442 | expressed sequence AI593442 | 45 | 55 | 10 | 24 | 3 | 18 | 11 | 38 | 18 | 128 | 11 |
| 920 | 0.30 | 0.15 | 2.00 | 0.05 | 0.98277 | Wdr24 | WD repeat domain 24 | 642 | 755 | 891 | 852 | 844 | 947 | 863 | 1189 | 1081 | 1112 | 950 |
| 36 | -2.92 | 1.46 | -2.00 | 0.05 | 0.98404 | Mfsd4a | major facilitator superfamily domain conta | 113 | 166 | 109 | 0 | 3 | 3 | 6 | 0 | 0 | 0 | 1 |
| 4261 | -0.13 | 0.07 | -1.99 | 0.05 | 0.9856 | 2310022B05Ril | RIKEN cDNA 2310022B05 gene | 5689 | 5540 | 5699 | 4022 | 4336 | 3132 | 4159 | 3512 | 3775 | 4132 | 2874 |
| 3155 | 0.14 | 0.07 | 1.99 | 0.05 | 0.9856 | Dusp3 | dual specificity phosphatase 3 (vaccinia v | 4493 | 4448 | 4252 | 2798 | 2536 | 2112 | 2790 | 3236 | 2696 | 3066 | 2284 |
| 2040 | 0.21 | 0.10 | 1.99 | 0.05 | 0.9856 | Cebpz | CCAAT/enhancer binding protein zeta | 2045 | 1791 | 1797 | 1725 | 2003 | 2112 | 1954 | 2255 | 2113 | 2009 | 2630 |
| 2133 | 0.22 | 0.11 | 1.99 | 0.05 | 0.9856 | Zfp622 | zinc finger protein 622 | 1717 | 2157 | 1533 | 2139 | 1725 | 2540 | 1879 | 3178 | 1776 | 2160 | 2664 |
| 2544 | -0.16 | 0.08 | -1.99 | 0.05 | 0.9856 | Pnn | pinin | 1805 | 2165 | 2013 | 1998 | 1964 | 5686 | 1829 | 1888 | 1704 | 1497 | 5435 |
| 1148 | 0.24 | 0.12 | 2.00 | 0.05 | 0.9856 | Nsmce3 | NSE3 homolog, SMC5-SMC6 complex cc | 859 | 1303 | 902 | 1272 | 1085 | 1009 | 1003 | 1630 | 1137 | 1319 | 1110 |
| 781 | -0.30 | 0.15 | -1.99 | 0.05 | 0.9856 | Slc35e2 | solute carrier family 35, member E2 | 608 | 514 | 673 | 864 | 790 | 1219 | 844 | 550 | 673 | 726 | 1133 |
| 880 | 0.33 | 0.17 | 1.99 | 0.05 | 0.9856 | Rfwd3 | ring finger and WD repeat domain 3 | 944 | 818 | 713 | 911 | 800 | 815 | 665 | 988 | 1057 | 1086 | 886 |
| 311 | -0.41 | 0.21 | -1.99 | 0.05 | 0.9856 | Il17re | interleukin 17 receptor E | 238 | 210 | 281 | 427 | 382 | 347 | 381 | 341 | 271 | 290 | 253 |
| 103 | -0.75 | 0.38 | -1.99 | 0.05 | 0.9856 | Gm47484 | | 94 | 130 | 87 | 151 | 114 | 158 | 97 | 56 | 102 | 62 | 87 |
| 143 | -0.87 | 0.44 | -1.99 | 0.05 | 0.9856 | Fstl5 | follistatin-like 5 | 261 | 668 | 235 | 67 | 77 | 50 | 63 | 50 | 45 | 15 | 39 |
| 6 | -3.87 | 1.94 | -1.99 | 0.05 | 0.9856 | Gm45242 | | 3 | 0 | 14 | 4 | 13 | 8 | 15 | 0 | 13 | 0 | 0 |
| 23 | -1.85 | 0.93 | -1.99 | 0.05 | 0.9856 | Pabpc4l | poly(A) binding protein, cytoplasmic 4-like | 57 | 64 | 48 | 30 | 13 | 14 | 12 | 0 | 5 | 6 | 8 |
| 14 | -2.68 | 1.35 | -1.99 | 0.05 | 0.9856 | 6820408C15Ril | RIKEN cDNA 6820408C15 gene | 15 | 12 | 36 | 32 | 6 | 17 | 12 | 0 | 10 | 0 | 9 |
| 4 | -6.21 | 3.12 | -1.99 | 0.05 | 0.9856 | Gm43201 | | 0 | 5 | 0 | 0 | 9 | 9 | 3 | 0 | 0 | 0 | 15 |
| 5 | -3.98 | 2.00 | -1.99 | 0.05 | 0.9856 | Gm3550 | | 2 | 11 | 22 | 7 | 8 | 2 | 1 | 0 | 0 | 0 | 0 |
| 15 | 3.90 | 1.96 | 1.99 | 0.05 | 0.9856 | Gm37249 | | 9 | 11 | 0 | 0 | 0 | 63 | 0 | 14 | 7 | 0 | 65 |
| 730 | 0.26 | 0.13 | 1.99 | 0.05 | 0.98573 | E130309D02Ri | RIKEN cDNA E130309D02 gene | 709 | 856 | 662 | 691 | 682 | 558 | 707 | 857 | 770 | 876 | 666 |
| 4956 | -0.15 | 0.07 | -1.99 | 0.05 | 0.98867 | Ncor2 | nuclear receptor co-repressor 2 | 5182 | 4989 | 5326 | 5189 | 5358 | 4432 | 5460 | 4282 | 5357 | 5224 | 3713 |
| 1035 | 0.31 | 0.15 | 1.99 | 0.05 | 0.98867 | Pfdn1 | prefoldin 1 | 1223 | 1033 | 1496 | 736 | 998 | 807 | 875 | 1116 | 1021 | 1224 | 854 |

| | | | | | | | | | | | | | | | | | |
|---|---|---|---|---|---|---|---|---|---|---|---|---|---|---|---|---|---|
| 613 | 0.28 | 0.14 | 1.99 | 0.05 | 0.98867 | Wdr46 | WD repeat domain 46 | 830 | 698 | 734 | 472 | 522 | 515 | 513 | 614 | 635 | 678 | 535 |
| 623 | 0.45 | 0.23 | 1.99 | 0.05 | 0.98867 | Plat | plasminogen activator, tissue | 2348 | 1462 | 1647 | 172 | 131 | 123 | 164 | 255 | 212 | 176 | 157 |
| 32 | -2.10 | 1.06 | -1.99 | 0.05 | 0.98867 | Fam89a | family with sequence similarity 89, membe | 85 | 74 | 117 | 25 | 13 | 9 | 9 | 0 | 14 | 0 | 5 |
| 787 | -3.39 | 1.71 | -1.99 | 0.05 | 0.98867 | Angptl2 | angiopoietin-like 2 | 3455 | 2577 | 2592 | 13 | 7 | 6 | 0 | 0 | 0 | 0 | 6 |
| 2117 | -0.16 | 0.08 | -1.99 | 0.05 | 0.98985 | Mllt6 | myeloid/lymphoid or mixed-lineage leuke | 2559 | 2193 | 2582 | 1857 | 2132 | 2538 | 1907 | 1712 | 1857 | 1672 | 2281 |
| 2489 | 0.17 | 0.08 | 1.99 | 0.05 | 0.9912 | As3mt | arsenic (+3 oxidation state) methyltransfe | 1839 | 2044 | 2088 | 2674 | 2615 | 2224 | 2568 | 3115 | 2788 | 3048 | 2380 |
| 3736 | -0.23 | 0.12 | -1.98 | 0.05 | 0.99142 | Ncoa2 | nuclear receptor coactivator 2 | 3689 | 3305 | 3085 | 4155 | 3974 | 4073 | 4561 | 3304 | 4274 | 3151 | 3528 |
| 2970 | 0.20 | 0.10 | 1.98 | 0.05 | 0.99142 | Neu1 | neuraminidase 1 | 1434 | 1550 | 1617 | 3271 | 3207 | 2896 | 3624 | 4238 | 3501 | 4334 | 3000 |
| 718 | 0.36 | 0.18 | 1.98 | 0.05 | 0.99142 | Med7 | mediator complex subunit 7 | 700 | 679 | 581 | 572 | 677 | 841 | 517 | 974 | 771 | 745 | 838 |
| 263 | -0.57 | 0.29 | -1.98 | 0.05 | 0.99142 | Borcs5 | BLOC-1 related complex subunit 5 | 290 | 234 | 396 | 270 | 349 | 205 | 341 | 109 | 278 | 188 | 231 |
| 36 | -1.55 | 0.78 | -1.98 | 0.05 | 0.99142 | Gm16401 | | 43 | 33 | 46 | 58 | 45 | 38 | 49 | 7 | 54 | 4 | 21 |
| 16 | -2.90 | 1.47 | -1.98 | 0.05 | 0.99142 | 5730420D15Rik | | 0 | 0 | 20 | 32 | 18 | 22 | 40 | 9 | 28 | 0 | 5 |
| 19 | -2.06 | 1.04 | -1.98 | 0.05 | 0.99142 | Pcp4 | Purkinje cell protein 4 | 13 | 21 | 60 | 19 | 19 | 21 | 25 | 7 | 4 | 0 | 24 |
| 3 | -6.34 | 3.20 | -1.98 | 0.05 | 0.99142 | Cnga4 | cyclic nucleotide gated channel alpha 4 | 0 | 6 | 0 | 0 | 3 | 13 | 8 | 0 | 0 | 0 | 6 |
| 895 | -0.38 | 0.19 | -1.98 | 0.05 | 0.99235 | Ino80 | INO80 complex subunit | 1070 | 656 | 885 | 890 | 884 | 1298 | 931 | 480 | 839 | 627 | 1291 |
| 1229 | -0.21 | 0.11 | -1.98 | 0.05 | 0.9931 | Tmem104 | transmembrane protein 104 | 1042 | 888 | 1080 | 1420 | 1527 | 1233 | 1436 | 1191 | 1423 | 1342 | 932 |
| 7997 | 0.12 | 0.06 | 1.98 | 0.05 | 0.99358 | Pum2 | pumilio RNA-binding family member 2 | 7893 | 7841 | 7257 | 7747 | 7604 | 7356 | 8452 | 8836 | 8783 | 8381 | 7820 |
| 35 | -2.05 | 1.03 | -1.98 | 0.05 | 0.99358 | Ldb3 | LIM domain binding 3 | 153 | 27 | 42 | 9 | 7 | 70 | 26 | 0 | 6 | 0 | 47 |
| 8 | -3.49 | 1.76 | -1.98 | 0.05 | 0.99358 | Gm10812 | | 6 | 0 | 0 | 12 | 23 | 6 | 13 | 0 | 3 | 28 | 0 |
| 356 | -4.71 | 2.38 | -1.98 | 0.05 | 0.99358 | Prkch | protein kinase C, eta | 1361 | 1231 | 1235 | 0 | 32 | 9 | 10 | 37 | 0 | 0 | 0 |
| 5504 | 0.20 | 0.10 | 1.98 | 0.05 | 0.99517 | Macroh2a1 | macroH2A.1 histone | 5426 | 5864 | 6457 | 4725 | 4839 | 5466 | 4859 | 6403 | 4813 | 5843 | 5844 |
| 80 | -1.73 | 0.87 | -1.98 | 0.05 | 0.99598 | Dab1 | disabled 1 | 302 | 181 | 205 | 37 | 20 | 44 | 23 | 0 | 7 | 59 | 6 |
| 81 | -1.40 | 0.71 | -1.98 | 0.05 | 0.99598 | Snx20 | sorting nexin 20 | 365 | 239 | 210 | 14 | 9 | 7 | 22 | 0 | 3 | 17 | 4 |
| 128 | -0.78 | 0.39 | -1.98 | 0.05 | 0.9966 | Frs3 | fibroblast growth factor receptor substrate | 99 | 224 | 141 | 189 | 112 | 145 | 150 | 63 | 91 | 67 | 132 |
| 40 | -1.26 | 0.64 | -1.98 | 0.05 | 0.99784 | Gm16794 | | 14 | 22 | 49 | 38 | 41 | 107 | 40 | 6 | 23 | 14 | 88 |
| 74 | -2.69 | 1.36 | -1.98 | 0.05 | 0.99784 | Phf11d | PHD finger protein 11D | 348 | 112 | 342 | 0 | 3 | 5 | 0 | 0 | 0 | 0 | 0 |
| 398 | 0.45 | 0.23 | 1.98 | 0.05 | 0.99844 | Rrp36 | ribosomal RNA processing 36 homolog (S | 431 | 426 | 367 | 291 | 365 | 348 | 321 | 616 | 368 | 433 | 414 |
| 154 | -3.80 | 1.93 | -1.97 | 0.05 | 0.99844 | Ggt1 | gamma-glutamyltransferase 1 | 208 | 740 | 685 | 26 | 17 | 3 | 0 | 0 | 0 | 0 | 13 |
| 5018585 | 0.21 | 0.69 | 0.30 | 0.76 | 1 | Ins2 | insulin II | 253447 | 268115 | 621793 | 6671989 | 7361267 | 4380689 | 6490272 | 10569108 | 7147986 | 7141738 | 4298029 |
| 2027032 | 0.23 | 0.72 | 0.32 | 0.75 | 1 | Ins1 | insulin I | 94961 | 100933 | 254397 | 2753665 | 3034748 | 1566568 | 2594784 | 4470561 | 2993671 | 2868680 | 1564378 |
| 578565 | 0.10 | 0.34 | 0.29 | 0.77 | 1 | Iapp | islet amyloid polypeptide | 57556 | 81741 | 102368 | 822932 | 771889 | 592422 | 767363 | 941290 | 780210 | 804300 | 642143 |
| 521715 | 0.06 | 0.19 | 0.30 | 0.76 | 1 | Chga | chromogranin A | 218706 | 261112 | 339847 | 650124 | 663529 | 503350 | 584099 | 825346 | 565489 | 659244 | 468021 |
| 224366 | 0.03 | 0.33 | 0.10 | 0.92 | 1 | mt-Co1 | cytochrome c oxidase subunit I | 164865 | 211597 | 122399 | 239412 | 169963 | 234212 | 336436 | 137738 | 272688 | 331293 | 247429 |
| 266160 | 0.12 | 0.08 | 1.49 | 0.14 | 1 | Scg2 | secretogranin II | 60134 | 91836 | 85075 | 421331 | 309163 | 248320 | 313656 | 452916 | 342383 | 311317 | 291628 |
| 206275 | 0.08 | 0.06 | 1.31 | 0.19 | 1 | Cpe | carboxypeptidase E | 153153 | 169274 | 182590 | 228566 | 227446 | 166273 | 235181 | 250676 | 228336 | 248522 | 179003 |
| 214413 | 0.00 | 0.07 | 0.01 | 0.99 | 1 | Pcsk2 | proprotein convertase subtilisin/kexin type | 238272 | 297812 | 287148 | 215568 | 192299 | 144071 | 213703 | 236365 | 183857 | 208252 | 141193 |
| 136459 | -0.11 | 0.35 | -0.32 | 0.75 | 1 | mt-Cytb | cytochrome b | 94223 | 132854 | 74743 | 152870 | 116376 | 148633 | 203380 | 59314 | 178788 | 192492 | 147378 |
| 101514 | -0.19 | 0.41 | -0.48 | 0.63 | 1 | mt-Nd5 | NADH dehydrogenase subunit 5 | 68683 | 97346 | 49678 | 118108 | 83717 | 118814 | 155908 | 28212 | 133708 | 147357 | 115120 |
| 106152 | 0.04 | 0.21 | 0.18 | 0.85 | 1 | G6pc2 | glucose-6-phosphatase, catalytic, 2 | 6341 | 6008 | 13593 | 136681 | 160462 | 108610 | 155122 | 117870 | 175070 | 168601 | 119316 |
| 106298 | 0.14 | 0.13 | 1.10 | 0.27 | 1 | Ero1lb | ERO1-like beta (S. cerevisiae) | 8094 | 9909 | 16028 | 135197 | 138589 | 131475 | 134692 | 155186 | 147403 | 147666 | 145041 |
| 1569790 | -1.02 | 1.68 | -0.61 | 0.54 | 1 | Gcg | glucagon | 4725010 | 6369833 | 5558150 | 80255 | 111012 | 93139 | 126172 | 39561 | 69536 | 52906 | 42112 |
| 93113 | 0.60 | 0.58 | 1.04 | 0.30 | 1 | | | 29594 | 44490 | 86110 | 75638 | 69731 | 58198 | 109092 | 82345 | 71927 | 331181 | 65937 |
| 69574 | -0.11 | 0.33 | -0.32 | 0.75 | 1 | mt-Nd1 | NADH dehydrogenase subunit 1 | 61075 | 72092 | 42695 | 75086 | 57035 | 68753 | 102994 | 28332 | 85435 | 101886 | 69930 |
| 109544 | -0.04 | 0.04 | -1.07 | 0.28 | 1 | Eef1a1 | eukaryotic translation elongation factor 1 | 181127 | 167726 | 176595 | 93705 | 94386 | 59698 | 97524 | 88334 | 91151 | 93439 | 61300 |
| 68115 | -0.07 | 0.08 | -0.91 | 0.36 | 1 | Acly | ATP citrate lyase | 27223 | 29400 | 32034 | 86147 | 90110 | 67130 | 94663 | 72064 | 89600 | 95243 | 65648 |
| 70898 | -0.03 | 0.05 | -0.54 | 0.59 | 1 | Scg3 | secretogranin III | 28663 | 37179 | 34943 | 96099 | 87686 | 72389 | 86740 | 95309 | 87661 | 79309 | 73896 |
| 53930 | -0.16 | 0.53 | -0.31 | 0.76 | 1 | mt-Nd2 | NADH dehydrogenase subunit 2 | 42275 | 57149 | 30473 | 59510 | 46121 | 53424 | 82824 | 15386 | 72184 | 81968 | 51920 |
| 67095 | 0.09 | 0.08 | 1.10 | 0.27 | 1 | Psap | prosaposin | 74564 | 52325 | 59157 | 71385 | 73305 | 46266 | 76582 | 81340 | 73720 | 81554 | 47848 |
| 79403 | 0.00 | 0.04 | 0.07 | 0.94 | 1 | Eef2 | eukaryotic translation elongation factor 2 | 109542 | 108951 | 110531 | 73784 | 73490 | 48462 | 75924 | 76540 | 71847 | 76782 | 47577 |
| 67004 | -0.10 | 0.10 | -1.01 | 0.31 | 1 | Chgb | chromogranin B | 46755 | 69901 | 75178 | 75290 | 72715 | 60952 | 72721 | 73558 | 67447 | 64351 | 58182 |
| 42485 | -0.18 | 0.62 | -0.28 | 0.78 | 1 | mt-Nd4 | NADH dehydrogenase subunit 4 | 29087 | 45173 | 20078 | 46871 | 34022 | 49433 | 65473 | 8898 | 56609 | 65741 | 45953 |
| 46027 | -0.01 | 0.22 | -0.06 | 0.95 | 1 | Slc2a2 | solute carrier family 2 (facilitated glucose | 2452 | 2546 | 5833 | 61680 | 69189 | 51994 | 64545 | 51646 | 75998 | 65694 | 54716 |
| 47658 | -0.04 | 0.05 | -0.89 | 0.38 | 1 | Atp2a3 | ATPase, Ca++ transporting, ubiquitous | 13721 | 14797 | 14840 | 64300 | 68274 | 46889 | 64297 | 59620 | 68027 | 65520 | 43951 |
| 49534 | 0.07 | 0.07 | 0.98 | 0.33 | 1 | Cd164 | CD164 antigen | 23659 | 28206 | 27594 | 61194 | 57920 | 45143 | 62737 | 58492 | 63023 | 68154 | 48750 |
| 57610 | 0.13 | 0.07 | 1.72 | 0.08 | 1 | Hsp90b1 | heat shock protein 90, beta (Grp94), mem | 54494 | 45832 | 44366 | 62208 | 56537 | 53472 | 61691 | 72852 | 58826 | 61784 | 61645 |
| 65636 | 0.04 | 0.08 | 0.47 | 0.64 | 1 | Peg3 | paternally expressed 3 | 38359 | 53389 | 42678 | 56807 | 59124 | 112968 | 61360 | 69482 | 62963 | 54670 | 110197 |
| 50669 | 0.06 | 0.07 | 0.84 | 0.40 | 1 | P4hb | prolyl 4-hydroxylase, beta polypeptide | 31406 | 33508 | 37223 | 62936 | 58834 | 39782 | 60971 | 71808 | 57112 | 62013 | 41759 |
| 44045 | 0.01 | 0.08 | 0.13 | 0.89 | 1 | Syt13 | synaptotagmin XIII | 15823 | 17131 | 23030 | 51389 | 63340 | 39470 | 59271 | 55071 | 61649 | 59705 | 38617 |
| 43120 | 0.02 | 0.08 | 0.30 | 0.76 | 1 | Hyou1 | hypoxia up-regulated 1 | 16337 | 14866 | 20465 | 51546 | 57074 | 42669 | 58302 | 53469 | 54709 | 62192 | 42691 |
| 50824 | 0.14 | 0.07 | 1.88 | 0.06 | 1 | Pdia3 | protein disulfide isomerase associated 3 | 36130 | 36991 | 38088 | 59940 | 54223 | 41708 | 57221 | 73637 | 54075 | 60346 | 46708 |

| | | | | | | | | | | | | | | | | | |
|---|---|---|---|---|---|---|---|---|---|---|---|---|---|---|---|---|---|
| 35770 | -0.02 | 0.11 | -0.15 | 0.88 | 1 | Pcsk1 | proprotein convertase subtilisin/kexin type | 2931 | 3158 | 4830 | 48989 | 48438 | 42920 | 51857 | 45029 | 50450 | 47728 | 47134 |
| 41931 | 0.02 | 0.07 | 0.32 | 0.75 | 1 | Calr | calreticulin | 30614 | 25069 | 29581 | 49182 | 50141 | 35822 | 51200 | 55256 | 50042 | 49003 | 35334 |
| 36052 | 0.00 | 0.19 | 0.00 | 1.00 | 1 | Glp1r | glucagon-like peptide 1 receptor | 2044 | 2380 | 4539 | 51927 | 55496 | 35509 | 50366 | 50147 | 58032 | 51453 | 34682 |
| 45064 | 0.00 | 0.06 | 0.00 | 1.00 | 1 | Maged1 | melanoma antigen, family D, 1 | 29670 | 33332 | 34591 | 50711 | 49239 | 49563 | 49465 | 53727 | 47387 | 48822 | 49192 |
| 37082 | 0.05 | 0.11 | 0.50 | 0.62 | 1 | Scg5 | secretogranin V | 32831 | 35311 | 30278 | 47500 | 28253 | 28264 | 48581 | 46437 | 28367 | 49261 | 32817 |
| 39779 | 0.05 | 0.05 | 1.03 | 0.30 | 1 | Eif4g2 | eukaryotic translation initiation factor 4, ga | 35130 | 34942 | 34919 | 43103 | 41202 | 34254 | 45203 | 43663 | 44006 | 43145 | 38003 |
| 41295 | 0.04 | 0.05 | 0.84 | 0.40 | 1 | Slc30a8 | solute carrier family 30 (zinc transporter), | 16443 | 21713 | 21753 | 51147 | 51988 | 48086 | 43301 | 51679 | 54834 | 44566 | 48734 |
| 29752 | 0.05 | 0.10 | 0.55 | 0.58 | 1 | Papss2 | 3'-phosphoadenosine 5'-phosphosulfate s | 2913 | 2940 | 4366 | 40972 | 40853 | 30842 | 42962 | 40906 | 43585 | 43664 | 33266 |
| 28761 | 0.07 | 0.11 | 0.60 | 0.55 | 1 | Gng12 | guanine nucleotide binding protein (G pro | 5190 | 5799 | 6439 | 37727 | 32494 | 33340 | 42438 | 32390 | 38288 | 45046 | 37216 |
| 37963 | -0.13 | 0.09 | -1.40 | 0.16 | 1 | Abcc8 | ATP-binding cassette, sub-family C (CFTF | 22609 | 31456 | 25840 | 42957 | 41773 | 50426 | 41822 | 36914 | 40538 | 42241 | 41014 |
| 48060 | 0.05 | 0.07 | 0.80 | 0.42 | 1 | Hsp90ab1 | heat shock protein 90 alpha (cytosolic), cl | 55357 | 54755 | 56100 | 44793 | 42842 | 49045 | 41153 | 52358 | 43841 | 40852 | 47560 |
| 35323 | 0.04 | 0.04 | 0.89 | 0.37 | 1 | Canx | calnexin | 25159 | 25367 | 26936 | 41721 | 40223 | 31181 | 40533 | 42053 | 41192 | 41350 | 32844 |
| 30764 | 0.00 | 0.09 | -0.03 | 0.97 | 1 | Spock2 | sparc/osteonectin, cwcv and kazal-like do | 4596 | 4644 | 6812 | 43071 | 45710 | 31668 | 40489 | 47516 | 45286 | 38709 | 29906 |
| 30648 | -0.07 | 0.07 | -0.89 | 0.37 | 1 | Gfpt1 | glutamine fructose-6-phosphate transami | 16190 | 17566 | 18393 | 36005 | 37748 | 31912 | 39963 | 30704 | 38726 | 38120 | 31797 |
| 32701 | -0.10 | 0.07 | -1.46 | 0.15 | 1 | Ptprn2 | protein tyrosine phosphatase, receptor typ | 21032 | 26514 | 23983 | 41368 | 38765 | 29461 | 39142 | 36643 | 38220 | 38122 | 26461 |
| 26697 | 0.03 | 0.10 | 0.26 | 0.79 | 1 | Chic1 | cysteine-rich hydrophobic domain 1 | 7188 | 12303 | 9943 | 36325 | 30134 | 29020 | 35073 | 39489 | 32205 | 31044 | 30945 |
| 29264 | 0.03 | 0.07 | 0.39 | 0.70 | 1 | Prkacb | protein kinase, cAMP dependent, catalytic | 17190 | 21548 | 19520 | 35268 | 33195 | 29062 | 33198 | 32174 | 36019 | 33661 | 31071 |
| 37745 | -0.04 | 0.07 | -0.57 | 0.57 | 1 | Aplp1 | amyloid beta (A4) precursor-like protein 1 | 56863 | 65837 | 63968 | 30853 | 30677 | 21618 | 32529 | 33641 | 26927 | 31639 | 20641 |
| 30471 | 0.05 | 0.05 | 0.87 | 0.39 | 1 | Prkar1a | protein kinase, cAMP dependent regulato | 31127 | 29620 | 32819 | 30168 | 31434 | 24998 | 32475 | 32802 | 31078 | 31697 | 26960 |
| 24667 | 0.25 | 0.25 | 1.01 | 0.31 | 1 | Mafa | v-maf musculoaponeurotic fibrosarcoma o | 1197 | 971 | 2395 | 31510 | 34597 | 23604 | 32291 | 36518 | 36443 | 44067 | 27747 |
| 28182 | 0.05 | 0.05 | 1.08 | 0.28 | 1 | Cltc | clathrin, heavy polypeptide (Hc) | 21178 | 22053 | 22939 | 29813 | 33382 | 24510 | 32096 | 30819 | 35750 | 32010 | 25454 |
| 31578 | 0.12 | 0.07 | 1.68 | 0.09 | 1 | Calm1 | calmodulin 1 | 33220 | 31995 | 36226 | 29674 | 31844 | 24197 | 31994 | 37813 | 31854 | 32010 | 26528 |
| 26801 | 0.07 | 0.06 | 1.20 | 0.23 | 1 | Lamp1 | lysosomal-associated membrane protein | 22834 | 21406 | 21479 | 30889 | 28479 | 20452 | 31894 | 28534 | 30806 | 36076 | 21962 |
| 23499 | 0.22 | 0.21 | 1.02 | 0.31 | 1 | Prlr | prolactin receptor | 1458 | 1295 | 2838 | 28692 | 32106 | 24597 | 31668 | 37429 | 36857 | 32273 | 29277 |
| 35835 | -0.11 | 0.09 | -1.19 | 0.24 | 1 | Resp18 | regulated endocrine-specific protein 18 | 41746 | 53948 | 68288 | 31070 | 34365 | 22572 | 31554 | 32114 | 30785 | 25088 | 22658 |
| 23695 | 0.06 | 0.11 | 0.51 | 0.61 | 1 | Hadh | hydroxyacyl-Coenzyme A dehydrogenase | 3561 | 4606 | 6249 | 31931 | 32195 | 25702 | 30974 | 31463 | 33631 | 33167 | 27165 |
| 24677 | 0.08 | 0.07 | 1.23 | 0.22 | 1 | Pdia6 | protein disulfide isomerase associated 6 | 13076 | 12012 | 12811 | 31308 | 27527 | 24278 | 30183 | 33985 | 27176 | 33349 | 25744 |
| 24747 | 0.02 | 0.07 | 0.27 | 0.79 | 1 | Ptprn | protein tyrosine phosphatase, receptor typ | 14464 | 17718 | 15386 | 30803 | 26239 | 24467 | 29791 | 30581 | 27002 | 33054 | 22708 |
| 23857 | 0.01 | 0.04 | 0.26 | 0.79 | 1 | Surf4 | surfeit gene 4 | 13260 | 13010 | 15411 | 27102 | 31644 | 21561 | 29709 | 27995 | 30929 | 29860 | 21944 |
| 24679 | 0.06 | 0.05 | 1.09 | 0.28 | 1 | Mlec | malectin | 13577 | 14043 | 15560 | 28128 | 30907 | 23221 | 29693 | 30943 | 30585 | 31028 | 23783 |
| 23397 | 0.01 | 0.06 | 0.16 | 0.87 | 1 | Lman1 | lectin, mannose-binding, 1 | 8663 | 8708 | 10980 | 27918 | 29967 | 26529 | 29559 | 26716 | 31397 | 30718 | 26213 |
| 24875 | -0.02 | 0.07 | -0.31 | 0.75 | 1 | Hdlbp | high density lipoprotein (HDL) binding pro | 14439 | 13277 | 14885 | 25207 | 33091 | 28701 | 29088 | 22103 | 34314 | 28493 | 30025 |
| 24910 | 0.18 | 0.09 | 1.95 | 0.05 | 1 | Pkm | pyruvate kinase, muscle | 18400 | 15306 | 17721 | 26413 | 27064 | 21747 | 29030 | 35941 | 26577 | 31953 | 23858 |
| 38335 | 0.20 | 0.24 | 0.87 | 0.39 | 1 | Pcsk1n | proprotein convertase subtilisin/kexin type | 50791 | 57365 | 72044 | 29121 | 33553 | 18908 | 28919 | 50171 | 27904 | 34274 | 18633 |
| 24610 | 0.13 | 0.08 | 1.58 | 0.11 | 1 | Rpn2 | ribophorin II | 14403 | 13591 | 13954 | 31256 | 28433 | 19844 | 28908 | 40481 | 29700 | 29668 | 20474 |
| 23460 | -0.01 | 0.08 | -0.14 | 0.88 | 1 | Disp2 | dispatched RND transporter family memb | 10207 | 13773 | 13431 | 29667 | 29632 | 22758 | 28533 | 27678 | 31399 | 29631 | 21352 |
| 34096 | -0.03 | 0.05 | -0.51 | 0.61 | 1 | Itm2b | integral membrane protein 2B | 58532 | 54282 | 60499 | 25472 | 26828 | 21056 | 28393 | 23655 | 25859 | 29377 | 21104 |
| 26125 | 0.05 | 0.04 | 1.25 | 0.21 | 1 | Copa | coatomer protein complex subunit alpha | 17981 | 20393 | 18227 | 29833 | 27591 | 27523 | 28380 | 32026 | 28557 | 29302 | 27563 |
| 22151 | -0.03 | 0.07 | -0.44 | 0.66 | 1 | Srpr | signal recognition particle receptor ('dock | 9865 | 11186 | 12644 | 28376 | 29224 | 20190 | 28366 | 25584 | 28830 | 29337 | 20055 |
| 22556 | 0.03 | 0.07 | 0.36 | 0.72 | 1 | Sphkap | SPHK1 interactor, AKAP domain containi | 13529 | 15819 | 14793 | 25989 | 25415 | 21483 | 28152 | 24601 | 27775 | 28674 | 21881 |
| 30808 | 0.14 | 0.07 | 1.91 | 0.06 | 1 | Rpl4 | ribosomal protein L4 | 44538 | 39019 | 46398 | 25820 | 26264 | 19880 | 27740 | 30738 | 26682 | 28791 | 23023 |
| 26149 | 0.12 | 0.08 | 1.51 | 0.13 | 1 | Atp5a1 | ATP synthase, H+ transporting, mitochond | 26367 | 23715 | 26006 | 27498 | 26908 | 19230 | 27583 | 34558 | 26638 | 27852 | 21285 |
| 26858 | 0.11 | 0.07 | 1.72 | 0.08 | 1 | Tgoln1 | trans-golgi network protein | 20321 | 23896 | 24491 | 30239 | 29610 | 21277 | 27542 | 37042 | 30932 | 26583 | 23503 |
| 20427 | 0.08 | 0.14 | 0.56 | 0.58 | 1 | Nkx6-1 | NK6 homeobox 1 | 1784 | 2096 | 3337 | 27679 | 28891 | 21525 | 27465 | 32625 | 27945 | 31282 | 20071 |
| 20042 | 0.07 | 0.06 | 1.27 | 0.20 | 1 | Mia3 | melanoma inhibitory activity 3 | 7689 | 6936 | 8573 | 23021 | 23700 | 23506 | 25836 | 24090 | 26615 | 26422 | 24070 |
| 17713 | 0.09 | 0.14 | 0.69 | 0.49 | 1 | Ucn3 | urocortin 3 | 939 | 1120 | 1720 | 25601 | 24406 | 16861 | 25622 | 24977 | 25930 | 29447 | 18215 |
| 20103 | 0.02 | 0.08 | 0.24 | 0.81 | 1 | Prnp | prion protein | 10172 | 11722 | 12475 | 25368 | 23880 | 18195 | 25453 | 22652 | 25517 | 26179 | 19515 |
| 21577 | 0.10 | 0.06 | 1.70 | 0.09 | 1 | Serp1 | stress-associated endoplasmic reticulum | 10050 | 10656 | 11627 | 22924 | 29608 | 21432 | 24940 | 23881 | 31389 | 27463 | 23371 |
| 18061 | -0.04 | 0.07 | -0.62 | 0.54 | 1 | Cyfip2 | cytoplasmic FMR1 interacting protein 2 | 6137 | 7129 | 6534 | 26838 | 23280 | 15737 | 24931 | 22660 | 25071 | 25456 | 14893 |
| 19415 | 0.19 | 0.34 | 0.56 | 0.58 | 1 | Sytl4 | synaptotagmin-like 4 | 919 | 823 | 2210 | 24236 | 26263 | 23417 | 24308 | 28777 | 27715 | 26384 | 28512 |
| 20459 | -0.10 | 0.09 | -1.18 | 0.24 | 1 | Flnb | filamin, beta | 10125 | 11336 | 11519 | 23609 | 28896 | 22463 | 24121 | 19442 | 30514 | 22477 | 20545 |
| 19090 | 0.06 | 0.05 | 1.27 | 0.20 | 1 | Spcs3 | signal peptidase complex subunit 3 homo | 8106 | 8293 | 9552 | 22446 | 25765 | 18015 | 23948 | 24002 | 25639 | 24373 | 19850 |
| 19968 | -0.04 | 0.09 | -0.50 | 0.62 | 1 | Fndc3a | fibronectin type III domain containing 3A | 12584 | 14656 | 13926 | 22659 | 23989 | 19941 | 23863 | 19098 | 25989 | 23122 | 19825 |
| 16799 | 0.07 | 0.13 | 0.57 | 0.57 | 1 | Bace2 | beta-site APP-cleaving enzyme 2 | 1663 | 1353 | 2658 | 23660 | 25113 | 14751 | 23774 | 25560 | 25101 | 25888 | 15273 |
| 18165 | -0.01 | 0.06 | -0.23 | 0.82 | 1 | Uggt1 | UDP-glucose glycoprotein glucosyltransfe | 7417 | 7227 | 8158 | 23296 | 23173 | 18726 | 23704 | 23845 | 23940 | 21486 | 18843 |
| 21945 | 0.04 | 0.09 | 0.45 | 0.65 | 1 | Cplx2 | complexin 2 | 19742 | 24196 | 22494 | 21222 | 23347 | 17418 | 23596 | 28587 | 22126 | 22647 | 16019 |
| 19063 | 0.02 | 0.07 | 0.28 | 0.78 | 1 | Fkbp9 | FK506 binding protein 9 | 16431 | 14203 | 15466 | 21585 | 21000 | 15077 | 23492 | 24730 | 19661 | 22912 | 15140 |
| 19021 | -0.10 | 0.11 | -0.91 | 0.37 | 1 | Usp9x | ubiquitin specific peptidase 9, X chromos | 10959 | 14594 | 12273 | 22353 | 23244 | 19811 | 23181 | 17423 | 25001 | 20845 | 19541 |
| 25430 | 0.07 | 0.10 | 0.78 | 0.43 | 1 | Selenop | selenoprotein P | 32077 | 24589 | 27537 | 25472 | 26043 | 20507 | 23168 | 28923 | 26722 | 23561 | 21132 |
| 19291 | 0.05 | 0.04 | 1.04 | 0.30 | 1 | Dnajc3 | DnaJ heat shock protein family (Hsp40) m | 10596 | 10461 | 11654 | 22215 | 23092 | 20007 | 23026 | 22236 | 24149 | 23232 | 21538 |

| | | | | | | | | | | | | | | | | | |
|---|---|---|---|---|---|---|---|---|---|---|---|---|---|---|---|---|---|
| 23164 | 0.10 | 0.08 | 1.23 | 0.22 | 1 | Atp5b | ATP synthase, H+ transporting mitochond | 27501 | 23112 | 28063 | 23006 | 21944 | 16939 | 22905 | 26986 | 21779 | 24840 | 17728 |
| 19037 | -0.01 | 0.06 | -0.11 | 0.91 | 1 | Ptprf | protein tyrosine phosphatase, receptor typ | 12527 | 14011 | 13175 | 22531 | 20962 | 18442 | 22900 | 23148 | 21647 | 23241 | 16827 |
| 19352 | 0.01 | 0.06 | 0.11 | 0.91 | 1 | Rap1gap2 | RAP1 GTPase activating protein 2 | 13155 | 13572 | 15125 | 20786 | 23086 | 18619 | 22677 | 22798 | 23492 | 22276 | 17288 |
| 19737 | 0.18 | 0.11 | 1.59 | 0.11 | 1 | Dynll2 | dynein light chain LC8-type 2 | 11165 | 12613 | 14533 | 22581 | 22035 | 16322 | 22428 | 32461 | 21855 | 24117 | 16998 |
| 20128 | -0.06 | 0.07 | -0.85 | 0.39 | 1 | Twsg1 | twisted gastrulation BMP signaling modul | 15291 | 15889 | 15408 | 24118 | 24749 | 18142 | 22335 | 19554 | 25018 | 22567 | 18335 |
| 21738 | -0.09 | 0.06 | -1.35 | 0.18 | 1 | Dync1h1 | dynein cytoplasmic 1 heavy chain 1 | 22512 | 20795 | 24510 | 20699 | 23900 | 21474 | 22001 | 21303 | 23962 | 18672 | 19286 |
| 18863 | -0.04 | 0.08 | -0.45 | 0.65 | 1 | Dst | dystonin | 10593 | 12821 | 12747 | 18920 | 20770 | 25506 | 21658 | 18004 | 22544 | 20617 | 23312 |
| 21310 | -0.04 | 0.14 | -0.29 | 0.77 | 1 | Il1r1 | interleukin 1 receptor, type I | 6001 | 13484 | 7861 | 27575 | 26276 | 29832 | 21653 | 23783 | 27871 | 21696 | 28384 |
| 18787 | 0.02 | 0.10 | 0.17 | 0.87 | 1 | Ubr4 | ubiquitin protein ligase E3 component n-r | 9367 | 11474 | 9789 | 17819 | 20940 | 27483 | 21554 | 19542 | 23835 | 20350 | 24502 |
| 17507 | 0.11 | 0.10 | 1.03 | 0.30 | 1 | Syp | synaptophysin | 4749 | 5816 | 7152 | 22949 | 24038 | 15558 | 21492 | 29323 | 22627 | 22751 | 16124 |
| 19806 | 0.10 | 0.09 | 1.19 | 0.23 | 1 | Tmbim6 | transmembrane BAX inhibitor motif contai | 18622 | 18294 | 20176 | 20935 | 20814 | 13883 | 21468 | 27104 | 20188 | 22494 | 13885 |
| 17428 | 0.00 | 0.06 | -0.01 | 0.99 | 1 | Ddost | dolichyl-di-phosphooligosaccharide-prote | 13568 | 12221 | 12565 | 20309 | 20667 | 14109 | 21443 | 22335 | 19300 | 21638 | 13550 |
| 17503 | 0.01 | 0.07 | 0.15 | 0.88 | 1 | Ids | iduronate 2-sulfatase | 11827 | 13658 | 14031 | 19962 | 20460 | 14416 | 21274 | 19186 | 21895 | 21654 | 14165 |
| 18323 | 0.08 | 0.06 | 1.27 | 0.20 | 1 | Serinc1 | serine incorporator 1 | 12218 | 12554 | 13179 | 20940 | 20451 | 16932 | 21257 | 23720 | 20336 | 20801 | 19162 |
| 18829 | -0.05 | 0.05 | -0.98 | 0.33 | 1 | Sez6l2 | seizure related 6 homolog like 2 | 16204 | 19243 | 18587 | 22371 | 20732 | 13492 | 21188 | 21816 | 20595 | 20274 | 12620 |
| 17359 | -0.07 | 0.07 | -1.00 | 0.32 | 1 | Ddc | dopa decarboxylase | 6095 | 7528 | 9442 | 22635 | 25795 | 16831 | 20899 | 21261 | 23601 | 20221 | 16640 |
| 17129 | 0.06 | 0.07 | 0.86 | 0.39 | 1 | Saraf | store-operated calcium entry-associated r | 8853 | 9439 | 10884 | 20503 | 20736 | 15831 | 20788 | 22274 | 20132 | 22474 | 16505 |
| 14192 | 0.12 | 0.11 | 1.10 | 0.27 | 1 | Ndufs2 | NADH:ubiquinone oxidoreductase core s | 5908 | 5322 | 7269 | 11153 | 23686 | 10414 | 20621 | 14277 | 23600 | 23728 | 10138 |
| 16336 | -0.02 | 0.11 | -0.17 | 0.87 | 1 | Tmod2 | tropomodulin 2 | 6729 | 9979 | 8380 | 19671 | 20268 | 17142 | 20555 | 17637 | 23705 | 18534 | 17098 |
| 15607 | 0.10 | 0.08 | 1.28 | 0.20 | 1 | Emb | embigin | 5703 | 6518 | 7618 | 18407 | 20702 | 13882 | 20500 | 19487 | 21639 | 21330 | 15890 |
| 15920 | -0.06 | 0.11 | -0.58 | 0.56 | 1 | Slc7a14 | solute carrier family 7 (cationic amino acio | 5771 | 7995 | 8204 | 22275 | 17740 | 17808 | 20432 | 17695 | 21023 | 18803 | 17377 |
| 17795 | 0.00 | 0.05 | -0.01 | 0.99 | 1 | Tm9sf3 | transmembrane 9 superfamily member 3 | 14062 | 13699 | 14675 | 19708 | 20205 | 16468 | 20360 | 18411 | 20967 | 19261 | 17931 |
| 15933 | 0.14 | 0.10 | 1.35 | 0.18 | 1 | Cd63 | CD63 antigen | 8487 | 8297 | 9907 | 20208 | 17934 | 11999 | 20271 | 24747 | 17818 | 22988 | 12603 |
| 15855 | 0.04 | 0.05 | 0.67 | 0.50 | 1 | Gns | glucosamine (N-acetyl)-6-sulfatase | 11458 | 10400 | 10148 | 19694 | 16736 | 13598 | 20199 | 20732 | 18070 | 19988 | 13382 |
| 25721 | 0.04 | 0.04 | 0.95 | 0.34 | 1 | Ddx5 | DEAD (Asp-Glu-Ala-Asp) box polypeptide | 24257 | 22443 | 21277 | 17914 | 19215 | 48504 | 20091 | 18844 | 19350 | 20501 | 50538 |
| 21678 | -0.11 | 0.07 | -1.52 | 0.13 | 1 | Nedd4 | neural precursor cell expressed, develop | 32197 | 31100 | 27807 | 20592 | 18431 | 17354 | 20074 | 16665 | 18945 | 18159 | 17131 |
| 17378 | -0.03 | 0.07 | -0.49 | 0.62 | 1 | Prrc2b | proline-rich coiled-coil 2B | 14240 | 15052 | 12537 | 20888 | 19301 | 15182 | 19847 | 20653 | 20858 | 19470 | 13132 |
| 17111 | 0.05 | 0.04 | 1.15 | 0.25 | 1 | Slc38a10 | solute carrier family 38, member 10 | 11669 | 11631 | 11599 | 20682 | 20216 | 14644 | 19755 | 21609 | 21051 | 20827 | 14532 |
| 61378 | -0.05 | 0.08 | -0.61 | 0.54 | 1 | Ttr | transthyretin | 145122 | 161074 | 204566 | 21551 | 21614 | 20701 | 19718 | 20280 | 20621 | 20667 | 19245 |
| 16769 | -0.11 | 0.07 | -1.61 | 0.11 | 1 | Prpf8 | pre-mRNA processing factor 8 | 14005 | 15047 | 14413 | 18960 | 19645 | 14938 | 19491 | 16035 | 19750 | 18356 | 13818 |
| 14936 | -0.03 | 0.06 | -0.43 | 0.66 | 1 | Itpkb | inositol 1,4,5-trisphosphate 3-kinase B | 4705 | 4251 | 5184 | 19920 | 20439 | 16013 | 19436 | 19902 | 20616 | 17685 | 16147 |
| 18648 | -0.04 | 0.06 | -0.68 | 0.50 | 1 | Itm2c | integral membrane protein 2C | 21443 | 23380 | 24187 | 19460 | 18693 | 11581 | 19407 | 19595 | 17219 | 18419 | 11741 |
| 19126 | -0.05 | 0.07 | -0.74 | 0.46 | 1 | Nisch | nischarin | 13348 | 15605 | 14110 | 19512 | 20999 | 25757 | 19112 | 17572 | 20699 | 20054 | 23615 |
| 16245 | 0.06 | 0.06 | 1.07 | 0.28 | 1 | Tram1 | translocating chain-associating membran | 7913 | 7976 | 9678 | 18485 | 21342 | 16141 | 19000 | 17917 | 21834 | 21005 | 17407 |
| 15728 | -0.04 | 0.10 | -0.37 | 0.71 | 1 | Dnajb9 | DnaJ heat shock protein family (Hsp40) m | 6394 | 5258 | 9197 | 13867 | 24649 | 18608 | 18964 | 10484 | 25944 | 20556 | 19089 |
| 14902 | 0.12 | 0.08 | 1.46 | 0.15 | 1 | Magt1 | magnesium transporter 1 | 6076 | 6618 | 7206 | 18755 | 17311 | 14119 | 18927 | 22021 | 17697 | 18987 | 16203 |
| 15356 | 0.07 | 0.06 | 1.13 | 0.26 | 1 | Tmem30a | transmembrane protein 30A | 10958 | 11035 | 10824 | 17645 | 16287 | 13748 | 18923 | 17075 | 17640 | 19022 | 15756 |
| 16438 | 0.08 | 0.06 | 1.25 | 0.21 | 1 | Copb2 | coatomer protein complex, subunit beta 2 | 11787 | 10906 | 13560 | 16819 | 18918 | 16142 | 18464 | 18994 | 19093 | 17934 | 18205 |
| 16441 | 0.00 | 0.06 | 0.07 | 0.94 | 1 | Ssr3 | signal sequence receptor, gamma | 12698 | 11382 | 13134 | 15781 | 20681 | 16850 | 18353 | 15101 | 21035 | 17634 | 18207 |
| 23327 | 0.01 | 0.10 | 0.05 | 0.96 | 1 | Cst3 | cystatin C | 43135 | 35321 | 45630 | 18854 | 20295 | 8539 | 18194 | 21653 | 17215 | 19650 | 8114 |
| 14847 | 0.06 | 0.06 | 1.09 | 0.28 | 1 | Trp53inp2 | transformation related protein 53 inducibl | 5272 | 5984 | 6325 | 19450 | 19334 | 14366 | 18134 | 20096 | 19822 | 19982 | 14556 |
| 14602 | -0.02 | 0.06 | -0.30 | 0.77 | 1 | Ptbp3 | polypyrimidine tract binding protein 3 | 9287 | 10323 | 9534 | 17234 | 16810 | 14302 | 17888 | 16682 | 17021 | 16138 | 15406 |
| 16942 | -0.12 | 0.09 | -1.26 | 0.21 | 1 | Gnao1 | guanine nucleotide binding protein, alpha | 15640 | 20483 | 17710 | 16696 | 16588 | 17866 | 17853 | 14272 | 17033 | 16657 | 15566 |
| 18749 | -0.05 | 0.06 | -0.81 | 0.42 | 1 | Ctnnb1 | catenin (cadherin associated protein), bet | 21917 | 22366 | 22284 | 19068 | 19329 | 14971 | 17683 | 16324 | 18818 | 18262 | 15215 |
| 11730 | -0.07 | 0.10 | -0.65 | 0.51 | 1 | Rnase4 | ribonuclease, RNase A family 4 | 9754 | 9503 | 8949 | 13453 | 12980 | 7555 | 17642 | 10648 | 13227 | 17582 | 7735 |
| 18894 | 0.10 | 0.08 | 1.25 | 0.21 | 1 | Bsg | basigin | 21745 | 23587 | 26048 | 18152 | 17736 | 12115 | 17631 | 22275 | 16972 | 18720 | 12850 |
| 14693 | 0.01 | 0.09 | 0.11 | 0.91 | 1 | Cdh1 | cadherin 1 | 7188 | 9499 | 10613 | 18447 | 17289 | 13568 | 17584 | 18781 | 17752 | 17264 | 13639 |
| 12691 | 0.14 | 0.10 | 1.37 | 0.17 | 1 | Phactr1 | phosphatase and actin regulator 1 | 2100 | 1887 | 2693 | 14888 | 16884 | 13750 | 17527 | 20258 | 18934 | 17064 | 13614 |
| 15839 | 0.07 | 0.09 | 0.72 | 0.47 | 1 | Mrfap1 | Morf4 family associated protein 1 | 11809 | 13358 | 15428 | 17674 | 17584 | 12553 | 17486 | 19904 | 16335 | 19008 | 13088 |
| 15832 | -0.03 | 0.07 | -0.39 | 0.69 | 1 | Wdr6 | WD repeat domain 6 | 12595 | 16595 | 14706 | 18565 | 17383 | 12243 | 17444 | 19564 | 15841 | 17303 | 11908 |
| 15661 | 0.00 | 0.05 | -0.01 | 0.99 | 1 | Csnk1a1 | casein kinase 1, alpha 1 | 11899 | 13688 | 11416 | 17839 | 16746 | 15649 | 17439 | 17576 | 17179 | 16736 | 16104 |
| 13817 | 0.00 | 0.07 | -0.05 | 0.96 | 1 | Uso1 | USO1 vesicle docking factor | 6427 | 6343 | 8133 | 14772 | 17730 | 15524 | 17427 | 13338 | 18896 | 16123 | 17276 |
| 14296 | 0.05 | 0.05 | 0.96 | 0.34 | 1 | Nckap1 | NCK-associated protein 1 | 9065 | 9311 | 8990 | 17358 | 15925 | 13147 | 17424 | 16784 | 17377 | 18370 | 13510 |
| 14651 | 0.04 | 0.08 | 0.52 | 0.60 | 1 | Ndfip1 | Nedd4 family interacting protein 1 | 12383 | 12905 | 13918 | 15762 | 15126 | 11925 | 17316 | 14659 | 15294 | 18743 | 13128 |
| 15040 | -0.01 | 0.07 | -0.09 | 0.93 | 1 | Plxnb2 | plexin B2 | 13473 | 11401 | 11824 | 17747 | 17100 | 12556 | 17032 | 17700 | 17768 | 16708 | 12130 |
| 15884 | 0.04 | 0.07 | 0.63 | 0.53 | 1 | Ddx3x | DEAD/H (Asp-Glu-Ala-Asp/His) box polyp | 14501 | 13875 | 13039 | 16742 | 16838 | 15122 | 16946 | 15868 | 18628 | 16263 | 16900 |
| 20276 | 0.14 | 0.13 | 1.13 | 0.26 | 1 | Fth1 | ferritin heavy polypeptide 1 | 37279 | 24902 | 28219 | 17811 | 17019 | 10974 | 16870 | 23919 | 16835 | 17809 | 11400 |
| 14015 | -0.10 | 0.08 | -1.27 | 0.21 | 1 | Sec31a | Sec31 homolog A (S. cerevisiae) | 9733 | 8261 | 9748 | 14413 | 17719 | 16092 | 16796 | 11415 | 17711 | 17321 | 14951 |
| 16363 | -0.03 | 0.06 | -0.52 | 0.61 | 1 | Kcnk16 | potassium channel, subfamily K, member | 11006 | 13961 | 14885 | 19484 | 20147 | 14459 | 16755 | 19238 | 18853 | 16923 | 14287 |
| 14733 | 0.15 | 0.11 | 1.27 | 0.20 | 1 | Rpn1 | ribophorin I | 10734 | 7691 | 10548 | 16968 | 16140 | 13344 | 16753 | 21133 | 16562 | 17473 | 14716 |

| | | | | | | | | | | | | | | | | | |
|---|---|---|---|---|---|---|---|---|---|---|---|---|---|---|---|---|---|
| 17116 | -0.05 | 0.06 | -0.82 | 0.41 | 1 | Slc16a10 | solute carrier family 16 (monocarboxylic a | 15332 | 20352 | 17061 | 18853 | 17932 | 15376 | 16735 | 17988 | 18328 | 16078 | 14244 |
| 15275 | 0.05 | 0.08 | 0.59 | 0.55 | 1 | Golga4 | golgi autoantigen, golgin subfamily a, 4 | 5630 | 6325 | 7894 | 16677 | 17199 | 22222 | 16712 | 18120 | 17536 | 16360 | 23349 |
| 16761 | 0.05 | 0.09 | 0.55 | 0.58 | 1 | Pnmal2 | PNMA-like 2 | 14237 | 20412 | 19375 | 15570 | 18606 | 13109 | 16666 | 18898 | 17671 | 16415 | 13412 |
| 14540 | -0.04 | 0.08 | -0.51 | 0.61 | 1 | Tm4sf4 | transmembrane 4 superfamily member 4 | 8257 | 11405 | 10862 | 16911 | 16488 | 15611 | 16644 | 16907 | 15303 | 16046 | 15509 |
| 13412 | -0.03 | 0.07 | -0.47 | 0.64 | 1 | Manf | mesencephalic astrocyte-derived neurotr | 6789 | 5847 | 8146 | 13974 | 18321 | 14921 | 16629 | 11705 | 18539 | 17649 | 15014 |
| 12379 | -0.02 | 0.08 | -0.24 | 0.81 | 1 | Nell1 | NEL-like 1 | 4737 | 5567 | 5969 | 16009 | 17207 | 10520 | 16586 | 17111 | 15029 | 17242 | 10190 |
| 11660 | -0.04 | 0.07 | -0.51 | 0.61 | 1 | Slc4a7 | solute carrier family 4, sodium bicarbonat | 2366 | 2438 | 2767 | 15882 | 14328 | 14408 | 16498 | 14885 | 15557 | 14966 | 14162 |
| 13787 | -0.06 | 0.08 | -0.77 | 0.44 | 1 | Abhd2 | abhydrolase domain containing 2 | 6575 | 7695 | 7188 | 18820 | 17672 | 13775 | 16279 | 14813 | 18711 | 16822 | 13301 |
| 15970 | 0.06 | 0.07 | 0.89 | 0.37 | 1 | Gnai2 | guanine nucleotide binding protein (G pro | 19016 | 16284 | 17246 | 16156 | 16655 | 10970 | 16123 | 18521 | 16283 | 18010 | 10407 |
| 13297 | -0.07 | 0.05 | -1.43 | 0.15 | 1 | Ogdh | oxoglutarate (alpha-ketoglutarate) dehydr | 10867 | 10113 | 10832 | 16427 | 16098 | 9984 | 16086 | 16037 | 15289 | 15213 | 9325 |
| 16122 | 0.02 | 0.07 | 0.25 | 0.80 | 1 | Ddb1 | damage specific DNA binding protein 1 | 19023 | 16777 | 18557 | 15519 | 17396 | 11929 | 16074 | 17990 | 16823 | 16234 | 11022 |
| 15101 | 0.04 | 0.07 | 0.57 | 0.57 | 1 | Eif4g1 | eukaryotic translation initiation factor 4, ga | 13074 | 10781 | 10648 | 15277 | 16291 | 17283 | 16060 | 16750 | 16859 | 15931 | 17157 |
| 13785 | 0.07 | 0.06 | 1.20 | 0.23 | 1 | Ssr1 | signal sequence receptor, alpha | 9515 | 8557 | 9790 | 12473 | 17322 | 14584 | 16060 | 13409 | 17540 | 16787 | 15597 |
| 13046 | 0.06 | 0.04 | 1.39 | 0.16 | 1 | Cyb5b | cytochrome b5 type B | 5333 | 5283 | 5552 | 16238 | 17143 | 12967 | 16030 | 16237 | 17611 | 17296 | 13814 |
| 14089 | 0.09 | 0.06 | 1.42 | 0.15 | 1 | Aldoa | aldolase A, fructose-bisphosphate | 12893 | 11315 | 12495 | 14999 | 14983 | 11112 | 16003 | 16914 | 15103 | 18087 | 11073 |
| 11200 | 0.12 | 0.12 | 0.95 | 0.34 | 1 | Tmem215 | transmembrane protein 215 | 728 | 542 | 1093 | 14320 | 15775 | 11940 | 15974 | 15171 | 16804 | 17377 | 13475 |
| 13851 | 0.00 | 0.06 | -0.03 | 0.98 | 1 | Tmem59 | transmembrane protein 59 | 11527 | 11873 | 13294 | 15543 | 15827 | 10661 | 15940 | 14680 | 15780 | 16013 | 11220 |
| 14102 | 0.04 | 0.07 | 0.58 | 0.56 | 1 | Os9 | amplified in osteosarcoma | 10370 | 11094 | 12080 | 16498 | 16280 | 11252 | 15913 | 18298 | 15354 | 16343 | 11639 |
| 13441 | -0.10 | 0.10 | -0.96 | 0.34 | 1 | Rab1a | RAB1A, member RAS oncogene family | 10574 | 11149 | 11890 | 13750 | 16175 | 13106 | 15840 | 10294 | 16250 | 14770 | 14052 |
| 15641 | 0.08 | 0.07 | 1.08 | 0.28 | 1 | Ywhae | tyrosine 3-monooxygenase/tryptophan 5- | 17344 | 15810 | 19136 | 14441 | 15879 | 12149 | 15813 | 17384 | 15912 | 15596 | 12588 |
| 18749 | 0.00 | 0.07 | 0.01 | 0.99 | 1 | Slc7a2 | solute carrier family 7 (cationic amino acid | 21784 | 27412 | 25334 | 18038 | 14008 | 18044 | 15770 | 18268 | 15555 | 14823 | 17197 |
| 14849 | -0.01 | 0.06 | -0.15 | 0.88 | 1 | Copg1 | coatomer protein complex, subunit gamm | 11620 | 11178 | 12681 | 15075 | 16918 | 16378 | 15756 | 14921 | 15800 | 17254 | 15762 |
| 12072 | 0.05 | 0.12 | 0.38 | 0.70 | 1 | Frzb | frizzled-related protein | 2666 | 4421 | 4362 | 16492 | 16516 | 11026 | 15756 | 16852 | 16026 | 17312 | 11366 |
| 14950 | -0.09 | 0.07 | -1.27 | 0.20 | 1 | Tspan7 | tetraspanin 7 | 14570 | 17060 | 18275 | 15721 | 16270 | 11336 | 15718 | 13782 | 15650 | 15434 | 10639 |
| 12589 | 0.09 | 0.09 | 1.00 | 0.32 | 1 | Fam174b | family with sequence similarity 174, meml | 5667 | 6482 | 7931 | 13717 | 16152 | 11760 | 15712 | 16740 | 16430 | 16058 | 11828 |
| 14002 | 0.05 | 0.05 | 0.86 | 0.39 | 1 | Glg1 | golgi apparatus protein 1 | 12942 | 12609 | 12469 | 14933 | 15819 | 10368 | 15702 | 16733 | 16030 | 16593 | 9818 |
| 15937 | 0.04 | 0.07 | 0.55 | 0.58 | 1 | Ppp1cb | protein phosphatase 1 catalytic subunit b | 16818 | 18603 | 17085 | 17184 | 14974 | 12945 | 15634 | 15124 | 15932 | 16767 | 14238 |
| 13539 | 0.10 | 0.06 | 1.60 | 0.11 | 1 | Spcs2 | signal peptidase complex subunit 2 homo | 7008 | 6289 | 8129 | 14952 | 17373 | 13752 | 15605 | 16619 | 17198 | 16611 | 15391 |
| 11810 | -0.05 | 0.14 | -0.37 | 0.71 | 1 | Gjd2 | gap junction protein, delta 2 | 1094 | 1555 | 2280 | 15873 | 15869 | 16169 | 15595 | 14433 | 15956 | 14978 | 16113 |
| 16830 | 0.10 | 0.07 | 1.35 | 0.18 | 1 | Pabpc1 | poly(A) binding protein, cytoplasmic 1 | 22976 | 19235 | 19097 | 15578 | 17339 | 11413 | 15441 | 16434 | 18213 | 17368 | 12041 |
| 12286 | 0.03 | 0.06 | 0.54 | 0.59 | 1 | Stxbp1 | syntaxin binding protein 1 | 6376 | 7069 | 7137 | 14717 | 15209 | 11199 | 15379 | 16348 | 15725 | 15220 | 10763 |
| 14630 | 0.00 | 0.07 | 0.00 | 1.00 | 1 | Sec61a1 | Sec61 alpha 1 subunit (S. cerevisiae) | 9929 | 8787 | 10625 | 13257 | 20210 | 17174 | 15261 | 13562 | 19325 | 16069 | 16734 |
| 12332 | 0.05 | 0.10 | 0.52 | 0.60 | 1 | Arfgef3 | ARFGEF family member 3 | 4085 | 6214 | 5783 | 13923 | 15206 | 14128 | 15121 | 16796 | 17266 | 13423 | 13703 |
| 15126 | 0.07 | 0.06 | 1.24 | 0.22 | 1 | Sqstm1 | sequestosome 1 | 13686 | 12727 | 14769 | 15057 | 17492 | 13418 | 15064 | 17401 | 17028 | 15942 | 13802 |
| 18179 | -0.08 | 0.06 | -1.34 | 0.18 | 1 | Lgals3bp | lectin, galactoside-binding, soluble, 3 bin | 27475 | 30278 | 30836 | 16045 | 15554 | 10478 | 15028 | 14642 | 14534 | 15542 | 9561 |
| 12934 | 0.05 | 0.08 | 0.62 | 0.53 | 1 | Atp6v1b2 | ATPase, H+ transporting, lysosomal V1 su | 8527 | 9956 | 10025 | 14445 | 15339 | 11188 | 15012 | 14137 | 15044 | 16690 | 11914 |
| 14134 | 0.00 | 0.07 | 0.02 | 0.98 | 1 | Golgb1 | golgi autoantigen, golgin subfamily b, ma | 6921 | 7920 | 7934 | 13179 | 16622 | 21494 | 14978 | 14206 | 17129 | 13608 | 21479 |
| 13176 | 0.01 | 0.05 | 0.27 | 0.79 | 1 | Lman2 | lectin, mannose-binding 2 | 8286 | 8410 | 9168 | 14358 | 17814 | 12246 | 14913 | 14810 | 17105 | 15389 | 12441 |
| 13558 | 0.03 | 0.07 | 0.44 | 0.66 | 1 | Uba1 | ubiquitin-like modifier activating enzyme | 12823 | 11633 | 12080 | 14566 | 15611 | 10418 | 14896 | 16760 | 15436 | 15192 | 9720 |
| 12491 | 0.06 | 0.06 | 1.06 | 0.29 | 1 | Ptpn11 | protein tyrosine phosphatase, non-recept | 7691 | 7887 | 8169 | 13580 | 15150 | 11973 | 14845 | 15472 | 14699 | 15930 | 12002 |
| 11930 | -0.09 | 0.12 | -0.78 | 0.43 | 1 | Cmtm4 | CKLF-like MARVEL transmembrane dom | 7384 | 11631 | 9320 | 14325 | 14198 | 9875 | 14790 | 11938 | 14296 | 14031 | 9444 |
| 17626 | 0.14 | 0.12 | 1.16 | 0.25 | 1 | Rpl8 | ribosomal protein L8 | 28529 | 24583 | 30732 | 13713 | 14077 | 9512 | 14715 | 20049 | 12585 | 15334 | 10057 |
| 12959 | 0.01 | 0.06 | 0.13 | 0.90 | 1 | Etnk1 | ethanolamine kinase 1 | 8324 | 9074 | 9053 | 15040 | 14899 | 13252 | 14713 | 13740 | 15211 | 15413 | 13824 |
| 15040 | 0.07 | 0.05 | 1.49 | 0.14 | 1 | Eif3a | eukaryotic translation initiation factor 3, su | 12036 | 12866 | 13709 | 15243 | 16047 | 15901 | 14689 | 16446 | 16521 | 14571 | 17414 |
| 12647 | -0.08 | 0.04 | -1.94 | 0.05 | 1 | Nfe2l1 | nuclear factor, erythroid derived 2,-like 1 | 10766 | 10147 | 10756 | 14428 | 14952 | 11243 | 14649 | 12964 | 14460 | 14228 | 10527 |
| 16337 | 0.02 | 0.07 | 0.30 | 0.77 | 1 | Tpt1 | tumor protein, translationally-controlled 1 | 25437 | 25175 | 25889 | 13402 | 13592 | 9562 | 14630 | 12099 | 14044 | 16046 | 9836 |
| 11774 | -0.07 | 0.10 | -0.66 | 0.51 | 1 | Kif1a | kinesin family member 1A | 5319 | 7781 | 6161 | 14323 | 14696 | 12723 | 14630 | 13270 | 14788 | 14661 | 11167 |
| 9770 | 0.00 | 0.11 | 0.04 | 0.96 | 1 | Ntrk2 | neurotrophic tyrosine kinase, receptor, typ | 4220 | 2420 | 3188 | 14036 | 12114 | 8128 | 14615 | 12500 | 13199 | 14250 | 8797 |
| 13704 | -0.04 | 0.08 | -0.45 | 0.65 | 1 | Stt3a | STT3, subunit of the oligosaccharyltransfe | 10153 | 9844 | 11872 | 13170 | 17894 | 14385 | 14544 | 11027 | 17814 | 15246 | 14790 |
| 10171 | -0.03 | 0.08 | -0.38 | 0.70 | 1 | Pde5a | phosphodiesterase 5A, cGMP-specific | 1573 | 1183 | 2034 | 12758 | 16125 | 10626 | 14533 | 12895 | 17129 | 12658 | 10369 |
| 17742 | 0.10 | 0.11 | 0.84 | 0.40 | 1 | Rps3 | ribosomal protein S3 | 31913 | 26482 | 30405 | 13892 | 13683 | 9118 | 14463 | 19113 | 12246 | 14367 | 9475 |
| 15373 | 0.08 | 0.13 | 0.61 | 0.54 | 1 | mt-Rnr2 | | 19199 | 23700 | 18660 | 12956 | 11272 | 13714 | 14406 | 10790 | 15888 | 14041 | 14479 |
| 14255 | 0.08 | 0.06 | 1.35 | 0.18 | 1 | Chd4 | chromodomain helicase DNA binding pro | 12705 | 14358 | 11701 | 16195 | 13850 | 12854 | 14340 | 19025 | 14073 | 15024 | 12687 |
| 11269 | 0.14 | 0.07 | 1.95 | 0.05 | 1 | Tspan13 | tetraspanin 13 | 5352 | 5658 | 6045 | 14683 | 12905 | 9079 | 14326 | 15748 | 13989 | 15668 | 10510 |
| 13285 | 0.04 | 0.08 | 0.44 | 0.66 | 1 | Tle5 | TLE family member 5, transcriptional mod | 10950 | 12038 | 13319 | 14941 | 14857 | 9938 | 14293 | 15257 | 14124 | 16914 | 9500 |
| 12657 | 0.06 | 0.07 | 0.90 | 0.37 | 1 | Lamp2 | lysosomal-associated membrane protein | 12882 | 10885 | 10671 | 14616 | 13227 | 9471 | 14160 | 13774 | 14042 | 14714 | 10791 |
| 11985 | 0.02 | 0.05 | 0.40 | 0.69 | 1 | Sort1 | sortilin 1 | 8404 | 9069 | 9216 | 14260 | 13875 | 9842 | 14094 | 13933 | 14862 | 14715 | 9566 |
| 13630 | 0.06 | 0.07 | 0.93 | 0.35 | 1 | Cyb5r3 | cytochrome b5 reductase 3 | 11535 | 12281 | 12422 | 15577 | 15176 | 10760 | 14064 | 17999 | 14355 | 14528 | 11227 |
| 12624 | -0.16 | 0.09 | -1.86 | 0.06 | 1 | Map1b | microtubule-associated protein 1B | 10642 | 14851 | 12305 | 13728 | 13804 | 11669 | 14032 | 12714 | 13410 | 12094 | 9617 |
| 12217 | -0.02 | 0.07 | -0.33 | 0.74 | 1 | Arhgef12 | Rho guanine nucleotide exchange factor | 9236 | 9739 | 9182 | 14048 | 13889 | 11498 | 13983 | 12600 | 15486 | 14108 | 10615 |

| | | | | | | | | | | | | | | | | | | |
|---|---|---|---|---|---|---|---|---|---|---|---|---|---|---|---|---|---|---|
| 12288 | 0.14 | 0.08 | 1.84 | 0.07 | 1 | Mtch1 | mitochondrial carrier 1 | 8682 | 8538 | 9308 | 13653 | 13799 | 10115 | 13929 | 17036 | 13536 | 15869 | 10700 |
| 17134 | 0.00 | 0.05 | 0.06 | 0.95 | 1 | Hspa8 | heat shock protein 8 | 18555 | 16367 | 22504 | 11879 | 18402 | 21294 | 13890 | 12359 | 18457 | 13315 | 21448 |
| 13601 | -0.05 | 0.10 | -0.46 | 0.65 | 1 | Ddx6 | DEAD (Asp-Glu-Ala-Asp) box polypeptide | 13351 | 15926 | 14202 | 12923 | 13428 | 13629 | 13850 | 10583 | 15083 | 13379 | 13262 |
| 10973 | -0.15 | 0.13 | -1.22 | 0.22 | 1 | Unc80 | unc-80, NALCN activator | 6131 | 9359 | 6769 | 11760 | 11443 | 14887 | 13828 | 10757 | 12444 | 10940 | 12390 |
| 23878 | -0.01 | 0.08 | -0.17 | 0.87 | 1 | Actb | actin, beta | 57700 | 44607 | 42674 | 13881 | 13803 | 17784 | 13791 | 14110 | 14222 | 13921 | 16162 |
| 10874 | 0.09 | 0.06 | 1.45 | 0.15 | 1 | Tmem65 | transmembrane protein 65 | 3044 | 3658 | 3609 | 13711 | 13457 | 12080 | 13664 | 15262 | 14103 | 13525 | 13504 |
| 12561 | -0.05 | 0.09 | -0.55 | 0.58 | 1 | Iqgap1 | IQ motif containing GTPase activating pro | 12234 | 8624 | 11065 | 13277 | 15020 | 12028 | 13644 | 12845 | 15374 | 12662 | 11395 |
| 10219 | -0.03 | 0.10 | -0.28 | 0.78 | 1 | Atp8b1 | ATPase, class I, type 8B, member 1 | 3007 | 3983 | 4067 | 12877 | 12320 | 12325 | 13627 | 11873 | 13163 | 13053 | 12120 |
| 11483 | -0.02 | 0.09 | -0.19 | 0.85 | 1 | Slc38a4 | solute carrier family 38, member 4 | 5340 | 7010 | 6416 | 13182 | 12822 | 14472 | 13622 | 11650 | 13547 | 13650 | 14606 |
| 10942 | 0.05 | 0.07 | 0.75 | 0.45 | 1 | Madd | MAP-kinase activating death domain | 3235 | 3802 | 3876 | 14570 | 13757 | 11573 | 13613 | 16844 | 14530 | 13660 | 10899 |
| 11310 | 0.14 | 0.11 | 1.30 | 0.20 | 1 | Nucb2 | nucleobindin 2 | 3892 | 5340 | 5722 | 15532 | 12829 | 10064 | 13540 | 18994 | 13314 | 13632 | 11548 |
| 14525 | 0.11 | 0.06 | 1.84 | 0.07 | 1 | Rrbp1 | ribosome binding protein 1 | 13569 | 12526 | 13054 | 12584 | 15330 | 16620 | 13512 | 13898 | 15962 | 15995 | 16724 |
| 11381 | -0.04 | 0.05 | -0.77 | 0.44 | 1 | Ganab | alpha glucosidase 2 alpha neutral subuni | 9194 | 8308 | 8733 | 13115 | 12717 | 10863 | 13481 | 13433 | 12328 | 12754 | 10264 |
| 12388 | -0.04 | 0.05 | -0.71 | 0.48 | 1 | Gtf2i | general transcription factor II I | 11235 | 12117 | 11720 | 12870 | 13680 | 11230 | 13443 | 12737 | 13069 | 13575 | 10589 |
| 12473 | -0.04 | 0.10 | -0.44 | 0.66 | 1 | Arcn1 | archain 1 | 9417 | 9557 | 9715 | 12689 | 15423 | 13311 | 13433 | 9960 | 16127 | 13440 | 14133 |
| 12155 | -0.05 | 0.06 | -0.83 | 0.41 | 1 | Syt7 | synaptotagmin VII | 8961 | 10698 | 10455 | 13677 | 13782 | 11675 | 13399 | 14035 | 13977 | 12313 | 10733 |
| 11341 | 0.02 | 0.07 | 0.23 | 0.82 | 1 | Selenof | selenoprotein F | 8925 | 8634 | 9414 | 13517 | 11856 | 9987 | 13358 | 13160 | 12637 | 12252 | 11012 |
| 11403 | 0.06 | 0.06 | 0.95 | 0.34 | 1 | Copb1 | coatomer protein complex, subunit beta 1 | 8199 | 8393 | 8073 | 11678 | 13568 | 10806 | 13349 | 12279 | 13728 | 13782 | 11576 |
| 12148 | 0.03 | 0.05 | 0.55 | 0.58 | 1 | Kif1b | kinesin family member 1B | 7127 | 8008 | 7596 | 15122 | 12937 | 13441 | 13309 | 16195 | 13753 | 12834 | 13307 |
| 12308 | 0.03 | 0.05 | 0.64 | 0.52 | 1 | Rap1b | RAS related protein 1b | 12004 | 12334 | 11686 | 12848 | 11723 | 11379 | 13246 | 12921 | 12068 | 13118 | 12064 |
| 12044 | -0.10 | 0.08 | -1.21 | 0.23 | 1 | Huwe1 | HECT, UBA and WWE domain containing | 9055 | 9983 | 8964 | 12059 | 12928 | 15814 | 13203 | 11234 | 13648 | 11670 | 13920 |
| 11781 | 0.05 | 0.07 | 0.70 | 0.49 | 1 | Sdha | succinate dehydrogenase complex, subu | 10398 | 9540 | 10890 | 12704 | 12524 | 10116 | 13195 | 14358 | 12312 | 13026 | 10530 |
| 11480 | -0.02 | 0.11 | -0.19 | 0.85 | 1 | Lrp11 | low density lipoprotein receptor-related pr | 6137 | 8582 | 9008 | 14078 | 13991 | 10465 | 13155 | 12227 | 13454 | 15018 | 10165 |
| 12926 | 0.00 | 0.08 | 0.04 | 0.97 | 1 | Pja2 | praja ring finger ubiquitin ligase 2 | 10939 | 13319 | 12633 | 12972 | 14027 | 12491 | 13109 | 11837 | 13964 | 13874 | 13019 |
| 9928 | 0.09 | 0.07 | 1.25 | 0.21 | 1 | Asah1 | N-acylsphingosine amidohydrolase 1 | 5707 | 4380 | 4965 | 12260 | 11960 | 8291 | 13062 | 11226 | 12591 | 15346 | 9416 |
| 13658 | -0.04 | 0.08 | -0.49 | 0.62 | 1 | Sdf4 | stromal cell derived factor 4 | 17537 | 17400 | 19628 | 13721 | 12879 | 8889 | 13034 | 14305 | 11699 | 12050 | 9100 |
| 9186 | 0.12 | 0.16 | 0.74 | 0.46 | 1 | Fam151a | family with sequence simliarity 151, meml | 606 | 535 | 978 | 12462 | 12519 | 9468 | 12998 | 16546 | 12124 | 12838 | 9968 |
| 15785 | 0.05 | 0.06 | 0.83 | 0.41 | 1 | Hnrnpa2b1 | heterogeneous nuclear ribonucleoprotein | 17563 | 14741 | 16524 | 12174 | 11583 | 24621 | 12958 | 13021 | 12477 | 12545 | 25425 |
| 10514 | -0.02 | 0.09 | -0.26 | 0.80 | 1 | Marchf6 | membrane associated ring-CH-type finge | 6838 | 8763 | 7182 | 12049 | 12157 | 9690 | 12958 | 10892 | 12501 | 12919 | 9704 |
| 11184 | 0.12 | 0.07 | 1.82 | 0.07 | 1 | Vps35 | VPS35 retromer complex component | 6556 | 6689 | 6257 | 13671 | 11803 | 11279 | 12929 | 13942 | 13328 | 12731 | 13844 |
| 9471 | -0.07 | 0.08 | -0.83 | 0.41 | 1 | Ldlr | low density lipoprotein receptor | 3802 | 4886 | 4753 | 12770 | 11889 | 8911 | 12881 | 12332 | 11282 | 12082 | 8595 |
| 10279 | 0.04 | 0.10 | 0.39 | 0.70 | 1 | Slc2a5 | solute carrier family 2 (facilitated glucose | 3502 | 5204 | 4778 | 13440 | 12690 | 10093 | 12863 | 13791 | 13109 | 13712 | 9882 |
| 11383 | -0.17 | 0.09 | -1.90 | 0.06 | 1 | Tmed8 | transmembrane p24 trafficking protein 8 | 11350 | 14376 | 13823 | 11131 | 11223 | 10202 | 12856 | 9816 | 10457 | 10584 | 9392 |
| 11384 | 0.01 | 0.06 | 0.26 | 0.80 | 1 | Tmed10 | transmembrane p24 trafficking protein 10 | 9652 | 9899 | 10188 | 12151 | 13511 | 8949 | 12853 | 12128 | 13148 | 13871 | 8870 |
| 10321 | 0.04 | 0.09 | 0.46 | 0.64 | 1 | Vapb | vesicle-associated membrane protein, as | 5876 | 6155 | 7240 | 11458 | 12256 | 9932 | 12844 | 12628 | 11488 | 13507 | 10142 |
| 14649 | 0.02 | 0.06 | 0.34 | 0.74 | 1 | Rplp0 | ribosomal protein, large, P0 | 24317 | 19715 | 23621 | 12274 | 12628 | 8719 | 12814 | 12741 | 12386 | 12948 | 8980 |
| 12067 | 0.02 | 0.07 | 0.22 | 0.83 | 1 | Snd1 | staphylococcal nuclease and tudor doma | 11013 | 13790 | 12031 | 12653 | 12185 | 9897 | 12773 | 14551 | 12017 | 12506 | 9326 |
| 9964 | -0.03 | 0.10 | -0.32 | 0.75 | 1 | Abcd3 | ATP-binding cassette, sub-family D (ALD) | 4829 | 5812 | 6584 | 11803 | 11993 | 10181 | 12771 | 10778 | 11588 | 12137 | 11130 |
| 12970 | 0.00 | 0.06 | -0.07 | 0.95 | 1 | Hnrnpk | heterogeneous nuclear ribonucleoprotein | 14520 | 15359 | 15377 | 12398 | 11652 | 12008 | 12714 | 11449 | 12536 | 12489 | 12170 |
| 12154 | -0.07 | 0.08 | -0.93 | 0.35 | 1 | Laptm4a | lysosomal-associated protein transmemb | 14327 | 11642 | 13688 | 13052 | 12450 | 9940 | 12691 | 10451 | 12858 | 12547 | 10049 |
| 9779 | -0.04 | 0.08 | -0.55 | 0.58 | 1 | Vdr | vitamin D (1,25-dihydroxyvitamin D3) rece | 3085 | 4427 | 4032 | 13723 | 12624 | 9720 | 12681 | 12946 | 12835 | 12157 | 9340 |
| 10868 | -0.02 | 0.05 | -0.53 | 0.59 | 1 | Arf1 | ADP-ribosylation factor 1 | 8192 | 8109 | 8393 | 11700 | 12750 | 10740 | 12636 | 11318 | 12784 | 12350 | 10579 |
| 12128 | -0.14 | 0.13 | -1.06 | 0.29 | 1 | Pcbp2 | poly(rC) binding protein 2 | 13358 | 13834 | 13757 | 12230 | 13720 | 9482 | 12618 | 7834 | 14135 | 13749 | 8688 |
| 12562 | -0.01 | 0.06 | -0.17 | 0.86 | 1 | Gnb1 | guanine nucleotide binding protein (G prc | 15404 | 15124 | 16490 | 11816 | 12390 | 8995 | 12616 | 12721 | 11983 | 11291 | 9357 |
| 11712 | 0.03 | 0.06 | 0.47 | 0.64 | 1 | Tnpo1 | transportin 1 | 7719 | 8286 | 8313 | 13063 | 12387 | 13697 | 12584 | 12102 | 13724 | 12662 | 14293 |
| 9831 | 0.12 | 0.09 | 1.43 | 0.15 | 1 | Zdhhc2 | zinc finger, DHHC domain containing 2 | 2182 | 2645 | 2854 | 13235 | 11252 | 11046 | 12553 | 14127 | 12962 | 13342 | 11942 |
| 12400 | -0.08 | 0.07 | -1.16 | 0.24 | 1 | Wnk1 | WNK lysine deficient protein kinase 1 | 14850 | 14883 | 13726 | 11630 | 12272 | 11353 | 12504 | 11251 | 12915 | 11098 | 9918 |
| 13672 | 0.04 | 0.08 | 0.57 | 0.57 | 1 | Tsc22d1 | TSC22 domain family, member 1 | 14903 | 15045 | 16575 | 13075 | 12476 | 13055 | 12469 | 15301 | 12104 | 12046 | 13348 |
| 13112 | -0.07 | 0.07 | -1.03 | 0.30 | 1 | Nucb1 | nucleobindin 1 | 16064 | 17643 | 18011 | 13999 | 12351 | 8487 | 12426 | 13541 | 11300 | 12568 | 7843 |
| 14807 | 0.08 | 0.10 | 0.83 | 0.40 | 1 | Son | Son DNA binding protein | 10358 | 13842 | 10437 | 11573 | 13180 | 25892 | 12378 | 14620 | 13643 | 12901 | 24058 |
| 11377 | 0.06 | 0.09 | 0.68 | 0.49 | 1 | Ranbp2 | RAN binding protein 2 | 7960 | 8725 | 8291 | 11097 | 12224 | 13230 | 12313 | 11478 | 13831 | 10600 | 15397 |
| 12574 | 0.07 | 0.06 | 1.20 | 0.23 | 1 | Slc25a3 | solute carrier family 25 (mitochondrial car | 13430 | 13217 | 13857 | 11595 | 12168 | 11656 | 12312 | 13496 | 12181 | 12667 | 11735 |
| 10162 | 0.01 | 0.10 | 0.14 | 0.89 | 1 | Atp1b1 | ATPase, Na+/K+ transporting, beta 1 poly | 6098 | 6908 | 7711 | 12142 | 10507 | 10378 | 12298 | 11077 | 11289 | 12801 | 10572 |
| 10511 | 0.04 | 0.06 | 0.71 | 0.48 | 1 | 4833439L19Rik | RIKEN cDNA 4833439L19 gene | 4815 | 5043 | 5941 | 13236 | 13445 | 10175 | 12293 | 13169 | 13925 | 13379 | 10200 |
| 11228 | 0.08 | 0.04 | 1.90 | 0.06 | 1 | Tm9sf2 | transmembrane 9 superfamily member 2 | 10322 | 9415 | 10230 | 11631 | 12019 | 9531 | 12275 | 11809 | 12824 | 13096 | 10358 |
| 10080 | 0.02 | 0.07 | 0.26 | 0.79 | 1 | Wls | wntless WNT ligand secretion mediator | 5386 | 4427 | 4425 | 12973 | 12506 | 10278 | 12223 | 12143 | 13243 | 12825 | 10455 |
| 10287 | 0.05 | 0.06 | 0.80 | 0.42 | 1 | Hepacam2 | HEPACAM family member 2 | 3481 | 3860 | 3877 | 12724 | 12900 | 12334 | 12170 | 12087 | 14031 | 12918 | 12776 |
| 11752 | 0.02 | 0.07 | 0.29 | 0.77 | 1 | Calm3 | calmodulin 3 | 12654 | 12137 | 14496 | 11586 | 12654 | 8289 | 12158 | 12770 | 11384 | 12718 | 8423 |
| 9979 | 0.00 | 0.08 | 0.04 | 0.97 | 1 | Rcan3 | regulator of calcineurin 3 | 6421 | 8512 | 7861 | 11729 | 11327 | 8346 | 12101 | 11484 | 11498 | 11747 | 8741 |
| 11474 | 0.03 | 0.04 | 0.80 | 0.43 | 1 | Dcaf7 | DDB1 and CUL4 associated factor 7 | 9036 | 10002 | 9160 | 13570 | 12505 | 10352 | 12051 | 13808 | 12608 | 12530 | 10591 |

| | | | | | | | | | | | | | | | | | | |
|---|---|---|---|---|---|---|---|---|---|---|---|---|---|---|---|---|---|---|
| 10097 | -0.06 | 0.08 | -0.86 | 0.39 | 1 | Tmx4 | thioredoxin-related transmembrane prote | 8462 | 9534 | 9913 | 10863 | 11091 | 8619 | 11976 | 9756 | 11013 | 11265 | 8578 |
| 11342 | 0.05 | 0.06 | 0.88 | 0.38 | 1 | Kidins220 | kinase D-interacting substrate 220 | 6483 | 7987 | 7075 | 13318 | 11054 | 14417 | 11971 | 13976 | 12364 | 11890 | 14232 |
| 10545 | -0.09 | 0.13 | -0.66 | 0.51 | 1 | Map2 | microtubule-associated protein 2 | 5834 | 9000 | 6801 | 12440 | 10445 | 13702 | 11887 | 8963 | 11340 | 10837 | 14750 |
| 11735 | 0.07 | 0.06 | 1.34 | 0.18 | 1 | Morf4l2 | mortality factor 4 like 2 | 8095 | 8033 | 9317 | 10784 | 12858 | 14952 | 11864 | 11485 | 12586 | 12805 | 16311 |
| 10485 | -0.05 | 0.06 | -0.84 | 0.40 | 1 | Creb3l2 | cAMP responsive element binding proteir | 8887 | 9103 | 10388 | 9827 | 12434 | 10231 | 11748 | 10159 | 12389 | 10843 | 9331 |
| 10979 | 0.10 | 0.08 | 1.36 | 0.17 | 1 | Eif5 | eukaryotic translation initiation factor 5 | 8832 | 8725 | 9071 | 10728 | 12404 | 10522 | 11727 | 13470 | 12830 | 10880 | 11575 |
| 9696 | 0.06 | 0.08 | 0.77 | 0.44 | 1 | Marchf8 | membrane associated ring-CH-type finge | 7042 | 7316 | 7987 | 10233 | 10473 | 8753 | 11690 | 9784 | 11124 | 13145 | 9115 |
| 9618 | 0.05 | 0.07 | 0.68 | 0.50 | 1 | Arfgef1 | ADP-ribosylation factor guanine nucleotic | 6355 | 6864 | 6438 | 10761 | 10405 | 9456 | 11667 | 10875 | 12519 | 10918 | 9538 |
| 14989 | -0.02 | 0.10 | -0.19 | 0.85 | 1 | H2-D1 | histocompatibility 2, D region locus 1 | 38444 | 18574 | 26867 | 9274 | 12061 | 7515 | 11666 | 8770 | 11617 | 12972 | 7115 |
| 10172 | 0.03 | 0.05 | 0.52 | 0.61 | 1 | Atp6ap1 | ATPase, H+ transporting, lysosomal acce | 8926 | 8999 | 8911 | 11741 | 10793 | 7850 | 11628 | 12361 | 11379 | 11817 | 7491 |
| 10438 | 0.14 | 0.11 | 1.30 | 0.19 | 1 | Ap1s2 | adaptor-related protein complex 1, sigma | 5294 | 7170 | 6959 | 10452 | 10062 | 13131 | 11616 | 10959 | 10990 | 13199 | 14990 |
| 11875 | 0.15 | 0.10 | 1.41 | 0.16 | 1 | Akr1a1 | aldo-keto reductase family 1, member A1 | 13135 | 10967 | 13342 | 11594 | 11942 | 8938 | 11550 | 15574 | 11439 | 13006 | 9137 |
| 10663 | 0.14 | 0.10 | 1.46 | 0.14 | 1 | Tmed3 | transmembrane p24 trafficking protein 3 | 7978 | 9183 | 9776 | 12250 | 11139 | 7769 | 11506 | 16404 | 11290 | 11767 | 8233 |
| 10708 | 0.02 | 0.10 | 0.16 | 0.87 | 1 | Gprasp1 | G protein-coupled receptor associated so | 9102 | 11120 | 10960 | 9125 | 9655 | 12844 | 11455 | 10262 | 9443 | 10813 | 13011 |
| 9861 | 0.12 | 0.08 | 1.54 | 0.12 | 1 | Eid1 | EP300 interacting inhibitor of differentiatic | 7427 | 7870 | 8572 | 10721 | 11166 | 7330 | 11431 | 12846 | 10927 | 11894 | 8287 |
| 9258 | 0.00 | 0.10 | 0.02 | 0.99 | 1 | Rab27a | RAB27A, member RAS oncogene family | 4817 | 6359 | 6674 | 11056 | 10160 | 9370 | 11380 | 10448 | 10914 | 11269 | 9396 |
| 10697 | -0.01 | 0.09 | -0.06 | 0.95 | 1 | Rtn3 | reticulon 3 | 11185 | 11899 | 12161 | 11036 | 10731 | 8173 | 11368 | 8985 | 11293 | 12034 | 8802 |
| 10925 | 0.05 | 0.07 | 0.66 | 0.51 | 1 | Cnot6l | CCR4-NOT transcription complex, subuni | 10035 | 12785 | 11127 | 10720 | 11113 | 9137 | 11367 | 12079 | 11589 | 10985 | 9244 |
| 8778 | 0.04 | 0.09 | 0.51 | 0.61 | 1 | Itgav | integrin alpha V | 4141 | 4675 | 4161 | 13034 | 9995 | 7102 | 11340 | 11088 | 11744 | 11019 | 8254 |
| 11083 | 0.00 | 0.05 | 0.01 | 0.99 | 1 | Arf3 | ADP-ribosylation factor 3 | 10857 | 11719 | 12693 | 11315 | 11858 | 8749 | 11324 | 11824 | 11848 | 11354 | 8366 |
| 9835 | -0.07 | 0.06 | -1.18 | 0.24 | 1 | Ahcyl1 | S-adenosylhomocysteine hydrolase-like ' | 6385 | 6497 | 6551 | 10754 | 10611 | 12761 | 11320 | 9175 | 10594 | 10739 | 12802 |
| 10822 | -0.03 | 0.06 | -0.48 | 0.63 | 1 | Nbr1 | NBR1, autophagy cargo receptor | 7707 | 9079 | 9033 | 11147 | 12719 | 11887 | 11274 | 11288 | 12834 | 10234 | 11835 |
| 9716 | -0.05 | 0.06 | -0.94 | 0.35 | 1 | Aco2 | aconitase 2, mitochondrial | 8067 | 7331 | 7575 | 11466 | 11253 | 8738 | 11270 | 10825 | 11252 | 10506 | 8593 |
| 8757 | 0.08 | 0.10 | 0.80 | 0.42 | 1 | Wnt4 | wingless-type MMTV integration site famil | 5606 | 6179 | 5763 | 8945 | 10025 | 7932 | 11258 | 10603 | 9531 | 12873 | 7617 |
| 13985 | 0.06 | 0.08 | 0.74 | 0.46 | 1 | Hnrnpu | heterogeneous nuclear ribonucleoproteir | 16612 | 15117 | 15216 | 13053 | 11519 | 16408 | 11255 | 13849 | 11555 | 11174 | 18079 |
| 11056 | 0.11 | 0.07 | 1.53 | 0.13 | 1 | Gdi2 | guanosine diphosphate (GDP) dissociatic | 13982 | 11328 | 11310 | 10489 | 11054 | 8263 | 11220 | 11515 | 11157 | 11721 | 9571 |
| 9706 | 0.09 | 0.07 | 1.31 | 0.19 | 1 | Sec62 | SEC62 homolog (S. cerevisiae) | 6989 | 7547 | 7349 | 9790 | 10456 | 9602 | 11171 | 12248 | 10686 | 11032 | 9897 |
| 9721 | -0.02 | 0.07 | -0.21 | 0.84 | 1 | Rab7 | RAB7, member RAS oncogene family | 7576 | 7743 | 7684 | 11283 | 11089 | 8591 | 11168 | 9607 | 11391 | 12324 | 8474 |
| 10266 | -0.04 | 0.07 | -0.59 | 0.56 | 1 | Xbp1 | X-box binding protein 1 | 9290 | 7981 | 8674 | 9603 | 12568 | 10729 | 11155 | 8827 | 11869 | 11791 | 10441 |
| 8674 | 0.02 | 0.11 | 0.22 | 0.83 | 1 | Insm1 | insulinoma-associated 1 | 2562 | 3557 | 4014 | 11169 | 12198 | 7690 | 11149 | 10733 | 12722 | 12221 | 7399 |
| 8860 | 0.01 | 0.07 | 0.16 | 0.87 | 1 | Rab3d | RAB3D, member RAS oncogene family | 3299 | 3993 | 4280 | 11842 | 12275 | 7486 | 11136 | 12889 | 11632 | 11229 | 7402 |
| 10645 | 0.07 | 0.08 | 0.96 | 0.34 | 1 | Tcf25 | transcription factor 25 (basic helix-loop-he | 7822 | 8166 | 8597 | 9540 | 10519 | 14021 | 11127 | 11543 | 10359 | 11785 | 13615 |
| 9921 | -0.01 | 0.09 | -0.08 | 0.93 | 1 | Srp72 | signal recognition particle 72 | 6064 | 6939 | 6946 | 10095 | 12317 | 11183 | 11054 | 9148 | 12253 | 11168 | 11968 |
| 10470 | 0.05 | 0.07 | 0.74 | 0.46 | 1 | Pafah1b1 | platelet-activating factor acetylhydrolase, | 8359 | 9526 | 8165 | 11589 | 10724 | 10457 | 11038 | 11073 | 11311 | 10873 | 12050 |
| 9564 | 0.00 | 0.09 | -0.04 | 0.97 | 1 | Scarb2 | scavenger receptor class B, member 2 | 9876 | 6988 | 8067 | 10890 | 10990 | 7321 | 11009 | 10118 | 11486 | 10913 | 7545 |
| 8340 | 0.00 | 0.13 | 0.00 | 1.00 | 1 | Ghr | growth hormone receptor | 2800 | 4112 | 3447 | 10454 | 10565 | 8772 | 10956 | 8403 | 11058 | 11343 | 9834 |
| 11355 | 0.16 | 0.08 | 1.92 | 0.06 | 1 | Cnbp | cellular nucleic acid binding protein | 12827 | 12352 | 12660 | 10902 | 10784 | 8408 | 10947 | 14325 | 10518 | 11425 | 9750 |
| 9034 | 0.01 | 0.11 | 0.13 | 0.90 | 1 | Gng4 | guanine nucleotide binding protein (G prc | 4083 | 5547 | 5484 | 9559 | 11167 | 10241 | 10947 | 9287 | 11580 | 11213 | 10264 |
| 8613 | 0.01 | 0.07 | 0.08 | 0.94 | 1 | Cacna2d1 | calcium channel, voltage-dependent, alpl | 5212 | 5927 | 5467 | 9508 | 9778 | 8819 | 10914 | 9456 | 10172 | 10190 | 9298 |
| 13465 | 0.00 | 0.09 | 0.05 | 0.96 | 1 | Ctsb | cathepsin B | 27481 | 19744 | 21587 | 10670 | 10959 | 7047 | 10874 | 11592 | 9966 | 11188 | 7006 |
| 9961 | -0.04 | 0.09 | -0.42 | 0.68 | 1 | Hipk3 | homeodomain interacting protein kinase : | 7565 | 9215 | 7761 | 11270 | 10841 | 10041 | 10867 | 9972 | 12111 | 10396 | 9537 |
| 9793 | 0.05 | 0.08 | 0.61 | 0.54 | 1 | Adgrg1 | adhesion G protein-coupled receptor G1 | 6993 | 8260 | 8457 | 11427 | 10962 | 7752 | 10862 | 12555 | 11280 | 12184 | 6985 |
| 14021 | 0.05 | 0.07 | 0.76 | 0.45 | 1 | Rack1 | receptor for activated C kinase 1 | 26570 | 22930 | 24679 | 10244 | 10585 | 7656 | 10831 | 11465 | 9719 | 11521 | 8031 |
| 12106 | 0.17 | 0.11 | 1.59 | 0.11 | 1 | Ncl | nucleolin | 16765 | 13284 | 13192 | 10436 | 10834 | 10197 | 10824 | 14328 | 11551 | 10741 | 11009 |
| 7636 | 0.08 | 0.09 | 0.85 | 0.40 | 1 | Creg1 | cellular repressor of E1A-stimulated gene | 3127 | 2975 | 3674 | 9898 | 9439 | 6040 | 10816 | 9142 | 9941 | 11898 | 7046 |
| 9421 | 0.02 | 0.09 | 0.22 | 0.83 | 1 | Sesn3 | sestrin 3 | 7971 | 10301 | 9005 | 9980 | 8924 | 8172 | 10789 | 10542 | 9610 | 10529 | 7804 |
| 10016 | 0.05 | 0.08 | 0.55 | 0.58 | 1 | Ccnd1 | cyclin D1 | 10370 | 10001 | 11826 | 7780 | 11468 | 8372 | 10738 | 8595 | 11300 | 11710 | 8012 |
| 9796 | 0.05 | 0.08 | 0.65 | 0.52 | 1 | Pax6 | paired box 6 | 5099 | 6240 | 6288 | 9922 | 10174 | 13459 | 10736 | 11005 | 10462 | 10705 | 13662 |
| 9020 | 0.00 | 0.07 | 0.01 | 0.99 | 1 | Ubr3 | ubiquitin protein ligase E3 component n-r | 5986 | 5890 | 6440 | 10690 | 9952 | 9187 | 10586 | 9499 | 11157 | 10504 | 9329 |
| 9548 | -0.11 | 0.16 | -0.69 | 0.49 | 1 | Il6st | interleukin 6 signal transducer | 17919 | 14058 | 14061 | 5256 | 9582 | 5015 | 10570 | 4022 | 10351 | 9387 | 4803 |
| 7486 | -0.13 | 0.15 | -0.92 | 0.36 | 1 | Lrrn1 | leucine rich repeat protein 1, neuronal | 3739 | 4595 | 3987 | 9322 | 9398 | 7284 | 10538 | 5999 | 9975 | 10512 | 6993 |
| 9913 | -0.03 | 0.09 | -0.32 | 0.75 | 1 | Aff4 | AF4/FMR2 family, member 4 | 6897 | 8318 | 6577 | 11427 | 10623 | 11520 | 10518 | 9932 | 11864 | 10257 | 11108 |
| 9175 | -0.09 | 0.11 | -0.84 | 0.40 | 1 | Sec16a | SEC16 homolog A, endoplasmic reticulur | 5798 | 6403 | 6325 | 9383 | 11597 | 10860 | 10502 | 7357 | 12772 | 10003 | 9924 |
| 9023 | 0.05 | 0.08 | 0.63 | 0.53 | 1 | Sel1l | sel-1 suppressor of lin-12-like (C. elegans | 6832 | 8335 | 6969 | 9688 | 9911 | 7744 | 10499 | 9747 | 10687 | 10958 | 7888 |
| 12839 | 0.00 | 0.05 | 0.07 | 0.95 | 1 | Eif4b | eukaryotic translation initiation factor 4B | 20061 | 21040 | 19314 | 10427 | 10361 | 9081 | 10490 | 10896 | 10473 | 9959 | 9129 |
| 7911 | 0.06 | 0.07 | 0.76 | 0.45 | 1 | Atp2c1 | ATPase, Ca++-sequestering | 3908 | 3823 | 4494 | 9227 | 9132 | 7872 | 10456 | 9962 | 9940 | 9954 | 8256 |
| 12158 | 0.14 | 0.12 | 1.17 | 0.24 | 1 | Anxa5 | annexin A5 | 15369 | 14762 | 15938 | 11928 | 9477 | 9609 | 10451 | 15819 | 8961 | 10820 | 10599 |
| 9294 | 0.06 | 0.07 | 0.92 | 0.36 | 1 | Gabarap | gamma-aminobutyric acid receptor assoc | 8121 | 7918 | 8708 | 10010 | 10055 | 7264 | 10445 | 11495 | 9890 | 11144 | 7183 |
| 10212 | -0.04 | 0.08 | -0.52 | 0.60 | 1 | Vegfa | vascular endothelial growth factor A | 4224 | 4781 | 5091 | 10248 | 10270 | 18941 | 10438 | 9294 | 10251 | 10878 | 17921 |
| 8922 | 0.00 | 0.05 | 0.04 | 0.97 | 1 | Amfr | autocrine motility factor receptor | 7003 | 7030 | 7411 | 10412 | 10216 | 7263 | 10435 | 10189 | 9932 | 10927 | 7324 |

| | | | | | | | |
|---|---|---|---|---|---|---|---|
| 15066 | -0.10 | 0.07 | -1.43 | 0.15 | 1 | Sptbn1 | spectrin beta, non-erythrocytic 1 |
| 7958 | -0.06 | 0.09 | -0.61 | 0.54 | 1 | Ppm1l | protein phosphatase 1 (formerly 2C)-like |
| 12864 | -0.06 | 0.07 | -0.92 | 0.36 | 1 | Itgb1 | integrin beta 1 (fibronectin receptor beta) |
| 9341 | 0.03 | 0.06 | 0.44 | 0.66 | 1 | Tmed7 | transmembrane p24 trafficking protein 7 |
| 9340 | -0.04 | 0.06 | -0.68 | 0.50 | 1 | Ccni | cyclin I |
| 9768 | 0.10 | 0.07 | 1.39 | 0.16 | 1 | Tnks2 | tankyrase, TRF1-interacting ankyrin-relate |
| 8516 | 0.05 | 0.10 | 0.49 | 0.62 | 1 | Golph3 | golgi phosphoprotein 3 |
| 11243 | 0.09 | 0.08 | 1.25 | 0.21 | 1 | Rps2 | ribosomal protein S2 |
| 9660 | 0.01 | 0.06 | 0.25 | 0.81 | 1 | Eif5a | eukaryotic translation initiation factor 5A |
| 8480 | 0.07 | 0.09 | 0.81 | 0.42 | 1 | Gaa | glucosidase, alpha, acid |
| 8929 | 0.07 | 0.07 | 0.96 | 0.34 | 1 | Ap3d1 | adaptor-related protein complex 3, delta 1 |
| 7705 | -0.01 | 0.11 | -0.05 | 0.96 | 1 | Ppp1r1a | protein phosphatase 1, regulatory inhibito |
| 9815 | -0.05 | 0.05 | -0.96 | 0.34 | 1 | Trip12 | thyroid hormone receptor interactor 12 |
| 9439 | 0.16 | 0.08 | 1.87 | 0.06 | 1 | Nrd1 | nardilysin, N-arginine dibasic convertase, |
| 10578 | 0.08 | 0.06 | 1.36 | 0.17 | 1 | Csde1 | cold shock domain containing E1, RNA bi |
| 10254 | 0.03 | 0.05 | 0.58 | 0.56 | 1 | Kpnb1 | karyopherin (importin) beta 1 |
| 11960 | -0.08 | 0.09 | -0.86 | 0.39 | 1 | Rps14 | ribosomal protein S14 |
| 8548 | 0.02 | 0.05 | 0.51 | 0.61 | 1 | Fam20b | family with sequence similarity 20, memb |
| 10592 | -0.12 | 0.10 | -1.19 | 0.23 | 1 | Slc7a8 | solute carrier family 7 (cationic amino aci |
| 7829 | 0.02 | 0.09 | 0.22 | 0.83 | 1 | Yipf6 | Yip1 domain family, member 6 |
| 9049 | -0.01 | 0.06 | -0.27 | 0.79 | 1 | Tbl1x | transducin (beta)-like 1 X-linked |
| 8287 | -0.10 | 0.12 | -0.88 | 0.38 | 1 | Ern1 | endoplasmic reticulum (ER) to nucleus si |
| 8458 | 0.11 | 0.08 | 1.34 | 0.18 | 1 | Tmem248 | transmembrane protein 248 |
| 8670 | 0.05 | 0.07 | 0.78 | 0.44 | 1 | Sec63 | SEC63-like (S. cerevisiae) |
| 8707 | -0.04 | 0.10 | -0.36 | 0.72 | 1 | Epb41l3 | erythrocyte membrane protein band 4.1 li |
| 9398 | 0.00 | 0.05 | -0.07 | 0.94 | 1 | Fam120a | family with sequence similarity 120, meml |
| 7293 | 0.02 | 0.06 | 0.34 | 0.73 | 1 | Arl8b | ADP-ribosylation factor-like 8B |
| 7846 | -0.12 | 0.08 | -1.62 | 0.10 | 1 | Sh3pxd2a | SH3 and PX domains 2A |
| 8298 | 0.00 | 0.09 | 0.01 | 0.99 | 1 | Utrn | utrophin |
| 7948 | -0.04 | 0.10 | -0.35 | 0.73 | 1 | Tmed4 | transmembrane p24 trafficking protein 4 |
| 9300 | 0.06 | 0.07 | 0.88 | 0.38 | 1 | Ubl3 | ubiquitin-like 3 |
| 8591 | 0.02 | 0.11 | 0.19 | 0.85 | 1 | Prepl | prolyl endopeptidase-like |
| 8262 | -0.02 | 0.09 | -0.19 | 0.85 | 1 | Zfp106 | zinc finger protein 106 |
| 8317 | 0.11 | 0.09 | 1.31 | 0.19 | 1 | Smim14 | small integral membrane protein 14 |
| 10129 | -0.03 | 0.06 | -0.62 | 0.53 | 1 | Cdc42 | cell division cycle 42 |
| 8908 | 0.02 | 0.09 | 0.22 | 0.82 | 1 | Arf4 | ADP-ribosylation factor 4 |
| 7803 | -0.07 | 0.14 | -0.49 | 0.63 | 1 | Pclo | piccolo (presynaptic cytomatrix protein) |
| 9081 | 0.16 | 0.15 | 1.13 | 0.26 | 1 | Tnr | tenascin R |
| 6838 | 0.20 | 0.12 | 1.66 | 0.10 | 1 | Atf5 | activating transcription factor 5 |
| 9418 | 0.07 | 0.09 | 0.82 | 0.41 | 1 | Vcp | valosin containing protein |
| 8031 | -0.03 | 0.06 | -0.43 | 0.67 | 1 | Pi4ka | phosphatidylinositol 4-kinase alpha |
| 8151 | -0.03 | 0.11 | -0.28 | 0.78 | 1 | Hspa13 | heat shock protein 70 family, member 13 |
| 9552 | 0.06 | 0.06 | 0.95 | 0.34 | 1 | Ncor1 | nuclear receptor co-repressor 1 |
| 8128 | -0.01 | 0.09 | -0.10 | 0.92 | 1 | Akap11 | A kinase (PRKA) anchor protein 11 |
| 9172 | -0.06 | 0.10 | -0.57 | 0.57 | 1 | Ube2d3 | ubiquitin-conjugating enzyme E2D 3 |
| 9745 | -0.02 | 0.09 | -0.22 | 0.83 | 1 | Clstn1 | calsyntenin 1 |
| 8180 | -0.01 | 0.07 | -0.13 | 0.89 | 1 | Myo6 | myosin VI |
| 8233 | 0.10 | 0.08 | 1.24 | 0.22 | 1 | Mdh1 | malate dehydrogenase 1, NAD (soluble) |
| 10730 | 0.00 | 0.08 | 0.04 | 0.97 | 1 | Spint2 | serine protease inhibitor, Kunitz type 2 |
| 7818 | -0.05 | 0.09 | -0.59 | 0.56 | 1 | Mia2 | MIA SH3 domain ER export factor 2 |
| 8540 | 0.02 | 0.07 | 0.26 | 0.80 | 1 | Tspan3 | tetraspanin 3 |
| 7014 | 0.04 | 0.08 | 0.50 | 0.61 | 1 | Entpd3 | ectonucleoside triphosphate diphosphoh |
| 9462 | -0.12 | 0.08 | -1.44 | 0.15 | 1 | Prrc2a | proline-rich coiled-coil 2A |
| 8934 | 0.01 | 0.10 | 0.12 | 0.91 | 1 | Fkbp4 | FK506 binding protein 4 |
| 6911 | -0.03 | 0.08 | -0.39 | 0.70 | 1 | Nbas | neuroblastoma amplified sequence |
| 5513 | -0.04 | 0.17 | -0.25 | 0.80 | 1 | Spock1 | sparc/osteonectin, cwcv and kazal-like do |
| 8166 | -0.10 | 0.09 | -1.09 | 0.28 | 1 | Gbf1 | golgi-specific brefeldin A-resistance facto |
| 8894 | 0.07 | 0.09 | 0.78 | 0.43 | 1 | Sstr3 | somatostatin receptor 3 |
| 6863 | 0.03 | 0.10 | 0.32 | 0.75 | 1 | Pdx1 | pancreatic and duodenal homeobox 1 |
| 7814 | -0.07 | 0.10 | -0.69 | 0.49 | 1 | Lrba | LPS-responsive beige-like anchor |

| | | | | | | | | | | | |
|---|---|---|---|---|---|---|---|---|---|---|---|
| 15066 | 31542 | 29083 | 28823 | 9773 | 9138 | 10133 | 10426 | 8960 | 9695 | 8451 | 9700 |
| 7958 | 5136 | 5977 | 6540 | 9556 | 9103 | 6579 | 10359 | 8463 | 9620 | 9885 | 6317 |
| 12864 | 24235 | 20782 | 20496 | 10352 | 9600 | 8582 | 10352 | 9168 | 9677 | 9291 | 8968 |
| 9341 | 7800 | 8067 | 8375 | 10233 | 10249 | 8087 | 10346 | 9461 | 10918 | 10596 | 8615 |
| 9340 | 10635 | 10744 | 10738 | 9660 | 9609 | 6203 | 10342 | 8583 | 10225 | 9999 | 6001 |
| 9768 | 8716 | 10591 | 8958 | 9735 | 9431 | 8645 | 10327 | 11601 | 10627 | 9765 | 9053 |
| 8516 | 5448 | 6909 | 5776 | 9160 | 9983 | 7728 | 10274 | 9134 | 10355 | 10956 | 7957 |
| 11243 | 19696 | 17009 | 14389 | 10165 | 7686 | 6713 | 10268 | 11624 | 7389 | 11951 | 6787 |
| 9660 | 10724 | 8340 | 10321 | 9484 | 10527 | 7957 | 10256 | 9619 | 10402 | 10694 | 7937 |
| 8480 | 6066 | 5937 | 6745 | 10269 | 9546 | 6024 | 10243 | 11099 | 9417 | 12152 | 5788 |
| 8929 | 5956 | 5925 | 5161 | 9564 | 9746 | 10092 | 10233 | 10989 | 10572 | 10463 | 9518 |
| 7705 | 705 | 814 | 1240 | 10522 | 11010 | 9276 | 10189 | 9829 | 11749 | 10419 | 9003 |
| 9815 | 9416 | 9893 | 9955 | 10326 | 10213 | 9340 | 10171 | 9273 | 10129 | 10088 | 9159 |
| 9439 | 6892 | 6502 | 6716 | 9798 | 9896 | 9630 | 10168 | 13240 | 10502 | 10388 | 10098 |
| 10578 | 9688 | 10118 | 9824 | 10860 | 10975 | 10181 | 10152 | 10791 | 11971 | 10310 | 11486 |
| 10254 | 12006 | 11585 | 11838 | 10064 | 10067 | 8012 | 10134 | 9718 | 11003 | 10093 | 8272 |
| 11960 | 20556 | 19840 | 23054 | 9657 | 9265 | 5890 | 10129 | 10121 | 7721 | 9825 | 5497 |
| 8548 | 6457 | 6980 | 6620 | 10294 | 9496 | 6816 | 10094 | 10386 | 9887 | 9970 | 7030 |
| 10592 | 12258 | 13582 | 13590 | 10971 | 11664 | 7304 | 10049 | 7965 | 12147 | 10346 | 6637 |
| 7829 | 4555 | 5166 | 5510 | 9226 | 8626 | 7360 | 10032 | 9427 | 9441 | 8432 | 8344 |
| 9049 | 6612 | 7561 | 7365 | 10248 | 10635 | 8258 | 10012 | 10289 | 11161 | 9332 | 8068 |
| 8287 | 5605 | 7903 | 7471 | 9433 | 9236 | 7556 | 9994 | 7860 | 10069 | 9377 | 6657 |
| 8458 | 3786 | 3928 | 4340 | 8741 | 11769 | 8444 | 9952 | 10054 | 11977 | 11961 | 8082 |
| 8670 | 5521 | 5598 | 6158 | 8815 | 10458 | 9174 | 9918 | 10360 | 10427 | 9229 | 9711 |
| 8707 | 3418 | 4612 | 3833 | 11294 | 10400 | 10954 | 9903 | 9361 | 10364 | 11174 | 10469 |
| 9398 | 9181 | 9588 | 8757 | 9573 | 9386 | 9116 | 9886 | 9758 | 9762 | 9712 | 8661 |
| 7293 | 3851 | 4009 | 3856 | 8707 | 8611 | 6807 | 9865 | 8968 | 9245 | 9569 | 6734 |
| 7846 | 3343 | 2633 | 3670 | 8248 | 12234 | 9397 | 9854 | 6873 | 12378 | 9673 | 8000 |
| 8298 | 7065 | 7196 | 6479 | 9045 | 7233 | 9187 | 9852 | 7590 | 8796 | 9516 | 9318 |
| 7948 | 5674 | 6667 | 7308 | 8619 | 9111 | 6725 | 9840 | 7621 | 8939 | 10262 | 6658 |
| 9300 | 9008 | 9574 | 9606 | 9786 | 9490 | 7052 | 9833 | 11668 | 9115 | 9917 | 7253 |
| 8591 | 4290 | 5770 | 5519 | 8496 | 10669 | 10097 | 9810 | 8292 | 11728 | 9077 | 10748 |
| 8262 | 5666 | 5451 | 4819 | 8974 | 10080 | 8831 | 9807 | 8536 | 10528 | 9078 | 9113 |
| 8317 | 4906 | 5786 | 5792 | 8811 | 8935 | 8469 | 9795 | 10211 | 9317 | 10806 | 8663 |
| 10129 | 13904 | 12330 | 12946 | 9627 | 9966 | 7216 | 9789 | 9575 | 9257 | 9454 | 7354 |
| 8908 | 9287 | 8863 | 8220 | 8128 | 9998 | 7624 | 9777 | 7408 | 10123 | 10396 | 8160 |
| 7803 | 1024 | 1660 | 1794 | 8954 | 9064 | 13867 | 9769 | 8270 | 9997 | 9007 | 12426 |
| 9081 | 7867 | 12207 | 9222 | 7727 | 6358 | 9498 | 9763 | 7862 | 8146 | 10253 | 10987 |
| 6838 | 2103 | 1895 | 2670 | 8538 | 8720 | 4472 | 9754 | 11969 | 9582 | 11067 | 4446 |
| 9418 | 8844 | 7936 | 8922 | 8467 | 11447 | 8363 | 9750 | 11038 | 10986 | 9148 | 8701 |
| 8031 | 5746 | 5795 | 6528 | 9338 | 9304 | 7020 | 9745 | 8745 | 9796 | 9603 | 6724 |
| 8151 | 3844 | 3550 | 4422 | 7941 | 12662 | 8826 | 9723 | 6467 | 12521 | 9360 | 10339 |
| 9552 | 7167 | 8252 | 8530 | 9954 | 9864 | 10094 | 9715 | 11269 | 11049 | 9412 | 9761 |
| 8128 | 5811 | 7026 | 6067 | 8998 | 8563 | 8105 | 9695 | 8855 | 9185 | 7874 | 9230 |
| 9172 | 8817 | 9726 | 9598 | 9045 | 9201 | 9066 | 9692 | 6978 | 9843 | 9883 | 9040 |
| 9745 | 8697 | 8147 | 9161 | 11431 | 11990 | 7595 | 9690 | 12799 | 11464 | 9208 | 7014 |
| 8180 | 5099 | 5873 | 6090 | 10003 | 9105 | 7837 | 9673 | 9250 | 9288 | 9737 | 8032 |
| 8233 | 5709 | 5061 | 5937 | 9674 | 9195 | 7224 | 9668 | 10384 | 9110 | 10280 | 8318 |
| 10730 | 13030 | 14440 | 16858 | 9656 | 10022 | 7433 | 9635 | 10618 | 9714 | 9485 | 7131 |
| 7818 | 4757 | 4463 | 5319 | 9071 | 9295 | 8302 | 9590 | 7072 | 9698 | 9833 | 8600 |
| 8540 | 8027 | 7816 | 7892 | 9182 | 8933 | 7075 | 9589 | 8339 | 9239 | 10910 | 6935 |
| 7014 | 1664 | 1748 | 2060 | 8584 | 9013 | 8154 | 9584 | 8609 | 9173 | 10051 | 8510 |
| 9462 | 10659 | 9752 | 9939 | 9695 | 9756 | 9241 | 9583 | 7743 | 10443 | 9229 | 8043 |
| 8934 | 8791 | 6679 | 9862 | 8598 | 9625 | 8590 | 9575 | 9582 | 8524 | 9636 | 8814 |
| 6911 | 2695 | 3129 | 3080 | 8783 | 8520 | 7121 | 9557 | 7941 | 8515 | 9269 | 7409 |
| 5513 | 1480 | 2274 | 1929 | 6799 | 8535 | 3070 | 9540 | 5666 | 8798 | 9418 | 3131 |
| 8166 | 5153 | 5452 | 5853 | 8496 | 11167 | 8624 | 9535 | 7148 | 11003 | 9539 | 7851 |
| 8894 | 3932 | 6069 | 4784 | 10220 | 10006 | 10763 | 9528 | 11041 | 10466 | 10084 | 10937 |
| 6863 | 868 | 1365 | 1286 | 10502 | 9122 | 6463 | 9520 | 10186 | 9827 | 9673 | 6684 |
| 7814 | 6016 | 7681 | 6545 | 8284 | 8951 | 6907 | 9496 | 7519 | 9481 | 8568 | 6512 |

| | | | | | | | | | | | | | | | | | |
|---|---|---|---|---|---|---|---|---|---|---|---|---|---|---|---|---|---|
| 8492 | -0.03 | 0.08 | -0.45 | 0.65 | 1 | Ddr1 | discoidin domain receptor family, membe | 6754 | 8674 | 7261 | 10160 | 9566 | 6552 | 9492 | 9443 | 9300 | 9928 | 6286 |
| 8300 | -0.01 | 0.07 | -0.21 | 0.84 | 1 | Gorasp2 | golgi reassembly stacking protein 2 | 5545 | 5191 | 6060 | 8391 | 11100 | 8386 | 9485 | 7651 | 11314 | 9936 | 8241 |
| 9038 | -0.06 | 0.05 | -1.19 | 0.23 | 1 | Numa1 | nuclear mitotic apparatus protein 1 | 8181 | 8537 | 8506 | 9282 | 9874 | 9235 | 9483 | 9014 | 9893 | 9000 | 8419 |
| 7687 | -0.15 | 0.08 | -1.80 | 0.07 | 1 | Slc43a2 | solute carrier family 43, member 2 | 4921 | 4771 | 5957 | 9138 | 9823 | 7699 | 9468 | 9432 | 9161 | 7691 | 6493 |
| 8258 | 0.06 | 0.06 | 1.05 | 0.29 | 1 | Jak1 | Janus kinase 1 | 7316 | 6258 | 6862 | 8594 | 8826 | 7537 | 9463 | 9592 | 8673 | 9838 | 7876 |
| 9352 | 0.01 | 0.09 | 0.13 | 0.90 | 1 | Rad21 | RAD21 cohesin complex component | 10195 | 10399 | 9700 | 9156 | 9255 | 8277 | 9427 | 7649 | 10022 | 9903 | 8886 |
| 6272 | -0.28 | 0.15 | -1.79 | 0.07 | 1 | Pla2g2f | phospholipase A2, group IIF | 3477 | 4660 | 5372 | 7623 | 7633 | 5652 | 9401 | 6482 | 5751 | 8518 | 4422 |
| 7635 | 0.03 | 0.07 | 0.37 | 0.71 | 1 | Ghitm | growth hormone inducible transmembran | 5614 | 5121 | 5675 | 8952 | 8541 | 6640 | 9396 | 7999 | 8807 | 9843 | 7394 |
| 8925 | 0.08 | 0.06 | 1.44 | 0.15 | 1 | Dstn | destrin | 10504 | 9149 | 9128 | 8991 | 8507 | 6884 | 9391 | 9655 | 8967 | 9492 | 7503 |
| 9920 | 0.09 | 0.08 | 1.13 | 0.26 | 1 | Caprin1 | cell cycle associated protein 1 | 10823 | 11494 | 9529 | 9127 | 9007 | 9896 | 9368 | 8686 | 10101 | 10600 | 10490 |
| 8375 | -0.04 | 0.06 | -0.62 | 0.54 | 1 | G3bp2 | GTPase activating protein (SH3 domain) I | 7138 | 8158 | 7496 | 8994 | 9047 | 7777 | 9354 | 8571 | 9079 | 8486 | 8022 |
| 9200 | -0.01 | 0.08 | -0.09 | 0.93 | 1 | Pdia4 | protein disulfide isomerase associated 4 | 9234 | 6779 | 8216 | 8364 | 11581 | 9299 | 9320 | 7972 | 10744 | 10154 | 9532 |
| 8863 | 0.05 | 0.06 | 0.92 | 0.36 | 1 | Pdxdc1 | pyridoxal-dependent decarboxylase dom | 6391 | 6128 | 6753 | 9564 | 10251 | 9297 | 9288 | 9641 | 9888 | 10512 | 9781 |
| 8392 | 0.03 | 0.08 | 0.39 | 0.69 | 1 | Gabarapl1 | gamma-aminobutyric acid (GABA) A rece | 8371 | 8374 | 8973 | 8833 | 8317 | 6395 | 9287 | 9813 | 8081 | 9747 | 6120 |
| 9911 | -0.10 | 0.09 | -1.05 | 0.29 | 1 | Macf1 | microtubule-actin crosslinking factor 1 | 8026 | 8054 | 8376 | 9902 | 10369 | 14094 | 9244 | 7869 | 11776 | 8584 | 12733 |
| 11424 | 0.17 | 0.12 | 1.42 | 0.16 | 1 | Hsp90aa1 | heat shock protein 90, alpha (cytosolic), c | 9288 | 8806 | 9885 | 10368 | 10147 | 16173 | 9211 | 15059 | 10424 | 8843 | 17464 |
| 8669 | 0.05 | 0.06 | 0.73 | 0.46 | 1 | Ubr5 | ubiquitin protein ligase E3 component n-r | 7248 | 7804 | 7022 | 8545 | 9160 | 9162 | 9202 | 8980 | 9506 | 9243 | 9486 |
| 11121 | 0.10 | 0.08 | 1.26 | 0.21 | 1 | Rps4x | ribosomal protein S4, X-linked | 22059 | 19102 | 19451 | 7752 | 7785 | 5253 | 9178 | 9274 | 7588 | 8706 | 6183 |
| 6681 | 0.07 | 0.10 | 0.69 | 0.49 | 1 | Pcx | pyruvate carboxylase | 1168 | 1048 | 1402 | 9557 | 9103 | 6154 | 9165 | 10547 | 8916 | 10359 | 6069 |
| 7107 | 0.01 | 0.08 | 0.14 | 0.89 | 1 | Rnf11 | ring finger protein 11 | 5565 | 6397 | 5992 | 8210 | 7471 | 5314 | 9127 | 7887 | 7648 | 8671 | 5897 |
| 8367 | 0.09 | 0.06 | 1.39 | 0.16 | 1 | Atp2b1 | ATPase, Ca++ transporting, plasma mem | 5770 | 6492 | 6377 | 9085 | 8440 | 8918 | 9116 | 10344 | 9381 | 9010 | 9103 |
| 6715 | 0.03 | 0.10 | 0.29 | 0.77 | 1 | 9330182L06Rik | RIKEN cDNA 9330182L06 gene | 2641 | 3250 | 2816 | 8363 | 8271 | 6584 | 9116 | 7582 | 8904 | 8950 | 7385 |
| 9062 | -0.04 | 0.10 | -0.37 | 0.71 | 1 | Txndc5 | thioredoxin domain containing 5 | 13849 | 8811 | 9129 | 8399 | 9450 | 7406 | 9113 | 8181 | 8753 | 9306 | 7285 |
| 7913 | 0.08 | 0.06 | 1.33 | 0.18 | 1 | Irak1 | interleukin-1 receptor-associated kinase 1 | 4876 | 5342 | 5115 | 9383 | 8603 | 7666 | 9105 | 10589 | 9145 | 9748 | 7469 |
| 8225 | -0.02 | 0.07 | -0.25 | 0.80 | 1 | Trim44 | tripartite motif-containing 44 | 7289 | 8176 | 7980 | 8334 | 8644 | 7651 | 9104 | 7725 | 9103 | 8563 | 7904 |
| 8297 | -0.03 | 0.06 | -0.42 | 0.67 | 1 | Herc2 | HECT and RLD domain containing E3 ub | 5298 | 4570 | 5982 | 9092 | 9969 | 9908 | 9039 | 8439 | 10289 | 9313 | 9372 |
| 10482 | 0.10 | 0.12 | 0.84 | 0.40 | 1 | Ptms | parathymosin | 13075 | 12140 | 13524 | 10045 | 10804 | 6684 | 9031 | 14263 | 9107 | 10222 | 6408 |
| 7161 | 0.02 | 0.07 | 0.22 | 0.83 | 1 | Enpp5 | ectonucleotide pyrophosphatase/phosph | 3562 | 3461 | 3453 | 9197 | 8519 | 7264 | 9015 | 8077 | 9343 | 9130 | 7751 |
| 7882 | 0.12 | 0.08 | 1.44 | 0.15 | 1 | Degs1 | delta(4)-desaturase, sphingolipid 1 | 4255 | 4076 | 3905 | 11714 | 8444 | 6572 | 8955 | 12362 | 9023 | 10191 | 7202 |
| 9100 | -0.07 | 0.07 | -0.92 | 0.36 | 1 | Papola | poly (A) polymerase alpha | 7886 | 8462 | 8952 | 9231 | 10556 | 9482 | 8949 | 7772 | 10253 | 9099 | 9457 |
| 7810 | 0.12 | 0.07 | 1.72 | 0.09 | 1 | Tpd52 | tumor protein D52 | 4701 | 5422 | 5427 | 9200 | 8661 | 6923 | 8934 | 9622 | 9064 | 10128 | 7833 |
| 8665 | -0.04 | 0.07 | -0.61 | 0.54 | 1 | Gdi1 | guanosine diphosphate (GDP) dissociatio | 8172 | 8743 | 9277 | 8291 | 8772 | 9039 | 8929 | 8930 | 8444 | 7641 | 9078 |
| 11024 | 0.11 | 0.08 | 1.48 | 0.14 | 1 | Rpl7 | ribosomal protein L7 | 20726 | 18714 | 18602 | 7798 | 8054 | 5657 | 8896 | 9897 | 7906 | 8768 | 6250 |
| 8123 | 0.00 | 0.06 | -0.01 | 0.99 | 1 | 9530068E07Rik | RIKEN cDNA 9530068E07 gene | 8013 | 7809 | 8626 | 8291 | 8745 | 6482 | 8896 | 8488 | 8617 | 9183 | 6206 |
| 6929 | 0.00 | 0.09 | -0.03 | 0.98 | 1 | Gosr2 | golgi SNAP receptor complex member 2 | 4526 | 4187 | 4230 | 6631 | 9023 | 7165 | 8857 | 6367 | 8631 | 8837 | 7759 |
| 5565 | 0.06 | 0.11 | 0.50 | 0.61 | 1 | Ckb | creatine kinase, brain | 2535 | 1856 | 1931 | 7956 | 5408 | 4582 | 8856 | 8631 | 5454 | 9463 | 4547 |
| 9077 | -0.12 | 0.11 | -1.10 | 0.27 | 1 | Hap1 | huntingtin-associated protein 1 | 5503 | 7913 | 8479 | 8915 | 8612 | 14270 | 8844 | 9088 | 7693 | 7887 | 12645 |
| 8420 | 0.09 | 0.06 | 1.40 | 0.16 | 1 | Hspa4 | heat shock protein 4 | 8098 | 8125 | 7996 | 7895 | 8705 | 7717 | 8844 | 8118 | 9014 | 9080 | 9032 |
| 11086 | 0.08 | 0.12 | 0.62 | 0.54 | 1 | Rps9 | ribosomal protein S9 | 20578 | 17164 | 19982 | 8247 | 8255 | 5786 | 8826 | 11566 | 7494 | 8217 | 5836 |
| 10587 | -0.13 | 0.10 | -1.26 | 0.21 | 1 | Rimbp2 | RIMS binding protein 2 | 4532 | 7630 | 5825 | 11109 | 10515 | 21168 | 8821 | 8934 | 10761 | 8233 | 18931 |
| 9224 | -0.04 | 0.07 | -0.61 | 0.54 | 1 | Cds2 | CDP-diacylglycerol synthase (phosphatid | 10794 | 12149 | 11475 | 9105 | 8746 | 7212 | 8798 | 10035 | 8820 | 7848 | 6486 |
| 7086 | -0.01 | 0.08 | -0.12 | 0.90 | 1 | Pde3b | phosphodiesterase 3B, cGMP-inhibited | 4408 | 5399 | 4865 | 8142 | 7779 | 7071 | 8786 | 7645 | 8368 | 8114 | 7370 |
| 8066 | 0.10 | 0.08 | 1.32 | 0.19 | 1 | Hook1 | hook microtubule tethering protein 1 | 3383 | 4146 | 4196 | 8587 | 8918 | 10787 | 8747 | 8797 | 9599 | 8939 | 12621 |
| 6949 | 0.04 | 0.07 | 0.58 | 0.56 | 1 | Clptm1l | CLPTM1-like | 4597 | 3445 | 4213 | 8015 | 8127 | 6761 | 8745 | 7763 | 8385 | 9244 | 7143 |
| 6658 | 0.03 | 0.07 | 0.48 | 0.63 | 1 | Edem2 | ER degradation enhancer, mannosidase | 3891 | 4194 | 3435 | 8748 | 7022 | 5921 | 8728 | 8855 | 7080 | 9650 | 5712 |
| 8598 | 0.08 | 0.07 | 1.15 | 0.25 | 1 | Eif3c | eukaryotic translation initiation factor 3, su | 9187 | 8282 | 8643 | 8594 | 8996 | 7012 | 8709 | 9973 | 8723 | 9042 | 7420 |
| 6619 | 0.05 | 0.12 | 0.37 | 0.71 | 1 | Efcab1 | EF-hand calcium binding domain 1 | 3162 | 4195 | 3849 | 8236 | 7223 | 6242 | 8709 | 6888 | 7968 | 8995 | 7342 |
| 6111 | -0.07 | 0.11 | -0.66 | 0.51 | 1 | Casc4 | cancer susceptibility candidate 4 | 2700 | 2937 | 3109 | 7465 | 7874 | 5893 | 8700 | 5821 | 8339 | 8263 | 6123 |
| 7006 | 0.02 | 0.06 | 0.33 | 0.74 | 1 | Peg13 | | 4676 | 4267 | 4470 | 7970 | 7594 | 7314 | 8696 | 7937 | 7714 | 9269 | 7153 |
| 6909 | -0.16 | 0.15 | -1.13 | 0.26 | 1 | Slc24a2 | solute carrier family 24 (sodium/potassiur | 1615 | 2897 | 2401 | 9784 | 8064 | 9990 | 8672 | 6670 | 8802 | 7652 | 9450 |
| 8212 | 0.04 | 0.08 | 0.51 | 0.61 | 1 | Anapc5 | anaphase-promoting complex subunit 5 | 6166 | 4912 | 4969 | 8413 | 8636 | 11015 | 8629 | 9406 | 8661 | 8970 | 10552 |
| 9359 | 0.18 | 0.25 | 0.72 | 0.47 | 1 | Rpl41 | ribosomal protein L41 | 10914 | 10466 | 13080 | 7813 | 8474 | 6697 | 8592 | 14610 | 7624 | 8168 | 6512 |
| 7386 | -0.10 | 0.07 | -1.46 | 0.14 | 1 | Fasn | fatty acid synthase | 6433 | 6455 | 6716 | 8578 | 9124 | 5511 | 8560 | 7275 | 9143 | 8669 | 4787 |
| 9539 | 0.13 | 0.09 | 1.51 | 0.13 | 1 | Rpl10 | ribosomal protein L10 | 13535 | 12254 | 13539 | 8077 | 8122 | 6508 | 8557 | 10500 | 8090 | 8745 | 7007 |
| 7482 | -0.05 | 0.10 | -0.54 | 0.59 | 1 | Fbxl16 | F-box and leucine-rich repeat protein 16 | 3972 | 5241 | 4890 | 8837 | 8735 | 8609 | 8554 | 7688 | 8732 | 9304 | 7739 |
| 7119 | 0.00 | 0.09 | 0.05 | 0.96 | 1 | Prps1 | phosphoribosyl pyrophosphate synthetas | 4505 | 4922 | 5501 | 8074 | 8150 | 6897 | 8554 | 7516 | 7849 | 8849 | 7496 |
| 7569 | -0.01 | 0.13 | -0.06 | 0.95 | 1 | Arhgap5 | Rho GTPase activating protein 5 | 4545 | 6869 | 4809 | 9441 | 8213 | 7544 | 8531 | 7159 | 9158 | 8318 | 8675 |
| 11545 | -0.02 | 0.09 | -0.26 | 0.79 | 1 | Serinc3 | serine incorporator 3 | 25406 | 18331 | 18559 | 9126 | 8179 | 6790 | 8526 | 8700 | 8400 | 8411 | 6571 |
| 8628 | 0.11 | 0.09 | 1.19 | 0.23 | 1 | Serbp1 | serpine1 mRNA binding protein 1 | 11108 | 10106 | 10027 | 7660 | 7570 | 6870 | 8515 | 10041 | 7901 | 7768 | 7344 |
| 6572 | 0.10 | 0.09 | 1.10 | 0.27 | 1 | Sirpa | signal-regulatory protein alpha | 4253 | 2811 | 3008 | 8093 | 7837 | 5508 | 8511 | 8322 | 8425 | 9837 | 5690 |

| | | | | | | | | | | | | | | | | | |
|---|---|---|---|---|---|---|---|---|---|---|---|---|---|---|---|---|---|
| 7774 | 0.12 | 0.07 | 1.84 | 0.07 | 1 | Hexa | hexosaminidase A | 6874 | 6801 | 6852 | 9065 | 8321 | 5159 | 8506 | 10670 | 8885 | 8991 | 5389 |
| 8107 | 0.04 | 0.07 | 0.64 | 0.52 | 1 | Psmd2 | proteasome (prosome, macropain) 26S su | 7550 | 6374 | 7644 | 8245 | 9096 | 7420 | 8497 | 9165 | 9156 | 8877 | 7152 |
| 7112 | -0.12 | 0.10 | -1.26 | 0.21 | 1 | Dsp | desmoplakin | 3432 | 4987 | 5319 | 9097 | 9296 | 6657 | 8485 | 7791 | 9455 | 7864 | 5850 |
| 7617 | 0.05 | 0.06 | 0.91 | 0.36 | 1 | Smarca4 | SWI/SNF related, matrix associated, actin | 6010 | 6142 | 6156 | 8244 | 8314 | 7013 | 8474 | 9369 | 8594 | 8778 | 6693 |
| 7620 | -0.04 | 0.08 | -0.52 | 0.60 | 1 | Fkbp1a | FK506 binding protein 1a | 7236 | 7537 | 8503 | 7896 | 8121 | 6226 | 8459 | 7091 | 7858 | 8548 | 6341 |
| 7753 | 0.00 | 0.06 | -0.04 | 0.97 | 1 | Rnf10 | ring finger protein 10 | 7367 | 6961 | 6757 | 8141 | 8909 | 6570 | 8455 | 8174 | 8380 | 9261 | 6314 |
| 7886 | 0.03 | 0.07 | 0.38 | 0.71 | 1 | Tax1bp1 | Tax1 (human T cell leukemia virus type I) | 7454 | 7273 | 7247 | 7647 | 8348 | 7690 | 8400 | 7120 | 8746 | 8482 | 8341 |
| 7319 | 0.08 | 0.08 | 1.00 | 0.32 | 1 | Fam91a1 | family with sequence similarity 91, membe | 4667 | 5600 | 4716 | 7815 | 8247 | 7430 | 8357 | 8244 | 9019 | 8729 | 7680 |
| 8005 | 0.11 | 0.08 | 1.40 | 0.16 | 1 | St13 | suppression of tumorigenicity 13 | 6785 | 7006 | 7839 | 8094 | 8277 | 7243 | 8347 | 9892 | 8531 | 8428 | 7611 |
| 8108 | -0.05 | 0.07 | -0.68 | 0.49 | 1 | Snrnp200 | small nuclear ribonucleoprotein 200 (U5) | 8854 | 9044 | 8288 | 8064 | 8751 | 6916 | 8343 | 7243 | 8648 | 7918 | 7117 |
| 7181 | 0.08 | 0.08 | 1.08 | 0.28 | 1 | Napa | N-ethylmaleimide sensitive fusion protein | 4268 | 4539 | 4699 | 7896 | 7454 | 8067 | 8342 | 8779 | 7633 | 9278 | 8031 |
| 6561 | -0.08 | 0.10 | -0.82 | 0.41 | 1 | Tspan5 | tetraspanin 5 | 2870 | 4313 | 3704 | 8633 | 8632 | 5933 | 8339 | 7861 | 8144 | 8148 | 5594 |
| 7593 | -0.14 | 0.11 | -1.32 | 0.19 | 1 | Zfp516 | zinc finger protein 516 | 4821 | 5573 | 5370 | 8145 | 9583 | 9377 | 8338 | 6184 | 10117 | 7759 | 8255 |
| 6483 | 0.03 | 0.08 | 0.35 | 0.73 | 1 | Aars | alanyl-tRNA synthetase | 3216 | 3704 | 3278 | 7459 | 8585 | 5806 | 8337 | 8617 | 8537 | 8199 | 5576 |
| 5752 | 0.06 | 0.07 | 0.80 | 0.42 | 1 | Tmem229b | transmembrane protein 229B | 1514 | 1485 | 1617 | 7873 | 7091 | 5439 | 8317 | 7949 | 8206 | 8281 | 5501 |
| 7779 | -0.02 | 0.06 | -0.39 | 0.69 | 1 | Dcaf8 | DDB1 and CUL4 associated factor 8 | 5595 | 6347 | 5968 | 8047 | 8736 | 9024 | 8295 | 8212 | 8377 | 8130 | 8835 |
| 7116 | -0.06 | 0.07 | -0.90 | 0.37 | 1 | Nipal3 | NIPA-like domain containing 3 | 3787 | 4411 | 4450 | 9195 | 8778 | 7144 | 8295 | 8384 | 9401 | 8123 | 6312 |
| 7105 | 0.03 | 0.08 | 0.30 | 0.76 | 1 | Sorbs2 | sorbin and SH3 domain containing 2 | 4549 | 4530 | 4784 | 7165 | 8210 | 8132 | 8274 | 6446 | 9428 | 8372 | 8270 |
| 7101 | 0.13 | 0.09 | 1.45 | 0.15 | 1 | Wsb2 | WD repeat and SOCS box-containing 2 | 3715 | 4169 | 4508 | 8458 | 8876 | 5657 | 8274 | 9994 | 8433 | 10177 | 5853 |
| 7055 | 0.04 | 0.11 | 0.37 | 0.71 | 1 | Kras | Kirsten rat sarcoma viral oncogene homo | 4266 | 6460 | 5104 | 8573 | 7152 | 6553 | 8270 | 7739 | 7748 | 8428 | 7315 |
| 6731 | -0.02 | 0.10 | -0.20 | 0.84 | 1 | Tspyl4 | TSPY-like 4 | 2668 | 3712 | 3639 | 7971 | 8138 | 7848 | 8269 | 7359 | 7997 | 7945 | 8498 |
| 8459 | 0.04 | 0.06 | 0.81 | 0.42 | 1 | BC005537 | cDNA sequence BC005537 | 7718 | 7739 | 7476 | 8436 | 8113 | 9720 | 8248 | 8058 | 8811 | 8840 | 9888 |
| 7575 | -0.05 | 0.10 | -0.50 | 0.62 | 1 | Atf6 | activating transcription factor 6 | 4308 | 4254 | 4290 | 8335 | 10311 | 8831 | 8241 | 7016 | 10768 | 7900 | 9071 |
| 7264 | 0.00 | 0.09 | -0.02 | 0.98 | 1 | Ssr4 | signal sequence receptor, delta | 4573 | 5485 | 6375 | 8784 | 8129 | 6565 | 8234 | 8462 | 8408 | 8657 | 6234 |
| 6822 | -0.04 | 0.06 | -0.58 | 0.56 | 1 | Fam102a | family with sequence similarity 102, meml | 4367 | 5133 | 4708 | 8272 | 8226 | 6001 | 8216 | 8475 | 8045 | 8088 | 5512 |
| 7337 | 0.08 | 0.07 | 1.16 | 0.25 | 1 | Deptor | DEP domain containing MTOR-interacting | 4833 | 5940 | 5516 | 8197 | 7737 | 7039 | 8199 | 9469 | 8369 | 8585 | 6820 |
| 6688 | 0.05 | 0.05 | 0.86 | 0.39 | 1 | Lbh | limb-bud and heart | 4318 | 4205 | 4236 | 8439 | 7802 | 5482 | 8185 | 8777 | 7841 | 8621 | 5662 |
| 6246 | 0.08 | 0.13 | 0.65 | 0.52 | 1 | Gpr158 | G protein-coupled receptor 158 | 378 | 494 | 664 | 10520 | 8207 | 5777 | 8174 | 9248 | 9745 | 9265 | 6236 |
| 6801 | 0.21 | 0.12 | 1.79 | 0.07 | 1 | Bex2 | brain expressed X-linked 2 | 3374 | 4349 | 4766 | 7549 | 7422 | 5608 | 8171 | 10559 | 7840 | 8965 | 6210 |
| 7331 | -0.03 | 0.07 | -0.44 | 0.66 | 1 | Pdcd6ip | programmed cell death 6 interacting prote | 6979 | 6155 | 6591 | 7471 | 8455 | 6688 | 8170 | 6977 | 8907 | 7399 | 6854 |
| 7831 | 0.09 | 0.10 | 0.90 | 0.37 | 1 | H13 | histocompatibility 13 | 5652 | 4995 | 5598 | 7552 | 9941 | 8259 | 8166 | 10155 | 9538 | 8209 | 8074 |
| 6437 | 0.03 | 0.09 | 0.38 | 0.70 | 1 | Gpd2 | glycerol phosphate dehydrogenase 2, mit | 3417 | 4615 | 3511 | 8053 | 5899 | 7230 | 8150 | 8284 | 6813 | 7618 | 7222 |
| 8687 | 0.08 | 0.08 | 0.99 | 0.32 | 1 | Hnrnpab | heterogeneous nuclear ribonucleoprotein | 11053 | 8835 | 11014 | 8070 | 8526 | 6673 | 8149 | 8793 | 8115 | 9390 | 6938 |
| 7457 | 0.03 | 0.08 | 0.38 | 0.70 | 1 | Cand1 | cullin associated and neddylation disasso | 6411 | 7703 | 7327 | 7416 | 7696 | 6709 | 8147 | 7955 | 7675 | 8241 | 6745 |
| 11064 | -0.03 | 0.07 | -0.41 | 0.68 | 1 | Myh9 | myosin, heavy polypeptide 9, non-muscle | 23273 | 19668 | 19424 | 7333 | 8053 | 6389 | 8141 | 7000 | 8387 | 8018 | 6020 |
| 7082 | 0.03 | 0.09 | 0.33 | 0.74 | 1 | Rab6a | RAB6A, member RAS oncogene family | 4215 | 4975 | 4082 | 8413 | 7429 | 8014 | 8137 | 7250 | 7977 | 8594 | 8810 |
| 6958 | -0.04 | 0.09 | -0.45 | 0.65 | 1 | Robo2 | roundabout guidance receptor 2 | 5431 | 4859 | 4485 | 7977 | 7314 | 7894 | 8135 | 6725 | 8340 | 7276 | 8103 |
| 7103 | 0.07 | 0.08 | 0.89 | 0.37 | 1 | Dpp8 | dipeptidylpeptidase 8 | 6457 | 6669 | 6109 | 7215 | 6752 | 6669 | 8128 | 6906 | 7820 | 8466 | 6938 |
| 7058 | 0.04 | 0.05 | 0.79 | 0.43 | 1 | Tmed9 | transmembrane p24 trafficking protein 9 | 4295 | 3263 | 4893 | 6629 | 9356 | 8039 | 8103 | 6813 | 9344 | 8805 | 8100 |
| 7696 | 0.05 | 0.07 | 0.71 | 0.48 | 1 | Stag2 | stromal antigen 2 | 7212 | 7593 | 6880 | 8157 | 8000 | 6784 | 8081 | 7987 | 7847 | 8190 | 7925 |
| 11977 | 0.02 | 0.13 | 0.12 | 0.91 | 1 | H2-K1 | histocompatibility 2, K1, K region | 21029 | 16639 | 14777 | 11760 | 8777 | 10751 | 8074 | 12871 | 8541 | 8163 | 10367 |
| 9952 | 0.00 | 0.09 | -0.01 | 0.99 | 1 | Castor2 | cytosolic arginine sensor for mTORC1 sul | 14210 | 16442 | 16435 | 7586 | 7775 | 7662 | 8061 | 9243 | 7885 | 7633 | 6535 |
| 6267 | 0.18 | 0.11 | 1.66 | 0.10 | 1 | Ogdhl | oxoglutarate dehydrogenase-like | 2550 | 3688 | 3176 | 7156 | 7566 | 5062 | 8057 | 9197 | 8378 | 8698 | 5410 |
| 6391 | 0.10 | 0.11 | 0.86 | 0.39 | 1 | Ffar1 | free fatty acid receptor 1 | 979 | 1372 | 1693 | 8205 | 8661 | 7045 | 8045 | 8716 | 9776 | 8834 | 6972 |
| 6280 | -0.02 | 0.06 | -0.28 | 0.78 | 1 | Atp6v1a | ATPase, H+ transporting, lysosomal V1 su | 3583 | 3176 | 3427 | 7709 | 8068 | 5814 | 8038 | 7416 | 8232 | 7683 | 5929 |
| 6166 | -0.01 | 0.08 | -0.10 | 0.92 | 1 | Pik3c2a | phosphatidylinositol-4-phosphate 3-kinas | 3117 | 3480 | 3073 | 6420 | 7056 | 7649 | 8031 | 6757 | 6846 | 7351 | 8045 |
| 6481 | 0.01 | 0.07 | 0.13 | 0.89 | 1 | Isoc1 | isochorismatase domain containing 1 | 4984 | 5301 | 4978 | 7487 | 6856 | 5639 | 8026 | 7110 | 7061 | 7502 | 6343 |
| 7550 | -0.04 | 0.07 | -0.60 | 0.55 | 1 | Golim4 | golgi integral membrane protein 4 | 5552 | 6492 | 6271 | 8229 | 7797 | 8776 | 8010 | 7210 | 8291 | 7284 | 9144 |
| 6926 | -0.08 | 0.08 | -0.99 | 0.32 | 1 | Cadps | Ca2+-dependent secretion activator | 3674 | 5127 | 4230 | 8485 | 8272 | 7757 | 7969 | 7666 | 8330 | 7205 | 7468 |
| 6229 | 0.04 | 0.07 | 0.50 | 0.62 | 1 | Naga | N-acetyl galactosaminidase, alpha | 4615 | 4070 | 4831 | 6077 | 7851 | 5312 | 7959 | 6114 | 7400 | 8537 | 5757 |
| 7509 | -0.03 | 0.06 | -0.52 | 0.61 | 1 | Ap2b1 | adaptor-related protein complex 2, beta 1 | 8154 | 6895 | 7709 | 7563 | 8812 | 5952 | 7956 | 7539 | 8567 | 7582 | 5875 |
| 7233 | -0.02 | 0.06 | -0.35 | 0.73 | 1 | Igf2r | insulin-like growth factor 2 receptor | 6181 | 5928 | 5904 | 8355 | 8202 | 6416 | 7955 | 8113 | 8796 | 7869 | 5842 |
| 7084 | 0.02 | 0.09 | 0.20 | 0.84 | 1 | Clcn3 | chloride channel, voltage-sensitive 3 | 5295 | 7640 | 5892 | 7839 | 7239 | 6354 | 7944 | 7958 | 7274 | 7764 | 6728 |
| 6745 | 0.01 | 0.07 | 0.11 | 0.91 | 1 | Golga3 | golgi autoantigen, golgin subfamily a, 3 | 4230 | 4268 | 4441 | 6788 | 8023 | 7783 | 7942 | 6823 | 8108 | 8331 | 7454 |
| 7548 | -0.04 | 0.09 | -0.45 | 0.65 | 1 | Dennd4c | DENN/MADD domain containing 4C | 2590 | 2899 | 2805 | 10855 | 8878 | 10290 | 7939 | 8601 | 10107 | 8388 | 9676 |
| 6947 | 0.10 | 0.10 | 0.96 | 0.34 | 1 | Dap | death-associated protein | 4564 | 5964 | 6416 | 7460 | 7839 | 5450 | 7934 | 8621 | 7964 | 8629 | 5573 |
| 6264 | 0.03 | 0.09 | 0.38 | 0.70 | 1 | Tvp23b | trans-golgi network vesicle protein 23B | 3716 | 3954 | 4546 | 7167 | 7057 | 5880 | 7933 | 6722 | 7591 | 7951 | 6392 |
| 6326 | -0.06 | 0.10 | -0.65 | 0.51 | 1 | Wfs1 | wolframin ER transmembrane glycoprotei | 1827 | 1891 | 2913 | 6934 | 9652 | 7597 | 7927 | 6050 | 9220 | 8375 | 7206 |
| 5767 | -0.04 | 0.08 | -0.47 | 0.64 | 1 | Arfgef2 | ADP-ribosylation factor guanine nucleotic | 3076 | 3136 | 3284 | 6722 | 6545 | 6122 | 7918 | 6221 | 7249 | 7325 | 5833 |
| 7661 | 0.11 | 0.07 | 1.70 | 0.09 | 1 | Slc25a4 | solute carrier family 25 (mitochondrial car | 7647 | 7917 | 7889 | 7697 | 7686 | 5906 | 7914 | 8985 | 7556 | 8760 | 6311 |

| | | | | | | | | | | | | | | | | | |
|---|---|---|---|---|---|---|---|---|---|---|---|---|---|---|---|---|---|
| 8117 | -0.06 | 0.08 | -0.85 | 0.39 | 1 | Ctsl | cathepsin L | 12121 | 10732 | 9990 | 8052 | 7600 | 5346 | 7912 | 6643 | 7524 | 7850 | 5516 |
| 5957 | 0.03 | 0.09 | 0.28 | 0.78 | 1 | Ociad2 | OCIA domain containing 2 | 2634 | 3201 | 2958 | 7420 | 6862 | 5984 | 7882 | 6794 | 7336 | 8110 | 6352 |
| 10753 | 0.12 | 0.07 | 1.77 | 0.08 | 1 | Sptan1 | spectrin alpha, non-erythrocytic 1 | 16821 | 15641 | 16337 | 8130 | 7375 | 9899 | 7872 | 9695 | 7910 | 8495 | 10106 |
| 6929 | -0.01 | 0.06 | -0.22 | 0.83 | 1 | Ostc | oligosaccharyltransferase complex subur | 4376 | 4390 | 5477 | 7011 | 9519 | 6793 | 7858 | 7549 | 9011 | 7350 | 6886 |
| 4643 | 0.19 | 0.12 | 1.56 | 0.12 | 1 | Cntnap2 | contactin associated protein-like 2 | 400 | 282 | 585 | 5249 | 6732 | 3354 | 7850 | 5182 | 8654 | 8705 | 4082 |
| 6522 | -0.01 | 0.11 | -0.11 | 0.91 | 1 | Edem3 | ER degradation enhancer, mannosidase | 3318 | 3667 | 3191 | 7128 | 8393 | 7495 | 7832 | 5861 | 8754 | 8285 | 7812 |
| 7439 | 0.03 | 0.06 | 0.55 | 0.58 | 1 | Alkbh5 | alkB homolog 5, RNA demethylase | 6006 | 7032 | 5955 | 8229 | 7971 | 6985 | 7826 | 8630 | 8066 | 8379 | 6751 |
| 8173 | -0.08 | 0.08 | -0.98 | 0.33 | 1 | Arhgdia | Rho GDP dissociation inhibitor (GDI) alph | 11607 | 9505 | 9967 | 7509 | 8953 | 5896 | 7802 | 6578 | 8303 | 8137 | 5650 |
| 8992 | 0.04 | 0.08 | 0.56 | 0.58 | 1 | Calm2 | calmodulin 2 | 12794 | 11694 | 13258 | 8040 | 7921 | 6345 | 7783 | 9066 | 7373 | 8156 | 6479 |
| 7069 | -0.09 | 0.07 | -1.37 | 0.17 | 1 | Sec23b | SEC23 homolog B, COPII coat complex c | 4034 | 3507 | 5048 | 6306 | 10439 | 9000 | 7783 | 5678 | 10048 | 6932 | 8983 |
| 6565 | -0.01 | 0.09 | -0.06 | 0.95 | 1 | Arl1 | ADP-ribosylation factor-like 1 | 5322 | 4954 | 5710 | 6862 | 7734 | 5763 | 7757 | 5663 | 7578 | 8642 | 6232 |
| 6188 | -0.07 | 0.11 | -0.63 | 0.53 | 1 | Abat | 4-aminobutyrate aminotransferase | 2796 | 3037 | 2812 | 8482 | 7345 | 6777 | 7755 | 6292 | 7530 | 8885 | 6363 |
| 6406 | 0.04 | 0.06 | 0.67 | 0.51 | 1 | Atp6v0a1 | ATPase, H+ transporting, lysosomal V0 su | 4250 | 3841 | 3951 | 7793 | 7250 | 6001 | 7736 | 8067 | 7698 | 7991 | 5882 |
| 6454 | 0.06 | 0.10 | 0.55 | 0.58 | 1 | Pank3 | pantothenate kinase 3 | 4004 | 4949 | 3619 | 7430 | 7253 | 6271 | 7719 | 6941 | 8196 | 7710 | 6897 |
| 8279 | -0.05 | 0.08 | -0.58 | 0.56 | 1 | Actn4 | actinin alpha 4 | 12319 | 10284 | 9483 | 7768 | 8019 | 6401 | 7718 | 7877 | 7419 | 7873 | 5910 |
| 6280 | -0.06 | 0.08 | -0.73 | 0.47 | 1 | Braf | Braf transforming gene | 3364 | 3483 | 3777 | 7664 | 7140 | 7286 | 7696 | 6330 | 7731 | 7409 | 7201 |
| 7171 | 0.02 | 0.08 | 0.22 | 0.83 | 1 | Usp34 | ubiquitin specific peptidase 34 | 6041 | 6665 | 6166 | 6835 | 7126 | 8158 | 7691 | 6552 | 7659 | 7494 | 8493 |
| 7200 | -0.05 | 0.07 | -0.63 | 0.53 | 1 | Hectd1 | HECT domain E3 ubiquitin protein ligase | 6402 | 6142 | 6597 | 7760 | 7399 | 7623 | 7688 | 6603 | 8263 | 7120 | 7605 |
| 5279 | -0.06 | 0.09 | -0.70 | 0.48 | 1 | Adra2a | adrenergic receptor, alpha 2a | 690 | 675 | 912 | 7425 | 7852 | 5520 | 7681 | 6781 | 7385 | 7983 | 5171 |
| 6478 | -0.07 | 0.14 | -0.51 | 0.61 | 1 | Rab3gap2 | RAB3 GTPase activating protein subunit 2 | 3227 | 4036 | 3994 | 5999 | 7985 | 8974 | 7675 | 4623 | 7989 | 8174 | 8584 |
| 7417 | 0.08 | 0.07 | 1.20 | 0.23 | 1 | Cpd | carboxypeptidase D | 8475 | 8691 | 8443 | 6877 | 6478 | 6220 | 7659 | 7000 | 7492 | 7372 | 6880 |
| 6407 | 0.04 | 0.07 | 0.52 | 0.60 | 1 | Tollip | toll interacting protein | 4384 | 4968 | 4514 | 6934 | 7221 | 6174 | 7636 | 7456 | 7211 | 7547 | 6428 |
| 4608 | 0.12 | 0.12 | 1.07 | 0.28 | 1 | Adh1 | alcohol dehydrogenase 1 (class I) | 1236 | 1123 | 1636 | 5100 | 6880 | 2798 | 7635 | 5593 | 6788 | 8734 | 3162 |
| 7997 | 0.11 | 0.11 | 0.97 | 0.33 | 1 | Man1a | mannosidase 1, alpha | 10308 | 8391 | 6842 | 9443 | 7555 | 5511 | 7625 | 9552 | 8451 | 8255 | 6036 |
| 6965 | -0.06 | 0.05 | -1.24 | 0.22 | 1 | Ssr2 | signal sequence receptor, beta | 6222 | 5228 | 6923 | 6658 | 8818 | 6660 | 7608 | 5990 | 8435 | 7567 | 6507 |
| 7869 | -0.03 | 0.07 | -0.45 | 0.65 | 1 | Slc3a2 | solute carrier family 3 (activators of dibasi | 8778 | 9447 | 9407 | 7894 | 8911 | 5426 | 7592 | 7052 | 8541 | 8081 | 5434 |
| 7392 | 0.13 | 0.09 | 1.43 | 0.15 | 1 | Eif3b | eukaryotic translation initiation factor 3, su | 7194 | 5972 | 6979 | 6718 | 8495 | 6398 | 7584 | 9066 | 8448 | 8050 | 6407 |
| 6208 | -0.17 | 0.14 | -1.20 | 0.23 | 1 | Nbea | neurobeachin | 4314 | 6363 | 5426 | 6336 | 6391 | 7237 | 7583 | 5007 | 7248 | 5647 | 6741 |
| 6866 | 0.08 | 0.06 | 1.46 | 0.15 | 1 | Ppp2r5c | protein phosphatase 2, regulatory subuni | 5262 | 4924 | 5440 | 7726 | 7668 | 6173 | 7561 | 8108 | 8044 | 7856 | 6766 |
| 9250 | 0.03 | 0.07 | 0.43 | 0.67 | 1 | Trim35 | tripartite motif-containing 35 | 14856 | 16015 | 14765 | 6645 | 6992 | 6607 | 7552 | 6761 | 7049 | 7511 | 7001 |
| 8954 | 0.09 | 0.09 | 1.01 | 0.31 | 1 | Eef1b2 | eukaryotic translation elongation factor 1 | 15585 | 14112 | 15433 | 6968 | 6521 | 4771 | 7550 | 8523 | 6167 | 7705 | 5155 |
| 8083 | 0.05 | 0.05 | 0.98 | 0.33 | 1 | Hp1bp3 | heterochromatin protein 1, binding proteir | 7834 | 8384 | 8423 | 7880 | 8128 | 8021 | 7537 | 8725 | 8044 | 7614 | 8327 |
| 7636 | 0.07 | 0.06 | 1.20 | 0.23 | 1 | Sdcbp | syndecan binding protein | 9404 | 8933 | 8790 | 7684 | 7308 | 5277 | 7520 | 7316 | 7659 | 8003 | 6103 |
| 6078 | 0.11 | 0.12 | 0.88 | 0.38 | 1 | Rims3 | regulating synaptic membrane exocytosis | 474 | 355 | 606 | 7941 | 7424 | 8599 | 7518 | 8416 | 7975 | 8374 | 9178 |
| 6982 | 0.00 | 0.06 | -0.06 | 0.95 | 1 | Specc1l | sperm antigen with calponin homology ar | 5455 | 6147 | 5236 | 8439 | 6978 | 7115 | 7511 | 8084 | 7682 | 7195 | 6965 |
| 5612 | 0.02 | 0.12 | 0.20 | 0.84 | 1 | Gch1 | GTP cyclohydrolase 1 | 2529 | 3224 | 3390 | 6746 | 6164 | 5679 | 7500 | 6364 | 6131 | 7868 | 6138 |
| 6590 | 0.02 | 0.07 | 0.34 | 0.74 | 1 | Arpc5 | actin related protein 2/3 complex, subunit | 6663 | 6328 | 6695 | 6763 | 6960 | 5007 | 7480 | 6135 | 7373 | 7746 | 5344 |
| 7028 | -0.08 | 0.08 | -0.92 | 0.36 | 1 | Herpud1 | homocysteine-inducible, endoplasmic ret | 4959 | 5793 | 5182 | 6981 | 7338 | 9788 | 7477 | 6363 | 7454 | 7357 | 8621 |
| 6545 | 0.04 | 0.06 | 0.71 | 0.48 | 1 | Smap2 | small ArfGAP 2 | 4173 | 4664 | 4828 | 7715 | 7617 | 5983 | 7461 | 7744 | 7899 | 7605 | 6305 |
| 7319 | 0.12 | 0.07 | 1.68 | 0.09 | 1 | Cct8 | chaperonin containing Tcp1, subunit 8 (th | 7828 | 6374 | 8243 | 6448 | 7179 | 6737 | 7459 | 6644 | 7335 | 8262 | 8001 |
| 7523 | -0.01 | 0.07 | -0.16 | 0.87 | 1 | Klhl7 | kelch-like 7 | 6037 | 7768 | 6704 | 8313 | 8375 | 7063 | 7456 | 8331 | 8628 | 6634 | 7440 |
| 7853 | -0.10 | 0.11 | -0.93 | 0.35 | 1 | Pcbp1 | poly(rC) binding protein 1 | 9189 | 9622 | 10035 | 7589 | 7689 | 6914 | 7455 | 5488 | 7672 | 8369 | 6360 |
| 7447 | -0.06 | 0.08 | -0.71 | 0.48 | 1 | Atp9a | ATPase, class II, type 9A | 6198 | 7092 | 7451 | 8784 | 9293 | 5510 | 7447 | 10039 | 8035 | 7240 | 4827 |
| 6665 | 0.08 | 0.08 | 1.09 | 0.27 | 1 | Sacm1l | SAC1 suppressor of actin mutations 1-like | 4120 | 4914 | 4342 | 7201 | 7642 | 6806 | 7447 | 7518 | 8096 | 8213 | 7021 |
| 6989 | -0.19 | 0.12 | -1.61 | 0.11 | 1 | Ubb | ubiquitin B | 6131 | 6031 | 6237 | 7198 | 7413 | 8890 | 7445 | 4586 | 7763 | 7019 | 8168 |
| 6923 | -0.08 | 0.08 | -1.06 | 0.29 | 1 | Rere | arginine glutamic acid dipeptide (RE) rep | 4853 | 6013 | 5615 | 8056 | 7849 | 7291 | 7430 | 7227 | 8306 | 7203 | 6314 |
| 5965 | 0.03 | 0.07 | 0.50 | 0.62 | 1 | Elovl5 | ELOVL family member 5, elongation of lor | 3157 | 3189 | 3092 | 7554 | 6879 | 5926 | 7426 | 6759 | 7504 | 7681 | 6453 |
| 5607 | 0.02 | 0.10 | 0.23 | 0.81 | 1 | Sdf2l1 | stromal cell-derived factor 2-like 1 | 2297 | 1913 | 2138 | 7740 | 6804 | 5435 | 7425 | 8018 | 5980 | 8231 | 5695 |
| 8184 | 0.04 | 0.06 | 0.57 | 0.57 | 1 | Rhoa | ras homolog family member A | 10804 | 9713 | 10051 | 7746 | 8275 | 5931 | 7421 | 7890 | 7904 | 8147 | 6139 |
| 6460 | -0.03 | 0.08 | -0.44 | 0.66 | 1 | 2610507B11Ril | RIKEN cDNA 2610507B11 gene | 5997 | 4473 | 5163 | 6329 | 6848 | 7485 | 7418 | 6534 | 7105 | 6658 | 7053 |
| 6956 | 0.11 | 0.08 | 1.33 | 0.18 | 1 | Larp1 | La ribonucleoprotein domain family, mem | 4803 | 5153 | 4987 | 7600 | 8067 | 6404 | 7403 | 9991 | 8298 | 7488 | 6319 |
| 6802 | -0.12 | 0.10 | -1.16 | 0.25 | 1 | Akap13 | A kinase (PRKA) anchor protein 13 | 3891 | 3501 | 4160 | 8200 | 9477 | 7630 | 7386 | 6969 | 10477 | 6380 | 6750 |
| 7214 | 0.02 | 0.09 | 0.28 | 0.78 | 1 | Hid1 | HID1 domain containing | 3502 | 4256 | 4874 | 6888 | 10392 | 8506 | 7382 | 8216 | 9922 | 7632 | 7783 |
| 6402 | -0.14 | 0.13 | -1.05 | 0.29 | 1 | Sec24d | Sec24 related gene family, member D (S. | 4755 | 3583 | 4397 | 5219 | 9529 | 7848 | 7339 | 3973 | 9738 | 7204 | 6835 |
| 6841 | 0.15 | 0.10 | 1.50 | 0.13 | 1 | Adi1 | acireductone dioxygenase 1 | 3173 | 5458 | 4586 | 8077 | 9414 | 4384 | 7337 | 10712 | 10011 | 7359 | 4746 |
| 7073 | 0.05 | 0.07 | 0.71 | 0.48 | 1 | H1f0 | H1.0 linker histone | 5346 | 5965 | 6532 | 7537 | 8483 | 6149 | 7325 | 7622 | 8319 | 8313 | 6216 |
| 7049 | 0.13 | 0.08 | 1.61 | 0.11 | 1 | Nars | asparaginyl-tRNA synthetase | 4392 | 4445 | 5027 | 5857 | 8793 | 8539 | 7325 | 6837 | 9232 | 8120 | 8971 |
| 6742 | -0.09 | 0.06 | -1.48 | 0.14 | 1 | Vamp2 | vesicle-associated membrane protein 2 | 4307 | 4903 | 4844 | 8327 | 7621 | 7705 | 7323 | 7332 | 7598 | 7191 | 7013 |
| 6831 | 0.07 | 0.08 | 0.87 | 0.39 | 1 | Lin7c | lin-7 homolog C (C. elegans) | 5679 | 6864 | 5875 | 7043 | 7179 | 6185 | 7311 | 7707 | 7368 | 6797 | 7135 |
| 7113 | 0.05 | 0.08 | 0.62 | 0.53 | 1 | Atrx | ATRX, chromatin remodeler | 6550 | 6522 | 6101 | 5720 | 6326 | 9669 | 7290 | 6442 | 7032 | 6349 | 10244 |

| | | | | | | | | | | | | | | | | | | |
|---|---|---|---|---|---|---|---|---|---|---|---|---|---|---|---|---|---|---|
| 7305 | -0.09 | 0.06 | -1.58 | 0.11 | 1 | Chd3 | chromodomain helicase DNA binding pro | 7540 | 7902 | 7176 | 7004 | 7058 | 8450 | 7289 | 6506 | 6849 | 6882 | 7697 |
| 5731 | -0.03 | 0.12 | -0.28 | 0.78 | 1 | Azin1 | antizyme inhibitor 1 | 4274 | 5469 | 4136 | 5816 | 5966 | 5809 | 7287 | 5341 | 5914 | 7006 | 6026 |
| 8207 | -0.05 | 0.06 | -0.83 | 0.41 | 1 | Brd2 | bromodomain containing 2 | 9007 | 8450 | 8454 | 8119 | 7634 | 9782 | 7281 | 7525 | 7680 | 7482 | 8860 |
| 7141 | 0.04 | 0.06 | 0.63 | 0.53 | 1 | Dhx9 | DEAH (Asp-Glu-Ala-His) box polypeptide | 6882 | 6847 | 6576 | 6701 | 7516 | 7223 | 7273 | 6977 | 7823 | 7056 | 7673 |
| 6670 | -0.02 | 0.06 | -0.30 | 0.76 | 1 | Por | P450 (cytochrome) oxidoreductase | 4886 | 4992 | 4715 | 7914 | 8241 | 6126 | 7252 | 8153 | 7668 | 7710 | 5713 |
| 6358 | -0.07 | 0.09 | -0.79 | 0.43 | 1 | C1galt1 | core 1 synthase, glycoprotein-N-acetylgal | 5012 | 4886 | 4804 | 7626 | 7340 | 6091 | 7239 | 6020 | 7697 | 6695 | 6526 |
| 6359 | 0.02 | 0.06 | 0.30 | 0.77 | 1 | Ilrun | inflammation and lipid regulator with UBA | 5904 | 6387 | 6163 | 6496 | 6810 | 5045 | 7235 | 7025 | 6974 | 6768 | 5142 |
| 5796 | -0.02 | 0.08 | -0.20 | 0.84 | 1 | Rab3gap1 | RAB3 GTPase activating protein subunit 1 | 3328 | 3897 | 3999 | 6447 | 7189 | 5587 | 7226 | 6401 | 6995 | 6761 | 5929 |
| 7660 | 0.02 | 0.07 | 0.31 | 0.76 | 1 | Smarcc2 | SWI/SNF related, matrix associated, actin | 5790 | 6258 | 5824 | 8458 | 8029 | 9292 | 7223 | 8281 | 8388 | 8383 | 8333 |
| 6543 | -0.03 | 0.11 | -0.27 | 0.78 | 1 | Abca1 | ATP-binding cassette, sub-family A (ABC1 | 7139 | 5991 | 5320 | 7811 | 6658 | 5336 | 7196 | 7606 | 7640 | 6375 | 4901 |
| 6850 | -0.03 | 0.07 | -0.50 | 0.61 | 1 | Taok2 | TAO kinase 2 | 4507 | 5045 | 5394 | 7246 | 7019 | 9131 | 7189 | 7170 | 7142 | 6986 | 8521 |
| 8228 | 0.16 | 0.10 | 1.60 | 0.11 | 1 | Rpsa | ribosomal protein SA | 13956 | 11897 | 14284 | 6039 | 6234 | 4280 | 7183 | 8061 | 5984 | 7868 | 4725 |
| 6417 | 0.02 | 0.08 | 0.29 | 0.77 | 1 | Tmem106b | transmembrane protein 106B | 6198 | 6103 | 6130 | 6476 | 6685 | 5555 | 7142 | 5735 | 6967 | 7628 | 5963 |
| 6521 | -0.02 | 0.06 | -0.28 | 0.78 | 1 | Kdelr2 | KDEL (Lys-Asp-Glu-Leu) endoplasmic ret | 4537 | 4136 | 5536 | 6560 | 8859 | 6325 | 7128 | 6040 | 9016 | 7172 | 6418 |
| 6577 | 0.10 | 0.09 | 1.16 | 0.25 | 1 | Ecpas | Ecm29 proteasome adaptor and scaffold | 5267 | 4551 | 4399 | 6757 | 7436 | 6692 | 7123 | 8528 | 7480 | 7176 | 6943 |
| 5827 | -0.01 | 0.11 | -0.05 | 0.96 | 1 | Megf9 | multiple EGF-like-domains 9 | 3249 | 4227 | 3522 | 7165 | 6835 | 5510 | 7123 | 5942 | 7618 | 7160 | 5740 |
| 10495 | -0.01 | 0.09 | -0.09 | 0.93 | 1 | Srrm2 | serine/arginine repetitive matrix 2 | 6916 | 7767 | 6584 | 8003 | 8173 | 24743 | 7122 | 7803 | 9073 | 7173 | 22092 |
| 7239 | 0.06 | 0.09 | 0.67 | 0.51 | 1 | Cct7 | chaperonin containing Tcp1, subunit 7 (et | 8621 | 6876 | 8837 | 6561 | 7253 | 6178 | 7120 | 7679 | 6866 | 7361 | 6276 |
| 6337 | 0.09 | 0.06 | 1.45 | 0.15 | 1 | Ap1b1 | adaptor protein complex AP-1, beta 1 sub | 5026 | 4821 | 4494 | 6933 | 7438 | 5286 | 7114 | 7770 | 7621 | 7798 | 5403 |
| 5859 | -0.03 | 0.09 | -0.37 | 0.71 | 1 | Alg2 | asparagine-linked glycosylation 2 (alpha- | 2981 | 3599 | 3806 | 6474 | 7738 | 5971 | 7112 | 5936 | 8083 | 7024 | 5725 |
| 6579 | 0.10 | 0.10 | 1.00 | 0.32 | 1 | Ppp1r37 | protein phosphatase 1, regulatory subuni | 4578 | 4990 | 5516 | 6696 | 7211 | 6548 | 7108 | 8919 | 6983 | 7628 | 6196 |
| 7101 | -0.16 | 0.12 | -1.35 | 0.18 | 1 | Baiap3 | BAI1-associated protein 3 | 6541 | 7955 | 8076 | 5126 | 8452 | 8814 | 7106 | 5268 | 7225 | 6538 | 7013 |
| 6030 | 0.09 | 0.06 | 1.50 | 0.13 | 1 | Setd3 | SET domain containing 3 | 4342 | 4156 | 4334 | 6879 | 6708 | 5261 | 7100 | 7580 | 6912 | 7166 | 5892 |
| 8075 | -0.02 | 0.33 | -0.06 | 0.95 | 1 | Enpp2 | ectonucleotide pyrophosphatase/phosphc | 19099 | 11121 | 5399 | 6897 | 7551 | 5286 | 7091 | 6940 | 6771 | 6845 | 5825 |
| 6899 | 0.07 | 0.10 | 0.67 | 0.50 | 1 | Mycbp2 | MYC binding protein 2, E3 ubiquitin prote | 5600 | 4827 | 5861 | 6433 | 6727 | 8889 | 7088 | 8349 | 7024 | 6393 | 8697 |
| 6983 | -0.04 | 0.10 | -0.43 | 0.67 | 1 | Ubqln2 | ubiquilin 2 | 7979 | 9967 | 7985 | 6512 | 6934 | 5257 | 7086 | 6104 | 6783 | 7216 | 4991 |
| 5801 | -0.03 | 0.09 | -0.31 | 0.75 | 1 | Pofut2 | protein O-fucosyltransferase 2 | 3440 | 2805 | 3874 | 7192 | 7191 | 5592 | 7083 | 6333 | 7182 | 7560 | 5558 |
| 6254 | 0.15 | 0.10 | 1.58 | 0.11 | 1 | Prdx6 | peroxiredoxin 6 | 4075 | 4319 | 4971 | 6799 | 7144 | 5186 | 7078 | 8820 | 6872 | 7824 | 5703 |
| 5689 | -0.12 | 0.13 | -0.88 | 0.38 | 1 | Emilin1 | elastin microfibril interfacer 1 | 2833 | 1547 | 2250 | 7497 | 8275 | 6170 | 7070 | 6661 | 7607 | 7278 | 5393 |
| 8841 | 0.03 | 0.08 | 0.39 | 0.70 | 1 | Sf3b1 | splicing factor 3b, subunit 1 | 8640 | 9203 | 7813 | 7235 | 6513 | 14306 | 7069 | 7996 | 6875 | 6008 | 15589 |
| 5826 | -0.03 | 0.11 | -0.26 | 0.79 | 1 | Gnl3l | guanine nucleotide binding protein-like 3 | 3895 | 4864 | 4651 | 6327 | 6697 | 5496 | 7063 | 5306 | 7277 | 6780 | 5732 |
| 5226 | -0.04 | 0.11 | -0.41 | 0.68 | 1 | Prune2 | prune homolog 2 | 1488 | 2047 | 1818 | 6974 | 6451 | 5992 | 7063 | 5910 | 7084 | 6444 | 6221 |
| 6748 | 0.08 | 0.09 | 0.81 | 0.42 | 1 | Mapk1 | mitogen-activated protein kinase 1 | 6515 | 6862 | 6843 | 6347 | 6642 | 6172 | 7062 | 5890 | 7105 | 8469 | 6319 |
| 6972 | 0.09 | 0.10 | 0.86 | 0.39 | 1 | Cox4i1 | cytochrome c oxidase subunit 4I1 | 9097 | 7699 | 9081 | 6642 | 6536 | 4306 | 7059 | 7974 | 5909 | 8017 | 4373 |
| 6954 | 0.02 | 0.11 | 0.16 | 0.87 | 1 | Cbx6 | chromobox 6 | 7663 | 7469 | 7590 | 6593 | 6815 | 6106 | 7059 | 8895 | 6607 | 6123 | 5571 |
| 7755 | 0.09 | 0.11 | 0.82 | 0.41 | 1 | Cfl1 | cofilin 1, non-muscle | 12420 | 8876 | 10656 | 6329 | 6769 | 5727 | 7057 | 8099 | 6551 | 6681 | 6139 |
| 8151 | 0.10 | 0.11 | 0.89 | 0.37 | 1 | Atf4 | activating transcription factor 4 | 6118 | 5211 | 6674 | 9027 | 9113 | 9400 | 7053 | 10589 | 9182 | 7323 | 9974 |
| 6412 | 0.07 | 0.08 | 0.82 | 0.41 | 1 | Vps37a | vacuolar protein sorting 37A | 3220 | 3461 | 3741 | 6250 | 6756 | 9307 | 7050 | 7568 | 7072 | 6417 | 9690 |
| 5801 | 0.01 | 0.09 | 0.15 | 0.88 | 1 | Ttll7 | tubulin tyrosine ligase-like family, membe | 3093 | 3795 | 3644 | 6756 | 6339 | 6361 | 7046 | 6615 | 7224 | 6247 | 6694 |
| 7035 | 0.13 | 0.10 | 1.22 | 0.22 | 1 | Krt7 | keratin 7 | 3864 | 4179 | 5122 | 7119 | 8259 | 8302 | 7041 | 9797 | 7744 | 7616 | 8348 |
| 6563 | -0.07 | 0.06 | -1.14 | 0.25 | 1 | Rbbp4 | retinoblastoma binding protein 4, chroma | 7756 | 7270 | 7469 | 5959 | 6832 | 5649 | 7038 | 5364 | 6495 | 6755 | 5602 |
| 5357 | 0.05 | 0.11 | 0.47 | 0.64 | 1 | Stk32a | serine/threonine kinase 32A | 2462 | 3196 | 3080 | 5667 | 6649 | 5314 | 7038 | 6294 | 6754 | 6968 | 5501 |
| 5112 | -0.04 | 0.12 | -0.34 | 0.74 | 1 | Slc7a5 | solute carrier family 7 (cationic amino acic | 2562 | 3086 | 2827 | 5712 | 7506 | 3902 | 7034 | 5174 | 7612 | 7081 | 3736 |
| 6212 | 0.04 | 0.08 | 0.46 | 0.65 | 1 | Sgpl1 | sphingosine phosphate lyase 1 | 4551 | 4632 | 4756 | 7685 | 7385 | 4728 | 7023 | 6521 | 8254 | 7888 | 4906 |
| 6988 | 0.10 | 0.09 | 1.18 | 0.24 | 1 | 2900097C17Rik | | 6029 | 5949 | 7220 | 6582 | 8004 | 6193 | 7021 | 8656 | 7640 | 7151 | 6421 |
| 6274 | 0.08 | 0.08 | 0.97 | 0.33 | 1 | Uhmk1 | U2AF homology motif (UHM) kinase 1 | 5552 | 5796 | 4806 | 6225 | 6747 | 5721 | 7000 | 6903 | 7438 | 6938 | 5883 |
| 6481 | 0.01 | 0.06 | 0.27 | 0.79 | 1 | Mapk8ip2 | mitogen-activated protein kinase 8 interac | 4526 | 5479 | 5318 | 7847 | 7253 | 5697 | 6997 | 8146 | 7593 | 6867 | 5571 |
| 6110 | 0.16 | 0.14 | 1.18 | 0.24 | 1 | Oxct1 | 3-oxoacid CoA transferase 1 | 4815 | 3424 | 3444 | 7781 | 6455 | 4973 | 6989 | 9960 | 6739 | 6600 | 6034 |
| 5518 | 0.01 | 0.11 | 0.07 | 0.94 | 1 | Slitrk6 | SLIT and NTRK-like family, member 6 | 2038 | 2828 | 2330 | 7487 | 5413 | 6828 | 6987 | 6355 | 5837 | 6999 | 7599 |
| 6638 | 0.06 | 0.06 | 1.12 | 0.26 | 1 | Scaf11 | SR-related CTD-associated factor 11 | 5739 | 5996 | 5689 | 6942 | 6530 | 6749 | 6981 | 6937 | 7288 | 7052 | 7119 |
| 7232 | -0.03 | 0.06 | -0.56 | 0.58 | 1 | Ndrg1 | N-myc downstream regulated gene 1 | 9873 | 10125 | 9888 | 7078 | 6502 | 4630 | 6971 | 6569 | 6743 | 6359 | 4817 |
| 6319 | -0.05 | 0.07 | -0.76 | 0.45 | 1 | Ociad1 | OCIA domain containing 1 | 4496 | 4616 | 4810 | 7040 | 7507 | 6785 | 6964 | 5943 | 7274 | 6999 | 7071 |
| 6057 | -0.13 | 0.10 | -1.29 | 0.20 | 1 | Srebf2 | sterol regulatory element binding factor 2 | 5199 | 4663 | 4139 | 7118 | 7836 | 5509 | 6963 | 5850 | 7445 | 6931 | 4975 |
| 6673 | -0.03 | 0.09 | -0.41 | 0.68 | 1 | Cnot1 | CCR4-NOT transcription complex, subuni | 5817 | 5773 | 5422 | 6912 | 7171 | 7445 | 6958 | 5814 | 7656 | 6770 | 7671 |
| 5997 | 0.04 | 0.08 | 0.52 | 0.60 | 1 | Derl1 | Der1-like domain family, member 1 | 3917 | 2849 | 3442 | 6623 | 7680 | 6239 | 6944 | 6754 | 7658 | 7302 | 6561 |
| 5388 | -0.02 | 0.11 | -0.15 | 0.88 | 1 | Sdk2 | sidekick cell adhesion molecule 2 | 834 | 897 | 1244 | 7297 | 7366 | 6547 | 6944 | 6494 | 8146 | 7953 | 5542 |
| 8559 | 0.09 | 0.12 | 0.80 | 0.42 | 1 | Rps5 | ribosomal protein S5 | 16957 | 13497 | 15665 | 5820 | 6233 | 4223 | 6941 | 8126 | 5575 | 6688 | 4429 |
| 6529 | -0.03 | 0.08 | -0.39 | 0.70 | 1 | Sec24c | Sec24 related gene family, member C (S. | 5129 | 5634 | 5387 | 6288 | 7626 | 7272 | 6940 | 6160 | 7805 | 6807 | 6766 |
| 5785 | 0.04 | 0.10 | 0.36 | 0.72 | 1 | Esrp1 | epithelial splicing regulatory protein 1 | 2916 | 3516 | 3526 | 6375 | 7667 | 5514 | 6937 | 5868 | 7828 | 7767 | 5727 |
| 6766 | 0.04 | 0.06 | 0.67 | 0.50 | 1 | Ppp2ca | protein phosphatase 2 (formerly 2A), cata | 5787 | 6198 | 5467 | 7113 | 7447 | 6582 | 6928 | 7182 | 7309 | 7287 | 7122 |

| | | | | | | | | | | | | | | | | | |
|---|---|---|---|---|---|---|---|---|---|---|---|---|---|---|---|---|---|
| 6770 | -0.06 | 0.08 | -0.79 | 0.43 | 1 | Ttc39b | tetratricopeptide repeat domain 39B | 5880 | 7564 | 7061 | 7013 | 6961 | 6655 | 6918 | 6459 | 7080 | 6471 | 6408 |
| 6134 | -0.02 | 0.08 | -0.26 | 0.80 | 1 | Pcyox1 | prenylcysteine oxidase 1 | 4809 | 4312 | 4399 | 7256 | 8264 | 4691 | 6901 | 6861 | 8609 | 6908 | 4469 |
| 5843 | 0.02 | 0.09 | 0.25 | 0.81 | 1 | Synrg | synergin, gamma | 3930 | 4728 | 3871 | 6693 | 6288 | 5728 | 6896 | 6699 | 7446 | 6791 | 5200 |
| 6274 | 0.01 | 0.08 | 0.13 | 0.90 | 1 | Cep170b | centrosomal protein 170B | 4359 | 4538 | 4770 | 6876 | 7698 | 5967 | 6888 | 6917 | 7961 | 7933 | 5113 |
| 6499 | -0.06 | 0.07 | -0.82 | 0.41 | 1 | Sec22b | SEC22 homolog B, vesicle trafficking prot | 4824 | 5516 | 5940 | 6575 | 8113 | 6629 | 6882 | 6007 | 7510 | 6480 | 7017 |
| 6163 | 0.07 | 0.07 | 1.01 | 0.31 | 1 | Gnai3 | guanine nucleotide binding protein (G prc | 5502 | 5675 | 4970 | 6600 | 6290 | 5444 | 6875 | 6429 | 6355 | 7123 | 6531 |
| 7818 | -0.01 | 0.08 | -0.16 | 0.88 | 1 | Pde4dip | phosphodiesterase 4D interacting protein | 3907 | 4848 | 4153 | 9254 | 8404 | 12315 | 6872 | 8014 | 8517 | 7834 | 11879 |
| 6830 | 0.03 | 0.06 | 0.50 | 0.61 | 1 | Ctbp1 | C-terminal binding protein 1 | 6980 | 6693 | 6617 | 7135 | 7384 | 5723 | 6867 | 7688 | 6923 | 7366 | 5757 |
| 6545 | 0.04 | 0.08 | 0.50 | 0.61 | 1 | Yme1l1 | YME1-like 1 (S. cerevisiae) | 4364 | 4832 | 4760 | 7337 | 7354 | 7057 | 6837 | 6350 | 7728 | 7757 | 7624 |
| 6180 | 0.03 | 0.06 | 0.40 | 0.69 | 1 | Rab10 | RAB10, member RAS oncogene family | 5860 | 5943 | 6048 | 6412 | 6697 | 4940 | 6828 | 6033 | 7071 | 6818 | 5335 |
| 6353 | -0.07 | 0.07 | -1.05 | 0.29 | 1 | Mapk3 | mitogen-activated protein kinase 3 | 6210 | 6555 | 6130 | 6697 | 7437 | 5159 | 6827 | 6138 | 7092 | 6766 | 4866 |
| 7222 | 0.01 | 0.07 | 0.11 | 0.91 | 1 | Hnrnpa3 | heterogeneous nuclear ribonucleoprotein | 8080 | 7315 | 8004 | 6504 | 6914 | 7677 | 6820 | 7105 | 6462 | 6386 | 8170 |
| 7329 | 0.03 | 0.07 | 0.44 | 0.66 | 1 | Hnrnpul2 | heterogeneous nuclear ribonucleoprotein | 7540 | 7398 | 7670 | 6944 | 7306 | 7604 | 6818 | 7706 | 7890 | 6331 | 7408 |
| 4904 | -0.04 | 0.08 | -0.54 | 0.59 | 1 | Ret | ret proto-oncogene | 1647 | 2038 | 1916 | 6532 | 6940 | 4218 | 6818 | 6398 | 6840 | 6534 | 4067 |
| 5163 | 0.17 | 0.16 | 1.02 | 0.31 | 1 | Vstm2l | V-set and transmembrane domain contair | 585 | 684 | 1141 | 6349 | 7951 | 4594 | 6808 | 7112 | 7231 | 9454 | 4888 |
| 5650 | -0.02 | 0.11 | -0.23 | 0.81 | 1 | Prkar1b | protein kinase, cAMP dependent regulatc | 4113 | 4622 | 5694 | 6399 | 6123 | 4753 | 6807 | 6810 | 5997 | 5947 | 4888 |
| 5746 | -0.07 | 0.09 | -0.80 | 0.42 | 1 | Insig1 | insulin induced gene 1 | 1815 | 1652 | 1989 | 8757 | 7884 | 6235 | 6798 | 7459 | 7451 | 6755 | 6417 |
| 9196 | 0.07 | 0.09 | 0.73 | 0.47 | 1 | Rsrp1 | arginine/serine rich protein 1 | 4756 | 5421 | 6675 | 9314 | 7715 | 17513 | 6792 | 8952 | 8266 | 7831 | 17916 |
| 8450 | -0.09 | 0.08 | -1.18 | 0.24 | 1 | H3f3b | H3.3 histone B | 12985 | 10736 | 12058 | 7100 | 6889 | 8642 | 6792 | 5839 | 6600 | 7385 | 7919 |
| 5355 | 0.06 | 0.07 | 0.82 | 0.41 | 1 | Erlec1 | endoplasmic reticulum lectin 1 | 4163 | 4282 | 4068 | 6407 | 5616 | 4018 | 6786 | 6306 | 5887 | 6877 | 4495 |
| 5456 | 0.03 | 0.07 | 0.37 | 0.71 | 1 | Sqle | squalene epoxidase | 1386 | 1955 | 1730 | 7384 | 7064 | 5965 | 6780 | 7635 | 7410 | 6460 | 6243 |
| 6122 | 0.05 | 0.06 | 0.82 | 0.41 | 1 | Ccny | cyclin Y | 5364 | 5560 | 5752 | 6579 | 6602 | 4945 | 6772 | 6612 | 7286 | 6874 | 4992 |
| 10804 | -0.04 | 0.11 | -0.37 | 0.71 | 1 | Ddx17 | DEAD (Asp-Glu-Ala-Asp) box polypeptide | 8382 | 8835 | 8125 | 6128 | 8356 | 25938 | 6768 | 5441 | 8790 | 6447 | 25640 |
| 5939 | -0.11 | 0.09 | -1.20 | 0.23 | 1 | Akap9 | A kinase (PRKA) anchor protein (yotiao) 9 | 3938 | 4215 | 4875 | 5812 | 6415 | 8014 | 6765 | 4867 | 6875 | 5612 | 7945 |
| 5133 | 0.02 | 0.08 | 0.27 | 0.79 | 1 | Suco | SUN domain containing ossification facto | 2989 | 3257 | 2833 | 5718 | 5734 | 5329 | 6758 | 5672 | 6143 | 6345 | 5682 |
| 6969 | 0.09 | 0.08 | 1.22 | 0.22 | 1 | Actr2 | ARP2 actin-related protein 2 | 8161 | 6748 | 7031 | 6701 | 6756 | 6248 | 6749 | 6727 | 7840 | 6894 | 6803 |
| 6557 | 0.02 | 0.09 | 0.18 | 0.86 | 1 | Itfg1 | integrin alpha FG-GAP repeat containing | 6441 | 7253 | 6807 | 7178 | 6870 | 4912 | 6748 | 5941 | 6962 | 7703 | 5316 |
| 6226 | 0.11 | 0.07 | 1.52 | 0.13 | 1 | Pafah1b2 | platelet-activating factor acetylhydrolase, | 5826 | 6720 | 6117 | 6154 | 5905 | 5100 | 6746 | 7068 | 6283 | 7251 | 5319 |
| 6284 | 0.03 | 0.07 | 0.38 | 0.70 | 1 | Cul3 | cullin 3 | 4995 | 5618 | 5410 | 6948 | 6652 | 5976 | 6741 | 6664 | 7209 | 6241 | 6676 |
| 5682 | 0.11 | 0.08 | 1.37 | 0.17 | 1 | Tshz1 | teashirt zinc finger family member 1 | 2832 | 3478 | 3161 | 6506 | 6534 | 5704 | 6738 | 7247 | 7235 | 7397 | 5674 |
| 6187 | 0.04 | 0.09 | 0.44 | 0.66 | 1 | Prrc1 | proline-rich coiled-coil 1 | 4030 | 4636 | 4374 | 5854 | 7536 | 7063 | 6721 | 6993 | 7131 | 6735 | 6988 |
| 5732 | -0.08 | 0.07 | -1.15 | 0.25 | 1 | Mapre2 | microtubule-associated protein, RP/EB fa | 4758 | 4919 | 4449 | 6676 | 6388 | 5407 | 6717 | 5962 | 6544 | 5845 | 5384 |
| 5668 | -0.11 | 0.11 | -0.99 | 0.32 | 1 | Sgsm1 | small G protein signaling modulator 1 | 4644 | 5753 | 4329 | 6734 | 5721 | 5481 | 6706 | 5697 | 6442 | 6404 | 4442 |
| 5285 | 0.09 | 0.07 | 1.30 | 0.19 | 1 | Gga2 | golgi associated, gamma adaptin ear con | 2592 | 2617 | 2879 | 6326 | 6141 | 5016 | 6704 | 6980 | 6549 | 7191 | 5140 |
| 6584 | 0.13 | 0.08 | 1.72 | 0.09 | 1 | Reep5 | receptor accessory protein 5 | 6258 | 5999 | 6598 | 7314 | 6950 | 4573 | 6684 | 8579 | 7161 | 7476 | 4826 |
| 7880 | 0.00 | 0.12 | 0.03 | 0.97 | 1 | Rplp1 | ribosomal protein, large, P1 | 11776 | 10775 | 14886 | 6364 | 6862 | 4579 | 6682 | 7546 | 5808 | 7270 | 4131 |
| 6366 | -0.02 | 0.06 | -0.36 | 0.72 | 1 | Zfr | zinc finger RNA binding protein | 5618 | 6078 | 5376 | 6491 | 6217 | 7263 | 6675 | 6563 | 6141 | 6005 | 7596 |
| 6828 | -0.01 | 0.07 | -0.19 | 0.85 | 1 | Supt6 | SPT6, histone chaperone and transcriptic | 5970 | 6601 | 6138 | 6748 | 7597 | 7312 | 6674 | 7196 | 7058 | 7057 | 6758 |
| 5712 | -0.13 | 0.11 | -1.17 | 0.24 | 1 | Osbp | oxysterol binding protein | 3337 | 3608 | 3815 | 5928 | 7261 | 7222 | 6668 | 4507 | 7520 | 6559 | 6406 |
| 7411 | 0.16 | 0.12 | 1.35 | 0.18 | 1 | Rpl3 | ribosomal protein L3 | 11807 | 9142 | 12068 | 5193 | 6438 | 4686 | 6662 | 7708 | 6122 | 6752 | 4940 |
| 6259 | 0.02 | 0.09 | 0.24 | 0.81 | 1 | Atxn10 | ataxin 10 | 6676 | 7547 | 7818 | 6213 | 5942 | 4438 | 6662 | 6981 | 5684 | 6140 | 4745 |
| 9197 | 0.09 | 0.06 | 1.52 | 0.13 | 1 | Rbm39 | RNA binding motif protein 39 | 6391 | 8744 | 6694 | 7724 | 6645 | 17359 | 6659 | 8466 | 7022 | 6792 | 18670 |
| 6502 | -0.14 | 0.08 | -1.72 | 0.08 | 1 | Mbtps1 | membrane-bound transcription factor pep | 7852 | 8315 | 7685 | 6762 | 6158 | 5405 | 6658 | 5237 | 6012 | 6114 | 5319 |
| 6502 | -0.13 | 0.07 | -1.75 | 0.08 | 1 | Il6ra | interleukin 6 receptor, alpha | 4476 | 5101 | 5061 | 7078 | 6713 | 9210 | 6657 | 6679 | 6988 | 5364 | 8197 |
| 5556 | 0.01 | 0.08 | 0.10 | 0.92 | 1 | Mbd2 | methyl-CpG binding domain protein 2 | 3801 | 3369 | 4100 | 6066 | 6802 | 5300 | 6644 | 5065 | 6935 | 7028 | 6006 |
| 5114 | -0.06 | 0.08 | -0.78 | 0.44 | 1 | Abca5 | ATP-binding cassette, sub-family A (ABC | 2500 | 2814 | 2432 | 6083 | 5723 | 6374 | 6622 | 5833 | 6195 | 5543 | 6139 |
| 6056 | -0.04 | 0.09 | -0.40 | 0.69 | 1 | Tor1aip2 | torsin A interacting protein 2 | 4168 | 5424 | 4973 | 7283 | 6411 | 6067 | 6617 | 6029 | 6964 | 6142 | 6540 |
| 5974 | -0.07 | 0.09 | -0.75 | 0.45 | 1 | Kcnj11 | potassium inwardly rectifying channel, su | 4709 | 6568 | 5722 | 6900 | 6331 | 5038 | 6616 | 6655 | 6048 | 6579 | 4545 |
| 7707 | 0.07 | 0.08 | 0.94 | 0.35 | 1 | Ybx1 | Y box protein 1 | 12163 | 9301 | 10717 | 6911 | 6354 | 5692 | 6609 | 6892 | 6871 | 7527 | 5736 |
| 5382 | 0.04 | 0.10 | 0.43 | 0.67 | 1 | Slc16a12 | solute carrier family 16 (monocarboxylic a | 3647 | 4367 | 3916 | 6043 | 5638 | 5038 | 6608 | 5574 | 5846 | 6842 | 5680 |
| 5977 | 0.12 | 0.09 | 1.32 | 0.19 | 1 | Akt1 | thymoma viral proto-oncogene 1 | 5819 | 4658 | 5351 | 5756 | 6788 | 4720 | 6597 | 7635 | 6853 | 6813 | 4757 |
| 6383 | -0.14 | 0.09 | -1.53 | 0.13 | 1 | Fndc3b | fibronectin type III domain containing 3B | 6150 | 4753 | 5938 | 6854 | 8325 | 6154 | 6575 | 5264 | 8087 | 6407 | 5704 |
| 6191 | 0.11 | 0.09 | 1.30 | 0.19 | 1 | Trabd | TraB domain containing | 2801 | 2792 | 3331 | 5491 | 8830 | 7708 | 6567 | 7133 | 8237 | 7240 | 7975 |
| 6105 | 0.00 | 0.07 | -0.04 | 0.97 | 1 | Impad1 | inositol monophosphatase domain contai | 5881 | 6377 | 6502 | 6777 | 6043 | 4862 | 6562 | 6692 | 6342 | 6049 | 5066 |
| 5363 | -0.07 | 0.10 | -0.69 | 0.49 | 1 | Qdpr | quinoid dihydropteridine reductase | 3622 | 3604 | 4315 | 6448 | 6027 | 5228 | 6561 | 6673 | 5432 | 6014 | 5064 |
| 5308 | -0.01 | 0.07 | -0.18 | 0.85 | 1 | Abca3 | ATP-binding cassette, sub-family A (ABC | 3484 | 3241 | 3929 | 6097 | 6340 | 4931 | 6543 | 5522 | 6905 | 6617 | 4775 |
| 6217 | 0.08 | 0.07 | 1.04 | 0.30 | 1 | Pdcd4 | programmed cell death 4 | 6308 | 6260 | 6925 | 5263 | 6261 | 5773 | 6530 | 5961 | 6084 | 6706 | 6319 |
| 5761 | 0.04 | 0.08 | 0.45 | 0.65 | 1 | Nr1d2 | nuclear receptor subfamily 1, group D, me | 4273 | 4346 | 4345 | 6342 | 5493 | 6486 | 6528 | 5543 | 6239 | 6551 | 7223 |
| 5885 | -0.04 | 0.08 | -0.52 | 0.60 | 1 | Ap2a2 | adaptor-related protein complex 2, alpha | 6064 | 5118 | 5251 | 6203 | 6889 | 4799 | 6512 | 6581 | 6148 | 6805 | 4359 |
| 4012 | 0.04 | 0.09 | 0.51 | 0.61 | 1 | Sorl1 | sortilin-related receptor, LDLR class A rep | 1411 | 1344 | 1298 | 5702 | 4453 | 3037 | 6503 | 6073 | 5221 | 6160 | 2924 |

| | | | | | | | | | | | | | | | | | |
|---|---|---|---|---|---|---|---|---|---|---|---|---|---|---|---|---|---|
| 5638 | 0.02 | 0.08 | 0.29 | 0.77 | 1 | Rnf14 | ring finger protein 14 | 3608 | 4117 | 3866 | 5948 | 7110 | 5480 | 6501 | 6722 | 6675 | 6139 | 5856 |
| 4963 | -0.14 | 0.09 | -1.52 | 0.13 | 1 | Plekhb2 | pleckstrin homology domain containing, fa | 4429 | 4134 | 4647 | 4940 | 5906 | 4277 | 6499 | 3995 | 5715 | 6073 | 3975 |
| 5800 | 0.11 | 0.08 | 1.36 | 0.17 | 1 | Pkp4 | plakophilin 4 | 3294 | 3661 | 3574 | 6553 | 6432 | 6045 | 6496 | 8367 | 6243 | 6738 | 6396 |
| 6112 | 0.03 | 0.07 | 0.35 | 0.72 | 1 | Mib1 | mindbomb E3 ubiquitin protein ligase 1 | 4224 | 4043 | 4491 | 6585 | 6356 | 7525 | 6494 | 5948 | 7337 | 6503 | 7725 |
| 6349 | 0.01 | 0.07 | 0.11 | 0.92 | 1 | Supt5 | suppressor of Ty 5, DSIF elongation facto | 4786 | 4719 | 5410 | 5943 | 7555 | 7406 | 6492 | 6610 | 7667 | 5899 | 7351 |
| 6401 | 0.07 | 0.07 | 1.05 | 0.29 | 1 | Tpr | translocated promoter region, nuclear bas | 6435 | 6022 | 5665 | 5603 | 6070 | 7333 | 6487 | 5817 | 6717 | 6429 | 7830 |
| 5325 | -0.03 | 0.07 | -0.50 | 0.62 | 1 | Arhgap35 | Rho GTPase activating protein 35 | 3456 | 3713 | 3848 | 6666 | 6279 | 4647 | 6486 | 6041 | 6357 | 6358 | 4723 |
| 7067 | 0.17 | 0.14 | 1.18 | 0.24 | 1 | Ptma | prothymosin alpha | 9374 | 7394 | 8536 | 6071 | 6223 | 5750 | 6481 | 9699 | 6367 | 5952 | 5884 |
| 6179 | 0.01 | 0.10 | 0.08 | 0.94 | 1 | B4galt6 | UDP-Gal:betaGlcNAc beta 1,4-galactosyl | 5744 | 7632 | 7168 | 5934 | 6118 | 4998 | 6480 | 7227 | 6244 | 5790 | 4639 |
| 5363 | -0.06 | 0.09 | -0.64 | 0.52 | 1 | Ywhaq | tyrosine 3-monooxygenase/tryptophan 5- | 4769 | 4554 | 4716 | 6048 | 5665 | 4745 | 6474 | 4942 | 5400 | 6434 | 5247 |
| 5507 | 0.04 | 0.09 | 0.41 | 0.68 | 1 | Usp33 | ubiquitin specific peptidase 33 | 3033 | 2645 | 3253 | 5803 | 5973 | 7194 | 6471 | 6187 | 6678 | 5458 | 7888 |
| 6229 | 0.03 | 0.09 | 0.31 | 0.75 | 1 | Paip2 | polyadenylate-binding protein-interacting | 5614 | 6304 | 5709 | 6436 | 6267 | 6013 | 6470 | 5549 | 6665 | 7046 | 6441 |
| 5041 | -0.18 | 0.12 | -1.57 | 0.12 | 1 | Arhgef17 | Rho guanine nucleotide exchange factor | 5685 | 5262 | 4449 | 4533 | 4817 | 5353 | 6467 | 3471 | 4345 | 6440 | 4628 |
| 7469 | 0.03 | 0.06 | 0.45 | 0.65 | 1 | Ivns1abp | influenza virus NS1A binding protein | 8023 | 8219 | 9379 | 5891 | 6562 | 9055 | 6462 | 5969 | 6768 | 6336 | 9491 |
| 5480 | -0.03 | 0.10 | -0.31 | 0.76 | 1 | Mtmr6 | myotubularin related protein 6 | 4483 | 4165 | 4422 | 5711 | 6076 | 5513 | 6459 | 4531 | 6974 | 6415 | 5536 |
| 5338 | -0.08 | 0.12 | -0.71 | 0.48 | 1 | Scnn1b | sodium channel, nonvoltage-gated 1 beta | 1136 | 1962 | 1825 | 7683 | 7309 | 6244 | 6434 | 6941 | 7121 | 6703 | 5360 |
| 5441 | -0.14 | 0.10 | -1.41 | 0.16 | 1 | Sipa1l3 | signal-induced proliferation-associated 1 | 3927 | 4629 | 4419 | 7375 | 5855 | 4896 | 6423 | 5824 | 6791 | 5170 | 4538 |
| 5507 | -0.18 | 0.11 | -1.61 | 0.11 | 1 | Txnrd1 | thioredoxin reductase 1 | 4941 | 5707 | 5695 | 5988 | 5905 | 5169 | 6420 | 4175 | 5844 | 6045 | 4687 |
| 4667 | -0.04 | 0.10 | -0.42 | 0.67 | 1 | Chst11 | carbohydrate sulfotransferase 11 | 1455 | 1225 | 2010 | 6014 | 7039 | 4105 | 6420 | 5147 | 6962 | 7003 | 3959 |
| 6621 | 0.11 | 0.08 | 1.39 | 0.16 | 1 | Myo18a | myosin XVIIIA | 5632 | 6475 | 6096 | 7252 | 6396 | 6140 | 6415 | 8858 | 6549 | 6977 | 6035 |
| 7388 | -0.11 | 0.13 | -0.87 | 0.39 | 1 | Mbnl2 | muscleblind like splicing factor 2 | 9981 | 13161 | 9558 | 6321 | 5740 | 6777 | 6410 | 4523 | 6387 | 5834 | 6576 |
| 6126 | -0.04 | 0.13 | -0.34 | 0.74 | 1 | Esyt1 | extended synaptotagmin-like protein 1 | 6372 | 4152 | 4264 | 7008 | 7971 | 5298 | 6408 | 6760 | 7436 | 6644 | 5069 |
| 5482 | -0.01 | 0.07 | -0.10 | 0.92 | 1 | Gtf3c1 | general transcription factor III C 1 | 4298 | 4371 | 4742 | 5510 | 6219 | 5341 | 6402 | 5778 | 6343 | 6363 | 4938 |
| 4853 | 0.06 | 0.09 | 0.65 | 0.52 | 1 | Cln6 | ceroid-lipofuscinosis, neuronal 6 | 1897 | 2609 | 2217 | 6412 | 6333 | 3729 | 6393 | 6534 | 6530 | 6791 | 3937 |
| 5968 | 0.09 | 0.05 | 1.67 | 0.09 | 1 | Elp3 | elongator acetyltransferase complex subu | 4020 | 4253 | 4862 | 6091 | 7268 | 5697 | 6383 | 6606 | 7543 | 6769 | 6159 |
| 5884 | -0.09 | 0.07 | -1.22 | 0.22 | 1 | Fam20c | family with sequence similarity 20, membe | 4736 | 5439 | 5506 | 6535 | 6491 | 5846 | 6377 | 6706 | 5723 | 6102 | 5267 |
| 6020 | 0.02 | 0.08 | 0.28 | 0.78 | 1 | Srp68 | signal recognition particle 68 | 3691 | 4195 | 4664 | 5862 | 7393 | 6958 | 6376 | 5663 | 7954 | 6359 | 7105 |
| 5427 | -0.03 | 0.08 | -0.40 | 0.69 | 1 | Arsb | arylsulfatase B | 5958 | 6159 | 5260 | 6260 | 4943 | 3873 | 6367 | 5370 | 5214 | 6360 | 3938 |
| 8216 | 0.09 | 0.14 | 0.65 | 0.51 | 1 | Rps15 | ribosomal protein S15 | 14141 | 12523 | 15567 | 5934 | 6643 | 4224 | 6341 | 8989 | 5308 | 6479 | 4229 |
| 5627 | 0.10 | 0.10 | 1.04 | 0.30 | 1 | Fam135a | family with sequence similarity 135, meml | 2240 | 2796 | 2907 | 6263 | 6175 | 7218 | 6340 | 5945 | 6840 | 7127 | 8051 |
| 7303 | 0.03 | 0.05 | 0.49 | 0.62 | 1 | Rmnd5a | required for meiotic nuclear division 5 hor | 8950 | 10598 | 9396 | 6854 | 6947 | 5322 | 6332 | 7145 | 6916 | 6412 | 5459 |
| 5746 | 0.08 | 0.07 | 1.13 | 0.26 | 1 | Atxn7l3b | ataxin 7-like 3B | 5111 | 5244 | 5166 | 5911 | 6323 | 4552 | 6324 | 7304 | 6132 | 6460 | 4680 |
| 5634 | 0.09 | 0.08 | 1.01 | 0.31 | 1 | Slc33a1 | solute carrier family 33 (acetyl-CoA transp | 2044 | 2581 | 3085 | 6343 | 7406 | 6271 | 6324 | 6559 | 7664 | 6765 | 6931 |
| 5449 | 0.23 | 0.12 | 1.93 | 0.05 | 1 | Spcs1 | signal peptidase complex subunit 1 homo | 3224 | 2687 | 3716 | 6159 | 5888 | 4807 | 6296 | 8212 | 6347 | 7171 | 5433 |
| 6174 | 0.05 | 0.08 | 0.59 | 0.55 | 1 | Cd47 | CD47 antigen (Rh-related antigen, integri | 8261 | 6484 | 6750 | 4920 | 6007 | 5590 | 6295 | 4839 | 6707 | 6313 | 5754 |
| 5829 | 0.04 | 0.08 | 0.53 | 0.59 | 1 | Pea15a | phosphoprotein enriched in astrocytes 15 | 6729 | 5292 | 5819 | 5636 | 6265 | 4670 | 6281 | 6205 | 6272 | 5893 | 5060 |
| 5637 | 0.02 | 0.07 | 0.36 | 0.72 | 1 | Desi1 | desumoylating isopeptidase 1 | 3520 | 3912 | 4015 | 6637 | 6562 | 5612 | 6276 | 6452 | 6385 | 6832 | 5809 |
| 6023 | 0.05 | 0.08 | 0.65 | 0.52 | 1 | Ttc3 | tetratricopeptide repeat domain 3 | 4797 | 4961 | 5219 | 5530 | 6216 | 7150 | 6268 | 6658 | 6113 | 5688 | 7651 |
| 4635 | 0.17 | 0.15 | 1.14 | 0.26 | 1 | Gad1 | glutamate decarboxylase 1 | 284 | 261 | 481 | 5014 | 5425 | 6882 | 6258 | 6448 | 6108 | 6766 | 7056 |
| 5153 | 0.00 | 0.10 | -0.01 | 0.99 | 1 | Rapgef4 | Rap guanine nucleotide exchange factor | 2999 | 4085 | 4023 | 5582 | 5656 | 5297 | 6245 | 5794 | 5937 | 5639 | 5423 |
| 4829 | -0.03 | 0.13 | -0.26 | 0.79 | 1 | Pnmal1 | PNMA-like 1 | 2982 | 3703 | 3470 | 5612 | 5358 | 4537 | 6245 | 4361 | 5964 | 5746 | 5144 |
| 6346 | -0.05 | 0.07 | -0.69 | 0.49 | 1 | Ppp1ca | protein phosphatase 1 catalytic subunit al | 8570 | 6849 | 8018 | 5922 | 6563 | 4772 | 6242 | 5429 | 6086 | 6846 | 4510 |
| 5727 | 0.02 | 0.06 | 0.40 | 0.69 | 1 | Pten | phosphatase and tensin homolog | 5630 | 5698 | 5054 | 6093 | 5783 | 5029 | 6240 | 6169 | 5934 | 5959 | 5408 |
| 5200 | -0.04 | 0.06 | -0.68 | 0.49 | 1 | Ube3c | ubiquitin protein ligase E3C | 2231 | 2119 | 2348 | 6318 | 7412 | 5655 | 6233 | 5786 | 7159 | 6362 | 5576 |
| 5393 | 0.01 | 0.10 | 0.05 | 0.96 | 1 | Slc39a9 | solute carrier family 39 (zinc transporter), | 3542 | 4558 | 3754 | 5945 | 6025 | 5498 | 6226 | 5465 | 6273 | 6568 | 5475 |
| 4949 | -0.10 | 0.08 | -1.28 | 0.20 | 1 | Rps6ka2 | ribosomal protein S6 kinase, polypeptide | 2055 | 2434 | 2440 | 6817 | 6567 | 4924 | 6224 | 5916 | 6453 | 6379 | 4230 |
| 5548 | 0.07 | 0.09 | 0.81 | 0.42 | 1 | Bex3 | brain expressed X-linked 3 | 4213 | 3997 | 4869 | 5995 | 5939 | 5217 | 6211 | 6895 | 5791 | 6455 | 5442 |
| 6508 | 0.07 | 0.08 | 0.85 | 0.40 | 1 | Purb | purine rich element binding protein B | 6629 | 7215 | 6434 | 6683 | 6599 | 5489 | 6207 | 7960 | 6953 | 5682 | 5733 |
| 5867 | -0.02 | 0.07 | -0.27 | 0.79 | 1 | Eif4ebp2 | eukaryotic translation initiation factor 4E b | 6745 | 6481 | 6078 | 6266 | 5997 | 4192 | 6201 | 5812 | 6573 | 6462 | 3725 |
| 4905 | 0.02 | 0.08 | 0.25 | 0.80 | 1 | Tlk1 | tousled-like kinase 1 | 2882 | 3141 | 2981 | 6021 | 5207 | 4906 | 6197 | 5568 | 5586 | 6467 | 4999 |
| 6928 | -0.03 | 0.07 | -0.36 | 0.72 | 1 | Pgrmc1 | progesterone receptor membrane compo | 9197 | 9950 | 10200 | 6450 | 6224 | 4771 | 6196 | 6034 | 5736 | 6774 | 4678 |
| 6379 | -0.15 | 0.10 | -1.50 | 0.13 | 1 | Birc6 | baculoviral IAP repeat-containing 6 | 5798 | 5887 | 5318 | 5549 | 6584 | 9595 | 6185 | 4845 | 6635 | 5020 | 8752 |
| 5731 | 0.10 | 0.07 | 1.38 | 0.17 | 1 | Tmx1 | thioredoxin-related transmembrane prote | 4550 | 5278 | 5512 | 6133 | 6112 | 4631 | 6180 | 6274 | 6672 | 6442 | 5255 |
| 4482 | -0.24 | 0.15 | -1.63 | 0.10 | 1 | Cdhr1 | cadherin-related family member 1 | 727 | 1051 | 1459 | 6303 | 6889 | 5489 | 6176 | 4046 | 6644 | 5360 | 5159 |
| 5169 | 0.02 | 0.09 | 0.20 | 0.84 | 1 | Aak1 | AP2 associated kinase 1 | 3288 | 3611 | 3514 | 5655 | 5099 | 6174 | 6164 | 5395 | 6233 | 5840 | 5890 |
| 6866 | 0.07 | 0.11 | 0.65 | 0.51 | 1 | Rps21 | ribosomal protein S21 | 10418 | 9327 | 11729 | 5613 | 5572 | 3983 | 6157 | 6613 | 5330 | 7119 | 3664 |
| 5192 | -0.03 | 0.13 | -0.21 | 0.83 | 1 | Dmxl2 | Dmx-like 2 | 1976 | 2994 | 2537 | 5350 | 5560 | 7870 | 6155 | 5195 | 6512 | 5029 | 7931 |
| 7651 | -0.07 | 0.08 | -0.93 | 0.35 | 1 | Cpq | carboxypeptidase Q | 13578 | 15117 | 13834 | 5829 | 5700 | 3745 | 6146 | 5074 | 5781 | 5441 | 3920 |
| 5234 | 0.03 | 0.09 | 0.34 | 0.73 | 1 | Clock | circadian locomotor output cycles kaput | 3221 | 3630 | 3541 | 5592 | 5622 | 5962 | 6143 | 5544 | 6565 | 5412 | 6340 |
| 5953 | 0.08 | 0.07 | 1.19 | 0.23 | 1 | M6pr | mannose-6-phosphate receptor, cation de | 6101 | 5539 | 5793 | 6354 | 6109 | 4809 | 6140 | 6160 | 6706 | 6328 | 5450 |

| | | | | | | | | | | | | | | | | | | |
|---|---|---|---|---|---|---|---|---|---|---|---|---|---|---|---|---|---|---|
| 5431 | -0.02 | 0.07 | -0.34 | 0.73 | 1 | Mtpn | myotrophin | 6111 | 5107 | 5329 | 5201 | 5768 | 4642 | 6140 | 4665 | 6048 | 6108 | 4626 |
| 5573 | -0.05 | 0.08 | -0.59 | 0.55 | 1 | Hadha | hydroxyacyl-CoA dehydrogenase trifuncti | 5689 | 5777 | 5772 | 5676 | 5881 | 4724 | 6130 | 4825 | 6122 | 5722 | 4991 |
| 4674 | -0.13 | 0.10 | -1.26 | 0.21 | 1 | Myo5a | myosin VA | 3859 | 3320 | 3115 | 5923 | 5373 | 4144 | 6124 | 4833 | 5605 | 4837 | 4284 |
| 5945 | -0.01 | 0.09 | -0.10 | 0.92 | 1 | Kif21a | kinesin family member 21A | 4160 | 6288 | 5264 | 6418 | 5758 | 6583 | 6122 | 6718 | 6038 | 5525 | 6526 |
| 5054 | 0.03 | 0.06 | 0.57 | 0.57 | 1 | Paip2b | poly(A) binding protein interacting protein | 3004 | 2929 | 3347 | 5880 | 6418 | 4516 | 6116 | 6037 | 6539 | 5934 | 4874 |
| 8928 | -0.07 | 0.07 | -1.05 | 0.30 | 1 | Hnrnph1 | heterogeneous nuclear ribonucleoprotein | 8619 | 9093 | 8663 | 5494 | 6124 | 18763 | 6113 | 5108 | 5886 | 5669 | 18672 |
| 5671 | 0.01 | 0.10 | 0.11 | 0.92 | 1 | Dnajc5 | DnaJ heat shock protein family (Hsp40) m | 4859 | 5321 | 4509 | 5450 | 6269 | 5916 | 6109 | 5174 | 6786 | 5867 | 6121 |
| 5779 | 0.06 | 0.07 | 0.89 | 0.37 | 1 | Etf1 | eukaryotic translation termination factor 1 | 5235 | 5799 | 5030 | 6441 | 5781 | 4961 | 6104 | 6040 | 6318 | 6390 | 5472 |
| 5486 | 0.09 | 0.07 | 1.18 | 0.24 | 1 | Trip11 | thyroid hormone receptor interactor 11 | 3327 | 3693 | 3517 | 5712 | 6098 | 6206 | 6104 | 5997 | 7007 | 5879 | 6804 |
| 5570 | 0.04 | 0.08 | 0.46 | 0.65 | 1 | Grsf1 | G-rich RNA sequence binding factor 1 | 4490 | 5123 | 4339 | 6532 | 6175 | 4622 | 6101 | 5932 | 6157 | 6609 | 5195 |
| 4672 | -0.02 | 0.11 | -0.18 | 0.86 | 1 | Slc22a23 | solute carrier family 22, member 23 | 818 | 1076 | 965 | 7435 | 6258 | 4702 | 6090 | 5709 | 6944 | 6729 | 4663 |
| 5628 | 0.09 | 0.10 | 0.85 | 0.39 | 1 | Rangap1 | RAN GTPase activating protein 1 | 4201 | 3587 | 4935 | 5708 | 6538 | 5564 | 6087 | 7007 | 6098 | 6091 | 6094 |
| 6372 | -0.10 | 0.05 | -1.81 | 0.07 | 1 | Ppp2r1a | protein phosphatase 2, regulatory subuni | 8298 | 7992 | 8274 | 6187 | 6481 | 4769 | 6079 | 5758 | 6156 | 5599 | 4495 |
| 6135 | -0.13 | 0.10 | -1.24 | 0.21 | 1 | Cap1 | CAP, adenylate cyclase-associated prote | 9760 | 7707 | 7296 | 5811 | 5710 | 4657 | 6079 | 4238 | 5782 | 5938 | 4511 |
| 5289 | 0.02 | 0.07 | 0.28 | 0.78 | 1 | Nt5c2 | 5'-nucleotidase, cytosolic II | 3348 | 3620 | 3755 | 5934 | 5864 | 5706 | 6069 | 6282 | 5735 | 6148 | 5723 |
| 4805 | -0.01 | 0.08 | -0.15 | 0.88 | 1 | Prkca | protein kinase C, alpha | 2092 | 2335 | 2277 | 6135 | 6061 | 4894 | 6068 | 5544 | 6922 | 5459 | 5069 |
| 6810 | 0.06 | 0.10 | 0.64 | 0.52 | 1 | Pak3 | p21 (RAC1) activated kinase 3 | 5836 | 7931 | 7913 | 5654 | 6534 | 7692 | 6066 | 7125 | 6583 | 5338 | 8236 |
| 5974 | -0.13 | 0.09 | -1.40 | 0.16 | 1 | Plekha1 | pleckstrin homology domain containing, fa | 5709 | 7453 | 6709 | 6081 | 5279 | 6505 | 6066 | 5142 | 5297 | 5326 | 6152 |
| 4962 | -0.01 | 0.07 | -0.09 | 0.93 | 1 | Nudcd3 | NudC domain containing 3 | 5014 | 4130 | 4567 | 5071 | 5253 | 4097 | 6066 | 5442 | 5587 | 5447 | 3910 |
| 6441 | 0.08 | 0.07 | 1.14 | 0.25 | 1 | Actr3 | ARP3 actin-related protein 3 | 9723 | 7956 | 8415 | 5901 | 5628 | 4202 | 6054 | 5663 | 6182 | 6174 | 4950 |
| 5512 | 0.06 | 0.11 | 0.52 | 0.60 | 1 | Cltb | clathrin, light polypeptide (Lcb) | 4352 | 3973 | 5292 | 6040 | 5959 | 4985 | 6045 | 7160 | 5619 | 6098 | 5112 |
| 5296 | 0.04 | 0.07 | 0.53 | 0.60 | 1 | Scamp1 | secretory carrier membrane protein 1 | 4002 | 4493 | 4286 | 5980 | 5908 | 4532 | 6045 | 5824 | 6169 | 6344 | 4675 |
| 4789 | -0.04 | 0.10 | -0.44 | 0.66 | 1 | Ergic1 | endoplasmic reticulum-golgi intermediate | 3976 | 3988 | 4021 | 5987 | 4743 | 3862 | 6035 | 4854 | 5250 | 6303 | 3661 |
| 6105 | -0.09 | 0.06 | -1.53 | 0.13 | 1 | Eif4g3 | eukaryotic translation initiation factor 4 ga | 5377 | 5314 | 5400 | 6329 | 6651 | 7340 | 6034 | 5946 | 6225 | 6108 | 6430 |
| 5525 | -0.08 | 0.08 | -1.04 | 0.30 | 1 | Arpc1a | actin related protein 2/3 complex, subunit | 5898 | 6257 | 5996 | 5658 | 5910 | 4313 | 6033 | 4818 | 5663 | 6133 | 4099 |
| 4808 | 0.19 | 0.14 | 1.33 | 0.18 | 1 | Rasgrf1 | RAS protein-specific guanine nucleotide- | 324 | 571 | 690 | 5371 | 6712 | 5799 | 6031 | 6597 | 7863 | 6111 | 6820 |
| 5745 | 0.03 | 0.09 | 0.38 | 0.70 | 1 | Usp47 | ubiquitin specific peptidase 47 | 5043 | 5612 | 4825 | 6194 | 5786 | 5573 | 6025 | 5962 | 6551 | 5322 | 6296 |
| 5954 | 0.10 | 0.08 | 1.31 | 0.19 | 1 | Prkaa1 | protein kinase, AMP-activated, alpha 1 ca | 4107 | 5574 | 5135 | 6845 | 6328 | 5293 | 6021 | 6929 | 6748 | 6655 | 5857 |
| 5842 | 0.11 | 0.07 | 1.67 | 0.09 | 1 | Scfd1 | Sec1 family domain containing 1 | 3465 | 3264 | 3889 | 5464 | 7122 | 7077 | 6013 | 5454 | 7546 | 6532 | 8431 |
| 6232 | 0.08 | 0.06 | 1.22 | 0.22 | 1 | Cct2 | chaperonin containing Tcp1, subunit 2 (be | 9116 | 7756 | 8375 | 4877 | 5618 | 4636 | 6006 | 5397 | 5558 | 5941 | 5276 |
| 4517 | 0.05 | 0.07 | 0.73 | 0.47 | 1 | Srebf1 | sterol regulatory element binding transcrip | 2725 | 2145 | 2240 | 5624 | 4851 | 4365 | 5997 | 5808 | 5066 | 6581 | 4287 |
| 5809 | -0.02 | 0.09 | -0.21 | 0.84 | 1 | Preb | prolactin regulatory element binding | 3022 | 2823 | 3187 | 5108 | 7675 | 8892 | 5996 | 5023 | 7659 | 6291 | 8221 |
| 5473 | 0.03 | 0.09 | 0.35 | 0.73 | 1 | Cope | coatomer protein complex, subunit epsilo | 3691 | 3873 | 4279 | 6059 | 5337 | 6554 | 5994 | 6291 | 5517 | 6301 | 6312 |
| 5450 | 0.13 | 0.12 | 1.05 | 0.29 | 1 | Edem1 | ER degradation enhancer, mannosidase | 4737 | 3132 | 2928 | 5674 | 6225 | 5591 | 5994 | 6737 | 6356 | 6733 | 5850 |
| 5321 | 0.14 | 0.09 | 1.54 | 0.12 | 1 | Fem1b | fem 1 homolog b | 4176 | 5502 | 4532 | 5747 | 5052 | 4259 | 5992 | 7100 | 5296 | 6210 | 4663 |
| 4970 | -0.06 | 0.07 | -0.83 | 0.40 | 1 | Rfk | riboflavin kinase | 2765 | 2924 | 3400 | 6053 | 6223 | 4956 | 5990 | 5328 | 6542 | 5872 | 4620 |
| 5990 | 0.05 | 0.07 | 0.70 | 0.48 | 1 | Ankrd17 | ankyrin repeat domain 17 | 4648 | 5293 | 5408 | 5746 | 6169 | 6980 | 5989 | 6137 | 6483 | 6242 | 6795 |
| 5686 | 0.01 | 0.08 | 0.12 | 0.91 | 1 | Gnaq | guanine nucleotide binding protein, alpha | 5678 | 6607 | 6451 | 5316 | 5729 | 4814 | 5982 | 5744 | 5813 | 5481 | 4936 |
| 5365 | 0.00 | 0.09 | 0.00 | 1.00 | 1 | Neurod1 | neurogenic differentiation 1 | 2475 | 3216 | 3014 | 6010 | 6014 | 7139 | 5981 | 5544 | 5938 | 6290 | 7400 |
| 3722 | -0.01 | 0.17 | -0.03 | 0.97 | 1 | Hapln4 | hyaluronan and proteoglycan link protein | 353 | 293 | 750 | 4588 | 6258 | 2919 | 5978 | 4412 | 6096 | 6492 | 2800 |
| 6179 | 0.09 | 0.09 | 1.07 | 0.29 | 1 | Dctn1 | dynactin 1 | 5705 | 5561 | 5914 | 6087 | 6869 | 5611 | 5977 | 7802 | 6637 | 6294 | 5515 |
| 5511 | -0.05 | 0.08 | -0.59 | 0.56 | 1 | Fbxo21 | F-box protein 21 | 4202 | 5464 | 4793 | 5838 | 6141 | 5489 | 5972 | 5841 | 6237 | 5474 | 5170 |
| 5304 | -0.08 | 0.12 | -0.71 | 0.48 | 1 | Arfgap3 | ADP-ribosylation factor GTPase activating | 2246 | 2670 | 3071 | 5641 | 7506 | 6606 | 5972 | 4238 | 7653 | 5738 | 7006 |
| 6518 | -0.01 | 0.07 | -0.10 | 0.92 | 1 | Ptbp1 | polypyrimidine tract binding protein 1 | 8003 | 7706 | 7506 | 6216 | 5689 | 6405 | 5971 | 5573 | 5772 | 6648 | 6205 |
| 6369 | -0.05 | 0.07 | -0.71 | 0.48 | 1 | Hnrnpl | heterogeneous nuclear ribonucleoprotein | 6198 | 6093 | 5590 | 6672 | 6339 | 7539 | 5969 | 5793 | 6248 | 6109 | 7504 |
| 7779 | 0.02 | 0.12 | 0.19 | 0.85 | 1 | Tln1 | talin 1 | 15762 | 11052 | 10983 | 5379 | 6786 | 5577 | 5960 | 5878 | 7065 | 5818 | 5313 |
| 5248 | -0.05 | 0.06 | -0.88 | 0.38 | 1 | Stat3 | signal transducer and activator of transcri | 5775 | 4883 | 5003 | 4977 | 5646 | 4842 | 5959 | 4877 | 5562 | 5622 | 4581 |
| 5204 | 0.12 | 0.09 | 1.30 | 0.19 | 1 | Nipal1 | NIPA-like domain containing 1 | 1176 | 1733 | 1545 | 6144 | 6374 | 6827 | 5954 | 7204 | 6816 | 6466 | 7003 |
| 5306 | -0.07 | 0.09 | -0.73 | 0.47 | 1 | B230219D22Ri | RIKEN cDNA B230219D22 gene | 5558 | 5590 | 5372 | 5511 | 5253 | 4707 | 5946 | 4435 | 5823 | 5304 | 4865 |
| 5273 | 0.05 | 0.07 | 0.76 | 0.45 | 1 | Atp6v0d1 | ATPase, H+ transporting, lysosomal V0 su | 3990 | 4001 | 4566 | 5677 | 6472 | 4242 | 5935 | 6116 | 5993 | 6531 | 4484 |
| 5177 | -0.13 | 0.12 | -1.13 | 0.26 | 1 | Zfhx2 | zinc finger homeobox 2 | 1706 | 2933 | 2541 | 6823 | 6364 | 6883 | 5935 | 6167 | 6538 | 5639 | 5412 |
| 5051 | 0.01 | 0.08 | 0.08 | 0.94 | 1 | Ap2a1 | adaptor-related protein complex 2, alpha | 4064 | 3066 | 4116 | 5623 | 6049 | 4414 | 5931 | 5562 | 5990 | 6645 | 4099 |
| 4800 | -0.08 | 0.10 | -0.79 | 0.43 | 1 | Grk3 | G protein-coupled receptor kinase 3 | 2173 | 2736 | 2119 | 5645 | 5056 | 6881 | 5931 | 5047 | 5508 | 5146 | 6563 |
| 5686 | -0.01 | 0.11 | -0.05 | 0.96 | 1 | Fxyd6 | FXYD domain-containing ion transport reg | 4030 | 4968 | 6170 | 5470 | 6175 | 6183 | 5929 | 6114 | 5700 | 5888 | 5920 |
| 4872 | -0.12 | 0.07 | -1.61 | 0.11 | 1 | Nfasc | neurofascin | 3929 | 4182 | 3680 | 5922 | 5457 | 4474 | 5929 | 5446 | 5409 | 4808 | 4350 |
| 6075 | -0.14 | 0.07 | -1.89 | 0.06 | 1 | Lasp1 | LIM and SH3 protein 1 | 8601 | 8264 | 9773 | 5471 | 5467 | 4160 | 5927 | 5001 | 5037 | 5385 | 3736 |
| 4393 | 0.03 | 0.12 | 0.22 | 0.83 | 1 | Nectin3 | nectin cell adhesion molecule 3 | 1859 | 2176 | 2304 | 5177 | 4724 | 4937 | 5925 | 4361 | 5355 | 6215 | 5284 |
| 6191 | 0.00 | 0.06 | -0.05 | 0.96 | 1 | Rac1 | Rac family small GTPase 1 | 8788 | 8286 | 8250 | 5487 | 5748 | 4248 | 5894 | 5223 | 6043 | 6033 | 4104 |
| 6802 | 0.08 | 0.07 | 1.16 | 0.25 | 1 | Rdx | radixin | 9606 | 9762 | 8868 | 5413 | 5997 | 5379 | 5892 | 6028 | 5816 | 6140 | 5916 |
| 5354 | 0.04 | 0.06 | 0.63 | 0.53 | 1 | Vdac1 | voltage-dependent anion channel 1 | 3957 | 4451 | 3911 | 6191 | 5535 | 5381 | 5891 | 6506 | 5934 | 5687 | 5454 |

| | | | | | | | | | | | | | | | | | | |
|---|---|---|---|---|---|---|---|---|---|---|---|---|---|---|---|---|---|---|
| 5214 | -0.10 | 0.08 | -1.15 | 0.25 | 1 | Glyr1 | glyoxylate reductase 1 homolog (Arabido | 4470 | 4223 | 4582 | 5234 | 5630 | 5963 | 5886 | 4255 | 6078 | 5300 | 5731 |
| 4853 | -0.04 | 0.07 | -0.53 | 0.59 | 1 | Pip4k2c | phosphatidylinositol-5-phosphate 4-kinas | 3510 | 3548 | 3609 | 5810 | 5201 | 4754 | 5882 | 5191 | 5179 | 5919 | 4780 |
| 5818 | -0.10 | 0.16 | -0.63 | 0.53 | 1 | Epm2aip1 | EPM2A (laforin) interacting protein 1 | 3026 | 4937 | 3654 | 5908 | 5493 | 9644 | 5880 | 3934 | 6341 | 5353 | 9832 |
| 5598 | 0.16 | 0.11 | 1.47 | 0.14 | 1 | Stard10 | START domain containing 10 | 4459 | 4872 | 5566 | 5681 | 6158 | 4203 | 5862 | 8030 | 5866 | 6570 | 4311 |
| 6871 | 0.25 | 0.13 | 1.93 | 0.05 | 1 | Rpl6 | ribosomal protein L6 | 10183 | 10379 | 10680 | 4885 | 5180 | 4132 | 5852 | 8564 | 5251 | 5903 | 4573 |
| 5989 | 0.04 | 0.06 | 0.61 | 0.54 | 1 | Arpc2 | actin related protein 2/3 complex, subunit | 8486 | 7092 | 7286 | 5446 | 5478 | 4427 | 5850 | 5468 | 5393 | 6500 | 4454 |
| 5951 | 0.01 | 0.10 | 0.07 | 0.94 | 1 | Picalm | phosphatidylinositol binding clathrin asse | 7460 | 7090 | 5639 | 5579 | 5467 | 5667 | 5850 | 4843 | 5832 | 5849 | 6180 |
| 5761 | -0.01 | 0.06 | -0.16 | 0.87 | 1 | Hnrnpul1 | heterogeneous nuclear ribonucleoprotein | 6124 | 5831 | 6133 | 5607 | 5944 | 5317 | 5838 | 5231 | 6358 | 5631 | 5357 |
| 6606 | 0.10 | 0.09 | 1.15 | 0.25 | 1 | Rps8 | ribosomal protein S8 | 11175 | 9399 | 10306 | 5155 | 5321 | 3805 | 5836 | 6658 | 5248 | 5769 | 3995 |
| 6028 | -0.09 | 0.12 | -0.79 | 0.43 | 1 | Ywhag | tyrosine 3-monooxygenase/tryptophan 5- | 6361 | 7945 | 6885 | 6279 | 5922 | 5257 | 5827 | 4508 | 6241 | 5747 | 5341 |
| 6348 | -0.01 | 0.06 | -0.19 | 0.85 | 1 | Cbx5 | chromobox 5 | 6355 | 6770 | 6513 | 7251 | 6300 | 5853 | 5824 | 6709 | 6243 | 5831 | 6177 |
| 5194 | -0.05 | 0.08 | -0.66 | 0.51 | 1 | Kdelr1 | KDEL (Lys-Asp-Glu-Leu) endoplasmic ret | 4302 | 4356 | 4527 | 5891 | 6034 | 4601 | 5817 | 4828 | 6159 | 6241 | 4376 |
| 5332 | 0.05 | 0.09 | 0.51 | 0.61 | 1 | Ppp3cb | protein phosphatase 3, catalytic subunit, I | 4700 | 5232 | 4974 | 5466 | 5430 | 4741 | 5814 | 6937 | 5385 | 5276 | 4699 |
| 4824 | -0.01 | 0.06 | -0.16 | 0.87 | 1 | Ndrg3 | N-myc downstream regulated gene 3 | 3956 | 3570 | 3565 | 5297 | 5292 | 4630 | 5814 | 4970 | 5520 | 5960 | 4487 |
| 6368 | 0.12 | 0.11 | 1.01 | 0.31 | 1 | Tmem176b | transmembrane protein 176B | 10027 | 6892 | 7829 | 5892 | 5670 | 4277 | 5813 | 7294 | 5757 | 6318 | 4274 |
| 5610 | 0.02 | 0.06 | 0.39 | 0.70 | 1 | Polr2m | polymerase (RNA) II (DNA directed) polyp | 6565 | 6406 | 6663 | 5840 | 5352 | 3901 | 5806 | 5539 | 5612 | 5906 | 4121 |
| 4543 | -0.02 | 0.10 | -0.21 | 0.84 | 1 | Slc8a1 | solute carrier family 8 (sodium/calcium ex | 3627 | 4145 | 3711 | 4672 | 4925 | 3999 | 5796 | 4850 | 5447 | 4815 | 3982 |
| 4654 | -0.06 | 0.11 | -0.59 | 0.56 | 1 | Avl9 | AVL9 cell migration associated | 3313 | 4435 | 3241 | 5634 | 4737 | 4433 | 5792 | 5003 | 5248 | 4815 | 4542 |
| 3156 | 0.18 | 0.16 | 1.09 | 0.28 | 1 | Pdyn | prodynorphin | 929 | 1147 | 761 | 4025 | 2527 | 2304 | 5791 | 5496 | 2801 | 6823 | 2108 |
| 10701 | 0.03 | 0.10 | 0.25 | 0.80 | 1 | Tmsb4x | thymosin, beta 4, X chromosome | 33049 | 21345 | 23476 | 5123 | 4694 | 4216 | 5769 | 5613 | 4575 | 5208 | 4640 |
| 4916 | -0.13 | 0.08 | -1.59 | 0.11 | 1 | Slc12a7 | solute carrier family 12, member 7 | 2534 | 2749 | 2871 | 6420 | 6357 | 5325 | 5769 | 5838 | 6103 | 5943 | 4161 |
| 5017 | -0.05 | 0.09 | -0.52 | 0.60 | 1 | Tbc1d9b | TBC1 domain family, member 9B | 5102 | 5776 | 4895 | 5252 | 5162 | 3795 | 5755 | 5394 | 5703 | 4911 | 3442 |
| 5075 | -0.01 | 0.06 | -0.25 | 0.81 | 1 | Rab18 | RAB18, member RAS oncogene family | 4376 | 3956 | 4427 | 4966 | 5842 | 5067 | 5752 | 4676 | 5907 | 5714 | 5139 |
| 5056 | -0.06 | 0.13 | -0.48 | 0.63 | 1 | St18 | suppression of tumorigenicity 18 | 2451 | 3762 | 2829 | 6126 | 4926 | 6974 | 5749 | 4742 | 5610 | 5240 | 7203 |
| 5014 | -0.02 | 0.08 | -0.22 | 0.83 | 1 | Txnl1 | thioredoxin-like 1 | 3362 | 3225 | 3564 | 5215 | 6336 | 5302 | 5744 | 4494 | 6472 | 5626 | 5811 |
| 4967 | 0.04 | 0.11 | 0.35 | 0.72 | 1 | Ube2j1 | ubiquitin-conjugating enzyme E2J 1 | 3591 | 3924 | 3579 | 5223 | 5648 | 4851 | 5741 | 4500 | 6388 | 5991 | 5204 |
| 4375 | -0.05 | 0.09 | -0.56 | 0.57 | 1 | Dhdds | dehydrodolichyl diphosphate synthase | 2786 | 2846 | 3170 | 4298 | 5479 | 4477 | 5739 | 4069 | 5442 | 5658 | 4162 |
| 5701 | 0.01 | 0.07 | 0.13 | 0.89 | 1 | Anp32e | acidic (leucine-rich) nuclear phosphoprot | 6594 | 5554 | 6780 | 5528 | 5707 | 4869 | 5735 | 5388 | 5647 | 5677 | 5234 |
| 5397 | -0.01 | 0.09 | -0.15 | 0.88 | 1 | Mcf2l | mcf.2 transforming sequence-like | 4018 | 4529 | 4823 | 4901 | 6395 | 6113 | 5730 | 5972 | 5856 | 5617 | 5413 |
| 4408 | -0.07 | 0.10 | -0.72 | 0.47 | 1 | Scly | selenocysteine lyase | 1473 | 1829 | 2002 | 5679 | 5743 | 4976 | 5727 | 4735 | 5859 | 5742 | 4725 |
| 5879 | -0.04 | 0.17 | -0.23 | 0.82 | 1 | Gm38070 | | 364 | 1096 | 1151 | 7967 | 8974 | 8572 | 5725 | 7042 | 8659 | 5242 | 9876 |
| 5072 | 0.06 | 0.11 | 0.53 | 0.60 | 1 | Nkx2-2 | NK2 homeobox 2 | 3148 | 3477 | 3479 | 5731 | 5775 | 5066 | 5722 | 5488 | 5901 | 7385 | 4623 |
| 5746 | 0.08 | 0.07 | 1.14 | 0.25 | 1 | Stt3b | STT3, subunit of the oligosaccharyltransfe | 6716 | 5985 | 6811 | 4730 | 6215 | 4570 | 5721 | 4828 | 6725 | 6205 | 4706 |
| 6272 | 0.08 | 0.07 | 1.15 | 0.25 | 1 | Hnrnpc | heterogeneous nuclear ribonucleoprotein | 7612 | 6092 | 8065 | 5053 | 6066 | 6116 | 5720 | 5609 | 6020 | 5794 | 6849 |
| 5033 | 0.09 | 0.08 | 1.09 | 0.28 | 1 | Scrn1 | secernin 1 | 3113 | 4163 | 3865 | 5713 | 5641 | 4302 | 5716 | 6758 | 5887 | 5674 | 4535 |
| 5630 | -0.04 | 0.09 | -0.48 | 0.63 | 1 | Zmynd11 | zinc finger, MYND domain containing 11 | 4557 | 4881 | 5393 | 6049 | 5836 | 6246 | 5713 | 4830 | 6361 | 5812 | 6250 |
| 5148 | -0.01 | 0.08 | -0.11 | 0.91 | 1 | Ahcyl2 | S-adenosylhomocysteine hydrolase-like 2 | 3722 | 3778 | 3974 | 5772 | 6422 | 4673 | 5713 | 6634 | 6311 | 5277 | 4353 |
| 6609 | -0.04 | 0.07 | -0.62 | 0.53 | 1 | Srsf6 | serine and arginine-rich splicing factor 6 | 5905 | 6751 | 6022 | 5589 | 5754 | 10312 | 5712 | 5281 | 5598 | 5604 | 10166 |
| 4637 | 0.12 | 0.11 | 1.08 | 0.28 | 1 | Pgap1 | post-GPI attachment to proteins 1 | 2412 | 3150 | 3175 | 5307 | 4459 | 4778 | 5704 | 5566 | 5473 | 5660 | 5326 |
| 6075 | 0.04 | 0.06 | 0.62 | 0.54 | 1 | Hmgcr | 3-hydroxy-3-methylglutaryl-Coenzyme A r | 2477 | 3186 | 2651 | 8690 | 6663 | 7881 | 5696 | 8332 | 6677 | 6453 | 8117 |
| 5745 | 0.01 | 0.11 | 0.14 | 0.89 | 1 | Rabac1 | Rab acceptor 1 (prenylated) | 5552 | 5108 | 6507 | 6624 | 6506 | 4008 | 5696 | 7065 | 5757 | 6579 | 3789 |
| 4431 | -0.21 | 0.16 | -1.34 | 0.18 | 1 | Casr | calcium-sensing receptor | 2241 | 3157 | 2736 | 6443 | 5273 | 4432 | 5691 | 3567 | 5799 | 5251 | 4146 |
| 4288 | 0.01 | 0.12 | 0.11 | 0.91 | 1 | Kcnmb2 | potassium large conductance calcium-act | 1781 | 2582 | 2483 | 5137 | 5085 | 4149 | 5690 | 5328 | 5344 | 5409 | 4182 |
| 4994 | 0.06 | 0.09 | 0.63 | 0.53 | 1 | Npepps | aminopeptidase puromycin sensitive | 4041 | 4340 | 4160 | 4987 | 5291 | 4756 | 5684 | 6508 | 5223 | 5244 | 4700 |
| 5862 | -0.06 | 0.12 | -0.53 | 0.60 | 1 | Lrrc59 | leucine rich repeat containing 59 | 4849 | 4361 | 4029 | 5777 | 7832 | 6874 | 5683 | 4975 | 7550 | 6133 | 6418 |
| 5309 | 0.03 | 0.08 | 0.31 | 0.76 | 1 | Nf2 | neurofibromin 2 | 4421 | 3743 | 4174 | 5462 | 5984 | 5701 | 5677 | 6370 | 5906 | 5513 | 5453 |
| 5694 | -0.04 | 0.07 | -0.53 | 0.60 | 1 | Herc1 | HECT and RLD domain containing E3 ub | 4634 | 4728 | 4633 | 5131 | 5656 | 8195 | 5675 | 5146 | 5993 | 5032 | 7814 |
| 5008 | 0.03 | 0.09 | 0.39 | 0.70 | 1 | Sec11c | SEC11 homolog C, signal peptidase com | 2413 | 2399 | 2504 | 7072 | 5861 | 5029 | 5672 | 7155 | 5648 | 5781 | 5550 |
| 4889 | 0.05 | 0.06 | 0.85 | 0.40 | 1 | Mthfd1 | methylenetetrahydrofolate dehydrogenas | 2436 | 2402 | 2545 | 5957 | 5565 | 5581 | 5672 | 5720 | 6045 | 6084 | 5778 |
| 5517 | 0.05 | 0.09 | 0.54 | 0.59 | 1 | Mapk8ip1 | mitogen-activated protein kinase 8 intera | 4288 | 4957 | 4870 | 6168 | 6077 | 4876 | 5669 | 7736 | 5803 | 5465 | 4774 |
| 5007 | -0.07 | 0.10 | -0.74 | 0.46 | 1 | Cdh2 | cadherin 2 | 3610 | 4994 | 4055 | 6231 | 5453 | 4437 | 5664 | 5205 | 5876 | 5072 | 4475 |
| 6473 | 0.00 | 0.07 | 0.03 | 0.98 | 1 | Eef1g | eukaryotic translation elongation factor 1 | 10062 | 9898 | 10657 | 5354 | 5237 | 4002 | 5655 | 5624 | 4893 | 5910 | 3908 |
| 5838 | 0.00 | 0.10 | -0.02 | 0.98 | 1 | Zcchc18 | zinc finger, CCHC domain containing 18 | 6646 | 7441 | 7594 | 4895 | 5214 | 5492 | 5654 | 4550 | 4981 | 6133 | 5617 |
| 5013 | -0.04 | 0.07 | -0.51 | 0.61 | 1 | Nomo1 | nodal modulator 1 | 4616 | 4479 | 4571 | 6102 | 5738 | 3458 | 5646 | 5884 | 5227 | 6161 | 3262 |
| 5085 | 0.05 | 0.08 | 0.65 | 0.52 | 1 | Slc44a1 | solute carrier family 44, member 1 | 4677 | 3914 | 3739 | 5612 | 5515 | 4645 | 5644 | 6097 | 5619 | 5670 | 4802 |
| 5631 | 0.08 | 0.09 | 0.90 | 0.37 | 1 | Parva | parvin, alpha | 5294 | 4568 | 4607 | 6244 | 6531 | 4634 | 5634 | 6634 | 7075 | 6041 | 4678 |
| 4944 | 0.14 | 0.09 | 1.51 | 0.13 | 1 | Enah | ENAH actin regulator | 3182 | 3894 | 3637 | 5080 | 5098 | 4945 | 5634 | 5933 | 5377 | 6398 | 5209 |
| 4814 | 0.11 | 0.13 | 0.91 | 0.36 | 1 | Slc35b1 | solute carrier family 35, member B1 | 1823 | 1856 | 2425 | 5404 | 5932 | 5536 | 5632 | 6941 | 5214 | 6594 | 5602 |
| 5661 | 0.03 | 0.07 | 0.47 | 0.64 | 1 | Ip6k1 | inositol hexaphosphate kinase 1 | 5485 | 5572 | 5033 | 5549 | 5637 | 6014 | 5631 | 5607 | 5784 | 6268 | 5694 |
| 5466 | -0.12 | 0.10 | -1.21 | 0.22 | 1 | Bptf | bromodomain PHD finger transcription fa | 5522 | 5603 | 4912 | 6076 | 5238 | 5980 | 5631 | 5729 | 5802 | 4526 | 5110 |

| | | | | | | | | | | | | | | | | | | |
|---|---|---|---|---|---|---|---|---|---|---|---|---|---|---|---|---|---|---|
| 5309 | -0.05 | 0.07 | -0.77 | 0.44 | 1 | Golga2 | golgi autoantigen, golgin subfamily a, 2 | 4543 | 4602 | 4287 | 5627 | 5687 | 5952 | 5625 | 5137 | 5738 | 5749 | 5459 |
| 8554 | -0.13 | 0.08 | -1.63 | 0.10 | 1 | Jund | jun D proto-oncogene | 17162 | 13810 | 14224 | 4881 | 5530 | 9695 | 5622 | 5140 | 4541 | 5526 | 7961 |
| 4705 | -0.07 | 0.09 | -0.72 | 0.47 | 1 | Snap25 | synaptosomal-associated protein 25 | 2520 | 2728 | 2684 | 5123 | 4734 | 6846 | 5621 | 4226 | 4847 | 5248 | 7183 |
| 5632 | -0.09 | 0.08 | -1.08 | 0.28 | 1 | Baz1b | bromodomain adjacent to zinc finger dom | 5322 | 5052 | 4714 | 5964 | 5567 | 6983 | 5611 | 5058 | 5874 | 5163 | 6648 |
| 5507 | 0.10 | 0.10 | 1.04 | 0.30 | 1 | Metap2 | methionine aminopeptidase 2 | 5950 | 5472 | 6085 | 4819 | 5228 | 5108 | 5597 | 6410 | 5282 | 5008 | 5616 |
| 4753 | -0.05 | 0.07 | -0.65 | 0.51 | 1 | Zmpste24 | zinc metallopeptidase, STE24 | 3332 | 3491 | 3646 | 5156 | 5986 | 4517 | 5597 | 4792 | 5993 | 5080 | 4688 |
| 4344 | -0.08 | 0.09 | -0.95 | 0.34 | 1 | Aldh18a1 | aldehyde dehydrogenase 18 family, mem | 3325 | 3364 | 3419 | 4420 | 5730 | 3603 | 5590 | 4233 | 5557 | 5299 | 3247 |
| 3422 | -0.08 | 0.20 | -0.40 | 0.69 | 1 | Ptprt | protein tyrosine phosphatase, receptor typ | 2322 | 2975 | 2182 | 2956 | 2881 | 4065 | 5586 | 2288 | 3181 | 5531 | 3673 |
| 5595 | 0.09 | 0.11 | 0.80 | 0.43 | 1 | Celf1 | CUGBP, Elav-like family member 1 | 4807 | 5766 | 5865 | 4662 | 5224 | 6416 | 5585 | 6444 | 5352 | 5201 | 6218 |
| 6595 | 0.02 | 0.08 | 0.23 | 0.82 | 1 | Smarca2 | SWI/SNF related, matrix associated, actin | 8309 | 9772 | 8276 | 5508 | 5690 | 6193 | 5579 | 5591 | 5832 | 5906 | 5892 |
| 5474 | -0.10 | 0.10 | -0.97 | 0.33 | 1 | Tent5c | terminal nucleotidyltransferase 5C | 5389 | 6452 | 4238 | 6918 | 5100 | 5273 | 5575 | 6179 | 5699 | 4692 | 4703 |
| 5279 | 0.13 | 0.09 | 1.46 | 0.14 | 1 | Nap1l1 | nucleosome assembly protein 1-like 1 | 5612 | 4569 | 4931 | 4835 | 5186 | 4987 | 5560 | 5679 | 5903 | 5157 | 5648 |
| 5359 | 0.11 | 0.08 | 1.39 | 0.16 | 1 | Dnajb11 | DnaJ heat shock protein family (Hsp40) m | 3514 | 2540 | 3475 | 5239 | 6866 | 6126 | 5559 | 5565 | 7102 | 6357 | 6612 |
| 5080 | 0.14 | 0.11 | 1.23 | 0.22 | 1 | Prkcsh | protein kinase C substrate 80K-H | 3840 | 3156 | 3579 | 5968 | 5638 | 4313 | 5541 | 7794 | 5446 | 6102 | 4502 |
| 4606 | -0.03 | 0.09 | -0.34 | 0.73 | 1 | Acadsb | acyl-Coenzyme A dehydrogenase, short/b | 3444 | 4235 | 3255 | 5151 | 4751 | 4642 | 5532 | 5297 | 4755 | 4611 | 4988 |
| 4273 | -0.05 | 0.10 | -0.51 | 0.61 | 1 | Oxr1 | oxidation resistance 1 | 2759 | 2617 | 2976 | 4974 | 4374 | 4762 | 5531 | 4171 | 4707 | 5111 | 5019 |
| 4956 | 0.02 | 0.11 | 0.19 | 0.85 | 1 | Klhdc10 | kelch domain containing 10 | 3849 | 5643 | 4380 | 4944 | 5383 | 4198 | 5526 | 6283 | 5372 | 4927 | 4013 |
| 5097 | -0.01 | 0.07 | -0.08 | 0.93 | 1 | Mlf2 | myeloid leukemia factor 2 | 4773 | 4500 | 5117 | 5391 | 5636 | 4324 | 5524 | 5310 | 5261 | 5902 | 4327 |
| 5170 | 0.07 | 0.08 | 0.90 | 0.37 | 1 | Chmp4b | charged multivesicular body protein 4B | 5736 | 4484 | 5286 | 4912 | 5602 | 4174 | 5521 | 5465 | 5104 | 5929 | 4662 |
| 5903 | -0.03 | 0.07 | -0.41 | 0.68 | 1 | Abr | active BCR-related gene | 5862 | 6658 | 6245 | 6438 | 5814 | 5527 | 5520 | 5870 | 6345 | 5744 | 4909 |
| 4827 | 0.10 | 0.08 | 1.33 | 0.18 | 1 | Derl2 | Der1-like domain family, member 2 | 2097 | 2064 | 2236 | 4592 | 6594 | 5830 | 5513 | 5134 | 6771 | 5868 | 6397 |
| 4632 | -0.10 | 0.06 | -1.74 | 0.08 | 1 | Kif13b | kinesin family member 13B | 2490 | 2642 | 2672 | 5285 | 5656 | 5906 | 5513 | 5266 | 5243 | 4999 | 5279 |
| 4987 | 0.09 | 0.08 | 1.10 | 0.27 | 1 | Pgrmc2 | progesterone receptor membrane compo | 3593 | 3718 | 4126 | 5410 | 5919 | 4210 | 5510 | 5574 | 5759 | 6579 | 4455 |
| 5835 | 0.00 | 0.06 | -0.03 | 0.97 | 1 | Plekha6 | pleckstrin homology domain containing, fa | 3929 | 4525 | 4526 | 6013 | 6358 | 7805 | 5509 | 6384 | 6443 | 5655 | 7038 |
| 5282 | -0.16 | 0.09 | -1.74 | 0.08 | 1 | Zswim8 | zinc finger SWIM-type containing 8 | 3885 | 4963 | 4566 | 5859 | 5785 | 6427 | 5499 | 4871 | 5707 | 5266 | 5275 |
| 4649 | -0.11 | 0.10 | -1.09 | 0.28 | 1 | Acox1 | acyl-Coenzyme A oxidase 1, palmitoyl | 3433 | 4414 | 4186 | 5253 | 5198 | 4352 | 5498 | 4346 | 5349 | 4832 | 4274 |
| 5208 | 0.04 | 0.08 | 0.44 | 0.66 | 1 | Rundc3a | RUN domain containing 3A | 3019 | 3947 | 3596 | 5525 | 5810 | 6246 | 5488 | 6334 | 5688 | 5580 | 6059 |
| 5391 | 0.01 | 0.09 | 0.17 | 0.87 | 1 | Erbin | Erbb2 interacting protein | 5683 | 6315 | 5089 | 5340 | 4965 | 5219 | 5485 | 5008 | 5557 | 5012 | 5626 |
| 4631 | -0.13 | 0.10 | -1.22 | 0.22 | 1 | Tulp4 | tubby like protein 4 | 3230 | 3960 | 3256 | 5374 | 5224 | 5045 | 5477 | 4360 | 5216 | 5360 | 4436 |
| 6414 | 0.09 | 0.08 | 1.03 | 0.31 | 1 | Sfpq | splicing factor proline/glutamine rich (poly | 6512 | 6533 | 6425 | 5849 | 5045 | 8336 | 5476 | 6958 | 5021 | 5352 | 9049 |
| 5106 | -0.01 | 0.07 | -0.17 | 0.86 | 1 | Trpc4ap | transient receptor potential cation channe | 4929 | 4267 | 4687 | 5186 | 5524 | 5029 | 5467 | 5493 | 5291 | 5680 | 4616 |
| 6153 | -0.13 | 0.10 | -1.27 | 0.21 | 1 | Tns3 | tensin 3 | 8531 | 7466 | 6479 | 6046 | 5573 | 6553 | 5457 | 5890 | 5462 | 4955 | 5277 |
| 4547 | 0.26 | 0.32 | 0.82 | 0.41 | 1 | Trpm5 | transient receptor potential cation channe | 257 | 158 | 571 | 5471 | 6622 | 4950 | 5442 | 7475 | 6715 | 6123 | 6230 |
| 4858 | 0.05 | 0.08 | 0.66 | 0.51 | 1 | Col4a3bp | collagen, type IV, alpha 3 (Goodpasture a | 3519 | 3807 | 3664 | 5594 | 4569 | 5203 | 5438 | 6220 | 4888 | 4942 | 5594 |
| 4652 | -0.06 | 0.14 | -0.39 | 0.69 | 1 | R3hdm2 | R3H domain containing 2 | 3888 | 4303 | 4365 | 4835 | 5441 | 3816 | 5438 | 3425 | 5774 | 6251 | 3636 |
| 5539 | 0.02 | 0.08 | 0.20 | 0.84 | 1 | Pttg1ip | pituitary tumor-transforming 1 interacting p | 6643 | 5766 | 6602 | 5246 | 6023 | 4152 | 5434 | 6036 | 5474 | 5329 | 4219 |
| 4716 | 0.03 | 0.10 | 0.24 | 0.81 | 1 | Pcmtd1 | protein-L-isoaspartate (D-aspartate) O-me | 3390 | 4356 | 3397 | 5338 | 4863 | 4605 | 5415 | 4781 | 5501 | 5326 | 4905 |
| 6198 | -0.05 | 0.08 | -0.60 | 0.55 | 1 | Anxa6 | annexin A6 | 10826 | 9007 | 9670 | 5241 | 5062 | 3934 | 5413 | 5417 | 4912 | 4910 | 3789 |
| 5046 | -0.05 | 0.10 | -0.48 | 0.63 | 1 | Lnpep | leucyl/cystinyl aminopeptidase | 4573 | 4045 | 4093 | 5368 | 5388 | 5559 | 5411 | 5135 | 5989 | 4432 | 5518 |
| 6509 | 0.11 | 0.10 | 1.00 | 0.31 | 1 | Tcp1 | t-complex protein 1 | 9042 | 8379 | 8786 | 5622 | 5764 | 5023 | 5407 | 7402 | 5250 | 5353 | 5571 |
| 5803 | -0.07 | 0.08 | -0.89 | 0.37 | 1 | Rpl9 | ribosomal protein L9 | 9778 | 9113 | 10454 | 3871 | 4781 | 3682 | 5398 | 4003 | 4378 | 4641 | 3737 |
| 4717 | 0.04 | 0.07 | 0.50 | 0.61 | 1 | Gsk3b | glycogen synthase kinase 3 beta | 3983 | 3826 | 3685 | 4897 | 4980 | 4646 | 5394 | 5619 | 5244 | 5291 | 4324 |
| 6018 | 0.12 | 0.11 | 1.16 | 0.25 | 1 | Hnrnpm | heterogeneous nuclear ribonucleoprotein | 7238 | 5965 | 7443 | 4711 | 5070 | 6708 | 5392 | 6381 | 5208 | 5313 | 6766 |
| 4325 | -0.11 | 0.10 | -1.12 | 0.26 | 1 | Pcnx | pecanex homolog | 3793 | 3736 | 3209 | 4767 | 4304 | 4690 | 5387 | 3732 | 4738 | 4920 | 4303 |
| 5816 | 0.08 | 0.14 | 0.61 | 0.54 | 1 | Sod1 | superoxide dismutase 1, soluble | 7413 | 6144 | 8389 | 4893 | 5326 | 4781 | 5385 | 6822 | 4595 | 5438 | 4786 |
| 5524 | -0.06 | 0.07 | -0.83 | 0.41 | 1 | Bag6 | BCL2-associated athanogene 6 | 5673 | 6094 | 5847 | 5513 | 6015 | 5062 | 5385 | 5248 | 5698 | 5628 | 4604 |
| 5136 | -0.02 | 0.08 | -0.26 | 0.79 | 1 | Csnk1d | casein kinase 1, delta | 4961 | 5471 | 4481 | 5780 | 5081 | 4686 | 5385 | 5101 | 4975 | 5970 | 4600 |
| 4879 | -0.01 | 0.08 | -0.16 | 0.87 | 1 | Ubqln1 | ubiquilin 1 | 4202 | 4064 | 3514 | 5237 | 5752 | 4657 | 5383 | 5012 | 5606 | 5695 | 4550 |
| 4743 | -0.05 | 0.07 | -0.70 | 0.48 | 1 | Bcr | BCR activator of RhoGEF and GTPase | 3657 | 3937 | 3344 | 5556 | 5228 | 4745 | 5383 | 5632 | 5168 | 5470 | 4058 |
| 5045 | 0.06 | 0.10 | 0.59 | 0.56 | 1 | Slc25a39 | solute carrier family 25, member 39 | 4466 | 5196 | 5619 | 5607 | 5221 | 3421 | 5379 | 5984 | 5207 | 6112 | 3286 |
| 4826 | 0.00 | 0.07 | 0.02 | 0.98 | 1 | Tusc3 | tumor suppressor candidate 3 | 3911 | 4292 | 4353 | 5380 | 5279 | 4248 | 5375 | 5350 | 4912 | 5419 | 4568 |
| 4578 | -0.02 | 0.07 | -0.22 | 0.83 | 1 | Cmip | c-Maf inducing protein | 2916 | 3459 | 3445 | 5601 | 5322 | 4034 | 5372 | 5897 | 5194 | 5361 | 3756 |
| 4675 | -0.09 | 0.12 | -0.76 | 0.45 | 1 | Dennd11 | DENN domain containing 11 | 3830 | 4665 | 3817 | 5160 | 4737 | 4888 | 5367 | 3989 | 5305 | 4852 | 4811 |
| 6451 | 0.02 | 0.08 | 0.25 | 0.80 | 1 | Ctnna1 | catenin (cadherin associated protein), alp | 9829 | 8219 | 8812 | 5540 | 5826 | 5163 | 5365 | 5066 | 5843 | 5603 | 5698 |
| 4305 | -0.01 | 0.08 | -0.17 | 0.87 | 1 | Vwa8 | von Willebrand factor A domain containin | 2550 | 2763 | 2772 | 5221 | 4844 | 4243 | 5357 | 5313 | 5483 | 5008 | 3805 |
| 4508 | 0.06 | 0.06 | 0.89 | 0.37 | 1 | Sgpp1 | sphingosine-1-phosphate phosphatase 1 | 3577 | 3568 | 3573 | 5311 | 4725 | 3674 | 5355 | 5297 | 5055 | 5612 | 3839 |
| 6149 | 0.05 | 0.12 | 0.42 | 0.67 | 1 | Zbtb20 | zinc finger and BTB domain containing 20 | 7085 | 8001 | 7614 | 5032 | 5611 | 5966 | 5348 | 7020 | 6340 | 4655 | 4966 |
| 4679 | -0.01 | 0.09 | -0.13 | 0.90 | 1 | Srpk2 | serine/arginine-rich protein specific kinas | 2509 | 3062 | 2776 | 4874 | 4978 | 6381 | 5343 | 4793 | 5142 | 4758 | 6849 |
| 4764 | 0.01 | 0.07 | 0.15 | 0.88 | 1 | Tmem167b | transmembrane protein 167B | 3127 | 3488 | 3348 | 5402 | 5654 | 4753 | 5341 | 4971 | 5818 | 5356 | 5140 |
| 4896 | 0.11 | 0.10 | 1.12 | 0.26 | 1 | Atp5g3 | ATP synthase, H+ transporting, mitochond | 5142 | 4253 | 5140 | 4819 | 5008 | 3718 | 5334 | 6141 | 4880 | 5536 | 3885 |

| | | | | | | | | | | | | | | | | | |
|---|---|---|---|---|---|---|---|---|---|---|---|---|---|---|---|---|---|
| 4412 | -0.06 | 0.07 | -0.91 | 0.36 | 1 | Znfx1 | zinc finger, NFX1-type containing 1 | 3761 | 4132 | 3975 | 4989 | 4900 | 3471 | 5332 | 5006 | 4863 | 4866 | 3232 |
| 4273 | -0.13 | 0.08 | -1.56 | 0.12 | 1 | Rab22a | RAB22A, member RAS oncogene family | 2795 | 2918 | 2627 | 5135 | 5160 | 4586 | 5331 | 4236 | 5166 | 4712 | 4341 |
| 5273 | 0.05 | 0.07 | 0.74 | 0.46 | 1 | Spop | speckle-type BTB/POZ protein | 5718 | 5402 | 5595 | 5163 | 5202 | 4589 | 5329 | 5811 | 5218 | 5072 | 4906 |
| 4371 | 0.07 | 0.09 | 0.87 | 0.39 | 1 | Fam234a | family with sequence similarity 234, meml | 2067 | 2204 | 2282 | 5494 | 5255 | 4018 | 5329 | 6455 | 5491 | 5838 | 3651 |
| 4633 | 0.04 | 0.14 | 0.31 | 0.76 | 1 | Vil1 | villin 1 | 540 | 927 | 1247 | 6494 | 6396 | 5515 | 5325 | 7012 | 6652 | 5360 | 5501 |
| 5117 | 0.02 | 0.09 | 0.25 | 0.80 | 1 | Ipo7 | importin 7 | 4508 | 4697 | 3899 | 5144 | 5625 | 5318 | 5323 | 5070 | 5892 | 5078 | 5730 |
| 4698 | 0.03 | 0.08 | 0.41 | 0.68 | 1 | Pitpnb | phosphatidylinositol transfer protein, beta | 3989 | 3735 | 4385 | 4772 | 5428 | 3996 | 5319 | 5500 | 5321 | 5519 | 3708 |
| 4554 | 0.13 | 0.09 | 1.53 | 0.13 | 1 | Gde1 | glycerophosphodiester phosphodiesteras | 2304 | 2308 | 2467 | 5763 | 5088 | 4345 | 5311 | 6067 | 5187 | 6518 | 4735 |
| 4644 | 0.12 | 0.11 | 1.10 | 0.27 | 1 | Cad | carbamoyl-phosphate synthetase 2, aspa | 2331 | 1510 | 2579 | 5267 | 6208 | 4546 | 5306 | 5729 | 6625 | 6416 | 4571 |
| 4821 | -0.05 | 0.11 | -0.43 | 0.67 | 1 | Dicer1 | dicer 1, ribonuclease type III | 3258 | 4036 | 3061 | 5202 | 4983 | 6146 | 5303 | 4473 | 5354 | 4710 | 6503 |
| 4907 | 0.07 | 0.08 | 0.85 | 0.39 | 1 | Inpp4a | inositol polyphosphate-4-phosphatase, ty | 1765 | 2161 | 1982 | 5917 | 4730 | 7582 | 5302 | 5772 | 5350 | 5896 | 7516 |
| 6050 | 0.12 | 0.08 | 1.53 | 0.13 | 1 | Tapbp | TAP binding protein | 8117 | 6791 | 7622 | 5450 | 6058 | 4226 | 5299 | 6342 | 6321 | 6102 | 4228 |
| 4652 | 0.20 | 0.11 | 1.85 | 0.06 | 1 | Tent5a | terminal nucleotidyltransferase 5A | 2411 | 3158 | 2367 | 6154 | 3831 | 4603 | 5299 | 8679 | 4323 | 5303 | 5048 |
| 4217 | -0.08 | 0.11 | -0.68 | 0.50 | 1 | Cxxc4 | CXXC finger 4 | 2103 | 2521 | 2507 | 4849 | 4945 | 5002 | 5299 | 3882 | 5416 | 4978 | 4879 |
| 5139 | -0.03 | 0.08 | -0.40 | 0.69 | 1 | Nek9 | NIMA (never in mitosis gene a)-related ex | 5924 | 5551 | 4820 | 5054 | 5255 | 4720 | 5294 | 4828 | 5210 | 5257 | 4614 |
| 4842 | 0.01 | 0.09 | 0.13 | 0.90 | 1 | Fbxo3 | F-box protein 3 | 3920 | 4818 | 3711 | 5386 | 4570 | 5113 | 5285 | 4803 | 4915 | 5365 | 5379 |
| 5017 | 0.04 | 0.06 | 0.59 | 0.56 | 1 | Retreg2 | reticulophagy regulator family member 2 | 4615 | 4929 | 4760 | 5333 | 5551 | 3979 | 5282 | 5961 | 5715 | 5147 | 3920 |
| 5576 | -0.13 | 0.09 | -1.48 | 0.14 | 1 | Epcam | epithelial cell adhesion molecule | 5579 | 6892 | 8273 | 5552 | 5952 | 4482 | 5281 | 5022 | 5113 | 4861 | 4332 |
| 5386 | -0.09 | 0.09 | -0.96 | 0.34 | 1 | Ubap2l | ubiquitin-associated protein 2-like | 5341 | 5014 | 4514 | 5365 | 5675 | 6554 | 5281 | 4453 | 5765 | 5406 | 5871 |
| 5382 | -0.01 | 0.07 | -0.13 | 0.89 | 1 | Ssrp1 | structure specific recognition protein 1 | 6720 | 5511 | 5993 | 4977 | 5554 | 4750 | 5281 | 5029 | 5391 | 5213 | 4785 |
| 4673 | 0.13 | 0.10 | 1.33 | 0.18 | 1 | Lars | leucyl-tRNA synthetase | 3058 | 2842 | 3196 | 5015 | 5262 | 4636 | 5272 | 6635 | 5285 | 5205 | 4995 |
| 4763 | 0.06 | 0.09 | 0.66 | 0.51 | 1 | Pdpk1 | 3-phosphoinositide dependent protein kir | 3245 | 3867 | 3236 | 4941 | 4475 | 5961 | 5269 | 5446 | 5305 | 5067 | 5580 |
| 5348 | 0.09 | 0.07 | 1.24 | 0.21 | 1 | Usp48 | ubiquitin specific peptidase 48 | 3134 | 3354 | 3800 | 4782 | 5475 | 8086 | 5265 | 5691 | 5815 | 5530 | 7895 |
| 4158 | 0.12 | 0.07 | 1.70 | 0.09 | 1 | Wdr7 | WD repeat domain 7 | 2054 | 1949 | 2193 | 5166 | 4915 | 3615 | 5244 | 5196 | 5647 | 5858 | 3898 |
| 4791 | 0.01 | 0.11 | 0.08 | 0.94 | 1 | Atf2 | activating transcription factor 2 | 3189 | 4583 | 4194 | 5182 | 5369 | 4503 | 5243 | 4712 | 5910 | 5076 | 4735 |
| 4646 | 0.04 | 0.11 | 0.33 | 0.74 | 1 | Zfp871 | zinc finger protein 871 | 3005 | 3457 | 4067 | 4351 | 4481 | 5941 | 5243 | 5123 | 5438 | 4391 | 5604 |
| 5139 | 0.04 | 0.08 | 0.49 | 0.63 | 1 | Pbrm1 | polybromo 1 | 4477 | 5290 | 4626 | 5363 | 5244 | 4854 | 5242 | 6457 | 5537 | 4714 | 4729 |
| 5031 | 0.14 | 0.08 | 1.79 | 0.07 | 1 | Psmd1 | proteasome (prosome, macropain) 26S su | 4971 | 4033 | 5200 | 5026 | 5230 | 4056 | 5236 | 5661 | 5550 | 5947 | 4426 |
| 3854 | 0.14 | 0.18 | 0.79 | 0.43 | 1 | Enho | energy homeostasis associated | 511 | 736 | 1052 | 5109 | 5481 | 3180 | 5232 | 7159 | 4878 | 5795 | 3263 |
| 5497 | -0.02 | 0.06 | -0.27 | 0.79 | 1 | Ehmt2 | euchromatic histone lysine N-methyltrans | 6100 | 5664 | 5724 | 5167 | 5760 | 5467 | 5228 | 5270 | 5230 | 5577 | 5283 |
| 4038 | 0.01 | 0.08 | 0.12 | 0.90 | 1 | 6430548M08Ri | RIKEN cDNA 6430548M08 gene | 2029 | 1777 | 1914 | 4283 | 5622 | 4122 | 5224 | 4016 | 5786 | 5433 | 4208 |
| 4600 | -0.11 | 0.06 | -1.75 | 0.08 | 1 | Ankrd40 | ankyrin repeat domain 40 | 3390 | 3525 | 4144 | 5721 | 5940 | 3709 | 5222 | 5146 | 5400 | 4757 | 3647 |
| 4907 | -0.06 | 0.09 | -0.73 | 0.46 | 1 | Uqcrc2 | ubiquinol cytochrome c reductase core pr | 5514 | 5477 | 5419 | 4660 | 5081 | 4218 | 5221 | 3918 | 4740 | 5516 | 4220 |
| 6757 | 0.01 | 0.09 | 0.08 | 0.94 | 1 | Luc7l2 | LUC7-like 2 (S. cerevisiae) | 5093 | 6441 | 5261 | 5464 | 5941 | 12097 | 5219 | 5258 | 5981 | 5467 | 12110 |
| 4331 | -0.07 | 0.11 | -0.67 | 0.50 | 1 | Mtmr7 | myotubularin related protein 7 | 2466 | 3125 | 2819 | 5598 | 4817 | 4532 | 5208 | 4208 | 5072 | 5150 | 4648 |
| 4241 | -0.03 | 0.07 | -0.41 | 0.68 | 1 | Ncoa1 | nuclear receptor coactivator 1 | 3233 | 3382 | 3000 | 4706 | 4479 | 4324 | 5205 | 4565 | 4764 | 4851 | 4144 |
| 4802 | 0.03 | 0.06 | 0.51 | 0.61 | 1 | Mtmr3 | myotubularin related protein 3 | 4447 | 4229 | 3865 | 5390 | 4779 | 4532 | 5204 | 5310 | 4868 | 5744 | 4459 |
| 4472 | -0.05 | 0.10 | -0.49 | 0.63 | 1 | Tmtc3 | transmembrane and tetratricopeptide repe | 1707 | 1903 | 1863 | 5612 | 5377 | 5947 | 5200 | 4458 | 5940 | 4820 | 6366 |
| 3947 | -0.18 | 0.12 | -1.43 | 0.15 | 1 | Ell2 | elongation factor RNA polymerase II 2 | 2485 | 2951 | 2259 | 5102 | 4108 | 4577 | 5199 | 3520 | 4405 | 4463 | 4344 |
| 4732 | 0.09 | 0.10 | 0.97 | 0.33 | 1 | Atp6v1h | ATPase, H+ transporting, lysosomal V1 su | 3946 | 3825 | 4310 | 4894 | 5366 | 3882 | 5198 | 6290 | 4961 | 5203 | 4178 |
| 5113 | -0.02 | 0.07 | -0.24 | 0.81 | 1 | Atp6v0b | ATPase, H+ transporting, lysosomal V0 su | 2876 | 3122 | 3269 | 6170 | 5957 | 6320 | 5194 | 5888 | 5322 | 5698 | 6428 |
| 6073 | -0.09 | 0.07 | -1.24 | 0.22 | 1 | Gnb2 | guanine nucleotide binding protein (G prc | 8789 | 7746 | 9163 | 5023 | 5916 | 5030 | 5193 | 4668 | 5184 | 5683 | 4404 |
| 5304 | 0.13 | 0.09 | 1.44 | 0.15 | 1 | Rtcb | RNA 2',3'-cyclic phosphate and 5'-OH liga | 5682 | 5547 | 6087 | 5114 | 5080 | 4205 | 5192 | 6315 | 4845 | 5500 | 4783 |
| 3927 | -0.10 | 0.12 | -0.89 | 0.37 | 1 | Vps13b | vacuolar protein sorting 13B | 3235 | 3125 | 2898 | 3917 | 4015 | 4450 | 5190 | 3266 | 4493 | 3960 | 4648 |
| 4724 | 0.15 | 0.09 | 1.63 | 0.10 | 1 | Chmp5 | charged multivesicular body protein 5 | 4207 | 4081 | 4433 | 4510 | 4889 | 4032 | 5183 | 5953 | 4803 | 5235 | 4634 |
| 4768 | 0.14 | 0.09 | 1.61 | 0.11 | 1 | Efr3a | EFR3 homolog A | 4861 | 4784 | 4353 | 4758 | 4678 | 3685 | 5174 | 6037 | 5218 | 4789 | 4112 |
| 4718 | -0.06 | 0.09 | -0.64 | 0.52 | 1 | Dnaja4 | DnaJ heat shock protein family (Hsp40) m | 2498 | 3155 | 3559 | 5125 | 5262 | 6195 | 5172 | 5035 | 5412 | 4757 | 5730 |
| 4695 | -0.05 | 0.10 | -0.49 | 0.62 | 1 | Dnm2 | dynamin 2 | 3834 | 3504 | 3716 | 4755 | 4709 | 5948 | 5167 | 3835 | 5065 | 5767 | 5342 |
| 4729 | 0.11 | 0.09 | 1.22 | 0.22 | 1 | Bpnt1 | bisphosphate 3'-nucleotidase 1 | 1790 | 2226 | 2401 | 6555 | 5667 | 4590 | 5163 | 6131 | 6168 | 6198 | 5128 |
| 3884 | 0.00 | 0.12 | 0.03 | 0.98 | 1 | Mboat2 | membrane bound O-acyltransferase dom | 987 | 1244 | 1596 | 4892 | 4772 | 4545 | 5154 | 4479 | 5573 | 5072 | 4409 |
| 4972 | 0.07 | 0.08 | 0.85 | 0.39 | 1 | Ppp6r1 | protein phosphatase 6, regulatory subuni | 4576 | 3766 | 4247 | 4866 | 5145 | 5370 | 5153 | 5814 | 5116 | 5262 | 5380 |
| 4986 | 0.13 | 0.08 | 1.78 | 0.08 | 1 | Gna11 | guanine nucleotide binding protein, alpha | 4305 | 4471 | 4927 | 5127 | 5369 | 3925 | 5147 | 6001 | 5577 | 6023 | 3979 |
| 5163 | 0.09 | 0.11 | 0.84 | 0.40 | 1 | Psmb4 | proteasome (prosome, macropain) subun | 5967 | 4923 | 6266 | 4831 | 5264 | 3948 | 5143 | 6148 | 4792 | 5400 | 4110 |
| 6412 | 0.06 | 0.07 | 0.86 | 0.39 | 1 | Eif4h | eukaryotic translation initiation factor 4H | 8945 | 8178 | 8876 | 5349 | 5720 | 5607 | 5142 | 5649 | 5874 | 5880 | 5306 |
| 5487 | 0.09 | 0.09 | 1.03 | 0.30 | 1 | Nr3c1 | nuclear receptor subfamily 3, group C, me | 7118 | 6258 | 5377 | 5378 | 4772 | 4853 | 5134 | 6163 | 4982 | 5021 | 5299 |
| 5014 | 0.07 | 0.08 | 0.83 | 0.41 | 1 | Rab37 | RAB37, member RAS oncogene family | 2249 | 3299 | 3116 | 6357 | 5802 | 5392 | 5134 | 7116 | 5689 | 5284 | 5713 |
| 6077 | 0.04 | 0.08 | 0.43 | 0.67 | 1 | Rpl7a | ribosomal protein L7A | 9632 | 8234 | 10198 | 5263 | 4892 | 3881 | 5133 | 5136 | 4968 | 5418 | 4097 |
| 4908 | 0.00 | 0.08 | 0.06 | 0.95 | 1 | Clint1 | clathrin interactor 1 | 3837 | 3944 | 4245 | 4672 | 5721 | 5394 | 5128 | 4342 | 6150 | 5318 | 5241 |
| 4277 | -0.19 | 0.10 | -1.84 | 0.07 | 1 | Ubr2 | ubiquitin protein ligase E3 component n-r | 2732 | 2928 | 2757 | 4930 | 5115 | 5376 | 5125 | 3638 | 5237 | 4596 | 4617 |
| 4148 | 0.03 | 0.10 | 0.27 | 0.79 | 1 | Man2b2 | mannosidase 2, alpha B2 | 4245 | 4290 | 3601 | 4008 | 4477 | 2980 | 5122 | 4282 | 4949 | 4758 | 2914 |

| | | | | | | | | | | | | | | | | | | |
|---|---|---|---|---|---|---|---|---|---|---|---|---|---|---|---|---|---|---|
| 4233 | -0.05 | 0.09 | -0.60 | 0.55 | 1 | Ncald | neurocalcin delta | 3958 | 4418 | 4194 | 4341 | 4295 | 3588 | 5121 | 4209 | 3963 | 4721 | 3755 |
| 5413 | -0.05 | 0.07 | -0.65 | 0.52 | 1 | Anxa4 | annexin A4 | 5449 | 5351 | 6059 | 5068 | 5639 | 5788 | 5116 | 4463 | 5883 | 4695 | 6034 |
| 4282 | 0.10 | 0.06 | 1.74 | 0.08 | 1 | Efcab14 | EF-hand calcium binding domain 14 | 3162 | 3000 | 2739 | 4761 | 4786 | 3759 | 5113 | 5049 | 5075 | 5495 | 4157 |
| 5194 | 0.11 | 0.09 | 1.24 | 0.21 | 1 | Ptpa | protein phosphatase 2 protein activator | 5630 | 5168 | 5745 | 4949 | 5336 | 4042 | 5112 | 6437 | 5290 | 5391 | 4031 |
| 4919 | 0.04 | 0.09 | 0.38 | 0.70 | 1 | Sympk | symplekin | 4960 | 4071 | 4413 | 5331 | 5462 | 4102 | 5107 | 6117 | 5393 | 5352 | 3801 |
| 5063 | 0.05 | 0.08 | 0.62 | 0.54 | 1 | Ampd2 | adenosine monophosphate deaminase 2 | 2935 | 2649 | 3193 | 4997 | 5952 | 7030 | 5106 | 4749 | 5964 | 6158 | 6958 |
| 4356 | 0.06 | 0.10 | 0.59 | 0.55 | 1 | Rragc | Ras-related GTP binding C | 3458 | 3818 | 3930 | 5062 | 4544 | 3296 | 5106 | 4450 | 4692 | 5771 | 3786 |
| 5836 | 0.08 | 0.07 | 1.08 | 0.28 | 1 | Rbbp7 | retinoblastoma binding protein 7, chroma | 8103 | 7397 | 8212 | 4836 | 5280 | 4492 | 5105 | 5562 | 5007 | 5208 | 4992 |
| 4333 | -0.15 | 0.10 | -1.47 | 0.14 | 1 | Ralgapb | Ral GTPase activating protein, beta subu | 3770 | 3712 | 3833 | 4671 | 4678 | 4604 | 5103 | 3596 | 5188 | 4166 | 4342 |
| 4330 | 0.07 | 0.09 | 0.78 | 0.44 | 1 | Mfn2 | mitofusin 2 | 3412 | 3350 | 3102 | 4386 | 5245 | 3615 | 5101 | 5516 | 4913 | 5544 | 3444 |
| 5442 | 0.04 | 0.09 | 0.51 | 0.61 | 1 | Msi2 | musashi RNA-binding protein 2 | 3596 | 4617 | 4630 | 5808 | 6416 | 5838 | 5096 | 5328 | 6989 | 5702 | 5845 |
| 5496 | -0.19 | 0.10 | -1.97 | 0.05 | 1 | Dag1 | dystroglycan 1 | 8787 | 8021 | 7050 | 5386 | 5087 | 3919 | 5094 | 3977 | 5006 | 4785 | 3342 |
| 4273 | -0.02 | 0.09 | -0.23 | 0.82 | 1 | Aga | aspartylglucosaminidase | 2976 | 3492 | 3641 | 5504 | 4480 | 3544 | 5084 | 4986 | 4898 | 4767 | 3633 |
| 4445 | 0.10 | 0.08 | 1.25 | 0.21 | 1 | Jade1 | jade family PHD finger 1 | 2633 | 2990 | 2589 | 5601 | 4685 | 4253 | 5079 | 6591 | 5152 | 4721 | 4598 |
| 6188 | -0.01 | 0.11 | -0.10 | 0.92 | 1 | Hsph1 | heat shock 105kDa/110kDa protein 1 | 2763 | 4209 | 3648 | 4672 | 5677 | 13685 | 5078 | 5638 | 5484 | 4708 | 12512 |
| 4573 | 0.14 | 0.08 | 1.65 | 0.10 | 1 | Ddx24 | DEAD (Asp-Glu-Ala-Asp) box polypeptide | 3540 | 3764 | 3897 | 4506 | 5098 | 3931 | 5076 | 5890 | 5222 | 5013 | 4366 |
| 4121 | -0.08 | 0.12 | -0.60 | 0.55 | 1 | Jpt2 | Jupiter microtubule associated homolog 2 | 3022 | 3055 | 2853 | 3963 | 5390 | 4210 | 5069 | 3230 | 5329 | 4926 | 4283 |
| 3470 | -0.07 | 0.18 | -0.39 | 0.70 | 1 | Car10 | carbonic anhydrase 10 | 149 | 170 | 364 | 5112 | 5167 | 3833 | 5067 | 4248 | 5185 | 4497 | 4376 |
| 4984 | -0.08 | 0.09 | -0.89 | 0.37 | 1 | Slc25a5 | solute carrier family 25 (mitochondrial car | 6296 | 6305 | 6626 | 4632 | 5003 | 3581 | 5066 | 3667 | 5013 | 5013 | 3620 |
| 5018 | 0.12 | 0.07 | 1.65 | 0.10 | 1 | Skp1a | S-phase kinase-associated protein 1A | 5557 | 5216 | 5360 | 4729 | 5113 | 3853 | 5059 | 5701 | 5015 | 5284 | 4311 |
| 4454 | 0.02 | 0.10 | 0.19 | 0.85 | 1 | Atad1 | ATPase family, AAA domain containing 1 | 3414 | 3476 | 3634 | 4936 | 4655 | 4444 | 5055 | 4126 | 5530 | 4792 | 4938 |
| 5088 | 0.24 | 0.16 | 1.54 | 0.12 | 1 | Ctsd | cathepsin D | 5057 | 4331 | 5332 | 5297 | 5179 | 3140 | 5028 | 8848 | 5012 | 5388 | 3356 |
| 5180 | 0.11 | 0.07 | 1.47 | 0.14 | 1 | Top1 | topoisomerase (DNA) I | 5688 | 5174 | 4859 | 4958 | 4348 | 5525 | 5024 | 5905 | 4694 | 5088 | 5722 |
| 4689 | 0.10 | 0.07 | 1.49 | 0.14 | 1 | Atp6v1c1 | ATPase, H+ transporting, lysosomal V1 su | 3346 | 3613 | 3432 | 4905 | 5037 | 4921 | 5009 | 5821 | 5439 | 5031 | 5024 |
| 4312 | 0.01 | 0.08 | 0.08 | 0.94 | 1 | Syt14 | synaptotagmin XIV | 2091 | 2795 | 2414 | 5234 | 4584 | 5169 | 5009 | 5363 | 5117 | 4431 | 5224 |
| 4541 | 0.06 | 0.11 | 0.58 | 0.56 | 1 | Cerk | ceramide kinase | 2877 | 2132 | 2460 | 5832 | 5310 | 4570 | 5006 | 6034 | 5700 | 5673 | 4352 |
| 4283 | 0.06 | 0.06 | 1.04 | 0.30 | 1 | Cog6 | component of oligomeric golgi complex 6 | 2798 | 2410 | 2922 | 4491 | 4783 | 4772 | 4998 | 4491 | 5242 | 5254 | 4956 |
| 5004 | 0.03 | 0.08 | 0.39 | 0.70 | 1 | Ipo5 | importin 5 | 6144 | 5196 | 5792 | 4900 | 5011 | 3870 | 4987 | 5289 | 5148 | 4641 | 4069 |
| 3995 | -0.03 | 0.08 | -0.39 | 0.70 | 1 | Slc25a44 | solute carrier family 25, member 44 | 2108 | 2359 | 2297 | 4663 | 5142 | 3964 | 4986 | 4460 | 5371 | 4861 | 3730 |
| 4520 | 0.05 | 0.07 | 0.78 | 0.44 | 1 | Stard7 | START domain containing 7 | 3312 | 3383 | 3551 | 5172 | 5127 | 4102 | 4984 | 5677 | 5004 | 5146 | 4261 |
| 5056 | 0.03 | 0.06 | 0.39 | 0.69 | 1 | Araf | Araf proto-oncogene, serine/threonine kir | 2274 | 2187 | 2315 | 5122 | 5219 | 8930 | 4980 | 4872 | 5787 | 5139 | 8793 |
| 5011 | 0.10 | 0.09 | 1.09 | 0.28 | 1 | Rock1 | Rho-associated coiled-coil containing pro | 4634 | 4109 | 4669 | 4461 | 4809 | 5868 | 4978 | 5431 | 5338 | 4379 | 6448 |
| 4818 | -0.09 | 0.07 | -1.26 | 0.21 | 1 | Ash1l | ASH1 like histone lysine methyltransferas | 4325 | 4756 | 4947 | 4510 | 4963 | 5628 | 4978 | 4402 | 4975 | 4297 | 5219 |
| 4100 | 0.06 | 0.07 | 0.90 | 0.37 | 1 | Stx7 | syntaxin 7 | 3284 | 2804 | 2581 | 4623 | 4379 | 3873 | 4978 | 4759 | 4401 | 5087 | 4328 |
| 5393 | -0.06 | 0.06 | -0.96 | 0.34 | 1 | Ncstn | nicastrin | 6324 | 7198 | 6202 | 6015 | 5055 | 4155 | 4974 | 5473 | 4971 | 5230 | 3732 |
| 4751 | 0.06 | 0.09 | 0.62 | 0.53 | 1 | Mdh2 | malate dehydrogenase 2, NAD (mitochon | 5415 | 4621 | 5344 | 4342 | 5065 | 3714 | 4973 | 5320 | 4518 | 5143 | 3807 |
| 4147 | 0.04 | 0.09 | 0.42 | 0.67 | 1 | Man2b1 | mannosidase 2, alpha B1 | 6747 | 3885 | 4555 | 3376 | 4398 | 2123 | 4973 | 3843 | 4262 | 5522 | 1929 |
| 4261 | 0.15 | 0.11 | 1.44 | 0.15 | 1 | Igf1r | insulin-like growth factor I receptor | 1939 | 2721 | 1950 | 4827 | 4617 | 4557 | 4972 | 6152 | 5432 | 5400 | 4304 |
| 4736 | 0.05 | 0.10 | 0.46 | 0.64 | 1 | H1f2 | H1.2 linker histone, cluster member | 2279 | 3351 | 3652 | 5110 | 5927 | 5060 | 4968 | 5852 | 5645 | 5047 | 5210 |
| 4442 | -0.06 | 0.10 | -0.60 | 0.55 | 1 | Cdc42se2 | CDC42 small effector 2 | 5028 | 4195 | 4296 | 4501 | 4964 | 3575 | 4967 | 3486 | 5175 | 4980 | 3696 |
| 5141 | 0.13 | 0.08 | 1.74 | 0.08 | 1 | Ppm1b | protein phosphatase 1B, magnesium dep | 3390 | 3602 | 3638 | 6104 | 6038 | 4789 | 4961 | 7250 | 6030 | 5597 | 5148 |
| 4879 | 0.01 | 0.07 | 0.22 | 0.83 | 1 | Pdha1 | pyruvate dehydrogenase E1 alpha 1 | 5246 | 4637 | 4964 | 5095 | 4919 | 4352 | 4960 | 4999 | 4905 | 4966 | 4623 |
| 4515 | 0.12 | 0.08 | 1.42 | 0.16 | 1 | Strap | serine/threonine kinase receptor associat | 4254 | 3983 | 3576 | 4377 | 5024 | 3832 | 4954 | 4886 | 5360 | 5075 | 4343 |
| 4227 | 0.14 | 0.10 | 1.42 | 0.16 | 1 | Gdap1 | ganglioside-induced differentiation-assoc | 2153 | 2904 | 2352 | 4945 | 4263 | 4487 | 4954 | 5132 | 4905 | 5437 | 4970 |
| 3831 | -0.03 | 0.10 | -0.27 | 0.78 | 1 | Lmbrd2 | LMBR1 domain containing 2 | 2159 | 2623 | 2432 | 4442 | 4396 | 3852 | 4952 | 3941 | 4799 | 4566 | 3979 |
| 4791 | 0.12 | 0.06 | 1.92 | 0.06 | 1 | Sumo3 | small ubiquitin-like modifier 3 | 4437 | 4609 | 4864 | 4636 | 5105 | 3912 | 4949 | 5186 | 5375 | 5483 | 4143 |
| 4738 | -0.02 | 0.08 | -0.23 | 0.82 | 1 | Tacc1 | transforming, acidic coiled-coil containing | 5345 | 6117 | 5429 | 4386 | 4462 | 3944 | 4943 | 4378 | 4532 | 4534 | 4047 |
| 4904 | 0.11 | 0.14 | 0.80 | 0.42 | 1 | Mtdh | metadherin | 6220 | 4552 | 4116 | 4474 | 4969 | 4417 | 4942 | 5717 | 5480 | 4354 | 4702 |
| 4299 | 0.07 | 0.08 | 0.85 | 0.40 | 1 | Chpf | chondroitin polymerizing factor | 2923 | 3320 | 3045 | 4951 | 4863 | 3710 | 4941 | 5697 | 4586 | 5611 | 3647 |
| 4012 | 0.05 | 0.10 | 0.47 | 0.64 | 1 | Dnajb14 | DnaJ heat shock protein family (Hsp40) m | 2557 | 2670 | 2263 | 4515 | 4700 | 3844 | 4936 | 5427 | 5086 | 4204 | 3926 |
| 4055 | 0.04 | 0.10 | 0.36 | 0.72 | 1 | Dpp4 | dipeptidylpeptidase 4 | 2484 | 2907 | 2732 | 4720 | 4445 | 3912 | 4934 | 4042 | 4973 | 5170 | 4285 |
| 4780 | 0.21 | 0.14 | 1.44 | 0.15 | 1 | Cox7a2l | cytochrome c oxidase subunit 7A2 like | 3651 | 4220 | 5055 | 4925 | 5325 | 3096 | 4932 | 7986 | 5217 | 4468 | 3705 |
| 4555 | -0.12 | 0.10 | -1.29 | 0.20 | 1 | Abca2 | ATP-binding cassette, sub-family A (ABC1 | 3416 | 3662 | 3768 | 5678 | 4776 | 5074 | 4926 | 4456 | 5050 | 5138 | 4155 |
| 4902 | -0.12 | 0.09 | -1.35 | 0.18 | 1 | Usp19 | ubiquitin specific peptidase 19 | 4302 | 3905 | 3607 | 4964 | 5641 | 6455 | 4924 | 4544 | 5343 | 4571 | 5663 |
| 5031 | 0.21 | 0.13 | 1.60 | 0.11 | 1 | Rrp1 | ribosomal RNA processing 1 homolog (S. | 4968 | 4684 | 5259 | 4308 | 4720 | 4715 | 4923 | 6985 | 4414 | 5045 | 5318 |
| 4857 | -0.03 | 0.10 | -0.31 | 0.76 | 1 | Ap1g1 | adaptor protein complex AP-1, gamma 1 s | 4299 | 5851 | 5041 | 4604 | 4981 | 4820 | 4923 | 4396 | 4843 | 4881 | 4787 |
| 4298 | -0.07 | 0.12 | -0.57 | 0.57 | 1 | Alg10b | asparagine-linked glycosylation 10B (alph | 3756 | 4750 | 3878 | 4713 | 4474 | 3774 | 4923 | 3889 | 4942 | 4043 | 4136 |
| 4340 | 0.01 | 0.09 | 0.17 | 0.87 | 1 | Atp6v0e2 | ATPase, H+ transporting, lysosomal V0 su | 2984 | 3515 | 4162 | 4516 | 5532 | 3510 | 4920 | 4734 | 5306 | 4968 | 3598 |
| 4022 | 0.02 | 0.08 | 0.22 | 0.83 | 1 | Arl5a | ADP-ribosylation factor-like 5A | 3553 | 2862 | 3049 | 4476 | 4317 | 3605 | 4920 | 4061 | 4652 | 4600 | 4144 |
| 4781 | -0.03 | 0.09 | -0.37 | 0.72 | 1 | Nptn | neuroplastin | 5544 | 4835 | 5100 | 4881 | 4874 | 4079 | 4907 | 3934 | 4885 | 5154 | 4403 |

| | | | | | | | | | | | | | | | | | | |
|---|---|---|---|---|---|---|---|---|---|---|---|---|---|---|---|---|---|---|
| 4853 | 0.03 | 0.06 | 0.56 | 0.57 | 1 | Zc3h14 | zinc finger CCCH type containing 14 | 4664 | 4552 | 4483 | 5106 | 4819 | 4782 | 4899 | 5150 | 5067 | 4726 | 5137 |
| 4384 | -0.10 | 0.09 | -1.06 | 0.29 | 1 | Epb41l4b | erythrocyte membrane protein band 4.1 li | 3267 | 3825 | 3831 | 5324 | 4875 | 4179 | 4897 | 4443 | 4547 | 5192 | 3840 |
| 4520 | -0.03 | 0.08 | -0.38 | 0.70 | 1 | Kdm5c | lysine (K)-specific demethylase 5C | 3529 | 3982 | 4064 | 4731 | 4873 | 4750 | 4896 | 4475 | 5233 | 4958 | 4227 |
| 4017 | 0.00 | 0.10 | -0.05 | 0.96 | 1 | Nptxr | neuronal pentraxin receptor | 2315 | 2332 | 2999 | 5200 | 4737 | 3434 | 4886 | 5122 | 5051 | 4829 | 3279 |
| 5095 | -0.03 | 0.07 | -0.41 | 0.68 | 1 | Atl3 | atlastin GTPase 3 | 7084 | 6887 | 6020 | 4649 | 4459 | 4245 | 4885 | 4281 | 4732 | 4540 | 4265 |
| 4663 | 0.02 | 0.08 | 0.20 | 0.84 | 1 | Zmynd8 | zinc finger, MYND-type containing 8 | 2379 | 3295 | 3148 | 5682 | 5209 | 5335 | 4885 | 5394 | 5805 | 5048 | 5112 |
| 3863 | -0.16 | 0.10 | -1.68 | 0.09 | 1 | Erlin2 | ER lipid raft associated 2 | 3462 | 3843 | 3767 | 4253 | 4145 | 3357 | 4885 | 3407 | 3967 | 4086 | 3324 |
| 4726 | 0.05 | 0.06 | 0.77 | 0.44 | 1 | Ywhaz | tyrosine 3-monooxygenase/tryptophan 5- | 4492 | 4219 | 4192 | 5221 | 5083 | 4039 | 4877 | 5220 | 5012 | 5382 | 4249 |
| 4614 | -0.05 | 0.07 | -0.74 | 0.46 | 1 | Uaca | uveal autoantigen with coiled-coil domair | 3251 | 3239 | 3770 | 5002 | 5135 | 5584 | 4877 | 4820 | 4935 | 4922 | 5223 |
| 4174 | 0.12 | 0.10 | 1.21 | 0.23 | 1 | Cyb561 | cytochrome b-561 | 2101 | 2064 | 2747 | 5037 | 5393 | 3375 | 4871 | 6043 | 5380 | 5392 | 3505 |
| 4865 | 0.12 | 0.13 | 0.99 | 0.32 | 1 | Idh2 | isocitrate dehydrogenase 2 (NADP+), mit | 3225 | 2909 | 3831 | 5410 | 4915 | 5646 | 4860 | 6431 | 4600 | 5848 | 5839 |
| 4740 | 0.11 | 0.07 | 1.59 | 0.11 | 1 | Gramd3 | GRAM domain containing 3 | 2033 | 2651 | 2103 | 6041 | 5217 | 5735 | 4855 | 6094 | 5487 | 5648 | 6277 |
| 4520 | 0.11 | 0.10 | 1.12 | 0.26 | 1 | Snx5 | sorting nexin 5 | 4871 | 3362 | 3663 | 4710 | 4721 | 3919 | 4855 | 4964 | 5201 | 5169 | 4288 |
| 4684 | -0.06 | 0.13 | -0.46 | 0.64 | 1 | Cacna1d | calcium channel, voltage-dependent, L ty | 2052 | 3019 | 2128 | 5312 | 5042 | 7517 | 4854 | 4163 | 5256 | 5289 | 6893 |
| 5862 | 0.15 | 0.10 | 1.46 | 0.14 | 1 | Rps24 | ribosomal protein S24 | 9532 | 9336 | 9596 | 3983 | 3867 | 4368 | 4853 | 5510 | 4133 | 4466 | 4842 |
| 4551 | 0.08 | 0.08 | 1.11 | 0.27 | 1 | Acbd5 | acyl-Coenzyme A binding domain contair | 3343 | 3481 | 3592 | 4911 | 4953 | 4511 | 4852 | 5680 | 5011 | 4573 | 5155 |
| 3981 | 0.07 | 0.10 | 0.72 | 0.47 | 1 | Dad1 | defender against cell death 1 | 2780 | 2792 | 3127 | 4592 | 4203 | 3425 | 4847 | 4801 | 4369 | 5525 | 3335 |
| 4984 | -0.04 | 0.10 | -0.46 | 0.65 | 1 | Klhl9 | kelch-like 9 | 4548 | 5789 | 5346 | 5048 | 4720 | 5213 | 4836 | 4240 | 4621 | 4613 | 5846 |
| 4743 | -0.02 | 0.08 | -0.29 | 0.77 | 1 | Capza2 | capping protein (actin filament) muscle Z- | 7032 | 5489 | 5715 | 4346 | 4230 | 3720 | 4836 | 4016 | 4478 | 4400 | 3914 |
| 4971 | 0.09 | 0.08 | 1.09 | 0.28 | 1 | Shc1 | src homology 2 domain-containing transf | 3128 | 2896 | 3304 | 5198 | 6645 | 5368 | 4834 | 5406 | 6416 | 5849 | 5634 |
| 4740 | -0.06 | 0.08 | -0.77 | 0.44 | 1 | Bmpr1a | bone morphogenetic protein receptor, typ | 5508 | 5807 | 5304 | 4625 | 4981 | 3749 | 4829 | 4411 | 4746 | 4130 | 4046 |
| 5002 | 0.05 | 0.10 | 0.45 | 0.65 | 1 | Aamp | angio-associated migratory protein | 4948 | 4755 | 6073 | 4654 | 5218 | 4599 | 4827 | 5858 | 4782 | 4735 | 4569 |
| 4326 | -0.13 | 0.10 | -1.31 | 0.19 | 1 | Coa5 | cytochrome C oxidase assembly factor 5 | 4649 | 4886 | 4608 | 5334 | 4130 | 3228 | 4827 | 3951 | 4529 | 4297 | 3143 |
| 5361 | -0.06 | 0.09 | -0.70 | 0.48 | 1 | Wdr26 | WD repeat domain 26 | 6439 | 6914 | 5780 | 5536 | 5011 | 4964 | 4824 | 5235 | 5200 | 4041 | 5023 |
| 4782 | 0.07 | 0.07 | 0.99 | 0.32 | 1 | Copz1 | coatomer protein complex, subunit zeta 1 | 4300 | 4212 | 4994 | 4496 | 5509 | 4279 | 4824 | 5048 | 5377 | 5309 | 4256 |
| 3741 | 0.10 | 0.10 | 0.99 | 0.32 | 1 | Agl | amylo-1,6-glucosidase, 4-alpha-glucanot | 2142 | 2543 | 2076 | 4321 | 4202 | 3324 | 4824 | 4400 | 4812 | 4678 | 3823 |
| 8528 | -0.01 | 0.15 | -0.04 | 0.96 | 1 | Btg2 | BTG anti-proliferation factor 2 | 19842 | 8982 | 10324 | 3969 | 5136 | 13737 | 4822 | 4014 | 5197 | 4973 | 12817 |
| 6151 | 0.13 | 0.09 | 1.41 | 0.16 | 1 | Gc | vitamin D binding protein | 10503 | 12338 | 12791 | 3885 | 4766 | 1882 | 4819 | 4591 | 5188 | 4786 | 2109 |
| 4172 | 0.04 | 0.10 | 0.36 | 0.72 | 1 | Pnpla8 | patatin-like phospholipase domain contai | 3399 | 3821 | 3445 | 4208 | 4498 | 3852 | 4817 | 3765 | 4748 | 5142 | 4193 |
| 4030 | 0.02 | 0.10 | 0.20 | 0.84 | 1 | Pebp1 | phosphatidylethanolamine binding protei | 3259 | 2857 | 3845 | 4542 | 4389 | 3305 | 4813 | 4946 | 4357 | 4759 | 3259 |
| 4110 | -0.09 | 0.07 | -1.19 | 0.24 | 1 | Smim7 | small integral membrane protein 7 | 3800 | 3576 | 4362 | 4270 | 4768 | 3406 | 4809 | 3719 | 4506 | 4620 | 3380 |
| 4818 | -0.08 | 0.08 | -1.02 | 0.31 | 1 | Neo1 | neogenin | 6323 | 6184 | 6048 | 4236 | 4428 | 4214 | 4804 | 3660 | 4628 | 4501 | 3969 |
| 4015 | -0.16 | 0.12 | -1.39 | 0.16 | 1 | Sik3 | SIK family kinase 3 | 3161 | 3121 | 2996 | 4253 | 4587 | 4696 | 4804 | 3141 | 4658 | 4942 | 3804 |
| 4415 | -0.09 | 0.07 | -1.31 | 0.19 | 1 | Mon2 | MON2 homolog, regulator of endosome t | 3382 | 2782 | 3151 | 4569 | 5405 | 5504 | 4800 | 4409 | 4970 | 4624 | 4970 |
| 4781 | 0.02 | 0.08 | 0.24 | 0.81 | 1 | 4932438A13Ril | RIKEN cDNA 4932438A13 gene | 3362 | 3321 | 3438 | 3932 | 4634 | 7725 | 4784 | 4038 | 5276 | 4264 | 7811 |
| 4099 | -0.08 | 0.11 | -0.73 | 0.47 | 1 | Scpep1 | serine carboxypeptidase 1 | 4639 | 3462 | 3261 | 5618 | 3940 | 2991 | 4779 | 4734 | 3656 | 5063 | 2951 |
| 4156 | 0.20 | 0.15 | 1.37 | 0.17 | 1 | Fkbp2 | FK506 binding protein 2 | 2284 | 2126 | 2439 | 4781 | 4735 | 3608 | 4774 | 7667 | 4395 | 5163 | 3748 |
| 3325 | -0.11 | 0.14 | -0.81 | 0.42 | 1 | Tmem163 | transmembrane protein 163 | 860 | 408 | 1010 | 3982 | 5274 | 3598 | 4771 | 3089 | 4961 | 5311 | 3306 |
| 4172 | 0.15 | 0.09 | 1.69 | 0.09 | 1 | Rcbtb1 | regulator of chromosome condensation (F | 2316 | 2806 | 2853 | 4453 | 4609 | 4135 | 4769 | 4799 | 5194 | 5343 | 4612 |
| 6496 | -0.05 | 0.08 | -0.66 | 0.51 | 1 | Mprip | myosin phosphatase Rho interacting prot | 10807 | 8944 | 8862 | 4748 | 5011 | 7304 | 4766 | 4533 | 4877 | 4798 | 6808 |
| 4175 | -0.09 | 0.11 | -0.78 | 0.44 | 1 | Wbp2 | WW domain binding protein 2 | 4040 | 5229 | 4699 | 4522 | 4301 | 2861 | 4766 | 3755 | 4373 | 4721 | 2661 |
| 4754 | 0.12 | 0.08 | 1.47 | 0.14 | 1 | Ro60 | Ro60, Y RNA binding protein | 3310 | 4354 | 3936 | 5562 | 4279 | 4843 | 4763 | 6211 | 5086 | 4488 | 5466 |
| 4442 | 0.00 | 0.10 | 0.02 | 0.99 | 1 | Psme4 | proteasome (prosome, macropain) activa | 2206 | 2102 | 1958 | 4991 | 5822 | 5712 | 4757 | 4805 | 6004 | 4824 | 5686 |
| 4360 | 0.21 | 0.16 | 1.27 | 0.20 | 1 | Ufc1 | ubiquitin-fold modifier conjugating enzym | 2515 | 2870 | 3389 | 4777 | 4623 | 3721 | 4757 | 8354 | 4692 | 4347 | 3920 |
| 5195 | 0.10 | 0.08 | 1.37 | 0.17 | 1 | Rad23b | RAD23 homolog B, nucleotide excision re | 5240 | 4891 | 5482 | 4784 | 5726 | 4785 | 4752 | 5636 | 5349 | 5342 | 5158 |
| 4384 | 0.07 | 0.07 | 1.10 | 0.27 | 1 | Agpat3 | 1-acylglycerol-3-phosphate O-acyltransfe | 2727 | 2747 | 2815 | 4737 | 4987 | 4970 | 4748 | 4962 | 5041 | 5534 | 4950 |
| 3934 | 0.12 | 0.14 | 0.81 | 0.42 | 1 | Syndig1l | synapse differentiation inducing 1 like | 779 | 1421 | 1282 | 5241 | 4921 | 4178 | 4741 | 6111 | 4505 | 5499 | 4592 |
| 3455 | -0.02 | 0.09 | -0.17 | 0.87 | 1 | Nrxn1 | neurexin I | 1575 | 1671 | 1730 | 3618 | 4115 | 4136 | 4739 | 3585 | 4187 | 4493 | 4151 |
| 8928 | -0.09 | 0.11 | -0.89 | 0.37 | 1 | Clu | clusterin | 18432 | 19280 | 24447 | 5463 | 5167 | 3103 | 4734 | 5890 | 4586 | 4524 | 2580 |
| 4335 | -0.01 | 0.06 | -0.13 | 0.90 | 1 | Ireb2 | iron responsive element binding protein 2 | 2430 | 2576 | 2679 | 5014 | 5102 | 5187 | 4733 | 4904 | 5153 | 4439 | 5472 |
| 4908 | 0.02 | 0.08 | 0.23 | 0.82 | 1 | Txlna | taxilin alpha | 4540 | 4485 | 4364 | 3994 | 4958 | 6482 | 4732 | 4230 | 5287 | 4356 | 6561 |
| 4410 | 0.12 | 0.13 | 0.91 | 0.36 | 1 | Pycr2 | pyrroline-5-carboxylate reductase family, | 2346 | 2406 | 2954 | 4688 | 5014 | 5071 | 4730 | 6832 | 4550 | 4785 | 5138 |
| 4760 | -0.03 | 0.07 | -0.47 | 0.64 | 1 | Ube4b | ubiquitination factor E4B | 3986 | 4776 | 4531 | 4946 | 5479 | 4586 | 4728 | 4908 | 5527 | 4758 | 4138 |
| 4466 | 0.04 | 0.10 | 0.41 | 0.68 | 1 | Tnks | tankyrase, TRF1-interacting ankyrin-relate | 3300 | 3681 | 3781 | 4765 | 4367 | 5040 | 4728 | 4391 | 5491 | 4933 | 4651 |
| 3961 | -0.06 | 0.08 | -0.83 | 0.41 | 1 | Gnptg | N-acetylglucosamine-1-phosphotransfera | 2466 | 2651 | 2737 | 5024 | 4639 | 3894 | 4726 | 4400 | 4409 | 4494 | 4135 |
| 5557 | 0.00 | 0.08 | -0.06 | 0.96 | 1 | Mxd4 | Max dimerization protein 4 | 7039 | 6577 | 6634 | 4754 | 5442 | 5569 | 4720 | 4926 | 5550 | 4841 | 5080 |
| 7469 | -0.04 | 0.10 | -0.38 | 0.70 | 1 | Ly6e | lymphocyte antigen 6 complex, locus E | 18224 | 14539 | 15651 | 4940 | 4266 | 3114 | 4711 | 5314 | 3974 | 4547 | 2881 |
| 4231 | -0.17 | 0.10 | -1.61 | 0.11 | 1 | Cdc42bpa | CDC42 binding protein kinase alpha | 4721 | 3758 | 3899 | 4616 | 4392 | 4319 | 4709 | 3514 | 4680 | 3972 | 3957 |
| 4288 | -0.11 | 0.09 | -1.18 | 0.24 | 1 | Pak2 | p21 (RAC1) activated kinase 2 | 5636 | 5434 | 4986 | 3882 | 4287 | 3320 | 4707 | 3388 | 4183 | 3867 | 3477 |
| 5465 | -0.01 | 0.08 | -0.07 | 0.95 | 1 | Sf1 | splicing factor 1 | 5756 | 6056 | 5681 | 4549 | 5291 | 6897 | 4703 | 4625 | 5519 | 4926 | 6113 |

| | | | | | | | | | | | | | | | | | | |
|---|---|---|---|---|---|---|---|---|---|---|---|---|---|---|---|---|---|---|
| 4590 | -0.07 | 0.11 | -0.62 | 0.54 | 1 | Dazap2 | DAZ associated protein 2 | 6198 | 4961 | 4646 | 4672 | 4772 | 3555 | 4696 | 3834 | 5063 | 4709 | 3380 |
| 4800 | -0.05 | 0.06 | -0.81 | 0.42 | 1 | Kdm1a | lysine (K)-specific demethylase 1A | 3447 | 4160 | 4565 | 5636 | 5451 | 4875 | 4695 | 5590 | 5403 | 4468 | 4507 |
| 4557 | -0.02 | 0.08 | -0.20 | 0.84 | 1 | Trp53bp1 | transformation related protein 53 binding | 3941 | 3929 | 3970 | 4075 | 4445 | 6031 | 4694 | 3807 | 4977 | 4401 | 5852 |
| 4255 | 0.12 | 0.08 | 1.51 | 0.13 | 1 | Nt5c3 | 5'-nucleotidase, cytosolic III | 1610 | 2005 | 2218 | 4877 | 5240 | 4826 | 4693 | 5106 | 5719 | 5161 | 5351 |
| 4846 | -0.01 | 0.08 | -0.17 | 0.86 | 1 | Nsd1 | nuclear receptor-binding SET-domain prc | 4154 | 4976 | 4248 | 4712 | 4207 | 6467 | 4692 | 4421 | 4572 | 4534 | 6316 |
| 4640 | 0.08 | 0.10 | 0.77 | 0.44 | 1 | Ap3b2 | adaptor-related protein complex 3, beta 2 | 2488 | 2743 | 3369 | 5224 | 5145 | 5586 | 4689 | 6335 | 5353 | 4758 | 5349 |
| 4623 | 0.05 | 0.11 | 0.43 | 0.66 | 1 | Hdac6 | histone deacetylase 6 | 2225 | 3090 | 2956 | 5256 | 4824 | 6135 | 4686 | 6733 | 4793 | 4683 | 5467 |
| 4087 | 0.18 | 0.12 | 1.55 | 0.12 | 1 | 0610040J01Rik | RIKEN cDNA 0610040J01 gene | 1701 | 1761 | 2257 | 4979 | 4514 | 4131 | 4681 | 6589 | 5156 | 4731 | 4457 |
| 4409 | -0.02 | 0.08 | -0.19 | 0.85 | 1 | Adipor1 | adiponectin receptor 1 | 4434 | 4204 | 4338 | 4579 | 5043 | 3554 | 4679 | 4039 | 5378 | 4534 | 3717 |
| 4582 | 0.11 | 0.10 | 1.16 | 0.25 | 1 | Erp44 | endoplasmic reticulum protein 44 | 3629 | 3119 | 3140 | 4485 | 5918 | 4429 | 4677 | 5172 | 5799 | 5165 | 4865 |
| 4070 | -0.01 | 0.09 | -0.16 | 0.88 | 1 | Serpina10 | serine (or cysteine) peptidase inhibitor, cl | 2444 | 3293 | 3012 | 4852 | 4526 | 4026 | 4668 | 4879 | 4631 | 4723 | 3721 |
| 4635 | 0.04 | 0.08 | 0.48 | 0.63 | 1 | Rab1b | RAB1B, member RAS oncogene family | 4827 | 4665 | 5117 | 4800 | 4854 | 3607 | 4665 | 5102 | 4556 | 5262 | 3533 |
| 3941 | 0.03 | 0.10 | 0.28 | 0.78 | 1 | Vps13c | vacuolar protein sorting 13C | 2088 | 2322 | 1575 | 4281 | 4222 | 5341 | 4663 | 4514 | 4307 | 4682 | 5353 |
| 3887 | -0.13 | 0.09 | -1.47 | 0.14 | 1 | Cox8a | cytochrome c oxidase subunit 8A | 3348 | 3282 | 3832 | 4412 | 4523 | 3238 | 4656 | 3521 | 4373 | 4704 | 2872 |
| 4249 | 0.08 | 0.10 | 0.79 | 0.43 | 1 | 4930402H24Rik | RIKEN cDNA 4930402H24 gene | 2911 | 2825 | 2861 | 5032 | 4304 | 4471 | 4652 | 6356 | 4492 | 4439 | 4402 |
| 4231 | -0.04 | 0.08 | -0.44 | 0.66 | 1 | Taf1 | TATA-box binding protein associated fact | 3239 | 3098 | 3335 | 3762 | 4651 | 5559 | 4649 | 3652 | 4877 | 4010 | 5709 |
| 4411 | 0.10 | 0.07 | 1.39 | 0.16 | 1 | Ube2b | ubiquitin-conjugating enzyme E2B | 4097 | 4223 | 4194 | 4114 | 4308 | 4324 | 4648 | 4545 | 4328 | 4838 | 4907 |
| 5507 | -0.08 | 0.08 | -1.09 | 0.28 | 1 | Nono | non-POU-domain-containing, octamer bir | 7110 | 7433 | 6941 | 4675 | 5473 | 5325 | 4645 | 4232 | 4933 | 4600 | 5213 |
| 4476 | 0.02 | 0.06 | 0.26 | 0.79 | 1 | Sun1 | Sad1 and UNC84 domain containing 1 | 3794 | 3547 | 3767 | 4770 | 4754 | 4795 | 4638 | 4655 | 5001 | 4727 | 4785 |
| 6445 | 0.01 | 0.06 | 0.23 | 0.82 | 1 | Glud1 | glutamate dehydrogenase 1 | 12154 | 11477 | 12100 | 4548 | 4396 | 3925 | 4636 | 4374 | 4371 | 4830 | 4082 |
| 4673 | -0.03 | 0.07 | -0.34 | 0.73 | 1 | Ano6 | anoctamin 6 | 5493 | 5397 | 4439 | 5213 | 4198 | 4156 | 4636 | 4741 | 4499 | 4803 | 3830 |
| 4216 | -0.08 | 0.08 | -1.02 | 0.31 | 1 | Fbxw11 | F-box and WD-40 domain protein 11 | 4257 | 4130 | 4089 | 4447 | 4637 | 3680 | 4636 | 3925 | 4818 | 4175 | 3582 |
| 3970 | 0.14 | 0.09 | 1.64 | 0.10 | 1 | Dynlt3 | dynein light chain Tctex-type 3 | 2238 | 1975 | 2281 | 4748 | 4623 | 3720 | 4636 | 5112 | 4895 | 4781 | 4657 |
| 3863 | 0.10 | 0.09 | 1.14 | 0.25 | 1 | Lmbrd1 | LMBR1 domain containing 1 | 3081 | 3044 | 2752 | 4133 | 3940 | 3502 | 4635 | 5138 | 4394 | 4176 | 3696 |
| 4241 | 0.09 | 0.07 | 1.31 | 0.19 | 1 | Hprt | hypoxanthine guanine phosphoribosyl tra | 4417 | 3788 | 4265 | 4405 | 4188 | 3334 | 4632 | 4458 | 4380 | 4956 | 3832 |
| 4736 | 0.06 | 0.11 | 0.52 | 0.60 | 1 | Mat1a | methionine adenosyltransferase I, alpha | 4576 | 6823 | 6137 | 4050 | 4711 | 3498 | 4630 | 4956 | 4796 | 4251 | 3665 |
| 3561 | -0.13 | 0.12 | -1.12 | 0.26 | 1 | Samd4b | sterile alpha motif domain containing 4B | 2572 | 2993 | 2380 | 4050 | 4416 | 3196 | 4626 | 3512 | 4563 | 4011 | 2849 |
| 4580 | 0.07 | 0.07 | 1.01 | 0.31 | 1 | Chtop | chromatin target of PRMT1 | 4175 | 4157 | 4150 | 4244 | 4716 | 4887 | 4621 | 5049 | 4539 | 5007 | 4842 |
| 4289 | -0.08 | 0.07 | -1.09 | 0.28 | 1 | Ndst1 | N-deacetylase/N-sulfotransferase (hepara | 4825 | 4529 | 4810 | 4586 | 4291 | 3439 | 4614 | 4635 | 4263 | 4146 | 3039 |
| 4450 | 0.09 | 0.10 | 0.95 | 0.34 | 1 | Spint1 | serine protease inhibitor, Kunitz type 1 | 2893 | 3369 | 4176 | 4728 | 5512 | 3697 | 4613 | 5804 | 5544 | 5206 | 3413 |
| 4511 | 0.00 | 0.08 | -0.01 | 0.99 | 1 | Nsd3 | nuclear receptor binding SET domain pro | 3367 | 3237 | 3014 | 4702 | 4454 | 6293 | 4610 | 4588 | 5053 | 4437 | 5864 |
| 4371 | 0.04 | 0.07 | 0.54 | 0.59 | 1 | Epn1 | epsin 1 | 4302 | 4185 | 4386 | 4329 | 4933 | 3507 | 4610 | 4791 | 4607 | 4892 | 3535 |
| 4842 | -0.01 | 0.09 | -0.16 | 0.87 | 1 | Setd5 | SET domain containing 5 | 4083 | 4475 | 4252 | 4752 | 5002 | 5985 | 4606 | 4283 | 5547 | 4877 | 5398 |
| 4452 | 0.02 | 0.07 | 0.25 | 0.81 | 1 | Senp6 | SUMO/sentrin specific peptidase 6 | 3245 | 3701 | 3705 | 4341 | 4909 | 5173 | 4599 | 4455 | 4890 | 4405 | 5548 |
| 4132 | 0.10 | 0.10 | 1.04 | 0.30 | 1 | Hsd17b12 | hydroxysteroid (17-beta) dehydrogenase | 2758 | 2144 | 2761 | 4811 | 5411 | 3412 | 4599 | 4437 | 6274 | 4896 | 3946 |
| 4631 | 0.02 | 0.10 | 0.17 | 0.86 | 1 | Chd8 | chromodomain helicase DNA binding pro | 4072 | 4306 | 4447 | 4481 | 4755 | 5053 | 4596 | 5642 | 5080 | 3909 | 4597 |
| 4403 | -0.04 | 0.10 | -0.35 | 0.72 | 1 | Ppp6r3 | protein phosphatase 6, regulatory subuni | 4534 | 3818 | 3889 | 4246 | 4444 | 5007 | 4595 | 3797 | 5164 | 3976 | 4963 |
| 4550 | -0.03 | 0.08 | -0.37 | 0.71 | 1 | Mageh1 | melanoma antigen, family H, 1 | 3826 | 4065 | 4611 | 4619 | 4508 | 5206 | 4590 | 4436 | 4311 | 4287 | 5590 |
| 4518 | -0.05 | 0.07 | -0.71 | 0.48 | 1 | Tspyl1 | testis-specific protein, Y-encoded-like 1 | 4703 | 5488 | 5096 | 4667 | 4100 | 4172 | 4588 | 4302 | 3983 | 4379 | 4215 |
| 4750 | -0.05 | 0.06 | -0.81 | 0.42 | 1 | Anxa7 | annexin A7 | 5063 | 4716 | 5438 | 4344 | 5416 | 4472 | 4587 | 4124 | 5287 | 4490 | 4311 |
| 4693 | 0.12 | 0.10 | 1.12 | 0.26 | 1 | Fkbp8 | FK506 binding protein 8 | 4673 | 4170 | 4951 | 4675 | 4657 | 4196 | 4587 | 5876 | 4448 | 5147 | 4245 |
| 4678 | 0.14 | 0.08 | 1.87 | 0.06 | 1 | Znrf2 | zinc and ring finger 2 | 4489 | 5235 | 5244 | 4466 | 4832 | 3469 | 4582 | 5058 | 5035 | 5286 | 3761 |
| 4914 | 0.03 | 0.11 | 0.29 | 0.77 | 1 | Rps27a | ribosomal protein S27A | 8109 | 6928 | 8001 | 3558 | 4219 | 3016 | 4577 | 4720 | 3924 | 3895 | 3102 |
| 6194 | -0.10 | 0.06 | -1.62 | 0.10 | 1 | Clic4 | chloride intracellular channel 4 (mitochon | 11644 | 11182 | 11524 | 4804 | 4965 | 3106 | 4571 | 4500 | 4797 | 4198 | 2847 |
| 4424 | 0.13 | 0.15 | 0.90 | 0.37 | 1 | Creld2 | cysteine-rich with EGF-like domains 2 | 3239 | 2485 | 2638 | 5658 | 4415 | 4401 | 4568 | 7489 | 3995 | 5379 | 4397 |
| 5185 | -0.10 | 0.09 | -1.11 | 0.27 | 1 | Pld3 | phospholipase D family, member 3 | 7375 | 7952 | 7667 | 4738 | 4907 | 3380 | 4562 | 3772 | 4649 | 5008 | 3021 |
| 4026 | -0.22 | 0.13 | -1.68 | 0.09 | 1 | Trrap | transformation/transcription domain-asso | 3750 | 4012 | 3535 | 4279 | 4207 | 4642 | 4562 | 2797 | 4197 | 4510 | 3800 |
| 4497 | 0.01 | 0.07 | 0.16 | 0.87 | 1 | Sh3glb1 | SH3-domain GRB2-like B1 (endophilin) | 5453 | 4573 | 5034 | 4368 | 4707 | 3490 | 4558 | 4017 | 4790 | 4908 | 3571 |
| 3917 | -0.03 | 0.09 | -0.28 | 0.78 | 1 | Golt1b | golgi transport 1B | 2817 | 2821 | 2969 | 4281 | 4823 | 3699 | 4556 | 3748 | 4564 | 5182 | 3632 |
| 4312 | 0.11 | 0.06 | 1.66 | 0.10 | 1 | Aftph | aftiphilin | 3644 | 4057 | 3562 | 4486 | 4359 | 4032 | 4554 | 4860 | 4547 | 4788 | 4548 |
| 4456 | -0.12 | 0.11 | -1.05 | 0.29 | 1 | Nav2 | neuron navigator 2 | 1739 | 2464 | 2910 | 4981 | 5597 | 6761 | 4551 | 4248 | 5457 | 4736 | 5573 |
| 4095 | -0.17 | 0.09 | -1.85 | 0.06 | 1 | Gpc4 | glypican 4 | 4076 | 4699 | 4549 | 4155 | 4378 | 3698 | 4550 | 3375 | 4120 | 4007 | 3434 |
| 4592 | -0.07 | 0.07 | -1.11 | 0.27 | 1 | Tcaf1 | TRPM8 channel-associated factor 1 | 4302 | 4886 | 4191 | 5027 | 4668 | 4794 | 4548 | 4475 | 4379 | 4482 | 4762 |
| 3005 | -0.17 | 0.12 | -1.38 | 0.17 | 1 | Slc9a7 | solute carrier family 9 (sodium/hydrogen e | 732 | 664 | 900 | 4453 | 3847 | 3398 | 4548 | 3165 | 4172 | 3875 | 3298 |
| 4274 | 0.13 | 0.12 | 1.04 | 0.30 | 1 | Clip1 | CAP-GLY domain containing linker protei | 3605 | 2750 | 3699 | 4618 | 4314 | 4159 | 4545 | 5717 | 4732 | 4911 | 3966 |
| 5111 | -0.08 | 0.12 | -0.65 | 0.52 | 1 | Celf3 | CUGBP, Elav-like family member 3 | 3022 | 4323 | 4190 | 4554 | 4891 | 9140 | 4544 | 4108 | 4635 | 4914 | 7902 |
| 5710 | 0.17 | 0.15 | 1.08 | 0.28 | 1 | Rps20 | ribosomal protein S20 | 9905 | 8323 | 10790 | 4191 | 4294 | 2757 | 4540 | 6762 | 4152 | 4247 | 2851 |
| 4642 | 0.01 | 0.08 | 0.11 | 0.91 | 1 | Pcm1 | pericentriolar material 1 | 3300 | 4470 | 3656 | 5155 | 4298 | 5761 | 4539 | 4889 | 4546 | 4434 | 6018 |
| 3616 | 0.00 | 0.08 | -0.06 | 0.96 | 1 | Prkce | protein kinase C, epsilon | 2203 | 2641 | 2565 | 4337 | 4262 | 3047 | 4536 | 4734 | 4126 | 4338 | 2981 |
| 3872 | 0.12 | 0.13 | 0.96 | 0.34 | 1 | Asns | asparagine synthetase | 1525 | 2591 | 2559 | 4498 | 4938 | 3178 | 4534 | 5398 | 5293 | 4781 | 3301 |

| | | | | | | | | | | | | | | | | | |
|---|---|---|---|---|---|---|---|---|---|---|---|---|---|---|---|---|---|
| 4155 | -0.18 | 0.11 | -1.66 | 0.10 | 1 | Tecr | trans-2,3-enoyl-CoA reductase | 3146 | 3430 | 3785 | 4738 | 4977 | 4481 | 4531 | 3485 | 4413 | 4878 | 3847 |
| 5347 | 0.17 | 0.09 | 1.89 | 0.06 | 1 | Sh3bgrl | SH3-binding domain glutamic acid-rich pı | 9097 | 7676 | 7458 | 4567 | 4131 | 3098 | 4525 | 5539 | 4236 | 4788 | 3700 |
| 3533 | -0.11 | 0.11 | -1.04 | 0.30 | 1 | Kif5c | kinesin family member 5C | 2376 | 1905 | 2619 | 4043 | 3834 | 4200 | 4525 | 3858 | 3580 | 3953 | 3971 |
| 4466 | 0.08 | 0.07 | 1.08 | 0.28 | 1 | Ei24 | etoposide induced 2.4 mRNA | 4342 | 4293 | 4602 | 4153 | 5029 | 3706 | 4520 | 4774 | 5147 | 5062 | 3498 |
| 4285 | -0.14 | 0.10 | -1.47 | 0.14 | 1 | Rph3al | rabphilin 3A-like (without C2 domains) | 3033 | 4373 | 4342 | 4928 | 4892 | 4242 | 4514 | 4098 | 4735 | 4131 | 3847 |
| 4286 | 0.17 | 0.10 | 1.80 | 0.07 | 1 | Atp5pb | ATP synthase peripheral stalk-membrane | 4338 | 3482 | 4351 | 4251 | 4238 | 3438 | 4511 | 5222 | 4380 | 5044 | 3886 |
| 3913 | 0.19 | 0.10 | 1.78 | 0.08 | 1 | Atp1b3 | ATPase, Na+/K+ transporting, beta 3 poly | 4184 | 3246 | 3282 | 4212 | 3654 | 2769 | 4505 | 5417 | 4140 | 4484 | 3147 |
| 3548 | 0.03 | 0.08 | 0.34 | 0.74 | 1 | Scp2 | sterol carrier protein 2, liver | 2809 | 2549 | 2717 | 3895 | 3663 | 3291 | 4503 | 4044 | 3536 | 4359 | 3660 |
| 4087 | -0.01 | 0.10 | -0.06 | 0.96 | 1 | Akt3 | thymoma viral proto-oncogene 3 | 3292 | 3210 | 3296 | 4736 | 4711 | 3658 | 4501 | 3709 | 4960 | 4895 | 3989 |
| 4020 | -0.06 | 0.09 | -0.67 | 0.50 | 1 | Emc7 | ER membrane protein complex subunit 7 | 3539 | 3739 | 3996 | 4579 | 4274 | 3461 | 4501 | 3761 | 4480 | 4312 | 3574 |
| 5497 | -0.03 | 0.07 | -0.45 | 0.65 | 1 | Rpl23 | ribosomal protein L23 | 9955 | 9171 | 10882 | 3728 | 4173 | 3038 | 4500 | 3711 | 4114 | 3978 | 3217 |
| 4338 | 0.06 | 0.10 | 0.65 | 0.51 | 1 | Ints3 | integrator complex subunit 3 | 2712 | 3099 | 2897 | 4634 | 5295 | 4636 | 4492 | 4536 | 5329 | 5621 | 4466 |
| 4796 | -0.06 | 0.10 | -0.66 | 0.51 | 1 | Isl1 | ISL1 transcription factor, LIM/homeodoma | 4462 | 5041 | 5317 | 4293 | 3696 | 6811 | 4490 | 3690 | 3825 | 4131 | 6995 |
| 3448 | 0.02 | 0.11 | 0.19 | 0.85 | 1 | Klhl32 | kelch-like 32 | 840 | 817 | 975 | 4698 | 4035 | 4275 | 4489 | 4796 | 4364 | 3801 | 4840 |
| 4296 | -0.01 | 0.07 | -0.12 | 0.90 | 1 | Atg13 | autophagy related 13 | 2799 | 3326 | 3703 | 4738 | 5270 | 4192 | 4488 | 4792 | 5868 | 4282 | 3801 |
| 3968 | -0.01 | 0.06 | -0.17 | 0.87 | 1 | Vps11 | VPS11, CORVET/HOPS core subunit | 3535 | 3220 | 3437 | 4036 | 4532 | 3727 | 4487 | 4075 | 4454 | 4499 | 3651 |
| 4651 | 0.05 | 0.07 | 0.77 | 0.44 | 1 | Cct6a | chaperonin containing Tcp1, subunit 6a (: | 5460 | 5303 | 6197 | 3673 | 4499 | 4144 | 4484 | 3940 | 4600 | 4338 | 4520 |
| 4310 | -0.11 | 0.07 | -1.43 | 0.15 | 1 | Mef2d | myocyte enhancer factor 2D | 4105 | 4471 | 4592 | 4496 | 4578 | 4174 | 4480 | 3866 | 4721 | 4191 | 3733 |
| 3429 | 0.16 | 0.13 | 1.18 | 0.24 | 1 | Gmpr | guanosine monophosphate reductase | 1735 | 1474 | 1591 | 3418 | 4214 | 3434 | 4477 | 5457 | 3801 | 4548 | 3571 |
| 3639 | -0.08 | 0.09 | -0.89 | 0.37 | 1 | Mfsd1 | major facilitator superfamily domain conta | 3460 | 2826 | 3022 | 4266 | 4323 | 2731 | 4476 | 3452 | 4393 | 4270 | 2807 |
| 3118 | -0.08 | 0.17 | -0.50 | 0.61 | 1 | Akr1c14 | aldo-keto reductase family 1, member C1 | 696 | 1371 | 1213 | 4601 | 3778 | 3162 | 4476 | 4035 | 3577 | 4100 | 3286 |
| 4705 | -0.05 | 0.07 | -0.69 | 0.49 | 1 | Gpi1 | glucose-6-phosphate isomerase 1 | 7147 | 6471 | 6015 | 3987 | 4137 | 3728 | 4474 | 3520 | 3833 | 4671 | 3777 |
| 4365 | 0.02 | 0.07 | 0.22 | 0.83 | 1 | Slc30a5 | solute carrier family 30 (zinc transporter), | 2862 | 2922 | 3103 | 4356 | 5734 | 4918 | 4472 | 4610 | 5401 | 4546 | 5088 |
| 4752 | -0.09 | 0.08 | -1.09 | 0.28 | 1 | Rps17 | ribosomal protein S17 | 7848 | 7413 | 8567 | 3436 | 3826 | 2966 | 4471 | 3210 | 3805 | 3698 | 3028 |
| 5170 | 0.06 | 0.08 | 0.69 | 0.49 | 1 | Capns1 | calpain, small subunit 1 | 7821 | 6273 | 7111 | 4768 | 4518 | 3705 | 4454 | 4459 | 4506 | 5311 | 3941 |
| 4622 | 0.00 | 0.06 | 0.00 | 1.00 | 1 | Usp7 | ubiquitin specific peptidase 7 | 4676 | 4441 | 4361 | 4210 | 4630 | 5393 | 4453 | 4188 | 4626 | 4466 | 5404 |
| 4034 | -0.05 | 0.09 | -0.54 | 0.59 | 1 | Nsf | N-ethylmaleimide sensitive fusion protein | 3186 | 3415 | 3310 | 4288 | 5095 | 3686 | 4449 | 3721 | 4984 | 4647 | 3593 |
| 3307 | 0.07 | 0.12 | 0.60 | 0.55 | 1 | Fam210b | family with sequence similarity 210, meml | 1518 | 1709 | 1830 | 3832 | 5179 | 1833 | 4448 | 3371 | 5649 | 4936 | 2070 |
| 3158 | 0.08 | 0.07 | 1.16 | 0.25 | 1 | Lrrc10b | leucine rich repeat containing 10B | 1246 | 1036 | 1038 | 4016 | 3721 | 3029 | 4448 | 3936 | 3943 | 5157 | 3172 |
| 3895 | 0.06 | 0.09 | 0.64 | 0.52 | 1 | Cyld | CYLD lysine 63 deubiquitinase | 3166 | 3500 | 3072 | 3805 | 4332 | 3641 | 4446 | 3875 | 4590 | 4382 | 4040 |
| 4161 | 0.07 | 0.10 | 0.67 | 0.50 | 1 | Sub1 | SUB1 homolog, transcriptional regulator | 4781 | 3075 | 3835 | 4266 | 4084 | 3821 | 4445 | 4068 | 4242 | 4691 | 4462 |
| 3863 | 0.08 | 0.07 | 1.24 | 0.21 | 1 | Mindy2 | MINDY lysine 48 deubiquitinase 2 | 3415 | 3435 | 2963 | 4112 | 4015 | 3308 | 4445 | 4250 | 4268 | 4638 | 3641 |
| 4596 | -0.05 | 0.08 | -0.59 | 0.55 | 1 | Anapc1 | anaphase promoting complex subunit 1 | 5225 | 5088 | 4679 | 4270 | 4493 | 4854 | 4443 | 4032 | 4662 | 4182 | 4623 |
| 4128 | 0.02 | 0.06 | 0.26 | 0.80 | 1 | Fxr2 | fragile X mental retardation, autosomal hc | 3166 | 3097 | 3611 | 4066 | 4782 | 4385 | 4440 | 4244 | 4568 | 4613 | 4430 |
| 4120 | 0.01 | 0.08 | 0.11 | 0.91 | 1 | Galnt1 | polypeptide N-acetylgalactosaminyltransf | 5030 | 4354 | 4376 | 4232 | 3905 | 3152 | 4440 | 3718 | 4366 | 4524 | 3220 |
| 4163 | -0.08 | 0.10 | -0.80 | 0.42 | 1 | Oga | O-GlcNAcase | 3363 | 2695 | 3611 | 4668 | 4299 | 5169 | 4438 | 4437 | 4192 | 4054 | 4871 |
| 4620 | 0.09 | 0.12 | 0.73 | 0.46 | 1 | Prdx1 | peroxiredoxin 1 | 6351 | 4557 | 6017 | 4547 | 4338 | 3086 | 4437 | 5472 | 4443 | 4323 | 3250 |
| 3427 | -0.04 | 0.07 | -0.61 | 0.54 | 1 | Sumf1 | sulfatase modifying factor 1 | 2906 | 2878 | 2838 | 3809 | 3833 | 2688 | 4434 | 3703 | 3683 | 4278 | 2651 |
| 3970 | 0.00 | 0.10 | -0.04 | 0.96 | 1 | Slc25a51 | solute carrier family 25, member 51 | 3087 | 3298 | 2873 | 4275 | 4354 | 4159 | 4431 | 3660 | 4954 | 4276 | 4304 |
| 3948 | -0.01 | 0.07 | -0.08 | 0.93 | 1 | Zzef1 | zinc finger, ZZ-type with EF hand domain | 2860 | 2934 | 3067 | 3991 | 4003 | 4909 | 4428 | 3877 | 4318 | 4368 | 4675 |
| 4333 | -0.09 | 0.08 | -1.17 | 0.24 | 1 | Lrpap1 | low density lipoprotein receptor-related pı | 4948 | 4961 | 5008 | 4300 | 4699 | 3483 | 4427 | 4493 | 3853 | 4032 | 3461 |
| 4247 | 0.02 | 0.09 | 0.24 | 0.81 | 1 | Sec23ip | Sec23 interacting protein | 2689 | 2781 | 2629 | 4431 | 5395 | 4895 | 4427 | 4306 | 5707 | 4633 | 4819 |
| 4243 | 0.08 | 0.07 | 1.18 | 0.24 | 1 | Fam129b | family with sequence similarity 129, meml | 4138 | 3641 | 4296 | 4169 | 4747 | 3446 | 4423 | 4716 | 5150 | 4387 | 3554 |
| 2648 | -0.20 | 0.16 | -1.24 | 0.21 | 1 | Cdk14 | cyclin-dependent kinase 14 | 1901 | 1778 | 1184 | 2856 | 3013 | 2722 | 4421 | 1981 | 2990 | 3711 | 2577 |
| 4414 | -0.04 | 0.09 | -0.47 | 0.64 | 1 | Zwint | ZW10 interactor | 5385 | 5476 | 5398 | 4430 | 3757 | 3794 | 4407 | 3580 | 3830 | 4309 | 4185 |
| 4016 | 0.04 | 0.08 | 0.54 | 0.59 | 1 | Slc30a9 | solute carrier family 30 (zinc transporter), | 2765 | 2635 | 3100 | 4375 | 4376 | 4438 | 4394 | 4671 | 4322 | 4636 | 4469 |
| 4294 | 0.14 | 0.08 | 1.74 | 0.08 | 1 | Chmp3 | charged multivesicular body protein 3 | 4991 | 4384 | 5045 | 4057 | 4203 | 3005 | 4393 | 4616 | 4152 | 4823 | 3563 |
| 3743 | -0.08 | 0.10 | -0.80 | 0.43 | 1 | Dnajc14 | DnaJ heat shock protein family (Hsp40) m | 3366 | 3253 | 3553 | 3871 | 4076 | 3540 | 4393 | 3016 | 4387 | 4189 | 3530 |
| 4369 | -0.10 | 0.08 | -1.30 | 0.19 | 1 | Tom1l2 | target of myb1-like 2 (chicken) | 3740 | 4275 | 4351 | 4746 | 4627 | 4678 | 4392 | 3932 | 4769 | 4033 | 4514 |
| 3958 | 0.13 | 0.09 | 1.42 | 0.15 | 1 | Polr2b | polymerase (RNA) II (DNA directed) polyp | 3377 | 3637 | 3475 | 3669 | 4493 | 3297 | 4392 | 4311 | 4413 | 4534 | 3942 |
| 3844 | 0.05 | 0.08 | 0.67 | 0.50 | 1 | Ppp3r1 | protein phosphatase 3, regulatory subuni | 2686 | 3010 | 3219 | 3999 | 4372 | 3619 | 4392 | 4449 | 4334 | 4453 | 3757 |
| 4722 | 0.00 | 0.09 | 0.04 | 0.97 | 1 | Tmpo | thymopoietin | 5721 | 5035 | 4764 | 4605 | 4077 | 5088 | 4386 | 3757 | 4483 | 4781 | 5241 |
| 3832 | -0.02 | 0.11 | -0.14 | 0.89 | 1 | Cab39 | calcium binding protein 39 | 3721 | 3938 | 3030 | 3975 | 3754 | 3727 | 4386 | 3670 | 4252 | 3686 | 4014 |
| 4387 | 0.00 | 0.07 | -0.06 | 0.95 | 1 | Vdac2 | voltage-dependent anion channel 2 | 5637 | 5069 | 5749 | 3848 | 4481 | 3217 | 4385 | 3498 | 4333 | 4623 | 3422 |
| 4174 | 0.06 | 0.08 | 0.74 | 0.46 | 1 | Sepsecs | Sep (O-phosphoserine) tRNA:Sec (selen | 1324 | 1665 | 1607 | 4144 | 4591 | 7038 | 4384 | 4368 | 4846 | 4307 | 7643 |
| 4672 | -0.14 | 0.08 | -1.78 | 0.08 | 1 | Rpl17 | ribosomal protein L17 | 8035 | 7517 | 8122 | 3455 | 3844 | 2869 | 4379 | 2986 | 3806 | 3614 | 2763 |
| 4083 | 0.08 | 0.10 | 0.79 | 0.43 | 1 | Rab21 | RAB21, member RAS oncogene family | 3744 | 4276 | 4054 | 4134 | 4067 | 3330 | 4376 | 3870 | 4366 | 5290 | 3403 |
| 3761 | -0.04 | 0.09 | -0.40 | 0.69 | 1 | Rhobtb1 | Rho-related BTB domain containing 1 | 1593 | 1926 | 1706 | 4897 | 4512 | 4518 | 4374 | 4230 | 5070 | 4369 | 4179 |
| 4255 | -0.02 | 0.07 | -0.32 | 0.75 | 1 | Ube2z | ubiquitin-conjugating enzyme E2Z | 4549 | 4838 | 4275 | 4239 | 4539 | 3559 | 4373 | 4135 | 4261 | 4475 | 3563 |
| 2589 | 0.11 | 0.18 | 0.63 | 0.53 | 1 | Tcf24 | transcription factor 24 | 427 | 409 | 735 | 3384 | 3449 | 1483 | 4371 | 3648 | 3922 | 5289 | 1363 |

| | | | | | | | | | | | | | | | | | | |
|---|---|---|---|---|---|---|---|---|---|---|---|---|---|---|---|---|---|---|
| 3619 | -0.02 | 0.12 | -0.17 | 0.86 | 1 | Eaf1 | ELL associated factor 1 | 2336 | 2950 | 2491 | 4028 | 3893 | 3842 | 4367 | 3349 | 4545 | 4072 | 3942 |
| 4626 | 0.06 | 0.10 | 0.62 | 0.54 | 1 | Cenpb | centromere protein B | 6288 | 5356 | 5861 | 4478 | 4563 | 2862 | 4366 | 5405 | 4145 | 4730 | 2827 |
| 4930 | 0.06 | 0.12 | 0.51 | 0.61 | 1 | Mbnl1 | muscleblind like splicing factor 1 | 8018 | 4904 | 5206 | 4155 | 4074 | 5040 | 4362 | 3766 | 4985 | 4682 | 5033 |
| 3877 | -0.02 | 0.09 | -0.18 | 0.86 | 1 | Ckap5 | cytoskeleton associated protein 5 | 3524 | 3134 | 2946 | 4278 | 4015 | 3950 | 4360 | 4663 | 4138 | 3907 | 3732 |
| 3881 | 0.05 | 0.07 | 0.65 | 0.52 | 1 | Slc35f5 | solute carrier family 35, member F5 | 2094 | 2263 | 2208 | 4635 | 4862 | 3885 | 4359 | 4482 | 5182 | 4900 | 3817 |
| 4556 | 0.22 | 0.13 | 1.67 | 0.09 | 1 | Park7 | Parkinson disease (autosomal recessive, | 4433 | 4148 | 5332 | 4463 | 4246 | 3575 | 4357 | 6539 | 4325 | 4682 | 4019 |
| 4302 | -0.02 | 0.07 | -0.28 | 0.78 | 1 | Dnmt3a | DNA methyltransferase 3A | 2832 | 2657 | 2698 | 4308 | 4209 | 6865 | 4357 | 4090 | 4610 | 4172 | 6529 |
| 3953 | 0.05 | 0.07 | 0.63 | 0.53 | 1 | Ube2q1 | ubiquitin-conjugating enzyme E2Q family | 3460 | 3503 | 3047 | 3977 | 4166 | 3971 | 4356 | 4143 | 4327 | 4520 | 4015 |
| 4922 | 0.15 | 0.11 | 1.33 | 0.18 | 1 | Klf9 | Kruppel-like factor 9 | 4580 | 4164 | 3549 | 5335 | 3791 | 6427 | 4355 | 6747 | 4486 | 4613 | 6092 |
| 3890 | 0.05 | 0.11 | 0.45 | 0.65 | 1 | Atp13a3 | ATPase type 13A3 | 2936 | 2986 | 2389 | 4225 | 4060 | 4309 | 4355 | 3696 | 4722 | 4647 | 4465 |
| 3833 | -0.16 | 0.09 | -1.83 | 0.07 | 1 | Tmem131 | transmembrane protein 131 | 3815 | 3576 | 3895 | 4204 | 4295 | 3388 | 4352 | 3209 | 4425 | 4106 | 2900 |
| 3632 | 0.06 | 0.11 | 0.54 | 0.59 | 1 | Phyh | phytanoyl-CoA hydroxylase | 2142 | 2694 | 2922 | 3919 | 3987 | 3509 | 4349 | 3896 | 4104 | 4561 | 3871 |
| 3832 | -0.04 | 0.08 | -0.53 | 0.60 | 1 | Ppt1 | palmitoyl-protein thioesterase 1 | 6365 | 4176 | 3884 | 3435 | 3393 | 2848 | 4347 | 3252 | 3030 | 4543 | 2876 |
| 4707 | 0.04 | 0.07 | 0.62 | 0.53 | 1 | Maco1 | macoilin 1 | 4013 | 5136 | 4109 | 4936 | 4667 | 4996 | 4345 | 5358 | 4734 | 4177 | 5308 |
| 3713 | -0.10 | 0.12 | -0.86 | 0.39 | 1 | Sdc3 | syndecan 3 | 6019 | 3622 | 3813 | 3924 | 3475 | 2379 | 4340 | 3189 | 3838 | 4130 | 2112 |
| 3630 | 0.06 | 0.11 | 0.55 | 0.58 | 1 | Bcar1 | breast cancer anti-estrogen resistance 1 | 1933 | 2571 | 2408 | 4031 | 3979 | 3796 | 4338 | 4316 | 4273 | 4685 | 3602 |
| 4763 | -0.01 | 0.08 | -0.17 | 0.87 | 1 | Qsox1 | quiescin Q6 sulfhydryl oxidase 1 | 3298 | 5255 | 3714 | 6751 | 4765 | 4104 | 4334 | 7670 | 4423 | 4446 | 3634 |
| 3352 | -0.09 | 0.18 | -0.48 | 0.63 | 1 | Ubxn2a | UBX domain protein 2A | 1977 | 2801 | 2139 | 3809 | 4005 | 3232 | 4333 | 2571 | 4466 | 4473 | 3069 |
| 4194 | 0.12 | 0.09 | 1.25 | 0.21 | 1 | Ktn1 | kinectin 1 | 3298 | 2839 | 3564 | 4070 | 4256 | 4832 | 4329 | 5028 | 4538 | 4284 | 5094 |
| 3836 | 0.08 | 0.09 | 0.96 | 0.34 | 1 | Cdipt | CDP-diacylglycerol--inositol 3-phosphatic | 2819 | 2651 | 3214 | 4480 | 4358 | 3033 | 4324 | 4626 | 4668 | 5087 | 2935 |
| 3806 | 0.10 | 0.11 | 0.87 | 0.39 | 1 | Cdc73 | cell division cycle 73, Paf1/RNA polymera | 2639 | 3190 | 3312 | 3601 | 3998 | 3850 | 4324 | 3449 | 4751 | 4422 | 4332 |
| 3479 | -0.03 | 0.14 | -0.22 | 0.82 | 1 | Slc35e1 | solute carrier family 35, member E1 | 2405 | 2585 | 2251 | 3234 | 4493 | 3593 | 4318 | 2869 | 4414 | 4598 | 3505 |
| 4073 | -0.04 | 0.08 | -0.50 | 0.61 | 1 | Zfp652 | zinc finger protein 652 | 3660 | 3941 | 4036 | 3935 | 4739 | 3789 | 4315 | 3930 | 4881 | 4027 | 3547 |
| 3583 | -0.20 | 0.13 | -1.49 | 0.14 | 1 | Phf2 | PHD finger protein 2 | 3581 | 3427 | 3035 | 4086 | 4052 | 3149 | 4314 | 2659 | 4235 | 4226 | 2654 |
| 3175 | -0.16 | 0.11 | -1.46 | 0.14 | 1 | Kcnn3 | potassium intermediate/small conductanc | 2180 | 2274 | 1771 | 3918 | 3571 | 3353 | 4311 | 3123 | 3828 | 3811 | 2785 |
| 4491 | -0.03 | 0.08 | -0.37 | 0.71 | 1 | Sf3b3 | splicing factor 3b, subunit 3 | 5413 | 5282 | 4903 | 4578 | 4229 | 3931 | 4309 | 4020 | 4532 | 4615 | 3588 |
| 4292 | -0.15 | 0.10 | -1.51 | 0.13 | 1 | Gse1 | genetic suppressor element 1, coiled-coil | 2982 | 3672 | 3323 | 4866 | 4975 | 5438 | 4301 | 3821 | 4835 | 4346 | 4648 |
| 4046 | -0.13 | 0.08 | -1.72 | 0.08 | 1 | Nfic | nuclear factor I/C | 5217 | 4873 | 4665 | 3727 | 4407 | 3094 | 4297 | 3563 | 3792 | 4231 | 2642 |
| 4886 | -0.08 | 0.10 | -0.83 | 0.41 | 1 | Vwa5b2 | von Willebrand factor A domain containin | 3541 | 5104 | 4206 | 5055 | 4602 | 7250 | 4295 | 5295 | 4258 | 4566 | 5578 |
| 4169 | 0.02 | 0.07 | 0.23 | 0.82 | 1 | Cyhr1 | cysteine and histidine rich 1 | 4036 | 4190 | 3978 | 3959 | 4188 | 4289 | 4287 | 4246 | 3851 | 4557 | 4281 |
| 5192 | -0.13 | 0.07 | -1.74 | 0.08 | 1 | Pkd1 | polycystin 1, transient receptor poteintial ( | 7470 | 6092 | 5732 | 3841 | 3491 | 8324 | 4285 | 3294 | 3491 | 4258 | 6833 |
| 5179 | 0.07 | 0.07 | 1.03 | 0.30 | 1 | Septin7 | septin 7 | 8277 | 7000 | 7023 | 4035 | 4000 | 4580 | 4284 | 3930 | 4470 | 4412 | 4959 |
| 3996 | 0.03 | 0.10 | 0.28 | 0.78 | 1 | Lrrc58 | leucine rich repeat containing 58 | 2827 | 4181 | 3337 | 4335 | 3978 | 4033 | 4284 | 4426 | 4428 | 4069 | 4054 |
| 3724 | 0.14 | 0.08 | 1.74 | 0.08 | 1 | Cfap36 | cilia and flagella associated protein 36 | 2744 | 2674 | 3063 | 3685 | 3822 | 3665 | 4282 | 4068 | 4098 | 4606 | 4257 |
| 3984 | 0.10 | 0.08 | 1.24 | 0.21 | 1 | Fh1 | fumarate hydratase 1 | 2721 | 2592 | 2629 | 4334 | 4160 | 4472 | 4281 | 4241 | 4033 | 5157 | 5203 |
| 3978 | -0.12 | 0.10 | -1.21 | 0.23 | 1 | Itpr3 | inositol 1,4,5-triphosphate receptor 3 | 3451 | 3362 | 3144 | 4238 | 4307 | 4746 | 4279 | 3557 | 4473 | 4399 | 3807 |
| 4009 | -0.15 | 0.10 | -1.42 | 0.16 | 1 | Med13 | mediator complex subunit 13 | 3858 | 4683 | 3735 | 3875 | 3823 | 4756 | 4275 | 3639 | 4008 | 3363 | 4089 |
| 3716 | -0.10 | 0.10 | -0.92 | 0.36 | 1 | Rraga | Ras-related GTP binding A | 3280 | 3387 | 3233 | 3802 | 4302 | 3602 | 4274 | 3002 | 4182 | 4160 | 3653 |
| 4173 | 0.06 | 0.08 | 0.76 | 0.45 | 1 | Zbtb7a | zinc finger and BTB domain containing 7a | 4135 | 4300 | 4083 | 4233 | 4341 | 3427 | 4273 | 4835 | 4782 | 4368 | 3125 |
| 4864 | 0.15 | 0.11 | 1.35 | 0.18 | 1 | Hdgf | heparin binding growth factor | 5870 | 4963 | 5949 | 4967 | 4506 | 3592 | 4272 | 6155 | 4523 | 4909 | 3795 |
| 4564 | -0.04 | 0.08 | -0.50 | 0.62 | 1 | Ik | IK cytokine | 4580 | 4659 | 5345 | 3680 | 4643 | 5371 | 4272 | 3437 | 4719 | 3773 | 5725 |
| 3800 | 0.03 | 0.07 | 0.41 | 0.68 | 1 | Fbxl5 | F-box and leucine-rich repeat protein 5 | 2907 | 2851 | 2963 | 4089 | 4381 | 3636 | 4272 | 4559 | 4231 | 4059 | 3847 |
| 4337 | 0.05 | 0.10 | 0.57 | 0.57 | 1 | G3bp1 | GTPase activating protein (SH3 domain) I | 5075 | 4373 | 4420 | 3935 | 4414 | 3991 | 4270 | 4682 | 4388 | 3877 | 4282 |
| 3950 | -0.11 | 0.07 | -1.56 | 0.12 | 1 | Kcnh2 | potassium voltage-gated channel, subfam | 2958 | 3106 | 3572 | 4663 | 4966 | 3680 | 4269 | 4218 | 4332 | 4351 | 3333 |
| 5242 | 1.09 | 0.64 | 1.71 | 0.09 | 1 | Lars2 | leucyl-tRNA synthetase, mitochondrial | 1760 | 2586 | 5172 | 3509 | 2775 | 2787 | 4265 | 4723 | 3826 | 22184 | 4076 |
| 4643 | -0.03 | 0.09 | -0.33 | 0.74 | 1 | Pkn1 | protein kinase N1 | 5536 | 4464 | 4874 | 4835 | 4883 | 4270 | 4263 | 4481 | 4879 | 4723 | 3859 |
| 3535 | -0.02 | 0.08 | -0.25 | 0.80 | 1 | Smpd1 | sphingomyelin phosphodiesterase 1, acic | 2666 | 2953 | 2528 | 4246 | 4057 | 2876 | 4262 | 3972 | 4080 | 4544 | 2702 |
| 4441 | 0.04 | 0.10 | 0.36 | 0.72 | 1 | Rnf187 | ring finger protein 187 | 5196 | 4729 | 5671 | 4202 | 4574 | 3340 | 4260 | 5222 | 4333 | 4132 | 3190 |
| 4248 | -0.01 | 0.09 | -0.12 | 0.90 | 1 | Arl6ip1 | ADP-ribosylation factor-like 6 interacting p | 5781 | 4807 | 4759 | 4176 | 4288 | 3067 | 4259 | 4062 | 4278 | 3941 | 3306 |
| 3858 | 0.02 | 0.07 | 0.23 | 0.82 | 1 | Ube2o | ubiquitin-conjugating enzyme E2O | 1935 | 2373 | 2023 | 5067 | 4286 | 4338 | 4258 | 4976 | 4759 | 4377 | 4048 |
| 3503 | -0.03 | 0.09 | -0.38 | 0.71 | 1 | Tug1 | | 2944 | 3058 | 2624 | 3707 | 3647 | 3527 | 4257 | 3985 | 3618 | 3449 | 3711 |
| 4813 | 0.10 | 0.10 | 1.05 | 0.29 | 1 | Phip | pleckstrin homology domain interacting p | 4115 | 4631 | 4275 | 4394 | 3923 | 6635 | 4252 | 5522 | 4447 | 3658 | 7096 |
| 3527 | 0.05 | 0.08 | 0.56 | 0.57 | 1 | Itprid2 | ITPR interacting domain containing 2 | 2310 | 2364 | 1992 | 4421 | 3884 | 3276 | 4252 | 3982 | 4301 | 4510 | 3507 |
| 3925 | 0.14 | 0.08 | 1.65 | 0.10 | 1 | Nus1 | NUS1 dehydrodolichyl diphosphate synth | 2782 | 2499 | 2688 | 4313 | 4228 | 3957 | 4251 | 3981 | 4872 | 5087 | 4522 |
| 3619 | 0.06 | 0.10 | 0.61 | 0.54 | 1 | Yipf2 | Yip1 domain family, member 2 | 1540 | 1602 | 2040 | 4141 | 4566 | 4013 | 4250 | 4068 | 4453 | 4874 | 4261 |
| 3449 | 0.01 | 0.14 | 0.10 | 0.92 | 1 | Nol4 | nucleolar protein 4 | 2139 | 3058 | 1930 | 3860 | 3874 | 3363 | 4249 | 3627 | 3886 | 4506 | 3451 |
| 3567 | -0.04 | 0.10 | -0.35 | 0.73 | 1 | Bcl2l2 | BCL2-like 2 | 2249 | 2328 | 2289 | 4024 | 3744 | 4332 | 4248 | 3161 | 4106 | 4271 | 4480 |
| 4438 | -0.20 | 0.14 | -1.46 | 0.15 | 1 | Nfat5 | nuclear factor of activated T cells 5 | 4283 | 4767 | 3831 | 4333 | 3997 | 6648 | 4245 | 3373 | 4808 | 2956 | 5575 |
| 2938 | -0.19 | 0.12 | -1.61 | 0.11 | 1 | Glis3 | GLIS family zinc finger 3 | 1149 | 1231 | 1394 | 3854 | 4153 | 2947 | 4243 | 2687 | 4225 | 3475 | 2965 |
| 4146 | 0.02 | 0.09 | 0.28 | 0.78 | 1 | Pja1 | praja ring finger ubiquitin ligase 1 | 3787 | 4718 | 4512 | 4049 | 4098 | 3770 | 4242 | 4035 | 4073 | 4291 | 4029 |

| | | | | | | | | | | | | | | | | | | |
|---|---|---|---|---|---|---|---|---|---|---|---|---|---|---|---|---|---|---|
| 4042 | -0.02 | 0.07 | -0.27 | 0.79 | 1 | Vps4b | vacuolar protein sorting 4B | 3513 | 3579 | 3697 | 3962 | 4475 | 4265 | 4241 | 3668 | 4403 | 4473 | 4187 |
| 4361 | 0.14 | 0.11 | 1.28 | 0.20 | 1 | Ralbp1 | ralA binding protein 1 | 4678 | 4614 | 4459 | 3898 | 4378 | 3647 | 4239 | 5699 | 4772 | 3974 | 3610 |
| 4806 | 0.03 | 0.07 | 0.47 | 0.64 | 1 | Btg1 | BTG anti-proliferation factor 1 | 5551 | 3728 | 4105 | 3376 | 3707 | 8193 | 4238 | 3400 | 3727 | 4489 | 8351 |
| 3852 | -0.13 | 0.10 | -1.28 | 0.20 | 1 | Prkaca | protein kinase, cAMP dependent, catalytic | 3389 | 3483 | 3812 | 4283 | 4654 | 3357 | 4234 | 3107 | 4544 | 4222 | 3284 |
| 4178 | -0.02 | 0.06 | -0.39 | 0.70 | 1 | Ubtf | upstream binding transcription factor, RNA | 4596 | 3968 | 4330 | 4020 | 4200 | 4223 | 4233 | 4071 | 4009 | 4155 | 4155 |
| 4213 | 0.12 | 0.08 | 1.55 | 0.12 | 1 | Api5 | apoptosis inhibitor 5 | 4187 | 3950 | 3927 | 4017 | 4170 | 3979 | 4230 | 3812 | 4675 | 4915 | 4485 |
| 3538 | -0.07 | 0.10 | -0.74 | 0.46 | 1 | Mgat3 | mannoside acetylglucosaminyltransferase | 3263 | 3134 | 3054 | 3686 | 4154 | 2995 | 4228 | 3064 | 3921 | 4516 | 2906 |
| 4233 | 0.04 | 0.10 | 0.38 | 0.71 | 1 | Mkrn1 | makorin, ring finger protein, 1 | 3965 | 4345 | 4773 | 4010 | 4442 | 3783 | 4225 | 5200 | 4210 | 4140 | 3472 |
| 3244 | 0.08 | 0.07 | 1.24 | 0.22 | 1 | Snx30 | sorting nexin family member 30 | 1925 | 1643 | 1883 | 3397 | 3777 | 3310 | 4223 | 3763 | 4008 | 4200 | 3559 |
| 3708 | -0.12 | 0.09 | -1.30 | 0.20 | 1 | Setd7 | SET domain containing (lysine methyltran | 2941 | 3176 | 2763 | 4394 | 4169 | 3841 | 4221 | 3612 | 4414 | 3671 | 3589 |
| 4708 | 0.09 | 0.10 | 0.84 | 0.40 | 1 | Ptp4a2 | protein tyrosine phosphatase 4a2 | 7333 | 5465 | 5872 | 4455 | 4421 | 3008 | 4220 | 4855 | 4265 | 4470 | 3424 |
| 3938 | -0.01 | 0.06 | -0.10 | 0.92 | 1 | Dlst | dihydrolipoamide S-succinyltransferase (I | 3922 | 3665 | 3791 | 3881 | 3891 | 4011 | 4218 | 3755 | 4011 | 3994 | 4174 |
| 4364 | 0.03 | 0.08 | 0.41 | 0.68 | 1 | Zfp148 | zinc finger protein 148 | 4426 | 4631 | 3928 | 4222 | 4306 | 4570 | 4215 | 4112 | 4421 | 4655 | 4522 |
| 4361 | 0.08 | 0.08 | 1.04 | 0.30 | 1 | Tpi1 | triosephosphate isomerase 1 | 5263 | 4632 | 5224 | 4141 | 4578 | 3045 | 4215 | 4613 | 4167 | 4695 | 3400 |
| 4058 | 0.12 | 0.11 | 1.13 | 0.26 | 1 | Cdk5rap3 | CDK5 regulatory subunit associated prote | 2835 | 2738 | 3187 | 4201 | 4428 | 4315 | 4215 | 5598 | 4212 | 4574 | 4332 |
| 4256 | 0.13 | 0.10 | 1.33 | 0.18 | 1 | Bzw1 | basic leucine zipper and W2 domains 1 | 4731 | 3678 | 3615 | 4111 | 4307 | 4006 | 4212 | 4504 | 4564 | 4379 | 4708 |
| 4383 | 0.05 | 0.13 | 0.38 | 0.70 | 1 | Zdbf2 | zinc finger, DBF-type containing 2 | 3830 | 6411 | 5076 | 4004 | 3499 | 4392 | 4210 | 4073 | 4289 | 3559 | 4865 |
| 4111 | 0.03 | 0.09 | 0.39 | 0.70 | 1 | Scaf1 | SR-related CTD-associated factor 1 | 3648 | 3224 | 3696 | 4197 | 4482 | 4231 | 4204 | 4891 | 4206 | 4396 | 4045 |
| 3669 | 0.14 | 0.14 | 0.98 | 0.32 | 1 | Guk1 | guanylate kinase 1 | 1925 | 1807 | 2852 | 4330 | 4237 | 3309 | 4202 | 5188 | 4130 | 4775 | 3609 |
| 3854 | -0.02 | 0.08 | -0.19 | 0.85 | 1 | Trappc11 | trafficking protein particle complex 11 | 2394 | 2860 | 2868 | 4001 | 4451 | 4568 | 4201 | 3794 | 4663 | 4139 | 4458 |
| 3895 | 0.01 | 0.06 | 0.22 | 0.82 | 1 | Immt | inner membrane protein, mitochondrial | 3451 | 2966 | 3120 | 3867 | 4014 | 4505 | 4194 | 3972 | 4131 | 4173 | 4449 |
| 3528 | -0.12 | 0.09 | -1.25 | 0.21 | 1 | Tm9sf4 | transmembrane 9 superfamily protein me | 2945 | 3018 | 3270 | 4088 | 3807 | 3290 | 4192 | 3186 | 3841 | 3936 | 3230 |
| 3334 | -0.06 | 0.10 | -0.62 | 0.53 | 1 | Cobll1 | Cobl-like 1 | 2031 | 2371 | 2040 | 4139 | 3736 | 3351 | 4192 | 3572 | 4194 | 4070 | 2975 |
| 4573 | 0.07 | 0.13 | 0.49 | 0.62 | 1 | Fmn2 | formin 2 | 1660 | 4845 | 2449 | 7002 | 4404 | 4690 | 4191 | 6831 | 5033 | 4271 | 4926 |
| 4444 | -0.06 | 0.10 | -0.60 | 0.55 | 1 | Bclaf1 | BCL2-associated transcription factor 1 | 4002 | 4073 | 4170 | 3145 | 3938 | 7317 | 4191 | 2838 | 4253 | 3555 | 7406 |
| 3997 | -0.01 | 0.07 | -0.16 | 0.87 | 1 | Aldh9a1 | aldehyde dehydrogenase 9, subfamily A1 | 2926 | 2994 | 3286 | 4684 | 5210 | 3397 | 4189 | 4408 | 4846 | 4546 | 3477 |
| 3673 | -0.14 | 0.08 | -1.83 | 0.07 | 1 | Rpa1 | replication protein A1 | 3348 | 2946 | 2806 | 4533 | 4242 | 3459 | 4185 | 3918 | 3826 | 3941 | 3198 |
| 3768 | 0.06 | 0.15 | 0.38 | 0.70 | 1 | Rhpn2 | rhophilin, Rho GTPase binding protein 2 | 1503 | 2606 | 2675 | 5086 | 4432 | 3364 | 4184 | 4569 | 4402 | 5078 | 3550 |
| 3798 | -0.01 | 0.09 | -0.14 | 0.89 | 1 | Gspt1 | G1 to S phase transition 1 | 3680 | 3484 | 3252 | 3653 | 4121 | 3778 | 4183 | 3208 | 4281 | 4217 | 3925 |
| 3668 | 0.14 | 0.09 | 1.48 | 0.14 | 1 | Cmas | cytidine monophospho-N-acetylneuramin | 2098 | 2255 | 2089 | 3717 | 4560 | 3701 | 4175 | 4209 | 4448 | 5022 | 4070 |
| 3915 | -0.02 | 0.07 | -0.28 | 0.78 | 1 | Cdk16 | cyclin-dependent kinase 16 | 2805 | 2448 | 2788 | 4255 | 4445 | 4771 | 4173 | 4305 | 4277 | 4303 | 4498 |
| 3752 | -0.04 | 0.08 | -0.53 | 0.60 | 1 | Gak | cyclin G associated kinase | 2823 | 2641 | 2564 | 4075 | 4749 | 3855 | 4173 | 4057 | 4304 | 4359 | 3667 |
| 4101 | 0.15 | 0.11 | 1.43 | 0.15 | 1 | Afdn | afadin, adherens junction formation factor | 3568 | 3167 | 3972 | 3774 | 3790 | 4583 | 4170 | 4928 | 4525 | 4337 | 4294 |
| 4120 | 0.04 | 0.07 | 0.50 | 0.61 | 1 | Ythdf3 | YTH N6-methyladenosine RNA binding p | 3262 | 4337 | 3314 | 4614 | 3837 | 4370 | 4166 | 4686 | 4034 | 4069 | 4628 |
| 4036 | 0.17 | 0.11 | 1.51 | 0.13 | 1 | Zranb1 | zinc finger, RAN-binding domain containi | 2347 | 2615 | 2616 | 5261 | 4399 | 3460 | 4166 | 6776 | 4422 | 4106 | 4230 |
| 3927 | 0.12 | 0.10 | 1.19 | 0.23 | 1 | Ube2r2 | ubiquitin-conjugating enzyme E2R 2 | 4371 | 4272 | 4661 | 3818 | 3688 | 2546 | 4166 | 5105 | 3878 | 4154 | 2537 |
| 5555 | 0.10 | 0.09 | 1.11 | 0.27 | 1 | Npm1 | nucleophosmin 1 | 11511 | 8699 | 9904 | 3501 | 3672 | 3635 | 4165 | 4097 | 3941 | 3761 | 4217 |
| 3618 | 0.02 | 0.09 | 0.18 | 0.85 | 1 | Coro2b | coronin, actin binding protein, 2B | 1694 | 2175 | 2409 | 4327 | 4596 | 3587 | 4165 | 4343 | 4621 | 4149 | 3729 |
| 3928 | 0.01 | 0.08 | 0.15 | 0.88 | 1 | Tsnax | translin-associated factor X | 4221 | 3891 | 4228 | 3513 | 4079 | 3636 | 4162 | 4063 | 3908 | 3768 | 3739 |
| 4221 | -0.03 | 0.11 | -0.29 | 0.77 | 1 | Zfp664 | zinc finger protein 664 | 4249 | 4512 | 4063 | 4078 | 4130 | 4582 | 4159 | 3183 | 4328 | 4552 | 4592 |
| 4302 | 0.09 | 0.10 | 0.91 | 0.36 | 1 | Prdx2 | peroxiredoxin 2 | 6029 | 5019 | 6241 | 3834 | 3809 | 2760 | 4158 | 4539 | 3845 | 4195 | 2894 |
| 3990 | 0.05 | 0.08 | 0.64 | 0.52 | 1 | Ccdc47 | coiled-coil domain containing 47 | 3269 | 3473 | 3085 | 3806 | 4323 | 4455 | 4158 | 4236 | 4325 | 4203 | 4556 |
| 4075 | -0.04 | 0.07 | -0.55 | 0.58 | 1 | Actr1b | ARP1 actin-related protein 1B, centractin | 3989 | 3739 | 4634 | 3987 | 4403 | 3908 | 4157 | 3922 | 4108 | 4005 | 3975 |
| 4035 | -0.08 | 0.06 | -1.26 | 0.21 | 1 | Cul4a | cullin 4A | 4107 | 3671 | 4152 | 4050 | 4313 | 4127 | 4156 | 3627 | 4186 | 4108 | 3891 |
| 4025 | -0.06 | 0.07 | -0.91 | 0.36 | 1 | Ppp2r5a | protein phosphatase 2, regulatory subuni | 4938 | 4779 | 4526 | 3512 | 3817 | 3879 | 4154 | 3729 | 3644 | 3729 | 3568 |
| 4087 | 0.00 | 0.08 | 0.04 | 0.96 | 1 | Arih1 | ariadne RBR E3 ubiquitin protein ligase 1 | 3782 | 3771 | 3680 | 4079 | 3883 | 4704 | 4152 | 3612 | 4359 | 4012 | 4921 |
| 4980 | -0.03 | 0.09 | -0.31 | 0.75 | 1 | Ip6k2 | inositol hexaphosphate kinase 2 | 2964 | 3406 | 3815 | 4489 | 4155 | 9669 | 4151 | 4672 | 3786 | 4030 | 9642 |
| 4208 | -0.09 | 0.09 | -0.96 | 0.34 | 1 | Fip1l1 | FIP1 like 1 (S. cerevisiae) | 3399 | 4256 | 4038 | 4494 | 4414 | 4685 | 4149 | 3907 | 4649 | 3429 | 4865 |
| 4131 | 0.05 | 0.08 | 0.63 | 0.53 | 1 | Dcaf11 | DDB1 and CUL4 associated factor 11 | 3269 | 3805 | 3672 | 4000 | 4617 | 4264 | 4149 | 4823 | 4508 | 4244 | 4093 |
| 3521 | 0.00 | 0.08 | -0.01 | 0.99 | 1 | Srprb | signal recognition particle receptor, B sub | 2374 | 2156 | 2680 | 3273 | 4705 | 3626 | 4147 | 3074 | 4603 | 4381 | 3714 |
| 4379 | -0.05 | 0.08 | -0.56 | 0.58 | 1 | Atp5c1 | ATP synthase, H+ transporting, mitochond | 6510 | 5851 | 6005 | 4151 | 3820 | 3078 | 4130 | 3720 | 4023 | 3587 | 3294 |
| 3723 | -0.14 | 0.08 | -1.87 | 0.06 | 1 | Slc29a1 | solute carrier family 29 (nucleoside transp | 3401 | 3540 | 4003 | 3461 | 4600 | 3565 | 4130 | 3276 | 4066 | 3662 | 3254 |
| 4164 | -0.02 | 0.06 | -0.32 | 0.75 | 1 | Ppm1a | protein phosphatase 1A, magnesium dep | 4657 | 4675 | 4247 | 4018 | 4220 | 3860 | 4127 | 4086 | 3981 | 4122 | 3814 |
| 4196 | 0.13 | 0.08 | 1.54 | 0.12 | 1 | Ibtk | inhibitor of Bruton agammaglobulinemia t | 3177 | 2962 | 3288 | 3772 | 5040 | 4652 | 4124 | 4544 | 5471 | 4604 | 4521 |
| 3332 | 0.01 | 0.14 | 0.07 | 0.94 | 1 | Atm | ataxia telangiectasia mutated | 1564 | 2356 | 1347 | 4467 | 3714 | 3415 | 4119 | 3724 | 3873 | 4148 | 3927 |
| 4746 | 0.03 | 0.14 | 0.23 | 0.82 | 1 | mt-Rnr1 | | 5328 | 8330 | 5614 | 4199 | 3757 | 4225 | 4115 | 3645 | 4857 | 4005 | 4131 |
| 4523 | -0.05 | 0.10 | -0.53 | 0.60 | 1 | Ctdsp2 | CTD (carboxy-terminal domain, RNA poly | 8019 | 5516 | 6413 | 3394 | 4344 | 3220 | 4103 | 2916 | 4475 | 4428 | 2923 |
| 3956 | 0.11 | 0.08 | 1.39 | 0.16 | 1 | Csnk1g2 | casein kinase 1, gamma 2 | 3770 | 3668 | 3659 | 3605 | 4276 | 3571 | 4103 | 4384 | 4159 | 4736 | 3580 |
| 3686 | -0.23 | 0.14 | -1.67 | 0.09 | 1 | Vat1l | vesicle amine transport protein 1 like | 4567 | 6245 | 4701 | 3090 | 3463 | 2886 | 4101 | 2638 | 2999 | 3157 | 2704 |
| 3420 | -0.01 | 0.10 | -0.14 | 0.89 | 1 | Blcap | bladder cancer associated protein | 1918 | 2285 | 1976 | 4022 | 3554 | 4106 | 4101 | 3655 | 3438 | 4638 | 3921 |

| | | | | | | | | | | | | | | | | | |
|---|---|---|---|---|---|---|---|---|---|---|---|---|---|---|---|---|---|
| 7961 | -0.05 | 0.13 | -0.38 | 0.71 | 1 | Rian | | 3442 | 4789 | 3982 | 4066 | 4417 | 25730 | 4100 | 3097 | 4569 | 4510 | 24871 |
| 3574 | -0.04 | 0.07 | -0.62 | 0.54 | 1 | Mgrn1 | mahogunin, ring finger 1 | 3386 | 3021 | 3141 | 3706 | 3754 | 3521 | 4100 | 3544 | 3982 | 3983 | 3172 |
| 3959 | 0.07 | 0.13 | 0.56 | 0.58 | 1 | Tbc1d15 | TBC1 domain family, member 15 | 2597 | 2992 | 2644 | 3770 | 5002 | 4306 | 4098 | 3337 | 5288 | 4629 | 4886 |
| 3744 | -0.07 | 0.06 | -1.15 | 0.25 | 1 | Dpp9 | dipeptidylpeptidase 9 | 2598 | 2539 | 2863 | 4236 | 4786 | 3880 | 4098 | 4146 | 4421 | 3887 | 3728 |
| 3667 | 0.00 | 0.09 | -0.03 | 0.98 | 1 | Ate1 | arginyltransferase 1 | 3046 | 3546 | 3387 | 3727 | 3760 | 3610 | 4092 | 3426 | 4102 | 3934 | 3706 |
| 3405 | -0.06 | 0.09 | -0.61 | 0.55 | 1 | Osbpl2 | oxysterol binding protein-like 2 | 2008 | 2663 | 2333 | 4158 | 4080 | 3211 | 4092 | 3779 | 4010 | 3945 | 3181 |
| 5264 | 0.03 | 0.07 | 0.45 | 0.66 | 1 | Ewsr1 | Ewing sarcoma breakpoint region 1 | 4494 | 5064 | 4910 | 4789 | 4444 | 8284 | 4091 | 4808 | 4401 | 4624 | 7998 |
| 3629 | 0.16 | 0.12 | 1.33 | 0.18 | 1 | Lhfpl4 | lipoma HMGIC fusion partner-like protein | 1711 | 2584 | 2152 | 3988 | 4465 | 3107 | 4091 | 5569 | 4303 | 4853 | 3090 |
| 4853 | 0.14 | 0.12 | 1.14 | 0.26 | 1 | Ppp1r9b | protein phosphatase 1, regulatory subuni | 7517 | 6598 | 6350 | 4237 | 3793 | 3390 | 4088 | 6097 | 3729 | 4307 | 3272 |
| 3701 | -0.01 | 0.09 | -0.11 | 0.91 | 1 | Tle3 | transducin-like enhancer of split 3 | 2202 | 1754 | 1924 | 4504 | 4351 | 4528 | 4088 | 4434 | 4402 | 4342 | 4177 |
| 3825 | 0.08 | 0.10 | 0.84 | 0.40 | 1 | Acat1 | acetyl-Coenzyme A acetyltransferase 1 | 4199 | 3848 | 4903 | 3644 | 3664 | 2800 | 4087 | 3879 | 3713 | 4088 | 3246 |
| 3537 | 0.05 | 0.08 | 0.62 | 0.54 | 1 | Tsc2 | TSC complex subunit 2 | 3380 | 3052 | 2688 | 3738 | 3471 | 3299 | 4086 | 4237 | 3710 | 4267 | 2981 |
| 4345 | -0.11 | 0.08 | -1.42 | 0.16 | 1 | Dhx15 | DEAH (Asp-Glu-Ala-His) box polypeptide | 5144 | 5135 | 4612 | 3783 | 4084 | 5062 | 4084 | 3556 | 3651 | 3550 | 5131 |
| 3433 | 0.18 | 0.11 | 1.61 | 0.11 | 1 | Cabp7 | calcium binding protein 7 | 1105 | 1582 | 1554 | 4332 | 3656 | 3581 | 4083 | 5247 | 4380 | 4540 | 3700 |
| 4647 | 0.14 | 0.11 | 1.27 | 0.20 | 1 | Tmem176a | transmembrane protein 176A | 7767 | 5706 | 6448 | 4060 | 3776 | 2884 | 4075 | 5084 | 4025 | 4341 | 2948 |
| 3164 | 0.07 | 0.11 | 0.58 | 0.56 | 1 | Msmo1 | methylsterol monoxygenase 1 | 804 | 931 | 1033 | 3787 | 4017 | 3756 | 4074 | 4751 | 3680 | 3871 | 4097 |
| 3763 | 0.00 | 0.09 | 0.04 | 0.97 | 1 | Tmem214 | transmembrane protein 214 | 2229 | 2434 | 2150 | 3683 | 4462 | 5065 | 4072 | 3731 | 4535 | 4124 | 4906 |
| 3694 | -0.03 | 0.11 | -0.22 | 0.82 | 1 | Morf4l1 | mortality factor 4 like 1 | 3087 | 2951 | 3834 | 3247 | 4248 | 3955 | 4065 | 4020 | 4158 | 3107 | 3960 |
| 4466 | 0.08 | 0.09 | 0.91 | 0.36 | 1 | Hmgn3 | high mobility group nucleosomal binding | 3023 | 3986 | 4030 | 3856 | 4516 | 6056 | 4064 | 4451 | 4631 | 3996 | 6514 |
| 4270 | 0.03 | 0.08 | 0.32 | 0.75 | 1 | Cnot7 | CCR4-NOT transcription complex, subuni | 4673 | 4736 | 5197 | 3848 | 3759 | 4352 | 4061 | 3493 | 4221 | 4288 | 4340 |
| 3669 | 0.07 | 0.09 | 0.81 | 0.42 | 1 | Gmfb | glia maturation factor, beta | 3827 | 3459 | 3274 | 3675 | 3586 | 3179 | 4059 | 3254 | 3692 | 4607 | 3749 |
| 4838 | -0.01 | 0.07 | -0.13 | 0.90 | 1 | Add1 | adducin 1 (alpha) | 8005 | 6836 | 7092 | 4212 | 4168 | 3238 | 4058 | 3934 | 4172 | 4302 | 3196 |
| 4502 | 0.16 | 0.11 | 1.38 | 0.17 | 1 | Cdc37 | cell division cycle 37 | 5083 | 4679 | 5878 | 4141 | 4117 | 3689 | 4049 | 5474 | 4132 | 4486 | 3795 |
| 3348 | 0.09 | 0.08 | 1.12 | 0.26 | 1 | Sar1b | secretion associated Ras related GTPase | 1877 | 1788 | 2076 | 4016 | 3994 | 3025 | 4049 | 4317 | 3994 | 4276 | 3415 |
| 3837 | -0.04 | 0.11 | -0.39 | 0.70 | 1 | Apc | APC, WNT signaling pathway regulator | 3203 | 3709 | 3029 | 3982 | 4070 | 4260 | 4046 | 3335 | 4462 | 3846 | 4262 |
| 3449 | 0.07 | 0.10 | 0.70 | 0.48 | 1 | Kcnma1 | potassium large conductance calcium-act | 1334 | 2097 | 1480 | 4467 | 3991 | 3595 | 4043 | 4625 | 4459 | 4141 | 3704 |
| 2783 | 0.11 | 0.08 | 1.29 | 0.20 | 1 | Grin1 | glutamate receptor, ionotropic, NMDA1 (z | 329 | 304 | 357 | 3589 | 3743 | 2884 | 4042 | 3681 | 3954 | 4602 | 3127 |
| 3830 | -0.02 | 0.08 | -0.29 | 0.77 | 1 | Mapk14 | mitogen-activated protein kinase 14 | 4106 | 3486 | 3436 | 3877 | 4157 | 3585 | 4038 | 3773 | 4202 | 4117 | 3355 |
| 5239 | 0.28 | 0.28 | 1.01 | 0.31 | 1 | Rplp2 | ribosomal protein, large P2 | 9111 | 7361 | 9534 | 3385 | 3939 | 2662 | 4036 | 6988 | 3531 | 4379 | 2697 |
| 3737 | 0.06 | 0.08 | 0.78 | 0.43 | 1 | Ppm1g | protein phosphatase 1G (formerly 2C), ma | 3999 | 3600 | 3988 | 3138 | 4143 | 3125 | 4033 | 3881 | 4065 | 4092 | 3044 |
| 4276 | -0.05 | 0.07 | -0.73 | 0.46 | 1 | Hnrnpd | heterogeneous nuclear ribonucleoprotein | 5415 | 5390 | 5108 | 3681 | 4135 | 3980 | 4032 | 3651 | 3845 | 3877 | 3924 |
| 3673 | 0.04 | 0.08 | 0.53 | 0.60 | 1 | Tef | thyrotroph embryonic factor | 3054 | 2601 | 2621 | 4052 | 3789 | 3935 | 4032 | 4158 | 3503 | 4443 | 4217 |
| 3767 | 0.01 | 0.09 | 0.09 | 0.93 | 1 | Actr10 | ARP10 actin-related protein 10 | 3532 | 3660 | 4077 | 3662 | 4100 | 3232 | 4030 | 3398 | 3942 | 4582 | 3228 |
| 5120 | 0.22 | 0.13 | 1.73 | 0.08 | 1 | Rpl14 | ribosomal protein L14 | 8549 | 7869 | 9582 | 3457 | 3561 | 2854 | 4029 | 5467 | 3733 | 4208 | 3006 |
| 3773 | 0.09 | 0.07 | 1.25 | 0.21 | 1 | Slc25a36 | solute carrier family 25, member 36 | 1981 | 2443 | 2215 | 4463 | 3762 | 4667 | 4025 | 4738 | 4216 | 4145 | 4847 |
| 3408 | -0.07 | 0.11 | -0.63 | 0.53 | 1 | Epb41 | erythrocyte membrane protein band 4.1 | 1899 | 1818 | 1715 | 4043 | 3741 | 4592 | 4025 | 3145 | 4457 | 3854 | 4196 |
| 4221 | 0.01 | 0.07 | 0.20 | 0.84 | 1 | Cttn | cortactin | 4721 | 4053 | 5088 | 3671 | 4578 | 3966 | 4015 | 3997 | 4210 | 4198 | 3937 |
| 3693 | 0.08 | 0.09 | 0.93 | 0.35 | 1 | Lrp10 | low-density lipoprotein receptor-related p | 3933 | 3486 | 3349 | 3851 | 4039 | 2499 | 4015 | 4716 | 3794 | 4508 | 2432 |
| 3442 | -0.06 | 0.10 | -0.65 | 0.51 | 1 | Kcnb1 | potassium voltage gated channel, Shab-r | 2539 | 2773 | 2272 | 4922 | 3186 | 3423 | 4015 | 3868 | 3591 | 3983 | 3287 |
| 3494 | -0.06 | 0.08 | -0.79 | 0.43 | 1 | Tbcd | tubulin-specific chaperone d | 2094 | 1959 | 2249 | 4578 | 4441 | 3398 | 4013 | 4457 | 4126 | 3826 | 3296 |
| 3324 | 0.13 | 0.09 | 1.33 | 0.18 | 1 | Crebrf | CREB3 regulatory factor | 2715 | 2122 | 2906 | 3452 | 3739 | 2597 | 4010 | 3938 | 4426 | 3797 | 2867 |
| 3296 | -0.17 | 0.11 | -1.49 | 0.14 | 1 | Paxip1 | PAX interacting (with transcription-activati | 1327 | 1439 | 1279 | 3924 | 4019 | 5074 | 4008 | 2902 | 4087 | 3819 | 4376 |
| 3546 | 0.02 | 0.08 | 0.28 | 0.78 | 1 | Daam1 | dishevelled associated activator of morph | 2154 | 2538 | 2226 | 3932 | 3678 | 4313 | 4007 | 4188 | 3886 | 4010 | 4069 |
| 3432 | 0.15 | 0.07 | 1.97 | 0.05 | 1 | Leprotl1 | leptin receptor overlapping transcript-like | 2400 | 2597 | 2425 | 3971 | 3537 | 2902 | 4007 | 4057 | 4205 | 4319 | 3328 |
| 3505 | -0.01 | 0.08 | -0.09 | 0.93 | 1 | Rab5a | RAB5A, member RAS oncogene family | 3019 | 3086 | 3134 | 3644 | 3761 | 3299 | 3999 | 3302 | 3931 | 3775 | 3608 |
| 4322 | 0.13 | 0.13 | 1.01 | 0.31 | 1 | Hip1r | huntingtin interacting protein 1 related | 3374 | 2892 | 3601 | 4445 | 4970 | 4543 | 3997 | 6125 | 5212 | 4199 | 4181 |
| 3953 | 0.09 | 0.08 | 1.14 | 0.25 | 1 | Gpat4 | glycerol-3-phosphate acyltransferase 4 | 3548 | 3617 | 3901 | 3915 | 4148 | 3617 | 3995 | 4821 | 4187 | 3965 | 3770 |
| 4453 | 0.11 | 0.10 | 1.06 | 0.29 | 1 | Trim28 | tripartite motif-containing 28 | 5925 | 4799 | 5343 | 3960 | 4024 | 3849 | 3993 | 4976 | 4106 | 4251 | 3752 |
| 3564 | 0.01 | 0.07 | 0.18 | 0.85 | 1 | Fam114a1 | family with sequence similarity 114, meml | 2714 | 3069 | 2848 | 3853 | 3945 | 3437 | 3992 | 3974 | 3925 | 3840 | 3610 |
| 3315 | 0.05 | 0.09 | 0.52 | 0.61 | 1 | Hacd3 | 3-hydroxyacyl-CoA dehydratase 3 | 2766 | 3010 | 2889 | 3576 | 3295 | 2827 | 3992 | 3413 | 3675 | 4033 | 2985 |
| 3550 | 0.07 | 0.08 | 0.92 | 0.36 | 1 | Ppme1 | protein phosphatase methylesterase 1 | 2735 | 2842 | 3174 | 3526 | 3943 | 3339 | 3988 | 3771 | 3979 | 4132 | 3622 |
| 3222 | -0.01 | 0.07 | -0.15 | 0.88 | 1 | Abcg1 | ATP binding cassette subfamily G membe | 3342 | 2938 | 3319 | 3227 | 3486 | 2245 | 3984 | 3385 | 3625 | 3788 | 2099 |
| 4083 | -0.01 | 0.08 | -0.12 | 0.91 | 1 | Emc10 | ER membrane protein complex subunit 1( | 5068 | 4725 | 5172 | 4187 | 4087 | 2740 | 3982 | 4239 | 3727 | 4333 | 2652 |
| 3675 | 0.13 | 0.11 | 1.15 | 0.25 | 1 | Ica1 | islet cell autoantigen 1 | 2020 | 2784 | 2710 | 3724 | 4181 | 3852 | 3980 | 4532 | 3997 | 4340 | 4306 |
| 3232 | 0.07 | 0.11 | 0.70 | 0.48 | 1 | Ormdl3 | ORM1-like 3 (S. cerevisiae) | 2113 | 2035 | 2334 | 3583 | 3534 | 2997 | 3976 | 4226 | 3525 | 4464 | 2764 |
| 4504 | 0.06 | 0.11 | 0.59 | 0.56 | 1 | Kmt2c | lysine (K)-specific methyltransferase 2C | 3891 | 3828 | 3858 | 4404 | 3872 | 6493 | 3975 | 4690 | 5039 | 4081 | 5415 |
| 4048 | 0.05 | 0.08 | 0.60 | 0.55 | 1 | Nek7 | NIMA (never in mitosis gene a)-related ex | 4807 | 4231 | 4310 | 4308 | 3915 | 3156 | 3971 | 3918 | 4269 | 4177 | 3461 |
| 3564 | -0.08 | 0.10 | -0.83 | 0.40 | 1 | Dhx40 | DEAH (Asp-Glu-Ala-His) box polypeptide | 2515 | 3069 | 3565 | 3449 | 4460 | 3562 | 3971 | 3290 | 4248 | 3574 | 3504 |
| 3320 | 0.05 | 0.16 | 0.29 | 0.77 | 1 | B830017H08Rik | | 188 | 182 | 363 | 4218 | 4878 | 4631 | 3969 | 4348 | 4319 | 4825 | 4600 |
| 4162 | 0.12 | 0.07 | 1.74 | 0.08 | 1 | Zfp706 | zinc finger protein 706 | 4672 | 4419 | 4213 | 4332 | 4210 | 3056 | 3965 | 4548 | 4502 | 4591 | 3269 |

| | | | | | | | | | | | | | | | | | |
|---|---|---|---|---|---|---|---|---|---|---|---|---|---|---|---|---|---|
| 3948 | 0.18 | 0.10 | 1.87 | 0.06 | 1 | Limd1 | LIM domains containing 1 | 3816 | 4118 | 3808 | 3970 | 3875 | 2901 | 3965 | 5670 | 4000 | 4315 | 2985 |
| 3877 | 0.06 | 0.08 | 0.77 | 0.44 | 1 | Ufl1 | UFM1 specific ligase 1 | 2018 | 1755 | 2378 | 3052 | 4941 | 5887 | 3963 | 3048 | 4981 | 4334 | 6291 |
| 4425 | 0.00 | 0.07 | -0.03 | 0.98 | 1 | Rpl28 | ribosomal protein L28 | 7541 | 6101 | 7715 | 3420 | 3765 | 2521 | 3958 | 3432 | 3633 | 4070 | 2518 |
| 4343 | 0.15 | 0.08 | 1.91 | 0.06 | 1 | Clta | clathrin, light polypeptide (Lca) | 5654 | 5099 | 5357 | 3839 | 3996 | 3204 | 3955 | 4532 | 3979 | 4794 | 3361 |
| 4110 | 0.16 | 0.09 | 1.88 | 0.06 | 1 | Set | SET nuclear oncogene | 4248 | 4423 | 4471 | 3375 | 4186 | 3610 | 3953 | 4551 | 4391 | 4298 | 3704 |
| 4631 | -0.15 | 0.12 | -1.30 | 0.20 | 1 | Prrc2c | proline-rich coiled-coil 2C | 4294 | 4576 | 4445 | 4347 | 3871 | 7776 | 3949 | 3039 | 4472 | 3903 | 6271 |
| 3197 | 0.06 | 0.10 | 0.56 | 0.57 | 1 | Adam9 | a disintegrin and metallopeptidase domai | 2257 | 1466 | 1868 | 3233 | 3140 | 4090 | 3948 | 2754 | 3680 | 4069 | 4666 |
| 3666 | -0.12 | 0.13 | -0.92 | 0.36 | 1 | Sgms2 | sphingomyelin synthase 2 | 2285 | 3656 | 3211 | 4216 | 4153 | 3871 | 3945 | 3429 | 4514 | 3580 | 3471 |
| 5022 | 0.02 | 0.07 | 0.23 | 0.82 | 1 | Srsf1 | serine and arginine-rich splicing factor 1 | 5556 | 5467 | 5085 | 3348 | 3760 | 8362 | 3940 | 3418 | 3940 | 3730 | 8640 |
| 3865 | -0.04 | 0.11 | -0.39 | 0.70 | 1 | Dync2h1 | dynein cytoplasmic 2 heavy chain 1 | 2725 | 3914 | 3362 | 3584 | 3550 | 5391 | 3939 | 3845 | 3947 | 3099 | 5154 |
| 3312 | 0.08 | 0.12 | 0.70 | 0.49 | 1 | Tlcd4 | TLC domain containing 4 | 963 | 1554 | 1475 | 4122 | 4372 | 3331 | 3939 | 4158 | 4809 | 4135 | 3578 |
| 2861 | -0.04 | 0.11 | -0.38 | 0.70 | 1 | Mroh1 | maestro heat-like repeat family member 1 | 1188 | 1260 | 1410 | 3721 | 3190 | 3146 | 3937 | 3392 | 3299 | 3909 | 3015 |
| 5165 | 0.02 | 0.09 | 0.25 | 0.80 | 1 | Rpl18 | ribosomal protein L18 | 8551 | 8267 | 8568 | 4413 | 4125 | 3105 | 3935 | 4779 | 3530 | 4279 | 3260 |
| 3621 | -0.01 | 0.08 | -0.19 | 0.85 | 1 | Ubr1 | ubiquitin protein ligase E3 component n-r | 2961 | 3104 | 3191 | 3728 | 3842 | 3852 | 3930 | 3322 | 3996 | 3907 | 4001 |
| 2908 | -0.17 | 0.11 | -1.62 | 0.11 | 1 | Atrn | attractin | 2588 | 1850 | 2468 | 2993 | 3534 | 2850 | 3930 | 2463 | 3503 | 2841 | 2964 |
| 3127 | -0.09 | 0.11 | -0.81 | 0.42 | 1 | Tmco3 | transmembrane and coiled-coil domains | 1754 | 1851 | 2050 | 4071 | 3822 | 2997 | 3929 | 3114 | 3617 | 4103 | 3092 |
| 3914 | 0.01 | 0.10 | 0.13 | 0.90 | 1 | Acbd3 | acyl-Coenzyme A binding domain contair | 3537 | 4205 | 3420 | 3934 | 3666 | 4335 | 3928 | 3500 | 3638 | 4391 | 4503 |
| 4534 | -0.10 | 0.07 | -1.55 | 0.12 | 1 | Vat1 | vesicle amine transport 1 | 8621 | 7677 | 7751 | 3650 | 3454 | 2358 | 3927 | 3155 | 3085 | 3869 | 2324 |
| 3492 | -0.08 | 0.15 | -0.51 | 0.61 | 1 | Slc30a7 | solute carrier family 30 (zinc transporter), | 2046 | 2196 | 2028 | 3291 | 4758 | 4401 | 3927 | 2471 | 4712 | 3898 | 4686 |
| 4525 | -0.11 | 0.08 | -1.42 | 0.15 | 1 | Gstz1 | glutathione transferase zeta 1 (maleylace | 3149 | 3930 | 3734 | 4732 | 3879 | 7710 | 3924 | 4076 | 3753 | 3818 | 7071 |
| 3961 | 0.01 | 0.13 | 0.11 | 0.91 | 1 | Pik3r1 | phosphoinositide-3-kinase regulatory sub | 3497 | 3169 | 2555 | 5468 | 3841 | 3817 | 3922 | 5917 | 3825 | 3578 | 3976 |
| 3626 | 0.01 | 0.07 | 0.17 | 0.87 | 1 | Nploc4 | NPL4 homolog, ubiquitin recognition facto | 2708 | 2823 | 2967 | 3846 | 4138 | 3697 | 3922 | 3602 | 4544 | 3975 | 3660 |
| 3948 | 0.13 | 0.12 | 1.14 | 0.25 | 1 | Glb1l2 | galactosidase, beta 1-like 2 | 1896 | 2800 | 1754 | 4926 | 4906 | 3898 | 3920 | 5370 | 5078 | 4815 | 4068 |
| 3698 | -0.18 | 0.16 | -1.14 | 0.25 | 1 | Hcfc1 | host cell factor C1 | 4447 | 4270 | 3282 | 3731 | 4016 | 3522 | 3918 | 2989 | 4565 | 2857 | 3079 |
| 3625 | -0.10 | 0.08 | -1.25 | 0.21 | 1 | Dlg1 | discs large MAGUK scaffold protein 1 | 4040 | 3782 | 3865 | 3619 | 3729 | 3314 | 3918 | 3130 | 3615 | 3372 | 3487 |
| 3661 | 0.13 | 0.09 | 1.55 | 0.12 | 1 | Zyg11b | zyg-ll family member B, cell cycle regulato | 2895 | 3512 | 3167 | 3615 | 3720 | 3364 | 3917 | 4574 | 3926 | 3890 | 3694 |
| 3327 | 0.05 | 0.10 | 0.49 | 0.62 | 1 | Sec61b | Sec61 beta subunit | 1997 | 2200 | 2455 | 3386 | 4185 | 3234 | 3915 | 3793 | 3848 | 4378 | 3202 |
| 3604 | -0.02 | 0.11 | -0.23 | 0.82 | 1 | Cnpy2 | canopy FGF signaling regulator 2 | 3424 | 3449 | 4103 | 3762 | 4151 | 2623 | 3911 | 4525 | 3738 | 3309 | 2645 |
| 5781 | -0.04 | 0.08 | -0.53 | 0.60 | 1 | Nktr | natural killer tumor recognition sequence | 2566 | 2928 | 2800 | 3681 | 4085 | 16775 | 3907 | 3466 | 4346 | 3804 | 15236 |
| 3922 | -0.06 | 0.07 | -0.74 | 0.46 | 1 | Ergic3 | ERGIC and golgi 3 | 4789 | 4367 | 4845 | 3950 | 4004 | 2999 | 3907 | 4031 | 3648 | 3665 | 2944 |
| 3810 | 0.05 | 0.09 | 0.49 | 0.63 | 1 | Pccb | propionyl Coenzyme A carboxylase, beta | 3008 | 3262 | 3487 | 3987 | 4450 | 3436 | 3907 | 4998 | 4072 | 4030 | 3276 |
| 3904 | 0.14 | 0.09 | 1.59 | 0.11 | 1 | Diaph1 | diaphanous related formin 1 | 5207 | 4394 | 4893 | 3674 | 3390 | 2543 | 3906 | 4475 | 3672 | 4034 | 2756 |
| 4714 | -0.14 | 0.14 | -1.02 | 0.31 | 1 | Mlxipl | MLX interacting protein-like | 380 | 516 | 516 | 3472 | 3933 | 15572 | 3901 | 2683 | 3981 | 4007 | 12891 |
| 3207 | 0.08 | 0.14 | 0.54 | 0.59 | 1 | Ank2 | ankyrin 2, brain | 1259 | 2047 | 1486 | 3533 | 3117 | 4330 | 3901 | 3626 | 3798 | 4014 | 4171 |
| 3252 | 0.04 | 0.07 | 0.66 | 0.51 | 1 | Bmi1 | Bmi1 polycomb ring finger oncogene | 2239 | 2310 | 2028 | 4038 | 3749 | 2713 | 3898 | 3951 | 3838 | 4079 | 2927 |
| 2976 | 0.16 | 0.09 | 1.74 | 0.08 | 1 | Glrb | glycine receptor, beta subunit | 1999 | 2242 | 2015 | 3501 | 3282 | 1853 | 3898 | 3589 | 3985 | 4218 | 2159 |
| 4641 | 0.02 | 0.07 | 0.27 | 0.79 | 1 | Hnrnpa0 | heterogeneous nuclear ribonucleoprotei | 6766 | 6719 | 6517 | 3954 | 4458 | 3137 | 3894 | 4173 | 3959 | 4154 | 3325 |
| 3678 | -0.07 | 0.08 | -0.83 | 0.40 | 1 | Wipi2 | WD repeat domain, phosphoinositide inte | 2998 | 3532 | 3305 | 3806 | 4408 | 3563 | 3892 | 4141 | 3827 | 3529 | 3461 |
| 3761 | -0.02 | 0.07 | -0.22 | 0.83 | 1 | Gcn1 | GCN1 activator of EIF2AK4 | 4242 | 3627 | 3998 | 3332 | 3942 | 3664 | 3890 | 3605 | 3852 | 3891 | 3322 |
| 3174 | -0.09 | 0.10 | -0.92 | 0.36 | 1 | Dhx32 | DEAH (Asp-Glu-Ala-His) box polypeptide | 1499 | 1832 | 1719 | 4068 | 3930 | 3535 | 3890 | 3209 | 3898 | 3817 | 3520 |
| 3777 | 0.10 | 0.12 | 0.82 | 0.41 | 1 | H2-Q2 | histocompatibility 2, Q region locus 2 | 2779 | 3789 | 3614 | 4266 | 3691 | 3210 | 3889 | 5436 | 3330 | 4132 | 3413 |
| 4401 | 0.04 | 0.08 | 0.46 | 0.64 | 1 | Eif4a1 | eukaryotic translation initiation factor 4A1 | 5793 | 5505 | 5531 | 3318 | 4332 | 4054 | 3887 | 3457 | 4083 | 4104 | 4349 |
| 3704 | -0.03 | 0.08 | -0.41 | 0.69 | 1 | Yy1 | YY1 transcription factor | 4143 | 4309 | 3970 | 3636 | 3847 | 2973 | 3886 | 3559 | 3799 | 3430 | 3186 |
| 4156 | -0.06 | 0.10 | -0.60 | 0.55 | 1 | Capzb | capping protein (actin filament) muscle Z- | 6152 | 4831 | 5142 | 3875 | 4151 | 3166 | 3885 | 4152 | 3664 | 3750 | 2946 |
| 3924 | 0.07 | 0.10 | 0.69 | 0.49 | 1 | Pck2 | phosphoenolpyruvate carboxykinase 2 (n | 2575 | 3476 | 3915 | 4676 | 4182 | 3423 | 3884 | 5197 | 4311 | 4072 | 3459 |
| 3778 | -0.04 | 0.12 | -0.32 | 0.75 | 1 | Usp8 | ubiquitin specific peptidase 8 | 3706 | 3399 | 3474 | 3822 | 3892 | 4009 | 3880 | 2746 | 4356 | 4418 | 3852 |
| 3573 | -0.06 | 0.10 | -0.57 | 0.57 | 1 | Agap1 | ArfGAP with GTPase domain, ankyrin rep | 3639 | 3110 | 2970 | 3868 | 4186 | 3106 | 3877 | 3353 | 4440 | 3951 | 2806 |
| 4448 | 0.00 | 0.12 | 0.01 | 0.99 | 1 | Rpl26 | ribosomal protein L26 | 6974 | 6511 | 8572 | 3201 | 3708 | 2620 | 3875 | 4257 | 3341 | 3419 | 2449 |
| 3522 | 0.03 | 0.10 | 0.28 | 0.78 | 1 | Bmp1 | bone morphogenetic protein 1 | 2648 | 1868 | 2175 | 3534 | 4143 | 4299 | 3874 | 3538 | 4667 | 4038 | 3953 |
| 3834 | -0.05 | 0.09 | -0.57 | 0.57 | 1 | Cdkn1b | cyclin-dependent kinase inhibitor 1B | 3408 | 3310 | 3823 | 3695 | 3970 | 4492 | 3873 | 2968 | 4130 | 3975 | 4523 |
| 3665 | 0.06 | 0.08 | 0.75 | 0.45 | 1 | Opa1 | OPA1, mitochondrial dynamin like GTPas | 3010 | 3420 | 3304 | 3539 | 3865 | 3699 | 3873 | 3641 | 4163 | 3905 | 3898 |
| 3588 | 0.12 | 0.09 | 1.41 | 0.16 | 1 | Dld | dihydrolipoamide dehydrogenase | 2937 | 2489 | 2671 | 2951 | 4235 | 3988 | 3871 | 3559 | 4290 | 3984 | 4497 |
| 3313 | 0.01 | 0.10 | 0.12 | 0.90 | 1 | Pigq | phosphatidylinositol glycan anchor biosy | 2367 | 2642 | 2990 | 3776 | 3568 | 2919 | 3866 | 4358 | 3618 | 3453 | 2889 |
| 3550 | 0.02 | 0.08 | 0.24 | 0.81 | 1 | Trio | triple functional domain (PTPRF interactir | 2438 | 2283 | 2366 | 3911 | 3751 | 4350 | 3864 | 3678 | 4369 | 3990 | 4052 |
| 3541 | -0.04 | 0.09 | -0.46 | 0.64 | 1 | Klhl24 | kelch-like 24 | 3786 | 2976 | 4211 | 3671 | 3698 | 2952 | 3862 | 3207 | 3692 | 3923 | 2970 |
| 3728 | 0.14 | 0.09 | 1.52 | 0.13 | 1 | Tmem127 | transmembrane protein 127 | 3103 | 3122 | 3206 | 3951 | 3800 | 3370 | 3861 | 5140 | 3990 | 3885 | 3578 |
| 3133 | -0.10 | 0.08 | -1.27 | 0.21 | 1 | AU040320 | expressed sequence AU040320 | 2110 | 2209 | 1921 | 3981 | 3862 | 2904 | 3860 | 3441 | 3814 | 3702 | 2664 |
| 3769 | 0.09 | 0.10 | 0.95 | 0.34 | 1 | Tgfbr1 | transforming growth factor, beta receptor | 3816 | 3242 | 3491 | 3894 | 3684 | 3499 | 3859 | 4634 | 4120 | 3875 | 3346 |
| 2912 | -0.10 | 0.08 | -1.25 | 0.21 | 1 | Nipsnap2 | nipsnap homolog 2 | 1748 | 1786 | 1791 | 3242 | 3637 | 3095 | 3859 | 3307 | 3428 | 3189 | 2948 |
| 3757 | 0.08 | 0.11 | 0.77 | 0.44 | 1 | Npdc1 | neural proliferation, differentiation and co | 3119 | 3668 | 3648 | 3977 | 3656 | 3418 | 3858 | 5219 | 3730 | 3840 | 3197 |

| | | | | | | | | | | | | | | | | | |
|---|---|---|---|---|---|---|---|---|---|---|---|---|---|---|---|---|---|
| 3482 | -0.18 | 0.14 | -1.25 | 0.21 | 1 | Nup98 | nucleoporin 98 | 2788 | 3113 | 2578 | 4254 | 3756 | 3922 | 3857 | 2899 | 4488 | 3040 | 3606 |
| 3397 | 0.05 | 0.13 | 0.35 | 0.73 | 1 | Cbs | cystathionine beta-synthase | 1645 | 2757 | 2316 | 3806 | 3625 | 3765 | 3857 | 4291 | 3384 | 3967 | 3957 |
| 4341 | 0.01 | 0.08 | 0.16 | 0.88 | 1 | Hdac5 | histone deacetylase 5 | 4477 | 5031 | 3753 | 5104 | 4086 | 4078 | 3853 | 5341 | 3888 | 4440 | 3703 |
| 3542 | 0.05 | 0.10 | 0.52 | 0.61 | 1 | Psmc2 | proteasome (prosome, macropain) 26S su | 3125 | 2796 | 3309 | 3539 | 3999 | 3221 | 3853 | 4319 | 3493 | 3792 | 3514 |
| 3342 | -0.06 | 0.08 | -0.73 | 0.47 | 1 | Impdh1 | inosine monophosphate dehydrogenase | 2247 | 1706 | 2580 | 3550 | 4168 | 3814 | 3852 | 3122 | 4356 | 3688 | 3680 |
| 3241 | -0.18 | 0.14 | -1.33 | 0.18 | 1 | Usp22 | ubiquitin specific peptidase 22 | 2971 | 3678 | 2659 | 3069 | 3852 | 3243 | 3852 | 2652 | 3428 | 3336 | 2908 |
| 3277 | -0.04 | 0.09 | -0.39 | 0.69 | 1 | Prps2 | phosphoribosyl pyrophosphate synthetas | 3727 | 3183 | 3167 | 3416 | 3550 | 2373 | 3847 | 3173 | 3806 | 3325 | 2478 |
| 4102 | 0.02 | 0.09 | 0.26 | 0.80 | 1 | Zbtb4 | zinc finger and BTB domain containing 4 | 5471 | 5586 | 4700 | 4049 | 3246 | 3383 | 3844 | 3795 | 3512 | 4535 | 3001 |
| 3546 | -0.08 | 0.11 | -0.70 | 0.49 | 1 | Man1b1 | mannosidase, alpha, class 1B, member 1 | 4852 | 3446 | 3441 | 3676 | 3679 | 2833 | 3842 | 3703 | 3280 | 3389 | 2867 |
| 3491 | -0.08 | 0.10 | -0.75 | 0.45 | 1 | Wasl | Wiskott-Aldrich syndrome-like (human) | 3023 | 3482 | 3180 | 4087 | 3422 | 3395 | 3838 | 3286 | 3973 | 3453 | 3265 |
| 3138 | 0.10 | 0.12 | 0.81 | 0.42 | 1 | Iars | isoleucine-tRNA synthetase | 2029 | 2111 | 1880 | 2807 | 3567 | 3538 | 3837 | 3135 | 4385 | 3893 | 3336 |
| 3671 | 0.02 | 0.08 | 0.28 | 0.78 | 1 | Zzz3 | zinc finger, ZZ domain containing 3 | 3155 | 3422 | 3698 | 3739 | 3496 | 3865 | 3835 | 3662 | 3885 | 3723 | 3903 |
| 2698 | 0.12 | 0.15 | 0.80 | 0.42 | 1 | B3galt5 | UDP-Gal:betaGlcNAc beta 1,3-galactosyl | 895 | 934 | 1393 | 2976 | 3773 | 2099 | 3832 | 2652 | 4037 | 4598 | 2489 |
| 3311 | 0.04 | 0.12 | 0.36 | 0.72 | 1 | Rpl30 | ribosomal protein L30 | 6184 | 4025 | 4960 | 1912 | 2413 | 2446 | 3830 | 2563 | 2282 | 3192 | 2610 |
| 3819 | 0.10 | 0.08 | 1.23 | 0.22 | 1 | Uqcrc1 | ubiquinol-cytochrome c reductase core pr | 4232 | 3639 | 4154 | 3588 | 4222 | 2860 | 3829 | 4024 | 3898 | 4453 | 3113 |
| 3794 | 0.07 | 0.08 | 0.77 | 0.44 | 1 | Rnf145 | ring finger protein 145 | 3225 | 3332 | 3160 | 3475 | 4239 | 4102 | 3829 | 3819 | 4510 | 3719 | 4327 |
| 4191 | 0.08 | 0.09 | 0.91 | 0.36 | 1 | Cct3 | chaperonin containing Tcp1, subunit 3 (ga | 5706 | 4505 | 5966 | 2947 | 3882 | 3914 | 3828 | 3505 | 3745 | 3571 | 4533 |
| 3072 | 0.07 | 0.10 | 0.68 | 0.50 | 1 | Bhlhe41 | basic helix-loop-helix family, member e41 | 1937 | 2071 | 1502 | 3461 | 3277 | 3274 | 3825 | 3768 | 3612 | 3592 | 3472 |
| 3455 | 0.00 | 0.08 | -0.04 | 0.97 | 1 | Mtss1 | MTSS I-BAR domain containing 1 | 3011 | 3274 | 2444 | 3679 | 3229 | 3947 | 3818 | 3644 | 3479 | 3573 | 3906 |
| 3310 | 0.02 | 0.08 | 0.18 | 0.86 | 1 | Wars | tryptophanyl-tRNA synthetase | 2327 | 2100 | 2279 | 3358 | 4409 | 3160 | 3816 | 3282 | 4461 | 4188 | 3031 |
| 3300 | -0.01 | 0.07 | -0.12 | 0.91 | 1 | Stam2 | signal transducing adaptor molecule (SH3 | 2376 | 2424 | 2470 | 3834 | 3796 | 3131 | 3815 | 3470 | 3824 | 3778 | 3385 |
| 4066 | 0.11 | 0.07 | 1.68 | 0.09 | 1 | Cd2ap | CD2-associated protein | 3940 | 4614 | 3654 | 3987 | 3620 | 4192 | 3812 | 4621 | 3959 | 3951 | 4370 |
| 3820 | 0.14 | 0.12 | 1.17 | 0.24 | 1 | Bag1 | BCL2-associated athanogene 1 | 4143 | 3724 | 4616 | 3727 | 3718 | 2700 | 3809 | 5044 | 3515 | 4280 | 2741 |
| 3751 | 0.16 | 0.11 | 1.45 | 0.15 | 1 | Phf20l1 | PHD finger protein 20-like 1 | 1669 | 1855 | 1703 | 3200 | 3065 | 6990 | 3806 | 4544 | 3420 | 3448 | 7565 |
| 3395 | 0.03 | 0.08 | 0.46 | 0.65 | 1 | Suds3 | suppressor of defective silencing 3 homol | 2819 | 2820 | 3170 | 3489 | 3649 | 3164 | 3805 | 3664 | 3561 | 3836 | 3371 |
| 3193 | 0.10 | 0.08 | 1.18 | 0.24 | 1 | Bccip | BRCA2 and CDKN1A interacting protein | 2319 | 2140 | 2531 | 3361 | 3349 | 3065 | 3804 | 3845 | 3639 | 3891 | 3178 |
| 3570 | -0.03 | 0.12 | -0.27 | 0.78 | 1 | Rps6ka3 | ribosomal protein S6 kinase polypeptide 3 | 4565 | 4262 | 3746 | 3406 | 3286 | 3016 | 3801 | 2870 | 3873 | 3229 | 3216 |
| 4028 | 0.01 | 0.10 | 0.05 | 0.96 | 1 | Ssb | Sjogren syndrome antigen B | 4706 | 3724 | 4419 | 3487 | 3505 | 4772 | 3799 | 2772 | 4041 | 3759 | 5320 |
| 3048 | 0.02 | 0.12 | 0.12 | 0.90 | 1 | Steap2 | six transmembrane epithelial antigen of p | 1487 | 2120 | 1864 | 3847 | 3128 | 3210 | 3792 | 3434 | 3511 | 3809 | 3322 |
| 3030 | 0.02 | 0.10 | 0.16 | 0.88 | 1 | Mrs2 | MRS2 magnesium transporter | 1779 | 1766 | 1553 | 3068 | 3422 | 3750 | 3791 | 2886 | 3759 | 3818 | 3736 |
| 3721 | 0.07 | 0.08 | 0.90 | 0.37 | 1 | Vapa | vesicle-associated membrane protein, as | 4100 | 3830 | 3638 | 3191 | 3693 | 3649 | 3790 | 3485 | 4039 | 3855 | 3659 |
| 5270 | 0.15 | 0.08 | 1.89 | 0.06 | 1 | Marcks | myristoylated alanine rich protein kinase ( | 8261 | 8380 | 8755 | 4176 | 3716 | 3722 | 3789 | 5025 | 4212 | 3786 | 4153 |
| 4168 | 0.15 | 0.15 | 0.95 | 0.34 | 1 | Rpl21 | ribosomal protein L21 | 6071 | 5319 | 7196 | 3048 | 3710 | 2443 | 3788 | 4747 | 3452 | 3080 | 3000 |
| 3656 | 0.06 | 0.08 | 0.75 | 0.46 | 1 | Ptpra | protein tyrosine phosphatase, receptor typ | 4592 | 3455 | 3650 | 3375 | 3565 | 3220 | 3788 | 3569 | 3782 | 4127 | 3096 |
| 3263 | -0.04 | 0.10 | -0.37 | 0.71 | 1 | Patj | PATJ, crumbs cell polarity complex compo | 2669 | 3198 | 3162 | 3176 | 3758 | 2866 | 3787 | 3300 | 4064 | 3099 | 2816 |
| 3473 | -0.20 | 0.10 | -1.94 | 0.05 | 1 | Sptbn2 | spectrin beta, non-erythrocytic 2 | 2641 | 2615 | 3399 | 4570 | 4280 | 3173 | 3782 | 3879 | 3541 | 3663 | 2662 |
| 4197 | 0.10 | 0.08 | 1.23 | 0.22 | 1 | Wdr1 | WD repeat domain 1 | 5135 | 4543 | 4656 | 3994 | 4361 | 3253 | 3781 | 3960 | 4503 | 4340 | 3643 |
| 4513 | -0.01 | 0.11 | -0.05 | 0.96 | 1 | Slc38a2 | solute carrier family 38, member 2 | 4199 | 3720 | 3355 | 4881 | 3969 | 6643 | 3780 | 4408 | 4609 | 3643 | 6438 |
| 4629 | 0.14 | 0.08 | 1.65 | 0.10 | 1 | Msl1 | male specific lethal 1 | 4471 | 5251 | 4520 | 5059 | 4250 | 4308 | 3773 | 6283 | 4368 | 4327 | 4311 |
| 3341 | -0.05 | 0.10 | -0.48 | 0.63 | 1 | Slc35b4 | solute carrier family 35, member B4 | 1737 | 1778 | 1418 | 3936 | 3791 | 4694 | 3773 | 3414 | 4077 | 3702 | 4432 |
| 3218 | 0.00 | 0.11 | 0.00 | 1.00 | 1 | Usp24 | ubiquitin specific peptidase 24 | 3325 | 2646 | 2863 | 3087 | 2969 | 3487 | 3771 | 3331 | 3661 | 3002 | 3253 |
| 3289 | 0.01 | 0.11 | 0.09 | 0.93 | 1 | Ppp6c | protein phosphatase 6, catalytic subunit | 2945 | 3264 | 2937 | 3249 | 3446 | 3004 | 3770 | 2871 | 3863 | 3640 | 3187 |
| 3115 | -0.02 | 0.13 | -0.16 | 0.87 | 1 | Rnf115 | ring finger protein 115 | 1995 | 2353 | 2153 | 3685 | 3567 | 2962 | 3769 | 2730 | 3958 | 3864 | 3231 |
| 3938 | 0.06 | 0.08 | 0.67 | 0.50 | 1 | Agfg1 | ArfGAP with FG repeats 1 | 3099 | 3114 | 3175 | 4106 | 3474 | 5293 | 3766 | 4630 | 3729 | 3608 | 5326 |
| 3712 | -0.06 | 0.08 | -0.77 | 0.44 | 1 | Pink1 | PTEN induced putative kinase 1 | 4157 | 4674 | 4351 | 3626 | 3868 | 2844 | 3762 | 3854 | 3740 | 3193 | 2759 |
| 3962 | 0.04 | 0.09 | 0.42 | 0.67 | 1 | Spag9 | sperm associated antigen 9 | 3761 | 3930 | 3618 | 3158 | 3587 | 5415 | 3761 | 3085 | 3895 | 3855 | 5513 |
| 3429 | 0.21 | 0.11 | 1.87 | 0.06 | 1 | Rab3a | RAB3A, member RAS oncogene family | 1592 | 1439 | 2016 | 3500 | 4534 | 3387 | 3753 | 4906 | 4380 | 4610 | 3606 |
| 3817 | -0.08 | 0.07 | -1.14 | 0.26 | 1 | Cry2 | cryptochrome 2 (photolyase-like) | 2856 | 3010 | 2915 | 3499 | 3397 | 6507 | 3751 | 3218 | 3516 | 3507 | 5805 |
| 4893 | -0.02 | 0.08 | -0.23 | 0.82 | 1 | Rps18 | ribosomal protein S18 | 7918 | 8877 | 8655 | 3834 | 3551 | 3135 | 3745 | 4142 | 3404 | 3305 | 3263 |
| 4196 | -0.04 | 0.10 | -0.39 | 0.69 | 1 | Kmt2e | lysine (K)-specific methyltransferase 2E | 4538 | 5155 | 4041 | 3873 | 3782 | 5101 | 3743 | 3459 | 3991 | 4066 | 4412 |
| 3374 | 0.12 | 0.07 | 1.77 | 0.08 | 1 | Mettl9 | methyltransferase like 9 | 2837 | 2723 | 3066 | 3436 | 3576 | 2883 | 3741 | 3728 | 4110 | 4017 | 2994 |
| 3091 | -0.05 | 0.07 | -0.76 | 0.45 | 1 | Xpot | exportin, tRNA (nuclear export receptor fo | 2230 | 2154 | 2236 | 3231 | 3815 | 3164 | 3738 | 3234 | 3494 | 3666 | 3040 |
| 4139 | -0.02 | 0.10 | -0.23 | 0.82 | 1 | Megf8 | multiple EGF-like-domains 8 | 5292 | 5023 | 4751 | 3808 | 3975 | 3783 | 3737 | 4487 | 3471 | 3912 | 3295 |
| 3535 | -0.01 | 0.08 | -0.16 | 0.88 | 1 | Spryd3 | SPRY domain containing 3 | 3388 | 3209 | 3762 | 3491 | 3948 | 3150 | 3737 | 3902 | 3708 | 3545 | 3047 |
| 3822 | 0.00 | 0.11 | -0.01 | 0.99 | 1 | Xpo7 | exportin 7 | 3715 | 4109 | 3875 | 4039 | 3912 | 3488 | 3729 | 3082 | 4107 | 4213 | 3774 |
| 3620 | 0.11 | 0.08 | 1.40 | 0.16 | 1 | Selenow | selenoprotein W | 4359 | 3698 | 4171 | 3348 | 3813 | 2332 | 3729 | 3806 | 3764 | 4457 | 2343 |
| 3773 | 0.00 | 0.07 | -0.06 | 0.95 | 1 | Nap1l4 | nucleosome assembly protein 1-like 4 | 3699 | 3634 | 4171 | 3332 | 4090 | 3879 | 3726 | 3396 | 3842 | 3840 | 3888 |
| 3610 | 0.14 | 0.08 | 1.80 | 0.07 | 1 | Prpf40a | pre-mRNA processing factor 40A | 2899 | 2973 | 2500 | 3665 | 3660 | 3842 | 3721 | 4533 | 3939 | 3923 | 4051 |
| 3743 | 0.04 | 0.08 | 0.48 | 0.63 | 1 | Rnf20 | ring finger protein 20 | 2823 | 3051 | 2685 | 3294 | 3736 | 5341 | 3720 | 3535 | 3953 | 3641 | 5387 |
| 3739 | 0.02 | 0.08 | 0.27 | 0.79 | 1 | Dgkd | diacylglycerol kinase, delta | 3321 | 2723 | 3082 | 3954 | 3873 | 4336 | 3717 | 3998 | 3989 | 4061 | 4072 |

| | | | | | | | | | | | | | | | | | | |
|---|---|---|---|---|---|---|---|---|---|---|---|---|---|---|---|---|---|---|
| 2963 | 0.05 | 0.07 | 0.64 | 0.52 | 1 | Rcan2 | regulator of calcineurin 2 | 1971 | 1796 | 1858 | 3553 | 3480 | 2488 | 3716 | 3926 | 3672 | 3683 | 2446 |
| 4306 | 0.11 | 0.10 | 1.12 | 0.26 | 1 | Krt18 | keratin 18 | 3714 | 3857 | 5809 | 2999 | 5224 | 4528 | 3714 | 4019 | 5050 | 4084 | 4361 |
| 3413 | 0.06 | 0.07 | 0.91 | 0.36 | 1 | Msantd4 | Myb/SANT-like DNA-binding domain cont | 3260 | 3521 | 3120 | 3671 | 3178 | 2980 | 3709 | 3766 | 3284 | 3769 | 3281 |
| 3617 | 0.09 | 0.12 | 0.79 | 0.43 | 1 | Ndufs1 | NADH:ubiquinone oxidoreductase core s | 3465 | 3101 | 3403 | 3658 | 3409 | 3634 | 3706 | 4488 | 3721 | 3102 | 4104 |
| 3930 | 0.07 | 0.10 | 0.76 | 0.45 | 1 | Esyt2 | extended synaptotagmin-like protein 2 | 5256 | 4406 | 4274 | 3683 | 3327 | 3566 | 3705 | 3513 | 4085 | 3567 | 3849 |
| 3599 | 0.09 | 0.08 | 1.08 | 0.28 | 1 | Fgfr1op2 | FGFR1 oncogene partner 2 | 3270 | 3129 | 2904 | 3942 | 3691 | 3358 | 3705 | 4149 | 4097 | 3607 | 3742 |
| 3226 | -0.12 | 0.12 | -0.96 | 0.34 | 1 | Lats1 | large tumor suppressor | 2316 | 2924 | 2422 | 3266 | 3436 | 4043 | 3705 | 2695 | 3554 | 3267 | 3854 |
| 3659 | -0.10 | 0.10 | -1.00 | 0.32 | 1 | Ctso | cathepsin O | 3795 | 4749 | 4365 | 3573 | 3640 | 3222 | 3704 | 2989 | 3634 | 3321 | 3251 |
| 3126 | -0.06 | 0.12 | -0.49 | 0.63 | 1 | Exoc6b | exocyst complex component 6B | 2551 | 3390 | 2436 | 3222 | 3241 | 3086 | 3704 | 3369 | 3420 | 3100 | 2871 |
| 4018 | -0.09 | 0.07 | -1.25 | 0.21 | 1 | Cdc42bpb | CDC42 binding protein kinase beta | 5366 | 4792 | 5497 | 3888 | 3860 | 3237 | 3701 | 3491 | 3684 | 3667 | 3014 |
| 3219 | 0.07 | 0.09 | 0.78 | 0.44 | 1 | Atp6v1g1 | ATPase, H+ transporting, lysosomal V1 su | 2547 | 2445 | 2634 | 3460 | 3797 | 2650 | 3698 | 3883 | 3398 | 3799 | 3100 |
| 3532 | 0.05 | 0.10 | 0.48 | 0.63 | 1 | Clcn4 | chloride channel, voltage-sensitive 4 | 2976 | 3032 | 2936 | 3541 | 3948 | 3502 | 3695 | 4492 | 3806 | 3364 | 3560 |
| 3289 | 0.02 | 0.09 | 0.22 | 0.82 | 1 | Slc37a3 | solute carrier family 37 (glycerol-3-phospl | 2780 | 2153 | 2560 | 3629 | 3705 | 3208 | 3695 | 3611 | 3423 | 4046 | 3366 |
| 3939 | -0.04 | 0.07 | -0.60 | 0.55 | 1 | Fxr1 | fragile X mental retardation gene 1, autos | 4509 | 4560 | 4575 | 3360 | 3509 | 4471 | 3694 | 3151 | 3630 | 3229 | 4641 |
| 3830 | 0.04 | 0.10 | 0.45 | 0.65 | 1 | Ago2 | argonaute RISC catalytic subunit 2 | 3100 | 3661 | 3116 | 4293 | 3437 | 4415 | 3693 | 5213 | 3858 | 3510 | 3833 |
| 3299 | 0.02 | 0.11 | 0.15 | 0.88 | 1 | Cul1 | cullin 1 | 2902 | 2572 | 2340 | 3461 | 3416 | 3550 | 3691 | 2779 | 3969 | 3714 | 3893 |
| 3141 | 0.03 | 0.09 | 0.33 | 0.74 | 1 | Cog7 | component of oligomeric golgi complex 7 | 2207 | 2326 | 2293 | 3702 | 3508 | 2747 | 3691 | 4075 | 3468 | 4063 | 2473 |
| 3636 | -0.04 | 0.08 | -0.45 | 0.65 | 1 | Papss1 | 3'-phosphoadenosine 5'-phosphosulfate s | 3139 | 3286 | 3537 | 3775 | 3906 | 3825 | 3690 | 3183 | 3891 | 3875 | 3893 |
| 3231 | 0.10 | 0.10 | 1.00 | 0.32 | 1 | Washc5 | WASH complex subunit 5 | 2405 | 2250 | 1922 | 3762 | 3633 | 2915 | 3689 | 3425 | 3922 | 4336 | 3281 |
| 3574 | -0.12 | 0.07 | -1.67 | 0.09 | 1 | Oaz1 | ornithine decarboxylase antizyme 1 | 3723 | 3319 | 4072 | 3499 | 3980 | 3542 | 3686 | 3392 | 3459 | 3565 | 3077 |
| 2913 | -0.03 | 0.11 | -0.24 | 0.81 | 1 | Slc44a3 | solute carrier family 44, member 3 | 1143 | 1714 | 1596 | 3863 | 3855 | 2554 | 3686 | 3602 | 3895 | 3399 | 2738 |
| 3769 | 0.08 | 0.09 | 0.87 | 0.38 | 1 | Trak1 | trafficking protein, kinesin binding 1 | 3339 | 3114 | 3425 | 3581 | 3683 | 4395 | 3685 | 3776 | 3760 | 4685 | 4020 |
| 3766 | 0.05 | 0.07 | 0.75 | 0.45 | 1 | Erap1 | endoplasmic reticulum aminopeptidase 1 | 4127 | 4162 | 4627 | 3436 | 3871 | 3016 | 3685 | 3470 | 4244 | 3573 | 3220 |
| 3107 | -0.07 | 0.08 | -0.88 | 0.38 | 1 | Hbp1 | high mobility group box transcription facto | 3071 | 2023 | 3443 | 1974 | 3839 | 3683 | 3685 | 2059 | 3609 | 3239 | 3555 |
| 4009 | -0.02 | 0.12 | -0.20 | 0.84 | 1 | Cd200 | CD200 antigen | 4829 | 5499 | 5378 | 2928 | 4452 | 3261 | 3684 | 3040 | 3974 | 3980 | 3070 |
| 4492 | -0.11 | 0.07 | -1.51 | 0.13 | 1 | Fus | fused in sarcoma | 5458 | 5094 | 6220 | 3587 | 3637 | 6012 | 3683 | 3069 | 3640 | 3482 | 5528 |
| 3176 | 0.15 | 0.08 | 1.84 | 0.07 | 1 | Lonp2 | lon peptidase 2, peroxisomal | 2726 | 2642 | 2827 | 2888 | 3241 | 2882 | 3680 | 3310 | 3499 | 4043 | 3199 |
| 3650 | -0.03 | 0.10 | -0.34 | 0.73 | 1 | Lrp6 | low density lipoprotein receptor-related pı | 4435 | 3985 | 3794 | 3026 | 3824 | 3603 | 3677 | 3179 | 3896 | 3515 | 3214 |
| 4054 | -0.01 | 0.08 | -0.14 | 0.89 | 1 | Ugt2b34 | UDP glucuronosyltransferase 2 family, po | 3656 | 4686 | 4202 | 5095 | 4233 | 3150 | 3676 | 4374 | 4530 | 3631 | 3364 |
| 3561 | 0.08 | 0.12 | 0.61 | 0.54 | 1 | Tex261 | testis expressed gene 261 | 3137 | 2236 | 3550 | 3771 | 4113 | 3133 | 3676 | 4268 | 4070 | 4266 | 2949 |
| 3266 | -0.07 | 0.15 | -0.48 | 0.63 | 1 | Gclm | glutamate-cysteine ligase, modifier subur | 2885 | 3880 | 3604 | 3495 | 3293 | 2624 | 3675 | 2476 | 3446 | 3980 | 2568 |
| 2492 | 0.06 | 0.11 | 0.54 | 0.59 | 1 | Serpini1 | serine (or cysteine) peptidase inhibitor, cl | 1179 | 1236 | 1210 | 3172 | 2979 | 1866 | 3673 | 3130 | 2829 | 4008 | 2133 |
| 3946 | 0.02 | 0.08 | 0.21 | 0.83 | 1 | Eif3h | eukaryotic translation initiation factor 3, su | 6161 | 5668 | 6131 | 3141 | 3233 | 2541 | 3671 | 3063 | 3206 | 4167 | 2420 |
| 3668 | 0.06 | 0.07 | 0.82 | 0.41 | 1 | Prpf19 | pre-mRNA processing factor 19 | 4027 | 3109 | 3891 | 3171 | 3762 | 3786 | 3671 | 3427 | 3897 | 3784 | 3828 |
| 3573 | 0.01 | 0.08 | 0.17 | 0.87 | 1 | Clasp2 | CLIP associating protein 2 | 3149 | 3265 | 3151 | 4337 | 3511 | 3297 | 3669 | 3731 | 3926 | 3922 | 3342 |
| 3432 | 0.03 | 0.09 | 0.39 | 0.70 | 1 | Rnpepl1 | arginyl aminopeptidase (aminopeptidase | 2713 | 2952 | 2993 | 3454 | 3771 | 3451 | 3668 | 4194 | 3818 | 3573 | 3162 |
| 3502 | 0.10 | 0.09 | 1.10 | 0.27 | 1 | Uri1 | URI1, prefoldin-like chaperone | 3009 | 3496 | 3106 | 3762 | 3866 | 2696 | 3666 | 3806 | 3704 | 4312 | 3103 |
| 4071 | 0.08 | 0.09 | 0.84 | 0.40 | 1 | Ubn1 | ubinuclein 1 | 3457 | 3909 | 3314 | 4309 | 3665 | 4934 | 3665 | 5430 | 3793 | 3677 | 4628 |
| 3501 | 0.07 | 0.11 | 0.60 | 0.55 | 1 | Map3k2 | mitogen-activated protein kinase kinase k | 3143 | 4116 | 3119 | 3337 | 3138 | 3588 | 3665 | 3691 | 3632 | 3094 | 3992 |
| 2920 | -0.01 | 0.10 | -0.14 | 0.89 | 1 | Tcta | T cell leukemia translocation altered gene | 1381 | 1332 | 1719 | 3454 | 3845 | 2986 | 3663 | 3734 | 3384 | 3431 | 3196 |
| 3798 | 0.05 | 0.09 | 0.57 | 0.57 | 1 | Prpf6 | pre-mRNA splicing factor 6 | 2831 | 3395 | 3553 | 3997 | 4196 | 3827 | 3660 | 4912 | 3984 | 3496 | 3929 |
| 3681 | 0.13 | 0.12 | 1.05 | 0.29 | 1 | Cirbp | cold inducible RNA binding protein | 2751 | 3194 | 3303 | 3816 | 3480 | 3903 | 3659 | 5297 | 3424 | 3940 | 3721 |
| 3281 | 0.16 | 0.10 | 1.53 | 0.13 | 1 | Usp16 | ubiquitin specific peptidase 16 | 1421 | 1430 | 1348 | 2960 | 4015 | 4467 | 3659 | 4014 | 4017 | 3735 | 5028 |
| 3771 | -0.02 | 0.10 | -0.20 | 0.84 | 1 | Chd9 | chromodomain helicase DNA binding pro | 3730 | 3570 | 3422 | 3273 | 3628 | 4936 | 3658 | 2870 | 4149 | 3648 | 4603 |
| 4158 | 0.11 | 0.12 | 0.92 | 0.36 | 1 | Ptov1 | prostate tumor over expressed gene 1 | 4220 | 4460 | 4765 | 3776 | 3807 | 4297 | 3655 | 4628 | 3310 | 4598 | 4226 |
| 4137 | 0.08 | 0.07 | 1.13 | 0.26 | 1 | Kdm2a | lysine (K)-specific demethylase 2A | 3589 | 4333 | 3962 | 4306 | 3859 | 4512 | 3654 | 5040 | 4049 | 3653 | 4555 |
| 3478 | 0.01 | 0.08 | 0.16 | 0.87 | 1 | Capn7 | calpain 7 | 2251 | 2632 | 2443 | 3616 | 3818 | 4295 | 3652 | 3501 | 3997 | 3531 | 4528 |
| 2789 | -0.08 | 0.12 | -0.66 | 0.51 | 1 | Asah2 | N-acylsphingosine amidohydrolase 2 | 1612 | 2063 | 1577 | 3623 | 3138 | 2646 | 3652 | 3034 | 3638 | 3296 | 2397 |
| 3595 | 0.14 | 0.08 | 1.73 | 0.08 | 1 | Swi5 | SWI5 recombination repair homolog (yea | 3468 | 3701 | 3762 | 3518 | 3314 | 3116 | 3643 | 3859 | 3524 | 4258 | 3377 |
| 2724 | 0.11 | 0.10 | 1.14 | 0.25 | 1 | Jag2 | jagged 2 | 1072 | 1144 | 1459 | 3450 | 3879 | 1667 | 3643 | 3998 | 3901 | 3967 | 1783 |
| 3054 | 0.00 | 0.13 | -0.03 | 0.97 | 1 | Sgpp2 | sphingosine-1-phosphate phosphotase 2 | 1393 | 1885 | 1911 | 4159 | 2972 | 3475 | 3641 | 3743 | 3198 | 3811 | 3402 |
| 3708 | 0.09 | 0.11 | 0.86 | 0.39 | 1 | Rcn2 | reticulocalbin 2 | 3210 | 3178 | 4024 | 3699 | 4197 | 3155 | 3640 | 4760 | 4106 | 3353 | 3462 |
| 3449 | -0.07 | 0.08 | -0.83 | 0.41 | 1 | Mtcl1 | microtubule crosslinking factor 1 | 1505 | 2333 | 2058 | 4451 | 4103 | 4170 | 3640 | 4637 | 3752 | 3336 | 3951 |
| 4658 | 0.04 | 0.09 | 0.45 | 0.65 | 1 | Rpl18a | ribosomal protein L18A | 8644 | 7161 | 9040 | 3464 | 3533 | 2386 | 3635 | 3927 | 3524 | 3408 | 2515 |
| 3128 | 0.07 | 0.12 | 0.60 | 0.55 | 1 | Cox6c | cytochrome c oxidase subunit 6C | 2960 | 2697 | 3155 | 3059 | 3392 | 2396 | 3635 | 4087 | 3040 | 3523 | 2469 |
| 3728 | -0.02 | 0.13 | -0.12 | 0.91 | 1 | Sppl2a | signal peptide peptidase like 2A | 4427 | 3783 | 4029 | 3788 | 3635 | 3300 | 3634 | 2586 | 4311 | 4164 | 3351 |
| 2579 | -0.03 | 0.11 | -0.31 | 0.75 | 1 | Tenm3 | teneurin transmembrane protein 3 | 969 | 954 | 809 | 3669 | 2994 | 2672 | 3633 | 2859 | 3234 | 3934 | 2645 |
| 4014 | -0.09 | 0.08 | -1.07 | 0.28 | 1 | St6gal1 | beta galactoside alpha 2,6 sialyltransfera | 6925 | 6618 | 5160 | 4092 | 3180 | 2275 | 3627 | 3603 | 2824 | 3412 | 2437 |
| 4284 | 0.03 | 0.09 | 0.35 | 0.72 | 1 | Baz2a | bromodomain adjacent to zinc finger dom | 4118 | 4626 | 3955 | 4535 | 4341 | 4537 | 3623 | 4723 | 4689 | 3936 | 4046 |
| 4489 | 0.21 | 0.12 | 1.76 | 0.08 | 1 | Eif3f | eukaryotic translation initiation factor 3, su | 7558 | 6388 | 7729 | 3334 | 3464 | 2328 | 3622 | 4627 | 3323 | 4600 | 2407 |

| | | | | | | | | | | | | | | | | | | |
|---|---|---|---|---|---|---|---|---|---|---|---|---|---|---|---|---|---|---|
| 4472 | 0.00 | 0.07 | 0.02 | 0.98 | 1 | Hnrnpdl | heterogeneous nuclear ribonucleoprotein | 4101 | 3855 | 4135 | 3525 | 3685 | 7640 | 3620 | 3209 | 3606 | 3951 | 7868 |
| 2947 | 0.00 | 0.10 | -0.05 | 0.96 | 1 | Tead1 | TEA domain family member 1 | 2741 | 2498 | 2921 | 2795 | 3004 | 2741 | 3616 | 3260 | 3194 | 3078 | 2569 |
| 3279 | 0.05 | 0.09 | 0.53 | 0.60 | 1 | Chid1 | chitinase domain containing 1 | 2328 | 2558 | 2508 | 3800 | 3367 | 3242 | 3614 | 4548 | 3357 | 3744 | 2997 |
| 3726 | 0.02 | 0.10 | 0.17 | 0.86 | 1 | Mga | MAX gene associated | 3197 | 4061 | 3332 | 3210 | 3214 | 5030 | 3613 | 3376 | 3647 | 3062 | 5242 |
| 2997 | 0.05 | 0.07 | 0.68 | 0.50 | 1 | Tes | testis derived transcript | 1301 | 1192 | 1329 | 3662 | 3992 | 3065 | 3613 | 3554 | 3939 | 4025 | 3293 |
| 3540 | 0.05 | 0.11 | 0.46 | 0.64 | 1 | Cops2 | COP9 signalosome subunit 2 | 3256 | 3492 | 2870 | 3524 | 3918 | 3345 | 3612 | 3209 | 4035 | 3779 | 3898 |
| 4214 | -0.13 | 0.08 | -1.60 | 0.11 | 1 | Ltbp3 | latent transforming growth factor beta binc | 7397 | 6255 | 6404 | 3735 | 3513 | 2849 | 3610 | 3609 | 3040 | 3497 | 2443 |
| 3536 | 0.00 | 0.11 | 0.02 | 0.99 | 1 | Cited2 | Cbp/p300-interacting transactivator, with ( | 2830 | 2383 | 2896 | 3145 | 2959 | 5624 | 3608 | 2713 | 2890 | 4376 | 5473 |
| 2922 | -0.05 | 0.11 | -0.49 | 0.62 | 1 | Snx13 | sorting nexin 13 | 2294 | 2408 | 2292 | 3250 | 2922 | 3034 | 3606 | 2726 | 3359 | 3013 | 3239 |
| 3839 | 0.11 | 0.07 | 1.57 | 0.12 | 1 | Dnaja2 | DnaJ heat shock protein family (Hsp40) m | 4211 | 4200 | 4289 | 3302 | 3790 | 3495 | 3601 | 3836 | 3832 | 3614 | 4064 |
| 3226 | -0.15 | 0.10 | -1.47 | 0.14 | 1 | Trim25 | tripartite motif-containing 25 | 2878 | 3127 | 3022 | 4034 | 3583 | 2713 | 3601 | 2876 | 3662 | 3470 | 2524 |
| 3763 | 0.21 | 0.16 | 1.32 | 0.19 | 1 | Chchd2 | coiled-coil-helix-coiled-coil-helix domain | 3377 | 3247 | 4413 | 3401 | 3785 | 3216 | 3600 | 5687 | 3483 | 3747 | 3438 |
| 3647 | 0.07 | 0.11 | 0.61 | 0.54 | 1 | Psmb1 | proteasome (prosome, macropain) subun | 4626 | 3969 | 5108 | 3196 | 3463 | 2656 | 3598 | 4025 | 3341 | 3336 | 2802 |
| 3303 | 0.04 | 0.11 | 0.36 | 0.72 | 1 | Tmem33 | transmembrane protein 33 | 2303 | 2799 | 2362 | 3552 | 3728 | 3352 | 3592 | 3119 | 3804 | 4122 | 3601 |
| 3583 | -0.08 | 0.11 | -0.68 | 0.50 | 1 | Meis2 | Meis homeobox 2 | 3855 | 4765 | 4141 | 3010 | 3619 | 3450 | 3591 | 2939 | 3770 | 3175 | 3103 |
| 3353 | -0.11 | 0.14 | -0.77 | 0.44 | 1 | Ahi1 | Abelson helper integration site 1 | 2721 | 4326 | 3373 | 2972 | 3460 | 3613 | 3589 | 3465 | 3197 | 2549 | 3613 |
| 4190 | -0.05 | 0.10 | -0.51 | 0.61 | 1 | Arid1a | AT rich interactive domain 1A (SWI-like) | 4281 | 4736 | 3932 | 4792 | 4453 | 3977 | 3585 | 4938 | 4318 | 3733 | 3345 |
| 3159 | -0.10 | 0.13 | -0.78 | 0.43 | 1 | Zdhhc5 | zinc finger, DHHC domain containing 5 | 3180 | 3522 | 2772 | 3254 | 3468 | 2776 | 3585 | 2579 | 3743 | 3180 | 2691 |
| 2984 | 0.02 | 0.10 | 0.24 | 0.81 | 1 | Vmac | vimentin-type intermediate filament assoc | 2219 | 2229 | 2608 | 3174 | 3766 | 2295 | 3581 | 3503 | 3203 | 3796 | 2444 |
| 3190 | -0.08 | 0.08 | -0.94 | 0.35 | 1 | Arfip2 | ADP-ribosylation factor interacting proteir | 1924 | 1791 | 2233 | 3054 | 4054 | 4263 | 3579 | 2676 | 3980 | 3710 | 3823 |
| 4034 | -0.13 | 0.11 | -1.15 | 0.25 | 1 | Srrm1 | serine/arginine repetitive matrix 1 | 4033 | 4523 | 4079 | 4161 | 4092 | 4738 | 3577 | 2981 | 4253 | 3626 | 4309 |
| 3692 | -0.02 | 0.09 | -0.22 | 0.83 | 1 | Pfdn5 | prefoldin 5 | 5028 | 5270 | 5640 | 3228 | 3241 | 2340 | 3576 | 3420 | 2996 | 3710 | 2163 |
| 4335 | 0.11 | 0.12 | 0.86 | 0.39 | 1 | Rpl37a | ribosomal protein L37a | 6029 | 5837 | 8300 | 3335 | 3837 | 2526 | 3573 | 4221 | 3420 | 3944 | 2665 |
| 3155 | 0.08 | 0.10 | 0.79 | 0.43 | 1 | Grina | glutamate receptor, ionotropic, N-methyl D | 2533 | 1875 | 2277 | 3769 | 3365 | 2904 | 3572 | 3815 | 3537 | 3994 | 3065 |
| 3543 | 0.11 | 0.07 | 1.54 | 0.12 | 1 | Gapvd1 | GTPase activating protein and VPS9 dom | 3625 | 3236 | 3169 | 3320 | 3265 | 3751 | 3570 | 3810 | 3736 | 3855 | 3633 |
| 3201 | -0.03 | 0.10 | -0.24 | 0.81 | 1 | Dcaf6 | DDB1 and CUL4 associated factor 6 | 2399 | 2824 | 2170 | 3562 | 3497 | 3410 | 3569 | 3259 | 3782 | 3327 | 3415 |
| 2758 | -0.07 | 0.10 | -0.71 | 0.48 | 1 | Fam171b | family with sequence similarity 171, meml | 1837 | 2102 | 1706 | 3649 | 3364 | 2117 | 3569 | 3246 | 3465 | 3055 | 2229 |
| 3502 | 0.01 | 0.08 | 0.09 | 0.93 | 1 | Armcx2 | armadillo repeat containing, X-linked 2 | 3395 | 3789 | 3691 | 3209 | 3671 | 3331 | 3567 | 3395 | 3930 | 3174 | 3373 |
| 4155 | 0.20 | 0.13 | 1.52 | 0.13 | 1 | Mt1 | metallothionein 1 | 1730 | 2744 | 2676 | 4624 | 4531 | 5011 | 3565 | 7539 | 4479 | 3902 | 4905 |
| 3249 | -0.03 | 0.09 | -0.28 | 0.78 | 1 | Taok3 | TAO kinase 3 | 3215 | 3407 | 2955 | 3350 | 3303 | 2994 | 3561 | 3137 | 3571 | 3069 | 3170 |
| 3102 | 0.01 | 0.11 | 0.05 | 0.96 | 1 | Uck2 | uridine-cytidine kinase 2 | 1754 | 2514 | 1938 | 3419 | 4024 | 2929 | 3561 | 3746 | 3674 | 3518 | 3043 |
| 3120 | 0.04 | 0.10 | 0.35 | 0.73 | 1 | Dyrk2 | dual-specificity tyrosine-(Y)-phosphorylati | 2218 | 2188 | 2029 | 3475 | 3540 | 3152 | 3560 | 4378 | 3494 | 3184 | 3097 |
| 3469 | 0.05 | 0.12 | 0.42 | 0.68 | 1 | Wapl | WAPL cohesin release factor | 2665 | 3213 | 2814 | 3374 | 3226 | 4308 | 3558 | 2889 | 3822 | 3712 | 4577 |
| 3321 | 0.02 | 0.08 | 0.21 | 0.83 | 1 | Man1a2 | mannosidase, alpha, class 1A, member 2 | 2822 | 2864 | 2532 | 3678 | 3664 | 3154 | 3558 | 3699 | 4018 | 3555 | 2991 |
| 3639 | 0.12 | 0.09 | 1.37 | 0.17 | 1 | Usp4 | ubiquitin specific peptidase 4 (proto-onco | 3362 | 2737 | 3789 | 3427 | 3883 | 3576 | 3556 | 3995 | 4142 | 3922 | 3640 |
| 3321 | 0.02 | 0.09 | 0.19 | 0.85 | 1 | Sdhc | succinate dehydrogenase complex, subu | 2537 | 2603 | 2830 | 4270 | 3704 | 2723 | 3554 | 4050 | 3577 | 3571 | 3107 |
| 3301 | 0.12 | 0.13 | 0.93 | 0.35 | 1 | Ndufb11 | NADH:ubiquinone oxidoreductase subun | 3155 | 3141 | 3877 | 3236 | 3239 | 2397 | 3554 | 4520 | 3159 | 3702 | 2332 |
| 3553 | 0.07 | 0.09 | 0.77 | 0.44 | 1 | Far1 | fatty acyl CoA reductase 1 | 3746 | 2935 | 3188 | 3088 | 3352 | 4191 | 3552 | 3004 | 3852 | 3321 | 4853 |
| 3674 | 0.12 | 0.08 | 1.53 | 0.13 | 1 | Ddx42 | DEAD (Asp-Glu-Ala-Asp) box polypeptide | 3301 | 3271 | 3156 | 3330 | 3696 | 4160 | 3548 | 3685 | 4254 | 3680 | 4333 |
| 2928 | 0.01 | 0.08 | 0.19 | 0.85 | 1 | Tmem245 | transmembrane protein 245 | 2663 | 2308 | 2255 | 2863 | 2840 | 3205 | 3544 | 3040 | 3125 | 3254 | 3108 |
| 3358 | -0.02 | 0.11 | -0.17 | 0.87 | 1 | Snx2 | sorting nexin 2 | 3713 | 3356 | 3116 | 3220 | 3388 | 3281 | 3542 | 2713 | 3582 | 3267 | 3759 |
| 3198 | 0.02 | 0.07 | 0.34 | 0.73 | 1 | Mapk9 | mitogen-activated protein kinase 9 | 2595 | 2439 | 2866 | 3337 | 3705 | 2931 | 3542 | 3638 | 3742 | 3541 | 2842 |
| 3553 | 0.06 | 0.06 | 0.95 | 0.34 | 1 | Tnpo3 | transportin 3 | 3219 | 3356 | 3303 | 3602 | 3688 | 3460 | 3540 | 3580 | 3891 | 3632 | 3813 |
| 4687 | 0.15 | 0.10 | 1.53 | 0.13 | 1 | Rpl13 | ribosomal protein L13 | 8869 | 8258 | 8961 | 3085 | 3179 | 2254 | 3539 | 4197 | 3181 | 3565 | 2471 |
| 3444 | -0.08 | 0.13 | -0.62 | 0.54 | 1 | Tmem184b | transmembrane protein 184b | 3926 | 3784 | 3667 | 3517 | 3522 | 2937 | 3535 | 2509 | 3625 | 4280 | 2584 |
| 3062 | 0.05 | 0.11 | 0.47 | 0.64 | 1 | Srp14 | signal recognition particle 14 | 2451 | 2095 | 2991 | 3278 | 3465 | 2569 | 3532 | 3440 | 3278 | 3922 | 2667 |
| 3424 | -0.03 | 0.07 | -0.45 | 0.65 | 1 | Rnf167 | ring finger protein 167 | 2813 | 3118 | 3528 | 3856 | 3912 | 2980 | 3530 | 3543 | 3774 | 3567 | 3037 |
| 3047 | 0.12 | 0.11 | 1.08 | 0.28 | 1 | Lrrfip2 | leucine rich repeat (in FLII) interacting prc | 1348 | 911 | 1344 | 3278 | 3667 | 3879 | 3530 | 3827 | 3865 | 3665 | 4203 |
| 3941 | 0.11 | 0.14 | 0.79 | 0.43 | 1 | Rpl12 | ribosomal protein L12 | 6278 | 4846 | 6646 | 3203 | 2969 | 2580 | 3528 | 4251 | 3029 | 3406 | 2612 |
| 3508 | 0.07 | 0.08 | 0.97 | 0.33 | 1 | Vps41 | VPS41 HOPS complex subunit | 3619 | 3637 | 3671 | 3567 | 3389 | 2987 | 3528 | 3946 | 3792 | 3297 | 3152 |
| 3229 | -0.03 | 0.13 | -0.21 | 0.83 | 1 | Unc13a | unc-13 homolog A | 1178 | 1934 | 1684 | 3762 | 3330 | 4963 | 3527 | 3652 | 3365 | 3770 | 4352 |
| 3067 | -0.10 | 0.08 | -1.33 | 0.18 | 1 | Ccdc6 | coiled-coil domain containing 6 | 1599 | 1694 | 1596 | 3798 | 3985 | 3619 | 3527 | 3408 | 3908 | 3296 | 3304 |
| 3636 | 0.08 | 0.07 | 1.23 | 0.22 | 1 | Tbrg1 | transforming growth factor beta regulated | 3505 | 3490 | 3720 | 3429 | 3910 | 3363 | 3526 | 3785 | 4205 | 3506 | 3555 |
| 3525 | 0.11 | 0.08 | 1.37 | 0.17 | 1 | Tmem263 | transmembrane protein 263 | 2624 | 3552 | 2956 | 3886 | 3369 | 3485 | 3522 | 4085 | 3696 | 3801 | 3797 |
| 3110 | 0.02 | 0.09 | 0.21 | 0.83 | 1 | Cog1 | component of oligomeric golgi complex 1 | 1824 | 1639 | 1746 | 3595 | 3278 | 4010 | 3522 | 3426 | 3921 | 3775 | 3478 |
| 3534 | 0.04 | 0.09 | 0.48 | 0.63 | 1 | Eif3l | eukaryotic translation initiation factor 3, su | 3900 | 3928 | 3382 | 3293 | 3772 | 3033 | 3519 | 3583 | 3782 | 3739 | 2947 |
| 4022 | 0.10 | 0.11 | 0.91 | 0.36 | 1 | Nop53 | NOP53 ribosome biogenesis factor | 6356 | 5387 | 6156 | 3181 | 3364 | 2607 | 3513 | 4323 | 3109 | 3601 | 2648 |
| 3511 | -0.12 | 0.10 | -1.21 | 0.23 | 1 | Pkn2 | protein kinase N2 | 3310 | 3502 | 2982 | 4131 | 3924 | 3480 | 3513 | 3302 | 3834 | 3062 | 3588 |
| 2982 | -0.02 | 0.09 | -0.21 | 0.84 | 1 | Yars | tyrosyl-tRNA synthetase | 2152 | 2283 | 2451 | 2982 | 3794 | 2775 | 3513 | 3067 | 3429 | 3496 | 2861 |
| 2442 | 0.02 | 0.15 | 0.10 | 0.92 | 1 | Eno2 | enolase 2, gamma neuronal | 1303 | 1625 | 1375 | 3158 | 2216 | 2245 | 3513 | 3653 | 2055 | 3622 | 2093 |

| | | | | | | | | | | | | | | | | | |
|---|---|---|---|---|---|---|---|---|---|---|---|---|---|---|---|---|---|
| 3454 | -0.08 | 0.11 | -0.69 | 0.49 | 1 | Clptm1 | cleft lip and palate associated transmemb | 3417 | 3489 | 3330 | 4090 | 3841 | 2794 | 3510 | 2983 | 3782 | 4075 | 2677 |
| 2854 | 0.03 | 0.09 | 0.32 | 0.75 | 1 | Tbc1d9 | TBC1 domain family, member 9 | 2136 | 2206 | 2105 | 3098 | 3347 | 2356 | 3508 | 3698 | 3514 | 3198 | 2231 |
| 2568 | 0.06 | 0.10 | 0.59 | 0.55 | 1 | Nceh1 | neutral cholesterol ester hydrolase 1 | 1515 | 1502 | 1753 | 2795 | 3324 | 1957 | 3507 | 3155 | 3213 | 3255 | 2278 |
| 3680 | 0.03 | 0.09 | 0.36 | 0.72 | 1 | Mybbp1a | MYB binding protein (P160) 1a | 4592 | 4153 | 4298 | 3266 | 3751 | 2948 | 3505 | 3088 | 4065 | 4171 | 2643 |
| 2572 | 0.24 | 0.13 | 1.88 | 0.06 | 1 | Prss53 | protease, serine 53 | 116 | 152 | 181 | 3198 | 3422 | 2618 | 3505 | 4455 | 3787 | 3988 | 2868 |
| 2860 | 0.00 | 0.13 | 0.02 | 0.98 | 1 | Qsox2 | quiescin Q6 sulfhydryl oxidase 2 | 1474 | 1927 | 1944 | 3530 | 3218 | 2779 | 3504 | 3106 | 3379 | 3936 | 2664 |
| 3424 | -0.01 | 0.10 | -0.14 | 0.89 | 1 | Ralgapa1 | Ral GTPase activating protein, alpha subu | 2813 | 2992 | 3055 | 3482 | 3600 | 3843 | 3501 | 2870 | 4083 | 3385 | 4043 |
| 3350 | 0.17 | 0.15 | 1.12 | 0.26 | 1 | Selenom | selenoprotein M | 2573 | 2148 | 3017 | 3588 | 3814 | 2747 | 3500 | 5347 | 3667 | 3572 | 2880 |
| 3042 | 0.04 | 0.11 | 0.40 | 0.69 | 1 | Fzd3 | frizzled class receptor 3 | 1174 | 2066 | 1565 | 4105 | 3245 | 3264 | 3500 | 4124 | 3508 | 3458 | 3453 |
| 3307 | -0.05 | 0.08 | -0.65 | 0.52 | 1 | Cul7 | cullin 7 | 3317 | 3141 | 3202 | 3854 | 3560 | 2698 | 3495 | 3787 | 3602 | 3086 | 2633 |
| 3279 | -0.14 | 0.12 | -1.17 | 0.24 | 1 | Syne1 | spectrin repeat containing, nuclear envelo | 3261 | 3900 | 3326 | 2604 | 2563 | 4845 | 3495 | 2352 | 2620 | 3057 | 4046 |
| 3082 | 0.08 | 0.09 | 0.94 | 0.35 | 1 | Zc3h15 | zinc finger CCCH-type containing 15 | 2352 | 2657 | 2649 | 3012 | 3081 | 3149 | 3495 | 3334 | 3366 | 3260 | 3551 |
| 3760 | 0.04 | 0.07 | 0.53 | 0.60 | 1 | Smc3 | structural maintenance of chromosomes 3 | 3831 | 3479 | 3751 | 3447 | 3849 | 4162 | 3494 | 3367 | 4091 | 3608 | 4285 |
| 3429 | -0.06 | 0.07 | -0.85 | 0.40 | 1 | Chp1 | calcineurin-like EF hand protein 1 | 3825 | 3619 | 3800 | 3607 | 3730 | 2643 | 3494 | 3368 | 3525 | 3692 | 2412 |
| 6393 | 0.12 | 0.06 | 1.86 | 0.06 | 1 | Aplp2 | amyloid beta (A4) precursor-like protein 2 | 15748 | 13363 | 14037 | 3342 | 3208 | 2964 | 3491 | 3653 | 3612 | 3940 | 2960 |
| 3659 | -0.13 | 0.09 | -1.43 | 0.15 | 1 | Atp2b4 | ATPase, Ca++ transporting, plasma mem | 4795 | 5509 | 4337 | 3652 | 3169 | 3102 | 3482 | 2977 | 3201 | 2979 | 3041 |
| 4567 | 0.00 | 0.08 | -0.04 | 0.97 | 1 | Dnaja1 | DnaJ heat shock protein family (Hsp40) m | 3103 | 4005 | 3473 | 3661 | 3381 | 9329 | 3480 | 3455 | 3732 | 3317 | 9296 |
| 2741 | -0.02 | 0.09 | -0.20 | 0.84 | 1 | Ttyh2 | tweety family member 2 | 853 | 778 | 905 | 3759 | 3602 | 3003 | 3480 | 3354 | 3832 | 3889 | 2699 |
| 3443 | 0.12 | 0.08 | 1.57 | 0.12 | 1 | Hif1a | hypoxia inducible factor 1, alpha subunit | 3528 | 4174 | 3169 | 3519 | 3219 | 2716 | 3477 | 3887 | 3569 | 3536 | 3078 |
| 2949 | -0.11 | 0.14 | -0.81 | 0.42 | 1 | Suox | sulfite oxidase | 1915 | 2794 | 2559 | 3301 | 3551 | 2741 | 3477 | 2673 | 3359 | 3457 | 2612 |
| 4040 | 0.04 | 0.08 | 0.49 | 0.63 | 1 | Rps7 | ribosomal protein S7 | 7640 | 6570 | 7206 | 2705 | 3048 | 2190 | 3474 | 2943 | 2892 | 3264 | 2511 |
| 4320 | -0.02 | 0.08 | -0.33 | 0.75 | 1 | Acin1 | apoptotic chromatin condensation induce | 3518 | 3752 | 3631 | 4210 | 3667 | 7208 | 3473 | 4345 | 3806 | 3295 | 6620 |
| 3159 | 0.07 | 0.15 | 0.45 | 0.65 | 1 | Fnip1 | folliculin interacting protein 1 | 1573 | 1410 | 1478 | 3576 | 3534 | 4070 | 3473 | 4692 | 3353 | 2670 | 4921 |
| 3483 | 0.04 | 0.09 | 0.43 | 0.66 | 1 | Rufy3 | RUN and FYVE domain containing 3 | 2205 | 2312 | 2586 | 3137 | 3715 | 5103 | 3472 | 3462 | 3395 | 3769 | 5152 |
| 3811 | -0.01 | 0.08 | -0.14 | 0.89 | 1 | Zfp638 | zinc finger protein 638 | 3063 | 3917 | 3111 | 3954 | 3251 | 5244 | 3468 | 4287 | 3385 | 2920 | 5325 |
| 2941 | 0.09 | 0.12 | 0.77 | 0.44 | 1 | Kpna3 | karyopherin (importin) alpha 3 | 2200 | 2959 | 2110 | 2909 | 2890 | 2863 | 3468 | 3166 | 3167 | 3675 | 2944 |
| 3633 | 0.01 | 0.10 | 0.14 | 0.89 | 1 | Sec14l1 | SEC14-like lipid binding 1 | 3947 | 3399 | 3341 | 4337 | 3550 | 3222 | 3467 | 3954 | 3901 | 3648 | 3198 |
| 2952 | 0.01 | 0.10 | 0.14 | 0.88 | 1 | Ccdc127 | coiled-coil domain containing 127 | 1830 | 1973 | 1976 | 3153 | 3366 | 3275 | 3466 | 3207 | 3809 | 2906 | 3515 |
| 5224 | 0.06 | 0.15 | 0.42 | 0.67 | 1 | Mcl1 | myeloid cell leukemia sequence 1 | 10659 | 7658 | 7725 | 4190 | 3660 | 3986 | 3465 | 5292 | 3515 | 3234 | 4076 |
| 3724 | 0.05 | 0.08 | 0.64 | 0.52 | 1 | Stk11 | serine/threonine kinase 11 | 3682 | 3531 | 3824 | 3248 | 3992 | 4012 | 3465 | 3877 | 3690 | 3804 | 3843 |
| 3484 | -0.08 | 0.12 | -0.68 | 0.49 | 1 | Tab2 | TGF-beta activated kinase 1/MAP3K7 bin | 3750 | 4166 | 3495 | 3718 | 3247 | 3408 | 3465 | 2718 | 3683 | 3538 | 3140 |
| 3275 | 0.01 | 0.09 | 0.08 | 0.94 | 1 | Klhdc2 | kelch domain containing 2 | 2779 | 3083 | 2732 | 3424 | 3560 | 3236 | 3465 | 3100 | 3664 | 3422 | 3562 |
| 3484 | 0.10 | 0.06 | 1.66 | 0.10 | 1 | Sdhd | succinate dehydrogenase complex, subu | 3415 | 3356 | 3478 | 3297 | 3710 | 3070 | 3464 | 3587 | 3835 | 3822 | 3289 |
| 3359 | -0.14 | 0.09 | -1.54 | 0.12 | 1 | Man2a2 | mannosidase 2, alpha 2 | 4219 | 2933 | 3945 | 3293 | 3712 | 3052 | 3464 | 2893 | 3502 | 3381 | 2552 |
| 3312 | 0.04 | 0.12 | 0.31 | 0.76 | 1 | Aff1 | AF4/FMR2 family, member 1 | 2475 | 2221 | 1862 | 4325 | 3390 | 3556 | 3463 | 3742 | 3992 | 4258 | 3151 |
| 3302 | 0.04 | 0.09 | 0.49 | 0.62 | 1 | Scap | SREBF chaperone | 2798 | 2464 | 2922 | 3663 | 3814 | 2871 | 3463 | 3828 | 3855 | 4008 | 2639 |
| 3591 | 0.04 | 0.08 | 0.49 | 0.62 | 1 | Pkdcc | protein kinase domain containing, cytopla | 2712 | 3105 | 2643 | 4096 | 4206 | 3551 | 3462 | 4087 | 4140 | 3916 | 3579 |
| 3383 | 0.01 | 0.08 | 0.20 | 0.84 | 1 | Kars | lysyl-tRNA synthetase | 3755 | 3289 | 3837 | 3254 | 3539 | 2831 | 3461 | 2904 | 3848 | 3466 | 3030 |
| 3233 | 0.10 | 0.11 | 0.88 | 0.38 | 1 | Micu2 | mitochondrial calcium uptake 2 | 3264 | 2796 | 3415 | 3154 | 3302 | 2684 | 3461 | 4029 | 3074 | 3420 | 2967 |
| 4218 | -0.05 | 0.12 | -0.39 | 0.70 | 1 | Top2b | topoisomerase (DNA) II beta | 7309 | 6641 | 7032 | 2865 | 3093 | 3475 | 3460 | 3155 | 3452 | 2407 | 3510 |
| 3839 | -0.17 | 0.15 | -1.14 | 0.25 | 1 | Fubp1 | far upstream element (FUSE) binding pro | 4245 | 5367 | 4558 | 2745 | 3220 | 5276 | 3460 | 2061 | 3341 | 2790 | 5169 |
| 3061 | -0.08 | 0.09 | -1.00 | 0.32 | 1 | Jmjd8 | jumonji domain containing 8 | 2436 | 2746 | 2714 | 3481 | 3471 | 2858 | 3460 | 3196 | 3047 | 3531 | 2736 |
| 3278 | 0.00 | 0.09 | 0.02 | 0.98 | 1 | Tbc1d1 | TBC1 domain family, member 1 | 2847 | 2858 | 2794 | 3521 | 3433 | 3337 | 3459 | 2933 | 3894 | 3536 | 3445 |
| 2878 | 0.24 | 0.18 | 1.32 | 0.19 | 1 | Uqcc2 | ubiquinol-cytochrome c reductase comple | 1447 | 1604 | 2232 | 2945 | 3146 | 2381 | 3455 | 4944 | 3032 | 4172 | 2295 |
| 2837 | 0.07 | 0.10 | 0.65 | 0.52 | 1 | Tceal8 | transcription elongation factor A (SII)-like | 1933 | 2265 | 2166 | 2823 | 3129 | 2738 | 3455 | 3196 | 3071 | 3238 | 3194 |
| 3492 | 0.05 | 0.06 | 0.77 | 0.44 | 1 | Adipor2 | adiponectin receptor 2 | 3269 | 3666 | 3280 | 3797 | 3440 | 3167 | 3452 | 4068 | 3628 | 3460 | 3181 |
| 3234 | -0.12 | 0.13 | -0.95 | 0.34 | 1 | Dnm1l | dynamin 1-like | 2940 | 2671 | 2650 | 3067 | 3683 | 3889 | 3452 | 2200 | 3595 | 3156 | 4273 |
| 3157 | -0.05 | 0.12 | -0.46 | 0.64 | 1 | Slain2 | SLAIN motif family, member 2 | 2686 | 2753 | 2321 | 3454 | 3416 | 3399 | 3445 | 2637 | 3567 | 3677 | 3374 |
| 3288 | 0.02 | 0.10 | 0.19 | 0.85 | 1 | Pdzd8 | PDZ domain containing 8 | 2928 | 3170 | 2646 | 3730 | 3649 | 2798 | 3444 | 3338 | 4051 | 3610 | 2806 |
| 3394 | -0.05 | 0.11 | -0.45 | 0.65 | 1 | Kdm5a | lysine (K)-specific demethylase 5A | 3644 | 4069 | 3671 | 2874 | 3420 | 3433 | 3442 | 3035 | 3479 | 2651 | 3619 |
| 3124 | 0.11 | 0.15 | 0.69 | 0.49 | 1 | Sptbn4 | spectrin beta, non-erythrocytic 4 | 1050 | 1015 | 1699 | 2897 | 2983 | 5570 | 3440 | 3502 | 3315 | 3636 | 5255 |
| 3547 | -0.01 | 0.08 | -0.18 | 0.86 | 1 | Pds5a | PDS5 cohesin associated factor A | 4172 | 3518 | 3847 | 3340 | 3492 | 3509 | 3439 | 2886 | 3595 | 3471 | 3747 |
| 4062 | 0.00 | 0.09 | 0.00 | 1.00 | 1 | Rps25 | ribosomal protein S25 | 7080 | 6936 | 6752 | 3247 | 3030 | 2249 | 3436 | 2742 | 3329 | 3529 | 2346 |
| 3553 | 0.05 | 0.08 | 0.64 | 0.53 | 1 | Vps13d | vacuolar protein sorting 13D | 3004 | 2958 | 3284 | 3245 | 3404 | 4606 | 3436 | 3305 | 4039 | 3424 | 4383 |
| 3492 | -0.05 | 0.09 | -0.60 | 0.55 | 1 | Flii | flightless I actin binding protein | 4168 | 3182 | 3604 | 3531 | 3747 | 3236 | 3436 | 3173 | 3546 | 3855 | 2939 |
| 3015 | -0.03 | 0.10 | -0.32 | 0.75 | 1 | Mbp | myelin basic protein | 2836 | 3191 | 3051 | 2959 | 3405 | 2377 | 3432 | 2872 | 3174 | 3536 | 2336 |
| 3600 | 0.13 | 0.07 | 1.81 | 0.07 | 1 | Puf60 | poly-U binding splicing factor 60 | 4092 | 3423 | 4113 | 3175 | 3459 | 3286 | 3431 | 3634 | 3589 | 3796 | 3599 |
| 3326 | -0.04 | 0.12 | -0.35 | 0.73 | 1 | Actn1 | actinin, alpha 1 | 4563 | 3222 | 3074 | 3526 | 3236 | 2827 | 3431 | 3397 | 3534 | 3328 | 2447 |
| 3191 | -0.05 | 0.07 | -0.77 | 0.44 | 1 | Ipo9 | importin 9 | 3149 | 2659 | 3100 | 3253 | 3483 | 3155 | 3430 | 2904 | 3417 | 3509 | 3042 |
| 3620 | 0.03 | 0.07 | 0.43 | 0.67 | 1 | Tprgl | transformation related protein 63 regulate | 4232 | 4180 | 4171 | 3665 | 3621 | 2748 | 3428 | 3801 | 3445 | 3795 | 2732 |

| | | | | | | | | | | | | | | | | | | |
|---|---|---|---|---|---|---|---|---|---|---|---|---|---|---|---|---|---|---|
| 3401 | -0.06 | 0.09 | -0.64 | 0.52 | 1 | Vezf1 | vascular endothelial zinc finger 1 | 3638 | 4265 | 3392 | 3565 | 3143 | 3189 | 3427 | 3212 | 3459 | 3294 | 2826 |
| 3297 | 0.13 | 0.08 | 1.70 | 0.09 | 1 | Rab11a | RAB11A, member RAS oncogene family | 3311 | 3470 | 3499 | 3454 | 3088 | 2468 | 3425 | 3658 | 3623 | 3393 | 2879 |
| 3659 | -0.02 | 0.09 | -0.17 | 0.86 | 1 | Cpn1 | carboxypeptidase N, polypeptide 1 | 4011 | 4823 | 5191 | 3420 | 3390 | 2951 | 3424 | 3214 | 3437 | 3328 | 3058 |
| 3097 | -0.07 | 0.10 | -0.75 | 0.45 | 1 | Xrn1 | 5'-3' exoribonuclease 1 | 2702 | 3020 | 2952 | 3155 | 3035 | 3389 | 3422 | 2999 | 3403 | 2692 | 3296 |
| 2910 | 0.10 | 0.12 | 0.89 | 0.38 | 1 | C2cd2l | C2 calcium-dependent domain containing | 1298 | 1502 | 1613 | 2698 | 4371 | 2849 | 3422 | 3334 | 4127 | 3922 | 2871 |
| 3771 | -0.15 | 0.10 | -1.59 | 0.11 | 1 | 2900026A02Ril | RIKEN cDNA 2900026A02 gene | 5204 | 5621 | 5060 | 3533 | 3417 | 3097 | 3419 | 2687 | 3451 | 3120 | 2869 |
| 2584 | -0.06 | 0.12 | -0.46 | 0.65 | 1 | Dach1 | dachshund family transcription factor 1 | 1168 | 1430 | 1668 | 3400 | 2880 | 2615 | 3415 | 3011 | 3263 | 2939 | 2633 |
| 3433 | -0.12 | 0.12 | -0.96 | 0.34 | 1 | Ep300 | E1A binding protein p300 | 3258 | 3981 | 3440 | 3805 | 3231 | 3623 | 3414 | 2878 | 3916 | 3008 | 3205 |
| 3199 | -0.08 | 0.07 | -1.13 | 0.26 | 1 | Supt16 | SPT16, facilitates chromatin remodeling s | 3235 | 3037 | 3125 | 3384 | 3128 | 3324 | 3414 | 3167 | 3140 | 3042 | 3194 |
| 2969 | 0.03 | 0.08 | 0.43 | 0.66 | 1 | Jade2 | jade family PHD finger 2 | 2693 | 3094 | 2232 | 3496 | 2914 | 2358 | 3411 | 3571 | 3109 | 3419 | 2364 |
| 2858 | -0.12 | 0.08 | -1.52 | 0.13 | 1 | Nr1d1 | nuclear receptor subfamily 1, group D, me | 4023 | 2350 | 2981 | 1961 | 2863 | 3260 | 3409 | 1891 | 2609 | 3277 | 2812 |
| 2873 | 0.06 | 0.13 | 0.44 | 0.66 | 1 | Fam174a | family with sequence similarity 174, meml | 1950 | 1720 | 2221 | 3544 | 3064 | 2521 | 3408 | 3081 | 3097 | 4400 | 2593 |
| 2789 | -0.06 | 0.10 | -0.59 | 0.55 | 1 | Trappc10 | trafficking protein particle complex 10 | 2314 | 1669 | 1811 | 2975 | 3287 | 3002 | 3408 | 2730 | 3482 | 3233 | 2765 |
| 2847 | -0.11 | 0.08 | -1.34 | 0.18 | 1 | Cops7a | COP9 signalosome subunit 7A | 2569 | 2239 | 2551 | 2895 | 3170 | 2986 | 3406 | 2979 | 2691 | 2996 | 2831 |
| 2974 | 0.02 | 0.11 | 0.15 | 0.88 | 1 | Vps39 | VPS39 HOPS complex subunit | 2105 | 2573 | 2121 | 3229 | 3049 | 3183 | 3405 | 3040 | 3154 | 3867 | 2984 |
| 3374 | -0.06 | 0.09 | -0.64 | 0.52 | 1 | Rassf3 | Ras association (RalGDS/AF-6) domain fa | 4548 | 4037 | 4109 | 3497 | 3089 | 2472 | 3400 | 3265 | 3306 | 2963 | 2429 |
| 2384 | -0.15 | 0.16 | -0.95 | 0.34 | 1 | Kcnip1 | Kv channel-interacting protein 1 | 495 | 850 | 739 | 3281 | 3484 | 2508 | 3400 | 2578 | 3543 | 3098 | 2251 |
| 3112 | 0.01 | 0.10 | 0.14 | 0.89 | 1 | Atp6v0a2 | ATPase, H+ transporting, lysosomal V0 su | 2658 | 2040 | 2241 | 3559 | 3415 | 3199 | 3399 | 3732 | 3439 | 3439 | 3108 |
| 2635 | -0.04 | 0.09 | -0.46 | 0.65 | 1 | Adcy9 | adenylate cyclase 9 | 1407 | 1133 | 1162 | 3357 | 3002 | 3067 | 3398 | 3097 | 3362 | 3278 | 2726 |
| 3659 | -0.19 | 0.14 | -1.39 | 0.17 | 1 | Smg1 | SMG1 homolog, phosphatidylinositol 3-ki | 3587 | 3422 | 2901 | 2780 | 3053 | 6782 | 3394 | 2164 | 3317 | 2419 | 6432 |
| 3500 | 0.02 | 0.08 | 0.30 | 0.76 | 1 | Hsbp1 | heat shock factor binding protein 1 | 3343 | 3549 | 3939 | 3579 | 3792 | 2954 | 3392 | 3916 | 3541 | 3583 | 2912 |
| 3345 | 0.09 | 0.08 | 1.12 | 0.26 | 1 | Rlim | ring finger protein, LIM domain interacting | 2622 | 2725 | 2793 | 3065 | 3577 | 3822 | 3388 | 3074 | 3735 | 3616 | 4376 |
| 2644 | -0.01 | 0.12 | -0.11 | 0.91 | 1 | Tbc1d8b | TBC1 domain family, member 8B | 1526 | 1613 | 1361 | 2686 | 3501 | 2789 | 3385 | 2711 | 3390 | 3218 | 2906 |
| 3556 | -0.24 | 0.14 | -1.67 | 0.09 | 1 | Col6a6 | collagen, type VI, alpha 6 | 2383 | 1632 | 1676 | 3223 | 4421 | 7029 | 3384 | 2644 | 3827 | 2910 | 5982 |
| 3158 | -0.02 | 0.08 | -0.29 | 0.77 | 1 | Lonp1 | lon peptidase 1, mitochondrial | 3073 | 3041 | 3072 | 3038 | 3330 | 3128 | 3378 | 3492 | 3045 | 3134 | 3002 |
| 2995 | -0.01 | 0.08 | -0.15 | 0.88 | 1 | Bcap31 | B cell receptor associated protein 31 | 3280 | 2853 | 3200 | 3174 | 2962 | 2341 | 3378 | 2791 | 3042 | 3496 | 2429 |
| 2795 | -0.16 | 0.18 | -0.90 | 0.37 | 1 | Ky | kyphoscoliosis peptidase | 610 | 410 | 1015 | 4185 | 4083 | 3461 | 3378 | 3481 | 3981 | 3144 | 2999 |
| 2795 | -0.09 | 0.13 | -0.73 | 0.47 | 1 | Psd3 | pleckstrin and Sec7 domain containing 3 | 3132 | 2961 | 2399 | 2880 | 2446 | 2809 | 3376 | 2172 | 2825 | 2710 | 3034 |
| 2662 | -0.01 | 0.10 | -0.13 | 0.90 | 1 | Mmut | methylmalonyl-Coenzyme A mutase | 2131 | 1880 | 1853 | 3022 | 3004 | 2397 | 3375 | 3094 | 3120 | 2784 | 2624 |
| 3364 | -0.01 | 0.09 | -0.07 | 0.94 | 1 | Dnajc13 | DnaJ heat shock protein family (Hsp40) m | 2436 | 2394 | 2495 | 3825 | 3448 | 4219 | 3374 | 3240 | 3856 | 3520 | 4197 |
| 3025 | -0.11 | 0.12 | -0.91 | 0.36 | 1 | Ago1 | argonaute RISC catalytic subunit 1 | 3123 | 3442 | 2781 | 2868 | 3212 | 2929 | 3372 | 2482 | 2974 | 3514 | 2583 |
| 2614 | 0.02 | 0.10 | 0.23 | 0.82 | 1 | Ergic2 | ERGIC and golgi 2 | 1564 | 1367 | 1450 | 2838 | 2927 | 2942 | 3371 | 2731 | 3119 | 2899 | 3542 |
| 3168 | -0.04 | 0.06 | -0.61 | 0.54 | 1 | Rab5b | RAB5B, member RAS oncogene family | 3310 | 3366 | 3402 | 3250 | 3348 | 2572 | 3369 | 3156 | 3257 | 3357 | 2456 |
| 3177 | 0.01 | 0.10 | 0.08 | 0.94 | 1 | Shoc2 | Shoc2, leucine rich repeat scaffold protei | 2539 | 2669 | 2282 | 3745 | 3393 | 3193 | 3366 | 3133 | 3673 | 3728 | 3227 |
| 3033 | 0.05 | 0.12 | 0.42 | 0.67 | 1 | Sil1 | endoplasmic reticulum chaperone SIL1 h | 2230 | 2695 | 2720 | 3358 | 3380 | 2472 | 3366 | 4217 | 2952 | 3562 | 2410 |
| 3157 | 0.06 | 0.09 | 0.61 | 0.54 | 1 | Ube4a | ubiquitination factor E4A | 1938 | 1761 | 1640 | 3301 | 3365 | 4402 | 3365 | 3633 | 3717 | 3203 | 4406 |
| 3416 | 0.10 | 0.12 | 0.81 | 0.42 | 1 | Tsn | translin | 3645 | 3555 | 3052 | 3137 | 3611 | 3051 | 3364 | 4382 | 3436 | 3151 | 3192 |
| 2915 | 0.02 | 0.09 | 0.22 | 0.82 | 1 | Timm29 | translocase of inner mitochondrial membr | 2201 | 2387 | 2444 | 3099 | 3361 | 2636 | 3362 | 3071 | 3161 | 3419 | 2929 |
| 3885 | 0.15 | 0.15 | 1.02 | 0.31 | 1 | Rpl11 | ribosomal protein L11 | 5394 | 4632 | 7152 | 2763 | 3399 | 2616 | 3361 | 4150 | 3057 | 3189 | 3017 |
| 2948 | -0.02 | 0.11 | -0.19 | 0.85 | 1 | Ppat | phosphoribosyl pyrophosphate amidotrar | 1215 | 2013 | 1381 | 4072 | 2588 | 3964 | 3360 | 3816 | 2825 | 2856 | 4341 |
| 3232 | -0.05 | 0.09 | -0.54 | 0.59 | 1 | Zfand3 | zinc finger, AN1-type domain 3 | 3994 | 3955 | 3935 | 2832 | 3184 | 2639 | 3358 | 2483 | 3258 | 3422 | 2488 |
| 3305 | -0.09 | 0.10 | -0.98 | 0.33 | 1 | Stat5b | signal transducer and activator of transcri | 3135 | 2816 | 2976 | 4097 | 3747 | 2943 | 3354 | 3309 | 3835 | 3507 | 2637 |
| 2895 | -0.06 | 0.13 | -0.46 | 0.65 | 1 | Necap1 | NECAP endocytosis associated 1 | 1820 | 2144 | 1958 | 3226 | 3483 | 3137 | 3353 | 2520 | 3260 | 3787 | 3157 |
| 3410 | 0.14 | 0.09 | 1.45 | 0.15 | 1 | Pfkm | phosphofructokinase, muscle | 2903 | 3014 | 3417 | 3247 | 3772 | 3032 | 3352 | 4258 | 3653 | 3769 | 3090 |
| 3805 | 0.17 | 0.10 | 1.77 | 0.08 | 1 | Crk | v-crk avian sarcoma virus CT10 oncogen | 4454 | 4429 | 3401 | 3607 | 3310 | 3618 | 3351 | 4431 | 3718 | 3424 | 4111 |
| 3750 | 0.04 | 0.10 | 0.42 | 0.68 | 1 | Tnrc6a | trinucleotide repeat containing 6a | 3472 | 3821 | 3236 | 2978 | 2670 | 6322 | 3351 | 3275 | 3304 | 3006 | 5815 |
| 3283 | -0.02 | 0.09 | -0.28 | 0.78 | 1 | Khdrbs1 | KH domain containing, RNA binding, sigr | 4921 | 4421 | 4259 | 2634 | 2950 | 2446 | 3351 | 2816 | 2983 | 2829 | 2500 |
| 3174 | 0.11 | 0.07 | 1.57 | 0.12 | 1 | Hdac2 | histone deacetylase 2 | 2738 | 2559 | 2641 | 3163 | 3270 | 3181 | 3346 | 3624 | 3247 | 3556 | 3596 |
| 2308 | -0.05 | 0.13 | -0.41 | 0.68 | 1 | Reps2 | RALBP1 associated Eps domain containi | 1072 | 930 | 732 | 3210 | 3024 | 2051 | 3345 | 2697 | 3156 | 2836 | 2339 |
| 3808 | 0.02 | 0.11 | 0.14 | 0.89 | 1 | Nav1 | neuron navigator 1 | 4088 | 3817 | 4008 | 4068 | 3887 | 3554 | 3341 | 4596 | 4203 | 3076 | 3248 |
| 2982 | 0.00 | 0.06 | 0.03 | 0.98 | 1 | Gcnt2 | glucosaminyl (N-acetyl) transferase 2, I-br | 2553 | 2774 | 2272 | 3464 | 2968 | 2824 | 3341 | 3474 | 2965 | 3267 | 2897 |
| 2434 | -0.05 | 0.12 | -0.42 | 0.68 | 1 | Syngr1 | synaptogyrin 1 | 1469 | 1629 | 1757 | 2807 | 2825 | 2133 | 3339 | 2777 | 2586 | 3554 | 1896 |
| 3031 | -0.06 | 0.12 | -0.48 | 0.63 | 1 | Exoc4 | exocyst complex component 4 | 3192 | 3553 | 2623 | 3060 | 3240 | 2580 | 3338 | 3166 | 3352 | 2731 | 2501 |
| 3168 | 0.10 | 0.08 | 1.16 | 0.24 | 1 | Amigo1 | adhesion molecule with Ig like domain 1 | 2287 | 2676 | 2164 | 3676 | 3401 | 2946 | 3336 | 3864 | 3541 | 4052 | 2905 |
| 3072 | 0.06 | 0.07 | 0.87 | 0.39 | 1 | Wdr11 | WD repeat domain 11 | 2165 | 2193 | 2088 | 3497 | 3387 | 3146 | 3336 | 3407 | 3786 | 3605 | 3181 |
| 3166 | 0.06 | 0.08 | 0.76 | 0.45 | 1 | Cxxc5 | CXXC finger 5 | 2831 | 3031 | 3173 | 3367 | 3279 | 2646 | 3335 | 3632 | 3256 | 3577 | 2704 |
| 2987 | 0.14 | 0.09 | 1.59 | 0.11 | 1 | Stx12 | syntaxin 12 | 2721 | 2799 | 2533 | 2681 | 3006 | 2797 | 3333 | 3231 | 3055 | 3580 | 3117 |
| 3646 | 0.18 | 0.12 | 1.55 | 0.12 | 1 | Pura | purine rich element binding protein A | 3402 | 4425 | 3845 | 3389 | 3326 | 3125 | 3329 | 5306 | 3432 | 3352 | 3180 |
| 2785 | -0.12 | 0.12 | -1.00 | 0.32 | 1 | Ficd | FIC domain containing | 1086 | 1341 | 1655 | 3104 | 4474 | 2908 | 3328 | 2394 | 4219 | 3243 | 2881 |
| 3147 | 0.01 | 0.11 | 0.07 | 0.95 | 1 | Psmc1 | protease (prosome, macropain) 26S subu | 3276 | 2824 | 3145 | 3160 | 3444 | 2729 | 3326 | 3732 | 3355 | 2822 | 2808 |

| | | | | | | | | | | | | | | | | | | |
|---|---|---|---|---|---|---|---|---|---|---|---|---|---|---|---|---|---|---|
| 3335 | -0.10 | 0.07 | -1.48 | 0.14 | 1 | Stk25 | serine/threonine kinase 25 (yeast) | 3077 | 2715 | 3248 | 3171 | 3393 | 4401 | 3320 | 2719 | 3386 | 3068 | 4187 |
| 2620 | 0.03 | 0.07 | 0.44 | 0.66 | 1 | Tmed2 | transmembrane p24 trafficking protein 2 | 1762 | 1648 | 1801 | 2866 | 3058 | 2423 | 3320 | 2901 | 3097 | 3482 | 2464 |
| 2817 | -0.12 | 0.10 | -1.20 | 0.23 | 1 | Arhgef11 | Rho guanine nucleotide exchange factor | 2027 | 2655 | 2169 | 3433 | 2975 | 2860 | 3316 | 2990 | 3130 | 2621 | 2813 |
| 3262 | 0.12 | 0.08 | 1.65 | 0.10 | 1 | Aldh3a2 | aldehyde dehydrogenase family 3, subfar | 2436 | 2346 | 2245 | 3800 | 3515 | 3171 | 3315 | 3972 | 3948 | 3836 | 3298 |
| 3129 | -0.06 | 0.09 | -0.72 | 0.47 | 1 | Ylpm1 | YLP motif containing 1 | 2738 | 3294 | 2853 | 3263 | 2930 | 3541 | 3313 | 3315 | 3175 | 2826 | 3172 |
| 2903 | -0.01 | 0.10 | -0.09 | 0.93 | 1 | Nf1 | neurofibromin 1 | 2671 | 2789 | 2930 | 2798 | 2884 | 2785 | 3312 | 2623 | 3341 | 3286 | 2515 |
| 3235 | 0.01 | 0.07 | 0.18 | 0.86 | 1 | Mtss2 | MTSS I-BAR domain containing 2 | 1610 | 1967 | 1557 | 4750 | 3320 | 3817 | 3309 | 4466 | 3565 | 3415 | 3809 |
| 3631 | -0.14 | 0.10 | -1.34 | 0.18 | 1 | Dennd5a | DENN/MADD domain containing 5A | 6165 | 5204 | 4871 | 3064 | 3101 | 2948 | 3308 | 2792 | 3206 | 2653 | 2633 |
| 2554 | -0.06 | 0.08 | -0.81 | 0.42 | 1 | Pip5k1b | phosphatidylinositol-4-phosphate 5-kinas | 772 | 639 | 864 | 3183 | 3772 | 2936 | 3308 | 2917 | 3443 | 3367 | 2891 |
| 3138 | 0.03 | 0.07 | 0.42 | 0.68 | 1 | Slc25a1 | solute carrier family 25 (mitochondrial car | 2337 | 2218 | 2489 | 3712 | 3872 | 2710 | 3307 | 3735 | 3708 | 3613 | 2813 |
| 3267 | -0.05 | 0.08 | -0.63 | 0.53 | 1 | Eif3d | eukaryotic translation initiation factor 3, su | 3520 | 3025 | 3124 | 3295 | 3566 | 3188 | 3306 | 3144 | 3369 | 3224 | 3174 |
| 3260 | -0.03 | 0.09 | -0.35 | 0.73 | 1 | Ep400 | E1A binding protein p400 | 3372 | 2992 | 3427 | 3011 | 2979 | 3892 | 3306 | 2740 | 3391 | 3474 | 3273 |
| 2904 | -0.08 | 0.12 | -0.68 | 0.49 | 1 | Pcbd1 | pterin 4 alpha carbinolamine dehydratase | 2448 | 2971 | 3096 | 3637 | 2814 | 2340 | 3306 | 2757 | 2962 | 3243 | 2372 |
| 3703 | 0.04 | 0.14 | 0.32 | 0.75 | 1 | Maged2 | melanoma antigen, family D, 2 | 4667 | 3794 | 5325 | 3713 | 3230 | 3012 | 3305 | 4196 | 3092 | 3321 | 3076 |
| 3817 | 0.10 | 0.11 | 0.93 | 0.35 | 1 | Nipbl | NIPBL cohesin loading factor | 3973 | 3885 | 3172 | 3380 | 3721 | 4539 | 3304 | 4326 | 3725 | 3376 | 4589 |
| 2674 | -0.10 | 0.15 | -0.66 | 0.51 | 1 | Fam120c | family with sequence similarity 120, meml | 1974 | 2511 | 2360 | 2666 | 3016 | 2646 | 3304 | 2061 | 3258 | 3109 | 2513 |
| 3216 | 0.13 | 0.09 | 1.33 | 0.18 | 1 | Mcfd2 | multiple coagulation factor deficiency 2 | 2188 | 2621 | 2822 | 2947 | 4052 | 3006 | 3302 | 3398 | 3995 | 3594 | 3457 |
| 3466 | 0.08 | 0.08 | 0.96 | 0.34 | 1 | Rabl6 | RAB, member RAS oncogene family-like | 3046 | 2834 | 3316 | 3185 | 3517 | 4097 | 3301 | 3709 | 3720 | 3625 | 3774 |
| 3374 | -0.02 | 0.09 | -0.27 | 0.79 | 1 | Bbx | bobby sox HMG box containing | 2686 | 3177 | 3274 | 3926 | 3053 | 3828 | 3300 | 3427 | 3427 | 3100 | 3913 |
| 3128 | 0.02 | 0.08 | 0.19 | 0.85 | 1 | Exoc5 | exocyst complex component 5 | 1833 | 2202 | 1999 | 3516 | 3997 | 3303 | 3300 | 3618 | 3803 | 3205 | 3634 |
| 2872 | 0.17 | 0.11 | 1.55 | 0.12 | 1 | Ube2w | ubiquitin-conjugating enzyme E2W (putat | 1851 | 2251 | 2335 | 3257 | 2847 | 2470 | 3299 | 3575 | 3183 | 3591 | 2933 |
| 3294 | -0.02 | 0.11 | -0.18 | 0.85 | 1 | Seh1l | SEH1-like (S. cerevisiae | 3186 | 3123 | 3194 | 3369 | 3743 | 3008 | 3298 | 2664 | 3924 | 3670 | 3052 |
| 3566 | 0.03 | 0.08 | 0.34 | 0.73 | 1 | Cmtm6 | CKLF-like MARVEL transmembrane dom | 5178 | 4339 | 5182 | 3035 | 3272 | 2507 | 3297 | 3348 | 3181 | 3532 | 2350 |
| 3168 | 0.11 | 0.10 | 1.07 | 0.28 | 1 | Plpbp | pyridoxal phosphate binding protein | 2195 | 2103 | 2339 | 2770 | 3009 | 4521 | 3296 | 3001 | 3075 | 4180 | 4363 |
| 3359 | -0.13 | 0.10 | -1.33 | 0.18 | 1 | Zmiz1 | zinc finger, MIZ-type containing 1 | 3945 | 3435 | 3519 | 3235 | 3766 | 3288 | 3293 | 2640 | 3729 | 3233 | 2866 |
| 4128 | -0.20 | 0.11 | -1.75 | 0.08 | 1 | Adcy6 | adenylate cyclase 6 | 5830 | 6279 | 5970 | 3888 | 3110 | 4345 | 3289 | 2550 | 3219 | 3319 | 3605 |
| 3489 | 0.00 | 0.10 | 0.02 | 0.98 | 1 | Impact | impact, RWD domain protein | 2805 | 3538 | 2756 | 3472 | 3230 | 4617 | 3287 | 3032 | 3402 | 3371 | 4870 |
| 3111 | 0.06 | 0.08 | 0.73 | 0.47 | 1 | Tor1b | torsin family 1, member B | 2356 | 2489 | 2382 | 2829 | 3327 | 3796 | 3285 | 3362 | 3368 | 3353 | 3681 |
| 2947 | 0.02 | 0.08 | 0.19 | 0.85 | 1 | Dhx30 | DEAH (Asp-Glu-Ala-His) box polypeptide | 2432 | 2318 | 2552 | 3107 | 3065 | 3028 | 3277 | 3366 | 3275 | 3263 | 2741 |
| 3242 | 0.16 | 0.14 | 1.11 | 0.27 | 1 | Ndufv1 | NADH:ubiquinone oxidoreductase core s | 3196 | 2782 | 3543 | 3174 | 3210 | 2674 | 3276 | 4509 | 2905 | 3421 | 2970 |
| 2833 | -0.04 | 0.09 | -0.41 | 0.68 | 1 | Foxa2 | forkhead box A2 | 2110 | 2400 | 2314 | 3059 | 3027 | 2956 | 3273 | 2936 | 3025 | 3321 | 2748 |
| 3498 | -0.04 | 0.09 | -0.38 | 0.71 | 1 | Ppig | peptidyl-prolyl isomerase G (cyclophilin G | 2949 | 3828 | 3308 | 3629 | 3482 | 3976 | 3272 | 3127 | 3716 | 3156 | 4032 |
| 3036 | 0.02 | 0.11 | 0.15 | 0.88 | 1 | Naa50 | N(alpha)-acetyltransferase 50, NatE catal | 3262 | 3024 | 2968 | 2917 | 3008 | 2811 | 3268 | 3327 | 3227 | 2584 | 3001 |
| 3362 | 0.03 | 0.11 | 0.26 | 0.79 | 1 | Cic | capicua transcriptional repressor | 4042 | 3347 | 3269 | 3116 | 3652 | 2916 | 3267 | 3537 | 3705 | 3697 | 2430 |
| 3550 | 0.08 | 0.09 | 0.88 | 0.38 | 1 | Mecp2 | methyl CpG binding protein 2 | 3546 | 4536 | 3634 | 3277 | 3220 | 3510 | 3258 | 3817 | 3691 | 3184 | 3372 |
| 3338 | 0.00 | 0.08 | -0.01 | 0.99 | 1 | Cacfd1 | calcium channel flower domain containing | 2431 | 2840 | 2712 | 3429 | 3900 | 3796 | 3258 | 3500 | 3650 | 3519 | 3677 |
| 4117 | -0.04 | 0.08 | -0.47 | 0.64 | 1 | Rpl22 | ribosomal protein L22 | 7935 | 6922 | 7678 | 2929 | 3147 | 2199 | 3257 | 3141 | 2888 | 2991 | 2199 |
| 3397 | -0.07 | 0.10 | -0.71 | 0.48 | 1 | Rab5c | RAB5C, member RAS oncogene family | 4130 | 3530 | 3822 | 3437 | 3576 | 2956 | 3257 | 2619 | 3617 | 3470 | 2948 |
| 2770 | 0.14 | 0.11 | 1.24 | 0.21 | 1 | Mars1 | methionine-tRNA synthetase 1 | 1768 | 1571 | 1659 | 2915 | 3688 | 2145 | 3256 | 4084 | 3922 | 3474 | 1989 |
| 2728 | -0.14 | 0.10 | -1.39 | 0.16 | 1 | Syap1 | synapse associated protein 1 | 2404 | 2127 | 2277 | 3121 | 3070 | 2686 | 3256 | 2333 | 2802 | 3078 | 2848 |
| 2750 | -0.16 | 0.12 | -1.27 | 0.20 | 1 | Lrrc41 | leucine rich repeat containing 41 | 2222 | 3035 | 2217 | 3201 | 2944 | 2627 | 3255 | 2500 | 2884 | 2977 | 2383 |
| 3766 | 0.03 | 0.12 | 0.24 | 0.81 | 1 | Anp32a | acidic (leucine-rich) nuclear phosphoprot | 5760 | 5480 | 6507 | 3039 | 2840 | 2590 | 3254 | 3598 | 2593 | 2910 | 2859 |
| 3138 | 0.07 | 0.09 | 0.76 | 0.45 | 1 | Agtpbp1 | ATP/GTP binding protein 1 | 1887 | 2209 | 2565 | 3658 | 3501 | 3151 | 3253 | 4035 | 3953 | 3026 | 3277 |
| 2922 | -0.02 | 0.10 | -0.16 | 0.87 | 1 | Abhd16a | abhydrolase domain containing 16A | 2288 | 1963 | 2405 | 3309 | 3222 | 2990 | 3252 | 3384 | 3112 | 3648 | 2572 |
| 2868 | -0.07 | 0.12 | -0.56 | 0.57 | 1 | Snx4 | sorting nexin 4 | 2302 | 2392 | 2163 | 3214 | 3128 | 3017 | 3249 | 2316 | 3252 | 3384 | 3127 |
| 4146 | 0.23 | 0.19 | 1.20 | 0.23 | 1 | Rps12 | ribosomal protein S12 | 7128 | 5572 | 7984 | 2690 | 3155 | 2274 | 3248 | 5220 | 3172 | 2861 | 2305 |
| 2781 | -0.04 | 0.13 | -0.34 | 0.74 | 1 | Syvn1 | synovial apoptosis inhibitor 1, synoviolin | 1693 | 1172 | 1428 | 2409 | 3441 | 4152 | 3248 | 1929 | 3588 | 4072 | 3460 |
| 3170 | -0.02 | 0.11 | -0.17 | 0.87 | 1 | Usp5 | ubiquitin specific peptidase 5 (isopeptida: | 3596 | 2933 | 3408 | 3085 | 3624 | 2603 | 3247 | 3683 | 3066 | 3031 | 2592 |
| 2721 | -0.01 | 0.11 | -0.12 | 0.91 | 1 | Tor3a | torsin family 3, member A | 1677 | 1623 | 1424 | 3526 | 3392 | 2461 | 3247 | 2801 | 3553 | 3822 | 2401 |
| 3321 | -0.03 | 0.09 | -0.29 | 0.77 | 1 | Pum1 | pumilio RNA-binding family member 1 | 3129 | 3241 | 2809 | 3505 | 3396 | 3641 | 3246 | 3783 | 3490 | 3076 | 3219 |
| 3791 | 0.08 | 0.07 | 1.05 | 0.30 | 1 | Cct4 | chaperonin containing Tcp1, subunit 4 (de | 5566 | 5296 | 5141 | 3204 | 2850 | 3193 | 3244 | 3402 | 2872 | 3225 | 3711 |
| 2961 | -0.08 | 0.12 | -0.66 | 0.51 | 1 | Atp5j | ATP synthase, H+ transporting, mitochon | 3154 | 3034 | 3108 | 3311 | 3036 | 2370 | 3244 | 2311 | 2937 | 3431 | 2637 |
| 2528 | -0.05 | 0.13 | -0.36 | 0.72 | 1 | Gm572 | predicted gene 572 | 697 | 1038 | 997 | 3222 | 3065 | 3217 | 3244 | 2884 | 3019 | 3337 | 3084 |
| 3168 | -0.10 | 0.11 | -0.92 | 0.36 | 1 | Tox4 | TOX high mobility group box family memb | 2823 | 3472 | 3181 | 3222 | 3307 | 3304 | 3243 | 2891 | 3691 | 2538 | 3178 |
| 2764 | 0.10 | 0.17 | 0.61 | 0.54 | 1 | Dbp | D site albumin promoter binding protein | 2479 | 1715 | 2533 | 2531 | 2726 | 2925 | 3242 | 3557 | 2196 | 3458 | 3045 |
| 3309 | 0.04 | 0.11 | 0.35 | 0.73 | 1 | Dctn2 | dynactin 2 | 3795 | 2856 | 3555 | 2762 | 3120 | 3833 | 3239 | 3318 | 3169 | 2865 | 3883 |
| 3054 | -0.12 | 0.14 | -0.86 | 0.39 | 1 | Ankfy1 | ankyrin repeat and FYVE domain contain | 4154 | 3473 | 3513 | 2850 | 2781 | 2824 | 3238 | 2703 | 3262 | 2109 | 2690 |
| 3359 | 0.14 | 0.12 | 1.12 | 0.26 | 1 | Pdap1 | PDGFA associated protein 1 | 3388 | 3025 | 3647 | 2993 | 3649 | 2909 | 3237 | 4389 | 3487 | 3272 | 2951 |
| 3091 | 0.00 | 0.10 | -0.02 | 0.98 | 1 | Htatsf1 | HIV TAT specific factor 1 | 3048 | 2933 | 2818 | 2989 | 3092 | 3263 | 3237 | 2514 | 3505 | 3130 | 3471 |
| 4232 | 0.07 | 0.08 | 0.87 | 0.39 | 1 | Tardbp | TAR DNA binding protein | 4363 | 4807 | 4680 | 2727 | 3029 | 7039 | 3235 | 3192 | 3127 | 3228 | 7126 |

| | | | | | | | | | | | | | | | | | | |
|---|---|---|---|---|---|---|---|---|---|---|---|---|---|---|---|---|---|---|
| 2823 | -0.06 | 0.09 | -0.65 | 0.51 | 1 | Gpr107 | G protein-coupled receptor 107 | 2720 | 2581 | 2707 | 2855 | 2987 | 2649 | 3235 | 2490 | 2840 | 3555 | 2435 |
| 2969 | -0.03 | 0.11 | -0.26 | 0.79 | 1 | Zmym4 | zinc finger, MYM-type 4 | 2418 | 3111 | 3042 | 2950 | 3158 | 2820 | 3233 | 2675 | 3267 | 3064 | 2924 |
| 3190 | 0.20 | 0.13 | 1.54 | 0.12 | 1 | Map2k2 | mitogen-activated protein kinase kinase 2 | 2667 | 2336 | 2899 | 3257 | 3187 | 2933 | 3229 | 4716 | 3381 | 3435 | 3051 |
| 3571 | 0.01 | 0.08 | 0.09 | 0.92 | 1 | Ctsa | cathepsin A | 4245 | 3699 | 4183 | 3671 | 3704 | 2927 | 3228 | 3493 | 3683 | 3576 | 2869 |
| 2903 | -0.04 | 0.09 | -0.40 | 0.69 | 1 | Vps26b | VPS26 retromer complex component B | 2817 | 2981 | 2554 | 3233 | 2836 | 2647 | 3228 | 2753 | 3117 | 3187 | 2575 |
| 3250 | -0.21 | 0.13 | -1.65 | 0.10 | 1 | Smad5 | SMAD family member 5 | 3645 | 4158 | 3692 | 3597 | 3088 | 3079 | 3227 | 2253 | 3031 | 3027 | 2948 |
| 8981 | -0.09 | 0.09 | -0.92 | 0.36 | 1 | Igfbp7 | insulin-like growth factor binding protein 7 | 25929 | 20637 | 23782 | 3867 | 3958 | 3596 | 3226 | 3558 | 3550 | 3336 | 3349 |
| 3116 | 0.05 | 0.09 | 0.60 | 0.55 | 1 | Slc25a12 | solute carrier family 25 (mitochondrial car | 2809 | 2617 | 2814 | 3359 | 3547 | 2644 | 3223 | 3608 | 3161 | 3743 | 2748 |
| 3182 | -0.03 | 0.08 | -0.35 | 0.73 | 1 | Map7 | microtubule-associated protein 7 | 2000 | 2399 | 2551 | 3571 | 3570 | 3803 | 3221 | 3390 | 3459 | 3352 | 3682 |
| 3097 | -0.09 | 0.13 | -0.69 | 0.49 | 1 | Paics | phosphoribosylaminoimidazole carboxyla | 4284 | 3136 | 3608 | 3122 | 2869 | 2657 | 3221 | 2555 | 3267 | 2516 | 2833 |
| 3853 | 0.06 | 0.07 | 0.80 | 0.42 | 1 | Dync1i2 | dynein cytoplasmic 1 intermediate chain 2 | 5067 | 4868 | 5432 | 3299 | 3179 | 3537 | 3220 | 3449 | 3184 | 3404 | 3747 |
| 3418 | 0.08 | 0.09 | 0.97 | 0.33 | 1 | Nucks1 | nuclear casein kinase and cyclin-depend | 3438 | 3270 | 3340 | 3708 | 3557 | 2898 | 3218 | 3696 | 4034 | 3234 | 3202 |
| 3089 | 0.00 | 0.12 | -0.02 | 0.99 | 1 | Socs7 | suppressor of cytokine signaling 7 | 2700 | 2841 | 2614 | 3540 | 3115 | 2942 | 3216 | 4539 | 3026 | 2713 | 2732 |
| 3285 | -0.07 | 0.10 | -0.70 | 0.48 | 1 | Tob1 | transducer of ErbB-2.1 | 2492 | 2968 | 2975 | 3189 | 3649 | 4146 | 3214 | 2898 | 3679 | 3357 | 3572 |
| 3278 | 0.04 | 0.08 | 0.52 | 0.60 | 1 | Ddx46 | DEAD (Asp-Glu-Ala-Asp) box polypeptide | 3011 | 2907 | 2586 | 3075 | 3279 | 3984 | 3208 | 3012 | 3160 | 3665 | 4171 |
| 3242 | 0.11 | 0.12 | 0.93 | 0.35 | 1 | Mvp | major vault protein | 4100 | 3278 | 3500 | 2966 | 3087 | 2582 | 3208 | 4151 | 2956 | 3327 | 2507 |
| 2770 | 0.01 | 0.12 | 0.06 | 0.96 | 1 | Pitpnm3 | PITPNM family member 3 | 594 | 580 | 949 | 3303 | 3602 | 4013 | 3207 | 2983 | 3851 | 3527 | 3864 |
| 3153 | 0.12 | 0.13 | 0.96 | 0.34 | 1 | Vipr1 | vasoactive intestinal peptide receptor 1 | 2829 | 4052 | 2569 | 2954 | 2623 | 3323 | 3204 | 3133 | 3044 | 3603 | 3350 |
| 3466 | 0.04 | 0.10 | 0.36 | 0.72 | 1 | Rps23 | ribosomal protein S23 | 5849 | 5111 | 6785 | 2389 | 2706 | 1813 | 3201 | 2745 | 2693 | 2857 | 1977 |
| 3324 | -0.03 | 0.10 | -0.26 | 0.79 | 1 | Lsm14a | LSM14A mRNA processing body assemb | 4244 | 4150 | 3655 | 3173 | 2942 | 3057 | 3201 | 2826 | 3431 | 2880 | 3004 |
| 3088 | -0.01 | 0.13 | -0.07 | 0.95 | 1 | Dmxl1 | Dmx-like 1 | 3203 | 2797 | 2493 | 2651 | 2919 | 3965 | 3200 | 2354 | 3498 | 2713 | 4181 |
| 2914 | 0.10 | 0.11 | 0.88 | 0.38 | 1 | Fem1a | fem 1 homolog a | 2099 | 2566 | 1690 | 3306 | 3132 | 2738 | 3200 | 3623 | 3430 | 3654 | 2614 |
| 2698 | 0.12 | 0.09 | 1.29 | 0.20 | 1 | Gpsm1 | G-protein signalling modulator 1 (AGS3-li | 1204 | 983 | 1275 | 2097 | 3327 | 3996 | 3199 | 2362 | 3480 | 3692 | 4066 |
| 2247 | 0.05 | 0.18 | 0.26 | 0.79 | 1 | Gpr45 | G protein-coupled receptor 45 | 281 | 536 | 599 | 3172 | 3107 | 2004 | 3198 | 3019 | 3221 | 3440 | 2134 |
| 3000 | 0.07 | 0.09 | 0.76 | 0.45 | 1 | Glmp | glycosylated lysosomal membrane protein | 2656 | 3025 | 2965 | 3143 | 3181 | 2286 | 3196 | 3375 | 3460 | 3648 | 2070 |
| 2771 | 0.08 | 0.13 | 0.59 | 0.56 | 1 | Hif1an | hypoxia-inducible factor 1, alpha subunit | 2077 | 2254 | 2577 | 2824 | 2750 | 2646 | 3196 | 3839 | 2664 | 3112 | 2540 |
| 2984 | -0.19 | 0.13 | -1.48 | 0.14 | 1 | Tor1aip1 | torsin A interacting protein 1 | 3306 | 2778 | 2799 | 3685 | 3387 | 2509 | 3194 | 2424 | 3419 | 2802 | 2522 |
| 2780 | 0.13 | 0.15 | 0.85 | 0.39 | 1 | Tssc4 | tumor-suppressing subchromosomal tran | 824 | 1219 | 1412 | 3138 | 3920 | 2631 | 3190 | 4829 | 3540 | 3297 | 2580 |
| 3296 | 0.01 | 0.08 | 0.17 | 0.86 | 1 | Stip1 | stress-induced phosphoprotein 1 | 3836 | 3018 | 4153 | 2384 | 3292 | 3744 | 3189 | 2725 | 3085 | 3094 | 3738 |
| 3251 | 0.01 | 0.09 | 0.12 | 0.91 | 1 | Hspa4l | heat shock protein 4 like | 2485 | 2925 | 3116 | 3027 | 3690 | 3634 | 3189 | 2860 | 3966 | 3142 | 3724 |
| 3275 | 0.17 | 0.09 | 1.94 | 0.05 | 1 | Matr3 | matrin 3 | 3130 | 3508 | 3291 | 2850 | 2713 | 3525 | 3186 | 3413 | 2936 | 3273 | 4204 |
| 3017 | 0.00 | 0.09 | 0.04 | 0.97 | 1 | Rnf181 | ring finger protein 181 | 2152 | 1986 | 2547 | 2465 | 3999 | 3629 | 3185 | 2595 | 3670 | 3447 | 3512 |
| 3259 | -0.03 | 0.08 | -0.45 | 0.65 | 1 | Appbp2 | amyloid beta precursor protein (cytoplasm | 3552 | 3815 | 3330 | 3458 | 2987 | 3109 | 3181 | 3171 | 3223 | 2887 | 3133 |
| 2975 | -0.08 | 0.15 | -0.56 | 0.57 | 1 | Gck | glucokinase | 1376 | 1878 | 2249 | 2868 | 3029 | 4858 | 3180 | 2639 | 2621 | 3102 | 4926 |
| 2854 | -0.03 | 0.12 | -0.22 | 0.82 | 1 | Tcn2 | transcobalamin 2 | 2801 | 2718 | 2798 | 3131 | 3262 | 2012 | 3180 | 2343 | 3327 | 3842 | 1981 |
| 2803 | -0.06 | 0.08 | -0.80 | 0.42 | 1 | Lmo7 | LIM domain only 7 | 1749 | 2197 | 1933 | 3199 | 3086 | 3281 | 3178 | 3228 | 2916 | 2947 | 3114 |
| 2781 | -0.06 | 0.11 | -0.52 | 0.60 | 1 | Kat2b | K(lysine) acetyltransferase 2B | 2116 | 1909 | 1864 | 3452 | 2825 | 3130 | 3178 | 2705 | 3117 | 2866 | 3427 |
| 3006 | 0.01 | 0.11 | 0.08 | 0.93 | 1 | Twf1 | twinfilin actin binding protein 1 | 2341 | 2579 | 2590 | 3522 | 3186 | 2845 | 3176 | 2697 | 3628 | 3543 | 2964 |
| 3537 | 0.03 | 0.11 | 0.24 | 0.81 | 1 | Grk2 | G protein-coupled receptor kinase 2 | 5044 | 3801 | 3757 | 3724 | 3292 | 2811 | 3175 | 3361 | 3470 | 3723 | 2749 |
| 3169 | 0.02 | 0.09 | 0.26 | 0.80 | 1 | Pcyt1a | phosphate cytidylyltransferase 1, choline, | 3414 | 2969 | 2859 | 3357 | 3432 | 2728 | 3173 | 3499 | 3213 | 3546 | 2672 |
| 2707 | -0.09 | 0.13 | -0.67 | 0.50 | 1 | Eea1 | early endosome antigen 1 | 1981 | 1810 | 1602 | 3063 | 2893 | 3381 | 3173 | 3045 | 2935 | 2237 | 3653 |
| 3053 | 0.11 | 0.09 | 1.32 | 0.19 | 1 | Pcmt1 | protein-L-isoaspartate (D-aspartate) O-me | 3176 | 3077 | 3316 | 2682 | 3105 | 2581 | 3171 | 3418 | 3178 | 3116 | 2761 |
| 14158 | 0.04 | 0.29 | 0.12 | 0.90 | 1 | Malat1 | | 2832 | 4152 | 3119 | 1751 | 1348 | 73451 | 3168 | 1711 | 2087 | 2960 | 59165 |
| 2812 | -0.04 | 0.08 | -0.50 | 0.61 | 1 | 2510009E07Ril | RIKEN cDNA 2510009E07 gene | 3237 | 2893 | 2841 | 3075 | 2668 | 2229 | 3168 | 2691 | 2841 | 3059 | 2225 |
| 3031 | -0.02 | 0.08 | -0.20 | 0.84 | 1 | Rabgap1 | RAB GTPase activating protein 1 | 2919 | 2917 | 2690 | 3295 | 2944 | 3068 | 3167 | 2820 | 3037 | 3290 | 3192 |
| 2851 | -0.10 | 0.08 | -1.23 | 0.22 | 1 | Brk1 | BRICK1, SCAR/WAVE actin-nucleating co | 3091 | 2936 | 3346 | 2937 | 3122 | 2139 | 3166 | 2677 | 2760 | 3147 | 2040 |
| 3417 | 0.09 | 0.11 | 0.79 | 0.43 | 1 | Rpl15 | ribosomal protein L15 | 4962 | 4394 | 5795 | 2764 | 2702 | 2266 | 3164 | 2960 | 2672 | 3364 | 2547 |
| 2838 | -0.15 | 0.10 | -1.50 | 0.13 | 1 | Ctsf | cathepsin F | 2563 | 3004 | 2702 | 3041 | 3287 | 2564 | 3164 | 2640 | 2805 | 3209 | 2236 |
| 3684 | 0.06 | 0.07 | 0.90 | 0.37 | 1 | Smc6 | structural maintenance of chromosomes 6 | 4443 | 4611 | 4413 | 3177 | 3061 | 3834 | 3161 | 3048 | 3349 | 3449 | 3982 |
| 2773 | -0.14 | 0.10 | -1.41 | 0.16 | 1 | Lmtk2 | lemur tyrosine kinase 2 | 2080 | 1881 | 1789 | 3439 | 3359 | 2996 | 3160 | 2879 | 3421 | 3063 | 2440 |
| 3456 | -0.06 | 0.07 | -0.83 | 0.41 | 1 | Nudt4 | nudix (nucleoside diphosphate linked mo | 5173 | 4552 | 4655 | 2922 | 3238 | 2737 | 3157 | 2836 | 2992 | 3049 | 2709 |
| 3099 | 0.09 | 0.12 | 0.76 | 0.45 | 1 | Anapc2 | anaphase promoting complex subunit 2 | 3929 | 3304 | 3387 | 2678 | 3043 | 2466 | 3152 | 3788 | 2888 | 3002 | 2446 |
| 3451 | -0.08 | 0.09 | -0.89 | 0.37 | 1 | Slk | STE20-like kinase | 3951 | 3864 | 3994 | 2858 | 3020 | 4361 | 3147 | 2399 | 2717 | 3344 | 4300 |
| 3126 | 0.06 | 0.07 | 0.85 | 0.40 | 1 | Keap1 | kelch-like ECH-associated protein 1 | 2661 | 2606 | 2994 | 2938 | 3562 | 3147 | 3145 | 3371 | 3619 | 3296 | 3049 |
| 2485 | 0.12 | 0.13 | 0.94 | 0.35 | 1 | Ndufc2 | NADH:ubiquinone oxidoreductase subun | 1955 | 1849 | 2232 | 2386 | 2443 | 2262 | 3143 | 3031 | 2399 | 3098 | 2538 |
| 3018 | 0.05 | 0.10 | 0.47 | 0.64 | 1 | Psma2 | proteasome subunit alpha 2 | 3621 | 3006 | 3745 | 2866 | 3030 | 2221 | 3142 | 3215 | 2732 | 3080 | 2541 |
| 2967 | 0.03 | 0.11 | 0.24 | 0.81 | 1 | Cdc37l1 | cell division cycle 37-like 1 | 1723 | 1830 | 2080 | 2779 | 2978 | 4386 | 3142 | 2405 | 3136 | 3315 | 4858 |
| 2961 | 0.07 | 0.08 | 0.94 | 0.35 | 1 | Smarca5 | SWI/SNF related, matrix associated, actin | 2786 | 2845 | 2581 | 3007 | 3011 | 2723 | 3142 | 3184 | 3161 | 3013 | 3118 |
| 2993 | 0.06 | 0.09 | 0.71 | 0.48 | 1 | Uqcrfs1 | ubiquinol-cytochrome c reductase, Rieske | 3653 | 3350 | 3609 | 2857 | 2816 | 2138 | 3140 | 2903 | 2689 | 3214 | 2555 |
| 4175 | 0.11 | 0.17 | 0.63 | 0.53 | 1 | Grn | granulin | 8887 | 5126 | 6469 | 3431 | 3269 | 2236 | 3139 | 4729 | 3169 | 3492 | 1973 |

| | | | | | | | | | | | | | | | | | | |
|---|---|---|---|---|---|---|---|---|---|---|---|---|---|---|---|---|---|---|
| 3742 | 0.08 | 0.07 | 1.02 | 0.31 | 1 | Rps26 | ribosomal protein S26 | 6824 | 5906 | 6591 | 2872 | 2768 | 1884 | 3139 | 2827 | 2860 | 3321 | 2168 |
| 3218 | 0.07 | 0.11 | 0.61 | 0.54 | 1 | Foxo1 | forkhead box O1 | 2957 | 3353 | 2970 | 3279 | 3371 | 2846 | 3136 | 4609 | 3175 | 2978 | 2721 |
| 2868 | 0.05 | 0.11 | 0.48 | 0.63 | 1 | Mtch2 | mitochondrial carrier 2 | 2231 | 2394 | 2545 | 3142 | 2799 | 2874 | 3135 | 2669 | 3152 | 3558 | 3048 |
| 2667 | 0.02 | 0.15 | 0.15 | 0.88 | 1 | Fnip2 | folliculin interacting protein 2 | 1636 | 1936 | 1606 | 3089 | 2928 | 2798 | 3131 | 2328 | 3694 | 3504 | 2692 |
| 3228 | 0.12 | 0.07 | 1.57 | 0.12 | 1 | Cebpg | CCAAT/enhancer binding protein (C/EBP | 2157 | 3067 | 2509 | 3810 | 3182 | 3208 | 3130 | 4035 | 3470 | 3310 | 3632 |
| 2663 | 0.01 | 0.10 | 0.11 | 0.92 | 1 | Phkb | phosphorylase kinase beta | 1598 | 2110 | 1943 | 2969 | 3130 | 2553 | 3128 | 3064 | 3161 | 2983 | 2650 |
| 3059 | -0.14 | 0.09 | -1.52 | 0.13 | 1 | Rapgef1 | Rap guanine nucleotide exchange factor | 4174 | 3095 | 3765 | 2977 | 2982 | 2743 | 3126 | 2651 | 2921 | 2844 | 2375 |
| 2924 | 0.03 | 0.09 | 0.39 | 0.70 | 1 | Sod2 | superoxide dismutase 2, mitochondrial | 2657 | 2798 | 2529 | 3019 | 3087 | 2722 | 3126 | 3230 | 3249 | 2787 | 2960 |
| 3573 | -0.04 | 0.09 | -0.42 | 0.68 | 1 | Smc1a | structural maintenance of chromosomes 1 | 5246 | 4307 | 5329 | 3048 | 2995 | 3193 | 3124 | 2980 | 2965 | 2893 | 3224 |
| 4158 | 0.02 | 0.08 | 0.22 | 0.82 | 1 | Amotl1 | angiomotin-like 1 | 7246 | 8009 | 7894 | 2972 | 3071 | 2059 | 3122 | 3373 | 2926 | 2950 | 2112 |
| 3079 | 0.06 | 0.09 | 0.71 | 0.48 | 1 | Mkln1 | muskelin 1, intracellular mediator contain | 2895 | 2892 | 2856 | 2977 | 2948 | 3293 | 3122 | 3352 | 3379 | 2799 | 3360 |
| 2907 | 0.04 | 0.13 | 0.33 | 0.75 | 1 | Mogs | mannosyl-oligosaccharide glucosidase | 1963 | 2451 | 2094 | 2876 | 3470 | 3073 | 3122 | 2660 | 3544 | 3618 | 3101 |
| 2613 | -0.17 | 0.11 | -1.54 | 0.12 | 1 | Adam22 | a disintegrin and metallopeptidase domai | 1247 | 1695 | 1643 | 2872 | 3052 | 3693 | 3121 | 2772 | 2976 | 2308 | 3361 |
| 3540 | 0.17 | 0.10 | 1.76 | 0.08 | 1 | D5Ertd579e | DNA segment, Chr 5, ERATO Doi 579, ex | 2027 | 2343 | 2086 | 4551 | 2915 | 4718 | 3118 | 5093 | 3465 | 3354 | 5270 |
| 3275 | 0.19 | 0.11 | 1.69 | 0.09 | 1 | Ggnbp2 | gametogenetin binding protein 2 | 2723 | 2891 | 2578 | 3096 | 3119 | 3591 | 3118 | 4749 | 3225 | 3250 | 3691 |
| 2926 | -0.04 | 0.10 | -0.37 | 0.71 | 1 | Mff | mitochondrial fission factor | 2481 | 2404 | 2429 | 3384 | 3049 | 3034 | 3118 | 2673 | 3269 | 3030 | 3322 |
| 2707 | 0.12 | 0.11 | 1.08 | 0.28 | 1 | Ptpn4 | protein tyrosine phosphatase, non-recept | 1807 | 2387 | 1857 | 2792 | 2877 | 2577 | 3117 | 3398 | 3251 | 2861 | 2857 |
| 2689 | 0.00 | 0.09 | 0.01 | 0.99 | 1 | Psme3 | proteaseome (prosome, macropain) activ | 2379 | 2299 | 2014 | 2888 | 2769 | 2654 | 3115 | 2670 | 3081 | 3218 | 2490 |
| 2488 | -0.05 | 0.12 | -0.45 | 0.65 | 1 | Ksr2 | kinase suppressor of ras 2 | 1752 | 2020 | 2340 | 2230 | 3031 | 2434 | 3115 | 2522 | 3121 | 2699 | 2106 |
| 2784 | -0.04 | 0.14 | -0.30 | 0.76 | 1 | Rprd1b | regulation of nuclear pre-mRNA domain c | 2159 | 2544 | 1982 | 3211 | 3409 | 2423 | 3114 | 2379 | 3413 | 3452 | 2537 |
| 2601 | 0.10 | 0.08 | 1.29 | 0.20 | 1 | Fam117a | family with sequence similarity 117, meml | 1275 | 1345 | 1403 | 2791 | 3032 | 2926 | 3111 | 3220 | 3207 | 3300 | 2999 |
| 3394 | 0.28 | 0.32 | 0.86 | 0.39 | 1 | Atp5e | ATP synthase, H+ transporting, mitochon | 3696 | 3040 | 4447 | 3068 | 3126 | 2132 | 3110 | 6798 | 2875 | 2826 | 2214 |
| 2778 | -0.05 | 0.10 | -0.57 | 0.57 | 1 | Ykt6 | YKT6 v-SNARE homolog (S. cerevisiae) | 2523 | 2510 | 2213 | 2939 | 2954 | 2870 | 3110 | 2926 | 2858 | 2532 | 3125 |
| 1910 | 0.03 | 0.17 | 0.15 | 0.88 | 1 | Tmem130 | transmembrane protein 130 | 2873 | 2272 | 2151 | 1317 | 1463 | 899 | 3110 | 1899 | 1241 | 2952 | 835 |
| 3267 | -0.07 | 0.09 | -0.73 | 0.46 | 1 | Pitpna | phosphatidylinositol transfer protein, alph | 4858 | 3662 | 4371 | 2855 | 3022 | 2803 | 3109 | 2694 | 3198 | 2710 | 2651 |
| 5478 | -0.09 | 0.16 | -0.59 | 0.55 | 1 | Rgs2 | regulator of G-protein signaling 2 | 5050 | 4635 | 4167 | 4695 | 1834 | 13950 | 3108 | 4156 | 1883 | 2632 | 14148 |
| 2843 | -0.15 | 0.12 | -1.24 | 0.21 | 1 | Mdm1 | transformed mouse 3T3 cell double minut | 992 | 1044 | 988 | 2738 | 2347 | 6629 | 3107 | 2031 | 2770 | 2485 | 6139 |
| 3334 | 0.08 | 0.09 | 0.90 | 0.37 | 1 | Snx3 | sorting nexin 3 | 4075 | 3775 | 4219 | 3121 | 3289 | 2444 | 3106 | 3103 | 3076 | 3792 | 2679 |
| 3306 | 0.00 | 0.09 | 0.05 | 0.96 | 1 | Cdk4 | cyclin-dependent kinase 4 | 3454 | 2751 | 2898 | 3170 | 3312 | 4033 | 3106 | 3298 | 3121 | 3425 | 3796 |
| 2991 | -0.01 | 0.07 | -0.16 | 0.87 | 1 | Mark2 | MAP/microtubule affinity regulating kinase | 2564 | 2699 | 2896 | 3107 | 3320 | 2882 | 3104 | 3273 | 3167 | 3216 | 2674 |
| 3006 | -0.15 | 0.08 | -1.95 | 0.05 | 1 | Septin9 | septin 9 | 4242 | 3609 | 4078 | 2661 | 2984 | 2348 | 3103 | 2149 | 2863 | 2929 | 2103 |
| 2805 | -0.01 | 0.07 | -0.19 | 0.85 | 1 | Virma | vir like m6A methyltransferase associated | 2730 | 2627 | 2679 | 2873 | 2810 | 2681 | 3103 | 2677 | 2855 | 3021 | 2800 |
| 2450 | 0.02 | 0.09 | 0.18 | 0.86 | 1 | Enpp4 | ectonucleotide pyrophosphatase/phosph | 2237 | 2152 | 2273 | 2493 | 2684 | 1839 | 3103 | 2756 | 2702 | 2759 | 1948 |
| 2927 | 0.06 | 0.11 | 0.58 | 0.56 | 1 | Pom121 | nuclear pore membrane protein 121 | 2231 | 2880 | 2234 | 3387 | 3213 | 2382 | 3101 | 3442 | 3478 | 3720 | 2129 |
| 2641 | -0.05 | 0.12 | -0.40 | 0.69 | 1 | Mid2 | midline 2 | 1633 | 1869 | 1965 | 3551 | 2767 | 2585 | 3098 | 2690 | 3282 | 2925 | 2682 |
| 2961 | -0.06 | 0.10 | -0.63 | 0.53 | 1 | Ap3b1 | adaptor-related protein complex 3, beta 1 | 3052 | 2831 | 2955 | 2961 | 3271 | 2765 | 3095 | 2345 | 3404 | 3126 | 2763 |
| 3255 | 0.01 | 0.14 | 0.06 | 0.95 | 1 | Abce1 | ATP-binding cassette, sub-family E (OABI | 2668 | 2637 | 2356 | 2997 | 3673 | 4145 | 3091 | 2237 | 3776 | 3599 | 4621 |
| 3156 | 0.02 | 0.11 | 0.15 | 0.88 | 1 | Otud5 | OTU domain containing 5 | 2772 | 2994 | 2711 | 2867 | 3399 | 3648 | 3091 | 2502 | 3388 | 3435 | 3910 |
| 3151 | 0.00 | 0.13 | -0.02 | 0.98 | 1 | Cbarp | calcium channel, voltage-dependent, beta | 3028 | 3688 | 3533 | 3082 | 3129 | 2892 | 3091 | 2594 | 2994 | 3813 | 2816 |
| 3271 | 0.10 | 0.11 | 0.89 | 0.38 | 1 | Psmd3 | proteasome (prosome, macropain) 26S su | 3585 | 2979 | 3145 | 3395 | 3282 | 2884 | 3090 | 4152 | 3426 | 3223 | 2817 |
| 2825 | 0.07 | 0.07 | 0.97 | 0.33 | 1 | Ptgr2 | prostaglandin reductase 2 | 2506 | 2558 | 2342 | 2781 | 2925 | 2754 | 3090 | 3017 | 2951 | 3169 | 2984 |
| 2604 | -0.02 | 0.11 | -0.22 | 0.82 | 1 | Gatad2a | GATA zinc finger domain containing 2A | 2406 | 2347 | 2151 | 2508 | 3076 | 2259 | 3090 | 2338 | 3142 | 3202 | 2127 |
| 3158 | 0.00 | 0.09 | 0.01 | 0.99 | 1 | Ndel1 | nudE neurodevelopment protein 1 like 1 | 2612 | 2502 | 2539 | 3355 | 3431 | 3647 | 3088 | 2895 | 3415 | 3486 | 3769 |
| 3084 | 0.13 | 0.09 | 1.38 | 0.17 | 1 | Nmt1 | N-myristoyltransferase 1 | 3315 | 3080 | 3252 | 2559 | 3264 | 2693 | 3086 | 3339 | 3217 | 3462 | 2657 |
| 3166 | 0.07 | 0.09 | 0.74 | 0.46 | 1 | Smad4 | SMAD family member 4 | 3535 | 4266 | 3343 | 2916 | 2809 | 2720 | 3085 | 3573 | 2940 | 2987 | 2647 |
| 3082 | 0.12 | 0.07 | 1.63 | 0.10 | 1 | Tmx3 | thioredoxin-related transmembrane prote | 2969 | 2739 | 2799 | 2914 | 3281 | 2908 | 3083 | 3296 | 3300 | 3291 | 3326 |
| 2914 | -0.10 | 0.10 | -0.95 | 0.34 | 1 | Htt | huntingtin | 2514 | 2325 | 2440 | 2963 | 2932 | 3806 | 3083 | 2277 | 3163 | 3182 | 3370 |
| 2607 | -0.08 | 0.10 | -0.84 | 0.40 | 1 | Atp6v1f | ATPase, H+ transporting, lysosomal V1 su | 2259 | 2262 | 2769 | 2801 | 3058 | 2072 | 3077 | 2698 | 2644 | 3076 | 1961 |
| 3633 | 0.09 | 0.09 | 0.93 | 0.35 | 1 | Rnh1 | ribonuclease/angiogenin inhibitor 1 | 4202 | 4017 | 4365 | 3348 | 3876 | 2991 | 3075 | 3899 | 3450 | 3667 | 3074 |
| 3170 | 0.04 | 0.08 | 0.48 | 0.63 | 1 | Abcf1 | ATP-binding cassette, sub-family F (GCN: | 3565 | 3288 | 3241 | 3019 | 3040 | 3083 | 3074 | 3238 | 3352 | 3121 | 2851 |
| 3073 | 0.21 | 0.14 | 1.45 | 0.15 | 1 | Drap1 | Dr1 associated protein 1 (negative cofact | 2744 | 2545 | 2978 | 2942 | 3238 | 2533 | 3074 | 4703 | 2971 | 3393 | 2685 |
| 2816 | 0.14 | 0.08 | 1.73 | 0.08 | 1 | Tmem50b | transmembrane protein 50B | 2083 | 1946 | 2453 | 2910 | 3329 | 2361 | 3073 | 2891 | 3637 | 3471 | 2822 |
| 2744 | -0.12 | 0.11 | -1.04 | 0.30 | 1 | Fmnl2 | formin-like 2 | 2044 | 1573 | 2145 | 2710 | 3533 | 3269 | 3073 | 1890 | 3552 | 2942 | 3454 |
| 3028 | 0.09 | 0.08 | 1.07 | 0.28 | 1 | Sucla2 | succinate-Coenzyme A ligase, ADP-formi | 2702 | 2505 | 2892 | 3103 | 3093 | 2961 | 3068 | 3344 | 3198 | 3062 | 3380 |
| 2870 | 0.01 | 0.13 | 0.04 | 0.97 | 1 | Ccar1 | cell division cycle and apoptosis regulato | 2235 | 2317 | 2102 | 2292 | 2736 | 4289 | 3068 | 2703 | 3064 | 2215 | 4544 |
| 2995 | 0.07 | 0.12 | 0.58 | 0.56 | 1 | Mgat2 | mannoside acetylglucosaminyltransferas | 2574 | 2532 | 2229 | 2686 | 3687 | 3061 | 3067 | 2727 | 3686 | 3439 | 3261 |
| 2913 | 0.21 | 0.12 | 1.79 | 0.07 | 1 | Rora | RAR-related orphan receptor alpha | 1941 | 2372 | 2559 | 3178 | 2954 | 2419 | 3065 | 4372 | 3245 | 3415 | 2527 |
| 2968 | -0.03 | 0.09 | -0.30 | 0.76 | 1 | Plpp5 | phospholipid phosphatase 5 | 1039 | 1171 | 1425 | 3259 | 3758 | 4549 | 3061 | 2829 | 3559 | 3352 | 4644 |
| 2974 | -0.17 | 0.10 | -1.66 | 0.10 | 1 | Smg7 | Smg-7 homolog, nonsense mediated mR | 2416 | 2929 | 2393 | 3419 | 3144 | 3576 | 3060 | 2655 | 3142 | 2637 | 3342 |
| 2709 | 0.24 | 0.16 | 1.47 | 0.14 | 1 | Fkbp1b | FK506 binding protein 1b | 220 | 190 | 397 | 3243 | 4333 | 2778 | 3060 | 4405 | 4008 | 4118 | 3045 |

| | | | | | | | | | | | | | | | | | | |
|---|---|---|---|---|---|---|---|---|---|---|---|---|---|---|---|---|---|---|
| 2836 | -0.13 | 0.09 | -1.49 | 0.14 | 1 | Lztr1 | leucine-zipper-like transcriptional regulat | 2605 | 2422 | 2835 | 2999 | 3065 | 3057 | 3059 | 2389 | 3082 | 3085 | 2601 |
| 3363 | -0.02 | 0.11 | -0.20 | 0.84 | 1 | Kif1c | kinesin family member 1C | 5366 | 3878 | 4399 | 3253 | 3155 | 2281 | 3057 | 3322 | 3085 | 3042 | 2155 |
| 3099 | 0.02 | 0.10 | 0.15 | 0.88 | 1 | Nub1 | negative regulator of ubiquitin-like proteir | 3082 | 3560 | 2984 | 2908 | 3412 | 2795 | 3056 | 2900 | 3338 | 3221 | 2835 |
| 3333 | 0.01 | 0.08 | 0.15 | 0.88 | 1 | Cnn3 | calponin 3, acidic | 3977 | 3414 | 3295 | 3304 | 2811 | 3744 | 3053 | 2925 | 2821 | 3477 | 3844 |
| 3036 | 0.08 | 0.12 | 0.69 | 0.49 | 1 | Rock2 | Rho-associated coiled-coil containing prc | 3636 | 3559 | 3122 | 2304 | 2770 | 3102 | 3052 | 3012 | 3226 | 2589 | 3028 |
| 2069 | 0.04 | 0.18 | 0.24 | 0.81 | 1 | Aldh1l2 | aldehyde dehydrogenase 1 family, memb | 415 | 824 | 646 | 3172 | 2495 | 1335 | 3050 | 3493 | 2746 | 3518 | 1066 |
| 2900 | 0.06 | 0.11 | 0.54 | 0.59 | 1 | Coa3 | cytochrome C oxidase assembly factor 3 | 2072 | 2088 | 2678 | 3552 | 3444 | 2231 | 3049 | 3961 | 3269 | 3246 | 2309 |
| 2512 | 0.03 | 0.07 | 0.46 | 0.65 | 1 | Vldlr | very low density lipoprotein receptor | 1116 | 1042 | 1069 | 2693 | 2815 | 3502 | 3048 | 2761 | 2838 | 3012 | 3741 |
| 2787 | 0.02 | 0.12 | 0.20 | 0.84 | 1 | Atpif1 | ATPase inhibitory factor 1 | 2835 | 2645 | 3229 | 2723 | 2869 | 2243 | 3047 | 3388 | 2506 | 2845 | 2327 |
| 3033 | 0.08 | 0.13 | 0.63 | 0.53 | 1 | Micos10 | mitochondrial contact site and cristae orgi | 2841 | 2809 | 2979 | 3044 | 3172 | 2689 | 3043 | 4173 | 2726 | 3380 | 2510 |
| 3399 | -0.08 | 0.09 | -0.81 | 0.42 | 1 | Atxn2l | ataxin 2-like | 2248 | 2380 | 2546 | 3237 | 3260 | 6025 | 3042 | 2631 | 3612 | 3075 | 5333 |
| 3378 | -0.01 | 0.09 | -0.12 | 0.90 | 1 | Ppp4r3b | protein phosphatase 4 regulatory subunit | 3372 | 3377 | 3376 | 3392 | 3386 | 3712 | 3041 | 2829 | 3386 | 3215 | 4076 |
| 2882 | -0.10 | 0.10 | -1.01 | 0.31 | 1 | Mier1 | MEIR1 treanscription regulator | 2594 | 3266 | 2895 | 3379 | 2536 | 2947 | 3038 | 2713 | 2610 | 2733 | 2990 |
| 3186 | 0.13 | 0.11 | 1.22 | 0.22 | 1 | Ist1 | increased sodium tolerance 1 homolog (y | 3217 | 3075 | 3187 | 2952 | 2655 | 3541 | 3036 | 3872 | 2685 | 2992 | 3837 |
| 3234 | 0.03 | 0.09 | 0.36 | 0.72 | 1 | Zdhhc20 | zinc finger, DHHC domain containing 20 | 2823 | 3701 | 2863 | 3692 | 3026 | 3225 | 3035 | 3370 | 3101 | 3433 | 3307 |
| 3030 | -0.07 | 0.13 | -0.56 | 0.57 | 1 | Spg20 | spastic paraplegia 20, spartin (Troyer syn | 3149 | 3571 | 3770 | 2576 | 3346 | 2689 | 3030 | 2048 | 3625 | 2928 | 2592 |
| 2974 | -0.01 | 0.10 | -0.13 | 0.90 | 1 | Ssbp3 | single-stranded DNA binding protein 3 | 2982 | 2911 | 3177 | 3069 | 3292 | 2450 | 3029 | 3579 | 2881 | 3098 | 2251 |
| 2882 | -0.08 | 0.09 | -0.91 | 0.36 | 1 | Grb2 | growth factor receptor bound protein 2 | 3717 | 3111 | 2897 | 2705 | 3072 | 2504 | 3029 | 2586 | 2826 | 2829 | 2428 |
| 2855 | -0.06 | 0.10 | -0.61 | 0.54 | 1 | Foxp4 | forkhead box P4 | 2665 | 2910 | 3276 | 2845 | 3181 | 2415 | 3029 | 2814 | 2964 | 3243 | 2064 |
| 3276 | -0.02 | 0.09 | -0.19 | 0.85 | 1 | Myo10 | myosin X | 2621 | 3293 | 2806 | 4175 | 3482 | 3034 | 3028 | 3924 | 3548 | 3571 | 2557 |
| 3067 | 0.01 | 0.10 | 0.11 | 0.92 | 1 | Rnf6 | ring finger protein (C3H2C3 type) 6 | 2186 | 2174 | 1923 | 2969 | 3242 | 4401 | 3028 | 2790 | 3406 | 2961 | 4657 |
| 3144 | 0.09 | 0.09 | 0.98 | 0.33 | 1 | Hexim1 | hexamethylene bis-acetamide inducible ' | 3209 | 3511 | 3131 | 3343 | 2826 | 2780 | 3027 | 3092 | 3200 | 3674 | 2790 |
| 2980 | 0.13 | 0.11 | 1.20 | 0.23 | 1 | Psmc5 | protease (prosome, macropain) 26S subu | 3062 | 2370 | 3068 | 2648 | 3252 | 2669 | 3027 | 3566 | 3113 | 3187 | 2820 |
| 2553 | 0.06 | 0.11 | 0.50 | 0.62 | 1 | Chrnb2 | cholinergic receptor, nicotinic, beta polyp | 1112 | 1633 | 1475 | 3010 | 3251 | 2362 | 3027 | 3371 | 3316 | 3301 | 2223 |
| 2500 | 0.01 | 0.08 | 0.08 | 0.94 | 1 | Dnajc1 | DnaJ heat shock protein family (Hsp40) n | 1940 | 1750 | 1792 | 2555 | 2405 | 3011 | 3027 | 2492 | 2604 | 2917 | 3012 |
| 2334 | 0.00 | 0.11 | -0.02 | 0.99 | 1 | Tbc1d22b | TBC1 domain family, member 22B | 986 | 934 | 1075 | 2584 | 2902 | 2830 | 3024 | 3020 | 3117 | 2458 | 2747 |
| 2560 | -0.05 | 0.16 | -0.30 | 0.77 | 1 | Erich3 | glutamate rich 3 | 847 | 1672 | 1037 | 3390 | 2756 | 3364 | 3023 | 2801 | 3095 | 2811 | 3362 |
| 2608 | -0.12 | 0.08 | -1.53 | 0.13 | 1 | Fam32a | family with sequence similarity 32, memb | 2782 | 2531 | 2721 | 2549 | 2691 | 2482 | 3022 | 2236 | 2739 | 2738 | 2193 |
| 3464 | 0.16 | 0.11 | 1.44 | 0.15 | 1 | Mapre1 | microtubule-associated protein, RP/EB fa | 4684 | 4199 | 3763 | 3075 | 3108 | 2807 | 3021 | 3984 | 3468 | 2988 | 3003 |
| 2624 | 0.20 | 0.12 | 1.68 | 0.09 | 1 | Ccdc91 | coiled-coil domain containing 91 | 1917 | 1745 | 2096 | 2336 | 3169 | 2237 | 3019 | 3573 | 2999 | 3321 | 2449 |
| 2982 | 0.00 | 0.09 | 0.01 | 0.99 | 1 | Parp1 | poly (ADP-ribose) polymerase family, mei | 3130 | 2535 | 2889 | 2681 | 3256 | 3150 | 3016 | 2587 | 3539 | 2878 | 3139 |
| 3158 | 0.06 | 0.10 | 0.62 | 0.53 | 1 | Map1lc3b | microtubule-associated protein 1 light cha | 3343 | 2725 | 3156 | 3248 | 2959 | 3251 | 3015 | 3573 | 3165 | 3292 | 3015 |
| 3108 | -0.11 | 0.08 | -1.40 | 0.16 | 1 | Actr1a | ARP1 actin-related protein 1A, centractin | 4181 | 3647 | 3901 | 2992 | 3122 | 2528 | 3015 | 2589 | 2795 | 2821 | 2592 |
| 2775 | 0.18 | 0.16 | 1.11 | 0.27 | 1 | Cox6a1 | cytochrome c oxidase subunit 6A1 | 2877 | 2252 | 3185 | 2789 | 2702 | 1866 | 3015 | 4102 | 2569 | 3174 | 1989 |
| 3197 | -0.09 | 0.10 | -0.97 | 0.33 | 1 | Lpp | LIM domain containing preferred transloc | 3714 | 4286 | 3604 | 2918 | 3190 | 3029 | 3014 | 2879 | 3159 | 2878 | 2502 |
| 2895 | -0.03 | 0.11 | -0.30 | 0.77 | 1 | Sec13 | SEC13 homolog, nuclear pore and COPII | 2601 | 1840 | 2327 | 3148 | 3038 | 3407 | 3014 | 2465 | 3029 | 3549 | 3422 |
| 3045 | 0.01 | 0.08 | 0.15 | 0.88 | 1 | Sypl | synaptophysin-like protein | 3500 | 3757 | 3675 | 2790 | 2801 | 2632 | 3012 | 2653 | 2923 | 2991 | 2757 |
| 2615 | 0.00 | 0.08 | -0.06 | 0.95 | 1 | Stau2 | staufen double-stranded RNA binding prc | 2117 | 1805 | 2060 | 2868 | 3213 | 2315 | 3012 | 2603 | 3266 | 3078 | 2422 |
| 2192 | 0.00 | 0.15 | 0.01 | 0.99 | 1 | Ush2a | usherin | 227 | 260 | 416 | 2444 | 2613 | 3479 | 3011 | 2216 | 3109 | 2857 | 3484 |
| 3566 | -0.13 | 0.09 | -1.47 | 0.14 | 1 | Rps16 | ribosomal protein S16 | 6374 | 5865 | 6781 | 2950 | 2627 | 1971 | 3009 | 2407 | 2644 | 2693 | 1900 |
| 2649 | -0.14 | 0.11 | -1.26 | 0.21 | 1 | Kpna1 | karyopherin (importin) alpha 1 | 1854 | 2510 | 1898 | 3103 | 2962 | 2921 | 3009 | 2624 | 2973 | 2565 | 2725 |
| 3282 | 0.14 | 0.11 | 1.26 | 0.21 | 1 | Ginm1 | glycoprotein integral membrane 1 | 2668 | 2655 | 3465 | 3889 | 3532 | 2591 | 3008 | 3817 | 3782 | 3887 | 2807 |
| 3022 | -0.11 | 0.12 | -0.90 | 0.37 | 1 | Wac | WW domain containing adaptor with coile | 2359 | 3335 | 2585 | 3190 | 3130 | 3630 | 3008 | 2609 | 3294 | 2787 | 3317 |
| 2805 | 0.07 | 0.09 | 0.70 | 0.48 | 1 | Zhx1 | zinc fingers and homeoboxes 1 | 2255 | 2245 | 2554 | 3008 | 2902 | 2691 | 3008 | 2781 | 3186 | 3504 | 2725 |
| 2686 | 0.03 | 0.08 | 0.32 | 0.75 | 1 | Trappc9 | trafficking protein particle complex 9 | 2449 | 2360 | 2087 | 3285 | 2904 | 2027 | 3007 | 3441 | 2685 | 3184 | 2122 |
| 2551 | 0.07 | 0.09 | 0.83 | 0.41 | 1 | Anks1 | ankyrin repeat and SAM domain containii | 1702 | 1387 | 1297 | 2690 | 2800 | 3038 | 3007 | 2692 | 2982 | 3422 | 3045 |
| 2447 | -0.08 | 0.15 | -0.54 | 0.59 | 1 | Arhgap24 | Rho GTPase activating protein 24 | 1796 | 1965 | 2082 | 2577 | 2846 | 2332 | 3007 | 1798 | 3033 | 3201 | 2279 |
| 3574 | 0.07 | 0.08 | 0.78 | 0.44 | 1 | Sbno1 | strawberry notch 1 | 2784 | 2641 | 2539 | 3367 | 3373 | 5658 | 3006 | 3313 | 3656 | 3513 | 5465 |
| 2846 | -0.03 | 0.14 | -0.21 | 0.83 | 1 | Gabrb3 | gamma-aminobutyric acid (GABA) A rece | 2249 | 3451 | 2985 | 3117 | 3151 | 2171 | 3005 | 2467 | 3384 | 3108 | 2215 |
| 2828 | -0.03 | 0.08 | -0.44 | 0.66 | 1 | Cbfa2t3 | CBFA2/RUNX1 translocation partner 3 | 3115 | 3087 | 3064 | 2861 | 3043 | 2089 | 3003 | 2855 | 3141 | 3015 | 1831 |
| 2940 | -0.03 | 0.10 | -0.29 | 0.77 | 1 | Irgq | immunity-related GTPase family, Q | 2616 | 2930 | 2593 | 3208 | 3344 | 2654 | 3001 | 2789 | 3467 | 3221 | 2514 |
| 2914 | -0.04 | 0.09 | -0.46 | 0.65 | 1 | Tmem50a | transmembrane protein 50A | 3322 | 2746 | 3359 | 3138 | 3119 | 2232 | 2999 | 2954 | 2968 | 2928 | 2287 |
| 2735 | -0.02 | 0.10 | -0.19 | 0.85 | 1 | Ide | insulin degrading enzyme | 2999 | 1966 | 2335 | 2533 | 3050 | 2855 | 2999 | 2316 | 3160 | 3113 | 2761 |
| 2723 | 0.09 | 0.09 | 1.04 | 0.30 | 1 | Abcb10 | ATP-binding cassette, sub-family B (MDR | 1483 | 1441 | 1825 | 3128 | 3280 | 2793 | 2999 | 3176 | 3422 | 3516 | 2892 |
| 2829 | -0.01 | 0.09 | -0.12 | 0.91 | 1 | Rnf103 | ring finger protein 103 | 2266 | 2509 | 2485 | 3283 | 2900 | 2781 | 2998 | 3213 | 3242 | 2510 | 2937 |
| 2740 | -0.13 | 0.11 | -1.11 | 0.26 | 1 | Tfg | Trk-fused gene | 2363 | 2255 | 2538 | 2552 | 3054 | 3314 | 2995 | 1945 | 3314 | 2593 | 3219 |
| 2719 | 0.12 | 0.07 | 1.79 | 0.07 | 1 | Sfxn1 | sideroflexin 1 | 2228 | 2226 | 1942 | 2885 | 2700 | 2669 | 2992 | 3154 | 3033 | 3297 | 2782 |
| 3082 | -0.09 | 0.10 | -0.95 | 0.34 | 1 | Ilf3 | interleukin enhancer binding factor 3 | 2986 | 3486 | 2936 | 2825 | 3069 | 3747 | 2991 | 2557 | 2716 | 3028 | 3557 |
| 2789 | -0.05 | 0.11 | -0.47 | 0.64 | 1 | Map3k7 | mitogen-activated protein kinase kinase k | 2352 | 2522 | 2815 | 2921 | 2840 | 2896 | 2987 | 2641 | 3199 | 2343 | 3160 |
| 2819 | 0.02 | 0.12 | 0.17 | 0.87 | 1 | Fis1 | fission, mitochondrial 1 | 3087 | 2432 | 3265 | 2779 | 3078 | 2203 | 2986 | 3283 | 2670 | 3022 | 2205 |

| | | | | | | | | | | | | | | | | | | |
|---|---|---|---|---|---|---|---|---|---|---|---|---|---|---|---|---|---|---|
| 2778 | -0.09 | 0.14 | -0.64 | 0.52 | 1 | Tmem123 | transmembrane protein 123 | 5027 | 3296 | 3065 | 2202 | 2625 | 2101 | 2984 | 1926 | 2811 | 2422 | 2104 |
| 2875 | 0.04 | 0.09 | 0.38 | 0.70 | 1 | Tspan31 | tetraspanin 31 | 2758 | 2340 | 2761 | 3232 | 3314 | 2200 | 2983 | 3432 | 3100 | 3267 | 2233 |
| 3562 | -0.02 | 0.09 | -0.20 | 0.84 | 1 | Pbx1 | pre B cell leukemia homeobox 1 | 6208 | 6122 | 5214 | 2797 | 2767 | 2357 | 2982 | 2834 | 2905 | 2563 | 2433 |
| 3240 | 0.08 | 0.08 | 0.99 | 0.32 | 1 | Khdrbs3 | KH domain containing, RNA binding, sigr | 3300 | 3562 | 4120 | 3158 | 3321 | 2551 | 2982 | 3097 | 3446 | 3323 | 2777 |
| 3107 | 0.07 | 0.09 | 0.70 | 0.48 | 1 | Ctcf | CCCTC-binding factor | 3676 | 3283 | 2732 | 2883 | 2928 | 3166 | 2980 | 2848 | 3108 | 3276 | 3297 |
| 2905 | -0.08 | 0.11 | -0.77 | 0.44 | 1 | Irf6 | interferon regulatory factor 6 | 2487 | 3462 | 3182 | 3214 | 2665 | 2860 | 2980 | 3209 | 2392 | 2657 | 2843 |
| 3010 | 0.07 | 0.08 | 0.85 | 0.40 | 1 | Snx17 | sorting nexin 17 | 2952 | 3123 | 3509 | 2801 | 3188 | 2540 | 2978 | 2981 | 3072 | 3131 | 2837 |
| 2683 | 0.01 | 0.10 | 0.12 | 0.90 | 1 | Comt | catechol-O-methyltransferase | 1881 | 1631 | 1984 | 3239 | 3308 | 2410 | 2978 | 2679 | 3408 | 3505 | 2495 |
| 2851 | 0.07 | 0.09 | 0.75 | 0.45 | 1 | Cep104 | centrosomal protein 104 | 2319 | 2154 | 1982 | 3083 | 3162 | 2922 | 2976 | 3574 | 3223 | 3085 | 2876 |
| 3078 | -0.02 | 0.11 | -0.19 | 0.85 | 1 | Maz | MYC-associated zinc finger protein (purin | 3233 | 2714 | 3098 | 3050 | 3058 | 3413 | 2975 | 3469 | 3077 | 2804 | 2964 |
| 2631 | -0.02 | 0.10 | -0.17 | 0.87 | 1 | Ltn1 | listerin E3 ubiquitin protein ligase 1 | 2187 | 2035 | 1914 | 2699 | 2830 | 2969 | 2972 | 2908 | 3046 | 2517 | 2863 |
| 2897 | -0.09 | 0.10 | -0.92 | 0.36 | 1 | Nr2c2 | nuclear receptor subfamily 2, group C, me | 3151 | 3459 | 3225 | 2646 | 2603 | 3145 | 2969 | 2280 | 2820 | 2664 | 2908 |
| 2974 | 0.03 | 0.10 | 0.28 | 0.78 | 1 | Prdx5 | peroxiredoxin 5 | 3834 | 3062 | 3607 | 2977 | 2899 | 2146 | 2968 | 3239 | 2611 | 3145 | 2223 |
| 2160 | -0.07 | 0.10 | -0.70 | 0.48 | 1 | Prkar2b | protein kinase, cAMP dependent regulato | 797 | 859 | 898 | 2703 | 2625 | 2556 | 2968 | 2391 | 2828 | 2567 | 2570 |
| 2909 | 0.09 | 0.11 | 0.84 | 0.40 | 1 | 5031439G07Ril | RIKEN cDNA 5031439G07 gene | 3889 | 3377 | 2616 | 3061 | 2659 | 2044 | 2964 | 3245 | 3047 | 2916 | 2182 |
| 2557 | -0.08 | 0.08 | -1.01 | 0.31 | 1 | Acp2 | acid phosphatase 2, lysosomal | 2650 | 2628 | 2419 | 2646 | 2518 | 2387 | 2964 | 2650 | 2477 | 2476 | 2317 |
| 2859 | 0.07 | 0.10 | 0.71 | 0.48 | 1 | Ppfia1 | protein tyrosine phosphatase, receptor typ | 2378 | 1846 | 2141 | 2618 | 3175 | 3500 | 2961 | 3071 | 3426 | 2799 | 3532 |
| 3262 | 0.00 | 0.11 | -0.01 | 0.99 | 1 | U2af2 | U2 small nuclear ribonucleoprotein auxili | 4095 | 3257 | 4376 | 2460 | 3001 | 3723 | 2960 | 3028 | 2992 | 2682 | 3302 |
| 2814 | 0.07 | 0.12 | 0.58 | 0.56 | 1 | Togaram1 | TOG array regulator of axonemal microtul | 1959 | 2128 | 1768 | 3124 | 2759 | 3395 | 2957 | 2533 | 3143 | 3612 | 3580 |
| 2680 | -0.07 | 0.10 | -0.77 | 0.44 | 1 | Lypla2 | lysophospholipase 2 | 2262 | 2311 | 2733 | 2772 | 3141 | 2477 | 2957 | 2259 | 3176 | 3004 | 2392 |
| 3139 | -0.01 | 0.10 | -0.05 | 0.96 | 1 | Cpeb4 | cytoplasmic polyadenylation element binc | 2098 | 2514 | 2446 | 3252 | 3189 | 4382 | 2956 | 2719 | 3618 | 3210 | 4141 |
| 3204 | -0.19 | 0.14 | -1.32 | 0.19 | 1 | Zfp207 | zinc finger protein 207 | 3154 | 3881 | 2846 | 2711 | 2882 | 4807 | 2952 | 2062 | 2896 | 2243 | 4804 |
| 2429 | -0.03 | 0.11 | -0.28 | 0.78 | 1 | Pde8a | phosphodiesterase 8A | 1814 | 1668 | 1964 | 2399 | 2831 | 2507 | 2948 | 2004 | 2949 | 3331 | 2306 |
| 2416 | -0.06 | 0.12 | -0.52 | 0.60 | 1 | Bcl2l13 | BCL2-like 13 (apoptosis facilitator) | 1697 | 2153 | 1876 | 2781 | 2777 | 2171 | 2948 | 2271 | 2815 | 2701 | 2380 |
| 3028 | -0.11 | 0.09 | -1.14 | 0.25 | 1 | Sp3 | trans-acting transcription factor 3 | 3715 | 3736 | 3494 | 2914 | 2665 | 3055 | 2947 | 2870 | 2630 | 2250 | 3038 |
| 2985 | 0.07 | 0.08 | 0.85 | 0.39 | 1 | Gapdh | glyceraldehyde-3-phosphate dehydroger | 3752 | 3128 | 3533 | 2549 | 2806 | 2654 | 2944 | 2987 | 2796 | 2981 | 2704 |
| 2979 | 0.10 | 0.10 | 0.99 | 0.32 | 1 | Psmb7 | proteasome (prosome, macropain) subun | 3487 | 3328 | 3452 | 2749 | 2907 | 2259 | 2942 | 3466 | 2641 | 3125 | 2407 |
| 2819 | 0.02 | 0.10 | 0.18 | 0.86 | 1 | Yipf3 | Yip1 domain family, member 3 | 2610 | 2761 | 2785 | 2901 | 3069 | 2394 | 2939 | 2699 | 3091 | 3554 | 2199 |
| 2332 | -0.09 | 0.12 | -0.77 | 0.44 | 1 | Ildr1 | immunoglobulin-like domain containing re | 992 | 1430 | 1483 | 3055 | 2994 | 2204 | 2937 | 2618 | 3250 | 2709 | 1983 |
| 3157 | -0.01 | 0.08 | -0.12 | 0.90 | 1 | Gna13 | guanine nucleotide binding protein, alpha | 3226 | 3689 | 2851 | 4052 | 2687 | 2898 | 2936 | 3605 | 2961 | 2876 | 2941 |
| 5410 | -0.09 | 0.10 | -0.85 | 0.40 | 1 | Srsf5 | serine and arginine-rich splicing factor 5 | 3438 | 2559 | 2921 | 2514 | 2784 | 17274 | 2935 | 2037 | 3133 | 2443 | 17470 |
| 2673 | -0.05 | 0.10 | -0.48 | 0.63 | 1 | Faf2 | Fas associated factor family member 2 | 2054 | 2306 | 2017 | 3065 | 2920 | 2773 | 2935 | 2739 | 3345 | 2660 | 2584 |
| 3004 | -0.07 | 0.13 | -0.53 | 0.60 | 1 | Zmiz2 | zinc finger, MIZ-type containing 2 | 3240 | 2743 | 2233 | 3418 | 3509 | 2835 | 2929 | 2975 | 3259 | 3219 | 2680 |
| 2528 | 0.03 | 0.12 | 0.27 | 0.78 | 1 | Grk5 | G protein-coupled receptor kinase 5 | 1652 | 1238 | 1270 | 3306 | 2530 | 2925 | 2929 | 2786 | 3154 | 3147 | 2869 |
| 3296 | -0.16 | 0.11 | -1.47 | 0.14 | 1 | Pcdh1 | protocadherin 1 | 3696 | 4484 | 3944 | 3628 | 3505 | 2707 | 2928 | 2623 | 3076 | 3127 | 2536 |
| 2998 | 0.22 | 0.12 | 1.87 | 0.06 | 1 | Anapc4 | anaphase promoting complex subunit 4 | 2352 | 1994 | 2803 | 2934 | 3006 | 3080 | 2928 | 3831 | 3285 | 3247 | 3514 |
| 2548 | 0.08 | 0.11 | 0.70 | 0.48 | 1 | Gusb | glucuronidase, beta | 3321 | 2107 | 2307 | 2634 | 2590 | 1704 | 2928 | 2860 | 2823 | 3026 | 1726 |
| 3220 | -0.11 | 0.12 | -0.94 | 0.35 | 1 | Mast3 | microtubule associated serine/threonine k | 2875 | 3601 | 3115 | 3407 | 3560 | 3531 | 2927 | 2991 | 3087 | 3547 | 2775 |
| 3127 | -0.12 | 0.10 | -1.21 | 0.23 | 1 | Cs | citrate synthase | 5049 | 4037 | 3978 | 2626 | 3308 | 2235 | 2925 | 2278 | 2737 | 3179 | 2042 |
| 2298 | -0.03 | 0.11 | -0.31 | 0.76 | 1 | Abhd15 | abhydrolase domain containing 15 | 812 | 1127 | 929 | 3338 | 2629 | 2472 | 2924 | 2801 | 2931 | 2933 | 2382 |
| 2979 | -0.09 | 0.09 | -1.05 | 0.29 | 1 | Tnpo2 | transportin 2 (importin 3, karyopherin beta | 2626 | 2641 | 2935 | 2906 | 3241 | 3585 | 2923 | 2431 | 3288 | 2988 | 3203 |
| 2651 | 0.02 | 0.11 | 0.21 | 0.83 | 1 | Atl2 | atlastin GTPase 2 | 1707 | 2268 | 1571 | 2878 | 2760 | 3144 | 2923 | 3001 | 2638 | 3113 | 3157 |
| 2609 | 0.07 | 0.12 | 0.59 | 0.56 | 1 | 1500011B03Ril | RIKEN cDNA 2610524H06 gene | 2112 | 2400 | 2593 | 2510 | 3159 | 1926 | 2923 | 3261 | 2784 | 3162 | 1873 |
| 2511 | -0.12 | 0.16 | -0.73 | 0.47 | 1 | Reep6 | receptor accessory protein 6 | 909 | 1540 | 1344 | 3466 | 3599 | 2390 | 2922 | 2535 | 3616 | 3300 | 2002 |
| 3804 | -0.06 | 0.10 | -0.55 | 0.58 | 1 | Pfn1 | profilin 1 | 7641 | 5573 | 6241 | 3044 | 3123 | 2331 | 2919 | 2910 | 3072 | 2760 | 2230 |
| 2713 | 0.00 | 0.10 | 0.02 | 0.98 | 1 | Tsc1 | TSC complex subunit 1 | 1851 | 2010 | 1909 | 3545 | 2618 | 2969 | 2918 | 2989 | 3142 | 2767 | 3127 |
| 4046 | 0.28 | 0.15 | 1.89 | 0.06 | 1 | Rbm25 | RNA binding motif protein 25 | 3065 | 3401 | 3645 | 3432 | 3275 | 5970 | 2917 | 5984 | 3369 | 3267 | 6180 |
| 3580 | 0.11 | 0.09 | 1.28 | 0.20 | 1 | Capn2 | calpain 2 | 5772 | 4711 | 4997 | 2940 | 2994 | 2638 | 2916 | 3187 | 3391 | 3117 | 2716 |
| 3220 | 0.02 | 0.09 | 0.27 | 0.79 | 1 | Baz2b | bromodomain adjacent to zinc finger dom | 1923 | 2541 | 2027 | 3227 | 2863 | 5459 | 2915 | 3278 | 3169 | 3037 | 4980 |
| 2687 | 0.00 | 0.09 | 0.00 | 1.00 | 1 | Arel1 | apoptosis resistant E3 ubiquitin protein lig | 2182 | 1672 | 1988 | 2847 | 3177 | 2916 | 2914 | 2734 | 3172 | 2857 | 3100 |
| 2783 | -0.07 | 0.13 | -0.54 | 0.59 | 1 | Slc48a1 | solute carrier family 48 (heme transporter | 3246 | 3511 | 3163 | 2869 | 2738 | 2008 | 2908 | 2206 | 2687 | 3502 | 1777 |
| 2406 | -0.04 | 0.10 | -0.39 | 0.69 | 1 | Krtcap2 | keratinocyte associated protein 2 | 1745 | 1900 | 1774 | 2791 | 2917 | 2019 | 2908 | 2557 | 2645 | 3334 | 1875 |
| 2168 | 0.01 | 0.11 | 0.11 | 0.92 | 1 | Dlg3 | discs large MAGUK scaffold protein 3 | 1168 | 896 | 1067 | 2567 | 2345 | 2517 | 2905 | 2759 | 2513 | 2459 | 2651 |
| 3042 | 0.12 | 0.08 | 1.50 | 0.13 | 1 | Calcoco1 | calcium binding and coiled coil domain 1 | 2896 | 3446 | 3126 | 3514 | 2843 | 2243 | 2903 | 3720 | 2959 | 3399 | 2413 |
| 3212 | 0.08 | 0.09 | 0.86 | 0.39 | 1 | P4ha1 | procollagen-proline, 2-oxoglutarate 4-dio | 3640 | 3423 | 3973 | 3082 | 3090 | 2750 | 2899 | 3456 | 2936 | 3120 | 2959 |
| 4215 | 0.01 | 0.13 | 0.08 | 0.94 | 1 | Kmt2a | lysine (K)-specific methyltransferase 2A | 5353 | 5765 | 3973 | 3179 | 2669 | 7183 | 2898 | 4085 | 2925 | 2644 | 5685 |
| 3748 | 0.05 | 0.09 | 0.59 | 0.56 | 1 | Safb | scaffold attachment factor B | 2931 | 3372 | 3356 | 3574 | 3251 | 5892 | 2898 | 4218 | 3200 | 3162 | 5377 |
| 2631 | 0.12 | 0.12 | 0.98 | 0.33 | 1 | Chmp1a | charged multivesicular body protein 1A | 2780 | 2456 | 2999 | 2216 | 2666 | 2164 | 2898 | 3136 | 2704 | 2718 | 2197 |
| 2419 | 0.13 | 0.11 | 1.20 | 0.23 | 1 | Ppp2r3a | protein phosphatase 2, regulatory subuni | 1609 | 2061 | 1554 | 2612 | 2498 | 2196 | 2898 | 3235 | 2867 | 2726 | 2353 |
| 2740 | 0.11 | 0.08 | 1.31 | 0.19 | 1 | Ift81 | intraflagellar transport 81 | 1500 | 1894 | 1464 | 3087 | 2863 | 3308 | 2893 | 3243 | 3023 | 3152 | 3714 |

| | | | | | | | | | | | | | | | | | | |
|---|---|---|---|---|---|---|---|---|---|---|---|---|---|---|---|---|---|---|
| 2604 | 0.11 | 0.08 | 1.40 | 0.16 | 1 | Ncbp1 | nuclear cap binding protein subunit 1 | 1860 | 1704 | 1903 | 2539 | 2963 | 2762 | 2893 | 3008 | 2907 | 3140 | 2966 |
| 2610 | 0.10 | 0.09 | 1.11 | 0.26 | 1 | Ube2e3 | ubiquitin-conjugating enzyme E2E 3 | 2365 | 2751 | 2455 | 2589 | 2624 | 2086 | 2892 | 3127 | 2611 | 2924 | 2288 |
| 2534 | 0.13 | 0.13 | 0.99 | 0.32 | 1 | Laptm4b | lysosomal-associated protein transmemb | 2400 | 2004 | 2619 | 2662 | 2485 | 1898 | 2889 | 3198 | 2388 | 3334 | 1998 |
| 2714 | 0.14 | 0.09 | 1.64 | 0.10 | 1 | Esd | esterase D/formylglutathione hydrolase | 2503 | 2851 | 2687 | 2581 | 2696 | 2202 | 2888 | 3184 | 2867 | 2969 | 2425 |
| 2706 | -0.05 | 0.10 | -0.47 | 0.64 | 1 | Rab8a | RAB8A, member RAS oncogene family | 2416 | 2064 | 2381 | 3019 | 3193 | 2521 | 2884 | 2381 | 3321 | 2960 | 2630 |
| 2599 | 0.07 | 0.10 | 0.78 | 0.44 | 1 | Snx19 | sorting nexin 19 | 2044 | 2199 | 2121 | 2778 | 2829 | 2319 | 2884 | 2743 | 2817 | 3462 | 2396 |
| 2446 | -0.15 | 0.18 | -0.85 | 0.39 | 1 | Ap3m1 | adaptor-related protein complex 3, mu 1 s | 1913 | 2134 | 2030 | 2269 | 2910 | 2774 | 2881 | 1383 | 2814 | 2717 | 3081 |
| 2825 | 0.10 | 0.07 | 1.46 | 0.15 | 1 | Psmd6 | proteasome (prosome, macropain) 26S su | 2447 | 2303 | 2619 | 2618 | 2955 | 2984 | 2880 | 3037 | 3047 | 3028 | 3162 |
| 2777 | 0.13 | 0.13 | 1.05 | 0.29 | 1 | Naa15 | N(alpha)-acetyltransferase 15, NatA auxil | 2300 | 2195 | 2150 | 2663 | 2554 | 3260 | 2877 | 3758 | 2805 | 2409 | 3570 |
| 3726 | 0.14 | 0.14 | 0.96 | 0.34 | 1 | Myl6 | myosin, light polypeptide 6, alkali, smootl | 5781 | 4414 | 5203 | 2674 | 2791 | 3871 | 2876 | 3996 | 2637 | 2892 | 3853 |
| 3206 | 0.13 | 0.13 | 1.04 | 0.30 | 1 | Marf1 | meiosis regulator and mRNA stability 1 | 2892 | 3011 | 2617 | 2991 | 3534 | 3310 | 2874 | 4445 | 3318 | 2991 | 3286 |
| 2973 | 0.16 | 0.11 | 1.45 | 0.15 | 1 | Ddx41 | DEAD (Asp-Glu-Ala-Asp) box polypeptide | 2598 | 2406 | 2899 | 3040 | 3059 | 2719 | 2874 | 3922 | 2979 | 3251 | 2954 |
| 2783 | 0.02 | 0.09 | 0.24 | 0.81 | 1 | Ift57 | intraflagellar transport 57 | 1590 | 1992 | 1887 | 3271 | 3320 | 3018 | 2874 | 2886 | 3502 | 3042 | 3231 |
| 2754 | 0.01 | 0.09 | 0.08 | 0.93 | 1 | Abcf3 | ATP-binding cassette, sub-family F (GCN: | 2044 | 2367 | 2432 | 2498 | 3219 | 3113 | 2873 | 2718 | 3049 | 2786 | 3198 |
| 3716 | -0.02 | 0.08 | -0.21 | 0.83 | 1 | Irf2bp2 | interferon regulatory factor 2 binding prote | 3773 | 3099 | 3408 | 2861 | 3193 | 6610 | 2872 | 2873 | 3061 | 3191 | 5929 |
| 2830 | 0.03 | 0.08 | 0.32 | 0.75 | 1 | Parp6 | poly (ADP-ribose) polymerase family, mer | 2199 | 2484 | 2557 | 2844 | 3016 | 3084 | 2872 | 3326 | 3020 | 2625 | 3101 |
| 2429 | -0.12 | 0.15 | -0.82 | 0.41 | 1 | Tmem184a | transmembrane protein 184a | 1108 | 1771 | 2023 | 2803 | 3091 | 2595 | 2872 | 2677 | 2794 | 2831 | 2151 |
| 2917 | 0.07 | 0.08 | 0.90 | 0.37 | 1 | Tkt | transketolase | 3927 | 3450 | 3312 | 2716 | 2729 | 2065 | 2871 | 2955 | 3043 | 3068 | 1949 |
| 2720 | -0.03 | 0.11 | -0.31 | 0.75 | 1 | Med25 | mediator complex subunit 25 | 1999 | 2282 | 2112 | 2747 | 2813 | 3476 | 2870 | 2376 | 2853 | 3220 | 3167 |
| 2538 | -0.05 | 0.08 | -0.56 | 0.57 | 1 | Xpo6 | exportin 6 | 2490 | 2214 | 2007 | 2505 | 2470 | 2940 | 2870 | 2363 | 2596 | 2861 | 2603 |
| 2786 | -0.07 | 0.11 | -0.64 | 0.52 | 1 | Syncrip | synaptotagmin binding, cytoplasmic RNA | 3171 | 3277 | 2820 | 2570 | 2571 | 2919 | 2868 | 2413 | 2714 | 2160 | 3163 |
| 2512 | -0.14 | 0.10 | -1.40 | 0.16 | 1 | Pomt2 | protein-O-mannosyltransferase 2 | 1966 | 2088 | 2272 | 2542 | 3007 | 2734 | 2867 | 2088 | 2913 | 2782 | 2378 |
| 3008 | -0.19 | 0.10 | -1.85 | 0.06 | 1 | Hook3 | hook microtubule tethering protein 3 | 2712 | 2759 | 2467 | 2873 | 2551 | 5082 | 2865 | 2080 | 2647 | 2437 | 4620 |
| 2354 | -0.20 | 0.11 | -1.82 | 0.07 | 1 | Slc23a2 | solute carrier family 23 (nucleobase trans | 2197 | 1884 | 2065 | 2626 | 2457 | 2598 | 2862 | 2052 | 2395 | 2083 | 2679 |
| 2778 | 0.01 | 0.08 | 0.16 | 0.87 | 1 | Fyttd1 | forty-two-three domain containing 1 | 2219 | 2578 | 2385 | 3109 | 2627 | 3052 | 2858 | 3096 | 2910 | 2781 | 2948 |
| 2858 | 0.14 | 0.08 | 1.68 | 0.09 | 1 | Armcx4 | armadillo repeat containing, X-linked 4 | 2093 | 2442 | 2287 | 2996 | 2647 | 3184 | 2856 | 3545 | 3081 | 2713 | 3596 |
| 2705 | 0.12 | 0.08 | 1.46 | 0.14 | 1 | Drg1 | developmentally regulated GTP binding p | 2386 | 2199 | 2678 | 2507 | 3013 | 2429 | 2856 | 2717 | 2976 | 3229 | 2769 |
| 2359 | -0.04 | 0.14 | -0.32 | 0.75 | 1 | Dnajc6 | DnaJ heat shock protein family (Hsp40) m | 1688 | 2243 | 1999 | 2584 | 2705 | 2037 | 2855 | 2048 | 2807 | 2795 | 2190 |
| 3152 | 0.09 | 0.08 | 1.17 | 0.24 | 1 | Hnrnpr | heterogeneous nuclear ribonucleoprotein | 3281 | 3393 | 3654 | 2918 | 2768 | 3239 | 2850 | 2951 | 2880 | 3076 | 3660 |
| 2331 | 0.05 | 0.10 | 0.55 | 0.59 | 1 | Srxn1 | sulfiredoxin 1 homolog (S. cerevisiae) | 1795 | 1708 | 1752 | 2208 | 3062 | 1930 | 2848 | 2458 | 3007 | 2733 | 2145 |
| 3186 | 0.01 | 0.08 | 0.09 | 0.93 | 1 | Fkbp5 | FK506 binding protein 5 | 4326 | 4281 | 4653 | 2495 | 2626 | 2897 | 2847 | 2537 | 2620 | 3010 | 2755 |
| 2729 | 0.01 | 0.08 | 0.14 | 0.89 | 1 | Gigyf2 | GRB10 interacting GYF protein 2 | 1802 | 1770 | 1962 | 2974 | 2859 | 3523 | 2847 | 2989 | 2839 | 2952 | 3505 |
| 2577 | -0.07 | 0.10 | -0.70 | 0.49 | 1 | Kifbp | kinesin family binding protein | 1996 | 2455 | 1999 | 2781 | 2900 | 2675 | 2847 | 2716 | 2989 | 2485 | 2505 |
| 2711 | 0.07 | 0.09 | 0.81 | 0.42 | 1 | Snap47 | synaptosomal-associated protein, 47 | 2513 | 2400 | 2787 | 2583 | 2955 | 2419 | 2846 | 3055 | 2812 | 2803 | 2653 |
| 3103 | -0.02 | 0.08 | -0.19 | 0.85 | 1 | Rbbp6 | retinoblastoma binding protein 6, ubiquitin | 2772 | 3150 | 2788 | 3152 | 2563 | 4197 | 2844 | 2790 | 2863 | 2628 | 4388 |
| 2860 | -0.05 | 0.12 | -0.39 | 0.69 | 1 | Abi1 | abl-interactor 1 | 3410 | 2796 | 2815 | 2785 | 2793 | 2922 | 2844 | 1983 | 3057 | 3015 | 3043 |
| 2535 | -0.21 | 0.13 | -1.52 | 0.13 | 1 | Tox3 | TOX high mobility group box family memb | 2037 | 3067 | 2391 | 2771 | 2853 | 2450 | 2844 | 2168 | 2732 | 2446 | 2123 |
| 2358 | -0.01 | 0.11 | -0.08 | 0.94 | 1 | Rab28 | RAB28, member RAS oncogene family | 1983 | 2304 | 2157 | 2395 | 2612 | 1964 | 2840 | 2397 | 2568 | 2429 | 2291 |
| 2645 | 0.00 | 0.09 | 0.05 | 0.96 | 1 | Lrrc8d | leucine rich repeat containing 8D | 1888 | 2250 | 2089 | 2956 | 2991 | 2623 | 2839 | 2820 | 3165 | 3012 | 2468 |
| 3346 | 0.01 | 0.09 | 0.10 | 0.92 | 1 | Ankrd11 | ankyrin repeat domain 11 | 3817 | 4117 | 3745 | 2859 | 2592 | 4298 | 2838 | 3359 | 2821 | 2593 | 3770 |
| 3026 | 0.00 | 0.09 | -0.06 | 0.96 | 1 | Pfkl | phosphofructokinase, liver, B-type | 3371 | 3101 | 3372 | 3230 | 3086 | 2571 | 2837 | 3315 | 2884 | 3104 | 2417 |
| 2713 | -0.06 | 0.15 | -0.36 | 0.72 | 1 | Ric8b | RIC8 guanine nucleotide exchange factor | 1778 | 2770 | 2164 | 3119 | 2513 | 3305 | 2834 | 2187 | 2933 | 2677 | 3563 |
| 3370 | -0.02 | 0.07 | -0.23 | 0.82 | 1 | Ddx39b | DEAD (Asp-Glu-Ala-Asp) box polypeptide | 3658 | 2783 | 3512 | 2427 | 2621 | 5842 | 2832 | 2362 | 2686 | 2941 | 5403 |
| 3316 | 0.17 | 0.10 | 1.69 | 0.09 | 1 | Dnajc7 | DnaJ heat shock protein family (Hsp40) m | 3677 | 3372 | 3518 | 2780 | 2828 | 3764 | 2831 | 3757 | 2869 | 3017 | 4066 |
| 3005 | -0.02 | 0.13 | -0.16 | 0.87 | 1 | Anp32b | acidic (leucine-rich) nuclear phosphoprot | 4412 | 3518 | 3818 | 2814 | 2822 | 2259 | 2831 | 3230 | 2816 | 2281 | 2254 |
| 2970 | 0.09 | 0.11 | 0.79 | 0.43 | 1 | Arfgap1 | ADP-ribosylation factor GTPase activating | 2346 | 1757 | 2281 | 2404 | 2884 | 4788 | 2831 | 3104 | 2874 | 3106 | 4295 |
| 2746 | 0.14 | 0.11 | 1.26 | 0.21 | 1 | Gnl1 | guanine nucleotide binding protein-like 1 | 2645 | 2196 | 2753 | 2613 | 3040 | 2219 | 2831 | 3569 | 3030 | 3138 | 2167 |
| 2539 | 0.07 | 0.09 | 0.80 | 0.43 | 1 | Scamp2 | secretory carrier membrane protein 2 | 2258 | 2044 | 2086 | 2371 | 2773 | 2494 | 2831 | 2654 | 2987 | 3188 | 2243 |
| 2396 | 0.06 | 0.10 | 0.60 | 0.55 | 1 | Strbp | spermatid perinuclear RNA binding prote | 1510 | 1547 | 1276 | 2700 | 2648 | 2603 | 2827 | 2863 | 3032 | 2562 | 2783 |
| 2192 | -0.14 | 0.08 | -1.66 | 0.10 | 1 | Ndufa1 | NADH:ubiquinone oxidoreductase subun | 1854 | 1677 | 1881 | 2563 | 2710 | 1740 | 2826 | 2184 | 2379 | 2516 | 1780 |
| 2725 | 0.02 | 0.16 | 0.10 | 0.92 | 1 | Atp5j2 | ATP synthase, H+ transporting, mitochond | 2784 | 2518 | 3777 | 2779 | 2755 | 2037 | 2825 | 3418 | 2574 | 2210 | 2302 |
| 3908 | -0.10 | 0.10 | -1.05 | 0.29 | 1 | Pitpnm2 | phosphatidylinositol transfer protein, mem | 6284 | 7526 | 5323 | 3324 | 2860 | 3361 | 2824 | 2973 | 2966 | 2579 | 2972 |
| 3191 | -0.16 | 0.14 | -1.10 | 0.27 | 1 | Atp8b2 | ATPase, class I, type 8B, member 2 | 4725 | 3885 | 3934 | 2050 | 3311 | 3663 | 2824 | 1566 | 3163 | 2820 | 3162 |
| 2488 | -0.01 | 0.09 | -0.10 | 0.92 | 1 | Pard3 | par-3 family cell polarity regulator | 1454 | 1313 | 1431 | 3239 | 2971 | 2576 | 2824 | 3335 | 2927 | 2845 | 2452 |
| 2812 | 0.04 | 0.13 | 0.32 | 0.75 | 1 | Vars | valyl-tRNA synthetase | 3273 | 2802 | 3043 | 2604 | 2908 | 2359 | 2823 | 3576 | 2539 | 2897 | 2113 |
| 2683 | 0.02 | 0.10 | 0.22 | 0.83 | 1 | Psen1 | presenilin 1 | 2502 | 2758 | 2871 | 2758 | 2863 | 2162 | 2823 | 2451 | 2878 | 3194 | 2251 |
| 2420 | 0.16 | 0.08 | 1.91 | 0.06 | 1 | Rhoq | ras homolog family member Q | 2393 | 2042 | 1822 | 2598 | 2341 | 1899 | 2823 | 2801 | 2645 | 2947 | 2308 |
| 2946 | 0.17 | 0.10 | 1.76 | 0.08 | 1 | U2surp | U2 snRNP-associated SURP domain con | 2636 | 2311 | 2527 | 2622 | 2744 | 3549 | 2822 | 3520 | 2983 | 2923 | 3764 |
| 2891 | 0.07 | 0.09 | 0.77 | 0.44 | 1 | Arhgap1 | Rho GTPase activating protein 1 | 3100 | 3172 | 3134 | 2657 | 2725 | 2696 | 2821 | 2492 | 3235 | 3075 | 2690 |
| 2620 | -0.06 | 0.12 | -0.49 | 0.63 | 1 | Xpo5 | exportin 5 | 2204 | 2295 | 2146 | 2786 | 3042 | 2626 | 2820 | 2199 | 2889 | 3314 | 2495 |

| | | | | | | | | | | | | | | | | | |
|---|---|---|---|---|---|---|---|---|---|---|---|---|---|---|---|---|---|
| 2436 | -0.04 | 0.10 | -0.45 | 0.66 | 1 | Atp13a1 | ATPase type 13A1 | 1611 | 1277 | 1674 | 2413 | 2603 | 3473 | 2820 | 2558 | 2643 | 2509 | 3214 |
| 1868 | -0.13 | 0.16 | -0.80 | 0.42 | 1 | Pxylp1 | 2-phosphoxylose phosphatase 1 | 983 | 1488 | 857 | 2637 | 2171 | 1368 | 2820 | 2178 | 2361 | 2505 | 1184 |
| 3395 | 0.05 | 0.10 | 0.50 | 0.62 | 1 | Tagln2 | transgelin 2 | 5582 | 4372 | 4500 | 2859 | 3025 | 2554 | 2818 | 2838 | 3183 | 2937 | 2679 |
| 2462 | -0.03 | 0.09 | -0.30 | 0.77 | 1 | Pias2 | protein inhibitor of activated STAT 2 | 1816 | 1996 | 2013 | 2752 | 2768 | 2373 | 2818 | 2655 | 3066 | 2473 | 2352 |
| 2828 | 0.10 | 0.10 | 1.01 | 0.31 | 1 | Sec11a | SEC11 homolog A, signal peptidase com | 2723 | 2085 | 2972 | 2451 | 3233 | 2784 | 2817 | 3040 | 3124 | 2883 | 2990 |
| 3213 | -0.01 | 0.09 | -0.12 | 0.91 | 1 | Pcmtd2 | protein-L-isoaspartate (D-aspartate) O-me | 3018 | 3720 | 3513 | 3264 | 3066 | 3459 | 2813 | 2837 | 2965 | 3161 | 3527 |
| 3133 | 0.02 | 0.11 | 0.22 | 0.82 | 1 | Prdm2 | PR domain containing 2, with ZNF domai | 2551 | 3150 | 2635 | 3346 | 3200 | 3599 | 2813 | 3450 | 3667 | 3139 | 2906 |
| 2760 | 0.14 | 0.13 | 1.08 | 0.28 | 1 | Ddrgk1 | DDRGK domain containing 1 | 1929 | 1823 | 2057 | 2877 | 2807 | 3152 | 2813 | 4104 | 2748 | 2995 | 3054 |
| 3209 | 0.18 | 0.12 | 1.55 | 0.12 | 1 | Eef1d | eukaryotic translation elongation factor 1 | 4409 | 3839 | 4599 | 2674 | 2662 | 2312 | 2812 | 3724 | 2747 | 3201 | 2318 |
| 2553 | 0.07 | 0.09 | 0.73 | 0.47 | 1 | Gne | glucosamine (UDP-N-acetyl)-2-epimeras | 1863 | 1826 | 1719 | 2505 | 2931 | 2837 | 2810 | 2917 | 3026 | 2982 | 2671 |
| 2832 | 0.13 | 0.11 | 1.17 | 0.24 | 1 | Spty2d1 | SPT2 chromatin protein domain containin | 2410 | 2406 | 2354 | 2882 | 2863 | 2875 | 2809 | 2542 | 3342 | 3628 | 3044 |
| 2714 | -0.10 | 0.09 | -1.17 | 0.24 | 1 | Mast4 | microtubule associated serine/threonine k | 2275 | 2281 | 2500 | 2837 | 3258 | 2876 | 2806 | 2411 | 3372 | 2745 | 2491 |
| 3129 | 0.02 | 0.10 | 0.23 | 0.82 | 1 | Rnf4 | ring finger protein 4 | 3298 | 3368 | 3175 | 2791 | 3390 | 3203 | 2805 | 2661 | 3517 | 2987 | 3225 |
| 2891 | 0.00 | 0.11 | 0.01 | 0.99 | 1 | Itch | itchy, E3 ubiquitin protein ligase | 2680 | 2779 | 2452 | 2668 | 3154 | 3293 | 2804 | 2562 | 3451 | 2586 | 3374 |
| 2829 | 0.03 | 0.12 | 0.27 | 0.79 | 1 | Gmppa | GDP-mannose pyrophosphorylase A | 1910 | 1998 | 2113 | 2290 | 3310 | 4069 | 2804 | 2544 | 2885 | 3532 | 3659 |
| 2787 | 0.06 | 0.09 | 0.64 | 0.52 | 1 | Adk | adenosine kinase | 2744 | 3079 | 3426 | 2932 | 2687 | 2062 | 2803 | 3019 | 2884 | 2894 | 2122 |
| 2706 | 0.04 | 0.11 | 0.33 | 0.74 | 1 | Unc45a | unc-45 myosin chaperone A | 2310 | 2050 | 2138 | 2837 | 3060 | 2766 | 2803 | 3564 | 2867 | 2778 | 2596 |
| 2703 | -0.16 | 0.09 | -1.82 | 0.07 | 1 | Apbb1 | amyloid beta (A4) precursor protein-bindi | 2495 | 2652 | 3015 | 3145 | 2934 | 2498 | 2803 | 2816 | 2554 | 2624 | 2201 |
| 3061 | 0.06 | 0.10 | 0.60 | 0.55 | 1 | Poldip3 | polymerase (DNA-directed), delta interact | 2625 | 3095 | 2789 | 3164 | 3136 | 3166 | 2799 | 3865 | 3013 | 3312 | 2709 |
| 2898 | 0.00 | 0.09 | 0.03 | 0.98 | 1 | Cul4b | cullin 4B | 2832 | 2668 | 2634 | 3307 | 2978 | 2782 | 2799 | 2838 | 3124 | 3046 | 2868 |
| 2688 | 0.00 | 0.11 | -0.03 | 0.98 | 1 | Cluh | clustered mitochondria (cluA/CLU1) homo | 2137 | 1600 | 1869 | 3129 | 3290 | 2734 | 2799 | 2884 | 3125 | 3467 | 2536 |
| 2516 | 0.16 | 0.10 | 1.71 | 0.09 | 1 | Faf1 | Fas-associated factor 1 | 2374 | 1752 | 2425 | 2216 | 2866 | 2112 | 2798 | 2801 | 2830 | 3140 | 2364 |
| 2635 | -0.05 | 0.09 | -0.51 | 0.61 | 1 | Egln1 | egl-9 family hypoxia-inducible factor 1 | 2865 | 2826 | 2562 | 2802 | 3070 | 1895 | 2796 | 2562 | 3002 | 2646 | 1959 |
| 2960 | 0.05 | 0.10 | 0.47 | 0.64 | 1 | Hdac11 | histone deacetylase 11 | 3082 | 3324 | 3668 | 3178 | 2675 | 2421 | 2795 | 3188 | 2626 | 3004 | 2596 |
| 2753 | -0.02 | 0.07 | -0.22 | 0.82 | 1 | Paip1 | polyadenylate binding protein-interacting | 2801 | 2971 | 2860 | 2926 | 2811 | 2347 | 2795 | 2756 | 2751 | 2991 | 2272 |
| 2589 | -0.09 | 0.13 | -0.70 | 0.48 | 1 | Rwdd4a | RWD domain containing 4A | 1337 | 1629 | 1745 | 2940 | 3482 | 3029 | 2794 | 2136 | 3018 | 3026 | 3339 |
| 2903 | -0.01 | 0.11 | -0.11 | 0.91 | 1 | Helz | helicase with zinc finger domain | 2432 | 2384 | 2517 | 2885 | 3464 | 3189 | 2793 | 3503 | 3421 | 2438 | 2909 |
| 2568 | 0.03 | 0.11 | 0.26 | 0.79 | 1 | Vps35l | VPS35 endosomal protein sorting factor li | 2379 | 2486 | 2113 | 2701 | 2921 | 2129 | 2793 | 2375 | 2975 | 3012 | 2370 |
| 2336 | -0.09 | 0.13 | -0.71 | 0.48 | 1 | Fech | ferrochelatase | 1990 | 2141 | 2198 | 2596 | 2425 | 2137 | 2792 | 1917 | 2666 | 2871 | 1960 |
| 2319 | 0.09 | 0.07 | 1.26 | 0.21 | 1 | Gmppb | GDP-mannose pyrophosphorylase B | 1054 | 908 | 1201 | 2170 | 2972 | 2887 | 2792 | 2376 | 3068 | 2992 | 3087 |
| 3044 | -0.03 | 0.15 | -0.17 | 0.86 | 1 | Hmgcs1 | 3-hydroxy-3-methylglutaryl-Coenzyme A s | 1860 | 2144 | 1846 | 4057 | 3147 | 3727 | 2791 | 4583 | 3100 | 1939 | 4291 |
| 2891 | 0.14 | 0.09 | 1.55 | 0.12 | 1 | Ank3 | ankyrin 3, epithelial | 2511 | 3396 | 2625 | 3104 | 2467 | 2696 | 2788 | 3695 | 2860 | 3030 | 2628 |
| 2884 | 0.02 | 0.08 | 0.21 | 0.83 | 1 | Zfp91 | zinc finger protein 91 | 2478 | 2767 | 2445 | 2954 | 3163 | 3041 | 2788 | 2966 | 2978 | 2921 | 3223 |
| 3023 | 0.19 | 0.11 | 1.66 | 0.10 | 1 | Arf5 | ADP-ribosylation factor 5 | 3346 | 3079 | 3487 | 2825 | 3056 | 2228 | 2785 | 3865 | 2891 | 3301 | 2393 |
| 2944 | 0.04 | 0.09 | 0.46 | 0.64 | 1 | Prr12 | proline rich 12 | 3050 | 3607 | 3078 | 3318 | 2633 | 2374 | 2785 | 3550 | 3023 | 2884 | 2081 |
| 2557 | -0.16 | 0.10 | -1.53 | 0.13 | 1 | Ppp2r5b | protein phosphatase 2, regulatory subuni | 2274 | 2145 | 2571 | 2741 | 3037 | 2536 | 2784 | 2104 | 2790 | 2988 | 2156 |
| 3103 | 0.06 | 0.09 | 0.65 | 0.52 | 1 | Stx5a | syntaxin 5A | 1970 | 1778 | 2313 | 2086 | 3348 | 5621 | 2783 | 2190 | 3538 | 2999 | 5503 |
| 3036 | -0.02 | 0.11 | -0.21 | 0.83 | 1 | Gatad2b | GATA zinc finger domain containing 2B | 3239 | 4039 | 3565 | 2711 | 2793 | 3063 | 2783 | 2764 | 3243 | 2589 | 2609 |
| 2344 | -0.04 | 0.09 | -0.38 | 0.70 | 1 | Smdt1 | single-pass membrane protein with aspar | 2123 | 2110 | 2544 | 2430 | 2566 | 1840 | 2783 | 2468 | 2511 | 2653 | 1760 |
| 3315 | -0.01 | 0.12 | -0.12 | 0.90 | 1 | Taok1 | TAO kinase 1 | 3339 | 3572 | 3116 | 2892 | 2887 | 4668 | 2782 | 2550 | 3536 | 2371 | 4752 |
| 2515 | 0.19 | 0.10 | 1.88 | 0.06 | 1 | Abcb7 | ATP-binding cassette, sub-family B (MDR | 2049 | 1791 | 2181 | 2796 | 2549 | 1978 | 2781 | 2774 | 3051 | 3491 | 2229 |
| 2513 | -0.15 | 0.15 | -1.03 | 0.30 | 1 | Ipo8 | importin 8 | 2609 | 2548 | 2402 | 2451 | 2682 | 2592 | 2780 | 1590 | 2899 | 2553 | 2542 |
| 2438 | 0.03 | 0.09 | 0.37 | 0.71 | 1 | Dgcr2 | DiGeorge syndrome critical region gene 2 | 2323 | 2267 | 1963 | 2479 | 2449 | 2267 | 2780 | 2974 | 2498 | 2790 | 2027 |
| 3049 | 0.12 | 0.10 | 1.25 | 0.21 | 1 | Gpbp1 | GC-rich promoter binding protein 1 | 2550 | 3515 | 3089 | 2887 | 2865 | 3159 | 2779 | 3130 | 3015 | 3143 | 3409 |
| 2749 | -0.05 | 0.12 | -0.38 | 0.70 | 1 | Cdip1 | cell death inducing Trp53 target 1 | 2491 | 3385 | 2631 | 2481 | 2815 | 2962 | 2779 | 2548 | 2830 | 2586 | 2732 |
| 2254 | -0.14 | 0.17 | -0.85 | 0.39 | 1 | Ndfip2 | Nedd4 family interacting protein 2 | 1406 | 1708 | 1477 | 2694 | 2341 | 2745 | 2779 | 1624 | 2316 | 2990 | 2720 |
| 2731 | 0.17 | 0.11 | 1.56 | 0.12 | 1 | Vti1b | vesicle transport through interaction with t | 2265 | 1944 | 2401 | 2680 | 3092 | 2489 | 2778 | 3597 | 3065 | 3048 | 2687 |
| 2592 | -0.13 | 0.11 | -1.22 | 0.22 | 1 | Dhx36 | DEAH (Asp-Glu-Ala-His) box polypeptide | 2117 | 2423 | 2753 | 2787 | 2670 | 2838 | 2778 | 2285 | 2408 | 2532 | 2922 |
| 3545 | -0.03 | 0.10 | -0.35 | 0.73 | 1 | Prpf4b | pre-mRNA processing factor 4B | 1822 | 2208 | 1634 | 3186 | 3025 | 7917 | 2777 | 2869 | 2927 | 3014 | 7617 |
| 2578 | -0.04 | 0.08 | -0.46 | 0.64 | 1 | Denn2b | DENN domain containing 2B | 2311 | 2070 | 2355 | 2758 | 2743 | 2671 | 2776 | 2640 | 2950 | 2489 | 2591 |
| 2449 | 0.18 | 0.11 | 1.62 | 0.11 | 1 | Aktip | thymoma viral proto-oncogene 1 interacti | 1759 | 1887 | 1986 | 2359 | 2696 | 2161 | 2771 | 2975 | 2647 | 3382 | 2316 |
| 2470 | -0.03 | 0.11 | -0.24 | 0.81 | 1 | Tcf20 | transcription factor 20 | 2203 | 1949 | 2077 | 2872 | 2508 | 2411 | 2770 | 2265 | 3012 | 2575 | 2534 |
| 2284 | 0.02 | 0.10 | 0.22 | 0.83 | 1 | Vkorc1l1 | vitamin K epoxide reductase complex, sul | 1548 | 1560 | 1641 | 2348 | 2838 | 2146 | 2770 | 2200 | 3109 | 2744 | 2223 |
| 3197 | -0.08 | 0.15 | -0.56 | 0.57 | 1 | Rfx6 | regulatory factor X, 6 | 1385 | 2208 | 1994 | 3389 | 2791 | 6371 | 2769 | 2397 | 3476 | 2764 | 5625 |
| 2532 | -0.05 | 0.09 | -0.52 | 0.61 | 1 | C77080 | expressed sequence C77080 | 1396 | 1749 | 1840 | 3350 | 2991 | 2447 | 2769 | 3554 | 3074 | 2549 | 2129 |
| 2867 | -0.06 | 0.09 | -0.69 | 0.49 | 1 | Hectd4 | HECT domain E3 ubiquitin protein ligase | 2622 | 2251 | 2222 | 2717 | 2989 | 4034 | 2766 | 2708 | 2804 | 2841 | 3578 |
| 2504 | 0.05 | 0.10 | 0.45 | 0.65 | 1 | Bckdk | branched chain ketoacid dehydrogenase | 1863 | 2044 | 2086 | 2852 | 2839 | 2106 | 2764 | 3170 | 2556 | 3250 | 2019 |
| 2793 | -0.01 | 0.10 | -0.13 | 0.89 | 1 | Reep3 | receptor accessory protein 3 | 3593 | 3221 | 3399 | 2659 | 2722 | 2159 | 2761 | 2786 | 2994 | 2277 | 2157 |
| 2658 | -0.12 | 0.09 | -1.42 | 0.16 | 1 | Hibadh | 3-hydroxyisobutyrate dehydrogenase | 2587 | 2840 | 2660 | 3089 | 2963 | 2168 | 2761 | 2688 | 2779 | 2890 | 1807 |
| 2818 | 0.04 | 0.09 | 0.47 | 0.64 | 1 | Atraid | all-trans retinoic acid induced differentiati | 2293 | 2593 | 2520 | 2973 | 3330 | 2576 | 2760 | 2987 | 3001 | 3251 | 2718 |

| | | | | | | | | | | | | | | | | | | |
|---|---|---|---|---|---|---|---|---|---|---|---|---|---|---|---|---|---|---|
| 2599 | 0.01 | 0.08 | 0.10 | 0.92 | 1 | Trim41 | tripartite motif-containing 41 | 1854 | 1925 | 1938 | 2510 | 2712 | 3438 | 2760 | 2500 | 2876 | 2818 | 3261 |
| 3126 | -0.03 | 0.11 | -0.26 | 0.79 | 1 | Phldb2 | pleckstrin homology like domain, family B | 2551 | 2546 | 2631 | 4071 | 2713 | 3950 | 2759 | 3561 | 2902 | 2736 | 3969 |
| 2704 | -0.07 | 0.09 | -0.78 | 0.43 | 1 | Gosr1 | golgi SNAP receptor complex member 1 | 2461 | 2729 | 2420 | 2903 | 2902 | 2768 | 2758 | 2558 | 3020 | 2621 | 2604 |
| 2641 | 0.03 | 0.14 | 0.19 | 0.85 | 1 | Fuca1 | fucosidase, alpha-L- 1, tissue | 2665 | 1987 | 2423 | 2905 | 3111 | 2083 | 2758 | 3707 | 2722 | 2673 | 2021 |
| 3221 | 0.08 | 0.15 | 0.55 | 0.58 | 1 | Cflar | CASP8 and FADD-like apoptosis regulato | 3342 | 3187 | 3197 | 1981 | 3960 | 3866 | 2757 | 2219 | 3731 | 3233 | 3965 |
| 2680 | -0.08 | 0.11 | -0.76 | 0.45 | 1 | Tmc4 | transmembrane channel-like gene family | 1656 | 2131 | 2440 | 2655 | 2855 | 3696 | 2756 | 2805 | 2730 | 2452 | 3308 |
| 2264 | 0.16 | 0.15 | 1.08 | 0.28 | 1 | Mns1 | meiosis-specific nuclear structural protein | 552 | 672 | 903 | 2528 | 2429 | 3035 | 2756 | 2870 | 2655 | 2898 | 3610 |
| 2612 | -0.11 | 0.07 | -1.47 | 0.14 | 1 | Mink1 | misshapen-like kinase 1 (zebrafish) | 2471 | 2375 | 2439 | 2875 | 2880 | 2605 | 2755 | 2506 | 2765 | 2732 | 2326 |
| 2568 | 0.06 | 0.10 | 0.63 | 0.53 | 1 | Copg2 | coatomer protein complex, subunit gamm | 2280 | 2066 | 2128 | 2990 | 2597 | 2277 | 2755 | 3474 | 2517 | 2826 | 2337 |
| 2329 | -0.02 | 0.18 | -0.12 | 0.90 | 1 | Bhlha15 | basic helix-loop-helix family, member a15 | 511 | 894 | 793 | 2227 | 3593 | 3177 | 2755 | 2263 | 3910 | 2514 | 2982 |
| 2325 | 0.04 | 0.14 | 0.27 | 0.79 | 1 | Tmem164 | transmembrane protein 164 | 1283 | 807 | 1134 | 2949 | 3028 | 2303 | 2753 | 3264 | 2846 | 2803 | 2402 |
| 2900 | -0.10 | 0.10 | -1.09 | 0.28 | 1 | Ubfd1 | ubiquitin family domain containing 1 | 2373 | 2683 | 3034 | 2626 | 3258 | 3680 | 2752 | 2172 | 3227 | 2680 | 3415 |
| 2369 | 0.03 | 0.11 | 0.25 | 0.80 | 1 | Cisd2 | CDGSH iron sulfur domain 2 | 1772 | 1620 | 1893 | 2691 | 2403 | 2441 | 2751 | 2940 | 2674 | 2388 | 2484 |
| 2906 | -0.03 | 0.08 | -0.38 | 0.70 | 1 | Map2k1 | mitogen-activated protein kinase kinase 1 | 4054 | 3769 | 3815 | 2465 | 2715 | 2316 | 2750 | 2301 | 2630 | 3004 | 2145 |
| 2705 | 0.13 | 0.15 | 0.85 | 0.39 | 1 | Cdc42bpg | CDC42 binding protein kinase gamma (D | 1719 | 1749 | 1951 | 2843 | 3069 | 2834 | 2750 | 4561 | 3139 | 2588 | 2550 |
| 2588 | -0.17 | 0.10 | -1.69 | 0.09 | 1 | Ptpn1 | protein tyrosine phosphatase, non-recept | 2678 | 2250 | 2298 | 2975 | 2846 | 2691 | 2750 | 2582 | 2658 | 2250 | 2489 |
| 2343 | 0.01 | 0.14 | 0.06 | 0.95 | 1 | Cacna1c | calcium channel, voltage-dependent, L ty | 1244 | 1574 | 1173 | 2535 | 1991 | 3674 | 2744 | 2208 | 2600 | 2660 | 3369 |
| 2746 | 0.00 | 0.11 | 0.04 | 0.97 | 1 | Eif2ak3 | eukaryotic translation initiation factor 2 alp | 2020 | 1804 | 2090 | 2332 | 3344 | 3658 | 2743 | 1996 | 3667 | 3000 | 3550 |
| 2930 | 0.05 | 0.08 | 0.62 | 0.54 | 1 | Thoc2 | THO complex 2 | 2702 | 2852 | 2707 | 2777 | 2504 | 3751 | 2742 | 2786 | 2886 | 2561 | 3962 |
| 2918 | -0.05 | 0.12 | -0.46 | 0.64 | 1 | Ap2m1 | adaptor-related protein complex 2, mu 1 s | 3346 | 3052 | 3134 | 2789 | 3120 | 2798 | 2741 | 2068 | 3044 | 3145 | 2864 |
| 2843 | -0.09 | 0.10 | -0.87 | 0.38 | 1 | Ncam1 | neural cell adhesion molecule 1 | 2592 | 3292 | 3487 | 2808 | 2979 | 2763 | 2740 | 2896 | 2613 | 2768 | 2337 |
| 2634 | 0.05 | 0.11 | 0.44 | 0.66 | 1 | Egln2 | egl-9 family hypoxia-inducible factor 2 | 2286 | 2343 | 2588 | 2875 | 2699 | 2325 | 2740 | 3617 | 2690 | 2517 | 2293 |
| 2569 | 0.05 | 0.09 | 0.56 | 0.57 | 1 | 2510039O18Ril | RIKEN cDNA 2510039O18 gene | 2209 | 2034 | 2367 | 2796 | 2927 | 2159 | 2739 | 2916 | 2677 | 3192 | 2239 |
| 2780 | 0.05 | 0.11 | 0.48 | 0.63 | 1 | Pfkfb2 | 6-phosphofructo-2-kinase/fructose-2,6-bip | 523 | 598 | 558 | 3120 | 2648 | 5608 | 2737 | 2574 | 3154 | 2884 | 6175 |
| 2789 | 0.18 | 0.11 | 1.62 | 0.11 | 1 | Tmem250-ps | transmembrane protein 250, pseudogene | 2645 | 3173 | 3238 | 2596 | 2759 | 1979 | 2736 | 3649 | 2802 | 3060 | 2036 |
| 2508 | -0.06 | 0.08 | -0.77 | 0.44 | 1 | Ppp1r10 | protein phosphatase 1, regulatory subuni | 1891 | 1639 | 1942 | 2612 | 3125 | 2827 | 2736 | 2702 | 2864 | 2679 | 2569 |
| 2750 | 0.08 | 0.07 | 1.10 | 0.27 | 1 | Tsr1 | TSR1 20S rRNA accumulation | 2132 | 2291 | 2180 | 2958 | 2700 | 3106 | 2735 | 2867 | 2941 | 2986 | 3353 |
| 1976 | 0.18 | 0.15 | 1.20 | 0.23 | 1 | Dcaf12l1 | DDB1 and CUL4 associated factor 12-like | 679 | 685 | 872 | 2552 | 2244 | 1598 | 2735 | 3421 | 2423 | 2762 | 1763 |
| 3670 | 0.01 | 0.12 | 0.10 | 0.92 | 1 | Krt8 | keratin 8 | 4338 | 4723 | 5760 | 2858 | 3340 | 3773 | 2734 | 3812 | 3187 | 2391 | 3457 |
| 2648 | 0.09 | 0.12 | 0.74 | 0.46 | 1 | Nsun2 | NOL1/NOP2/Sun domain family member | 3110 | 2149 | 2616 | 2337 | 2796 | 2450 | 2732 | 3027 | 2798 | 2575 | 2540 |
| 2205 | 0.11 | 0.09 | 1.13 | 0.26 | 1 | Tmem8 | transmembrane protein 8 | 791 | 600 | 875 | 2360 | 2837 | 2663 | 2732 | 2712 | 3147 | 2727 | 2808 |
| 2547 | 0.01 | 0.09 | 0.12 | 0.90 | 1 | Bap1 | Brca1 associated protein 1 | 2330 | 2292 | 2440 | 2364 | 2897 | 2444 | 2731 | 2895 | 2803 | 2504 | 2321 |
| 2753 | 0.05 | 0.12 | 0.38 | 0.70 | 1 | Qrich1 | glutamine-rich 1 | 2399 | 2451 | 2178 | 2430 | 2972 | 3307 | 2730 | 2304 | 3377 | 2852 | 3288 |
| 2590 | -0.02 | 0.10 | -0.15 | 0.88 | 1 | Tbl2 | transducin (beta)-like 2 | 1512 | 1561 | 1518 | 2611 | 3359 | 3309 | 2729 | 2645 | 3453 | 2590 | 3201 |
| 2228 | -0.09 | 0.12 | -0.70 | 0.48 | 1 | Slc7a1 | solute carrier family 7 (cationic amino acid | 1881 | 2257 | 2021 | 2335 | 2568 | 1809 | 2729 | 1998 | 2717 | 2476 | 1720 |
| 3656 | 0.06 | 0.11 | 0.56 | 0.57 | 1 | Map4k4 | mitogen-activated protein kinase kinase k | 5571 | 4865 | 5697 | 2961 | 2707 | 3385 | 2727 | 3594 | 2942 | 2754 | 3012 |
| 2306 | -0.02 | 0.09 | -0.19 | 0.85 | 1 | Ero1l | ERO1-like (S. cerevisiae) | 1682 | 1615 | 1584 | 2517 | 2828 | 2240 | 2725 | 2316 | 2684 | 2738 | 2438 |
| 2731 | 0.02 | 0.08 | 0.27 | 0.79 | 1 | Ankrd13c | ankyrin repeat domain 13c | 2243 | 2502 | 2386 | 2816 | 2999 | 2822 | 2724 | 2683 | 2832 | 2898 | 3134 |
| 2550 | 0.06 | 0.09 | 0.68 | 0.50 | 1 | Larp4 | La ribonucleoprotein domain family, mem | 1567 | 1973 | 1638 | 2875 | 2653 | 2925 | 2724 | 2826 | 3091 | 2610 | 3171 |
| 2517 | 0.05 | 0.12 | 0.40 | 0.69 | 1 | Acap2 | ArfGAP with coiled-coil, ankyrin repeat an | 2461 | 2110 | 2054 | 2344 | 2318 | 2994 | 2724 | 2764 | 2719 | 2197 | 3006 |
| 2699 | 0.07 | 0.09 | 0.83 | 0.40 | 1 | Ndufv2 | NADH:ubiquinone oxidoreductase core s | 2796 | 2471 | 2719 | 2816 | 2867 | 2178 | 2720 | 3169 | 2807 | 2807 | 2341 |
| 2381 | 0.02 | 0.09 | 0.21 | 0.84 | 1 | Apmap | adipocyte plasma membrane associated | 2477 | 2619 | 2577 | 2388 | 2398 | 1728 | 2720 | 2428 | 2380 | 2538 | 1935 |
| 2739 | 0.07 | 0.10 | 0.78 | 0.44 | 1 | Cdc5l | cell division cycle 5-like (S. pombe) | 2418 | 2376 | 2278 | 2940 | 2852 | 2704 | 2719 | 3295 | 2864 | 2517 | 3170 |
| 2582 | -0.01 | 0.08 | -0.12 | 0.90 | 1 | Dlat | dihydrolipoamide S-acetyltransferase (E2 | 2587 | 2435 | 2521 | 2742 | 2747 | 2237 | 2719 | 2496 | 2780 | 3002 | 2141 |
| 3148 | 0.09 | 0.11 | 0.81 | 0.42 | 1 | Tmem63a | transmembrane protein 63a | 2243 | 2283 | 2553 | 3424 | 3304 | 3924 | 2718 | 4106 | 3045 | 3084 | 3944 |
| 2648 | 0.14 | 0.15 | 0.96 | 0.34 | 1 | Aagab | alpha- and gamma-adaptin binding prote | 2088 | 1652 | 2244 | 2911 | 2735 | 2637 | 2718 | 3734 | 2518 | 2934 | 2960 |
| 2397 | -0.16 | 0.12 | -1.40 | 0.16 | 1 | Mtor | mechanistic target of rapamycin kinase | 2068 | 1522 | 1737 | 3033 | 2712 | 2643 | 2716 | 2409 | 2714 | 2468 | 2348 |
| 2364 | -0.04 | 0.11 | -0.39 | 0.69 | 1 | BC004004 | cDNA sequence BC004004 | 1933 | 1746 | 1662 | 2907 | 2661 | 2194 | 2716 | 2981 | 2748 | 2315 | 2143 |
| 2615 | 0.11 | 0.11 | 1.02 | 0.31 | 1 | Ypel5 | yippee like 5 | 2478 | 3075 | 3026 | 2298 | 2763 | 1959 | 2715 | 2624 | 2784 | 2798 | 2249 |
| 1945 | 0.04 | 0.11 | 0.41 | 0.68 | 1 | Robo1 | roundabout guidance receptor 1 | 413 | 486 | 448 | 2241 | 2588 | 2318 | 2715 | 2384 | 2737 | 2804 | 2259 |
| 2793 | 0.13 | 0.10 | 1.26 | 0.21 | 1 | Gcc2 | GRIP and coiled-coil domain containing 2 | 2185 | 2368 | 2764 | 2321 | 2797 | 3326 | 2709 | 3066 | 3133 | 2410 | 3643 |
| 2314 | -0.21 | 0.13 | -1.62 | 0.11 | 1 | Synj1 | synaptojanin 1 | 1917 | 1944 | 1844 | 2503 | 2367 | 3009 | 2709 | 1696 | 2630 | 2397 | 2439 |
| 2902 | 0.11 | 0.08 | 1.50 | 0.13 | 1 | Pnrc2 | proline-rich nuclear receptor coactivator 2 | 2889 | 3218 | 3364 | 2822 | 2700 | 2557 | 2706 | 3072 | 2860 | 2898 | 2838 |
| 2789 | 0.05 | 0.08 | 0.72 | 0.47 | 1 | Ivd | isovaleryl coenzyme A dehydrogenase | 2367 | 2388 | 2293 | 2762 | 2286 | 3834 | 2706 | 2826 | 2378 | 2808 | 4033 |
| 2422 | 0.10 | 0.09 | 1.13 | 0.26 | 1 | Creb3 | cAMP responsive element binding protein | 2095 | 2052 | 2140 | 2142 | 2662 | 2343 | 2703 | 2550 | 2657 | 2775 | 2526 |
| 2404 | -0.09 | 0.13 | -0.71 | 0.48 | 1 | Rptor | regulatory associated protein of MTOR, co | 2077 | 2614 | 2141 | 2649 | 2626 | 2145 | 2702 | 2018 | 2730 | 2701 | 2040 |
| 2834 | -0.03 | 0.09 | -0.32 | 0.75 | 1 | Vamp3 | vesicle-associated membrane protein 3 | 3218 | 3446 | 3321 | 2778 | 2653 | 2554 | 2698 | 3038 | 2496 | 2293 | 2682 |
| 2642 | -0.03 | 0.12 | -0.27 | 0.78 | 1 | Zc3hav1 | zinc finger CCCH type, antiviral 1 | 2974 | 3400 | 2787 | 2612 | 2591 | 2176 | 2698 | 2127 | 2916 | 2473 | 2312 |
| 2089 | 0.04 | 0.14 | 0.30 | 0.77 | 1 | Mthfd2 | methylenetetrahydrofolate dehydrogenas | 821 | 1100 | 929 | 2334 | 2942 | 1945 | 2697 | 2350 | 2881 | 3019 | 1962 |
| 2502 | 0.14 | 0.10 | 1.40 | 0.16 | 1 | Ipmk | inositol polyphosphate multikinase | 1835 | 2344 | 2075 | 2707 | 2474 | 2260 | 2696 | 2820 | 2864 | 2997 | 2449 |

| | | | | | | | | | | | | | | | | | |
|---|---|---|---|---|---|---|---|---|---|---|---|---|---|---|---|---|---|
| 2289 | 0.04 | 0.09 | 0.42 | 0.68 | 1 | Letm1 | leucine zipper-EF-hand containing transm | 1835 | 1478 | 1653 | 2381 | 2665 | 2240 | 2695 | 2592 | 2817 | 2586 | 2232 |
| 2675 | 0.05 | 0.10 | 0.50 | 0.61 | 1 | Azi2 | 5-azacytidine induced gene 2 | 2087 | 2504 | 2237 | 2791 | 2870 | 2733 | 2693 | 2874 | 3107 | 2383 | 3150 |
| 2616 | -0.11 | 0.19 | -0.55 | 0.58 | 1 | Lgalsl | lectin, galactoside binding-like | 1870 | 1519 | 1478 | 2737 | 3858 | 2923 | 2692 | 1694 | 4350 | 2997 | 2657 |
| 2781 | 0.13 | 0.08 | 1.57 | 0.12 | 1 | Pdhb | pyruvate dehydrogenase (lipoamide) beta | 2103 | 2492 | 2596 | 2662 | 2770 | 3062 | 2687 | 3011 | 2972 | 2914 | 3324 |
| 2670 | 0.11 | 0.07 | 1.48 | 0.14 | 1 | Dgke | diacylglycerol kinase, epsilon | 1060 | 1388 | 1092 | 3517 | 2785 | 3413 | 2687 | 4122 | 2936 | 2817 | 3552 |
| 2387 | 0.05 | 0.11 | 0.48 | 0.63 | 1 | Otud7b | OTU domain containing 7B | 1544 | 1622 | 1702 | 2970 | 2288 | 2557 | 2687 | 3015 | 2869 | 2433 | 2573 |
| 2289 | 0.05 | 0.09 | 0.54 | 0.59 | 1 | Elp1 | elongator complex protein 1 | 1748 | 1574 | 1796 | 2146 | 2581 | 2461 | 2685 | 2524 | 2614 | 2396 | 2651 |
| 2635 | -0.04 | 0.11 | -0.32 | 0.75 | 1 | Aqr | aquarius | 2184 | 1788 | 2113 | 2861 | 3431 | 2598 | 2683 | 2546 | 3536 | 2490 | 2754 |
| 2432 | 0.09 | 0.09 | 0.99 | 0.32 | 1 | Fbxl17 | F-box and leucine-rich repeat protein 17 | 1874 | 2228 | 1903 | 2611 | 2717 | 2044 | 2683 | 2830 | 2828 | 2908 | 2126 |
| 2793 | 0.21 | 0.15 | 1.42 | 0.16 | 1 | Trir | telomerase RNA component interacting R | 2853 | 2268 | 2981 | 2469 | 2871 | 2408 | 2682 | 3834 | 2601 | 3388 | 2373 |
| 2430 | 0.04 | 0.07 | 0.49 | 0.63 | 1 | Cask | calcium/calmodulin-dependent serine pro | 2401 | 2483 | 2368 | 2633 | 2392 | 1910 | 2682 | 2637 | 2446 | 2820 | 1961 |
| 2568 | 0.07 | 0.09 | 0.78 | 0.43 | 1 | Kdm5b | lysine (K)-specific demethylase 5B | 2608 | 2611 | 2639 | 2322 | 2374 | 2575 | 2681 | 2506 | 2820 | 2669 | 2442 |
| 2216 | -0.01 | 0.13 | -0.07 | 0.94 | 1 | Lcorl | ligand dependent nuclear receptor corepr | 1475 | 1865 | 1535 | 2703 | 2055 | 2366 | 2681 | 2162 | 2517 | 2324 | 2699 |
| 2606 | -0.01 | 0.12 | -0.10 | 0.92 | 1 | Zmym2 | zinc finger, MYM-type 2 | 2137 | 2559 | 1870 | 2590 | 2528 | 3280 | 2680 | 2286 | 2743 | 2512 | 3483 |
| 2952 | -0.01 | 0.11 | -0.14 | 0.89 | 1 | Spen | spen family transcription repressor | 2913 | 3310 | 2699 | 3611 | 2580 | 2931 | 2679 | 3725 | 2950 | 2744 | 2326 |
| 2723 | 0.13 | 0.09 | 1.40 | 0.16 | 1 | Arfgap2 | ADP-ribosylation factor GTPase activating | 2447 | 2546 | 2552 | 2407 | 2729 | 2895 | 2679 | 2969 | 2824 | 3134 | 2773 |
| 2832 | 0.02 | 0.11 | 0.15 | 0.88 | 1 | AW549877 | expressed sequence AW549877 | 1658 | 1773 | 1597 | 3122 | 2460 | 4654 | 2677 | 2367 | 2865 | 2871 | 5105 |
| 2734 | -0.21 | 0.11 | -1.88 | 0.06 | 1 | Etl4 | enhancer trap locus 4 | 2049 | 3382 | 2884 | 3548 | 2946 | 2565 | 2677 | 2641 | 2561 | 2487 | 2339 |
| 2521 | -0.11 | 0.12 | -0.90 | 0.37 | 1 | Traf3 | TNF receptor-associated factor 3 | 2355 | 2166 | 2350 | 2571 | 2688 | 2833 | 2674 | 1868 | 2819 | 2984 | 2426 |
| 3695 | -0.06 | 0.11 | -0.51 | 0.61 | 1 | Myo1c | myosin IC | 7647 | 6173 | 6461 | 2777 | 2847 | 2030 | 2673 | 2361 | 3275 | 2575 | 1824 |
| 2516 | 0.00 | 0.11 | 0.01 | 0.99 | 1 | Hagh | hydroxyacyl glutathione hydrolase | 1804 | 1789 | 2382 | 2675 | 3245 | 2258 | 2673 | 2231 | 3014 | 3160 | 2449 |
| 3210 | -0.14 | 0.11 | -1.21 | 0.23 | 1 | Abca7 | ATP-binding cassette, sub-family A (ABC1 | 1923 | 2906 | 2593 | 3315 | 3597 | 5187 | 2671 | 3093 | 3068 | 2959 | 3995 |
| 2413 | -0.06 | 0.12 | -0.51 | 0.61 | 1 | Dennd6a | DENN/MADD domain containing 6A | 2482 | 3048 | 2647 | 2411 | 2159 | 2151 | 2671 | 2010 | 2338 | 2395 | 2234 |
| 2964 | 0.02 | 0.12 | 0.15 | 0.88 | 1 | Abcc5 | ATP-binding cassette, sub-family C (CFTR | 1886 | 2095 | 2167 | 2945 | 2177 | 5406 | 2669 | 2727 | 3018 | 2294 | 5217 |
| 2150 | -0.12 | 0.11 | -1.04 | 0.30 | 1 | Ppp2r1b | protein phosphatase 2, regulatory subuni | 1595 | 1882 | 1733 | 2462 | 2369 | 2105 | 2668 | 2096 | 2340 | 2102 | 2299 |
| 3131 | -0.08 | 0.09 | -0.91 | 0.36 | 1 | Phf3 | PHD finger protein 3 | 2724 | 2656 | 2744 | 3594 | 3316 | 3994 | 2667 | 3186 | 3224 | 2722 | 3615 |
| 2518 | 0.06 | 0.08 | 0.78 | 0.44 | 1 | Gtf3c4 | general transcription factor IIIC, polypepti | 1871 | 2059 | 1658 | 2892 | 2691 | 2574 | 2667 | 3077 | 2738 | 2822 | 2645 |
| 2517 | 0.11 | 0.14 | 0.78 | 0.44 | 1 | 9930021J03Rik | RIKEN cDNA 9930021J03 gene | 1829 | 2347 | 1955 | 2511 | 2331 | 2887 | 2665 | 2299 | 3095 | 2955 | 2816 |
| 2349 | -0.08 | 0.13 | -0.61 | 0.54 | 1 | Rnf13 | ring finger protein 13 | 2920 | 2296 | 2038 | 2566 | 2398 | 1928 | 2665 | 1901 | 2462 | 2512 | 2147 |
| 2800 | 0.06 | 0.10 | 0.59 | 0.55 | 1 | Rnf40 | ring finger protein 40 | 2376 | 2620 | 2721 | 2637 | 2976 | 3043 | 2663 | 3110 | 2630 | 2984 | 3040 |
| 2494 | 0.10 | 0.10 | 1.01 | 0.31 | 1 | Fam8a1 | family with sequence similarity 8, member | 2481 | 2575 | 2429 | 2482 | 2200 | 2292 | 2662 | 2290 | 2631 | 2849 | 2537 |
| 2430 | 0.09 | 0.07 | 1.21 | 0.23 | 1 | Slc39a6 | solute carrier family 39 (metal ion transpo | 1988 | 1991 | 1910 | 2565 | 2723 | 2140 | 2661 | 2827 | 2957 | 2731 | 2232 |
| 2918 | 0.09 | 0.12 | 0.82 | 0.41 | 1 | Stub1 | STIP1 homology and U-Box containing pr | 2967 | 2918 | 3140 | 2945 | 2633 | 2889 | 2660 | 3715 | 2469 | 2869 | 2898 |
| 2584 | 0.05 | 0.13 | 0.35 | 0.72 | 1 | Trib1 | tribbles pseudokinase 1 | 2919 | 1521 | 1932 | 1951 | 1861 | 4362 | 2660 | 2176 | 1930 | 2360 | 4752 |
| 2783 | -0.09 | 0.09 | -1.03 | 0.30 | 1 | Mta1 | metastasis associated 1 | 3012 | 3053 | 3447 | 3069 | 2747 | 2401 | 2658 | 2549 | 2761 | 2686 | 2231 |
| 2641 | -0.04 | 0.13 | -0.33 | 0.74 | 1 | Lpin1 | lipin 1 | 1736 | 1369 | 2036 | 2437 | 4349 | 2559 | 2658 | 2624 | 4700 | 2603 | 1975 |
| 2525 | -0.03 | 0.07 | -0.37 | 0.71 | 1 | Pds5b | PDS5 cohesin associated factor B | 2998 | 2413 | 2991 | 2128 | 2459 | 2525 | 2656 | 2025 | 2376 | 2669 | 2535 |
| 3035 | -0.14 | 0.12 | -1.13 | 0.26 | 1 | Ppfibp1 | PTPRF interacting protein, binding protein | 4906 | 4558 | 5031 | 2504 | 2160 | 2541 | 2655 | 1863 | 2579 | 2078 | 2509 |
| 2815 | 0.06 | 0.13 | 0.42 | 0.67 | 1 | Adgrl1 | adhesion G protein-coupled receptor L1 | 1839 | 1410 | 1932 | 2435 | 2649 | 5138 | 2654 | 3150 | 2897 | 2628 | 4235 |
| 2516 | -0.06 | 0.08 | -0.80 | 0.42 | 1 | Gsk3a | glycogen synthase kinase 3 alpha | 2453 | 2125 | 2475 | 2596 | 2936 | 2339 | 2652 | 2265 | 2813 | 2775 | 2251 |
| 3302 | 0.06 | 0.07 | 0.81 | 0.42 | 1 | Ppp4r3a | protein phosphatase 4 regulatory subunit | 3473 | 3791 | 3166 | 3319 | 2787 | 3911 | 2651 | 3239 | 2956 | 2794 | 4234 |
| 2637 | -0.09 | 0.09 | -0.93 | 0.35 | 1 | Git1 | GIT ArfGAP 1 | 3304 | 2927 | 2909 | 2669 | 2836 | 2041 | 2651 | 2293 | 2883 | 2718 | 1779 |
| 2637 | 0.02 | 0.07 | 0.30 | 0.76 | 1 | Kpna6 | karyopherin (importin) alpha 6 | 2058 | 2380 | 2171 | 2957 | 2813 | 2682 | 2651 | 3031 | 3036 | 2552 | 2672 |
| 2463 | -0.10 | 0.12 | -0.81 | 0.42 | 1 | Ccdc186 | coiled-coil domain containing 186 | 1917 | 2386 | 2206 | 2525 | 2500 | 2902 | 2649 | 2089 | 2656 | 2078 | 3190 |
| 2462 | 0.03 | 0.09 | 0.37 | 0.71 | 1 | Smarcc1 | SWI/SNF related, matrix associated, actin | 2450 | 2255 | 2085 | 2386 | 2813 | 2183 | 2649 | 2607 | 2796 | 2610 | 2247 |
| 2708 | -0.08 | 0.08 | -0.95 | 0.34 | 1 | Akt2 | thymoma viral proto-oncogene 2 | 2664 | 2864 | 2465 | 3261 | 2778 | 2504 | 2648 | 2887 | 2765 | 2764 | 2189 |
| 2627 | 0.07 | 0.08 | 0.87 | 0.39 | 1 | Med28 | mediator complex subunit 28 | 3056 | 3132 | 3202 | 2546 | 2433 | 1876 | 2645 | 2782 | 2522 | 2848 | 1858 |
| 3228 | -0.05 | 0.10 | -0.53 | 0.60 | 1 | Ppib | peptidylprolyl isomerase B | 5034 | 4127 | 4447 | 2313 | 3191 | 3018 | 2644 | 2471 | 2850 | 2468 | 2943 |
| 2677 | 0.17 | 0.10 | 1.70 | 0.09 | 1 | Cyp51 | cytochrome P450, family 51 | 1300 | 1382 | 1208 | 3762 | 2805 | 2839 | 2643 | 3976 | 2870 | 3139 | 3526 |
| 19887 | 0.05 | 0.23 | 0.24 | 0.81 | 1 | Gsn | gelsolin | 96097 | 41854 | 60536 | 2848 | 2869 | 1605 | 2642 | 2808 | 2701 | 2961 | 1831 |
| 2541 | 0.05 | 0.08 | 0.59 | 0.56 | 1 | Vps33a | VPS33A CORVET/HOPS core subunit | 2410 | 2455 | 2047 | 2827 | 2763 | 2101 | 2642 | 2821 | 2750 | 2968 | 2166 |
| 2949 | -0.01 | 0.10 | -0.08 | 0.94 | 1 | Insr | insulin receptor | 2906 | 3242 | 3392 | 3344 | 2664 | 2805 | 2641 | 3188 | 3223 | 2525 | 2503 |
| 2542 | 0.04 | 0.11 | 0.35 | 0.72 | 1 | Slc37a4 | solute carrier family 37 (glucose-6-phospl | 933 | 929 | 1348 | 2904 | 2854 | 3799 | 2641 | 2887 | 2892 | 2807 | 3967 |
| 2406 | 0.18 | 0.12 | 1.52 | 0.13 | 1 | Pithd1 | PITH (C-terminal proteasome-interacting | 1819 | 1948 | 2227 | 2276 | 2538 | 2129 | 2641 | 2937 | 2537 | 3200 | 2219 |
| 2419 | -0.06 | 0.12 | -0.55 | 0.58 | 1 | Zdhhc8 | zinc finger, DHHC domain containing 8 | 1869 | 2068 | 1952 | 2607 | 2787 | 2525 | 2640 | 2114 | 2931 | 2870 | 2244 |
| 2728 | -0.09 | 0.09 | -0.90 | 0.37 | 1 | Idh3g | isocitrate dehydrogenase 3 (NAD+), gamı | 2607 | 2447 | 2629 | 2925 | 2920 | 2998 | 2639 | 2508 | 2500 | 2982 | 2855 |
| 2858 | -0.12 | 0.08 | -1.53 | 0.13 | 1 | Szrd1 | SUZ RNA binding domain containing 1 | 3891 | 3809 | 3999 | 2530 | 2818 | 2286 | 2637 | 2135 | 2734 | 2570 | 2025 |
| 2426 | -0.03 | 0.08 | -0.39 | 0.69 | 1 | Ncbp2 | nuclear cap binding protein subunit 2 | 2566 | 2469 | 2421 | 2414 | 2416 | 2262 | 2633 | 2300 | 2590 | 2360 | 2254 |
| 2495 | -0.16 | 0.10 | -1.63 | 0.10 | 1 | Dop1b | DOP1 leucine zipper like protein B | 1989 | 2073 | 2031 | 2969 | 2785 | 2891 | 2631 | 2384 | 2609 | 2825 | 2263 |
| 2437 | 0.11 | 0.11 | 0.96 | 0.34 | 1 | Abhd13 | abhydrolase domain containing 13 | 1477 | 2247 | 1711 | 2752 | 2488 | 2428 | 2631 | 2904 | 2808 | 2701 | 2661 |

| | | | | | | | | | | | | | | | | | |
|---|---|---|---|---|---|---|---|---|---|---|---|---|---|---|---|---|---|
| 2724 | 0.04 | 0.09 | 0.42 | 0.67 | 1 | Serf2 | small EDRK-rich factor 2 | 2607 | 2322 | 2604 | 2763 | 2694 | 2970 | 2630 | 3068 | 2558 | 2660 | 3088 |
| 2460 | 0.01 | 0.10 | 0.05 | 0.96 | 1 | Tomm20 | translocase of outer mitochondrial membr | 2243 | 2862 | 2487 | 2474 | 2718 | 1899 | 2630 | 2420 | 2796 | 2598 | 1934 |
| 2312 | -0.06 | 0.10 | -0.63 | 0.53 | 1 | Cers6 | ceramide synthase 6 | 1959 | 2215 | 1801 | 2685 | 2382 | 2235 | 2630 | 2347 | 2617 | 2583 | 1972 |
| 2251 | -0.28 | 0.15 | -1.79 | 0.07 | 1 | Nalcn | sodium leak channel, non-selective | 2006 | 2200 | 2221 | 2341 | 2496 | 2505 | 2630 | 1356 | 2254 | 2380 | 2374 |
| 2526 | 0.21 | 0.13 | 1.70 | 0.09 | 1 | Suclg1 | succinate-CoA ligase, GDP-forming, alph | 2149 | 1958 | 2601 | 2207 | 2672 | 2257 | 2629 | 3340 | 2911 | 2575 | 2487 |
| 2498 | 0.04 | 0.11 | 0.37 | 0.71 | 1 | Ddi2 | DNA-damage inducible protein 2 | 2114 | 2270 | 1727 | 2552 | 2487 | 2876 | 2629 | 2635 | 2861 | 2462 | 2863 |
| 2915 | 0.03 | 0.08 | 0.43 | 0.67 | 1 | Nfx1 | nuclear transcription factor, X-box binding | 2469 | 2435 | 2561 | 2661 | 2746 | 4133 | 2628 | 3040 | 2775 | 2512 | 4103 |
| 2389 | 0.04 | 0.12 | 0.32 | 0.75 | 1 | Mydgf | myeloid derived growth factor | 1698 | 1588 | 1972 | 2734 | 2822 | 2161 | 2628 | 3297 | 2880 | 2188 | 2310 |
| 2582 | -0.10 | 0.14 | -0.72 | 0.47 | 1 | Tasor2 | transcription activation suppressor family | 1828 | 2173 | 2061 | 2364 | 2531 | 3965 | 2627 | 1732 | 2832 | 2436 | 3853 |
| 2331 | -0.09 | 0.10 | -0.89 | 0.38 | 1 | Atp6v0e | ATPase, H+ transporting, lysosomal V0 su | 1999 | 1819 | 2156 | 2710 | 2723 | 2046 | 2627 | 2065 | 2862 | 2682 | 1950 |
| 2037 | -0.08 | 0.11 | -0.75 | 0.45 | 1 | Tmem147 | transmembrane protein 147 | 1283 | 1036 | 1402 | 2610 | 2366 | 2001 | 2627 | 2333 | 2350 | 2512 | 1885 |
| 2189 | -0.18 | 0.14 | -1.23 | 0.22 | 1 | C2cd3 | C2 calcium-dependent domain containing | 1778 | 1829 | 1753 | 2154 | 2434 | 2673 | 2626 | 1599 | 2769 | 2337 | 2127 |
| 2936 | 0.04 | 0.10 | 0.39 | 0.70 | 1 | Ankhd1 | ankyrin repeat and KH domain containing | 2186 | 1961 | 2138 | 2615 | 3074 | 4568 | 2625 | 2732 | 3564 | 2562 | 4271 |
| 2920 | 0.04 | 0.10 | 0.39 | 0.70 | 1 | Rpl37 | ribosomal protein L37 | 4454 | 4480 | 5114 | 2039 | 2477 | 1795 | 2623 | 2521 | 2348 | 2364 | 1901 |
| 2961 | -0.04 | 0.08 | -0.54 | 0.59 | 1 | Tnrc18 | trinucleotide repeat containing 18 | 4121 | 3832 | 4222 | 2521 | 2518 | 2685 | 2622 | 2515 | 2581 | 2700 | 2254 |
| 2242 | -0.09 | 0.10 | -0.87 | 0.38 | 1 | Ost4 | oligosaccharyltransferase complex subur | 1844 | 1591 | 2113 | 2346 | 2744 | 2090 | 2622 | 1814 | 2821 | 2634 | 2046 |
| 2839 | -0.09 | 0.11 | -0.82 | 0.41 | 1 | Cep170 | centrosomal protein 170 | 2459 | 2325 | 2288 | 3118 | 2581 | 4127 | 2620 | 2331 | 2867 | 2623 | 3886 |
| 2668 | -0.02 | 0.12 | -0.14 | 0.88 | 1 | Rsf1 | remodeling and spacing factor 1 | 2255 | 2829 | 2921 | 2414 | 2774 | 2887 | 2618 | 3086 | 2652 | 2114 | 2801 |
| 2288 | 0.00 | 0.07 | 0.06 | 0.95 | 1 | Ttc39a | tetratricopeptide repeat domain 39A | 955 | 1080 | 1199 | 2780 | 3011 | 2534 | 2618 | 2845 | 3091 | 2518 | 2538 |
| 2257 | 0.05 | 0.09 | 0.58 | 0.56 | 1 | Txndc15 | thioredoxin domain containing 15 | 1881 | 1640 | 1804 | 2470 | 2609 | 1859 | 2617 | 2881 | 2501 | 2742 | 1828 |
| 2005 | 0.00 | 0.11 | 0.02 | 0.99 | 1 | Creb3l1 | cAMP responsive element binding proteir | 947 | 1117 | 1415 | 2181 | 2811 | 1632 | 2616 | 2439 | 2887 | 2532 | 1482 |
| 2370 | 0.03 | 0.10 | 0.26 | 0.80 | 1 | Clpx | caseinolytic mitochondrial matrix peptidas | 1958 | 2190 | 1844 | 2430 | 2290 | 2576 | 2614 | 2760 | 2220 | 2199 | 2989 |
| 3366 | 0.11 | 0.09 | 1.26 | 0.21 | 1 | Srsf3 | serine and arginine-rich splicing factor 3 | 4329 | 3315 | 3609 | 2176 | 2509 | 5144 | 2610 | 2688 | 2420 | 2834 | 5390 |
| 2381 | -0.06 | 0.11 | -0.55 | 0.58 | 1 | Ap3s2 | adaptor-related protein complex 3, sigma | 2035 | 2192 | 2040 | 2452 | 2866 | 2237 | 2609 | 2212 | 2993 | 2263 | 2296 |
| 2304 | 0.21 | 0.11 | 1.88 | 0.06 | 1 | Gnaz | guanine nucleotide binding protein, alpha | 1068 | 1342 | 1211 | 2436 | 2481 | 2497 | 2608 | 3505 | 2789 | 2915 | 2490 |
| 2226 | 0.14 | 0.12 | 1.12 | 0.26 | 1 | Cdc16 | CDC16 cell division cycle 16 | 1971 | 1891 | 1904 | 1908 | 2094 | 2314 | 2607 | 2570 | 2318 | 2149 | 2759 |
| 3032 | 0.22 | 0.12 | 1.87 | 0.06 | 1 | Sltm | SAFB-like, transcription modulator | 2274 | 2511 | 2415 | 2589 | 2372 | 4509 | 2606 | 3991 | 2535 | 2723 | 4823 |
| 2859 | 0.08 | 0.11 | 0.72 | 0.47 | 1 | Pnrc1 | proline-rich nuclear receptor coactivator 1 | 4115 | 3074 | 3763 | 2085 | 2739 | 2426 | 2606 | 1815 | 2845 | 3509 | 2470 |
| 4280 | -0.06 | 0.14 | -0.44 | 0.66 | 1 | Tubb5 | tubulin, beta 5 class I | 10813 | 6825 | 7327 | 3238 | 2820 | 2598 | 2605 | 3180 | 2404 | 2897 | 2375 |
| 2836 | -0.15 | 0.10 | -1.52 | 0.13 | 1 | Shisa5 | shisa family member 5 | 5533 | 3910 | 4274 | 2442 | 2389 | 1713 | 2605 | 2059 | 2365 | 2459 | 1449 |
| 2194 | 0.04 | 0.08 | 0.48 | 0.63 | 1 | Pigk | phosphatidylinositol glycan anchor biosyı | 1858 | 1606 | 1782 | 2323 | 2391 | 2010 | 2605 | 2340 | 2331 | 2705 | 2185 |
| 3022 | 0.00 | 0.15 | -0.02 | 0.98 | 1 | Dio1 | deiodinase, iodothyronine, type I | 480 | 1250 | 1515 | 5218 | 4692 | 2641 | 2603 | 4412 | 4897 | 2970 | 2565 |
| 2642 | 0.03 | 0.07 | 0.40 | 0.69 | 1 | Chmp7 | charged multivesicular body protein 7 | 2480 | 2479 | 2522 | 2702 | 2959 | 2416 | 2603 | 2937 | 2819 | 2671 | 2471 |
| 2572 | -0.04 | 0.11 | -0.32 | 0.75 | 1 | Eml1 | echinoderm microtubule associated prote | 3439 | 2972 | 2448 | 2397 | 2781 | 2062 | 2603 | 2513 | 2555 | 2507 | 2019 |
| 2473 | -0.11 | 0.08 | -1.36 | 0.17 | 1 | Unc119b | unc-119 lipid binding chaperone B | 3037 | 3249 | 3008 | 2451 | 2299 | 1957 | 2603 | 2441 | 2048 | 2197 | 1917 |
| 2674 | -0.08 | 0.10 | -0.86 | 0.39 | 1 | Zfp24 | zinc finger protein 24 | 2621 | 3329 | 2838 | 2790 | 2574 | 2610 | 2601 | 2803 | 2600 | 1986 | 2659 |
| 2666 | -0.10 | 0.12 | -0.84 | 0.40 | 1 | Ankrd44 | ankyrin repeat domain 44 | 2128 | 2008 | 2055 | 3557 | 2315 | 3515 | 2601 | 3045 | 2390 | 2420 | 3298 |
| 2391 | 0.13 | 0.09 | 1.43 | 0.15 | 1 | BC031181 | cDNA sequence BC031181 | 1788 | 1716 | 1850 | 2509 | 2647 | 2258 | 2600 | 2976 | 2570 | 2965 | 2425 |
| 2170 | 0.08 | 0.09 | 0.87 | 0.38 | 1 | Gm15459 | | 3746 | 3172 | 617 | 2407 | 372 | 2628 | 2599 | 2516 | 424 | 2610 | 2783 |
| 2109 | 0.15 | 0.13 | 1.14 | 0.25 | 1 | Pde4b | phosphodiesterase 4B, cAMP specific | 2485 | 1283 | 1455 | 2138 | 2238 | 1560 | 2598 | 2170 | 2519 | 2867 | 1891 |
| 2591 | 0.08 | 0.11 | 0.71 | 0.48 | 1 | H6pd | hexose-6-phosphate dehydrogenase (glu | 4228 | 2926 | 2717 | 2354 | 2407 | 1611 | 2597 | 2398 | 2564 | 3189 | 1509 |
| 2485 | 0.07 | 0.09 | 0.77 | 0.44 | 1 | Vps13a | vacuolar protein sorting 13A | 1962 | 2245 | 2087 | 2393 | 2389 | 2884 | 2597 | 2732 | 2373 | 2557 | 3113 |
| 2143 | 0.06 | 0.10 | 0.59 | 0.55 | 1 | Nampt | nicotinamide phosphoribosyltransferase | 2144 | 1757 | 2081 | 2086 | 2221 | 1753 | 2594 | 2155 | 2375 | 2288 | 2120 |
| 2473 | -0.14 | 0.11 | -1.24 | 0.21 | 1 | Ube2k | ubiquitin-conjugating enzyme E2K | 2192 | 2180 | 2092 | 2795 | 3075 | 2416 | 2591 | 2060 | 2850 | 2476 | 2483 |
| 2452 | 0.01 | 0.10 | 0.10 | 0.92 | 1 | Tmco1 | transmembrane and coiled-coil domains | 2119 | 1939 | 2022 | 3191 | 2517 | 2127 | 2591 | 2652 | 2833 | 2768 | 2211 |
| 5440 | -0.10 | 0.11 | -0.97 | 0.33 | 1 | Dnajb1 | DnaJ heat shock protein family (Hsp40) n | 5485 | 6017 | 6266 | 2825 | 2251 | 14275 | 2590 | 2975 | 2376 | 1936 | 12849 |
| 3626 | 0.09 | 0.06 | 1.32 | 0.19 | 1 | Srsf2 | serine and arginine-rich splicing factor 2 | 4177 | 3345 | 4118 | 2286 | 2681 | 6136 | 2589 | 2395 | 2720 | 2903 | 6531 |
| 3201 | 0.07 | 0.11 | 0.64 | 0.52 | 1 | Zfp445 | zinc finger protein 445 | 2358 | 2956 | 2389 | 3328 | 2735 | 4781 | 2589 | 2750 | 3316 | 2894 | 5110 |
| 2983 | 0.06 | 0.10 | 0.60 | 0.55 | 1 | Ankrd10 | ankyrin repeat domain 10 | 964 | 1097 | 1140 | 2980 | 2458 | 6477 | 2589 | 3353 | 2687 | 2387 | 6685 |
| 2215 | 0.00 | 0.12 | 0.02 | 0.98 | 1 | Stxbp5 | syntaxin binding protein 5 (tomosyn) | 1719 | 2079 | 1952 | 2365 | 2148 | 2175 | 2588 | 2388 | 2720 | 2289 | 1948 |
| 2509 | -0.10 | 0.08 | -1.21 | 0.23 | 1 | Agap3 | ArfGAP with GTPase domain, ankyrin rep | 2602 | 2542 | 2526 | 2432 | 2890 | 2398 | 2587 | 2352 | 2523 | 2628 | 2122 |
| 3045 | 0.16 | 0.17 | 0.94 | 0.35 | 1 | C1qa | complement component 1, q subcompone | 7140 | 3859 | 5360 | 2233 | 1595 | 1700 | 2585 | 2844 | 1723 | 2486 | 1967 |
| 2084 | 0.02 | 0.12 | 0.18 | 0.86 | 1 | Polr3k | polymerase (RNA) III (DNA directed) poly | 1592 | 1648 | 1890 | 1905 | 2420 | 1948 | 2583 | 2275 | 2183 | 2163 | 2315 |
| 3080 | 0.04 | 0.08 | 0.53 | 0.59 | 1 | Rsrc2 | arginine/serine-rich coiled-coil 2 | 2128 | 2357 | 1958 | 2487 | 2367 | 5969 | 2582 | 2513 | 2554 | 2433 | 6533 |
| 1806 | 0.03 | 0.12 | 0.21 | 0.83 | 1 | Igfals | insulin-like growth factor binding protein, | 452 | 559 | 597 | 2643 | 2416 | 1456 | 2581 | 2794 | 2503 | 2096 | 1775 |
| 2347 | 0.05 | 0.12 | 0.45 | 0.66 | 1 | Tmbim4 | transmembrane BAX inhibitor motif contai | 1513 | 1834 | 1784 | 2591 | 2897 | 2081 | 2580 | 2822 | 2453 | 3142 | 2122 |
| 2394 | -0.01 | 0.10 | -0.05 | 0.96 | 1 | Armc1 | armadillo repeat containing 1 | 1796 | 2309 | 2051 | 2572 | 2752 | 2207 | 2579 | 2511 | 2781 | 2297 | 2479 |
| 2328 | 0.17 | 0.09 | 1.81 | 0.07 | 1 | Fam126b | family with sequence similarity 126, meml | 1579 | 1857 | 1794 | 2346 | 2570 | 2113 | 2579 | 2847 | 2764 | 2630 | 2525 |
| 4080 | 0.03 | 0.12 | 0.23 | 0.82 | 1 | Kif12 | kinesin family member 12 | 1170 | 1556 | 1939 | 2629 | 2625 | 12500 | 2578 | 2597 | 2860 | 2782 | 11644 |
| 2902 | 0.16 | 0.13 | 1.22 | 0.22 | 1 | Dpp3 | dipeptidylpeptidase 3 | 2137 | 1560 | 2162 | 2350 | 3884 | 3617 | 2577 | 3369 | 3578 | 2974 | 3719 |

| | | | | | | | | | | | | | | | | | |
|---|---|---|---|---|---|---|---|---|---|---|---|---|---|---|---|---|---|
| 2137 | 0.05 | 0.16 | 0.31 | 0.75 | 1 | Ccdc88a | coiled coil domain containing 88A | 1985 | 2133 | 1590 | 1820 | 1534 | 2925 | 2577 | 1858 | 2275 | 2102 | 2708 |
| 2370 | 0.16 | 0.09 | 1.72 | 0.09 | 1 | Ndufa4 | Ndufa4, mitochondrial complex associate | 1863 | 1724 | 1642 | 2735 | 2674 | 1843 | 2576 | 2891 | 2696 | 3385 | 2038 |
| 2111 | 0.01 | 0.10 | 0.14 | 0.89 | 1 | Vma21 | VMA21 vacuolar H+-ATPase homolog (S. | 1855 | 1743 | 1727 | 2311 | 2152 | 1866 | 2576 | 1926 | 2422 | 2579 | 2060 |
| 2210 | 0.05 | 0.07 | 0.73 | 0.47 | 1 | Dpy19l4 | dpy-19-like 4 (C. elegans) | 1728 | 1565 | 1655 | 2386 | 2433 | 2102 | 2575 | 2463 | 2441 | 2803 | 2163 |
| 1960 | -0.05 | 0.12 | -0.39 | 0.69 | 1 | Mettl7a1 | methyltransferase like 7A1 | 1177 | 1064 | 891 | 2551 | 2278 | 1946 | 2575 | 1906 | 2379 | 2835 | 1955 |
| 2046 | -0.15 | 0.13 | -1.20 | 0.23 | 1 | Pmpcb | peptidase (mitochondrial processing) beta | 1884 | 2039 | 1933 | 1987 | 2119 | 2074 | 2574 | 1615 | 2193 | 2027 | 2056 |
| 2633 | 0.18 | 0.11 | 1.61 | 0.11 | 1 | Serpinb6a | serine (or cysteine) peptidase inhibitor, cl | 3656 | 2473 | 2544 | 2372 | 2522 | 2060 | 2573 | 2628 | 2571 | 2968 | 2596 |
| 2037 | 0.14 | 0.12 | 1.16 | 0.25 | 1 | Hspa12a | heat shock protein 12A | 630 | 611 | 472 | 2659 | 2200 | 2412 | 2573 | 2292 | 2762 | 3019 | 2777 |
| 2549 | 0.12 | 0.10 | 1.20 | 0.23 | 1 | Cse1l | chromosome segregation 1-like (S. cerev | 2540 | 2692 | 1927 | 2515 | 2480 | 2422 | 2572 | 2698 | 2556 | 3010 | 2623 |
| 2234 | -0.06 | 0.14 | -0.45 | 0.65 | 1 | Mapk10 | mitogen-activated protein kinase 10 | 738 | 1309 | 1267 | 2466 | 2949 | 2851 | 2571 | 2689 | 2948 | 2338 | 2446 |
| 2287 | 0.12 | 0.12 | 0.97 | 0.33 | 1 | Krcc1 | lysine-rich coiled-coil 1 | 2126 | 2459 | 2251 | 2114 | 2123 | 1973 | 2570 | 2801 | 2261 | 2174 | 2301 |
| 2248 | 0.11 | 0.09 | 1.28 | 0.20 | 1 | Sptlc1 | serine palmitoyltransferase, long chain ba | 1612 | 1478 | 1699 | 2485 | 2589 | 1932 | 2570 | 2573 | 2565 | 3059 | 2170 |
| 1952 | -0.14 | 0.10 | -1.45 | 0.15 | 1 | Tm2d2 | TM2 domain containing 2 | 1464 | 1397 | 1724 | 2248 | 2439 | 1613 | 2570 | 1943 | 2166 | 2322 | 1581 |
| 2717 | 0.04 | 0.09 | 0.40 | 0.69 | 1 | Eif4a3 | eukaryotic translation initiation factor 4A3 | 3170 | 2997 | 2790 | 2724 | 2662 | 2348 | 2567 | 2644 | 2728 | 3064 | 2186 |
| 2424 | -0.02 | 0.08 | -0.28 | 0.78 | 1 | Kif3b | kinesin family member 3B | 2124 | 1827 | 1996 | 2704 | 2408 | 2766 | 2567 | 2544 | 2450 | 2632 | 2647 |
| 2232 | -0.10 | 0.09 | -1.14 | 0.25 | 1 | Usp31 | ubiquitin specific peptidase 31 | 1088 | 1401 | 1194 | 2816 | 2137 | 3312 | 2567 | 2623 | 2187 | 2253 | 2974 |
| 2351 | -0.05 | 0.11 | -0.40 | 0.69 | 1 | Flcn | folliculin | 991 | 739 | 1163 | 2477 | 2742 | 3874 | 2566 | 2124 | 2721 | 2933 | 3536 |
| 2426 | 0.05 | 0.09 | 0.51 | 0.61 | 1 | Map9 | microtubule-associated protein 9 | 1680 | 2099 | 1925 | 2714 | 2382 | 2660 | 2563 | 2704 | 2632 | 2402 | 2927 |
| 2388 | 0.01 | 0.11 | 0.11 | 0.92 | 1 | Mtmr2 | myotubularin related protein 2 | 2008 | 1976 | 2177 | 2604 | 2544 | 2268 | 2563 | 2063 | 3047 | 2644 | 2373 |
| 2361 | 0.03 | 0.11 | 0.30 | 0.77 | 1 | Myt1 | myelin transcription factor 1 | 1681 | 1497 | 1929 | 1955 | 2362 | 3483 | 2562 | 2363 | 2641 | 2170 | 3326 |
| 2231 | 0.08 | 0.13 | 0.61 | 0.54 | 1 | Ttc37 | tetratricopeptide repeat domain 37 | 1597 | 1780 | 2115 | 1801 | 2459 | 2414 | 2562 | 1790 | 2790 | 2462 | 2773 |
| 2543 | 0.09 | 0.10 | 0.86 | 0.39 | 1 | Atic | 5-aminoimidazole-4-carboxamide ribonu | 2702 | 2400 | 2974 | 2483 | 2467 | 2129 | 2561 | 2928 | 2684 | 2492 | 2158 |
| 2004 | 0.16 | 0.11 | 1.53 | 0.13 | 1 | Skap1 | src family associated phosphoprotein 1 | 1005 | 1160 | 1171 | 2304 | 2344 | 1629 | 2560 | 2857 | 2395 | 2745 | 1879 |
| 2705 | 0.11 | 0.16 | 0.72 | 0.47 | 1 | Uqcrh | ubiquinol-cytochrome c reductase hinge p | 3619 | 3020 | 3926 | 2432 | 2453 | 1750 | 2558 | 3527 | 2249 | 2330 | 1893 |
| 2336 | 0.04 | 0.10 | 0.43 | 0.67 | 1 | Cops4 | COP9 signalosome subunit 4 | 2635 | 2570 | 2552 | 1973 | 2386 | 1946 | 2558 | 2339 | 2342 | 2319 | 2079 |
| 2307 | -0.06 | 0.15 | -0.41 | 0.68 | 1 | Foxj3 | forkhead box J3 | 1806 | 2251 | 2016 | 2371 | 2420 | 2483 | 2557 | 1722 | 2685 | 2671 | 2395 |
| 3980 | 0.00 | 0.12 | 0.00 | 1.00 | 1 | Heg1 | heart development protein with EGF-like d | 8317 | 7490 | 6399 | 3101 | 2580 | 2486 | 2556 | 3707 | 2515 | 2362 | 2264 |
| 2252 | -0.05 | 0.12 | -0.44 | 0.66 | 1 | Fitm2 | fat storage-inducing transmembrane prote | 1122 | 1763 | 1407 | 2738 | 2472 | 2559 | 2556 | 3316 | 2461 | 1895 | 2481 |
| 2405 | 0.03 | 0.10 | 0.34 | 0.74 | 1 | Herpud2 | HERPUD family member 2 | 1837 | 2201 | 1891 | 2428 | 2726 | 2432 | 2555 | 2581 | 2530 | 2803 | 2470 |
| 2055 | 0.02 | 0.15 | 0.12 | 0.91 | 1 | Cpeb3 | cytoplasmic polyadenylation element bind | 1646 | 1741 | 1567 | 1800 | 2493 | 1915 | 2555 | 1929 | 2922 | 2084 | 1949 |
| 2564 | 0.08 | 0.10 | 0.75 | 0.45 | 1 | Spg7 | SPG7, paraplegin matrix AAA peptidase s | 2382 | 2441 | 2522 | 2612 | 2528 | 2435 | 2554 | 3162 | 2413 | 2820 | 2333 |
| 2467 | -0.01 | 0.12 | -0.11 | 0.91 | 1 | Dpf2 | D4, zinc and double PHD fingers family 2 | 1841 | 2313 | 1960 | 2502 | 2364 | 3163 | 2553 | 2138 | 2645 | 2729 | 2934 |
| 2453 | 0.03 | 0.12 | 0.28 | 0.78 | 1 | Rnf114 | ring finger protein 114 | 2261 | 2967 | 2221 | 2358 | 2364 | 2379 | 2551 | 2396 | 2263 | 2776 | 2449 |
| 2057 | 0.07 | 0.16 | 0.46 | 0.64 | 1 | Klc2 | kinesin light chain 2 | 1303 | 988 | 1768 | 1933 | 2376 | 2222 | 2551 | 2482 | 2153 | 2704 | 2147 |
| 2361 | 0.07 | 0.08 | 0.87 | 0.39 | 1 | Sfmbt1 | Scm-like with four mbt domains 1 | 1168 | 1295 | 1184 | 2863 | 2785 | 2702 | 2550 | 2914 | 3093 | 2673 | 2741 |
| 2188 | -0.01 | 0.14 | -0.09 | 0.93 | 1 | Rrm2b | ribonucleotide reductase M2 B (TP53 indu | 1887 | 2350 | 2126 | 2332 | 2045 | 1955 | 2550 | 2213 | 2588 | 1872 | 2153 |
| 2636 | -0.17 | 0.10 | -1.64 | 0.10 | 1 | Dlg4 | discs large MAGUK scaffold protein 4 | 2346 | 2986 | 2867 | 2848 | 2502 | 3121 | 2549 | 2266 | 2465 | 2452 | 2588 |
| 2562 | -0.04 | 0.07 | -0.59 | 0.56 | 1 | Arl6ip5 | ADP-ribosylation factor-like 6 interacting p | 2649 | 2728 | 2816 | 2662 | 2860 | 2057 | 2549 | 2558 | 2801 | 2550 | 1954 |
| 2687 | 0.15 | 0.09 | 1.79 | 0.07 | 1 | Heca | hdc homolog, cell cycle regulator | 2497 | 2800 | 2463 | 2970 | 2442 | 2344 | 2547 | 3067 | 3028 | 3060 | 2337 |
| 2712 | 0.05 | 0.12 | 0.44 | 0.66 | 1 | Uba2 | ubiquitin-like modifier activating enzyme 2 | 3985 | 2904 | 3435 | 2356 | 2356 | 2330 | 2545 | 2601 | 2557 | 2185 | 2576 |
| 2661 | 0.00 | 0.08 | -0.04 | 0.97 | 1 | Kctd20 | potassium channel tetramerisation domai | 2871 | 2994 | 2651 | 2444 | 2261 | 3137 | 2545 | 2435 | 2447 | 2291 | 3192 |
| 2215 | 0.17 | 0.10 | 1.60 | 0.11 | 1 | Miga1 | mitoguardin 1 | 1330 | 1410 | 1106 | 2475 | 2443 | 2213 | 2545 | 2498 | 2696 | 2919 | 2731 |
| 1919 | -0.11 | 0.14 | -0.79 | 0.43 | 1 | Atp8a1 | ATPase, aminophospholipid transporter ( | 1066 | 1121 | 944 | 2305 | 2397 | 2083 | 2544 | 1793 | 2577 | 2065 | 2218 |
| 2759 | 0.00 | 0.10 | -0.02 | 0.98 | 1 | Wdfy3 | WD repeat and FYVE domain containing | 3268 | 2753 | 2481 | 2324 | 2415 | 3729 | 2542 | 2592 | 2547 | 2514 | 3181 |
| 2340 | -0.12 | 0.12 | -0.97 | 0.33 | 1 | Bcl9 | B cell CLL/lymphoma 9 | 1398 | 1728 | 1265 | 2827 | 2585 | 3190 | 2542 | 2288 | 2814 | 2383 | 2716 |
| 2866 | 0.17 | 0.09 | 1.97 | 0.05 | 1 | Snx1 | sorting nexin 1 | 3369 | 3041 | 3453 | 2327 | 2709 | 2614 | 2541 | 2993 | 2725 | 2961 | 2787 |
| 2785 | 0.11 | 0.08 | 1.34 | 0.18 | 1 | Dars | aspartyl-tRNA synthetase | 3168 | 3074 | 3148 | 2667 | 2395 | 2589 | 2541 | 2993 | 2549 | 2513 | 2994 |
| 2019 | 0.00 | 0.10 | -0.02 | 0.99 | 1 | Ina | internexin neuronal intermediate filament | 1068 | 1121 | 923 | 2487 | 2410 | 2105 | 2540 | 2815 | 2463 | 2192 | 2086 |
| 2688 | -0.01 | 0.12 | -0.11 | 0.91 | 1 | Trp53inp1 | transformation related protein 53 inducibl | 4050 | 2941 | 2947 | 1585 | 2066 | 3628 | 2538 | 1364 | 2293 | 2474 | 3682 |
| 2412 | 0.17 | 0.09 | 1.80 | 0.07 | 1 | Rtl5 | retrotransposon Gag like 5 | 1135 | 1279 | 1279 | 2766 | 2396 | 3061 | 2538 | 3305 | 2615 | 2966 | 3188 |
| 2664 | 0.05 | 0.14 | 0.34 | 0.73 | 1 | Taldo1 | transaldolase 1 | 3290 | 2586 | 3380 | 2398 | 2806 | 2099 | 2537 | 3260 | 2425 | 2538 | 1983 |
| 2084 | -0.23 | 0.14 | -1.67 | 0.09 | 1 | Cldnd1 | claudin domain containing 1 | 1981 | 2201 | 2032 | 2201 | 2259 | 2015 | 2536 | 1444 | 2168 | 2244 | 1843 |
| 2838 | 0.13 | 0.15 | 0.86 | 0.39 | 1 | Chmp2a | charged multivesicular body protein 2A | 3155 | 2882 | 3803 | 2588 | 2661 | 2396 | 2535 | 3667 | 2348 | 2794 | 2388 |
| 2133 | 0.00 | 0.11 | -0.02 | 0.99 | 1 | Idh3a | isocitrate dehydrogenase 3 (NAD+) alpha | 2251 | 1932 | 2270 | 2100 | 2163 | 1736 | 2533 | 2267 | 2356 | 2020 | 1834 |
| 3411 | -0.02 | 0.12 | -0.19 | 0.85 | 1 | Map4 | microtubule-associated protein 4 | 7091 | 5612 | 5076 | 2969 | 2394 | 2045 | 2530 | 2664 | 2736 | 2553 | 1854 |
| 2788 | -0.02 | 0.11 | -0.16 | 0.88 | 1 | Erp29 | endoplasmic reticulum protein 29 | 2738 | 2202 | 2616 | 2922 | 3470 | 2714 | 2530 | 2638 | 3112 | 2889 | 2840 |
| 2339 | 0.05 | 0.09 | 0.57 | 0.57 | 1 | Pik3ca | phosphatidylinositol-4,5-bisphosphate 3-k | 2237 | 1872 | 1789 | 2332 | 2280 | 2601 | 2530 | 2418 | 2610 | 2458 | 2599 |
| 2221 | -0.03 | 0.09 | -0.33 | 0.74 | 1 | Pgm3 | phosphoglucomutase 3 | 1412 | 1328 | 1756 | 1799 | 2761 | 2981 | 2530 | 1798 | 2838 | 2298 | 2932 |
| 2751 | -0.05 | 0.07 | -0.69 | 0.49 | 1 | Ywhah | tyrosine 3-monooxygenase/tryptophan 5- | 3322 | 3558 | 3373 | 2801 | 2452 | 2386 | 2528 | 2711 | 2341 | 2521 | 2269 |
| 2684 | 0.19 | 0.15 | 1.26 | 0.21 | 1 | Atp5d | ATP synthase, H+ transporting, mitochond | 2996 | 2795 | 3440 | 2395 | 2532 | 1977 | 2528 | 3690 | 2340 | 2805 | 2023 |

| | | | | | | | | | | | | | | | | | | |
|---|---|---|---|---|---|---|---|---|---|---|---|---|---|---|---|---|---|---|
| 2745 | -0.06 | 0.09 | -0.67 | 0.50 | 1 | Ldb1 | LIM domain binding 1 | 2597 | 2489 | 2647 | 2822 | 2438 | 3729 | 2526 | 2402 | 2483 | 2790 | 3278 |
| 2391 | -0.10 | 0.10 | -0.95 | 0.34 | 1 | Snx27 | sorting nexin family member 27 | 2331 | 2165 | 1961 | 2533 | 2541 | 2647 | 2525 | 2198 | 2581 | 2194 | 2623 |
| 2187 | 0.12 | 0.10 | 1.19 | 0.24 | 1 | Extl2 | exostosin-like glycosyltransferase 2 | 1545 | 1438 | 1531 | 2219 | 2587 | 2049 | 2524 | 2892 | 2489 | 2567 | 2222 |
| 3178 | 0.09 | 0.12 | 0.76 | 0.45 | 1 | Rpl23a | ribosomal protein L23A | 6158 | 5057 | 6275 | 2096 | 2256 | 1593 | 2523 | 2721 | 2474 | 2160 | 1645 |
| 2795 | 0.05 | 0.09 | 0.61 | 0.54 | 1 | Brd1 | bromodomain containing 1 | 1979 | 2348 | 1833 | 3135 | 2723 | 3729 | 2523 | 3135 | 2867 | 3013 | 3460 |
| 2617 | -0.03 | 0.09 | -0.31 | 0.76 | 1 | Pcnx3 | pecanex homolog 3 | 2533 | 2300 | 2406 | 2513 | 2817 | 3045 | 2521 | 2699 | 2858 | 2523 | 2574 |
| 2368 | 0.08 | 0.10 | 0.79 | 0.43 | 1 | Tmem94 | transmembrane protein 94 | 1760 | 2097 | 1768 | 2706 | 2235 | 2428 | 2520 | 2955 | 2625 | 2869 | 2087 |
| 2246 | 0.07 | 0.10 | 0.75 | 0.46 | 1 | Usp45 | ubiquitin specific petidase 45 | 1312 | 1594 | 1114 | 2520 | 2079 | 2990 | 2520 | 2603 | 2445 | 2487 | 3046 |
| 2526 | -0.16 | 0.14 | -1.15 | 0.25 | 1 | Fto | fat mass and obesity associated | 3176 | 2646 | 2978 | 2908 | 2624 | 1883 | 2519 | 1855 | 2592 | 2946 | 1662 |
| 2507 | -0.07 | 0.09 | -0.78 | 0.43 | 1 | Ints1 | integrator complex subunit 1 | 2704 | 2539 | 2544 | 2626 | 2532 | 2440 | 2519 | 2222 | 2650 | 2607 | 2192 |
| 2324 | 0.01 | 0.09 | 0.13 | 0.90 | 1 | Icmt | isoprenylcysteine carboxyl methyltransfer | 2684 | 2338 | 2254 | 2044 | 2422 | 2138 | 2514 | 2194 | 2393 | 2402 | 2178 |
| 2954 | -0.07 | 0.13 | -0.55 | 0.58 | 1 | Kmt2d | lysine (K)-specific methyltransferase 2D | 3011 | 3173 | 2825 | 2333 | 2445 | 4977 | 2512 | 2909 | 2379 | 2436 | 3493 |
| 2389 | -0.11 | 0.14 | -0.83 | 0.40 | 1 | Soga1 | suppressor of glucose, autophagy associ | 2117 | 2243 | 1906 | 2485 | 3141 | 2183 | 2512 | 2687 | 3142 | 2169 | 1697 |
| 2682 | 0.12 | 0.10 | 1.20 | 0.23 | 1 | Armc10 | armadillo repeat containing 10 | 1833 | 2852 | 2421 | 3049 | 2682 | 2499 | 2511 | 3443 | 2692 | 2710 | 2810 |
| 2336 | 0.01 | 0.08 | 0.12 | 0.90 | 1 | Inpp5a | inositol polyphosphate-5-phosphatase A | 2238 | 2010 | 2088 | 2479 | 2450 | 2213 | 2511 | 2494 | 2565 | 2369 | 2281 |
| 2012 | -0.05 | 0.08 | -0.63 | 0.53 | 1 | Pip4p1 | phosphatidylinositol-4,5-bisphosphate 4-| | 1328 | 1151 | 1375 | 2226 | 2334 | 2225 | 2511 | 1961 | 2323 | 2485 | 2217 |
| 3026 | 0.02 | 0.14 | 0.18 | 0.86 | 1 | Gpx1 | glutathione peroxidase 1 | 5876 | 3922 | 4322 | 2667 | 2580 | 1757 | 2509 | 2988 | 2326 | 2462 | 1879 |
| 2360 | 0.03 | 0.08 | 0.38 | 0.70 | 1 | Tpst2 | protein-tyrosine sulfotransferase 2 | 2161 | 2157 | 2206 | 2516 | 2628 | 1953 | 2508 | 2546 | 2525 | 2820 | 1939 |
| 2337 | 0.05 | 0.11 | 0.47 | 0.63 | 1 | Ptdss1 | phosphatidylserine synthase 1 | 2578 | 2188 | 2022 | 2247 | 2622 | 1923 | 2506 | 2694 | 2289 | 2446 | 2189 |
| 2683 | 0.08 | 0.12 | 0.73 | 0.47 | 1 | Tmem39a | transmembrane protein 39a | 1362 | 1697 | 1589 | 2339 | 3110 | 4166 | 2504 | 2429 | 3365 | 2843 | 4107 |
| 2347 | 0.07 | 0.09 | 0.72 | 0.47 | 1 | Aggf1 | angiogenic factor with G patch and FHA d | 2108 | 1985 | 1946 | 2493 | 2193 | 2459 | 2503 | 2760 | 2367 | 2257 | 2742 |
| 3583 | 0.01 | 0.14 | 0.08 | 0.94 | 1 | Ogt | O-linked N-acetylglucosamine (GlcNAc) t | 1953 | 4010 | 1854 | 3250 | 1977 | 8119 | 2501 | 3332 | 2532 | 2018 | 7866 |
| 2543 | 0.14 | 0.09 | 1.60 | 0.11 | 1 | Tmem167 | transmembrane protein 167 | 1754 | 1873 | 1987 | 2581 | 2699 | 2815 | 2501 | 2913 | 3351 | 2386 | 3113 |
| 2439 | -0.09 | 0.11 | -0.79 | 0.43 | 1 | Hs6st1 | heparan sulfate 6-O-sulfotransferase 1 | 2828 | 3070 | 2893 | 2236 | 2450 | 2080 | 2500 | 1863 | 2478 | 2561 | 1865 |
| 2399 | 0.04 | 0.09 | 0.50 | 0.62 | 1 | Usp15 | ubiquitin specific peptidase 15 | 1869 | 1764 | 1766 | 2117 | 2567 | 3159 | 2500 | 2235 | 2777 | 2425 | 3215 |
| 2210 | 0.16 | 0.09 | 1.82 | 0.07 | 1 | Aida | axin interactor, dorsalization associated | 1609 | 1420 | 1393 | 2508 | 2516 | 1865 | 2499 | 2830 | 2514 | 3086 | 2073 |
| 2086 | 0.07 | 0.14 | 0.50 | 0.62 | 1 | Pdzd4 | PDZ domain containing 4 | 1429 | 1326 | 1586 | 2315 | 2549 | 1686 | 2499 | 3105 | 2143 | 2623 | 1682 |
| 2571 | 0.08 | 0.11 | 0.67 | 0.50 | 1 | Psmc3 | proteasome (prosome, macropain) 26S su | 3808 | 2641 | 2709 | 2182 | 2570 | 2040 | 2498 | 2618 | 2285 | 2853 | 2081 |
| 2422 | 0.08 | 0.10 | 0.82 | 0.41 | 1 | Plaa | phospholipase A2, activating protein | 1807 | 1963 | 1850 | 2402 | 2604 | 2682 | 2497 | 2411 | 2965 | 2330 | 3127 |
| 2315 | 0.00 | 0.12 | -0.02 | 0.99 | 1 | Appl1 | adaptor protein, phosphotyrosine interact | 2667 | 2679 | 2610 | 1885 | 2060 | 2314 | 2497 | 1963 | 2377 | 1887 | 2527 |
| 2503 | -0.01 | 0.10 | -0.13 | 0.90 | 1 | Adar | adenosine deaminase, RNA-specific | 2774 | 2674 | 2381 | 2597 | 2391 | 2376 | 2496 | 2324 | 2385 | 3005 | 2126 |
| 2459 | 0.00 | 0.11 | 0.02 | 0.98 | 1 | Coro1c | coronin, actin binding protein 1C | 3070 | 2944 | 2478 | 2494 | 2316 | 1968 | 2496 | 2724 | 2432 | 2058 | 2065 |
| 2454 | 0.02 | 0.11 | 0.21 | 0.84 | 1 | Fbxo38 | F-box protein 38 | 2251 | 1861 | 2128 | 2482 | 2511 | 2805 | 2495 | 2647 | 2715 | 2196 | 2905 |
| 2344 | 0.12 | 0.11 | 1.09 | 0.28 | 1 | Ap1s1 | adaptor protein complex AP-1, sigma 1 | 2007 | 1747 | 2250 | 2442 | 2678 | 1851 | 2493 | 3107 | 2606 | 2639 | 1960 |
| 2812 | -0.04 | 0.09 | -0.41 | 0.68 | 1 | Atp11a | ATPase, class VI, type 11A | 3197 | 2972 | 2743 | 2703 | 2322 | 3678 | 2490 | 2302 | 2555 | 2646 | 3326 |
| 2448 | -0.08 | 0.13 | -0.64 | 0.52 | 1 | 1110004F10Rik | RIKEN cDNA 1110004F10 gene | 2956 | 3092 | 2771 | 1949 | 2615 | 2242 | 2489 | 1738 | 2397 | 2539 | 2136 |
| 2094 | -0.05 | 0.14 | -0.34 | 0.73 | 1 | Rab39b | RAB39B, member RAS oncogene family | 893 | 1144 | 1154 | 2168 | 2843 | 2535 | 2489 | 1796 | 3018 | 2253 | 2747 |
| 2579 | 0.04 | 0.13 | 0.30 | 0.76 | 1 | Rb1cc1 | RB1-inducible coiled-coil 1 | 1949 | 2115 | 1730 | 2296 | 2975 | 3378 | 2487 | 2201 | 3184 | 2749 | 3301 |
| 2517 | 0.06 | 0.10 | 0.65 | 0.51 | 1 | Txndc11 | thioredoxin domain containing 11 | 2144 | 1724 | 1917 | 2052 | 2876 | 3330 | 2487 | 2260 | 2893 | 2939 | 3065 |
| 2604 | 0.08 | 0.13 | 0.66 | 0.51 | 1 | Fau | Finkel-Biskis-Reilly murine sarcoma virus | 4149 | 3475 | 4325 | 1827 | 2237 | 1536 | 2486 | 2558 | 2064 | 2531 | 1450 |
| 2186 | 0.01 | 0.11 | 0.13 | 0.90 | 1 | Pik3r4 | phosphoinositide-3-kinase regulatory sub | 1672 | 1874 | 1538 | 2270 | 2426 | 2219 | 2483 | 2844 | 2479 | 2209 | 2032 |
| 3007 | -0.02 | 0.09 | -0.17 | 0.86 | 1 | Cggbp1 | CGG triplet repeat binding protein 1 | 4248 | 4332 | 3851 | 3028 | 2425 | 2461 | 2480 | 2655 | 2807 | 2443 | 2346 |
| 2586 | 0.04 | 0.07 | 0.54 | 0.59 | 1 | Ube2l3 | ubiquitin-conjugating enzyme E2L 3 | 3064 | 3258 | 3124 | 2401 | 2617 | 1855 | 2480 | 2595 | 2619 | 2602 | 1827 |
| 2491 | 0.21 | 0.16 | 1.28 | 0.20 | 1 | Cox6b1 | cytochrome c oxidase, subunit 6B1 | 2943 | 2181 | 3105 | 2290 | 2239 | 1798 | 2479 | 3538 | 2399 | 2710 | 1714 |
| 2442 | 0.13 | 0.09 | 1.45 | 0.15 | 1 | Slc25a46 | solute carrier family 25, member 46 | 1718 | 2260 | 1882 | 2585 | 2709 | 2244 | 2478 | 2965 | 2889 | 2728 | 2402 |
| 1784 | -0.11 | 0.09 | -1.15 | 0.25 | 1 | Prkag2 | protein kinase, AMP-activated, gamma 2 | | 1135 | 812 | 839 | 2084 | 2094 | 2084 | 2476 | 1825 | 1946 | 2218 | 2109 |
| 2415 | 0.08 | 0.14 | 0.60 | 0.55 | 1 | Sh3d19 | SH3 domain protein D19 | 2218 | 1831 | 1699 | 2736 | 2478 | 2378 | 2475 | 3512 | 2596 | 2304 | 2337 |
| 2033 | -0.05 | 0.12 | -0.41 | 0.68 | 1 | Ndufv3 | NADH:ubiquinone oxidoreductase core s | 1621 | 1452 | 1955 | 2257 | 2416 | 1680 | 2475 | 2558 | 2159 | 2150 | 1642 |
| 1813 | 0.06 | 0.16 | 0.37 | 0.71 | 1 | Chrm3 | cholinergic receptor, muscarinic 3, cardia | 372 | 187 | 248 | 2416 | 2343 | 2152 | 2471 | 2379 | 2642 | 2397 | 2340 |
| 2738 | -0.19 | 0.12 | -1.64 | 0.10 | 1 | Myh10 | myosin, heavy polypeptide 10, non-muscl | 4824 | 3528 | 4517 | 2358 | 2193 | 2185 | 2469 | 2156 | 2063 | 1878 | 1945 |
| 2604 | -0.06 | 0.09 | -0.68 | 0.50 | 1 | Psmd7 | proteasome (prosome, macropain) 26S su | 2718 | 2842 | 2756 | 2841 | 2736 | 2320 | 2469 | 3032 | 2307 | 2378 | 2245 |
| 2217 | 0.01 | 0.09 | 0.11 | 0.92 | 1 | Cog5 | component of oligomeric golgi complex 5 | 2086 | 1839 | 2059 | 2223 | 2332 | 2128 | 2469 | 1924 | 2354 | 2680 | 2296 |
| 2202 | -0.05 | 0.07 | -0.63 | 0.53 | 1 | Ppp2r5e | protein phosphatase 2, regulatory subuni | 2261 | 1954 | 2094 | 2075 | 2337 | 2221 | 2469 | 2076 | 2324 | 2320 | 2091 |
| 2285 | 0.03 | 0.09 | 0.35 | 0.73 | 1 | Ufsp2 | UFM1-specific peptidase 2 | 1611 | 1512 | 1533 | 2190 | 2633 | 2816 | 2468 | 2087 | 2688 | 2534 | 3062 |
| 3354 | 0.11 | 0.14 | 0.76 | 0.45 | 1 | Luc7l3 | LUC7-like 3 (S. cerevisiae) | 2304 | 3404 | 2979 | 2758 | 2135 | 6007 | 2467 | 3932 | 1916 | 2304 | 6692 |
| 2512 | -0.13 | 0.11 | -1.20 | 0.23 | 1 | Cep350 | centrosomal protein 350 | 2583 | 2479 | 2092 | 2322 | 2138 | 3754 | 2467 | 1997 | 2276 | 1940 | 3584 |
| 2232 | -0.12 | 0.10 | -1.30 | 0.19 | 1 | Rhbdd2 | rhomboid domain containing 2 | 1754 | 2035 | 2134 | 2498 | 2526 | 2220 | 2467 | 2238 | 2241 | 2445 | 1991 |
| 2130 | -0.12 | 0.11 | -1.07 | 0.28 | 1 | Ppp1r13b | protein phosphatase 1, regulatory subuni | 1956 | 2434 | 1994 | 2258 | 2098 | 2050 | 2467 | 1972 | 2022 | 2340 | 1835 |
| 2812 | 0.05 | 0.09 | 0.51 | 0.61 | 1 | Psma7 | proteasome subunit alpha 7 | 3423 | 3029 | 3800 | 2572 | 2659 | 2475 | 2466 | 2794 | 2594 | 2473 | 2649 |
| 2356 | -0.02 | 0.10 | -0.20 | 0.84 | 1 | Fundc2 | FUN14 domain containing 2 | 2685 | 2781 | 2615 | 2563 | 2374 | 1599 | 2466 | 2094 | 2365 | 2650 | 1723 |

| | | | | | | | | | | | | | | | | | |
|---|---|---|---|---|---|---|---|---|---|---|---|---|---|---|---|---|---|
| 2303 | -0.02 | 0.09 | -0.28 | 0.78 | 1 | Nelfb | negative elongation factor complex memb | 2364 | 1962 | 2218 | 2225 | 2306 | 2483 | 2466 | 2414 | 2220 | 2330 | 2340 |
| 2348 | -0.08 | 0.10 | -0.79 | 0.43 | 1 | Wbp11 | WW domain binding protein 11 | 2191 | 2487 | 2237 | 2707 | 2379 | 2166 | 2465 | 2173 | 2496 | 2481 | 2046 |
| 2640 | 0.00 | 0.10 | -0.03 | 0.97 | 1 | Fam3c | family with sequence similarity 3, member | 3820 | 3991 | 4266 | 2427 | 2030 | 1603 | 2462 | 2374 | 2089 | 2206 | 1771 |
| 2469 | -0.09 | 0.11 | -0.79 | 0.43 | 1 | Senp5 | SUMO/sentrin specific peptidase 5 | 2214 | 2289 | 2036 | 2408 | 2832 | 2901 | 2462 | 2458 | 2709 | 2044 | 2807 |
| 2517 | -0.04 | 0.09 | -0.45 | 0.65 | 1 | Sin3a | transcriptional regulator, SIN3A (yeast) | 2506 | 2794 | 2448 | 2461 | 2731 | 2458 | 2461 | 2387 | 2566 | 2568 | 2309 |
| 2486 | 0.04 | 0.08 | 0.45 | 0.65 | 1 | Kmt5b | lysine methyltransferase 5B | 2401 | 2464 | 2282 | 2366 | 2186 | 2971 | 2461 | 2723 | 2324 | 2349 | 2823 |
| 2216 | -0.01 | 0.10 | -0.09 | 0.93 | 1 | Max | Max protein | 1842 | 1837 | 2288 | 2340 | 2320 | 2111 | 2461 | 2464 | 2259 | 2338 | 2110 |
| 2246 | -0.12 | 0.15 | -0.80 | 0.42 | 1 | Gatad1 | GATA zinc finger domain containing 1 | 2484 | 2744 | 2295 | 2205 | 2506 | 1728 | 2460 | 1632 | 2281 | 2794 | 1574 |
| 2246 | -0.02 | 0.09 | -0.21 | 0.84 | 1 | Rasa1 | RAS p21 protein activator 1 | 2027 | 1694 | 1963 | 2100 | 2289 | 2713 | 2457 | 1884 | 2601 | 2420 | 2560 |
| 2199 | -0.06 | 0.10 | -0.57 | 0.57 | 1 | Wdr45b | WD repeat domain 45B | 1981 | 2151 | 2031 | 2089 | 2553 | 2103 | 2457 | 1961 | 2346 | 2317 | 2198 |
| 2091 | 0.00 | 0.09 | -0.02 | 0.98 | 1 | Hectd3 | HECT domain E3 ubiquitin protein ligase | 2034 | 1541 | 1503 | 2153 | 2274 | 2063 | 2457 | 2202 | 2145 | 2662 | 1970 |
| 2586 | 0.16 | 0.11 | 1.39 | 0.17 | 1 | Slmap | sarcolemma associated protein | 3271 | 2291 | 2407 | 2264 | 2488 | 2485 | 2456 | 2740 | 2747 | 2726 | 2572 |
| 2357 | -0.02 | 0.11 | -0.22 | 0.82 | 1 | Psmd12 | proteasome (prosome, macropain) 26S su | 2364 | 2240 | 2180 | 2252 | 2545 | 2392 | 2456 | 2404 | 2655 | 2004 | 2436 |
| 2753 | -0.12 | 0.10 | -1.14 | 0.26 | 1 | Mast2 | microtubule associated serine/threonine k | 3777 | 3072 | 3859 | 2398 | 2963 | 2320 | 2454 | 2293 | 2686 | 2701 | 1763 |
| 2418 | 0.02 | 0.10 | 0.21 | 0.83 | 1 | Eif3i | eukaryotic translation initiation factor 3, su | 3367 | 2959 | 3403 | 1994 | 2294 | 1648 | 2452 | 2246 | 1953 | 2517 | 1763 |
| 2655 | 0.09 | 0.13 | 0.69 | 0.49 | 1 | Hint1 | histidine triad nucleotide binding protein 1 | 3513 | 3069 | 4200 | 2413 | 2337 | 1702 | 2451 | 2761 | 2254 | 2829 | 1680 |
| 2225 | 0.18 | 0.12 | 1.52 | 0.13 | 1 | Aig1 | androgen-induced 1 | 1653 | 1515 | 1941 | 2570 | 2249 | 1811 | 2449 | 3057 | 2571 | 2567 | 2089 |
| 2159 | 0.06 | 0.12 | 0.52 | 0.61 | 1 | Gsr | glutathione reductase | 1814 | 1849 | 1730 | 1998 | 2639 | 1880 | 2449 | 2098 | 2432 | 2956 | 1904 |
| 2426 | -0.05 | 0.09 | -0.51 | 0.61 | 1 | Cpne3 | copine III | 3019 | 2507 | 2447 | 2528 | 2341 | 2199 | 2448 | 2380 | 2393 | 2286 | 2136 |
| 2237 | -0.15 | 0.11 | -1.39 | 0.17 | 1 | B4galt3 | UDP-Gal:betaGlcNAc beta 1,4-galactosyl | 1962 | 2368 | 2515 | 2332 | 2309 | 2243 | 2448 | 1970 | 2172 | 2203 | 2089 |
| 2110 | -0.09 | 0.11 | -0.77 | 0.44 | 1 | Rnf2 | ring finger protein 2 | 1765 | 1726 | 1858 | 2563 | 2084 | 2107 | 2448 | 1966 | 2176 | 2280 | 2238 |
| 2614 | 0.07 | 0.10 | 0.71 | 0.48 | 1 | Scyl1 | SCY1-like 1 (S. cerevisiae) | 2193 | 1825 | 2139 | 2608 | 2917 | 3051 | 2447 | 2502 | 3085 | 3072 | 2914 |
| 2256 | 0.08 | 0.10 | 0.78 | 0.43 | 1 | Tmem183a | transmembrane protein 183A | 2095 | 1911 | 2008 | 2235 | 2256 | 2219 | 2447 | 2705 | 2290 | 2277 | 2377 |
| 2186 | 0.03 | 0.09 | 0.40 | 0.69 | 1 | Pi4k2a | phosphatidylinositol 4-kinase type 2 alpha | 1688 | 1602 | 1537 | 2214 | 2549 | 2274 | 2447 | 2241 | 2456 | 2736 | 2299 |
| 2887 | -0.02 | 0.10 | -0.24 | 0.81 | 1 | Kpna4 | karyopherin (importin) alpha 4 | 3349 | 4458 | 2838 | 2906 | 2405 | 2889 | 2446 | 2867 | 2490 | 2293 | 2820 |
| 2810 | 0.05 | 0.10 | 0.55 | 0.58 | 1 | Ddx23 | DEAD (Asp-Glu-Ala-Asp) box polypeptide | 2910 | 3074 | 2861 | 2648 | 2941 | 2790 | 2446 | 2666 | 2649 | 3053 | 2869 |
| 2154 | -0.13 | 0.19 | -0.70 | 0.49 | 1 | Syn1 | synapsin I | 1527 | 2098 | 1896 | 2669 | 2560 | 1804 | 2446 | 1493 | 2666 | 2811 | 1720 |
| 2589 | -0.03 | 0.10 | -0.25 | 0.80 | 1 | Btrc | beta-transducin repeat containing protein | 2039 | 2693 | 2429 | 2694 | 2423 | 3224 | 2445 | 2577 | 2418 | 2678 | 2857 |
| 2453 | 0.12 | 0.10 | 1.17 | 0.24 | 1 | Tm9sf1 | transmembrane 9 superfamily member 1 | 2000 | 1560 | 1783 | 2626 | 2736 | 2552 | 2445 | 2748 | 2754 | 3149 | 2635 |
| 2018 | 0.03 | 0.10 | 0.27 | 0.79 | 1 | Uvrag | UV radiation resistance associated gene | 1945 | 1295 | 1941 | 1923 | 2165 | 1894 | 2444 | 1831 | 2367 | 2391 | 2008 |
| 2366 | -0.07 | 0.10 | -0.71 | 0.48 | 1 | Skiv2l | superkiller viralicidic activity 2-like (S. cere | 2497 | 2149 | 2174 | 2512 | 2317 | 2547 | 2443 | 2633 | 2037 | 2480 | 2234 |
| 2344 | -0.04 | 0.08 | -0.51 | 0.61 | 1 | Mat2b | methionine adenosyltransferase II, beta | 2283 | 2088 | 2471 | 2320 | 2536 | 2314 | 2443 | 2336 | 2259 | 2504 | 2229 |
| 2484 | -0.17 | 0.11 | -1.59 | 0.11 | 1 | Gprasp2 | G protein-coupled receptor associated so | 2403 | 2935 | 2752 | 2037 | 2528 | 3199 | 2442 | 2180 | 2195 | 1954 | 2697 |
| 2377 | 0.03 | 0.08 | 0.33 | 0.74 | 1 | Aacs | acetoacetyl-CoA synthetase | 1665 | 1834 | 1697 | 2560 | 2605 | 2778 | 2442 | 2687 | 2636 | 2651 | 2592 |
| 2584 | -0.21 | 0.11 | -1.85 | 0.06 | 1 | Inppl1 | inositol polyphosphate phosphatase-like | 2508 | 2878 | 2505 | 2951 | 2670 | 2963 | 2441 | 2092 | 2561 | 2489 | 2372 |
| 1686 | 0.01 | 0.18 | 0.06 | 0.95 | 1 | Tmem150c | transmembrane protein 150C | 425 | 624 | 565 | 2041 | 2280 | 1678 | 2440 | 1623 | 2419 | 2554 | 1898 |
| 2361 | -0.01 | 0.11 | -0.11 | 0.91 | 1 | Mzt1 | mitotic spindle organizing protein 1 | 2035 | 2281 | 2181 | 2629 | 2681 | 2039 | 2439 | 2367 | 2707 | 2123 | 2486 |
| 2333 | 0.08 | 0.11 | 0.77 | 0.44 | 1 | Rbm47 | RNA binding motif protein 47 | 1165 | 1354 | 1542 | 2508 | 2611 | 2947 | 2438 | 2701 | 2952 | 2845 | 2605 |
| 2253 | 0.10 | 0.14 | 0.75 | 0.45 | 1 | Mob1b | MOB kinase activator 1B | 1024 | 1864 | 1217 | 2788 | 2451 | 2260 | 2438 | 3088 | 2905 | 2669 | 2079 |
| 2342 | 0.08 | 0.13 | 0.61 | 0.54 | 1 | Mbd3 | methyl-CpG binding domain protein 3 | 2660 | 1983 | 2736 | 2037 | 2502 | 1948 | 2437 | 2772 | 2291 | 2590 | 1803 |
| 2540 | 0.02 | 0.12 | 0.17 | 0.86 | 1 | Ybx3 | Y box protein 3 | 3511 | 2525 | 3557 | 2262 | 2492 | 1912 | 2436 | 2638 | 2392 | 2431 | 1789 |
| 1769 | -0.22 | 0.12 | -1.79 | 0.07 | 1 | Upb1 | ureidopropionase, beta | 987 | 1191 | 1220 | 2108 | 2035 | 2060 | 2436 | 1806 | 1999 | 1681 | 1933 |
| 2060 | 0.21 | 0.14 | 1.55 | 0.12 | 1 | Tanc2 | tetratricopeptide repeat, ankyrin repeat ar | 2072 | 1655 | 1781 | 2016 | 1760 | 1710 | 2435 | 2986 | 2165 | 2360 | 1721 |
| 2720 | 0.05 | 0.08 | 0.60 | 0.55 | 1 | Gabpb2 | GA repeat binding protein, beta 2 | 3042 | 3084 | 3002 | 2480 | 2257 | 3043 | 2434 | 2703 | 2333 | 2298 | 3243 |
| 1859 | 0.01 | 0.14 | 0.09 | 0.93 | 1 | Fhdc1 | FH2 domain containing 1 | 202 | 163 | 308 | 2595 | 2834 | 1948 | 2434 | 2677 | 3142 | 2082 | 2060 |
| 2474 | 0.01 | 0.08 | 0.16 | 0.87 | 1 | Tmem259 | transmembrane protein 259 | 2364 | 2230 | 2317 | 2339 | 2633 | 2700 | 2433 | 2519 | 2369 | 2666 | 2641 |
| 2282 | 0.02 | 0.08 | 0.19 | 0.85 | 1 | Txndc16 | thioredoxin domain containing 16 | 2094 | 2257 | 1954 | 2636 | 2430 | 1877 | 2433 | 2587 | 2394 | 2338 | 2107 |
| 2786 | 0.08 | 0.10 | 0.85 | 0.39 | 1 | Npepl1 | aminopeptidase-like 1 | 1493 | 1603 | 1774 | 2943 | 2775 | 4444 | 2432 | 3246 | 2713 | 2881 | 4345 |
| 2249 | -0.01 | 0.09 | -0.13 | 0.89 | 1 | Eif4e | eukaryotic translation initiation factor 4E | 2179 | 2190 | 2313 | 2165 | 2439 | 2032 | 2431 | 1959 | 2391 | 2450 | 2189 |
| 2008 | 0.20 | 0.14 | 1.43 | 0.15 | 1 | Kl | klotho | 1935 | 2734 | 1791 | 2130 | 1388 | 1366 | 2431 | 2308 | 1715 | 2655 | 1635 |
| 2367 | 0.02 | 0.09 | 0.21 | 0.83 | 1 | Angel2 | angel homolog 2 | 1699 | 1964 | 1714 | 2846 | 2555 | 2439 | 2430 | 2641 | 2649 | 2330 | 2773 |
| 1965 | -0.11 | 0.16 | -0.71 | 0.48 | 1 | Ube2q2 | ubiquitin-conjugating enzyme E2Q family | 1233 | 1357 | 1631 | 2244 | 2320 | 2001 | 2430 | 1496 | 2195 | 2572 | 2138 |
| 2220 | -0.08 | 0.09 | -0.95 | 0.34 | 1 | Hipk2 | homeodomain interacting protein kinase 2 | 1839 | 2063 | 2062 | 2495 | 2467 | 2079 | 2429 | 2582 | 2513 | 2015 | 1872 |
| 2369 | 0.12 | 0.10 | 1.16 | 0.25 | 1 | Ankle2 | ankyrin repeat and LEM domain containir | 1784 | 1674 | 1620 | 2424 | 2893 | 2322 | 2428 | 2805 | 3068 | 2474 | 2562 |
| 2276 | 0.15 | 0.09 | 1.75 | 0.08 | 1 | Parn | poly(A)-specific ribonuclease (deadenyla | 1524 | 1275 | 1636 | 2304 | 2799 | 2236 | 2425 | 2720 | 2795 | 2822 | 2499 |
| 2317 | 0.10 | 0.09 | 1.12 | 0.26 | 1 | Gm16286 | thioredoxin-like 4A | 2174 | 2265 | 2255 | 2259 | 2173 | 2207 | 2424 | 2714 | 2277 | 2548 | 2194 |
| 2807 | -0.06 | 0.10 | -0.59 | 0.56 | 1 | Trim56 | tripartite motif-containing 56 | 3728 | 3510 | 3745 | 2834 | 2589 | 2295 | 2423 | 2732 | 2682 | 2079 | 2260 |
| 2448 | -0.01 | 0.09 | -0.15 | 0.88 | 1 | Prpf18 | pre-mRNA processing factor 18 | 2442 | 2809 | 2779 | 2396 | 2428 | 2244 | 2422 | 2166 | 2599 | 2391 | 2251 |
| 2224 | -0.04 | 0.09 | -0.42 | 0.67 | 1 | Atp7a | ATPase, Cu++ transporting, alpha polype | 1194 | 1363 | 1310 | 2707 | 2517 | 2769 | 2422 | 2352 | 2680 | 2200 | 2948 |
| 2882 | -0.04 | 0.12 | -0.30 | 0.76 | 1 | Sp1 | trans-acting transcription factor 1 | 3197 | 3096 | 2823 | 2865 | 2979 | 3156 | 2421 | 2266 | 2923 | 2862 | 3117 |

| | | | | | | | | | | | | | | | | | |
|---|---|---|---|---|---|---|---|---|---|---|---|---|---|---|---|---|---|
| 2561 | 0.12 | 0.09 | 1.36 | 0.17 | 1 | Mark3 | MAP/microtubule affinity regulating kinase | 2085 | 2334 | 2329 | 2319 | 2308 | 3251 | 2421 | 2451 | 2617 | 2709 | 3345 |
| 2395 | 0.03 | 0.11 | 0.32 | 0.75 | 1 | Exoc3 | exocyst complex component 3 | 2643 | 2475 | 2494 | 2210 | 2399 | 2198 | 2421 | 1861 | 2486 | 2714 | 2442 |
| 2324 | -0.06 | 0.09 | -0.64 | 0.52 | 1 | Lpcat3 | lysophosphatidylcholine acyltransferase 3 | 1504 | 1614 | 1766 | 3065 | 2835 | 2236 | 2421 | 2615 | 2772 | 2595 | 2147 |
| 2102 | -0.07 | 0.14 | -0.54 | 0.59 | 1 | Osbpl8 | oxysterol binding protein-like 8 | 1606 | 1677 | 1373 | 2554 | 2243 | 2255 | 2421 | 1806 | 2550 | 2263 | 2377 |
| 2131 | -0.04 | 0.11 | -0.36 | 0.72 | 1 | Micu1 | mitochondrial calcium uptake 1 | 2026 | 2049 | 1959 | 2383 | 2466 | 1588 | 2419 | 1914 | 2309 | 2525 | 1799 |
| 2071 | 0.09 | 0.12 | 0.74 | 0.46 | 1 | Asic1 | acid-sensing (proton-gated) ion channel 1 | 613 | 971 | 1017 | 2661 | 2620 | 2042 | 2418 | 2734 | 3162 | 2362 | 2176 |
| 2483 | 0.01 | 0.09 | 0.09 | 0.92 | 1 | Tbc1d20 | TBC1 domain family, member 20 | 2377 | 2544 | 2330 | 2793 | 2336 | 2428 | 2417 | 3110 | 2172 | 2422 | 2383 |
| 2199 | -0.07 | 0.12 | -0.61 | 0.54 | 1 | Zfp9 | zinc finger protein 9 | 1263 | 1300 | 1732 | 2132 | 2271 | 3323 | 2417 | 1982 | 2767 | 1850 | 3151 |
| 2077 | -0.09 | 0.12 | -0.74 | 0.46 | 1 | Tbc1d23 | TBC1 domain family, member 23 | 1989 | 1743 | 1484 | 1812 | 1842 | 3010 | 2417 | 2094 | 1710 | 1789 | 2955 |
| 2369 | 0.09 | 0.10 | 0.92 | 0.36 | 1 | Cops3 | COP9 signalosome subunit 3 | 1919 | 2237 | 2027 | 2404 | 2355 | 2445 | 2416 | 2297 | 2473 | 2745 | 2741 |
| 2031 | -0.17 | 0.15 | -1.11 | 0.27 | 1 | Scn9a | sodium channel, voltage-gated, type IX, a | 1395 | 1966 | 1863 | 1922 | 2070 | 2606 | 2416 | 1541 | 2018 | 1924 | 2618 |
| 2515 | 0.10 | 0.10 | 0.92 | 0.36 | 1 | Cdk8 | cyclin-dependent kinase 8 | 958 | 1277 | 1193 | 2404 | 2456 | 4529 | 2415 | 2553 | 2881 | 2602 | 4400 |
| 2345 | 0.02 | 0.10 | 0.23 | 0.82 | 1 | Flywch1 | FLYWCH-type zinc finger 1 | 2340 | 2255 | 2314 | 2467 | 2341 | 2126 | 2415 | 2785 | 2084 | 2565 | 2102 |
| 2319 | -0.04 | 0.10 | -0.37 | 0.71 | 1 | Got2 | glutamatic-oxaloacetic transaminase 2, m | 2581 | 2503 | 2403 | 2428 | 2574 | 1674 | 2415 | 2138 | 2480 | 2759 | 1559 |
| 2502 | -0.07 | 0.09 | -0.82 | 0.41 | 1 | Wdr13 | WD repeat domain 13 | 2289 | 1874 | 1915 | 2391 | 2303 | 3897 | 2414 | 2237 | 2328 | 2234 | 3643 |
| 2400 | -0.08 | 0.10 | -0.80 | 0.43 | 1 | Ankrd13a | ankyrin repeat domain 13a | 3295 | 2501 | 2659 | 2248 | 2404 | 2146 | 2412 | 2038 | 2349 | 2210 | 2134 |
| 2033 | -0.08 | 0.10 | -0.81 | 0.42 | 1 | Dip2c | disco interacting protein 2 homolog C | 1576 | 1517 | 1596 | 2212 | 2294 | 2148 | 2411 | 1851 | 2518 | 2230 | 2014 |
| 2218 | 0.05 | 0.15 | 0.36 | 0.72 | 1 | Mgat1 | mannoside acetylglucosaminyltransferase | 3462 | 2496 | 2966 | 1770 | 1877 | 1563 | 2410 | 2407 | 2079 | 1824 | 1540 |
| 1971 | -0.06 | 0.10 | -0.61 | 0.54 | 1 | Vmp1 | vacuole membrane protein 1 | 1549 | 1323 | 1634 | 2208 | 2446 | 1729 | 2410 | 1980 | 2502 | 1984 | 1918 |
| 2206 | -0.06 | 0.10 | -0.60 | 0.55 | 1 | Gpr108 | G protein-coupled receptor 108 | 1700 | 1471 | 1565 | 2526 | 2592 | 2423 | 2408 | 2636 | 2322 | 2438 | 2183 |
| 2743 | 0.10 | 0.13 | 0.73 | 0.46 | 1 | Acadvl | acyl-Coenzyme A dehydrogenase, very lo | 3568 | 2932 | 4125 | 2678 | 2293 | 2048 | 2407 | 3018 | 2383 | 2626 | 2092 |
| 2274 | -0.01 | 0.11 | -0.07 | 0.94 | 1 | Arid2 | AT rich interactive domain 2 (ARID, RFX-l | 2156 | 1795 | 1678 | 2320 | 2060 | 2973 | 2407 | 2224 | 2435 | 2423 | 2546 |
| 2293 | 0.07 | 0.09 | 0.80 | 0.43 | 1 | Agpat1 | 1-acylglycerol-3-phosphate O-acyltransfe | 1845 | 1804 | 1984 | 2308 | 2846 | 1997 | 2406 | 2630 | 2567 | 2784 | 2056 |
| 1989 | 0.18 | 0.09 | 1.95 | 0.05 | 1 | Galk2 | galactokinase 2 | 1191 | 1309 | 1211 | 2262 | 2023 | 1809 | 2405 | 2848 | 2396 | 2456 | 1964 |
| 3575 | 0.04 | 0.08 | 0.46 | 0.64 | 1 | Srsf11 | serine and arginine-rich splicing factor 11 | 2367 | 3081 | 2571 | 2808 | 2649 | 7695 | 2404 | 3366 | 2698 | 2271 | 7412 |
| 2157 | 0.12 | 0.12 | 0.95 | 0.34 | 1 | Isg20 | interferon-stimulated protein | 1217 | 1602 | 1732 | 2283 | 2756 | 1796 | 2404 | 2482 | 2612 | 2767 | 2070 |
| 2374 | 0.08 | 0.08 | 0.93 | 0.35 | 1 | Cacybp | calcyclin binding protein | 2490 | 2348 | 2327 | 2139 | 2140 | 2532 | 2402 | 2368 | 2110 | 2357 | 2907 |
| 2133 | 0.02 | 0.09 | 0.21 | 0.84 | 1 | Pmpca | peptidase (mitochondrial processing) alph | 1795 | 1756 | 1875 | 2005 | 2075 | 2477 | 2401 | 2004 | 2175 | 2290 | 2606 |
| 2210 | 0.04 | 0.15 | 0.26 | 0.79 | 1 | Slc30a4 | solute carrier family 30 (zinc transporter), | 1870 | 2727 | 1528 | 2540 | 2329 | 1732 | 2400 | 2435 | 2637 | 2143 | 1965 |
| 2156 | -0.15 | 0.16 | -0.93 | 0.35 | 1 | Heatr5b | HEAT repeat containing 5B | 2104 | 1916 | 1815 | 2237 | 2461 | 2228 | 2398 | 1368 | 2588 | 2396 | 2206 |
| 2089 | 0.02 | 0.12 | 0.19 | 0.85 | 1 | Sos2 | SOS Ras/Rho guanine nucleotide exchar | 2016 | 2175 | 2069 | 1897 | 2141 | 1831 | 2396 | 1756 | 2558 | 2392 | 1749 |
| 2254 | 0.09 | 0.14 | 0.63 | 0.53 | 1 | Rexo2 | RNA exonuclease 2 | 2449 | 1603 | 2006 | 2324 | 2433 | 1955 | 2395 | 2779 | 2315 | 2365 | 2175 |
| 1906 | -0.07 | 0.12 | -0.53 | 0.60 | 1 | Sgsm2 | small G protein signaling modulator 2 | 1594 | 1284 | 1172 | 1834 | 2089 | 2301 | 2394 | 1598 | 2218 | 2593 | 1892 |
| 2246 | 0.03 | 0.12 | 0.27 | 0.79 | 1 | Tesk1 | testis specific protein kinase 1 | 1591 | 2006 | 1663 | 2512 | 2638 | 2029 | 2393 | 2500 | 2565 | 2999 | 1814 |
| 2081 | -0.15 | 0.08 | -1.83 | 0.07 | 1 | Asna1 | arsA arsenite transporter, ATP-binding, ho | 1686 | 1711 | 1692 | 2574 | 2205 | 2189 | 2393 | 2163 | 2050 | 2164 | 2061 |
| 1399 | 0.28 | 0.28 | 1.00 | 0.32 | 1 | Cox6a2 | cytochrome c oxidase subunit 6A2 | 73 | 32 | 118 | 1223 | 1453 | 1867 | 2392 | 1829 | 1354 | 2865 | 2188 |
| 2214 | 0.04 | 0.15 | 0.30 | 0.77 | 1 | Mfsd14a | major facilitator superfamily domain conta | 1738 | 2230 | 1616 | 2344 | 2183 | 2321 | 2391 | 1899 | 2505 | 2765 | 2356 |
| 2340 | -0.01 | 0.08 | -0.09 | 0.93 | 1 | Mtfr1l | mitochondrial fission regulator 1-like | 2366 | 2439 | 2603 | 2243 | 2607 | 1938 | 2389 | 2353 | 2541 | 2437 | 1825 |
| 2328 | 0.07 | 0.11 | 0.68 | 0.50 | 1 | Slf2 | SMC5-SMC6 complex localization factor 2 | 1784 | 2329 | 2146 | 2312 | 2018 | 2716 | 2385 | 2322 | 2356 | 2416 | 2821 |
| 2325 | 0.23 | 0.13 | 1.85 | 0.06 | 1 | Arpc5l | actin related protein 2/3 complex, subunit | 1832 | 1712 | 2125 | 2058 | 2408 | 2296 | 2385 | 2944 | 2325 | 3014 | 2476 |
| 2252 | -0.11 | 0.12 | -0.87 | 0.38 | 1 | Kat7 | K(lysine) acetyltransferase 7 | 2460 | 2610 | 2512 | 2159 | 2030 | 2321 | 2383 | 1661 | 2365 | 2063 | 2202 |
| 2233 | -0.04 | 0.08 | -0.48 | 0.63 | 1 | Farp1 | FERM, RhoGEF (Arhgef) and pleckstrin d | 2599 | 2442 | 2220 | 2218 | 2042 | 2125 | 2383 | 2300 | 2109 | 2200 | 1926 |
| 2230 | -0.01 | 0.10 | -0.13 | 0.90 | 1 | Rp2 | retinitis pigmentosa 2 homolog | 2106 | 1940 | 2125 | 2416 | 2314 | 2101 | 2383 | 2000 | 2465 | 2366 | 2316 |
| 2462 | -0.14 | 0.13 | -1.08 | 0.28 | 1 | Cers2 | ceramide synthase 2 | 3113 | 2484 | 2963 | 2451 | 2389 | 2412 | 2382 | 1590 | 2558 | 2556 | 2184 |
| 2291 | 0.02 | 0.11 | 0.19 | 0.85 | 1 | Arpp19 | cAMP-regulated phosphoprotein 19 | 2692 | 2721 | 2850 | 2058 | 2361 | 1634 | 2380 | 2484 | 2093 | 2109 | 1818 |
| 2251 | 0.06 | 0.10 | 0.62 | 0.54 | 1 | Stk4 | serine/threonine kinase 4 | 2667 | 2374 | 2105 | 1987 | 2333 | 1924 | 2380 | 2226 | 2411 | 2266 | 2084 |
| 2448 | 0.00 | 0.09 | -0.05 | 0.96 | 1 | Flot2 | flotillin 2 | 3037 | 2896 | 3046 | 2158 | 2572 | 1857 | 2379 | 2453 | 2348 | 2514 | 1671 |
| 2823 | -0.02 | 0.10 | -0.24 | 0.81 | 1 | Adam10 | a disintegrin and metallopeptidase domai | 4406 | 3386 | 4057 | 2472 | 2525 | 2269 | 2378 | 2048 | 2547 | 2750 | 2215 |
| 2167 | 0.04 | 0.11 | 0.36 | 0.72 | 1 | Hccs | holocytochrome c synthetase | 1885 | 2123 | 1993 | 2269 | 2051 | 2102 | 2378 | 2010 | 2390 | 2388 | 2244 |
| 1908 | -0.10 | 0.12 | -0.78 | 0.44 | 1 | Mfsd6 | major facilitator superfamily domain conta | 1758 | 1754 | 1800 | 2127 | 1716 | 1871 | 2378 | 1671 | 1878 | 2347 | 1690 |
| 2650 | -0.16 | 0.16 | -0.98 | 0.33 | 1 | Nfkb1 | nuclear factor of kappa light polypeptide g | 3808 | 2554 | 2597 | 2742 | 2494 | 3021 | 2377 | 2176 | 2674 | 1893 | 2814 |
| 2201 | -0.05 | 0.10 | -0.54 | 0.59 | 1 | Camta2 | calmodulin binding transcription activator | 1929 | 1636 | 1960 | 2349 | 2405 | 2347 | 2377 | 2132 | 2607 | 2523 | 1943 |
| 1944 | -0.02 | 0.11 | -0.18 | 0.85 | 1 | Aldh5a1 | aldhehyde dehydrogenase family 5, subfa | 1251 | 1554 | 1401 | 2440 | 1858 | 1984 | 2376 | 2337 | 2080 | 2190 | 1917 |
| 2881 | 0.19 | 0.14 | 1.37 | 0.17 | 1 | Rps13 | ribosomal protein S13 | 4055 | 3709 | 5189 | 2075 | 2418 | 1866 | 2375 | 3019 | 2239 | 2767 | 1976 |
| 2154 | 0.00 | 0.12 | -0.02 | 0.99 | 1 | Mtmr4 | myotubularin related protein 4 | 1620 | 1517 | 1485 | 1806 | 2560 | 2814 | 2375 | 1817 | 2659 | 2445 | 2595 |
| 2874 | -0.15 | 0.09 | -1.58 | 0.11 | 1 | Zfp740 | zinc finger protein 740 | 4079 | 3697 | 3736 | 2774 | 2607 | 2802 | 2373 | 2195 | 2430 | 2168 | 2748 |
| 2006 | -0.10 | 0.15 | -0.64 | 0.52 | 1 | Cntfr | ciliary neurotrophic factor receptor | 351 | 312 | 492 | 3185 | 2551 | 2702 | 2373 | 2619 | 2545 | 2472 | 2469 |
| 2452 | 0.00 | 0.13 | -0.04 | 0.97 | 1 | Akirin1 | akirin 1 | 1648 | 2413 | 1385 | 3127 | 2816 | 2497 | 2372 | 2818 | 2761 | 2482 | 2651 |
| 1648 | -0.06 | 0.13 | -0.48 | 0.63 | 1 | Focad | focadhesin | 915 | 985 | 826 | 2059 | 1828 | 1622 | 2372 | 2084 | 2162 | 1702 | 1571 |
| 2797 | -0.02 | 0.09 | -0.21 | 0.84 | 1 | Hmgn1 | high mobility group nucleosomal binding | 4240 | 3668 | 4052 | 2319 | 2447 | 2331 | 2371 | 2490 | 2261 | 2222 | 2372 |

| | | | | | | | | | | | | | | | | | | |
|---|---|---|---|---|---|---|---|---|---|---|---|---|---|---|---|---|---|---|
| 1698 | 0.07 | 0.11 | 0.62 | 0.54 | 1 | Stk26 | serine/threonine kinase 26 | 390 | 396 | 354 | 1791 | 2080 | 2295 | 2371 | 1846 | 2069 | 2386 | 2698 |
| 2399 | -0.04 | 0.08 | -0.54 | 0.59 | 1 | Dsg2 | desmoglein 2 | 3148 | 3099 | 3108 | 2165 | 2317 | 1763 | 2369 | 2129 | 2369 | 2395 | 1527 |
| 2090 | -0.20 | 0.13 | -1.62 | 0.11 | 1 | Wipi1 | WD repeat domain, phosphoinositide inte | 1235 | 1317 | 1333 | 2587 | 2815 | 2418 | 2368 | 1770 | 2758 | 2396 | 1989 |
| 2429 | 0.06 | 0.08 | 0.70 | 0.49 | 1 | Axin1 | axin 1 | 2213 | 2310 | 2039 | 2594 | 2548 | 2351 | 2367 | 2896 | 2676 | 2397 | 2326 |
| 2371 | -0.07 | 0.11 | -0.69 | 0.49 | 1 | Cdc23 | CDC23 cell division cycle 23 | 2104 | 1960 | 2149 | 2525 | 2648 | 2628 | 2367 | 2166 | 2163 | 2714 | 2661 |
| 2202 | -0.09 | 0.12 | -0.76 | 0.45 | 1 | Nbeal1 | neurobeachin like 1 | 1519 | 1350 | 1526 | 2159 | 2175 | 3453 | 2365 | 1690 | 2617 | 1926 | 3438 |
| 2541 | -0.10 | 0.11 | -0.91 | 0.36 | 1 | Brd8 | bromodomain containing 8 | 1841 | 2151 | 2012 | 2319 | 2266 | 4389 | 2363 | 1818 | 2363 | 2295 | 4136 |
| 2079 | -0.04 | 0.09 | -0.43 | 0.66 | 1 | Slc35g2 | solute carrier family 35, member G2 | 1451 | 1794 | 1776 | 2368 | 2389 | 1933 | 2362 | 2175 | 2446 | 2239 | 1934 |
| 2205 | 0.01 | 0.14 | 0.09 | 0.93 | 1 | Slc36a4 | solute carrier family 36 (proton/amino acid | 1683 | 2415 | 1599 | 2512 | 2437 | 1941 | 2361 | 2169 | 2655 | 2545 | 1934 |
| 2052 | -0.14 | 0.19 | -0.74 | 0.46 | 1 | Tmem59l | transmembrane protein 59-like | 3841 | 3542 | 4692 | 1335 | 1309 | 516 | 2360 | 1045 | 1216 | 2209 | 513 |
| 2732 | 0.04 | 0.09 | 0.38 | 0.70 | 1 | Chka | choline kinase alpha | 1391 | 2194 | 1509 | 2725 | 2304 | 4939 | 2359 | 3165 | 2332 | 2253 | 4885 |
| 2408 | -0.11 | 0.12 | -0.88 | 0.38 | 1 | Cldn4 | claudin 4 | 2659 | 2551 | 3102 | 2833 | 2352 | 1879 | 2359 | 2530 | 2010 | 2508 | 1704 |
| 2646 | -0.07 | 0.08 | -0.87 | 0.39 | 1 | Fbxw8 | F-box and WD-40 domain protein 8 | 3771 | 3139 | 3950 | 2241 | 2641 | 2101 | 2358 | 2131 | 2340 | 2481 | 1955 |
| 2332 | -0.10 | 0.13 | -0.75 | 0.45 | 1 | Phf8 | PHD finger protein 8 | 2724 | 2781 | 2533 | 2013 | 2581 | 2133 | 2358 | 1659 | 2691 | 2177 | 2007 |
| 2009 | 0.03 | 0.11 | 0.27 | 0.79 | 1 | Unc50 | unc-50 homolog | 1573 | 1434 | 1615 | 2011 | 2160 | 2088 | 2355 | 1759 | 2295 | 2714 | 2093 |
| 2133 | -0.11 | 0.10 | -1.07 | 0.29 | 1 | Gga1 | golgi associated, gamma adaptin ear con | 2327 | 1931 | 2165 | 2306 | 2009 | 2171 | 2354 | 2000 | 2233 | 2100 | 1873 |
| 1986 | -0.05 | 0.09 | -0.58 | 0.56 | 1 | Def8 | differentially expressed in FDCP 8 | 1467 | 1648 | 1649 | 2325 | 2244 | 1748 | 2354 | 2483 | 2245 | 2093 | 1591 |
| 2139 | 0.01 | 0.08 | 0.13 | 0.90 | 1 | Samm50 | SAMM50 sorting and assembly machiner | 2559 | 1969 | 2282 | 2012 | 2030 | 1935 | 2353 | 2033 | 2081 | 2364 | 1914 |
| 2365 | 0.21 | 0.13 | 1.57 | 0.12 | 1 | Denr | density-regulated protein | 1903 | 2006 | 2199 | 2267 | 2511 | 2041 | 2352 | 3614 | 2463 | 2445 | 2213 |
| 2530 | -0.02 | 0.09 | -0.17 | 0.86 | 1 | Cul5 | cullin 5 | 2059 | 2509 | 2108 | 2874 | 2729 | 2682 | 2351 | 2570 | 2576 | 2432 | 2940 |
| 2239 | -0.04 | 0.08 | -0.43 | 0.67 | 1 | Pi4kb | phosphatidylinositol 4-kinase beta | 1771 | 1883 | 1806 | 2542 | 2481 | 2313 | 2351 | 2446 | 2609 | 2388 | 2039 |
| 2553 | 0.12 | 0.09 | 1.25 | 0.21 | 1 | Ccnt1 | cyclin T1 | 1651 | 1949 | 2076 | 2986 | 2518 | 2891 | 2349 | 3396 | 2756 | 2594 | 2914 |
| 2492 | 0.09 | 0.07 | 1.29 | 0.20 | 1 | Ensa | endosulfine alpha | 2771 | 2594 | 2526 | 2343 | 2298 | 2451 | 2349 | 2350 | 2463 | 2568 | 2695 |
| 2261 | -0.06 | 0.09 | -0.66 | 0.51 | 1 | Iars2 | isoleucine-tRNA synthetase 2, mitochond | 2317 | 2051 | 2171 | 2517 | 2330 | 2166 | 2349 | 2165 | 2214 | 2244 | 2347 |
| 2829 | -0.10 | 0.11 | -0.87 | 0.38 | 1 | Mef2a | myocyte enhancer factor 2A | 4869 | 4329 | 3689 | 2608 | 2194 | 2284 | 2348 | 1975 | 2344 | 2255 | 2223 |
| 2325 | -0.09 | 0.15 | -0.64 | 0.52 | 1 | Fam13b | family with sequence similarity 13, membe | 2267 | 2588 | 2140 | 2342 | 2069 | 2811 | 2347 | 1580 | 2259 | 2474 | 2694 |
| 2216 | -0.11 | 0.11 | -0.98 | 0.32 | 1 | Rims2 | regulating synaptic membrane exocytosis | 1583 | 2196 | 2103 | 2800 | 2101 | 2352 | 2347 | 2385 | 2339 | 1986 | 2183 |
| 2432 | 0.20 | 0.13 | 1.52 | 0.13 | 1 | Edf1 | endothelial differentiation-related factor 1 | 2626 | 2251 | 2818 | 2241 | 2392 | 1893 | 2345 | 3035 | 2191 | 2831 | 2126 |
| 2396 | 0.04 | 0.09 | 0.43 | 0.67 | 1 | Psma1 | proteasome subunit alpha 1 | 2817 | 2014 | 2683 | 2145 | 2313 | 2450 | 2344 | 1894 | 2441 | 2606 | 2653 |
| 2457 | -0.17 | 0.09 | -1.84 | 0.07 | 1 | Tubb4b | tubulin, beta 4B class IVB | 2539 | 2375 | 2176 | 2493 | 2515 | 3195 | 2340 | 2105 | 2141 | 2410 | 2736 |
| 2644 | 0.08 | 0.11 | 0.68 | 0.50 | 1 | Endod1 | endonuclease domain containing 1 | 4163 | 3698 | 4506 | 2313 | 2038 | 1400 | 2339 | 2635 | 2397 | 2257 | 1332 |
| 2358 | 0.15 | 0.10 | 1.56 | 0.12 | 1 | Hbs1l | Hbs1-like (S. cerevisiae) | 1678 | 2199 | 1836 | 2245 | 2393 | 2598 | 2339 | 2619 | 2615 | 2501 | 2919 |
| 2301 | -0.02 | 0.11 | -0.17 | 0.87 | 1 | Washc2 | WASH complex subunit 2` | 2400 | 1681 | 1828 | 2444 | 2264 | 2684 | 2337 | 2187 | 2141 | 2565 | 2776 |
| 2645 | 0.23 | 0.14 | 1.65 | 0.10 | 1 | Acsl5 | acyl-CoA synthetase long-chain family me | 3967 | 2813 | 2844 | 1858 | 2737 | 2065 | 2335 | 2714 | 2892 | 2607 | 2259 |
| 1747 | -0.04 | 0.12 | -0.37 | 0.71 | 1 | Negr1 | neuronal growth regulator 1 | 1089 | 1229 | 1342 | 1944 | 2022 | 1578 | 2335 | 1834 | 2200 | 2179 | 1466 |
| 2525 | 0.00 | 0.10 | 0.01 | 0.99 | 1 | Wdr33 | WD repeat domain 33 | 2456 | 2678 | 2205 | 2653 | 2577 | 2616 | 2334 | 3135 | 2505 | 2139 | 2478 |
| 2461 | 0.12 | 0.13 | 0.97 | 0.33 | 1 | Ppia | peptidylprolyl isomerase A | 2570 | 2290 | 2578 | 2292 | 2632 | 2101 | 2334 | 3107 | 2305 | 2853 | 2007 |
| 2212 | -0.08 | 0.08 | -1.07 | 0.29 | 1 | Parp8 | poly (ADP-ribose) polymerase family, mer | 1893 | 1779 | 1718 | 2276 | 2157 | 2977 | 2334 | 2134 | 2171 | 2109 | 2781 |
| 2113 | -0.07 | 0.12 | -0.58 | 0.56 | 1 | Atg2b | autophagy related 2B | 1506 | 1415 | 1579 | 2115 | 2170 | 2977 | 2333 | 1833 | 2754 | 2074 | 2491 |
| 2112 | -0.14 | 0.10 | -1.37 | 0.17 | 1 | Insig2 | insulin induced gene 2 | 1310 | 1482 | 1655 | 2878 | 2397 | 2249 | 2333 | 2282 | 2304 | 2143 | 2193 |
| 2147 | 0.00 | 0.09 | 0.05 | 0.96 | 1 | Slc36a1 | solute carrier family 36 (proton/amino acid | 1040 | 1052 | 945 | 2766 | 2486 | 2699 | 2331 | 2551 | 2537 | 2538 | 2670 |
| 2290 | 0.10 | 0.14 | 0.69 | 0.49 | 1 | Upf3b | UPF3 regulator of nonsense transcripts h | 992 | 1501 | 1421 | 2016 | 2005 | 3786 | 2330 | 2545 | 2164 | 1930 | 4501 |
| 2220 | -0.10 | 0.11 | -0.94 | 0.35 | 1 | Dnaja3 | DnaJ heat shock protein family (Hsp40) m | 1797 | 1822 | 1921 | 2335 | 2834 | 2257 | 2330 | 1959 | 2478 | 2617 | 2069 |
| 2785 | 0.00 | 0.10 | 0.00 | 1.00 | 1 | Asph | aspartate-beta-hydroxylase | 3950 | 4119 | 4134 | 2395 | 2326 | 2141 | 2328 | 2859 | 2203 | 2024 | 2156 |
| 2049 | 0.08 | 0.12 | 0.65 | 0.52 | 1 | Pcgf5 | polycomb group ring finger 5 | 1374 | 1639 | 1744 | 2182 | 2179 | 1944 | 2327 | 2142 | 2329 | 2736 | 1943 |
| 2368 | 0.05 | 0.12 | 0.42 | 0.67 | 1 | Setx | senataxin | 2507 | 2655 | 2244 | 2079 | 2220 | 2541 | 2326 | 1994 | 2639 | 2478 | 2370 |
| 3020 | -0.03 | 0.13 | -0.24 | 0.81 | 1 | Pbxip1 | pre B cell leukemia transcription factor int | 6138 | 4984 | 4815 | 2070 | 2214 | 2100 | 2325 | 2680 | 2111 | 1999 | 1782 |
| 2353 | 0.15 | 0.08 | 1.87 | 0.06 | 1 | Smu1 | smu-1 suppressor of mec-8 and unc-52 h | 2589 | 2278 | 2599 | 2091 | 2316 | 2004 | 2325 | 2524 | 2465 | 2500 | 2185 |
| 2317 | 0.17 | 0.11 | 1.57 | 0.12 | 1 | Marchf5 | membrane associated ring-CH-type finge | 1813 | 2078 | 2009 | 2435 | 2192 | 2259 | 2325 | 2381 | 2560 | 2995 | 2435 |
| 2212 | 0.00 | 0.11 | 0.02 | 0.98 | 1 | Nacc1 | nucleus accumbens associated 1, BEN a | 2251 | 2192 | 1905 | 2208 | 2347 | 2081 | 2325 | 2484 | 2494 | 2231 | 1818 |
| 2337 | 0.06 | 0.12 | 0.53 | 0.60 | 1 | Zbtb41 | zinc finger and BTB domain containing 4 | 1627 | 2420 | 1796 | 2622 | 2333 | 2423 | 2324 | 2614 | 2930 | 2177 | 2436 |
| 2330 | -0.16 | 0.15 | -1.05 | 0.30 | 1 | Med12 | mediator complex subunit 12 | 1720 | 2522 | 1956 | 2301 | 2386 | 3252 | 2322 | 1746 | 2615 | 2084 | 2722 |
| 3226 | -0.01 | 0.13 | -0.07 | 0.95 | 1 | Hnrnpa1 | heterogeneous nuclear ribonucleoprotein | 5270 | 4026 | 4830 | 1949 | 2224 | 4231 | 2321 | 2412 | 2249 | 1770 | 4206 |
| 2487 | -0.17 | 0.09 | -1.86 | 0.06 | 1 | Pamr1 | peptidase domain containing associated | 2796 | 3591 | 3196 | 2374 | 2502 | 2211 | 2321 | 2081 | 2301 | 1940 | 2048 |
| 2334 | 0.02 | 0.12 | 0.14 | 0.89 | 1 | Chuk | conserved helix-loop-helix ubiquitous kin | 1731 | 1185 | 1828 | 2278 | 2324 | 3428 | 2321 | 2150 | 2334 | 2367 | 3726 |
| 2430 | 0.05 | 0.12 | 0.41 | 0.68 | 1 | Strn3 | striatin, calmodulin binding protein 3 | 2408 | 2302 | 2205 | 2619 | 2121 | 2646 | 2320 | 3155 | 2272 | 1987 | 2695 |
| 2492 | -0.09 | 0.12 | -0.77 | 0.44 | 1 | Cacul1 | CDK2 associated, cullin domain 1 | 1997 | 2251 | 2259 | 3110 | 2803 | 2516 | 2318 | 2463 | 3150 | 1936 | 2609 |
| 1937 | -0.12 | 0.14 | -0.84 | 0.40 | 1 | B630019K06Rik | | 1118 | 1762 | 1668 | 2026 | 2252 | 2087 | 2318 | 2249 | 2172 | 1879 | 1772 |
| 1802 | -0.14 | 0.15 | -0.92 | 0.36 | 1 | Faxc | failed axon connections homolog | 848 | 1205 | 1157 | 2533 | 2187 | 1678 | 2317 | 1744 | 2387 | 2128 | 1638 |
| 2434 | 0.12 | 0.13 | 0.92 | 0.36 | 1 | Ran | RAN, member RAS oncogene family | 2998 | 2486 | 3115 | 2202 | 2278 | 1914 | 2315 | 2833 | 2053 | 2547 | 2030 |

| | | | | | | | | | | | | | | | | | |
|---|---|---|---|---|---|---|---|---|---|---|---|---|---|---|---|---|---|
| 2321 | 0.09 | 0.08 | 1.12 | 0.26 | 1 | Med22 | mediator complex subunit 22 | 2197 | 2157 | 2245 | 2337 | 2508 | 1986 | 2315 | 2778 | 2528 | 2467 | 2008 |
| 2531 | -0.13 | 0.08 | -1.53 | 0.13 | 1 | Plxnb1 | plexin B1 | 2931 | 2763 | 2835 | 2084 | 2470 | 3241 | 2313 | 1936 | 2360 | 2164 | 2742 |
| 2290 | 0.17 | 0.13 | 1.32 | 0.19 | 1 | Polr2g | polymerase (RNA) II (DNA directed) polyp | 2056 | 2032 | 2532 | 2212 | 2355 | 1825 | 2313 | 3032 | 2245 | 2702 | 1891 |
| 2044 | -0.13 | 0.10 | -1.31 | 0.19 | 1 | Auh | AU RNA binding protein/enoyl-coenzyme | 1416 | 1731 | 1774 | 2504 | 2283 | 2055 | 2312 | 2416 | 2181 | 1824 | 1991 |
| 2283 | -0.07 | 0.12 | -0.60 | 0.55 | 1 | Ubxn7 | UBX domain protein 7 | 2121 | 2668 | 2219 | 2125 | 2274 | 2553 | 2310 | 1901 | 2334 | 2124 | 2484 |
| 2248 | 0.01 | 0.10 | 0.08 | 0.93 | 1 | Foxk2 | forkhead box K2 | 1790 | 2272 | 1913 | 2453 | 2261 | 2331 | 2310 | 2261 | 2472 | 2382 | 2282 |
| 2084 | -0.06 | 0.11 | -0.58 | 0.56 | 1 | Abhd12 | abhydrolase domain containing 12 | 2234 | 1879 | 2140 | 2333 | 2275 | 1581 | 2309 | 2351 | 2437 | 1955 | 1435 |
| 2309 | 0.14 | 0.12 | 1.15 | 0.25 | 1 | Ppp2cb | protein phosphatase 2 (formerly 2A), cata | 2294 | 2118 | 2386 | 2362 | 2430 | 1744 | 2307 | 3171 | 2402 | 2291 | 1897 |
| 2195 | -0.11 | 0.14 | -0.75 | 0.45 | 1 | Mlxip | MLX interacting protein | 1804 | 1666 | 1460 | 2669 | 2175 | 2809 | 2307 | 2023 | 2675 | 1958 | 2598 |
| 2926 | -0.09 | 0.08 | -1.19 | 0.24 | 1 | Mid1ip1 | Mid1 interacting protein 1 (gastrulation sp | 3961 | 5088 | 4443 | 2792 | 2339 | 2221 | 2306 | 2467 | 2216 | 2330 | 2019 |
| 2068 | 0.04 | 0.12 | 0.31 | 0.75 | 1 | Arl3 | ADP-ribosylation factor-like 3 | 1797 | 2192 | 2373 | 2009 | 1896 | 1876 | 2305 | 2117 | 2083 | 2164 | 1939 |
| 2092 | 0.05 | 0.09 | 0.54 | 0.59 | 1 | Mtmr9 | myotubularin related protein 9 | 2119 | 2313 | 2187 | 2028 | 2137 | 1582 | 2304 | 2143 | 2403 | 2204 | 1595 |
| 2774 | -0.02 | 0.10 | -0.22 | 0.82 | 1 | Mtus1 | mitochondrial tumor suppressor 1 | 3111 | 3673 | 3087 | 3110 | 2227 | 2798 | 2303 | 2505 | 2293 | 2642 | 2766 |
| 2521 | 0.04 | 0.12 | 0.36 | 0.72 | 1 | Stag1 | stromal antigen 1 | 3722 | 2881 | 3197 | 2287 | 2382 | 1868 | 2303 | 2582 | 2424 | 1987 | 2095 |
| 2214 | -0.16 | 0.09 | -1.73 | 0.08 | 1 | Atf7ip | activating transcription factor 7 interacting | 2784 | 2611 | 2606 | 2379 | 2244 | 1696 | 2302 | 2148 | 2232 | 1877 | 1480 |
| 1611 | -0.02 | 0.15 | -0.12 | 0.91 | 1 | Adora1 | adenosine A1 receptor | 288 | 502 | 417 | 2338 | 1932 | 1704 | 2302 | 2403 | 2153 | 2142 | 1541 |
| 2274 | 0.00 | 0.08 | 0.03 | 0.98 | 1 | Adprh | ADP-ribosylarginine hydrolase | 2648 | 2801 | 2685 | 2222 | 2213 | 1680 | 2301 | 2139 | 2208 | 2498 | 1617 |
| 1972 | -0.08 | 0.20 | -0.38 | 0.70 | 1 | Tmem184c | transmembrane protein 184C | 1214 | 1251 | 1059 | 2487 | 2367 | 2090 | 2300 | 1303 | 2585 | 2789 | 2243 |
| 1878 | -0.05 | 0.09 | -0.55 | 0.58 | 1 | Rassf4 | Ras association (RalGDS/AF-6) domain fa | 1802 | 1430 | 1464 | 1992 | 2174 | 1673 | 2300 | 1888 | 2173 | 2067 | 1693 |
| 2134 | 0.03 | 0.08 | 0.38 | 0.70 | 1 | Sirt2 | sirtuin 2 | 2151 | 1969 | 1763 | 2145 | 2101 | 2155 | 2299 | 2214 | 2149 | 2446 | 2085 |
| 2585 | 0.00 | 0.12 | -0.01 | 0.99 | 1 | Fads1 | fatty acid desaturase 1 | 3359 | 3373 | 2792 | 2769 | 2653 | 1707 | 2298 | 2389 | 2555 | 2921 | 1616 |
| 2092 | -0.03 | 0.14 | -0.21 | 0.83 | 1 | Dcaf1 | DDB1 and CUL4 associated factor 1 | 1913 | 2244 | 1909 | 1983 | 2154 | 2104 | 2298 | 1602 | 2513 | 2178 | 2109 |
| 1838 | 0.00 | 0.13 | 0.00 | 1.00 | 1 | Phka1 | phosphorylase kinase alpha 1 | 1096 | 1285 | 1481 | 1866 | 1986 | 2020 | 2296 | 1966 | 2054 | 2344 | 1822 |
| 2578 | -0.10 | 0.10 | -0.93 | 0.35 | 1 | Brd4 | bromodomain containing 4 | 3022 | 3303 | 2717 | 2652 | 2443 | 2580 | 2294 | 2623 | 2561 | 2092 | 2069 |
| 2173 | 0.08 | 0.13 | 0.63 | 0.53 | 1 | Rbx1 | ring-box 1 | 1753 | 2064 | 2552 | 2015 | 2245 | 1980 | 2294 | 2332 | 2140 | 2333 | 2191 |
| 2144 | -0.06 | 0.09 | -0.68 | 0.49 | 1 | Mcur1 | mitochondrial calcium uniporter regulator | 2075 | 2233 | 2314 | 2312 | 2261 | 1777 | 2293 | 2051 | 2358 | 2245 | 1665 |
| 2069 | 0.07 | 0.10 | 0.64 | 0.53 | 1 | Cox7a2 | cytochrome c oxidase subunit 7A2 | 2164 | 2232 | 1844 | 2113 | 2141 | 1496 | 2293 | 2651 | 2196 | 2145 | 1487 |
| 2203 | 0.02 | 0.08 | 0.25 | 0.80 | 1 | Gtpbp1 | GTP binding protein 1 | 1820 | 1869 | 1986 | 2337 | 2413 | 2169 | 2292 | 2271 | 2316 | 2568 | 2197 |
| 2180 | 0.12 | 0.11 | 1.14 | 0.25 | 1 | Amz2 | archaelysin family metallopeptidase 2 | 1605 | 1942 | 2083 | 2328 | 2098 | 2055 | 2292 | 2741 | 2318 | 2240 | 2276 |
| 3900 | -0.02 | 0.09 | -0.18 | 0.86 | 1 | Kctd12 | potassium channel tetramerisation domai | 10783 | 7811 | 7196 | 1671 | 1868 | 2813 | 2291 | 1820 | 1858 | 2215 | 2576 |
| 2456 | 0.09 | 0.11 | 0.81 | 0.42 | 1 | Cant1 | calcium activated nucleotidase 1 | 2140 | 2242 | 1923 | 2419 | 2853 | 2439 | 2291 | 3103 | 2706 | 2580 | 2319 |
| 2188 | 0.09 | 0.11 | 0.82 | 0.41 | 1 | Mcrip1 | MAPK regulated corepressor interacting p | 1924 | 2231 | 2362 | 2212 | 2301 | 1660 | 2291 | 2759 | 2220 | 2486 | 1623 |
| 2115 | 0.07 | 0.10 | 0.74 | 0.46 | 1 | Pdcl | phosducin-like | 1657 | 1592 | 1678 | 1853 | 2650 | 2156 | 2291 | 2065 | 2676 | 2258 | 2392 |
| 2112 | 0.02 | 0.09 | 0.25 | 0.81 | 1 | Odc1 | ornithine decarboxylase, structural 1 | 3082 | 2249 | 1925 | 1698 | 1392 | 2526 | 2291 | 1824 | 1294 | 2471 | 2479 |
| 2254 | 0.04 | 0.14 | 0.25 | 0.80 | 1 | Eif3k | eukaryotic translation initiation factor 3, su | 3508 | 2809 | 3190 | 1857 | 1960 | 1440 | 2290 | 2462 | 1556 | 2170 | 1551 |
| 3224 | 0.13 | 0.11 | 1.09 | 0.27 | 1 | Rpl27a | ribosomal protein L27A | 5234 | 5097 | 6334 | 2531 | 2215 | 1957 | 2289 | 2706 | 2209 | 2692 | 2195 |
| 2343 | 0.05 | 0.11 | 0.48 | 0.63 | 1 | Psmb6 | proteasome (prosome, macropain) subun | 3052 | 2137 | 3053 | 2017 | 2352 | 1933 | 2289 | 2163 | 2211 | 2695 | 1873 |
| 1962 | 0.19 | 0.12 | 1.50 | 0.13 | 1 | Podxl2 | podocalyxin-like 2 | 1015 | 1418 | 1210 | 2316 | 2222 | 1506 | 2288 | 3206 | 2161 | 2616 | 1629 |
| 2433 | -0.08 | 0.11 | -0.74 | 0.46 | 1 | Arhgap21 | Rho GTPase activating protein 21 | 2608 | 2268 | 2205 | 2453 | 2180 | 3224 | 2287 | 1938 | 2510 | 2253 | 2836 |
| 2392 | -0.05 | 0.09 | -0.50 | 0.62 | 1 | Cabin1 | calcineurin binding protein 1 | 2152 | 2248 | 2373 | 2487 | 2331 | 2837 | 2287 | 2441 | 2254 | 2501 | 2398 |
| 1749 | 0.00 | 0.11 | -0.02 | 0.98 | 1 | Tmem30b | transmembrane protein 30B | 1027 | 1331 | 1080 | 2408 | 2117 | 1110 | 2287 | 2209 | 2127 | 2409 | 1130 |
| 2362 | 0.06 | 0.12 | 0.50 | 0.61 | 1 | Dbnl | drebrin-like | 3248 | 2506 | 2963 | 2133 | 2228 | 1806 | 2285 | 2635 | 2217 | 2061 | 1898 |
| 1997 | -0.21 | 0.13 | -1.56 | 0.12 | 1 | R3hdm1 | R3H domain containing 1 | 1410 | 1367 | 1615 | 2089 | 2349 | 2607 | 2285 | 1404 | 2600 | 1854 | 2391 |
| 2164 | 0.03 | 0.09 | 0.27 | 0.79 | 1 | Rgp1 | RAB6A GEF compex partner 1 | 2047 | 1982 | 2298 | 1989 | 2135 | 2250 | 2283 | 2193 | 2375 | 1981 | 2266 |
| 2074 | -0.03 | 0.14 | -0.23 | 0.82 | 1 | Bicd1 | BICD cargo adaptor 1 | 1080 | 1106 | 1191 | 2471 | 2153 | 2878 | 2283 | 1731 | 2554 | 2517 | 2847 |
| 1733 | -0.22 | 0.16 | -1.36 | 0.17 | 1 | Dnajc12 | DnaJ heat shock protein family (Hsp40) m | 1412 | 1967 | 1869 | 1871 | 1856 | 1451 | 2283 | 1367 | 1717 | 1765 | 1502 |
| 2271 | -0.12 | 0.11 | -1.12 | 0.26 | 1 | Wdtc1 | WD and tetratricopeptide repeats 1 | 3045 | 3106 | 2454 | 2367 | 2378 | 1436 | 2281 | 2148 | 2131 | 2446 | 1187 |
| 2079 | -0.01 | 0.12 | -0.12 | 0.91 | 1 | Dnajb12 | DnaJ heat shock protein family (Hsp40) m | 1867 | 1537 | 1755 | 2430 | 2524 | 1625 | 2281 | 1878 | 2599 | 2719 | 1652 |
| 2291 | 0.07 | 0.13 | 0.52 | 0.60 | 1 | Pomgnt2 | protein O-linked mannose beta 1,4-N-ace | 1929 | 2443 | 2349 | 2552 | 2311 | 1833 | 2279 | 2809 | 2114 | 2893 | 1693 |
| 2165 | 0.14 | 0.10 | 1.50 | 0.13 | 1 | Fuca2 | fucosidase, alpha-L- 2, plasma | 1952 | 2139 | 2381 | 2249 | 2220 | 1499 | 2279 | 2556 | 2348 | 2531 | 1665 |
| 2162 | 0.01 | 0.09 | 0.06 | 0.96 | 1 | Pigt | phosphatidylinositol glycan anchor biosyn | 2450 | 2234 | 2247 | 2224 | 2414 | 1472 | 2275 | 2110 | 2243 | 2664 | 1447 |
| 2273 | 0.04 | 0.15 | 0.25 | 0.80 | 1 | Atf6b | activating transcription factor 6 beta | 2336 | 1366 | 1890 | 2152 | 2776 | 2407 | 2274 | 2520 | 2536 | 2255 | 2490 |
| 2006 | -0.21 | 0.12 | -1.69 | 0.09 | 1 | Epb41l1 | erythrocyte membrane protein band 4.1 li | 2220 | 2308 | 1896 | 2199 | 2349 | 1561 | 2274 | 1618 | 2094 | 2116 | 1426 |
| 2375 | 0.09 | 0.11 | 0.79 | 0.43 | 1 | Lsr | lipolysis stimulated lipoprotein receptor | 2523 | 2817 | 2985 | 2178 | 2261 | 1864 | 2272 | 2932 | 2232 | 2299 | 1767 |
| 2226 | -0.03 | 0.12 | -0.29 | 0.77 | 1 | Acsl4 | acyl-CoA synthetase long-chain family me | 2368 | 1951 | 1996 | 2669 | 2302 | 1966 | 2271 | 2439 | 2370 | 2062 | 2092 |
| 2024 | 0.02 | 0.10 | 0.16 | 0.87 | 1 | Casd1 | CAS1 domain containing 1 | 1928 | 2030 | 2051 | 2207 | 1851 | 1767 | 2271 | 1980 | 2015 | 2190 | 1971 |
| 2040 | 0.14 | 0.10 | 1.43 | 0.15 | 1 | Ppip5k2 | diphosphoinositol pentakisphosphate kin | 1446 | 1475 | 1612 | 1686 | 2261 | 2308 | 2269 | 2012 | 2425 | 2349 | 2593 |
| 2565 | 0.10 | 0.09 | 1.16 | 0.25 | 1 | Zfp260 | zinc finger protein 260 | 2566 | 2736 | 2635 | 2156 | 2480 | 2830 | 2268 | 2204 | 2495 | 2454 | 3389 |
| 1831 | 0.00 | 0.10 | 0.00 | 1.00 | 1 | Tmem135 | transmembrane protein 135 | 1457 | 1438 | 1354 | 1900 | 2121 | 1676 | 2268 | 2022 | 2186 | 1935 | 1788 |
| 2740 | -0.05 | 0.11 | -0.44 | 0.66 | 1 | Jmjd1c | jumonji domain containing 1C | 3069 | 3492 | 2724 | 2435 | 2387 | 3496 | 2267 | 2103 | 2489 | 2204 | 3470 |

| | | | | | | | | | | | | | | | | | | |
|---|---|---|---|---|---|---|---|---|---|---|---|---|---|---|---|---|---|---|
| 2298 | 0.13 | 0.11 | 1.20 | 0.23 | 1 | Arl2bp | ADP-ribosylation factor-like 2 binding pro | 1741 | 1568 | 1874 | 2682 | 2592 | 2068 | 2266 | 3039 | 2593 | 2643 | 2214 |
| 2312 | -0.07 | 0.09 | -0.74 | 0.46 | 1 | Gripap1 | GRIP1 associated protein 1 | 1463 | 1639 | 1859 | 2091 | 2446 | 3642 | 2265 | 2212 | 2311 | 1914 | 3595 |
| 2263 | 0.11 | 0.10 | 1.15 | 0.25 | 1 | Afg3l1 | AFG3-like AAA ATPase 1 | 1970 | 1931 | 2147 | 2392 | 2294 | 2071 | 2265 | 2815 | 2422 | 2633 | 1960 |
| 2238 | 0.02 | 0.08 | 0.27 | 0.79 | 1 | Ranbp3 | RAN binding protein 3 | 1949 | 1761 | 1979 | 2309 | 2211 | 2605 | 2265 | 2250 | 2250 | 2485 | 2554 |
| 2014 | 0.01 | 0.10 | 0.14 | 0.89 | 1 | Clns1a | chloride channel, nucleotide-sensitive, 1A | 1659 | 1863 | 2218 | 2019 | 2245 | 1641 | 2265 | 2248 | 2228 | 2097 | 1678 |
| 2738 | -0.05 | 0.12 | -0.40 | 0.69 | 1 | Xiap | X-linked inhibitor of apoptosis | 4123 | 3846 | 4057 | 2347 | 2202 | 2346 | 2264 | 1730 | 2616 | 2288 | 2302 |
| 4188 | 0.21 | 0.11 | 1.90 | 0.06 | 1 | Snrnp70 | small nuclear ribonucleoprotein 70 (U1) | 3180 | 2904 | 3087 | 1591 | 2143 | 11932 | 2263 | 2332 | 2177 | 2836 | 11622 |
| 2397 | -0.09 | 0.12 | -0.75 | 0.46 | 1 | Tm7sf3 | transmembrane 7 superfamily member 3 | 2619 | 2956 | 2553 | 2905 | 2263 | 1920 | 2263 | 3148 | 2088 | 1748 | 1907 |
| 2077 | -0.06 | 0.11 | -0.54 | 0.59 | 1 | Umps | uridine monophosphate synthetase | 1706 | 1888 | 2080 | 2385 | 2165 | 1953 | 2263 | 2014 | 2101 | 2252 | 2043 |
| 2264 | -0.11 | 0.11 | -1.05 | 0.30 | 1 | Arpc3 | actin related protein 2/3 complex, subunit | 3769 | 2557 | 2826 | 1974 | 2113 | 1838 | 2262 | 1922 | 2036 | 2002 | 1607 |
| 1576 | -0.26 | 0.16 | -1.70 | 0.09 | 1 | Lgi1 | leucine-rich repeat LGI family, member 1 | 753 | 1145 | 753 | 2111 | 1795 | 1773 | 2262 | 1953 | 1349 | 2176 | 1261 |
| 2368 | 0.08 | 0.11 | 0.71 | 0.48 | 1 | Psmd8 | proteasome (prosome, macropain) 26S su | 3213 | 2293 | 2782 | 2162 | 2305 | 1905 | 2261 | 2468 | 2363 | 2427 | 1867 |
| 2273 | -0.02 | 0.12 | -0.19 | 0.85 | 1 | Trappc6b | trafficking protein particle complex 6B | 1965 | 2124 | 1924 | 2739 | 2450 | 2133 | 2261 | 2063 | 2479 | 2367 | 2494 |
| 2222 | 0.04 | 0.11 | 0.38 | 0.70 | 1 | Etfa | electron transferring flavoprotein, alpha p | 2585 | 2260 | 2676 | 2254 | 2085 | 1755 | 2261 | 2261 | 2052 | 2062 | 2194 |
| 1971 | -0.03 | 0.11 | -0.26 | 0.80 | 1 | Thap12 | THAP domain containing 12 | 2177 | 1650 | 1772 | 2199 | 1886 | 1795 | 2260 | 1887 | 2117 | 2000 | 1938 |
| 2492 | 0.02 | 0.09 | 0.26 | 0.80 | 1 | Kansl1 | KAT8 regulatory NSL complex subunit 1 | 2605 | 2579 | 2486 | 2401 | 2109 | 3061 | 2259 | 2461 | 2446 | 2263 | 2742 |
| 2157 | 0.11 | 0.13 | 0.89 | 0.37 | 1 | Ppip5k1 | diphosphoinositol pentakisphosphate kin | 1050 | 1211 | 1438 | 2294 | 2142 | 2962 | 2258 | 2631 | 2450 | 2656 | 2632 |
| 1957 | 0.16 | 0.13 | 1.29 | 0.20 | 1 | Ffar2 | free fatty acid receptor 2 | 1277 | 1478 | 1513 | 2255 | 2233 | 1265 | 2258 | 3290 | 2347 | 2407 | 1207 |
| 2212 | 0.03 | 0.12 | 0.25 | 0.81 | 1 | Fam98a | family with sequence similarity 98, membe | 1326 | 1668 | 1768 | 2120 | 2208 | 3075 | 2257 | 1917 | 2468 | 2268 | 3260 |
| 2000 | 0.04 | 0.12 | 0.38 | 0.71 | 1 | Zfp687 | zinc finger protein 687 | 1286 | 1436 | 1337 | 2067 | 2080 | 2414 | 2257 | 2012 | 2648 | 1985 | 2482 |
| 2444 | -0.07 | 0.12 | -0.60 | 0.55 | 1 | Cbl | Casitas B-lineage lymphoma | 3266 | 3037 | 2599 | 2418 | 1952 | 2609 | 2255 | 2022 | 2346 | 2164 | 2215 |
| 2259 | -0.08 | 0.09 | -0.90 | 0.37 | 1 | Kdm4a | lysine (K)-specific demethylase 4A | 2351 | 1968 | 2294 | 2098 | 2287 | 2741 | 2255 | 2114 | 2208 | 2203 | 2326 |
| 1661 | 0.28 | 0.16 | 1.80 | 0.07 | 1 | Fer1l6 | fer-1-like 6 (C. elegans) | 169 | 130 | 175 | 2057 | 1539 | 2104 | 2254 | 1854 | 2075 | 2742 | 3172 |
| 2106 | 0.08 | 0.08 | 0.95 | 0.34 | 1 | Fktn | fukutin | 1814 | 1629 | 1741 | 2039 | 2361 | 2093 | 2253 | 2381 | 2435 | 2288 | 2133 |
| 1676 | -0.10 | 0.13 | -0.77 | 0.44 | 1 | Filip1 | filamin A interacting protein 1 | 878 | 1007 | 1146 | 1787 | 1870 | 2047 | 2252 | 1665 | 2145 | 1674 | 1962 |
| 2147 | -0.01 | 0.11 | -0.08 | 0.94 | 1 | Wwp2 | WW domain containing E3 ubiquitin prote | 1981 | 1929 | 1730 | 2332 | 2338 | 2047 | 2250 | 2757 | 2291 | 2210 | 1751 |
| 2216 | -0.01 | 0.11 | -0.14 | 0.89 | 1 | Rnf220 | ring finger protein 220 | 2082 | 2004 | 2315 | 2585 | 2266 | 1922 | 2249 | 2564 | 2126 | 2429 | 1834 |
| 2353 | 0.03 | 0.08 | 0.38 | 0.71 | 1 | Ccdc117 | coiled-coil domain containing 117 | 1854 | 2190 | 1959 | 2451 | 2496 | 2629 | 2248 | 2755 | 2584 | 2183 | 2534 |
| 1847 | 0.15 | 0.12 | 1.24 | 0.21 | 1 | Kat14 | lysine acetyltransferase 14 | 1348 | 1216 | 1359 | 1936 | 2242 | 1348 | 2248 | 2639 | 2035 | 2465 | 1481 |
| 2355 | 0.04 | 0.11 | 0.35 | 0.73 | 1 | Oip5os1 | | 3694 | 2941 | 2846 | 1847 | 1994 | 2030 | 2244 | 2195 | 2131 | 1926 | 2050 |
| 2258 | 0.00 | 0.12 | -0.03 | 0.98 | 1 | Nsd2 | nuclear receptor binding SET domain pro | 2213 | 1851 | 1921 | 2317 | 1838 | 3092 | 2243 | 2150 | 2350 | 2101 | 2764 |
| 2547 | -0.16 | 0.13 | -1.22 | 0.22 | 1 | Setd2 | SET domain containing 2 | 3153 | 3197 | 2502 | 2666 | 2235 | 2947 | 2241 | 2009 | 2368 | 1801 | 2901 |
| 2197 | 0.14 | 0.08 | 1.70 | 0.09 | 1 | Dr1 | down-regulator of transcription 1 | 1985 | 2491 | 2024 | 2318 | 2083 | 1757 | 2240 | 2607 | 2234 | 2404 | 2024 |
| 2271 | 0.02 | 0.08 | 0.28 | 0.78 | 1 | Kcmf1 | potassium channel modulatory factor 1 | 2193 | 2386 | 2024 | 2641 | 2300 | 1954 | 2239 | 2420 | 2303 | 2369 | 2153 |
| 2156 | 0.05 | 0.09 | 0.59 | 0.55 | 1 | Ube2g2 | ubiquitin-conjugating enzyme E2G 2 | 1708 | 1624 | 1796 | 2170 | 2559 | 2161 | 2239 | 2077 | 2451 | 2494 | 2442 |
| 1987 | -0.14 | 0.08 | -1.64 | 0.10 | 1 | Psmd11 | proteasome (prosome, macropain) 26S su | 1772 | 1883 | 2003 | 2015 | 2193 | 2035 | 2239 | 1872 | 2030 | 1939 | 1872 |
| 2179 | 0.00 | 0.09 | -0.02 | 0.99 | 1 | Zkscan1 | zinc finger with KRAB and SCAN domains | 1726 | 1757 | 1850 | 2038 | 2629 | 2417 | 2238 | 1928 | 2520 | 2392 | 2477 |
| 2054 | -0.14 | 0.09 | -1.51 | 0.13 | 1 | Prxl2c | peroxiredoxin like 2C | 2181 | 2563 | 2247 | 2191 | 2181 | 1556 | 2238 | 1975 | 2113 | 1951 | 1399 |
| 2439 | -0.08 | 0.13 | -0.61 | 0.54 | 1 | Golga5 | golgi autoantigen, golgin subfamily a, 5 | 1854 | 1999 | 1785 | 2646 | 2703 | 3277 | 2237 | 1900 | 2864 | 2361 | 3207 |
| 2338 | -0.09 | 0.14 | -0.66 | 0.51 | 1 | Fry | FRY microtubule binding protein | 1668 | 2231 | 1705 | 3022 | 2707 | 2429 | 2237 | 2192 | 2932 | 2420 | 2174 |
| 1929 | -0.05 | 0.11 | -0.40 | 0.69 | 1 | Sobp | sine oculis binding protein | 1507 | 1329 | 1691 | 2244 | 2353 | 1617 | 2235 | 2499 | 2284 | 2002 | 1458 |
| 1720 | 0.08 | 0.11 | 0.76 | 0.45 | 1 | Galc | galactosylceramidase | 1212 | 1224 | 1278 | 1946 | 1942 | 1308 | 2235 | 1858 | 2055 | 2230 | 1635 |
| 2321 | 0.05 | 0.10 | 0.50 | 0.61 | 1 | Vbp1 | von Hippel-Lindau binding protein 1 | 2322 | 2307 | 2635 | 2347 | 2452 | 1960 | 2234 | 2182 | 2316 | 2534 | 2246 |
| 2315 | -0.09 | 0.11 | -0.81 | 0.42 | 1 | Susd6 | sushi domain containing 6 | 3013 | 2512 | 2463 | 2430 | 2143 | 2195 | 2234 | 2186 | 2262 | 1993 | 2030 |
| 2278 | 0.07 | 0.09 | 0.79 | 0.43 | 1 | N4bp2l2 | NEDD4 binding protein 2-like 2 | 1915 | 2023 | 1938 | 2067 | 2086 | 2933 | 2234 | 2485 | 2164 | 2007 | 3203 |
| 2060 | 0.10 | 0.10 | 1.00 | 0.32 | 1 | Txndc12 | thioredoxin domain containing 12 (endop | 1839 | 1875 | 2033 | 2002 | 2297 | 1630 | 2234 | 2497 | 2155 | 2378 | 1722 |
| 2162 | -0.19 | 0.11 | -1.71 | 0.09 | 1 | Nup210 | nucleoporin 210 | 2781 | 2017 | 2218 | 2210 | 2293 | 2187 | 2233 | 1761 | 2132 | 1926 | 2020 |
| 1995 | 0.19 | 0.15 | 1.31 | 0.19 | 1 | Cyp4v3 | cytochrome P450, family 4, subfamily v, p | 1837 | 2383 | 2400 | 1778 | 1874 | 1281 | 2233 | 2056 | 1977 | 2610 | 1519 |
| 1897 | -0.08 | 0.10 | -0.83 | 0.41 | 1 | Tmcc2 | transmembrane and coiled-coil domains 2 | 2088 | 2145 | 2010 | 2029 | 1974 | 1287 | 2233 | 1697 | 1958 | 2206 | 1240 |
| 3127 | 0.10 | 0.11 | 0.91 | 0.36 | 1 | Tpm4 | tropomyosin 4 | 6823 | 4368 | 5252 | 2117 | 2397 | 1915 | 2232 | 2112 | 2391 | 2638 | 2153 |
| 2251 | 0.01 | 0.10 | 0.08 | 0.94 | 1 | Ext2 | exostosin glycosyltransferase 2 | 2979 | 2599 | 2772 | 2177 | 2264 | 1530 | 2232 | 2176 | 2391 | 1987 | 1652 |
| 2088 | -0.16 | 0.10 | -1.54 | 0.12 | 1 | Nup214 | nucleoporin 214 | 2050 | 1997 | 1951 | 2186 | 2450 | 2054 | 2231 | 1898 | 2254 | 2326 | 1572 |
| 2792 | 0.13 | 0.11 | 1.23 | 0.22 | 1 | Ubn2 | ubinuclein 2 | 1743 | 1914 | 1912 | 2238 | 2552 | 5197 | 2230 | 3056 | 2593 | 2593 | 4682 |
| 2322 | 0.03 | 0.12 | 0.30 | 0.77 | 1 | Ninj1 | ninjurin 1 | 1995 | 2183 | 1803 | 2633 | 3010 | 1811 | 2230 | 2738 | 2512 | 2726 | 1908 |
| 1835 | 0.02 | 0.12 | 0.18 | 0.86 | 1 | Frk | fyn-related kinase | 1391 | 1900 | 1648 | 2195 | 1750 | 1409 | 2230 | 2200 | 1866 | 2071 | 1520 |
| 1817 | 0.19 | 0.11 | 1.78 | 0.08 | 1 | Bnip3 | BCL2/adenovirus E1B interacting protein | 1125 | 1174 | 1222 | 1856 | 2027 | 1593 | 2230 | 2417 | 2095 | 2286 | 1960 |
| 2158 | -0.04 | 0.09 | -0.40 | 0.69 | 1 | Rabgap1l | RAB GTPase activating protein 1-like | 1947 | 2015 | 2013 | 2084 | 2201 | 2478 | 2228 | 1915 | 2460 | 2036 | 2365 |
| 2081 | 0.02 | 0.13 | 0.14 | 0.89 | 1 | Rap1a | RAS-related protein 1a | 2835 | 2109 | 2093 | 2047 | 1819 | 1792 | 2228 | 1682 | 2277 | 2012 | 1999 |
| 1965 | -0.06 | 0.10 | -0.57 | 0.57 | 1 | Utp6 | UTP6 small subunit processome compon | 1975 | 2095 | 1998 | 1806 | 2104 | 1797 | 2228 | 1694 | 1952 | 2174 | 1793 |
| 1926 | -0.01 | 0.10 | -0.07 | 0.94 | 1 | Rnf7 | ring finger protein 7 | 1537 | 1392 | 1688 | 2144 | 2339 | 1613 | 2228 | 2041 | 2074 | 2369 | 1758 |

| | | | | | | | | | | | | | | | | | | |
|---|---|---|---|---|---|---|---|---|---|---|---|---|---|---|---|---|---|---|
| 2290 | 0.08 | 0.12 | 0.67 | 0.50 | 1 | Trak2 | trafficking protein, kinesin binding 2 | 2799 | 2208 | 2280 | 2162 | 2323 | 1966 | 2225 | 2757 | 2389 | 2155 | 1923 |
| 2237 | 0.05 | 0.08 | 0.61 | 0.54 | 1 | Tecpr1 | tectonin beta-propeller repeat containing | 2331 | 2341 | 2430 | 2366 | 2146 | 1857 | 2225 | 2435 | 2137 | 2428 | 1909 |
| 2172 | -0.03 | 0.09 | -0.31 | 0.76 | 1 | Smyd2 | SET and MYND domain containing 2 | 1863 | 1801 | 2019 | 2240 | 2544 | 2178 | 2225 | 2375 | 2277 | 2371 | 1995 |
| 2098 | 0.16 | 0.10 | 1.59 | 0.11 | 1 | Phf20 | PHD finger protein 20 | 2315 | 2157 | 1857 | 2117 | 1797 | 1735 | 2225 | 2819 | 1997 | 2226 | 1834 |
| 1706 | 0.03 | 0.13 | 0.20 | 0.84 | 1 | Akr1c19 | aldo-keto reductase family 1, member C1 | 1706 | 1365 | 1608 | 1609 | 1520 | 1631 | 2223 | 1693 | 1391 | 2174 | 1848 |
| 2015 | 0.03 | 0.11 | 0.25 | 0.80 | 1 | Tenm2 | teneurin transmembrane protein 2 | 1662 | 1946 | 1521 | 2166 | 2312 | 1716 | 2222 | 2354 | 2425 | 2214 | 1625 |
| 2154 | 0.03 | 0.10 | 0.30 | 0.77 | 1 | Carmil3 | capping protein regulator and myosin 1 li | 333 | 370 | 357 | 1803 | 1878 | 5429 | 2221 | 2121 | 1905 | 2168 | 5113 |
| 2132 | -0.09 | 0.11 | -0.83 | 0.41 | 1 | Frs2 | fibroblast growth factor receptor substrate | 1937 | 2441 | 1913 | 2325 | 2059 | 2241 | 2221 | 2162 | 2241 | 1713 | 2202 |
| 2506 | 0.08 | 0.07 | 1.16 | 0.25 | 1 | Wdr82 | WD repeat domain containing 82 | 2939 | 3319 | 2681 | 2350 | 1972 | 2508 | 2220 | 2603 | 2140 | 2320 | 2513 |
| 2291 | -0.04 | 0.12 | -0.33 | 0.74 | 1 | Ufd1 | ubiquitin recognition factor in ER-associa | 2033 | 1708 | 2246 | 2135 | 2689 | 2717 | 2220 | 2552 | 2398 | 1982 | 2522 |
| 1903 | -0.07 | 0.13 | -0.52 | 0.60 | 1 | Kdm4c | lysine (K)-specific demethylase 4C | 1576 | 1733 | 1778 | 1851 | 1907 | 2103 | 2220 | 1506 | 2170 | 1911 | 2179 |
| 2569 | -0.12 | 0.11 | -1.04 | 0.30 | 1 | Nlgn2 | neuroligin 2 | 2732 | 3417 | 2841 | 2170 | 2124 | 3596 | 2218 | 2014 | 1981 | 2335 | 2832 |
| 1971 | -0.11 | 0.12 | -0.94 | 0.35 | 1 | Slc9a8 | solute carrier family 9 (sodium/hydrogen e | 1325 | 1874 | 1380 | 2342 | 2098 | 2233 | 2217 | 2027 | 2182 | 2071 | 1933 |
| 2840 | -0.05 | 0.13 | -0.39 | 0.70 | 1 | Add3 | adducin 3 (gamma) | 5737 | 4790 | 3570 | 2162 | 2315 | 2012 | 2216 | 1948 | 2035 | 2319 | 2139 |
| 2374 | -0.12 | 0.09 | -1.37 | 0.17 | 1 | Rbfox2 | RNA binding protein, fox-1 homolog (C. e | 2571 | 2878 | 2781 | 2345 | 1921 | 2835 | 2216 | 2201 | 1933 | 1847 | 2588 |
| 2235 | -0.08 | 0.09 | -0.84 | 0.40 | 1 | Odf2 | outer dense fiber of sperm tails 2 | 1950 | 1763 | 1771 | 2140 | 2177 | 3285 | 2216 | 2124 | 2218 | 1919 | 3026 |
| 2036 | -0.21 | 0.15 | -1.41 | 0.16 | 1 | Wiz | widely-interspaced zinc finger motifs | 2014 | 2308 | 1931 | 2178 | 2493 | 1735 | 2213 | 1517 | 2262 | 2319 | 1428 |
| 2009 | -0.12 | 0.09 | -1.37 | 0.17 | 1 | Wrnip1 | Werner helicase interacting protein 1 | 1927 | 1942 | 1820 | 1946 | 2173 | 2221 | 2213 | 1883 | 2076 | 1969 | 1926 |
| 1862 | 0.06 | 0.12 | 0.46 | 0.65 | 1 | Lyrm9 | LYR motif containing 9 | 1167 | 1746 | 1443 | 2144 | 1889 | 1631 | 2213 | 2529 | 2079 | 1870 | 1766 |
| 2289 | 0.17 | 0.12 | 1.36 | 0.17 | 1 | Ubxn6 | UBX domain protein 6 | 1974 | 2197 | 2125 | 2319 | 2244 | 2051 | 2211 | 3221 | 2135 | 2722 | 1982 |
| 2258 | 0.03 | 0.09 | 0.28 | 0.78 | 1 | Tmem109 | transmembrane protein 109 | 2735 | 2364 | 2745 | 2197 | 2304 | 1700 | 2211 | 2401 | 2114 | 2378 | 1688 |
| 1915 | -0.02 | 0.09 | -0.17 | 0.86 | 1 | Usp32 | ubiquitin specific peptidase 32 | 1570 | 1372 | 1365 | 2078 | 1947 | 2188 | 2210 | 1819 | 2190 | 2234 | 2094 |
| 1903 | -0.01 | 0.10 | -0.09 | 0.93 | 1 | Vps29 | VPS29 retromer complex component | 1742 | 1793 | 1662 | 1906 | 1886 | 1894 | 2210 | 1758 | 2055 | 2035 | 1990 |
| 2298 | 0.03 | 0.12 | 0.28 | 0.78 | 1 | Msl3 | MSL complex subunit 3 | 2786 | 2439 | 2406 | 2371 | 2175 | 1964 | 2209 | 2216 | 2608 | 1915 | 2184 |
| 2073 | -0.07 | 0.09 | -0.79 | 0.43 | 1 | Scamp5 | secretory carrier membrane protein 5 | 1471 | 1511 | 1694 | 2347 | 2317 | 2406 | 2209 | 2266 | 2075 | 2154 | 2353 |
| 2256 | 0.03 | 0.13 | 0.22 | 0.82 | 1 | Ctdsp1 | CTD (carboxy-terminal domain, RNA poly | 2949 | 2695 | 2933 | 2158 | 1975 | 1658 | 2208 | 2778 | 1955 | 1892 | 1613 |
| 2285 | 0.04 | 0.09 | 0.43 | 0.67 | 1 | Cpsf3 | cleavage and polyadenylation specificity | 2769 | 2277 | 2598 | 2071 | 2078 | 2266 | 2206 | 2183 | 2126 | 2490 | 2075 |
| 2131 | 0.04 | 0.10 | 0.35 | 0.73 | 1 | Tnk2 | tyrosine kinase, non-receptor, 2 | 2144 | 2343 | 2294 | 1899 | 2185 | 1914 | 2205 | 2253 | 2218 | 2294 | 1693 |
| 1827 | 0.00 | 0.11 | 0.03 | 0.98 | 1 | Akr1c12 | aldo-keto reductase family 1, member C1 | 640 | 866 | 789 | 2708 | 2255 | 1757 | 2205 | 2591 | 2032 | 2172 | 2083 |
| 1986 | 0.09 | 0.11 | 0.83 | 0.41 | 1 | Galnt11 | polypeptide N-acetylgalactosaminyltransf | 1223 | 1435 | 1378 | 1827 | 2376 | 2214 | 2204 | 2138 | 2387 | 2464 | 2201 |
| 1538 | 0.03 | 0.15 | 0.20 | 0.84 | 1 | Cryl1 | crystallin, lambda 1 | 713 | 575 | 678 | 1722 | 1936 | 1516 | 2202 | 2405 | 1636 | 2124 | 1411 |
| 2239 | 0.07 | 0.12 | 0.58 | 0.56 | 1 | Sae1 | SUMO1 activating enzyme subunit 1 | 2666 | 1898 | 2680 | 2219 | 2267 | 1798 | 2201 | 2419 | 2304 | 2290 | 1885 |
| 2026 | 0.15 | 0.11 | 1.45 | 0.15 | 1 | Tmed5 | transmembrane p24 trafficking protein 5 | 2145 | 1791 | 1564 | 2073 | 2184 | 1500 | 2201 | 2147 | 2407 | 2512 | 1768 |
| 2284 | 0.07 | 0.11 | 0.63 | 0.53 | 1 | Cnst | consortin, connexin sorting protein | 1675 | 1778 | 1556 | 2732 | 2407 | 2481 | 2200 | 2275 | 2570 | 2710 | 2739 |
| 2516 | 0.08 | 0.11 | 0.67 | 0.51 | 1 | Nup50 | nucleoporin 50 | 2195 | 2617 | 2423 | 2938 | 2726 | 2092 | 2199 | 2942 | 2413 | 3008 | 2121 |
| 2509 | -0.06 | 0.13 | -0.47 | 0.64 | 1 | Foxo3 | forkhead box O3 | 2665 | 3786 | 2394 | 2473 | 2364 | 2533 | 2197 | 2474 | 2326 | 2415 | 1970 |
| 1968 | -0.01 | 0.09 | -0.07 | 0.95 | 1 | Atp6v0c | ATPase, H+ transporting, lysosomal V0 su | 1408 | 1538 | 1190 | 2551 | 1909 | 2088 | 2197 | 2575 | 1735 | 2513 | 1948 |
| 2093 | -0.08 | 0.09 | -0.84 | 0.40 | 1 | Cnpy3 | canopy FGF signaling regulator 3 | 2120 | 1641 | 2230 | 2024 | 2493 | 2017 | 2196 | 1694 | 2276 | 2261 | 2076 |
| 1732 | 0.05 | 0.11 | 0.46 | 0.65 | 1 | Chm | choroidermia (RAB escort protein 1) | 1422 | 1438 | 1545 | 1634 | 1848 | 1534 | 2196 | 1673 | 1940 | 2110 | 1713 |
| 2213 | 0.07 | 0.12 | 0.61 | 0.54 | 1 | Suz12 | SUZ12 polycomb repressive complex 2 su | 2177 | 2172 | 1605 | 2347 | 1916 | 2521 | 2195 | 1996 | 2275 | 2474 | 2669 |
| 2178 | -0.01 | 0.14 | -0.10 | 0.92 | 1 | Btaf1 | B-TFIID TATA-box binding protein associ | 1736 | 1997 | 1440 | 2307 | 1798 | 3139 | 2195 | 1937 | 2305 | 1852 | 3257 |
| 1487 | 0.15 | 0.12 | 1.23 | 0.22 | 1 | Vgf | VGF nerve growth factor inducible | 1021 | 929 | 768 | 1697 | 1617 | 898 | 2195 | 1740 | 1643 | 2851 | 994 |
| 2342 | 0.12 | 0.12 | 1.07 | 0.28 | 1 | Zfyve27 | zinc finger, FYVE domain containing 27 | 1801 | 1870 | 2012 | 1970 | 2250 | 3252 | 2194 | 2801 | 2320 | 2249 | 3046 |
| 1344 | -0.19 | 0.14 | -1.37 | 0.17 | 1 | Met | met proto-oncogene | 1928 | 1577 | 1199 | 1393 | 886 | 996 | 2194 | 1116 | 977 | 1571 | 942 |
| 1922 | 0.01 | 0.11 | 0.12 | 0.90 | 1 | Trmt6 | tRNA methyltransferase 6 | 1226 | 1133 | 1404 | 1897 | 2486 | 2061 | 2193 | 2104 | 2544 | 1743 | 2352 |
| 2329 | -0.23 | 0.13 | -1.77 | 0.08 | 1 | Camsap2 | calmodulin regulated spectrin-associated | 2383 | 3038 | 2362 | 2682 | 2313 | 2439 | 2192 | 1746 | 2208 | 1948 | 2309 |
| 2431 | 0.05 | 0.11 | 0.43 | 0.67 | 1 | Tut7 | terminal uridylyl transferase 7 | 2410 | 1969 | 2205 | 2142 | 2239 | 3305 | 2190 | 2266 | 2458 | 1918 | 3643 |
| 2382 | 0.15 | 0.13 | 1.13 | 0.26 | 1 | Zmat2 | zinc finger, matrin type 2 | 2918 | 2411 | 2814 | 2071 | 2433 | 1866 | 2190 | 2971 | 2348 | 2160 | 2019 |
| 3075 | -0.07 | 0.11 | -0.60 | 0.55 | 1 | Ski | ski sarcoma viral oncogene homolog (avi | 5579 | 4837 | 4555 | 2759 | 2559 | 2106 | 2188 | 2450 | 2434 | 2597 | 1762 |
| 2542 | 0.03 | 0.11 | 0.30 | 0.76 | 1 | Phc2 | polyhomeotic 2 | 3230 | 3356 | 3254 | 2375 | 2370 | 1975 | 2188 | 2961 | 2410 | 1991 | 1851 |
| 2048 | 0.15 | 0.11 | 1.36 | 0.17 | 1 | Pitrm1 | pitrilysin metallepetidase 1 | 1549 | 1339 | 1452 | 2353 | 1892 | 2204 | 2187 | 2541 | 2399 | 2203 | 2405 |
| 2424 | 0.14 | 0.10 | 1.36 | 0.17 | 1 | Upf2 | UPF2 regulator of nonsense transcripts h | 2428 | 2689 | 2374 | 2329 | 1873 | 2747 | 2186 | 2951 | 1970 | 2420 | 2699 |
| 1704 | 0.16 | 0.17 | 0.98 | 0.33 | 1 | Xk | X-linked Kx blood group | 382 | 683 | 583 | 2124 | 1941 | 1792 | 2185 | 2285 | 2495 | 2496 | 1780 |
| 2268 | -0.17 | 0.11 | -1.48 | 0.14 | 1 | Vcpip1 | valosin containing protein (p97)/p47 com | 2225 | 2370 | 2064 | 2795 | 2236 | 2454 | 2184 | 2248 | 2289 | 1716 | 2372 |
| 1845 | 0.01 | 0.12 | 0.04 | 0.97 | 1 | 5730455P16Ril | RIKEN cDNA 5730455P16 gene | 1322 | 1466 | 1293 | 1923 | 2011 | 1966 | 2184 | 1668 | 2128 | 2142 | 2190 |
| 2508 | -0.01 | 0.08 | -0.07 | 0.95 | 1 | Jade3 | jade family PHD finger 3 | 2212 | 2357 | 2607 | 2513 | 2262 | 3253 | 2183 | 2504 | 2330 | 2048 | 3325 |
| 1707 | 0.24 | 0.15 | 1.67 | 0.09 | 1 | Fam83g | family with sequence similarity 83, memb | 394 | 349 | 520 | 2233 | 2262 | 1271 | 2180 | 3126 | 2546 | 2590 | 1309 |
| 2624 | 0.09 | 0.11 | 0.80 | 0.42 | 1 | Ube2h | ubiquitin-conjugating enzyme E2H | 2665 | 3381 | 3018 | 2608 | 2651 | 2134 | 2179 | 3202 | 2201 | 2603 | 2227 |
| 2022 | 0.07 | 0.10 | 0.72 | 0.47 | 1 | Glod4 | glyoxalase domain containing 4 | 2371 | 2198 | 2244 | 1868 | 1917 | 1557 | 2179 | 2311 | 1962 | 2066 | 1568 |
| 2760 | -0.16 | 0.13 | -1.25 | 0.21 | 1 | Lmna | lamin A | 4593 | 3701 | 4166 | 1953 | 2836 | 2425 | 2178 | 1400 | 2613 | 2255 | 2239 |

| | | | | | | | | | | | | | | | | | | |
|---|---|---|---|---|---|---|---|---|---|---|---|---|---|---|---|---|---|---|
| 2362 | -0.04 | 0.08 | -0.46 | 0.65 | 1 | Rab11b | RAB11B, member RAS oncogene family | 2197 | 2547 | 2181 | 2745 | 2393 | 2337 | 2177 | 2667 | 2158 | 2424 | 2153 |
| 2304 | 0.11 | 0.11 | 1.02 | 0.31 | 1 | Rif1 | replication timing regulatory factor 1 | 1738 | 1411 | 1950 | 1961 | 2033 | 3499 | 2175 | 2004 | 2679 | 1976 | 3920 |
| 2127 | 0.04 | 0.08 | 0.41 | 0.68 | 1 | Cdk13 | cyclin-dependent kinase 13 | 1832 | 1932 | 1944 | 1986 | 2254 | 2321 | 2175 | 1992 | 2272 | 2312 | 2377 |
| 1796 | 0.09 | 0.12 | 0.70 | 0.48 | 1 | Trappc13 | trafficking protein particle complex 13 | 1103 | 1059 | 1000 | 1836 | 2003 | 2002 | 2175 | 1584 | 2229 | 2356 | 2408 |
| 1783 | 0.10 | 0.15 | 0.67 | 0.50 | 1 | Cnih1 | cornichon family AMPA receptor auxiliary | 1501 | 1530 | 1843 | 1748 | 1851 | 1314 | 2175 | 2489 | 2099 | 1665 | 1402 |
| 2190 | -0.01 | 0.11 | -0.11 | 0.92 | 1 | Urgcp | upregulator of cell proliferation | 1836 | 2438 | 1940 | 2357 | 2365 | 2079 | 2173 | 2208 | 2382 | 2376 | 1938 |
| 2106 | 0.09 | 0.15 | 0.62 | 0.54 | 1 | Plxna2 | plexin A2 | 1968 | 2352 | 2606 | 1924 | 1578 | 2201 | 2172 | 1949 | 1937 | 2371 | 2104 |
| 2137 | 0.16 | 0.12 | 1.36 | 0.17 | 1 | Chmp4c | charged multivesicular body protein 4C | 1059 | 1222 | 1314 | 2601 | 2219 | 2431 | 2171 | 2375 | 2692 | 2809 | 2619 |
| 2034 | -0.05 | 0.08 | -0.61 | 0.54 | 1 | Atg4b | autophagy related 4B, cysteine peptidase | 1995 | 1947 | 2147 | 2122 | 2032 | 1963 | 2171 | 1912 | 2055 | 2026 | 2008 |
| 1922 | -0.04 | 0.12 | -0.35 | 0.72 | 1 | Otud6b | OTU domain containing 6B | 1609 | 1798 | 1654 | 1941 | 2175 | 1866 | 2171 | 1766 | 2264 | 1810 | 2093 |
| 2315 | -0.06 | 0.10 | -0.58 | 0.56 | 1 | Mapkapk2 | MAP kinase-activated protein kinase 2 | 3877 | 2730 | 2954 | 1884 | 2313 | 1719 | 2170 | 1821 | 2128 | 2342 | 1528 |
| 2169 | 0.11 | 0.13 | 0.84 | 0.40 | 1 | Rab33b | RAB33B, member RAS oncogene family | 1545 | 2134 | 1988 | 2263 | 2255 | 2065 | 2170 | 2351 | 2232 | 2750 | 2108 |
| 2139 | 0.00 | 0.08 | 0.01 | 0.99 | 1 | Stx6 | syntaxin 6 | 2319 | 2415 | 2406 | 2181 | 2037 | 1811 | 2170 | 2096 | 2185 | 2000 | 1910 |
| 2312 | 0.12 | 0.13 | 0.92 | 0.36 | 1 | Brd3 | bromodomain containing 3 | 2168 | 2376 | 2313 | 1996 | 2259 | 2454 | 2169 | 2936 | 2503 | 1969 | 2285 |
| 2160 | 0.09 | 0.12 | 0.76 | 0.45 | 1 | Tmem242 | transmembrane protein 242 | 2046 | 1785 | 2427 | 2431 | 2420 | 1483 | 2169 | 2686 | 2370 | 2280 | 1660 |
| 2551 | 0.16 | 0.11 | 1.52 | 0.13 | 1 | Bod1l | biorientation of chromosomes in cell divis | 2086 | 2378 | 2316 | 2118 | 2261 | 3502 | 2168 | 2833 | 2723 | 1984 | 3695 |
| 2415 | 0.08 | 0.11 | 0.73 | 0.47 | 1 | Sowahc | sosondowah ankyrin repeat domain famil | 2285 | 2318 | 1794 | 2280 | 2402 | 2950 | 2168 | 2252 | 2544 | 2514 | 3059 |
| 1927 | 0.26 | 0.14 | 1.85 | 0.06 | 1 | Vps8 | VPS8 CORVET complex subunit | 1363 | 1038 | 1175 | 1988 | 1944 | 1890 | 2167 | 3071 | 2235 | 2268 | 2059 |
| 2239 | 0.03 | 0.08 | 0.41 | 0.68 | 1 | Drosha | drosha, ribonuclease type III | 2056 | 2260 | 2172 | 2450 | 2341 | 2018 | 2166 | 2316 | 2532 | 2223 | 2095 |
| 2142 | 0.04 | 0.12 | 0.30 | 0.77 | 1 | Nans | N-acetylneuraminic acid synthase (sialic a | 1763 | 1276 | 1992 | 2295 | 2567 | 2090 | 2166 | 2002 | 2586 | 2691 | 2133 |
| 1956 | 0.05 | 0.10 | 0.53 | 0.60 | 1 | Ttc33 | tetratricopeptide repeat domain 33 | 1633 | 1795 | 1845 | 2068 | 2158 | 1581 | 2166 | 2319 | 1929 | 2307 | 1714 |
| 2139 | 0.06 | 0.09 | 0.70 | 0.48 | 1 | 1810013L24Rik | RIKEN cDNA 1810013L24 gene | 1866 | 2110 | 2171 | 2055 | 1948 | 2326 | 2164 | 2099 | 2119 | 2173 | 2501 |
| 2092 | -0.08 | 0.11 | -0.70 | 0.49 | 1 | Crebl2 | cAMP responsive element binding proteir | 1806 | 2219 | 2135 | 2283 | 2159 | 2049 | 2164 | 1782 | 2188 | 2085 | 2140 |
| 2246 | -0.12 | 0.09 | -1.30 | 0.19 | 1 | Stk24 | serine/threonine kinase 24 | 2642 | 2654 | 2711 | 2493 | 2292 | 1747 | 2162 | 1974 | 2172 | 2286 | 1572 |
| 2167 | -0.09 | 0.12 | -0.76 | 0.45 | 1 | Atxn7l3 | ataxin 7-like 3 | 2315 | 2528 | 2521 | 2263 | 1975 | 2081 | 2162 | 1685 | 2155 | 2190 | 1963 |
| 2005 | 0.16 | 0.22 | 0.74 | 0.46 | 1 | Mien1 | migration and invasion enhancer 1 | 1440 | 1301 | 1943 | 2406 | 2070 | 1441 | 2162 | 3926 | 1968 | 1816 | 1584 |
| 2285 | 0.06 | 0.12 | 0.50 | 0.61 | 1 | Cfdp1 | craniofacial development protein 1 | 3046 | 2673 | 3030 | 1896 | 2141 | 1831 | 2161 | 2436 | 2098 | 1854 | 1968 |
| 1963 | -0.06 | 0.13 | -0.51 | 0.61 | 1 | Hivep2 | human immunodeficiency virus type I enh | 1746 | 1856 | 1186 | 2031 | 1896 | 2497 | 2161 | 2067 | 1885 | 1803 | 2463 |
| 2212 | -0.11 | 0.12 | -0.88 | 0.38 | 1 | Tut4 | terminal uridylyl transferase 4 | 1591 | 2447 | 2281 | 2192 | 2340 | 2626 | 2159 | 2025 | 2131 | 1955 | 2578 |
| 2035 | 0.03 | 0.10 | 0.26 | 0.80 | 1 | Abraxas2 | BRISC complex subunit | 1297 | 1242 | 1065 | 2610 | 2343 | 2201 | 2159 | 2406 | 2410 | 2246 | 2408 |
| 2578 | 0.06 | 0.11 | 0.53 | 0.60 | 1 | Sf3a1 | splicing factor 3a, subunit 1 | 3125 | 3370 | 3067 | 2333 | 2648 | 2028 | 2158 | 3015 | 2384 | 2330 | 1897 |
| 1988 | 0.03 | 0.11 | 0.31 | 0.76 | 1 | Chchd3 | coiled-coil-helix-coiled-coil-helix domain | 2226 | 1838 | 2245 | 2058 | 1867 | 1601 | 2158 | 1747 | 2119 | 2214 | 1788 |
| 1983 | 0.07 | 0.14 | 0.46 | 0.64 | 1 | Map3k10 | mitogen-activated protein kinase kinase k | 1673 | 1260 | 1507 | 1978 | 2416 | 1896 | 2158 | 2733 | 2119 | 2410 | 1662 |
| 1951 | -0.07 | 0.10 | -0.69 | 0.49 | 1 | Ptpn23 | protein tyrosine phosphatase, non-recept | 1371 | 1524 | 1559 | 1962 | 2318 | 2263 | 2158 | 1886 | 2497 | 2007 | 1916 |
| 1943 | 0.10 | 0.12 | 0.80 | 0.42 | 1 | Cldn12 | claudin 12 | 1389 | 1697 | 1677 | 2326 | 1841 | 1697 | 2158 | 2426 | 1900 | 2463 | 1797 |
| 1886 | 0.08 | 0.10 | 0.73 | 0.47 | 1 | Ttc7b | tetratricopeptide repeat domain 7B | 1440 | 1321 | 1689 | 2190 | 2226 | 1366 | 2155 | 2144 | 2242 | 2509 | 1463 |
| 1882 | -0.03 | 0.10 | -0.27 | 0.79 | 1 | Lss | lanosterol synthase | 817 | 926 | 1025 | 2046 | 2254 | 2567 | 2155 | 1879 | 2580 | 1922 | 2533 |
| 2557 | 0.06 | 0.13 | 0.43 | 0.67 | 1 | Itgb5 | integrin beta 5 | 5498 | 3535 | 4322 | 2076 | 1889 | 1095 | 2153 | 2421 | 1945 | 2121 | 1071 |
| 2286 | -0.18 | 0.10 | -1.75 | 0.08 | 1 | Sri | sorcin | 3703 | 3256 | 3621 | 1986 | 1972 | 1648 | 2152 | 1659 | 1760 | 1631 | 1762 |
| 1923 | -0.05 | 0.09 | -0.55 | 0.59 | 1 | Med14 | mediator complex subunit 14 | 1869 | 1797 | 1873 | 1975 | 1754 | 2072 | 2152 | 1953 | 1896 | 1972 | 1843 |
| 2228 | -0.12 | 0.12 | -1.02 | 0.31 | 1 | Strn4 | striatin, calmodulin binding protein 4 | 2429 | 1797 | 2342 | 2451 | 2227 | 2512 | 2151 | 1943 | 2391 | 2161 | 2107 |
| 2141 | -0.09 | 0.09 | -0.99 | 0.32 | 1 | Ppp5c | protein phosphatase 5, catalytic subunit | 2133 | 1838 | 2124 | 2252 | 2447 | 2146 | 2150 | 1929 | 2170 | 2221 | 2146 |
| 2734 | -0.07 | 0.12 | -0.56 | 0.57 | 1 | Srrt | serrate RNA effector molecule homolog (/ | 2916 | 2082 | 2656 | 2430 | 2754 | 4132 | 2149 | 2399 | 2329 | 2154 | 4073 |
| 2053 | -0.01 | 0.09 | -0.08 | 0.94 | 1 | Arl8a | ADP-ribosylation factor-like 8A | 1793 | 1705 | 1878 | 2286 | 2174 | 1997 | 2149 | 2052 | 2407 | 2300 | 1843 |
| 1838 | -0.02 | 0.11 | -0.17 | 0.86 | 1 | Usp46 | ubiquitin specific peptidase 46 | 1285 | 1423 | 1421 | 2441 | 2051 | 1468 | 2149 | 1963 | 2200 | 2335 | 1482 |
| 1906 | -0.06 | 0.11 | -0.52 | 0.60 | 1 | Naa35 | N(alpha)-acetyltransferase 35, NatC auxil | 1452 | 1494 | 1533 | 1915 | 2118 | 2202 | 2148 | 1555 | 2107 | 2137 | 2310 |
| 2030 | 0.11 | 0.19 | 0.58 | 0.56 | 1 | Uqcrq | ubiquinol-cytochrome c reductase, compl | 1377 | 1175 | 1666 | 2364 | 2483 | 1634 | 2147 | 3674 | 2177 | 2027 | 1608 |
| 1911 | -0.01 | 0.12 | -0.05 | 0.96 | 1 | Lamtor1 | late endosomal/lysosomal adaptor, MAPK | 1687 | 1879 | 2035 | 2098 | 2116 | 1325 | 2147 | 2266 | 1895 | 2360 | 1214 |
| 2257 | 0.00 | 0.10 | -0.05 | 0.96 | 1 | Cr1l | complement component (3b/4b) receptor | 2967 | 2588 | 2560 | 2230 | 2136 | 1857 | 2146 | 1946 | 2303 | 2150 | 1942 |
| 2224 | -0.04 | 0.10 | -0.44 | 0.66 | 1 | Asxl2 | additional sex combs like 2, transcriptiona | 1292 | 1645 | 1489 | 2490 | 2389 | 3150 | 2146 | 2098 | 2536 | 2185 | 3042 |
| 2068 | -0.15 | 0.11 | -1.44 | 0.15 | 1 | Ctdnep1 | CTD nuclear envelope phosphatase 1 | 2286 | 2488 | 2182 | 2205 | 2309 | 1664 | 2146 | 1808 | 2234 | 1811 | 1612 |
| 1930 | -0.22 | 0.12 | -1.77 | 0.08 | 1 | Cnot2 | CCR4-NOT transcription complex, subuni | 2123 | 2073 | 1797 | 1909 | 1943 | 2162 | 2145 | 1389 | 1812 | 1712 | 2166 |
| 2650 | -0.06 | 0.10 | -0.56 | 0.58 | 1 | Tnrc6b | trinucleotide repeat containing 6b | 2800 | 3323 | 2988 | 2588 | 2881 | 2581 | 2142 | 2886 | 2603 | 1863 | 2498 |
| 2444 | 0.01 | 0.10 | 0.08 | 0.94 | 1 | Eif3g | eukaryotic translation initiation factor 3, su | 2913 | 2566 | 3180 | 1977 | 2800 | 2202 | 2142 | 2215 | 2367 | 2155 | 2364 |
| 2010 | 0.02 | 0.10 | 0.24 | 0.81 | 1 | Uba6 | ubiquitin-like modifier activating enzyme 6 | 1523 | 1977 | 1641 | 2263 | 1902 | 2104 | 2142 | 2142 | 2109 | 1959 | 2347 |
| 1884 | -0.08 | 0.09 | -0.84 | 0.40 | 1 | Ankib1 | ankyrin repeat and IBR domain containing | 1754 | 1699 | 1625 | 1963 | 1976 | 1950 | 2142 | 1988 | 1773 | 1823 | 2035 |
| 1922 | -0.01 | 0.13 | -0.08 | 0.94 | 1 | Fam171a1 | family with sequence similarity 171, meml | 1945 | 1666 | 1967 | 2177 | 1869 | 1565 | 2141 | 2420 | 1751 | 2282 | 1354 |
| 2165 | 0.16 | 0.12 | 1.38 | 0.17 | 1 | Irf2bp1 | interferon regulatory factor 2 binding prote | 2076 | 2308 | 2458 | 2314 | 2029 | 1512 | 2140 | 2793 | 1970 | 2513 | 1705 |
| 2236 | 0.05 | 0.10 | 0.46 | 0.65 | 1 | Slc25a11 | solute carrier family 25 (mitochondrial car | 2347 | 2374 | 2492 | 2368 | 2322 | 1680 | 2139 | 2544 | 2158 | 2617 | 1559 |
| 2014 | -0.01 | 0.12 | -0.05 | 0.96 | 1 | Otub1 | OTU domain, ubiquitin aldehyde binding | 2208 | 2013 | 2215 | 1859 | 2089 | 1774 | 2138 | 2345 | 2025 | 1854 | 1630 |

| | | | | | | | | | | | | | | | | | |
|---|---|---|---|---|---|---|---|---|---|---|---|---|---|---|---|---|---|
| 1967 | -0.06 | 0.09 | -0.69 | 0.49 | 1 | Cdc123 | cell division cycle 123 | 1916 | 1753 | 1989 | 1971 | 2172 | 1877 | 2138 | 1774 | 1984 | 2033 | 2032 |
| 1951 | 0.01 | 0.14 | 0.10 | 0.92 | 1 | Adarb1 | adenosine deaminase, RNA-specific, B1 | 1129 | 1095 | 1405 | 2471 | 1864 | 2390 | 2138 | 2389 | 2394 | 1933 | 2252 |
| 1889 | 0.13 | 0.12 | 1.07 | 0.29 | 1 | Ctnnal1 | catenin (cadherin associated protein), alp | 1331 | 1249 | 1397 | 1771 | 1628 | 2433 | 2137 | 1615 | 1865 | 2425 | 2929 |
| 1831 | -0.07 | 0.09 | -0.79 | 0.43 | 1 | Snx12 | sorting nexin 12 | 1841 | 1547 | 1737 | 2016 | 2010 | 1543 | 2137 | 1887 | 2007 | 1866 | 1546 |
| 1955 | 0.02 | 0.10 | 0.25 | 0.81 | 1 | Ift52 | intraflagellar transport 52 | 1488 | 1820 | 1785 | 2107 | 2157 | 1739 | 2136 | 2192 | 2131 | 2217 | 1737 |
| 2278 | -0.01 | 0.12 | -0.06 | 0.95 | 1 | Slc4a2 | solute carrier family 4 (anion exchanger), | 2580 | 2186 | 2555 | 2505 | 2163 | 2053 | 2134 | 2362 | 2215 | 2531 | 1770 |
| 2263 | 0.02 | 0.09 | 0.25 | 0.80 | 1 | Dock7 | dedicator of cytokinesis 7 | 2449 | 2591 | 2303 | 1947 | 2113 | 2519 | 2134 | 1978 | 2288 | 2170 | 2402 |
| 1710 | 0.01 | 0.13 | 0.07 | 0.95 | 1 | Isca1 | iron-sulfur cluster assembly 1 | 1368 | 1413 | 1336 | 1576 | 1695 | 1909 | 2134 | 1417 | 1908 | 2159 | 1892 |
| 2178 | 0.16 | 0.09 | 1.67 | 0.09 | 1 | Slc25a25 | solute carrier family 25 (mitochondrial car | 1527 | 1896 | 1800 | 2500 | 1979 | 2243 | 2133 | 2735 | 2220 | 2235 | 2692 |
| 1869 | 0.06 | 0.12 | 0.51 | 0.61 | 1 | C1galt1c1 | C1GALT1-specific chaperone 1 | 1360 | 1625 | 1749 | 2163 | 2032 | 1441 | 2133 | 2284 | 1932 | 2239 | 1603 |
| 2201 | 0.08 | 0.17 | 0.47 | 0.64 | 1 | Hsd17b4 | hydroxysteroid (17-beta) dehydrogenase | 3151 | 2027 | 3004 | 1970 | 2106 | 1560 | 2132 | 2595 | 1744 | 2425 | 1501 |
| 2137 | 0.06 | 0.11 | 0.60 | 0.55 | 1 | Fbh1 | F-box DNA helicase 1 | 2460 | 2258 | 2060 | 1930 | 1771 | 2336 | 2132 | 1909 | 2100 | 1869 | 2683 |
| 2128 | 0.11 | 0.15 | 0.74 | 0.46 | 1 | Ift80 | intraflagellar transport 80 | 1574 | 2195 | 1971 | 2032 | 2175 | 2157 | 2132 | 2075 | 2095 | 2765 | 2230 |
| 1907 | -0.03 | 0.09 | -0.35 | 0.72 | 1 | Hs2st1 | heparan sulfate 2-O-sulfotransferase 1 | 1977 | 1940 | 1921 | 2123 | 1848 | 1562 | 2129 | 1871 | 2017 | 2021 | 1569 |
| 2282 | -0.12 | 0.11 | -1.09 | 0.27 | 1 | Kdm7a | lysine (K)-specific demethylase 7A | 2684 | 2888 | 2221 | 2199 | 1988 | 2711 | 2127 | 2026 | 2099 | 1770 | 2393 |
| 2600 | 0.05 | 0.10 | 0.49 | 0.62 | 1 | Cpsf6 | cleavage and polyadenylation specific fac | 2388 | 2751 | 2273 | 2161 | 2120 | 4012 | 2126 | 2034 | 2219 | 2352 | 4165 |
| 1910 | 0.07 | 0.16 | 0.42 | 0.67 | 1 | Rnf217 | ring finger protein 217 | 893 | 1354 | 1230 | 2153 | 1985 | 2293 | 2126 | 1797 | 2262 | 2593 | 2321 |
| 2415 | 0.20 | 0.10 | 1.94 | 0.05 | 1 | Tfrc | transferrin receptor | 1102 | 1370 | 1266 | 2381 | 2053 | 3971 | 2125 | 2345 | 2344 | 2585 | 5028 |
| 2218 | -0.03 | 0.09 | -0.38 | 0.71 | 1 | Pex6 | peroxisomal biogenesis factor 6 | 2342 | 2537 | 2390 | 2136 | 2356 | 2038 | 2125 | 2297 | 2072 | 2230 | 1876 |
| 1926 | -0.05 | 0.09 | -0.62 | 0.54 | 1 | Eya3 | EYA transcriptional coactivator and phosp | 1481 | 1534 | 1574 | 2114 | 2179 | 2030 | 2124 | 2241 | 2107 | 1837 | 1964 |
| 2877 | 0.18 | 0.11 | 1.69 | 0.09 | 1 | Cotl1 | coactosin-like 1 (Dictyostelium) | 4912 | 4391 | 4515 | 2354 | 2270 | 1575 | 2123 | 2993 | 2186 | 2611 | 1713 |
| 2211 | -0.14 | 0.10 | -1.45 | 0.15 | 1 | Pex19 | peroxisomal biogenesis factor 19 | 2441 | 2647 | 2445 | 2353 | 2268 | 2072 | 2123 | 1797 | 2194 | 2076 | 1910 |
| 1900 | -0.06 | 0.10 | -0.66 | 0.51 | 1 | Xpr1 | xenotropic and polytropic retrovirus recep | 1765 | 1778 | 1584 | 1925 | 1982 | 2033 | 2123 | 1879 | 2006 | 1792 | 2033 |
| 2147 | 0.08 | 0.13 | 0.58 | 0.56 | 1 | Ecd | ecdysoneless cell cycle regulator | 1762 | 1719 | 1712 | 2100 | 2695 | 2032 | 2121 | 2994 | 2343 | 2063 | 2082 |
| 1713 | 0.09 | 0.10 | 0.89 | 0.37 | 1 | Ggact | gamma-glutamylamine cyclotransferase | 940 | 1016 | 1006 | 1847 | 2188 | 1568 | 2120 | 2092 | 2131 | 2138 | 1800 |
| 2716 | 0.13 | 0.08 | 1.68 | 0.09 | 1 | Ppp3ca | protein phosphatase 3, catalytic subunit, a | 4155 | 4323 | 3658 | 2442 | 2200 | 1692 | 2117 | 2692 | 2230 | 2563 | 1807 |
| 1835 | 0.11 | 0.11 | 1.00 | 0.32 | 1 | Vps50 | VPS50 EARP/GARPII complex subunit | 1219 | 1322 | 1264 | 1872 | 1940 | 1933 | 2115 | 1829 | 2275 | 2022 | 2397 |
| 1613 | -0.02 | 0.15 | -0.13 | 0.89 | 1 | Ppm1h | protein phosphatase 1H (PP2C domain c | 1013 | 535 | 894 | 1939 | 1833 | 1811 | 2115 | 1786 | 1977 | 2018 | 1823 |
| 2119 | -0.08 | 0.12 | -0.70 | 0.48 | 1 | Cux1 | cut-like homeobox 1 | 2677 | 2406 | 2340 | 2039 | 2114 | 1882 | 2113 | 1639 | 2374 | 1968 | 1760 |
| 1994 | 0.03 | 0.12 | 0.27 | 0.79 | 1 | Tmem185a | transmembrane protein 185A | 1525 | 1979 | 1626 | 2211 | 2161 | 1836 | 2112 | 1907 | 2212 | 2235 | 2124 |
| 1979 | -0.06 | 0.10 | -0.60 | 0.55 | 1 | Nol6 | nucleolar protein family 6 (RNA-associate | 2255 | 1751 | 2295 | 1635 | 2425 | 1696 | 2112 | 1452 | 2553 | 1942 | 1654 |
| 2389 | -0.07 | 0.09 | -0.75 | 0.45 | 1 | Rnf44 | ring finger protein 44 | 2849 | 2904 | 2690 | 2460 | 2328 | 2232 | 2109 | 2170 | 2368 | 2264 | 1906 |
| 2155 | -0.02 | 0.09 | -0.20 | 0.84 | 1 | Cd82 | CD82 antigen | 2904 | 2774 | 2549 | 2186 | 2078 | 1433 | 2109 | 1926 | 2154 | 2078 | 1511 |
| 1839 | 0.20 | 0.15 | 1.40 | 0.16 | 1 | Ostm1 | osteopetrosis associated transmembrane | 1366 | 884 | 985 | 2215 | 1917 | 1676 | 2109 | 2602 | 2268 | 2183 | 2023 |
| 1958 | -0.12 | 0.12 | -1.02 | 0.31 | 1 | Agtrap | angiotensin II, type I receptor-associated p | 2418 | 2240 | 2483 | 2077 | 2054 | 1270 | 2108 | 1974 | 1516 | 2139 | 1256 |
| 1904 | 0.08 | 0.09 | 0.91 | 0.36 | 1 | Pdcd6 | programmed cell death 6 | 1733 | 1648 | 1833 | 1820 | 1910 | 1811 | 2108 | 1855 | 1999 | 2169 | 2058 |
| 1853 | -0.05 | 0.10 | -0.55 | 0.58 | 1 | Nt5dc3 | 5'-nucleotidase domain containing 3 | 1319 | 1506 | 1143 | 2084 | 1987 | 2188 | 2108 | 1958 | 1953 | 2110 | 2030 |
| 2548 | 0.10 | 0.11 | 0.96 | 0.34 | 1 | Psma6 | proteasome subunit alpha 6 | 3607 | 2966 | 3418 | 2306 | 2405 | 1893 | 2107 | 2611 | 2340 | 2254 | 2122 |
| 2203 | -0.01 | 0.09 | -0.10 | 0.92 | 1 | Cep120 | centrosomal protein 120 | 1680 | 1989 | 1583 | 2531 | 2476 | 2413 | 2107 | 2406 | 2378 | 2174 | 2499 |
| 1630 | -0.02 | 0.14 | -0.15 | 0.88 | 1 | Bag4 | BCL2-associated athanogene 4 | 889 | 1147 | 1069 | 1843 | 1756 | 1716 | 2107 | 2053 | 2114 | 1812 | 1426 |
| 2317 | 0.13 | 0.15 | 0.86 | 0.39 | 1 | Ankrd12 | ankyrin repeat domain 12 | 1486 | 1318 | 1402 | 2044 | 1924 | 4010 | 2106 | 2797 | 2261 | 1604 | 4534 |
| 2255 | -0.13 | 0.11 | -1.18 | 0.24 | 1 | Ahctf1 | AT hook containing transcription factor 1 | 2411 | 2264 | 2038 | 2477 | 2230 | 2618 | 2105 | 2026 | 2122 | 1803 | 2714 |
| 2102 | 0.01 | 0.14 | 0.08 | 0.94 | 1 | Ndufb10 | NADH:ubiquinone oxidoreductase subun | 1906 | 2165 | 2545 | 2144 | 2210 | 1718 | 2104 | 2074 | 2094 | 2655 | 1502 |
| 1839 | 0.03 | 0.14 | 0.18 | 0.86 | 1 | Defb1 | defensin beta 1 | 268 | 410 | 559 | 2998 | 2729 | 1571 | 2104 | 2551 | 3157 | 2311 | 1567 |
| 1824 | 0.14 | 0.11 | 1.30 | 0.19 | 1 | Uap1l1 | UDP-N-acteylglucosamine pyrophosphor | 1015 | 1125 | 1129 | 2138 | 1908 | 1811 | 2104 | 2742 | 2120 | 2119 | 1848 |
| 2322 | -0.03 | 0.14 | -0.20 | 0.84 | 1 | Rc3h1 | RING CCCH (C3H) domains 1 | 2240 | 2742 | 2044 | 2176 | 2152 | 2938 | 2103 | 1734 | 2437 | 2244 | 2735 |
| 2182 | 0.00 | 0.09 | -0.03 | 0.98 | 1 | Pex5l | peroxisomal biogenesis factor 5-like | 1182 | 1553 | 1298 | 2681 | 1975 | 3222 | 2103 | 2786 | 2140 | 1864 | 3203 |
| 2135 | -0.06 | 0.09 | -0.63 | 0.53 | 1 | Maea | macrophage erythroblast attacher | 2456 | 2484 | 2414 | 1974 | 2145 | 1990 | 2103 | 1789 | 1982 | 2195 | 1950 |
| 2365 | -0.13 | 0.12 | -1.08 | 0.28 | 1 | Cbfb | core binding factor beta | 3464 | 3757 | 3474 | 2180 | 1934 | 1787 | 2102 | 1536 | 2093 | 1923 | 1762 |
| 2088 | -0.12 | 0.18 | -0.65 | 0.51 | 1 | St8sia3 | ST8 alpha-N-acetyl-neuraminide alpha-2 | 2734 | 3871 | 3403 | 1910 | 1454 | 1285 | 2100 | 1467 | 1259 | 2147 | 1334 |
| 1984 | 0.02 | 0.10 | 0.17 | 0.86 | 1 | Rundc1 | RUN domain containing 1 | 1476 | 2032 | 1499 | 2140 | 1951 | 2144 | 2100 | 2423 | 1813 | 2115 | 2133 |
| 1943 | -0.04 | 0.13 | -0.35 | 0.73 | 1 | Atp2b2 | ATPase, Ca++ transporting, plasma mem | 794 | 1291 | 984 | 2358 | 2404 | 2395 | 2099 | 2300 | 2683 | 1780 | 2283 |
| 1957 | -0.15 | 0.14 | -1.05 | 0.29 | 1 | Brpf3 | bromodomain and PHD finger containing | 1736 | 1991 | 1668 | 2350 | 1861 | 2172 | 2098 | 1517 | 2072 | 1932 | 2127 |
| 1433 | -0.13 | 0.17 | -0.77 | 0.44 | 1 | Kcnh1 | potassium voltage-gated channel, subfam | 160 | 265 | 348 | 2250 | 2184 | 1292 | 2097 | 1784 | 2276 | 1895 | 1209 |
| 1873 | 0.12 | 0.10 | 1.20 | 0.23 | 1 | Snph | syntaphilin | 915 | 806 | 952 | 2222 | 2120 | 2162 | 2096 | 2413 | 2329 | 2423 | 2161 |
| 1791 | -0.10 | 0.12 | -0.83 | 0.41 | 1 | Tmem63b | transmembrane protein 63b | 1492 | 1129 | 1654 | 1950 | 2072 | 1865 | 2094 | 1956 | 2114 | 1695 | 1680 |
| 1755 | -0.07 | 0.13 | -0.52 | 0.60 | 1 | Cdcp1 | CUB domain containing protein 1 | 1339 | 1149 | 1648 | 1750 | 2430 | 1385 | 2094 | 1621 | 2273 | 2522 | 1094 |
| 2710 | 0.17 | 0.13 | 1.34 | 0.18 | 1 | Zranb2 | zinc finger, RAN-binding domain containi | 2042 | 2152 | 2037 | 2129 | 1973 | 4843 | 2093 | 1790 | 2413 | 2758 | 5581 |
| 2018 | 0.14 | 0.08 | 1.70 | 0.09 | 1 | Trim23 | tripartite motif-containing 23 | 1649 | 1738 | 1885 | 2000 | 2066 | 1895 | 2093 | 2258 | 2297 | 2127 | 2194 |
| 2088 | -0.10 | 0.12 | -0.84 | 0.40 | 1 | Appl2 | adaptor protein, phosphotyrosine interact | 1180 | 1167 | 1145 | 1854 | 1749 | 4291 | 2091 | 1368 | 1965 | 1865 | 4293 |

| | | | | | | | | | | | | | | | | | | |
|---|---|---|---|---|---|---|---|---|---|---|---|---|---|---|---|---|---|---|
| 1110 | 0.17 | 0.28 | 0.60 | 0.55 | 1 | Gm44850 | | 53 | 24 | 114 | 828 | 1234 | 1462 | 2090 | 862 | 1271 | 2445 | 1821 |
| 2085 | -0.04 | 0.12 | -0.33 | 0.74 | 1 | Trim33 | tripartite motif-containing 33 | 1696 | 1941 | 1599 | 2304 | 2079 | 2480 | 2089 | 1932 | 2384 | 1830 | 2596 |
| 1903 | -0.01 | 0.11 | -0.08 | 0.94 | 1 | Slc35a4 | solute carrier family 35, member A4 | 1917 | 1838 | 1795 | 1758 | 2032 | 1826 | 2089 | 1760 | 2317 | 1817 | 1781 |
| 1877 | -0.18 | 0.18 | -0.97 | 0.33 | 1 | Lyst | lysosomal trafficking regulator | 1932 | 1392 | 1409 | 1941 | 1640 | 2690 | 2089 | 1099 | 2153 | 1714 | 2586 |
| 2021 | -0.14 | 0.09 | -1.51 | 0.13 | 1 | Ift172 | intraflagellar transport 172 | 1406 | 1548 | 1262 | 2584 | 2408 | 2357 | 2087 | 2365 | 2068 | 2048 | 2103 |
| 1895 | -0.11 | 0.14 | -0.78 | 0.44 | 1 | Tmem258 | transmembrane protein 258 | 1265 | 1351 | 1447 | 2035 | 2157 | 2392 | 2086 | 1441 | 2211 | 2320 | 2145 |
| 2357 | -0.01 | 0.12 | -0.09 | 0.93 | 1 | Fbxo11 | F-box protein 11 | 2201 | 2837 | 2286 | 2509 | 2183 | 2571 | 2085 | 1958 | 2351 | 2310 | 2632 |
| 1965 | -0.03 | 0.12 | -0.22 | 0.82 | 1 | Zfp275 | zinc finger protein 275 | 1515 | 2093 | 1491 | 2298 | 1708 | 2273 | 2085 | 1879 | 1894 | 2102 | 2276 |
| 1966 | 0.05 | 0.08 | 0.57 | 0.57 | 1 | Fam76a | family with sequence similarity 76, memb | 1876 | 1801 | 1705 | 2214 | 2028 | 1677 | 2082 | 2179 | 2144 | 2125 | 1791 |
| 1802 | 0.13 | 0.13 | 0.95 | 0.34 | 1 | Necab2 | N-terminal EF-hand calcium binding prote | 1060 | 1301 | 1180 | 1868 | 1986 | 1779 | 2082 | 2781 | 2061 | 2096 | 1629 |
| 2200 | 0.15 | 0.12 | 1.32 | 0.19 | 1 | Pdcd11 | programmed cell death 11 | 1941 | 1842 | 2108 | 2067 | 2187 | 2302 | 2080 | 1970 | 2467 | 2920 | 2314 |
| 1992 | -0.26 | 0.17 | -1.49 | 0.14 | 1 | Golph3l | golgi phosphoprotein 3-like | 1845 | 1670 | 2035 | 1651 | 2873 | 2185 | 2080 | 963 | 2592 | 1955 | 2061 |
| 1734 | 0.02 | 0.12 | 0.13 | 0.90 | 1 | Tdrd7 | tudor domain containing 7 | 1695 | 1485 | 1315 | 1644 | 1948 | 1587 | 2080 | 1938 | 2013 | 1799 | 1565 |
| 1975 | 0.11 | 0.14 | 0.76 | 0.45 | 1 | Metap1 | methionyl aminopeptidase 1 | 1450 | 1969 | 1759 | 1601 | 1962 | 2316 | 2079 | 2379 | 2082 | 1913 | 2221 |
| 2123 | -0.21 | 0.14 | -1.49 | 0.14 | 1 | Ncoa6 | nuclear receptor coactivator 6 | 1476 | 1871 | 1439 | 2905 | 2410 | 2592 | 2078 | 1808 | 2378 | 2043 | 2348 |
| 2310 | 0.00 | 0.11 | -0.03 | 0.98 | 1 | Ythdf1 | YTH N6-methyladenosine RNA binding p | 2184 | 2521 | 2181 | 2437 | 2168 | 2598 | 2077 | 2026 | 2478 | 2155 | 2584 |
| 2212 | -0.07 | 0.16 | -0.45 | 0.65 | 1 | Septin11 | septin 11 | 3040 | 2138 | 2250 | 2035 | 2364 | 2164 | 2077 | 1886 | 2506 | 1635 | 2233 |
| 2501 | 0.06 | 0.10 | 0.66 | 0.51 | 1 | Myef2 | myelin basic protein expression factor 2, r | 2008 | 2185 | 2304 | 2235 | 2186 | 3822 | 2074 | 2133 | 2260 | 2535 | 3772 |
| 2068 | -0.12 | 0.12 | -1.04 | 0.30 | 1 | Mapre3 | microtubule-associated protein, RP/EB fa | 1929 | 2476 | 2214 | 1813 | 2065 | 2461 | 2074 | 1693 | 2007 | 1888 | 2126 |
| 1972 | -0.09 | 0.11 | -0.86 | 0.39 | 1 | Rbsn | rabenosyn, RAB effector | 2088 | 1799 | 1717 | 1833 | 2199 | 2183 | 2072 | 1558 | 2060 | 2124 | 2060 |
| 2242 | 0.19 | 0.10 | 1.80 | 0.07 | 1 | Chd7 | chromodomain helicase DNA binding pro | 869 | 1063 | 836 | 2165 | 2187 | 3822 | 2071 | 2926 | 2485 | 2079 | 4153 |
| 2211 | 0.17 | 0.14 | 1.22 | 0.22 | 1 | Ndufa13 | NADH:ubiquinone oxidoreductase subun | 2372 | 2150 | 2801 | 2154 | 2248 | 1500 | 2071 | 2914 | 2108 | 2455 | 1544 |
| 2766 | 0.05 | 0.14 | 0.38 | 0.70 | 1 | Plec | plectin | 5049 | 3920 | 4519 | 2160 | 2169 | 1839 | 2070 | 2799 | 2174 | 2224 | 1501 |
| 1969 | 0.16 | 0.17 | 0.97 | 0.33 | 1 | Glrx5 | glutaredoxin 5 | 1157 | 981 | 1447 | 2105 | 2416 | 1922 | 2070 | 3099 | 2048 | 2487 | 1928 |
| 1556 | -0.07 | 0.21 | -0.31 | 0.75 | 1 | Gabbr2 | gamma-aminobutyric acid (GABA) B rece | 305 | 359 | 376 | 2495 | 2051 | 1569 | 2070 | 1218 | 2594 | 2311 | 1765 |
| 2541 | -0.22 | 0.14 | -1.55 | 0.12 | 1 | Ggcx | gamma-glutamyl carboxylase | 1682 | 2062 | 1979 | 2465 | 2465 | 4943 | 2069 | 1510 | 2298 | 2200 | 4279 |
| 2270 | -0.08 | 0.13 | -0.63 | 0.53 | 1 | Eif2a | eukaryotic translation initiation factor 2A | 2909 | 3044 | 2818 | 1957 | 2257 | 2016 | 2069 | 1484 | 2130 | 2210 | 2073 |
| 2116 | -0.02 | 0.11 | -0.22 | 0.83 | 1 | Usp20 | ubiquitin specific peptidase 20 | 1680 | 1560 | 1996 | 2309 | 2511 | 2203 | 2069 | 2518 | 2329 | 2176 | 1931 |
| 2221 | 0.03 | 0.09 | 0.33 | 0.74 | 1 | Dock9 | dedicator of cytokinesis 9 | 2131 | 2224 | 2333 | 2174 | 2042 | 2497 | 2067 | 2207 | 2373 | 1927 | 2455 |
| 2055 | -0.01 | 0.11 | -0.04 | 0.96 | 1 | Tgfbrap1 | transforming growth factor, beta receptor a | 1611 | 1511 | 1377 | 2157 | 2154 | 2696 | 2066 | 1876 | 2452 | 2179 | 2525 |
| 1751 | -0.21 | 0.11 | -1.93 | 0.05 | 1 | Maml3 | mastermind like transcriptional coactivato | 1180 | 1300 | 1344 | 2378 | 1809 | 2042 | 2066 | 1873 | 1902 | 1691 | 1679 |
| 1967 | 0.04 | 0.09 | 0.45 | 0.66 | 1 | Dhrs7 | dehydrogenase/reductase (SDR family) m | 1686 | 1738 | 1655 | 2278 | 2163 | 1649 | 2061 | 2591 | 2050 | 2026 | 1736 |
| 1860 | -0.16 | 0.10 | -1.54 | 0.12 | 1 | Naxe | NAD(P)HX epimerase | 1948 | 2069 | 2398 | 1958 | 1928 | 1464 | 2061 | 1758 | 1717 | 1817 | 1345 |
| 2316 | -0.03 | 0.13 | -0.27 | 0.79 | 1 | F11r | F11 receptor | 3455 | 3089 | 3999 | 1805 | 2077 | 1595 | 2060 | 2296 | 1919 | 1767 | 1412 |
| 2055 | 0.10 | 0.11 | 0.90 | 0.37 | 1 | Pgm1 | phosphoglucomutase 1 | 2203 | 2168 | 2324 | 2024 | 2032 | 1572 | 2060 | 2483 | 2129 | 1855 | 1755 |
| 1933 | -0.15 | 0.11 | -1.31 | 0.19 | 1 | Glt8d1 | glycosyltransferase 8 domain containing | 1317 | 1496 | 1566 | 2226 | 1940 | 2600 | 2060 | 1664 | 2005 | 1738 | 2646 |
| 1744 | 0.15 | 0.13 | 1.16 | 0.24 | 1 | Cisd1 | CDGSH iron sulfur domain 1 | 1294 | 1332 | 1426 | 1902 | 1831 | 1346 | 2060 | 2684 | 1883 | 1986 | 1444 |
| 2524 | 0.11 | 0.15 | 0.74 | 0.46 | 1 | Sdc4 | syndecan 4 | 4561 | 3136 | 3399 | 2477 | 2206 | 1179 | 2059 | 2753 | 2694 | 2204 | 1098 |
| 1857 | -0.24 | 0.12 | -1.93 | 0.05 | 1 | Psmf1 | proteasome (prosome, macropain) inhibit | 1212 | 1533 | 1703 | 2232 | 2702 | 1671 | 2059 | 1518 | 2361 | 1864 | 1570 |
| 2455 | -0.04 | 0.08 | -0.51 | 0.61 | 1 | Rab12 | RAB12, member RAS oncogene family | 3643 | 4060 | 3561 | 2008 | 1797 | 2120 | 2057 | 1878 | 1726 | 2004 | 2154 |
| 1906 | -0.05 | 0.10 | -0.55 | 0.58 | 1 | Armc8 | armadillo repeat containing 8 | 1473 | 1302 | 1547 | 1999 | 2270 | 2137 | 2057 | 1899 | 2324 | 1757 | 2204 |
| 1859 | -0.10 | 0.13 | -0.76 | 0.45 | 1 | Fam160a2 | family with sequence similarity 160, meml | 1150 | 1669 | 1468 | 2040 | 1955 | 2314 | 2056 | 1666 | 2076 | 1909 | 2148 |
| 2722 | -0.22 | 0.14 | -1.56 | 0.12 | 1 | Cpt1a | carnitine palmitoyltransferase 1a, liver | 3650 | 3775 | 4159 | 3311 | 2845 | 1682 | 2055 | 2354 | 2952 | 1538 | 1621 |
| 2773 | 0.15 | 0.12 | 1.31 | 0.19 | 1 | Xpo1 | exportin 1 | 3426 | 2730 | 2342 | 2413 | 1838 | 4133 | 2054 | 2442 | 2300 | 2249 | 4571 |
| 2150 | -0.03 | 0.13 | -0.20 | 0.84 | 1 | Agps | alkylglycerone phosphate synthase | 2611 | 2319 | 2189 | 1954 | 2366 | 1930 | 2054 | 1557 | 2179 | 2452 | 2034 |
| 2065 | 0.14 | 0.14 | 0.99 | 0.32 | 1 | Gabarapl2 | gamma-aminobutyric acid (GABA) A rece | 2074 | 1827 | 2235 | 2122 | 2049 | 1638 | 2054 | 2878 | 2140 | 1825 | 1876 |
| 1893 | -0.21 | 0.12 | -1.80 | 0.07 | 1 | Pcgf3 | polycomb group ring finger 3 | 1688 | 2404 | 1868 | 2197 | 2100 | 1631 | 2054 | 1783 | 2019 | 1601 | 1480 |
| 2378 | -0.14 | 0.16 | -0.88 | 0.38 | 1 | Kctd12b | potassium channel tetramerisation domai | 3440 | 3851 | 3696 | 1801 | 2029 | 2015 | 2053 | 1176 | 2170 | 1997 | 1935 |
| 2008 | 0.07 | 0.08 | 0.86 | 0.39 | 1 | Abi2 | abl-interactor 2 | 1567 | 1913 | 1705 | 2391 | 2082 | 1689 | 2053 | 2518 | 2322 | 2241 | 1607 |
| 1993 | 0.00 | 0.10 | -0.02 | 0.98 | 1 | Rad23a | RAD23 homolog A, nucleotide excision re | 2110 | 1803 | 2466 | 2032 | 2051 | 1617 | 2053 | 1893 | 2328 | 2013 | 1558 |
| 1862 | 0.06 | 0.13 | 0.44 | 0.66 | 1 | Taf9b | TATA-box binding protein associated fact | 1470 | 1739 | 1639 | 2024 | 2069 | 1527 | 2053 | 1659 | 2226 | 2284 | 1797 |
| 1444 | -0.08 | 0.14 | -0.61 | 0.54 | 1 | Ubxn10 | UBX domain protein 10 | 912 | 914 | 1249 | 1364 | 1790 | 1376 | 2053 | 1382 | 1763 | 1855 | 1223 |
| 2082 | -0.04 | 0.12 | -0.30 | 0.76 | 1 | Ryk | receptor-like tyrosine kinase | 2644 | 2256 | 2244 | 2156 | 2079 | 1680 | 2052 | 2249 | 2179 | 1769 | 1596 |
| 2249 | -0.03 | 0.10 | -0.33 | 0.74 | 1 | Fbxl3 | F-box and leucine-rich repeat protein 3 | 2515 | 2549 | 2934 | 2071 | 1845 | 2459 | 2051 | 1884 | 1815 | 1870 | 2743 |
| 2044 | 0.13 | 0.10 | 1.33 | 0.18 | 1 | Cpsf1 | cleavage and polyadenylation specific fac | 2225 | 1737 | 2084 | 1904 | 1900 | 1990 | 2051 | 2219 | 2179 | 2213 | 1985 |
| 1935 | 0.09 | 0.11 | 0.80 | 0.42 | 1 | Crocc | ciliary rootlet coiled-coil, rootletin | 928 | 857 | 976 | 2070 | 1964 | 2930 | 2051 | 2172 | 2057 | 2576 | 2707 |
| 1817 | -0.15 | 0.13 | -1.18 | 0.24 | 1 | Tmem106c | transmembrane protein 106C | 1758 | 2270 | 2292 | 1771 | 1947 | 1419 | 2049 | 1470 | 1997 | 1708 | 1307 |
| 2290 | 0.22 | 0.11 | 1.96 | 0.05 | 1 | Fnta | farnesyltransferase, CAAX box, alpha | 2183 | 2425 | 2385 | 2204 | 2061 | 2037 | 2048 | 3218 | 2179 | 2231 | 2224 |
| 1994 | 0.09 | 0.12 | 0.69 | 0.49 | 1 | Terf2 | telomeric repeat binding factor 2 | 1035 | 1138 | 1310 | 2463 | 2000 | 2418 | 2048 | 2925 | 2231 | 1966 | 2396 |
| 1966 | -0.18 | 0.17 | -1.06 | 0.29 | 1 | Fam183b | family with sequence similarity 183, meml | 1451 | 2421 | 2225 | 2258 | 1984 | 1958 | 2046 | 1534 | 1825 | 1882 | 2048 |

| | | | | | | | | | | | | | | | | | | |
|---|---|---|---|---|---|---|---|---|---|---|---|---|---|---|---|---|---|---|
| 1836 | -0.02 | 0.11 | -0.15 | 0.88 | 1 | Eif2ak1 | eukaryotic translation initiation factor 2 alp | 1638 | 1401 | 1618 | 1854 | 2164 | 1756 | 2046 | 2051 | 2160 | 1743 | 1769 |
| 1796 | 0.10 | 0.13 | 0.78 | 0.44 | 1 | Pla2g15 | phospholipase A2, group XV | 1951 | 1498 | 1765 | 2020 | 1888 | 1098 | 2046 | 2355 | 1795 | 2070 | 1270 |
| 1720 | -0.07 | 0.13 | -0.50 | 0.62 | 1 | Slc39a3 | solute carrier family 39 (zinc transporter), | 1670 | 1943 | 1801 | 1600 | 1831 | 1430 | 2046 | 1744 | 1456 | 1981 | 1423 |
| 1576 | -0.04 | 0.12 | -0.37 | 0.71 | 1 | Abcc4 | ATP-binding cassette, sub-family C (CFTF | 1194 | 973 | 897 | 1965 | 1886 | 1373 | 2046 | 1751 | 2024 | 1783 | 1441 |
| 1830 | -0.13 | 0.10 | -1.27 | 0.20 | 1 | Pgd | phosphogluconate dehydrogenase | 2522 | 1934 | 2067 | 1664 | 1735 | 1687 | 2045 | 1639 | 1549 | 1609 | 1680 |
| 2316 | -0.03 | 0.10 | -0.33 | 0.74 | 1 | Tpp2 | tripeptidyl peptidase II | 2250 | 2382 | 2729 | 2170 | 2322 | 2562 | 2044 | 1835 | 2346 | 1902 | 2937 |
| 1841 | -0.02 | 0.13 | -0.16 | 0.87 | 1 | Klhl21 | kelch-like 21 | 1222 | 1723 | 1309 | 2265 | 1912 | 1839 | 2044 | 1950 | 1943 | 2365 | 1684 |
| 1458 | 0.00 | 0.16 | 0.03 | 0.98 | 1 | Itgb8 | integrin beta 8 | 879 | 1291 | 957 | 1795 | 1274 | 1373 | 2044 | 1654 | 1501 | 1622 | 1645 |
| 2153 | -0.17 | 0.10 | -1.62 | 0.11 | 1 | Zer1 | zyg-11 related, cell cycle regulator | 2001 | 2490 | 2193 | 2020 | 2390 | 2552 | 2042 | 1915 | 2077 | 1901 | 2108 |
| 2241 | -0.05 | 0.11 | -0.42 | 0.67 | 1 | Rubcn | RUN domain and cysteine-rich domain cc | 2395 | 2619 | 2172 | 2627 | 1956 | 2273 | 2041 | 2073 | 2105 | 2297 | 2096 |
| 2171 | -0.19 | 0.11 | -1.65 | 0.10 | 1 | Atxn1l | ataxin 1-like | 2466 | 2608 | 2142 | 2534 | 2277 | 2033 | 2040 | 2027 | 2251 | 1707 | 1794 |
| 2120 | -0.03 | 0.11 | -0.25 | 0.80 | 1 | Kansl3 | KAT8 regulatory NSL complex subunit 3 | 2075 | 1803 | 1636 | 2507 | 2073 | 2355 | 2040 | 2500 | 1900 | 2364 | 2069 |
| 1964 | -0.17 | 0.13 | -1.33 | 0.18 | 1 | Setd1b | SET domain containing 1B | 1959 | 2198 | 2125 | 1851 | 2191 | 1981 | 2040 | 1665 | 2289 | 1878 | 1431 |
| 2420 | 0.14 | 0.10 | 1.52 | 0.13 | 1 | Msl2 | MSL complex subunit 2 | 2763 | 3307 | 2426 | 2143 | 2072 | 2357 | 2038 | 2412 | 2258 | 2365 | 2478 |
| 2097 | -0.03 | 0.12 | -0.22 | 0.83 | 1 | Usp12 | ubiquitin specific peptidase 12 | 1962 | 2041 | 1967 | 2383 | 2134 | 2064 | 2038 | 1814 | 2088 | 2498 | 2081 |
| 2040 | -0.02 | 0.12 | -0.15 | 0.88 | 1 | Slc25a23 | solute carrier family 25 (mitochondrial car | 1769 | 2206 | 1970 | 2222 | 2121 | 1899 | 2038 | 2026 | 2130 | 2405 | 1657 |
| 1770 | -0.18 | 0.11 | -1.74 | 0.08 | 1 | Heatr5a | HEAT repeat containing 5A | 1937 | 1327 | 1560 | 1717 | 2196 | 1827 | 2038 | 1372 | 1902 | 1886 | 1712 |
| 2346 | 0.08 | 0.12 | 0.67 | 0.50 | 1 | Upf1 | UPF1 regulator of nonsense transcripts ho | 2213 | 2306 | 2251 | 2083 | 2636 | 2502 | 2035 | 2717 | 2161 | 2410 | 2488 |
| 2131 | 0.01 | 0.08 | 0.11 | 0.91 | 1 | Cd2bp2 | CD2 cytoplasmic tail binding protein 2 | 2273 | 2241 | 2250 | 1949 | 2194 | 2134 | 2033 | 2165 | 2006 | 2044 | 2148 |
| 1803 | 0.05 | 0.09 | 0.51 | 0.61 | 1 | Rchy1 | ring finger and CHY zinc finger domain cc | 1300 | 1413 | 1223 | 1851 | 1797 | 2123 | 2033 | 1913 | 1859 | 1911 | 2409 |
| 1795 | 0.08 | 0.10 | 0.80 | 0.42 | 1 | Synj2bp | synaptojanin 2 binding protein | 1663 | 1782 | 1721 | 1908 | 1629 | 1534 | 2033 | 1907 | 1880 | 1932 | 1758 |
| 1822 | -0.03 | 0.10 | -0.29 | 0.78 | 1 | Uhrf1bp1l | UHRF1 (ICBP90) binding protein 1-like | 1400 | 1089 | 1086 | 1756 | 1876 | 2620 | 2031 | 1471 | 1945 | 2102 | 2664 |
| 2267 | 0.16 | 0.16 | 1.00 | 0.32 | 1 | Nudc | nudC nuclear distribution protein | 2516 | 2156 | 2483 | 1959 | 2342 | 2018 | 2030 | 3227 | 2010 | 2199 | 1997 |
| 2171 | 0.02 | 0.10 | 0.22 | 0.82 | 1 | Cgn | cingulin | 1780 | 2040 | 2148 | 2668 | 2139 | 2033 | 2029 | 2830 | 2272 | 1966 | 1979 |
| 2026 | -0.02 | 0.12 | -0.17 | 0.86 | 1 | Osbpl9 | oxysterol binding protein-like 9 | 2771 | 2398 | 2203 | 2022 | 1672 | 1787 | 2029 | 2236 | 1858 | 1709 | 1604 |
| 1895 | -0.13 | 0.12 | -1.07 | 0.29 | 1 | Arhgap32 | Rho GTPase activating protein 32 | 1989 | 2198 | 2098 | 2200 | 1727 | 1639 | 2029 | 1700 | 1642 | 2167 | 1456 |
| 2350 | 0.08 | 0.08 | 1.05 | 0.29 | 1 | Gpbp1l1 | GC-rich promoter binding protein 1-like 1 | 2347 | 2687 | 2736 | 2184 | 2105 | 2469 | 2027 | 2433 | 2240 | 2177 | 2448 |
| 2280 | 0.00 | 0.10 | -0.02 | 0.98 | 1 | Plekhg3 | pleckstrin homology domain containing, fa | 2281 | 2415 | 1976 | 2114 | 2153 | 2925 | 2026 | 2457 | 2131 | 1863 | 2738 |
| 2160 | 0.02 | 0.10 | 0.15 | 0.88 | 1 | Prmt1 | protein arginine N-methyltransferase 1 | 3072 | 2309 | 3173 | 1687 | 1877 | 1980 | 2026 | 1803 | 2089 | 1888 | 1861 |
| 1604 | -0.20 | 0.15 | -1.35 | 0.18 | 1 | Soat1 | sterol O-acyltransferase 1 | 1736 | 971 | 1261 | 2002 | 1736 | 1503 | 2025 | 1694 | 1394 | 2061 | 1256 |
| 2068 | -0.03 | 0.09 | -0.33 | 0.75 | 1 | Scaf8 | SR-related CTD-associated factor 8 | 2065 | 2114 | 2033 | 2156 | 1970 | 2194 | 2024 | 2358 | 1998 | 1959 | 1873 |
| 1651 | -0.02 | 0.11 | -0.22 | 0.83 | 1 | Alg11 | asparagine-linked glycosylation 11 (alpha | 1054 | 1328 | 1051 | 2028 | 1853 | 1538 | 2024 | 1848 | 1941 | 1878 | 1620 |
| 1590 | -0.03 | 0.12 | -0.24 | 0.81 | 1 | Pcbp3 | poly(rC) binding protein 3 | 704 | 923 | 928 | 1890 | 1826 | 1802 | 2024 | 1800 | 1965 | 1906 | 1727 |
| 2581 | -0.02 | 0.11 | -0.22 | 0.83 | 1 | Pkd2 | polycystin 2, transient receptor potential c | 3649 | 2975 | 3021 | 2350 | 1993 | 3126 | 2022 | 2173 | 2119 | 2200 | 2766 |
| 2332 | 0.18 | 0.13 | 1.34 | 0.18 | 1 | Ubxn1 | UBX domain protein 1 | 2929 | 2980 | 3343 | 1749 | 2089 | 1799 | 2022 | 2801 | 2144 | 1981 | 1818 |
| 1959 | 0.16 | 0.17 | 0.94 | 0.35 | 1 | B3gat3 | beta-1,3-glucuronyltransferase 3 (glucuro | 1447 | 1272 | 1864 | 2080 | 2222 | 1660 | 2021 | 2962 | 1857 | 2443 | 1719 |
| 1603 | -0.03 | 0.13 | -0.21 | 0.83 | 1 | Dusp18 | dual specificity phosphatase 18 | 697 | 757 | 601 | 2113 | 1682 | 2065 | 2021 | 1748 | 2092 | 1793 | 2066 |
| 1951 | -0.10 | 0.10 | -1.06 | 0.29 | 1 | Clip2 | CAP-GLY domain containing linker protei | 2225 | 2394 | 2411 | 1949 | 1954 | 1539 | 2020 | 1820 | 1705 | 2084 | 1364 |
| 1976 | 0.09 | 0.10 | 0.89 | 0.38 | 1 | Cops5 | COP9 signalosome subunit 5 | 1943 | 1769 | 2074 | 1923 | 1825 | 1951 | 2019 | 1883 | 1870 | 2293 | 2187 |
| 1713 | -0.06 | 0.10 | -0.59 | 0.56 | 1 | Ube2e1 | ubiquitin-conjugating enzyme E2E 1 | 1673 | 1387 | 1596 | 1893 | 1776 | 1549 | 2019 | 1530 | 1868 | 1941 | 1607 |
| 2270 | 0.12 | 0.11 | 1.09 | 0.28 | 1 | Nop58 | NOP58 ribonucleoprotein | 2657 | 2691 | 2144 | 1713 | 1986 | 2633 | 2018 | 1903 | 2172 | 2053 | 2996 |
| 2405 | 0.12 | 0.14 | 0.90 | 0.37 | 1 | Ush1c | USH1 protein network component harmo | 1299 | 2087 | 1940 | 2152 | 2190 | 3826 | 2017 | 3021 | 2160 | 2209 | 3551 |
| 2673 | -0.02 | 0.08 | -0.25 | 0.80 | 1 | Ets2 | E26 avian leukemia oncogene 2, 3' doma | 4113 | 4198 | 3954 | 2366 | 2237 | 2004 | 2016 | 2255 | 2352 | 2105 | 1805 |
| 1693 | 0.02 | 0.10 | 0.26 | 0.79 | 1 | Mmgt1 | membrane magnesium transporter 1 | 1383 | 1441 | 1288 | 1778 | 1917 | 1485 | 2016 | 1844 | 2025 | 1958 | 1493 |
| 2197 | 0.04 | 0.08 | 0.45 | 0.65 | 1 | Dcaf5 | DDB1 and CUL4 associated factor 5 | 2083 | 2337 | 2252 | 2242 | 2124 | 2250 | 2015 | 2429 | 2144 | 2210 | 2082 |
| 2155 | -0.01 | 0.08 | -0.09 | 0.93 | 1 | Arhgef16 | Rho guanine nucleotide exchange factor | 1192 | 1665 | 1527 | 2667 | 2287 | 2706 | 2015 | 2752 | 2208 | 1987 | 2697 |
| 1346 | -0.23 | 0.12 | -1.88 | 0.06 | 1 | Fbxo32 | F-box protein 32 | 1008 | 555 | 1005 | 869 | 1869 | 1821 | 2015 | 706 | 1588 | 1900 | 1474 |
| 1804 | 0.05 | 0.11 | 0.48 | 0.63 | 1 | Gid4 | GID complex subunit 4, VID24 homolog | 1969 | 1916 | 1975 | 1595 | 1771 | 1464 | 2013 | 1890 | 2090 | 1823 | 1340 |
| 1466 | 0.15 | 0.15 | 1.01 | 0.31 | 1 | Arxes2 | adipocyte-related X-chromosome express | 755 | 1197 | 911 | 1722 | 1549 | 1012 | 2013 | 2109 | 1751 | 1896 | 1211 |
| 2074 | 0.11 | 0.15 | 0.71 | 0.48 | 1 | Usp35 | ubiquitin specific peptidase 35 | 577 | 889 | 785 | 2486 | 1941 | 3478 | 2011 | 3076 | 1723 | 2456 | 3388 |
| 2515 | 0.18 | 0.11 | 1.71 | 0.09 | 1 | Ddx21 | DEAD (Asp-Glu-Ala-Asp) box polypeptide | 3165 | 2849 | 2323 | 2399 | 2085 | 2573 | 2007 | 2741 | 2336 | 2505 | 2678 |
| 2048 | -0.17 | 0.12 | -1.43 | 0.15 | 1 | Cbx7 | chromobox 7 | 1939 | 1859 | 1671 | 2039 | 2130 | 2816 | 2007 | 1493 | 2003 | 1962 | 2613 |
| 2480 | 0.11 | 0.10 | 1.07 | 0.28 | 1 | Sart1 | squamous cell carcinoma antigen recogn | 3066 | 2572 | 2794 | 2205 | 2263 | 2564 | 2005 | 2087 | 2290 | 2709 | 2724 |
| 2034 | -0.11 | 0.11 | -1.05 | 0.29 | 1 | Prkar2a | protein kinase, cAMP dependent regulato | 2050 | 1991 | 1941 | 2198 | 2322 | 1985 | 2004 | 2100 | 2167 | 1642 | 1973 |
| 1705 | -0.19 | 0.14 | -1.31 | | 1 | Atxn2 | ataxin 2 | 1307 | 1406 | 1176 | 2017 | 1834 | 2044 | 2004 | 1323 | 1933 | 1826 | 1880 |
| 2316 | -0.06 | 0.09 | -0.71 | 0.48 | 1 | Dync1li2 | dynein, cytoplasmic 1 light intermediate c | 2430 | 2589 | 2938 | 1957 | 2003 | 2969 | 2003 | 1716 | 2046 | 1922 | 2906 |
| 1769 | 0.10 | 0.13 | 0.77 | 0.44 | 1 | Ak1 | adenylate kinase 1 | 1497 | 1264 | 1389 | 1757 | 2147 | 1478 | 2003 | 2465 | 1919 | 1933 | 1604 |
| 1354 | -0.14 | 0.20 | -0.71 | 0.48 | 1 | Rspo4 | R-spondin 4 | 243 | 399 | 468 | 1694 | 1891 | 1640 | 2003 | 1394 | 1668 | 2017 | 1477 |
| 1941 | 0.02 | 0.10 | 0.17 | 0.87 | 1 | C2cd5 | C2 calcium-dependent domain containing | 1659 | 1553 | 1465 | 1905 | 1804 | 2577 | 2002 | 1796 | 1806 | 2376 | 2404 |
| 2101 | -0.15 | 0.21 | -0.70 | 0.48 | 1 | Cdon | cell adhesion molecule-related/down-reg | 4436 | 2240 | 1925 | 2012 | 1800 | 1825 | 2001 | 1734 | 2030 | 1658 | 1453 |

| | | | | | | | | | | | | | | | | | | |
|---|---|---|---|---|---|---|---|---|---|---|---|---|---|---|---|---|---|---|
| 2204 | 0.07 | 0.15 | 0.46 | 0.65 | 1 | Vamp8 | vesicle-associated membrane protein 8 | 2302 | 2166 | 2591 | 2115 | 2461 | 1752 | 1999 | 3106 | 2097 | 2035 | 1620 |
| 1851 | -0.05 | 0.13 | -0.34 | 0.74 | 1 | Usp36 | ubiquitin specific peptidase 36 | 1154 | 1406 | 1414 | 1587 | 1975 | 2743 | 1999 | 1996 | 1950 | 1485 | 2655 |
| 2932 | 0.15 | 0.09 | 1.64 | 0.10 | 1 | Hexb | hexosaminidase B | 5882 | 4931 | 6233 | 1886 | 2151 | 1131 | 1998 | 2274 | 2270 | 2379 | 1121 |
| 2088 | 0.19 | 0.10 | 1.79 | 0.07 | 1 | Celf4 | CUGBP, Elav-like family member 4 | 1638 | 2175 | 1838 | 2098 | 2048 | 1928 | 1998 | 2628 | 2243 | 2447 | 1928 |
| 1773 | -0.07 | 0.12 | -0.54 | 0.59 | 1 | Lrp12 | low density lipoprotein-related protein 12 | 911 | 1264 | 1114 | 2070 | 2178 | 2051 | 1997 | 1886 | 1888 | 2105 | 2039 |
| 2514 | 0.02 | 0.12 | 0.16 | 0.87 | 1 | Fam168b | family with sequence similarity 168, meml | 3920 | 3241 | 3817 | 1927 | 2136 | 2226 | 1996 | 2434 | 1914 | 2013 | 2030 |
| 2289 | -0.22 | 0.12 | -1.88 | 0.06 | 1 | Extl3 | exostosin-like glycosyltransferase 3 | 3503 | 3240 | 3203 | 2473 | 2038 | 1698 | 1996 | 1935 | 2027 | 1541 | 1527 |
| 2198 | 0.08 | 0.09 | 0.88 | 0.38 | 1 | Tmf1 | TATA element modulatory factor 1 | 1438 | 1629 | 1225 | 2609 | 2089 | 2985 | 1996 | 2742 | 2245 | 2085 | 3137 |
| 2156 | 0.04 | 0.13 | 0.28 | 0.78 | 1 | Dnajb2 | DnaJ heat shock protein family (Hsp40) m | 1677 | 1705 | 2153 | 1689 | 2485 | 2861 | 1996 | 2383 | 2074 | 1946 | 2747 |
| 1961 | 0.12 | 0.10 | 1.21 | 0.22 | 1 | Odr4 | odr4 GPCR localization factor homolog | 1040 | 1149 | 1094 | 1763 | 2087 | 2910 | 1996 | 1897 | 2195 | 2183 | 3253 |
| 1645 | 0.18 | 0.13 | 1.42 | 0.16 | 1 | Mthfd1l | methylenetetrahydrofolate dehydrogenas | 681 | 508 | 754 | 1604 | 2112 | 1874 | 1996 | 2288 | 2306 | 1989 | 1981 |
| 2047 | -0.10 | 0.10 | -1.06 | 0.29 | 1 | Dynlrb1 | dynein light chain roadblock-type 1 | 2421 | 2446 | 2962 | 2003 | 2072 | 1543 | 1995 | 1742 | 2020 | 1677 | 1637 |
| 2065 | 0.04 | 0.14 | 0.26 | 0.79 | 1 | Atrnl1 | attractin like 1 | 2826 | 2646 | 2032 | 1722 | 2100 | 1681 | 1992 | 1595 | 2277 | 2101 | 1738 |
| 2336 | -0.07 | 0.11 | -0.65 | 0.52 | 1 | Zfand5 | zinc finger, AN1-type domain 5 | 2871 | 2952 | 2997 | 2188 | 1810 | 2647 | 1991 | 2278 | 1925 | 1557 | 2479 |
| 1648 | 0.09 | 0.09 | 1.02 | 0.31 | 1 | Fhod3 | formin homology 2 domain containing 3 | 799 | 736 | 780 | 2036 | 2034 | 1565 | 1989 | 2170 | 2380 | 2092 | 1546 |
| 2342 | 0.03 | 0.12 | 0.26 | 0.80 | 1 | Cyfip1 | cytoplasmic FMR1 interacting protein 1 | 4319 | 3016 | 3270 | 2044 | 1794 | 1670 | 1988 | 1754 | 2141 | 1960 | 1809 |
| 2220 | 0.02 | 0.11 | 0.23 | 0.82 | 1 | Mdm2 | transformed mouse 3T3 cell double minut | 1977 | 2127 | 1671 | 2449 | 2228 | 2589 | 1988 | 2202 | 2349 | 2177 | 2666 |
| 2185 | -0.01 | 0.11 | -0.06 | 0.95 | 1 | Fam168a | family with sequence similarity 168, meml | 2615 | 2471 | 3058 | 2159 | 1866 | 1943 | 1987 | 2102 | 2015 | 2020 | 1795 |
| 1588 | -0.03 | 0.12 | -0.27 | 0.79 | 1 | Tmem38a | transmembrane protein 38A | 1192 | 1235 | 1436 | 1506 | 2080 | 1323 | 1986 | 1449 | 1839 | 2052 | 1370 |
| 2242 | -0.02 | 0.16 | -0.14 | 0.89 | 1 | N4bp1 | NEDD4 binding protein 1 | 2103 | 2264 | 2055 | 2546 | 2232 | 2406 | 1985 | 1676 | 2551 | 2657 | 2183 |
| 2017 | -0.10 | 0.12 | -0.87 | 0.38 | 1 | Crtc1 | CREB regulated transcription coactivator | 1772 | 2036 | 1540 | 2200 | 2223 | 2319 | 1985 | 1928 | 2122 | 2125 | 1936 |
| 2127 | -0.12 | 0.10 | -1.20 | 0.23 | 1 | Sin3b | transcriptional regulator, SIN3B (yeast) | 2141 | 2307 | 2273 | 2177 | 2082 | 2430 | 1984 | 1740 | 1844 | 2106 | 2309 |
| 1818 | -0.14 | 0.16 | -0.88 | 0.38 | 1 | Foxk1 | forkhead box K1 | 2283 | 2151 | 2105 | 1609 | 1810 | 1557 | 1984 | 1115 | 1910 | 2282 | 1189 |
| 2726 | 0.00 | 0.12 | 0.01 | 0.99 | 1 | Wasf2 | WAS protein family, member 2 | 5509 | 3881 | 4416 | 2065 | 2083 | 1874 | 1983 | 1752 | 2117 | 2664 | 1645 |
| 2762 | -0.01 | 0.07 | -0.10 | 0.92 | 1 | Tcf4 | transcription factor 4 | 5407 | 5160 | 4666 | 2032 | 1763 | 1812 | 1982 | 2033 | 1804 | 1993 | 1727 |
| 1780 | 0.15 | 0.15 | 1.02 | 0.31 | 1 | Armc5 | armadillo repeat containing 5 | 1032 | 781 | 975 | 2249 | 1966 | 1731 | 1981 | 2777 | 1919 | 2400 | 1772 |
| 2465 | 0.17 | 0.10 | 1.75 | 0.08 | 1 | Nemf | nuclear export mediator factor | 1438 | 1339 | 1492 | 1946 | 2460 | 4388 | 1980 | 2520 | 2541 | 2097 | 4919 |
| 2440 | -0.03 | 0.11 | -0.27 | 0.79 | 1 | Thra | thyroid hormone receptor alpha | 4837 | 4163 | 4639 | 1586 | 1837 | 1275 | 1975 | 1897 | 1481 | 1896 | 1256 |
| 2202 | 0.12 | 0.13 | 0.96 | 0.34 | 1 | Plcg1 | phospholipase C, gamma 1 | 2541 | 1925 | 2407 | 1894 | 1829 | 2570 | 1974 | 1623 | 2242 | 2731 | 2485 |
| 2136 | -0.03 | 0.09 | -0.30 | 0.76 | 1 | Rpl7l1 | ribosomal protein L7-like 1 | 2387 | 2244 | 2336 | 2278 | 2200 | 1895 | 1972 | 2094 | 2214 | 1968 | 1911 |
| 1768 | -0.04 | 0.11 | -0.34 | 0.73 | 1 | Mapk6 | mitogen-activated protein kinase 6 | 1601 | 1937 | 1734 | 1708 | 1833 | 1659 | 1972 | 1961 | 1826 | 1598 | 1620 |
| 1835 | 0.04 | 0.10 | 0.41 | 0.68 | 1 | Mfsd11 | major facilitator superfamily domain conta | 973 | 1112 | 1245 | 1984 | 2244 | 2070 | 1970 | 2221 | 2490 | 1647 | 2230 |
| 1987 | -0.12 | 0.12 | -0.99 | 0.32 | 1 | Ubap2 | ubiquitin-associated protein 2 | 2095 | 1983 | 1965 | 2051 | 2085 | 2095 | 1968 | 1614 | 2122 | 2178 | 1705 |
| 1507 | 0.10 | 0.12 | 0.88 | 0.38 | 1 | Mlx | MAX-like protein X | 1228 | 968 | 1201 | 1520 | 1486 | 1385 | 1968 | 1434 | 1647 | 2026 | 1714 |
| 2073 | 0.22 | 0.15 | 1.47 | 0.14 | 1 | Fbxo9 | f-box protein 9 | 1325 | 1265 | 1627 | 2409 | 2104 | 2066 | 1967 | 3322 | 2150 | 2405 | 2161 |
| 2020 | -0.11 | 0.08 | -1.38 | 0.17 | 1 | Pcif1 | PDX1 C-terminal inhibiting factor 1 | 2346 | 2615 | 2405 | 2072 | 1967 | 1695 | 1967 | 2030 | 1922 | 1735 | 1464 |
| 1744 | -0.15 | 0.14 | -1.07 | 0.28 | 1 | Naa60 | N(alpha)-acetyltransferase 60, NatF catal | 1534 | 1279 | 1467 | 1978 | 2020 | 1817 | 1966 | 1243 | 2001 | 2214 | 1664 |
| 2412 | -0.05 | 0.11 | -0.40 | 0.69 | 1 | Zfp266 | zinc finger protein 266 | 2430 | 2926 | 3013 | 2731 | 2033 | 2515 | 1965 | 2065 | 2264 | 2049 | 2541 |
| 2347 | -0.10 | 0.11 | -0.95 | 0.34 | 1 | Chd6 | chromodomain helicase DNA binding pro | 2178 | 3280 | 2253 | 2241 | 2193 | 2954 | 1965 | 2456 | 1912 | 1793 | 2594 |
| 1880 | -0.01 | 0.11 | -0.10 | 0.92 | 1 | Tasor | transcription activation suppressor | 1365 | 1222 | 1272 | 1785 | 1711 | 2959 | 1965 | 1650 | 2075 | 1631 | 3048 |
| 2058 | 0.00 | 0.13 | 0.02 | 0.98 | 1 | Retreg3 | reticulophagy regulator family member 3 | 2177 | 1752 | 2211 | 1966 | 2186 | 2118 | 1964 | 2272 | 2099 | 1650 | 2241 |
| 2048 | 0.01 | 0.12 | 0.06 | 0.95 | 1 | Psmc4 | proteasome (prosome, macropain) 26S su | 2430 | 1741 | 2245 | 1952 | 2025 | 2096 | 1963 | 2005 | 2135 | 1834 | 2099 |
| 2016 | 0.15 | 0.13 | 1.19 | 0.23 | 1 | Eps8 | epidermal growth factor receptor pathway | 999 | 1173 | 1057 | 2317 | 2642 | 2040 | 1963 | 2107 | 3180 | 2236 | 2467 |
| 1973 | -0.05 | 0.13 | -0.36 | 0.72 | 1 | Smchd1 | SMC hinge domain containing 1 | 2689 | 2286 | 1890 | 1764 | 2011 | 1812 | 1963 | 1793 | 1934 | 1662 | 1899 |
| 1803 | -0.12 | 0.10 | -1.24 | 0.21 | 1 | Trip4 | thyroid hormone receptor interactor 4 | 1741 | 1942 | 1874 | 1904 | 1762 | 1805 | 1963 | 1825 | 1669 | 1551 | 1802 |
| 1595 | 0.14 | 0.10 | 1.33 | 0.18 | 1 | 0610010F05Ril | RIKEN cDNA 0610010F05 gene | 1107 | 816 | 1127 | 1620 | 1805 | 1507 | 1963 | 1786 | 1861 | 2285 | 1667 |
| 1426 | 0.01 | 0.17 | 0.08 | 0.93 | 1 | Dcx | doublecortin | 589 | 973 | 715 | 1548 | 1368 | 1792 | 1963 | 1691 | 1538 | 1649 | 1859 |
| 1952 | -0.12 | 0.11 | -1.08 | 0.28 | 1 | Exoc2 | exocyst complex component 2 | 2093 | 1871 | 1966 | 2294 | 1978 | 1850 | 1962 | 1702 | 1962 | 1981 | 1809 |
| 1938 | -0.07 | 0.08 | -0.91 | 0.37 | 1 | Nt5dc2 | 5'-nucleotidase domain containing 2 | 2044 | 2115 | 2237 | 1904 | 2013 | 1762 | 1962 | 1802 | 1928 | 1843 | 1710 |
| 1996 | 0.00 | 0.13 | 0.03 | 0.98 | 1 | Nolc1 | nucleolar and coiled-body phosphoprotei | 1840 | 1863 | 1606 | 1946 | 2099 | 2286 | 1960 | 1563 | 2280 | 2235 | 2277 |
| 1924 | -0.08 | 0.08 | -0.95 | 0.34 | 1 | Rbm15b | RNA binding motif protein 15B | 1563 | 1770 | 1640 | 2262 | 2143 | 1954 | 1960 | 2009 | 2118 | 1980 | 1765 |
| 1514 | -0.07 | 0.16 | -0.44 | 0.66 | 1 | Gdap2 | ganglioside-induced differentiation-assoc | 909 | 1041 | 954 | 1519 | 1940 | 1602 | 1960 | 1252 | 2070 | 1604 | 1804 |
| 2674 | 0.01 | 0.11 | 0.11 | 0.91 | 1 | Rps15a | ribosomal protein S15A | 5510 | 4475 | 5488 | 1764 | 1877 | 1319 | 1959 | 2090 | 1804 | 1913 | 1211 |
| 1940 | 0.02 | 0.13 | 0.19 | 0.85 | 1 | Nudt3 | nudix (nucleotide diphosphate linked moi | 1811 | 1776 | 2496 | 1880 | 2036 | 1658 | 1959 | 2344 | 2119 | 1676 | 1583 |
| 1870 | -0.09 | 0.13 | -0.68 | 0.50 | 1 | Sos1 | SOS Ras/Rac guanine nucleotide exchar | 1592 | 2142 | 1508 | 2040 | 1970 | 1942 | 1959 | 1704 | 2048 | 1922 | 1737 |
| 1736 | 0.01 | 0.14 | 0.08 | 0.94 | 1 | Samd8 | sterile alpha motif domain containing 8 | 1029 | 1672 | 1341 | 2073 | 1768 | 1701 | 1959 | 1897 | 2011 | 1731 | 1911 |
| 1919 | -0.08 | 0.14 | -0.60 | 0.55 | 1 | Fyco1 | FYVE and coiled-coil domain containing ' | 1645 | 1224 | 1191 | 2335 | 1978 | 2480 | 1957 | 1676 | 2099 | 2176 | 2344 |
| 1866 | -0.12 | 0.15 | -0.76 | 0.44 | 1 | Hsd17b11 | hydroxysteroid (17-beta) dehydrogenase | 2331 | 2189 | 2152 | 2016 | 1975 | 1230 | 1956 | 1337 | 1632 | 2428 | 1275 |
| 2309 | 0.05 | 0.10 | 0.53 | 0.60 | 1 | Sfr1 | SWI5 dependent recombination repair 1 | 3303 | 2924 | 2925 | 2004 | 2127 | 1864 | 1954 | 1756 | 2148 | 2388 | 2002 |
| 1886 | 0.07 | 0.08 | 0.85 | 0.40 | 1 | Brox | BRO1 domain and CAAX motif containing | 1542 | 1593 | 1586 | 1945 | 1899 | 2019 | 1952 | 1906 | 1980 | 2158 | 2172 |

| | | | | | | | | | | | | | | | | | |
|---|---|---|---|---|---|---|---|---|---|---|---|---|---|---|---|---|---|
| 1850 | -0.06 | 0.16 | -0.37 | 0.71 | 1 | Tmem64 | transmembrane protein 64 | 2163 | 2178 | 1883 | 1958 | 1765 | 1503 | 1952 | 1269 | 2014 | 2194 | 1477 |
| 1663 | 0.03 | 0.10 | 0.28 | 0.78 | 1 | Abcf2 | ATP-binding cassette, sub-family F (GCN2 | 1975 | 1545 | 1815 | 1540 | 1518 | 1421 | 1950 | 1600 | 1700 | 1772 | 1452 |
| 2118 | 0.02 | 0.07 | 0.25 | 0.80 | 1 | Fmr1 | fragile X mental retardation 1 | 2004 | 2150 | 2096 | 1995 | 2026 | 2498 | 1949 | 1987 | 2030 | 2030 | 2532 |
| 2040 | -0.07 | 0.09 | -0.75 | 0.45 | 1 | Zfand6 | zinc finger, AN1-type domain 6 | 2132 | 1754 | 1893 | 1878 | 2176 | 2511 | 1949 | 1745 | 1996 | 1948 | 2456 |
| 1984 | -0.17 | 0.11 | -1.52 | 0.13 | 1 | Eci2 | enoyl-Coenzyme A delta isomerase 2 | 2507 | 2648 | 2553 | 1744 | 1734 | 2024 | 1949 | 1587 | 1751 | 1374 | 1949 |
| 2377 | -0.07 | 0.13 | -0.53 | 0.60 | 1 | Sft2d2 | SFT2 domain containing 2 | 3570 | 3161 | 2755 | 2492 | 1995 | 2084 | 1948 | 2409 | 2109 | 1799 | 1824 |
| 1776 | 0.02 | 0.10 | 0.16 | 0.87 | 1 | Slc27a4 | solute carrier family 27 (fatty acid transpor | 1484 | 1585 | 1599 | 1947 | 2089 | 1386 | 1948 | 1961 | 1978 | 2252 | 1309 |
| 2306 | -0.15 | 0.13 | -1.16 | 0.25 | 1 | Ccnt2 | cyclin T2 | 1326 | 1637 | 1461 | 1658 | 1607 | 5722 | 1946 | 1307 | 1785 | 1519 | 5403 |
| 2231 | 0.11 | 0.14 | 0.79 | 0.43 | 1 | Bnip3l | BCL2/adenovirus E1B interacting protein | 3196 | 3231 | 3529 | 1779 | 1852 | 1417 | 1946 | 2470 | 1994 | 1524 | 1608 |
| 1623 | -0.17 | 0.10 | -1.69 | 0.09 | 1 | Klhl22 | kelch-like 22 | 1516 | 1548 | 1346 | 1775 | 1557 | 1835 | 1946 | 1453 | 1514 | 1783 | 1584 |
| 2130 | -0.17 | 0.10 | -1.60 | 0.11 | 1 | Shroom3 | shroom family member 3 | 1805 | 2562 | 2144 | 2646 | 2370 | 1894 | 1945 | 2732 | 2233 | 1713 | 1384 |
| 1642 | -0.15 | 0.15 | -1.06 | 0.29 | 1 | Ube2g1 | ubiquitin-conjugating enzyme E2G 1 | 1597 | 2192 | 1509 | 1722 | 1685 | 1393 | 1945 | 1430 | 1571 | 1562 | 1453 |
| 2321 | -0.02 | 0.11 | -0.21 | 0.83 | 1 | Klf13 | Kruppel-like factor 13 | 3215 | 3568 | 2787 | 2623 | 1891 | 1516 | 1943 | 2926 | 1857 | 1845 | 1355 |
| 2334 | -0.03 | 0.11 | -0.31 | 0.76 | 1 | Qars | glutaminyl-tRNA synthetase | 3653 | 2968 | 2933 | 2101 | 2117 | 1994 | 1941 | 2011 | 2077 | 1893 | 1987 |
| 1934 | 0.02 | 0.14 | 0.14 | 0.89 | 1 | Eftud2 | elongation factor Tu GTP binding domain | 2827 | 2261 | 1984 | 1735 | 1861 | 1515 | 1941 | 2170 | 1802 | 1663 | 1513 |
| 1838 | -0.10 | 0.13 | -0.75 | 0.45 | 1 | Evi5 | ecotropic viral integration site 5 | 2209 | 2164 | 1772 | 1896 | 1799 | 1650 | 1941 | 1456 | 1895 | 1650 | 1787 |
| 2439 | 0.15 | 0.12 | 1.23 | 0.22 | 1 | Tspyl2 | TSPY-like 2 | 1307 | 1414 | 1204 | 1525 | 1884 | 5688 | 1940 | 2118 | 2104 | 1868 | 5771 |
| 2184 | 0.03 | 0.08 | 0.37 | 0.71 | 1 | Elk3 | ELK3, member of ETS oncogene family | 3131 | 3032 | 3246 | 1820 | 1918 | 1547 | 1940 | 1709 | 2107 | 1994 | 1581 |
| 1664 | 0.07 | 0.09 | 0.76 | 0.45 | 1 | Uqcc1 | ubiquinol-cytochrome c reductase comple | 1213 | 1227 | 1348 | 1707 | 1938 | 1504 | 1940 | 1857 | 1900 | 2051 | 1618 |
| 2087 | 0.05 | 0.10 | 0.55 | 0.58 | 1 | Hnrnpll | heterogeneous nuclear ribonucleoprotein | 2087 | 2273 | 1966 | 2223 | 1769 | 2223 | 1939 | 1913 | 1924 | 2112 | 2523 |
| 1832 | -0.04 | 0.09 | -0.44 | 0.66 | 1 | Atg3 | autophagy related 3 | 2231 | 1780 | 1903 | 1460 | 1854 | 1950 | 1939 | 1368 | 1703 | 1975 | 1986 |
| 1715 | 0.08 | 0.09 | 0.89 | 0.37 | 1 | Bach1 | BTB and CNC homology 1, basic leucine | 1585 | 1254 | 1244 | 1680 | 1655 | 1902 | 1939 | 1774 | 1803 | 1872 | 2157 |
| 1540 | -0.13 | 0.14 | -0.98 | 0.33 | 1 | Acad11 | acyl-Coenzyme A dehydrogenase family, | 1004 | 1135 | 1208 | 1457 | 2012 | 1692 | 1939 | 1223 | 1901 | 1790 | 1575 |
| 2246 | 0.03 | 0.14 | 0.19 | 0.85 | 1 | Dock11 | dedicator of cytokinesis 11 | 2726 | 4306 | 2394 | 2015 | 1981 | 1633 | 1938 | 2104 | 1913 | 2000 | 1698 |
| 1977 | 0.02 | 0.13 | 0.17 | 0.87 | 1 | Ankrd33b | ankyrin repeat domain 33B | 1397 | 1097 | 1164 | 2388 | 2081 | 2546 | 1938 | 1905 | 2176 | 2499 | 2557 |
| 1780 | 0.15 | 0.10 | 1.47 | 0.14 | 1 | Gm20300 | | 1258 | 1530 | 1209 | 1820 | 1793 | 1827 | 1938 | 2113 | 2070 | 1917 | 2102 |
| 2156 | -0.08 | 0.11 | -0.74 | 0.46 | 1 | Ilk | integrin linked kinase | 3309 | 2558 | 2758 | 1665 | 2218 | 1910 | 1937 | 1479 | 2028 | 2127 | 1723 |
| 1963 | 0.13 | 0.10 | 1.24 | 0.22 | 1 | Eny2 | ENY2 transcription and export complex 2 | 1696 | 1873 | 2091 | 1768 | 1811 | 2096 | 1937 | 1803 | 2087 | 2051 | 2386 |
| 1497 | 0.16 | 0.14 | 1.20 | 0.23 | 1 | Idh1 | isocitrate dehydrogenase 1 (NADP+), solu | 1679 | 1038 | 1219 | 1483 | 1298 | 1237 | 1937 | 1782 | 1633 | 1753 | 1412 |
| 2152 | -0.10 | 0.11 | -0.88 | 0.38 | 1 | Cacnb3 | calcium channel, voltage-dependent, beta | 2113 | 3147 | 2791 | 2063 | 2073 | 2001 | 1936 | 1864 | 2191 | 1748 | 1750 |
| 2143 | 0.24 | 0.21 | 1.16 | 0.25 | 1 | Sem1 | SEM1, 26S proteasome complex subunit | 2657 | 2397 | 2982 | 1799 | 1803 | 1457 | 1936 | 3490 | 1526 | 1959 | 1572 |
| 2143 | 0.09 | 0.09 | 1.08 | 0.28 | 1 | Slu7 | SLU7 splicing factor homolog (S. cerevisi | 2659 | 2259 | 2483 | 1947 | 2097 | 1842 | 1936 | 2066 | 2254 | 2011 | 2020 |
| 2015 | 0.06 | 0.10 | 0.60 | 0.55 | 1 | Vps4a | vacuolar protein sorting 4A | 2014 | 2107 | 2110 | 2012 | 1986 | 1850 | 1936 | 1968 | 2151 | 2321 | 1710 |
| 1888 | -0.04 | 0.12 | -0.31 | 0.75 | 1 | Pxk | PX domain containing serine/threonine ki | 1222 | 902 | 1305 | 2090 | 2451 | 2247 | 1936 | 1577 | 2626 | 2234 | 2179 |
| 1839 | -0.13 | 0.12 | -1.08 | 0.28 | 1 | Amigo2 | adhesion molecule with Ig like domain 2 | 1209 | 1613 | 1581 | 1881 | 2049 | 2407 | 1936 | 1736 | 1778 | 1913 | 2123 |
| 1614 | 0.02 | 0.15 | 0.15 | 0.88 | 1 | Dcun1d4 | DCN1, defective in cullin neddylation 1, d | 1060 | 1538 | 1236 | 1964 | 1724 | 1291 | 1936 | 1696 | 1719 | 2268 | 1328 |
| 2222 | 0.02 | 0.11 | 0.15 | 0.88 | 1 | Oaz2 | ornithine decarboxylase antizyme 2 | 3328 | 3198 | 4057 | 1727 | 1881 | 1353 | 1935 | 2027 | 1748 | 1780 | 1403 |
| 1898 | -0.07 | 0.12 | -0.61 | 0.54 | 1 | Unc13b | unc-13 homolog B | 716 | 1163 | 935 | 2385 | 1877 | 3084 | 1935 | 2045 | 1994 | 1857 | 2883 |
| 2001 | 0.05 | 0.08 | 0.64 | 0.52 | 1 | Dnajc8 | DnaJ heat shock protein family (Hsp40) m | 2511 | 2106 | 2288 | 1770 | 1929 | 1789 | 1933 | 1791 | 1942 | 1990 | 1961 |
| 1686 | 0.08 | 0.13 | 0.58 | 0.56 | 1 | Chmp2b | charged multivesicular body protein 2B | 2159 | 2236 | 2184 | 1409 | 1382 | 1140 | 1933 | 1394 | 1623 | 1690 | 1397 |
| 2152 | 0.05 | 0.13 | 0.38 | 0.70 | 1 | Rpl36 | ribosomal protein L36 | 3185 | 2852 | 3352 | 1657 | 1952 | 1462 | 1931 | 2266 | 1765 | 1873 | 1374 |
| 2034 | -0.04 | 0.11 | -0.37 | 0.71 | 1 | Ctr9 | CTR9 homolog, Paf1/RNA polymerase II c | 2375 | 2684 | 2187 | 1755 | 1855 | 2117 | 1931 | 2067 | 1770 | 1580 | 2054 |
| 1704 | 0.12 | 0.15 | 0.76 | 0.44 | 1 | Pcsk9 | proprotein convertase subtilisin/kexin type | 140 | 341 | 301 | 2653 | 2229 | 1756 | 1931 | 2822 | 2820 | 1951 | 1803 |
| 2023 | -0.07 | 0.10 | -0.71 | 0.48 | 1 | Eif1a | eukaryotic translation initiation factor 1A | 2058 | 2666 | 2211 | 2124 | 1772 | 2003 | 1930 | 2061 | 1859 | 1526 | 2040 |
| 1739 | 0.11 | 0.11 | 1.00 | 0.32 | 1 | Eif2b5 | eukaryotic translation initiation factor 2B, s | 1305 | 1151 | 1385 | 1773 | 2069 | 1598 | 1930 | 2189 | 1901 | 2106 | 1728 |
| 1683 | -0.02 | 0.13 | -0.16 | 0.87 | 1 | Slc35a5 | solute carrier family 35, member A5 | 1216 | 1654 | 1366 | 1909 | 1685 | 1679 | 1930 | 1642 | 1730 | 1986 | 1723 |
| 3013 | 0.05 | 0.08 | 0.63 | 0.53 | 1 | Luc7l | Luc7-like | 2103 | 2860 | 2621 | 2458 | 2043 | 6161 | 1929 | 2838 | 2096 | 1893 | 6139 |
| 1792 | 0.00 | 0.10 | -0.01 | 0.99 | 1 | Men1 | multiple endocrine neoplasia 1 | 1249 | 1284 | 1394 | 2010 | 2160 | 1793 | 1929 | 1827 | 2055 | 2187 | 1820 |
| 2018 | 0.06 | 0.09 | 0.67 | 0.50 | 1 | Ndufa6 | NADH:ubiquinone oxidoreductase subun | 2595 | 2536 | 2887 | 1568 | 1847 | 1613 | 1927 | 1782 | 1860 | 1868 | 1719 |
| 1747 | 0.04 | 0.11 | 0.41 | 0.68 | 1 | Rrn3 | RRN3 RNA polymerase I transcription fac | 1567 | 1380 | 1246 | 1876 | 2004 | 1587 | 1927 | 1840 | 2144 | 2027 | 1618 |
| 1745 | 0.01 | 0.09 | 0.06 | 0.95 | 1 | Zadh2 | zinc binding alcohol dehydrogenase, don | 1857 | 2010 | 1982 | 1697 | 1667 | 1369 | 1927 | 1763 | 1692 | 1843 | 1383 |
| 1529 | 0.03 | 0.13 | 0.22 | 0.82 | 1 | Tigar | Trp53 induced glycolysis regulatory phos | 588 | 813 | 549 | 1980 | 1851 | 1604 | 1927 | 1847 | 1973 | 2070 | 1615 |
| 2494 | 0.15 | 0.12 | 1.28 | 0.20 | 1 | Brwd1 | bromodomain and WD repeat domain cor | 1603 | 1493 | 1602 | 1873 | 2382 | 4761 | 1926 | 2336 | 2173 | 2727 | 4556 |
| 2682 | 0.08 | 0.15 | 0.53 | 0.60 | 1 | Arap1 | ArfGAP with RhoGAP domain, ankyrin rep | 4169 | 3301 | 3450 | 2269 | 2563 | 2265 | 1925 | 2955 | 2336 | 2174 | 2090 |
| 1826 | 0.01 | 0.11 | 0.11 | 0.91 | 1 | Traf6 | TNF receptor-associated factor 6 | 1540 | 1920 | 1321 | 1977 | 1873 | 1848 | 1925 | 1912 | 1863 | 1991 | 1916 |
| 1857 | -0.11 | 0.11 | -0.98 | 0.33 | 1 | H3f3a | H3.3 histone A | 2325 | 2525 | 2584 | 1539 | 1829 | 1451 | 1924 | 1413 | 1705 | 1654 | 1476 |
| 1513 | 0.01 | 0.13 | 0.05 | 0.96 | 1 | Ctnnd2 | catenin (cadherin associated protein), del | 1266 | 1463 | 1207 | 1504 | 1487 | 1412 | 1924 | 1771 | 1721 | 1618 | 1266 |
| 1671 | -0.01 | 0.10 | -0.10 | 0.92 | 1 | Cnnm2 | cyclin M2 | 883 | 960 | 1196 | 1980 | 2239 | 1531 | 1923 | 2146 | 2343 | 1681 | 1500 |
| 2372 | 0.11 | 0.12 | 0.88 | 0.38 | 1 | Tbc1d22a | TBC1 domain family, member 22a | 3125 | 4888 | 2692 | 2079 | 1892 | 1453 | 1921 | 2680 | 1899 | 1923 | 1541 |
| 1927 | -0.04 | 0.11 | -0.34 | 0.74 | 1 | Cbx1 | chromobox 1 | 1961 | 1867 | 2321 | 1743 | 1736 | 2234 | 1921 | 1841 | 1681 | 1870 | 2022 |

| | | | | | | | | | | | | | | | | | | |
|---|---|---|---|---|---|---|---|---|---|---|---|---|---|---|---|---|---|---|
| 1644 | -0.17 | 0.14 | -1.23 | 0.22 | 1 | Tfdp2 | transcription factor Dp 2 | 1480 | 2186 | 1548 | 1960 | 1539 | 1412 | 1920 | 1654 | 1557 | 1346 | 1483 |
| 1908 | 0.19 | 0.14 | 1.30 | 0.19 | 1 | Rogdi | rogdi homolog | 1428 | 1469 | 1610 | 1771 | 1896 | 2103 | 1919 | 2712 | 1780 | 2227 | 2075 |
| 2165 | -0.18 | 0.11 | -1.65 | 0.10 | 1 | Nup153 | nucleoporin 153 | 2416 | 2467 | 2248 | 2187 | 2261 | 2488 | 1918 | 1622 | 2092 | 1739 | 2379 |
| 1996 | -0.11 | 0.10 | -1.10 | 0.27 | 1 | Dipk2a | divergent protein kinase domain 2A | 2181 | 3087 | 2687 | 2192 | 1793 | 1366 | 1918 | 1950 | 1910 | 1605 | 1263 |
| 2086 | -0.02 | 0.09 | -0.18 | 0.86 | 1 | Tmem234 | transmembrane protein 234 | 2263 | 1970 | 2361 | 2024 | 2138 | 2136 | 1917 | 2008 | 2040 | 1814 | 2272 |
| 1666 | 0.16 | 0.13 | 1.27 | 0.20 | 1 | Echdc1 | enoyl Coenzyme A hydratase domain cor | 1046 | 1169 | 1055 | 1844 | 1833 | 1511 | 1917 | 1673 | 2118 | 2356 | 1799 |
| 2641 | 0.17 | 0.15 | 1.18 | 0.24 | 1 | Map7d1 | MAP7 domain containing 1 | 3565 | 2557 | 2759 | 2458 | 2240 | 2886 | 1915 | 3009 | 2271 | 2507 | 2890 |
| 1999 | -0.01 | 0.11 | -0.07 | 0.94 | 1 | Bub3 | BUB3 mitotic checkpoint protein | 2371 | 2396 | 2363 | 1866 | 1959 | 1702 | 1915 | 2131 | 1629 | 1781 | 1876 |
| 1907 | 0.11 | 0.14 | 0.76 | 0.45 | 1 | Ubl5 | ubiquitin-like 5 | 1936 | 1859 | 2393 | 1676 | 1955 | 1548 | 1915 | 2372 | 1776 | 2071 | 1470 |
| 1860 | -0.01 | 0.13 | -0.07 | 0.95 | 1 | Hivep1 | human immunodeficiency virus type I enh | 1945 | 2491 | 1869 | 1743 | 1524 | 1928 | 1914 | 1544 | 1787 | 1740 | 1975 |
| 1804 | 0.07 | 0.12 | 0.59 | 0.55 | 1 | Rint1 | RAD50 interactor 1 | 945 | 804 | 995 | 1372 | 2286 | 2797 | 1914 | 1708 | 2419 | 1694 | 2910 |
| 1709 | -0.19 | 0.10 | -1.88 | 0.06 | 1 | Smpd3 | sphingomyelin phosphodiesterase 3, neu | 2160 | 2019 | 1869 | 1708 | 1849 | 1311 | 1914 | 1781 | 1514 | 1613 | 1065 |
| 2577 | 0.02 | 0.15 | 0.11 | 0.92 | 1 | Rpl35a | ribosomal protein L35A | 3256 | 4621 | 5036 | 2388 | 1802 | 1501 | 1913 | 2899 | 1735 | 1502 | 1696 |
| 1984 | -0.12 | 0.12 | -0.96 | 0.34 | 1 | Ptch1 | patched 1 | 3483 | 2382 | 2285 | 1870 | 1872 | 1439 | 1913 | 1461 | 1682 | 2024 | 1410 |
| 2097 | -0.08 | 0.16 | -0.47 | 0.64 | 1 | Irs2 | insulin receptor substrate 2 | 3270 | 1525 | 2285 | 1320 | 2032 | 3073 | 1912 | 1562 | 1921 | 1667 | 2504 |
| 1750 | -0.17 | 0.10 | -1.64 | 0.10 | 1 | Slc22a17 | solute carrier family 22 (organic cation tra | 1122 | 1195 | 1246 | 1672 | 1718 | 3016 | 1912 | 1395 | 1624 | 1757 | 2589 |
| 1490 | -0.12 | 0.09 | -1.33 | 0.18 | 1 | Unc5a | unc-5 netrin receptor A | 1219 | 1172 | 1135 | 1703 | 1568 | 1513 | 1911 | 1578 | 1455 | 1784 | 1358 |
| 2005 | 0.16 | 0.13 | 1.19 | 0.23 | 1 | Dnajc21 | DnaJ heat shock protein family (Hsp40) m | 1122 | 1263 | 1386 | 1780 | 2276 | 2689 | 1909 | 2367 | 1893 | 2392 | 2983 |
| 1824 | -0.05 | 0.09 | -0.52 | 0.60 | 1 | Fundc1 | FUN14 domain containing 1 | 1608 | 1868 | 1822 | 2116 | 2079 | 1430 | 1909 | 1849 | 2019 | 1787 | 1573 |
| 2280 | -0.05 | 0.11 | -0.45 | 0.66 | 1 | Stau1 | staufen double-stranded RNA binding pro | 2558 | 2847 | 2545 | 2206 | 2161 | 2422 | 1908 | 1756 | 2088 | 2097 | 2490 |
| 1827 | 0.01 | 0.12 | 0.06 | 0.95 | 1 | Spg21 | SPG21, maspardin | 1870 | 1861 | 2180 | 1802 | 2003 | 1358 | 1908 | 1499 | 1981 | 2188 | 1450 |
| 1636 | 0.10 | 0.10 | 1.00 | 0.32 | 1 | Pdxk | pyridoxal (pyridoxine, vitamin B6) kinase | 1649 | 1639 | 1512 | 1641 | 1533 | 1293 | 1908 | 1691 | 1846 | 1942 | 1338 |
| 2670 | 0.16 | 0.11 | 1.41 | 0.16 | 1 | Klc1 | kinesin light chain 1 | 2955 | 2489 | 2851 | 2109 | 1913 | 4132 | 1907 | 2376 | 1976 | 2552 | 4108 |
| 1730 | -0.04 | 0.12 | -0.30 | 0.76 | 1 | Fbxo33 | F-box protein 33 | 1326 | 1662 | 1155 | 2006 | 1928 | 1706 | 1907 | 1841 | 1937 | 1729 | 1830 |
| 2382 | -0.02 | 0.10 | -0.18 | 0.86 | 1 | Pcf11 | PCF11 cleavage and polyadenylation fac | 2323 | 2513 | 2298 | 2254 | 2166 | 3274 | 1906 | 1949 | 2080 | 2239 | 3197 |
| 1917 | 0.15 | 0.17 | 0.87 | 0.39 | 1 | 2210016F16Rik | RIKEN cDNA 2210016F16 gene | 1556 | 1648 | 1914 | 2049 | 2034 | 1478 | 1905 | 3209 | 1781 | 2130 | 1384 |
| 1788 | -0.15 | 0.11 | -1.30 | 0.20 | 1 | Fbrsl1 | fibrosin-like 1 | 1176 | 1209 | 1304 | 2088 | 2066 | 2335 | 1905 | 1542 | 1984 | 2048 | 2014 |
| 1772 | -0.13 | 0.14 | -0.92 | 0.36 | 1 | Whrn | whirlin | 500 | 516 | 406 | 2654 | 2313 | 2570 | 1905 | 2187 | 2420 | 1741 | 2278 |
| 1958 | 0.19 | 0.14 | 1.30 | 0.20 | 1 | Zfp292 | zinc finger protein 292 | 1900 | 1501 | 1410 | 1312 | 1539 | 3233 | 1903 | 1579 | 2253 | 1933 | 2978 |
| 1769 | -0.05 | 0.08 | -0.60 | 0.55 | 1 | Rtl6 | retrotransposon Gag like 6 | 1407 | 1665 | 1317 | 2510 | 1846 | 1398 | 1902 | 2375 | 1793 | 1933 | 1309 |
| 2456 | -0.09 | 0.11 | -0.82 | 0.41 | 1 | Zscan26 | zinc finger and SCAN domain containing | 1616 | 1550 | 1803 | 1833 | 2206 | 5296 | 1901 | 1379 | 2240 | 1930 | 5268 |
| 1872 | -0.06 | 0.11 | -0.59 | 0.56 | 1 | Fam160b1 | family with sequence similarity 160, meml | 1547 | 1788 | 1377 | 2148 | 1863 | 2204 | 1901 | 1752 | 1827 | 1972 | 2217 |
| 1727 | -0.18 | 0.13 | -1.37 | 0.17 | 1 | Clasp1 | CLIP associating protein 1 | 1823 | 1928 | 1587 | 2006 | 1608 | 1739 | 1901 | 1440 | 1772 | 1861 | 1333 |
| 1709 | -0.03 | 0.12 | -0.23 | 0.82 | 1 | Uggt2 | UDP-glucose glycoprotein glucosyltransfe | 1505 | 1551 | 1737 | 1815 | 1819 | 1532 | 1901 | 1703 | 1580 | 2051 | 1603 |
| 1685 | -0.01 | 0.16 | -0.06 | 0.96 | 1 | Abcb9 | ATP-binding cassette, sub-family B (MDR | 822 | 1264 | 976 | 1758 | 1972 | 2128 | 1901 | 1847 | 1857 | 2218 | 1787 |
| 1614 | -0.09 | 0.10 | -0.90 | 0.37 | 1 | Rai1 | retinoic acid induced 1 | 1228 | 1212 | 1250 | 1646 | 1762 | 1940 | 1901 | 1774 | 1620 | 1641 | 1785 |
| 1845 | 0.07 | 0.10 | 0.74 | 0.46 | 1 | Impa1 | inositol (myo)-1(or 4)-monophosphatase | 1964 | 2037 | 1853 | 1846 | 1773 | 1536 | 1900 | 1793 | 1840 | 1875 | 1877 |
| 1723 | 0.13 | 0.13 | 1.02 | 0.31 | 1 | Uqcrb | ubiquinol-cytochrome c reductase binding | 1650 | 1642 | 1874 | 1595 | 1610 | 1484 | 1900 | 2103 | 1624 | 1830 | 1646 |
| 1716 | 0.08 | 0.12 | 0.67 | 0.51 | 1 | Cars | cysteinyl-tRNA synthetase | 824 | 1046 | 1018 | 1629 | 2080 | 2173 | 1900 | 2099 | 2098 | 1896 | 2112 |
| 1707 | 0.11 | 0.12 | 0.90 | 0.37 | 1 | Xrcc5 | X-ray repair complementing defective rep | 1186 | 1376 | 1530 | 1852 | 1872 | 1464 | 1900 | 1790 | 1954 | 2003 | 1847 |
| 2031 | 0.18 | 0.15 | 1.18 | 0.24 | 1 | Cc2d1b | coiled-coil and C2 domain containing 1B | 1543 | 1168 | 1362 | 2091 | 1961 | 2617 | 1899 | 2946 | 1975 | 2069 | 2709 |
| 1583 | -0.06 | 0.13 | -0.43 | 0.67 | 1 | Apba1 | amyloid beta (A4) precursor protein bindi | 752 | 690 | 978 | 2139 | 1863 | 1721 | 1898 | 1908 | 2010 | 1900 | 1552 |
| 2961 | 0.08 | 0.10 | 0.77 | 0.44 | 1 | Rbm5 | RNA binding motif protein 5 | 1972 | 2533 | 2249 | 2089 | 1805 | 7007 | 1896 | 2063 | 2009 | 2209 | 6738 |
| 2268 | 0.15 | 0.15 | 1.05 | 0.29 | 1 | Man2c1 | mannosidase, alpha, class 2C, member 1 | 2086 | 2043 | 2350 | 1784 | 2190 | 2937 | 1896 | 2806 | 1981 | 2206 | 2669 |
| 2178 | 0.02 | 0.09 | 0.28 | 0.78 | 1 | Nup88 | nucleoporin 88 | 2127 | 2269 | 2114 | 1938 | 2278 | 2538 | 1896 | 1982 | 2206 | 1975 | 2640 |
| 1979 | 0.02 | 0.14 | 0.11 | 0.91 | 1 | Fcho2 | FCH domain only 2 | 2406 | 1967 | 1953 | 1724 | 1673 | 2341 | 1894 | 1269 | 1965 | 2244 | 2335 |
| 1755 | -0.14 | 0.10 | -1.33 | 0.18 | 1 | Magee1 | melanoma antigen, family E, 1 | 1538 | 2045 | 1745 | 2241 | 1916 | 1286 | 1894 | 1876 | 1675 | 1877 | 1211 |
| 1719 | -0.24 | 0.14 | -1.71 | 0.09 | 1 | Wdr37 | WD repeat domain 37 | 1339 | 1735 | 1427 | 1939 | 1902 | 2030 | 1894 | 1360 | 1806 | 1436 | 2045 |
| 1506 | -0.03 | 0.12 | -0.23 | 0.82 | 1 | Rhbdd1 | rhomboid domain containing 1 | 1029 | 1053 | 1114 | 1568 | 1694 | 1565 | 1894 | 1466 | 1917 | 1932 | 1338 |
| 1585 | 0.06 | 0.10 | 0.57 | 0.57 | 1 | Sdr39u1 | short chain dehydrogenase/reductase far | 1198 | 1313 | 1223 | 1738 | 1725 | 1368 | 1893 | 1695 | 1836 | 1982 | 1468 |
| 1507 | 0.16 | 0.14 | 1.13 | 0.26 | 1 | Cmpk2 | cytidine monophosphate (UMP-CMP) kin | 1194 | 988 | 1130 | 1723 | 1453 | 1188 | 1893 | 1997 | 1361 | 2244 | 1406 |
| 1965 | -0.09 | 0.11 | -0.82 | 0.41 | 1 | Sbds | SBDS ribosome maturation factor | 2398 | 2014 | 2300 | 1971 | 1835 | 1970 | 1892 | 1671 | 1860 | 1570 | 2130 |
| 1798 | 0.07 | 0.12 | 0.57 | 0.57 | 1 | Bcl7b | B cell CLL/lymphoma 7B | 1732 | 1939 | 1778 | 1650 | 2003 | 1446 | 1892 | 1850 | 1811 | 2203 | 1472 |
| 2025 | -0.03 | 0.11 | -0.29 | 0.77 | 1 | Elavl1 | ELAV (embryonic lethal, abnormal vision) | 2389 | 2462 | 2124 | 1824 | 2046 | 1973 | 1891 | 1788 | 2140 | 1698 | 1945 |
| 1218 | 0.08 | 0.14 | 0.55 | 0.58 | 1 | Mreg | melanoregulin | 221 | 151 | 198 | 1799 | 1419 | 1159 | 1891 | 1663 | 1656 | 1919 | 1324 |
| 2050 | -0.15 | 0.11 | -1.32 | 0.19 | 1 | Phc3 | polyhomeotic 3 | 2252 | 2303 | 1941 | 2037 | 1845 | 2688 | 1890 | 1652 | 1988 | 1614 | 2340 |
| 1915 | 0.05 | 0.11 | 0.44 | 0.66 | 1 | Cwc15 | CWC15 spliceosome-associated protein | 1911 | 1709 | 2160 | 1933 | 2018 | 1671 | 1890 | 2263 | 1926 | 1891 | 1698 |
| 1907 | -0.01 | 0.09 | -0.08 | 0.94 | 1 | Eif1b | eukaryotic translation initiation factor 1B | 2254 | 2509 | 2512 | 1898 | 1707 | 1391 | 1890 | 1825 | 1705 | 1779 | 1513 |
| 1878 | 0.03 | 0.12 | 0.25 | 0.80 | 1 | Hdgfl3 | HDGF like 3 | 1990 | 2336 | 1926 | 1757 | 1970 | 1520 | 1889 | 1631 | 1926 | 2016 | 1697 |
| 1749 | 0.07 | 0.10 | 0.73 | 0.47 | 1 | Cbfa2t2 | CBFA2/RUNX1 translocation partner 2 | 1521 | 1703 | 1701 | 1708 | 1646 | 1732 | 1889 | 1795 | 1810 | 1984 | 1753 |

| | | | | | | | | | | | | | | | | | | |
|---|---|---|---|---|---|---|---|---|---|---|---|---|---|---|---|---|---|---|
| 1553 | 0.15 | 0.11 | 1.32 | 0.19 | 1 | Hopx | HOP homeobox | 608 | 524 | 647 | 1810 | 1896 | 1676 | 1889 | 2230 | 1940 | 2033 | 1834 |
| 2352 | 0.02 | 0.20 | 0.11 | 0.91 | 1 | Muc4 | mucin 4 | 55 | 128 | 173 | 2596 | 2345 | 6076 | 1888 | 1988 | 3452 | 2159 | 5015 |
| 1948 | 0.08 | 0.09 | 0.83 | 0.41 | 1 | Pbx2 | pre B cell leukemia homeobox 2 | 1571 | 1809 | 1655 | 1977 | 2120 | 1997 | 1888 | 2045 | 2104 | 2203 | 2063 |
| 1775 | 0.01 | 0.13 | 0.06 | 0.95 | 1 | Slc35a3 | solute carrier family 35 (UDP-N-acetylgluc | 1678 | 1693 | 1687 | 2084 | 1594 | 1652 | 1888 | 1573 | 2083 | 1838 | 1758 |
| 2026 | 0.00 | 0.13 | 0.01 | 0.99 | 1 | Fbxw5 | F-box and WD-40 domain protein 5 | 1527 | 1759 | 1640 | 2198 | 2242 | 2357 | 1886 | 2068 | 2022 | 2526 | 2065 |
| 1829 | 0.23 | 0.12 | 1.87 | 0.06 | 1 | Ssx2ip | synovial sarcoma, X 2 interacting protein | 865 | 1393 | 939 | 2071 | 2085 | 1734 | 1886 | 2595 | 2217 | 2284 | 2054 |
| 1743 | -0.06 | 0.11 | -0.58 | 0.56 | 1 | Aco1 | aconitase 1 | 1821 | 1706 | 1718 | 1717 | 1869 | 1637 | 1886 | 1802 | 1962 | 1516 | 1535 |
| 1511 | -0.15 | 0.17 | -0.89 | 0.37 | 1 | Clstn3 | calsyntenin 3 | 1508 | 1959 | 1706 | 1515 | 1533 | 1028 | 1886 | 1576 | 1121 | 1922 | 866 |
| 2010 | -0.01 | 0.10 | -0.15 | 0.88 | 1 | Ddx47 | DEAD (Asp-Glu-Ala-Asp) box polypeptide | 1991 | 2220 | 1930 | 2033 | 2033 | 2057 | 1885 | 2366 | 1952 | 1794 | 1853 |
| 1883 | -0.10 | 0.12 | -0.82 | 0.41 | 1 | Ambra1 | autophagy/beclin 1 regulator 1 | 1839 | 1535 | 1857 | 2150 | 2003 | 1941 | 1885 | 1663 | 2228 | 1727 | 1883 |
| 1653 | 0.03 | 0.10 | 0.31 | 0.76 | 1 | Cab39l | calcium binding protein 39-like | 1650 | 1646 | 1626 | 1735 | 1763 | 1204 | 1885 | 1595 | 1780 | 1900 | 1398 |
| 2044 | 0.01 | 0.09 | 0.13 | 0.90 | 1 | Bsdc1 | BSD domain containing 1 | 1791 | 2014 | 2239 | 1932 | 2374 | 2013 | 1884 | 2100 | 2104 | 1978 | 2055 |
| 2013 | -0.02 | 0.11 | -0.18 | 0.86 | 1 | Hnrnph2 | heterogeneous nuclear ribonucleoprotein | 2080 | 1938 | 1770 | 1850 | 2155 | 2321 | 1884 | 1939 | 1916 | 1738 | 2558 |
| 1307 | 0.09 | 0.15 | 0.61 | 0.54 | 1 | Elovl6 | ELOVL family member 6, elongation of lo | 572 | 692 | 743 | 1372 | 1408 | 1333 | 1884 | 1663 | 1639 | 1660 | 1413 |
| 1901 | 0.10 | 0.10 | 0.96 | 0.33 | 1 | Hltf | helicase-like transcription factor | 1655 | 1602 | 1627 | 1771 | 1874 | 2199 | 1882 | 1806 | 2282 | 1774 | 2441 |
| 1272 | 0.19 | 0.14 | 1.37 | 0.17 | 1 | Slc4a10 | solute carrier family 4, sodium bicarbonat | 228 | 178 | 247 | 1529 | 1355 | 1491 | 1882 | 1747 | 1686 | 1887 | 1764 |
| 1411 | -0.02 | 0.14 | -0.12 | 0.90 | 1 | Pbx3 | pre B cell leukemia homeobox 3 | 633 | 700 | 668 | 1866 | 1711 | 1335 | 1880 | 1469 | 1735 | 2172 | 1358 |
| 1920 | 0.05 | 0.12 | 0.39 | 0.70 | 1 | Eloc | elongin C | 1932 | 1992 | 2131 | 1723 | 2024 | 1758 | 1879 | 2178 | 2110 | 1480 | 1914 |
| 1855 | -0.02 | 0.12 | -0.15 | 0.88 | 1 | Gnl2 | guanine nucleotide binding protein-like 2 | 1543 | 1517 | 1392 | 1693 | 1954 | 2446 | 1878 | 1621 | 2075 | 1546 | 2741 |
| 1798 | 0.07 | 0.12 | 0.60 | 0.55 | 1 | Ptk2 | PTK2 protein tyrosine kinase 2 | 2608 | 1644 | 2388 | 1449 | 1704 | 1391 | 1877 | 1710 | 1718 | 1874 | 1411 |
| 2433 | -0.02 | 0.10 | -0.25 | 0.81 | 1 | Mafg | v-maf musculoaponeurotic fibrosarcoma c | 1886 | 1782 | 1724 | 2297 | 2024 | 4641 | 1876 | 2200 | 1865 | 2165 | 4300 |
| 1852 | 0.06 | 0.11 | 0.52 | 0.60 | 1 | Abhd8 | abhydrolase domain containing 8 | 1703 | 1854 | 1715 | 1897 | 2107 | 1494 | 1876 | 2391 | 1827 | 2029 | 1482 |
| 2111 | 0.11 | 0.10 | 1.11 | 0.27 | 1 | Rc3h2 | ring finger and CCCH-type zinc finger dor | 1416 | 1510 | 1433 | 1947 | 2028 | 3163 | 1874 | 2294 | 2195 | 1713 | 3644 |
| 1929 | 0.04 | 0.12 | 0.39 | 0.70 | 1 | Nup205 | nucleoporin 205 | 2346 | 2493 | 1967 | 1613 | 1872 | 1733 | 1874 | 1676 | 1890 | 1987 | 1766 |
| 1928 | 0.04 | 0.12 | 0.36 | 0.72 | 1 | Cyth2 | cytohesin 2 | 1561 | 1838 | 1805 | 1833 | 2207 | 1949 | 1874 | 2442 | 2092 | 1698 | 1909 |
| 1724 | -0.02 | 0.12 | -0.15 | 0.88 | 1 | Sf3b6 | splicing factor 3B, subunit 6 | 1635 | 1496 | 1807 | 1713 | 2043 | 1427 | 1874 | 1568 | 2162 | 1515 | 1722 |
| 1687 | -0.08 | 0.11 | -0.71 | 0.48 | 1 | Pskh1 | protein serine kinase H1 | 1648 | 1735 | 1506 | 1825 | 1824 | 1505 | 1874 | 1500 | 1843 | 1743 | 1550 |
| 2016 | 0.01 | 0.10 | 0.10 | 0.92 | 1 | Vps52 | VPS52 GARP complex subunit | 2189 | 2314 | 2241 | 1758 | 2036 | 2020 | 1873 | 1764 | 2217 | 1962 | 1806 |
| 1893 | -0.07 | 0.11 | -0.66 | 0.51 | 1 | Rbm3 | RNA binding motif (RNP1, RRM) protein 3 | 2816 | 1644 | 1881 | 1448 | 1693 | 2357 | 1873 | 1281 | 1548 | 1829 | 2448 |
| 1985 | 0.31 | 0.21 | 1.45 | 0.15 | 1 | Atp5h | ATP synthase, H+ transporting, mitochonc | 2137 | 1780 | 2747 | 1658 | 1902 | 1281 | 1871 | 3349 | 1907 | 1734 | 1473 |
| 1935 | 0.15 | 0.11 | 1.30 | 0.19 | 1 | Psmc6 | proteasome (prosome, macropain) 26S su | 2453 | 1885 | 1987 | 1706 | 1709 | 1807 | 1871 | 1822 | 1978 | 1784 | 2281 |
| 1907 | 0.13 | 0.10 | 1.26 | 0.21 | 1 | Bcas2 | breast carcinoma amplified sequence 2 | 1552 | 1899 | 1917 | 1952 | 1711 | 1926 | 1871 | 2109 | 1949 | 1922 | 2165 |
| 1727 | -0.11 | 0.13 | -0.84 | 0.40 | 1 | Pts | 6-pyruvoyl-tetrahydropterin synthase | 1203 | 1434 | 1416 | 1712 | 1703 | 2433 | 1871 | 1323 | 1796 | 1713 | 2388 |
| 1694 | 0.05 | 0.13 | 0.43 | 0.66 | 1 | Mrpl10 | mitochondrial ribosomal protein L10 | 1251 | 1611 | 1758 | 1738 | 1781 | 1474 | 1871 | 2068 | 1777 | 1803 | 1504 |
| 941 | -0.02 | 0.18 | -0.12 | 0.91 | 1 | Rab6b | RAB6B, member RAS oncogene family | 1318 | 459 | 220 | 528 | 172 | 1713 | 1871 | 611 | 167 | 1817 | 1476 |
| 2130 | 0.26 | 0.20 | 1.27 | 0.20 | 1 | Rpl19 | ribosomal protein L19 | 2907 | 2324 | 3872 | 1605 | 1643 | 1369 | 1870 | 2844 | 1691 | 1775 | 1525 |
| 1911 | -0.06 | 0.12 | -0.54 | 0.59 | 1 | Ankrd52 | ankyrin repeat domain 52 | 2030 | 2104 | 1953 | 1834 | 1706 | 2234 | 1870 | 1936 | 1981 | 1638 | 1731 |
| 1969 | -0.12 | 0.13 | -0.94 | 0.35 | 1 | Dyrk1a | dual-specificity tyrosine-(Y)-phosphorylati | 2255 | 2042 | 1706 | 2282 | 1827 | 2188 | 1869 | 1745 | 1988 | 1856 | 1898 |
| 1831 | -0.10 | 0.11 | -0.86 | 0.39 | 1 | R3hdm4 | R3H domain containing 4 | 1958 | 1743 | 2033 | 1695 | 2020 | 1857 | 1869 | 1814 | 1850 | 1454 | 1848 |
| 1897 | 0.25 | 0.14 | 1.73 | 0.08 | 1 | Bex1 | brain expressed X-linked 1 | 2172 | 1923 | 2315 | 1586 | 1806 | 1335 | 1868 | 2435 | 1691 | 2263 | 1474 |
| 2478 | -0.14 | 0.11 | -1.28 | 0.20 | 1 | Rap1gap | Rap1 GTPase-activating protein | 2835 | 4234 | 4064 | 2209 | 2185 | 2200 | 1867 | 2023 | 1886 | 1875 | 1878 |
| 1688 | -0.15 | 0.10 | -1.58 | 0.11 | 1 | Brap | BRCA1 associated protein | 1607 | 1536 | 1640 | 1579 | 2015 | 1796 | 1867 | 1400 | 1810 | 1557 | 1761 |
| 2314 | -0.16 | 0.13 | -1.22 | 0.22 | 1 | Rev3l | REV3 like, DNA directed polymerase zeta | 2021 | 2341 | 1794 | 2387 | 2092 | 3816 | 1866 | 1707 | 2171 | 1738 | 3522 |
| 2068 | 0.04 | 0.09 | 0.40 | 0.69 | 1 | Sgta | small glutamine-rich tetratricopeptide rep | 2076 | 2217 | 2280 | 2164 | 2096 | 1833 | 1866 | 2479 | 2116 | 1906 | 1712 |
| 1929 | 0.04 | 0.10 | 0.36 | 0.72 | 1 | Nsfl1c | NSFL1 (p97) cofactor (p47) | 1985 | 1649 | 2193 | 1436 | 1936 | 2390 | 1866 | 1744 | 1818 | 1778 | 2425 |
| 2129 | 0.05 | 0.11 | 0.46 | 0.64 | 1 | Rars | arginyl-tRNA synthetase | 2442 | 2140 | 2304 | 1815 | 2244 | 2173 | 1864 | 1887 | 2066 | 1825 | 2662 |
| 1991 | 0.08 | 0.13 | 0.62 | 0.53 | 1 | Tapt1 | transmembrane anterior posterior transfor | 1633 | 1663 | 1660 | 2055 | 2124 | 2167 | 1864 | 1756 | 2002 | 2549 | 2427 |
| 1848 | 0.08 | 0.11 | 0.74 | 0.46 | 1 | Cog4 | component of oligomeric golgi complex 4 | 1674 | 1542 | 1539 | 1780 | 2055 | 1864 | 1863 | 2258 | 2040 | 1785 | 1930 |
| 1729 | 0.14 | 0.18 | 0.78 | 0.43 | 1 | Cldn7 | claudin 7 | 767 | 1035 | 1657 | 1834 | 2330 | 1405 | 1863 | 2504 | 2073 | 2057 | 1493 |
| 1681 | -0.13 | 0.13 | -1.06 | 0.29 | 1 | Dnal1 | dynein, axonemal, light chain 1 | 1554 | 1679 | 1617 | 1744 | 1615 | 1896 | 1859 | 1369 | 1788 | 1425 | 1946 |
| 2337 | 0.26 | 0.19 | 1.40 | 0.16 | 1 | Clic1 | chloride intracellular channel 1 | 3815 | 2665 | 3223 | 1859 | 2016 | 1477 | 1858 | 3269 | 1763 | 2084 | 1675 |
| 1755 | 0.11 | 0.11 | 0.95 | 0.34 | 1 | Vps53 | VPS53 GARP complex subunit | 1360 | 1626 | 1426 | 1668 | 1966 | 1668 | 1858 | 2183 | 1975 | 1891 | 1684 |
| 2333 | 0.05 | 0.11 | 0.43 | 0.67 | 1 | Ahsa1 | AHA1, activator of heat shock protein ATF | 2709 | 2507 | 2922 | 1745 | 2204 | 2879 | 1857 | 2229 | 2113 | 1917 | 2582 |
| 1724 | -0.02 | 0.13 | -0.13 | 0.90 | 1 | Prr13 | proline rich 13 | 1973 | 2062 | 1700 | 1599 | 1911 | 1298 | 1856 | 1482 | 1854 | 1823 | 1401 |
| 1944 | -0.06 | 0.13 | -0.49 | 0.62 | 1 | Pitpnm1 | phosphatidylinositol transfer protein, mem | 1686 | 1890 | 1528 | 2000 | 1693 | 2802 | 1855 | 1559 | 1819 | 2094 | 2458 |
| 1579 | 0.08 | 0.13 | 0.57 | 0.57 | 1 | Glce | glucuronyl C5-epimerase | 960 | 780 | 1139 | 1823 | 1656 | 1694 | 1853 | 1678 | 2110 | 1707 | 1968 |
| 1916 | 0.23 | 0.13 | 1.73 | 0.08 | 1 | Knop1 | lysine rich nucleolar protein 1 | 1399 | 1414 | 1548 | 2023 | 1888 | 1898 | 1851 | 2989 | 2053 | 1950 | 2061 |
| 1815 | -0.02 | 0.09 | -0.25 | 0.80 | 1 | Cstf2t | cleavage stimulation factor, 3' pre-RNA su | 1507 | 1749 | 1775 | 1956 | 1813 | 1903 | 1850 | 1910 | 1808 | 1747 | 1947 |
| 1678 | 0.06 | 0.10 | 0.63 | 0.53 | 1 | Atmin | ATM interactor | 1432 | 1735 | 1367 | 1873 | 1820 | 1259 | 1850 | 2200 | 1849 | 1718 | 1356 |
| 1641 | 0.12 | 0.09 | 1.34 | 0.18 | 1 | Pik3r3 | phosphoinositide-3-kinase regulatory sub | 1018 | 1122 | 1077 | 1701 | 1900 | 1654 | 1850 | 1930 | 1970 | 1909 | 1924 |

| | | | | | | | | | | | | | | | | | | |
|---|---|---|---|---|---|---|---|---|---|---|---|---|---|---|---|---|---|---|
| 2137 | 0.06 | 0.15 | 0.40 | 0.69 | 1 | Ncaph2 | non-SMC condensin II complex, subunit H | 2980 | 2268 | 2409 | 2114 | 1878 | 1908 | 1849 | 2522 | 1985 | 1694 | 1897 |
| 2344 | -0.02 | 0.11 | -0.15 | 0.88 | 1 | Mta2 | metastasis-associated gene family, memb | 3487 | 3081 | 3200 | 2173 | 2023 | 2003 | 1848 | 2242 | 1838 | 1732 | 2161 |
| 1255 | -0.26 | 0.22 | -1.17 | 0.24 | 1 | Galr1 | galanin receptor 1 | 70 | 74 | 222 | 1663 | 2057 | 1705 | 1848 | 1257 | 1914 | 1484 | 1514 |
| 1994 | 0.10 | 0.14 | 0.74 | 0.46 | 1 | Lsm6 | LSM6 homolog, U6 small nuclear RNA ar | 2177 | 2957 | 3075 | 1572 | 1869 | 1314 | 1847 | 2163 | 1809 | 1810 | 1340 |
| 1564 | 0.02 | 0.11 | 0.19 | 0.85 | 1 | Lrfn3 | leucine rich repeat and fibronectin type III | 1034 | 1298 | 1130 | 2126 | 1736 | 1069 | 1847 | 1981 | 1881 | 2138 | 963 |
| 2976 | 0.05 | 0.11 | 0.41 | 0.68 | 1 | Aldh2 | aldehyde dehydrogenase 2, mitochondria | 7859 | 5570 | 5734 | 1841 | 1617 | 1348 | 1845 | 2094 | 1539 | 1964 | 1324 |
| 2034 | 0.25 | 0.14 | 1.74 | 0.08 | 1 | Raly | hnRNP-associated with lethal yellow | 3740 | 2470 | 3161 | 1352 | 1430 | 1277 | 1845 | 2011 | 1404 | 2342 | 1341 |
| 1908 | -0.09 | 0.10 | -0.84 | 0.40 | 1 | Phrf1 | PHD and ring finger domains 1 | 1627 | 1738 | 1559 | 1995 | 2121 | 2311 | 1845 | 1941 | 2148 | 1651 | 2053 |
| 1801 | -0.22 | 0.14 | -1.61 | 0.11 | 1 | Fam117b | family with sequence similarity 117, meml | 2201 | 2528 | 2445 | 1582 | 1802 | 1551 | 1845 | 1439 | 1756 | 1132 | 1524 |
| 1776 | 0.08 | 0.13 | 0.58 | 0.56 | 1 | Tmem165 | transmembrane protein 165 | 1521 | 1630 | 1506 | 1721 | 1891 | 1762 | 1845 | 1508 | 1912 | 2321 | 1915 |
| 1833 | -0.03 | 0.10 | -0.29 | 0.77 | 1 | Hgs | HGF-regulated tyrosine kinase substrate | 1741 | 1804 | 1810 | 1621 | 2088 | 1927 | 1844 | 1624 | 2162 | 1744 | 1802 |
| 1890 | -0.04 | 0.11 | -0.39 | 0.70 | 1 | Tmem87b | transmembrane protein 87B | 2056 | 1876 | 1640 | 1659 | 1800 | 2396 | 1843 | 1537 | 1839 | 1622 | 2524 |
| 1864 | -0.17 | 0.10 | -1.73 | 0.08 | 1 | Phf12 | PHD finger protein 12 | 1832 | 1664 | 1919 | 1848 | 1941 | 2366 | 1843 | 1477 | 1986 | 1748 | 1880 |
| 1736 | 0.02 | 0.13 | 0.12 | 0.90 | 1 | Gpr27 | G protein-coupled receptor 27 | 944 | 1260 | 1304 | 2086 | 2271 | 1564 | 1843 | 2421 | 1782 | 1863 | 1755 |
| 1551 | 0.12 | 0.14 | 0.86 | 0.39 | 1 | Micos13 | mitochondrial contact site and cristae orga | 1250 | 1316 | 1406 | 1483 | 1590 | 1341 | 1842 | 2134 | 1736 | 1654 | 1314 |
| 2245 | 0.07 | 0.10 | 0.68 | 0.50 | 1 | Abl1 | c-abl oncogene 1, non-receptor tyrosine k | 2714 | 2405 | 2366 | 2148 | 2106 | 2303 | 1841 | 2337 | 2162 | 2130 | 2179 |
| 1746 | 0.04 | 0.09 | 0.46 | 0.65 | 1 | Arih2 | ariadne RBR E3 ubiquitin protein ligase 2 | 1717 | 1462 | 1585 | 1563 | 1856 | 1850 | 1841 | 1641 | 1686 | 1930 | 2080 |
| 1616 | -0.12 | 0.09 | -1.26 | 0.21 | 1 | Txn2 | thioredoxin 2 | 1566 | 1554 | 1675 | 1588 | 1832 | 1499 | 1841 | 1489 | 1804 | 1501 | 1429 |
| 1830 | -0.10 | 0.09 | -1.14 | 0.25 | 1 | Sesn2 | sestrin 2 | 2316 | 2413 | 2509 | 1757 | 1746 | 1316 | 1840 | 1773 | 1590 | 1726 | 1139 |
| 1676 | -0.12 | 0.13 | -0.96 | 0.34 | 1 | Zbtb7b | zinc finger and BTB domain containing 7E | 1404 | 1056 | 1185 | 1848 | 2330 | 1708 | 1840 | 1768 | 1982 | 1637 | 1676 |
| 1890 | -0.03 | 0.10 | -0.34 | 0.73 | 1 | Dctn4 | dynactin 4 | 2046 | 2027 | 1991 | 1932 | 1605 | 2066 | 1839 | 1884 | 1714 | 1571 | 2112 |
| 1877 | 0.02 | 0.09 | 0.25 | 0.80 | 1 | Asb1 | ankyrin repeat and SOCS box-containing | 1840 | 2157 | 1651 | 1902 | 1757 | 1945 | 1839 | 1959 | 1796 | 1926 | 1880 |
| 1875 | -0.11 | 0.10 | -1.14 | 0.26 | 1 | Fbxw2 | F-box and WD-40 domain protein 2 | 2204 | 2101 | 2251 | 1731 | 1936 | 1800 | 1839 | 1647 | 1898 | 1439 | 1782 |
| 1559 | 0.13 | 0.11 | 1.21 | 0.23 | 1 | Prkd1 | protein kinase D1 | 1442 | 1070 | 1583 | 1539 | 1690 | 1160 | 1839 | 1618 | 2054 | 1805 | 1347 |
| 1927 | 0.02 | 0.18 | 0.09 | 0.93 | 1 | Cndp2 | CNDP dipeptidase 2 (metallopeptidase M | 2801 | 1628 | 2690 | 1687 | 2011 | 1483 | 1838 | 2223 | 1745 | 1667 | 1426 |
| 1783 | -0.03 | 0.12 | -0.27 | 0.78 | 1 | Plcxd2 | phosphatidylinositol-specific phospholipa | 830 | 799 | 1280 | 1453 | 1968 | 3251 | 1838 | 1636 | 2153 | 1584 | 2819 |
| 1934 | -0.04 | 0.10 | -0.38 | 0.70 | 1 | Fkbp15 | FK506 binding protein 15 | 2204 | 1963 | 2077 | 1989 | 1856 | 1920 | 1837 | 1669 | 1993 | 1954 | 1807 |
| 2186 | 0.23 | 0.15 | 1.57 | 0.12 | 1 | Cul9 | cullin 9 | 1834 | 1736 | 1665 | 2288 | 2082 | 2407 | 1836 | 3240 | 2212 | 2683 | 2065 |
| 2301 | -0.11 | 0.09 | -1.23 | 0.22 | 1 | Mapk8ip3 | mitogen-activated protein kinase 8 interac | 848 | 814 | 1192 | 1661 | 2064 | 6087 | 1835 | 1419 | 2091 | 1702 | 5592 |
| 1613 | -0.06 | 0.12 | -0.50 | 0.62 | 1 | Camk2n2 | calcium/calmodulin-dependent protein kin | 1224 | 1372 | 1424 | 1998 | 1825 | 1286 | 1835 | 1807 | 1836 | 2084 | 1047 |
| 1807 | 0.10 | 0.10 | 0.95 | 0.34 | 1 | Slc25a32 | solute carrier family 25, member 32 | 1167 | 1422 | 1358 | 2004 | 1669 | 2168 | 1834 | 2073 | 1813 | 1788 | 2582 |
| 2334 | 0.10 | 0.12 | 0.79 | 0.43 | 1 | Mt2 | metallothionein 2 | 1109 | 2321 | 1505 | 2973 | 2316 | 2984 | 1833 | 3005 | 2563 | 2293 | 2772 |
| 2146 | -0.18 | 0.12 | -1.44 | 0.15 | 1 | Kmt2b | lysine (K)-specific methyltransferase 2B | 2059 | 2613 | 2163 | 2353 | 2164 | 2576 | 1833 | 1700 | 2062 | 1950 | 2138 |
| 1783 | -0.17 | 0.16 | -1.08 | 0.28 | 1 | Rbl2 | RB transcriptional corepressor like 2 | 2370 | 1940 | 1821 | 1800 | 2035 | 1477 | 1833 | 1550 | 1990 | 1278 | 1514 |
| 1599 | 0.08 | 0.12 | 0.67 | 0.51 | 1 | Chmp1b | charged multivesicular body protein 1B | 1543 | 1537 | 1826 | 1500 | 1446 | 1381 | 1833 | 1448 | 1607 | 1845 | 1620 |
| 1936 | 0.04 | 0.10 | 0.37 | 0.71 | 1 | Zhx3 | zinc fingers and homeoboxes 3 | 1666 | 1809 | 1916 | 1965 | 1942 | 2112 | 1832 | 2219 | 1915 | 2018 | 1903 |
| 1672 | 0.01 | 0.11 | 0.07 | 0.94 | 1 | Plpp6 | phospholipid phosphatase 6 | 1107 | 1285 | 1293 | 2145 | 1881 | 1487 | 1832 | 1812 | 1987 | 2025 | 1537 |
| 1648 | -0.11 | 0.13 | -0.83 | 0.41 | 1 | Cfap97 | cilia and flagella associated protein 97 | 1241 | 1577 | 1361 | 1910 | 1774 | 1713 | 1832 | 1456 | 1828 | 1579 | 1856 |
| 1762 | 0.06 | 0.11 | 0.54 | 0.59 | 1 | Ap1ar | adaptor-related protein complex 1 associ | 1346 | 1236 | 1100 | 1983 | 2044 | 1834 | 1831 | 1903 | 2016 | 2165 | 1928 |
| 1756 | 0.05 | 0.12 | 0.43 | 0.67 | 1 | Pikfyve | phosphoinositide kinase, FYVE type zinc | 769 | 833 | 799 | 1454 | 1946 | 3100 | 1831 | 1467 | 2069 | 2012 | 3032 |
| 1752 | 0.07 | 0.13 | 0.56 | 0.58 | 1 | Cdkl2 | cyclin-dependent kinase-like 2 (CDC2-rel | 871 | 1226 | 1162 | 2083 | 2219 | 1702 | 1831 | 1892 | 2091 | 2168 | 2029 |
| 2056 | 0.00 | 0.11 | -0.02 | 0.98 | 1 | Crybg3 | beta-gamma crystallin domain containing | 1740 | 1667 | 1895 | 2396 | 1889 | 2554 | 1830 | 1945 | 2125 | 1999 | 2578 |
| 1711 | -0.05 | 0.14 | -0.36 | 0.72 | 1 | Fubp3 | far upstream element (FUSE) binding pro | 1251 | 1632 | 1464 | 1490 | 1716 | 2308 | 1830 | 1335 | 1635 | 1803 | 2361 |
| 1730 | -0.12 | 0.09 | -1.39 | 0.16 | 1 | Zfyve9 | zinc finger, FYVE domain containing 9 | 1959 | 1787 | 2051 | 1693 | 1870 | 1515 | 1829 | 1686 | 1685 | 1561 | 1398 |
| 1616 | -0.16 | 0.09 | -1.72 | 0.09 | 1 | Stxbp6 | syntaxin binding protein 6 (amisyn) | 1113 | 1396 | 1410 | 2044 | 1982 | 1478 | 1825 | 1742 | 1856 | 1497 | 1439 |
| 2160 | 0.15 | 0.11 | 1.41 | 0.16 | 1 | Zfp568 | zinc finger protein 568 | 1801 | 1960 | 1657 | 1723 | 1658 | 3601 | 1824 | 1872 | 2149 | 2026 | 3491 |
| 1896 | -0.07 | 0.10 | -0.69 | 0.49 | 1 | Trim8 | tripartite motif-containing 8 | 2056 | 2324 | 2190 | 1924 | 1925 | 1617 | 1823 | 2204 | 1751 | 1557 | 1485 |
| 1540 | -0.10 | 0.09 | -1.09 | 0.27 | 1 | Ddx19b | DEAD (Asp-Glu-Ala-Asp) box polypeptide | 1111 | 1119 | 1220 | 1795 | 1697 | 1669 | 1823 | 1580 | 1685 | 1588 | 1650 |
| 1420 | -0.13 | 0.16 | -0.84 | 0.40 | 1 | Dagla | diacylglycerol lipase, alpha | 645 | 944 | 801 | 1876 | 1669 | 1542 | 1823 | 1391 | 1845 | 1789 | 1291 |
| 2134 | -0.15 | 0.08 | -1.79 | 0.07 | 1 | Foxp1 | forkhead box P1 | 3144 | 3355 | 3256 | 1923 | 1699 | 1767 | 1822 | 1812 | 1618 | 1499 | 1579 |
| 2411 | -0.01 | 0.16 | -0.05 | 0.96 | 1 | Ppp1r15b | protein phosphatase 1, regulatory subuni | 3083 | 2620 | 2257 | 2943 | 2224 | 2337 | 1821 | 2676 | 2414 | 1861 | 2281 |
| 1640 | 0.05 | 0.13 | 0.39 | 0.70 | 1 | Zfyve21 | zinc finger, FYVE domain containing 21 | 1002 | 807 | 1005 | 1841 | 1859 | 1896 | 1821 | 1473 | 1861 | 2442 | 2032 |
| 1776 | 0.02 | 0.11 | 0.15 | 0.88 | 1 | Kif16b | kinesin family member 16B | 1567 | 1491 | 1341 | 2103 | 1986 | 1608 | 1820 | 2087 | 2157 | 1825 | 1550 |
| 1909 | -0.09 | 0.12 | -0.77 | 0.44 | 1 | Cherp | calcium homeostasis endoplasmic reticul | 2126 | 1478 | 2003 | 1687 | 2171 | 2249 | 1818 | 1548 | 2097 | 2070 | 1752 |
| 2444 | -0.07 | 0.08 | -0.92 | 0.36 | 1 | Rps19 | ribosomal protein S19 | 4472 | 3743 | 4498 | 1756 | 2086 | 1605 | 1817 | 1707 | 1923 | 1843 | 1440 |
| 2197 | 0.06 | 0.10 | 0.56 | 0.57 | 1 | Phb2 | prohibitin 2 | 3148 | 2624 | 2846 | 1928 | 2013 | 1838 | 1817 | 1865 | 2006 | 2325 | 1760 |
| 1792 | 0.01 | 0.12 | 0.08 | 0.94 | 1 | Entr1 | endosome associated trafficking regulato | 1140 | 969 | 1395 | 1811 | 1829 | 2647 | 1816 | 1937 | 1774 | 1896 | 2495 |
| 1768 | -0.17 | 0.12 | -1.38 | 0.17 | 1 | Cd99l2 | CD99 antigen-like 2 | 2236 | 2431 | 2522 | 1625 | 1722 | 1301 | 1816 | 1287 | 1560 | 1826 | 1125 |
| 1428 | 0.00 | 0.10 | -0.04 | 0.97 | 1 | Mnt | max binding protein | 1597 | 934 | 1356 | 1043 | 1717 | 1307 | 1816 | 1054 | 1733 | 2004 | 1145 |
| 1606 | 0.13 | 0.12 | 1.12 | 0.26 | 1 | Sgsm3 | small G protein signaling modulator 3 | 829 | 771 | 882 | 1711 | 1726 | 2004 | 1815 | 2195 | 1764 | 2000 | 1969 |

| | | | | | | | | | | | | | | | | | | |
|---|---|---|---|---|---|---|---|---|---|---|---|---|---|---|---|---|---|---|
| 2108 | -0.12 | 0.09 | -1.35 | 0.18 | 1 | Spin1 | spindlin 1 | 3505 | 3431 | 3380 | 1831 | 1650 | 1430 | 1814 | 1544 | 1667 | 1512 | 1427 |
| 1653 | -0.04 | 0.14 | -0.25 | 0.81 | 1 | Slc25a17 | solute carrier family 25 (mitochondrial car | 2041 | 1602 | 1840 | 1296 | 1671 | 1571 | 1813 | 989 | 1515 | 2192 | 1654 |
| 1919 | 0.21 | 0.13 | 1.55 | 0.12 | 1 | Phf14 | PHD finger protein 14 | 1867 | 1700 | 1627 | 1702 | 1848 | 2000 | 1811 | 2616 | 1906 | 1797 | 2232 |
| 1860 | 0.03 | 0.09 | 0.36 | 0.72 | 1 | Gcc1 | golgi coiled coil 1 | 1209 | 1448 | 1308 | 2181 | 1872 | 2302 | 1811 | 2195 | 2148 | 1776 | 2213 |
| 1844 | 0.11 | 0.11 | 0.97 | 0.33 | 1 | Map3k4 | mitogen-activated protein kinase kinase k | 1506 | 1862 | 1518 | 1936 | 1812 | 1848 | 1811 | 1968 | 2071 | 2201 | 1755 |
| 1945 | -0.10 | 0.11 | -0.95 | 0.34 | 1 | Rbck1 | RanBP-type and C3HC4-type zinc finger | 2378 | 2122 | 2361 | 1810 | 1757 | 2123 | 1809 | 1433 | 1631 | 2003 | 1969 |
| 1664 | 0.15 | 0.10 | 1.45 | 0.15 | 1 | Mfn1 | mitofusin 1 | 1485 | 1488 | 1464 | 1473 | 1863 | 1443 | 1808 | 1875 | 1860 | 1931 | 1612 |
| 1661 | 0.22 | 0.14 | 1.58 | 0.11 | 1 | Rps4l | | 1453 | 1224 | 1909 | 1242 | 1807 | 1494 | 1808 | 1710 | 1760 | 2228 | 1637 |
| 1837 | -0.07 | 0.10 | -0.67 | 0.50 | 1 | Tbc1d10b | TBC1 domain family, member 10b | 1648 | 2269 | 1734 | 2114 | 1895 | 1605 | 1807 | 2175 | 1660 | 1913 | 1387 |
| 2234 | -0.06 | 0.10 | -0.59 | 0.55 | 1 | Dnajb4 | DnaJ heat shock protein family (Hsp40) m | 2489 | 2550 | 2041 | 1993 | 1860 | 3216 | 1806 | 1907 | 1781 | 1608 | 3326 |
| 1964 | -0.22 | 0.13 | -1.76 | 0.08 | 1 | Aldh6a1 | aldehyde dehydrogenase family 6, subfar | 2673 | 3481 | 2842 | 1908 | 1478 | 1610 | 1806 | 1327 | 1390 | 1604 | 1480 |
| 1917 | -0.14 | 0.10 | -1.38 | 0.17 | 1 | Ptk2b | PTK2 protein tyrosine kinase 2 beta | 3086 | 3422 | 2573 | 1768 | 1543 | 1192 | 1806 | 1629 | 1570 | 1445 | 1058 |
| 1822 | 0.15 | 0.14 | 1.09 | 0.27 | 1 | Plcl2 | phospholipase C-like 2 | 1615 | 1595 | 1184 | 1960 | 1706 | 1860 | 1806 | 2869 | 1514 | 1958 | 1980 |
| 1716 | -0.08 | 0.11 | -0.72 | 0.47 | 1 | Fads6 | fatty acid desaturase domain family, mem | 1170 | 1289 | 1559 | 1822 | 2341 | 1675 | 1806 | 2133 | 1929 | 1619 | 1534 |
| 1710 | 0.14 | 0.10 | 1.38 | 0.17 | 1 | Arhgap12 | Rho GTPase activating protein 12 | 1506 | 1495 | 1376 | 1816 | 1358 | 1892 | 1806 | 1841 | 1628 | 1819 | 2274 |
| 2059 | -0.04 | 0.13 | -0.33 | 0.74 | 1 | Arid1b | AT rich interactive domain 1B (SWI-like) | 2702 | 1951 | 2381 | 2174 | 2059 | 1858 | 1803 | 1841 | 2096 | 2155 | 1631 |
| 1681 | 0.04 | 0.09 | 0.46 | 0.65 | 1 | Rnf185 | ring finger protein 185 | 983 | 1175 | 1188 | 1889 | 1972 | 1795 | 1803 | 1910 | 2142 | 1887 | 1750 |
| 2227 | 0.11 | 0.10 | 1.12 | 0.26 | 1 | Pa2g4 | proliferation-associated 2G4 | 3480 | 2799 | 3211 | 1742 | 1881 | 1791 | 1802 | 2054 | 2050 | 1838 | 1851 |
| 1812 | 0.06 | 0.13 | 0.47 | 0.64 | 1 | Syt5 | synaptotagmin V | 1434 | 1829 | 1893 | 2086 | 1842 | 1516 | 1801 | 1956 | 1775 | 2194 | 1606 |
| 2818 | -0.07 | 0.15 | -0.51 | 0.61 | 1 | Eml5 | echinoderm microtubule associated prote | 919 | 1282 | 1067 | 1885 | 1445 | 8630 | 1800 | 1430 | 1857 | 1380 | 9300 |
| 2180 | 0.01 | 0.11 | 0.13 | 0.89 | 1 | Tnrc6c | trinucleotide repeat containing 6C | 2165 | 2549 | 2435 | 1965 | 1788 | 2848 | 1800 | 2391 | 1859 | 1690 | 2487 |
| 1810 | 0.09 | 0.10 | 0.95 | 0.34 | 1 | Cpox | coproporphyrinogen oxidase | 1088 | 1142 | 1217 | 1728 | 2330 | 2123 | 1800 | 1958 | 2381 | 2087 | 2058 |
| 1542 | 0.01 | 0.14 | 0.04 | 0.97 | 1 | Mapt | microtubule-associated protein tau | 511 | 546 | 801 | 1617 | 1779 | 2373 | 1799 | 1877 | 1847 | 1717 | 2100 |
| 1518 | 0.09 | 0.12 | 0.79 | 0.43 | 1 | Satb1 | special AT-rich sequence binding protein | 635 | 629 | 577 | 1785 | 1506 | 2081 | 1799 | 2330 | 1539 | 1735 | 2081 |
| 2604 | 0.05 | 0.14 | 0.32 | 0.75 | 1 | Rpl34 | ribosomal protein L34 | 3856 | 4462 | 4555 | 2213 | 1723 | 1947 | 1797 | 2944 | 1590 | 1660 | 1893 |
| 2000 | 0.02 | 0.09 | 0.20 | 0.84 | 1 | Elf2 | E74-like factor 2 | 2460 | 2433 | 2102 | 1914 | 1740 | 2010 | 1797 | 1826 | 1897 | 1783 | 2040 |
| 1757 | 0.02 | 0.13 | 0.13 | 0.90 | 1 | Ccz1 | CCZ1 vacuolar protein trafficking and bio | 1732 | 1309 | 1539 | 1937 | 2012 | 1581 | 1797 | 1593 | 2022 | 1861 | 1946 |
| 1494 | -0.03 | 0.11 | -0.29 | 0.77 | 1 | Tbc1d8 | TBC1 domain family, member 8 | 1095 | 1199 | 1035 | 1909 | 1591 | 1356 | 1797 | 1574 | 1755 | 1663 | 1463 |
| 1734 | 0.07 | 0.13 | 0.52 | 0.60 | 1 | Dnajc18 | DnaJ heat shock protein family (Hsp40) m | 1707 | 1809 | 1904 | 1631 | 1760 | 1461 | 1796 | 2226 | 1700 | 1584 | 1497 |
| 1946 | -0.04 | 0.10 | -0.43 | 0.67 | 1 | Mpdz | multiple PDZ domain protein | 1896 | 1946 | 1719 | 2103 | 2019 | 2130 | 1795 | 1876 | 2108 | 1761 | 2056 |
| 1717 | -0.03 | 0.10 | -0.32 | 0.75 | 1 | Pip5k1a | phosphatidylinositol-4-phosphate 5-kinas | 1620 | 1447 | 1502 | 1935 | 1784 | 1725 | 1795 | 1905 | 1766 | 1638 | 1769 |
| 1647 | -0.01 | 0.09 | -0.10 | 0.92 | 1 | Gorasp1 | golgi reassembly stacking protein 1 | 1237 | 1308 | 1317 | 1718 | 2003 | 1633 | 1795 | 1795 | 1913 | 1836 | 1565 |
| 1541 | -0.11 | 0.14 | -0.81 | 0.42 | 1 | Csgalnact2 | chondroitin sulfate N-acetylgalactosaminy | 1700 | 1687 | 1451 | 1310 | 1789 | 1418 | 1795 | 1368 | 1545 | 1369 | 1518 |
| 1435 | 0.18 | 0.10 | 1.84 | 0.07 | 1 | Rwdd1 | RWD domain containing 1 | 903 | 779 | 812 | 1650 | 1505 | 1305 | 1795 | 1812 | 1740 | 1875 | 1612 |
| 1737 | 0.02 | 0.10 | 0.23 | 0.82 | 1 | Napg | N-ethylmaleimide sensitive fusion protein | 1451 | 1320 | 1486 | 1593 | 1908 | 2054 | 1793 | 1663 | 1638 | 1887 | 2314 |
| 1946 | 0.05 | 0.11 | 0.46 | 0.65 | 1 | Atp13a2 | ATPase type 13A2 | 2070 | 2486 | 2315 | 1862 | 1732 | 1710 | 1792 | 2328 | 1853 | 1767 | 1488 |
| 1752 | 0.07 | 0.08 | 0.82 | 0.41 | 1 | Hmox2 | heme oxygenase 2 | 1550 | 1558 | 1441 | 2090 | 1745 | 1565 | 1792 | 2081 | 1884 | 1924 | 1645 |
| 1894 | -0.17 | 0.11 | -1.51 | 0.13 | 1 | Stx16 | syntaxin 16 | 1899 | 2170 | 2231 | 1777 | 1858 | 2260 | 1788 | 1387 | 1993 | 1557 | 1911 |
| 1874 | -0.08 | 0.08 | -1.08 | 0.28 | 1 | Ccar2 | cell cycle activator and apoptosis regulato | 1694 | 1937 | 1961 | 1919 | 1926 | 2094 | 1788 | 1818 | 1833 | 1686 | 1958 |
| 1753 | -0.13 | 0.10 | -1.24 | 0.22 | 1 | Atp1b2 | ATPase, Na+/K+ transporting, beta 2 poly | 2545 | 2369 | 2186 | 1665 | 1720 | 1151 | 1788 | 1448 | 1715 | 1736 | 960 |
| 1269 | 0.19 | 0.18 | 1.06 | 0.29 | 1 | Abracl | ABRA C-terminal like | 647 | 476 | 774 | 1250 | 1425 | 1210 | 1788 | 1884 | 1484 | 1659 | 1367 |
| 1736 | 0.08 | 0.09 | 0.84 | 0.40 | 1 | Oxsr1 | oxidative-stress responsive 1 | 1458 | 1238 | 1386 | 1765 | 1817 | 1931 | 1787 | 1681 | 1949 | 2062 | 2021 |
| 1663 | -0.11 | 0.10 | -1.12 | 0.26 | 1 | Nedd8 | neural precursor cell expressed, develop | 1972 | 1721 | 2232 | 1675 | 1685 | 1281 | 1787 | 1446 | 1567 | 1690 | 1232 |
| 1567 | 0.00 | 0.10 | -0.02 | 0.99 | 1 | Nlk | nemo like kinase | 890 | 794 | 821 | 1831 | 1847 | 1899 | 1787 | 1628 | 1830 | 1993 | 1924 |
| 1549 | -0.20 | 0.21 | -0.96 | 0.34 | 1 | Exph5 | exophilin 5 | 570 | 411 | 648 | 1791 | 1849 | 2594 | 1787 | 821 | 2385 | 1946 | 2235 |
| 1546 | -0.03 | 0.12 | -0.27 | 0.79 | 1 | Ccdc149 | coiled-coil domain containing 149 | 1084 | 1066 | 1063 | 1646 | 1832 | 1692 | 1787 | 1569 | 2132 | 1547 | 1593 |
| 1847 | 0.02 | 0.13 | 0.14 | 0.89 | 1 | Mib2 | mindbomb E3 ubiquitin protein ligase 2 | 1747 | 1411 | 1882 | 1757 | 1892 | 2170 | 1786 | 2038 | 1667 | 1895 | 2070 |
| 1678 | 0.02 | 0.10 | 0.25 | 0.80 | 1 | Vipas39 | VPS33B interacting protein, apical-basola | 1682 | 1544 | 1687 | 1465 | 1755 | 1707 | 1786 | 1478 | 1712 | 1995 | 1653 |
| 2617 | -0.12 | 0.17 | -0.73 | 0.47 | 1 | Rgs4 | regulator of G-protein signaling 4 | 5655 | 6552 | 6368 | 961 | 1478 | 1077 | 1785 | 980 | 1416 | 1680 | 836 |
| 2288 | 0.05 | 0.10 | 0.56 | 0.58 | 1 | Chordc1 | cysteine and histidine-rich domain (CHOI | 1791 | 2302 | 1868 | 1884 | 1801 | 3841 | 1784 | 2073 | 1895 | 1571 | 4357 |
| 1866 | -0.04 | 0.15 | -0.31 | 0.76 | 1 | Aldh7a1 | aldehyde dehydrogenase family 7, memb | 2076 | 2493 | 2707 | 1735 | 1907 | 1306 | 1784 | 1857 | 1385 | 1946 | 1325 |
| 1591 | -0.11 | 0.11 | -1.03 | 0.30 | 1 | Sumo1 | small ubiquitin-like modifier 1 | 1586 | 1704 | 1615 | 1602 | 1795 | 1374 | 1784 | 1397 | 1622 | 1565 | 1460 |
| 1617 | 0.04 | 0.09 | 0.45 | 0.65 | 1 | Cox7b | cytochrome c oxidase subunit 7B | 1419 | 1239 | 1533 | 1727 | 1724 | 1461 | 1783 | 1681 | 1819 | 1886 | 1517 |
| 1602 | -0.01 | 0.13 | -0.08 | 0.94 | 1 | Ngrn | neugrin, neurite outgrowth associated | 1199 | 1224 | 1514 | 1539 | 1995 | 1575 | 1783 | 1881 | 1555 | 1699 | 1663 |
| 1548 | -0.31 | 0.17 | -1.85 | 0.06 | 1 | Pcdh15 | protocadherin 15 | 1088 | 1820 | 1379 | 1713 | 1460 | 2062 | 1782 | 1191 | 1251 | 1382 | 1895 |
| 1929 | 0.13 | 0.15 | 0.84 | 0.40 | 1 | Cuedc1 | CUE domain containing 1 | 1995 | 2270 | 1745 | 1799 | 1486 | 2101 | 1781 | 2844 | 1275 | 1816 | 2106 |
| 3457 | 0.12 | 0.10 | 1.22 | 0.22 | 1 | Srsf7 | serine and arginine-rich splicing factor 7 | 2759 | 2783 | 2259 | 1438 | 1603 | 9850 | 1780 | 1923 | 1543 | 1877 | 10216 |
| 1964 | 0.00 | 0.12 | 0.00 | 1.00 | 1 | Stxbp2 | syntaxin binding protein 2 | 1603 | 1755 | 1492 | 1954 | 1846 | 2772 | 1780 | 2387 | 1820 | 1501 | 2690 |
| 2088 | 0.10 | 0.12 | 0.82 | 0.41 | 1 | Hapln1 | hyaluronan and proteoglycan link protein | 3220 | 3732 | 3740 | 1656 | 1566 | 927 | 1779 | 1584 | 1740 | 2036 | 986 |
| 1857 | -0.06 | 0.17 | -0.37 | 0.71 | 1 | Ier5 | immediate early response 5 | 3081 | 1586 | 1921 | 1764 | 2159 | 1374 | 1779 | 1673 | 1962 | 1779 | 1347 |

| | | | | | | | | | | | | | | | | | | |
|---|---|---|---|---|---|---|---|---|---|---|---|---|---|---|---|---|---|---|
| 1800 | 0.16 | 0.16 | 0.99 | 0.32 | 1 | 1810037I17Rik | RIKEN cDNA 1810037I17 gene | 1791 | 1317 | 1862 | 2072 | 1753 | 1372 | 1779 | 2584 | 1819 | 2017 | 1433 |
| 1713 | 0.07 | 0.12 | 0.62 | 0.53 | 1 | Vps28 | vacuolar protein sorting 28 | 1748 | 1634 | 1963 | 1541 | 1630 | 1631 | 1779 | 1923 | 1628 | 1757 | 1608 |
| 1504 | -0.10 | 0.11 | -0.97 | 0.33 | 1 | Shprh | SNF2 histone linker PHD RING helicase | 1267 | 1164 | 1345 | 1455 | 1495 | 1865 | 1779 | 1236 | 1559 | 1516 | 1858 |
| 1739 | 0.03 | 0.11 | 0.29 | 0.78 | 1 | Cetn3 | centrin 3 | 2135 | 1589 | 2217 | 1528 | 1745 | 1463 | 1778 | 1400 | 1674 | 1794 | 1802 |
| 1836 | 0.05 | 0.12 | 0.48 | 0.63 | 1 | Sfxn2 | sideroflexin 2 | 895 | 750 | 801 | 2166 | 1756 | 2987 | 1777 | 2032 | 1711 | 2137 | 3190 |
| 2074 | -0.10 | 0.11 | -0.85 | 0.40 | 1 | Adss | adenylosuccinate synthetase, non muscle | 3366 | 3302 | 3050 | 1836 | 1623 | 1536 | 1776 | 1438 | 1699 | 1503 | 1684 |
| 2026 | -0.06 | 0.15 | -0.39 | 0.70 | 1 | Lims1 | LIM and senescent cell antigen-like doma | 3796 | 2965 | 2763 | 1931 | 1484 | 1338 | 1776 | 1317 | 1842 | 1646 | 1430 |
| 1457 | 0.04 | 0.17 | 0.25 | 0.80 | 1 | Sh3kbp1 | SH3-domain kinase binding protein 1 | 1445 | 1593 | 1315 | 1523 | 1300 | 1175 | 1776 | 1164 | 1785 | 1553 | 1398 |
| 1643 | 0.08 | 0.17 | 0.48 | 0.63 | 1 | Fam20a | family with sequence similarity 20, memb | 1048 | 1534 | 1131 | 2010 | 1598 | 1614 | 1774 | 1540 | 1841 | 2291 | 1694 |
| 2070 | 0.14 | 0.11 | 1.26 | 0.21 | 1 | Psmd4 | proteasome (prosome, macropain) 26S s | 2477 | 2039 | 2335 | 1701 | 2158 | 1948 | 1773 | 2142 | 1992 | 2213 | 1989 |
| 2004 | 0.30 | 0.17 | 1.76 | 0.08 | 1 | Mphosph8 | M-phase phosphoprotein 8 | 1470 | 1512 | 1351 | 1936 | 1848 | 2281 | 1773 | 3523 | 2113 | 1635 | 2596 |
| 1454 | -0.15 | 0.11 | -1.38 | 0.17 | 1 | Gstm1 | glutathione S-transferase, mu 1 | 2805 | 1839 | 1811 | 1102 | 1212 | 974 | 1773 | 965 | 1161 | 1387 | 967 |
| 1449 | -0.02 | 0.13 | -0.11 | 0.91 | 1 | Copz2 | coatomer protein complex, subunit zeta 2 | 1119 | 1024 | 1222 | 1697 | 1517 | 1270 | 1773 | 1831 | 1617 | 1864 | 1003 |
| 1402 | 0.10 | 0.14 | 0.72 | 0.47 | 1 | Ctnnbip1 | catenin beta interacting protein 1 | 847 | 998 | 1066 | 1346 | 1815 | 1096 | 1773 | 1938 | 1661 | 1726 | 1153 |
| 1743 | -0.04 | 0.13 | -0.31 | 0.76 | 1 | Usp29 | ubiquitin specific peptidase 29 | 1140 | 1706 | 1402 | 2245 | 1664 | 1917 | 1772 | 1760 | 1754 | 1890 | 1925 |
| 1528 | 0.12 | 0.13 | 0.91 | 0.36 | 1 | Lypla1 | lysophospholipase 1 | 1249 | 1397 | 1249 | 1735 | 1487 | 1225 | 1772 | 1493 | 1733 | 1884 | 1583 |
| 1936 | 0.00 | 0.10 | 0.01 | 0.99 | 1 | Morc2a | microrchidia 2A | 1770 | 1653 | 2246 | 1876 | 1980 | 2191 | 1771 | 1927 | 2084 | 1826 | 1975 |
| 1840 | -0.01 | 0.10 | -0.11 | 0.91 | 1 | Nhlrc2 | NHL repeat containing 2 | 2029 | 1985 | 1945 | 1769 | 1704 | 1916 | 1771 | 1511 | 1797 | 1963 | 1850 |
| 1720 | 0.05 | 0.12 | 0.41 | 0.68 | 1 | Fam49b | family with sequence similarity 49, memb | 3016 | 1911 | 2005 | 1453 | 1352 | 1333 | 1770 | 1402 | 1597 | 1629 | 1446 |
| 1708 | -0.09 | 0.12 | -0.75 | 0.45 | 1 | Mapk1ip1l | mitogen-activated protein kinase 1 intera | 1484 | 1935 | 1719 | 1602 | 1843 | 1806 | 1770 | 1707 | 1842 | 1359 | 1723 |
| 1713 | -0.04 | 0.10 | -0.42 | 0.67 | 1 | Ezh1 | enhancer of zeste 1 polycomb repressive | 1678 | 1287 | 1641 | 1500 | 1538 | 2454 | 1769 | 1571 | 1560 | 1638 | 2208 |
| 2139 | -0.03 | 0.09 | -0.33 | 0.74 | 1 | Smurf2 | SMAD specific E3 ubiquitin protein ligase | 2685 | 3066 | 2828 | 1888 | 1903 | 1980 | 1766 | 2107 | 1806 | 1602 | 1899 |
| 1946 | -0.15 | 0.10 | -1.49 | 0.14 | 1 | Gprin1 | G protein-regulated inducer of neurite out | 2411 | 3142 | 2381 | 1928 | 1821 | 1528 | 1766 | 2026 | 1641 | 1507 | 1251 |
| 1937 | -0.13 | 0.12 | -1.05 | 0.29 | 1 | Plod3 | procollagen-lysine, 2-oxoglutarate 5-diox | 2972 | 2038 | 2942 | 1844 | 1896 | 1453 | 1766 | 1607 | 1744 | 1752 | 1295 |
| 1484 | 0.19 | 0.11 | 1.73 | 0.08 | 1 | Dnajc24 | DnaJ heat shock protein family (Hsp40) m | 317 | 404 | 355 | 1785 | 1785 | 1788 | 1765 | 2133 | 1803 | 2088 | 2099 |
| 1717 | 0.10 | 0.15 | 0.65 | 0.51 | 1 | Atp5md | ATP synthase membrane subunit DAPIT | 1640 | 1497 | 1970 | 1558 | 1845 | 1487 | 1763 | 2259 | 1667 | 1714 | 1488 |
| 1790 | -0.03 | 0.10 | -0.30 | 0.76 | 1 | Cog3 | component of oligomeric golgi complex 3 | 1151 | 1109 | 1276 | 1506 | 1933 | 2896 | 1762 | 1391 | 2053 | 1547 | 3062 |
| 1631 | 0.02 | 0.10 | 0.21 | 0.83 | 1 | Spg11 | SPG11, spatacsin vesicle trafficking asso | 1343 | 1277 | 1106 | 1826 | 1636 | 1833 | 1762 | 1644 | 1812 | 1762 | 1943 |
| 1562 | 0.03 | 0.11 | 0.27 | 0.79 | 1 | Tsg101 | tumor susceptibility gene 101 | 1501 | 1408 | 1588 | 1567 | 1639 | 1308 | 1762 | 1380 | 1693 | 1886 | 1448 |
| 2218 | -0.09 | 0.12 | -0.78 | 0.43 | 1 | Khsrp | KH-type splicing regulatory protein | 2052 | 2158 | 2051 | 2334 | 2292 | 3010 | 1761 | 1932 | 2449 | 1861 | 2493 |
| 1797 | 0.11 | 0.11 | 1.05 | 0.29 | 1 | Cpt2 | carnitine palmitoyltransferase 2 | 1415 | 1762 | 1705 | 2159 | 1719 | 1483 | 1760 | 2442 | 1820 | 2043 | 1461 |
| 1647 | 0.04 | 0.13 | 0.30 | 0.76 | 1 | Anapc13 | anaphase promoting complex subunit 13 | 1305 | 1428 | 1567 | 1830 | 1847 | 1369 | 1760 | 2017 | 1570 | 2054 | 1370 |
| 1957 | 0.20 | 0.17 | 1.21 | 0.23 | 1 | Gadd45g | growth arrest and DNA-damage-inducible | 1301 | 1043 | 1316 | 1567 | 1699 | 3325 | 1759 | 2504 | 1455 | 2061 | 3492 |
| 1825 | 0.05 | 0.11 | 0.46 | 0.65 | 1 | Ulk2 | unc-51 like kinase 2 | 1741 | 1348 | 1549 | 1796 | 1938 | 2093 | 1759 | 1974 | 2056 | 1834 | 1981 |
| 1536 | -0.14 | 0.12 | -1.12 | 0.26 | 1 | Med23 | mediator complex subunit 23 | 1597 | 1282 | 1438 | 1834 | 1372 | 1625 | 1757 | 1496 | 1461 | 1699 | 1339 |
| 1937 | -0.08 | 0.12 | -0.67 | 0.50 | 1 | Tap2 | transporter 2, ATP-binding cassette, sub-f | 3514 | 2414 | 3159 | 1693 | 1608 | 1197 | 1756 | 1607 | 1637 | 1689 | 1037 |
| 1506 | -0.07 | 0.12 | -0.64 | 0.53 | 1 | Gid8 | GID complex subunit 8 | 1402 | 1283 | 1135 | 1549 | 1790 | 1446 | 1756 | 1671 | 1635 | 1502 | 1397 |
| 1856 | 0.23 | 0.15 | 1.50 | 0.13 | 1 | Sdhb | succinate dehydrogenase complex, subu | 2182 | 1579 | 2418 | 1655 | 1630 | 1523 | 1755 | 2250 | 1797 | 1896 | 1730 |
| 1450 | -0.10 | 0.11 | -0.90 | 0.37 | 1 | Marchf4 | membrane associated ring-CH-type finge | 610 | 798 | 706 | 2050 | 1809 | 1543 | 1755 | 1732 | 1745 | 1788 | 1420 |
| 1692 | -0.19 | 0.18 | -1.06 | 0.29 | 1 | Limch1 | LIM and calponin homology domains 1 | 891 | 559 | 1005 | 2485 | 2445 | 1877 | 1754 | 2046 | 2289 | 1927 | 1334 |
| 1584 | 0.00 | 0.13 | -0.01 | 0.99 | 1 | Stk35 | serine/threonine kinase 35 | 889 | 1274 | 1130 | 2245 | 1354 | 1715 | 1754 | 2251 | 1585 | 1539 | 1690 |
| 1895 | 0.12 | 0.12 | 1.00 | 0.32 | 1 | Trappc3 | trafficking protein particle complex 3 | 1991 | 1848 | 2115 | 1877 | 1981 | 1503 | 1752 | 2411 | 1807 | 2045 | 1520 |
| 1648 | 0.04 | 0.10 | 0.38 | 0.70 | 1 | Ncdn | neurochondrin | 1447 | 1602 | 1618 | 1687 | 1672 | 1515 | 1752 | 1718 | 1671 | 1977 | 1466 |
| 1596 | -0.01 | 0.10 | -0.12 | 0.90 | 1 | Fam3a | family with sequence similarity 3, membe | 1338 | 1401 | 1557 | 1663 | 1824 | 1416 | 1752 | 1486 | 1852 | 1819 | 1448 |
| 1711 | -0.09 | 0.10 | -0.82 | 0.41 | 1 | Uap1 | UDP-N-acetylglucosamine pyrophosphor | 1961 | 1269 | 1907 | 1224 | 2123 | 1915 | 1751 | 1067 | 1867 | 1678 | 2055 |
| 1633 | -0.01 | 0.11 | -0.08 | 0.94 | 1 | Cers5 | ceramide synthase 5 | 1722 | 1772 | 1789 | 1422 | 1718 | 1457 | 1750 | 1347 | 1704 | 1897 | 1385 |
| 1590 | 0.08 | 0.11 | 0.76 | 0.45 | 1 | Nudt16l1 | nudix (nucleoside diphosphate linked mo | 1633 | 1611 | 1775 | 1582 | 1478 | 1251 | 1749 | 1848 | 1609 | 1623 | 1335 |
| 2062 | 0.11 | 0.19 | 0.58 | 0.56 | 1 | Pomp | proteasome maturation protein | 2578 | 2377 | 2882 | 1929 | 1785 | 1582 | 1748 | 2916 | 1855 | 1256 | 1772 |
| 1905 | -0.20 | 0.15 | -1.34 | 0.18 | 1 | Svil | supervillin | 2637 | 2068 | 2043 | 1608 | 2129 | 2079 | 1748 | 1313 | 2052 | 1439 | 1838 |
| 1606 | 0.03 | 0.09 | 0.30 | 0.76 | 1 | Reps1 | RalBP1 associated Eps domain containin | 1097 | 976 | 1026 | 1918 | 1720 | 1828 | 1748 | 1856 | 1768 | 1854 | 1879 |
| 1742 | 0.08 | 0.10 | 0.77 | 0.44 | 1 | Snx14 | sorting nexin 14 | 1368 | 1294 | 1254 | 1747 | 1986 | 1931 | 1747 | 2061 | 1901 | 1752 | 2124 |
| 1726 | 0.06 | 0.12 | 0.47 | 0.64 | 1 | Slc49a4 | solute carrier family 49 member 4 | 1838 | 1946 | 1901 | 1965 | 1433 | 1333 | 1747 | 2338 | 1535 | 1682 | 1267 |
| 1244 | 0.00 | 0.13 | 0.02 | 0.98 | 1 | Scrt1 | scratch family zinc finger 1 | 604 | 691 | 644 | 1672 | 1473 | 987 | 1747 | 1371 | 1620 | 1837 | 1041 |
| 1947 | 0.13 | 0.15 | 0.84 | 0.40 | 1 | Nfatc3 | nuclear factor of activated T cells, cytopla | 2198 | 1901 | 1661 | 2053 | 1713 | 1963 | 1746 | 1687 | 2450 | 2042 | 2002 |
| 1482 | -0.14 | 0.12 | -1.22 | 0.22 | 1 | Gpalpp1 | GPALPP motifs containing 1 | 1772 | 1871 | 1559 | 1495 | 1415 | 1191 | 1746 | 1216 | 1361 | 1459 | 1218 |
| 1733 | 0.07 | 0.13 | 0.55 | 0.58 | 1 | Fam149b | family with sequence similarity 149, meml | 1404 | 1175 | 1488 | 2083 | 1842 | 1637 | 1745 | 1751 | 1903 | 2177 | 1858 |
| 1535 | 0.06 | 0.10 | 0.56 | 0.58 | 1 | Ric3 | RIC3 acetylcholine receptor chaperone | 1001 | 964 | 1068 | 1695 | 1611 | 1734 | 1745 | 1543 | 1815 | 1855 | 1855 |
| 2790 | -0.21 | 0.11 | -1.96 | 0.05 | 1 | Bmpr2 | bone morphogenetic protein receptor, typ | 5926 | 6520 | 5800 | 1607 | 1619 | 1685 | 1744 | 1445 | 1571 | 1213 | 1557 |
| 1684 | -0.02 | 0.13 | -0.15 | 0.88 | 1 | Zmat3 | zinc finger matrin type 3 | 859 | 1492 | 1195 | 2184 | 1750 | 1825 | 1744 | 2153 | 2153 | 1508 | 1661 |
| 1556 | 0.06 | 0.15 | 0.40 | 0.69 | 1 | Ap1m2 | adaptor protein complex AP-1, mu 2 subu | 1033 | 1591 | 1430 | 1502 | 1625 | 1514 | 1744 | 1817 | 1508 | 1812 | 1546 |

| | | | | | | | | | | | | | | | | | | |
|---|---|---|---|---|---|---|---|---|---|---|---|---|---|---|---|---|---|---|
| 1726 | 0.04 | 0.11 | 0.31 | 0.76 | 1 | Wdr41 | WD repeat domain 41 | 1415 | 1221 | 1334 | 2044 | 1982 | 1642 | 1743 | 2126 | 1786 | 2030 | 1662 |
| 1663 | 0.02 | 0.09 | 0.22 | 0.83 | 1 | Dram2 | DNA-damage regulated autophagy modu | 1579 | 1506 | 1759 | 1725 | 1917 | 1311 | 1743 | 1711 | 1916 | 1611 | 1514 |
| 1602 | 0.12 | 0.13 | 0.93 | 0.35 | 1 | Opa3 | optic atrophy 3 | 1162 | 1385 | 1207 | 1741 | 1740 | 1325 | 1742 | 2545 | 1701 | 1864 | 1209 |
| 1725 | 0.00 | 0.14 | 0.00 | 1.00 | 1 | Arhgef18 | rho/rac guanine nucleotide exchange fac | 1722 | 1588 | 1397 | 1771 | 1850 | 1735 | 1741 | 2257 | 1878 | 1524 | 1508 |
| 2039 | -0.07 | 0.10 | -0.69 | 0.49 | 1 | Tpp1 | tripeptidyl peptidase I | 2779 | 2099 | 2449 | 1600 | 1966 | 2430 | 1740 | 1492 | 1870 | 1839 | 2167 |
| 1966 | -0.11 | 0.09 | -1.22 | 0.22 | 1 | Pitpnc1 | phosphatidylinositol transfer protein, cyto | 2826 | 3149 | 3403 | 1687 | 1700 | 1224 | 1740 | 1595 | 1608 | 1614 | 1078 |
| 1791 | 0.11 | 0.13 | 0.82 | 0.41 | 1 | Rrp7a | ribosomal RNA processing 7 homolog A ( | 1759 | 1877 | 1945 | 1680 | 1795 | 1562 | 1739 | 2179 | 1555 | 2067 | 1544 |
| 1728 | -0.02 | 0.11 | -0.14 | 0.89 | 1 | Smcr8 | Smith-Magenis syndrome chromosome re | 1658 | 1557 | 1592 | 1813 | 1564 | 2024 | 1739 | 1976 | 1684 | 1569 | 1825 |
| 1530 | 0.06 | 0.14 | 0.43 | 0.66 | 1 | Galnt14 | polypeptide N-acetylgalactosaminyltransf | 166 | 323 | 264 | 1990 | 1988 | 2146 | 1738 | 2328 | 1893 | 1917 | 2080 |
| 2117 | 0.18 | 0.10 | 1.73 | 0.08 | 1 | Cdk11b | cyclin-dependent kinase 11B | 1892 | 1744 | 1998 | 1859 | 1851 | 2905 | 1737 | 2250 | 2252 | 2052 | 2750 |
| 2009 | -0.05 | 0.10 | -0.45 | 0.65 | 1 | Bcl2l1 | BCL2-like 1 | 2207 | 2869 | 2388 | 2349 | 1905 | 1399 | 1737 | 2655 | 1664 | 1559 | 1370 |
| 1919 | -0.07 | 0.10 | -0.67 | 0.50 | 1 | Edc4 | enhancer of mRNA decapping 4 | 1819 | 1855 | 1921 | 2010 | 1882 | 2314 | 1737 | 1701 | 1982 | 1857 | 2029 |
| 1905 | 0.10 | 0.12 | 0.86 | 0.39 | 1 | Carhsp1 | calcium regulated heat stable protein 1 | 2841 | 2147 | 2482 | 1518 | 1891 | 1367 | 1736 | 1914 | 1955 | 1735 | 1373 |
| 1649 | -0.02 | 0.16 | -0.12 | 0.90 | 1 | Anapc16 | anaphase promoting complex subunit 16 | 1322 | 831 | 1260 | 1718 | 2177 | 1807 | 1736 | 1959 | 1752 | 1693 | 1882 |
| 1735 | 0.03 | 0.11 | 0.28 | 0.78 | 1 | Rmdn3 | regulator of microtubule dynamics 3 | 1312 | 1358 | 1463 | 1840 | 1999 | 1816 | 1735 | 1844 | 1915 | 2147 | 1657 |
| 1699 | -0.07 | 0.15 | -0.45 | 0.65 | 1 | 2610002M06Ri | RIKEN cDNA 2610002M06 gene | 2080 | 2392 | 1974 | 1576 | 1644 | 1292 | 1735 | 1335 | 2034 | 1368 | 1254 |
| 1794 | -0.01 | 0.10 | -0.13 | 0.89 | 1 | Ilf2 | interleukin enhancer binding factor 2 | 2196 | 1617 | 2065 | 1515 | 1681 | 2027 | 1734 | 1521 | 1570 | 1873 | 1932 |
| 1724 | -0.11 | 0.12 | -0.93 | 0.35 | 1 | Mturn | maturin, neural progenitor differentiation r | 2172 | 2411 | 2017 | 2045 | 1633 | 1079 | 1733 | 1636 | 1617 | 1371 | 1250 |
| 1540 | -0.03 | 0.11 | -0.24 | 0.81 | 1 | Akt1s1 | AKT1 substrate 1 (proline-rich) | 1597 | 1121 | 1354 | 1572 | 1811 | 1378 | 1733 | 1571 | 1552 | 1838 | 1411 |
| 1812 | 0.01 | 0.11 | 0.11 | 0.91 | 1 | Pggt1b | protein geranylgeranyltransferase type I, I | 1858 | 2024 | 2058 | 1533 | 1537 | 2133 | 1731 | 1404 | 1757 | 1543 | 2358 |
| 1866 | 0.01 | 0.14 | 0.04 | 0.96 | 1 | Ptpn9 | protein tyrosine phosphatase, non-recept | 2412 | 2563 | 2147 | 1712 | 1623 | 1563 | 1730 | 1546 | 1618 | 2322 | 1294 |
| 1844 | -0.03 | 0.11 | -0.26 | 0.79 | 1 | Srsf9 | serine and arginine-rich splicing factor 9 | 2080 | 1956 | 2074 | 1684 | 2128 | 1622 | 1730 | 1515 | 1813 | 1926 | 1760 |
| 1438 | -0.03 | 0.12 | -0.27 | 0.79 | 1 | Ndufb8 | NADH:ubiquinone oxidoreductase subun | 1173 | 1102 | 1293 | 1771 | 1528 | 1172 | 1730 | 1736 | 1628 | 1480 | 1205 |
| 1901 | 0.08 | 0.10 | 0.81 | 0.42 | 1 | Mical2 | microtubule associated monooxygenase, | 1392 | 1546 | 1400 | 1805 | 2008 | 2527 | 1729 | 2239 | 1925 | 1875 | 2462 |
| 1804 | 0.12 | 0.08 | 1.47 | 0.14 | 1 | Golga7 | golgi autoantigen, golgin subfamily a, 7 | 1669 | 1850 | 1670 | 1753 | 1711 | 1840 | 1729 | 1946 | 1858 | 1838 | 1981 |
| 1594 | 0.11 | 0.12 | 0.93 | 0.35 | 1 | Slc35c2 | solute carrier family 35, member C2 | 1085 | 1350 | 1278 | 1491 | 1958 | 1475 | 1729 | 1836 | 1902 | 1839 | 1590 |
| 1591 | -0.02 | 0.11 | -0.23 | 0.82 | 1 | Abhd14b | abhydrolase domain containing 14b | 1485 | 1558 | 1583 | 1587 | 1760 | 1384 | 1729 | 1592 | 1715 | 1926 | 1182 |
| 1451 | -0.01 | 0.11 | -0.11 | 0.91 | 1 | Kdm2b | lysine (K)-specific demethylase 2B | 872 | 908 | 762 | 1729 | 1694 | 1575 | 1729 | 1878 | 1811 | 1532 | 1468 |
| 1885 | 0.25 | 0.14 | 1.82 | 0.07 | 1 | Eif2s2 | eukaryotic translation initiation factor 2, su | 1866 | 2187 | 1745 | 1449 | 1996 | 1646 | 1728 | 2207 | 1982 | 1891 | 2037 |
| 1858 | 0.09 | 0.09 | 0.90 | 0.37 | 1 | Rbm27 | RNA binding motif protein 27 | 1676 | 1838 | 1556 | 1960 | 1695 | 2083 | 1728 | 2111 | 1941 | 1942 | 1905 |
| 1713 | -0.03 | 0.11 | -0.30 | 0.77 | 1 | Tiprl | TIP41, TOR signalling pathway regulator- | 1814 | 1937 | 1704 | 1526 | 1848 | 1667 | 1728 | 1463 | 1677 | 1767 | 1711 |
| 1276 | -0.07 | 0.13 | -0.50 | 0.61 | 1 | Zfp346 | zinc finger protein 346 | 809 | 513 | 616 | 1292 | 1490 | 1655 | 1728 | 1109 | 1585 | 1436 | 1800 |
| 2179 | -0.21 | 0.15 | -1.38 | 0.17 | 1 | Kdm6b | KDM1 lysine (K)-specific demethylase 6B | 3079 | 3338 | 2777 | 2044 | 2199 | 1916 | 1727 | 1453 | 2077 | 1948 | 1410 |
| 1923 | -0.05 | 0.12 | -0.40 | 0.69 | 1 | Bcor | BCL6 interacting corepressor | 1985 | 2222 | 1810 | 2219 | 2017 | 1673 | 1726 | 2582 | 1878 | 1555 | 1482 |
| 1848 | 0.00 | 0.11 | 0.00 | 1.00 | 1 | Cops8 | COP9 signalosome subunit 8 | 1889 | 2091 | 1927 | 1876 | 1916 | 1691 | 1726 | 1601 | 1947 | 2012 | 1658 |
| 1494 | 0.06 | 0.15 | 0.40 | 0.69 | 1 | Gtf2h5 | general transcription factor IIH, polypeptic | 1760 | 1252 | 1650 | 1486 | 1383 | 1191 | 1726 | 1763 | 1542 | 1405 | 1279 |
| 1757 | -0.04 | 0.11 | -0.36 | 0.72 | 1 | Rbm7 | RNA binding motif protein 7 | 2006 | 2218 | 2010 | 1753 | 1522 | 1637 | 1724 | 1398 | 1571 | 1695 | 1792 |
| 1718 | 0.08 | 0.09 | 0.84 | 0.40 | 1 | Trim27 | tripartite motif-containing 27 | 1499 | 1781 | 1688 | 1827 | 1634 | 1576 | 1724 | 2038 | 1825 | 1805 | 1507 |
| 1654 | 0.04 | 0.13 | 0.36 | 0.72 | 1 | Ugp2 | UDP-glucose pyrophosphorylase 2 | 2191 | 1724 | 1760 | 1648 | 1622 | 1181 | 1724 | 1337 | 1808 | 1767 | 1436 |
| 1611 | -0.13 | 0.14 | -0.90 | 0.37 | 1 | Snap23 | synaptosomal-associated protein 23 | 1920 | 1095 | 1399 | 1406 | 1846 | 1899 | 1724 | 1014 | 1805 | 1723 | 1896 |
| 1590 | 0.10 | 0.11 | 0.91 | 0.36 | 1 | Sqor | sulfide quinone oxidoreductase | 1331 | 1159 | 1318 | 1713 | 1917 | 1279 | 1724 | 1930 | 1759 | 1805 | 1555 |
| 1671 | 0.01 | 0.13 | 0.12 | 0.91 | 1 | Habp4 | hyaluronic acid binding protein 4 | 1383 | 1030 | 1208 | 1804 | 1943 | 1834 | 1723 | 1475 | 2256 | 1926 | 1802 |
| 1606 | -0.04 | 0.11 | -0.41 | 0.68 | 1 | Parg | poly (ADP-ribose) glycohydrolase | 1485 | 1231 | 1428 | 1444 | 1863 | 1801 | 1723 | 1214 | 1763 | 1762 | 1949 |
| 1619 | -0.06 | 0.09 | -0.69 | 0.49 | 1 | Gpatch2 | G patch domain containing 2 | 910 | 966 | 1131 | 1837 | 1853 | 2145 | 1721 | 1740 | 1758 | 1646 | 2098 |
| 1529 | 0.01 | 0.12 | 0.06 | 0.95 | 1 | Dhx8 | DEAH (Asp-Glu-Ala-His) box polypeptide | 1289 | 1277 | 1241 | 1521 | 1624 | 1607 | 1721 | 1828 | 1691 | 1703 | 1321 |
| 1451 | 0.05 | 0.16 | 0.33 | 0.74 | 1 | Dusp5 | dual specificity phosphatase 5 | 1534 | 842 | 1162 | 1151 | 1784 | 1500 | 1721 | 1571 | 1728 | 1541 | 1432 |
| 1420 | 0.21 | 0.16 | 1.31 | 0.19 | 1 | Fbxo2 | F-box protein 2 | 509 | 574 | 750 | 1768 | 1630 | 1206 | 1721 | 2574 | 1797 | 1836 | 1250 |
| 1558 | 0.05 | 0.11 | 0.48 | 0.63 | 1 | Sra1 | steroid receptor RNA activator 1 | 1569 | 1390 | 1593 | 1522 | 1543 | 1394 | 1720 | 1780 | 1472 | 1739 | 1421 |
| 1807 | -0.03 | 0.12 | -0.22 | 0.83 | 1 | Cxxc1 | CXXC finger 1 (PHD domain) | 2113 | 1827 | 1931 | 1593 | 1735 | 2030 | 1719 | 1841 | 1866 | 1462 | 1759 |
| 1788 | -0.06 | 0.12 | -0.49 | 0.63 | 1 | Tbc1d14 | TBC1 domain family, member 14 | 1961 | 1673 | 1718 | 1916 | 1713 | 1938 | 1719 | 1454 | 1881 | 1763 | 1930 |
| 1625 | -0.03 | 0.08 | -0.39 | 0.70 | 1 | Ndufa10 | NADH:ubiquinone oxidoreductase subun | 1736 | 1445 | 1676 | 1606 | 1700 | 1555 | 1718 | 1533 | 1720 | 1680 | 1504 |
| 1568 | -0.02 | 0.12 | -0.16 | 0.88 | 1 | Ift46 | intraflagellar transport 46 | 1701 | 2089 | 1601 | 1723 | 1316 | 1153 | 1718 | 2121 | 1241 | 1538 | 1048 |
| 2412 | 0.02 | 0.08 | 0.28 | 0.78 | 1 | Prpf38b | PRP38 pre-mRNA processing factor 38 (y | 2175 | 2638 | 2274 | 1929 | 1600 | 4395 | 1717 | 1858 | 1631 | 1814 | 4501 |
| 1892 | 0.14 | 0.16 | 0.90 | 0.37 | 1 | Eif6 | eukaryotic translation initiation factor 6 | 2236 | 2044 | 2284 | 1693 | 1856 | 1450 | 1717 | 2650 | 1662 | 1838 | 1384 |
| 1681 | 0.13 | 0.13 | 1.03 | 0.30 | 1 | Gps1 | G protein pathway suppressor 1 | 1381 | 1140 | 1513 | 1586 | 1824 | 1773 | 1717 | 2121 | 1842 | 1923 | 1668 |
| 1524 | 0.01 | 0.13 | 0.09 | 0.93 | 1 | Arhgef5 | Rho guanine nucleotide exchange factor | 1029 | 1019 | 1026 | 1551 | 1472 | 2077 | 1717 | 1298 | 1812 | 1811 | 1954 |
| 1465 | 0.11 | 0.14 | 0.77 | 0.44 | 1 | 1700019D03Ril | RIKEN cDNA 1700019D03 gene | 933 | 951 | 1274 | 1657 | 1645 | 1210 | 1716 | 1972 | 1649 | 1878 | 1234 |
| 1969 | 0.10 | 0.10 | 0.94 | 0.35 | 1 | Mob4 | MOB family member 4, phocein | 1627 | 2216 | 1918 | 2149 | 2007 | 1833 | 1715 | 2051 | 2030 | 2042 | 2066 |
| 1637 | -0.20 | 0.13 | -1.60 | 0.11 | 1 | Ankrd46 | ankyrin repeat domain 46 | 1783 | 1835 | 1997 | 1779 | 1477 | 1656 | 1714 | 1220 | 1485 | 1619 | 1443 |
| 1488 | 0.06 | 0.10 | 0.61 | 0.54 | 1 | Pip4p2 | phosphatidylinositol-4,5-bisphosphate 4-( | 1227 | 1127 | 1253 | 1527 | 1688 | 1319 | 1712 | 1549 | 1626 | 1955 | 1389 |

| | | | | | | | | | | | | | | | | | | |
|---|---|---|---|---|---|---|---|---|---|---|---|---|---|---|---|---|---|---|
| 1916 | 0.07 | 0.14 | 0.52 | 0.60 | 1 | C2cd4b | C2 calcium-dependent domain containing | 1089 | 1011 | 1513 | 1088 | 1467 | 4308 | 1711 | 1126 | 1549 | 1919 | 4300 |
| 1896 | 0.02 | 0.14 | 0.15 | 0.88 | 1 | Trpm7 | transient receptor potential cation channe | 2200 | 2179 | 1993 | 1651 | 1350 | 2427 | 1711 | 1477 | 1819 | 1310 | 2738 |
| 1614 | -0.09 | 0.10 | -0.93 | 0.35 | 1 | Arpin | actin-related protein 2/3 complex inhibitor | 1631 | 1614 | 1741 | 1709 | 1744 | 1418 | 1711 | 1393 | 1721 | 1668 | 1407 |
| 1486 | 0.13 | 0.15 | 0.87 | 0.38 | 1 | Bag5 | BCL2-associated athanogene 5 | 1427 | 1679 | 1540 | 1308 | 1280 | 1270 | 1711 | 1923 | 1361 | 1407 | 1441 |
| 1298 | 0.04 | 0.13 | 0.27 | 0.79 | 1 | Ttc19 | tetratricopeptide repeat domain 19 | 761 | 755 | 710 | 1238 | 1413 | 1599 | 1711 | 1495 | 1493 | 1346 | 1764 |
| 1923 | 0.15 | 0.10 | 1.47 | 0.14 | 1 | Nck1 | non-catalytic region of tyrosine kinase ada | 2042 | 1945 | 1582 | 2007 | 1646 | 2047 | 1710 | 2236 | 1884 | 1917 | 2143 |
| 1635 | 0.01 | 0.10 | 0.11 | 0.91 | 1 | Spire1 | spire type actin nucleation factor 1 | 1560 | 1178 | 1458 | 1764 | 1863 | 1528 | 1710 | 1691 | 1836 | 1851 | 1551 |
| 1725 | -0.11 | 0.13 | -0.83 | 0.41 | 1 | Tlk2 | tousled-like kinase 2 (Arabidopsis) | 1444 | 1724 | 1703 | 1742 | 1785 | 2016 | 1709 | 1311 | 1825 | 1452 | 2263 |
| 1617 | 0.02 | 0.10 | 0.18 | 0.86 | 1 | Dph3 | diphthamine biosynthesis 3 | 1412 | 1383 | 1618 | 1631 | 1814 | 1493 | 1709 | 1784 | 1836 | 1489 | 1620 |
| 1800 | -0.05 | 0.10 | -0.48 | 0.63 | 1 | Pfdn2 | prefoldin 2 | 1964 | 1832 | 2295 | 1747 | 1908 | 1615 | 1708 | 1744 | 1649 | 1641 | 1696 |
| 1632 | 0.03 | 0.10 | 0.32 | 0.75 | 1 | Scrn3 | secernin 3 | 1292 | 1563 | 1441 | 1659 | 1627 | 1756 | 1708 | 1709 | 1712 | 1604 | 1884 |
| 1340 | 0.22 | 0.16 | 1.39 | 0.17 | 1 | Ica1l | islet cell autoantigen 1-like | 124 | 125 | 253 | 1724 | 1981 | 1177 | 1708 | 1899 | 2417 | 1852 | 1486 |
| 1776 | 0.09 | 0.13 | 0.68 | 0.49 | 1 | Cdadc1 | cytidine and dCMP deaminase domain co | 1724 | 1251 | 1719 | 1901 | 1723 | 1850 | 1707 | 1916 | 1748 | 1998 | 1994 |
| 1754 | -0.16 | 0.12 | -1.27 | 0.20 | 1 | Dgkz | diacylglycerol kinase zeta | 2505 | 2077 | 2206 | 1461 | 1577 | 1861 | 1707 | 1614 | 1500 | 1377 | 1409 |
| 2057 | 0.15 | 0.15 | 1.03 | 0.30 | 1 | Snrpb | small nuclear ribonucleoprotein B | 3173 | 2679 | 2741 | 1465 | 1883 | 1583 | 1706 | 2324 | 1767 | 1740 | 1567 |
| 1832 | 0.04 | 0.11 | 0.38 | 0.70 | 1 | Rcor1 | REST corepressor 1 | 1921 | 1880 | 1838 | 1715 | 1729 | 1997 | 1706 | 1986 | 2009 | 1551 | 1821 |
| 1697 | -0.15 | 0.09 | -1.74 | 0.08 | 1 | Mcm3ap | minichromosome maintenance complex c | 1661 | 1608 | 1619 | 1832 | 1896 | 1819 | 1706 | 1674 | 1664 | 1507 | 1681 |
| 1658 | 0.08 | 0.10 | 0.76 | 0.44 | 1 | Pex2 | peroxisomal biogenesis factor 2 | 1259 | 1091 | 1274 | 1566 | 2030 | 1798 | 1706 | 1573 | 2108 | 1699 | 2134 |
| 1886 | -0.04 | 0.12 | -0.30 | 0.76 | 1 | Dvl3 | dishevelled segment polarity protein 3 | 1043 | 1526 | 1371 | 2100 | 2278 | 2452 | 1705 | 1934 | 2410 | 1891 | 2032 |
| 1819 | 0.16 | 0.08 | 1.93 | 0.05 | 1 | Uba3 | ubiquitin-like modifier activating enzyme 3 | 1669 | 1569 | 1611 | 1758 | 1646 | 2062 | 1705 | 1975 | 1807 | 1931 | 2280 |
| 1349 | 0.09 | 0.11 | 0.78 | 0.43 | 1 | Tmem218 | transmembrane protein 218 | 691 | 794 | 694 | 1505 | 1582 | 1339 | 1705 | 1868 | 1642 | 1663 | 1358 |
| 1257 | -0.11 | 0.14 | -0.78 | 0.43 | 1 | Gcdh | glutaryl-Coenzyme A dehydrogenase | 814 | 884 | 979 | 1491 | 1323 | 1271 | 1705 | 1315 | 1398 | 1284 | 1362 |
| 1555 | -0.06 | 0.13 | -0.46 | 0.65 | 1 | Cdr2 | cerebellar degeneration-related 2 | 648 | 1072 | 931 | 1960 | 1698 | 2017 | 1704 | 1652 | 1779 | 1579 | 2070 |
| 1179 | 0.23 | 0.16 | 1.49 | 0.14 | 1 | Plppr4 | phospholipid phosphatase related 4 | 262 | 243 | 393 | 1115 | 1625 | 1113 | 1704 | 1565 | 1764 | 2027 | 1163 |
| 1957 | 0.13 | 0.13 | 0.99 | 0.32 | 1 | Mtarc2 | mitochondrial amidoxime reducing compo | 2045 | 2139 | 2165 | 2275 | 1996 | 1295 | 1703 | 2710 | 1801 | 1767 | 1636 |
| 1743 | -0.02 | 0.14 | -0.12 | 0.90 | 1 | Fdft1 | farnesyl diphosphate farnesyl transferase | 1137 | 1077 | 1326 | 2276 | 2096 | 1788 | 1703 | 1672 | 1993 | 2257 | 1849 |
| 1575 | 0.02 | 0.13 | 0.18 | 0.86 | 1 | Rnf5 | ring finger protein 5 | 1215 | 1373 | 1515 | 1755 | 1646 | 1440 | 1703 | 1667 | 1572 | 2066 | 1378 |
| 1540 | -0.18 | 0.13 | -1.33 | 0.18 | 1 | Abcd1 | ATP-binding cassette, sub-family D (ALD) | 1692 | 1632 | 1536 | 1579 | 1743 | 1358 | 1703 | 1121 | 1637 | 1735 | 1203 |
| 1457 | 0.10 | 0.23 | 0.44 | 0.66 | 1 | Mrln | myoregulin | 762 | 908 | 1067 | 1536 | 1540 | 1504 | 1703 | 2821 | 1147 | 1589 | 1445 |
| 1287 | -0.28 | 0.16 | -1.79 | 0.07 | 1 | Lyplal1 | lysophospholipase-like 1 | 1005 | 1177 | 1227 | 1236 | 1430 | 1512 | 1702 | 880 | 1307 | 1283 | 1400 |
| 1953 | -0.08 | 0.08 | -0.91 | 0.36 | 1 | Fam234b | family with sequence similarity 234, meml | 1866 | 2365 | 2313 | 2211 | 1889 | 1874 | 1701 | 1967 | 1859 | 1658 | 1785 |
| 1841 | 0.02 | 0.09 | 0.23 | 0.82 | 1 | Sec24b | Sec24 related gene family, member B (S. | 1683 | 1810 | 1769 | 1994 | 1705 | 2054 | 1701 | 1915 | 1777 | 1879 | 1967 |
| 1702 | 0.04 | 0.10 | 0.39 | 0.70 | 1 | Exoc7 | exocyst complex component 7 | 1675 | 1344 | 1777 | 1535 | 1952 | 1679 | 1701 | 1672 | 1915 | 1932 | 1538 |
| 2310 | 0.05 | 0.09 | 0.57 | 0.57 | 1 | Mau2 | MAU2 sister chromatid cohesion factor | 1996 | 1991 | 1896 | 1857 | 2045 | 4069 | 1700 | 2077 | 2039 | 1872 | 3865 |
| 1812 | -0.24 | 0.18 | -1.32 | 0.19 | 1 | Elf1 | E74-like factor 1 | 2507 | 1903 | 1942 | 1621 | 2034 | 1898 | 1700 | 869 | 1961 | 1743 | 1751 |
| 1582 | -0.04 | 0.14 | -0.29 | 0.77 | 1 | Cdk5r2 | cyclin-dependent kinase 5, regulatory sub | 1031 | 1757 | 1199 | 1918 | 1903 | 1264 | 1699 | 1863 | 1854 | 1789 | 1127 |
| 1222 | 0.02 | 0.13 | 0.15 | 0.88 | 1 | B3gnt3 | UDP-GlcNAc:betaGal beta-1,3-N-acetylgl | 614 | 492 | 843 | 1372 | 1711 | 888 | 1699 | 1346 | 1771 | 1857 | 843 |
| 2116 | 0.13 | 0.09 | 1.42 | 0.16 | 1 | Ythdc1 | YTH domain containing 1 | 2032 | 2167 | 1715 | 1757 | 1483 | 3443 | 1698 | 1853 | 1772 | 1913 | 3444 |
| 1790 | 0.19 | 0.12 | 1.57 | 0.12 | 1 | Prelid1 | PRELI domain containing 1 | 2132 | 2087 | 2062 | 1464 | 1695 | 1379 | 1698 | 2220 | 1667 | 1792 | 1493 |
| 1575 | -0.05 | 0.13 | -0.35 | 0.72 | 1 | Ddhd2 | DDHD domain containing 2 | 1097 | 1283 | 1104 | 1741 | 1563 | 2036 | 1698 | 1359 | 1729 | 1850 | 1863 |
| 1542 | -0.11 | 0.10 | -1.13 | 0.26 | 1 | Ppa1 | pyrophosphatase (inorganic) 1 | 1959 | 1446 | 1701 | 1504 | 1526 | 1411 | 1697 | 1204 | 1467 | 1705 | 1339 |
| 1881 | 0.17 | 0.10 | 1.62 | 0.11 | 1 | Etfdh | electron transferring flavoprotein, dehydro | 1808 | 1716 | 1893 | 1824 | 1951 | 1712 | 1696 | 1814 | 2003 | 2303 | 1973 |
| 1853 | 0.05 | 0.12 | 0.38 | 0.70 | 1 | Thumpd1 | THUMP domain containing 1 | 1943 | 2415 | 2252 | 1921 | 1582 | 1593 | 1696 | 1602 | 1784 | 1848 | 1749 |
| 1938 | 0.00 | 0.11 | 0.04 | 0.97 | 1 | Zfp869 | zinc finger protein 869 | 1781 | 1827 | 1827 | 1834 | 2162 | 2212 | 1693 | 1598 | 1997 | 1869 | 2514 |
| 1722 | 0.03 | 0.13 | 0.20 | 0.85 | 1 | Acvr2a | activin receptor IIA | 1286 | 1885 | 1474 | 2388 | 1542 | 1524 | 1693 | 1853 | 1807 | 1854 | 1634 |
| 1939 | -0.01 | 0.11 | -0.13 | 0.90 | 1 | Trappc8 | trafficking protein particle complex 8 | 2016 | 2185 | 2075 | 2009 | 1750 | 2123 | 1692 | 1804 | 2128 | 1758 | 1793 |
| 1898 | -0.04 | 0.17 | -0.26 | 0.79 | 1 | Ezr | ezrin | 4153 | 2331 | 2395 | 1516 | 1686 | 1213 | 1692 | 1453 | 1770 | 1465 | 1206 |
| 1646 | 0.10 | 0.15 | 0.69 | 0.49 | 1 | Xpo4 | exportin 4 | 1417 | 1674 | 1460 | 1346 | 1525 | 1983 | 1691 | 1404 | 2042 | 1609 | 1951 |
| 1750 | 0.16 | 0.14 | 1.14 | 0.25 | 1 | Acaa1a | acetyl-Coenzyme A acyltransferase 1A | 2019 | 1660 | 1868 | 1224 | 1844 | 1741 | 1690 | 1886 | 1822 | 1944 | 1554 |
| 2081 | 0.05 | 0.09 | 0.57 | 0.57 | 1 | Gramd1a | GRAM domain containing 1A | 1517 | 1252 | 1433 | 1747 | 1659 | 4111 | 1689 | 1688 | 1852 | 1796 | 4153 |
| 1608 | -0.03 | 0.10 | -0.36 | 0.72 | 1 | Vps37c | vacuolar protein sorting 37C | 1143 | 1399 | 1562 | 1716 | 1882 | 1577 | 1689 | 1811 | 1859 | 1529 | 1522 |
| 1474 | -0.07 | 0.15 | -0.44 | 0.66 | 1 | Iscu | iron-sulfur cluster assembly enzyme | 1409 | 1235 | 1107 | 1742 | 1539 | 1379 | 1688 | 1183 | 1672 | 1902 | 1358 |
| 1387 | 0.08 | 0.15 | 0.54 | 0.59 | 1 | Gmds | GDP-mannose 4, 6-dehydratase | 905 | 868 | 963 | 1919 | 1348 | 1123 | 1688 | 1796 | 1339 | 2096 | 1211 |
| 1614 | -0.06 | 0.14 | -0.39 | 0.69 | 1 | Fam219a | family with sequence similarity 219, meml | 1142 | 1007 | 1356 | 1529 | 1536 | 2530 | 1687 | 1777 | 1593 | 1325 | 2269 |
| 1777 | 0.31 | 0.16 | 1.93 | 0.05 | 1 | Ndufb5 | NADH:ubiquinone oxidoreductase subun | 1638 | 1476 | 1963 | 1635 | 1784 | 1331 | 1686 | 2764 | 1853 | 1830 | 1588 |
| 1594 | -0.03 | 0.09 | -0.31 | 0.76 | 1 | Dda1 | DET1 and DDB1 associated 1 | 1461 | 1432 | 1769 | 1555 | 1814 | 1451 | 1686 | 1562 | 1698 | 1637 | 1470 |
| 2075 | 0.09 | 0.12 | 0.79 | 0.43 | 1 | Dcaf10 | DDB1 and CUL4 associated factor 10 | 1552 | 1828 | 1635 | 2342 | 2281 | 2345 | 1685 | 2198 | 2468 | 2272 | 2217 |
| 1779 | -0.06 | 0.10 | -0.60 | 0.55 | 1 | Tsr2 | TSR2 20S rRNA accumulation | 1979 | 1883 | 2409 | 1770 | 1900 | 1438 | 1685 | 1701 | 1709 | 1718 | 1384 |
| 1775 | 0.01 | 0.10 | 0.12 | 0.91 | 1 | Nagk | N-acetylglucosamine kinase | 1575 | 1615 | 1701 | 1907 | 1897 | 1797 | 1685 | 1755 | 1784 | 2025 | 1782 |
| 1615 | -0.05 | 0.13 | -0.42 | 0.67 | 1 | Usp30 | ubiquitin specific peptidase 30 | 1075 | 1679 | 1387 | 1995 | 1780 | 1504 | 1685 | 1753 | 1586 | 1796 | 1529 |

| | | | | | | | | | | | | | | | | | | |
|---|---|---|---|---|---|---|---|---|---|---|---|---|---|---|---|---|---|---|
| 1581 | 0.14 | 0.14 | 1.05 | 0.30 | 1 | Ankrd54 | ankyrin repeat domain 54 | 961 | 739 | 951 | 1469 | 1852 | 2019 | 1685 | 2048 | 1634 | 1887 | 2148 |
| 1518 | -0.02 | 0.16 | -0.11 | 0.91 | 1 | Faah | fatty acid amide hydrolase | 1046 | 1043 | 1611 | 1613 | 1656 | 1581 | 1685 | 1686 | 1563 | 1817 | 1397 |
| 1747 | 0.35 | 0.21 | 1.70 | 0.09 | 1 | Zcrb1 | zinc finger CCHC-type and RNA binding | 1523 | 1358 | 1763 | 1436 | 1408 | 1817 | 1683 | 2964 | 1483 | 1628 | 2157 |
| 1609 | 0.16 | 0.12 | 1.29 | 0.20 | 1 | Tapbpl | TAP binding protein-like | 1347 | 1249 | 1555 | 1402 | 1829 | 1497 | 1683 | 1981 | 1978 | 1544 | 1640 |
| 2476 | 0.11 | 0.09 | 1.22 | 0.22 | 1 | Gas5 | | 3858 | 3840 | 4300 | 1391 | 1529 | 2706 | 1681 | 1530 | 1606 | 1736 | 3052 |
| 1738 | -0.01 | 0.14 | -0.06 | 0.95 | 1 | Neurl4 | neuralized E3 ubiquitin protein ligase 4 | 2145 | 1435 | 1907 | 1416 | 1442 | 2365 | 1680 | 1625 | 1617 | 1525 | 1961 |
| 1670 | 0.01 | 0.13 | 0.05 | 0.96 | 1 | Zfp398 | zinc finger protein 398 | 952 | 1234 | 957 | 1750 | 1668 | 2515 | 1680 | 1490 | 1866 | 1799 | 2462 |
| 1588 | 0.00 | 0.09 | 0.00 | 1.00 | 1 | Ccndbp1 | cyclin D-type binding-protein 1 | 1249 | 1388 | 1473 | 1633 | 1813 | 1551 | 1680 | 1686 | 1851 | 1618 | 1529 |
| 1762 | 0.06 | 0.12 | 0.50 | 0.62 | 1 | Rnf41 | ring finger protein 41 | 1795 | 1735 | 1719 | 1778 | 1785 | 1656 | 1678 | 1964 | 1542 | 2123 | 1602 |
| 1641 | 0.02 | 0.10 | 0.22 | 0.83 | 1 | Elof1 | ELF1 homolog, elongation factor 1 | 1408 | 1331 | 1605 | 1672 | 2048 | 1421 | 1678 | 1791 | 1752 | 1866 | 1481 |
| 1613 | -0.15 | 0.15 | -0.97 | 0.33 | 1 | Ncoa3 | nuclear receptor coactivator 3 | 2475 | 1877 | 1803 | 1356 | 1433 | 1620 | 1678 | 1120 | 1684 | 1276 | 1422 |
| 1589 | 0.09 | 0.10 | 0.95 | 0.34 | 1 | Scaper | S phase cyclin A-associated protein in the | 1533 | 1496 | 1588 | 1421 | 1562 | 1563 | 1678 | 1586 | 1723 | 1539 | 1789 |
| 2026 | 0.05 | 0.15 | 0.34 | 0.73 | 1 | Txn1 | thioredoxin 1 | 3359 | 2493 | 2841 | 1817 | 1789 | 1352 | 1677 | 2330 | 1512 | 1807 | 1307 |
| 1612 | -0.02 | 0.14 | -0.14 | 0.89 | 1 | Mettl26 | methyltransferase like 26 | 1038 | 1432 | 1528 | 1853 | 1895 | 1509 | 1676 | 1609 | 1700 | 1937 | 1555 |
| 1607 | -0.04 | 0.11 | -0.33 | 0.74 | 1 | Glb1 | galactosidase, beta 1 | 1561 | 1364 | 1704 | 1903 | 1937 | 1098 | 1676 | 1629 | 1788 | 1891 | 1120 |
| 1404 | 0.06 | 0.15 | 0.41 | 0.69 | 1 | Trmt2b | TRM2 tRNA methyltransferase 2B | 1161 | 1187 | 1238 | 1563 | 1434 | 1145 | 1676 | 1601 | 1676 | 1242 | 1518 |
| 1948 | -0.06 | 0.10 | -0.54 | 0.59 | 1 | Coro1b | coronin, actin binding protein 1B | 2794 | 2376 | 2384 | 1739 | 1833 | 1811 | 1675 | 1518 | 1732 | 1902 | 1667 |
| 1571 | 0.05 | 0.10 | 0.47 | 0.64 | 1 | Wdr36 | WD repeat domain 36 | 1716 | 1298 | 1411 | 1597 | 1626 | 1424 | 1675 | 1539 | 1739 | 1698 | 1562 |
| 1406 | -0.04 | 0.13 | -0.29 | 0.78 | 1 | Naip2 | NLR family, apoptosis inhibitory protein 2 | 1021 | 888 | 850 | 1688 | 1642 | 1390 | 1675 | 1303 | 1614 | 2080 | 1316 |
| 1816 | 0.06 | 0.10 | 0.59 | 0.56 | 1 | Sdf2 | stromal cell derived factor 2 | 1859 | 1996 | 2209 | 1827 | 1761 | 1539 | 1674 | 2070 | 1755 | 1779 | 1502 |
| 1714 | 0.12 | 0.10 | 1.13 | 0.26 | 1 | Ak3 | adenylate kinase 3 | 2557 | 2164 | 1848 | 1539 | 1487 | 1183 | 1674 | 1705 | 1703 | 1801 | 1191 |
| 1513 | -0.08 | 0.14 | -0.57 | 0.57 | 1 | Nenf | neuron derived neurotrophic factor | 1793 | 1912 | 2002 | 1409 | 1601 | 899 | 1674 | 1817 | 1323 | 1427 | 788 |
| 1503 | -0.02 | 0.10 | -0.18 | 0.86 | 1 | Ttpal | tocopherol (alpha) transfer protein-like | 1019 | 1126 | 865 | 1810 | 1668 | 1651 | 1674 | 1638 | 1581 | 1796 | 1701 |
| 1933 | 0.15 | 0.13 | 1.12 | 0.26 | 1 | Syf2 | SYF2 homolog, RNA splicing factor (S. ce | 2264 | 1891 | 2323 | 1735 | 1927 | 1680 | 1672 | 2347 | 1901 | 1726 | 1797 |
| 1681 | 0.13 | 0.10 | 1.32 | 0.19 | 1 | Wtap | Wilms tumour 1-associating protein | 1575 | 1640 | 1418 | 1706 | 1366 | 1877 | 1672 | 1620 | 1623 | 1848 | 2144 |
| 2070 | -0.04 | 0.12 | -0.35 | 0.72 | 1 | Dido1 | death inducer-obliterator 1 | 1960 | 1710 | 1662 | 2094 | 2083 | 3040 | 1671 | 1897 | 1943 | 2038 | 2671 |
| 1719 | 0.09 | 0.12 | 0.74 | 0.46 | 1 | H2az1 | H2A.Z variant histone 1 | 1829 | 1648 | 1934 | 1557 | 1861 | 1456 | 1671 | 2056 | 1735 | 1615 | 1548 |
| 1718 | -0.16 | 0.14 | -1.14 | 0.25 | 1 | Srsf10 | serine and arginine-rich splicing factor 10 | 1705 | 1707 | 1542 | 1197 | 1430 | 2906 | 1671 | 1053 | 1371 | 1162 | 3152 |
| 1626 | 0.00 | 0.10 | 0.04 | 0.97 | 1 | Ctps2 | cytidine 5'-triphosphate synthase 2 | 1683 | 1541 | 1525 | 1779 | 1585 | 1519 | 1671 | 1490 | 1673 | 1879 | 1543 |
| 1464 | -0.11 | 0.14 | -0.83 | 0.41 | 1 | Cacnb2 | calcium channel, voltage-dependent, beta | 1085 | 1327 | 1343 | 1557 | 1622 | 1562 | 1671 | 1247 | 1437 | 1726 | 1525 |
| 1934 | -0.04 | 0.08 | -0.44 | 0.66 | 1 | Uhrf2 | ubiquitin-like, containing PHD and RING f | 1783 | 2193 | 1908 | 1887 | 1550 | 2669 | 1670 | 1812 | 1563 | 1557 | 2677 |
| 1740 | -0.01 | 0.11 | -0.05 | 0.96 | 1 | Rpl38 | ribosomal protein L38 | 2495 | 2376 | 2695 | 1487 | 1500 | 1128 | 1669 | 1688 | 1369 | 1660 | 1071 |
| 1424 | 0.01 | 0.17 | 0.06 | 0.95 | 1 | Gm30648 | | 545 | 666 | 372 | 1931 | 1814 | 1620 | 1669 | 1716 | 1981 | 1609 | 1743 |
| 1789 | 0.06 | 0.10 | 0.61 | 0.54 | 1 | Rbm18 | RNA binding motif protein 18 | 1654 | 1603 | 1808 | 1953 | 1772 | 1748 | 1668 | 2143 | 1858 | 1741 | 1728 |
| 1973 | 0.05 | 0.09 | 0.58 | 0.56 | 1 | Eif2s3y | eukaryotic translation initiation factor 2, su | 1531 | 1397 | 1696 | 1627 | 1630 | 3400 | 1667 | 1463 | 1833 | 1758 | 3701 |
| 1577 | 0.00 | 0.09 | 0.01 | 0.99 | 1 | Gba | glucosidase, beta, acid | 1828 | 1896 | 1651 | 1607 | 1507 | 1211 | 1667 | 1622 | 1571 | 1538 | 1250 |
| 1962 | 0.03 | 0.11 | 0.30 | 0.77 | 1 | Arnt | aryl hydrocarbon receptor nuclear translo | 1782 | 1853 | 1546 | 2264 | 1823 | 2386 | 1666 | 2194 | 1880 | 2027 | 2162 |
| 1600 | 0.00 | 0.11 | 0.04 | 0.97 | 1 | Mtfr1 | mitochondrial fission regulator 1 | 1178 | 1392 | 1175 | 1905 | 1810 | 1536 | 1666 | 1693 | 1806 | 1940 | 1496 |
| 1862 | 0.12 | 0.12 | 1.05 | 0.29 | 1 | Apeh | acylpeptide hydrolase | 1606 | 1846 | 1962 | 1916 | 1897 | 1702 | 1665 | 2567 | 1933 | 1721 | 1670 |
| 1738 | 0.05 | 0.12 | 0.42 | 0.67 | 1 | Stk16 | serine/threonine kinase 16 | 1595 | 1205 | 1650 | 1692 | 1915 | 1930 | 1665 | 1682 | 1757 | 2083 | 1943 |
| 1683 | -0.14 | 0.12 | -1.16 | 0.25 | 1 | Ralgds | ral guanine nucleotide dissociation stimul | 1905 | 1890 | 1590 | 2076 | 1591 | 1561 | 1665 | 1484 | 1639 | 1721 | 1389 |
| 1369 | -0.16 | 0.11 | -1.46 | 0.14 | 1 | Pdk1 | pyruvate dehydrogenase kinase, isoenzy | 1397 | 1200 | 1265 | 1520 | 1522 | 1209 | 1665 | 1382 | 1301 | 1345 | 1248 |
| 1562 | -0.12 | 0.12 | -1.06 | 0.29 | 1 | Pptc7 | PTC7 protein phosphatase homolog | 1428 | 1730 | 1069 | 2050 | 1529 | 1532 | 1664 | 1756 | 1554 | 1456 | 1410 |
| 1526 | 0.11 | 0.09 | 1.16 | 0.25 | 1 | Syn2 | synapsin II | 692 | 606 | 823 | 1553 | 2050 | 1803 | 1664 | 1657 | 2046 | 1960 | 1929 |
| 1673 | 0.11 | 0.14 | 0.79 | 0.43 | 1 | Imp3 | IMP3, U3 small nucleolar ribonucleoprote | 1677 | 1822 | 2097 | 1455 | 1762 | 1265 | 1663 | 1776 | 1501 | 2097 | 1290 |
| 1313 | -0.29 | 0.20 | -1.45 | 0.15 | 1 | Rab3b | RAB3B, member RAS oncogene family | 1148 | 1658 | 1685 | 1198 | 1364 | 1222 | 1663 | 941 | 1135 | 1546 | 885 |
| 1877 | -0.06 | 0.15 | -0.41 | 0.69 | 1 | Zcchc2 | zinc finger, CCHC domain containing 2 | 1159 | 913 | 837 | 2189 | 1715 | 3510 | 1662 | 1622 | 1971 | 1771 | 3295 |
| 2234 | 0.09 | 0.10 | 0.86 | 0.39 | 1 | Btf3 | basic transcription factor 3 | 3689 | 3147 | 3618 | 1781 | 1826 | 1568 | 1658 | 2027 | 1914 | 1829 | 1512 |
| 2214 | 0.30 | 0.18 | 1.67 | 0.09 | 1 | Rpl10a | ribosomal protein L10A | 3481 | 2840 | 3922 | 1414 | 1512 | 1727 | 1658 | 2555 | 1487 | 1923 | 1829 |
| 1641 | 0.08 | 0.16 | 0.48 | 0.63 | 1 | Igsf8 | immunoglobulin superfamily, member 8 | 2156 | 1591 | 1578 | 1610 | 1645 | 1253 | 1657 | 2160 | 1759 | 1426 | 1215 |
| 1624 | -0.10 | 0.14 | -0.70 | 0.48 | 1 | Cdv3 | carnitine deficiency-associated gene expr | 1911 | 1985 | 1487 | 1534 | 1548 | 1713 | 1657 | 1243 | 1660 | 1407 | 1716 |
| 1446 | 0.08 | 0.18 | 0.45 | 0.66 | 1 | Fras1 | Fraser extracellular matrix complex subur | 702 | 327 | 424 | 1550 | 1600 | 2214 | 1657 | 1490 | 1910 | 1712 | 2318 |
| 1295 | -0.05 | 0.14 | -0.37 | 0.71 | 1 | Foxred2 | FAD-dependent oxidoreductase domain c | 465 | 460 | 328 | 1571 | 1627 | 1756 | 1657 | 1644 | 1640 | 1450 | 1645 |
| 1643 | -0.11 | 0.10 | -1.04 | 0.30 | 1 | Fam172a | family with sequence similarity 172, meml | 2447 | 2288 | 2144 | 1532 | 1464 | 1161 | 1655 | 1250 | 1452 | 1486 | 1190 |
| 1172 | 0.15 | 0.12 | 1.32 | 0.19 | 1 | Phyhip | phytanoyl-CoA hydroxylase interacting pr | 359 | 392 | 391 | 1427 | 1501 | 939 | 1654 | 1713 | 1791 | 1814 | 913 |
| 1625 | 0.12 | 0.10 | 1.17 | 0.24 | 1 | Fbxl14 | F-box and leucine-rich repeat protein 14 | 1702 | 1535 | 1312 | 1642 | 1603 | 1455 | 1653 | 1922 | 1655 | 1998 | 1399 |
| 1542 | -0.03 | 0.13 | -0.25 | 0.80 | 1 | Tmem178b | transmembrane protein 178B | 1113 | 1410 | 1592 | 1725 | 1478 | 1614 | 1653 | 1658 | 1837 | 1428 | 1451 |
| 1732 | -0.01 | 0.19 | -0.04 | 0.97 | 1 | Vps18 | VPS18 CORVET/HOPS core subunit | 1569 | 1183 | 1253 | 2397 | 2045 | 1358 | 1652 | 2966 | 1736 | 1667 | 1225 |
| 1713 | 0.14 | 0.10 | 1.44 | 0.15 | 1 | Sephs2 | selenophosphate synthetase 2 | 1085 | 1275 | 1187 | 2161 | 2004 | 1455 | 1652 | 2579 | 2040 | 1756 | 1646 |
| 1562 | 0.06 | 0.16 | 0.36 | 0.72 | 1 | Kbtbd7 | kelch repeat and BTB (POZ) domain conta | 1282 | 1994 | 1716 | 1625 | 1346 | 1347 | 1652 | 1520 | 1573 | 1845 | 1283 |

| | | | | | | | | | | | | | | | | | | |
|---|---|---|---|---|---|---|---|---|---|---|---|---|---|---|---|---|---|---|
| 1639 | 0.00 | 0.19 | 0.02 | 0.98 | 1 | Nup93 | nucleoporin 93 | 758 | 344 | 515 | 1905 | 1917 | 2739 | 1651 | 1913 | 1817 | 2047 | 2423 |
| 1887 | -0.08 | 0.17 | -0.47 | 0.63 | 1 | Rbm26 | RNA binding motif protein 26 | 1392 | 1472 | 1122 | 1747 | 1668 | 3545 | 1650 | 1142 | 1936 | 1758 | 3330 |
| 1806 | 0.10 | 0.10 | 1.01 | 0.31 | 1 | Smg6 | Smg-6 homolog, nonsense mediated mR | 2003 | 2114 | 2050 | 1629 | 1618 | 1697 | 1650 | 1698 | 1826 | 1994 | 1585 |
| 1687 | 0.05 | 0.10 | 0.54 | 0.59 | 1 | Pik3c3 | phosphatidylinositol 3-kinase catalytic sul | 1434 | 1581 | 1538 | 1864 | 1882 | 1441 | 1650 | 2233 | 1850 | 1734 | 1347 |
| 1658 | 0.20 | 0.13 | 1.56 | 0.12 | 1 | Jtb | jumping translocation breakpoint | 1225 | 1150 | 1357 | 1836 | 1698 | 1541 | 1650 | 2472 | 1833 | 1754 | 1720 |
| 1360 | 0.05 | 0.15 | 0.34 | 0.73 | 1 | Akr1c13 | aldo-keto reductase family 1, member C1 | 701 | 601 | 788 | 1710 | 1790 | 1199 | 1650 | 1964 | 1693 | 1409 | 1456 |
| 1792 | -0.03 | 0.10 | -0.32 | 0.75 | 1 | Zc3h4 | zinc finger CCCH-type containing 4 | 1953 | 1926 | 1595 | 1862 | 1604 | 2094 | 1649 | 1691 | 1704 | 1693 | 1936 |
| 1625 | 0.26 | 0.14 | 1.90 | 0.06 | 1 | Smap1 | small ArfGAP 1 | 1349 | 1442 | 1233 | 1280 | 1737 | 1624 | 1649 | 2102 | 1814 | 1930 | 1720 |
| 1548 | -0.02 | 0.11 | -0.15 | 0.88 | 1 | Cox5b | cytochrome c oxidase subunit 5B | 1436 | 1477 | 1829 | 1578 | 1703 | 1248 | 1649 | 1708 | 1596 | 1605 | 1204 |
| 1734 | -0.07 | 0.10 | -0.77 | 0.44 | 1 | Marveld1 | MARVEL (membrane-associating) domai | 3089 | 2487 | 2501 | 1436 | 1593 | 959 | 1648 | 1437 | 1302 | 1689 | 933 |
| 1602 | -0.03 | 0.09 | -0.32 | 0.75 | 1 | Epg5 | ectopic P-granules autophagy protein 5 h | 1416 | 1276 | 1418 | 1590 | 1813 | 1767 | 1648 | 1501 | 1824 | 1771 | 1597 |
| 1373 | 0.16 | 0.11 | 1.40 | 0.16 | 1 | Tmem9b | TMEM9 domain family, member B | 986 | 963 | 1116 | 1349 | 1466 | 1226 | 1648 | 1557 | 1579 | 1901 | 1316 |
| 1384 | -0.10 | 0.09 | -1.08 | 0.28 | 1 | Gm47163 | | 808 | 981 | 850 | 2045 | 1647 | 1167 | 1647 | 1818 | 1602 | 1552 | 1102 |
| 2282 | -0.03 | 0.14 | -0.18 | 0.86 | 1 | Zim1 | zinc finger, imprinted 1 | 2070 | 3243 | 1938 | 2072 | 1201 | 4223 | 1646 | 1651 | 1590 | 1556 | 3906 |
| 1694 | 0.18 | 0.09 | 1.95 | 0.05 | 1 | Cyth1 | cytohesin 1 | 1697 | 1405 | 1709 | 1460 | 1563 | 1817 | 1645 | 1761 | 1720 | 1870 | 1987 |
| 1515 | -0.01 | 0.11 | -0.09 | 0.93 | 1 | Mpp5 | membrane protein, palmitoylated 5 (MAG | 1397 | 1477 | 1277 | 1893 | 1308 | 1461 | 1645 | 1598 | 1561 | 1573 | 1480 |
| 1519 | -0.03 | 0.14 | -0.18 | 0.85 | 1 | Rnf123 | ring finger protein 123 | 1337 | 1229 | 1043 | 1476 | 1671 | 1809 | 1644 | 1280 | 1850 | 1789 | 1585 |
| 1982 | 0.09 | 0.10 | 0.93 | 0.35 | 1 | Snx6 | sorting nexin 6 | 2300 | 2405 | 2268 | 1848 | 1814 | 1868 | 1643 | 2237 | 1799 | 1808 | 1809 |
| 1508 | -0.16 | 0.13 | -1.18 | 0.24 | 1 | Npnt | nephronectin | 594 | 720 | 865 | 2379 | 1749 | 1837 | 1642 | 1831 | 1793 | 1550 | 1625 |
| 1982 | 0.05 | 0.11 | 0.52 | 0.60 | 1 | Fam120b | family with sequence similarity 120, meml | 1621 | 1908 | 1773 | 1548 | 1959 | 2933 | 1641 | 1605 | 2015 | 1690 | 3109 |
| 1688 | -0.06 | 0.13 | -0.47 | 0.64 | 1 | Wdr48 | WD repeat domain 48 | 1730 | 1709 | 1412 | 1771 | 1559 | 2024 | 1641 | 1302 | 1690 | 1687 | 2045 |
| 1554 | 0.01 | 0.15 | 0.06 | 0.95 | 1 | Cox14 | cytochrome c oxidase assembly protein 1 | 1636 | 1401 | 2155 | 1516 | 1497 | 1282 | 1641 | 1674 | 1374 | 1605 | 1308 |
| 1544 | 0.13 | 0.10 | 1.26 | 0.21 | 1 | Siae | sialic acid acetylesterase | 1481 | 1395 | 1537 | 1684 | 1443 | 1230 | 1641 | 1881 | 1690 | 1684 | 1317 |
| 1718 | 0.16 | 0.12 | 1.31 | 0.19 | 1 | Commd7 | COMM domain containing 7 | 1466 | 1962 | 1844 | 1550 | 1665 | 1578 | 1640 | 1949 | 1727 | 1882 | 1632 |
| 1634 | -0.05 | 0.13 | -0.40 | 0.69 | 1 | Mtrex | Mtr4 exosome RNA helicase | 1798 | 1774 | 2030 | 1399 | 1363 | 1863 | 1640 | 1109 | 1653 | 1432 | 1915 |
| 1537 | -0.14 | 0.14 | -0.97 | 0.33 | 1 | Carm1 | coactivator-associated arginine methyltra | 2026 | 1512 | 2042 | 1422 | 1463 | 1332 | 1639 | 975 | 1640 | 1761 | 1100 |
| 1838 | 0.12 | 0.11 | 1.10 | 0.27 | 1 | Smg5 | Smg-5 homolog, nonsense mediated mR | 1722 | 1648 | 1758 | 1926 | 1559 | 2132 | 1638 | 2141 | 1771 | 1788 | 2136 |
| 1545 | -0.07 | 0.09 | -0.70 | 0.48 | 1 | Cyc1 | cytochrome c-1 | 1506 | 1324 | 1471 | 1456 | 1677 | 1713 | 1637 | 1306 | 1759 | 1546 | 1597 |
| 1666 | -0.12 | 0.16 | -0.72 | 0.47 | 1 | Crbn | cereblon | 1748 | 1572 | 1319 | 2007 | 1902 | 1565 | 1636 | 1307 | 1715 | 1694 | 1858 |
| 1637 | -0.06 | 0.11 | -0.54 | 0.59 | 1 | Smad2 | SMAD family member 2 | 1818 | 1658 | 1965 | 1591 | 1597 | 1574 | 1636 | 1320 | 1466 | 1727 | 1659 |
| 1198 | -0.06 | 0.14 | -0.45 | 0.65 | 1 | Slc25a16 | solute carrier family 25 (mitochondrial car | 583 | 688 | 749 | 1520 | 1475 | 1081 | 1636 | 1462 | 1540 | 1220 | 1227 |
| 1629 | -0.03 | 0.13 | -0.22 | 0.82 | 1 | Mdc1 | mediator of DNA damage checkpoint 1 | 1271 | 1290 | 1243 | 1869 | 1565 | 2046 | 1635 | 1965 | 1808 | 1318 | 1905 |
| 2006 | -0.13 | 0.14 | -0.94 | 0.35 | 1 | Csk | c-src tyrosine kinase | 3519 | 3149 | 2822 | 1780 | 1774 | 1361 | 1633 | 1207 | 1737 | 1872 | 1218 |
| 1744 | -0.07 | 0.12 | -0.57 | 0.57 | 1 | Pex5 | peroxisomal biogenesis factor 5 | 1609 | 2011 | 1950 | 1941 | 1603 | 1773 | 1632 | 2030 | 1508 | 1695 | 1434 |
| 1311 | 0.09 | 0.14 | 0.63 | 0.53 | 1 | Prkdc | protein kinase, DNA activated, catalytic pc | 768 | 865 | 594 | 1228 | 1303 | 1759 | 1632 | 1481 | 1506 | 1477 | 1806 |
| 1812 | -0.02 | 0.10 | -0.16 | 0.87 | 1 | Tcf12 | transcription factor 12 | 2493 | 2195 | 2239 | 1434 | 1483 | 1975 | 1631 | 1391 | 1573 | 1404 | 2117 |
| 1580 | 0.21 | 0.13 | 1.68 | 0.09 | 1 | Wdfy1 | WD repeat and FYVE domain containing | 1574 | 1069 | 1533 | 1082 | 1922 | 1522 | 1631 | 1548 | 1885 | 2103 | 1513 |
| 1408 | 0.14 | 0.13 | 1.07 | 0.28 | 1 | Clpb | ClpB caseinolytic peptidase B | 1212 | 1440 | 1281 | 1357 | 1339 | 1136 | 1631 | 1775 | 1699 | 1530 | 1087 |
| 1831 | -0.13 | 0.12 | -1.05 | 0.30 | 1 | Unc79 | unc-79 homolog | 1929 | 2172 | 1976 | 1392 | 1676 | 2638 | 1630 | 1181 | 1625 | 1512 | 2409 |
| 1549 | 0.04 | 0.10 | 0.39 | 0.70 | 1 | Trp53bp2 | transformation related protein 53 binding | 1418 | 1602 | 1654 | 1581 | 1477 | 1404 | 1630 | 1675 | 1531 | 1691 | 1379 |
| 1496 | 0.11 | 0.11 | 1.00 | 0.32 | 1 | Pla2g12a | phospholipase A2, group XIIA | 1366 | 1437 | 1670 | 1452 | 1654 | 1038 | 1630 | 1658 | 1647 | 1790 | 1113 |
| 1939 | 0.20 | 0.16 | 1.25 | 0.21 | 1 | Rbm8a | RNA binding motif protein 8a | 2072 | 2332 | 2525 | 1579 | 1739 | 1717 | 1629 | 2637 | 1592 | 1684 | 1819 |
| 1590 | -0.09 | 0.12 | -0.73 | 0.47 | 1 | Med24 | mediator complex subunit 24 | 1531 | 1337 | 1597 | 1552 | 1751 | 1778 | 1629 | 1759 | 1534 | 1504 | 1519 |
| 1478 | 0.08 | 0.15 | 0.55 | 0.58 | 1 | Ppp1r16a | protein phosphatase 1, regulatory subuni | 1526 | 1225 | 1301 | 1321 | 1524 | 1458 | 1629 | 1929 | 1460 | 1465 | 1426 |
| 1474 | -0.08 | 0.11 | -0.74 | 0.46 | 1 | Ogfod1 | 2-oxoglutarate and iron-dependent oxyge | 1292 | 1174 | 1397 | 1411 | 1717 | 1572 | 1629 | 1149 | 1668 | 1729 | 1481 |
| 1282 | 0.00 | 0.09 | 0.00 | 1.00 | 1 | Klhl2 | kelch-like 2, Mayven | 1023 | 852 | 1008 | 1325 | 1435 | 1227 | 1629 | 1337 | 1506 | 1515 | 1246 |
| 1689 | -0.21 | 0.14 | -1.51 | 0.13 | 1 | Leprot | leptin receptor overlapping transcript | 3034 | 2210 | 2274 | 1609 | 1404 | 1290 | 1628 | 1109 | 1384 | 1305 | 1332 |
| 1663 | -0.03 | 0.13 | -0.21 | 0.84 | 1 | Tfip11 | tuftelin interacting protein 11 | 1738 | 2066 | 1726 | 1675 | 1628 | 1466 | 1628 | 1615 | 1889 | 1709 | 1148 |
| 1513 | -0.09 | 0.11 | -0.85 | 0.40 | 1 | Mrpl17 | mitochondrial ribosomal protein L17 | 1408 | 1518 | 1677 | 1725 | 1746 | 1149 | 1628 | 1492 | 1606 | 1378 | 1318 |
| 1090 | -0.15 | 0.17 | -0.91 | 0.37 | 1 | Gnpda1 | glucosamine-6-phosphate deaminase 1 | 468 | 391 | 550 | 1551 | 1276 | 1095 | 1628 | 994 | 1431 | 1495 | 1108 |
| 1426 | -0.08 | 0.15 | -0.49 | 0.62 | 1 | Lztfl1 | leucine zipper transcription factor-like 1 | 1005 | 1006 | 1006 | 1288 | 1384 | 2154 | 1626 | 918 | 1481 | 1681 | 2142 |
| 1314 | 0.08 | 0.15 | 0.56 | 0.58 | 1 | Nxt2 | nuclear transport factor 2-like export facto | 787 | 879 | 1104 | 1528 | 1466 | 1055 | 1626 | 1328 | 1647 | 1841 | 1198 |
| 1594 | -0.37 | 0.30 | -1.24 | 0.21 | 1 | Rasgrf2 | RAS protein-specific guanine nucleotide- | 2515 | 4338 | 3106 | 742 | 1272 | 454 | 1625 | 601 | 928 | 1713 | 243 |
| 1749 | 0.08 | 0.12 | 0.72 | 0.47 | 1 | Sptlc2 | serine palmitoyltransferase, long chain ba | 2183 | 2072 | 2012 | 1385 | 1655 | 1629 | 1624 | 1840 | 1437 | 1698 | 1706 |
| 1573 | -0.08 | 0.17 | -0.47 | 0.64 | 1 | Scyl2 | SCY1-like 2 (S. cerevisiae) | 1235 | 1980 | 1479 | 1779 | 1362 | 1722 | 1624 | 1288 | 1787 | 1369 | 1673 |
| 2642 | 0.02 | 0.10 | 0.23 | 0.82 | 1 | Sdc2 | syndecan 2 | 5274 | 5527 | 5651 | 1901 | 1797 | 944 | 1623 | 1730 | 1689 | 1953 | 975 |
| 1796 | 0.10 | 0.11 | 0.89 | 0.37 | 1 | Nap1l5 | nucleosome assembly protein 1-like 5 | 1778 | 1702 | 1512 | 1912 | 1209 | 2350 | 1623 | 1961 | 1282 | 1694 | 2738 |
| 1692 | 0.06 | 0.21 | 0.28 | 0.78 | 1 | Sec16b | SEC16 homolog B (S. cerevisiae) | 353 | 401 | 168 | 2227 | 1715 | 3132 | 1622 | 1647 | 1917 | 2109 | 3325 |
| 1353 | 0.10 | 0.19 | 0.49 | 0.62 | 1 | Rassf8 | Ras association (RalGDS/AF-6) domain fa | 1593 | 777 | 1250 | 1318 | 1301 | 1235 | 1622 | 1585 | 1575 | 1297 | 1325 |
| 2045 | 0.09 | 0.12 | 0.69 | 0.49 | 1 | Tra2b | transformer 2 beta | 2262 | 2515 | 1785 | 1620 | 1519 | 2982 | 1621 | 1488 | 1677 | 1923 | 3102 |

| | | | | | | | | | | | | | | | | | | |
|---|---|---|---|---|---|---|---|---|---|---|---|---|---|---|---|---|---|---|
| 1871 | 0.14 | 0.11 | 1.27 | 0.20 | 1 | Blmh | bleomycin hydrolase | 2406 | 2044 | 2119 | 1806 | 1660 | 1567 | 1621 | 2113 | 1632 | 1887 | 1722 |
| 1479 | 0.08 | 0.14 | 0.55 | 0.59 | 1 | Gpx4 | glutathione peroxidase 4 | 1518 | 1604 | 1501 | 1273 | 1620 | 1140 | 1621 | 1866 | 1492 | 1517 | 1119 |
| 1443 | 0.15 | 0.14 | 1.05 | 0.29 | 1 | Crot | carnitine O-octanoyltransferase | 1097 | 829 | 1172 | 1646 | 1368 | 1410 | 1621 | 1559 | 1546 | 1937 | 1690 |
| 1914 | 0.03 | 0.11 | 0.24 | 0.81 | 1 | Lrp3 | low density lipoprotein receptor-related pı | 2173 | 2602 | 2119 | 2084 | 1657 | 1641 | 1620 | 1979 | 1978 | 1837 | 1371 |
| 1692 | -0.13 | 0.16 | -0.85 | 0.39 | 1 | Cacna1a | calcium channel, voltage-dependent, P/Q | 686 | 1251 | 865 | 1733 | 1344 | 3680 | 1620 | 1530 | 1331 | 1565 | 3005 |
| 1647 | 0.17 | 0.11 | 1.49 | 0.14 | 1 | Rgmb | repulsive guidance molecule family meml | 1440 | 1508 | 1412 | 1809 | 1640 | 1383 | 1620 | 2411 | 1733 | 1695 | 1467 |
| 1647 | -0.04 | 0.14 | -0.31 | 0.76 | 1 | Orc3 | origin recognition complex, subunit 3 | 1543 | 1858 | 1327 | 1528 | 1783 | 1854 | 1620 | 1372 | 1686 | 1583 | 1964 |
| 1998 | 0.04 | 0.13 | 0.28 | 0.78 | 1 | Nfxl1 | nuclear transcription factor, X-box binding | 1239 | 932 | 1291 | 1241 | 2235 | 4106 | 1617 | 1380 | 2506 | 1459 | 3975 |
| 1631 | -0.14 | 0.12 | -1.12 | 0.26 | 1 | Als2 | alsin Rho guanine nucleotide exchange f | 1561 | 2120 | 1785 | 1813 | 1415 | 1700 | 1617 | 1457 | 1593 | 1410 | 1472 |
| 1592 | -0.09 | 0.14 | -0.68 | 0.49 | 1 | Ube3b | ubiquitin protein ligase E3B | 2115 | 1495 | 1553 | 1595 | 1601 | 1556 | 1617 | 1637 | 1569 | 1456 | 1316 |
| 1392 | 0.00 | 0.10 | 0.05 | 0.96 | 1 | Shc2 | SHC (Src homology 2 domain containing) | 1205 | 1030 | 1227 | 1370 | 1416 | 1518 | 1617 | 1457 | 1489 | 1568 | 1420 |
| 1328 | -0.07 | 0.17 | -0.38 | 0.70 | 1 | Tmod1 | tropomodulin 1 | 519 | 306 | 402 | 1897 | 1624 | 1703 | 1617 | 1639 | 1752 | 1426 | 1720 |
| 1515 | 0.08 | 0.11 | 0.70 | 0.48 | 1 | Ankrd28 | ankyrin repeat domain 28 | 1588 | 1464 | 1163 | 1484 | 1453 | 1512 | 1616 | 1563 | 1609 | 1629 | 1582 |
| 1586 | -0.25 | 0.14 | -1.75 | 0.08 | 1 | Polr1a | polymerase (RNA) I polypeptide A | 1724 | 1497 | 1815 | 1438 | 2140 | 1515 | 1615 | 935 | 1765 | 1434 | 1571 |
| 1415 | 0.06 | 0.16 | 0.37 | 0.71 | 1 | Cep290 | centrosomal protein 290 | 486 | 654 | 563 | 1607 | 1480 | 2005 | 1615 | 1362 | 1989 | 1334 | 2465 |
| 1585 | -0.03 | 0.12 | -0.25 | 0.80 | 1 | Zfp598 | zinc finger protein 598 | 1404 | 937 | 1553 | 1249 | 1748 | 2291 | 1614 | 1441 | 1636 | 1543 | 2025 |
| 1639 | 0.17 | 0.15 | 1.12 | 0.26 | 1 | Cetn2 | centrin 2 | 1686 | 1843 | 2105 | 1482 | 1587 | 1124 | 1612 | 2243 | 1478 | 1672 | 1196 |
| 1817 | -0.07 | 0.14 | -0.54 | 0.59 | 1 | Ranbp9 | RAN binding protein 9 | 1878 | 2009 | 1633 | 2241 | 1692 | 1890 | 1610 | 1537 | 1762 | 1727 | 2006 |
| 1587 | -0.04 | 0.10 | -0.42 | 0.67 | 1 | Abcc1 | ATP-binding cassette, sub-family C (CFTF | 1684 | 1460 | 1389 | 1744 | 1736 | 1462 | 1610 | 1642 | 1662 | 1696 | 1372 |
| 1475 | -0.02 | 0.13 | -0.13 | 0.90 | 1 | Manba | mannosidase, beta A, lysosomal | 1721 | 1547 | 1456 | 1746 | 1267 | 1155 | 1610 | 1892 | 1371 | 1363 | 1100 |
| 1580 | 0.26 | 0.20 | 1.30 | 0.19 | 1 | Ndufa2 | NADH:ubiquinone oxidoreductase subun | 1514 | 1327 | 1789 | 1394 | 1714 | 1061 | 1609 | 2602 | 1437 | 1739 | 1194 |
| 1960 | 0.13 | 0.12 | 1.04 | 0.30 | 1 | Lzts2 | leucine zipper, putative tumor suppressor | 2098 | 2314 | 2283 | 2115 | 1565 | 1783 | 1608 | 2615 | 1652 | 1792 | 1731 |
| 1511 | -0.05 | 0.12 | -0.45 | 0.65 | 1 | Wdr35 | WD repeat domain 35 | 1221 | 1503 | 1297 | 1722 | 1416 | 1653 | 1608 | 1630 | 1525 | 1230 | 1819 |
| 1935 | 0.02 | 0.09 | 0.16 | 0.87 | 1 | Idh3b | isocitrate dehydrogenase 3 (NAD+) beta | 2262 | 2036 | 2206 | 1542 | 1738 | 2448 | 1607 | 1605 | 1506 | 1784 | 2547 |
| 1653 | 0.09 | 0.14 | 0.63 | 0.53 | 1 | Ccdc92 | coiled-coil domain containing 92 | 1127 | 1743 | 1564 | 1829 | 1814 | 1352 | 1606 | 2518 | 1598 | 1714 | 1319 |
| 1462 | -0.17 | 0.12 | -1.48 | 0.14 | 1 | Chpf2 | chondroitin polymerizing factor 2 | 2022 | 1312 | 1392 | 1443 | 1537 | 1421 | 1606 | 1243 | 1320 | 1564 | 1229 |
| 1464 | -0.12 | 0.12 | -0.99 | 0.32 | 1 | Dixdc1 | DIX domain containing 1 | 1436 | 1216 | 1394 | 1887 | 1496 | 1286 | 1605 | 1596 | 1431 | 1601 | 1157 |
| 1307 | -0.10 | 0.12 | -0.86 | 0.39 | 1 | Tk2 | thymidine kinase 2, mitochondrial | 916 | 1022 | 1074 | 1536 | 1553 | 1186 | 1605 | 1266 | 1491 | 1635 | 1098 |
| 1650 | -0.21 | 0.15 | -1.36 | 0.17 | 1 | Tmcc3 | transmembrane and coiled coil domains 3 | 2316 | 1983 | 1882 | 1812 | 1540 | 1454 | 1603 | 1457 | 1620 | 1101 | 1377 |
| 1623 | -0.23 | 0.19 | -1.21 | 0.22 | 1 | A330076H08Rik | | 455 | 594 | 647 | 1859 | 1335 | 3800 | 1603 | 1018 | 1794 | 1234 | 3519 |
| 1527 | 0.05 | 0.13 | 0.42 | 0.68 | 1 | Spata13 | spermatogenesis associated 13 | 2202 | 1946 | 2054 | 1108 | 1506 | 941 | 1603 | 1408 | 1670 | 1564 | 793 |
| 2033 | 0.20 | 0.13 | 1.53 | 0.13 | 1 | Use1 | unconventional SNARE in the ER 1 homc | 3155 | 2883 | 3126 | 1427 | 1645 | 1439 | 1602 | 2191 | 1554 | 1868 | 1471 |
| 1735 | 0.02 | 0.10 | 0.20 | 0.84 | 1 | Mmadhc | methylmalonic aciduria (cobalamin deficie | 1779 | 1716 | 1878 | 1685 | 1871 | 1648 | 1602 | 1759 | 1800 | 1479 | 1872 |
| 1560 | 0.10 | 0.12 | 0.91 | 0.36 | 1 | Coasy | Coenzyme A synthase | 1359 | 1087 | 1252 | 1518 | 1833 | 1526 | 1602 | 1865 | 1745 | 1892 | 1476 |
| 1462 | -0.04 | 0.12 | -0.31 | 0.76 | 1 | Suclg2 | succinate-Coenzyme A ligase, GDP-formi | 1662 | 1649 | 1872 | 1523 | 1302 | 1089 | 1602 | 1281 | 1459 | 1586 | 1058 |
| 1402 | -0.04 | 0.10 | -0.43 | 0.67 | 1 | BC029722 | | 1226 | 1309 | 1469 | 1483 | 1643 | 1090 | 1602 | 1460 | 1486 | 1493 | 1165 |
| 1371 | 0.00 | 0.13 | -0.02 | 0.98 | 1 | Tceal3 | transcription elongation factor A (SII)-like | 860 | 957 | 1003 | 1616 | 1378 | 1530 | 1602 | 1712 | 1361 | 1749 | 1319 |
| 1779 | 0.07 | 0.10 | 0.73 | 0.46 | 1 | Mysm1 | myb-like, SWIRM and MPN domains 1 | 1119 | 1154 | 1045 | 1518 | 1530 | 3272 | 1599 | 1577 | 1787 | 1520 | 3443 |
| 1666 | -0.03 | 0.10 | -0.32 | 0.75 | 1 | Tjp2 | tight junction protein 2 | 1436 | 1545 | 1614 | 1919 | 1845 | 1563 | 1599 | 2023 | 1856 | 1425 | 1502 |
| 1653 | 0.02 | 0.16 | 0.10 | 0.92 | 1 | Matn2 | matrilin 2 | 4531 | 2015 | 2070 | 1085 | 1143 | 931 | 1599 | 1318 | 926 | 1573 | 995 |
| 1612 | -0.23 | 0.14 | -1.65 | 0.10 | 1 | Ncoa5 | nuclear receptor coactivator 5 | 1627 | 1822 | 1809 | 1676 | 1505 | 1929 | 1599 | 1103 | 1323 | 1622 | 1717 |
| 1608 | 0.17 | 0.10 | 1.78 | 0.08 | 1 | Ube2d2a | ubiquitin-conjugating enzyme E2D 2A | 1351 | 1281 | 1414 | 1745 | 1540 | 1520 | 1599 | 1922 | 1732 | 1905 | 1673 |
| 1685 | -0.06 | 0.12 | -0.49 | 0.62 | 1 | Lgals8 | lectin, galactose binding, soluble 8 | 1999 | 1913 | 1829 | 1578 | 1499 | 1834 | 1598 | 1223 | 1667 | 1592 | 1796 |
| 1557 | 0.06 | 0.12 | 0.51 | 0.61 | 1 | Med16 | mediator complex subunit 16 | 1348 | 1168 | 1357 | 1750 | 1822 | 1287 | 1598 | 2116 | 1704 | 1743 | 1233 |
| 1788 | 0.07 | 0.16 | 0.46 | 0.65 | 1 | Ckap4 | cytoskeleton-associated protein 4 | 2586 | 1811 | 1544 | 1639 | 2003 | 1434 | 1597 | 1901 | 1798 | 1863 | 1489 |
| 1684 | -0.01 | 0.13 | -0.06 | 0.95 | 1 | Pogk | pogo transposable element with KRAB dc | 1795 | 1659 | 1428 | 2016 | 1532 | 1707 | 1597 | 1685 | 1846 | 1711 | 1551 |
| 1455 | 0.13 | 0.10 | 1.23 | 0.22 | 1 | Ankmy2 | ankyrin repeat and MYND domain contair | 1524 | 1377 | 1300 | 1331 | 1448 | 1261 | 1596 | 1571 | 1710 | 1622 | 1267 |
| 1625 | -0.02 | 0.11 | -0.15 | 0.88 | 1 | Pes1 | pescadillo ribosomal biogenesis factor 1 | 1863 | 1608 | 1707 | 1529 | 1783 | 1466 | 1595 | 1302 | 1864 | 1596 | 1560 |
| 1581 | 0.02 | 0.13 | 0.17 | 0.87 | 1 | BC005624 | cDNA sequence BC005624 | 1823 | 1335 | 1599 | 1486 | 1464 | 1733 | 1595 | 1707 | 1600 | 1410 | 1636 |
| 1464 | 0.02 | 0.10 | 0.17 | 0.87 | 1 | Qpctl | glutaminyl-peptide cyclotransferase-like | 881 | 889 | 1054 | 1701 | 1803 | 1508 | 1595 | 1785 | 1685 | 1691 | 1514 |
| 1758 | -0.07 | 0.10 | -0.70 | 0.49 | 1 | Atp9b | ATPase, class II, type 9B | 2000 | 1674 | 1902 | 1736 | 1589 | 2125 | 1594 | 1442 | 1672 | 1559 | 2044 |
| 1331 | -0.14 | 0.17 | -0.78 | 0.43 | 1 | Ttc8 | tetratricopeptide repeat domain 8 | 1120 | 1282 | 1173 | 1334 | 1315 | 1512 | 1594 | 879 | 1250 | 1673 | 1506 |
| 1235 | -0.16 | 0.11 | -1.44 | 0.15 | 1 | Usp6nl | USP6 N-terminal like | 1089 | 818 | 802 | 1294 | 1428 | 1434 | 1594 | 1197 | 1362 | 1349 | 1212 |
| 1206 | 0.32 | 0.18 | 1.76 | 0.08 | 1 | Rgs7bp | regulator of G-protein signalling 7 binding | 1444 | 1310 | 860 | 1065 | 1031 | 673 | 1594 | 1431 | 1603 | 1377 | 882 |
| 1650 | -0.07 | 0.11 | -0.68 | 0.50 | 1 | Dync1li1 | dynein cytoplasmic 1 light intermediate ch | 1898 | 2074 | 1874 | 1595 | 1599 | 1510 | 1593 | 1636 | 1624 | 1271 | 1477 |
| 1713 | 0.12 | 0.12 | 0.98 | 0.33 | 1 | Vps26a | VPS26 retromer complex component A | 1969 | 1577 | 1795 | 1838 | 1505 | 1531 | 1592 | 2023 | 1660 | 1736 | 1618 |
| 1624 | 0.05 | 0.13 | 0.41 | 0.68 | 1 | Zmym5 | zinc finger, MYM-type 5 | 1271 | 1609 | 1628 | 1387 | 1373 | 2188 | 1592 | 1367 | 1699 | 1453 | 2299 |
| 1442 | 0.01 | 0.13 | 0.08 | 0.94 | 1 | Wdr81 | WD repeat domain 81 | 1220 | 1142 | 1387 | 1435 | 1428 | 1546 | 1592 | 1214 | 1614 | 1891 | 1391 |
| 1360 | 0.02 | 0.17 | 0.12 | 0.90 | 1 | Hlcs | holocarboxylase synthetase (biotin- [prop | 1058 | 760 | 818 | 1651 | 1567 | 1266 | 1592 | 2087 | 1624 | 1447 | 1092 |
| 1798 | -0.01 | 0.11 | -0.10 | 0.92 | 1 | Pdlim5 | PDZ and LIM domain 5 | 2742 | 2471 | 2317 | 1518 | 1520 | 1511 | 1591 | 1694 | 1631 | 1407 | 1372 |

| | | | | | | | | | | | | | | | | | | |
|---|---|---|---|---|---|---|---|---|---|---|---|---|---|---|---|---|---|---|
| 1556 | 0.07 | 0.11 | 0.67 | 0.51 | 1 | Psmd14 | proteasome (prosome, macropain) 26S su | 1910 | 1513 | 1822 | 1239 | 1620 | 1327 | 1591 | 1395 | 1410 | 1900 | 1386 |
| 1650 | -0.07 | 0.09 | -0.72 | 0.47 | 1 | Ikbkg | inhibitor of kappaB kinase gamma | 1395 | 1651 | 1449 | 1707 | 1744 | 1935 | 1590 | 1682 | 1505 | 1560 | 1931 |
| 1494 | 0.11 | 0.14 | 0.79 | 0.43 | 1 | Mocs2 | molybdenum cofactor synthesis 2 | 946 | 1073 | 1350 | 1485 | 1552 | 1631 | 1590 | 2008 | 1581 | 1474 | 1740 |
| 1438 | 0.02 | 0.14 | 0.16 | 0.88 | 1 | Arhgap44 | Rho GTPase activating protein 44 | 1196 | 990 | 1554 | 1583 | 1669 | 1092 | 1590 | 1500 | 1689 | 1993 | 963 |
| 1667 | -0.01 | 0.11 | -0.10 | 0.92 | 1 | Washc4 | WASH complex subunit 4 | 1589 | 1537 | 1758 | 1624 | 1556 | 1974 | 1589 | 1660 | 1809 | 1345 | 1893 |
| 1665 | 0.09 | 0.10 | 0.82 | 0.41 | 1 | Bod1 | biorientation of chromosomes in cell divis | 1595 | 1657 | 1598 | 1491 | 1834 | 1614 | 1589 | 1505 | 1813 | 1785 | 1830 |
| 1486 | 0.03 | 0.12 | 0.22 | 0.83 | 1 | Nol8 | nucleolar protein 8 | 706 | 715 | 1110 | 1224 | 2013 | 2003 | 1589 | 1170 | 2011 | 1637 | 2171 |
| 1866 | -0.22 | 0.11 | -1.95 | 0.05 | 1 | Clip3 | CAP-GLY domain containing linker protei | 2748 | 2993 | 2745 | 1815 | 1789 | 1265 | 1588 | 1344 | 1747 | 1422 | 1065 |
| 1602 | -0.07 | 0.13 | -0.58 | 0.56 | 1 | 1810030O07Ril | RIKEN cDNA 1810030O07 gene | 1639 | 1584 | 1702 | 1660 | 1783 | 1461 | 1588 | 1204 | 1848 | 1608 | 1549 |
| 1468 | 0.02 | 0.11 | 0.22 | 0.83 | 1 | Ube2m | ubiquitin-conjugating enzyme E2M | 1358 | 1170 | 1180 | 1386 | 1778 | 1411 | 1588 | 1612 | 1667 | 1684 | 1312 |
| 1868 | 0.12 | 0.12 | 1.05 | 0.29 | 1 | Rufy1 | RUN and FYVE domain containing 1 | 1772 | 1824 | 2095 | 1883 | 1941 | 1696 | 1587 | 2360 | 1808 | 1793 | 1787 |
| 1730 | 0.06 | 0.11 | 0.52 | 0.60 | 1 | Ric8a | RIC8 guanine nucleotide exchange factor | 1937 | 1737 | 1891 | 1577 | 1934 | 1505 | 1587 | 1917 | 1706 | 1662 | 1574 |
| 1377 | -0.10 | 0.12 | -0.82 | 0.41 | 1 | Eif2b2 | eukaryotic translation initiation factor 2B, s | 1322 | 1361 | 1289 | 1560 | 1421 | 1220 | 1587 | 1490 | 1401 | 1182 | 1318 |
| 1481 | 0.09 | 0.10 | 0.88 | 0.38 | 1 | Pin1 | protein (peptidyl-prolyl cis/trans isomeras | 1362 | 1160 | 1410 | 1391 | 1601 | 1410 | 1586 | 1599 | 1585 | 1807 | 1385 |
| 2509 | 0.07 | 0.11 | 0.61 | 0.54 | 1 | Arglu1 | arginine and glutamate rich 1 | 1422 | 2037 | 1605 | 1735 | 1458 | 6362 | 1585 | 1995 | 1390 | 1770 | 6240 |
| 1673 | -0.07 | 0.12 | -0.57 | 0.57 | 1 | Plekha5 | pleckstrin homology domain containing, fa | 1376 | 1854 | 1462 | 1837 | 1448 | 2157 | 1585 | 1479 | 1623 | 1480 | 2106 |
| 1390 | 0.02 | 0.10 | 0.23 | 0.82 | 1 | Naglu | alpha-N-acetylglucosaminidase (Sanfilipp | 1464 | 1224 | 1253 | 1539 | 1530 | 978 | 1585 | 1519 | 1415 | 1740 | 1043 |
| 1670 | -0.04 | 0.10 | -0.37 | 0.71 | 1 | Zfp451 | zinc finger protein 451 | 1621 | 1833 | 1577 | 1627 | 1383 | 2150 | 1584 | 1422 | 1477 | 1486 | 2213 |
| 1919 | 0.17 | 0.11 | 1.58 | 0.11 | 1 | Eno1 | enolase 1, alpha non-neuron | 2215 | 2338 | 1757 | 1974 | 1708 | 1670 | 1583 | 2458 | 1690 | 1837 | 1877 |
| 1419 | 0.02 | 0.10 | 0.25 | 0.80 | 1 | Cstf2 | cleavage stimulation factor, 3' pre-RNA su | 1602 | 1311 | 1422 | 1136 | 1412 | 1463 | 1583 | 1261 | 1300 | 1645 | 1477 |
| 2281 | 0.01 | 0.12 | 0.04 | 0.97 | 1 | Safb2 | scaffold attachment factor B2 | 1740 | 1549 | 1596 | 1833 | 1693 | 5134 | 1581 | 2155 | 1668 | 1524 | 4618 |
| 1387 | 0.06 | 0.11 | 0.55 | 0.58 | 1 | Yipf1 | Yip1 domain family, member 1 | 1259 | 1100 | 1220 | 1455 | 1494 | 1188 | 1581 | 1712 | 1456 | 1541 | 1247 |
| 1873 | 0.07 | 0.12 | 0.55 | 0.58 | 1 | Gopc | golgi associated PDZ and coiled-coil moti | 1952 | 1898 | 1463 | 1721 | 1792 | 2375 | 1580 | 1726 | 1781 | 1824 | 2492 |
| 1701 | -0.13 | 0.14 | -0.89 | 0.37 | 1 | Rpl31 | ribosomal protein L31 | 2405 | 2529 | 2718 | 1317 | 1565 | 1282 | 1579 | 929 | 1682 | 1450 | 1252 |
| 1526 | -0.06 | 0.18 | -0.34 | 0.73 | 1 | Nbeal2 | neurobeachin-like 2 | 1798 | 1351 | 1858 | 1063 | 1200 | 2280 | 1579 | 1559 | 1135 | 1226 | 1734 |
| 1463 | -0.12 | 0.12 | -1.00 | 0.32 | 1 | Klhl42 | kelch-like 42 | 1295 | 1365 | 1493 | 1428 | 1625 | 1567 | 1579 | 1133 | 1589 | 1597 | 1419 |
| 1528 | -0.12 | 0.12 | -1.02 | 0.31 | 1 | Lratd2 | LRAT domain containing 1 | 1169 | 1604 | 1416 | 1848 | 1690 | 1465 | 1578 | 1444 | 1748 | 1448 | 1402 |
| 1444 | -0.06 | 0.13 | -0.43 | 0.67 | 1 | Lias | lipoic acid synthetase | 1259 | 1360 | 1612 | 1631 | 1464 | 1265 | 1578 | 1629 | 1420 | 1273 | 1392 |
| 1756 | 0.17 | 0.12 | 1.45 | 0.15 | 1 | Irf2bpl | interferon regulatory factor 2 binding prote | 2234 | 2287 | 2333 | 1409 | 1677 | 1142 | 1576 | 1774 | 1701 | 2098 | 1081 |
| 1445 | 0.14 | 0.13 | 1.08 | 0.28 | 1 | Timm17a | translocase of inner mitochondrial membr | 1196 | 1233 | 1451 | 1340 | 1476 | 1327 | 1576 | 1784 | 1631 | 1407 | 1473 |
| 1184 | 0.22 | 0.17 | 1.28 | 0.20 | 1 | Frmd5 | FERM domain containing 5 | 208 | 97 | 261 | 1204 | 1489 | 1453 | 1576 | 1367 | 1918 | 1899 | 1551 |
| 2051 | -0.03 | 0.11 | -0.24 | 0.81 | 1 | Crkl | v-crk avian sarcoma virus CT10 oncogene | 2605 | 2798 | 2410 | 2332 | 1939 | 1591 | 1575 | 2385 | 1634 | 1747 | 1549 |
| 1261 | 0.09 | 0.14 | 0.62 | 0.53 | 1 | Fkbp11 | FK506 binding protein 11 | 379 | 661 | 498 | 1790 | 1373 | 1165 | 1574 | 2314 | 1441 | 1544 | 1127 |
| 1157 | -0.08 | 0.19 | -0.43 | 0.67 | 1 | Rundc3b | RUN domain containing 3B | 867 | 1027 | 911 | 1236 | 1201 | 1079 | 1574 | 828 | 1451 | 1396 | 1153 |
| 1934 | 0.15 | 0.12 | 1.26 | 0.21 | 1 | Ncoa7 | nuclear receptor coactivator 7 | 1807 | 2398 | 2286 | 1779 | 1951 | 1688 | 1573 | 1796 | 2495 | 1812 | 1685 |
| 1424 | 0.11 | 0.13 | 0.81 | 0.42 | 1 | Pnpo | pyridoxine 5'-phosphate oxidase | 842 | 1014 | 963 | 1399 | 1805 | 1390 | 1573 | 1688 | 1717 | 1958 | 1317 |
| 1616 | 0.01 | 0.09 | 0.16 | 0.87 | 1 | Babam1 | BRISC and BRCA1 A complex member 1 | 1778 | 1818 | 1837 | 1687 | 1625 | 1250 | 1572 | 1826 | 1589 | 1553 | 1241 |
| 1558 | -0.05 | 0.14 | -0.37 | 0.71 | 1 | Med15 | mediator complex subunit 15 | 1601 | 1630 | 1604 | 1348 | 1517 | 1806 | 1572 | 1084 | 1591 | 1814 | 1569 |
| 1744 | 0.18 | 0.10 | 1.77 | 0.08 | 1 | Lemd2 | LEM domain containing 2 | 1564 | 1363 | 1541 | 1660 | 1631 | 2024 | 1569 | 1813 | 1691 | 2098 | 2232 |
| 1635 | -0.02 | 0.09 | -0.25 | 0.80 | 1 | Mta3 | metastasis associated 3 | 1673 | 1622 | 1649 | 1671 | 1693 | 1640 | 1569 | 1751 | 1678 | 1450 | 1593 |
| 1347 | 0.11 | 0.11 | 1.03 | 0.30 | 1 | Clcn7 | chloride channel, voltage-sensitive 7 | 1120 | 959 | 942 | 1531 | 1228 | 1355 | 1569 | 1681 | 1432 | 1541 | 1461 |
| 1434 | -0.05 | 0.14 | -0.39 | 0.70 | 1 | Tnik | TRAF2 and NCK interacting kinase | 879 | 714 | 899 | 1922 | 1617 | 1648 | 1568 | 1577 | 1803 | 1392 | 1756 |
| 1569 | -0.03 | 0.10 | -0.32 | 0.75 | 1 | Lrrc1 | leucine rich repeat containing 1 | 1477 | 1460 | 1473 | 1781 | 1750 | 1391 | 1567 | 1916 | 1637 | 1440 | 1362 |
| 1486 | -0.14 | 0.09 | -1.44 | 0.15 | 1 | Sh2b1 | SH2B adaptor protein 1 | 1660 | 1444 | 1520 | 1298 | 1437 | 1814 | 1567 | 1102 | 1310 | 1380 | 1815 |
| 1268 | 0.20 | 0.17 | 1.21 | 0.23 | 1 | Jph3 | junctophilin 3 | 714 | 891 | 865 | 1322 | 1308 | 1095 | 1567 | 2016 | 1245 | 1794 | 1133 |
| 1776 | -0.02 | 0.11 | -0.18 | 0.86 | 1 | Tial1 | Tia1 cytotoxic granule-associated RNA bi | 1472 | 1301 | 1399 | 1375 | 1509 | 3241 | 1566 | 1479 | 1595 | 1234 | 3369 |
| 1420 | 0.10 | 0.11 | 0.93 | 0.35 | 1 | Gpr180 | G protein-coupled receptor 180 | 1134 | 1115 | 1122 | 1209 | 1580 | 1556 | 1564 | 1367 | 1589 | 1778 | 1605 |
| 1827 | 0.16 | 0.13 | 1.26 | 0.21 | 1 | Adh5 | alcohol dehydrogenase 5 (class III), chi p | 2708 | 1986 | 2716 | 1558 | 1536 | 1341 | 1563 | 1805 | 1566 | 1704 | 1618 |
| 1652 | 0.00 | 0.12 | -0.01 | 1.00 | 1 | Pla2g6 | phospholipase A2, group VI | 1292 | 1833 | 1632 | 1758 | 1537 | 1854 | 1562 | 1908 | 1538 | 1686 | 1576 |
| 1381 | -0.11 | 0.11 | -1.06 | 0.29 | 1 | Ext1 | exostosin glycosyltransferase 1 | 1917 | 1186 | 1715 | 1070 | 1712 | 1052 | 1562 | 923 | 1474 | 1488 | 1091 |
| 2111 | -0.05 | 0.12 | -0.40 | 0.69 | 1 | Rxra | retinoid X receptor alpha | 4238 | 4221 | 4333 | 1387 | 1272 | 1053 | 1561 | 1666 | 1215 | 1368 | 910 |
| 1792 | 0.22 | 0.18 | 1.23 | 0.22 | 1 | Zc3h13 | zinc finger CCCH type containing 13 | 1742 | 1525 | 1968 | 1614 | 1393 | 2111 | 1561 | 2529 | 1645 | 1426 | 2197 |
| 1792 | -0.04 | 0.15 | -0.24 | 0.81 | 1 | Kcnh6 | potassium voltage-gated channel, subfan | 1313 | 1325 | 1379 | 1608 | 1338 | 3517 | 1561 | 1302 | 1294 | 2070 | 3003 |
| 1574 | -0.20 | 0.10 | -1.89 | 0.06 | 1 | Mgat5 | mannoside acetylglucosaminyltransferase | 2219 | 1695 | 2056 | 1402 | 1790 | 1291 | 1561 | 1200 | 1647 | 1368 | 1081 |
| 1508 | -0.04 | 0.13 | -0.34 | 0.73 | 1 | Phc1 | polyhomeotic 1 | 1488 | 1843 | 1673 | 1396 | 1431 | 1471 | 1561 | 1528 | 1619 | 1389 | 1188 |
| 1594 | 0.10 | 0.16 | 0.61 | 0.54 | 1 | Hars | histidyl-tRNA synthetase | 1605 | 1137 | 1224 | 1390 | 1926 | 1696 | 1560 | 1738 | 1999 | 1450 | 1805 |
| 1708 | 0.06 | 0.10 | 0.62 | 0.53 | 1 | Socs5 | suppressor of cytokine signaling 5 | 2098 | 2097 | 1766 | 1805 | 1456 | 1448 | 1559 | 1996 | 1538 | 1622 | 1402 |
| 1595 | -0.07 | 0.13 | -0.57 | 0.57 | 1 | Ngly1 | N-glycanase 1 | 1879 | 1469 | 1727 | 1737 | 1491 | 1604 | 1558 | 1497 | 1623 | 1340 | 1623 |
| 1706 | 0.04 | 0.10 | 0.34 | 0.73 | 1 | Med13l | mediator complex subunit 13-like | 1739 | 1541 | 1851 | 1541 | 1622 | 2034 | 1557 | 1653 | 1790 | 1686 | 1756 |
| 1390 | -0.09 | 0.12 | -0.75 | 0.45 | 1 | Zbtb14 | zinc finger and BTB domain containing 14 | 984 | 1109 | 1136 | 1743 | 1630 | 1301 | 1557 | 1349 | 1501 | 1676 | 1305 |

| | | | | | | | | | | | | | | | | | | |
|---|---|---|---|---|---|---|---|---|---|---|---|---|---|---|---|---|---|---|
| 1566 | 0.10 | 0.11 | 0.86 | 0.39 | 1 | Gatd1 | glutamine amidotransferase like class 1 d | 1196 | 1305 | 1334 | 1406 | 1910 | 1604 | 1556 | 1709 | 1846 | 1820 | 1539 |
| 1594 | 0.05 | 0.09 | 0.51 | 0.61 | 1 | Pias1 | protein inhibitor of activated STAT 1 | 1730 | 1904 | 1589 | 1510 | 1354 | 1640 | 1555 | 1615 | 1513 | 1586 | 1539 |
| 1446 | 0.12 | 0.11 | 1.13 | 0.26 | 1 | Tti1 | TELO2 interacting protein 1 | 1158 | 1123 | 1218 | 1477 | 1449 | 1450 | 1555 | 1776 | 1458 | 1772 | 1471 |
| 1946 | 0.06 | 0.09 | 0.70 | 0.48 | 1 | Cpsf7 | cleavage and polyadenylation specific fac | 1824 | 1856 | 1996 | 1563 | 1470 | 3145 | 1553 | 1763 | 1635 | 1507 | 3097 |
| 1175 | -0.11 | 0.12 | -0.94 | 0.35 | 1 | Pcca | propionyl-Coenzyme A carboxylase, alph | 914 | 860 | 905 | 1427 | 1220 | 1129 | 1553 | 1130 | 1268 | 1378 | 1139 |
| 1378 | 0.04 | 0.12 | 0.31 | 0.75 | 1 | Cog2 | component of oligomeric golgi complex 2 | 842 | 1044 | 999 | 1406 | 1524 | 1571 | 1552 | 1712 | 1483 | 1483 | 1543 |
| 1629 | 0.01 | 0.10 | 0.12 | 0.90 | 1 | Smarcal1 | SWI/SNF related matrix associated, actin | 1315 | 1639 | 1476 | 1816 | 1813 | 1546 | 1551 | 1800 | 1590 | 1739 | 1637 |
| 1369 | -0.21 | 0.16 | -1.28 | 0.20 | 1 | Herc3 | hect domain and RLD 3 | 1281 | 959 | 1441 | 1693 | 1444 | 1369 | 1551 | 995 | 1566 | 1440 | 1320 |
| 1783 | -0.03 | 0.15 | -0.19 | 0.85 | 1 | Rbm6 | RNA binding motif protein 6 | 1830 | 1678 | 1513 | 1573 | 1582 | 2601 | 1550 | 1099 | 1850 | 1624 | 2713 |
| 1519 | 0.12 | 0.11 | 1.05 | 0.29 | 1 | Pigs | phosphatidylinositol glycan anchor biosyı | 1861 | 1538 | 1697 | 1468 | 1444 | 1074 | 1550 | 1810 | 1497 | 1731 | 1037 |
| 1348 | -0.12 | 0.13 | -0.88 | 0.38 | 1 | Ankrd27 | ankyrin repeat domain 27 (VPS9 domain) | 1325 | 1390 | 1279 | 1472 | 1223 | 1394 | 1550 | 1435 | 1382 | 1086 | 1296 |
| 1658 | -0.03 | 0.10 | -0.33 | 0.74 | 1 | Rnf216 | ring finger protein 216 | 1758 | 1763 | 1777 | 1494 | 1676 | 1827 | 1549 | 1583 | 1784 | 1432 | 1589 |
| 1649 | 0.00 | 0.09 | -0.04 | 0.97 | 1 | Fbxo28 | F-box protein 28 | 1723 | 1769 | 1652 | 1856 | 1534 | 1569 | 1549 | 1774 | 1568 | 1614 | 1531 |
| 1781 | -0.15 | 0.13 | -1.12 | 0.26 | 1 | Cdc42se1 | CDC42 small effector 1 | 2437 | 2958 | 2398 | 1601 | 1639 | 1386 | 1548 | 1647 | 1642 | 1061 | 1274 |
| 1504 | 0.11 | 0.12 | 0.89 | 0.38 | 1 | Mrps23 | mitochondrial ribosomal protein S23 | 1238 | 1276 | 1393 | 1466 | 1792 | 1266 | 1547 | 2001 | 1619 | 1662 | 1286 |
| 1966 | -0.04 | 0.10 | -0.41 | 0.68 | 1 | Tuba1a | tubulin, alpha 1A | 2998 | 2593 | 2794 | 1414 | 1669 | 2090 | 1546 | 1259 | 1726 | 1673 | 1866 |
| 1536 | -0.18 | 0.17 | -1.04 | 0.30 | 1 | Ebna1bp2 | EBNA1 binding protein 2 | 2231 | 2070 | 2413 | 986 | 1430 | 1447 | 1546 | 1245 | 1232 | 880 | 1416 |
| 1331 | 0.02 | 0.10 | 0.23 | 0.82 | 1 | Yif1a | Yip1 interacting factor homolog A (S. cere | 1226 | 1027 | 1119 | 1172 | 1588 | 1273 | 1546 | 1216 | 1504 | 1713 | 1251 |
| 1223 | -0.08 | 0.12 | -0.67 | 0.50 | 1 | Tmem231 | transmembrane protein 231 | 829 | 737 | 844 | 1546 | 1539 | 1042 | 1546 | 1573 | 1608 | 1224 | 970 |
| 1894 | -0.11 | 0.15 | -0.74 | 0.46 | 1 | Chd2 | chromodomain helicase DNA binding pro | 2394 | 2180 | 2226 | 1770 | 1509 | 2456 | 1545 | 1182 | 1849 | 1605 | 2118 |
| 1765 | -0.10 | 0.15 | -0.67 | 0.50 | 1 | Zcchc14 | zinc finger, CCHC domain containing 14 | 2432 | 2310 | 1594 | 1703 | 1699 | 1804 | 1545 | 1506 | 1581 | 1798 | 1446 |
| 1752 | -0.20 | 0.19 | -1.06 | 0.29 | 1 | Lgi2 | leucine-rich repeat LGI family, member 2 | 2877 | 1939 | 2113 | 1863 | 2092 | 1073 | 1544 | 1386 | 2137 | 1264 | 979 |
| 1303 | 0.11 | 0.15 | 0.75 | 0.46 | 1 | Hcn2 | hyperpolarization-activated, cyclic nucleo | 743 | 908 | 885 | 1453 | 1597 | 1034 | 1544 | 1652 | 1530 | 2030 | 960 |
| 1451 | -0.23 | 0.18 | -1.25 | 0.21 | 1 | Fkrp | fukutin related protein | 1298 | 1769 | 1403 | 1663 | 1684 | 1296 | 1543 | 1001 | 1608 | 1598 | 1098 |
| 1409 | 0.13 | 0.14 | 0.93 | 0.35 | 1 | Gfm2 | G elongation factor, mitochondrial 2 | 828 | 1011 | 911 | 1175 | 1479 | 1889 | 1543 | 1690 | 1461 | 1450 | 2062 |
| 1325 | -0.26 | 0.14 | -1.89 | 0.06 | 1 | Slc4a8 | solute carrier family 4 (anion exchanger), | 905 | 1177 | 1177 | 1542 | 1406 | 1643 | 1543 | 1305 | 1517 | 1029 | 1326 |
| 1806 | 0.13 | 0.12 | 1.07 | 0.28 | 1 | Ptges3 | prostaglandin E synthase 3 | 1984 | 2247 | 2161 | 1588 | 1654 | 1653 | 1542 | 1818 | 1540 | 2013 | 1667 |
| 1356 | 0.01 | 0.11 | 0.11 | 0.92 | 1 | Zfp422 | zinc finger protein 422 | 1120 | 1186 | 1059 | 1382 | 1409 | 1412 | 1542 | 1654 | 1364 | 1437 | 1351 |
| 1570 | 0.02 | 0.09 | 0.18 | 0.86 | 1 | Klhdc3 | kelch domain containing 3 | 1886 | 1692 | 2022 | 1296 | 1557 | 1414 | 1541 | 1412 | 1594 | 1494 | 1365 |
| 1524 | 0.03 | 0.10 | 0.24 | 0.81 | 1 | Ripk4 | receptor-interacting serine-threonine kina | 1064 | 1449 | 1363 | 1741 | 1694 | 1412 | 1541 | 1763 | 1666 | 1660 | 1407 |
| 1583 | -0.01 | 0.09 | -0.07 | 0.94 | 1 | Pak4 | p21 (RAC1) activated kinase 4 | 1198 | 1464 | 1369 | 1756 | 1652 | 1748 | 1539 | 1930 | 1636 | 1477 | 1641 |
| 1579 | -0.16 | 0.10 | -1.66 | 0.10 | 1 | Ascc2 | activating signal cointegrator 1 complex s | 1975 | 1951 | 1951 | 1415 | 1519 | 1581 | 1539 | 1160 | 1419 | 1356 | 1499 |
| 1152 | -0.16 | 0.12 | -1.33 | 0.18 | 1 | Naxd | NAD(P)HX dehydratase | 1256 | 865 | 931 | 942 | 1377 | 1231 | 1539 | 940 | 1198 | 1311 | 1081 |
| 1797 | -0.04 | 0.14 | -0.29 | 0.77 | 1 | Nedd4l | neural precursor cell expressed, developı | 2163 | 2412 | 2393 | 1482 | 1560 | 1908 | 1538 | 1881 | 1395 | 1617 | 1421 |
| 1460 | 0.01 | 0.16 | 0.06 | 0.95 | 1 | Hcfc2 | host cell factor C2 | 1092 | 1288 | 879 | 1335 | 1530 | 1972 | 1538 | 1248 | 1632 | 1530 | 2014 |
| 1260 | -0.18 | 0.17 | -1.02 | 0.31 | 1 | Igsf1 | immunoglobulin superfamily, member 1 | 557 | 721 | 671 | 1182 | 1746 | 1797 | 1538 | 936 | 1439 | 1327 | 1939 |
| 1710 | 0.05 | 0.13 | 0.37 | 0.71 | 1 | Tbca | tubulin cofactor A | 2173 | 2071 | 2762 | 1494 | 1583 | 1199 | 1537 | 1601 | 1645 | 1275 | 1471 |
| 1651 | -0.07 | 0.08 | -0.84 | 0.40 | 1 | Zc3h18 | zinc finger CCCH-type containing 18 | 1766 | 1655 | 1839 | 1456 | 1630 | 1983 | 1537 | 1440 | 1512 | 1483 | 1858 |
| 1533 | -0.11 | 0.10 | -1.14 | 0.25 | 1 | Lrch1 | leucine-rich repeats and calponin homolc | 1930 | 1655 | 1805 | 1394 | 1424 | 1595 | 1537 | 1336 | 1210 | 1561 | 1411 |
| 1440 | -0.11 | 0.10 | -1.08 | 0.28 | 1 | Bud31 | BUD31 homolog | 1488 | 1330 | 1415 | 1348 | 1623 | 1522 | 1537 | 1311 | 1512 | 1255 | 1496 |
| 1802 | 0.03 | 0.15 | 0.20 | 0.84 | 1 | Syt11 | synaptotagmin XI | 1873 | 2394 | 2110 | 2054 | 1624 | 1350 | 1536 | 2566 | 1783 | 1119 | 1416 |
| 1952 | 0.21 | 0.17 | 1.26 | 0.21 | 1 | Aopep | aminopeptidase O | 1015 | 653 | 994 | 1866 | 1733 | 3610 | 1535 | 2333 | 1754 | 1931 | 4047 |
| 1525 | -0.16 | 0.17 | -0.93 | 0.35 | 1 | Ggps1 | geranylgeranyl diphosphate synthase 1 | 1808 | 1742 | 1416 | 1494 | 1381 | 1762 | 1535 | 884 | 1380 | 1605 | 1764 |
| 1497 | -0.09 | 0.12 | -0.81 | 0.42 | 1 | Ppp2r2d | protein phosphatase 2, regulatory subuni | 1390 | 1451 | 1533 | 1535 | 1563 | 1587 | 1535 | 1338 | 1757 | 1226 | 1551 |
| 1478 | 0.22 | 0.14 | 1.51 | 0.13 | 1 | Larp1b | La ribonucleoprotein domain family, mem | 669 | 650 | 695 | 1644 | 1584 | 1789 | 1535 | 2361 | 1795 | 1407 | 2126 |
| 1379 | 0.02 | 0.14 | 0.16 | 0.88 | 1 | Pip4k2a | phosphatidylinositol-5-phosphate 4-kinas | 2021 | 1185 | 1196 | 1347 | 1257 | 1172 | 1535 | 1116 | 1498 | 1705 | 1137 |
| 1355 | 0.05 | 0.13 | 0.35 | 0.73 | 1 | Rasal2 | RAS protein activator like 2 | 1249 | 1206 | 1630 | 1116 | 1333 | 1335 | 1535 | 1137 | 1366 | 1631 | 1369 |
| 1793 | 0.08 | 0.11 | 0.76 | 0.45 | 1 | Aebp2 | AE binding protein 2 | 2133 | 2240 | 1790 | 1601 | 1448 | 2024 | 1534 | 1700 | 1734 | 1497 | 2027 |
| 1444 | 0.08 | 0.13 | 0.63 | 0.53 | 1 | Gpd1l | glycerol-3-phosphate dehydrogenase 1-li | 1965 | 1653 | 1630 | 1085 | 1523 | 1053 | 1534 | 1406 | 1499 | 1395 | 1140 |
| 1380 | 0.13 | 0.12 | 1.10 | 0.27 | 1 | Ift88 | intraflagellar transport 88 | 884 | 769 | 894 | 1384 | 1483 | 1644 | 1534 | 1795 | 1492 | 1632 | 1675 |
| 1365 | 0.08 | 0.13 | 0.60 | 0.55 | 1 | Nsmce4a | NSE4 homolog A, SMC5-SMC6 complex | 1792 | 1602 | 1726 | 1131 | 1075 | 1100 | 1533 | 1292 | 1253 | 1261 | 1249 |
| 1686 | -0.12 | 0.09 | -1.23 | 0.22 | 1 | Zfp512b | zinc finger protein 512B | 1119 | 1291 | 1208 | 1995 | 1617 | 2629 | 1532 | 1938 | 1530 | 1382 | 2301 |
| 1636 | -0.06 | 0.12 | -0.46 | 0.65 | 1 | Senp3 | SUMO/sentrin specific peptidase 3 | 1870 | 1352 | 1794 | 1564 | 1621 | 1858 | 1532 | 1167 | 1745 | 1606 | 1892 |
| 1408 | 0.19 | 0.11 | 1.68 | 0.09 | 1 | Ctnnbl1 | catenin, beta like 1 | 1277 | 1079 | 1217 | 1386 | 1362 | 1292 | 1532 | 1758 | 1473 | 1565 | 1553 |
| 2082 | 0.24 | 0.13 | 1.77 | 0.08 | 1 | Colgalt1 | collagen beta(1-O)galactosyltransferase 1 | 3639 | 2456 | 2945 | 1514 | 1781 | 1541 | 1531 | 2025 | 1794 | 1897 | 1775 |
| 1258 | -0.07 | 0.14 | -0.52 | 0.60 | 1 | Zdhhc24 | zinc finger, DHHC domain containing 24 | 848 | 756 | 1031 | 1563 | 1476 | 1182 | 1531 | 1556 | 1304 | 1368 | 1220 |
| 1459 | -0.19 | 0.16 | -1.22 | 0.22 | 1 | Slc31a1 | solute carrier family 31, member 1 | 1186 | 996 | 1169 | 1729 | 1909 | 1554 | 1530 | 1084 | 1923 | 1302 | 1668 |
| 1309 | -0.14 | 0.13 | -1.08 | 0.28 | 1 | Hpcal1 | hippocalcin-like 1 | 1344 | 1170 | 1241 | 1455 | 1507 | 1039 | 1530 | 1588 | 1228 | 1489 | 811 |
| 1541 | 0.26 | 0.16 | 1.64 | 0.10 | 1 | Btbd9 | BTB (POZ) domain containing 9 | 1480 | 1254 | 1298 | 1494 | 1536 | 1251 | 1529 | 2402 | 1406 | 2007 | 1298 |
| 1607 | 0.07 | 0.16 | 0.41 | 0.68 | 1 | Farsb | phenylalanyl-tRNA synthetase, beta subu | 1758 | 1666 | 1521 | 1315 | 1820 | 1524 | 1528 | 1096 | 1988 | 1761 | 1697 |

| | | | | | | | | | | | | | | | | | | |
|---|---|---|---|---|---|---|---|---|---|---|---|---|---|---|---|---|---|---|
| 1558 | 0.24 | 0.19 | 1.25 | 0.21 | 1 | Lamtor2 | late endosomal/lysosomal adaptor, MAPK | 1459 | 1319 | 2139 | 1299 | 1659 | 1110 | 1528 | 2297 | 1600 | 1537 | 1186 |
| 1654 | 0.28 | 0.15 | 1.90 | 0.06 | 1 | Ap2s1 | adaptor-related protein complex 2, sigma | 2011 | 1853 | 1744 | 1415 | 1577 | 1102 | 1527 | 2403 | 1480 | 1846 | 1233 |
| 1483 | 0.00 | 0.10 | 0.01 | 0.99 | 1 | Hdac3 | histone deacetylase 3 | 1429 | 1455 | 1537 | 1328 | 1505 | 1588 | 1527 | 1501 | 1391 | 1534 | 1516 |
| 2198 | -0.12 | 0.12 | -0.96 | 0.34 | 1 | Myadm | myeloid-associated differentiation marker | 4973 | 3800 | 3876 | 1518 | 1676 | 1254 | 1526 | 1134 | 1562 | 1635 | 1220 |
| 1845 | 0.03 | 0.09 | 0.37 | 0.71 | 1 | Casc3 | cancer susceptibility candidate 3 | 1632 | 2087 | 1970 | 1821 | 1625 | 2270 | 1526 | 1800 | 1858 | 1596 | 2113 |
| 1787 | -0.14 | 0.12 | -1.17 | 0.24 | 1 | Srsf4 | serine and arginine-rich splicing factor 4 | 2035 | 1842 | 1752 | 1785 | 1666 | 2348 | 1526 | 1280 | 1537 | 1629 | 2256 |
| 1322 | 0.16 | 0.14 | 1.21 | 0.23 | 1 | Amd1 | S-adenosylmethionine decarboxylase 1 | 798 | 673 | 802 | 1609 | 1528 | 1123 | 1526 | 1557 | 1462 | 2034 | 1431 |
| 1733 | 0.17 | 0.12 | 1.42 | 0.16 | 1 | Stk40 | serine/threonine kinase 40 | 1778 | 2565 | 1710 | 1602 | 1508 | 1447 | 1525 | 2089 | 1662 | 1770 | 1408 |
| 1636 | 0.13 | 0.11 | 1.23 | 0.22 | 1 | Ssu72 | Ssu72 RNA polymerase II CTD phosphat | 1583 | 1347 | 1440 | 1595 | 1705 | 1670 | 1524 | 1704 | 1904 | 1917 | 1605 |
| 1478 | 0.04 | 0.13 | 0.29 | 0.77 | 1 | Slc35b2 | solute carrier family 35, member B2 | 1659 | 1441 | 1206 | 1618 | 1671 | 1091 | 1524 | 1581 | 1719 | 1584 | 1159 |
| 1461 | 0.04 | 0.09 | 0.48 | 0.63 | 1 | Grpel1 | GrpE-like 1, mitochondrial | 1360 | 1363 | 1433 | 1396 | 1662 | 1291 | 1522 | 1398 | 1689 | 1568 | 1389 |
| 1426 | -0.04 | 0.12 | -0.31 | 0.76 | 1 | Pacs1 | phosphofurin acidic cluster sorting proteir | 1711 | 1386 | 1374 | 1451 | 1344 | 1346 | 1522 | 1190 | 1521 | 1620 | 1225 |
| 1564 | -0.01 | 0.09 | -0.10 | 0.92 | 1 | Usf2 | upstream transcription factor 2 | 1571 | 1675 | 1596 | 1679 | 1658 | 1336 | 1520 | 1614 | 1582 | 1703 | 1267 |
| 1545 | 0.02 | 0.11 | 0.18 | 0.85 | 1 | Lsm14b | LSM family member 14B | 1337 | 1487 | 1577 | 1565 | 1601 | 1568 | 1520 | 1410 | 1622 | 1720 | 1593 |
| 1619 | 0.17 | 0.13 | 1.34 | 0.18 | 1 | Prxl2a | peroxiredoxin like 2A | 1549 | 2051 | 2074 | 1600 | 1538 | 1073 | 1519 | 1793 | 1496 | 1718 | 1398 |
| 1511 | -0.02 | 0.11 | -0.22 | 0.83 | 1 | Tmub2 | transmembrane and ubiquitin-like domair | 1337 | 1332 | 1138 | 1799 | 1708 | 1430 | 1519 | 1812 | 1646 | 1520 | 1382 |
| 1438 | 0.03 | 0.15 | 0.22 | 0.83 | 1 | Marveld3 | MARVEL (membrane-associating) domai | 567 | 931 | 802 | 1746 | 1537 | 1906 | 1519 | 1599 | 1679 | 1735 | 1796 |
| 1396 | 0.16 | 0.13 | 1.29 | 0.20 | 1 | Echs1 | enoyl Coenzyme A hydratase, short chair | 1694 | 1351 | 1566 | 1267 | 1246 | 1057 | 1519 | 1621 | 1376 | 1408 | 1254 |
| 1333 | -0.12 | 0.14 | -0.85 | 0.40 | 1 | Zfp251 | zinc finger protein 251 | 1029 | 1123 | 1131 | 1680 | 1387 | 1336 | 1519 | 1190 | 1528 | 1275 | 1465 |
| 1878 | 0.08 | 0.10 | 0.75 | 0.46 | 1 | Ndufa9 | NADH:ubiquinone oxidoreductase subun | 2370 | 2545 | 2280 | 1841 | 1751 | 1418 | 1518 | 2082 | 1952 | 1552 | 1354 |
| 1648 | 0.12 | 0.10 | 1.20 | 0.23 | 1 | Glo1 | glyoxalase 1 | 1862 | 2096 | 1901 | 1542 | 1536 | 1265 | 1518 | 1935 | 1613 | 1425 | 1434 |
| 1472 | -0.05 | 0.18 | -0.27 | 0.79 | 1 | Tomm7 | translocase of outer mitochondrial membr | 1833 | 1578 | 2129 | 1291 | 1539 | 1066 | 1518 | 1796 | 1254 | 1108 | 1080 |
| 1386 | -0.10 | 0.13 | -0.74 | 0.46 | 1 | Coq9 | coenzyme Q9 | 1049 | 989 | 1372 | 1350 | 1505 | 1714 | 1518 | 1077 | 1388 | 1566 | 1717 |
| 2169 | 0.03 | 0.13 | 0.24 | 0.81 | 1 | Rragb | Ras-related GTP binding B | 3400 | 3842 | 4247 | 1562 | 1465 | 1560 | 1516 | 1869 | 1279 | 1516 | 1599 |
| 2072 | -0.10 | 0.10 | -1.06 | 0.29 | 1 | Bambi | BMP and activin membrane-bound inhibit | 2782 | 3538 | 2916 | 1580 | 1469 | 2416 | 1516 | 1669 | 1222 | 1333 | 2348 |
| 1713 | -0.23 | 0.14 | -1.63 | 0.10 | 1 | Btbd7 | BTB (POZ) domain containing 7 | 2538 | 2594 | 2062 | 1476 | 1456 | 1839 | 1515 | 1048 | 1499 | 1275 | 1537 |
| 1594 | 0.07 | 0.11 | 0.61 | 0.55 | 1 | Rabggtb | Rab geranylgeranyl transferase, b subuni | 1748 | 1587 | 1262 | 1519 | 1336 | 1958 | 1515 | 1416 | 1536 | 1664 | 1995 |
| 1582 | -0.02 | 0.09 | -0.25 | 0.80 | 1 | Bbs2 | Bardet-Biedl syndrome 2 (human) | 1170 | 1363 | 1342 | 1515 | 1447 | 2341 | 1515 | 1475 | 1553 | 1399 | 2278 |
| 1456 | -0.19 | 0.11 | -1.79 | 0.07 | 1 | Gle1 | GLE1 RNA export mediator (yeast) | 1807 | 1692 | 1835 | 1434 | 1480 | 1269 | 1515 | 1279 | 1191 | 1226 | 1293 |
| 2385 | -0.13 | 0.25 | -0.53 | 0.60 | 1 | Msln | mesothelin | 788 | 323 | 682 | 1919 | 4094 | 5493 | 1514 | 1859 | 3896 | 1529 | 4140 |
| 1028 | -0.10 | 0.13 | -0.73 | 0.47 | 1 | Angel1 | angel homolog 1 | 512 | 519 | 503 | 1239 | 1403 | 897 | 1514 | 1021 | 1462 | 1345 | 891 |
| 1735 | 0.01 | 0.10 | 0.13 | 0.90 | 1 | Ash2l | ASH2 like histone lysine methyltransferas | 1689 | 1769 | 1724 | 1694 | 1752 | 1964 | 1513 | 1704 | 1951 | 1493 | 1833 |
| 1609 | 0.05 | 0.10 | 0.51 | 0.61 | 1 | Zmym3 | zinc finger, MYM-type 3 | 1586 | 1855 | 1326 | 1676 | 1300 | 1881 | 1512 | 1736 | 1498 | 1480 | 1845 |
| 1537 | 0.11 | 0.11 | 0.98 | 0.33 | 1 | 8030462N17Ril | RIKEN cDNA 8030462N17 gene | 1257 | 1675 | 1512 | 1532 | 1391 | 1560 | 1512 | 1618 | 1598 | 1608 | 1638 |
| 1512 | -0.10 | 0.13 | -0.79 | 0.43 | 1 | Bckdha | branched chain ketoacid dehydrogenase | 1684 | 1795 | 1904 | 1620 | 1593 | 1047 | 1512 | 1600 | 1323 | 1702 | 852 |
| 1544 | -0.16 | 0.12 | -1.31 | 0.19 | 1 | Prkci | protein kinase C, iota | 1916 | 1862 | 1723 | 1445 | 1628 | 1462 | 1511 | 1134 | 1376 | 1644 | 1288 |
| 1509 | -0.01 | 0.10 | -0.15 | 0.88 | 1 | Dap3 | death associated protein 3 | 1646 | 1360 | 1511 | 1250 | 1387 | 1895 | 1511 | 1148 | 1271 | 1674 | 1941 |
| 1465 | 0.04 | 0.11 | 0.42 | 0.68 | 1 | Socs4 | suppressor of cytokine signaling 4 | 1678 | 1794 | 1405 | 1477 | 1240 | 1309 | 1511 | 1593 | 1320 | 1310 | 1482 |
| 1446 | 0.00 | 0.13 | 0.04 | 0.97 | 1 | Taz | tafazzin | 843 | 896 | 925 | 1387 | 1315 | 2366 | 1511 | 1660 | 1304 | 1225 | 2477 |
| 1322 | 0.03 | 0.13 | 0.20 | 0.84 | 1 | Gpd1 | glycerol-3-phosphate dehydrogenase 1 (s | 879 | 1056 | 1012 | 1373 | 1570 | 1272 | 1510 | 1804 | 1388 | 1421 | 1253 |
| 1565 | -0.21 | 0.12 | -1.70 | 0.09 | 1 | Ccser2 | coiled-coil serine rich 2 | 2105 | 2280 | 1999 | 1455 | 1450 | 1387 | 1509 | 1044 | 1253 | 1313 | 1420 |
| 1982 | -0.03 | 0.15 | -0.18 | 0.86 | 1 | Ppp1r12a | protein phosphatase 1, regulatory subuni | 2609 | 2283 | 2108 | 1585 | 1412 | 2953 | 1508 | 1685 | 1649 | 1101 | 2907 |
| 1398 | -0.11 | 0.13 | -0.89 | 0.38 | 1 | Trp53i13 | transformation related protein 53 inducibl | 1109 | 931 | 1068 | 1749 | 1596 | 1514 | 1508 | 1767 | 1406 | 1416 | 1312 |
| 1434 | -0.03 | 0.14 | -0.22 | 0.82 | 1 | Atg4d | autophagy related 4D, cysteine peptidase | 977 | 1294 | 1145 | 1743 | 1521 | 1456 | 1507 | 1603 | 1479 | 1864 | 1187 |
| 1773 | 0.08 | 0.11 | 0.73 | 0.47 | 1 | Ppp1r12c | protein phosphatase 1, regulatory subuni | 2311 | 2129 | 2152 | 1336 | 1543 | 1893 | 1506 | 1696 | 1369 | 1669 | 1893 |
| 1769 | 0.14 | 0.10 | 1.32 | 0.19 | 1 | Btbd1 | BTB (POZ) domain containing 1 | 2449 | 2725 | 2698 | 1406 | 1277 | 1323 | 1506 | 1573 | 1640 | 1529 | 1335 |
| 1482 | 0.18 | 0.10 | 1.84 | 0.07 | 1 | Tfam | transcription factor A, mitochondrial | 1574 | 1522 | 1476 | 1370 | 1321 | 1279 | 1506 | 1742 | 1415 | 1754 | 1338 |
| 1409 | 0.13 | 0.11 | 1.12 | 0.26 | 1 | Prpsap1 | phosphoribosyl pyrophosphate synthetas | 1393 | 1221 | 1117 | 1342 | 1140 | 1645 | 1506 | 1645 | 1276 | 1727 | 1487 |
| 2110 | 0.09 | 0.13 | 0.69 | 0.49 | 1 | Tpm3 | tropomyosin 3, gamma | 4248 | 3198 | 3231 | 1324 | 1549 | 1715 | 1505 | 1669 | 1634 | 1448 | 1689 |
| 1499 | 0.11 | 0.11 | 0.99 | 0.32 | 1 | Igip | IgA inducing protein | 1374 | 1565 | 1580 | 1633 | 1288 | 1320 | 1505 | 1814 | 1428 | 1614 | 1365 |
| 1545 | -0.02 | 0.16 | -0.15 | 0.88 | 1 | Pacs2 | phosphofurin acidic cluster sorting proteir | 2439 | 1411 | 1421 | 1484 | 1426 | 1495 | 1504 | 1499 | 1509 | 1508 | 1299 |
| 1392 | -0.05 | 0.10 | -0.54 | 0.59 | 1 | Zfp667 | zinc finger protein 667 | 741 | 916 | 847 | 1886 | 1524 | 1596 | 1504 | 1856 | 1609 | 1320 | 1510 |
| 1319 | 0.10 | 0.15 | 0.66 | 0.51 | 1 | Gas2 | growth arrest specific 2 | 782 | 893 | 779 | 1764 | 1212 | 1317 | 1504 | 1547 | 1200 | 2084 | 1424 |
| 1572 | -0.03 | 0.10 | -0.28 | 0.78 | 1 | Trap1 | TNF receptor-associated protein 1 | 1701 | 1386 | 1514 | 1603 | 1568 | 1730 | 1503 | 1564 | 1455 | 1523 | 1741 |
| 1555 | 0.07 | 0.13 | 0.55 | 0.59 | 1 | Map3k5 | mitogen-activated protein kinase kinase k | 1424 | 1519 | 1540 | 1751 | 1305 | 1584 | 1503 | 2070 | 1447 | 1526 | 1439 |
| 1517 | 0.02 | 0.11 | 0.14 | 0.89 | 1 | Hmgxb3 | HMG box domain containing 3 | 1128 | 1326 | 1328 | 1834 | 1547 | 1529 | 1503 | 1736 | 1641 | 1729 | 1383 |
| 1509 | 0.21 | 0.11 | 1.97 | 0.05 | 1 | Ccdc25 | coiled-coil domain containing 25 | 1005 | 1307 | 1112 | 1669 | 1539 | 1374 | 1502 | 2257 | 1713 | 1512 | 1604 |
| 1364 | -0.02 | 0.13 | -0.13 | 0.89 | 1 | Pcyt1b | phosphate cytidylyltransferase 1, choline, | 661 | 845 | 1057 | 1623 | 1586 | 1538 | 1502 | 1548 | 1656 | 1477 | 1506 |
| 1878 | 0.04 | 0.11 | 0.40 | 0.69 | 1 | Itsn2 | intersectin 2 | 2497 | 1977 | 1972 | 1601 | 1381 | 2537 | 1500 | 1682 | 1467 | 1532 | 2508 |
| 1606 | 0.04 | 0.12 | 0.36 | 0.72 | 1 | Nipa2 | non imprinted in Prader-Willi/Angelman s | 1363 | 1536 | 1221 | 1813 | 1945 | 1415 | 1500 | 1850 | 1812 | 1824 | 1392 |

| | | | | | | | | | | | | | | | | | | |
|---|---|---|---|---|---|---|---|---|---|---|---|---|---|---|---|---|---|---|
| 1529 | -0.03 | 0.13 | -0.23 | 0.82 | 1 | Tex2 | testis expressed gene 2 | 1208 | 1256 | 1578 | 1455 | 1771 | 1692 | 1500 | 1141 | 1942 | 1647 | 1629 |
| 1655 | 0.08 | 0.12 | 0.62 | 0.53 | 1 | Ftsj3 | FtsJ RNA methyltransferase homolog 3 (E | 2054 | 1800 | 1596 | 1276 | 1671 | 1767 | 1499 | 1387 | 1665 | 1718 | 1773 |
| 1648 | -0.16 | 0.10 | -1.54 | 0.12 | 1 | Fbrs | fibrosin | 1654 | 1511 | 2083 | 1519 | 1665 | 2103 | 1499 | 1208 | 1749 | 1374 | 1765 |
| 1503 | -0.13 | 0.12 | -1.12 | 0.26 | 1 | Eif4enif1 | eukaryotic translation initiation factor 4E r | 1556 | 1749 | 1438 | 1642 | 1401 | 1627 | 1499 | 1284 | 1482 | 1238 | 1622 |
| 1459 | -0.11 | 0.13 | -0.85 | 0.40 | 1 | Runx1t1 | RUNX1 translocation partner 1 | 1240 | 1811 | 1438 | 1443 | 1316 | 1748 | 1499 | 1400 | 1445 | 1198 | 1515 |
| 1353 | -0.13 | 0.14 | -0.96 | 0.34 | 1 | Tmem14c | transmembrane protein 14C | 1367 | 1121 | 1197 | 1628 | 1548 | 1187 | 1499 | 1585 | 1210 | 1283 | 1256 |
| 1243 | 0.07 | 0.11 | 0.66 | 0.51 | 1 | Vegfb | vascular endothelial growth factor B | 781 | 781 | 634 | 1555 | 1539 | 990 | 1499 | 1659 | 1487 | 1736 | 1013 |
| 1580 | 0.04 | 0.17 | 0.26 | 0.80 | 1 | Ehd1 | EH-domain containing 1 | 3055 | 1885 | 1658 | 1127 | 1573 | 1125 | 1498 | 1269 | 1588 | 1447 | 1155 |
| 1452 | -0.06 | 0.14 | -0.40 | 0.69 | 1 | Cyp20a1 | cytochrome P450, family 20, subfamily a, | 1383 | 1065 | 1335 | 1874 | 1743 | 1092 | 1498 | 1382 | 1765 | 1732 | 1103 |
| 1240 | 0.08 | 0.13 | 0.57 | 0.57 | 1 | Slc35d3 | solute carrier family 35, member D3 | 427 | 596 | 516 | 1403 | 1586 | 1410 | 1497 | 1556 | 1537 | 1632 | 1485 |
| 1124 | 0.10 | 0.14 | 0.74 | 0.46 | 1 | Cux2 | cut-like homeobox 2 | 617 | 697 | 854 | 1225 | 1316 | 871 | 1497 | 1424 | 1502 | 1465 | 898 |
| 1819 | 0.02 | 0.10 | 0.18 | 0.86 | 1 | Rgl1 | ral guanine nucleotide dissociation stimul | 3451 | 2619 | 2650 | 1432 | 1371 | 1290 | 1496 | 1420 | 1358 | 1714 | 1207 |
| 1431 | 0.06 | 0.14 | 0.45 | 0.65 | 1 | Fam155a | family with sequence similarity 155, meml | 943 | 1433 | 1045 | 1534 | 1491 | 1510 | 1496 | 1481 | 1654 | 1548 | 1605 |
| 1442 | 0.11 | 0.11 | 1.04 | 0.30 | 1 | Ppp1r7 | protein phosphatase 1, regulatory subuni | 1908 | 1682 | 1459 | 1244 | 1362 | 1106 | 1494 | 1327 | 1468 | 1513 | 1303 |
| 1318 | -0.03 | 0.11 | -0.25 | 0.80 | 1 | Orai3 | ORAI calcium release-activated calcium r | 1174 | 1180 | 1164 | 1481 | 1515 | 1049 | 1494 | 1680 | 1405 | 1301 | 1053 |
| 2277 | 0.09 | 0.13 | 0.68 | 0.50 | 1 | Zc3h7a | zinc finger CCCH type containing 7 A | 1023 | 1157 | 1211 | 1122 | 1672 | 6203 | 1493 | 1623 | 1727 | 1225 | 6596 |
| 1367 | 0.04 | 0.11 | 0.34 | 0.74 | 1 | Tex264 | testis expressed gene 264 | 1129 | 1098 | 982 | 1483 | 1632 | 1212 | 1493 | 1734 | 1477 | 1614 | 1180 |
| 1435 | -0.03 | 0.13 | -0.23 | 0.82 | 1 | Uty | ubiquitously transcribed tetratricopeptide | 876 | 900 | 769 | 1711 | 1431 | 2041 | 1491 | 1278 | 1524 | 1642 | 2123 |
| 1377 | 0.03 | 0.10 | 0.34 | 0.73 | 1 | Bmt2 | base methyltransferase of 25S rRNA 2 | 1515 | 1430 | 1305 | 1326 | 1205 | 1357 | 1491 | 1526 | 1149 | 1422 | 1423 |
| 1736 | 0.06 | 0.11 | 0.58 | 0.56 | 1 | Vwa5a | von Willebrand factor A domain containin | 2920 | 2234 | 2781 | 1228 | 1526 | 1236 | 1490 | 1427 | 1470 | 1391 | 1395 |
| 1664 | 0.12 | 0.11 | 1.13 | 0.26 | 1 | Arsa | arylsulfatase A | 1730 | 1681 | 1845 | 1836 | 1612 | 1301 | 1490 | 1974 | 1677 | 1814 | 1339 |
| 1403 | 0.17 | 0.13 | 1.29 | 0.20 | 1 | Elavl4 | ELAV like RNA binding protein 4 | 671 | 918 | 892 | 1399 | 1388 | 1830 | 1490 | 1657 | 1752 | 1638 | 1793 |
| 1396 | 0.17 | 0.14 | 1.26 | 0.21 | 1 | Jak2 | Janus kinase 2 | 1844 | 1562 | 1037 | 1239 | 1122 | 1274 | 1490 | 1248 | 1362 | 1829 | 1354 |
| 1722 | 0.02 | 0.09 | 0.23 | 0.82 | 1 | Mbd1 | methyl-CpG binding domain protein 1 | 1667 | 1882 | 1991 | 1401 | 1558 | 2221 | 1489 | 1446 | 1644 | 1511 | 2136 |
| 1655 | 0.09 | 0.10 | 0.89 | 0.38 | 1 | Srpk1 | serine/arginine-rich protein specific kinas | 1158 | 1098 | 1185 | 1577 | 1508 | 2647 | 1489 | 1731 | 1677 | 1671 | 2468 |
| 1152 | -0.20 | 0.10 | -1.92 | 0.05 | 1 | Pycr1 | pyrroline-5-carboxylate reductase 1 | 795 | 855 | 779 | 1644 | 1441 | 877 | 1489 | 1438 | 1210 | 1427 | 721 |
| 1760 | 0.20 | 0.15 | 1.33 | 0.18 | 1 | D8Ertd738e | DNA segment, Chr 8, ERATO Doi 738, ex | 1647 | 1511 | 1982 | 1640 | 1951 | 1539 | 1488 | 2389 | 1797 | 1879 | 1532 |
| 1176 | 0.14 | 0.15 | 0.93 | 0.35 | 1 | Ptprr | protein tyrosine phosphatase, receptor typ | 608 | 666 | 854 | 1368 | 1351 | 933 | 1488 | 1387 | 1439 | 1789 | 1051 |
| 1592 | -0.01 | 0.11 | -0.07 | 0.95 | 1 | Rab35 | RAB35, member RAS oncogene family | 2139 | 2270 | 1926 | 1409 | 1373 | 1316 | 1487 | 1424 | 1309 | 1680 | 1183 |
| 1194 | -0.07 | 0.19 | -0.36 | 0.72 | 1 | Tmem255a | transmembrane protein 255A | 588 | 758 | 397 | 1841 | 1252 | 1283 | 1486 | 1474 | 1672 | 1282 | 1106 |
| 2521 | -0.29 | 0.17 | -1.72 | 0.09 | 1 | C8b | complement component 8, beta polypepti | 6136 | 6647 | 6559 | 1015 | 916 | 1169 | 1485 | 767 | 674 | 1382 | 981 |
| 1557 | -0.01 | 0.11 | -0.06 | 0.95 | 1 | Dcaf12 | DDB1 and CUL4 associated factor 12 | 1914 | 1435 | 1648 | 1400 | 1722 | 1461 | 1485 | 1264 | 1564 | 1654 | 1582 |
| 1696 | 0.06 | 0.14 | 0.42 | 0.68 | 1 | Eps8l2 | EPS8-like 2 | 1484 | 1562 | 1887 | 1694 | 1471 | 2092 | 1484 | 1534 | 1817 | 1875 | 1753 |
| 1596 | 0.06 | 0.11 | 0.57 | 0.57 | 1 | Zkscan8 | zinc finger with KRAB and SCAN domains | 1362 | 1535 | 1372 | 1588 | 1290 | 2175 | 1484 | 1510 | 1583 | 1667 | 1987 |
| 1427 | 0.02 | 0.11 | 0.17 | 0.87 | 1 | Nphp1 | nephronophthisis 1 (juvenile) homolog (h | 1025 | 1183 | 1248 | 1611 | 1550 | 1430 | 1484 | 1790 | 1447 | 1392 | 1537 |
| 1279 | -0.17 | 0.13 | -1.26 | 0.21 | 1 | Slc11a2 | solute carrier family 11 (proton-coupled d | 1165 | 1025 | 1347 | 1154 | 1350 | 1546 | 1484 | 848 | 1192 | 1441 | 1513 |
| 1367 | -0.04 | 0.15 | -0.28 | 0.78 | 1 | Ccdc92b | coiled-coil domain containing 92B | 1126 | 1072 | 1372 | 1665 | 1436 | 1189 | 1483 | 1322 | 1496 | 1863 | 1013 |
| 1359 | 0.11 | 0.12 | 0.91 | 0.36 | 1 | Dcun1d3 | DCN1, defective in cullin neddylation 1, d | 754 | 1027 | 774 | 1604 | 1350 | 1527 | 1483 | 1532 | 1654 | 1617 | 1626 |
| 1626 | -0.03 | 0.09 | -0.33 | 0.74 | 1 | Rps27l | ribosomal protein S27-like | 1929 | 1843 | 2263 | 1545 | 1745 | 1224 | 1482 | 1466 | 1703 | 1405 | 1287 |
| 1599 | -0.03 | 0.16 | -0.17 | 0.86 | 1 | Qser1 | glutamine and serine rich 1 | 1802 | 2155 | 1625 | 1513 | 1386 | 1680 | 1482 | 1163 | 1714 | 1510 | 1554 |
| 1591 | -0.18 | 0.14 | -1.25 | 0.21 | 1 | Tmbim1 | transmembrane BAX inhibitor motif contai | 2520 | 1897 | 2563 | 1410 | 1587 | 1095 | 1482 | 1532 | 1341 | 1180 | 897 |
| 1294 | 0.17 | 0.11 | 1.50 | 0.13 | 1 | Tm2d1 | TM2 domain containing 1 | 935 | 995 | 1068 | 1378 | 1363 | 1061 | 1482 | 1452 | 1491 | 1771 | 1236 |
| 1249 | 0.15 | 0.19 | 0.80 | 0.43 | 1 | Egf | epidermal growth factor | 601 | 1319 | 862 | 1504 | 1161 | 1040 | 1482 | 1680 | 1413 | 1597 | 1086 |
| 1525 | -0.33 | 0.18 | -1.79 | 0.07 | 1 | Dop1a | DOP1 leucine zipper like protein A | 1367 | 1079 | 1148 | 1592 | 1667 | 2440 | 1481 | 753 | 1458 | 1466 | 2326 |
| 1284 | -0.09 | 0.14 | -0.65 | 0.51 | 1 | Strn | striatin, calmodulin binding protein | 1234 | 967 | 1116 | 1352 | 1224 | 1521 | 1481 | 1276 | 1364 | 1055 | 1538 |
| 1983 | -0.04 | 0.14 | -0.31 | 0.75 | 1 | Tns2 | tensin 2 | 3973 | 3189 | 2623 | 1670 | 1658 | 1269 | 1480 | 1455 | 1662 | 1704 | 1126 |
| 1674 | 0.00 | 0.13 | -0.02 | 0.98 | 1 | Trappc12 | trafficking protein particle complex 12 | 1382 | 1694 | 1583 | 1761 | 2028 | 1593 | 1480 | 1464 | 2345 | 1489 | 1597 |
| 1326 | 0.12 | 0.13 | 0.90 | 0.37 | 1 | Surf1 | surfeit gene 1 | 1205 | 1061 | 1381 | 1264 | 1351 | 1161 | 1480 | 1554 | 1321 | 1503 | 1300 |
| 1224 | 0.00 | 0.10 | 0.05 | 0.96 | 1 | E2f6 | E2F transcription factor 6 | 773 | 739 | 922 | 1385 | 1530 | 1104 | 1480 | 1457 | 1572 | 1437 | 1065 |
| 1548 | 0.03 | 0.10 | 0.33 | 0.74 | 1 | Polr2e | polymerase (RNA) II (DNA directed) polyp | 2257 | 1675 | 2034 | 1319 | 1522 | 1144 | 1479 | 1469 | 1457 | 1530 | 1138 |
| 1383 | 0.15 | 0.10 | 1.55 | 0.12 | 1 | Rpe | ribulose-5-phosphate-3-epimerase | 1324 | 1112 | 1115 | 1452 | 1328 | 1275 | 1479 | 1565 | 1560 | 1608 | 1399 |
| 1204 | -0.08 | 0.11 | -0.76 | 0.45 | 1 | Rtl8c | retrotransposon Gag like 8C | 1167 | 1133 | 1172 | 1171 | 1545 | 823 | 1479 | 1251 | 1376 | 1412 | 719 |
| 1249 | 0.00 | 0.13 | 0.03 | 0.98 | 1 | Ror2 | receptor tyrosine kinase-like orphan recep | 820 | 627 | 676 | 1619 | 1589 | 1075 | 1478 | 1328 | 1662 | 1830 | 1031 |
| 1746 | 0.03 | 0.11 | 0.28 | 0.78 | 1 | Ice1 | interactor of little elongation complex ELL | 1711 | 1608 | 1431 | 1651 | 1392 | 2688 | 1477 | 1464 | 1660 | 1676 | 2448 |
| 1469 | 0.04 | 0.09 | 0.46 | 0.64 | 1 | Ptpn21 | protein tyrosine phosphatase, non-recept | 1257 | 1380 | 1229 | 1764 | 1487 | 1336 | 1477 | 1710 | 1597 | 1544 | 1373 |
| 1460 | -0.02 | 0.10 | -0.17 | 0.87 | 1 | Wdr61 | WD repeat domain 61 | 1662 | 1561 | 1715 | 1311 | 1422 | 1378 | 1477 | 1178 | 1468 | 1435 | 1457 |
| 1360 | -0.05 | 0.12 | -0.42 | 0.68 | 1 | Ypel3 | yippee like 3 | 1533 | 1260 | 1689 | 1172 | 1298 | 1393 | 1477 | 1288 | 1303 | 1200 | 1343 |
| 1803 | 0.09 | 0.17 | 0.50 | 0.62 | 1 | Gatm | glycine amidinotransferase (L-arginine:gl | 2558 | 3616 | 2835 | 1323 | 1176 | 1265 | 1476 | 1179 | 1788 | 1346 | 1273 |
| 1592 | -0.07 | 0.15 | -0.48 | 0.63 | 1 | Ddx54 | DEAD (Asp-Glu-Ala-Asp) box polypeptide | 2181 | 1749 | 1442 | 1567 | 1792 | 1381 | 1475 | 1425 | 1648 | 1544 | 1307 |
| 1526 | -0.10 | 0.15 | -0.65 | 0.51 | 1 | Sbf1 | SET binding factor 1 | 2178 | 1400 | 1549 | 1455 | 1319 | 1794 | 1474 | 1387 | 1361 | 1168 | 1705 |

| | | | | | | | | | | | | | | | | | | |
|---|---|---|---|---|---|---|---|---|---|---|---|---|---|---|---|---|---|---|
| 1447 | -0.02 | 0.13 | -0.16 | 0.88 | 1 | Ccdc97 | coiled-coil domain containing 97 | 1598 | 1405 | 1454 | 1216 | 1604 | 1484 | 1474 | 1500 | 1611 | 1256 | 1317 |
| 1399 | 0.19 | 0.16 | 1.21 | 0.23 | 1 | Trappc2l | trafficking protein particle complex 2-like | 1150 | 991 | 1266 | 1602 | 1381 | 1112 | 1474 | 2121 | 1370 | 1727 | 1197 |
| 1676 | 0.05 | 0.14 | 0.35 | 0.73 | 1 | Gart | phosphoribosylglycinamide formyltransfe | 2294 | 1460 | 1833 | 1399 | 1609 | 1837 | 1473 | 1610 | 1379 | 1640 | 1906 |
| 1508 | 0.00 | 0.11 | 0.04 | 0.97 | 1 | Ptar1 | protein prenyltransferase alpha subunit re | 1022 | 1040 | 985 | 1455 | 1546 | 2312 | 1472 | 1525 | 1785 | 1351 | 2094 |
| 1491 | 0.06 | 0.11 | 0.60 | 0.55 | 1 | Ola1 | Obg-like ATPase 1 | 1816 | 1643 | 1860 | 1444 | 1315 | 1190 | 1472 | 1327 | 1512 | 1421 | 1401 |
| 1875 | 0.16 | 0.13 | 1.20 | 0.23 | 1 | Ikbip | IKBKB interacting protein | 2569 | 2663 | 2820 | 1331 | 1655 | 1468 | 1470 | 1808 | 1441 | 1939 | 1457 |
| 1495 | -0.02 | 0.12 | -0.17 | 0.87 | 1 | Phactr4 | phosphatase and actin regulator 4 | 1247 | 1334 | 1489 | 1744 | 1604 | 1393 | 1469 | 1455 | 1727 | 1757 | 1222 |
| 1055 | -0.10 | 0.16 | -0.60 | 0.55 | 1 | Fam169a | family with sequence similarity 169, meml | 540 | 572 | 578 | 1520 | 1116 | 1028 | 1469 | 1017 | 1198 | 1485 | 1077 |
| 1459 | -0.15 | 0.16 | -0.98 | 0.33 | 1 | Rybp | RING1 and YY1 binding protein | 1373 | 1819 | 1981 | 1446 | 1552 | 1254 | 1468 | 1072 | 1410 | 1562 | 1111 |
| 1376 | 0.03 | 0.12 | 0.27 | 0.79 | 1 | Kifap3 | kinesin-associated protein 3 | 1384 | 1252 | 1101 | 1455 | 1451 | 1244 | 1467 | 1636 | 1557 | 1468 | 1119 |
| 1616 | 0.22 | 0.16 | 1.39 | 0.17 | 1 | Rtf1 | RTF1, Paf1/RNA polymerase II complex c | 2001 | 1721 | 1573 | 1334 | 1528 | 1401 | 1466 | 2234 | 1376 | 1622 | 1523 |
| 1518 | -0.13 | 0.14 | -0.93 | 0.35 | 1 | Marchf7 | membrane associated ring-CH-type finge | 1498 | 1147 | 1218 | 999 | 1294 | 2903 | 1466 | 994 | 1318 | 1012 | 2850 |
| 1461 | -0.05 | 0.14 | -0.36 | 0.72 | 1 | Ralgapa2 | Ral GTPase activating protein, alpha subu | 1079 | 1613 | 1378 | 1538 | 1425 | 1659 | 1465 | 1368 | 1695 | 1217 | 1637 |
| 1459 | 0.05 | 0.13 | 0.38 | 0.70 | 1 | Ndufa7 | NADH:ubiquinone oxidoreductase subun | 1317 | 1332 | 1634 | 1635 | 1400 | 1261 | 1465 | 1865 | 1426 | 1488 | 1223 |
| 1420 | -0.08 | 0.12 | -0.73 | 0.47 | 1 | Ipp | IAP promoted placental gene | 1322 | 1624 | 1260 | 1414 | 1525 | 1461 | 1465 | 1441 | 1415 | 1392 | 1296 |
| 1247 | -0.12 | 0.12 | -1.01 | 0.31 | 1 | Vopp1 | vesicular, overexpressed in cancer, prosu | 1730 | 1529 | 1961 | 1032 | 1251 | 701 | 1465 | 980 | 1155 | 1197 | 711 |
| 1244 | -0.01 | 0.14 | -0.04 | 0.97 | 1 | Chd5 | chromodomain helicase DNA binding pro | 681 | 816 | 850 | 1273 | 1407 | 1532 | 1464 | 1345 | 1456 | 1571 | 1284 |
| 1620 | 0.06 | 0.11 | 0.56 | 0.57 | 1 | Rbpms | RNA binding protein gene with multiple sp | 1957 | 1887 | 1577 | 1522 | 1396 | 1685 | 1463 | 1506 | 1417 | 1820 | 1592 |
| 1435 | 0.07 | 0.09 | 0.73 | 0.47 | 1 | Hadhb | hydroxyacyl-CoA dehydrogenase trifuncti | 1280 | 1513 | 1255 | 1517 | 1417 | 1325 | 1463 | 1695 | 1477 | 1556 | 1293 |
| 1219 | -0.09 | 0.19 | -0.48 | 0.63 | 1 | Dph6 | diphthamine biosynthesis 6 | 1066 | 1437 | 1065 | 1174 | 1331 | 1101 | 1463 | 1003 | 1426 | 992 | 1354 |
| 1918 | 0.08 | 0.11 | 0.71 | 0.48 | 1 | Hmgb1 | high mobility group box 1 | 2302 | 2119 | 2429 | 1614 | 1931 | 1901 | 1462 | 1795 | 1984 | 1381 | 2180 |
| 1672 | 0.10 | 0.14 | 0.69 | 0.49 | 1 | Abl2 | v-abl Abelson murine leukemia viral onco | 1685 | 1612 | 1451 | 1885 | 1531 | 1673 | 1462 | 2418 | 1628 | 1597 | 1452 |
| 1480 | -0.25 | 0.15 | -1.66 | 0.10 | 1 | Tmem260 | transmembrane protein 260 | 1487 | 1517 | 1653 | 1588 | 1625 | 1595 | 1462 | 937 | 1637 | 1360 | 1422 |
| 1296 | 0.18 | 0.13 | 1.33 | 0.18 | 1 | Dhrs4 | dehydrogenase/reductase (SDR family) m | 642 | 463 | 607 | 1337 | 1373 | 1695 | 1461 | 1495 | 1438 | 2065 | 1675 |
| 1502 | 0.04 | 0.10 | 0.39 | 0.70 | 1 | Poglut1 | protein O-glucosyltransferase 1 | 1838 | 1727 | 1558 | 1403 | 1537 | 1227 | 1460 | 1539 | 1425 | 1489 | 1324 |
| 1405 | 0.01 | 0.15 | 0.09 | 0.93 | 1 | Pelp1 | proline, glutamic acid and leucine rich pro | 1568 | 1402 | 1282 | 1486 | 1384 | 1207 | 1460 | 1915 | 1416 | 1255 | 1078 |
| 1258 | -0.12 | 0.15 | -0.79 | 0.43 | 1 | Eml6 | echinoderm microtubule associated prote | 647 | 598 | 762 | 1074 | 1416 | 2119 | 1460 | 779 | 1605 | 1231 | 2151 |
| 1647 | 0.26 | 0.13 | 1.95 | 0.05 | 1 | Anks4b | ankyrin repeat and sterile alpha motif dom | 462 | 770 | 864 | 2331 | 1871 | 1581 | 1459 | 3103 | 2233 | 1703 | 1737 |
| 8475 | -0.23 | 0.36 | -0.66 | 0.51 | 1 | Meg3 | | 689 | 1481 | 906 | 753 | 1189 | 44722 | 1458 | 708 | 1183 | 976 | 39156 |
| 1539 | -0.01 | 0.14 | -0.07 | 0.95 | 1 | Tmem134 | transmembrane protein 134 | 1529 | 1323 | 1714 | 1556 | 1702 | 1469 | 1456 | 1573 | 1397 | 1918 | 1292 |
| 1432 | 0.03 | 0.13 | 0.23 | 0.82 | 1 | Krr1 | KRR1, small subunit (SSU) processome c | 1310 | 1269 | 1732 | 1332 | 1307 | 1554 | 1456 | 1447 | 1334 | 1314 | 1695 |
| 1320 | 0.15 | 0.12 | 1.34 | 0.18 | 1 | Nelfe | negative elongation factor complex memb | 1257 | 1127 | 1377 | 1182 | 1375 | 1097 | 1456 | 1495 | 1390 | 1465 | 1305 |
| 2047 | 0.10 | 0.13 | 0.72 | 0.47 | 1 | Ttc14 | tetratricopeptide repeat domain 14 | 773 | 872 | 598 | 1347 | 1233 | 5762 | 1455 | 1196 | 1410 | 1717 | 6157 |
| 1685 | 0.13 | 0.12 | 1.07 | 0.29 | 1 | Mesd | mesoderm development LRP chaperone | 2429 | 1976 | 2294 | 1325 | 1676 | 1173 | 1455 | 1711 | 1648 | 1767 | 1080 |
| 1449 | -0.02 | 0.10 | -0.18 | 0.86 | 1 | Kdm4b | lysine (K)-specific demethylase 4B | 1473 | 1202 | 1381 | 1444 | 1561 | 1517 | 1455 | 1498 | 1503 | 1546 | 1361 |
| 1328 | 0.02 | 0.15 | 0.14 | 0.89 | 1 | Myt1l | myelin transcription factor 1-like | 872 | 1196 | 654 | 1180 | 992 | 2320 | 1455 | 1327 | 1042 | 1428 | 2144 |
| 1326 | 0.25 | 0.17 | 1.47 | 0.14 | 1 | Tmprss4 | transmembrane protease, serine 4 | 78 | 172 | 148 | 2072 | 1609 | 1407 | 1455 | 2016 | 1930 | 1922 | 1775 |
| 1523 | 0.07 | 0.11 | 0.60 | 0.55 | 1 | Ptprm | protein tyrosine phosphatase, receptor typ | 1990 | 1618 | 1673 | 1503 | 1625 | 984 | 1454 | 1498 | 1796 | 1684 | 934 |
| 1499 | 0.08 | 0.12 | 0.67 | 0.50 | 1 | Vps54 | VPS54 GARP complex subunit | 1386 | 1282 | 1545 | 1502 | 1369 | 1620 | 1454 | 1374 | 1613 | 1396 | 1947 |
| 1380 | 0.00 | 0.09 | 0.05 | 0.96 | 1 | BC017158 | cDNA sequence BC017158 | 782 | 719 | 737 | 1456 | 1437 | 2134 | 1454 | 1548 | 1448 | 1483 | 1985 |
| 1660 | 0.06 | 0.13 | 0.44 | 0.66 | 1 | Rtf2 | replication termination factor 2 | 2144 | 1782 | 1937 | 1608 | 1572 | 1432 | 1453 | 1836 | 1722 | 1359 | 1411 |
| 1544 | 0.25 | 0.15 | 1.63 | 0.10 | 1 | Ndufb3 | NADH:ubiquinone oxidoreductase subun | 1424 | 1246 | 1763 | 1572 | 1545 | 1163 | 1453 | 2017 | 1484 | 1890 | 1424 |
| 1496 | 0.06 | 0.13 | 0.47 | 0.64 | 1 | Psmb2 | proteasome (prosome, macropain) subun | 1590 | 1376 | 1282 | 1284 | 1763 | 1473 | 1453 | 1362 | 1717 | 1681 | 1472 |
| 1450 | -0.17 | 0.14 | -1.22 | 0.22 | 1 | Tanc1 | tetratricopeptide repeat, ankyrin repeat ar | 2314 | 2204 | 2086 | 1590 | 978 | 938 | 1453 | 1181 | 1171 | 1262 | 771 |
| 1376 | 0.03 | 0.13 | 0.25 | 0.80 | 1 | Armcx1 | armadillo repeat containing, X-linked 1 | 1137 | 1220 | 981 | 1333 | 1169 | 1839 | 1452 | 1184 | 1340 | 1293 | 2191 |
| 1334 | 0.00 | 0.14 | 0.02 | 0.99 | 1 | Ndufs6 | NADH:ubiquinone oxidoreductase core s | 1212 | 1231 | 1614 | 1420 | 1500 | 927 | 1452 | 1679 | 1382 | 1334 | 921 |
| 1709 | 0.22 | 0.16 | 1.39 | 0.17 | 1 | Phf1 | PHD finger protein 1 | 1191 | 1291 | 1436 | 1217 | 1391 | 2967 | 1451 | 2085 | 1538 | 1550 | 2682 |
| 1580 | 0.14 | 0.13 | 1.04 | 0.30 | 1 | Ppp4c | protein phosphatase 4, catalytic subunit | 1948 | 1518 | 1622 | 1322 | 1679 | 1391 | 1450 | 1747 | 1420 | 1778 | 1500 |
| 1547 | 0.13 | 0.11 | 1.20 | 0.23 | 1 | Arl4a | ADP-ribosylation factor-like 4A | 1685 | 1375 | 1413 | 1371 | 1345 | 1827 | 1450 | 1628 | 1492 | 1432 | 1999 |
| 1540 | -0.04 | 0.12 | -0.38 | 0.70 | 1 | St3gal1 | ST3 beta-galactoside alpha-2,3-sialyltran | 2434 | 1731 | 1966 | 1484 | 1324 | 1226 | 1450 | 1331 | 1291 | 1468 | 1233 |
| 1194 | -0.18 | 0.13 | -1.39 | 0.16 | 1 | Etv5 | ets variant 5 | 741 | 607 | 571 | 1828 | 1365 | 1329 | 1450 | 1425 | 1152 | 1365 | 1298 |
| 1166 | -0.13 | 0.16 | -0.79 | 0.43 | 1 | Ehbp1 | EH domain binding protein 1 | 1512 | 1090 | 1049 | 1181 | 1014 | 1168 | 1450 | 1200 | 1014 | 949 | 1199 |
| 1091 | -0.09 | 0.13 | -0.68 | 0.50 | 1 | Dhcr24 | 24-dehydrocholesterol reductase | 735 | 797 | 546 | 1443 | 1252 | 987 | 1450 | 1247 | 1221 | 1306 | 1020 |
| 1835 | 0.18 | 0.12 | 1.54 | 0.12 | 1 | Rnf146 | ring finger protein 146 | 1801 | 1672 | 1943 | 1757 | 1698 | 2038 | 1449 | 2184 | 1901 | 1723 | 2019 |
| 1332 | 0.04 | 0.16 | 0.23 | 0.82 | 1 | Slc25a22 | solute carrier family 25 (mitochondrial car | 966 | 1130 | 1329 | 1293 | 1476 | 1295 | 1449 | 1015 | 1672 | 1653 | 1372 |
| 1185 | 0.07 | 0.13 | 0.53 | 0.60 | 1 | Acaca | acetyl-Coenzyme A carboxylase alpha | 913 | 828 | 979 | 1314 | 1289 | 972 | 1449 | 1623 | 1423 | 1239 | 1005 |
| 1544 | 0.13 | 0.09 | 1.56 | 0.12 | 1 | Fbxo22 | F-box protein 22 | 1430 | 1654 | 1430 | 1628 | 1473 | 1397 | 1447 | 1755 | 1693 | 1538 | 1534 |
| 1428 | 0.21 | 0.11 | 1.89 | 0.06 | 1 | Hsdl1 | hydroxysteroid dehydrogenase like 1 | 1424 | 1396 | 1343 | 1496 | 1234 | 1151 | 1446 | 1870 | 1437 | 1767 | 1148 |
| 3121 | 0.06 | 0.16 | 0.37 | 0.71 | 1 | Ctsz | cathepsin Z | 8471 | 7209 | 6942 | 1750 | 1528 | 974 | 1445 | 2321 | 1416 | 1247 | 1034 |
| 1261 | -0.08 | 0.15 | -0.55 | 0.58 | 1 | Abitram | actin binding transcription modulator | 718 | 998 | 769 | 1546 | 1176 | 1702 | 1445 | 1244 | 1455 | 1220 | 1603 |

| | | | | | | | | | | | | | | | | | | |
|---|---|---|---|---|---|---|---|---|---|---|---|---|---|---|---|---|---|---|
| 1610 | -0.04 | 0.16 | -0.27 | 0.79 | 1 | Zfp612 | zinc finger protein 612 | 741 | 1418 | 1060 | 1561 | 1470 | 2892 | 1444 | 1371 | 1512 | 1447 | 2798 |
| 1477 | -0.03 | 0.14 | -0.22 | 0.83 | 1 | Ncln | nicalin | 2027 | 1354 | 1642 | 1588 | 1448 | 1177 | 1444 | 1248 | 1454 | 1614 | 1253 |
| 2577 | -0.11 | 0.17 | -0.64 | 0.52 | 1 | Tpm1 | tropomyosin 1, alpha | 7715 | 5124 | 4745 | 1453 | 1321 | 1363 | 1443 | 1621 | 1122 | 1024 | 1417 |
| 1342 | 0.05 | 0.13 | 0.37 | 0.71 | 1 | Tmem19 | transmembrane protein 19 | 1249 | 973 | 892 | 1678 | 1384 | 1210 | 1443 | 1535 | 1515 | 1704 | 1180 |
| 1294 | 0.01 | 0.14 | 0.04 | 0.97 | 1 | Dpagt1 | dolichyl-phosphate (UDP-N-acetylglucosa | 994 | 721 | 884 | 1032 | 1780 | 1530 | 1443 | 965 | 1670 | 1761 | 1449 |
| 1174 | 0.07 | 0.14 | 0.55 | 0.58 | 1 | Six4 | sine oculis-related homeobox 4 | 453 | 414 | 431 | 1048 | 1126 | 2128 | 1443 | 1356 | 1300 | 1419 | 1799 |
| 1068 | -0.02 | 0.14 | -0.16 | 0.88 | 1 | Ptprs | protein tyrosine phosphatase, receptor typ | 1371 | 755 | 1155 | 584 | 1089 | 1129 | 1443 | 542 | 1176 | 1497 | 1008 |
| 1549 | -0.05 | 0.11 | -0.41 | 0.68 | 1 | Sphk2 | sphingosine kinase 2 | 1269 | 1245 | 1318 | 1846 | 1761 | 1664 | 1442 | 1778 | 1518 | 1668 | 1527 |
| 1405 | -0.04 | 0.11 | -0.34 | 0.73 | 1 | Prkag1 | protein kinase, AMP-activated, gamma 1 r | 1821 | 1612 | 1750 | 1111 | 1395 | 1263 | 1442 | 1247 | 1191 | 1322 | 1303 |
| 1391 | 0.20 | 0.20 | 1.00 | 0.32 | 1 | Timm13 | translocase of inner mitochondrial membr | 1293 | 1150 | 1598 | 1332 | 1403 | 1024 | 1442 | 2288 | 1344 | 1337 | 1094 |
| 2188 | 0.08 | 0.09 | 0.95 | 0.34 | 1 | Tnfaip1 | tumor necrosis factor, alpha-induced prot | 3370 | 3440 | 3711 | 1574 | 1896 | 1677 | 1441 | 1621 | 1912 | 1720 | 1711 |
| 1866 | -0.07 | 0.10 | -0.71 | 0.48 | 1 | Slc40a1 | solute carrier family 40 (iron-regulated tra | 1897 | 2852 | 1937 | 2531 | 1497 | 1622 | 1441 | 2353 | 1481 | 1211 | 1704 |
| 1847 | -0.15 | 0.10 | -1.54 | 0.12 | 1 | Fermt2 | fermitin family member 2 | 2900 | 3229 | 2958 | 1555 | 1231 | 1673 | 1441 | 1243 | 1209 | 1278 | 1595 |
| 1378 | -0.20 | 0.15 | -1.35 | 0.18 | 1 | Dlg5 | discs large MAGUK scaffold protein 5 | 1206 | 1627 | 1376 | 1285 | 1431 | 1696 | 1441 | 993 | 1446 | 1177 | 1483 |
| 1435 | -0.16 | 0.17 | -0.96 | 0.33 | 1 | Zfyve1 | zinc finger, FYVE domain containing 1 | 1295 | 1268 | 1478 | 1633 | 1411 | 1676 | 1440 | 977 | 1376 | 1685 | 1547 |
| 1483 | -0.12 | 0.11 | -1.13 | 0.26 | 1 | Gnpat | glyceronephosphate O-acyltransferase | 1600 | 1842 | 1534 | 1519 | 1474 | 1472 | 1439 | 1382 | 1161 | 1507 | 1385 |
| 1482 | -0.08 | 0.11 | -0.75 | 0.45 | 1 | Tsc22d4 | TSC22 domain family, member 4 | 2239 | 2247 | 2406 | 1234 | 1292 | 883 | 1439 | 1328 | 1145 | 1225 | 867 |
| 1371 | 0.11 | 0.12 | 0.90 | 0.37 | 1 | Coro7 | coronin 7 | 1156 | 905 | 942 | 1463 | 1298 | 1614 | 1439 | 1651 | 1551 | 1580 | 1481 |
| 1896 | 0.12 | 0.11 | 1.17 | 0.24 | 1 | Hspe1 | heat shock protein 1 (chaperonin 10) | 1690 | 1590 | 1797 | 1399 | 1265 | 3468 | 1438 | 1427 | 1359 | 1732 | 3690 |
| 1354 | 0.04 | 0.14 | 0.27 | 0.79 | 1 | Taf10 | TATA-box binding protein associated fact | 1251 | 1288 | 1319 | 1316 | 1603 | 1079 | 1438 | 1785 | 1240 | 1475 | 1105 |
| 1214 | -0.04 | 0.18 | -0.22 | 0.83 | 1 | Sh3gl2 | SH3-domain GRB2-like 2 | 672 | 818 | 1110 | 1226 | 1685 | 1070 | 1438 | 877 | 1764 | 1532 | 1163 |
| 1403 | -0.06 | 0.12 | -0.47 | 0.64 | 1 | Ppp6r2 | protein phosphatase 6, regulatory subuni | 1241 | 969 | 1169 | 1341 | 1213 | 2205 | 1437 | 1240 | 1375 | 1456 | 1792 |
| 1098 | -0.21 | 0.17 | -1.20 | 0.23 | 1 | Cds1 | CDP-diacylglycerol synthase 1 | 327 | 635 | 529 | 1371 | 1590 | 1259 | 1437 | 1447 | 1250 | 1199 | 1038 |
| 1075 | 0.03 | 0.12 | 0.28 | 0.78 | 1 | 9130023H24Ril | RIKEN cDNA 9130023H24 gene | 462 | 533 | 499 | 1497 | 1148 | 1037 | 1437 | 1485 | 1192 | 1383 | 1150 |
| 1330 | 0.10 | 0.14 | 0.73 | 0.46 | 1 | Uqcr11 | ubiquinol-cytochrome c reductase, compl | 1181 | 1237 | 1465 | 1467 | 1243 | 1027 | 1436 | 1724 | 1291 | 1507 | 1057 |
| 1732 | 0.13 | 0.16 | 0.85 | 0.39 | 1 | Paf1 | Paf1, RNA polymerase II complex compon | 2023 | 1665 | 2017 | 1402 | 1630 | 1897 | 1434 | 2100 | 1494 | 1398 | 1986 |
| 1611 | 0.10 | 0.10 | 1.06 | 0.29 | 1 | Noc2l | NOC2 like nucleolar associated transcript | 1555 | 1426 | 1525 | 1498 | 1415 | 2026 | 1434 | 1729 | 1467 | 1525 | 2119 |
| 2210 | -0.09 | 0.14 | -0.64 | 0.52 | 1 | Dusp11 | dual specificity phosphatase 11 (RNA/RN | 2323 | 2038 | 1977 | 1641 | 2061 | 4113 | 1433 | 1294 | 2048 | 1488 | 3895 |
| 2072 | 0.02 | 0.11 | 0.19 | 0.85 | 1 | Cnot6 | CCR4-NOT transcription complex, subuni | 2677 | 2721 | 2798 | 1754 | 1708 | 2387 | 1433 | 1596 | 1975 | 1651 | 2091 |
| 1747 | -0.11 | 0.11 | -1.01 | 0.31 | 1 | Rbm33 | RNA binding motif protein 33 | 1612 | 1815 | 1599 | 1574 | 1409 | 2975 | 1433 | 1388 | 1584 | 1179 | 2648 |
| 1345 | 0.05 | 0.16 | 0.33 | 0.74 | 1 | Rab3ip | RAB3A interacting protein | 1066 | 1423 | 1355 | 1350 | 1382 | 1195 | 1433 | 1413 | 1187 | 1787 | 1201 |
| 1643 | 0.06 | 0.08 | 0.67 | 0.50 | 1 | Arf2 | ADP-ribosylation factor 2 | 2322 | 2256 | 2173 | 1462 | 1381 | 1280 | 1432 | 1432 | 1457 | 1495 | 1385 |
| 2017 | 0.05 | 0.11 | 0.47 | 0.64 | 1 | Ndn | necdin | 3550 | 3522 | 3666 | 1598 | 1485 | 1122 | 1430 | 1335 | 1481 | 1573 | 1422 |
| 1487 | 0.04 | 0.11 | 0.37 | 0.71 | 1 | B4galt5 | UDP-Gal:betaGlcNAc beta 1,4-galactosyl | 1309 | 1389 | 1057 | 1732 | 1608 | 1430 | 1430 | 1865 | 1590 | 1561 | 1387 |
| 1427 | -0.20 | 0.12 | -1.73 | 0.08 | 1 | Amph | amphiphysin | 1368 | 1781 | 1524 | 1659 | 1427 | 1382 | 1430 | 1424 | 1069 | 1391 | 1244 |
| 1334 | 0.17 | 0.14 | 1.20 | 0.23 | 1 | Jagn1 | jagunal homolog 1 | 800 | 1212 | 983 | 1322 | 1516 | 1232 | 1430 | 1592 | 1595 | 1543 | 1443 |
| 1542 | 0.15 | 0.12 | 1.32 | 0.19 | 1 | Stard4 | StAR-related lipid transfer (START) doma | 1205 | 1450 | 1166 | 1708 | 1603 | 1488 | 1429 | 1673 | 1703 | 1910 | 1631 |
| 1540 | -0.21 | 0.13 | -1.63 | 0.10 | 1 | Pim3 | proviral integration site 3 | 2228 | 1489 | 1640 | 1396 | 1641 | 1733 | 1429 | 1146 | 1222 | 1520 | 1502 |
| 1453 | -0.22 | 0.15 | -1.44 | 0.15 | 1 | Wnk4 | WNK lysine deficient protein kinase 4 | 1414 | 2071 | 2003 | 1301 | 1266 | 1617 | 1429 | 1445 | 1146 | 972 | 1322 |
| 1359 | 0.08 | 0.16 | 0.54 | 0.59 | 1 | Zrsr1 | zinc finger (CCCH type), RNA binding mo | 963 | 1357 | 823 | 1248 | 1321 | 1743 | 1429 | 1404 | 1395 | 1544 | 1725 |
| 1484 | 0.18 | 0.14 | 1.35 | 0.18 | 1 | Rab11fip3 | RAB11 family interacting protein 3 (class | 1511 | 1358 | 1283 | 1175 | 1476 | 1625 | 1428 | 1797 | 1398 | 1626 | 1647 |
| 1455 | -0.10 | 0.14 | -0.73 | 0.46 | 1 | Ubap1 | ubiquitin-associated protein 1 | 1531 | 1483 | 1260 | 1578 | 1641 | 1425 | 1427 | 1186 | 1572 | 1386 | 1518 |
| 1243 | 0.15 | 0.14 | 1.07 | 0.28 | 1 | Tbc1d24 | TBC1 domain family, member 24 | 544 | 535 | 550 | 1540 | 978 | 1785 | 1427 | 1462 | 1337 | 1690 | 1819 |
| 1344 | 0.01 | 0.13 | 0.07 | 0.95 | 1 | Ciao1 | cytosolic iron-sulfur protein assembly 1 | 1275 | 1050 | 1234 | 1458 | 1552 | 1169 | 1426 | 1644 | 1423 | 1248 | 1304 |
| 1315 | -0.11 | 0.16 | -0.64 | 0.52 | 1 | 1600012H06Ril | RIKEN cDNA 1600012H06 gene | 1160 | 1064 | 1064 | 1561 | 1464 | 1310 | 1426 | 951 | 1420 | 1623 | 1423 |
| 1674 | -0.04 | 0.10 | -0.45 | 0.65 | 1 | Brip1os | | 2852 | 2506 | 2594 | 1399 | 1489 | 1000 | 1425 | 1482 | 1360 | 1342 | 970 |
| 1664 | -0.18 | 0.11 | -1.66 | 0.10 | 1 | Nadk | NAD kinase | 2538 | 1959 | 2171 | 1394 | 1608 | 1763 | 1425 | 1254 | 1438 | 1296 | 1458 |
| 1514 | 0.04 | 0.13 | 0.34 | 0.73 | 1 | Coq2 | coenzyme Q2 4-hydroxybenzoate polypre | 1224 | 1383 | 1556 | 1495 | 1337 | 1904 | 1425 | 1792 | 1430 | 1380 | 1730 |
| 1242 | 0.13 | 0.14 | 0.97 | 0.33 | 1 | Slc7a4 | solute carrier family 7 (cationic amino aci | 578 | 680 | 620 | 1406 | 1550 | 1184 | 1425 | 1815 | 1486 | 1851 | 1065 |
| 1237 | -0.07 | 0.10 | -0.69 | 0.49 | 1 | Aar2 | AAR2 splicing factor homolog | 1319 | 1367 | 1246 | 1376 | 1207 | 959 | 1425 | 1235 | 1250 | 1262 | 960 |
| 1688 | 0.16 | 0.11 | 1.41 | 0.16 | 1 | Plod1 | procollagen-lysine, 2-oxoglutarate 5-diox | 2563 | 2177 | 2368 | 1581 | 1435 | 947 | 1424 | 1786 | 1480 | 1815 | 987 |
| 1427 | -0.02 | 0.12 | -0.18 | 0.86 | 1 | Ubqln4 | ubiquilin 4 | 1318 | 1294 | 1182 | 1356 | 1704 | 1510 | 1424 | 1381 | 1586 | 1313 | 1629 |
| 1368 | -0.07 | 0.12 | -0.63 | 0.53 | 1 | Poc1b | POC1 centriolar protein B | 1407 | 1677 | 1468 | 1373 | 1536 | 1034 | 1424 | 1478 | 1522 | 1154 | 972 |
| 1300 | -0.07 | 0.17 | -0.42 | 0.67 | 1 | Zfp654 | zinc finger protein 654 | 834 | 995 | 689 | 1469 | 1348 | 1746 | 1424 | 1342 | 1506 | 998 | 1950 |
| 1227 | -0.11 | 0.13 | -0.80 | 0.42 | 1 | Pex3 | peroxisomal biogenesis factor 3 | 939 | 1005 | 955 | 1365 | 1373 | 1318 | 1424 | 1022 | 1435 | 1292 | 1363 |
| 1568 | 0.09 | 0.10 | 0.87 | 0.38 | 1 | Dele1 | DAP3 binding cell death enhancer 1 | 1409 | 1550 | 1409 | 1376 | 1642 | 1808 | 1423 | 1579 | 1640 | 1524 | 1889 |
| 1393 | -0.07 | 0.10 | -0.69 | 0.49 | 1 | Pex26 | peroxisomal biogenesis factor 26 | 826 | 891 | 944 | 1600 | 1487 | 1943 | 1423 | 1455 | 1495 | 1253 | 2002 |
| 1338 | -0.01 | 0.13 | -0.06 | 0.95 | 1 | Tmem9 | transmembrane protein 9 | 1371 | 1152 | 1259 | 1586 | 1387 | 1061 | 1423 | 1408 | 1266 | 1775 | 1030 |
| 1217 | -0.11 | 0.17 | -0.64 | 0.52 | 1 | Rcor3 | REST corepressor 3 | 985 | 1123 | 1031 | 1021 | 1186 | 1642 | 1423 | 774 | 1202 | 1288 | 1708 |
| 1407 | -0.02 | 0.13 | -0.15 | 0.88 | 1 | Poldip2 | polymerase (DNA-directed), delta interact | 1209 | 1151 | 1571 | 1461 | 1551 | 1383 | 1422 | 1501 | 1368 | 1580 | 1285 |

| | | | | | | | | | | | | | | | | | |
|---|---|---|---|---|---|---|---|---|---|---|---|---|---|---|---|---|---|
| 4578 | 0.12 | 0.14 | 0.85 | 0.39 | 1 | B2m | beta-2 microglobulin | 20239 | 9550 | 11493 | 1266 | 1021 | 579 | 1421 | 988 | 1309 | 1860 | 635 |
| 1324 | -0.04 | 0.10 | -0.41 | 0.68 | 1 | Sppl3 | signal peptide peptidase 3 | 1471 | 1415 | 1476 | 1241 | 1315 | 1201 | 1421 | 1320 | 1295 | 1235 | 1179 |
| 1549 | 0.14 | 0.11 | 1.32 | 0.19 | 1 | Rnf168 | ring finger protein 168 | 1307 | 1742 | 1531 | 1607 | 1474 | 1389 | 1420 | 2125 | 1512 | 1466 | 1469 |
| 1466 | -0.07 | 0.09 | -0.79 | 0.43 | 1 | Gtf2h1 | general transcription factor II H, polypepti | 1561 | 1592 | 1457 | 1621 | 1385 | 1483 | 1419 | 1446 | 1322 | 1301 | 1543 |
| 2020 | 0.21 | 0.12 | 1.67 | 0.10 | 1 | Hnrnph3 | heterogeneous nuclear ribonucleoprotein | 2384 | 2559 | 2576 | 1222 | 1491 | 2794 | 1418 | 1853 | 1515 | 1680 | 2731 |
| 1480 | -0.04 | 0.13 | -0.29 | 0.77 | 1 | Cfap20 | cilia and flagella associated protein 20 | 1106 | 1281 | 1037 | 1189 | 1374 | 2543 | 1418 | 1105 | 1390 | 1417 | 2418 |
| 1245 | 0.20 | 0.12 | 1.68 | 0.09 | 1 | Plk3 | polo like kinase 3 | 261 | 204 | 249 | 1395 | 1608 | 1602 | 1418 | 1491 | 1639 | 2000 | 1833 |
| 1614 | -0.11 | 0.13 | -0.81 | 0.42 | 1 | Ulk1 | unc-51 like kinase 1 | 1461 | 1607 | 1766 | 2179 | 1538 | 1585 | 1416 | 1691 | 1635 | 1532 | 1344 |
| 1581 | 0.17 | 0.14 | 1.19 | 0.23 | 1 | Sart3 | squamous cell carcinoma antigen recogn | 1533 | 1824 | 1800 | 1311 | 1381 | 1631 | 1416 | 2034 | 1562 | 1409 | 1490 |
| 1565 | 0.02 | 0.11 | 0.15 | 0.88 | 1 | Gdap1l1 | ganglioside-induced differentiation-assoc | 1215 | 1293 | 1203 | 1708 | 1630 | 1938 | 1416 | 1897 | 1606 | 1285 | 2019 |
| 1453 | -0.06 | 0.11 | -0.60 | 0.55 | 1 | Flot1 | flotillin 1 | 2146 | 1718 | 1966 | 1326 | 1303 | 1144 | 1416 | 1340 | 1253 | 1275 | 1092 |
| 1451 | -0.02 | 0.11 | -0.15 | 0.88 | 1 | Csnk2a2 | casein kinase 2, alpha prime polypeptide | 1642 | 1433 | 1340 | 1400 | 1279 | 1710 | 1416 | 1512 | 1300 | 1238 | 1687 |
| 1687 | -0.22 | 0.13 | -1.68 | 0.09 | 1 | Arhgef7 | Rho guanine nucleotide exchange factor | 2937 | 2492 | 2140 | 1463 | 1336 | 1680 | 1415 | 1255 | 1188 | 1030 | 1616 |
| 1485 | 0.07 | 0.10 | 0.70 | 0.49 | 1 | Tsen34 | tRNA splicing endonuclease subunit 34 | 1910 | 1718 | 2168 | 1221 | 1381 | 1123 | 1415 | 1378 | 1380 | 1504 | 1139 |
| 1298 | -0.12 | 0.14 | -0.89 | 0.37 | 1 | Sel1l3 | sel-1 suppressor of lin-12-like 3 (C. elega | 890 | 1299 | 1334 | 1582 | 1325 | 1270 | 1415 | 1530 | 1282 | 1276 | 1075 |
| 1203 | 0.03 | 0.14 | 0.19 | 0.85 | 1 | Atp5g1 | ATP synthase, H+ transporting, mitochon | 881 | 914 | 1170 | 1292 | 1293 | 1095 | 1415 | 1315 | 1200 | 1445 | 1213 |
| 1193 | 0.10 | 0.12 | 0.84 | 0.40 | 1 | Ripply3 | ripply transcriptional repressor 3 | 1060 | 945 | 1115 | 1372 | 1297 | 777 | 1415 | 1367 | 1304 | 1486 | 982 |
| 1161 | -0.09 | 0.13 | -0.68 | 0.50 | 1 | Mtmr10 | myotubularin related protein 10 | 940 | 1214 | 969 | 1390 | 1158 | 1024 | 1414 | 1169 | 1229 | 1262 | 1004 |
| 1347 | -0.06 | 0.12 | -0.53 | 0.59 | 1 | Rbm28 | RNA binding motif protein 28 | 1501 | 1022 | 1478 | 1159 | 1278 | 1675 | 1413 | 1161 | 1216 | 1240 | 1668 |
| 1215 | 0.04 | 0.11 | 0.34 | 0.73 | 1 | Pdia5 | protein disulfide isomerase associated 5 | 838 | 820 | 990 | 1120 | 1403 | 1357 | 1413 | 1295 | 1341 | 1350 | 1432 |
| 1685 | -0.10 | 0.09 | -1.08 | 0.28 | 1 | Rprd2 | regulation of nuclear pre-mRNA domain c | 1780 | 1893 | 1818 | 1651 | 1466 | 2215 | 1412 | 1542 | 1476 | 1282 | 1994 |
| 1286 | -0.03 | 0.10 | -0.27 | 0.79 | 1 | Fam98b | family with sequence similarity 98, memb | 1319 | 1307 | 1369 | 1175 | 1252 | 1283 | 1412 | 1097 | 1258 | 1360 | 1314 |
| 1131 | 0.11 | 0.17 | 0.67 | 0.50 | 1 | Maneal | mannosidase, endo-alpha-like | 622 | 987 | 754 | 1176 | 1221 | 1024 | 1412 | 1415 | 1250 | 1511 | 1067 |
| 1570 | 0.09 | 0.13 | 0.72 | 0.47 | 1 | Gigyf1 | GRB10 interacting GYF protein 1 | 1377 | 1285 | 1434 | 1602 | 1389 | 1992 | 1411 | 1906 | 1394 | 1606 | 1872 |
| 1327 | 0.21 | 0.11 | 1.80 | 0.07 | 1 | Atg5 | autophagy related 5 | 1115 | 867 | 932 | 1357 | 1437 | 1227 | 1410 | 1542 | 1682 | 1520 | 1507 |
| 1301 | -0.11 | 0.11 | -0.99 | 0.32 | 1 | Kremen1 | kringle containing transmembrane proteir | 2034 | 1695 | 1872 | 1147 | 1059 | 897 | 1410 | 1162 | 1045 | 1225 | 760 |
| 1274 | -0.06 | 0.17 | -0.34 | 0.74 | 1 | Casz1 | castor zinc finger 1 | 672 | 1056 | 1032 | 1187 | 1419 | 1700 | 1410 | 1557 | 1572 | 1137 | 1270 |
| 1178 | 0.00 | 0.16 | -0.01 | 0.99 | 1 | Gnptab | N-acetylglucosamine-1-phosphate transf | 1504 | 1095 | 894 | 1148 | 1057 | 1133 | 1410 | 1203 | 1350 | 1222 | 944 |
| 1385 | 0.12 | 0.12 | 1.04 | 0.30 | 1 | Skap2 | src family associated phosphoprotein 2 | 1934 | 1434 | 1542 | 1314 | 1226 | 1013 | 1409 | 1439 | 1346 | 1392 | 1184 |
| 2026 | 0.04 | 0.12 | 0.32 | 0.75 | 1 | Dek | DEK oncogene (DNA binding) | 3875 | 3281 | 2907 | 1564 | 1522 | 1538 | 1407 | 1610 | 1607 | 1476 | 1499 |
| 1729 | 0.00 | 0.09 | 0.02 | 0.98 | 1 | Zfp146 | zinc finger protein 146 | 1812 | 1885 | 1931 | 1676 | 1453 | 2149 | 1407 | 1530 | 1499 | 1466 | 2209 |
| 1580 | -0.02 | 0.11 | -0.14 | 0.89 | 1 | Taf15 | TATA-box binding protein associated fact | 1640 | 1472 | 1521 | 1294 | 1605 | 2113 | 1407 | 1390 | 1613 | 1305 | 2017 |
| 1129 | 0.09 | 0.15 | 0.63 | 0.53 | 1 | Trim9 | tripartite motif-containing 9 | 649 | 727 | 502 | 967 | 950 | 1829 | 1407 | 1234 | 1065 | 1404 | 1684 |
| 1480 | 0.07 | 0.12 | 0.60 | 0.55 | 1 | Rb1 | RB transcriptional corepressor 1 | 1541 | 1544 | 1266 | 1382 | 1498 | 1522 | 1406 | 1563 | 1675 | 1372 | 1507 |
| 1476 | 0.09 | 0.12 | 0.77 | 0.44 | 1 | Pgp | phosphoglycolate phosphatase | 1193 | 1639 | 1273 | 1665 | 1545 | 1209 | 1406 | 1903 | 1570 | 1761 | 1072 |
| 1465 | -0.23 | 0.17 | -1.30 | 0.19 | 1 | Golga1 | golgi autoantigen, golgin subfamily a, 1 | 757 | 918 | 954 | 1083 | 1573 | 3001 | 1406 | 745 | 1516 | 1070 | 3088 |
| 1405 | -0.13 | 0.15 | -0.84 | 0.40 | 1 | Cplane1 | ciliogenesis and planar polarity effector 1 | 974 | 1307 | 1022 | 1525 | 1411 | 2004 | 1405 | 1112 | 1581 | 1293 | 1819 |
| 1307 | 0.01 | 0.13 | 0.10 | 0.92 | 1 | Hgsnat | heparan-alpha-glucosaminide N-acetyltra | 1796 | 1173 | 1415 | 1319 | 1319 | 895 | 1403 | 1057 | 1497 | 1601 | 899 |
| 1463 | -0.22 | 0.17 | -1.24 | 0.21 | 1 | Fbxl20 | F-box and leucine-rich repeat protein 20 | 1824 | 1473 | 1594 | 1627 | 1438 | 1514 | 1401 | 1061 | 1715 | 1047 | 1399 |
| 1367 | 0.09 | 0.15 | 0.58 | 0.56 | 1 | Pawr | PRKC, apoptosis, WT1, regulator | 1374 | 1503 | 1500 | 1224 | 1260 | 1254 | 1401 | 1102 | 1396 | 1814 | 1207 |
| 1298 | 0.13 | 0.11 | 1.11 | 0.27 | 1 | Cul2 | cullin 2 | 966 | 1144 | 901 | 1325 | 1357 | 1303 | 1401 | 1463 | 1509 | 1408 | 1501 |
| 1693 | -0.01 | 0.10 | -0.14 | 0.89 | 1 | Tpcn1 | two pore channel 1 | 2306 | 2098 | 2059 | 1507 | 1449 | 1752 | 1400 | 1614 | 1374 | 1334 | 1734 |
| 1263 | 0.06 | 0.12 | 0.53 | 0.60 | 1 | Uhrf1bp1 | UHRF1 (ICBP90) binding protein 1 | 716 | 1048 | 815 | 1510 | 1289 | 1332 | 1400 | 1585 | 1415 | 1497 | 1291 |
| 1215 | -0.01 | 0.14 | -0.09 | 0.93 | 1 | Tcea1 | transcription elongation factor A (SII) 1 | 1264 | 1100 | 1050 | 1301 | 1151 | 1147 | 1400 | 1097 | 1279 | 1166 | 1408 |
| 1467 | 0.16 | 0.14 | 1.20 | 0.23 | 1 | Cdc14b | CDC14 cell division cycle 14B | 788 | 743 | 908 | 1344 | 2201 | 1526 | 1399 | 1859 | 2329 | 1358 | 1684 |
| 1707 | 0.09 | 0.08 | 1.10 | 0.27 | 1 | Sugt1 | SGT1, suppressor of G2 allele of SKP1 (S | 2458 | 2603 | 2507 | 1364 | 1356 | 1314 | 1398 | 1442 | 1474 | 1432 | 1433 |
| 1458 | 0.16 | 0.17 | 0.95 | 0.34 | 1 | Psma3 | proteasome subunit alpha 3 | 1507 | 1313 | 1640 | 1159 | 1552 | 1365 | 1398 | 1840 | 1324 | 1274 | 1666 |
| 1451 | 0.22 | 0.15 | 1.46 | 0.15 | 1 | Xrcc6 | X-ray repair complementing defective rep | 1327 | 1145 | 1264 | 875 | 1346 | 2051 | 1398 | 1051 | 1395 | 2051 | 2060 |
| 1424 | -0.07 | 0.12 | -0.54 | 0.59 | 1 | Cox15 | cytochrome c oxidase assembly protein 1 | 1247 | 1248 | 1156 | 1683 | 1727 | 1323 | 1398 | 1542 | 1473 | 1723 | 1149 |
| 1466 | 0.02 | 0.12 | 0.20 | 0.84 | 1 | Stk39 | serine/threonine kinase 39 | 2028 | 1773 | 1874 | 1351 | 1473 | 987 | 1397 | 1354 | 1253 | 1381 | 1255 |
| 1452 | -0.18 | 0.11 | -1.69 | 0.09 | 1 | Rreb1 | ras responsive element binding protein 1 | 1444 | 1681 | 1378 | 1936 | 1539 | 1215 | 1397 | 1644 | 1505 | 1217 | 1014 |
| 1369 | 0.00 | 0.12 | 0.01 | 0.99 | 1 | Lta4h | leukotriene A4 hydrolase | 1986 | 1303 | 1792 | 1118 | 1269 | 1205 | 1397 | 1226 | 1261 | 1413 | 1092 |
| 1330 | 0.07 | 0.13 | 0.54 | 0.59 | 1 | Tbck | TBC1 domain containing kinase | 1165 | 1219 | 1005 | 1302 | 1258 | 1513 | 1397 | 1182 | 1422 | 1726 | 1436 |
| 1873 | -0.18 | 0.12 | -1.50 | 0.13 | 1 | Sh3bp4 | SH3-domain binding protein 4 | 3133 | 3405 | 3897 | 1616 | 1261 | 1111 | 1396 | 1346 | 1330 | 1184 | 926 |
| 1622 | -0.09 | 0.12 | -0.73 | 0.47 | 1 | Snx9 | sorting nexin 9 | 2949 | 2090 | 2343 | 1410 | 1346 | 1219 | 1396 | 1110 | 1351 | 1437 | 1190 |
| 1286 | 0.12 | 0.14 | 0.81 | 0.42 | 1 | Nipal2 | NIPA-like domain containing 2 | 555 | 876 | 895 | 1451 | 1483 | 1316 | 1396 | 1764 | 1656 | 1503 | 1248 |
| 1601 | 0.11 | 0.09 | 1.22 | 0.22 | 1 | Zfx | zinc finger protein X-linked | 1610 | 1822 | 1800 | 1532 | 1409 | 1614 | 1395 | 1625 | 1597 | 1376 | 1834 |
| 1371 | -0.23 | 0.15 | -1.54 | 0.12 | 1 | Stard13 | StAR-related lipid transfer (START) doma | 1284 | 1540 | 1501 | 1309 | 1266 | 1796 | 1395 | 843 | 1204 | 1198 | 1750 |
| 1202 | -0.05 | 0.12 | -0.46 | 0.65 | 1 | Meis3 | Meis homeobox 3 | 751 | 849 | 852 | 1189 | 1264 | 1643 | 1395 | 1306 | 1293 | 1163 | 1522 |
| 1810 | -0.03 | 0.11 | -0.24 | 0.81 | 1 | Pim2 | proviral integration site 2 | 2348 | 2660 | 2498 | 1603 | 1515 | 1733 | 1394 | 1851 | 1300 | 1405 | 1603 |

| | | | | | | | | | | | | | | | | | | |
|---|---|---|---|---|---|---|---|---|---|---|---|---|---|---|---|---|---|---|
| 1777 | 0.28 | 0.16 | 1.74 | 0.08 | 1 | Eml2 | echinoderm microtubule associated prote | 2299 | 2277 | 2544 | 1305 | 1580 | 1287 | 1394 | 2266 | 1502 | 1842 | 1249 |
| 1294 | -0.07 | 0.14 | -0.53 | 0.60 | 1 | Ncbp3 | nuclear cap binding subunit 3 | 1313 | 1401 | 1163 | 1196 | 1305 | 1401 | 1394 | 1416 | 1200 | 1023 | 1424 |
| 1150 | 0.00 | 0.13 | 0.02 | 0.99 | 1 | Lamc2 | laminin, gamma 2 | 1106 | 1098 | 1220 | 1213 | 1086 | 922 | 1394 | 1340 | 1186 | 1153 | 934 |
| 1654 | 0.22 | 0.14 | 1.60 | 0.11 | 1 | Arid4b | AT rich interactive domain 4B (RBP1-like) | 1494 | 1638 | 1430 | 1384 | 1445 | 2034 | 1393 | 2158 | 1600 | 1279 | 2339 |
| 1341 | 0.11 | 0.11 | 1.02 | 0.31 | 1 | Rad17 | RAD17 checkpoint clamp loader compon | 1054 | 1160 | 1076 | 1309 | 1441 | 1357 | 1391 | 1316 | 1629 | 1419 | 1596 |
| 1295 | 0.02 | 0.15 | 0.11 | 0.92 | 1 | Hras | Harvey rat sarcoma virus oncogene | 1244 | 1270 | 1728 | 1082 | 1410 | 1084 | 1391 | 1499 | 1364 | 1202 | 976 |
| 1688 | 0.05 | 0.19 | 0.27 | 0.78 | 1 | Zbtb33 | zinc finger and BTB domain containing 33 | 1604 | 2202 | 1821 | 1924 | 1459 | 1605 | 1390 | 1265 | 1562 | 2070 | 1671 |
| 1464 | 0.06 | 0.13 | 0.48 | 0.63 | 1 | Smarcd2 | SWI/SNF related, matrix associated, actin | 2006 | 1511 | 1467 | 1357 | 1360 | 1323 | 1390 | 1664 | 1326 | 1394 | 1305 |
| 1438 | -0.07 | 0.14 | -0.47 | 0.64 | 1 | Kat6b | K(lysine) acetyltransferase 6B | 1032 | 1353 | 1088 | 1341 | 1502 | 2117 | 1390 | 1314 | 1635 | 1360 | 1683 |
| 1423 | 0.04 | 0.14 | 0.27 | 0.79 | 1 | Cdk12 | cyclin-dependent kinase 12 | 1594 | 1306 | 1140 | 1231 | 1352 | 1756 | 1390 | 1189 | 1579 | 1285 | 1834 |
| 1311 | -0.15 | 0.11 | -1.27 | 0.20 | 1 | Ppard | peroxisome proliferator activator receptor | 1423 | 1428 | 1706 | 1403 | 1339 | 1052 | 1390 | 1121 | 1340 | 1146 | 1067 |
| 1545 | -0.11 | 0.11 | -0.98 | 0.33 | 1 | Limk2 | LIM motif-containing protein kinase 2 | 1753 | 1923 | 2209 | 1286 | 1353 | 1755 | 1389 | 1278 | 1238 | 1336 | 1474 |
| 1521 | 0.03 | 0.11 | 0.26 | 0.80 | 1 | Actr8 | ARP8 actin-related protein 8 | 1151 | 1559 | 1413 | 1481 | 1525 | 1838 | 1389 | 1462 | 1800 | 1324 | 1787 |
| 1474 | 0.12 | 0.12 | 0.96 | 0.34 | 1 | Wdr43 | WD repeat domain 43 | 1525 | 1242 | 1330 | 1321 | 1328 | 1767 | 1389 | 1658 | 1341 | 1338 | 1979 |
| 1335 | 0.19 | 0.10 | 1.97 | 0.05 | 1 | Stx3 | syntaxin 3 | 842 | 925 | 747 | 1477 | 1307 | 1516 | 1389 | 1660 | 1591 | 1532 | 1699 |
| 1141 | -0.03 | 0.13 | -0.27 | 0.79 | 1 | Fam45a | family with sequence similarity 45, membe | 885 | 954 | 949 | 1177 | 1129 | 1241 | 1389 | 1020 | 1249 | 1208 | 1352 |
| 1104 | -0.09 | 0.14 | -0.65 | 0.52 | 1 | Kat2a | K(lysine) acetyltransferase 2A | 1187 | 1048 | 1041 | 918 | 1219 | 1051 | 1389 | 963 | 1284 | 1121 | 923 |
| 1227 | -0.22 | 0.14 | -1.59 | 0.11 | 1 | Osbpl5 | oxysterol binding protein-like 5 | 1182 | 878 | 862 | 1391 | 1548 | 1358 | 1387 | 1058 | 1358 | 1266 | 1215 |
| 1439 | -0.04 | 0.13 | -0.31 | 0.76 | 1 | Zfp110 | zinc finger protein 110 | 1451 | 1885 | 1384 | 1315 | 1403 | 1519 | 1386 | 1250 | 1299 | 1333 | 1605 |
| 1421 | 0.07 | 0.11 | 0.67 | 0.50 | 1 | Tomm34 | translocase of outer mitochondrial membr | 1468 | 1341 | 1531 | 1257 | 1503 | 1351 | 1386 | 1506 | 1452 | 1597 | 1241 |
| 1189 | 0.03 | 0.12 | 0.30 | 0.76 | 1 | Dhx29 | DEAH (Asp-Glu-Ala-His) box polypeptide | 985 | 663 | 962 | 1131 | 1446 | 1192 | 1386 | 1033 | 1631 | 1337 | 1315 |
| 1127 | 0.03 | 0.19 | 0.15 | 0.88 | 1 | Elovl2 | elongation of very long chain fatty acids (I | 440 | 661 | 659 | 1349 | 1470 | 1050 | 1386 | 1086 | 1561 | 1687 | 1046 |
| 1622 | 0.12 | 0.11 | 1.11 | 0.27 | 1 | Ssbp4 | single stranded DNA binding protein 4 | 1470 | 1621 | 1713 | 1588 | 1562 | 1703 | 1384 | 1983 | 1508 | 1477 | 1835 |
| 1578 | 0.01 | 0.11 | 0.05 | 0.96 | 1 | Tnfrsf1a | tumor necrosis factor receptor superfamily | 3512 | 2315 | 2489 | 1139 | 1384 | 581 | 1384 | 1074 | 1326 | 1570 | 579 |
| 1465 | -0.06 | 0.12 | -0.49 | 0.62 | 1 | Phf21a | PHD finger protein 21A | 1508 | 1447 | 1293 | 1188 | 1298 | 2227 | 1384 | 1274 | 1128 | 1542 | 1831 |
| 1448 | -0.10 | 0.15 | -0.65 | 0.52 | 1 | Prr14l | proline rich 14-like | 1177 | 1479 | 1102 | 1641 | 1532 | 1752 | 1384 | 1259 | 1583 | 1604 | 1419 |
| 1225 | -0.07 | 0.17 | -0.43 | 0.67 | 1 | Fam210a | family with sequence similarity 210, meml | 989 | 745 | 588 | 1539 | 1298 | 1489 | 1384 | 1138 | 1416 | 1346 | 1537 |
| 1436 | -0.06 | 0.15 | -0.43 | 0.66 | 1 | Dock8 | dedicator of cytokinesis 8 | 2877 | 1679 | 1990 | 1162 | 1313 | 833 | 1383 | 812 | 1370 | 1490 | 884 |
| 1349 | 0.05 | 0.11 | 0.51 | 0.61 | 1 | Fgd4 | FYVE, RhoGEF and PH domain containin | 1197 | 1247 | 1216 | 1452 | 1247 | 1394 | 1383 | 1607 | 1292 | 1260 | 1546 |
| 1216 | -0.09 | 0.16 | -0.59 | 0.55 | 1 | B3galnt1 | UDP-GalNAc:betaGlcNAc beta 1,3-galact | 1177 | 1119 | 943 | 1406 | 1341 | 1093 | 1383 | 935 | 1313 | 1490 | 1172 |
| 1455 | -0.03 | 0.16 | -0.19 | 0.85 | 1 | Clcn5 | chloride channel, voltage-sensitive 5 | 1706 | 2386 | 2304 | 1312 | 1120 | 1043 | 1382 | 1111 | 1213 | 1422 | 1010 |
| 1211 | 0.04 | 0.14 | 0.26 | 0.79 | 1 | C330007P06Ri | RIKEN cDNA C330007P06 gene | 812 | 962 | 805 | 1294 | 1329 | 1289 | 1382 | 1182 | 1620 | 1252 | 1393 |
| 1369 | -0.04 | 0.12 | -0.32 | 0.75 | 1 | Dstyk | dual serine/threonine and tyrosine proteir | 1568 | 1575 | 1504 | 1275 | 1223 | 1394 | 1380 | 1497 | 1226 | 1100 | 1320 |
| 1252 | 0.05 | 0.12 | 0.40 | 0.69 | 1 | Hivep3 | human immunodeficiency virus type I enh | 828 | 1166 | 848 | 1756 | 1166 | 1085 | 1380 | 1736 | 1337 | 1163 | 1303 |
| 1097 | 0.20 | 0.15 | 1.31 | 0.19 | 1 | Cd59a | CD59a antigen | 1129 | 1085 | 1185 | 1133 | 997 | 573 | 1380 | 1354 | 1020 | 1432 | 778 |
| 1501 | 0.22 | 0.14 | 1.51 | 0.13 | 1 | D1Ertd622e | DNA segment, Chr 1, ERATO Doi 622, ex | 846 | 893 | 920 | 2033 | 1505 | 1448 | 1379 | 2670 | 1588 | 1691 | 1532 |
| 1190 | 0.00 | 0.16 | -0.03 | 0.98 | 1 | Tmem41b | transmembrane protein 41B | 546 | 790 | 616 | 1235 | 1505 | 1458 | 1379 | 1174 | 1516 | 1418 | 1452 |
| 1085 | 0.03 | 0.16 | 0.17 | 0.86 | 1 | Tmtc2 | transmembrane and tetratricopeptide repe | 950 | 822 | 961 | 1196 | 996 | 980 | 1379 | 894 | 1208 | 1399 | 1152 |
| 1666 | 0.24 | 0.13 | 1.76 | 0.08 | 1 | Fam207a | family with sequence similarity 207, meml | 1454 | 1585 | 1461 | 1881 | 1563 | 1509 | 1378 | 2045 | 1696 | 2241 | 1515 |
| 1511 | -0.25 | 0.15 | -1.68 | 0.09 | 1 | Fbxo42 | F-box protein 42 | 1738 | 2250 | 1560 | 1657 | 1297 | 1707 | 1378 | 1052 | 1315 | 1231 | 1440 |
| 1447 | -0.09 | 0.11 | -0.86 | 0.39 | 1 | Pnpla2 | patatin-like phospholipase domain contai | 1931 | 1741 | 1827 | 1449 | 1376 | 1141 | 1378 | 1420 | 1323 | 1401 | 924 |
| 1127 | -0.27 | 0.18 | -1.52 | 0.13 | 1 | Ttc17 | tetratricopeptide repeat domain 17 | 1185 | 808 | 1039 | 1347 | 1099 | 1268 | 1378 | 812 | 1243 | 951 | 1263 |
| 1765 | 0.22 | 0.16 | 1.36 | 0.17 | 1 | Las1l | LAS1-like (S. cerevisiae) | 1439 | 1625 | 1633 | 1463 | 1934 | 1996 | 1377 | 2586 | 1798 | 1396 | 2170 |
| 1497 | -0.16 | 0.15 | -1.09 | 0.27 | 1 | Xrn2 | 5'-3' exoribonuclease 2 | 2142 | 1721 | 1476 | 1265 | 1502 | 1735 | 1377 | 1111 | 1443 | 1240 | 1459 |
| 1470 | 0.05 | 0.13 | 0.41 | 0.69 | 1 | Nol7 | nucleolar protein 7 | 1701 | 1832 | 1693 | 1160 | 1305 | 1513 | 1377 | 1052 | 1386 | 1623 | 1523 |
| 1189 | -0.01 | 0.14 | -0.06 | 0.95 | 1 | Gpr89 | G protein-coupled receptor 89 | 848 | 1020 | 987 | 1034 | 1405 | 1309 | 1375 | 1047 | 1476 | 1320 | 1251 |
| 1429 | -0.15 | 0.11 | -1.30 | 0.19 | 1 | Cnot3 | CCR4-NOT transcription complex, subuni | 1585 | 1347 | 1583 | 1238 | 1327 | 1948 | 1374 | 1197 | 1324 | 1056 | 1738 |
| 1122 | 0.05 | 0.12 | 0.41 | 0.68 | 1 | Tafa5 | TAFA chemokine like family member 5 | 643 | 488 | 710 | 1335 | 1426 | 1016 | 1374 | 1369 | 1578 | 1391 | 1014 |
| 1869 | 0.11 | 0.16 | 0.66 | 0.51 | 1 | Ccdc50 | coiled-coil domain containing 50 | 3553 | 2184 | 1822 | 1170 | 1200 | 2560 | 1373 | 1206 | 1497 | 1387 | 2605 |
| 1429 | -0.36 | 0.19 | -1.91 | 0.06 | 1 | Syne2 | spectrin repeat containing, nuclear envelo | 1972 | 1812 | 1664 | 912 | 1210 | 2288 | 1373 | 650 | 1332 | 853 | 1649 |
| 2384 | -0.14 | 0.08 | -1.91 | 0.06 | 1 | Gipr | gastric inhibitory polypeptide receptor | 387 | 377 | 407 | 1259 | 1362 | 9039 | 1372 | 1159 | 1206 | 1200 | 8453 |
| 1475 | 0.13 | 0.15 | 0.86 | 0.39 | 1 | Fam222b | family with sequence similarity 222, meml | 1090 | 1397 | 1339 | 1583 | 1455 | 1518 | 1372 | 1459 | 1567 | 1946 | 1497 |
| 1324 | -0.14 | 0.12 | -1.17 | 0.24 | 1 | Zfp827 | zinc finger protein 827 | 1000 | 1466 | 1235 | 1508 | 1403 | 1404 | 1372 | 1493 | 1225 | 1315 | 1139 |
| 1172 | 0.03 | 0.14 | 0.19 | 0.85 | 1 | Got1 | glutamic-oxaloacetic transaminase 1, solu | 1417 | 1005 | 1027 | 1112 | 1248 | 926 | 1372 | 1404 | 1126 | 1401 | 848 |
| 1584 | 0.18 | 0.13 | 1.36 | 0.17 | 1 | Tuba4a | tubulin, alpha 4A | 1658 | 1875 | 1658 | 1516 | 1421 | 1363 | 1371 | 2310 | 1442 | 1428 | 1384 |
| 1502 | 0.11 | 0.12 | 0.98 | 0.33 | 1 | Nod1 | nucleotide-binding oligomerization doma | 1273 | 1262 | 1137 | 1458 | 1279 | 2077 | 1371 | 1815 | 1453 | 1302 | 2094 |
| 1233 | 0.05 | 0.12 | 0.37 | 0.71 | 1 | 2810013P06Rik | | 1004 | 1092 | 1226 | 1320 | 1301 | 1059 | 1371 | 1285 | 1266 | 1377 | 1261 |
| 1804 | 0.12 | 0.15 | 0.80 | 0.42 | 1 | Cdc27 | cell division cycle 27 | 1606 | 1749 | 1306 | 1830 | 1848 | 2248 | 1370 | 1689 | 2134 | 1662 | 2403 |
| 1348 | 0.06 | 0.18 | 0.31 | 0.76 | 1 | Plcb4 | phospholipase C, beta 4 | 1516 | 1609 | 1280 | 1190 | 1134 | 1395 | 1369 | 853 | 1510 | 1521 | 1453 |
| 1347 | 0.13 | 0.11 | 1.14 | 0.26 | 1 | Ppp1r11 | protein phosphatase 1, regulatory inhibito | 1539 | 1397 | 1485 | 1191 | 1393 | 1000 | 1369 | 1557 | 1350 | 1529 | 1004 |

| | | | | | | | | | | | | | | | | | |
|---|---|---|---|---|---|---|---|---|---|---|---|---|---|---|---|---|---|
| 1383 | 0.00 | 0.12 | 0.00 | 1.00 | 1 | Bicd2 | BICD cargo adaptor 2 | 1596 | 1415 | 1346 | 1493 | 1087 | 1491 | 1368 | 1506 | 1255 | 1212 | 1439 |
| 1375 | -0.03 | 0.16 | -0.16 | 0.87 | 1 | Prdx3 | peroxiredoxin 3 | 1452 | 1240 | 1174 | 1601 | 1465 | 1223 | 1368 | 1124 | 1680 | 1606 | 1194 |
| 1101 | -0.05 | 0.13 | -0.42 | 0.67 | 1 | Bpgm | 2,3-bisphosphoglycerate mutase | 732 | 743 | 862 | 1280 | 1298 | 1035 | 1368 | 1033 | 1303 | 1297 | 1159 |
| 1279 | -0.11 | 0.15 | -0.71 | 0.47 | 1 | Ermp1 | endoplasmic reticulum metallopeptidase | 1749 | 1367 | 1385 | 1294 | 1247 | 1044 | 1367 | 960 | 1508 | 1172 | 980 |
| 1370 | 0.07 | 0.13 | 0.55 | 0.58 | 1 | Tbcb | tubulin folding cofactor B | 1651 | 1401 | 1622 | 1219 | 1375 | 1086 | 1366 | 1529 | 1288 | 1564 | 971 |
| 1250 | 0.20 | 0.13 | 1.56 | 0.12 | 1 | Slc26a2 | solute carrier family 26 (sulfate transporte | 1033 | 1026 | 1083 | 1210 | 1139 | 1215 | 1366 | 1552 | 1548 | 1246 | 1332 |
| 1111 | 0.03 | 0.17 | 0.15 | 0.88 | 1 | Dpm3 | dolichyl-phosphate mannosyltransferase | 873 | 673 | 885 | 1285 | 1241 | 934 | 1366 | 1661 | 1140 | 1310 | 854 |
| 1998 | -0.07 | 0.11 | -0.60 | 0.55 | 1 | Snrk | SNF related kinase | 3350 | 3528 | 3359 | 1794 | 1355 | 1478 | 1365 | 1842 | 1279 | 1285 | 1343 |
| 1528 | -0.03 | 0.13 | -0.27 | 0.79 | 1 | Iws1 | IWS1, SUPT6 interacting protein | 1266 | 1071 | 1225 | 1584 | 1622 | 2087 | 1364 | 1177 | 1616 | 1588 | 2205 |
| 1444 | 0.14 | 0.13 | 1.06 | 0.29 | 1 | Vps26c | VPS26 endosomal protein sorting factor C | 1404 | 1351 | 1287 | 1520 | 1364 | 1354 | 1364 | 1564 | 1268 | 1989 | 1425 |
| 1232 | 0.01 | 0.11 | 0.11 | 0.91 | 1 | Gnb5 | guanine nucleotide binding protein (G pro | 1306 | 1264 | 1315 | 1146 | 1336 | 975 | 1364 | 1065 | 1303 | 1383 | 1094 |
| 13687 | -0.03 | 0.08 | -0.34 | 0.73 | 1 | Col4a1 | collagen, type IV, alpha 1 | 47013 | 42421 | 50681 | 1148 | 1569 | 1207 | 1363 | 1218 | 1258 | 1329 | 1350 |
| 2490 | 0.11 | 0.15 | 0.72 | 0.47 | 1 | Fgl2 | fibrinogen-like protein 2 | 6627 | 4883 | 4429 | 1329 | 1731 | 1049 | 1362 | 1240 | 1786 | 1859 | 1094 |
| 1416 | 0.00 | 0.11 | 0.00 | 1.00 | 1 | Sap18 | Sin3-associated polypeptide 18 | 1472 | 1523 | 1898 | 1333 | 1467 | 1182 | 1362 | 1466 | 1367 | 1300 | 1205 |
| 1339 | 0.26 | 0.25 | 1.07 | 0.29 | 1 | Romo1 | reactive oxygen species modulator 1 | 1010 | 838 | 1397 | 1215 | 1469 | 1079 | 1362 | 2600 | 1292 | 1468 | 996 |
| 1307 | -0.09 | 0.18 | -0.51 | 0.61 | 1 | Rapgef6 | Rap guanine nucleotide exchange factor | 1623 | 1247 | 1214 | 1252 | 1278 | 1358 | 1361 | 739 | 1484 | 1453 | 1369 |
| 1211 | 0.17 | 0.16 | 1.04 | 0.30 | 1 | Ier3ip1 | immediate early response 3 interacting pr | 1054 | 1002 | 1190 | 1324 | 1113 | 953 | 1361 | 1701 | 1333 | 1131 | 1165 |
| 1541 | -0.25 | 0.13 | -1.95 | 0.05 | 1 | Zfp609 | zinc finger protein 609 | 1272 | 1466 | 1409 | 1579 | 1831 | 2176 | 1359 | 1184 | 1780 | 1161 | 1734 |
| 1364 | 0.09 | 0.14 | 0.68 | 0.50 | 1 | Zfyve16 | zinc finger, FYVE domain containing 16 | 1233 | 1502 | 1012 | 1318 | 1249 | 1529 | 1359 | 1318 | 1467 | 1385 | 1633 |
| 1647 | 0.04 | 0.11 | 0.39 | 0.69 | 1 | Arf6 | ADP-ribosylation factor 6 | 2068 | 2341 | 2011 | 1595 | 1510 | 1290 | 1358 | 1647 | 1360 | 1674 | 1262 |
| 1630 | -0.09 | 0.13 | -0.65 | 0.51 | 1 | Tmtc1 | transmembrane and tetratricopeptide repe | 3374 | 3236 | 2696 | 1160 | 1022 | 904 | 1358 | 870 | 1134 | 1301 | 876 |
| 1310 | -0.01 | 0.10 | -0.06 | 0.95 | 1 | Mboat7 | membrane bound O-acyltransferase dom | 1522 | 1322 | 1482 | 1204 | 1313 | 1166 | 1358 | 1226 | 1342 | 1431 | 1045 |
| 1159 | 0.09 | 0.16 | 0.56 | 0.57 | 1 | Dner | delta/notch-like EGF repeat containing | 975 | 626 | 722 | 1591 | 1325 | 775 | 1358 | 1650 | 1359 | 1538 | 831 |
| 1573 | 0.06 | 0.17 | 0.32 | 0.75 | 1 | Celf2 | CUGBP, Elav-like family member 2 | 3257 | 2229 | 2128 | 1175 | 1248 | 952 | 1357 | 1630 | 1299 | 1153 | 871 |
| 1427 | -0.06 | 0.10 | -0.60 | 0.55 | 1 | Josd1 | Josephin domain containing 1 | 1239 | 1638 | 1485 | 1507 | 1539 | 1372 | 1357 | 1622 | 1394 | 1253 | 1297 |
| 1260 | -0.07 | 0.11 | -0.70 | 0.49 | 1 | Strip1 | striatin interacting protein 1 | 1448 | 1253 | 1313 | 1142 | 1265 | 1289 | 1357 | 1269 | 1140 | 1226 | 1156 |
| 1462 | -0.08 | 0.13 | -0.63 | 0.53 | 1 | Tep1 | telomerase associated protein 1 | 1586 | 1365 | 1426 | 1726 | 1362 | 1570 | 1356 | 1501 | 1326 | 1564 | 1298 |
| 1398 | 0.09 | 0.13 | 0.69 | 0.49 | 1 | Srm | spermidine synthase | 996 | 881 | 958 | 1106 | 1828 | 1800 | 1355 | 1308 | 1911 | 1350 | 1888 |
| 1269 | -0.16 | 0.16 | -1.01 | 0.31 | 1 | Lman2l | lectin, mannose-binding 2-like | 1039 | 993 | 1463 | 1578 | 1336 | 1253 | 1355 | 1418 | 1182 | 1119 | 1224 |
| 1237 | 0.04 | 0.14 | 0.28 | 0.78 | 1 | Itpr2 | inositol 1,4,5-triphosphate receptor 2 | 1165 | 913 | 846 | 1349 | 1343 | 1232 | 1355 | 1386 | 1416 | 1249 | 1357 |
| 1502 | 0.21 | 0.14 | 1.47 | 0.14 | 1 | Mettl1 | methyltransferase like 1 | 993 | 1001 | 1456 | 1299 | 1645 | 1786 | 1354 | 1929 | 1643 | 1483 | 1930 |
| 1287 | 0.13 | 0.13 | 0.99 | 0.32 | 1 | Kiz | kizuna centrosomal protein | 1331 | 1692 | 1231 | 1141 | 1237 | 995 | 1354 | 1389 | 1298 | 1369 | 1118 |
| 1481 | 0.19 | 0.12 | 1.56 | 0.12 | 1 | Dcaf13 | DDB1 and CUL4 associated factor 13 | 1132 | 1252 | 1121 | 1393 | 1187 | 2019 | 1352 | 1760 | 1131 | 1713 | 2235 |
| 1327 | -0.04 | 0.11 | -0.33 | 0.74 | 1 | Zbtb38 | zinc finger and BTB domain containing 38 | 1531 | 1333 | 1275 | 1318 | 1204 | 1421 | 1352 | 1279 | 1244 | 1144 | 1496 |
| 1141 | -0.10 | 0.11 | -0.92 | 0.36 | 1 | Pknox1 | Pbx/knotted 1 homeobox | 1025 | 962 | 1024 | 1177 | 1178 | 1225 | 1352 | 981 | 1130 | 1280 | 1222 |
| 1100 | 0.35 | 0.18 | 1.89 | 0.06 | 1 | Cib2 | calcium and integrin binding family memb | 367 | 464 | 419 | 1224 | 1225 | 884 | 1351 | 2435 | 1339 | 1297 | 1095 |
| 938 | 0.29 | 0.19 | 1.53 | 0.13 | 1 | Dlgap1 | DLG associated protein 1 | 96 | 90 | 160 | 1154 | 1091 | 862 | 1350 | 1555 | 1397 | 1606 | 954 |
| 1218 | -0.17 | 0.11 | -1.46 | 0.15 | 1 | Galns | galactosamine (N-acetyl)-6-sulfate sulfata | 1307 | 1360 | 1304 | 1149 | 1455 | 1038 | 1349 | 1135 | 1262 | 1095 | 945 |
| 1211 | -0.06 | 0.18 | -0.34 | 0.73 | 1 | H2bc4 | H2B clustered histone 4 | 526 | 560 | 817 | 1428 | 1395 | 1627 | 1349 | 1098 | 1218 | 1569 | 1729 |
| 1426 | 0.02 | 0.14 | 0.11 | 0.91 | 1 | Ercc6l2 | excision repair cross-complementing rode | 1284 | 1176 | 974 | 1464 | 1610 | 1663 | 1348 | 1306 | 1713 | 1419 | 1731 |
| 1198 | -0.16 | 0.14 | -1.16 | 0.25 | 1 | Slc35a2 | solute carrier family 35 (UDP-galactose tr | 849 | 937 | 780 | 1326 | 1360 | 1560 | 1348 | 1009 | 1389 | 1239 | 1377 |
| 1322 | 0.02 | 0.10 | 0.20 | 0.84 | 1 | Relch | RAB11 binding and LisH domain, coiled-c | 961 | 812 | 864 | 1166 | 1036 | 2347 | 1347 | 1082 | 1125 | 1364 | 2435 |
| 2010 | 0.05 | 0.15 | 0.35 | 0.73 | 1 | Mob1a | MOB kinase activator 1A | 4511 | 3106 | 3157 | 1342 | 1401 | 1481 | 1346 | 1448 | 1563 | 1213 | 1542 |
| 1264 | 0.02 | 0.15 | 0.12 | 0.90 | 1 | Thada | thyroid adenoma associated | 1214 | 779 | 1017 | 1419 | 1358 | 1276 | 1346 | 1320 | 1595 | 1346 | 1232 |
| 1206 | 0.06 | 0.14 | 0.44 | 0.66 | 1 | Exoc8 | exocyst complex component 8 | 782 | 1038 | 857 | 1224 | 1408 | 1183 | 1346 | 1667 | 1317 | 1278 | 1169 |
| 1348 | -0.11 | 0.15 | -0.76 | 0.45 | 1 | Rnf19b | ring finger protein 19B | 1563 | 1667 | 1239 | 1307 | 1470 | 1259 | 1345 | 1064 | 1419 | 1347 | 1150 |
| 1116 | -0.16 | 0.12 | -1.28 | 0.20 | 1 | Cln5 | ceroid-lipofuscinosis, neuronal 5 | 886 | 998 | 925 | 1357 | 1240 | 1044 | 1345 | 1188 | 1350 | 1025 | 917 |
| 877 | -0.21 | 0.15 | -1.46 | 0.15 | 1 | Pcdhga5 | protocadherin gamma subfamily A, 5 | 632 | 560 | 538 | 1078 | 1178 | 677 | 1345 | 777 | 1106 | 1052 | 699 |
| 2120 | -0.09 | 0.10 | -0.94 | 0.35 | 1 | Srek1 | splicing regulatory glutamine/lysine-rich p | 1389 | 1959 | 1535 | 1625 | 1396 | 5178 | 1344 | 1442 | 1475 | 1209 | 4761 |
| 1575 | -0.02 | 0.14 | -0.18 | 0.86 | 1 | Kbtbd2 | kelch repeat and BTB (POZ) domain conta | 1722 | 1918 | 1828 | 1500 | 1247 | 1895 | 1344 | 1105 | 1566 | 1414 | 1790 |
| 1244 | 0.09 | 0.11 | 0.84 | 0.40 | 1 | 5031425E22Rik | | 1043 | 1134 | 954 | 1279 | 1159 | 1305 | 1344 | 1500 | 1055 | 1430 | 1484 |
| 1157 | 0.20 | 0.12 | 1.64 | 0.10 | 1 | A530058N18Rik | | 245 | 280 | 319 | 1261 | 1192 | 1725 | 1344 | 1570 | 1482 | 1363 | 1951 |
| 1460 | 0.08 | 0.15 | 0.56 | 0.58 | 1 | Ppil2 | peptidylprolyl isomerase (cyclophilin)-like | 1936 | 1375 | 1613 | 1194 | 1339 | 1531 | 1343 | 1549 | 1319 | 1163 | 1700 |
| 1182 | 0.03 | 0.14 | 0.24 | 0.81 | 1 | G6pdx | glucose-6-phosphate dehydrogenase X-l | 1553 | 1200 | 935 | 1147 | 1115 | 986 | 1343 | 995 | 1247 | 1481 | 1002 |
| 1541 | -0.09 | 0.16 | -0.55 | 0.58 | 1 | Secisbp2l | SECIS binding protein 2-like | 2311 | 1938 | 1888 | 1373 | 1337 | 1498 | 1342 | 1242 | 1540 | 967 | 1513 |
| 1486 | -0.10 | 0.11 | -0.91 | 0.36 | 1 | Eri3 | exoribonuclease 3 | 1905 | 1752 | 1674 | 1455 | 1410 | 1479 | 1342 | 1368 | 1324 | 1426 | 1213 |
| 1399 | 0.05 | 0.13 | 0.41 | 0.68 | 1 | Epc2 | enhancer of polycomb homolog 2 | 1499 | 1890 | 1424 | 1240 | 1208 | 1390 | 1341 | 1324 | 1385 | 1127 | 1566 |
| 1265 | 0.04 | 0.11 | 0.38 | 0.70 | 1 | 1600014C10Ril | RIKEN cDNA 1600014C10 gene | 1297 | 1395 | 1290 | 1248 | 1204 | 1098 | 1341 | 1441 | 1308 | 1238 | 1059 |
| 1268 | -0.26 | 0.16 | -1.65 | 0.10 | 1 | Acvr1b | activin A receptor, type 1B | 1341 | 1357 | 1098 | 1655 | 1302 | 1239 | 1340 | 1039 | 1409 | 1177 | 994 |
| 1162 | -0.08 | 0.13 | -0.63 | 0.53 | 1 | Ercc3 | excision repair cross-complementing rode | 1239 | 1019 | 979 | 1329 | 1197 | 1052 | 1339 | 980 | 1225 | 1261 | 1166 |

| | | | | | | | | | | | | | | | | | | |
|---|---|---|---|---|---|---|---|---|---|---|---|---|---|---|---|---|---|---|
| 1522 | -0.16 | 0.10 | -1.54 | 0.12 | 1 | Zfp646 | zinc finger protein 646 | 1632 | 1857 | 1724 | 1522 | 1581 | 1642 | 1338 | 1229 | 1426 | 1313 | 1484 |
| 1332 | -0.06 | 0.15 | -0.42 | 0.67 | 1 | Yeats4 | YEATS domain containing 4 | 1510 | 1373 | 1353 | 1298 | 1396 | 1276 | 1338 | 1360 | 1360 | 936 | 1450 |
| 1176 | -0.10 | 0.13 | -0.76 | 0.45 | 1 | D430042O09Ri | RIKEN cDNA D430042O09 gene | 1038 | 1092 | 920 | 1079 | 1203 | 1473 | 1338 | 960 | 1151 | 1106 | 1574 |
| 1173 | -0.21 | 0.19 | -1.14 | 0.25 | 1 | Nup160 | nucleoporin 160 | 1486 | 1053 | 1036 | 1169 | 1319 | 1111 | 1338 | 631 | 1179 | 1440 | 1139 |
| 1654 | -0.03 | 0.13 | -0.27 | 0.79 | 1 | Nsmaf | neutral sphingomyelinase (N-SMase) acti | 1660 | 1639 | 1455 | 1572 | 1361 | 2549 | 1337 | 1245 | 1598 | 1378 | 2397 |
| 1330 | 0.03 | 0.11 | 0.32 | 0.75 | 1 | Lancl1 | LanC (bacterial lantibiotic synthetase com | 986 | 976 | 1085 | 1428 | 1553 | 1401 | 1337 | 1550 | 1367 | 1356 | 1586 |
| 1435 | -0.02 | 0.12 | -0.16 | 0.88 | 1 | Gpatch8 | G patch domain containing 8 | 1404 | 1079 | 1472 | 1297 | 1333 | 1986 | 1336 | 1199 | 1546 | 1270 | 1860 |
| 1256 | 0.15 | 0.11 | 1.35 | 0.18 | 1 | Prelid3b | PRELI domain containing 3B | 1293 | 1270 | 1316 | 1176 | 1147 | 1051 | 1336 | 1495 | 1278 | 1359 | 1101 |
| 867 | -0.03 | 0.15 | -0.22 | 0.83 | 1 | Bag2 | BCL2-associated athanogene 2 | 281 | 329 | 262 | 1160 | 1044 | 860 | 1336 | 1091 | 1200 | 1069 | 909 |
| 2094 | -0.11 | 0.11 | -1.02 | 0.31 | 1 | Mical3 | microtubule associated monooxygenase, | 1799 | 2534 | 2171 | 2395 | 1952 | 2944 | 1335 | 2297 | 1920 | 1306 | 2378 |
| 1355 | 0.04 | 0.12 | 0.37 | 0.71 | 1 | Usp37 | ubiquitin specific peptidase 37 | 958 | 1137 | 846 | 1392 | 1395 | 1759 | 1335 | 1466 | 1493 | 1256 | 1866 |
| 1227 | 0.02 | 0.13 | 0.19 | 0.85 | 1 | Fam204a | family with sequence similarity 204, meml | 999 | 921 | 1242 | 1217 | 1288 | 1268 | 1335 | 1065 | 1299 | 1386 | 1481 |
| 1397 | -0.01 | 0.11 | -0.06 | 0.95 | 1 | Rnpep | arginyl aminopeptidase (aminopeptidase | 1808 | 1492 | 1731 | 1261 | 1408 | 1185 | 1334 | 1364 | 1300 | 1258 | 1230 |
| 1454 | 0.25 | 0.15 | 1.65 | 0.10 | 1 | Ndufa8 | NADH:ubiquinone oxidoreductase subun | 1614 | 1572 | 2010 | 1291 | 1293 | 992 | 1333 | 1939 | 1389 | 1499 | 1061 |
| 1429 | 0.18 | 0.18 | 1.00 | 0.32 | 1 | Ndufs3 | NADH:ubiquinone oxidoreductase core s | 1038 | 1124 | 1502 | 1357 | 1472 | 1467 | 1333 | 2172 | 1316 | 1377 | 1563 |
| 1195 | 0.16 | 0.15 | 1.08 | 0.28 | 1 | Lrfn4 | leucine rich repeat and fibronectin type III | 579 | 738 | 669 | 1296 | 1220 | 1404 | 1333 | 1812 | 1227 | 1556 | 1315 |
| 1184 | -0.03 | 0.16 | -0.19 | 0.85 | 1 | Cdc25a | cell division cycle 25A | 904 | 1290 | 956 | 1248 | 1308 | 1106 | 1333 | 1045 | 1357 | 1325 | 1152 |
| 1155 | -0.05 | 0.14 | -0.33 | 0.74 | 1 | Lactb2 | lactamase, beta 2 | 563 | 508 | 653 | 1332 | 1349 | 1513 | 1333 | 944 | 1584 | 1319 | 1601 |
| 1334 | 0.04 | 0.12 | 0.34 | 0.73 | 1 | Lclat1 | lysocardiolipin acyltransferase 1 | 1514 | 1530 | 1402 | 1323 | 1306 | 1072 | 1332 | 1166 | 1420 | 1533 | 1076 |
| 1486 | 0.00 | 0.11 | 0.04 | 0.97 | 1 | Csnk1g3 | casein kinase 1, gamma 3 | 1447 | 1682 | 1455 | 1635 | 1454 | 1431 | 1330 | 1736 | 1408 | 1146 | 1620 |
| 1354 | 0.02 | 0.13 | 0.15 | 0.88 | 1 | 4931406C07Ril | RIKEN cDNA 4931406C07 gene | 1473 | 1686 | 1269 | 1334 | 1223 | 1303 | 1330 | 1193 | 1286 | 1586 | 1211 |
| 1285 | -0.02 | 0.14 | -0.12 | 0.90 | 1 | Urod | uroporphyrinogen decarboxylase | 1422 | 1216 | 1484 | 1238 | 1453 | 1022 | 1330 | 1357 | 1077 | 1440 | 1091 |
| 1197 | -0.05 | 0.15 | -0.34 | 0.73 | 1 | Ptpru | protein tyrosine phosphatase, receptor typ | 1087 | 1188 | 779 | 1207 | 1115 | 1515 | 1330 | 1188 | 1247 | 1327 | 1187 |
| 1381 | -0.14 | 0.17 | -0.85 | 0.40 | 1 | Hnrnpf | heterogeneous nuclear ribonucleoprotein | 2056 | 1683 | 1374 | 1159 | 1295 | 1490 | 1329 | 849 | 1397 | 1184 | 1377 |
| 1316 | 0.06 | 0.13 | 0.49 | 0.63 | 1 | Tada2b | transcriptional adaptor 2B | 1005 | 1163 | 875 | 1512 | 1466 | 1291 | 1329 | 1424 | 1649 | 1365 | 1397 |
| 1262 | 0.14 | 0.16 | 0.86 | 0.39 | 1 | Alg1 | asparagine-linked glycosylation 1 (beta-1 | 694 | 712 | 844 | 1631 | 1118 | 1445 | 1329 | 1903 | 1286 | 1569 | 1345 |
| 1104 | 0.01 | 0.14 | 0.04 | 0.97 | 1 | Ptpdc1 | protein tyrosine phosphatase domain con | 586 | 807 | 449 | 1529 | 1145 | 1146 | 1329 | 1474 | 1325 | 1284 | 1068 |
| 1487 | 0.05 | 0.10 | 0.52 | 0.61 | 1 | Nrbp1 | nuclear receptor binding protein 1 | 1860 | 1853 | 1754 | 1123 | 1227 | 1682 | 1328 | 1294 | 1274 | 1273 | 1689 |
| 1270 | -0.08 | 0.14 | -0.55 | 0.58 | 1 | Sc5d | sterol-C5-desaturase | 793 | 1251 | 1022 | 1312 | 1387 | 1564 | 1328 | 1186 | 1295 | 1231 | 1598 |
| 1276 | 0.09 | 0.14 | 0.63 | 0.53 | 1 | Rnmt | RNA (guanine-7-) methyltransferase | 1176 | 1273 | 1186 | 1013 | 1220 | 1479 | 1327 | 1404 | 1241 | 1113 | 1603 |
| 1264 | 0.10 | 0.10 | 1.02 | 0.31 | 1 | 2610001J05Rik | RIKEN cDNA 2610001J05 gene | 1351 | 1222 | 1371 | 1220 | 1318 | 936 | 1326 | 1432 | 1376 | 1395 | 957 |
| 991 | 0.16 | 0.11 | 1.40 | 0.16 | 1 | Hs3st6 | heparan sulfate (glucosamine) 3-O-sulfot | 271 | 253 | 228 | 1223 | 1236 | 1016 | 1326 | 1455 | 1238 | 1430 | 1227 |
| 1752 | 0.06 | 0.12 | 0.52 | 0.60 | 1 | Midn | midnolin | 3087 | 2627 | 2590 | 1332 | 1426 | 1279 | 1325 | 1583 | 1491 | 1288 | 1247 |
| 1413 | -0.06 | 0.10 | -0.65 | 0.52 | 1 | Patz1 | POZ (BTB) and AT hook containing zinc fi | 1470 | 1958 | 1552 | 1572 | 1320 | 1188 | 1325 | 1425 | 1284 | 1242 | 1209 |
| 1268 | 0.17 | 0.13 | 1.29 | 0.20 | 1 | Chn1 | chimerin 1 | 538 | 813 | 816 | 1472 | 1496 | 1263 | 1325 | 1584 | 1652 | 1570 | 1419 |
| 1680 | -0.12 | 0.10 | -1.18 | 0.24 | 1 | Efhd2 | EF hand domain containing 2 | 4173 | 2912 | 3083 | 1041 | 1189 | 778 | 1324 | 1025 | 940 | 1262 | 749 |
| 1454 | 0.01 | 0.14 | 0.04 | 0.96 | 1 | Tob2 | transducer of ERBB2, 2 | 2700 | 2100 | 2206 | 920 | 1158 | 1044 | 1324 | 818 | 1446 | 1376 | 900 |
| 1291 | -0.02 | 0.10 | -0.21 | 0.83 | 1 | Mlh3 | mutL homolog 3 | 1234 | 1178 | 1227 | 1331 | 1250 | 1416 | 1324 | 1294 | 1400 | 1269 | 1277 |
| 4908 | -0.14 | 0.11 | -1.25 | 0.21 | 1 | Junb | jun B proto-oncogene | 19964 | 8072 | 9381 | 728 | 973 | 5662 | 1323 | 692 | 791 | 1329 | 5076 |
| 1459 | -0.08 | 0.14 | -0.61 | 0.54 | 1 | Ccnd3 | cyclin D3 | 2254 | 2128 | 1791 | 1350 | 1239 | 1174 | 1323 | 1213 | 1387 | 1045 | 1147 |
| 1387 | 0.14 | 0.12 | 1.20 | 0.23 | 1 | Marchf9 | membrane associated ring-CH-type finge | 842 | 998 | 855 | 1754 | 1666 | 1230 | 1323 | 1958 | 1529 | 1766 | 1333 |
| 1330 | 0.19 | 0.15 | 1.29 | 0.20 | 1 | Tma7 | translational machinery associated 7 | 1052 | 1037 | 1553 | 1201 | 1503 | 1116 | 1322 | 1643 | 1452 | 1295 | 1455 |
| 1674 | -0.04 | 0.11 | -0.34 | 0.73 | 1 | Ptp4a3 | protein tyrosine phosphatase 4a3 | 2992 | 2307 | 2655 | 1410 | 1487 | 1090 | 1320 | 1317 | 1424 | 1301 | 1114 |
| 1617 | 0.11 | 0.13 | 0.86 | 0.39 | 1 | Tpd52l2 | tumor protein D52-like 2 | 2100 | 1730 | 2035 | 1417 | 1501 | 1500 | 1320 | 1764 | 1442 | 1452 | 1525 |
| 1222 | -0.05 | 0.15 | -0.36 | 0.72 | 1 | Tbc1d19 | TBC1 domain family, member 19 | 758 | 872 | 853 | 1546 | 1299 | 1407 | 1320 | 1073 | 1568 | 1349 | 1396 |
| 1848 | -0.17 | 0.15 | -1.10 | 0.27 | 1 | Per1 | period circadian clock 1 | 2677 | 2283 | 1584 | 1897 | 1165 | 2989 | 1319 | 1542 | 1208 | 1211 | 2448 |
| 1363 | -0.13 | 0.17 | -0.74 | 0.46 | 1 | Tmem131l | transmembrane 131 like | 1704 | 1740 | 1553 | 1476 | 938 | 1483 | 1319 | 1252 | 1238 | 845 | 1439 |
| 1242 | 0.28 | 0.16 | 1.77 | 0.08 | 1 | Hes6 | hairy and enhancer of split 6 | 783 | 703 | 864 | 1322 | 1255 | 1185 | 1319 | 2059 | 1354 | 1404 | 1412 |
| 1132 | -0.17 | 0.14 | -1.16 | 0.25 | 1 | Pigg | phosphatidylinositol glycan anchor biosyı | 820 | 638 | 789 | 1378 | 1459 | 1232 | 1319 | 1186 | 1373 | 992 | 1262 |
| 1387 | 0.04 | 0.11 | 0.31 | 0.75 | 1 | Usp38 | ubiquitin specific peptidase 38 | 1005 | 1436 | 994 | 1697 | 1486 | 1326 | 1318 | 1793 | 1465 | 1212 | 1522 |
| 1371 | 0.03 | 0.12 | 0.28 | 0.78 | 1 | Sde2 | SDE2 telomere maintenance homolog (S | 1455 | 1412 | 1490 | 1443 | 1287 | 1231 | 1318 | 1702 | 1331 | 1279 | 1130 |
| 1316 | 0.04 | 0.11 | 0.39 | 0.70 | 1 | Rsbn1l | round spermatid basic protein 1-like | 1256 | 1406 | 1156 | 1351 | 1075 | 1496 | 1317 | 1574 | 1150 | 1131 | 1568 |
| 1119 | 0.14 | 0.13 | 1.06 | 0.29 | 1 | Gramd1c | GRAM domain containing 1C | 406 | 464 | 588 | 1335 | 1335 | 1169 | 1317 | 1451 | 1494 | 1552 | 1203 |
| 1526 | -0.02 | 0.14 | -0.15 | 0.88 | 1 | Tubb2a | tubulin, beta 2A class IIA | 1634 | 1577 | 1632 | 1762 | 1372 | 1543 | 1316 | 1967 | 1305 | 1208 | 1468 |
| 1359 | -0.01 | 0.11 | -0.09 | 0.93 | 1 | Mllt10 | myeloid/lymphoid or mixed-lineage leuke | 1238 | 1322 | 1246 | 1254 | 1205 | 1808 | 1316 | 1170 | 1420 | 1145 | 1822 |
| 1328 | 0.01 | 0.14 | 0.05 | 0.96 | 1 | Senp2 | SUMO/sentrin specific peptidase 2 | 954 | 1286 | 1006 | 1428 | 1400 | 1522 | 1316 | 1188 | 1480 | 1383 | 1647 |
| 1210 | 0.01 | 0.12 | 0.11 | 0.91 | 1 | Get4 | golgi to ER traffic protein 4 | 1003 | 986 | 1086 | 1157 | 1335 | 1285 | 1316 | 1374 | 1302 | 1119 | 1347 |
| 1607 | 0.09 | 0.08 | 1.10 | 0.27 | 1 | Tmem87a | transmembrane protein 87A | 1598 | 1760 | 1742 | 1355 | 1411 | 2013 | 1315 | 1453 | 1583 | 1373 | 2078 |
| 1444 | 0.10 | 0.14 | 0.70 | 0.49 | 1 | Mospd2 | motile sperm domain containing 2 | 1128 | 932 | 864 | 1486 | 1363 | 2066 | 1315 | 1666 | 1491 | 1212 | 2362 |
| 1376 | 0.02 | 0.13 | 0.13 | 0.90 | 1 | Inpp5b | inositol polyphosphate-5-phosphatase B | 1276 | 1518 | 1387 | 1297 | 1567 | 1264 | 1315 | 1247 | 1532 | 1539 | 1195 |

| | | | | | | | | | | | | | | | | | |
|---|---|---|---|---|---|---|---|---|---|---|---|---|---|---|---|---|---|
| 1276 | -0.08 | 0.11 | -0.73 | 0.47 | 1 | Slc4a3 | solute carrier family 4 (anion exchanger), | 1551 | 1349 | 1527 | 915 | 967 | 1752 | 1315 | 854 | 1038 | 1104 | 1663 |
| 1247 | -0.15 | 0.10 | -1.49 | 0.14 | 1 | Parl | presenilin associated, rhomboid-like | 1338 | 1225 | 1389 | 1288 | 1306 | 1233 | 1315 | 1153 | 1271 | 1060 | 1139 |
| 1428 | -0.19 | 0.11 | -1.72 | 0.09 | 1 | Ptprj | protein tyrosine phosphatase, receptor typ | 1819 | 1568 | 1459 | 1510 | 1436 | 1538 | 1314 | 1288 | 1203 | 1252 | 1325 |
| 1227 | 0.03 | 0.11 | 0.25 | 0.80 | 1 | Tomm40l | translocase of outer mitochondrial membr | 1026 | 949 | 1105 | 1371 | 1305 | 1171 | 1314 | 1395 | 1295 | 1385 | 1183 |
| 1205 | 0.05 | 0.13 | 0.43 | 0.67 | 1 | Zc3h10 | zinc finger CCCH type containing 10 | 852 | 947 | 1033 | 1281 | 1192 | 1314 | 1314 | 1571 | 1285 | 1189 | 1278 |
| 1173 | 0.01 | 0.16 | 0.06 | 0.95 | 1 | Micu3 | mitochondrial calcium uptake family, mem | 654 | 734 | 919 | 1360 | 1162 | 1413 | 1314 | 1047 | 1393 | 1291 | 1612 |
| 1321 | 0.10 | 0.15 | 0.70 | 0.48 | 1 | Sh3bp5l | SH3 binding domain protein 5 like | 1054 | 1243 | 1363 | 1226 | 1353 | 1340 | 1313 | 1316 | 1320 | 1723 | 1276 |
| 1943 | -0.09 | 0.13 | -0.70 | 0.48 | 1 | Ddit4 | DNA-damage-inducible transcript 4 | 2074 | 2064 | 1829 | 891 | 982 | 4573 | 1312 | 856 | 773 | 1283 | 4734 |
| 1469 | -0.10 | 0.15 | -0.62 | 0.54 | 1 | Ifrd1 | interferon-related developmental regulato | 1384 | 2032 | 1549 | 1225 | 1444 | 1740 | 1310 | 1280 | 1401 | 925 | 1873 |
| 1548 | 0.03 | 0.13 | 0.19 | 0.85 | 1 | Psma4 | proteasome subunit alpha 4 | 2135 | 1862 | 2020 | 1382 | 1482 | 1265 | 1309 | 1110 | 1347 | 1618 | 1499 |
| 1200 | -0.14 | 0.13 | -1.07 | 0.29 | 1 | Brpf1 | bromodomain and PHD finger containing | 1031 | 1105 | 1004 | 1300 | 1142 | 1517 | 1308 | 1295 | 1180 | 895 | 1429 |
| 1199 | -0.04 | 0.12 | -0.30 | 0.77 | 1 | Clec16a | C-type lectin domain family 16, member A | 1170 | 836 | 1064 | 1223 | 1554 | 1036 | 1306 | 1259 | 1438 | 1345 | 958 |
| 1353 | -0.09 | 0.12 | -0.76 | 0.45 | 1 | Slc35c1 | solute carrier family 35, member C1 | 931 | 1114 | 976 | 1633 | 1626 | 1560 | 1305 | 1360 | 1422 | 1461 | 1498 |
| 1214 | -0.21 | 0.17 | -1.24 | 0.21 | 1 | Prep | prolyl endopeptidase | 1280 | 1304 | 1104 | 1360 | 1404 | 1101 | 1305 | 861 | 1483 | 1154 | 996 |
| 1190 | -0.24 | 0.16 | -1.52 | 0.13 | 1 | Eml4 | echinoderm microtubule associated prote | 1558 | 1096 | 1256 | 1332 | 1039 | 1298 | 1305 | 1042 | 1038 | 906 | 1225 |
| 1581 | 0.00 | 0.13 | -0.02 | 0.98 | 1 | Ints6 | integrator complex subunit 6 | 1234 | 1466 | 1049 | 1580 | 1545 | 2375 | 1304 | 1621 | 1557 | 1199 | 2464 |
| 1474 | 0.16 | 0.13 | 1.18 | 0.24 | 1 | Sugp1 | SURP and G patch domain containing 1 | 1001 | 951 | 1335 | 1008 | 1487 | 2378 | 1302 | 1500 | 1523 | 1282 | 2445 |
| 1364 | 0.32 | 0.20 | 1.59 | 0.11 | 1 | Ndufb7 | NADH:ubiquinone oxidoreductase subun | 1411 | 1169 | 1679 | 1275 | 1222 | 937 | 1302 | 2213 | 1164 | 1511 | 1119 |
| 991 | 0.02 | 0.15 | 0.13 | 0.89 | 1 | Aqp11 | aquaporin 11 | 198 | 190 | 247 | 1258 | 1377 | 1158 | 1302 | 1401 | 1152 | 1507 | 1113 |
| 828 | -0.20 | 0.22 | -0.92 | 0.36 | 1 | Aff2 | AF4/FMR2 family, member 2 | 268 | 431 | 284 | 989 | 1070 | 976 | 1302 | 828 | 1224 | 943 | 796 |
| 1406 | 0.17 | 0.15 | 1.09 | 0.28 | 1 | Fam216a | family with sequence similarity 216, meml | 867 | 1293 | 1378 | 1419 | 1434 | 1472 | 1301 | 1679 | 1357 | 1628 | 1636 |
| 1321 | -0.09 | 0.16 | -0.53 | 0.60 | 1 | Irf2 | interferon regulatory factor 2 | 1808 | 1662 | 1563 | 1138 | 1370 | 1055 | 1301 | 790 | 1223 | 1459 | 1166 |
| 1533 | -0.02 | 0.11 | -0.14 | 0.89 | 1 | Mettl16 | methyltransferase like 16 | 1631 | 1809 | 1857 | 1551 | 1419 | 1518 | 1300 | 1601 | 1490 | 1058 | 1625 |
| 1246 | -0.05 | 0.13 | -0.42 | 0.67 | 1 | Nudt5 | nudix (nucleoside diphosphate linked mo | 1147 | 873 | 959 | 1350 | 1502 | 1298 | 1300 | 1083 | 1444 | 1395 | 1355 |
| 1221 | -0.08 | 0.15 | -0.52 | 0.60 | 1 | Sp4 | trans-acting transcription factor 4 | 1315 | 1898 | 1593 | 1107 | 1010 | 1013 | 1300 | 1096 | 1043 | 1106 | 951 |
| 1525 | -0.03 | 0.14 | -0.22 | 0.82 | 1 | Snap29 | synaptosomal-associated protein 29 | 1586 | 1865 | 1535 | 1607 | 1672 | 1384 | 1298 | 1255 | 1663 | 1319 | 1587 |
| 1786 | 0.09 | 0.14 | 0.64 | 0.52 | 1 | Cnot10 | CCR4-NOT transcription complex, subuni | 1894 | 1538 | 1785 | 1428 | 1596 | 2713 | 1297 | 1854 | 1582 | 1295 | 2667 |
| 1542 | 0.04 | 0.11 | 0.39 | 0.69 | 1 | Gxylt1 | glucoside xylosyltransferase 1 | 1713 | 2066 | 1587 | 1584 | 1346 | 1490 | 1297 | 1554 | 1384 | 1560 | 1379 |
| 1262 | -0.13 | 0.12 | -1.03 | 0.30 | 1 | Pou2f1 | POU domain, class 2, transcription factor | 1094 | 1478 | 1210 | 1364 | 1276 | 1322 | 1297 | 1305 | 1355 | 1119 | 1060 |
| 1240 | -0.03 | 0.12 | -0.25 | 0.80 | 1 | Gzf1 | GDNF-inducible zinc finger protein 1 | 871 | 1021 | 859 | 1402 | 1207 | 1586 | 1297 | 1413 | 1377 | 1028 | 1582 |
| 1801 | 0.08 | 0.14 | 0.56 | 0.58 | 1 | Gm38394 | zinc finger, BED type containing 6 | 1943 | 2109 | 1726 | 1131 | 1143 | 3297 | 1296 | 999 | 1398 | 1435 | 3335 |
| 1571 | -0.20 | 0.15 | -1.27 | 0.20 | 1 | Triobp | TRIO and F-actin binding protein | 2243 | 2390 | 2384 | 1336 | 1659 | 1166 | 1296 | 1667 | 1137 | 1005 | 1003 |
| 1407 | 0.07 | 0.11 | 0.67 | 0.50 | 1 | Pum3 | pumilio RNA-binding family member 3 | 1750 | 1489 | 1531 | 1183 | 1372 | 1363 | 1296 | 1251 | 1329 | 1304 | 1612 |
| 1389 | -0.03 | 0.12 | -0.23 | 0.82 | 1 | Scrib | scribbled planar cell polarity | 1893 | 1319 | 1431 | 1170 | 1322 | 1580 | 1296 | 1121 | 1344 | 1345 | 1453 |
| 1185 | 0.05 | 0.12 | 0.43 | 0.67 | 1 | Gatd3a | glutamine amidotransferase like class 1 d | 1038 | 923 | 1185 | 1149 | 1284 | 1128 | 1296 | 1375 | 1298 | 1279 | 1082 |
| 1487 | -0.04 | 0.13 | -0.28 | 0.78 | 1 | Foxj2 | forkhead box J2 | 1337 | 1860 | 1301 | 1778 | 1380 | 1571 | 1295 | 1512 | 1328 | 1604 | 1388 |
| 1404 | -0.17 | 0.14 | -1.17 | 0.24 | 1 | Exosc10 | exosome component 10 | 1581 | 1334 | 1617 | 1341 | 1405 | 1675 | 1294 | 1024 | 1521 | 952 | 1696 |
| 1290 | 0.10 | 0.11 | 0.95 | 0.34 | 1 | Ap3m2 | adaptor-related protein complex 3, mu 2 s | 1138 | 1157 | 1267 | 1412 | 1350 | 1087 | 1293 | 1412 | 1364 | 1401 | 1312 |
| 1234 | 0.22 | 0.12 | 1.86 | 0.06 | 1 | Kif19a | kinesin family member 19A | 497 | 438 | 460 | 1347 | 1383 | 1600 | 1292 | 1793 | 1513 | 1622 | 1629 |
| 1155 | -0.02 | 0.13 | -0.13 | 0.89 | 1 | Prune1 | prune exopolyphosphatase | 1226 | 1337 | 1091 | 1226 | 1117 | 866 | 1292 | 1259 | 1120 | 1475 | 697 |
| 1268 | 0.30 | 0.15 | 1.95 | 0.05 | 1 | Banp | BTG3 associated nuclear protein | 564 | 495 | 711 | 956 | 1278 | 2011 | 1291 | 1610 | 1429 | 1457 | 2145 |
| 1195 | 0.15 | 0.14 | 1.08 | 0.28 | 1 | Cript | cysteine-rich PDZ-binding protein | 1356 | 1254 | 1208 | 947 | 1179 | 1011 | 1291 | 1402 | 1221 | 1190 | 1083 |
| 1411 | -0.05 | 0.14 | -0.37 | 0.71 | 1 | Plekha3 | pleckstrin homology domain-containing, f | 1648 | 1502 | 1456 | 1513 | 1506 | 1235 | 1290 | 1119 | 1538 | 1466 | 1249 |
| 1223 | 0.17 | 0.18 | 0.96 | 0.34 | 1 | Topors | topoisomerase I binding, arginine/serine- | 1379 | 979 | 1052 | 942 | 1116 | 1397 | 1290 | 1621 | 1200 | 1173 | 1306 |
| 1186 | 0.07 | 0.15 | 0.45 | 0.65 | 1 | Trpc1 | transient receptor potential cation channe | 667 | 694 | 417 | 1424 | 1273 | 1516 | 1290 | 1593 | 1337 | 1328 | 1504 |
| 1101 | -0.02 | 0.14 | -0.11 | 0.91 | 1 | Naa20 | N(alpha)-acetyltransferase 20, NatB catal | 764 | 810 | 784 | 1017 | 1226 | 1358 | 1290 | 886 | 1260 | 1355 | 1361 |
| 1348 | 0.21 | 0.14 | 1.49 | 0.14 | 1 | Ddx19a | DEAD (Asp-Glu-Ala-Asp) box polypeptide | 1433 | 1206 | 1482 | 1119 | 1369 | 1186 | 1289 | 1719 | 1428 | 1363 | 1231 |
| 1569 | 0.00 | 0.12 | -0.04 | 0.97 | 1 | Lpcat1 | lysophosphatidylcholine acyltransferase 1 | 2735 | 2182 | 2620 | 1299 | 1272 | 1010 | 1288 | 1400 | 1203 | 1234 | 1013 |
| 1316 | -0.04 | 0.14 | -0.30 | 0.76 | 1 | Gpkow | G patch domain and KOW motifs | 1363 | 1223 | 1060 | 1414 | 1339 | 1448 | 1288 | 1207 | 1474 | 1188 | 1470 |
| 1108 | -0.04 | 0.15 | -0.27 | 0.79 | 1 | Fam122a | family with sequence similarity 122, meml | 839 | 759 | 677 | 1222 | 1373 | 1126 | 1288 | 949 | 1321 | 1452 | 1184 |
| 1507 | 0.18 | 0.20 | 0.90 | 0.37 | 1 | Cops9 | COP9 signalosome subunit 9 | 1694 | 1452 | 1949 | 1494 | 1449 | 1070 | 1287 | 2454 | 1283 | 1427 | 1025 |
| 1388 | -0.10 | 0.11 | -0.95 | 0.34 | 1 | Pcnt | pericentrin (kendrin) | 1595 | 1756 | 1442 | 1472 | 1392 | 1261 | 1287 | 1343 | 1293 | 1368 | 1057 |
| 1070 | -0.09 | 0.12 | -0.79 | 0.43 | 1 | Sec22c | SEC22 homolog C, vesicle trafficking prot | 737 | 902 | 867 | 1189 | 1322 | 974 | 1287 | 1295 | 1206 | 1123 | 870 |
| 1265 | -0.10 | 0.13 | -0.78 | 0.44 | 1 | Arhgef28 | Rho guanine nucleotide exchange factor | 981 | 1068 | 968 | 1897 | 1141 | 1338 | 1286 | 1609 | 1248 | 1301 | 1081 |
| 1248 | 0.21 | 0.15 | 1.43 | 0.15 | 1 | Ddx49 | DEAD (Asp-Glu-Ala-Asp) box polypeptide | 1239 | 1045 | 1250 | 836 | 1316 | 1318 | 1286 | 1338 | 1220 | 1516 | 1362 |
| 1125 | 0.21 | 0.17 | 1.29 | 0.20 | 1 | Rassf10 | Ras association (RalGDS/AF-6) domain fa | 248 | 269 | 430 | 1306 | 1401 | 1271 | 1286 | 1828 | 1683 | 1404 | 1244 |
| 1103 | -0.06 | 0.14 | -0.43 | 0.66 | 1 | Mrps14 | mitochondrial ribosomal protein S14 | 911 | 862 | 978 | 1140 | 1332 | 1008 | 1285 | 871 | 1352 | 1403 | 991 |
| 1529 | 0.15 | 0.14 | 1.09 | 0.27 | 1 | Eif2s1 | eukaryotic translation initiation factor 2, su | 1945 | 1698 | 1502 | 1396 | 1420 | 1414 | 1284 | 1225 | 1528 | 1713 | 1696 |
| 1289 | 0.13 | 0.13 | 0.98 | 0.33 | 1 | Mlycd | malonyl-CoA decarboxylase | 909 | 1058 | 1052 | 1450 | 1438 | 1132 | 1284 | 1939 | 1325 | 1437 | 1149 |
| 1092 | -0.10 | 0.15 | -0.69 | 0.49 | 1 | 4632415L05Rik | | 917 | 1216 | 924 | 1306 | 1143 | 921 | 1284 | 989 | 1183 | 1222 | 911 |

| | | | | | | | | | | | | | | | | | |
|---|---|---|---|---|---|---|---|---|---|---|---|---|---|---|---|---|---|
| 1361 | 0.12 | 0.13 | 0.98 | 0.33 | 1 | Vps51 | VPS51 GARP complex subunit | 1232 | 1202 | 1554 | 1208 | 1399 | 1374 | 1283 | 1584 | 1376 | 1398 | 1367 |
| 1248 | 0.03 | 0.11 | 0.29 | 0.78 | 1 | Prkx | protein kinase, X-linked | 1453 | 1363 | 1220 | 1226 | 1314 | 968 | 1283 | 1170 | 1294 | 1443 | 998 |
| 1366 | 0.02 | 0.12 | 0.14 | 0.89 | 1 | Ubl4a | ubiquitin-like 4A | 1301 | 1372 | 1767 | 1336 | 1380 | 1265 | 1282 | 1456 | 1365 | 1213 | 1291 |
| 1333 | -0.10 | 0.12 | -0.86 | 0.39 | 1 | Csnk2b | casein kinase 2, beta polypeptide | 1655 | 1456 | 1910 | 1126 | 1513 | 1079 | 1282 | 1148 | 1386 | 975 | 1130 |
| 1256 | 0.12 | 0.13 | 0.90 | 0.37 | 1 | Pmm2 | phosphomannomutase 2 | 1119 | 956 | 1012 | 1071 | 1333 | 1477 | 1282 | 1491 | 1384 | 1301 | 1391 |
| 1183 | 0.11 | 0.19 | 0.57 | 0.57 | 1 | Ttc7 | tetratricopeptide repeat domain 7 | 1708 | 797 | 1137 | 1044 | 1254 | 946 | 1282 | 1404 | 1148 | 1319 | 978 |
| 982 | 0.02 | 0.17 | 0.13 | 0.90 | 1 | Hfe | homeostatic iron regulator | 1385 | 1073 | 1247 | 850 | 697 | 673 | 1282 | 885 | 660 | 1396 | 657 |
| 1466 | 0.07 | 0.13 | 0.53 | 0.60 | 1 | Imp4 | IMP4, U3 small nucleolar ribonucleoprote | 1571 | 1465 | 1593 | 1193 | 1626 | 1511 | 1281 | 1501 | 1653 | 1139 | 1590 |
| 1255 | 0.09 | 0.12 | 0.77 | 0.44 | 1 | Pfn2 | profilin 2 | 1052 | 1312 | 1074 | 1336 | 1323 | 1084 | 1281 | 1295 | 1308 | 1399 | 1342 |
| 1163 | -0.09 | 0.12 | -0.79 | 0.43 | 1 | Ap3s1 | adaptor-related protein complex 3, sigma | 878 | 920 | 968 | 1566 | 1286 | 1063 | 1281 | 1247 | 1323 | 1141 | 1125 |
| 1057 | 0.10 | 0.18 | 0.54 | 0.59 | 1 | F8a | factor 8-associated gene A | 428 | 479 | 686 | 1273 | 1224 | 1063 | 1281 | 1544 | 1144 | 1428 | 1072 |
| 990 | -0.02 | 0.13 | -0.13 | 0.90 | 1 | Mllt11 | myeloid/lymphoid or mixed-lineage leuke | 745 | 862 | 727 | 1043 | 1032 | 950 | 1281 | 1033 | 962 | 1231 | 1022 |
| 1670 | -0.13 | 0.10 | -1.24 | 0.22 | 1 | Ripor1 | RHO family interacting cell polarization re | 2318 | 2781 | 2532 | 1683 | 1339 | 1305 | 1280 | 1425 | 1308 | 1346 | 1048 |
| 1410 | -0.03 | 0.33 | -0.10 | 0.92 | 1 | Bc1 | | 2036 | 2291 | 3922 | 662 | 1212 | 390 | 1280 | 1870 | 834 | 688 | 323 |
| 1556 | 0.02 | 0.14 | 0.18 | 0.86 | 1 | Bahd1 | bromo adjacent homology domain contai | 1869 | 2110 | 1877 | 1722 | 1240 | 1316 | 1279 | 1630 | 1463 | 1588 | 1018 |
| 1108 | 0.03 | 0.11 | 0.25 | 0.80 | 1 | Eif2d | eukaryotic translation initiation factor 2D | 967 | 869 | 971 | 1014 | 1075 | 1284 | 1279 | 1081 | 1145 | 1143 | 1367 |
| 1259 | -0.14 | 0.14 | -0.98 | 0.33 | 1 | Inafm2 | InaF motif containing 2 | 1146 | 1513 | 1415 | 1471 | 1263 | 1084 | 1278 | 1156 | 1382 | 1287 | 851 |
| 1235 | 0.09 | 0.18 | 0.49 | 0.62 | 1 | Scand1 | SCAN domain-containing 1 | 1068 | 1264 | 1238 | 1358 | 1241 | 913 | 1278 | 1641 | 944 | 1763 | 878 |
| 1449 | -0.11 | 0.15 | -0.73 | 0.47 | 1 | Rnf38 | ring finger protein 38 | 1314 | 1699 | 1512 | 1608 | 1350 | 1714 | 1277 | 1101 | 1454 | 1467 | 1448 |
| 1399 | 0.07 | 0.13 | 0.58 | 0.57 | 1 | Wbp4 | WW domain binding protein 4 | 1433 | 1432 | 1463 | 1245 | 1348 | 1515 | 1277 | 1588 | 1183 | 1510 | 1396 |
| 1391 | -0.07 | 0.10 | -0.71 | 0.48 | 1 | Ahdc1 | AT hook, DNA binding motif, containing 1 | 1804 | 1872 | 1713 | 1330 | 1166 | 1303 | 1276 | 1381 | 1153 | 1198 | 1111 |
| 1338 | 0.07 | 0.11 | 0.60 | 0.55 | 1 | Prmt5 | protein arginine N-methyltransferase 5 | 1648 | 1564 | 1683 | 1206 | 1190 | 1117 | 1276 | 1453 | 1260 | 1291 | 1035 |
| 1246 | -0.02 | 0.15 | -0.12 | 0.91 | 1 | Hsdl2 | hydroxysteroid dehydrogenase like 2 | 1429 | 1420 | 1377 | 1215 | 1330 | 942 | 1276 | 890 | 1512 | 1239 | 1075 |
| 1092 | 0.20 | 0.14 | 1.41 | 0.16 | 1 | Ccpg1 | cell cycle progression 1 | 747 | 887 | 888 | 1090 | 1121 | 930 | 1276 | 1369 | 1157 | 1494 | 1048 |
| 1294 | -0.10 | 0.19 | -0.54 | 0.59 | 1 | Fgd2 | FYVE, RhoGEF and PH domain containin | 1309 | 1118 | 1193 | 1487 | 1280 | 1407 | 1275 | 1864 | 1142 | 916 | 1246 |
| 1590 | 0.16 | 0.09 | 1.74 | 0.08 | 1 | Zfp330 | zinc finger protein 330 | 1044 | 996 | 1041 | 1418 | 1458 | 2688 | 1274 | 1519 | 1661 | 1575 | 2814 |
| 1502 | 0.08 | 0.17 | 0.49 | 0.62 | 1 | Ilkap | integrin-linked kinase-associated serine/t | 1237 | 1218 | 1570 | 1119 | 1524 | 2161 | 1274 | 1827 | 1274 | 1132 | 2184 |
| 1398 | 0.10 | 0.11 | 0.96 | 0.34 | 1 | Crnkl1 | crooked neck pre-mRNA splicing factor 1 | 1284 | 1244 | 1486 | 1247 | 1445 | 1513 | 1274 | 1488 | 1653 | 1213 | 1534 |
| 1200 | -0.26 | 0.17 | -1.58 | 0.11 | 1 | Dennd1a | DENN/MADD domain containing 1A | 1237 | 1672 | 1236 | 1417 | 1257 | 978 | 1274 | 921 | 1198 | 1265 | 741 |
| 992 | 0.01 | 0.19 | 0.06 | 0.95 | 1 | Mras | muscle and microspikes RAS | 702 | 758 | 648 | 1114 | 1093 | 875 | 1274 | 761 | 1144 | 1597 | 950 |
| 1478 | -0.03 | 0.12 | -0.25 | 0.80 | 1 | Crebbp | CREB binding protein | 1801 | 1875 | 1643 | 1372 | 1142 | 1775 | 1273 | 1243 | 1335 | 1347 | 1457 |
| 1253 | -0.26 | 0.19 | -1.38 | 0.17 | 1 | Zfp142 | zinc finger protein 142 | 866 | 1009 | 935 | 1374 | 1294 | 2036 | 1273 | 761 | 1382 | 1313 | 1546 |
| 1092 | 0.10 | 0.15 | 0.68 | 0.50 | 1 | S100a1 | S100 calcium binding protein A1 | 758 | 615 | 910 | 1390 | 1314 | 747 | 1273 | 1452 | 1313 | 1314 | 922 |
| 1085 | 0.05 | 0.11 | 0.47 | 0.64 | 1 | Trappc1 | trafficking protein particle complex 1 | 944 | 996 | 943 | 1145 | 1064 | 960 | 1273 | 1330 | 1101 | 1185 | 992 |
| 1173 | 0.13 | 0.18 | 0.72 | 0.47 | 1 | Ndufa11 | NADH:ubiquinone oxidoreductase subun | 900 | 708 | 1078 | 1199 | 1353 | 1030 | 1272 | 1764 | 1330 | 1322 | 942 |
| 1419 | 0.27 | 0.17 | 1.53 | 0.13 | 1 | Rpia | ribose 5-phosphate isomerase A | 1507 | 1159 | 1223 | 1134 | 1636 | 1290 | 1270 | 1799 | 1727 | 1230 | 1630 |
| 1331 | -0.04 | 0.12 | -0.34 | 0.74 | 1 | Ythdf2 | YTH N6-methyladenosine RNA binding p | 1437 | 1542 | 1560 | 1274 | 1297 | 1266 | 1270 | 1395 | 1319 | 961 | 1321 |
| 1275 | 0.03 | 0.13 | 0.24 | 0.81 | 1 | Naa25 | N(alpha)-acetyltransferase 25, NatB auxil | 1052 | 1163 | 926 | 1409 | 1298 | 1402 | 1270 | 1175 | 1311 | 1565 | 1457 |
| 1211 | -0.05 | 0.14 | -0.31 | 0.75 | 1 | Lig3 | ligase III, DNA, ATP-dependent | 1080 | 1020 | 854 | 1042 | 1266 | 1696 | 1270 | 963 | 1369 | 1233 | 1524 |
| 1195 | 0.23 | 0.21 | 1.11 | 0.27 | 1 | Ubc | ubiquitin C | 1035 | 663 | 1190 | 661 | 1086 | 1791 | 1270 | 1271 | 1158 | 1006 | 2010 |
| 1432 | 0.00 | 0.11 | 0.03 | 0.98 | 1 | Usp10 | ubiquitin specific peptidase 10 | 1471 | 1578 | 1233 | 1466 | 1371 | 1629 | 1269 | 1366 | 1344 | 1466 | 1561 |
| 1077 | -0.08 | 0.14 | -0.57 | 0.57 | 1 | Gbe1 | glucan (1,4-alpha-), branching enzyme 1 | 966 | 998 | 837 | 1280 | 1084 | 1013 | 1269 | 1109 | 1336 | 1055 | 897 |
| 1386 | -0.04 | 0.12 | -0.35 | 0.72 | 1 | Rictor | RPTOR independent companion of MTOF | 1148 | 1358 | 1274 | 1217 | 1155 | 2221 | 1268 | 1138 | 1217 | 1316 | 1935 |
| 1299 | 0.15 | 0.13 | 1.14 | 0.25 | 1 | Tmem115 | transmembrane protein 115 | 1398 | 1210 | 1281 | 1406 | 1259 | 990 | 1268 | 1746 | 1268 | 1391 | 1070 |
| 1236 | 0.10 | 0.11 | 0.95 | 0.34 | 1 | Ccnc | cyclin C | 966 | 1063 | 966 | 1379 | 1206 | 1264 | 1268 | 1303 | 1367 | 1314 | 1503 |
| 1231 | 0.06 | 0.12 | 0.51 | 0.61 | 1 | Polr3b | polymerase (RNA) III (DNA directed) poly | 1313 | 1197 | 992 | 1211 | 1334 | 1095 | 1268 | 1362 | 1213 | 1378 | 1177 |
| 1196 | 0.18 | 0.16 | 1.11 | 0.27 | 1 | Rab25 | RAB25, member RAS oncogene family | 851 | 1084 | 1168 | 1116 | 1274 | 1029 | 1268 | 1752 | 1227 | 1329 | 1062 |
| 1162 | 0.02 | 0.12 | 0.19 | 0.85 | 1 | Yrdc | yrdC domain containing (E.coli) | 640 | 718 | 765 | 1147 | 1550 | 1325 | 1268 | 1127 | 1503 | 1426 | 1315 |
| 2355 | -0.19 | 0.14 | -1.41 | 0.16 | 1 | Gm2a | GM2 ganglioside activator protein | 6723 | 5283 | 4507 | 1334 | 1532 | 857 | 1267 | 1175 | 1222 | 1274 | 727 |
| 1188 | -0.06 | 0.14 | -0.44 | 0.66 | 1 | Grb7 | growth factor receptor bound protein 7 | 1140 | 1065 | 1654 | 983 | 1270 | 1178 | 1266 | 1072 | 1261 | 1176 | 997 |
| 1069 | 0.05 | 0.17 | 0.31 | 0.76 | 1 | Snx25 | sorting nexin 25 | 1105 | 675 | 843 | 1099 | 1183 | 953 | 1266 | 1373 | 1117 | 1090 | 1055 |
| 1064 | -0.03 | 0.16 | -0.21 | 0.83 | 1 | Chchd10 | coiled-coil-helix-coiled-coil-helix domain | 596 | 737 | 789 | 1317 | 1331 | 911 | 1266 | 1031 | 1364 | 1520 | 839 |
| 1207 | -0.07 | 0.13 | -0.50 | 0.62 | 1 | Tbc1d5 | TBC1 domain family, member 5 | 1389 | 1178 | 963 | 1379 | 1273 | 1056 | 1265 | 1126 | 1190 | 1453 | 1009 |
| 1421 | 0.13 | 0.10 | 1.24 | 0.21 | 1 | Rtraf | RNA transcription, translation and transpo | 1786 | 1749 | 1785 | 1195 | 1441 | 1031 | 1264 | 1482 | 1431 | 1310 | 1155 |
| 1131 | -0.21 | 0.18 | -1.16 | 0.25 | 1 | Slc25a53 | solute carrier family 25, member 53 | 1410 | 1660 | 1635 | 881 | 978 | 1004 | 1264 | 767 | 1013 | 748 | 1077 |
| 1037 | 0.20 | 0.16 | 1.23 | 0.22 | 1 | Map7d2 | MAP7 domain containing 2 | 741 | 489 | 635 | 1223 | 979 | 970 | 1264 | 1508 | 1259 | 1351 | 988 |
| 1344 | -0.05 | 0.14 | -0.36 | 0.72 | 1 | Dnajc11 | DnaJ heat shock protein family (Hsp40) m | 1647 | 1275 | 1363 | 1154 | 1424 | 1464 | 1263 | 923 | 1499 | 1224 | 1541 |
| 945 | -0.18 | 0.20 | -0.92 | 0.36 | 1 | Rnf150 | ring finger protein 150 | 841 | 620 | 498 | 1078 | 1264 | 896 | 1263 | 1072 | 1202 | 846 | 816 |
| 1488 | -0.10 | 0.13 | -0.82 | 0.41 | 1 | Gabpa | GA repeat binding protein, alpha | 2091 | 1764 | 1891 | 1409 | 1327 | 1489 | 1262 | 1045 | 1432 | 1208 | 1449 |
| 1381 | -0.04 | 0.15 | -0.24 | 0.81 | 1 | Lap3 | leucine aminopeptidase 3 | 1835 | 1946 | 2007 | 1179 | 1338 | 972 | 1262 | 959 | 1122 | 1533 | 1039 |

| | | | | | | | | | | | | | | | | | | |
|---|---|---|---|---|---|---|---|---|---|---|---|---|---|---|---|---|---|---|
| 1375 | 0.06 | 0.15 | 0.42 | 0.68 | 1 | Kmt5a | lysine methyltransferase 5A | 1315 | 1590 | 1534 | 1553 | 1088 | 1282 | 1262 | 1936 | 1296 | 1137 | 1129 |
| 1360 | 0.11 | 0.13 | 0.81 | 0.42 | 1 | Plrg1 | pleiotropic regulator 1 | 1556 | 1505 | 1685 | 1189 | 1250 | 1205 | 1262 | 1540 | 1138 | 1444 | 1184 |
| 1128 | -0.11 | 0.11 | -1.03 | 0.30 | 1 | Xpnpep1 | X-prolyl aminopeptidase (aminopeptidase | 1100 | 899 | 1110 | 1113 | 1178 | 1281 | 1262 | 1046 | 1187 | 1030 | 1201 |
| 1315 | -0.13 | 0.11 | -1.16 | 0.25 | 1 | Nvl | nuclear VCP-like | 1192 | 975 | 1246 | 1297 | 1342 | 1860 | 1260 | 994 | 1397 | 1208 | 1688 |
| 1505 | -0.01 | 0.15 | -0.05 | 0.96 | 1 | Hmbox1 | homeobox containing 1 | 1820 | 1745 | 1694 | 1210 | 1324 | 1864 | 1258 | 1017 | 1725 | 1202 | 1699 |
| 1281 | 0.00 | 0.13 | 0.02 | 0.98 | 1 | Psmd5 | proteasome (prosome, macropain) 26S su | 1257 | 1690 | 1591 | 1161 | 1169 | 1173 | 1258 | 1181 | 1201 | 1090 | 1318 |
| 1127 | 0.12 | 0.14 | 0.90 | 0.37 | 1 | Atp8a2 | ATPase, aminophospholipid transporter-l | 734 | 1042 | 818 | 1322 | 1191 | 921 | 1258 | 1287 | 1456 | 1383 | 983 |
| 1077 | -0.03 | 0.14 | -0.20 | 0.84 | 1 | Iqce | IQ motif containing E | 1011 | 1014 | 895 | 1039 | 991 | 1221 | 1257 | 914 | 1192 | 1065 | 1246 |
| 544 | -0.39 | 0.29 | -1.34 | 0.18 | 1 | Rec8 | REC8 meiotic recombination protein | 377 | 135 | 51 | 494 | 235 | 821 | 1257 | 492 | 71 | 1269 | 784 |
| 1364 | -0.11 | 0.16 | -0.72 | 0.47 | 1 | Sec24a | Sec24 related gene family, member A (S. | 885 | 1219 | 792 | 1489 | 1312 | 2262 | 1256 | 1281 | 1545 | 1023 | 1944 |
| 1354 | -0.02 | 0.14 | -0.18 | 0.86 | 1 | Cpsf2 | cleavage and polyadenylation specific fac | 1352 | 1252 | 1302 | 1364 | 1706 | 1226 | 1256 | 1462 | 1491 | 1110 | 1377 |
| 1334 | 0.07 | 0.12 | 0.57 | 0.57 | 1 | Ints4 | integrator complex subunit 4 | 1323 | 1036 | 1086 | 1155 | 1404 | 1657 | 1256 | 1095 | 1497 | 1559 | 1610 |
| 1254 | 0.06 | 0.15 | 0.40 | 0.69 | 1 | Uqcr10 | ubiquinol-cytochrome c reductase, compl | 1187 | 1311 | 1577 | 1252 | 1301 | 952 | 1256 | 1454 | 1099 | 1400 | 1004 |
| 1110 | 0.07 | 0.14 | 0.51 | 0.61 | 1 | Trappc4 | trafficking protein particle complex 4 | 973 | 1052 | 1128 | 1077 | 1148 | 927 | 1256 | 1397 | 1217 | 1005 | 1027 |
| 909 | 0.01 | 0.19 | 0.06 | 0.95 | 1 | Sytl1 | synaptotagmin-like 1 | 359 | 397 | 414 | 852 | 962 | 1315 | 1256 | 982 | 786 | 1483 | 1192 |
| 1353 | 0.02 | 0.17 | 0.14 | 0.89 | 1 | Zbtb40 | zinc finger and BTB domain containing 40 | 601 | 716 | 826 | 1633 | 1522 | 1854 | 1255 | 2224 | 1251 | 1148 | 1853 |
| 1325 | 0.15 | 0.17 | 0.86 | 0.39 | 1 | Lsm4 | LSM4 homolog, U6 small nuclear RNA ar | 1760 | 1329 | 1803 | 1056 | 1386 | 889 | 1255 | 1551 | 1059 | 1562 | 926 |
| 1038 | 0.06 | 0.13 | 0.48 | 0.63 | 1 | Mkrn2 | makorin, ring finger protein, 2 | 1023 | 886 | 949 | 996 | 975 | 970 | 1255 | 1159 | 1170 | 1020 | 1018 |
| 1033 | -0.11 | 0.20 | -0.55 | 0.59 | 1 | Amer2 | APC membrane recruitment 2 | 626 | 933 | 722 | 1131 | 1157 | 1161 | 1254 | 825 | 1225 | 1347 | 983 |
| 1251 | 0.26 | 0.16 | 1.60 | 0.11 | 1 | Ndufb6 | NADH:ubiquinone oxidoreductase subun | 1362 | 1192 | 1553 | 1165 | 1075 | 889 | 1253 | 1544 | 1067 | 1624 | 1039 |
| 1496 | -0.11 | 0.12 | -0.94 | 0.35 | 1 | Pan3 | PAN3 poly(A) specific ribonuclease subu | 1567 | 1317 | 1301 | 1100 | 1357 | 2652 | 1252 | 966 | 1115 | 1404 | 2425 |
| 1263 | -0.05 | 0.10 | -0.52 | 0.60 | 1 | Dctn6 | dynactin 6 | 1242 | 1154 | 1374 | 1202 | 1397 | 1300 | 1251 | 1145 | 1294 | 1112 | 1424 |
| 1074 | 0.12 | 0.15 | 0.78 | 0.43 | 1 | Tor2a | torsin family 2, member A | 880 | 831 | 864 | 1028 | 1186 | 948 | 1251 | 1564 | 1137 | 1137 | 986 |
| 984 | 0.03 | 0.14 | 0.24 | 0.81 | 1 | Large2 | LARGE xylosyl- and glucuronyltransferas | 515 | 489 | 472 | 1117 | 1189 | 1013 | 1251 | 1426 | 1197 | 1359 | 794 |
| 1445 | 0.24 | 0.12 | 1.96 | 0.05 | 1 | Cspp1 | centrosome and spindle pole associated | 557 | 504 | 429 | 1053 | 906 | 3629 | 1250 | 1375 | 1274 | 1355 | 3560 |
| 1665 | 0.06 | 0.14 | 0.44 | 0.66 | 1 | Skil | SKI-like | 2711 | 2096 | 1745 | 1516 | 1328 | 1650 | 1249 | 1432 | 1328 | 1447 | 1812 |
| 1455 | -0.05 | 0.12 | -0.39 | 0.69 | 1 | Smarcad1 | SWI/SNF-related, matrix-associated actin- | 1235 | 1247 | 1019 | 1565 | 1433 | 2095 | 1249 | 1330 | 1335 | 1381 | 2116 |
| 1265 | -0.09 | 0.15 | -0.60 | 0.55 | 1 | Slc9a6 | solute carrier family 9 (sodium/hydrogen e | 1134 | 1214 | 1066 | 1630 | 1417 | 1130 | 1248 | 1124 | 1252 | 1530 | 1168 |
| 1166 | 0.07 | 0.12 | 0.61 | 0.54 | 1 | Scoc | short coiled-coil protein | 978 | 987 | 1169 | 1223 | 1193 | 1063 | 1248 | 1296 | 1188 | 1266 | 1211 |
| 990 | -0.24 | 0.13 | -1.85 | 0.06 | 1 | Sik2 | salt inducible kinase 2 | 815 | 731 | 815 | 1077 | 961 | 1329 | 1248 | 909 | 927 | 877 | 1203 |
| 1419 | 0.02 | 0.12 | 0.21 | 0.84 | 1 | Akirin2 | akirin 2 | 1537 | 1689 | 1704 | 1455 | 1232 | 1357 | 1247 | 1229 | 1219 | 1399 | 1543 |
| 1314 | 0.04 | 0.11 | 0.35 | 0.73 | 1 | Zbtb11 | zinc finger and BTB domain containing 11 | 1011 | 1156 | 957 | 1546 | 1106 | 1702 | 1247 | 1371 | 1357 | 1211 | 1795 |
| 1260 | -0.09 | 0.13 | -0.71 | 0.48 | 1 | Fgd6 | FYVE, RhoGEF and PH domain containin | 561 | 651 | 606 | 1494 | 1693 | 1762 | 1247 | 1196 | 1639 | 1255 | 1752 |
| 1164 | 0.07 | 0.14 | 0.51 | 0.61 | 1 | Mief1 | mitochondrial elongation factor 1 | 987 | 789 | 814 | 999 | 1353 | 1391 | 1247 | 1350 | 1245 | 1405 | 1222 |
| 1128 | 0.25 | 0.17 | 1.47 | 0.14 | 1 | Cdc34 | cell division cycle 34 | 708 | 850 | 876 | 1087 | 1200 | 988 | 1247 | 1681 | 1125 | 1613 | 1029 |
| 990 | -0.07 | 0.15 | -0.44 | 0.66 | 1 | Tceal1 | transcription elongation factor A (SII)-like | 817 | 1033 | 951 | 1027 | 962 | 904 | 1247 | 1036 | 880 | 1112 | 923 |
| 1251 | -0.10 | 0.14 | -0.70 | 0.48 | 1 | G2e3 | G2/M-phase specific E3 ubiquitin ligase | 674 | 1171 | 894 | 1552 | 1098 | 1817 | 1246 | 1265 | 1230 | 1090 | 1725 |
| 1561 | 0.08 | 0.14 | 0.60 | 0.55 | 1 | Mrpl24 | mitochondrial ribosomal protein L24 | 1646 | 1562 | 1644 | 1171 | 1355 | 2226 | 1245 | 1649 | 1286 | 1135 | 2256 |
| 1421 | 0.02 | 0.16 | 0.13 | 0.90 | 1 | Med1 | mediator complex subunit 1 | 1534 | 1345 | 1145 | 1261 | 1228 | 2087 | 1245 | 1362 | 1540 | 1119 | 1769 |
| 1525 | -0.14 | 0.13 | -1.13 | 0.26 | 1 | Stk38 | serine/threonine kinase 38 | 1869 | 2089 | 1566 | 1491 | 1302 | 1816 | 1243 | 1599 | 1025 | 1007 | 1771 |
| 1415 | 0.10 | 0.13 | 0.78 | 0.43 | 1 | Pepd | peptidase D | 1982 | 1438 | 1953 | 1294 | 1306 | 1080 | 1243 | 1279 | 1328 | 1358 | 1302 |
| 1170 | 0.11 | 0.19 | 0.58 | 0.56 | 1 | H2aj | H2J.A histone | 783 | 806 | 1165 | 1288 | 1374 | 946 | 1243 | 1799 | 1100 | 1383 | 979 |
| 1464 | -0.05 | 0.11 | -0.46 | 0.65 | 1 | Asxl1 | additional sex combs like 1 | 2070 | 1865 | 1993 | 1234 | 1261 | 1442 | 1242 | 1226 | 1306 | 1319 | 1149 |
| 1420 | -0.28 | 0.18 | -1.56 | 0.12 | 1 | Arnt2 | aryl hydrocarbon receptor nuclear translo | 2105 | 3097 | 2428 | 1133 | 1169 | 847 | 1242 | 726 | 1088 | 976 | 810 |
| 1122 | -0.06 | 0.13 | -0.46 | 0.64 | 1 | Brd7 | bromodomain containing 7 | 1073 | 927 | 920 | 1097 | 1015 | 1452 | 1242 | 882 | 1190 | 1222 | 1320 |
| 2116 | 0.12 | 0.13 | 0.87 | 0.38 | 1 | Mdm4 | transformed mouse 3T3 cell double minut | 1645 | 1582 | 1363 | 1105 | 1025 | 5993 | 1241 | 1431 | 1346 | 1100 | 5439 |
| 1619 | -0.19 | 0.21 | -0.94 | 0.35 | 1 | Arhgef9 | CDC42 guanine nucleotide exchange fac | 2080 | 3054 | 2435 | 1434 | 1227 | 1549 | 1241 | 744 | 1169 | 1426 | 1448 |
| 1344 | 0.18 | 0.12 | 1.47 | 0.14 | 1 | Osbpl1a | oxysterol binding protein-like 1A | 1377 | 1677 | 1449 | 1251 | 1273 | 1043 | 1241 | 1434 | 1273 | 1641 | 1130 |
| 1273 | -0.23 | 0.14 | -1.62 | 0.11 | 1 | Mbd5 | methyl-CpG binding domain protein 5 | 1452 | 1698 | 1435 | 1398 | 1055 | 1399 | 1241 | 911 | 1136 | 1072 | 1207 |
| 1209 | -0.10 | 0.13 | -0.81 | 0.42 | 1 | Gfm1 | G elongation factor, mitochondrial 1 | 1331 | 1294 | 1209 | 1146 | 1325 | 1184 | 1241 | 943 | 1323 | 1105 | 1197 |
| 1094 | -0.04 | 0.12 | -0.29 | 0.77 | 1 | Ccm2 | cerebral cavernous malformation 2 | 1000 | 689 | 913 | 1212 | 1303 | 1009 | 1241 | 1029 | 1292 | 1302 | 1044 |
| 1366 | -0.05 | 0.14 | -0.36 | 0.72 | 1 | Dlgap4 | DLG associated protein 4 | 1627 | 1407 | 1562 | 1301 | 1278 | 1487 | 1240 | 1526 | 1298 | 1146 | 1157 |
| 1344 | -0.32 | 0.19 | -1.70 | 0.09 | 1 | Lrch3 | leucine-rich repeats and calponin homolo | 1731 | 1468 | 1641 | 1327 | 1117 | 1759 | 1240 | 860 | 1310 | 715 | 1617 |
| 1119 | -0.07 | 0.10 | -0.70 | 0.48 | 1 | Pi4k2b | phosphatidylinositol 4-kinase type 2 beta | 661 | 738 | 707 | 1361 | 1460 | 1166 | 1240 | 1293 | 1409 | 1112 | 1158 |
| 1023 | -0.03 | 0.17 | -0.20 | 0.84 | 1 | Snrpn | small nuclear ribonucleoprotein N | 900 | 823 | 1204 | 1168 | 1020 | 782 | 1240 | 1031 | 1000 | 1278 | 801 |
| 1408 | -0.03 | 0.13 | -0.25 | 0.80 | 1 | Emg1 | EMG1 N1-specific pseudouridine methylt | 1857 | 1539 | 2150 | 1222 | 1272 | 1271 | 1239 | 974 | 1270 | 1477 | 1220 |
| 1317 | 0.02 | 0.11 | 0.18 | 0.86 | 1 | Shtn1 | shootin 1 | 1494 | 1222 | 1496 | 1218 | 1319 | 1317 | 1239 | 1172 | 1339 | 1471 | 1203 |
| 1317 | 0.06 | 0.13 | 0.44 | 0.66 | 1 | Zfp68 | zinc finger protein 68 | 1190 | 1758 | 1295 | 1312 | 1109 | 1350 | 1239 | 1344 | 1337 | 1067 | 1486 |
| 1216 | -0.04 | 0.13 | -0.33 | 0.74 | 1 | Mrpl50 | mitochondrial ribosomal protein L50 | 1156 | 1316 | 1182 | 1300 | 1236 | 1157 | 1239 | 1029 | 1185 | 1356 | 1220 |
| 1153 | 0.08 | 0.16 | 0.51 | 0.61 | 1 | Kctd13 | potassium channel tetramerisation domai | 691 | 874 | 1015 | 1247 | 1265 | 1139 | 1239 | 1461 | 1271 | 1484 | 996 |

| | | | | | | | | | | | | | | | | | | |
|---|---|---|---|---|---|---|---|---|---|---|---|---|---|---|---|---|---|---|
| 1186 | -0.07 | 0.13 | -0.57 | 0.57 | 1 | Eml3 | echinoderm microtubule associated prote | 1317 | 1125 | 1053 | 1142 | 1161 | 1358 | 1238 | 1261 | 1129 | 1048 | 1212 |
| 929 | -0.12 | 0.11 | -1.09 | 0.27 | 1 | Cd276 | CD276 antigen | 1031 | 938 | 682 | 1111 | 933 | 660 | 1238 | 1102 | 788 | 1112 | 624 |
| 1547 | 0.02 | 0.12 | 0.15 | 0.88 | 1 | Brf1 | BRF1, RNA polymerase III transcription in | 1482 | 1658 | 1216 | 1205 | 1299 | 2555 | 1236 | 1315 | 1205 | 1292 | 2558 |
| 1497 | -0.13 | 0.19 | -0.70 | 0.49 | 1 | Btbd2 | BTB (POZ) domain containing 2 | 1737 | 2464 | 1960 | 1399 | 1536 | 1214 | 1236 | 1017 | 1163 | 1626 | 1119 |
| 1374 | -0.13 | 0.17 | -0.77 | 0.44 | 1 | Sytl5 | synaptotagmin-like 5 | 1742 | 2787 | 2113 | 1069 | 1060 | 1056 | 1235 | 898 | 1109 | 899 | 1142 |
| 1217 | -0.18 | 0.14 | -1.28 | 0.20 | 1 | Parp4 | poly (ADP-ribose) polymerase family, mer | 1560 | 1191 | 1291 | 1276 | 1196 | 1226 | 1235 | 849 | 1173 | 1304 | 1088 |
| 1137 | -0.12 | 0.15 | -0.84 | 0.40 | 1 | Bsn | bassoon | 813 | 763 | 757 | 1332 | 1379 | 1332 | 1235 | 999 | 1436 | 1413 | 1046 |
| 1302 | -0.03 | 0.13 | -0.23 | 0.82 | 1 | Cep63 | centrosomal protein 63 | 1161 | 1250 | 1254 | 1175 | 1309 | 1634 | 1234 | 1447 | 1129 | 991 | 1741 |
| 1298 | -0.09 | 0.15 | -0.60 | 0.55 | 1 | Spred2 | sprouty-related EVH1 domain containing | 1678 | 2221 | 1874 | 1233 | 960 | 960 | 1233 | 979 | 1090 | 1231 | 821 |
| 1604 | -0.25 | 0.17 | -1.47 | 0.14 | 1 | Lrp5 | low density lipoprotein receptor-related pı | 2540 | 2148 | 2577 | 1808 | 1504 | 997 | 1232 | 1617 | 1332 | 1291 | 601 |
| 1395 | 0.18 | 0.13 | 1.44 | 0.15 | 1 | Tmem168 | transmembrane protein 168 | 1832 | 1370 | 1345 | 1416 | 1233 | 1166 | 1232 | 1530 | 1381 | 1551 | 1284 |
| 1152 | -0.10 | 0.18 | -0.56 | 0.57 | 1 | Map4k3 | mitogen-activated protein kinase kinase k | 1248 | 1327 | 986 | 1028 | 890 | 1590 | 1232 | 772 | 1166 | 1064 | 1372 |
| 1284 | -0.25 | 0.14 | -1.79 | 0.07 | 1 | Ptpn13 | protein tyrosine phosphatase, non-recept | 2051 | 1616 | 1653 | 1317 | 936 | 1319 | 1231 | 1065 | 990 | 1007 | 935 |
| 1450 | 0.03 | 0.13 | 0.21 | 0.83 | 1 | Zfp592 | zinc finger protein 592 | 1515 | 1744 | 1433 | 1685 | 1236 | 1434 | 1230 | 1437 | 1390 | 1604 | 1240 |
| 1203 | 0.16 | 0.15 | 1.07 | 0.29 | 1 | Dis3l | DIS3 like exosome 3'-5' exoribonuclease | 1136 | 1216 | 828 | 1125 | 1289 | 1090 | 1230 | 1416 | 1346 | 1465 | 1087 |
| 1102 | -0.02 | 0.13 | -0.13 | 0.89 | 1 | Wdr47 | WD repeat domain 47 | 763 | 882 | 707 | 1066 | 1112 | 1487 | 1230 | 1273 | 940 | 1137 | 1527 |
| 991 | 0.00 | 0.16 | -0.02 | 0.98 | 1 | Nectin1 | nectin cell adhesion molecule 1 | 562 | 529 | 589 | 1589 | 1024 | 751 | 1230 | 1548 | 1190 | 1219 | 671 |
| 810 | -0.15 | 0.21 | -0.70 | 0.48 | 1 | Dtnbos | | 95 | 189 | 128 | 1275 | 1102 | 874 | 1230 | 913 | 1083 | 1208 | 807 |
| 1103 | 0.01 | 0.15 | 0.09 | 0.93 | 1 | Mtif2 | mitochondrial translational initiation factor | 939 | 885 | 989 | 978 | 961 | 1436 | 1229 | 987 | 1129 | 912 | 1689 |
| 1047 | 0.18 | 0.12 | 1.45 | 0.15 | 1 | Fbxl4 | F-box and leucine-rich repeat protein 4 | 747 | 736 | 745 | 1180 | 992 | 964 | 1229 | 1135 | 1236 | 1372 | 1175 |
| 1313 | 0.03 | 0.10 | 0.26 | 0.79 | 1 | Smim13 | small integral membrane protein 13 | 1531 | 1567 | 1366 | 1414 | 1278 | 1024 | 1228 | 1454 | 1310 | 1207 | 1061 |
| 1404 | 0.06 | 0.16 | 0.40 | 0.69 | 1 | Mepce | methylphosphate capping enzyme | 1637 | 1361 | 1640 | 1328 | 1433 | 1285 | 1226 | 1776 | 1213 | 1209 | 1332 |
| 1229 | 0.01 | 0.12 | 0.10 | 0.92 | 1 | Fam192a | family with sequence similarity 192, meml | 1536 | 1267 | 1293 | 932 | 1313 | 1218 | 1226 | 937 | 1260 | 1349 | 1189 |
| 1180 | 0.03 | 0.12 | 0.20 | 0.84 | 1 | Rnf214 | ring finger protein 214 | 968 | 923 | 913 | 1370 | 1104 | 1349 | 1226 | 1212 | 1377 | 1311 | 1230 |
| 1169 | 0.05 | 0.13 | 0.41 | 0.68 | 1 | Zrsr2 | zinc finger (CCCH type), RNA binding mc | 1121 | 1154 | 1323 | 960 | 1262 | 1106 | 1226 | 1113 | 1154 | 1115 | 1327 |
| 1398 | -0.15 | 0.15 | -1.02 | 0.31 | 1 | Zfp523 | zinc finger protein 523 | 1174 | 1567 | 1430 | 1255 | 1503 | 1926 | 1225 | 938 | 1430 | 1260 | 1672 |
| 1263 | -0.04 | 0.14 | -0.32 | 0.75 | 1 | Orc2 | origin recognition complex, subunit 2 | 1317 | 1179 | 1097 | 1231 | 1477 | 1289 | 1225 | 1104 | 1471 | 1130 | 1374 |
| 1214 | -0.11 | 0.13 | -0.81 | 0.42 | 1 | Fam131a | family with sequence similarity 131, meml | 1180 | 1479 | 1213 | 1399 | 1050 | 1204 | 1225 | 1578 | 870 | 1054 | 1100 |
| 1641 | -0.09 | 0.14 | -0.68 | 0.50 | 1 | Lix1l | Lix1-like | 3105 | 3440 | 2571 | 1652 | 814 | 926 | 1224 | 1330 | 806 | 1383 | 800 |
| 1186 | 0.19 | 0.11 | 1.78 | 0.07 | 1 | C1d | C1D nuclear receptor co-repressor | 1380 | 1057 | 1240 | 1019 | 1075 | 1064 | 1224 | 1229 | 1303 | 1350 | 1110 |
| 1147 | -0.09 | 0.16 | -0.55 | 0.58 | 1 | Tor4a | torsin family 4, member A | 1518 | 1207 | 1405 | 1245 | 1244 | 631 | 1224 | 1401 | 1246 | 951 | 548 |
| 1047 | -0.17 | 0.15 | -1.12 | 0.26 | 1 | Mbtps2 | membrane-bound transcription factor pep | 1014 | 885 | 1150 | 1063 | 1131 | 1047 | 1223 | 872 | 1251 | 797 | 1081 |
| 990 | -0.08 | 0.16 | -0.51 | 0.61 | 1 | Samd10 | sterile alpha motif domain containing 10 | 765 | 756 | 746 | 1133 | 1170 | 876 | 1223 | 801 | 1264 | 1361 | 797 |
| 1075 | -0.11 | 0.14 | -0.78 | 0.44 | 1 | Kat5 | K(lysine) acetyltransferase 5 | 1071 | 1024 | 977 | 1020 | 1112 | 1172 | 1222 | 781 | 1158 | 1123 | 1167 |
| 1024 | 0.03 | 0.17 | 0.19 | 0.85 | 1 | Phyhipl | phytanoyl-CoA hydroxylase interacting pr | 792 | 1201 | 911 | 999 | 892 | 1025 | 1222 | 1016 | 1037 | 1182 | 986 |
| 1932 | -0.17 | 0.14 | -1.19 | 0.24 | 1 | Kctd10 | potassium channel tetramerisation domai | 4839 | 3868 | 3392 | 1378 | 1256 | 985 | 1221 | 900 | 1204 | 1265 | 940 |
| 1653 | -0.01 | 0.13 | -0.06 | 0.95 | 1 | Maf1 | MAF1 homolog, negative regulator of RN/ | 2717 | 2343 | 2787 | 1136 | 1518 | 1307 | 1221 | 1235 | 1136 | 1538 | 1241 |
| 1347 | -0.04 | 0.16 | -0.24 | 0.81 | 1 | Dcp2 | decapping mRNA 2 | 2284 | 2176 | 1771 | 939 | 986 | 1210 | 1221 | 1242 | 1019 | 833 | 1139 |
| 1273 | 0.13 | 0.14 | 0.87 | 0.38 | 1 | Pqbp1 | polyglutamine binding protein 1 | 1197 | 1046 | 1271 | 1111 | 1463 | 1225 | 1221 | 1543 | 1280 | 1499 | 1153 |
| 1269 | 0.11 | 0.11 | 0.93 | 0.35 | 1 | Ddx56 | DEAD (Asp-Glu-Ala-Asp) box polypeptide | 1288 | 1261 | 1277 | 1161 | 1300 | 1185 | 1221 | 1147 | 1438 | 1501 | 1175 |
| 1177 | 0.04 | 0.15 | 0.30 | 0.77 | 1 | Thoc3 | THO complex 3 | 1111 | 1102 | 1333 | 1035 | 1327 | 1040 | 1221 | 1470 | 1187 | 1175 | 951 |
| 1445 | -0.10 | 0.11 | -0.98 | 0.33 | 1 | Evl | Ena-vasodilator stimulated phosphoprote | 2039 | 2019 | 2089 | 1349 | 1360 | 1114 | 1220 | 1369 | 1113 | 1190 | 1029 |
| 1295 | 0.04 | 0.13 | 0.27 | 0.78 | 1 | Zfp12 | zinc finger protein 12 | 952 | 1196 | 1014 | 1280 | 1247 | 1719 | 1220 | 1102 | 1372 | 1342 | 1795 |
| 1249 | -0.03 | 0.12 | -0.21 | 0.83 | 1 | Rbbp5 | retinoblastoma binding protein 5, histone | 1295 | 1345 | 1178 | 1154 | 1268 | 1341 | 1220 | 1216 | 1110 | 1068 | 1547 |
| 1377 | -0.08 | 0.20 | -0.39 | 0.69 | 1 | Bicral | BRD4 interacting chromatin remodeling c | 1639 | 1530 | 1295 | 1427 | 1324 | 1483 | 1219 | 834 | 1666 | 1391 | 1343 |
| 1128 | 0.01 | 0.11 | 0.09 | 0.93 | 1 | Mccc1 | methylcrotonoyl-Coenzyme A carboxylas | 855 | 1152 | 707 | 1295 | 958 | 1349 | 1219 | 1425 | 991 | 1067 | 1393 |
| 1417 | -0.09 | 0.14 | -0.66 | 0.51 | 1 | Cops6 | COP9 signalosome subunit 6 | 1793 | 1698 | 1738 | 1423 | 1628 | 1073 | 1218 | 1069 | 1453 | 1434 | 1061 |
| 1051 | 0.05 | 0.13 | 0.36 | 0.72 | 1 | Bex4 | brain expressed X-linked 4 | 854 | 923 | 1005 | 1086 | 1092 | 911 | 1218 | 1042 | 1156 | 1360 | 910 |
| 1280 | 0.20 | 0.14 | 1.46 | 0.14 | 1 | Fbxo8 | F-box protein 8 | 931 | 1078 | 1446 | 1270 | 1367 | 1093 | 1217 | 1386 | 1517 | 1569 | 1204 |
| 1257 | -0.01 | 0.17 | -0.06 | 0.95 | 1 | Rps6kb1 | ribosomal protein S6 kinase, polypeptide | 902 | 1109 | 831 | 940 | 1132 | 2241 | 1217 | 892 | 1292 | 1134 | 2132 |
| 1195 | -0.08 | 0.12 | -0.63 | 0.53 | 1 | Rsrc1 | arginine/serine-rich coiled-coil 1 | 1428 | 1029 | 1159 | 1193 | 1217 | 1254 | 1217 | 1061 | 1255 | 1267 | 1061 |
| 953 | -0.12 | 0.12 | -0.99 | 0.32 | 1 | L3mbtl2 | L3MBTL2 polycomb repressive complex ´ | 782 | 666 | 684 | 1282 | 1018 | 837 | 1217 | 1102 | 974 | 1110 | 809 |
| 1402 | 0.04 | 0.12 | 0.36 | 0.72 | 1 | Rfc1 | replication factor C (activator 1) 1 | 1617 | 1397 | 1420 | 1501 | 1182 | 1521 | 1216 | 1423 | 1285 | 1386 | 1470 |
| 1266 | 0.03 | 0.10 | 0.29 | 0.77 | 1 | Elmo3 | engulfment and cell motility 3 | 876 | 909 | 846 | 1407 | 1338 | 1638 | 1216 | 1418 | 1340 | 1340 | 1595 |
| 1194 | 0.16 | 0.13 | 1.29 | 0.20 | 1 | Bet1l | Bet1 golgi vesicular membrane trafficking | 848 | 898 | 1057 | 1114 | 1202 | 1348 | 1216 | 1460 | 1235 | 1301 | 1460 |
| 802 | 0.05 | 0.26 | 0.17 | 0.86 | 1 | Gm27033 | | 56 | 104 | 153 | 1169 | 1037 | 768 | 1216 | 985 | 1114 | 1416 | 807 |
| 1595 | -0.05 | 0.09 | -0.61 | 0.54 | 1 | Emc1 | ER membrane protein complex subunit 1 | 1374 | 2028 | 2008 | 1644 | 1776 | 1543 | 1215 | 1695 | 1627 | 1188 | 1445 |
| 1030 | -0.05 | 0.18 | -0.30 | 0.76 | 1 | Vac14 | Vac14 homolog (S. cerevisiae) | 950 | 566 | 950 | 1077 | 1024 | 1205 | 1215 | 1143 | 947 | 1152 | 1103 |
| 1379 | 0.18 | 0.12 | 1.49 | 0.14 | 1 | Lysmd3 | LysM, putative peptidoglycan-binding, do | 1096 | 1093 | 1112 | 1187 | 1220 | 1946 | 1213 | 1200 | 1324 | 1698 | 2078 |
| 1343 | -0.01 | 0.13 | -0.08 | 0.94 | 1 | Terf2ip | telomeric repeat binding factor 2, interacti | 1247 | 1712 | 1276 | 1632 | 1243 | 1215 | 1213 | 1354 | 1350 | 1427 | 1101 |

| | | | | | | | | | | | | | | | | | |
|---|---|---|---|---|---|---|---|---|---|---|---|---|---|---|---|---|---|
| 1267 | -0.10 | 0.15 | -0.69 | 0.49 | 1 | Nkiras2 | NFKB inhibitor interacting Ras-like protei | 1319 | 1114 | 1524 | 1300 | 1526 | 1077 | 1213 | 900 | 1544 | 1463 | 959 |
| 1442 | -0.08 | 0.11 | -0.71 | 0.48 | 1 | Zdhhc17 | zinc finger, DHHC domain containing 17 | 1199 | 1260 | 1227 | 1252 | 1274 | 2437 | 1212 | 1025 | 1159 | 1215 | 2608 |
| 1186 | -0.20 | 0.13 | -1.56 | 0.12 | 1 | Sema4d | sema domain, immunoglobulin domain (I | 1559 | 1351 | 1377 | 1266 | 1208 | 976 | 1211 | 901 | 1055 | 1318 | 827 |
| 1127 | 0.04 | 0.13 | 0.32 | 0.75 | 1 | Coq8a | coenzyme Q8A | 744 | 780 | 971 | 1297 | 1481 | 854 | 1211 | 1584 | 1418 | 1315 | 746 |
| 1416 | -0.09 | 0.13 | -0.67 | 0.50 | 1 | Mpzl1 | myelin protein zero-like 1 | 1732 | 1919 | 2136 | 894 | 1341 | 1577 | 1210 | 850 | 1130 | 1112 | 1670 |
| 1216 | -0.14 | 0.11 | -1.32 | 0.19 | 1 | Usf1 | upstream transcription factor 1 | 1422 | 1166 | 1372 | 1171 | 1393 | 1160 | 1210 | 1086 | 1294 | 1097 | 1001 |
| 1077 | 0.12 | 0.11 | 1.07 | 0.29 | 1 | Elac1 | elaC ribonuclease Z 1 | 1040 | 1011 | 884 | 1041 | 991 | 1041 | 1210 | 1145 | 1067 | 1173 | 1249 |
| 1981 | -0.19 | 0.28 | -0.68 | 0.50 | 1 | Cd44 | CD44 antigen | 3034 | 1996 | 2178 | 1370 | 3527 | 1570 | 1209 | 1171 | 3444 | 976 | 1312 |
| 1469 | -0.02 | 0.11 | -0.16 | 0.87 | 1 | Epn2 | epsin 2 | 2322 | 2566 | 2372 | 1052 | 1003 | 1214 | 1209 | 1121 | 1071 | 1052 | 1173 |
| 1211 | 0.08 | 0.13 | 0.62 | 0.53 | 1 | Tbc1d13 | TBC1 domain family, member 13 | 1066 | 1065 | 952 | 1083 | 1419 | 1265 | 1209 | 1360 | 1437 | 1271 | 1196 |
| 1034 | 0.18 | 0.17 | 1.05 | 0.29 | 1 | Mtx2 | metaxin 2 | 892 | 989 | 1209 | 747 | 998 | 946 | 1209 | 1096 | 1010 | 1221 | 1063 |
| 926 | -0.02 | 0.14 | -0.15 | 0.88 | 1 | Rars2 | arginyl-tRNA synthetase 2, mitochondrial | 617 | 576 | 594 | 795 | 1014 | 1210 | 1209 | 852 | 1035 | 951 | 1331 |
| 1509 | -0.09 | 0.13 | -0.69 | 0.49 | 1 | Dhx38 | DEAH (Asp-Glu-Ala-His) box polypeptide | 2013 | 1677 | 1891 | 1433 | 1694 | 1347 | 1208 | 1356 | 1472 | 1206 | 1300 |
| 1303 | 0.09 | 0.14 | 0.65 | 0.52 | 1 | Aup1 | ancient ubiquitous protein 1 | 1412 | 1066 | 1180 | 1206 | 1280 | 1475 | 1208 | 1540 | 1255 | 1193 | 1520 |
| 1148 | 0.06 | 0.15 | 0.42 | 0.67 | 1 | Vta1 | vesicle (multivesicular body) trafficking 1 | 1233 | 1118 | 1191 | 1094 | 1126 | 1002 | 1208 | 1478 | 976 | 1166 | 1031 |
| 1100 | 0.15 | 0.12 | 1.21 | 0.23 | 1 | Mmgt2 | membrane magnesium transporter 2 | 917 | 970 | 915 | 1168 | 1185 | 811 | 1208 | 1536 | 1326 | 1305 | 759 |
| 1239 | 0.01 | 0.16 | 0.06 | 0.95 | 1 | Commd6 | COMM domain containing 6 | 1136 | 1177 | 1193 | 1277 | 1399 | 1145 | 1207 | 928 | 1325 | 1439 | 1401 |
| 1168 | -0.10 | 0.10 | -0.95 | 0.34 | 1 | Slc39a14 | solute carrier family 39 (zinc transporter), | 1229 | 1259 | 1204 | 1275 | 1115 | 1132 | 1207 | 1281 | 986 | 1101 | 1056 |
| 1161 | -0.07 | 0.13 | -0.52 | 0.60 | 1 | Spag7 | sperm associated antigen 7 | 1486 | 1140 | 1510 | 1165 | 1193 | 875 | 1207 | 1052 | 1098 | 1043 | 1005 |
| 970 | 0.08 | 0.15 | 0.51 | 0.61 | 1 | Mrgpre | MAS-related GPR, member E | 347 | 612 | 539 | 1334 | 1249 | 687 | 1207 | 1415 | 1348 | 1150 | 786 |
| 1396 | -0.13 | 0.15 | -0.87 | 0.38 | 1 | Ppil4 | peptidylprolyl isomerase (cyclophilin)-like | 1277 | 1525 | 1276 | 1420 | 1306 | 1929 | 1206 | 941 | 1414 | 1104 | 1963 |
| 1391 | 0.16 | 0.11 | 1.46 | 0.14 | 1 | Ikbkb | inhibitor of kappaB kinase beta | 1671 | 1140 | 1387 | 903 | 1004 | 2216 | 1206 | 1039 | 1226 | 1396 | 2112 |
| 1292 | 0.07 | 0.14 | 0.51 | 0.61 | 1 | Lancl2 | LanC (bacterial lantibiotic synthetase com | 1398 | 1532 | 1358 | 1249 | 1317 | 1024 | 1206 | 1749 | 1222 | 1199 | 954 |
| 1350 | -0.11 | 0.12 | -0.87 | 0.38 | 1 | Rapgef2 | Rap guanine nucleotide exchange factor | 1864 | 1658 | 1559 | 1164 | 1384 | 1307 | 1205 | 975 | 1292 | 1231 | 1209 |
| 1212 | 0.31 | 0.20 | 1.53 | 0.13 | 1 | Bscl2 | Berardinelli-Seip congenital lipodystroph | 1172 | 946 | 1106 | 1177 | 1188 | 871 | 1205 | 2237 | 1177 | 1337 | 921 |
| 1198 | 0.17 | 0.18 | 0.95 | 0.34 | 1 | Polr3d | polymerase (RNA) III (DNA directed) poly | 888 | 1019 | 1270 | 1024 | 1037 | 1457 | 1205 | 1305 | 1006 | 1467 | 1505 |
| 1133 | -0.16 | 0.16 | -0.96 | 0.34 | 1 | Slc9a1 | solute carrier family 9 (sodium/hydrogen e | 1465 | 1389 | 1148 | 1112 | 1213 | 923 | 1205 | 765 | 1182 | 1118 | 940 |
| 1095 | -0.07 | 0.13 | -0.60 | 0.55 | 1 | Grb14 | growth factor receptor bound protein 14 | 1254 | 1331 | 1296 | 971 | 1086 | 928 | 1205 | 1115 | 927 | 1047 | 889 |
| 1427 | 0.09 | 0.11 | 0.83 | 0.40 | 1 | Eif1ax | eukaryotic translation initiation factor 1A, | 2322 | 1903 | 1868 | 1274 | 1152 | 1019 | 1204 | 1263 | 1149 | 1427 | 1115 |
| 1159 | 0.01 | 0.19 | 0.06 | 0.95 | 1 | Ano8 | anoctamin 8 | 896 | 690 | 800 | 1359 | 1132 | 1445 | 1204 | 1799 | 1185 | 1065 | 1172 |
| 1556 | -0.09 | 0.13 | -0.73 | 0.47 | 1 | Map3k3 | mitogen-activated protein kinase kinase k | 2533 | 2192 | 2090 | 1053 | 1376 | 1688 | 1203 | 1029 | 1325 | 1082 | 1543 |
| 1228 | 0.04 | 0.12 | 0.34 | 0.73 | 1 | Dact2 | dishevelled-binding antagonist of beta-ca | 1007 | 1015 | 966 | 1402 | 1287 | 1281 | 1203 | 1644 | 1300 | 1206 | 1199 |
| 1167 | 0.08 | 0.15 | 0.55 | 0.58 | 1 | Vps36 | vacuolar protein sorting 36 | 1164 | 945 | 1125 | 1223 | 1133 | 1099 | 1203 | 1425 | 1344 | 1009 | 1168 |
| 1112 | -0.09 | 0.11 | -0.85 | 0.40 | 1 | Fbxo25 | F-box protein 25 | 900 | 933 | 918 | 1357 | 1236 | 1107 | 1203 | 1101 | 1145 | 1171 | 1165 |
| 1093 | -0.19 | 0.12 | -1.61 | 0.11 | 1 | Acss2 | acyl-CoA synthetase short-chain family m | 434 | 609 | 721 | 1225 | 1642 | 1385 | 1203 | 1119 | 1484 | 1006 | 1189 |
| 1035 | -0.03 | 0.15 | -0.21 | 0.83 | 1 | Fsd1l | fibronectin type III and SPRY domain cont | 665 | 1018 | 809 | 1103 | 997 | 1177 | 1203 | 1207 | 977 | 980 | 1254 |
| 1375 | -0.02 | 0.12 | -0.17 | 0.86 | 1 | Ap1m1 | adaptor-related protein complex AP-1, mu | 1669 | 1709 | 1409 | 1164 | 1103 | 1770 | 1202 | 1110 | 1222 | 1284 | 1477 |
| 1258 | -0.12 | 0.12 | -0.95 | 0.34 | 1 | Shroom2 | shroom family member 2 | 1384 | 1338 | 1810 | 1242 | 1231 | 1160 | 1202 | 1208 | 1201 | 1046 | 1017 |
| 1231 | 0.04 | 0.10 | 0.41 | 0.68 | 1 | Ogfod2 | 2-oxoglutarate and iron-dependent oxyge | 980 | 1001 | 1116 | 1158 | 1401 | 1383 | 1202 | 1186 | 1293 | 1287 | 1534 |
| 1111 | -0.18 | 0.12 | -1.44 | 0.15 | 1 | Dync2li1 | dynein cytoplasmic 2 light intermediate ch | 1279 | 1276 | 1296 | 1160 | 1178 | 912 | 1202 | 901 | 961 | 1063 | 989 |
| 686 | 0.28 | 0.22 | 1.26 | 0.21 | 1 | Nrcam | neuronal cell adhesion molecule | 467 | 506 | 486 | 509 | 656 | 425 | 1202 | 708 | 826 | 1231 | 534 |
| 1469 | 0.09 | 0.11 | 0.77 | 0.44 | 1 | Ogfr | opioid growth factor receptor | 1971 | 1860 | 2043 | 1244 | 1341 | 1198 | 1200 | 1542 | 1368 | 1184 | 1211 |
| 1151 | 0.10 | 0.13 | 0.79 | 0.43 | 1 | Rtn2 | reticulon 2 (Z-band associated protein) | 932 | 1155 | 1034 | 1310 | 1145 | 907 | 1200 | 1490 | 1230 | 1456 | 803 |
| 1031 | -0.13 | 0.13 | -1.02 | 0.31 | 1 | Gm44250 | | 1095 | 979 | 886 | 1183 | 1127 | 872 | 1200 | 933 | 1130 | 1079 | 854 |
| 1531 | -0.08 | 0.11 | -0.72 | 0.47 | 1 | Cnot8 | CCR4-NOT transcription complex, subuni | 1466 | 1646 | 1348 | 1284 | 1315 | 2513 | 1199 | 1263 | 1013 | 1248 | 2549 |
| 1059 | -0.09 | 0.12 | -0.80 | 0.43 | 1 | Sema4b | sema domain, immunoglobulin domain (I | 1060 | 963 | 923 | 1052 | 1244 | 994 | 1199 | 919 | 1134 | 1182 | 976 |
| 1006 | -0.09 | 0.15 | -0.61 | 0.54 | 1 | Egfr | epidermal growth factor receptor | 1280 | 764 | 812 | 962 | 1191 | 885 | 1199 | 897 | 1103 | 1110 | 860 |
| 1639 | -0.13 | 0.12 | -1.11 | 0.26 | 1 | Ppp1r15a | protein phosphatase 1, regulatory subuni | 2923 | 1837 | 2125 | 825 | 1251 | 2519 | 1197 | 739 | 1050 | 1191 | 2368 |
| 1075 | 0.07 | 0.11 | 0.64 | 0.52 | 1 | Tbc1d30 | TBC1 domain family, member 30 | 771 | 749 | 700 | 1198 | 1128 | 1164 | 1197 | 1201 | 1299 | 1283 | 1133 |
| 1034 | 0.15 | 0.14 | 1.03 | 0.30 | 1 | Mnx1 | motor neuron and pancreas homeobox 1 | 71 | 69 | 80 | 1654 | 1270 | 1177 | 1197 | 1656 | 1273 | 1541 | 1383 |
| 1018 | 0.03 | 0.12 | 0.26 | 0.80 | 1 | Ddx31 | DEAD/H (Asp-Glu-Ala-Asp/His) box polyp | 334 | 337 | 371 | 1318 | 1157 | 1344 | 1197 | 1379 | 1200 | 1105 | 1455 |
| 1404 | 0.10 | 0.10 | 1.02 | 0.31 | 1 | Fam104a | family with sequence similarity 104, meml | 1531 | 1720 | 1476 | 1352 | 1314 | 1298 | 1196 | 1526 | 1445 | 1351 | 1234 |
| 1147 | -0.08 | 0.13 | -0.66 | 0.51 | 1 | Fkbp3 | FK506 binding protein 3 | 1325 | 1225 | 1312 | 1009 | 918 | 1364 | 1196 | 969 | 929 | 906 | 1468 |
| 1537 | 0.15 | 0.11 | 1.39 | 0.17 | 1 | Tbc1d17 | TBC1 domain family, member 17 | 2051 | 1665 | 1918 | 1207 | 1336 | 1612 | 1195 | 1411 | 1358 | 1510 | 1640 |
| 1392 | -0.18 | 0.14 | -1.23 | 0.22 | 1 | Mdn1 | midasin AAA ATPase 1 | 1508 | 1166 | 1224 | 851 | 1388 | 2639 | 1195 | 949 | 1078 | 953 | 2361 |
| 1250 | -0.10 | 0.11 | -0.88 | 0.38 | 1 | Hnf4a | hepatic nuclear factor 4, alpha | 1122 | 1580 | 1554 | 1391 | 1222 | 1097 | 1195 | 1365 | 1152 | 1012 | 1060 |
| 1154 | -0.03 | 0.10 | -0.26 | 0.80 | 1 | Decr2 | 2-4-dienoyl-Coenzyme A reductase 2, pe | 930 | 1157 | 1117 | 1301 | 1299 | 985 | 1195 | 1351 | 1298 | 1124 | 939 |
| 1391 | -0.04 | 0.19 | -0.20 | 0.84 | 1 | Ptpn6 | protein tyrosine phosphatase, non-recept | 2068 | 1004 | 1426 | 1437 | 1441 | 1363 | 1194 | 1123 | 1488 | 1452 | 1310 |
| 1366 | -0.02 | 0.15 | -0.10 | 0.92 | 1 | Pip4k2b | phosphatidylinositol-5-phosphate 4-kinas | 2354 | 1582 | 1568 | 1270 | 1250 | 1064 | 1194 | 1184 | 1304 | 1229 | 1023 |
| 1132 | 0.10 | 0.13 | 0.75 | 0.46 | 1 | Ofd1 | OFD1, centriole and centriolar satellite pr | 686 | 885 | 792 | 1028 | 1232 | 1417 | 1193 | 1284 | 1322 | 1101 | 1516 |

| | | | | | | | | | | | | | | | | | | |
|---|---|---|---|---|---|---|---|---|---|---|---|---|---|---|---|---|---|---|
| 1121 | 0.03 | 0.15 | 0.19 | 0.85 | 1 | Mrpl36 | mitochondrial ribosomal protein L36 | 1011 | 1096 | 1415 | 1132 | 1263 | 757 | 1193 | 1188 | 1165 | 1412 | 701 |
| 949 | -0.13 | 0.15 | -0.84 | 0.40 | 1 | Tmem199 | transmembrane protein 199 | 595 | 783 | 571 | 948 | 1060 | 1230 | 1193 | 1042 | 962 | 961 | 1093 |
| 1247 | -0.04 | 0.10 | -0.45 | 0.65 | 1 | Kctd3 | potassium channel tetramerisation domai | 1126 | 1186 | 1241 | 1210 | 1273 | 1490 | 1192 | 1237 | 1218 | 1202 | 1343 |
| 1215 | 0.11 | 0.17 | 0.64 | 0.52 | 1 | Ugcg | UDP-glucose ceramide glucosyltransfera | 1398 | 1100 | 832 | 1268 | 1125 | 1236 | 1192 | 1581 | 1334 | 1184 | 1117 |
| 1190 | 0.15 | 0.13 | 1.22 | 0.22 | 1 | Nuak1 | NUAK family, SNF1-like kinase, 1 | 1360 | 936 | 1356 | 966 | 1372 | 923 | 1192 | 1152 | 1333 | 1588 | 913 |
| 1130 | 0.05 | 0.15 | 0.31 | 0.75 | 1 | Npat | nuclear protein in the AT region | 822 | 861 | 744 | 1208 | 1235 | 1276 | 1192 | 977 | 1410 | 1347 | 1358 |
| 1597 | 0.19 | 0.11 | 1.69 | 0.09 | 1 | Bms1 | BMS1, ribosome biogenesis factor | 1724 | 1715 | 1811 | 1441 | 1295 | 1822 | 1191 | 1820 | 1376 | 1340 | 2034 |
| 1038 | -0.03 | 0.16 | -0.17 | 0.86 | 1 | Intu | inturned planar cell polarity protein | 531 | 495 | 536 | 1201 | 1211 | 1341 | 1191 | 1065 | 1602 | 943 | 1305 |
| 1294 | 0.14 | 0.11 | 1.24 | 0.21 | 1 | Map2k4 | mitogen-activated protein kinase kinase 4 | 1306 | 1355 | 1128 | 1280 | 1202 | 1292 | 1190 | 1393 | 1256 | 1548 | 1281 |
| 1134 | -0.03 | 0.15 | -0.18 | 0.86 | 1 | Dnajc9 | DnaJ heat shock protein family (Hsp40) m | 1275 | 890 | 1052 | 1136 | 1369 | 986 | 1190 | 1293 | 1061 | 1219 | 1007 |
| 1124 | 0.10 | 0.17 | 0.60 | 0.55 | 1 | Ccdc124 | coiled-coil domain containing 124 | 1067 | 987 | 1250 | 1072 | 1123 | 984 | 1190 | 1513 | 1021 | 1095 | 1066 |
| 1887 | -0.19 | 0.16 | -1.17 | 0.24 | 1 | Bicc1 | BicC family RNA binding protein 1 | 4556 | 3110 | 3509 | 1594 | 1061 | 1256 | 1187 | 980 | 1069 | 1337 | 1098 |
| 1114 | 0.10 | 0.19 | 0.51 | 0.61 | 1 | Slc50a1 | solute carrier family 50 (sugar transporter | 764 | 678 | 1116 | 1030 | 1139 | 1348 | 1187 | 1396 | 1068 | 1279 | 1244 |
| 1245 | -0.15 | 0.11 | -1.35 | 0.18 | 1 | Kank1 | KN motif and ankyrin repeat domains 1 | 1644 | 1435 | 1982 | 1023 | 1140 | 1183 | 1186 | 933 | 1115 | 996 | 1058 |
| 1198 | 0.15 | 0.11 | 1.33 | 0.18 | 1 | Nifk | nucleolar protein interacting with the FHA | 1390 | 1278 | 1208 | 1041 | 1184 | 993 | 1186 | 1286 | 1092 | 1354 | 1169 |
| 1181 | 0.12 | 0.10 | 1.12 | 0.26 | 1 | Pgam5 | phosphoglycerate mutase family member | 1117 | 1099 | 1136 | 1131 | 1243 | 1067 | 1186 | 1342 | 1201 | 1313 | 1157 |
| 1059 | -0.31 | 0.18 | -1.75 | 0.08 | 1 | Hspb8 | heat shock protein 8 | 1553 | 1341 | 1221 | 1317 | 1000 | 512 | 1186 | 1322 | 817 | 1100 | 274 |
| 1365 | 0.10 | 0.12 | 0.86 | 0.39 | 1 | Phf23 | PHD finger protein 23 | 1422 | 1665 | 1346 | 1478 | 1277 | 1129 | 1185 | 1844 | 1175 | 1392 | 1102 |
| 1345 | -0.04 | 0.12 | -0.37 | 0.71 | 1 | Pcdhgc3 | protocadherin gamma subfamily C, 3 | 2451 | 2046 | 1835 | 1190 | 1184 | 726 | 1185 | 1064 | 1218 | 1204 | 686 |
| 1235 | 0.39 | 0.21 | 1.82 | 0.07 | 1 | Cuta | cutA divalent cation tolerance homolog | 1142 | 966 | 1212 | 981 | 1280 | 952 | 1185 | 2234 | 1232 | 1323 | 1080 |
| 1312 | -0.10 | 0.12 | -0.87 | 0.38 | 1 | Trim26 | tripartite motif-containing 26 | 1438 | 1494 | 1449 | 1189 | 1131 | 1707 | 1183 | 1025 | 1007 | 1324 | 1482 |
| 1180 | -0.10 | 0.10 | -0.99 | 0.32 | 1 | Rnf139 | ring finger protein 139 | 1086 | 1137 | 1182 | 1396 | 1295 | 1079 | 1182 | 1249 | 1189 | 1127 | 1054 |
| 1180 | 0.03 | 0.15 | 0.18 | 0.86 | 1 | Heatr1 | HEAT repeat containing 1 | 1266 | 794 | 1021 | 1127 | 1304 | 1293 | 1182 | 1261 | 1349 | 1155 | 1222 |
| 936 | -0.24 | 0.14 | -1.77 | 0.08 | 1 | Phlpp2 | PH domain and leucine rich repeat protei | 842 | 964 | 784 | 1095 | 803 | 1109 | 1182 | 839 | 828 | 938 | 906 |
| 1333 | 0.07 | 0.16 | 0.43 | 0.67 | 1 | Cdk9 | cyclin-dependent kinase 9 (CDC2-relatec | 1494 | 1282 | 1536 | 1152 | 1333 | 1385 | 1180 | 1629 | 1119 | 1168 | 1382 |
| 1169 | 0.14 | 0.18 | 0.77 | 0.44 | 1 | Snf8 | SNF8, ESCRT-II complex subunit, homolc | 1196 | 958 | 1252 | 1261 | 1022 | 1019 | 1180 | 1700 | 1054 | 1143 | 1078 |
| 1139 | -0.03 | 0.13 | -0.24 | 0.81 | 1 | Ech1 | enoyl coenzyme A hydratase 1, peroxisor | 1573 | 1215 | 1502 | 1044 | 942 | 1003 | 1180 | 1104 | 924 | 1047 | 993 |
| 1121 | -0.08 | 0.15 | -0.52 | 0.60 | 1 | Dtnb | dystrobrevin, beta | 511 | 666 | 725 | 1346 | 1174 | 1657 | 1180 | 1029 | 1291 | 1219 | 1537 |
| 1117 | -0.11 | 0.11 | -1.00 | 0.32 | 1 | Tspan33 | tetraspanin 33 | 891 | 938 | 998 | 1408 | 1353 | 975 | 1180 | 1415 | 1150 | 1063 | 918 |
| 914 | 0.00 | 0.18 | -0.01 | 0.99 | 1 | Fndc10 | fibronectin type III domain containing 10 | 451 | 333 | 518 | 1263 | 1185 | 732 | 1180 | 1349 | 1050 | 1316 | 677 |
| 1438 | -0.11 | 0.13 | -0.86 | 0.39 | 1 | Sbf2 | SET binding factor 2 | 2190 | 1894 | 1798 | 1397 | 1098 | 1485 | 1179 | 1083 | 1206 | 1030 | 1453 |
| 1222 | -0.20 | 0.11 | -1.78 | 0.07 | 1 | Sesn1 | sestrin 1 | 1526 | 1248 | 1540 | 1018 | 1313 | 1352 | 1178 | 880 | 1085 | 943 | 1361 |
| 1539 | 0.10 | 0.14 | 0.74 | 0.46 | 1 | Map3k11 | mitogen-activated protein kinase kinase k | 1730 | 2288 | 1852 | 1380 | 1484 | 1275 | 1177 | 1712 | 1281 | 1570 | 1185 |
| 1450 | -0.18 | 0.14 | -1.26 | 0.21 | 1 | Pcsk6 | proprotein convertase subtilisin/kexin type | 2719 | 1920 | 2388 | 1442 | 1384 | 730 | 1177 | 1125 | 1225 | 1191 | 652 |
| 1083 | -0.14 | 0.20 | -0.71 | 0.48 | 1 | Prkab1 | protein kinase, AMP-activated, beta 1 nor | 866 | 1001 | 998 | 1236 | 1155 | 1145 | 1177 | 699 | 1167 | 1444 | 1029 |
| 810 | 0.12 | 0.14 | 0.91 | 0.36 | 1 | Ppm1e | protein phosphatase 1E (PP2C domain c | 370 | 299 | 380 | 911 | 989 | 705 | 1177 | 1003 | 1167 | 1043 | 865 |
| 693 | 0.15 | 0.21 | 0.74 | 0.46 | 1 | St8sia5 | ST8 alpha-N-acetyl-neuraminide alpha-2 | 94 | 101 | 47 | 956 | 765 | 626 | 1177 | 818 | 974 | 1320 | 749 |
| 1140 | 0.05 | 0.12 | 0.40 | 0.69 | 1 | Yaf2 | YY1 associated factor 2 | 1260 | 1232 | 1228 | 1164 | 1056 | 938 | 1176 | 994 | 1197 | 1280 | 1010 |
| 1114 | -0.07 | 0.14 | -0.53 | 0.59 | 1 | Pgpep1 | pyroglutamyl-peptidase I | 1238 | 1246 | 1309 | 1159 | 1197 | 802 | 1176 | 1360 | 1025 | 933 | 808 |
| 1072 | -0.04 | 0.10 | -0.41 | 0.68 | 1 | Cstb | cystatin B | 1202 | 1129 | 1111 | 1196 | 1065 | 791 | 1176 | 1130 | 1076 | 1190 | 730 |
| 1357 | 0.04 | 0.20 | 0.22 | 0.83 | 1 | Slc39a1 | solute carrier family 39 (zinc transporter), | 2460 | 1420 | 2168 | 1247 | 1119 | 815 | 1175 | 1570 | 1017 | 1159 | 781 |
| 913 | 0.06 | 0.18 | 0.34 | 0.73 | 1 | Rfesd | Rieske (Fe-S) domain containing | 509 | 724 | 730 | 884 | 1089 | 813 | 1175 | 1001 | 993 | 1221 | 901 |
| 1287 | 0.04 | 0.11 | 0.42 | 0.68 | 1 | Ints13 | integrator complex subunit 13 | 1178 | 1085 | 1068 | 1262 | 1122 | 1766 | 1174 | 1163 | 1296 | 1184 | 1858 |
| 1496 | -0.01 | 0.18 | -0.03 | 0.98 | 1 | Cxadr | coxsackie virus and adenovirus receptor | 1564 | 1710 | 1863 | 1510 | 1195 | 1774 | 1173 | 1032 | 1146 | 1696 | 1793 |
| 1385 | 0.03 | 0.10 | 0.27 | 0.79 | 1 | Trim24 | tripartite motif-containing 24 | 1283 | 1606 | 1321 | 1484 | 1300 | 1494 | 1173 | 1552 | 1462 | 1104 | 1449 |
| 1036 | -0.09 | 0.12 | -0.69 | 0.49 | 1 | Acsl1 | acyl-CoA synthetase long-chain family me | 996 | 974 | 1153 | 1219 | 1079 | 777 | 1173 | 1071 | 1067 | 1176 | 707 |
| 1323 | 0.15 | 0.13 | 1.18 | 0.24 | 1 | Mul1 | mitochondrial ubiquitin ligase activator of | 1288 | 1163 | 1171 | 1512 | 1395 | 1097 | 1172 | 1710 | 1367 | 1520 | 1162 |
| 1247 | 0.03 | 0.17 | 0.21 | 0.84 | 1 | Kdm6a | lysine (K)-specific demethylase 6A | 1323 | 1396 | 1078 | 993 | 1142 | 1604 | 1172 | 942 | 1456 | 1114 | 1494 |
| 1189 | -0.15 | 0.13 | -1.14 | 0.25 | 1 | Rfx5 | regulatory factor X, 5 (influences HLA clas | 906 | 855 | 1145 | 1308 | 1294 | 1569 | 1172 | 1273 | 1161 | 993 | 1405 |
| 1150 | 0.09 | 0.13 | 0.70 | 0.48 | 1 | Acbd6 | acyl-Coenzyme A binding domain contair | 1178 | 1082 | 1321 | 1189 | 1150 | 877 | 1172 | 1410 | 1206 | 1145 | 916 |
| 1128 | -0.13 | 0.13 | -0.96 | 0.34 | 1 | Emd | emerin | 1183 | 918 | 1026 | 1111 | 1255 | 1287 | 1172 | 843 | 1326 | 1164 | 1121 |
| 1032 | 0.00 | 0.13 | 0.00 | 1.00 | 1 | Pex14 | peroxisomal biogenesis factor 14 | 909 | 870 | 1127 | 972 | 1139 | 921 | 1172 | 1109 | 1139 | 1215 | 777 |
| 1013 | 0.00 | 0.16 | 0.03 | 0.98 | 1 | Pdzd11 | PDZ domain containing 11 | 704 | 696 | 749 | 983 | 1184 | 1140 | 1172 | 1347 | 1148 | 871 | 1150 |
| 895 | -0.17 | 0.11 | -1.54 | 0.12 | 1 | Sgk3 | serum/glucocorticoid regulated kinase 3 | 818 | 698 | 716 | 950 | 893 | 1016 | 1172 | 875 | 775 | 1010 | 918 |
| 1166 | 0.15 | 0.10 | 1.54 | 0.12 | 1 | Spast | spastin | 1234 | 1338 | 961 | 1244 | 939 | 1039 | 1171 | 1473 | 1046 | 1310 | 1068 |
| 1063 | -0.03 | 0.12 | -0.22 | 0.83 | 1 | Pcnx4 | pecanex homolog 4 | 903 | 1004 | 936 | 1361 | 1011 | 907 | 1171 | 1438 | 1050 | 1069 | 840 |
| 1430 | -0.16 | 0.10 | -1.64 | 0.10 | 1 | Ift122 | intraflagellar transport 122 | 1561 | 1852 | 1614 | 1543 | 1280 | 1661 | 1170 | 1201 | 1184 | 1100 | 1566 |
| 878 | 0.16 | 0.25 | 0.64 | 0.52 | 1 | Gm17767 | | 54 | 63 | 161 | 943 | 1214 | 1060 | 1170 | 937 | 1423 | 1623 | 1005 |
| 1072 | 0.09 | 0.12 | 0.75 | 0.46 | 1 | Ccser1 | coiled-coil serine rich 1 | 602 | 686 | 638 | 1281 | 987 | 1347 | 1169 | 1409 | 1165 | 1085 | 1425 |
| 1046 | -0.13 | 0.14 | -0.87 | 0.38 | 1 | Cipc | CLOCK interacting protein, circadian | 854 | 930 | 778 | 1141 | 1068 | 1283 | 1169 | 827 | 1077 | 1173 | 1207 |

| | | | | | | | | | | | | | | | | | |
|---|---|---|---|---|---|---|---|---|---|---|---|---|---|---|---|---|---|
| 1392 | -0.03 | 0.13 | -0.24 | 0.81 | 1 | Cast | calpastatin | 2926 | 1947 | 2244 | 963 | 1001 | 986 | 1168 | 722 | 1085 | 1164 | 1105 |
| 1065 | 0.02 | 0.12 | 0.18 | 0.86 | 1 | P3h1 | prolyl 3-hydroxylase 1 | 1223 | 995 | 1136 | 881 | 1051 | 1063 | 1168 | 1091 | 1010 | 1042 | 1060 |
| 930 | -0.16 | 0.13 | -1.23 | 0.22 | 1 | Pdpr | pyruvate dehydrogenase phosphatase re | 850 | 827 | 672 | 1181 | 1062 | 746 | 1168 | 929 | 1069 | 1072 | 653 |
| 1631 | -0.04 | 0.12 | -0.34 | 0.74 | 1 | Dtx3 | deltex 3, E3 ubiquitin ligase | 1226 | 1486 | 1410 | 1562 | 1206 | 3127 | 1167 | 1758 | 1190 | 1068 | 2738 |
| 1322 | -0.08 | 0.17 | -0.47 | 0.64 | 1 | Wbp1l | WW domain binding protein 1 like | 1923 | 1618 | 1477 | 1362 | 1184 | 1170 | 1167 | 1481 | 1169 | 849 | 1147 |
| 1609 | -0.05 | 0.16 | -0.29 | 0.77 | 1 | Hgfac | hepatocyte growth factor activator | 1035 | 2650 | 1906 | 2017 | 1130 | 1839 | 1166 | 2066 | 1148 | 1141 | 1595 |
| 1362 | 0.10 | 0.15 | 0.66 | 0.51 | 1 | Gstp1 | glutathione S-transferase, pi 1 | 2405 | 2282 | 2566 | 975 | 915 | 684 | 1166 | 1229 | 807 | 1154 | 804 |
| 1168 | -0.08 | 0.12 | -0.71 | 0.48 | 1 | Thap11 | THAP domain containing 11 | 1255 | 1400 | 1433 | 1274 | 1131 | 921 | 1166 | 1342 | 1033 | 1023 | 865 |
| 1059 | 0.14 | 0.19 | 0.76 | 0.45 | 1 | Mpp6 | membrane protein, palmitoylated 6 (MAG | 1140 | 836 | 610 | 1172 | 1273 | 706 | 1166 | 1123 | 1423 | 1352 | 844 |
| 1418 | -0.07 | 0.14 | -0.47 | 0.64 | 1 | Iffo1 | intermediate filament family orphan 1 | 943 | 890 | 1142 | 1157 | 1138 | 2981 | 1165 | 1276 | 922 | 1095 | 2887 |
| 1351 | 0.12 | 0.10 | 1.17 | 0.24 | 1 | Pcnp | PEST proteolytic signal containing nuclea | 1091 | 1367 | 1411 | 1490 | 1344 | 1278 | 1165 | 1461 | 1563 | 1329 | 1359 |
| 1225 | -0.03 | 0.12 | -0.29 | 0.77 | 1 | Mcmbp | minichromosome maintenance complex b | 1616 | 1462 | 1349 | 1205 | 1135 | 1072 | 1165 | 995 | 1195 | 1134 | 1143 |
| 1036 | 0.07 | 0.11 | 0.62 | 0.54 | 1 | Slc17a5 | solute carrier family 17 (anion/sugar trans | 589 | 778 | 658 | 1482 | 1159 | 781 | 1165 | 1602 | 1160 | 1122 | 906 |
| 977 | -0.04 | 0.16 | -0.27 | 0.79 | 1 | Gclc | glutamate-cysteine ligase, catalytic subur | 917 | 951 | 779 | 838 | 986 | 1117 | 1165 | 804 | 1085 | 889 | 1212 |
| 1211 | -0.23 | 0.15 | -1.60 | 0.11 | 1 | Sh3gl1 | SH3-domain GRB2-like 1 | 1409 | 1140 | 1257 | 1144 | 1204 | 1552 | 1164 | 714 | 995 | 1145 | 1593 |
| 1190 | 0.13 | 0.13 | 1.01 | 0.31 | 1 | Katna1 | katanin p60 (ATPase-containing) subunit | 1096 | 1022 | 1143 | 1238 | 1117 | 1161 | 1164 | 1409 | 1109 | 1155 | 1482 |
| 1169 | 0.11 | 0.12 | 0.88 | 0.38 | 1 | Ldah | lipid droplet associated hydrolase | 1069 | 1048 | 1290 | 1058 | 1177 | 1148 | 1164 | 1309 | 1175 | 1112 | 1307 |
| 1118 | 0.14 | 0.15 | 0.90 | 0.37 | 1 | Ccnh | cyclin H | 839 | 881 | 901 | 1146 | 1056 | 1229 | 1164 | 992 | 1186 | 1574 | 1333 |
| 1028 | -0.23 | 0.12 | -1.85 | 0.06 | 1 | Zfp746 | zinc finger protein 746 | 825 | 829 | 846 | 980 | 1075 | 1489 | 1164 | 801 | 937 | 872 | 1486 |
| 1203 | 0.06 | 0.21 | 0.31 | 0.76 | 1 | Psen2 | presenilin 2 | 1098 | 833 | 1146 | 1350 | 1338 | 1082 | 1162 | 1923 | 1230 | 974 | 1100 |
| 965 | -0.14 | 0.16 | -0.84 | 0.40 | 1 | Elp4 | elongator acetyltransferase complex subu | 797 | 839 | 717 | 1041 | 1195 | 925 | 1162 | 752 | 1128 | 1105 | 957 |
| 1177 | 0.04 | 0.13 | 0.31 | 0.76 | 1 | Pak1ip1 | PAK1 interacting protein 1 | 1307 | 1076 | 1321 | 1069 | 1125 | 1206 | 1161 | 1276 | 1057 | 1103 | 1250 |
| 717 | -0.08 | 0.26 | -0.32 | 0.75 | 1 | Shisa2 | shisa family member 2 | 188 | 360 | 325 | 936 | 770 | 737 | 1161 | 730 | 783 | 1216 | 680 |
| 1331 | 0.24 | 0.18 | 1.33 | 0.18 | 1 | Rpl39 | ribosomal protein L39 | 2010 | 1573 | 2273 | 1138 | 947 | 772 | 1159 | 1617 | 1105 | 1211 | 842 |
| 1386 | 0.06 | 0.15 | 0.41 | 0.68 | 1 | Syngr2 | synaptogyrin 2 | 2124 | 1482 | 1723 | 1123 | 1526 | 1049 | 1158 | 1386 | 1228 | 1394 | 1056 |
| 1273 | 0.08 | 0.13 | 0.60 | 0.55 | 1 | Cnep1r1 | CTD nuclear envelope phosphatase 1 reg | 966 | 1282 | 1241 | 1367 | 1239 | 1352 | 1158 | 1506 | 1123 | 1322 | 1448 |
| 1151 | -0.15 | 0.10 | -1.40 | 0.16 | 1 | Socs6 | suppressor of cytokine signaling 6 | 1355 | 1317 | 1269 | 1182 | 1190 | 1055 | 1158 | 1124 | 1040 | 970 | 1006 |
| 1122 | -0.05 | 0.15 | -0.35 | 0.73 | 1 | Ap5m1 | adaptor-related protein complex 5, mu 1 s | 783 | 878 | 801 | 1171 | 1196 | 1482 | 1158 | 874 | 1204 | 1258 | 1539 |
| 936 | 0.18 | 0.20 | 0.93 | 0.35 | 1 | Rnf227 | | 700 | 738 | 608 | 1140 | 823 | 688 | 1158 | 1836 | 919 | 855 | 826 |
| 2170 | -0.26 | 0.17 | -1.57 | 0.12 | 1 | Dpp10 | dipeptidylpeptidase 10 | 5023 | 6726 | 5138 | 803 | 931 | 911 | 1157 | 774 | 847 | 670 | 893 |
| 2111 | 0.03 | 0.19 | 0.14 | 0.89 | 1 | Zfp36l2 | zinc finger protein 36, C3H type-like 2 | 6092 | 3430 | 3602 | 1299 | 1448 | 1101 | 1156 | 1378 | 1425 | 1148 | 1139 |
| 771 | 0.22 | 0.28 | 0.78 | 0.43 | 1 | Kcna2 | potassium voltage-gated channel, shaker | 627 | 664 | 874 | 922 | 406 | 445 | 1156 | 1145 | 679 | 1090 | 470 |
| 2361 | -0.04 | 0.09 | -0.44 | 0.66 | 1 | Grb10 | growth factor receptor bound protein 10 | 5959 | 5702 | 5891 | 1040 | 943 | 1132 | 1155 | 1040 | 906 | 1159 | 1046 |
| 1285 | 0.14 | 0.16 | 0.91 | 0.36 | 1 | Stim2 | stromal interaction molecule 2 | 1355 | 1021 | 1138 | 1278 | 1228 | 1385 | 1155 | 1400 | 1557 | 1074 | 1551 |
| 1435 | -0.05 | 0.12 | -0.45 | 0.65 | 1 | Zbtb44 | zinc finger and BTB domain containing 44 | 1519 | 1405 | 1489 | 1379 | 1361 | 1879 | 1154 | 1222 | 1429 | 1052 | 1897 |
| 1059 | -0.07 | 0.14 | -0.49 | 0.62 | 1 | Gphn | gephyrin | 664 | 555 | 615 | 1280 | 1148 | 1433 | 1154 | 991 | 1339 | 1190 | 1279 |
| 1201 | 0.02 | 0.14 | 0.15 | 0.88 | 1 | Stox2 | storkhead box 2 | 845 | 1039 | 832 | 1028 | 1091 | 1970 | 1153 | 1127 | 1320 | 1007 | 1798 |
| 1119 | 0.09 | 0.11 | 0.84 | 0.40 | 1 | Fam92a | family with sequence similarity 92, membe | 879 | 872 | 1007 | 1168 | 1160 | 1141 | 1153 | 1272 | 1165 | 1240 | 1247 |
| 1277 | 0.21 | 0.13 | 1.62 | 0.11 | 1 | Nom1 | nucleolar protein with MIF4G domain 1 | 1165 | 1109 | 1098 | 1233 | 1124 | 1435 | 1152 | 1719 | 1187 | 1266 | 1554 |
| 1254 | -0.10 | 0.12 | -0.87 | 0.38 | 1 | Dut | deoxyuridine triphosphatase | 1721 | 1405 | 1500 | 1302 | 1232 | 1063 | 1152 | 1153 | 1094 | 1126 | 1048 |
| 1207 | 0.08 | 0.15 | 0.52 | 0.60 | 1 | Rbmx | RNA binding motif protein, X chromosome | 909 | 1082 | 1019 | 1012 | 1026 | 1803 | 1152 | 926 | 1090 | 1401 | 1853 |
| 1190 | 0.12 | 0.14 | 0.86 | 0.39 | 1 | Rnf121 | ring finger protein 121 | 1351 | 1170 | 1042 | 1012 | 1306 | 1091 | 1152 | 1333 | 1147 | 1262 | 1219 |
| 1084 | -0.03 | 0.13 | -0.21 | 0.83 | 1 | Dpp7 | dipeptidylpeptidase 7 | 830 | 808 | 846 | 1281 | 1042 | 1288 | 1152 | 1222 | 1229 | 980 | 1246 |
| 1069 | 0.11 | 0.16 | 0.69 | 0.49 | 1 | Cbr1 | carbonyl reductase 1 | 1077 | 968 | 1045 | 901 | 1156 | 964 | 1152 | 1347 | 966 | 1181 | 1002 |
| 1254 | 0.06 | 0.14 | 0.40 | 0.69 | 1 | Zfp60 | zinc finger protein 60 | 976 | 1439 | 1017 | 1350 | 1334 | 1254 | 1151 | 1258 | 1309 | 1260 | 1450 |
| 970 | -0.03 | 0.13 | -0.25 | 0.80 | 1 | Dgkh | diacylglycerol kinase, eta | 864 | 710 | 742 | 973 | 1038 | 1067 | 1151 | 980 | 1146 | 960 | 1042 |
| 1822 | -0.08 | 0.13 | -0.61 | 0.54 | 1 | Arhgef1 | Rho guanine nucleotide exchange factor | 2642 | 2270 | 2334 | 1467 | 1381 | 2607 | 1150 | 1467 | 1299 | 1106 | 2319 |
| 1192 | 0.06 | 0.16 | 0.35 | 0.73 | 1 | Babam2 | BRISC and BRCA1 A complex member 2 | 1329 | 1073 | 1140 | 1244 | 1308 | 996 | 1150 | 1384 | 1333 | 969 | 1186 |
| 1082 | -0.02 | 0.18 | -0.14 | 0.89 | 1 | Mrpl1 | mitochondrial ribosomal protein L1 | 860 | 1041 | 1213 | 1097 | 1036 | 1139 | 1149 | 920 | 1011 | 1386 | 1051 |
| 1066 | 0.15 | 0.13 | 1.13 | 0.26 | 1 | Tmx2 | thioredoxin-related transmembrane prote | 818 | 879 | 949 | 1050 | 1216 | 893 | 1149 | 1428 | 1164 | 1108 | 1075 |
| 1049 | -0.03 | 0.14 | -0.21 | 0.84 | 1 | Nmd3 | NMD3 ribosome export adaptor | 983 | 1118 | 1016 | 961 | 1092 | 1042 | 1149 | 1189 | 1020 | 873 | 1096 |
| 1308 | -0.06 | 0.13 | -0.47 | 0.64 | 1 | Map4k5 | mitogen-activated protein kinase kinase k | 1329 | 1088 | 1081 | 1323 | 1134 | 1988 | 1148 | 1238 | 1231 | 1091 | 1736 |
| 1561 | 0.14 | 0.13 | 1.08 | 0.28 | 1 | Crebzf | CREB/ATF bZIP transcription factor | 1146 | 1337 | 1168 | 1155 | 1129 | 2979 | 1147 | 1143 | 1114 | 1498 | 3355 |
| 1219 | -0.17 | 0.13 | -1.26 | 0.21 | 1 | Atg16l1 | autophagy related 16-like 1 (S. cerevisiae | 886 | 1008 | 830 | 1317 | 1164 | 2019 | 1147 | 953 | 1205 | 1060 | 1817 |
| 1217 | 0.06 | 0.12 | 0.45 | 0.65 | 1 | Kif2a | kinesin family member 2A | 1153 | 1339 | 1224 | 1265 | 1090 | 1242 | 1147 | 1141 | 1303 | 1327 | 1158 |
| 1123 | -0.13 | 0.13 | -1.03 | 0.30 | 1 | Carmil1 | capping protein regulator and myosin 1 lin | 1096 | 1118 | 1321 | 1143 | 1049 | 1263 | 1147 | 1011 | 1204 | 903 | 1098 |
| 1069 | 0.10 | 0.17 | 0.57 | 0.57 | 1 | Fam50a | family with sequence similarity 50, membe | 1106 | 946 | 1127 | 928 | 900 | 1174 | 1147 | 1278 | 1083 | 864 | 1206 |
| 945 | -0.15 | 0.14 | -1.05 | 0.30 | 1 | Ramac | RNA guanine-7 methyltransferase activat | 866 | 813 | 1022 | 1043 | 958 | 898 | 1147 | 798 | 950 | 949 | 951 |
| 1231 | 0.06 | 0.12 | 0.54 | 0.59 | 1 | Sf3a3 | splicing factor 3a, subunit 3 | 1473 | 1360 | 1506 | 1147 | 1070 | 1122 | 1146 | 964 | 1201 | 1235 | 1318 |
| 1188 | 0.07 | 0.16 | 0.40 | 0.69 | 1 | Rprd1a | regulation of nuclear pre-mRNA domain c | 760 | 1259 | 1123 | 1058 | 1147 | 1502 | 1146 | 1072 | 1275 | 1206 | 1521 |

| | | | | | | | | | | | | | | | | | | |
|---|---|---|---|---|---|---|---|---|---|---|---|---|---|---|---|---|---|---|
| 1126 | 0.08 | 0.11 | 0.71 | 0.48 | 1 | Deaf1 | DEAF1, transcription factor | 829 | 913 | 837 | 1137 | 1304 | 1181 | 1146 | 1357 | 1260 | 1266 | 1160 |
| 1103 | 0.18 | 0.16 | 1.13 | 0.26 | 1 | Ift74 | intraflagellar transport 74 | 795 | 985 | 819 | 929 | 1015 | 1384 | 1146 | 937 | 1270 | 1276 | 1578 |
| 846 | 0.09 | 0.16 | 0.56 | 0.57 | 1 | Zswim5 | zinc finger SWIM-type containing 5 | 437 | 532 | 382 | 907 | 897 | 885 | 1146 | 1185 | 1002 | 1153 | 777 |
| 1204 | 0.09 | 0.14 | 0.65 | 0.52 | 1 | Mon1b | MON1 homolog B, secretory traffciking as | 1061 | 1178 | 1006 | 1249 | 1285 | 1154 | 1145 | 1156 | 1409 | 1511 | 1095 |
| 1176 | -0.03 | 0.13 | -0.24 | 0.81 | 1 | Frmd4a | FERM domain containing 4A | 1466 | 1487 | 1612 | 1100 | 881 | 1108 | 1145 | 921 | 1026 | 1065 | 1130 |
| 1046 | 0.09 | 0.12 | 0.79 | 0.43 | 1 | Smim19 | small integral membrane protein 19 | 635 | 752 | 670 | 1449 | 1167 | 835 | 1145 | 1365 | 1155 | 1334 | 1001 |
| 1240 | -0.05 | 0.14 | -0.37 | 0.71 | 1 | Rgs9 | regulator of G-protein signaling 9 | 918 | 1323 | 847 | 1097 | 1013 | 2117 | 1144 | 1061 | 975 | 1103 | 2045 |
| 1188 | -0.12 | 0.14 | -0.89 | 0.37 | 1 | Vhl | von Hippel-Lindau tumor suppressor | 1418 | 1479 | 1317 | 1294 | 1309 | 867 | 1144 | 1388 | 1116 | 871 | 869 |
| 1180 | 0.12 | 0.15 | 0.84 | 0.40 | 1 | Anapc11 | anaphase promoting complex subunit 11 | 1660 | 1259 | 1650 | 1075 | 1077 | 751 | 1144 | 1281 | 1112 | 1056 | 913 |
| 1102 | 0.03 | 0.17 | 0.17 | 0.86 | 1 | Entpd7 | ectonucleoside triphosphate diphosphohy | 687 | 953 | 626 | 1170 | 1072 | 1480 | 1144 | 1298 | 1400 | 884 | 1406 |
| 854 | -0.03 | 0.15 | -0.17 | 0.86 | 1 | Fam135b | family with sequence similarity 135, meml | 181 | 273 | 207 | 1217 | 973 | 1072 | 1143 | 1119 | 1118 | 931 | 1163 |
| 1132 | -0.01 | 0.15 | -0.07 | 0.94 | 1 | Utp20 | UTP20 small subunit processome compo | 1410 | 880 | 1021 | 1048 | 1115 | 1277 | 1142 | 952 | 1320 | 1121 | 1164 |
| 1231 | -0.08 | 0.12 | -0.71 | 0.48 | 1 | Prcc | papillary renal cell carcinoma (translocati | 1295 | 1415 | 1583 | 1171 | 1274 | 1160 | 1141 | 1250 | 1201 | 889 | 1163 |
| 959 | -0.22 | 0.14 | -1.61 | 0.11 | 1 | Tlr2 | toll-like receptor 2 | 1177 | 795 | 1066 | 887 | 1204 | 783 | 1141 | 606 | 1118 | 1033 | 734 |
| 1187 | -0.09 | 0.16 | -0.53 | 0.60 | 1 | Pkia | protein kinase inhibitor, alpha | 1735 | 1875 | 1423 | 1145 | 932 | 922 | 1140 | 853 | 1045 | 882 | 1102 |
| 1180 | -0.11 | 0.15 | -0.75 | 0.46 | 1 | Scamp3 | secretory carrier membrane protein 3 | 997 | 1024 | 1168 | 1331 | 1416 | 1185 | 1140 | 864 | 1379 | 1302 | 1173 |
| 1016 | -0.09 | 0.12 | -0.80 | 0.42 | 1 | Taf5l | TATA-box binding protein associated fact | 874 | 966 | 1015 | 1103 | 1174 | 890 | 1140 | 996 | 1020 | 1073 | 930 |
| 1074 | 0.05 | 0.14 | 0.37 | 0.71 | 1 | Med12l | mediator complex subunit 12-like | 595 | 739 | 703 | 1122 | 984 | 1580 | 1139 | 1073 | 1213 | 1195 | 1469 |
| 2188 | 0.03 | 0.11 | 0.25 | 0.80 | 1 | Sox4 | SRY (sex determining region Y)-box 4 | 2913 | 3307 | 3114 | 1308 | 946 | 3862 | 1138 | 1368 | 914 | 1150 | 4049 |
| 1396 | 0.02 | 0.10 | 0.22 | 0.83 | 1 | Hmg20a | high mobility group 20A | 1562 | 1754 | 1698 | 1168 | 1169 | 1661 | 1138 | 1084 | 1256 | 1208 | 1660 |
| 1273 | 0.12 | 0.11 | 1.11 | 0.27 | 1 | Asb13 | ankyrin repeat and SOCS box-containing | 830 | 959 | 923 | 1371 | 1350 | 1556 | 1138 | 1468 | 1512 | 1447 | 1446 |
| 1115 | -0.21 | 0.15 | -1.45 | 0.15 | 1 | Scyl3 | SCY1-like 3 (S. cerevisiae) | 695 | 980 | 705 | 1407 | 1215 | 1551 | 1138 | 956 | 1123 | 1048 | 1451 |
| 1110 | 0.11 | 0.14 | 0.76 | 0.45 | 1 | Ndufs8 | NADH:ubiquinone oxidoreductase core s | 1176 | 874 | 1268 | 1088 | 1088 | 954 | 1138 | 1191 | 1156 | 1343 | 929 |
| 1087 | -0.05 | 0.17 | -0.31 | 0.76 | 1 | Tmem161a | transmembrane protein 161A | 905 | 800 | 975 | 1388 | 948 | 1248 | 1138 | 1228 | 1239 | 909 | 1182 |
| 1020 | -0.25 | 0.15 | -1.73 | 0.08 | 1 | Cnnm3 | cyclin M3 | 1262 | 1183 | 878 | 1107 | 1092 | 953 | 1138 | 820 | 993 | 1021 | 770 |
| 1207 | 0.00 | 0.13 | -0.02 | 0.98 | 1 | Kctd2 | potassium channel tetramerisation domai | 932 | 1261 | 1368 | 1222 | 1351 | 1157 | 1137 | 1317 | 1170 | 1155 | 1204 |
| 1188 | 0.15 | 0.15 | 1.05 | 0.30 | 1 | Rab11fip2 | RAB11 family interacting protein 2 (class | 1115 | 1439 | 1032 | 1238 | 926 | 1209 | 1137 | 1108 | 1234 | 1368 | 1265 |
| 1025 | 0.19 | 0.11 | 1.64 | 0.10 | 1 | Cop1 | COP1, E3 ubiquitin ligase | 924 | 700 | 810 | 995 | 1004 | 1000 | 1137 | 1127 | 1251 | 1190 | 1134 |
| 974 | 0.14 | 0.16 | 0.84 | 0.40 | 1 | Cln3 | ceroid lipofuscinosis, neuronal 3, juvenile | 492 | 687 | 735 | 1104 | 1091 | 831 | 1137 | 1376 | 1164 | 1309 | 785 |
| 1407 | 0.14 | 0.10 | 1.40 | 0.16 | 1 | Aimp1 | aminoacyl tRNA synthetase complex-inte | 1825 | 1678 | 1830 | 1228 | 1309 | 1163 | 1136 | 1368 | 1362 | 1329 | 1250 |
| 1318 | 0.09 | 0.12 | 0.72 | 0.47 | 1 | Wipf2 | WAS/WASL interacting protein family, me | 1284 | 1536 | 1171 | 1336 | 1358 | 1244 | 1136 | 1627 | 1425 | 1193 | 1186 |
| 957 | -0.20 | 0.14 | -1.43 | 0.15 | 1 | Tmem219 | transmembrane protein 219 | 825 | 591 | 814 | 1099 | 1186 | 997 | 1136 | 773 | 1139 | 972 | 995 |
| 1081 | -0.17 | 0.10 | -1.67 | 0.10 | 1 | Itfg2 | integrin alpha FG-GAP repeat containing | 1267 | 1097 | 1217 | 1042 | 1061 | 1164 | 1135 | 947 | 981 | 938 | 1039 |
| 939 | 0.08 | 0.19 | 0.40 | 0.69 | 1 | Acsl6 | acyl-CoA synthetase long-chain family me | 390 | 569 | 676 | 1220 | 1037 | 817 | 1135 | 1413 | 1050 | 1260 | 765 |
| 1163 | -0.10 | 0.14 | -0.75 | 0.46 | 1 | Capn5 | calpain 5 | 1616 | 1418 | 1188 | 1181 | 1036 | 1068 | 1134 | 1082 | 1101 | 1204 | 770 |
| 1303 | -0.02 | 0.12 | -0.20 | 0.85 | 1 | Trmt2a | TRM2 tRNA methyltransferase 2A | 983 | 863 | 1189 | 1119 | 1493 | 1985 | 1133 | 1313 | 1267 | 1131 | 1854 |
| 1183 | 0.09 | 0.13 | 0.71 | 0.48 | 1 | Dmwd | dystrophia myotonica-containing WD repe | 1241 | 1515 | 1161 | 1063 | 897 | 1295 | 1133 | 1417 | 870 | 1182 | 1240 |
| 1105 | -0.21 | 0.14 | -1.53 | 0.12 | 1 | Exd2 | exonuclease 3'-5' domain containing 2 | 874 | 974 | 800 | 1114 | 1384 | 1467 | 1133 | 929 | 1121 | 1043 | 1316 |
| 1017 | 0.04 | 0.15 | 0.30 | 0.77 | 1 | Timm17b | translocase of inner mitochondrial membr | 1059 | 780 | 997 | 913 | 1239 | 819 | 1133 | 1169 | 1028 | 1293 | 755 |
| 1321 | -0.11 | 0.16 | -0.70 | 0.49 | 1 | Tpst1 | protein-tyrosine sulfotransferase 1 | 2463 | 1853 | 2162 | 1071 | 1148 | 803 | 1132 | 1321 | 955 | 978 | 645 |
| 1185 | 0.00 | 0.13 | -0.04 | 0.97 | 1 | Emc3 | ER membrane protein complex subunit 3 | 1549 | 1276 | 1308 | 1064 | 1146 | 1111 | 1132 | 969 | 1326 | 1065 | 1093 |
| 1485 | -0.08 | 0.13 | -0.62 | 0.54 | 1 | Nfe2l2 | nuclear factor, erythroid derived 2, like 2 | 2718 | 2695 | 2067 | 1039 | 1179 | 1199 | 1131 | 929 | 1100 | 1191 | 1081 |
| 1198 | 0.17 | 0.13 | 1.29 | 0.20 | 1 | Esf1 | ESF1 nucleolar pre-rRNA processing pro | 1344 | 1303 | 1102 | 1018 | 1093 | 1190 | 1131 | 1343 | 1256 | 1052 | 1343 |
| 1140 | -0.12 | 0.15 | -0.79 | 0.43 | 1 | Scmh1 | sex comb on midleg homolog 1 | 1082 | 1244 | 1116 | 1412 | 1100 | 1096 | 1131 | 958 | 1286 | 1063 | 1049 |
| 1040 | -0.01 | 0.12 | -0.05 | 0.96 | 1 | Nfs1 | nitrogen fixation gene 1 (S. cerevisiae) | 824 | 925 | 895 | 1085 | 1111 | 1078 | 1131 | 1184 | 983 | 1167 | 1061 |
| 1333 | 0.24 | 0.14 | 1.73 | 0.08 | 1 | Stk38l | serine/threonine kinase 38 like | 1034 | 1410 | 1003 | 1267 | 1143 | 1549 | 1130 | 2074 | 1163 | 1311 | 1577 |
| 1215 | 0.11 | 0.12 | 0.85 | 0.39 | 1 | Gtf2a1 | general transcription factor II A, 1 | 1096 | 1086 | 1011 | 1161 | 1335 | 1264 | 1130 | 1105 | 1322 | 1418 | 1433 |
| 1032 | 0.02 | 0.17 | 0.13 | 0.90 | 1 | Ccdc167 | coiled-coil domain containing 167 | 514 | 684 | 545 | 1071 | 1174 | 1380 | 1130 | 947 | 1061 | 1405 | 1439 |
| 768 | 0.04 | 0.20 | 0.18 | 0.86 | 1 | Ccl28 | chemokine (C-C motif) ligand 28 | 256 | 160 | 322 | 643 | 876 | 1060 | 1130 | 439 | 1151 | 1297 | 1115 |
| 1175 | -0.04 | 0.13 | -0.34 | 0.74 | 1 | Auts2 | autism susceptibility candidate 2 | 731 | 1021 | 899 | 1212 | 1410 | 1454 | 1129 | 1264 | 1508 | 1051 | 1241 |
| 1091 | 0.00 | 0.11 | -0.04 | 0.97 | 1 | Nat10 | N-acetyltransferase 10 | 873 | 777 | 981 | 1053 | 1153 | 1357 | 1129 | 1120 | 1117 | 1052 | 1388 |
| 1544 | -0.10 | 0.13 | -0.81 | 0.42 | 1 | Rbm10 | RNA binding motif protein 10 | 1572 | 1192 | 1579 | 1285 | 1456 | 2707 | 1128 | 1181 | 1413 | 1085 | 2391 |
| 1375 | 0.05 | 0.13 | 0.40 | 0.69 | 1 | Fam53c | family with sequence similarity 53, memb | 1264 | 1642 | 1362 | 1467 | 1228 | 1530 | 1128 | 1256 | 1354 | 1463 | 1436 |
| 1035 | -0.13 | 0.18 | -0.74 | 0.46 | 1 | Hdac4 | histone deacetylase 4 | 793 | 1081 | 840 | 889 | 1272 | 1229 | 1128 | 1113 | 1158 | 764 | 1120 |
| 845 | -0.13 | 0.23 | -0.57 | 0.57 | 1 | Alkbh8 | alkB homolog 8, tRNA methyltransferase | 598 | 814 | 616 | 830 | 892 | 930 | 1128 | 605 | 1173 | 747 | 965 |
| 1150 | 0.08 | 0.14 | 0.57 | 0.57 | 1 | Anapc7 | anaphase promoting complex subunit 7 | 1150 | 1249 | 1317 | 1176 | 1000 | 1014 | 1127 | 1530 | 1032 | 1094 | 965 |
| 1148 | 0.09 | 0.16 | 0.52 | 0.60 | 1 | Ciao2a | cytosolic iron-sulfur assembly component | 1372 | 1017 | 1265 | 1154 | 1251 | 836 | 1127 | 1454 | 1063 | 1113 | 979 |
| 1045 | 0.00 | 0.15 | -0.03 | 0.98 | 1 | Commd8 | COMM domain containing 8 | 1209 | 797 | 874 | 1189 | 1044 | 958 | 1127 | 1069 | 1060 | 1097 | 1070 |
| 1007 | -0.04 | 0.17 | -0.22 | 0.82 | 1 | Galnt18 | polypeptide N-acetylgalactosaminyltransf | 875 | 595 | 700 | 1125 | 999 | 1256 | 1127 | 1284 | 884 | 1009 | 1223 |
| 977 | -0.04 | 0.12 | -0.33 | 0.74 | 1 | Cbx4 | chromobox 4 | 993 | 895 | 1054 | 821 | 1010 | 992 | 1127 | 759 | 1072 | 1104 | 920 |

| | | | | | | | | | | | | | | | | | |
|---|---|---|---|---|---|---|---|---|---|---|---|---|---|---|---|---|---|
| 934 | 0.02 | 0.20 | 0.12 | 0.90 | 1 | Zbtb18 | zinc finger and BTB domain containing 18 | 556 | 817 | 421 | 1020 | 919 | 1154 | 1127 | 883 | 1042 | 1188 | 1153 |
| 1391 | -0.04 | 0.12 | -0.29 | 0.77 | 1 | Arpc4 | actin related protein 2/3 complex, subunit | 2528 | 1891 | 2088 | 1065 | 1285 | 972 | 1126 | 1118 | 1212 | 1137 | 882 |
| 1215 | 0.11 | 0.11 | 0.94 | 0.35 | 1 | Nae1 | NEDD8 activating enzyme E1 subunit 1 | 1167 | 1068 | 1120 | 1004 | 1330 | 1379 | 1126 | 1230 | 1302 | 1233 | 1412 |
| 1141 | 0.23 | 0.14 | 1.67 | 0.09 | 1 | Eif2ak2 | eukaryotic translation initiation factor 2-al | 1294 | 1268 | 1260 | 1049 | 904 | 933 | 1126 | 1373 | 1278 | 1046 | 1017 |
| 1091 | -0.23 | 0.17 | -1.39 | 0.16 | 1 | Jazf1 | JAZF zinc finger 1 | 992 | 1227 | 1259 | 1208 | 1190 | 1067 | 1126 | 729 | 1170 | 1070 | 967 |
| 972 | 0.06 | 0.18 | 0.33 | 0.74 | 1 | Zxdb | zinc finger, X-linked, duplicated B | 951 | 1016 | 738 | 994 | 953 | 823 | 1126 | 750 | 1016 | 1391 | 935 |
| 1124 | 0.02 | 0.12 | 0.17 | 0.86 | 1 | Zfp92 | zinc finger protein 92 | 377 | 562 | 460 | 1355 | 1233 | 1740 | 1125 | 1279 | 1403 | 1135 | 1693 |
| 1112 | -0.05 | 0.12 | -0.38 | 0.70 | 1 | Prpf4 | pre-mRNA processing factor 4 | 859 | 995 | 858 | 1183 | 1001 | 1541 | 1125 | 1060 | 1165 | 1043 | 1401 |
| 1362 | 0.07 | 0.10 | 0.65 | 0.51 | 1 | Bdp1 | B double prime 1, subunit of RNA polyme | 1136 | 1356 | 1093 | 1295 | 1131 | 2042 | 1124 | 1324 | 1303 | 1144 | 2038 |
| 1338 | 0.05 | 0.15 | 0.31 | 0.76 | 1 | Creld1 | cysteine-rich with EGF-like domains 1 | 1585 | 1594 | 1706 | 1278 | 1409 | 1001 | 1124 | 1685 | 1100 | 1305 | 936 |
| 896 | -0.20 | 0.20 | -0.99 | 0.32 | 1 | Synpr | synaptoporin | 305 | 350 | 449 | 1597 | 1118 | 853 | 1124 | 1155 | 874 | 1302 | 732 |
| 1234 | 0.13 | 0.11 | 1.22 | 0.22 | 1 | Crem | cAMP responsive element modulator | 758 | 1222 | 799 | 1525 | 1218 | 1271 | 1123 | 1704 | 1338 | 1114 | 1503 |
| 1215 | -0.07 | 0.13 | -0.51 | 0.61 | 1 | Tatdn2 | TatD DNase domain containing 2 | 1472 | 1281 | 1301 | 1090 | 1332 | 1221 | 1123 | 1169 | 1153 | 949 | 1277 |
| 1186 | -0.08 | 0.10 | -0.76 | 0.44 | 1 | Atg2a | autophagy related 2A | 1411 | 1474 | 1399 | 1112 | 1218 | 1044 | 1123 | 1125 | 1161 | 1024 | 957 |
| 1234 | -0.01 | 0.16 | -0.05 | 0.96 | 1 | Rap1gds1 | RAP1, GTP-GDP dissociation stimulator 1 | 1857 | 1170 | 1297 | 1283 | 1086 | 1144 | 1122 | 1147 | 1269 | 1170 | 1035 |
| 1280 | 0.17 | 0.11 | 1.60 | 0.11 | 1 | Pdrg1 | p53 and DNA damage regulated 1 | 1128 | 1052 | 1322 | 992 | 1265 | 1600 | 1121 | 1273 | 1285 | 1194 | 1846 |
| 1129 | 0.03 | 0.12 | 0.26 | 0.79 | 1 | Taf6 | TATA-box binding protein associated fact | 955 | 864 | 1066 | 1222 | 1242 | 1121 | 1121 | 1081 | 1295 | 1333 | 1116 |
| 1207 | 0.03 | 0.12 | 0.25 | 0.81 | 1 | Armcx3 | armadillo repeat containing, X-linked 3 | 895 | 1038 | 962 | 1182 | 1469 | 1376 | 1120 | 1210 | 1342 | 1225 | 1462 |
| 1071 | -0.22 | 0.15 | -1.47 | 0.14 | 1 | Gtf2e2 | general transcription factor II E, polypeptic | 1046 | 1100 | 1239 | 1118 | 1057 | 1199 | 1120 | 713 | 985 | 1041 | 1163 |
| 1030 | -0.02 | 0.14 | -0.18 | 0.86 | 1 | Zfp719 | zinc finger protein 719 | 708 | 860 | 874 | 1111 | 1022 | 1217 | 1120 | 918 | 1136 | 1046 | 1321 |
| 1001 | 0.15 | 0.17 | 0.86 | 0.39 | 1 | Ebag9 | estrogen receptor-binding fragment-asso | 725 | 925 | 663 | 1061 | 908 | 1043 | 1120 | 984 | 1347 | 1158 | 1075 |
| 1246 | 0.04 | 0.12 | 0.38 | 0.71 | 1 | Etv3 | ets variant 3 | 1585 | 1515 | 1625 | 1097 | 922 | 1292 | 1119 | 1073 | 1146 | 1072 | 1260 |
| 869 | 0.20 | 0.14 | 1.43 | 0.15 | 1 | St8sia1 | ST8 alpha-N-acetyl-neuraminide alpha-2 | 185 | 211 | 175 | 1201 | 1091 | 781 | 1119 | 1155 | 1441 | 1239 | 958 |
| 1560 | 0.31 | 0.22 | 1.42 | 0.16 | 1 | Rpl35 | ribosomal protein L35 | 2536 | 2254 | 2789 | 1014 | 1248 | 829 | 1118 | 2204 | 1181 | 1052 | 931 |
| 1199 | 0.03 | 0.12 | 0.27 | 0.79 | 1 | Pef1 | penta-EF hand domain containing 1 | 1289 | 1310 | 1289 | 1131 | 1245 | 1098 | 1118 | 1067 | 1173 | 1390 | 1080 |
| 1151 | -0.02 | 0.14 | -0.12 | 0.90 | 1 | Ssh1 | slingshot protein phosphatase 1 | 1169 | 1100 | 1189 | 1233 | 1246 | 1010 | 1118 | 1511 | 1168 | 970 | 943 |
| 1069 | -0.12 | 0.18 | -0.66 | 0.51 | 1 | Stxbp3 | syntaxin binding protein 3 | 1325 | 1212 | 1117 | 833 | 1117 | 1133 | 1118 | 842 | 1116 | 702 | 1245 |
| 942 | -0.02 | 0.13 | -0.14 | 0.89 | 1 | Fig4 | FIG4 phosphoinositide 5-phosphatase | 998 | 926 | 740 | 920 | 1056 | 777 | 1118 | 975 | 1040 | 1091 | 725 |
| 1158 | -0.08 | 0.15 | -0.53 | 0.60 | 1 | Cc2d2a | coiled-coil and C2 domain containing 2A | 971 | 1255 | 872 | 1071 | 1205 | 1553 | 1117 | 966 | 1075 | 1143 | 1507 |
| 1150 | 0.11 | 0.13 | 0.85 | 0.40 | 1 | Mvb12a | multivesicular body subunit 12A | 884 | 1054 | 1200 | 1055 | 1291 | 1113 | 1117 | 1329 | 1200 | 1274 | 1130 |
| 1288 | -0.09 | 0.15 | -0.61 | 0.54 | 1 | Dazap1 | DAZ associated protein 1 | 1325 | 1325 | 1283 | 1199 | 1202 | 1741 | 1116 | 864 | 1056 | 1416 | 1643 |
| 1376 | 0.05 | 0.15 | 0.29 | 0.77 | 1 | Parp12 | poly (ADP-ribose) polymerase family, me | 1613 | 1393 | 1548 | 1252 | 1433 | 1403 | 1114 | 1372 | 1112 | 1580 | 1317 |
| 1034 | 0.06 | 0.11 | 0.56 | 0.58 | 1 | 2410131K14Ril | RIKEN cDNA 2410131K14 gene | 740 | 767 | 789 | 1138 | 1256 | 926 | 1114 | 1232 | 1213 | 1305 | 897 |
| 956 | 0.18 | 0.14 | 1.33 | 0.19 | 1 | Pgk1 | phosphoglycerate kinase 1 | 912 | 673 | 768 | 892 | 973 | 854 | 1114 | 1216 | 983 | 1170 | 960 |
| 1199 | 0.07 | 0.12 | 0.55 | 0.58 | 1 | St6galnac6 | ST6 (alpha-N-acetyl-neuraminyl-2,3-beta | 1526 | 1528 | 1130 | 1168 | 1116 | 979 | 1113 | 1416 | 1014 | 1190 | 1008 |
| 1048 | 0.07 | 0.12 | 0.55 | 0.58 | 1 | AI597479 | expressed sequence AI597479 | 1152 | 1252 | 1044 | 990 | 854 | 992 | 1113 | 1174 | 975 | 1015 | 967 |
| 981 | -0.06 | 0.14 | -0.42 | 0.67 | 1 | Relt | RELT tumor necrosis factor receptor | 840 | 679 | 827 | 1175 | 1177 | 851 | 1113 | 1110 | 1201 | 926 | 896 |
| 955 | 0.06 | 0.13 | 0.49 | 0.63 | 1 | Glrx | glutaredoxin | 401 | 454 | 610 | 970 | 1341 | 1011 | 1113 | 1086 | 1239 | 1143 | 1136 |
| 1286 | -0.11 | 0.12 | -0.89 | 0.37 | 1 | Eps15l1 | epidermal growth factor receptor pathway | 1189 | 1419 | 1218 | 1066 | 1160 | 2039 | 1112 | 999 | 979 | 1179 | 1787 |
| 864 | -0.09 | 0.12 | -0.74 | 0.46 | 1 | Atg4c | autophagy related 4C, cysteine peptidase | 639 | 560 | 673 | 871 | 1025 | 918 | 1112 | 832 | 1077 | 978 | 819 |
| 1361 | -0.09 | 0.12 | -0.71 | 0.48 | 1 | Ttc9c | tetratricopeptide repeat domain 9C | 1504 | 1714 | 1694 | 1417 | 1311 | 1349 | 1111 | 1060 | 1227 | 1233 | 1356 |
| 1300 | -0.11 | 0.14 | -0.76 | 0.45 | 1 | Ifnar1 | interferon (alpha and beta) receptor 1 | 1671 | 1575 | 1511 | 1161 | 1115 | 1555 | 1111 | 886 | 1336 | 974 | 1401 |
| 1205 | 0.19 | 0.17 | 1.16 | 0.25 | 1 | Ndufs7 | NADH:ubiquinone oxidoreductase core s | 1124 | 959 | 1211 | 1156 | 1267 | 1098 | 1111 | 1753 | 1193 | 1279 | 1103 |
| 1135 | 0.01 | 0.12 | 0.06 | 0.95 | 1 | Gpam | glycerol-3-phosphate acyltransferase, mit | 1325 | 1087 | 1181 | 1086 | 1138 | 1097 | 1111 | 1127 | 961 | 1191 | 1186 |
| 1131 | 0.05 | 0.12 | 0.43 | 0.67 | 1 | Tent4b | terminal nucleotidyltransferase 4B | 927 | 952 | 748 | 1381 | 950 | 1397 | 1111 | 1270 | 1170 | 1103 | 1434 |
| 1033 | 0.10 | 0.17 | 0.60 | 0.55 | 1 | Mindy3 | MINDY lysine 48 deubiquitinase 3 | 1062 | 780 | 1002 | 992 | 1113 | 904 | 1111 | 1281 | 1184 | 907 | 1027 |
| 800 | -0.17 | 0.14 | -1.21 | 0.23 | 1 | Slc38a7 | solute carrier family 38, member 7 | 754 | 660 | 645 | 823 | 773 | 863 | 1111 | 731 | 843 | 899 | 698 |
| 1324 | 0.19 | 0.15 | 1.26 | 0.21 | 1 | Rsbn1 | rosbin, round spermatid basic protein 1 | 1173 | 1928 | 1148 | 1119 | 1025 | 1570 | 1110 | 1552 | 1078 | 1302 | 1564 |
| 1177 | -0.15 | 0.18 | -0.85 | 0.40 | 1 | Traf7 | TNF receptor-associated factor 7 | 1770 | 1303 | 1337 | 847 | 991 | 1595 | 1110 | 999 | 1050 | 735 | 1206 |
| 1122 | -0.01 | 0.12 | -0.07 | 0.94 | 1 | Rnf34 | ring finger protein 34 | 994 | 904 | 1048 | 1217 | 1128 | 1245 | 1110 | 1091 | 1071 | 1175 | 1354 |
| 1081 | 0.11 | 0.14 | 0.74 | 0.46 | 1 | Clpp | caseinolytic mitochondrial matrix peptidas | 890 | 751 | 1081 | 943 | 1358 | 1017 | 1110 | 1330 | 1333 | 1091 | 987 |
| 1029 | 0.00 | 0.10 | -0.01 | 0.99 | 1 | Agpat5 | 1-acylglycerol-3-phosphate O-acyltransfe | 758 | 831 | 807 | 1252 | 1155 | 948 | 1110 | 1237 | 1229 | 1038 | 959 |
| 3105 | 0.21 | 0.43 | 0.49 | 0.62 | 1 | Nr4a1 | nuclear receptor subfamily 4, group A, me | 9873 | 3993 | 3426 | 1690 | 1062 | 4377 | 1108 | 2363 | 1474 | 1096 | 3688 |
| 1334 | -0.09 | 0.15 | -0.59 | 0.56 | 1 | Tmem43 | transmembrane protein 43 | 2520 | 1715 | 1833 | 1055 | 1406 | 869 | 1108 | 1135 | 1157 | 1109 | 770 |
| 1138 | 0.09 | 0.11 | 0.82 | 0.41 | 1 | Emc2 | ER membrane protein complex subunit 2 | 1215 | 906 | 1173 | 987 | 1274 | 1102 | 1108 | 1096 | 1307 | 1137 | 1212 |
| 1037 | -0.05 | 0.16 | -0.32 | 0.75 | 1 | Ipo4 | importin 4 | 1082 | 724 | 933 | 1170 | 1040 | 1090 | 1108 | 1193 | 1027 | 1086 | 960 |
| 1924 | -0.06 | 0.11 | -0.51 | 0.61 | 1 | Rnf213 | ring finger protein 213 | 4064 | 3415 | 3437 | 979 | 964 | 2191 | 1106 | 833 | 1138 | 1002 | 2032 |
| 1353 | -0.13 | 0.19 | -0.69 | 0.49 | 1 | Synpo | synaptopodin | 2256 | 1943 | 1871 | 1292 | 1364 | 754 | 1106 | 838 | 1172 | 1673 | 616 |
| 1321 | 0.18 | 0.11 | 1.64 | 0.10 | 1 | Gtf2b | general transcription factor IIB | 1115 | 1047 | 1187 | 1274 | 1429 | 1434 | 1106 | 1327 | 1459 | 1450 | 1700 |
| 1041 | -0.19 | 0.12 | -1.55 | 0.12 | 1 | Dnajc16 | DnaJ heat shock protein family (Hsp40) m | 830 | 809 | 706 | 1427 | 1181 | 1138 | 1106 | 1145 | 1062 | 1124 | 924 |

| | | | | | | | | | | | | | | | | | | |
|---|---|---|---|---|---|---|---|---|---|---|---|---|---|---|---|---|---|---|
| 1209 | 0.01 | 0.15 | 0.07 | 0.95 | 1 | Gon4l | gon-4-like (C.elegans) | 1253 | 883 | 1355 | 901 | 1368 | 1559 | 1105 | 1199 | 1355 | 1036 | 1279 |
| 1157 | 0.12 | 0.12 | 0.94 | 0.35 | 1 | Orc4 | origin recognition complex, subunit 4 | 892 | 945 | 998 | 856 | 1075 | 1711 | 1105 | 1014 | 1287 | 1006 | 1838 |
| 1488 | -0.31 | 0.19 | -1.62 | 0.11 | 1 | Sidt2 | SID1 transmembrane family, member 2 | 2600 | 1934 | 1903 | 1324 | 1220 | 1831 | 1104 | 1013 | 1242 | 702 | 1494 |
| 1195 | 0.21 | 0.17 | 1.21 | 0.23 | 1 | Eif4e3 | eukaryotic translation initiation factor 4E r | 1246 | 1101 | 1369 | 1088 | 1102 | 1054 | 1104 | 1730 | 1176 | 1023 | 1158 |
| 1192 | 0.15 | 0.16 | 0.97 | 0.33 | 1 | Cnot11 | CCR4-NOT transcription complex, subuni | 842 | 1031 | 812 | 1322 | 1337 | 1120 | 1104 | 1682 | 1294 | 1626 | 942 |
| 1324 | 0.12 | 0.17 | 0.70 | 0.48 | 1 | Camsap1 | calmodulin regulated spectrin-associated | 1107 | 1753 | 922 | 1206 | 1253 | 1616 | 1103 | 1402 | 1351 | 1264 | 1581 |
| 1180 | -0.03 | 0.16 | -0.20 | 0.84 | 1 | Snx10 | sorting nexin 10 | 1223 | 1055 | 1171 | 1521 | 1186 | 1004 | 1103 | 1530 | 1189 | 984 | 1013 |
| 1121 | -0.16 | 0.15 | -1.13 | 0.26 | 1 | Zfp426 | zinc finger protein 426 | 1108 | 1221 | 1270 | 1278 | 893 | 1332 | 1103 | 979 | 1038 | 832 | 1274 |
| 1084 | 0.11 | 0.13 | 0.90 | 0.37 | 1 | E2f4 | E2F transcription factor 4 | 1037 | 805 | 1029 | 798 | 1176 | 1248 | 1103 | 757 | 1366 | 1315 | 1289 |
| 993 | 0.01 | 0.16 | 0.04 | 0.97 | 1 | Ctns | cystinosis, nephropathic | 553 | 577 | 682 | 1233 | 990 | 1212 | 1103 | 959 | 1189 | 1320 | 1107 |
| 977 | 0.02 | 0.15 | 0.16 | 0.87 | 1 | Dym | dymeclin | 949 | 1073 | 854 | 922 | 1077 | 789 | 1103 | 879 | 1132 | 1209 | 760 |
| 777 | -0.04 | 0.15 | -0.30 | 0.77 | 1 | Srrm4 | serine/arginine repetitive matrix 4 | 320 | 270 | 427 | 906 | 1040 | 754 | 1103 | 751 | 1135 | 1099 | 745 |
| 1266 | 0.09 | 0.10 | 0.89 | 0.37 | 1 | Brix1 | BRX1, biogenesis of ribosomes | 1301 | 1372 | 1461 | 1143 | 1232 | 1272 | 1102 | 1327 | 1321 | 1130 | 1265 |
| 1024 | -0.09 | 0.19 | -0.46 | 0.65 | 1 | Pcid2 | PCI domain containing 2 | 969 | 1240 | 715 | 971 | 1163 | 1043 | 1102 | 1128 | 1129 | 845 | 954 |
| 973 | 0.01 | 0.17 | 0.04 | 0.97 | 1 | 2610301B20Ril | RIKEN cDNA 2610301B20 gene | 810 | 829 | 810 | 1020 | 1049 | 924 | 1102 | 707 | 1140 | 1166 | 1143 |
| 1839 | 0.00 | 0.12 | 0.00 | 1.00 | 1 | Tnks1bp1 | tankyrase 1 binding protein 1 | 3890 | 3545 | 3808 | 1182 | 995 | 1205 | 1101 | 966 | 1097 | 1346 | 1095 |
| 1326 | 0.15 | 0.14 | 1.11 | 0.27 | 1 | Dctn5 | dynactin 5 | 1483 | 1244 | 1080 | 1161 | 1258 | 1591 | 1101 | 1400 | 1283 | 1492 | 1490 |
| 1263 | 0.09 | 0.18 | 0.52 | 0.60 | 1 | Elob | elongin B | 1425 | 1021 | 1766 | 1253 | 1378 | 971 | 1101 | 1356 | 1138 | 1409 | 1076 |
| 1017 | 0.07 | 0.15 | 0.49 | 0.62 | 1 | Dffa | DNA fragmentation factor, alpha subunit | 727 | 896 | 862 | 1017 | 1039 | 1080 | 1100 | 959 | 1120 | 1310 | 1074 |
| 1012 | 0.02 | 0.22 | 0.07 | 0.94 | 1 | Zc3h12c | zinc finger CCCH type containing 12C | 746 | 984 | 819 | 850 | 858 | 1407 | 1100 | 572 | 1092 | 1096 | 1606 |
| 942 | 0.01 | 0.13 | 0.10 | 0.92 | 1 | Tarsl2 | threonyl-tRNA synthetase-like 2 | 640 | 514 | 682 | 1061 | 1004 | 1076 | 1100 | 1003 | 1013 | 1188 | 1088 |
| 1154 | -0.02 | 0.12 | -0.18 | 0.86 | 1 | Atf1 | activating transcription factor 1 | 1322 | 1354 | 1171 | 1119 | 1225 | 1015 | 1099 | 1084 | 1220 | 1083 | 1005 |
| 1147 | 0.01 | 0.12 | 0.09 | 0.93 | 1 | Hdgfl2 | HDGF like 2 | 1282 | 1068 | 1287 | 837 | 1060 | 1485 | 1099 | 972 | 1133 | 902 | 1492 |
| 1441 | 0.09 | 0.18 | 0.49 | 0.62 | 1 | Smc4 | structural maintenance of chromosomes 4 | 2708 | 2117 | 1397 | 1125 | 1109 | 1325 | 1097 | 1264 | 1102 | 1063 | 1547 |
| 1288 | 0.04 | 0.13 | 0.28 | 0.78 | 1 | Elmo2 | engulfment and cell motility 2 | 1962 | 1785 | 1316 | 1189 | 1076 | 1111 | 1097 | 1357 | 1052 | 1233 | 987 |
| 1197 | 0.06 | 0.16 | 0.38 | 0.71 | 1 | Cstf3 | cleavage stimulation factor, 3' pre-RNA, s | 1214 | 1075 | 854 | 928 | 1271 | 1611 | 1097 | 962 | 1302 | 1173 | 1682 |
| 1195 | 0.23 | 0.16 | 1.45 | 0.15 | 1 | Cystm1 | cysteine-rich transmembrane module con | 1182 | 1555 | 1499 | 1117 | 1014 | 844 | 1097 | 1238 | 1185 | 1578 | 841 |
| 1140 | 0.00 | 0.15 | 0.02 | 0.98 | 1 | Ttbk2 | tau tubulin kinase 2 | 1023 | 1101 | 935 | 1133 | 1023 | 1485 | 1097 | 872 | 1201 | 1276 | 1391 |
| 1055 | 0.05 | 0.15 | 0.36 | 0.72 | 1 | Epb41l5 | erythrocyte membrane protein band 4.1 li | 921 | 1246 | 1124 | 1034 | 902 | 1042 | 1097 | 933 | 1005 | 1123 | 1174 |
| 1039 | 0.01 | 0.13 | 0.11 | 0.91 | 1 | Tmem68 | transmembrane protein 68 | 774 | 901 | 896 | 1162 | 1127 | 1022 | 1097 | 971 | 1210 | 1118 | 1156 |
| 987 | 0.05 | 0.12 | 0.40 | 0.69 | 1 | Nck2 | non-catalytic region of tyrosine kinase ada | 824 | 761 | 896 | 1084 | 956 | 977 | 1097 | 1087 | 1000 | 1087 | 1081 |
| 1055 | 0.04 | 0.17 | 0.23 | 0.81 | 1 | Mpnd | MPN domain containing | 1293 | 1007 | 1233 | 912 | 1075 | 892 | 1095 | 1340 | 969 | 943 | 846 |
| 1296 | -0.18 | 0.19 | -0.94 | 0.35 | 1 | Atf7 | activating transcription factor 7 | 1452 | 1597 | 1399 | 1494 | 1173 | 1438 | 1094 | 748 | 1217 | 1257 | 1383 |
| 884 | -0.01 | 0.16 | -0.06 | 0.95 | 1 | 1110065P20Ril | RIKEN cDNA 1110065P20 gene | 526 | 603 | 695 | 977 | 1097 | 790 | 1094 | 1071 | 894 | 1225 | 753 |
| 1232 | 0.12 | 0.14 | 0.82 | 0.41 | 1 | Ubxn2b | UBX domain protein 2B | 766 | 1036 | 1044 | 1571 | 1298 | 1203 | 1093 | 1323 | 1402 | 1436 | 1378 |
| 1203 | -0.06 | 0.25 | -0.24 | 0.81 | 1 | Dmpk | dystrophia myotonica-protein kinase | 1248 | 574 | 535 | 1439 | 1424 | 1582 | 1093 | 1427 | 1239 | 1140 | 1528 |
| 1202 | 0.09 | 0.15 | 0.61 | 0.54 | 1 | Wbp1 | WW domain binding protein 1 | 974 | 1052 | 1148 | 1329 | 1108 | 1336 | 1093 | 1229 | 1130 | 1543 | 1284 |
| 631 | 0.02 | 0.19 | 0.09 | 0.93 | 1 | Akain1 | A kinase (PRKA) anchor inhibitor 1 | 116 | 148 | 134 | 845 | 819 | 527 | 1093 | 789 | 770 | 1029 | 669 |
| 1062 | -0.14 | 0.13 | -1.07 | 0.28 | 1 | Zfp335 | zinc finger protein 335 | 1048 | 763 | 905 | 990 | 1097 | 1513 | 1092 | 741 | 1064 | 1225 | 1245 |
| 1022 | 0.00 | 0.13 | -0.02 | 0.99 | 1 | Alg5 | asparagine-linked glycosylation 5 (dolich | 987 | 803 | 995 | 1078 | 1165 | 903 | 1092 | 1200 | 1020 | 1055 | 944 |
| 979 | -0.10 | 0.12 | -0.79 | 0.43 | 1 | Plekhm2 | pleckstrin homology domain containing, fa | 1200 | 1131 | 1051 | 877 | 856 | 998 | 1092 | 875 | 895 | 853 | 935 |
| 928 | -0.13 | 0.15 | -0.90 | 0.37 | 1 | Fntb | farnesyltransferase, CAAX box, beta | 731 | 839 | 822 | 1073 | 1023 | 877 | 1092 | 833 | 1129 | 1064 | 720 |
| 749 | 0.10 | 0.19 | 0.51 | 0.61 | 1 | Bckdhb | branched chain ketoacid dehydrogenase | 518 | 358 | 329 | 908 | 583 | 813 | 1092 | 1256 | 608 | 915 | 855 |
| 1224 | -0.05 | 0.13 | -0.38 | 0.70 | 1 | Zfp512 | zinc finger protein 512 | 1295 | 1617 | 1004 | 1262 | 1091 | 1404 | 1090 | 1237 | 1046 | 987 | 1437 |
| 1098 | 0.02 | 0.14 | 0.14 | 0.89 | 1 | Sdhaf2 | succinate dehydrogenase complex asser | 975 | 1215 | 1030 | 1008 | 1277 | 1024 | 1090 | 1023 | 1208 | 1039 | 1184 |
| 1074 | 0.18 | 0.13 | 1.37 | 0.17 | 1 | Taf11 | TATA-box binding protein associated fact | 748 | 804 | 804 | 1160 | 956 | 1220 | 1090 | 1243 | 980 | 1372 | 1436 |
| 1002 | 0.12 | 0.17 | 0.70 | 0.48 | 1 | Nkiras1 | NFKB inhibitor interacting Ras-like protei | 735 | 480 | 760 | 1001 | 945 | 1308 | 1090 | 1186 | 958 | 1271 | 1289 |
| 1069 | -0.03 | 0.12 | -0.21 | 0.83 | 1 | Atxn7 | ataxin 7 | 836 | 956 | 815 | 946 | 862 | 1737 | 1088 | 988 | 972 | 906 | 1653 |
| 1036 | -0.24 | 0.21 | -1.15 | 0.25 | 1 | Slc41a1 | solute carrier family 41, member 1 | 1416 | 1733 | 1322 | 961 | 1023 | 680 | 1088 | 591 | 1109 | 768 | 702 |
| 1014 | -0.06 | 0.15 | -0.40 | 0.69 | 1 | Ralb | v-ral simian leukemia viral oncogene B | 1484 | 1255 | 1396 | 796 | 980 | 738 | 1088 | 944 | 892 | 772 | 804 |
| 1013 | -0.14 | 0.13 | -1.05 | 0.29 | 1 | Bcas3 | breast carcinoma amplified sequence 3 | 1090 | 1054 | 1190 | 1204 | 1088 | 713 | 1088 | 948 | 933 | 1158 | 675 |
| 955 | 0.01 | 0.13 | 0.07 | 0.94 | 1 | Asb8 | ankyrin repeat and SOCS box-containing | 707 | 759 | 754 | 1093 | 1161 | 797 | 1088 | 1241 | 1053 | 951 | 906 |
| 1356 | -0.19 | 0.13 | -1.50 | 0.13 | 1 | Tfe3 | transcription factor E3 | 1896 | 1768 | 1834 | 1229 | 1361 | 1336 | 1087 | 926 | 1240 | 1175 | 1063 |
| 1230 | 0.10 | 0.11 | 0.99 | 0.32 | 1 | Mrpl9 | mitochondrial ribosomal protein L9 | 1287 | 1436 | 1379 | 1375 | 1044 | 1046 | 1087 | 1368 | 1155 | 1166 | 1191 |
| 1056 | -0.04 | 0.14 | -0.29 | 0.78 | 1 | Sephs1 | selenophosphate synthetase 1 | 1416 | 1314 | 1118 | 910 | 931 | 1013 | 1087 | 840 | 1086 | 948 | 950 |
| 971 | 0.14 | 0.17 | 0.85 | 0.39 | 1 | Hmgcll1 | 3-hydroxymethyl-3-methylglutaryl-Coenzy | 408 | 708 | 608 | 1202 | 1030 | 949 | 1087 | 1150 | 1204 | 1127 | 1209 |
| 1501 | -0.08 | 0.14 | -0.54 | 0.59 | 1 | Raf1 | v-raf-leukemia viral oncogene 1 | 2296 | 1615 | 1776 | 1089 | 1057 | 2278 | 1086 | 794 | 1187 | 1086 | 2246 |
| 1145 | 0.17 | 0.16 | 1.07 | 0.28 | 1 | Nsrp1 | nuclear speckle regulatory protein 1 | 1081 | 972 | 1169 | 952 | 1091 | 1276 | 1086 | 1463 | 1179 | 952 | 1375 |
| 962 | 0.00 | 0.18 | -0.01 | 1.00 | 1 | Chst12 | carbohydrate sulfotransferase 12 | 1084 | 512 | 840 | 1010 | 1173 | 782 | 1086 | 981 | 1298 | 1103 | 716 |
| 1141 | -0.12 | 0.21 | -0.55 | 0.58 | 1 | Invs | inversin | 982 | 1136 | 974 | 949 | 1415 | 1447 | 1085 | 635 | 1301 | 1338 | 1294 |

| | | | | | | | | | | | | | | | | | | |
|---|---|---|---|---|---|---|---|---|---|---|---|---|---|---|---|---|---|---|
| 1263 | 0.02 | 0.18 | 0.12 | 0.90 | 1 | Dhx16 | DEAH (Asp-Glu-Ala-His) box polypeptide | 1568 | 1063 | 997 | 1034 | 1503 | 1483 | 1084 | 1177 | 1330 | 1343 | 1306 |
| 1027 | -0.08 | 0.15 | -0.52 | 0.60 | 1 | Dcun1d2 | DCN1, defective in cullin neddylation 1, d | 637 | 840 | 692 | 1196 | 1126 | 1262 | 1084 | 911 | 1221 | 947 | 1379 |
| 926 | -0.07 | 0.16 | -0.43 | 0.67 | 1 | Gpatch11 | G patch domain containing 11 | 924 | 1061 | 924 | 842 | 782 | 1019 | 1084 | 686 | 856 | 1011 | 1002 |
| 1141 | -0.24 | 0.16 | -1.52 | 0.13 | 1 | Psd | pleckstrin and Sec7 domain containing | 860 | 1099 | 918 | 853 | 1047 | 2282 | 1083 | 729 | 893 | 967 | 1820 |
| 1447 | -0.14 | 0.13 | -1.11 | 0.27 | 1 | Nptx1 | neuronal pentraxin 1 | 2401 | 3253 | 2356 | 1097 | 857 | 1102 | 1082 | 934 | 736 | 1036 | 1058 |
| 1173 | 0.07 | 0.16 | 0.47 | 0.64 | 1 | Zfp322a | zinc finger protein 322A | 1080 | 1153 | 1030 | 1396 | 1098 | 1121 | 1082 | 1009 | 1217 | 1258 | 1459 |
| 1155 | -0.22 | 0.23 | -0.93 | 0.35 | 1 | Rufy2 | RUN and FYVE domain-containing 2 | 982 | 912 | 664 | 1243 | 1143 | 1909 | 1082 | 570 | 1094 | 1355 | 1753 |
| 1149 | 0.04 | 0.13 | 0.30 | 0.76 | 1 | Zxdc | ZXD family zinc finger C | 1246 | 1007 | 1047 | 1165 | 1015 | 1354 | 1082 | 1280 | 1081 | 1117 | 1246 |
| 1099 | 0.14 | 0.18 | 0.77 | 0.44 | 1 | Cdk5 | cyclin-dependent kinase 5 | 902 | 1088 | 953 | 925 | 1128 | 1197 | 1082 | 1557 | 952 | 1202 | 1108 |
| 940 | -0.09 | 0.12 | -0.79 | 0.43 | 1 | Rxylt1 | ribitol xylosyltransferase 1 | 1035 | 969 | 904 | 1080 | 999 | 687 | 1082 | 995 | 834 | 1028 | 731 |
| 1815 | 0.04 | 0.18 | 0.22 | 0.82 | 1 | Ablim1 | actin-binding LIM protein 1 | 3659 | 2596 | 2160 | 1602 | 1316 | 1697 | 1081 | 1522 | 1278 | 1524 | 1527 |
| 1222 | -0.08 | 0.10 | -0.86 | 0.39 | 1 | Smarcb1 | SWI/SNF related, matrix associated, actin | 1701 | 1383 | 1844 | 943 | 1146 | 1198 | 1081 | 839 | 1086 | 1038 | 1177 |
| 1165 | 0.21 | 0.13 | 1.60 | 0.11 | 1 | Cyb5a | cytochrome b5 type A (microsomal) | 1339 | 1026 | 1051 | 915 | 1178 | 1168 | 1081 | 931 | 1362 | 1413 | 1355 |
| 948 | -0.08 | 0.15 | -0.50 | 0.62 | 1 | Myo5b | myosin VB | 625 | 831 | 800 | 1108 | 1073 | 895 | 1081 | 1164 | 1261 | 751 | 838 |
| 1012 | -0.09 | 0.15 | -0.57 | 0.57 | 1 | Snrpd3 | small nuclear ribonucleoprotein D3 | 1456 | 1355 | 1419 | 737 | 952 | 776 | 1080 | 594 | 970 | 998 | 791 |
| 888 | 0.00 | 0.15 | 0.03 | 0.98 | 1 | Ubxn8 | UBX domain protein 8 | 864 | 747 | 711 | 940 | 971 | 742 | 1080 | 1089 | 948 | 859 | 823 |
| 1059 | -0.04 | 0.14 | -0.29 | 0.77 | 1 | Spns1 | spinster homolog 1 | 1101 | 874 | 1036 | 1047 | 1173 | 1083 | 1079 | 1199 | 963 | 1027 | 1069 |
| 928 | -0.06 | 0.13 | -0.43 | 0.67 | 1 | Slc10a7 | solute carrier family 10 (sodium/bile acid | 757 | 597 | 902 | 779 | 1234 | 992 | 1078 | 884 | 988 | 991 | 1011 |
| 1221 | 0.07 | 0.14 | 0.52 | 0.60 | 1 | Ing2 | inhibitor of growth family, member 2 | 819 | 1049 | 996 | 1447 | 1299 | 1280 | 1077 | 1665 | 1408 | 860 | 1532 |
| 1029 | -0.14 | 0.11 | -1.28 | 0.20 | 1 | Ints10 | integrator complex subunit 10 | 837 | 844 | 770 | 1115 | 1035 | 1412 | 1077 | 992 | 984 | 903 | 1356 |
| 1273 | 0.01 | 0.10 | 0.13 | 0.90 | 1 | Zfp397 | zinc finger protein 397 | 1135 | 1470 | 1145 | 1369 | 984 | 1647 | 1076 | 1330 | 983 | 1024 | 1841 |
| 1189 | 0.12 | 0.14 | 0.84 | 0.40 | 1 | Tomm40 | translocase of outer mitochondrial membr | 1344 | 1166 | 1375 | 887 | 1281 | 1168 | 1076 | 1181 | 1261 | 1350 | 990 |
| 1136 | 0.11 | 0.16 | 0.71 | 0.48 | 1 | Dennd2d | DENN/MADD domain containing 2D | 461 | 580 | 428 | 917 | 1164 | 2144 | 1076 | 1169 | 1324 | 962 | 2275 |
| 1045 | 0.20 | 0.12 | 1.70 | 0.09 | 1 | Exosc9 | exosome component 9 | 1200 | 896 | 907 | 983 | 989 | 902 | 1076 | 1199 | 1040 | 1282 | 1024 |
| 904 | -0.07 | 0.15 | -0.47 | 0.64 | 1 | Gnat2 | guanine nucleotide binding protein, alpha | 254 | 241 | 388 | 1090 | 1205 | 1270 | 1076 | 1036 | 1030 | 1029 | 1325 |
| 1353 | 0.07 | 0.12 | 0.60 | 0.55 | 1 | Txndc9 | thioredoxin domain containing 9 | 1566 | 1587 | 1625 | 1151 | 1381 | 1318 | 1075 | 1371 | 1157 | 1163 | 1495 |
| 1127 | -0.04 | 0.15 | -0.26 | 0.79 | 1 | Slc12a6 | solute carrier family 12, member 6 | 1380 | 1369 | 1192 | 1028 | 855 | 1341 | 1075 | 1229 | 978 | 840 | 1111 |
| 969 | 0.02 | 0.17 | 0.12 | 0.90 | 1 | Pdk2 | pyruvate dehydrogenase kinase, isoenzy | 914 | 748 | 1077 | 1144 | 1013 | 680 | 1075 | 1324 | 973 | 1082 | 633 |
| 1105 | 0.06 | 0.11 | 0.56 | 0.58 | 1 | Ndufs4 | NADH:ubiquinone oxidoreductase core s | 1527 | 1378 | 1362 | 908 | 1057 | 828 | 1074 | 1026 | 1072 | 1055 | 867 |
| 1021 | -0.06 | 0.14 | -0.44 | 0.66 | 1 | B230354K17Rik | | 1167 | 1111 | 1120 | 1062 | 917 | 941 | 1074 | 1133 | 861 | 846 | 1000 |
| 955 | 0.06 | 0.15 | 0.39 | 0.70 | 1 | 1110032A03Ril | RIKEN cDNA 1110032A03 gene | 620 | 791 | 653 | 957 | 1185 | 919 | 1074 | 1007 | 1147 | 1211 | 940 |
| 1143 | 0.03 | 0.13 | 0.25 | 0.80 | 1 | Rilpl2 | Rab interacting lysosomal protein-like 2 | 1422 | 1258 | 1427 | 1189 | 1084 | 829 | 1073 | 1335 | 1021 | 1108 | 825 |
| 1063 | -0.11 | 0.15 | -0.70 | 0.48 | 1 | Tuft1 | tuftelin 1 | 925 | 995 | 1112 | 1238 | 895 | 1283 | 1073 | 916 | 953 | 1087 | 1211 |
| 1036 | -0.14 | 0.15 | -0.95 | 0.34 | 1 | Snx15 | sorting nexin 15 | 883 | 1152 | 1022 | 915 | 1229 | 1155 | 1073 | 1127 | 968 | 841 | 1035 |
| 1032 | 0.06 | 0.20 | 0.30 | 0.76 | 1 | Atp5mpl | ATP synthase membrane subunit 6.8PL | 1295 | 1043 | 1328 | 1005 | 952 | 712 | 1073 | 1457 | 932 | 799 | 753 |
| 1236 | 0.13 | 0.15 | 0.84 | 0.40 | 1 | Rbm43 | RNA binding motif protein 43 | 1539 | 1650 | 1554 | 1200 | 976 | 975 | 1072 | 950 | 1183 | 1436 | 1062 |
| 1095 | 0.04 | 0.14 | 0.31 | 0.76 | 1 | Ado | 2-aminoethanethiol (cysteamine) dioxyge | 999 | 1015 | 974 | 1389 | 1076 | 898 | 1072 | 1652 | 1092 | 978 | 907 |
| 994 | -0.08 | 0.11 | -0.70 | 0.49 | 1 | Fopnl | Fgfr1op N-terminal like | 1215 | 939 | 1151 | 916 | 976 | 948 | 1072 | 906 | 919 | 1050 | 839 |
| 1300 | 0.06 | 0.16 | 0.35 | 0.72 | 1 | Nab1 | Ngfi-A binding protein 1 | 2394 | 2361 | 1624 | 1005 | 1126 | 660 | 1071 | 1380 | 948 | 1012 | 715 |
| 1076 | -0.17 | 0.14 | -1.24 | 0.21 | 1 | Wdr75 | WD repeat domain 75 | 1067 | 956 | 938 | 1141 | 1079 | 1357 | 1071 | 850 | 959 | 872 | 1540 |
| 917 | -0.02 | 0.16 | -0.16 | 0.88 | 1 | Atp5k | ATP synthase, H+ transporting, mitochon | 840 | 891 | 1083 | 849 | 1025 | 716 | 1071 | 1084 | 868 | 1003 | 654 |
| 2584 | -0.29 | 0.16 | -1.84 | 0.07 | 1 | Spp1 | secreted phosphoprotein 1 | 6957 | 5377 | 10796 | 656 | 956 | 139 | 1070 | 584 | 1112 | 690 | 90 |
| 1286 | -0.09 | 0.14 | -0.66 | 0.51 | 1 | Stat6 | signal transducer and activator of transcri | 3055 | 2026 | 2127 | 843 | 920 | 762 | 1070 | 833 | 939 | 836 | 735 |
| 994 | 0.03 | 0.13 | 0.21 | 0.83 | 1 | Rhobtb2 | Rho-related BTB domain containing 2 | 1090 | 1208 | 1142 | 965 | 834 | 838 | 1070 | 995 | 919 | 1108 | 770 |
| 743 | -0.09 | 0.14 | -0.65 | 0.51 | 1 | Pstpip2 | proline-serine-threonine phosphatase-int | 252 | 261 | 253 | 1021 | 978 | 749 | 1070 | 802 | 961 | 1097 | 728 |
| 879 | 0.07 | 0.13 | 0.51 | 0.61 | 1 | Acox3 | acyl-Coenzyme A oxidase 3, pristanoyl | 951 | 744 | 738 | 901 | 795 | 770 | 1069 | 1022 | 957 | 976 | 741 |
| 1211 | 0.15 | 0.16 | 0.91 | 0.37 | 1 | Elmod2 | ELMO/CED-12 domain containing 2 | 720 | 709 | 652 | 806 | 1464 | 2030 | 1068 | 1090 | 1466 | 1130 | 2190 |
| 1133 | 0.02 | 0.19 | 0.09 | 0.93 | 1 | Cdk2ap2 | CDK2-associated protein 2 | 1004 | 807 | 860 | 800 | 1563 | 1434 | 1068 | 809 | 1237 | 1627 | 1258 |
| 1001 | -0.07 | 0.11 | -0.61 | 0.54 | 1 | Sigmar1 | sigma non-opioid intracellular receptor 1 | 910 | 920 | 904 | 1223 | 1069 | 871 | 1068 | 1125 | 1140 | 997 | 786 |
| 1224 | -0.09 | 0.11 | -0.80 | 0.42 | 1 | Nipsnap1 | nipsnap homolog 1 | 1078 | 1171 | 1283 | 1504 | 1453 | 1098 | 1067 | 1371 | 1329 | 1132 | 979 |
| 1213 | 0.10 | 0.15 | 0.66 | 0.51 | 1 | Vps16 | VSP16 CORVET/HOPS core subunit | 1211 | 1206 | 1110 | 964 | 1123 | 1602 | 1067 | 1400 | 964 | 1200 | 1493 |
| 1013 | -0.06 | 0.13 | -0.42 | 0.68 | 1 | Fbxo45 | F-box protein 45 | 805 | 957 | 825 | 1157 | 1080 | 1059 | 1067 | 930 | 1192 | 1034 | 1041 |
| 974 | -0.02 | 0.14 | -0.13 | 0.90 | 1 | Spopl | speckle-type BTB/POZ protein-like | 815 | 947 | 835 | 873 | 1040 | 1103 | 1067 | 1001 | 1043 | 878 | 1117 |
| 820 | 0.00 | 0.14 | 0.03 | 0.98 | 1 | Gm15706 | | 426 | 611 | 402 | 1046 | 937 | 729 | 1067 | 1123 | 966 | 966 | 741 |
| 1357 | 0.10 | 0.16 | 0.62 | 0.54 | 1 | Ints8 | integrator complex subunit 8 | 1222 | 1871 | 998 | 1349 | 1055 | 1799 | 1066 | 1306 | 1375 | 1081 | 1805 |
| 1137 | -0.12 | 0.15 | -0.85 | 0.40 | 1 | Syngap1 | synaptic Ras GTPase activating protein 1 | 1166 | 1036 | 911 | 1052 | 1075 | 1738 | 1066 | 863 | 1152 | 1115 | 1327 |
| 976 | 0.00 | 0.12 | -0.04 | 0.97 | 1 | Gpatch2l | G patch domain containing 2 like | 1261 | 1102 | 991 | 783 | 909 | 939 | 1066 | 839 | 826 | 1079 | 944 |
| 948 | -0.03 | 0.15 | -0.22 | 0.83 | 1 | Bicdl1 | BICD family like cargo adaptor 1 | 571 | 564 | 559 | 929 | 984 | 1431 | 1066 | 757 | 999 | 1265 | 1305 |
| 1288 | 0.18 | 0.11 | 1.71 | 0.09 | 1 | Esco1 | establishment of sister chromatid cohesio | 854 | 1242 | 926 | 1204 | 1063 | 1884 | 1065 | 1434 | 1268 | 1118 | 2110 |
| 1076 | -0.11 | 0.20 | -0.57 | 0.57 | 1 | Pigp | phosphatidylinositol glycan anchor biosy | 868 | 874 | 1036 | 1168 | 1179 | 1243 | 1065 | 668 | 1199 | 1391 | 1140 |

| | | | | | | | | | | | | | | | | | |
|---|---|---|---|---|---|---|---|---|---|---|---|---|---|---|---|---|---|
| 1050 | 0.12 | 0.14 | 0.87 | 0.38 | 1 | Stard3nl | STARD3 N-terminal like | 1167 | 817 | 883 | 1079 | 1049 | 970 | 1065 | 1198 | 1143 | 1143 | 1038 |
| 1038 | 0.17 | 0.13 | 1.33 | 0.18 | 1 | Ttc13 | tetratricopeptide repeat domain 13 | 951 | 708 | 699 | 1012 | 1070 | 1116 | 1065 | 1182 | 1167 | 1293 | 1150 |
| 968 | -0.06 | 0.13 | -0.45 | 0.65 | 1 | Kif3a | kinesin family member 3A | 842 | 832 | 1023 | 1048 | 914 | 1035 | 1065 | 998 | 933 | 993 | 971 |
| 1061 | -0.12 | 0.14 | -0.85 | 0.40 | 1 | Cand2 | cullin-associated and neddylation-dissoci | 953 | 1396 | 961 | 1265 | 1172 | 835 | 1064 | 1171 | 1210 | 933 | 717 |
| 1808 | 0.11 | 0.12 | 0.90 | 0.37 | 1 | Akap8 | A kinase (PRKA) anchor protein 8 | 1589 | 1441 | 1727 | 1408 | 1165 | 3741 | 1063 | 1394 | 1309 | 1313 | 3741 |
| 1553 | -0.13 | 0.12 | -1.15 | 0.25 | 1 | Nxf1 | nuclear RNA export factor 1 | 1282 | 1227 | 1212 | 911 | 1077 | 3818 | 1063 | 907 | 990 | 795 | 3803 |
| 1255 | -0.02 | 0.14 | -0.16 | 0.87 | 1 | Cfl2 | cofilin 2, muscle | 1474 | 1443 | 1305 | 1204 | 1194 | 1360 | 1063 | 941 | 1300 | 1194 | 1322 |
| 1251 | -0.08 | 0.14 | -0.59 | 0.55 | 1 | Slc12a2 | solute carrier family 12, member 2 | 2303 | 1841 | 1913 | 1001 | 1019 | 872 | 1063 | 1054 | 1090 | 862 | 744 |
| 1214 | -0.14 | 0.12 | -1.15 | 0.25 | 1 | Fam193a | family with sequence homology 193, men | 1296 | 1482 | 1273 | 1124 | 1081 | 1627 | 1063 | 960 | 999 | 1134 | 1309 |
| 1147 | 0.00 | 0.15 | 0.03 | 0.97 | 1 | Ppp2r2a | protein phosphatase 2, regulatory subuni | 1424 | 1807 | 1915 | 894 | 879 | 880 | 1063 | 905 | 1102 | 845 | 900 |
| 1077 | 0.16 | 0.13 | 1.17 | 0.24 | 1 | Gadd45gip1 | growth arrest and DNA-damage-inducible | 713 | 714 | 843 | 1182 | 1266 | 1019 | 1063 | 1476 | 1168 | 1311 | 1095 |
| 950 | -0.18 | 0.15 | -1.24 | 0.21 | 1 | Nabp2 | nucleic acid binding protein 2 | 912 | 907 | 927 | 1049 | 1105 | 866 | 1063 | 730 | 1149 | 904 | 836 |
| 1244 | 0.18 | 0.13 | 1.36 | 0.17 | 1 | Eva1b | eva-1 homolog B (C. elegans) | 1152 | 1094 | 1357 | 1168 | 1226 | 1272 | 1062 | 1514 | 1199 | 1276 | 1361 |
| 1081 | 0.10 | 0.14 | 0.69 | 0.49 | 1 | Meaf6 | MYST/Esa1-associated factor 6 | 1132 | 937 | 1185 | 923 | 1115 | 1059 | 1062 | 1077 | 967 | 1399 | 1032 |
| 993 | -0.01 | 0.12 | -0.11 | 0.91 | 1 | Zdhhc21 | zinc finger, DHHC domain containing 21 | 1068 | 968 | 835 | 908 | 960 | 1116 | 1062 | 905 | 984 | 936 | 1184 |
| 1117 | -0.26 | 0.18 | -1.43 | 0.15 | 1 | Gltp | glycolipid transfer protein | 2053 | 1233 | 1312 | 1189 | 1296 | 651 | 1061 | 816 | 1082 | 997 | 601 |
| 961 | 0.06 | 0.16 | 0.36 | 0.72 | 1 | Me2 | malic enzyme 2, NAD(+)-dependent, mito | 1068 | 665 | 691 | 1094 | 1045 | 798 | 1061 | 1096 | 1040 | 1143 | 874 |
| 905 | -0.08 | 0.14 | -0.60 | 0.55 | 1 | Pdzk1ip1 | PDZK1 interacting protein 1 | 738 | 760 | 767 | 1008 | 989 | 874 | 1061 | 1087 | 1094 | 900 | 682 |
| 1143 | -0.05 | 0.18 | -0.30 | 0.77 | 1 | Zfp655 | zinc finger protein 655 | 1310 | 1462 | 1124 | 1149 | 980 | 1190 | 1060 | 776 | 1142 | 887 | 1493 |
| 1118 | -0.03 | 0.16 | -0.22 | 0.83 | 1 | Nelfcd | negative elongation factor complex memt | 1374 | 1219 | 1464 | 1004 | 998 | 1095 | 1059 | 1235 | 1087 | 868 | 892 |
| 1101 | 0.23 | 0.14 | 1.61 | 0.11 | 1 | Mrpl48 | mitochondrial ribosomal protein L48 | 1337 | 939 | 1153 | 1044 | 1006 | 883 | 1059 | 1288 | 1005 | 1306 | 1096 |
| 1012 | 0.13 | 0.17 | 0.75 | 0.45 | 1 | Dkc1 | dyskeratosis congenita 1, dyskerin | 1244 | 1081 | 1049 | 898 | 938 | 812 | 1059 | 1156 | 1066 | 775 | 1051 |
| 984 | 0.12 | 0.19 | 0.60 | 0.55 | 1 | Mrpl28 | mitochondrial ribosomal protein L28 | 858 | 1019 | 813 | 901 | 1127 | 765 | 1059 | 1590 | 899 | 927 | 862 |
| 977 | -0.09 | 0.20 | -0.46 | 0.65 | 1 | 4933421O10Rik | | 480 | 977 | 632 | 1139 | 1154 | 1123 | 1059 | 1023 | 985 | 1150 | 1030 |
| 782 | 0.02 | 0.12 | 0.18 | 0.86 | 1 | Borcs7 | BLOC-1 related complex subunit 7 | 589 | 526 | 568 | 837 | 841 | 703 | 1059 | 904 | 876 | 958 | 736 |
| 1289 | -0.08 | 0.16 | -0.49 | 0.63 | 1 | Sap130 | Sin3A associated protein | 1365 | 1135 | 1298 | 1155 | 1598 | 1511 | 1058 | 1172 | 1571 | 904 | 1411 |
| 1241 | -0.16 | 0.16 | -1.04 | 0.30 | 1 | Krit1 | KRIT1, ankyrin repeat containing | 770 | 878 | 829 | 1051 | 1159 | 2482 | 1058 | 689 | 1034 | 1027 | 2678 |
| 1206 | 0.03 | 0.12 | 0.28 | 0.78 | 1 | Smarce1 | SWI/SNF related, matrix associated, actin | 1250 | 1376 | 1231 | 1034 | 1200 | 1327 | 1058 | 1029 | 1076 | 1091 | 1589 |
| 1046 | -0.08 | 0.14 | -0.52 | 0.60 | 1 | Slc31a2 | solute carrier family 31, member 2 | 1155 | 986 | 1408 | 1090 | 1110 | 829 | 1058 | 1109 | 1040 | 878 | 843 |
| 874 | -0.15 | 0.14 | -1.08 | 0.28 | 1 | Snap91 | synaptosomal-associated protein 91 | 673 | 652 | 753 | 1024 | 1011 | 867 | 1058 | 755 | 1096 | 851 | 877 |
| 1405 | 0.28 | 0.18 | 1.59 | 0.11 | 1 | Psme2 | proteasome (prosome, macropain) activa | 2026 | 1583 | 2071 | 1119 | 1230 | 992 | 1057 | 1828 | 1155 | 1226 | 1172 |
| 1206 | 0.12 | 0.11 | 1.08 | 0.28 | 1 | Rpap1 | RNA polymerase II associated protein 1 | 1160 | 1232 | 1082 | 1264 | 1156 | 1213 | 1057 | 1402 | 1244 | 1313 | 1145 |
| 939 | -0.05 | 0.18 | -0.28 | 0.78 | 1 | Hace1 | HECT domain and ankyrin repeat contain | 749 | 750 | 750 | 1090 | 923 | 1014 | 1057 | 674 | 1149 | 1024 | 1143 |
| 1020 | 0.08 | 0.19 | 0.42 | 0.68 | 1 | Ckmt1 | creatine kinase, mitochondrial 1, ubiquito | 625 | 978 | 1100 | 1115 | 1227 | 695 | 1056 | 1663 | 993 | 1096 | 673 |
| 844 | 0.08 | 0.15 | 0.50 | 0.62 | 1 | Cpm | carboxypeptidase M | 486 | 406 | 442 | 1039 | 1059 | 681 | 1056 | 1347 | 1131 | 1018 | 619 |
| 722 | -0.19 | 0.23 | -0.81 | 0.42 | 1 | Akr7a5 | aldo-keto reductase family 7, member A5 | 629 | 658 | 778 | 710 | 714 | 626 | 1056 | 1013 | 675 | 604 | 483 |
| 996 | 0.06 | 0.14 | 0.40 | 0.69 | 1 | Src | Rous sarcoma oncogene | 1244 | 1062 | 1033 | 1139 | 777 | 756 | 1055 | 1235 | 893 | 1067 | 700 |
| 1036 | -0.11 | 0.13 | -0.82 | 0.41 | 1 | Snapin | SNAP-associated protein | 1196 | 1210 | 990 | 1041 | 1027 | 1023 | 1054 | 861 | 1031 | 965 | 993 |
| 964 | -0.07 | 0.13 | -0.57 | 0.57 | 1 | Mrpl2 | mitochondrial ribosomal protein L2 | 856 | 869 | 1085 | 980 | 1023 | 939 | 1054 | 1010 | 950 | 1018 | 824 |
| 1503 | 0.01 | 0.12 | 0.05 | 0.96 | 1 | Psme1 | proteasome (prosome, macropain) activa | 3060 | 2283 | 3050 | 978 | 1126 | 902 | 1053 | 1008 | 957 | 1204 | 913 |
| 885 | -0.23 | 0.19 | -1.21 | 0.23 | 1 | Wrn | Werner syndrome RecQ like helicase | 975 | 898 | 814 | 1082 | 881 | 798 | 1053 | 848 | 934 | 598 | 857 |
| 802 | -0.15 | 0.22 | -0.66 | 0.51 | 1 | Pde10a | phosphodiesterase 10A | 495 | 407 | 475 | 530 | 1250 | 1045 | 1053 | 465 | 1378 | 943 | 782 |
| 1257 | 0.01 | 0.17 | 0.07 | 0.94 | 1 | Tfdp1 | transcription factor Dp 1 | 1504 | 1025 | 1097 | 1161 | 1112 | 1736 | 1052 | 864 | 1395 | 1217 | 1658 |
| 1185 | 0.25 | 0.17 | 1.46 | 0.14 | 1 | Ascc1 | activating signal cointegrator 1 complex s | 1078 | 885 | 1019 | 1205 | 1298 | 1004 | 1052 | 1873 | 1282 | 1324 | 1014 |
| 1079 | 0.09 | 0.16 | 0.55 | 0.58 | 1 | Ube2n | ubiquitin-conjugating enzyme E2N | 1335 | 1565 | 1234 | 881 | 925 | 890 | 1052 | 970 | 1218 | 924 | 878 |
| 1070 | 0.13 | 0.13 | 1.04 | 0.30 | 1 | Mrps12 | mitochondrial ribosomal protein S12 | 697 | 653 | 888 | 1081 | 1422 | 1005 | 1052 | 1072 | 1318 | 1347 | 1234 |
| 1052 | 0.04 | 0.17 | 0.26 | 0.79 | 1 | Cnot4 | CCR4-NOT transcription complex, subuni | 1003 | 1334 | 1022 | 908 | 903 | 1183 | 1052 | 1068 | 1166 | 978 | 950 |
| 985 | -0.03 | 0.20 | -0.16 | 0.87 | 1 | Epm2a | epilepsy, progressive myoclonic epilepsy | 945 | 1004 | 1074 | 1083 | 1009 | 762 | 1052 | 1498 | 911 | 690 | 810 |
| 1471 | 0.17 | 0.16 | 1.11 | 0.27 | 1 | Xab2 | XPA binding protein 2 | 1690 | 1653 | 1696 | 1259 | 1401 | 1525 | 1051 | 1491 | 1164 | 1702 | 1547 |
| 1223 | -0.21 | 0.11 | -1.88 | 0.06 | 1 | Trim2 | tripartite motif-containing 2 | 2155 | 2199 | 2350 | 1047 | 925 | 613 | 1051 | 814 | 816 | 900 | 578 |
| 1110 | 0.18 | 0.19 | 0.93 | 0.35 | 1 | Inafm1 | InaF motif containing 1 | 1084 | 1151 | 1283 | 1027 | 1041 | 914 | 1051 | 1452 | 852 | 1493 | 863 |
| 959 | -0.25 | 0.19 | -1.36 | 0.17 | 1 | Gpcpd1 | glycerophosphocholine phosphodiestera | 1106 | 682 | 913 | 1026 | 771 | 1364 | 1051 | 590 | 867 | 828 | 1350 |
| 642 | 0.00 | 0.23 | -0.02 | 0.99 | 1 | Rtkn2 | rhotekin 2 | 347 | 526 | 168 | 1112 | 267 | 509 | 1051 | 1118 | 205 | 1242 | 514 |
| 1695 | 0.00 | 0.15 | 0.02 | 0.98 | 1 | Lima1 | LIM domain and actin binding 1 | 4123 | 3237 | 2844 | 1222 | 928 | 1023 | 1050 | 1226 | 1081 | 909 | 997 |
| 1355 | 0.05 | 0.15 | 0.35 | 0.72 | 1 | Dnmt1 | DNA methyltransferase (cytosine-5) 1 | 2255 | 1614 | 1457 | 1175 | 1083 | 1406 | 1049 | 1194 | 1184 | 1144 | 1350 |
| 909 | -0.15 | 0.13 | -1.20 | 0.23 | 1 | Sall2 | spalt like transcription factor 2 | 1133 | 1211 | 942 | 1027 | 854 | 592 | 1049 | 836 | 843 | 1020 | 491 |
| 1189 | 0.11 | 0.13 | 0.86 | 0.39 | 1 | Sprtn | SprT-like N-terminal domain | 1095 | 1179 | 1137 | 1054 | 1080 | 1484 | 1048 | 1354 | 1025 | 1229 | 1395 |
| 1024 | 0.14 | 0.13 | 1.05 | 0.29 | 1 | Hax1 | HCLS1 associated X-1 | 774 | 699 | 1034 | 868 | 1111 | 1144 | 1048 | 1155 | 1262 | 1041 | 1127 |
| 901 | -0.06 | 0.20 | -0.32 | 0.75 | 1 | Abhd5 | abhydrolase domain containing 5 | 873 | 888 | 845 | 892 | 883 | 875 | 1048 | 1243 | 967 | 654 | 742 |
| 1266 | -0.17 | 0.17 | -1.04 | 0.30 | 1 | S100pbp | S100P binding protein | 1402 | 1709 | 1511 | 1207 | 1047 | 1625 | 1047 | 757 | 983 | 1194 | 1446 |

| | | | | | | | | | | | | | | | | | |
|---|---|---|---|---|---|---|---|---|---|---|---|---|---|---|---|---|---|
| 1200 | 0.05 | 0.19 | 0.24 | 0.81 | 1 | Npr1 | natriuretic peptide receptor 1 | 2269 | 1406 | 1334 | 788 | 1283 | 933 | 1047 | 970 | 1226 | 992 | 950 |
| 1058 | 0.16 | 0.16 | 1.03 | 0.30 | 1 | Commd3 | COMM domain containing 3 | 858 | 1023 | 1164 | 995 | 931 | 1067 | 1047 | 1379 | 957 | 1023 | 1196 |
| 807 | -0.02 | 0.23 | -0.09 | 0.93 | 1 | Erc2 | ELKS/RAB6-interacting/CAST family mem | 200 | 425 | 319 | 787 | 737 | 1440 | 1047 | 894 | 923 | 826 | 1283 |
| 1197 | 0.15 | 0.13 | 1.22 | 0.22 | 1 | Thoc7 | THO complex 7 | 1368 | 1468 | 1486 | 854 | 1041 | 1243 | 1046 | 891 | 1244 | 1185 | 1343 |
| 1117 | -0.18 | 0.14 | -1.29 | 0.20 | 1 | Zhx2 | zinc fingers and homeoboxes 2 | 1203 | 1290 | 1054 | 1163 | 1202 | 1220 | 1046 | 855 | 1043 | 967 | 1240 |
| 1009 | 0.21 | 0.11 | 1.82 | 0.07 | 1 | Dnajc27 | DnaJ heat shock protein family (Hsp40) m | 1085 | 953 | 854 | 885 | 991 | 884 | 1046 | 1071 | 1049 | 1288 | 994 |
| 967 | -0.01 | 0.13 | -0.09 | 0.93 | 1 | Mettl4 | methyltransferase like 4 | 1034 | 1111 | 1043 | 839 | 881 | 975 | 1046 | 860 | 947 | 884 | 1017 |
| 882 | 0.02 | 0.16 | 0.15 | 0.88 | 1 | Kantr | | 511 | 607 | 551 | 721 | 942 | 1285 | 1046 | 760 | 1058 | 984 | 1241 |
| 1020 | 0.13 | 0.13 | 0.97 | 0.33 | 1 | Tusc2 | tumor suppressor 2, mitochondrial calciur | 775 | 792 | 925 | 1102 | 1220 | 806 | 1045 | 1419 | 1189 | 1000 | 952 |
| 947 | -0.21 | 0.13 | -1.60 | 0.11 | 1 | Plekhm3 | pleckstrin homology domain containing, fa | 935 | 875 | 821 | 1081 | 1129 | 921 | 1045 | 981 | 1006 | 795 | 827 |
| 874 | -0.26 | 0.17 | -1.51 | 0.13 | 1 | Cep83os | | 812 | 1109 | 1036 | 901 | 903 | 768 | 1045 | 755 | 922 | 622 | 737 |
| 1558 | 0.08 | 0.15 | 0.54 | 0.59 | 1 | Hmgn2 | high mobility group nucleosomal binding | 1379 | 1689 | 2333 | 1573 | 1493 | 1566 | 1044 | 1995 | 1424 | 1028 | 1614 |
| 1178 | 0.08 | 0.22 | 0.38 | 0.70 | 1 | Itsn1 | intersectin 1 (SH3 domain protein 1A) | 1924 | 1104 | 1210 | 1033 | 1082 | 1062 | 1044 | 1476 | 1214 | 943 | 872 |
| 1157 | 0.10 | 0.15 | 0.65 | 0.51 | 1 | Smndc1 | survival motor neuron domain containing | 1176 | 1095 | 1033 | 984 | 1187 | 1330 | 1044 | 1218 | 1364 | 900 | 1393 |
| 1039 | 0.06 | 0.11 | 0.58 | 0.56 | 1 | Tada3 | transcriptional adaptor 3 | 1013 | 1122 | 1018 | 976 | 952 | 1079 | 1044 | 1065 | 986 | 1086 | 1088 |
| 976 | 0.17 | 0.10 | 1.67 | 0.09 | 1 | Phlpp1 | PH domain and leucine rich repeat protei | 1248 | 1091 | 956 | 863 | 826 | 758 | 1044 | 981 | 948 | 1202 | 817 |
| 1119 | 0.15 | 0.14 | 1.01 | 0.31 | 1 | Nr2f6 | nuclear receptor subfamily 2, group F, me | 1202 | 1160 | 1083 | 1044 | 1165 | 913 | 1043 | 1360 | 1129 | 1416 | 797 |
| 924 | -0.09 | 0.16 | -0.57 | 0.57 | 1 | Ypel2 | yippee like 2 | 597 | 376 | 632 | 824 | 1053 | 1435 | 1043 | 710 | 1180 | 781 | 1531 |
| 1188 | -0.08 | 0.15 | -0.57 | 0.57 | 1 | Tomm22 | translocase of outer mitochondrial membr | 1777 | 1409 | 1620 | 1203 | 1036 | 964 | 1042 | 893 | 1191 | 974 | 964 |
| 1097 | -0.12 | 0.15 | -0.82 | 0.41 | 1 | Zfp866 | zinc finger protein 866 | 894 | 1040 | 978 | 1353 | 989 | 1385 | 1042 | 916 | 1134 | 966 | 1375 |
| 1065 | 0.03 | 0.14 | 0.25 | 0.80 | 1 | Uimc1 | ubiquitin interaction motif containing 1 | 1156 | 1022 | 977 | 800 | 767 | 1623 | 1042 | 752 | 768 | 1283 | 1521 |
| 895 | 0.01 | 0.12 | 0.05 | 0.96 | 1 | Selenoo | selenoprotein O | 662 | 576 | 698 | 820 | 961 | 1128 | 1042 | 880 | 980 | 1122 | 974 |
| 1059 | 0.05 | 0.13 | 0.37 | 0.71 | 1 | Coq10a | coenzyme Q10A | 622 | 740 | 791 | 1322 | 1143 | 1164 | 1040 | 1116 | 1173 | 1164 | 1372 |
| 1410 | -0.05 | 0.14 | -0.33 | 0.74 | 1 | Fads2 | fatty acid desaturase 2 | 2335 | 2347 | 2701 | 1203 | 1018 | 870 | 1039 | 849 | 1183 | 1005 | 966 |
| 1184 | -0.08 | 0.20 | -0.41 | 0.68 | 1 | Zkscan17 | zinc finger with KRAB and SCAN domain: | 1602 | 1300 | 1201 | 1431 | 1215 | 877 | 1039 | 805 | 1369 | 1342 | 845 |
| 1085 | -0.09 | 0.16 | -0.59 | 0.56 | 1 | Szt2 | SZT2 subunit of KICSTOR complex | 1121 | 892 | 826 | 1027 | 834 | 1852 | 1039 | 836 | 1083 | 929 | 1491 |
| 3845 | -0.30 | 0.22 | -1.39 | 0.16 | 1 | Hspa1a | heat shock protein 1A | 5844 | 6602 | 8382 | 821 | 888 | 8393 | 1038 | 392 | 666 | 1155 | 8114 |
| 1183 | -0.06 | 0.11 | -0.60 | 0.55 | 1 | H2az2 | H2A.Z histone variant 2 | 1458 | 1633 | 1627 | 1147 | 1108 | 935 | 1038 | 1152 | 1033 | 1078 | 804 |
| 1083 | -0.01 | 0.11 | -0.14 | 0.89 | 1 | Fbxo31 | F-box protein 31 | 1204 | 1340 | 1318 | 1039 | 1109 | 855 | 1038 | 981 | 1128 | 1094 | 808 |
| 1072 | 0.05 | 0.16 | 0.29 | 0.77 | 1 | Ints2 | integrator complex subunit 2 | 560 | 607 | 572 | 911 | 1086 | 1866 | 1038 | 750 | 1325 | 1050 | 2027 |
| 844 | -0.14 | 0.17 | -0.85 | 0.40 | 1 | Eif5a2 | eukaryotic translation initiation factor 5A2 | 758 | 692 | 614 | 972 | 1173 | 625 | 1038 | 693 | 1085 | 972 | 667 |
| 811 | -0.16 | 0.19 | -0.88 | 0.38 | 1 | Mdp1 | magnesium-dependent phosphatase 1 | 746 | 508 | 779 | 856 | 923 | 830 | 1038 | 838 | 807 | 674 | 925 |
| 1305 | 0.05 | 0.16 | 0.33 | 0.74 | 1 | Pofut1 | protein O-fucosyltransferase 1 | 1871 | 1785 | 1476 | 1123 | 1172 | 1164 | 1037 | 1199 | 1207 | 1432 | 889 |
| 1019 | -0.06 | 0.14 | -0.40 | 0.69 | 1 | Csnk1g1 | casein kinase 1, gamma 1 | 887 | 959 | 832 | 1002 | 927 | 1384 | 1037 | 799 | 1078 | 1002 | 1299 |
| 944 | -0.04 | 0.13 | -0.34 | 0.74 | 1 | D630045J12Ril | RIKEN cDNA D630045J12 gene | 532 | 713 | 509 | 1234 | 1034 | 1045 | 1037 | 1415 | 1018 | 846 | 997 |
| 923 | 0.05 | 0.16 | 0.33 | 0.74 | 1 | Dpy30 | dpy-30, histone methyltransferase comple | 990 | 975 | 1011 | 698 | 988 | 810 | 1037 | 1047 | 902 | 899 | 799 |
| 1031 | 0.00 | 0.12 | 0.01 | 0.99 | 1 | Gtf2a2 | general transcription factor II A, 2 | 870 | 772 | 947 | 825 | 935 | 1535 | 1036 | 816 | 972 | 884 | 1751 |
| 972 | 0.05 | 0.11 | 0.45 | 0.65 | 1 | Ehd3 | EH-domain containing 3 | 951 | 1085 | 955 | 1037 | 982 | 715 | 1036 | 1207 | 1026 | 1015 | 688 |
| 824 | -0.02 | 0.13 | -0.15 | 0.88 | 1 | Grip1 | glutamate receptor interacting protein 1 | 130 | 179 | 135 | 1084 | 983 | 1235 | 1036 | 1156 | 1032 | 931 | 1164 |
| 1183 | 0.06 | 0.13 | 0.48 | 0.63 | 1 | Slc66a2 | solute carrier family 66 member 2 | 1337 | 1586 | 1510 | 1097 | 1206 | 856 | 1035 | 1247 | 988 | 1238 | 910 |
| 1103 | -0.01 | 0.12 | -0.11 | 0.91 | 1 | Apbb2 | amyloid beta (A4) precursor protein-bindi | 1576 | 1494 | 1628 | 864 | 860 | 974 | 1035 | 930 | 801 | 1000 | 967 |
| 1010 | -0.18 | 0.15 | -1.19 | 0.23 | 1 | Wdr59 | WD repeat domain 59 | 740 | 908 | 705 | 1259 | 1102 | 1265 | 1035 | 943 | 1216 | 943 | 999 |
| 932 | -0.10 | 0.13 | -0.77 | 0.44 | 1 | Rpp14 | ribonuclease P 14 subunit | 821 | 847 | 852 | 1224 | 942 | 820 | 1035 | 960 | 885 | 967 | 901 |
| 904 | -0.12 | 0.15 | -0.79 | 0.43 | 1 | Nop10 | NOP10 ribonucleoprotein | 722 | 684 | 860 | 935 | 1090 | 904 | 1035 | 723 | 1074 | 1170 | 746 |
| 1462 | -0.11 | 0.12 | -0.92 | 0.36 | 1 | Trafd1 | TRAF type zinc finger domain containing | 2302 | 1891 | 1907 | 1090 | 1233 | 1817 | 1034 | 936 | 1133 | 1032 | 1704 |
| 1455 | 0.01 | 0.15 | 0.08 | 0.94 | 1 | Vasp | vasodilator-stimulated phosphoprotein | 2687 | 2362 | 2116 | 1251 | 1021 | 1067 | 1034 | 951 | 1046 | 1493 | 980 |
| 1133 | 0.11 | 0.13 | 0.84 | 0.40 | 1 | Stx18 | syntaxin 18 | 897 | 726 | 983 | 1138 | 1270 | 1288 | 1034 | 1073 | 1277 | 1409 | 1364 |
| 947 | -0.16 | 0.14 | -1.14 | 0.25 | 1 | Med9 | mediator complex subunit 9 | 1132 | 1068 | 958 | 974 | 1083 | 739 | 1034 | 752 | 859 | 1117 | 708 |
| 1338 | 0.16 | 0.18 | 0.89 | 0.38 | 1 | Fastk | Fas-activated serine/threonine kinase | 1020 | 939 | 1198 | 1109 | 1307 | 2090 | 1033 | 1686 | 1084 | 1332 | 1922 |
| 1268 | 0.09 | 0.12 | 0.72 | 0.47 | 1 | Ino80d | INO80 complex subunit D | 1066 | 1281 | 947 | 1089 | 1021 | 2093 | 1033 | 1294 | 1108 | 1152 | 1863 |
| 1185 | 0.11 | 0.11 | 1.03 | 0.30 | 1 | D6Wsu163e | DNA segment, Chr 6, Wayne State Univer | 939 | 1140 | 1114 | 1119 | 1334 | 1248 | 1033 | 1225 | 1296 | 1233 | 1354 |
| 1093 | -0.10 | 0.14 | -0.71 | 0.48 | 1 | Dcp1a | decapping mRNA 1A | 951 | 969 | 881 | 1085 | 1178 | 1481 | 1033 | 868 | 1222 | 1103 | 1255 |
| 1017 | -0.15 | 0.15 | -1.00 | 0.32 | 1 | Erc1 | ELKS/RAB6-interacting/CAST family mem | 1244 | 1078 | 1076 | 847 | 1075 | 1119 | 1033 | 636 | 948 | 1137 | 998 |
| 895 | -0.03 | 0.13 | -0.22 | 0.82 | 1 | Mrpl58 | mitochondrial ribosomal protein L58 | 751 | 681 | 844 | 939 | 998 | 848 | 1033 | 1069 | 944 | 982 | 758 |
| 1028 | 0.20 | 0.19 | 1.06 | 0.29 | 1 | Btbd17 | BTB (POZ) domain containing 17 | 467 | 474 | 659 | 1231 | 1331 | 918 | 1032 | 1792 | 1072 | 1307 | 1027 |
| 1028 | 0.02 | 0.13 | 0.16 | 0.87 | 1 | Stk11ip | serine/threonine kinase 11 interacting pro | 901 | 792 | 1071 | 1047 | 985 | 1172 | 1032 | 1026 | 979 | 1048 | 1255 |
| 1027 | -0.11 | 0.14 | -0.79 | 0.43 | 1 | Tubgcp4 | tubulin, gamma complex associated prote | 1021 | 914 | 1038 | 914 | 1002 | 1355 | 1032 | 689 | 1163 | 916 | 1258 |
| 988 | 0.06 | 0.10 | 0.62 | 0.53 | 1 | Shb | src homology 2 domain-containing transfo | 708 | 808 | 735 | 1150 | 1021 | 1009 | 1032 | 1249 | 1086 | 1048 | 1022 |
| 972 | -0.07 | 0.15 | -0.49 | 0.63 | 1 | Cdc26 | cell division cycle 26 | 1017 | 980 | 1171 | 972 | 829 | 1019 | 1032 | 952 | 914 | 756 | 1053 |
| 910 | 0.02 | 0.19 | 0.12 | 0.91 | 1 | Klhl8 | kelch-like 8 | 506 | 378 | 511 | 1187 | 1033 | 1001 | 1032 | 1369 | 882 | 1221 | 891 |

| | | | | | | | | | | | | | | | | | | |
|---|---|---|---|---|---|---|---|---|---|---|---|---|---|---|---|---|---|---|
| 789 | -0.36 | 0.19 | -1.96 | 0.05 | 1 | Nipsnap3b | nipsnap homolog 3B | 866 | 609 | 829 | 809 | 924 | 795 | 1032 | 543 | 1011 | 590 | 668 |
| 1217 | -0.22 | 0.17 | -1.30 | 0.19 | 1 | Plekha2 | pleckstrin homology domain-containing, f | 2933 | 1877 | 1820 | 1265 | 793 | 573 | 1031 | 914 | 766 | 884 | 529 |
| 1153 | -0.10 | 0.13 | -0.82 | 0.41 | 1 | Asrgl1 | asparaginase like 1 | 1496 | 1536 | 1437 | 1046 | 1259 | 915 | 1031 | 1143 | 957 | 1009 | 850 |
| 1008 | -0.18 | 0.12 | -1.54 | 0.12 | 1 | Fzr1 | fizzy and cell division cycle 20 related 1 | 1202 | 1287 | 1182 | 1030 | 937 | 940 | 1031 | 803 | 838 | 909 | 927 |
| 1138 | 0.13 | 0.18 | 0.73 | 0.47 | 1 | Crtc3 | CREB regulated transcription coactivator | 1479 | 983 | 1110 | 1283 | 1000 | 925 | 1029 | 1345 | 1359 | 1209 | 800 |
| 996 | 0.00 | 0.11 | -0.01 | 0.99 | 1 | Ttc9 | tetratricopeptide repeat domain 9 | 966 | 1026 | 1070 | 1155 | 1040 | 719 | 1029 | 1113 | 1013 | 1132 | 693 |
| 962 | -0.06 | 0.17 | -0.38 | 0.70 | 1 | Peli1 | pellino 1 | 1273 | 1143 | 749 | 1047 | 796 | 915 | 1029 | 1182 | 784 | 756 | 908 |
| 1247 | 0.09 | 0.12 | 0.75 | 0.45 | 1 | Ripk1 | receptor (TNFRSF)-interacting serine-thre | 1544 | 1470 | 1175 | 1245 | 1102 | 1242 | 1027 | 1279 | 1144 | 1137 | 1357 |
| 951 | -0.08 | 0.18 | -0.46 | 0.64 | 1 | Bcl6b | B cell CLL/lymphoma 6, member B | 595 | 766 | 852 | 1144 | 1004 | 1066 | 1027 | 925 | 943 | 1238 | 907 |
| 986 | 0.00 | 0.17 | 0.00 | 1.00 | 1 | Tctn3 | tectonic family member 3 | 456 | 538 | 624 | 859 | 1168 | 1565 | 1026 | 765 | 1079 | 1260 | 1510 |
| 970 | -0.21 | 0.13 | -1.60 | 0.11 | 1 | Mtr | 5-methyltetrahydrofolate-homocysteine m | 1388 | 1315 | 1242 | 810 | 879 | 891 | 1026 | 847 | 754 | 739 | 778 |
| 955 | 0.07 | 0.13 | 0.51 | 0.61 | 1 | Mtap | methylthioadenosine phosphorylase | 962 | 1114 | 931 | 893 | 914 | 817 | 1026 | 998 | 1035 | 1068 | 746 |
| 907 | -0.07 | 0.14 | -0.50 | 0.61 | 1 | 2310030G06Ril | RIKEN cDNA 2310030G06 gene | 669 | 612 | 768 | 1067 | 1036 | 921 | 1026 | 804 | 1071 | 1038 | 967 |
| 871 | -0.08 | 0.13 | -0.63 | 0.53 | 1 | Sys1 | SYS1 Golgi-localized integral membrane | 747 | 542 | 605 | 1009 | 1042 | 866 | 1026 | 925 | 921 | 1068 | 828 |
| 1321 | 0.08 | 0.13 | 0.64 | 0.52 | 1 | Uspl1 | ubiquitin specific peptidase like 1 | 1196 | 1252 | 991 | 1479 | 1146 | 1749 | 1024 | 1668 | 1102 | 1219 | 1707 |
| 1102 | 0.16 | 0.10 | 1.55 | 0.12 | 1 | Tmem222 | transmembrane protein 222 | 1027 | 1082 | 1059 | 1146 | 1121 | 933 | 1024 | 1329 | 1201 | 1229 | 970 |
| 1004 | 0.05 | 0.15 | 0.29 | 0.77 | 1 | Mcoln1 | mucolipin 1 | 546 | 533 | 742 | 1002 | 1039 | 1506 | 1024 | 1220 | 1086 | 1081 | 1266 |
| 899 | 0.18 | 0.17 | 1.02 | 0.31 | 1 | Trub1 | TruB pseudouridine (psi) synthase family | 567 | 541 | 690 | 885 | 781 | 1115 | 1024 | 1167 | 823 | 1126 | 1175 |
| 707 | 0.30 | 0.17 | 1.74 | 0.08 | 1 | Esrrg | estrogen-related receptor gamma | 260 | 265 | 204 | 901 | 610 | 632 | 1024 | 1291 | 897 | 875 | 812 |
| 1168 | -0.15 | 0.12 | -1.31 | 0.19 | 1 | Trim3 | tripartite motif-containing 3 | 1053 | 1157 | 1077 | 1192 | 1140 | 1662 | 1022 | 907 | 985 | 1048 | 1600 |
| 1012 | -0.01 | 0.11 | -0.08 | 0.94 | 1 | Zfp810 | zinc finger protein 810 | 674 | 581 | 769 | 928 | 1032 | 1557 | 1022 | 897 | 1025 | 953 | 1692 |
| 1629 | 0.14 | 0.14 | 1.00 | 0.32 | 1 | Tmod3 | tropomodulin 3 | 3651 | 2842 | 2894 | 1006 | 961 | 1067 | 1021 | 1360 | 978 | 1032 | 1106 |
| 1382 | 0.17 | 0.13 | 1.30 | 0.19 | 1 | Luzp1 | leucine zipper protein 1 | 2057 | 2512 | 2028 | 1083 | 909 | 1010 | 1021 | 1404 | 1160 | 1148 | 867 |
| 1072 | -0.06 | 0.13 | -0.46 | 0.65 | 1 | P3h4 | prolyl 3-hydroxylase family member 4 (no | 1658 | 1279 | 1227 | 975 | 1017 | 855 | 1021 | 996 | 819 | 1177 | 767 |
| 1031 | 0.11 | 0.16 | 0.67 | 0.50 | 1 | Mpc2 | mitochondrial pyruvate carrier 2 | 1035 | 1048 | 1344 | 924 | 1091 | 777 | 1021 | 1294 | 951 | 957 | 898 |
| 1022 | -0.18 | 0.13 | -1.43 | 0.15 | 1 | Herc4 | hect domain and RLD 4 | 1017 | 1014 | 1140 | 827 | 1109 | 1310 | 1021 | 753 | 992 | 747 | 1316 |
| 823 | 0.19 | 0.17 | 1.14 | 0.25 | 1 | Hint3 | histidine triad nucleotide binding protein 3 | 479 | 570 | 513 | 807 | 801 | 850 | 1021 | 933 | 964 | 1310 | 805 |
| 1111 | 0.10 | 0.10 | 0.92 | 0.36 | 1 | Wdr5 | WD repeat domain 5 | 1387 | 1172 | 1239 | 957 | 903 | 1199 | 1020 | 994 | 1042 | 1126 | 1184 |
| 3059 | 0.14 | 0.17 | 0.85 | 0.40 | 1 | Calu | calumenin | 9631 | 5555 | 6311 | 1194 | 1237 | 2357 | 1019 | 1427 | 1300 | 1168 | 2446 |
| 1117 | 0.17 | 0.12 | 1.43 | 0.15 | 1 | Lsg1 | large 60S subunit nuclear export GTPase | 848 | 917 | 1028 | 935 | 1085 | 1441 | 1019 | 1192 | 1116 | 1209 | 1492 |
| 1104 | 0.07 | 0.12 | 0.60 | 0.55 | 1 | Gtf3c3 | general transcription factor IIIC, polypepti | 790 | 976 | 971 | 1249 | 1238 | 1079 | 1019 | 1075 | 1373 | 1175 | 1196 |
| 1063 | -0.24 | 0.15 | -1.61 | 0.11 | 1 | Wdr90 | WD repeat domain 90 | 682 | 609 | 497 | 1242 | 916 | 2188 | 1019 | 811 | 923 | 931 | 1881 |
| 1013 | -0.01 | 0.13 | -0.09 | 0.93 | 1 | Rit1 | Ras-like without CAAX 1 | 1207 | 1359 | 1207 | 1020 | 891 | 753 | 1019 | 1139 | 760 | 1028 | 754 |
| 993 | -0.09 | 0.12 | -0.76 | 0.45 | 1 | Ift22 | intraflagellar transport 22 | 978 | 1115 | 1106 | 999 | 1017 | 938 | 1019 | 1118 | 922 | 855 | 850 |
| 912 | 0.04 | 0.18 | 0.22 | 0.83 | 1 | 0610009B22Ril | RIKEN cDNA 0610009B22 gene | 851 | 927 | 754 | 923 | 922 | 823 | 1019 | 677 | 1039 | 1162 | 935 |
| 813 | -0.03 | 0.14 | -0.21 | 0.84 | 1 | Hs1bp3 | HCLS1 binding protein 3 | 746 | 711 | 829 | 804 | 815 | 729 | 1019 | 795 | 909 | 836 | 750 |
| 1641 | -0.30 | 0.16 | -1.93 | 0.05 | 1 | Slco1a6 | solute carrier organic anion transporter fa | 43 | 260 | 140 | 4389 | 1470 | 2793 | 1018 | 3532 | 1351 | 723 | 2330 |
| 1103 | -0.05 | 0.10 | -0.46 | 0.64 | 1 | Ing5 | inhibitor of growth family, member 5 | 1031 | 1245 | 1145 | 1188 | 1164 | 1056 | 1018 | 1081 | 1192 | 1005 | 1003 |
| 1009 | -0.01 | 0.18 | -0.03 | 0.97 | 1 | Tmtc4 | transmembrane and tetratricopeptide repe | 927 | 661 | 819 | 1105 | 1245 | 980 | 1018 | 1273 | 1261 | 858 | 948 |
| 867 | 0.33 | 0.18 | 1.83 | 0.07 | 1 | Akap6 | A kinase (PRKA) anchor protein 6 | 334 | 350 | 219 | 931 | 852 | 1036 | 1018 | 1083 | 1345 | 1112 | 1253 |
| 722 | 0.06 | 0.15 | 0.40 | 0.69 | 1 | Fam110b | family with sequence similarity 110, meml | 480 | 455 | 498 | 633 | 810 | 730 | 1018 | 794 | 831 | 1009 | 688 |
| 961 | 0.06 | 0.13 | 0.49 | 0.63 | 1 | Tmem126b | transmembrane protein 126B | 899 | 932 | 854 | 1129 | 836 | 886 | 1017 | 977 | 1012 | 1000 | 1031 |
| 1380 | 0.13 | 0.15 | 0.86 | 0.39 | 1 | Eps15 | epidermal growth factor receptor pathway | 2927 | 1633 | 1969 | 928 | 1058 | 1129 | 1016 | 1015 | 1076 | 1197 | 1231 |
| 1155 | -0.07 | 0.15 | -0.44 | 0.66 | 1 | Dhx57 | DEAH (Asp-Glu-Ala-Asp/His) box polypep | 1105 | 1483 | 1146 | 865 | 1023 | 1692 | 1016 | 1088 | 871 | 911 | 1504 |
| 1127 | -0.09 | 0.13 | -0.71 | 0.48 | 1 | Rxrb | retinoid X receptor beta | 1310 | 1063 | 1290 | 1048 | 1154 | 1275 | 1016 | 901 | 1275 | 951 | 1112 |
| 978 | 0.16 | 0.20 | 0.81 | 0.42 | 1 | Rex1bd | required for excision 1-B domain containi | 845 | 841 | 997 | 938 | 997 | 826 | 1016 | 1559 | 850 | 1091 | 803 |
| 931 | 0.00 | 0.12 | 0.01 | 0.99 | 1 | Triap1 | TP53 regulated inhibitor of apoptosis 1 | 776 | 731 | 839 | 916 | 1139 | 877 | 1016 | 1097 | 1055 | 938 | 857 |
| 1285 | 0.21 | 0.13 | 1.58 | 0.11 | 1 | Dnttip2 | deoxynucleotidyltransferase, terminal, inte | 1344 | 1760 | 1218 | 1161 | 1054 | 1281 | 1015 | 1693 | 1165 | 983 | 1457 |
| 1096 | -0.19 | 0.11 | -1.74 | 0.08 | 1 | Mark4 | MAP/microtubule affinity regulating kinase | 1182 | 1239 | 1403 | 1163 | 1085 | 1126 | 1014 | 1061 | 977 | 866 | 941 |
| 1064 | 0.28 | 0.18 | 1.50 | 0.13 | 1 | Mrps21 | mitochondrial ribosomal protein S21 | 1069 | 767 | 1066 | 1112 | 1050 | 797 | 1014 | 1598 | 1024 | 1235 | 969 |
| 920 | 0.07 | 0.15 | 0.48 | 0.63 | 1 | Kif13a | kinesin family member 13A | 1657 | 1240 | 1303 | 513 | 559 | 835 | 1014 | 613 | 704 | 828 | 850 |
| 782 | -0.10 | 0.24 | -0.40 | 0.69 | 1 | D630039A03Ri | RIKEN cDNA D630039A03 gene | 347 | 518 | 642 | 936 | 765 | 939 | 1014 | 652 | 764 | 1068 | 958 |
| 987 | 0.19 | 0.14 | 1.34 | 0.18 | 1 | Lrif1 | ligand dependent nuclear receptor intera | 599 | 704 | 717 | 890 | 921 | 1289 | 1013 | 1235 | 1084 | 904 | 1495 |
| 1908 | -0.09 | 0.11 | -0.80 | 0.42 | 1 | Myo1b | myosin IB | 3837 | 4693 | 4288 | 1175 | 967 | 1055 | 1012 | 1076 | 1023 | 1050 | 818 |
| 1433 | -0.25 | 0.19 | -1.27 | 0.20 | 1 | Tspan9 | tetraspanin 9 | 3427 | 2986 | 3182 | 868 | 673 | 754 | 1012 | 437 | 726 | 1055 | 644 |
| 1019 | -0.07 | 0.16 | -0.41 | 0.68 | 1 | Cc2d1a | coiled-coil and C2 domain containing 1A | 1108 | 883 | 884 | 1104 | 1021 | 1123 | 1012 | 1104 | 1166 | 878 | 926 |
| 1136 | 0.08 | 0.13 | 0.63 | 0.53 | 1 | Ano10 | anoctamin 10 | 1301 | 1426 | 1034 | 1264 | 1113 | 823 | 1011 | 1413 | 993 | 1311 | 805 |
| 1120 | 0.14 | 0.13 | 1.03 | 0.30 | 1 | Msh2 | mutS homolog 2 | 1841 | 1266 | 1327 | 971 | 910 | 866 | 1011 | 1093 | 923 | 1096 | 1020 |
| 989 | -0.09 | 0.18 | -0.52 | 0.60 | 1 | Sgms1 | sphingomyelin synthase 1 | 1344 | 1340 | 1110 | 809 | 869 | 937 | 1011 | 589 | 839 | 1168 | 863 |
| 892 | -0.09 | 0.14 | -0.69 | 0.49 | 1 | Map3k12 | mitogen-activated protein kinase kinase k | 815 | 923 | 870 | 896 | 883 | 924 | 1011 | 931 | 828 | 757 | 975 |

| | | | | | | | | | | | | | | | | | | |
|---|---|---|---|---|---|---|---|---|---|---|---|---|---|---|---|---|---|---|
| 1339 | -0.09 | 0.11 | -0.85 | 0.39 | 1 | Nras | neuroblastoma ras oncogene | 2032 | 2056 | 1846 | 1338 | 1011 | 1184 | 1010 | 1177 | 995 | 989 | 1089 |
| 1052 | 0.05 | 0.16 | 0.31 | 0.76 | 1 | Slc39a13 | solute carrier family 39 (metal ion transpo | 989 | 791 | 857 | 909 | 1036 | 1387 | 1010 | 697 | 1054 | 1401 | 1437 |
| 1194 | 0.07 | 0.16 | 0.43 | 0.67 | 1 | Plcb1 | phospholipase C, beta 1 | 1511 | 1878 | 1506 | 1123 | 988 | 906 | 1009 | 880 | 1169 | 1234 | 933 |
| 1097 | 0.04 | 0.11 | 0.34 | 0.73 | 1 | Kdm5d | lysine (K)-specific demethylase 5D | 902 | 989 | 791 | 966 | 891 | 1771 | 1009 | 1137 | 890 | 956 | 1765 |
| 1073 | 0.00 | 0.17 | 0.01 | 0.99 | 1 | Gm4419 | | 239 | 531 | 643 | 1329 | 1356 | 1447 | 1009 | 1236 | 1632 | 906 | 1472 |
| 1435 | 0.21 | 0.15 | 1.40 | 0.16 | 1 | Tra2a | transformer 2 alpha | 1706 | 1517 | 1281 | 657 | 716 | 3013 | 1007 | 1004 | 953 | 887 | 3045 |
| 1063 | -0.08 | 0.15 | -0.54 | 0.59 | 1 | Slc45a4 | solute carrier family 45, member 4 | 1175 | 1180 | 1056 | 962 | 1227 | 1046 | 1007 | 969 | 1125 | 1139 | 809 |
| 986 | -0.18 | 0.15 | -1.18 | 0.24 | 1 | Blzf1 | basic leucine zipper nuclear factor 1 | 805 | 1000 | 824 | 992 | 1109 | 1237 | 1007 | 855 | 1119 | 724 | 1176 |
| 947 | -0.17 | 0.14 | -1.21 | 0.23 | 1 | Slc39a10 | solute carrier family 39 (zinc transporter), | 1178 | 1041 | 1082 | 797 | 937 | 1002 | 1007 | 584 | 882 | 926 | 976 |
| 815 | 0.05 | 0.18 | 0.28 | 0.78 | 1 | Nudt7 | nudix (nucleoside diphosphate linked mo | 554 | 419 | 636 | 845 | 855 | 905 | 1007 | 1083 | 967 | 811 | 879 |
| 1211 | -0.22 | 0.16 | -1.37 | 0.17 | 1 | Zfp280d | zinc finger protein 280D | 927 | 1325 | 1140 | 1255 | 1146 | 1869 | 1006 | 755 | 1053 | 917 | 1925 |
| 1050 | -0.16 | 0.15 | -1.07 | 0.28 | 1 | Mthfr | methylenetetrahydrofolate reductase | 831 | 816 | 897 | 1056 | 941 | 1751 | 1006 | 1164 | 854 | 750 | 1483 |
| 920 | 0.06 | 0.13 | 0.43 | 0.67 | 1 | Slc6a8 | solute carrier family 6 (neurotransmitter tra | 1008 | 1002 | 913 | 869 | 799 | 842 | 1006 | 920 | 951 | 1079 | 726 |
| 1254 | -0.23 | 0.13 | -1.76 | 0.08 | 1 | 2700081O15Ril | RIKEN cDNA 2700081O15 gene | 2098 | 2400 | 1948 | 1160 | 854 | 946 | 1005 | 836 | 822 | 1007 | 720 |
| 1004 | -0.01 | 0.13 | -0.07 | 0.94 | 1 | Trappc5 | trafficking protein particle complex 5 | 1041 | 1169 | 938 | 953 | 1060 | 931 | 1005 | 1139 | 893 | 1019 | 896 |
| 968 | 0.08 | 0.13 | 0.59 | 0.55 | 1 | Mxi1 | MAX interactor 1, dimerization protein | 1182 | 1212 | 1102 | 932 | 920 | 626 | 1005 | 877 | 1122 | 993 | 675 |
| 914 | 0.22 | 0.16 | 1.36 | 0.17 | 1 | Memo1 | mediator of cell motility 1 | 743 | 710 | 759 | 878 | 746 | 990 | 1005 | 1295 | 884 | 947 | 1093 |
| 1091 | 0.05 | 0.13 | 0.43 | 0.66 | 1 | Pgghg | protein glucosylgalactosylhydroxylysine g | 1111 | 1209 | 1063 | 1076 | 1122 | 1012 | 1004 | 1235 | 997 | 1225 | 946 |
| 1022 | -0.07 | 0.14 | -0.47 | 0.64 | 1 | Akap10 | A kinase (PRKA) anchor protein 10 | 1038 | 1254 | 854 | 1043 | 989 | 1094 | 1004 | 1086 | 974 | 824 | 1079 |
| 1005 | 0.10 | 0.11 | 0.98 | 0.33 | 1 | Pwp1 | PWP1 homolog, endonuclein | 1072 | 947 | 991 | 914 | 1007 | 953 | 1004 | 1029 | 1000 | 1051 | 1087 |
| 857 | -0.19 | 0.19 | -1.00 | 0.32 | 1 | Rabgef1 | RAB guanine nucleotide exchange factor | 682 | 690 | 676 | 1089 | 808 | 1004 | 1004 | 911 | 921 | 578 | 1060 |
| 1127 | -0.07 | 0.17 | -0.42 | 0.67 | 1 | Nol11 | nucleolar protein 11 | 1260 | 1202 | 1050 | 1426 | 1092 | 1041 | 1003 | 882 | 1060 | 1226 | 1159 |
| 1191 | 0.08 | 0.12 | 0.65 | 0.51 | 1 | Fam118a | family with sequence similarity 118, meml | 1123 | 1633 | 1229 | 1076 | 1040 | 1308 | 1002 | 1323 | 1035 | 1039 | 1290 |
| 1171 | 0.14 | 0.14 | 0.99 | 0.32 | 1 | Mms19 | MMS19 cytosolic iron-sulfur assembly cor | 1120 | 1171 | 1189 | 943 | 1008 | 1550 | 1002 | 1292 | 1235 | 949 | 1417 |
| 1071 | -0.21 | 0.17 | -1.23 | 0.22 | 1 | Camk2d | calcium/calmodulin-dependent protein kir | 2103 | 1850 | 1546 | 745 | 917 | 671 | 1002 | 531 | 817 | 1056 | 542 |
| 1200 | -0.16 | 0.17 | -0.90 | 0.37 | 1 | Epc1 | enhancer of polycomb homolog 1 | 1375 | 1419 | 1118 | 1148 | 989 | 1686 | 1001 | 673 | 910 | 1099 | 1779 |
| 1191 | 0.10 | 0.12 | 0.81 | 0.42 | 1 | Crtc2 | CREB regulated transcription coactivator | 1189 | 1274 | 1321 | 935 | 1248 | 1330 | 1001 | 1194 | 1235 | 1158 | 1213 |
| 1166 | -0.18 | 0.14 | -1.22 | 0.22 | 1 | Tle1 | transducin-like enhancer of split 1 | 1505 | 1534 | 1400 | 936 | 977 | 1543 | 1001 | 713 | 882 | 1113 | 1224 |
| 977 | 0.14 | 0.14 | 1.06 | 0.29 | 1 | Spryd7 | SPRY domain containing 7 | 830 | 861 | 957 | 984 | 1040 | 823 | 1000 | 877 | 1148 | 1194 | 1036 |
| 1143 | 0.14 | 0.15 | 0.96 | 0.34 | 1 | Mafk | v-maf musculoaponeurotic fibrosarcoma ( | 1431 | 1378 | 1359 | 915 | 1051 | 1013 | 999 | 1402 | 1041 | 1039 | 945 |
| 965 | 0.00 | 0.19 | 0.00 | 1.00 | 1 | Mtx3 | metaxin 3 | 670 | 884 | 662 | 634 | 962 | 1623 | 999 | 847 | 891 | 876 | 1567 |
| 916 | -0.02 | 0.14 | -0.11 | 0.91 | 1 | Acot11 | acyl-CoA thioesterase 11 | 368 | 378 | 377 | 958 | 1162 | 1374 | 999 | 875 | 1338 | 974 | 1269 |
| 859 | -0.03 | 0.17 | -0.17 | 0.86 | 1 | Phf21b | PHD finger protein 21B | 456 | 423 | 501 | 953 | 737 | 1386 | 999 | 725 | 906 | 1078 | 1279 |
| 1158 | -0.22 | 0.14 | -1.54 | 0.12 | 1 | Srf | serum response factor | 1436 | 1700 | 1273 | 1111 | 1030 | 1352 | 998 | 775 | 943 | 985 | 1132 |
| 1113 | -0.22 | 0.17 | -1.29 | 0.20 | 1 | Sav1 | salvador family WW domain containing 1 | 1561 | 1821 | 1295 | 1050 | 882 | 1135 | 998 | 651 | 859 | 992 | 995 |
| 1064 | -0.10 | 0.16 | -0.61 | 0.54 | 1 | Dip2b | disco interacting protein 2 homolog B | 1223 | 1046 | 1255 | 989 | 864 | 1386 | 998 | 1131 | 855 | 774 | 1179 |
| 905 | -0.23 | 0.15 | -1.58 | 0.11 | 1 | Sox12 | SRY (sex determining region Y)-box 12 | 1008 | 1202 | 826 | 1258 | 936 | 589 | 998 | 827 | 845 | 893 | 576 |
| 877 | 0.00 | 0.14 | -0.01 | 0.99 | 1 | Mfsd13a | major facilitator superfamily domain conta | 342 | 303 | 430 | 1183 | 1142 | 967 | 998 | 1150 | 1078 | 1029 | 1018 |
| 1131 | 0.02 | 0.22 | 0.10 | 0.92 | 1 | Ak2 | adenylate kinase 2 | 1760 | 1056 | 1167 | 922 | 1132 | 1143 | 997 | 1401 | 906 | 925 | 1035 |
| 994 | -0.01 | 0.12 | -0.11 | 0.92 | 1 | Utp18 | UTP18 small subunit processome compo | 899 | 770 | 1022 | 989 | 1061 | 1081 | 997 | 889 | 1012 | 978 | 1242 |
| 1002 | 0.14 | 0.15 | 0.95 | 0.34 | 1 | Stambp | STAM binding protein | 931 | 1244 | 910 | 1026 | 903 | 858 | 996 | 1039 | 1202 | 972 | 946 |
| 981 | -0.13 | 0.12 | -1.07 | 0.29 | 1 | Kif3c | kinesin family member 3C | 1271 | 1274 | 1278 | 1003 | 924 | 711 | 996 | 1059 | 780 | 867 | 629 |
| 901 | -0.19 | 0.14 | -1.35 | 0.18 | 1 | Zfp324 | zinc finger protein 324 | 556 | 566 | 641 | 780 | 1143 | 1411 | 996 | 655 | 1137 | 791 | 1235 |
| 731 | 0.00 | 0.17 | 0.01 | 0.99 | 1 | Pdk3 | pyruvate dehydrogenase kinase, isoenzy | 638 | 600 | 660 | 889 | 705 | 503 | 996 | 674 | 821 | 942 | 612 |
| 700 | 0.23 | 0.25 | 0.91 | 0.36 | 1 | Eya1 | EYA transcriptional coactivator and phosp | 341 | 627 | 476 | 695 | 623 | 569 | 996 | 717 | 874 | 1140 | 645 |
| 875 | 0.03 | 0.14 | 0.25 | 0.81 | 1 | Eif1ad | eukaryotic translation initiation factor 1A c | 833 | 796 | 719 | 903 | 884 | 802 | 995 | 744 | 944 | 1123 | 876 |
| 865 | 0.15 | 0.19 | 0.82 | 0.41 | 1 | Hcn4 | hyperpolarization-activated, cyclic nucleo | 173 | 375 | 191 | 1162 | 1086 | 866 | 995 | 1483 | 1247 | 1122 | 811 |
| 850 | -0.01 | 0.17 | -0.04 | 0.96 | 1 | Fbxl2 | F-box and leucine-rich repeat protein 2 | 254 | 297 | 269 | 1076 | 814 | 1387 | 995 | 907 | 773 | 1256 | 1319 |
| 1183 | -0.08 | 0.18 | -0.44 | 0.66 | 1 | Grk6 | G protein-coupled receptor kinase 6 | 2100 | 1440 | 1216 | 1095 | 1161 | 983 | 994 | 1156 | 939 | 994 | 938 |
| 910 | -0.19 | 0.16 | -1.21 | 0.23 | 1 | Manea | mannosidase, endo-alpha | 1347 | 1097 | 923 | 935 | 890 | 727 | 994 | 633 | 834 | 877 | 754 |
| 1271 | 0.00 | 0.17 | 0.03 | 0.98 | 1 | Enc1 | ectodermal-neural cortex 1 | 2760 | 2090 | 1545 | 1224 | 882 | 656 | 993 | 1395 | 690 | 1039 | 708 |
| 1022 | 0.11 | 0.12 | 0.87 | 0.38 | 1 | Phf10 | PHD finger protein 10 | 1007 | 1083 | 923 | 915 | 908 | 1139 | 993 | 893 | 1052 | 1036 | 1292 |
| 873 | -0.06 | 0.13 | -0.44 | 0.66 | 1 | Sdad1 | SDA1 domain containing 1 | 962 | 802 | 705 | 672 | 787 | 1201 | 993 | 782 | 721 | 855 | 1121 |
| 1090 | -0.11 | 0.15 | -0.71 | 0.48 | 1 | Golm1 | golgi membrane protein 1 | 1288 | 1033 | 1297 | 1245 | 1043 | 1056 | 992 | 1026 | 977 | 889 | 1142 |
| 852 | -0.04 | 0.20 | -0.19 | 0.85 | 1 | Rab11fip1 | RAB11 family interacting protein 1 (class | 736 | 998 | 545 | 829 | 898 | 877 | 992 | 839 | 954 | 899 | 807 |
| 1313 | 0.14 | 0.14 | 1.01 | 0.31 | 1 | Lurap1l | leucine rich adaptor protein 1-like | 1680 | 1473 | 2050 | 1246 | 1248 | 915 | 991 | 1170 | 1262 | 1350 | 1052 |
| 1076 | -0.25 | 0.19 | -1.26 | 0.21 | 1 | Rnf111 | ring finger 111 | 941 | 1154 | 958 | 1231 | 989 | 1522 | 991 | 677 | 1196 | 782 | 1396 |
| 974 | -0.03 | 0.16 | -0.20 | 0.84 | 1 | Gcsh | glycine cleavage system protein H (aminc | 964 | 1186 | 1578 | 808 | 998 | 740 | 991 | 898 | 905 | 903 | 740 |
| 1406 | 0.11 | 0.15 | 0.73 | 0.46 | 1 | Rab24 | RAB24, member RAS oncogene family | 824 | 759 | 1064 | 1595 | 1470 | 2098 | 990 | 1664 | 1503 | 1121 | 2376 |
| 973 | 0.02 | 0.14 | 0.14 | 0.89 | 1 | Mpp7 | membrane protein, palmitoylated 7 (MAG | 644 | 762 | 567 | 1215 | 928 | 1221 | 990 | 1067 | 1141 | 1060 | 1103 |

| | | | | | | | | | | | | | | | | | | |
|---|---|---|---|---|---|---|---|---|---|---|---|---|---|---|---|---|---|---|
| 944 | 0.12 | 0.12 | 0.99 | 0.32 | 1 | Stam | signal transducing adaptor molecule (SH3 | 955 | 827 | 748 | 857 | 893 | 1031 | 990 | 988 | 1016 | 1074 | 1001 |
| 1406 | -0.17 | 0.12 | -1.45 | 0.15 | 1 | Ubp1 | upstream binding protein 1 | 1038 | 948 | 1065 | 1202 | 1284 | 3066 | 989 | 898 | 1134 | 1024 | 2817 |
| 1189 | 0.03 | 0.13 | 0.23 | 0.82 | 1 | 2210016L21Rik | RIKEN cDNA 2210016L21 gene | 1383 | 1325 | 1343 | 1141 | 1096 | 1237 | 989 | 1344 | 1024 | 974 | 1226 |
| 997 | -0.03 | 0.15 | -0.19 | 0.85 | 1 | St7l | suppression of tumorigenicity 7-like | 781 | 968 | 784 | 1076 | 903 | 1296 | 989 | 823 | 999 | 1077 | 1271 |
| 983 | 0.38 | 0.20 | 1.94 | 0.05 | 1 | Pank1 | pantothenate kinase 1 | 647 | 521 | 737 | 1005 | 981 | 871 | 989 | 1697 | 1006 | 1356 | 999 |
| 973 | -0.01 | 0.11 | -0.10 | 0.92 | 1 | Pfdn6 | prefoldin subunit 6 | 1231 | 1105 | 1259 | 819 | 926 | 833 | 989 | 882 | 1003 | 907 | 753 |
| 971 | 0.04 | 0.15 | 0.25 | 0.80 | 1 | Ddx25 | DEAD (Asp-Glu-Ala-Asp) box polypeptide | 577 | 631 | 551 | 1242 | 1297 | 873 | 989 | 1240 | 1056 | 1350 | 879 |
| 826 | -0.02 | 0.12 | -0.20 | 0.84 | 1 | Zfp799 | zinc finger protein 799 | 888 | 711 | 737 | 739 | 824 | 856 | 989 | 751 | 774 | 887 | 934 |
| 1342 | 0.01 | 0.18 | 0.03 | 0.98 | 1 | Chd1 | chromodomain helicase DNA binding pro | 1259 | 1901 | 985 | 1438 | 1081 | 1814 | 988 | 1264 | 1357 | 881 | 1797 |
| 1326 | 0.05 | 0.12 | 0.45 | 0.65 | 1 | Phtf1 | putative homeodomain transcription facto | 1157 | 1426 | 1281 | 1663 | 1105 | 1543 | 988 | 1515 | 1250 | 1166 | 1495 |
| 1114 | 0.15 | 0.14 | 1.07 | 0.28 | 1 | Ppp2r5d | protein phosphatase 2, regulatory subuni | 1341 | 1008 | 1203 | 1099 | 1074 | 928 | 988 | 1200 | 1228 | 1351 | 839 |
| 997 | -0.06 | 0.13 | -0.46 | 0.65 | 1 | Bcl7a | B cell CLL/lymphoma 7A | 709 | 766 | 525 | 1185 | 949 | 1478 | 988 | 1082 | 930 | 1128 | 1231 |
| 812 | 0.02 | 0.17 | 0.15 | 0.88 | 1 | Ndrg4 | N-myc downstream regulated gene 4 | 847 | 780 | 896 | 782 | 707 | 699 | 988 | 718 | 1015 | 766 | 737 |
| 1077 | 0.12 | 0.15 | 0.82 | 0.41 | 1 | Ddx52 | DEAD (Asp-Glu-Ala-Asp) box polypeptide | 1222 | 973 | 869 | 1036 | 1077 | 1088 | 987 | 957 | 1176 | 1114 | 1342 |
| 1045 | 0.11 | 0.16 | 0.64 | 0.52 | 1 | Pno1 | partner of NOB1 homolog | 858 | 886 | 1001 | 1101 | 1191 | 923 | 987 | 814 | 1308 | 1276 | 1149 |
| 996 | 0.02 | 0.16 | 0.14 | 0.89 | 1 | Arl5b | ADP-ribosylation factor-like 5B | 1192 | 1168 | 917 | 994 | 855 | 967 | 987 | 1121 | 987 | 739 | 1024 |
| 980 | 0.08 | 0.12 | 0.68 | 0.50 | 1 | Abcb8 | ATP-binding cassette, sub-family B (MDR | 949 | 1010 | 866 | 1032 | 975 | 855 | 987 | 1159 | 1045 | 1123 | 775 |
| 887 | -0.19 | 0.18 | -1.01 | 0.31 | 1 | Tnfrsf21 | tumor necrosis factor receptor superfamily | 852 | 700 | 496 | 1076 | 867 | 1160 | 987 | 687 | 869 | 875 | 1192 |
| 884 | 0.01 | 0.23 | 0.04 | 0.97 | 1 | Ang | angiogenin, ribonuclease, RNase A famil | 462 | 911 | 1080 | 908 | 885 | 792 | 987 | 1210 | 991 | 836 | 669 |
| 863 | -0.12 | 0.13 | -0.92 | 0.36 | 1 | Pacsin1 | protein kinase C and casein kinase subst | 841 | 696 | 822 | 1007 | 947 | 779 | 987 | 937 | 843 | 877 | 762 |
| 704 | 0.44 | 0.29 | 1.49 | 0.14 | 1 | Gm12892 | | 273 | 151 | 234 | 932 | 569 | 468 | 987 | 1659 | 529 | 1310 | 633 |
| 683 | -0.04 | 0.20 | -0.19 | 0.85 | 1 | Derl3 | Der1-like domain family, member 3 | 411 | 368 | 326 | 1119 | 510 | 631 | 987 | 806 | 548 | 1177 | 624 |
| 1200 | 0.01 | 0.13 | 0.12 | 0.91 | 1 | Pwwp3a | PWWP domain containing 3A, DNA repai | 1196 | 1073 | 1238 | 938 | 903 | 2004 | 986 | 1106 | 865 | 912 | 1980 |
| 1161 | 0.06 | 0.15 | 0.39 | 0.70 | 1 | Ppp1r21 | protein phosphatase 1, regulatory subuni | 1301 | 1026 | 1313 | 1172 | 1075 | 1228 | 986 | 1018 | 1207 | 1354 | 1096 |
| 949 | 0.23 | 0.17 | 1.39 | 0.16 | 1 | Dnajb6 | DnaJ heat shock protein family (Hsp40) n | 835 | 771 | 1030 | 831 | 713 | 1068 | 986 | 1018 | 1119 | 1007 | 1062 |
| 1064 | 0.24 | 0.17 | 1.41 | 0.16 | 1 | Siah2 | siah E3 ubiquitin protein ligase 2 | 591 | 489 | 447 | 1108 | 1226 | 1351 | 985 | 1620 | 1358 | 1131 | 1399 |
| 986 | 0.01 | 0.15 | 0.04 | 0.97 | 1 | Pigv | phosphatidylinositol glycan anchor biosyı | 550 | 834 | 866 | 1237 | 1044 | 1024 | 985 | 1259 | 980 | 1032 | 1032 |
| 923 | 0.00 | 0.15 | 0.00 | 1.00 | 1 | Mxd1 | MAX dimerization protein 1 | 971 | 885 | 995 | 712 | 942 | 997 | 985 | 651 | 1184 | 839 | 998 |
| 879 | 0.09 | 0.13 | 0.68 | 0.50 | 1 | Lrrc61 | leucine rich repeat containing 61 | 796 | 872 | 791 | 850 | 946 | 715 | 985 | 876 | 1022 | 1054 | 762 |
| 965 | 0.06 | 0.16 | 0.40 | 0.69 | 1 | Tdrkh | tudor and KH domain containing protein | 672 | 931 | 989 | 1026 | 953 | 957 | 984 | 1130 | 967 | 1115 | 893 |
| 1127 | -0.06 | 0.12 | -0.50 | 0.62 | 1 | Dalrd3 | DALR anticodon binding domain containi | 734 | 869 | 848 | 1169 | 950 | 2016 | 983 | 1038 | 942 | 1077 | 1768 |
| 987 | 0.08 | 0.14 | 0.62 | 0.54 | 1 | Cntln | centlein, centrosomal protein | 824 | 921 | 999 | 830 | 987 | 1129 | 983 | 789 | 1071 | 1097 | 1227 |
| 957 | 0.02 | 0.16 | 0.14 | 0.89 | 1 | Parp10 | poly (ADP-ribose) polymerase family, mei | 1018 | 892 | 1060 | 746 | 972 | 1053 | 983 | 1044 | 862 | 810 | 1082 |
| 952 | -0.02 | 0.13 | -0.17 | 0.87 | 1 | Cbll1 | Casitas B-lineage lymphoma-like 1 | 818 | 986 | 679 | 1133 | 935 | 979 | 983 | 1006 | 987 | 1052 | 911 |
| 1477 | -0.23 | 0.14 | -1.63 | 0.10 | 1 | Zfp395 | zinc finger protein 395 | 3102 | 2488 | 3675 | 824 | 940 | 1012 | 982 | 745 | 878 | 957 | 643 |
| 1268 | -0.16 | 0.18 | -0.89 | 0.37 | 1 | Rasd1 | RAS, dexamethasone-induced 1 | 1568 | 961 | 1675 | 445 | 1053 | 2661 | 982 | 315 | 950 | 1131 | 2205 |
| 1081 | -0.14 | 0.13 | -1.09 | 0.28 | 1 | Kdm3a | lysine (K)-specific demethylase 3A | 916 | 975 | 957 | 1130 | 999 | 1602 | 982 | 937 | 1072 | 768 | 1555 |
| 944 | -0.08 | 0.14 | -0.59 | 0.55 | 1 | Dzip3 | DAZ interacting protein 3, zinc finger | 838 | 1042 | 815 | 814 | 838 | 1325 | 982 | 827 | 853 | 809 | 1243 |
| 805 | 0.26 | 0.16 | 1.67 | 0.09 | 1 | Abhd6 | abhydrolase domain containing 6 | 354 | 350 | 425 | 1033 | 768 | 732 | 981 | 1230 | 1002 | 1077 | 902 |
| 1018 | -0.16 | 0.15 | -1.07 | 0.28 | 1 | Usp3 | ubiquitin specific peptidase 3 | 1255 | 1433 | 1188 | 768 | 1075 | 1050 | 980 | 775 | 922 | 811 | 944 |
| 953 | -0.11 | 0.14 | -0.77 | 0.44 | 1 | Uck1 | uridine-cytidine kinase 1 | 1021 | 877 | 1100 | 963 | 903 | 1035 | 980 | 966 | 827 | 967 | 842 |
| 930 | -0.15 | 0.13 | -1.22 | 0.22 | 1 | Trmt112 | tRNA methyltransferase 11-2 | 979 | 1029 | 900 | 1010 | 1042 | 822 | 980 | 1029 | 865 | 782 | 791 |
| 888 | -0.02 | 0.13 | -0.15 | 0.88 | 1 | Galnt12 | polypeptide N-acetylgalactosaminyltransf | 668 | 741 | 741 | 1078 | 876 | 890 | 980 | 1066 | 909 | 1036 | 779 |
| 872 | -0.01 | 0.15 | -0.05 | 0.96 | 1 | Chchd4 | coiled-coil-helix-coiled-coil-helix domain | 770 | 909 | 731 | 850 | 947 | 811 | 980 | 1081 | 940 | 795 | 780 |
| 794 | 0.13 | 0.17 | 0.76 | 0.45 | 1 | Slc5a6 | solute carrier family 5 (sodium-dependen | 454 | 378 | 466 | 866 | 937 | 729 | 980 | 1132 | 941 | 1247 | 609 |
| 943 | -0.06 | 0.15 | -0.43 | 0.67 | 1 | Nol9 | nucleolar protein 9 | 900 | 1184 | 797 | 920 | 907 | 1022 | 979 | 899 | 824 | 1045 | 901 |
| 890 | -0.03 | 0.13 | -0.23 | 0.82 | 1 | Ndufb1-ps | | 752 | 913 | 1028 | 998 | 930 | 680 | 979 | 1002 | 954 | 857 | 699 |
| 1033 | 0.29 | 0.23 | 1.29 | 0.20 | 1 | Rps28 | ribosomal protein S28 | 1422 | 1305 | 1740 | 680 | 760 | 655 | 978 | 1425 | 881 | 793 | 721 |
| 1335 | 0.03 | 0.11 | 0.27 | 0.79 | 1 | Pacsin2 | protein kinase C and casein kinase subst | 2578 | 1932 | 2230 | 707 | 1053 | 1195 | 977 | 724 | 1041 | 1088 | 1157 |
| 1090 | -0.17 | 0.18 | -0.93 | 0.35 | 1 | Kcnk1 | potassium channel, subfamily K, member | 998 | 1327 | 1558 | 1360 | 1062 | 915 | 977 | 801 | 1051 | 1065 | 882 |
| 1050 | -0.06 | 0.19 | -0.30 | 0.76 | 1 | Utp4 | UTP4 small subunit processome compon | 945 | 1127 | 874 | 1199 | 1047 | 1164 | 977 | 759 | 1149 | 1240 | 1070 |
| 1043 | 0.06 | 0.13 | 0.46 | 0.65 | 1 | Dcun1d5 | DCN1, defective in cullin neddylation 1, d | 1023 | 888 | 760 | 843 | 754 | 1727 | 977 | 948 | 836 | 833 | 1885 |
| 1976 | -0.17 | 0.14 | -1.15 | 0.25 | 1 | Pnisr | PNN interacting serine/arginine-rich | 647 | 996 | 766 | 1003 | 938 | 7036 | 976 | 752 | 864 | 896 | 6862 |
| 1114 | 0.01 | 0.11 | 0.05 | 0.96 | 1 | Tcerg1 | transcription elongation regulator 1 (CA15 | 1235 | 1043 | 1133 | 924 | 967 | 1554 | 976 | 973 | 1033 | 900 | 1512 |
| 961 | -0.07 | 0.11 | -0.63 | 0.53 | 1 | Dgcr8 | DGCR8, microprocessor complex subunit | 1087 | 1015 | 1004 | 948 | 882 | 1013 | 976 | 916 | 881 | 855 | 989 |
| 756 | -0.25 | 0.14 | -1.83 | 0.07 | 1 | Pon3 | paraoxonase 3 | 1034 | 703 | 719 | 922 | 795 | 501 | 976 | 679 | 690 | 850 | 450 |
| 1033 | -0.07 | 0.14 | -0.52 | 0.61 | 1 | Acsf3 | acyl-CoA synthetase family member 3 | 928 | 975 | 1036 | 1182 | 1159 | 1003 | 975 | 883 | 1060 | 1130 | 1034 |
| 891 | -0.21 | 0.13 | -1.59 | 0.11 | 1 | Endov | endonuclease V | 959 | 940 | 864 | 874 | 817 | 1095 | 975 | 649 | 782 | 766 | 1076 |
| 1320 | -0.15 | 0.13 | -1.18 | 0.24 | 1 | Trp53i11 | transformation related protein 53 inducibl | 2234 | 2577 | 2330 | 1075 | 941 | 880 | 974 | 1058 | 689 | 894 | 873 |
| 1062 | -0.24 | 0.16 | -1.53 | 0.13 | 1 | Top3b | topoisomerase (DNA) III beta | 1120 | 987 | 958 | 964 | 1007 | 1692 | 974 | 595 | 884 | 1056 | 1441 |

| | | | | | | | | | | | | | | | | | |
|---|---|---|---|---|---|---|---|---|---|---|---|---|---|---|---|---|---|
| 1037 | -0.10 | 0.15 | -0.64 | 0.52 | 1 | Vdac3 | voltage-dependent anion channel 3 | 1295 | 1030 | 1273 | 1208 | 1057 | 806 | 974 | 993 | 877 | 1039 | 857 |
| 1011 | 0.05 | 0.13 | 0.39 | 0.70 | 1 | Oard1 | O-acyl-ADP-ribose deacylase 1 | 897 | 989 | 909 | 992 | 1161 | 972 | 974 | 982 | 1061 | 1150 | 1040 |
| 986 | -0.11 | 0.13 | -0.83 | 0.41 | 1 | Wdr18 | WD repeat domain 18 | 1235 | 1022 | 988 | 896 | 1118 | 953 | 974 | 907 | 897 | 994 | 860 |
| 872 | -0.47 | 0.25 | -1.84 | 0.07 | 1 | Pgbd5 | piggyBac transposable element derived 5 | 828 | 742 | 985 | 1119 | 1013 | 903 | 974 | 402 | 728 | 1141 | 757 |
| 992 | -0.12 | 0.14 | -0.91 | 0.36 | 1 | Mttp | microsomal triglyceride transfer protein | 1141 | 1332 | 1159 | 975 | 1106 | 752 | 973 | 849 | 869 | 969 | 785 |
| 968 | -0.06 | 0.14 | -0.39 | 0.69 | 1 | Tecpr2 | tectonin beta-propeller repeat containing | 611 | 547 | 813 | 1033 | 1130 | 1300 | 973 | 1077 | 1084 | 1003 | 1077 |
| 1069 | -0.05 | 0.12 | -0.39 | 0.70 | 1 | Zfp710 | zinc finger protein 710 | 1545 | 1387 | 1323 | 1018 | 912 | 895 | 972 | 918 | 842 | 1164 | 785 |
| 955 | 0.03 | 0.13 | 0.20 | 0.84 | 1 | Rdh14 | retinol dehydrogenase 14 (all-trans and 9 | 861 | 880 | 762 | 1002 | 1081 | 911 | 972 | 911 | 1080 | 1025 | 1021 |
| 745 | -0.33 | 0.19 | -1.68 | 0.09 | 1 | Zfp365 | zinc finger protein 365 | 1085 | 1358 | 825 | 900 | 348 | 540 | 972 | 731 | 315 | 674 | 444 |
| 726 | 0.01 | 0.18 | 0.08 | 0.94 | 1 | Pdp2 | pyruvate dehyrogenase phosphatase cata | 395 | 436 | 489 | 677 | 849 | 823 | 972 | 965 | 817 | 772 | 796 |
| 1169 | -0.02 | 0.14 | -0.12 | 0.90 | 1 | Dpysl2 | dihydropyrimidinase-like 2 | 1295 | 1297 | 1187 | 1033 | 1153 | 1414 | 970 | 883 | 1112 | 1242 | 1271 |
| 1004 | 0.21 | 0.11 | 1.90 | 0.06 | 1 | Dnajc2 | DnaJ heat shock protein family (Hsp40) m | 988 | 1047 | 963 | 907 | 861 | 985 | 970 | 1030 | 971 | 1051 | 1266 |
| 1429 | 0.10 | 0.16 | 0.63 | 0.53 | 1 | Trim39 | tripartite motif-containing 39 | 688 | 1092 | 803 | 945 | 1019 | 3633 | 969 | 1067 | 1133 | 1155 | 3214 |
| 956 | 0.03 | 0.19 | 0.17 | 0.87 | 1 | Prrg2 | proline-rich Gla (G-carboxyglutamic acid) | 685 | 540 | 809 | 1208 | 1075 | 925 | 969 | 1475 | 1023 | 916 | 892 |
| 935 | -0.26 | 0.15 | -1.76 | 0.08 | 1 | Snapc1 | small nuclear RNA activating complex, po | 859 | 861 | 776 | 946 | 951 | 1316 | 969 | 637 | 775 | 800 | 1395 |
| 1159 | 0.02 | 0.12 | 0.17 | 0.87 | 1 | Ric1 | RAB6A GEF complex partner 1 | 1457 | 1321 | 1373 | 1000 | 853 | 1449 | 968 | 1112 | 891 | 867 | 1463 |
| 1009 | -0.21 | 0.18 | -1.20 | 0.23 | 1 | Bend7 | BEN domain containing 7 | 1211 | 1772 | 1173 | 927 | 796 | 1048 | 968 | 736 | 903 | 768 | 801 |
| 991 | -0.20 | 0.15 | -1.34 | 0.18 | 1 | Gale | galactose-4-epimerase, UDP | 467 | 390 | 573 | 1069 | 1168 | 1795 | 968 | 687 | 1095 | 976 | 1716 |
| 944 | -0.30 | 0.21 | -1.40 | 0.16 | 1 | Tent4a | terminal nucleotidyltransferase 4A | 807 | 1462 | 821 | 1122 | 1011 | 931 | 968 | 719 | 942 | 940 | 663 |
| 928 | -0.08 | 0.19 | -0.44 | 0.66 | 1 | Nek1 | NIMA (never in mitosis gene a)-related ex | 1002 | 701 | 799 | 854 | 897 | 1175 | 968 | 516 | 1096 | 894 | 1312 |
| 917 | -0.15 | 0.15 | -0.96 | 0.34 | 1 | Tmem170b | transmembrane protein 170B | 758 | 837 | 772 | 1063 | 901 | 1114 | 968 | 714 | 961 | 880 | 1124 |
| 1786 | -0.13 | 0.18 | -0.72 | 0.47 | 1 | Csrp1 | cysteine and glycine-rich protein 1 | 6486 | 3429 | 3163 | 993 | 899 | 565 | 966 | 911 | 818 | 908 | 506 |
| 1146 | -0.05 | 0.16 | -0.28 | 0.78 | 1 | Zfp651 | zinc finger protein 651 | 2005 | 1593 | 1750 | 809 | 1042 | 825 | 966 | 586 | 1071 | 1197 | 766 |
| 957 | -0.01 | 0.14 | -0.08 | 0.94 | 1 | Cttnbp2 | cortactin binding protein 2 | 241 | 155 | 258 | 1010 | 1118 | 1839 | 966 | 843 | 1185 | 1074 | 1836 |
| 901 | 0.13 | 0.15 | 0.83 | 0.41 | 1 | Pop5 | processing of precursor 5, ribonuclease P | 675 | 571 | 848 | 927 | 1033 | 816 | 966 | 1130 | 980 | 1105 | 863 |
| 1348 | 0.14 | 0.14 | 1.02 | 0.31 | 1 | Pfkp | phosphofructokinase, platelet | 2778 | 1861 | 2016 | 1049 | 1040 | 824 | 965 | 1113 | 1085 | 1191 | 904 |
| 1187 | 0.18 | 0.16 | 1.11 | 0.27 | 1 | Dvl1 | dishevelled segment polarity protein 1 | 982 | 693 | 1079 | 985 | 872 | 2075 | 965 | 1153 | 1067 | 1113 | 2076 |
| 1134 | -0.12 | 0.15 | -0.83 | 0.41 | 1 | Baiap2l1 | BAI1-associated protein 2-like 1 | 1300 | 1546 | 1414 | 922 | 1038 | 1333 | 965 | 703 | 906 | 993 | 1356 |
| 1067 | -0.05 | 0.13 | -0.35 | 0.72 | 1 | Slc4a1ap | solute carrier family 4 (anion exchanger), | 898 | 1101 | 1221 | 1079 | 1043 | 1224 | 965 | 974 | 917 | 962 | 1351 |
| 916 | -0.17 | 0.18 | -0.92 | 0.36 | 1 | Fam83h | family with sequence similarity 83, membe | 468 | 1002 | 788 | 1310 | 854 | 1024 | 965 | 1014 | 839 | 857 | 956 |
| 1285 | 0.08 | 0.12 | 0.61 | 0.54 | 1 | Sntb2 | syntrophin, basic 2 | 2463 | 1932 | 1732 | 981 | 872 | 1081 | 964 | 1053 | 953 | 1141 | 965 |
| 939 | -0.10 | 0.19 | -0.54 | 0.59 | 1 | AI661453 | expressed sequence AI661453 | 1005 | 1091 | 1024 | 969 | 1102 | 636 | 964 | 762 | 894 | 1380 | 503 |
| 848 | -0.26 | 0.22 | -1.15 | 0.25 | 1 | Ndufa12 | NADH:ubiquinone oxidoreductase subun | 838 | 837 | 918 | 782 | 1083 | 783 | 964 | 400 | 1012 | 1020 | 695 |
| 782 | 0.08 | 0.14 | 0.56 | 0.58 | 1 | Ttc21b | tetratricopeptide repeat domain 21B | 798 | 796 | 719 | 656 | 702 | 746 | 964 | 777 | 790 | 836 | 816 |
| 1399 | -0.05 | 0.14 | -0.33 | 0.74 | 1 | Sec23a | SEC23 homolog A, COPII coat complex c | 1930 | 1762 | 1318 | 1442 | 1050 | 1829 | 963 | 1353 | 964 | 1077 | 1696 |
| 1253 | 0.09 | 0.13 | 0.68 | 0.49 | 1 | Phax | phosphorylated adaptor for RNA export | 1369 | 1431 | 1443 | 1378 | 1266 | 1030 | 963 | 1337 | 1130 | 1197 | 1236 |
| 1052 | -0.03 | 0.11 | -0.29 | 0.77 | 1 | Vti1a | vesicle transport through interaction with t | 1281 | 1169 | 1398 | 1016 | 1030 | 891 | 963 | 892 | 1139 | 925 | 871 |
| 986 | -0.01 | 0.14 | -0.06 | 0.95 | 1 | Trnt1 | tRNA nucleotidyl transferase, CCA-adding | 1005 | 1017 | 1133 | 847 | 945 | 1075 | 963 | 743 | 822 | 1045 | 1252 |
| 726 | 0.17 | 0.13 | 1.31 | 0.19 | 1 | Cgrrf1 | cell growth regulator with ring finger doma | 556 | 399 | 453 | 782 | 718 | 641 | 963 | 826 | 813 | 1036 | 798 |
| 1202 | 0.09 | 0.10 | 0.87 | 0.39 | 1 | Nop56 | NOP56 ribonucleoprotein | 1278 | 1206 | 1357 | 985 | 1102 | 1477 | 961 | 1060 | 1191 | 936 | 1663 |
| 1136 | 0.01 | 0.17 | 0.05 | 0.96 | 1 | Tubgcp2 | tubulin, gamma complex associated prote | 1452 | 1162 | 1396 | 1155 | 1137 | 962 | 961 | 1432 | 962 | 979 | 898 |
| 1060 | -0.20 | 0.16 | -1.22 | 0.22 | 1 | Wdfy2 | WD repeat and FYVE domain containing | 1122 | 1346 | 1562 | 808 | 998 | 1307 | 961 | 586 | 1027 | 899 | 1048 |
| 1025 | 0.00 | 0.14 | -0.03 | 0.98 | 1 | Slc25a28 | solute carrier family 25, member 28 | 592 | 647 | 817 | 863 | 1145 | 1636 | 961 | 824 | 993 | 1082 | 1714 |
| 1008 | 0.15 | 0.14 | 1.10 | 0.27 | 1 | Rp9 | retinitis pigmentosa 9 (human) | 1154 | 1350 | 1290 | 782 | 942 | 762 | 961 | 1043 | 1031 | 1050 | 730 |
| 977 | 0.14 | 0.14 | 1.02 | 0.31 | 1 | Vrk3 | vaccinia related kinase 3 | 823 | 1083 | 1005 | 835 | 962 | 958 | 961 | 1116 | 1051 | 1027 | 926 |
| 919 | 0.09 | 0.11 | 0.81 | 0.42 | 1 | Ralgps2 | Ral GEF with PH domain and SH3 bindin | 1079 | 863 | 877 | 807 | 890 | 870 | 961 | 933 | 866 | 1060 | 904 |
| 898 | -0.05 | 0.15 | -0.35 | 0.72 | 1 | Ccdc93 | coiled-coil domain containing 93 | 982 | 802 | 762 | 905 | 800 | 1065 | 961 | 769 | 934 | 781 | 1122 |
| 849 | 0.05 | 0.15 | 0.36 | 0.72 | 1 | Rpgr | retinitis pigmentosa GTPase regulator | 518 | 718 | 668 | 883 | 869 | 932 | 961 | 935 | 897 | 962 | 990 |
| 835 | -0.16 | 0.24 | -0.67 | 0.50 | 1 | Trpm3 | transient receptor potential cation channe | 147 | 327 | 277 | 908 | 1118 | 1442 | 961 | 666 | 1293 | 836 | 1205 |
| 1050 | 0.07 | 0.19 | 0.36 | 0.72 | 1 | Foxn3 | forkhead box N3 | 1148 | 1023 | 1004 | 999 | 826 | 1290 | 960 | 1453 | 890 | 746 | 1214 |
| 854 | 0.00 | 0.15 | 0.02 | 0.98 | 1 | Gm45606 | | 462 | 725 | 686 | 1235 | 908 | 647 | 960 | 1177 | 1001 | 991 | 605 |
| 685 | 0.13 | 0.18 | 0.73 | 0.47 | 1 | Fam217b | family with sequence similarity 217, meml | 275 | 167 | 232 | 862 | 830 | 630 | 960 | 1062 | 920 | 936 | 662 |
| 1145 | 0.22 | 0.15 | 1.52 | 0.13 | 1 | Polr1c | polymerase (RNA) I polypeptide C | 1432 | 1425 | 1209 | 1142 | 880 | 921 | 959 | 1659 | 912 | 1096 | 966 |
| 926 | -0.24 | 0.16 | -1.49 | 0.14 | 1 | Ammecr1l | AMME chromosomal region gene 1-like | 989 | 1325 | 904 | 811 | 916 | 1076 | 959 | 792 | 773 | 650 | 986 |
| 810 | -0.13 | 0.15 | -0.85 | 0.39 | 1 | 1110051M20Ri | RIKEN cDNA 1110051M20 gene | 643 | 852 | 842 | 873 | 822 | 776 | 959 | 820 | 782 | 782 | 757 |
| 1321 | 0.01 | 0.15 | 0.07 | 0.95 | 1 | Dynll1 | dynein light chain LC8-type 1 | 2519 | 2011 | 1916 | 843 | 1222 | 1006 | 958 | 977 | 1141 | 1052 | 882 |
| 1085 | 0.01 | 0.15 | 0.10 | 0.92 | 1 | Oxa1l | oxidase assembly 1-like | 1495 | 1303 | 1183 | 950 | 1075 | 969 | 958 | 1163 | 1056 | 906 | 881 |
| 1016 | 0.13 | 0.22 | 0.57 | 0.57 | 1 | P4ha2 | procollagen-proline, 2-oxoglutarate 4-dio | 1119 | 643 | 1180 | 1073 | 1097 | 812 | 958 | 1400 | 1121 | 871 | 905 |
| 988 | 0.03 | 0.16 | 0.19 | 0.85 | 1 | Dcbld2 | discoidin, CUB and LCCL domain contair | 1354 | 911 | 1008 | 965 | 853 | 951 | 957 | 704 | 1024 | 1152 | 985 |
| 1024 | -0.09 | 0.18 | -0.51 | 0.61 | 1 | Crtap | cartilage associated protein | 1796 | 1174 | 1032 | 1062 | 1023 | 697 | 956 | 757 | 907 | 1112 | 752 |

| | | | | | | | | | | | | | | | | | | |
|---|---|---|---|---|---|---|---|---|---|---|---|---|---|---|---|---|---|---|
| 895 | -0.14 | 0.14 | -1.02 | 0.31 | 1 | Dusp22 | dual specificity phosphatase 22 | 899 | 814 | 846 | 1065 | 992 | 806 | 956 | 931 | 1037 | 755 | 743 |
| 889 | 0.22 | 0.20 | 1.15 | 0.25 | 1 | Fuom | fucose mutarotase | 370 | 442 | 703 | 960 | 1131 | 765 | 956 | 1471 | 1083 | 1016 | 883 |
| 855 | -0.17 | 0.12 | -1.38 | 0.17 | 1 | Immp1l | IMP1 inner mitochondrial membrane pept | 610 | 653 | 676 | 1069 | 1136 | 800 | 956 | 949 | 972 | 764 | 820 |
| 1266 | -0.05 | 0.17 | -0.29 | 0.77 | 1 | Abhd17a | abhydrolase domain containing 17A | 1813 | 1334 | 1878 | 1340 | 1336 | 897 | 955 | 1393 | 1140 | 1019 | 821 |
| 815 | 0.07 | 0.14 | 0.46 | 0.64 | 1 | Pecr | peroxisomal trans-2-enoyl-CoA reductase | 473 | 571 | 611 | 943 | 879 | 791 | 955 | 1033 | 880 | 1023 | 805 |
| 1133 | -0.03 | 0.15 | -0.22 | 0.83 | 1 | Nfyb | nuclear transcription factor-Y beta | 1332 | 1498 | 1394 | 1053 | 948 | 1197 | 954 | 750 | 1030 | 1045 | 1263 |
| 1055 | 0.03 | 0.16 | 0.21 | 0.84 | 1 | Arl14ep | ADP-ribosylation factor-like 14 effector pr | 897 | 1491 | 1096 | 1046 | 958 | 1051 | 954 | 900 | 1002 | 979 | 1227 |
| 1301 | -0.01 | 0.12 | -0.11 | 0.91 | 1 | Fem1c | fem 1 homolog c | 1935 | 2148 | 1726 | 1144 | 1172 | 1003 | 953 | 1084 | 1139 | 1058 | 948 |
| 973 | 0.23 | 0.17 | 1.33 | 0.18 | 1 | Otulin | OTU deubiquitinase with linear linkage sp | 1064 | 926 | 886 | 754 | 1026 | 871 | 953 | 1279 | 903 | 1069 | 974 |
| 859 | 0.06 | 0.14 | 0.44 | 0.66 | 1 | Ss18l2 | SS18, nBAF chromatin remodeling compl | 484 | 452 | 729 | 796 | 966 | 1093 | 953 | 842 | 1098 | 952 | 1087 |
| 968 | -0.25 | 0.18 | -1.38 | 0.17 | 1 | Cdkn2aip | CDKN2A interacting protein | 1126 | 1012 | 1312 | 839 | 1017 | 1074 | 952 | 906 | 770 | 558 | 1083 |
| 859 | 0.02 | 0.13 | 0.18 | 0.85 | 1 | Nol12 | nucleolar protein 12 | 1147 | 966 | 1052 | 693 | 731 | 751 | 952 | 813 | 805 | 784 | 753 |
| 826 | -0.22 | 0.15 | -1.49 | 0.14 | 1 | Ell | elongation factor RNA polymerase II | 768 | 976 | 863 | 959 | 724 | 851 | 952 | 804 | 728 | 809 | 655 |
| 803 | -0.02 | 0.14 | -0.13 | 0.90 | 1 | Srp54c | signal recognition particle 54C | 404 | 387 | 574 | 644 | 1111 | 1058 | 952 | 673 | 1119 | 989 | 920 |
| 895 | -0.22 | 0.14 | -1.55 | 0.12 | 1 | Sf3a2 | splicing factor 3a, subunit 2 | 1063 | 1141 | 1275 | 843 | 870 | 744 | 951 | 646 | 910 | 809 | 591 |
| 845 | -0.01 | 0.15 | -0.08 | 0.94 | 1 | Zdhhc6 | zinc finger, DHHC domain containing 6 | 752 | 745 | 632 | 950 | 746 | 946 | 951 | 726 | 812 | 1077 | 958 |
| 1500 | 0.08 | 0.11 | 0.71 | 0.48 | 1 | Itga6 | integrin alpha 6 | 2642 | 2898 | 3242 | 916 | 939 | 947 | 950 | 963 | 964 | 1069 | 968 |
| 1108 | -0.14 | 0.21 | -0.65 | 0.52 | 1 | Zfp62 | zinc finger protein 62 | 1040 | 1201 | 812 | 1294 | 1004 | 1480 | 950 | 690 | 1124 | 867 | 1728 |
| 1065 | -0.02 | 0.12 | -0.21 | 0.83 | 1 | Smad1 | SMAD family member 1 | 1108 | 1383 | 1111 | 1119 | 940 | 1093 | 950 | 1020 | 1027 | 994 | 976 |
| 1160 | -0.19 | 0.13 | -1.42 | 0.15 | 1 | Pwwp3b | PWWP domain containing 3B | 1132 | 1686 | 1343 | 1330 | 891 | 1403 | 949 | 930 | 898 | 788 | 1405 |
| 1057 | 0.11 | 0.11 | 0.92 | 0.36 | 1 | Alyref | Aly/REF export factor | 1100 | 1173 | 1291 | 862 | 921 | 1139 | 949 | 876 | 998 | 975 | 1344 |
| 1457 | -0.08 | 0.17 | -0.50 | 0.61 | 1 | 4931406P16Ril | RIKEN cDNA 4931406P16 gene | 2802 | 2781 | 2218 | 1167 | 937 | 1187 | 948 | 763 | 1104 | 1085 | 1039 |
| 912 | 0.23 | 0.16 | 1.43 | 0.15 | 1 | Mrps34 | mitochondrial ribosomal protein S34 | 556 | 829 | 745 | 831 | 1016 | 840 | 948 | 1197 | 1046 | 1076 | 947 |
| 895 | 0.02 | 0.15 | 0.10 | 0.92 | 1 | Cenpc1 | centromere protein C1 | 918 | 641 | 762 | 855 | 894 | 1052 | 948 | 1001 | 867 | 851 | 1058 |
| 846 | -0.12 | 0.19 | -0.62 | 0.54 | 1 | Zkscan16 | zinc finger with KRAB and SCAN domains | 347 | 540 | 680 | 1114 | 876 | 1071 | 948 | 1082 | 818 | 746 | 1087 |
| 812 | -0.16 | 0.13 | -1.19 | 0.23 | 1 | Triqk | triple QxxK/R motif containing | 1126 | 1121 | 903 | 796 | 730 | 587 | 948 | 633 | 654 | 859 | 579 |
| 959 | 0.13 | 0.12 | 1.10 | 0.27 | 1 | Atr | ataxia telangiectasia and Rad3 related | 1021 | 898 | 921 | 854 | 841 | 1041 | 947 | 870 | 1066 | 960 | 1132 |
| 869 | -0.10 | 0.16 | -0.64 | 0.52 | 1 | Polr3f | polymerase (RNA) III (DNA directed) poly | 862 | 927 | 718 | 908 | 847 | 923 | 947 | 679 | 729 | 929 | 1086 |
| 854 | -0.29 | 0.27 | -1.07 | 0.28 | 1 | Dapl1 | death associated protein-like 1 | 166 | 278 | 182 | 1147 | 1256 | 1451 | 947 | 509 | 1299 | 930 | 1226 |
| 822 | -0.08 | 0.20 | -0.38 | 0.70 | 1 | Nudcd1 | NudC domain containing 1 | 836 | 770 | 812 | 551 | 925 | 937 | 947 | 414 | 812 | 1021 | 1018 |
| 790 | 0.09 | 0.16 | 0.59 | 0.56 | 1 | Aasdhppt | aminoadipate-semialdehyde dehydrogen | 636 | 508 | 611 | 654 | 746 | 1019 | 947 | 899 | 710 | 863 | 1093 |
| 1281 | -0.09 | 0.10 | -0.81 | 0.42 | 1 | Usp11 | ubiquitin specific peptidase 11 | 2190 | 2361 | 1967 | 933 | 965 | 1056 | 946 | 918 | 897 | 907 | 951 |
| 1033 | 0.15 | 0.16 | 0.96 | 0.34 | 1 | Scfd2 | Sec1 family domain containing 2 | 854 | 857 | 723 | 1015 | 1184 | 1067 | 946 | 1504 | 1085 | 1052 | 1079 |
| 871 | 0.01 | 0.21 | 0.05 | 0.96 | 1 | Klf5 | Kruppel-like factor 5 | 184 | 298 | 406 | 893 | 982 | 1548 | 946 | 905 | 994 | 1050 | 1380 |
| 1253 | 0.12 | 0.16 | 0.77 | 0.44 | 1 | Lbr | lamin B receptor | 2052 | 1466 | 1459 | 1144 | 920 | 1202 | 945 | 928 | 1173 | 1198 | 1301 |
| 1025 | -0.02 | 0.15 | -0.12 | 0.90 | 1 | Rnf31 | ring finger protein 31 | 1009 | 989 | 1044 | 1041 | 923 | 1231 | 945 | 908 | 884 | 1264 | 1036 |
| 976 | 0.15 | 0.16 | 0.98 | 0.33 | 1 | Pex13 | peroxisomal biogenesis factor 13 | 891 | 905 | 794 | 1128 | 777 | 997 | 945 | 1445 | 994 | 844 | 1022 |
| 1216 | 0.14 | 0.17 | 0.79 | 0.43 | 1 | P2rx4 | purinergic receptor P2X, ligand-gated ion | 1572 | 1170 | 1176 | 1284 | 1050 | 1206 | 944 | 1440 | 1051 | 1419 | 1062 |
| 950 | -0.13 | 0.13 | -0.99 | 0.32 | 1 | Dedd | death effector domain-containing | 844 | 1171 | 1014 | 950 | 878 | 1096 | 944 | 879 | 831 | 795 | 1050 |
| 911 | 0.16 | 0.15 | 1.10 | 0.27 | 1 | Mpv17 | MpV17 mitochondrial inner membrane pr | 675 | 704 | 882 | 905 | 849 | 962 | 944 | 1120 | 1024 | 1032 | 927 |
| 877 | -0.06 | 0.13 | -0.46 | 0.64 | 1 | Yif1b | Yip1 interacting factor homolog B (S. cere | 776 | 775 | 763 | 1018 | 1052 | 718 | 944 | 1120 | 891 | 943 | 653 |
| 780 | -0.12 | 0.15 | -0.78 | 0.44 | 1 | Rnf157 | ring finger protein 157 | 644 | 676 | 592 | 1028 | 945 | 563 | 944 | 988 | 965 | 671 | 567 |
| 1147 | -0.03 | 0.11 | -0.31 | 0.75 | 1 | Mex3c | mex3 RNA binding family member C | 1442 | 1825 | 1512 | 1238 | 891 | 900 | 942 | 1145 | 902 | 828 | 993 |
| 901 | 0.10 | 0.16 | 0.61 | 0.54 | 1 | Pde1c | phosphodiesterase 1C | 775 | 717 | 664 | 654 | 707 | 1402 | 941 | 563 | 838 | 1002 | 1650 |
| 974 | 0.19 | 0.13 | 1.52 | 0.13 | 1 | Mak16 | MAK16 homolog | 854 | 959 | 884 | 922 | 837 | 1035 | 940 | 1174 | 849 | 1106 | 1152 |
| 944 | -0.04 | 0.12 | -0.35 | 0.73 | 1 | Ppp1r8 | protein phosphatase 1, regulatory subuni | 1029 | 1081 | 879 | 1035 | 862 | 917 | 940 | 852 | 853 | 935 | 1002 |
| 998 | 0.04 | 0.12 | 0.30 | 0.76 | 1 | Nuak2 | NUAK family, SNF1-like kinase, 2 | 902 | 910 | 829 | 1184 | 953 | 1036 | 939 | 1256 | 989 | 1007 | 969 |
| 825 | -0.08 | 0.14 | -0.59 | 0.56 | 1 | Mipep | mitochondrial intermediate peptidase | 810 | 948 | 739 | 958 | 856 | 648 | 939 | 777 | 857 | 840 | 704 |
| 1287 | 0.20 | 0.13 | 1.53 | 0.13 | 1 | Nsa2 | NSA2 ribosome biogenesis homolog | 1981 | 2176 | 1837 | 941 | 1113 | 811 | 938 | 1025 | 1109 | 1114 | 1112 |
| 1092 | 0.23 | 0.15 | 1.50 | 0.13 | 1 | Gnl3 | guanine nucleotide binding protein-like 3 | 796 | 925 | 635 | 809 | 902 | 1817 | 938 | 943 | 1120 | 1104 | 2019 |
| 1025 | 0.17 | 0.11 | 1.54 | 0.12 | 1 | 2610008E11Ril | RIKEN cDNA 2610008E11 gene | 977 | 935 | 877 | 811 | 880 | 1368 | 938 | 982 | 893 | 1105 | 1506 |
| 929 | -0.02 | 0.17 | -0.12 | 0.90 | 1 | Tbc1d32 | TBC1 domain family, member 32 | 891 | 992 | 865 | 1048 | 939 | 838 | 938 | 728 | 1087 | 960 | 935 |
| 851 | -0.08 | 0.15 | -0.51 | 0.61 | 1 | Ccdc43 | coiled-coil domain containing 43 | 841 | 987 | 759 | 898 | 946 | 708 | 938 | 737 | 867 | 911 | 774 |
| 830 | 0.03 | 0.11 | 0.27 | 0.78 | 1 | Acad8 | acyl-Coenzyme A dehydrogenase family, | 873 | 716 | 666 | 791 | 669 | 1007 | 938 | 781 | 717 | 982 | 992 |
| 1080 | 0.01 | 0.12 | 0.08 | 0.94 | 1 | Cdkn2aipnl | CDKN2A interacting protein N-terminal lik | 1203 | 1331 | 1249 | 1194 | 1082 | 812 | 937 | 1145 | 1013 | 1170 | 745 |
| 929 | 0.03 | 0.14 | 0.18 | 0.86 | 1 | Zfp120 | zinc finger protein 120 | 806 | 797 | 983 | 950 | 1044 | 856 | 937 | 1101 | 906 | 871 | 970 |
| 909 | -0.14 | 0.14 | -1.01 | 0.31 | 1 | Zfp358 | zinc finger protein 358 | 874 | 892 | 1043 | 870 | 1178 | 800 | 937 | 978 | 856 | 822 | 751 |
| 951 | 0.07 | 0.15 | 0.48 | 0.63 | 1 | Rspry1 | ring finger and SPRY domain containing | 941 | 856 | 662 | 906 | 776 | 1303 | 936 | 913 | 1013 | 867 | 1286 |
| 888 | 0.17 | 0.13 | 1.31 | 0.19 | 1 | Exosc7 | exosome component 7 | 804 | 543 | 757 | 715 | 843 | 1129 | 936 | 914 | 960 | 972 | 1198 |
| 1649 | 0.03 | 0.12 | 0.28 | 0.78 | 1 | Khdc4 | KH domain containing 4, pre-mRNA splic | 1077 | 1601 | 1356 | 1093 | 869 | 4177 | 935 | 1130 | 1116 | 822 | 3962 |

| | | | | | | | | | | | | | | | | | | |
|---|---|---|---|---|---|---|---|---|---|---|---|---|---|---|---|---|---|---|
| 1116 | 0.00 | 0.13 | 0.00 | 1.00 | 1 | Acvr1 | activin A receptor, type 1 | 1567 | 1292 | 1579 | 934 | 1022 | 1017 | 935 | 907 | 1069 | 1114 | 844 |
| 1063 | 0.01 | 0.16 | 0.09 | 0.93 | 1 | Mrpl3 | mitochondrial ribosomal protein L3 | 1114 | 816 | 1042 | 1161 | 1036 | 1183 | 935 | 890 | 1068 | 1270 | 1175 |
| 1012 | -0.24 | 0.16 | -1.50 | 0.13 | 1 | Nr3c2 | nuclear receptor subfamily 3, group C, me | 1320 | 1830 | 1445 | 1118 | 817 | 677 | 935 | 731 | 783 | 904 | 571 |
| 914 | -0.04 | 0.14 | -0.26 | 0.80 | 1 | Zdhhc7 | zinc finger, DHHC domain containing 7 | 963 | 1013 | 747 | 873 | 959 | 933 | 935 | 1002 | 831 | 925 | 870 |
| 878 | -0.11 | 0.12 | -0.95 | 0.34 | 1 | C9orf72 | C9orf72, member of C9orf72-SMCR8 con | 903 | 900 | 785 | 989 | 840 | 912 | 935 | 798 | 828 | 908 | 863 |
| 1040 | 0.11 | 0.12 | 0.92 | 0.36 | 1 | Rab8b | RAB8B, member RAS oncogene family | 2236 | 1502 | 1609 | 689 | 642 | 668 | 934 | 706 | 807 | 997 | 653 |
| 982 | -0.22 | 0.16 | -1.39 | 0.17 | 1 | Rfx3 | regulatory factor X, 3 (influences HLA clas | 710 | 669 | 882 | 1048 | 962 | 1607 | 934 | 716 | 1134 | 717 | 1424 |
| 905 | 0.14 | 0.12 | 1.14 | 0.25 | 1 | Xpc | xeroderma pigmentosum, complementati | 837 | 669 | 809 | 862 | 914 | 926 | 934 | 1045 | 1000 | 974 | 987 |
| 888 | -0.08 | 0.13 | -0.57 | 0.57 | 1 | Yes1 | YES proto-oncogene 1, Src family tyrosin | 821 | 1010 | 894 | 971 | 803 | 907 | 933 | 879 | 825 | 949 | 777 |
| 872 | -0.05 | 0.13 | -0.42 | 0.68 | 1 | Gm12258 | predicted gene 12258 | 613 | 531 | 525 | 732 | 765 | 1579 | 933 | 699 | 819 | 742 | 1651 |
| 851 | 0.01 | 0.24 | 0.03 | 0.97 | 1 | Nabp1 | nucleic acid binding protein 1 | 446 | 322 | 187 | 858 | 878 | 1453 | 933 | 716 | 970 | 792 | 1811 |
| 1102 | 0.05 | 0.16 | 0.32 | 0.75 | 1 | Zfp318 | zinc finger protein 318 | 1075 | 1341 | 877 | 945 | 1002 | 1485 | 932 | 943 | 1096 | 1067 | 1362 |
| 1074 | -0.10 | 0.15 | -0.66 | 0.51 | 1 | Gm37494 | | 1066 | 980 | 776 | 957 | 815 | 1852 | 932 | 953 | 756 | 669 | 2054 |
| 1236 | -0.05 | 0.17 | -0.28 | 0.78 | 1 | Grik5 | glutamate receptor, ionotropic, kainate 5 ( | 1475 | 1203 | 1707 | 890 | 1020 | 1884 | 931 | 1157 | 882 | 817 | 1627 |
| 1142 | 0.06 | 0.11 | 0.50 | 0.61 | 1 | Nr1h2 | nuclear receptor subfamily 1, group H, me | 1410 | 1281 | 1277 | 802 | 1019 | 1460 | 931 | 901 | 924 | 1039 | 1513 |
| 1040 | 0.00 | 0.17 | 0.02 | 0.98 | 1 | Dmtf1 | cyclin D binding myb-like transcription fac | 763 | 882 | 998 | 845 | 825 | 1805 | 931 | 707 | 876 | 1104 | 1700 |
| 1003 | -0.01 | 0.15 | -0.05 | 0.96 | 1 | Tcp11l2 | t-complex 11 (mouse) like 2 | 1511 | 1246 | 1600 | 689 | 1127 | 596 | 931 | 562 | 1071 | 1038 | 666 |
| 887 | -0.09 | 0.14 | -0.66 | 0.51 | 1 | Cep57 | centrosomal protein 57 | 975 | 737 | 829 | 901 | 822 | 1063 | 931 | 748 | 900 | 764 | 1089 |
| 707 | -0.27 | 0.24 | -1.16 | 0.25 | 1 | Csmd2 | CUB and Sushi multiple domains 2 | 635 | 587 | 349 | 589 | 783 | 1097 | 931 | 451 | 757 | 690 | 904 |
| 996 | 0.06 | 0.18 | 0.35 | 0.73 | 1 | Czib | CXXC motif containing zinc binding prote | 920 | 774 | 986 | 869 | 920 | 1338 | 929 | 1269 | 917 | 783 | 1252 |
| 954 | 0.10 | 0.13 | 0.80 | 0.43 | 1 | Rngtt | RNA guanylyltransferase and 5'-phospha | 906 | 1069 | 1005 | 850 | 901 | 946 | 929 | 931 | 1034 | 1016 | 909 |
| 926 | 0.10 | 0.15 | 0.67 | 0.50 | 1 | Mea1 | male enhanced antigen 1 | 933 | 876 | 1035 | 905 | 931 | 761 | 929 | 1243 | 934 | 864 | 775 |
| 681 | 0.22 | 0.17 | 1.33 | 0.18 | 1 | Dusp4 | dual specificity phosphatase 4 | 362 | 238 | 216 | 816 | 791 | 541 | 929 | 1045 | 971 | 975 | 601 |
| 1084 | 0.00 | 0.18 | 0.01 | 0.99 | 1 | Usf3 | upstream transcription factor family memb | 931 | 1086 | 822 | 1020 | 990 | 1606 | 928 | 736 | 1152 | 1083 | 1575 |
| 1065 | 0.28 | 0.16 | 1.73 | 0.08 | 1 | Farsa | phenylalanyl-tRNA synthetase, alpha sub | 1072 | 757 | 1187 | 648 | 914 | 1523 | 928 | 1060 | 986 | 1131 | 1514 |
| 992 | 0.04 | 0.12 | 0.36 | 0.72 | 1 | Pias3 | protein inhibitor of activated STAT 3 | 978 | 1197 | 1067 | 1007 | 886 | 959 | 928 | 998 | 1015 | 991 | 891 |
| 903 | -0.05 | 0.11 | -0.50 | 0.62 | 1 | Zfp790 | zinc finger protein 790 | 786 | 814 | 711 | 864 | 862 | 1237 | 927 | 852 | 856 | 857 | 1174 |
| 624 | -0.08 | 0.28 | -0.28 | 0.78 | 1 | Slc6a19 | solute carrier family 6 (neurotransmitter tr | 28 | 47 | 86 | 977 | 744 | 766 | 927 | 1073 | 793 | 781 | 646 |
| 1235 | -0.01 | 0.17 | -0.05 | 0.96 | 1 | Mrtfb | myocardin related transcription factor B | 1918 | 1817 | 1837 | 909 | 827 | 1348 | 926 | 701 | 879 | 1282 | 1137 |
| 1039 | 0.13 | 0.17 | 0.73 | 0.46 | 1 | Zcchc17 | zinc finger, CCHC domain containing 17 | 1195 | 794 | 1072 | 951 | 1250 | 895 | 926 | 1198 | 1114 | 982 | 1055 |
| 1030 | 0.09 | 0.16 | 0.53 | 0.59 | 1 | Washc1 | WASH complex subunit 1 | 1152 | 1053 | 1203 | 1084 | 820 | 997 | 926 | 1270 | 980 | 787 | 1051 |
| 909 | 0.05 | 0.12 | 0.41 | 0.68 | 1 | Pigu | phosphatidylinositol glycan anchor biosyı | 1016 | 914 | 885 | 810 | 951 | 844 | 926 | 890 | 966 | 1003 | 799 |
| 848 | -0.01 | 0.15 | -0.07 | 0.95 | 1 | Emc4 | ER membrane protein complex subunit 4 | 659 | 554 | 685 | 1115 | 872 | 816 | 926 | 912 | 963 | 967 | 856 |
| 828 | -0.05 | 0.19 | -0.29 | 0.77 | 1 | Pde6d | phosphodiesterase 6D, cGMP-specific, ro | 679 | 671 | 921 | 1034 | 827 | 680 | 926 | 679 | 954 | 1039 | 693 |
| 1142 | 0.14 | 0.19 | 0.75 | 0.45 | 1 | Usp1 | ubiquitin specific peptidase 1 | 1441 | 1266 | 824 | 1065 | 1106 | 1199 | 925 | 1207 | 1287 | 1021 | 1219 |
| 1116 | 0.02 | 0.13 | 0.18 | 0.86 | 1 | Utp3 | UTP3 small subunit processome compon | 1659 | 1516 | 1351 | 988 | 1028 | 902 | 925 | 1091 | 944 | 924 | 948 |
| 1081 | 0.13 | 0.14 | 0.93 | 0.35 | 1 | Il13ra1 | interleukin 13 receptor, alpha 1 | 1623 | 1303 | 1323 | 909 | 1074 | 735 | 925 | 938 | 1259 | 994 | 803 |
| 1016 | 0.03 | 0.14 | 0.19 | 0.85 | 1 | Drg2 | developmentally regulated GTP binding p | 1236 | 1056 | 1083 | 821 | 1055 | 1068 | 925 | 1014 | 940 | 878 | 1096 |
| 807 | 0.00 | 0.14 | 0.00 | 1.00 | 1 | Mrpl57 | mitochondrial ribosomal protein L57 | 745 | 682 | 800 | 901 | 840 | 647 | 925 | 721 | 925 | 1041 | 652 |
| 2586 | -0.04 | 0.11 | -0.34 | 0.74 | 1 | Clk1 | CDC-like kinase 1 | 2086 | 1718 | 1997 | 1002 | 950 | 8295 | 924 | 767 | 1078 | 902 | 8724 |
| 1028 | -0.01 | 0.15 | -0.06 | 0.95 | 1 | Wdsub1 | WD repeat, SAM and U-box domain conta | 898 | 849 | 870 | 1089 | 894 | 1449 | 924 | 942 | 806 | 1222 | 1362 |
| 985 | -0.02 | 0.12 | -0.17 | 0.86 | 1 | Zfp672 | zinc finger protein 672 | 816 | 1053 | 809 | 969 | 885 | 1344 | 924 | 1025 | 878 | 933 | 1203 |
| 843 | 0.04 | 0.21 | 0.20 | 0.84 | 1 | Ikzf5 | IKAROS family zinc finger 5 | 834 | 1051 | 829 | 657 | 674 | 966 | 924 | 1001 | 825 | 596 | 911 |
| 832 | -0.24 | 0.22 | -1.11 | 0.27 | 1 | Tat | tyrosine aminotransferase | 226 | 393 | 378 | 1215 | 722 | 1559 | 924 | 892 | 592 | 894 | 1355 |
| 1047 | -0.14 | 0.15 | -0.94 | 0.35 | 1 | Rexo1 | REX1, RNA exonuclease 1 | 1022 | 1100 | 1025 | 923 | 1133 | 1418 | 923 | 761 | 930 | 1087 | 1197 |
| 1384 | 0.08 | 0.10 | 0.75 | 0.45 | 1 | Nufip2 | nuclear fragile X mental retardation protei | 1407 | 1612 | 1281 | 1238 | 958 | 2205 | 922 | 1388 | 1017 | 938 | 2258 |
| 989 | 0.07 | 0.17 | 0.37 | 0.71 | 1 | Ctdp1 | CTD (carboxy-terminal domain, RNA poly | 794 | 1128 | 576 | 1075 | 1099 | 985 | 922 | 1255 | 1039 | 1039 | 969 |
| 964 | -0.10 | 0.14 | -0.72 | 0.47 | 1 | Slain1 | SLAIN motif family, member 1 | 1013 | 912 | 827 | 1170 | 1065 | 910 | 922 | 991 | 1090 | 886 | 817 |
| 1078 | 0.06 | 0.15 | 0.43 | 0.67 | 1 | Hook2 | hook microtubule tethering protein 2 | 666 | 772 | 756 | 1002 | 948 | 1846 | 921 | 1374 | 977 | 775 | 1816 |
| 1060 | 0.21 | 0.14 | 1.49 | 0.14 | 1 | Actl6a | actin-like 6A | 1108 | 923 | 1222 | 824 | 997 | 1182 | 921 | 1176 | 1057 | 1020 | 1232 |
| 849 | -0.04 | 0.17 | -0.21 | 0.83 | 1 | Hps3 | HPS3, biogenesis of lysosomal organelle | 636 | 540 | 475 | 838 | 1045 | 1054 | 921 | 668 | 878 | 1118 | 1169 |
| 1209 | 0.12 | 0.17 | 0.73 | 0.46 | 1 | Rpl36a | ribosomal protein L36A | 2003 | 1576 | 2612 | 891 | 991 | 604 | 920 | 1138 | 919 | 1006 | 639 |
| 1181 | 0.11 | 0.18 | 0.62 | 0.54 | 1 | Zfp219 | zinc finger protein 219 | 1211 | 1552 | 1469 | 1003 | 1057 | 1236 | 920 | 965 | 1148 | 1417 | 1014 |
| 918 | -0.07 | 0.15 | -0.49 | 0.62 | 1 | Pign | phosphatidylinositol glycan anchor biosyı | 1255 | 1038 | 1234 | 654 | 808 | 965 | 920 | 491 | 921 | 867 | 948 |
| 898 | -0.05 | 0.14 | -0.38 | 0.70 | 1 | Armh3 | armadillo-like helical domain containing 3 | 788 | 836 | 841 | 974 | 961 | 899 | 920 | 916 | 967 | 1067 | 706 |
| 1053 | 0.09 | 0.15 | 0.65 | 0.52 | 1 | Aurkaip1 | aurora kinase A interacting protein 1 | 927 | 987 | 997 | 1129 | 1187 | 954 | 919 | 1402 | 961 | 1110 | 1011 |
| 1023 | -0.08 | 0.14 | -0.56 | 0.58 | 1 | Cuedc2 | CUE domain containing 2 | 1453 | 1300 | 1551 | 927 | 973 | 751 | 919 | 1022 | 881 | 741 | 740 |
| 847 | -0.25 | 0.20 | -1.26 | 0.21 | 1 | Mtmr1 | myotubularin related protein 1 | 825 | 940 | 945 | 1038 | 713 | 899 | 919 | 623 | 853 | 605 | 963 |
| 790 | -0.11 | 0.22 | -0.51 | 0.61 | 1 | Gem | GTP binding protein (gene overexpresse | 1301 | 521 | 631 | 382 | 859 | 1043 | 919 | 293 | 888 | 828 | 1023 |
| 1240 | 0.06 | 0.17 | 0.32 | 0.75 | 1 | Pcna | proliferating cell nuclear antigen | 2175 | 1360 | 1363 | 1019 | 1028 | 1313 | 918 | 955 | 1144 | 979 | 1383 |

| | | | | | | | | | | | | | | | | | |
|---|---|---|---|---|---|---|---|---|---|---|---|---|---|---|---|---|---|
| 1355 | -0.13 | 0.15 | -0.84 | 0.40 | 1 | Adam15 | a disintegrin and metallopeptidase domai | 3683 | 2543 | 2541 | 830 | 854 | 604 | 917 | 890 | 748 | 730 | 560 |
| 1087 | 0.23 | 0.16 | 1.44 | 0.15 | 1 | Arfrp1 | ADP-ribosylation factor related protein 1 | 1102 | 869 | 1324 | 812 | 982 | 1307 | 917 | 1190 | 994 | 1105 | 1351 |
| 933 | -0.07 | 0.15 | -0.48 | 0.63 | 1 | Polr3a | polymerase (RNA) III (DNA directed) poly | 666 | 624 | 943 | 996 | 1039 | 1150 | 917 | 759 | 967 | 1002 | 1205 |
| 1196 | 0.05 | 0.14 | 0.34 | 0.73 | 1 | Slc30a1 | solute carrier family 30 (zinc transporter), | 1337 | 1613 | 1419 | 1334 | 1022 | 1007 | 916 | 1587 | 1112 | 996 | 810 |
| 1039 | -0.13 | 0.17 | -0.75 | 0.46 | 1 | Tmem28 | transmembrane protein 28 | 1255 | 1989 | 1225 | 1013 | 949 | 739 | 916 | 907 | 768 | 1038 | 628 |
| 1028 | -0.20 | 0.14 | -1.44 | 0.15 | 1 | Limd2 | LIM domain containing 2 | 1577 | 1390 | 1406 | 1182 | 928 | 697 | 916 | 920 | 831 | 782 | 681 |
| 785 | 0.10 | 0.18 | 0.53 | 0.60 | 1 | Gfod2 | glucose-fructose oxidoreductase domain | 511 | 582 | 660 | 775 | 918 | 707 | 916 | 1140 | 749 | 952 | 726 |
| 1053 | 0.20 | 0.12 | 1.72 | 0.09 | 1 | Ttc1 | tetratricopeptide repeat domain 1 | 1204 | 1032 | 1247 | 975 | 934 | 941 | 915 | 1126 | 1100 | 1082 | 1024 |
| 932 | -0.10 | 0.13 | -0.73 | 0.46 | 1 | Polr3c | polymerase (RNA) III (DNA directed) poly | 1084 | 965 | 1078 | 975 | 978 | 817 | 915 | 862 | 1002 | 737 | 844 |
| 854 | 0.13 | 0.15 | 0.85 | 0.39 | 1 | Snrnp27 | small nuclear ribonucleoprotein 27 (U4/U | 827 | 780 | 992 | 689 | 851 | 794 | 915 | 952 | 803 | 948 | 839 |
| 1342 | 0.07 | 0.16 | 0.44 | 0.66 | 1 | Dctn3 | dynactin 3 | 1126 | 1307 | 1710 | 1467 | 1432 | 1409 | 914 | 1229 | 1494 | 1376 | 1302 |
| 909 | 0.09 | 0.14 | 0.64 | 0.52 | 1 | Zbtb22 | zinc finger and BTB domain containing 22 | 921 | 823 | 1113 | 827 | 896 | 815 | 914 | 977 | 954 | 1020 | 741 |
| 872 | 0.17 | 0.18 | 0.97 | 0.33 | 1 | Wdr83os | WD repeat domain 83 opposite strand | 1049 | 733 | 986 | 868 | 825 | 556 | 914 | 1214 | 854 | 1073 | 517 |
| 842 | -0.05 | 0.13 | -0.36 | 0.72 | 1 | AI837181 | expressed sequence AI837181 | 1084 | 1005 | 1062 | 802 | 654 | 740 | 914 | 729 | 677 | 867 | 732 |
| 755 | 0.10 | 0.17 | 0.61 | 0.54 | 1 | Brsk2 | BR serine/threonine kinase 2 | 263 | 221 | 291 | 838 | 752 | 1125 | 914 | 1062 | 826 | 809 | 1200 |
| 1116 | 0.21 | 0.15 | 1.35 | 0.18 | 1 | Ttc4 | tetratricopeptide repeat domain 4 | 986 | 672 | 1066 | 943 | 1049 | 1560 | 913 | 1150 | 1120 | 1234 | 1579 |
| 912 | 0.00 | 0.17 | 0.00 | 1.00 | 1 | 2810004N23Ril | RIKEN cDNA 2810004N23 gene | 989 | 897 | 854 | 826 | 881 | 1018 | 913 | 985 | 1027 | 662 | 976 |
| 869 | -0.07 | 0.18 | -0.41 | 0.68 | 1 | Gyg | glycogenin | 981 | 820 | 804 | 1040 | 848 | 753 | 913 | 709 | 1105 | 871 | 713 |
| 816 | -0.13 | 0.17 | -0.75 | 0.45 | 1 | Eef2k | eukaryotic elongation factor-2 kinase | 924 | 697 | 792 | 768 | 995 | 701 | 913 | 840 | 987 | 916 | 446 |
| 1015 | -0.02 | 0.14 | -0.17 | 0.86 | 1 | Rab5if | RAB5 interacting factor | 1343 | 1268 | 1146 | 1035 | 942 | 831 | 912 | 821 | 960 | 1122 | 779 |
| 993 | -0.01 | 0.13 | -0.09 | 0.93 | 1 | Cnot9 | CCR4-NOT transcription complex, subuni | 1071 | 1059 | 1115 | 935 | 964 | 1035 | 912 | 832 | 1151 | 877 | 972 |
| 1045 | 0.14 | 0.12 | 1.11 | 0.27 | 1 | Zfp386 | zinc finger protein 386 (Kruppel-like) | 993 | 918 | 1024 | 1037 | 1012 | 1101 | 911 | 947 | 1074 | 1109 | 1372 |
| 1273 | -0.16 | 0.18 | -0.91 | 0.36 | 1 | Fhl1 | four and a half LIM domains 1 | 1983 | 1194 | 1092 | 920 | 846 | 2538 | 910 | 828 | 856 | 772 | 2064 |
| 1103 | -0.04 | 0.16 | -0.27 | 0.79 | 1 | Dedd2 | death effector domain-containing DNA bir | 1083 | 900 | 1134 | 1246 | 865 | 1575 | 910 | 1134 | 965 | 872 | 1454 |
| 1014 | -0.20 | 0.15 | -1.30 | 0.19 | 1 | Zfp608 | zinc finger protein 608 | 921 | 910 | 1016 | 896 | 965 | 1653 | 910 | 587 | 971 | 929 | 1391 |
| 825 | -0.17 | 0.16 | -1.07 | 0.28 | 1 | Higd2a | HIG1 domain family, member 2A | 781 | 738 | 1160 | 833 | 910 | 728 | 910 | 675 | 816 | 752 | 766 |
| 771 | 0.02 | 0.16 | 0.12 | 0.91 | 1 | Timm8b | translocase of inner mitochondrial membr | 595 | 684 | 469 | 908 | 800 | 733 | 910 | 747 | 898 | 983 | 757 |
| 826 | -0.03 | 0.13 | -0.21 | 0.84 | 1 | Ndufa3 | NADH:ubiquinone oxidoreductase subun | 754 | 764 | 963 | 769 | 868 | 783 | 909 | 817 | 878 | 854 | 722 |
| 1233 | -0.29 | 0.17 | -1.75 | 0.08 | 1 | Dnpep | aspartyl aminopeptidase | 1855 | 1680 | 1930 | 1033 | 1079 | 1385 | 908 | 545 | 1007 | 877 | 1263 |
| 1136 | 0.15 | 0.14 | 1.04 | 0.30 | 1 | Polr1d | polymerase (RNA) I polypeptide D | 1186 | 1467 | 1452 | 1171 | 971 | 906 | 908 | 1495 | 921 | 1096 | 920 |
| 995 | -0.17 | 0.16 | -1.11 | 0.27 | 1 | Eif4e2 | eukaryotic translation initiation factor 4E r | 1211 | 1196 | 1058 | 991 | 951 | 1096 | 908 | 657 | 944 | 822 | 1116 |
| 963 | 0.04 | 0.11 | 0.38 | 0.70 | 1 | Cyb5r4 | cytochrome b5 reductase 4 | 715 | 831 | 872 | 901 | 1118 | 1106 | 908 | 991 | 1088 | 913 | 1156 |
| 846 | -0.06 | 0.15 | -0.41 | 0.68 | 1 | Erlin1 | ER lipid raft associated 1 | 897 | 842 | 707 | 940 | 770 | 892 | 907 | 713 | 894 | 801 | 945 |
| 834 | -0.02 | 0.13 | -0.13 | 0.90 | 1 | Nup62 | nucleoporin 62 | 775 | 902 | 676 | 909 | 782 | 826 | 907 | 836 | 685 | 947 | 929 |
| 782 | -0.14 | 0.15 | -0.93 | 0.35 | 1 | Fer | fer (fms/fps related) protein kinase | 847 | 637 | 814 | 795 | 765 | 826 | 907 | 592 | 833 | 715 | 868 |
| 957 | 0.04 | 0.11 | 0.37 | 0.71 | 1 | Psma5 | proteasome subunit alpha 5 | 1546 | 1233 | 1315 | 708 | 825 | 737 | 906 | 733 | 879 | 899 | 750 |
| 635 | 0.31 | 0.23 | 1.40 | 0.16 | 1 | Lrrc24 | leucine rich repeat containing 24 | 258 | 400 | 417 | 661 | 685 | 395 | 906 | 979 | 746 | 992 | 542 |
| 883 | -0.21 | 0.16 | -1.36 | 0.17 | 1 | Ccng2 | cyclin G2 | 1315 | 627 | 1088 | 723 | 921 | 1020 | 905 | 585 | 744 | 859 | 924 |
| 799 | -0.05 | 0.13 | -0.39 | 0.69 | 1 | Mrpl42 | mitochondrial ribosomal protein L42 | 667 | 720 | 726 | 749 | 926 | 815 | 905 | 708 | 930 | 857 | 784 |
| 895 | 0.17 | 0.13 | 1.31 | 0.19 | 1 | Macroh2a2 | macroH2A.2 histone | 616 | 830 | 807 | 927 | 851 | 887 | 904 | 1058 | 969 | 979 | 1017 |
| 875 | 0.10 | 0.15 | 0.64 | 0.52 | 1 | Sgsh | N-sulfoglucosamine sulfohydrolase (sulfa | 1098 | 1059 | 848 | 820 | 794 | 620 | 904 | 965 | 810 | 1193 | 509 |
| 854 | 0.11 | 0.17 | 0.63 | 0.53 | 1 | Ttc26 | tetratricopeptide repeat domain 26 | 404 | 434 | 485 | 929 | 1117 | 927 | 904 | 775 | 1083 | 1264 | 1070 |
| 1014 | -0.11 | 0.15 | -0.73 | 0.47 | 1 | Vps72 | vacuolar protein sorting 72 | 1139 | 1188 | 1342 | 954 | 1023 | 996 | 903 | 897 | 720 | 962 | 1026 |
| 814 | -0.16 | 0.13 | -1.25 | 0.21 | 1 | Map6 | microtubule-associated protein 6 | 715 | 672 | 613 | 810 | 814 | 1128 | 903 | 755 | 739 | 696 | 1106 |
| 680 | -0.12 | 0.13 | -0.93 | 0.35 | 1 | Gatc | glutamyl-tRNA(Gln) amidotransferase, su | 601 | 553 | 614 | 747 | 756 | 576 | 903 | 744 | 725 | 734 | 530 |
| 1112 | -0.12 | 0.17 | -0.69 | 0.49 | 1 | Scaf4 | SR-related CTD-associated factor 4 | 787 | 1011 | 853 | 1350 | 1319 | 1443 | 901 | 878 | 1267 | 1128 | 1299 |
| 953 | 0.12 | 0.12 | 1.01 | 0.31 | 1 | Ddx18 | DEAD (Asp-Glu-Ala-Asp) box polypeptide | 887 | 824 | 790 | 1013 | 971 | 943 | 901 | 1067 | 1013 | 976 | 1100 |
| 887 | 0.18 | 0.14 | 1.23 | 0.22 | 1 | Nudt9 | nudix (nucleoside diphosphate linked mo | 1023 | 769 | 900 | 916 | 862 | 635 | 901 | 1113 | 876 | 1027 | 730 |
| 850 | 0.09 | 0.18 | 0.52 | 0.60 | 1 | P4htm | prolyl 4-hydroxylase, transmembrane (en | 655 | 453 | 506 | 813 | 867 | 1172 | 901 | 1153 | 847 | 836 | 1145 |
| 794 | 0.06 | 0.13 | 0.44 | 0.66 | 1 | Mrrf | mitochondrial ribosome recycling factor | 714 | 659 | 740 | 899 | 828 | 625 | 901 | 818 | 960 | 866 | 726 |
| 700 | -0.05 | 0.21 | -0.22 | 0.83 | 1 | Zdhhc22 | zinc finger, DHHC-type containing 22 | 78 | 74 | 211 | 929 | 1312 | 527 | 901 | 1028 | 1382 | 828 | 433 |
| 976 | -0.04 | 0.13 | -0.32 | 0.75 | 1 | Micall1 | microtubule associated monooxygenase, | 1194 | 1209 | 1274 | 841 | 802 | 1045 | 900 | 911 | 832 | 885 | 848 |
| 909 | -0.01 | 0.16 | -0.09 | 0.93 | 1 | Dpm2 | dolichol-phosphate (beta-D) mannosyltra | 697 | 791 | 785 | 897 | 1049 | 1024 | 900 | 746 | 1059 | 1157 | 890 |
| 839 | 0.00 | 0.15 | -0.01 | 0.99 | 1 | Mrpl20 | mitochondrial ribosomal protein L20 | 843 | 690 | 821 | 779 | 1053 | 715 | 900 | 858 | 789 | 1010 | 767 |
| 703 | 0.01 | 0.19 | 0.04 | 0.97 | 1 | Clba1 | clathrin binding box of aftiphilin containin | 310 | 480 | 499 | 851 | 846 | 584 | 900 | 1108 | 804 | 837 | 519 |
| 1054 | -0.04 | 0.18 | -0.23 | 0.82 | 1 | Ehmt1 | euchromatic histone methyltransferase 1 | 1241 | 1162 | 1130 | 740 | 1133 | 1333 | 899 | 1079 | 823 | 848 | 1207 |
| 856 | -0.15 | 0.23 | -0.64 | 0.52 | 1 | Sptb | spectrin beta, erythrocytic | 1179 | 1354 | 1452 | 830 | 654 | 429 | 899 | 1076 | 686 | 461 | 398 |
| 778 | -0.09 | 0.19 | -0.46 | 0.65 | 1 | Ppp1r26 | protein phosphatase 1, regulatory subuni | 715 | 704 | 621 | 1010 | 621 | 838 | 899 | 630 | 820 | 884 | 811 |
| 1114 | -0.27 | 0.23 | -1.17 | 0.24 | 1 | Sun2 | Sad1 and UNC84 domain containing 2 | 2943 | 1607 | 1834 | 937 | 845 | 551 | 898 | 874 | 683 | 527 | 552 |
| 1192 | 0.02 | 0.29 | 0.05 | 0.96 | 1 | Nupr1 | nuclear protein transcription regulator 1 | 989 | 3038 | 3583 | 765 | 858 | 223 | 897 | 718 | 935 | 873 | 237 |

| | | | | | | | | | | | | | | | | | | |
|---|---|---|---|---|---|---|---|---|---|---|---|---|---|---|---|---|---|---|
| 974 | -0.10 | 0.18 | -0.56 | 0.58 | 1 | Rai2 | retinoic acid induced 2 | 1262 | 1902 | 1673 | 705 | 680 | 753 | 897 | 697 | 725 | 692 | 723 |
| 930 | 0.14 | 0.13 | 1.06 | 0.29 | 1 | Dapk3 | death-associated protein kinase 3 | 837 | 844 | 948 | 757 | 964 | 1008 | 897 | 991 | 925 | 1027 | 1031 |
| 889 | -0.03 | 0.20 | -0.16 | 0.87 | 1 | Bbip1 | BBSome interacting protein 1 | 770 | 751 | 768 | 1041 | 840 | 981 | 897 | 584 | 1023 | 1100 | 1027 |
| 789 | 0.06 | 0.18 | 0.33 | 0.74 | 1 | Lrrc42 | leucine rich repeat containing 42 | 641 | 530 | 803 | 820 | 770 | 796 | 897 | 969 | 796 | 863 | 789 |
| 1138 | 0.09 | 0.12 | 0.75 | 0.45 | 1 | Acaa2 | acetyl-Coenzyme A acyltransferase 2 (mit | 1618 | 1594 | 1792 | 978 | 1161 | 617 | 896 | 1089 | 1021 | 1081 | 671 |
| 1137 | 0.08 | 0.18 | 0.48 | 0.63 | 1 | Fam214a | family with sequence similarity 214, meml | 1117 | 722 | 943 | 1006 | 1002 | 1863 | 896 | 1226 | 1035 | 936 | 1763 |
| 868 | -0.18 | 0.17 | -1.07 | 0.29 | 1 | 2410002F23Rik | RIKEN cDNA 2410002F23 gene | 710 | 722 | 962 | 729 | 793 | 1391 | 896 | 841 | 670 | 641 | 1187 |
| 866 | -0.07 | 0.14 | -0.48 | 0.63 | 1 | Nhlrc3 | NHL repeat containing 3 | 685 | 845 | 873 | 765 | 872 | 1112 | 896 | 746 | 905 | 797 | 1034 |
| 760 | -0.10 | 0.17 | -0.58 | 0.56 | 1 | Slc17a9 | solute carrier family 17, member 9 | 341 | 350 | 354 | 1088 | 797 | 979 | 896 | 1206 | 702 | 748 | 902 |
| 678 | -0.16 | 0.18 | -0.89 | 0.38 | 1 | Mrpl39 | mitochondrial ribosomal protein L39 | 583 | 538 | 610 | 733 | 689 | 703 | 896 | 663 | 796 | 558 | 695 |
| 716 | -0.02 | 0.16 | -0.14 | 0.89 | 1 | Rft1 | RFT1 homolog | 763 | 796 | 636 | 589 | 750 | 634 | 895 | 643 | 718 | 810 | 639 |
| 1361 | -0.31 | 0.17 | -1.82 | 0.07 | 1 | Hjurp | RIKEN cDNA A730008H23 gene | 1517 | 1925 | 1356 | 1414 | 774 | 2517 | 894 | 1254 | 771 | 506 | 2044 |
| 1035 | 0.10 | 0.14 | 0.76 | 0.45 | 1 | Mrps17 | mitochondrial ribosomal protein S17 | 1217 | 1137 | 1065 | 891 | 971 | 1076 | 894 | 1156 | 846 | 1077 | 1054 |
| 851 | 0.17 | 0.20 | 0.86 | 0.39 | 1 | Mios | meiosis regulator for oocyte development | 960 | 518 | 575 | 784 | 765 | 1015 | 894 | 1061 | 934 | 863 | 991 |
| 1191 | -0.07 | 0.11 | -0.62 | 0.53 | 1 | Tbcel | tubulin folding cofactor E-like | 1979 | 2049 | 2089 | 915 | 963 | 794 | 893 | 978 | 953 | 775 | 711 |
| 978 | -0.20 | 0.14 | -1.45 | 0.15 | 1 | Serpinb9 | serine (or cysteine) peptidase inhibitor, cl | 1929 | 1753 | 1667 | 753 | 687 | 581 | 893 | 611 | 627 | 625 | 637 |
| 891 | -0.18 | 0.17 | -1.07 | 0.28 | 1 | Ganc | glucosidase, alpha; neutral C | 719 | 849 | 832 | 713 | 1106 | 1209 | 893 | 557 | 988 | 799 | 1139 |
| 787 | 0.13 | 0.14 | 0.91 | 0.36 | 1 | Mfsd5 | major facilitator superfamily domain conta | 926 | 693 | 833 | 738 | 770 | 538 | 893 | 900 | 822 | 1046 | 496 |
| 728 | -0.09 | 0.15 | -0.64 | 0.52 | 1 | Sgtb | small glutamine-rich tetratricopeptide rep | 599 | 763 | 633 | 811 | 712 | 691 | 893 | 761 | 761 | 689 | 691 |
| 1142 | -0.10 | 0.15 | -0.70 | 0.48 | 1 | Polg | polymerase (DNA directed), gamma | 1710 | 1319 | 1303 | 1111 | 1008 | 1256 | 892 | 964 | 987 | 867 | 1145 |
| 1067 | -0.06 | 0.13 | -0.45 | 0.65 | 1 | Smurf1 | SMAD specific E3 ubiquitin protein ligase | 1980 | 1848 | 1345 | 856 | 824 | 762 | 892 | 781 | 773 | 1000 | 673 |
| 990 | -0.12 | 0.11 | -1.09 | 0.28 | 1 | Slc27a1 | solute carrier family 27 (fatty acid transpo | 1259 | 1169 | 1201 | 880 | 888 | 1128 | 892 | 865 | 793 | 878 | 935 |
| 871 | -0.04 | 0.15 | -0.28 | 0.78 | 1 | Tmem159 | transmembrane protein 159 | 1032 | 739 | 968 | 946 | 937 | 691 | 892 | 900 | 1018 | 802 | 657 |
| 852 | -0.04 | 0.16 | -0.25 | 0.80 | 1 | Setbp1 | SET binding protein 1 | 936 | 870 | 992 | 959 | 656 | 827 | 892 | 916 | 771 | 851 | 701 |
| 829 | 0.03 | 0.13 | 0.22 | 0.82 | 1 | Tmem186 | transmembrane protein 186 | 565 | 676 | 659 | 816 | 866 | 986 | 892 | 879 | 911 | 761 | 1105 |
| 789 | -0.16 | 0.19 | -0.84 | 0.40 | 1 | Ccdc115 | coiled-coil domain containing 115 | 838 | 593 | 989 | 787 | 912 | 724 | 892 | 775 | 664 | 680 | 827 |
| 906 | 0.00 | 0.13 | -0.04 | 0.97 | 1 | AW209491 | expressed sequence AW209491 | 618 | 804 | 721 | 1088 | 879 | 1062 | 891 | 933 | 994 | 851 | 1127 |
| 882 | -0.04 | 0.18 | -0.20 | 0.84 | 1 | Wdr3 | WD repeat domain 3 | 1062 | 986 | 698 | 896 | 702 | 1020 | 891 | 609 | 813 | 926 | 1093 |
| 861 | 0.13 | 0.13 | 0.98 | 0.33 | 1 | Phospho2 | phosphatase, orphan 2 | 658 | 869 | 689 | 995 | 783 | 807 | 891 | 975 | 924 | 916 | 961 |
| 1788 | -0.19 | 0.12 | -1.66 | 0.10 | 1 | Pls3 | plastin 3 (T-isoform) | 5357 | 4185 | 4368 | 771 | 832 | 578 | 890 | 726 | 800 | 683 | 477 |
| 918 | 0.00 | 0.14 | 0.01 | 0.99 | 1 | Sccpdh | saccharopine dehydrogenase (putative) | 803 | 883 | 843 | 970 | 1085 | 842 | 890 | 867 | 914 | 1010 | 988 |
| 674 | -0.07 | 0.14 | -0.47 | 0.64 | 1 | Rab39 | RAB39, member RAS oncogene family | 146 | 174 | 98 | 1090 | 630 | 971 | 890 | 1101 | 664 | 732 | 917 |
| 1006 | 0.01 | 0.15 | 0.06 | 0.95 | 1 | Map4k2 | mitogen-activated protein kinase kinase k | 738 | 668 | 653 | 974 | 956 | 1717 | 889 | 923 | 932 | 1221 | 1399 |
| 871 | 0.14 | 0.17 | 0.82 | 0.41 | 1 | Hdhd5 | haloacid dehalogenase like hydrolase dc | 392 | 336 | 445 | 555 | 589 | 2036 | 889 | 568 | 763 | 1009 | 1997 |
| 854 | -0.15 | 0.15 | -0.95 | 0.34 | 1 | Bphl | biphenyl hydrolase-like (serine hydrolase | 1036 | 1081 | 921 | 852 | 843 | 742 | 889 | 641 | 768 | 987 | 639 |
| 776 | 0.12 | 0.17 | 0.71 | 0.48 | 1 | Mtmr11 | myotubularin related protein 11 | 640 | 612 | 748 | 690 | 684 | 882 | 889 | 899 | 882 | 836 | 780 |
| 1255 | 0.07 | 0.14 | 0.49 | 0.62 | 1 | Clk3 | CDC-like kinase 3 | 1363 | 1031 | 1056 | 956 | 1022 | 2237 | 888 | 1097 | 989 | 1010 | 2153 |
| 986 | -0.09 | 0.14 | -0.66 | 0.51 | 1 | Zfp462 | zinc finger protein 462 | 763 | 1010 | 958 | 1279 | 946 | 1085 | 888 | 945 | 983 | 913 | 1072 |
| 871 | -0.11 | 0.16 | -0.69 | 0.49 | 1 | Aspscr1 | alveolar soft part sarcoma chromosome re | 806 | 750 | 888 | 969 | 838 | 1002 | 888 | 1016 | 846 | 647 | 938 |
| 866 | -0.01 | 0.15 | -0.04 | 0.97 | 1 | Gemin5 | gem nuclear organelle associated protein | 906 | 748 | 792 | 864 | 912 | 879 | 888 | 1048 | 878 | 804 | 806 |
| 991 | -0.22 | 0.17 | -1.31 | 0.19 | 1 | Arhgap6 | Rho GTPase activating protein 6 | 1763 | 1722 | 1420 | 968 | 662 | 702 | 887 | 584 | 676 | 971 | 551 |
| 878 | -0.04 | 0.18 | -0.21 | 0.83 | 1 | Lrrc40 | leucine rich repeat containing 40 | 1002 | 981 | 759 | 735 | 1086 | 794 | 887 | 666 | 950 | 967 | 827 |
| 739 | -0.13 | 0.24 | -0.52 | 0.60 | 1 | Thsd4 | thrombospondin, type I, domain containin | 823 | 609 | 453 | 689 | 946 | 707 | 887 | 465 | 1040 | 846 | 660 |
| 1006 | 0.14 | 0.17 | 0.81 | 0.42 | 1 | Nit1 | nitrilase 1 | 658 | 1245 | 788 | 930 | 956 | 1201 | 886 | 1227 | 896 | 1059 | 1215 |
| 971 | -0.04 | 0.11 | -0.35 | 0.72 | 1 | Mphosph9 | M-phase phosphoprotein 9 | 654 | 856 | 685 | 981 | 931 | 1507 | 886 | 947 | 933 | 845 | 1460 |
| 765 | -0.18 | 0.12 | -1.45 | 0.15 | 1 | Rhot2 | ras homolog family member T2 | 633 | 619 | 685 | 708 | 859 | 982 | 886 | 646 | 745 | 716 | 934 |
| 1269 | 0.06 | 0.11 | 0.50 | 0.61 | 1 | Ppid | peptidylprolyl isomerase D (cyclophilin D) | 1017 | 1306 | 1259 | 871 | 1014 | 2253 | 885 | 987 | 994 | 823 | 2547 |
| 1026 | 0.08 | 0.15 | 0.51 | 0.61 | 1 | Ift140 | intraflagellar transport 140 | 921 | 960 | 1052 | 947 | 1134 | 1092 | 885 | 1271 | 1093 | 1070 | 864 |
| 1020 | 0.17 | 0.16 | 1.04 | 0.30 | 1 | Vrk1 | vaccinia related kinase 1 | 1015 | 697 | 693 | 1031 | 956 | 1282 | 885 | 1166 | 1052 | 1048 | 1392 |
| 1007 | 0.01 | 0.23 | 0.06 | 0.95 | 1 | Dennd5b | DENN/MADD domain containing 5B | 1428 | 1878 | 1043 | 621 | 921 | 926 | 885 | 718 | 1006 | 854 | 797 |
| 908 | -0.17 | 0.18 | -0.95 | 0.34 | 1 | Sdk1 | sidekick cell adhesion molecule 1 | 1202 | 1537 | 1401 | 604 | 892 | 713 | 885 | 599 | 818 | 752 | 587 |
| 883 | 0.03 | 0.12 | 0.27 | 0.79 | 1 | Coq5 | coenzyme Q5 methyltransferase | 850 | 892 | 890 | 879 | 944 | 795 | 885 | 1014 | 873 | 893 | 804 |
| 986 | -0.03 | 0.19 | -0.15 | 0.88 | 1 | Cep85 | centrosomal protein 85 | 643 | 861 | 681 | 987 | 1319 | 1187 | 884 | 762 | 1267 | 1070 | 1185 |
| 718 | 0.09 | 0.15 | 0.60 | 0.55 | 1 | Tmem192 | transmembrane protein 192 | 461 | 459 | 586 | 876 | 876 | 478 | 884 | 1003 | 864 | 863 | 553 |
| 1074 | 0.06 | 0.11 | 0.54 | 0.59 | 1 | Inpp5k | inositol polyphosphate 5-phosphatase K | 1538 | 1524 | 1261 | 911 | 800 | 1081 | 883 | 919 | 891 | 988 | 1017 |
| 1004 | 0.01 | 0.15 | 0.09 | 0.93 | 1 | Ss18 | SS18, nBAF chromatin remodeling compl | 1429 | 1016 | 1147 | 811 | 835 | 1200 | 883 | 962 | 838 | 904 | 1016 |
| 970 | 0.02 | 0.11 | 0.21 | 0.84 | 1 | Chic2 | cysteine-rich hydrophobic domain 2 | 1142 | 1008 | 1206 | 845 | 971 | 921 | 883 | 750 | 1008 | 953 | 985 |
| 940 | -0.11 | 0.16 | -0.72 | 0.47 | 1 | Klhl28 | kelch-like 28 | 728 | 805 | 748 | 1034 | 954 | 1287 | 882 | 730 | 1073 | 772 | 1328 |
| 658 | 0.22 | 0.18 | 1.22 | 0.22 | 1 | Acot13 | acyl-CoA thioesterase 13 | 380 | 446 | 475 | 639 | 748 | 492 | 882 | 921 | 753 | 884 | 622 |
| 954 | -0.05 | 0.16 | -0.32 | 0.75 | 1 | Cwc25 | CWC25 spliceosome-associated protein | 863 | 921 | 1025 | 1026 | 1024 | 944 | 881 | 1208 | 1087 | 640 | 870 |

| | | | | | | | | | | | | | | | | | | |
|---|---|---|---|---|---|---|---|---|---|---|---|---|---|---|---|---|---|---|
| 921 | 0.18 | 0.16 | 1.14 | 0.25 | 1 | Brcc3 | BRCA1/BRCA2-containing complex, subu | 765 | 704 | 536 | 990 | 893 | 1047 | 881 | 936 | 1119 | 1171 | 1086 |
| 682 | -0.05 | 0.17 | -0.32 | 0.75 | 1 | Nipa1 | non imprinted in Prader-Willi/Angelman s | 640 | 598 | 499 | 780 | 632 | 643 | 881 | 557 | 775 | 884 | 613 |
| 1436 | -0.27 | 0.20 | -1.36 | 0.17 | 1 | Samhd1 | SAM domain and HD domain, 1 | 4059 | 2592 | 2528 | 835 | 1025 | 862 | 880 | 649 | 853 | 594 | 918 |
| 1229 | -0.01 | 0.15 | -0.08 | 0.93 | 1 | Exoc6 | exocyst complex component 6 | 2062 | 2673 | 2081 | 734 | 883 | 868 | 880 | 739 | 914 | 784 | 902 |
| 1178 | 0.09 | 0.13 | 0.75 | 0.46 | 1 | Gm49336 | nucleoporin like 1 | 1137 | 1086 | 1145 | 1190 | 936 | 1636 | 880 | 1101 | 1116 | 971 | 1759 |
| 889 | 0.07 | 0.15 | 0.47 | 0.64 | 1 | Sncb | synuclein, beta | 500 | 707 | 743 | 1018 | 1066 | 866 | 880 | 1119 | 904 | 1000 | 980 |
| 857 | -0.12 | 0.16 | -0.75 | 0.45 | 1 | Btbd10 | BTB (POZ) domain containing 10 | 721 | 961 | 794 | 911 | 889 | 918 | 880 | 711 | 775 | 778 | 1087 |
| 605 | -0.17 | 0.17 | -1.03 | 0.30 | 1 | Tcp11l1 | t-complex 11 like 1 | 398 | 397 | 440 | 696 | 627 | 667 | 880 | 536 | 618 | 788 | 607 |
| 917 | 0.09 | 0.16 | 0.61 | 0.54 | 1 | Zw10 | zw10 kinetochore protein | 900 | 923 | 919 | 620 | 894 | 1177 | 879 | 890 | 949 | 769 | 1164 |
| 894 | -0.15 | 0.14 | -1.09 | 0.28 | 1 | Eif2b1 | eukaryotic translation initiation factor 2B, : | 1021 | 1069 | 964 | 863 | 972 | 855 | 879 | 672 | 807 | 824 | 914 |
| 658 | 0.02 | 0.15 | 0.12 | 0.90 | 1 | Tubb3 | tubulin, beta 3 class III | 175 | 174 | 165 | 972 | 810 | 670 | 879 | 823 | 774 | 1095 | 698 |
| 1019 | 0.07 | 0.16 | 0.43 | 0.67 | 1 | Hsf1 | heat shock factor 1 | 1066 | 665 | 1114 | 765 | 1109 | 1323 | 878 | 723 | 1010 | 1313 | 1243 |
| 924 | -0.06 | 0.13 | -0.43 | 0.66 | 1 | Tysnd1 | trypsin domain containing 1 | 686 | 713 | 737 | 960 | 1117 | 1141 | 878 | 827 | 985 | 1039 | 1080 |
| 1158 | 0.12 | 0.16 | 0.76 | 0.45 | 1 | Exoc1 | exocyst complex component 1 | 1240 | 1343 | 1165 | 964 | 912 | 1544 | 877 | 1444 | 927 | 790 | 1529 |
| 1155 | 0.05 | 0.16 | 0.33 | 0.74 | 1 | Tcf3 | transcription factor 3 | 2033 | 1498 | 1568 | 801 | 919 | 1160 | 877 | 1000 | 992 | 822 | 1038 |
| 865 | 0.10 | 0.14 | 0.68 | 0.50 | 1 | Tmem256 | transmembrane protein 256 | 664 | 627 | 722 | 1075 | 983 | 669 | 877 | 1283 | 990 | 975 | 650 |
| 819 | -0.03 | 0.14 | -0.18 | 0.85 | 1 | Lamtor5 | late endosomal/lysosomal adaptor, MAPK | 956 | 931 | 939 | 817 | 757 | 668 | 877 | 668 | 868 | 804 | 720 |
| 985 | -0.12 | 0.12 | -1.03 | 0.30 | 1 | Dip2a | disco interacting protein 2 homolog A | 1457 | 1344 | 1235 | 998 | 918 | 752 | 876 | 830 | 830 | 871 | 723 |
| 881 | -0.25 | 0.17 | -1.44 | 0.15 | 1 | Zfp629 | zinc finger protein 629 | 820 | 1069 | 899 | 1061 | 775 | 986 | 876 | 1104 | 687 | 511 | 899 |
| 810 | 0.00 | 0.15 | 0.03 | 0.98 | 1 | Btbd6 | BTB (POZ) domain containing 6 | 750 | 727 | 758 | 873 | 897 | 661 | 876 | 740 | 922 | 1122 | 590 |
| 743 | -0.03 | 0.18 | -0.17 | 0.86 | 1 | Slc16a9 | solute carrier family 16 (monocarboxylic a | 683 | 332 | 531 | 828 | 868 | 762 | 876 | 736 | 822 | 958 | 772 |
| 741 | -0.01 | 0.15 | -0.05 | 0.96 | 1 | Rdh11 | retinol dehydrogenase 11 | 637 | 502 | 497 | 949 | 817 | 632 | 876 | 789 | 905 | 862 | 685 |
| 627 | 0.21 | 0.14 | 1.52 | 0.13 | 1 | Acp1 | acid phosphatase 1, soluble | 1082 | 676 | 337 | 496 | 283 | 548 | 876 | 550 | 313 | 1112 | 623 |
| 1080 | -0.04 | 0.23 | -0.19 | 0.85 | 1 | Rad54l2 | RAD54 like 2 (S. cerevisiae) | 1241 | 750 | 729 | 1263 | 1094 | 1415 | 875 | 1075 | 1216 | 830 | 1389 |
| 902 | -0.04 | 0.16 | -0.24 | 0.81 | 1 | Champ1 | chromosome alignment maintaining phos | 853 | 943 | 812 | 803 | 960 | 1057 | 875 | 1077 | 741 | 815 | 989 |
| 1330 | -0.23 | 0.17 | -1.30 | 0.19 | 1 | Spon2 | spondin 2, extracellular matrix protein | 2134 | 2138 | 2160 | 1319 | 1481 | 692 | 874 | 1417 | 1025 | 916 | 475 |
| 892 | -0.21 | 0.16 | -1.32 | 0.19 | 1 | Lcor | ligand dependent nuclear receptor corepr | 717 | 850 | 727 | 1009 | 833 | 1289 | 874 | 1003 | 824 | 547 | 1142 |
| 885 | -0.07 | 0.14 | -0.49 | 0.62 | 1 | Ccdc181 | coiled-coil domain containing 181 | 905 | 893 | 893 | 901 | 955 | 864 | 874 | 699 | 906 | 982 | 858 |
| 824 | -0.05 | 0.20 | -0.24 | 0.81 | 1 | Eif2s3x | eukaryotic translation initiation factor 2, su | 739 | 957 | 895 | 693 | 886 | 811 | 874 | 584 | 1133 | 747 | 748 |
| 586 | -0.18 | 0.25 | -0.70 | 0.48 | 1 | Atp4a | ATPase, H+/K+ exchanging, gastric, alpha | 168 | 336 | 293 | 1040 | 527 | 526 | 874 | 1000 | 531 | 763 | 389 |
| 1303 | -0.08 | 0.15 | -0.58 | 0.56 | 1 | Vcl | vinculin | 3832 | 2595 | 2504 | 655 | 546 | 722 | 873 | 543 | 695 | 696 | 671 |
| 1062 | 0.19 | 0.17 | 1.11 | 0.26 | 1 | Coq10b | coenzyme Q10B | 976 | 845 | 602 | 1275 | 956 | 1182 | 873 | 1555 | 929 | 1021 | 1466 |
| 1029 | 0.03 | 0.16 | 0.21 | 0.84 | 1 | St3gal5 | ST3 beta-galactoside alpha-2,3-sialyltran | 1331 | 1047 | 1051 | 1072 | 1075 | 864 | 873 | 1143 | 946 | 1128 | 791 |
| 1016 | 0.03 | 0.14 | 0.19 | 0.85 | 1 | Cog8 | component of oligomeric golgi complex 8 | 704 | 811 | 696 | 1099 | 1264 | 1215 | 873 | 1047 | 1132 | 1135 | 1194 |
| 975 | -0.10 | 0.22 | -0.44 | 0.66 | 1 | Dcdc2a | doublecortin domain containing 2a | 833 | 679 | 1277 | 1354 | 862 | 1015 | 873 | 1054 | 1012 | 900 | 870 |
| 950 | -0.10 | 0.19 | -0.55 | 0.58 | 1 | Fut8 | fucosyltransferase 8 | 1657 | 928 | 1026 | 1011 | 888 | 734 | 873 | 694 | 809 | 1108 | 724 |
| 939 | -0.06 | 0.14 | -0.39 | 0.69 | 1 | Cep164 | centrosomal protein 164 | 737 | 716 | 733 | 958 | 1011 | 1302 | 873 | 1057 | 1012 | 697 | 1238 |
| 935 | -0.25 | 0.13 | -1.88 | 0.06 | 1 | Nfrkb | nuclear factor related to kappa B binding | 811 | 930 | 977 | 880 | 789 | 1558 | 873 | 668 | 814 | 645 | 1338 |
| 912 | -0.08 | 0.16 | -0.50 | 0.62 | 1 | Nprl3 | nitrogen permease regulator-like 3 | 432 | 440 | 546 | 913 | 912 | 1721 | 873 | 1039 | 737 | 799 | 1617 |
| 869 | -0.02 | 0.13 | -0.12 | 0.90 | 1 | Lgr4 | leucine-rich repeat-containing G protein-c | 795 | 839 | 938 | 854 | 917 | 855 | 873 | 743 | 1039 | 935 | 768 |
| 838 | -0.05 | 0.14 | -0.37 | 0.71 | 1 | Zscan29 | zinc finger SCAN domains 29 | 796 | 808 | 682 | 908 | 782 | 945 | 873 | 963 | 701 | 692 | 1071 |
| 672 | -0.02 | 0.18 | -0.13 | 0.90 | 1 | Morn4 | MORN repeat containing 4 | 251 | 350 | 359 | 801 | 986 | 587 | 873 | 821 | 793 | 1002 | 571 |
| 1136 | 0.12 | 0.18 | 0.67 | 0.51 | 1 | Stx4a | syntaxin 4A (placental) | 1793 | 1170 | 1233 | 915 | 1044 | 1150 | 872 | 1186 | 931 | 1009 | 1191 |
| 845 | 0.29 | 0.25 | 1.17 | 0.24 | 1 | Ndufc1 | NADH:ubiquinone oxidoreductase subun | 764 | 488 | 749 | 864 | 914 | 597 | 872 | 1629 | 913 | 828 | 675 |
| 940 | -0.05 | 0.13 | -0.37 | 0.71 | 1 | Rae1 | ribonucleic acid export 1 | 950 | 843 | 901 | 928 | 948 | 1134 | 871 | 761 | 941 | 1007 | 1055 |
| 954 | -0.14 | 0.15 | -0.93 | 0.35 | 1 | Ddhd1 | DDHD domain containing 1 | 1286 | 1520 | 1250 | 766 | 667 | 1084 | 870 | 619 | 754 | 748 | 930 |
| 873 | 0.01 | 0.16 | 0.04 | 0.97 | 1 | Rps6ka4 | ribosomal protein S6 kinase, polypeptide | 1277 | 814 | 945 | 897 | 891 | 619 | 870 | 951 | 808 | 902 | 631 |
| 852 | -0.10 | 0.18 | -0.55 | 0.58 | 1 | Chpt1 | choline phosphotransferase 1 | 1330 | 826 | 905 | 716 | 942 | 745 | 870 | 773 | 896 | 716 | 658 |
| 846 | 0.18 | 0.16 | 1.14 | 0.25 | 1 | Cfap298 | cilia and flagella associate protien 298 | 723 | 833 | 922 | 746 | 843 | 742 | 870 | 926 | 774 | 1010 | 917 |
| 1173 | 0.09 | 0.17 | 0.53 | 0.60 | 1 | Bin1 | bridging integrator 1 | 1211 | 859 | 983 | 930 | 868 | 2197 | 869 | 1226 | 831 | 989 | 1936 |
| 706 | 0.04 | 0.15 | 0.24 | 0.81 | 1 | Lnpk | lunapark, ER junction formation factor | 448 | 524 | 440 | 708 | 816 | 746 | 869 | 737 | 903 | 837 | 743 |
| 661 | -0.12 | 0.19 | -0.66 | 0.51 | 1 | Celf6 | CUGBP, Elav-like family member 6 | 471 | 562 | 493 | 770 | 751 | 604 | 869 | 591 | 589 | 881 | 691 |
| 1025 | 0.00 | 0.12 | -0.02 | 0.98 | 1 | Patl1 | protein associated with topoisomerase II l | 1232 | 1171 | 1098 | 964 | 938 | 1124 | 868 | 978 | 946 | 957 | 996 |
| 925 | 0.10 | 0.11 | 0.91 | 0.36 | 1 | Lrrc14 | leucine rich repeat containing 14 | 851 | 811 | 835 | 960 | 930 | 943 | 868 | 1008 | 994 | 1024 | 946 |
| 877 | 0.05 | 0.14 | 0.32 | 0.75 | 1 | Tmem107 | transmembrane protein 107 | 783 | 776 | 733 | 937 | 984 | 825 | 868 | 890 | 823 | 1095 | 932 |
| 784 | -0.14 | 0.17 | -0.85 | 0.40 | 1 | Tmem8b | transmembrane protein 8B | 893 | 773 | 595 | 719 | 892 | 864 | 868 | 785 | 755 | 708 | 772 |
| 639 | -0.13 | 0.16 | -0.80 | 0.42 | 1 | Zdhhc23 | zinc finger, DHHC domain containing 23 | 422 | 251 | 306 | 619 | 937 | 718 | 868 | 601 | 770 | 952 | 585 |
| 1359 | 0.01 | 0.17 | 0.06 | 0.96 | 1 | Pogz | pogo transposable element with ZNF dom | 1413 | 1557 | 1590 | 1337 | 975 | 2104 | 867 | 1030 | 1099 | 1289 | 1691 |
| 1037 | 0.10 | 0.10 | 0.99 | 0.32 | 1 | Numb | NUMB endocytic adaptor protein | 1399 | 1376 | 1343 | 924 | 822 | 900 | 867 | 918 | 926 | 987 | 942 |
| 970 | 0.01 | 0.11 | 0.13 | 0.89 | 1 | Tstd2 | thiosulfate sulfurtransferase (rhodanese)- | 899 | 1104 | 870 | 1066 | 954 | 987 | 867 | 1066 | 939 | 826 | 1094 |

| | | | | | | | | | | | | | | | | | |
|---|---|---|---|---|---|---|---|---|---|---|---|---|---|---|---|---|---|
| 938 | 0.05 | 0.13 | 0.35 | 0.73 | 1 | Banf1 | BAF nuclear assembly factor 1 | 1383 | 1027 | 1213 | 735 | 1020 | 671 | 867 | 879 | 980 | 895 | 644 |
| 1067 | -0.14 | 0.13 | -1.05 | 0.29 | 1 | Setdb1 | SET domain, bifurcated 1 | 1069 | 939 | 1007 | 915 | 1092 | 1666 | 866 | 777 | 804 | 956 | 1641 |
| 1025 | 0.14 | 0.17 | 0.81 | 0.42 | 1 | Map1s | microtubule-associated protein 1S | 1116 | 1111 | 1192 | 881 | 948 | 1035 | 866 | 1343 | 843 | 998 | 944 |
| 782 | -0.22 | 0.20 | -1.09 | 0.27 | 1 | Alg9 | asparagine-linked glycosylation 9 (alpha | 804 | 787 | 720 | 962 | 896 | 650 | 866 | 507 | 792 | 960 | 656 |
| 1020 | 0.24 | 0.16 | 1.45 | 0.15 | 1 | Naa40 | N(alpha)-acetyltransferase 40, NatD catal | 796 | 618 | 710 | 1027 | 924 | 1391 | 865 | 1313 | 1064 | 1253 | 1262 |
| 961 | 0.10 | 0.14 | 0.71 | 0.48 | 1 | Rps6kc1 | ribosomal protein S6 kinase polypeptide | 934 | 950 | 1083 | 971 | 877 | 952 | 865 | 1197 | 984 | 831 | 932 |
| 946 | 0.02 | 0.12 | 0.16 | 0.87 | 1 | Zpr1 | ZPR1 zinc finger | 1206 | 1075 | 1148 | 837 | 760 | 1004 | 865 | 757 | 883 | 947 | 923 |
| 802 | 0.25 | 0.21 | 1.23 | 0.22 | 1 | Slc35e4 | solute carrier family 35, member E4 | 585 | 454 | 593 | 876 | 827 | 645 | 865 | 1445 | 908 | 1063 | 563 |
| 780 | 0.17 | 0.12 | 1.45 | 0.15 | 1 | Ptcd2 | pentatricopeptide repeat domain 2 | 812 | 654 | 580 | 782 | 708 | 710 | 865 | 827 | 773 | 1065 | 802 |
| 734 | 0.14 | 0.18 | 0.78 | 0.44 | 1 | Hmbs | hydroxymethylbilane synthase | 613 | 541 | 455 | 692 | 761 | 756 | 865 | 1053 | 818 | 763 | 752 |
| 943 | -0.01 | 0.17 | -0.03 | 0.97 | 1 | Kcnc3 | potassium voltage gated channel, Shaw-r | 503 | 717 | 757 | 848 | 803 | 1746 | 864 | 1035 | 756 | 877 | 1471 |
| 769 | 0.01 | 0.13 | 0.04 | 0.97 | 1 | Chrac1 | chromatin accessibility complex 1 | 809 | 615 | 784 | 796 | 767 | 691 | 864 | 787 | 785 | 900 | 666 |
| 767 | -0.14 | 0.15 | -0.94 | 0.35 | 1 | Stxbp4 | syntaxin binding protein 4 | 776 | 881 | 716 | 711 | 777 | 830 | 864 | 620 | 752 | 693 | 819 |
| 923 | 0.19 | 0.14 | 1.39 | 0.16 | 1 | Vps37b | vacuolar protein sorting 37B | 1066 | 1080 | 853 | 1040 | 767 | 681 | 863 | 999 | 1054 | 1006 | 747 |
| 779 | -0.10 | 0.16 | -0.64 | 0.52 | 1 | Gtf3c6 | general transcription factor IIIC, polypepti | 897 | 1005 | 918 | 701 | 789 | 621 | 863 | 573 | 745 | 848 | 611 |
| 742 | 0.06 | 0.12 | 0.50 | 0.62 | 1 | Lmf1 | lipase maturation factor 1 | 631 | 594 | 540 | 904 | 756 | 611 | 863 | 838 | 800 | 971 | 657 |
| 1144 | -0.03 | 0.17 | -0.20 | 0.84 | 1 | Frmd4b | FERM domain containing 4B | 2187 | 1258 | 1663 | 721 | 1289 | 914 | 862 | 751 | 1205 | 949 | 789 |
| 983 | -0.28 | 0.17 | -1.62 | 0.11 | 1 | Gtpbp2 | GTP binding protein 2 | 924 | 579 | 844 | 680 | 706 | 2460 | 862 | 429 | 864 | 723 | 1743 |
| 907 | -0.14 | 0.14 | -1.01 | 0.31 | 1 | Pphln1 | periphilin 1 | 1056 | 939 | 947 | 856 | 908 | 1052 | 862 | 686 | 938 | 720 | 1017 |
| 866 | -0.07 | 0.13 | -0.53 | 0.60 | 1 | Trim32 | tripartite motif-containing 32 | 1058 | 947 | 936 | 1001 | 766 | 742 | 862 | 893 | 779 | 818 | 720 |
| 864 | 0.07 | 0.16 | 0.43 | 0.67 | 1 | Ctps | cytidine 5'-triphosphate synthase | 994 | 815 | 749 | 703 | 892 | 932 | 862 | 950 | 927 | 908 | 768 |
| 666 | 0.23 | 0.20 | 1.15 | 0.25 | 1 | B3galt6 | UDP-Gal:betaGal beta 1,3-galactosyltran: | 485 | 608 | 640 | 579 | 648 | 489 | 860 | 899 | 693 | 868 | 558 |
| 628 | -0.05 | 0.21 | -0.25 | 0.80 | 1 | Sntg1 | syntrophin, gamma 1 | 152 | 134 | 244 | 783 | 659 | 931 | 860 | 672 | 773 | 773 | 925 |
| 999 | 0.01 | 0.12 | 0.06 | 0.95 | 1 | Lemd3 | LEM domain containing 3 | 913 | 956 | 943 | 1125 | 881 | 1212 | 859 | 1142 | 920 | 818 | 1224 |
| 929 | 0.13 | 0.20 | 0.66 | 0.51 | 1 | Urm1 | ubiquitin related modifier 1 | 1021 | 887 | 1040 | 726 | 981 | 896 | 858 | 1162 | 932 | 652 | 1065 |
| 859 | -0.15 | 0.13 | -1.17 | 0.24 | 1 | Ruvbl2 | RuvB-like protein 2 | 1049 | 1026 | 1208 | 768 | 814 | 799 | 858 | 679 | 775 | 648 | 824 |
| 510 | 0.31 | 0.18 | 1.68 | 0.09 | 1 | Ust | uronyl-2-sulfotransferase | 429 | 204 | 209 | 489 | 333 | 493 | 858 | 555 | 547 | 862 | 631 |
| 1033 | 0.12 | 0.14 | 0.86 | 0.39 | 1 | Cryzl1 | crystallin, zeta (quinone reductase)-like 1 | 1069 | 1053 | 997 | 859 | 994 | 1232 | 857 | 1134 | 877 | 909 | 1376 |
| 1005 | -0.08 | 0.14 | -0.61 | 0.54 | 1 | Tnip1 | TNFAIP3 interacting protein 1 | 1605 | 1405 | 1358 | 871 | 827 | 872 | 857 | 774 | 893 | 930 | 665 |
| 903 | 0.12 | 0.18 | 0.68 | 0.50 | 1 | Gtf3a | general transcription factor III A | 745 | 882 | 787 | 855 | 968 | 882 | 857 | 1409 | 879 | 781 | 891 |
| 708 | 0.16 | 0.17 | 0.91 | 0.36 | 1 | Harbi1 | harbinger transposase derived 1 | 429 | 514 | 359 | 642 | 660 | 921 | 857 | 842 | 849 | 763 | 960 |
| 694 | -0.17 | 0.15 | -1.12 | 0.26 | 1 | 1110008P14Ril | RIKEN cDNA 1110008P14 gene | 498 | 475 | 653 | 740 | 885 | 698 | 856 | 686 | 716 | 835 | 595 |
| 689 | -0.16 | 0.16 | -0.98 | 0.33 | 1 | Parp16 | poly (ADP-ribose) polymerase family, mer | 404 | 476 | 506 | 725 | 893 | 781 | 856 | 719 | 778 | 849 | 591 |
| 524 | 0.23 | 0.44 | 0.54 | 0.59 | 1 | Angptl7 | angiopoietin-like 7 | 292 | 24 | 53 | 598 | 692 | 334 | 856 | 814 | 842 | 935 | 328 |
| 1187 | -0.21 | 0.17 | -1.27 | 0.20 | 1 | Myo9b | myosin IXb | 2153 | 1231 | 1390 | 857 | 710 | 2025 | 855 | 750 | 638 | 687 | 1765 |
| 833 | -0.07 | 0.14 | -0.55 | 0.58 | 1 | Rnf138 | ring finger protein 138 | 878 | 1009 | 954 | 844 | 711 | 832 | 855 | 751 | 816 | 781 | 729 |
| 948 | 0.03 | 0.16 | 0.21 | 0.83 | 1 | Mrpl30 | mitochondrial ribosomal protein L30 | 1195 | 1114 | 988 | 870 | 1031 | 765 | 854 | 722 | 1039 | 1002 | 850 |
| 861 | 0.18 | 0.17 | 1.06 | 0.29 | 1 | Fibp | fibroblast growth factor (acidic) intracellul: | 878 | 660 | 791 | 763 | 771 | 964 | 854 | 1140 | 853 | 841 | 956 |
| 839 | 0.11 | 0.12 | 0.86 | 0.39 | 1 | Arl6 | ADP-ribosylation factor-like 6 | 873 | 873 | 913 | 885 | 771 | 656 | 854 | 962 | 903 | 805 | 734 |
| 640 | -0.26 | 0.20 | -1.29 | 0.20 | 1 | Cdh22 | cadherin 22 | 389 | 639 | 444 | 702 | 864 | 608 | 854 | 722 | 572 | 724 | 521 |
| 836 | -0.17 | 0.14 | -1.21 | 0.23 | 1 | Ttf1 | transcription termination factor, RNA poly | 795 | 683 | 787 | 803 | 746 | 1260 | 853 | 726 | 771 | 616 | 1152 |
| 828 | -0.16 | 0.17 | -0.93 | 0.35 | 1 | Dusp26 | dual specificity phosphatase 26 (putative) | 1396 | 1095 | 1533 | 574 | 756 | 496 | 853 | 696 | 597 | 717 | 394 |
| 860 | -0.11 | 0.15 | -0.76 | 0.45 | 1 | Kdm1b | lysine (K)-specific demethylase 1B | 779 | 775 | 977 | 963 | 1020 | 762 | 852 | 1022 | 879 | 681 | 753 |
| 798 | -0.21 | 0.13 | -1.53 | 0.13 | 1 | Srr | serine racemase | 759 | 634 | 737 | 820 | 1019 | 864 | 852 | 630 | 922 | 725 | 820 |
| 731 | -0.06 | 0.15 | -0.40 | 0.69 | 1 | Bola3 | bolA-like 3 (E. coli) | 701 | 600 | 741 | 761 | 685 | 754 | 852 | 633 | 843 | 769 | 698 |
| 938 | -0.04 | 0.13 | -0.27 | 0.79 | 1 | Zfp507 | zinc finger protein 507 | 915 | 772 | 824 | 876 | 831 | 1412 | 851 | 746 | 920 | 958 | 1211 |
| 863 | -0.10 | 0.17 | -0.58 | 0.56 | 1 | LTO1 | ABCE maturation factor | 936 | 875 | 828 | 870 | 892 | 905 | 851 | 590 | 784 | 1003 | 963 |
| 986 | 0.30 | 0.16 | 1.82 | 0.07 | 1 | Jkamp | JNK1/MAPK8-associated membrane prot | 964 | 799 | 761 | 934 | 860 | 1081 | 849 | 1456 | 974 | 1094 | 1074 |
| 873 | 0.01 | 0.18 | 0.04 | 0.97 | 1 | Gramd1b | GRAM domain containing 1B | 859 | 917 | 984 | 994 | 771 | 770 | 849 | 1207 | 819 | 614 | 816 |
| 1105 | 0.05 | 0.13 | 0.36 | 0.72 | 1 | Sike1 | suppressor of IKBKE 1 | 1113 | 1014 | 1047 | 1006 | 1147 | 1434 | 848 | 1153 | 1039 | 1001 | 1356 |
| 982 | -0.12 | 0.12 | -0.94 | 0.35 | 1 | Map2k7 | mitogen-activated protein kinase kinase 7 | 985 | 1103 | 986 | 855 | 1004 | 1313 | 848 | 762 | 871 | 849 | 1226 |
| 942 | -0.18 | 0.14 | -1.29 | 0.20 | 1 | Trerf1 | transcriptional regulating factor 1 | 1204 | 1235 | 1003 | 1052 | 868 | 905 | 848 | 899 | 810 | 666 | 877 |
| 853 | 0.09 | 0.11 | 0.76 | 0.45 | 1 | Zfp367 | zinc finger protein 367 | 784 | 681 | 788 | 859 | 877 | 872 | 848 | 909 | 953 | 940 | 873 |
| 830 | -0.15 | 0.18 | -0.80 | 0.43 | 1 | Mrpl49 | mitochondrial ribosomal protein L49 | 860 | 832 | 963 | 958 | 881 | 694 | 848 | 979 | 902 | 524 | 687 |
| 1451 | -0.20 | 0.19 | -1.06 | 0.29 | 1 | Mat2a | methionine adenosyltransferase II, alpha | 2617 | 1854 | 1583 | 1241 | 874 | 2240 | 847 | 1029 | 733 | 698 | 2242 |
| 868 | 0.24 | 0.17 | 1.41 | 0.16 | 1 | Fars2 | phenylalanine-tRNA synthetase 2 (mitoch | 813 | 909 | 1013 | 804 | 806 | 647 | 847 | 1225 | 800 | 989 | 691 |
| 845 | -0.09 | 0.17 | -0.52 | 0.60 | 1 | Capn15 | calpain 15 | 718 | 893 | 825 | 626 | 777 | 1315 | 847 | 774 | 801 | 680 | 1034 |
| 814 | 0.29 | 0.27 | 1.06 | 0.29 | 1 | Ndufb9 | NADH:ubiquinone oxidoreductase subun | 626 | 513 | 804 | 744 | 850 | 646 | 847 | 1683 | 867 | 733 | 643 |
| 1142 | 0.10 | 0.13 | 0.76 | 0.45 | 1 | Rnf19a | ring finger protein 19A | 1564 | 1791 | 1463 | 853 | 971 | 1079 | 846 | 1022 | 963 | 989 | 1024 |
| 850 | -0.10 | 0.13 | -0.74 | 0.46 | 1 | Mrps2 | mitochondrial ribosomal protein S2 | 690 | 944 | 767 | 954 | 939 | 855 | 846 | 816 | 934 | 760 | 846 |

| | | | | | | | | | | | | | | | | | | |
|---|---|---|---|---|---|---|---|---|---|---|---|---|---|---|---|---|---|---|
| 763 | 0.35 | 0.24 | 1.46 | 0.15 | 1 | Cabp1 | calcium binding protein 1 | 93 | 128 | 181 | 917 | 743 | 997 | 846 | 1426 | 929 | 1224 | 914 |
| 986 | -0.04 | 0.18 | -0.25 | 0.80 | 1 | Alas1 | aminolevulinic acid synthase 1 | 1651 | 1175 | 1392 | 1020 | 718 | 770 | 845 | 1033 | 752 | 853 | 638 |
| 766 | -0.07 | 0.22 | -0.31 | 0.76 | 1 | Gprin3 | GPRIN family member 3 | 418 | 644 | 347 | 875 | 875 | 986 | 845 | 690 | 997 | 706 | 1036 |
| 1251 | 0.01 | 0.11 | 0.07 | 0.94 | 1 | Rbm4b | RNA binding motif protein 4B | 1074 | 1469 | 1034 | 1371 | 860 | 2030 | 844 | 1243 | 955 | 872 | 2003 |
| 889 | 0.08 | 0.15 | 0.57 | 0.57 | 1 | Dhx34 | DEAH (Asp-Glu-Ala-His) box polypeptide | 825 | 1087 | 865 | 742 | 753 | 1062 | 844 | 915 | 713 | 940 | 1037 |
| 700 | -0.29 | 0.17 | -1.70 | 0.09 | 1 | Pnkd | paroxysmal nonkinesiogenic dyskinesia | 703 | 722 | 588 | 895 | 783 | 588 | 844 | 567 | 725 | 851 | 435 |
| 1218 | 0.11 | 0.13 | 0.84 | 0.40 | 1 | Irf1 | interferon regulatory factor 1 | 2138 | 1741 | 2047 | 766 | 785 | 1206 | 843 | 776 | 798 | 1121 | 1177 |
| 847 | 0.12 | 0.17 | 0.70 | 0.48 | 1 | Zfp597 | zinc finger protein 597 | 746 | 700 | 776 | 901 | 662 | 987 | 843 | 708 | 882 | 1090 | 1024 |
| 679 | -0.06 | 0.21 | -0.27 | 0.78 | 1 | Fzd7 | frizzled class receptor 7 | 879 | 433 | 654 | 818 | 762 | 376 | 843 | 730 | 886 | 732 | 355 |
| 977 | 0.12 | 0.10 | 1.14 | 0.25 | 1 | Arl6ip4 | ADP-ribosylation factor-like 6 interacting p | 1155 | 1188 | 1091 | 892 | 818 | 956 | 842 | 977 | 870 | 978 | 985 |
| 905 | 0.09 | 0.13 | 0.72 | 0.47 | 1 | Rnps1 | RNA binding protein with serine rich dom | 1355 | 1103 | 845 | 710 | 636 | 1048 | 842 | 853 | 689 | 922 | 957 |
| 879 | -0.02 | 0.15 | -0.13 | 0.90 | 1 | Prps1l3 | phosphoribosyl pyrophosphate synthetas | 995 | 1250 | 910 | 867 | 798 | 774 | 842 | 813 | 887 | 692 | 845 |
| 858 | -0.03 | 0.15 | -0.23 | 0.81 | 1 | Dhcr7 | 7-dehydrocholesterol reductase | 599 | 521 | 660 | 1114 | 1109 | 788 | 842 | 984 | 1180 | 966 | 672 |
| 735 | 0.09 | 0.21 | 0.43 | 0.67 | 1 | Aifm1 | apoptosis-inducing factor, mitochondrion- | 741 | 526 | 535 | 525 | 880 | 816 | 842 | 796 | 797 | 643 | 985 |
| 721 | 0.20 | 0.14 | 1.39 | 0.16 | 1 | Mast1 | microtubule associated serine/threonine k | 477 | 363 | 407 | 818 | 765 | 682 | 842 | 1015 | 900 | 947 | 715 |
| 1030 | 0.01 | 0.12 | 0.13 | 0.90 | 1 | Desi2 | desumoylating isopeptidase 2 | 981 | 1350 | 964 | 1128 | 986 | 1042 | 841 | 1132 | 1026 | 873 | 1001 |
| 921 | 0.14 | 0.15 | 0.96 | 0.34 | 1 | C1qbp | complement component 1, q subcompone | 1376 | 927 | 1070 | 758 | 890 | 706 | 841 | 927 | 864 | 1091 | 677 |
| 763 | 0.03 | 0.15 | 0.17 | 0.86 | 1 | Zfp553 | zinc finger protein 553 | 666 | 882 | 735 | 814 | 696 | 670 | 841 | 799 | 892 | 779 | 619 |
| 930 | 0.01 | 0.16 | 0.05 | 0.96 | 1 | Ctif | CBP80/20-dependent translation initiatior | 1229 | 1100 | 1075 | 712 | 1175 | 672 | 840 | 814 | 1132 | 877 | 604 |
| 873 | 0.08 | 0.20 | 0.41 | 0.68 | 1 | Smug1 | single-strand selective monofunctional ur | 481 | 442 | 812 | 1126 | 982 | 883 | 840 | 973 | 1007 | 1051 | 1010 |
| 872 | 0.05 | 0.13 | 0.36 | 0.72 | 1 | Mrpl51 | mitochondrial ribosomal protein L51 | 699 | 908 | 990 | 914 | 1069 | 637 | 840 | 979 | 933 | 896 | 728 |
| 829 | 0.07 | 0.13 | 0.55 | 0.58 | 1 | Crcp | calcitonin gene-related peptide-receptor c | 760 | 778 | 726 | 831 | 822 | 845 | 840 | 1001 | 925 | 768 | 827 |
| 819 | 0.14 | 0.13 | 1.13 | 0.26 | 1 | Flad1 | flavin adenine dinucleotide synthetase 1 | 620 | 530 | 582 | 799 | 897 | 919 | 840 | 889 | 1016 | 1052 | 869 |
| 724 | 0.04 | 0.20 | 0.22 | 0.83 | 1 | Cryz | crystallin, zeta | 720 | 846 | 837 | 723 | 662 | 501 | 840 | 604 | 639 | 1061 | 533 |
| 999 | 0.04 | 0.12 | 0.34 | 0.73 | 1 | Evi5l | ecotropic viral integration site 5 like | 1079 | 1110 | 1175 | 905 | 984 | 1035 | 839 | 1054 | 1010 | 860 | 942 |
| 961 | 0.11 | 0.12 | 0.86 | 0.39 | 1 | Smyd3 | SET and MYND domain containing 3 | 877 | 1001 | 897 | 1026 | 1067 | 813 | 839 | 1216 | 997 | 1029 | 808 |
| 924 | 0.16 | 0.18 | 0.91 | 0.36 | 1 | Ccdc9b | coiled-coil domain containing 9B | 786 | 735 | 873 | 865 | 793 | 1173 | 839 | 1305 | 809 | 873 | 1116 |
| 858 | -0.18 | 0.16 | -1.13 | 0.26 | 1 | Fbxo10 | F-box protein 10 | 806 | 1007 | 1030 | 936 | 840 | 858 | 839 | 831 | 1021 | 685 | 586 |
| 3052 | -0.21 | 0.20 | -1.08 | 0.28 | 1 | Hspa1b | heat shock protein 1B | 3918 | 3946 | 6257 | 527 | 593 | 7767 | 838 | 302 | 606 | 735 | 8077 |
| 2349 | -0.19 | 0.25 | -0.75 | 0.45 | 1 | Errfi1 | ERBB receptor feedback inhibitor 1 | 7200 | 2696 | 2491 | 933 | 778 | 4436 | 838 | 845 | 647 | 666 | 4309 |
| 878 | 0.18 | 0.17 | 1.03 | 0.30 | 1 | Elp5 | elongator acetyltransferase complex subu | 848 | 892 | 870 | 783 | 938 | 717 | 838 | 703 | 1069 | 1239 | 754 |
| 871 | 0.06 | 0.16 | 0.36 | 0.72 | 1 | Mllt3 | myeloid/lymphoid or mixed-lineage leuke | 1056 | 973 | 1027 | 644 | 1006 | 711 | 838 | 612 | 864 | 980 | 873 |
| 816 | 0.03 | 0.17 | 0.17 | 0.86 | 1 | Adsl | adenylosuccinate lyase | 1124 | 778 | 783 | 604 | 804 | 885 | 838 | 817 | 773 | 769 | 800 |
| 665 | -0.12 | 0.16 | -0.77 | 0.44 | 1 | Pygo1 | pygopus 1 | 821 | 771 | 538 | 845 | 667 | 365 | 838 | 846 | 608 | 640 | 373 |
| 1429 | -0.08 | 0.12 | -0.68 | 0.49 | 1 | Arrdc3 | arrestin domain containing 3 | 3037 | 2139 | 2270 | 802 | 722 | 1877 | 837 | 726 | 726 | 738 | 1841 |
| 903 | -0.03 | 0.18 | -0.19 | 0.85 | 1 | Mier3 | MIER family member 3 | 728 | 1063 | 782 | 919 | 649 | 1255 | 837 | 1087 | 758 | 505 | 1347 |
| 897 | 0.28 | 0.14 | 1.95 | 0.05 | 1 | Fam219b | family with sequence similarity 219, meml | 1048 | 843 | 941 | 763 | 807 | 771 | 837 | 1103 | 888 | 896 | 969 |
| 886 | -0.06 | 0.19 | -0.31 | 0.76 | 1 | Cep192 | centrosomal protein 192 | 885 | 635 | 597 | 961 | 925 | 1163 | 837 | 794 | 1087 | 748 | 1111 |
| 1238 | -0.01 | 0.14 | -0.10 | 0.92 | 1 | Hip1 | huntingtin interacting protein 1 | 2771 | 2143 | 2149 | 902 | 684 | 853 | 836 | 677 | 761 | 1047 | 791 |
| 1106 | -0.10 | 0.13 | -0.75 | 0.45 | 1 | Fbf1 | Fas (TNFRSF6) binding factor 1 | 1410 | 1459 | 1577 | 749 | 1020 | 1423 | 836 | 875 | 831 | 831 | 1155 |
| 854 | -0.25 | 0.17 | -1.52 | 0.13 | 1 | Taf2 | TATA-box binding protein associated fact | 761 | 615 | 761 | 775 | 1036 | 1238 | 836 | 460 | 868 | 823 | 1219 |
| 847 | 0.04 | 0.12 | 0.32 | 0.75 | 1 | Uckl1 | uridine-cytidine kinase 1-like 1 | 595 | 654 | 474 | 811 | 701 | 1409 | 836 | 894 | 669 | 942 | 1332 |
| 748 | 0.11 | 0.17 | 0.64 | 0.52 | 1 | Eef1akmt2 | EEF1A lysine methyltransferase 2 | 588 | 635 | 677 | 824 | 697 | 675 | 836 | 1091 | 808 | 699 | 701 |
| 714 | -0.15 | 0.14 | -1.04 | 0.30 | 1 | Urb1 | URB1 ribosome biogenesis 1 homolog (S | 554 | 501 | 589 | 769 | 710 | 947 | 836 | 591 | 767 | 782 | 808 |
| 1250 | -0.01 | 0.13 | -0.07 | 0.94 | 1 | Rpl29 | ribosomal protein L29 | 1701 | 1492 | 2369 | 982 | 1336 | 953 | 835 | 1003 | 1284 | 775 | 1013 |
| 978 | 0.11 | 0.13 | 0.85 | 0.39 | 1 | Tbk1 | TANK-binding kinase 1 | 836 | 713 | 793 | 929 | 890 | 1421 | 835 | 1045 | 973 | 984 | 1335 |
| 957 | 0.21 | 0.25 | 0.87 | 0.38 | 1 | Hspbp1 | HSPA (heat shock 70kDa) binding proteir | 1070 | 817 | 1151 | 742 | 1027 | 836 | 835 | 1566 | 905 | 752 | 829 |
| 648 | 0.00 | 0.20 | -0.02 | 0.98 | 1 | Mif | macrophage migration inhibitory factor (g | 583 | 464 | 681 | 760 | 541 | 561 | 835 | 733 | 631 | 831 | 507 |
| 941 | 0.07 | 0.14 | 0.49 | 0.62 | 1 | Prmt2 | protein arginine N-methyltransferase 2 | 1048 | 1295 | 1182 | 778 | 903 | 817 | 834 | 945 | 815 | 922 | 813 |
| 761 | 0.10 | 0.12 | 0.82 | 0.41 | 1 | Prpsap2 | phosphoribosyl pyrophosphate synthetas | 692 | 656 | 706 | 728 | 760 | 726 | 834 | 754 | 791 | 894 | 825 |
| 631 | 0.02 | 0.22 | 0.09 | 0.93 | 1 | Alad | aminolevulinate, delta-, dehydratase | 1242 | 627 | 725 | 347 | 516 | 474 | 834 | 592 | 351 | 799 | 439 |
| 974 | -0.09 | 0.22 | -0.41 | 0.68 | 1 | Mynn | myoneurin | 822 | 1250 | 1062 | 936 | 855 | 1240 | 833 | 519 | 936 | 884 | 1379 |
| 911 | 0.13 | 0.14 | 0.93 | 0.35 | 1 | Hmgcl | 3-hydroxy-3-methylglutaryl-Coenzyme A l | 1051 | 773 | 1058 | 764 | 917 | 894 | 833 | 958 | 903 | 1015 | 856 |
| 906 | 0.07 | 0.18 | 0.38 | 0.70 | 1 | Pank2 | pantothenate kinase 2 | 1108 | 913 | 743 | 983 | 768 | 915 | 833 | 1160 | 771 | 707 | 1064 |
| 855 | -0.11 | 0.15 | -0.73 | 0.47 | 1 | Zfp316 | zinc finger protein 316 | 831 | 975 | 837 | 1042 | 846 | 796 | 833 | 740 | 827 | 934 | 746 |
| 811 | 0.06 | 0.17 | 0.32 | 0.75 | 1 | Pag1 | phosphoprotein associated with glycosph | 542 | 493 | 342 | 1014 | 818 | 1044 | 833 | 842 | 955 | 1005 | 1036 |
| 807 | -0.25 | 0.20 | -1.30 | 0.19 | 1 | Aph1a | aph1 homolog A, gamma secretase subu | 847 | 792 | 1045 | 900 | 873 | 731 | 833 | 461 | 837 | 850 | 706 |
| 1174 | -0.03 | 0.12 | -0.27 | 0.78 | 1 | Btbd3 | BTB (POZ) domain containing 3 | 1764 | 1958 | 1752 | 979 | 842 | 1105 | 832 | 904 | 964 | 734 | 1074 |
| 1115 | -0.03 | 0.16 | -0.21 | 0.84 | 1 | Map3k1 | mitogen-activated protein kinase kinase k | 1659 | 1237 | 1091 | 1099 | 846 | 1422 | 832 | 963 | 899 | 854 | 1366 |
| 968 | 0.02 | 0.11 | 0.21 | 0.83 | 1 | Stoml2 | stomatin (Epb7.2)-like 2 | 1200 | 968 | 1164 | 820 | 996 | 980 | 832 | 831 | 990 | 890 | 976 |

| | | | | | | | | | | | | | | | | | |
|---|---|---|---|---|---|---|---|---|---|---|---|---|---|---|---|---|---|
| 930 | 0.12 | 0.18 | 0.64 | 0.53 | 1 | Tbl3 | transducin (beta)-like 3 | 860 | 682 | 854 | 1002 | 944 | 959 | 832 | 1321 | 910 | 1046 | 815 |
| 893 | 0.20 | 0.13 | 1.59 | 0.11 | 1 | Adnp2 | ADNP homeobox 2 | 851 | 889 | 855 | 923 | 699 | 910 | 832 | 1123 | 818 | 943 | 982 |
| 801 | 0.04 | 0.18 | 0.23 | 0.82 | 1 | Ghdc | GH3 domain containing | 507 | 760 | 718 | 883 | 1005 | 645 | 832 | 1039 | 775 | 985 | 659 |
| 780 | 0.16 | 0.15 | 1.10 | 0.27 | 1 | Txnl4a | thioredoxin-like 4A | 645 | 537 | 733 | 789 | 928 | 609 | 832 | 962 | 940 | 809 | 792 |
| 715 | 0.00 | 0.18 | 0.01 | 0.99 | 1 | Mppe1 | metallophosphoesterase 1 | 643 | 584 | 657 | 645 | 682 | 828 | 832 | 847 | 703 | 589 | 853 |
| 707 | 0.06 | 0.13 | 0.48 | 0.63 | 1 | Bmyc | brain expressed myelocytomatosis oncog | 587 | 623 | 533 | 796 | 803 | 527 | 832 | 887 | 748 | 849 | 591 |
| 668 | -0.17 | 0.16 | -1.07 | 0.29 | 1 | Ubb-ps | | 731 | 677 | 749 | 782 | 778 | 296 | 832 | 640 | 779 | 877 | 210 |
| 1003 | -0.09 | 0.13 | -0.66 | 0.51 | 1 | Pfas | phosphoribosylformylglycinamidine synth | 1004 | 875 | 963 | 968 | 1146 | 1284 | 831 | 867 | 1027 | 929 | 1140 |
| 993 | 0.06 | 0.16 | 0.37 | 0.71 | 1 | Sap30bp | SAP30 binding protein | 1314 | 949 | 1132 | 789 | 1182 | 895 | 831 | 981 | 986 | 1002 | 857 |
| 938 | 0.14 | 0.17 | 0.83 | 0.41 | 1 | Phf5a | PHD finger protein 5A | 1247 | 917 | 1027 | 803 | 920 | 833 | 831 | 1142 | 907 | 810 | 882 |
| 892 | 0.14 | 0.13 | 1.04 | 0.30 | 1 | Taf3 | TATA-box binding protein associated fact | 1126 | 1017 | 942 | 855 | 753 | 757 | 831 | 1081 | 839 | 877 | 738 |
| 880 | 0.18 | 0.14 | 1.28 | 0.20 | 1 | Ovca2 | candidate tumor suppressor in ovarian ca | 637 | 682 | 762 | 801 | 1057 | 878 | 831 | 993 | 1031 | 1126 | 880 |
| 683 | -0.06 | 0.23 | -0.26 | 0.79 | 1 | Dusp28 | dual specificity phosphatase 28 | 343 | 501 | 534 | 814 | 877 | 595 | 831 | 523 | 886 | 1007 | 596 |
| 983 | -0.22 | 0.16 | -1.39 | 0.17 | 1 | St14 | suppression of tumorigenicity 14 (colon ca | 1579 | 1322 | 1706 | 831 | 936 | 724 | 830 | 879 | 705 | 808 | 491 |
| 961 | -0.18 | 0.14 | -1.33 | 0.18 | 1 | Stx17 | syntaxin 17 | 923 | 1119 | 1108 | 1075 | 892 | 1116 | 830 | 1017 | 746 | 626 | 1120 |
| 918 | 0.04 | 0.15 | 0.23 | 0.81 | 1 | Lamtor3 | late endosomal/lysosomal adaptor, MAPK | 724 | 659 | 831 | 992 | 765 | 1317 | 830 | 937 | 966 | 797 | 1285 |
| 899 | -0.12 | 0.18 | -0.67 | 0.51 | 1 | Mark1 | MAP/microtubule affinity regulating kinase | 978 | 690 | 774 | 1168 | 1006 | 871 | 830 | 893 | 972 | 887 | 818 |
| 791 | -0.08 | 0.14 | -0.58 | 0.56 | 1 | Rcan1 | regulator of calcineurin 1 | 947 | 880 | 761 | 866 | 785 | 669 | 830 | 730 | 805 | 721 | 706 |
| 747 | 0.13 | 0.16 | 0.81 | 0.42 | 1 | Prmt9 | protein arginine methyltransferase 9 | 618 | 768 | 531 | 649 | 730 | 786 | 830 | 870 | 671 | 894 | 873 |
| 736 | -0.34 | 0.21 | -1.65 | 0.10 | 1 | Fancm | Fanconi anemia, complementation group | 554 | 528 | 435 | 766 | 886 | 1128 | 830 | 430 | 779 | 612 | 1144 |
| 1052 | -0.05 | 0.17 | -0.27 | 0.79 | 1 | Myo9a | myosin IXa | 1444 | 1541 | 1246 | 894 | 712 | 1266 | 829 | 684 | 913 | 665 | 1376 |
| 960 | -0.07 | 0.13 | -0.56 | 0.58 | 1 | Snrpa | small nuclear ribonucleoprotein polypepti | 1084 | 1015 | 1053 | 779 | 1047 | 1150 | 829 | 816 | 831 | 898 | 1055 |
| 854 | -0.14 | 0.13 | -1.08 | 0.28 | 1 | Gucd1 | guanylyl cyclase domain containing 1 | 841 | 805 | 750 | 878 | 911 | 1049 | 829 | 716 | 773 | 913 | 928 |
| 853 | 0.15 | 0.13 | 1.17 | 0.24 | 1 | Utp15 | UTP15 small subunit processome compo | 896 | 758 | 740 | 802 | 785 | 903 | 829 | 889 | 943 | 840 | 1003 |
| 717 | -0.06 | 0.17 | -0.34 | 0.73 | 1 | Fahd1 | fumarylacetoacetate hydrolase domain cc | 547 | 753 | 776 | 770 | 788 | 583 | 829 | 675 | 740 | 822 | 603 |
| 716 | -0.02 | 0.16 | -0.14 | 0.89 | 1 | 5730409E04Ril | RIKEN cDNA 5730409E04Rik gene | 837 | 833 | 850 | 621 | 668 | 587 | 829 | 755 | 631 | 608 | 654 |
| 1173 | 0.04 | 0.12 | 0.29 | 0.78 | 1 | Shfl | shiftless antiviral inhibitor of ribosomal fra | 667 | 737 | 750 | 909 | 932 | 2746 | 828 | 1083 | 855 | 936 | 2457 |
| 1028 | -0.03 | 0.16 | -0.16 | 0.88 | 1 | Magoh | mago homolog, exon junction complex cc | 1308 | 1248 | 1597 | 854 | 996 | 910 | 828 | 634 | 851 | 1012 | 1074 |
| 996 | 0.10 | 0.12 | 0.83 | 0.40 | 1 | Cerkl | ceramide kinase-like | 524 | 662 | 550 | 1022 | 912 | 1675 | 828 | 1161 | 818 | 1007 | 1790 |
| 959 | -0.26 | 0.14 | -1.84 | 0.07 | 1 | Ranbp10 | RAN binding protein 10 | 927 | 954 | 985 | 910 | 966 | 1494 | 828 | 616 | 906 | 809 | 1159 |
| 845 | 0.33 | 0.21 | 1.63 | 0.10 | 1 | Kpna2 | karyopherin (importin) alpha 2 | 420 | 431 | 304 | 814 | 1130 | 806 | 828 | 1453 | 1225 | 911 | 967 |
| 784 | -0.09 | 0.12 | -0.77 | 0.44 | 1 | Mrpl44 | mitochondrial ribosomal protein L44 | 854 | 857 | 779 | 879 | 750 | 708 | 828 | 769 | 775 | 764 | 660 |
| 1140 | 0.16 | 0.15 | 1.03 | 0.30 | 1 | Cnppd1 | cyclin Pas1/PHO80 domain containing 1 | 1334 | 1143 | 1123 | 935 | 1150 | 1323 | 827 | 1153 | 986 | 1114 | 1455 |
| 1014 | 0.08 | 0.12 | 0.71 | 0.48 | 1 | Cwf19l2 | CWF19-like 2, cell cycle control (S. pomb | 902 | 1172 | 1065 | 914 | 894 | 1271 | 827 | 953 | 942 | 945 | 1271 |
| 962 | 0.22 | 0.12 | 1.80 | 0.07 | 1 | Diaph2 | diaphanous related formin 2 | 1010 | 964 | 869 | 1058 | 894 | 789 | 827 | 1157 | 1005 | 1124 | 884 |
| 856 | -0.13 | 0.16 | -0.82 | 0.41 | 1 | Asb7 | ankyrin repeat and SOCS box-containing | 852 | 947 | 900 | 929 | 874 | 864 | 827 | 718 | 1055 | 665 | 786 |
| 827 | -0.05 | 0.16 | -0.29 | 0.77 | 1 | Glrx3 | glutaredoxin 3 | 1145 | 906 | 1104 | 709 | 842 | 641 | 827 | 622 | 708 | 717 | 878 |
| 774 | -0.01 | 0.14 | -0.06 | 0.95 | 1 | Pbdc1 | polysaccharide biosynthesis domain cont | 686 | 736 | 704 | 726 | 796 | 844 | 827 | 842 | 693 | 715 | 941 |
| 768 | -0.25 | 0.16 | -1.57 | 0.12 | 1 | Rbm45 | RNA binding motif protein 45 | 778 | 677 | 758 | 761 | 876 | 896 | 827 | 492 | 870 | 701 | 812 |
| 952 | -0.11 | 0.20 | -0.57 | 0.57 | 1 | Nup188 | nucleoporin 188 | 1229 | 763 | 868 | 1075 | 989 | 1060 | 826 | 890 | 1059 | 836 | 880 |
| 742 | -0.09 | 0.15 | -0.58 | 0.56 | 1 | Slc7a6 | solute carrier family 7 (cationic amino acid | 791 | 524 | 573 | 735 | 705 | 963 | 826 | 652 | 746 | 712 | 931 |
| 700 | -0.05 | 0.19 | -0.28 | 0.78 | 1 | Arhgap26 | Rho GTPase activating protein 26 | 498 | 473 | 409 | 805 | 703 | 882 | 826 | 703 | 942 | 620 | 843 |
| 1069 | -0.13 | 0.17 | -0.78 | 0.43 | 1 | Ggh | gamma-glutamyl hydrolase | 1606 | 2576 | 2115 | 788 | 702 | 543 | 825 | 655 | 713 | 743 | 494 |
| 912 | -0.14 | 0.16 | -0.90 | 0.37 | 1 | Dennd3 | DENN/MADD domain containing 3 | 1506 | 1320 | 1447 | 596 | 767 | 839 | 825 | 715 | 770 | 630 | 613 |
| 897 | 0.04 | 0.13 | 0.34 | 0.73 | 1 | Ebp | phenylalkylamine Ca2+ antagonist (emop | 918 | 844 | 828 | 966 | 1023 | 762 | 825 | 987 | 959 | 1010 | 743 |
| 885 | -0.08 | 0.18 | -0.45 | 0.65 | 1 | Nfatc2 | nuclear factor of activated T cells, cytopla: | 913 | 753 | 733 | 960 | 838 | 1151 | 825 | 781 | 1073 | 750 | 961 |
| 846 | 0.00 | 0.13 | -0.01 | 0.99 | 1 | Cfap410 | cilia and flagella associated protein 410 | 714 | 689 | 671 | 813 | 919 | 1063 | 825 | 959 | 876 | 738 | 1043 |
| 841 | 0.08 | 0.16 | 0.54 | 0.59 | 1 | Nudt19 | nudix (nucleoside diphosphate linked mo | 859 | 749 | 881 | 1032 | 854 | 581 | 825 | 1054 | 793 | 962 | 663 |
| 826 | 0.06 | 0.18 | 0.33 | 0.74 | 1 | Larp7 | La ribonucleoprotein domain family, mem | 1161 | 717 | 1133 | 753 | 687 | 713 | 825 | 783 | 807 | 708 | 799 |
| 825 | -0.24 | 0.18 | -1.34 | 0.18 | 1 | Pced1a | PC-esterase domain containing 1A | 564 | 647 | 449 | 1107 | 786 | 1289 | 825 | 628 | 773 | 770 | 1235 |
| 920 | -0.34 | 0.22 | -1.56 | 0.12 | 1 | Ephx1 | epoxide hydrolase 1, microsomal | 2219 | 1468 | 1325 | 720 | 808 | 485 | 824 | 358 | 778 | 693 | 444 |
| 904 | -0.06 | 0.13 | -0.47 | 0.64 | 1 | Lrig2 | leucine-rich repeats and immunoglobulin | 985 | 1001 | 849 | 894 | 775 | 1124 | 824 | 781 | 805 | 724 | 1187 |
| 851 | 0.07 | 0.17 | 0.42 | 0.67 | 1 | Ddx58 | DEAD (Asp-Glu-Ala-Asp) box polypeptide | 1395 | 1400 | 1195 | 513 | 667 | 626 | 824 | 717 | 704 | 674 | 644 |
| 798 | -0.15 | 0.15 | -0.98 | 0.33 | 1 | Heatr6 | HEAT repeat containing 6 | 714 | 741 | 800 | 794 | 885 | 919 | 824 | 900 | 778 | 585 | 843 |
| 680 | -0.26 | 0.23 | -1.14 | 0.25 | 1 | Pgm2 | phosphoglucomutase 2 | 589 | 563 | 577 | 826 | 725 | 729 | 824 | 372 | 774 | 791 | 715 |
| 867 | 0.00 | 0.17 | -0.01 | 0.99 | 1 | Pkhd1 | polycystic kidney and hepatic disease 1 | 1404 | 1292 | 1678 | 430 | 647 | 693 | 823 | 511 | 774 | 638 | 643 |
| 850 | 0.07 | 0.13 | 0.49 | 0.62 | 1 | Armc9 | armadillo repeat containing 9 | 640 | 632 | 782 | 814 | 850 | 1090 | 822 | 943 | 858 | 904 | 1016 |
| 828 | 0.07 | 0.14 | 0.48 | 0.63 | 1 | Zfp777 | zinc finger protein 777 | 779 | 832 | 797 | 863 | 778 | 806 | 822 | 767 | 820 | 1006 | 837 |
| 958 | 0.05 | 0.14 | 0.37 | 0.71 | 1 | Mapk8 | mitogen-activated protein kinase 8 | 935 | 984 | 1106 | 803 | 838 | 1210 | 821 | 960 | 901 | 666 | 1318 |
| 917 | -0.05 | 0.17 | -0.27 | 0.79 | 1 | Igbp1 | immunoglobulin (CD79A) binding protein | 1415 | 1231 | 1225 | 802 | 685 | 823 | 821 | 524 | 746 | 984 | 829 |

| | | | | | | | | | | | | | | | | | | |
|---|---|---|---|---|---|---|---|---|---|---|---|---|---|---|---|---|---|---|
| 862 | -0.34 | 0.23 | -1.49 | 0.14 | 1 | D230025D16Ri | RIKEN cDNA D230025D16 gene | 961 | 1179 | 1001 | 849 | 807 | 1041 | 821 | 389 | 710 | 877 | 844 |
| 827 | 0.03 | 0.23 | 0.15 | 0.88 | 1 | Crb3 | crumbs family member 3 | 415 | 487 | 721 | 869 | 1078 | 889 | 821 | 708 | 1033 | 1349 | 721 |
| 809 | 0.12 | 0.19 | 0.61 | 0.54 | 1 | Nbn | nibrin | 597 | 590 | 452 | 875 | 857 | 911 | 821 | 675 | 1041 | 915 | 1161 |
| 778 | 0.11 | 0.20 | 0.55 | 0.58 | 1 | Zfhx3 | zinc finger homeobox 3 | 945 | 1078 | 832 | 523 | 603 | 805 | 821 | 850 | 730 | 699 | 670 |
| 774 | 0.07 | 0.21 | 0.32 | 0.75 | 1 | Prkaa2 | protein kinase, AMP-activated, alpha 2 ca | 1002 | 1277 | 1048 | 653 | 511 | 535 | 821 | 557 | 614 | 994 | 501 |
| 752 | -0.01 | 0.14 | -0.06 | 0.95 | 1 | Washc3 | WASH complex subunit 3 | 753 | 765 | 629 | 859 | 724 | 672 | 821 | 708 | 795 | 869 | 679 |
| 609 | 0.01 | 0.19 | 0.07 | 0.95 | 1 | Tmem126a | transmembrane protein 126A | 415 | 445 | 417 | 688 | 642 | 551 | 821 | 518 | 766 | 756 | 677 |
| 856 | -0.06 | 0.15 | -0.42 | 0.67 | 1 | Ubl7 | ubiquitin-like 7 (bone marrow stromal cell | 942 | 1057 | 1116 | 949 | 808 | 628 | 820 | 857 | 746 | 949 | 543 |
| 792 | 0.06 | 0.19 | 0.33 | 0.74 | 1 | Efl1 | elongation factor like GTPase 1 | 862 | 857 | 811 | 612 | 747 | 814 | 820 | 478 | 819 | 1025 | 862 |
| 659 | -0.05 | 0.17 | -0.31 | 0.76 | 1 | Pdp1 | pyruvate dehyrogenase phosphatase cata | 550 | 392 | 520 | 762 | 629 | 733 | 820 | 652 | 677 | 678 | 839 |
| 972 | 0.08 | 0.13 | 0.57 | 0.57 | 1 | Gstt1 | glutathione S-transferase, theta 1 | 1778 | 1599 | 1326 | 770 | 742 | 550 | 819 | 905 | 678 | 1019 | 507 |
| 821 | -0.21 | 0.16 | -1.33 | 0.18 | 1 | Etv6 | ets variant 6 | 1492 | 1053 | 1025 | 670 | 752 | 696 | 819 | 674 | 611 | 600 | 639 |
| 818 | -0.05 | 0.17 | -0.27 | 0.79 | 1 | Zfand2a | zinc finger, AN1-type domain 2A | 725 | 816 | 903 | 796 | 762 | 940 | 819 | 571 | 851 | 916 | 899 |
| 765 | 0.17 | 0.14 | 1.18 | 0.24 | 1 | Cdkl5 | cyclin-dependent kinase-like 5 | 624 | 604 | 586 | 677 | 729 | 882 | 819 | 700 | 922 | 991 | 881 |
| 702 | 0.20 | 0.20 | 0.96 | 0.34 | 1 | Ndufaf7 | NADH:ubiquinone oxidoreductase compl | 656 | 749 | 451 | 504 | 634 | 784 | 819 | 721 | 728 | 739 | 935 |
| 1101 | 0.24 | 0.14 | 1.73 | 0.08 | 1 | Pygo2 | pygopus 2 | 1297 | 1124 | 1200 | 906 | 1010 | 1157 | 818 | 1257 | 989 | 1087 | 1264 |
| 764 | 0.00 | 0.12 | -0.02 | 0.99 | 1 | Psmb3 | proteasome (prosome, macropain) subun | 999 | 724 | 864 | 675 | 788 | 632 | 818 | 660 | 809 | 782 | 653 |
| 968 | -0.09 | 0.19 | -0.49 | 0.62 | 1 | Acap3 | ArfGAP with coiled-coil, ankyrin repeat an | 1370 | 1193 | 856 | 1058 | 930 | 903 | 817 | 773 | 889 | 1095 | 759 |
| 925 | -0.08 | 0.12 | -0.66 | 0.51 | 1 | Ube2j2 | ubiquitin-conjugating enzyme E2J 2 | 771 | 915 | 829 | 1127 | 1017 | 970 | 817 | 1065 | 866 | 778 | 1020 |
| 792 | -0.03 | 0.19 | -0.18 | 0.86 | 1 | Idua | iduronidase, alpha-L | 675 | 639 | 756 | 893 | 786 | 837 | 817 | 754 | 658 | 1166 | 733 |
| 667 | -0.09 | 0.13 | -0.67 | 0.50 | 1 | Pgap3 | post-GPI attachment to proteins 3 | 442 | 481 | 425 | 798 | 781 | 683 | 817 | 825 | 761 | 743 | 587 |
| 850 | -0.03 | 0.12 | -0.23 | 0.82 | 1 | Glrx2 | glutaredoxin 2 (thioltransferase) | 903 | 999 | 709 | 854 | 668 | 1057 | 816 | 953 | 631 | 747 | 1011 |
| 850 | -0.13 | 0.14 | -0.93 | 0.35 | 1 | Tor1a | torsin family 1, member A (torsin A) | 973 | 774 | 953 | 892 | 934 | 823 | 816 | 680 | 811 | 849 | 844 |
| 779 | 0.01 | 0.13 | 0.05 | 0.96 | 1 | Tm2d3 | TM2 domain containing 3 | 640 | 694 | 661 | 894 | 918 | 653 | 816 | 798 | 854 | 969 | 672 |
| 738 | -0.06 | 0.15 | -0.42 | 0.67 | 1 | Mblac2 | metallo-beta-lactamase domain containin | 600 | 796 | 642 | 833 | 810 | 652 | 816 | 732 | 722 | 875 | 642 |
| 715 | 0.27 | 0.22 | 1.23 | 0.22 | 1 | Prorsd1 | prolyl-tRNA synthetase domain containin | 450 | 676 | 633 | 625 | 657 | 669 | 816 | 666 | 813 | 1092 | 773 |
| 972 | 0.20 | 0.16 | 1.25 | 0.21 | 1 | Prpf38a | PRP38 pre-mRNA processing factor 38 (y | 1211 | 1142 | 842 | 952 | 830 | 860 | 815 | 1356 | 759 | 945 | 979 |
| 917 | -0.21 | 0.13 | -1.66 | 0.10 | 1 | Arvcf | armadillo repeat gene deleted in velocarc | 1713 | 1545 | 1675 | 623 | 618 | 709 | 815 | 510 | 558 | 645 | 675 |
| 806 | 0.21 | 0.16 | 1.28 | 0.20 | 1 | Mphosph10 | M-phase phosphoprotein 10 (U3 small nu | 510 | 690 | 624 | 771 | 809 | 840 | 815 | 1105 | 857 | 708 | 1131 |
| 682 | 0.00 | 0.19 | 0.03 | 0.98 | 1 | Dsc2 | desmocollin 2 | 299 | 261 | 334 | 985 | 739 | 745 | 815 | 690 | 1023 | 859 | 752 |
| 1128 | -0.11 | 0.13 | -0.84 | 0.40 | 1 | Hdac1 | histone deacetylase 1 | 1169 | 1340 | 1443 | 1069 | 1373 | 1148 | 814 | 951 | 1040 | 859 | 1202 |
| 908 | 0.15 | 0.16 | 0.94 | 0.35 | 1 | Loxl2 | lysyl oxidase-like 2 | 1251 | 931 | 1063 | 815 | 703 | 849 | 814 | 706 | 1056 | 855 | 945 |
| 893 | 0.16 | 0.14 | 1.17 | 0.24 | 1 | Enoph1 | enolase-phosphatase 1 | 705 | 843 | 853 | 835 | 989 | 868 | 814 | 1043 | 926 | 1052 | 900 |
| 829 | 0.01 | 0.17 | 0.09 | 0.93 | 1 | Ncapd3 | non-SMC condensin II complex, subunit D | 616 | 623 | 551 | 823 | 856 | 1143 | 814 | 657 | 853 | 1128 | 1056 |
| 827 | 0.09 | 0.12 | 0.75 | 0.45 | 1 | 2310011J03Rik | RIKEN cDNA 2310011J03 gene | 796 | 885 | 831 | 815 | 913 | 643 | 814 | 914 | 876 | 930 | 674 |
| 825 | -0.05 | 0.16 | -0.34 | 0.73 | 1 | Klhl26 | kelch-like 26 | 866 | 835 | 849 | 899 | 696 | 911 | 813 | 714 | 764 | 987 | 741 |
| 789 | -0.32 | 0.26 | -1.19 | 0.23 | 1 | Bop1 | block of proliferation 1 | 1006 | 612 | 982 | 726 | 838 | 854 | 813 | 250 | 856 | 942 | 798 |
| 892 | -0.11 | 0.16 | -0.70 | 0.48 | 1 | Bri3bp | Bri3 binding protein | 1070 | 1068 | 1151 | 1020 | 903 | 634 | 812 | 1161 | 721 | 719 | 558 |
| 1025 | 0.13 | 0.15 | 0.85 | 0.40 | 1 | Rfc2 | replication factor C (activator 1) 2 | 1240 | 1202 | 909 | 942 | 946 | 1084 | 811 | 1191 | 951 | 982 | 1014 |
| 780 | -0.11 | 0.15 | -0.74 | 0.46 | 1 | Pgs1 | phosphatidylglycerophosphate synthase | 779 | 706 | 723 | 758 | 791 | 922 | 811 | 572 | 651 | 891 | 977 |
| 655 | 0.08 | 0.24 | 0.31 | 0.76 | 1 | Ehhadh | enoyl-Coenzyme A, hydratase/3-hydroxya | 284 | 385 | 497 | 907 | 679 | 500 | 811 | 664 | 958 | 1085 | 441 |
| 948 | -0.07 | 0.14 | -0.52 | 0.61 | 1 | Ccnk | cyclin K | 1071 | 1097 | 868 | 1059 | 948 | 975 | 810 | 882 | 894 | 961 | 860 |
| 908 | -0.02 | 0.15 | -0.15 | 0.88 | 1 | Plekhh2 | pleckstrin homology domain containing, fa | 1491 | 1169 | 963 | 831 | 704 | 850 | 810 | 630 | 763 | 949 | 824 |
| 847 | 0.13 | 0.14 | 0.89 | 0.37 | 1 | Acad9 | acyl-Coenzyme A dehydrogenase family, | 1193 | 792 | 989 | 654 | 780 | 782 | 810 | 830 | 747 | 980 | 755 |
| 732 | 0.13 | 0.19 | 0.65 | 0.52 | 1 | Fbxw4 | F-box and WD-40 domain protein 4 | 794 | 694 | 894 | 571 | 681 | 653 | 810 | 921 | 634 | 737 | 663 |
| 1298 | 0.09 | 0.16 | 0.60 | 0.55 | 1 | Arhgef3 | Rho guanine nucleotide exchange factor | 1997 | 1758 | 1955 | 1090 | 912 | 1303 | 809 | 1189 | 1003 | 730 | 1532 |
| 1110 | -0.19 | 0.19 | -1.00 | 0.32 | 1 | Tgfbr3 | transforming growth factor, beta receptor I | 2837 | 1709 | 1583 | 950 | 738 | 740 | 808 | 573 | 779 | 811 | 684 |
| 881 | -0.21 | 0.17 | -1.25 | 0.21 | 1 | Ctdspl2 | CTD (carboxy-terminal domain, RNA poly | 1103 | 925 | 957 | 916 | 835 | 1008 | 808 | 620 | 856 | 634 | 1032 |
| 822 | 0.17 | 0.17 | 1.01 | 0.31 | 1 | Decr1 | 2,4-dienoyl CoA reductase 1, mitochondri | 1126 | 723 | 806 | 855 | 815 | 538 | 808 | 981 | 828 | 898 | 664 |
| 813 | -0.41 | 0.21 | -1.92 | 0.05 | 1 | Rab23 | RAB23, member RAS oncogene family | 908 | 832 | 667 | 1187 | 821 | 910 | 808 | 549 | 758 | 625 | 877 |
| 790 | -0.11 | 0.15 | -0.74 | 0.46 | 1 | Sbk1 | SH3-binding kinase 1 | 588 | 780 | 582 | 1045 | 1020 | 615 | 808 | 1042 | 744 | 903 | 563 |
| 757 | 0.16 | 0.15 | 1.05 | 0.29 | 1 | Reep2 | receptor accessory protein 2 | 653 | 823 | 724 | 689 | 686 | 708 | 808 | 921 | 748 | 837 | 727 |
| 753 | -0.09 | 0.15 | -0.58 | 0.56 | 1 | Ptpn2 | protein tyrosine phosphatase, non-recept | 665 | 807 | 713 | 657 | 698 | 967 | 808 | 591 | 672 | 706 | 1001 |
| 933 | 0.07 | 0.15 | 0.47 | 0.64 | 1 | Ccdc71 | coiled-coil domain containing 71 | 1058 | 1081 | 1130 | 964 | 820 | 804 | 807 | 936 | 789 | 1130 | 746 |
| 742 | -0.15 | 0.25 | -0.59 | 0.55 | 1 | Zfp174 | zinc finger protein 174 | 353 | 577 | 612 | 919 | 851 | 889 | 807 | 470 | 850 | 933 | 899 |
| 1146 | -0.07 | 0.14 | -0.51 | 0.61 | 1 | Zkscan3 | zinc finger with KRAB and SCAN domains | 867 | 894 | 1075 | 724 | 626 | 2849 | 806 | 675 | 716 | 661 | 2712 |
| 881 | 0.27 | 0.18 | 1.47 | 0.14 | 1 | Eef2kmt | eukaryotic elongation factor 2 lysine meth | 878 | 742 | 904 | 732 | 814 | 893 | 806 | 1262 | 847 | 942 | 870 |
| 855 | -0.05 | 0.20 | -0.28 | 0.78 | 1 | Nup155 | nucleoporin 155 | 902 | 864 | 878 | 811 | 645 | 1177 | 806 | 789 | 927 | 518 | 1092 |
| 845 | 0.12 | 0.13 | 0.91 | 0.36 | 1 | Aptx | aprataxin | 770 | 918 | 840 | 782 | 825 | 822 | 806 | 951 | 981 | 819 | 777 |
| 729 | -0.25 | 0.16 | -1.63 | 0.10 | 1 | Mavs | mitochondrial antiviral signaling protein | 987 | 902 | 875 | 691 | 649 | 708 | 806 | 448 | 622 | 653 | 681 |

| | | | | | | | | | | | | | | | | | |
|---|---|---|---|---|---|---|---|---|---|---|---|---|---|---|---|---|---|
| 959 | -0.20 | 0.13 | -1.50 | 0.13 | 1 | Ino80e | INO80 complex subunit E | 1186 | 1074 | 1055 | 981 | 976 | 1106 | 805 | 781 | 836 | 831 | 925 |
| 813 | 0.07 | 0.16 | 0.43 | 0.67 | 1 | Itgb1bp1 | integrin beta 1 binding protein 1 | 910 | 710 | 1018 | 762 | 808 | 698 | 805 | 848 | 729 | 940 | 709 |
| 788 | 0.00 | 0.15 | -0.03 | 0.98 | 1 | Pex1 | peroxisomal biogenesis factor 1 | 434 | 366 | 371 | 780 | 897 | 1253 | 805 | 638 | 854 | 1001 | 1273 |
| 754 | -0.35 | 0.21 | -1.70 | 0.09 | 1 | Ccdc157 | coiled-coil domain containing 157 | 836 | 719 | 810 | 861 | 571 | 1075 | 805 | 741 | 568 | 420 | 883 |
| 708 | -0.18 | 0.16 | -1.11 | 0.27 | 1 | Arhgap18 | Rho GTPase activating protein 18 | 1107 | 974 | 902 | 737 | 531 | 494 | 805 | 505 | 498 | 684 | 552 |
| 696 | -0.14 | 0.16 | -0.86 | 0.39 | 1 | Ctnnd1 | catenin (cadherin associated protein), del | 735 | 730 | 797 | 655 | 694 | 664 | 805 | 502 | 785 | 645 | 645 |
| 909 | 0.14 | 0.20 | 0.70 | 0.48 | 1 | Tcof1 | treacle ribosome biogenesis factor 1 | 1504 | 1239 | 1137 | 607 | 772 | 705 | 804 | 1105 | 748 | 764 | 612 |
| 761 | -0.05 | 0.16 | -0.33 | 0.74 | 1 | Prkcz | protein kinase C, zeta | 791 | 652 | 786 | 780 | 833 | 702 | 804 | 907 | 827 | 658 | 627 |
| 743 | 0.06 | 0.16 | 0.36 | 0.72 | 1 | Manbal | mannosidase, beta A, lysosomal-like | 740 | 662 | 865 | 681 | 865 | 535 | 804 | 870 | 751 | 899 | 498 |
| 888 | 0.04 | 0.17 | 0.25 | 0.80 | 1 | Mrpl38 | mitochondrial ribosomal protein L38 | 825 | 879 | 1235 | 898 | 897 | 771 | 802 | 1038 | 780 | 837 | 807 |
| 723 | -0.15 | 0.14 | -1.06 | 0.29 | 1 | Fiz1 | Flt3 interacting zinc finger protein 1 | 749 | 705 | 846 | 866 | 739 | 572 | 802 | 759 | 661 | 720 | 534 |
| 713 | 0.05 | 0.18 | 0.30 | 0.77 | 1 | Zfp738 | zinc finger protein 738 | 461 | 403 | 504 | 560 | 577 | 1229 | 802 | 711 | 656 | 587 | 1347 |
| 1568 | 0.03 | 0.17 | 0.20 | 0.85 | 1 | Lgmn | legumain | 5023 | 2725 | 3437 | 920 | 844 | 402 | 801 | 836 | 931 | 942 | 383 |
| 1064 | 0.17 | 0.70 | 0.24 | 0.81 | 1 | Olfm4 | olfactomedin 4 | 4 | 72 | 78 | 731 | 967 | 3075 | 801 | 768 | 898 | 1101 | 3205 |
| 996 | 0.01 | 0.14 | 0.10 | 0.92 | 1 | Eapp | E2F-associated phosphoprotein | 1193 | 1480 | 1217 | 942 | 873 | 888 | 801 | 805 | 817 | 837 | 1097 |
| 860 | 0.11 | 0.16 | 0.72 | 0.47 | 1 | Tubgcp5 | tubulin, gamma complex associated prote | 688 | 498 | 707 | 789 | 826 | 1206 | 801 | 701 | 888 | 1143 | 1213 |
| 838 | 0.01 | 0.12 | 0.09 | 0.93 | 1 | Nicn1 | nicolin 1 | 1039 | 1047 | 806 | 850 | 786 | 710 | 801 | 882 | 739 | 800 | 752 |
| 764 | 0.03 | 0.15 | 0.17 | 0.87 | 1 | Gtf2f2 | general transcription factor IIF, polypeptid | 891 | 698 | 652 | 703 | 799 | 754 | 801 | 702 | 908 | 763 | 739 |
| 638 | 0.21 | 0.14 | 1.53 | 0.13 | 1 | Susd1 | sushi domain containing 1 | 294 | 288 | 273 | 650 | 690 | 718 | 801 | 878 | 772 | 894 | 763 |
| 1000 | 0.10 | 0.14 | 0.67 | 0.50 | 1 | Ppp4r1 | protein phosphatase 4, regulatory subuni | 1141 | 1168 | 1145 | 935 | 726 | 1201 | 800 | 1155 | 846 | 799 | 1083 |
| 978 | -0.09 | 0.16 | -0.56 | 0.58 | 1 | Zfp384 | zinc finger protein 384 | 1016 | 979 | 884 | 931 | 860 | 1439 | 800 | 629 | 913 | 842 | 1462 |
| 974 | 0.11 | 0.16 | 0.66 | 0.51 | 1 | Emc8 | ER membrane protein complex subunit 8 | 1052 | 1441 | 1400 | 747 | 846 | 894 | 800 | 893 | 817 | 987 | 836 |
| 880 | 0.05 | 0.19 | 0.24 | 0.81 | 1 | Pls1 | plastin 1 (I-isoform) | 962 | 1636 | 984 | 822 | 837 | 560 | 800 | 759 | 825 | 853 | 647 |
| 973 | -0.04 | 0.16 | -0.25 | 0.80 | 1 | Zfp180 | zinc finger protein 180 | 1133 | 916 | 976 | 838 | 1111 | 1146 | 799 | 941 | 948 | 730 | 1166 |
| 860 | 0.02 | 0.13 | 0.18 | 0.85 | 1 | Mrpl4 | mitochondrial ribosomal protein L4 | 784 | 744 | 870 | 913 | 914 | 875 | 799 | 862 | 859 | 983 | 858 |
| 1177 | -0.25 | 0.18 | -1.37 | 0.17 | 1 | Slco2a1 | solute carrier organic anion transporter fa | 3202 | 2346 | 2284 | 773 | 625 | 584 | 798 | 810 | 510 | 492 | 525 |
| 990 | 0.13 | 0.12 | 1.10 | 0.27 | 1 | Zfp329 | zinc finger protein 329 | 691 | 1002 | 704 | 1078 | 704 | 1490 | 798 | 1040 | 884 | 875 | 1619 |
| 984 | 0.11 | 0.15 | 0.78 | 0.43 | 1 | Hsf2 | heat shock factor 2 | 678 | 731 | 564 | 937 | 1006 | 1500 | 798 | 1068 | 857 | 1068 | 1613 |
| 767 | -0.01 | 0.17 | -0.08 | 0.93 | 1 | Uchl5 | ubiquitin carboxyl-terminal esterase L5 | 835 | 808 | 663 | 642 | 886 | 751 | 798 | 792 | 649 | 754 | 859 |
| 765 | -0.05 | 0.15 | -0.34 | 0.74 | 1 | Tmem181a | transmembrane protein 181A | 522 | 642 | 656 | 927 | 793 | 835 | 798 | 795 | 859 | 884 | 705 |
| 726 | -0.04 | 0.13 | -0.30 | 0.76 | 1 | Exosc3 | exosome component 3 | 709 | 773 | 669 | 729 | 740 | 687 | 798 | 795 | 708 | 701 | 674 |
| 709 | 0.04 | 0.19 | 0.24 | 0.81 | 1 | Cyp2j6 | cytochrome P450, family 2, subfamily j, pc | 755 | 824 | 852 | 673 | 679 | 491 | 798 | 503 | 839 | 815 | 575 |
| 868 | 0.06 | 0.11 | 0.58 | 0.56 | 1 | Mob2 | MOB kinase activator 2 | 939 | 918 | 976 | 773 | 832 | 885 | 797 | 790 | 855 | 891 | 898 |
| 756 | 0.08 | 0.15 | 0.51 | 0.61 | 1 | Apopt1 | apoptogenic, mitochondrial 1 | 570 | 561 | 835 | 781 | 817 | 691 | 797 | 867 | 928 | 760 | 711 |
| 667 | -0.09 | 0.20 | -0.43 | 0.67 | 1 | Thrb | thyroid hormone receptor beta | 840 | 961 | 756 | 633 | 458 | 582 | 797 | 439 | 540 | 638 | 692 |
| 618 | 0.00 | 0.30 | 0.01 | 1.00 | 1 | Galr3 | galanin receptor 3 | 57 | 127 | 203 | 833 | 903 | 692 | 797 | 715 | 769 | 998 | 701 |
| 1230 | -0.04 | 0.20 | -0.22 | 0.82 | 1 | Ppp1r12b | protein phosphatase 1, regulatory subuni | 3132 | 1632 | 1464 | 691 | 612 | 1674 | 796 | 725 | 695 | 662 | 1444 |
| 859 | -0.03 | 0.18 | -0.17 | 0.87 | 1 | Ncoa4 | nuclear receptor coactivator 4 | 902 | 872 | 566 | 869 | 897 | 1008 | 796 | 718 | 832 | 831 | 1154 |
| 804 | 0.02 | 0.17 | 0.14 | 0.89 | 1 | Ltbr | lymphotoxin B receptor | 1360 | 1288 | 1282 | 542 | 700 | 354 | 796 | 784 | 688 | 784 | 268 |
| 760 | -0.08 | 0.18 | -0.48 | 0.63 | 1 | Emc6 | ER membrane protein complex subunit 6 | 607 | 898 | 923 | 810 | 763 | 675 | 796 | 808 | 715 | 821 | 550 |
| 741 | -0.05 | 0.15 | -0.35 | 0.72 | 1 | Dolk | dolichol kinase | 543 | 645 | 708 | 937 | 830 | 615 | 796 | 905 | 748 | 859 | 563 |
| 730 | 0.15 | 0.13 | 1.19 | 0.24 | 1 | Dusp10 | dual specificity phosphatase 10 | 752 | 619 | 592 | 588 | 644 | 838 | 796 | 629 | 705 | 826 | 1047 |
| 728 | 0.11 | 0.15 | 0.72 | 0.47 | 1 | Lrrc73 | leucine rich repeat containing 73 | 449 | 488 | 395 | 894 | 766 | 729 | 796 | 1166 | 705 | 912 | 705 |
| 852 | -0.03 | 0.18 | -0.17 | 0.87 | 1 | Tbl1xr1 | transducin (beta)-like 1X-linked receptor · | 1622 | 875 | 961 | 695 | 818 | 684 | 795 | 699 | 711 | 734 | 777 |
| 716 | -0.06 | 0.17 | -0.33 | 0.74 | 1 | Cox7c | cytochrome c oxidase subunit 7C | 669 | 572 | 925 | 616 | 781 | 729 | 795 | 733 | 717 | 603 | 739 |
| 1082 | -0.25 | 0.18 | -1.38 | 0.17 | 1 | Mknk2 | MAP kinase-interacting serine/threonine I | 2774 | 1681 | 1645 | 799 | 745 | 811 | 794 | 652 | 748 | 676 | 577 |
| 970 | 0.00 | 0.16 | 0.00 | 1.00 | 1 | Tet3 | tet methylcytosine dioxygenase 3 | 828 | 939 | 893 | 725 | 752 | 1784 | 794 | 623 | 807 | 976 | 1544 |
| 869 | 0.08 | 0.17 | 0.47 | 0.64 | 1 | Nelfa | negative elongation factor complex memb | 798 | 950 | 813 | 782 | 789 | 1036 | 794 | 651 | 874 | 1009 | 1066 |
| 860 | 0.05 | 0.23 | 0.21 | 0.83 | 1 | Faap20 | Fanconi anemia core complex associatec | 1132 | 951 | 1115 | 768 | 864 | 596 | 794 | 1324 | 649 | 737 | 529 |
| 885 | 0.23 | 0.14 | 1.68 | 0.09 | 1 | Jarid2 | jumonji, AT rich interactive domain 2 | 968 | 829 | 886 | 677 | 691 | 1120 | 793 | 879 | 929 | 924 | 1045 |
| 879 | -0.11 | 0.16 | -0.74 | 0.46 | 1 | Pold3 | polymerase (DNA-directed), delta 3, acce | 1192 | 1079 | 935 | 726 | 817 | 992 | 793 | 723 | 676 | 613 | 1121 |
| 681 | 0.08 | 0.16 | 0.52 | 0.60 | 1 | Dgcr6 | DiGeorge syndrome critical region gene 6 | 529 | 613 | 589 | 707 | 706 | 575 | 793 | 988 | 666 | 750 | 580 |
| 1081 | -0.06 | 0.21 | -0.29 | 0.77 | 1 | Rrm1 | ribonucleotide reductase M1 | 2086 | 1473 | 1114 | 1105 | 1015 | 779 | 792 | 1053 | 903 | 760 | 813 |
| 974 | 0.06 | 0.14 | 0.43 | 0.66 | 1 | N4bp2 | NEDD4 binding protein 2 | 1081 | 1227 | 1151 | 678 | 764 | 1350 | 792 | 704 | 925 | 845 | 1194 |
| 919 | 0.01 | 0.14 | 0.05 | 0.96 | 1 | Zfp282 | zinc finger protein 282 | 878 | 933 | 876 | 1005 | 840 | 1068 | 792 | 860 | 1035 | 796 | 1022 |
| 800 | 0.22 | 0.18 | 1.24 | 0.21 | 1 | Snx8 | sorting nexin 8 | 674 | 496 | 617 | 838 | 809 | 794 | 792 | 1198 | 841 | 890 | 854 |
| 758 | -0.25 | 0.16 | -1.58 | 0.12 | 1 | Akr1e1 | aldo-keto reductase family 1, member E1 | 1032 | 803 | 926 | 720 | 649 | 841 | 792 | 525 | 599 | 559 | 889 |
| 896 | -0.05 | 0.14 | -0.39 | 0.70 | 1 | Pcyt2 | phosphate cytidylyltransferase 2, ethanol: | 488 | 428 | 524 | 964 | 827 | 1703 | 791 | 885 | 816 | 782 | 1649 |
| 879 | -0.15 | 0.18 | -0.86 | 0.39 | 1 | Acadm | acyl-Coenzyme A dehydrogenase, mediu | 1349 | 1113 | 1243 | 674 | 794 | 847 | 791 | 485 | 907 | 568 | 897 |
| 818 | 0.02 | 0.16 | 0.09 | 0.92 | 1 | Ipo13 | importin 13 | 839 | 793 | 659 | 884 | 845 | 806 | 791 | 949 | 823 | 662 | 943 |

| | | | | | | | | | | | | | | | | | | |
|---|---|---|---|---|---|---|---|---|---|---|---|---|---|---|---|---|---|---|
| 766 | -0.03 | 0.13 | -0.22 | 0.83 | 1 | Gpatch1 | G patch domain containing 1 | 721 | 642 | 558 | 817 | 716 | 967 | 791 | 781 | 779 | 743 | 909 |
| 708 | -0.13 | 0.14 | -0.95 | 0.34 | 1 | Tmem129 | transmembrane protein 129 | 547 | 614 | 749 | 660 | 878 | 753 | 791 | 690 | 752 | 699 | 659 |
| 619 | -0.11 | 0.19 | -0.56 | 0.57 | 1 | Lcmt2 | leucine carboxyl methyltransferase 2 | 374 | 347 | 589 | 774 | 728 | 567 | 791 | 668 | 683 | 720 | 573 |
| 850 | 0.05 | 0.14 | 0.38 | 0.71 | 1 | Rnf149 | ring finger protein 149 | 927 | 968 | 817 | 932 | 859 | 664 | 790 | 1115 | 727 | 784 | 762 |
| 780 | -0.28 | 0.16 | -1.77 | 0.08 | 1 | Mrpl43 | mitochondrial ribosomal protein L43 | 927 | 985 | 898 | 845 | 948 | 597 | 790 | 671 | 752 | 546 | 627 |
| 730 | -0.14 | 0.16 | -0.88 | 0.38 | 1 | Zbtb1 | zinc finger and BTB domain containing 1 | 715 | 1043 | 791 | 685 | 636 | 757 | 790 | 708 | 604 | 568 | 731 |
| 1030 | 0.10 | 0.13 | 0.74 | 0.46 | 1 | Ago3 | argonaute RISC catalytic subunit 3 | 625 | 605 | 590 | 688 | 729 | 2538 | 789 | 908 | 785 | 832 | 2239 |
| 896 | -0.23 | 0.20 | -1.17 | 0.24 | 1 | Gnpda2 | glucosamine-6-phosphate deaminase 2 | 1023 | 889 | 900 | 1089 | 911 | 988 | 789 | 565 | 834 | 890 | 973 |
| 808 | 0.04 | 0.14 | 0.29 | 0.77 | 1 | Arhgap17 | Rho GTPase activating protein 17 | 1192 | 932 | 984 | 637 | 605 | 829 | 789 | 738 | 696 | 674 | 810 |
| 803 | 0.17 | 0.13 | 1.36 | 0.17 | 1 | Spsb3 | splA/ryanodine receptor domain and SOC | 904 | 874 | 682 | 766 | 678 | 752 | 789 | 982 | 717 | 942 | 749 |
| 771 | -0.06 | 0.18 | -0.31 | 0.76 | 1 | Acd | adrenocortical dysplasia | 915 | 852 | 772 | 636 | 790 | 784 | 789 | 551 | 632 | 1042 | 719 |
| 725 | -0.22 | 0.18 | -1.20 | 0.23 | 1 | Bahcc1 | BAH domain and coiled-coil containing 1 | 860 | 1176 | 991 | 632 | 617 | 610 | 789 | 608 | 672 | 559 | 459 |
| 660 | -0.03 | 0.16 | -0.19 | 0.85 | 1 | Pgam1 | phosphoglycerate mutase 1 | 559 | 482 | 563 | 612 | 851 | 600 | 789 | 738 | 676 | 869 | 520 |
| 885 | -0.06 | 0.15 | -0.41 | 0.68 | 1 | Cep68 | centrosomal protein 68 | 1262 | 1098 | 926 | 898 | 688 | 930 | 788 | 801 | 815 | 772 | 755 |
| 722 | 0.09 | 0.14 | 0.66 | 0.51 | 1 | Tom1l1 | target of myb1-like 1 (chicken) | 491 | 546 | 485 | 747 | 714 | 858 | 788 | 701 | 789 | 841 | 987 |
| 855 | -0.02 | 0.19 | -0.13 | 0.90 | 1 | Tmem209 | transmembrane protein 209 | 1087 | 1140 | 954 | 823 | 818 | 705 | 787 | 576 | 960 | 730 | 830 |
| 764 | -0.01 | 0.21 | -0.03 | 0.97 | 1 | Zfp729a | zinc finger protein 729a | 446 | 627 | 473 | 757 | 810 | 1051 | 787 | 520 | 849 | 860 | 1227 |
| 533 | -0.28 | 0.19 | -1.47 | 0.14 | 1 | Nr2c1 | nuclear receptor subfamily 2, group C, me | 478 | 402 | 386 | 565 | 523 | 647 | 787 | 379 | 588 | 563 | 548 |
| 1009 | 0.17 | 0.14 | 1.23 | 0.22 | 1 | Ube2f | ubiquitin-conjugating enzyme E2F (putati | 993 | 1179 | 897 | 937 | 1023 | 1043 | 786 | 1132 | 1034 | 989 | 1089 |
| 891 | 0.05 | 0.13 | 0.38 | 0.70 | 1 | Smarcd1 | SWI/SNF related, matrix associated, actin | 909 | 1029 | 1057 | 856 | 848 | 854 | 786 | 804 | 889 | 952 | 816 |
| 858 | 0.08 | 0.14 | 0.57 | 0.57 | 1 | Zfp518a | zinc finger protein 518A | 733 | 758 | 645 | 995 | 806 | 964 | 786 | 1028 | 934 | 737 | 1054 |
| 836 | -0.16 | 0.17 | -0.95 | 0.34 | 1 | Depdc5 | DEP domain containing 5 | 917 | 798 | 707 | 973 | 893 | 921 | 786 | 693 | 912 | 738 | 860 |
| 818 | 0.06 | 0.13 | 0.47 | 0.64 | 1 | Fbxo44 | F-box protein 44 | 514 | 714 | 640 | 838 | 862 | 997 | 786 | 918 | 802 | 846 | 1078 |
| 812 | -0.01 | 0.15 | -0.04 | 0.97 | 1 | Osbpl3 | oxysterol binding protein-like 3 | 891 | 1069 | 657 | 976 | 583 | 849 | 786 | 852 | 735 | 717 | 823 |
| 772 | 0.21 | 0.19 | 1.14 | 0.25 | 1 | Ndufa5 | NADH:ubiquinone oxidoreductase subun | 821 | 757 | 985 | 671 | 707 | 568 | 786 | 1075 | 768 | 750 | 600 |
| 947 | 0.16 | 0.12 | 1.39 | 0.16 | 1 | Rbm22 | RNA binding motif protein 22 | 1190 | 1317 | 1138 | 776 | 817 | 822 | 784 | 849 | 889 | 917 | 921 |
| 754 | -0.11 | 0.14 | -0.77 | 0.44 | 1 | Natd1 | N-acetyltransferase domain containing 1 | 789 | 715 | 845 | 868 | 774 | 647 | 784 | 728 | 733 | 850 | 556 |
| 736 | -0.09 | 0.14 | -0.62 | 0.54 | 1 | Ggt7 | gamma-glutamyltransferase 7 | 194 | 156 | 219 | 897 | 886 | 1314 | 784 | 773 | 848 | 821 | 1203 |
| 733 | -0.21 | 0.16 | -1.32 | 0.19 | 1 | Gba2 | glucosidase beta 2 | 672 | 675 | 620 | 652 | 826 | 995 | 784 | 497 | 709 | 746 | 886 |
| 712 | 0.09 | 0.16 | 0.59 | 0.55 | 1 | Orc5 | origin recognition complex, subunit 5 | 650 | 566 | 604 | 611 | 810 | 708 | 784 | 821 | 815 | 663 | 795 |
| 1375 | -0.05 | 0.15 | -0.35 | 0.73 | 1 | Ppp4r1l-ps | | 326 | 458 | 329 | 728 | 878 | 5012 | 783 | 901 | 888 | 661 | 4158 |
| 874 | 0.13 | 0.13 | 0.95 | 0.34 | 1 | Zfp809 | zinc finger protein 809 | 745 | 807 | 820 | 884 | 741 | 1041 | 783 | 1007 | 726 | 846 | 1213 |
| 830 | 0.19 | 0.13 | 1.52 | 0.13 | 1 | Thumpd3 | THUMP domain containing 3 | 881 | 753 | 791 | 691 | 678 | 983 | 783 | 766 | 827 | 1012 | 962 |
| 779 | 0.03 | 0.14 | 0.23 | 0.82 | 1 | Sec22a | SEC22 homolog A, vesicle trafficking prot | 788 | 670 | 872 | 766 | 863 | 664 | 783 | 659 | 867 | 948 | 689 |
| 778 | 0.06 | 0.12 | 0.45 | 0.65 | 1 | Tab1 | TGF-beta activated kinase 1/MAP3K7 bin | 742 | 712 | 783 | 719 | 788 | 808 | 783 | 825 | 775 | 849 | 770 |
| 772 | 0.19 | 0.18 | 1.10 | 0.27 | 1 | Esrra | estrogen related receptor, alpha | 745 | 758 | 910 | 628 | 817 | 592 | 783 | 914 | 758 | 1024 | 562 |
| 762 | 0.11 | 0.15 | 0.70 | 0.48 | 1 | Mypop | Myb-related transcription factor, partner o | 517 | 661 | 597 | 823 | 850 | 700 | 783 | 1169 | 863 | 733 | 689 |
| 925 | -0.01 | 0.15 | -0.04 | 0.97 | 1 | Per2 | period circadian clock 2 | 413 | 663 | 448 | 890 | 709 | 1956 | 782 | 816 | 714 | 828 | 1952 |
| 851 | 0.06 | 0.13 | 0.44 | 0.66 | 1 | Dhdh | dihydrodiol dehydrogenase (dimeric) | 691 | 928 | 700 | 1056 | 875 | 746 | 782 | 984 | 911 | 952 | 736 |
| 798 | -0.02 | 0.16 | -0.12 | 0.90 | 1 | Lcmt1 | leucine carboxyl methyltransferase 1 | 915 | 1102 | 1163 | 608 | 718 | 707 | 782 | 723 | 658 | 625 | 778 |
| 709 | 0.15 | 0.17 | 0.94 | 0.35 | 1 | Pcsk2os1 | | 328 | 486 | 530 | 811 | 734 | 714 | 782 | 979 | 902 | 793 | 738 |
| 825 | -0.02 | 0.22 | -0.08 | 0.93 | 1 | Fam222a | family with sequence similarity 222, meml | 569 | 760 | 602 | 1077 | 938 | 796 | 781 | 701 | 927 | 1211 | 719 |
| 732 | -0.13 | 0.13 | -0.98 | 0.33 | 1 | Gas8 | growth arrest specific 8 | 700 | 490 | 635 | 728 | 804 | 942 | 781 | 642 | 750 | 714 | 872 |
| 4767 | -0.26 | 0.29 | -0.90 | 0.37 | 1 | Flna | filamin, alpha | 23715 | 11969 | 11327 | 744 | 668 | 774 | 780 | 551 | 761 | 482 | 664 |
| 798 | -0.03 | 0.13 | -0.23 | 0.82 | 1 | Osgep | O-sialoglycoprotein endopeptidase | 935 | 811 | 856 | 840 | 729 | 764 | 780 | 784 | 702 | 853 | 719 |
| 671 | -0.05 | 0.16 | -0.30 | 0.77 | 1 | Tmem62 | transmembrane protein 62 | 454 | 495 | 503 | 787 | 655 | 791 | 780 | 612 | 749 | 808 | 751 |
| 950 | 0.11 | 0.14 | 0.76 | 0.44 | 1 | Noct | nocturnin | 406 | 415 | 447 | 700 | 727 | 2254 | 778 | 838 | 666 | 942 | 2277 |
| 914 | -0.02 | 0.18 | -0.11 | 0.91 | 1 | Tsc22d2 | TSC22 domain family, member 2 | 969 | 875 | 676 | 1054 | 766 | 1205 | 778 | 1090 | 870 | 676 | 1094 |
| 851 | -0.17 | 0.13 | -1.28 | 0.20 | 1 | Pigo | phosphatidylinositol glycan anchor biosyı | 696 | 830 | 659 | 959 | 941 | 1126 | 778 | 806 | 886 | 720 | 955 |
| 846 | 0.01 | 0.12 | 0.10 | 0.92 | 1 | Tmem70 | transmembrane protein 70 | 797 | 918 | 779 | 941 | 892 | 787 | 778 | 849 | 923 | 817 | 829 |
| 748 | 0.06 | 0.19 | 0.31 | 0.76 | 1 | Cib1 | calcium and integrin binding 1 (calmyrin) | 743 | 547 | 930 | 829 | 804 | 545 | 778 | 871 | 738 | 779 | 658 |
| 1156 | 0.08 | 0.22 | 0.37 | 0.71 | 1 | Parp14 | poly (ADP-ribose) polymerase family, mei | 1871 | 1972 | 2445 | 684 | 876 | 728 | 777 | 1115 | 1016 | 435 | 801 |
| 792 | 0.08 | 0.18 | 0.45 | 0.66 | 1 | Polr3e | polymerase (RNA) III (DNA directed) poly | 677 | 393 | 680 | 707 | 725 | 1168 | 777 | 667 | 748 | 1065 | 1103 |
| 1490 | 0.18 | 0.19 | 0.99 | 0.32 | 1 | Nfkbia | nuclear factor of kappa light polypeptide g | 3861 | 2494 | 2382 | 625 | 830 | 1424 | 776 | 1008 | 882 | 867 | 1236 |
| 813 | 0.25 | 0.17 | 1.48 | 0.14 | 1 | Prxl2b | peroxiredoxin like 2B | 812 | 844 | 736 | 833 | 783 | 565 | 776 | 1339 | 750 | 810 | 691 |
| 791 | -0.10 | 0.14 | -0.75 | 0.46 | 1 | Zc2hc1a | zinc finger, C2HC-type containing 1A | 984 | 977 | 945 | 748 | 723 | 751 | 776 | 761 | 752 | 591 | 694 |
| 533 | 0.49 | 0.28 | 1.76 | 0.08 | 1 | mt-Nd6 | NADH dehydrogenase subunit 6 | 225 | 285 | 230 | 376 | 325 | 656 | 776 | 482 | 570 | 1211 | 724 |
| 1070 | 0.02 | 0.16 | 0.15 | 0.88 | 1 | Zfp644 | zinc finger protein 644 | 1217 | 1354 | 1142 | 879 | 985 | 1331 | 775 | 1015 | 1065 | 616 | 1395 |
| 766 | 0.07 | 0.16 | 0.46 | 0.64 | 1 | Mtif3 | mitochondrial translational initiation factor | 619 | 652 | 574 | 760 | 868 | 793 | 775 | 1039 | 720 | 842 | 786 |
| 718 | 0.01 | 0.25 | 0.03 | 0.98 | 1 | Tbc1d4 | TBC1 domain family, member 4 | 1059 | 418 | 594 | 754 | 843 | 529 | 775 | 872 | 792 | 761 | 496 |

| | | | | | | | | | | | | | | | | | | |
|---|---|---|---|---|---|---|---|---|---|---|---|---|---|---|---|---|---|---|
| 919 | 0.14 | 0.13 | 1.05 | 0.29 | 1 | Zfp788 | zinc finger protein 788 | 820 | 1138 | 1003 | 787 | 874 | 944 | 774 | 972 | 862 | 760 | 1172 |
| 841 | 0.02 | 0.16 | 0.14 | 0.89 | 1 | Commd9 | COMM domain containing 9 | 823 | 832 | 1005 | 733 | 879 | 883 | 774 | 799 | 723 | 1003 | 794 |
| 811 | 0.10 | 0.12 | 0.87 | 0.38 | 1 | Hmces | 5-hydroxymethylcytosine (hmC) binding, I | 801 | 824 | 769 | 795 | 720 | 853 | 774 | 796 | 743 | 869 | 971 |
| 791 | -0.32 | 0.18 | -1.80 | 0.07 | 1 | Marchf2 | membrane associated ring-CH-type finge | 1277 | 936 | 1196 | 818 | 718 | 631 | 774 | 704 | 624 | 493 | 527 |
| 688 | -0.11 | 0.19 | -0.58 | 0.56 | 1 | Tmem170 | transmembrane protein 170 | 424 | 324 | 390 | 908 | 864 | 781 | 774 | 936 | 887 | 590 | 692 |
| 664 | 0.13 | 0.20 | 0.68 | 0.50 | 1 | Asl | argininosuccinate lyase | 671 | 560 | 688 | 677 | 490 | 619 | 774 | 872 | 540 | 822 | 590 |
| 657 | 0.08 | 0.32 | 0.26 | 0.80 | 1 | Abcb4 | ATP-binding cassette, sub-family B (MDR | 41 | 239 | 174 | 983 | 781 | 750 | 774 | 1016 | 851 | 784 | 839 |
| 602 | 0.13 | 0.17 | 0.80 | 0.43 | 1 | Peli3 | pellino 3 | 324 | 354 | 330 | 597 | 683 | 615 | 774 | 604 | 774 | 922 | 643 |
| 1126 | 0.17 | 0.17 | 1.01 | 0.31 | 1 | Lmf2 | lipase maturation factor 2 | 1940 | 1417 | 1885 | 966 | 915 | 682 | 773 | 1128 | 858 | 1124 | 695 |
| 879 | -0.18 | 0.14 | -1.28 | 0.20 | 1 | Ssh3 | slingshot protein phosphatase 3 | 786 | 904 | 880 | 670 | 840 | 1505 | 773 | 560 | 745 | 745 | 1260 |
| 786 | -0.01 | 0.15 | -0.07 | 0.94 | 1 | Pramef8 | PRAME family member 8 | 911 | 724 | 696 | 800 | 763 | 826 | 773 | 665 | 837 | 858 | 793 |
| 751 | -0.02 | 0.15 | -0.13 | 0.90 | 1 | Adrm1 | adhesion regulating molecule 1 | 558 | 489 | 527 | 688 | 906 | 1013 | 773 | 839 | 837 | 784 | 845 |
| 642 | 0.03 | 0.22 | 0.15 | 0.88 | 1 | Ube2d1 | ubiquitin-conjugating enzyme E2D 1 | 647 | 737 | 829 | 468 | 591 | 564 | 773 | 731 | 523 | 679 | 516 |
| 933 | 0.12 | 0.14 | 0.88 | 0.38 | 1 | Tubgcp6 | tubulin, gamma complex associated prote | 893 | 882 | 855 | 902 | 892 | 1099 | 772 | 1083 | 1003 | 943 | 941 |
| 804 | 0.08 | 0.16 | 0.50 | 0.61 | 1 | Gemin7 | gem nuclear organelle associated protein | 751 | 755 | 920 | 817 | 868 | 639 | 772 | 1128 | 825 | 774 | 597 |
| 792 | -0.24 | 0.14 | -1.68 | 0.09 | 1 | Fbxo34 | F-box protein 34 | 811 | 760 | 762 | 783 | 1024 | 870 | 772 | 688 | 791 | 764 | 682 |
| 770 | -0.12 | 0.17 | -0.67 | 0.50 | 1 | Mre11a | MRE11A homolog A, double strand break | 979 | 981 | 757 | 753 | 706 | 761 | 772 | 776 | 759 | 536 | 695 |
| 1205 | -0.39 | 0.21 | -1.87 | 0.06 | 1 | Ankrd50 | ankyrin repeat domain 50 | 2279 | 2651 | 2330 | 923 | 781 | 907 | 771 | 392 | 766 | 711 | 742 |
| 914 | -0.04 | 0.17 | -0.23 | 0.82 | 1 | Rcbtb2 | regulator of chromosome condensation (F | 1198 | 958 | 1151 | 934 | 896 | 814 | 771 | 1037 | 844 | 657 | 798 |
| 756 | 0.10 | 0.18 | 0.54 | 0.59 | 1 | Plch1 | phospholipase C, eta 1 | 268 | 231 | 202 | 934 | 841 | 1125 | 771 | 736 | 1113 | 908 | 1187 |
| 749 | -0.03 | 0.17 | -0.20 | 0.84 | 1 | Farp2 | FERM, RhoGEF and pleckstrin domain pr | 540 | 539 | 531 | 898 | 869 | 803 | 771 | 653 | 965 | 930 | 740 |
| 685 | -0.08 | 0.15 | -0.51 | 0.61 | 1 | Haghl | hydroxyacylglutathione hydrolase-like | 498 | 441 | 387 | 712 | 672 | 1031 | 771 | 843 | 580 | 656 | 946 |
| 829 | 0.31 | 0.19 | 1.62 | 0.11 | 1 | Mrps7 | mitchondrial ribosomal protein S7 | 909 | 751 | 707 | 616 | 887 | 734 | 770 | 1166 | 947 | 859 | 774 |
| 739 | -0.08 | 0.16 | -0.50 | 0.62 | 1 | Dnm3 | dynamin 3 | 661 | 686 | 615 | 1118 | 745 | 550 | 770 | 1014 | 738 | 647 | 586 |
| 604 | 0.05 | 0.16 | 0.34 | 0.73 | 1 | Nars2 | asparaginyl-tRNA synthetase 2 (mitochor | 450 | 306 | 348 | 521 | 538 | 898 | 770 | 525 | 693 | 653 | 939 |
| 1092 | -0.17 | 0.16 | -1.08 | 0.28 | 1 | Neurl1a | neuralized E3 ubiquitin protein ligase 1A | 1587 | 1458 | 1825 | 1124 | 877 | 1009 | 769 | 761 | 795 | 800 | 1005 |
| 1090 | 0.04 | 0.11 | 0.42 | 0.68 | 1 | Cep70 | centrosomal protein 70 | 795 | 1094 | 1026 | 943 | 834 | 1900 | 769 | 1002 | 882 | 725 | 2022 |
| 1057 | -0.16 | 0.14 | -1.12 | 0.26 | 1 | Sh3glb2 | SH3-domain GRB2-like endophilin B2 | 1079 | 858 | 1283 | 813 | 858 | 1980 | 769 | 725 | 693 | 741 | 1832 |
| 807 | -0.02 | 0.14 | -0.11 | 0.92 | 1 | Ntan1 | N-terminal Asn amidase | 1090 | 954 | 871 | 733 | 830 | 648 | 769 | 666 | 791 | 929 | 594 |
| 698 | -0.05 | 0.16 | -0.30 | 0.76 | 1 | Pklr | pyruvate kinase liver and red blood cell | 523 | 735 | 595 | 770 | 774 | 614 | 769 | 929 | 814 | 653 | 501 |
| 626 | -0.09 | 0.15 | -0.60 | 0.55 | 1 | Riox2 | ribosomal oxygenase 2 | 616 | 519 | 663 | 527 | 595 | 736 | 769 | 512 | 621 | 589 | 742 |
| 620 | 0.01 | 0.14 | 0.09 | 0.93 | 1 | Fzd8 | frizzled class receptor 8 | 503 | 580 | 440 | 741 | 695 | 434 | 769 | 818 | 712 | 681 | 448 |
| 990 | 0.11 | 0.14 | 0.76 | 0.45 | 1 | Zfp410 | zinc finger protein 410 | 778 | 1032 | 1005 | 963 | 818 | 1301 | 768 | 1172 | 738 | 792 | 1523 |
| 902 | 0.17 | 0.13 | 1.31 | 0.19 | 1 | Grtp1 | GH regulated TBC protein 1 | 438 | 646 | 552 | 934 | 967 | 1199 | 768 | 1214 | 1105 | 712 | 1391 |
| 793 | 0.05 | 0.14 | 0.39 | 0.70 | 1 | 3830406C13Ril | RIKEN cDNA 3830406C13 gene | 691 | 803 | 747 | 726 | 942 | 745 | 768 | 777 | 953 | 850 | 722 |
| 786 | 0.07 | 0.14 | 0.51 | 0.61 | 1 | Srp19 | signal recognition particle 19 | 613 | 497 | 491 | 588 | 810 | 1278 | 768 | 658 | 861 | 770 | 1312 |
| 719 | -0.22 | 0.18 | -1.23 | 0.22 | 1 | Kbtbd4 | kelch repeat and BTB (POZ) domain conta | 762 | 704 | 721 | 820 | 812 | 662 | 768 | 504 | 637 | 826 | 688 |
| 693 | -0.19 | 0.17 | -1.13 | 0.26 | 1 | Zscan25 | zinc finger and SCAN domain containing | 522 | 491 | 666 | 782 | 834 | 784 | 768 | 788 | 611 | 647 | 732 |
| 974 | 0.34 | 0.18 | 1.91 | 0.06 | 1 | Pgls | 6-phosphogluconolactonase | 1166 | 991 | 1104 | 869 | 886 | 745 | 767 | 1441 | 875 | 1025 | 845 |
| 921 | -0.09 | 0.14 | -0.69 | 0.49 | 1 | Mgat4b | mannoside acetylglucosaminyltransferase | 1647 | 1480 | 1799 | 708 | 645 | 566 | 767 | 593 | 739 | 635 | 552 |
| 876 | -0.01 | 0.19 | -0.07 | 0.95 | 1 | Ttc5 | tetratricopeptide repeat domain 5 | 851 | 525 | 701 | 846 | 909 | 1259 | 767 | 941 | 756 | 742 | 1339 |
| 790 | 0.18 | 0.12 | 1.53 | 0.13 | 1 | Bcap29 | B cell receptor associated protein 29 | 811 | 765 | 850 | 691 | 822 | 656 | 767 | 864 | 894 | 869 | 698 |
| 718 | -0.28 | 0.20 | -1.39 | 0.16 | 1 | 1110059E24Ril | RIKEN cDNA 1110059E24 gene | 1066 | 745 | 830 | 787 | 768 | 562 | 767 | 524 | 634 | 524 | 685 |
| 839 | -0.14 | 0.13 | -1.12 | 0.26 | 1 | Hinfp | histone H4 transcription factor | 711 | 805 | 709 | 662 | 714 | 1522 | 766 | 631 | 653 | 632 | 1425 |
| 774 | -0.11 | 0.13 | -0.81 | 0.42 | 1 | Dzank1 | double zinc ribbon and ankyrin repeat do | 747 | 830 | 768 | 776 | 746 | 903 | 766 | 833 | 686 | 599 | 862 |
| 690 | 0.07 | 0.19 | 0.39 | 0.70 | 1 | Wdpcp | WD repeat containing planar cell polarity | 389 | 641 | 581 | 887 | 706 | 574 | 766 | 752 | 752 | 826 | 716 |
| 983 | -0.28 | 0.17 | -1.63 | 0.10 | 1 | Rfx7 | regulatory factor X, 7 | 1418 | 1442 | 1298 | 803 | 878 | 1198 | 765 | 504 | 866 | 719 | 922 |
| 610 | -0.20 | 0.21 | -0.97 | 0.33 | 1 | Mlph | melanophilin | 171 | 223 | 232 | 1366 | 632 | 494 | 765 | 1077 | 726 | 640 | 388 |
| 436 | 0.02 | 0.26 | 0.08 | 0.93 | 1 | Stk32b | serine/threonine kinase 32B | 245 | 189 | 280 | 292 | 712 | 254 | 765 | 225 | 699 | 806 | 323 |
| 1170 | 0.09 | 0.15 | 0.61 | 0.54 | 1 | Rela | v-rel reticuloendotheliosis viral oncogene | 1975 | 1722 | 1665 | 1097 | 775 | 1009 | 764 | 944 | 919 | 1036 | 967 |
| 952 | -0.08 | 0.14 | -0.60 | 0.55 | 1 | Dtx3l | deltex 3-like, E3 ubiquitin ligase | 1352 | 1572 | 1346 | 859 | 783 | 777 | 764 | 723 | 846 | 814 | 632 |
| 920 | 0.15 | 0.17 | 0.86 | 0.39 | 1 | Gtf3c5 | general transcription factor IIIC, polypepti | 872 | 1019 | 935 | 740 | 1091 | 876 | 764 | 787 | 990 | 1087 | 958 |
| 902 | -0.05 | 0.16 | -0.33 | 0.74 | 1 | Zfp865 | zinc finger protein 865 | 806 | 1078 | 710 | 950 | 844 | 1203 | 764 | 801 | 907 | 853 | 1010 |
| 884 | -0.09 | 0.18 | -0.53 | 0.60 | 1 | Capn10 | calpain 10 | 661 | 711 | 921 | 1070 | 901 | 1091 | 764 | 679 | 1042 | 869 | 1015 |
| 847 | -0.03 | 0.16 | -0.22 | 0.83 | 1 | Tbccd1 | TBCC domain containing 1 | 479 | 682 | 751 | 855 | 1023 | 1103 | 764 | 685 | 1056 | 842 | 1077 |
| 787 | 0.06 | 0.13 | 0.44 | 0.66 | 1 | Mrps5 | mitochondrial ribosomal protein S5 | 690 | 654 | 864 | 669 | 860 | 869 | 764 | 806 | 928 | 683 | 872 |
| 630 | -0.23 | 0.17 | -1.32 | 0.19 | 1 | Ccdc85c | coiled-coil domain containing 85C | 791 | 856 | 884 | 527 | 618 | 465 | 764 | 417 | 598 | 603 | 408 |
| 888 | 0.03 | 0.16 | 0.19 | 0.85 | 1 | Rabif | RAB interacting factor | 752 | 979 | 973 | 1003 | 847 | 872 | 763 | 752 | 1097 | 751 | 980 |
| 751 | 0.12 | 0.18 | 0.68 | 0.50 | 1 | Svop | SV2 related protein | 755 | 763 | 910 | 770 | 637 | 614 | 763 | 1018 | 664 | 690 | 678 |
| 810 | -0.05 | 0.14 | -0.38 | 0.70 | 1 | B3gnt9 | UDP-GlcNAc:betaGal beta-1,3-N-acetylgl | 1260 | 957 | 972 | 744 | 806 | 603 | 762 | 648 | 776 | 746 | 634 |

| | | | | | | | | | | | | | | | | | | |
|---|---|---|---|---|---|---|---|---|---|---|---|---|---|---|---|---|---|---|
| 628 | -0.18 | 0.17 | -1.02 | 0.31 | 1 | Plekhh1 | pleckstrin homology domain containing, fa | 275 | 222 | 171 | 923 | 774 | 849 | 762 | 688 | 719 | 783 | 738 |
| 1610 | 0.16 | 0.18 | 0.89 | 0.37 | 1 | Gria2 | glutamate receptor, ionotropic, AMPA2 (a | 2302 | 3945 | 2809 | 764 | 991 | 1633 | 761 | 887 | 1188 | 915 | 1510 |
| 957 | 0.18 | 0.17 | 1.07 | 0.29 | 1 | Plekhm1 | pleckstrin homology domain containing, fa | 1028 | 1015 | 987 | 1001 | 917 | 796 | 761 | 1438 | 796 | 963 | 821 |
| 768 | -0.22 | 0.17 | -1.25 | 0.21 | 1 | Cdk7 | cyclin-dependent kinase 7 | 717 | 805 | 674 | 773 | 778 | 1022 | 761 | 551 | 820 | 551 | 989 |
| 712 | 0.04 | 0.16 | 0.27 | 0.79 | 1 | Srek1ip1 | splicing regulatory glutamine/lysine-rich p | 478 | 648 | 597 | 733 | 708 | 806 | 761 | 826 | 669 | 827 | 780 |
| 594 | -0.06 | 0.23 | -0.25 | 0.80 | 1 | 6330403L08Rik | | 486 | 538 | 469 | 828 | 427 | 571 | 761 | 621 | 766 | 582 | 485 |
| 1041 | 0.03 | 0.14 | 0.25 | 0.80 | 1 | Msh6 | mutS homolog 6 | 1289 | 1419 | 1157 | 1115 | 852 | 1007 | 760 | 1284 | 847 | 823 | 893 |
| 889 | -0.12 | 0.13 | -0.94 | 0.35 | 1 | Zfp263 | zinc finger protein 263 | 764 | 810 | 785 | 810 | 871 | 1376 | 760 | 746 | 683 | 711 | 1462 |
| 758 | -0.08 | 0.16 | -0.54 | 0.59 | 1 | Iqcb1 | IQ calmodulin-binding motif containing 1 | 621 | 890 | 725 | 729 | 697 | 932 | 760 | 737 | 731 | 550 | 965 |
| 722 | -0.04 | 0.16 | -0.25 | 0.80 | 1 | Ciao3 | cytosolic iron-sulfur assembly component | 627 | 559 | 608 | 752 | 698 | 890 | 760 | 532 | 781 | 841 | 897 |
| 643 | 0.19 | 0.18 | 1.05 | 0.29 | 1 | Fbxo6 | F-box protein 6 | 642 | 473 | 597 | 635 | 524 | 576 | 760 | 849 | 633 | 836 | 545 |
| 583 | -0.01 | 0.20 | -0.06 | 0.95 | 1 | Mfsd8 | major facilitator superfamily domain conta | 476 | 545 | 345 | 596 | 525 | 661 | 760 | 485 | 643 | 765 | 614 |
| 981 | -0.09 | 0.13 | -0.66 | 0.51 | 1 | Camk1 | calcium/calmodulin-dependent protein kir | 1829 | 1599 | 1772 | 831 | 678 | 608 | 759 | 738 | 645 | 786 | 550 |
| 923 | 0.12 | 0.15 | 0.80 | 0.42 | 1 | Thoc5 | THO complex 5 | 1055 | 856 | 1003 | 948 | 772 | 994 | 759 | 974 | 984 | 835 | 975 |
| 765 | -0.05 | 0.15 | -0.32 | 0.75 | 1 | Tkfc | triokinase, FMN cyclase | 693 | 764 | 559 | 813 | 901 | 777 | 759 | 855 | 806 | 775 | 713 |
| 752 | 0.01 | 0.16 | 0.07 | 0.94 | 1 | Cept1 | choline/ethanolaminephosphotransferase | 905 | 727 | 801 | 705 | 679 | 762 | 759 | 692 | 850 | 604 | 789 |
| 751 | -0.08 | 0.15 | -0.52 | 0.60 | 1 | Arl15 | ADP-ribosylation factor-like 15 | 951 | 804 | 1237 | 642 | 725 | 569 | 759 | 524 | 732 | 750 | 566 |
| 698 | 0.03 | 0.15 | 0.23 | 0.82 | 1 | Rpap2 | RNA polymerase II associated protein 2 | 720 | 695 | 659 | 635 | 813 | 560 | 759 | 635 | 731 | 878 | 590 |
| 676 | -0.27 | 0.14 | -1.88 | 0.06 | 1 | Fam241b | family with sequence similarity 241, meml | 709 | 905 | 845 | 704 | 723 | 538 | 759 | 621 | 573 | 568 | 487 |
| 674 | -0.09 | 0.19 | -0.47 | 0.64 | 1 | Plekhj1 | pleckstrin homology domain containing, fa | 739 | 511 | 1038 | 557 | 596 | 705 | 759 | 422 | 667 | 743 | 675 |
| 608 | -0.08 | 0.21 | -0.39 | 0.70 | 1 | Lrsam1 | leucine rich repeat and sterile alpha motif | 634 | 407 | 498 | 528 | 746 | 629 | 758 | 665 | 580 | 535 | 708 |
| 889 | 0.07 | 0.24 | 0.28 | 0.78 | 1 | Tox | thymocyte selection-associated high mob | 965 | 1757 | 1683 | 747 | 536 | 566 | 757 | 887 | 487 | 836 | 553 |
| 861 | 0.04 | 0.15 | 0.27 | 0.79 | 1 | Gipc1 | GIPC PDZ domain containing family, mem | 1008 | 1153 | 1165 | 760 | 902 | 606 | 757 | 747 | 804 | 969 | 597 |
| 800 | 0.09 | 0.13 | 0.68 | 0.50 | 1 | Tprkb | Tp53rk binding protein | 937 | 1032 | 1052 | 628 | 640 | 773 | 757 | 725 | 682 | 714 | 863 |
| 789 | -0.03 | 0.19 | -0.17 | 0.87 | 1 | Hps1 | HPS1, biogenesis of lysosomal organelle | 816 | 519 | 500 | 911 | 894 | 893 | 757 | 880 | 795 | 842 | 875 |
| 782 | -0.08 | 0.16 | -0.50 | 0.62 | 1 | Cstf1 | cleavage stimulation factor, 3' pre-RNA, s | 587 | 937 | 858 | 747 | 832 | 858 | 757 | 759 | 769 | 724 | 771 |
| 738 | 0.05 | 0.19 | 0.25 | 0.80 | 1 | Homez | homeodomain leucine zipper-encoding g | 727 | 803 | 732 | 821 | 806 | 501 | 757 | 652 | 656 | 1043 | 624 |
| 736 | -0.30 | 0.19 | -1.59 | 0.11 | 1 | Iqsec2 | IQ motif and Sec7 domain 2 | 1071 | 740 | 892 | 812 | 793 | 593 | 757 | 481 | 862 | 604 | 489 |
| 946 | -0.11 | 0.15 | -0.70 | 0.48 | 1 | Prr14 | proline rich 14 | 969 | 1204 | 984 | 995 | 772 | 1248 | 756 | 679 | 808 | 827 | 1168 |
| 906 | -0.22 | 0.16 | -1.42 | 0.16 | 1 | Tubgcp3 | tubulin, gamma complex associated prote | 894 | 803 | 695 | 766 | 800 | 1712 | 756 | 510 | 694 | 756 | 1579 |
| 729 | 0.14 | 0.23 | 0.63 | 0.53 | 1 | Tmem198 | transmembrane protein 198 | 410 | 549 | 694 | 669 | 750 | 852 | 756 | 799 | 622 | 1023 | 891 |
| 640 | 0.26 | 0.15 | 1.76 | 0.08 | 1 | AA414768 | expressed sequence AA414768 | 446 | 420 | 351 | 628 | 694 | 576 | 756 | 739 | 881 | 851 | 696 |
| 821 | -0.08 | 0.15 | -0.52 | 0.60 | 1 | Zfp574 | zinc finger protein 574 | 754 | 735 | 768 | 1041 | 821 | 862 | 755 | 779 | 803 | 871 | 837 |
| 540 | 0.23 | 0.29 | 0.79 | 0.43 | 1 | Gm45847 | | 166 | 294 | 261 | 715 | 414 | 503 | 755 | 728 | 439 | 1101 | 566 |
| 863 | -0.28 | 0.22 | -1.25 | 0.21 | 1 | Ppl | periplakin | 1670 | 1292 | 1100 | 782 | 720 | 716 | 754 | 795 | 635 | 403 | 631 |
| 777 | 0.20 | 0.17 | 1.17 | 0.24 | 1 | Ltv1 | LTV1 ribosome biogenesis factor | 642 | 656 | 532 | 628 | 834 | 891 | 754 | 696 | 775 | 997 | 1136 |
| 846 | 0.21 | 0.14 | 1.44 | 0.15 | 1 | Wdr12 | WD repeat domain 12 | 585 | 547 | 530 | 769 | 979 | 1058 | 753 | 1038 | 980 | 974 | 1093 |
| 749 | -0.15 | 0.15 | -1.05 | 0.29 | 1 | Nadk2 | NAD kinase 2, mitochondrial | 653 | 970 | 736 | 824 | 731 | 794 | 753 | 690 | 670 | 697 | 726 |
| 651 | -0.18 | 0.13 | -1.37 | 0.17 | 1 | Gss | glutathione synthetase | 730 | 593 | 677 | 687 | 764 | 540 | 753 | 658 | 613 | 693 | 459 |
| 1018 | 0.20 | 0.17 | 1.20 | 0.23 | 1 | Edrf1 | erythroid differentiation regulatory factor 1 | 685 | 762 | 516 | 1031 | 1042 | 1492 | 752 | 1090 | 1196 | 1003 | 1627 |
| 842 | -0.17 | 0.13 | -1.30 | 0.19 | 1 | Magi1 | membrane associated guanylate kinase, | 475 | 727 | 633 | 1007 | 926 | 1234 | 752 | 955 | 949 | 568 | 1040 |
| 782 | -0.01 | 0.14 | -0.08 | 0.94 | 1 | Gca | grancalcin | 643 | 773 | 831 | 692 | 871 | 870 | 752 | 633 | 880 | 784 | 869 |
| 521 | -0.19 | 0.24 | -0.79 | 0.43 | 1 | Taar1 | trace amine-associated receptor 1 | 81 | 72 | 158 | 833 | 728 | 565 | 752 | 553 | 728 | 611 | 648 |
| 1009 | -0.09 | 0.13 | -0.66 | 0.51 | 1 | Ostf1 | osteoclast stimulating factor 1 | 2381 | 1677 | 1772 | 634 | 766 | 564 | 751 | 635 | 613 | 747 | 561 |
| 668 | -0.40 | 0.26 | -1.52 | 0.13 | 1 | Slc25a42 | solute carrier family 25, member 42 | 551 | 315 | 398 | 789 | 792 | 1013 | 751 | 273 | 736 | 782 | 949 |
| 1373 | 0.07 | 0.18 | 0.40 | 0.69 | 1 | Akap8l | A kinase (PRKA) anchor protein 8-like | 600 | 988 | 615 | 738 | 675 | 4507 | 750 | 739 | 783 | 930 | 3773 |
| 1082 | -0.25 | 0.15 | -1.65 | 0.10 | 1 | Abhd4 | abhydrolase domain containing 4 | 1721 | 1609 | 1706 | 910 | 1178 | 882 | 750 | 692 | 1042 | 746 | 664 |
| 1051 | 0.03 | 0.13 | 0.25 | 0.80 | 1 | Ube2i | ubiquitin-conjugating enzyme E2I | 1143 | 1072 | 1145 | 655 | 837 | 1854 | 749 | 634 | 946 | 799 | 1729 |
| 1008 | -0.23 | 0.17 | -1.35 | 0.18 | 1 | Rap2a | RAS related protein 2a | 1975 | 1890 | 1587 | 970 | 652 | 684 | 749 | 670 | 780 | 556 | 571 |
| 974 | 0.16 | 0.12 | 1.36 | 0.17 | 1 | Zfp319 | zinc finger protein 319 | 1081 | 1300 | 998 | 1090 | 785 | 839 | 749 | 1183 | 874 | 987 | 835 |
| 834 | 0.13 | 0.21 | 0.59 | 0.55 | 1 | Tnfaip8 | tumor necrosis factor, alpha-induced prot | 1558 | 791 | 873 | 883 | 748 | 466 | 749 | 915 | 846 | 845 | 505 |
| 828 | 0.05 | 0.16 | 0.34 | 0.73 | 1 | B3galnt2 | UDP-GalNAc:betaGlcNAc beta 1,3-galact | 966 | 643 | 936 | 700 | 698 | 1046 | 749 | 543 | 816 | 956 | 1058 |
| 813 | 0.12 | 0.15 | 0.80 | 0.42 | 1 | Psenen | presenilin enhancer gamma secretase su | 800 | 787 | 806 | 884 | 813 | 669 | 749 | 1026 | 801 | 994 | 607 |
| 1521 | 0.15 | 0.16 | 0.95 | 0.34 | 1 | Capn1 | calpain 1 | 1031 | 1729 | 2022 | 1743 | 1877 | 1325 | 748 | 2195 | 1853 | 1121 | 1087 |
| 839 | -0.12 | 0.14 | -0.84 | 0.40 | 1 | Hspa14 | heat shock protein 14 | 1166 | 1013 | 1041 | 765 | 702 | 908 | 748 | 605 | 779 | 636 | 869 |
| 774 | 0.25 | 0.19 | 1.30 | 0.19 | 1 | Ube2a | ubiquitin-conjugating enzyme E2A | 553 | 564 | 730 | 849 | 710 | 726 | 748 | 1050 | 743 | 1096 | 744 |
| 735 | -0.21 | 0.16 | -1.29 | 0.20 | 1 | Hps5 | HPS5, biogenesis of lysosomal organelle | 786 | 461 | 777 | 589 | 626 | 1227 | 748 | 412 | 587 | 643 | 1232 |
| 734 | -0.02 | 0.15 | -0.12 | 0.91 | 1 | Snrnp40 | small nuclear ribonucleoprotein 40 (U5) | 930 | 679 | 758 | 613 | 794 | 724 | 748 | 715 | 738 | 648 | 726 |
| 607 | -0.16 | 0.18 | -0.92 | 0.36 | 1 | Sec61g | SEC61, gamma subunit | 347 | 418 | 499 | 577 | 823 | 710 | 748 | 551 | 629 | 602 | 775 |
| 89113 | -0.72 | 1.16 | -0.62 | 0.54 | 1 | Ctrb1 | chymotrypsinogen B1 | 143105 | 429687 | 380840 | 15515 | 1449 | 84 | 747 | 1733 | 790 | 415 | 5884 |

| | | | | | | | | | | | | | | | | | | |
|---|---|---|---|---|---|---|---|---|---|---|---|---|---|---|---|---|---|---|
| 790 | 0.06 | 0.22 | 0.26 | 0.79 | 1 | Mrps26 | mitochondrial ribosomal protein S26 | 798 | 792 | 704 | 816 | 813 | 707 | 747 | 1274 | 778 | 509 | 757 |
| 783 | 0.02 | 0.15 | 0.11 | 0.91 | 1 | 6030458C11Ril | RIKEN cDNA 6030458C11 gene | 637 | 753 | 699 | 829 | 721 | 945 | 747 | 695 | 722 | 887 | 980 |
| 554 | 0.17 | 0.21 | 0.79 | 0.43 | 1 | Slc1a4 | solute carrier family 1 (glutamate/neutral a | 242 | 339 | 350 | 484 | 669 | 531 | 747 | 586 | 722 | 835 | 584 |
| 551 | 0.27 | 0.31 | 0.89 | 0.37 | 1 | Kcnf1 | potassium voltage-gated channel, subfam | 12 | 54 | 31 | 805 | 477 | 694 | 747 | 831 | 624 | 975 | 815 |
| 771 | -0.01 | 0.13 | -0.08 | 0.93 | 1 | Mccc2 | methylcrotonoyl-Coenzyme A carboxylase | 661 | 713 | 671 | 824 | 921 | 737 | 746 | 909 | 797 | 697 | 800 |
| 747 | -0.20 | 0.15 | -1.32 | 0.19 | 1 | Bbs1 | Bardet-Biedl syndrome 1 (human) | 773 | 778 | 617 | 908 | 750 | 836 | 746 | 650 | 730 | 756 | 676 |
| 895 | -0.10 | 0.15 | -0.65 | 0.51 | 1 | Pkig | protein kinase inhibitor, gamma | 1630 | 1263 | 1299 | 811 | 752 | 601 | 745 | 798 | 746 | 696 | 503 |
| 824 | -0.02 | 0.18 | -0.13 | 0.90 | 1 | Bcl10 | B cell leukemia/lymphoma 10 | 1148 | 1297 | 886 | 808 | 697 | 626 | 745 | 601 | 628 | 969 | 655 |
| 749 | -0.07 | 0.18 | -0.39 | 0.70 | 1 | Shmt2 | serine hydroxymethyltransferase 2 (mitoc | 1375 | 893 | 833 | 611 | 723 | 559 | 745 | 699 | 637 | 645 | 523 |
| 713 | 0.01 | 0.15 | 0.07 | 0.94 | 1 | C030006K11Ril | RIKEN cDNA C030006K11 gene | 518 | 609 | 726 | 829 | 755 | 641 | 745 | 952 | 774 | 714 | 583 |
| 660 | -0.02 | 0.24 | -0.10 | 0.92 | 1 | Nrde2 | nrde-2 necessary for RNA interference, do | 643 | 341 | 735 | 593 | 838 | 617 | 745 | 842 | 800 | 641 | 466 |
| 596 | 0.04 | 0.21 | 0.21 | 0.83 | 1 | Pigh | phosphatidylinositol glycan anchor biosyn | 366 | 254 | 436 | 650 | 684 | 626 | 745 | 824 | 747 | 589 | 631 |
| 846 | -0.14 | 0.18 | -0.78 | 0.43 | 1 | Senp7 | SUMO1/sentrin specific peptidase 7 | 999 | 995 | 1128 | 661 | 603 | 1198 | 744 | 412 | 707 | 690 | 1165 |
| 842 | 1.15 | 0.77 | 1.50 | 0.13 | 1 | Gm23935 | | 196 | 266 | 924 | 515 | 530 | 411 | 744 | 1170 | 375 | 3551 | 582 |
| 795 | 0.01 | 0.16 | 0.08 | 0.94 | 1 | Prpf31 | pre-mRNA processing factor 31 | 969 | 985 | 1015 | 596 | 731 | 797 | 743 | 507 | 845 | 809 | 752 |
| 785 | -0.05 | 0.19 | -0.24 | 0.81 | 1 | Recql | RecQ protein-like | 965 | 689 | 902 | 656 | 857 | 833 | 743 | 839 | 687 | 594 | 866 |
| 736 | 0.11 | 0.15 | 0.71 | 0.48 | 1 | Ppm1k | protein phosphatase 1K (PP2C domain co | 532 | 566 | 404 | 856 | 713 | 868 | 743 | 770 | 824 | 895 | 922 |
| 710 | -0.03 | 0.13 | -0.21 | 0.83 | 1 | Mnat1 | menage a trois 1 | 799 | 788 | 701 | 767 | 652 | 631 | 743 | 690 | 641 | 669 | 731 |
| 973 | -0.21 | 0.15 | -1.45 | 0.15 | 1 | Arhgap23 | Rho GTPase activating protein 23 | 1774 | 1723 | 1563 | 906 | 713 | 648 | 742 | 920 | 593 | 591 | 525 |
| 789 | -0.20 | 0.17 | -1.15 | 0.25 | 1 | Mapkbp1 | mitogen-activated protein kinase binding | 780 | 759 | 648 | 868 | 779 | 1089 | 742 | 569 | 819 | 770 | 860 |
| 783 | 0.07 | 0.16 | 0.47 | 0.64 | 1 | Polr3gl | polymerase (RNA) III (DNA directed) poly | 731 | 826 | 839 | 636 | 710 | 943 | 742 | 906 | 686 | 707 | 884 |
| 658 | 0.00 | 0.17 | -0.02 | 0.98 | 1 | Ercc2 | excision repair cross-complementing rode | 692 | 629 | 544 | 529 | 505 | 943 | 742 | 671 | 593 | 558 | 831 |
| 876 | 0.07 | 0.14 | 0.52 | 0.60 | 1 | Nbdy | negative regulator of P-body association | 1063 | 1085 | 1236 | 758 | 807 | 739 | 741 | 959 | 756 | 708 | 787 |
| 567 | 0.22 | 0.22 | 1.03 | 0.30 | 1 | Dapp1 | dual adaptor for phosphotyrosine and 3-p | 538 | 452 | 375 | 439 | 520 | 544 | 741 | 390 | 750 | 769 | 724 |
| 1126 | 0.00 | 0.13 | 0.03 | 0.97 | 1 | Ankrd24 | ankyrin repeat domain 24 | 347 | 375 | 450 | 852 | 750 | 3459 | 740 | 887 | 868 | 752 | 2903 |
| 867 | -0.03 | 0.17 | -0.20 | 0.84 | 1 | Dnttip1 | deoxynucleotidyltransferase, terminal, inte | 996 | 778 | 1060 | 799 | 830 | 1034 | 740 | 890 | 721 | 848 | 844 |
| 838 | 0.15 | 0.16 | 0.93 | 0.35 | 1 | Rgs3 | regulator of G-protein signaling 3 | 1036 | 772 | 855 | 787 | 753 | 806 | 740 | 728 | 719 | 1091 | 933 |
| 809 | -0.07 | 0.16 | -0.46 | 0.64 | 1 | Usp53 | ubiquitin specific peptidase 53 | 885 | 1116 | 1009 | 555 | 720 | 1013 | 740 | 640 | 664 | 641 | 918 |
| 791 | -0.02 | 0.15 | -0.14 | 0.89 | 1 | Ttll5 | tubulin tyrosine ligase-like family, membe | 741 | 831 | 840 | 811 | 856 | 749 | 740 | 689 | 970 | 823 | 653 |
| 781 | 0.17 | 0.12 | 1.38 | 0.17 | 1 | Phf13 | PHD finger protein 13 | 785 | 918 | 674 | 922 | 702 | 541 | 740 | 1126 | 773 | 863 | 547 |
| 737 | 0.15 | 0.16 | 0.93 | 0.35 | 1 | Rrp12 | ribosomal RNA processing 12 homolog (S | 603 | 486 | 550 | 603 | 1053 | 684 | 740 | 761 | 975 | 934 | 713 |
| 710 | 0.23 | 0.17 | 1.36 | 0.17 | 1 | Atg7 | autophagy related 7 | 883 | 689 | 655 | 551 | 754 | 524 | 740 | 820 | 862 | 796 | 537 |
| 579 | 0.14 | 0.19 | 0.74 | 0.46 | 1 | Gm11942 | | 1755 | 1377 | 187 | 656 | 82 | 43 | 740 | 593 | 88 | 778 | 71 |
| 982 | -0.11 | 0.13 | -0.86 | 0.39 | 1 | Git2 | GIT ArfGAP 2 | 1374 | 1343 | 1189 | 908 | 716 | 1213 | 739 | 701 | 725 | 732 | 1165 |
| 920 | 0.21 | 0.20 | 1.06 | 0.29 | 1 | Chchd7 | coiled-coil-helix-coiled-coil-helix domain | 687 | 679 | 862 | 964 | 734 | 1216 | 739 | 1413 | 888 | 729 | 1203 |
| 1196 | -0.18 | 0.16 | -1.08 | 0.28 | 1 | Adam17 | a disintegrin and metallopeptidase domai | 2189 | 1766 | 2062 | 950 | 1005 | 1058 | 738 | 609 | 941 | 715 | 1118 |
| 972 | -0.03 | 0.15 | -0.18 | 0.86 | 1 | Foxo4 | forkhead box O4 | 1295 | 1253 | 1431 | 940 | 956 | 752 | 738 | 931 | 768 | 924 | 702 |
| 921 | -0.19 | 0.17 | -1.15 | 0.25 | 1 | Npr2 | natriuretic peptide receptor 2 | 1816 | 1810 | 1705 | 499 | 556 | 767 | 738 | 520 | 424 | 730 | 569 |
| 728 | -0.11 | 0.16 | -0.70 | 0.49 | 1 | Zfp407 | zinc finger protein 407 | 925 | 957 | 999 | 600 | 572 | 751 | 738 | 503 | 662 | 681 | 618 |
| 711 | -0.18 | 0.16 | -1.13 | 0.26 | 1 | Tmem246 | transmembrane protein 246 | 1003 | 857 | 876 | 792 | 750 | 428 | 738 | 819 | 561 | 596 | 403 |
| 699 | 0.13 | 0.17 | 0.77 | 0.44 | 1 | Tab3 | TGF-beta activated kinase 1/MAP3K7 bin | 634 | 804 | 632 | 662 | 564 | 702 | 738 | 987 | 633 | 662 | 673 |
| 553 | -0.26 | 0.20 | -1.28 | 0.20 | 1 | Fzd6 | frizzled class receptor 6 | 528 | 458 | 759 | 575 | 631 | 440 | 738 | 524 | 469 | 532 | 433 |
| 1173 | 0.07 | 0.15 | 0.47 | 0.64 | 1 | H2-Q4 | histocompatibility 2, Q region locus 4 | 2758 | 1759 | 2288 | 592 | 824 | 837 | 737 | 789 | 733 | 784 | 803 |
| 899 | 0.11 | 0.17 | 0.66 | 0.51 | 1 | Cdc40 | cell division cycle 40 | 1158 | 926 | 721 | 834 | 800 | 1037 | 737 | 852 | 880 | 827 | 1115 |
| 830 | 0.20 | 0.20 | 0.98 | 0.33 | 1 | Lamtor4 | late endosomal/lysosomal adaptor, MAPK | 750 | 724 | 1018 | 824 | 768 | 747 | 737 | 1249 | 764 | 777 | 776 |
| 768 | 0.17 | 0.22 | 0.79 | 0.43 | 1 | Bloc1s6 | biogenesis of lysosomal organelles comp | 967 | 692 | 539 | 682 | 802 | 718 | 737 | 957 | 913 | 687 | 749 |
| 739 | 0.11 | 0.12 | 0.91 | 0.36 | 1 | Nkap | NFKB activating protein | 742 | 790 | 668 | 732 | 709 | 666 | 737 | 818 | 697 | 875 | 699 |
| 729 | 0.17 | 0.13 | 1.29 | 0.20 | 1 | Prmt7 | protein arginine N-methyltransferase 7 | 919 | 769 | 701 | 611 | 610 | 691 | 737 | 645 | 745 | 864 | 723 |
| 759 | 0.02 | 0.14 | 0.14 | 0.89 | 1 | Uros | uroporphyrinogen III synthase | 432 | 467 | 451 | 826 | 698 | 1201 | 736 | 813 | 834 | 617 | 1273 |
| 531 | 0.27 | 0.20 | 1.36 | 0.17 | 1 | B3galt2 | UDP-Gal:betaGlcNAc beta 1,3-galactosyl | 197 | 272 | 245 | 610 | 519 | 474 | 736 | 662 | 685 | 804 | 641 |
| 657 | -0.06 | 0.14 | -0.43 | 0.67 | 1 | Ppa2 | pyrophosphatase (inorganic) 2 | 741 | 584 | 711 | 715 | 698 | 505 | 735 | 577 | 678 | 716 | 561 |
| 816 | 0.17 | 0.12 | 1.42 | 0.16 | 1 | Ints5 | integrator complex subunit 5 | 633 | 669 | 745 | 933 | 897 | 682 | 734 | 998 | 1066 | 918 | 698 |
| 737 | 0.14 | 0.19 | 0.75 | 0.45 | 1 | Fra10ac1 | FRA10AC1 homolog (human) | 598 | 461 | 674 | 572 | 636 | 1076 | 734 | 825 | 624 | 627 | 1284 |
| 867 | 0.00 | 0.14 | -0.02 | 0.98 | 1 | Zfp800 | zinc finger protein 800 | 676 | 839 | 613 | 1077 | 733 | 1190 | 733 | 913 | 892 | 735 | 1137 |
| 650 | 0.24 | 0.15 | 1.58 | 0.11 | 1 | Lrrc57 | leucine rich repeat containing 57 | 624 | 690 | 638 | 555 | 593 | 506 | 733 | 665 | 754 | 801 | 591 |
| 648 | -0.06 | 0.19 | -0.30 | 0.77 | 1 | Gtpbp8 | GTP-binding protein 8 (putative) | 438 | 636 | 573 | 582 | 709 | 769 | 733 | 557 | 685 | 693 | 755 |
| 917 | -0.02 | 0.15 | -0.15 | 0.88 | 1 | Rab11fip5 | RAB11 family interacting protein 5 (class | 1924 | 1380 | 1441 | 816 | 662 | 447 | 732 | 716 | 692 | 907 | 373 |
| 841 | -0.02 | 0.13 | -0.13 | 0.90 | 1 | Tdrd3 | tudor domain containing 3 | 1078 | 1065 | 894 | 750 | 774 | 868 | 732 | 744 | 756 | 702 | 891 |
| 780 | 0.11 | 0.12 | 0.85 | 0.39 | 1 | Spout1 | SPOUT domain containing methyltransfer | 833 | 765 | 776 | 626 | 805 | 828 | 732 | 721 | 797 | 815 | 879 |
| 759 | 0.03 | 0.21 | 0.15 | 0.88 | 1 | Adck2 | aarF domain containing kinase 2 | 723 | 626 | 898 | 626 | 841 | 828 | 732 | 848 | 555 | 858 | 813 |

| | | | | | | | | | | | | | | | | | | |
|---|---|---|---|---|---|---|---|---|---|---|---|---|---|---|---|---|---|---|
| 563 | -0.04 | 0.20 | -0.22 | 0.82 | 1 | Poll | polymerase (DNA directed), lambda | 468 | 478 | 380 | 542 | 636 | 546 | 732 | 803 | 553 | 574 | 480 |
| 850 | -0.06 | 0.16 | -0.37 | 0.71 | 1 | Tmem51 | transmembrane protein 51 | 1090 | 1112 | 1255 | 819 | 856 | 587 | 731 | 670 | 743 | 951 | 534 |
| 828 | -0.07 | 0.20 | -0.33 | 0.74 | 1 | Zfp326 | zinc finger protein 326 | 1064 | 755 | 944 | 768 | 610 | 1120 | 731 | 795 | 672 | 495 | 1158 |
| 827 | -0.10 | 0.15 | -0.68 | 0.50 | 1 | Cblb | Casitas B-lineage lymphoma b | 1038 | 820 | 921 | 784 | 829 | 910 | 731 | 567 | 854 | 754 | 890 |
| 750 | 0.08 | 0.20 | 0.41 | 0.68 | 1 | Rgs7 | regulator of G protein signaling 7 | 518 | 792 | 712 | 819 | 773 | 705 | 731 | 757 | 682 | 1020 | 745 |
| 735 | 0.04 | 0.15 | 0.28 | 0.78 | 1 | Tmem11 | transmembrane protein 11 | 755 | 685 | 838 | 761 | 695 | 668 | 731 | 876 | 748 | 683 | 644 |
| 698 | -0.02 | 0.16 | -0.14 | 0.89 | 1 | Cmtr2 | cap methyltransferase 2 | 536 | 630 | 492 | 857 | 729 | 713 | 731 | 780 | 822 | 587 | 798 |
| 654 | -0.30 | 0.21 | -1.41 | 0.16 | 1 | Septin3 | septin 3 | 1043 | 949 | 730 | 638 | 684 | 434 | 731 | 542 | 530 | 412 | 502 |
| 626 | -0.25 | 0.17 | -1.49 | 0.14 | 1 | Magi2 | membrane associated guanylate kinase, | 512 | 623 | 691 | 782 | 729 | 517 | 731 | 541 | 595 | 667 | 495 |
| 473 | -0.06 | 0.23 | -0.26 | 0.79 | 1 | Pcdhb22 | protocadherin beta 22 | 478 | 512 | 281 | 482 | 482 | 331 | 731 | 619 | 411 | 461 | 413 |
| 1062 | 0.27 | 0.19 | 1.43 | 0.15 | 1 | Ccdc9 | coiled-coil domain containing 9 | 1197 | 1299 | 1570 | 807 | 797 | 1137 | 730 | 1352 | 817 | 913 | 1067 |
| 861 | 0.03 | 0.14 | 0.19 | 0.85 | 1 | Timmdc1 | translocase of inner mitochondrial membr | 803 | 774 | 1057 | 871 | 924 | 860 | 730 | 753 | 901 | 909 | 885 |
| 856 | 0.01 | 0.16 | 0.05 | 0.96 | 1 | Cpped1 | calcineurin-like phosphoesterase domain | 1132 | 867 | 1010 | 905 | 898 | 650 | 730 | 746 | 920 | 934 | 626 |
| 852 | 0.14 | 0.14 | 0.95 | 0.34 | 1 | Nop14 | NOP14 nucleolar protein | 846 | 851 | 1031 | 733 | 851 | 855 | 730 | 823 | 778 | 963 | 913 |
| 834 | 0.00 | 0.13 | -0.02 | 0.99 | 1 | Atg12 | autophagy related 12 | 844 | 874 | 1056 | 705 | 757 | 1019 | 730 | 797 | 754 | 680 | 964 |
| 794 | -0.01 | 0.12 | -0.05 | 0.96 | 1 | Hsd17b10 | hydroxysteroid (17-beta) dehydrogenase | 1316 | 1101 | 1270 | 639 | 662 | 500 | 730 | 680 | 628 | 701 | 506 |
| 781 | -0.07 | 0.15 | -0.49 | 0.62 | 1 | Parp9 | poly (ADP-ribose) polymerase family, mer | 1261 | 1111 | 1367 | 566 | 652 | 541 | 730 | 542 | 715 | 527 | 578 |
| 695 | 0.13 | 0.14 | 0.89 | 0.37 | 1 | Kdsr | 3-ketodihydrosphingosine reductase | 640 | 561 | 650 | 717 | 710 | 606 | 730 | 887 | 790 | 773 | 586 |
| 871 | -0.01 | 0.17 | -0.04 | 0.96 | 1 | Ccdc59 | coiled-coil domain containing 59 | 842 | 940 | 1025 | 714 | 608 | 1281 | 729 | 816 | 598 | 542 | 1484 |
| 847 | 0.12 | 0.21 | 0.57 | 0.57 | 1 | Pdgfa | platelet derived growth factor, alpha | 852 | 815 | 786 | 815 | 740 | 957 | 729 | 1375 | 610 | 710 | 933 |
| 770 | 0.06 | 0.13 | 0.45 | 0.65 | 1 | Hpf1 | histone PARylation factor 1 | 876 | 786 | 920 | 645 | 782 | 729 | 729 | 773 | 762 | 670 | 797 |
| 756 | 0.02 | 0.15 | 0.11 | 0.91 | 1 | Inpp5e | inositol polyphosphate-5-phosphatase E | 812 | 790 | 779 | 659 | 652 | 915 | 729 | 819 | 622 | 790 | 744 |
| 695 | -0.30 | 0.19 | -1.57 | 0.12 | 1 | Snrpd2 | small nuclear ribonucleoprotein D2 | 975 | 925 | 872 | 684 | 648 | 612 | 729 | 365 | 620 | 630 | 580 |
| 822 | -0.06 | 0.12 | -0.48 | 0.63 | 1 | Trmt10c | tRNA methyltransferase 10C, mitochondri | 800 | 926 | 897 | 880 | 862 | 805 | 728 | 785 | 756 | 714 | 891 |
| 767 | -0.02 | 0.14 | -0.15 | 0.88 | 1 | Elac2 | elaC ribonuclease Z 2 | 691 | 602 | 603 | 782 | 874 | 908 | 728 | 798 | 845 | 692 | 912 |
| 737 | 0.13 | 0.15 | 0.86 | 0.39 | 1 | Ring1 | ring finger protein 1 | 709 | 694 | 660 | 559 | 690 | 920 | 728 | 768 | 636 | 801 | 944 |
| 668 | 0.20 | 0.16 | 1.21 | 0.23 | 1 | Fam185a | family with sequence similarity 185, meml | 542 | 358 | 506 | 603 | 675 | 752 | 728 | 711 | 634 | 988 | 856 |
| 2512 | -0.12 | 0.13 | -0.92 | 0.36 | 1 | Ier2 | immediate early response 2 | 7965 | 4334 | 4994 | 452 | 521 | 3909 | 727 | 543 | 447 | 641 | 3097 |
| 1141 | -0.19 | 0.15 | -1.30 | 0.19 | 1 | Tmsb10 | thymosin, beta 10 | 3012 | 2025 | 2275 | 653 | 847 | 566 | 727 | 592 | 757 | 634 | 469 |
| 1069 | -0.11 | 0.16 | -0.73 | 0.47 | 1 | Asap1 | ArfGAP with SH3 domain, ankyrin repeat | 2201 | 1522 | 1876 | 664 | 972 | 815 | 727 | 517 | 957 | 819 | 687 |
| 830 | 0.10 | 0.13 | 0.76 | 0.45 | 1 | Eif2b4 | eukaryotic translation initiation factor 2B, : | 723 | 805 | 740 | 866 | 799 | 912 | 727 | 1079 | 753 | 811 | 913 |
| 825 | 0.23 | 0.17 | 1.32 | 0.19 | 1 | Gpaa1 | GPI anchor attachment protein 1 | 655 | 598 | 687 | 838 | 740 | 986 | 727 | 1103 | 758 | 1039 | 943 |
| 874 | -0.07 | 0.16 | -0.45 | 0.66 | 1 | Pnpla6 | patatin-like phospholipase domain contai | 814 | 912 | 810 | 574 | 870 | 1417 | 726 | 562 | 754 | 659 | 1513 |
| 657 | -0.10 | 0.14 | -0.67 | 0.50 | 1 | Pex11b | peroxisomal biogenesis factor 11 beta | 490 | 526 | 644 | 664 | 874 | 609 | 726 | 688 | 738 | 741 | 522 |
| 543 | -0.33 | 0.19 | -1.71 | 0.09 | 1 | Ptges2 | prostaglandin E synthase 2 | 395 | 555 | 493 | 664 | 572 | 561 | 726 | 440 | 540 | 544 | 478 |
| 961 | 0.07 | 0.13 | 0.52 | 0.60 | 1 | Senp1 | SUMO1/sentrin specific peptidase 1 | 1061 | 1011 | 928 | 836 | 918 | 1228 | 725 | 958 | 867 | 826 | 1218 |
| 878 | 0.11 | 0.14 | 0.81 | 0.42 | 1 | Mtmr12 | myotubularin related protein 12 | 1115 | 1458 | 1025 | 766 | 534 | 897 | 725 | 927 | 671 | 667 | 872 |
| 820 | 0.20 | 0.22 | 0.88 | 0.38 | 1 | Ngdn | neuroguidin, EIF4E binding protein | 847 | 736 | 845 | 586 | 844 | 910 | 725 | 1160 | 712 | 638 | 1022 |
| 817 | -0.22 | 0.18 | -1.25 | 0.21 | 1 | Pdcl3 | phosducin-like 3 | 1436 | 1218 | 1104 | 764 | 695 | 629 | 725 | 490 | 654 | 572 | 704 |
| 748 | -0.04 | 0.15 | -0.27 | 0.79 | 1 | Eef1aknmt | EEF1A lysine and N-terminal methyltrans | 698 | 764 | 718 | 774 | 715 | 850 | 725 | 703 | 778 | 844 | 655 |
| 741 | 0.07 | 0.16 | 0.41 | 0.68 | 1 | Dtd1 | D-tyrosyl-tRNA deacylase 1 | 918 | 950 | 1250 | 536 | 692 | 516 | 725 | 677 | 643 | 687 | 560 |
| 609 | -0.41 | 0.22 | -1.90 | 0.06 | 1 | Mcm2 | minichromosome maintenance complex c | 1127 | 687 | 731 | 671 | 501 | 483 | 725 | 405 | 562 | 441 | 368 |
| 1144 | 0.12 | 0.15 | 0.80 | 0.42 | 1 | Emsy | EMSY, BRCA2-interacting transcriptional | 1183 | 1363 | 1187 | 939 | 963 | 1660 | 724 | 860 | 1025 | 1068 | 1613 |
| 824 | -0.12 | 0.16 | -0.75 | 0.45 | 1 | Klhl11 | kelch-like 11 | 734 | 1201 | 686 | 914 | 806 | 889 | 724 | 969 | 771 | 555 | 817 |
| 767 | 0.12 | 0.18 | 0.64 | 0.52 | 1 | Nfu1 | NFU1 iron-sulfur cluster scaffold | 1021 | 753 | 937 | 768 | 620 | 634 | 724 | 893 | 809 | 644 | 638 |
| 766 | -0.01 | 0.15 | -0.08 | 0.93 | 1 | Pdik1l | PDLIM1 interacting kinase 1 like | 541 | 830 | 737 | 873 | 811 | 756 | 724 | 986 | 656 | 723 | 792 |
| 729 | 0.06 | 0.15 | 0.39 | 0.70 | 1 | Zfp113 | zinc finger protein 113 | 710 | 693 | 842 | 634 | 656 | 820 | 724 | 794 | 698 | 712 | 736 |
| 722 | 0.05 | 0.18 | 0.28 | 0.78 | 1 | Tbrg4 | transforming growth factor beta regulated | 803 | 572 | 746 | 611 | 653 | 887 | 724 | 820 | 722 | 611 | 792 |
| 693 | -0.08 | 0.16 | -0.49 | 0.63 | 1 | Tmem185b | transmembrane protein 185B | 730 | 888 | 908 | 708 | 646 | 540 | 724 | 560 | 657 | 726 | 534 |
| 404 | -0.23 | 0.29 | -0.81 | 0.42 | 1 | Lancl3 | LanC lantibiotic synthetase component C | 88 | 28 | 69 | 505 | 473 | 612 | 724 | 484 | 410 | 422 | 623 |
| 972 | -0.43 | 0.24 | -1.79 | 0.07 | 1 | S100a11 | S100 calcium binding protein A11 | 1736 | 2126 | 1309 | 942 | 893 | 627 | 723 | 402 | 732 | 602 | 604 |
| 723 | 0.03 | 0.14 | 0.21 | 0.83 | 1 | Snip1 | Smad nuclear interacting protein 1 | 728 | 775 | 780 | 719 | 642 | 721 | 723 | 723 | 644 | 814 | 685 |
| 626 | -0.21 | 0.17 | -1.26 | 0.21 | 1 | Mlst8 | MTOR associated protein, LST8 homolog | 519 | 559 | 537 | 650 | 645 | 791 | 723 | 590 | 637 | 456 | 774 |
| 770 | 0.22 | 0.15 | 1.45 | 0.15 | 1 | Polr2c | polymerase (RNA) II (DNA directed) polyp | 867 | 837 | 886 | 583 | 723 | 683 | 722 | 825 | 679 | 922 | 738 |
| 759 | 0.19 | 0.15 | 1.28 | 0.20 | 1 | B230118H07Ri | RIKEN cDNA B230118H07 gene | 705 | 677 | 924 | 760 | 823 | 531 | 722 | 836 | 811 | 835 | 721 |
| 789 | -0.01 | 0.18 | -0.07 | 0.94 | 1 | Thap2 | THAP domain containing, apoptosis asso | 680 | 885 | 663 | 906 | 715 | 909 | 721 | 646 | 905 | 788 | 858 |
| 769 | 0.23 | 0.20 | 1.16 | 0.25 | 1 | Zbtb6 | zinc finger and BTB domain containing 6 | 603 | 968 | 714 | 627 | 556 | 949 | 721 | 660 | 836 | 742 | 1088 |
| 752 | -0.01 | 0.15 | -0.07 | 0.94 | 1 | Gnpnat1 | glucosamine-phosphate N-acetyltransfera | 598 | 660 | 659 | 690 | 821 | 942 | 721 | 565 | 815 | 799 | 996 |
| 550 | 0.03 | 0.17 | 0.15 | 0.88 | 1 | Prob1 | proline rich basic protein 1 | 253 | 268 | 178 | 746 | 651 | 509 | 721 | 956 | 540 | 698 | 529 |
| 1026 | -0.23 | 0.30 | -0.76 | 0.45 | 1 | Dock10 | dedicator of cytokinesis 10 | 1746 | 925 | 833 | 1301 | 863 | 1314 | 720 | 1090 | 866 | 489 | 1133 |

| | | | | | | | | | | | | | | | | | | |
|---|---|---|---|---|---|---|---|---|---|---|---|---|---|---|---|---|---|---|
| 972 | 0.31 | 0.20 | 1.54 | 0.12 | 1 | Ccdc85b | coiled-coil domain containing 85B | 1057 | 1114 | 1093 | 964 | 798 | 772 | 720 | 1590 | 765 | 1112 | 711 |
| 829 | 0.03 | 0.19 | 0.18 | 0.86 | 1 | Syne4 | spectrin repeat containing, nuclear envelo | 389 | 546 | 474 | 677 | 716 | 1671 | 720 | 1066 | 600 | 598 | 1663 |
| 1824 | 0.17 | 0.15 | 1.17 | 0.24 | 1 | Ccnl2 | cyclin L2 | 924 | 969 | 692 | 597 | 621 | 6637 | 719 | 706 | 848 | 710 | 6637 |
| 914 | 0.05 | 0.14 | 0.33 | 0.74 | 1 | Trmt1 | tRNA methyltransferase 1 | 1155 | 1112 | 1337 | 629 | 613 | 1236 | 719 | 651 | 645 | 793 | 1168 |
| 902 | 0.02 | 0.19 | 0.08 | 0.94 | 1 | Mindy1 | MINDY lysine 48 deubiquitinase 1 | 1139 | 1140 | 1265 | 839 | 793 | 789 | 719 | 784 | 780 | 1095 | 579 |
| 898 | -0.15 | 0.17 | -0.89 | 0.37 | 1 | Ing1 | inhibitor of growth family, member 1 | 1336 | 1700 | 1341 | 618 | 852 | 704 | 719 | 636 | 772 | 542 | 663 |
| 861 | -0.01 | 0.14 | -0.07 | 0.94 | 1 | Cactin | cactin, spliceosome C complex subunit | 1014 | 927 | 1190 | 789 | 852 | 824 | 719 | 877 | 798 | 706 | 779 |
| 629 | 0.19 | 0.24 | 0.77 | 0.44 | 1 | Plut | | 32 | 37 | 78 | 977 | 899 | 582 | 719 | 1244 | 852 | 808 | 689 |
| 840 | 0.25 | 0.21 | 1.21 | 0.23 | 1 | Frg1 | FSHD region gene 1 | 1149 | 1021 | 1137 | 533 | 708 | 745 | 718 | 1088 | 720 | 680 | 742 |
| 654 | 0.03 | 0.15 | 0.21 | 0.83 | 1 | Ints14 | integrator complex subunit 14 | 731 | 633 | 730 | 571 | 621 | 611 | 718 | 547 | 743 | 607 | 683 |
| 1000 | 0.05 | 0.14 | 0.34 | 0.73 | 1 | Bag3 | BCL2-associated athanogene 3 | 1596 | 1685 | 1288 | 963 | 727 | 741 | 717 | 944 | 684 | 976 | 676 |
| 749 | 0.07 | 0.16 | 0.45 | 0.66 | 1 | Gtf2ird1 | general transcription factor II I repeat dom | 760 | 780 | 562 | 737 | 722 | 795 | 717 | 1021 | 674 | 710 | 759 |
| 670 | -0.02 | 0.19 | -0.12 | 0.91 | 1 | Slc46a3 | solute carrier family 46, member 3 | 713 | 509 | 539 | 686 | 843 | 585 | 717 | 641 | 852 | 598 | 686 |
| 108064 | -1.08 | 1.14 | -0.95 | 0.34 | 1 | Pnlip | pancreatic lipase | 149417 | 563096 | 450050 | 18616 | 1300 | 81 | 716 | 1305 | 736 | 281 | 3106 |
| 785 | 0.16 | 0.15 | 1.09 | 0.27 | 1 | Gskip | GSK3B interacting protein | 717 | 781 | 627 | 820 | 853 | 692 | 716 | 810 | 925 | 893 | 805 |
| 731 | 0.04 | 0.17 | 0.22 | 0.83 | 1 | Synj2 | synaptojanin 2 | 940 | 851 | 742 | 659 | 599 | 735 | 716 | 883 | 576 | 591 | 747 |
| 699 | -0.01 | 0.18 | -0.07 | 0.94 | 1 | Kctd11 | potassium channel tetramerisation domai | 589 | 551 | 736 | 946 | 714 | 548 | 716 | 723 | 772 | 823 | 568 |
| 630 | -0.04 | 0.23 | -0.19 | 0.85 | 1 | Zbtb5 | zinc finger and BTB domain containing 5 | 532 | 459 | 493 | 596 | 639 | 806 | 716 | 802 | 760 | 415 | 718 |
| 1125 | -0.02 | 0.19 | -0.08 | 0.94 | 1 | Smc5 | structural maintenance of chromosomes 5 | 1372 | 1306 | 1360 | 879 | 762 | 1822 | 715 | 1126 | 824 | 489 | 1718 |
| 978 | -0.11 | 0.14 | -0.78 | 0.43 | 1 | Trmt1l | tRNA methyltransferase 1 like | 798 | 1089 | 822 | 1037 | 897 | 1519 | 715 | 825 | 953 | 620 | 1488 |
| 979 | 0.08 | 0.15 | 0.57 | 0.57 | 1 | Smim15 | small integral membrane protein 15 | 1547 | 1312 | 1322 | 827 | 889 | 762 | 714 | 967 | 917 | 783 | 726 |
| 859 | 0.14 | 0.14 | 0.99 | 0.32 | 1 | Tcim | transcriptional and immune response reg | 1140 | 1343 | 1396 | 656 | 753 | 518 | 714 | 655 | 950 | 746 | 574 |
| 704 | 0.22 | 0.13 | 1.61 | 0.11 | 1 | Gmcl1 | germ cell-less, spermatogenesis associat | 573 | 590 | 524 | 625 | 655 | 803 | 714 | 675 | 736 | 858 | 993 |
| 690 | -0.01 | 0.19 | -0.05 | 0.96 | 1 | Rpusd1 | RNA pseudouridylate synthase domain co | 411 | 343 | 468 | 727 | 796 | 966 | 714 | 953 | 676 | 678 | 863 |
| 675 | 0.15 | 0.17 | 0.84 | 0.40 | 1 | Bbs7 | Bardet-Biedl syndrome 7 (human) | 441 | 504 | 613 | 582 | 591 | 905 | 714 | 658 | 844 | 711 | 858 |
| 645 | 0.16 | 0.17 | 0.93 | 0.35 | 1 | Tctn2 | tectonic family member 2 | 561 | 474 | 606 | 631 | 621 | 613 | 714 | 855 | 675 | 738 | 611 |
| 899 | 0.01 | 0.13 | 0.06 | 0.95 | 1 | Phf6 | PHD finger protein 6 | 1137 | 1358 | 1196 | 804 | 816 | 753 | 713 | 747 | 776 | 693 | 894 |
| 786 | 0.12 | 0.18 | 0.65 | 0.51 | 1 | Exosc4 | exosome component 4 | 760 | 724 | 859 | 770 | 967 | 585 | 713 | 1015 | 719 | 885 | 649 |
| 633 | 0.21 | 0.17 | 1.24 | 0.22 | 1 | Slc35f1 | solute carrier family 35, member F1 | 283 | 226 | 332 | 741 | 784 | 560 | 713 | 935 | 996 | 900 | 496 |
| 989 | -0.24 | 0.18 | -1.32 | 0.19 | 1 | Ifi27 | interferon, alpha-inducible protein 27 | 1847 | 1338 | 1751 | 721 | 939 | 800 | 712 | 436 | 1020 | 714 | 606 |
| 729 | 0.12 | 0.13 | 0.91 | 0.36 | 1 | Enox2 | ecto-NOX disulfide-thiol exchanger 2 | 675 | 768 | 770 | 691 | 718 | 661 | 711 | 810 | 838 | 729 | 649 |
| 704 | 0.15 | 0.19 | 0.81 | 0.42 | 1 | Srd5a3 | steroid 5 alpha-reductase 3 | 551 | 386 | 492 | 763 | 787 | 716 | 711 | 1048 | 836 | 796 | 656 |
| 695 | 0.09 | 0.25 | 0.36 | 0.72 | 1 | Usp54 | ubiquitin specific peptidase 54 | 642 | 863 | 633 | 528 | 556 | 878 | 711 | 431 | 882 | 723 | 798 |
| 615 | -0.04 | 0.14 | -0.27 | 0.79 | 1 | Dtd2 | D-tyrosyl-tRNA deacylase 2 | 546 | 438 | 477 | 738 | 673 | 566 | 711 | 685 | 684 | 690 | 557 |
| 972 | 0.03 | 0.16 | 0.20 | 0.84 | 1 | Hcfc1r1 | host cell factor C1 regulator 1 (XPO1-dep | 1138 | 1198 | 1256 | 922 | 1101 | 748 | 710 | 1127 | 1034 | 876 | 579 |
| 708 | 0.16 | 0.19 | 0.82 | 0.41 | 1 | 1810058I24Rik | | 921 | 590 | 915 | 621 | 701 | 505 | 710 | 869 | 751 | 688 | 519 |
| 823 | 0.09 | 0.14 | 0.68 | 0.50 | 1 | Rab31 | RAB31, member RAS oncogene family | 1925 | 1449 | 1235 | 521 | 525 | 412 | 709 | 573 | 554 | 645 | 505 |
| 818 | 0.08 | 0.17 | 0.45 | 0.65 | 1 | Spr | sepiapterin reductase | 1115 | 992 | 972 | 794 | 865 | 500 | 709 | 1045 | 815 | 689 | 503 |
| 720 | -0.07 | 0.16 | -0.45 | 0.65 | 1 | Pias4 | protein inhibitor of activated STAT 4 | 641 | 601 | 619 | 964 | 701 | 735 | 709 | 860 | 743 | 614 | 738 |
| 664 | 0.07 | 0.18 | 0.39 | 0.70 | 1 | Sap30l | SAP30-like | 770 | 577 | 554 | 774 | 561 | 576 | 709 | 944 | 601 | 733 | 508 |
| 1197 | -0.01 | 0.16 | -0.06 | 0.95 | 1 | B4galt1 | UDP-Gal:betaGlcNAc beta 1,4- galactosy | 3068 | 2624 | 1880 | 775 | 760 | 565 | 708 | 821 | 697 | 687 | 584 |
| 877 | -0.13 | 0.19 | -0.71 | 0.48 | 1 | Cep250 | centrosomal protein 250 | 1009 | 735 | 839 | 870 | 817 | 1289 | 708 | 667 | 956 | 583 | 1176 |
| 707 | 0.18 | 0.18 | 0.99 | 0.32 | 1 | Rrp1b | ribosomal RNA processing 1 homolog B ( | 571 | 702 | 440 | 541 | 561 | 1057 | 708 | 843 | 593 | 766 | 995 |
| 1098 | 0.17 | 0.15 | 1.12 | 0.26 | 1 | Ugdh | UDP-glucose dehydrogenase | 1934 | 1569 | 1613 | 760 | 962 | 857 | 706 | 935 | 999 | 858 | 881 |
| 977 | -0.17 | 0.13 | -1.26 | 0.21 | 1 | Plxna1 | plexin A1 | 1910 | 1685 | 1515 | 752 | 757 | 758 | 706 | 622 | 581 | 786 | 675 |
| 961 | -0.05 | 0.12 | -0.39 | 0.69 | 1 | Gtf2f1 | general transcription factor IIF, polypeptid | 1369 | 1133 | 1281 | 726 | 868 | 1161 | 706 | 733 | 769 | 757 | 1064 |
| 798 | -0.17 | 0.15 | -1.15 | 0.25 | 1 | Dhx33 | DEAH (Asp-Glu-Ala-His) box polypeptide | 891 | 848 | 803 | 536 | 654 | 1394 | 706 | 428 | 544 | 714 | 1265 |
| 908 | 0.09 | 0.16 | 0.55 | 0.58 | 1 | Zcchc8 | zinc finger, CCHC domain containing 8 | 1033 | 706 | 902 | 779 | 735 | 1341 | 705 | 641 | 918 | 889 | 1342 |
| 700 | 0.03 | 0.17 | 0.19 | 0.85 | 1 | Gm7072 | predicted gene 7072 | 839 | 749 | 659 | 608 | 663 | 713 | 705 | 520 | 808 | 670 | 768 |
| 631 | -0.11 | 0.22 | -0.52 | 0.60 | 1 | Zfp938 | zinc finger protein 938 | 574 | 550 | 726 | 670 | 515 | 724 | 705 | 397 | 591 | 671 | 820 |
| 249 | -0.28 | 1.20 | -0.23 | 0.82 | 1 | Itih1 | inter-alpha trypsin inhibitor, heavy chain 1 | 137 | 90 | 67 | 60 | 36 | 404 | 705 | 0 | 61 | 763 | 413 |
| 904 | -0.10 | 0.14 | -0.71 | 0.48 | 1 | Gabbr1 | gamma-aminobutyric acid (GABA) B rece | 906 | 904 | 1248 | 645 | 716 | 1523 | 704 | 689 | 731 | 574 | 1304 |
| 815 | 0.05 | 0.17 | 0.28 | 0.78 | 1 | Edc3 | enhancer of mRNA decapping 3 | 992 | 749 | 782 | 846 | 792 | 811 | 704 | 743 | 873 | 956 | 715 |
| 792 | -0.09 | 0.12 | -0.74 | 0.46 | 1 | Arl2 | ADP-ribosylation factor-like 2 | 899 | 892 | 854 | 816 | 762 | 841 | 704 | 772 | 662 | 672 | 834 |
| 722 | 0.00 | 0.20 | -0.02 | 0.98 | 1 | Zfp943 | zinc finger prtoein 943 | 637 | 864 | 751 | 624 | 626 | 865 | 703 | 466 | 754 | 630 | 1026 |
| 634 | -0.02 | 0.20 | -0.08 | 0.93 | 1 | Nqo2 | N-ribosyldihydronicotinamide quinone re | 775 | 591 | 646 | 736 | 638 | 440 | 703 | 724 | 642 | 509 | 571 |
| 581 | -0.12 | 0.18 | -0.66 | 0.51 | 1 | Vamp7-ps | | 520 | 514 | 507 | 621 | 663 | 536 | 703 | 431 | 663 | 611 | 624 |
| 859 | 0.19 | 0.13 | 1.43 | 0.15 | 1 | Ercc5 | excision repair cross-complementing rode | 1005 | 978 | 859 | 728 | 605 | 1050 | 702 | 898 | 594 | 880 | 1147 |
| 767 | 0.01 | 0.14 | 0.08 | 0.94 | 1 | Soga3 | SOGA family member 3 | 717 | 697 | 666 | 929 | 629 | 917 | 702 | 783 | 732 | 769 | 893 |
| 731 | -0.04 | 0.12 | -0.33 | 0.74 | 1 | Rad51d | RAD51 paralog D | 736 | 729 | 758 | 779 | 725 | 741 | 702 | 725 | 757 | 665 | 721 |

| | | | | | | | | | | | | | | | | | |
|---|---|---|---|---|---|---|---|---|---|---|---|---|---|---|---|---|---|
| 722 | -0.17 | 0.22 | -0.76 | 0.45 | 1 | Osbpl6 | oxysterol binding protein-like 6 | 733 | 1066 | 885 | 748 | 674 | 653 | 702 | 429 | 704 | 766 | 578 |
| 695 | -0.01 | 0.19 | -0.05 | 0.96 | 1 | Zfp952 | zinc finger protein 952 | 620 | 822 | 681 | 627 | 594 | 822 | 702 | 595 | 784 | 489 | 903 |
| 673 | 0.14 | 0.16 | 0.89 | 0.37 | 1 | Ndufaf4 | NADH:ubiquinone oxidoreductase compl | 544 | 629 | 531 | 607 | 744 | 656 | 702 | 665 | 730 | 757 | 835 |
| 861 | 0.07 | 0.16 | 0.45 | 0.65 | 1 | Klc4 | kinesin light chain 4 | 896 | 993 | 1023 | 787 | 770 | 942 | 701 | 801 | 659 | 960 | 947 |
| 583 | 0.09 | 0.15 | 0.61 | 0.54 | 1 | Tgfa | transforming growth factor alpha | 645 | 522 | 481 | 636 | 576 | 369 | 701 | 571 | 652 | 891 | 367 |
| 918 | -0.26 | 0.16 | -1.68 | 0.09 | 1 | Rarg | retinoic acid receptor, gamma | 1906 | 1708 | 1765 | 735 | 610 | 522 | 700 | 687 | 599 | 474 | 396 |
| 723 | 0.07 | 0.15 | 0.43 | 0.66 | 1 | Cwc27 | CWC27 spliceosome-associated protein | 615 | 639 | 482 | 641 | 749 | 941 | 700 | 743 | 683 | 751 | 1005 |
| 712 | 0.10 | 0.16 | 0.62 | 0.54 | 1 | Gstm5 | glutathione S-transferase, mu 5 | 824 | 929 | 822 | 656 | 696 | 459 | 700 | 842 | 583 | 872 | 448 |
| 599 | -0.06 | 0.16 | -0.39 | 0.70 | 1 | B9d1 | B9 protein domain 1 | 534 | 453 | 662 | 606 | 645 | 575 | 700 | 615 | 590 | 675 | 537 |
| 538 | -0.05 | 0.19 | -0.27 | 0.78 | 1 | Slc18a1 | solute carrier family 18 (vesicular monoar | 154 | 155 | 243 | 655 | 732 | 614 | 700 | 465 | 890 | 715 | 599 |
| 794 | -0.14 | 0.13 | -1.07 | 0.29 | 1 | Xrcc1 | X-ray repair complementing defective rep | 945 | 747 | 1028 | 693 | 822 | 920 | 699 | 539 | 783 | 662 | 895 |
| 761 | -0.16 | 0.17 | -0.96 | 0.34 | 1 | Letmd1 | LETM1 domain containing 1 | 853 | 863 | 802 | 590 | 771 | 1000 | 699 | 476 | 761 | 541 | 1010 |
| 691 | 0.09 | 0.26 | 0.35 | 0.72 | 1 | Pigc | phosphatidylinositol glycan anchor biosyı | 709 | 519 | 814 | 577 | 568 | 838 | 699 | 986 | 558 | 492 | 837 |
| 641 | 0.06 | 0.18 | 0.32 | 0.75 | 1 | Pdcd10 | programmed cell death 10 | 743 | 713 | 650 | 571 | 507 | 641 | 699 | 677 | 555 | 495 | 806 |
| 602 | -0.15 | 0.26 | -0.59 | 0.55 | 1 | Kdf1 | keratinocyte differentiation factor 1 | 461 | 591 | 625 | 638 | 683 | 558 | 699 | 319 | 654 | 774 | 617 |
| 582 | -0.24 | 0.20 | -1.19 | 0.24 | 1 | Pus1 | pseudouridine synthase 1 | 558 | 675 | 560 | 551 | 618 | 609 | 699 | 648 | 524 | 392 | 564 |
| 576 | -0.44 | 0.29 | -1.54 | 0.12 | 1 | Atg9a | autophagy related 9A | 572 | 592 | 560 | 497 | 705 | 704 | 699 | 200 | 677 | 630 | 505 |
| 862 | 0.20 | 0.20 | 0.98 | 0.33 | 1 | Zfp444 | zinc finger protein 444 | 933 | 1090 | 1174 | 550 | 906 | 786 | 698 | 944 | 675 | 869 | 852 |
| 728 | 0.03 | 0.17 | 0.20 | 0.85 | 1 | Cadm4 | cell adhesion molecule 4 | 1064 | 726 | 824 | 743 | 708 | 496 | 698 | 835 | 613 | 832 | 467 |
| 643 | -0.12 | 0.23 | -0.53 | 0.60 | 1 | Pex12 | peroxisomal biogenesis factor 12 | 394 | 623 | 473 | 777 | 735 | 700 | 698 | 480 | 760 | 824 | 609 |
| 538 | -0.14 | 0.22 | -0.61 | 0.54 | 1 | Foxo6 | forkhead box O6 | 313 | 419 | 277 | 740 | 667 | 457 | 698 | 480 | 617 | 830 | 415 |
| 534 | -0.18 | 0.21 | -0.87 | 0.39 | 1 | Nsun3 | NOL1/NOP2/Sun domain family member | 397 | 601 | 434 | 583 | 582 | 492 | 698 | 553 | 611 | 432 | 489 |
| 883 | 0.26 | 0.17 | 1.49 | 0.14 | 1 | Cdk5rap2 | CDK5 regulatory subunit associated prote | 1159 | 883 | 869 | 768 | 807 | 814 | 696 | 1071 | 776 | 1051 | 824 |
| 810 | 0.25 | 0.19 | 1.32 | 0.19 | 1 | Rps29 | ribosomal protein S29 | 1008 | 734 | 1463 | 571 | 728 | 604 | 696 | 877 | 921 | 706 | 606 |
| 629 | -0.13 | 0.16 | -0.79 | 0.43 | 1 | Btd | biotinidase | 969 | 830 | 735 | 603 | 511 | 481 | 696 | 615 | 569 | 533 | 382 |
| 553 | 0.03 | 0.22 | 0.12 | 0.90 | 1 | Cmbl | carboxymethylenebutenolidase-like (Pseu | 329 | 374 | 350 | 699 | 576 | 511 | 696 | 481 | 576 | 903 | 590 |
| 1053 | 0.07 | 0.15 | 0.45 | 0.65 | 1 | Smpdl3a | sphingomyelin phosphodiesterase, acid-l | 2104 | 1894 | 1726 | 806 | 801 | 542 | 695 | 934 | 883 | 696 | 505 |
| 765 | 0.06 | 0.14 | 0.42 | 0.67 | 1 | Bri3 | brain protein I3 | 795 | 809 | 954 | 766 | 854 | 550 | 695 | 789 | 793 | 882 | 531 |
| 698 | 0.04 | 0.15 | 0.29 | 0.77 | 1 | Polr1b | polymerase (RNA) I polypeptide B | 571 | 593 | 541 | 735 | 723 | 786 | 695 | 618 | 820 | 746 | 855 |
| 693 | 0.05 | 0.14 | 0.40 | 0.69 | 1 | Camsap3 | calmodulin regulated spectrin-associated | 668 | 660 | 693 | 719 | 628 | 703 | 695 | 744 | 691 | 787 | 636 |
| 812 | 0.20 | 0.20 | 0.97 | 0.33 | 1 | Smpd2 | sphingomyelin phosphodiesterase 2, neu | 558 | 429 | 640 | 805 | 767 | 1149 | 694 | 1192 | 795 | 755 | 1146 |
| 738 | 0.31 | 0.17 | 1.78 | 0.07 | 1 | Pmvk | phosphomevalonate kinase | 626 | 624 | 759 | 651 | 627 | 756 | 694 | 1017 | 708 | 795 | 857 |
| 624 | 0.11 | 0.21 | 0.50 | 0.62 | 1 | Lyrm2 | LYR motif containing 2 | 560 | 683 | 639 | 699 | 492 | 517 | 694 | 498 | 669 | 791 | 617 |
| 1054 | 0.13 | 0.11 | 1.12 | 0.26 | 1 | Fam76b | family with sequence similarity 76, memb | 1126 | 1095 | 1043 | 744 | 668 | 1909 | 693 | 884 | 738 | 729 | 1962 |
| 702 | -0.08 | 0.13 | -0.62 | 0.54 | 1 | Akap1 | A kinase (PRKA) anchor protein 1 | 865 | 794 | 699 | 770 | 592 | 708 | 693 | 680 | 627 | 653 | 642 |
| 1435 | 0.11 | 0.15 | 0.73 | 0.46 | 1 | Clk4 | CDC like kinase 4 | 619 | 641 | 769 | 744 | 621 | 4507 | 692 | 864 | 799 | 560 | 4971 |
| 1133 | 0.17 | 0.19 | 0.87 | 0.38 | 1 | Pml | promyelocytic leukemia | 1737 | 1466 | 1420 | 1006 | 956 | 1013 | 692 | 1061 | 1019 | 1284 | 807 |
| 983 | -0.10 | 0.17 | -0.60 | 0.55 | 1 | Wnk2 | WNK lysine deficient protein kinase 2 | 1820 | 1553 | 1567 | 938 | 657 | 744 | 692 | 688 | 709 | 860 | 585 |
| 835 | 0.08 | 0.18 | 0.44 | 0.66 | 1 | Arid4a | AT rich interactive domain 4A (RBP1-like) | 1038 | 1218 | 1148 | 529 | 622 | 945 | 692 | 732 | 714 | 483 | 1059 |
| 766 | -0.06 | 0.17 | -0.36 | 0.72 | 1 | Agbl5 | ATP/GTP binding protein-like 5 | 482 | 547 | 487 | 800 | 628 | 1407 | 692 | 955 | 679 | 492 | 1259 |
| 714 | 0.15 | 0.16 | 0.90 | 0.37 | 1 | Poc5 | POC5 centriolar protein | 739 | 591 | 628 | 602 | 763 | 732 | 692 | 741 | 626 | 957 | 787 |
| 618 | 0.20 | 0.15 | 1.30 | 0.19 | 1 | Pomk | protein-O-mannose kinase | 701 | 677 | 721 | 543 | 482 | 471 | 692 | 674 | 604 | 714 | 515 |
| 554 | -0.02 | 0.20 | -0.09 | 0.93 | 1 | Chd1l | chromodomain helicase DNA binding pro | 514 | 406 | 440 | 495 | 610 | 592 | 692 | 697 | 553 | 504 | 592 |
| 837 | 0.06 | 0.18 | 0.34 | 0.73 | 1 | Mosmo | modulator of smoothened | 681 | 777 | 750 | 858 | 668 | 1220 | 691 | 629 | 797 | 930 | 1203 |
| 766 | 0.19 | 0.19 | 0.98 | 0.33 | 1 | Srbd1 | S1 RNA binding domain 1 | 1074 | 888 | 758 | 496 | 803 | 691 | 691 | 759 | 806 | 742 | 718 |
| 890 | -0.16 | 0.19 | -0.81 | 0.42 | 1 | Camkk2 | calcium/calmodulin-dependent protein kir | 641 | 503 | 794 | 1058 | 940 | 1498 | 690 | 966 | 822 | 757 | 1118 |
| 750 | 0.12 | 0.17 | 0.70 | 0.48 | 1 | Tasp1 | taspase, threonine aspartase 1 | 525 | 435 | 666 | 834 | 742 | 912 | 690 | 889 | 850 | 808 | 898 |
| 699 | 0.00 | 0.15 | 0.02 | 0.99 | 1 | Pus10 | pseudouridylate synthase 10 | 577 | 630 | 691 | 802 | 671 | 723 | 690 | 853 | 672 | 600 | 781 |
| 967 | 0.00 | 0.14 | -0.01 | 0.99 | 1 | Bcl9l | B cell CLL/lymphoma 9-like | 1651 | 1510 | 1349 | 824 | 776 | 776 | 689 | 817 | 811 | 711 | 722 |
| 788 | -0.05 | 0.15 | -0.34 | 0.74 | 1 | Sgcb | sarcoglycan, beta (dystrophin-associated | 1028 | 994 | 898 | 852 | 727 | 659 | 689 | 633 | 794 | 737 | 657 |
| 771 | -0.07 | 0.15 | -0.50 | 0.62 | 1 | Kalrn | kalirin, RhoGEF kinase | 769 | 816 | 773 | 733 | 838 | 877 | 689 | 580 | 844 | 741 | 821 |
| 686 | 0.03 | 0.12 | 0.24 | 0.81 | 1 | Spata5 | spermatogenesis associated 5 | 750 | 707 | 655 | 595 | 601 | 801 | 689 | 630 | 593 | 676 | 846 |
| 730 | 0.07 | 0.18 | 0.37 | 0.71 | 1 | Hacd2 | 3-hydroxyacyl-CoA dehydratase 2 | 763 | 1105 | 749 | 684 | 737 | 532 | 688 | 688 | 870 | 645 | 568 |
| 562 | -0.04 | 0.17 | -0.25 | 0.80 | 1 | A130010J15Rik | RIKEN cDNA A130010J15 gene | 386 | 453 | 469 | 599 | 602 | 575 | 688 | 737 | 539 | 634 | 501 |
| 654 | -0.29 | 0.18 | -1.64 | 0.10 | 1 | Stard3 | START domain containing 3 | 685 | 719 | 707 | 777 | 739 | 590 | 687 | 443 | 683 | 626 | 540 |
| 634 | -0.06 | 0.22 | -0.28 | 0.78 | 1 | Nacc2 | nucleus accumbens associated 2, BEN ar | 1107 | 670 | 756 | 649 | 463 | 482 | 687 | 718 | 495 | 470 | 476 |
| 764 | 0.07 | 0.16 | 0.42 | 0.68 | 1 | Zmynd19 | zinc finger, MYND domain containing 19 | 726 | 924 | 645 | 847 | 682 | 751 | 686 | 870 | 597 | 971 | 705 |
| 702 | 0.04 | 0.15 | 0.23 | 0.81 | 1 | Prdm4 | PR domain containing 4 | 779 | 778 | 814 | 600 | 588 | 766 | 686 | 595 | 774 | 593 | 746 |
| 622 | 0.02 | 0.16 | 0.11 | 0.91 | 1 | R3hcc1 | R3H domain and coiled-coil containing 1 | 577 | 556 | 529 | 586 | 727 | 569 | 686 | 768 | 686 | 623 | 531 |
| 607 | -0.11 | 0.15 | -0.75 | 0.46 | 1 | Ecsit | ECSIT signalling integrator | 620 | 575 | 687 | 614 | 595 | 587 | 686 | 547 | 565 | 679 | 520 |

| | | | | | | | | | | | | | | | | | | |
|---|---|---|---|---|---|---|---|---|---|---|---|---|---|---|---|---|---|---|
| 844 | -0.17 | 0.16 | -1.07 | 0.28 | 1 | Ints7 | integrator complex subunit 7 | 783 | 787 | 778 | 998 | 976 | 1019 | 685 | 731 | 778 | 830 | 918 |
| 699 | -0.10 | 0.20 | -0.51 | 0.61 | 1 | Dnal4 | dynein, axonemal, light chain 4 | 692 | 675 | 696 | 763 | 797 | 647 | 685 | 434 | 812 | 773 | 712 |
| 579 | 0.12 | 0.18 | 0.67 | 0.50 | 1 | Inpp5j | inositol polyphosphate 5-phosphatase J | 235 | 344 | 295 | 670 | 643 | 617 | 685 | 751 | 827 | 755 | 541 |
| 736 | 0.11 | 0.26 | 0.42 | 0.68 | 1 | Hnf1a | HNF1 homeobox A | 372 | 665 | 792 | 779 | 841 | 709 | 684 | 600 | 867 | 1105 | 679 |
| 723 | -0.08 | 0.15 | -0.49 | 0.62 | 1 | Zbtb24 | zinc finger and BTB domain containing 24 | 573 | 665 | 706 | 796 | 697 | 896 | 684 | 616 | 641 | 746 | 927 |
| 710 | -0.05 | 0.15 | -0.34 | 0.73 | 1 | Zbtb39 | zinc finger and BTB domain containing 39 | 644 | 662 | 504 | 827 | 748 | 784 | 684 | 907 | 685 | 672 | 692 |
| 824 | -0.07 | 0.25 | -0.26 | 0.79 | 1 | Nrep | neuronal regeneration related protein | 1167 | 1171 | 1623 | 649 | 508 | 725 | 682 | 825 | 572 | 349 | 792 |
| 804 | -0.20 | 0.18 | -1.10 | 0.27 | 1 | Stard5 | StAR-related lipid transfer (START) doma | 1240 | 1379 | 1270 | 600 | 576 | 779 | 682 | 685 | 414 | 675 | 542 |
| 748 | -0.12 | 0.16 | -0.73 | 0.47 | 1 | Ttll12 | tubulin tyrosine ligase-like family, membe | 1075 | 1057 | 806 | 726 | 823 | 519 | 682 | 686 | 739 | 663 | 455 |
| 730 | 0.00 | 0.15 | 0.02 | 0.99 | 1 | Plscr1 | phospholipid scramblase 1 | 782 | 695 | 813 | 699 | 804 | 670 | 682 | 589 | 914 | 647 | 735 |
| 725 | -0.21 | 0.15 | -1.38 | 0.17 | 1 | Tex10 | testis expressed gene 10 | 845 | 748 | 796 | 670 | 815 | 812 | 682 | 582 | 614 | 535 | 878 |
| 724 | 0.04 | 0.16 | 0.28 | 0.78 | 1 | Gpr161 | G protein-coupled receptor 161 | 524 | 793 | 478 | 1055 | 550 | 707 | 681 | 1309 | 623 | 553 | 686 |
| 630 | -0.08 | 0.18 | -0.42 | 0.68 | 1 | Fn3k | fructosamine 3 kinase | 279 | 445 | 373 | 998 | 561 | 727 | 681 | 1062 | 668 | 529 | 606 |
| 1033 | -0.03 | 0.15 | -0.19 | 0.85 | 1 | Pxn | paxillin | 2532 | 2058 | 1878 | 697 | 576 | 512 | 680 | 577 | 740 | 674 | 442 |
| 848 | 0.02 | 0.20 | 0.11 | 0.91 | 1 | Dhrs1 | dehydrogenase/reductase (SDR family) n | 1117 | 850 | 1028 | 969 | 738 | 731 | 680 | 643 | 876 | 1001 | 691 |
| 833 | 0.15 | 0.13 | 1.16 | 0.25 | 1 | Rpl27 | ribosomal protein L27 | 1154 | 1089 | 1280 | 586 | 748 | 664 | 680 | 782 | 752 | 707 | 721 |
| 764 | -0.03 | 0.16 | -0.19 | 0.85 | 1 | Lmnb2 | lamin B2 | 853 | 631 | 828 | 701 | 682 | 1019 | 680 | 755 | 647 | 644 | 962 |
| 800 | 0.12 | 0.16 | 0.71 | 0.48 | 1 | Polr2f | polymerase (RNA) II (DNA directed) polyp | 1121 | 845 | 1077 | 673 | 802 | 613 | 679 | 875 | 690 | 768 | 661 |
| 739 | 0.09 | 0.13 | 0.69 | 0.49 | 1 | Fam133b | family with sequence similarity 133, meml | 892 | 697 | 638 | 651 | 583 | 941 | 679 | 741 | 569 | 770 | 963 |
| 720 | 0.19 | 0.18 | 1.08 | 0.28 | 1 | Polb | polymerase (DNA directed), beta | 649 | 621 | 698 | 539 | 718 | 839 | 679 | 857 | 625 | 678 | 1017 |
| 709 | -0.11 | 0.14 | -0.78 | 0.44 | 1 | Nkapd1 | NKAP domain containing 1 | 974 | 963 | 873 | 527 | 539 | 833 | 679 | 572 | 444 | 551 | 840 |
| 699 | 0.05 | 0.14 | 0.36 | 0.72 | 1 | Rpf1 | ribosome production factor 1 homolog | 595 | 767 | 741 | 655 | 702 | 709 | 679 | 739 | 659 | 723 | 726 |
| 689 | 0.13 | 0.17 | 0.77 | 0.44 | 1 | Bysl | bystin-like | 491 | 608 | 556 | 857 | 707 | 560 | 679 | 1171 | 652 | 635 | 667 |
| 876 | -0.07 | 0.17 | -0.45 | 0.66 | 1 | Chfr | checkpoint with forkhead and ring finger d | 854 | 797 | 683 | 859 | 873 | 1351 | 678 | 962 | 790 | 627 | 1166 |
| 745 | -0.15 | 0.16 | -0.89 | 0.37 | 1 | Tmem128 | transmembrane protein 128 | 788 | 710 | 646 | 843 | 929 | 731 | 678 | 693 | 746 | 681 | 748 |
| 662 | 0.11 | 0.19 | 0.56 | 0.58 | 1 | Naa38 | N(alpha)-acetyltransferase 38, NatC auxil | 658 | 478 | 684 | 691 | 742 | 522 | 678 | 869 | 625 | 746 | 584 |
| 888 | 0.18 | 0.21 | 0.87 | 0.39 | 1 | Topbp1 | topoisomerase (DNA) II binding protein 1 | 1494 | 1070 | 804 | 909 | 690 | 729 | 677 | 955 | 817 | 692 | 931 |
| 749 | -0.02 | 0.15 | -0.15 | 0.88 | 1 | Daxx | Fas death domain-associated protein | 929 | 800 | 668 | 778 | 731 | 756 | 677 | 696 | 739 | 752 | 709 |
| 699 | 0.08 | 0.15 | 0.51 | 0.61 | 1 | Il17ra | interleukin 17 receptor A | 972 | 769 | 729 | 687 | 507 | 675 | 677 | 636 | 675 | 735 | 630 |
| 1608 | 0.10 | 0.14 | 0.67 | 0.50 | 1 | Leng8 | leukocyte receptor cluster (LRC) member | 717 | 704 | 515 | 642 | 663 | 6275 | 676 | 808 | 754 | 766 | 5163 |
| 976 | 0.11 | 0.18 | 0.63 | 0.53 | 1 | Map1lc3a | microtubule-associated protein 1 light cha | 1305 | 1164 | 1736 | 807 | 955 | 703 | 676 | 1066 | 826 | 848 | 651 |
| 825 | -0.17 | 0.18 | -0.94 | 0.35 | 1 | Zfp949 | zinc finger protein 949 | 781 | 1012 | 979 | 714 | 606 | 1273 | 676 | 714 | 648 | 380 | 1289 |
| 739 | 0.05 | 0.20 | 0.26 | 0.79 | 1 | Plppr2 | phospholipid phosphatase related 2 | 404 | 350 | 617 | 701 | 849 | 1122 | 676 | 810 | 687 | 842 | 1070 |
| 721 | -0.11 | 0.21 | -0.54 | 0.59 | 1 | Phkg2 | phosphorylase kinase, gamma 2 (testis) | 682 | 749 | 814 | 557 | 706 | 987 | 676 | 820 | 561 | 424 | 952 |
| 671 | 0.15 | 0.26 | 0.58 | 0.56 | 1 | Gm5860 | | 92 | 175 | 74 | 968 | 1078 | 603 | 676 | 928 | 1200 | 1041 | 545 |
| 649 | -0.10 | 0.13 | -0.72 | 0.47 | 1 | Ndufab1 | NADH:ubiquinone oxidoreductase subun | 751 | 536 | 661 | 582 | 610 | 809 | 676 | 541 | 554 | 639 | 776 |
| 795 | 0.01 | 0.15 | 0.09 | 0.93 | 1 | Ssbp2 | single-stranded DNA binding protein 2 | 1042 | 907 | 1113 | 661 | 812 | 664 | 675 | 535 | 799 | 851 | 680 |
| 776 | 0.01 | 0.15 | 0.06 | 0.95 | 1 | Eif2ak4 | eukaryotic translation initiation factor 2 alp | 737 | 754 | 805 | 782 | 662 | 1003 | 675 | 689 | 709 | 837 | 878 |
| 742 | -0.18 | 0.16 | -1.12 | 0.26 | 1 | Pyroxd1 | pyridine nucleotide-disulphide oxidoredu | 740 | 870 | 707 | 794 | 612 | 980 | 675 | 538 | 553 | 598 | 1093 |
| 662 | -0.07 | 0.18 | -0.42 | 0.67 | 1 | Vps45 | vacuolar protein sorting 45 | 839 | 764 | 633 | 550 | 705 | 653 | 675 | 726 | 550 | 611 | 576 |
| 649 | 0.16 | 0.14 | 1.16 | 0.24 | 1 | Mfap1a | microfibrillar-associated protein 1A | 613 | 515 | 599 | 601 | 609 | 669 | 675 | 629 | 764 | 663 | 805 |
| 1885 | 0.01 | 0.11 | 0.05 | 0.96 | 1 | Oxtr | oxytocin receptor | 4110 | 7195 | 3715 | 980 | 520 | 653 | 674 | 1126 | 562 | 607 | 589 |
| 773 | 0.15 | 0.16 | 0.93 | 0.35 | 1 | Nap1l2 | nucleosome assembly protein 1-like 2 | 551 | 1067 | 845 | 829 | 646 | 707 | 674 | 974 | 743 | 707 | 764 |
| 743 | -0.20 | 0.13 | -1.58 | 0.11 | 1 | Rbm14 | RNA binding motif protein 14 | 653 | 768 | 621 | 687 | 622 | 1295 | 674 | 674 | 542 | 522 | 1120 |
| 699 | 0.32 | 0.22 | 1.43 | 0.15 | 1 | Akr1b3 | aldo-keto reductase family 1, member B3 | 639 | 577 | 722 | 682 | 602 | 555 | 674 | 1267 | 658 | 724 | 583 |
| 575 | -0.08 | 0.14 | -0.61 | 0.54 | 1 | Rcc1l | reculator of chromosome condensation 1 | 604 | 486 | 657 | 511 | 636 | 535 | 674 | 445 | 604 | 623 | 553 |
| 553 | -0.34 | 0.27 | -1.24 | 0.21 | 1 | Kcnab3 | potassium voltage-gated channel, shaker | 82 | 33 | 108 | 842 | 819 | 862 | 674 | 403 | 793 | 732 | 732 |
| 507 | -0.05 | 0.22 | -0.23 | 0.82 | 1 | Stat5a | signal transducer and activator of transcri | 485 | 308 | 641 | 475 | 504 | 453 | 674 | 491 | 645 | 478 | 426 |
| 973 | -0.21 | 0.14 | -1.51 | 0.13 | 1 | Gprc5c | G protein-coupled receptor, family C, grou | 1420 | 1843 | 1661 | 967 | 984 | 452 | 673 | 880 | 752 | 737 | 331 |
| 803 | 0.04 | 0.19 | 0.19 | 0.85 | 1 | Orc6 | origin recognition complex, subunit 6 | 643 | 426 | 563 | 922 | 858 | 1087 | 673 | 896 | 716 | 858 | 1185 |
| 783 | -0.04 | 0.21 | -0.20 | 0.84 | 1 | Ddx27 | DEAD (Asp-Glu-Ala-Asp) box polypeptide | 1008 | 904 | 1110 | 541 | 638 | 953 | 673 | 750 | 612 | 390 | 1034 |
| 747 | 0.12 | 0.16 | 0.76 | 0.44 | 1 | AU022252 | expressed sequence AU022252 | 635 | 637 | 853 | 849 | 695 | 700 | 673 | 907 | 762 | 756 | 752 |
| 727 | 0.07 | 0.15 | 0.47 | 0.63 | 1 | Klhl15 | kelch-like 15 | 472 | 651 | 693 | 835 | 719 | 774 | 673 | 980 | 725 | 638 | 836 |
| 797 | -0.04 | 0.17 | -0.21 | 0.83 | 1 | Plekhh3 | pleckstrin homology domain containing, fa | 1054 | 1067 | 1014 | 873 | 691 | 580 | 672 | 1067 | 692 | 574 | 485 |
| 610 | -0.19 | 0.20 | -0.95 | 0.34 | 1 | Rabl3 | RAB, member RAS oncogene family-like | 541 | 614 | 520 | 580 | 689 | 724 | 672 | 388 | 622 | 657 | 700 |
| 612 | -0.22 | 0.19 | -1.14 | 0.25 | 1 | Mob3c | MOB kinase activator 3C | 602 | 630 | 466 | 674 | 659 | 684 | 671 | 780 | 503 | 428 | 639 |
| 604 | -0.12 | 0.20 | -0.59 | 0.56 | 1 | Psat1 | phosphoserine aminotransferase 1 | 1161 | 1106 | 895 | 543 | 397 | 221 | 671 | 481 | 532 | 425 | 209 |
| 597 | 0.11 | 0.15 | 0.73 | 0.47 | 1 | Nubp1 | nucleotide binding protein 1 | 651 | 374 | 469 | 479 | 585 | 684 | 671 | 506 | 569 | 902 | 674 |
| 871 | -0.14 | 0.15 | -0.97 | 0.33 | 1 | Gmeb2 | glucocorticoid modulatory element bindin | 988 | 948 | 896 | 850 | 944 | 1073 | 670 | 690 | 812 | 689 | 1018 |
| 737 | -0.01 | 0.12 | -0.10 | 0.92 | 1 | Bbs9 | Bardet-Biedl syndrome 9 (human) | 649 | 795 | 803 | 781 | 723 | 763 | 670 | 828 | 751 | 599 | 746 |

| | | | | | | | | | | | | | | | | | | |
|---|---|---|---|---|---|---|---|---|---|---|---|---|---|---|---|---|---|---|
| 736 | 0.13 | 0.14 | 0.96 | 0.34 | 1 | Fam53a | family with sequence similarity 53, membe | 658 | 660 | 687 | 702 | 708 | 829 | 670 | 877 | 792 | 746 | 764 |
| 680 | 0.14 | 0.15 | 0.98 | 0.33 | 1 | Rab40c | Rab40C, member RAS oncogene family | 818 | 779 | 793 | 508 | 611 | 630 | 670 | 657 | 570 | 694 | 749 |
| 795 | -0.23 | 0.17 | -1.33 | 0.18 | 1 | Etfrf1 | electron transfer flavoprotein regulatory fa | 890 | 1053 | 931 | 890 | 856 | 757 | 669 | 543 | 741 | 732 | 680 |
| 782 | 0.26 | 0.15 | 1.73 | 0.08 | 1 | Tsga10 | testis specific 10 | 269 | 396 | 296 | 705 | 575 | 1553 | 669 | 1068 | 656 | 738 | 1684 |
| 717 | -0.23 | 0.26 | -0.86 | 0.39 | 1 | Pold2 | polymerase (DNA directed), delta 2, regul | 731 | 500 | 862 | 586 | 1047 | 713 | 669 | 231 | 959 | 791 | 803 |
| 712 | -0.32 | 0.23 | -1.43 | 0.15 | 1 | Pid1 | phosphotyrosine interaction domain conta | 2118 | 696 | 975 | 594 | 497 | 434 | 669 | 348 | 399 | 739 | 365 |
| 649 | 0.11 | 0.14 | 0.78 | 0.44 | 1 | Timm50 | translocase of inner mitochondrial membr | 744 | 543 | 633 | 640 | 644 | 560 | 669 | 688 | 694 | 697 | 629 |
| 597 | -0.05 | 0.15 | -0.30 | 0.76 | 1 | Stox1 | storkhead box 1 | 362 | 426 | 390 | 689 | 662 | 713 | 669 | 556 | 682 | 670 | 745 |
| 554 | -0.17 | 0.19 | -0.87 | 0.39 | 1 | L2hgdh | L-2-hydroxyglutarate dehydrogenase | 451 | 500 | 647 | 590 | 587 | 526 | 669 | 430 | 552 | 647 | 496 |
| 965 | -0.13 | 0.15 | -0.85 | 0.40 | 1 | Isyna1 | myo-inositol 1-phosphate synthase A1 | 1766 | 1663 | 1643 | 656 | 637 | 939 | 668 | 604 | 540 | 811 | 689 |
| 945 | 0.05 | 0.18 | 0.28 | 0.78 | 1 | Plbd2 | phospholipase B domain containing 2 | 1984 | 1203 | 1321 | 777 | 697 | 705 | 668 | 702 | 660 | 1054 | 628 |
| 844 | 0.02 | 0.15 | 0.15 | 0.88 | 1 | Nosip | nitric oxide synthase interacting protein | 1144 | 1048 | 1151 | 672 | 871 | 740 | 668 | 813 | 735 | 779 | 664 |
| 711 | -0.14 | 0.22 | -0.62 | 0.54 | 1 | Supt7l | SPT7-like, STAGA complex gamma subur | 750 | 634 | 862 | 786 | 746 | 678 | 668 | 389 | 825 | 782 | 706 |
| 690 | 0.15 | 0.17 | 0.87 | 0.38 | 1 | Zbtb37 | zinc finger and BTB domain containing 37 | 650 | 687 | 533 | 515 | 473 | 1060 | 668 | 476 | 616 | 755 | 1160 |
| 614 | 0.01 | 0.15 | 0.08 | 0.94 | 1 | Cenpj | centromere protein J | 555 | 391 | 480 | 464 | 695 | 821 | 668 | 475 | 720 | 600 | 880 |
| 605 | -0.07 | 0.16 | -0.44 | 0.66 | 1 | Golga7b | golgi autoantigen, golgin subfamily a, 7B | 531 | 734 | 593 | 631 | 732 | 412 | 668 | 666 | 661 | 663 | 358 |
| 544 | 0.23 | 0.26 | 0.91 | 0.36 | 1 | Crb1 | crumbs family member 1, photoreceptor m | 35 | 126 | 57 | 850 | 628 | 519 | 668 | 879 | 815 | 795 | 607 |
| 467 | 0.27 | 0.17 | 1.60 | 0.11 | 1 | Xrcc3 | X-ray repair complementing defective rep | 138 | 138 | 110 | 541 | 450 | 495 | 668 | 592 | 666 | 781 | 556 |
| 805 | 0.15 | 0.18 | 0.84 | 0.40 | 1 | Twf2 | twinfilin actin binding protein 2 | 1202 | 919 | 877 | 743 | 740 | 602 | 667 | 1055 | 660 | 775 | 619 |
| 741 | 0.07 | 0.14 | 0.49 | 0.62 | 1 | Ranbp6 | RAN binding protein 6 | 824 | 783 | 619 | 769 | 528 | 931 | 667 | 876 | 591 | 631 | 930 |
| 723 | 0.03 | 0.14 | 0.19 | 0.85 | 1 | Ints11 | integrator complex subunit 11 | 914 | 744 | 730 | 621 | 717 | 746 | 667 | 674 | 628 | 805 | 706 |
| 855 | 0.10 | 0.12 | 0.88 | 0.38 | 1 | Hars2 | histidyl-tRNA synthetase 2 | 864 | 822 | 899 | 668 | 705 | 1227 | 666 | 739 | 739 | 654 | 1421 |
| 555 | -0.15 | 0.21 | -0.69 | 0.49 | 1 | Gys1 | glycogen synthase 1, muscle | 514 | 851 | 613 | 522 | 548 | 417 | 666 | 534 | 604 | 492 | 340 |
| 550 | -0.01 | 0.20 | -0.06 | 0.95 | 1 | Ankrd26 | ankyrin repeat domain 26 | 516 | 630 | 466 | 403 | 453 | 725 | 666 | 510 | 492 | 538 | 653 |
| 518 | 0.01 | 0.18 | 0.04 | 0.97 | 1 | Osbpl7 | oxysterol binding protein-like 7 | 436 | 503 | 375 | 464 | 520 | 530 | 666 | 587 | 529 | 627 | 455 |
| 794 | 0.05 | 0.16 | 0.28 | 0.78 | 1 | Fut11 | fucosyltransferase 11 | 1194 | 1101 | 1071 | 628 | 737 | 603 | 665 | 794 | 820 | 576 | 549 |
| 677 | -0.12 | 0.14 | -0.89 | 0.38 | 1 | Riok2 | RIO kinase 2 | 768 | 749 | 753 | 671 | 655 | 702 | 665 | 529 | 611 | 589 | 761 |
| 605 | -0.10 | 0.20 | -0.52 | 0.60 | 1 | Tti2 | TELO2 interacting protein 2 | 503 | 569 | 566 | 450 | 709 | 772 | 664 | 577 | 681 | 433 | 732 |
| 565 | 0.03 | 0.16 | 0.17 | 0.86 | 1 | Nt5m | 5',3'-nucleotidase, mitochondrial | 393 | 332 | 469 | 526 | 705 | 601 | 664 | 671 | 631 | 658 | 570 |
| 878 | 0.02 | 0.14 | 0.14 | 0.89 | 1 | Samd9l | sterile alpha motif domain containing 9-lik | 1936 | 1470 | 1622 | 543 | 529 | 544 | 663 | 477 | 500 | 802 | 567 |
| 870 | 0.39 | 0.21 | 1.87 | 0.06 | 1 | Slc25a14 | solute carrier family 25 (mitochondrial car | 644 | 889 | 919 | 694 | 913 | 806 | 663 | 1139 | 884 | 1234 | 785 |
| 808 | -0.17 | 0.21 | -0.83 | 0.40 | 1 | Dact1 | dishevelled-binding antagonist of beta-ca | 1715 | 1299 | 1130 | 501 | 630 | 687 | 663 | 335 | 633 | 518 | 773 |
| 769 | -0.14 | 0.15 | -0.93 | 0.35 | 1 | Nup133 | nucleoporin 133 | 1162 | 1010 | 1118 | 726 | 759 | 569 | 663 | 687 | 621 | 535 | 609 |
| 747 | -0.07 | 0.16 | -0.46 | 0.64 | 1 | Zfp280b | zinc finger protein 280B | 975 | 840 | 738 | 706 | 735 | 791 | 663 | 726 | 627 | 580 | 837 |
| 742 | -0.23 | 0.16 | -1.42 | 0.16 | 1 | Zfyve26 | zinc finger, FYVE domain containing 26 | 726 | 831 | 670 | 845 | 649 | 1049 | 663 | 568 | 705 | 549 | 908 |
| 496 | -0.19 | 0.16 | -1.18 | 0.24 | 1 | Ldlrad4 | low density lipoprotein receptor class A d | 619 | 348 | 612 | 417 | 558 | 425 | 663 | 307 | 494 | 571 | 443 |
| 1162 | -0.30 | 0.16 | -1.95 | 0.05 | 1 | Lrig1 | leucine-rich repeats and immunoglobulin | 2138 | 2693 | 2564 | 757 | 517 | 1029 | 662 | 599 | 562 | 398 | 863 |
| 902 | -0.10 | 0.20 | -0.48 | 0.63 | 1 | Supt20 | SPT20 SAGA complex component | 1126 | 1255 | 888 | 478 | 448 | 1889 | 662 | 650 | 493 | 385 | 1643 |
| 695 | 0.21 | 0.17 | 1.27 | 0.21 | 1 | Uqcc3 | ubiquinol-cytochrome c reductase comple | 660 | 779 | 818 | 623 | 729 | 478 | 662 | 780 | 689 | 909 | 518 |
| 657 | -0.24 | 0.32 | -0.75 | 0.45 | 1 | Atxn1 | ataxin 1 | 449 | 694 | 557 | 670 | 481 | 1119 | 662 | 231 | 533 | 724 | 1111 |
| 577 | 0.01 | 0.16 | 0.08 | 0.94 | 1 | Mrpl41 | mitochondrial ribosomal protein L41 | 651 | 612 | 583 | 513 | 613 | 448 | 662 | 491 | 547 | 739 | 483 |
| 924 | -0.23 | 0.23 | -1.02 | 0.31 | 1 | Tiparp | TCDD-inducible poly(ADP-ribose) polyme | 1846 | 1206 | 948 | 824 | 705 | 1067 | 661 | 604 | 587 | 496 | 1215 |
| 775 | -0.20 | 0.24 | -0.84 | 0.40 | 1 | Scai | suppressor of cancer cell invasion | 444 | 596 | 506 | 572 | 686 | 1690 | 661 | 342 | 503 | 761 | 1761 |
| 673 | 0.14 | 0.15 | 0.92 | 0.36 | 1 | N4bp2l1 | NEDD4 binding protein 2-like 1 | 631 | 603 | 619 | 643 | 520 | 820 | 661 | 754 | 649 | 603 | 895 |
| 651 | 0.06 | 0.21 | 0.30 | 0.76 | 1 | Rnf215 | ring finger protein 215 | 569 | 475 | 658 | 621 | 628 | 745 | 661 | 674 | 501 | 934 | 693 |
| 643 | 0.19 | 0.20 | 0.94 | 0.35 | 1 | Cep76 | centrosomal protein 76 | 411 | 561 | 451 | 850 | 536 | 609 | 661 | 633 | 707 | 920 | 737 |
| 894 | -0.02 | 0.19 | -0.10 | 0.92 | 1 | Brd9 | bromodomain containing 9 | 1029 | 686 | 884 | 794 | 707 | 1480 | 660 | 824 | 738 | 577 | 1451 |
| 757 | 0.34 | 0.18 | 1.93 | 0.05 | 1 | Wdr34 | WD repeat domain 34 | 701 | 933 | 806 | 570 | 672 | 682 | 660 | 1023 | 705 | 830 | 747 |
| 644 | 0.17 | 0.25 | 0.67 | 0.50 | 1 | Racgap1 | Rac GTPase-activating protein 1 | 770 | 395 | 333 | 593 | 706 | 681 | 660 | 827 | 727 | 690 | 707 |
| 643 | 0.22 | 0.18 | 1.25 | 0.21 | 1 | Zfp65 | zinc finger protein 65 | 532 | 539 | 535 | 528 | 678 | 644 | 660 | 520 | 785 | 936 | 718 |
| 628 | 0.00 | 0.14 | 0.00 | 1.00 | 1 | Gng5 | guanine nucleotide binding protein (G pro | 713 | 574 | 705 | 543 | 696 | 548 | 660 | 476 | 697 | 728 | 564 |
| 626 | -0.14 | 0.16 | -0.87 | 0.38 | 1 | Pcgf2 | polycomb group ring finger 2 | 735 | 803 | 893 | 493 | 603 | 583 | 660 | 519 | 512 | 562 | 528 |
| 620 | 0.06 | 0.21 | 0.27 | 0.78 | 1 | Phb | prohibitin | 798 | 595 | 736 | 507 | 629 | 508 | 660 | 731 | 638 | 460 | 555 |
| 547 | -0.30 | 0.17 | -1.75 | 0.08 | 1 | Cep97 | centrosomal protein 97 | 522 | 463 | 399 | 480 | 589 | 826 | 660 | 373 | 524 | 513 | 666 |
| 811 | -0.15 | 0.14 | -1.09 | 0.28 | 1 | Sirt1 | sirtuin 1 | 969 | 1130 | 815 | 803 | 702 | 1001 | 659 | 678 | 680 | 594 | 893 |
| 778 | 0.12 | 0.16 | 0.76 | 0.45 | 1 | Nudcd2 | NudC domain containing 2 | 826 | 863 | 760 | 696 | 834 | 735 | 659 | 929 | 706 | 705 | 850 |
| 746 | -0.08 | 0.22 | -0.36 | 0.72 | 1 | Btf3l4 | basic transcription factor 3-like 4 | 638 | 951 | 695 | 813 | 824 | 747 | 659 | 496 | 812 | 743 | 828 |
| 668 | 0.25 | 0.23 | 1.09 | 0.28 | 1 | Twnk | twinkle mtDNA helicase | 681 | 466 | 776 | 492 | 663 | 678 | 659 | 925 | 630 | 616 | 766 |
| 638 | 0.15 | 0.18 | 0.82 | 0.41 | 1 | Mrpl11 | mitochondrial ribosomal protein L11 | 684 | 530 | 644 | 615 | 660 | 505 | 659 | 893 | 681 | 634 | 516 |
| 601 | -0.09 | 0.22 | -0.41 | 0.68 | 1 | Lgals12 | lectin, galactose binding, soluble 12 | 92 | 145 | 217 | 569 | 645 | 1315 | 659 | 568 | 636 | 589 | 1181 |

| | | | | | | | | | | | | | | | | | | |
|---|---|---|---|---|---|---|---|---|---|---|---|---|---|---|---|---|---|---|
| 528 | 0.06 | 0.18 | 0.32 | 0.75 | 1 | Zfp27 | zinc finger protein 27 | 450 | 277 | 403 | 543 | 471 | 621 | 659 | 541 | 583 | 583 | 673 |
| 936 | 0.10 | 0.16 | 0.61 | 0.54 | 1 | Klf7 | Kruppel-like factor 7 (ubiquitous) | 1641 | 1536 | 1604 | 633 | 498 | 888 | 658 | 849 | 570 | 638 | 782 |
| 670 | 0.17 | 0.17 | 1.00 | 0.32 | 1 | Cep19 | centrosomal protein 19 | 511 | 418 | 499 | 887 | 668 | 586 | 658 | 780 | 756 | 880 | 732 |
| 606 | -0.17 | 0.18 | -0.96 | 0.33 | 1 | Tmem175 | transmembrane protein 175 | 649 | 488 | 628 | 725 | 616 | 588 | 658 | 468 | 640 | 598 | 606 |
| 606 | -0.06 | 0.17 | -0.33 | 0.74 | 1 | Rrp8 | ribosomal RNA processing 8, methyltrans | 533 | 729 | 433 | 715 | 591 | 569 | 658 | 717 | 638 | 501 | 580 |
| 571 | 0.19 | 0.22 | 0.88 | 0.38 | 1 | Alg6 | asparagine-linked glycosylation 6 (alpha- | 344 | 578 | 382 | 529 | 521 | 616 | 658 | 583 | 637 | 769 | 662 |
| 767 | -0.06 | 0.16 | -0.37 | 0.71 | 1 | Psmd9 | proteasome (prosome, macropain) 26S su | 1084 | 884 | 1053 | 782 | 702 | 617 | 657 | 831 | 635 | 608 | 580 |
| 697 | 0.16 | 0.19 | 0.86 | 0.39 | 1 | Wdr77 | WD repeat domain 77 | 874 | 778 | 841 | 455 | 658 | 681 | 657 | 809 | 636 | 631 | 651 |
| 672 | -0.21 | 0.16 | -1.37 | 0.17 | 1 | Pcbp4 | poly(rC) binding protein 4 | 1038 | 737 | 893 | 692 | 622 | 561 | 657 | 567 | 497 | 609 | 514 |
| 661 | 0.04 | 0.14 | 0.28 | 0.78 | 1 | Gtf2h3 | general transcription factor IIH, polypeptic | 565 | 644 | 476 | 782 | 579 | 746 | 657 | 695 | 688 | 653 | 788 |
| 645 | -0.07 | 0.19 | -0.38 | 0.71 | 1 | Pwp2 | PWP2 periodic tryptophan protein homolo | 825 | 847 | 685 | 507 | 676 | 580 | 657 | 739 | 542 | 526 | 512 |
| 639 | 0.09 | 0.18 | 0.51 | 0.61 | 1 | Eif2b3 | eukaryotic translation initiation factor 2B, s | 552 | 483 | 586 | 632 | 716 | 609 | 657 | 741 | 537 | 730 | 788 |
| 565 | -0.17 | 0.18 | -0.93 | 0.35 | 1 | Zdhhc14 | zinc finger, DHHC domain containing 14 | 768 | 758 | 716 | 553 | 453 | 440 | 657 | 401 | 546 | 556 | 372 |
| 806 | 0.09 | 0.14 | 0.69 | 0.49 | 1 | Sestd1 | SEC14 and spectrin domains 1 | 1116 | 878 | 969 | 668 | 599 | 913 | 656 | 653 | 672 | 647 | 1093 |
| 679 | -0.04 | 0.15 | -0.28 | 0.78 | 1 | Nudt18 | nudix (nucleoside diphosphate linked mo | 731 | 808 | 815 | 745 | 703 | 474 | 656 | 715 | 637 | 787 | 400 |
| 755 | 0.13 | 0.15 | 0.88 | 0.38 | 1 | Llgl1 | LLGL1 scribble cell polarity complex com | 1101 | 895 | 890 | 631 | 612 | 686 | 655 | 774 | 695 | 754 | 609 |
| 711 | 0.11 | 0.16 | 0.66 | 0.51 | 1 | Pnpt1 | polyribonucleotide nucleotidyltransferase | 656 | 597 | 653 | 537 | 684 | 967 | 655 | 548 | 635 | 926 | 963 |
| 647 | -0.03 | 0.17 | -0.19 | 0.85 | 1 | Cd274 | CD274 antigen | 844 | 971 | 1081 | 514 | 508 | 451 | 655 | 567 | 571 | 569 | 389 |
| 546 | -0.17 | 0.22 | -0.81 | 0.42 | 1 | Get1 | guided entry of tail-anchored proteins fact | 477 | 650 | 498 | 487 | 581 | 597 | 655 | 380 | 585 | 544 | 552 |
| 524 | -0.38 | 0.20 | -1.90 | 0.06 | 1 | Coq7 | demethyl-Q 7 | 558 | 556 | 545 | 589 | 644 | 433 | 655 | 323 | 470 | 568 | 423 |
| 726 | -0.05 | 0.13 | -0.35 | 0.72 | 1 | Kri1 | KRI1 homolog | 645 | 595 | 574 | 690 | 742 | 1045 | 654 | 619 | 691 | 707 | 1021 |
| 632 | 0.00 | 0.16 | 0.01 | 0.99 | 1 | Gpn1 | GPN-loop GTPase 1 | 711 | 546 | 814 | 508 | 624 | 653 | 654 | 524 | 525 | 714 | 682 |
| 584 | -0.05 | 0.18 | -0.29 | 0.77 | 1 | Lrrc20 | leucine rich repeat containing 20 | 492 | 476 | 422 | 722 | 690 | 484 | 654 | 777 | 745 | 506 | 453 |
| 553 | -0.03 | 0.20 | -0.14 | 0.89 | 1 | Flrt1 | fibronectin leucine rich transmembrane pr | 245 | 231 | 301 | 782 | 675 | 531 | 654 | 950 | 745 | 545 | 429 |
| 442 | 0.07 | 0.27 | 0.26 | 0.79 | 1 | Cnnm1 | cyclin M1 | 27 | 19 | 64 | 575 | 678 | 411 | 654 | 710 | 714 | 599 | 414 |
| 797 | -0.20 | 0.18 | -1.09 | 0.28 | 1 | H2-T22 | histocompatibility 2, T region locus 22 | 1023 | 1437 | 686 | 979 | 679 | 703 | 653 | 747 | 626 | 515 | 716 |
| 693 | -0.16 | 0.18 | -0.90 | 0.37 | 1 | Noc3l | NOC3 like DNA replication regulator | 585 | 715 | 566 | 855 | 649 | 866 | 653 | 529 | 670 | 587 | 941 |
| 691 | -0.09 | 0.16 | -0.57 | 0.57 | 1 | Kti12 | KTI12 homolog, chromatin associated | 581 | 924 | 705 | 742 | 714 | 665 | 653 | 674 | 574 | 604 | 761 |
| 680 | -0.14 | 0.20 | -0.71 | 0.48 | 1 | Scn8a | sodium channel, voltage-gated, type VIII, | 397 | 491 | 507 | 712 | 541 | 1255 | 653 | 785 | 613 | 384 | 1137 |
| 659 | -0.08 | 0.23 | -0.36 | 0.72 | 1 | Rnft1 | ring finger protein, transmembrane 1 | 957 | 567 | 547 | 575 | 719 | 691 | 653 | 440 | 751 | 540 | 806 |
| 656 | -0.23 | 0.15 | -1.53 | 0.13 | 1 | Ticam1 | toll-like receptor adaptor molecule 1 | 652 | 686 | 759 | 651 | 799 | 653 | 653 | 461 | 694 | 603 | 605 |
| 386 | -0.45 | 0.23 | -1.97 | 0.05 | 1 | Lrfn1 | leucine rich repeat and fibronectin type III | 292 | 360 | 229 | 420 | 507 | 361 | 653 | 371 | 395 | 424 | 240 |
| 853 | 0.03 | 0.16 | 0.18 | 0.86 | 1 | Atg16l2 | autophagy related 16-like 2 (S. cerevisiae | 583 | 426 | 472 | 580 | 661 | 2123 | 652 | 754 | 640 | 638 | 1850 |
| 836 | 0.05 | 0.19 | 0.28 | 0.78 | 1 | Csnk1e | casein kinase 1, epsilon | 1277 | 1089 | 1415 | 694 | 497 | 820 | 652 | 820 | 568 | 613 | 745 |
| 823 | -0.10 | 0.19 | -0.52 | 0.61 | 1 | Slc25a20 | solute carrier family 25 (mitochondrial car | 752 | 1146 | 1031 | 882 | 987 | 665 | 652 | 620 | 839 | 818 | 657 |
| 793 | 0.08 | 0.16 | 0.48 | 0.63 | 1 | Myd88 | myeloid differentiation primary response g | 1404 | 946 | 1024 | 726 | 706 | 521 | 652 | 732 | 669 | 755 | 592 |
| 675 | -0.02 | 0.17 | -0.13 | 0.90 | 1 | Brat1 | BRCA1-associated ATM activator 1 | 646 | 660 | 650 | 801 | 628 | 666 | 652 | 852 | 715 | 502 | 657 |
| 647 | 0.09 | 0.17 | 0.55 | 0.58 | 1 | Nip7 | NIP7, nucleolar pre-rRNA processing pro | 643 | 687 | 814 | 611 | 635 | 511 | 652 | 743 | 603 | 573 | 645 |
| 606 | 0.09 | 0.15 | 0.58 | 0.56 | 1 | Hibch | 3-hydroxyisobutyryl-Coenzyme A hydrola | 580 | 610 | 588 | 586 | 507 | 621 | 652 | 643 | 570 | 730 | 575 |
| 578 | 0.14 | 0.14 | 0.98 | 0.33 | 1 | Gstm4 | glutathione S-transferase, mu 4 | 433 | 478 | 386 | 698 | 576 | 478 | 652 | 737 | 563 | 795 | 558 |
| 515 | -0.16 | 0.22 | -0.74 | 0.46 | 1 | Tjp3 | tight junction protein 3 | 182 | 354 | 395 | 556 | 739 | 549 | 652 | 589 | 663 | 546 | 447 |
| 899 | -0.01 | 0.18 | -0.04 | 0.96 | 1 | Prickle1 | prickle planar cell polarity protein 1 | 1534 | 1286 | 1188 | 539 | 868 | 878 | 651 | 485 | 785 | 742 | 929 |
| 865 | -0.12 | 0.12 | -0.95 | 0.34 | 1 | D10Wsu102e | DNA segment, Chr 10, Wayne State Unive | 1271 | 1754 | 1359 | 753 | 568 | 695 | 651 | 721 | 587 | 522 | 631 |
| 839 | -0.21 | 0.17 | -1.20 | 0.23 | 1 | Fchsd2 | FCH and double SH3 domains 2 | 1464 | 1226 | 1175 | 872 | 614 | 744 | 651 | 594 | 689 | 581 | 619 |
| 808 | 0.31 | 0.26 | 1.20 | 0.23 | 1 | Stx8 | syntaxin 8 | 779 | 719 | 769 | 864 | 781 | 566 | 651 | 1748 | 671 | 645 | 695 |
| 774 | 0.08 | 0.18 | 0.44 | 0.66 | 1 | Mrpl13 | mitochondrial ribosomal protein L13 | 904 | 856 | 1117 | 797 | 734 | 567 | 651 | 879 | 624 | 674 | 711 |
| 771 | 0.05 | 0.16 | 0.29 | 0.77 | 1 | H1f10 | H1.10 linker histone | 806 | 984 | 956 | 796 | 921 | 474 | 651 | 894 | 692 | 778 | 532 |
| 769 | 0.10 | 0.15 | 0.64 | 0.52 | 1 | Caml | calcium modulating ligand | 611 | 815 | 793 | 707 | 804 | 856 | 651 | 713 | 732 | 819 | 963 |
| 712 | 0.02 | 0.13 | 0.14 | 0.89 | 1 | Tmc6 | transmembrane channel-like gene family | 1093 | 1066 | 779 | 649 | 571 | 556 | 651 | 716 | 511 | 639 | 603 |
| 618 | -0.01 | 0.19 | -0.05 | 0.96 | 1 | Nat9 | N-acetyltransferase 9 (GCN5-related, puta | 685 | 474 | 710 | 438 | 710 | 701 | 651 | 629 | 580 | 526 | 701 |
| 606 | -0.23 | 0.19 | -1.20 | 0.23 | 1 | Snu13 | SNU13 homolog, small nuclear ribonucle | 692 | 747 | 830 | 514 | 673 | 523 | 651 | 336 | 644 | 504 | 551 |
| 588 | 0.02 | 0.17 | 0.10 | 0.92 | 1 | Twistnb | twist basic helix-loop-helix transcription fa | 709 | 623 | 642 | 561 | 457 | 570 | 651 | 665 | 500 | 526 | 566 |
| 428 | 0.31 | 0.21 | 1.50 | 0.13 | 1 | Kcnj12 | potassium inwardly-rectifying channel, su | 67 | 41 | 70 | 495 | 513 | 353 | 651 | 724 | 727 | 685 | 377 |
| 1065 | 0.10 | 0.13 | 0.75 | 0.45 | 1 | 4833420G17Ril | RIKEN cDNA 4833420G17 gene | 767 | 811 | 994 | 845 | 843 | 2128 | 650 | 842 | 840 | 871 | 2125 |
| 727 | 0.10 | 0.17 | 0.60 | 0.55 | 1 | Coq8b | coenzyme Q8B | 639 | 573 | 638 | 519 | 719 | 1112 | 650 | 747 | 612 | 787 | 996 |
| 714 | -0.09 | 0.15 | -0.61 | 0.54 | 1 | Mrps18a | mitochondrial ribosomal protein S18A | 657 | 592 | 801 | 672 | 1001 | 686 | 650 | 660 | 747 | 681 | 703 |
| 649 | -0.13 | 0.17 | -0.74 | 0.46 | 1 | Stmn3 | stathmin-like 3 | 759 | 850 | 833 | 820 | 622 | 369 | 650 | 702 | 463 | 711 | 364 |
| 740 | -0.14 | 0.14 | -0.97 | 0.33 | 1 | Nup107 | nucleoporin 107 | 953 | 972 | 947 | 655 | 757 | 690 | 649 | 565 | 603 | 589 | 759 |
| 641 | 0.05 | 0.20 | 0.23 | 0.82 | 1 | Tmem201 | transmembrane protein 201 | 819 | 631 | 789 | 568 | 558 | 550 | 649 | 585 | 490 | 953 | 456 |
| 621 | 0.21 | 0.16 | 1.30 | 0.19 | 1 | Gstm2 | glutathione S-transferase, mu 2 | 1019 | 724 | 658 | 522 | 565 | 316 | 649 | 521 | 575 | 858 | 419 |

| | | | | | | | | | | | | | | | | | |
|---|---|---|---|---|---|---|---|---|---|---|---|---|---|---|---|---|---|
| 469 | -0.15 | 0.23 | -0.66 | 0.51 | 1 | Casp9 | caspase 9 | 686 | 603 | 599 | 383 | 311 | 405 | 649 | 400 | 366 | 346 | 412 |
| 1020 | 0.02 | 0.17 | 0.11 | 0.91 | 1 | Anks3 | ankyrin repeat and sterile alpha motif don | 908 | 771 | 1064 | 822 | 937 | 1827 | 648 | 950 | 716 | 772 | 1801 |
| 860 | 0.13 | 0.16 | 0.85 | 0.40 | 1 | Eri1 | exoribonuclease 1 | 1579 | 1307 | 1013 | 741 | 603 | 658 | 648 | 796 | 658 | 648 | 807 |
| 736 | -0.15 | 0.19 | -0.77 | 0.44 | 1 | Mcts1 | malignant T cell amplified sequence 1 | 639 | 693 | 653 | 470 | 565 | 1515 | 648 | 342 | 530 | 675 | 1366 |
| 672 | -0.14 | 0.17 | -0.80 | 0.43 | 1 | Pigm | phosphatidylinositol glycan anchor biosyı | 641 | 574 | 581 | 584 | 673 | 970 | 648 | 550 | 590 | 455 | 1125 |
| 580 | -0.16 | 0.17 | -0.94 | 0.35 | 1 | Hspb6 | heat shock protein, alpha-crystallin-relate | 244 | 210 | 272 | 805 | 902 | 632 | 648 | 577 | 746 | 752 | 598 |
| 573 | 0.21 | 0.42 | 0.49 | 0.62 | 1 | Rps18-ps6 | | 1814 | 46 | 1842 | 23 | 607 | 7 | 648 | 65 | 501 | 748 | 1 |
| 686 | 0.09 | 0.19 | 0.47 | 0.64 | 1 | Haus2 | HAUS augmin-like complex, subunit 2 | 663 | 858 | 634 | 734 | 741 | 502 | 647 | 624 | 645 | 804 | 691 |
| 673 | 0.11 | 0.18 | 0.60 | 0.55 | 1 | Brsk1 | BR serine/threonine kinase 1 | 376 | 297 | 505 | 571 | 753 | 1065 | 647 | 762 | 718 | 761 | 944 |
| 598 | -0.11 | 0.17 | -0.64 | 0.52 | 1 | Nova1 | NOVA alternative splicing regulator 1 | 651 | 485 | 666 | 598 | 489 | 747 | 647 | 527 | 524 | 531 | 717 |
| 544 | -0.11 | 0.15 | -0.75 | 0.46 | 1 | Pyurf | Pigy upstream reading frame | 498 | 571 | 579 | 549 | 639 | 421 | 647 | 510 | 581 | 599 | 393 |
| 704 | -0.14 | 0.15 | -0.94 | 0.35 | 1 | Necap2 | NECAP endocytosis associated 2 | 1661 | 927 | 1026 | 416 | 377 | 702 | 646 | 293 | 369 | 643 | 686 |
| 603 | -0.28 | 0.18 | -1.51 | 0.13 | 1 | Chmp6 | charged multivesicular body protein 6 | 757 | 646 | 628 | 703 | 597 | 572 | 646 | 403 | 552 | 536 | 594 |
| 589 | 0.08 | 0.16 | 0.48 | 0.63 | 1 | Sdhaf4 | succinate dehydrogenase complex asser | 589 | 685 | 621 | 581 | 620 | 384 | 646 | 753 | 560 | 596 | 439 |
| 511 | 0.08 | 0.18 | 0.45 | 0.65 | 1 | Tmem268 | transmembrane protein 268 | 417 | 448 | 343 | 539 | 523 | 428 | 646 | 740 | 572 | 542 | 421 |
| 897 | 0.25 | 0.15 | 1.64 | 0.10 | 1 | Thoc1 | THO complex 1 | 745 | 828 | 803 | 684 | 642 | 1440 | 645 | 1064 | 739 | 620 | 1655 |
| 749 | 0.06 | 0.20 | 0.31 | 0.76 | 1 | Maoa | monoamine oxidase A | 1096 | 1295 | 899 | 593 | 748 | 440 | 645 | 527 | 687 | 800 | 511 |
| 711 | -0.11 | 0.16 | -0.65 | 0.52 | 1 | Zbtb10 | zinc finger and BTB domain containing 1( | 752 | 989 | 692 | 712 | 637 | 790 | 645 | 642 | 681 | 483 | 800 |
| 705 | -0.07 | 0.14 | -0.46 | 0.65 | 1 | 2010315B03Rik | RIKEN cDNA 2010315B03 gene | 467 | 676 | 651 | 756 | 680 | 946 | 645 | 685 | 752 | 517 | 980 |
| 702 | -0.19 | 0.20 | -0.96 | 0.34 | 1 | Cables2 | CDK5 and Abl enzyme substrate 2 | 722 | 583 | 662 | 998 | 687 | 735 | 645 | 869 | 678 | 559 | 588 |
| 671 | 0.14 | 0.18 | 0.79 | 0.43 | 1 | Bloc1s5 | biogenesis of lysosomal organelles comp | 667 | 902 | 805 | 576 | 658 | 478 | 645 | 869 | 609 | 724 | 453 |
| 574 | 0.01 | 0.19 | 0.06 | 0.95 | 1 | Rom1 | rod outer segment membrane protein 1 | 493 | 530 | 451 | 540 | 719 | 485 | 645 | 675 | 561 | 784 | 424 |
| 570 | -0.28 | 0.15 | -1.81 | 0.07 | 1 | Dhx37 | DEAH (Asp-Glu-Ala-His) box polypeptide | 571 | 641 | 604 | 586 | 581 | 623 | 645 | 583 | 478 | 437 | 516 |
| 540 | -0.09 | 0.20 | -0.44 | 0.66 | 1 | Traf3ip2 | TRAF3 interacting protein 2 | 505 | 611 | 519 | 459 | 563 | 546 | 645 | 426 | 519 | 675 | 475 |
| 1010 | -0.09 | 0.25 | -0.35 | 0.72 | 1 | Cryba2 | crystallin, beta A2 | 1490 | 2512 | 2339 | 431 | 662 | 713 | 644 | 703 | 405 | 600 | 610 |
| 1006 | -0.20 | 0.16 | -1.27 | 0.20 | 1 | Slc9a3r1 | solute carrier family 9 (sodium/hydrogen e | 1829 | 1811 | 1657 | 900 | 895 | 658 | 644 | 649 | 804 | 564 | 658 |
| 819 | 0.03 | 0.19 | 0.18 | 0.86 | 1 | Ptcd3 | pentatricopeptide repeat domain 3 | 904 | 941 | 713 | 611 | 761 | 1145 | 644 | 496 | 777 | 746 | 1275 |
| 751 | -0.01 | 0.18 | -0.03 | 0.98 | 1 | Entpd5 | ectonucleoside triphosphate diphosphohy | 1230 | 1220 | 931 | 524 | 680 | 598 | 644 | 628 | 637 | 504 | 669 |
| 635 | 0.17 | 0.18 | 0.93 | 0.35 | 1 | Rnf126 | ring finger protein 126 | 669 | 534 | 458 | 484 | 675 | 711 | 644 | 714 | 649 | 665 | 782 |
| 618 | -0.24 | 0.17 | -1.47 | 0.14 | 1 | Cyb5d1 | cytochrome b5 domain containing 1 | 642 | 663 | 759 | 674 | 606 | 631 | 644 | 525 | 609 | 422 | 622 |
| 711 | 0.15 | 0.18 | 0.83 | 0.40 | 1 | Crmp1 | collapsin response mediator protein 1 | 619 | 838 | 698 | 702 | 643 | 686 | 643 | 816 | 563 | 906 | 704 |
| 613 | -0.16 | 0.13 | -1.25 | 0.21 | 1 | Tufm | Tu translation elongation factor, mitochon | 811 | 699 | 666 | 586 | 598 | 588 | 643 | 524 | 553 | 571 | 508 |
| 776 | -0.24 | 0.18 | -1.29 | 0.20 | 1 | Igsf9b | immunoglobulin superfamily, member 9B | 326 | 652 | 475 | 808 | 638 | 1710 | 642 | 580 | 697 | 464 | 1546 |
| 751 | -0.04 | 0.15 | -0.28 | 0.78 | 1 | Abhd17b | abhydrolase domain containing 17B | 921 | 992 | 731 | 877 | 590 | 743 | 642 | 770 | 643 | 544 | 804 |
| 744 | 0.08 | 0.16 | 0.51 | 0.61 | 1 | Ssbp1 | single-stranded DNA binding protein 1 | 486 | 573 | 645 | 475 | 654 | 1350 | 642 | 490 | 700 | 641 | 1531 |
| 699 | 0.04 | 0.20 | 0.19 | 0.85 | 1 | 1110038F14Rik | RIKEN cDNA 1110038F14 gene | 660 | 536 | 795 | 636 | 786 | 757 | 642 | 906 | 629 | 647 | 701 |
| 641 | 0.13 | 0.21 | 0.64 | 0.52 | 1 | Ints12 | integrator complex subunit 12 | 503 | 505 | 867 | 561 | 595 | 664 | 642 | 534 | 665 | 811 | 708 |
| 519 | -0.07 | 0.22 | -0.32 | 0.75 | 1 | Lyrm4 | LYR motif containing 4 | 554 | 402 | 685 | 530 | 504 | 404 | 642 | 541 | 627 | 407 | 410 |
| 902 | 0.21 | 0.16 | 1.36 | 0.17 | 1 | St6galnac5 | ST6 (alpha-N-acetyl-neuraminyl-2,3-beta | 1264 | 1792 | 1458 | 810 | 495 | 533 | 641 | 1101 | 636 | 599 | 591 |
| 745 | 0.16 | 0.17 | 0.95 | 0.34 | 1 | Tubg1 | tubulin, gamma 1 | 966 | 669 | 868 | 614 | 806 | 636 | 641 | 816 | 777 | 714 | 686 |
| 641 | -0.14 | 0.16 | -0.87 | 0.38 | 1 | Mtrr | 5-methyltetrahydrofolate-homocysteine m | 638 | 661 | 692 | 563 | 772 | 667 | 641 | 465 | 801 | 567 | 584 |
| 463 | -0.14 | 0.20 | -0.70 | 0.48 | 1 | Gm13394 | | 929 | 690 | 77 | 630 | 84 | 395 | 641 | 487 | 61 | 638 | 461 |
| 706 | -0.28 | 0.22 | -1.25 | 0.21 | 1 | Hlf | hepatic leukemia factor | 1140 | 1348 | 1502 | 422 | 571 | 426 | 640 | 264 | 563 | 444 | 448 |
| 626 | -0.10 | 0.22 | -0.44 | 0.66 | 1 | AW554918 | expressed sequence AW554918 | 665 | 681 | 897 | 526 | 528 | 670 | 640 | 346 | 474 | 707 | 750 |
| 557 | -0.17 | 0.17 | -1.00 | 0.32 | 1 | Zfp862-ps | | 548 | 568 | 565 | 549 | 500 | 662 | 640 | 400 | 566 | 519 | 605 |
| 1086 | 0.06 | 0.18 | 0.33 | 0.74 | 1 | Sfswap | splicing factor SWAP | 935 | 701 | 816 | 962 | 797 | 2323 | 639 | 1050 | 712 | 848 | 2161 |
| 1025 | -0.02 | 0.23 | -0.09 | 0.93 | 1 | Ifi30 | interferon gamma inducible protein 30 | 2373 | 1456 | 1260 | 949 | 834 | 652 | 639 | 1088 | 651 | 826 | 554 |
| 808 | 0.06 | 0.20 | 0.31 | 0.75 | 1 | Gab2 | growth factor receptor bound protein 2-as | 1691 | 1127 | 1335 | 640 | 407 | 615 | 639 | 418 | 590 | 791 | 640 |
| 685 | 0.13 | 0.16 | 0.83 | 0.41 | 1 | C2cd4c | C2 calcium-dependent domain containing | 383 | 609 | 431 | 822 | 901 | 571 | 639 | 907 | 855 | 746 | 676 |
| 684 | 0.10 | 0.20 | 0.52 | 0.60 | 1 | Ccs | copper chaperone for superoxide dismuta | 971 | 655 | 766 | 575 | 638 | 628 | 639 | 792 | 611 | 555 | 697 |
| 674 | 0.21 | 0.21 | 1.02 | 0.31 | 1 | Smim22 | small integral membrane protein 22 | 495 | 597 | 645 | 692 | 615 | 626 | 639 | 1170 | 761 | 680 | 499 |
| 669 | 0.14 | 0.14 | 1.05 | 0.29 | 1 | Isca2 | iron-sulfur cluster assembly 2 | 546 | 483 | 549 | 713 | 665 | 725 | 639 | 730 | 734 | 699 | 872 |
| 583 | -0.19 | 0.16 | -1.22 | 0.22 | 1 | Map2k5 | mitogen-activated protein kinase kinase 5 | 668 | 709 | 734 | 628 | 528 | 500 | 639 | 466 | 501 | 550 | 489 |
| 505 | -0.36 | 0.23 | -1.54 | 0.12 | 1 | Lingo1 | leucine rich repeat and Ig domain contain | 464 | 662 | 448 | 608 | 586 | 362 | 639 | 418 | 566 | 594 | 206 |
| 811 | -0.03 | 0.15 | -0.18 | 0.86 | 1 | Rnf166 | ring finger protein 166 | 864 | 758 | 870 | 792 | 974 | 850 | 638 | 838 | 827 | 694 | 814 |
| 693 | 0.27 | 0.17 | 1.63 | 0.10 | 1 | Helq | helicase, POLQ-like | 368 | 300 | 276 | 669 | 651 | 1089 | 638 | 839 | 730 | 1027 | 1035 |
| 681 | -0.05 | 0.15 | -0.31 | 0.76 | 1 | Plekha8 | pleckstrin homology domain containing, fa | 585 | 687 | 502 | 772 | 706 | 788 | 638 | 718 | 635 | 757 | 701 |
| 599 | 0.00 | 0.17 | 0.03 | 0.98 | 1 | Mpdu1 | mannose-P-dolichol utilization defect 1 | 669 | 396 | 554 | 555 | 655 | 632 | 638 | 608 | 667 | 648 | 563 |
| 589 | 0.01 | 0.17 | 0.07 | 0.95 | 1 | Fastkd2 | FAST kinase domains 2 | 450 | 583 | 420 | 562 | 613 | 682 | 638 | 640 | 531 | 672 | 683 |
| 570 | -0.07 | 0.18 | -0.39 | 0.69 | 1 | Ffar3 | free fatty acid receptor 3 | 63 | 101 | 140 | 753 | 822 | 806 | 638 | 644 | 875 | 514 | 911 |

| | | | | | | | | | | | | | | | | | | |
|---|---|---|---|---|---|---|---|---|---|---|---|---|---|---|---|---|---|---|
| 911 | -0.15 | 0.16 | -0.93 | 0.35 | 1 | Csf1 | colony stimulating factor 1 (macrophage) | 2675 | 1493 | 1614 | 543 | 600 | 423 | 637 | 416 | 554 | 711 | 358 |
| 721 | 0.06 | 0.13 | 0.46 | 0.64 | 1 | Zfp362 | zinc finger protein 362 | 878 | 878 | 885 | 648 | 652 | 653 | 637 | 680 | 758 | 599 | 666 |
| 647 | 0.22 | 0.18 | 1.23 | 0.22 | 1 | Alkbh6 | alkB homolog 6 | 471 | 585 | 558 | 509 | 611 | 792 | 637 | 748 | 632 | 818 | 758 |
| 619 | 0.10 | 0.20 | 0.49 | 0.63 | 1 | Fadd | Fas (TNFRSF6)-associated via death dom | 635 | 686 | 530 | 635 | 655 | 396 | 637 | 962 | 561 | 795 | 315 |
| 563 | -0.27 | 0.16 | -1.74 | 0.08 | 1 | Ormdl2 | ORM1-like 2 (S. cerevisiae) | 517 | 458 | 637 | 637 | 682 | 552 | 637 | 542 | 574 | 455 | 505 |
| 544 | -0.17 | 0.19 | -0.87 | 0.39 | 1 | Ap4s1 | adaptor-related protein complex AP-4, sig | 705 | 405 | 427 | 602 | 632 | 483 | 637 | 454 | 584 | 571 | 485 |
| 520 | 0.00 | 0.26 | 0.01 | 0.99 | 1 | Dnah9 | dynein, axonemal, heavy chain 9 | 494 | 611 | 354 | 340 | 439 | 735 | 637 | 433 | 547 | 451 | 675 |
| 832 | -0.12 | 0.20 | -0.57 | 0.57 | 1 | Rab36 | RAB36, member RAS oncogene family | 625 | 975 | 867 | 790 | 672 | 1429 | 636 | 520 | 731 | 763 | 1147 |
| 676 | 0.14 | 0.15 | 0.94 | 0.35 | 1 | Tbpl1 | TATA box binding protein-like 1 | 577 | 527 | 564 | 754 | 624 | 731 | 636 | 820 | 798 | 639 | 771 |
| 634 | -0.13 | 0.14 | -0.93 | 0.35 | 1 | Atpsckmt | ATP synthase C subunit lysine N-methyltr | 414 | 452 | 432 | 679 | 591 | 1041 | 636 | 539 | 549 | 603 | 1035 |
| 747 | 0.17 | 0.18 | 0.93 | 0.35 | 1 | D3Ertd751e | DNA segment, Chr 3, ERATO Doi 751, ex | 204 | 480 | 187 | 998 | 475 | 1369 | 635 | 941 | 673 | 627 | 1634 |
| 745 | -0.11 | 0.17 | -0.65 | 0.52 | 1 | Tshz3 | teashirt zinc finger family member 3 | 656 | 866 | 786 | 817 | 717 | 863 | 635 | 630 | 857 | 479 | 893 |
| 733 | -0.03 | 0.16 | -0.22 | 0.83 | 1 | Dnajc15 | DnaJ heat shock protein family (Hsp40) m | 842 | 826 | 997 | 754 | 769 | 579 | 635 | 621 | 637 | 692 | 708 |
| 680 | 0.09 | 0.21 | 0.43 | 0.67 | 1 | Eci1 | enoyl-Coenzyme A delta isomerase 1 | 1050 | 640 | 962 | 578 | 633 | 493 | 635 | 761 | 701 | 501 | 522 |
| 572 | -0.12 | 0.20 | -0.59 | 0.56 | 1 | Slc9a9 | solute carrier family 9 (sodium/hydrogen e | 746 | 561 | 504 | 486 | 714 | 511 | 635 | 543 | 626 | 487 | 484 |
| 510 | 0.14 | 0.18 | 0.80 | 0.42 | 1 | Ppp2r3c | protein phosphatase 2, regulatory subuni | 414 | 357 | 395 | 445 | 430 | 617 | 635 | 597 | 546 | 540 | 639 |
| 1056 | -0.17 | 0.13 | -1.29 | 0.20 | 1 | Efnb1 | ephrin B1 | 2189 | 2259 | 2064 | 775 | 696 | 591 | 634 | 621 | 549 | 707 | 528 |
| 662 | -0.13 | 0.14 | -0.90 | 0.37 | 1 | Snx21 | sorting nexin family member 21 | 726 | 867 | 881 | 571 | 593 | 708 | 634 | 541 | 593 | 493 | 674 |
| 648 | -0.13 | 0.15 | -0.83 | 0.41 | 1 | Atg14 | autophagy related 14 | 771 | 528 | 960 | 494 | 678 | 738 | 634 | 450 | 553 | 608 | 720 |
| 641 | -0.06 | 0.14 | -0.43 | 0.66 | 1 | Slx4 | SLX4 structure-specific endonuclease sul | 589 | 710 | 627 | 604 | 550 | 832 | 634 | 561 | 550 | 616 | 778 |
| 709 | -0.30 | 0.19 | -1.58 | 0.11 | 1 | Lnx2 | ligand of numb-protein X 2 | 782 | 1006 | 749 | 753 | 786 | 725 | 633 | 492 | 757 | 489 | 624 |
| 545 | -0.02 | 0.20 | -0.12 | 0.90 | 1 | Htatip2 | HIV-1 Tat interactive protein 2 | 626 | 585 | 636 | 503 | 529 | 423 | 633 | 716 | 467 | 464 | 417 |
| 952 | -0.05 | 0.14 | -0.37 | 0.71 | 1 | Fnbp4 | formin binding protein 4 | 767 | 741 | 754 | 659 | 626 | 2244 | 632 | 496 | 738 | 613 | 2206 |
| 755 | 0.11 | 0.14 | 0.77 | 0.44 | 1 | Zfp317 | zinc finger protein 317 | 600 | 777 | 599 | 633 | 541 | 1253 | 632 | 823 | 595 | 595 | 1256 |
| 729 | -0.12 | 0.14 | -0.80 | 0.42 | 1 | Mcc | mutated in colorectal cancers | 548 | 576 | 703 | 691 | 650 | 1221 | 632 | 608 | 712 | 486 | 1191 |
| 648 | 0.09 | 0.19 | 0.46 | 0.64 | 1 | Lig4 | ligase IV, DNA, ATP-dependent | 377 | 600 | 552 | 741 | 645 | 653 | 632 | 1082 | 596 | 544 | 706 |
| 821 | 0.08 | 0.18 | 0.44 | 0.66 | 1 | Bcat2 | branched chain aminotransferase 2, mito | 709 | 663 | 723 | 494 | 1004 | 1287 | 631 | 579 | 843 | 849 | 1251 |
| 728 | -0.07 | 0.16 | -0.42 | 0.67 | 1 | Map11 | microtubule associated protein 11 | 564 | 519 | 528 | 629 | 761 | 1307 | 631 | 682 | 756 | 670 | 959 |
| 718 | 0.15 | 0.15 | 0.99 | 0.32 | 1 | Gtf2ird2 | GTF2I repeat domain containing 2 | 568 | 526 | 494 | 781 | 739 | 845 | 631 | 838 | 897 | 705 | 872 |
| 704 | 0.01 | 0.19 | 0.06 | 0.95 | 1 | Ccdc137 | coiled-coil domain containing 137 | 565 | 885 | 738 | 536 | 675 | 937 | 631 | 536 | 688 | 675 | 882 |
| 666 | -0.07 | 0.16 | -0.48 | 0.63 | 1 | Alg12 | asparagine-linked glycosylation 12 (alpha | 353 | 536 | 483 | 651 | 806 | 939 | 631 | 667 | 708 | 520 | 1032 |
| 610 | -0.28 | 0.22 | -1.28 | 0.20 | 1 | Bcl2l11 | BCL2-like 11 (apoptosis facilitator) | 944 | 586 | 953 | 532 | 522 | 593 | 631 | 331 | 351 | 659 | 611 |
| 805 | 0.11 | 0.18 | 0.60 | 0.55 | 1 | Adprhl2 | ADP-ribosylhydrolase like 2 | 862 | 750 | 843 | 890 | 752 | 801 | 630 | 894 | 665 | 979 | 793 |
| 784 | -0.01 | 0.14 | -0.07 | 0.94 | 1 | H2ax | H2A.X variant histone | 855 | 985 | 843 | 821 | 890 | 642 | 630 | 886 | 720 | 723 | 630 |
| 773 | 0.25 | 0.16 | 1.57 | 0.12 | 1 | Fam184a | family with sequence similarity 184, meml | 330 | 308 | 307 | 529 | 653 | 1737 | 630 | 595 | 864 | 899 | 1649 |
| 690 | 0.26 | 0.21 | 1.23 | 0.22 | 1 | Znhit2 | zinc finger, HIT domain containing 2 | 434 | 568 | 550 | 766 | 738 | 569 | 630 | 1038 | 719 | 1085 | 490 |
| 669 | 0.19 | 0.17 | 1.07 | 0.29 | 1 | Cntrob | centrobin, centrosomal BRCA2 interacting | 370 | 297 | 431 | 517 | 568 | 1284 | 630 | 677 | 633 | 795 | 1163 |
| 634 | -0.11 | 0.15 | -0.76 | 0.45 | 1 | Dis3l2 | DIS3 like 3'-5' exoribonuclease 2 | 673 | 599 | 769 | 574 | 698 | 659 | 630 | 543 | 579 | 701 | 548 |
| 617 | -0.02 | 0.18 | -0.09 | 0.93 | 1 | Sarm1 | sterile alpha and HEAT/Armadillo motif cc | 473 | 348 | 480 | 779 | 695 | 643 | 630 | 707 | 689 | 772 | 567 |
| 608 | -0.23 | 0.18 | -1.25 | 0.21 | 1 | Ccdc66 | coiled-coil domain containing 66 | 467 | 346 | 451 | 582 | 636 | 1051 | 630 | 540 | 581 | 395 | 1004 |
| 571 | -0.05 | 0.16 | -0.29 | 0.77 | 1 | Aph1b | aph1 homolog B, gamma secretase subu | 655 | 569 | 641 | 594 | 541 | 479 | 630 | 465 | 630 | 546 | 530 |
| 489 | 0.20 | 0.24 | 0.83 | 0.41 | 1 | Tmprss2 | transmembrane protease, serine 2 | 127 | 156 | 191 | 461 | 480 | 719 | 630 | 799 | 516 | 608 | 694 |
| 1242 | -0.23 | 0.21 | -1.10 | 0.27 | 1 | Clic5 | chloride intracellular channel 5 | 2973 | 3220 | 2871 | 810 | 625 | 430 | 629 | 497 | 668 | 367 | 570 |
| 724 | -0.03 | 0.21 | -0.12 | 0.90 | 1 | Vezt | vezatin, adherens junctions transmembra | 718 | 720 | 549 | 774 | 639 | 959 | 629 | 462 | 704 | 840 | 968 |
| 625 | -0.06 | 0.16 | -0.35 | 0.72 | 1 | Ndufaf3 | NADH:ubiquinone oxidoreductase compl | 522 | 557 | 765 | 579 | 743 | 612 | 629 | 636 | 659 | 644 | 524 |
| 606 | -0.27 | 0.15 | -1.74 | 0.08 | 1 | Polr2j | polymerase (RNA) II (DNA directed) polyp | 573 | 456 | 630 | 696 | 755 | 651 | 629 | 548 | 680 | 492 | 555 |
| 576 | -0.24 | 0.17 | -1.39 | 0.16 | 1 | Tmem86a | transmembrane protein 86A | 975 | 566 | 913 | 519 | 583 | 356 | 629 | 451 | 493 | 586 | 270 |
| 794 | 0.03 | 0.16 | 0.21 | 0.84 | 1 | Rnf25 | ring finger protein 25 | 622 | 592 | 634 | 775 | 658 | 1338 | 628 | 905 | 706 | 535 | 1342 |
| 755 | -0.05 | 0.14 | -0.37 | 0.71 | 1 | Eed | embryonic ectoderm development | 860 | 823 | 908 | 743 | 660 | 865 | 628 | 631 | 713 | 573 | 895 |
| 658 | 0.06 | 0.18 | 0.32 | 0.75 | 1 | Tada1 | transcriptional adaptor 1 | 744 | 733 | 693 | 572 | 527 | 760 | 628 | 690 | 689 | 474 | 729 |
| 641 | -0.05 | 0.15 | -0.36 | 0.72 | 1 | Zfp267 | zinc finger protein 267 | 578 | 524 | 499 | 649 | 676 | 825 | 628 | 586 | 724 | 589 | 778 |
| 565 | 0.18 | 0.18 | 0.97 | 0.33 | 1 | Sft2d3 | SFT2 domain containing 3 | 359 | 326 | 458 | 568 | 680 | 487 | 628 | 846 | 801 | 590 | 474 |
| 494 | -0.24 | 0.17 | -1.46 | 0.15 | 1 | Eid2b | EP300 interacting inhibitor of differentiatio | 488 | 520 | 533 | 600 | 512 | 380 | 628 | 454 | 427 | 518 | 374 |
| 815 | 0.03 | 0.20 | 0.13 | 0.90 | 1 | Nup85 | nucleoporin 85 | 1202 | 886 | 1071 | 596 | 817 | 842 | 627 | 834 | 736 | 526 | 824 |
| 729 | 0.06 | 0.18 | 0.31 | 0.76 | 1 | Tut1 | terminal uridylyl transferase 1, U6 snRNA | 528 | 521 | 746 | 711 | 706 | 1029 | 627 | 790 | 630 | 763 | 974 |
| 713 | 0.14 | 0.19 | 0.70 | 0.48 | 1 | Tpgs1 | tubulin polyglutamylase complex subunit | 640 | 616 | 897 | 725 | 820 | 540 | 627 | 802 | 680 | 938 | 555 |
| 707 | 0.21 | 0.14 | 1.53 | 0.13 | 1 | Rdh13 | retinol dehydrogenase 13 (all-trans and 9 | 524 | 713 | 587 | 647 | 631 | 844 | 627 | 868 | 657 | 676 | 1008 |
| 663 | -0.01 | 0.14 | -0.10 | 0.92 | 1 | Nob1 | NIN1/RPN12 binding protein 1 homolog | 760 | 602 | 607 | 579 | 623 | 840 | 627 | 528 | 588 | 742 | 795 |
| 661 | -0.13 | 0.15 | -0.87 | 0.38 | 1 | Tceanc2 | transcription elongation factor A (SII) N-te | 728 | 754 | 685 | 719 | 630 | 686 | 627 | 706 | 648 | 495 | 592 |
| 494 | -0.23 | 0.19 | -1.26 | 0.21 | 1 | Slc52a2 | solute carrier protein 52, member 2 | 418 | 294 | 416 | 642 | 518 | 543 | 627 | 490 | 493 | 500 | 497 |

| | | | | | | | | | | | | | | | | | |
|---|---|---|---|---|---|---|---|---|---|---|---|---|---|---|---|---|---|
| 492 | 0.00 | 0.22 | 0.01 | 0.99 | 1 | Cycs | cytochrome c, somatic | 384 | 335 | 445 | 435 | 631 | 439 | 627 | 536 | 553 | 416 | 608 |
| 1498 | -0.08 | 0.14 | -0.54 | 0.59 | 1 | H2-T23 | histocompatibility 2, T region locus 23 | 2843 | 3875 | 2892 | 1148 | 858 | 864 | 626 | 1280 | 723 | 688 | 676 |
| 644 | -0.35 | 0.22 | -1.63 | 0.10 | 1 | Wwox | WW domain-containing oxidoreductase | 782 | 873 | 686 | 695 | 744 | 582 | 626 | 372 | 561 | 684 | 475 |
| 854 | -0.13 | 0.16 | -0.79 | 0.43 | 1 | Lpin2 | lipin 2 | 1254 | 1523 | 1229 | 692 | 515 | 990 | 625 | 707 | 618 | 434 | 802 |
| 777 | 0.25 | 0.18 | 1.38 | 0.17 | 1 | Gm17018 | deleted in primary ciliary dyskinesia | 820 | 725 | 810 | 799 | 747 | 642 | 625 | 1169 | 707 | 754 | 752 |
| 763 | 0.09 | 0.17 | 0.49 | 0.63 | 1 | Timm44 | translocase of inner mitochondrial membr | 723 | 896 | 921 | 487 | 630 | 1144 | 625 | 643 | 611 | 717 | 994 |
| 718 | 0.13 | 0.15 | 0.90 | 0.37 | 1 | Ino80c | INO80 complex subunit C | 1007 | 1071 | 864 | 709 | 648 | 402 | 625 | 696 | 675 | 647 | 553 |
| 690 | -0.08 | 0.15 | -0.53 | 0.59 | 1 | Rab34 | RAB34, member RAS oncogene family | 965 | 770 | 924 | 645 | 627 | 630 | 625 | 538 | 531 | 730 | 608 |
| 545 | -0.01 | 0.18 | -0.08 | 0.94 | 1 | Rtn4ip1 | reticulon 4 interacting protein 1 | 353 | 333 | 397 | 552 | 576 | 707 | 625 | 680 | 568 | 464 | 740 |
| 1590 | 0.22 | 0.13 | 1.76 | 0.08 | 1 | Qk | quaking | 4372 | 3567 | 3706 | 502 | 462 | 1120 | 624 | 487 | 694 | 708 | 1245 |
| 1372 | 0.08 | 0.14 | 0.56 | 0.57 | 1 | Rflnb | refilin B | 2847 | 3381 | 3456 | 893 | 677 | 440 | 624 | 1008 | 613 | 643 | 516 |
| 782 | -0.13 | 0.16 | -0.83 | 0.41 | 1 | Obsl1 | obscurin-like 1 | 968 | 919 | 986 | 915 | 714 | 746 | 624 | 688 | 668 | 729 | 645 |
| 764 | 0.17 | 0.18 | 0.95 | 0.34 | 1 | Znhit1 | zinc finger, HIT domain containing 1 | 692 | 720 | 894 | 665 | 795 | 776 | 624 | 1045 | 768 | 599 | 825 |
| 722 | 0.09 | 0.19 | 0.46 | 0.65 | 1 | Snrpb2 | U2 small nuclear ribonucleoprotein B | 1036 | 822 | 731 | 670 | 598 | 702 | 624 | 850 | 686 | 527 | 699 |
| 617 | 0.12 | 0.14 | 0.86 | 0.39 | 1 | Lmbr1 | limb region 1 | 638 | 507 | 599 | 526 | 596 | 671 | 624 | 631 | 651 | 714 | 626 |
| 2910 | -0.03 | 0.13 | -0.21 | 0.83 | 1 | Neat1 | | 414 | 532 | 400 | 732 | 614 | 14410 | 623 | 579 | 753 | 688 | 12264 |
| 660 | 0.23 | 0.16 | 1.48 | 0.14 | 1 | Kctd9 | potassium channel tetramerisation domai | 485 | 518 | 406 | 543 | 589 | 957 | 623 | 658 | 712 | 809 | 958 |
| 653 | -0.05 | 0.15 | -0.31 | 0.76 | 1 | Phka2 | phosphorylase kinase alpha 2 | 681 | 626 | 606 | 583 | 714 | 766 | 623 | 658 | 699 | 611 | 620 |
| 515 | 0.01 | 0.21 | 0.03 | 0.98 | 1 | Zfp605 | zinc finger protein 605 | 500 | 466 | 342 | 519 | 586 | 448 | 623 | 595 | 672 | 457 | 456 |
| 506 | -0.06 | 0.19 | -0.31 | 0.76 | 1 | Tec | tec protein tyrosine kinase | 598 | 366 | 332 | 587 | 572 | 392 | 623 | 473 | 463 | 705 | 461 |
| 632 | -0.14 | 0.28 | -0.52 | 0.60 | 1 | Pnkp | polynucleotide kinase 3'- phosphatase | 645 | 360 | 766 | 626 | 683 | 779 | 622 | 768 | 642 | 365 | 693 |
| 748 | -0.05 | 0.18 | -0.30 | 0.77 | 1 | Rsu1 | Ras suppressor protein 1 | 1117 | 1108 | 1194 | 588 | 668 | 554 | 621 | 424 | 663 | 783 | 512 |
| 717 | -0.04 | 0.17 | -0.23 | 0.82 | 1 | Cenpv | centromere protein V | 559 | 565 | 720 | 805 | 857 | 787 | 621 | 643 | 726 | 828 | 780 |
| 689 | 0.18 | 0.19 | 0.94 | 0.35 | 1 | Ears2 | glutamyl-tRNA synthetase 2, mitochondria | 455 | 384 | 751 | 507 | 674 | 1045 | 621 | 687 | 704 | 705 | 1046 |
| 670 | 0.07 | 0.19 | 0.39 | 0.70 | 1 | Pou6f1 | POU domain, class 6, transcription factor | 704 | 560 | 500 | 679 | 581 | 845 | 621 | 524 | 757 | 857 | 744 |
| 561 | -0.03 | 0.20 | -0.13 | 0.90 | 1 | Dnajc19 | DnaJ heat shock protein family (Hsp40) m | 494 | 402 | 662 | 635 | 562 | 511 | 621 | 616 | 582 | 523 | 565 |
| 544 | -0.27 | 0.20 | -1.30 | 0.19 | 1 | Tmed10-ps | | 563 | 476 | 385 | 669 | 721 | 457 | 621 | 454 | 665 | 649 | 330 |
| 507 | -0.06 | 0.26 | -0.23 | 0.82 | 1 | Coprs | coordinator of PRMT5, differentiation stim | 480 | 303 | 485 | 586 | 605 | 399 | 621 | 679 | 536 | 372 | 507 |
| 660 | -0.17 | 0.20 | -0.85 | 0.40 | 1 | Fam189b | family with sequence similarity 189, meml | 879 | 591 | 630 | 820 | 633 | 663 | 620 | 572 | 678 | 555 | 622 |
| 627 | 0.07 | 0.23 | 0.30 | 0.76 | 1 | Snx11 | sorting nexin 11 | 663 | 572 | 420 | 489 | 588 | 870 | 620 | 818 | 599 | 470 | 793 |
| 563 | -0.17 | 0.21 | -0.80 | 0.42 | 1 | Thnsl1 | threonine synthase-like 1 (bacterial) | 333 | 684 | 449 | 635 | 543 | 668 | 620 | 655 | 550 | 363 | 694 |
| 549 | -0.34 | 0.19 | -1.79 | 0.07 | 1 | Atxn3 | ataxin 3 | 760 | 466 | 529 | 614 | 490 | 656 | 620 | 416 | 403 | 446 | 639 |
| 761 | -0.03 | 0.17 | -0.17 | 0.86 | 1 | Wwp1 | WW domain containing E3 ubiquitin prote | 1428 | 898 | 1259 | 598 | 557 | 646 | 619 | 562 | 608 | 551 | 651 |
| 739 | 0.01 | 0.18 | 0.08 | 0.94 | 1 | Mrpl37 | mitochondrial ribosomal protein L37 | 718 | 582 | 677 | 621 | 698 | 1106 | 619 | 799 | 529 | 598 | 1186 |
| 700 | 0.20 | 0.16 | 1.23 | 0.22 | 1 | Lrrcc1 | leucine rich repeat and coiled-coil domair | 791 | 647 | 660 | 622 | 577 | 794 | 619 | 863 | 670 | 650 | 802 |
| 664 | 0.17 | 0.18 | 0.90 | 0.37 | 1 | A430005L14Ril | RIKEN cDNA A430005L14 gene | 573 | 702 | 542 | 592 | 696 | 653 | 619 | 998 | 628 | 638 | 665 |
| 595 | 0.01 | 0.22 | 0.03 | 0.97 | 1 | Dtnbp1 | dystrobrevin binding protein 1 | 769 | 507 | 532 | 734 | 572 | 435 | 619 | 790 | 662 | 504 | 420 |
| 538 | 0.08 | 0.18 | 0.42 | 0.67 | 1 | Mrps28 | mitochondrial ribosomal protein S28 | 488 | 391 | 577 | 530 | 593 | 440 | 619 | 659 | 579 | 567 | 479 |
| 461 | -0.05 | 0.22 | -0.24 | 0.81 | 1 | Slc25a40 | solute carrier family 25, member 40 | 207 | 325 | 309 | 457 | 473 | 598 | 619 | 526 | 450 | 471 | 638 |
| 660 | 0.22 | 0.24 | 0.90 | 0.37 | 1 | Tipin | timeless interacting protein | 889 | 456 | 810 | 684 | 635 | 437 | 618 | 852 | 662 | 641 | 574 |
| 658 | -0.31 | 0.20 | -1.54 | 0.12 | 1 | Eme2 | essential meiotic structure-specific endon | 420 | 607 | 469 | 749 | 660 | 1146 | 618 | 417 | 673 | 528 | 946 |
| 630 | 0.07 | 0.19 | 0.37 | 0.71 | 1 | Adal | adenosine deaminase-like | 415 | 463 | 367 | 684 | 688 | 768 | 618 | 533 | 666 | 791 | 939 |
| 619 | 0.18 | 0.15 | 1.18 | 0.24 | 1 | N6amt1 | N-6 adenine-specific DNA methyltransfer | 405 | 408 | 541 | 543 | 657 | 742 | 618 | 594 | 677 | 741 | 889 |
| 576 | 0.27 | 0.18 | 1.48 | 0.14 | 1 | Phgdh | 3-phosphoglycerate dehydrogenase | 481 | 586 | 443 | 607 | 645 | 298 | 618 | 1014 | 630 | 643 | 367 |
| 961 | 0.07 | 0.22 | 0.33 | 0.74 | 1 | Sdccag8 | serologically defined colon cancer antige | 1012 | 1010 | 720 | 982 | 1113 | 1082 | 617 | 977 | 769 | 944 | 1340 |
| 757 | 0.14 | 0.18 | 0.75 | 0.46 | 1 | Clcn6 | chloride channel, voltage-sensitive 6 | 843 | 615 | 747 | 808 | 616 | 871 | 617 | 711 | 716 | 930 | 850 |
| 730 | -0.15 | 0.15 | -0.99 | 0.32 | 1 | Krba1 | KRAB-A domain containing 1 | 562 | 465 | 620 | 726 | 652 | 1332 | 617 | 576 | 618 | 560 | 1301 |
| 705 | -0.24 | 0.12 | -1.91 | 0.06 | 1 | Septin10 | septin 10 | 992 | 1103 | 1032 | 635 | 615 | 638 | 617 | 510 | 515 | 528 | 574 |
| 674 | -0.07 | 0.20 | -0.34 | 0.73 | 1 | Usp40 | ubiquitin specific peptidase 40 | 511 | 546 | 643 | 625 | 545 | 1109 | 617 | 782 | 527 | 413 | 1097 |
| 623 | -0.25 | 0.13 | -1.91 | 0.06 | 1 | Hmg20b | high mobility group 20B | 873 | 649 | 855 | 507 | 632 | 664 | 617 | 395 | 542 | 528 | 586 |
| 604 | -0.18 | 0.25 | -0.72 | 0.47 | 1 | Urb2 | URB2 ribosome biogenesis 2 homolog (S | 787 | 599 | 537 | 776 | 442 | 669 | 617 | 510 | 583 | 383 | 738 |
| 575 | -0.03 | 0.18 | -0.14 | 0.89 | 1 | Dpy19l3 | dpy-19-like 3 (C. elegans) | 452 | 529 | 548 | 633 | 666 | 477 | 617 | 801 | 546 | 678 | 381 |
| 1250 | -0.14 | 0.21 | -0.64 | 0.52 | 1 | Unc93b1 | unc-93 homolog B1, TLR signaling regula | 3994 | 2275 | 2331 | 635 | 698 | 751 | 616 | 760 | 552 | 522 | 620 |
| 683 | -0.29 | 0.21 | -1.40 | 0.16 | 1 | Dus3l | dihydrouridine synthase 3-like (S. cerevis | 639 | 543 | 494 | 535 | 584 | 1401 | 616 | 476 | 537 | 325 | 1363 |
| 679 | 0.25 | 0.23 | 1.06 | 0.29 | 1 | Miga2 | mitoguardin 2 | 658 | 760 | 642 | 392 | 674 | 799 | 616 | 886 | 676 | 573 | 791 |
| 646 | -0.04 | 0.16 | -0.28 | 0.78 | 1 | Six5 | sine oculis-related homeobox 5 | 628 | 538 | 537 | 554 | 728 | 859 | 616 | 652 | 610 | 658 | 727 |
| 967 | -0.11 | 0.14 | -0.77 | 0.44 | 1 | Jam3 | junction adhesion molecule 3 | 1921 | 1938 | 2013 | 709 | 569 | 574 | 614 | 612 | 489 | 670 | 526 |
| 822 | 0.10 | 0.19 | 0.55 | 0.58 | 1 | Sirt7 | sirtuin 7 | 922 | 634 | 639 | 628 | 582 | 1498 | 614 | 789 | 659 | 542 | 1532 |
| 701 | -0.12 | 0.16 | -0.77 | 0.44 | 1 | Mrtfa | myocardin related transcription factor A | 1052 | 971 | 877 | 677 | 559 | 656 | 614 | 484 | 636 | 595 | 585 |
| 673 | -0.07 | 0.17 | -0.40 | 0.69 | 1 | Traf2 | TNF receptor-associated factor 2 | 995 | 759 | 713 | 599 | 713 | 598 | 614 | 615 | 587 | 589 | 617 |

| | | | | | | | | | | | | | | | | | |
|---|---|---|---|---|---|---|---|---|---|---|---|---|---|---|---|---|---|
| 619 | -0.08 | 0.18 | -0.45 | 0.65 | 1 | Pus7 | pseudouridylate synthase 7 | 755 | 509 | 534 | 493 | 615 | 844 | 614 | 409 | 610 | 772 | 653 |
| 576 | 0.00 | 0.15 | -0.01 | 0.99 | 1 | Atg101 | autophagy related 101 | 776 | 543 | 561 | 570 | 543 | 489 | 614 | 503 | 524 | 733 | 482 |
| 568 | -0.24 | 0.18 | -1.31 | 0.19 | 1 | Wdr19 | WD repeat domain 19 | 643 | 539 | 548 | 559 | 507 | 751 | 614 | 350 | 562 | 492 | 681 |
| 626 | 0.06 | 0.20 | 0.29 | 0.77 | 1 | Pspc1 | paraspeckle protein 1 | 838 | 544 | 812 | 451 | 551 | 687 | 613 | 598 | 550 | 457 | 784 |
| 611 | 0.11 | 0.13 | 0.84 | 0.40 | 1 | Zfp64 | zinc finger protein 64 | 596 | 624 | 595 | 642 | 540 | 567 | 613 | 672 | 625 | 657 | 591 |
| 605 | 0.02 | 0.17 | 0.11 | 0.91 | 1 | Ttyh1 | tweety family member 1 | 496 | 669 | 542 | 812 | 450 | 580 | 613 | 825 | 518 | 495 | 652 |
| 857 | -0.06 | 0.19 | -0.34 | 0.74 | 1 | Morc3 | microrchidia 3 | 1087 | 1059 | 709 | 730 | 576 | 1406 | 612 | 816 | 574 | 438 | 1416 |
| 726 | 0.28 | 0.16 | 1.82 | 0.07 | 1 | Dus1l | dihydrouridine synthase 1-like (S. cerevis | 645 | 404 | 549 | 410 | 683 | 1232 | 612 | 631 | 762 | 725 | 1336 |
| 711 | 0.12 | 0.21 | 0.55 | 0.58 | 1 | Papolg | poly(A) polymerase gamma | 687 | 893 | 805 | 607 | 564 | 810 | 612 | 463 | 615 | 876 | 884 |
| 656 | -0.10 | 0.20 | -0.47 | 0.64 | 1 | Arl13b | ADP-ribosylation factor-like 13B | 897 | 854 | 670 | 549 | 586 | 715 | 612 | 410 | 686 | 489 | 745 |
| 594 | -0.15 | 0.21 | -0.73 | 0.47 | 1 | Bcorl1 | BCL6 co-repressor-like 1 | 636 | 754 | 583 | 717 | 518 | 537 | 612 | 699 | 649 | 357 | 469 |
| 571 | 0.05 | 0.16 | 0.30 | 0.76 | 1 | Gla | galactosidase, alpha | 448 | 413 | 453 | 653 | 718 | 455 | 612 | 796 | 684 | 605 | 447 |
| 543 | 0.21 | 0.17 | 1.19 | 0.24 | 1 | Cyb561d1 | cytochrome b-561 domain containing 1 | 282 | 423 | 319 | 572 | 486 | 630 | 612 | 701 | 588 | 698 | 664 |
| 1144 | -0.04 | 0.17 | -0.25 | 0.80 | 1 | Fnbp1 | formin binding protein 1 | 3429 | 2097 | 2072 | 620 | 513 | 793 | 611 | 509 | 631 | 596 | 711 |
| 746 | 0.26 | 0.18 | 1.43 | 0.15 | 1 | Zkscan7 | zinc finger with KRAB and SCAN domains | 415 | 542 | 436 | 486 | 517 | 1559 | 611 | 670 | 557 | 857 | 1559 |
| 711 | -0.09 | 0.22 | -0.39 | 0.70 | 1 | Casp3 | caspase 3 | 1226 | 851 | 857 | 770 | 676 | 463 | 611 | 821 | 560 | 468 | 516 |
| 700 | -0.01 | 0.21 | -0.06 | 0.95 | 1 | Cntrl | centriolin | 848 | 539 | 454 | 697 | 395 | 1292 | 611 | 529 | 587 | 576 | 1174 |
| 596 | -0.23 | 0.17 | -1.38 | 0.17 | 1 | Atpaf2 | ATP synthase mitochondrial F1 complex a | 598 | 627 | 534 | 589 | 705 | 681 | 611 | 454 | 632 | 541 | 584 |
| 591 | 0.05 | 0.26 | 0.21 | 0.84 | 1 | Apex1 | apurinic/apyrimidinic endonuclease 1 | 975 | 531 | 932 | 366 | 654 | 402 | 611 | 619 | 407 | 551 | 454 |
| 566 | 0.22 | 0.24 | 0.92 | 0.36 | 1 | Thyn1 | thymocyte nuclear protein 1 | 462 | 565 | 575 | 449 | 340 | 743 | 610 | 697 | 358 | 679 | 751 |
| 435 | -0.14 | 0.16 | -0.89 | 0.37 | 1 | Gpr19 | G protein-coupled receptor 19 | 427 | 365 | 350 | 448 | 419 | 441 | 610 | 429 | 372 | 489 | 439 |
| 691 | 0.16 | 0.13 | 1.23 | 0.22 | 1 | Brms1l | breast cancer metastasis-suppressor 1-lik | 825 | 714 | 682 | 589 | 563 | 768 | 609 | 649 | 560 | 774 | 861 |
| 581 | -0.02 | 0.14 | -0.17 | 0.87 | 1 | Tmem237 | transmembrane protein 237 | 599 | 627 | 687 | 542 | 599 | 505 | 609 | 608 | 618 | 518 | 479 |
| 565 | 0.03 | 0.16 | 0.18 | 0.86 | 1 | Ercc4 | excision repair cross-complementing rode | 732 | 692 | 500 | 484 | 462 | 570 | 609 | 456 | 489 | 617 | 606 |
| 526 | -0.02 | 0.14 | -0.12 | 0.91 | 1 | Zfyve19 | zinc finger, FYVE domain containing 19 | 589 | 442 | 466 | 395 | 506 | 648 | 609 | 407 | 466 | 625 | 637 |
| 456 | -0.08 | 0.24 | -0.34 | 0.74 | 1 | Tmem267 | transmembrane protein 267 | 345 | 345 | 330 | 447 | 524 | 454 | 609 | 272 | 574 | 625 | 494 |
| 761 | 0.21 | 0.13 | 1.66 | 0.10 | 1 | Ippk | inositol 1,3,4,5,6-pentakisphosphate 2-kir | 650 | 850 | 775 | 648 | 705 | 865 | 608 | 769 | 846 | 675 | 977 |
| 705 | -0.31 | 0.17 | -1.78 | 0.07 | 1 | Arhgef10 | Rho guanine nucleotide exchange factor | 1719 | 1249 | 1357 | 396 | 349 | 563 | 608 | 346 | 372 | 366 | 432 |
| 644 | 0.06 | 0.15 | 0.42 | 0.67 | 1 | Ttl | tubulin tyrosine ligase | 786 | 619 | 645 | 671 | 654 | 528 | 608 | 651 | 674 | 657 | 586 |
| 533 | 0.10 | 0.16 | 0.63 | 0.53 | 1 | Slc41a3 | solute carrier family 41, member 3 | 408 | 427 | 387 | 670 | 611 | 331 | 608 | 875 | 590 | 629 | 331 |
| 605 | -0.14 | 0.13 | -1.09 | 0.27 | 1 | Mier2 | MIER family member 2 | 539 | 596 | 539 | 620 | 537 | 853 | 607 | 600 | 515 | 506 | 738 |
| 551 | 0.01 | 0.16 | 0.06 | 0.95 | 1 | Grhpr | glyoxylate reductase/hydroxypyruvate rec | 541 | 618 | 582 | 549 | 518 | 474 | 607 | 525 | 538 | 647 | 458 |
| 517 | 0.08 | 0.18 | 0.44 | 0.66 | 1 | AI413582 | expressed sequence AI413582 | 337 | 498 | 337 | 548 | 534 | 508 | 607 | 600 | 595 | 558 | 570 |
| 528 | -0.04 | 0.14 | -0.29 | 0.77 | 1 | Coa7 | cytochrome c oxidase assembly factor 7 | 399 | 410 | 427 | 628 | 606 | 482 | 606 | 575 | 615 | 533 | 525 |
| 751 | 0.15 | 0.16 | 0.89 | 0.37 | 1 | Ppp1cc | protein phosphatase 1 catalytic subunit ga | 788 | 768 | 811 | 672 | 642 | 877 | 605 | 920 | 730 | 555 | 898 |
| 683 | 0.04 | 0.16 | 0.27 | 0.79 | 1 | Atad3a | ATPase family, AAA domain containing 3A | 818 | 532 | 659 | 612 | 582 | 910 | 605 | 592 | 587 | 800 | 811 |
| 682 | 0.25 | 0.16 | 1.52 | 0.13 | 1 | Polrmt | polymerase (RNA) mitochondrial (DNA di | 667 | 660 | 506 | 573 | 632 | 766 | 605 | 743 | 599 | 908 | 842 |
| 664 | -0.24 | 0.29 | -0.85 | 0.40 | 1 | Oxnad1 | oxidoreductase NAD-binding domain con | 483 | 728 | 482 | 782 | 846 | 773 | 605 | 329 | 860 | 551 | 861 |
| 628 | -0.09 | 0.20 | -0.43 | 0.67 | 1 | Mospd3 | motile sperm domain containing 3 | 636 | 788 | 754 | 628 | 761 | 445 | 605 | 528 | 536 | 801 | 430 |
| 798 | 0.06 | 0.20 | 0.31 | 0.76 | 1 | Zdhhc18 | zinc finger, DHHC domain containing 18 | 1190 | 1095 | 1249 | 679 | 633 | 624 | 604 | 943 | 527 | 707 | 524 |
| 717 | -0.10 | 0.17 | -0.57 | 0.57 | 1 | Ing3 | inhibitor of growth family, member 3 | 914 | 1028 | 933 | 643 | 558 | 764 | 604 | 577 | 648 | 404 | 812 |
| 636 | -0.15 | 0.17 | -0.84 | 0.40 | 1 | Dguok | deoxyguanosine kinase | 743 | 514 | 545 | 586 | 647 | 888 | 604 | 494 | 642 | 510 | 828 |
| 601 | -0.02 | 0.14 | -0.15 | 0.88 | 1 | Thap7 | THAP domain containing 7 | 500 | 559 | 463 | 733 | 688 | 541 | 604 | 685 | 604 | 678 | 556 |
| 806 | 0.05 | 0.15 | 0.31 | 0.76 | 1 | Cdkn1a | cyclin-dependent kinase inhibitor 1A (P21 | 1120 | 1108 | 894 | 698 | 667 | 856 | 603 | 802 | 704 | 581 | 828 |
| 774 | 0.16 | 0.14 | 1.14 | 0.26 | 1 | Rexo4 | REX4, 3'-5' exonuclease | 752 | 853 | 805 | 565 | 543 | 1177 | 603 | 753 | 637 | 554 | 1270 |
| 731 | -0.17 | 0.18 | -0.92 | 0.36 | 1 | Irf3 | interferon regulatory factor 3 | 809 | 708 | 875 | 600 | 545 | 1199 | 603 | 636 | 460 | 406 | 1199 |
| 687 | 0.08 | 0.24 | 0.33 | 0.74 | 1 | Cyb5r1 | cytochrome b5 reductase 1 | 696 | 503 | 698 | 719 | 750 | 659 | 603 | 1052 | 745 | 477 | 660 |
| 648 | 0.19 | 0.25 | 0.74 | 0.46 | 1 | Vsnl1 | visinin-like 1 | 119 | 317 | 325 | 920 | 956 | 560 | 603 | 972 | 751 | 884 | 720 |
| 589 | 0.18 | 0.16 | 1.09 | 0.28 | 1 | Chchd1 | coiled-coil-helix-coiled-coil-helix domain | 606 | 555 | 697 | 567 | 538 | 462 | 603 | 645 | 551 | 694 | 556 |
| 569 | -0.02 | 0.18 | -0.08 | 0.93 | 1 | Lnx1 | ligand of numb-protein X 1 | 269 | 399 | 422 | 621 | 652 | 719 | 603 | 533 | 644 | 653 | 739 |
| 718 | -0.17 | 0.14 | -1.18 | 0.24 | 1 | Irgm1 | immunity-related GTPase family M memb | 1300 | 1133 | 1200 | 540 | 679 | 435 | 602 | 417 | 581 | 599 | 412 |
| 653 | 0.09 | 0.14 | 0.65 | 0.52 | 1 | Pycrl | pyrroline-5-carboxylate reductase-like | 588 | 654 | 836 | 538 | 674 | 648 | 602 | 624 | 754 | 532 | 730 |
| 645 | -0.14 | 0.15 | -0.93 | 0.35 | 1 | Cbx3 | chromobox 3 | 784 | 698 | 835 | 586 | 692 | 620 | 602 | 535 | 601 | 455 | 686 |
| 593 | 0.26 | 0.14 | 1.88 | 0.06 | 1 | Eid2 | EP300 interacting inhibitor of differentiatio | 394 | 459 | 417 | 637 | 585 | 557 | 602 | 848 | 717 | 724 | 584 |
| 515 | -0.09 | 0.18 | -0.50 | 0.62 | 1 | Rcl1 | RNA terminal phosphate cyclase-like 1 | 574 | 437 | 594 | 366 | 643 | 472 | 602 | 381 | 550 | 683 | 365 |
| 694 | -0.11 | 0.16 | -0.69 | 0.49 | 1 | Dgkq | diacylglycerol kinase, theta | 583 | 607 | 618 | 693 | 604 | 1165 | 601 | 564 | 673 | 656 | 874 |
| 691 | -0.06 | 0.14 | -0.46 | 0.65 | 1 | H2bc21 | H2B clustered histone 21 | 622 | 872 | 857 | 799 | 644 | 631 | 601 | 737 | 716 | 489 | 632 |
| 684 | 0.12 | 0.13 | 0.89 | 0.37 | 1 | Rnf113a2 | ring finger protein 113A2 | 823 | 799 | 900 | 578 | 562 | 658 | 601 | 649 | 591 | 641 | 721 |
| 667 | 0.21 | 0.15 | 1.44 | 0.15 | 1 | Nudt21 | nudix (nucleoside diphosphate linked mo | 691 | 661 | 744 | 667 | 552 | 600 | 601 | 830 | 674 | 681 | 632 |
| 712 | -0.14 | 0.13 | -1.13 | 0.26 | 1 | Gtf2e1 | general transcription factor II E, polypeptic | 595 | 906 | 640 | 908 | 734 | 743 | 600 | 832 | 695 | 533 | 647 |

| | | | | | | | | | | | | | | | | | | |
|---|---|---|---|---|---|---|---|---|---|---|---|---|---|---|---|---|---|---|
| 674 | 0.31 | 0.16 | 1.91 | 0.06 | 1 | Lsm8 | LSM8 homolog, U6 small nuclear RNA as | 798 | 728 | 874 | 545 | 625 | 469 | 600 | 827 | 622 | 690 | 635 |
| 635 | -0.11 | 0.16 | -0.68 | 0.50 | 1 | Snapc4 | small nuclear RNA activating complex, pc | 380 | 492 | 352 | 561 | 567 | 1234 | 600 | 644 | 495 | 451 | 1211 |
| 586 | 0.00 | 0.20 | -0.02 | 0.99 | 1 | Mmachc | methylmalonic aciduria cblC type, with hc | 473 | 561 | 605 | 656 | 602 | 538 | 600 | 462 | 677 | 746 | 525 |
| 489 | 0.08 | 0.19 | 0.42 | 0.68 | 1 | Dnajc30 | DnaJ heat shock protein family (Hsp40) m | 332 | 424 | 369 | 491 | 603 | 370 | 600 | 483 | 604 | 701 | 397 |
| 913 | -0.02 | 0.13 | -0.16 | 0.87 | 1 | Syt9 | synaptotagmin IX | 1666 | 1851 | 1445 | 731 | 678 | 546 | 599 | 704 | 646 | 662 | 510 |
| 865 | 0.06 | 0.18 | 0.36 | 0.72 | 1 | Socs2 | suppressor of cytokine signaling 2 | 534 | 753 | 607 | 1048 | 850 | 1173 | 599 | 1415 | 828 | 509 | 1204 |
| 700 | 0.08 | 0.16 | 0.48 | 0.63 | 1 | Nek4 | NIMA (never in mitosis gene a)-related ex | 514 | 657 | 551 | 881 | 647 | 795 | 599 | 694 | 745 | 792 | 821 |
| 672 | -0.24 | 0.17 | -1.38 | 0.17 | 1 | Slc29a3 | solute carrier family 29 (nucleoside transp | 1449 | 826 | 1034 | 499 | 696 | 401 | 599 | 349 | 556 | 631 | 356 |
| 642 | 0.03 | 0.20 | 0.15 | 0.88 | 1 | Thap4 | THAP domain containing 4 | 614 | 686 | 744 | 698 | 608 | 571 | 599 | 463 | 641 | 777 | 661 |
| 546 | -0.11 | 0.24 | -0.49 | 0.63 | 1 | Kcnb2 | potassium voltage gated channel, Shab-r | 199 | 376 | 239 | 934 | 546 | 650 | 599 | 520 | 716 | 537 | 684 |
| 526 | -0.02 | 0.19 | -0.09 | 0.93 | 1 | Tmem101 | transmembrane protein 101 | 422 | 423 | 552 | 574 | 591 | 440 | 599 | 692 | 481 | 564 | 444 |
| 759 | -0.26 | 0.18 | -1.45 | 0.15 | 1 | Nin | ninein | 1506 | 1056 | 1167 | 607 | 680 | 600 | 598 | 377 | 546 | 690 | 516 |
| 687 | 0.19 | 0.18 | 1.08 | 0.28 | 1 | Cln8 | CLN8 transmembrane ER and ERGIC prc | 928 | 734 | 832 | 738 | 520 | 498 | 598 | 729 | 546 | 863 | 573 |
| 672 | 0.20 | 0.22 | 0.94 | 0.35 | 1 | Pde12 | phosphodiesterase 12 | 750 | 802 | 550 | 663 | 565 | 626 | 598 | 806 | 842 | 468 | 725 |
| 597 | -0.05 | 0.16 | -0.33 | 0.74 | 1 | Supv3l1 | suppressor of var1, 3-like 1 (S. cerevisiae | 593 | 400 | 492 | 595 | 714 | 678 | 598 | 565 | 597 | 674 | 661 |
| 422 | -0.24 | 0.20 | -1.16 | 0.25 | 1 | Bcs1l | BCS1-like (yeast) | 473 | 347 | 449 | 396 | 404 | 433 | 598 | 342 | 450 | 350 | 398 |
| 702 | -0.02 | 0.14 | -0.16 | 0.87 | 1 | Nop2 | NOP2 nucleolar protein | 939 | 850 | 869 | 572 | 608 | 778 | 597 | 614 | 636 | 523 | 735 |
| 700 | -0.06 | 0.18 | -0.31 | 0.75 | 1 | Gm26782 | | 262 | 731 | 582 | 875 | 825 | 765 | 597 | 1015 | 814 | 432 | 805 |
| 688 | -0.11 | 0.13 | -0.82 | 0.41 | 1 | Zbtb21 | zinc finger and BTB domain containing 2 | 591 | 761 | 667 | 797 | 610 | 882 | 597 | 663 | 662 | 550 | 787 |
| 678 | 0.04 | 0.14 | 0.26 | 0.80 | 1 | Zfp513 | zinc finger protein 513 | 640 | 654 | 669 | 710 | 573 | 838 | 597 | 667 | 681 | 577 | 857 |
| 629 | -0.10 | 0.15 | -0.66 | 0.51 | 1 | Asf1a | anti-silencing function 1A histone chaperc | 791 | 655 | 697 | 721 | 614 | 539 | 597 | 584 | 602 | 605 | 514 |
| 765 | -0.02 | 0.17 | -0.11 | 0.91 | 1 | Sfxn3 | sideroflexin 3 | 1370 | 981 | 926 | 635 | 663 | 686 | 596 | 539 | 701 | 621 | 698 |
| 724 | -0.02 | 0.20 | -0.08 | 0.93 | 1 | Ap4m1 | adaptor-related protein complex AP-4, mu | 824 | 956 | 813 | 654 | 517 | 923 | 596 | 476 | 484 | 859 | 857 |
| 670 | 0.01 | 0.15 | 0.03 | 0.97 | 1 | Map2k3 | mitogen-activated protein kinase kinase 3 | 1041 | 771 | 840 | 630 | 563 | 569 | 596 | 604 | 615 | 573 | 573 |
| 599 | 0.18 | 0.18 | 0.97 | 0.33 | 1 | Mrpl12 | mitochondrial ribosomal protein L12 | 558 | 543 | 672 | 563 | 638 | 457 | 596 | 814 | 562 | 676 | 504 |
| 534 | 0.00 | 0.15 | 0.00 | 1.00 | 1 | Bzw2 | basic leucine zipper and W2 domains 2 | 812 | 544 | 577 | 388 | 461 | 526 | 596 | 374 | 464 | 544 | 590 |
| 678 | -0.08 | 0.20 | -0.40 | 0.69 | 1 | Asb6 | ankyrin repeat and SOCS box-containing | 415 | 695 | 555 | 744 | 762 | 896 | 595 | 608 | 659 | 770 | 763 |
| 576 | -0.06 | 0.15 | -0.42 | 0.67 | 1 | Nhsl2 | NHS-like 2 | 1014 | 760 | 845 | 493 | 442 | 375 | 595 | 463 | 485 | 489 | 371 |
| 866 | -0.19 | 0.23 | -0.83 | 0.41 | 1 | S100a10 | S100 calcium binding protein A10 (calpac | 2229 | 1258 | 1411 | 586 | 581 | 674 | 594 | 514 | 491 | 379 | 806 |
| 768 | -0.09 | 0.12 | -0.72 | 0.47 | 1 | Plscr3 | phospholipid scramblase 3 | 1243 | 1272 | 1117 | 694 | 598 | 589 | 594 | 655 | 526 | 631 | 523 |
| 675 | -0.30 | 0.21 | -1.47 | 0.14 | 1 | Ralgps1 | Ral GEF with PH domain and SH3 bindin | 453 | 479 | 471 | 762 | 611 | 1347 | 594 | 408 | 740 | 475 | 1086 |
| 579 | -0.05 | 0.19 | -0.25 | 0.81 | 1 | Zfp758 | zinc finger protein 758 | 434 | 707 | 572 | 628 | 489 | 646 | 594 | 554 | 595 | 465 | 682 |
| 536 | -0.10 | 0.16 | -0.63 | 0.53 | 1 | Pcbd2 | pterin 4 alpha carbinolamine dehydratase | 441 | 548 | 501 | 666 | 548 | 473 | 594 | 524 | 636 | 542 | 423 |
| 667 | 0.09 | 0.19 | 0.46 | 0.65 | 1 | Usp42 | ubiquitin specific peptidase 42 | 674 | 687 | 567 | 547 | 604 | 870 | 593 | 440 | 777 | 672 | 907 |
| 640 | 0.16 | 0.14 | 1.15 | 0.25 | 1 | Mcrs1 | microspherule protein 1 | 664 | 624 | 614 | 543 | 645 | 642 | 593 | 722 | 664 | 684 | 645 |
| 519 | -0.04 | 0.16 | -0.27 | 0.78 | 1 | Zfp526 | zinc finger protein 526 | 630 | 501 | 542 | 457 | 483 | 515 | 593 | 499 | 539 | 537 | 411 |
| 867 | -0.08 | 0.15 | -0.53 | 0.60 | 1 | Zdhhc1 | zinc finger, DHHC domain containing 1 | 761 | 795 | 750 | 673 | 700 | 1777 | 592 | 546 | 639 | 726 | 1577 |
| 863 | 0.11 | 0.16 | 0.70 | 0.48 | 1 | Jmjd6 | jumonji domain containing 6 | 852 | 873 | 938 | 740 | 597 | 1358 | 592 | 921 | 714 | 527 | 1377 |
| 719 | 0.09 | 0.12 | 0.73 | 0.46 | 1 | Pdcd7 | programmed cell death 7 | 733 | 844 | 798 | 754 | 621 | 713 | 592 | 780 | 719 | 626 | 724 |
| 681 | 0.25 | 0.22 | 1.14 | 0.25 | 1 | Eipr1 | EARP complex and GARP complex intera | 378 | 685 | 487 | 620 | 847 | 632 | 592 | 1080 | 723 | 826 | 621 |
| 612 | -0.03 | 0.23 | -0.15 | 0.88 | 1 | Tmem208 | transmembrane protein 208 | 502 | 421 | 704 | 760 | 582 | 636 | 592 | 736 | 547 | 488 | 766 |
| 687 | 0.21 | 0.19 | 1.14 | 0.25 | 1 | Cep83 | centrosomal protein 83 | 642 | 357 | 506 | 604 | 732 | 861 | 591 | 611 | 963 | 721 | 969 |
| 680 | -0.06 | 0.19 | -0.31 | 0.76 | 1 | Plxna3 | plexin A3 | 625 | 883 | 576 | 638 | 610 | 937 | 591 | 638 | 589 | 723 | 671 |
| 567 | -0.21 | 0.20 | -1.04 | 0.30 | 1 | Mrps31 | mitochondrial ribosomal protein S31 | 627 | 604 | 757 | 661 | 446 | 573 | 591 | 429 | 443 | 475 | 631 |
| 561 | -0.35 | 0.25 | -1.42 | 0.15 | 1 | 2700049A03Ril | RIKEN cDNA 2700049A03 gene | 511 | 745 | 597 | 650 | 531 | 605 | 591 | 377 | 581 | 298 | 685 |
| 544 | -0.06 | 0.20 | -0.33 | 0.74 | 1 | Gm10524 | | 459 | 411 | 380 | 700 | 645 | 466 | 591 | 824 | 602 | 462 | 439 |
| 743 | 0.21 | 0.16 | 1.29 | 0.20 | 1 | Mterf3 | mitochondrial transcription termination fac | 663 | 517 | 657 | 537 | 633 | 1159 | 590 | 554 | 608 | 921 | 1329 |
| 738 | 0.12 | 0.22 | 0.55 | 0.58 | 1 | Cpeb2 | cytoplasmic polyadenylation element binc | 696 | 1114 | 523 | 852 | 512 | 823 | 590 | 964 | 844 | 488 | 709 |
| 669 | -0.04 | 0.16 | -0.26 | 0.80 | 1 | Mthfsd | methenyltetrahydrofolate synthetase dom | 508 | 488 | 410 | 684 | 498 | 1223 | 590 | 624 | 573 | 459 | 1306 |
| 635 | 0.17 | 0.14 | 1.24 | 0.21 | 1 | Mrpl15 | mitochondrial ribosomal protein L15 | 827 | 650 | 770 | 469 | 528 | 637 | 590 | 477 | 680 | 678 | 674 |
| 448 | -0.32 | 0.25 | -1.29 | 0.20 | 1 | E2f5 | E2F transcription factor 5 | 420 | 346 | 386 | 572 | 391 | 536 | 590 | 380 | 468 | 292 | 547 |
| 1091 | -0.14 | 0.11 | -1.25 | 0.21 | 1 | Gadd45b | growth arrest and DNA-damage-inducible | 2817 | 2088 | 2357 | 391 | 449 | 1077 | 589 | 392 | 422 | 513 | 908 |
| 684 | -0.03 | 0.16 | -0.20 | 0.84 | 1 | Zfp35 | zinc finger protein 35 | 794 | 739 | 824 | 747 | 684 | 588 | 589 | 773 | 617 | 540 | 626 |
| 631 | 0.03 | 0.17 | 0.18 | 0.86 | 1 | Samd1 | sterile alpha motif domain containing 1 | 660 | 620 | 626 | 617 | 610 | 666 | 589 | 799 | 527 | 614 | 610 |
| 624 | 0.04 | 0.18 | 0.22 | 0.82 | 1 | Prkab2 | protein kinase, AMP-activated, beta 2 nor | 423 | 464 | 299 | 548 | 371 | 1264 | 589 | 613 | 453 | 421 | 1416 |
| 568 | -0.05 | 0.19 | -0.28 | 0.78 | 1 | Fam214b | family with sequence similarity 214, meml | 449 | 429 | 487 | 463 | 758 | 693 | 589 | 666 | 547 | 542 | 629 |
| 553 | 0.00 | 0.24 | -0.01 | 0.99 | 1 | Syt2 | synaptotagmin II | 201 | 250 | 228 | 817 | 542 | 686 | 589 | 1042 | 583 | 343 | 798 |
| 491 | 0.09 | 0.24 | 0.40 | 0.69 | 1 | Dlgap3 | DLG associated protein 3 | 88 | 192 | 230 | 543 | 650 | 565 | 589 | 685 | 743 | 533 | 583 |
| 801 | 0.10 | 0.17 | 0.57 | 0.57 | 1 | Foxn2 | forkhead box N2 | 1035 | 1228 | 938 | 793 | 584 | 759 | 588 | 613 | 785 | 688 | 802 |
| 693 | 0.08 | 0.15 | 0.52 | 0.60 | 1 | Rmnd5b | required for meiotic nuclear division 5 hor | 802 | 717 | 669 | 699 | 696 | 653 | 588 | 700 | 687 | 802 | 614 |

| | | | | | | | | | | | | | | | | | | |
|---|---|---|---|---|---|---|---|---|---|---|---|---|---|---|---|---|---|---|
| 644 | 0.01 | 0.19 | 0.07 | 0.95 | 1 | Tank | TRAF family member-associated Nf-kapp | 851 | 642 | 518 | 538 | 645 | 752 | 588 | 523 | 646 | 593 | 792 |
| 581 | -0.05 | 0.18 | -0.28 | 0.78 | 1 | Ttc27 | tetratricopeptide repeat domain 27 | 824 | 517 | 637 | 578 | 627 | 455 | 588 | 598 | 587 | 544 | 438 |
| 429 | -0.11 | 0.30 | -0.36 | 0.72 | 1 | 5330434G04Rik | | 93 | 215 | 204 | 378 | 567 | 617 | 588 | 585 | 427 | 388 | 653 |
| 822 | 0.06 | 0.19 | 0.30 | 0.77 | 1 | Rlf | rearranged L-myc fusion sequence | 771 | 799 | 510 | 691 | 715 | 1426 | 587 | 612 | 769 | 703 | 1454 |
| 813 | -0.14 | 0.15 | -0.95 | 0.34 | 1 | Nectin2 | nectin cell adhesion molecule 2 | 1471 | 1515 | 1257 | 791 | 584 | 480 | 587 | 805 | 485 | 583 | 384 |
| 535 | -0.24 | 0.23 | -1.04 | 0.30 | 1 | Zfp748 | zinc finger protein 748 | 506 | 524 | 541 | 705 | 563 | 473 | 587 | 377 | 551 | 428 | 634 |
| 535 | -0.18 | 0.24 | -0.76 | 0.45 | 1 | Cdh4 | cadherin 4 | 283 | 514 | 349 | 539 | 845 | 530 | 587 | 538 | 787 | 535 | 381 |
| 787 | 0.11 | 0.13 | 0.86 | 0.39 | 1 | Ddx55 | DEAD (Asp-Glu-Ala-Asp) box polypeptide | 614 | 729 | 673 | 629 | 682 | 1313 | 586 | 816 | 675 | 626 | 1312 |
| 616 | -0.12 | 0.19 | -0.63 | 0.53 | 1 | 1700123O20Ril | RIKEN cDNA 1700123O20 gene | 696 | 614 | 558 | 788 | 623 | 564 | 586 | 575 | 607 | 495 | 672 |
| 602 | -0.07 | 0.24 | -0.28 | 0.78 | 1 | Fpgs | folylpolyglutamyl synthetase | 579 | 364 | 601 | 523 | 733 | 773 | 586 | 781 | 606 | 459 | 616 |
| 541 | -0.17 | 0.16 | -1.10 | 0.27 | 1 | Zfp277 | zinc finger protein 277 | 693 | 508 | 689 | 438 | 507 | 622 | 586 | 422 | 525 | 487 | 468 |
| 481 | 0.19 | 0.18 | 1.09 | 0.28 | 1 | Afg1l | AFG1 like ATPase | 281 | 194 | 181 | 474 | 595 | 499 | 586 | 529 | 706 | 715 | 529 |
| 458 | -0.06 | 0.16 | -0.39 | 0.69 | 1 | Tmem60 | transmembrane protein 60 | 371 | 306 | 376 | 510 | 508 | 434 | 586 | 481 | 526 | 493 | 446 |
| 309 | -0.11 | 0.37 | -0.30 | 0.77 | 1 | Vwa5b1 | von Willebrand factor A domain containin | 66 | 93 | 30 | 301 | 279 | 512 | 586 | 181 | 354 | 484 | 510 |
| 1052 | -0.05 | 0.15 | -0.30 | 0.76 | 1 | Gria3 | glutamate receptor, ionotropic, AMPA3 (a | 2109 | 2086 | 1611 | 562 | 458 | 1252 | 585 | 414 | 439 | 620 | 1439 |
| 927 | 0.04 | 0.12 | 0.32 | 0.75 | 1 | Ciapin1 | cytokine induced apoptosis inhibitor 1 | 571 | 843 | 904 | 935 | 1163 | 1219 | 585 | 996 | 1100 | 634 | 1248 |
| 847 | -0.21 | 0.15 | -1.43 | 0.15 | 1 | Rin2 | Ras and Rab interactor 2 | 2402 | 1510 | 1913 | 495 | 419 | 368 | 585 | 346 | 408 | 544 | 325 |
| 643 | -0.15 | 0.14 | -1.04 | 0.30 | 1 | Anxa11 | annexin A11 | 808 | 789 | 780 | 647 | 728 | 505 | 585 | 557 | 689 | 558 | 430 |
| 613 | 0.20 | 0.21 | 0.96 | 0.34 | 1 | Fam173a | family with sequence similarity 173, meml | 372 | 288 | 475 | 480 | 579 | 1016 | 585 | 719 | 579 | 741 | 905 |
| 612 | -0.09 | 0.17 | -0.49 | 0.63 | 1 | Ptpn3 | protein tyrosine phosphatase, non-recept | 807 | 855 | 493 | 782 | 498 | 502 | 585 | 561 | 494 | 666 | 488 |
| 611 | -0.03 | 0.17 | -0.20 | 0.84 | 1 | Mrps9 | mitochondrial ribosomal protein S9 | 701 | 721 | 509 | 511 | 569 | 748 | 585 | 558 | 527 | 502 | 784 |
| 528 | 0.09 | 0.15 | 0.64 | 0.52 | 1 | Zswim1 | zinc finger SWIM-type containing 1 | 305 | 397 | 307 | 605 | 569 | 560 | 585 | 667 | 623 | 613 | 574 |
| 509 | 0.05 | 0.23 | 0.23 | 0.82 | 1 | Prr15l | proline rich 15-like | 209 | 386 | 262 | 489 | 706 | 542 | 585 | 641 | 605 | 638 | 533 |
| 694 | -0.04 | 0.19 | -0.19 | 0.85 | 1 | Ercc6 | excision repair cross-complementing rode | 323 | 516 | 338 | 715 | 595 | 1423 | 584 | 569 | 724 | 609 | 1241 |
| 670 | 0.15 | 0.18 | 0.84 | 0.40 | 1 | Psmb5 | proteasome (prosome, macropain) subun | 641 | 568 | 899 | 670 | 681 | 561 | 584 | 787 | 687 | 707 | 585 |
| 557 | -0.13 | 0.21 | -0.60 | 0.55 | 1 | Ccdc166 | coiled-coil domain containing 166 | 453 | 406 | 365 | 664 | 750 | 538 | 584 | 535 | 488 | 844 | 504 |
| 811 | -0.18 | 0.14 | -1.31 | 0.19 | 1 | Gspt2 | G1 to S phase transition 2 | 967 | 1369 | 1257 | 725 | 690 | 829 | 583 | 693 | 504 | 554 | 745 |
| 751 | -0.03 | 0.24 | -0.13 | 0.90 | 1 | Zfp131 | zinc finger protein 131 | 793 | 856 | 562 | 864 | 590 | 1003 | 583 | 445 | 667 | 769 | 1133 |
| 663 | 0.15 | 0.16 | 0.91 | 0.36 | 1 | Usp39 | ubiquitin specific peptidase 39 | 695 | 674 | 575 | 643 | 712 | 596 | 583 | 763 | 609 | 694 | 748 |
| 550 | -0.19 | 0.14 | -1.31 | 0.19 | 1 | Mal2 | mal, T cell differentiation protein 2 | 469 | 676 | 604 | 720 | 676 | 306 | 583 | 642 | 584 | 536 | 258 |
| 860 | -0.10 | 0.29 | -0.33 | 0.74 | 1 | Iqgap2 | IQ motif containing GTPase activating pro | 884 | 421 | 1058 | 452 | 1911 | 581 | 582 | 385 | 2195 | 375 | 620 |
| 674 | -0.03 | 0.18 | -0.15 | 0.88 | 1 | Ccdc12 | coiled-coil domain containing 12 | 822 | 662 | 767 | 627 | 704 | 691 | 582 | 784 | 576 | 533 | 666 |
| 640 | 0.03 | 0.18 | 0.14 | 0.89 | 1 | Cby1 | chibby family member 1, beta catenin anta | 752 | 703 | 697 | 648 | 707 | 484 | 582 | 809 | 525 | 598 | 538 |
| 561 | 0.11 | 0.21 | 0.51 | 0.61 | 1 | Dbt | dihydrolipoamide branched chain transac | 700 | 755 | 521 | 361 | 528 | 544 | 582 | 470 | 495 | 684 | 528 |
| 1499 | -0.13 | 0.18 | -0.72 | 0.47 | 1 | Agrn | agrin | 4938 | 2914 | 4854 | 446 | 425 | 510 | 581 | 441 | 478 | 576 | 326 |
| 1000 | 0.04 | 0.28 | 0.15 | 0.88 | 1 | Lrrfip1 | leucine rich repeat (in FLII) interacting prc | 1765 | 1012 | 1273 | 1393 | 553 | 920 | 581 | 1227 | 576 | 714 | 983 |
| 746 | 0.05 | 0.26 | 0.21 | 0.84 | 1 | Grpel2 | GrpE-like 2, mitochondrial | 613 | 673 | 727 | 661 | 612 | 1129 | 581 | 1017 | 796 | 312 | 1083 |
| 601 | 0.19 | 0.23 | 0.82 | 0.41 | 1 | Rabggta | Rab geranylgeranyl transferase, a subuni | 591 | 355 | 536 | 509 | 599 | 724 | 581 | 833 | 584 | 634 | 667 |
| 590 | 0.06 | 0.24 | 0.25 | 0.80 | 1 | Rbfa | ribosome binding factor A | 680 | 537 | 695 | 586 | 532 | 527 | 581 | 811 | 600 | 388 | 554 |
| 553 | 0.22 | 0.16 | 1.33 | 0.18 | 1 | Sh3rf1 | SH3 domain containing ring finger 1 | 733 | 678 | 664 | 429 | 408 | 435 | 581 | 618 | 516 | 578 | 443 |
| 441 | -0.21 | 0.25 | -0.84 | 0.40 | 1 | Mrps11 | mitochondrial ribosomal protein S11 | 448 | 489 | 364 | 341 | 456 | 524 | 581 | 257 | 495 | 473 | 421 |
| 633 | 0.18 | 0.19 | 0.93 | 0.35 | 1 | Tmem223 | transmembrane protein 223 | 649 | 479 | 656 | 733 | 589 | 519 | 580 | 878 | 674 | 648 | 555 |
| 633 | -0.10 | 0.14 | -0.72 | 0.47 | 1 | Asb3 | ankyrin repeat and SOCS box-containing | 624 | 803 | 876 | 685 | 662 | 485 | 580 | 550 | 687 | 520 | 486 |
| 626 | 0.28 | 0.24 | 1.18 | 0.24 | 1 | Adgrb3 | adhesion G protein-coupled receptor B3 | 487 | 647 | 361 | 629 | 675 | 546 | 580 | 562 | 763 | 948 | 691 |
| 618 | 0.08 | 0.16 | 0.48 | 0.63 | 1 | Best3 | bestrophin 3 | 288 | 404 | 383 | 852 | 547 | 819 | 580 | 857 | 715 | 599 | 758 |
| 485 | -0.32 | 0.24 | -1.33 | 0.19 | 1 | Acad12 | acyl-Coenzyme A dehydrogenase family, | 319 | 544 | 535 | 473 | 582 | 549 | 580 | 355 | 445 | 519 | 435 |
| 458 | -0.07 | 0.17 | -0.39 | 0.70 | 1 | Tnfrsf23 | tumor necrosis factor receptor superfamily | 146 | 183 | 165 | 515 | 699 | 531 | 580 | 545 | 630 | 520 | 522 |
| 744 | 0.06 | 0.23 | 0.25 | 0.81 | 1 | Kazn | kazrin, periplakin interacting protein | 909 | 1320 | 1125 | 688 | 564 | 503 | 579 | 576 | 535 | 965 | 416 |
| 640 | 0.10 | 0.19 | 0.51 | 0.61 | 1 | Ogfod3 | 2-oxoglutarate and iron-dependent oxyge | 890 | 707 | 836 | 606 | 599 | 408 | 579 | 693 | 521 | 845 | 360 |
| 640 | 0.02 | 0.15 | 0.13 | 0.90 | 1 | Ppm1d | protein phosphatase 1D magnesium-dep | 887 | 704 | 782 | 593 | 530 | 607 | 578 | 571 | 468 | 649 | 668 |
| 599 | -0.06 | 0.16 | -0.34 | 0.74 | 1 | Dcun1d1 | DCN1, defective in cullin neddylation 1, d | 511 | 751 | 550 | 625 | 516 | 717 | 578 | 634 | 612 | 483 | 612 |
| 482 | -0.16 | 0.19 | -0.86 | 0.39 | 1 | Trmt12 | tRNA methyltranferase 12 | 386 | 295 | 307 | 626 | 601 | 476 | 578 | 584 | 449 | 483 | 517 |
| 774 | -0.01 | 0.15 | -0.08 | 0.94 | 1 | Glis2 | GLIS family zinc finger 2 | 1590 | 1247 | 1094 | 558 | 559 | 608 | 577 | 563 | 552 | 545 | 622 |
| 724 | 0.13 | 0.17 | 0.77 | 0.44 | 1 | Ankra2 | ankyrin repeat, family A (RFXANK-like), 2 | 722 | 607 | 689 | 747 | 555 | 946 | 577 | 611 | 637 | 795 | 1075 |
| 576 | -0.39 | 0.21 | -1.89 | 0.06 | 1 | Flvcr1 | feline leukemia virus subgroup C cellular | 643 | 564 | 565 | 710 | 490 | 810 | 577 | 378 | 553 | 421 | 624 |
| 366 | -0.11 | 0.21 | -0.49 | 0.62 | 1 | Scnn1g | sodium channel, nonvoltage-gated 1 gam | 98 | 111 | 99 | 528 | 530 | 308 | 577 | 365 | 559 | 506 | 350 |
| 802 | -0.10 | 0.14 | -0.74 | 0.46 | 1 | Atad2b | ATPase family, AAA domain containing 2I | 825 | 858 | 776 | 710 | 709 | 1313 | 576 | 589 | 702 | 590 | 1176 |
| 708 | 0.17 | 0.17 | 1.01 | 0.31 | 1 | Thop1 | thimet oligopeptidase 1 | 903 | 716 | 667 | 688 | 747 | 572 | 576 | 810 | 727 | 773 | 613 |
| 598 | 0.04 | 0.19 | 0.21 | 0.84 | 1 | Itpa | inosine triphosphatase (nucleoside tripho | 506 | 512 | 515 | 406 | 789 | 730 | 576 | 516 | 770 | 527 | 734 |
| 522 | -0.36 | 0.21 | -1.71 | 0.09 | 1 | Zbtb43 | zinc finger and BTB domain containing 43 | 522 | 511 | 465 | 554 | 365 | 891 | 576 | 359 | 437 | 323 | 744 |

| | | | | | | | | | | | | | | | | | | |
|---|---|---|---|---|---|---|---|---|---|---|---|---|---|---|---|---|---|---|
| 520 | -0.02 | 0.24 | -0.07 | 0.95 | 1 | Wdr74 | WD repeat domain 74 | 423 | 411 | 631 | 357 | 595 | 636 | 576 | 601 | 453 | 495 | 544 |
| 700 | -0.09 | 0.17 | -0.53 | 0.59 | 1 | Arl6ip6 | ADP-ribosylation factor-like 6 interacting p | 789 | 891 | 932 | 683 | 564 | 762 | 575 | 615 | 584 | 397 | 905 |
| 617 | 0.05 | 0.18 | 0.25 | 0.80 | 1 | Tmem160 | transmembrane protein 160 | 666 | 565 | 583 | 658 | 696 | 496 | 575 | 868 | 641 | 605 | 435 |
| 576 | -0.20 | 0.25 | -0.82 | 0.41 | 1 | Xxylt1 | xyloside xylosyltransferase 1 | 931 | 608 | 610 | 586 | 622 | 420 | 575 | 283 | 511 | 717 | 477 |
| 520 | 0.17 | 0.20 | 0.83 | 0.41 | 1 | Gm5113 | | 301 | 424 | 332 | 598 | 561 | 462 | 575 | 514 | 660 | 761 | 529 |
| 513 | -0.09 | 0.21 | -0.45 | 0.65 | 1 | Sp2 | Sp2 transcription factor | 665 | 521 | 373 | 463 | 601 | 457 | 575 | 429 | 560 | 630 | 374 |
| 484 | -0.14 | 0.20 | -0.72 | 0.47 | 1 | 4933404O12Rik | | 309 | 324 | 419 | 630 | 567 | 450 | 575 | 425 | 627 | 616 | 379 |
| 406 | 0.13 | 0.21 | 0.64 | 0.52 | 1 | Trpc4 | transient receptor potential cation channe | 55 | 77 | 54 | 712 | 388 | 396 | 575 | 629 | 565 | 517 | 498 |
| 682 | 0.03 | 0.16 | 0.20 | 0.84 | 1 | H2-M3 | histocompatibility 2, M region locus 3 | 856 | 731 | 944 | 682 | 565 | 631 | 574 | 618 | 609 | 696 | 592 |
| 670 | -0.06 | 0.15 | -0.41 | 0.69 | 1 | Rgl3 | ral guanine nucleotide dissociation stimul | 585 | 605 | 777 | 448 | 575 | 1164 | 574 | 414 | 640 | 508 | 1086 |
| 524 | -0.11 | 0.15 | -0.75 | 0.45 | 1 | Zfp961 | zinc finger protein 961 | 523 | 488 | 486 | 570 | 491 | 578 | 574 | 555 | 492 | 480 | 522 |
| 600 | 0.06 | 0.16 | 0.38 | 0.71 | 1 | Dhps | deoxyhypusine synthase | 645 | 540 | 713 | 576 | 606 | 545 | 573 | 637 | 530 | 605 | 626 |
| 589 | -0.20 | 0.21 | -0.94 | 0.35 | 1 | Crat | carnitine acetyltransferase | 1005 | 772 | 815 | 454 | 424 | 602 | 573 | 264 | 472 | 652 | 448 |
| 563 | -0.28 | 0.18 | -1.52 | 0.13 | 1 | Mr1 | major histocompatibility complex, class I-r | 1078 | 1157 | 815 | 501 | 400 | 253 | 573 | 370 | 267 | 514 | 262 |
| 500 | 0.16 | 0.21 | 0.78 | 0.44 | 1 | Atrip | ATR interacting protein | 373 | 336 | 327 | 284 | 471 | 823 | 573 | 481 | 465 | 603 | 767 |
| 457 | 0.03 | 0.19 | 0.14 | 0.88 | 1 | Ttll1 | tubulin tyrosine ligase-like 1 | 403 | 460 | 367 | 422 | 479 | 397 | 573 | 577 | 477 | 511 | 357 |
| 866 | -0.18 | 0.16 | -1.11 | 0.27 | 1 | Usp25 | ubiquitin specific peptidase 25 | 1964 | 1600 | 1406 | 651 | 610 | 573 | 572 | 564 | 396 | 607 | 584 |
| 807 | -0.07 | 0.14 | -0.47 | 0.64 | 1 | Napepld | N-acyl phosphatidylethanolamine phosph | 622 | 1081 | 687 | 1144 | 644 | 977 | 571 | 1013 | 764 | 505 | 869 |
| 620 | -0.17 | 0.17 | -1.00 | 0.32 | 1 | Anks6 | ankyrin repeat and sterile alpha motif dom | 635 | 633 | 428 | 613 | 525 | 1027 | 571 | 502 | 540 | 524 | 819 |
| 562 | -0.12 | 0.15 | -0.79 | 0.43 | 1 | Timm22 | translocase of inner mitochondrial membr | 617 | 633 | 722 | 586 | 580 | 457 | 571 | 570 | 478 | 533 | 437 |
| 431 | -0.25 | 0.18 | -1.38 | 0.17 | 1 | Rsph3b | radial spoke 3B homolog (Chlamydomon | 341 | 402 | 295 | 550 | 473 | 427 | 571 | 395 | 417 | 438 | 435 |
| 661 | -0.13 | 0.27 | -0.50 | 0.62 | 1 | Lyve1 | lymphatic vessel endothelial hyaluronan r | 2412 | 785 | 749 | 428 | 424 | 320 | 570 | 477 | 300 | 475 | 326 |
| 609 | -0.23 | 0.17 | -1.37 | 0.17 | 1 | Elmod1 | ELMO/CED-12 domain containing 1 | 690 | 815 | 1047 | 601 | 585 | 461 | 570 | 466 | 664 | 408 | 389 |
| 863 | 0.09 | 0.15 | 0.59 | 0.55 | 1 | Mbtd1 | mbt domain containing 1 | 1168 | 1020 | 1051 | 704 | 561 | 1207 | 569 | 577 | 723 | 688 | 1227 |
| 694 | -0.17 | 0.16 | -1.05 | 0.29 | 1 | Prpf3 | pre-mRNA processing factor 3 | 595 | 802 | 620 | 561 | 496 | 1329 | 569 | 578 | 499 | 368 | 1221 |
| 668 | -0.13 | 0.16 | -0.86 | 0.39 | 1 | Gps2 | G protein pathway suppressor 2 | 754 | 681 | 747 | 625 | 689 | 828 | 569 | 656 | 650 | 488 | 664 |
| 658 | 0.01 | 0.16 | 0.09 | 0.93 | 1 | Mllt1 | myeloid/lymphoid or mixed-lineage leuke | 1033 | 708 | 762 | 585 | 591 | 606 | 569 | 536 | 618 | 612 | 613 |
| 624 | 0.01 | 0.35 | 0.03 | 0.98 | 1 | Gcgr | glucagon receptor | 15 | 10 | 55 | 691 | 582 | 1485 | 569 | 773 | 607 | 464 | 1615 |
| 538 | -0.01 | 0.19 | -0.04 | 0.97 | 1 | Lin52 | lin-52 homolog (C. elegans) | 504 | 485 | 537 | 513 | 645 | 467 | 569 | 545 | 462 | 720 | 467 |
| 497 | 0.06 | 0.25 | 0.24 | 0.81 | 1 | Trappc2 | trafficking protein particle complex 2 | 272 | 395 | 200 | 529 | 644 | 510 | 569 | 484 | 594 | 652 | 617 |
| 374 | -0.31 | 0.25 | -1.20 | 0.23 | 1 | Zfp874a | zinc finger protein 874a | 539 | 342 | 344 | 365 | 273 | 384 | 569 | 181 | 302 | 380 | 439 |
| 926 | 0.17 | 0.24 | 0.71 | 0.48 | 1 | Psmb8 | proteasome (prosome, macropain) subun | 3519 | 1384 | 2107 | 242 | 420 | 317 | 568 | 452 | 424 | 397 | 354 |
| 717 | -0.08 | 0.17 | -0.49 | 0.63 | 1 | Plcb3 | phospholipase C, beta 3 | 1115 | 909 | 981 | 730 | 600 | 604 | 568 | 627 | 518 | 732 | 508 |
| 743 | 0.22 | 0.33 | 0.66 | 0.51 | 1 | Ncapd2 | non-SMC condensin I complex, subunit D | 1562 | 728 | 715 | 553 | 614 | 657 | 567 | 1015 | 745 | 412 | 608 |
| 636 | -0.07 | 0.17 | -0.42 | 0.67 | 1 | Kctd5 | potassium channel tetramerisation domai | 814 | 898 | 687 | 545 | 676 | 565 | 567 | 531 | 601 | 576 | 532 |
| 540 | 0.16 | 0.19 | 0.83 | 0.41 | 1 | Uxs1 | UDP-glucuronate decarboxylase 1 | 607 | 370 | 368 | 515 | 477 | 621 | 567 | 612 | 559 | 564 | 677 |
| 442 | -0.03 | 0.17 | -0.20 | 0.84 | 1 | 1500009L16Rik | RIKEN cDNA 1500009L16 gene | 234 | 199 | 249 | 579 | 569 | 405 | 567 | 547 | 533 | 519 | 457 |
| 722 | -0.12 | 0.18 | -0.68 | 0.50 | 1 | Casp2 | caspase 2 | 829 | 1141 | 1089 | 461 | 741 | 768 | 566 | 509 | 714 | 403 | 725 |
| 689 | -0.02 | 0.15 | -0.13 | 0.90 | 1 | Rgs17 | regulator of G-protein signaling 17 | 724 | 1011 | 776 | 644 | 609 | 720 | 566 | 773 | 508 | 571 | 677 |
| 434 | 0.23 | 0.28 | 0.83 | 0.41 | 1 | Pfdn4 | prefoldin 4 | 462 | 437 | 297 | 362 | 335 | 359 | 566 | 809 | 370 | 359 | 418 |
| 893 | -0.07 | 0.16 | -0.42 | 0.67 | 1 | Mknk1 | MAP kinase-interacting serine/threonine k | 948 | 827 | 882 | 439 | 602 | 2090 | 565 | 339 | 642 | 613 | 1875 |
| 703 | 0.06 | 0.17 | 0.38 | 0.71 | 1 | Ddx10 | DEAD (Asp-Glu-Ala-Asp) box polypeptide | 898 | 860 | 744 | 605 | 690 | 683 | 565 | 682 | 506 | 715 | 787 |
| 693 | 0.25 | 0.17 | 1.46 | 0.15 | 1 | Rasa2 | RAS p21 protein activator 2 | 539 | 635 | 487 | 688 | 562 | 904 | 565 | 1093 | 640 | 692 | 813 |
| 544 | 0.11 | 0.19 | 0.57 | 0.57 | 1 | Minpp1 | multiple inositol polyphosphate histidine p | 701 | 548 | 439 | 523 | 599 | 380 | 565 | 599 | 645 | 564 | 415 |
| 538 | 0.12 | 0.21 | 0.59 | 0.56 | 1 | Txlng | taxilin gamma | 365 | 243 | 332 | 431 | 517 | 874 | 565 | 543 | 705 | 415 | 928 |
| 511 | -0.01 | 0.26 | -0.06 | 0.96 | 1 | Atg10 | autophagy related 10 | 292 | 440 | 260 | 459 | 705 | 594 | 565 | 416 | 638 | 620 | 628 |
| 500 | 0.02 | 0.20 | 0.08 | 0.93 | 1 | Gkap1 | G kinase anchoring protein 1 | 445 | 401 | 448 | 449 | 462 | 613 | 565 | 655 | 400 | 519 | 541 |
| 458 | -0.01 | 0.19 | -0.05 | 0.96 | 1 | 2210408I21Rik | RIKEN cDNA 2210408I21 gene | 353 | 419 | 388 | 452 | 455 | 470 | 565 | 380 | 482 | 560 | 510 |
| 704 | -0.20 | 0.17 | -1.21 | 0.23 | 1 | Lrch2 | leucine-rich repeats and calponin homolo | 672 | 1211 | 844 | 844 | 616 | 674 | 564 | 612 | 568 | 582 | 553 |
| 624 | -0.01 | 0.16 | -0.09 | 0.93 | 1 | Ccdc174 | coiled-coil domain containing 174 | 550 | 629 | 641 | 598 | 544 | 799 | 564 | 627 | 468 | 498 | 949 |
| 617 | -0.10 | 0.21 | -0.48 | 0.63 | 1 | Cep126 | centrosomal protein 126 | 489 | 626 | 441 | 638 | 652 | 839 | 564 | 445 | 526 | 741 | 823 |
| 589 | -0.21 | 0.16 | -1.30 | 0.19 | 1 | Zfp212 | Zinc finger protein 212 | 573 | 504 | 704 | 651 | 618 | 675 | 564 | 453 | 551 | 509 | 673 |
| 700 | 0.13 | 0.14 | 0.88 | 0.38 | 1 | Foxred1 | FAD-dependent oxidoreductase domain c | 398 | 455 | 509 | 501 | 686 | 1317 | 563 | 684 | 694 | 600 | 1292 |
| 641 | 0.08 | 0.22 | 0.35 | 0.73 | 1 | Pigyl | phosphatidylinositol glycan anchor biosyn | 601 | 583 | 828 | 689 | 685 | 499 | 563 | 982 | 563 | 583 | 478 |
| 593 | -0.05 | 0.17 | -0.29 | 0.77 | 1 | Vars2 | valyl-tRNA synthetase 2, mitochondrial | 691 | 646 | 695 | 488 | 462 | 773 | 563 | 612 | 418 | 464 | 711 |
| 591 | 0.09 | 0.20 | 0.43 | 0.66 | 1 | Cdkn2c | cyclin dependent kinase inhibitor 2C | 690 | 536 | 375 | 850 | 524 | 435 | 563 | 813 | 586 | 701 | 424 |
| 412 | -0.30 | 0.25 | -1.20 | 0.23 | 1 | Iqcc | IQ motif containing C | 304 | 408 | 348 | 333 | 427 | 590 | 563 | 383 | 404 | 285 | 489 |
| 790 | -0.01 | 0.18 | -0.06 | 0.95 | 1 | Prpf40b | pre-mRNA processing factor 40B | 910 | 756 | 876 | 555 | 574 | 1401 | 562 | 678 | 419 | 622 | 1339 |
| 702 | 0.23 | 0.23 | 1.00 | 0.32 | 1 | Rbpj | recombination signal binding protein for i | 1310 | 931 | 872 | 395 | 531 | 641 | 562 | 812 | 505 | 576 | 592 |
| 698 | 0.08 | 0.15 | 0.53 | 0.60 | 1 | Cep131 | centrosomal protein 131 | 698 | 854 | 658 | 666 | 567 | 874 | 562 | 753 | 576 | 728 | 738 |

| | | | | | | | | | | | | | | | | | | |
|---|---|---|---|---|---|---|---|---|---|---|---|---|---|---|---|---|---|---|
| 666 | -0.21 | 0.18 | -1.22 | 0.22 | 1 | Efr3b | EFR3 homolog B | 856 | 1215 | 1066 | 546 | 609 | 529 | 562 | 478 | 419 | 501 | 541 |
| 613 | 0.17 | 0.14 | 1.20 | 0.23 | 1 | Tmem230 | transmembrane protein 230 | 547 | 532 | 627 | 591 | 557 | 663 | 562 | 689 | 692 | 621 | 664 |
| 601 | -0.08 | 0.19 | -0.41 | 0.68 | 1 | Ccp110 | centriolar coiled coil protein 110 | 403 | 439 | 483 | 460 | 348 | 1380 | 562 | 416 | 434 | 471 | 1218 |
| 476 | -0.14 | 0.17 | -0.86 | 0.39 | 1 | Orai2 | ORAI calcium release-activated calcium r | 478 | 455 | 470 | 529 | 498 | 420 | 562 | 388 | 523 | 497 | 412 |
| 588 | 0.04 | 0.18 | 0.21 | 0.83 | 1 | Toe1 | target of EGR1, member 1 (nuclear) | 743 | 548 | 587 | 442 | 631 | 633 | 561 | 582 | 504 | 553 | 680 |
| 486 | -0.03 | 0.18 | -0.14 | 0.89 | 1 | Pdhx | pyruvate dehydrogenase complex, compo | 401 | 394 | 504 | 475 | 454 | 549 | 561 | 411 | 538 | 581 | 483 |
| 741 | 0.09 | 0.14 | 0.62 | 0.54 | 1 | Hikeshi | heat shock protein nuclear import factor | 861 | 871 | 937 | 726 | 774 | 604 | 560 | 740 | 695 | 734 | 648 |
| 666 | 0.16 | 0.17 | 0.95 | 0.34 | 1 | Ranbp1 | RAN binding protein 1 | 854 | 654 | 798 | 671 | 594 | 542 | 560 | 763 | 589 | 705 | 598 |
| 665 | -0.17 | 0.20 | -0.84 | 0.40 | 1 | Camk2g | calcium/calmodulin-dependent protein kir | 960 | 623 | 779 | 702 | 460 | 915 | 560 | 538 | 519 | 519 | 740 |
| 657 | -0.03 | 0.17 | -0.18 | 0.86 | 1 | St6galnac4 | ST6 (alpha-N-acetyl-neuraminyl-2,3-beta | 1120 | 771 | 998 | 630 | 596 | 403 | 560 | 633 | 579 | 563 | 378 |
| 605 | -0.11 | 0.18 | -0.58 | 0.56 | 1 | Arrb2 | arrestin, beta 2 | 932 | 487 | 732 | 389 | 534 | 886 | 560 | 446 | 493 | 468 | 735 |
| 550 | 0.02 | 0.18 | 0.09 | 0.93 | 1 | Pard3b | par-3 family cell polarity regulator beta | 687 | 495 | 469 | 516 | 552 | 554 | 560 | 470 | 638 | 577 | 527 |
| 510 | -0.10 | 0.19 | -0.53 | 0.59 | 1 | 2810021J22Rik | RIKEN cDNA 2810021J22 gene | 403 | 411 | 403 | 548 | 575 | 574 | 560 | 594 | 486 | 392 | 666 |
| 497 | 0.00 | 0.19 | 0.01 | 0.99 | 1 | Fam160a1 | family with sequence similarity 160, meml | 256 | 257 | 261 | 672 | 550 | 534 | 560 | 819 | 573 | 587 | 398 |
| 672 | 0.06 | 0.17 | 0.37 | 0.71 | 1 | Snrpd1 | small nuclear ribonucleoprotein D1 | 1147 | 940 | 835 | 639 | 496 | 494 | 559 | 714 | 515 | 517 | 537 |
| 643 | 0.10 | 0.15 | 0.63 | 0.53 | 1 | Lsm12 | LSM12 homolog | 525 | 575 | 570 | 859 | 577 | 625 | 559 | 777 | 688 | 622 | 692 |
| 621 | -0.13 | 0.16 | -0.80 | 0.43 | 1 | Tmem98 | transmembrane protein 98 | 1029 | 912 | 902 | 505 | 629 | 385 | 559 | 575 | 514 | 478 | 339 |
| 595 | -0.19 | 0.19 | -1.00 | 0.32 | 1 | Erg28 | ergosterol biosynthesis 28 | 699 | 684 | 644 | 569 | 679 | 587 | 559 | 389 | 518 | 634 | 585 |
| 592 | -0.06 | 0.21 | -0.27 | 0.79 | 1 | Arhgef37 | Rho guanine nucleotide exchange factor | 332 | 354 | 220 | 888 | 816 | 594 | 559 | 876 | 737 | 542 | 591 |
| 590 | 0.17 | 0.16 | 1.06 | 0.29 | 1 | Thtpa | thiamine triphosphatase | 526 | 574 | 533 | 545 | 648 | 532 | 559 | 541 | 673 | 721 | 636 |
| 572 | 0.02 | 0.17 | 0.12 | 0.90 | 1 | Rpgrip1l | Rpgrip1-like | 632 | 707 | 449 | 534 | 524 | 621 | 559 | 560 | 558 | 540 | 608 |
| 458 | -0.02 | 0.16 | -0.11 | 0.91 | 1 | Tbc1d31 | TBC1 domain family, member 31 | 390 | 303 | 342 | 415 | 427 | 606 | 559 | 351 | 473 | 577 | 593 |
| 413 | -0.50 | 0.29 | -1.72 | 0.09 | 1 | Shld2 | shieldin complex subunit 2 | 432 | 477 | 404 | 310 | 461 | 532 | 558 | 180 | 443 | 259 | 484 |
| 699 | 0.07 | 0.18 | 0.41 | 0.68 | 1 | Dcps | decapping enzyme, scavenger | 921 | 712 | 778 | 622 | 708 | 687 | 557 | 740 | 730 | 537 | 700 |
| 697 | 0.21 | 0.18 | 1.19 | 0.23 | 1 | Scml4 | Scm polycomb group protein like 4 | 616 | 754 | 747 | 698 | 601 | 717 | 557 | 752 | 628 | 887 | 707 |
| 630 | 0.02 | 0.15 | 0.11 | 0.91 | 1 | Rce1 | Ras converting CAAX endopeptidase 1 | 593 | 560 | 678 | 595 | 697 | 693 | 557 | 695 | 610 | 626 | 630 |
| 536 | -0.09 | 0.25 | -0.37 | 0.71 | 1 | Alms1 | ALMS1, centrosome and basal body assc | 639 | 558 | 415 | 574 | 370 | 714 | 557 | 409 | 575 | 354 | 730 |
| 1029 | 0.09 | 0.28 | 0.31 | 0.76 | 1 | Crip1 | cysteine-rich protein 1 (intestinal) | 3548 | 1884 | 1874 | 462 | 474 | 403 | 556 | 936 | 367 | 480 | 337 |
| 612 | 0.13 | 0.15 | 0.90 | 0.37 | 1 | Tbc1d10a | TBC1 domain family, member 10a | 597 | 475 | 602 | 487 | 641 | 731 | 556 | 542 | 597 | 735 | 770 |
| 684 | 0.06 | 0.16 | 0.38 | 0.71 | 1 | Tsta3 | tissue specific transplantation antigen P3 | 675 | 697 | 985 | 512 | 737 | 742 | 555 | 597 | 665 | 698 | 660 |
| 677 | -0.24 | 0.15 | -1.55 | 0.12 | 1 | Repin1 | replication initiator 1 | 654 | 814 | 749 | 782 | 699 | 797 | 555 | 525 | 614 | 568 | 684 |
| 629 | 0.34 | 0.19 | 1.77 | 0.08 | 1 | Mrpl19 | mitochondrial ribosomal protein L19 | 584 | 696 | 668 | 469 | 621 | 541 | 555 | 831 | 616 | 781 | 560 |
| 670 | -0.15 | 0.18 | -0.84 | 0.40 | 1 | Stat2 | signal transducer and activator of transcri | 849 | 563 | 896 | 514 | 616 | 954 | 554 | 404 | 737 | 413 | 865 |
| 641 | 0.04 | 0.13 | 0.34 | 0.73 | 1 | Zfp84 | zinc finger protein 84 | 613 | 715 | 561 | 709 | 549 | 724 | 554 | 694 | 546 | 572 | 811 |
| 622 | 0.18 | 0.20 | 0.86 | 0.39 | 1 | Rbm19 | RNA binding motif protein 19 | 827 | 623 | 701 | 576 | 523 | 528 | 554 | 874 | 525 | 632 | 478 |
| 614 | -0.07 | 0.18 | -0.40 | 0.69 | 1 | Dnlz | DNL-type zinc finger | 636 | 679 | 661 | 614 | 664 | 597 | 554 | 752 | 565 | 607 | 426 |
| 581 | 0.08 | 0.19 | 0.41 | 0.68 | 1 | Acads | acyl-Coenzyme A dehydrogenase, short c | 730 | 885 | 700 | 494 | 544 | 377 | 554 | 733 | 465 | 462 | 442 |
| 560 | -0.02 | 0.22 | -0.08 | 0.94 | 1 | Swsap1 | SWIM type zinc finger 7 associated protei | 270 | 456 | 438 | 482 | 899 | 604 | 554 | 532 | 703 | 623 | 602 |
| 544 | -0.22 | 0.19 | -1.14 | 0.25 | 1 | Dcakd | dephospho-CoA kinase domain containir | 693 | 630 | 915 | 458 | 588 | 408 | 554 | 312 | 459 | 569 | 400 |
| 521 | -0.12 | 0.22 | -0.56 | 0.58 | 1 | Stat1 | signal transducer and activator of transcri | 933 | 594 | 935 | 395 | 429 | 341 | 554 | 390 | 467 | 321 | 378 |
| 514 | -0.07 | 0.25 | -0.26 | 0.79 | 1 | F8 | coagulation factor VIII | 294 | 365 | 332 | 640 | 517 | 650 | 554 | 330 | 607 | 607 | 759 |
| 513 | -0.05 | 0.24 | -0.19 | 0.85 | 1 | 2810402E24Rik | | 491 | 655 | 624 | 452 | 517 | 402 | 554 | 753 | 496 | 323 | 374 |
| 496 | -0.09 | 0.15 | -0.61 | 0.54 | 1 | Limk1 | LIM-domain containing, protein kinase | 653 | 468 | 563 | 398 | 560 | 434 | 554 | 403 | 528 | 523 | 375 |
| 1188 | -0.18 | 0.21 | -0.89 | 0.37 | 1 | Prex1 | phosphatidylinositol-3,4,5-trisphosphate-c | 3263 | 2618 | 2971 | 542 | 718 | 397 | 553 | 527 | 816 | 339 | 323 |
| 702 | 0.08 | 0.21 | 0.40 | 0.69 | 1 | Hat1 | histone aminotransferase 1 | 1225 | 841 | 825 | 616 | 664 | 500 | 553 | 812 | 492 | 617 | 576 |
| 652 | -0.21 | 0.19 | -1.11 | 0.27 | 1 | Unk | unkempt family zinc finger | 679 | 547 | 504 | 675 | 631 | 1016 | 553 | 431 | 582 | 493 | 1066 |
| 623 | -0.10 | 0.19 | -0.52 | 0.61 | 1 | Rhbdl3 | rhomboid like 3 | 850 | 677 | 709 | 592 | 537 | 683 | 553 | 366 | 565 | 652 | 668 |
| 570 | -0.18 | 0.18 | -0.99 | 0.32 | 1 | Fam57b | family with sequence similarity 57, membe | 607 | 946 | 735 | 520 | 478 | 562 | 553 | 484 | 492 | 437 | 460 |
| 409 | 0.10 | 0.29 | 0.34 | 0.74 | 1 | Fancl | Fanconi anemia, complementation group | 551 | 450 | 538 | 257 | 352 | 292 | 553 | 418 | 458 | 273 | 356 |
| 396 | 0.00 | 0.48 | 0.00 | 1.00 | 1 | Gm30173 | | 8 | 31 | 103 | 533 | 542 | 484 | 553 | 416 | 533 | 670 | 485 |
| 701 | 0.29 | 0.17 | 1.68 | 0.09 | 1 | Smyd4 | SET and MYND domain containing 4 | 539 | 602 | 505 | 786 | 696 | 678 | 552 | 1108 | 652 | 858 | 734 |
| 513 | -0.01 | 0.26 | -0.04 | 0.97 | 1 | Zfp945 | zinc finger protein 945 | 547 | 698 | 595 | 382 | 352 | 592 | 552 | 577 | 371 | 283 | 699 |
| 740 | 0.01 | 0.17 | 0.08 | 0.94 | 1 | Parp2 | poly (ADP-ribose) polymerase family, mer | 773 | 734 | 794 | 598 | 742 | 1037 | 551 | 796 | 608 | 603 | 900 |
| 699 | 0.10 | 0.22 | 0.46 | 0.65 | 1 | Nasp | nuclear autoantigenic sperm protein (hist | 1208 | 684 | 822 | 604 | 483 | 775 | 551 | 728 | 570 | 496 | 774 |
| 598 | 0.05 | 0.22 | 0.24 | 0.81 | 1 | Bud23 | BUD23, rRNA methyltransferase and ribo | 492 | 598 | 730 | 600 | 563 | 612 | 551 | 510 | 460 | 738 | 723 |
| 575 | 0.11 | 0.16 | 0.73 | 0.46 | 1 | Bivm | basic, immunoglobulin-like variable motif | 510 | 557 | 472 | 519 | 607 | 621 | 551 | 630 | 644 | 636 | 576 |
| 696 | -0.11 | 0.20 | -0.55 | 0.58 | 1 | Dot1l | DOT1-like, histone H3 methyltransferase | 582 | 665 | 524 | 481 | 435 | 1675 | 550 | 351 | 537 | 559 | 1302 |
| 537 | -0.38 | 0.26 | -1.45 | 0.15 | 1 | Gpr155 | G protein-coupled receptor 155 | 673 | 363 | 350 | 618 | 597 | 783 | 550 | 453 | 504 | 343 | 674 |
| 527 | 0.16 | 0.21 | 0.78 | 0.44 | 1 | AI987944 | expressed sequence AI987944 | 344 | 481 | 500 | 566 | 490 | 502 | 550 | 684 | 445 | 659 | 579 |
| 482 | 0.04 | 0.16 | 0.24 | 0.81 | 1 | Pkp2 | plakophilin 2 | 350 | 247 | 342 | 362 | 573 | 659 | 550 | 353 | 552 | 687 | 622 |

| | | | | | | | | | | | | | | | | | | |
|---|---|---|---|---|---|---|---|---|---|---|---|---|---|---|---|---|---|---|
| 730 | 0.16 | 0.16 | 0.99 | 0.32 | 1 | Nedd9 | neural precursor cell expressed, developı | 1500 | 1348 | 1276 | 466 | 395 | 424 | 549 | 424 | 455 | 735 | 461 |
| 684 | 0.14 | 0.20 | 0.70 | 0.48 | 1 | Galnt7 | polypeptide N-acetylgalactosaminyltransf | 871 | 808 | 828 | 578 | 664 | 582 | 549 | 928 | 633 | 529 | 558 |
| 600 | 0.07 | 0.17 | 0.40 | 0.69 | 1 | Abhd10 | abhydrolase domain containing 10 | 612 | 530 | 639 | 676 | 544 | 583 | 549 | 578 | 588 | 697 | 605 |
| 528 | -0.07 | 0.19 | -0.36 | 0.72 | 1 | B230217C12Rik | | 522 | 561 | 557 | 586 | 515 | 478 | 549 | 395 | 559 | 590 | 496 |
| 499 | 0.15 | 0.21 | 0.73 | 0.47 | 1 | Dexi | dexamethasone-induced transcript | 348 | 453 | 460 | 403 | 605 | 449 | 549 | 569 | 521 | 663 | 467 |
| 709 | -0.03 | 0.18 | -0.15 | 0.88 | 1 | Dbi | diazepam binding inhibitor | 1221 | 878 | 1395 | 569 | 656 | 391 | 548 | 648 | 651 | 504 | 343 |
| 704 | 0.21 | 0.18 | 1.17 | 0.24 | 1 | Umad1 | UMAP1-MVP12 associated (UMA) domai | 635 | 851 | 799 | 792 | 591 | 614 | 548 | 737 | 648 | 846 | 679 |
| 618 | 0.11 | 0.15 | 0.70 | 0.49 | 1 | Nap1l3 | nucleosome assembly protein 1-like 3 | 676 | 788 | 696 | 641 | 475 | 563 | 548 | 751 | 457 | 600 | 604 |
| 577 | 0.05 | 0.16 | 0.33 | 0.74 | 1 | Tdp2 | tyrosyl-DNA phosphodiesterase 2 | 589 | 572 | 549 | 518 | 395 | 825 | 548 | 600 | 456 | 487 | 802 |
| 542 | 0.13 | 0.16 | 0.79 | 0.43 | 1 | Trub2 | TruB pseudouridine (psi) synthase family | 493 | 324 | 516 | 456 | 530 | 685 | 548 | 526 | 628 | 555 | 705 |
| 453 | -0.29 | 0.18 | -1.65 | 0.10 | 1 | Mmp24 | matrix metallopeptidase 24 | 320 | 430 | 360 | 569 | 519 | 500 | 548 | 401 | 450 | 475 | 415 |
| 809 | -0.21 | 0.18 | -1.17 | 0.24 | 1 | Znrf1 | zinc and ring finger 1 | 1263 | 1401 | 1110 | 518 | 648 | 1047 | 547 | 405 | 543 | 586 | 833 |
| 592 | -0.11 | 0.20 | -0.55 | 0.58 | 1 | Net1 | neuroepithelial cell transforming gene 1 | 772 | 613 | 562 | 651 | 668 | 505 | 547 | 524 | 667 | 471 | 532 |
| 407 | -0.07 | 0.18 | -0.37 | 0.71 | 1 | Smim26 | small integral membrane protein 26 | 426 | 430 | 416 | 439 | 369 | 288 | 547 | 472 | 408 | 432 | 257 |
| 395 | -0.35 | 0.22 | -1.54 | 0.12 | 1 | D130017N08Rik | | 315 | 260 | 314 | 421 | 424 | 517 | 547 | 219 | 396 | 523 | 408 |
| 720 | 0.04 | 0.22 | 0.19 | 0.85 | 1 | Gpr119 | G-protein coupled receptor 119 | 754 | 1594 | 1070 | 544 | 445 | 675 | 546 | 499 | 572 | 474 | 743 |
| 592 | -0.12 | 0.15 | -0.84 | 0.40 | 1 | Fbxl18 | F-box and leucine-rich repeat protein 18 | 462 | 579 | 441 | 635 | 655 | 788 | 546 | 597 | 587 | 510 | 713 |
| 527 | -0.08 | 0.29 | -0.27 | 0.79 | 1 | Pex11a | peroxisomal biogenesis factor 11 alpha | 220 | 499 | 497 | 544 | 490 | 778 | 546 | 361 | 514 | 621 | 730 |
| 485 | -0.05 | 0.19 | -0.26 | 0.79 | 1 | Cdk18 | cyclin-dependent kinase 18 | 339 | 341 | 355 | 430 | 657 | 546 | 546 | 390 | 667 | 591 | 471 |
| 619 | -0.06 | 0.24 | -0.25 | 0.80 | 1 | Tars2 | threonyl-tRNA synthetase 2, mitochondria | 660 | 573 | 650 | 432 | 554 | 995 | 545 | 685 | 322 | 572 | 818 |
| 564 | -0.17 | 0.19 | -0.89 | 0.37 | 1 | Dnaaf5 | dynein, axonemal assembly factor 5 | 724 | 563 | 565 | 599 | 609 | 542 | 545 | 407 | 669 | 532 | 453 |
| 563 | -0.32 | 0.20 | -1.59 | 0.11 | 1 | Bbs4 | Bardet-Biedl syndrome 4 (human) | 627 | 671 | 674 | 569 | 541 | 662 | 545 | 305 | 449 | 513 | 635 |
| 507 | 0.12 | 0.18 | 0.68 | 0.49 | 1 | Gm5617 | predicted gene 5617 | 445 | 446 | 592 | 563 | 518 | 327 | 545 | 667 | 533 | 605 | 337 |
| 503 | 0.02 | 0.18 | 0.10 | 0.92 | 1 | Cyb5d2 | cytochrome b5 domain containing 2 | 311 | 406 | 361 | 508 | 660 | 506 | 545 | 579 | 526 | 572 | 561 |
| 745 | 0.03 | 0.15 | 0.17 | 0.87 | 1 | Rflna | refilin A | 1138 | 1679 | 1244 | 563 | 577 | 363 | 544 | 664 | 525 | 480 | 416 |
| 731 | -0.09 | 0.14 | -0.59 | 0.55 | 1 | Yeats2 | YEATS domain containing 2 | 783 | 707 | 710 | 649 | 711 | 1106 | 544 | 589 | 591 | 614 | 1040 |
| 569 | 0.01 | 0.14 | 0.07 | 0.95 | 1 | Efcab2 | EF-hand calcium binding domain 2 | 592 | 685 | 541 | 577 | 529 | 560 | 544 | 587 | 530 | 486 | 628 |
| 546 | -0.07 | 0.17 | -0.39 | 0.69 | 1 | Zc2hc1c | zinc finger, C2HC-type containing 1C | 293 | 511 | 446 | 652 | 626 | 609 | 544 | 701 | 530 | 537 | 560 |
| 396 | -0.23 | 0.26 | -0.86 | 0.39 | 1 | H2bc6 | H2B clustered histone 6 | 140 | 216 | 181 | 621 | 415 | 473 | 544 | 330 | 501 | 377 | 555 |
| 853 | -0.08 | 0.18 | -0.42 | 0.68 | 1 | Dbn1 | drebrin 1 | 2124 | 1567 | 1937 | 515 | 410 | 461 | 543 | 572 | 409 | 420 | 425 |
| 807 | -0.15 | 0.20 | -0.79 | 0.43 | 1 | Ssh2 | slingshot protein phosphatase 2 | 1780 | 1129 | 1319 | 691 | 462 | 738 | 543 | 428 | 489 | 553 | 748 |
| 672 | 0.14 | 0.20 | 0.73 | 0.47 | 1 | Me1 | malic enzyme 1, NADP(+)-dependent, cyt | 889 | 742 | 1000 | 756 | 536 | 418 | 543 | 689 | 641 | 775 | 407 |
| 605 | 0.04 | 0.19 | 0.21 | 0.83 | 1 | Zfr2 | zinc finger RNA binding protein 2 | 354 | 577 | 513 | 635 | 506 | 915 | 543 | 739 | 553 | 613 | 711 |
| 591 | 0.03 | 0.14 | 0.20 | 0.84 | 1 | R3hcc1l | R3H domain and coiled-coil containing 1 | 676 | 751 | 681 | 537 | 514 | 577 | 543 | 494 | 529 | 554 | 645 |
| 579 | 0.01 | 0.18 | 0.03 | 0.97 | 1 | Idnk | idnK gluconokinase homolog (E. coli) | 598 | 535 | 612 | 571 | 615 | 573 | 543 | 719 | 496 | 497 | 610 |
| 492 | -0.03 | 0.17 | -0.20 | 0.84 | 1 | Isy1 | ISY1 splicing factor homolog | 632 | 632 | 570 | 434 | 435 | 396 | 543 | 412 | 353 | 565 | 443 |
| 825 | -0.19 | 0.26 | -0.70 | 0.48 | 1 | Plin2 | perilipin 2 | 1884 | 1594 | 1904 | 691 | 420 | 320 | 542 | 621 | 432 | 255 | 412 |
| 680 | 0.34 | 0.20 | 1.66 | 0.10 | 1 | Mtpap | mitochondrial poly(A) polymerase | 767 | 585 | 752 | 523 | 542 | 781 | 542 | 922 | 619 | 688 | 755 |
| 675 | -0.08 | 0.17 | -0.49 | 0.62 | 1 | Nrf1 | nuclear respiratory factor 1 | 869 | 714 | 715 | 715 | 646 | 738 | 542 | 590 | 675 | 547 | 677 |
| 617 | 0.27 | 0.16 | 1.70 | 0.09 | 1 | Gm48226 | | 685 | 806 | 784 | 551 | 508 | 426 | 542 | 729 | 550 | 741 | 460 |
| 583 | 0.02 | 0.22 | 0.09 | 0.93 | 1 | Msl3l2 | MSL3 like 2 | 478 | 662 | 536 | 596 | 633 | 579 | 542 | 484 | 558 | 792 | 556 |
| 538 | -0.03 | 0.22 | -0.12 | 0.90 | 1 | Siah1a | siah E3 ubiquitin protein ligase 1A | 646 | 644 | 465 | 603 | 472 | 474 | 542 | 374 | 468 | 705 | 528 |
| 638 | 0.03 | 0.15 | 0.17 | 0.86 | 1 | Heatr3 | HEAT repeat containing 3 | 833 | 723 | 597 | 555 | 495 | 813 | 541 | 480 | 532 | 573 | 881 |
| 599 | -0.10 | 0.13 | -0.77 | 0.44 | 1 | Xpnpep3 | X-prolyl aminopeptidase 3, mitochondrial | 549 | 531 | 626 | 606 | 591 | 795 | 541 | 523 | 591 | 515 | 728 |
| 561 | 0.01 | 0.19 | 0.08 | 0.94 | 1 | Wdr44 | WD repeat domain 44 | 688 | 669 | 559 | 516 | 524 | 528 | 541 | 385 | 599 | 549 | 616 |
| 535 | 0.07 | 0.17 | 0.40 | 0.69 | 1 | 2310061I04Rik | RIKEN cDNA 2310061I04 gene | 392 | 404 | 344 | 468 | 575 | 735 | 541 | 461 | 581 | 670 | 718 |
| 523 | -0.11 | 0.21 | -0.50 | 0.62 | 1 | Ilvbl | ilvB (bacterial acetolactate synthase)-like | 933 | 426 | 658 | 484 | 441 | 466 | 541 | 437 | 476 | 511 | 381 |
| 497 | 0.15 | 0.18 | 0.82 | 0.41 | 1 | Ptpmt1 | protein tyrosine phosphatase, mitochondı | 517 | 465 | 477 | 463 | 522 | 368 | 541 | 681 | 479 | 567 | 384 |
| 489 | -0.01 | 0.16 | -0.03 | 0.97 | 1 | T2 | brachyury 2 | 100 | 123 | 123 | 640 | 620 | 707 | 540 | 633 | 665 | 452 | 777 |
| 471 | 0.27 | 0.20 | 1.37 | 0.17 | 1 | Adck1 | aarF domain containing kinase 1 | 388 | 410 | 388 | 396 | 449 | 418 | 540 | 692 | 435 | 569 | 501 |
| 693 | 0.13 | 0.17 | 0.77 | 0.44 | 1 | Upf3a | UPF3 regulator of nonsense transcripts h | 754 | 646 | 672 | 741 | 559 | 795 | 538 | 826 | 521 | 636 | 932 |
| 626 | 0.05 | 0.16 | 0.31 | 0.76 | 1 | Kxd1 | KxDL motif containing 1 | 860 | 859 | 798 | 459 | 617 | 528 | 538 | 503 | 630 | 620 | 471 |
| 615 | -0.06 | 0.15 | -0.41 | 0.68 | 1 | Cep89 | centrosomal protein 89 | 618 | 524 | 487 | 531 | 632 | 909 | 538 | 507 | 541 | 536 | 940 |
| 560 | 0.10 | 0.23 | 0.44 | 0.66 | 1 | Ssna1 | SS nuclear autoantigen 1 | 746 | 613 | 661 | 564 | 457 | 429 | 538 | 768 | 535 | 367 | 487 |
| 715 | 0.23 | 0.16 | 1.48 | 0.14 | 1 | Ahsa2 | AHA1, activator of heat shock protein ATF | 486 | 523 | 482 | 462 | 384 | 1651 | 537 | 532 | 546 | 645 | 1621 |
| 589 | 0.04 | 0.22 | 0.16 | 0.87 | 1 | Zdhhc13 | zinc finger, DHHC domain containing 13 | 658 | 813 | 525 | 506 | 598 | 561 | 537 | 450 | 559 | 760 | 515 |
| 636 | 0.11 | 0.14 | 0.79 | 0.43 | 1 | Dusp7 | dual specificity phosphatase 7 | 650 | 724 | 749 | 625 | 645 | 542 | 536 | 587 | 669 | 659 | 610 |
| 633 | -0.04 | 0.20 | -0.22 | 0.83 | 1 | Gpr142 | G protein-coupled receptor 142 | 503 | 726 | 766 | 590 | 734 | 675 | 536 | 565 | 566 | 705 | 599 |
| 533 | 0.09 | 0.18 | 0.52 | 0.60 | 1 | Trp53rka | transformation related protein 53 regulati | 468 | 542 | 520 | 608 | 471 | 472 | 536 | 752 | 451 | 484 | 560 |
| 513 | -0.04 | 0.19 | -0.19 | 0.85 | 1 | Tnk1 | tyrosine kinase, non-receptor, 1 | 360 | 417 | 583 | 498 | 519 | 617 | 536 | 527 | 530 | 546 | 507 |

| | | | | | | | | | | | | | | | | | |
|---|---|---|---|---|---|---|---|---|---|---|---|---|---|---|---|---|---|
| 456 | -0.07 | 0.20 | -0.37 | 0.71 | 1 | Tmem243 | transmembrane protein 243, mitochondria | 352 | 298 | 313 | 591 | 491 | 440 | 536 | 404 | 521 | 683 | 387 |
| 707 | 0.06 | 0.17 | 0.37 | 0.71 | 1 | Ppp1r14b | protein phosphatase 1, regulatory inhibitc | 938 | 830 | 954 | 630 | 750 | 548 | 535 | 615 | 706 | 790 | 484 |
| 641 | -0.23 | 0.20 | -1.15 | 0.25 | 1 | Cep162 | centrosomal protein 162 | 560 | 454 | 471 | 634 | 662 | 1134 | 535 | 360 | 656 | 522 | 1068 |
| 522 | -0.08 | 0.21 | -0.39 | 0.69 | 1 | Elf3 | E74-like factor 3 | 301 | 325 | 577 | 639 | 613 | 549 | 535 | 490 | 591 | 599 | 525 |
| 478 | -0.30 | 0.27 | -1.13 | 0.26 | 1 | Taf7 | TATA-box binding protein associated fact | 537 | 382 | 487 | 405 | 543 | 567 | 535 | 326 | 430 | 243 | 797 |
| 758 | -0.25 | 0.18 | -1.35 | 0.18 | 1 | Zfp715 | zinc finger protein 715 | 740 | 662 | 942 | 756 | 542 | 1402 | 534 | 525 | 516 | 473 | 1246 |
| 696 | -0.13 | 0.16 | -0.83 | 0.41 | 1 | Zfp46 | zinc finger protein 46 | 1035 | 954 | 1214 | 624 | 571 | 593 | 534 | 465 | 559 | 612 | 490 |
| 670 | 0.32 | 0.17 | 1.86 | 0.06 | 1 | Taf8 | TATA-box binding protein associated fact | 766 | 538 | 661 | 479 | 648 | 760 | 534 | 689 | 703 | 828 | 762 |
| 621 | 0.35 | 0.23 | 1.52 | 0.13 | 1 | Hint2 | histidine triad nucleotide binding protein 2 | 500 | 560 | 644 | 614 | 559 | 494 | 534 | 1110 | 595 | 787 | 433 |
| 573 | 0.24 | 0.17 | 1.38 | 0.17 | 1 | Mtg1 | mitochondrial ribosome-associated GTPa | 485 | 396 | 586 | 529 | 570 | 586 | 534 | 667 | 630 | 734 | 589 |
| 508 | 0.17 | 0.19 | 0.89 | 0.37 | 1 | Aven | apoptosis, caspase activation inhibitor | 470 | 435 | 476 | 446 | 622 | 366 | 534 | 715 | 599 | 591 | 339 |
| 500 | 0.05 | 0.19 | 0.25 | 0.80 | 1 | Mmab | methylmalonic aciduria (cobalamin deficie | 329 | 337 | 371 | 333 | 602 | 728 | 534 | 344 | 629 | 563 | 728 |
| 472 | -0.04 | 0.23 | -0.17 | 0.87 | 1 | Gdpd1 | glycerophosphodiester phosphodiesteras | 324 | 219 | 377 | 651 | 556 | 408 | 534 | 451 | 551 | 711 | 413 |
| 820 | -0.12 | 0.19 | -0.67 | 0.50 | 1 | Maml1 | mastermind like transcriptional coactivato | 988 | 1385 | 1098 | 918 | 748 | 697 | 532 | 821 | 554 | 760 | 520 |
| 748 | 0.03 | 0.24 | 0.12 | 0.91 | 1 | Slc20a1 | solute carrier family 20, member 1 | 649 | 423 | 469 | 753 | 854 | 1185 | 532 | 947 | 611 | 701 | 1108 |
| 595 | 0.11 | 0.14 | 0.83 | 0.41 | 1 | Rab9 | RAB9, member RAS oncogene family | 754 | 767 | 632 | 556 | 522 | 498 | 532 | 564 | 541 | 621 | 555 |
| 537 | 0.24 | 0.18 | 1.35 | 0.18 | 1 | Mb21d2 | Mab-21 domain containing 2 | 891 | 580 | 598 | 398 | 397 | 402 | 532 | 364 | 525 | 792 | 429 |
| 631 | 0.02 | 0.15 | 0.10 | 0.92 | 1 | Taf4 | TATA-box binding protein associated fact | 740 | 608 | 536 | 489 | 515 | 1005 | 531 | 507 | 556 | 562 | 891 |
| 557 | -0.12 | 0.16 | -0.73 | 0.46 | 1 | Gtpbp3 | GTP binding protein 3 | 679 | 588 | 689 | 398 | 520 | 714 | 531 | 411 | 385 | 500 | 715 |
| 535 | -0.17 | 0.17 | -1.02 | 0.31 | 1 | Kcnk12 | potassium channel, subfamily K, member | 476 | 648 | 683 | 607 | 541 | 468 | 531 | 619 | 473 | 466 | 370 |
| 521 | -0.17 | 0.22 | -0.76 | 0.45 | 1 | Gpr85 | G protein-coupled receptor 85 | 586 | 817 | 856 | 495 | 365 | 446 | 531 | 352 | 399 | 430 | 455 |
| 714 | 0.11 | 0.14 | 0.79 | 0.43 | 1 | Riok1 | RIO kinase 1 | 638 | 718 | 644 | 632 | 664 | 972 | 530 | 735 | 575 | 622 | 1125 |
| 595 | 0.23 | 0.15 | 1.56 | 0.12 | 1 | Mettl3 | methyltransferase like 3 | 565 | 440 | 491 | 447 | 430 | 941 | 530 | 561 | 579 | 564 | 997 |
| 591 | 0.09 | 0.17 | 0.51 | 0.61 | 1 | Zbed3 | zinc finger, BED type containing 3 | 695 | 697 | 765 | 685 | 436 | 455 | 530 | 684 | 529 | 578 | 448 |
| 580 | 0.05 | 0.19 | 0.27 | 0.79 | 1 | Osgin2 | oxidative stress induced growth inhibitor f | 457 | 747 | 604 | 693 | 397 | 638 | 530 | 563 | 519 | 556 | 670 |
| 533 | 0.15 | 0.17 | 0.84 | 0.40 | 1 | Slc25a15 | solute carrier family 25 (mitochondrial car | 422 | 455 | 460 | 561 | 487 | 561 | 530 | 663 | 586 | 672 | 466 |
| 513 | 0.24 | 0.18 | 1.34 | 0.18 | 1 | 2300009A05Ril | RIKEN cDNA 2300009A05 gene | 349 | 404 | 338 | 544 | 501 | 500 | 530 | 592 | 636 | 777 | 474 |
| 607 | -0.22 | 0.19 | -1.19 | 0.23 | 1 | Nop9 | NOP9 nucleolar protein | 833 | 771 | 653 | 556 | 723 | 573 | 529 | 434 | 501 | 540 | 570 |
| 598 | 0.00 | 0.17 | 0.01 | 0.99 | 1 | Mon1a | MON1 homolog A, secretory traffciking as | 405 | 562 | 513 | 599 | 754 | 675 | 529 | 622 | 630 | 652 | 636 |
| 595 | -0.08 | 0.16 | -0.52 | 0.60 | 1 | Smg9 | smg-9 homolog, nonsense mediated mRI | 593 | 504 | 577 | 592 | 660 | 713 | 529 | 459 | 606 | 596 | 713 |
| 534 | 0.10 | 0.21 | 0.49 | 0.63 | 1 | Slc2a1 | solute carrier family 2 (facilitated glucose | 824 | 488 | 441 | 440 | 620 | 400 | 529 | 535 | 557 | 607 | 437 |
| 528 | -0.14 | 0.18 | -0.76 | 0.44 | 1 | Recql5 | RecQ protein-like 5 | 640 | 577 | 465 | 416 | 522 | 686 | 529 | 353 | 452 | 506 | 666 |
| 485 | 0.12 | 0.23 | 0.54 | 0.59 | 1 | B4galt7 | xylosylprotein beta1,4-galactosyltransfera | 498 | 454 | 504 | 271 | 465 | 620 | 529 | 474 | 398 | 580 | 544 |
| 422 | -0.25 | 0.20 | -1.27 | 0.20 | 1 | Bend3 | BEN domain containing 3 | 431 | 333 | 482 | 482 | 493 | 341 | 529 | 409 | 490 | 339 | 316 |
| 421 | 0.10 | 0.26 | 0.39 | 0.70 | 1 | Tstd3 | thiosulfate sulfurtransferase (rhodanese)- | 498 | 252 | 444 | 459 | 404 | 276 | 529 | 593 | 405 | 444 | 322 |
| 744 | 0.02 | 0.13 | 0.15 | 0.88 | 1 | Lrrc49 | leucine rich repeat containing 49 | 1054 | 1228 | 972 | 709 | 582 | 631 | 528 | 657 | 611 | 547 | 661 |
| 706 | -0.07 | 0.17 | -0.43 | 0.67 | 1 | Ttll4 | tubulin tyrosine ligase-like family, membe | 611 | 636 | 624 | 483 | 464 | 1593 | 528 | 386 | 589 | 497 | 1351 |
| 569 | -0.20 | 0.19 | -1.08 | 0.28 | 1 | 1500015A07Rik | | 676 | 720 | 739 | 609 | 562 | 500 | 528 | 357 | 558 | 502 | 505 |
| 418 | -0.02 | 0.21 | -0.11 | 0.91 | 1 | Sh3yl1 | Sh3 domain YSC-like 1 | 483 | 572 | 372 | 360 | 372 | 334 | 528 | 385 | 382 | 526 | 288 |
| 803 | -0.03 | 0.17 | -0.16 | 0.87 | 1 | Ccdc82 | coiled-coil domain containing 82 | 1096 | 1077 | 792 | 707 | 662 | 1053 | 527 | 620 | 609 | 583 | 1111 |
| 718 | 0.13 | 0.17 | 0.78 | 0.43 | 1 | Zfp787 | zinc finger protein 787 | 800 | 795 | 896 | 655 | 695 | 705 | 527 | 677 | 620 | 818 | 711 |
| 656 | -0.27 | 0.27 | -1.03 | 0.30 | 1 | Elmo1 | engulfment and cell motility 1 | 1070 | 603 | 662 | 822 | 629 | 678 | 527 | 444 | 732 | 514 | 533 |
| 566 | 0.15 | 0.16 | 0.95 | 0.34 | 1 | Aen | apoptosis enhancing nuclease | 481 | 340 | 406 | 543 | 581 | 721 | 527 | 607 | 645 | 675 | 701 |
| 484 | 0.27 | 0.26 | 1.02 | 0.31 | 1 | Eepd1 | endonuclease/exonuclease/phosphatase | 336 | 313 | 466 | 659 | 350 | 354 | 527 | 840 | 563 | 553 | 360 |
| 419 | 0.13 | 0.23 | 0.56 | 0.58 | 1 | Trpm2 | transient receptor potential cation channe | 281 | 138 | 99 | 323 | 280 | 881 | 527 | 451 | 341 | 455 | 832 |
| 685 | 0.23 | 0.14 | 1.70 | 0.09 | 1 | Med10 | mediator complex subunit 10 | 791 | 835 | 765 | 593 | 524 | 720 | 526 | 624 | 598 | 737 | 817 |
| 606 | 0.01 | 0.20 | 0.05 | 0.96 | 1 | Ctbs | chitobiase | 606 | 719 | 571 | 737 | 600 | 527 | 526 | 477 | 640 | 638 | 631 |
| 573 | 0.27 | 0.19 | 1.39 | 0.16 | 1 | Slf1 | SMC5-SMC6 complex localization factor | 565 | 788 | 593 | 424 | 483 | 530 | 526 | 656 | 458 | 657 | 617 |
| 509 | -0.03 | 0.22 | -0.14 | 0.89 | 1 | Nsdhl | NAD(P) dependent steroid dehydrogenas | 519 | 446 | 344 | 587 | 619 | 438 | 526 | 429 | 679 | 522 | 494 |
| 509 | 0.26 | 0.19 | 1.32 | 0.19 | 1 | Spidr | scaffolding protein involved in DNA repair | 440 | 577 | 486 | 444 | 510 | 381 | 526 | 655 | 598 | 443 | 537 |
| 436 | -0.23 | 0.20 | -1.11 | 0.27 | 1 | Ube2e2 | ubiquitin-conjugating enzyme E2E 2 | 477 | 385 | 488 | 475 | 450 | 398 | 526 | 364 | 549 | 346 | 340 |
| 861 | 0.18 | 0.16 | 1.11 | 0.27 | 1 | Guf1 | GUF1 homolog, GTPase | 556 | 364 | 403 | 489 | 393 | 2456 | 525 | 561 | 517 | 515 | 2695 |
| 682 | 0.02 | 0.14 | 0.17 | 0.86 | 1 | Hmgxb4 | HMG box domain containing 4 | 823 | 726 | 743 | 596 | 590 | 887 | 525 | 578 | 633 | 617 | 784 |
| 658 | 0.05 | 0.17 | 0.30 | 0.77 | 1 | Elovl1 | elongation of very long chain fatty acids (I | 988 | 822 | 775 | 604 | 610 | 538 | 525 | 527 | 702 | 631 | 512 |
| 588 | -0.13 | 0.16 | -0.84 | 0.40 | 1 | Aldoc | aldolase C, fructose-bisphosphate | 712 | 926 | 895 | 524 | 596 | 412 | 525 | 462 | 597 | 410 | 413 |
| 548 | 0.05 | 0.15 | 0.35 | 0.72 | 1 | Mettl6 | methyltransferase like 6 | 533 | 574 | 531 | 548 | 490 | 587 | 525 | 662 | 463 | 502 | 616 |
| 518 | 0.25 | 0.30 | 0.82 | 0.41 | 1 | Rhbdd3 | rhomboid domain containing 3 | 328 | 221 | 443 | 542 | 425 | 658 | 525 | 918 | 431 | 553 | 660 |
| 485 | -0.05 | 0.17 | -0.30 | 0.77 | 1 | Nif3l1 | Ngg1 interacting factor 3-like 1 (S. pombe | 366 | 422 | 451 | 485 | 511 | 553 | 525 | 436 | 577 | 424 | 587 |
| 418 | -0.07 | 0.30 | -0.24 | 0.81 | 1 | Rab2b | RAB2B, member RAS oncogene family | 672 | 439 | 422 | 277 | 272 | 423 | 525 | 133 | 290 | 734 | 412 |
| 872 | 0.03 | 0.19 | 0.17 | 0.87 | 1 | Erf | Ets2 repressor factor | 1143 | 1604 | 1456 | 926 | 681 | 541 | 524 | 795 | 583 | 849 | 488 |

| | | | | | | | | | | | | | | | | | | |
|---|---|---|---|---|---|---|---|---|---|---|---|---|---|---|---|---|---|---|
| 737 | 0.06 | 0.18 | 0.31 | 0.75 | 1 | Pcsk7 | proprotein convertase subtilisin/kexin type | 781 | 976 | 968 | 664 | 573 | 887 | 524 | 562 | 539 | 764 | 874 |
| 639 | -0.17 | 0.22 | -0.77 | 0.44 | 1 | Nfyc | nuclear transcription factor-Y gamma | 598 | 496 | 434 | 647 | 482 | 1247 | 524 | 411 | 620 | 426 | 1143 |
| 533 | -0.04 | 0.19 | -0.22 | 0.82 | 1 | St3gal3 | ST3 beta-galactoside alpha-2,3-sialyltran | 666 | 591 | 626 | 537 | 449 | 492 | 524 | 651 | 403 | 529 | 391 |
| 512 | -0.12 | 0.16 | -0.77 | 0.44 | 1 | Zfp994 | zinc finger protein 994 | 522 | 463 | 556 | 543 | 490 | 569 | 524 | 439 | 527 | 460 | 538 |
| 1037 | -0.16 | 0.14 | -1.15 | 0.25 | 1 | Ints6l | integrator complex subunit 6 like | 428 | 349 | 321 | 523 | 522 | 3731 | 523 | 375 | 499 | 484 | 3656 |
| 598 | -0.20 | 0.19 | -1.03 | 0.30 | 1 | Capsl | calcyphosine-like | 620 | 1001 | 1167 | 528 | 516 | 456 | 523 | 475 | 456 | 393 | 445 |
| 570 | 0.06 | 0.19 | 0.30 | 0.76 | 1 | Ago4 | argonaute RISC catalytic subunit 4 | 760 | 723 | 500 | 550 | 456 | 549 | 523 | 781 | 438 | 513 | 475 |
| 562 | -0.21 | 0.23 | -0.92 | 0.36 | 1 | Zfp821 | zinc finger protein 821 | 614 | 814 | 720 | 473 | 520 | 608 | 523 | 667 | 403 | 277 | 563 |
| 480 | -0.04 | 0.20 | -0.20 | 0.85 | 1 | Znhit6 | zinc finger, HIT type 6 | 654 | 561 | 546 | 304 | 423 | 529 | 523 | 272 | 483 | 457 | 523 |
| 1250 | 0.10 | 0.15 | 0.66 | 0.51 | 1 | Prpf39 | pre-mRNA processing factor 39 | 742 | 836 | 775 | 771 | 418 | 3883 | 522 | 717 | 592 | 526 | 3964 |
| 622 | 0.24 | 0.19 | 1.26 | 0.21 | 1 | Mad1l1 | MAD1 mitotic arrest deficient 1-like 1 | 787 | 554 | 617 | 648 | 540 | 533 | 522 | 751 | 674 | 574 | 643 |
| 611 | -0.12 | 0.18 | -0.65 | 0.51 | 1 | Tmem18 | transmembrane protein 18 | 560 | 543 | 554 | 638 | 721 | 734 | 522 | 453 | 588 | 569 | 843 |
| 593 | 0.16 | 0.17 | 0.94 | 0.35 | 1 | Slc12a9 | solute carrier family 12 (potassium/chloric | 525 | 451 | 581 | 555 | 548 | 727 | 522 | 648 | 544 | 689 | 730 |
| 415 | 0.00 | 0.17 | -0.02 | 0.99 | 1 | Polr3g | polymerase (RNA) III (DNA directed) poly | 145 | 139 | 130 | 584 | 460 | 505 | 522 | 520 | 479 | 638 | 442 |
| 385 | 0.04 | 0.18 | 0.25 | 0.81 | 1 | Dhtkd1 | dehydrogenase E1 and transketolase dor | 191 | 180 | 215 | 334 | 564 | 378 | 522 | 365 | 565 | 533 | 385 |
| 612 | 0.01 | 0.19 | 0.05 | 0.96 | 1 | Tpra1 | transmembrane protein, adipocyte assoc | 498 | 455 | 607 | 550 | 617 | 910 | 521 | 618 | 492 | 684 | 781 |
| 542 | -0.11 | 0.20 | -0.55 | 0.58 | 1 | Rpap3 | RNA polymerase II associated protein 3 | 688 | 456 | 528 | 582 | 638 | 477 | 521 | 407 | 515 | 602 | 552 |
| 515 | 0.15 | 0.17 | 0.86 | 0.39 | 1 | Mrpl54 | mitochondrial ribosomal protein L54 | 402 | 410 | 566 | 515 | 610 | 391 | 521 | 485 | 629 | 657 | 473 |
| 476 | 0.13 | 0.15 | 0.82 | 0.41 | 1 | Pctp | phosphatidylcholine transfer protein | 311 | 252 | 367 | 480 | 623 | 441 | 521 | 554 | 613 | 585 | 490 |
| 438 | 0.11 | 0.22 | 0.52 | 0.60 | 1 | Mtm1 | X-linked myotubular myopathy gene 1 | 508 | 259 | 283 | 430 | 348 | 518 | 521 | 428 | 452 | 456 | 613 |
| 772 | 0.09 | 0.13 | 0.67 | 0.50 | 1 | Nfya | nuclear transcription factor-Y alpha | 1200 | 1067 | 1058 | 526 | 474 | 997 | 520 | 492 | 536 | 654 | 967 |
| 661 | 0.42 | 0.22 | 1.89 | 0.06 | 1 | Bad | BCL2-associated agonist of cell death | 623 | 668 | 966 | 511 | 616 | 484 | 520 | 900 | 616 | 853 | 510 |
| 597 | -0.09 | 0.17 | -0.52 | 0.60 | 1 | Usp21 | ubiquitin specific peptidase 21 | 609 | 668 | 738 | 574 | 447 | 816 | 520 | 525 | 417 | 560 | 688 |
| 596 | 0.16 | 0.18 | 0.88 | 0.38 | 1 | Stk3 | serine/threonine kinase 3 | 1062 | 772 | 943 | 511 | 344 | 406 | 520 | 446 | 483 | 625 | 446 |
| 546 | 0.23 | 0.21 | 1.12 | 0.26 | 1 | Cox19 | cytochrome c oxidase assembly protein 1 | 468 | 529 | 606 | 596 | 428 | 473 | 520 | 765 | 439 | 657 | 530 |
| 442 | -0.26 | 0.20 | -1.31 | 0.19 | 1 | Bud13 | BUD13 homolog | 454 | 356 | 351 | 466 | 563 | 463 | 520 | 301 | 521 | 428 | 443 |
| 640 | -0.09 | 0.14 | -0.61 | 0.54 | 1 | Tctn1 | tectonic family member 1 | 692 | 736 | 718 | 516 | 563 | 941 | 519 | 567 | 549 | 473 | 763 |
| 618 | 0.16 | 0.18 | 0.89 | 0.37 | 1 | Usp51 | ubiquitin specific protease 51 | 281 | 484 | 277 | 808 | 465 | 975 | 519 | 693 | 614 | 683 | 1004 |
| 613 | 0.28 | 0.20 | 1.36 | 0.17 | 1 | Swt1 | SWT1 RNA endoribonuclease homolog (: | 530 | 668 | 501 | 435 | 559 | 770 | 519 | 697 | 760 | 477 | 827 |
| 608 | -0.24 | 0.15 | -1.60 | 0.11 | 1 | Klf12 | Kruppel-like factor 12 | 818 | 848 | 761 | 661 | 519 | 615 | 519 | 468 | 520 | 435 | 523 |
| 601 | 0.00 | 0.23 | -0.02 | 0.99 | 1 | Med8 | mediator complex subunit 8 | 441 | 625 | 382 | 639 | 647 | 776 | 519 | 437 | 719 | 577 | 852 |
| 593 | 0.15 | 0.19 | 0.79 | 0.43 | 1 | Rbm15 | RNA binding motif protein 15 | 715 | 681 | 478 | 494 | 285 | 983 | 519 | 527 | 465 | 514 | 865 |
| 580 | -0.11 | 0.24 | -0.45 | 0.65 | 1 | Dtx2 | deltex 2, E3 ubiquitin ligase | 482 | 348 | 307 | 868 | 581 | 759 | 519 | 561 | 684 | 536 | 733 |
| 568 | 0.08 | 0.18 | 0.44 | 0.66 | 1 | Cptp | ceramide-1-phosphate transfer protein | 654 | 673 | 511 | 539 | 649 | 429 | 519 | 688 | 563 | 594 | 429 |
| 532 | 0.13 | 0.15 | 0.87 | 0.38 | 1 | Zfp37 | zinc finger protein 37 | 497 | 496 | 510 | 504 | 512 | 538 | 519 | 506 | 646 | 574 | 546 |
| 483 | 0.01 | 0.19 | 0.04 | 0.97 | 1 | Ccdc112 | coiled-coil domain containing 112 | 382 | 554 | 342 | 463 | 473 | 553 | 519 | 534 | 476 | 475 | 538 |
| 388 | -0.02 | 0.17 | -0.12 | 0.90 | 1 | Ulk4 | unc-51-like kinase 4 | 231 | 183 | 299 | 348 | 541 | 367 | 519 | 336 | 617 | 511 | 318 |
| 789 | -0.03 | 0.15 | -0.21 | 0.83 | 1 | Zcchc7 | zinc finger, CCHC domain containing 7 | 527 | 579 | 631 | 456 | 573 | 1975 | 518 | 568 | 565 | 403 | 1884 |
| 525 | -0.03 | 0.17 | -0.19 | 0.85 | 1 | Hdac9 | histone deacetylase 9 | 394 | 551 | 398 | 824 | 474 | 422 | 518 | 738 | 605 | 496 | 353 |
| 512 | -0.07 | 0.19 | -0.38 | 0.70 | 1 | Zfp637 | zinc finger protein 637 | 486 | 640 | 457 | 508 | 608 | 440 | 518 | 524 | 457 | 538 | 456 |
| 458 | -0.17 | 0.17 | -0.97 | 0.33 | 1 | Irak1bp1 | interleukin-1 receptor-associated kinase | 450 | 492 | 558 | 508 | 485 | 369 | 518 | 466 | 402 | 406 | 389 |
| 426 | -0.14 | 0.22 | -0.64 | 0.52 | 1 | Meak7 | MTOR associated protein , eak-7 homolo | 351 | 279 | 409 | 538 | 401 | 446 | 518 | 422 | 545 | 395 | 381 |
| 391 | 0.09 | 0.19 | 0.44 | 0.66 | 1 | Bdh1 | 3-hydroxybutyrate dehydrogenase, type 1 | 178 | 104 | 105 | 462 | 495 | 418 | 518 | 470 | 461 | 611 | 480 |
| 285 | -0.32 | 0.33 | -0.99 | 0.32 | 1 | Rhd | Rh blood group, D antigen | 92 | 191 | 195 | 369 | 332 | 262 | 518 | 272 | 347 | 277 | 283 |
| 1233 | -0.09 | 0.15 | -0.59 | 0.55 | 1 | Ifih1 | interferon induced with helicase C domai | 2931 | 3409 | 3042 | 578 | 651 | 397 | 517 | 718 | 552 | 393 | 378 |
| 647 | -0.06 | 0.17 | -0.35 | 0.73 | 1 | Tmem132a | transmembrane protein 132A | 897 | 875 | 943 | 593 | 572 | 549 | 517 | 678 | 630 | 453 | 412 |
| 581 | -0.04 | 0.20 | -0.18 | 0.86 | 1 | Nubp2 | nucleotide binding protein 2 | 892 | 621 | 837 | 318 | 633 | 603 | 517 | 452 | 568 | 401 | 551 |
| 453 | 0.15 | 0.17 | 0.87 | 0.39 | 1 | Syt3 | synaptotagmin III | 250 | 230 | 231 | 516 | 583 | 391 | 517 | 539 | 615 | 742 | 375 |
| 619 | -0.08 | 0.20 | -0.41 | 0.68 | 1 | Dnmbp | dynamin binding protein | 1072 | 775 | 680 | 565 | 490 | 632 | 516 | 422 | 579 | 492 | 592 |
| 546 | 0.03 | 0.17 | 0.21 | 0.84 | 1 | Rbm34 | RNA binding motif protein 34 | 650 | 509 | 615 | 407 | 535 | 616 | 516 | 338 | 601 | 625 | 589 |
| 538 | 0.04 | 0.17 | 0.24 | 0.81 | 1 | Fbxo16 | F-box protein 16 | 286 | 431 | 468 | 535 | 703 | 573 | 516 | 606 | 659 | 457 | 685 |
| 512 | -0.24 | 0.19 | -1.24 | 0.21 | 1 | Arl10 | ADP-ribosylation factor-like 10 | 522 | 602 | 442 | 646 | 554 | 477 | 516 | 560 | 563 | 353 | 397 |
| 431 | -0.25 | 0.18 | -1.39 | 0.16 | 1 | Tbc1d12 | TBC1D12: TBC1 domain family, member | 550 | 437 | 475 | 394 | 350 | 511 | 516 | 292 | 284 | 498 | 434 |
| 757 | -0.34 | 0.21 | -1.62 | 0.11 | 1 | Abca8b | ATP-binding cassette, sub-family A (ABC | 1527 | 1351 | 1349 | 684 | 462 | 620 | 515 | 375 | 533 | 332 | 573 |
| 659 | -0.04 | 0.21 | -0.18 | 0.86 | 1 | Med20 | mediator complex subunit 20 | 615 | 700 | 686 | 515 | 413 | 1215 | 515 | 593 | 559 | 314 | 1122 |
| 651 | 0.20 | 0.21 | 0.96 | 0.34 | 1 | Myg1 | melanocyte proliferating gene 1 | 821 | 787 | 803 | 470 | 614 | 606 | 515 | 804 | 466 | 609 | 668 |
| 555 | -0.17 | 0.19 | -0.90 | 0.37 | 1 | Rida | reactive intermediate imine deaminase A | 589 | 598 | 720 | 589 | 609 | 494 | 515 | 710 | 487 | 419 | 377 |
| 661 | -0.23 | 0.21 | -1.10 | 0.27 | 1 | Cers4 | ceramide synthase 4 | 1126 | 1057 | 1153 | 594 | 428 | 572 | 514 | 646 | 411 | 313 | 451 |
| 571 | -0.04 | 0.19 | -0.22 | 0.83 | 1 | Crls1 | cardiolipin synthase 1 | 528 | 428 | 414 | 624 | 649 | 702 | 514 | 623 | 641 | 466 | 693 |
| 471 | 0.14 | 0.22 | 0.64 | 0.52 | 1 | BC003965 | cDNA sequence BC003965 | 426 | 502 | 491 | 330 | 432 | 511 | 514 | 584 | 400 | 506 | 483 |

| | | | | | | | | | | | | | | | | | | |
|---|---|---|---|---|---|---|---|---|---|---|---|---|---|---|---|---|---|---|
| 456 | 0.23 | 0.17 | 1.38 | 0.17 | 1 | Rnf170 | ring finger protein 170 | 339 | 214 | 347 | 347 | 475 | 569 | 514 | 469 | 523 | 593 | 628 |
| 614 | 0.09 | 0.21 | 0.43 | 0.67 | 1 | Zdhhc4 | zinc finger, DHHC domain containing 4 | 682 | 517 | 741 | 466 | 615 | 754 | 513 | 711 | 490 | 574 | 686 |
| 558 | -0.09 | 0.14 | -0.61 | 0.54 | 1 | 2310057M21Ri | RIKEN cDNA 2310057M21 gene | 452 | 583 | 486 | 558 | 588 | 717 | 513 | 546 | 503 | 491 | 706 |
| 553 | 0.00 | 0.20 | 0.02 | 0.98 | 1 | Xylt2 | xylosyltransferase II | 823 | 716 | 724 | 488 | 516 | 362 | 513 | 701 | 439 | 510 | 293 |
| 547 | -0.13 | 0.17 | -0.76 | 0.45 | 1 | Ptrh2 | peptidyl-tRNA hydrolase 2 | 616 | 576 | 658 | 520 | 626 | 515 | 513 | 585 | 513 | 402 | 490 |
| 536 | 0.01 | 0.17 | 0.06 | 0.96 | 1 | Mycbp | MYC binding protein | 751 | 687 | 757 | 382 | 431 | 511 | 513 | 341 | 431 | 599 | 497 |
| 477 | 0.18 | 0.16 | 1.13 | 0.26 | 1 | Narf | nuclear prelamin A recognition factor | 625 | 380 | 440 | 313 | 473 | 484 | 513 | 370 | 520 | 620 | 513 |
| 459 | -0.31 | 0.26 | -1.18 | 0.24 | 1 | Fam136a | family with sequence similarity 136, meml | 452 | 519 | 406 | 570 | 509 | 417 | 513 | 533 | 476 | 219 | 436 |
| 450 | 0.07 | 0.23 | 0.30 | 0.76 | 1 | Caap1 | caspase activity and apoptosis inhibitor 1 | 315 | 434 | 321 | 396 | 498 | 482 | 513 | 367 | 497 | 549 | 578 |
| 448 | -0.14 | 0.19 | -0.75 | 0.45 | 1 | Rtl8b | retrotransposon Gag like 8B | 514 | 404 | 579 | 443 | 498 | 341 | 513 | 307 | 507 | 422 | 396 |
| 317 | -0.42 | 0.27 | -1.53 | 0.13 | 1 | Mpz | myelin protein zero | 198 | 251 | 263 | 320 | 489 | 235 | 513 | 217 | 405 | 452 | 143 |
| 1134 | -0.04 | 0.17 | -0.25 | 0.80 | 1 | Fam193b | family with sequence similarity 193, meml | 482 | 494 | 407 | 325 | 433 | 4433 | 512 | 475 | 378 | 390 | 4141 |
| 684 | 0.00 | 0.22 | 0.01 | 1.00 | 1 | Mapkap1 | mitogen-activated protein kinase associat | 957 | 562 | 734 | 718 | 675 | 709 | 512 | 554 | 721 | 777 | 605 |
| 573 | 0.14 | 0.20 | 0.71 | 0.48 | 1 | Ten1 | TEN1 telomerase capping complex subur | 544 | 560 | 640 | 613 | 607 | 424 | 512 | 877 | 501 | 541 | 483 |
| 507 | 0.02 | 0.20 | 0.12 | 0.90 | 1 | Mcu | mitochondrial calcium uniporter | 532 | 519 | 493 | 473 | 495 | 503 | 512 | 377 | 550 | 697 | 433 |
| 481 | 0.05 | 0.21 | 0.25 | 0.80 | 1 | Agpat2 | 1-acylglycerol-3-phosphate O-acyltransfe | 469 | 420 | 508 | 615 | 441 | 333 | 512 | 595 | 411 | 648 | 334 |
| 460 | -0.36 | 0.23 | -1.55 | 0.12 | 1 | Lrrn2 | leucine rich repeat protein 2, neuronal | 414 | 336 | 280 | 788 | 542 | 399 | 512 | 728 | 352 | 407 | 307 |
| 749 | -0.02 | 0.21 | -0.10 | 0.92 | 1 | Tyk2 | tyrosine kinase 2 | 831 | 627 | 638 | 778 | 749 | 1074 | 511 | 619 | 846 | 650 | 917 |
| 629 | 0.18 | 0.18 | 0.99 | 0.32 | 1 | Rmc1 | regulator of MON1-CCZ1 | 492 | 394 | 416 | 530 | 611 | 1008 | 511 | 623 | 606 | 826 | 898 |
| 610 | 0.10 | 0.15 | 0.68 | 0.50 | 1 | Gadd45a | growth arrest and DNA-damage-inducible | 930 | 837 | 715 | 542 | 462 | 516 | 511 | 499 | 440 | 645 | 618 |
| 530 | -0.27 | 0.19 | -1.43 | 0.15 | 1 | Cep78 | centrosomal protein 78 | 577 | 549 | 432 | 467 | 524 | 804 | 511 | 397 | 440 | 346 | 777 |
| 507 | -0.05 | 0.17 | -0.31 | 0.76 | 1 | Hps4 | HPS4, biogenesis of lysosomal organelle | 501 | 395 | 447 | 392 | 422 | 826 | 511 | 468 | 376 | 399 | 843 |
| 499 | 0.07 | 0.16 | 0.45 | 0.65 | 1 | Kansl2 | KAT8 regulatory NSL complex subunit 2 | 384 | 350 | 497 | 427 | 515 | 628 | 511 | 467 | 542 | 585 | 585 |
| 495 | 0.05 | 0.17 | 0.27 | 0.79 | 1 | Kctd7 | potassium channel tetramerisation domai | 301 | 438 | 372 | 511 | 537 | 565 | 511 | 602 | 616 | 475 | 512 |
| 746 | 0.23 | 0.19 | 1.25 | 0.21 | 1 | Slbp | stem-loop binding protein | 1301 | 1043 | 1011 | 589 | 451 | 688 | 510 | 726 | 664 | 498 | 719 |
| 677 | 0.15 | 0.17 | 0.89 | 0.37 | 1 | Noa1 | nitric oxide associated 1 | 625 | 764 | 676 | 719 | 694 | 624 | 510 | 928 | 577 | 666 | 666 |
| 525 | 0.14 | 0.19 | 0.75 | 0.45 | 1 | Zswim4 | zinc finger SWIM-type containing 4 | 612 | 499 | 651 | 546 | 425 | 418 | 510 | 637 | 524 | 550 | 400 |
| 512 | 0.13 | 0.18 | 0.74 | 0.46 | 1 | Aatf | apoptosis antagonizing transcription facto | 575 | 371 | 513 | 462 | 448 | 573 | 510 | 495 | 584 | 496 | 607 |
| 464 | 0.18 | 0.19 | 0.98 | 0.33 | 1 | Dgkg | diacylglycerol kinase, gamma | 92 | 83 | 100 | 653 | 565 | 527 | 510 | 710 | 589 | 524 | 750 |
| 422 | 0.05 | 0.24 | 0.19 | 0.85 | 1 | Morc4 | microrchidia 4 | 320 | 437 | 299 | 466 | 474 | 320 | 510 | 345 | 472 | 607 | 393 |
| 587 | -0.01 | 0.16 | -0.08 | 0.94 | 1 | Katnal1 | katanin p60 subunit A-like 1 | 577 | 739 | 771 | 563 | 534 | 573 | 509 | 636 | 535 | 386 | 636 |
| 539 | -0.11 | 0.17 | -0.66 | 0.51 | 1 | Gorab | golgin, RAB6-interacting | 500 | 488 | 604 | 541 | 521 | 690 | 509 | 545 | 515 | 509 | 512 |
| 720 | 0.08 | 0.16 | 0.50 | 0.62 | 1 | Arhgap39 | Rho GTPase activating protein 39 | 613 | 503 | 602 | 598 | 662 | 1276 | 508 | 708 | 608 | 598 | 1247 |
| 628 | 0.07 | 0.20 | 0.33 | 0.74 | 1 | Ctbp2 | C-terminal binding protein 2 | 1003 | 692 | 845 | 457 | 447 | 729 | 508 | 626 | 446 | 441 | 710 |
| 588 | 0.01 | 0.14 | 0.04 | 0.97 | 1 | Commd10 | COMM domain containing 10 | 733 | 760 | 767 | 555 | 571 | 467 | 508 | 505 | 525 | 594 | 485 |
| 564 | 0.19 | 0.20 | 0.92 | 0.36 | 1 | Ppp4r4 | protein phosphatase 4, regulatory subuni | 484 | 587 | 523 | 546 | 383 | 734 | 508 | 454 | 569 | 665 | 749 |
| 532 | 0.29 | 0.21 | 1.38 | 0.17 | 1 | Isoc2a | isochorismatase domain containing 2a | 510 | 344 | 287 | 557 | 473 | 575 | 508 | 821 | 522 | 585 | 673 |
| 846 | -0.03 | 0.24 | -0.12 | 0.91 | 1 | Tet2 | tet methylcytosine dioxygenase 2 | 510 | 622 | 420 | 445 | 402 | 2769 | 506 | 314 | 618 | 502 | 2195 |
| 714 | -0.18 | 0.16 | -1.14 | 0.26 | 1 | Rell1 | RELT-like 1 | 1033 | 893 | 876 | 606 | 521 | 1013 | 506 | 409 | 452 | 519 | 1025 |
| 714 | 0.01 | 0.20 | 0.03 | 0.98 | 1 | F3 | coagulation factor III | 1962 | 1130 | 883 | 469 | 403 | 543 | 506 | 573 | 350 | 571 | 461 |
| 533 | 0.36 | 0.28 | 1.27 | 0.20 | 1 | Bola2 | bolA-like 2 (E. coli) | 619 | 571 | 689 | 347 | 547 | 320 | 506 | 929 | 438 | 515 | 376 |
| 518 | -0.13 | 0.21 | -0.63 | 0.53 | 1 | 3110082I17Rik | RIKEN cDNA 3110082I17 gene | 575 | 698 | 704 | 350 | 570 | 510 | 506 | 415 | 561 | 317 | 488 |
| 497 | -0.13 | 0.16 | -0.79 | 0.43 | 1 | Oxsm | 3-oxoacyl-ACP synthase, mitochondrial | 510 | 406 | 667 | 486 | 554 | 477 | 506 | 414 | 530 | 483 | 434 |
| 455 | -0.18 | 0.19 | -0.96 | 0.34 | 1 | Dhrs7b | dehydrogenase/reductase (SDR family) n | 381 | 325 | 457 | 591 | 554 | 399 | 506 | 483 | 421 | 487 | 401 |
| 420 | -0.04 | 0.26 | -0.14 | 0.89 | 1 | Maip1 | matrix AAA peptidase interacting protein ' | 486 | 385 | 430 | 343 | 436 | 392 | 506 | 563 | 324 | 331 | 422 |
| 816 | -0.01 | 0.13 | -0.08 | 0.94 | 1 | Zeb1 | zinc finger E-box binding homeobox 1 | 2023 | 1302 | 1464 | 335 | 374 | 894 | 505 | 383 | 346 | 480 | 865 |
| 592 | 0.02 | 0.15 | 0.16 | 0.88 | 1 | Mapk1ip1 | mitogen-activated protein kinase 1 intera | 359 | 272 | 344 | 502 | 473 | 1239 | 504 | 445 | 484 | 551 | 1339 |
| 525 | 0.18 | 0.17 | 1.09 | 0.27 | 1 | Tsfm | Ts translation elongation factor, mitochon | 502 | 401 | 469 | 377 | 603 | 581 | 504 | 469 | 560 | 696 | 612 |
| 500 | -0.10 | 0.25 | -0.42 | 0.67 | 1 | 1810008I18Rik | | 203 | 368 | 254 | 649 | 745 | 544 | 504 | 434 | 599 | 614 | 589 |
| 456 | -0.15 | 0.16 | -0.90 | 0.37 | 1 | Slc2a8 | solute carrier family 2, (facilitated glucose | 405 | 292 | 394 | 513 | 543 | 498 | 504 | 405 | 507 | 517 | 440 |
| 421 | -0.31 | 0.27 | -1.17 | 0.24 | 1 | Hdhd2 | haloacid dehalogenase-like hydrolase dc | 408 | 524 | 480 | 396 | 481 | 389 | 504 | 203 | 460 | 469 | 322 |
| 1166 | -0.11 | 0.20 | -0.54 | 0.59 | 1 | Ifngr1 | interferon gamma receptor 1 | 3787 | 2488 | 2306 | 740 | 440 | 528 | 503 | 636 | 439 | 457 | 507 |
| 578 | -0.24 | 0.17 | -1.36 | 0.17 | 1 | Rorc | RAR-related orphan receptor gamma | 406 | 733 | 741 | 554 | 714 | 643 | 503 | 505 | 652 | 410 | 496 |
| 490 | 0.00 | 0.20 | -0.01 | 0.99 | 1 | Mto1 | mitochondrial tRNA translation optimizatio | 369 | 466 | 407 | 408 | 528 | 639 | 503 | 415 | 457 | 582 | 620 |
| 489 | -0.13 | 0.20 | -0.63 | 0.53 | 1 | Il17rc | interleukin 17 receptor C | 728 | 607 | 623 | 471 | 357 | 438 | 503 | 514 | 344 | 510 | 284 |
| 439 | -0.04 | 0.19 | -0.23 | 0.82 | 1 | Mrps25 | mitochondrial ribosomal protein S25 | 522 | 477 | 450 | 329 | 414 | 475 | 503 | 416 | 390 | 357 | 498 |
| 405 | -0.10 | 0.17 | -0.57 | 0.57 | 1 | Pkib | protein kinase inhibitor beta, cAMP deper | 247 | 226 | 289 | 527 | 554 | 336 | 503 | 489 | 507 | 421 | 360 |
| 684 | 0.29 | 0.17 | 1.67 | 0.09 | 1 | Klhdc4 | kelch domain containing 4 | 718 | 584 | 562 | 646 | 604 | 806 | 502 | 846 | 679 | 702 | 873 |
| 638 | 0.18 | 0.17 | 1.06 | 0.29 | 1 | Faap100 | Fanconi anemia core complex associatec | 820 | 939 | 942 | 513 | 556 | 443 | 502 | 645 | 537 | 701 | 421 |
| 611 | -0.07 | 0.17 | -0.40 | 0.69 | 1 | Rev1 | REV1, DNA directed polymerase | 521 | 520 | 582 | 501 | 393 | 1221 | 502 | 429 | 522 | 397 | 1132 |

| | | | | | | | | | | | | | | | | | | |
|---|---|---|---|---|---|---|---|---|---|---|---|---|---|---|---|---|---|---|
| 600 | -0.08 | 0.17 | -0.50 | 0.62 | 1 | Mfsd10 | major facilitator superfamily domain conta | 701 | 492 | 599 | 527 | 472 | 966 | 502 | 386 | 527 | 522 | 910 |
| 538 | -0.07 | 0.20 | -0.37 | 0.71 | 1 | Raph1 | Ras association (RalGDS/AF-6) and plecl | 978 | 781 | 571 | 452 | 461 | 428 | 502 | 497 | 468 | 412 | 372 |
| 496 | 0.19 | 0.20 | 0.97 | 0.33 | 1 | Mrm1 | mitochondrial rRNA methyltransferase 1 | 452 | 405 | 413 | 442 | 586 | 421 | 502 | 726 | 556 | 484 | 472 |
| 491 | 0.03 | 0.21 | 0.14 | 0.89 | 1 | Elmod3 | ELMO/CED-12 domain containing 3 | 416 | 324 | 325 | 591 | 576 | 429 | 502 | 626 | 495 | 790 | 328 |
| 481 | 0.22 | 0.17 | 1.27 | 0.20 | 1 | 2310039H08Ril | RIKEN cDNA 2310039H08 gene | 364 | 474 | 409 | 499 | 475 | 375 | 502 | 660 | 505 | 644 | 381 |
| 643 | 0.09 | 0.19 | 0.47 | 0.64 | 1 | Zcchc9 | zinc finger, CCHC domain containing 9 | 593 | 738 | 459 | 548 | 655 | 868 | 501 | 697 | 639 | 572 | 805 |
| 643 | 0.09 | 0.16 | 0.58 | 0.56 | 1 | Zfp281 | zinc finger protein 281 | 747 | 813 | 526 | 625 | 550 | 742 | 501 | 674 | 551 | 608 | 737 |
| 471 | 0.18 | 0.23 | 0.80 | 0.42 | 1 | Nfkbil1 | nuclear factor of kappa light polypeptide g | 438 | 516 | 565 | 347 | 461 | 398 | 501 | 655 | 450 | 447 | 407 |
| 409 | -0.23 | 0.24 | -0.96 | 0.34 | 1 | Rsph9 | radial spoke head 9 homolog (Chlamydor | 382 | 255 | 380 | 527 | 500 | 346 | 501 | 549 | 436 | 337 | 286 |
| 409 | 0.09 | 0.24 | 0.36 | 0.72 | 1 | Paqr9 | progestin and adipoQ receptor family mer | 288 | 412 | 287 | 408 | 400 | 370 | 501 | 642 | 437 | 314 | 438 |
| 405 | -0.01 | 0.24 | -0.06 | 0.95 | 1 | Pmm1 | phosphomannomutase 1 | 540 | 403 | 599 | 294 | 460 | 240 | 501 | 425 | 356 | 344 | 294 |
| 229 | -0.23 | 0.28 | -0.80 | 0.42 | 1 | 4931431B13Rik | | 0 | 1 | 0 | 134 | 292 | 384 | 501 | 86 | 228 | 514 | 384 |
| 1199 | 0.09 | 0.17 | 0.52 | 0.61 | 1 | Arpc1b | actin related protein 2/3 complex, subunit | 3954 | 2244 | 2657 | 526 | 532 | 509 | 500 | 444 | 469 | 750 | 602 |
| 931 | 0.19 | 0.20 | 0.93 | 0.35 | 1 | Zfc3h1 | zinc finger, C3H1-type containing | 688 | 434 | 321 | 538 | 529 | 2747 | 500 | 711 | 549 | 693 | 2530 |
| 612 | 0.21 | 0.18 | 1.16 | 0.25 | 1 | Sf3b5 | splicing factor 3b, subunit 5 | 815 | 748 | 953 | 488 | 530 | 423 | 500 | 631 | 518 | 720 | 408 |
| 588 | -0.25 | 0.24 | -1.02 | 0.31 | 1 | Plcg2 | phospholipase C, gamma 2 | 1728 | 754 | 768 | 487 | 418 | 328 | 500 | 330 | 450 | 457 | 249 |
| 471 | -0.03 | 0.16 | -0.18 | 0.86 | 1 | Tmem179b | transmembrane protein 179B | 476 | 387 | 460 | 403 | 590 | 462 | 500 | 421 | 510 | 470 | 506 |
| 446 | 0.08 | 0.23 | 0.36 | 0.72 | 1 | Neil2 | nei like 2 (E. coli) | 247 | 148 | 118 | 519 | 377 | 737 | 500 | 447 | 504 | 462 | 848 |
| 738 | 0.19 | 0.13 | 1.48 | 0.14 | 1 | Rnf169 | ring finger protein 169 | 876 | 1018 | 730 | 621 | 445 | 1024 | 499 | 656 | 523 | 634 | 1092 |
| 572 | 0.02 | 0.17 | 0.13 | 0.89 | 1 | Cwc22 | CWC22 spliceosome-associated protein | 466 | 423 | 568 | 654 | 640 | 608 | 499 | 557 | 596 | 594 | 692 |
| 462 | 0.08 | 0.17 | 0.49 | 0.63 | 1 | Dusp12 | dual specificity phosphatase 12 | 595 | 392 | 514 | 302 | 439 | 485 | 499 | 301 | 411 | 509 | 634 |
| 754 | 0.07 | 0.18 | 0.41 | 0.68 | 1 | U2af1 | U2 small nuclear ribonucleoprotein auxili | 891 | 719 | 833 | 558 | 699 | 1093 | 498 | 704 | 621 | 495 | 1187 |
| 731 | 0.01 | 0.14 | 0.08 | 0.94 | 1 | Pprc1 | peroxisome proliferative activated recepto | 670 | 882 | 711 | 914 | 625 | 850 | 498 | 904 | 653 | 529 | 802 |
| 557 | 0.01 | 0.15 | 0.07 | 0.95 | 1 | Slc7a6os | solute carrier family 7, member 6 opposite | 605 | 497 | 600 | 361 | 432 | 944 | 498 | 410 | 484 | 451 | 849 |
| 521 | -0.15 | 0.21 | -0.71 | 0.48 | 1 | Taf12 | TATA-box binding protein associated fact | 590 | 606 | 502 | 570 | 545 | 498 | 498 | 579 | 518 | 304 | 526 |
| 435 | -0.17 | 0.19 | -0.86 | 0.39 | 1 | Marchf3 | membrane associated ring-CH-type finge | 243 | 175 | 189 | 515 | 557 | 615 | 498 | 389 | 399 | 549 | 656 |
| 398 | -0.07 | 0.22 | -0.33 | 0.74 | 1 | Fam187b | family with sequence similarity 187, meml | 232 | 349 | 293 | 464 | 390 | 440 | 498 | 370 | 366 | 517 | 453 |
| 1166 | -0.14 | 0.12 | -1.11 | 0.27 | 1 | Upp1 | uridine phosphorylase 1 | 2409 | 3021 | 2938 | 579 | 671 | 588 | 497 | 524 | 517 | 461 | 623 |
| 1164 | 0.37 | 0.25 | 1.49 | 0.14 | 1 | Fam129a | family with sequence similarity 129, meml | 4062 | 2135 | 2628 | 655 | 279 | 331 | 497 | 759 | 487 | 511 | 456 |
| 511 | 0.07 | 0.24 | 0.30 | 0.76 | 1 | Fut10 | fucosyltransferase 10 | 849 | 454 | 644 | 518 | 471 | 283 | 497 | 313 | 576 | 659 | 359 |
| 499 | 0.26 | 0.23 | 1.13 | 0.26 | 1 | Fen1 | flap structure specific endonuclease 1 | 791 | 409 | 369 | 420 | 495 | 370 | 497 | 492 | 622 | 612 | 414 |
| 442 | -0.06 | 0.20 | -0.28 | 0.78 | 1 | F2rl1 | coagulation factor II (thrombin) receptor-li | 280 | 382 | 415 | 508 | 447 | 478 | 497 | 423 | 455 | 508 | 471 |
| 357 | -0.21 | 0.24 | -0.85 | 0.39 | 1 | Osbpl10 | oxysterol binding protein-like 10 | 211 | 120 | 191 | 508 | 470 | 315 | 497 | 285 | 424 | 608 | 293 |
| 1108 | -0.06 | 0.16 | -0.35 | 0.73 | 1 | Scarb1 | scavenger receptor class B, member 1 | 2436 | 3079 | 3031 | 390 | 544 | 423 | 496 | 549 | 458 | 394 | 389 |
| 598 | 0.07 | 0.21 | 0.33 | 0.74 | 1 | Zfp334 | zinc finger protein 334 | 528 | 810 | 706 | 545 | 492 | 649 | 496 | 853 | 563 | 487 | 448 |
| 585 | -0.19 | 0.24 | -0.78 | 0.43 | 1 | Pak1 | p21 (RAC1) activated kinase 1 | 952 | 927 | 760 | 626 | 579 | 292 | 496 | 744 | 516 | 290 | 256 |
| 541 | 0.18 | 0.20 | 0.88 | 0.38 | 1 | Nefh | neurofilament, heavy polypeptide | 38 | 131 | 136 | 787 | 1027 | 283 | 496 | 1081 | 1156 | 535 | 283 |
| 478 | -0.05 | 0.23 | -0.21 | 0.83 | 1 | Mplkip | M-phase specific PLK1 intereacting prote | 359 | 429 | 511 | 493 | 412 | 578 | 496 | 335 | 422 | 480 | 744 |
| 473 | -0.02 | 0.21 | -0.10 | 0.92 | 1 | Agbl3 | ATP/GTP binding protein-like 3 | 335 | 406 | 403 | 478 | 509 | 547 | 496 | 571 | 335 | 528 | 594 |
| 466 | 0.05 | 0.22 | 0.23 | 0.82 | 1 | Zfp87 | zinc finger protein 87 | 422 | 308 | 413 | 476 | 490 | 494 | 496 | 618 | 490 | 386 | 537 |
| 460 | -0.03 | 0.15 | -0.18 | 0.86 | 1 | Tmem117 | transmembrane protein 117 | 266 | 303 | 283 | 576 | 568 | 486 | 496 | 542 | 564 | 459 | 521 |
| 615 | -0.04 | 0.16 | -0.27 | 0.78 | 1 | Adgra3 | adhesion G protein-coupled receptor A3 | 935 | 813 | 933 | 626 | 583 | 358 | 495 | 586 | 603 | 517 | 316 |
| 594 | 0.19 | 0.17 | 1.10 | 0.27 | 1 | Zfp830 | zinc finger protein 830 | 577 | 508 | 705 | 531 | 526 | 668 | 495 | 641 | 578 | 662 | 638 |
| 584 | 0.22 | 0.19 | 1.21 | 0.23 | 1 | Rad50 | RAD50 double strand break repair protei | 820 | 550 | 614 | 581 | 396 | 573 | 495 | 624 | 490 | 678 | 598 |
| 476 | -0.34 | 0.28 | -1.23 | 0.22 | 1 | Fbxo46 | F-box protein 46 | 758 | 481 | 685 | 460 | 475 | 418 | 494 | 488 | 383 | 225 | 371 |
| 451 | -0.31 | 0.27 | -1.16 | 0.25 | 1 | Pola1 | polymerase (DNA directed), alpha 1 | 1087 | 703 | 603 | 321 | 287 | 341 | 494 | 317 | 296 | 201 | 308 |
| 432 | -0.21 | 0.21 | -0.98 | 0.33 | 1 | Pxmp4 | peroxisomal membrane protein 4 | 504 | 413 | 484 | 427 | 486 | 375 | 494 | 483 | 495 | 322 | 268 |
| 421 | 0.00 | 0.25 | 0.00 | 1.00 | 1 | Thap6 | | 232 | 331 | 394 | 403 | 390 | 520 | 494 | 490 | 519 | 291 | 566 |
| 410 | 0.07 | 0.19 | 0.37 | 0.71 | 1 | Tmem42 | transmembrane protein 42 | 189 | 277 | 257 | 506 | 516 | 331 | 494 | 621 | 472 | 443 | 401 |
| 734 | -0.30 | 0.17 | -1.74 | 0.08 | 1 | Sh2b3 | SH2B adaptor protein 3 | 1723 | 1428 | 1433 | 543 | 419 | 468 | 493 | 476 | 353 | 327 | 411 |
| 580 | -0.34 | 0.18 | -1.84 | 0.07 | 1 | Mov10 | Mov10 RISC complex RNA helicase | 837 | 995 | 956 | 560 | 505 | 444 | 493 | 372 | 352 | 513 | 352 |
| 560 | 0.14 | 0.18 | 0.75 | 0.46 | 1 | Pus3 | pseudouridine synthase 3 | 446 | 526 | 390 | 514 | 554 | 740 | 493 | 621 | 603 | 654 | 621 |
| 492 | 0.06 | 0.18 | 0.35 | 0.73 | 1 | Mrpl46 | mitochondrial ribosomal protein L46 | 438 | 462 | 455 | 451 | 545 | 493 | 493 | 490 | 478 | 641 | 467 |
| 470 | 0.14 | 0.29 | 0.49 | 0.63 | 1 | Mtg2 | mitochondrial ribosome associated GTPa | 407 | 363 | 484 | 284 | 445 | 666 | 493 | 665 | 364 | 416 | 586 |
| 429 | -0.28 | 0.25 | -1.11 | 0.27 | 1 | Mfap1b | microfibrillar-associated protein 1B | 258 | 440 | 429 | 515 | 463 | 487 | 493 | 283 | 445 | 395 | 511 |
| 1139 | -0.09 | 0.15 | -0.57 | 0.57 | 1 | Socs3 | suppressor of cytokine signaling 3 | 4236 | 1756 | 2200 | 271 | 394 | 1077 | 492 | 224 | 384 | 549 | 945 |
| 663 | 0.08 | 0.13 | 0.60 | 0.55 | 1 | Parp11 | poly (ADP-ribose) polymerase family, mer | 691 | 947 | 794 | 679 | 546 | 651 | 492 | 701 | 534 | 581 | 680 |
| 521 | 0.12 | 0.20 | 0.61 | 0.54 | 1 | Ccdc32 | coiled-coil domain containing 32 | 635 | 619 | 457 | 596 | 431 | 378 | 492 | 520 | 426 | 802 | 371 |
| 369 | 0.16 | 0.28 | 0.60 | 0.55 | 1 | Arhgap36 | Rho GTPase activating protein 36 | 297 | 189 | 164 | 185 | 412 | 533 | 492 | 300 | 424 | 365 | 699 |
| 579 | 0.29 | 0.16 | 1.84 | 0.07 | 1 | Mcm9 | minichromosome maintenance 9 homolog | 581 | 530 | 448 | 541 | 526 | 598 | 491 | 715 | 530 | 715 | 698 |

| | | | | | | | | | | | | | | | | | | |
|---|---|---|---|---|---|---|---|---|---|---|---|---|---|---|---|---|---|---|
| 545 | -0.09 | 0.18 | -0.49 | 0.63 | 1 | Pot1a | protection of telomeres 1A | 684 | 465 | 550 | 592 | 549 | 573 | 491 | 475 | 466 | 593 | 556 |
| 479 | 0.03 | 0.17 | 0.17 | 0.86 | 1 | Erh | ERH mRNA splicing and mitosis factor | 465 | 372 | 559 | 361 | 476 | 604 | 491 | 438 | 513 | 480 | 515 |
| 705 | -0.13 | 0.17 | -0.75 | 0.45 | 1 | Snhg8 | | 1133 | 1341 | 1531 | 435 | 434 | 609 | 490 | 466 | 418 | 450 | 453 |
| 553 | -0.09 | 0.15 | -0.63 | 0.53 | 1 | Tmem67 | transmembrane protein 67 | 516 | 519 | 564 | 553 | 668 | 604 | 490 | 455 | 647 | 495 | 576 |
| 544 | -0.08 | 0.17 | -0.44 | 0.66 | 1 | Chchd6 | coiled-coil-helix-coiled-coil-helix domain | 502 | 598 | 528 | 621 | 673 | 435 | 490 | 754 | 530 | 465 | 387 |
| 490 | 0.11 | 0.17 | 0.63 | 0.53 | 1 | 0610012G03Ril | RIKEN cDNA 0610012G03 gene | 446 | 454 | 456 | 507 | 586 | 366 | 490 | 491 | 529 | 623 | 441 |
| 480 | 0.26 | 0.20 | 1.29 | 0.20 | 1 | Fam118b | family with sequence similarity 118, meml | 383 | 384 | 321 | 437 | 392 | 592 | 490 | 690 | 568 | 446 | 577 |
| 460 | -0.15 | 0.17 | -0.89 | 0.37 | 1 | Zkscan5 | zinc finger with KRAB and SCAN domains | 414 | 426 | 547 | 460 | 484 | 493 | 490 | 374 | 453 | 478 | 438 |
| 622 | -0.05 | 0.19 | -0.27 | 0.78 | 1 | Nfatc1 | nuclear factor of activated T cells, cytopla: | 1121 | 794 | 717 | 523 | 551 | 573 | 489 | 516 | 546 | 586 | 430 |
| 575 | 0.01 | 0.16 | 0.09 | 0.93 | 1 | Psmd10 | proteasome (prosome, macropain) 26S su | 742 | 732 | 865 | 509 | 501 | 478 | 489 | 521 | 599 | 388 | 506 |
| 489 | 0.03 | 0.16 | 0.19 | 0.85 | 1 | Tbp | TATA box binding protein | 392 | 380 | 374 | 413 | 394 | 820 | 489 | 431 | 443 | 518 | 721 |
| 438 | 0.15 | 0.19 | 0.81 | 0.42 | 1 | Iba57 | IBA57 homolog, iron-sulfur cluster assem | 263 | 332 | 382 | 494 | 468 | 369 | 489 | 590 | 468 | 501 | 459 |
| 387 | 0.17 | 0.32 | 0.54 | 0.59 | 1 | Ormdl1 | ORM1-like 1 (S. cerevisiae) | 336 | 428 | 293 | 291 | 393 | 304 | 489 | 664 | 462 | 251 | 350 |
| 605 | 0.20 | 0.17 | 1.19 | 0.24 | 1 | Nxn | nucleoredoxin | 1059 | 788 | 812 | 572 | 497 | 308 | 488 | 580 | 512 | 662 | 381 |
| 502 | 0.00 | 0.19 | 0.02 | 0.99 | 1 | Mrpl21 | mitochondrial ribosomal protein L21 | 423 | 625 | 627 | 515 | 473 | 439 | 488 | 569 | 426 | 402 | 541 |
| 484 | 0.13 | 0.23 | 0.56 | 0.57 | 1 | Tmem25 | transmembrane protein 25 | 247 | 406 | 459 | 468 | 520 | 544 | 488 | 492 | 507 | 613 | 584 |
| 473 | -0.35 | 0.24 | -1.47 | 0.14 | 1 | Rbbp8 | retinoblastoma binding protein 8, endonu | 573 | 355 | 549 | 577 | 454 | 539 | 488 | 276 | 425 | 416 | 553 |
| 429 | 0.08 | 0.27 | 0.28 | 0.78 | 1 | Prdm10 | PR domain containing 10 | 282 | 365 | 294 | 464 | 383 | 494 | 488 | 276 | 499 | 614 | 566 |
| 413 | 0.05 | 0.24 | 0.20 | 0.84 | 1 | Gm7160 | | 138 | 212 | 238 | 463 | 367 | 606 | 488 | 516 | 353 | 438 | 724 |
| 662 | -0.11 | 0.13 | -0.82 | 0.41 | 1 | Cnksr3 | Cnksr family member 3 | 883 | 1228 | 927 | 600 | 567 | 544 | 487 | 612 | 489 | 466 | 478 |
| 624 | -0.09 | 0.18 | -0.48 | 0.63 | 1 | Alkbh1 | alkB homolog 1, histone H2A dioxygenas | 479 | 654 | 584 | 527 | 395 | 1150 | 487 | 562 | 355 | 339 | 1330 |
| 542 | -0.39 | 0.22 | -1.75 | 0.08 | 1 | Spsb1 | splA/ryanodine receptor domain and SOC | 1126 | 954 | 1040 | 335 | 416 | 336 | 487 | 306 | 421 | 386 | 152 |
| 508 | 0.09 | 0.18 | 0.50 | 0.62 | 1 | Foxa3 | forkhead box A3 | 440 | 653 | 658 | 531 | 539 | 300 | 487 | 642 | 479 | 529 | 327 |
| 504 | 0.10 | 0.22 | 0.43 | 0.66 | 1 | Pgm2l1 | phosphoglucomutase 2-like 1 | 863 | 763 | 644 | 357 | 444 | 282 | 487 | 619 | 354 | 419 | 308 |
| 492 | -0.01 | 0.17 | -0.06 | 0.95 | 1 | Gtf2h2 | general transcription factor II H, polypepti | 425 | 481 | 514 | 407 | 410 | 693 | 487 | 388 | 421 | 473 | 711 |
| 481 | 0.23 | 0.22 | 1.04 | 0.30 | 1 | Tmem205 | transmembrane protein 205 | 446 | 400 | 459 | 438 | 469 | 419 | 487 | 777 | 439 | 537 | 416 |
| 370 | -0.04 | 0.22 | -0.20 | 0.84 | 1 | Chrd | chordin | 389 | 289 | 267 | 340 | 441 | 309 | 487 | 492 | 326 | 475 | 260 |
| 695 | 0.17 | 0.15 | 1.11 | 0.27 | 1 | Gpr137 | G protein-coupled receptor 137 | 535 | 612 | 685 | 571 | 646 | 1036 | 486 | 644 | 585 | 652 | 1190 |
| 626 | 0.24 | 0.20 | 1.20 | 0.23 | 1 | Snrpe | small nuclear ribonucleoprotein E | 1046 | 736 | 850 | 532 | 483 | 447 | 486 | 747 | 496 | 524 | 537 |
| 605 | -0.02 | 0.22 | -0.10 | 0.92 | 1 | Ptpn12 | protein tyrosine phosphatase, non-recept | 927 | 906 | 692 | 625 | 359 | 624 | 486 | 399 | 543 | 433 | 667 |
| 594 | -0.31 | 0.18 | -1.73 | 0.08 | 1 | Trmt9b | tRNA methyltransferase 9B | 546 | 706 | 676 | 672 | 564 | 835 | 486 | 402 | 473 | 501 | 677 |
| 647 | -0.20 | 0.26 | -0.79 | 0.43 | 1 | Casp8ap2 | caspase 8 associated protein 2 | 782 | 643 | 605 | 706 | 592 | 891 | 485 | 331 | 715 | 448 | 922 |
| 523 | 0.27 | 0.25 | 1.10 | 0.27 | 1 | Asphd1 | aspartate beta-hydroxylase domain conta | 336 | 409 | 424 | 435 | 400 | 749 | 485 | 905 | 482 | 428 | 703 |
| 521 | -0.05 | 0.22 | -0.23 | 0.82 | 1 | Mtmr14 | myotubularin related protein 14 | 845 | 600 | 469 | 400 | 567 | 489 | 485 | 383 | 508 | 473 | 514 |
| 415 | -0.24 | 0.19 | -1.24 | 0.21 | 1 | Rsad1 | radical S-adenosyl methionine domain cc | 376 | 372 | 499 | 429 | 400 | 475 | 485 | 327 | 379 | 374 | 450 |
| 365 | -0.14 | 0.29 | -0.47 | 0.64 | 1 | B3glct | beta-3-glucosyltransferase | 398 | 318 | 259 | 403 | 425 | 284 | 485 | 345 | 546 | 256 | 300 |
| 718 | -0.27 | 0.14 | -1.92 | 0.06 | 1 | Ppm1f | protein phosphatase 1F (PP2C domain cc | 1791 | 1664 | 1432 | 382 | 370 | 414 | 484 | 371 | 294 | 340 | 359 |
| 532 | 0.03 | 0.19 | 0.15 | 0.88 | 1 | Mrpl18 | mitochondrial ribosomal protein L18 | 785 | 544 | 548 | 525 | 498 | 462 | 484 | 576 | 530 | 465 | 441 |
| 519 | -0.22 | 0.22 | -1.00 | 0.32 | 1 | Sdhaf1 | succinate dehydrogenase complex asser | 445 | 460 | 533 | 640 | 733 | 436 | 484 | 336 | 640 | 582 | 414 |
| 509 | 0.17 | 0.23 | 0.73 | 0.46 | 1 | 3110040N11Ril | RIKEN cDNA 3110040N11 gene | 537 | 512 | 554 | 384 | 522 | 485 | 484 | 728 | 443 | 501 | 453 |
| 711 | 0.17 | 0.18 | 0.94 | 0.35 | 1 | Gas2l1 | growth arrest-specific 2 like 1 | 921 | 1014 | 954 | 573 | 669 | 593 | 483 | 637 | 613 | 800 | 564 |
| 665 | 0.04 | 0.14 | 0.29 | 0.77 | 1 | C2cd2 | C2 calcium-dependent domain containing | 1197 | 1134 | 1087 | 543 | 471 | 421 | 483 | 562 | 518 | 480 | 414 |
| 466 | -0.18 | 0.23 | -0.76 | 0.45 | 1 | Snapc2 | small nuclear RNA activating complex, pc | 583 | 519 | 508 | 481 | 538 | 347 | 483 | 630 | 429 | 313 | 298 |
| 397 | 0.14 | 0.33 | 0.43 | 0.66 | 1 | Gm281 | predicted gene 281 | 30 | 3 | 25 | 381 | 435 | 704 | 483 | 355 | 434 | 662 | 853 |
| 590 | -0.33 | 0.22 | -1.51 | 0.13 | 1 | Ptcd1 | pentatricopeptide repeat domain 1 | 715 | 740 | 625 | 708 | 601 | 664 | 482 | 346 | 578 | 492 | 534 |
| 479 | -0.01 | 0.19 | -0.04 | 0.97 | 1 | Cbwd1 | COBW domain containing 1 | 529 | 583 | 565 | 413 | 439 | 457 | 482 | 328 | 472 | 519 | 476 |
| 473 | -0.19 | 0.25 | -0.78 | 0.44 | 1 | Malt1 | MALT1 paracaspase | 731 | 594 | 410 | 496 | 351 | 519 | 482 | 271 | 427 | 377 | 547 |
| 448 | 0.25 | 0.24 | 1.03 | 0.30 | 1 | Mrpl34 | mitochondrial ribosomal protein L34 | 440 | 405 | 506 | 398 | 434 | 298 | 482 | 723 | 370 | 551 | 318 |
| 444 | 0.09 | 0.18 | 0.50 | 0.62 | 1 | Acyp1 | acylphosphatase 1, erythrocyte (common | 391 | 361 | 467 | 464 | 411 | 417 | 482 | 514 | 437 | 509 | 430 |
| 335 | -0.46 | 0.25 | -1.83 | 0.07 | 1 | Rab9b | RAB9B, member RAS oncogene family | 223 | 293 | 349 | 392 | 490 | 283 | 482 | 232 | 317 | 313 | 315 |
| 656 | -0.21 | 0.16 | -1.37 | 0.17 | 1 | Zfp868 | zinc finger protein 868 | 631 | 695 | 833 | 516 | 683 | 993 | 481 | 495 | 547 | 335 | 1009 |
| 563 | -0.21 | 0.24 | -0.85 | 0.39 | 1 | Map3k21 | mitogen-activated protein kinase kinase k | 997 | 871 | 902 | 360 | 453 | 511 | 481 | 191 | 527 | 466 | 436 |
| 555 | 0.10 | 0.25 | 0.39 | 0.69 | 1 | Glt1d1 | glycosyltransferase 1 domain containing | 90 | 190 | 302 | 554 | 527 | 1102 | 481 | 607 | 682 | 465 | 1104 |
| 489 | 0.25 | 0.21 | 1.18 | 0.24 | 1 | Nrtn | neurturin | 166 | 164 | 163 | 669 | 662 | 399 | 481 | 1008 | 558 | 654 | 450 |
| 472 | 0.19 | 0.20 | 0.98 | 0.33 | 1 | Nr2c2ap | nuclear receptor 2C2-associated protein | 558 | 438 | 381 | 392 | 455 | 451 | 481 | 611 | 471 | 464 | 489 |
| 399 | -0.16 | 0.23 | -0.72 | 0.47 | 1 | Susd4 | sushi domain containing 4 | 309 | 290 | 161 | 471 | 421 | 552 | 481 | 409 | 443 | 425 | 425 |
| 396 | -0.27 | 0.28 | -0.97 | 0.33 | 1 | Pcdhb14 | protocadherin beta 14 | 445 | 371 | 270 | 526 | 401 | 375 | 481 | 232 | 476 | 433 | 343 |
| 512 | 0.16 | 0.19 | 0.86 | 0.39 | 1 | Camkmt | calmodulin-lysine N-methyltransferase | 371 | 552 | 448 | 534 | 564 | 436 | 480 | 509 | 621 | 639 | 480 |
| 435 | -0.08 | 0.20 | -0.40 | 0.69 | 1 | D130043K22Ri | RIKEN cDNA D130043K22 gene | 130 | 188 | 195 | 689 | 443 | 571 | 480 | 594 | 529 | 362 | 600 |
| 447 | -0.28 | 0.28 | -1.01 | 0.31 | 1 | Mansc1 | MANSC domain containing 1 | 539 | 715 | 584 | 628 | 333 | 283 | 479 | 246 | 375 | 424 | 313 |

| | | | | | | | | | | | | | | | | | | |
|---|---|---|---|---|---|---|---|---|---|---|---|---|---|---|---|---|---|---|
| 414 | 0.02 | 0.20 | 0.09 | 0.93 | 1 | Ap1s3 | adaptor-related protein complex AP-1, sig | 365 | 346 | 407 | 445 | 351 | 428 | 479 | 378 | 422 | 553 | 383 |
| 702 | -0.04 | 0.16 | -0.25 | 0.80 | 1 | Elk4 | ELK4, member of ETS oncogene family | 908 | 892 | 922 | 481 | 590 | 984 | 478 | 569 | 432 | 506 | 959 |
| 456 | -0.32 | 0.25 | -1.31 | 0.19 | 1 | Mettl27 | methyltransferase like 27 | 561 | 966 | 552 | 450 | 332 | 363 | 478 | 392 | 354 | 219 | 344 |
| 444 | -0.04 | 0.20 | -0.22 | 0.82 | 1 | Thap1 | THAP domain containing, apoptosis asso | 371 | 419 | 431 | 464 | 534 | 379 | 478 | 525 | 523 | 323 | 439 |
| 371 | 0.02 | 0.26 | 0.07 | 0.95 | 1 | Sult1c2 | sulfotransferase family, cytosolic, 1C, mer | 104 | 248 | 167 | 555 | 403 | 311 | 478 | 600 | 387 | 576 | 253 |
| 307 | -0.45 | 0.28 | -1.60 | 0.11 | 1 | Arxes1 | adipocyte-related X-chromosome express | 109 | 253 | 206 | 474 | 392 | 267 | 478 | 358 | 379 | 222 | 241 |
| 537 | 0.10 | 0.16 | 0.67 | 0.50 | 1 | Smpd4 | sphingomyelin phosphodiesterase 4 | 546 | 527 | 474 | 434 | 437 | 778 | 477 | 548 | 464 | 541 | 686 |
| 535 | 0.04 | 0.22 | 0.17 | 0.86 | 1 | Fdx1 | ferredoxin 1 | 502 | 793 | 637 | 518 | 420 | 542 | 477 | 395 | 542 | 569 | 491 |
| 529 | 0.29 | 0.17 | 1.70 | 0.09 | 1 | Dnajc4 | DnaJ heat shock protein family (Hsp40) m | 408 | 551 | 470 | 474 | 546 | 467 | 477 | 758 | 571 | 569 | 525 |
| 434 | -0.07 | 0.21 | -0.33 | 0.74 | 1 | Msto1 | misato 1, mitochondrial distribution and m | 500 | 480 | 619 | 441 | 390 | 318 | 477 | 497 | 319 | 413 | 319 |
| 430 | 0.10 | 0.22 | 0.46 | 0.65 | 1 | Hemk1 | HemK methyltransferase family member 1 | 263 | 213 | 124 | 471 | 254 | 775 | 477 | 660 | 209 | 621 | 658 |
| 426 | 0.16 | 0.24 | 0.68 | 0.50 | 1 | Nr1h4 | nuclear receptor subfamily 1, group H, me | 277 | 296 | 399 | 472 | 327 | 477 | 477 | 549 | 360 | 531 | 524 |
| 516 | 0.02 | 0.16 | 0.11 | 0.91 | 1 | Ythdc2 | YTH domain containing 2 | 536 | 357 | 385 | 450 | 466 | 794 | 476 | 417 | 493 | 529 | 772 |
| 510 | -0.16 | 0.23 | -0.69 | 0.49 | 1 | Pwwp2a | PWWP domain containing 2A | 462 | 483 | 554 | 508 | 426 | 713 | 476 | 528 | 450 | 246 | 764 |
| 426 | -0.01 | 0.18 | -0.08 | 0.94 | 1 | Kbtbd3 | kelch repeat and BTB (POZ) domain conta | 417 | 284 | 303 | 418 | 335 | 599 | 476 | 312 | 346 | 560 | 635 |
| 725 | 0.00 | 0.25 | 0.00 | 1.00 | 1 | Nefm | neurofilament, medium polypeptide | 1294 | 1847 | 1723 | 216 | 471 | 407 | 475 | 306 | 467 | 415 | 359 |
| 557 | 0.08 | 0.16 | 0.48 | 0.63 | 1 | Tmem80 | transmembrane protein 80 | 388 | 538 | 449 | 627 | 508 | 702 | 475 | 652 | 450 | 586 | 755 |
| 482 | -0.22 | 0.23 | -0.93 | 0.35 | 1 | 2310015A10Rik | | 354 | 757 | 424 | 557 | 471 | 525 | 475 | 418 | 504 | 410 | 402 |
| 434 | 0.21 | 0.28 | 0.77 | 0.44 | 1 | Tchp | trichoplein, keratin filament binding | 343 | 269 | 335 | 326 | 318 | 679 | 475 | 678 | 338 | 451 | 558 |
| 426 | -0.02 | 0.19 | -0.10 | 0.92 | 1 | Cdkal1 | CDK5 regulatory subunit associated prote | 365 | 308 | 445 | 418 | 456 | 443 | 475 | 439 | 465 | 352 | 524 |
| 289 | 0.03 | 0.28 | 0.10 | 0.92 | 1 | AW011738 | | 225 | 260 | 269 | 261 | 308 | 137 | 475 | 248 | 478 | 403 | 118 |
| 640 | 0.03 | 0.21 | 0.13 | 0.89 | 1 | Poglut3 | protein O-glucosyltransferase 3 | 1426 | 994 | 909 | 383 | 527 | 458 | 474 | 454 | 498 | 393 | 524 |
| 574 | -0.26 | 0.24 | -1.10 | 0.27 | 1 | Cyb561a3 | cytochrome b561 family, member A3 | 1005 | 582 | 743 | 610 | 476 | 593 | 474 | 313 | 548 | 471 | 500 |
| 552 | 0.04 | 0.22 | 0.16 | 0.87 | 1 | Gtf3c2 | general transcription factor IIIC, polypepti | 456 | 337 | 234 | 561 | 524 | 936 | 474 | 455 | 573 | 605 | 921 |
| 435 | -0.09 | 0.23 | -0.40 | 0.69 | 1 | Cbr4 | carbonyl reductase 4 | 311 | 422 | 424 | 471 | 409 | 508 | 474 | 304 | 491 | 468 | 500 |
| 380 | 0.09 | 0.24 | 0.39 | 0.70 | 1 | 1700052K11Rik | | 352 | 405 | 260 | 471 | 284 | 309 | 474 | 327 | 372 | 599 | 331 |
| 526 | -0.01 | 0.15 | -0.07 | 0.95 | 1 | Gm50240 | | 381 | 563 | 438 | 709 | 480 | 553 | 473 | 620 | 510 | 448 | 611 |
| 525 | 0.09 | 0.21 | 0.41 | 0.68 | 1 | Slc35f6 | solute carrier family 35, member F6 | 659 | 607 | 655 | 465 | 455 | 466 | 473 | 644 | 546 | 345 | 463 |
| 475 | -0.01 | 0.18 | -0.03 | 0.98 | 1 | Vps33b | vacuolar protein sorting 33B | 345 | 292 | 407 | 403 | 444 | 774 | 473 | 447 | 513 | 368 | 758 |
| 461 | -0.05 | 0.19 | -0.24 | 0.81 | 1 | Med6 | mediator complex subunit 6 | 456 | 571 | 496 | 392 | 414 | 513 | 473 | 489 | 417 | 322 | 527 |
| 404 | 0.12 | 0.20 | 0.62 | 0.54 | 1 | D630024D03Rik | | 100 | 145 | 117 | 398 | 471 | 603 | 473 | 560 | 463 | 429 | 686 |
| 734 | -0.08 | 0.21 | -0.38 | 0.71 | 1 | Dusp16 | dual specificity phosphatase 16 | 1096 | 2218 | 1104 | 646 | 351 | 434 | 472 | 460 | 407 | 515 | 372 |
| 578 | 0.21 | 0.26 | 0.81 | 0.42 | 1 | Mcat | malonyl CoA:ACP acyltransferase (mitoch | 472 | 619 | 588 | 478 | 607 | 585 | 472 | 546 | 521 | 974 | 492 |
| 562 | 0.02 | 0.23 | 0.09 | 0.93 | 1 | Ube2s | ubiquitin-conjugating enzyme E2S | 572 | 443 | 633 | 562 | 588 | 609 | 472 | 719 | 472 | 381 | 727 |
| 545 | 0.10 | 0.21 | 0.50 | 0.62 | 1 | Slx4ip | SLX4 interacting protein | 478 | 686 | 548 | 437 | 583 | 578 | 472 | 450 | 555 | 623 | 586 |
| 520 | 0.15 | 0.19 | 0.78 | 0.43 | 1 | Katnb1 | katanin p80 (WD40-containing) subunit B | 703 | 676 | 653 | 322 | 463 | 501 | 472 | 431 | 507 | 567 | 432 |
| 465 | 0.15 | 0.18 | 0.82 | 0.41 | 1 | Ap5s1 | adaptor-related protein 5 complex, sigma | 395 | 429 | 462 | 389 | 502 | 451 | 472 | 584 | 556 | 425 | 455 |
| 459 | -0.27 | 0.17 | -1.58 | 0.11 | 1 | Kctd15 | potassium channel tetramerisation domai | 362 | 382 | 273 | 420 | 373 | 956 | 472 | 324 | 351 | 394 | 739 |
| 413 | -0.11 | 0.21 | -0.50 | 0.62 | 1 | Mapk7 | mitogen-activated protein kinase 7 | 624 | 566 | 500 | 375 | 282 | 347 | 472 | 241 | 338 | 462 | 332 |
| 381 | 0.03 | 0.26 | 0.11 | 0.91 | 1 | Ppp1r3d | protein phosphatase 1, regulatory subuni | 212 | 96 | 100 | 428 | 553 | 407 | 472 | 523 | 414 | 571 | 411 |
| 307 | -0.01 | 0.27 | -0.03 | 0.98 | 1 | Fancf | Fanconi anemia, complementation group | 168 | 91 | 200 | 351 | 319 | 321 | 472 | 370 | 332 | 451 | 302 |
| 636 | 0.15 | 0.20 | 0.76 | 0.44 | 1 | Mrps30 | mitochondrial ribosomal protein S30 | 689 | 611 | 883 | 667 | 589 | 559 | 471 | 710 | 562 | 689 | 570 |
| 563 | 0.16 | 0.22 | 0.73 | 0.46 | 1 | Nsmce1 | NSE1 homolog, SMC5-SMC6 complex cc | 602 | 709 | 775 | 629 | 356 | 467 | 471 | 772 | 485 | 533 | 391 |
| 456 | -0.15 | 0.22 | -0.67 | 0.50 | 1 | Slirp | SRA stem-loop interacting RNA binding p | 512 | 389 | 511 | 543 | 324 | 567 | 471 | 418 | 408 | 450 | 422 |
| 437 | 0.12 | 0.21 | 0.55 | 0.58 | 1 | Tmub1 | transmembrane and ubiquitin-like domair | 347 | 399 | 503 | 384 | 490 | 361 | 471 | 504 | 431 | 576 | 345 |
| 325 | 0.00 | 0.23 | -0.01 | 0.99 | 1 | Mfsd6l | major facilitator superfamily domain conta | 117 | 161 | 145 | 393 | 375 | 314 | 471 | 572 | 382 | 397 | 252 |
| 653 | 0.03 | 0.14 | 0.26 | 0.80 | 1 | Tspan14 | tetraspanin 14 | 1347 | 1108 | 1229 | 455 | 474 | 333 | 470 | 431 | 479 | 478 | 378 |
| 550 | -0.05 | 0.16 | -0.35 | 0.72 | 1 | Ccdc136 | coiled-coil domain containing 136 | 638 | 740 | 603 | 504 | 520 | 576 | 470 | 538 | 529 | 410 | 519 |
| 500 | 0.10 | 0.21 | 0.46 | 0.65 | 1 | Sirt3 | sirtuin 3 | 368 | 488 | 593 | 476 | 423 | 586 | 470 | 621 | 409 | 478 | 590 |
| 478 | 0.04 | 0.18 | 0.24 | 0.81 | 1 | Dyrk1b | dual-specificity tyrosine-(Y)-phosphorylati | 408 | 326 | 382 | 542 | 482 | 543 | 470 | 542 | 457 | 596 | 509 |
| 478 | 0.18 | 0.24 | 0.75 | 0.46 | 1 | Xkr7 | X-linked Kx blood group related 7 | 349 | 626 | 483 | 487 | 351 | 456 | 470 | 599 | 462 | 622 | 349 |
| 300 | -0.15 | 0.38 | -0.39 | 0.70 | 1 | Gm13736 | | 279 | 3 | 301 | 21 | 458 | 303 | 470 | 5 | 450 | 618 | 393 |
| 1278 | 0.03 | 0.16 | 0.17 | 0.87 | 1 | Amotl2 | angiomotin-like 2 | 3706 | 2897 | 3082 | 549 | 611 | 517 | 469 | 590 | 566 | 667 | 401 |
| 825 | 0.20 | 0.26 | 0.78 | 0.43 | 1 | Cebpb | CCAAT/enhancer binding protein (C/EBP | 2581 | 1003 | 1170 | 459 | 529 | 522 | 469 | 674 | 442 | 741 | 487 |
| 811 | -0.02 | 0.15 | -0.14 | 0.89 | 1 | Trp53 | transformation related protein 53 | 1453 | 1260 | 1584 | 461 | 654 | 766 | 469 | 541 | 586 | 509 | 640 |
| 580 | -0.26 | 0.20 | -1.31 | 0.19 | 1 | Per3 | period circadian clock 3 | 346 | 446 | 466 | 349 | 381 | 1530 | 469 | 284 | 397 | 295 | 1418 |
| 570 | 0.22 | 0.17 | 1.30 | 0.19 | 1 | Gins4 | GINS complex subunit 4 (Sld5 homolog) | 802 | 618 | 958 | 382 | 542 | 398 | 469 | 447 | 527 | 693 | 433 |
| 516 | 0.20 | 0.21 | 0.98 | 0.33 | 1 | Cluap1 | clusterin associated protein 1 | 614 | 496 | 518 | 500 | 435 | 470 | 469 | 679 | 520 | 392 | 582 |
| 497 | -0.06 | 0.17 | -0.38 | 0.71 | 1 | Zfp709 | zinc finger protein 709 | 529 | 450 | 447 | 578 | 490 | 525 | 469 | 462 | 435 | 562 | 522 |
| 435 | -0.30 | 0.22 | -1.39 | 0.16 | 1 | Gpt2 | glutamic pyruvate transaminase (alanine | 524 | 599 | 816 | 309 | 396 | 389 | 469 | 301 | 418 | 281 | 281 |

| | | | | | | | | | | | | | | | | | | |
|---|---|---|---|---|---|---|---|---|---|---|---|---|---|---|---|---|---|---|
| 374 | 0.43 | 0.38 | 1.15 | 0.25 | 1 | Eef1akmt3 | EEF1A lysine methyltransferase 3 | 30 | 9 | 92 | 326 | 568 | 356 | 469 | 480 | 633 | 729 | 428 |
| 834 | -0.09 | 0.18 | -0.54 | 0.59 | 1 | Myl12a | myosin, light chain 12A, regulatory, non-s | 1905 | 1650 | 1898 | 518 | 415 | 510 | 468 | 358 | 455 | 577 | 420 |
| 661 | -0.29 | 0.19 | -1.56 | 0.12 | 1 | Pmepa1 | prostate transmembrane protein, androge | 1832 | 1071 | 1687 | 343 | 404 | 267 | 468 | 343 | 323 | 323 | 211 |
| 619 | 0.08 | 0.15 | 0.51 | 0.61 | 1 | Prkcd | protein kinase C, delta | 1212 | 846 | 889 | 449 | 408 | 559 | 468 | 436 | 465 | 527 | 556 |
| 446 | -0.01 | 0.20 | -0.03 | 0.97 | 1 | Amdhd2 | amidohydrolase domain containing 2 | 386 | 403 | 427 | 501 | 413 | 460 | 468 | 591 | 354 | 505 | 403 |
| 434 | -0.01 | 0.29 | -0.05 | 0.96 | 1 | Arhgdig | Rho GDP dissociation inhibitor (GDI) gam | 149 | 344 | 442 | 635 | 572 | 277 | 468 | 681 | 401 | 489 | 316 |
| 421 | -0.19 | 0.20 | -0.94 | 0.35 | 1 | Mapkapk3 | mitogen-activated protein kinase-activate | 507 | 297 | 416 | 437 | 353 | 553 | 468 | 321 | 367 | 413 | 500 |
| 350 | -0.38 | 0.23 | -1.62 | 0.11 | 1 | Hddc3 | HD domain containing 3 | 345 | 210 | 349 | 371 | 528 | 308 | 468 | 334 | 397 | 258 | 280 |
| 669 | 0.01 | 0.14 | 0.07 | 0.94 | 1 | Ccdc184 | coiled-coil domain containing 184 | 836 | 1316 | 866 | 754 | 444 | 500 | 467 | 716 | 408 | 508 | 542 |
| 630 | 0.07 | 0.16 | 0.43 | 0.67 | 1 | Whamm | WAS protein homolog associated with act | 473 | 499 | 610 | 550 | 679 | 923 | 467 | 487 | 681 | 647 | 909 |
| 540 | -0.12 | 0.22 | -0.57 | 0.57 | 1 | Sf3b4 | splicing factor 3b, subunit 4 | 800 | 759 | 637 | 572 | 497 | 412 | 467 | 324 | 485 | 562 | 426 |
| 532 | 0.07 | 0.23 | 0.31 | 0.76 | 1 | Zscan21 | zinc finger and SCAN domain containing | 480 | 473 | 576 | 566 | 462 | 590 | 467 | 830 | 457 | 406 | 541 |
| 502 | 0.01 | 0.25 | 0.05 | 0.96 | 1 | Ubac2 | ubiquitin associated domain containing 2 | 821 | 625 | 680 | 302 | 526 | 398 | 467 | 556 | 442 | 368 | 334 |
| 465 | 0.04 | 0.17 | 0.26 | 0.79 | 1 | Nt5dc1 | 5'-nucleotidase domain containing 1 | 374 | 409 | 354 | 523 | 480 | 484 | 467 | 470 | 471 | 595 | 482 |
| 416 | 0.14 | 0.25 | 0.56 | 0.58 | 1 | Otud7a | OTU domain containing 7A | 82 | 57 | 131 | 523 | 593 | 451 | 467 | 567 | 579 | 721 | 400 |
| 376 | -0.05 | 0.19 | -0.25 | 0.80 | 1 | Prorp | protein only RNase P catalytic subunit | 404 | 349 | 362 | 344 | 426 | 297 | 467 | 305 | 489 | 404 | 290 |
| 364 | -0.21 | 0.19 | -1.10 | 0.27 | 1 | Mettl21a | methyltransferase like 21A | 340 | 324 | 287 | 341 | 359 | 477 | 467 | 286 | 373 | 340 | 415 |
| 579 | -0.25 | 0.18 | -1.36 | 0.17 | 1 | Nova2 | NOVA alternative splicing regulator 2 | 993 | 1438 | 806 | 503 | 361 | 368 | 466 | 468 | 360 | 283 | 319 |
| 534 | 0.05 | 0.17 | 0.31 | 0.76 | 1 | Fhad1 | forkhead-associated (FHA) phosphopepti | 269 | 479 | 425 | 614 | 536 | 686 | 466 | 760 | 495 | 484 | 664 |
| 523 | -0.04 | 0.17 | -0.22 | 0.83 | 1 | Wdr73 | WD repeat domain 73 | 591 | 523 | 630 | 552 | 531 | 482 | 466 | 524 | 440 | 479 | 536 |
| 412 | -0.18 | 0.17 | -1.03 | 0.30 | 1 | Itpk1 | inositol 1,3,4-triphosphate 5/6 kinase | 643 | 411 | 518 | 357 | 352 | 384 | 466 | 270 | 341 | 469 | 320 |
| 345 | -0.29 | 0.24 | -1.23 | 0.22 | 1 | Krt222 | keratin 222 | 245 | 209 | 325 | 393 | 443 | 354 | 466 | 393 | 378 | 249 | 336 |
| 700 | -0.03 | 0.16 | -0.17 | 0.87 | 1 | Impdh2 | inosine monophosphate dehydrogenase | 1309 | 1149 | 1079 | 581 | 493 | 555 | 465 | 614 | 399 | 495 | 564 |
| 476 | 0.06 | 0.18 | 0.32 | 0.75 | 1 | Tsr3 | TSR3 20S rRNA accumulation | 286 | 370 | 426 | 489 | 452 | 632 | 465 | 526 | 453 | 482 | 660 |
| 471 | 0.22 | 0.22 | 1.00 | 0.32 | 1 | Mpi | mannose phosphate isomerase | 371 | 281 | 473 | 507 | 407 | 486 | 465 | 612 | 606 | 459 | 516 |
| 466 | -0.09 | 0.20 | -0.44 | 0.66 | 1 | Abt1 | activator of basal transcription 1 | 522 | 418 | 585 | 536 | 386 | 469 | 465 | 393 | 442 | 473 | 443 |
| 459 | 0.05 | 0.20 | 0.25 | 0.80 | 1 | Fggy | FGGY carbohydrate kinase domain conta | 387 | 466 | 426 | 374 | 438 | 559 | 465 | 511 | 342 | 429 | 651 |
| 392 | 0.04 | 0.24 | 0.18 | 0.86 | 1 | Efhc2 | EF-hand domain (C-terminal) containing 2 | 119 | 273 | 180 | 486 | 482 | 406 | 465 | 556 | 410 | 466 | 471 |
| 335 | -0.24 | 0.23 | -1.05 | 0.30 | 1 | Tafa1 | TAFA chemokine like family member 1 | 68 | 51 | 55 | 622 | 421 | 400 | 465 | 496 | 386 | 322 | 397 |
| 1177 | 0.28 | 0.27 | 1.06 | 0.29 | 1 | Fosl2 | fos-like antigen 2 | 3881 | 2098 | 1820 | 790 | 562 | 473 | 464 | 1140 | 622 | 602 | 500 |
| 552 | 0.23 | 0.20 | 1.16 | 0.24 | 1 | Rbm12 | RNA binding motif protein 12 | 489 | 797 | 604 | 593 | 352 | 513 | 464 | 751 | 479 | 589 | 445 |
| 522 | 0.02 | 0.15 | 0.10 | 0.92 | 1 | Rsph3a | radial spoke 3A homolog (Chlamydomon | 382 | 361 | 444 | 522 | 741 | 543 | 464 | 557 | 699 | 522 | 504 |
| 513 | 0.22 | 0.17 | 1.34 | 0.18 | 1 | Mrpl45 | mitochondrial ribosomal protein L45 | 636 | 591 | 588 | 473 | 466 | 356 | 464 | 657 | 450 | 542 | 415 |
| 460 | -0.33 | 0.18 | -1.86 | 0.06 | 1 | Serpina7 | serine (or cysteine) peptidase inhibitor, cl | 389 | 670 | 443 | 637 | 546 | 304 | 464 | 556 | 474 | 377 | 196 |
| 444 | -0.08 | 0.20 | -0.41 | 0.68 | 1 | Cnksr1 | connector enhancer of kinase suppressor | 260 | 465 | 308 | 472 | 470 | 566 | 464 | 529 | 494 | 372 | 484 |
| 416 | -0.06 | 0.19 | -0.34 | 0.74 | 1 | Snx7 | sorting nexin 7 | 402 | 402 | 584 | 382 | 423 | 359 | 464 | 351 | 444 | 367 | 399 |
| 522 | 0.06 | 0.20 | 0.29 | 0.77 | 1 | Faim | Fas apoptotic inhibitory molecule | 569 | 493 | 508 | 533 | 496 | 546 | 463 | 674 | 521 | 384 | 560 |
| 463 | 0.30 | 0.17 | 1.78 | 0.07 | 1 | Rbm48 | RNA binding motif protein 48 | 342 | 300 | 258 | 412 | 447 | 560 | 463 | 528 | 578 | 617 | 584 |
| 431 | 0.10 | 0.17 | 0.60 | 0.55 | 1 | Mpc1 | mitochondrial pyruvate carrier 1 | 311 | 331 | 218 | 445 | 417 | 546 | 463 | 534 | 412 | 515 | 549 |
| 536 | 0.12 | 0.15 | 0.80 | 0.42 | 1 | Lyar | Ly1 antibody reactive clone | 716 | 677 | 623 | 471 | 445 | 478 | 462 | 590 | 439 | 491 | 506 |
| 511 | 0.12 | 0.25 | 0.49 | 0.62 | 1 | Med21 | mediator complex subunit 21 | 672 | 553 | 695 | 392 | 362 | 506 | 462 | 254 | 389 | 743 | 593 |
| 474 | 0.13 | 0.19 | 0.68 | 0.50 | 1 | Macrod1 | mono-ADP ribosylhydrolase 1 | 304 | 340 | 476 | 499 | 558 | 442 | 462 | 583 | 503 | 587 | 464 |
| 461 | 0.39 | 0.25 | 1.57 | 0.12 | 1 | Macrod2 | mono-ADP ribosylhydrolase 2 | 218 | 302 | 306 | 489 | 431 | 436 | 462 | 758 | 357 | 687 | 630 |
| 562 | -0.02 | 0.16 | -0.15 | 0.88 | 1 | Zscan22 | zinc finger and SCAN domain containing | 548 | 747 | 477 | 647 | 522 | 599 | 461 | 592 | 563 | 429 | 601 |
| 540 | -0.09 | 0.28 | -0.32 | 0.75 | 1 | Dsel | dermatan sulfate epimerase-like | 951 | 585 | 609 | 482 | 456 | 505 | 461 | 226 | 647 | 594 | 424 |
| 440 | -0.28 | 0.23 | -1.24 | 0.22 | 1 | Zc3h6 | zinc finger CCCH type containing 6 | 475 | 347 | 433 | 529 | 395 | 572 | 461 | 482 | 311 | 314 | 524 |
| 426 | -0.19 | 0.27 | -0.69 | 0.49 | 1 | Trmt61b | | 199 | 303 | 281 | 350 | 602 | 624 | 461 | 240 | 613 | 356 | 655 |
| 413 | -0.33 | 0.21 | -1.54 | 0.12 | 1 | Ift27 | intraflagellar transport 27 | 459 | 564 | 493 | 434 | 431 | 343 | 461 | 454 | 373 | 241 | 285 |
| 810 | 0.09 | 0.18 | 0.48 | 0.63 | 1 | Slc39a11 | solute carrier family 39 (metal ion transpo | 1417 | 1511 | 1844 | 583 | 570 | 390 | 460 | 703 | 451 | 598 | 387 |
| 628 | 0.09 | 0.16 | 0.55 | 0.58 | 1 | Ifi35 | interferon-induced protein 35 | 1078 | 842 | 960 | 420 | 457 | 623 | 460 | 518 | 509 | 441 | 597 |
| 568 | -0.02 | 0.14 | -0.11 | 0.92 | 1 | Ccdc191 | coiled-coil domain containing 191 | 543 | 625 | 496 | 499 | 457 | 881 | 460 | 564 | 390 | 457 | 876 |
| 473 | 0.10 | 0.20 | 0.49 | 0.62 | 1 | Eefsec | eukaryotic elongation factor, selenocystei | 593 | 547 | 536 | 408 | 426 | 377 | 460 | 379 | 431 | 710 | 334 |
| 450 | 0.19 | 0.18 | 1.07 | 0.29 | 1 | Tmco6 | transmembrane and coiled-coil domains 6 | 305 | 301 | 304 | 310 | 464 | 666 | 460 | 402 | 532 | 497 | 704 |
| 263 | -0.34 | 0.25 | -1.37 | 0.17 | 1 | Cgref1 | cell growth regulator with EF hand domai | 65 | 99 | 98 | 410 | 317 | 287 | 460 | 345 | 286 | 270 | 255 |
| 842 | -0.12 | 0.20 | -0.61 | 0.54 | 1 | Helz2 | helicase with zinc finger 2, transcriptional | 1597 | 1635 | 1357 | 657 | 583 | 730 | 459 | 431 | 684 | 457 | 667 |
| 566 | 0.12 | 0.17 | 0.70 | 0.49 | 1 | Casp8 | caspase 8 | 1078 | 893 | 899 | 508 | 401 | 257 | 459 | 550 | 423 | 435 | 329 |
| 501 | 0.01 | 0.16 | 0.06 | 0.95 | 1 | Gabpb1 | GA repeat binding protein, beta 1 | 673 | 661 | 677 | 344 | 401 | 542 | 459 | 405 | 410 | 383 | 554 |
| 453 | -0.26 | 0.20 | -1.28 | 0.20 | 1 | Dbr1 | debranching RNA lariats 1 | 293 | 338 | 416 | 609 | 494 | 572 | 459 | 359 | 468 | 410 | 568 |
| 431 | 0.11 | 0.25 | 0.43 | 0.67 | 1 | Brd3os | bromodomain containing 3, opposite stra | 328 | 384 | 561 | 400 | 455 | 355 | 459 | 564 | 371 | 496 | 365 |
| 833 | -0.11 | 0.19 | -0.57 | 0.57 | 1 | Paxbp1 | PAX3 and PAX7 binding protein 1 | 492 | 588 | 501 | 564 | 403 | 2430 | 458 | 335 | 433 | 509 | 2455 |

| | | | | | | | | | | | | | | | | | | |
|---|---|---|---|---|---|---|---|---|---|---|---|---|---|---|---|---|---|---|
| 772 | -0.20 | 0.21 | -0.93 | 0.35 | 1 | Cyth3 | cytohesin 3 | 1736 | 1405 | 1996 | 440 | 429 | 453 | 458 | 484 | 421 | 263 | 402 |
| 756 | 0.21 | 0.18 | 1.15 | 0.25 | 1 | Zgpat | zinc finger, CCCH-type with G patch doma | 589 | 545 | 624 | 426 | 538 | 1728 | 458 | 715 | 569 | 473 | 1658 |
| 716 | -0.24 | 0.18 | -1.33 | 0.18 | 1 | Nrip1 | nuclear receptor interacting protein 1 | 1418 | 1291 | 1113 | 501 | 538 | 694 | 458 | 367 | 529 | 403 | 560 |
| 496 | 0.04 | 0.18 | 0.25 | 0.81 | 1 | Rnaseh1 | ribonuclease H1 | 355 | 402 | 426 | 448 | 525 | 678 | 458 | 452 | 463 | 595 | 654 |
| 480 | -0.02 | 0.17 | -0.10 | 0.92 | 1 | Oma1 | OMA1 zinc metallopeptidase | 424 | 410 | 567 | 492 | 485 | 512 | 458 | 444 | 498 | 519 | 467 |
| 471 | -0.06 | 0.20 | -0.32 | 0.75 | 1 | Otoa | otoancorin | 79 | 99 | 116 | 565 | 967 | 489 | 458 | 708 | 887 | 453 | 361 |
| 408 | 0.16 | 0.26 | 0.60 | 0.55 | 1 | Tmem69 | transmembrane protein 69 | 295 | 446 | 477 | 332 | 345 | 411 | 458 | 404 | 323 | 511 | 489 |
| 698 | -0.10 | 0.14 | -0.73 | 0.47 | 1 | Uvssa | UV stimulated scaffold protein A | 387 | 471 | 417 | 670 | 601 | 1558 | 457 | 664 | 544 | 385 | 1520 |
| 634 | 0.10 | 0.16 | 0.65 | 0.51 | 1 | Rilpl1 | Rab interacting lysosomal protein-like 1 | 848 | 864 | 880 | 596 | 563 | 499 | 457 | 555 | 578 | 639 | 498 |
| 618 | -0.16 | 0.18 | -0.91 | 0.36 | 1 | Slc15a4 | solute carrier family 15, member 4 | 940 | 940 | 823 | 664 | 429 | 612 | 457 | 495 | 468 | 367 | 599 |
| 583 | -0.07 | 0.19 | -0.37 | 0.71 | 1 | Numbl | numb-like | 875 | 722 | 561 | 662 | 448 | 603 | 457 | 482 | 401 | 554 | 645 |
| 575 | 0.29 | 0.16 | 1.73 | 0.08 | 1 | Mtx1 | metaxin 1 | 655 | 601 | 533 | 411 | 530 | 654 | 457 | 584 | 611 | 538 | 754 |
| 486 | 0.04 | 0.17 | 0.25 | 0.80 | 1 | Ccdc90b | coiled-coil domain containing 90B | 626 | 414 | 616 | 363 | 509 | 483 | 457 | 355 | 442 | 593 | 492 |
| 453 | 0.25 | 0.21 | 1.18 | 0.24 | 1 | Fahd2a | fumarylacetoacetate hydrolase domain cc | 374 | 362 | 401 | 425 | 422 | 440 | 457 | 711 | 398 | 489 | 505 |
| 294 | -0.11 | 0.31 | -0.37 | 0.71 | 1 | Map3k9 | mitogen-activated protein kinase kinase k | 201 | 261 | 125 | 410 | 273 | 259 | 457 | 273 | 406 | 300 | 266 |
| 269 | -0.31 | 0.27 | -1.17 | 0.24 | 1 | Fmn1 | formin 1 | 265 | 236 | 143 | 304 | 364 | 152 | 457 | 368 | 185 | 346 | 140 |
| 805 | 0.22 | 0.15 | 1.42 | 0.15 | 1 | Tbce | tubulin-specific chaperone E | 397 | 461 | 397 | 483 | 482 | 2182 | 456 | 562 | 490 | 685 | 2257 |
| 472 | 0.16 | 0.23 | 0.67 | 0.50 | 1 | Coil | coilin | 286 | 449 | 493 | 408 | 437 | 581 | 456 | 426 | 476 | 598 | 583 |
| 467 | -0.09 | 0.16 | -0.53 | 0.59 | 1 | Ubiad1 | UbiA prenyltransferase domain containin | 551 | 570 | 466 | 455 | 484 | 431 | 456 | 443 | 472 | 446 | 364 |
| 351 | -0.14 | 0.26 | -0.54 | 0.59 | 1 | Mrps18b | mitochondrial ribosomal protein S18B | 243 | 280 | 288 | 220 | 455 | 476 | 456 | 288 | 447 | 285 | 422 |
| 630 | 0.39 | 0.23 | 1.72 | 0.09 | 1 | Dvl2 | dishevelled segment polarity protein 2 | 635 | 544 | 655 | 491 | 628 | 627 | 455 | 903 | 511 | 772 | 710 |
| 582 | 0.08 | 0.14 | 0.60 | 0.55 | 1 | Vps9d1 | VPS9 domain containing 1 | 571 | 528 | 605 | 525 | 541 | 757 | 455 | 536 | 529 | 528 | 828 |
| 525 | -0.21 | 0.22 | -0.95 | 0.34 | 1 | Dcaf17 | DDB1 and CUL4 associated factor 17 | 433 | 483 | 379 | 621 | 522 | 752 | 455 | 425 | 522 | 308 | 878 |
| 502 | -0.05 | 0.18 | -0.31 | 0.76 | 1 | Utp11 | UTP11 small subunit processome compo | 657 | 681 | 595 | 373 | 325 | 672 | 455 | 447 | 325 | 331 | 659 |
| 478 | 0.12 | 0.21 | 0.60 | 0.55 | 1 | Shpk | sedoheptulokinase | 298 | 463 | 474 | 473 | 620 | 382 | 455 | 680 | 481 | 493 | 441 |
| 437 | 0.00 | 0.19 | -0.02 | 0.98 | 1 | Nsmce2 | NSE2/MMS21 homolog, SMC5-SMC6 co | 594 | 592 | 541 | 385 | 343 | 364 | 455 | 333 | 432 | 361 | 412 |
| 437 | 0.02 | 0.20 | 0.09 | 0.93 | 1 | Zbtb26 | zinc finger and BTB domain containing 26 | 257 | 246 | 320 | 403 | 456 | 652 | 455 | 512 | 494 | 316 | 700 |
| 428 | 0.05 | 0.20 | 0.23 | 0.82 | 1 | Ighmbp2 | immunoglobulin mu binding protein 2 | 346 | 386 | 493 | 311 | 474 | 483 | 455 | 463 | 450 | 388 | 463 |
| 397 | -0.13 | 0.29 | -0.47 | 0.64 | 1 | Acvr1c | activin A receptor, type IC | 360 | 139 | 294 | 403 | 483 | 531 | 455 | 401 | 508 | 299 | 497 |
| 375 | -0.08 | 0.17 | -0.46 | 0.65 | 1 | Ddx28 | DEAD (Asp-Glu-Ala-Asp) box polypeptide | 322 | 265 | 280 | 437 | 508 | 281 | 455 | 390 | 416 | 473 | 299 |
| 941 | 0.03 | 0.20 | 0.16 | 0.87 | 1 | Lipa | lysosomal acid lipase A | 2931 | 1850 | 1625 | 494 | 595 | 390 | 454 | 476 | 554 | 629 | 357 |
| 511 | 0.12 | 0.15 | 0.76 | 0.45 | 1 | Med19 | mediator complex subunit 19 | 700 | 660 | 737 | 433 | 404 | 400 | 454 | 485 | 419 | 483 | 444 |
| 483 | 0.09 | 0.19 | 0.49 | 0.62 | 1 | Acsl3 | acyl-CoA synthetase long-chain family me | 402 | 251 | 330 | 511 | 586 | 536 | 454 | 473 | 647 | 542 | 580 |
| 469 | -0.02 | 0.24 | -0.10 | 0.92 | 1 | 4632404H12Rik | | 284 | 413 | 405 | 626 | 313 | 677 | 454 | 463 | 410 | 515 | 594 |
| 466 | 0.12 | 0.17 | 0.73 | 0.46 | 1 | Lzts3 | leucine zipper, putative tumor suppressor | 493 | 417 | 457 | 453 | 397 | 488 | 454 | 507 | 378 | 514 | 562 |
| 410 | -0.36 | 0.21 | -1.67 | 0.09 | 1 | Fam163a | family with sequence similarity 163, meml | 517 | 301 | 673 | 205 | 745 | 279 | 454 | 139 | 569 | 413 | 213 |
| 398 | -0.16 | 0.21 | -0.78 | 0.44 | 1 | Cdca4 | cell division cycle associated 4 | 578 | 446 | 459 | 511 | 293 | 286 | 454 | 357 | 297 | 386 | 310 |
| 365 | -0.37 | 0.29 | -1.28 | 0.20 | 1 | Pcdhga7 | protocadherin gamma subfamily A, 7 | 287 | 365 | 262 | 460 | 455 | 372 | 454 | 207 | 486 | 308 | 360 |
| 630 | -0.20 | 0.20 | -1.00 | 0.32 | 1 | Clk2 | CDC-like kinase 2 | 455 | 504 | 430 | 473 | 367 | 1643 | 453 | 272 | 431 | 401 | 1507 |
| 533 | 0.35 | 0.22 | 1.61 | 0.11 | 1 | Pdcd2 | programmed cell death 2 | 445 | 554 | 497 | 483 | 487 | 459 | 453 | 949 | 431 | 542 | 565 |
| 506 | -0.03 | 0.14 | -0.20 | 0.84 | 1 | Snx24 | sorting nexing 24 | 594 | 584 | 653 | 452 | 507 | 473 | 453 | 480 | 494 | 434 | 440 |
| 478 | 0.15 | 0.31 | 0.47 | 0.64 | 1 | Rsad2 | radical S-adenosyl methionine domain cc | 1298 | 680 | 1225 | 202 | 127 | 167 | 453 | 155 | 212 | 585 | 159 |
| 463 | 0.09 | 0.19 | 0.48 | 0.63 | 1 | Setd1a | SET domain containing 1A | 453 | 410 | 279 | 529 | 399 | 538 | 453 | 505 | 510 | 529 | 484 |
| 1006 | 0.15 | 0.27 | 0.55 | 0.58 | 1 | Bhlhe40 | basic helix-loop-helix family, member e40 | 3217 | 1595 | 1696 | 736 | 340 | 651 | 452 | 823 | 437 | 483 | 637 |
| 563 | 0.28 | 0.20 | 1.36 | 0.17 | 1 | Adpgk | ADP-dependent glucokinase | 545 | 364 | 328 | 413 | 420 | 981 | 452 | 488 | 617 | 486 | 1096 |
| 447 | -0.04 | 0.20 | -0.18 | 0.86 | 1 | Kctd21 | potassium channel tetramerisation domai | 388 | 246 | 367 | 501 | 536 | 494 | 452 | 494 | 453 | 501 | 485 |
| 393 | -0.15 | 0.24 | -0.62 | 0.53 | 1 | Cops7b | COP9 signalosome subunit 7B | 380 | 267 | 310 | 389 | 456 | 467 | 452 | 443 | 390 | 277 | 489 |
| 373 | -0.18 | 0.18 | -0.97 | 0.33 | 1 | BC052040 | cDNA sequence BC052040 | 446 | 446 | 455 | 322 | 383 | 309 | 452 | 312 | 339 | 330 | 306 |
| 338 | -0.03 | 0.27 | -0.10 | 0.92 | 1 | Plcxd3 | phosphatidylinositol-specific phospholipa | 150 | 256 | 90 | 495 | 245 | 463 | 452 | 408 | 364 | 318 | 473 |
| 1125 | 0.17 | 0.25 | 0.68 | 0.50 | 1 | Depp1 | DEPP1 autophagy regulator | 2060 | 2879 | 2891 | 361 | 347 | 1006 | 450 | 462 | 268 | 684 | 971 |
| 569 | 0.04 | 0.20 | 0.20 | 0.84 | 1 | Mphosph6 | M phase phosphoprotein 6 | 820 | 558 | 694 | 564 | 442 | 613 | 450 | 554 | 536 | 441 | 590 |
| 471 | 0.18 | 0.21 | 0.84 | 0.40 | 1 | Chst10 | carbohydrate sulfotransferase 10 | 740 | 644 | 471 | 471 | 373 | 260 | 450 | 359 | 409 | 675 | 331 |
| 436 | -0.18 | 0.19 | -0.96 | 0.34 | 1 | Srrm3 | serine/arginine repetitive matrix 3 | 233 | 318 | 263 | 417 | 417 | 844 | 450 | 351 | 451 | 361 | 695 |
| 431 | -0.31 | 0.17 | -1.80 | 0.07 | 1 | Gemin6 | gem nuclear organelle associated proteir | 371 | 449 | 429 | 590 | 478 | 418 | 450 | 385 | 414 | 417 | 341 |
| 347 | 0.21 | 0.32 | 0.66 | 0.51 | 1 | Abhd3 | abhydrolase domain containing 3 | 74 | 229 | 150 | 357 | 329 | 393 | 450 | 542 | 526 | 353 | 410 |
| 261 | -0.26 | 0.33 | -0.78 | 0.43 | 1 | Sgcd | sarcoglycan, delta (dystrophin-associatec | 166 | 138 | 114 | 407 | 319 | 178 | 449 | 212 | 442 | 238 | 206 |
| 754 | 0.02 | 0.28 | 0.07 | 0.95 | 1 | Sema3c | sema domain, immunoglobulin domain (I | 2838 | 951 | 1477 | 456 | 347 | 261 | 448 | 469 | 439 | 367 | 246 |
| 549 | -0.26 | 0.20 | -1.32 | 0.19 | 1 | Kansl1l | KAT8 regulatory NSL complex subunit 1-l | 617 | 627 | 553 | 672 | 378 | 810 | 448 | 387 | 411 | 383 | 750 |
| 539 | 0.16 | 0.26 | 0.62 | 0.54 | 1 | Spsb2 | splA/ryanodine receptor domain and SOC | 466 | 450 | 588 | 674 | 465 | 465 | 448 | 1013 | 433 | 486 | 445 |
| 492 | -0.20 | 0.22 | -0.88 | 0.38 | 1 | Zfp111 | zinc finger protein 111 | 547 | 405 | 623 | 538 | 405 | 664 | 448 | 475 | 459 | 372 | 473 |

| | | | | | | | | | | | | | | | | | | |
|---|---|---|---|---|---|---|---|---|---|---|---|---|---|---|---|---|---|---|
| 471 | 0.05 | 0.17 | 0.31 | 0.76 | 1 | 4933431E20Rik | | 496 | 461 | 551 | 483 | 450 | 422 | 448 | 546 | 456 | 399 | 474 |
| 463 | 0.32 | 0.22 | 1.46 | 0.14 | 1 | Nprl2 | NPR2 like, GATOR1 complex subunit | 334 | 380 | 494 | 295 | 431 | 572 | 448 | 560 | 491 | 466 | 628 |
| 459 | -0.28 | 0.15 | -1.79 | 0.07 | 1 | Spindoc | spindlin interactor and repressor of chrom | 565 | 472 | 535 | 440 | 479 | 532 | 448 | 339 | 372 | 377 | 490 |
| 429 | 0.12 | 0.17 | 0.72 | 0.47 | 1 | Zc3h3 | zinc finger CCCH type containing 3 | 77 | 94 | 86 | 499 | 465 | 735 | 448 | 516 | 606 | 487 | 706 |
| 363 | -0.12 | 0.27 | -0.44 | 0.66 | 1 | Tctex1d2 | Tctex1 domain containing 2 | 434 | 331 | 672 | 272 | 294 | 318 | 448 | 293 | 343 | 256 | 330 |
| 277 | -0.38 | 0.30 | -1.29 | 0.20 | 1 | Gm15163 | | 428 | 254 | 257 | 274 | 217 | 183 | 448 | 134 | 156 | 569 | 129 |
| 675 | -0.22 | 0.17 | -1.26 | 0.21 | 1 | Sema4a | sema domain, immunoglobulin domain (Ig | 1368 | 1377 | 1266 | 515 | 424 | 445 | 447 | 451 | 445 | 294 | 392 |
| 586 | 0.11 | 0.19 | 0.57 | 0.57 | 1 | Unc5b | unc-5 netrin receptor B | 1069 | 669 | 1087 | 480 | 489 | 308 | 447 | 511 | 520 | 587 | 282 |
| 529 | 0.28 | 0.25 | 1.09 | 0.28 | 1 | Psmg1 | proteasome (prosome, macropain) assem | 706 | 386 | 625 | 444 | 547 | 421 | 447 | 729 | 450 | 573 | 494 |
| 491 | 0.10 | 0.17 | 0.60 | 0.55 | 1 | Mrpl40 | mitochondrial ribosomal protein L40 | 462 | 479 | 413 | 490 | 582 | 429 | 447 | 547 | 545 | 581 | 426 |
| 487 | 0.03 | 0.18 | 0.18 | 0.86 | 1 | Commd2 | COMM domain containing 2 | 619 | 547 | 571 | 484 | 458 | 395 | 447 | 387 | 406 | 535 | 511 |
| 480 | 0.05 | 0.19 | 0.25 | 0.80 | 1 | Mpped2 | metallophosphoesterase domain containi | 560 | 605 | 552 | 621 | 351 | 331 | 447 | 579 | 468 | 480 | 286 |
| 456 | 0.22 | 0.19 | 1.21 | 0.22 | 1 | Lipo3 | lipase, member O3 | 351 | 344 | 287 | 527 | 485 | 399 | 447 | 489 | 647 | 573 | 463 |
| 453 | -0.03 | 0.20 | -0.16 | 0.87 | 1 | Ints9 | integrator complex subunit 9 | 703 | 701 | 449 | 409 | 349 | 380 | 447 | 474 | 356 | 325 | 395 |
| 427 | -0.27 | 0.17 | -1.66 | 0.10 | 1 | Zmat1 | zinc finger, matrin type 1 | 420 | 469 | 410 | 506 | 380 | 521 | 447 | 444 | 325 | 314 | 459 |
| 320 | -0.26 | 0.22 | -1.17 | 0.24 | 1 | Tmem121b | transmembrane protein 121B | 184 | 149 | 136 | 501 | 394 | 328 | 447 | 382 | 308 | 316 | 378 |
| 554 | 0.08 | 0.21 | 0.39 | 0.69 | 1 | Mrps24 | mitochondrial ribosomal protein S24 | 682 | 425 | 761 | 578 | 619 | 414 | 446 | 573 | 601 | 540 | 454 |
| 520 | -0.14 | 0.20 | -0.66 | 0.51 | 1 | Exosc2 | exosome component 2 | 749 | 456 | 658 | 488 | 524 | 560 | 446 | 461 | 466 | 376 | 535 |
| 486 | -0.03 | 0.18 | -0.18 | 0.86 | 1 | AI467606 | expressed sequence AI467606 | 641 | 578 | 660 | 375 | 591 | 348 | 446 | 462 | 456 | 442 | 342 |
| 484 | 0.02 | 0.17 | 0.11 | 0.91 | 1 | Mcee | methylmalonyl CoA epimerase | 644 | 569 | 612 | 497 | 423 | 373 | 446 | 424 | 429 | 424 | 482 |
| 463 | -0.14 | 0.20 | -0.70 | 0.49 | 1 | Cdkl1 | cyclin-dependent kinase-like 1 (CDC2-rel | 218 | 392 | 325 | 726 | 376 | 617 | 446 | 634 | 384 | 344 | 626 |
| 432 | 0.00 | 0.19 | 0.00 | 1.00 | 1 | Mrps27 | mitochondrial ribosomal protein S27 | 508 | 361 | 641 | 308 | 478 | 397 | 446 | 394 | 418 | 438 | 357 |
| 430 | -0.14 | 0.23 | -0.59 | 0.55 | 1 | Rnf24 | ring finger protein 24 | 342 | 482 | 279 | 522 | 419 | 523 | 446 | 349 | 496 | 399 | 475 |
| 527 | 0.01 | 0.15 | 0.07 | 0.94 | 1 | Tmcc1 | transmembrane and coiled coil domains 1 | 746 | 751 | 597 | 444 | 434 | 524 | 445 | 445 | 428 | 474 | 514 |
| 497 | -0.25 | 0.23 | -1.09 | 0.27 | 1 | Nagpa | N-acetylglucosamine-1-phosphodiester a | 891 | 661 | 730 | 481 | 433 | 348 | 445 | 235 | 454 | 493 | 297 |
| 483 | 0.12 | 0.23 | 0.51 | 0.61 | 1 | Tinf2 | Terf1 (TRF1)-interacting nuclear factor 2 | 494 | 546 | 603 | 438 | 328 | 534 | 445 | 656 | 321 | 397 | 549 |
| 449 | 0.25 | 0.23 | 1.10 | 0.27 | 1 | Bloc1s3 | biogenesis of lysosomal organelles comp | 353 | 384 | 455 | 444 | 435 | 350 | 445 | 694 | 475 | 618 | 285 |
| 395 | -0.31 | 0.24 | -1.31 | 0.19 | 1 | Cacna1b | calcium channel, voltage-dependent, N ty | 279 | 454 | 264 | 431 | 427 | 522 | 445 | 446 | 229 | 420 | 422 |
| 662 | -0.03 | 0.16 | -0.16 | 0.87 | 1 | Trim11 | tripartite motif-containing 11 | 373 | 602 | 536 | 600 | 698 | 1173 | 444 | 584 | 565 | 519 | 1188 |
| 569 | 0.06 | 0.16 | 0.40 | 0.69 | 1 | Zfp160 | zinc finger protein 160 | 456 | 608 | 534 | 452 | 449 | 897 | 444 | 466 | 419 | 455 | 1078 |
| 539 | 0.14 | 0.16 | 0.85 | 0.40 | 1 | Senp8 | SUMO/sentrin specific peptidase 8 | 496 | 680 | 594 | 538 | 458 | 529 | 444 | 567 | 585 | 386 | 655 |
| 443 | -0.08 | 0.22 | -0.37 | 0.71 | 1 | Prmt3 | protein arginine N-methyltransferase 3 | 519 | 318 | 303 | 607 | 365 | 493 | 444 | 469 | 351 | 551 | 447 |
| 361 | 0.19 | 0.24 | 0.78 | 0.43 | 1 | Lmx1b | LIM homeobox transcription factor 1 beta | 172 | 138 | 112 | 406 | 353 | 447 | 444 | 653 | 449 | 394 | 399 |
| 658 | 0.14 | 0.22 | 0.64 | 0.52 | 1 | Zmym6 | zinc finger, MYM-type 6 | 330 | 413 | 283 | 361 | 410 | 1789 | 443 | 343 | 414 | 650 | 1803 |
| 495 | -0.35 | 0.28 | -1.27 | 0.20 | 1 | Isg20l2 | interferon stimulated exonuclease gene 2 | 691 | 749 | 620 | 324 | 485 | 622 | 443 | 324 | 486 | 182 | 520 |
| 478 | 0.16 | 0.23 | 0.71 | 0.48 | 1 | Fbxo4 | F-box protein 4 | 540 | 407 | 528 | 459 | 423 | 452 | 443 | 690 | 432 | 406 | 478 |
| 457 | 0.17 | 0.21 | 0.81 | 0.42 | 1 | Msrb2 | methionine sulfoxide reductase B2 | 395 | 397 | 441 | 425 | 521 | 397 | 443 | 611 | 400 | 483 | 513 |
| 667 | 0.22 | 0.14 | 1.56 | 0.12 | 1 | Zfp26 | zinc finger protein 26 | 728 | 694 | 677 | 476 | 460 | 1069 | 442 | 496 | 598 | 574 | 1126 |
| 451 | 0.23 | 0.16 | 1.45 | 0.15 | 1 | 4930430F08Rik | RIKEN cDNA 4930430F08 gene | 430 | 361 | 397 | 411 | 410 | 470 | 442 | 528 | 440 | 483 | 592 |
| 436 | 0.10 | 0.22 | 0.46 | 0.65 | 1 | Zfp13 | zinc finger protein 13 | 426 | 450 | 493 | 459 | 385 | 358 | 442 | 307 | 485 | 524 | 464 |
| 409 | 0.08 | 0.17 | 0.47 | 0.64 | 1 | Rfxank | regulatory factor X-associated ankyrin-co | 479 | 334 | 508 | 338 | 401 | 369 | 442 | 364 | 419 | 433 | 417 |
| 342 | 0.11 | 0.20 | 0.53 | 0.59 | 1 | D730003I15Rik | | 216 | 256 | 193 | 368 | 315 | 366 | 442 | 434 | 431 | 388 | 350 |
| 491 | -0.05 | 0.20 | -0.28 | 0.78 | 1 | Armt1 | acidic residue methyltransferase 1 | 401 | 522 | 360 | 546 | 542 | 544 | 441 | 689 | 473 | 341 | 540 |
| 464 | -0.11 | 0.19 | -0.59 | 0.55 | 1 | Fance | Fanconi anemia, complementation group | 607 | 428 | 654 | 364 | 419 | 559 | 441 | 409 | 346 | 385 | 494 |
| 460 | -0.07 | 0.18 | -0.37 | 0.71 | 1 | Tdp1 | tyrosyl-DNA phosphodiesterase 1 | 656 | 392 | 532 | 359 | 511 | 468 | 441 | 370 | 390 | 442 | 496 |
| 445 | 0.18 | 0.26 | 0.69 | 0.49 | 1 | Them6 | thioesterase superfamily member 6 | 317 | 176 | 234 | 558 | 535 | 344 | 441 | 651 | 517 | 853 | 269 |
| 378 | 0.11 | 0.17 | 0.68 | 0.50 | 1 | Zbtb8os | zinc finger and BTB domain containing 8 | 323 | 295 | 317 | 316 | 382 | 413 | 441 | 382 | 408 | 457 | 425 |
| 362 | -0.01 | 0.23 | -0.05 | 0.96 | 1 | Gm15663 | | 269 | 439 | 348 | 369 | 301 | 347 | 441 | 477 | 337 | 325 | 328 |
| 359 | 0.00 | 0.18 | -0.01 | 0.99 | 1 | Zfp617 | zinc finger protein 617 | 287 | 314 | 312 | 351 | 353 | 371 | 441 | 350 | 354 | 442 | 370 |
| 251 | -0.11 | 0.33 | -0.33 | 0.74 | 1 | 5930430L01Rik | | 400 | 101 | 268 | 218 | 223 | 148 | 441 | 215 | 153 | 437 | 153 |
| 700 | -0.16 | 0.22 | -0.72 | 0.47 | 1 | Phactr2 | phosphatase and actin regulator 2 | 1651 | 1327 | 1268 | 624 | 362 | 406 | 440 | 373 | 484 | 380 | 385 |
| 475 | -0.37 | 0.26 | -1.45 | 0.15 | 1 | Pigb | phosphatidylinositol glycan anchor biosyn | 412 | 514 | 487 | 461 | 548 | 682 | 440 | 210 | 459 | 456 | 560 |
| 456 | -0.10 | 0.20 | -0.47 | 0.64 | 1 | Irs1 | insulin receptor substrate 1 | 377 | 270 | 274 | 456 | 456 | 776 | 440 | 519 | 419 | 375 | 655 |
| 455 | -0.01 | 0.18 | -0.04 | 0.97 | 1 | Med30 | mediator complex subunit 30 | 521 | 366 | 449 | 490 | 501 | 397 | 440 | 454 | 457 | 559 | 369 |
| 444 | 0.02 | 0.22 | 0.07 | 0.94 | 1 | Zfp946 | zinc finger protein 946 | 456 | 506 | 467 | 373 | 325 | 572 | 440 | 305 | 495 | 339 | 611 |
| 749 | -0.23 | 0.22 | -1.02 | 0.31 | 1 | Tap1 | transporter 1, ATP-binding cassette, sub-f | 2304 | 1360 | 1294 | 388 | 542 | 407 | 439 | 379 | 440 | 331 | 353 |
| 517 | 0.05 | 0.20 | 0.26 | 0.79 | 1 | Tgds | TDP-glucose 4,6-dehydratase | 605 | 457 | 352 | 548 | 465 | 639 | 439 | 548 | 451 | 471 | 715 |
| 507 | 0.13 | 0.19 | 0.69 | 0.49 | 1 | Rffl | ring finger and FYVE like domain containi | 492 | 371 | 414 | 508 | 497 | 603 | 439 | 540 | 529 | 662 | 518 |
| 438 | 0.11 | 0.27 | 0.40 | 0.69 | 1 | Mfsd12 | major facilitator superfamily domain conta | 588 | 493 | 559 | 246 | 524 | 339 | 439 | 498 | 420 | 325 | 383 |
| 403 | -0.44 | 0.27 | -1.67 | 0.09 | 1 | Dnajc25 | DnaJ heat shock protein family (Hsp40) m | 454 | 288 | 362 | 331 | 425 | 651 | 439 | 129 | 397 | 335 | 623 |

| | | | | | | | | | | | | | | | | | | |
|---|---|---|---|---|---|---|---|---|---|---|---|---|---|---|---|---|---|---|
| 366 | 0.19 | 0.23 | 0.82 | 0.41 | 1 | Ppcs | phosphopantothenoylcysteine synthetase | 268 | 294 | 368 | 386 | 339 | 277 | 439 | 570 | 385 | 376 | 327 |
| 554 | 0.02 | 0.23 | 0.08 | 0.94 | 1 | Lin54 | lin-54 homolog (C. elegans) | 611 | 649 | 357 | 594 | 578 | 617 | 438 | 523 | 588 | 501 | 636 |
| 546 | 0.17 | 0.20 | 0.85 | 0.40 | 1 | Zbtb17 | zinc finger and BTB domain containing 17 | 550 | 664 | 554 | 470 | 430 | 661 | 438 | 497 | 437 | 711 | 596 |
| 538 | 0.04 | 0.15 | 0.28 | 0.78 | 1 | Pex16 | peroxisomal biogenesis factor 16 | 489 | 509 | 503 | 511 | 564 | 662 | 438 | 510 | 499 | 510 | 727 |
| 520 | 0.21 | 0.19 | 1.09 | 0.28 | 1 | Psmg2 | proteasome (prosome, macropain) assem | 437 | 649 | 537 | 399 | 462 | 593 | 438 | 657 | 461 | 491 | 590 |
| 414 | 0.07 | 0.16 | 0.40 | 0.69 | 1 | Ppif | peptidylprolyl isomerase F (cyclophilin F) | 498 | 352 | 451 | 333 | 402 | 408 | 438 | 356 | 420 | 524 | 367 |
| 323 | -0.08 | 0.24 | -0.33 | 0.74 | 1 | Bspry | B-box and SPRY domain containing | 186 | 302 | 280 | 377 | 335 | 273 | 438 | 367 | 361 | 416 | 220 |
| 633 | 0.39 | 0.21 | 1.82 | 0.07 | 1 | Fbl | fibrillarin | 788 | 693 | 1024 | 398 | 505 | 593 | 437 | 783 | 581 | 452 | 709 |
| 620 | -0.05 | 0.28 | -0.17 | 0.86 | 1 | Pigr | polymeric immunoglobulin receptor | 600 | 1335 | 1666 | 588 | 347 | 276 | 437 | 520 | 358 | 392 | 306 |
| 541 | -0.06 | 0.17 | -0.38 | 0.71 | 1 | Snx33 | sorting nexin 33 | 1048 | 971 | 868 | 479 | 367 | 278 | 436 | 434 | 284 | 486 | 294 |
| 514 | -0.18 | 0.20 | -0.87 | 0.38 | 1 | Znrd1 | zinc ribbon domain containing 1 | 612 | 552 | 524 | 586 | 558 | 517 | 436 | 358 | 529 | 496 | 481 |
| 504 | 0.16 | 0.23 | 0.70 | 0.48 | 1 | Nop16 | NOP16 nucleolar protein | 676 | 565 | 444 | 384 | 524 | 470 | 436 | 643 | 475 | 439 | 482 |
| 503 | -0.02 | 0.23 | -0.08 | 0.94 | 1 | Mus81 | MUS81 structure-specific endonuclease s | 436 | 436 | 472 | 391 | 530 | 746 | 436 | 428 | 333 | 620 | 703 |
| 467 | 0.18 | 0.29 | 0.64 | 0.52 | 1 | Dusp19 | dual specificity phosphatase 19 | 380 | 585 | 392 | 463 | 421 | 445 | 436 | 280 | 559 | 596 | 582 |
| 465 | -0.10 | 0.21 | -0.47 | 0.64 | 1 | Eral1 | Era (G-protein)-like 1 (E. coli) | 532 | 555 | 549 | 385 | 412 | 549 | 436 | 278 | 355 | 532 | 530 |
| 421 | -0.20 | 0.18 | -1.06 | 0.29 | 1 | Fbxl15 | F-box and leucine-rich repeat protein 15 | 314 | 398 | 325 | 621 | 445 | 412 | 436 | 510 | 356 | 500 | 316 |
| 416 | 0.10 | 0.18 | 0.55 | 0.58 | 1 | Tmem102 | transmembrane protein 102 | 284 | 405 | 363 | 418 | 451 | 388 | 436 | 446 | 556 | 429 | 396 |
| 400 | -0.08 | 0.23 | -0.35 | 0.72 | 1 | 4931428F04Rik | RIKEN cDNA 4931428F04 gene | 281 | 340 | 367 | 275 | 519 | 525 | 436 | 284 | 556 | 367 | 455 |
| 524 | 0.24 | 0.18 | 1.36 | 0.17 | 1 | Sclt1 | sodium channel and clathrin linker 1 | 463 | 346 | 344 | 557 | 411 | 708 | 435 | 613 | 482 | 560 | 844 |
| 443 | 0.06 | 0.16 | 0.35 | 0.72 | 1 | Exosc1 | exosome component 1 | 423 | 466 | 394 | 384 | 431 | 511 | 435 | 462 | 451 | 424 | 491 |
| 440 | 0.07 | 0.18 | 0.38 | 0.70 | 1 | Epha7 | Eph receptor A7 | 396 | 508 | 295 | 502 | 377 | 446 | 435 | 672 | 391 | 412 | 407 |
| 427 | -0.44 | 0.24 | -1.80 | 0.07 | 1 | Mrpl27 | mitochondrial ribosomal protein L27 | 503 | 461 | 526 | 495 | 489 | 408 | 435 | 304 | 457 | 210 | 413 |
| 518 | 0.10 | 0.27 | 0.35 | 0.72 | 1 | Dennd4b | DENN/MADD domain containing 4B | 762 | 418 | 594 | 606 | 287 | 576 | 434 | 459 | 452 | 616 | 494 |
| 515 | 0.12 | 0.25 | 0.49 | 0.62 | 1 | Dmap1 | DNA methyltransferase 1-associated prot | 474 | 605 | 696 | 431 | 542 | 438 | 434 | 747 | 361 | 459 | 473 |
| 480 | 0.13 | 0.28 | 0.48 | 0.63 | 1 | Tspan17 | tetraspanin 17 | 565 | 374 | 635 | 344 | 490 | 508 | 434 | 657 | 391 | 474 | 406 |
| 620 | -0.06 | 0.13 | -0.41 | 0.68 | 1 | Mapk15 | mitogen-activated protein kinase 15 | 861 | 842 | 780 | 436 | 389 | 959 | 433 | 444 | 348 | 438 | 895 |
| 559 | 0.02 | 0.18 | 0.12 | 0.91 | 1 | Ndst2 | N-deacetylase/N-sulfotransferase (hepara | 639 | 604 | 704 | 538 | 460 | 664 | 433 | 455 | 535 | 573 | 547 |
| 541 | 0.02 | 0.19 | 0.08 | 0.94 | 1 | Tmem251 | transmembrane protein 251 | 355 | 372 | 415 | 724 | 673 | 580 | 433 | 589 | 558 | 618 | 640 |
| 524 | -0.04 | 0.21 | -0.19 | 0.85 | 1 | Ess2 | ess-2 splicing factor | 534 | 762 | 549 | 532 | 422 | 610 | 433 | 381 | 439 | 572 | 530 |
| 524 | 0.32 | 0.20 | 1.61 | 0.11 | 1 | Bax | BCL2-associated X protein | 445 | 541 | 662 | 355 | 495 | 557 | 433 | 634 | 528 | 528 | 585 |
| 510 | -0.09 | 0.24 | -0.37 | 0.71 | 1 | Rbbp9 | retinoblastoma binding protein 9, serine h | 746 | 798 | 776 | 475 | 421 | 352 | 433 | 254 | 542 | 408 | 398 |
| 706 | -0.22 | 0.17 | -1.26 | 0.21 | 1 | Ttyh3 | tweety family member 3 | 1809 | 1288 | 1350 | 505 | 444 | 399 | 432 | 356 | 430 | 404 | 346 |
| 451 | 0.09 | 0.20 | 0.46 | 0.65 | 1 | Rfng | RFNG O-fucosylpeptide 3-beta-N-acetylg | 453 | 380 | 463 | 355 | 470 | 522 | 432 | 535 | 451 | 385 | 511 |
| 443 | 0.15 | 0.21 | 0.72 | 0.47 | 1 | Gal3st1 | galactose-3-O-sulfotransferase 1 | 223 | 321 | 269 | 431 | 487 | 580 | 432 | 433 | 486 | 603 | 613 |
| 435 | -0.07 | 0.17 | -0.39 | 0.69 | 1 | Mrps33 | mitochondrial ribosomal protein S33 | 378 | 419 | 430 | 461 | 482 | 440 | 432 | 389 | 532 | 379 | 444 |
| 435 | 0.20 | 0.21 | 0.97 | 0.33 | 1 | Mrpl55 | mitochondrial ribosomal protein L55 | 341 | 378 | 471 | 441 | 397 | 386 | 432 | 650 | 473 | 447 | 368 |
| 389 | -0.18 | 0.22 | -0.83 | 0.40 | 1 | Carf | calcium response factor | 250 | 205 | 234 | 348 | 325 | 758 | 432 | 198 | 355 | 386 | 791 |
| 330 | -0.34 | 0.25 | -1.35 | 0.18 | 1 | Thnsl2 | threonine synthase-like 2 (bacterial) | 373 | 322 | 274 | 385 | 416 | 260 | 432 | 357 | 350 | 218 | 244 |
| 274 | 0.08 | 0.24 | 0.32 | 0.75 | 1 | Lrrtm2 | leucine rich repeat transmembrane neuro | 108 | 121 | 118 | 263 | 348 | 255 | 432 | 269 | 442 | 397 | 264 |
| 527 | 0.02 | 0.17 | 0.10 | 0.92 | 1 | Dmac2 | distal membrane arm assembly complex 2 | 585 | 558 | 562 | 450 | 517 | 641 | 431 | 462 | 598 | 435 | 555 |
| 465 | 0.10 | 0.19 | 0.53 | 0.59 | 1 | Rara | retinoic acid receptor, alpha | 825 | 575 | 481 | 450 | 310 | 341 | 431 | 414 | 344 | 643 | 302 |
| 448 | -0.22 | 0.26 | -0.87 | 0.39 | 1 | Mmaa | methylmalonic aciduria (cobalamin deficie | 289 | 316 | 418 | 587 | 370 | 673 | 431 | 291 | 384 | 410 | 757 |
| 395 | 0.01 | 0.30 | 0.02 | 0.98 | 1 | Isoc2b | isochorismatase domain containing 2b | 188 | 126 | 378 | 413 | 504 | 465 | 431 | 580 | 537 | 323 | 399 |
| 326 | 0.11 | 0.48 | 0.24 | 0.81 | 1 | Cib3 | calcium and integrin binding family memb | 29 | 5 | 61 | 283 | 306 | 703 | 431 | 475 | 313 | 307 | 676 |
| 54256 | -1.04 | 1.17 | -0.89 | 0.37 | 1 | Cpb1 | carboxypeptidase B1 (tissue) | 75608 | 294145 | 214113 | 9625 | 587 | 38 | 430 | 379 | 403 | 202 | 1288 |
| 510 | -0.30 | 0.23 | -1.30 | 0.19 | 1 | Adprm | ADP-ribose/CDP-alcohol diphosphatase, | 547 | 512 | 531 | 662 | 435 | 670 | 430 | 363 | 494 | 312 | 654 |
| 404 | -0.12 | 0.20 | -0.61 | 0.54 | 1 | C1qtnf4 | C1q and tumor necrosis factor related pro | 496 | 381 | 630 | 401 | 386 | 314 | 430 | 351 | 352 | 424 | 283 |
| 383 | -0.01 | 0.20 | -0.03 | 0.97 | 1 | Saysd1 | SAYSVFN motif domain containing 1 | 315 | 321 | 314 | 340 | 491 | 374 | 430 | 300 | 425 | 455 | 452 |
| 1060 | 0.02 | 0.17 | 0.12 | 0.90 | 1 | Ccnl1 | cyclin L1 | 1446 | 967 | 814 | 461 | 376 | 2900 | 429 | 394 | 440 | 438 | 2991 |
| 617 | -0.26 | 0.21 | -1.26 | 0.21 | 1 | Nfia | nuclear factor I/A | 1645 | 1292 | 1148 | 432 | 306 | 306 | 429 | 236 | 353 | 343 | 290 |
| 505 | -0.07 | 0.18 | -0.39 | 0.69 | 1 | Tsen54 | tRNA splicing endonuclease subunit 54 | 595 | 511 | 548 | 403 | 571 | 589 | 429 | 348 | 472 | 532 | 552 |
| 499 | -0.01 | 0.22 | -0.06 | 0.95 | 1 | Yod1 | YOD1 deubiquitinase | 444 | 374 | 387 | 408 | 397 | 911 | 429 | 482 | 474 | 280 | 899 |
| 495 | -0.08 | 0.21 | -0.37 | 0.71 | 1 | Fn3krp | fructosamine 3 kinase related protein | 462 | 582 | 579 | 520 | 494 | 517 | 429 | 349 | 501 | 558 | 453 |
| 488 | 0.24 | 0.20 | 1.19 | 0.23 | 1 | Iah1 | isoamyl acetate-hydrolyzing esterase 1 h | 713 | 613 | 500 | 437 | 350 | 386 | 429 | 719 | 357 | 469 | 399 |
| 467 | 0.01 | 0.26 | 0.04 | 0.97 | 1 | Dhx35 | DEAH (Asp-Glu-Ala-His) box polypeptide | 542 | 515 | 510 | 442 | 387 | 485 | 429 | 286 | 413 | 726 | 400 |
| 460 | -0.08 | 0.17 | -0.46 | 0.64 | 1 | Mrps15 | mitochondrial ribosomal protein S15 | 540 | 426 | 559 | 446 | 413 | 526 | 429 | 400 | 415 | 420 | 487 |
| 445 | 0.02 | 0.21 | 0.10 | 0.92 | 1 | Tmem203 | transmembrane protein 203 | 304 | 307 | 348 | 511 | 598 | 412 | 429 | 543 | 436 | 639 | 372 |
| 418 | 0.07 | 0.16 | 0.44 | 0.66 | 1 | Lsm1 | LSM1 homolog, mRNA degradation asso | 389 | 387 | 394 | 382 | 418 | 438 | 429 | 426 | 409 | 502 | 418 |
| 394 | -0.38 | 0.21 | -1.82 | 0.07 | 1 | Slc18b1 | solute carrier family 18, subfamily B, mem | 409 | 396 | 476 | 490 | 439 | 354 | 429 | 280 | 487 | 321 | 257 |
| 1159 | -0.33 | 0.18 | -1.88 | 0.06 | 1 | Slc44a2 | solute carrier family 44, member 2 | 3611 | 3087 | 2877 | 596 | 278 | 477 | 428 | 400 | 276 | 327 | 388 |

| | | | | | | | | | | | | | | | | | | |
|---|---|---|---|---|---|---|---|---|---|---|---|---|---|---|---|---|---|---|
| 627 | 0.02 | 0.16 | 0.10 | 0.92 | 1 | Efemp2 | epidermal growth factor-containing fibulin | 1347 | 1082 | 1033 | 510 | 391 | 370 | 428 | 460 | 379 | 550 | 348 |
| 539 | -0.06 | 0.17 | -0.36 | 0.72 | 1 | Wdr4 | WD repeat domain 4 | 490 | 567 | 470 | 396 | 455 | 983 | 428 | 454 | 347 | 465 | 877 |
| 509 | 0.39 | 0.20 | 1.93 | 0.05 | 1 | Homer1 | homer scaffolding protein 1 | 455 | 303 | 395 | 359 | 438 | 729 | 428 | 634 | 529 | 505 | 827 |
| 452 | -0.17 | 0.22 | -0.77 | 0.44 | 1 | Tpt1-ps3 | | 736 | 762 | 642 | 396 | 431 | 249 | 428 | 244 | 415 | 407 | 264 |
| 423 | 0.22 | 0.22 | 1.03 | 0.30 | 1 | Fan1 | FANCD2/FANCI-associated nuclease 1 | 294 | 400 | 322 | 473 | 254 | 540 | 428 | 517 | 363 | 549 | 511 |
| 409 | 0.18 | 0.19 | 0.93 | 0.35 | 1 | Nudt14 | nudix (nucleoside diphosphate linked mo | 295 | 344 | 366 | 428 | 452 | 314 | 428 | 624 | 453 | 486 | 304 |
| 357 | -0.11 | 0.23 | -0.49 | 0.63 | 1 | Ggct | gamma-glutamyl cyclotransferase | 481 | 395 | 423 | 341 | 256 | 344 | 428 | 345 | 292 | 252 | 365 |
| 329 | -0.18 | 0.22 | -0.80 | 0.42 | 1 | Gm20208 | | 196 | 277 | 230 | 371 | 343 | 401 | 428 | 341 | 369 | 265 | 402 |
| 52906 | -0.82 | 1.15 | -0.72 | 0.47 | 1 | Cpa1 | carboxypeptidase A1, pancreatic | 69036 | 284899 | 215580 | 8586 | 792 | 30 | 427 | 570 | 350 | 296 | 1402 |
| 603 | -0.03 | 0.18 | -0.14 | 0.89 | 1 | Sfi1 | Sfi1 homolog, spindle assembly associate | 244 | 310 | 319 | 599 | 474 | 1522 | 427 | 553 | 532 | 531 | 1117 |
| 475 | 0.16 | 0.23 | 0.70 | 0.48 | 1 | Unc119 | unc-119 lipid binding chaperone | 643 | 574 | 529 | 366 | 470 | 350 | 427 | 650 | 385 | 533 | 298 |
| 451 | 0.10 | 0.20 | 0.50 | 0.62 | 1 | Trim21 | tripartite motif-containing 21 | 482 | 493 | 427 | 450 | 454 | 371 | 427 | 656 | 376 | 430 | 392 |
| 405 | 0.12 | 0.24 | 0.52 | 0.60 | 1 | Polr2i | polymerase (RNA) II (DNA directed) polyp | 383 | 437 | 463 | 346 | 385 | 334 | 427 | 608 | 444 | 331 | 294 |
| 398 | -0.02 | 0.19 | -0.12 | 0.91 | 1 | Hsd17b7 | hydroxysteroid (17-beta) dehydrogenase | 377 | 356 | 360 | 375 | 431 | 421 | 427 | 447 | 460 | 319 | 406 |
| 599 | -0.48 | 0.28 | -1.73 | 0.08 | 1 | Slc12a4 | solute carrier family 12, member 4 | 1923 | 1107 | 1255 | 294 | 318 | 274 | 426 | 399 | 186 | 280 | 128 |
| 546 | 0.01 | 0.21 | 0.04 | 0.97 | 1 | Ppp1r3e | protein phosphatase 1, regulatory subuni | 303 | 611 | 507 | 683 | 718 | 488 | 426 | 606 | 604 | 603 | 460 |
| 510 | -0.07 | 0.18 | -0.38 | 0.71 | 1 | Plagl2 | pleiomorphic adenoma gene-like 2 | 558 | 647 | 603 | 519 | 464 | 538 | 426 | 366 | 471 | 505 | 514 |
| 469 | -0.02 | 0.18 | -0.13 | 0.89 | 1 | Surf2 | surfeit gene 2 | 358 | 266 | 301 | 452 | 441 | 798 | 426 | 337 | 499 | 466 | 808 |
| 448 | 0.13 | 0.22 | 0.62 | 0.54 | 1 | Mettl25 | methyltransferase like 25 | 383 | 463 | 534 | 451 | 365 | 445 | 426 | 577 | 358 | 497 | 431 |
| 437 | 0.25 | 0.22 | 1.13 | 0.26 | 1 | Ccdc86 | coiled-coil domain containing 86 | 529 | 400 | 481 | 382 | 333 | 412 | 426 | 580 | 446 | 363 | 457 |
| 781 | -0.12 | 0.23 | -0.52 | 0.60 | 1 | Dusp8 | dual specificity phosphatase 8 | 1235 | 2741 | 1167 | 464 | 422 | 421 | 425 | 678 | 304 | 435 | 292 |
| 510 | 0.10 | 0.20 | 0.52 | 0.60 | 1 | Nln | neurolysin (metallopeptidase M3 family) | 496 | 521 | 578 | 700 | 427 | 389 | 425 | 690 | 538 | 392 | 458 |
| 457 | 0.35 | 0.33 | 1.07 | 0.28 | 1 | Gm10736 | | 451 | 359 | 605 | 341 | 454 | 324 | 425 | 942 | 397 | 422 | 313 |
| 437 | -0.03 | 0.18 | -0.17 | 0.87 | 1 | Pigx | phosphatidylinositol glycan anchor biosyı | 559 | 415 | 646 | 382 | 447 | 359 | 425 | 419 | 418 | 408 | 332 |
| 429 | -0.12 | 0.23 | -0.53 | 0.60 | 1 | Samd12 | sterile alpha motif domain containing 12 | 566 | 569 | 465 | 622 | 338 | 263 | 425 | 335 | 408 | 470 | 261 |
| 425 | -0.09 | 0.20 | -0.44 | 0.66 | 1 | Efna5 | ephrin A5 | 560 | 627 | 523 | 536 | 343 | 204 | 425 | 489 | 287 | 508 | 172 |
| 325 | 0.17 | 0.25 | 0.68 | 0.50 | 1 | Cited4 | Cbp/p300-interacting transactivator, with | 147 | 182 | 162 | 298 | 366 | 348 | 425 | 555 | 317 | 430 | 345 |
| 273 | -0.18 | 0.48 | -0.38 | 0.70 | 1 | Gm10263 | | 790 | 41 | 730 | 10 | 277 | 0 | 425 | 2 | 289 | 438 | 0 |
| 1020 | 0.14 | 0.60 | 0.24 | 0.81 | 1 | Top2a | topoisomerase (DNA) II alpha | 3329 | 892 | 708 | 918 | 904 | 706 | 424 | 1178 | 733 | 602 | 827 |
| 773 | -0.35 | 0.20 | -1.75 | 0.08 | 1 | Ccdc162 | coiled-coil domain containing 162 | 228 | 187 | 164 | 689 | 507 | 2810 | 424 | 396 | 482 | 368 | 2244 |
| 571 | 0.09 | 0.21 | 0.44 | 0.66 | 1 | Lig1 | ligase I, DNA, ATP-dependent | 1260 | 718 | 786 | 421 | 454 | 390 | 424 | 505 | 510 | 502 | 316 |
| 512 | 0.28 | 0.22 | 1.28 | 0.20 | 1 | Cebpzos | CCAAT/enhancer binding protein (C/EBP | 420 | 424 | 783 | 535 | 575 | 303 | 424 | 606 | 582 | 495 | 490 |
| 500 | 0.01 | 0.18 | 0.05 | 0.96 | 1 | Msrb1 | methionine sulfoxide reductase B1 | 835 | 684 | 790 | 338 | 525 | 292 | 424 | 315 | 557 | 478 | 266 |
| 451 | 0.29 | 0.22 | 1.32 | 0.19 | 1 | Utp14a | UTP14A small subunit processome comp | 576 | 406 | 475 | 248 | 398 | 529 | 424 | 494 | 387 | 439 | 590 |
| 625 | -0.12 | 0.20 | -0.60 | 0.55 | 1 | Gramd4 | GRAM domain containing 4 | 1172 | 847 | 917 | 600 | 525 | 493 | 423 | 525 | 431 | 524 | 414 |
| 533 | -0.13 | 0.27 | -0.47 | 0.64 | 1 | Mlh1 | mutL homolog 1 | 488 | 461 | 368 | 332 | 496 | 1098 | 423 | 196 | 547 | 450 | 1005 |
| 411 | -0.30 | 0.24 | -1.29 | 0.20 | 1 | Ocln | occludin | 183 | 310 | 451 | 607 | 487 | 456 | 423 | 380 | 473 | 336 | 416 |
| 364 | 0.22 | 0.23 | 0.98 | 0.33 | 1 | Zfp930 | zinc finger protein 930 | 311 | 320 | 289 | 326 | 259 | 416 | 423 | 519 | 341 | 326 | 478 |
| 320 | -0.06 | 0.24 | -0.27 | 0.79 | 1 | Pard6g | par-6 family cell polarity regulator gamma | 311 | 220 | 198 | 475 | 265 | 274 | 423 | 299 | 294 | 426 | 333 |
| 48209 | -0.77 | 1.18 | -0.66 | 0.51 | 1 | 2210010C04Ril | RIKEN cDNA 2210010C04 gene | 63841 | 244693 | 207812 | 9253 | 728 | 24 | 422 | 992 | 446 | 197 | 1890 |
| 847 | 0.01 | 0.22 | 0.06 | 0.95 | 1 | St3gal2 | ST3 beta-galactoside alpha-2,3-sialyltran | 2151 | 1507 | 1781 | 530 | 563 | 381 | 422 | 462 | 408 | 760 | 354 |
| 528 | -0.24 | 0.24 | -0.97 | 0.33 | 1 | Daglb | diacylglycerol lipase, beta | 830 | 519 | 717 | 552 | 562 | 459 | 422 | 299 | 395 | 574 | 478 |
| 492 | 0.04 | 0.27 | 0.16 | 0.87 | 1 | Slc25a19 | solute carrier family 25 (mitochondrial thia | 498 | 346 | 418 | 470 | 519 | 615 | 422 | 578 | 638 | 272 | 634 |
| 490 | 0.09 | 0.19 | 0.47 | 0.64 | 1 | Vamp1 | vesicle-associated membrane protein 1 | 380 | 596 | 492 | 390 | 481 | 613 | 422 | 443 | 479 | 484 | 610 |
| 488 | 0.27 | 0.18 | 1.48 | 0.14 | 1 | Fbxw9 | F-box and WD-40 domain protein 9 | 496 | 529 | 548 | 290 | 309 | 725 | 422 | 422 | 400 | 444 | 788 |
| 454 | 0.07 | 0.22 | 0.33 | 0.74 | 1 | Maff | v-maf musculoaponeurotic fibrosarcoma | 851 | 656 | 504 | 340 | 255 | 428 | 422 | 253 | 334 | 573 | 381 |
| 478 | -0.21 | 0.20 | -1.04 | 0.30 | 1 | Zfp74 | zinc finger protein 74 | 522 | 477 | 558 | 580 | 465 | 511 | 421 | 478 | 460 | 286 | 502 |
| 466 | -0.05 | 0.21 | -0.25 | 0.80 | 1 | Gm7964 | | 510 | 525 | 666 | 500 | 421 | 395 | 421 | 320 | 456 | 465 | 444 |
| 445 | 0.01 | 0.16 | 0.06 | 0.95 | 1 | Card6 | caspase recruitment domain family, meml | 396 | 461 | 466 | 449 | 530 | 375 | 421 | 526 | 487 | 422 | 357 |
| 436 | -0.16 | 0.21 | -0.77 | 0.44 | 1 | Gpr137b | G protein-coupled receptor 137B | 576 | 688 | 420 | 382 | 385 | 452 | 421 | 424 | 384 | 298 | 363 |
| 411 | 0.14 | 0.24 | 0.57 | 0.57 | 1 | Ndufaf8 | NADH:ubiquinone oxidoreductase compl | 348 | 343 | 433 | 513 | 402 | 259 | 421 | 468 | 390 | 684 | 255 |
| 392 | 0.21 | 0.20 | 1.05 | 0.29 | 1 | Ccnq | cyclin Q | 349 | 321 | 363 | 375 | 400 | 312 | 421 | 548 | 374 | 544 | 311 |
| 354 | -0.19 | 0.25 | -0.76 | 0.45 | 1 | Zfp955b | zinc finger protein 955B | 334 | 405 | 307 | 367 | 304 | 412 | 421 | 212 | 277 | 442 | 409 |
| 258 | 0.20 | 0.26 | 0.77 | 0.44 | 1 | Elovl4 | elongation of very long chain fatty acids (I | 75 | 76 | 89 | 384 | 239 | 183 | 421 | 349 | 356 | 416 | 250 |
| 976 | -0.14 | 0.33 | -0.41 | 0.68 | 1 | Sik1 | salt inducible kinase 1 | 4164 | 2421 | 1675 | 197 | 394 | 269 | 420 | 226 | 552 | 249 | 167 |
| 475 | 0.09 | 0.17 | 0.51 | 0.61 | 1 | Setd6 | SET domain containing 6 | 298 | 331 | 417 | 488 | 459 | 651 | 420 | 565 | 495 | 389 | 711 |
| 462 | -0.15 | 0.22 | -0.67 | 0.50 | 1 | Mrps16 | mitochondrial ribosomal protein S16 | 519 | 343 | 644 | 459 | 507 | 464 | 420 | 288 | 487 | 546 | 399 |
| 518 | 0.33 | 0.26 | 1.29 | 0.20 | 1 | 2310033P09Ril | RIKEN cDNA 2310033P09 gene | 578 | 647 | 630 | 391 | 408 | 440 | 419 | 883 | 356 | 508 | 437 |
| 396 | 0.13 | 0.21 | 0.63 | 0.53 | 1 | Rad1 | RAD1 checkpoint DNA exonuclease | 422 | 446 | 338 | 301 | 381 | 404 | 419 | 315 | 448 | 455 | 431 |
| 569 | 0.17 | 0.20 | 0.87 | 0.38 | 1 | Ddit3 | DNA-damage inducible transcript 3 | 381 | 331 | 378 | 360 | 566 | 1114 | 418 | 569 | 438 | 531 | 1169 |

| | | | | | | | | | | | | | | | | | |
|---|---|---|---|---|---|---|---|---|---|---|---|---|---|---|---|---|---|
| 550 | -0.30 | 0.19 | -1.58 | 0.12 | 1 | Nrbp2 | nuclear receptor binding protein 2 | 575 | 454 | 431 | 542 | 573 | 1005 | 418 | 383 | 454 | 421 | 791 |
| 509 | 0.13 | 0.24 | 0.56 | 0.57 | 1 | Ophn1 | oligophrenin 1 | 625 | 567 | 626 | 472 | 387 | 509 | 418 | 709 | 497 | 323 | 464 |
| 493 | 0.01 | 0.17 | 0.07 | 0.95 | 1 | Zfp956 | zinc finger protein 956 | 469 | 617 | 470 | 415 | 436 | 646 | 418 | 504 | 401 | 366 | 683 |
| 460 | 0.11 | 0.21 | 0.51 | 0.61 | 1 | Nfkbib | nuclear factor of kappa light polypeptide g | 586 | 378 | 369 | 433 | 527 | 415 | 418 | 432 | 528 | 498 | 477 |
| 457 | 0.04 | 0.22 | 0.16 | 0.87 | 1 | Sult4a1 | sulfotransferase family 4A, member 1 | 457 | 747 | 754 | 317 | 428 | 352 | 418 | 375 | 425 | 389 | 364 |
| 417 | -0.04 | 0.17 | -0.22 | 0.83 | 1 | Mrpl32 | mitochondrial ribosomal protein L32 | 353 | 414 | 397 | 441 | 391 | 472 | 418 | 410 | 375 | 358 | 554 |
| 409 | -0.34 | 0.23 | -1.51 | 0.13 | 1 | Tnfrsf11b | tumor necrosis factor receptor superfamily | 241 | 220 | 244 | 776 | 602 | 320 | 418 | 452 | 573 | 415 | 235 |
| 280 | -0.32 | 0.28 | -1.16 | 0.25 | 1 | A030001D20Rik | | 194 | 276 | 280 | 305 | 307 | 260 | 418 | 288 | 301 | 191 | 260 |
| 254 | -0.19 | 0.29 | -0.67 | 0.50 | 1 | Pcdhgb2 | protocadherin gamma subfamily B, 2 | 182 | 247 | 233 | 302 | 290 | 140 | 418 | 196 | 338 | 285 | 159 |
| 579 | -0.18 | 0.23 | -0.81 | 0.42 | 1 | Klc3 | kinesin light chain 3 | 728 | 743 | 974 | 523 | 558 | 564 | 417 | 686 | 439 | 295 | 437 |
| 554 | 0.07 | 0.14 | 0.46 | 0.65 | 1 | Specc1 | sperm antigen with calponin homology ar | 989 | 863 | 856 | 407 | 407 | 419 | 417 | 383 | 450 | 464 | 434 |
| 527 | -0.02 | 0.16 | -0.14 | 0.89 | 1 | Pole3 | polymerase (DNA directed), epsilon 3 (p1 | 887 | 637 | 806 | 374 | 529 | 428 | 417 | 367 | 428 | 497 | 430 |
| 408 | -0.01 | 0.26 | -0.05 | 0.96 | 1 | Zfp408 | zinc finger protein 408 | 374 | 393 | 474 | 279 | 303 | 620 | 417 | 447 | 252 | 286 | 643 |
| 397 | -0.06 | 0.18 | -0.31 | 0.76 | 1 | Kcnd1 | potassium voltage-gated channel, Shal-re | 317 | 391 | 314 | 450 | 405 | 421 | 417 | 519 | 388 | 417 | 326 |
| 363 | -0.01 | 0.22 | -0.04 | 0.97 | 1 | Armcx5 | armadillo repeat containing, X-linked 5 | 270 | 158 | 190 | 414 | 302 | 562 | 417 | 320 | 411 | 428 | 520 |
| 673 | 0.20 | 0.24 | 0.86 | 0.39 | 1 | Zfp950 | zinc finger protein 950 | 275 | 335 | 196 | 270 | 284 | 2171 | 416 | 586 | 287 | 331 | 2254 |
| 630 | 0.06 | 0.28 | 0.22 | 0.82 | 1 | Mfhas1 | malignant fibrous histiocytoma amplified s | 1422 | 1000 | 771 | 565 | 358 | 456 | 416 | 802 | 475 | 359 | 304 |
| 606 | 0.02 | 0.25 | 0.07 | 0.94 | 1 | Rfxap | regulatory factor X-associated protein | 699 | 870 | 642 | 706 | 563 | 533 | 416 | 444 | 481 | 754 | 563 |
| 499 | 0.22 | 0.28 | 0.80 | 0.42 | 1 | Trps1 | transcriptional repressor GATA binding 1 | 719 | 499 | 435 | 342 | 344 | 686 | 416 | 715 | 387 | 346 | 598 |
| 375 | 0.02 | 0.25 | 0.07 | 0.94 | 1 | Gm10762 | | 186 | 329 | 360 | 460 | 468 | 279 | 416 | 507 | 359 | 470 | 294 |
| 555 | -0.10 | 0.19 | -0.51 | 0.61 | 1 | Rtca | RNA 3'-terminal phosphate cyclase | 649 | 585 | 699 | 567 | 658 | 527 | 415 | 505 | 459 | 555 | 488 |
| 516 | 0.08 | 0.23 | 0.34 | 0.73 | 1 | Mtln | | 446 | 408 | 480 | 560 | 725 | 419 | 415 | 750 | 551 | 392 | 529 |
| 509 | 0.02 | 0.17 | 0.15 | 0.88 | 1 | Mxra7 | matrix-remodelling associated 7 | 1317 | 764 | 1137 | 276 | 357 | 142 | 415 | 309 | 317 | 407 | 156 |
| 477 | -0.02 | 0.27 | -0.07 | 0.95 | 1 | Ppdpf | pancreatic progenitor cell differentiation a | 827 | 509 | 828 | 376 | 435 | 313 | 415 | 582 | 343 | 366 | 254 |
| 437 | 0.17 | 0.23 | 0.73 | 0.47 | 1 | Gm44950 | | 420 | 381 | 491 | 401 | 453 | 365 | 415 | 374 | 402 | 721 | 388 |
| 401 | 0.14 | 0.25 | 0.55 | 0.58 | 1 | Eef1akmt1 | EEF1A alpha lysine methyltransferase 1 | 401 | 360 | 458 | 345 | 418 | 336 | 415 | 609 | 330 | 398 | 347 |
| 1176 | -0.15 | 0.22 | -0.67 | 0.50 | 1 | Nfkbiz | nuclear factor of kappa light polypeptide g | 2467 | 1098 | 729 | 246 | 272 | 3486 | 414 | 166 | 264 | 422 | 3373 |
| 694 | -0.13 | 0.23 | -0.55 | 0.58 | 1 | Reck | reversion-inducing-cysteine-rich protein w | 1868 | 1246 | 1436 | 526 | 352 | 308 | 414 | 584 | 300 | 340 | 264 |
| 572 | 0.15 | 0.18 | 0.84 | 0.40 | 1 | Fam160b2 | family with sequence similarity 160, meml | 454 | 532 | 459 | 632 | 547 | 711 | 414 | 794 | 501 | 593 | 657 |
| 353 | 0.38 | 0.21 | 1.77 | 0.08 | 1 | Foxp2 | forkhead box P2 | 357 | 524 | 168 | 400 | 194 | 227 | 414 | 491 | 269 | 564 | 276 |
| 630 | -0.15 | 0.26 | -0.57 | 0.57 | 1 | Lmnb1 | lamin B1 | 1724 | 917 | 758 | 437 | 548 | 457 | 413 | 410 | 439 | 371 | 452 |
| 519 | -0.24 | 0.28 | -0.88 | 0.38 | 1 | Xkr6 | X-linked Kx blood group related 6 | 331 | 554 | 493 | 388 | 434 | 1046 | 413 | 199 | 368 | 428 | 1051 |
| 509 | 0.06 | 0.17 | 0.38 | 0.71 | 1 | Zfp639 | zinc finger protein 639 | 456 | 631 | 468 | 480 | 299 | 797 | 413 | 451 | 385 | 402 | 814 |
| 429 | -0.02 | 0.20 | -0.11 | 0.91 | 1 | Otud3 | OTU domain containing 3 | 253 | 450 | 272 | 480 | 477 | 514 | 413 | 507 | 508 | 386 | 461 |
| 416 | 0.37 | 0.25 | 1.47 | 0.14 | 1 | Ddx60 | DEAD (Asp-Glu-Ala-Asp) box polypeptide | 273 | 356 | 333 | 459 | 287 | 424 | 413 | 378 | 558 | 578 | 517 |
| 353 | -0.05 | 0.21 | -0.22 | 0.83 | 1 | Tsen2 | tRNA splicing endonuclease subunit 2 | 341 | 226 | 243 | 314 | 407 | 422 | 413 | 273 | 454 | 359 | 430 |
| 344 | -0.28 | 0.27 | -1.05 | 0.30 | 1 | Wscd1 | WSC domain containing 1 | 325 | 286 | 458 | 358 | 376 | 340 | 413 | 358 | 220 | 281 | 367 |
| 309 | -0.25 | 0.21 | -1.20 | 0.23 | 1 | Efhc1 | EF-hand domain (C-terminal) containing | 286 | 264 | 299 | 284 | 279 | 412 | 413 | 250 | 284 | 274 | 354 |
| 758 | -0.06 | 0.28 | -0.24 | 0.81 | 1 | Naaa | N-acylethanolamine acid amidase | 2663 | 1269 | 947 | 454 | 504 | 357 | 412 | 368 | 522 | 573 | 269 |
| 470 | 0.06 | 0.22 | 0.27 | 0.79 | 1 | Nmral1 | NmrA-like family domain containing 1 | 440 | 495 | 457 | 389 | 555 | 487 | 412 | 308 | 519 | 559 | 551 |
| 354 | 0.06 | 0.21 | 0.30 | 0.76 | 1 | Slx1b | SLX1 structure-specific endonuclease sul | 221 | 169 | 259 | 307 | 340 | 536 | 412 | 353 | 351 | 483 | 457 |
| 569 | -0.02 | 0.19 | -0.13 | 0.89 | 1 | Qtrt2 | queuine tRNA-ribosyltransferase accessc | 479 | 562 | 562 | 446 | 571 | 912 | 411 | 339 | 551 | 500 | 928 |
| 501 | 0.00 | 0.16 | 0.00 | 1.00 | 1 | Rbms2 | RNA binding motif, single stranded intera | 1050 | 880 | 856 | 306 | 314 | 325 | 411 | 314 | 355 | 434 | 264 |
| 465 | -0.18 | 0.20 | -0.88 | 0.38 | 1 | Ppfia3 | protein tyrosine phosphatase, receptor typ | 535 | 476 | 645 | 487 | 423 | 512 | 411 | 436 | 413 | 443 | 332 |
| 460 | 0.34 | 0.22 | 1.58 | 0.11 | 1 | Tmem151b | transmembrane protein 151B | 333 | 602 | 483 | 493 | 408 | 298 | 411 | 618 | 416 | 609 | 395 |
| 394 | -0.33 | 0.22 | -1.49 | 0.14 | 1 | Med27 | mediator complex subunit 27 | 447 | 309 | 357 | 492 | 404 | 477 | 411 | 268 | 385 | 424 | 366 |
| 1096 | 0.13 | 0.13 | 0.98 | 0.33 | 1 | Cx3cl1 | chemokine (C-X3-C motif) ligand 1 | 2261 | 2862 | 3079 | 562 | 536 | 344 | 410 | 526 | 543 | 520 | 409 |
| 767 | 0.07 | 0.20 | 0.36 | 0.72 | 1 | Myo1e | myosin IE | 1798 | 1505 | 1785 | 431 | 408 | 361 | 410 | 636 | 408 | 380 | 310 |
| 693 | 0.12 | 0.19 | 0.64 | 0.52 | 1 | Tspo | translocator protein | 1839 | 1134 | 1496 | 347 | 499 | 250 | 410 | 489 | 401 | 508 | 250 |
| 452 | -0.05 | 0.24 | -0.21 | 0.83 | 1 | Rnaseh2a | ribonuclease H2, large subunit | 469 | 592 | 573 | 400 | 458 | 396 | 410 | 634 | 376 | 254 | 408 |
| 405 | 0.19 | 0.21 | 0.92 | 0.36 | 1 | Cyp4f39 | cytochrome P450, family 4, subfamily f, pc | 68 | 71 | 56 | 663 | 571 | 346 | 410 | 648 | 619 | 657 | 351 |
| 404 | -0.04 | 0.20 | -0.21 | 0.83 | 1 | Cenpx | centromere protein X | 511 | 359 | 561 | 422 | 377 | 315 | 410 | 380 | 427 | 401 | 285 |
| 381 | -0.15 | 0.21 | -0.72 | 0.47 | 1 | Tnfrsf11a | tumor necrosis factor receptor superfamily | 335 | 274 | 320 | 477 | 558 | 279 | 410 | 350 | 534 | 348 | 308 |
| 358 | 0.03 | 0.22 | 0.14 | 0.89 | 1 | Med11 | mediator complex subunit 11 | 390 | 347 | 319 | 342 | 363 | 307 | 410 | 502 | 339 | 323 | 301 |
| 657 | -0.11 | 0.19 | -0.58 | 0.56 | 1 | Dis3 | DIS3 homolog, exosome endoribonuclea | 896 | 791 | 735 | 603 | 507 | 932 | 409 | 415 | 500 | 394 | 1047 |
| 586 | 0.06 | 0.24 | 0.23 | 0.82 | 1 | Resf1 | retroelement silencing factor 1 | 973 | 736 | 577 | 412 | 285 | 953 | 409 | 336 | 478 | 327 | 958 |
| 496 | 0.19 | 0.20 | 0.94 | 0.35 | 1 | Ctu1 | cytosolic thiouridylase subunit 1 | 514 | 465 | 518 | 548 | 404 | 483 | 409 | 731 | 449 | 456 | 484 |
| 424 | -0.09 | 0.20 | -0.46 | 0.64 | 1 | Sema6b | sema domain, transmembrane domain (T | 680 | 596 | 644 | 473 | 356 | 194 | 409 | 343 | 399 | 339 | 227 |
| 347 | -0.24 | 0.20 | -1.24 | 0.22 | 1 | 4933431K14Rik | | 143 | 213 | 179 | 517 | 506 | 349 | 409 | 407 | 436 | 286 | 375 |
| 573 | 0.11 | 0.16 | 0.67 | 0.50 | 1 | Polr3h | polymerase (RNA) III (DNA directed) poly | 589 | 756 | 642 | 602 | 582 | 494 | 408 | 627 | 531 | 553 | 519 |

| | | | | | | | | | | | | | | | | | | |
|---|---|---|---|---|---|---|---|---|---|---|---|---|---|---|---|---|---|---|
| 521 | -0.07 | 0.21 | -0.33 | 0.74 | 1 | Acp6 | acid phosphatase 6, lysophosphatidic | 440 | 362 | 541 | 410 | 492 | 934 | 408 | 290 | 434 | 600 | 817 |
| 410 | 0.02 | 0.18 | 0.09 | 0.93 | 1 | Nt5c | 5',3'-nucleotidase, cytosolic | 580 | 389 | 624 | 326 | 423 | 298 | 408 | 331 | 367 | 419 | 343 |
| 374 | 0.11 | 0.19 | 0.57 | 0.57 | 1 | Dus2 | dihydrouridine synthase 2 | 322 | 334 | 363 | 294 | 405 | 385 | 408 | 345 | 381 | 401 | 480 |
| 623 | 0.09 | 0.21 | 0.44 | 0.66 | 1 | Mtf2 | metal response element binding transcrip | 746 | 935 | 574 | 659 | 376 | 788 | 407 | 803 | 518 | 327 | 718 |
| 557 | 0.10 | 0.18 | 0.58 | 0.56 | 1 | Rrs1 | ribosome biogenesis regulator 1 | 626 | 706 | 669 | 580 | 536 | 470 | 407 | 510 | 532 | 618 | 474 |
| 490 | -0.22 | 0.19 | -1.15 | 0.25 | 1 | B4galt4 | UDP-Gal:betaGlcNAc beta 1,4-galactosyl | 793 | 948 | 1030 | 381 | 358 | 269 | 407 | 308 | 348 | 259 | 294 |
| 468 | -0.07 | 0.18 | -0.42 | 0.68 | 1 | Plekhf2 | pleckstrin homology domain containing, fa | 812 | 704 | 750 | 407 | 338 | 324 | 407 | 343 | 414 | 328 | 315 |
| 465 | 0.21 | 0.19 | 1.10 | 0.27 | 1 | Kctd6 | potassium channel tetramerisation domai | 316 | 304 | 265 | 528 | 418 | 600 | 407 | 670 | 508 | 383 | 721 |
| 463 | 0.13 | 0.19 | 0.70 | 0.48 | 1 | Zfp213 | zinc finger protein 213 | 492 | 495 | 442 | 513 | 398 | 412 | 407 | 706 | 376 | 439 | 415 |
| 11498 | -0.46 | 1.27 | -0.36 | 0.72 | 1 | Cd74 | CD74 antigen (invariant polypeptide of m | 61807 | 27355 | 36301 | 93 | 125 | 40 | 406 | 109 | 194 | 0 | 52 |
| 693 | 0.01 | 0.19 | 0.03 | 0.97 | 1 | Cramp1l | cramped chromatin regulator homolog 1 | 625 | 819 | 682 | 470 | 647 | 1295 | 406 | 561 | 705 | 416 | 997 |
| 674 | 0.11 | 0.28 | 0.38 | 0.71 | 1 | Il11ra1 | interleukin 11 receptor, alpha chain 1 | 1502 | 890 | 1293 | 465 | 464 | 431 | 406 | 811 | 392 | 402 | 357 |
| 501 | 0.01 | 0.16 | 0.06 | 0.95 | 1 | Wee1 | WEE 1 homolog 1 (S. pombe) | 796 | 799 | 682 | 492 | 386 | 330 | 406 | 432 | 445 | 407 | 332 |
| 399 | -0.07 | 0.21 | -0.34 | 0.73 | 1 | Slc8a2 | solute carrier family 8 (sodium/calcium ex | 571 | 522 | 413 | 370 | 419 | 288 | 406 | 320 | 421 | 328 | 329 |
| 337 | 0.04 | 0.26 | 0.15 | 0.88 | 1 | Zfp958 | zinc finger protein 958 | 369 | 327 | 182 | 322 | 282 | 386 | 406 | 254 | 353 | 353 | 473 |
| 285 | -0.09 | 0.44 | -0.21 | 0.84 | 1 | Gm48837 | | 51 | 10 | 76 | 17 | 358 | 684 | 406 | 6 | 361 | 474 | 692 |
| 534 | -0.05 | 0.22 | -0.21 | 0.83 | 1 | Zfp157 | zinc finger protein 157 | 536 | 768 | 535 | 508 | 382 | 773 | 405 | 333 | 492 | 437 | 711 |
| 507 | -0.14 | 0.22 | -0.65 | 0.52 | 1 | Frrs1l | ferric-chelate reductase 1 like | 1019 | 1095 | 906 | 400 | 261 | 254 | 405 | 397 | 366 | 325 | 150 |
| 495 | -0.11 | 0.19 | -0.59 | 0.55 | 1 | Kin | Kin17 DNA and RNA binding protein | 432 | 574 | 599 | 417 | 454 | 690 | 405 | 295 | 456 | 370 | 755 |
| 421 | -0.24 | 0.18 | -1.37 | 0.17 | 1 | 1110002L01Rik | | 391 | 365 | 357 | 411 | 477 | 620 | 405 | 335 | 433 | 394 | 441 |
| 328 | -0.32 | 0.20 | -1.60 | 0.11 | 1 | Armcx6 | armadillo repeat containing, X-linked 6 | 379 | 354 | 350 | 343 | 312 | 337 | 405 | 220 | 308 | 326 | 270 |
| 290 | 0.03 | 0.26 | 0.13 | 0.89 | 1 | Gm49492 | | 224 | 232 | 184 | 348 | 313 | 172 | 405 | 293 | 242 | 581 | 193 |
| 288 | -0.17 | 0.23 | -0.72 | 0.47 | 1 | Klf15 | Kruppel-like factor 15 | 248 | 194 | 222 | 304 | 267 | 345 | 405 | 322 | 205 | 300 | 353 |
| 609 | -0.34 | 0.23 | -1.46 | 0.14 | 1 | Rusc2 | RUN and SH3 domain containing 2 | 1356 | 967 | 1114 | 492 | 405 | 503 | 404 | 346 | 384 | 225 | 501 |
| 603 | -0.25 | 0.14 | -1.77 | 0.08 | 1 | Retreg1 | reticulophagy regulator 1 | 911 | 1261 | 1209 | 487 | 439 | 435 | 404 | 395 | 372 | 334 | 385 |
| 467 | 0.09 | 0.23 | 0.39 | 0.70 | 1 | Zfp628 | zinc finger protein 628 | 539 | 399 | 427 | 460 | 500 | 447 | 404 | 688 | 404 | 448 | 418 |
| 460 | -0.12 | 0.18 | -0.68 | 0.49 | 1 | Pard6b | par-6 family cell polarity regulator beta | 422 | 491 | 513 | 536 | 572 | 381 | 404 | 459 | 453 | 492 | 331 |
| 448 | 0.23 | 0.20 | 1.18 | 0.24 | 1 | Wdr70 | WD repeat domain 70 | 550 | 529 | 445 | 286 | 388 | 494 | 404 | 456 | 447 | 394 | 532 |
| 432 | 0.16 | 0.24 | 0.66 | 0.51 | 1 | Mvk | mevalonate kinase | 371 | 421 | 454 | 536 | 367 | 331 | 404 | 591 | 334 | 630 | 315 |
| 400 | 0.22 | 0.23 | 0.95 | 0.34 | 1 | Zfp563 | zinc finger protein 563 | 286 | 331 | 321 | 400 | 316 | 449 | 404 | 564 | 377 | 290 | 666 |
| 255 | -0.47 | 0.29 | -1.63 | 0.10 | 1 | Pdss2 | prenyl (solanesyl) diphosphate synthase, | 221 | 240 | 224 | 288 | 251 | 287 | 404 | 173 | 262 | 174 | 284 |
| 59825 | -0.84 | 1.11 | -0.76 | 0.45 | 1 | Cela1 | chymotrypsin-like elastase family, membe | 69385 | 301018 | 270952 | 11418 | 936 | 68 | 403 | 969 | 448 | 249 | 2229 |
| 560 | -0.04 | 0.17 | -0.25 | 0.81 | 1 | Wdr20 | WD repeat domain 20 | 571 | 594 | 671 | 536 | 570 | 676 | 403 | 392 | 577 | 453 | 723 |
| 474 | -0.04 | 0.22 | -0.17 | 0.87 | 1 | Trim62 | tripartite motif-containing 62 | 589 | 843 | 836 | 354 | 359 | 374 | 403 | 359 | 333 | 457 | 311 |
| 440 | -0.19 | 0.22 | -0.88 | 0.38 | 1 | Utp25 | UTP25 small subunit processome compo | 407 | 515 | 357 | 393 | 448 | 621 | 403 | 414 | 386 | 242 | 649 |
| 402 | 0.02 | 0.20 | 0.09 | 0.93 | 1 | Dmac1 | distal membrane arm assembly complex | 470 | 486 | 393 | 342 | 430 | 346 | 403 | 489 | 355 | 365 | 344 |
| 377 | 0.16 | 0.17 | 0.95 | 0.34 | 1 | Rrp9 | RRP9, small subunit (SSU) processome c | 480 | 292 | 351 | 271 | 339 | 410 | 403 | 322 | 321 | 493 | 466 |
| 337 | -0.12 | 0.24 | -0.49 | 0.62 | 1 | Sac3d1 | SAC3 domain containing 1 | 261 | 408 | 368 | 325 | 348 | 314 | 403 | 262 | 334 | 337 | 346 |
| 519 | 0.03 | 0.22 | 0.14 | 0.89 | 1 | Rfx1 | regulatory factor X, 1 (influences HLA clas | 774 | 594 | 520 | 436 | 471 | 575 | 402 | 426 | 473 | 622 | 420 |
| 474 | 0.16 | 0.23 | 0.69 | 0.49 | 1 | Dxo | decapping exoribonuclease | 467 | 253 | 343 | 393 | 604 | 572 | 402 | 513 | 492 | 591 | 587 |
| 444 | -0.12 | 0.19 | -0.61 | 0.54 | 1 | Klhl36 | kelch-like 36 | 596 | 488 | 430 | 531 | 408 | 403 | 402 | 495 | 350 | 453 | 329 |
| 379 | 0.08 | 0.19 | 0.41 | 0.68 | 1 | Ntmt1 | N-terminal Xaa-Pro-Lys N-methyltransfera | 412 | 286 | 418 | 296 | 364 | 419 | 402 | 315 | 351 | 517 | 390 |
| 513 | 0.21 | 0.19 | 1.14 | 0.26 | 1 | Prkrip1 | Prkr interacting protein 1 (IL11 inducible) | 516 | 668 | 581 | 410 | 503 | 478 | 401 | 606 | 428 | 532 | 520 |
| 493 | 0.22 | 0.27 | 0.82 | 0.41 | 1 | Dse | dermatan sulfate epimerase | 1358 | 659 | 739 | 298 | 297 | 255 | 401 | 383 | 441 | 278 | 318 |
| 425 | 0.02 | 0.22 | 0.11 | 0.91 | 1 | Gtpbp6 | GTP binding protein 6 (putative) | 520 | 425 | 566 | 284 | 331 | 555 | 401 | 392 | 264 | 372 | 563 |
| 390 | -0.03 | 0.20 | -0.16 | 0.87 | 1 | Amacr | alpha-methylacyl-CoA racemase | 277 | 395 | 498 | 413 | 429 | 332 | 401 | 363 | 449 | 380 | 350 |
| 607 | -0.08 | 0.27 | -0.32 | 0.75 | 1 | Pan2 | PAN2 poly(A) specific ribonuclease subur | 528 | 329 | 333 | 352 | 328 | 1832 | 400 | 412 | 419 | 224 | 1524 |
| 450 | -0.02 | 0.18 | -0.12 | 0.90 | 1 | Dph7 | diphthamine biosynethesis 7 | 365 | 450 | 439 | 358 | 473 | 643 | 400 | 388 | 396 | 456 | 585 |
| 434 | -0.19 | 0.27 | -0.70 | 0.48 | 1 | Fez1 | fasciculation and elongation protein zeta | 503 | 504 | 573 | 333 | 579 | 363 | 400 | 372 | 581 | 191 | 372 |
| 390 | 0.20 | 0.26 | 0.74 | 0.46 | 1 | 2510002D24Ril | RIKEN cDNA 2510002D24 gene | 296 | 222 | 339 | 367 | 416 | 412 | 400 | 609 | 343 | 375 | 509 |
| 389 | -0.31 | 0.37 | -0.82 | 0.41 | 1 | Uprt | uracil phosphoribosyltransferase | 403 | 554 | 439 | 337 | 396 | 389 | 400 | 528 | 382 | 99 | 349 |
| 352 | 0.20 | 0.28 | 0.71 | 0.48 | 1 | Pparg | peroxisome proliferator activated receptor | 170 | 216 | 289 | 352 | 390 | 295 | 400 | 400 | 557 | 607 | 200 |
| 351 | -0.02 | 0.27 | -0.08 | 0.94 | 1 | Lysmd4 | LysM, putative peptidoglycan-binding, do | 207 | 236 | 373 | 278 | 315 | 559 | 400 | 282 | 395 | 446 | 377 |
| 350 | -0.25 | 0.20 | -1.27 | 0.21 | 1 | Ccdc148 | coiled-coil domain containing 148 | 246 | 338 | 311 | 343 | 347 | 506 | 400 | 314 | 315 | 264 | 462 |
| 501 | -0.10 | 0.18 | -0.58 | 0.56 | 1 | Zswim9 | zinc finger SWIM-type containing 9 | 630 | 623 | 618 | 483 | 480 | 520 | 399 | 523 | 454 | 325 | 460 |
| 467 | 0.05 | 0.19 | 0.28 | 0.78 | 1 | Cox20 | cytochrome c oxidase assembly protein 2 | 567 | 424 | 414 | 492 | 485 | 453 | 399 | 488 | 417 | 477 | 527 |
| 443 | 0.04 | 0.18 | 0.23 | 0.82 | 1 | 2610021A01Rik | | 378 | 427 | 419 | 457 | 343 | 598 | 399 | 502 | 414 | 335 | 601 |
| 434 | -0.30 | 0.22 | -1.39 | 0.17 | 1 | Zfp606 | zinc finger protein 606 | 444 | 421 | 518 | 403 | 411 | 612 | 399 | 238 | 282 | 379 | 672 |
| 425 | -0.08 | 0.18 | -0.42 | 0.68 | 1 | Zcchc3 | zinc finger, CCHC domain containing 3 | 596 | 606 | 414 | 396 | 384 | 383 | 399 | 441 | 333 | 408 | 317 |
| 397 | -0.03 | 0.23 | -0.14 | 0.89 | 1 | Mpv17l | Mpv17 transgene, kidney disease mutant | 595 | 740 | 595 | 303 | 282 | 251 | 399 | 283 | 353 | 258 | 304 |

| | | | | | | | | | | | | | | | | | |
|---|---|---|---|---|---|---|---|---|---|---|---|---|---|---|---|---|---|
| 300 | 0.04 | 0.25 | 0.16 | 0.87 | 1 | Zc3h12b | zinc finger CCCH-type containing 12B | 245 | 313 | 239 | 303 | 241 | 298 | 399 | 260 | 338 | 350 | 315 |
| 485 | 0.20 | 0.16 | 1.27 | 0.20 | 1 | Fcf1 | FCF1 rRNA processing protein | 715 | 548 | 607 | 398 | 382 | 437 | 398 | 437 | 473 | 475 | 468 |
| 480 | 0.31 | 0.25 | 1.26 | 0.21 | 1 | Paxx | non-homologous end joining factor | 415 | 195 | 382 | 345 | 482 | 715 | 398 | 562 | 438 | 585 | 758 |
| 438 | 0.10 | 0.25 | 0.39 | 0.69 | 1 | Ppie | peptidylprolyl isomerase E (cyclophilin E) | 706 | 590 | 798 | 258 | 307 | 356 | 398 | 445 | 316 | 283 | 358 |
| 424 | 0.07 | 0.24 | 0.30 | 0.77 | 1 | Bcl7c | B cell CLL/lymphoma 7C | 723 | 632 | 482 | 308 | 369 | 299 | 398 | 439 | 429 | 300 | 280 |
| 362 | -0.35 | 0.22 | -1.60 | 0.11 | 1 | Pcdhga6 | protocadherin gamma subfamily A, 6 | 305 | 465 | 243 | 543 | 423 | 312 | 398 | 352 | 379 | 300 | 259 |
| 517 | -0.24 | 0.25 | -0.98 | 0.33 | 1 | Rhog | ras homolog family member G | 1181 | 924 | 757 | 505 | 359 | 265 | 397 | 226 | 323 | 466 | 280 |
| 477 | -0.26 | 0.22 | -1.16 | 0.25 | 1 | Ttc22 | tetratricopeptide repeat domain 22 | 576 | 543 | 788 | 486 | 473 | 466 | 397 | 353 | 306 | 457 | 408 |
| 440 | -0.17 | 0.29 | -0.59 | 0.55 | 1 | Cep295 | centrosomal protein 295 | 493 | 335 | 223 | 463 | 376 | 764 | 397 | 356 | 435 | 267 | 734 |
| 424 | -0.12 | 0.23 | -0.54 | 0.59 | 1 | Ulk3 | unc-51-like kinase 3 | 283 | 320 | 277 | 291 | 335 | 997 | 397 | 238 | 361 | 425 | 736 |
| 408 | 0.21 | 0.24 | 0.88 | 0.38 | 1 | Zfp729b | zinc finger protein 729b | 369 | 344 | 324 | 420 | 351 | 404 | 397 | 357 | 358 | 733 | 434 |
| 335 | 0.39 | 0.29 | 1.35 | 0.18 | 1 | Rnaseh2b | ribonuclease H2, subunit B | 310 | 157 | 312 | 312 | 266 | 291 | 397 | 507 | 336 | 433 | 361 |
| 199 | -0.27 | 0.38 | -0.71 | 0.48 | 1 | Nanos1 | nanos C2HC-type zinc finger 1 | 195 | 266 | 192 | 185 | 115 | 167 | 397 | 112 | 147 | 216 | 199 |
| 581 | -0.07 | 0.17 | -0.41 | 0.68 | 1 | Arhgef25 | Rho guanine nucleotide exchange factor | 945 | 980 | 941 | 382 | 440 | 576 | 396 | 368 | 320 | 439 | 607 |
| 439 | -0.20 | 0.20 | -1.01 | 0.31 | 1 | Prdm15 | PR domain containing 15 | 378 | 323 | 404 | 412 | 388 | 783 | 396 | 304 | 470 | 289 | 682 |
| 419 | -0.27 | 0.23 | -1.18 | 0.24 | 1 | Zfp141 | zinc finger protein 141 | 457 | 380 | 321 | 525 | 445 | 518 | 396 | 379 | 437 | 286 | 466 |
| 362 | -0.24 | 0.17 | -1.42 | 0.16 | 1 | Lactb | lactamase, beta | 333 | 372 | 367 | 399 | 387 | 392 | 396 | 313 | 361 | 343 | 317 |
| 357 | 0.07 | 0.23 | 0.32 | 0.75 | 1 | Fam122b | family with sequence similarity 122, meml | 336 | 375 | 347 | 322 | 342 | 322 | 396 | 469 | 442 | 286 | 286 |
| 327 | -0.43 | 0.24 | -1.76 | 0.08 | 1 | Nudt12 | nudix (nucleoside diphosphate linked mo | 395 | 334 | 401 | 328 | 363 | 315 | 396 | 159 | 361 | 263 | 286 |
| 59223 | -1.15 | 1.15 | -1.00 | 0.32 | 1 | Cel | carboxyl ester lipase | 82350 | 329593 | 225203 | 11467 | 583 | 38 | 395 | 392 | 347 | 196 | 890 |
| 28783 | -1.33 | 1.10 | -1.21 | 0.23 | 1 | Pnliprp1 | pancreatic lipase related protein 1 | 34320 | 144864 | 131186 | 4085 | 370 | 24 | 395 | 368 | 186 | 91 | 728 |
| 457 | 0.33 | 0.28 | 1.19 | 0.23 | 1 | Sacs | sacsin | 431 | 317 | 439 | 405 | 194 | 772 | 395 | 483 | 448 | 415 | 729 |
| 527 | -0.21 | 0.27 | -0.81 | 0.42 | 1 | Zfp780b | zinc finger protein 780B | 601 | 464 | 427 | 527 | 605 | 746 | 394 | 254 | 562 | 469 | 752 |
| 443 | 0.00 | 0.17 | -0.02 | 0.98 | 1 | Rmi1 | RecQ mediated genome instability 1 | 305 | 280 | 306 | 429 | 496 | 669 | 394 | 469 | 389 | 426 | 710 |
| 384 | -0.53 | 0.33 | -1.59 | 0.11 | 1 | Amn1 | antagonist of mitotic exit network 1 | 370 | 468 | 292 | 386 | 354 | 669 | 394 | 120 | 331 | 332 | 507 |
| 378 | -0.32 | 0.22 | -1.45 | 0.15 | 1 | Grin2c | glutamate receptor, ionotropic, NMDA2C | 121 | 123 | 106 | 555 | 353 | 818 | 394 | 492 | 347 | 241 | 608 |
| 532 | -0.06 | 0.17 | -0.35 | 0.73 | 1 | Med17 | mediator complex subunit 17 | 622 | 598 | 584 | 477 | 546 | 647 | 393 | 512 | 444 | 393 | 632 |
| 468 | -0.09 | 0.21 | -0.42 | 0.68 | 1 | Ino80b | INO80 complex subunit B | 757 | 553 | 612 | 424 | 385 | 435 | 393 | 323 | 349 | 574 | 339 |
| 466 | -0.16 | 0.20 | -0.83 | 0.41 | 1 | Ppwd1 | peptidylprolyl isomerase domain and WD | 421 | 330 | 417 | 544 | 442 | 710 | 393 | 374 | 425 | 434 | 639 |
| 445 | -0.11 | 0.21 | -0.54 | 0.59 | 1 | Plekhg5 | pleckstrin homology domain containing, fa | 715 | 649 | 680 | 407 | 330 | 330 | 393 | 344 | 351 | 486 | 214 |
| 413 | -0.23 | 0.19 | -1.21 | 0.23 | 1 | Adck5 | aarF domain containing kinase 5 | 393 | 195 | 376 | 313 | 398 | 811 | 393 | 234 | 339 | 371 | 725 |
| 398 | 0.11 | 0.32 | 0.35 | 0.73 | 1 | Cks1b | CDC28 protein kinase 1b | 556 | 252 | 336 | 364 | 462 | 342 | 393 | 459 | 510 | 255 | 447 |
| 597 | -0.04 | 0.19 | -0.23 | 0.82 | 1 | Ctc1 | CTS telomere maintenance complex com | 931 | 927 | 875 | 659 | 420 | 471 | 392 | 609 | 489 | 447 | 344 |
| 417 | 0.17 | 0.20 | 0.87 | 0.39 | 1 | Rnf219 | ring finger protein 219 | 475 | 584 | 473 | 309 | 343 | 393 | 392 | 388 | 458 | 344 | 432 |
| 357 | 0.13 | 0.22 | 0.57 | 0.57 | 1 | Rps6ka5 | ribosomal protein S6 kinase, polypeptide | 329 | 403 | 231 | 468 | 284 | 275 | 392 | 563 | 349 | 281 | 354 |
| 307 | -0.27 | 0.26 | -1.03 | 0.30 | 1 | Eef1e1 | eukaryotic translation elongation factor 1 | 352 | 334 | 340 | 237 | 334 | 319 | 392 | 269 | 277 | 185 | 333 |
| 573 | 0.13 | 0.16 | 0.78 | 0.43 | 1 | Vgll4 | vestigial like family member 4 | 1365 | 1175 | 1173 | 311 | 280 | 259 | 391 | 393 | 294 | 348 | 311 |
| 416 | 0.18 | 0.20 | 0.91 | 0.36 | 1 | Ifrd2 | interferon-related developmental regulato | 501 | 318 | 361 | 309 | 379 | 526 | 391 | 429 | 452 | 444 | 463 |
| 331 | -0.18 | 0.27 | -0.65 | 0.52 | 1 | Carmil2 | capping protein regulator and myosin 1 lii | 336 | 389 | 234 | 305 | 347 | 366 | 391 | 228 | 328 | 453 | 260 |
| 310 | -0.30 | 0.23 | -1.34 | 0.18 | 1 | Slc10a3 | solute carrier family 10 (sodium/bile acid c | 270 | 229 | 242 | 415 | 401 | 264 | 391 | 264 | 424 | 277 | 234 |
| 647 | -0.11 | 0.19 | -0.62 | 0.54 | 1 | Nudt11 | nudix (nucleoside diphosphate linked mo | 1538 | 1841 | 1491 | 291 | 251 | 231 | 390 | 308 | 285 | 323 | 170 |
| 511 | 0.19 | 0.20 | 0.97 | 0.33 | 1 | Akap7 | A kinase (PRKA) anchor protein 7 | 563 | 559 | 717 | 464 | 435 | 478 | 390 | 575 | 386 | 506 | 552 |
| 460 | 0.00 | 0.21 | -0.02 | 0.99 | 1 | Borcs6 | BLOC-1 related complex subunit 6 | 661 | 630 | 550 | 450 | 473 | 292 | 390 | 572 | 331 | 407 | 308 |
| 456 | 0.20 | 0.23 | 0.88 | 0.38 | 1 | Lage3 | L antigen family, member 3 | 502 | 391 | 524 | 516 | 420 | 344 | 390 | 688 | 438 | 389 | 419 |
| 453 | 0.00 | 0.19 | -0.02 | 0.99 | 1 | Msh3 | mutS homolog 3 | 552 | 629 | 661 | 337 | 396 | 446 | 390 | 282 | 384 | 447 | 464 |
| 449 | -0.12 | 0.25 | -0.47 | 0.64 | 1 | Proser1 | proline and serine rich 1 | 480 | 399 | 483 | 374 | 382 | 695 | 390 | 198 | 431 | 532 | 575 |
| 447 | 0.24 | 0.26 | 0.92 | 0.36 | 1 | C87436 | expressed sequence C87436 | 372 | 491 | 472 | 330 | 398 | 491 | 390 | 660 | 501 | 258 | 551 |
| 445 | 0.15 | 0.23 | 0.66 | 0.51 | 1 | Cyp4f13 | cytochrome P450, family 4, subfamily f, pc | 450 | 509 | 388 | 341 | 352 | 598 | 390 | 616 | 389 | 322 | 543 |
| 422 | 0.11 | 0.19 | 0.57 | 0.57 | 1 | Oscp1 | organic solute carrier partner 1 | 295 | 414 | 454 | 461 | 485 | 338 | 390 | 564 | 450 | 468 | 328 |
| 412 | 0.00 | 0.21 | 0.01 | 0.99 | 1 | Nufip1 | nuclear fragile X mental retardation protei | 457 | 543 | 538 | 368 | 269 | 464 | 390 | 288 | 374 | 330 | 512 |
| 806 | -0.03 | 0.20 | -0.14 | 0.89 | 1 | Sat1 | spermidine/spermine N1-acetyl transferas | 2123 | 1380 | 1830 | 289 | 246 | 927 | 389 | 374 | 246 | 368 | 692 |
| 618 | 0.17 | 0.19 | 0.85 | 0.40 | 1 | Mob3a | MOB kinase activator 3A | 1151 | 1031 | 1148 | 460 | 363 | 408 | 389 | 622 | 388 | 480 | 354 |
| 593 | 0.06 | 0.19 | 0.31 | 0.76 | 1 | Rap2c | RAP2C, member of RAS oncogene family | 1061 | 1104 | 572 | 560 | 365 | 536 | 389 | 649 | 378 | 442 | 465 |
| 513 | -0.03 | 0.21 | -0.15 | 0.88 | 1 | Crybg1 | crystallin beta-gamma domain containing | 927 | 873 | 491 | 633 | 314 | 380 | 389 | 539 | 367 | 348 | 385 |
| 502 | -0.23 | 0.25 | -0.94 | 0.35 | 1 | Brwd3 | bromodomain and WD repeat domain co | 587 | 734 | 506 | 363 | 387 | 800 | 389 | 257 | 424 | 233 | 843 |
| 335 | -0.16 | 0.26 | -0.62 | 0.54 | 1 | Jrk | jerky | 303 | 434 | 286 | 385 | 402 | 229 | 389 | 387 | 366 | 214 | 287 |
| 333 | -0.28 | 0.19 | -1.49 | 0.14 | 1 | Snx29 | sorting nexin 29 | 456 | 363 | 315 | 323 | 323 | 350 | 389 | 284 | 217 | 307 | 341 |
| 325 | -0.16 | 0.21 | -0.77 | 0.44 | 1 | Mad2l2 | MAD2 mitotic arrest deficient-like 2 | 253 | 219 | 382 | 329 | 390 | 330 | 389 | 298 | 333 | 346 | 307 |
| 610 | 0.35 | 0.19 | 1.81 | 0.07 | 1 | Bin3 | bridging integrator 3 | 893 | 705 | 896 | 537 | 472 | 456 | 388 | 591 | 515 | 683 | 576 |
| 571 | 0.04 | 0.20 | 0.19 | 0.85 | 1 | Tjap1 | tight junction associated protein 1 | 669 | 652 | 593 | 486 | 550 | 731 | 388 | 469 | 420 | 604 | 719 |

| | | | | | | | | | | | | | | | | | |
|---|---|---|---|---|---|---|---|---|---|---|---|---|---|---|---|---|---|
| 462 | -0.37 | 0.36 | -1.02 | 0.31 | 1 | Ltbp2 | latent transforming growth factor beta binc | 1975 | 362 | 308 | 219 | 269 | 484 | 388 | 134 | 241 | 289 | 412 |
| 448 | 0.36 | 0.22 | 1.66 | 0.10 | 1 | 4933434E20Ril | RIKEN cDNA 4933434E20 gene | 281 | 200 | 340 | 457 | 488 | 473 | 388 | 627 | 510 | 545 | 623 |
| 382 | -0.02 | 0.19 | -0.11 | 0.91 | 1 | Zkscan14 | zinc finger with KRAB and SCAN domains | 380 | 397 | 425 | 341 | 395 | 388 | 388 | 400 | 313 | 392 | 389 |
| 378 | 0.08 | 0.23 | 0.35 | 0.73 | 1 | Tcaim | T cell activation inhibitor, mitochondrial | 305 | 379 | 226 | 369 | 309 | 500 | 388 | 357 | 392 | 304 | 624 |
| 315 | -0.13 | 0.23 | -0.57 | 0.57 | 1 | Tpd52l1 | tumor protein D52-like 1 | 206 | 260 | 339 | 383 | 376 | 221 | 388 | 396 | 453 | 255 | 191 |
| 619 | -0.15 | 0.29 | -0.51 | 0.61 | 1 | Stx2 | syntaxin 2 | 1181 | 1094 | 1244 | 584 | 379 | 359 | 387 | 221 | 446 | 577 | 339 |
| 489 | 0.01 | 0.18 | 0.07 | 0.94 | 1 | Atox1 | antioxidant 1 copper chaperone | 959 | 615 | 800 | 400 | 394 | 313 | 387 | 364 | 400 | 407 | 337 |
| 475 | -0.14 | 0.18 | -0.79 | 0.43 | 1 | Ipo11 | importin 11 | 813 | 745 | 651 | 312 | 430 | 454 | 387 | 294 | 360 | 376 | 404 |
| 420 | -0.29 | 0.23 | -1.27 | 0.20 | 1 | Aldh16a1 | aldehyde dehydrogenase 16 family, mem | 900 | 535 | 601 | 380 | 329 | 325 | 387 | 308 | 274 | 252 | 324 |
| 377 | 0.20 | 0.25 | 0.78 | 0.44 | 1 | 2410021H03Rik | | 14 | 14 | 30 | 275 | 302 | 993 | 387 | 365 | 422 | 389 | 952 |
| 371 | -0.05 | 0.18 | -0.29 | 0.77 | 1 | Magef1 | | 438 | 443 | 440 | 326 | 352 | 336 | 387 | 277 | 326 | 362 | 391 |
| 369 | -0.10 | 0.24 | -0.43 | 0.67 | 1 | 9330151L19Rik | | 238 | 286 | 296 | 336 | 375 | 571 | 387 | 248 | 324 | 478 | 520 |
| 365 | -0.07 | 0.30 | -0.22 | 0.83 | 1 | Jrkl | Jrk-like | 380 | 437 | 283 | 368 | 251 | 467 | 387 | 324 | 392 | 183 | 538 |
| 319 | 0.27 | 0.30 | 0.89 | 0.37 | 1 | Alkbh7 | alkB homolog 7 | 223 | 340 | 296 | 228 | 293 | 270 | 387 | 516 | 297 | 412 | 242 |
| 547 | 0.08 | 0.17 | 0.44 | 0.66 | 1 | Tigd2 | tigger transposable element derived 2 | 578 | 588 | 599 | 600 | 578 | 515 | 386 | 543 | 537 | 540 | 553 |
| 477 | -0.28 | 0.24 | -1.17 | 0.24 | 1 | Cry1 | cryptochrome 1 (photolyase-like) | 617 | 439 | 683 | 525 | 598 | 386 | 386 | 258 | 619 | 397 | 344 |
| 470 | -0.14 | 0.17 | -0.84 | 0.40 | 1 | Pisd | phosphatidylserine decarboxylase | 614 | 513 | 535 | 345 | 462 | 638 | 386 | 315 | 392 | 340 | 632 |
| 405 | 0.10 | 0.25 | 0.40 | 0.69 | 1 | Alg3 | asparagine-linked glycosylation 3 (alpha- | 208 | 286 | 360 | 448 | 512 | 396 | 386 | 385 | 425 | 609 | 437 |
| 379 | 0.26 | 0.22 | 1.15 | 0.25 | 1 | Trmt5 | TRM5 tRNA methyltransferase 5 | 305 | 246 | 313 | 424 | 301 | 394 | 386 | 569 | 367 | 415 | 450 |
| 375 | 0.26 | 0.20 | 1.28 | 0.20 | 1 | Acbd4 | acyl-Coenzyme A binding domain contair | 511 | 537 | 419 | 317 | 252 | 237 | 386 | 418 | 291 | 536 | 224 |
| 335 | -0.13 | 0.28 | -0.47 | 0.64 | 1 | Dph1 | diphthamide biosynthesis 1 | 190 | 290 | 273 | 333 | 394 | 407 | 386 | 204 | 398 | 402 | 407 |
| 237 | 0.05 | 0.38 | 0.14 | 0.89 | 1 | Tmem144 | transmembrane protein 144 | 29 | 79 | 87 | 333 | 198 | 247 | 386 | 429 | 283 | 317 | 214 |
| 556 | 0.08 | 0.21 | 0.37 | 0.71 | 1 | Nup54 | nucleoporin 54 | 775 | 675 | 714 | 480 | 404 | 664 | 385 | 403 | 481 | 613 | 520 |
| 486 | -0.19 | 0.17 | -1.07 | 0.29 | 1 | Odf2l | outer dense fiber of sperm tails 2-like | 323 | 301 | 365 | 417 | 343 | 1143 | 385 | 356 | 339 | 285 | 1088 |
| 398 | 0.11 | 0.22 | 0.51 | 0.61 | 1 | Mrps10 | mitochondrial ribosomal protein S10 | 393 | 292 | 375 | 354 | 338 | 513 | 385 | 476 | 285 | 406 | 563 |
| 386 | 0.17 | 0.21 | 0.78 | 0.43 | 1 | Kyat1 | kynurenine aminotransferase 1 | 225 | 325 | 294 | 332 | 380 | 505 | 385 | 507 | 358 | 429 | 503 |
| 382 | 0.07 | 0.25 | 0.30 | 0.77 | 1 | Il1rap | interleukin 1 receptor accessory protein | 618 | 506 | 545 | 381 | 263 | 219 | 385 | 265 | 323 | 341 | 359 |
| 369 | -0.12 | 0.27 | -0.45 | 0.65 | 1 | Med31 | mediator complex subunit 31 | 388 | 396 | 325 | 303 | 388 | 451 | 385 | 403 | 410 | 200 | 409 |
| 368 | -0.05 | 0.24 | -0.23 | 0.82 | 1 | Arl16 | ADP-ribosylation factor-like 16 | 372 | 342 | 484 | 319 | 355 | 374 | 385 | 210 | 340 | 482 | 388 |
| 489 | 0.15 | 0.17 | 0.89 | 0.37 | 1 | Fam110a | family with sequence similarity 110, meml | 552 | 549 | 488 | 596 | 484 | 340 | 384 | 586 | 503 | 497 | 402 |
| 425 | 0.27 | 0.23 | 1.18 | 0.24 | 1 | Tpk1 | thiamine pyrophosphokinase | 362 | 648 | 354 | 415 | 323 | 382 | 384 | 407 | 410 | 523 | 462 |
| 416 | 0.09 | 0.24 | 0.38 | 0.70 | 1 | Gm9856 | | 245 | 488 | 442 | 419 | 423 | 417 | 384 | 474 | 348 | 420 | 516 |
| 391 | -0.11 | 0.24 | -0.44 | 0.66 | 1 | Morn2 | MORN repeat containing 2 | 362 | 332 | 491 | 398 | 467 | 363 | 384 | 497 | 318 | 417 | 276 |
| 384 | 0.04 | 0.18 | 0.20 | 0.84 | 1 | Gan | giant axonal neuropathy | 456 | 338 | 421 | 376 | 329 | 397 | 384 | 387 | 330 | 398 | 411 |
| 354 | 0.07 | 0.21 | 0.35 | 0.73 | 1 | Ccdc58 | coiled-coil domain containing 58 | 336 | 329 | 329 | 298 | 301 | 423 | 384 | 271 | 311 | 484 | 425 |
| 302 | -0.11 | 0.26 | -0.43 | 0.67 | 1 | 4833438C02Rik | | 221 | 309 | 179 | 304 | 385 | 288 | 384 | 272 | 330 | 345 | 308 |
| 455 | 0.18 | 0.22 | 0.80 | 0.42 | 1 | Trappc6a | trafficking protein particle complex 6A | 472 | 443 | 661 | 465 | 387 | 378 | 383 | 528 | 390 | 495 | 408 |
| 455 | 0.18 | 0.16 | 1.14 | 0.25 | 1 | Zfp768 | zinc finger protein 768 | 508 | 610 | 637 | 406 | 375 | 358 | 383 | 461 | 422 | 433 | 409 |
| 414 | 0.24 | 0.21 | 1.13 | 0.26 | 1 | Epb41l4a | erythrocyte membrane protein band 4.1 li | 253 | 329 | 442 | 424 | 374 | 429 | 383 | 389 | 535 | 464 | 530 |
| 404 | 0.01 | 0.21 | 0.05 | 0.96 | 1 | Lurap1 | leucine rich adaptor protein 1 | 293 | 566 | 414 | 494 | 403 | 306 | 383 | 451 | 364 | 457 | 312 |
| 402 | -0.03 | 0.22 | -0.13 | 0.89 | 1 | Rpp25l | ribonuclease P/MRP 25 subunit-like | 380 | 401 | 525 | 421 | 403 | 351 | 383 | 556 | 379 | 357 | 270 |
| 400 | 0.14 | 0.24 | 0.59 | 0.55 | 1 | Zfp933 | zinc finger protein 933 | 260 | 308 | 398 | 469 | 272 | 518 | 383 | 456 | 408 | 406 | 524 |
| 376 | 0.17 | 0.25 | 0.66 | 0.51 | 1 | Uchl3 | ubiquitin carboxyl-terminal esterase L3 (u | 306 | 377 | 330 | 337 | 352 | 383 | 383 | 294 | 453 | 583 | 334 |
| 333 | -0.06 | 0.21 | -0.26 | 0.79 | 1 | Tpgs2 | tubulin polyglutamylase complex subunit | 212 | 272 | 333 | 298 | 372 | 396 | 383 | 293 | 362 | 368 | 370 |
| 319 | -0.05 | 0.22 | -0.25 | 0.80 | 1 | Psph | phosphoserine phosphatase | 338 | 225 | 346 | 302 | 402 | 249 | 383 | 363 | 328 | 316 | 259 |
| 484 | 0.11 | 0.16 | 0.67 | 0.50 | 1 | Pik3cb | phosphatidylinositol-4,5-bisphosphate 3-l | 808 | 593 | 692 | 361 | 452 | 358 | 382 | 388 | 460 | 453 | 374 |
| 473 | -0.17 | 0.20 | -0.84 | 0.40 | 1 | Meis1 | Meis homeobox 1 | 573 | 541 | 468 | 395 | 439 | 717 | 382 | 454 | 339 | 403 | 489 |
| 417 | 0.04 | 0.18 | 0.21 | 0.83 | 1 | Baiap2 | brain-specific angiogenesis inhibitor 1-as | 601 | 476 | 415 | 340 | 217 | 593 | 382 | 367 | 256 | 319 | 619 |
| 359 | 0.42 | 0.27 | 1.55 | 0.12 | 1 | Srp54a | signal recognition particle 54A | 182 | 93 | 264 | 257 | 437 | 411 | 382 | 504 | 506 | 434 | 479 |
| 473 | 0.08 | 0.17 | 0.45 | 0.65 | 1 | Snx16 | sorting nexin 16 | 571 | 568 | 574 | 459 | 425 | 431 | 381 | 473 | 370 | 417 | 538 |
| 446 | -0.02 | 0.18 | -0.14 | 0.89 | 1 | Mocs3 | molybdenum cofactor synthesis 3 | 440 | 418 | 413 | 490 | 568 | 401 | 381 | 492 | 474 | 439 | 394 |
| 409 | 0.09 | 0.17 | 0.55 | 0.58 | 1 | Cnih4 | cornichon family AMPA receptor auxiliary | 730 | 492 | 624 | 279 | 318 | 302 | 381 | 271 | 300 | 444 | 363 |
| 557 | 0.08 | 0.19 | 0.42 | 0.68 | 1 | Sarnp | SAP domain containing ribonucleoproteir | 822 | 707 | 738 | 447 | 500 | 545 | 380 | 579 | 408 | 422 | 574 |
| 552 | -0.38 | 0.30 | -1.26 | 0.21 | 1 | Nfil3 | nuclear factor, interleukin 3, regulated | 1083 | 545 | 719 | 583 | 398 | 742 | 380 | 407 | 410 | 222 | 582 |
| 503 | -0.01 | 0.16 | -0.06 | 0.95 | 1 | Kat8 | K(lysine) acetyltransferase 8 | 568 | 468 | 645 | 417 | 477 | 642 | 380 | 345 | 440 | 441 | 707 |
| 415 | -0.10 | 0.19 | -0.51 | 0.61 | 1 | Snrpf | small nuclear ribonucleoprotein polypepti | 577 | 495 | 708 | 236 | 483 | 358 | 380 | 307 | 369 | 330 | 323 |
| 346 | -0.15 | 0.22 | -0.69 | 0.49 | 1 | Etaa1 | Ewing tumor-associated antigen 1 | 371 | 388 | 355 | 307 | 353 | 365 | 380 | 309 | 302 | 243 | 433 |
| 45206 | -0.45 | 1.14 | -0.39 | 0.70 | 1 | Prss2 | protease, serine 2 | 59729 | 236238 | 188700 | 7833 | 755 | 25 | 379 | 1045 | 403 | 289 | 1869 |
| 633 | -0.14 | 0.24 | -0.58 | 0.56 | 1 | Prkd2 | protein kinase D2 | 1253 | 1537 | 1150 | 312 | 458 | 420 | 379 | 345 | 314 | 515 | 280 |
| 571 | -0.02 | 0.20 | -0.10 | 0.92 | 1 | Fuz | fuzzy planar cell polarity protein | 409 | 433 | 334 | 505 | 483 | 1217 | 379 | 666 | 355 | 416 | 1079 |

| | | | | | | | | | | | | | | | | | | |
|---|---|---|---|---|---|---|---|---|---|---|---|---|---|---|---|---|---|---|
| 440 | -0.11 | 0.28 | -0.39 | 0.69 | 1 | Supt4a | SPT4A, DSIF elongation factor subunit | 724 | 461 | 488 | 413 | 452 | 390 | 379 | 494 | 474 | 249 | 313 |
| 373 | -0.38 | 0.22 | -1.75 | 0.08 | 1 | Dcaf15 | DDB1 and CUL4 associated factor 15 | 493 | 413 | 450 | 392 | 296 | 477 | 379 | 326 | 245 | 213 | 417 |
| 352 | 0.03 | 0.21 | 0.16 | 0.88 | 1 | Ctu2 | cytosolic thiouridylase subunit 2 | 479 | 221 | 342 | 222 | 318 | 466 | 379 | 244 | 231 | 483 | 492 |
| 348 | 0.23 | 0.27 | 0.84 | 0.40 | 1 | Ccdc126 | coiled-coil domain containing 126 | 298 | 487 | 434 | 297 | 242 | 272 | 379 | 372 | 384 | 433 | 231 |
| 335 | 0.25 | 0.23 | 1.10 | 0.27 | 1 | Mkrn2os | makorin, ring finger protein 2, opposite str | 165 | 255 | 299 | 366 | 374 | 243 | 379 | 399 | 442 | 425 | 339 |
| 321 | 0.06 | 0.26 | 0.24 | 0.81 | 1 | Tubg2 | tubulin, gamma 2 | 182 | 241 | 348 | 240 | 399 | 332 | 379 | 236 | 394 | 437 | 344 |
| 154 | 0.28 | 0.40 | 0.71 | 0.48 | 1 | Klhl33 | kelch-like 33 | 280 | 143 | 113 | 83 | 35 | 60 | 379 | 133 | 67 | 350 | 54 |
| 441 | -0.20 | 0.22 | -0.93 | 0.35 | 1 | Nol10 | nucleolar protein 10 | 497 | 350 | 527 | 450 | 523 | 498 | 378 | 381 | 447 | 274 | 527 |
| 422 | 0.51 | 0.34 | 1.51 | 0.13 | 1 | Cdh7 | cadherin 7, type 2 | 31 | 89 | 50 | 480 | 435 | 569 | 378 | 414 | 573 | 855 | 765 |
| 360 | -0.17 | 0.19 | -0.94 | 0.35 | 1 | Otub2 | OTU domain, ubiquitin aldehyde binding | 230 | 300 | 230 | 402 | 455 | 459 | 378 | 434 | 351 | 319 | 404 |
| 299 | -0.01 | 0.26 | -0.04 | 0.96 | 1 | Trmt11 | tRNA methyltransferase 11 | 298 | 334 | 258 | 308 | 210 | 315 | 378 | 250 | 320 | 246 | 375 |
| 285 | -0.43 | 0.32 | -1.34 | 0.18 | 1 | Sybu | syntabulin (syntaxin-interacting) | 333 | 458 | 342 | 225 | 308 | 237 | 378 | 134 | 319 | 189 | 216 |
| 518 | -0.23 | 0.32 | -0.74 | 0.46 | 1 | Cnpy4 | canopy FGF signaling regulator 4 | 1047 | 872 | 986 | 402 | 362 | 331 | 377 | 582 | 288 | 155 | 299 |
| 456 | -0.14 | 0.19 | -0.71 | 0.48 | 1 | Sertad2 | SERTA domain containing 2 | 927 | 731 | 730 | 390 | 310 | 282 | 377 | 319 | 340 | 420 | 193 |
| 438 | -0.24 | 0.20 | -1.23 | 0.22 | 1 | Simc1 | SUMO-interacting motifs containing 1 | 369 | 513 | 404 | 375 | 446 | 714 | 377 | 307 | 428 | 294 | 591 |
| 415 | -0.13 | 0.18 | -0.77 | 0.44 | 1 | D2hgdh | D-2-hydroxyglutarate dehydrogenase | 264 | 450 | 271 | 505 | 318 | 692 | 377 | 445 | 342 | 341 | 558 |
| 411 | -0.13 | 0.18 | -0.74 | 0.46 | 1 | Katnbl1 | katanin p80 subunit B like 1 | 399 | 468 | 436 | 443 | 449 | 416 | 377 | 367 | 329 | 394 | 448 |
| 335 | 0.36 | 0.36 | 0.97 | 0.33 | 1 | Emid1 | EMI domain containing 1 | 409 | 224 | 367 | 384 | 193 | 216 | 377 | 628 | 180 | 386 | 323 |
| 645 | -0.07 | 0.20 | -0.33 | 0.74 | 1 | Rab10os | | 540 | 574 | 494 | 430 | 444 | 1590 | 376 | 589 | 409 | 273 | 1377 |
| 516 | 0.11 | 0.19 | 0.62 | 0.54 | 1 | Apba3 | amyloid beta (A4) precursor protein-bindi | 544 | 654 | 680 | 341 | 490 | 636 | 376 | 368 | 532 | 488 | 572 |
| 442 | 0.01 | 0.30 | 0.05 | 0.96 | 1 | Ldhd | lactate dehydrogenase D | 347 | 343 | 458 | 446 | 370 | 640 | 376 | 291 | 381 | 721 | 485 |
| 410 | -0.42 | 0.24 | -1.75 | 0.08 | 1 | Snrpc | U1 small nuclear ribonucleoprotein C | 529 | 572 | 542 | 366 | 413 | 461 | 376 | 168 | 308 | 326 | 447 |
| 407 | 0.08 | 0.25 | 0.34 | 0.73 | 1 | Cdkn2d | cyclin dependent kinase inhibitor 2D | 504 | 342 | 281 | 352 | 512 | 365 | 376 | 428 | 360 | 587 | 369 |
| 371 | 0.06 | 0.25 | 0.24 | 0.81 | 1 | Mettl8 | methyltransferase like 8 | 432 | 261 | 336 | 467 | 318 | 339 | 376 | 395 | 369 | 352 | 442 |
| 280 | 0.02 | 0.22 | 0.07 | 0.94 | 1 | Tefm | transcription elongation factor, mitochond | 270 | 197 | 321 | 248 | 276 | 243 | 376 | 265 | 284 | 335 | 261 |
| 625 | -0.23 | 0.20 | -1.13 | 0.26 | 1 | Adgb | androglobin | 406 | 461 | 425 | 401 | 659 | 1556 | 375 | 308 | 463 | 413 | 1405 |
| 482 | 0.06 | 0.32 | 0.19 | 0.85 | 1 | Hbegf | heparin-binding EGF-like growth factor | 537 | 333 | 507 | 831 | 436 | 279 | 375 | 841 | 352 | 505 | 301 |
| 461 | 0.22 | 0.23 | 0.92 | 0.36 | 1 | Ccdc134 | coiled-coil domain containing 134 | 378 | 267 | 369 | 248 | 337 | 951 | 375 | 459 | 355 | 313 | 1020 |
| 409 | -0.35 | 0.19 | -1.81 | 0.07 | 1 | Zfp354c | zinc finger protein 354C | 641 | 684 | 603 | 345 | 360 | 358 | 375 | 357 | 293 | 236 | 253 |
| 529 | -0.17 | 0.28 | -0.62 | 0.54 | 1 | Ifnar2 | interferon (alpha and beta) receptor 2 | 1359 | 878 | 1209 | 284 | 370 | 229 | 374 | 380 | 350 | 174 | 209 |
| 523 | 0.34 | 0.21 | 1.64 | 0.10 | 1 | Naa16 | N(alpha)-acetyltransferase 16, NatA auxil | 345 | 375 | 544 | 459 | 375 | 784 | 374 | 660 | 483 | 442 | 913 |
| 394 | 0.09 | 0.21 | 0.44 | 0.66 | 1 | Bco1 | beta-carotene oxygenase 1 | 225 | 395 | 300 | 458 | 430 | 363 | 374 | 692 | 358 | 366 | 377 |
| 380 | -0.01 | 0.24 | -0.03 | 0.98 | 1 | Zfp974 | zinc finger protein 974 | 215 | 489 | 250 | 395 | 368 | 475 | 374 | 404 | 333 | 385 | 488 |
| 374 | -0.30 | 0.25 | -1.22 | 0.22 | 1 | Mocs1 | molybdenum cofactor synthesis 1 | 484 | 515 | 680 | 386 | 288 | 292 | 374 | 200 | 309 | 318 | 269 |
| 650 | -0.15 | 0.17 | -0.85 | 0.40 | 1 | Atp5l | ATP synthase, H+ transporting, mitochonc | 613 | 808 | 695 | 564 | 455 | 1226 | 373 | 364 | 424 | 411 | 1214 |
| 515 | -0.34 | 0.21 | -1.59 | 0.11 | 1 | Gpc1 | glypican 1 | 1341 | 766 | 1137 | 394 | 365 | 219 | 373 | 257 | 347 | 270 | 192 |
| 431 | 0.10 | 0.23 | 0.45 | 0.65 | 1 | Lins1 | lines homolog 1 | 525 | 326 | 439 | 441 | 316 | 538 | 373 | 386 | 449 | 505 | 445 |
| 412 | -0.10 | 0.24 | -0.42 | 0.68 | 1 | Mfsd4b4 | | 243 | 394 | 340 | 260 | 377 | 819 | 373 | 260 | 413 | 274 | 781 |
| 364 | 0.06 | 0.26 | 0.24 | 0.81 | 1 | Wdr55 | WD repeat domain 55 | 552 | 357 | 478 | 307 | 320 | 247 | 373 | 185 | 311 | 536 | 334 |
| 314 | -0.33 | 0.21 | -1.58 | 0.11 | 1 | Mpv17l2 | MPV17 mitochondrial membrane protein- | 402 | 341 | 310 | 433 | 319 | 219 | 373 | 323 | 303 | 255 | 178 |
| 312 | -0.14 | 0.24 | -0.59 | 0.56 | 1 | Ocel1 | occludin/ELL domain containing 1 | 277 | 313 | 336 | 266 | 289 | 377 | 373 | 183 | 333 | 321 | 363 |
| 293 | -0.04 | 0.29 | -0.14 | 0.89 | 1 | Pde7b | phosphodiesterase 7B | 526 | 533 | 372 | 138 | 203 | 196 | 373 | 156 | 185 | 361 | 182 |
| 545 | 0.02 | 0.19 | 0.09 | 0.93 | 1 | Dtymk | deoxythymidylate kinase | 678 | 541 | 530 | 431 | 510 | 797 | 372 | 367 | 469 | 522 | 779 |
| 498 | -0.18 | 0.21 | -0.85 | 0.40 | 1 | Cars2 | cysteinyl-tRNA synthetase 2 (mitochondri | 333 | 352 | 286 | 375 | 356 | 1219 | 372 | 242 | 398 | 285 | 1265 |
| 460 | 0.15 | 0.17 | 0.91 | 0.36 | 1 | Zfp81 | zinc finger protein 81 | 489 | 530 | 406 | 498 | 419 | 433 | 372 | 509 | 417 | 435 | 556 |
| 377 | 0.16 | 0.28 | 0.57 | 0.57 | 1 | Mzt2 | mitotic spindle organizing protein 2 | 404 | 258 | 466 | 383 | 403 | 265 | 372 | 532 | 308 | 493 | 268 |
| 364 | -0.15 | 0.23 | -0.65 | 0.52 | 1 | Snx32 | sorting nexin 32 | 89 | 131 | 74 | 295 | 282 | 963 | 372 | 313 | 278 | 231 | 977 |
| 358 | 0.01 | 0.23 | 0.05 | 0.96 | 1 | Tmem189 | transmembrane protein 189 | 417 | 282 | 324 | 479 | 294 | 305 | 372 | 465 | 327 | 407 | 270 |
| 346 | -0.05 | 0.20 | -0.26 | 0.80 | 1 | Dusp23 | dual specificity phosphatase 23 | 151 | 209 | 190 | 457 | 545 | 287 | 372 | 534 | 409 | 325 | 326 |
| 314 | -0.51 | 0.28 | -1.82 | 0.07 | 1 | Disp1 | dispatched RND transporter family memb | 554 | 514 | 378 | 300 | 227 | 275 | 372 | 116 | 210 | 291 | 216 |
| 286 | 0.18 | 0.21 | 0.84 | 0.40 | 1 | Rnf182 | ring finger protein 182 | 48 | 64 | 56 | 351 | 325 | 348 | 372 | 371 | 339 | 429 | 443 |
| 460 | 0.14 | 0.26 | 0.52 | 0.61 | 1 | Kctd18 | potassium channel tetramerisation domai | 491 | 355 | 431 | 361 | 358 | 690 | 371 | 604 | 279 | 327 | 795 |
| 441 | 0.01 | 0.21 | 0.05 | 0.96 | 1 | Slc30a6 | solute carrier family 30 (zinc transporter), | 281 | 295 | 254 | 553 | 419 | 667 | 371 | 386 | 534 | 424 | 669 |
| 409 | -0.02 | 0.21 | -0.09 | 0.93 | 1 | Osgin1 | oxidative stress induced growth inhibitor ' | 432 | 384 | 500 | 431 | 380 | 417 | 371 | 498 | 413 | 309 | 370 |
| 403 | 0.25 | 0.19 | 1.34 | 0.18 | 1 | C330018D20Ri | RIKEN cDNA C330018D20 gene | 369 | 489 | 375 | 384 | 322 | 375 | 371 | 510 | 308 | 466 | 464 |
| 392 | 0.01 | 0.25 | 0.06 | 0.96 | 1 | Cd3eap | CD3E antigen, epsilon polypeptide assoc | 566 | 542 | 563 | 257 | 384 | 298 | 371 | 463 | 261 | 336 | 275 |
| 384 | 0.12 | 0.18 | 0.68 | 0.50 | 1 | Cda | cytidine deaminase | 125 | 164 | 235 | 397 | 551 | 452 | 371 | 523 | 563 | 398 | 441 |
| 382 | -0.12 | 0.18 | -0.69 | 0.49 | 1 | Zfpl1 | zinc finger like protein 1 | 305 | 273 | 402 | 298 | 414 | 573 | 371 | 232 | 380 | 343 | 606 |
| 381 | 0.00 | 0.25 | 0.00 | 1.00 | 1 | Endog | endonuclease G | 256 | 149 | 342 | 426 | 534 | 399 | 371 | 485 | 494 | 336 | 403 |
| 313 | -0.04 | 0.24 | -0.15 | 0.88 | 1 | Napb | N-ethylmaleimide sensitive fusion protein | 208 | 189 | 188 | 219 | 275 | 582 | 371 | 291 | 301 | 222 | 597 |

| | | | | | | | | | | | | | | | | | | |
|---|---|---|---|---|---|---|---|---|---|---|---|---|---|---|---|---|---|---|
| 300 | -0.15 | 0.25 | -0.60 | 0.55 | 1 | Zfp689 | zinc finger protein 689 | 224 | 341 | 335 | 360 | 320 | 213 | 371 | 257 | 320 | 377 | 187 |
| 532 | -0.35 | 0.18 | -1.95 | 0.05 | 1 | Stxbp5l | syntaxin binding protein 5-like | 605 | 981 | 843 | 550 | 356 | 636 | 370 | 407 | 379 | 223 | 501 |
| 387 | -0.11 | 0.19 | -0.57 | 0.57 | 1 | Fcsk | fucose kinase | 405 | 315 | 435 | 381 | 417 | 439 | 370 | 353 | 452 | 296 | 398 |
| 368 | 0.13 | 0.20 | 0.67 | 0.50 | 1 | BC024978 | cDNA sequence BC024978 | 178 | 282 | 234 | 354 | 414 | 462 | 370 | 470 | 406 | 371 | 514 |
| 344 | -0.09 | 0.18 | -0.49 | 0.62 | 1 | Pop7 | processing of precursor 7, ribonuclease F | 374 | 335 | 398 | 380 | 357 | 270 | 370 | 372 | 328 | 354 | 246 |
| 573 | -0.26 | 0.18 | -1.45 | 0.15 | 1 | Ptbp2 | polypyrimidine tract binding protein 2 | 545 | 616 | 451 | 347 | 294 | 1494 | 368 | 227 | 244 | 346 | 1374 |
| 556 | 0.00 | 0.16 | -0.02 | 0.98 | 1 | Sugp2 | SURP and G patch domain containing 2 | 458 | 531 | 414 | 329 | 273 | 1361 | 368 | 377 | 286 | 289 | 1428 |
| 481 | 0.00 | 0.18 | -0.01 | 0.99 | 1 | Pank4 | pantothenate kinase 4 | 501 | 450 | 481 | 260 | 455 | 846 | 368 | 284 | 447 | 332 | 870 |
| 465 | -0.26 | 0.22 | -1.16 | 0.25 | 1 | Arsk | arylsulfatase K | 922 | 665 | 600 | 365 | 523 | 339 | 368 | 280 | 381 | 367 | 308 |
| 458 | -0.10 | 0.28 | -0.38 | 0.70 | 1 | Gpn3 | GPN-loop GTPase 3 | 768 | 464 | 504 | 453 | 418 | 435 | 368 | 215 | 489 | 433 | 488 |
| 417 | 0.07 | 0.28 | 0.25 | 0.80 | 1 | Tmem97 | transmembrane protein 97 | 345 | 937 | 685 | 347 | 266 | 278 | 368 | 466 | 323 | 299 | 274 |
| 381 | -0.29 | 0.28 | -1.05 | 0.29 | 1 | Zfp882 | zinc finger protein 882 | 352 | 518 | 430 | 490 | 338 | 392 | 368 | 214 | 283 | 402 | 399 |
| 258 | -0.25 | 0.29 | -0.86 | 0.39 | 1 | Nol3 | nucleolar protein 3 (apoptosis repressor v | 156 | 216 | 236 | 395 | 259 | 146 | 368 | 454 | 273 | 250 | 89 |
| 1018 | -0.30 | 0.21 | -1.39 | 0.16 | 1 | Tbc1d2b | TBC1 domain family, member 2B | 3652 | 2330 | 2205 | 368 | 479 | 424 | 367 | 214 | 414 | 393 | 346 |
| 750 | 0.08 | 0.17 | 0.51 | 0.61 | 1 | Echdc2 | enoyl Coenzyme A hydratase domain cor | 408 | 406 | 349 | 502 | 513 | 2166 | 367 | 604 | 484 | 455 | 1997 |
| 736 | 0.04 | 0.22 | 0.17 | 0.86 | 1 | Jpt1 | Jupiter microtubule associated homolog 1 | 2297 | 1285 | 1276 | 428 | 476 | 323 | 367 | 436 | 432 | 442 | 335 |
| 567 | -0.06 | 0.24 | -0.23 | 0.82 | 1 | Mgat4a | mannoside acetylglucosaminyltransferas | 1176 | 702 | 842 | 621 | 449 | 355 | 367 | 498 | 451 | 459 | 321 |
| 497 | 0.05 | 0.19 | 0.24 | 0.81 | 1 | Wdr92 | WD repeat domain 92 | 367 | 440 | 553 | 614 | 502 | 529 | 367 | 476 | 592 | 374 | 654 |
| 472 | -0.17 | 0.19 | -0.87 | 0.38 | 1 | Dck | deoxycytidine kinase | 884 | 660 | 696 | 444 | 394 | 350 | 367 | 293 | 338 | 425 | 340 |
| 371 | -0.41 | 0.28 | -1.49 | 0.14 | 1 | Hcn3 | hyperpolarization-activated, cyclic nucleo | 359 | 421 | 447 | 319 | 296 | 614 | 367 | 264 | 346 | 135 | 515 |
| 312 | -0.31 | 0.24 | -1.27 | 0.21 | 1 | Upp2 | uridine phosphorylase 2 | 251 | 174 | 227 | 457 | 443 | 278 | 367 | 363 | 302 | 274 | 291 |
| 300 | 0.17 | 0.28 | 0.60 | 0.55 | 1 | St7 | suppression of tumorigenicity 7 | 274 | 139 | 234 | 388 | 244 | 226 | 367 | 390 | 338 | 501 | 197 |
| 529 | -0.21 | 0.17 | -1.18 | 0.24 | 1 | Ppp2r3d | protein phosphatase 2 (formerly 2A), regu | 518 | 439 | 524 | 453 | 540 | 967 | 366 | 425 | 420 | 327 | 841 |
| 359 | 0.06 | 0.22 | 0.27 | 0.79 | 1 | Cdk2ap1 | CDK2 (cyclin-dependent kinase 2)-assoc | 209 | 204 | 334 | 307 | 309 | 585 | 366 | 338 | 332 | 348 | 620 |
| 330 | 0.06 | 0.24 | 0.26 | 0.80 | 1 | Snn | stannin | 457 | 402 | 282 | 396 | 284 | 191 | 366 | 375 | 342 | 273 | 258 |
| 644 | -0.08 | 0.16 | -0.48 | 0.63 | 1 | Cdk17 | cyclin-dependent kinase 17 | 835 | 1182 | 1000 | 483 | 342 | 936 | 365 | 436 | 424 | 337 | 742 |
| 519 | 0.21 | 0.21 | 0.98 | 0.33 | 1 | Inpp1 | inositol polyphosphate-1-phosphatase | 522 | 634 | 774 | 484 | 460 | 451 | 365 | 560 | 417 | 574 | 468 |
| 377 | 0.08 | 0.22 | 0.37 | 0.71 | 1 | Bnip1 | BCL2/adenovirus E1B interacting protein | 387 | 256 | 428 | 305 | 408 | 421 | 365 | 401 | 423 | 298 | 457 |
| 324 | -0.33 | 0.20 | -1.66 | 0.10 | 1 | Gpn2 | GPN-loop GTPase 2 | 299 | 382 | 352 | 317 | 358 | 367 | 365 | 255 | 291 | 254 | 328 |
| 648 | -0.47 | 0.25 | -1.92 | 0.05 | 1 | Gng2 | guanine nucleotide binding protein (G pro | 1601 | 1492 | 1330 | 462 | 456 | 286 | 364 | 176 | 374 | 349 | 241 |
| 542 | 0.33 | 0.23 | 1.42 | 0.15 | 1 | Aprt | adenine phosphoribosyl transferase | 835 | 559 | 701 | 496 | 421 | 410 | 364 | 728 | 535 | 529 | 379 |
| 467 | -0.04 | 0.16 | -0.24 | 0.81 | 1 | Zup1 | zinc finger containing ubiquitin peptidase | 436 | 440 | 356 | 377 | 344 | 888 | 364 | 349 | 361 | 332 | 895 |
| 448 | 0.09 | 0.18 | 0.53 | 0.60 | 1 | Aldh4a1 | aldehyde dehydrogenase 4 family, memb | 627 | 668 | 556 | 395 | 414 | 313 | 364 | 505 | 408 | 341 | 340 |
| 363 | 0.10 | 0.21 | 0.47 | 0.64 | 1 | Wrap73 | WD repeat containing, antisense to Trp73 | 335 | 274 | 369 | 386 | 354 | 344 | 363 | 337 | 382 | 510 | 340 |
| 507 | -0.11 | 0.19 | -0.59 | 0.55 | 1 | Spry2 | sprouty RTK signaling antagonist 2 | 1201 | 924 | 820 | 323 | 351 | 314 | 362 | 276 | 258 | 457 | 292 |
| 419 | 0.35 | 0.23 | 1.52 | 0.13 | 1 | Exosc5 | exosome component 5 | 484 | 401 | 493 | 286 | 323 | 453 | 362 | 551 | 339 | 453 | 458 |
| 406 | 0.18 | 0.23 | 0.77 | 0.44 | 1 | Rpf2 | ribosome production factor 2 homolog | 565 | 399 | 460 | 299 | 381 | 384 | 362 | 507 | 321 | 379 | 409 |
| 383 | -0.04 | 0.20 | -0.22 | 0.83 | 1 | Tex9 | testis expressed gene 9 | 275 | 343 | 368 | 363 | 338 | 565 | 362 | 292 | 347 | 372 | 587 |
| 377 | 0.16 | 0.20 | 0.78 | 0.43 | 1 | Tmem38b | transmembrane protein 38B | 538 | 366 | 488 | 273 | 400 | 276 | 362 | 394 | 413 | 337 | 304 |
| 336 | -0.25 | 0.29 | -0.87 | 0.38 | 1 | Polr2h | polymerase (RNA) II (DNA directed) polyp | 450 | 258 | 549 | 339 | 291 | 334 | 362 | 334 | 249 | 238 | 289 |
| 317 | -0.39 | 0.25 | -1.54 | 0.12 | 1 | Tmem241 | transmembrane protein 241 | 387 | 317 | 448 | 339 | 352 | 258 | 362 | 221 | 387 | 179 | 235 |
| 265 | -0.11 | 0.27 | -0.41 | 0.68 | 1 | Actr3b | ARP3 actin-related protein 3B | 196 | 95 | 113 | 288 | 415 | 249 | 362 | 316 | 353 | 287 | 240 |
| 257 | -0.42 | 0.34 | -1.24 | 0.22 | 1 | Wdr31 | WD repeat domain 31 | 165 | 238 | 285 | 311 | 235 | 292 | 362 | 314 | 247 | 120 | 264 |
| 412 | 0.02 | 0.20 | 0.11 | 0.91 | 1 | Pafah1b3 | platelet-activating factor acetylhydrolase, | 641 | 572 | 619 | 363 | 301 | 299 | 361 | 382 | 292 | 453 | 245 |
| 366 | -0.03 | 0.22 | -0.14 | 0.89 | 1 | Cisd3 | CDGSH iron sulfur domain 3 | 299 | 304 | 533 | 316 | 420 | 358 | 361 | 402 | 436 | 305 | 293 |
| 313 | -0.26 | 0.27 | -0.97 | 0.33 | 1 | Brf2 | BRF2, RNA polymerase III transcription in | 484 | 480 | 339 | 324 | 247 | 240 | 361 | 158 | 273 | 290 | 248 |
| 308 | -0.02 | 0.27 | -0.08 | 0.93 | 1 | Top3a | topoisomerase (DNA) III alpha | 366 | 327 | 320 | 203 | 298 | 341 | 361 | 372 | 242 | 269 | 292 |
| 217 | -0.30 | 0.32 | -0.93 | 0.35 | 1 | Fam107a | family with sequence similarity 107, meml | 159 | 149 | 149 | 171 | 175 | 352 | 361 | 93 | 193 | 298 | 292 |
| 473 | 0.01 | 0.21 | 0.06 | 0.95 | 1 | Trpm4 | transient receptor potential cation channe | 782 | 481 | 644 | 404 | 512 | 364 | 360 | 447 | 407 | 437 | 361 |
| 467 | -0.05 | 0.22 | -0.21 | 0.83 | 1 | Prkra | protein kinase, interferon inducible doubl | 753 | 776 | 645 | 366 | 403 | 358 | 360 | 529 | 347 | 270 | 325 |
| 451 | -0.20 | 0.24 | -0.84 | 0.40 | 1 | Mmd | monocyte to macrophage differentiation-a | 704 | 585 | 653 | 430 | 364 | 441 | 360 | 381 | 348 | 204 | 495 |
| 439 | -0.35 | 0.29 | -1.21 | 0.23 | 1 | Pgap2 | post-GPI attachment to proteins 2 | 812 | 494 | 636 | 383 | 447 | 418 | 360 | 367 | 345 | 175 | 394 |
| 420 | -0.05 | 0.21 | -0.26 | 0.80 | 1 | Tomm5 | translocase of outer mitochondrial membr | 521 | 414 | 436 | 450 | 488 | 350 | 360 | 334 | 402 | 488 | 373 |
| 391 | 0.06 | 0.25 | 0.25 | 0.80 | 1 | Thap3 | THAP domain containing, apoptosis asso | 358 | 378 | 531 | 390 | 329 | 402 | 360 | 367 | 294 | 480 | 409 |
| 315 | -0.07 | 0.24 | -0.29 | 0.77 | 1 | Fgfbp3 | fibroblast growth factor binding protein 3 | 187 | 256 | 304 | 422 | 378 | 226 | 360 | 446 | 292 | 397 | 198 |
| 678 | 0.11 | 0.14 | 0.77 | 0.44 | 1 | Cttnbp2nl | CTTNBP2 N-terminal like | 1778 | 1596 | 1533 | 310 | 281 | 282 | 359 | 305 | 340 | 362 | 316 |
| 467 | 0.33 | 0.23 | 1.42 | 0.15 | 1 | Zfp839 | zinc finger protein 839 | 402 | 367 | 419 | 439 | 280 | 707 | 359 | 644 | 373 | 508 | 635 |
| 429 | -0.21 | 0.19 | -1.13 | 0.26 | 1 | Abhd14a | abhydrolase domain containing 14A | 585 | 716 | 736 | 346 | 353 | 372 | 359 | 366 | 297 | 238 | 349 |
| 355 | 0.11 | 0.23 | 0.49 | 0.62 | 1 | Mrpl14 | mitochondrial ribosomal protein L14 | 414 | 330 | 531 | 270 | 370 | 258 | 359 | 319 | 316 | 497 | 244 |
| 424 | 0.21 | 0.24 | 0.87 | 0.39 | 1 | Shkbp1 | Sh3kbp1 binding protein 1 | 510 | 460 | 643 | 335 | 316 | 402 | 358 | 498 | 402 | 443 | 301 |

| | | | | | | | | | | | | | | | | | | |
|---|---|---|---|---|---|---|---|---|---|---|---|---|---|---|---|---|---|---|
| 421 | 0.24 | 0.22 | 1.10 | 0.27 | 1 | Ptdss2 | phosphatidylserine synthase 2 | 487 | 443 | 479 | 386 | 230 | 514 | 358 | 407 | 299 | 514 | 514 |
| 416 | -0.27 | 0.21 | -1.30 | 0.19 | 1 | Wdr83 | WD repeat domain containing 83 | 556 | 401 | 492 | 389 | 418 | 525 | 358 | 334 | 311 | 258 | 530 |
| 414 | 0.22 | 0.22 | 1.03 | 0.30 | 1 | Yae1d1 | Yae1 domain containing 1 | 340 | 593 | 327 | 370 | 334 | 443 | 358 | 575 | 432 | 321 | 461 |
| 400 | 0.00 | 0.23 | 0.00 | 1.00 | 1 | Ss18l1 | SS18, nBAF chromatin remodeling compl | 435 | 292 | 397 | 369 | 320 | 608 | 358 | 425 | 377 | 317 | 501 |
| 398 | -0.65 | 0.37 | -1.76 | 0.08 | 1 | Dcp1b | decapping mRNA 1B | 436 | 327 | 284 | 319 | 257 | 936 | 358 | 89 | 312 | 139 | 923 |
| 391 | 0.25 | 0.24 | 1.05 | 0.29 | 1 | Cox18 | cytochrome c oxidase assembly protein 1 | 275 | 216 | 394 | 353 | 464 | 391 | 358 | 455 | 400 | 581 | 417 |
| 346 | 0.24 | 0.25 | 0.96 | 0.33 | 1 | Polr2l | polymerase (RNA) II (DNA directed) polyp | 302 | 340 | 399 | 284 | 314 | 290 | 358 | 505 | 355 | 424 | 235 |
| 338 | -0.22 | 0.21 | -1.04 | 0.30 | 1 | Pacrg | PARK2 co-regulated | 263 | 374 | 251 | 407 | 421 | 339 | 358 | 310 | 325 | 370 | 298 |
| 467 | 0.17 | 0.29 | 0.58 | 0.56 | 1 | Mrpl52 | mitochondrial ribosomal protein L52 | 605 | 529 | 911 | 341 | 406 | 344 | 357 | 638 | 308 | 357 | 343 |
| 451 | 0.26 | 0.16 | 1.61 | 0.11 | 1 | Ftsj1 | FtsJ RNA methyltransferase homolog 1 (E | 514 | 413 | 560 | 299 | 473 | 450 | 357 | 364 | 597 | 473 | 462 |
| 382 | -0.07 | 0.32 | -0.22 | 0.83 | 1 | Zfp420 | zinc finger protein 420 | 190 | 555 | 289 | 333 | 316 | 616 | 357 | 270 | 265 | 404 | 612 |
| 370 | 0.02 | 0.21 | 0.07 | 0.94 | 1 | 2610044O15Ril | RIKEN cDNA 2610044O15 gene | 419 | 455 | 511 | 299 | 341 | 331 | 357 | 361 | 345 | 406 | 248 |
| 423 | -0.08 | 0.18 | -0.44 | 0.66 | 1 | Sgk2 | serum/glucocorticoid regulated kinase 2 | 414 | 533 | 584 | 412 | 509 | 320 | 356 | 483 | 456 | 264 | 324 |
| 372 | -0.08 | 0.18 | -0.42 | 0.68 | 1 | B330016D10Rik | | 275 | 379 | 348 | 477 | 426 | 334 | 356 | 419 | 344 | 358 | 379 |
| 348 | -0.02 | 0.19 | -0.12 | 0.91 | 1 | Pomt1 | protein-O-mannosyltransferase 1 | 326 | 224 | 287 | 293 | 335 | 531 | 356 | 327 | 360 | 305 | 479 |
| 343 | 0.07 | 0.21 | 0.34 | 0.73 | 1 | Amigo3 | adhesion molecule with Ig like domain 3 | 152 | 104 | 122 | 406 | 418 | 470 | 356 | 377 | 501 | 444 | 419 |
| 330 | 0.04 | 0.18 | 0.22 | 0.82 | 1 | Zfp770 | zinc finger protein 770 | 282 | 292 | 285 | 333 | 307 | 371 | 356 | 363 | 292 | 370 | 384 |
| 234 | -0.15 | 0.25 | -0.58 | 0.56 | 1 | Cpeb1os1 | | 226 | 88 | 175 | 255 | 295 | 186 | 356 | 188 | 274 | 325 | 203 |
| 48810 | -1.02 | 1.12 | -0.91 | 0.36 | 1 | Cela2a | chymotrypsin-like elastase family, membe | 61280 | 265700 | 197896 | 9065 | 488 | 32 | 355 | 665 | 286 | 155 | 985 |
| 496 | -0.03 | 0.20 | -0.13 | 0.90 | 1 | Spata6 | spermatogenesis associated 6 | 767 | 702 | 658 | 548 | 438 | 337 | 355 | 570 | 371 | 344 | 365 |
| 493 | 0.03 | 0.29 | 0.11 | 0.91 | 1 | Fbxl6 | F-box and leucine-rich repeat protein 6 | 638 | 342 | 493 | 350 | 438 | 793 | 355 | 528 | 347 | 267 | 868 |
| 444 | -0.15 | 0.19 | -0.81 | 0.42 | 1 | Fgd1 | FYVE, RhoGEF and PH domain containin | 593 | 574 | 575 | 456 | 433 | 405 | 355 | 405 | 470 | 310 | 310 |
| 429 | -0.02 | 0.18 | -0.13 | 0.90 | 1 | Rhbdl2 | rhomboid like 2 | 194 | 319 | 332 | 465 | 515 | 615 | 355 | 401 | 487 | 374 | 668 |
| 426 | 0.19 | 0.23 | 0.80 | 0.42 | 1 | Rpusd4 | RNA pseudouridylate synthase domain c | 464 | 402 | 457 | 482 | 375 | 347 | 355 | 662 | 455 | 316 | 375 |
| 389 | 0.00 | 0.18 | -0.01 | 0.99 | 1 | Zfp446 | zinc finger protein 446 | 266 | 277 | 315 | 312 | 333 | 730 | 355 | 301 | 414 | 343 | 633 |
| 378 | -0.06 | 0.20 | -0.32 | 0.75 | 1 | Zfp143 | zinc finger protein 143 | 595 | 433 | 471 | 291 | 354 | 346 | 355 | 204 | 356 | 370 | 384 |
| 339 | 0.04 | 0.26 | 0.15 | 0.88 | 1 | Mcph1 | microcephaly, primary autosomal recessiv | 359 | 216 | 206 | 262 | 418 | 427 | 355 | 295 | 461 | 319 | 412 |
| 320 | -0.13 | 0.26 | -0.51 | 0.61 | 1 | Gdpgp1 | GDP-D-glucose phosphorylase 1 | 106 | 199 | 220 | 519 | 434 | 251 | 355 | 324 | 511 | 386 | 212 |
| 455 | -0.01 | 0.18 | -0.04 | 0.97 | 1 | Rsl24d1 | ribosomal L24 domain containing 1 | 531 | 591 | 671 | 458 | 399 | 392 | 354 | 512 | 411 | 377 | 312 |
| 435 | -0.04 | 0.18 | -0.24 | 0.81 | 1 | Ublcp1 | ubiquitin-like domain containing CTD phc | 334 | 395 | 310 | 438 | 339 | 758 | 354 | 411 | 397 | 278 | 766 |
| 426 | 0.15 | 0.22 | 0.66 | 0.51 | 1 | Pnpla7 | patatin-like phospholipase domain contai | 683 | 419 | 507 | 362 | 302 | 445 | 354 | 465 | 347 | 415 | 384 |
| 409 | -0.07 | 0.21 | -0.36 | 0.72 | 1 | Pcdhgb6 | protocadherin gamma subfamily B, 6 | 559 | 478 | 489 | 384 | 483 | 277 | 354 | 368 | 496 | 416 | 194 |
| 366 | 0.04 | 0.23 | 0.19 | 0.85 | 1 | Ska2 | spindle and kinetochore associated comp | 381 | 465 | 348 | 318 | 339 | 369 | 354 | 257 | 374 | 323 | 496 |
| 452 | 0.22 | 0.21 | 1.07 | 0.28 | 1 | Taf1b | TATA-box binding protein associated fact | 459 | 482 | 453 | 475 | 419 | 388 | 353 | 682 | 451 | 326 | 481 |
| 401 | 0.14 | 0.24 | 0.59 | 0.56 | 1 | Cyp4f17 | cytochrome P450, family 4, subfamily f, pc | 295 | 329 | 254 | 359 | 378 | 590 | 353 | 292 | 429 | 571 | 561 |
| 386 | -0.06 | 0.20 | -0.29 | 0.77 | 1 | Cep135 | centrosomal protein 135 | 233 | 400 | 248 | 438 | 361 | 555 | 353 | 374 | 328 | 354 | 602 |
| 349 | 0.00 | 0.26 | -0.01 | 0.99 | 1 | Ppcdc | phosphopantothenoylcysteine decarboxy | 319 | 466 | 372 | 260 | 260 | 474 | 353 | 382 | 343 | 241 | 369 |
| 331 | 0.10 | 0.24 | 0.40 | 0.69 | 1 | Atg4a | autophagy related 4A, cysteine peptidase | 299 | 246 | 258 | 330 | 358 | 309 | 353 | 387 | 254 | 540 | 308 |
| 331 | 0.09 | 0.20 | 0.44 | 0.66 | 1 | Alg8 | asparagine-linked glycosylation 8 (alpha- | 360 | 286 | 215 | 263 | 305 | 418 | 353 | 279 | 302 | 365 | 494 |
| 304 | -0.20 | 0.28 | -0.71 | 0.48 | 1 | Pcdhgc4 | protocadherin gamma subfamily C, 4 | 296 | 308 | 299 | 402 | 335 | 217 | 353 | 182 | 298 | 380 | 270 |
| 277 | -0.06 | 0.36 | -0.18 | 0.86 | 1 | Bcl6 | B cell leukemia/lymphoma 6 | 1144 | 286 | 474 | 37 | 158 | 59 | 353 | 28 | 202 | 245 | 66 |
| 225 | -0.01 | 0.23 | -0.05 | 0.96 | 1 | Gm50353 | | 29 | 24 | 26 | 280 | 241 | 331 | 353 | 262 | 294 | 313 | 320 |
| 606 | -0.07 | 0.15 | -0.48 | 0.63 | 1 | Rassf1 | Ras association (RalGDS/AF-6) domain fa | 1001 | 961 | 908 | 432 | 438 | 715 | 352 | 392 | 368 | 380 | 718 |
| 560 | 0.15 | 0.15 | 1.03 | 0.30 | 1 | Tgs1 | trimethylguanosine synthase 1 | 797 | 879 | 714 | 426 | 323 | 682 | 352 | 477 | 321 | 442 | 742 |
| 485 | 0.05 | 0.27 | 0.17 | 0.86 | 1 | B930095G15Rik | | 354 | 462 | 463 | 410 | 391 | 833 | 352 | 703 | 492 | 216 | 661 |
| 448 | 0.14 | 0.18 | 0.80 | 0.43 | 1 | Ubac1 | ubiquitin associated domain containing 1 | 468 | 464 | 494 | 389 | 427 | 499 | 352 | 483 | 384 | 465 | 499 |
| 430 | 0.21 | 0.22 | 0.98 | 0.32 | 1 | Ntpcr | nucleoside-triphosphatase, cancer-relate | 561 | 512 | 642 | 408 | 376 | 255 | 352 | 498 | 329 | 518 | 284 |
| 270 | -0.20 | 0.31 | -0.64 | 0.52 | 1 | Gm6658 | | 123 | 288 | 220 | 353 | 303 | 241 | 352 | 247 | 285 | 356 | 198 |
| 261 | -0.39 | 0.26 | -1.50 | 0.13 | 1 | Cutal | cutA divalent cation tolerance homolog-lil | 209 | 198 | 243 | 411 | 254 | 237 | 352 | 321 | 273 | 206 | 166 |
| 537 | 0.15 | 0.24 | 0.60 | 0.55 | 1 | Hus1 | HUS1 checkpoint clamp component | 516 | 763 | 434 | 487 | 565 | 593 | 351 | 470 | 462 | 598 | 671 |
| 404 | -0.32 | 0.26 | -1.25 | 0.21 | 1 | Syde2 | synapse defective 1, Rho GTPase, homol | 420 | 708 | 671 | 494 | 307 | 327 | 351 | 235 | 329 | 300 | 302 |
| 393 | 0.24 | 0.23 | 1.05 | 0.29 | 1 | Ppil3 | peptidylprolyl isomerase (cyclophilin)-like | 570 | 390 | 450 | 331 | 352 | 290 | 351 | 536 | 357 | 412 | 281 |
| 352 | 0.01 | 0.23 | 0.02 | 0.98 | 1 | Gfer | growth factor, augmenter of liver regenera | 376 | 442 | 397 | 328 | 336 | 288 | 351 | 514 | 278 | 282 | 274 |
| 349 | 0.31 | 0.23 | 1.37 | 0.17 | 1 | Gm49338 | | 299 | 213 | 211 | 305 | 323 | 415 | 351 | 512 | 420 | 374 | 416 |
| 629 | -0.07 | 0.26 | -0.28 | 0.78 | 1 | Mcm7 | minichromosome maintenance complex c | 1559 | 784 | 937 | 464 | 549 | 494 | 350 | 448 | 389 | 421 | 519 |
| 472 | -0.10 | 0.25 | -0.38 | 0.70 | 1 | Ddx51 | DEAD (Asp-Glu-Ala-Asp) box polypeptide | 560 | 547 | 414 | 510 | 353 | 663 | 350 | 272 | 452 | 341 | 727 |
| 452 | 0.08 | 0.28 | 0.29 | 0.77 | 1 | Tent2 | terminal nucleotidyltransferase 2 | 680 | 490 | 428 | 242 | 351 | 691 | 350 | 331 | 433 | 211 | 765 |
| 451 | 0.32 | 0.19 | 1.69 | 0.09 | 1 | Pdcd2l | programmed cell death 2-like | 370 | 514 | 463 | 403 | 489 | 363 | 350 | 644 | 468 | 491 | 410 |
| 423 | -0.25 | 0.18 | -1.33 | 0.18 | 1 | Polr2d | polymerase (RNA) II (DNA directed) polyp | 559 | 533 | 553 | 493 | 428 | 361 | 350 | 364 | 366 | 287 | 353 |
| 409 | -0.19 | 0.27 | -0.70 | 0.48 | 1 | Suv39h1 | suppressor of variegation 3-9 1 | 667 | 627 | 477 | 338 | 384 | 374 | 350 | 320 | 426 | 187 | 352 |

| | | | | | | | | | | | | | | | | | | |
|---|---|---|---|---|---|---|---|---|---|---|---|---|---|---|---|---|---|---|
| 372 | 0.22 | 0.21 | 1.01 | 0.31 | 1 | Zfp2 | zinc finger protein 2 | 371 | 383 | 327 | 326 | 336 | 360 | 350 | 389 | 265 | 492 | 486 |
| 357 | -0.08 | 0.19 | -0.40 | 0.69 | 1 | Zfp763 | zinc finger protein 763 | 348 | 465 | 363 | 453 | 303 | 316 | 350 | 338 | 342 | 312 | 341 |
| 353 | 0.08 | 0.19 | 0.44 | 0.66 | 1 | Klhl23 | kelch-like 23 | 430 | 491 | 350 | 383 | 291 | 250 | 350 | 402 | 291 | 310 | 337 |
| 315 | -0.24 | 0.23 | -1.03 | 0.30 | 1 | Arid3b | AT rich interactive domain 3B (BRIGHT-lil | 142 | 151 | 141 | 347 | 399 | 505 | 350 | 218 | 334 | 282 | 598 |
| 274 | 0.02 | 0.20 | 0.10 | 0.92 | 1 | E130311K13Ril | RIKEN cDNA E130311K13 gene | 257 | 215 | 203 | 281 | 289 | 243 | 350 | 319 | 247 | 316 | 294 |
| 263 | 0.09 | 0.32 | 0.28 | 0.78 | 1 | Gm50431 | | 233 | 138 | 171 | 280 | 267 | 246 | 350 | 353 | 395 | 193 | 268 |
| 562 | -0.12 | 0.17 | -0.72 | 0.47 | 1 | Lrrc45 | leucine rich repeat containing 45 | 417 | 482 | 447 | 426 | 432 | 1319 | 349 | 481 | 316 | 340 | 1171 |
| 490 | 0.20 | 0.21 | 0.95 | 0.34 | 1 | Arrdc1 | arrestin domain containing 1 | 536 | 524 | 594 | 376 | 353 | 671 | 349 | 594 | 353 | 376 | 659 |
| 430 | 0.08 | 0.18 | 0.46 | 0.64 | 1 | Gmpr2 | guanosine monophosphate reductase 2 | 580 | 627 | 593 | 288 | 356 | 430 | 349 | 402 | 325 | 357 | 420 |
| 391 | 0.11 | 0.18 | 0.62 | 0.54 | 1 | Jmjd4 | jumonji domain containing 4 | 395 | 424 | 310 | 412 | 339 | 422 | 349 | 488 | 405 | 341 | 413 |
| 351 | -0.40 | 0.25 | -1.60 | 0.11 | 1 | Spice1 | spindle and centriole associated protein 1 | 387 | 291 | 396 | 319 | 341 | 563 | 349 | 276 | 323 | 162 | 453 |
| 319 | -0.09 | 0.21 | -0.41 | 0.68 | 1 | Nsun4 | NOL1/NOP2/Sun domain family, member | 359 | 202 | 268 | 326 | 291 | 416 | 349 | 297 | 294 | 313 | 394 |
| 401 | 0.10 | 0.25 | 0.41 | 0.68 | 1 | Stoml1 | stomatin-like 1 | 569 | 334 | 504 | 350 | 277 | 478 | 348 | 447 | 320 | 312 | 467 |
| 388 | 0.07 | 0.21 | 0.34 | 0.74 | 1 | Terf1 | telomeric repeat binding factor 1 | 420 | 377 | 531 | 309 | 349 | 424 | 348 | 419 | 327 | 294 | 469 |
| 339 | 0.06 | 0.23 | 0.26 | 0.79 | 1 | Ubox5 | U box domain containing 5 | 172 | 273 | 243 | 374 | 370 | 392 | 348 | 292 | 454 | 386 | 420 |
| 845 | -0.25 | 0.18 | -1.38 | 0.17 | 1 | Cish | cytokine inducible SH2-containing proteir | 320 | 1033 | 465 | 1252 | 304 | 2112 | 347 | 1033 | 310 | 236 | 1882 |
| 492 | 0.02 | 0.18 | 0.10 | 0.92 | 1 | Klf16 | Kruppel-like factor 16 | 613 | 869 | 468 | 592 | 432 | 353 | 347 | 568 | 421 | 420 | 332 |
| 477 | -0.02 | 0.17 | -0.12 | 0.90 | 1 | Zfp61 | zinc finger protein 61 | 340 | 400 | 470 | 393 | 484 | 787 | 347 | 456 | 387 | 312 | 875 |
| 472 | 0.27 | 0.19 | 1.40 | 0.16 | 1 | Smn1 | survival motor neuron 1 | 661 | 560 | 540 | 389 | 387 | 429 | 347 | 579 | 406 | 415 | 474 |
| 467 | 0.24 | 0.19 | 1.29 | 0.20 | 1 | Tlr3 | toll-like receptor 3 | 1158 | 660 | 696 | 288 | 308 | 231 | 347 | 309 | 354 | 550 | 233 |
| 370 | -0.23 | 0.24 | -0.97 | 0.33 | 1 | Zfp229 | zinc finger protein 229 | 474 | 464 | 485 | 337 | 349 | 374 | 347 | 330 | 289 | 182 | 437 |
| 362 | 0.17 | 0.34 | 0.49 | 0.63 | 1 | Dgka | diacylglycerol kinase, alpha | 548 | 364 | 533 | 232 | 163 | 493 | 347 | 412 | 320 | 222 | 344 |
| 361 | 0.02 | 0.23 | 0.09 | 0.93 | 1 | Pex10 | peroxisomal biogenesis factor 10 | 236 | 346 | 333 | 252 | 317 | 614 | 347 | 290 | 330 | 340 | 564 |
| 335 | -0.22 | 0.22 | -0.96 | 0.33 | 1 | Eif3j1 | eukaryotic translation initiation factor 3, su | 220 | 342 | 388 | 360 | 362 | 393 | 347 | 291 | 289 | 278 | 416 |
| 275 | -0.31 | 0.25 | -1.20 | 0.23 | 1 | Lpin3 | lipin 3 | 247 | 229 | 226 | 354 | 228 | 351 | 347 | 188 | 261 | 249 | 346 |
| 271 | -0.23 | 0.34 | -0.65 | 0.51 | 1 | Zfp941 | zinc finger protein 941 | 166 | 287 | 265 | 294 | 254 | 312 | 347 | 141 | 249 | 363 | 299 |
| 482 | 0.04 | 0.23 | 0.15 | 0.88 | 1 | Slc25a38 | solute carrier family 25, member 38 | 495 | 382 | 404 | 588 | 455 | 603 | 346 | 518 | 438 | 583 | 491 |
| 377 | -0.03 | 0.22 | -0.13 | 0.89 | 1 | Ikzf4 | IKAROS family zinc finger 4 | 292 | 657 | 374 | 457 | 258 | 369 | 346 | 418 | 303 | 316 | 351 |
| 369 | 0.04 | 0.23 | 0.18 | 0.86 | 1 | Taf1c | TATA-box binding protein associated fact | 317 | 497 | 308 | 362 | 259 | 459 | 346 | 577 | 261 | 274 | 403 |
| 333 | 0.13 | 0.19 | 0.71 | 0.48 | 1 | Pax6os1 | | 112 | 131 | 135 | 450 | 421 | 340 | 346 | 570 | 469 | 330 | 359 |
| 329 | 0.09 | 0.26 | 0.33 | 0.74 | 1 | Clp1 | CLP1, cleavage and polyadenylation fact | 371 | 178 | 274 | 323 | 383 | 304 | 346 | 329 | 442 | 298 | 370 |
| 282 | 0.01 | 0.30 | 0.05 | 0.96 | 1 | Bloc1s4 | biogenesis of lysosomal organelles comp | 355 | 214 | 263 | 263 | 272 | 248 | 346 | 418 | 279 | 231 | 215 |
| 111 | -0.41 | 0.57 | -0.73 | 0.46 | 1 | Gm13453 | | 251 | 5 | 12 | 15 | 4 | 95 | 346 | 8 | 4 | 432 | 45 |
| 645 | 0.03 | 0.17 | 0.15 | 0.88 | 1 | Srgap2 | SLIT-ROBO Rho GTPase activating prote | 1773 | 1314 | 1670 | 230 | 256 | 332 | 345 | 282 | 282 | 361 | 255 |
| 594 | -0.17 | 0.30 | -0.57 | 0.57 | 1 | Mcm4 | minichromosome maintenance complex c | 1718 | 841 | 640 | 553 | 458 | 393 | 345 | 343 | 413 | 464 | 367 |
| 422 | -0.16 | 0.28 | -0.58 | 0.56 | 1 | Prr3 | proline-rich polypeptide 3 | 393 | 470 | 434 | 451 | 361 | 560 | 345 | 679 | 321 | 188 | 439 |
| 358 | 0.11 | 0.17 | 0.64 | 0.52 | 1 | Yars2 | tyrosyl-tRNA synthetase 2 (mitochondrial) | 314 | 324 | 378 | 296 | 375 | 385 | 345 | 298 | 408 | 371 | 444 |
| 342 | 0.30 | 0.28 | 1.05 | 0.29 | 1 | Cox10 | heme A:farnesyltransferase cytochrome c | 363 | 200 | 316 | 228 | 391 | 342 | 345 | 502 | 376 | 402 | 303 |
| 327 | 0.09 | 0.25 | 0.35 | 0.72 | 1 | Itpkc | inositol 1,4,5-trisphosphate 3-kinase C | 383 | 317 | 388 | 290 | 241 | 331 | 345 | 357 | 270 | 234 | 439 |
| 645 | -0.28 | 0.28 | -0.99 | 0.32 | 1 | Sv2b | synaptic vesicle glycoprotein 2 b | 1548 | 2099 | 1333 | 348 | 182 | 247 | 344 | 402 | 136 | 131 | 320 |
| 373 | 0.01 | 0.20 | 0.05 | 0.96 | 1 | Rap2b | RAP2B, member of RAS oncogene family | 663 | 437 | 546 | 317 | 316 | 232 | 344 | 300 | 274 | 447 | 222 |
| 335 | -0.05 | 0.24 | -0.20 | 0.84 | 1 | Tmem238 | transmembrane protein 238 | 310 | 215 | 378 | 363 | 387 | 317 | 344 | 430 | 364 | 310 | 268 |
| 710 | -0.10 | 0.19 | -0.53 | 0.60 | 1 | Iqsec1 | IQ motif and Sec7 domain 1 | 1901 | 1974 | 1623 | 319 | 309 | 200 | 343 | 445 | 270 | 267 | 154 |
| 521 | 0.28 | 0.26 | 1.05 | 0.30 | 1 | Lpar1 | lysophosphatidic acid receptor 1 | 1489 | 693 | 916 | 333 | 285 | 227 | 343 | 511 | 314 | 353 | 264 |
| 440 | 0.03 | 0.25 | 0.11 | 0.91 | 1 | Fam162a | family with sequence similarity 162, meml | 631 | 514 | 449 | 458 | 442 | 349 | 343 | 285 | 442 | 541 | 385 |
| 360 | -0.36 | 0.21 | -1.67 | 0.09 | 1 | Tfcp2 | transcription factor CP2 | 313 | 235 | 364 | 276 | 356 | 746 | 343 | 270 | 308 | 198 | 552 |
| 356 | -0.11 | 0.25 | -0.42 | 0.68 | 1 | Tada2a | transcriptional adaptor 2A | 283 | 166 | 300 | 396 | 293 | 596 | 343 | 299 | 274 | 380 | 586 |
| 328 | -0.14 | 0.31 | -0.46 | 0.65 | 1 | Ripk2 | receptor (TNFRSF)-interacting serine-thre | 403 | 457 | 330 | 393 | 236 | 297 | 343 | 166 | 318 | 314 | 350 |
| 290 | 0.06 | 0.25 | 0.23 | 0.82 | 1 | Smim4 | small integral membrane protein 4 | 242 | 240 | 313 | 270 | 296 | 260 | 343 | 415 | 282 | 268 | 260 |
| 526 | -0.06 | 0.21 | -0.28 | 0.78 | 1 | Camta1 | calmodulin binding transcription activator | 734 | 633 | 580 | 406 | 460 | 729 | 342 | 322 | 337 | 502 | 736 |
| 501 | -0.48 | 0.27 | -1.80 | 0.07 | 1 | Sbno2 | strawberry notch 2 | 1022 | 670 | 758 | 375 | 342 | 708 | 342 | 177 | 406 | 209 | 507 |
| 463 | 0.03 | 0.23 | 0.12 | 0.90 | 1 | Otulinl | OTU deubiquitinase with linear linkage sp | 704 | 418 | 746 | 378 | 353 | 500 | 342 | 283 | 348 | 493 | 522 |
| 447 | 0.22 | 0.16 | 1.37 | 0.17 | 1 | Tdrp | testis development related protein | 648 | 871 | 588 | 443 | 326 | 199 | 342 | 467 | 371 | 390 | 271 |
| 389 | -0.22 | 0.21 | -1.02 | 0.31 | 1 | Zfp626 | zinc finger protein 626 | 362 | 460 | 441 | 377 | 411 | 466 | 342 | 249 | 402 | 254 | 516 |
| 378 | 0.13 | 0.24 | 0.52 | 0.60 | 1 | Zfp518b | zinc finger protein 518B | 281 | 378 | 312 | 509 | 352 | 329 | 342 | 336 | 486 | 493 | 341 |
| 358 | 0.24 | 0.18 | 1.36 | 0.17 | 1 | Pafah2 | platelet-activating factor acetylhydrolase 2 | 323 | 341 | 298 | 357 | 290 | 374 | 342 | 418 | 414 | 349 | 428 |
| 288 | -0.20 | 0.27 | -0.74 | 0.46 | 1 | Usp28 | ubiquitin specific peptidase 28 | 337 | 305 | 320 | 283 | 194 | 358 | 342 | 280 | 213 | 175 | 365 |
| 272 | -0.02 | 0.25 | -0.10 | 0.92 | 1 | Gm16536 | | 211 | 186 | 116 | 293 | 296 | 309 | 342 | 405 | 242 | 272 | 319 |
| 742 | -0.16 | 0.20 | -0.81 | 0.42 | 1 | Evpl | envoplakin | 1288 | 1590 | 1389 | 796 | 485 | 435 | 341 | 708 | 430 | 318 | 378 |
| 470 | -0.09 | 0.22 | -0.43 | 0.67 | 1 | Nudt16 | nudix (nucleoside diphosphate linked mo | 625 | 569 | 606 | 501 | 513 | 380 | 341 | 476 | 392 | 287 | 474 |

| | | | | | | | | | | | | | | | | | | |
|---|---|---|---|---|---|---|---|---|---|---|---|---|---|---|---|---|---|---|
| 402 | -0.06 | 0.21 | -0.31 | 0.76 | 1 | Ankrd13b | ankyrin repeat domain 13b | 421 | 319 | 472 | 343 | 337 | 626 | 341 | 377 | 372 | 265 | 544 |
| 368 | -0.08 | 0.26 | -0.32 | 0.75 | 1 | Snapc5 | small nuclear RNA activating complex, pc | 483 | 317 | 481 | 465 | 330 | 278 | 341 | 266 | 362 | 437 | 288 |
| 256 | -0.29 | 0.31 | -0.93 | 0.35 | 1 | Zcchc12 | zinc finger, CCHC domain containing 12 | 170 | 365 | 191 | 236 | 224 | 337 | 341 | 307 | 191 | 252 | 198 |
| 517 | 0.26 | 0.26 | 1.03 | 0.31 | 1 | Tuba1b | tubulin, alpha 1B | 1203 | 565 | 877 | 405 | 363 | 270 | 340 | 531 | 405 | 434 | 297 |
| 516 | -0.04 | 0.18 | -0.23 | 0.82 | 1 | Zfp467 | zinc finger protein 467 | 820 | 1026 | 771 | 453 | 369 | 392 | 340 | 372 | 325 | 441 | 372 |
| 438 | -0.32 | 0.28 | -1.16 | 0.25 | 1 | Fam53b | family with sequence similarity 53, membe | 872 | 770 | 773 | 375 | 348 | 260 | 340 | 251 | 392 | 157 | 277 |
| 394 | 0.13 | 0.25 | 0.54 | 0.59 | 1 | Gatb | glutamyl-tRNA(Gln) amidotransferase, su | 373 | 306 | 430 | 371 | 446 | 376 | 340 | 480 | 319 | 524 | 365 |
| 393 | -0.03 | 0.22 | -0.13 | 0.90 | 1 | Taco1 | translational activator of mitochondrially e | 277 | 374 | 321 | 431 | 487 | 435 | 340 | 426 | 337 | 483 | 409 |
| 316 | 0.03 | 0.19 | 0.15 | 0.88 | 1 | Slc25a13 | solute carrier family 25 (mitochondrial car | 298 | 308 | 296 | 438 | 315 | 193 | 340 | 424 | 301 | 340 | 226 |
| 220 | -0.43 | 0.33 | -1.33 | 0.18 | 1 | Gk5 | glycerol kinase 5 (putative) | 220 | 125 | 177 | 196 | 227 | 300 | 340 | 79 | 308 | 216 | 234 |
| 468 | 0.08 | 0.30 | 0.25 | 0.80 | 1 | Atad2 | ATPase family, AAA domain containing 2 | 784 | 320 | 373 | 392 | 446 | 586 | 339 | 308 | 568 | 362 | 673 |
| 416 | 0.17 | 0.16 | 1.05 | 0.30 | 1 | Strada | STE20-related kinase adaptor alpha | 437 | 472 | 438 | 313 | 360 | 511 | 339 | 395 | 379 | 367 | 567 |
| 393 | -0.40 | 0.23 | -1.71 | 0.09 | 1 | Cinp | cyclin-dependent kinase 2 interacting pro | 522 | 476 | 634 | 328 | 404 | 451 | 339 | 329 | 344 | 167 | 333 |
| 362 | 0.16 | 0.21 | 0.75 | 0.45 | 1 | Ubtd2 | ubiquitin domain containing 2 | 238 | 238 | 207 | 501 | 436 | 281 | 339 | 533 | 558 | 358 | 298 |
| 345 | -0.11 | 0.26 | -0.40 | 0.69 | 1 | 1110018N20Rik | | 148 | 269 | 243 | 367 | 426 | 496 | 339 | 393 | 266 | 408 | 439 |
| 321 | -0.18 | 0.31 | -0.59 | 0.56 | 1 | Vash1 | vasohibin 1 | 322 | 177 | 277 | 377 | 352 | 388 | 339 | 475 | 290 | 225 | 304 |
| 2635 | -0.08 | 0.18 | -0.46 | 0.65 | 1 | Col6a3 | collagen, type VI, alpha 3 | 9971 | 7549 | 8504 | 503 | 327 | 355 | 338 | 349 | 390 | 282 | 413 |
| 489 | -0.15 | 0.31 | -0.49 | 0.63 | 1 | Fmo1 | flavin containing monooxygenase 1 | 1312 | 518 | 662 | 318 | 508 | 327 | 338 | 247 | 312 | 310 | 525 |
| 444 | 0.19 | 0.25 | 0.77 | 0.44 | 1 | Tnip2 | TNFAIP3 interacting protein 2 | 571 | 340 | 515 | 375 | 296 | 609 | 338 | 485 | 274 | 438 | 647 |
| 391 | -0.18 | 0.29 | -0.61 | 0.54 | 1 | Tango6 | transport and golgi organization 6 | 619 | 279 | 499 | 434 | 376 | 341 | 338 | 228 | 378 | 553 | 261 |
| 388 | -0.01 | 0.17 | -0.07 | 0.94 | 1 | Wdr91 | WD repeat domain 91 | 619 | 579 | 541 | 310 | 340 | 278 | 338 | 367 | 297 | 349 | 251 |
| 376 | 0.04 | 0.20 | 0.21 | 0.83 | 1 | Dhodh | dihydroorotate dehydrogenase | 425 | 367 | 417 | 319 | 359 | 422 | 338 | 351 | 327 | 471 | 339 |
| 367 | -0.17 | 0.25 | -0.69 | 0.49 | 1 | Atl1 | atlastin GTPase 1 | 210 | 229 | 292 | 501 | 379 | 524 | 338 | 288 | 302 | 420 | 555 |
| 342 | -0.04 | 0.24 | -0.16 | 0.87 | 1 | BC048403 | cDNA sequence BC048403 | 349 | 313 | 236 | 408 | 332 | 378 | 338 | 334 | 440 | 289 | 349 |
| 222 | -0.14 | 0.28 | -0.49 | 0.63 | 1 | Acyp2 | acylphosphatase 2, muscle type | 159 | 109 | 165 | 271 | 215 | 209 | 338 | 189 | 198 | 408 | 181 |
| 858 | 0.03 | 0.16 | 0.16 | 0.88 | 1 | Arl4d | ADP-ribosylation factor-like 4D | 1777 | 1913 | 1995 | 291 | 298 | 935 | 337 | 390 | 302 | 259 | 944 |
| 836 | -0.02 | 0.17 | -0.13 | 0.90 | 1 | Phldb1 | pleckstrin homology like domain, family B | 2714 | 1823 | 1866 | 375 | 292 | 396 | 337 | 373 | 261 | 407 | 349 |
| 491 | -0.15 | 0.17 | -0.86 | 0.39 | 1 | A430033K04Ril | RIKEN cDNA A430033K04 gene | 425 | 564 | 522 | 649 | 430 | 630 | 337 | 524 | 380 | 321 | 624 |
| 388 | 0.23 | 0.23 | 1.00 | 0.32 | 1 | Bmp2k | BMP2 inducible kinase | 992 | 485 | 414 | 292 | 212 | 260 | 337 | 328 | 259 | 350 | 340 |
| 382 | -0.09 | 0.25 | -0.35 | 0.73 | 1 | 5530601H04Rik | | 444 | 357 | 567 | 307 | 291 | 508 | 337 | 242 | 223 | 422 | 506 |
| 320 | -0.04 | 0.23 | -0.17 | 0.86 | 1 | Ankrd49 | ankyrin repeat domain 49 | 359 | 206 | 235 | 337 | 315 | 384 | 337 | 292 | 328 | 290 | 438 |
| 310 | -0.31 | 0.24 | -1.25 | 0.21 | 1 | Naf1 | nuclear assembly factor 1 ribonucleoprote | 437 | 466 | 369 | 310 | 299 | 239 | 337 | 256 | 274 | 174 | 251 |
| 648 | -0.03 | 0.28 | -0.09 | 0.93 | 1 | Sh3bgrl3 | SH3 domain binding glutamic acid-rich pr | 1935 | 1172 | 1583 | 345 | 361 | 192 | 336 | 468 | 306 | 196 | 233 |
| 474 | -0.05 | 0.18 | -0.30 | 0.77 | 1 | Inha | inhibin alpha | 194 | 206 | 217 | 256 | 269 | 1592 | 336 | 299 | 290 | 301 | 1250 |
| 393 | 0.25 | 0.22 | 1.15 | 0.25 | 1 | Ccdc34 | coiled-coil domain containing 34 | 553 | 472 | 442 | 253 | 345 | 366 | 336 | 451 | 321 | 433 | 348 |
| 379 | -0.31 | 0.22 | -1.40 | 0.16 | 1 | Srfbp1 | serum response factor binding protein 1 | 392 | 475 | 321 | 447 | 349 | 516 | 336 | 249 | 320 | 295 | 474 |
| 372 | -0.18 | 0.18 | -0.99 | 0.32 | 1 | Trmt10b | tRNA methyltransferase 10B | 362 | 405 | 408 | 304 | 424 | 483 | 336 | 312 | 321 | 289 | 448 |
| 346 | 0.16 | 0.22 | 0.76 | 0.45 | 1 | 4632428C04Rik | | 187 | 249 | 228 | 345 | 449 | 334 | 336 | 547 | 481 | 289 | 357 |
| 433 | 0.00 | 0.31 | 0.00 | 1.00 | 1 | Zdhhc16 | zinc finger, DHHC domain containing 16 | 492 | 343 | 375 | 348 | 419 | 653 | 335 | 616 | 299 | 236 | 648 |
| 406 | 0.04 | 0.17 | 0.25 | 0.80 | 1 | Cnrip1 | cannabinoid receptor interacting protein 1 | 584 | 512 | 562 | 354 | 335 | 361 | 335 | 389 | 355 | 366 | 319 |
| 366 | 0.06 | 0.19 | 0.28 | 0.78 | 1 | Fgf9 | fibroblast growth factor 9 | 389 | 467 | 427 | 346 | 305 | 353 | 335 | 448 | 290 | 354 | 313 |
| 362 | 0.36 | 0.28 | 1.25 | 0.21 | 1 | Scel | sciellin | 168 | 288 | 143 | 331 | 381 | 445 | 335 | 363 | 513 | 535 | 484 |
| 293 | 0.01 | 0.21 | 0.03 | 0.98 | 1 | Bola1 | bolA-like 1 (E. coli) | 243 | 218 | 259 | 296 | 366 | 243 | 335 | 246 | 348 | 434 | 237 |
| 403 | 0.12 | 0.20 | 0.58 | 0.56 | 1 | Aimp2 | aminoacyl tRNA synthetase complex-inte | 528 | 477 | 438 | 314 | 409 | 377 | 334 | 433 | 341 | 424 | 363 |
| 362 | -0.18 | 0.21 | -0.85 | 0.39 | 1 | Aasdh | aminoadipate-semialdehyde dehydrogen | 208 | 250 | 265 | 259 | 305 | 780 | 334 | 260 | 227 | 252 | 839 |
| 333 | 0.10 | 0.24 | 0.42 | 0.67 | 1 | Zscan12 | zinc finger and SCAN domain containing | 344 | 297 | 292 | 270 | 323 | 382 | 334 | 404 | 328 | 242 | 441 |
| 330 | 0.10 | 0.22 | 0.48 | 0.63 | 1 | Dohh | deoxyhypusine hydroxylase/monooxyger | 281 | 389 | 322 | 369 | 282 | 266 | 334 | 527 | 311 | 263 | 283 |
| 330 | 0.15 | 0.25 | 0.60 | 0.55 | 1 | Mipol1 | mirror-image polydactyly 1 | 250 | 266 | 248 | 307 | 291 | 411 | 334 | 212 | 391 | 453 | 462 |
| 317 | 0.06 | 0.28 | 0.22 | 0.83 | 1 | Thoc6 | THO complex 6 | 473 | 274 | 549 | 200 | 262 | 291 | 334 | 315 | 231 | 273 | 286 |
| 312 | -0.21 | 0.20 | -1.09 | 0.27 | 1 | Slc16a6 | solute carrier family 16 (monocarboxylic a | 307 | 292 | 256 | 318 | 352 | 380 | 334 | 316 | 311 | 265 | 300 |
| 303 | -0.18 | 0.22 | -0.82 | 0.41 | 1 | Atp5g2 | ATP synthase, H+ transporting, mitochond | 300 | 341 | 298 | 256 | 334 | 349 | 334 | 220 | 335 | 281 | 288 |
| 1003 | 0.20 | 0.18 | 1.10 | 0.27 | 1 | Sash1 | SAM and SH3 domain containing 1 | 2979 | 2501 | 2785 | 444 | 297 | 209 | 333 | 474 | 455 | 340 | 216 |
| 384 | 0.16 | 0.21 | 0.74 | 0.46 | 1 | 2810025M15Rik | | 535 | 497 | 608 | 367 | 296 | 226 | 333 | 466 | 309 | 314 | 274 |
| 340 | 0.15 | 0.25 | 0.61 | 0.54 | 1 | 1700008J07Rik | | 230 | 462 | 193 | 381 | 361 | 275 | 333 | 436 | 371 | 379 | 317 |
| 322 | 0.00 | 0.27 | 0.01 | 0.99 | 1 | Snupn | snurportin 1 | 304 | 337 | 291 | 279 | 333 | 343 | 333 | 383 | 378 | 187 | 372 |
| 305 | 0.18 | 0.25 | 0.72 | 0.47 | 1 | Pet100 | PET100 homolog | 274 | 215 | 354 | 272 | 319 | 254 | 333 | 423 | 321 | 325 | 264 |
| 851 | 0.13 | 0.66 | 0.19 | 0.85 | 1 | Mki67 | antigen identified by monoclonal antibody | 2762 | 734 | 482 | 577 | 918 | 752 | 332 | 725 | 801 | 420 | 863 |
| 596 | 0.06 | 0.23 | 0.24 | 0.81 | 1 | Cdk19 | cyclin-dependent kinase 19 | 1222 | 951 | 973 | 375 | 495 | 467 | 332 | 549 | 405 | 307 | 481 |
| 371 | 0.06 | 0.23 | 0.28 | 0.78 | 1 | Atpaf1 | ATP synthase mitochondrial F1 complex a | 273 | 255 | 221 | 411 | 454 | 422 | 332 | 329 | 413 | 546 | 424 |
| 332 | 0.14 | 0.30 | 0.47 | 0.64 | 1 | Ube2d-ps | | 290 | 326 | 194 | 211 | 421 | 391 | 332 | 382 | 426 | 307 | 369 |

| | | | | | | | | | | | | | | | | | |
|---|---|---|---|---|---|---|---|---|---|---|---|---|---|---|---|---|---|
| 329 | 0.09 | 0.23 | 0.38 | 0.71 | 1 | Rps19bp1 | ribosomal protein S19 binding protein 1 | 309 | 380 | 417 | 331 | 257 | 297 | 332 | 390 | 256 | 295 | 359 |
| 316 | 0.14 | 0.21 | 0.66 | 0.51 | 1 | Aars2 | alanyl-tRNA synthetase 2, mitochondrial | 202 | 190 | 274 | 344 | 325 | 331 | 332 | 293 | 392 | 388 | 408 |
| 310 | 0.16 | 0.29 | 0.56 | 0.58 | 1 | Zfp438 | zinc finger protein 438 | 164 | 319 | 239 | 242 | 320 | 378 | 332 | 312 | 387 | 402 | 317 |
| 269 | -0.19 | 0.36 | -0.54 | 0.59 | 1 | Zfp28 | zinc finger protein 28 | 332 | 168 | 261 | 182 | 205 | 475 | 332 | 290 | 197 | 131 | 387 |
| 268 | -0.34 | 0.29 | -1.17 | 0.24 | 1 | Gm49636 | | 177 | 82 | 121 | 322 | 398 | 347 | 332 | 153 | 415 | 307 | 291 |
| 438 | 0.05 | 0.25 | 0.22 | 0.83 | 1 | Zfp983 | zinc finger protein 983 | 304 | 364 | 246 | 431 | 328 | 822 | 331 | 262 | 383 | 483 | 862 |
| 386 | -0.12 | 0.18 | -0.66 | 0.51 | 1 | Fchsd1 | FCH and double SH3 domains 1 | 685 | 524 | 542 | 335 | 308 | 319 | 331 | 258 | 261 | 363 | 321 |
| 375 | 0.12 | 0.34 | 0.37 | 0.71 | 1 | Cdk10 | cyclin-dependent kinase 10 | 386 | 264 | 294 | 268 | 267 | 671 | 331 | 570 | 289 | 193 | 594 |
| 368 | -0.10 | 0.21 | -0.49 | 0.63 | 1 | Polm | polymerase (DNA directed), mu | 316 | 318 | 295 | 296 | 315 | 662 | 331 | 198 | 333 | 346 | 640 |
| 351 | -0.01 | 0.24 | -0.04 | 0.97 | 1 | Smim11 | small integral membrane protein 11 | 530 | 362 | 550 | 290 | 349 | 248 | 331 | 373 | 307 | 258 | 262 |
| 341 | 0.11 | 0.19 | 0.57 | 0.57 | 1 | Tspyl5 | testis-specific protein, Y-encoded-like 5 | 428 | 387 | 344 | 310 | 315 | 287 | 331 | 279 | 347 | 398 | 320 |
| 331 | -0.20 | 0.26 | -0.79 | 0.43 | 1 | Myo7a | myosin VIIA | 907 | 400 | 349 | 148 | 173 | 456 | 331 | 171 | 188 | 232 | 287 |
| 269 | -0.04 | 0.28 | -0.13 | 0.89 | 1 | Slc25a35 | solute carrier family 25, member 35 | 154 | 198 | 115 | 250 | 314 | 363 | 331 | 212 | 339 | 294 | 388 |
| 552 | 0.22 | 0.22 | 1.01 | 0.31 | 1 | Elf4 | E74-like factor 4 (ets domain transcription | 1288 | 1108 | 933 | 362 | 370 | 194 | 330 | 356 | 556 | 358 | 212 |
| 469 | 0.13 | 0.25 | 0.53 | 0.60 | 1 | Srgap1 | SLIT-ROBO Rho GTPase activating prote | 434 | 627 | 667 | 502 | 361 | 406 | 330 | 792 | 315 | 345 | 380 |
| 421 | 0.33 | 0.23 | 1.41 | 0.16 | 1 | Selenoh | selenoprotein H | 589 | 416 | 566 | 316 | 401 | 302 | 330 | 560 | 411 | 450 | 292 |
| 372 | 0.16 | 0.22 | 0.73 | 0.47 | 1 | Card19 | caspase recruitment domain family, meml | 348 | 499 | 446 | 387 | 278 | 314 | 330 | 504 | 286 | 425 | 279 |
| 368 | 0.03 | 0.22 | 0.14 | 0.89 | 1 | Cyb561d2 | cytochrome b-561 domain containing 2 | 364 | 360 | 306 | 377 | 491 | 295 | 330 | 329 | 416 | 470 | 311 |
| 368 | -0.11 | 0.22 | -0.51 | 0.61 | 1 | Elk1 | ELK1, member of ETS oncogene family | 479 | 322 | 372 | 276 | 378 | 487 | 330 | 199 | 396 | 295 | 512 |
| 237 | -0.06 | 0.28 | -0.21 | 0.83 | 1 | Sdcbp2 | syndecan binding protein (syntenin) 2 | 61 | 115 | 54 | 326 | 182 | 334 | 330 | 453 | 165 | 196 | 385 |
| 463 | -0.40 | 0.33 | -1.19 | 0.23 | 1 | Nfkb2 | nuclear factor of kappa light polypeptide g | 1440 | 901 | 857 | 226 | 273 | 251 | 329 | 262 | 205 | 100 | 250 |
| 402 | -0.15 | 0.35 | -0.44 | 0.66 | 1 | 1700017B05Ril | RIKEN cDNA 1700017B05 gene | 1219 | 500 | 508 | 331 | 315 | 186 | 329 | 359 | 332 | 189 | 155 |
| 380 | 0.14 | 0.20 | 0.68 | 0.50 | 1 | Wdr25 | WD repeat domain 25 | 211 | 294 | 265 | 446 | 418 | 420 | 329 | 621 | 367 | 415 | 392 |
| 346 | 0.13 | 0.34 | 0.37 | 0.71 | 1 | Cmc1 | COX assembly mitochondrial protein 1 | 443 | 296 | 375 | 358 | 349 | 234 | 329 | 605 | 285 | 198 | 334 |
| 288 | -0.31 | 0.28 | -1.09 | 0.27 | 1 | Plpp7 | phospholipid phosphatase 7 (inactive) | 173 | 282 | 249 | 416 | 315 | 306 | 329 | 210 | 299 | 361 | 228 |
| 253 | -0.08 | 0.24 | -0.31 | 0.76 | 1 | Etohd2 | | 208 | 289 | 138 | 304 | 216 | 260 | 329 | 260 | 225 | 268 | 288 |
| 235 | 0.20 | 0.31 | 0.63 | 0.53 | 1 | Adam32 | a disintegrin and metallopeptidase domai | 45 | 35 | 77 | 321 | 254 | 215 | 329 | 271 | 321 | 470 | 247 |
| 568 | -0.30 | 0.22 | -1.38 | 0.17 | 1 | Gnb4 | guanine nucleotide binding protein (G prc | 1582 | 1179 | 1304 | 288 | 319 | 247 | 328 | 315 | 199 | 346 | 142 |
| 444 | -0.18 | 0.18 | -0.97 | 0.33 | 1 | Mettl14 | methyltransferase like 14 | 800 | 645 | 700 | 341 | 266 | 515 | 328 | 248 | 297 | 272 | 468 |
| 407 | -0.04 | 0.19 | -0.21 | 0.83 | 1 | Zfp90 | zinc finger protein 90 | 444 | 514 | 537 | 330 | 420 | 435 | 328 | 371 | 311 | 371 | 412 |
| 403 | -0.32 | 0.23 | -1.38 | 0.17 | 1 | Scarf2 | scavenger receptor class F, member 2 | 1167 | 622 | 811 | 224 | 213 | 216 | 328 | 153 | 174 | 392 | 131 |
| 358 | -0.03 | 0.20 | -0.17 | 0.87 | 1 | Zfp771 | zinc finger protein 771 | 330 | 377 | 401 | 357 | 403 | 339 | 328 | 412 | 342 | 385 | 266 |
| 333 | -0.07 | 0.20 | -0.33 | 0.74 | 1 | Phykpl | 5-phosphohydroxy-L-lysine phospholyase | 263 | 282 | 277 | 354 | 261 | 516 | 328 | 293 | 280 | 357 | 449 |
| 314 | -0.04 | 0.29 | -0.13 | 0.89 | 1 | Yju2 | YJU2 splicing factor | 357 | 278 | 403 | 341 | 256 | 291 | 328 | 430 | 251 | 201 | 321 |
| 523 | 0.16 | 0.31 | 0.52 | 0.60 | 1 | Psd4 | pleckstrin and Sec7 domain containing 4 | 498 | 406 | 257 | 726 | 711 | 366 | 327 | 994 | 630 | 502 | 338 |
| 397 | -0.25 | 0.22 | -1.13 | 0.26 | 1 | B4galt2 | UDP-Gal:betaGlcNAc beta 1,4- galactosy | 696 | 490 | 489 | 416 | 403 | 315 | 327 | 272 | 358 | 316 | 284 |
| 386 | 0.31 | 0.26 | 1.20 | 0.23 | 1 | Rassf7 | Ras association (RalGDS/AF-6) domain fa | 255 | 363 | 346 | 332 | 433 | 360 | 327 | 505 | 397 | 609 | 319 |
| 381 | 0.32 | 0.31 | 1.02 | 0.31 | 1 | Crlf2 | cytokine receptor-like factor 2 | 458 | 417 | 512 | 326 | 231 | 325 | 327 | 693 | 258 | 376 | 264 |
| 355 | 0.21 | 0.24 | 0.88 | 0.38 | 1 | Prim1 | DNA primase, p49 subunit | 420 | 317 | 392 | 316 | 301 | 336 | 327 | 502 | 365 | 303 | 323 |
| 244 | -0.38 | 0.31 | -1.22 | 0.22 | 1 | Oser1 | oxidative stress responsive serine rich 1 | 186 | 255 | 229 | 228 | 275 | 296 | 327 | 114 | 239 | 234 | 306 |
| 210 | -0.12 | 0.27 | -0.44 | 0.66 | 1 | Degs2 | delta(4)-desaturase, sphingolipid 2 | 80 | 126 | 57 | 346 | 172 | 206 | 327 | 339 | 207 | 316 | 129 |
| 378 | 0.18 | 0.21 | 0.84 | 0.40 | 1 | Ccdc30 | coiled-coil domain containing 30 | 159 | 202 | 249 | 481 | 318 | 547 | 326 | 491 | 415 | 376 | 595 |
| 362 | 0.09 | 0.20 | 0.44 | 0.66 | 1 | Eif4ebp1 | eukaryotic translation initiation factor 4E b | 399 | 487 | 366 | 344 | 419 | 232 | 326 | 378 | 388 | 403 | 237 |
| 351 | -0.24 | 0.24 | -0.99 | 0.32 | 1 | Kdelr3 | KDEL (Lys-Asp-Glu-Leu) endoplasmic ret | 663 | 788 | 673 | 222 | 247 | 154 | 326 | 170 | 175 | 278 | 169 |
| 344 | -0.20 | 0.24 | -0.86 | 0.39 | 1 | Cacnb1 | calcium channel, voltage-dependent, beta | 389 | 402 | 426 | 336 | 278 | 423 | 326 | 183 | 304 | 335 | 378 |
| 336 | -0.09 | 0.18 | -0.52 | 0.61 | 1 | Tmem177 | transmembrane protein 177 | 346 | 433 | 366 | 343 | 359 | 292 | 326 | 298 | 334 | 300 | 302 |
| 305 | -0.20 | 0.26 | -0.76 | 0.45 | 1 | Rnf135 | ring finger protein 135 | 260 | 285 | 309 | 370 | 285 | 334 | 326 | 301 | 337 | 165 | 379 |
| 266 | 0.14 | 0.25 | 0.53 | 0.59 | 1 | 1300002E11Rik | | 200 | 176 | 185 | 180 | 242 | 390 | 326 | 315 | 249 | 250 | 416 |
| 233 | -0.24 | 0.28 | -0.87 | 0.38 | 1 | E130317F20Rik | | 188 | 182 | 159 | 301 | 301 | 184 | 326 | 192 | 261 | 213 | 257 |
| 559 | -0.25 | 0.18 | -1.38 | 0.17 | 1 | Sox13 | SRY (sex determining region Y)-box 13 | 852 | 1056 | 985 | 419 | 439 | 603 | 325 | 396 | 326 | 362 | 388 |
| 469 | 0.37 | 0.22 | 1.71 | 0.09 | 1 | 0610030E20Ril | RIKEN cDNA 0610030E20 gene | 462 | 389 | 404 | 314 | 290 | 822 | 325 | 557 | 312 | 488 | 794 |
| 445 | -0.05 | 0.20 | -0.24 | 0.81 | 1 | Atp11c | ATPase, class VI, type 11C | 1192 | 680 | 564 | 287 | 230 | 407 | 325 | 234 | 252 | 312 | 412 |
| 382 | 0.14 | 0.20 | 0.71 | 0.48 | 1 | Dera | deoxyribose-phosphate aldolase (putative | 478 | 386 | 383 | 327 | 390 | 360 | 325 | 344 | 483 | 359 | 367 |
| 354 | 0.01 | 0.21 | 0.07 | 0.95 | 1 | Itgb3bp | integrin beta 3 binding protein (beta3-enc | 421 | 403 | 379 | 356 | 298 | 352 | 325 | 476 | 291 | 280 | 315 |
| 354 | -0.05 | 0.25 | -0.19 | 0.85 | 1 | Slc25a10 | solute carrier family 25 (mitochondrial car | 516 | 321 | 455 | 277 | 435 | 264 | 325 | 177 | 349 | 448 | 324 |
| 345 | 0.05 | 0.33 | 0.16 | 0.88 | 1 | Zfp560 | zinc finger protein 560 | 373 | 253 | 458 | 375 | 309 | 297 | 325 | 512 | 403 | 180 | 305 |
| 327 | 0.26 | 0.26 | 0.98 | 0.33 | 1 | Mri1 | methylthioribose-1-phosphate isomerase | 380 | 287 | 413 | 193 | 282 | 363 | 325 | 402 | 289 | 294 | 375 |
| 263 | -0.07 | 0.23 | -0.30 | 0.77 | 1 | Zbtb25 | zinc finger and BTB domain containing 25 | 292 | 268 | 217 | 237 | 279 | 247 | 325 | 234 | 306 | 243 | 247 |
| 921 | 0.30 | 0.26 | 1.15 | 0.25 | 1 | F13a1 | coagulation factor XIII, A1 subunit | 3979 | 1592 | 1928 | 237 | 399 | 233 | 324 | 412 | 407 | 331 | 286 |
| 544 | -0.12 | 0.19 | -0.62 | 0.53 | 1 | Eogt | EGF domain-specific O-linked N-acetylglu | 1103 | 741 | 1331 | 284 | 359 | 504 | 324 | 312 | 306 | 273 | 449 |

| | | | | | | | | | | | | | | | | | | |
|---|---|---|---|---|---|---|---|---|---|---|---|---|---|---|---|---|---|---|
| 361 | -0.03 | 0.30 | -0.09 | 0.93 | 1 | Rdh10 | retinol dehydrogenase 10 (all-trans) | 665 | 453 | 338 | 323 | 227 | 395 | 324 | 423 | 252 | 180 | 394 |
| 352 | 0.03 | 0.27 | 0.10 | 0.92 | 1 | Panx2 | pannexin 2 | 229 | 428 | 475 | 396 | 345 | 293 | 324 | 394 | 311 | 428 | 253 |
| 317 | -0.10 | 0.23 | -0.45 | 0.65 | 1 | Ebpl | emopamil binding protein-like | 344 | 440 | 455 | 287 | 301 | 248 | 324 | 214 | 299 | 345 | 227 |
| 284 | 0.00 | 0.26 | 0.00 | 1.00 | 1 | Cdk5r1 | cyclin-dependent kinase 5, regulatory sub | 285 | 184 | 316 | 320 | 288 | 230 | 324 | 387 | 308 | 280 | 200 |
| 271 | -0.07 | 0.31 | -0.21 | 0.83 | 1 | Ankrd39 | ankyrin repeat domain 39 | 185 | 219 | 239 | 299 | 167 | 385 | 324 | 321 | 243 | 156 | 440 |
| 1756 | 0.01 | 0.24 | 0.05 | 0.96 | 1 | Thbd | thrombomodulin | 7111 | 4572 | 4645 | 678 | 445 | 96 | 323 | 554 | 384 | 374 | 133 |
| 421 | 0.32 | 0.21 | 1.49 | 0.14 | 1 | Enox1 | ecto-NOX disulfide-thiol exchanger 1 | 339 | 255 | 342 | 396 | 474 | 454 | 323 | 533 | 503 | 556 | 452 |
| 335 | -0.48 | 0.27 | -1.78 | 0.08 | 1 | Mad2l1 | MAD2 mitotic arrest deficient-like 1 | 752 | 450 | 387 | 295 | 382 | 228 | 323 | 211 | 226 | 209 | 221 |
| 307 | -0.40 | 0.24 | -1.62 | 0.11 | 1 | Fdxr | ferredoxin reductase | 311 | 371 | 445 | 259 | 302 | 366 | 323 | 203 | 199 | 188 | 406 |
| 280 | -0.36 | 0.21 | -1.70 | 0.09 | 1 | Ccpg1os | cell cycle progression 1, opposite strand | 351 | 250 | 277 | 349 | 283 | 284 | 323 | 215 | 239 | 270 | 243 |
| 268 | 0.10 | 0.32 | 0.32 | 0.75 | 1 | Ccdc189 | coiled-coil domain containing 189 | 132 | 223 | 241 | 227 | 219 | 359 | 323 | 414 | 224 | 278 | 308 |
| 153 | -0.28 | 0.33 | -0.86 | 0.39 | 1 | Fabp3 | fatty acid binding protein 3, muscle and h | 60 | 47 | 78 | 202 | 161 | 142 | 323 | 167 | 205 | 188 | 114 |
| 352 | 0.11 | 0.22 | 0.51 | 0.61 | 1 | Tmem71 | transmembrane protein 71 | 337 | 489 | 311 | 388 | 262 | 356 | 322 | 324 | 343 | 395 | 349 |
| 349 | 0.42 | 0.29 | 1.48 | 0.14 | 1 | Rusc1 | RUN and SH3 domain containing 1 | 283 | 230 | 162 | 253 | 385 | 396 | 322 | 502 | 545 | 376 | 385 |
| 307 | -0.22 | 0.25 | -0.87 | 0.38 | 1 | Mecr | mitochondrial trans-2-enoyl-CoA reducta | 390 | 401 | 276 | 312 | 199 | 412 | 322 | 200 | 262 | 220 | 377 |
| 549 | 0.33 | 0.22 | 1.48 | 0.14 | 1 | Ppp1r3b | protein phosphatase 1, regulatory subuni | 554 | 926 | 654 | 743 | 245 | 421 | 321 | 958 | 333 | 422 | 457 |
| 511 | 0.24 | 0.29 | 0.84 | 0.40 | 1 | Ddx39 | DEAD (Asp-Glu-Ala-Asp) box polypeptide | 1021 | 537 | 617 | 307 | 338 | 597 | 321 | 540 | 321 | 265 | 760 |
| 440 | 0.10 | 0.22 | 0.46 | 0.64 | 1 | Mif4gd | MIF4G domain containing | 587 | 508 | 457 | 383 | 449 | 429 | 321 | 504 | 408 | 327 | 463 |
| 413 | 0.10 | 0.21 | 0.49 | 0.62 | 1 | Ppp3cc | protein phosphatase 3, catalytic subunit, | 637 | 580 | 383 | 350 | 280 | 458 | 321 | 278 | 324 | 380 | 547 |
| 407 | -0.35 | 0.24 | -1.44 | 0.15 | 1 | Slc35b3 | solute carrier family 35, member B3 | 359 | 272 | 398 | 361 | 340 | 845 | 321 | 166 | 278 | 345 | 786 |
| 314 | -0.14 | 0.21 | -0.69 | 0.49 | 1 | Bend4 | BEN domain containing 4 | 361 | 368 | 197 | 422 | 293 | 290 | 321 | 380 | 252 | 300 | 270 |
| 277 | 0.03 | 0.26 | 0.10 | 0.92 | 1 | Tmem108 | transmembrane protein 108 | 197 | 180 | 196 | 277 | 283 | 325 | 321 | 173 | 387 | 383 | 321 |
| 14076 | -1.00 | 1.17 | -0.86 | 0.39 | 1 | Clps | colipase, pancreatic | 18467 | 72879 | 59470 | 2556 | 157 | 15 | 320 | 141 | 143 | 112 | 576 |
| 406 | 0.14 | 0.23 | 0.60 | 0.55 | 1 | Stk19 | serine/threonine kinase 19 | 368 | 280 | 545 | 259 | 274 | 730 | 320 | 299 | 302 | 372 | 722 |
| 343 | 0.15 | 0.26 | 0.59 | 0.55 | 1 | Wars2 | tryptophanyl tRNA synthetase 2 (mitocho | 402 | 426 | 457 | 176 | 333 | 357 | 320 | 335 | 378 | 250 | 335 |
| 341 | 0.15 | 0.22 | 0.69 | 0.49 | 1 | Shq1 | SHQ1 homolog (S. cerevisiae) | 352 | 350 | 334 | 263 | 310 | 391 | 320 | 419 | 371 | 277 | 358 |
| 309 | -0.31 | 0.26 | -1.18 | 0.24 | 1 | Kctd1 | potassium channel tetramerisation domai | 324 | 321 | 477 | 334 | 291 | 309 | 320 | 236 | 204 | 336 | 243 |
| 289 | 0.48 | 0.50 | 0.96 | 0.34 | 1 | Asb11 | ankyrin repeat and SOCS box-containing | 3 | 24 | 54 | 382 | 271 | 314 | 320 | 482 | 464 | 540 | 325 |
| 261 | -0.01 | 0.28 | -0.04 | 0.97 | 1 | Fkbpl | FK506 binding protein-like | 202 | 307 | 190 | 214 | 256 | 293 | 320 | 299 | 297 | 205 | 284 |
| 362 | -0.12 | 0.32 | -0.38 | 0.70 | 1 | Zbtb7c | zinc finger and BTB domain containing 7( | 759 | 681 | 788 | 130 | 238 | 228 | 319 | 261 | 141 | 291 | 151 |
| 354 | 0.10 | 0.21 | 0.47 | 0.64 | 1 | Zmym1 | zinc finger, MYM domain containing 1 | 341 | 514 | 329 | 321 | 237 | 442 | 319 | 349 | 335 | 283 | 429 |
| 319 | 0.17 | 0.19 | 0.89 | 0.37 | 1 | Sms | spermine synthase | 359 | 325 | 379 | 233 | 248 | 357 | 319 | 286 | 296 | 350 | 356 |
| 310 | 0.08 | 0.21 | 0.38 | 0.71 | 1 | Tfpt | TCF3 (E2A) fusion partner | 262 | 300 | 310 | 254 | 289 | 367 | 319 | 343 | 267 | 281 | 416 |
| 303 | 0.36 | 0.25 | 1.43 | 0.15 | 1 | Tifa | TRAF-interacting protein with forkhead-as | 524 | 248 | 273 | 262 | 267 | 161 | 319 | 255 | 372 | 384 | 268 |
| 294 | -0.01 | 0.28 | -0.02 | 0.98 | 1 | Zfp955a | zinc finger protein 955A | 347 | 361 | 181 | 253 | 286 | 309 | 319 | 382 | 252 | 227 | 322 |
| 280 | -0.18 | 0.33 | -0.53 | 0.60 | 1 | Gtpbp10 | GTP-binding protein 10 (putative) | 302 | 376 | 235 | 319 | 257 | 250 | 319 | 129 | 263 | 334 | 299 |
| 232 | 0.06 | 0.33 | 0.17 | 0.86 | 1 | Smim12 | small integral membrane protein 12 | 377 | 216 | 273 | 153 | 190 | 182 | 319 | 270 | 217 | 165 | 186 |
| 227 | -0.01 | 0.26 | -0.04 | 0.97 | 1 | Hnf4g | hepatocyte nuclear factor 4, gamma | 108 | 89 | 153 | 270 | 263 | 221 | 319 | 220 | 350 | 243 | 266 |
| 376 | -0.27 | 0.18 | -1.47 | 0.14 | 1 | D030056L22Ril | RIKEN cDNA D030056L22 gene | 513 | 512 | 497 | 443 | 369 | 305 | 318 | 313 | 341 | 273 | 258 |
| 361 | -0.13 | 0.18 | -0.75 | 0.45 | 1 | Coq6 | coenzyme Q6 monooxygenase | 335 | 389 | 383 | 319 | 412 | 449 | 318 | 309 | 322 | 292 | 448 |
| 351 | -0.18 | 0.20 | -0.91 | 0.36 | 1 | Neil1 | nei endonuclease VIII-like 1 (E. coli) | 351 | 481 | 357 | 294 | 322 | 495 | 318 | 275 | 300 | 280 | 392 |
| 343 | 0.14 | 0.18 | 0.78 | 0.44 | 1 | Smim8 | small integral membrane protein 8 | 370 | 439 | 372 | 343 | 285 | 292 | 318 | 336 | 346 | 361 | 316 |
| 318 | -0.25 | 0.23 | -1.12 | 0.26 | 1 | Abhd11 | abhydrolase domain containing 11 | 339 | 399 | 428 | 224 | 233 | 474 | 318 | 205 | 195 | 206 | 478 |
| 316 | 0.04 | 0.18 | 0.20 | 0.84 | 1 | Serhl | serine hydrolase-like | 287 | 278 | 290 | 327 | 282 | 363 | 318 | 356 | 281 | 303 | 388 |
| 260 | 0.17 | 0.23 | 0.75 | 0.46 | 1 | Hhat | hedgehog acyltransferase | 189 | 190 | 120 | 279 | 260 | 245 | 318 | 408 | 310 | 299 | 241 |
| 245 | -0.37 | 0.34 | -1.11 | 0.27 | 1 | Gm20163 | | 75 | 111 | 136 | 390 | 366 | 255 | 318 | 138 | 337 | 305 | 259 |
| 481 | 0.12 | 0.23 | 0.55 | 0.58 | 1 | Josd2 | Josephin domain containing 2 | 449 | 582 | 599 | 397 | 354 | 679 | 317 | 637 | 297 | 337 | 638 |
| 367 | -0.41 | 0.25 | -1.64 | 0.10 | 1 | Slc4a4 | solute carrier family 4 (anion exchanger), | 629 | 629 | 751 | 268 | 220 | 345 | 317 | 176 | 270 | 157 | 279 |
| 319 | -0.03 | 0.21 | -0.14 | 0.89 | 1 | Erich1 | glutamate rich 1 | 287 | 314 | 313 | 297 | 318 | 372 | 317 | 248 | 281 | 390 | 368 |
| 4065 | 0.19 | 0.35 | 0.55 | 0.58 | 1 | Igfbp5 | insulin-like growth factor binding protein 5 | 22151 | 11124 | 8822 | 227 | 483 | 197 | 316 | 212 | 477 | 426 | 280 |
| 430 | -0.05 | 0.18 | -0.25 | 0.80 | 1 | Actl6b | actin-like 6B | 408 | 453 | 470 | 372 | 496 | 553 | 316 | 330 | 485 | 368 | 483 |
| 347 | -0.12 | 0.20 | -0.60 | 0.55 | 1 | Mthfd2l | methylenetetrahydrofolate dehydrogenas | 495 | 440 | 377 | 324 | 341 | 315 | 316 | 327 | 258 | 363 | 262 |
| 344 | -0.08 | 0.33 | -0.25 | 0.80 | 1 | Smyd5 | SET and MYND domain containing 5 | 464 | 226 | 261 | 205 | 322 | 562 | 316 | 94 | 452 | 352 | 528 |
| 264 | 0.01 | 0.27 | 0.03 | 0.98 | 1 | Dipk1b | divergent protein kinase domain 1B | 160 | 153 | 221 | 375 | 288 | 199 | 316 | 283 | 275 | 429 | 205 |
| 184 | -0.09 | 0.35 | -0.25 | 0.80 | 1 | Ndufs5-ps | | 269 | 134 | 108 | 213 | 105 | 125 | 316 | 85 | 123 | 386 | 159 |
| 529 | 0.12 | 0.20 | 0.64 | 0.52 | 1 | Cyb5rl | cytochrome b5 reductase-like | 406 | 609 | 500 | 597 | 496 | 685 | 315 | 573 | 492 | 528 | 613 |
| 384 | -0.08 | 0.21 | -0.40 | 0.69 | 1 | Rtel1 | regulator of telomere elongation helicase | 384 | 377 | 336 | 225 | 375 | 712 | 315 | 282 | 314 | 281 | 624 |
| 358 | 0.00 | 0.25 | 0.00 | 1.00 | 1 | D17H6S53E | DNA segment, Chr 17, human D6S53E | 376 | 386 | 340 | 425 | 335 | 325 | 315 | 547 | 376 | 223 | 295 |
| 340 | 0.02 | 0.28 | 0.08 | 0.94 | 1 | Snrnp35 | small nuclear ribonucleoprotein 35 (U11/ | 398 | 351 | 386 | 338 | 299 | 317 | 315 | 169 | 371 | 425 | 371 |
| 319 | 0.11 | 0.27 | 0.43 | 0.67 | 1 | 4930578M01Rik | | 119 | 107 | 151 | 305 | 370 | 525 | 315 | 518 | 304 | 349 | 442 |

| | | | | | | | | | | | | | | | | | |
|---|---|---|---|---|---|---|---|---|---|---|---|---|---|---|---|---|---|
| 304 | 0.04 | 0.27 | 0.15 | 0.88 | 1 | Krtcap3 | keratinocyte associated protein 3 | 176 | 242 | 308 | 261 | 364 | 350 | 315 | 380 | 246 | 304 | 397 |
| 239 | -0.08 | 0.25 | -0.32 | 0.75 | 1 | Slc39a7 | solute carrier family 39 (zinc transporter), | 130 | 89 | 213 | 222 | 298 | 277 | 315 | 215 | 389 | 245 | 229 |
| 497 | 0.03 | 0.21 | 0.15 | 0.88 | 1 | Kmt5c | lysine methyltransferase 5C | 399 | 466 | 372 | 344 | 301 | 1240 | 314 | 298 | 377 | 408 | 947 |
| 455 | 0.18 | 0.17 | 1.06 | 0.29 | 1 | Ctnna2 | catenin (cadherin associated protein), alp | 401 | 551 | 610 | 427 | 431 | 452 | 314 | 490 | 493 | 412 | 429 |
| 324 | 0.08 | 0.22 | 0.35 | 0.73 | 1 | Cyp4f16 | cytochrome P450, family 4, subfamily f, pc | 357 | 293 | 332 | 359 | 267 | 309 | 314 | 341 | 257 | 444 | 293 |
| 310 | 0.15 | 0.31 | 0.49 | 0.62 | 1 | Cd164l2 | CD164 sialomucin-like 2 | 299 | 301 | 533 | 236 | 306 | 224 | 314 | 306 | 210 | 435 | 246 |
| 294 | -0.31 | 0.23 | -1.38 | 0.17 | 1 | Stradb | STE20-related kinase adaptor beta | 312 | 290 | 266 | 324 | 294 | 371 | 314 | 197 | 297 | 227 | 341 |
| 268 | -0.14 | 0.28 | -0.51 | 0.61 | 1 | Gm6206 | | 311 | 191 | 244 | 303 | 300 | 220 | 314 | 148 | 314 | 352 | 253 |
| 217 | 0.06 | 0.27 | 0.23 | 0.82 | 1 | Ndufa12-ps | | 335 | 249 | 164 | 249 | 142 | 119 | 314 | 262 | 196 | 216 | 137 |
| 546 | 0.25 | 0.21 | 1.18 | 0.24 | 1 | Secisbp2 | SECIS binding protein 2 | 339 | 437 | 329 | 421 | 447 | 1101 | 313 | 465 | 416 | 573 | 1163 |
| 473 | 0.28 | 0.22 | 1.25 | 0.21 | 1 | Zkscan6 | zinc finger with KRAB and SCAN domains | 581 | 441 | 552 | 403 | 475 | 451 | 313 | 563 | 404 | 469 | 548 |
| 412 | 0.02 | 0.17 | 0.13 | 0.90 | 1 | Marveld2 | MARVEL (membrane-associating) domai | 327 | 472 | 302 | 542 | 391 | 453 | 313 | 556 | 349 | 335 | 492 |
| 372 | 0.14 | 0.24 | 0.57 | 0.57 | 1 | Pih1d1 | PIH1 domain containing 1 | 541 | 450 | 426 | 319 | 332 | 295 | 313 | 525 | 302 | 303 | 285 |
| 371 | 0.05 | 0.31 | 0.17 | 0.87 | 1 | Arid3a | AT rich interactive domain 3A (BRIGHT-lik | 631 | 296 | 226 | 390 | 335 | 371 | 313 | 262 | 387 | 497 | 369 |
| 309 | 0.08 | 0.27 | 0.31 | 0.76 | 1 | Anapc15 | anaphase promoting complex C subunit 1 | 271 | 343 | 227 | 205 | 351 | 380 | 313 | 343 | 323 | 336 | 312 |
| 278 | -0.44 | 0.27 | -1.64 | 0.10 | 1 | Det1 | de-etiolated homolog 1 (Arabidopsis) | 232 | 371 | 208 | 333 | 275 | 373 | 313 | 171 | 220 | 236 | 332 |
| 220 | -0.68 | 0.36 | -1.88 | 0.06 | 1 | Tmem154 | transmembrane protein 154 | 422 | 279 | 166 | 216 | 291 | 156 | 313 | 145 | 180 | 116 | 141 |
| 657 | 0.03 | 0.17 | 0.17 | 0.86 | 1 | Rgl2 | ral guanine nucleotide dissociation stimul | 815 | 875 | 1019 | 361 | 352 | 1286 | 312 | 372 | 329 | 416 | 1093 |
| 462 | 0.10 | 0.19 | 0.52 | 0.60 | 1 | Fbxl19 | F-box and leucine-rich repeat protein 19 | 540 | 477 | 511 | 386 | 357 | 687 | 312 | 425 | 351 | 443 | 595 |
| 444 | 0.23 | 0.18 | 1.28 | 0.20 | 1 | Oasl2 | 2'-5' oligoadenylate synthetase-like 2 | 922 | 606 | 943 | 291 | 387 | 112 | 312 | 291 | 494 | 398 | 129 |
| 364 | 0.02 | 0.24 | 0.08 | 0.94 | 1 | Pcgf6 | polycomb group ring finger 6 | 195 | 408 | 348 | 330 | 270 | 585 | 312 | 304 | 321 | 269 | 663 |
| 348 | -0.05 | 0.25 | -0.19 | 0.85 | 1 | Zfp688 | zinc finger protein 688 | 327 | 427 | 280 | 315 | 357 | 435 | 312 | 253 | 351 | 384 | 382 |
| 347 | -0.06 | 0.29 | -0.20 | 0.84 | 1 | Cc2d2b | coiled-coil and C2 domain containing 2B | 29 | 19 | 59 | 354 | 303 | 923 | 312 | 293 | 318 | 335 | 876 |
| 332 | 0.23 | 0.33 | 0.69 | 0.49 | 1 | Engase | endo-beta-N-acetylglucosaminidase | 460 | 360 | 560 | 217 | 229 | 281 | 312 | 480 | 287 | 193 | 279 |
| 325 | 0.18 | 0.25 | 0.74 | 0.46 | 1 | Spa17 | sperm autoantigenic protein 17 | 322 | 262 | 389 | 375 | 292 | 244 | 312 | 453 | 304 | 317 | 311 |
| 311 | -0.11 | 0.18 | -0.58 | 0.56 | 1 | Trim68 | tripartite motif-containing 68 | 254 | 208 | 238 | 209 | 234 | 653 | 312 | 196 | 228 | 272 | 615 |
| 296 | -0.11 | 0.27 | -0.40 | 0.69 | 1 | Bclaf3 | Bclaf1 and Thrap3 family member 3 | 266 | 143 | 240 | 285 | 243 | 489 | 312 | 195 | 215 | 434 | 433 |
| 279 | 0.00 | 0.24 | -0.01 | 0.99 | 1 | Nkrf | NF-kappaB repressing factor | 125 | 170 | 96 | 448 | 307 | 288 | 312 | 309 | 339 | 334 | 342 |
| 495 | -0.30 | 0.27 | -1.13 | 0.26 | 1 | Map1a | microtubule-associated protein 1 A | 1138 | 955 | 971 | 340 | 331 | 287 | 311 | 408 | 238 | 343 | 125 |
| 433 | -0.33 | 0.20 | -1.61 | 0.11 | 1 | Pgf | placental growth factor | 761 | 821 | 895 | 358 | 335 | 245 | 311 | 336 | 283 | 282 | 134 |
| 389 | 0.11 | 0.17 | 0.65 | 0.51 | 1 | 1110059G10Ril | RIKEN cDNA 1110059G10 gene | 432 | 480 | 565 | 308 | 354 | 369 | 311 | 320 | 370 | 313 | 456 |
| 366 | -0.04 | 0.33 | -0.13 | 0.89 | 1 | Ikzf2 | IKAROS family zinc finger 2 | 292 | 263 | 112 | 270 | 334 | 813 | 311 | 402 | 322 | 222 | 682 |
| 362 | -0.01 | 0.21 | -0.04 | 0.96 | 1 | Wrap53 | WD repeat containing, antisense to Trp53 | 395 | 458 | 465 | 263 | 345 | 424 | 311 | 279 | 292 | 388 | 365 |
| 328 | 0.08 | 0.21 | 0.36 | 0.72 | 1 | Dnajc28 | DnaJ heat shock protein family (Hsp40) m | 364 | 322 | 285 | 294 | 344 | 326 | 311 | 244 | 362 | 388 | 364 |
| 308 | 0.18 | 0.22 | 0.80 | 0.43 | 1 | Hira | histone cell cycle regulator | 346 | 207 | 284 | 261 | 317 | 309 | 311 | 322 | 344 | 285 | 403 |
| 166 | 0.25 | 0.41 | 0.59 | 0.55 | 1 | Nmnat2 | nicotinamide nucleotide adenylyltransfera | 64 | 146 | 19 | 212 | 93 | 120 | 311 | 287 | 159 | 316 | 100 |
| 440 | -0.31 | 0.21 | -1.44 | 0.15 | 1 | Sema4c | sema domain, immunoglobulin domain (Ig | 1222 | 713 | 839 | 249 | 335 | 245 | 310 | 180 | 326 | 249 | 180 |
| 364 | 0.12 | 0.24 | 0.49 | 0.62 | 1 | Rpl14-ps1 | | 660 | 669 | 729 | 144 | 309 | 179 | 310 | 188 | 309 | 304 | 204 |
| 312 | -0.05 | 0.31 | -0.18 | 0.86 | 1 | Zfp383 | zinc finger protein 383 | 370 | 346 | 444 | 307 | 192 | 337 | 310 | 344 | 300 | 155 | 324 |
| 286 | -0.03 | 0.24 | -0.12 | 0.90 | 1 | Mettl23 | methyltransferase like 23 | 220 | 307 | 306 | 274 | 330 | 244 | 310 | 410 | 261 | 247 | 237 |
| 407 | -0.32 | 0.26 | -1.23 | 0.22 | 1 | Sppl2b | signal peptide peptidase like 2B | 540 | 307 | 465 | 252 | 289 | 824 | 309 | 100 | 291 | 303 | 801 |
| 386 | -0.03 | 0.19 | -0.16 | 0.87 | 1 | Ndufb2 | NADH:ubiquinone oxidoreductase subun | 546 | 424 | 509 | 317 | 388 | 382 | 309 | 345 | 310 | 314 | 397 |
| 378 | 0.28 | 0.20 | 1.41 | 0.16 | 1 | Tmem143 | transmembrane protein 143 | 325 | 418 | 387 | 301 | 375 | 385 | 309 | 431 | 372 | 437 | 415 |
| 323 | 0.51 | 0.36 | 1.41 | 0.16 | 1 | Gm11769 | | 125 | 47 | 99 | 302 | 346 | 405 | 309 | 615 | 511 | 270 | 522 |
| 288 | 0.11 | 0.26 | 0.42 | 0.68 | 1 | Gm45330 | | 177 | 220 | 191 | 242 | 389 | 295 | 309 | 256 | 472 | 359 | 260 |
| 267 | 0.35 | 0.32 | 1.10 | 0.27 | 1 | Tfb2m | transcription factor B2, mitochondrial | 214 | 91 | 187 | 183 | 315 | 292 | 309 | 379 | 376 | 251 | 338 |
| 685 | -0.41 | 0.30 | -1.35 | 0.18 | 1 | Fgf1 | fibroblast growth factor 1 | 2539 | 1680 | 1157 | 432 | 175 | 307 | 308 | 314 | 152 | 121 | 345 |
| 425 | 0.01 | 0.19 | 0.06 | 0.95 | 1 | Nmi | N-myc (and STAT) interactor | 676 | 592 | 678 | 416 | 380 | 262 | 308 | 375 | 325 | 346 | 316 |
| 411 | 0.00 | 0.18 | 0.01 | 0.99 | 1 | Mtfp1 | mitochondrial fission process 1 | 540 | 628 | 695 | 405 | 382 | 242 | 308 | 390 | 363 | 276 | 295 |
| 401 | -0.37 | 0.19 | -1.92 | 0.05 | 1 | Plekha7 | pleckstrin homology domain containing, fa | 663 | 669 | 732 | 375 | 274 | 365 | 308 | 244 | 210 | 252 | 322 |
| 385 | 0.39 | 0.22 | 1.75 | 0.08 | 1 | Cwf19l1 | CWF19-like 1, cell cycle control (S. pombe | 337 | 475 | 415 | 324 | 268 | 411 | 308 | 346 | 445 | 453 | 450 |
| 343 | 0.11 | 0.25 | 0.44 | 0.66 | 1 | Exo5 | exonuclease 5 | 329 | 274 | 404 | 326 | 384 | 313 | 308 | 423 | 278 | 416 | 317 |
| 284 | 0.05 | 0.26 | 0.18 | 0.86 | 1 | Sgf29 | SAGA complex associated factor 29 | 227 | 181 | 320 | 221 | 331 | 331 | 308 | 351 | 276 | 273 | 307 |
| 208 | -0.17 | 0.37 | -0.46 | 0.65 | 1 | Gm2011 | | 190 | 186 | 112 | 279 | 146 | 215 | 308 | 343 | 176 | 103 | 226 |
| 788 | 0.07 | 0.16 | 0.43 | 0.66 | 1 | Dusp6 | dual specificity phosphatase 6 | 2564 | 1470 | 1951 | 235 | 342 | 415 | 307 | 222 | 330 | 437 | 393 |
| 447 | -0.13 | 0.23 | -0.56 | 0.57 | 1 | Nol4l | nucleolar protein 4-like | 563 | 766 | 481 | 481 | 384 | 466 | 307 | 326 | 421 | 371 | 353 |
| 421 | 0.33 | 0.20 | 1.66 | 0.10 | 1 | Zfp942 | zinc finger protein 942 | 516 | 508 | 332 | 304 | 328 | 516 | 307 | 404 | 363 | 459 | 592 |
| 397 | -0.28 | 0.16 | -1.71 | 0.09 | 1 | Sec61a2 | Sec61, alpha subunit 2 (S. cerevisiae) | 473 | 427 | 429 | 252 | 248 | 856 | 307 | 222 | 204 | 234 | 714 |
| 341 | 0.08 | 0.24 | 0.35 | 0.73 | 1 | Eri2 | exoribonuclease 2 | 350 | 309 | 250 | 382 | 282 | 408 | 307 | 256 | 366 | 398 | 446 |
| 336 | -0.02 | 0.19 | -0.12 | 0.90 | 1 | Tradd | TNFRSF1A-associated via death domain | 389 | 358 | 404 | 274 | 324 | 380 | 307 | 334 | 302 | 317 | 303 |

| | | | | | | | | | | | | | | | | | | |
|---|---|---|---|---|---|---|---|---|---|---|---|---|---|---|---|---|---|---|
| 330 | 0.07 | 0.28 | 0.23 | 0.82 | 1 | Ceacam1 | carcinoembryonic antigen-related cell adl | 475 | 634 | 479 | 299 | 207 | 172 | 307 | 312 | 132 | 397 | 218 |
| 254 | -0.39 | 0.26 | -1.51 | 0.13 | 1 | Jph4 | junctophilin 4 | 161 | 110 | 163 | 309 | 267 | 451 | 307 | 184 | 319 | 211 | 315 |
| 243 | -0.22 | 0.25 | -0.89 | 0.37 | 1 | Akip1 | A kinase (PRKA) interacting protein 1 | 243 | 308 | 251 | 305 | 202 | 188 | 307 | 224 | 170 | 310 | 159 |
| 544 | -0.13 | 0.23 | -0.56 | 0.57 | 1 | Cnp | 2',3'-cyclic nucleotide 3' phosphodiesteras | 1780 | 895 | 970 | 365 | 343 | 206 | 306 | 291 | 311 | 334 | 189 |
| 419 | -0.16 | 0.17 | -0.94 | 0.35 | 1 | Oaf | out at first homolog | 788 | 456 | 805 | 240 | 348 | 456 | 306 | 209 | 298 | 269 | 440 |
| 400 | -0.19 | 0.21 | -0.90 | 0.37 | 1 | Dzip1l | DAZ interacting protein 1-like | 426 | 342 | 311 | 417 | 355 | 701 | 306 | 313 | 344 | 312 | 575 |
| 369 | -0.29 | 0.25 | -1.17 | 0.24 | 1 | Chml | choroideremia-like | 447 | 533 | 430 | 451 | 322 | 375 | 306 | 207 | 317 | 264 | 407 |
| 306 | 0.16 | 0.20 | 0.79 | 0.43 | 1 | Coa6 | cytochrome c oxidase assembly factor 6 | 312 | 252 | 300 | 324 | 338 | 216 | 306 | 389 | 315 | 358 | 255 |
| 300 | -0.41 | 0.25 | -1.62 | 0.10 | 1 | Gpa33 | glycoprotein A33 (transmembrane) | 214 | 431 | 503 | 275 | 355 | 284 | 306 | 229 | 298 | 192 | 212 |
| 298 | -0.18 | 0.28 | -0.63 | 0.53 | 1 | Ap2m1-ps | | 336 | 434 | 267 | 283 | 332 | 269 | 306 | 191 | 342 | 289 | 236 |
| 396 | 0.04 | 0.20 | 0.20 | 0.84 | 1 | Ruvbl1 | RuvB-like protein 1 | 514 | 583 | 430 | 398 | 328 | 365 | 305 | 331 | 399 | 362 | 336 |
| 340 | 0.13 | 0.21 | 0.62 | 0.53 | 1 | Slc22a5 | solute carrier family 22 (organic cation tra | 431 | 343 | 364 | 304 | 285 | 331 | 305 | 257 | 274 | 453 | 389 |
| 340 | 0.07 | 0.24 | 0.28 | 0.78 | 1 | Med4 | mediator complex subunit 4 | 255 | 396 | 334 | 332 | 387 | 322 | 305 | 331 | 287 | 404 | 383 |
| 334 | 0.04 | 0.20 | 0.19 | 0.85 | 1 | Acot8 | acyl-CoA thioesterase 8 | 298 | 319 | 357 | 275 | 323 | 436 | 305 | 321 | 287 | 357 | 397 |
| 327 | -0.37 | 0.34 | -1.07 | 0.28 | 1 | Pms2 | PMS1 homolog2, mismatch repair system | 441 | 367 | 254 | 305 | 335 | 448 | 305 | 117 | 356 | 227 | 448 |
| 322 | 0.24 | 0.27 | 0.88 | 0.38 | 1 | Gsto1 | glutathione S-transferase omega 1 | 471 | 316 | 409 | 309 | 250 | 209 | 305 | 466 | 244 | 298 | 263 |
| 298 | -0.24 | 0.24 | -1.01 | 0.31 | 1 | Nrbf2 | nuclear receptor binding factor 2 | 301 | 260 | 297 | 371 | 267 | 366 | 305 | 218 | 328 | 285 | 281 |
| 289 | 0.26 | 0.39 | 0.68 | 0.50 | 1 | Shld1 | shieldin complex subunit 1 | 189 | 139 | 264 | 267 | 342 | 242 | 305 | 636 | 194 | 353 | 250 |
| 252 | 0.39 | 0.32 | 1.23 | 0.22 | 1 | Tnfrsf9 | tumor necrosis factor receptor superfamily | 28 | 37 | 8 | 278 | 278 | 297 | 305 | 469 | 347 | 353 | 373 |
| 240 | 0.11 | 0.29 | 0.38 | 0.70 | 1 | Zbtb11os1 | | 154 | 181 | 281 | 226 | 282 | 160 | 305 | 321 | 246 | 319 | 161 |
| 186 | 0.30 | 0.39 | 0.77 | 0.44 | 1 | Gm7292 | | 294 | 307 | 22 | 419 | 10 | 26 | 305 | 339 | 35 | 247 | 38 |
| 478 | 0.22 | 0.22 | 0.98 | 0.32 | 1 | Zfp276 | zinc finger protein (C2H2 type) 276 | 405 | 453 | 356 | 362 | 343 | 914 | 304 | 554 | 301 | 497 | 767 |
| 380 | 0.04 | 0.21 | 0.20 | 0.85 | 1 | Med26 | mediator complex subunit 26 | 396 | 438 | 397 | 438 | 369 | 346 | 304 | 310 | 409 | 404 | 368 |
| 347 | -0.09 | 0.21 | -0.45 | 0.65 | 1 | 2410004B18Ril | RIKEN cDNA 2410004B18 gene | 402 | 482 | 316 | 374 | 245 | 433 | 304 | 371 | 291 | 218 | 384 |
| 17658 | -0.04 | 0.40 | -0.10 | 0.92 | 1 | C3 | complement component 3 | 122973 | 45972 | 23216 | 298 | 295 | 154 | 303 | 280 | 234 | 327 | 184 |
| 501 | 0.10 | 0.30 | 0.35 | 0.73 | 1 | Meox1 | mesenchyme homeobox 1 | 932 | 849 | 1234 | 288 | 369 | 244 | 303 | 351 | 180 | 450 | 315 |
| 423 | -0.40 | 0.26 | -1.58 | 0.11 | 1 | Aox1 | aldehyde oxidase 1 | 371 | 294 | 319 | 424 | 355 | 1090 | 303 | 429 | 264 | 255 | 553 |
| 383 | 0.21 | 0.28 | 0.76 | 0.44 | 1 | Klhl25 | kelch-like 25 | 563 | 392 | 478 | 399 | 307 | 270 | 303 | 563 | 363 | 250 | 325 |
| 351 | 0.21 | 0.23 | 0.93 | 0.35 | 1 | Lad1 | ladinin | 100 | 164 | 129 | 586 | 408 | 246 | 303 | 940 | 410 | 345 | 231 |
| 291 | 0.18 | 0.25 | 0.70 | 0.48 | 1 | Klhdc1 | kelch domain containing 1 | 224 | 329 | 334 | 297 | 198 | 291 | 303 | 288 | 296 | 291 | 353 |
| 222 | -0.01 | 0.29 | -0.04 | 0.97 | 1 | Gm8121 | | 199 | 193 | 192 | 226 | 238 | 168 | 303 | 284 | 235 | 164 | 236 |
| 165 | -0.55 | 0.28 | -1.93 | 0.05 | 1 | Gm6733 | | 93 | 106 | 18 | 382 | 75 | 172 | 303 | 224 | 36 | 256 | 147 |
| 484 | 0.40 | 0.23 | 1.74 | 0.08 | 1 | Osbpl11 | oxysterol binding protein-like 11 | 668 | 533 | 667 | 416 | 265 | 519 | 302 | 578 | 335 | 444 | 597 |
| 452 | 0.02 | 0.26 | 0.07 | 0.94 | 1 | Fam126a | family with sequence similarity 126, meml | 1219 | 837 | 830 | 212 | 257 | 264 | 302 | 372 | 216 | 225 | 238 |
| 417 | 0.20 | 0.21 | 0.98 | 0.33 | 1 | Nudt13 | nudix (nucleoside diphosphate linked mo | 490 | 609 | 383 | 261 | 298 | 591 | 302 | 422 | 285 | 356 | 589 |
| 409 | -0.01 | 0.21 | -0.05 | 0.96 | 1 | Cdan1 | congenital dyserythropoietic anemia, type | 500 | 535 | 327 | 364 | 267 | 663 | 302 | 431 | 303 | 249 | 563 |
| 381 | 0.06 | 0.27 | 0.20 | 0.84 | 1 | Chac1 | ChaC, cation transport regulator 1 | 68 | 117 | 188 | 600 | 426 | 573 | 302 | 526 | 464 | 432 | 489 |
| 375 | -0.15 | 0.23 | -0.67 | 0.50 | 1 | Igtp | interferon gamma induced GTPase | 709 | 671 | 545 | 346 | 296 | 221 | 302 | 287 | 280 | 200 | 269 |
| 286 | 0.10 | 0.28 | 0.34 | 0.73 | 1 | 4931414P19Ril | RIKEN cDNA 4931414P19 gene | 346 | 266 | 338 | 207 | 278 | 278 | 302 | 377 | 238 | 227 | 292 |
| 445 | 0.07 | 0.21 | 0.34 | 0.73 | 1 | Mpp1 | membrane protein, palmitoylated | 1211 | 772 | 752 | 215 | 301 | 240 | 301 | 225 | 315 | 274 | 290 |
| 407 | -0.39 | 0.26 | -1.51 | 0.13 | 1 | Atxn7l1 | ataxin 7-like 1 | 687 | 547 | 716 | 366 | 261 | 472 | 301 | 214 | 283 | 153 | 476 |
| 346 | -0.08 | 0.17 | -0.46 | 0.64 | 1 | Spata7 | spermatogenesis associated 7 | 320 | 380 | 359 | 314 | 327 | 466 | 301 | 297 | 309 | 282 | 449 |
| 309 | 0.23 | 0.24 | 0.95 | 0.34 | 1 | Hps6 | HPS6, biogenesis of lysosomal organelle | 398 | 306 | 352 | 249 | 296 | 220 | 301 | 372 | 421 | 286 | 195 |
| 307 | -0.19 | 0.26 | -0.71 | 0.48 | 1 | Nr1h3 | nuclear receptor subfamily 1, group H, me | 607 | 323 | 345 | 312 | 280 | 225 | 301 | 308 | 244 | 241 | 189 |
| 289 | 0.12 | 0.32 | 0.36 | 0.72 | 1 | Aass | aminoadipate-semialdehyde synthase | 228 | 173 | 97 | 390 | 385 | 221 | 301 | 352 | 454 | 291 | 284 |
| 261 | 0.15 | 0.29 | 0.53 | 0.60 | 1 | Slc2a6 | solute carrier family 2 (facilitated glucose | 159 | 131 | 96 | 212 | 378 | 280 | 301 | 204 | 421 | 386 | 306 |
| 746 | -0.01 | 0.24 | -0.05 | 0.96 | 1 | Apold1 | apolipoprotein L domain containing 1 | 1920 | 2040 | 1940 | 347 | 336 | 176 | 300 | 394 | 168 | 340 | 245 |
| 355 | -0.10 | 0.22 | -0.45 | 0.66 | 1 | Cbx8 | chromobox 8 | 320 | 372 | 408 | 354 | 433 | 363 | 300 | 251 | 366 | 404 | 330 |
| 342 | 0.06 | 0.22 | 0.28 | 0.78 | 1 | 2010320M18Rik | | 273 | 365 | 379 | 392 | 381 | 258 | 300 | 507 | 309 | 368 | 229 |
| 332 | 0.02 | 0.29 | 0.08 | 0.93 | 1 | Leng1 | leukocyte receptor cluster (LRC) member | 412 | 436 | 368 | 277 | 197 | 410 | 300 | 423 | 117 | 334 | 382 |
| 268 | 0.02 | 0.24 | 0.08 | 0.94 | 1 | Hsbp1l1 | heat shock factor binding protein 1-like 1 | 317 | 272 | 407 | 219 | 262 | 180 | 300 | 197 | 248 | 370 | 176 |
| 243 | -0.38 | 0.22 | -1.70 | 0.09 | 1 | Zfp3 | zinc finger protein 3 | 309 | 336 | 262 | 223 | 269 | 206 | 300 | 192 | 190 | 219 | 165 |
| 367 | 0.08 | 0.28 | 0.30 | 0.77 | 1 | Zfp623 | zinc finger protein 623 | 388 | 449 | 514 | 360 | 318 | 315 | 299 | 294 | 285 | 541 | 278 |
| 296 | -0.16 | 0.21 | -0.78 | 0.44 | 1 | Uevld | UEV and lactate/malate dehyrogenase dc | 360 | 411 | 330 | 277 | 222 | 336 | 299 | 258 | 223 | 198 | 344 |
| 258 | 0.24 | 0.24 | 1.02 | 0.31 | 1 | Rnf186 | ring finger protein 186 | 82 | 99 | 115 | 272 | 276 | 309 | 299 | 440 | 314 | 283 | 346 |
| 223 | -0.26 | 0.28 | -0.92 | 0.36 | 1 | Dand5 | DAN domain family member 5, BMP antag | 184 | 151 | 189 | 224 | 294 | 213 | 299 | 139 | 194 | 368 | 193 |
| 220 | -0.77 | 0.41 | -1.90 | 0.06 | 1 | Timm10b | translocase of inner mitochondrial membr | 114 | 266 | 251 | 184 | 300 | 293 | 299 | 102 | 257 | 77 | 281 |
| 362 | 0.22 | 0.18 | 1.22 | 0.22 | 1 | Nmnat3 | nicotinamide nucleotide adenylyltransfera | 238 | 344 | 308 | 369 | 396 | 361 | 298 | 523 | 433 | 367 | 351 |
| 344 | -0.14 | 0.23 | -0.62 | 0.53 | 1 | Dmac2l | distal membrane arm assembly complex 2 | 291 | 438 | 367 | 320 | 323 | 444 | 298 | 227 | 338 | 229 | 508 |
| 339 | 0.25 | 0.27 | 0.95 | 0.34 | 1 | Plagl1 | pleiomorphic adenoma gene-like 1 | 313 | 328 | 170 | 341 | 329 | 365 | 298 | 416 | 434 | 292 | 447 |

| | | | | | | | | | | | | | | | | | | |
|---|---|---|---|---|---|---|---|---|---|---|---|---|---|---|---|---|---|---|
| 322 | 0.08 | 0.22 | 0.36 | 0.72 | 1 | Zfp944 | zinc finger protein 944 | 347 | 279 | 302 | 320 | 230 | 427 | 298 | 265 | 337 | 291 | 450 |
| 309 | -0.22 | 0.21 | -1.05 | 0.29 | 1 | Phf7 | PHD finger protein 7 | 281 | 274 | 306 | 395 | 285 | 388 | 298 | 272 | 301 | 237 | 367 |
| 279 | 0.08 | 0.25 | 0.31 | 0.76 | 1 | Timm9 | translocase of inner mitochondrial membr | 208 | 237 | 240 | 165 | 287 | 399 | 298 | 197 | 258 | 282 | 498 |
| 279 | -0.12 | 0.33 | -0.38 | 0.70 | 1 | Apip | APAF1 interacting protein | 428 | 374 | 324 | 214 | 263 | 225 | 298 | 246 | 240 | 125 | 328 |
| 272 | -0.10 | 0.23 | -0.42 | 0.68 | 1 | Mettl5 | methyltransferase like 5 | 306 | 287 | 231 | 267 | 232 | 316 | 298 | 190 | 256 | 238 | 373 |
| 631 | 0.11 | 0.19 | 0.61 | 0.54 | 1 | Plekhn1 | pleckstrin homology domain containing, fa | 475 | 502 | 446 | 477 | 413 | 1515 | 297 | 527 | 344 | 460 | 1489 |
| 399 | 0.18 | 0.20 | 0.93 | 0.35 | 1 | Mpg | N-methylpurine-DNA glycosylase | 551 | 439 | 502 | 378 | 288 | 395 | 297 | 409 | 370 | 331 | 425 |
| 332 | 0.01 | 0.34 | 0.02 | 0.99 | 1 | Zfp449 | zinc finger protein 449 | 296 | 498 | 230 | 335 | 340 | 331 | 297 | 205 | 281 | 483 | 359 |
| 203 | 0.09 | 0.31 | 0.28 | 0.78 | 1 | Hnmt | histamine N-methyltransferase | 267 | 186 | 208 | 230 | 139 | 102 | 297 | 326 | 139 | 225 | 117 |
| 492 | -0.30 | 0.39 | -0.75 | 0.45 | 1 | Gm6999 | | 12 | 2 | 16 | 259 | 271 | 2118 | 296 | 319 | 266 | 106 | 1748 |
| 385 | -0.29 | 0.23 | -1.25 | 0.21 | 1 | Rnpc3 | RNA-binding region (RNP1, RRM) contai | 261 | 202 | 200 | 272 | 410 | 929 | 296 | 173 | 284 | 313 | 898 |
| 343 | -0.15 | 0.19 | -0.78 | 0.44 | 1 | Pex11g | peroxisomal biogenesis factor 11 gamma | 142 | 169 | 238 | 240 | 477 | 676 | 296 | 232 | 430 | 238 | 639 |
| 333 | -0.46 | 0.29 | -1.59 | 0.11 | 1 | Mex3d | mex3 RNA binding family member D | 462 | 632 | 337 | 400 | 357 | 250 | 296 | 171 | 270 | 265 | 220 |
| 325 | 0.16 | 0.46 | 0.35 | 0.72 | 1 | E2f2 | E2F transcription factor 2 | 487 | 162 | 284 | 263 | 394 | 276 | 296 | 656 | 344 | 167 | 244 |
| 289 | 0.20 | 0.36 | 0.54 | 0.59 | 1 | 2610306M01Rik | | 279 | 231 | 374 | 270 | 230 | 251 | 296 | 541 | 179 | 312 | 222 |
| 233 | -0.30 | 0.24 | -1.27 | 0.20 | 1 | Pheta1 | PH domain containing endocytic traffickin | 301 | 342 | 244 | 194 | 158 | 281 | 296 | 177 | 135 | 205 | 232 |
| 633 | 0.05 | 0.19 | 0.27 | 0.78 | 1 | Fam43a | family with sequence similarity 43, membe | 1732 | 1311 | 1541 | 199 | 319 | 369 | 295 | 298 | 315 | 294 | 293 |
| 478 | 0.15 | 0.24 | 0.61 | 0.54 | 1 | Ifngr2 | interferon gamma receptor 2 | 1133 | 686 | 827 | 347 | 330 | 246 | 295 | 429 | 254 | 466 | 246 |
| 475 | 0.06 | 0.24 | 0.24 | 0.81 | 1 | Vav2 | vav 2 oncogene | 885 | 552 | 551 | 385 | 420 | 480 | 295 | 348 | 397 | 363 | 554 |
| 384 | -0.06 | 0.19 | -0.35 | 0.73 | 1 | Mrpl47 | mitochondrial ribosomal protein L47 | 398 | 447 | 444 | 410 | 380 | 408 | 295 | 453 | 360 | 258 | 367 |
| 312 | 0.02 | 0.18 | 0.10 | 0.92 | 1 | Pick1 | protein interacting with C kinase 1 | 214 | 211 | 213 | 315 | 249 | 534 | 295 | 294 | 280 | 298 | 533 |
| 295 | 0.09 | 0.25 | 0.35 | 0.72 | 1 | Zfp932 | zinc finger protein 932 | 414 | 278 | 164 | 182 | 224 | 437 | 295 | 185 | 204 | 362 | 495 |
| 280 | -0.21 | 0.37 | -0.56 | 0.57 | 1 | Clip4 | CAP-GLY domain containing linker protei | 220 | 142 | 127 | 385 | 255 | 418 | 295 | 487 | 203 | 134 | 415 |
| 268 | 0.08 | 0.27 | 0.31 | 0.76 | 1 | Zfp931 | zinc finger protein 931 | 286 | 240 | 280 | 317 | 203 | 230 | 295 | 334 | 273 | 205 | 289 |
| 240 | -0.25 | 0.38 | -0.66 | 0.51 | 1 | Serinc2 | serine incorporator 2 | 321 | 267 | 370 | 243 | 158 | 154 | 295 | 124 | 213 | 415 | 75 |
| 238 | 0.27 | 0.48 | 0.55 | 0.58 | 1 | Phactr3 | phosphatase and actin regulator 3 | 10 | 37 | 87 | 210 | 246 | 372 | 295 | 383 | 230 | 286 | 459 |
| 326 | 0.24 | 0.33 | 0.73 | 0.46 | 1 | Gm46430 | predicted gene, 46430 | 300 | 211 | 503 | 270 | 180 | 432 | 294 | 355 | 284 | 242 | 518 |
| 324 | 0.37 | 0.27 | 1.37 | 0.17 | 1 | 2810006K23Ril | RIKEN cDNA 2810006K23 gene | 184 | 398 | 303 | 242 | 352 | 270 | 294 | 360 | 500 | 294 | 369 |
| 260 | -0.25 | 0.27 | -0.93 | 0.35 | 1 | Slc38a6 | solute carrier family 38, member 6 | 107 | 69 | 73 | 299 | 231 | 577 | 294 | 190 | 278 | 197 | 541 |
| 242 | -0.22 | 0.27 | -0.82 | 0.41 | 1 | Stambpl1 | STAM binding protein like 1 | 287 | 317 | 369 | 220 | 201 | 188 | 294 | 260 | 170 | 207 | 146 |
| 205 | 0.25 | 0.29 | 0.86 | 0.39 | 1 | Agbl4 | ATP/GTP binding protein-like 4 | 65 | 104 | 82 | 226 | 187 | 204 | 294 | 360 | 204 | 250 | 279 |
| 672 | -0.03 | 0.26 | -0.11 | 0.91 | 1 | Fyn | Fyn proto-oncogene | 1905 | 1405 | 1805 | 284 | 284 | 272 | 293 | 430 | 179 | 300 | 232 |
| 552 | -0.17 | 0.22 | -0.80 | 0.43 | 1 | Ccdc8 | coiled-coil domain containing 8 | 808 | 949 | 1244 | 458 | 537 | 324 | 293 | 468 | 460 | 179 | 349 |
| 477 | 0.06 | 0.23 | 0.27 | 0.79 | 1 | Arhgef10l | Rho guanine nucleotide exchange factor | 780 | 647 | 748 | 444 | 450 | 290 | 293 | 561 | 463 | 354 | 217 |
| 447 | 0.15 | 0.16 | 0.97 | 0.33 | 1 | 9930104L06Rik | RIKEN cDNA 9930104L06 gene | 341 | 530 | 525 | 389 | 370 | 623 | 293 | 455 | 412 | 319 | 664 |
| 301 | -0.09 | 0.24 | -0.37 | 0.71 | 1 | Esrp2 | epithelial splicing regulatory protein 2 | 170 | 225 | 328 | 337 | 338 | 363 | 293 | 305 | 345 | 328 | 273 |
| 362 | 0.00 | 0.19 | -0.01 | 0.99 | 1 | Zfand2b | zinc finger, AN1 type domain 2B | 360 | 452 | 382 | 303 | 355 | 442 | 292 | 343 | 366 | 250 | 438 |
| 305 | 0.03 | 0.27 | 0.13 | 0.90 | 1 | 2700062C07Ril | RIKEN cDNA 2700062C07 gene | 233 | 319 | 382 | 226 | 294 | 374 | 292 | 389 | 301 | 204 | 336 |
| 607 | -0.04 | 0.18 | -0.22 | 0.82 | 1 | 1810026B05Rik | | 491 | 485 | 439 | 227 | 272 | 1751 | 291 | 275 | 194 | 258 | 2000 |
| 575 | 0.00 | 0.24 | -0.02 | 0.99 | 1 | Fez2 | fasciculation and elongation protein zeta | 1281 | 1194 | 1345 | 349 | 389 | 215 | 291 | 527 | 262 | 255 | 222 |
| 444 | 0.15 | 0.21 | 0.71 | 0.48 | 1 | 1700025G04Ril | RIKEN cDNA 1700025G04 gene | 1245 | 941 | 907 | 256 | 134 | 161 | 291 | 231 | 183 | 366 | 163 |
| 401 | -0.43 | 0.35 | -1.22 | 0.22 | 1 | Frmd8 | FERM domain containing 8 | 1151 | 675 | 807 | 249 | 236 | 236 | 291 | 308 | 202 | 95 | 158 |
| 314 | -0.17 | 0.28 | -0.61 | 0.54 | 1 | 4921524J17Rik | RIKEN cDNA 4921524J17 gene | 354 | 518 | 425 | 272 | 283 | 289 | 291 | 168 | 269 | 352 | 231 |
| 280 | -0.29 | 0.22 | -1.29 | 0.20 | 1 | B9d2 | B9 protein domain 2 | 281 | 309 | 396 | 281 | 343 | 239 | 291 | 249 | 216 | 255 | 217 |
| 265 | -0.12 | 0.26 | -0.46 | 0.64 | 1 | Mgst3 | microsomal glutathione S-transferase 3 | 381 | 472 | 462 | 172 | 253 | 127 | 291 | 178 | 195 | 240 | 142 |
| 456 | 0.12 | 0.18 | 0.63 | 0.53 | 1 | C1qtnf6 | C1q and tumor necrosis factor related pro | 913 | 705 | 815 | 214 | 343 | 393 | 290 | 271 | 274 | 352 | 445 |
| 314 | 0.13 | 0.22 | 0.59 | 0.56 | 1 | Arhgef19 | Rho guanine nucleotide exchange factor | 402 | 301 | 253 | 242 | 238 | 408 | 290 | 329 | 176 | 358 | 462 |
| 299 | 0.11 | 0.20 | 0.53 | 0.59 | 1 | Mrgbp | MRG/MORF4L binding protein | 298 | 257 | 244 | 169 | 208 | 552 | 290 | 224 | 233 | 274 | 543 |
| 268 | 0.00 | 0.31 | -0.01 | 1.00 | 1 | Tceanc | transcription elongation factor A (SII) N-te | 147 | 324 | 274 | 193 | 269 | 338 | 290 | 200 | 279 | 236 | 399 |
| 425 | 0.11 | 0.20 | 0.56 | 0.58 | 1 | Gsap | gamma-secretase activating protein | 440 | 411 | 316 | 432 | 369 | 599 | 289 | 455 | 372 | 353 | 639 |
| 350 | 0.10 | 0.21 | 0.50 | 0.62 | 1 | Pibf1 | progesterone immunomodulatory binding | 290 | 379 | 361 | 266 | 284 | 514 | 289 | 371 | 316 | 224 | 563 |
| 324 | -0.09 | 0.20 | -0.44 | 0.66 | 1 | Sall4 | spalt like transcription factor 4 | 232 | 375 | 362 | 438 | 284 | 323 | 289 | 416 | 324 | 249 | 273 |
| 321 | -0.01 | 0.27 | -0.02 | 0.98 | 1 | Kcnj6 | potassium inwardly-rectifying channel, su | 271 | 481 | 455 | 290 | 256 | 293 | 289 | 418 | 368 | 193 | 212 |
| 266 | 0.07 | 0.29 | 0.23 | 0.82 | 1 | B130055M24Rik | | 145 | 138 | 118 | 340 | 184 | 433 | 289 | 202 | 303 | 395 | 382 |
| 165 | -0.49 | 0.27 | -1.79 | 0.07 | 1 | Gm5069 | | 43 | 26 | 52 | 218 | 250 | 230 | 289 | 143 | 230 | 166 | 173 |
| 128 | 0.16 | 0.39 | 0.42 | 0.68 | 1 | Kcnh5 | potassium voltage-gated channel, subfam | 11 | 15 | 16 | 128 | 120 | 107 | 289 | 114 | 141 | 339 | 134 |
| 436 | -0.43 | 0.37 | -1.18 | 0.24 | 1 | Birc3 | baculoviral IAP repeat-containing 3 | 1143 | 750 | 654 | 341 | 313 | 309 | 288 | 81 | 331 | 290 | 297 |
| 280 | 0.03 | 0.27 | 0.12 | 0.90 | 1 | Gm9776 | | 208 | 310 | 319 | 341 | 256 | 216 | 288 | 225 | 408 | 289 | 220 |
| 262 | 0.04 | 0.26 | 0.17 | 0.87 | 1 | Pigf | phosphatidylinositol glycan anchor biosyr | 310 | 288 | 271 | 253 | 240 | 213 | 288 | 300 | 287 | 179 | 258 |
| 156 | -0.23 | 0.75 | -0.31 | 0.76 | 1 | Padi2 | peptidyl arginine deiminase, type II | 175 | 402 | 272 | 27 | 19 | 44 | 288 | 15 | 12 | 432 | 26 |

| | | | | | | | | | | | | | | | | | | |
|---|---|---|---|---|---|---|---|---|---|---|---|---|---|---|---|---|---|---|
| 702 | -0.15 | 0.23 | -0.67 | 0.50 | 1 | A1cf | APOBEC1 complementation factor | 957 | 2487 | 1372 | 500 | 329 | 441 | 286 | 305 | 352 | 304 | 391 |
| 603 | -0.04 | 0.18 | -0.23 | 0.82 | 1 | Irf9 | interferon regulatory factor 9 | 1286 | 1050 | 1188 | 315 | 433 | 549 | 286 | 344 | 315 | 341 | 528 |
| 528 | 0.07 | 0.23 | 0.29 | 0.77 | 1 | Clasrp | CLK4-associating serine/arginine rich prc | 364 | 315 | 415 | 363 | 221 | 1544 | 286 | 382 | 237 | 365 | 1313 |
| 403 | 0.31 | 0.22 | 1.41 | 0.16 | 1 | D11Wsu47e | DNA segment, Chr 11, Wayne State Unive | 294 | 402 | 296 | 341 | 337 | 562 | 286 | 623 | 300 | 398 | 587 |
| 382 | -0.13 | 0.18 | -0.72 | 0.47 | 1 | E4f1 | E4F transcription factor 1 | 310 | 276 | 321 | 214 | 290 | 946 | 286 | 189 | 298 | 255 | 818 |
| 338 | -0.11 | 0.17 | -0.66 | 0.51 | 1 | Ciao2b | cytosolic iron-sulfur assembly component | 404 | 374 | 451 | 298 | 338 | 373 | 286 | 292 | 297 | 267 | 339 |
| 327 | 0.02 | 0.34 | 0.06 | 0.95 | 1 | Arhgap11a | Rho GTPase activating protein 11A | 609 | 323 | 277 | 310 | 346 | 235 | 286 | 423 | 373 | 216 | 199 |
| 314 | -0.21 | 0.37 | -0.57 | 0.57 | 1 | Pde4d | phosphodiesterase 4D, cAMP specific | 513 | 960 | 674 | 123 | 99 | 195 | 286 | 202 | 127 | 135 | 134 |
| 278 | -0.25 | 0.28 | -0.92 | 0.36 | 1 | Retsat | retinol saturase (all trans retinol 13,14 rec | 382 | 349 | 341 | 298 | 295 | 196 | 286 | 202 | 332 | 158 | 215 |
| 236 | -0.36 | 0.27 | -1.30 | 0.19 | 1 | Zfp976 | zinc finger protein 976 | 210 | 193 | 184 | 256 | 252 | 318 | 286 | 197 | 231 | 135 | 336 |
| 488 | -0.06 | 0.23 | -0.25 | 0.80 | 1 | Pdgfrl | platelet-derived growth factor receptor-like | 631 | 735 | 692 | 642 | 306 | 456 | 285 | 608 | 288 | 278 | 449 |
| 328 | 0.20 | 0.22 | 0.90 | 0.37 | 1 | Fam57a | family with sequence similarity 57, membe | 289 | 301 | 312 | 380 | 314 | 271 | 285 | 345 | 397 | 466 | 251 |
| 327 | 0.15 | 0.21 | 0.69 | 0.49 | 1 | Txnrd2 | thioredoxin reductase 2 | 389 | 352 | 348 | 300 | 252 | 340 | 285 | 240 | 268 | 395 | 429 |
| 297 | -0.07 | 0.22 | -0.34 | 0.73 | 1 | Coq4 | coenzyme Q4 | 272 | 235 | 333 | 348 | 311 | 298 | 285 | 260 | 319 | 344 | 267 |
| 286 | 0.11 | 0.26 | 0.42 | 0.67 | 1 | Nphp4 | nephronophthisis 4 (juvenile) homolog (h | 170 | 154 | 185 | 185 | 298 | 521 | 285 | 187 | 340 | 417 | 406 |
| 267 | -0.26 | 0.22 | -1.22 | 0.22 | 1 | Mid1 | midline 1 | 397 | 343 | 251 | 264 | 260 | 241 | 285 | 231 | 192 | 299 | 173 |
| 483 | -0.25 | 0.25 | -1.01 | 0.31 | 1 | Cemip2 | cell migration inducing hyaluronidase 2 | 819 | 741 | 562 | 588 | 372 | 480 | 284 | 539 | 287 | 243 | 394 |
| 457 | 0.07 | 0.28 | 0.26 | 0.80 | 1 | Smox | spermine oxidase | 793 | 656 | 901 | 442 | 368 | 188 | 284 | 290 | 333 | 587 | 187 |
| 432 | -0.11 | 0.27 | -0.41 | 0.68 | 1 | Ddx20 | DEAD (Asp-Glu-Ala-Asp) box polypeptide | 597 | 549 | 328 | 541 | 438 | 437 | 284 | 388 | 351 | 339 | 500 |
| 342 | -0.16 | 0.21 | -0.77 | 0.44 | 1 | Spats2 | spermatogenesis associated, serine-rich | 629 | 608 | 695 | 236 | 236 | 199 | 284 | 203 | 273 | 261 | 138 |
| 275 | 0.17 | 0.23 | 0.76 | 0.45 | 1 | Med29 | mediator complex subunit 29 | 229 | 250 | 298 | 268 | 249 | 249 | 284 | 382 | 273 | 282 | 257 |
| 265 | -0.06 | 0.25 | -0.24 | 0.81 | 1 | Txnl4b | thioredoxin-like 4B | 227 | 204 | 227 | 382 | 268 | 222 | 284 | 243 | 307 | 260 | 287 |
| 200 | -0.54 | 0.32 | -1.69 | 0.09 | 1 | Pcdhgb1 | protocadherin gamma subfamily B, 1 | 113 | 155 | 138 | 250 | 362 | 158 | 284 | 126 | 320 | 151 | 140 |
| 402 | 0.08 | 0.19 | 0.44 | 0.66 | 1 | Pde7a | phosphodiesterase 7A | 552 | 485 | 452 | 319 | 311 | 525 | 283 | 322 | 339 | 377 | 456 |
| 382 | -0.21 | 0.24 | -0.88 | 0.38 | 1 | Ypel4 | yippee like 4 | 196 | 300 | 220 | 242 | 359 | 975 | 283 | 209 | 318 | 236 | 861 |
| 353 | 0.25 | 0.27 | 0.92 | 0.36 | 1 | Gm43737 | | 325 | 526 | 505 | 383 | 191 | 307 | 283 | 339 | 313 | 395 | 312 |
| 341 | 0.08 | 0.24 | 0.32 | 0.75 | 1 | Msra | methionine sulfoxide reductase A | 312 | 319 | 362 | 510 | 301 | 248 | 283 | 495 | 399 | 252 | 269 |
| 265 | -0.43 | 0.30 | -1.41 | 0.16 | 1 | Adamts17 | a disintegrin-like and metallopeptidase (re | 156 | 245 | 306 | 305 | 343 | 307 | 283 | 378 | 203 | 129 | 262 |
| 255 | -0.20 | 0.28 | -0.72 | 0.47 | 1 | Fam174c | family with sequence similarity 174, meml | 285 | 198 | 163 | 258 | 293 | 320 | 283 | 273 | 257 | 185 | 288 |
| 219 | 0.19 | 0.32 | 0.59 | 0.55 | 1 | C030014I23Rik | | 48 | 136 | 96 | 242 | 265 | 195 | 283 | 364 | 287 | 300 | 199 |
| 198 | 0.04 | 0.28 | 0.13 | 0.90 | 1 | Sowaha | sosondowah ankyrin repeat domain famil | 80 | 92 | 144 | 198 | 255 | 174 | 283 | 221 | 280 | 308 | 144 |
| 554 | 0.03 | 0.29 | 0.10 | 0.92 | 1 | Palm | paralemmin | 1676 | 874 | 1173 | 475 | 234 | 179 | 282 | 477 | 241 | 294 | 182 |
| 475 | -0.23 | 0.25 | -0.92 | 0.36 | 1 | Piga | phosphatidylinositol glycan anchor biosyr | 132 | 127 | 201 | 495 | 312 | 1345 | 282 | 316 | 347 | 200 | 1463 |
| 474 | 0.28 | 0.22 | 1.29 | 0.20 | 1 | Nt5c3b | 5'-nucleotidase, cytosolic IIIB | 600 | 508 | 480 | 498 | 404 | 460 | 282 | 571 | 472 | 441 | 495 |
| 349 | 0.35 | 0.25 | 1.37 | 0.17 | 1 | E130308A19Ril | RIKEN cDNA E130308A19 gene | 333 | 378 | 288 | 368 | 275 | 302 | 282 | 373 | 268 | 608 | 359 |
| 342 | 0.04 | 0.26 | 0.14 | 0.89 | 1 | Xylt1 | xylosyltransferase 1 | 650 | 580 | 637 | 270 | 200 | 167 | 282 | 170 | 303 | 346 | 152 |
| 314 | -0.22 | 0.23 | -0.98 | 0.33 | 1 | Bicra | BRD4 interacting chromatin remodeling c | 467 | 476 | 325 | 236 | 266 | 402 | 282 | 200 | 253 | 246 | 306 |
| 303 | 0.65 | 0.66 | 0.98 | 0.33 | 1 | Gm24187 | | 76 | 141 | 451 | 261 | 149 | 160 | 282 | 152 | 146 | 1305 | 211 |
| 277 | 0.03 | 0.22 | 0.15 | 0.88 | 1 | Pym1 | PYM homolog 1, exon junction complex a | 304 | 264 | 259 | 260 | 279 | 273 | 282 | 320 | 248 | 237 | 320 |
| 234 | -0.41 | 0.28 | -1.50 | 0.13 | 1 | Gm13456 | | 383 | 367 | 418 | 189 | 192 | 140 | 282 | 96 | 175 | 201 | 131 |
| 467 | -0.05 | 0.20 | -0.23 | 0.81 | 1 | Rabep2 | rabaptin, RAB GTPase binding effector pr | 571 | 553 | 575 | 499 | 442 | 529 | 281 | 454 | 363 | 344 | 524 |
| 409 | -0.16 | 0.27 | -0.60 | 0.55 | 1 | Rbl1 | RB transcriptional corepressor like 1 | 571 | 489 | 419 | 508 | 283 | 525 | 281 | 239 | 317 | 386 | 475 |
| 407 | -0.10 | 0.22 | -0.43 | 0.67 | 1 | Gmip | Gem-interacting protein | 534 | 415 | 507 | 341 | 343 | 574 | 281 | 367 | 247 | 252 | 613 |
| 397 | 0.07 | 0.23 | 0.29 | 0.77 | 1 | Il10rb | interleukin 10 receptor, beta | 848 | 600 | 788 | 323 | 233 | 199 | 281 | 359 | 283 | 243 | 203 |
| 281 | 0.16 | 0.25 | 0.64 | 0.52 | 1 | Ube2v2 | ubiquitin-conjugating enzyme E2 variant 2 | 298 | 270 | 367 | 279 | 181 | 276 | 281 | 307 | 288 | 264 | 276 |
| 251 | -0.28 | 0.26 | -1.08 | 0.28 | 1 | Gpr39 | G protein-coupled receptor 39 | 231 | 218 | 273 | 344 | 253 | 231 | 281 | 262 | 304 | 170 | 192 |
| 249 | 0.50 | 0.28 | 1.75 | 0.08 | 1 | Mfap3l | microfibrillar-associated protein 3-like | 191 | 193 | 291 | 252 | 211 | 115 | 281 | 271 | 295 | 447 | 189 |
| 795 | -0.22 | 0.21 | -1.03 | 0.30 | 1 | Prkd3 | protein kinase D3 | 2515 | 1892 | 1877 | 391 | 402 | 253 | 280 | 268 | 281 | 278 | 304 |
| 682 | -0.07 | 0.23 | -0.31 | 0.75 | 1 | Tnfsf10 | tumor necrosis factor (ligand) superfamily | 2087 | 1519 | 2174 | 155 | 280 | 152 | 280 | 109 | 381 | 206 | 161 |
| 346 | -0.09 | 0.24 | -0.35 | 0.72 | 1 | Zkscan2 | zinc finger with KRAB and SCAN domains | 225 | 468 | 408 | 493 | 267 | 370 | 280 | 357 | 336 | 290 | 316 |
| 249 | -0.05 | 0.25 | -0.21 | 0.84 | 1 | Rabepk | Rab9 effector protein with kelch motifs | 315 | 255 | 365 | 225 | 192 | 219 | 280 | 185 | 200 | 223 | 282 |
| 249 | 0.12 | 0.24 | 0.49 | 0.63 | 1 | Diras1 | DIRAS family, GTP-binding RAS-like 1 | 214 | 261 | 279 | 309 | 219 | 136 | 280 | 378 | 286 | 236 | 141 |
| 241 | -0.10 | 0.31 | -0.32 | 0.75 | 1 | P2ry12 | purinergic receptor P2Y, G-protein couple | 220 | 183 | 147 | 331 | 290 | 186 | 280 | 349 | 317 | 156 | 196 |
| 513 | -0.10 | 0.19 | -0.50 | 0.62 | 1 | Irgm2 | immunity-related GTPase family M memb | 1037 | 990 | 1194 | 350 | 430 | 203 | 279 | 246 | 347 | 337 | 229 |
| 389 | 0.16 | 0.20 | 0.81 | 0.42 | 1 | Xrcc4 | X-ray repair complementing defective rep | 482 | 428 | 511 | 382 | 368 | 314 | 279 | 402 | 383 | 291 | 433 |
| 368 | -0.01 | 0.22 | -0.05 | 0.96 | 1 | Wdcp | WD repeat and coiled coil containing | 228 | 341 | 257 | 302 | 424 | 636 | 279 | 356 | 355 | 332 | 542 |
| 345 | 0.06 | 0.23 | 0.28 | 0.78 | 1 | Dgat1 | diacylglycerol O-acyltransferase 1 | 297 | 327 | 510 | 222 | 329 | 475 | 279 | 238 | 278 | 346 | 491 |
| 316 | -0.28 | 0.21 | -1.29 | 0.20 | 1 | Lpcat4 | lysophosphatidylcholine acyltransferase 4 | 572 | 493 | 513 | 294 | 285 | 185 | 279 | 253 | 235 | 182 | 184 |
| 291 | -0.16 | 0.29 | -0.53 | 0.59 | 1 | Sumf2 | sulfatase modifying factor 2 | 483 | 369 | 351 | 311 | 209 | 246 | 279 | 353 | 264 | 178 | 161 |
| 252 | -0.01 | 0.24 | -0.06 | 0.95 | 1 | Tmppe | transmembrane protein with metallophos | 263 | 294 | 168 | 286 | 233 | 228 | 279 | 218 | 227 | 332 | 243 |

| | | | | | | | | | | | | | | | | | | |
|---|---|---|---|---|---|---|---|---|---|---|---|---|---|---|---|---|---|---|
| 251 | -0.12 | 0.27 | -0.45 | 0.65 | 1 | Zdhhc12 | zinc finger, DHHC domain containing 12 | 238 | 178 | 201 | 204 | 339 | 281 | 279 | 182 | 402 | 201 | 250 |
| 238 | -0.40 | 0.30 | -1.33 | 0.18 | 1 | Fndc5 | fibronectin type III domain containing 5 | 269 | 351 | 147 | 316 | 311 | 152 | 279 | 195 | 191 | 267 | 138 |
| 365 | 0.41 | 0.31 | 1.33 | 0.18 | 1 | Fah | fumarylacetoacetate hydrolase | 391 | 334 | 454 | 320 | 194 | 435 | 278 | 549 | 279 | 444 | 333 |
| 362 | 0.15 | 0.25 | 0.58 | 0.56 | 1 | Cmtr1 | cap methyltransferase 1 | 467 | 286 | 367 | 238 | 365 | 494 | 278 | 367 | 287 | 419 | 411 |
| 349 | -0.06 | 0.27 | -0.23 | 0.82 | 1 | Vps37d | vacuolar protein sorting 37D | 366 | 600 | 656 | 387 | 243 | 234 | 278 | 290 | 228 | 330 | 232 |
| 309 | -0.01 | 0.20 | -0.03 | 0.98 | 1 | Zfp707 | zinc finger protein 707 | 309 | 317 | 320 | 277 | 217 | 473 | 278 | 307 | 250 | 263 | 392 |
| 292 | -0.17 | 0.25 | -0.69 | 0.49 | 1 | Sertad4 | SERTA domain containing 4 | 472 | 313 | 286 | 375 | 248 | 238 | 278 | 247 | 215 | 278 | 267 |
| 328 | 0.23 | 0.19 | 1.19 | 0.23 | 1 | Zbtb12 | zinc finger and BTB domain containing 1 | 372 | 422 | 309 | 336 | 270 | 263 | 277 | 433 | 249 | 353 | 323 |
| 276 | -0.36 | 0.22 | -1.64 | 0.10 | 1 | Fam161a | family with sequence similarity 161, meml | 227 | 270 | 259 | 327 | 307 | 355 | 277 | 271 | 192 | 188 | 364 |
| 266 | 0.12 | 0.23 | 0.50 | 0.61 | 1 | Actr6 | ARP6 actin-related protein 6 | 281 | 328 | 275 | 189 | 243 | 272 | 277 | 213 | 293 | 274 | 281 |
| 460 | 0.33 | 0.27 | 1.22 | 0.22 | 1 | Nde1 | nudE neurodevelopment protein 1 | 888 | 435 | 559 | 415 | 351 | 359 | 276 | 529 | 378 | 457 | 415 |
| 365 | 0.19 | 0.20 | 0.93 | 0.35 | 1 | Ap5z1 | adaptor-related protein complex 5, zeta 1 | 283 | 268 | 272 | 307 | 214 | 728 | 276 | 344 | 325 | 280 | 720 |
| 330 | 0.21 | 0.22 | 0.91 | 0.36 | 1 | Tbc1d7 | TBC1 domain family, member 7 | 260 | 277 | 320 | 357 | 338 | 302 | 276 | 539 | 398 | 267 | 301 |
| 318 | -0.24 | 0.27 | -0.89 | 0.37 | 1 | Il3ra | interleukin 3 receptor, alpha chain | 473 | 328 | 497 | 277 | 283 | 352 | 276 | 269 | 152 | 292 | 303 |
| 310 | -0.20 | 0.26 | -0.75 | 0.45 | 1 | Gm20008 | | 265 | 378 | 185 | 358 | 298 | 451 | 276 | 233 | 297 | 252 | 418 |
| 277 | 0.03 | 0.25 | 0.12 | 0.91 | 1 | A730017L22Rik | | 238 | 172 | 186 | 414 | 269 | 252 | 276 | 293 | 331 | 366 | 248 |
| 208 | -0.21 | 0.30 | -0.70 | 0.48 | 1 | Kyat3 | kynurenine aminotransferase 3 | 141 | 170 | 159 | 213 | 296 | 187 | 276 | 137 | 244 | 215 | 248 |
| 463 | 0.16 | 0.18 | 0.92 | 0.36 | 1 | Trnau1ap | tRNA selenocysteine 1 associated proteir | 613 | 538 | 685 | 325 | 392 | 559 | 275 | 402 | 375 | 366 | 565 |
| 409 | -0.02 | 0.22 | -0.08 | 0.94 | 1 | Klhl20 | kelch-like 20 | 600 | 528 | 498 | 306 | 332 | 544 | 275 | 349 | 381 | 233 | 454 |
| 372 | -0.07 | 0.39 | -0.19 | 0.85 | 1 | Cep44 | centrosomal protein 44 | 408 | 384 | 341 | 235 | 275 | 670 | 275 | 426 | 284 | 81 | 711 |
| 335 | 0.05 | 0.29 | 0.16 | 0.87 | 1 | Rpa2 | replication protein A2 | 717 | 324 | 378 | 295 | 239 | 305 | 275 | 365 | 228 | 249 | 311 |
| 301 | 0.15 | 0.25 | 0.59 | 0.56 | 1 | Gm37305 | | 263 | 415 | 316 | 322 | 251 | 253 | 275 | 269 | 291 | 401 | 258 |
| 277 | 0.11 | 0.22 | 0.52 | 0.60 | 1 | Dtwd1 | DTW domain containing 1 | 220 | 175 | 250 | 325 | 278 | 273 | 275 | 300 | 303 | 304 | 341 |
| 251 | -0.05 | 0.25 | -0.19 | 0.85 | 1 | Wdr49 | WD repeat domain 49 | 312 | 193 | 325 | 98 | 306 | 309 | 275 | 101 | 287 | 274 | 284 |
| 245 | -0.09 | 0.24 | -0.37 | 0.71 | 1 | Tex30 | testis expressed 30 | 146 | 194 | 118 | 339 | 290 | 240 | 275 | 363 | 265 | 194 | 267 |
| 530 | 0.01 | 0.16 | 0.09 | 0.93 | 1 | Sorbs1 | sorbin and SH3 domain containing 1 | 1105 | 999 | 1090 | 339 | 329 | 369 | 274 | 343 | 318 | 280 | 386 |
| 425 | -0.07 | 0.28 | -0.24 | 0.81 | 1 | Sspn | sarcospan | 1001 | 642 | 638 | 321 | 362 | 249 | 274 | 203 | 461 | 263 | 255 |
| 388 | 0.08 | 0.20 | 0.39 | 0.69 | 1 | Tet1 | tet methylcytosine dioxygenase 1 | 410 | 518 | 532 | 340 | 256 | 514 | 274 | 353 | 323 | 318 | 428 |
| 362 | 0.19 | 0.24 | 0.79 | 0.43 | 1 | Pard6a | par-6 family cell polarity regulator alpha | 302 | 261 | 370 | 421 | 387 | 341 | 274 | 535 | 343 | 350 | 393 |
| 333 | 0.41 | 0.24 | 1.71 | 0.09 | 1 | Frrs1 | ferric-chelate reductase 1 | 426 | 280 | 301 | 251 | 298 | 309 | 274 | 453 | 325 | 435 | 309 |
| 314 | 0.39 | 0.23 | 1.71 | 0.09 | 1 | Taf13 | TATA-box binding protein associated fact | 306 | 262 | 268 | 343 | 193 | 325 | 274 | 465 | 273 | 322 | 426 |
| 296 | 0.07 | 0.22 | 0.35 | 0.73 | 1 | Msantd3 | Myb/SANT-like DNA-binding domain cont | 363 | 480 | 218 | 326 | 250 | 207 | 274 | 397 | 214 | 295 | 231 |
| 275 | 0.22 | 0.22 | 1.02 | 0.31 | 1 | Pkp3 | plakophilin 3 | 269 | 228 | 268 | 294 | 279 | 179 | 274 | 272 | 370 | 404 | 182 |
| 272 | -0.27 | 0.28 | -0.95 | 0.34 | 1 | Spag1 | sperm associated antigen 1 | 173 | 160 | 132 | 327 | 318 | 452 | 274 | 166 | 295 | 348 | 344 |
| 270 | 0.19 | 0.24 | 0.79 | 0.43 | 1 | Zswim7 | zinc finger SWIM-type containing 7 | 157 | 108 | 160 | 277 | 306 | 331 | 274 | 381 | 314 | 264 | 401 |
| 221 | -0.03 | 0.30 | -0.09 | 0.93 | 1 | Nqo1 | NAD(P)H dehydrogenase, quinone 1 | 154 | 83 | 129 | 354 | 256 | 145 | 274 | 212 | 276 | 398 | 153 |
| 170 | 0.16 | 0.63 | 0.25 | 0.80 | 1 | Sgcz | sarcoglycan zeta | 0 | 4 | 19 | 228 | 187 | 185 | 274 | 285 | 163 | 340 | 180 |
| 349 | 0.36 | 0.29 | 1.26 | 0.21 | 1 | Zfp30 | zinc finger protein 30 | 282 | 202 | 308 | 382 | 366 | 298 | 273 | 656 | 415 | 286 | 367 |
| 328 | 0.27 | 0.23 | 1.17 | 0.24 | 1 | Rtl8a | retrotransposon Gag like 8A | 384 | 355 | 387 | 243 | 372 | 242 | 273 | 386 | 288 | 366 | 311 |
| 285 | 0.16 | 0.19 | 0.82 | 0.41 | 1 | Wdr53 | WD repeat domain 53 | 274 | 324 | 211 | 329 | 216 | 277 | 273 | 386 | 249 | 259 | 333 |
| 241 | 0.11 | 0.27 | 0.41 | 0.68 | 1 | A530017D24Rik | | 120 | 207 | 182 | 250 | 219 | 276 | 273 | 323 | 313 | 188 | 305 |
| 229 | -0.41 | 0.30 | -1.37 | 0.17 | 1 | Tusc1 | tumor suppressor candidate 1 | 220 | 170 | 147 | 351 | 327 | 188 | 273 | 238 | 275 | 155 | 180 |
| 166 | -0.31 | 0.29 | -1.06 | 0.29 | 1 | Zfp791 | zinc finger protein 791 | 149 | 86 | 112 | 161 | 225 | 166 | 273 | 189 | 163 | 165 | 134 |
| 612 | 0.01 | 0.24 | 0.03 | 0.98 | 1 | Slc25a24 | solute carrier family 25 (mitochondrial car | 1686 | 1287 | 1516 | 333 | 303 | 202 | 272 | 444 | 284 | 197 | 209 |
| 393 | -0.37 | 0.24 | -1.53 | 0.13 | 1 | Adcy3 | adenylate cyclase 3 | 987 | 950 | 891 | 230 | 196 | 132 | 272 | 123 | 161 | 290 | 93 |
| 382 | 0.00 | 0.20 | -0.01 | 0.99 | 1 | Plgrkt | plasminogen receptor, C-terminal lysine t | 384 | 307 | 375 | 267 | 209 | 840 | 272 | 277 | 253 | 231 | 791 |
| 380 | 0.28 | 0.23 | 1.22 | 0.22 | 1 | Atp5o | ATP synthase, H+ transporting, mitochonc | 389 | 366 | 562 | 400 | 305 | 316 | 272 | 474 | 376 | 366 | 353 |
| 368 | -0.09 | 0.20 | -0.46 | 0.64 | 1 | Ccdc71l | coiled-coil domain containing 71 like | 794 | 651 | 580 | 340 | 228 | 204 | 272 | 272 | 248 | 276 | 182 |
| 350 | -0.06 | 0.24 | -0.25 | 0.81 | 1 | Spata2 | spermatogenesis associated 2 | 583 | 404 | 526 | 278 | 285 | 337 | 272 | 188 | 255 | 428 | 299 |
| 329 | 0.13 | 0.27 | 0.48 | 0.63 | 1 | Armc6 | armadillo repeat containing 6 | 423 | 393 | 459 | 252 | 239 | 341 | 272 | 206 | 224 | 468 | 346 |
| 296 | -0.22 | 0.34 | -0.64 | 0.52 | 1 | Zbtb8a | zinc finger and BTB domain containing 8a | 370 | 293 | 189 | 269 | 375 | 344 | 272 | 133 | 282 | 388 | 339 |
| 295 | 0.11 | 0.22 | 0.50 | 0.62 | 1 | Noc4l | NOC4 like | 399 | 367 | 289 | 271 | 246 | 264 | 272 | 220 | 308 | 301 | 311 |
| 253 | -0.25 | 0.29 | -0.86 | 0.39 | 1 | 1700096K18Rik | | 252 | 166 | 294 | 308 | 344 | 188 | 272 | 161 | 253 | 365 | 184 |
| 221 | -0.12 | 0.27 | -0.46 | 0.64 | 1 | Camkk1 | calcium/calmodulin-dependent protein kir | 152 | 128 | 204 | 236 | 207 | 298 | 272 | 236 | 206 | 191 | 304 |
| 556 | -0.06 | 0.21 | -0.28 | 0.78 | 1 | Zfp217 | zinc finger protein 217 | 1170 | 992 | 1238 | 349 | 372 | 388 | 271 | 417 | 368 | 240 | 306 |
| 527 | -0.26 | 0.19 | -1.38 | 0.17 | 1 | Ppox | protoporphyrinogen oxidase | 341 | 376 | 314 | 447 | 363 | 1466 | 271 | 304 | 250 | 309 | 1354 |
| 294 | -0.01 | 0.31 | -0.04 | 0.97 | 1 | Gstt2 | glutathione S-transferase, theta 2 | 490 | 407 | 386 | 224 | 176 | 293 | 271 | 401 | 152 | 167 | 268 |
| 293 | -0.48 | 0.35 | -1.37 | 0.17 | 1 | Qpct | glutaminyl-peptide cyclotransferase (gluta | 730 | 567 | 776 | 109 | 141 | 134 | 271 | 37 | 126 | 175 | 153 |
| 284 | 0.07 | 0.26 | 0.27 | 0.79 | 1 | Arsg | arylsulfatase G | 286 | 399 | 262 | 221 | 302 | 257 | 271 | 322 | 278 | 319 | 205 |
| 261 | 0.05 | 0.29 | 0.17 | 0.87 | 1 | Tfcp2l1 | transcription factor CP2-like 1 | 166 | 403 | 319 | 262 | 259 | 174 | 271 | 284 | 311 | 270 | 155 |

| | | | | | | | | | | | | | | | | | | |
|---|---|---|---|---|---|---|---|---|---|---|---|---|---|---|---|---|---|---|
| 233 | -0.20 | 0.27 | -0.72 | 0.47 | 1 | Gm11175 | | 151 | 192 | 143 | 236 | 291 | 305 | 271 | 180 | 203 | 313 | 280 |
| 231 | 0.15 | 0.31 | 0.50 | 0.62 | 1 | Rcor2 | REST corepressor 2 | 129 | 228 | 181 | 233 | 194 | 232 | 271 | 367 | 182 | 332 | 192 |
| 12838 | -1.21 | 1.24 | -0.97 | 0.33 | 1 | Try4 | trypsin 4 | 33629 | 48831 | 55005 | 2114 | 198 | 7 | 270 | 187 | 104 | 75 | 801 |
| 322 | -0.33 | 0.31 | -1.09 | 0.28 | 1 | 4930503L19Ril | RIKEN cDNA 4930503L19 gene | 503 | 501 | 435 | 140 | 161 | 548 | 270 | 161 | 142 | 95 | 585 |
| 317 | -0.20 | 0.25 | -0.78 | 0.44 | 1 | Tert | telomerase reverse transcriptase | 215 | 319 | 238 | 294 | 375 | 509 | 270 | 197 | 320 | 289 | 461 |
| 283 | -0.41 | 0.30 | -1.37 | 0.17 | 1 | Zfp979 | zinc finger protein 979 | 259 | 509 | 255 | 313 | 354 | 260 | 270 | 190 | 296 | 198 | 212 |
| 259 | -0.16 | 0.41 | -0.40 | 0.69 | 1 | AU041133 | expressed sequence AU041133 | 151 | 237 | 308 | 292 | 192 | 352 | 270 | 88 | 267 | 309 | 386 |
| 245 | 0.07 | 0.23 | 0.30 | 0.76 | 1 | Particl | | 236 | 255 | 218 | 250 | 236 | 199 | 270 | 300 | 225 | 340 | 166 |
| 391 | 0.21 | 0.25 | 0.84 | 0.40 | 1 | Zfp280c | zinc finger protein 280C | 305 | 452 | 273 | 297 | 398 | 565 | 269 | 339 | 377 | 417 | 608 |
| 371 | 0.16 | 0.24 | 0.68 | 0.49 | 1 | Gar1 | GAR1 ribonucleoprotein | 615 | 483 | 631 | 233 | 298 | 306 | 269 | 343 | 319 | 198 | 390 |
| 360 | 0.49 | 0.32 | 1.55 | 0.12 | 1 | Ccdc28b | coiled coil domain containing 28B | 465 | 427 | 370 | 167 | 205 | 492 | 269 | 562 | 229 | 277 | 491 |
| 327 | -0.08 | 0.23 | -0.35 | 0.72 | 1 | Fut4 | fucosyltransferase 4 | 649 | 465 | 632 | 251 | 253 | 148 | 269 | 246 | 274 | 321 | 90 |
| 326 | 0.24 | 0.27 | 0.88 | 0.38 | 1 | Mitd1 | MIT, microtubule interacting and transport | 269 | 341 | 457 | 282 | 204 | 404 | 269 | 400 | 240 | 294 | 428 |
| 320 | 0.17 | 0.22 | 0.77 | 0.44 | 1 | Trit1 | tRNA isopentenyltransferase 1 | 337 | 280 | 248 | 168 | 278 | 535 | 269 | 222 | 252 | 325 | 605 |
| 307 | -0.02 | 0.22 | -0.10 | 0.92 | 1 | Zfp764 | zinc finger protein 764 | 299 | 245 | 296 | 395 | 359 | 254 | 269 | 312 | 351 | 367 | 234 |
| 306 | 0.14 | 0.30 | 0.49 | 0.63 | 1 | Malsu1 | mitochondrial assembly of ribosomal larg | 326 | 285 | 359 | 213 | 333 | 320 | 269 | 455 | 246 | 254 | 304 |
| 297 | -0.24 | 0.22 | -1.09 | 0.28 | 1 | Zfp182 | zinc finger protein 182 | 255 | 244 | 317 | 218 | 269 | 553 | 269 | 233 | 203 | 179 | 525 |
| 261 | -0.03 | 0.28 | -0.09 | 0.93 | 1 | Rassf6 | Ras association (RalGDS/AF-6) domain fa | 227 | 229 | 271 | 216 | 262 | 303 | 269 | 130 | 304 | 247 | 412 |
| 234 | -0.34 | 0.27 | -1.24 | 0.21 | 1 | Fxn | frataxin | 184 | 187 | 181 | 278 | 304 | 269 | 269 | 138 | 235 | 264 | 268 |
| 156 | -0.12 | 0.37 | -0.33 | 0.74 | 1 | Gm6969 | | 341 | 37 | 32 | 59 | 39 | 282 | 269 | 24 | 54 | 282 | 300 |
| 478 | 0.22 | 0.21 | 1.04 | 0.30 | 1 | Commd4 | COMM domain containing 4 | 654 | 489 | 526 | 358 | 328 | 728 | 268 | 470 | 351 | 343 | 746 |
| 423 | -0.17 | 0.20 | -0.86 | 0.39 | 1 | Tle2 | transducin-like enhancer of split 2 | 190 | 191 | 300 | 291 | 375 | 1246 | 268 | 261 | 296 | 317 | 920 |
| 342 | -0.13 | 0.31 | -0.43 | 0.66 | 1 | Zcchc4 | zinc finger, CCHC domain containing 4 | 266 | 313 | 292 | 379 | 303 | 507 | 268 | 484 | 313 | 116 | 516 |
| 318 | -0.33 | 0.22 | -1.53 | 0.13 | 1 | Dleu2 | | 371 | 409 | 465 | 277 | 262 | 439 | 268 | 272 | 179 | 178 | 378 |
| 294 | 0.06 | 0.20 | 0.30 | 0.77 | 1 | Emc9 | ER membrane protein complex subunit 9 | 251 | 334 | 196 | 346 | 256 | 332 | 268 | 333 | 289 | 276 | 347 |
| 245 | -0.18 | 0.31 | -0.59 | 0.55 | 1 | Kcnq2 | potassium voltage-gated channel, subfam | 632 | 290 | 492 | 152 | 141 | 127 | 268 | 114 | 125 | 157 | 199 |
| 239 | -0.16 | 0.25 | -0.63 | 0.53 | 1 | Pfkfb4 | 6-phosphofructo-2-kinase/fructose-2,6-bip | 298 | 283 | 233 | 172 | 259 | 261 | 268 | 218 | 192 | 249 | 200 |
| 182 | 0.17 | 0.30 | 0.55 | 0.58 | 1 | Cbr3 | carbonyl reductase 3 | 261 | 101 | 203 | 151 | 192 | 85 | 268 | 209 | 170 | 256 | 111 |
| 409 | 0.28 | 0.34 | 0.82 | 0.41 | 1 | Ncapg2 | non-SMC condensin II complex, subunit G | 752 | 331 | 262 | 428 | 425 | 295 | 267 | 422 | 499 | 438 | 378 |
| 408 | 0.03 | 0.21 | 0.15 | 0.88 | 1 | Abcc10 | ATP-binding cassette, sub-family C (CFTF | 276 | 147 | 191 | 174 | 308 | 1251 | 267 | 229 | 277 | 296 | 1068 |
| 395 | 0.07 | 0.24 | 0.29 | 0.77 | 1 | Polh | polymerase (DNA directed), eta (RAD 30 | 590 | 452 | 405 | 431 | 352 | 364 | 267 | 403 | 394 | 356 | 328 |
| 345 | 0.19 | 0.20 | 0.94 | 0.34 | 1 | Borcs8 | BLOC-1 related complex subunit 8 | 416 | 352 | 435 | 258 | 315 | 371 | 267 | 339 | 280 | 374 | 384 |
| 314 | -0.05 | 0.18 | -0.30 | 0.77 | 1 | Lrrc56 | leucine rich repeat containing 56 | 252 | 222 | 262 | 302 | 331 | 487 | 267 | 276 | 338 | 276 | 438 |
| 284 | -0.04 | 0.24 | -0.16 | 0.88 | 1 | Cd320 | CD320 antigen | 343 | 302 | 332 | 312 | 284 | 220 | 267 | 352 | 269 | 201 | 238 |
| 255 | -0.01 | 0.24 | -0.05 | 0.96 | 1 | Rita1 | RBPJ interacting and tubulin associated 1 | 234 | 204 | 275 | 308 | 244 | 228 | 267 | 236 | 255 | 260 | 288 |
| 217 | -0.19 | 0.36 | -0.53 | 0.60 | 1 | Cblc | Casitas B-lineage lymphoma c | 285 | 388 | 238 | 200 | 105 | 213 | 267 | 94 | 125 | 291 | 181 |
| 216 | -0.05 | 0.27 | -0.18 | 0.85 | 1 | Tmem138 | transmembrane protein 138 | 217 | 131 | 190 | 234 | 203 | 230 | 267 | 221 | 269 | 196 | 215 |
| 661 | -0.34 | 0.20 | -1.69 | 0.09 | 1 | Cdc42ep4 | CDC42 effector protein (Rho GTPase bin | 1722 | 1638 | 1850 | 361 | 335 | 190 | 266 | 346 | 207 | 192 | 166 |
| 356 | -0.05 | 0.23 | -0.20 | 0.84 | 1 | Dcaf4 | DDB1 and CUL4 associated factor 4 | 381 | 524 | 359 | 436 | 263 | 389 | 266 | 374 | 380 | 202 | 347 |
| 352 | 0.24 | 0.23 | 1.02 | 0.31 | 1 | Alkbh3 | alkB homolog 3, alpha-ketoglutarate-dep | 385 | 300 | 354 | 276 | 307 | 460 | 266 | 454 | 308 | 319 | 439 |
| 273 | 0.15 | 0.42 | 0.36 | 0.72 | 1 | Ikzf3 | IKAROS family zinc finger 3 | 501 | 262 | 75 | 263 | 292 | 183 | 266 | 366 | 261 | 344 | 191 |
| 531 | -0.20 | 0.19 | -1.04 | 0.30 | 1 | Sharpin | SHANK-associated RH domain interactin | 634 | 637 | 600 | 403 | 473 | 972 | 265 | 350 | 359 | 255 | 895 |
| 454 | 0.07 | 0.19 | 0.35 | 0.72 | 1 | Gbp7 | guanylate binding protein 7 | 1271 | 888 | 822 | 184 | 173 | 366 | 265 | 165 | 234 | 255 | 372 |
| 417 | -0.11 | 0.19 | -0.58 | 0.56 | 1 | Ap4e1 | adaptor-related protein complex AP-4, ep | 468 | 413 | 375 | 278 | 352 | 810 | 265 | 249 | 274 | 274 | 831 |
| 371 | -0.26 | 0.23 | -1.11 | 0.27 | 1 | Polk | polymerase (DNA directed), kappa | 509 | 407 | 549 | 365 | 361 | 426 | 265 | 331 | 281 | 192 | 396 |
| 285 | -0.12 | 0.33 | -0.36 | 0.72 | 1 | Stc2 | stanniocalcin 2 | 859 | 348 | 585 | 172 | 211 | 62 | 265 | 236 | 199 | 140 | 55 |
| 267 | -0.01 | 0.26 | -0.03 | 0.97 | 1 | Ptrhd1 | peptidyl-tRNA hydrolase domain containi | 192 | 179 | 319 | 281 | 412 | 188 | 265 | 247 | 284 | 332 | 237 |
| 220 | -0.44 | 0.32 | -1.35 | 0.18 | 1 | Scrn2 | secernin 2 | 364 | 265 | 310 | 185 | 195 | 190 | 265 | 65 | 201 | 202 | 179 |
| 446 | -0.06 | 0.21 | -0.26 | 0.79 | 1 | Cdk5rap1 | CDK5 regulatory subunit associated prote | 383 | 427 | 839 | 402 | 618 | 405 | 264 | 373 | 481 | 372 | 340 |
| 365 | -0.08 | 0.23 | -0.37 | 0.71 | 1 | Sufu | SUFU negative regulator of hedgehog sig | 700 | 552 | 456 | 277 | 262 | 375 | 264 | 353 | 191 | 265 | 314 |
| 322 | -0.05 | 0.27 | -0.19 | 0.85 | 1 | Ap5b1 | adaptor-related protein complex 5, beta 1 | 302 | 340 | 297 | 439 | 354 | 253 | 264 | 308 | 316 | 468 | 198 |
| 320 | -0.23 | 0.26 | -0.90 | 0.37 | 1 | Inip | INTS3 and NABP interacting protein | 460 | 346 | 308 | 383 | 357 | 298 | 264 | 254 | 339 | 234 | 279 |
| 318 | -0.18 | 0.31 | -0.60 | 0.55 | 1 | Acer3 | alkaline ceramidase 3 | 688 | 373 | 434 | 275 | 262 | 259 | 264 | 212 | 326 | 153 | 250 |
| 314 | -0.01 | 0.29 | -0.04 | 0.97 | 1 | Btg3 | BTG anti-proliferation factor 3 | 276 | 249 | 372 | 275 | 335 | 406 | 264 | 452 | 231 | 219 | 380 |
| 288 | -0.39 | 0.36 | -1.06 | 0.29 | 1 | Helb | helicase (DNA) B | 327 | 240 | 204 | 336 | 307 | 408 | 264 | 115 | 370 | 193 | 400 |
| 263 | -0.09 | 0.26 | -0.36 | 0.72 | 1 | Tfb1m | transcription factor B1, mitochondrial | 222 | 178 | 163 | 242 | 372 | 325 | 264 | 276 | 283 | 218 | 353 |
| 246 | -0.04 | 0.27 | -0.15 | 0.88 | 1 | Ddx59 | DEAD (Asp-Glu-Ala-Asp) box polypeptide | 320 | 276 | 303 | 249 | 210 | 192 | 264 | 264 | 222 | 160 | 244 |
| 230 | 0.22 | 0.36 | 0.62 | 0.53 | 1 | Gm47204 | | 133 | 124 | 289 | 182 | 299 | 190 | 264 | 330 | 195 | 258 | 267 |
| 210 | 0.02 | 0.34 | 0.06 | 0.95 | 1 | Pradc1 | protease-associated domain containing 1 | 170 | 93 | 164 | 248 | 215 | 188 | 264 | 373 | 221 | 240 | 132 |
| 663 | -0.33 | 0.20 | -1.68 | 0.09 | 1 | Sipa1l2 | signal-induced proliferation-associated 1 | 1866 | 1676 | 1948 | 134 | 301 | 314 | 263 | 143 | 201 | 204 | 239 |

| | | | | | | | | | | | | | | | | | |
|---|---|---|---|---|---|---|---|---|---|---|---|---|---|---|---|---|---|
| 344 | -0.02 | 0.27 | -0.06 | 0.95 | 1 | Rasgef1b | RasGEF domain family, member 1B | 1005 | 554 | 399 | 248 | 175 | 239 | 263 | 226 | 206 | 204 | 262 |
| 332 | 0.34 | 0.21 | 1.68 | 0.09 | 1 | Mrto4 | mRNA turnover 4, ribosome maturation fa | 383 | 307 | 421 | 283 | 265 | 307 | 263 | 345 | 324 | 408 | 345 |
| 301 | 0.03 | 0.22 | 0.16 | 0.88 | 1 | Sco1 | SCO1 cytochrome c oxidase assembly pr | 300 | 282 | 334 | 327 | 314 | 276 | 263 | 397 | 287 | 252 | 278 |
| 301 | 0.22 | 0.25 | 0.86 | 0.39 | 1 | Pced1b | PC-esterase domain containing 1B | 223 | 262 | 282 | 333 | 220 | 351 | 263 | 506 | 278 | 280 | 310 |
| 282 | 0.18 | 0.30 | 0.60 | 0.55 | 1 | Cdpf1 | cysteine rich, DPF motif domain containin | 311 | 181 | 293 | 225 | 295 | 298 | 263 | 404 | 361 | 241 | 226 |
| 259 | -0.36 | 0.23 | -1.54 | 0.12 | 1 | B4galnt4 | beta-1,4-N-acetyl-galactosaminyl transfer | 390 | 430 | 328 | 231 | 238 | 224 | 263 | 171 | 189 | 164 | 223 |
| 711 | 0.04 | 0.31 | 0.12 | 0.90 | 1 | Mcm6 | minichromosome maintenance complex c | 1970 | 1402 | 899 | 739 | 355 | 427 | 262 | 605 | 441 | 300 | 424 |
| 488 | -0.03 | 0.28 | -0.11 | 0.91 | 1 | Akap17b | A kinase (PRKA) anchor protein 17B | 987 | 1338 | 911 | 242 | 249 | 309 | 262 | 209 | 236 | 421 | 202 |
| 389 | 0.08 | 0.20 | 0.38 | 0.70 | 1 | Trip10 | thyroid hormone receptor interactor 10 | 1060 | 701 | 877 | 169 | 188 | 182 | 262 | 167 | 257 | 215 | 200 |
| 375 | 0.41 | 0.22 | 1.89 | 0.06 | 1 | Akr1b10 | aldo-keto reductase family 1, member B1( | 389 | 318 | 397 | 304 | 304 | 414 | 262 | 411 | 289 | 430 | 605 |
| 346 | 0.30 | 0.28 | 1.05 | 0.29 | 1 | Actr5 | ARP5 actin-related protein 5 | 335 | 442 | 361 | 207 | 266 | 466 | 262 | 485 | 216 | 337 | 432 |
| 318 | 0.36 | 0.22 | 1.67 | 0.10 | 1 | Coa4 | cytochrome c oxidase assembly factor 4 | 225 | 255 | 277 | 292 | 351 | 294 | 262 | 400 | 386 | 457 | 301 |
| 290 | 0.21 | 0.28 | 0.76 | 0.45 | 1 | Apool | apolipoprotein O-like | 329 | 209 | 300 | 277 | 301 | 243 | 262 | 400 | 222 | 339 | 303 |
| 289 | -0.14 | 0.21 | -0.67 | 0.50 | 1 | Chchd5 | coiled-coil-helix-coiled-coil-helix domain | 298 | 316 | 365 | 341 | 391 | 178 | 262 | 291 | 278 | 267 | 198 |
| 287 | 0.15 | 0.21 | 0.71 | 0.48 | 1 | Trmt61a | tRNA methyltransferase 61A | 175 | 265 | 194 | 305 | 309 | 318 | 262 | 316 | 334 | 269 | 413 |
| 221 | 0.06 | 0.27 | 0.23 | 0.81 | 1 | Gm15417 | | 118 | 130 | 190 | 221 | 223 | 271 | 262 | 292 | 231 | 246 | 251 |
| 388 | 0.00 | 0.22 | 0.01 | 0.99 | 1 | Mrpl35 | mitochondrial ribosomal protein L35 | 341 | 625 | 502 | 347 | 469 | 330 | 261 | 334 | 366 | 287 | 405 |
| 367 | 0.14 | 0.22 | 0.63 | 0.53 | 1 | Prmt6 | protein arginine N-methyltransferase 6 | 366 | 451 | 503 | 311 | 290 | 428 | 261 | 439 | 250 | 298 | 438 |
| 300 | 0.35 | 0.26 | 1.34 | 0.18 | 1 | 6330403K07Ril | RIKEN cDNA 6330403K07 gene | 81 | 94 | 70 | 318 | 447 | 314 | 261 | 439 | 530 | 267 | 485 |
| 256 | 0.33 | 0.30 | 1.11 | 0.27 | 1 | Cyp2u1 | cytochrome P450, family 2, subfamily u, p | 95 | 102 | 48 | 232 | 285 | 363 | 261 | 386 | 370 | 291 | 379 |
| 232 | 0.29 | 0.28 | 1.03 | 0.31 | 1 | Ydjc | YdjC homolog (bacterial) | 154 | 125 | 149 | 193 | 237 | 260 | 261 | 359 | 199 | 362 | 256 |
| 165 | -0.32 | 0.26 | -1.23 | 0.22 | 1 | Setmar | SET domain without mariner transposase | 163 | 135 | 123 | 198 | 175 | 147 | 261 | 154 | 180 | 160 | 121 |
| 397 | -0.01 | 0.30 | -0.04 | 0.97 | 1 | Nedd1 | neural precursor cell expressed, develop | 575 | 659 | 598 | 279 | 370 | 329 | 260 | 444 | 334 | 134 | 383 |
| 353 | -0.06 | 0.26 | -0.21 | 0.83 | 1 | Plag1 | pleiomorphic adenoma gene 1 | 268 | 513 | 248 | 249 | 271 | 686 | 260 | 243 | 274 | 247 | 626 |
| 352 | 0.04 | 0.31 | 0.13 | 0.89 | 1 | Gm2415 | | 614 | 462 | 584 | 237 | 303 | 280 | 260 | 415 | 272 | 156 | 291 |
| 275 | 0.14 | 0.33 | 0.43 | 0.67 | 1 | Pigbos1 | Pigb opposite strand 1 | 231 | 148 | 351 | 215 | 339 | 279 | 260 | 395 | 331 | 169 | 311 |
| 236 | -0.03 | 0.27 | -0.09 | 0.93 | 1 | Commd1 | COMM domain containing 1 | 200 | 208 | 236 | 331 | 198 | 192 | 260 | 352 | 189 | 206 | 219 |
| 222 | 0.10 | 0.46 | 0.22 | 0.82 | 1 | Ndufaf6 | NADH:ubiquinone oxidoreductase compl | 155 | 77 | 192 | 151 | 292 | 274 | 260 | 401 | 228 | 107 | 309 |
| 215 | 0.17 | 0.30 | 0.56 | 0.57 | 1 | Acot6 | acyl-CoA thioesterase 6 | 165 | 121 | 192 | 201 | 237 | 169 | 260 | 202 | 191 | 450 | 180 |
| 169 | -0.06 | 0.46 | -0.12 | 0.90 | 1 | Gm33447 | | 5 | 4 | 22 | 288 | 175 | 202 | 260 | 286 | 203 | 202 | 209 |
| 396 | 0.17 | 0.25 | 0.70 | 0.49 | 1 | Agfg2 | ArfGAP with FG repeats 2 | 649 | 413 | 485 | 354 | 349 | 357 | 259 | 444 | 374 | 300 | 368 |
| 351 | -0.08 | 0.23 | -0.33 | 0.74 | 1 | Epdr1 | ependymin related protein 1 (zebrafish) | 587 | 542 | 498 | 414 | 242 | 232 | 259 | 392 | 223 | 215 | 255 |
| 335 | 0.35 | 0.27 | 1.28 | 0.20 | 1 | Nxt1 | NTF2-related export protein 1 | 406 | 295 | 428 | 347 | 263 | 249 | 259 | 402 | 248 | 473 | 320 |
| 332 | -0.39 | 0.21 | -1.85 | 0.06 | 1 | Pik3ip1 | phosphoinositide-3-kinase interacting pro | 782 | 442 | 829 | 168 | 208 | 276 | 259 | 140 | 175 | 185 | 188 |
| 317 | -0.01 | 0.24 | -0.03 | 0.97 | 1 | Rab40b | Rab40B, member RAS oncogene family | 398 | 315 | 323 | 355 | 382 | 221 | 259 | 421 | 295 | 326 | 190 |
| 281 | 0.31 | 0.21 | 1.48 | 0.14 | 1 | Nubpl | nucleotide binding protein-like | 259 | 321 | 253 | 277 | 274 | 200 | 259 | 323 | 290 | 380 | 258 |
| 276 | 0.02 | 0.22 | 0.09 | 0.93 | 1 | Leng9 | leukocyte receptor cluster (LRC) member | 327 | 239 | 272 | 261 | 267 | 308 | 259 | 294 | 267 | 233 | 314 |
| 270 | 0.15 | 0.28 | 0.52 | 0.60 | 1 | Nudt22 | nudix (nucleoside diphosphate linked mo | 193 | 250 | 297 | 205 | 305 | 282 | 259 | 381 | 278 | 295 | 218 |
| 238 | -0.20 | 0.21 | -0.93 | 0.35 | 1 | Rmnd1 | required for meiotic nuclear division 1 hor | 142 | 145 | 142 | 263 | 218 | 423 | 259 | 233 | 172 | 227 | 398 |
| 213 | 0.11 | 0.38 | 0.29 | 0.77 | 1 | Zfp937 | zinc finger protein 937 | 135 | 312 | 153 | 144 | 207 | 226 | 259 | 196 | 211 | 300 | 200 |
| 201 | -0.31 | 0.42 | -0.72 | 0.47 | 1 | Tsku | tsukushi, small leucine rich proteoglycan | 447 | 109 | 134 | 207 | 243 | 139 | 259 | 129 | 263 | 149 | 135 |
| 200 | 0.02 | 0.29 | 0.07 | 0.94 | 1 | Gm8234 | | 69 | 70 | 57 | 354 | 166 | 210 | 259 | 421 | 226 | 189 | 184 |
| 189 | -0.11 | 0.40 | -0.26 | 0.79 | 1 | Ttc30a1 | tetratricopeptide repeat domain 30A1 | 90 | 179 | 132 | 200 | 187 | 182 | 259 | 352 | 212 | 81 | 207 |
| 1091 | -0.15 | 0.25 | -0.63 | 0.53 | 1 | Slc9a3r2 | solute carrier family 9 (sodium/hydrogen e | 3338 | 2873 | 3876 | 252 | 299 | 175 | 258 | 402 | 187 | 215 | 123 |
| 377 | 0.17 | 0.29 | 0.58 | 0.56 | 1 | Gm15427 | | 760 | 578 | 653 | 300 | 304 | 146 | 258 | 491 | 232 | 222 | 199 |
| 370 | -0.06 | 0.23 | -0.28 | 0.78 | 1 | Ogfrl1 | opioid growth factor receptor-like 1 | 865 | 830 | 722 | 215 | 167 | 197 | 258 | 283 | 189 | 209 | 135 |
| 280 | -0.07 | 0.28 | -0.25 | 0.80 | 1 | Dyrk3 | dual-specificity tyrosine-(Y)-phosphorylati | 264 | 154 | 154 | 381 | 286 | 354 | 258 | 338 | 282 | 341 | 266 |
| 246 | -0.04 | 0.27 | -0.16 | 0.87 | 1 | H1f3 | H1.3 linker histone, cluster member | 85 | 54 | 58 | 349 | 265 | 391 | 258 | 308 | 252 | 233 | 456 |
| 221 | -0.43 | 0.25 | -1.72 | 0.09 | 1 | Bcdin3d | BCDIN3 domain containing | 305 | 254 | 254 | 267 | 214 | 194 | 258 | 156 | 196 | 155 | 180 |
| 477 | 0.09 | 0.20 | 0.42 | 0.67 | 1 | Fkbp14 | FK506 binding protein 14 | 1010 | 655 | 606 | 211 | 235 | 778 | 257 | 222 | 292 | 274 | 703 |
| 334 | 0.46 | 0.26 | 1.79 | 0.07 | 1 | Cutc | cutC copper transporter | 283 | 256 | 255 | 336 | 313 | 282 | 257 | 668 | 385 | 307 | 328 |
| 333 | 0.16 | 0.22 | 0.73 | 0.47 | 1 | Tango2 | transport and golgi organization 2 | 247 | 345 | 306 | 353 | 333 | 371 | 257 | 314 | 347 | 406 | 388 |
| 269 | 0.22 | 0.28 | 0.80 | 0.42 | 1 | Thg1l | tRNA-histidine guanylyltransferase 1-like | 311 | 238 | 266 | 216 | 273 | 234 | 257 | 418 | 233 | 258 | 250 |
| 163 | 0.29 | 0.31 | 0.95 | 0.34 | 1 | Gm45716 | | 41 | 49 | 61 | 172 | 152 | 163 | 257 | 251 | 219 | 234 | 199 |
| 440 | -0.11 | 0.26 | -0.43 | 0.67 | 1 | Hyal2 | hyaluronoglucosaminidase 2 | 631 | 498 | 603 | 525 | 493 | 341 | 256 | 346 | 458 | 401 | 286 |
| 402 | -0.02 | 0.19 | -0.09 | 0.93 | 1 | Ankzf1 | ankyrin repeat and zinc finger domain cor | 276 | 223 | 316 | 162 | 243 | 1191 | 256 | 141 | 284 | 265 | 1063 |
| 390 | 0.07 | 0.24 | 0.28 | 0.78 | 1 | Rfx2 | regulatory factor X, 2 (influences HLA clas | 478 | 319 | 339 | 326 | 352 | 633 | 256 | 351 | 354 | 370 | 511 |
| 324 | -0.05 | 0.22 | -0.24 | 0.81 | 1 | Ppil1 | peptidylprolyl isomerase (cyclophilin)-like | 392 | 338 | 366 | 393 | 276 | 331 | 256 | 351 | 283 | 247 | 327 |
| 302 | 0.39 | 0.24 | 1.61 | 0.11 | 1 | Hmox1 | heme oxygenase 1 | 629 | 441 | 382 | 233 | 202 | 123 | 256 | 357 | 279 | 252 | 166 |
| 266 | 0.21 | 0.29 | 0.71 | 0.48 | 1 | Krt10 | keratin 10 | 264 | 228 | 392 | 241 | 290 | 167 | 256 | 393 | 248 | 225 | 224 |

| | | | | | | | | | | | | | | | | | | |
|---|---|---|---|---|---|---|---|---|---|---|---|---|---|---|---|---|---|---|
| 259 | -0.44 | 0.37 | -1.19 | 0.23 | 1 | Ttc39aos1 | | 97 | 102 | 101 | 146 | 174 | 808 | 256 | 52 | 224 | 148 | 739 |
| 257 | -0.27 | 0.23 | -1.18 | 0.24 | 1 | Gm13421 | | 148 | 91 | 198 | 186 | 316 | 517 | 256 | 132 | 216 | 236 | 528 |
| 187 | -0.43 | 0.27 | -1.60 | 0.11 | 1 | Vcpkmt | valosin containing protein lysine (K) meth | 252 | 180 | 200 | 148 | 211 | 202 | 256 | 96 | 188 | 153 | 173 |
| 414 | 0.36 | 0.30 | 1.18 | 0.24 | 1 | Scnm1 | sodium channel modifier 1 | 471 | 317 | 472 | 273 | 365 | 582 | 255 | 572 | 287 | 450 | 508 |
| 357 | -0.11 | 0.19 | -0.61 | 0.54 | 1 | Bak1 | BCL2-antagonist/killer 1 | 793 | 641 | 586 | 273 | 236 | 229 | 255 | 236 | 213 | 228 | 239 |
| 324 | 0.21 | 0.27 | 0.78 | 0.43 | 1 | Riox1 | ribosomal oxygenase 1 | 534 | 367 | 504 | 295 | 275 | 167 | 255 | 446 | 253 | 277 | 187 |
| 319 | -0.08 | 0.21 | -0.40 | 0.69 | 1 | Lzic | leucine zipper and CTNNBIP1 domain co | 486 | 504 | 471 | 277 | 247 | 275 | 255 | 277 | 280 | 188 | 254 |
| 309 | -0.04 | 0.24 | -0.19 | 0.85 | 1 | Alg14 | asparagine-linked glycosylation 14 | 436 | 466 | 310 | 309 | 290 | 256 | 255 | 224 | 301 | 300 | 250 |
| 300 | 0.01 | 0.22 | 0.04 | 0.97 | 1 | Gm21926 | | 329 | 283 | 299 | 271 | 280 | 380 | 255 | 226 | 257 | 376 | 344 |
| 293 | -0.04 | 0.22 | -0.18 | 0.85 | 1 | Entpd6 | ectonucleoside triphosphate diphosphohy | 396 | 389 | 413 | 258 | 231 | 280 | 255 | 319 | 242 | 198 | 243 |
| 285 | -0.45 | 0.38 | -1.20 | 0.23 | 1 | Cdc25b | cell division cycle 25B | 735 | 343 | 399 | 245 | 196 | 269 | 255 | 217 | 200 | 88 | 193 |
| 264 | -0.16 | 0.25 | -0.64 | 0.52 | 1 | 9530077C05Ril | RIKEN cDNA 9530077C05 gene | 182 | 253 | 254 | 283 | 232 | 402 | 255 | 187 | 269 | 263 | 320 |
| 244 | 0.03 | 0.25 | 0.14 | 0.89 | 1 | Atn1 | atrophin 1 | 210 | 163 | 347 | 237 | 256 | 214 | 255 | 199 | 290 | 289 | 222 |
| 227 | -0.09 | 0.27 | -0.34 | 0.74 | 1 | Cdkl3 | cyclin-dependent kinase-like 3 | 179 | 121 | 244 | 208 | 244 | 294 | 255 | 181 | 193 | 287 | 289 |
| 218 | -0.22 | 0.31 | -0.72 | 0.47 | 1 | Mrm2 | mitochondrial rRNA methyltransferase 2 | 230 | 195 | 228 | 245 | 213 | 218 | 255 | 280 | 182 | 125 | 226 |
| 353 | -0.35 | 0.30 | -1.15 | 0.25 | 1 | Rps6ka1 | ribosomal protein S6 kinase polypeptide | 1296 | 703 | 714 | 117 | 174 | 123 | 254 | 161 | 145 | 109 | 87 |
| 318 | 0.13 | 0.28 | 0.46 | 0.64 | 1 | 1700066M21Ri | RIKEN cDNA 1700066M21 gene | 312 | 246 | 276 | 489 | 318 | 231 | 254 | 381 | 352 | 283 | 357 |
| 284 | -0.07 | 0.26 | -0.27 | 0.79 | 1 | Gmeb1 | glucocorticoid modulatory element bindin | 378 | 297 | 376 | 193 | 279 | 329 | 254 | 209 | 325 | 149 | 341 |
| 245 | 0.38 | 0.24 | 1.57 | 0.12 | 1 | Tprn | taperin | 201 | 307 | 229 | 228 | 173 | 192 | 254 | 351 | 241 | 309 | 213 |
| 227 | -0.32 | 0.35 | -0.91 | 0.36 | 1 | Gm9958 | | 240 | 285 | 253 | 300 | 220 | 176 | 254 | 101 | 191 | 305 | 176 |
| 211 | -0.20 | 0.27 | -0.71 | 0.48 | 1 | Fbxo17 | F-box protein 17 | 176 | 222 | 139 | 186 | 260 | 254 | 254 | 181 | 215 | 214 | 222 |
| 495 | -0.36 | 0.27 | -1.34 | 0.18 | 1 | Cdo1 | cysteine dioxygenase 1, cytosolic | 1184 | 1406 | 1636 | 172 | 168 | 96 | 253 | 204 | 107 | 139 | 84 |
| 304 | -0.17 | 0.39 | -0.43 | 0.66 | 1 | Tacc3 | transforming, acidic coiled-coil containing | 528 | 341 | 289 | 397 | 206 | 288 | 253 | 433 | 248 | 106 | 252 |
| 292 | 0.25 | 0.22 | 1.15 | 0.25 | 1 | 5830417I10Rik | | 177 | 255 | 186 | 254 | 271 | 409 | 253 | 388 | 268 | 318 | 433 |
| 285 | 0.35 | 0.27 | 1.30 | 0.19 | 1 | Spryd4 | SPRY domain containing 4 | 245 | 185 | 273 | 333 | 255 | 216 | 253 | 506 | 297 | 332 | 240 |
| 252 | 0.14 | 0.27 | 0.50 | 0.61 | 1 | Rimkla | ribosomal modification protein rimK-like fa | 80 | 143 | 89 | 280 | 151 | 505 | 253 | 235 | 254 | 247 | 539 |
| 243 | -0.20 | 0.24 | -0.82 | 0.41 | 1 | Zfp846 | zinc finger protein 846 | 184 | 205 | 243 | 210 | 233 | 372 | 253 | 147 | 237 | 180 | 414 |
| 446 | -0.47 | 0.32 | -1.49 | 0.14 | 1 | Zfp532 | zinc finger protein 532 | 821 | 843 | 793 | 383 | 305 | 460 | 252 | 111 | 352 | 215 | 376 |
| 427 | -0.22 | 0.24 | -0.90 | 0.37 | 1 | Capza1 | capping protein (actin filament) muscle Z- | 727 | 920 | 496 | 373 | 358 | 377 | 252 | 294 | 258 | 211 | 436 |
| 427 | -0.05 | 0.30 | -0.17 | 0.87 | 1 | Nostrin | nitric oxide synthase trafficker | 889 | 1183 | 916 | 134 | 304 | 168 | 252 | 170 | 249 | 308 | 120 |
| 414 | 0.29 | 0.25 | 1.18 | 0.24 | 1 | Fam98c | family with sequence similarity 98, membe | 480 | 303 | 402 | 363 | 362 | 544 | 252 | 502 | 363 | 370 | 611 |
| 314 | 0.09 | 0.27 | 0.32 | 0.75 | 1 | Rbmx2 | RNA binding motif protein, X-linked 2 | 297 | 279 | 373 | 286 | 352 | 311 | 252 | 429 | 295 | 183 | 400 |
| 280 | -0.11 | 0.22 | -0.50 | 0.62 | 1 | Cpsf4 | cleavage and polyadenylation specific fac | 455 | 324 | 394 | 212 | 172 | 363 | 252 | 199 | 204 | 192 | 315 |
| 261 | 0.01 | 0.25 | 0.05 | 0.96 | 1 | 2410022M11Rik | | 220 | 282 | 270 | 284 | 318 | 193 | 252 | 209 | 284 | 335 | 220 |
| 228 | 0.29 | 0.29 | 1.02 | 0.31 | 1 | Diras2 | DIRAS family, GTP-binding RAS-like 2 | 94 | 76 | 37 | 426 | 182 | 202 | 252 | 386 | 269 | 308 | 280 |
| 228 | 0.02 | 0.26 | 0.08 | 0.94 | 1 | 9030407P20Rik | | 112 | 105 | 106 | 310 | 263 | 252 | 252 | 234 | 238 | 401 | 242 |
| 182 | -0.22 | 0.36 | -0.61 | 0.54 | 1 | Tmem121 | transmembrane protein 121 | 71 | 121 | 105 | 235 | 250 | 150 | 252 | 173 | 208 | 353 | 85 |
| 541 | -0.23 | 0.23 | -0.99 | 0.32 | 1 | Spry4 | sprouty RTK signaling antagonist 4 | 1230 | 1501 | 1484 | 270 | 291 | 134 | 251 | 224 | 227 | 139 | 197 |
| 409 | -0.13 | 0.26 | -0.51 | 0.61 | 1 | Ccrl2 | chemokine (C-C motif) receptor-like 2 | 1148 | 632 | 762 | 284 | 278 | 216 | 251 | 263 | 269 | 189 | 212 |
| 371 | -0.02 | 0.23 | -0.07 | 0.94 | 1 | Slc12a5 | solute carrier family 12, member 5 | 163 | 194 | 210 | 374 | 387 | 777 | 251 | 275 | 362 | 389 | 694 |
| 332 | -0.05 | 0.21 | -0.24 | 0.81 | 1 | Zfp1 | zinc finger protein 1 | 394 | 465 | 450 | 290 | 233 | 394 | 251 | 233 | 270 | 188 | 482 |
| 309 | -0.20 | 0.23 | -0.86 | 0.39 | 1 | Irak2 | interleukin-1 receptor-associated kinase 2 | 567 | 505 | 591 | 269 | 232 | 172 | 251 | 155 | 204 | 272 | 176 |
| 269 | -0.16 | 0.23 | -0.71 | 0.48 | 1 | Kptn | kaptin | 348 | 338 | 387 | 202 | 298 | 242 | 251 | 151 | 246 | 222 | 280 |
| 248 | -0.40 | 0.32 | -1.26 | 0.21 | 1 | Dnajb5 | DnaJ heat shock protein family (Hsp40) m | 371 | 188 | 239 | 151 | 297 | 365 | 251 | 81 | 289 | 156 | 336 |
| 240 | -0.24 | 0.24 | -0.98 | 0.33 | 1 | Mgme1 | mitochondrial genome maintenance exor | 236 | 203 | 258 | 239 | 235 | 319 | 251 | 212 | 255 | 153 | 275 |
| 219 | -0.15 | 0.32 | -0.46 | 0.65 | 1 | B3galt4 | UDP-Gal:betaGlcNAc beta 1,3-galactosyl | 120 | 250 | 179 | 279 | 294 | 151 | 251 | 284 | 175 | 175 | 246 |
| 187 | -0.20 | 0.39 | -0.51 | 0.61 | 1 | Klhl3 | kelch-like 3 | 207 | 193 | 313 | 181 | 98 | 185 | 251 | 211 | 156 | 152 | 108 |
| 161 | -0.46 | 0.43 | -1.08 | 0.28 | 1 | Gm13375 | | 142 | 103 | 147 | 119 | 208 | 180 | 251 | 31 | 232 | 166 | 188 |
| 482 | -0.42 | 0.25 | -1.71 | 0.09 | 1 | Id2 | inhibitor of DNA binding 2 | 1391 | 994 | 1037 | 256 | 309 | 252 | 250 | 125 | 278 | 180 | 232 |
| 284 | -0.08 | 0.22 | -0.35 | 0.73 | 1 | Cdr2l | cerebellar degeneration-related protein 2 | 563 | 564 | 440 | 195 | 173 | 177 | 250 | 215 | 186 | 241 | 124 |
| 283 | 0.12 | 0.29 | 0.41 | 0.68 | 1 | Zfp511 | zinc finger protein 511 | 212 | 273 | 164 | 284 | 201 | 446 | 250 | 180 | 250 | 392 | 459 |
| 282 | 0.14 | 0.26 | 0.54 | 0.59 | 1 | Gpatch4 | G patch domain containing 4 | 358 | 496 | 383 | 225 | 149 | 262 | 250 | 305 | 227 | 175 | 271 |
| 238 | 0.12 | 0.28 | 0.43 | 0.67 | 1 | Gm14097 | | 121 | 79 | 85 | 284 | 343 | 225 | 250 | 211 | 440 | 332 | 245 |
| 203 | -0.24 | 0.33 | -0.73 | 0.46 | 1 | Polr2k | polymerase (RNA) II (DNA directed) polyp | 150 | 218 | 196 | 215 | 210 | 206 | 250 | 277 | 189 | 100 | 218 |
| 388 | -0.01 | 0.20 | -0.03 | 0.97 | 1 | Apbb3 | amyloid beta (A4) precursor protein-bindi | 261 | 320 | 309 | 253 | 289 | 952 | 249 | 348 | 242 | 259 | 787 |
| 377 | -0.25 | 0.29 | -0.88 | 0.38 | 1 | Fbxl12 | F-box and leucine-rich repeat protein 12 | 353 | 287 | 405 | 260 | 326 | 837 | 249 | 136 | 230 | 402 | 659 |
| 339 | -0.14 | 0.21 | -0.65 | 0.51 | 1 | Hgh1 | HGH1 homolog | 295 | 538 | 338 | 345 | 343 | 388 | 249 | 374 | 230 | 223 | 410 |
| 302 | -0.18 | 0.22 | -0.79 | 0.43 | 1 | Axin2 | axin 2 | 412 | 428 | 488 | 243 | 243 | 321 | 249 | 180 | 220 | 289 | 250 |
| 302 | 0.26 | 0.21 | 1.23 | 0.22 | 1 | Mtres1 | mitochondrial transcription rescue factor 1 | 293 | 373 | 373 | 307 | 207 | 276 | 249 | 320 | 266 | 314 | 339 |
| 294 | -0.01 | 0.27 | -0.04 | 0.97 | 1 | Cfap126 | cilia and flagella associated protein 126 | 278 | 420 | 348 | 308 | 268 | 270 | 249 | 209 | 328 | 335 | 220 |

| | | | | | | | | | | | | | | | | | | |
|---|---|---|---|---|---|---|---|---|---|---|---|---|---|---|---|---|---|---|
| 290 | 0.24 | 0.18 | 1.30 | 0.19 | 1 | H2aw | H2A.W histone | 184 | 242 | 214 | 295 | 236 | 386 | 249 | 358 | 293 | 268 | 463 |
| 286 | 0.17 | 0.32 | 0.52 | 0.60 | 1 | Sigirr | single immunoglobulin and toll-interleukin | 301 | 168 | 294 | 261 | 189 | 451 | 249 | 342 | 315 | 292 | 282 |
| 249 | -0.03 | 0.27 | -0.10 | 0.92 | 1 | 1700030K09Ril | RIKEN cDNA 1700030K09 gene | 293 | 264 | 214 | 158 | 249 | 347 | 249 | 209 | 271 | 176 | 313 |
| 247 | 0.23 | 0.26 | 0.90 | 0.37 | 1 | Kbtbd8 | kelch repeat and BTB (POZ) domain conta | 160 | 208 | 193 | 292 | 144 | 315 | 249 | 315 | 228 | 265 | 344 |
| 227 | 0.08 | 0.30 | 0.28 | 0.78 | 1 | Kcnu1 | potassium channel, subfamily U, member | 100 | 194 | 136 | 219 | 253 | 277 | 249 | 202 | 240 | 276 | 351 |
| 539 | 0.18 | 0.18 | 0.98 | 0.33 | 1 | Ftx | | 183 | 301 | 179 | 272 | 267 | 1670 | 248 | 314 | 306 | 269 | 1919 |
| 375 | 0.02 | 0.33 | 0.07 | 0.94 | 1 | Nr4a2 | nuclear receptor subfamily 4, group A, me | 856 | 490 | 335 | 346 | 259 | 366 | 248 | 379 | 336 | 229 | 278 |
| 317 | -0.08 | 0.22 | -0.34 | 0.73 | 1 | Wdr60 | WD repeat domain 60 | 294 | 255 | 372 | 347 | 311 | 406 | 248 | 266 | 277 | 290 | 417 |
| 309 | -0.03 | 0.26 | -0.10 | 0.92 | 1 | Stx1a | syntaxin 1A (brain) | 516 | 400 | 636 | 133 | 253 | 311 | 248 | 227 | 212 | 174 | 286 |
| 293 | 0.10 | 0.26 | 0.38 | 0.71 | 1 | Pdcd5 | programmed cell death 5 | 329 | 364 | 317 | 226 | 300 | 288 | 248 | 170 | 338 | 304 | 341 |
| 288 | -0.17 | 0.24 | -0.73 | 0.46 | 1 | Cnnm4 | cyclin M4 | 365 | 388 | 301 | 376 | 215 | 291 | 248 | 215 | 232 | 268 | 270 |
| 277 | -0.04 | 0.24 | -0.16 | 0.88 | 1 | 2310010J17Rik | | 297 | 277 | 360 | 339 | 244 | 238 | 248 | 228 | 322 | 270 | 227 |
| 274 | 0.29 | 0.21 | 1.41 | 0.16 | 1 | Tnfsf12 | tumor necrosis factor (ligand) superfamily | 312 | 336 | 274 | 254 | 239 | 188 | 248 | 314 | 270 | 383 | 195 |
| 244 | 0.16 | 0.33 | 0.48 | 0.63 | 1 | Adssl1 | adenylosuccinate synthetase like 1 | 348 | 442 | 168 | 193 | 257 | 118 | 248 | 249 | 279 | 245 | 136 |
| 215 | 0.26 | 0.26 | 0.98 | 0.33 | 1 | Nudt2 | nudix (nucleoside diphosphate linked mo | 227 | 173 | 251 | 168 | 233 | 139 | 248 | 231 | 193 | 278 | 218 |
| 208 | -0.04 | 0.30 | -0.14 | 0.89 | 1 | 1700037C18Ril | RIKEN cDNA 1700037C18 gene | 92 | 154 | 156 | 302 | 201 | 205 | 248 | 266 | 170 | 291 | 200 |
| 457 | -0.04 | 0.25 | -0.16 | 0.87 | 1 | Rrnad1 | ribosomal RNA adenine dimethylase dom | 292 | 316 | 221 | 208 | 347 | 1313 | 247 | 190 | 275 | 312 | 1304 |
| 351 | 0.20 | 0.21 | 0.97 | 0.33 | 1 | Fgfr1op | Fgfr1 oncogene partner | 387 | 287 | 414 | 249 | 269 | 536 | 247 | 322 | 326 | 249 | 582 |
| 345 | 0.00 | 0.21 | 0.02 | 0.98 | 1 | Aifm2 | apoptosis-inducing factor, mitochondrion- | 384 | 326 | 421 | 339 | 310 | 434 | 247 | 257 | 330 | 330 | 416 |
| 307 | 0.08 | 0.26 | 0.32 | 0.75 | 1 | Pole4 | polymerase (DNA-directed), epsilon 4 (p1 | 596 | 370 | 448 | 175 | 331 | 202 | 247 | 176 | 255 | 247 | 335 |
| 291 | 0.02 | 0.26 | 0.09 | 0.93 | 1 | Tatdn1 | TatD DNase domain containing 1 | 297 | 393 | 359 | 161 | 259 | 384 | 247 | 152 | 280 | 219 | 447 |
| 244 | -0.37 | 0.24 | -1.58 | 0.11 | 1 | Cfap74 | cilia and flagella associated protein 74 | 68 | 76 | 84 | 175 | 223 | 718 | 247 | 163 | 143 | 171 | 614 |
| 402 | -0.06 | 0.19 | -0.31 | 0.75 | 1 | Gm49980 | | 222 | 417 | 587 | 451 | 689 | 221 | 246 | 418 | 728 | 281 | 166 |
| 316 | -0.04 | 0.24 | -0.19 | 0.85 | 1 | Gtf2h4 | general transcription factor II H, polypepti | 408 | 465 | 401 | 257 | 303 | 305 | 246 | 349 | 200 | 296 | 248 |
| 296 | 0.07 | 0.23 | 0.32 | 0.75 | 1 | Smg8 | smg-8 homolog, nonsense mediated mRI | 505 | 366 | 280 | 292 | 216 | 272 | 246 | 286 | 253 | 305 | 239 |
| 289 | 0.01 | 0.30 | 0.04 | 0.97 | 1 | Primpol | primase and polymerase (DNA-directed) | 230 | 436 | 348 | 283 | 216 | 321 | 246 | 189 | 192 | 310 | 406 |
| 258 | -0.10 | 0.28 | -0.34 | 0.73 | 1 | Dglucy | D-glutamate cyclase | 393 | 321 | 306 | 297 | 132 | 265 | 246 | 181 | 164 | 321 | 212 |
| 244 | -0.25 | 0.29 | -0.86 | 0.39 | 1 | Cox11 | cytochrome c oxidase assembly protein 1 | 261 | 259 | 207 | 220 | 219 | 366 | 246 | 110 | 208 | 269 | 320 |
| 239 | -0.02 | 0.25 | -0.10 | 0.92 | 1 | C030034I22Rik | | 219 | 228 | 214 | 248 | 193 | 306 | 246 | 171 | 224 | 298 | 287 |
| 212 | -0.23 | 0.28 | -0.80 | 0.43 | 1 | Slc46a1 | solute carrier family 46, member 1 | 204 | 135 | 210 | 309 | 273 | 146 | 246 | 254 | 193 | 207 | 159 |
| 188 | -0.35 | 0.26 | -1.35 | 0.18 | 1 | Gm4285 | | 155 | 133 | 177 | 237 | 210 | 202 | 246 | 168 | 190 | 151 | 196 |
| 314 | 0.06 | 0.18 | 0.30 | 0.76 | 1 | Cir1 | corepressor interacting with RBPJ, 1 | 349 | 335 | 395 | 296 | 303 | 324 | 245 | 305 | 322 | 276 | 309 |
| 276 | -0.44 | 0.31 | -1.43 | 0.15 | 1 | Lmln | leishmanolysin-like (metallopeptidase M8 | 221 | 304 | 295 | 382 | 276 | 363 | 245 | 131 | 260 | 233 | 323 |
| 223 | 0.05 | 0.34 | 0.15 | 0.88 | 1 | Elp6 | elongator acetyltransferase complex subu | 124 | 192 | 219 | 307 | 194 | 199 | 245 | 204 | 196 | 348 | 225 |
| 10587 | -1.14 | 0.65 | -1.73 | 0.08 | 1 | Apoe | apolipoprotein E | 46824 | 25572 | 42517 | 149 | 390 | 206 | 244 | 60 | 246 | 18 | 228 |
| 502 | -0.23 | 0.28 | -0.80 | 0.42 | 1 | Scd1 | stearoyl-Coenzyme A desaturase 1 | 1462 | 1233 | 850 | 311 | 389 | 123 | 244 | 198 | 371 | 236 | 109 |
| 404 | 0.06 | 0.26 | 0.23 | 0.82 | 1 | Ppan | peter pan homolog | 329 | 410 | 347 | 331 | 347 | 736 | 244 | 589 | 273 | 246 | 595 |
| 228 | 0.08 | 0.28 | 0.29 | 0.77 | 1 | Pigw | phosphatidylinositol glycan anchor biosyr | 160 | 177 | 246 | 278 | 194 | 218 | 244 | 214 | 251 | 290 | 236 |
| 181 | -0.10 | 0.37 | -0.27 | 0.79 | 1 | Syngr3 | synaptogyrin 3 | 114 | 235 | 104 | 235 | 187 | 130 | 244 | 138 | 166 | 300 | 134 |
| 288 | -0.34 | 0.33 | -1.03 | 0.30 | 1 | Zfp641 | zinc finger protein 641 | 196 | 242 | 231 | 309 | 206 | 572 | 243 | 337 | 191 | 82 | 558 |
| 287 | 0.27 | 0.21 | 1.31 | 0.19 | 1 | Zfp59 | zinc finger protein 59 | 290 | 247 | 271 | 236 | 247 | 330 | 243 | 226 | 278 | 367 | 426 |
| 276 | -0.27 | 0.31 | -0.89 | 0.37 | 1 | Tmem106a | transmembrane protein 106A | 673 | 315 | 474 | 240 | 188 | 185 | 243 | 221 | 210 | 160 | 122 |
| 240 | -0.21 | 0.21 | -0.96 | 0.33 | 1 | Elovl7 | ELOVL family member 7, elongation of lor | 244 | 286 | 209 | 385 | 225 | 165 | 243 | 323 | 219 | 213 | 130 |
| 167 | 0.52 | 0.38 | 1.35 | 0.18 | 1 | Doc2b | double C2, beta | 86 | 91 | 32 | 115 | 141 | 171 | 243 | 288 | 187 | 188 | 290 |
| 1044 | -0.08 | 0.17 | -0.49 | 0.62 | 1 | Tia1 | cytotoxic granule-associated RNA binding | 516 | 658 | 527 | 353 | 245 | 4100 | 242 | 351 | 284 | 157 | 4047 |
| 275 | -0.09 | 0.28 | -0.31 | 0.76 | 1 | Zfp52 | zinc finger protein 52 | 337 | 375 | 239 | 238 | 247 | 329 | 242 | 149 | 261 | 240 | 365 |
| 269 | -0.47 | 0.25 | -1.89 | 0.06 | 1 | Dph2 | DPH2 homolog | 388 | 372 | 322 | 280 | 270 | 291 | 242 | 234 | 192 | 125 | 243 |
| 265 | 0.13 | 0.22 | 0.58 | 0.57 | 1 | Camk1g | calcium/calmodulin-dependent protein kir | 275 | 250 | 189 | 359 | 259 | 187 | 242 | 399 | 252 | 295 | 205 |
| 261 | -0.21 | 0.24 | -0.87 | 0.38 | 1 | Nme7 | NME/NM23 family member 7 | 441 | 361 | 359 | 166 | 266 | 238 | 242 | 161 | 182 | 281 | 173 |
| 246 | 0.15 | 0.25 | 0.60 | 0.55 | 1 | Pcdhga10 | protocadherin gamma subfamily A, 10 | 240 | 328 | 309 | 223 | 237 | 168 | 242 | 207 | 241 | 268 | 241 |
| 235 | 0.10 | 0.33 | 0.29 | 0.77 | 1 | Cmc2 | COX assembly mitochondrial protein 2 | 265 | 163 | 232 | 242 | 209 | 229 | 242 | 317 | 314 | 153 | 214 |
| 230 | 0.00 | 0.31 | 0.01 | 0.99 | 1 | Lhfpl5 | lipoma HMGIC fusion partner-like 5 | 160 | 159 | 234 | 348 | 179 | 216 | 242 | 336 | 208 | 250 | 194 |
| 204 | -0.05 | 0.37 | -0.13 | 0.90 | 1 | Psmc3ip | proteasome (prosome, macropain) 26S su | 282 | 307 | 112 | 288 | 51 | 228 | 242 | 117 | 86 | 258 | 273 |
| 186 | 0.06 | 0.26 | 0.22 | 0.82 | 1 | Tfap4 | transcription factor AP4 | 212 | 133 | 206 | 88 | 188 | 221 | 242 | 110 | 202 | 250 | 193 |
| 358 | -0.38 | 0.23 | -1.62 | 0.11 | 1 | Cdc14a | CDC14 cell division cycle 14A | 631 | 582 | 612 | 272 | 237 | 438 | 241 | 134 | 230 | 240 | 316 |
| 346 | -0.39 | 0.25 | -1.57 | 0.12 | 1 | Rnf138rt1 | ring finger protein 138, retrogene 1 | 591 | 601 | 713 | 106 | 204 | 543 | 241 | 93 | 154 | 178 | 385 |
| 323 | -0.13 | 0.22 | -0.58 | 0.56 | 1 | Slc25a37 | solute carrier family 25, member 37 | 251 | 194 | 319 | 242 | 357 | 631 | 241 | 231 | 244 | 298 | 546 |
| 316 | 0.12 | 0.31 | 0.40 | 0.69 | 1 | Grwd1 | glutamate-rich WD repeat containing 1 | 421 | 293 | 458 | 326 | 255 | 264 | 241 | 490 | 280 | 238 | 212 |
| 281 | 0.24 | 0.21 | 1.14 | 0.25 | 1 | Apex2 | apurinic/apyrimidinic endonuclease 2 | 257 | 367 | 176 | 288 | 196 | 331 | 241 | 366 | 246 | 274 | 350 |
| 268 | -0.52 | 0.30 | -1.74 | 0.08 | 1 | Stk36 | serine/threonine kinase 36 | 196 | 262 | 260 | 346 | 298 | 423 | 241 | 125 | 299 | 213 | 285 |

| | | | | | | | | | | | | | | | | | |
|---|---|---|---|---|---|---|---|---|---|---|---|---|---|---|---|---|---|
| 250 | 0.04 | 0.27 | 0.16 | 0.87 | 1 | Zfp959 | zinc finger protein 959 | 283 | 399 | 318 | 171 | 166 | 269 | 241 | 175 | 154 | 214 | 361 |
| 249 | -0.05 | 0.29 | -0.18 | 0.85 | 1 | Slc19a1 | solute carrier family 19 (folate transporter) | 303 | 297 | 296 | 299 | 162 | 235 | 241 | 207 | 288 | 225 | 181 |
| 578 | 0.25 | 0.24 | 1.01 | 0.31 | 1 | Gprc5b | G protein-coupled receptor, family C, grou | 1751 | 1391 | 1315 | 339 | 162 | 97 | 240 | 343 | 253 | 398 | 72 |
| 554 | -0.28 | 0.28 | -1.00 | 0.32 | 1 | Kank2 | KN motif and ankyrin repeat domains 2 | 2107 | 1129 | 1193 | 276 | 213 | 186 | 240 | 272 | 197 | 143 | 141 |
| 551 | 0.18 | 0.21 | 0.85 | 0.39 | 1 | Trib2 | tribbles pseudokinase 2 | 1351 | 1313 | 1440 | 237 | 155 | 283 | 240 | 222 | 178 | 322 | 322 |
| 347 | 0.34 | 0.25 | 1.38 | 0.17 | 1 | Ppp1r35 | protein phosphatase 1, regulatory subuni | 439 | 342 | 523 | 322 | 269 | 273 | 240 | 381 | 284 | 416 | 325 |
| 324 | -0.24 | 0.34 | -0.72 | 0.47 | 1 | Adgrv1 | adhesion G protein-coupled receptor V1 | 51 | 83 | 165 | 247 | 296 | 919 | 240 | 310 | 388 | 84 | 783 |
| 296 | 0.38 | 0.31 | 1.23 | 0.22 | 1 | Ccdc107 | coiled-coil domain containing 107 | 333 | 289 | 408 | 198 | 268 | 249 | 240 | 496 | 248 | 267 | 257 |
| 274 | -0.18 | 0.28 | -0.63 | 0.53 | 1 | Azin2 | antizyme inhibitor 2 | 334 | 201 | 246 | 395 | 302 | 240 | 240 | 302 | 262 | 294 | 193 |
| 253 | 0.41 | 0.34 | 1.20 | 0.23 | 1 | Lsm10 | U7 snRNP-specific Sm-like protein LSM1 | 230 | 216 | 353 | 174 | 220 | 195 | 240 | 470 | 293 | 220 | 174 |
| 223 | -0.12 | 0.29 | -0.41 | 0.68 | 1 | 2900052L18Rik | | 201 | 228 | 182 | 179 | 200 | 334 | 240 | 260 | 199 | 125 | 307 |
| 215 | -0.15 | 0.26 | -0.57 | 0.57 | 1 | Pde4a | phosphodiesterase 4A, cAMP specific | 342 | 289 | 229 | 154 | 210 | 189 | 240 | 187 | 147 | 174 | 206 |
| 198 | -0.37 | 0.35 | -1.03 | 0.30 | 1 | Dok5 | docking protein 5 | 84 | 203 | 171 | 292 | 237 | 178 | 240 | 181 | 285 | 97 | 213 |
| 176 | 0.29 | 0.34 | 0.87 | 0.38 | 1 | Dusp14 | dual specificity phosphatase 14 | 87 | 32 | 46 | 179 | 237 | 149 | 240 | 309 | 225 | 249 | 187 |
| 172 | 0.02 | 0.40 | 0.06 | 0.95 | 1 | Them4 | thioesterase superfamily member 4 | 172 | 117 | 86 | 182 | 177 | 156 | 240 | 264 | 242 | 104 | 153 |
| 408 | -0.10 | 0.26 | -0.37 | 0.71 | 1 | Gpr146 | G protein-coupled receptor 146 | 1299 | 923 | 798 | 140 | 189 | 171 | 239 | 206 | 116 | 294 | 115 |
| 339 | -0.05 | 0.28 | -0.19 | 0.85 | 1 | Spred3 | sprouty-related EVH1 domain containing | 341 | 352 | 301 | 242 | 259 | 641 | 239 | 146 | 233 | 381 | 589 |
| 313 | 0.30 | 0.19 | 1.58 | 0.11 | 1 | Mpst | mercaptopyruvate sulfurtransferase | 652 | 638 | 547 | 203 | 163 | 116 | 239 | 244 | 200 | 285 | 151 |
| 286 | 0.17 | 0.21 | 0.80 | 0.42 | 1 | 9130221H12Rik | | 358 | 269 | 299 | 270 | 282 | 254 | 239 | 313 | 298 | 287 | 274 |
| 260 | 0.25 | 0.25 | 1.00 | 0.32 | 1 | Acat2 | acetyl-Coenzyme A acetyltransferase 2 | 229 | 244 | 246 | 284 | 221 | 230 | 239 | 226 | 370 | 313 | 258 |
| 256 | -0.11 | 0.26 | -0.44 | 0.66 | 1 | Ndc1 | NDC1 transmembrane nucleoporin | 528 | 351 | 347 | 198 | 264 | 136 | 239 | 207 | 200 | 187 | 159 |
| 242 | 0.05 | 0.28 | 0.18 | 0.86 | 1 | Zfp414 | zinc finger protein 414 | 195 | 266 | 332 | 132 | 255 | 296 | 239 | 176 | 208 | 227 | 341 |
| 233 | 0.07 | 0.25 | 0.27 | 0.79 | 1 | Ogg1 | 8-oxoguanine DNA-glycosylase 1 | 175 | 230 | 218 | 194 | 214 | 306 | 239 | 220 | 278 | 256 | 235 |
| 204 | -0.10 | 0.32 | -0.31 | 0.75 | 1 | Ccdc122 | coiled-coil domain containing 122 | 117 | 166 | 70 | 324 | 207 | 211 | 239 | 175 | 185 | 326 | 222 |
| 1270 | 0.11 | 0.16 | 0.67 | 0.50 | 1 | Sgk1 | serum/glucocorticoid regulated kinase 1 | 4103 | 3925 | 3922 | 200 | 314 | 223 | 238 | 291 | 313 | 241 | 206 |
| 577 | 0.18 | 0.18 | 1.01 | 0.31 | 1 | Sh3bp5 | SH3-domain binding protein 5 (BTK-asso | 1971 | 1244 | 1453 | 200 | 219 | 113 | 238 | 224 | 239 | 345 | 103 |
| 373 | 0.06 | 0.22 | 0.27 | 0.78 | 1 | Rab11fip4 | RAB11 family interacting protein 4 (class | 753 | 918 | 567 | 290 | 239 | 139 | 238 | 358 | 165 | 258 | 178 |
| 320 | 0.16 | 0.30 | 0.55 | 0.58 | 1 | 1110012L19Rik | RIKEN cDNA 1110012L19 gene | 516 | 416 | 452 | 233 | 293 | 225 | 238 | 470 | 216 | 242 | 217 |
| 296 | 0.20 | 0.22 | 0.92 | 0.36 | 1 | Mrps22 | mitochondrial ribosomal protein S22 | 453 | 460 | 295 | 208 | 217 | 292 | 238 | 250 | 261 | 261 | 321 |
| 264 | -0.23 | 0.26 | -0.87 | 0.38 | 1 | Rad9a | RAD9 checkpoint clamp component A | 385 | 260 | 299 | 318 | 251 | 243 | 238 | 181 | 229 | 294 | 210 |
| 213 | -0.33 | 0.28 | -1.16 | 0.24 | 1 | Zranb3 | zinc finger, RAN-binding domain containi | 307 | 276 | 205 | 172 | 191 | 252 | 238 | 176 | 133 | 124 | 267 |
| 193 | 0.16 | 0.35 | 0.46 | 0.64 | 1 | BC037704 | | 100 | 116 | 245 | 184 | 166 | 195 | 238 | 227 | 206 | 247 | 199 |
| 177 | 0.03 | 0.34 | 0.10 | 0.92 | 1 | Pm20d2 | peptidase M20 domain containing 2 | 125 | 90 | 102 | 186 | 203 | 157 | 238 | 92 | 278 | 276 | 197 |
| 473 | -0.16 | 0.29 | -0.57 | 0.57 | 1 | Tmed6 | transmembrane p24 trafficking protein 6 | 393 | 1443 | 991 | 448 | 417 | 183 | 237 | 300 | 334 | 307 | 152 |
| 352 | 0.18 | 0.28 | 0.64 | 0.52 | 1 | Wdr5b | WD repeat domain 5B | 349 | 423 | 368 | 282 | 393 | 363 | 237 | 248 | 311 | 487 | 409 |
| 293 | 0.19 | 0.30 | 0.63 | 0.53 | 1 | 5730480H06Rik | PARK2 co-regulated-like | 160 | 243 | 273 | 273 | 232 | 467 | 237 | 396 | 202 | 354 | 385 |
| 293 | -0.18 | 0.21 | -0.82 | 0.41 | 1 | Lin37 | lin-37 homolog (C. elegans) | 343 | 349 | 339 | 261 | 302 | 353 | 237 | 282 | 214 | 192 | 346 |
| 269 | 0.06 | 0.24 | 0.25 | 0.81 | 1 | Lrrc28 | leucine rich repeat containing 28 | 262 | 184 | 236 | 267 | 389 | 230 | 237 | 301 | 331 | 268 | 254 |
| 216 | 0.12 | 0.26 | 0.46 | 0.65 | 1 | Crppa | CDP-L-ribitol pyrophosphorylase A | 301 | 315 | 301 | 142 | 121 | 206 | 237 | 154 | 170 | 220 | 210 |
| 214 | 0.36 | 0.32 | 1.12 | 0.26 | 1 | Xylb | xylulokinase homolog (H. influenzae) | 79 | 182 | 110 | 198 | 212 | 216 | 237 | 305 | 252 | 344 | 220 |
| 207 | -0.06 | 0.38 | -0.15 | 0.88 | 1 | Znrd1as | zinc ribbon domain containing 1, antisens | 142 | 243 | 269 | 133 | 194 | 238 | 237 | 260 | 207 | 91 | 263 |
| 354 | 0.31 | 0.25 | 1.24 | 0.21 | 1 | Dock5 | dedicator of cytokinesis 5 | 766 | 504 | 489 | 220 | 255 | 235 | 236 | 393 | 264 | 300 | 233 |
| 342 | -0.07 | 0.34 | -0.20 | 0.84 | 1 | Kcnip3 | Kv channel interacting protein 3, calsenili | 794 | 946 | 818 | 127 | 182 | 85 | 236 | 154 | 163 | 99 | 160 |
| 300 | -0.24 | 0.20 | -1.24 | 0.22 | 1 | Zfp39 | zinc finger protein 39 | 338 | 331 | 410 | 292 | 289 | 374 | 236 | 206 | 252 | 204 | 361 |
| 290 | 0.33 | 0.23 | 1.44 | 0.15 | 1 | L1cam | L1 cell adhesion molecule | 355 | 327 | 510 | 232 | 228 | 180 | 236 | 249 | 307 | 361 | 205 |
| 265 | 0.00 | 0.24 | -0.02 | 0.98 | 1 | Efcab15 | EF-hand calcium binding domain 15 | 188 | 317 | 311 | 277 | 335 | 183 | 236 | 375 | 345 | 155 | 193 |
| 263 | -0.18 | 0.35 | -0.51 | 0.61 | 1 | Tspoap1 | TSPO associated protein 1 | 258 | 399 | 273 | 95 | 221 | 506 | 236 | 173 | 130 | 197 | 405 |
| 223 | -0.15 | 0.29 | -0.52 | 0.61 | 1 | Gm26559 | | 97 | 182 | 200 | 339 | 265 | 212 | 236 | 211 | 238 | 259 | 219 |
| 214 | -0.04 | 0.36 | -0.10 | 0.92 | 1 | Ephx4 | epoxide hydrolase 4 | 225 | 315 | 335 | 173 | 177 | 146 | 236 | 312 | 108 | 151 | 176 |
| 211 | -0.10 | 0.30 | -0.34 | 0.73 | 1 | Prkn | parkin RBR E3 ubiquitin protein ligase | 116 | 85 | 137 | 263 | 181 | 350 | 236 | 221 | 260 | 236 | 232 |
| 200 | 0.24 | 0.27 | 0.90 | 0.37 | 1 | Sdhaf3 | succinate dehydrogenase complex asser | 158 | 126 | 209 | 157 | 203 | 185 | 236 | 215 | 203 | 308 | 196 |
| 196 | 0.05 | 0.26 | 0.19 | 0.85 | 1 | Uxt | ubiquitously expressed prefoldin like cha | 197 | 182 | 202 | 176 | 189 | 172 | 236 | 243 | 211 | 175 | 170 |
| 158 | 0.20 | 0.37 | 0.54 | 0.59 | 1 | Nckap5 | NCK-associated protein 5 | 180 | 53 | 77 | 187 | 146 | 101 | 236 | 130 | 244 | 238 | 151 |
| 400 | 0.04 | 0.25 | 0.16 | 0.87 | 1 | Tmem140 | transmembrane protein 140 | 1099 | 745 | 1000 | 174 | 212 | 151 | 235 | 270 | 166 | 192 | 160 |
| 375 | -0.02 | 0.17 | -0.10 | 0.92 | 1 | Mtrf1l | mitochondrial translational release factor | 359 | 557 | 495 | 333 | 315 | 464 | 235 | 335 | 307 | 200 | 524 |
| 320 | -0.43 | 0.26 | -1.66 | 0.10 | 1 | Nebl | nebulette | 484 | 766 | 523 | 259 | 244 | 251 | 235 | 151 | 152 | 171 | 283 |
| 316 | -0.32 | 0.32 | -0.99 | 0.32 | 1 | Ice2 | interactor of little elongation complex ELL | 340 | 416 | 406 | 76 | 209 | 748 | 235 | 96 | 164 | 128 | 658 |
| 282 | -0.08 | 0.28 | -0.30 | 0.77 | 1 | Snrpa1 | small nuclear ribonucleoprotein polypepti | 473 | 269 | 376 | 172 | 252 | 347 | 235 | 184 | 249 | 138 | 408 |
| 172 | 0.03 | 0.27 | 0.10 | 0.92 | 1 | | | 82 | 111 | 95 | 224 | 215 | 127 | 235 | 236 | 179 | 204 | 182 |
| 351 | -0.05 | 0.22 | -0.24 | 0.81 | 1 | Ikbke | inhibitor of kappaB kinase epsilon | 248 | 192 | 276 | 202 | 332 | 822 | 234 | 143 | 327 | 296 | 785 |

| | | | | | | | | | | | | | | | | | |
|---|---|---|---|---|---|---|---|---|---|---|---|---|---|---|---|---|---|
| 221 | -0.31 | 0.28 | -1.11 | 0.27 | 1 | H1f4 | H1.4 linker histone, cluster member | 30 | 78 | 98 | 280 | 365 | 344 | 234 | 221 | 268 | 167 | 348 |
| 406 | -0.04 | 0.29 | -0.15 | 0.88 | 1 | Htra2 | HtrA serine peptidase 2 | 859 | 451 | 526 | 352 | 332 | 409 | 233 | 305 | 336 | 216 | 443 |
| 356 | 0.09 | 0.22 | 0.40 | 0.69 | 1 | Rps6kb2 | ribosomal protein S6 kinase, polypeptide | 232 | 175 | 219 | 263 | 280 | 869 | 233 | 263 | 269 | 363 | 753 |
| 354 | 0.22 | 0.30 | 0.76 | 0.45 | 1 | Pold1 | polymerase (DNA directed), delta 1, catal | 858 | 385 | 459 | 284 | 232 | 246 | 233 | 393 | 195 | 341 | 267 |
| 341 | 0.24 | 0.23 | 1.05 | 0.30 | 1 | Rrp15 | ribosomal RNA processing 15 homolog (S | 582 | 417 | 518 | 189 | 248 | 373 | 233 | 310 | 309 | 242 | 336 |
| 277 | 0.48 | 0.30 | 1.57 | 0.12 | 1 | Abcd2 | ATP-binding cassette, sub-family D (ALD) | 280 | 130 | 253 | 296 | 246 | 223 | 233 | 425 | 303 | 356 | 308 |
| 253 | -0.21 | 0.29 | -0.73 | 0.46 | 1 | Zfp760 | zinc finger protein 760 | 230 | 314 | 346 | 274 | 159 | 328 | 233 | 163 | 199 | 153 | 384 |
| 229 | -0.01 | 0.28 | -0.04 | 0.97 | 1 | Trmt10a | tRNA methyltransferase 10A | 161 | 204 | 202 | 152 | 263 | 328 | 233 | 243 | 246 | 157 | 325 |
| 218 | 0.17 | 0.24 | 0.69 | 0.49 | 1 | Acad10 | acyl-Coenzyme A dehydrogenase family, | 199 | 153 | 88 | 230 | 179 | 281 | 233 | 247 | 199 | 295 | 297 |
| 84 | 0.80 | 0.76 | 1.06 | 0.29 | 1 | Mir6236 | | 13 | 10 | 61 | 16 | 35 | 68 | 233 | 108 | 108 | 222 | 53 |
| 919 | -0.18 | 0.28 | -0.62 | 0.53 | 1 | Pxdn | peroxidasin | 3037 | 2616 | 2809 | 233 | 68 | 337 | 232 | 202 | 84 | 144 | 345 |
| 322 | 0.02 | 0.25 | 0.07 | 0.94 | 1 | Sirt5 | sirtuin 5 | 289 | 302 | 336 | 364 | 387 | 305 | 232 | 460 | 334 | 327 | 208 |
| 289 | -0.18 | 0.22 | -0.81 | 0.42 | 1 | Gm11837 | predicted gene 11837 | 459 | 516 | 606 | 232 | 185 | 200 | 232 | 219 | 204 | 166 | 165 |
| 234 | 0.16 | 0.28 | 0.59 | 0.56 | 1 | Aamdc | adipogenesis associated Mth938 domain | 330 | 369 | 260 | 158 | 177 | 183 | 232 | 304 | 143 | 224 | 193 |
| 186 | 0.31 | 0.37 | 0.85 | 0.40 | 1 | Gm10941 | | 20 | 63 | 59 | 228 | 246 | 132 | 232 | 421 | 209 | 228 | 204 |
| 170 | -0.10 | 0.25 | -0.41 | 0.68 | 1 | Ctf1 | cardiotrophin 1 | 126 | 120 | 147 | 185 | 236 | 113 | 232 | 189 | 183 | 225 | 111 |
| 668 | -0.06 | 0.22 | -0.26 | 0.80 | 1 | AI506816 | | 1160 | 1684 | 1585 | 600 | 432 | 202 | 231 | 626 | 462 | 176 | 190 |
| 301 | 0.35 | 0.26 | 1.32 | 0.19 | 1 | Gpx2 | glutathione peroxidase 2 | 82 | 131 | 233 | 300 | 312 | 430 | 231 | 417 | 379 | 318 | 481 |
| 268 | -0.16 | 0.37 | -0.44 | 0.66 | 1 | Slc5a3 | solute carrier family 5 (inositol transporter | 253 | 396 | 404 | 292 | 103 | 362 | 231 | 338 | 172 | 103 | 292 |
| 251 | -0.27 | 0.30 | -0.91 | 0.37 | 1 | Rnf32 | ring finger protein 32 | 321 | 385 | 427 | 172 | 171 | 303 | 231 | 198 | 178 | 94 | 283 |
| 203 | -0.04 | 0.39 | -0.10 | 0.92 | 1 | Rps6kl1 | ribosomal protein S6 kinase-like 1 | 154 | 391 | 150 | 148 | 178 | 208 | 231 | 181 | 212 | 125 | 251 |
| 186 | 0.05 | 0.41 | 0.12 | 0.90 | 1 | Dcxr | dicarbonyl L-xylulose reductase | 251 | 225 | 204 | 152 | 147 | 112 | 231 | 381 | 145 | 103 | 100 |
| 179 | -0.28 | 0.40 | -0.71 | 0.48 | 1 | Gnmt | glycine N-methyltransferase | 88 | 261 | 242 | 232 | 151 | 134 | 231 | 214 | 112 | 130 | 168 |
| 176 | -0.33 | 0.47 | -0.70 | 0.49 | 1 | Dnaic1 | dynein, axonemal, intermediate chain 1 | 15 | 80 | 106 | 235 | 167 | 324 | 231 | 156 | 175 | 161 | 290 |
| 143 | -0.09 | 0.33 | -0.26 | 0.79 | 1 | C130013H08Rik | | 63 | 89 | 78 | 124 | 176 | 164 | 231 | 182 | 149 | 155 | 167 |
| 560 | 0.02 | 0.26 | 0.07 | 0.95 | 1 | Csf2ra | colony stimulating factor 2 receptor, alpha | 807 | 487 | 438 | 313 | 213 | 1572 | 230 | 243 | 321 | 255 | 1286 |
| 384 | -0.02 | 0.19 | -0.11 | 0.91 | 1 | Ccdc28a | coiled-coil domain containing 28A | 213 | 492 | 388 | 521 | 410 | 433 | 230 | 424 | 365 | 270 | 477 |
| 288 | 0.27 | 0.25 | 1.09 | 0.27 | 1 | Mettl18 | methyltransferase like 18 | 230 | 310 | 218 | 297 | 285 | 275 | 230 | 342 | 269 | 422 | 286 |
| 270 | -0.21 | 0.28 | -0.74 | 0.46 | 1 | Zbtb34 | zinc finger and BTB domain containing 34 | 293 | 394 | 329 | 288 | 199 | 312 | 230 | 226 | 273 | 113 | 309 |
| 266 | -0.25 | 0.33 | -0.75 | 0.45 | 1 | Osbp2 | oxysterol binding protein 2 | 419 | 706 | 432 | 160 | 140 | 194 | 230 | 196 | 104 | 245 | 98 |
| 218 | 0.02 | 0.27 | 0.08 | 0.93 | 1 | Tspan6 | tetraspanin 6 | 368 | 182 | 240 | 233 | 206 | 127 | 230 | 221 | 181 | 269 | 140 |
| 200 | -0.12 | 0.29 | -0.40 | 0.69 | 1 | Gemin8 | gem nuclear organelle associated protein | 223 | 213 | 174 | 159 | 229 | 205 | 230 | 117 | 222 | 189 | 242 |
| 189 | 0.20 | 0.31 | 0.63 | 0.53 | 1 | Tmem41a | transmembrane protein 41a | 214 | 229 | 249 | 155 | 127 | 120 | 230 | 217 | 157 | 286 | 94 |
| 184 | -0.16 | 0.33 | -0.47 | 0.63 | 1 | Gpr157 | G protein-coupled receptor 157 | 164 | 205 | 289 | 223 | 144 | 124 | 230 | 224 | 135 | 171 | 117 |
| 267 | 0.07 | 0.22 | 0.32 | 0.75 | 1 | Hs6st3 | heparan sulfate 6-O-sulfotransferase 3 | 243 | 289 | 258 | 270 | 236 | 313 | 229 | 317 | 271 | 269 | 240 |
| 263 | -0.15 | 0.42 | -0.35 | 0.73 | 1 | Zfat | zinc finger and AT hook domain containin | 218 | 370 | 198 | 276 | 300 | 290 | 229 | 104 | 355 | 346 | 204 |
| 246 | -0.14 | 0.23 | -0.61 | 0.54 | 1 | Bbs10 | Bardet-Biedl syndrome 10 (human) | 171 | 235 | 191 | 323 | 325 | 236 | 229 | 246 | 241 | 274 | 234 |
| 229 | 0.13 | 0.22 | 0.58 | 0.56 | 1 | Fastkd3 | FAST kinase domains 3 | 263 | 235 | 201 | 184 | 166 | 287 | 229 | 176 | 202 | 229 | 343 |
| 222 | -0.17 | 0.31 | -0.57 | 0.57 | 1 | Zfp593 | zinc finger protein 593 | 330 | 217 | 410 | 228 | 152 | 175 | 229 | 166 | 172 | 167 | 191 |
| 202 | -0.51 | 0.36 | -1.41 | 0.16 | 1 | Gm2115 | | 215 | 169 | 160 | 228 | 305 | 201 | 229 | 183 | 322 | 98 | 113 |
| 153 | -0.12 | 0.30 | -0.39 | 0.69 | 1 | Sugct | succinyl-CoA glutarate-CoA transferase | 130 | 71 | 103 | 168 | 173 | 145 | 229 | 177 | 116 | 236 | 138 |
| 312 | -0.02 | 0.23 | -0.07 | 0.94 | 1 | Zfp579 | zinc finger protein 579 | 251 | 244 | 291 | 270 | 231 | 641 | 228 | 231 | 288 | 285 | 469 |
| 311 | 0.05 | 0.24 | 0.22 | 0.83 | 1 | Hmga1 | high mobility group AT-hook 1 | 364 | 388 | 275 | 420 | 312 | 224 | 228 | 327 | 289 | 337 | 252 |
| 290 | 0.22 | 0.33 | 0.69 | 0.49 | 1 | Tnfrsf22 | tumor necrosis factor receptor superfamily | 202 | 149 | 322 | 363 | 249 | 326 | 228 | 395 | 270 | 370 | 317 |
| 258 | -0.20 | 0.29 | -0.68 | 0.49 | 1 | L3mbtl3 | L3MBTL3 histone methyl-lysine binding p | 553 | 417 | 486 | 225 | 133 | 156 | 228 | 210 | 128 | 126 | 176 |
| 254 | 0.31 | 0.31 | 1.02 | 0.31 | 1 | Reep4 | receptor accessory protein 4 | 305 | 226 | 271 | 146 | 258 | 262 | 228 | 359 | 236 | 211 | 290 |
| 247 | 0.04 | 0.24 | 0.18 | 0.86 | 1 | Ankrd13d | ankyrin repeat domain 13 family, member | 206 | 147 | 254 | 201 | 232 | 392 | 228 | 217 | 233 | 290 | 323 |
| 247 | -0.27 | 0.28 | -0.95 | 0.34 | 1 | Slc35d1 | solute carrier family 35 (UDP-glucuronic a | 275 | 253 | 205 | 267 | 243 | 337 | 228 | 144 | 269 | 183 | 316 |
| 242 | 0.04 | 0.28 | 0.14 | 0.89 | 1 | Fmc1 | formation of mitochondrial complex V ass | 269 | 290 | 341 | 202 | 246 | 184 | 228 | 310 | 239 | 142 | 209 |
| 238 | -0.24 | 0.23 | -1.04 | 0.30 | 1 | Tfeb | transcription factor EB | 424 | 411 | 382 | 176 | 198 | 156 | 228 | 185 | 130 | 185 | 142 |
| 176 | -0.03 | 0.37 | -0.09 | 0.93 | 1 | Tbc1d2 | TBC1 domain family, member 2 | 160 | 138 | 219 | 144 | 160 | 174 | 228 | 262 | 202 | 143 | 107 |
| 286 | -0.05 | 0.30 | -0.15 | 0.88 | 1 | Zfp40 | zinc finger protein 40 | 231 | 299 | 197 | 366 | 234 | 401 | 227 | 185 | 217 | 379 | 414 |
| 266 | -0.17 | 0.23 | -0.75 | 0.45 | 1 | Aaas | achalasia, adrenocortical insufficiency, al | 408 | 404 | 327 | 247 | 217 | 255 | 227 | 254 | 222 | 165 | 203 |
| 246 | -0.30 | 0.27 | -1.14 | 0.25 | 1 | Znhit3 | zinc finger, HIT type 3 | 267 | 207 | 368 | 264 | 276 | 250 | 227 | 140 | 217 | 220 | 268 |
| 168 | -0.61 | 0.35 | -1.75 | 0.08 | 1 | Ak5 | adenylate kinase 5 | 146 | 67 | 87 | 277 | 210 | 224 | 227 | 147 | 172 | 120 | 176 |
| 160 | -0.48 | 0.30 | -1.62 | 0.10 | 1 | Alkbh4 | alkB homolog 4, lysine demethylase | 154 | 187 | 170 | 186 | 130 | 186 | 227 | 158 | 133 | 115 | 116 |
| 424 | 0.24 | 0.21 | 1.16 | 0.25 | 1 | Map3k20 | mitogen-activated protein kinase kinase k | 886 | 898 | 935 | 226 | 190 | 244 | 226 | 205 | 213 | 339 | 305 |
| 402 | -0.05 | 0.33 | -0.16 | 0.87 | 1 | Ptges | prostaglandin E synthase | 958 | 663 | 546 | 188 | 170 | 570 | 226 | 288 | 189 | 100 | 525 |
| 334 | -0.59 | 0.33 | -1.77 | 0.08 | 1 | Gpr179 | G protein-coupled receptor 179 | 673 | 981 | 743 | 151 | 79 | 309 | 226 | 131 | 45 | 148 | 184 |
| 324 | -0.07 | 0.24 | -0.30 | 0.77 | 1 | Nhp2 | NHP2 ribonucleoprotein | 770 | 538 | 441 | 268 | 250 | 179 | 226 | 215 | 261 | 260 | 153 |

| | | | | | | | | | | | | | | | | | | |
|---|---|---|---|---|---|---|---|---|---|---|---|---|---|---|---|---|---|---|
| 318 | 0.21 | 0.21 | 1.01 | 0.31 | 1 | Lonrf1 | LON peptidase N-terminal domain and rir | 230 | 349 | 275 | 391 | 302 | 302 | 226 | 481 | 278 | 346 | 312 |
| 293 | 0.15 | 0.23 | 0.64 | 0.52 | 1 | Zscan2 | zinc finger and SCAN domain containing | 525 | 337 | 449 | 214 | 196 | 274 | 226 | 251 | 263 | 245 | 241 |
| 268 | 0.22 | 0.31 | 0.71 | 0.48 | 1 | Brdt | bromodomain, testis-specific | 174 | 271 | 338 | 283 | 228 | 263 | 226 | 275 | 241 | 402 | 249 |
| 260 | -0.16 | 0.41 | -0.39 | 0.70 | 1 | Hpse | heparanase | 778 | 301 | 117 | 246 | 202 | 176 | 226 | 141 | 196 | 318 | 154 |
| 247 | -0.09 | 0.30 | -0.31 | 0.75 | 1 | Snta1 | syntrophin, acidic 1 | 534 | 367 | 346 | 255 | 164 | 107 | 226 | 299 | 175 | 152 | 90 |
| 187 | 0.27 | 0.36 | 0.75 | 0.45 | 1 | Timm23 | translocase of inner mitochondrial membr | 126 | 98 | 174 | 154 | 170 | 196 | 226 | 290 | 264 | 129 | 231 |
| 169 | -0.40 | 0.40 | -1.00 | 0.32 | 1 | Gpr135 | G protein-coupled receptor 135 | 244 | 294 | 153 | 204 | 132 | 96 | 226 | 70 | 149 | 233 | 63 |
| 163 | 0.04 | 0.29 | 0.14 | 0.89 | 1 | Glt28d2 | glycosyltransferase 28 domain containing | 101 | 107 | 140 | 175 | 181 | 127 | 226 | 217 | 165 | 233 | 120 |
| 142 | 0.26 | 0.37 | 0.71 | 0.48 | 1 | Gm19461 | | 27 | 13 | 24 | 143 | 166 | 150 | 226 | 300 | 183 | 184 | 150 |
| 139 | 0.06 | 0.39 | 0.15 | 0.88 | 1 | 9230112J17Rik | | 37 | 88 | 41 | 143 | 163 | 136 | 226 | 163 | 216 | 175 | 140 |
| 495 | -0.12 | 0.28 | -0.43 | 0.67 | 1 | Ldlrap1 | low density lipoprotein receptor adaptor p | 1271 | 1153 | 1255 | 237 | 247 | 183 | 225 | 317 | 317 | 128 | 117 |
| 381 | 0.32 | 0.21 | 1.54 | 0.12 | 1 | Baz1a | bromodomain adjacent to zinc finger dom | 819 | 609 | 594 | 196 | 238 | 306 | 225 | 315 | 233 | 273 | 388 |
| 347 | -0.03 | 0.25 | -0.13 | 0.90 | 1 | Tle4 | transducin-like enhancer of split 4 | 803 | 527 | 601 | 206 | 255 | 243 | 225 | 163 | 230 | 372 | 188 |
| 227 | -0.15 | 0.31 | -0.47 | 0.64 | 1 | Nadsyn1 | NAD synthetase 1 | 210 | 132 | 140 | 212 | 243 | 374 | 225 | 285 | 189 | 143 | 341 |
| 221 | 0.15 | 0.32 | 0.48 | 0.63 | 1 | 9130401M01Ri | RIKEN cDNA 9130401M01 gene | 179 | 200 | 162 | 229 | 223 | 189 | 225 | 454 | 184 | 164 | 223 |
| 207 | 0.03 | 0.26 | 0.12 | 0.91 | 1 | Gm49602 | | 336 | 279 | 365 | 141 | 146 | 136 | 225 | 168 | 142 | 174 | 168 |
| 192 | 0.13 | 0.30 | 0.42 | 0.67 | 1 | 1500002C15Rik | | 167 | 236 | 135 | 272 | 127 | 149 | 225 | 200 | 231 | 210 | 166 |
| 177 | 0.25 | 0.65 | 0.39 | 0.69 | 1 | Meltf | melanotransferrin | 12 | 0 | 23 | 256 | 113 | 235 | 225 | 190 | 132 | 446 | 317 |
| 116 | 0.90 | 0.47 | 1.91 | 0.06 | 1 | Astn1 | astrotactin 1 | 59 | 91 | 77 | 61 | 53 | 51 | 225 | 171 | 113 | 267 | 106 |
| 457 | -0.18 | 0.25 | -0.71 | 0.48 | 1 | Wsb1 | WD repeat and SOCS box-containing 1 | 1227 | 933 | 885 | 219 | 105 | 493 | 224 | 190 | 132 | 122 | 498 |
| 281 | 0.10 | 0.22 | 0.46 | 0.64 | 1 | Gin1 | gypsy retrotransposon integrase 1 | 253 | 319 | 284 | 257 | 270 | 326 | 224 | 319 | 230 | 298 | 307 |
| 263 | -0.17 | 0.27 | -0.64 | 0.52 | 1 | Tma16 | translation machinery associated 16 | 355 | 390 | 433 | 126 | 269 | 293 | 224 | 180 | 210 | 142 | 270 |
| 257 | -0.05 | 0.27 | -0.19 | 0.85 | 1 | Zfp53 | zinc finger protein 53 | 283 | 199 | 231 | 262 | 226 | 341 | 224 | 147 | 273 | 232 | 413 |
| 248 | -0.03 | 0.24 | -0.11 | 0.91 | 1 | Ttf2 | transcription termination factor, RNA poly | 407 | 373 | 310 | 185 | 183 | 234 | 224 | 262 | 174 | 182 | 196 |
| 248 | -0.03 | 0.28 | -0.10 | 0.92 | 1 | Magohb | mago homolog B, exon junction complex | 246 | 329 | 362 | 279 | 246 | 162 | 224 | 233 | 165 | 269 | 211 |
| 238 | -0.01 | 0.35 | -0.03 | 0.98 | 1 | Prrt3 | proline-rich transmembrane protein 3 | 106 | 47 | 109 | 375 | 345 | 235 | 224 | 374 | 361 | 220 | 222 |
| 233 | -0.40 | 0.41 | -0.98 | 0.33 | 1 | Rad51ap2 | RAD51 associated protein 2 | 17 | 63 | 21 | 509 | 276 | 414 | 224 | 144 | 272 | 193 | 428 |
| 227 | -0.28 | 0.31 | -0.92 | 0.36 | 1 | Anapc10 | anaphase promoting complex subunit 10 | 333 | 417 | 280 | 169 | 158 | 253 | 224 | 123 | 196 | 117 | 231 |
| 184 | 0.15 | 0.27 | 0.54 | 0.59 | 1 | Gm42984 | | 124 | 208 | 160 | 223 | 152 | 129 | 224 | 273 | 161 | 224 | 151 |
| 165 | -0.30 | 0.33 | -0.92 | 0.36 | 1 | Amer3 | APC membrane recruitment 3 | 142 | 165 | 120 | 155 | 262 | 123 | 224 | 112 | 179 | 229 | 104 |
| 13361 | -1.19 | 1.25 | -0.96 | 0.34 | 1 | Try5 | trypsin 5 | 22990 | 56728 | 63223 | 1765 | 145 | 13 | 223 | 215 | 52 | 67 | 1553 |
| 390 | -0.21 | 0.20 | -1.06 | 0.29 | 1 | Il4ra | interleukin 4 receptor, alpha | 1198 | 781 | 1118 | 150 | 167 | 105 | 223 | 118 | 175 | 149 | 105 |
| 387 | 0.21 | 0.18 | 1.21 | 0.23 | 1 | Ramp2 | receptor (calcitonin) activity modifying pro | 1162 | 721 | 930 | 143 | 164 | 134 | 223 | 154 | 199 | 285 | 144 |
| 311 | 0.22 | 0.30 | 0.73 | 0.46 | 1 | Slc35e3 | solute carrier family 35, member E3 | 277 | 340 | 381 | 242 | 335 | 323 | 223 | 293 | 225 | 442 | 340 |
| 249 | 0.08 | 0.25 | 0.32 | 0.75 | 1 | C1qtnf12 | C1q and tumor necrosis factor related 12 | 202 | 349 | 227 | 316 | 186 | 231 | 223 | 284 | 286 | 234 | 196 |
| 242 | -0.16 | 0.25 | -0.65 | 0.51 | 1 | Pnck | pregnancy upregulated non-ubiquitously | 340 | 417 | 390 | 179 | 157 | 243 | 223 | 155 | 145 | 215 | 200 |
| 224 | -0.04 | 0.31 | -0.13 | 0.90 | 1 | Fancc | Fanconi anemia, complementation group | 446 | 230 | 268 | 158 | 156 | 242 | 223 | 190 | 178 | 124 | 253 |
| 221 | -0.07 | 0.25 | -0.30 | 0.77 | 1 | 2810410L24Rik | | 206 | 293 | 215 | 197 | 135 | 324 | 223 | 180 | 144 | 187 | 330 |
| 220 | -0.36 | 0.31 | -1.16 | 0.25 | 1 | A430018G15Rik | | 186 | 331 | 230 | 341 | 231 | 150 | 223 | 197 | 229 | 107 | 194 |
| 143 | -0.02 | 0.34 | -0.05 | 0.96 | 1 | Cd63-ps | | 91 | 58 | 89 | 201 | 148 | 105 | 223 | 120 | 176 | 206 | 158 |
| 432 | -0.10 | 0.23 | -0.43 | 0.67 | 1 | Pyroxd2 | pyridine nucleotide-disulphide oxidoredu | 216 | 235 | 269 | 351 | 192 | 1400 | 222 | 284 | 281 | 188 | 1110 |
| 298 | -0.15 | 0.24 | -0.61 | 0.54 | 1 | Prrt1 | proline-rich transmembrane protein 1 | 313 | 287 | 233 | 130 | 187 | 784 | 222 | 132 | 198 | 175 | 613 |
| 289 | -0.24 | 0.25 | -0.99 | 0.32 | 1 | Ppp2r2b | protein phosphatase 2, regulatory subuni | 667 | 784 | 520 | 151 | 161 | 126 | 222 | 124 | 146 | 170 | 112 |
| 283 | 0.19 | 0.23 | 0.79 | 0.43 | 1 | Rnft2 | ring finger protein, transmembrane 2 | 196 | 359 | 180 | 249 | 225 | 418 | 222 | 359 | 226 | 270 | 406 |
| 270 | -0.33 | 0.25 | -1.35 | 0.18 | 1 | Lca5 | Leber congenital amaurosis 5 (human) | 377 | 384 | 325 | 219 | 274 | 334 | 222 | 138 | 208 | 238 | 252 |
| 254 | 0.00 | 0.27 | 0.00 | 1.00 | 1 | Gm10033 | predicted gene 10033 | 293 | 277 | 255 | 233 | 181 | 336 | 222 | 231 | 231 | 137 | 405 |
| 246 | 0.32 | 0.27 | 1.19 | 0.23 | 1 | Zfp759 | zinc finger protein 759 | 216 | 164 | 245 | 224 | 195 | 290 | 222 | 326 | 222 | 319 | 287 |
| 235 | -0.18 | 0.28 | -0.66 | 0.51 | 1 | Cdk20 | cyclin-dependent kinase 20 | 228 | 243 | 293 | 193 | 212 | 333 | 222 | 110 | 249 | 232 | 267 |
| 206 | 0.05 | 0.25 | 0.18 | 0.86 | 1 | Gm20257 | | 149 | 141 | 166 | 285 | 172 | 216 | 222 | 251 | 227 | 223 | 217 |
| 199 | 0.00 | 0.28 | -0.02 | 0.99 | 1 | Inava | innate immunity activator | 59 | 76 | 80 | 262 | 305 | 172 | 222 | 410 | 260 | 216 | 126 |
| 172 | 0.06 | 0.29 | 0.22 | 0.83 | 1 | D830050J10Rik | | 124 | 130 | 161 | 167 | 206 | 123 | 222 | 153 | 177 | 286 | 139 |
| 423 | -0.07 | 0.27 | -0.27 | 0.78 | 1 | Rassf2 | Ras association (RalGDS/AF-6) domain fa | 1642 | 800 | 704 | 295 | 144 | 107 | 221 | 235 | 115 | 274 | 116 |
| 368 | -0.43 | 0.23 | -1.85 | 0.06 | 1 | Amt | aminomethyltransferase | 504 | 519 | 666 | 333 | 272 | 516 | 221 | 158 | 207 | 219 | 434 |
| 351 | 0.02 | 0.20 | 0.09 | 0.93 | 1 | Ndufs5 | NADH:ubiquinone oxidoreductase core s | 196 | 379 | 481 | 451 | 465 | 282 | 221 | 407 | 399 | 260 | 325 |
| 338 | -0.28 | 0.34 | -0.81 | 0.42 | 1 | Cilk1 | ciliogenesis associated kinase 1 | 818 | 524 | 492 | 187 | 281 | 346 | 221 | 211 | 277 | 102 | 262 |
| 325 | 0.06 | 0.21 | 0.29 | 0.77 | 1 | Ahcy | S-adenosylhomocysteine hydrolase | 396 | 434 | 643 | 162 | 301 | 337 | 221 | 169 | 360 | 196 | 355 |
| 307 | 0.25 | 0.24 | 1.06 | 0.29 | 1 | Rhno1 | RAD9-HUS1-RAD1 interacting nuclear or | 351 | 266 | 184 | 262 | 233 | 467 | 221 | 350 | 257 | 300 | 481 |
| 271 | 0.38 | 0.26 | 1.43 | 0.15 | 1 | Traf5 | TNF receptor-associated factor 5 | 466 | 297 | 298 | 172 | 274 | 172 | 221 | 315 | 266 | 294 | 208 |
| 228 | 0.11 | 0.26 | 0.41 | 0.68 | 1 | Zkscan4 | zinc finger with KRAB and SCAN domains | 188 | 205 | 250 | 232 | 219 | 220 | 221 | 254 | 191 | 314 | 210 |
| 160 | 0.21 | 0.45 | 0.46 | 0.65 | 1 | Plppr1 | phospholipid phosphatase related 1 | 23 | 72 | 92 | 174 | 176 | 152 | 221 | 200 | 190 | 313 | 142 |

| | | | | | | | | | | | | | | | | | |
|---|---|---|---|---|---|---|---|---|---|---|---|---|---|---|---|---|---|
| 155 | -0.25 | 0.43 | -0.58 | 0.56 | 1 | D030028A08Rik | | 104 | 82 | 80 | 158 | 162 | 206 | 221 | 44 | 195 | 268 | 184 |
| 377 | -0.54 | 0.29 | -1.90 | 0.06 | 1 | Prickle2 | prickle planar cell polarity protein 2 | 1120 | 886 | 852 | 169 | 226 | 143 | 220 | 110 | 226 | 95 | 100 |
| 328 | -0.07 | 0.25 | -0.29 | 0.77 | 1 | Rbm12b2 | RNA binding motif protein 12 B2 | 291 | 357 | 396 | 336 | 202 | 509 | 220 | 265 | 253 | 146 | 634 |
| 327 | 0.10 | 0.25 | 0.40 | 0.69 | 1 | Mks1 | Meckel syndrome, type 1 | 277 | 333 | 210 | 176 | 247 | 744 | 220 | 238 | 251 | 247 | 657 |
| 300 | -0.20 | 0.20 | -1.04 | 0.30 | 1 | Utp23 | UTP23 small subunit processome compo | 281 | 419 | 321 | 358 | 321 | 317 | 220 | 367 | 241 | 189 | 270 |
| 257 | -0.32 | 0.32 | -1.02 | 0.31 | 1 | Psmg4 | proteasome (prosome, macropain) assem | 283 | 259 | 491 | 269 | 314 | 192 | 220 | 141 | 191 | 304 | 166 |
| 256 | -0.04 | 0.42 | -0.09 | 0.93 | 1 | Nav3 | neuron navigator 3 | 165 | 252 | 116 | 368 | 233 | 277 | 220 | 510 | 321 | 68 | 287 |
| 244 | 0.06 | 0.24 | 0.25 | 0.80 | 1 | Slc19a2 | solute carrier family 19 (thiamine transpor | 281 | 328 | 235 | 215 | 273 | 187 | 220 | 282 | 227 | 255 | 180 |
| 196 | -0.25 | 0.30 | -0.83 | 0.41 | 1 | Kcnip4 | Kv channel interacting protein 4 | 211 | 260 | 259 | 146 | 178 | 231 | 220 | 184 | 145 | 192 | 132 |
| 176 | 0.11 | 0.27 | 0.40 | 0.69 | 1 | Lrrc27 | leucine rich repeat containing 27 | 115 | 65 | 88 | 138 | 210 | 243 | 220 | 204 | 183 | 224 | 246 |
| 540 | -0.27 | 0.22 | -1.25 | 0.21 | 1 | St3gal6 | ST3 beta-galactoside alpha-2,3-sialyltran | 1200 | 1405 | 1379 | 365 | 303 | 186 | 219 | 191 | 298 | 250 | 147 |
| 443 | -0.24 | 0.24 | -1.01 | 0.31 | 1 | Arhgef40 | Rho guanine nucleotide exchange factor | 1429 | 942 | 963 | 235 | 188 | 187 | 219 | 127 | 216 | 185 | 177 |
| 365 | 0.45 | 0.30 | 1.48 | 0.14 | 1 | Ldhb | lactate dehydrogenase B | 577 | 264 | 231 | 300 | 242 | 498 | 219 | 452 | 301 | 388 | 547 |
| 333 | 0.15 | 0.24 | 0.63 | 0.53 | 1 | Hyi | | 311 | 287 | 396 | 242 | 275 | 540 | 219 | 242 | 242 | 372 | 538 |
| 283 | 0.18 | 0.21 | 0.86 | 0.39 | 1 | 9330159F19Rik | RIKEN cDNA 9330159F19 gene | 301 | 356 | 333 | 209 | 188 | 391 | 219 | 231 | 248 | 260 | 374 |
| 213 | 0.02 | 0.28 | 0.09 | 0.93 | 1 | Ccne2 | cyclin E2 | 236 | 148 | 142 | 247 | 184 | 250 | 219 | 169 | 247 | 229 | 276 |
| 204 | 0.27 | 0.34 | 0.79 | 0.43 | 1 | Zfp458 | zinc finger protein 458 | 261 | 278 | 143 | 138 | 125 | 234 | 219 | 155 | 231 | 162 | 295 |
| 110 | -0.50 | 0.67 | -0.74 | 0.46 | 1 | BC023719 | | 160 | 0 | 183 | 7 | 204 | 0 | 219 | 0 | 193 | 241 | 1 |
| 500 | 0.08 | 0.20 | 0.39 | 0.70 | 1 | D430019H16Ri | RIKEN cDNA D430019H16 gene | 1049 | 1485 | 1313 | 168 | 216 | 201 | 218 | 227 | 192 | 210 | 221 |
| 322 | 0.35 | 0.25 | 1.41 | 0.16 | 1 | Snapc3 | small nuclear RNA activating complex, pc | 226 | 256 | 224 | 119 | 215 | 746 | 218 | 224 | 251 | 261 | 800 |
| 314 | -0.07 | 0.32 | -0.21 | 0.83 | 1 | Incenp | inner centromere protein | 682 | 334 | 412 | 254 | 320 | 243 | 218 | 328 | 189 | 227 | 249 |
| 312 | 0.09 | 0.22 | 0.39 | 0.69 | 1 | Mtfmt | mitochondrial methionyl-tRNA formyltrans | 212 | 250 | 259 | 325 | 301 | 496 | 218 | 331 | 325 | 330 | 385 |
| 270 | 0.37 | 0.22 | 1.66 | 0.10 | 1 | Gm45871 | predicted gene 45871 | 263 | 236 | 200 | 241 | 194 | 341 | 218 | 389 | 242 | 260 | 387 |
| 256 | -0.10 | 0.24 | -0.43 | 0.67 | 1 | 2900093K20Rik | | 160 | 293 | 204 | 333 | 312 | 266 | 218 | 236 | 276 | 240 | 281 |
| 256 | -0.03 | 0.27 | -0.10 | 0.92 | 1 | Glmn | glomulin, FKBP associated protein | 263 | 452 | 319 | 186 | 230 | 267 | 218 | 174 | 221 | 228 | 261 |
| 228 | -0.15 | 0.40 | -0.38 | 0.70 | 1 | Dgkb | diacylglycerol kinase, beta | 485 | 850 | 352 | 92 | 70 | 73 | 218 | 72 | 93 | 120 | 86 |
| 210 | 0.17 | 0.33 | 0.51 | 0.61 | 1 | Llph | LLP homolog, long-term synaptic facilitati | 202 | 239 | 178 | 140 | 232 | 195 | 218 | 331 | 196 | 160 | 220 |
| 209 | -0.01 | 0.25 | -0.05 | 0.96 | 1 | Mir22hg | | 175 | 123 | 117 | 168 | 235 | 314 | 218 | 141 | 207 | 265 | 334 |
| 207 | 0.19 | 0.24 | 0.78 | 0.43 | 1 | Rad51c | RAD51 paralog C | 174 | 222 | 160 | 171 | 152 | 263 | 218 | 197 | 179 | 243 | 293 |
| 195 | -0.44 | 0.32 | -1.40 | 0.16 | 1 | Zc4h2 | zinc finger, C4H2 domain containing | 359 | 328 | 366 | 110 | 170 | 137 | 218 | 138 | 137 | 89 | 97 |
| 189 | -0.11 | 0.39 | -0.28 | 0.78 | 1 | Spata2l | spermatogenesis associated 2-like | 236 | 76 | 142 | 217 | 232 | 179 | 218 | 276 | 157 | 173 | 175 |
| 188 | 0.25 | 0.36 | 0.68 | 0.50 | 1 | Syce2 | synaptonemal complex central element p | 196 | 111 | 173 | 150 | 232 | 119 | 218 | 353 | 175 | 215 | 120 |
| 357 | 0.32 | 0.25 | 1.28 | 0.20 | 1 | Ccdc130 | coiled-coil domain containing 130 | 210 | 316 | 196 | 339 | 322 | 571 | 217 | 395 | 311 | 456 | 594 |
| 347 | 0.12 | 0.22 | 0.55 | 0.58 | 1 | Lrwd1 | leucine-rich repeats and WD repeat doma | 444 | 486 | 389 | 270 | 327 | 390 | 217 | 357 | 303 | 283 | 348 |
| 237 | 0.18 | 0.24 | 0.75 | 0.45 | 1 | Cradd | CASP2 and RIPK1 domain containing ad | 290 | 246 | 338 | 158 | 249 | 184 | 217 | 235 | 269 | 264 | 153 |
| 221 | -0.12 | 0.32 | -0.38 | 0.71 | 1 | Zfp248 | zinc finger protein 248 | 203 | 169 | 195 | 305 | 203 | 235 | 217 | 220 | 275 | 122 | 287 |
| 214 | 0.12 | 0.28 | 0.44 | 0.66 | 1 | Gm49463 | | 247 | 112 | 236 | 214 | 217 | 192 | 217 | 222 | 243 | 245 | 208 |
| 203 | 0.16 | 0.29 | 0.57 | 0.57 | 1 | 4930431P19Rik | | 131 | 151 | 200 | 212 | 236 | 161 | 217 | 319 | 212 | 171 | 227 |
| 197 | -0.02 | 0.26 | -0.10 | 0.92 | 1 | Gm17251 | | 172 | 138 | 161 | 229 | 178 | 232 | 217 | 262 | 198 | 182 | 199 |
| 366 | 0.15 | 0.33 | 0.44 | 0.66 | 1 | Exog | endo/exonuclease (5'-3'), endonuclease ( | 420 | 356 | 292 | 305 | 363 | 502 | 216 | 556 | 337 | 165 | 519 |
| 359 | 0.29 | 0.33 | 0.89 | 0.37 | 1 | Arfip1 | ADP-ribosylation factor interacting proteir | 264 | 206 | 307 | 367 | 411 | 408 | 216 | 736 | 441 | 224 | 369 |
| 283 | -0.37 | 0.29 | -1.27 | 0.20 | 1 | Plp2 | proteolipid protein 2 | 565 | 449 | 532 | 194 | 232 | 216 | 216 | 97 | 196 | 283 | 131 |
| 260 | -0.35 | 0.27 | -1.31 | 0.19 | 1 | Dync1i1 | dynein cytoplasmic 1 intermediate chain ' | 408 | 593 | 502 | 183 | 200 | 155 | 216 | 190 | 144 | 107 | 160 |
| 255 | -0.05 | 0.27 | -0.20 | 0.84 | 1 | Gstcd | glutathione S-transferase, C-terminal dom | 389 | 358 | 391 | 146 | 226 | 260 | 216 | 134 | 220 | 287 | 183 |
| 380 | 0.29 | 0.23 | 1.26 | 0.21 | 1 | Wtip | WT1-interacting protein | 1002 | 558 | 701 | 260 | 246 | 140 | 215 | 280 | 281 | 308 | 185 |
| 322 | -0.19 | 0.23 | -0.80 | 0.42 | 1 | Tvp23a | trans-golgi network vesicle protein 23A | 646 | 556 | 535 | 235 | 259 | 249 | 215 | 221 | 242 | 146 | 237 |
| 302 | -0.13 | 0.34 | -0.39 | 0.70 | 1 | Tm7sf2 | transmembrane 7 superfamily member 2 | 104 | 38 | 107 | 390 | 398 | 613 | 215 | 277 | 362 | 331 | 490 |
| 286 | -0.05 | 0.37 | -0.15 | 0.88 | 1 | Tha1 | threonine aldolase 1 | 189 | 82 | 300 | 286 | 242 | 592 | 215 | 271 | 212 | 269 | 487 |
| 274 | 0.43 | 0.28 | 1.52 | 0.13 | 1 | Ttc32 | tetratricopeptide repeat domain 32 | 178 | 96 | 186 | 271 | 289 | 318 | 215 | 386 | 332 | 399 | 347 |
| 267 | 0.07 | 0.25 | 0.30 | 0.77 | 1 | Timm21 | translocase of inner mitochondrial membr | 438 | 336 | 277 | 137 | 172 | 405 | 215 | 180 | 206 | 161 | 412 |
| 260 | 0.06 | 0.23 | 0.27 | 0.79 | 1 | Zmat5 | zinc finger, matrin type 5 | 327 | 230 | 332 | 270 | 253 | 227 | 215 | 261 | 261 | 247 | 238 |
| 234 | -0.08 | 0.28 | -0.28 | 0.78 | 1 | Mvd | mevalonate (diphospho) decarboxylase | 276 | 155 | 165 | 248 | 286 | 259 | 215 | 184 | 274 | 290 | 226 |
| 232 | 0.22 | 0.25 | 0.88 | 0.38 | 1 | Mrpl22 | mitochondrial ribosomal protein L22 | 229 | 257 | 246 | 175 | 188 | 267 | 215 | 236 | 241 | 285 | 215 |
| 205 | -0.39 | 0.30 | -1.31 | 0.19 | 1 | Pisd-ps2 | | 196 | 189 | 98 | 123 | 98 | 621 | 215 | 118 | 106 | 125 | 361 |
| 170 | 0.38 | 0.40 | 0.97 | 0.33 | 1 | Mrps18c | mitochondrial ribosomal protein S18C | 150 | 107 | 231 | 105 | 135 | 154 | 215 | 272 | 162 | 182 | 161 |
| 159 | -0.47 | 0.37 | -1.28 | 0.20 | 1 | Caly | calcyon neuron-specific vesicular protein | 220 | 312 | 211 | 125 | 118 | 108 | 215 | 192 | 95 | 103 | 51 |
| 289 | -0.02 | 0.24 | -0.10 | 0.92 | 1 | Sirt6 | sirtuin 6 | 232 | 241 | 323 | 244 | 283 | 452 | 214 | 161 | 301 | 276 | 451 |
| 272 | -0.29 | 0.28 | -1.01 | 0.31 | 1 | Tarbp1 | TAR RNA binding protein 1 | 343 | 228 | 156 | 190 | 267 | 576 | 214 | 154 | 190 | 229 | 442 |
| 261 | 0.03 | 0.23 | 0.11 | 0.91 | 1 | Srrd | SRR1 domain containing | 281 | 328 | 267 | 296 | 192 | 281 | 214 | 279 | 235 | 166 | 329 |
| 228 | -0.31 | 0.32 | -0.95 | 0.34 | 1 | Ppp1r3c | protein phosphatase 1, regulatory subuni | 421 | 296 | 516 | 161 | 181 | 135 | 214 | 68 | 167 | 157 | 195 |

| | | | | | | | | | | | | | | | | | |
|---|---|---|---|---|---|---|---|---|---|---|---|---|---|---|---|---|---|
| 219 | -0.02 | 0.27 | -0.09 | 0.93 | 1 | Eif3j2 | eukaryotic translation initiation factor 3, su | 245 | 251 | 182 | 231 | 261 | 167 | 214 | 171 | 254 | 246 | 184 |
| 174 | -0.42 | 0.28 | -1.51 | 0.13 | 1 | Ntm | neurotrimin | 169 | 286 | 215 | 193 | 180 | 133 | 214 | 137 | 136 | 147 | 112 |
| 261 | 0.04 | 0.28 | 0.13 | 0.90 | 1 | Ndufaf1 | NADH:ubiquinone oxidoreductase compl | 230 | 225 | 402 | 152 | 300 | 320 | 213 | 238 | 244 | 139 | 410 |
| 255 | 0.02 | 0.29 | 0.06 | 0.96 | 1 | Tsen15 | tRNA splicing endonuclease subunit 15 | 313 | 385 | 283 | 283 | 176 | 234 | 213 | 160 | 191 | 304 | 261 |
| 243 | 0.10 | 0.21 | 0.47 | 0.64 | 1 | Lsm2 | LSM2 homolog, U6 small nuclear RNA ar | 295 | 313 | 286 | 202 | 218 | 223 | 213 | 258 | 222 | 227 | 211 |
| 239 | 0.42 | 0.34 | 1.23 | 0.22 | 1 | Mcoln3 | mucolipin 3 | 15 | 6 | 24 | 367 | 289 | 234 | 213 | 455 | 425 | 341 | 264 |
| 228 | -0.42 | 0.33 | -1.27 | 0.21 | 1 | Zfp677 | zinc finger protein 677 | 253 | 177 | 205 | 267 | 242 | 306 | 213 | 107 | 235 | 134 | 367 |
| 225 | 0.19 | 0.36 | 0.52 | 0.61 | 1 | Pcdhga9 | protocadherin gamma subfamily A, 9 | 68 | 257 | 213 | 253 | 317 | 147 | 213 | 251 | 274 | 259 | 225 |
| 221 | -0.42 | 0.27 | -1.53 | 0.13 | 1 | Mtrf1 | mitochondrial translational release factor | 209 | 195 | 276 | 245 | 286 | 240 | 213 | 107 | 260 | 180 | 215 |
| 144 | -0.28 | 0.39 | -0.70 | 0.49 | 1 | Tmem169 | transmembrane protein 169 | 25 | 59 | 41 | 128 | 227 | 221 | 213 | 126 | 250 | 120 | 176 |
| 302 | -0.19 | 0.28 | -0.67 | 0.50 | 1 | Dmrtc1a | DMRT-like family C1a | 478 | 742 | 603 | 193 | 225 | 168 | 212 | 127 | 165 | 236 | 172 |
| 295 | 0.41 | 0.26 | 1.57 | 0.12 | 1 | S100a13 | S100 calcium binding protein A13 | 522 | 380 | 472 | 222 | 217 | 150 | 212 | 399 | 253 | 237 | 187 |
| 291 | -0.06 | 0.31 | -0.18 | 0.85 | 1 | Amer1 | APC membrane recruitment 1 | 387 | 434 | 405 | 256 | 215 | 302 | 212 | 351 | 282 | 104 | 250 |
| 285 | 0.17 | 0.26 | 0.66 | 0.51 | 1 | Lrguk | leucine-rich repeats and guanylate kinase | 118 | 183 | 154 | 215 | 310 | 534 | 212 | 200 | 378 | 287 | 539 |
| 280 | 0.24 | 0.27 | 0.90 | 0.37 | 1 | Siva1 | SIVA1, apoptosis-inducing factor | 309 | 270 | 360 | 262 | 235 | 260 | 212 | 423 | 288 | 249 | 211 |
| 253 | -0.05 | 0.27 | -0.20 | 0.84 | 1 | Tmem35b | transmembrane protein 35B | 177 | 255 | 396 | 307 | 246 | 232 | 212 | 291 | 233 | 210 | 224 |
| 241 | 0.09 | 0.25 | 0.38 | 0.70 | 1 | Tlcd1 | TLC domain containing 1 | 219 | 270 | 181 | 244 | 220 | 274 | 212 | 341 | 259 | 218 | 209 |
| 200 | -0.30 | 0.33 | -0.91 | 0.36 | 1 | 3425401B19Ril | RIKEN cDNA 3425401B19 gene | 143 | 190 | 200 | 334 | 229 | 134 | 212 | 158 | 237 | 281 | 85 |
| 180 | 0.05 | 0.25 | 0.21 | 0.84 | 1 | Fdxacb1 | ferredoxin-fold anticodon binding domain | 157 | 181 | 111 | 187 | 140 | 213 | 212 | 192 | 155 | 179 | 254 |
| 480 | 0.02 | 0.24 | 0.10 | 0.92 | 1 | Zfas1 | | 842 | 819 | 1192 | 248 | 279 | 468 | 211 | 280 | 184 | 280 | 476 |
| 284 | 0.14 | 0.23 | 0.63 | 0.53 | 1 | Tmem216 | transmembrane protein 216 | 266 | 397 | 325 | 282 | 216 | 313 | 211 | 238 | 267 | 287 | 322 |
| 227 | -0.27 | 0.44 | -0.61 | 0.54 | 1 | Cip2a | cell proliferation regulating inhibitor of prc | 395 | 190 | 115 | 316 | 275 | 181 | 211 | 162 | 264 | 146 | 239 |
| 219 | 0.27 | 0.28 | 0.94 | 0.35 | 1 | Paqr3 | progestin and adipoQ receptor family mer | 127 | 209 | 170 | 228 | 217 | 207 | 211 | 228 | 241 | 348 | 225 |
| 198 | 0.04 | 0.33 | 0.12 | 0.90 | 1 | Gm15651 | | 34 | 21 | 48 | 266 | 312 | 236 | 211 | 377 | 239 | 232 | 205 |
| 189 | 0.09 | 0.30 | 0.29 | 0.77 | 1 | Apoo | apolipoprotein O | 155 | 216 | 134 | 200 | 216 | 141 | 211 | 158 | 241 | 215 | 193 |
| 181 | 0.06 | 0.31 | 0.19 | 0.85 | 1 | Dnaic2 | dynein, axonemal, intermediate chain 2 | 39 | 65 | 74 | 264 | 207 | 207 | 211 | 174 | 275 | 255 | 224 |
| 179 | -0.32 | 0.34 | -0.94 | 0.35 | 1 | Fam161b | family with sequence similarity 161, meml | 162 | 69 | 148 | 163 | 185 | 330 | 211 | 182 | 120 | 137 | 261 |
| 177 | -0.13 | 0.25 | -0.51 | 0.61 | 1 | Slc16a13 | solute carrier family 16 (monocarboxylic a | 194 | 214 | 137 | 221 | 164 | 141 | 211 | 170 | 169 | 183 | 142 |
| 175 | 0.37 | 0.32 | 1.16 | 0.24 | 1 | Nek5 | NIMA (never in mitosis gene a)-related ex | 78 | 104 | 152 | 141 | 177 | 171 | 211 | 242 | 196 | 231 | 228 |
| 163 | 0.23 | 0.28 | 0.81 | 0.42 | 1 | Nme4 | NME/NM23 nucleoside diphosphate kina | 113 | 160 | 125 | 154 | 157 | 117 | 211 | 229 | 197 | 216 | 117 |
| 343 | 0.31 | 0.27 | 1.16 | 0.25 | 1 | Taf1a | TATA-box binding protein associated fact | 392 | 307 | 336 | 186 | 230 | 604 | 210 | 377 | 177 | 263 | 697 |
| 278 | -0.33 | 0.31 | -1.05 | 0.30 | 1 | Fam167a | family with sequence similarity 167, meml | 258 | 459 | 244 | 312 | 304 | 353 | 210 | 181 | 310 | 224 | 208 |
| 223 | -0.17 | 0.26 | -0.67 | 0.50 | 1 | Ndufaf2 | NADH:ubiquinone oxidoreductase compl | 243 | 258 | 318 | 270 | 233 | 156 | 210 | 176 | 212 | 173 | 201 |
| 4531 | -0.82 | 1.32 | -0.62 | 0.53 | 1 | Fn1 | fibronectin 1 | 27068 | 10822 | 11656 | 0 | 4 | 2 | 209 | 6 | 2 | 75 | 0 |
| 233 | 0.10 | 0.31 | 0.32 | 0.75 | 1 | Zfp772 | zinc finger protein 772 | 139 | 305 | 277 | 232 | 163 | 292 | 209 | 263 | 180 | 243 | 261 |
| 202 | 0.11 | 0.29 | 0.38 | 0.70 | 1 | Rsph1 | radial spoke head 1 homolog (Chlamydor | 175 | 211 | 254 | 232 | 145 | 168 | 209 | 248 | 239 | 207 | 136 |
| 197 | 0.23 | 0.31 | 0.73 | 0.47 | 1 | Adora2b | adenosine A2b receptor | 55 | 26 | 49 | 139 | 273 | 326 | 209 | 246 | 265 | 171 | 408 |
| 194 | 0.08 | 0.39 | 0.21 | 0.83 | 1 | Jakmip1 | janus kinase and microtubule interacting | 18 | 29 | 74 | 242 | 214 | 318 | 209 | 285 | 194 | 233 | 317 |
| 185 | -0.29 | 0.40 | -0.72 | 0.47 | 1 | Zfp783 | | 208 | 286 | 127 | 67 | 71 | 440 | 209 | 65 | 86 | 102 | 372 |
| 172 | -0.33 | 0.33 | -0.98 | 0.33 | 1 | Cenpq | centromere protein Q | 226 | 185 | 159 | 192 | 148 | 185 | 209 | 144 | 185 | 94 | 160 |
| 163 | -0.38 | 0.34 | -1.10 | 0.27 | 1 | Mrm3 | mitochondrial rRNA methyltransferase 3 | 142 | 169 | 219 | 246 | 135 | 130 | 209 | 123 | 146 | 130 | 146 |
| 155 | -0.25 | 0.27 | -0.92 | 0.36 | 1 | Gm46620 | | 120 | 143 | 98 | 200 | 184 | 141 | 209 | 152 | 147 | 152 | 158 |
| 133 | -0.22 | 0.35 | -0.63 | 0.53 | 1 | Gm42809 | | 86 | 98 | 75 | 127 | 171 | 141 | 209 | 90 | 181 | 133 | 153 |
| 332 | -0.18 | 0.24 | -0.75 | 0.45 | 1 | Wasf3 | WAS protein family, member 3 | 616 | 536 | 521 | 284 | 195 | 363 | 208 | 189 | 244 | 165 | 329 |
| 316 | 0.11 | 0.25 | 0.46 | 0.65 | 1 | Gtdc1 | glycosyltransferase-like domain containin | 438 | 446 | 476 | 224 | 337 | 246 | 208 | 277 | 269 | 354 | 203 |
| 309 | 0.27 | 0.36 | 0.73 | 0.47 | 1 | Pbk | PDZ binding kinase | 179 | 162 | 87 | 472 | 405 | 232 | 208 | 762 | 296 | 243 | 354 |
| 297 | 0.13 | 0.24 | 0.55 | 0.58 | 1 | Taf6l | TATA-box binding protein associated fact | 379 | 361 | 314 | 280 | 314 | 255 | 208 | 336 | 238 | 301 | 285 |
| 222 | 0.06 | 0.23 | 0.25 | 0.80 | 1 | Cfap69 | cilia and flagella associated protein 69 | 147 | 187 | 95 | 215 | 129 | 439 | 208 | 229 | 161 | 173 | 464 |
| 214 | 0.14 | 0.22 | 0.62 | 0.54 | 1 | Zfp653 | zinc finger protein 653 | 205 | 206 | 207 | 192 | 152 | 277 | 208 | 199 | 184 | 231 | 292 |
| 200 | -0.60 | 0.51 | -1.18 | 0.24 | 1 | Rbis | ribosomal biogenesis factor | 202 | 222 | 111 | 136 | 208 | 365 | 208 | 103 | 184 | 32 | 431 |
| 194 | -0.10 | 0.34 | -0.30 | 0.76 | 1 | Wasf1 | WAS protein family, member 1 | 338 | 321 | 188 | 172 | 131 | 149 | 208 | 244 | 145 | 98 | 135 |
| 123 | 0.57 | 0.37 | 1.52 | 0.13 | 1 | Tmem35a | transmembrane protein 35A | 49 | 45 | 55 | 95 | 154 | 43 | 208 | 249 | 153 | 222 | 76 |
| 452 | 0.18 | 0.36 | 0.49 | 0.62 | 1 | Tnfaip3 | tumor necrosis factor, alpha-induced prot | 2238 | 621 | 612 | 95 | 140 | 299 | 207 | 232 | 129 | 183 | 221 |
| 391 | -0.27 | 0.30 | -0.90 | 0.37 | 1 | Kcnq1ot1 | | 172 | 223 | 126 | 119 | 198 | 1499 | 207 | 104 | 114 | 220 | 1319 |
| 309 | -0.10 | 0.24 | -0.44 | 0.66 | 1 | Hdac10 | histone deacetylase 10 | 281 | 239 | 242 | 290 | 239 | 622 | 207 | 313 | 201 | 171 | 596 |
| 294 | 0.03 | 0.27 | 0.11 | 0.91 | 1 | Casp6 | caspase 6 | 553 | 395 | 378 | 320 | 217 | 196 | 207 | 350 | 198 | 200 | 217 |
| 273 | 0.12 | 0.27 | 0.43 | 0.66 | 1 | Rcc1 | regulator of chromosome condensation 1 | 589 | 338 | 257 | 195 | 190 | 275 | 207 | 168 | 229 | 277 | 274 |
| 217 | -0.38 | 0.25 | -1.48 | 0.14 | 1 | Zfp119a | zinc finger protein 119a | 295 | 233 | 264 | 214 | 204 | 257 | 207 | 118 | 157 | 157 | 276 |
| 141 | -0.13 | 0.33 | -0.40 | 0.69 | 1 | Gm20712 | | 34 | 67 | 37 | 194 | 189 | 142 | 207 | 204 | 220 | 117 | 141 |
| 423 | -0.06 | 0.29 | -0.21 | 0.84 | 1 | 9130016M20Rik | | 718 | 1622 | 768 | 172 | 182 | 228 | 206 | 163 | 178 | 210 | 202 |

| | | | | | | | | | | | | | | | | | |
|---|---|---|---|---|---|---|---|---|---|---|---|---|---|---|---|---|---|
| 326 | -0.20 | 0.20 | -1.00 | 0.32 | 1 | Qtrt1 | queuine tRNA-ribosyltransferase catalytic | 208 | 200 | 327 | 305 | 315 | 697 | 206 | 222 | 300 | 196 | 606 |
| 251 | 0.20 | 0.25 | 0.79 | 0.43 | 1 | Rpp38 | ribonuclease P/MRP 38 subunit | 210 | 249 | 265 | 293 | 193 | 260 | 206 | 285 | 200 | 308 | 295 |
| 195 | -0.43 | 0.27 | -1.60 | 0.11 | 1 | Pqlc2 | PQ loop repeat containing 2 | 103 | 172 | 170 | 297 | 185 | 284 | 206 | 188 | 150 | 138 | 258 |
| 277 | -0.06 | 0.34 | -0.19 | 0.85 | 1 | Gcfc2 | GC-rich sequence DNA binding factor 2 | 268 | 178 | 127 | 182 | 330 | 562 | 204 | 162 | 345 | 216 | 473 |
| 259 | 0.08 | 0.24 | 0.33 | 0.74 | 1 | Pelo | pelota mRNA surveillance and ribosome | 268 | 208 | 273 | 243 | 300 | 277 | 204 | 314 | 282 | 220 | 261 |
| 235 | 0.00 | 0.27 | 0.01 | 0.99 | 1 | Usp2 | ubiquitin specific peptidase 2 | 341 | 304 | 267 | 235 | 177 | 204 | 204 | 323 | 157 | 225 | 147 |
| 231 | 0.05 | 0.21 | 0.26 | 0.80 | 1 | Fam92b | family with sequence similarity 92, membe | 220 | 284 | 228 | 224 | 165 | 294 | 204 | 238 | 173 | 206 | 306 |
| 230 | -0.08 | 0.26 | -0.29 | 0.77 | 1 | Zfp551 | zinc fingr protein 551 | 180 | 241 | 180 | 268 | 268 | 243 | 204 | 184 | 275 | 187 | 294 |
| 224 | -0.30 | 0.23 | -1.26 | 0.21 | 1 | Zfp94 | zinc finger protein 94 | 247 | 192 | 201 | 258 | 245 | 295 | 204 | 174 | 195 | 204 | 248 |
| 219 | -0.02 | 0.37 | -0.05 | 0.96 | 1 | Zfp93 | zinc finger protein 93 | 225 | 184 | 247 | 298 | 138 | 234 | 204 | 119 | 250 | 265 | 246 |
| 219 | 0.03 | 0.28 | 0.11 | 0.91 | 1 | Zcchc10 | zinc finger, CCHC domain containing 10 | 287 | 229 | 223 | 203 | 221 | 193 | 204 | 277 | 161 | 174 | 239 |
| 208 | -0.29 | 0.32 | -0.91 | 0.36 | 1 | Mars2 | methionine-tRNA synthetase 2 (mitochon | 226 | 199 | 118 | 259 | 232 | 256 | 204 | 262 | 188 | 125 | 215 |
| 141 | -0.05 | 0.33 | -0.15 | 0.88 | 1 | Rell2 | RELT-like 2 | 134 | 125 | 128 | 104 | 95 | 200 | 204 | 136 | 128 | 151 | 144 |
| 585 | -0.19 | 0.21 | -0.88 | 0.38 | 1 | Wipf1 | WAS/WASL interacting protein family, me | 2245 | 1300 | 1428 | 186 | 204 | 183 | 203 | 126 | 216 | 179 | 168 |
| 318 | -0.04 | 0.24 | -0.19 | 0.85 | 1 | Casp7 | caspase 7 | 605 | 608 | 565 | 183 | 270 | 213 | 203 | 266 | 222 | 201 | 161 |
| 290 | -0.17 | 0.22 | -0.77 | 0.44 | 1 | Arntl | aryl hydrocarbon receptor nuclear translo | 418 | 460 | 403 | 260 | 302 | 244 | 203 | 224 | 248 | 171 | 258 |
| 275 | 0.36 | 0.31 | 1.18 | 0.24 | 1 | Rel | reticuloendotheliosis oncogene | 631 | 329 | 257 | 207 | 198 | 181 | 203 | 334 | 215 | 227 | 243 |
| 267 | -0.37 | 0.31 | -1.20 | 0.23 | 1 | Lefty1 | left right determination factor 1 | 407 | 422 | 617 | 324 | 200 | 105 | 203 | 284 | 182 | 100 | 90 |
| 244 | 0.22 | 0.25 | 0.88 | 0.38 | 1 | Nim1k | NIM1 serine/threonine protein kinase | 300 | 299 | 200 | 214 | 245 | 202 | 203 | 272 | 197 | 308 | 249 |
| 229 | 0.19 | 0.27 | 0.71 | 0.48 | 1 | Rmdn2 | regulator of microtubule dynamics 2 | 167 | 253 | 181 | 262 | 206 | 221 | 203 | 251 | 247 | 341 | 188 |
| 223 | 0.04 | 0.27 | 0.16 | 0.87 | 1 | Arrdc2 | arrestin domain containing 2 | 296 | 250 | 261 | 105 | 163 | 370 | 203 | 167 | 164 | 209 | 268 |
| 217 | 0.12 | 0.27 | 0.44 | 0.66 | 1 | Zfp948 | zinc finger protein 948 | 272 | 155 | 162 | 195 | 112 | 353 | 203 | 200 | 148 | 185 | 398 |
| 197 | -0.48 | 0.29 | -1.70 | 0.09 | 1 | 2010204K13Rik | | 152 | 216 | 276 | 219 | 273 | 184 | 203 | 229 | 171 | 97 | 148 |
| 183 | 0.12 | 0.33 | 0.37 | 0.71 | 1 | Ptges3-ps | | 154 | 198 | 128 | 195 | 165 | 136 | 203 | 400 | 154 | 135 | 142 |
| 126 | 0.11 | 0.35 | 0.33 | 0.74 | 1 | Nacad | NAC alpha domain containing | 101 | 56 | 34 | 84 | 108 | 171 | 203 | 105 | 124 | 277 | 119 |
| 435 | -0.26 | 0.21 | -1.24 | 0.21 | 1 | Clmn | calmin | 970 | 907 | 1143 | 300 | 301 | 163 | 202 | 197 | 265 | 167 | 171 |
| 230 | -0.06 | 0.29 | -0.21 | 0.83 | 1 | Hdac8 | histone deacetylase 8 | 208 | 464 | 260 | 179 | 205 | 217 | 202 | 256 | 157 | 164 | 219 |
| 224 | -0.19 | 0.29 | -0.67 | 0.50 | 1 | Ajm1 | apical junction component 1 | 141 | 95 | 127 | 274 | 267 | 359 | 202 | 193 | 165 | 280 | 355 |
| 182 | 0.27 | 0.30 | 0.90 | 0.37 | 1 | Shc4 | SHC (Src homology 2 domain containing) | 118 | 145 | 121 | 214 | 154 | 153 | 202 | 182 | 172 | 352 | 187 |
| 154 | -0.11 | 0.28 | -0.38 | 0.70 | 1 | Zfp455 | zinc finger protein 455 | 157 | 131 | 116 | 163 | 203 | 107 | 202 | 131 | 188 | 164 | 131 |
| 146 | 0.09 | 0.35 | 0.25 | 0.80 | 1 | Pcdha11 | protocadherin alpha 10 | 33 | 87 | 56 | 215 | 137 | 143 | 202 | 195 | 152 | 213 | 172 |
| 134 | 0.56 | 0.41 | 1.36 | 0.17 | 1 | Gabra4 | gamma-aminobutyric acid (GABA) A rece | 135 | 204 | 120 | 136 | 42 | 31 | 202 | 232 | 107 | 238 | 31 |
| 134 | -0.16 | 0.41 | -0.39 | 0.70 | 1 | Gm50303 | | 6 | 14 | 28 | 155 | 172 | 213 | 202 | 185 | 192 | 139 | 163 |
| 411 | -0.37 | 0.21 | -1.72 | 0.08 | 1 | Nab2 | Ngfi-A binding protein 2 | 1071 | 1155 | 829 | 235 | 214 | 173 | 201 | 197 | 163 | 120 | 158 |
| 406 | -0.26 | 0.31 | -0.86 | 0.39 | 1 | Lrrc8b | leucine rich repeat containing 8 family, m | 1005 | 1049 | 1098 | 185 | 146 | 174 | 201 | 127 | 231 | 81 | 173 |
| 372 | 0.21 | 0.33 | 0.64 | 0.52 | 1 | Scn1b | sodium channel, voltage-gated, type I, be | 775 | 303 | 541 | 214 | 574 | 167 | 201 | 235 | 610 | 273 | 200 |
| 247 | -0.02 | 0.47 | -0.04 | 0.97 | 1 | Abhd18 | abhydrolase domain containing 18 | 145 | 215 | 95 | 99 | 161 | 718 | 201 | 49 | 215 | 250 | 571 |
| 231 | -0.18 | 0.33 | -0.56 | 0.58 | 1 | Fzd5 | frizzled class receptor 5 | 285 | 304 | 242 | 259 | 273 | 156 | 201 | 348 | 233 | 104 | 140 |
| 207 | 0.25 | 0.26 | 0.95 | 0.34 | 1 | Gstk1 | glutathione S-transferase kappa 1 | 219 | 260 | 178 | 182 | 197 | 149 | 201 | 321 | 181 | 238 | 154 |
| 199 | 0.07 | 0.30 | 0.22 | 0.82 | 1 | Siah1b | siah E3 ubiquitin protein ligase 1B | 196 | 215 | 200 | 172 | 208 | 181 | 201 | 285 | 217 | 122 | 195 |
| 195 | -0.14 | 0.29 | -0.49 | 0.63 | 1 | Rbm20 | RNA binding motif protein 20 | 148 | 173 | 164 | 159 | 191 | 289 | 201 | 151 | 185 | 112 | 374 |
| 150 | -0.45 | 0.32 | -1.40 | 0.16 | 1 | Map2k6 | mitogen-activated protein kinase kinase 6 | 232 | 170 | 236 | 147 | 141 | 99 | 201 | 98 | 124 | 90 | 108 |
| 551 | -0.12 | 0.20 | -0.58 | 0.56 | 1 | Fam81a | family with sequence similarity 81, membe | 1305 | 1511 | 1668 | 183 | 243 | 185 | 200 | 181 | 274 | 128 | 181 |
| 305 | -0.21 | 0.20 | -1.04 | 0.30 | 1 | Pcyox1l | prenylcysteine oxidase 1 like | 276 | 529 | 463 | 290 | 308 | 324 | 200 | 263 | 276 | 191 | 234 |
| 281 | -0.41 | 0.36 | -1.14 | 0.26 | 1 | Kifc2 | kinesin family member C2 | 82 | 155 | 122 | 221 | 205 | 931 | 200 | 75 | 233 | 179 | 691 |
| 281 | -0.20 | 0.43 | -0.48 | 0.63 | 1 | Cenpf | centromere protein F | 428 | 207 | 147 | 389 | 308 | 324 | 200 | 410 | 233 | 137 | 307 |
| 271 | 0.07 | 0.27 | 0.26 | 0.80 | 1 | Gpank1 | G patch domain and ankyrin repeats 1 | 220 | 262 | 368 | 405 | 225 | 219 | 200 | 356 | 270 | 213 | 248 |
| 263 | -0.05 | 0.27 | -0.18 | 0.85 | 1 | Kif9 | kinesin family member 9 | 206 | 170 | 210 | 308 | 330 | 333 | 200 | 228 | 267 | 346 | 290 |
| 237 | 0.01 | 0.33 | 0.02 | 0.98 | 1 | Mkks | McKusick-Kaufman syndrome | 223 | 169 | 337 | 236 | 193 | 293 | 200 | 290 | 196 | 130 | 344 |
| 203 | 0.21 | 0.32 | 0.64 | 0.52 | 1 | Rwdd3 | RWD domain containing 3 | 139 | 269 | 153 | 153 | 184 | 239 | 200 | 193 | 186 | 269 | 248 |
| 190 | -0.18 | 0.40 | -0.46 | 0.65 | 1 | Tmem145 | transmembrane protein 145 | 112 | 236 | 211 | 119 | 171 | 332 | 200 | 86 | 179 | 218 | 221 |
| 179 | -0.06 | 0.35 | -0.18 | 0.86 | 1 | Apc2 | APC regulator of WNT signaling pathway | 240 | 172 | 190 | 187 | 146 | 128 | 200 | 254 | 179 | 211 | 57 |
| 173 | -0.06 | 0.37 | -0.16 | 0.87 | 1 | Platr25 | | 79 | 195 | 150 | 195 | 116 | 213 | 200 | 234 | 113 | 113 | 291 |
| 169 | -0.13 | 0.32 | -0.42 | 0.68 | 1 | Herc6 | hect domain and RLD 6 | 172 | 184 | 228 | 158 | 135 | 168 | 200 | 169 | 198 | 108 | 139 |
| 273 | -0.28 | 0.21 | -1.34 | 0.18 | 1 | Mterf4 | mitochondrial transcription termination fac | 323 | 375 | 369 | 248 | 286 | 330 | 199 | 178 | 198 | 198 | 304 |
| 246 | 0.44 | 0.35 | 1.27 | 0.21 | 1 | Itga2 | integrin alpha 2 | 226 | 109 | 138 | 355 | 142 | 265 | 199 | 389 | 290 | 307 | 284 |
| 240 | -0.30 | 0.28 | -1.07 | 0.28 | 1 | Cep41 | centrosomal protein 41 | 389 | 383 | 408 | 228 | 150 | 225 | 199 | 108 | 168 | 188 | 195 |
| 213 | -0.44 | 0.40 | -1.09 | 0.27 | 1 | Dnah2 | dynein, axonemal, heavy chain 2 | 170 | 261 | 179 | 78 | 100 | 617 | 199 | 98 | 115 | 59 | 462 |
| 191 | -0.26 | 0.35 | -0.74 | 0.46 | 1 | E430018J23Rik | RIKEN cDNA E430018J23 gene | 227 | 179 | 231 | 249 | 210 | 138 | 199 | 258 | 149 | 97 | 168 |
| 181 | 0.04 | 0.31 | 0.13 | 0.90 | 1 | Gpr162 | G protein-coupled receptor 162 | 99 | 76 | 44 | 324 | 188 | 136 | 199 | 407 | 161 | 231 | 122 |

| | | | | | | | | | | | | | | | | | |
|---|---|---|---|---|---|---|---|---|---|---|---|---|---|---|---|---|---|
| 179 | 0.15 | 0.33 | 0.44 | 0.66 | 1 | 2310034G01Rik | | 148 | 136 | 176 | 157 | 187 | 167 | 199 | 283 | 201 | 121 | 195 |
| 153 | -0.04 | 0.31 | -0.12 | 0.91 | 1 | Ccdc173 | coiled-coil domain containing 173 | 73 | 106 | 91 | 143 | 110 | 255 | 199 | 140 | 117 | 148 | 305 |
| 597 | -0.14 | 0.31 | -0.45 | 0.65 | 1 | Fam107b | family with sequence similarity 107, meml | 2195 | 1497 | 1582 | 64 | 157 | 286 | 198 | 134 | 150 | 91 | 217 |
| 342 | 0.17 | 0.30 | 0.55 | 0.58 | 1 | Zfp235 | zinc finger protein 235 | 278 | 424 | 272 | 319 | 259 | 579 | 198 | 221 | 330 | 397 | 481 |
| 301 | 0.34 | 0.38 | 0.90 | 0.37 | 1 | Zc3hc1 | zinc finger, C3HC type 1 | 587 | 340 | 553 | 187 | 181 | 220 | 198 | 474 | 187 | 166 | 212 |
| 297 | 0.15 | 0.26 | 0.58 | 0.56 | 1 | Rpa3 | replication protein A3 | 532 | 371 | 487 | 181 | 296 | 218 | 198 | 263 | 214 | 218 | 283 |
| 295 | 0.29 | 0.22 | 1.27 | 0.20 | 1 | Tagap1 | T cell activation GTPase activating proteir | 321 | 345 | 249 | 233 | 237 | 383 | 198 | 313 | 224 | 330 | 409 |
| 279 | 0.19 | 0.26 | 0.72 | 0.47 | 1 | Zc3h11a | zinc finger CCCH type containing 11A | 282 | 176 | 223 | 150 | 236 | 547 | 198 | 258 | 199 | 213 | 581 |
| 258 | -0.52 | 0.29 | -1.79 | 0.07 | 1 | Rab20 | RAB20, member RAS oncogene family | 490 | 317 | 442 | 247 | 250 | 227 | 198 | 125 | 122 | 237 | 180 |
| 252 | -0.26 | 0.29 | -0.91 | 0.36 | 1 | Xpa | xeroderma pigmentosum, complementati | 300 | 339 | 356 | 193 | 242 | 299 | 198 | 218 | 159 | 100 | 361 |
| 239 | -0.10 | 0.26 | -0.37 | 0.71 | 1 | Ccdc51 | coiled-coil domain containing 51 | 196 | 259 | 196 | 301 | 295 | 215 | 198 | 343 | 211 | 149 | 260 |
| 227 | 0.31 | 0.23 | 1.34 | 0.18 | 1 | Rpp30 | ribonuclease P/MRP 30 subunit | 376 | 267 | 294 | 159 | 166 | 172 | 198 | 243 | 195 | 219 | 201 |
| 218 | 0.15 | 0.31 | 0.46 | 0.64 | 1 | Pcdhb20 | protocadherin beta 20 | 151 | 301 | 166 | 202 | 249 | 185 | 198 | 244 | 242 | 299 | 155 |
| 216 | 0.37 | 0.35 | 1.04 | 0.30 | 1 | Gm10095 | | 327 | 399 | 364 | 92 | 164 | 99 | 198 | 253 | 129 | 252 | 103 |
| 178 | 0.35 | 0.41 | 0.85 | 0.40 | 1 | Pbld2 | phenazine biosynthesis-like protein doma | 58 | 145 | 138 | 120 | 182 | 172 | 198 | 371 | 199 | 122 | 247 |
| 174 | -0.13 | 0.30 | -0.44 | 0.66 | 1 | Carnmt1 | carnosine N-methyltransferase 1 | 142 | 214 | 108 | 222 | 176 | 166 | 198 | 178 | 205 | 138 | 169 |
| 162 | 0.38 | 0.37 | 1.02 | 0.31 | 1 | Rasd2 | RASD family, member 2 | 63 | 28 | 95 | 143 | 236 | 97 | 198 | 279 | 274 | 296 | 74 |
| 353 | 0.19 | 0.30 | 0.62 | 0.53 | 1 | Cdhr2 | cadherin-related family member 2 | 356 | 901 | 777 | 171 | 240 | 245 | 197 | 328 | 258 | 223 | 187 |
| 348 | -0.01 | 0.21 | -0.04 | 0.96 | 1 | Metap1d | methionyl aminopeptidase type 1D (mitoc | 127 | 265 | 185 | 245 | 208 | 932 | 197 | 235 | 230 | 160 | 1047 |
| 286 | 0.12 | 0.27 | 0.46 | 0.65 | 1 | Haus7 | HAUS augmin-like complex, subunit 7 | 447 | 283 | 295 | 262 | 288 | 262 | 197 | 308 | 226 | 268 | 307 |
| 276 | 0.19 | 0.22 | 0.85 | 0.40 | 1 | Gpsm2 | G-protein signalling modulator 2 (AGS3-li | 554 | 472 | 362 | 233 | 175 | 166 | 197 | 207 | 230 | 249 | 190 |
| 246 | 0.17 | 0.23 | 0.71 | 0.48 | 1 | AV356131 | | 181 | 310 | 272 | 259 | 288 | 177 | 197 | 278 | 264 | 269 | 207 |
| 211 | 0.14 | 0.29 | 0.50 | 0.62 | 1 | Map10 | microtubule-associated protein 10 | 132 | 217 | 167 | 151 | 276 | 234 | 197 | 218 | 255 | 193 | 280 |
| 125 | -0.25 | 0.33 | -0.77 | 0.44 | 1 | Nxpe3 | neurexophilin and PC-esterase domain fa | 331 | 129 | 116 | 90 | 45 | 117 | 197 | 64 | 57 | 117 | 116 |
| 119 | 0.02 | 0.42 | 0.05 | 0.96 | 1 | 5033421B08Rik | | 3 | 13 | 8 | 132 | 134 | 167 | 197 | 112 | 140 | 169 | 234 |
| 296 | -0.01 | 0.38 | -0.01 | 0.99 | 1 | Prc1 | protein regulator of cytokinesis 1 | 516 | 244 | 191 | 323 | 316 | 309 | 196 | 361 | 254 | 194 | 346 |
| 290 | -0.01 | 0.32 | -0.05 | 0.96 | 1 | Rnf112 | ring finger protein 112 | 264 | 485 | 491 | 234 | 140 | 404 | 196 | 136 | 250 | 183 | 403 |
| 217 | 0.04 | 0.30 | 0.13 | 0.90 | 1 | Zbtb42 | zinc finger and BTB domain containing 42 | 273 | 268 | 195 | 260 | 251 | 116 | 196 | 185 | 186 | 289 | 170 |
| 215 | 0.37 | 0.37 | 0.99 | 0.32 | 1 | Rasl10b | RAS-like, family 10, member B | 285 | 209 | 267 | 209 | 133 | 107 | 196 | 133 | 223 | 501 | 104 |
| 211 | 0.19 | 0.28 | 0.66 | 0.51 | 1 | Nup43 | nucleoporin 43 | 301 | 358 | 294 | 107 | 154 | 184 | 196 | 165 | 204 | 166 | 187 |
| 124 | 0.70 | 0.51 | 1.39 | 0.17 | 1 | Cnr1 | cannabinoid receptor 1 (brain) | 73 | 8 | 31 | 116 | 71 | 109 | 196 | 255 | 127 | 232 | 144 |
| 535 | -0.42 | 0.24 | -1.74 | 0.08 | 1 | Stard8 | START domain containing 8 | 1648 | 1179 | 1439 | 252 | 246 | 225 | 195 | 114 | 239 | 157 | 195 |
| 296 | 0.07 | 0.30 | 0.24 | 0.81 | 1 | 1700037H04Ril | RIKEN cDNA 1700037H04 gene | 625 | 434 | 433 | 151 | 293 | 216 | 195 | 271 | 227 | 267 | 146 |
| 270 | 0.17 | 0.30 | 0.57 | 0.57 | 1 | Cep85l | centrosomal protein 85-like | 503 | 439 | 382 | 159 | 158 | 254 | 195 | 329 | 147 | 229 | 175 |
| 246 | 0.04 | 0.27 | 0.14 | 0.89 | 1 | Acy1 | aminoacylase 1 | 255 | 249 | 222 | 293 | 242 | 243 | 195 | 202 | 229 | 323 | 252 |
| 229 | -0.13 | 0.42 | -0.30 | 0.76 | 1 | Coro2a | coronin, actin binding protein 2A | 710 | 163 | 251 | 212 | 214 | 112 | 195 | 228 | 172 | 162 | 104 |
| 223 | -0.17 | 0.24 | -0.69 | 0.49 | 1 | Txnrd3 | thioredoxin reductase 3 | 350 | 324 | 320 | 226 | 186 | 161 | 195 | 187 | 213 | 144 | 142 |
| 149 | 0.36 | 0.43 | 0.84 | 0.40 | 1 | Fbxl21 | F-box and leucine-rich repeat protein 21 | 46 | 62 | 133 | 152 | 100 | 149 | 195 | 181 | 220 | 261 | 134 |
| 141 | -0.45 | 0.44 | -1.02 | 0.31 | 1 | Mblac1 | metallo-beta-lactamase domain containin | 122 | 166 | 145 | 212 | 125 | 96 | 195 | 165 | 188 | 46 | 85 |
| 680 | 0.07 | 0.40 | 0.18 | 0.86 | 1 | Fmnl3 | formin-like 3 | 2271 | 1679 | 2467 | 116 | 111 | 75 | 194 | 316 | 85 | 81 | 80 |
| 437 | -0.33 | 0.24 | -1.37 | 0.17 | 1 | Pisd-ps1 | | 344 | 333 | 154 | 152 | 142 | 1751 | 194 | 129 | 124 | 137 | 1351 |
| 437 | 0.05 | 0.39 | 0.14 | 0.89 | 1 | Smc2 | structural maintenance of chromosomes 2 | 1215 | 435 | 555 | 382 | 335 | 353 | 194 | 469 | 248 | 232 | 388 |
| 271 | -0.57 | 0.30 | -1.90 | 0.06 | 1 | Angpt2 | angiopoietin 2 | 565 | 521 | 377 | 222 | 295 | 193 | 194 | 98 | 181 | 176 | 155 |
| 265 | 0.02 | 0.28 | 0.06 | 0.95 | 1 | Cep95 | centrosomal protein 95 | 241 | 119 | 195 | 183 | 131 | 680 | 194 | 123 | 186 | 256 | 603 |
| 254 | 0.12 | 0.26 | 0.46 | 0.65 | 1 | Ndufaf5 | NADH:ubiquinone oxidoreductase compl | 278 | 272 | 209 | 269 | 292 | 218 | 194 | 255 | 230 | 307 | 268 |
| 223 | -0.02 | 0.23 | -0.09 | 0.93 | 1 | Npm3 | nucleoplasmin 3 | 383 | 241 | 349 | 175 | 190 | 188 | 194 | 162 | 193 | 176 | 206 |
| 205 | 0.15 | 0.30 | 0.48 | 0.63 | 1 | Mettl15 | methyltransferase like 15 | 198 | 224 | 270 | 180 | 169 | 188 | 194 | 162 | 175 | 313 | 178 |
| 194 | 0.17 | 0.26 | 0.63 | 0.53 | 1 | C530005A16Rik | | 118 | 154 | 135 | 321 | 183 | 134 | 194 | 261 | 228 | 207 | 202 |
| 153 | -0.06 | 0.27 | -0.21 | 0.83 | 1 | Trim13 | tripartite motif-containing 13 | 135 | 92 | 138 | 153 | 181 | 141 | 194 | 148 | 143 | 205 | 148 |
| 415 | -0.29 | 0.25 | -1.18 | 0.24 | 1 | Mylip | myosin regulatory light chain interacting p | 1047 | 838 | 936 | 227 | 338 | 204 | 193 | 190 | 236 | 133 | 221 |
| 367 | 0.01 | 0.30 | 0.04 | 0.97 | 1 | Slc25a30 | solute carrier family 25, member 30 | 698 | 696 | 587 | 307 | 273 | 223 | 193 | 198 | 308 | 395 | 152 |
| 277 | 0.28 | 0.24 | 1.18 | 0.24 | 1 | Mief2 | mitochondrial elongation factor 2 | 174 | 173 | 174 | 270 | 335 | 355 | 193 | 382 | 339 | 310 | 345 |
| 258 | -0.05 | 0.26 | -0.22 | 0.83 | 1 | Tmed1 | transmembrane p24 trafficking protein 1 | 486 | 308 | 452 | 234 | 207 | 179 | 193 | 200 | 225 | 169 | 190 |
| 252 | 0.02 | 0.27 | 0.09 | 0.93 | 1 | Myrip | myosin VIIA and Rab interacting protein | 392 | 269 | 292 | 255 | 248 | 204 | 193 | 205 | 302 | 196 | 216 |
| 232 | 0.32 | 0.21 | 1.55 | 0.12 | 1 | Vcp-rs | | 191 | 211 | 196 | 233 | 222 | 216 | 193 | 307 | 257 | 285 | 238 |
| 225 | -0.34 | 0.30 | -1.13 | 0.26 | 1 | Tle6 | transducin-like enhancer of split 6 | 243 | 340 | 381 | 202 | 220 | 217 | 193 | 258 | 118 | 140 | 159 |
| 215 | 0.09 | 0.32 | 0.29 | 0.77 | 1 | Supt3 | SPT3, SAGA and STAGA complex compo | 457 | 309 | 233 | 156 | 170 | 141 | 193 | 107 | 223 | 192 | 189 |
| 211 | -0.38 | 0.25 | -1.55 | 0.12 | 1 | Pcdhb18 | protocadherin beta 18 | 147 | 236 | 264 | 257 | 277 | 208 | 193 | 202 | 251 | 102 | 186 |
| 185 | 0.25 | 0.37 | 0.68 | 0.50 | 1 | Vegfc | vascular endothelial growth factor C | 487 | 296 | 326 | 72 | 55 | 107 | 193 | 147 | 51 | 192 | 108 |
| 180 | 0.05 | 0.28 | 0.17 | 0.87 | 1 | 1700001L05Rik | | 111 | 178 | 131 | 182 | 196 | 193 | 193 | 228 | 144 | 224 | 204 |

| | | | | | | | | | | | | | | | | | | |
|---|---|---|---|---|---|---|---|---|---|---|---|---|---|---|---|---|---|---|
| 173 | -0.29 | 0.30 | -0.99 | 0.32 | 1 | Vangl2 | VANGL planar cell polarity 2 | 238 | 243 | 201 | 182 | 143 | 150 | 193 | 102 | 164 | 182 | 105 |
| 160 | -0.08 | 0.32 | -0.26 | 0.79 | 1 | Septin5 | septin 5 | 116 | 130 | 104 | 144 | 222 | 163 | 193 | 105 | 201 | 207 | 175 |
| 126 | 0.10 | 0.50 | 0.19 | 0.85 | 1 | Dok7 | docking protein 7 | 9 | 32 | 45 | 138 | 107 | 183 | 193 | 100 | 150 | 211 | 217 |
| 108 | -0.48 | 0.35 | -1.35 | 0.18 | 1 | Pcdha2 | protocadherin alpha 2 | 67 | 53 | 49 | 182 | 89 | 130 | 193 | 79 | 83 | 142 | 120 |
| 90 | -0.52 | 0.47 | -1.11 | 0.27 | 1 | Gsdma | gasdermin A | 6 | 5 | 0 | 146 | 149 | 90 | 193 | 71 | 103 | 140 | 84 |
| 529 | -0.30 | 0.36 | -0.84 | 0.40 | 1 | Pim1 | proviral integration site 1 | 2847 | 1024 | 516 | 126 | 217 | 238 | 192 | 96 | 144 | 223 | 198 |
| 314 | 0.04 | 0.33 | 0.14 | 0.89 | 1 | Zfp51 | zinc finger protein 51 | 314 | 233 | 229 | 396 | 317 | 401 | 192 | 198 | 380 | 337 | 452 |
| 245 | -0.15 | 0.27 | -0.53 | 0.59 | 1 | Usp27x | ubiquitin specific peptidase 27, X chromo | 421 | 369 | 381 | 224 | 181 | 197 | 192 | 280 | 134 | 144 | 177 |
| 206 | -0.02 | 0.29 | -0.08 | 0.94 | 1 | Bbs5 | Bardet-Biedl syndrome 5 (human) | 193 | 230 | 189 | 194 | 155 | 287 | 192 | 158 | 245 | 130 | 295 |
| 201 | 0.18 | 0.37 | 0.49 | 0.62 | 1 | Kcnip2 | Kv channel-interacting protein 2 | 9 | 55 | 29 | 291 | 328 | 182 | 192 | 290 | 399 | 263 | 172 |
| 194 | 0.01 | 0.30 | 0.04 | 0.97 | 1 | Actr3-ps | | 206 | 239 | 283 | 150 | 215 | 138 | 192 | 136 | 169 | 278 | 130 |
| 150 | -0.40 | 0.28 | -1.46 | 0.14 | 1 | Gpx4-ps2 | | 129 | 119 | 213 | 147 | 219 | 121 | 192 | 114 | 149 | 157 | 91 |
| 133 | -0.23 | 0.33 | -0.69 | 0.49 | 1 | 1700086O06Rik | | 107 | 127 | 110 | 123 | 146 | 139 | 192 | 136 | 109 | 103 | 171 |
| 367 | -0.02 | 0.37 | -0.07 | 0.95 | 1 | Gm37459 | | 60 | 209 | 33 | 193 | 170 | 1308 | 191 | 284 | 99 | 250 | 1238 |
| 325 | -0.12 | 0.23 | -0.52 | 0.60 | 1 | Perp | PERP, TP53 apoptosis effector | 324 | 537 | 399 | 417 | 354 | 218 | 191 | 487 | 312 | 183 | 157 |
| 239 | -0.05 | 0.30 | -0.18 | 0.86 | 1 | Gm9794 | | 428 | 346 | 448 | 188 | 217 | 124 | 191 | 244 | 159 | 122 | 163 |
| 237 | 0.21 | 0.26 | 0.83 | 0.41 | 1 | H4c9 | H4 clustered histone 9 | 62 | 164 | 96 | 390 | 267 | 229 | 191 | 323 | 337 | 237 | 310 |
| 229 | -0.09 | 0.24 | -0.37 | 0.71 | 1 | Haus6 | HAUS augmin-like complex, subunit 6 | 432 | 281 | 320 | 204 | 164 | 198 | 191 | 151 | 154 | 228 | 191 |
| 211 | 0.01 | 0.38 | 0.02 | 0.98 | 1 | Nek10 | NIMA (never in mitosis gene a)- related ki | 27 | 104 | 63 | 186 | 112 | 550 | 191 | 317 | 158 | 102 | 510 |
| 179 | -0.27 | 0.46 | -0.59 | 0.56 | 1 | Dph5 | diphthamide biosynthesis 5 | 237 | 193 | 203 | 161 | 123 | 233 | 191 | 227 | 140 | 40 | 217 |
| 144 | -0.18 | 0.36 | -0.50 | 0.62 | 1 | Gng7 | guanine nucleotide binding protein (G prc | 198 | 151 | 193 | 133 | 122 | 99 | 191 | 202 | 102 | 117 | 72 |
| 320 | 0.03 | 0.23 | 0.13 | 0.90 | 1 | Poli | polymerase (DNA directed), iota | 503 | 522 | 401 | 212 | 249 | 389 | 190 | 181 | 249 | 243 | 384 |
| 303 | 0.28 | 0.25 | 1.09 | 0.27 | 1 | Jak3 | Janus kinase 3 | 604 | 511 | 589 | 148 | 114 | 309 | 190 | 215 | 191 | 211 | 254 |
| 241 | 0.08 | 0.29 | 0.28 | 0.78 | 1 | Nlrx1 | NLR family member X1 | 311 | 302 | 379 | 195 | 215 | 189 | 190 | 339 | 205 | 197 | 131 |
| 232 | -0.01 | 0.26 | -0.03 | 0.97 | 1 | Lrrc75a | leucine rich repeat containing 75A | 236 | 207 | 269 | 228 | 263 | 232 | 190 | 191 | 244 | 308 | 180 |
| 223 | -0.10 | 0.23 | -0.43 | 0.67 | 1 | Nhlrc1 | NHL repeat containing 1 | 157 | 280 | 249 | 333 | 217 | 167 | 190 | 293 | 271 | 144 | 151 |
| 130 | -0.22 | 0.41 | -0.54 | 0.59 | 1 | 1700113A16Rik | | 49 | 130 | 65 | 152 | 129 | 164 | 190 | 118 | 169 | 103 | 162 |
| 113 | -0.27 | 0.40 | -0.67 | 0.50 | 1 | Catspere2 | cation channel sperm associated auxiliar | 115 | 72 | 79 | 54 | 154 | 145 | 190 | 90 | 80 | 106 | 157 |
| 360 | 0.32 | 0.33 | 0.96 | 0.34 | 1 | Trim46 | tripartite motif-containing 46 | 336 | 380 | 503 | 192 | 262 | 611 | 189 | 437 | 161 | 281 | 611 |
| 196 | -0.17 | 0.40 | -0.44 | 0.66 | 1 | Aste1 | asteroid homolog 1 | 124 | 268 | 152 | 172 | 152 | 325 | 189 | 80 | 208 | 142 | 347 |
| 179 | -0.47 | 0.33 | -1.42 | 0.16 | 1 | Snhg3 | | 208 | 199 | 258 | 198 | 154 | 196 | 189 | 93 | 173 | 89 | 210 |
| 177 | 0.15 | 0.34 | 0.45 | 0.65 | 1 | Fhit | fragile histidine triad gene | 149 | 127 | 111 | 121 | 214 | 206 | 189 | 226 | 134 | 301 | 173 |
| 162 | -0.32 | 0.37 | -0.87 | 0.38 | 1 | Tubd1 | tubulin, delta 1 | 88 | 136 | 215 | 161 | 150 | 217 | 189 | 83 | 203 | 109 | 227 |
| 157 | -0.74 | 0.43 | -1.72 | 0.09 | 1 | Mfsd4b1 | major facilitator superfamily domain conta | 250 | 298 | 318 | 127 | 74 | 137 | 189 | 32 | 67 | 102 | 131 |
| 126 | -0.33 | 0.41 | -0.79 | 0.43 | 1 | Pcdhga8 | protocadherin gamma subfamily A, 8 | 118 | 68 | 91 | 106 | 199 | 127 | 189 | 151 | 163 | 70 | 102 |
| 779 | -0.32 | 0.33 | -0.98 | 0.33 | 1 | Myof | myoferlin | 3931 | 1913 | 1962 | 51 | 138 | 68 | 188 | 58 | 106 | 62 | 93 |
| 598 | -0.02 | 0.28 | -0.06 | 0.95 | 1 | Gm34759 | | 32 | 26 | 9 | 143 | 139 | 2838 | 188 | 238 | 126 | 117 | 2719 |
| 358 | -0.27 | 0.21 | -1.25 | 0.21 | 1 | Cpt1c | carnitine palmitoyltransferase 1c | 538 | 639 | 593 | 214 | 256 | 551 | 188 | 163 | 239 | 191 | 367 |
| 266 | -0.38 | 0.34 | -1.13 | 0.26 | 1 | Arhgap33 | Rho GTPase activating protein 33 | 168 | 294 | 229 | 333 | 310 | 451 | 188 | 140 | 220 | 285 | 311 |
| 259 | 0.02 | 0.24 | 0.08 | 0.94 | 1 | Dact3 | dishevelled-binding antagonist of beta-ca | 618 | 402 | 405 | 200 | 187 | 125 | 188 | 222 | 159 | 219 | 120 |
| 211 | 0.26 | 0.31 | 0.85 | 0.39 | 1 | Qrsl1 | glutaminyl-tRNA synthase (glutamine-hyc | 127 | 99 | 148 | 223 | 130 | 363 | 188 | 229 | 206 | 291 | 314 |
| 172 | 0.22 | 0.41 | 0.53 | 0.59 | 1 | Alox8 | arachidonate 8-lipoxygenase | 75 | 174 | 157 | 139 | 244 | 96 | 188 | 183 | 229 | 335 | 73 |
| 143 | 0.10 | 0.38 | 0.26 | 0.80 | 1 | Abtb2 | ankyrin repeat and BTB (POZ) domain co | 218 | 83 | 136 | 173 | 100 | 59 | 188 | 204 | 127 | 245 | 39 |
| 124 | -0.46 | 0.38 | -1.22 | 0.22 | 1 | D430020J02Rik | | 168 | 136 | 204 | 106 | 94 | 112 | 188 | 80 | 89 | 72 | 119 |
| 121 | -0.44 | 0.54 | -0.81 | 0.42 | 1 | Cdx4 | caudal type homeobox 4 | 18 | 164 | 73 | 193 | 167 | 64 | 188 | 145 | 101 | 170 | 43 |
| 71 | -0.45 | 0.63 | -0.70 | 0.48 | 1 | Gm3375 | | 252 | 12 | 3 | 17 | 10 | 6 | 188 | 3 | 5 | 270 | 8 |
| 290 | -0.06 | 0.25 | -0.23 | 0.82 | 1 | Acvr2b | activin receptor IIB | 308 | 352 | 335 | 322 | 253 | 348 | 187 | 349 | 268 | 139 | 332 |
| 247 | 0.06 | 0.32 | 0.18 | 0.86 | 1 | Porcn | porcupine O-acyltransferase | 173 | 286 | 166 | 222 | 236 | 389 | 187 | 211 | 374 | 185 | 285 |
| 215 | 0.18 | 0.23 | 0.79 | 0.43 | 1 | Cetn4 | centrin 4 | 256 | 270 | 287 | 235 | 179 | 132 | 187 | 250 | 224 | 179 | 172 |
| 213 | 0.19 | 0.34 | 0.56 | 0.58 | 1 | Bmf | BCL2 modifying factor | 639 | 299 | 421 | 51 | 89 | 160 | 187 | 108 | 139 | 138 | 115 |
| 196 | -0.02 | 0.29 | -0.08 | 0.94 | 1 | Gm11478 | | 335 | 296 | 330 | 134 | 154 | 125 | 187 | 213 | 114 | 161 | 111 |
| 172 | -0.08 | 0.26 | -0.29 | 0.77 | 1 | Fam71e1 | family with sequence similarity 71, membe | 94 | 129 | 88 | 242 | 128 | 265 | 187 | 193 | 154 | 201 | 210 |
| 147 | 0.05 | 0.33 | 0.14 | 0.89 | 1 | Pcdhgb7 | protocadherin gamma subfamily B, 7 | 227 | 169 | 138 | 137 | 110 | 90 | 187 | 152 | 190 | 151 | 63 |
| 606 | 0.01 | 0.39 | 0.02 | 0.98 | 1 | A230057D06Rik | | 97 | 114 | 28 | 97 | 112 | 2858 | 186 | 67 | 170 | 151 | 2788 |
| 415 | -0.30 | 0.22 | -1.37 | 0.17 | 1 | Ankrd16 | ankyrin repeat domain 16 | 132 | 186 | 215 | 194 | 171 | 1543 | 186 | 158 | 164 | 106 | 1506 |
| 295 | 0.15 | 0.26 | 0.58 | 0.56 | 1 | Pdlim7 | PDZ and LIM domain 7 | 720 | 351 | 491 | 149 | 166 | 298 | 186 | 148 | 220 | 307 | 214 |
| 259 | -0.35 | 0.42 | -0.84 | 0.40 | 1 | Otud1 | OTU domain containing 1 | 501 | 383 | 549 | 226 | 206 | 149 | 186 | 275 | 189 | 46 | 135 |
| 246 | -0.07 | 0.30 | -0.25 | 0.80 | 1 | Zbtb49 | zinc finger and BTB domain containing 49 | 209 | 305 | 187 | 214 | 258 | 341 | 186 | 330 | 134 | 180 | 363 |
| 237 | -0.21 | 0.24 | -0.89 | 0.37 | 1 | B3gntl1 | UDP-GlcNAc:betaGal beta-1,3-N-acetylgl | 203 | 143 | 270 | 215 | 250 | 418 | 186 | 154 | 222 | 202 | 345 |
| 233 | 0.07 | 0.25 | 0.26 | 0.79 | 1 | Zfp691 | zinc finger protein 691 | 285 | 266 | 337 | 177 | 232 | 227 | 186 | 243 | 191 | 232 | 190 |

| | | | | | | | | | | | | | | | | | | |
|---|---|---|---|---|---|---|---|---|---|---|---|---|---|---|---|---|---|---|
| 213 | 0.11 | 0.33 | 0.32 | 0.75 | 1 | Rab19 | RAB19, member RAS oncogene family | 190 | 158 | 88 | 237 | 308 | 185 | 186 | 280 | 250 | 251 | 208 |
| 205 | -0.24 | 0.23 | -1.05 | 0.29 | 1 | 2700046G09Rik | | 155 | 244 | 168 | 232 | 221 | 273 | 186 | 220 | 156 | 167 | 234 |
| 198 | 0.14 | 0.27 | 0.54 | 0.59 | 1 | Ankrd37 | ankyrin repeat domain 37 | 187 | 75 | 181 | 107 | 166 | 336 | 186 | 115 | 162 | 183 | 477 |
| 190 | 0.19 | 0.36 | 0.54 | 0.59 | 1 | Nupl2 | nucleoporin like 2 | 147 | 194 | 193 | 121 | 210 | 204 | 186 | 123 | 180 | 335 | 202 |
| 182 | 0.04 | 0.32 | 0.13 | 0.90 | 1 | Gm36543 | | 50 | 64 | 151 | 239 | 230 | 194 | 186 | 265 | 277 | 169 | 183 |
| 181 | -0.06 | 0.49 | -0.13 | 0.90 | 1 | Fam189a1 | family with sequence similarity 189, meml | 81 | 228 | 224 | 203 | 140 | 212 | 186 | 75 | 183 | 282 | 179 |
| 175 | -0.62 | 0.38 | -1.62 | 0.10 | 1 | Cmss1 | cms small ribosomal subunit 1 | 317 | 245 | 246 | 181 | 146 | 143 | 186 | 46 | 147 | 174 | 89 |
| 127 | -0.24 | 0.37 | -0.64 | 0.52 | 1 | Lrriq3 | leucine-rich repeats and IQ motif containi | 60 | 102 | 60 | 175 | 108 | 162 | 186 | 101 | 122 | 108 | 215 |
| 610 | 0.12 | 0.23 | 0.52 | 0.60 | 1 | Snrnp48 | small nuclear ribonucleoprotein 48 (U11/ | 318 | 444 | 311 | 263 | 193 | 2075 | 185 | 366 | 262 | 117 | 2177 |
| 348 | 0.23 | 0.24 | 0.96 | 0.34 | 1 | Zfp287 | zinc finger protein 287 | 331 | 403 | 306 | 232 | 300 | 600 | 185 | 397 | 301 | 238 | 534 |
| 262 | -0.21 | 0.32 | -0.66 | 0.51 | 1 | Rnd2 | Rho family GTPase 2 | 629 | 455 | 482 | 207 | 152 | 158 | 185 | 206 | 177 | 86 | 142 |
| 207 | 0.37 | 0.26 | 1.44 | 0.15 | 1 | 1810014B01Rik | | 178 | 176 | 173 | 197 | 194 | 176 | 185 | 345 | 185 | 251 | 211 |
| 176 | -0.01 | 0.29 | -0.04 | 0.97 | 1 | Gm44509 | | 215 | 166 | 125 | 183 | 190 | 156 | 185 | 131 | 197 | 194 | 189 |
| 105 | 0.25 | 0.36 | 0.69 | 0.49 | 1 | Nrip3 | nuclear receptor interacting protein 3 | 62 | 66 | 36 | 83 | 93 | 100 | 185 | 149 | 120 | 148 | 114 |
| 319 | -0.33 | 0.28 | -1.18 | 0.24 | 1 | Lipe | lipase, hormone sensitive | 758 | 683 | 780 | 134 | 165 | 223 | 184 | 75 | 158 | 210 | 136 |
| 238 | -0.07 | 0.33 | -0.22 | 0.83 | 1 | C630043F03Rik | | 125 | 234 | 170 | 207 | 192 | 443 | 184 | 400 | 110 | 135 | 416 |
| 208 | 0.25 | 0.29 | 0.86 | 0.39 | 1 | Rwdd2b | RWD domain containing 2B | 284 | 218 | 179 | 135 | 232 | 179 | 184 | 229 | 197 | 270 | 179 |
| 182 | -0.09 | 0.38 | -0.23 | 0.82 | 1 | Bid | BH3 interacting domain death agonist | 414 | 310 | 163 | 137 | 123 | 123 | 184 | 59 | 144 | 220 | 130 |
| 169 | -0.07 | 0.36 | -0.21 | 0.84 | 1 | Prim2 | DNA primase, p58 subunit | 374 | 232 | 239 | 96 | 122 | 126 | 184 | 115 | 140 | 79 | 156 |
| 137 | -0.44 | 0.52 | -0.85 | 0.40 | 1 | Gm7353 | | 33 | 67 | 54 | 183 | 168 | 222 | 184 | 34 | 161 | 200 | 198 |
| 551 | 0.00 | 0.17 | -0.02 | 0.98 | 1 | Cat | catalase | 1896 | 1476 | 1381 | 162 | 184 | 123 | 183 | 177 | 176 | 191 | 112 |
| 522 | -0.10 | 0.35 | -0.29 | 0.77 | 1 | Csrnp1 | cysteine-serine-rich nuclear protein 1 | 2056 | 1069 | 932 | 169 | 139 | 380 | 183 | 71 | 270 | 189 | 287 |
| 337 | 0.14 | 0.25 | 0.58 | 0.56 | 1 | Cntn3 | contactin 3 | 123 | 368 | 178 | 570 | 229 | 498 | 183 | 583 | 208 | 312 | 455 |
| 251 | -0.15 | 0.27 | -0.57 | 0.57 | 1 | Snrpg | small nuclear ribonucleoprotein polypepti | 419 | 389 | 417 | 185 | 219 | 217 | 183 | 262 | 166 | 153 | 153 |
| 213 | -0.08 | 0.26 | -0.32 | 0.75 | 1 | Sat2 | spermidine/spermine N1-acetyl transfera: | 118 | 203 | 253 | 280 | 240 | 205 | 183 | 282 | 194 | 152 | 238 |
| 212 | 0.15 | 0.31 | 0.47 | 0.63 | 1 | Ankrd42 | ankyrin repeat domain 42 | 190 | 145 | 286 | 170 | 211 | 241 | 183 | 155 | 172 | 317 | 263 |
| 183 | 0.03 | 0.26 | 0.13 | 0.90 | 1 | Gm42732 | | 99 | 88 | 151 | 158 | 192 | 306 | 183 | 190 | 188 | 194 | 268 |
| 179 | 0.07 | 0.31 | 0.23 | 0.82 | 1 | Sh3d21 | SH3 domain containing 21 | 204 | 222 | 195 | 128 | 172 | 170 | 183 | 249 | 150 | 152 | 147 |
| 177 | 0.31 | 0.26 | 1.18 | 0.24 | 1 | Hexim2 | hexamethylene bis-acetamide inducible 2 | 181 | 190 | 143 | 203 | 117 | 134 | 183 | 285 | 190 | 193 | 131 |
| 176 | 0.39 | 0.27 | 1.42 | 0.15 | 1 | Scp2-ps2 | | 185 | 241 | 178 | 133 | 128 | 134 | 183 | 204 | 206 | 180 | 165 |
| 148 | 0.50 | 0.30 | 1.68 | 0.09 | 1 | A330094K24Rik | | 81 | 64 | 82 | 151 | 167 | 85 | 183 | 268 | 170 | 234 | 144 |
| 250 | -0.30 | 0.24 | -1.24 | 0.21 | 1 | Xkr4 | X-linked Kx blood group related 4 | 331 | 400 | 394 | 176 | 229 | 309 | 182 | 176 | 212 | 109 | 237 |
| 245 | 0.22 | 0.23 | 0.94 | 0.35 | 1 | Dhfr | dihydrofolate reductase | 429 | 382 | 351 | 224 | 156 | 140 | 182 | 200 | 184 | 283 | 157 |
| 228 | 0.08 | 0.26 | 0.30 | 0.76 | 1 | Cyp7b1 | cytochrome P450, family 7, subfamily b, p | 238 | 333 | 292 | 247 | 211 | 145 | 182 | 373 | 172 | 156 | 163 |
| 212 | -0.46 | 0.31 | -1.46 | 0.14 | 1 | Slc38a9 | solute carrier family 38, member 9 | 249 | 295 | 179 | 174 | 179 | 391 | 182 | 122 | 177 | 89 | 299 |
| 208 | 0.04 | 0.25 | 0.17 | 0.87 | 1 | Cep112 | centrosomal protein 112 | 262 | 201 | 399 | 100 | 187 | 245 | 182 | 124 | 199 | 189 | 204 |
| 198 | -0.32 | 0.28 | -1.14 | 0.26 | 1 | Dok1 | docking protein 1 | 405 | 303 | 399 | 154 | 161 | 102 | 182 | 144 | 109 | 109 | 109 |
| 184 | -0.13 | 0.32 | -0.41 | 0.68 | 1 | Rpp21 | ribonuclease P 21 subunit | 234 | 106 | 271 | 191 | 186 | 178 | 182 | 163 | 164 | 178 | 170 |
| 179 | 0.00 | 0.29 | 0.00 | 1.00 | 1 | Vsig10l | V-set and immunoglobulin domain contai | 158 | 91 | 134 | 182 | 199 | 231 | 182 | 219 | 178 | 165 | 231 |
| 561 | -0.20 | 0.22 | -0.89 | 0.37 | 1 | Lmtk3 | lemur tyrosine kinase 3 | 336 | 397 | 489 | 218 | 197 | 2177 | 181 | 137 | 284 | 156 | 1596 |
| 208 | -0.22 | 0.36 | -0.61 | 0.54 | 1 | Pak6 | p21 (RAC1) activated kinase 6 | 409 | 596 | 396 | 90 | 50 | 168 | 181 | 112 | 59 | 111 | 119 |
| 196 | 0.13 | 0.26 | 0.49 | 0.62 | 1 | Tom1 | target of myb1 trafficking protein | 132 | 136 | 122 | 208 | 175 | 285 | 181 | 177 | 259 | 232 | 250 |
| 196 | 0.00 | 0.31 | 0.00 | 1.00 | 1 | Zfp697 | zinc finger protein 697 | 173 | 167 | 214 | 184 | 189 | 227 | 181 | 97 | 286 | 196 | 241 |
| 196 | 0.33 | 0.34 | 0.98 | 0.33 | 1 | Ly96 | lymphocyte antigen 96 | 146 | 173 | 188 | 166 | 236 | 130 | 181 | 400 | 162 | 211 | 159 |
| 188 | 0.25 | 0.30 | 0.83 | 0.41 | 1 | Lsm3 | LSM3 homolog, U6 small nuclear RNA ar | 311 | 223 | 304 | 109 | 136 | 139 | 181 | 215 | 147 | 161 | 143 |
| 172 | 0.14 | 0.27 | 0.50 | 0.62 | 1 | Hddc2 | HD domain containing 2 | 160 | 103 | 124 | 188 | 207 | 145 | 181 | 202 | 193 | 193 | 198 |
| 148 | -0.20 | 0.33 | -0.60 | 0.55 | 1 | Gm41584 | | 31 | 97 | 69 | 236 | 211 | 139 | 181 | 199 | 184 | 129 | 156 |
| 148 | 0.30 | 0.30 | 0.99 | 0.32 | 1 | Pars2 | prolyl-tRNA synthetase (mitochondrial)(pu | 119 | 135 | 131 | 152 | 116 | 108 | 181 | 158 | 137 | 250 | 144 |
| 124 | 0.01 | 0.43 | 0.01 | 0.99 | 1 | Ccdc160 | coiled-coil domain containing 160 | 91 | 63 | 98 | 124 | 105 | 140 | 181 | 192 | 155 | 71 | 139 |
| 284 | 0.41 | 0.24 | 1.70 | 0.09 | 1 | Telo2 | telomere maintenance 2 | 389 | 359 | 407 | 131 | 200 | 350 | 180 | 280 | 230 | 220 | 373 |
| 223 | -0.05 | 0.30 | -0.18 | 0.86 | 1 | Pycard | PYD and CARD domain containing | 602 | 413 | 430 | 157 | 106 | 75 | 180 | 83 | 157 | 165 | 88 |
| 209 | -0.02 | 0.37 | -0.04 | 0.97 | 1 | Nudt10 | nudix (nucleoside diphosphate linked mo | 386 | 559 | 336 | 72 | 132 | 125 | 180 | 140 | 140 | 144 | 84 |
| 167 | -0.20 | 0.28 | -0.72 | 0.47 | 1 | Zfp54 | zinc finger protein 54 | 144 | 101 | 126 | 186 | 197 | 210 | 180 | 129 | 151 | 151 | 268 |
| 146 | -0.18 | 0.31 | -0.56 | 0.57 | 1 | Utp14b | UTP14B small subunit processome comp | 133 | 117 | 76 | 149 | 161 | 184 | 180 | 151 | 130 | 113 | 212 |
| 127 | -0.03 | 0.42 | -0.07 | 0.95 | 1 | 5430403G16Rik | | 146 | 226 | 145 | 94 | 82 | 92 | 180 | 67 | 122 | 119 | 119 |
| 1033 | 0.17 | 0.24 | 0.74 | 0.46 | 1 | Esam | endothelial cell-specific adhesion molecu | 3076 | 2894 | 3391 | 293 | 240 | 214 | 179 | 228 | 252 | 407 | 188 |
| 490 | -0.38 | 0.23 | -1.67 | 0.10 | 1 | Kctd17 | potassium channel tetramerisation domai | 1413 | 1154 | 1303 | 172 | 263 | 239 | 179 | 116 | 162 | 227 | 160 |
| 468 | 0.15 | 0.29 | 0.52 | 0.60 | 1 | Dpy19l1 | dpy-19-like 1 (C. elegans) | 1633 | 854 | 799 | 221 | 238 | 236 | 179 | 241 | 233 | 187 | 322 |
| 247 | -0.30 | 0.22 | -1.39 | 0.17 | 1 | Ccdc120 | coiled-coil domain containing 120 | 317 | 385 | 386 | 258 | 207 | 259 | 179 | 171 | 180 | 175 | 201 |
| 241 | -0.08 | 0.30 | -0.27 | 0.78 | 1 | Mitf | melanogenesis associated transcription f: | 241 | 187 | 265 | 201 | 246 | 372 | 179 | 111 | 293 | 272 | 284 |

| | | | | | | | | | | | | | | | | | | |
|---|---|---|---|---|---|---|---|---|---|---|---|---|---|---|---|---|---|---|
| 237 | 0.25 | 0.25 | 1.01 | 0.31 | 1 | Mettl2 | methyltransferase like 2 | 389 | 359 | 249 | 158 | 109 | 310 | 179 | 191 | 190 | 174 | 303 |
| 224 | -0.05 | 0.29 | -0.17 | 0.86 | 1 | Zfp874b | zinc finger protein 874b | 290 | 266 | 346 | 188 | 201 | 214 | 179 | 249 | 189 | 104 | 234 |
| 223 | -0.76 | 0.39 | -1.95 | 0.05 | 1 | Asb4 | ankyrin repeat and SOCS box-containing | 359 | 644 | 577 | 146 | 147 | 77 | 179 | 65 | 52 | 129 | 74 |
| 221 | 0.06 | 0.25 | 0.22 | 0.82 | 1 | AW146154 | expressed sequence AW146154 | 164 | 200 | 162 | 202 | 201 | 353 | 179 | 154 | 238 | 220 | 359 |
| 203 | 0.07 | 0.27 | 0.27 | 0.79 | 1 | Ankrd35 | ankyrin repeat domain 35 | 106 | 155 | 86 | 252 | 243 | 244 | 179 | 257 | 183 | 258 | 275 |
| 190 | -0.07 | 0.33 | -0.22 | 0.82 | 1 | Rit2 | Ras-like without CAAX 2 | 136 | 193 | 176 | 151 | 221 | 259 | 179 | 241 | 121 | 211 | 198 |
| 180 | 0.09 | 0.29 | 0.30 | 0.76 | 1 | Gm9905 | | 133 | 177 | 134 | 251 | 196 | 129 | 179 | 166 | 220 | 222 | 174 |
| 120 | -0.35 | 0.37 | -0.96 | 0.34 | 1 | Gm13340 | | 102 | 90 | 56 | 157 | 129 | 135 | 179 | 70 | 139 | 157 | 109 |
| 119 | -0.62 | 0.40 | -1.56 | 0.12 | 1 | G730003C15Rik | | 65 | 137 | 156 | 121 | 159 | 114 | 179 | 107 | 85 | 112 | 70 |
| 404 | -0.17 | 0.30 | -0.56 | 0.57 | 1 | Magi3 | membrane associated guanylate kinase, | 1173 | 878 | 1081 | 186 | 146 | 178 | 178 | 162 | 233 | 88 | 146 |
| 347 | -0.30 | 0.24 | -1.26 | 0.21 | 1 | Snhg1 | | 420 | 426 | 595 | 144 | 158 | 733 | 178 | 92 | 157 | 108 | 801 |
| 255 | -0.22 | 0.22 | -0.96 | 0.34 | 1 | Gm48768 | | 263 | 446 | 391 | 272 | 278 | 195 | 178 | 206 | 198 | 178 | 199 |
| 235 | -0.07 | 0.25 | -0.27 | 0.79 | 1 | Gt(ROSA)26Sor | | 206 | 315 | 255 | 270 | 185 | 301 | 178 | 195 | 245 | 194 | 238 |
| 217 | 0.47 | 0.38 | 1.24 | 0.21 | 1 | Chp2 | calcineurin-like EF hand protein 2 | 474 | 193 | 282 | 198 | 181 | 57 | 178 | 357 | 164 | 201 | 104 |
| 171 | 0.18 | 0.34 | 0.54 | 0.59 | 1 | Cfap54 | cilia and flagella associated protein 54 | 128 | 157 | 124 | 155 | 112 | 242 | 178 | 304 | 167 | 129 | 187 |
| 170 | -0.01 | 0.25 | -0.04 | 0.97 | 1 | Lekr1 | leucine, glutamate and lysine rich 1 | 187 | 194 | 137 | 150 | 128 | 219 | 178 | 174 | 106 | 184 | 212 |
| 163 | -0.02 | 0.33 | -0.07 | 0.94 | 1 | Mterf2 | mitochondrial transcription termination fa | 165 | 71 | 103 | 200 | 161 | 196 | 178 | 196 | 178 | 169 | 179 |
| 150 | -0.28 | 0.30 | -0.94 | 0.35 | 1 | 4933406P04Rik | | 85 | 114 | 139 | 224 | 162 | 147 | 178 | 203 | 162 | 100 | 131 |
| 132 | 0.16 | 0.42 | 0.37 | 0.71 | 1 | Gm49654 | | 73 | 93 | 100 | 134 | 81 | 153 | 178 | 248 | 173 | 126 | 91 |
| 312 | 0.02 | 0.24 | 0.09 | 0.93 | 1 | Rpl13a | ribosomal protein L13A | 635 | 542 | 681 | 215 | 184 | 199 | 177 | 156 | 223 | 160 | 265 |
| 293 | -0.09 | 0.20 | -0.43 | 0.67 | 1 | Slc22a15 | solute carrier family 22 (organic anion/cat | 403 | 557 | 487 | 291 | 208 | 236 | 177 | 285 | 200 | 184 | 190 |
| 226 | 0.20 | 0.31 | 0.66 | 0.51 | 1 | Wdr76 | WD repeat domain 76 | 369 | 193 | 293 | 184 | 192 | 205 | 177 | 247 | 212 | 153 | 259 |
| 225 | -0.23 | 0.39 | -0.58 | 0.56 | 1 | Kcnq4 | potassium voltage-gated channel, subfam | 212 | 115 | 148 | 211 | 130 | 519 | 177 | 60 | 188 | 225 | 490 |
| 182 | -0.09 | 0.38 | -0.23 | 0.82 | 1 | 4921531C22Rik | | 94 | 149 | 182 | 242 | 235 | 157 | 177 | 136 | 177 | 314 | 143 |
| 153 | -0.17 | 0.30 | -0.56 | 0.58 | 1 | A830073O21Rik | | 112 | 191 | 109 | 157 | 94 | 248 | 177 | 141 | 104 | 122 | 233 |
| 395 | -0.04 | 0.24 | -0.17 | 0.87 | 1 | Phlda1 | pleckstrin homology like domain, family A | 1223 | 693 | 787 | 163 | 195 | 301 | 176 | 203 | 142 | 223 | 238 |
| 239 | 0.04 | 0.26 | 0.13 | 0.89 | 1 | Wdr45 | WD repeat domain 45 | 169 | 296 | 178 | 224 | 236 | 356 | 176 | 220 | 225 | 238 | 310 |
| 213 | -0.44 | 0.33 | -1.31 | 0.19 | 1 | Tyms | thymidylate synthase | 511 | 221 | 264 | 174 | 217 | 204 | 176 | 131 | 188 | 128 | 126 |
| 199 | 0.24 | 0.33 | 0.73 | 0.47 | 1 | Rps6ka6 | ribosomal protein S6 kinase polypeptide | 190 | 352 | 266 | 132 | 198 | 117 | 176 | 275 | 146 | 205 | 133 |
| 191 | 0.09 | 0.35 | 0.24 | 0.81 | 1 | Agk | acylglycerol kinase | 329 | 296 | 214 | 86 | 150 | 211 | 176 | 173 | 179 | 115 | 174 |
| 173 | 0.14 | 0.36 | 0.38 | 0.70 | 1 | 5830454E08Rik | | 94 | 165 | 190 | 165 | 230 | 125 | 176 | 214 | 147 | 263 | 133 |
| 430 | -0.31 | 0.22 | -1.39 | 0.17 | 1 | Aard | alanine and arginine rich domain contain | 1156 | 1087 | 1313 | 135 | 106 | 238 | 175 | 101 | 100 | 147 | 174 |
| 202 | -0.03 | 0.34 | -0.08 | 0.93 | 1 | Tmem120a | transmembrane protein 120A | 368 | 289 | 151 | 190 | 182 | 160 | 175 | 256 | 166 | 130 | 153 |
| 175 | -0.07 | 0.29 | -0.22 | 0.82 | 1 | Pam16 | presequence translocase-asssociated mo | 114 | 147 | 226 | 144 | 215 | 200 | 175 | 184 | 155 | 144 | 216 |
| 168 | -0.47 | 0.39 | -1.22 | 0.22 | 1 | Gm46603 | | 247 | 239 | 280 | 150 | 175 | 112 | 175 | 228 | 93 | 73 | 78 |
| 160 | -0.47 | 0.56 | -0.82 | 0.41 | 1 | Fcho1 | FCH domain only 1 | 419 | 134 | 160 | 112 | 125 | 197 | 175 | 102 | 138 | 23 | 175 |
| 151 | -0.55 | 0.48 | -1.14 | 0.25 | 1 | Gm14403 | | 172 | 266 | 100 | 130 | 156 | 183 | 175 | 70 | 153 | 50 | 209 |
| 142 | 0.28 | 0.29 | 0.97 | 0.33 | 1 | Snca | synuclein, alpha | 102 | 142 | 68 | 185 | 90 | 119 | 175 | 226 | 115 | 153 | 188 |
| 304 | 0.05 | 0.21 | 0.23 | 0.82 | 1 | Dnm1 | dynamin 1 | 840 | 530 | 624 | 150 | 179 | 153 | 174 | 148 | 158 | 242 | 145 |
| 238 | 0.33 | 0.22 | 1.51 | 0.13 | 1 | Ercc1 | excision repair cross-complementing rode | 317 | 248 | 257 | 165 | 161 | 304 | 174 | 233 | 213 | 209 | 338 |
| 186 | 0.02 | 0.37 | 0.06 | 0.95 | 1 | 8430429K09Rik | | 114 | 162 | 152 | 212 | 212 | 192 | 174 | 93 | 224 | 246 | 263 |
| 143 | -0.10 | 0.47 | -0.21 | 0.84 | 1 | Ccsap | centriole, cilia and spindle associated pro | 112 | 303 | 88 | 174 | 92 | 129 | 174 | 71 | 138 | 153 | 136 |
| 142 | -0.51 | 0.44 | -1.17 | 0.24 | 1 | Thumpd2 | THUMP domain containing 2 | 108 | 179 | 98 | 180 | 152 | 165 | 174 | 57 | 81 | 169 | 199 |
| 136 | 0.05 | 0.50 | 0.10 | 0.92 | 1 | 9330020H09Rik | | 29 | 134 | 104 | 159 | 100 | 122 | 174 | 297 | 106 | 73 | 196 |
| 131 | -0.17 | 0.33 | -0.50 | 0.61 | 1 | Pcdhga1 | protocadherin gamma subfamily A, 1 | 100 | 125 | 81 | 144 | 104 | 181 | 174 | 126 | 134 | 164 | 107 |
| 85 | -0.16 | 0.48 | -0.34 | 0.74 | 1 | 1810019D21Rik | | 14 | 51 | 14 | 123 | 52 | 106 | 174 | 117 | 57 | 149 | 78 |
| 490 | 0.65 | 0.39 | 1.66 | 0.10 | 1 | Col27a1 | collagen, type XXVII, alpha 1 | 394 | 323 | 332 | 67 | 26 | 1570 | 173 | 170 | 100 | 102 | 2130 |
| 368 | -0.02 | 0.29 | -0.08 | 0.93 | 1 | Apobec3 | apolipoprotein B mRNA editing enzyme, c | 1201 | 651 | 516 | 191 | 128 | 385 | 173 | 166 | 193 | 166 | 279 |
| 274 | 0.13 | 0.25 | 0.51 | 0.61 | 1 | Acot9 | acyl-CoA thioesterase 9 | 688 | 442 | 504 | 185 | 177 | 127 | 173 | 236 | 171 | 179 | 138 |
| 252 | 0.47 | 0.38 | 1.26 | 0.21 | 1 | Pnp | purine-nucleoside phosphorylase | 424 | 228 | 329 | 174 | 250 | 147 | 173 | 434 | 256 | 157 | 199 |
| 151 | 0.20 | 0.31 | 0.65 | 0.51 | 1 | Eva1c | eva-1 homolog C (C. elegans) | 88 | 116 | 79 | 182 | 138 | 123 | 173 | 255 | 177 | 243 | 82 |
| 151 | 0.16 | 0.43 | 0.36 | 0.72 | 1 | Gm10451 | | 80 | 184 | 152 | 115 | 78 | 199 | 173 | 218 | 68 | 126 | 264 |
| 147 | 0.28 | 0.35 | 0.79 | 0.43 | 1 | Strip2 | striatin interacting protein 2 | 95 | 123 | 89 | 144 | 143 | 132 | 173 | 103 | 220 | 198 | 203 |
| 135 | 0.03 | 0.39 | 0.07 | 0.94 | 1 | Edaradd | EDAR (ectodysplasin-A receptor)-associa | 38 | 74 | 69 | 172 | 89 | 216 | 173 | 177 | 101 | 198 | 174 |
| 99 | -0.23 | 0.37 | -0.62 | 0.53 | 1 | Depdc7 | DEP domain containing 7 | 88 | 96 | 56 | 112 | 95 | 69 | 173 | 98 | 108 | 155 | 40 |
| 358 | -0.04 | 0.24 | -0.15 | 0.88 | 1 | Uba7 | ubiquitin-like modifier activating enzyme | 801 | 705 | 739 | 139 | 185 | 380 | 172 | 136 | 183 | 242 | 254 |
| 231 | 0.17 | 0.25 | 0.66 | 0.51 | 1 | Pcgf1 | polycomb group ring finger 1 | 154 | 164 | 181 | 289 | 208 | 303 | 172 | 243 | 238 | 274 | 320 |
| 226 | 0.44 | 0.24 | 1.81 | 0.07 | 1 | Accs | 1-aminocyclopropane-1-carboxylate synt | 191 | 129 | 155 | 198 | 193 | 293 | 172 | 280 | 239 | 270 | 360 |
| 214 | -0.31 | 0.38 | -0.81 | 0.42 | 1 | Tmem39b | transmembrane protein 39b | 239 | 166 | 238 | 286 | 257 | 212 | 172 | 198 | 284 | 75 | 231 |
| 208 | -0.09 | 0.27 | -0.34 | 0.73 | 1 | Zdhhc15 | zinc finger, DHHC domain containing 15 | 194 | 179 | 211 | 269 | 225 | 209 | 172 | 171 | 242 | 164 | 251 |

| | | | | | | | | | | | | | | | | | | |
|---|---|---|---|---|---|---|---|---|---|---|---|---|---|---|---|---|---|---|
| 197 | -0.11 | 0.35 | -0.33 | 0.74 | 1 | Nle1 | notchless homolog 1 | 239 | 136 | 234 | 140 | 150 | 317 | 172 | 73 | 278 | 131 | 293 |
| 197 | -0.25 | 0.23 | -1.09 | 0.28 | 1 | Cenpo | centromere protein O | 190 | 189 | 183 | 216 | 213 | 268 | 172 | 159 | 173 | 165 | 237 |
| 149 | -0.55 | 0.32 | -1.73 | 0.08 | 1 | Mpzl3 | myelin protein zero-like 3 | 121 | 149 | 124 | 214 | 192 | 154 | 172 | 182 | 113 | 73 | 146 |
| 142 | -0.05 | 0.33 | -0.15 | 0.88 | 1 | Map3k19 | mitogen-activated protein kinase kinase k | 3 | 10 | 6 | 277 | 183 | 157 | 172 | 244 | 141 | 201 | 169 |
| 428 | -0.48 | 0.30 | -1.57 | 0.12 | 1 | Parm1 | prostate androgen-regulated mucin-like p | 1553 | 1140 | 740 | 242 | 137 | 201 | 171 | 107 | 168 | 124 | 129 |
| 268 | 0.11 | 0.26 | 0.42 | 0.67 | 1 | Mindy4 | MINDY lysine 48 deubiquitinase 4 | 324 | 311 | 389 | 225 | 238 | 291 | 171 | 337 | 209 | 191 | 259 |
| 249 | -0.20 | 0.41 | -0.49 | 0.63 | 1 | Tpx2 | TPX2, microtubule-associated | 747 | 169 | 237 | 224 | 256 | 182 | 171 | 189 | 171 | 220 | 169 |
| 200 | 0.26 | 0.29 | 0.89 | 0.37 | 1 | Ubtd1 | ubiquitin domain containing 1 | 315 | 361 | 233 | 140 | 165 | 95 | 171 | 254 | 135 | 245 | 92 |
| 200 | 0.00 | 0.31 | -0.01 | 1.00 | 1 | Polr1e | polymerase (RNA) I polypeptide E | 234 | 155 | 137 | 131 | 243 | 277 | 171 | 126 | 164 | 255 | 301 |
| 168 | -0.40 | 0.40 | -0.99 | 0.32 | 1 | Zfp119b | zinc finger protein 119b | 196 | 207 | 226 | 159 | 152 | 188 | 171 | 129 | 212 | 46 | 162 |
| 140 | -0.28 | 0.30 | -0.92 | 0.36 | 1 | Rbm41 | RNA binding motif protein 41 | 225 | 162 | 149 | 111 | 96 | 181 | 171 | 89 | 118 | 106 | 136 |
| 134 | -0.39 | 0.39 | -1.01 | 0.31 | 1 | Lrrc18 | leucine rich repeat containing 18 | 117 | 225 | 205 | 144 | 131 | 83 | 171 | 94 | 75 | 151 | 79 |
| 1834 | -0.23 | 0.23 | -0.99 | 0.32 | 1 | Cavin1 | caveolae associated 1 | 8558 | 5359 | 5588 | 97 | 77 | 31 | 170 | 112 | 69 | 73 | 38 |
| 255 | 0.03 | 0.31 | 0.09 | 0.93 | 1 | Naif1 | nuclear apoptosis inducing factor 1 | 406 | 352 | 233 | 288 | 239 | 197 | 170 | 192 | 271 | 280 | 176 |
| 235 | 0.01 | 0.28 | 0.05 | 0.96 | 1 | Rdm1 | RAD52 motif 1 | 304 | 389 | 511 | 120 | 239 | 165 | 170 | 125 | 189 | 209 | 167 |
| 216 | 0.18 | 0.33 | 0.55 | 0.58 | 1 | Tedc1 | tubulin epsilon and delta complex 1 | 214 | 117 | 144 | 306 | 164 | 247 | 170 | 286 | 177 | 314 | 238 |
| 209 | 0.41 | 0.32 | 1.27 | 0.21 | 1 | Tmem191c | transmembrane protein 191C | 55 | 73 | 120 | 132 | 207 | 409 | 170 | 185 | 244 | 323 | 385 |
| 202 | 0.15 | 0.27 | 0.55 | 0.58 | 1 | Chac2 | ChaC, cation transport regulator 2 | 192 | 268 | 279 | 263 | 140 | 139 | 170 | 222 | 200 | 178 | 173 |
| 190 | -0.05 | 0.29 | -0.16 | 0.87 | 1 | Zfp661 | zinc finger protein 661 | 170 | 179 | 211 | 207 | 179 | 208 | 170 | 220 | 202 | 102 | 240 |
| 189 | -0.07 | 0.29 | -0.24 | 0.81 | 1 | Cabcoco1 | ciliary associated calcium binding coiled- | 189 | 240 | 142 | 204 | 256 | 145 | 170 | 193 | 220 | 158 | 160 |
| 182 | -0.24 | 0.26 | -0.93 | 0.35 | 1 | Pttg1 | pituitary tumor-transforming gene 1 | 310 | 199 | 210 | 167 | 163 | 190 | 170 | 111 | 127 | 162 | 197 |
| 166 | -0.14 | 0.32 | -0.43 | 0.67 | 1 | Haus8 | 4HAUS augmin-like complex, subunit 8 | 454 | 242 | 206 | 102 | 127 | 87 | 170 | 118 | 77 | 128 | 113 |
| 157 | 0.26 | 0.29 | 0.91 | 0.36 | 1 | 6330418K02Rik | | 157 | 122 | 183 | 131 | 124 | 153 | 170 | 189 | 165 | 165 | 170 |
| 140 | 0.25 | 0.38 | 0.65 | 0.52 | 1 | Sult2b1 | sulfotransferase family, cytosolic, 2B, mer | 39 | 56 | 84 | 119 | 209 | 107 | 170 | 256 | 190 | 229 | 77 |
| 138 | -0.11 | 0.35 | -0.32 | 0.75 | 1 | Mtus2 | microtubule associated tumor suppressor | 232 | 185 | 126 | 106 | 64 | 173 | 170 | 129 | 77 | 79 | 182 |
| 109 | -0.36 | 0.33 | -1.10 | 0.27 | 1 | Gm15496 | | 99 | 51 | 104 | 84 | 112 | 156 | 170 | 52 | 87 | 144 | 137 |
| 108 | -0.01 | 0.42 | -0.02 | 0.99 | 1 | Lpar2 | lysophosphatidic acid receptor 2 | 96 | 84 | 111 | 81 | 102 | 80 | 170 | 168 | 95 | 153 | 44 |
| 637 | -0.21 | 0.19 | -1.13 | 0.26 | 1 | Mmp15 | matrix metallopeptidase 15 | 1782 | 1621 | 2136 | 247 | 216 | 154 | 169 | 193 | 184 | 187 | 118 |
| 480 | -0.36 | 0.28 | -1.29 | 0.20 | 1 | Asap2 | ArfGAP with SH3 domain, ankyrin repeat | 810 | 1308 | 816 | 429 | 435 | 278 | 169 | 316 | 352 | 107 | 256 |
| 263 | 0.06 | 0.24 | 0.25 | 0.81 | 1 | Tarbp2 | TARBP2, RISC loading complex RNA bin | 176 | 207 | 184 | 268 | 229 | 489 | 169 | 343 | 250 | 164 | 416 |
| 245 | -0.39 | 0.36 | -1.07 | 0.29 | 1 | Zfp668 | zinc finger protein 668 | 345 | 284 | 377 | 343 | 193 | 239 | 169 | 110 | 248 | 133 | 257 |
| 242 | -0.04 | 0.30 | -0.13 | 0.90 | 1 | Tcf7l2 | transcription factor 7 like 2, T cell specific, | 307 | 398 | 309 | 172 | 223 | 270 | 169 | 207 | 151 | 270 | 185 |
| 196 | 0.28 | 0.26 | 1.06 | 0.29 | 1 | Bloc1s2 | biogenesis of lysosomal organelles comp | 189 | 224 | 152 | 158 | 170 | 212 | 169 | 279 | 152 | 202 | 246 |
| 178 | -0.29 | 0.32 | -0.91 | 0.36 | 1 | 9230112E08Rik | | 95 | 76 | 140 | 170 | 155 | 423 | 169 | 94 | 181 | 194 | 257 |
| 167 | -0.52 | 0.32 | -1.65 | 0.10 | 1 | Ubxn11 | UBX domain protein 11 | 128 | 147 | 143 | 210 | 160 | 276 | 169 | 82 | 112 | 139 | 271 |
| 144 | -0.48 | 0.45 | -1.07 | 0.29 | 1 | Ndufb4 | NADH:ubiquinone oxidoreductase subun | 174 | 266 | 165 | 123 | 110 | 161 | 169 | 39 | 144 | 124 | 104 |
| 91 | -0.76 | 0.42 | -1.80 | 0.07 | 1 | Ablim2 | actin-binding LIM protein 2 | 100 | 63 | 86 | 57 | 115 | 128 | 169 | 38 | 85 | 46 | 110 |
| 490 | 0.24 | 0.23 | 1.06 | 0.29 | 1 | Tmem179 | transmembrane protein 179 | 1123 | 1120 | 1022 | 274 | 224 | 300 | 168 | 277 | 316 | 161 | 410 |
| 363 | -0.12 | 0.23 | -0.55 | 0.58 | 1 | Atat1 | alpha tubulin acetyltransferase 1 | 220 | 326 | 381 | 221 | 180 | 1049 | 168 | 200 | 152 | 176 | 916 |
| 259 | -0.23 | 0.29 | -0.80 | 0.42 | 1 | Dennd6b | DENN/MADD domain containing 6B | 106 | 68 | 94 | 99 | 176 | 946 | 168 | 143 | 121 | 97 | 836 |
| 230 | 0.18 | 0.35 | 0.53 | 0.60 | 1 | Rnaseh2c | ribonuclease H2, subunit C | 442 | 239 | 418 | 206 | 184 | 112 | 168 | 299 | 151 | 157 | 152 |
| 178 | 0.18 | 0.39 | 0.46 | 0.64 | 1 | Gmnn | geminin | 325 | 180 | 193 | 140 | 113 | 157 | 168 | 90 | 252 | 125 | 212 |
| 171 | -0.06 | 0.31 | -0.18 | 0.86 | 1 | Gm50107 | | 187 | 283 | 232 | 109 | 98 | 227 | 168 | 149 | 114 | 109 | 200 |
| 163 | 0.15 | 0.31 | 0.49 | 0.63 | 1 | 9230114K14Rik | | 134 | 140 | 134 | 157 | 159 | 161 | 168 | 263 | 140 | 128 | 207 |
| 135 | 0.04 | 0.46 | 0.09 | 0.93 | 1 | Vipr2 | vasoactive intestinal peptide receptor 2 | 282 | 143 | 162 | 208 | 55 | 29 | 168 | 171 | 144 | 100 | 26 |
| 134 | 0.07 | 0.35 | 0.19 | 0.85 | 1 | Paox | polyamine oxidase (exo-N4-amino) | 143 | 151 | 157 | 104 | 89 | 140 | 168 | 169 | 81 | 146 | 130 |
| 249 | 0.00 | 0.35 | -0.01 | 0.99 | 1 | Kcnab2 | potassium voltage-gated channel, shaker | 804 | 321 | 342 | 142 | 182 | 114 | 167 | 67 | 168 | 282 | 151 |
| 222 | 0.12 | 0.26 | 0.46 | 0.64 | 1 | Dll1 | delta like canonical Notch ligand 1 | 343 | 278 | 395 | 193 | 153 | 167 | 167 | 199 | 159 | 222 | 163 |
| 221 | 0.41 | 0.28 | 1.47 | 0.14 | 1 | Lxn | latexin | 426 | 438 | 282 | 102 | 152 | 131 | 167 | 181 | 162 | 188 | 202 |
| 207 | 0.17 | 0.32 | 0.53 | 0.59 | 1 | Gpatch3 | G patch domain containing 3 | 247 | 136 | 221 | 236 | 170 | 215 | 167 | 269 | 211 | 225 | 184 |
| 190 | -0.23 | 0.28 | -0.82 | 0.41 | 1 | Ppp1r13l | protein phosphatase 1, regulatory subuni | 186 | 181 | 221 | 192 | 272 | 178 | 167 | 118 | 178 | 185 | 207 |
| 181 | 0.16 | 0.28 | 0.55 | 0.58 | 1 | Gm46404 | | 195 | 253 | 194 | 138 | 158 | 176 | 167 | 156 | 230 | 157 | 172 |
| 177 | 0.12 | 0.25 | 0.48 | 0.63 | 1 | Dpf3 | D4, zinc and double PHD fingers, family 3 | 206 | 184 | 188 | 138 | 158 | 190 | 167 | 124 | 182 | 198 | 211 |
| 162 | -0.42 | 0.34 | -1.21 | 0.23 | 1 | Nsun6 | NOL1/NOP2/Sun domain family member | 190 | 199 | 175 | 193 | 142 | 188 | 167 | 74 | 164 | 115 | 178 |
| 135 | -0.69 | 0.36 | -1.93 | 0.05 | 1 | Cdyl2 | chromodomain protein, Y chromosome-lil | 245 | 144 | 246 | 130 | 96 | 125 | 167 | 45 | 81 | 100 | 109 |
| 62 | -0.59 | 0.79 | -0.75 | 0.45 | 1 | 1700030C10Rik | | 62 | 13 | 6 | 46 | 5 | 107 | 167 | 1 | 10 | 161 | 103 |
| 593 | 0.27 | 0.25 | 1.08 | 0.28 | 1 | Emp2 | epithelial membrane protein 2 | 1935 | 1464 | 1668 | 237 | 115 | 138 | 166 | 295 | 170 | 216 | 123 |
| 544 | 0.39 | 0.29 | 1.32 | 0.19 | 1 | Psmb10 | proteasome (prosome, macropain) subun | 1359 | 1012 | 1329 | 185 | 366 | 285 | 166 | 444 | 319 | 222 | 293 |
| 228 | -0.25 | 0.23 | -1.13 | 0.26 | 1 | Trim14 | tripartite motif-containing 14 | 345 | 287 | 365 | 177 | 249 | 228 | 166 | 136 | 212 | 164 | 177 |
| 224 | -0.05 | 0.24 | -0.22 | 0.83 | 1 | Nup37 | nucleoporin 37 | 219 | 247 | 244 | 248 | 232 | 240 | 166 | 218 | 257 | 134 | 257 |

| | | | | | | | | | | | | | | | | | |
|---|---|---|---|---|---|---|---|---|---|---|---|---|---|---|---|---|---|
| 192 | -0.19 | 0.28 | -0.66 | 0.51 | 1 | Pwwp2b | PWWP domain containing 2B | 263 | 228 | 303 | 239 | 181 | 101 | 166 | 199 | 172 | 189 | 66 |
| 171 | -0.24 | 0.26 | -0.89 | 0.37 | 1 | 1810010H24Ril | RIKEN cDNA 1810010H24 gene | 167 | 279 | 226 | 176 | 163 | 151 | 166 | 138 | 158 | 129 | 133 |
| 163 | -0.55 | 0.45 | -1.23 | 0.22 | 1 | Pkmyt1 | protein kinase, membrane associated tyrc | 178 | 158 | 120 | 221 | 110 | 271 | 166 | 39 | 115 | 158 | 253 |
| 131 | 0.09 | 0.37 | 0.24 | 0.81 | 1 | Gm5823 | | 101 | 108 | 87 | 139 | 155 | 49 | 166 | 213 | 153 | 249 | 24 |
| 117 | -0.64 | 0.49 | -1.30 | 0.19 | 1 | Lrp2bp | Lrp2 binding protein | 65 | 167 | 86 | 182 | 149 | 99 | 166 | 45 | 155 | 90 | 87 |
| 307 | 0.01 | 0.26 | 0.03 | 0.97 | 1 | Nepro | nucleolus and neural progenitor protein | 281 | 295 | 249 | 420 | 275 | 429 | 165 | 323 | 278 | 229 | 435 |
| 298 | 0.09 | 0.26 | 0.35 | 0.72 | 1 | Rbak | RB-associated KRAB zinc finger | 305 | 209 | 224 | 235 | 264 | 592 | 165 | 232 | 250 | 282 | 517 |
| 245 | 0.28 | 0.25 | 1.11 | 0.27 | 1 | Mbip | MAP3K12 binding inhibitory protein 1 | 228 | 266 | 196 | 164 | 188 | 390 | 165 | 255 | 155 | 249 | 438 |
| 204 | 0.08 | 0.33 | 0.25 | 0.80 | 1 | Sars2 | seryl-aminoacyl-tRNA synthetase 2 | 196 | 107 | 127 | 143 | 253 | 332 | 165 | 236 | 167 | 241 | 273 |
| 198 | -0.36 | 0.30 | -1.18 | 0.24 | 1 | Echdc3 | enoyl Coenzyme A hydratase domain cor | 79 | 203 | 120 | 370 | 169 | 314 | 165 | 196 | 227 | 128 | 209 |
| 195 | 0.33 | 0.37 | 0.88 | 0.38 | 1 | Pomc | pro-opiomelanocortin-alpha | 102 | 123 | 132 | 211 | 185 | 179 | 165 | 547 | 171 | 184 | 147 |
| 179 | -0.30 | 0.29 | -1.05 | 0.29 | 1 | Trim45 | tripartite motif-containing 45 | 249 | 333 | 295 | 129 | 134 | 173 | 165 | 145 | 106 | 134 | 106 |
| 171 | -0.04 | 0.30 | -0.15 | 0.88 | 1 | Dhrs11 | dehydrogenase/reductase (SDR family) n | 180 | 127 | 151 | 166 | 171 | 209 | 165 | 132 | 125 | 265 | 194 |
| 165 | 0.01 | 0.35 | 0.02 | 0.98 | 1 | 5430405H02Rik | | 94 | 143 | 226 | 78 | 216 | 206 | 165 | 117 | 242 | 95 | 237 |
| 145 | -0.02 | 0.47 | -0.03 | 0.97 | 1 | Ccne1 | cyclin E1 | 248 | 71 | 52 | 188 | 169 | 101 | 165 | 184 | 173 | 137 | 109 |
| 131 | 0.05 | 0.35 | 0.14 | 0.88 | 1 | Gm45447 | | 80 | 98 | 155 | 126 | 144 | 109 | 165 | 123 | 125 | 193 | 121 |
| 101 | 0.30 | 0.69 | 0.44 | 0.66 | 1 | Gm16315 | | 29 | 24 | 0 | 114 | 96 | 120 | 165 | 104 | 190 | 149 | 124 |
| 18422 | -0.89 | 1.16 | -0.76 | 0.45 | 1 | Cela3b | chymotrypsin-like elastase family, membe | 24064 | 88732 | 83701 | 3667 | 340 | 16 | 164 | 531 | 171 | 57 | 1204 |
| 273 | 0.08 | 0.28 | 0.27 | 0.78 | 1 | Zscan20 | zinc finger and SCAN domains 20 | 259 | 323 | 246 | 312 | 256 | 341 | 164 | 214 | 273 | 292 | 323 |
| 268 | -0.17 | 0.24 | -0.69 | 0.49 | 1 | Zbtb48 | zinc finger and BTB domain containing 48 | 145 | 259 | 203 | 187 | 208 | 649 | 164 | 196 | 208 | 99 | 624 |
| 241 | -0.17 | 0.23 | -0.76 | 0.45 | 1 | Aldh1b1 | aldehyde dehydrogenase 1 family, memb | 376 | 491 | 428 | 231 | 191 | 128 | 164 | 234 | 171 | 119 | 118 |
| 219 | 0.35 | 0.29 | 1.20 | 0.23 | 1 | Ermard | ER membrane associated RNA degradati | 169 | 165 | 91 | 126 | 179 | 418 | 164 | 224 | 194 | 214 | 463 |
| 185 | -0.30 | 0.33 | -0.89 | 0.37 | 1 | Ccnj | cyclin J | 207 | 224 | 184 | 200 | 166 | 245 | 164 | 90 | 153 | 222 | 177 |
| 175 | -0.16 | 0.29 | -0.56 | 0.58 | 1 | Nek3 | NIMA (never in mitosis gene a)-related ex | 100 | 320 | 110 | 242 | 119 | 202 | 164 | 229 | 95 | 142 | 197 |
| 169 | -0.19 | 0.28 | -0.67 | 0.50 | 1 | Zfp78 | zinc finger protein 78 | 142 | 135 | 197 | 155 | 170 | 246 | 164 | 102 | 194 | 178 | 176 |
| 134 | -0.42 | 0.43 | -0.97 | 0.33 | 1 | 9030612E09Rik | | 129 | 106 | 75 | 181 | 154 | 151 | 164 | 48 | 151 | 178 | 135 |
| 220 | -0.19 | 0.30 | -0.62 | 0.53 | 1 | Spef1 | sperm flagellar 1 | 420 | 466 | 353 | 117 | 152 | 181 | 163 | 79 | 110 | 152 | 226 |
| 186 | -0.06 | 0.27 | -0.24 | 0.81 | 1 | Tmem151a | transmembrane protein 151A | 329 | 331 | 233 | 147 | 159 | 115 | 163 | 139 | 149 | 198 | 88 |
| 185 | 0.03 | 0.35 | 0.07 | 0.94 | 1 | Ass1 | argininosuccinate synthetase 1 | 498 | 296 | 318 | 139 | 63 | 101 | 163 | 147 | 91 | 93 | 123 |
| 181 | -0.06 | 0.34 | -0.18 | 0.86 | 1 | Rnf180 | ring finger protein 180 | 175 | 240 | 213 | 153 | 198 | 178 | 163 | 93 | 184 | 224 | 173 |
| 175 | -0.07 | 0.36 | -0.19 | 0.85 | 1 | Gm17066 | | 76 | 185 | 110 | 149 | 101 | 373 | 163 | 101 | 121 | 151 | 390 |
| 171 | 0.23 | 0.26 | 0.91 | 0.36 | 1 | Phldb3 | pleckstrin homology like domain, family B | 241 | 207 | 160 | 152 | 133 | 128 | 163 | 207 | 159 | 218 | 113 |
| 142 | 0.60 | 0.34 | 1.78 | 0.08 | 1 | Gcat | glycine C-acetyltransferase (2-amino-3-ke | 109 | 114 | 112 | 113 | 114 | 88 | 163 | 275 | 118 | 223 | 128 |
| 112 | 0.69 | 0.45 | 1.53 | 0.13 | 1 | Inhba | inhibin beta-A | 305 | 61 | 71 | 36 | 44 | 82 | 163 | 139 | 61 | 184 | 83 |
| 266 | 0.10 | 0.29 | 0.36 | 0.72 | 1 | Rbmxl1 | RNA binding motif protein, X-linked like-1 | 334 | 303 | 449 | 207 | 266 | 250 | 162 | 173 | 196 | 314 | 271 |
| 238 | 0.09 | 0.29 | 0.31 | 0.75 | 1 | Irak4 | interleukin-1 receptor-associated kinase 4 | 470 | 329 | 308 | 171 | 178 | 216 | 162 | 260 | 163 | 213 | 151 |
| 200 | 0.24 | 0.31 | 0.78 | 0.43 | 1 | Acot2 | acyl-CoA thioesterase 2 | 263 | 278 | 337 | 144 | 183 | 115 | 162 | 219 | 126 | 243 | 131 |
| 185 | 0.21 | 0.26 | 0.81 | 0.42 | 1 | Timm10 | translocase of inner mitochondrial membr | 236 | 202 | 233 | 151 | 149 | 168 | 162 | 222 | 168 | 191 | 151 |
| 175 | 0.32 | 0.26 | 1.20 | 0.23 | 1 | Efna2 | ephrin A2 | 213 | 163 | 148 | 173 | 145 | 138 | 162 | 242 | 194 | 209 | 135 |
| 172 | 0.02 | 0.29 | 0.07 | 0.94 | 1 | Gm32031 | | 126 | 69 | 110 | 172 | 227 | 230 | 162 | 198 | 190 | 176 | 232 |
| 161 | -0.09 | 0.28 | -0.32 | 0.75 | 1 | Rnls | renalase, FAD-dependent amine oxidase | 163 | 148 | 111 | 163 | 215 | 155 | 162 | 166 | 157 | 185 | 146 |
| 160 | 0.00 | 0.27 | 0.00 | 1.00 | 1 | Nme6 | NME/NM23 nucleoside diphosphate kina | 120 | 94 | 101 | 164 | 233 | 163 | 162 | 171 | 195 | 188 | 163 |
| 410 | 0.08 | 0.25 | 0.34 | 0.73 | 1 | Hspa12b | heat shock protein 12B | 1048 | 839 | 1144 | 205 | 163 | 182 | 161 | 231 | 157 | 135 | 242 |
| 336 | 0.14 | 0.25 | 0.56 | 0.57 | 1 | Cyp39a1 | cytochrome P450, family 39, subfamily a, | 592 | 984 | 591 | 202 | 229 | 132 | 161 | 269 | 239 | 126 | 177 |
| 263 | 0.07 | 0.25 | 0.27 | 0.79 | 1 | Tdg | thymine DNA glycosylase | 197 | 204 | 222 | 261 | 316 | 369 | 161 | 196 | 323 | 222 | 426 |
| 239 | -0.16 | 0.25 | -0.64 | 0.52 | 1 | Sh2d5 | SH2 domain containing 5 | 107 | 191 | 269 | 288 | 275 | 381 | 161 | 231 | 270 | 185 | 272 |
| 227 | -0.16 | 0.25 | -0.65 | 0.52 | 1 | Zswim6 | zinc finger SWIM-type containing 6 | 493 | 408 | 349 | 157 | 167 | 175 | 161 | 137 | 157 | 117 | 181 |
| 215 | -0.35 | 0.32 | -1.09 | 0.27 | 1 | Rabl2 | RAB, member RAS oncogene family-like | 202 | 128 | 132 | 190 | 239 | 467 | 161 | 155 | 197 | 100 | 396 |
| 203 | -0.15 | 0.30 | -0.51 | 0.61 | 1 | Nat8f4 | N-acetyltransferase 8 (GCN5-related) fam | 237 | 265 | 345 | 207 | 184 | 162 | 161 | 192 | 234 | 82 | 162 |
| 189 | -0.07 | 0.30 | -0.24 | 0.81 | 1 | 2810001G20Rik | | 215 | 206 | 197 | 162 | 302 | 138 | 161 | 171 | 194 | 161 | 175 |
| 174 | -0.46 | 0.27 | -1.73 | 0.08 | 1 | Pcdhga11 | protocadherin gamma subfamily A, 11 | 192 | 221 | 287 | 225 | 175 | 139 | 161 | 143 | 172 | 106 | 95 |
| 171 | -0.34 | 0.30 | -1.13 | 0.26 | 1 | Prss36 | protease, serine 36 | 161 | 191 | 167 | 175 | 159 | 257 | 161 | 109 | 98 | 193 | 204 |
| 139 | 0.34 | 0.35 | 0.97 | 0.33 | 1 | A330008L17Rik | | 62 | 106 | 101 | 128 | 124 | 140 | 161 | 123 | 187 | 220 | 175 |
| 131 | -0.28 | 0.30 | -0.93 | 0.35 | 1 | Morn1 | MORN repeat containing 1 | 125 | 146 | 145 | 115 | 112 | 173 | 161 | 85 | 110 | 122 | 145 |
| 125 | 0.46 | 0.38 | 1.23 | 0.22 | 1 | Pold4 | polymerase (DNA-directed), delta 4 | 164 | 91 | 144 | 58 | 81 | 126 | 161 | 119 | 177 | 115 | 142 |
| 497 | -0.24 | 0.23 | -1.05 | 0.29 | 1 | Pld1 | phospholipase D1 | 1424 | 1343 | 1461 | 171 | 145 | 192 | 160 | 145 | 182 | 143 | 103 |
| 293 | -0.29 | 0.31 | -0.94 | 0.35 | 1 | Inpp4b | inositol polyphosphate-4-phosphatase, ty | 867 | 654 | 818 | 85 | 119 | 112 | 160 | 36 | 135 | 124 | 114 |
| 289 | 0.38 | 0.32 | 1.21 | 0.23 | 1 | Sertad1 | SERTA domain containing 1 | 823 | 415 | 574 | 142 | 132 | 163 | 160 | 250 | 189 | 134 | 197 |
| 276 | -0.03 | 0.36 | -0.10 | 0.92 | 1 | Krt80 | keratin 80 | 502 | 270 | 264 | 292 | 285 | 267 | 160 | 337 | 266 | 215 | 178 |
| 228 | -0.75 | 0.44 | -1.68 | 0.09 | 1 | Tnfrsf12a | tumor necrosis factor receptor superfamily | 411 | 407 | 531 | 225 | 162 | 158 | 160 | 32 | 122 | 180 | 117 |

| | | | | | | | | | | | | | | | | | | |
|---|---|---|---|---|---|---|---|---|---|---|---|---|---|---|---|---|---|---|
| 227 | 0.36 | 0.33 | 1.08 | 0.28 | 1 | Wdhd1 | WD repeat and HMG-box DNA binding pr | 263 | 209 | 103 | 256 | 147 | 293 | 160 | 302 | 254 | 180 | 328 |
| 224 | -0.09 | 0.33 | -0.29 | 0.78 | 1 | Plpp2 | phospholipid phosphatase 2 | 411 | 347 | 365 | 207 | 163 | 133 | 160 | 129 | 225 | 250 | 74 |
| 200 | 0.29 | 0.25 | 1.15 | 0.25 | 1 | Gm43254 | | 220 | 189 | 190 | 207 | 192 | 160 | 160 | 285 | 192 | 206 | 198 |
| 181 | 0.25 | 0.46 | 0.55 | 0.58 | 1 | Clybl | citrate lyase beta like | 219 | 176 | 363 | 68 | 175 | 156 | 160 | 282 | 161 | 137 | 92 |
| 173 | -0.07 | 0.31 | -0.21 | 0.83 | 1 | Trmo | tRNA methyltransferase O | 261 | 160 | 184 | 224 | 120 | 161 | 160 | 175 | 128 | 142 | 186 |
| 154 | 0.08 | 0.30 | 0.27 | 0.79 | 1 | Gm30551 | | 127 | 91 | 183 | 100 | 198 | 157 | 160 | 72 | 203 | 224 | 177 |
| 118 | 0.38 | 0.40 | 0.95 | 0.34 | 1 | Rpp25 | ribonuclease P/MRP 25 subunit | 67 | 58 | 32 | 137 | 104 | 94 | 160 | 266 | 157 | 100 | 126 |
| 93 | 0.21 | 0.57 | 0.37 | 0.71 | 1 | Eif5al3-ps | | 114 | 110 | 36 | 125 | 11 | 35 | 160 | 294 | 14 | 79 | 40 |
| 548 | -0.02 | 0.28 | -0.06 | 0.95 | 1 | Apod | apolipoprotein D | 2190 | 1111 | 1462 | 211 | 171 | 99 | 159 | 220 | 143 | 134 | 124 |
| 190 | 0.09 | 0.29 | 0.32 | 0.75 | 1 | Panx1 | pannexin 1 | 444 | 232 | 243 | 98 | 173 | 134 | 159 | 137 | 171 | 185 | 110 |
| 189 | 0.16 | 0.30 | 0.55 | 0.59 | 1 | Eda | ectodysplasin-A | 309 | 402 | 287 | 139 | 112 | 105 | 159 | 109 | 165 | 155 | 140 |
| 174 | -0.07 | 0.25 | -0.28 | 0.78 | 1 | | RIKEN cDNA A930007A09 gene | 115 | 171 | 179 | 226 | 177 | 180 | 159 | 193 | 190 | 128 | 201 |
| 171 | -0.40 | 0.32 | -1.28 | 0.20 | 1 | Sirt4 | sirtuin 4 | 122 | 86 | 135 | 153 | 112 | 430 | 159 | 81 | 140 | 93 | 373 |
| 159 | 0.04 | 0.40 | 0.10 | 0.92 | 1 | Tmem120b | transmembrane protein 120B | 157 | 71 | 80 | 131 | 165 | 247 | 159 | 251 | 86 | 165 | 242 |
| 150 | 0.12 | 0.50 | 0.24 | 0.81 | 1 | Lhx1 | LIM homeobox protein 1 | 19 | 71 | 17 | 184 | 210 | 196 | 159 | 122 | 227 | 269 | 176 |
| 149 | 0.01 | 0.31 | 0.02 | 0.98 | 1 | Gm14326 | predicted gene 14326 | 117 | 135 | 159 | 186 | 130 | 134 | 159 | 188 | 135 | 111 | 188 |
| 127 | -0.27 | 0.41 | -0.67 | 0.51 | 1 | Pcdhga2 | protocadherin gamma subfamily A, 2 | 78 | 98 | 78 | 177 | 114 | 165 | 159 | 73 | 114 | 236 | 108 |
| 125 | 0.00 | 0.36 | 0.00 | 1.00 | 1 | Nme5 | NME/NM23 family member 5 | 101 | 138 | 166 | 116 | 119 | 91 | 159 | 125 | 93 | 156 | 108 |
| 104 | -0.13 | 0.33 | -0.38 | 0.70 | 1 | C030015A19Rik | | 112 | 82 | 67 | 97 | 57 | 157 | 159 | 74 | 75 | 129 | 140 |
| 5206 | -1.05 | 0.74 | -1.41 | 0.16 | 1 | H2-Aa | histocompatibility 2, class II antigen A, alp | 29502 | 11115 | 16270 | 13 | 60 | 25 | 158 | 1 | 65 | 36 | 18 |
| 225 | 0.11 | 0.28 | 0.39 | 0.70 | 1 | Phtf2 | putative homeodomain transcription facto | 337 | 320 | 181 | 158 | 171 | 297 | 158 | 226 | 159 | 158 | 305 |
| 182 | -0.44 | 0.27 | -1.63 | 0.10 | 1 | Rnf208 | ring finger protein 208 | 281 | 306 | 262 | 165 | 164 | 167 | 158 | 147 | 102 | 153 | 90 |
| 177 | -0.52 | 0.34 | -1.54 | 0.12 | 1 | Nutf2 | nuclear transport factor 2 | 252 | 192 | 209 | 235 | 174 | 186 | 158 | 83 | 144 | 117 | 199 |
| 177 | -0.29 | 0.32 | -0.89 | 0.37 | 1 | Tatdn3 | TatD DNase domain containing 3 | 148 | 150 | 77 | 135 | 183 | 369 | 158 | 136 | 92 | 161 | 341 |
| 175 | 0.79 | 0.42 | 1.88 | 0.06 | 1 | Bicdl2 | BICD family like cargo adaptor 2 | 81 | 83 | 165 | 118 | 150 | 146 | 158 | 353 | 181 | 346 | 139 |
| 157 | -0.03 | 0.33 | -0.10 | 0.92 | 1 | Mrps36 | mitochondrial ribosomal protein S36 | 147 | 176 | 157 | 179 | 155 | 128 | 158 | 118 | 169 | 246 | 96 |
| 150 | -0.21 | 0.37 | -0.56 | 0.58 | 1 | Zfp850 | zinc finger protein 850 | 208 | 129 | 243 | 159 | 99 | 148 | 158 | 83 | 153 | 106 | 168 |
| 241 | -0.31 | 0.32 | -0.99 | 0.32 | 1 | Dclre1c | DNA cross-link repair 1C | 277 | 303 | 324 | 221 | 114 | 449 | 157 | 168 | 146 | 68 | 426 |
| 232 | -0.85 | 0.50 | -1.70 | 0.09 | 1 | Evc | EvC ciliary complex subunit 1 | 1013 | 518 | 541 | 38 | 72 | 54 | 157 | 79 | 23 | 27 | 36 |
| 222 | -0.13 | 0.28 | -0.44 | 0.66 | 1 | Dolpp1 | dolichyl pyrophosphate phosphatase 1 | 245 | 218 | 188 | 151 | 226 | 399 | 157 | 126 | 156 | 229 | 342 |
| 194 | -0.33 | 0.36 | -0.92 | 0.36 | 1 | Orai1 | ORAI calcium release-activated calcium r | 328 | 410 | 242 | 219 | 148 | 122 | 157 | 87 | 197 | 129 | 95 |
| 192 | 0.31 | 0.25 | 1.24 | 0.21 | 1 | Tcf19 | transcription factor 19 | 288 | 179 | 262 | 152 | 173 | 134 | 157 | 183 | 191 | 210 | 178 |
| 186 | -0.28 | 0.34 | -0.84 | 0.40 | 1 | Sh3bgr | SH3-binding domain glutamic acid-rich pi | 407 | 330 | 387 | 142 | 113 | 84 | 157 | 54 | 125 | 164 | 78 |
| 132 | -0.43 | 0.39 | -1.11 | 0.27 | 1 | Rsl1 | regulator of sex limited protein 1 | 285 | 108 | 137 | 147 | 110 | 105 | 157 | 56 | 83 | 133 | 137 |
| 123 | 0.21 | 0.49 | 0.42 | 0.67 | 1 | Asnsd1 | asparagine synthetase domain containing | 39 | 73 | 87 | 123 | 120 | 91 | 157 | 364 | 110 | 75 | 116 |
| 292 | 0.12 | 0.24 | 0.49 | 0.62 | 1 | Afap1 | actin filament associated protein 1 | 857 | 575 | 577 | 153 | 118 | 153 | 156 | 137 | 174 | 151 | 165 |
| 215 | 0.36 | 0.28 | 1.27 | 0.21 | 1 | Tmem141 | transmembrane protein 141 | 235 | 209 | 215 | 187 | 201 | 197 | 156 | 168 | 228 | 335 | 240 |
| 188 | 0.10 | 0.28 | 0.37 | 0.71 | 1 | Mad2l1bp | MAD2L1 binding protein | 213 | 197 | 345 | 137 | 223 | 112 | 156 | 226 | 217 | 152 | 92 |
| 179 | -0.09 | 0.27 | -0.31 | 0.75 | 1 | Gm42547 | | 121 | 192 | 187 | 156 | 173 | 270 | 156 | 177 | 129 | 158 | 244 |
| 172 | 0.04 | 0.29 | 0.13 | 0.90 | 1 | Zfp354b | zinc finger protein 354B | 119 | 152 | 193 | 165 | 165 | 218 | 156 | 241 | 169 | 146 | 171 |
| 122 | -0.08 | 0.37 | -0.20 | 0.84 | 1 | Srsf12 | serine and arginine-rich splicing factor 12 | 130 | 220 | 126 | 81 | 113 | 92 | 156 | 116 | 109 | 112 | 83 |
| 313 | -0.09 | 0.30 | -0.30 | 0.76 | 1 | Eno3 | enolase 3, beta muscle | 193 | 212 | 257 | 155 | 157 | 951 | 155 | 249 | 127 | 88 | 902 |
| 227 | 0.36 | 0.23 | 1.57 | 0.12 | 1 | Gemin2 | gem nuclear organelle associated proteir | 251 | 264 | 282 | 151 | 193 | 249 | 155 | 205 | 196 | 247 | 305 |
| 212 | -0.06 | 0.27 | -0.24 | 0.81 | 1 | Nphp3 | nephronophthisis 3 (adolescent) | 228 | 274 | 147 | 232 | 102 | 388 | 155 | 239 | 139 | 112 | 312 |
| 210 | -0.05 | 0.28 | -0.19 | 0.85 | 1 | Haus1 | HAUS augmin-like complex, subunit 1 | 335 | 309 | 263 | 263 | 130 | 173 | 155 | 158 | 144 | 187 | 196 |
| 202 | 0.27 | 0.30 | 0.92 | 0.36 | 1 | Zfp784 | zinc finger protein 784 | 248 | 177 | 209 | 192 | 151 | 216 | 155 | 256 | 189 | 267 | 161 |
| 201 | 0.19 | 0.26 | 0.76 | 0.45 | 1 | Fbxw17 | F-box and WD-40 domain protein 17 | 263 | 236 | 194 | 119 | 137 | 313 | 155 | 140 | 188 | 178 | 288 |
| 188 | 0.12 | 0.28 | 0.42 | 0.67 | 1 | Gcnt1 | glucosaminyl (N-acetyl) transferase 1, cor | 200 | 189 | 125 | 208 | 210 | 173 | 155 | 191 | 198 | 211 | 210 |
| 185 | -0.12 | 0.36 | -0.34 | 0.73 | 1 | Smim10l2a | small integral membrane protein 10 like 2 | 94 | 115 | 142 | 210 | 211 | 298 | 155 | 98 | 247 | 252 | 211 |
| 183 | 0.02 | 0.24 | 0.10 | 0.92 | 1 | Smim27 | small integral membrane protein 27 | 150 | 175 | 177 | 228 | 205 | 161 | 155 | 258 | 186 | 164 | 157 |
| 181 | 0.29 | 0.33 | 0.87 | 0.38 | 1 | Cep128 | centrosomal protein 128 | 372 | 266 | 223 | 104 | 116 | 133 | 155 | 232 | 138 | 130 | 126 |
| 176 | -0.02 | 0.27 | -0.08 | 0.94 | 1 | Nup35 | nucleoporin 35 | 183 | 119 | 138 | 168 | 187 | 243 | 155 | 175 | 151 | 187 | 232 |
| 165 | -0.10 | 0.38 | -0.27 | 0.79 | 1 | Srcap | Snf2-related CREBBP activator protein | 112 | 216 | 85 | 212 | 149 | 223 | 155 | 117 | 180 | 201 | 167 |
| 164 | 0.25 | 0.29 | 0.85 | 0.40 | 1 | Ethe1 | ethylmalonic encephalopathy 1 | 192 | 208 | 285 | 109 | 140 | 106 | 155 | 102 | 175 | 188 | 145 |
| 148 | -0.30 | 0.39 | -0.77 | 0.44 | 1 | Gm10699 | | 172 | 111 | 198 | 206 | 138 | 130 | 155 | 159 | 170 | 71 | 123 |
| 147 | -0.09 | 0.36 | -0.25 | 0.80 | 1 | Glyctk | glycerate kinase | 122 | 197 | 96 | 140 | 198 | 124 | 155 | 148 | 116 | 170 | 149 |
| 144 | -0.21 | 0.40 | -0.52 | 0.60 | 1 | Lipt1 | lipoyltransferase 1 | 106 | 135 | 156 | 197 | 128 | 150 | 155 | 73 | 125 | 193 | 166 |
| 140 | -0.31 | 0.45 | -0.70 | 0.48 | 1 | Angptl6 | angiopoietin-like 6 | 172 | 98 | 207 | 155 | 121 | 148 | 155 | 165 | 88 | 58 | 179 |
| 108 | -0.27 | 0.43 | -0.62 | 0.53 | 1 | 4930550C14Ril | RIKEN cDNA 4930550C14 gene | 15 | 10 | 28 | 164 | 135 | 156 | 155 | 98 | 85 | 156 | 189 |
| 373 | -0.19 | 0.30 | -0.66 | 0.51 | 1 | Arhgef6 | Rac/Cdc42 guanine nucleotide exchange | 1795 | 618 | 672 | 136 | 156 | 94 | 154 | 115 | 108 | 153 | 98 |

| | | | | | | | | | | | | | | | | | | |
|---|---|---|---|---|---|---|---|---|---|---|---|---|---|---|---|---|---|---|
| 273 | -0.11 | 0.28 | -0.38 | 0.71 | 1 | Trim7 | tripartite motif-containing 7 | 255 | 322 | 246 | 200 | 296 | 486 | 154 | 282 | 163 | 182 | 413 |
| 232 | 0.13 | 0.30 | 0.44 | 0.66 | 1 | Slc25a26 | solute carrier family 25 (mitochondrial car | 226 | 363 | 139 | 260 | 237 | 214 | 154 | 326 | 221 | 223 | 192 |
| 169 | -0.20 | 0.30 | -0.66 | 0.51 | 1 | Sdr42e1 | short chain dehydrogenase/reductase far | 152 | 319 | 182 | 187 | 134 | 148 | 154 | 251 | 110 | 85 | 137 |
| 162 | 0.08 | 0.30 | 0.25 | 0.80 | 1 | Gm9833 | | 113 | 165 | 156 | 179 | 147 | 161 | 154 | 271 | 119 | 133 | 179 |
| 138 | -0.34 | 0.44 | -0.78 | 0.44 | 1 | 4930579K19Rik | | 102 | 183 | 147 | 179 | 166 | 103 | 154 | 100 | 60 | 200 | 123 |
| 130 | -0.10 | 0.28 | -0.36 | 0.72 | 1 | Kcnn1 | potassium intermediate/small conductanc | 119 | 139 | 97 | 180 | 93 | 135 | 154 | 151 | 112 | 119 | 132 |
| 126 | 0.39 | 0.41 | 0.95 | 0.34 | 1 | Cox16 | cytochrome c oxidase assembly protein 1 | 64 | 43 | 135 | 97 | 98 | 125 | 154 | 72 | 212 | 213 | 178 |
| 317 | 0.42 | 0.28 | 1.50 | 0.13 | 1 | Crlf3 | cytokine receptor-like factor 3 | 734 | 491 | 441 | 253 | 146 | 224 | 153 | 291 | 222 | 319 | 208 |
| 209 | -0.05 | 0.27 | -0.17 | 0.86 | 1 | Cchcr1 | coiled-coil alpha-helical rod protein 1 | 243 | 224 | 253 | 187 | 159 | 290 | 153 | 209 | 120 | 134 | 330 |
| 207 | 0.28 | 0.39 | 0.72 | 0.47 | 1 | Gm44109 | | 38 | 59 | 75 | 148 | 77 | 626 | 153 | 282 | 105 | 142 | 577 |
| 167 | -0.10 | 0.26 | -0.36 | 0.72 | 1 | Ybey | ybeY metallopeptidase | 176 | 174 | 186 | 169 | 215 | 134 | 153 | 133 | 188 | 142 | 161 |
| 137 | -0.25 | 0.45 | -0.56 | 0.58 | 1 | Taf4b | TATA-box binding protein associated fact | 135 | 246 | 148 | 230 | 88 | 77 | 153 | 89 | 132 | 72 | 136 |
| 133 | -0.09 | 0.43 | -0.22 | 0.83 | 1 | Gm20633 | | 144 | 65 | 120 | 150 | 163 | 99 | 153 | 236 | 169 | 113 | 54 |
| 123 | 0.09 | 0.37 | 0.25 | 0.80 | 1 | Gm4013 | | 75 | 170 | 128 | 122 | 113 | 76 | 153 | 199 | 111 | 103 | 97 |
| 260 | 0.22 | 0.30 | 0.71 | 0.48 | 1 | Lmo4 | LIM domain only 4 | 957 | 441 | 738 | 53 | 84 | 62 | 152 | 68 | 104 | 93 | 115 |
| 255 | 0.54 | 0.30 | 1.76 | 0.08 | 1 | Ap1g2 | adaptor protein complex AP-1, gamma 2 s | 272 | 221 | 185 | 158 | 206 | 346 | 152 | 188 | 229 | 434 | 411 |
| 212 | 0.36 | 0.31 | 1.18 | 0.24 | 1 | Rps16-ps2 | | 413 | 375 | 397 | 154 | 109 | 81 | 152 | 115 | 142 | 274 | 121 |
| 207 | 0.44 | 0.32 | 1.38 | 0.17 | 1 | Dimt1 | DIM1 dimethyladenosine transferase 1-lik | 176 | 173 | 235 | 98 | 139 | 352 | 152 | 118 | 187 | 313 | 337 |
| 201 | 0.16 | 0.29 | 0.55 | 0.58 | 1 | Purg | purine-rich element binding protein G | 254 | 221 | 205 | 167 | 172 | 232 | 152 | 268 | 186 | 135 | 220 |
| 194 | 0.06 | 0.30 | 0.22 | 0.83 | 1 | Bok | BCL2-related ovarian killer | 472 | 232 | 413 | 144 | 119 | 83 | 152 | 143 | 123 | 171 | 86 |
| 175 | -0.01 | 0.29 | -0.02 | 0.98 | 1 | Tyw5 | tRNA-yW synthesizing protein 5 | 227 | 146 | 128 | 164 | 110 | 269 | 152 | 132 | 116 | 129 | 356 |
| 173 | -0.34 | 0.33 | -1.03 | 0.30 | 1 | Mbd4 | methyl-CpG binding domain protein 4 | 274 | 271 | 160 | 140 | 112 | 236 | 152 | 178 | 50 | 97 | 232 |
| 172 | -0.34 | 0.38 | -0.89 | 0.37 | 1 | Zfp239 | zinc finger protein 239 | 193 | 201 | 188 | 292 | 130 | 155 | 152 | 147 | 170 | 68 | 193 |
| 152 | 0.49 | 0.56 | 0.88 | 0.38 | 1 | Etfb | electron transferring flavoprotein, beta po | 97 | 57 | 128 | 88 | 159 | 134 | 152 | 495 | 144 | 113 | 104 |
| 145 | -0.65 | 0.41 | -1.58 | 0.12 | 1 | Gpr173 | G-protein coupled receptor 173 | 168 | 187 | 143 | 117 | 214 | 170 | 152 | 104 | 129 | 41 | 167 |
| 239 | 0.40 | 0.29 | 1.39 | 0.16 | 1 | Tepsin | TEPSIN, adaptor related protein complex | 120 | 203 | 184 | 223 | 127 | 464 | 151 | 233 | 257 | 240 | 433 |
| 189 | -0.19 | 0.36 | -0.53 | 0.60 | 1 | Gm14325 | predicted gene 14325 | 191 | 127 | 207 | 208 | 249 | 201 | 151 | 122 | 144 | 291 | 184 |
| 175 | 0.39 | 0.28 | 1.38 | 0.17 | 1 | Ranbp17 | RAN binding protein 17 | 141 | 99 | 134 | 95 | 181 | 259 | 151 | 171 | 170 | 251 | 275 |
| 161 | -0.44 | 0.46 | -0.96 | 0.34 | 1 | Med18 | mediator complex subunit 18 | 293 | 146 | 127 | 137 | 181 | 179 | 151 | 30 | 140 | 191 | 194 |
| 146 | 0.35 | 0.33 | 1.06 | 0.29 | 1 | Spata33 | spermatogenesis associated 33 | 99 | 111 | 85 | 122 | 177 | 116 | 151 | 282 | 161 | 174 | 128 |
| 142 | -0.31 | 0.27 | -1.12 | 0.26 | 1 | Tmc7 | transmembrane channel-like gene family | 58 | 83 | 48 | 286 | 146 | 186 | 151 | 177 | 125 | 144 | 157 |
| 126 | 0.31 | 0.48 | 0.66 | 0.51 | 1 | Rbks | ribokinase | 87 | 88 | 104 | 50 | 111 | 169 | 151 | 39 | 179 | 278 | 134 |
| 84 | 0.31 | 0.53 | 0.58 | 0.56 | 1 | Zfp125 | | 147 | 37 | 0 | 62 | 8 | 127 | 151 | 96 | 11 | 176 | 112 |
| 933 | -0.33 | 0.26 | -1.29 | 0.20 | 1 | Slc38a5 | solute carrier family 38, member 5 | 1995 | 3765 | 2976 | 277 | 260 | 171 | 150 | 108 | 250 | 165 | 144 |
| 284 | 0.12 | 0.50 | 0.25 | 0.81 | 1 | Rrm2 | ribonucleotide reductase M2 | 1077 | 261 | 196 | 186 | 289 | 140 | 150 | 248 | 235 | 155 | 182 |
| 214 | -0.04 | 0.23 | -0.18 | 0.86 | 1 | Armc7 | armadillo repeat containing 7 | 260 | 306 | 249 | 253 | 192 | 184 | 150 | 257 | 173 | 149 | 180 |
| 192 | 0.19 | 0.50 | 0.38 | 0.71 | 1 | Cdc20 | cell division cycle 20 | 654 | 176 | 118 | 107 | 211 | 94 | 150 | 153 | 206 | 122 | 126 |
| 189 | 0.35 | 0.25 | 1.40 | 0.16 | 1 | Rnf8 | ring finger protein 8 | 131 | 143 | 124 | 161 | 106 | 331 | 150 | 243 | 142 | 166 | 385 |
| 181 | 0.18 | 0.29 | 0.63 | 0.53 | 1 | Fbxo36 | F-box protein 36 | 197 | 185 | 149 | 159 | 226 | 150 | 150 | 219 | 172 | 205 | 181 |
| 107 | -0.17 | 0.46 | -0.38 | 0.70 | 1 | Slc35f2 | solute carrier family 35, member F2 | 14 | 10 | 23 | 195 | 78 | 169 | 150 | 161 | 135 | 76 | 163 |
| 254 | -0.45 | 0.23 | -1.95 | 0.05 | 1 | Traf4 | TNF receptor associated factor 4 | 516 | 546 | 494 | 207 | 130 | 230 | 149 | 122 | 121 | 120 | 155 |
| 223 | 0.08 | 0.34 | 0.24 | 0.81 | 1 | Paqr8 | progestin and adipoQ receptor family mer | 458 | 403 | 264 | 213 | 147 | 134 | 149 | 103 | 171 | 215 | 194 |
| 197 | 0.19 | 0.37 | 0.51 | 0.61 | 1 | Apcdd1 | adenomatosis polyposis coli down-regula | 616 | 304 | 328 | 116 | 93 | 73 | 149 | 204 | 106 | 97 | 79 |
| 161 | -0.43 | 0.34 | -1.25 | 0.21 | 1 | Trdmt1 | tRNA aspartic acid methyltransferase 1 | 257 | 254 | 157 | 181 | 158 | 147 | 149 | 87 | 152 | 103 | 130 |
| 147 | -0.33 | 0.43 | -0.78 | 0.43 | 1 | 5330413P13Rik | | 150 | 240 | 208 | 103 | 130 | 167 | 149 | 38 | 116 | 124 | 193 |
| 133 | -0.70 | 0.58 | -1.21 | 0.22 | 1 | Mrps6 | mitochondrial ribosomal protein S6 | 228 | 165 | 141 | 157 | 103 | 134 | 149 | 12 | 101 | 134 | 134 |
| 267 | -0.32 | 0.34 | -0.96 | 0.34 | 1 | Srgap3 | SLIT-ROBO Rho GTPase activating prote | 555 | 871 | 624 | 67 | 124 | 156 | 148 | 64 | 95 | 109 | 123 |
| 265 | 0.60 | 0.39 | 1.55 | 0.12 | 1 | Mrpl33 | mitochondrial ribosomal protein L33 | 238 | 152 | 263 | 214 | 177 | 378 | 148 | 563 | 245 | 222 | 319 |
| 249 | 0.27 | 0.30 | 0.90 | 0.37 | 1 | Trmt13 | tRNA methyltransferase 13 | 138 | 197 | 226 | 125 | 136 | 612 | 148 | 112 | 190 | 229 | 626 |
| 238 | -0.41 | 0.35 | -1.17 | 0.24 | 1 | Grasp | GRP1 (general receptor for phosphoinosi | 513 | 434 | 496 | 242 | 194 | 89 | 148 | 73 | 178 | 151 | 97 |
| 228 | 0.05 | 0.24 | 0.22 | 0.82 | 1 | Naa80 | N(alpha)-acetyltransferase 80, NatH catal | 264 | 291 | 376 | 211 | 192 | 219 | 148 | 243 | 215 | 139 | 205 |
| 211 | 0.17 | 0.25 | 0.68 | 0.50 | 1 | Xrcc2 | X-ray repair complementing defective rep | 196 | 271 | 192 | 249 | 177 | 214 | 148 | 225 | 213 | 233 | 200 |
| 181 | 0.35 | 0.29 | 1.19 | 0.24 | 1 | Pop4 | processing of precursor 4, ribonuclease F | 356 | 165 | 232 | 131 | 89 | 175 | 148 | 118 | 156 | 210 | 213 |
| 177 | -0.52 | 0.28 | -1.85 | 0.06 | 1 | Trim65 | tripartite motif-containing 65 | 251 | 319 | 230 | 127 | 134 | 271 | 148 | 94 | 119 | 88 | 163 |
| 166 | -0.08 | 0.31 | -0.25 | 0.80 | 1 | Nmnat1 | nicotinamide nucleotide adenylyltransfera | 196 | 146 | 151 | 179 | 204 | 150 | 148 | 138 | 127 | 213 | 179 |
| 164 | -0.29 | 0.41 | -0.70 | 0.49 | 1 | E430024I08Rik | | 93 | 273 | 160 | 174 | 139 | 230 | 148 | 87 | 90 | 157 | 257 |
| 157 | 0.27 | 0.30 | 0.88 | 0.38 | 1 | Hilpda | hypoxia inducible lipid droplet associated | 269 | 198 | 299 | 110 | 89 | 88 | 148 | 126 | 112 | 194 | 97 |
| 128 | -0.21 | 0.30 | -0.70 | 0.49 | 1 | Rpl38-ps2 | | 184 | 164 | 175 | 112 | 113 | 100 | 148 | 90 | 79 | 148 | 96 |
| 127 | -0.31 | 0.55 | -0.57 | 0.57 | 1 | 4833418N02Rik | | 100 | 116 | 42 | 151 | 129 | 194 | 148 | 34 | 185 | 126 | 171 |
| 125 | -0.20 | 0.35 | -0.57 | 0.57 | 1 | 4930451G09Rik | | 129 | 131 | 109 | 71 | 160 | 167 | 148 | 105 | 100 | 120 | 139 |

| | | | | | | | | | | | | | | | | | | |
|---|---|---|---|---|---|---|---|---|---|---|---|---|---|---|---|---|---|---|
| 368 | 0.09 | 0.43 | 0.21 | 0.83 | 1 | Pfkfb3 | 6-phosphofructo-2-kinase/fructose-2,6-bip | 1126 | 787 | 778 | 182 | 85 | 226 | 147 | 342 | 136 | 53 | 182 |
| 149 | 0.13 | 0.55 | 0.23 | 0.82 | 1 | Il1r2 | interleukin 1 receptor, type II | 57 | 7 | 48 | 239 | 187 | 152 | 147 | 293 | 189 | 169 | 145 |
| 114 | 0.13 | 0.45 | 0.28 | 0.78 | 1 | Sfxn4 | sideroflexin 4 | 110 | 145 | 57 | 96 | 56 | 150 | 147 | 48 | 89 | 171 | 183 |
| 102 | -0.13 | 0.43 | -0.30 | 0.76 | 1 | Bambi-ps1 | | 35 | 50 | 22 | 81 | 143 | 161 | 147 | 81 | 142 | 106 | 158 |
| 411 | 0.03 | 0.20 | 0.16 | 0.87 | 1 | Smad3 | SMAD family member 3 | 1416 | 1146 | 972 | 124 | 103 | 110 | 146 | 132 | 78 | 164 | 129 |
| 255 | -0.15 | 0.25 | -0.60 | 0.55 | 1 | Lacc1 | laccase domain containing 1 | 399 | 383 | 383 | 195 | 225 | 287 | 146 | 152 | 222 | 115 | 300 |
| 217 | 0.48 | 0.31 | 1.55 | 0.12 | 1 | Zfp658 | zinc finger protein 658 | 229 | 130 | 215 | 100 | 199 | 356 | 146 | 220 | 209 | 270 | 314 |
| 193 | 0.17 | 0.23 | 0.74 | 0.46 | 1 | Alyref2 | Aly/REF export factor 2 | 117 | 229 | 165 | 213 | 173 | 221 | 146 | 263 | 190 | 149 | 253 |
| 172 | 0.25 | 0.33 | 0.77 | 0.44 | 1 | Aplf | aprataxin and PNKP like factor | 228 | 135 | 168 | 148 | 161 | 163 | 146 | 221 | 189 | 120 | 208 |
| 137 | -0.43 | 0.40 | -1.09 | 0.28 | 1 | Gm49284 | | 75 | 90 | 103 | 91 | 137 | 292 | 146 | 75 | 48 | 121 | 325 |
| 114 | -0.15 | 0.52 | -0.29 | 0.77 | 1 | Ppp1r3f | protein phosphatase 1, regulatory subuni | 35 | 141 | 53 | 110 | 131 | 151 | 146 | 67 | 169 | 126 | 121 |
| 466 | -0.44 | 0.30 | -1.47 | 0.14 | 1 | St3gal4 | ST3 beta-galactoside alpha-2,3-sialyltran | 1621 | 886 | 1481 | 210 | 180 | 120 | 145 | 139 | 148 | 80 | 115 |
| 262 | -0.26 | 0.26 | -1.01 | 0.31 | 1 | Rem2 | rad and gem related GTP binding protein | 403 | 568 | 392 | 222 | 169 | 276 | 145 | 126 | 160 | 115 | 303 |
| 245 | -0.21 | 0.32 | -0.65 | 0.52 | 1 | Rassf5 | Ras association (RalGDS/AF-6) domain fa | 952 | 560 | 514 | 74 | 77 | 54 | 145 | 38 | 103 | 133 | 42 |
| 205 | -0.07 | 0.23 | -0.28 | 0.78 | 1 | Pcdhb17 | protocadherin beta 17 | 277 | 262 | 312 | 182 | 200 | 190 | 145 | 188 | 183 | 139 | 175 |
| 152 | 0.00 | 0.31 | 0.00 | 1.00 | 1 | Osgepl1 | O-sialoglycoprotein endopeptidase-like 1 | 97 | 102 | 64 | 88 | 121 | 366 | 145 | 97 | 126 | 146 | 320 |
| 136 | -0.78 | 0.50 | -1.57 | 0.12 | 1 | Bcl3 | B cell leukemia/lymphoma 3 | 344 | 203 | 246 | 81 | 69 | 126 | 145 | 9 | 64 | 104 | 108 |
| 120 | 0.03 | 0.31 | 0.11 | 0.91 | 1 | Dus4l | dihydrouridine synthase 4-like (S. cerevis | 142 | 106 | 107 | 80 | 72 | 187 | 145 | 98 | 86 | 102 | 200 |
| 261 | -0.39 | 0.28 | -1.42 | 0.16 | 1 | Ly75 | lymphocyte antigen 75 | 685 | 665 | 692 | 101 | 119 | 102 | 144 | 104 | 125 | 79 | 56 |
| 211 | -0.06 | 0.37 | -0.16 | 0.87 | 1 | Ccdc77 | coiled-coil domain containing 77 | 165 | 235 | 217 | 201 | 139 | 361 | 144 | 74 | 200 | 167 | 417 |
| 209 | -0.02 | 0.25 | -0.09 | 0.93 | 1 | Dcbld1 | discoidin, CUB and LCCL domain contair | 388 | 336 | 355 | 167 | 180 | 117 | 144 | 155 | 204 | 157 | 94 |
| 128 | -0.51 | 0.38 | -1.33 | 0.18 | 1 | 3222401L13Rik | | 123 | 158 | 85 | 172 | 109 | 184 | 144 | 117 | 122 | 52 | 142 |
| 118 | -0.33 | 0.42 | -0.77 | 0.44 | 1 | Ntrk1 | neurotrophic tyrosine kinase, receptor, typ | 194 | 52 | 96 | 112 | 126 | 126 | 144 | 42 | 154 | 125 | 125 |
| 105 | -0.23 | 0.49 | -0.47 | 0.64 | 1 | Gm26910 | | 87 | 72 | 87 | 117 | 74 | 132 | 144 | 29 | 101 | 165 | 142 |
| 97 | -0.03 | 0.40 | -0.08 | 0.93 | 1 | 1810044D09Rik | | 52 | 94 | 82 | 94 | 100 | 84 | 144 | 149 | 87 | 94 | 89 |
| 540 | -0.22 | 0.23 | -0.99 | 0.32 | 1 | Gucy1b1 | guanylate cyclase 1, soluble, beta 1 | 1944 | 1586 | 1564 | 102 | 76 | 140 | 143 | 115 | 86 | 97 | 89 |
| 540 | -0.46 | 0.26 | -1.81 | 0.07 | 1 | Rbms3 | RNA binding motif, single stranded intera | 1536 | 1524 | 1512 | 241 | 170 | 238 | 143 | 189 | 110 | 174 | 106 |
| 524 | -0.09 | 0.27 | -0.33 | 0.74 | 1 | Lyn | LYN proto-oncogene, Src family tyrosine k | 2213 | 1051 | 1248 | 196 | 186 | 119 | 143 | 144 | 154 | 158 | 148 |
| 324 | -0.19 | 0.30 | -0.63 | 0.53 | 1 | Itprip | inositol 1,4,5-triphosphate receptor intera | 898 | 688 | 743 | 176 | 192 | 130 | 143 | 226 | 100 | 185 | 83 |
| 215 | -0.20 | 0.28 | -0.70 | 0.48 | 1 | Rab13 | RAB13, member RAS oncogene family | 589 | 450 | 524 | 106 | 92 | 90 | 143 | 104 | 114 | 80 | 73 |
| 208 | -0.78 | 0.46 | -1.68 | 0.09 | 1 | Nek6 | NIMA (never in mitosis gene a)-related ex | 641 | 358 | 457 | 152 | 110 | 102 | 143 | 42 | 169 | 41 | 75 |
| 192 | 0.25 | 0.32 | 0.76 | 0.45 | 1 | Rmdn1 | regulator of microtubule dynamics 1 | 118 | 149 | 102 | 134 | 187 | 331 | 143 | 211 | 122 | 255 | 358 |
| 170 | -0.30 | 0.88 | -0.34 | 0.73 | 1 | Mgam | maltase-glucoamylase | 162 | 447 | 372 | 39 | 200 | 75 | 143 | 0 | 235 | 120 | 79 |
| 166 | -0.54 | 0.29 | -1.88 | 0.06 | 1 | Ctxn1 | cortexin 1 | 206 | 247 | 388 | 133 | 171 | 128 | 143 | 68 | 138 | 91 | 108 |
| 160 | 0.62 | 0.35 | 1.79 | 0.07 | 1 | Ppargc1a | peroxisome proliferative activated recepto | 245 | 244 | 179 | 139 | 46 | 96 | 143 | 263 | 93 | 216 | 95 |
| 135 | 0.02 | 0.37 | 0.07 | 0.95 | 1 | Alg13 | asparagine-linked glycosylation 13 | 133 | 125 | 72 | 122 | 101 | 224 | 143 | 149 | 164 | 111 | 145 |
| 115 | -0.35 | 0.44 | -0.78 | 0.43 | 1 | Dnmt3b | DNA methyltransferase 3B | 124 | 109 | 97 | 65 | 96 | 215 | 143 | 64 | 122 | 41 | 194 |
| 102 | 0.01 | 0.69 | 0.01 | 0.99 | 1 | Mt3 | metallothionein 3 | 35 | 0 | 20 | 151 | 116 | 116 | 143 | 166 | 126 | 158 | 89 |
| 86 | -0.19 | 0.47 | -0.41 | 0.68 | 1 | Gm28529 | | 82 | 76 | 127 | 79 | 72 | 59 | 143 | 41 | 98 | 77 | 97 |
| 327 | -0.06 | 0.23 | -0.25 | 0.80 | 1 | Tmem181b-ps | | 101 | 75 | 63 | 158 | 119 | 1311 | 142 | 109 | 114 | 198 | 1210 |
| 299 | 0.24 | 0.28 | 0.87 | 0.39 | 1 | Sap30 | sin3 associated polypeptide | 359 | 441 | 382 | 394 | 210 | 240 | 142 | 330 | 210 | 303 | 279 |
| 286 | -0.12 | 0.34 | -0.36 | 0.72 | 1 | Tmem161b | transmembrane protein 161B | 423 | 418 | 288 | 124 | 188 | 593 | 142 | 207 | 156 | 73 | 536 |
| 272 | 0.10 | 0.26 | 0.41 | 0.68 | 1 | Zdhhc9 | zinc finger, DHHC domain containing 9 | 657 | 457 | 480 | 186 | 186 | 159 | 142 | 218 | 165 | 175 | 169 |
| 203 | 0.26 | 0.34 | 0.76 | 0.45 | 1 | Ak6 | TATA-box binding protein associated fact | 285 | 263 | 200 | 131 | 242 | 150 | 142 | 138 | 197 | 313 | 169 |
| 159 | -0.16 | 0.28 | -0.57 | 0.57 | 1 | Lrrc3 | leucine rich repeat containing 3 | 289 | 310 | 269 | 132 | 112 | 69 | 142 | 145 | 100 | 137 | 46 |
| 137 | -0.43 | 0.35 | -1.23 | 0.22 | 1 | Prr7 | proline rich 7 (synaptic) | 162 | 132 | 215 | 116 | 185 | 126 | 142 | 72 | 85 | 160 | 112 |
| 134 | -0.28 | 0.31 | -0.92 | 0.36 | 1 | 3000002C10Rik | | 119 | 72 | 78 | 113 | 85 | 302 | 142 | 87 | 53 | 142 | 277 |
| 128 | -0.63 | 0.44 | -1.42 | 0.16 | 1 | Pcdhga3 | protocadherin gamma subfamily A, 3 | 69 | 147 | 82 | 156 | 178 | 168 | 142 | 127 | 209 | 40 | 90 |
| 128 | 0.34 | 0.34 | 0.99 | 0.32 | 1 | Ptrh1 | peptidyl-tRNA hydrolase 1 homolog | 88 | 72 | 135 | 77 | 146 | 134 | 142 | 136 | 125 | 198 | 155 |
| 117 | 0.13 | 0.42 | 0.32 | 0.75 | 1 | A930005H10Rik | | 97 | 79 | 131 | 85 | 131 | 107 | 142 | 170 | 98 | 81 | 162 |
| 351 | -0.11 | 0.27 | -0.42 | 0.67 | 1 | Smad7 | SMAD family member 7 | 760 | 743 | 1022 | 123 | 240 | 200 | 141 | 145 | 139 | 200 | 150 |
| 271 | 0.00 | 0.45 | -0.01 | 0.99 | 1 | Ezh2 | enhancer of zeste 2 polycomb repressive | 943 | 221 | 206 | 228 | 152 | 281 | 141 | 263 | 130 | 183 | 233 |
| 188 | -0.05 | 0.33 | -0.14 | 0.89 | 1 | Noxo1 | NADPH oxidase organizer 1 | 94 | 123 | 134 | 174 | 170 | 367 | 141 | 293 | 114 | 102 | 354 |
| 175 | 0.26 | 0.41 | 0.63 | 0.53 | 1 | Pik3r5 | phosphoinositide-3-kinase regulatory sub | 368 | 133 | 139 | 181 | 76 | 205 | 141 | 204 | 151 | 126 | 200 |
| 175 | 0.26 | 0.29 | 0.92 | 0.36 | 1 | Ovol2 | ovo like zinc finger 2 | 22 | 52 | 63 | 236 | 225 | 183 | 141 | 382 | 302 | 134 | 182 |
| 165 | -0.24 | 0.46 | -0.53 | 0.60 | 1 | Tmem229a | transmembrane protein 229A | 317 | 298 | 525 | 38 | 56 | 149 | 141 | 81 | 55 | 73 | 81 |
| 127 | -0.27 | 0.37 | -0.73 | 0.47 | 1 | Rpl31-ps8 | | 196 | 257 | 154 | 155 | 70 | 77 | 141 | 68 | 87 | 122 | 73 |
| 483 | -0.29 | 0.28 | -1.04 | 0.30 | 1 | Stom | stomatin | 1954 | 1227 | 1119 | 212 | 140 | 77 | 140 | 103 | 125 | 120 | 95 |
| 339 | -0.21 | 0.29 | -0.73 | 0.46 | 1 | Grhl1 | grainyhead like transcription factor 1 | 184 | 306 | 453 | 283 | 247 | 785 | 140 | 344 | 194 | 80 | 718 |
| 233 | 0.04 | 0.42 | 0.09 | 0.93 | 1 | Kntc1 | kinetochore associated 1 | 359 | 238 | 91 | 328 | 226 | 236 | 140 | 305 | 243 | 191 | 209 |

| | | | | | | | | | | | | | | | | | | |
|---|---|---|---|---|---|---|---|---|---|---|---|---|---|---|---|---|---|---|
| 225 | -0.27 | 0.33 | -0.83 | 0.41 | 1 | Clec11a | C-type lectin domain family 11, member a | 688 | 420 | 467 | 146 | 122 | 80 | 140 | 153 | 125 | 76 | 57 |
| 215 | 0.20 | 0.26 | 0.77 | 0.44 | 1 | Zfp775 | zinc finger protein 775 | 204 | 202 | 219 | 253 | 224 | 192 | 140 | 265 | 254 | 234 | 175 |
| 182 | -0.18 | 0.25 | -0.72 | 0.47 | 1 | Zfp619 | zinc finger protein 619 | 125 | 122 | 132 | 178 | 218 | 333 | 140 | 139 | 183 | 162 | 270 |
| 182 | 0.21 | 0.31 | 0.68 | 0.49 | 1 | Zfp995 | zinc finger protein 995 | 280 | 206 | 202 | 158 | 129 | 175 | 140 | 183 | 188 | 223 | 117 |
| 177 | 0.00 | 0.42 | 0.00 | 1.00 | 1 | Zfp870 | zinc finger protein 870 | 181 | 338 | 310 | 200 | 98 | 125 | 140 | 72 | 180 | 165 | 139 |
| 176 | -0.25 | 0.35 | -0.72 | 0.47 | 1 | Pms1 | PMS1 homolog 1, mismatch repair systen | 190 | 163 | 202 | 196 | 186 | 205 | 140 | 74 | 195 | 131 | 257 |
| 174 | -0.15 | 0.28 | -0.52 | 0.60 | 1 | Hspbap1 | Hspb associated protein 1 | 198 | 278 | 149 | 180 | 180 | 173 | 140 | 214 | 147 | 126 | 133 |
| 153 | -0.07 | 0.31 | -0.23 | 0.82 | 1 | Efcab7 | EF-hand calcium binding domain 7 | 88 | 75 | 179 | 139 | 138 | 265 | 140 | 111 | 169 | 125 | 260 |
| 147 | -0.18 | 0.31 | -0.59 | 0.56 | 1 | Gm10433 | | 44 | 108 | 53 | 224 | 250 | 132 | 140 | 219 | 206 | 121 | 117 |
| 145 | -0.24 | 0.28 | -0.88 | 0.38 | 1 | Gm8437 | | 154 | 135 | 122 | 131 | 122 | 256 | 140 | 92 | 125 | 134 | 189 |
| 142 | -0.28 | 0.40 | -0.70 | 0.49 | 1 | Ttc41 | tetratricopeptide repeat domain 41 | 88 | 45 | 126 | 95 | 71 | 419 | 140 | 49 | 100 | 148 | 278 |
| 346 | -0.34 | 0.22 | -1.60 | 0.11 | 1 | Zfpm1 | zinc finger protein, multitype 1 | 709 | 910 | 882 | 270 | 201 | 117 | 139 | 196 | 187 | 125 | 74 |
| 207 | -0.16 | 0.33 | -0.50 | 0.62 | 1 | Trim59 | tripartite motif-containing 59 | 412 | 239 | 247 | 243 | 192 | 154 | 139 | 175 | 155 | 142 | 175 |
| 191 | -0.52 | 0.43 | -1.22 | 0.22 | 1 | Lmcd1 | LIM and cysteine-rich domains 1 | 506 | 409 | 453 | 130 | 109 | 56 | 139 | 67 | 130 | 33 | 69 |
| 182 | 0.13 | 0.35 | 0.37 | 0.71 | 1 | Hic2 | hypermethylated in cancer 2 | 172 | 110 | 160 | 195 | 137 | 277 | 139 | 162 | 185 | 290 | 180 |
| 179 | 0.11 | 0.35 | 0.31 | 0.75 | 1 | Ufsp1 | UFM1-specific peptidase 1 | 143 | 209 | 181 | 234 | 161 | 153 | 139 | 178 | 130 | 292 | 146 |
| 162 | -0.30 | 0.32 | -0.94 | 0.35 | 1 | Rpusd2 | RNA pseudouridylate synthase domain co | 157 | 181 | 109 | 262 | 184 | 161 | 139 | 129 | 154 | 164 | 144 |
| 144 | -0.91 | 0.48 | -1.89 | 0.06 | 1 | Myorg | myogenesis regulating glycosidase (puta | 335 | 240 | 397 | 133 | 74 | 42 | 139 | 49 | 22 | 129 | 26 |
| 133 | 0.44 | 0.33 | 1.33 | 0.18 | 1 | Gm17435 | | 40 | 38 | 23 | 137 | 128 | 166 | 139 | 287 | 168 | 169 | 166 |
| 119 | 0.24 | 0.36 | 0.67 | 0.50 | 1 | Fam151b | family with sequence similarity 151, meml | 100 | 142 | 130 | 115 | 67 | 111 | 139 | 137 | 124 | 139 | 104 |
| 109 | 0.58 | 0.77 | 0.75 | 0.45 | 1 | Crhr1 | corticotropin releasing hormone receptor | 1 | 0 | 32 | 88 | 169 | 107 | 139 | 177 | 148 | 210 | 132 |
| 109 | 0.20 | 0.33 | 0.60 | 0.55 | 1 | Ttc34 | tetratricopeptide repeat domain 34 | 95 | 71 | 85 | 80 | 138 | 76 | 139 | 86 | 203 | 151 | 73 |
| 108 | 0.18 | 0.39 | 0.47 | 0.64 | 1 | Gp5 | glycoprotein 5 (platelet) | 8 | 26 | 21 | 155 | 126 | 85 | 139 | 270 | 171 | 119 | 74 |
| 624 | 0.46 | 0.26 | 1.77 | 0.08 | 1 | Shank3 | SH3 and multiple ankyrin repeat domains | 1969 | 1637 | 2074 | 102 | 60 | 219 | 138 | 158 | 116 | 187 | 208 |
| 429 | -0.44 | 0.28 | -1.53 | 0.13 | 1 | Zfp703 | zinc finger protein 703 | 1224 | 979 | 1122 | 242 | 164 | 243 | 138 | 170 | 139 | 70 | 222 |
| 189 | 0.07 | 0.33 | 0.21 | 0.83 | 1 | Hexdc | hexosaminidase (glycosyl hydrolase fami | 176 | 203 | 221 | 146 | 168 | 263 | 138 | 129 | 111 | 250 | 275 |
| 174 | -0.36 | 0.40 | -0.91 | 0.36 | 1 | Zfp41 | zinc finger protein 41 | 373 | 266 | 194 | 170 | 146 | 145 | 138 | 178 | 155 | 67 | 80 |
| 174 | 0.11 | 0.29 | 0.37 | 0.71 | 1 | A730020E08Rik | | 106 | 166 | 76 | 216 | 176 | 222 | 138 | 189 | 185 | 160 | 277 |
| 160 | 0.34 | 0.31 | 1.12 | 0.26 | 1 | Dhrs13 | dehydrogenase/reductase (SDR family) m | 161 | 188 | 134 | 139 | 142 | 127 | 138 | 284 | 141 | 198 | 112 |
| 156 | -0.20 | 0.36 | -0.56 | 0.57 | 1 | Psmg3 | proteasome (prosome, macropain) assem | 214 | 162 | 208 | 150 | 157 | 153 | 138 | 205 | 96 | 84 | 149 |
| 143 | -0.57 | 0.67 | -0.86 | 0.39 | 1 | Pusl1 | pseudouridylate synthase-like 1 | 147 | 108 | 156 | 112 | 113 | 283 | 138 | 196 | 87 | 6 | 222 |
| 130 | -0.50 | 0.29 | -1.72 | 0.09 | 1 | Gm31992 | | 124 | 127 | 168 | 172 | 151 | 131 | 138 | 123 | 97 | 98 | 100 |
| 126 | -0.51 | 0.53 | -0.97 | 0.33 | 1 | Gm4791 | predicted gene 4791 | 6 | 0 | 2 | 214 | 225 | 194 | 138 | 63 | 222 | 131 | 191 |
| 118 | -0.13 | 0.30 | -0.42 | 0.67 | 1 | BB218582 | | 95 | 119 | 74 | 119 | 69 | 204 | 138 | 97 | 74 | 119 | 192 |
| 103 | -0.08 | 0.41 | -0.19 | 0.85 | 1 | Gm3924 | | 47 | 16 | 44 | 80 | 191 | 121 | 138 | 87 | 164 | 73 | 175 |
| 101 | -0.30 | 0.33 | -0.91 | 0.36 | 1 | Gm12500 | predicted gene 12500 | 57 | 62 | 79 | 116 | 134 | 115 | 138 | 94 | 88 | 102 | 125 |
| 14596 | -0.28 | 1.09 | -0.26 | 0.79 | 1 | Rnase1 | ribonuclease, RNase A family, 1 (pancrea | 19137 | 80062 | 56965 | 2966 | 250 | 19 | 137 | 190 | 158 | 174 | 504 |
| 290 | 0.01 | 0.25 | 0.04 | 0.97 | 1 | Zbtb2 | zinc finger and BTB domain containing 2 | 491 | 521 | 383 | 284 | 201 | 277 | 137 | 287 | 159 | 207 | 247 |
| 198 | 0.02 | 0.30 | 0.08 | 0.93 | 1 | Stn1 | STN1, CST complex subunit | 482 | 367 | 383 | 72 | 131 | 126 | 137 | 103 | 84 | 178 | 114 |
| 181 | -0.10 | 0.43 | -0.24 | 0.81 | 1 | Sertad3 | SERTA domain containing 3 | 221 | 177 | 209 | 184 | 167 | 195 | 137 | 360 | 146 | 73 | 125 |
| 176 | -0.53 | 0.43 | -1.22 | 0.22 | 1 | Ccnyl1 | cyclin Y-like 1 | 362 | 289 | 183 | 144 | 175 | 172 | 137 | 37 | 109 | 161 | 172 |
| 175 | 0.22 | 0.33 | 0.66 | 0.51 | 1 | Gm16124 | | 166 | 211 | 250 | 116 | 208 | 134 | 137 | 246 | 173 | 187 | 102 |
| 172 | -0.35 | 0.31 | -1.13 | 0.26 | 1 | Hspb11 | heat shock protein family B (small), memb | 204 | 252 | 227 | 193 | 209 | 139 | 137 | 93 | 190 | 104 | 146 |
| 148 | -0.30 | 0.50 | -0.59 | 0.55 | 1 | 4930570G19Rik | | 139 | 90 | 169 | 119 | 132 | 226 | 137 | 21 | 114 | 225 | 256 |
| 125 | -0.32 | 0.30 | -1.09 | 0.28 | 1 | Igsf5 | immunoglobulin superfamily, member 5 | 108 | 130 | 103 | 161 | 144 | 132 | 137 | 134 | 136 | 82 | 110 |
| 118 | 0.43 | 0.36 | 1.19 | 0.23 | 1 | Tesk2 | testis-specific kinase 2 | 127 | 114 | 94 | 67 | 110 | 95 | 137 | 166 | 93 | 183 | 115 |
| 101 | -0.63 | 0.44 | -1.43 | 0.15 | 1 | Mettl22 | methyltransferase like 22 | 83 | 96 | 109 | 94 | 106 | 147 | 137 | 32 | 54 | 124 | 132 |
| 73 | -0.33 | 0.37 | -0.89 | 0.37 | 1 | Gm47962 | | 39 | 22 | 27 | 87 | 95 | 83 | 137 | 87 | 61 | 95 | 72 |
| 458 | 0.49 | 0.28 | 1.73 | 0.08 | 1 | C1qtnf1 | C1q and tumor necrosis factor related pro | 1742 | 880 | 954 | 221 | 143 | 98 | 136 | 291 | 203 | 254 | 115 |
| 170 | 0.03 | 0.28 | 0.10 | 0.92 | 1 | Ccdc159 | coiled-coil domain containing 159 | 106 | 96 | 128 | 85 | 122 | 445 | 136 | 97 | 112 | 161 | 386 |
| 164 | 0.05 | 0.36 | 0.13 | 0.90 | 1 | Gm12183 | | 228 | 133 | 97 | 89 | 146 | 277 | 136 | 136 | 116 | 100 | 344 |
| 148 | -0.28 | 0.31 | -0.90 | 0.37 | 1 | Sfxn5 | sideroflexin 5 | 236 | 232 | 155 | 133 | 110 | 168 | 136 | 144 | 98 | 75 | 137 |
| 136 | 0.55 | 0.45 | 1.22 | 0.22 | 1 | Pdxp | pyridoxal (pyridoxine, vitamin B6) phosph | 67 | 82 | 100 | 74 | 164 | 121 | 136 | 333 | 130 | 140 | 148 |
| 125 | 0.20 | 0.45 | 0.44 | 0.66 | 1 | Gm10698 | | 53 | 116 | 52 | 110 | 162 | 126 | 136 | 102 | 138 | 241 | 141 |
| 122 | 0.22 | 0.28 | 0.79 | 0.43 | 1 | Gdf11 | growth differentiation factor 11 | 124 | 141 | 108 | 145 | 118 | 52 | 136 | 176 | 121 | 148 | 72 |
| 106 | -0.12 | 0.43 | -0.28 | 0.78 | 1 | 4732491K20Rik | | 56 | 99 | 125 | 102 | 135 | 81 | 136 | 107 | 96 | 73 | 152 |
| 76 | -0.24 | 0.51 | -0.47 | 0.64 | 1 | Gm29054 | | 9 | 32 | 14 | 98 | 108 | 81 | 136 | 67 | 109 | 99 | 78 |
| 469 | 0.18 | 0.33 | 0.55 | 0.58 | 1 | Rasa3 | RAS p21 protein activator 3 | 2105 | 937 | 937 | 130 | 168 | 118 | 135 | 232 | 138 | 124 | 136 |
| 260 | -0.04 | 0.30 | -0.14 | 0.89 | 1 | Trmu | tRNA 5-methylaminomethyl-2-thiouridylat | 245 | 279 | 284 | 89 | 211 | 608 | 135 | 145 | 111 | 148 | 605 |
| 253 | 0.29 | 0.38 | 0.77 | 0.44 | 1 | Tuba1c | tubulin, alpha 1C | 681 | 241 | 256 | 203 | 166 | 205 | 135 | 306 | 166 | 231 | 191 |

| | | | | | | | | | | | | | | | | | | |
|---|---|---|---|---|---|---|---|---|---|---|---|---|---|---|---|---|---|---|
| 143 | -0.20 | 0.44 | -0.45 | 0.65 | 1 | Dach2 | dachshund family transcription factor 2 | 118 | 155 | 83 | 181 | 173 | 156 | 135 | 58 | 160 | 173 | 185 |
| 135 | 0.54 | 0.33 | 1.62 | 0.11 | 1 | Pcdhb11 | protocadherin beta 11 | 84 | 150 | 113 | 139 | 95 | 96 | 135 | 160 | 177 | 213 | 124 |
| 131 | 0.01 | 0.30 | 0.04 | 0.97 | 1 | Fbxl8 | F-box and leucine-rich repeat protein 8 | 226 | 126 | 148 | 101 | 127 | 104 | 135 | 120 | 110 | 165 | 84 |
| 127 | -0.17 | 0.32 | -0.53 | 0.60 | 1 | 1110019D14Rik | | 91 | 108 | 145 | 140 | 141 | 133 | 135 | 86 | 146 | 107 | 163 |
| 287 | -0.30 | 0.30 | -1.00 | 0.32 | 1 | Bcl11a | B cell CLL/lymphoma 11A (zinc finger pro | 731 | 637 | 482 | 190 | 173 | 221 | 134 | 96 | 182 | 143 | 163 |
| 248 | -0.27 | 0.23 | -1.18 | 0.24 | 1 | Col9a2 | collagen, type IX, alpha 2 | 369 | 534 | 643 | 197 | 184 | 134 | 134 | 155 | 133 | 116 | 127 |
| 222 | -0.23 | 0.43 | -0.54 | 0.59 | 1 | Ifit1 | interferon-induced protein with tetratricop | 484 | 423 | 556 | 81 | 188 | 91 | 134 | 24 | 230 | 155 | 73 |
| 182 | 0.05 | 0.49 | 0.11 | 0.92 | 1 | Izumo4 | IZUMO family member 4 | 152 | 99 | 53 | 78 | 157 | 500 | 134 | 226 | 138 | 67 | 397 |
| 174 | 0.09 | 0.29 | 0.32 | 0.75 | 1 | A230051N06Rik | | 102 | 165 | 207 | 154 | 135 | 275 | 134 | 196 | 150 | 130 | 262 |
| 158 | -0.26 | 0.48 | -0.55 | 0.58 | 1 | Hscb | HscB iron-sulfur cluster co-chaperone | 204 | 169 | 186 | 220 | 149 | 123 | 134 | 41 | 181 | 178 | 151 |
| 153 | 0.25 | 0.34 | 0.73 | 0.47 | 1 | H2ac6 | H2A clustered histone 6 | 77 | 70 | 123 | 150 | 199 | 166 | 134 | 257 | 151 | 156 | 198 |
| 149 | 0.00 | 0.44 | -0.01 | 0.99 | 1 | Comtd1 | catechol-O-methyltransferase domain cor | 151 | 69 | 174 | 216 | 184 | 96 | 134 | 225 | 174 | 84 | 138 |
| 146 | -0.30 | 0.49 | -0.61 | 0.54 | 1 | Pask | PAS domain containing serine/threonine | 196 | 101 | 153 | 137 | 164 | 190 | 134 | 202 | 122 | 39 | 170 |
| 129 | 0.03 | 0.31 | 0.10 | 0.92 | 1 | Atp6v1g2 | ATPase, H+ transporting, lysosomal V1 su | 55 | 48 | 88 | 148 | 129 | 194 | 134 | 116 | 168 | 147 | 196 |
| 119 | -0.13 | 0.37 | -0.36 | 0.72 | 1 | Rsph4a | radial spoke head 4 homolog A (Chlamyd | 45 | 96 | 78 | 210 | 95 | 124 | 134 | 218 | 74 | 121 | 113 |
| 655 | -0.42 | 0.24 | -1.78 | 0.07 | 1 | Mamld1 | mastermind-like domain containing 1 | 2219 | 1995 | 2270 | 74 | 77 | 128 | 133 | 57 | 89 | 58 | 103 |
| 261 | -0.22 | 0.24 | -0.91 | 0.36 | 1 | Klhl18 | kelch-like 18 | 607 | 493 | 432 | 125 | 107 | 362 | 133 | 127 | 105 | 86 | 289 |
| 220 | 0.35 | 0.31 | 1.12 | 0.26 | 1 | Naa10 | N(alpha)-acetyltransferase 10, NatA catal | 231 | 201 | 341 | 170 | 161 | 260 | 133 | 294 | 175 | 144 | 308 |
| 215 | 0.06 | 0.38 | 0.15 | 0.88 | 1 | Ect2 | ect2 oncogene | 301 | 208 | 222 | 242 | 256 | 160 | 133 | 214 | 216 | 109 | 302 |
| 207 | 0.38 | 0.27 | 1.38 | 0.17 | 1 | Cables1 | CDK5 and Abl enzyme substrate 1 | 284 | 317 | 398 | 144 | 131 | 143 | 133 | 251 | 182 | 125 | 174 |
| 180 | -0.23 | 0.30 | -0.79 | 0.43 | 1 | Dclre1b | DNA cross-link repair 1B | 403 | 307 | 246 | 123 | 118 | 176 | 133 | 158 | 77 | 117 | 123 |
| 180 | -0.33 | 0.27 | -1.23 | 0.22 | 1 | Hyls1 | HYLS1, centriolar and ciliogenesis assoc | 180 | 185 | 167 | 229 | 182 | 255 | 133 | 148 | 121 | 134 | 241 |
| 178 | 0.28 | 0.37 | 0.75 | 0.45 | 1 | 1810059H22Rik | | 66 | 182 | 161 | 218 | 210 | 157 | 133 | 178 | 207 | 264 | 184 |
| 176 | -0.12 | 0.34 | -0.35 | 0.73 | 1 | 3110056K07Rik | | 131 | 131 | 95 | 206 | 201 | 259 | 133 | 251 | 92 | 148 | 291 |
| 170 | -0.41 | 0.42 | -0.98 | 0.33 | 1 | Zfp7 | zinc finger protein 7 | 205 | 199 | 220 | 176 | 167 | 210 | 133 | 39 | 158 | 156 | 205 |
| 143 | 0.10 | 0.34 | 0.29 | 0.77 | 1 | D130020L05Rik | | 161 | 194 | 215 | 145 | 87 | 120 | 133 | 112 | 120 | 182 | 108 |
| 111 | -0.35 | 0.35 | -0.98 | 0.33 | 1 | Gm10177 | | 87 | 113 | 65 | 129 | 146 | 123 | 133 | 103 | 129 | 76 | 112 |
| 109 | -0.29 | 0.42 | -0.69 | 0.49 | 1 | Gsdme | gasdermin E | 104 | 142 | 76 | 105 | 124 | 109 | 133 | 169 | 102 | 50 | 88 |
| 4273 | -1.38 | 1.03 | -1.34 | 0.18 | 1 | H2-Ab1 | histocompatibility 2, class II antigen A, bet | 23279 | 10323 | 13088 | 46 | 35 | 12 | 132 | 0 | 63 | 19 | 11 |
| 216 | -0.15 | 0.38 | -0.39 | 0.69 | 1 | Akna | AT-hook transcription factor | 1047 | 398 | 371 | 64 | 26 | 67 | 132 | 44 | 58 | 142 | 28 |
| 211 | -0.09 | 0.25 | -0.37 | 0.71 | 1 | Lsm11 | U7 snRNP-specific Sm-like protein LSM1 | 431 | 498 | 300 | 158 | 133 | 137 | 132 | 126 | 108 | 144 | 148 |
| 198 | -0.53 | 0.35 | -1.51 | 0.13 | 1 | Hnf1aos1 | | 153 | 128 | 149 | 242 | 242 | 361 | 132 | 122 | 184 | 67 | 397 |
| 179 | 0.19 | 0.32 | 0.58 | 0.56 | 1 | Snrnp25 | small nuclear ribonucleoprotein 25 (U11/ | 329 | 161 | 220 | 171 | 177 | 100 | 132 | 196 | 198 | 191 | 92 |
| 159 | -0.36 | 0.35 | -1.02 | 0.31 | 1 | Kazald1 | Kazal-type serine peptidase inhibitor dom | 274 | 249 | 224 | 166 | 118 | 141 | 132 | 139 | 138 | 55 | 109 |
| 156 | -0.20 | 0.44 | -0.46 | 0.65 | 1 | A630052C17Rik | | 80 | 134 | 92 | 158 | 199 | 258 | 132 | 64 | 146 | 228 | 225 |
| 150 | -0.12 | 0.34 | -0.37 | 0.71 | 1 | Nat8f1 | N-acetyltransferase 8 (GCN5-related) fam | 101 | 193 | 219 | 179 | 129 | 155 | 132 | 130 | 141 | 152 | 118 |
| 148 | 0.09 | 0.47 | 0.18 | 0.85 | 1 | Diablo | diablo, IAP-binding mitochondrial protein | 93 | 29 | 133 | 88 | 171 | 311 | 132 | 180 | 96 | 137 | 258 |
| 143 | 0.39 | 0.30 | 1.30 | 0.19 | 1 | A830082N09Rik | | 100 | 96 | 142 | 124 | 97 | 182 | 132 | 137 | 168 | 175 | 221 |
| 125 | -0.06 | 0.29 | -0.20 | 0.84 | 1 | Tcea1-ps1 | | 203 | 131 | 142 | 117 | 105 | 105 | 132 | 103 | 122 | 133 | 86 |
| 110 | -0.14 | 0.45 | -0.32 | 0.75 | 1 | Gm49797 | | 88 | 93 | 134 | 104 | 133 | 86 | 132 | 215 | 86 | 62 | 79 |
| 107 | 0.16 | 0.46 | 0.36 | 0.72 | 1 | E230013L22Rik | | 65 | 117 | 124 | 114 | 73 | 87 | 132 | 80 | 75 | 176 | 129 |
| 97 | -0.41 | 0.50 | -0.82 | 0.41 | 1 | Pgbd1 | piggyBac transposable element derived 1 | 142 | 170 | 137 | 92 | 50 | 76 | 132 | 64 | 72 | 28 | 104 |
| 92 | 0.03 | 0.41 | 0.07 | 0.95 | 1 | 4921536K21Ril | RIKEN cDNA 4921536K21 gene | 24 | 34 | 32 | 82 | 129 | 106 | 132 | 129 | 141 | 63 | 140 |
| 79 | 0.02 | 0.56 | 0.04 | 0.97 | 1 | Gm36298 | | 58 | 78 | 64 | 74 | 40 | 83 | 132 | 140 | 67 | 31 | 107 |
| 489 | -0.32 | 0.23 | -1.43 | 0.15 | 1 | Gab1 | growth factor receptor bound protein 2-as | 1553 | 1327 | 1505 | 170 | 94 | 161 | 131 | 110 | 84 | 122 | 125 |
| 177 | -0.40 | 0.32 | -1.26 | 0.21 | 1 | Zscan18 | zinc finger and SCAN domain containing | 182 | 198 | 144 | 187 | 113 | 348 | 131 | 120 | 52 | 146 | 328 |
| 174 | 0.51 | 0.32 | 1.61 | 0.11 | 1 | Gm11423 | | 148 | 120 | 163 | 187 | 118 | 173 | 131 | 286 | 202 | 139 | 249 |
| 158 | -0.11 | 0.28 | -0.40 | 0.69 | 1 | B230322F03Rik | | 130 | 176 | 136 | 239 | 163 | 124 | 131 | 262 | 163 | 119 | 89 |
| 156 | -0.14 | 0.28 | -0.51 | 0.61 | 1 | C130021I20Rik | | 65 | 100 | 118 | 110 | 162 | 338 | 131 | 108 | 134 | 106 | 348 |
| 151 | 0.22 | 0.35 | 0.65 | 0.52 | 1 | Cyren | cell cycle regulator of NHEJ | 131 | 219 | 204 | 88 | 99 | 205 | 131 | 156 | 141 | 133 | 158 |
| 147 | -0.30 | 0.38 | -0.79 | 0.43 | 1 | Sdsl | serine dehydratase-like | 215 | 171 | 241 | 153 | 112 | 145 | 131 | 189 | 96 | 67 | 103 |
| 144 | -0.61 | 0.32 | -1.89 | 0.06 | 1 | Tnfaip8l3 | tumor necrosis factor, alpha-induced prot | 368 | 345 | 310 | 68 | 88 | 54 | 131 | 43 | 65 | 63 | 45 |
| 135 | -0.35 | 0.35 | -0.98 | 0.33 | 1 | Katnal2 | katanin p60 subunit A-like 2 | 157 | 162 | 138 | 177 | 140 | 128 | 131 | 70 | 127 | 152 | 108 |
| 93 | -0.19 | 0.41 | -0.46 | 0.65 | 1 | Gm12576 | | 67 | 103 | 92 | 80 | 71 | 122 | 131 | 64 | 59 | 107 | 133 |
| 223 | -0.08 | 0.40 | -0.20 | 0.84 | 1 | Rad18 | RAD18 E3 ubiquitin protein ligase | 511 | 436 | 439 | 128 | 149 | 123 | 130 | 221 | 133 | 46 | 132 |
| 200 | 0.05 | 0.31 | 0.16 | 0.87 | 1 | Pigl | phosphatidylinositol glycan anchor biosyr | 169 | 111 | 116 | 210 | 191 | 366 | 130 | 178 | 190 | 246 | 287 |
| 162 | -0.16 | 0.33 | -0.50 | 0.62 | 1 | Polg2 | polymerase (DNA directed), gamma 2, ac | 146 | 104 | 120 | 133 | 132 | 339 | 130 | 160 | 111 | 73 | 336 |
| 156 | -0.06 | 0.27 | -0.21 | 0.83 | 1 | Atxn7l2 | ataxin 7-like 2 | 126 | 153 | 180 | 132 | 123 | 265 | 130 | 153 | 134 | 112 | 210 |
| 152 | -0.21 | 0.27 | -0.76 | 0.45 | 1 | Mthfsl | 5, 10-methenyltetrahydrofolate synthetase | 144 | 200 | 190 | 144 | 192 | 148 | 130 | 123 | 138 | 139 | 127 |
| 123 | -0.55 | 0.43 | -1.27 | 0.20 | 1 | 1810062G17Ril | RIKEN cDNA 1810062G17 gene | 40 | 96 | 22 | 235 | 114 | 220 | 130 | 79 | 81 | 90 | 250 |

| | | | | | | | | | | | | | | | | | |
|---|---|---|---|---|---|---|---|---|---|---|---|---|---|---|---|---|---|
| 106 | -0.17 | 0.41 | -0.42 | 0.68 | 1 | Skida1 | SKI/DACH domain containing 1 | 73 | 80 | 35 | 124 | 152 | 105 | 130 | 82 | 106 | 178 | 102 |
| 97 | 0.04 | 0.35 | 0.12 | 0.90 | 1 | Gm5417 | | 68 | 73 | 85 | 73 | 110 | 102 | 130 | 116 | 93 | 116 | 97 |
| 92 | -0.18 | 0.37 | -0.48 | 0.63 | 1 | Rangrf | RAN guanine nucleotide release factor | 132 | 102 | 94 | 69 | 89 | 76 | 130 | 108 | 58 | 102 | 58 |
| 351 | -0.15 | 0.30 | -0.50 | 0.62 | 1 | Apaf1 | apoptotic peptidase activating factor 1 | 1091 | 549 | 667 | 188 | 221 | 272 | 129 | 119 | 209 | 188 | 224 |
| 261 | 0.02 | 0.35 | 0.06 | 0.95 | 1 | Zfp369 | zinc finger protein 369 | 258 | 297 | 140 | 209 | 202 | 544 | 129 | 202 | 256 | 104 | 531 |
| 229 | -0.29 | 0.40 | -0.73 | 0.47 | 1 | P3h2 | prolyl 3-hydroxylase 2 | 675 | 634 | 467 | 102 | 105 | 68 | 129 | 138 | 110 | 36 | 57 |
| 202 | -0.83 | 0.58 | -1.43 | 0.15 | 1 | Plscr4 | phospholipid scramblase 4 | 447 | 416 | 402 | 117 | 131 | 186 | 129 | 71 | 130 | 6 | 187 |
| 162 | -0.21 | 0.26 | -0.80 | 0.43 | 1 | Trpt1 | tRNA phosphotransferase 1 | 157 | 145 | 165 | 123 | 177 | 270 | 129 | 110 | 150 | 94 | 265 |
| 161 | -0.12 | 0.33 | -0.36 | 0.72 | 1 | Zfp595 | zinc finger protein 595 | 202 | 228 | 176 | 145 | 71 | 266 | 129 | 76 | 99 | 135 | 244 |
| 154 | 0.02 | 0.31 | 0.08 | 0.94 | 1 | Usp49 | ubiquitin specific peptidase 49 | 112 | 129 | 140 | 133 | 181 | 216 | 129 | 161 | 123 | 193 | 182 |
| 139 | -0.14 | 0.41 | -0.34 | 0.73 | 1 | Speg | SPEG complex locus | 376 | 236 | 293 | 25 | 95 | 82 | 129 | 27 | 104 | 75 | 84 |
| 137 | 0.32 | 0.42 | 0.77 | 0.44 | 1 | Gm8730 | | 189 | 173 | 281 | 116 | 93 | 19 | 129 | 256 | 118 | 120 | 15 |
| 112 | 0.36 | 0.52 | 0.69 | 0.49 | 1 | BC037032 | | 44 | 13 | 58 | 125 | 79 | 160 | 129 | 209 | 81 | 169 | 170 |
| 103 | -0.79 | 0.88 | -0.91 | 0.36 | 1 | H4c11 | H4 clustered histone 11 | 28 | 25 | 16 | 125 | 112 | 227 | 129 | 0 | 81 | 130 | 257 |
| 83 | -0.26 | 0.46 | -0.57 | 0.57 | 1 | Oaz1-ps | | 85 | 69 | 41 | 123 | 95 | 52 | 129 | 43 | 90 | 113 | 74 |
| 221 | -0.18 | 0.26 | -0.70 | 0.48 | 1 | Gpt | glutamic pyruvic transaminase, soluble | 252 | 334 | 277 | 217 | 131 | 333 | 128 | 125 | 137 | 109 | 383 |
| 187 | -0.38 | 0.26 | -1.46 | 0.15 | 1 | A130014A01Rik | | 172 | 327 | 226 | 237 | 148 | 224 | 128 | 177 | 118 | 71 | 225 |
| 151 | -0.07 | 0.39 | -0.19 | 0.85 | 1 | Hsd17b13 | hydroxysteroid (17-beta) dehydrogenase | 117 | 479 | 232 | 159 | 66 | 76 | 128 | 153 | 78 | 99 | 76 |
| 149 | 0.39 | 0.29 | 1.37 | 0.17 | 1 | 2810414N06Rik | | 66 | 91 | 125 | 169 | 157 | 134 | 128 | 199 | 197 | 200 | 170 |
| 140 | 0.25 | 0.35 | 0.71 | 0.48 | 1 | Gm10138 | | 127 | 96 | 52 | 150 | 158 | 137 | 128 | 165 | 189 | 196 | 139 |
| 136 | -0.47 | 0.54 | -0.88 | 0.38 | 1 | Zfp867 | zinc finger protein 867 | 215 | 148 | 342 | 96 | 105 | 104 | 128 | 96 | 88 | 18 | 153 |
| 123 | 0.23 | 0.51 | 0.44 | 0.66 | 1 | Gm50145 | | 18 | 48 | 54 | 174 | 119 | 145 | 128 | 75 | 161 | 267 | 166 |
| 113 | 0.07 | 0.39 | 0.17 | 0.86 | 1 | 1110020A21Rik | | 88 | 40 | 102 | 140 | 141 | 88 | 128 | 123 | 112 | 149 | 130 |
| 99 | 0.29 | 0.70 | 0.42 | 0.68 | 1 | Gm50144 | | 22 | 1 | 28 | 117 | 99 | 123 | 128 | 225 | 87 | 72 | 192 |
| 93 | 0.23 | 0.46 | 0.51 | 0.61 | 1 | Gm11789 | | 0 | 0 | 0 | 122 | 112 | 85 | 128 | 202 | 165 | 155 | 53 |
| 90 | 0.37 | 0.38 | 0.95 | 0.34 | 1 | Pemt | phosphatidylethanolamine N-methyltrans | 64 | 98 | 61 | 81 | 65 | 64 | 128 | 94 | 95 | 149 | 90 |
| 356 | 0.29 | 0.31 | 0.92 | 0.36 | 1 | Stk10 | serine/threonine kinase 10 | 951 | 643 | 707 | 225 | 258 | 91 | 127 | 326 | 239 | 272 | 75 |
| 210 | -0.28 | 0.31 | -0.90 | 0.37 | 1 | Rfc5 | replication factor C (activator 1) 5 | 412 | 274 | 253 | 200 | 172 | 250 | 127 | 166 | 127 | 107 | 224 |
| 169 | -0.11 | 0.26 | -0.42 | 0.68 | 1 | Zfp11 | zinc finger protein 11 | 133 | 154 | 140 | 243 | 168 | 206 | 127 | 188 | 160 | 135 | 200 |
| 161 | -0.11 | 0.33 | -0.34 | 0.73 | 1 | 5330438D12Rik | | 210 | 278 | 286 | 129 | 103 | 149 | 127 | 162 | 129 | 61 | 135 |
| 152 | -0.44 | 0.46 | -0.94 | 0.35 | 1 | Zfp975 | zinc finger protein 975 | 163 | 110 | 73 | 201 | 195 | 212 | 127 | 52 | 176 | 129 | 231 |
| 120 | -0.47 | 0.52 | -0.90 | 0.37 | 1 | Ppil6 | peptidylprolyl isomerase (cyclophilin)-like | 87 | 106 | 144 | 29 | 92 | 312 | 127 | 75 | 57 | 31 | 260 |
| 104 | -1.08 | 0.61 | -1.78 | 0.07 | 1 | Tpmt | thiopurine methyltransferase | 239 | 184 | 92 | 67 | 112 | 102 | 127 | 6 | 55 | 85 | 76 |
| 103 | -0.24 | 0.45 | -0.54 | 0.59 | 1 | Gm19590 | | 79 | 61 | 131 | 108 | 96 | 131 | 127 | 162 | 90 | 68 | 80 |
| 70 | -0.43 | 0.53 | -0.81 | 0.42 | 1 | AI427809 | | 49 | 10 | 22 | 97 | 103 | 39 | 127 | 23 | 95 | 160 | 43 |
| 448 | -0.62 | 0.43 | -1.45 | 0.15 | 1 | Gfra3 | glial cell line derived neurotrophic factor f | 1235 | 1449 | 1588 | 69 | 124 | 69 | 126 | 88 | 100 | 14 | 65 |
| 434 | 0.11 | 0.40 | 0.28 | 0.78 | 1 | Mcm5 | minichromosome maintenance complex c | 1583 | 552 | 661 | 353 | 256 | 215 | 126 | 313 | 248 | 250 | 221 |
| 219 | 0.27 | 0.33 | 0.82 | 0.41 | 1 | E130307A14Rik | | 248 | 323 | 235 | 164 | 171 | 270 | 126 | 123 | 206 | 265 | 274 |
| 183 | 0.03 | 0.27 | 0.11 | 0.91 | 1 | Dzip1 | DAZ interacting protein 1 | 226 | 238 | 196 | 111 | 142 | 303 | 126 | 133 | 168 | 112 | 259 |
| 179 | -0.39 | 0.45 | -0.86 | 0.39 | 1 | Nit2 | nitrilase family, member 2 | 276 | 257 | 381 | 164 | 104 | 180 | 126 | 236 | 95 | 39 | 115 |
| 138 | 0.40 | 0.46 | 0.88 | 0.38 | 1 | Zbed5 | zinc finger, BED type containing 5 | 163 | 126 | 261 | 64 | 67 | 155 | 126 | 83 | 68 | 273 | 138 |
| 138 | 0.22 | 0.42 | 0.52 | 0.60 | 1 | Fbxo27 | F-box protein 27 | 98 | 42 | 74 | 123 | 195 | 166 | 126 | 251 | 170 | 119 | 160 |
| 128 | 0.17 | 0.39 | 0.44 | 0.66 | 1 | Klhdc8a | kelch domain containing 8A | 75 | 60 | 64 | 130 | 215 | 85 | 126 | 284 | 147 | 135 | 82 |
| 106 | -0.27 | 0.44 | -0.61 | 0.54 | 1 | Ccdc14 | coiled-coil domain containing 14 | 84 | 143 | 91 | 88 | 76 | 170 | 126 | 59 | 128 | 62 | 135 |
| 105 | -0.12 | 0.49 | -0.24 | 0.81 | 1 | C820005J03Rik | | 83 | 123 | 45 | 151 | 101 | 100 | 126 | 49 | 120 | 128 | 134 |
| 98 | 0.13 | 0.45 | 0.30 | 0.77 | 1 | Gm30606 | | 74 | 55 | 28 | 102 | 122 | 75 | 126 | 217 | 66 | 120 | 98 |
| 92 | -0.32 | 0.41 | -0.77 | 0.44 | 1 | Ccdc142os | | 33 | 68 | 70 | 81 | 135 | 121 | 126 | 68 | 129 | 97 | 82 |
| 64 | -0.09 | 0.43 | -0.21 | 0.83 | 1 | A930015D03Rik | | 51 | 16 | 68 | 27 | 102 | 46 | 126 | 41 | 109 | 79 | 38 |
| 23842 | -0.47 | 1.14 | -0.41 | 0.68 | 1 | Ctrl | chymotrypsin-like | 30217 | 122450 | 102923 | 4578 | 447 | 19 | 125 | 316 | 222 | 153 | 808 |
| 247 | -0.21 | 0.28 | -0.74 | 0.46 | 1 | Ifitm2 | interferon induced transmembrane protei | 753 | 575 | 570 | 107 | 94 | 110 | 125 | 88 | 130 | 91 | 70 |
| 177 | 0.16 | 0.31 | 0.53 | 0.59 | 1 | 1700088E04Ril | RIKEN cDNA 1700088E04 gene | 96 | 168 | 193 | 167 | 206 | 215 | 125 | 178 | 184 | 211 | 200 |
| 156 | 0.16 | 0.29 | 0.56 | 0.58 | 1 | Tmem81 | transmembrane protein 81 | 190 | 148 | 247 | 74 | 161 | 182 | 125 | 134 | 154 | 115 | 182 |
| 149 | -0.25 | 0.34 | -0.73 | 0.47 | 1 | Ccdc57 | coiled-coil domain containing 57 | 172 | 144 | 101 | 116 | 145 | 263 | 125 | 88 | 74 | 182 | 233 |
| 147 | 0.10 | 0.37 | 0.26 | 0.80 | 1 | Cep57l1 | centrosomal protein 57-like 1 | 154 | 259 | 110 | 89 | 115 | 206 | 125 | 153 | 133 | 124 | 148 |
| 146 | -0.14 | 0.57 | -0.25 | 0.80 | 1 | Fstl3 | follistatin-like 3 | 385 | 128 | 241 | 96 | 139 | 79 | 125 | 193 | 122 | 31 | 62 |
| 145 | -0.26 | 0.42 | -0.62 | 0.53 | 1 | Eps8l1 | EPS8-like 1 | 196 | 50 | 132 | 73 | 67 | 369 | 125 | 24 | 71 | 207 | 286 |
| 137 | -0.52 | 0.32 | -1.62 | 0.11 | 1 | Serp2 | stress-associated endoplasmic reticulum | 145 | 262 | 279 | 113 | 136 | 102 | 125 | 88 | 111 | 55 | 85 |
| 136 | -0.24 | 0.37 | -0.65 | 0.52 | 1 | Gm2058 | | 123 | 228 | 98 | 201 | 139 | 112 | 125 | 97 | 149 | 128 | 95 |
| 134 | 0.15 | 0.30 | 0.49 | 0.62 | 1 | A430035B10Rik | | 149 | 114 | 127 | 137 | 136 | 109 | 125 | 130 | 141 | 207 | 101 |
| 130 | -0.01 | 0.29 | -0.04 | 0.97 | 1 | 9330162012Rik | | 179 | 167 | 178 | 116 | 114 | 95 | 125 | 151 | 102 | 122 | 79 |

| | | | | | | | | | | | | | | | | | | |
|---|---|---|---|---|---|---|---|---|---|---|---|---|---|---|---|---|---|---|
| 124 | -0.16 | 0.36 | -0.45 | 0.65 | 1 | Trib3 | tribbles pseudokinase 3 | 95 | 67 | 149 | 176 | 137 | 121 | 125 | 136 | 122 | 139 | 103 |
| 66 | -0.61 | 0.63 | -0.98 | 0.33 | 1 | E330040D14Rik | | 15 | 2 | 2 | 162 | 92 | 52 | 125 | 124 | 62 | 54 | 35 |
| 1152 | -1.44 | 1.01 | -1.42 | 0.15 | 1 | Egflam | EGF-like, fibronectin type III and laminin G | 4305 | 4227 | 3770 | 43 | 60 | 43 | 124 | 43 | 38 | 0 | 21 |
| 257 | -0.32 | 0.25 | -1.29 | 0.20 | 1 | Erbb3 | erb-b2 receptor tyrosine kinase 3 | 620 | 546 | 715 | 82 | 161 | 156 | 124 | 76 | 93 | 94 | 158 |
| 225 | -0.05 | 0.31 | -0.17 | 0.86 | 1 | Fam189a2 | family with sequence similarity 189, meml | 426 | 416 | 507 | 165 | 143 | 128 | 124 | 171 | 141 | 187 | 69 |
| 183 | -0.15 | 0.30 | -0.49 | 0.63 | 1 | Setdb2 | SET domain, bifurcated 2 | 280 | 182 | 204 | 204 | 142 | 243 | 124 | 165 | 124 | 138 | 214 |
| 175 | -0.27 | 0.37 | -0.74 | 0.46 | 1 | Slc2a9 | solute carrier family 2 (facilitated glucose | 340 | 332 | 185 | 162 | 189 | 110 | 124 | 83 | 118 | 155 | 129 |
| 153 | 0.57 | 0.34 | 1.69 | 0.09 | 1 | Abca4 | ATP-binding cassette, sub-family A (ABC1 | 94 | 42 | 78 | 172 | 163 | 124 | 124 | 229 | 223 | 258 | 171 |
| 149 | -0.04 | 0.26 | -0.17 | 0.87 | 1 | Zfp472 | zinc finger protein 472 | 172 | 154 | 185 | 104 | 128 | 225 | 124 | 122 | 106 | 128 | 198 |
| 127 | -0.47 | 0.31 | -1.51 | 0.13 | 1 | Zfp442 | zinc finger protein 442 | 126 | 126 | 158 | 100 | 141 | 204 | 124 | 70 | 130 | 62 | 162 |
| 113 | 0.24 | 0.35 | 0.68 | 0.50 | 1 | Zfp947 | zinc finger protein 947 | 85 | 58 | 116 | 114 | 133 | 85 | 124 | 115 | 123 | 140 | 154 |
| 226 | 0.47 | 0.51 | 0.93 | 0.35 | 1 | Ier3 | immediate early response 3 | 933 | 532 | 528 | 28 | 23 | 59 | 122 | 68 | 85 | 33 | 76 |
| 183 | -0.18 | 0.34 | -0.53 | 0.60 | 1 | Vrk2 | vaccinia related kinase 2 | 341 | 229 | 282 | 173 | 162 | 158 | 122 | 184 | 140 | 80 | 146 |
| 163 | -0.37 | 0.32 | -1.15 | 0.25 | 1 | Taf5 | TATA-box binding protein associated fact | 202 | 159 | 217 | 135 | 202 | 220 | 122 | 65 | 157 | 158 | 159 |
| 148 | -0.05 | 0.32 | -0.15 | 0.88 | 1 | Kcnh3 | potassium voltage-gated channel, subfam | 195 | 149 | 137 | 157 | 163 | 131 | 122 | 153 | 121 | 201 | 100 |
| 145 | 0.30 | 0.43 | 0.69 | 0.49 | 1 | Zfp607a | zinc finger protein 607A | 142 | 72 | 148 | 136 | 81 | 213 | 122 | 203 | 155 | 79 | 242 |
| 141 | -0.44 | 0.35 | -1.27 | 0.20 | 1 | Ift43 | intraflagellar transport 43 | 204 | 220 | 284 | 127 | 101 | 128 | 122 | 101 | 109 | 46 | 109 |
| 139 | -0.90 | 0.48 | -1.89 | 0.06 | 1 | 1810055G02Ril | RIKEN cDNA 1810055G02 gene | 222 | 212 | 180 | 109 | 133 | 210 | 122 | 65 | 114 | 18 | 145 |
| 138 | 0.19 | 0.35 | 0.55 | 0.58 | 1 | 5033417F24Rik | | 82 | 70 | 167 | 149 | 190 | 101 | 122 | 156 | 167 | 218 | 101 |
| 136 | -0.15 | 0.39 | -0.39 | 0.70 | 1 | Chadl | chondroadherin-like | 112 | 99 | 241 | 151 | 162 | 112 | 122 | 87 | 122 | 185 | 106 |
| 115 | 0.32 | 0.49 | 0.65 | 0.52 | 1 | Lsm5 | LSM5 homolog, U6 small nuclear RNA ar | 161 | 79 | 125 | 47 | 120 | 124 | 122 | 160 | 115 | 59 | 155 |
| 90 | -0.78 | 0.60 | -1.30 | 0.19 | 1 | Garnl3 | GTPase activating RANGAP domain-like | 203 | 124 | 140 | 29 | 67 | 101 | 122 | 3 | 69 | 73 | 61 |
| 86 | 0.07 | 0.44 | 0.15 | 0.88 | 1 | Trim66 | tripartite motif-containing 66 | 116 | 111 | 103 | 46 | 49 | 80 | 122 | 65 | 71 | 137 | 44 |
| 297 | 0.00 | 0.24 | 0.01 | 0.99 | 1 | Trim12c | tripartite motif-containing 12C | 781 | 619 | 740 | 153 | 155 | 133 | 121 | 156 | 127 | 113 | 169 |
| 273 | -0.06 | 0.33 | -0.20 | 0.84 | 1 | Tcf7l1 | transcription factor 7 like 1 (T cell specific, | 537 | 381 | 413 | 277 | 301 | 162 | 121 | 195 | 279 | 187 | 154 |
| 242 | -0.06 | 0.27 | -0.22 | 0.83 | 1 | Neurl1b | neuralized E3 ubiquitin protein ligase 1B | 518 | 542 | 296 | 225 | 142 | 178 | 121 | 176 | 113 | 187 | 166 |
| 238 | 0.28 | 0.46 | 0.61 | 0.54 | 1 | Fcer1g | Fc receptor, IgE, high affinity I, gamma po | 816 | 378 | 636 | 86 | 56 | 85 | 121 | 75 | 163 | 44 | 160 |
| 233 | 0.36 | 0.39 | 0.93 | 0.35 | 1 | Slc16a11 | solute carrier family 16 (monocarboxylic a | 263 | 261 | 309 | 116 | 69 | 452 | 121 | 290 | 73 | 95 | 515 |
| 222 | -0.28 | 0.50 | -0.56 | 0.57 | 1 | Map3k14 | mitogen-activated protein kinase kinase k | 569 | 273 | 343 | 123 | 153 | 278 | 121 | 131 | 187 | 26 | 238 |
| 187 | 0.13 | 0.59 | 0.22 | 0.83 | 1 | Kif20a | kinesin family member 20A | 417 | 151 | 106 | 233 | 127 | 177 | 121 | 257 | 184 | 40 | 242 |
| 147 | -0.05 | 0.31 | -0.15 | 0.88 | 1 | Zc3h8 | zinc finger CCCH type containing 8 | 155 | 80 | 162 | 96 | 110 | 286 | 121 | 60 | 151 | 135 | 263 |
| 138 | 0.28 | 0.36 | 0.76 | 0.44 | 1 | Zfp109 | zinc finger protein 109 | 154 | 204 | 125 | 110 | 105 | 129 | 121 | 78 | 141 | 173 | 184 |
| 136 | -0.18 | 0.29 | -0.62 | 0.54 | 1 | Serf1 | small EDRK-rich factor 1 | 148 | 152 | 166 | 153 | 149 | 121 | 121 | 98 | 152 | 130 | 105 |
| 131 | -0.46 | 0.33 | -1.36 | 0.17 | 1 | Pus7l | pseudouridylate synthase 7-like | 148 | 188 | 111 | 176 | 169 | 111 | 121 | 102 | 99 | 94 | 121 |
| 129 | -0.04 | 0.40 | -0.09 | 0.93 | 1 | 1810021B22Rik | | 85 | 76 | 60 | 129 | 170 | 172 | 121 | 64 | 176 | 200 | 163 |
| 127 | 0.20 | 0.37 | 0.53 | 0.59 | 1 | 2810030D12Rik | | 40 | 78 | 86 | 164 | 163 | 104 | 121 | 246 | 110 | 115 | 175 |
| 108 | 0.24 | 0.35 | 0.68 | 0.50 | 1 | C330013E15Rik | | 70 | 44 | 51 | 69 | 145 | 136 | 121 | 110 | 152 | 189 | 105 |
| 98 | -0.58 | 0.41 | -1.42 | 0.15 | 1 | 2010001A14Rik | | 50 | 91 | 157 | 77 | 144 | 122 | 121 | 48 | 100 | 81 | 83 |
| 88 | 0.29 | 0.41 | 0.70 | 0.48 | 1 | Ftl1-ps1 | ferritin light chain 1 | 44 | 25 | 58 | 81 | 106 | 70 | 121 | 159 | 127 | 102 | 73 |
| 329 | 0.26 | 0.29 | 0.88 | 0.38 | 1 | 2410006H16Rik | | 619 | 415 | 621 | 162 | 223 | 391 | 120 | 229 | 150 | 214 | 469 |
| 244 | -0.01 | 0.25 | -0.05 | 0.96 | 1 | Lmbr1l | limb region 1 like | 274 | 295 | 248 | 190 | 132 | 541 | 120 | 166 | 157 | 161 | 396 |
| 199 | 0.20 | 0.30 | 0.66 | 0.51 | 1 | Adamts6 | a disintegrin-like and metallopeptidase (re | 190 | 285 | 209 | 109 | 98 | 372 | 120 | 148 | 161 | 81 | 416 |
| 146 | -0.16 | 0.40 | -0.39 | 0.70 | 1 | Zgrf1 | zinc finger, GRF-type containing 1 | 174 | 197 | 113 | 139 | 152 | 167 | 120 | 79 | 177 | 81 | 210 |
| 141 | 0.10 | 0.32 | 0.32 | 0.75 | 1 | 5033430I15Rik | | 75 | 134 | 71 | 177 | 146 | 172 | 120 | 168 | 151 | 188 | 147 |
| 135 | -0.01 | 0.32 | -0.02 | 0.98 | 1 | Gm8292 | | 317 | 240 | 262 | 58 | 100 | 57 | 120 | 56 | 82 | 125 | 68 |
| 133 | 0.13 | 0.46 | 0.27 | 0.79 | 1 | Sema6c | sema domain, transmembrane domain (T | 283 | 190 | 212 | 101 | 58 | 69 | 120 | 238 | 68 | 81 | 44 |
| 127 | -0.18 | 0.29 | -0.60 | 0.55 | 1 | Tmem266 | transmembrane protein 266 | 144 | 228 | 132 | 130 | 116 | 104 | 120 | 147 | 83 | 100 | 95 |
| 123 | 0.18 | 0.41 | 0.43 | 0.67 | 1 | B230307C23Ril | RIKEN cDNA B230307C23 gene | 109 | 77 | 97 | 119 | 94 | 161 | 120 | 197 | 73 | 207 | 105 |
| 110 | 0.07 | 0.39 | 0.18 | 0.85 | 1 | H3c15 | H3 clustered histone 15 | 55 | 90 | 38 | 104 | 138 | 127 | 120 | 192 | 138 | 119 | 91 |
| 90 | -0.43 | 0.37 | -1.16 | 0.25 | 1 | Gm19605 | | 77 | 86 | 104 | 86 | 81 | 123 | 120 | 61 | 71 | 58 | 126 |
| 273 | -0.56 | 0.40 | -1.40 | 0.16 | 1 | Cdc42ep3 | CDC42 effector protein (Rho GTPase bin | 1015 | 720 | 604 | 107 | 85 | 79 | 119 | 32 | 98 | 46 | 97 |
| 159 | 0.12 | 0.31 | 0.37 | 0.71 | 1 | Gsto2 | glutathione S-transferase omega 2 | 111 | 152 | 134 | 153 | 240 | 131 | 119 | 253 | 190 | 109 | 151 |
| 128 | -0.61 | 0.33 | -1.83 | 0.07 | 1 | Nptx2 | neuronal pentraxin 2 | 403 | 254 | 179 | 79 | 77 | 68 | 119 | 37 | 53 | 77 | 58 |
| 125 | 0.34 | 0.36 | 0.93 | 0.35 | 1 | Tmem17 | transmembrane protein 17 | 128 | 79 | 138 | 116 | 124 | 93 | 119 | 156 | 95 | 204 | 124 |
| 112 | 0.05 | 0.40 | 0.12 | 0.90 | 1 | Gm7818 | | 153 | 172 | 58 | 106 | 96 | 98 | 119 | 95 | 102 | 102 | 133 |
| 111 | -0.06 | 0.32 | -0.18 | 0.86 | 1 | 9130019O22Ril | RIKEN cDNA 9130019O22 gene | 118 | 187 | 129 | 105 | 117 | 62 | 119 | 117 | 110 | 99 | 62 |
| 108 | -0.97 | 0.64 | -1.51 | 0.13 | 1 | H2-T10 | | 425 | 63 | 212 | 46 | 110 | 14 | 119 | 1 | 92 | 89 | 14 |
| 106 | -0.17 | 0.47 | -0.37 | 0.71 | 1 | Kcng3 | potassium voltage-gated channel, subfam | 9 | 2 | 18 | 134 | 100 | 230 | 119 | 74 | 116 | 124 | 245 |
| 313 | 0.32 | 0.26 | 1.25 | 0.21 | 1 | Dok4 | docking protein 4 | 786 | 734 | 917 | 111 | 97 | 120 | 118 | 195 | 128 | 115 | 127 |
| 250 | -0.04 | 0.30 | -0.14 | 0.89 | 1 | Saal1 | serum amyloid A-like 1 | 406 | 341 | 472 | 185 | 259 | 218 | 118 | 242 | 177 | 115 | 217 |

| | | | | | | | | | | | | | | | | | | |
|---|---|---|---|---|---|---|---|---|---|---|---|---|---|---|---|---|---|---|
| 234 | 0.14 | 0.27 | 0.53 | 0.60 | 1 | Glb1l | galactosidase, beta 1-like | 283 | 265 | 197 | 157 | 128 | 524 | 118 | 160 | 170 | 171 | 405 |
| 226 | -0.21 | 0.36 | -0.59 | 0.55 | 1 | Iffo2 | intermediate filament family orphan 2 | 568 | 529 | 497 | 126 | 123 | 104 | 118 | 137 | 116 | 43 | 126 |
| 153 | -0.53 | 0.38 | -1.38 | 0.17 | 1 | Zswim3 | zinc finger SWIM-type containing 3 | 190 | 186 | 176 | 146 | 232 | 150 | 118 | 63 | 147 | 76 | 194 |
| 120 | -0.26 | 0.39 | -0.67 | 0.50 | 1 | Sinhcaf | SIN3-HDAC complex associated factor | 105 | 172 | 216 | 103 | 108 | 120 | 118 | 54 | 103 | 119 | 106 |
| 120 | 0.03 | 0.33 | 0.10 | 0.92 | 1 | Lbhd1 | LBH domain containing 1 | 135 | 99 | 116 | 116 | 107 | 133 | 118 | 70 | 147 | 139 | 144 |
| 117 | 0.29 | 0.35 | 0.82 | 0.41 | 1 | Pih1d2 | PIH1 domain containing 2 | 44 | 76 | 61 | 127 | 78 | 186 | 118 | 158 | 144 | 143 | 155 |
| 101 | -0.48 | 0.47 | -1.03 | 0.30 | 1 | 0610009L18Rik | | 82 | 215 | 148 | 158 | 78 | 49 | 118 | 56 | 72 | 67 | 70 |
| 99 | 0.22 | 0.42 | 0.52 | 0.60 | 1 | Ung | uracil DNA glycosylase | 155 | 106 | 113 | 85 | 48 | 83 | 118 | 110 | 119 | 84 | 67 |
| 90 | 0.72 | 0.47 | 1.51 | 0.13 | 1 | Npas3 | neuronal PAS domain protein 3 | 27 | 65 | 61 | 55 | 65 | 78 | 118 | 141 | 100 | 146 | 132 |
| 87 | -1.15 | 0.65 | -1.77 | 0.08 | 1 | Nkapl | NFKB activating protein-like | 112 | 162 | 53 | 100 | 119 | 78 | 118 | 46 | 61 | 9 | 102 |
| 69 | -0.40 | 0.41 | -0.99 | 0.32 | 1 | Hdac1-ps | | 116 | 52 | 53 | 65 | 50 | 63 | 118 | 34 | 31 | 111 | 64 |
| 46 | 0.38 | 1.01 | 0.38 | 0.71 | 1 | Insyn2a | inhibitory synaptic factor 2A | 0 | 5 | 0 | 19 | 24 | 61 | 118 | 8 | 48 | 164 | 61 |
| 293 | 0.22 | 0.44 | 0.50 | 0.61 | 1 | Synpo2 | synaptopodin 2 | 1301 | 612 | 380 | 108 | 93 | 116 | 117 | 143 | 195 | 76 | 86 |
| 205 | 0.42 | 0.28 | 1.53 | 0.13 | 1 | Rfc3 | replication factor C (activator 1) 3 | 302 | 290 | 204 | 144 | 170 | 192 | 117 | 235 | 173 | 188 | 242 |
| 199 | 0.32 | 0.27 | 1.18 | 0.24 | 1 | Snhg4 | | 299 | 281 | 320 | 116 | 176 | 166 | 117 | 200 | 174 | 185 | 152 |
| 190 | -0.08 | 0.28 | -0.30 | 0.77 | 1 | Ercc8 | excision repaiross-complementing rodent | 261 | 204 | 249 | 160 | 218 | 216 | 117 | 153 | 152 | 160 | 199 |
| 182 | 0.02 | 0.28 | 0.06 | 0.95 | 1 | Ccdc61 | coiled-coil domain containing 61 | 232 | 198 | 263 | 155 | 144 | 233 | 117 | 181 | 115 | 122 | 243 |
| 166 | 0.00 | 0.33 | 0.00 | 1.00 | 1 | Rccd1 | RCC1 domain containing 1 | 119 | 157 | 79 | 74 | 103 | 423 | 117 | 89 | 92 | 103 | 465 |
| 157 | -0.29 | 0.35 | -0.81 | 0.42 | 1 | BC002059 | | 148 | 208 | 175 | 179 | 148 | 184 | 117 | 214 | 100 | 52 | 202 |
| 101 | -0.70 | 0.39 | -1.78 | 0.07 | 1 | Chrm4 | cholinergic receptor, muscarinic 4 | 88 | 224 | 91 | 133 | 93 | 82 | 117 | 109 | 42 | 91 | 40 |
| 92 | -0.34 | 0.45 | -0.76 | 0.45 | 1 | Atp7b | ATPase, Cu++ transporting, beta polypep | 112 | 95 | 138 | 137 | 60 | 59 | 117 | 116 | 86 | 48 | 48 |
| 76 | -0.61 | 0.54 | -1.14 | 0.25 | 1 | Samd13 | RIKEN cDNA 4930503B20 gene | 48 | 83 | 76 | 103 | 55 | 106 | 117 | 27 | 70 | 98 | 59 |
| 65 | -0.14 | 0.44 | -0.32 | 0.75 | 1 | Gm47889 | | 43 | 38 | 26 | 88 | 46 | 70 | 117 | 75 | 60 | 55 | 90 |
| 64 | -0.45 | 0.83 | -0.54 | 0.59 | 1 | Gm10642 | | 33 | 2 | 73 | 123 | 54 | 52 | 117 | 89 | 41 | 46 | 69 |
| 409 | -0.52 | 0.35 | -1.50 | 0.13 | 1 | Lratd1 | LRAT domain containing 1 | 1131 | 1047 | 1294 | 240 | 100 | 145 | 116 | 173 | 42 | 107 | 107 |
| 377 | 0.44 | 0.40 | 1.11 | 0.27 | 1 | Hmgb2 | high mobility group box 2 | 1542 | 338 | 480 | 92 | 211 | 366 | 116 | 188 | 200 | 200 | 413 |
| 283 | 0.37 | 0.53 | 0.69 | 0.49 | 1 | Slc14a2 | solute carrier family 14 (urea transporter), | 0 | 26 | 71 | 668 | 431 | 198 | 116 | 718 | 389 | 173 | 322 |
| 205 | 0.41 | 0.36 | 1.16 | 0.25 | 1 | Tmem150a | transmembrane protein 150A | 222 | 87 | 169 | 154 | 172 | 346 | 116 | 235 | 187 | 240 | 331 |
| 191 | -0.37 | 0.34 | -1.11 | 0.27 | 1 | Rbm12b1 | RNA binding motif protein 12 B1 | 156 | 260 | 214 | 245 | 133 | 301 | 116 | 202 | 138 | 43 | 292 |
| 167 | -0.55 | 0.49 | -1.14 | 0.26 | 1 | Prdm16 | PR domain containing 16 | 546 | 384 | 425 | 39 | 58 | 79 | 116 | 58 | 55 | 17 | 61 |
| 165 | -0.08 | 0.27 | -0.28 | 0.78 | 1 | Fam229b | family with sequence similarity 229, meml | 133 | 125 | 194 | 144 | 196 | 242 | 116 | 130 | 143 | 138 | 253 |
| 163 | -0.17 | 0.29 | -0.60 | 0.55 | 1 | Gm34220 | | 143 | 243 | 165 | 229 | 184 | 138 | 116 | 146 | 149 | 152 | 126 |
| 159 | -0.36 | 0.30 | -1.21 | 0.23 | 1 | Rpl37rt | ribosomal protein L37, retrotransposed | 332 | 284 | 256 | 122 | 142 | 108 | 116 | 102 | 129 | 98 | 60 |
| 149 | 0.05 | 0.33 | 0.14 | 0.89 | 1 | N4bp3 | NEDD4 binding protein 3 | 276 | 272 | 285 | 94 | 103 | 76 | 116 | 140 | 70 | 137 | 68 |
| 149 | 0.22 | 0.27 | 0.79 | 0.43 | 1 | Gm19412 | | 104 | 168 | 154 | 157 | 146 | 133 | 116 | 233 | 158 | 103 | 166 |
| 142 | 0.19 | 0.35 | 0.54 | 0.59 | 1 | Gm16845 | | 136 | 158 | 121 | 161 | 134 | 113 | 116 | 119 | 144 | 247 | 108 |
| 138 | 0.43 | 0.41 | 1.04 | 0.30 | 1 | Gm16740 | | 97 | 76 | 130 | 128 | 140 | 116 | 116 | 301 | 137 | 196 | 85 |
| 135 | 0.45 | 0.34 | 1.32 | 0.19 | 1 | Gm7658 | | 86 | 114 | 118 | 96 | 189 | 88 | 116 | 234 | 168 | 171 | 100 |
| 108 | -0.15 | 0.36 | -0.40 | 0.69 | 1 | Wdr89 | WD repeat domain 89 | 103 | 153 | 79 | 101 | 121 | 101 | 116 | 137 | 86 | 75 | 112 |
| 80 | -0.25 | 0.49 | -0.51 | 0.61 | 1 | Zfp273 | zinc finger protein 273 | 79 | 27 | 56 | 74 | 58 | 116 | 116 | 36 | 81 | 54 | 181 |
| 77 | -0.40 | 0.55 | -0.73 | 0.46 | 1 | Gpr62 | G protein-coupled receptor 62 | 23 | 25 | 4 | 170 | 96 | 56 | 116 | 167 | 110 | 54 | 27 |
| 13358 | 0.62 | 1.17 | 0.53 | 0.59 | 1 | Ighm | | 103671 | 39689 | 3361 | 0 | 4 | 3 | 115 | 43 | 2 | 40 | 10 |
| 1472 | -0.83 | 0.55 | -1.52 | 0.13 | 1 | Lcp1 | lymphocyte cytosolic protein 1 | 8821 | 3435 | 3442 | 67 | 89 | 31 | 115 | 8 | 84 | 68 | 26 |
| 303 | -0.11 | 0.30 | -0.38 | 0.71 | 1 | Rgs12 | regulator of G-protein signaling 12 | 722 | 579 | 568 | 275 | 225 | 147 | 115 | 241 | 168 | 153 | 135 |
| 271 | 0.28 | 0.29 | 0.96 | 0.33 | 1 | Galt | galactose-1-phosphate uridyl transferase | 450 | 306 | 382 | 181 | 214 | 341 | 115 | 221 | 217 | 250 | 302 |
| 266 | -0.07 | 0.29 | -0.25 | 0.80 | 1 | Sh2d3c | SH2 domain containing 3C | 565 | 633 | 686 | 172 | 154 | 94 | 115 | 88 | 150 | 162 | 106 |
| 218 | 0.43 | 0.44 | 0.96 | 0.34 | 1 | Exosc8 | exosome component 8 | 513 | 205 | 360 | 86 | 138 | 233 | 115 | 222 | 178 | 66 | 281 |
| 193 | 0.15 | 0.32 | 0.48 | 0.63 | 1 | Faap24 | Fanconi anemia core complex associatec | 240 | 240 | 195 | 242 | 192 | 139 | 115 | 193 | 175 | 252 | 142 |
| 145 | 0.00 | 0.32 | 0.01 | 1.00 | 1 | Tmem220 | transmembrane protein 220 | 139 | 116 | 150 | 197 | 135 | 142 | 115 | 146 | 178 | 166 | 107 |
| 137 | 0.25 | 0.42 | 0.60 | 0.55 | 1 | Gm379 | | 75 | 64 | 108 | 200 | 139 | 95 | 115 | 342 | 133 | 170 | 69 |
| 127 | -0.38 | 0.33 | -1.14 | 0.25 | 1 | Maats1 | MYCBP-associated, testis expressed 1 | 118 | 164 | 130 | 166 | 98 | 179 | 115 | 78 | 112 | 85 | 154 |
| 103 | -0.18 | 0.34 | -0.53 | 0.60 | 1 | Mcrip2 | MAPK regulated corepressor interacting p | 109 | 84 | 70 | 123 | 96 | 123 | 115 | 78 | 89 | 156 | 94 |
| 100 | 0.30 | 0.60 | 0.50 | 0.62 | 1 | 4930524O07Rik | | 3 | 20 | 53 | 45 | 188 | 127 | 115 | 82 | 181 | 137 | 148 |
| 77 | 0.03 | 0.47 | 0.07 | 0.95 | 1 | Glns-ps1 | | 87 | 8 | 78 | 30 | 166 | 25 | 115 | 44 | 109 | 161 | 19 |
| 74 | -0.73 | 0.54 | -1.33 | 0.18 | 1 | Gm9920 | | 82 | 144 | 34 | 70 | 48 | 114 | 115 | 49 | 54 | 28 | 71 |
| 59 | -0.84 | 0.63 | -1.32 | 0.19 | 1 | Gm7887 | | 69 | 17 | 16 | 40 | 47 | 123 | 115 | 3 | 45 | 67 | 111 |
| 468 | -0.36 | 0.33 | -1.08 | 0.28 | 1 | Itga9 | integrin alpha 9 | 1911 | 1003 | 1382 | 63 | 72 | 200 | 114 | 23 | 106 | 54 | 217 |
| 448 | -0.29 | 0.35 | -0.82 | 0.41 | 1 | Prcp | prolylcarboxypeptidase (angiotensinase ( | 1919 | 904 | 1036 | 213 | 159 | 98 | 114 | 74 | 141 | 149 | 122 |
| 203 | 0.09 | 0.27 | 0.34 | 0.73 | 1 | Letm2 | leucine zipper-EF-hand containing transn | 153 | 171 | 208 | 133 | 162 | 426 | 114 | 206 | 127 | 124 | 405 |
| 127 | -0.03 | 0.45 | -0.07 | 0.95 | 1 | Nat2 | N-acetyltransferase 2 (arylamine N-acetyl | 178 | 94 | 166 | 146 | 119 | 97 | 114 | 146 | 165 | 50 | 119 |

| | | | | | | | | | | | | | | | | | |
|---|---|---|---|---|---|---|---|---|---|---|---|---|---|---|---|---|---|
| 121 | -0.65 | 0.41 | -1.61 | 0.11 | 1 | Rgs19 | regulator of G-protein signaling 19 | 320 | 220 | 186 | 90 | 77 | 86 | 114 | 57 | 90 | 41 | 46 |
| 118 | -0.32 | 0.34 | -0.95 | 0.34 | 1 | A330035P11Rik | | 37 | 73 | 75 | 102 | 164 | 227 | 114 | 98 | 147 | 62 | 202 |
| 113 | 0.26 | 0.43 | 0.59 | 0.55 | 1 | 2610035D17Rik | | 332 | 202 | 202 | 21 | 42 | 57 | 114 | 50 | 53 | 138 | 36 |
| 91 | -0.02 | 0.42 | -0.04 | 0.97 | 1 | Zfp558 | zinc finger protein 558 | 109 | 154 | 103 | 98 | 30 | 81 | 114 | 71 | 47 | 98 | 91 |
| 86 | 0.04 | 0.56 | 0.07 | 0.94 | 1 | Atp8b5 | ATPase, class I, type 8B, member 5 | 72 | 58 | 75 | 28 | 60 | 167 | 114 | 75 | 85 | 27 | 184 |
| 538 | -0.06 | 0.36 | -0.16 | 0.88 | 1 | Cebpd | CCAAT/enhancer binding protein (C/EBP | 1816 | 828 | 929 | 341 | 146 | 561 | 113 | 277 | 112 | 157 | 632 |
| 234 | 0.23 | 0.37 | 0.61 | 0.54 | 1 | Ednra | endothelin receptor type A | 1035 | 621 | 501 | 26 | 29 | 28 | 113 | 64 | 47 | 89 | 18 |
| 188 | 0.14 | 0.30 | 0.45 | 0.65 | 1 | Bbc3 | BCL2 binding component 3 | 181 | 191 | 134 | 117 | 159 | 386 | 113 | 132 | 183 | 161 | 313 |
| 137 | -0.47 | 0.36 | -1.29 | 0.20 | 1 | Pstk | phosphoseryl-tRNA kinase | 151 | 186 | 101 | 137 | 101 | 243 | 113 | 56 | 86 | 77 | 254 |
| 135 | -0.17 | 0.35 | -0.48 | 0.63 | 1 | Il15ra | interleukin 15 receptor, alpha chain | 200 | 130 | 83 | 88 | 122 | 238 | 113 | 72 | 105 | 100 | 238 |
| 123 | -0.32 | 0.31 | -1.03 | 0.30 | 1 | Gm49748 | | 61 | 104 | 91 | 170 | 119 | 201 | 113 | 94 | 109 | 93 | 195 |
| 117 | -0.34 | 0.36 | -0.95 | 0.34 | 1 | Zfp184 | zinc finger protein 184 (Kruppel-like) | 116 | 93 | 124 | 151 | 111 | 151 | 113 | 137 | 75 | 68 | 148 |
| 99 | 0.15 | 0.41 | 0.37 | 0.71 | 1 | Gm38910 | | 118 | 146 | 112 | 83 | 89 | 35 | 113 | 120 | 54 | 182 | 34 |
| 81 | -0.67 | 0.74 | -0.91 | 0.37 | 1 | Hmga1b | high mobility group AT-hook 1B | 160 | 19 | 83 | 39 | 110 | 9 | 113 | 1 | 138 | 211 | 4 |
| 80 | -0.35 | 0.49 | -0.72 | 0.47 | 1 | Alb | albumin | 216 | 103 | 160 | 26 | 45 | 21 | 113 | 30 | 36 | 121 | 6 |
| 484 | -0.47 | 0.32 | -1.49 | 0.14 | 1 | Dll4 | delta like canonical Notch ligand 4 | 1351 | 1212 | 1578 | 107 | 105 | 299 | 112 | 54 | 120 | 36 | 347 |
| 255 | 0.30 | 0.31 | 0.98 | 0.33 | 1 | Tcirg1 | T cell, immune regulator 1, ATPase, H+ tra | 701 | 434 | 439 | 102 | 102 | 248 | 112 | 207 | 107 | 131 | 218 |
| 187 | 0.14 | 0.34 | 0.40 | 0.69 | 1 | Tbc1d25 | TBC1 domain family, member 25 | 299 | 338 | 190 | 158 | 163 | 143 | 112 | 221 | 143 | 90 | 203 |
| 156 | 0.41 | 0.42 | 0.98 | 0.33 | 1 | Trp53rkb | transformation related protein 53 regulatin | 183 | 107 | 231 | 143 | 116 | 131 | 112 | 215 | 105 | 252 | 118 |
| 152 | -0.33 | 0.40 | -0.83 | 0.41 | 1 | Pcdhb16 | protocadherin beta 16 | 208 | 210 | 148 | 151 | 186 | 153 | 112 | 181 | 161 | 52 | 107 |
| 132 | 0.03 | 0.44 | 0.08 | 0.94 | 1 | Cdc45 | cell division cycle 45 | 257 | 120 | 71 | 109 | 172 | 102 | 112 | 120 | 150 | 120 | 115 |
| 117 | 0.01 | 0.38 | 0.02 | 0.99 | 1 | Gm10076 | | 182 | 198 | 201 | 130 | 60 | 52 | 112 | 97 | 126 | 85 | 47 |
| 105 | -0.29 | 0.69 | -0.42 | 0.67 | 1 | Ptch2 | patched 2 | 314 | 41 | 33 | 115 | 105 | 99 | 112 | 89 | 133 | 41 | 70 |
| 95 | -0.94 | 0.72 | -1.30 | 0.19 | 1 | Nmrk1 | nicotinamide riboside kinase 1 | 152 | 153 | 90 | 62 | 93 | 142 | 112 | 70 | 65 | 3 | 108 |
| 266 | 0.32 | 0.44 | 0.72 | 0.47 | 1 | Alox5ap | arachidonate 5-lipoxygenase activating p | 1068 | 314 | 436 | 136 | 93 | 134 | 111 | 246 | 141 | 183 | 66 |
| 173 | -0.05 | 0.31 | -0.15 | 0.88 | 1 | E2f3 | E2F transcription factor 3 | 261 | 310 | 294 | 130 | 145 | 138 | 111 | 82 | 126 | 165 | 138 |
| 137 | -0.20 | 0.31 | -0.65 | 0.51 | 1 | Gnb1l | guanine nucleotide binding protein (G pro | 102 | 93 | 138 | 172 | 217 | 128 | 111 | 175 | 158 | 88 | 122 |
| 136 | 0.45 | 0.29 | 1.52 | 0.13 | 1 | Avpi1 | arginine vasopressin-induced 1 | 216 | 204 | 164 | 110 | 92 | 74 | 111 | 109 | 117 | 185 | 118 |
| 135 | 0.25 | 0.27 | 0.93 | 0.35 | 1 | Ropn1l | ropporin 1-like | 79 | 109 | 114 | 130 | 159 | 141 | 111 | 159 | 166 | 146 | 169 |
| 128 | 0.05 | 0.36 | 0.13 | 0.89 | 1 | 2210408F21Rik | | 128 | 147 | 165 | 145 | 141 | 81 | 111 | 145 | 88 | 109 | 147 |
| 126 | -0.03 | 0.29 | -0.11 | 0.92 | 1 | Kcnh8 | potassium voltage-gated channel, subfam | 103 | 152 | 161 | 99 | 120 | 160 | 111 | 114 | 123 | 91 | 153 |
| 103 | -0.20 | 0.36 | -0.55 | 0.58 | 1 | Psg16 | pregnancy specific glycoprotein 16 | 57 | 65 | 53 | 117 | 104 | 177 | 111 | 88 | 142 | 71 | 147 |
| 102 | -0.32 | 0.45 | -0.71 | 0.48 | 1 | Gm10705 | | 92 | 145 | 101 | 123 | 129 | 73 | 111 | 42 | 102 | 111 | 91 |
| 97 | -0.70 | 0.41 | -1.73 | 0.08 | 1 | Dgat2 | diacylglycerol O-acyltransferase 2 | 132 | 114 | 143 | 140 | 81 | 88 | 111 | 90 | 57 | 39 | 75 |
| 76 | 0.17 | 0.41 | 0.41 | 0.68 | 1 | Scnn1a | sodium channel, nonvoltage-gated 1 alph | 77 | 48 | 49 | 74 | 83 | 25 | 111 | 79 | 137 | 138 | 16 |
| 71 | -1.05 | 0.83 | -1.27 | 0.21 | 1 | Hist1h2al | | 61 | 0 | 39 | 23 | 145 | 36 | 111 | 0 | 121 | 213 | 26 |
| 1068 | -1.34 | 0.71 | -1.89 | 0.06 | 1 | Klf2 | Kruppel-like factor 2 (lung) | 4987 | 2895 | 3085 | 30 | 26 | 336 | 110 | 35 | 8 | 5 | 226 |
| 599 | -0.34 | 0.33 | -1.01 | 0.31 | 1 | Arrb1 | arrestin, beta 1 | 3013 | 1287 | 1564 | 74 | 112 | 78 | 110 | 70 | 121 | 137 | 20 |
| 319 | -0.04 | 0.24 | -0.14 | 0.89 | 1 | Prdm1 | PR domain containing 1, with ZNF domai | 1406 | 560 | 843 | 67 | 78 | 87 | 110 | 44 | 78 | 151 | 83 |
| 280 | 0.01 | 0.51 | 0.01 | 0.99 | 1 | Ccna2 | cyclin A2 | 882 | 269 | 191 | 275 | 220 | 239 | 110 | 374 | 147 | 137 | 238 |
| 201 | -0.14 | 0.41 | -0.33 | 0.74 | 1 | Zfp14 | zinc finger protein 14 | 179 | 249 | 287 | 231 | 134 | 321 | 110 | 75 | 220 | 182 | 224 |
| 141 | 0.08 | 0.33 | 0.23 | 0.82 | 1 | Ino80dos | | 86 | 99 | 126 | 168 | 89 | 233 | 110 | 188 | 102 | 95 | 259 |
| 139 | 0.28 | 0.29 | 0.97 | 0.33 | 1 | Fbxo41 | F-box protein 41 | 234 | 186 | 173 | 98 | 60 | 148 | 110 | 101 | 71 | 155 | 190 |
| 136 | -0.10 | 0.37 | -0.26 | 0.79 | 1 | Ubash3b | ubiquitin associated and SH3 domain cor | 330 | 140 | 224 | 60 | 127 | 96 | 110 | 68 | 107 | 183 | 46 |
| 130 | -0.09 | 0.35 | -0.25 | 0.80 | 1 | Xkr8 | X-linked Kx blood group related 8 | 225 | 177 | 175 | 142 | 95 | 81 | 110 | 134 | 144 | 90 | 52 |
| 129 | -0.42 | 0.40 | -1.04 | 0.30 | 1 | Radil | Ras association and DIL domains | 267 | 301 | 188 | 115 | 47 | 96 | 110 | 57 | 25 | 130 | 82 |
| 108 | 0.16 | 0.37 | 0.43 | 0.67 | 1 | Zfp108 | zinc finger protein 108 | 115 | 115 | 68 | 78 | 86 | 151 | 110 | 78 | 128 | 122 | 135 |
| 107 | -0.53 | 0.45 | -1.19 | 0.24 | 1 | Tigd5 | tigger transposable element derived 5 | 33 | 188 | 112 | 158 | 139 | 85 | 110 | 117 | 88 | 91 | 49 |
| 104 | 0.42 | 0.40 | 1.04 | 0.30 | 1 | AI182371 | expressed sequence AI182371 | 82 | 61 | 64 | 76 | 100 | 109 | 110 | 207 | 110 | 99 | 120 |
| 103 | -0.36 | 0.45 | -0.81 | 0.42 | 1 | Sp6 | trans-acting transcription factor 6 | 106 | 117 | 64 | 221 | 92 | 63 | 110 | 64 | 108 | 130 | 53 |
| 99 | -1.37 | 0.82 | -1.68 | 0.09 | 1 | H2-Bl | histocompatibility 2, blastocyst | 396 | 8 | 260 | 21 | 107 | 0 | 110 | 0 | 81 | 107 | 0 |
| 98 | -0.05 | 0.40 | -0.13 | 0.89 | 1 | Gm6560 | | 104 | 101 | 57 | 105 | 115 | 83 | 110 | 139 | 91 | 67 | 108 |
| 93 | -0.91 | 0.73 | -1.25 | 0.21 | 1 | Mrpl23 | mitochondrial ribosomal protein L23 | 88 | 125 | 113 | 74 | 87 | 144 | 110 | 2 | 110 | 45 | 127 |
| 92 | 0.17 | 0.50 | 0.35 | 0.73 | 1 | 5830444B04Rik | | 105 | 31 | 0 | 39 | 7 | 253 | 110 | 60 | 4 | 170 | 233 |
| 89 | -0.32 | 0.45 | -0.70 | 0.48 | 1 | Ttll13 | tubulin tyrosine ligase-like family, membe | 57 | 52 | 111 | 71 | 78 | 166 | 110 | 98 | 81 | 49 | 109 |
| 77 | -0.25 | 0.42 | -0.59 | 0.55 | 1 | BC024063 | cDNA sequence BC024063 | 60 | 47 | 58 | 90 | 56 | 110 | 110 | 43 | 80 | 91 | 100 |
| 75 | 0.23 | 0.41 | 0.56 | 0.58 | 1 | Stamos | | 76 | 29 | 88 | 59 | 67 | 50 | 110 | 52 | 99 | 120 | 69 |
| 61 | 0.38 | 0.55 | 0.70 | 0.49 | 1 | Rtraf-ps | | 184 | 20 | 36 | 13 | 37 | 37 | 110 | 45 | 21 | 121 | 46 |
| 1048 | 0.12 | 0.22 | 0.53 | 0.59 | 1 | Oat | ornithine aminotransferase | 4379 | 3037 | 3085 | 61 | 140 | 183 | 109 | 103 | 102 | 106 | 225 |
| 324 | -0.56 | 0.34 | -1.64 | 0.10 | 1 | AI480526 | | 28 | 61 | 52 | 86 | 98 | 1624 | 109 | 43 | 113 | 45 | 1308 |

| | | | | | | | | | | | | | | | | | | |
|---|---|---|---|---|---|---|---|---|---|---|---|---|---|---|---|---|---|---|
| 177 | -0.47 | 0.29 | -1.59 | 0.11 | 1 | Hectd2 | HECT domain E3 ubiquitin protein ligase | 324 | 284 | 361 | 125 | 180 | 150 | 109 | 58 | 124 | 104 | 130 |
| 146 | 0.15 | 0.42 | 0.36 | 0.72 | 1 | Lrrc36 | leucine rich repeat containing 36 | 33 | 99 | 35 | 144 | 106 | 329 | 109 | 95 | 111 | 179 | 367 |
| 144 | -0.33 | 0.63 | -0.52 | 0.60 | 1 | Dbf4 | DBF4 zinc finger | 293 | 98 | 135 | 238 | 80 | 166 | 109 | 178 | 140 | 21 | 131 |
| 133 | -0.14 | 0.33 | -0.44 | 0.66 | 1 | Gm15500 | | 225 | 182 | 359 | 80 | 91 | 81 | 109 | 54 | 123 | 80 | 81 |
| 129 | -0.40 | 0.48 | -0.84 | 0.40 | 1 | BC051142 | cDNA sequence BC051142 | 71 | 140 | 102 | 71 | 35 | 425 | 109 | 63 | 76 | 31 | 297 |
| 127 | -0.30 | 0.60 | -0.50 | 0.62 | 1 | Gm4924 | predicted gene 4924 | 108 | 267 | 121 | 103 | 74 | 186 | 109 | 16 | 117 | 72 | 227 |
| 106 | -0.06 | 0.31 | -0.20 | 0.85 | 1 | 9330102E08Rik | | 59 | 60 | 57 | 134 | 119 | 142 | 109 | 137 | 133 | 99 | 116 |
| 102 | 0.14 | 0.41 | 0.34 | 0.73 | 1 | Gm4202 | | 155 | 132 | 69 | 88 | 89 | 70 | 109 | 166 | 103 | 86 | 56 |
| 101 | 0.13 | 0.36 | 0.36 | 0.72 | 1 | Lyrm1 | LYR motif containing 1 | 78 | 99 | 51 | 116 | 98 | 92 | 109 | 174 | 113 | 80 | 100 |
| 98 | -0.10 | 0.42 | -0.23 | 0.82 | 1 | Gm10509 | | 65 | 62 | 62 | 112 | 141 | 91 | 109 | 82 | 64 | 171 | 123 |
| 85 | -0.30 | 0.42 | -0.73 | 0.47 | 1 | Gm8210 | | 176 | 166 | 38 | 117 | 35 | 34 | 109 | 57 | 26 | 143 | 28 |
| 654 | 0.07 | 0.20 | 0.33 | 0.75 | 1 | Rab3c | RAB3C, member RAS oncogene family | 2295 | 2193 | 1633 | 194 | 103 | 120 | 108 | 191 | 101 | 130 | 127 |
| 213 | 0.16 | 0.23 | 0.67 | 0.50 | 1 | Trmt44 | tRNA methyltransferase 44 | 219 | 296 | 270 | 195 | 150 | 293 | 108 | 233 | 164 | 133 | 286 |
| 205 | -0.02 | 0.35 | -0.06 | 0.95 | 1 | Ccdc88b | coiled-coil domain containing 88B | 701 | 328 | 330 | 94 | 126 | 119 | 108 | 139 | 109 | 131 | 72 |
| 185 | -0.28 | 0.39 | -0.73 | 0.46 | 1 | A530020G20Rik | | 75 | 76 | 34 | 109 | 112 | 622 | 108 | 64 | 120 | 63 | 652 |
| 172 | -0.69 | 0.38 | -1.81 | 0.07 | 1 | Sall1 | spalt like transcription factor 1 | 228 | 198 | 131 | 321 | 169 | 226 | 108 | 189 | 120 | 111 | 86 |
| 130 | -0.05 | 0.33 | -0.16 | 0.88 | 1 | 4930522L14Rik | zinc finger protein 431-like | 178 | 234 | 260 | 125 | 103 | 56 | 108 | 87 | 99 | 93 | 86 |
| 119 | 0.46 | 0.37 | 1.23 | 0.22 | 1 | C8g | complement component 8, gamma polype | 67 | 81 | 36 | 131 | 146 | 90 | 108 | 145 | 152 | 213 | 145 |
| 113 | -1.01 | 0.88 | -1.14 | 0.25 | 1 | Gm5165 | predicted gene 5165 | 193 | 159 | 49 | 148 | 102 | 159 | 108 | 0 | 60 | 106 | 156 |
| 97 | 0.58 | 0.77 | 0.75 | 0.45 | 1 | Mc5r | melanocortin 5 receptor | 0 | 4 | 26 | 177 | 85 | 36 | 108 | 298 | 122 | 156 | 50 |
| 92 | -0.09 | 0.55 | -0.16 | 0.87 | 1 | Zfp940 | zinc finger protein 940 | 45 | 108 | 39 | 73 | 108 | 102 | 108 | 173 | 101 | 27 | 133 |
| 81 | 0.06 | 0.40 | 0.16 | 0.88 | 1 | Gm14399 | predicted gene 14393 | 75 | 31 | 28 | 73 | 85 | 101 | 108 | 60 | 108 | 103 | 114 |
| 75 | -0.26 | 0.41 | -0.64 | 0.53 | 1 | Trp53cor1 | | 39 | 85 | 49 | 87 | 85 | 75 | 108 | 81 | 85 | 79 | 55 |
| 65 | -0.29 | 0.68 | -0.43 | 0.67 | 1 | Lmo1 | LIM domain only 1 | 73 | 32 | 10 | 56 | 96 | 63 | 108 | 15 | 118 | 73 | 74 |
| 57 | 0.33 | 0.45 | 0.74 | 0.46 | 1 | Olfm2 | olfactomedin 2 | 63 | 53 | 25 | 43 | 47 | 24 | 108 | 70 | 58 | 106 | 32 |
| 236 | 0.47 | 0.49 | 0.96 | 0.34 | 1 | Xlr3a | X-linked lymphocyte-regulated 3A | 23 | 248 | 39 | 179 | 99 | 714 | 107 | 199 | 174 | 197 | 617 |
| 217 | 0.01 | 0.33 | 0.02 | 0.99 | 1 | Rad52 | RAD52 homolog, DNA repair protein | 193 | 253 | 399 | 143 | 119 | 429 | 107 | 156 | 96 | 146 | 341 |
| 175 | 0.17 | 0.29 | 0.60 | 0.55 | 1 | Serac1 | serine active site containing 1 | 146 | 178 | 144 | 211 | 94 | 288 | 107 | 246 | 143 | 108 | 264 |
| 160 | 0.19 | 0.43 | 0.43 | 0.67 | 1 | Tmem29 | transmembrane protein 29 | 188 | 271 | 183 | 82 | 143 | 179 | 107 | 206 | 67 | 189 | 142 |
| 131 | -0.37 | 0.33 | -1.09 | 0.27 | 1 | Klhdc9 | kelch domain containing 9 | 110 | 145 | 163 | 214 | 151 | 95 | 107 | 180 | 111 | 119 | 49 |
| 120 | 0.08 | 0.36 | 0.23 | 0.82 | 1 | Churc1 | churchill domain containing 1 | 115 | 68 | 96 | 121 | 95 | 189 | 107 | 158 | 114 | 112 | 142 |
| 112 | 0.31 | 0.42 | 0.73 | 0.47 | 1 | Pcdhb19 | protocadherin beta 19 | 91 | 166 | 79 | 95 | 103 | 94 | 107 | 81 | 143 | 178 | 98 |
| 112 | -0.23 | 0.36 | -0.64 | 0.52 | 1 | Mfsd3 | major facilitator superfamily domain conta | 147 | 79 | 84 | 123 | 114 | 158 | 107 | 118 | 105 | 84 | 116 |
| 111 | -0.16 | 0.39 | -0.40 | 0.69 | 1 | BC051226 | | 58 | 80 | 127 | 144 | 128 | 116 | 107 | 183 | 97 | 63 | 122 |
| 99 | -0.20 | 0.37 | -0.54 | 0.59 | 1 | Ccdc103 | coiled-coil domain containing 103 | 97 | 98 | 127 | 117 | 78 | 101 | 107 | 86 | 107 | 109 | 58 |
| 86 | -0.30 | 0.53 | -0.56 | 0.58 | 1 | Wdr17 | WD repeat domain 17 | 80 | 144 | 89 | 90 | 46 | 107 | 107 | 24 | 73 | 86 | 104 |
| 85 | -0.61 | 0.44 | -1.40 | 0.16 | 1 | Gm49937 | | 42 | 95 | 58 | 123 | 113 | 99 | 107 | 49 | 96 | 54 | 95 |
| 85 | 0.36 | 0.40 | 0.90 | 0.37 | 1 | Actrt3 | actin related protein T3 | 34 | 17 | 34 | 85 | 130 | 52 | 107 | 124 | 181 | 82 | 83 |
| 83 | -0.51 | 0.40 | -1.28 | 0.20 | 1 | Zfp971 | zinc finger protein 971 | 56 | 116 | 75 | 92 | 75 | 114 | 107 | 67 | 66 | 50 | 95 |
| 646 | 0.12 | 0.23 | 0.51 | 0.61 | 1 | Upk3a | uroplakin 3A | 1950 | 2526 | 1869 | 96 | 103 | 58 | 106 | 145 | 66 | 119 | 73 |
| 209 | -0.09 | 0.33 | -0.28 | 0.78 | 1 | Shank2 | SH3 and multiple ankyrin repeat domains | 415 | 527 | 608 | 94 | 85 | 96 | 106 | 120 | 70 | 119 | 60 |
| 189 | 0.41 | 0.35 | 1.17 | 0.24 | 1 | Anln | anillin, actin binding protein | 373 | 235 | 176 | 146 | 154 | 148 | 106 | 176 | 222 | 140 | 201 |
| 153 | 0.20 | 0.33 | 0.61 | 0.54 | 1 | Dnph1 | 2'-deoxynucleoside 5'-phosphate N-hydro | 151 | 157 | 155 | 209 | 155 | 93 | 106 | 279 | 114 | 155 | 110 |
| 138 | -0.44 | 0.59 | -0.75 | 0.45 | 1 | Aspm | abnormal spindle microtubule assembly | 312 | 63 | 63 | 204 | 135 | 171 | 106 | 86 | 152 | 67 | 159 |
| 136 | -0.11 | 0.61 | -0.19 | 0.85 | 1 | Cdca3 | cell division cycle associated 3 | 311 | 62 | 73 | 121 | 198 | 109 | 106 | 242 | 84 | 85 | 102 |
| 119 | 0.18 | 0.34 | 0.54 | 0.59 | 1 | Adnp | activity-dependent neuroprotective protei | 101 | 121 | 130 | 134 | 76 | 134 | 106 | 133 | 138 | 120 | 114 |
| 116 | -0.33 | 0.57 | -0.57 | 0.57 | 1 | Svbp | small vasohibin binding protein | 311 | 107 | 246 | 96 | 64 | 71 | 106 | 94 | 95 | 19 | 63 |
| 105 | 0.05 | 0.51 | 0.10 | 0.92 | 1 | Mcm8 | minichromosome maintenance 8 homolo | 85 | 59 | 65 | 130 | 77 | 111 | 106 | 43 | 106 | 290 | 86 |
| 102 | -1.08 | 0.56 | -1.92 | 0.05 | 1 | Gm1673 | predicted gene 1673 | 106 | 210 | 145 | 94 | 135 | 90 | 106 | 96 | 53 | 9 | 84 |
| 91 | -0.40 | 0.43 | -0.92 | 0.36 | 1 | | | 46 | 42 | 18 | 125 | 159 | 110 | 106 | 89 | 97 | 134 | 72 |
| 88 | -0.49 | 0.46 | -1.05 | 0.29 | 1 | Capza1-ps1 | | 293 | 34 | 111 | 28 | 133 | 47 | 106 | 25 | 55 | 82 | 47 |
| 77 | -0.58 | 0.49 | -1.17 | 0.24 | 1 | A930037H05Rik | | 53 | 27 | 42 | 113 | 64 | 127 | 106 | 39 | 58 | 157 | 56 |
| 74 | 0.17 | 0.51 | 0.33 | 0.74 | 1 | Capn9 | calpain 9 | 7 | 18 | 22 | 71 | 78 | 97 | 106 | 165 | 65 | 85 | 103 |
| 69 | -0.75 | 0.87 | -0.86 | 0.39 | 1 | Plaat1 | phospholipase A and acyltransferase 1 | 100 | 155 | 142 | 28 | 65 | 16 | 106 | 0 | 53 | 84 | 12 |
| 58 | 0.00 | 0.44 | -0.01 | 0.99 | 1 | Gm7467 | | 45 | 36 | 21 | 54 | 52 | 54 | 106 | 94 | 43 | 79 | 51 |
| 279 | -1.39 | 1.00 | -1.39 | 0.16 | 1 | Gm609 | predicted gene 609 | 597 | 1042 | 738 | 45 | 110 | 224 | 105 | 0 | 56 | 64 | 86 |
| 264 | -0.08 | 0.29 | -0.28 | 0.78 | 1 | Pald1 | phosphatase domain containing, paladin | 819 | 516 | 630 | 150 | 141 | 79 | 105 | 148 | 140 | 121 | 56 |
| 253 | 0.25 | 0.44 | 0.58 | 0.56 | 1 | Efna1 | ephrin A1 | 837 | 721 | 777 | 18 | 60 | 30 | 105 | 82 | 47 | 71 | 32 |
| 153 | 0.29 | 0.31 | 0.92 | 0.36 | 1 | Nemp2 | nuclear envelope integral membrane prot | 205 | 175 | 166 | 96 | 118 | 183 | 105 | 162 | 84 | 140 | 247 |
| 139 | 0.21 | 0.47 | 0.44 | 0.66 | 1 | Nfatc2ip | nuclear factor of activated T cells, cytopla | 233 | 131 | 150 | 101 | 75 | 173 | 105 | 214 | 75 | 48 | 223 |

| | | | | | | | | | | | | | | | | | |
|---|---|---|---|---|---|---|---|---|---|---|---|---|---|---|---|---|---|
| 127 | 0.01 | 0.30 | 0.02 | 0.98 | 1 | Pla2g2c | phospholipase A2, group IIC | 55 | 129 | 151 | 190 | 153 | 81 | 105 | 180 | 169 | 100 | 84 |
| 122 | 0.52 | 0.38 | 1.38 | 0.17 | 1 | Gm12481 | | 152 | 116 | 64 | 106 | 98 | 76 | 105 | 249 | 92 | 206 | 75 |
| 97 | -0.08 | 0.33 | -0.25 | 0.81 | 1 | Gm3699 | | 130 | 157 | 105 | 121 | 67 | 49 | 105 | 137 | 61 | 97 | 41 |
| 91 | 0.04 | 0.41 | 0.09 | 0.93 | 1 | Zfp58 | zinc finger protein 58 | 107 | 53 | 81 | 84 | 73 | 113 | 105 | 126 | 60 | 82 | 117 |
| 56 | -0.56 | 1.05 | -0.53 | 0.59 | 1 | Gm11460 | | 0 | 45 | 1 | 52 | 124 | 30 | 105 | 71 | 42 | 124 | 18 |
| 1321 | -0.48 | 0.29 | -1.64 | 0.10 | 1 | Firre | | 68 | 246 | 82 | 163 | 97 | 6956 | 104 | 61 | 86 | 72 | 6593 |
| 229 | 0.18 | 0.37 | 0.49 | 0.62 | 1 | Nemp1 | nuclear envelope integral membrane prot | 461 | 295 | 450 | 140 | 158 | 194 | 104 | 118 | 147 | 310 | 140 |
| 216 | -0.44 | 0.36 | -1.21 | 0.23 | 1 | Nfkbie | nuclear factor of kappa light polypeptide g | 824 | 444 | 406 | 98 | 85 | 117 | 104 | 49 | 110 | 63 | 79 |
| 177 | -0.18 | 0.40 | -0.44 | 0.66 | 1 | Nxpe4 | neurexophilin and PC-esterase domain fa | 323 | 368 | 540 | 98 | 71 | 105 | 104 | 94 | 38 | 104 | 106 |
| 148 | 0.18 | 0.43 | 0.42 | 0.68 | 1 | Wdr78 | WD repeat domain 78 | 112 | 183 | 132 | 106 | 184 | 167 | 104 | 72 | 147 | 229 | 197 |
| 128 | -0.19 | 0.42 | -0.44 | 0.66 | 1 | 2310014F06Rik | | 168 | 233 | 124 | 100 | 152 | 108 | 104 | 60 | 86 | 128 | 148 |
| 123 | -0.59 | 0.37 | -1.62 | 0.10 | 1 | Prickle3 | prickle planar cell polarity protein 3 | 160 | 251 | 202 | 109 | 94 | 136 | 104 | 52 | 49 | 94 | 106 |
| 111 | 0.43 | 0.34 | 1.25 | 0.21 | 1 | Cenps | centromere protein S | 166 | 189 | 162 | 61 | 81 | 53 | 104 | 123 | 81 | 124 | 76 |
| 110 | -0.50 | 0.40 | -1.23 | 0.22 | 1 | Zfp963 | zinc finger protein 963 | 52 | 121 | 94 | 115 | 123 | 175 | 104 | 57 | 97 | 52 | 216 |
| 104 | -0.39 | 0.40 | -0.98 | 0.33 | 1 | Gm13420 | predicted gene 13420 | 95 | 162 | 100 | 72 | 98 | 170 | 104 | 117 | 65 | 66 | 93 |
| 93 | -0.50 | 0.52 | -0.96 | 0.34 | 1 | Gm50069 | | 115 | 127 | 105 | 77 | 126 | 74 | 104 | 20 | 67 | 107 | 97 |
| 78 | 0.16 | 0.51 | 0.31 | 0.76 | 1 | Celf5 | CUGBP, Elav-like family member 5 | 68 | 64 | 76 | 60 | 66 | 56 | 104 | 188 | 68 | 80 | 32 |
| 76 | 0.02 | 0.59 | 0.03 | 0.98 | 1 | Pcdhb21 | protocadherin beta 21 | 86 | 66 | 25 | 74 | 84 | 63 | 104 | 138 | 113 | 35 | 52 |
| 75 | -0.85 | 0.45 | -1.90 | 0.06 | 1 | Gfra4 | glial cell line derived neurotrophic factor f | 139 | 95 | 49 | 91 | 74 | 64 | 104 | 88 | 25 | 77 | 23 |
| 67 | -0.49 | 0.68 | -0.72 | 0.47 | 1 | Eno1b | enolase 1B, retrotransposed | 102 | 19 | 120 | 5 | 111 | 40 | 104 | 0 | 94 | 120 | 21 |
| 5289 | -0.97 | 0.57 | -1.70 | 0.09 | 1 | Timp3 | tissue inhibitor of metalloproteinase 3 | 22107 | 15372 | 20435 | 47 | 20 | 10 | 103 | 25 | 34 | 14 | 6 |
| 279 | 0.26 | 0.29 | 0.89 | 0.37 | 1 | Mettl17 | methyltransferase like 17 | 165 | 223 | 212 | 121 | 182 | 822 | 103 | 256 | 158 | 144 | 677 |
| 220 | -0.16 | 0.32 | -0.50 | 0.62 | 1 | Misp | mitotic spindle positioning | 431 | 515 | 650 | 60 | 160 | 105 | 103 | 72 | 185 | 75 | 66 |
| 191 | -0.10 | 0.28 | -0.35 | 0.72 | 1 | Gcnt4 | glucosaminyl (N-acetyl) transferase 4, cor | 417 | 482 | 448 | 118 | 73 | 94 | 103 | 124 | 71 | 81 | 88 |
| 188 | 0.37 | 0.36 | 1.01 | 0.31 | 1 | Lin9 | lin-9 homolog (C. elegans) | 271 | 219 | 126 | 165 | 182 | 181 | 103 | 269 | 203 | 182 | 169 |
| 147 | 0.08 | 0.31 | 0.27 | 0.79 | 1 | Miip | migration and invasion inhibitory protein | 96 | 90 | 105 | 84 | 78 | 401 | 103 | 104 | 125 | 72 | 361 |
| 145 | 0.26 | 0.37 | 0.69 | 0.49 | 1 | Zfp341 | zinc finger protein 341 | 163 | 122 | 129 | 125 | 100 | 218 | 103 | 239 | 117 | 135 | 144 |
| 115 | 0.24 | 0.41 | 0.58 | 0.56 | 1 | Cep152 | centrosomal protein 152 | 122 | 129 | 103 | 56 | 60 | 219 | 103 | 153 | 79 | 76 | 167 |
| 114 | -0.12 | 0.43 | -0.28 | 0.78 | 1 | C920006O11Rik | | 84 | 63 | 113 | 54 | 115 | 196 | 103 | 23 | 166 | 71 | 261 |
| 94 | -0.71 | 0.58 | -1.22 | 0.22 | 1 | Ccdc138 | coiled-coil domain containing 138 | 74 | 224 | 121 | 88 | 48 | 131 | 103 | 21 | 82 | 35 | 105 |
| 90 | 0.25 | 0.32 | 0.79 | 0.43 | 1 | Zfp429 | zinc finger protein 429 | 107 | 68 | 84 | 74 | 73 | 82 | 103 | 82 | 100 | 130 | 85 |
| 78 | -0.21 | 0.67 | -0.31 | 0.76 | 1 | Rbp3 | retinol binding protein 3, interstitial | 0 | 14 | 3 | 142 | 152 | 52 | 103 | 94 | 117 | 151 | 30 |
| 74 | -0.02 | 0.49 | -0.05 | 0.96 | 1 | 2810403D21Rik | | 64 | 76 | 72 | 82 | 54 | 60 | 103 | 51 | 105 | 49 | 94 |
| 1077 | -0.09 | 0.59 | -0.15 | 0.88 | 1 | Plpp1 | phospholipid phosphatase 1 | 3703 | 3561 | 4065 | 16 | 115 | 19 | 102 | 25 | 59 | 167 | 12 |
| 423 | 0.15 | 0.23 | 0.67 | 0.50 | 1 | Ahr | aryl-hydrocarbon receptor | 1102 | 1106 | 1088 | 151 | 206 | 190 | 102 | 195 | 197 | 146 | 174 |
| 420 | -0.08 | 0.25 | -0.32 | 0.75 | 1 | Hes1 | hes family bHLH transcription factor 1 | 1470 | 957 | 1190 | 123 | 163 | 121 | 102 | 92 | 122 | 122 | 152 |
| 198 | 0.26 | 0.52 | 0.49 | 0.62 | 1 | Nek2 | NIMA (never in mitosis gene a)-related ex | 347 | 155 | 65 | 249 | 213 | 164 | 102 | 324 | 244 | 135 | 177 |
| 192 | -0.12 | 0.48 | -0.24 | 0.81 | 1 | 1700029J07Rik | RIKEN cDNA 1700029J07 gene | 146 | 82 | 105 | 81 | 74 | 664 | 102 | 161 | 66 | 28 | 609 |
| 190 | 0.00 | 0.43 | 0.01 | 0.99 | 1 | Bub1b | BUB1B, mitotic checkpoint serine/threonir | 847 | 221 | 215 | 95 | 97 | 109 | 102 | 85 | 109 | 82 | 126 |
| 153 | 0.23 | 0.28 | 0.79 | 0.43 | 1 | Zfp964 | zinc finger protein 964 | 216 | 263 | 222 | 121 | 83 | 139 | 102 | 175 | 84 | 102 | 173 |
| 141 | 0.12 | 0.39 | 0.31 | 0.76 | 1 | Zfp354a | zinc finger protein 354A | 118 | 174 | 143 | 108 | 128 | 199 | 102 | 127 | 93 | 218 | 141 |
| 127 | -0.09 | 0.35 | -0.26 | 0.80 | 1 | C1rl | complement component 1, r subcompone | 148 | 206 | 143 | 178 | 113 | 51 | 102 | 241 | 106 | 62 | 46 |
| 122 | -0.81 | 0.44 | -1.85 | 0.06 | 1 | Zfp202 | zinc finger protein 202 | 146 | 270 | 154 | 161 | 75 | 134 | 102 | 57 | 71 | 26 | 144 |
| 112 | 0.08 | 0.52 | 0.16 | 0.87 | 1 | Dlgap5 | DLG associated protein 5 | 290 | 31 | 48 | 108 | 77 | 88 | 102 | 67 | 83 | 242 | 96 |
| 104 | 0.19 | 0.33 | 0.58 | 0.56 | 1 | Zfp428 | zinc finger protein 428 | 139 | 81 | 87 | 96 | 95 | 93 | 102 | 131 | 102 | 130 | 85 |
| 103 | 0.93 | 0.48 | 1.92 | 0.05 | 1 | Tmigd3 | transmembrane and immunoglobulin don | 9 | 1 | 19 | 50 | 128 | 114 | 102 | 132 | 171 | 252 | 151 |
| 53 | 0.43 | 0.70 | 0.61 | 0.54 | 1 | Dock3 | dedicator of cyto-kinesis 3 | 17 | 3 | 0 | 102 | 28 | 30 | 102 | 153 | 43 | 61 | 49 |
| 660 | -0.49 | 0.33 | -1.49 | 0.14 | 1 | Rhoc | ras homolog family member C | 2157 | 1423 | 2270 | 243 | 296 | 184 | 101 | 222 | 142 | 140 | 84 |
| 589 | -0.32 | 0.38 | -0.85 | 0.40 | 1 | Olfml2b | olfactomedin-like 2B | 2653 | 1736 | 1265 | 227 | 68 | 66 | 101 | 193 | 75 | 39 | 58 |
| 327 | -0.53 | 0.34 | -1.58 | 0.11 | 1 | Rras | related RAS viral (r-ras) oncogene | 1201 | 747 | 790 | 146 | 166 | 91 | 101 | 66 | 122 | 62 | 100 |
| 204 | 0.39 | 0.38 | 1.02 | 0.31 | 1 | Knstrn | kinetochore-localized astrin/SPAG5 bindi | 288 | 191 | 135 | 276 | 170 | 156 | 101 | 342 | 171 | 170 | 241 |
| 186 | -0.02 | 0.35 | -0.05 | 0.96 | 1 | Tmem198b | transmembrane protein 198b | 338 | 188 | 279 | 123 | 183 | 213 | 101 | 156 | 104 | 119 | 239 |
| 168 | -0.47 | 0.36 | -1.30 | 0.19 | 1 | Prr36 | proline rich 36 | 327 | 487 | 458 | 66 | 113 | 55 | 101 | 79 | 48 | 79 | 39 |
| 158 | 0.06 | 0.31 | 0.21 | 0.83 | 1 | Caprin2 | caprin family member 2 | 160 | 167 | 125 | 94 | 96 | 356 | 101 | 97 | 143 | 85 | 316 |
| 119 | 0.05 | 0.35 | 0.14 | 0.89 | 1 | Wdr54 | WD repeat domain 54 | 125 | 146 | 153 | 147 | 95 | 88 | 101 | 115 | 98 | 162 | 76 |
| 116 | 0.36 | 0.34 | 1.07 | 0.29 | 1 | 9330104G04Rik | | 75 | 134 | 136 | 116 | 94 | 96 | 101 | 140 | 107 | 121 | 156 |
| 113 | -0.33 | 0.34 | -0.97 | 0.33 | 1 | Ttc39c | tetratricopeptide repeat domain 39C | 191 | 135 | 165 | 130 | 88 | 95 | 101 | 76 | 69 | 113 | 76 |
| 106 | -0.07 | 0.35 | -0.19 | 0.85 | 1 | Cdnf | cerebral dopamine neurotrophic factor | 130 | 123 | 110 | 136 | 87 | 87 | 101 | 94 | 94 | 77 | 126 |
| 103 | 0.12 | 0.36 | 0.33 | 0.74 | 1 | Galnt9 | polypeptide N-acetylgalactosaminyltransf | 36 | 30 | 20 | 130 | 101 | 161 | 101 | 127 | 81 | 196 | 151 |
| 102 | 0.07 | 0.44 | 0.16 | 0.87 | 1 | 2810002D19Rik | | 134 | 178 | 113 | 82 | 57 | 99 | 101 | 41 | 101 | 100 | 115 |

| | | | | | | | | | | | | | | | | | | |
|---|---|---|---|---|---|---|---|---|---|---|---|---|---|---|---|---|---|---|
| 100 | -0.03 | 0.47 | -0.06 | 0.95 | 1 | H4c8 | H4 clustered histone 8 | 23 | 13 | 59 | 131 | 142 | 132 | 101 | 166 | 106 | 84 | 138 |
| 94 | -0.09 | 0.57 | -0.15 | 0.88 | 1 | 0610009E02Rik | | 17 | 70 | 96 | 127 | 100 | 106 | 101 | 42 | 124 | 121 | 131 |
| 91 | -0.04 | 0.56 | -0.08 | 0.94 | 1 | Gm41177 | | 0 | 10 | 10 | 203 | 73 | 123 | 101 | 137 | 135 | 80 | 132 |
| 89 | 0.51 | 0.52 | 0.97 | 0.33 | 1 | Ppp1r42 | protein phosphatase 1, regulatory subuni | 9 | 12 | 43 | 44 | 121 | 120 | 101 | 71 | 120 | 218 | 119 |
| 89 | -0.27 | 0.42 | -0.65 | 0.52 | 1 | Il17d | interleukin 17D | 141 | 79 | 144 | 88 | 73 | 62 | 101 | 44 | 55 | 133 | 57 |
| 87 | -0.05 | 0.46 | -0.12 | 0.91 | 1 | Epha8 | Eph receptor A8 | 42 | 109 | 70 | 89 | 75 | 90 | 101 | 173 | 47 | 86 | 74 |
| 520 | -0.04 | 0.31 | -0.13 | 0.90 | 1 | Plek | pleckstrin | 2559 | 1168 | 949 | 144 | 112 | 172 | 100 | 102 | 133 | 160 | 127 |
| 185 | -0.53 | 0.38 | -1.41 | 0.16 | 1 | Pvr | poliovirus receptor | 528 | 610 | 378 | 68 | 61 | 78 | 100 | 30 | 64 | 44 | 77 |
| 164 | 0.56 | 0.30 | 1.85 | 0.06 | 1 | Gm4204 | | 133 | 153 | 222 | 106 | 168 | 153 | 100 | 124 | 215 | 227 | 203 |
| 159 | -0.12 | 0.42 | -0.29 | 0.77 | 1 | Metrnl | meteorin, glial cell differentiation regulato | 706 | 260 | 271 | 66 | 75 | 37 | 100 | 61 | 83 | 45 | 47 |
| 150 | -0.20 | 0.30 | -0.67 | 0.50 | 1 | Gon7 | GON7 subunit of KEOPS complex | 151 | 164 | 173 | 175 | 173 | 171 | 100 | 174 | 103 | 103 | 158 |
| 140 | 0.10 | 0.33 | 0.31 | 0.76 | 1 | Gins3 | GINS complex subunit 3 (Psf3 homolog) | 93 | 214 | 127 | 119 | 156 | 149 | 100 | 137 | 133 | 97 | 214 |
| 134 | 0.25 | 0.30 | 0.86 | 0.39 | 1 | Fam241a | family with sequence similarity 241, meml | 133 | 182 | 177 | 123 | 112 | 112 | 100 | 141 | 130 | 158 | 107 |
| 134 | -0.11 | 0.45 | -0.25 | 0.80 | 1 | Klhl29 | kelch-like 29 | 274 | 101 | 204 | 173 | 104 | 84 | 100 | 108 | 171 | 79 | 79 |
| 118 | -0.27 | 0.37 | -0.72 | 0.47 | 1 | Rasal3 | RAS protein activator like 3 | 289 | 151 | 157 | 113 | 79 | 80 | 100 | 57 | 99 | 113 | 54 |
| 93 | 0.08 | 0.44 | 0.18 | 0.86 | 1 | Ddt | D-dopachrome tautomerase | 93 | 47 | 105 | 90 | 89 | 95 | 100 | 119 | 115 | 54 | 111 |
| 84 | -1.44 | 0.74 | -1.94 | 0.05 | 1 | 4732496C06Rik | | 42 | 17 | 30 | 93 | 133 | 212 | 100 | 16 | 130 | 4 | 149 |
| 81 | -0.69 | 0.51 | -1.34 | 0.18 | 1 | Grk4 | G protein-coupled receptor kinase 4 | 71 | 88 | 57 | 51 | 100 | 146 | 100 | 21 | 82 | 39 | 137 |
| 76 | -0.38 | 0.46 | -0.83 | 0.41 | 1 | 2510016D11Rik | | 43 | 69 | 79 | 86 | 50 | 132 | 100 | 65 | 52 | 85 | 73 |
| 73 | -0.45 | 0.40 | -1.12 | 0.26 | 1 | Hdhd3 | haloacid dehalogenase-like hydrolase dc | 109 | 58 | 88 | 68 | 91 | 56 | 100 | 59 | 41 | 84 | 47 |
| 5139 | -3.33 | 2.76 | -1.20 | 0.23 | 1 | Flt1 | FMS-like tyrosine kinase 1 | 17242 | 18869 | 20315 | 0 | 0 | 3 | 99 | 2 | 0 | 0 | 0 |
| 248 | 0.12 | 0.28 | 0.43 | 0.67 | 1 | Irf7 | interferon regulatory factor 7 | 698 | 309 | 911 | 61 | 139 | 88 | 99 | 85 | 150 | 128 | 63 |
| 211 | 0.20 | 0.36 | 0.57 | 0.57 | 1 | Lrrc8a | leucine rich repeat containing 8A VRAC s | 414 | 349 | 351 | 101 | 244 | 127 | 99 | 199 | 147 | 128 | 157 |
| 204 | 0.00 | 0.40 | 0.00 | 1.00 | 1 | Relb | avian reticuloendotheliosis viral (v-rel) on | 643 | 336 | 378 | 115 | 87 | 143 | 99 | 140 | 118 | 50 | 134 |
| 192 | 0.23 | 0.35 | 0.66 | 0.51 | 1 | Asap3 | ArfGAP with SH3 domain, ankyrin repeat | 468 | 396 | 398 | 116 | 66 | 107 | 99 | 174 | 129 | 77 | 82 |
| 174 | -0.15 | 0.33 | -0.44 | 0.66 | 1 | Arid5a | AT rich interactive domain 5A (MRF1-like) | 504 | 365 | 184 | 120 | 75 | 145 | 99 | 153 | 43 | 107 | 117 |
| 164 | 0.30 | 0.35 | 0.86 | 0.39 | 1 | Ndor1 | NADPH dependent diflavin oxidoreductas | 113 | 144 | 176 | 116 | 144 | 276 | 99 | 154 | 123 | 219 | 238 |
| 145 | -0.70 | 0.50 | -1.40 | 0.16 | 1 | Plk4 | polo like kinase 4 | 269 | 129 | 121 | 257 | 137 | 151 | 99 | 73 | 93 | 58 | 208 |
| 132 | 0.22 | 0.31 | 0.71 | 0.47 | 1 | Prdm11 | PR domain containing 11 | 240 | 153 | 178 | 86 | 90 | 131 | 99 | 132 | 92 | 103 | 143 |
| 114 | -0.17 | 0.43 | -0.41 | 0.69 | 1 | Zik1 | zinc finger protein interacting with K prote | 69 | 213 | 177 | 151 | 63 | 100 | 99 | 163 | 61 | 88 | 68 |
| 108 | 0.54 | 0.57 | 0.96 | 0.34 | 1 | Vkorc1 | vitamin K epoxide reductase complex, sul | 108 | 27 | 115 | 68 | 102 | 114 | 99 | 238 | 153 | 81 | 86 |
| 105 | 0.19 | 0.33 | 0.59 | 0.56 | 1 | Hsf4 | heat shock transcription factor 4 | 37 | 44 | 59 | 120 | 117 | 137 | 99 | 110 | 146 | 146 | 138 |
| 87 | -0.29 | 0.48 | -0.60 | 0.55 | 1 | Rbfox3 | RNA binding protein, fox-1 homolog (C. e | 58 | 145 | 197 | 65 | 73 | 67 | 99 | 88 | 60 | 53 | 56 |
| 81 | -1.33 | 0.74 | -1.81 | 0.07 | 1 | Nexmif | neurite extension and migration factor | 233 | 201 | 171 | 16 | 46 | 37 | 99 | 0 | 21 | 53 | 17 |
| 77 | 0.43 | 0.60 | 0.72 | 0.47 | 1 | S100z | S100 calcium binding protein, zeta | 14 | 17 | 0 | 97 | 59 | 104 | 99 | 120 | 103 | 94 | 143 |
| 556 | 0.05 | 0.84 | 0.06 | 0.95 | 1 | Gm49542 | | 0 | 13 | 14 | 124 | 71 | 2758 | 98 | 44 | 118 | 138 | 2735 |
| 224 | -1.10 | 0.69 | -1.60 | 0.11 | 1 | H2bu2 | H2B.U histone 2 | 536 | 690 | 581 | 60 | 94 | 170 | 98 | 0 | 76 | 62 | 93 |
| 160 | 0.33 | 0.38 | 0.87 | 0.39 | 1 | 4930481A15Rik | | 105 | 154 | 181 | 103 | 146 | 234 | 98 | 273 | 97 | 119 | 251 |
| 154 | -0.12 | 0.35 | -0.34 | 0.73 | 1 | Zfp105 | zinc finger protein 105 | 250 | 335 | 312 | 125 | 78 | 112 | 98 | 61 | 98 | 119 | 103 |
| 134 | -0.05 | 0.37 | -0.12 | 0.90 | 1 | Tcea2 | transcription elongation factor A (SII), 2 | 116 | 117 | 148 | 75 | 50 | 351 | 98 | 94 | 82 | 53 | 295 |
| 124 | -0.20 | 0.41 | -0.49 | 0.63 | 1 | Def6 | differentially expressed in FDCP 6 | 335 | 165 | 156 | 142 | 75 | 63 | 98 | 64 | 109 | 111 | 51 |
| 117 | 0.03 | 0.39 | 0.07 | 0.94 | 1 | Lyrm7 | LYR motif containing 7 | 88 | 187 | 117 | 73 | 101 | 172 | 98 | 153 | 92 | 89 | 119 |
| 104 | -0.39 | 0.43 | -0.92 | 0.36 | 1 | 1700007L15Rik | | 121 | 93 | 68 | 121 | 168 | 100 | 98 | 78 | 108 | 63 | 127 |
| 86 | -0.20 | 0.44 | -0.46 | 0.65 | 1 | 4833413G10Rik | | 68 | 104 | 63 | 83 | 95 | 97 | 98 | 63 | 99 | 125 | 51 |
| 84 | -0.55 | 0.40 | -1.36 | 0.17 | 1 | Tyw3 | tRNA-yW synthesizing protein 3 homolog | 124 | 100 | 122 | 99 | 57 | 90 | 98 | 68 | 46 | 43 | 81 |
| 79 | -0.73 | 0.83 | -0.87 | 0.38 | 1 | Hdx | highly divergent homeobox | 117 | 85 | 77 | 48 | 61 | 126 | 98 | 0 | 91 | 59 | 102 |
| 57 | -0.22 | 0.47 | -0.47 | 0.64 | 1 | Pcdha9 | protocadherin alpha 9 | 20 | 22 | 16 | 67 | 77 | 55 | 98 | 39 | 55 | 121 | 52 |
| 417 | -0.25 | 0.36 | -0.68 | 0.50 | 1 | Kit | KIT proto-oncogene receptor tyrosine kina | 1265 | 1441 | 870 | 137 | 84 | 196 | 97 | 198 | 66 | 30 | 198 |
| 254 | -0.06 | 0.39 | -0.16 | 0.87 | 1 | Cdkn1c | cyclin-dependent kinase inhibitor 1C (P57 | 736 | 604 | 847 | 67 | 58 | 81 | 97 | 119 | 36 | 100 | 48 |
| 244 | -0.53 | 0.29 | -1.82 | 0.07 | 1 | Susd2 | sushi domain containing 2 | 1351 | 474 | 456 | 51 | 43 | 46 | 97 | 23 | 34 | 68 | 41 |
| 240 | -0.10 | 0.25 | -0.39 | 0.70 | 1 | Tmeff1 | transmembrane protein with EGF-like anc | 534 | 621 | 443 | 202 | 123 | 123 | 97 | 162 | 119 | 102 | 118 |
| 235 | -0.23 | 0.34 | -0.68 | 0.50 | 1 | Plod2 | procollagen lysine, 2-oxoglutarate 5-diox | 659 | 480 | 406 | 215 | 99 | 146 | 97 | 197 | 88 | 76 | 117 |
| 224 | -0.11 | 0.27 | -0.42 | 0.67 | 1 | Tro | trophinin | 506 | 554 | 351 | 164 | 137 | 148 | 97 | 155 | 90 | 113 | 153 |
| 203 | 0.36 | 0.31 | 1.17 | 0.24 | 1 | Gsdmd | gasdermin D | 490 | 388 | 367 | 95 | 64 | 183 | 97 | 175 | 99 | 91 | 179 |
| 171 | -0.13 | 0.47 | -0.28 | 0.78 | 1 | Cdk1 | cyclin-dependent kinase 1 | 439 | 230 | 121 | 137 | 202 | 128 | 97 | 126 | 133 | 91 | 173 |
| 168 | -0.14 | 0.28 | -0.51 | 0.61 | 1 | A2ml1 | alpha-2-macroglobulin like 1 | 320 | 314 | 421 | 101 | 100 | 120 | 97 | 107 | 89 | 99 | 83 |
| 154 | -0.54 | 0.35 | -1.53 | 0.13 | 1 | Gstm7 | glutathione S-transferase, mu 7 | 303 | 380 | 336 | 131 | 77 | 87 | 97 | 105 | 59 | 33 | 82 |
| 119 | -0.63 | 0.40 | -1.57 | 0.12 | 1 | Tcp11 | t-complex protein 11 | 140 | 297 | 190 | 95 | 58 | 156 | 97 | 66 | 67 | 30 | 110 |
| 105 | 0.90 | 0.50 | 1.82 | 0.07 | 1 | Gm35315 | predicted gene, 35315 | 52 | 60 | 126 | 63 | 93 | 66 | 97 | 232 | 84 | 120 | 160 |
| 101 | -0.62 | 0.35 | -1.77 | 0.08 | 1 | Fastkd5 | FAST kinase domains 5 | 136 | 164 | 133 | 135 | 71 | 107 | 97 | 56 | 56 | 66 | 88 |

| | | | | | | | | | | | | | | | | | | |
|---|---|---|---|---|---|---|---|---|---|---|---|---|---|---|---|---|---|---|
| 83 | -0.67 | 0.49 | -1.35 | 0.18 | 1 | Zfp647 | zinc finger protein 647 | 160 | 75 | 121 | 63 | 89 | 70 | 97 | 13 | 60 | 124 | 47 |
| 63 | -0.92 | 0.94 | -0.98 | 0.32 | 1 | Gm12758 | | 21 | 54 | 62 | 70 | 86 | 68 | 97 | 0 | 99 | 35 | 98 |
| 56 | 0.13 | 0.62 | 0.21 | 0.83 | 1 | Rprml | reprimo-like | 10 | 0 | 4 | 86 | 69 | 36 | 97 | 66 | 57 | 116 | 75 |
| 395 | 0.47 | 0.29 | 1.62 | 0.11 | 1 | H2-DMa | histocompatibility 2, class II, locus DMa | 1669 | 853 | 717 | 116 | 117 | 117 | 96 | 146 | 128 | 264 | 126 |
| 336 | -0.46 | 0.39 | -1.18 | 0.24 | 1 | Agpat4 | 1-acylglycerol-3-phosphate O-acyltransfe | 860 | 1114 | 1123 | 37 | 112 | 98 | 96 | 88 | 71 | 33 | 59 |
| 134 | -0.20 | 0.30 | -0.69 | 0.49 | 1 | Gm17259 | | 104 | 119 | 137 | 163 | 229 | 113 | 96 | 127 | 179 | 91 | 113 |
| 124 | -0.54 | 0.39 | -1.40 | 0.16 | 1 | Rnf113a1 | ring finger protein 113A1 | 84 | 97 | 123 | 223 | 147 | 164 | 96 | 71 | 107 | 103 | 150 |
| 120 | -0.02 | 0.35 | -0.05 | 0.96 | 1 | Zfp128 | zinc finger protein 128 | 142 | 124 | 166 | 111 | 106 | 122 | 96 | 64 | 91 | 133 | 163 |
| 116 | 0.13 | 0.41 | 0.33 | 0.74 | 1 | Sp3os | | 116 | 171 | 109 | 67 | 154 | 98 | 96 | 148 | 145 | 81 | 89 |
| 103 | 0.02 | 0.32 | 0.05 | 0.96 | 1 | Trim67 | tripartite motif-containing 67 | 89 | 120 | 63 | 129 | 85 | 113 | 96 | 162 | 90 | 90 | 93 |
| 96 | -0.16 | 0.34 | -0.47 | 0.64 | 1 | Pin4 | protein (peptidyl-prolyl cis/trans isomeras | 100 | 92 | 145 | 75 | 125 | 84 | 96 | 72 | 79 | 99 | 84 |
| 86 | -0.62 | 0.44 | -1.42 | 0.15 | 1 | Gramd2 | GRAM domain containing 2 | 30 | 42 | 86 | 98 | 89 | 168 | 96 | 52 | 66 | 44 | 176 |
| 83 | -0.01 | 0.43 | -0.02 | 0.98 | 1 | Gm4430 | | 58 | 47 | 48 | 91 | 112 | 72 | 96 | 42 | 117 | 148 | 83 |
| 55 | -0.41 | 0.73 | -0.55 | 0.58 | 1 | Gm6483 | | 65 | 50 | 31 | 31 | 43 | 80 | 96 | 2 | 54 | 80 | 78 |
| 173 | 0.25 | 0.36 | 0.67 | 0.50 | 1 | Haus4 | HAUS augmin-like complex, subunit 4 | 319 | 199 | 227 | 221 | 112 | 106 | 95 | 156 | 156 | 196 | 120 |
| 158 | 0.05 | 0.49 | 0.10 | 0.92 | 1 | Abcd4 | ATP-binding cassette, sub-family D (ALD) | 351 | 151 | 152 | 87 | 81 | 280 | 95 | 170 | 84 | 41 | 246 |
| 152 | -0.35 | 0.45 | -0.78 | 0.44 | 1 | Pmf1 | polyamine-modulated factor 1 | 428 | 214 | 185 | 116 | 128 | 122 | 95 | 169 | 76 | 64 | 71 |
| 143 | -0.71 | 0.75 | -0.94 | 0.35 | 1 | Procr | protein C receptor, endothelial | 492 | 317 | 323 | 90 | 14 | 14 | 95 | 156 | 10 | 62 | 0 |
| 139 | 0.38 | 0.42 | 0.92 | 0.36 | 1 | Gm36738 | | 67 | 109 | 182 | 175 | 133 | 90 | 95 | 343 | 114 | 126 | 98 |
| 136 | -0.13 | 0.30 | -0.45 | 0.66 | 1 | 2310009B15Ril | RIKEN cDNA 2310009B15 gene | 214 | 201 | 167 | 149 | 120 | 112 | 95 | 124 | 102 | 91 | 116 |
| 128 | 0.17 | 0.29 | 0.60 | 0.55 | 1 | C330011M18Rik | | 100 | 128 | 140 | 106 | 154 | 135 | 95 | 140 | 163 | 126 | 119 |
| 126 | 0.35 | 0.35 | 1.01 | 0.31 | 1 | C2cd4a | C2 calcium-dependent domain containing | 39 | 40 | 26 | 116 | 107 | 240 | 95 | 93 | 134 | 182 | 318 |
| 120 | -0.06 | 0.43 | -0.15 | 0.88 | 1 | Prr15 | proline rich 15 | 256 | 183 | 262 | 71 | 84 | 57 | 95 | 147 | 58 | 70 | 40 |
| 86 | -0.01 | 0.53 | -0.01 | 0.99 | 1 | Gm45495 | | 39 | 64 | 90 | 75 | 106 | 77 | 95 | 181 | 98 | 28 | 95 |
| 80 | 0.00 | 0.50 | 0.01 | 0.99 | 1 | Hyal3 | hyaluronoglucosaminidase 3 | 46 | 100 | 86 | 43 | 95 | 87 | 95 | 51 | 135 | 88 | 58 |
| 80 | 0.35 | 0.53 | 0.66 | 0.51 | 1 | Xlr3b | X-linked lymphocyte-regulated 3B | 37 | 52 | 8 | 60 | 8 | 243 | 95 | 92 | 33 | 61 | 193 |
| 71 | -0.30 | 0.55 | -0.55 | 0.58 | 1 | Gm47882 | | 33 | 13 | 43 | 81 | 88 | 84 | 95 | 19 | 73 | 137 | 115 |
| 64 | -0.30 | 0.54 | -0.56 | 0.58 | 1 | C130046K22Rik | | 42 | 100 | 38 | 43 | 74 | 67 | 95 | 96 | 34 | 59 | 58 |
| 140 | 0.36 | 0.39 | 0.92 | 0.36 | 1 | Dct | dopachrome tautomerase | 34 | 109 | 55 | 152 | 187 | 151 | 94 | 271 | 149 | 196 | 140 |
| 118 | -0.14 | 0.42 | -0.33 | 0.74 | 1 | 1500026H17Rik | | 89 | 46 | 84 | 168 | 166 | 123 | 94 | 68 | 151 | 143 | 162 |
| 115 | -0.02 | 0.41 | -0.04 | 0.96 | 1 | Ppt2 | palmitoyl-protein thioesterase 2 | 235 | 159 | 144 | 53 | 119 | 102 | 94 | 102 | 93 | 61 | 98 |
| 111 | 0.33 | 0.46 | 0.71 | 0.48 | 1 | Gm2830 | | 148 | 243 | 190 | 47 | 95 | 41 | 94 | 152 | 58 | 79 | 69 |
| 108 | -0.02 | 0.34 | -0.06 | 0.95 | 1 | Cdc42ep5 | CDC42 effector protein (Rho GTPase bin | 213 | 177 | 185 | 80 | 95 | 36 | 94 | 112 | 70 | 98 | 29 |
| 99 | 0.29 | 0.48 | 0.61 | 0.54 | 1 | Tomm6os | | 27 | 33 | 79 | 61 | 136 | 156 | 94 | 123 | 98 | 173 | 113 |
| 96 | 0.20 | 0.36 | 0.55 | 0.59 | 1 | Pcdhga4 | protocadherin gamma subfamily A, 4 | 96 | 67 | 94 | 68 | 144 | 68 | 94 | 126 | 139 | 99 | 62 |
| 93 | -0.49 | 0.37 | -1.32 | 0.19 | 1 | Sv2c | synaptic vesicle glycoprotein 2c | 104 | 134 | 143 | 68 | 81 | 129 | 94 | 67 | 68 | 37 | 100 |
| 93 | 0.67 | 0.60 | 1.11 | 0.26 | 1 | Prelid3a | PRELI domain containing 3A | 51 | 89 | 125 | 87 | 21 | 90 | 94 | 162 | 89 | 156 | 59 |
| 92 | -0.65 | 0.63 | -1.04 | 0.30 | 1 | Gm5345 | | 213 | 104 | 245 | 50 | 69 | 24 | 94 | 7 | 25 | 162 | 22 |
| 88 | 0.11 | 0.39 | 0.29 | 0.78 | 1 | Zfp991 | zinc finger protein 991 | 53 | 98 | 105 | 90 | 86 | 70 | 94 | 79 | 101 | 106 | 81 |
| 81 | -0.60 | 0.62 | -0.98 | 0.33 | 1 | 8430432A02Rik | | 73 | 60 | 127 | 72 | 166 | 47 | 94 | 101 | 73 | 12 | 65 |
| 78 | -0.04 | 0.42 | -0.09 | 0.93 | 1 | A230072E10Rik | | 45 | 64 | 51 | 53 | 55 | 159 | 94 | 46 | 65 | 99 | 130 |
| 72 | -0.17 | 0.51 | -0.33 | 0.74 | 1 | Hadhb-ps | | 73 | 57 | 91 | 67 | 62 | 69 | 94 | 129 | 72 | 45 | 37 |
| 66 | -0.27 | 0.53 | -0.51 | 0.61 | 1 | Myo3a | myosin IIIA | 2 | 8 | 15 | 121 | 107 | 68 | 94 | 75 | 73 | 85 | 79 |
| 58 | -0.19 | 0.64 | -0.30 | 0.77 | 1 | Gm15728 | | 22 | 30 | 43 | 109 | 16 | 75 | 94 | 49 | 61 | 76 | 59 |
| 233 | -0.40 | 0.26 | -1.54 | 0.12 | 1 | Chsy1 | chondroitin sulfate synthase 1 | 943 | 421 | 587 | 58 | 82 | 115 | 93 | 39 | 64 | 66 | 98 |
| 230 | 0.19 | 0.38 | 0.50 | 0.61 | 1 | Ppm1m | protein phosphatase 1M | 826 | 439 | 396 | 89 | 130 | 93 | 93 | 162 | 118 | 76 | 107 |
| 186 | -0.02 | 0.36 | -0.05 | 0.96 | 1 | Igdcc4 | immunoglobulin superfamily, DCC subcla | 511 | 386 | 323 | 101 | 119 | 96 | 93 | 170 | 98 | 71 | 78 |
| 168 | -0.05 | 0.35 | -0.14 | 0.89 | 1 | Snhg5 | | 348 | 130 | 193 | 78 | 132 | 296 | 93 | 79 | 124 | 86 | 290 |
| 126 | 0.67 | 0.40 | 1.66 | 0.10 | 1 | Gm6563 | | 158 | 162 | 143 | 74 | 85 | 89 | 93 | 265 | 91 | 135 | 94 |
| 124 | 0.27 | 0.29 | 0.91 | 0.36 | 1 | Kcnn2 | potassium intermediate/small conductanc | 235 | 155 | 124 | 79 | 51 | 169 | 93 | 90 | 69 | 133 | 166 |
| 99 | 0.51 | 0.37 | 1.37 | 0.17 | 1 | Zfp953 | zinc finger protein 953 | 118 | 124 | 85 | 59 | 60 | 108 | 93 | 130 | 108 | 93 | 112 |
| 85 | -0.42 | 0.39 | -1.10 | 0.27 | 1 | Gm14261 | | 43 | 91 | 50 | 132 | 118 | 90 | 93 | 80 | 107 | 54 | 82 |
| 81 | 0.18 | 0.59 | 0.30 | 0.77 | 1 | Rps18-ps5 | | 178 | 35 | 247 | 13 | 104 | 12 | 93 | 56 | 100 | 44 | 8 |
| 71 | 0.50 | 0.49 | 1.02 | 0.31 | 1 | Mzf1 | myeloid zinc finger 1 | 49 | 30 | 32 | 45 | 36 | 119 | 93 | 116 | 77 | 58 | 127 |
| 59 | 0.12 | 0.58 | 0.20 | 0.84 | 1 | A930029G22Rik | | 29 | 25 | 67 | 68 | 40 | 47 | 93 | 103 | 70 | 43 | 60 |
| 652 | -0.36 | 0.48 | -0.77 | 0.44 | 1 | Col16a1 | collagen, type XVI, alpha 1 | 3083 | 961 | 1378 | 335 | 233 | 316 | 92 | 358 | 152 | 90 | 172 |
| 319 | -1.56 | 0.94 | -1.66 | 0.10 | 1 | Ncs1 | neuronal calcium sensor 1 | 1136 | 876 | 962 | 82 | 98 | 111 | 92 | 48 | 48 | 0 | 58 |
| 182 | 0.01 | 0.43 | 0.02 | 0.98 | 1 | Prss8 | protease, serine 8 (prostasin) | 274 | 475 | 316 | 152 | 97 | 126 | 92 | 49 | 125 | 158 | 133 |
| 137 | -0.35 | 0.38 | -0.90 | 0.37 | 1 | Cenpa | centromere protein A | 447 | 169 | 186 | 109 | 81 | 105 | 92 | 48 | 75 | 102 | 96 |
| 128 | -0.39 | 0.51 | -0.77 | 0.44 | 1 | Hunk | hormonally upregulated Neu-associated l | 170 | 285 | 263 | 74 | 122 | 77 | 92 | 67 | 147 | 15 | 94 |

| | | | | | | | | | | | | | | | | | | |
|---|---|---|---|---|---|---|---|---|---|---|---|---|---|---|---|---|---|---|
| 125 | 0.45 | 0.41 | 1.10 | 0.27 | 1 | Gm2423 | | 140 | 77 | 122 | 126 | 122 | 92 | 92 | 249 | 144 | 100 | 112 |
| 122 | -0.19 | 0.34 | -0.56 | 0.58 | 1 | H2bc8 | H2B clustered histone 8 | 45 | 58 | 74 | 157 | 125 | 242 | 92 | 97 | 190 | 76 | 181 |
| 118 | 0.12 | 0.36 | 0.32 | 0.75 | 1 | Kdm8 | lysine (K)-specific demethylase 8 | 195 | 90 | 126 | 70 | 82 | 181 | 92 | 108 | 62 | 98 | 192 |
| 103 | 0.43 | 0.47 | 0.91 | 0.36 | 1 | Grcc10 | gene rich cluster, C10 gene | 65 | 62 | 108 | 37 | 114 | 157 | 92 | 160 | 100 | 90 | 147 |
| 102 | 0.04 | 0.45 | 0.08 | 0.93 | 1 | Gm40466 | | 105 | 122 | 29 | 66 | 68 | 208 | 92 | 71 | 62 | 139 | 161 |
| 92 | 0.12 | 0.45 | 0.26 | 0.79 | 1 | Slc26a6 | solute carrier family 26, member 6 | 76 | 73 | 75 | 48 | 48 | 199 | 92 | 122 | 43 | 59 | 181 |
| 87 | -0.43 | 0.37 | -1.16 | 0.25 | 1 | Cldn23 | claudin 23 | 120 | 69 | 91 | 56 | 142 | 101 | 92 | 51 | 94 | 63 | 75 |
| 79 | -0.78 | 0.48 | -1.62 | 0.11 | 1 | Slco1a5 | solute carrier organic anion transporter fa | 15 | 28 | 53 | 124 | 127 | 107 | 92 | 38 | 162 | 32 | 85 |
| 508 | -0.18 | 0.33 | -0.54 | 0.59 | 1 | Plin3 | perilipin 3 | 1945 | 1322 | 1464 | 133 | 127 | 79 | 91 | 59 | 116 | 202 | 52 |
| 497 | -0.37 | 0.24 | -1.54 | 0.12 | 1 | Peak1 | pseudopodium-enriched atypical kinase 1 | 1982 | 1488 | 1275 | 91 | 92 | 136 | 91 | 76 | 86 | 76 | 75 |
| 173 | 0.15 | 0.35 | 0.44 | 0.66 | 1 | 2700097O09Ril | RIKEN cDNA 2700097O09 gene | 125 | 146 | 128 | 100 | 127 | 339 | 91 | 67 | 100 | 151 | 533 |
| 169 | 0.18 | 0.32 | 0.57 | 0.57 | 1 | Ubald2 | UBA-like domain containing 2 | 274 | 235 | 265 | 121 | 180 | 117 | 91 | 112 | 152 | 187 | 125 |
| 168 | -0.04 | 0.29 | -0.14 | 0.89 | 1 | Rad9b | RAD9 checkpoint clamp component B | 72 | 76 | 43 | 81 | 59 | 622 | 91 | 68 | 75 | 80 | 583 |
| 148 | -0.29 | 0.48 | -0.60 | 0.55 | 1 | Hebp1 | heme binding protein 1 | 543 | 316 | 273 | 84 | 42 | 43 | 91 | 143 | 28 | 35 | 31 |
| 133 | 0.22 | 0.43 | 0.51 | 0.61 | 1 | Fam221a | family with sequence similarity 221, meml | 144 | 174 | 190 | 115 | 81 | 147 | 91 | 169 | 73 | 198 | 82 |
| 126 | -0.10 | 0.41 | -0.25 | 0.81 | 1 | Gm15446 | | 131 | 72 | 79 | 97 | 224 | 156 | 91 | 90 | 119 | 121 | 204 |
| 125 | 0.56 | 0.39 | 1.42 | 0.16 | 1 | Lrrc29 | leucine rich repeat containing 29 | 47 | 111 | 88 | 121 | 98 | 149 | 91 | 188 | 119 | 210 | 147 |
| 119 | 0.40 | 0.52 | 0.78 | 0.44 | 1 | Kif22 | kinesin family member 22 | 331 | 129 | 89 | 95 | 73 | 68 | 91 | 188 | 92 | 53 | 100 |
| 108 | -0.15 | 0.43 | -0.36 | 0.72 | 1 | Shf | Src homology 2 domain containing F | 124 | 206 | 216 | 82 | 84 | 78 | 91 | 100 | 41 | 104 | 64 |
| 107 | -0.42 | 0.41 | -1.04 | 0.30 | 1 | Gm4366 | | 223 | 108 | 203 | 120 | 108 | 51 | 91 | 61 | 82 | 62 | 63 |
| 101 | 0.27 | 0.37 | 0.72 | 0.47 | 1 | Tmem53 | transmembrane protein 53 | 130 | 77 | 143 | 97 | 106 | 51 | 91 | 122 | 87 | 138 | 66 |
| 92 | 0.48 | 0.44 | 1.09 | 0.28 | 1 | Nxph3 | neurexophilin 3 | 30 | 46 | 71 | 158 | 84 | 40 | 91 | 133 | 121 | 169 | 68 |
| 85 | -0.46 | 0.39 | -1.17 | 0.24 | 1 | Gm9762 | | 97 | 99 | 115 | 124 | 73 | 75 | 91 | 52 | 78 | 64 | 68 |
| 70 | -0.12 | 0.55 | -0.23 | 0.82 | 1 | Gm2861 | | 0 | 0 | 0 | 90 | 113 | 82 | 91 | 149 | 138 | 40 | 62 |
| 69 | 0.00 | 0.50 | 0.00 | 1.00 | 1 | 4933440N22Rik | | 41 | 27 | 78 | 61 | 104 | 51 | 91 | 116 | 82 | 49 | 55 |
| 62 | 0.15 | 0.94 | 0.16 | 0.87 | 1 | Barhl1 | BarH like homeobox 1 | 0 | 1 | 32 | 55 | 94 | 43 | 91 | 17 | 83 | 174 | 89 |
| 45 | -0.63 | 0.56 | -1.13 | 0.26 | 1 | Shox2 | short stature homeobox 2 | 47 | 47 | 70 | 54 | 29 | 30 | 91 | 31 | 39 | 32 | 23 |
| 171 | 0.58 | 0.31 | 1.88 | 0.06 | 1 | Nckap5l | NCK-associated protein 5-like | 438 | 375 | 350 | 60 | 64 | 69 | 90 | 144 | 71 | 131 | 86 |
| 157 | 0.25 | 0.34 | 0.74 | 0.46 | 1 | Atad5 | ATPase family, AAA domain containing 5 | 261 | 191 | 162 | 134 | 123 | 161 | 90 | 118 | 185 | 113 | 189 |
| 154 | 0.24 | 0.56 | 0.43 | 0.67 | 1 | Foxm1 | forkhead box M1 | 353 | 106 | 82 | 136 | 158 | 142 | 90 | 258 | 150 | 79 | 145 |
| 118 | -0.05 | 0.36 | -0.14 | 0.89 | 1 | Csnk2a3 | | 129 | 80 | 116 | 109 | 158 | 128 | 90 | 76 | 201 | 113 | 98 |
| 97 | 0.15 | 0.35 | 0.44 | 0.66 | 1 | 1190007I07Rik | RIKEN cDNA 1190007I07 gene | 122 | 89 | 131 | 47 | 102 | 112 | 90 | 88 | 101 | 82 | 104 |
| 82 | -0.27 | 0.51 | -0.53 | 0.59 | 1 | Ssc4d | scavenger receptor cysteine rich family, 4 | 71 | 26 | 57 | 93 | 60 | 150 | 90 | 58 | 91 | 35 | 172 |
| 80 | -0.55 | 0.43 | -1.27 | 0.20 | 1 | Meig1 | meiosis expressed gene 1 | 51 | 38 | 27 | 135 | 114 | 107 | 90 | 53 | 73 | 73 | 115 |
| 68 | 0.27 | 0.56 | 0.48 | 0.63 | 1 | Rpl34-ps1 | ribosomal protein L34, pseudogene 1 | 238 | 19 | 179 | 15 | 49 | 7 | 90 | 16 | 70 | 46 | 19 |
| 58 | 0.17 | 0.57 | 0.30 | 0.77 | 1 | Paupar | | 46 | 53 | 35 | 56 | 20 | 76 | 90 | 83 | 47 | 28 | 109 |
| 45 | -0.22 | 0.58 | -0.38 | 0.71 | 1 | Pcsk2os2 | | 45 | 27 | 43 | 56 | 25 | 30 | 90 | 20 | 49 | 75 | 31 |
| 8963 | -0.97 | 1.15 | -0.85 | 0.40 | 1 | Col4a2 | collagen, type IV, alpha 2 | 32758 | 30209 | 35446 | 14 | 24 | 6 | 89 | 28 | 9 | 0 | 12 |
| 6833 | -2.13 | 1.53 | -1.39 | 0.17 | 1 | Plvap | plasmalemma vesicle associated protein | 23018 | 23576 | 28447 | 0 | 6 | 3 | 89 | 0 | 3 | 17 | 0 |
| 463 | 0.09 | 0.34 | 0.26 | 0.79 | 1 | Rhoj | ras homolog family member J | 1826 | 1168 | 1539 | 81 | 33 | 60 | 89 | 35 | 72 | 122 | 63 |
| 428 | -0.48 | 0.28 | -1.71 | 0.09 | 1 | Gpm6a | glycoprotein m6a | 1336 | 1328 | 1337 | 130 | 110 | 76 | 89 | 133 | 51 | 82 | 41 |
| 198 | 0.15 | 0.32 | 0.46 | 0.65 | 1 | Igf2bp2 | insulin-like growth factor 2 mRNA binding | 280 | 453 | 212 | 187 | 57 | 282 | 89 | 124 | 114 | 106 | 273 |
| 194 | 0.45 | 0.31 | 1.48 | 0.14 | 1 | Dpm1 | dolichol-phosphate (beta-D) mannosyltra | 86 | 74 | 59 | 80 | 73 | 681 | 89 | 102 | 109 | 192 | 588 |
| 174 | -0.14 | 0.32 | -0.44 | 0.66 | 1 | Snhg12 | | 226 | 293 | 242 | 61 | 81 | 373 | 89 | 45 | 74 | 93 | 341 |
| 157 | -0.35 | 0.53 | -0.66 | 0.51 | 1 | Plet1 | placenta expressed transcript 1 | 337 | 279 | 355 | 124 | 116 | 78 | 89 | 134 | 106 | 13 | 98 |
| 140 | -0.33 | 0.32 | -1.05 | 0.29 | 1 | Gm16023 | | 126 | 161 | 182 | 162 | 158 | 194 | 89 | 122 | 83 | 109 | 156 |
| 139 | -0.82 | 0.68 | -1.21 | 0.23 | 1 | Slc39a8 | solute carrier family 39 (metal ion transpo | 508 | 251 | 375 | 20 | 94 | 21 | 89 | 0 | 69 | 86 | 11 |
| 123 | -0.27 | 0.44 | -0.62 | 0.54 | 1 | Spc24 | SPC24, NDC80 kinetochore complex con | 220 | 63 | 110 | 136 | 154 | 132 | 89 | 80 | 121 | 157 | 95 |
| 120 | -0.02 | 0.45 | -0.05 | 0.96 | 1 | Tram1l1 | translocation associated membrane prote | 201 | 165 | 137 | 131 | 73 | 94 | 89 | 183 | 27 | 119 | 96 |
| 117 | -0.31 | 0.43 | -0.72 | 0.47 | 1 | Garem1 | GRB2 associated regulator of MAPK1 sub | 169 | 59 | 135 | 134 | 151 | 128 | 89 | 79 | 85 | 146 | 115 |
| 112 | -0.37 | 0.46 | -0.81 | 0.42 | 1 | 9030622O22Rik | | 193 | 157 | 219 | 63 | 95 | 122 | 89 | 86 | 25 | 73 | 112 |
| 107 | -0.51 | 0.46 | -1.12 | 0.26 | 1 | Gm5805 | | 262 | 162 | 150 | 119 | 94 | 50 | 89 | 28 | 77 | 95 | 53 |
| 98 | -0.32 | 0.39 | -0.83 | 0.41 | 1 | Fignl2 | fidgetin-like 2 | 124 | 112 | 169 | 83 | 75 | 130 | 89 | 102 | 64 | 52 | 83 |
| 91 | 0.22 | 0.51 | 0.43 | 0.67 | 1 | Samd15 | sterile alpha motif domain containing 15 | 66 | 105 | 157 | 58 | 62 | 101 | 89 | 119 | 51 | 119 | 72 |
| 76 | -0.33 | 0.47 | -0.71 | 0.48 | 1 | Cst6 | cystatin E/M | 24 | 51 | 28 | 127 | 123 | 63 | 89 | 101 | 98 | 32 | 101 |
| 67 | 0.33 | 0.45 | 0.72 | 0.47 | 1 | C130036L24Rik | | 74 | 74 | 54 | 53 | 43 | 53 | 89 | 105 | 79 | 55 | 56 |
| 64 | -0.02 | 0.78 | -0.02 | 0.98 | 1 | Gm9403 | | 10 | 15 | 49 | 19 | 86 | 106 | 89 | 136 | 79 | 12 | 108 |
| 58 | -0.45 | 0.79 | -0.57 | 0.57 | 1 | Gm41349 | | 23 | 0 | 21 | 123 | 71 | 55 | 89 | 72 | 83 | 54 | 44 |
| 390 | -0.03 | 0.40 | -0.08 | 0.93 | 1 | Stk17b | serine/threonine kinase 17b (apoptosis-ir | 2102 | 867 | 688 | 92 | 105 | 44 | 88 | 107 | 71 | 54 | 76 |
| 134 | 0.44 | 0.37 | 1.18 | 0.24 | 1 | Arv1 | ARV1 homolog, fatty acid homeostasis m | 85 | 76 | 117 | 99 | 89 | 254 | 88 | 114 | 122 | 213 | 220 |

| | | | | | | | | | | | | | | | | | | |
|---|---|---|---|---|---|---|---|---|---|---|---|---|---|---|---|---|---|---|
| 126 | -0.24 | 0.32 | -0.75 | 0.45 | 1 | Tedc2 | tubulin epsilon and delta complex 2 | 132 | 123 | 168 | 100 | 163 | 165 | 88 | 63 | 109 | 107 | 165 |
| 108 | 0.40 | 0.51 | 0.80 | 0.43 | 1 | Proc | protein C | 69 | 187 | 96 | 64 | 129 | 54 | 88 | 236 | 61 | 126 | 73 |
| 107 | -0.57 | 0.41 | -1.39 | 0.16 | 1 | Sumo2 | small ubiquitin-like modifier 2 | 85 | 153 | 197 | 81 | 114 | 136 | 88 | 41 | 113 | 33 | 138 |
| 94 | -0.11 | 0.44 | -0.25 | 0.80 | 1 | Gprc6a | G protein-coupled receptor, family C, grou | 58 | 29 | 74 | 154 | 107 | 99 | 88 | 82 | 115 | 97 | 129 |
| 89 | 0.38 | 0.47 | 0.82 | 0.41 | 1 | H4c12 | H4 clustered histone 12 | 15 | 55 | 27 | 100 | 57 | 140 | 88 | 158 | 64 | 153 | 123 |
| 85 | -0.55 | 0.41 | -1.35 | 0.18 | 1 | 6330415G19Rik | | 64 | 92 | 62 | 135 | 86 | 117 | 88 | 53 | 69 | 99 | 66 |
| 81 | 0.07 | 0.50 | 0.14 | 0.89 | 1 | Gm29650 | | 73 | 17 | 57 | 72 | 72 | 136 | 88 | 87 | 101 | 53 | 139 |
| 71 | -0.26 | 0.50 | -0.53 | 0.60 | 1 | A830019P07Rik | | 24 | 63 | 50 | 97 | 88 | 81 | 88 | 39 | 91 | 91 | 73 |
| 61 | -0.19 | 0.95 | -0.20 | 0.84 | 1 | Wdr27 | WD repeat domain 27 | 64 | 58 | 59 | 22 | 58 | 84 | 88 | 67 | 71 | 0 | 103 |
| 45 | -0.77 | 0.59 | -1.31 | 0.19 | 1 | 6030443J06Rik | | 13 | 19 | 8 | 43 | 68 | 84 | 88 | 15 | 37 | 59 | 62 |
| 39 | -0.42 | 0.75 | -0.56 | 0.57 | 1 | Hrh3 | histamine receptor H3 | 5 | 11 | 0 | 47 | 53 | 34 | 88 | 51 | 15 | 76 | 52 |
| 8902 | -1.37 | 1.02 | -1.34 | 0.18 | 1 | Hspg2 | perlecan (heparan sulfate proteoglycan 2 | 31279 | 31576 | 34790 | 60 | 27 | 19 | 87 | 0 | 49 | 18 | 16 |
| 984 | 0.88 | 1.61 | 0.54 | 0.59 | 1 | Laptm5 | lysosomal-associated protein transmemb | 5250 | 2566 | 2828 | 15 | 0 | 3 | 87 | 49 | 14 | 0 | 12 |
| 164 | -0.08 | 0.34 | -0.24 | 0.81 | 1 | Rnasel | ribonuclease L (2', 5'-oligoisoadenylate sy | 429 | 317 | 296 | 117 | 52 | 133 | 87 | 126 | 43 | 98 | 102 |
| 152 | 0.47 | 0.37 | 1.29 | 0.20 | 1 | Tpcn2 | two pore segment channel 2 | 221 | 197 | 197 | 70 | 77 | 199 | 87 | 57 | 96 | 214 | 261 |
| 146 | -0.53 | 0.40 | -1.33 | 0.18 | 1 | Airn | | 183 | 225 | 216 | 151 | 63 | 271 | 87 | 70 | 102 | 35 | 202 |
| 145 | 0.15 | 0.42 | 0.36 | 0.72 | 1 | Rad51 | RAD51 recombinase | 341 | 106 | 143 | 85 | 165 | 137 | 87 | 125 | 133 | 156 | 113 |
| 131 | -0.29 | 0.43 | -0.68 | 0.50 | 1 | Naprt | nicotinate phosphoribosyltransferase | 197 | 168 | 266 | 119 | 154 | 71 | 87 | 168 | 65 | 104 | 38 |
| 121 | 0.18 | 0.42 | 0.43 | 0.67 | 1 | Gm37254 | | 69 | 114 | 78 | 105 | 96 | 192 | 87 | 132 | 191 | 41 | 219 |
| 119 | -0.03 | 0.31 | -0.09 | 0.93 | 1 | Iqcg | IQ motif containing G | 123 | 122 | 186 | 109 | 132 | 113 | 87 | 76 | 115 | 109 | 135 |
| 113 | 0.25 | 0.36 | 0.69 | 0.49 | 1 | Ube2n-ps1 | | 82 | 172 | 103 | 96 | 127 | 92 | 87 | 153 | 92 | 122 | 117 |
| 106 | -0.18 | 0.41 | -0.45 | 0.65 | 1 | Fv1 | Friend virus susceptibility 1 | 102 | 141 | 185 | 111 | 58 | 130 | 87 | 111 | 51 | 55 | 139 |
| 90 | -0.57 | 0.41 | -1.39 | 0.17 | 1 | Ccl25 | chemokine (C-C motif) ligand 25 | 25 | 51 | 39 | 90 | 94 | 220 | 87 | 43 | 81 | 41 | 214 |
| 68 | 0.01 | 0.43 | 0.02 | 0.99 | 1 | H2bc9 | H2B clustered histone 9 | 15 | 6 | 18 | 76 | 80 | 109 | 87 | 66 | 86 | 121 | 88 |
| 68 | -0.35 | 0.47 | -0.74 | 0.46 | 1 | Ush1g | USH1 protein network component sans | 95 | 102 | 57 | 88 | 62 | 36 | 87 | 42 | 85 | 70 | 23 |
| 63 | -0.47 | 0.56 | -0.85 | 0.40 | 1 | 9230116N13Rik | | 26 | 67 | 56 | 55 | 67 | 89 | 87 | 53 | 23 | 49 | 122 |
| 57 | -0.74 | 0.71 | -1.05 | 0.29 | 1 | Mok | MOK protein kinase | 96 | 54 | 18 | 67 | 21 | 123 | 87 | 31 | 37 | 9 | 81 |
| 50 | -0.21 | 0.50 | -0.42 | 0.67 | 1 | Acta1 | actin, alpha 1, skeletal muscle | 37 | 13 | 28 | 50 | 67 | 46 | 87 | 53 | 85 | 46 | 32 |
| 972 | -0.53 | 0.38 | -1.38 | 0.17 | 1 | Arx | aristaless related homeobox | 3029 | 3761 | 3459 | 31 | 77 | 57 | 86 | 87 | 24 | 50 | 26 |
| 227 | 0.03 | 0.34 | 0.09 | 0.93 | 1 | Adora2a | adenosine A2a receptor | 668 | 597 | 580 | 101 | 43 | 91 | 86 | 92 | 40 | 95 | 105 |
| 124 | 0.39 | 0.38 | 1.02 | 0.31 | 1 | Rapgefl1 | Rap guanine nucleotide exchange factor | 118 | 197 | 138 | 118 | 62 | 134 | 86 | 146 | 136 | 143 | 87 |
| 107 | 0.51 | 0.38 | 1.35 | 0.18 | 1 | 5930420M18Rik | | 29 | 72 | 45 | 85 | 96 | 165 | 86 | 141 | 129 | 152 | 174 |
| 103 | 0.50 | 0.40 | 1.25 | 0.21 | 1 | Tnfrsf14 | tumor necrosis factor receptor superfamily | 153 | 145 | 135 | 53 | 60 | 91 | 86 | 152 | 93 | 77 | 89 |
| 89 | -1.13 | 0.83 | -1.37 | 0.17 | 1 | Rps27rt | ribosomal protein S27, retrogene | 196 | 174 | 197 | 61 | 54 | 52 | 86 | 0 | 74 | 24 | 54 |
| 86 | -0.17 | 0.47 | -0.35 | 0.72 | 1 | Zfp112 | zinc finger protein 112 | 72 | 119 | 140 | 45 | 70 | 120 | 86 | 90 | 75 | 32 | 98 |
| 58 | 0.20 | 0.48 | 0.41 | 0.68 | 1 | 3110070M22Rik | | 43 | 61 | 22 | 81 | 49 | 25 | 86 | 97 | 83 | 53 | 32 |
| 56 | -0.50 | 0.61 | -0.82 | 0.41 | 1 | Gm47469 | | 85 | 41 | 36 | 55 | 40 | 74 | 86 | 9 | 68 | 82 | 42 |
| 51 | -0.22 | 0.49 | -0.45 | 0.65 | 1 | Gm17509 | | 90 | 67 | 51 | 41 | 39 | 28 | 86 | 56 | 40 | 36 | 24 |
| 13016 | -0.90 | 0.71 | -1.26 | 0.21 | 1 | Col1a2 | collagen, type I, alpha 2 | 77783 | 26891 | 38232 | 46 | 26 | 17 | 85 | 2 | 36 | 31 | 24 |
| 318 | 0.61 | 0.34 | 1.77 | 0.08 | 1 | Pabpn1 | poly(A) binding protein, nuclear 1 | 139 | 177 | 151 | 132 | 142 | 1040 | 85 | 299 | 117 | 256 | 962 |
| 158 | -0.58 | 0.62 | -0.94 | 0.35 | 1 | Gm7967 | | 15 | 74 | 14 | 52 | 34 | 798 | 85 | 20 | 63 | 26 | 557 |
| 151 | 0.01 | 0.39 | 0.02 | 0.98 | 1 | Slfn9 | schlafen 9 | 206 | 127 | 194 | 157 | 134 | 187 | 85 | 133 | 147 | 183 | 106 |
| 148 | 0.27 | 0.50 | 0.55 | 0.58 | 1 | Cmtm8 | CKLF-like MARVEL transmembrane dom | 311 | 252 | 440 | 35 | 86 | 73 | 85 | 165 | 70 | 57 | 57 |
| 125 | -0.08 | 0.29 | -0.28 | 0.78 | 1 | 4933412E12Rik | | 219 | 253 | 232 | 81 | 86 | 92 | 85 | 75 | 94 | 80 | 76 |
| 108 | -0.50 | 0.49 | -1.02 | 0.31 | 1 | Aph1c | aph1 homolog C, gamma secretase subu | 174 | 204 | 152 | 186 | 61 | 40 | 85 | 64 | 35 | 158 | 24 |
| 100 | -0.11 | 0.47 | -0.23 | 0.82 | 1 | Map3k13 | mitogen-activated protein kinase kinase k | 178 | 254 | 189 | 41 | 38 | 85 | 85 | 70 | 61 | 63 | 34 |
| 86 | 0.15 | 0.45 | 0.34 | 0.73 | 1 | Gm48623 | | 67 | 72 | 98 | 78 | 76 | 84 | 85 | 162 | 69 | 102 | 51 |
| 78 | -0.21 | 0.39 | -0.53 | 0.60 | 1 | Eif4a-ps4 | | 93 | 78 | 138 | 65 | 87 | 61 | 85 | 52 | 54 | 81 | 69 |
| 73 | -0.10 | 0.42 | -0.23 | 0.82 | 1 | Pdss1 | prenyl (solanesyl) diphosphate synthase, | 84 | 52 | 57 | 41 | 74 | 109 | 85 | 32 | 88 | 54 | 127 |
| 70 | 0.39 | 0.52 | 0.75 | 0.45 | 1 | Gm7240 | | 88 | 51 | 23 | 30 | 84 | 69 | 85 | 54 | 99 | 103 | 85 |
| 64 | -0.09 | 0.54 | -0.16 | 0.87 | 1 | Gm12993 | | 57 | 99 | 74 | 81 | 44 | 35 | 85 | 29 | 60 | 108 | 35 |
| 62 | 0.43 | 0.40 | 1.10 | 0.27 | 1 | 4933439K11Rik | | 20 | 13 | 17 | 53 | 59 | 71 | 85 | 70 | 84 | 89 | 121 |
| 34 | 0.06 | 0.81 | 0.07 | 0.94 | 1 | Asb16 | ankyrin repeat and SOCS box-containing | 13 | 21 | 3 | 22 | 27 | 19 | 85 | 108 | 49 | 21 | 11 |
| 211 | -0.33 | 0.41 | -0.81 | 0.42 | 1 | Arhgap19 | Rho GTPase activating protein 19 | 657 | 381 | 457 | 158 | 127 | 85 | 84 | 49 | 146 | 77 | 100 |
| 185 | -0.37 | 0.41 | -0.89 | 0.37 | 1 | Chst14 | carbohydrate sulfotransferase 14 | 566 | 347 | 376 | 157 | 131 | 54 | 84 | 70 | 124 | 54 | 71 |
| 164 | -0.09 | 0.37 | -0.24 | 0.81 | 1 | Blvra | biliverdin reductase A | 414 | 327 | 284 | 112 | 58 | 145 | 84 | 46 | 91 | 107 | 131 |
| 161 | 0.50 | 0.36 | 1.40 | 0.16 | 1 | 1810034E14Rik | | 135 | 98 | 190 | 212 | 159 | 109 | 84 | 213 | 227 | 196 | 147 |
| 125 | 0.28 | 0.45 | 0.62 | 0.53 | 1 | Dlec1 | deleted in lung and esophageal cancer 1 | 75 | 123 | 110 | 117 | 85 | 182 | 84 | 210 | 40 | 143 | 202 |
| 118 | 0.52 | 0.39 | 1.32 | 0.19 | 1 | AI429214 | expressed sequence AI429214 | 106 | 94 | 72 | 86 | 103 | 135 | 84 | 187 | 65 | 162 | 204 |
| 116 | -0.10 | 0.43 | -0.25 | 0.81 | 1 | Pthlh | parathyroid hormone-like peptide | 158 | 95 | 111 | 137 | 118 | 129 | 84 | 166 | 101 | 55 | 123 |

| | | | | | | | | | | | | | | | | | |
|---|---|---|---|---|---|---|---|---|---|---|---|---|---|---|---|---|---|
| 110 | 0.17 | 0.40 | 0.42 | 0.68 | 1 | Gm20939 | predicted gene, 20939 | 57 | 83 | 99 | 118 | 129 | 128 | 84 | 73 | 124 | 169 | 150 |
| 107 | 0.62 | 0.36 | 1.74 | 0.08 | 1 | Cnbd2 | cyclic nucleotide binding domain containi | 59 | 125 | 74 | 95 | 79 | 88 | 84 | 235 | 85 | 121 | 127 |
| 100 | -0.39 | 0.39 | -1.00 | 0.32 | 1 | Gm10643 | | 98 | 86 | 132 | 125 | 146 | 86 | 84 | 134 | 78 | 55 | 71 |
| 90 | -0.34 | 0.58 | -0.59 | 0.55 | 1 | Tmem14a | transmembrane protein 14A | 81 | 43 | 160 | 110 | 65 | 98 | 84 | 57 | 138 | 23 | 128 |
| 87 | 0.00 | 0.55 | 0.01 | 0.99 | 1 | Gm4737 | predicted gene 4737 | 216 | 185 | 83 | 88 | 23 | 12 | 84 | 97 | 7 | 148 | 11 |
| 75 | -0.42 | 0.49 | -0.85 | 0.39 | 1 | Apoa2 | apolipoprotein A-II | 57 | 121 | 120 | 108 | 57 | 54 | 84 | 44 | 65 | 43 | 76 |
| 61 | 0.10 | 0.48 | 0.21 | 0.84 | 1 | Lipo2 | lipase, member O2 | 45 | 91 | 60 | 45 | 51 | 47 | 84 | 89 | 47 | 66 | 49 |
| 48 | -1.60 | 0.86 | -1.86 | 0.06 | 1 | Gm10734 | | 29 | 30 | 24 | 74 | 74 | 72 | 84 | 1 | 71 | 13 | 54 |
| 166 | 0.32 | 0.38 | 0.85 | 0.40 | 1 | Adat1 | adenosine deaminase, tRNA-specific 1 | 193 | 122 | 232 | 103 | 110 | 274 | 83 | 173 | 171 | 61 | 301 |
| 118 | 0.24 | 0.35 | 0.68 | 0.50 | 1 | B230369F24Rik | | 96 | 123 | 74 | 119 | 145 | 117 | 83 | 125 | 176 | 125 | 118 |
| 112 | 0.07 | 0.32 | 0.23 | 0.82 | 1 | Glcci1 | glucocorticoid induced transcript 1 | 75 | 141 | 49 | 121 | 59 | 205 | 83 | 119 | 72 | 73 | 229 |
| 111 | 0.10 | 0.39 | 0.25 | 0.80 | 1 | Cfap61 | cilia and flagella associated protein 61 | 108 | 129 | 92 | 102 | 125 | 111 | 83 | 189 | 87 | 126 | 74 |
| 86 | -0.34 | 0.50 | -0.68 | 0.49 | 1 | 2510046G10Rik | | 75 | 27 | 45 | 137 | 121 | 100 | 83 | 93 | 143 | 55 | 68 |
| 83 | -0.28 | 0.45 | -0.63 | 0.53 | 1 | Gm30025 | | 94 | 87 | 43 | 115 | 112 | 67 | 83 | 127 | 66 | 49 | 75 |
| 77 | -0.18 | 0.47 | -0.39 | 0.70 | 1 | Brinp2 | bone morphogenic protein/retinoic acid ir | 23 | 24 | 10 | 170 | 125 | 49 | 83 | 85 | 81 | 135 | 60 |
| 66 | -0.44 | 0.46 | -0.94 | 0.34 | 1 | Ccnf | cyclin F | 154 | 103 | 38 | 78 | 36 | 59 | 83 | 45 | 43 | 52 | 38 |
| 66 | -0.28 | 0.46 | -0.62 | 0.54 | 1 | Gm20632 | | 41 | 33 | 95 | 83 | 81 | 55 | 83 | 68 | 76 | 77 | 36 |
| 63 | -0.39 | 0.57 | -0.68 | 0.49 | 1 | Mastl | microtubule associated serine/threonine k | 62 | 61 | 50 | 40 | 78 | 74 | 83 | 16 | 79 | 33 | 119 |
| 61 | -0.39 | 0.49 | -0.81 | 0.42 | 1 | Usp13 | ubiquitin specific peptidase 13 (isopeptida | 80 | 55 | 51 | 57 | 63 | 59 | 83 | 29 | 77 | 91 | 25 |
| 49 | 0.24 | 0.57 | 0.42 | 0.67 | 1 | Gm10110 | | 154 | 14 | 14 | 29 | 24 | 31 | 83 | 43 | 28 | 95 | 28 |
| 2883 | -1.25 | 0.86 | -1.45 | 0.15 | 1 | Msn | moesin | 12590 | 9200 | 9692 | 19 | 35 | 23 | 82 | 0 | 30 | 17 | 27 |
| 660 | 0.15 | 0.34 | 0.44 | 0.66 | 1 | Fbln5 | fibulin 5 | 3306 | 1451 | 1868 | 91 | 96 | 39 | 82 | 103 | 115 | 50 | 60 |
| 295 | 0.30 | 0.25 | 1.21 | 0.23 | 1 | Bst2 | bone marrow stromal cell antigen 2 | 1066 | 638 | 932 | 39 | 96 | 54 | 82 | 56 | 105 | 115 | 59 |
| 246 | -0.31 | 0.27 | -1.15 | 0.25 | 1 | Slc15a2 | solute carrier family 15 (H+/peptide transp | 207 | 342 | 424 | 72 | 175 | 637 | 82 | 75 | 144 | 54 | 488 |
| 216 | 0.14 | 0.35 | 0.40 | 0.69 | 1 | 1600020E01Rik | | 112 | 89 | 118 | 109 | 45 | 720 | 82 | 64 | 96 | 91 | 853 |
| 210 | -0.06 | 0.64 | -0.10 | 0.92 | 1 | Kif11 | kinesin family member 11 | 801 | 242 | 131 | 186 | 182 | 123 | 82 | 210 | 147 | 36 | 167 |
| 206 | 0.30 | 0.40 | 0.74 | 0.46 | 1 | Blvrb | biliverdin reductase B (flavin reductase (N | 625 | 319 | 434 | 120 | 88 | 75 | 82 | 116 | 76 | 272 | 63 |
| 202 | 0.00 | 0.44 | 0.00 | 1.00 | 1 | Taf1d | TATA-box binding protein associated fact | 193 | 216 | 137 | 88 | 58 | 569 | 82 | 104 | 102 | 22 | 656 |
| 161 | 0.26 | 0.31 | 0.82 | 0.41 | 1 | Pot1b | protection of telomeres 1B | 213 | 273 | 117 | 110 | 93 | 261 | 82 | 127 | 98 | 144 | 257 |
| 159 | 0.11 | 0.35 | 0.31 | 0.76 | 1 | Zfp873 | zinc finger protein 873 | 213 | 302 | 176 | 102 | 75 | 277 | 82 | 86 | 134 | 95 | 204 |
| 121 | 0.35 | 0.39 | 0.89 | 0.37 | 1 | Zfp984 | zinc finger protein 984 | 206 | 168 | 177 | 71 | 109 | 77 | 82 | 124 | 59 | 125 | 131 |
| 112 | -0.52 | 0.38 | -1.36 | 0.17 | 1 | Oxld1 | oxidoreductase like domain containing 1 | 124 | 145 | 120 | 131 | 172 | 108 | 82 | 70 | 130 | 49 | 101 |
| 107 | -0.11 | 0.48 | -0.23 | 0.82 | 1 | Gm29170 | | 127 | 97 | 198 | 133 | 43 | 138 | 82 | 100 | 95 | 58 | 109 |
| 103 | 0.52 | 0.37 | 1.43 | 0.15 | 1 | Nudt1 | nudix (nucleoside diphosphate linked mo | 195 | 135 | 106 | 62 | 74 | 62 | 82 | 144 | 68 | 116 | 86 |
| 97 | -0.08 | 0.47 | -0.17 | 0.87 | 1 | Pdzrn4 | PDZ domain containing RING finger 4 | 161 | 268 | 232 | 43 | 30 | 52 | 82 | 30 | 39 | 52 | 77 |
| 96 | -0.30 | 0.45 | -0.67 | 0.50 | 1 | Gm14586 | | 168 | 213 | 195 | 67 | 55 | 57 | 82 | 73 | 75 | 23 | 50 |
| 84 | -0.37 | 0.44 | -0.85 | 0.40 | 1 | Srl | sarcalumenin | 110 | 160 | 90 | 64 | 105 | 63 | 82 | 50 | 82 | 84 | 35 |
| 84 | 0.02 | 0.60 | 0.04 | 0.97 | 1 | Gm34225 | | 125 | 179 | 55 | 32 | 103 | 57 | 82 | 39 | 123 | 85 | 41 |
| 68 | -0.72 | 0.48 | -1.49 | 0.14 | 1 | A530072M11Rik | | 31 | 36 | 72 | 81 | 79 | 116 | 82 | 56 | 53 | 23 | 118 |
| 54 | -0.17 | 0.48 | -0.35 | 0.72 | 1 | 1700018L02Rik | | 45 | 53 | 54 | 69 | 49 | 35 | 82 | 39 | 41 | 79 | 46 |
| 559 | -0.34 | 0.27 | -1.25 | 0.21 | 1 | Frmd6 | FERM domain containing 6 | 2509 | 1382 | 1545 | 130 | 96 | 87 | 81 | 82 | 62 | 94 | 79 |
| 437 | -1.11 | 0.87 | -1.28 | 0.20 | 1 | Slc1a5 | solute carrier family 1 (neutral amino acid | 1998 | 1229 | 1095 | 85 | 62 | 77 | 81 | 30 | 51 | 1 | 98 |
| 391 | -0.13 | 0.41 | -0.33 | 0.74 | 1 | Cxcl16 | chemokine (C-X-C motif) ligand 16 | 1479 | 851 | 1371 | 53 | 92 | 87 | 81 | 131 | 71 | 33 | 57 |
| 193 | -0.52 | 0.37 | -1.40 | 0.16 | 1 | Dpyd | dihydropyrimidine dehydrogenase | 448 | 412 | 289 | 165 | 126 | 203 | 81 | 58 | 115 | 88 | 137 |
| 183 | 0.26 | 0.43 | 0.61 | 0.54 | 1 | Ipcef1 | interaction protein for cytohesin exchange | 201 | 89 | 196 | 230 | 235 | 148 | 81 | 284 | 255 | 169 | 127 |
| 182 | 0.32 | 0.32 | 1.01 | 0.31 | 1 | Zfp169 | zinc finger protein 169 | 154 | 105 | 80 | 94 | 84 | 564 | 81 | 117 | 124 | 137 | 461 |
| 122 | 0.07 | 0.39 | 0.18 | 0.86 | 1 | Mctp1 | multiple C2 domains, transmembrane 1 | 432 | 241 | 168 | 51 | 49 | 66 | 81 | 57 | 78 | 57 | 58 |
| 118 | 0.49 | 0.53 | 0.93 | 0.35 | 1 | Rpp40 | ribonuclease P 40 subunit | 150 | 131 | 36 | 60 | 137 | 128 | 81 | 127 | 168 | 91 | 184 |
| 111 | -0.10 | 0.44 | -0.22 | 0.83 | 1 | Dmtn | dematin actin binding protein | 214 | 280 | 226 | 76 | 37 | 68 | 81 | 32 | 72 | 73 | 65 |
| 109 | -0.32 | 0.34 | -0.93 | 0.35 | 1 | Iqck | IQ motif containing K | 98 | 171 | 166 | 119 | 97 | 121 | 81 | 70 | 116 | 52 | 109 |
| 104 | -0.43 | 0.48 | -0.90 | 0.37 | 1 | Snhg6 | | 115 | 89 | 117 | 141 | 62 | 174 | 81 | 38 | 48 | 122 | 160 |
| 102 | 0.09 | 0.43 | 0.20 | 0.84 | 1 | Adat2 | adenosine deaminase, tRNA-specific 2 | 199 | 98 | 89 | 95 | 85 | 85 | 81 | 46 | 113 | 134 | 97 |
| 95 | -0.12 | 0.42 | -0.28 | 0.78 | 1 | Gm3854 | | 75 | 104 | 104 | 105 | 78 | 116 | 81 | 64 | 144 | 44 | 125 |
| 94 | 0.19 | 0.38 | 0.50 | 0.62 | 1 | Gm45769 | | 38 | 99 | 77 | 107 | 104 | 91 | 81 | 109 | 113 | 117 | 92 |
| 77 | -0.53 | 0.51 | -1.04 | 0.30 | 1 | Iglon5 | IgLON family member 5 | 118 | 112 | 130 | 92 | 43 | 69 | 81 | 20 | 49 | 68 | 68 |
| 75 | -0.17 | 0.46 | -0.38 | 0.70 | 1 | Oas1c | 2'-5' oligoadenylate synthetase 1C | 105 | 94 | 89 | 32 | 72 | 97 | 81 | 25 | 83 | 48 | 104 |
| 71 | -0.09 | 0.40 | -0.23 | 0.82 | 1 | Btbd19 | | 123 | 65 | 73 | 67 | 61 | 63 | 81 | 67 | 56 | 58 | 72 |
| 62 | -0.33 | 0.54 | -0.62 | 0.54 | 1 | Rpl38-ps1 | | 78 | 100 | 70 | 77 | 40 | 24 | 81 | 75 | 16 | 106 | 13 |
| 61 | -1.09 | 0.63 | -1.75 | 0.08 | 1 | Ankrd45 | ankyrin repeat domain 45 | 92 | 113 | 84 | 83 | 33 | 56 | 81 | 36 | 35 | 6 | 46 |
| 48 | 0.13 | 0.72 | 0.18 | 0.86 | 1 | Nudc-ps1 | | 124 | 9 | 20 | 24 | 3 | 52 | 81 | 7 | 15 | 142 | 50 |

| | | | | | | | | | | | | | | | | | | |
|---|---|---|---|---|---|---|---|---|---|---|---|---|---|---|---|---|---|---|
| 136 | -0.03 | 0.32 | -0.10 | 0.92 | 1 | Gm15545 | | 97 | 73 | 80 | 114 | 132 | 301 | 80 | 127 | 94 | 93 | 303 |
| 135 | -0.37 | 0.31 | -1.19 | 0.23 | 1 | Tekt2 | tektin 2 | 265 | 232 | 157 | 109 | 97 | 182 | 80 | 79 | 68 | 79 | 136 |
| 106 | -0.35 | 0.54 | -0.64 | 0.52 | 1 | Star | steroidogenic acute regulatory protein | 445 | 159 | 84 | 67 | 54 | 52 | 80 | 88 | 16 | 44 | 75 |
| 104 | 0.37 | 0.49 | 0.75 | 0.45 | 1 | 1700056E22Ril | RIKEN cDNA 1700056E22 gene | 74 | 75 | 143 | 96 | 112 | 72 | 80 | 83 | 80 | 251 | 82 |
| 104 | 0.11 | 0.38 | 0.29 | 0.77 | 1 | Idi1 | isopentenyl-diphosphate delta isomerase | 75 | 46 | 42 | 95 | 140 | 159 | 80 | 96 | 136 | 112 | 166 |
| 93 | -0.43 | 0.60 | -0.71 | 0.48 | 1 | Zbed4 | zinc finger, BED type containing 4 | 102 | 98 | 99 | 137 | 81 | 98 | 80 | 14 | 82 | 109 | 119 |
| 77 | -1.09 | 0.56 | -1.95 | 0.05 | 1 | Cthrc1 | collagen triple helix repeat containing 1 | 45 | 32 | 41 | 139 | 107 | 156 | 80 | 56 | 90 | 12 | 93 |
| 72 | -0.27 | 0.38 | -0.73 | 0.47 | 1 | Grin2d | glutamate receptor, ionotropic, NMDA2D | 121 | 72 | 95 | 67 | 54 | 72 | 80 | 56 | 43 | 73 | 55 |
| 69 | 0.06 | 0.53 | 0.10 | 0.92 | 1 | Tex26 | testis expressed 26 | 29 | 49 | 48 | 92 | 46 | 86 | 80 | 58 | 37 | 148 | 87 |
| 66 | -0.16 | 0.39 | -0.41 | 0.68 | 1 | Mesp2 | mesoderm posterior 2 | 63 | 60 | 43 | 107 | 65 | 45 | 80 | 74 | 77 | 72 | 39 |
| 62 | 0.40 | 0.41 | 0.96 | 0.33 | 1 | Gm14026 | | 43 | 57 | 41 | 67 | 49 | 36 | 80 | 79 | 80 | 116 | 39 |
| 52 | -0.31 | 0.53 | -0.59 | 0.55 | 1 | 3110009E18Ril | RIKEN cDNA 3110009E18 gene | 43 | 8 | 28 | 51 | 58 | 63 | 80 | 17 | 68 | 99 | 55 |
| 34 | 0.46 | 0.69 | 0.66 | 0.51 | 1 | Tma7-ps | | 28 | 29 | 2 | 35 | 4 | 13 | 80 | 115 | 3 | 44 | 25 |
| 1222 | -4.23 | 3.55 | -1.19 | 0.23 | 1 | Btnl9 | butyrophilin-like 9 | 4439 | 4525 | 4391 | 0 | 0 | 0 | 79 | 0 | 0 | 0 | 5 |
| 310 | -0.19 | 0.45 | -0.43 | 0.67 | 1 | Hspb1 | heat shock protein 1 | 686 | 815 | 1113 | 39 | 57 | 278 | 79 | 28 | 46 | 149 | 118 |
| 250 | 0.07 | 0.44 | 0.16 | 0.87 | 1 | Cit | citron | 657 | 874 | 518 | 36 | 77 | 152 | 79 | 74 | 47 | 160 | 78 |
| 166 | 0.23 | 0.36 | 0.64 | 0.52 | 1 | Tirap | toll-interleukin 1 receptor (TIR) domain-cc | 364 | 248 | 241 | 160 | 117 | 91 | 79 | 162 | 98 | 108 | 157 |
| 112 | 0.20 | 0.43 | 0.47 | 0.64 | 1 | Flywch2 | FLYWCH family member 2 | 92 | 163 | 160 | 88 | 127 | 87 | 79 | 114 | 79 | 167 | 79 |
| 111 | -0.07 | 0.35 | -0.19 | 0.85 | 1 | Nudt6 | nudix (nucleoside diphosphate linked mo | 128 | 79 | 79 | 118 | 93 | 187 | 79 | 103 | 68 | 112 | 180 |
| 111 | -0.37 | 0.60 | -0.61 | 0.54 | 1 | Nuf2 | NUF2, NDC80 kinetochore complex comp | 292 | 30 | 57 | 140 | 124 | 94 | 79 | 56 | 127 | 153 | 69 |
| 103 | 0.28 | 0.45 | 0.63 | 0.53 | 1 | Rbm4 | RNA binding motif protein 4 | 40 | 76 | 84 | 58 | 77 | 219 | 79 | 171 | 80 | 73 | 173 |
| 93 | 0.08 | 0.45 | 0.17 | 0.86 | 1 | Ccdc171 | coiled-coil domain containing 171 | 54 | 51 | 107 | 105 | 47 | 163 | 79 | 71 | 101 | 80 | 171 |
| 90 | 0.18 | 0.51 | 0.35 | 0.73 | 1 | Armc4 | armadillo repeat containing 4 | 22 | 11 | 60 | 122 | 112 | 111 | 79 | 136 | 96 | 125 | 112 |
| 89 | -0.33 | 0.44 | -0.75 | 0.45 | 1 | Apof | apolipoprotein F | 43 | 54 | 96 | 121 | 166 | 52 | 79 | 148 | 109 | 91 | 19 |
| 87 | -0.15 | 0.43 | -0.36 | 0.72 | 1 | Wnt5b | wingless-type MMTV integration site famil | 131 | 72 | 91 | 86 | 143 | 40 | 79 | 114 | 74 | 97 | 31 |
| 87 | 0.00 | 0.59 | 0.00 | 1.00 | 1 | Tbx1 | T-box 1 | 153 | 130 | 116 | 41 | 99 | 43 | 79 | 12 | 65 | 140 | 79 |
| 85 | -0.44 | 0.47 | -0.95 | 0.34 | 1 | Gm11696 | | 49 | 144 | 69 | 96 | 89 | 120 | 79 | 45 | 54 | 77 | 110 |
| 84 | -0.26 | 0.35 | -0.74 | 0.46 | 1 | Gm4673 | | 76 | 82 | 50 | 77 | 42 | 184 | 79 | 59 | 41 | 50 | 181 |
| 83 | -0.14 | 0.50 | -0.28 | 0.78 | 1 | Hacd1 | 3-hydroxyacyl-CoA dehydratase 1 | 45 | 120 | 167 | 81 | 85 | 62 | 79 | 53 | 59 | 95 | 65 |
| 81 | -0.12 | 0.39 | -0.30 | 0.77 | 1 | Immp2l | IMP2 inner mitochondrial membrane pept | 99 | 102 | 73 | 88 | 81 | 66 | 79 | 115 | 48 | 62 | 76 |
| 65 | 0.01 | 0.53 | 0.01 | 0.99 | 1 | Lin28a | lin-28 homolog A (C. elegans) | 3 | 0 | 0 | 88 | 105 | 84 | 79 | 114 | 71 | 90 | 87 |
| 57 | -0.89 | 0.88 | -1.01 | 0.31 | 1 | 4930525G20Rik | | 45 | 31 | 0 | 116 | 107 | 56 | 79 | 52 | 45 | 31 | 65 |
| 49 | -0.01 | 0.71 | -0.02 | 0.99 | 1 | Gm15910 | | 9 | 30 | 56 | 46 | 55 | 36 | 79 | 39 | 39 | 128 | 26 |
| 49 | -0.55 | 0.81 | -0.68 | 0.50 | 1 | Gm5898 | | 78 | 33 | 22 | 10 | 30 | 68 | 79 | 1 | 10 | 164 | 45 |
| 577 | -1.28 | 0.93 | -1.38 | 0.17 | 1 | Notch2 | notch 2 | 2700 | 1674 | 1617 | 70 | 43 | 49 | 78 | 0 | 38 | 41 | 37 |
| 300 | -0.46 | 0.62 | -0.73 | 0.46 | 1 | Ddah2 | dimethylarginine dimethylaminohydrolase | 1267 | 701 | 986 | 40 | 33 | 25 | 78 | 0 | 36 | 102 | 28 |
| 216 | -0.51 | 0.35 | -1.44 | 0.15 | 1 | Slc8b1 | solute carrier family 8 (sodium/lithium/calc | 974 | 371 | 443 | 102 | 90 | 70 | 78 | 65 | 61 | 89 | 39 |
| 172 | 0.23 | 0.38 | 0.61 | 0.54 | 1 | Inka2 | inka box actin regulator 2 | 247 | 253 | 246 | 197 | 110 | 148 | 78 | 130 | 106 | 211 | 170 |
| 164 | 0.26 | 0.51 | 0.51 | 0.61 | 1 | Hells | helicase, lymphoid specific | 473 | 145 | 113 | 130 | 122 | 148 | 78 | 217 | 151 | 142 | 86 |
| 138 | -0.19 | 0.39 | -0.47 | 0.64 | 1 | Uox | urate oxidase | 87 | 110 | 78 | 280 | 152 | 175 | 78 | 124 | 170 | 128 | 139 |
| 138 | 0.26 | 0.37 | 0.70 | 0.48 | 1 | Dnajc17 | DnaJ heat shock protein family (Hsp40) m | 162 | 124 | 156 | 112 | 122 | 182 | 78 | 170 | 194 | 82 | 137 |
| 133 | -0.32 | 0.48 | -0.68 | 0.50 | 1 | Ckap2l | cytoskeleton associated protein 2-like | 291 | 96 | 107 | 141 | 163 | 146 | 78 | 95 | 141 | 137 | 74 |
| 124 | -0.42 | 0.50 | -0.85 | 0.40 | 1 | A930017M01Rik | | 47 | 186 | 171 | 251 | 125 | 109 | 78 | 86 | 138 | 125 | 48 |
| 110 | -0.23 | 0.37 | -0.62 | 0.54 | 1 | Sft2d1 | SFT2 domain containing 1 | 161 | 105 | 112 | 73 | 37 | 263 | 78 | 35 | 52 | 77 | 215 |
| 106 | 0.35 | 0.42 | 0.83 | 0.41 | 1 | Gm26982 | | 114 | 129 | 127 | 74 | 74 | 115 | 78 | 51 | 93 | 179 | 128 |
| 105 | 0.15 | 0.36 | 0.43 | 0.67 | 1 | Lsm7 | LSM7 homolog, U6 small nuclear RNA ar | 118 | 67 | 154 | 54 | 80 | 170 | 78 | 78 | 83 | 63 | 206 |
| 87 | -0.18 | 0.54 | -0.33 | 0.74 | 1 | Mfsd9 | major facilitator superfamily domain conta | 70 | 171 | 75 | 113 | 93 | 55 | 78 | 38 | 101 | 125 | 38 |
| 78 | 0.38 | 0.48 | 0.78 | 0.44 | 1 | Zfp970 | zinc finger protein 970 | 81 | 43 | 43 | 85 | 42 | 92 | 78 | 111 | 71 | 46 | 167 |
| 74 | 0.05 | 0.46 | 0.11 | 0.91 | 1 | Gm8983 | | 105 | 101 | 118 | 46 | 107 | 8 | 78 | 37 | 100 | 104 | 8 |
| 68 | -0.44 | 0.49 | -0.92 | 0.36 | 1 | Zfp469 | zinc finger protein 469 | 135 | 30 | 83 | 55 | 85 | 65 | 78 | 39 | 85 | 45 | 43 |
| 53 | 0.68 | 0.71 | 0.96 | 0.34 | 1 | Gm15610 | | 69 | 81 | 0 | 83 | 3 | 25 | 78 | 126 | 14 | 79 | 20 |
| 51 | -0.73 | 0.85 | -0.86 | 0.39 | 1 | Gm32224 | | 17 | 1 | 0 | 84 | 101 | 62 | 78 | 36 | 56 | 73 | 51 |
| 49 | -0.14 | 0.62 | -0.22 | 0.83 | 1 | Gm14494 | | 21 | 66 | 58 | 28 | 67 | 35 | 78 | 30 | 48 | 55 | 50 |
| 44 | -0.01 | 0.71 | -0.01 | 0.99 | 1 | AI463170 | | 4 | 29 | 49 | 41 | 52 | 31 | 78 | 68 | 50 | 48 | 39 |
| 42 | 0.21 | 0.80 | 0.27 | 0.79 | 1 | Gm48529 | | 45 | 20 | 42 | 12 | 59 | 28 | 78 | 71 | 31 | 14 | 59 |
| 27 | -0.82 | 1.09 | -0.76 | 0.45 | 1 | Gm4864 | | 110 | 2 | 0 | 0 | 3 | 33 | 78 | 1 | 0 | 40 | 29 |
| 915 | -0.27 | 0.29 | -0.93 | 0.35 | 1 | Rgs11 | regulator of G-protein signaling 11 | 23 | 35 | 12 | 99 | 81 | 5125 | 77 | 41 | 88 | 80 | 4406 |
| 342 | 0.04 | 0.29 | 0.13 | 0.90 | 1 | Pygl | liver glycogen phosphorylase | 1038 | 1024 | 813 | 131 | 97 | 123 | 77 | 189 | 70 | 68 | 136 |
| 334 | 0.43 | 0.32 | 1.33 | 0.18 | 1 | Tgfb3 | transforming growth factor, beta 3 | 1505 | 733 | 719 | 108 | 57 | 43 | 77 | 131 | 65 | 194 | 41 |
| 285 | -0.19 | 0.39 | -0.50 | 0.62 | 1 | Fam111a | family with sequence similarity 111, meml | 1079 | 612 | 655 | 83 | 142 | 103 | 77 | 29 | 141 | 97 | 119 |

| | | | | | | | | | | | | | | | | | | |
|---|---|---|---|---|---|---|---|---|---|---|---|---|---|---|---|---|---|---|
| 284 | 0.18 | 0.36 | 0.51 | 0.61 | 1 | Palld | palladin, cytoskeletal associated protein | 1263 | 701 | 597 | 28 | 101 | 66 | 77 | 58 | 92 | 76 | 68 |
| 235 | -0.52 | 0.43 | -1.21 | 0.23 | 1 | Plp1 | proteolipid protein (myelin) 1 | 474 | 660 | 685 | 28 | 38 | 261 | 77 | 30 | 38 | 15 | 272 |
| 191 | 0.54 | 0.35 | 1.54 | 0.12 | 1 | Slc6a9 | solute carrier family 6 (neurotransmitter tr | 361 | 525 | 345 | 88 | 87 | 90 | 77 | 180 | 192 | 75 | 82 |
| 163 | -0.41 | 0.43 | -0.96 | 0.33 | 1 | Paqr7 | progestin and adipoQ receptor family mer | 445 | 288 | 506 | 109 | 103 | 30 | 77 | 93 | 89 | 26 | 31 |
| 161 | -0.45 | 0.39 | -1.13 | 0.26 | 1 | Zfp692 | zinc finger protein 692 | 84 | 75 | 82 | 49 | 51 | 667 | 77 | 29 | 75 | 26 | 556 |
| 117 | -0.12 | 0.41 | -0.29 | 0.77 | 1 | Frg2f1 | FSHD region gene 2 family member 1 | 135 | 82 | 69 | 147 | 148 | 147 | 77 | 93 | 114 | 140 | 140 |
| 109 | -0.40 | 0.43 | -0.93 | 0.35 | 1 | Mis18a | MIS18 kinetochore protein A | 266 | 239 | 190 | 43 | 98 | 64 | 77 | 23 | 54 | 80 | 64 |
| 103 | 0.35 | 0.43 | 0.82 | 0.41 | 1 | Zfp820 | zinc finger protein 820 | 191 | 169 | 141 | 45 | 77 | 72 | 77 | 147 | 67 | 75 | 70 |
| 103 | -0.52 | 0.46 | -1.12 | 0.26 | 1 | Zfp566 | zinc finger protein 566 | 125 | 146 | 168 | 78 | 132 | 91 | 77 | 32 | 102 | 31 | 147 |
| 102 | -0.78 | 0.42 | -1.87 | 0.06 | 1 | Gm5124 | | 160 | 285 | 287 | 58 | 78 | 36 | 77 | 43 | 29 | 48 | 23 |
| 102 | -0.06 | 0.35 | -0.17 | 0.87 | 1 | Csdc2 | cold shock domain containing C2, RNA b | 135 | 149 | 137 | 144 | 82 | 58 | 77 | 89 | 87 | 97 | 63 |
| 92 | -0.34 | 0.51 | -0.66 | 0.51 | 1 | Zfp934 | zinc finger protein 934 | 115 | 81 | 127 | 105 | 77 | 118 | 77 | 123 | 23 | 82 | 86 |
| 87 | -0.17 | 0.51 | -0.34 | 0.73 | 1 | Pcdhgb4 | protocadherin gamma subfamily B, 4 | 99 | 164 | 72 | 52 | 94 | 110 | 77 | 32 | 88 | 89 | 85 |
| 85 | -0.75 | 0.47 | -1.60 | 0.11 | 1 | Kcnk13 | potassium channel, subfamily K, member | 193 | 103 | 111 | 115 | 80 | 57 | 77 | 30 | 78 | 54 | 39 |
| 75 | 0.07 | 0.44 | 0.16 | 0.87 | 1 | Gm11868 | | 107 | 120 | 38 | 137 | 17 | 46 | 77 | 114 | 16 | 104 | 53 |
| 72 | -0.22 | 0.39 | -0.57 | 0.57 | 1 | Zbtb3 | zinc finger and BTB domain containing 3 | 65 | 70 | 101 | 67 | 78 | 77 | 77 | 53 | 67 | 85 | 55 |
| 72 | -0.37 | 0.53 | -0.70 | 0.48 | 1 | 5330439M10Rik | | 82 | 138 | 133 | 60 | 31 | 70 | 77 | 68 | 37 | 19 | 74 |
| 52 | -0.25 | 0.58 | -0.43 | 0.67 | 1 | Mkln1os | | 40 | 78 | 35 | 34 | 66 | 52 | 77 | 25 | 58 | 67 | 44 |
| 43 | -0.34 | 0.85 | -0.40 | 0.69 | 1 | Chil6 | chitinase-like 6 | 0 | 12 | 0 | 77 | 52 | 55 | 77 | 44 | 43 | 58 | 50 |
| 367 | 0.45 | 0.43 | 1.05 | 0.29 | 1 | Mcm3 | minichromosome maintenance complex c | 1520 | 687 | 433 | 234 | 125 | 149 | 76 | 195 | 232 | 258 | 133 |
| 140 | -0.43 | 0.43 | -1.01 | 0.31 | 1 | Ttc23 | tetratricopeptide repeat domain 23 | 340 | 206 | 251 | 108 | 66 | 159 | 76 | 25 | 65 | 93 | 154 |
| 134 | -0.45 | 0.52 | -0.85 | 0.39 | 1 | Slc25a27 | solute carrier family 25, member 27 | 147 | 127 | 70 | 56 | 44 | 431 | 76 | 9 | 62 | 54 | 398 |
| 109 | 0.25 | 0.43 | 0.58 | 0.56 | 1 | 1700112J16Rik | | 45 | 141 | 139 | 93 | 130 | 79 | 76 | 242 | 104 | 84 | 68 |
| 103 | -0.45 | 0.45 | -0.99 | 0.32 | 1 | Etfbkmt | electron transfer flavoprotein beta subunit | 209 | 136 | 116 | 87 | 111 | 103 | 76 | 29 | 91 | 111 | 67 |
| 103 | -0.47 | 0.36 | -1.31 | 0.19 | 1 | Luzp2 | leucine zipper protein 2 | 189 | 211 | 227 | 60 | 102 | 57 | 76 | 50 | 71 | 37 | 54 |
| 98 | -1.25 | 0.77 | -1.63 | 0.10 | 1 | Trim34a | tripartite motif-containing 34A | 248 | 163 | 155 | 72 | 60 | 111 | 76 | 0 | 39 | 40 | 117 |
| 93 | -0.34 | 0.54 | -0.64 | 0.52 | 1 | 1110025M09Rik | | 57 | 181 | 114 | 131 | 91 | 83 | 76 | 37 | 61 | 111 | 79 |
| 89 | 0.36 | 0.32 | 1.13 | 0.26 | 1 | Gm4890 | | 71 | 82 | 72 | 88 | 77 | 85 | 76 | 125 | 106 | 102 | 89 |
| 87 | 0.08 | 0.35 | 0.24 | 0.81 | 1 | Gm10644 | | 39 | 74 | 34 | 128 | 53 | 141 | 76 | 109 | 79 | 73 | 145 |
| 79 | 0.75 | 0.55 | 1.35 | 0.18 | 1 | Gm13835 | | 49 | 33 | 81 | 81 | 73 | 36 | 76 | 193 | 48 | 126 | 77 |
| 79 | -0.41 | 0.37 | -1.12 | 0.26 | 1 | Tube1 | tubulin, epsilon 1 | 95 | 129 | 92 | 91 | 52 | 99 | 76 | 51 | 52 | 54 | 80 |
| 71 | 0.40 | 0.50 | 0.80 | 0.42 | 1 | D830035M03Rik | | 47 | 36 | 132 | 73 | 60 | 35 | 76 | 100 | 100 | 67 | 57 |
| 65 | -0.62 | 0.51 | -1.20 | 0.23 | 1 | Gm5884 | | 46 | 78 | 50 | 105 | 93 | 47 | 76 | 78 | 59 | 18 | 65 |
| 62 | 0.25 | 0.49 | 0.50 | 0.62 | 1 | Zfp879 | zinc finger protein 879 | 79 | 61 | 68 | 44 | 26 | 79 | 76 | 83 | 55 | 54 | 59 |
| 62 | -0.32 | 0.50 | -0.64 | 0.52 | 1 | Gm16861 | | 20 | 36 | 11 | 173 | 63 | 46 | 76 | 66 | 84 | 55 | 46 |
| 61 | -0.35 | 0.52 | -0.67 | 0.50 | 1 | Gm527 | predicted gene 527 | 96 | 45 | 47 | 53 | 68 | 68 | 76 | 68 | 31 | 36 | 79 |
| 52 | 0.56 | 0.75 | 0.74 | 0.46 | 1 | 4930555F03Rik | | 10 | 22 | 0 | 42 | 43 | 62 | 76 | 78 | 88 | 102 | 54 |
| 50 | 0.24 | 0.82 | 0.30 | 0.77 | 1 | Gm7049 | | 69 | 55 | 22 | 32 | 11 | 33 | 76 | 143 | 25 | 84 | 2 |
| 44 | -0.52 | 0.68 | -0.76 | 0.45 | 1 | Gm45380 | | 13 | 17 | 67 | 43 | 52 | 47 | 76 | 12 | 50 | 73 | 39 |
| 4522 | -1.43 | 1.16 | -1.23 | 0.22 | 1 | Reg1 | regenerating islet-derived 1 | 7834 | 19939 | 20739 | 536 | 59 | 10 | 75 | 41 | 29 | 15 | 465 |
| 619 | -0.55 | 0.47 | -1.17 | 0.24 | 1 | Pmp22 | peripheral myelin protein 22 | 2588 | 1564 | 1746 | 158 | 105 | 161 | 75 | 20 | 153 | 50 | 192 |
| 300 | -0.71 | 0.42 | -1.68 | 0.09 | 1 | Ccdc88c | coiled-coil domain containing 88C | 906 | 934 | 890 | 67 | 62 | 139 | 75 | 38 | 89 | 13 | 87 |
| 170 | 0.32 | 0.39 | 0.81 | 0.42 | 1 | Dffb | DNA fragmentation factor, beta subunit | 80 | 57 | 68 | 58 | 69 | 573 | 75 | 110 | 122 | 40 | 618 |
| 159 | 0.19 | 0.41 | 0.46 | 0.65 | 1 | Ccdc40 | coiled-coil domain containing 40 | 135 | 209 | 291 | 154 | 157 | 145 | 75 | 114 | 123 | 206 | 138 |
| 133 | -0.45 | 0.51 | -0.88 | 0.38 | 1 | Dctpp1 | dCTP pyrophosphatase 1 | 268 | 158 | 178 | 158 | 129 | 112 | 75 | 30 | 97 | 167 | 94 |
| 129 | -0.04 | 0.30 | -0.14 | 0.89 | 1 | Smim24 | small integral membrane protein 24 | 182 | 176 | 234 | 113 | 132 | 102 | 75 | 103 | 115 | 72 | 119 |
| 128 | 0.54 | 0.47 | 1.15 | 0.25 | 1 | Myo19 | myosin XIX | 76 | 29 | 125 | 89 | 112 | 212 | 75 | 134 | 87 | 180 | 288 |
| 124 | 0.55 | 0.45 | 1.24 | 0.22 | 1 | Rfc4 | replication factor C (activator 1) 4 | 346 | 181 | 148 | 53 | 77 | 68 | 75 | 168 | 96 | 82 | 65 |
| 112 | 0.08 | 0.36 | 0.22 | 0.83 | 1 | Spin4 | spindlin family, member 4 | 181 | 120 | 162 | 91 | 115 | 82 | 75 | 59 | 122 | 139 | 80 |
| 110 | 0.45 | 0.41 | 1.09 | 0.28 | 1 | Mroh7 | maestro heat-like repeat family member 7 | 34 | 31 | 41 | 36 | 59 | 355 | 75 | 112 | 66 | 85 | 321 |
| 108 | 0.12 | 0.36 | 0.33 | 0.74 | 1 | Cbx2 | chromobox 2 | 170 | 107 | 154 | 83 | 89 | 116 | 75 | 111 | 108 | 90 | 81 |
| 105 | -0.18 | 0.39 | -0.47 | 0.64 | 1 | Blm | Bloom syndrome, RecQ like helicase | 207 | 117 | 147 | 126 | 65 | 100 | 75 | 74 | 74 | 80 | 92 |
| 102 | -0.06 | 0.44 | -0.13 | 0.89 | 1 | Gm11613 | | 94 | 77 | 94 | 123 | 134 | 99 | 75 | 87 | 63 | 157 | 118 |
| 85 | -0.46 | 0.36 | -1.30 | 0.19 | 1 | Gm10432 | | 107 | 184 | 107 | 108 | 73 | 50 | 75 | 105 | 46 | 55 | 29 |
| 78 | -0.02 | 0.35 | -0.05 | 0.96 | 1 | Gm14295 | predicted gene 14295 | 57 | 61 | 67 | 60 | 75 | 135 | 75 | 59 | 81 | 80 | 113 |
| 63 | -0.22 | 0.97 | -0.23 | 0.82 | 1 | C030029H02Rik | | 0 | 0 | 16 | 31 | 106 | 115 | 75 | 129 | 93 | 8 | 117 |
| 59 | -0.38 | 0.57 | -0.67 | 0.50 | 1 | Tnfsf9 | tumor necrosis factor (ligand) superfamily | 115 | 73 | 44 | 36 | 72 | 51 | 75 | 71 | 36 | 28 | 44 |
| 56 | -0.07 | 0.49 | -0.13 | 0.89 | 1 | Gm45486 | | 5 | 3 | 9 | 65 | 91 | 70 | 75 | 57 | 68 | 54 | 122 |
| 52 | -1.12 | 0.59 | -1.90 | 0.06 | 1 | Gm14305 | predicted gene 14432 | 63 | 7 | 43 | 62 | 82 | 83 | 75 | 14 | 46 | 43 | 60 |
| 43 | 0.84 | 0.58 | 1.44 | 0.15 | 1 | Gm3772 | | 0 | 0 | 2 | 31 | 30 | 36 | 75 | 53 | 68 | 116 | 66 |

| | | | | | | | | | | | | | | | | | | |
|---|---|---|---|---|---|---|---|---|---|---|---|---|---|---|---|---|---|---|
| 11585 | 0.40 | 0.47 | 0.85 | 0.40 | 1 | Sparc | secreted acidic cysteine rich glycoprotein | 52991 | 31242 | 42869 | 18 | 50 | 12 | 74 | 65 | 55 | 32 | 21 |
| 1447 | -0.30 | 0.23 | -1.29 | 0.20 | 1 | Edn3 | endothelin 3 | 3050 | 6875 | 5230 | 125 | 126 | 89 | 74 | 100 | 107 | 98 | 40 |
| 358 | 0.10 | 0.33 | 0.31 | 0.76 | 1 | Tubb6 | tubulin, beta 6 class V | 1645 | 907 | 740 | 79 | 68 | 92 | 74 | 75 | 108 | 62 | 86 |
| 222 | -0.19 | 0.30 | -0.65 | 0.52 | 1 | Spaca6 | sperm acrosome associated 6 | 74 | 37 | 65 | 70 | 71 | 1045 | 74 | 70 | 76 | 52 | 806 |
| 170 | -0.08 | 0.42 | -0.20 | 0.84 | 1 | Samd4 | sterile alpha motif domain containing 4 | 630 | 368 | 401 | 73 | 40 | 55 | 74 | 50 | 98 | 48 | 38 |
| 168 | -0.08 | 0.34 | -0.24 | 0.81 | 1 | Acsf2 | acyl-CoA synthetase family member 2 | 329 | 187 | 337 | 116 | 169 | 152 | 74 | 96 | 174 | 76 | 140 |
| 121 | 0.05 | 0.87 | 0.06 | 0.95 | 1 | Dlg2 | discs large MAGUK scaffold protein 2 | 406 | 240 | 183 | 78 | 72 | 24 | 74 | 112 | 104 | 0 | 42 |
| 112 | 0.24 | 0.74 | 0.32 | 0.75 | 1 | Plk1 | polo like kinase 1 | 379 | 74 | 81 | 86 | 81 | 57 | 74 | 249 | 64 | 18 | 66 |
| 110 | 0.44 | 0.41 | 1.06 | 0.29 | 1 | Ckap2 | cytoskeleton associated protein 2 | 98 | 88 | 80 | 95 | 146 | 82 | 74 | 174 | 193 | 62 | 120 |
| 109 | -0.27 | 0.42 | -0.63 | 0.53 | 1 | Gm5454 | | 229 | 360 | 237 | 40 | 53 | 32 | 74 | 70 | 29 | 48 | 26 |
| 100 | 0.14 | 0.37 | 0.39 | 0.70 | 1 | Suv39h2 | suppressor of variegation 3-9 2 | 76 | 107 | 113 | 145 | 95 | 68 | 74 | 173 | 71 | 90 | 89 |
| 81 | -0.16 | 0.38 | -0.43 | 0.67 | 1 | A730063M14Rik | | 67 | 58 | 67 | 112 | 99 | 86 | 74 | 103 | 62 | 89 | 79 |
| 77 | -0.04 | 0.43 | -0.09 | 0.93 | 1 | Gm10657 | | 51 | 68 | 56 | 117 | 93 | 56 | 74 | 73 | 65 | 131 | 61 |
| 74 | 0.24 | 0.37 | 0.65 | 0.51 | 1 | Rpsa-ps10 | | 101 | 123 | 89 | 69 | 50 | 35 | 74 | 107 | 54 | 72 | 42 |
| 67 | -0.97 | 0.52 | -1.88 | 0.06 | 1 | Rtl10 | | 47 | 107 | 107 | 95 | 55 | 83 | 74 | 41 | 61 | 14 | 59 |
| 61 | 0.70 | 0.60 | 1.18 | 0.24 | 1 | Zfp459 | zinc finger protein 459 | 27 | 7 | 54 | 25 | 118 | 39 | 74 | 89 | 64 | 113 | 65 |
| 57 | 0.65 | 0.52 | 1.26 | 0.21 | 1 | 4930577N17Rik | | 27 | 51 | 67 | 28 | 59 | 30 | 74 | 57 | 87 | 85 | 61 |
| 1100 | -1.71 | 1.13 | -1.51 | 0.13 | 1 | Klf4 | Kruppel-like factor 4 (gut) | 5277 | 2571 | 3090 | 43 | 17 | 538 | 73 | 0 | 40 | 4 | 444 |
| 395 | 0.02 | 0.28 | 0.08 | 0.94 | 1 | Fam102b | family with sequence similarity 102, meml | 1735 | 936 | 1007 | 93 | 73 | 86 | 73 | 72 | 93 | 116 | 61 |
| 259 | -0.62 | 0.52 | -1.18 | 0.24 | 1 | Gm49759 | | 71 | 64 | 36 | 69 | 50 | 1257 | 73 | 31 | 81 | 9 | 1108 |
| 181 | -0.29 | 0.34 | -0.86 | 0.39 | 1 | Pola2 | polymerase (DNA directed), alpha 2 | 252 | 296 | 233 | 163 | 193 | 248 | 73 | 127 | 170 | 71 | 170 |
| 163 | -0.22 | 0.37 | -0.60 | 0.55 | 1 | Dpysl5 | dihydropyrimidinase-like 5 | 467 | 520 | 425 | 37 | 49 | 45 | 73 | 61 | 46 | 33 | 35 |
| 140 | 0.00 | 0.63 | 0.00 | 1.00 | 1 | Ube2c | ubiquitin-conjugating enzyme E2C | 395 | 126 | 64 | 144 | 173 | 74 | 73 | 240 | 93 | 64 | 92 |
| 132 | -0.99 | 0.72 | -1.37 | 0.17 | 1 | Snhg14 | | 45 | 79 | 52 | 66 | 53 | 489 | 73 | 70 | 33 | 0 | 490 |
| 131 | -0.15 | 0.33 | -0.46 | 0.65 | 1 | Ppfia2 | protein tyrosine phosphatase, receptor ty | 236 | 222 | 306 | 73 | 80 | 127 | 73 | 65 | 113 | 45 | 101 |
| 114 | 0.31 | 0.33 | 0.95 | 0.34 | 1 | Zfp580 | zinc finger protein 580 | 155 | 135 | 150 | 74 | 137 | 80 | 73 | 102 | 134 | 125 | 87 |
| 111 | 0.00 | 0.41 | 0.00 | 1.00 | 1 | Dynlt1f | dynein light chain Tctex-type 1F | 147 | 212 | 157 | 64 | 85 | 136 | 73 | 56 | 99 | 112 | 81 |
| 110 | -0.36 | 0.38 | -0.97 | 0.33 | 1 | Mcts2 | malignant T cell amplified sequence 2 | 71 | 80 | 69 | 90 | 58 | 311 | 73 | 76 | 58 | 28 | 300 |
| 105 | 0.06 | 0.40 | 0.14 | 0.89 | 1 | Ntn3 | netrin 3 | 232 | 127 | 100 | 95 | 97 | 76 | 73 | 79 | 103 | 100 | 77 |
| 101 | -0.29 | 0.40 | -0.71 | 0.48 | 1 | Ephb6 | Eph receptor B6 | 361 | 166 | 195 | 18 | 52 | 80 | 73 | 31 | 37 | 44 | 56 |
| 98 | -0.38 | 0.49 | -0.78 | 0.43 | 1 | 2900005J15Rik | | 3 | 11 | 10 | 81 | 38 | 408 | 73 | 32 | 65 | 50 | 307 |
| 93 | -0.33 | 0.35 | -0.93 | 0.35 | 1 | BC025920 | | 124 | 179 | 86 | 110 | 47 | 111 | 73 | 88 | 26 | 59 | 115 |
| 89 | 0.26 | 0.47 | 0.55 | 0.58 | 1 | Bco2 | beta-carotene oxygenase 2 | 82 | 109 | 91 | 84 | 40 | 110 | 73 | 197 | 67 | 45 | 85 |
| 87 | 0.22 | 0.53 | 0.41 | 0.68 | 1 | Mrnip | MRN complex interacting protein | 88 | 125 | 197 | 47 | 47 | 74 | 73 | 23 | 72 | 122 | 84 |
| 84 | -1.20 | 0.68 | -1.76 | 0.08 | 1 | Nectin4 | nectin cell adhesion molecule 4 | 184 | 108 | 166 | 97 | 93 | 41 | 73 | 21 | 6 | 91 | 42 |
| 76 | -0.56 | 0.50 | -1.12 | 0.26 | 1 | Ccdc113 | coiled-coil domain containing 113 | 106 | 149 | 135 | 56 | 54 | 77 | 73 | 15 | 52 | 59 | 58 |
| 73 | -0.57 | 0.77 | -0.74 | 0.46 | 1 | Abhd1 | abhydrolase domain containing 1 | 164 | 36 | 105 | 17 | 95 | 47 | 73 | 0 | 72 | 174 | 21 |
| 68 | -0.36 | 0.54 | -0.66 | 0.51 | 1 | Pabpc5 | poly(A) binding protein, cytoplasmic 5 | 67 | 22 | 29 | 149 | 66 | 68 | 73 | 66 | 58 | 58 | 91 |
| 67 | -0.35 | 0.48 | -0.72 | 0.47 | 1 | 4930594M22Rik | | 30 | 34 | 68 | 102 | 84 | 72 | 73 | 41 | 89 | 97 | 47 |
| 3952 | -0.06 | 1.32 | -0.05 | 0.96 | 1 | Podxl | podocalyxin-like | 14503 | 14051 | 14799 | 1 | 0 | 7 | 72 | 13 | 9 | 12 | 2 |
| 243 | -0.52 | 0.40 | -1.31 | 0.19 | 1 | Slc16a7 | solute carrier family 16 (monocarboxylic a | 599 | 841 | 747 | 39 | 55 | 120 | 72 | 15 | 51 | 62 | 70 |
| 189 | -0.21 | 0.52 | -0.40 | 0.69 | 1 | Kif21b | kinesin family member 21B | 637 | 405 | 310 | 81 | 122 | 104 | 72 | 151 | 84 | 22 | 92 |
| 186 | -0.72 | 0.62 | -1.16 | 0.24 | 1 | Arhgap27 | Rho GTPase activating protein 27 | 783 | 482 | 581 | 33 | 20 | 6 | 72 | 6 | 39 | 10 | 12 |
| 102 | 0.05 | 0.37 | 0.13 | 0.90 | 1 | Zfp185 | zinc finger protein 185 | 293 | 177 | 94 | 65 | 53 | 76 | 72 | 70 | 47 | 115 | 60 |
| 91 | 0.51 | 0.62 | 0.83 | 0.41 | 1 | Gm26384 | | 86 | 47 | 108 | 85 | 107 | 35 | 72 | 293 | 55 | 50 | 68 |
| 91 | -0.12 | 0.43 | -0.29 | 0.77 | 1 | Gm16378 | | 106 | 67 | 94 | 97 | 114 | 94 | 72 | 108 | 134 | 49 | 67 |
| 90 | -0.46 | 0.51 | -0.90 | 0.37 | 1 | Slco4a1 | solute carrier organic anion transporter fa | 111 | 79 | 193 | 106 | 62 | 112 | 72 | 73 | 28 | 73 | 84 |
| 84 | -0.88 | 0.83 | -1.06 | 0.29 | 1 | Rab26os | | 223 | 162 | 193 | 36 | 49 | 53 | 72 | 20 | 70 | 0 | 50 |
| 82 | 0.19 | 0.35 | 0.54 | 0.59 | 1 | 4930481B07Rik | | 24 | 28 | 25 | 84 | 87 | 146 | 72 | 83 | 114 | 97 | 140 |
| 71 | -0.68 | 0.41 | -1.67 | 0.10 | 1 | Ggn | gametogenetin | 82 | 133 | 67 | 86 | 60 | 82 | 72 | 79 | 45 | 33 | 39 |
| 64 | -0.16 | 0.48 | -0.33 | 0.74 | 1 | Hcar2 | hydroxycarboxylic acid receptor 2 | 106 | 29 | 74 | 63 | 45 | 80 | 72 | 49 | 45 | 58 | 87 |
| 60 | 0.39 | 0.46 | 0.85 | 0.39 | 1 | Gm16589 | | 106 | 5 | 56 | 10 | 62 | 62 | 72 | 10 | 83 | 108 | 81 |
| 58 | 0.18 | 0.49 | 0.37 | 0.71 | 1 | Gm10275 | | 88 | 95 | 74 | 63 | 35 | 18 | 72 | 35 | 51 | 80 | 32 |
| 58 | -0.24 | 0.69 | -0.34 | 0.73 | 1 | 4930519K11Rik | | 9 | 100 | 65 | 101 | 40 | 46 | 72 | 44 | 55 | 67 | 40 |
| 52 | 0.16 | 0.50 | 0.32 | 0.75 | 1 | Tmem51os1 | | 91 | 14 | 10 | 23 | 18 | 98 | 72 | 27 | 17 | 67 | 138 |
| 51 | -0.29 | 0.63 | -0.46 | 0.65 | 1 | 9530051G07Rik | | 39 | 44 | 25 | 70 | 48 | 51 | 72 | 12 | 49 | 93 | 58 |
| 48 | -0.49 | 0.60 | -0.82 | 0.41 | 1 | Gm9910 | | 18 | 46 | 29 | 45 | 77 | 57 | 72 | 19 | 76 | 45 | 46 |
| 37 | -0.33 | 0.89 | -0.37 | 0.71 | 1 | Pkd2l2 | polycystic kidney disease 2-like 2 | 56 | 24 | 2 | 52 | 24 | 37 | 72 | 9 | 36 | 21 | 78 |
| 33 | -2.34 | 1.24 | -1.89 | 0.06 | 1 | Lrrc38 | leucine rich repeat containing 38 | 0 | 0 | 12 | 61 | 85 | 39 | 72 | 0 | 26 | 23 | 41 |
| 566 | -0.35 | 0.43 | -0.82 | 0.41 | 1 | Swap70 | SWA-70 protein | 2661 | 1614 | 1460 | 106 | 37 | 64 | 71 | 73 | 61 | 14 | 65 |

| | | | | | | | | | | | | | | | | | | |
|---|---|---|---|---|---|---|---|---|---|---|---|---|---|---|---|---|---|---|
| 421 | -0.59 | 0.36 | -1.62 | 0.11 | 1 | Dennd2a | DENN/MADD domain containing 2A | 1536 | 1383 | 1095 | 87 | 79 | 117 | 71 | 38 | 23 | 117 | 87 |
| 277 | 0.32 | 0.33 | 0.97 | 0.33 | 1 | Myc | myelocytomatosis oncogene | 946 | 538 | 314 | 93 | 62 | 347 | 71 | 74 | 85 | 171 | 343 |
| 177 | 0.05 | 0.41 | 0.13 | 0.90 | 1 | Tmco4 | transmembrane and coiled-coil domains 4 | 483 | 324 | 324 | 109 | 96 | 117 | 71 | 169 | 117 | 54 | 79 |
| 174 | 0.37 | 0.57 | 0.65 | 0.52 | 1 | Hmmr | hyaluronan mediated motility receptor (RI | 383 | 88 | 73 | 186 | 188 | 142 | 71 | 261 | 178 | 169 | 176 |
| 139 | 0.64 | 0.49 | 1.30 | 0.19 | 1 | Jpx | | 78 | 164 | 52 | 123 | 97 | 208 | 71 | 83 | 158 | 252 | 239 |
| 127 | 0.07 | 0.51 | 0.14 | 0.89 | 1 | Tonsl | tonsoku-like, DNA repair protein | 193 | 121 | 77 | 138 | 106 | 147 | 71 | 45 | 105 | 233 | 163 |
| 116 | -0.03 | 0.39 | -0.07 | 0.94 | 1 | Alkbh2 | alkB homolog 2, alpha-ketoglutarate-depe | 132 | 63 | 129 | 95 | 164 | 161 | 71 | 115 | 121 | 108 | 116 |
| 112 | -1.20 | 0.68 | -1.75 | 0.08 | 1 | Atp1a3 | ATPase, Na+/K+ transporting, alpha 3 pol | 468 | 221 | 176 | 55 | 90 | 40 | 71 | 41 | 47 | 3 | 21 |
| 109 | 0.36 | 0.49 | 0.73 | 0.46 | 1 | 6430571L13Rik | RIKEN cDNA 6430571L13 gene | 109 | 126 | 41 | 104 | 93 | 109 | 71 | 266 | 100 | 50 | 128 |
| 88 | 0.37 | 0.44 | 0.83 | 0.40 | 1 | Ciart | circadian associated repressor of transcri | 38 | 27 | 19 | 54 | 41 | 250 | 71 | 53 | 66 | 174 | 172 |
| 77 | -0.19 | 0.41 | -0.46 | 0.65 | 1 | C030037D09Rik | | 55 | 46 | 32 | 133 | 66 | 112 | 71 | 64 | 60 | 91 | 123 |
| 58 | -0.19 | 0.48 | -0.39 | 0.69 | 1 | Tph2 | tryptophan hydroxylase 2 | 41 | 40 | 30 | 98 | 57 | 55 | 71 | 54 | 89 | 44 | 56 |
| 54 | 0.27 | 0.56 | 0.48 | 0.63 | 1 | Golt1a | golgi transport 1A | 65 | 74 | 72 | 28 | 39 | 36 | 71 | 71 | 58 | 26 | 53 |
| 52 | 0.01 | 0.60 | 0.02 | 0.98 | 1 | 4930412F12Rik | | 25 | 42 | 21 | 53 | 79 | 23 | 71 | 34 | 50 | 148 | 23 |
| 48 | 0.11 | 0.63 | 0.18 | 0.86 | 1 | Myadml2 | myeloid-associated differentiation marker | 77 | 55 | 101 | 28 | 43 | 11 | 71 | 30 | 20 | 53 | 42 |
| 35 | -0.31 | 0.63 | -0.49 | 0.63 | 1 | Chrna4 | cholinergic receptor, nicotinic, alpha poly | 20 | 3 | 3 | 39 | 52 | 32 | 71 | 48 | 26 | 72 | 22 |
| 354 | 0.01 | 0.39 | 0.04 | 0.97 | 1 | Krt19 | keratin 19 | 1295 | 879 | 1176 | 33 | 153 | 24 | 70 | 57 | 148 | 35 | 28 |
| 342 | -1.76 | 1.36 | -1.30 | 0.19 | 1 | Adcy7 | adenylate cyclase 7 | 2048 | 793 | 804 | 7 | 13 | 6 | 70 | 0 | 14 | 0 | 11 |
| 148 | 0.86 | 0.56 | 1.52 | 0.13 | 1 | Sox5 | SRY (sex determining region Y)-box 5 | 451 | 357 | 266 | 14 | 28 | 115 | 70 | 105 | 94 | 64 | 62 |
| 127 | -0.62 | 0.41 | -1.50 | 0.13 | 1 | Shank1 | SH3 and multiple ankyrin repeat domains | 156 | 212 | 157 | 124 | 120 | 209 | 70 | 155 | 76 | 31 | 91 |
| 122 | 0.08 | 0.46 | 0.17 | 0.86 | 1 | Igsf9 | immunoglobulin superfamily, member 9 | 216 | 143 | 245 | 110 | 59 | 116 | 70 | 173 | 61 | 57 | 95 |
| 102 | -0.30 | 0.38 | -0.79 | 0.43 | 1 | Fgfr4 | fibroblast growth factor receptor 4 | 175 | 187 | 234 | 67 | 95 | 56 | 70 | 97 | 56 | 53 | 35 |
| 83 | 0.00 | 0.48 | 0.00 | 1.00 | 1 | 2810454H06Rik | | 126 | 124 | 144 | 39 | 61 | 78 | 70 | 56 | 84 | 22 | 105 |
| 78 | -0.87 | 0.48 | -1.82 | 0.07 | 1 | Gm49768 | | 56 | 77 | 113 | 94 | 129 | 84 | 70 | 36 | 46 | 32 | 122 |
| 78 | -0.34 | 0.42 | -0.82 | 0.41 | 1 | Saxo2 | stabilizer of axonemal microtubules 2 | 58 | 57 | 136 | 74 | 83 | 101 | 70 | 34 | 84 | 63 | 94 |
| 77 | -0.38 | 0.55 | -0.68 | 0.49 | 1 | Gm43909 | | 65 | 74 | 88 | 122 | 81 | 64 | 70 | 27 | 86 | 128 | 42 |
| 75 | 0.12 | 0.58 | 0.22 | 0.83 | 1 | Gm14410 | predicted gene 14410 | 57 | 128 | 31 | 46 | 90 | 84 | 70 | 44 | 82 | 93 | 98 |
| 57 | -0.62 | 0.51 | -1.23 | 0.22 | 1 | Slc22a4 | solute carrier family 22 (organic cation tra | 104 | 31 | 101 | 42 | 62 | 52 | 70 | 17 | 30 | 45 | 71 |
| 54 | -0.49 | 0.53 | -0.92 | 0.36 | 1 | Gm19719 | | 70 | 29 | 35 | 107 | 45 | 42 | 70 | 32 | 48 | 82 | 32 |
| 52 | -0.17 | 0.58 | -0.29 | 0.77 | 1 | Pir | pirin | 124 | 85 | 72 | 19 | 26 | 43 | 70 | 42 | 38 | 26 | 23 |
| 46 | -1.15 | 1.01 | -1.14 | 0.25 | 1 | Rptoros | | 20 | 4 | 17 | 71 | 77 | 85 | 70 | 71 | 24 | 0 | 69 |
| 29 | 0.42 | 1.15 | 0.36 | 0.72 | 1 | Grid2ip | glutamate receptor, ionotropic, delta 2 (Gr | 0 | 0 | 8 | 7 | 30 | 35 | 70 | 82 | 29 | 30 | 26 |
| 1619 | -2.44 | 1.30 | -1.87 | 0.06 | 1 | Csf1r | colony stimulating factor 1 receptor | 8688 | 3774 | 5183 | 25 | 23 | 14 | 69 | 0 | 20 | 0 | 13 |
| 1265 | -0.04 | 0.32 | -0.13 | 0.90 | 1 | Epb41l2 | erythrocyte membrane protein band 4.1 li | 5439 | 4033 | 3864 | 67 | 44 | 117 | 69 | 82 | 98 | 33 | 64 |
| 355 | -0.38 | 0.27 | -1.38 | 0.17 | 1 | Edil3 | EGF-like repeats and discoidin I-like dom | 688 | 1369 | 1120 | 154 | 125 | 48 | 69 | 175 | 83 | 36 | 37 |
| 155 | -0.20 | 0.36 | -0.56 | 0.58 | 1 | Slc28a2 | solute carrier family 28 (sodium-coupled r | 245 | 210 | 178 | 129 | 59 | 265 | 69 | 93 | 48 | 46 | 367 |
| 139 | 0.03 | 0.42 | 0.06 | 0.95 | 1 | Kif23 | kinesin family member 23 | 456 | 192 | 143 | 81 | 87 | 123 | 69 | 108 | 59 | 85 | 127 |
| 111 | 0.03 | 0.43 | 0.08 | 0.94 | 1 | Frem2 | Fras1 related extracellular matrix protein 2 | 229 | 91 | 121 | 85 | 106 | 128 | 69 | 85 | 133 | 72 | 106 |
| 96 | 0.00 | 0.47 | 0.00 | 1.00 | 1 | Gls2 | glutaminase 2 (liver, mitochondrial) | 68 | 214 | 184 | 76 | 89 | 63 | 69 | 50 | 79 | 94 | 68 |
| 95 | -0.27 | 0.42 | -0.63 | 0.53 | 1 | Kif17 | kinesin family member 17 | 92 | 123 | 102 | 59 | 109 | 160 | 69 | 50 | 129 | 40 | 118 |
| 92 | 0.49 | 0.42 | 1.17 | 0.24 | 1 | 9930014A18Rik | | 219 | 158 | 116 | 48 | 48 | 52 | 69 | 98 | 95 | 62 | 50 |
| 92 | -0.48 | 0.88 | -0.54 | 0.59 | 1 | Chek1 | checkpoint kinase 1 | 138 | 48 | 86 | 78 | 50 | 207 | 69 | 88 | 54 | 0 | 197 |
| 91 | 0.28 | 0.39 | 0.73 | 0.47 | 1 | Gm44899 | | 106 | 82 | 83 | 114 | 66 | 76 | 69 | 165 | 80 | 80 | 79 |
| 89 | 0.35 | 0.43 | 0.80 | 0.42 | 1 | Gm10130 | predicted gene 10130 | 136 | 116 | 104 | 46 | 67 | 91 | 69 | 120 | 81 | 49 | 97 |
| 87 | -0.04 | 0.39 | -0.11 | 0.92 | 1 | Onecut2 | one cut domain, family member 2 | 134 | 167 | 111 | 81 | 37 | 93 | 69 | 74 | 62 | 39 | 94 |
| 82 | -0.21 | 0.45 | -0.47 | 0.64 | 1 | Tmem79 | transmembrane protein 79 | 106 | 67 | 101 | 89 | 97 | 73 | 69 | 45 | 53 | 128 | 77 |
| 76 | 0.03 | 0.49 | 0.07 | 0.94 | 1 | Gm16174 | | 49 | 20 | 39 | 109 | 60 | 126 | 69 | 110 | 101 | 70 | 84 |
| 76 | -0.10 | 0.42 | -0.23 | 0.82 | 1 | Ccl27a | chemokine (C-C motif) ligand 27A | 74 | 110 | 77 | 67 | 33 | 130 | 69 | 43 | 50 | 62 | 118 |
| 70 | -0.14 | 0.42 | -0.32 | 0.75 | 1 | 9430085M18Rik | | 47 | 27 | 58 | 57 | 103 | 96 | 69 | 35 | 66 | 100 | 110 |
| 69 | -0.33 | 0.54 | -0.61 | 0.54 | 1 | E030042O20Rik | | 6 | 42 | 24 | 137 | 91 | 80 | 69 | 134 | 43 | 88 | 46 |
| 69 | 0.32 | 0.54 | 0.60 | 0.55 | 1 | 2610507I01Rik | | 49 | 66 | 43 | 44 | 29 | 116 | 69 | 136 | 18 | 76 | 111 |
| 68 | -0.38 | 0.40 | -0.94 | 0.35 | 1 | Rpl36a-ps2 | | 160 | 124 | 103 | 60 | 44 | 36 | 69 | 37 | 39 | 41 | 38 |
| 65 | 0.42 | 0.49 | 0.85 | 0.40 | 1 | Pcdhac2 | protocadherin alpha subfamily C, 2 | 35 | 47 | 29 | 64 | 54 | 52 | 69 | 75 | 70 | 179 | 37 |
| 59 | -0.10 | 0.62 | -0.17 | 0.87 | 1 | Gm28151 | | 88 | 60 | 37 | 58 | 21 | 96 | 69 | 51 | 49 | 14 | 110 |
| 57 | -0.04 | 0.50 | -0.08 | 0.93 | 1 | Fam166a | family with sequence similarity 166, meml | 17 | 14 | 10 | 61 | 80 | 75 | 69 | 30 | 89 | 111 | 66 |
| 46 | -0.19 | 0.70 | -0.27 | 0.79 | 1 | Glo1-ps | | 101 | 13 | 103 | 15 | 64 | 25 | 69 | 38 | 29 | 23 | 24 |
| 44 | -0.48 | 1.08 | -0.45 | 0.66 | 1 | Pcdha12 | protocadherin alpha 12 | 13 | 57 | 32 | 93 | 12 | 55 | 69 | 0 | 39 | 57 | 52 |
| 39 | -0.13 | 0.86 | -0.15 | 0.88 | 1 | 1700007K13Rik | RIKEN cDNA 1700007K13 gene | 25 | 32 | 33 | 17 | 54 | 27 | 69 | 87 | 4 | 54 | 26 |
| 37 | 0.92 | 0.71 | 1.30 | 0.20 | 1 | Cplx3 | complexin 3 | 9 | 11 | 0 | 28 | 15 | 22 | 69 | 111 | 41 | 70 | 25 |
| 34 | -0.36 | 0.66 | -0.55 | 0.58 | 1 | Gm4815 | | 38 | 28 | 34 | 16 | 22 | 46 | 69 | 9 | 40 | 49 | 20 |

| | | | | | | | | | | | | | | | | | |
|---|---|---|---|---|---|---|---|---|---|---|---|---|---|---|---|---|---|
| 891 | -1.21 | 0.66 | -1.84 | 0.07 | 1 | Akap12 | A kinase (PRKA) anchor protein (gravin) 1 | 3262 | 2856 | 2888 | 221 | 185 | 57 | 68 | 124 | 116 | 8 | 21 |
| 290 | -0.34 | 0.32 | -1.07 | 0.29 | 1 | Maml2 | mastermind like transcriptional coactivato | 952 | 941 | 739 | 82 | 73 | 87 | 68 | 81 | 73 | 28 | 67 |
| 186 | -0.21 | 0.51 | -0.40 | 0.69 | 1 | Snhg17 | | 119 | 153 | 144 | 50 | 46 | 623 | 68 | 68 | 73 | 8 | 696 |
| 165 | -0.42 | 0.39 | -1.08 | 0.28 | 1 | Plekha4 | pleckstrin homology domain containing, fa | 214 | 144 | 177 | 206 | 112 | 347 | 68 | 90 | 86 | 93 | 280 |
| 124 | 0.03 | 0.38 | 0.09 | 0.93 | 1 | 9430015G10Ril | RIKEN cDNA 9430015G10 gene | 161 | 93 | 91 | 65 | 98 | 303 | 68 | 89 | 103 | 75 | 213 |
| 104 | -0.65 | 0.47 | -1.37 | 0.17 | 1 | Gm42047 | | 22 | 65 | 59 | 120 | 57 | 303 | 68 | 85 | 61 | 13 | 298 |
| 101 | 0.64 | 0.41 | 1.54 | 0.12 | 1 | Ppih | peptidyl prolyl isomerase H | 138 | 82 | 128 | 78 | 47 | 107 | 68 | 98 | 152 | 85 | 128 |
| 86 | -0.07 | 0.49 | -0.14 | 0.89 | 1 | Rpl36al | ribosomal protein L36A-like | 58 | 73 | 33 | 46 | 129 | 161 | 68 | 65 | 109 | 53 | 153 |
| 72 | 0.06 | 0.39 | 0.17 | 0.87 | 1 | Dsn1 | DSN1 homolog, MIS12 kinetochore comp | 100 | 66 | 52 | 69 | 62 | 75 | 68 | 81 | 50 | 99 | 67 |
| 68 | 0.47 | 0.45 | 1.05 | 0.29 | 1 | Gm49477 | | 16 | 19 | 9 | 95 | 96 | 40 | 68 | 101 | 91 | 121 | 86 |
| 66 | 0.58 | 0.46 | 1.26 | 0.21 | 1 | Ano9 | anoctamin 9 | 23 | 33 | 14 | 43 | 40 | 109 | 68 | 94 | 35 | 135 | 137 |
| 62 | -0.79 | 0.51 | -1.55 | 0.12 | 1 | Pcdhb1 | protocadherin beta 1 | 51 | 99 | 105 | 65 | 40 | 89 | 68 | 56 | 28 | 18 | 62 |
| 58 | -0.48 | 0.78 | -0.61 | 0.54 | 1 | Cngb3 | cyclic nucleotide gated channel beta 3 | 0 | 20 | 44 | 57 | 102 | 78 | 68 | 93 | 100 | 30 | 44 |
| 51 | -0.21 | 0.50 | -0.41 | 0.68 | 1 | Gm43545 | | 52 | 83 | 39 | 37 | 63 | 37 | 68 | 49 | 43 | 54 | 32 |
| 47 | 0.98 | 0.70 | 1.40 | 0.16 | 1 | Fut1 | fucosyltransferase 1 | 29 | 69 | 74 | 14 | 53 | 2 | 68 | 59 | 65 | 81 | 8 |
| 46 | 0.01 | 1.10 | 0.01 | 1.00 | 1 | Flicr | | 0 | 0 | 31 | 81 | 52 | 27 | 68 | 39 | 56 | 121 | 31 |
| 599 | -0.23 | 0.34 | -0.70 | 0.49 | 1 | Notch4 | notch 4 | 2028 | 1840 | 2306 | 78 | 50 | 30 | 67 | 68 | 38 | 26 | 56 |
| 405 | 0.23 | 0.52 | 0.44 | 0.66 | 1 | Marcksl1 | MARCKS-like 1 | 1597 | 1006 | 1362 | 35 | 62 | 40 | 67 | 156 | 80 | 18 | 26 |
| 139 | 0.16 | 0.36 | 0.46 | 0.64 | 1 | Tmem86b | transmembrane protein 86B | 148 | 193 | 184 | 69 | 46 | 330 | 67 | 95 | 94 | 52 | 251 |
| 115 | -0.63 | 0.41 | -1.54 | 0.12 | 1 | Hmgb3 | high mobility group box 3 | 266 | 225 | 310 | 88 | 62 | 59 | 67 | 82 | 34 | 49 | 24 |
| 113 | 0.16 | 0.39 | 0.42 | 0.68 | 1 | 2610005L07Rik | | 112 | 169 | 87 | 76 | 42 | 225 | 67 | 46 | 54 | 99 | 267 |
| 104 | -0.12 | 0.54 | -0.21 | 0.83 | 1 | 1110038B12Rik | | 135 | 155 | 170 | 47 | 34 | 171 | 67 | 52 | 66 | 12 | 231 |
| 100 | -0.50 | 0.43 | -1.15 | 0.25 | 1 | Brca2 | breast cancer 2, early onset | 183 | 201 | 126 | 112 | 46 | 129 | 67 | 57 | 73 | 32 | 77 |
| 99 | -0.08 | 0.32 | -0.24 | 0.81 | 1 | Proser3 | proline and serine rich 3 | 62 | 77 | 77 | 88 | 112 | 184 | 67 | 95 | 84 | 76 | 165 |
| 97 | 0.45 | 0.48 | 0.94 | 0.35 | 1 | Eef1a1-ps1 | | 136 | 94 | 171 | 69 | 82 | 52 | 67 | 196 | 74 | 71 | 54 |
| 92 | -0.72 | 0.66 | -1.10 | 0.27 | 1 | Mapk11 | mitogen-activated protein kinase 11 | 259 | 164 | 253 | 75 | 18 | 44 | 67 | 6 | 41 | 45 | 40 |
| 87 | -0.65 | 0.50 | -1.30 | 0.20 | 1 | Styx | serine/threonine/tyrosine interaction prote | 99 | 45 | 82 | 95 | 75 | 194 | 67 | 80 | 32 | 28 | 165 |
| 82 | -0.41 | 0.52 | -0.79 | 0.43 | 1 | Gm15361 | | 121 | 92 | 88 | 92 | 99 | 57 | 67 | 29 | 85 | 143 | 27 |
| 62 | -0.29 | 0.92 | -0.31 | 0.75 | 1 | Abcc2 | ATP-binding cassette, sub-family C (CFTF | 0 | 20 | 0 | 85 | 79 | 133 | 67 | 35 | 50 | 66 | 153 |
| 62 | -0.65 | 0.48 | -1.36 | 0.17 | 1 | Rab43 | RAB43, member RAS oncogene family | 127 | 76 | 67 | 44 | 77 | 57 | 67 | 21 | 29 | 64 | 54 |
| 61 | -0.15 | 0.43 | -0.36 | 0.72 | 1 | Aqp7 | aquaporin 7 | 35 | 52 | 33 | 101 | 52 | 68 | 67 | 89 | 70 | 37 | 63 |
| 59 | -0.29 | 0.49 | -0.60 | 0.55 | 1 | Sgip1 | SH3-domain GRB2-like (endophilin) inter | 147 | 106 | 77 | 56 | 30 | 19 | 67 | 30 | 51 | 50 | 11 |
| 58 | 0.29 | 0.54 | 0.54 | 0.59 | 1 | Nthl1 | nth (endonuclease III)-like 1 (E.coli) | 48 | 57 | 121 | 36 | 32 | 52 | 67 | 64 | 48 | 62 | 53 |
| 55 | -0.37 | 0.47 | -0.79 | 0.43 | 1 | Zmiz1os1 | | 56 | 45 | 63 | 71 | 75 | 36 | 67 | 30 | 49 | 62 | 50 |
| 52 | -0.88 | 0.92 | -0.96 | 0.34 | 1 | Gm8909 | predicted gene 8909 | 180 | 15 | 174 | 12 | 49 | 0 | 67 | 0 | 58 | 17 | 3 |
| 49 | -0.56 | 0.54 | -1.03 | 0.30 | 1 | Oacyl | O-acyltransferase like | 14 | 14 | 16 | 80 | 74 | 70 | 67 | 20 | 48 | 72 | 71 |
| 35 | -0.13 | 0.58 | -0.23 | 0.82 | 1 | Gm5182 | | 15 | 15 | 6 | 33 | 47 | 33 | 67 | 23 | 38 | 70 | 37 |
| 3362 | -1.17 | 1.13 | -1.03 | 0.30 | 1 | Ahnak | AHNAK nucleoprotein (desmoyokin) | 17228 | 9565 | 10031 | 1 | 26 | 17 | 66 | 0 | 16 | 6 | 27 |
| 1105 | -1.26 | 0.80 | -1.58 | 0.11 | 1 | Atf3 | activating transcription factor 3 | 5796 | 2155 | 2522 | 39 | 47 | 933 | 66 | 0 | 40 | 39 | 520 |
| 241 | -0.02 | 0.36 | -0.05 | 0.96 | 1 | Zfp385a | zinc finger protein 385A | 836 | 542 | 665 | 106 | 81 | 46 | 66 | 138 | 47 | 76 | 46 |
| 231 | 0.63 | 0.39 | 1.61 | 0.11 | 1 | Palmd | palmdelphin | 766 | 672 | 733 | 24 | 23 | 38 | 66 | 71 | 59 | 45 | 46 |
| 88 | 0.52 | 0.41 | 1.26 | 0.21 | 1 | Il18 | interleukin 18 | 178 | 185 | 143 | 28 | 30 | 63 | 66 | 53 | 35 | 103 | 77 |
| 87 | 0.48 | 0.50 | 0.96 | 0.34 | 1 | 4933434E20Rik | | 21 | 25 | 53 | 81 | 67 | 148 | 66 | 108 | 59 | 155 | 169 |
| 83 | 0.42 | 0.52 | 0.80 | 0.42 | 1 | Adam5 | a disintegrin and metallopeptidase domai | 153 | 157 | 143 | 23 | 89 | 30 | 66 | 66 | 64 | 46 | 79 |
| 75 | -0.05 | 0.52 | -0.10 | 0.92 | 1 | 4430402I18Rik | RIKEN cDNA 4430402I18 gene | 45 | 95 | 59 | 56 | 118 | 64 | 66 | 136 | 79 | 27 | 84 |
| 70 | -0.08 | 0.60 | -0.13 | 0.90 | 1 | Gm11110 | | 86 | 90 | 56 | 64 | 79 | 49 | 66 | 147 | 60 | 15 | 59 |
| 64 | -0.58 | 0.54 | -1.07 | 0.28 | 1 | Eldr | | 34 | 19 | 30 | 73 | 73 | 122 | 66 | 14 | 66 | 48 | 157 |
| 62 | -0.54 | 0.55 | -0.97 | 0.33 | 1 | Erich6 | glutamate rich 6 | 11 | 33 | 45 | 80 | 66 | 103 | 66 | 67 | 35 | 22 | 152 |
| 61 | -0.09 | 0.55 | -0.16 | 0.87 | 1 | 4632411P08Rik | | 32 | 59 | 36 | 99 | 56 | 49 | 66 | 112 | 82 | 15 | 68 |
| 61 | 0.78 | 0.51 | 1.52 | 0.13 | 1 | Gm14167 | | 41 | 38 | 18 | 97 | 20 | 42 | 66 | 137 | 79 | 59 | 71 |
| 60 | 0.56 | 0.48 | 1.16 | 0.24 | 1 | Gm10209 | | 39 | 37 | 30 | 82 | 37 | 39 | 66 | 53 | 83 | 131 | 60 |
| 57 | 0.72 | 0.74 | 0.97 | 0.33 | 1 | Crlf1 | cytokine receptor-like factor 1 | 69 | 65 | 123 | 21 | 19 | 26 | 66 | 136 | 26 | 26 | 46 |
| 54 | -0.17 | 0.58 | -0.29 | 0.77 | 1 | Gm20275 | | 35 | 82 | 30 | 46 | 49 | 67 | 66 | 25 | 54 | 37 | 105 |
| 53 | 0.40 | 0.49 | 0.81 | 0.42 | 1 | Gm12751 | | 63 | 26 | 58 | 43 | 53 | 29 | 66 | 81 | 50 | 84 | 35 |
| 52 | 0.71 | 0.75 | 0.94 | 0.35 | 1 | Gm5499 | | 30 | 2 | 101 | 35 | 61 | 10 | 66 | 70 | 92 | 94 | 15 |
| 51 | -0.70 | 0.51 | -1.39 | 0.17 | 1 | Gab3 | growth factor receptor bound protein 2-as | 69 | 81 | 76 | 55 | 33 | 48 | 66 | 21 | 20 | 46 | 40 |
| 45 | -0.55 | 0.66 | -0.83 | 0.41 | 1 | Styk1 | serine/threonine/tyrosine kinase 1 | 16 | 60 | 61 | 43 | 26 | 75 | 66 | 20 | 40 | 37 | 47 |
| 33 | 0.52 | 0.91 | 0.57 | 0.57 | 1 | Gm49871 | | 23 | 2 | 55 | 4 | 34 | 43 | 66 | 35 | 29 | 26 | 45 |
| 27 | -0.61 | 0.82 | -0.75 | 0.45 | 1 | Gm9484 | | 5 | 16 | 39 | 25 | 30 | 23 | 66 | 21 | 16 | 24 | 29 |
| 540 | -0.22 | 0.37 | -0.60 | 0.55 | 1 | Rapgef5 | Rap guanine nucleotide exchange factor | 2041 | 1495 | 1309 | 223 | 232 | 74 | 65 | 120 | 202 | 113 | 61 |

| | | | | | | | | | | | | | | | | | |
|---|---|---|---|---|---|---|---|---|---|---|---|---|---|---|---|---|---|
| 233 | -0.13 | 0.60 | -0.21 | 0.83 | 1 | Plekhg2 | pleckstrin homology domain containing, fa | 908 | 650 | 584 | 60 | 22 | 59 | 65 | 9 | 101 | 30 | 73 |
| 178 | -0.38 | 0.41 | -0.91 | 0.36 | 1 | Ccdc163 | coiled-coil domain containing 163 | 223 | 282 | 183 | 139 | 98 | 380 | 65 | 76 | 122 | 26 | 369 |
| 146 | 0.68 | 0.63 | 1.08 | 0.28 | 1 | Fcgbp | Fc fragment of IgG binding protein | 44 | 220 | 690 | 47 | 100 | 30 | 65 | 207 | 86 | 90 | 29 |
| 128 | -0.29 | 0.86 | -0.34 | 0.73 | 1 | Armh4 | armadillo-like helical domain containing 4 | 644 | 407 | 226 | 4 | 3 | 12 | 65 | 0 | 20 | 21 | 7 |
| 112 | 0.46 | 0.41 | 1.13 | 0.26 | 1 | Gipc2 | GIPC PDZ domain containing family, mem | 168 | 154 | 126 | 57 | 115 | 91 | 65 | 118 | 176 | 95 | 71 |
| 78 | -0.02 | 0.41 | -0.04 | 0.97 | 1 | Esyt3 | extended synaptotagmin-like protein 3 | 85 | 99 | 73 | 81 | 91 | 51 | 65 | 126 | 86 | 68 | 29 |
| 77 | 0.11 | 0.48 | 0.23 | 0.82 | 1 | Ncmap | noncompact myelin associated protein | 63 | 73 | 122 | 95 | 46 | 67 | 65 | 83 | 116 | 35 | 81 |
| 56 | -0.75 | 0.88 | -0.85 | 0.40 | 1 | Zfp72 | zinc finger protein 72 | 154 | 81 | 102 | 42 | 34 | 35 | 65 | 37 | 33 | 0 | 37 |
| 43 | -1.42 | 1.29 | -1.10 | 0.27 | 1 | Cemip | cell migration inducing protein, hyalurona | 0 | 7 | 50 | 39 | 62 | 86 | 65 | 0 | 9 | 62 | 89 |
| 36 | -0.90 | 0.85 | -1.06 | 0.29 | 1 | Zfp85os | | 21 | 36 | 18 | 21 | 64 | 46 | 65 | 24 | 41 | 3 | 58 |
| 2669 | -0.35 | 0.26 | -1.38 | 0.17 | 1 | Thbs1 | thrombospondin 1 | 11250 | 9832 | 7976 | 53 | 34 | 23 | 64 | 19 | 46 | 36 | 29 |
| 337 | 0.40 | 0.46 | 0.87 | 0.39 | 1 | Cped1 | cadherin-like and PC-esterase domain co | 1644 | 605 | 518 | 206 | 36 | 117 | 64 | 198 | 76 | 71 | 170 |
| 253 | -0.39 | 0.61 | -0.64 | 0.52 | 1 | Tram2 | translocating chain-associating membran | 1086 | 609 | 549 | 48 | 60 | 126 | 64 | 94 | 53 | 4 | 86 |
| 219 | 0.38 | 0.38 | 0.99 | 0.32 | 1 | Timeless | timeless circadian clock 1 | 643 | 355 | 551 | 121 | 89 | 94 | 64 | 197 | 81 | 117 | 95 |
| 162 | 0.05 | 0.34 | 0.14 | 0.89 | 1 | Bend5 | BEN domain containing 5 | 380 | 340 | 221 | 113 | 95 | 141 | 64 | 112 | 93 | 67 | 158 |
| 142 | 0.36 | 0.72 | 0.50 | 0.62 | 1 | Gm49599 | | 8 | 44 | 24 | 6 | 13 | 683 | 64 | 13 | 40 | 49 | 621 |
| 124 | -0.12 | 0.53 | -0.22 | 0.82 | 1 | Aurka | aurora kinase A | 272 | 82 | 99 | 207 | 106 | 96 | 64 | 96 | 125 | 128 | 90 |
| 123 | -0.05 | 0.36 | -0.14 | 0.89 | 1 | Unkl | unkempt family like zinc finger | 305 | 161 | 263 | 86 | 104 | 62 | 64 | 56 | 92 | 95 | 67 |
| 119 | 0.27 | 0.38 | 0.70 | 0.48 | 1 | Zfp939 | zinc finger protein 939 | 151 | 64 | 166 | 60 | 102 | 221 | 64 | 90 | 97 | 119 | 177 |
| 107 | -0.01 | 0.43 | -0.02 | 0.98 | 1 | Btbd11 | BTB (POZ) domain containing 11 | 178 | 126 | 121 | 123 | 115 | 69 | 64 | 165 | 76 | 67 | 71 |
| 95 | 0.35 | 0.40 | 0.87 | 0.38 | 1 | Zfp382 | zinc finger protein 382 | 160 | 137 | 53 | 97 | 60 | 85 | 64 | 119 | 79 | 119 | 77 |
| 92 | -0.64 | 0.54 | -1.18 | 0.24 | 1 | Abraxas1 | BRCA1 A complex subunit | 130 | 178 | 144 | 31 | 41 | 192 | 64 | 8 | 23 | 64 | 133 |
| 89 | 0.11 | 0.39 | 0.28 | 0.78 | 1 | Itpka | inositol 1,4,5-trisphosphate 3-kinase A | 184 | 108 | 104 | 102 | 55 | 55 | 64 | 89 | 52 | 99 | 62 |
| 84 | 0.28 | 0.57 | 0.49 | 0.62 | 1 | Kifc1 | kinesin family member C1 | 141 | 89 | 34 | 50 | 88 | 99 | 64 | 129 | 101 | 49 | 85 |
| 84 | 0.07 | 0.38 | 0.18 | 0.85 | 1 | Gm15510 | | 27 | 85 | 87 | 84 | 105 | 94 | 64 | 116 | 112 | 58 | 87 |
| 74 | -0.07 | 0.43 | -0.17 | 0.87 | 1 | Tdo2 | tryptophan 2,3-dioxygenase | 144 | 111 | 56 | 75 | 77 | 40 | 64 | 73 | 60 | 66 | 44 |
| 73 | -1.45 | 1.06 | -1.37 | 0.17 | 1 | H2-T-ps | histocompatibility 2, T region locus, pseu | 365 | 0 | 222 | 15 | 43 | 4 | 64 | 0 | 52 | 35 | 5 |
| 63 | -0.32 | 0.61 | -0.52 | 0.60 | 1 | Gm16835 | | 5 | 71 | 15 | 82 | 102 | 73 | 64 | 108 | 61 | 57 | 53 |
| 62 | -1.95 | 1.10 | -1.78 | 0.08 | 1 | Tmsb15b1 | thymosin beta 15b1 | 100 | 156 | 214 | 6 | 30 | 41 | 64 | 0 | 38 | 0 | 27 |
| 61 | 0.12 | 0.64 | 0.19 | 0.85 | 1 | Etnk2 | ethanolamine kinase 2 | 66 | 110 | 27 | 71 | 58 | 36 | 64 | 20 | 70 | 102 | 53 |
| 59 | -0.97 | 0.61 | -1.57 | 0.12 | 1 | Lin7b | lin-7 homolog B (C. elegans) | 17 | 34 | 73 | 49 | 71 | 112 | 64 | 10 | 35 | 26 | 160 |
| 58 | -0.96 | 0.61 | -1.58 | 0.11 | 1 | Syce1 | synaptonemal complex central element p | 20 | 49 | 85 | 114 | 56 | 87 | 64 | 24 | 31 | 45 | 67 |
| 51 | -0.14 | 0.49 | -0.28 | 0.78 | 1 | 2500002B13Rik | | 32 | 29 | 25 | 83 | 46 | 50 | 64 | 63 | 24 | 90 | 55 |
| 50 | 0.03 | 0.50 | 0.06 | 0.95 | 1 | Gm48079 | | 32 | 58 | 39 | 39 | 51 | 54 | 64 | 36 | 67 | 54 | 57 |
| 50 | -1.42 | 0.83 | -1.71 | 0.09 | 1 | 9630028I04Rik | | 34 | 62 | 89 | 50 | 90 | 37 | 64 | 46 | 57 | 0 | 17 |
| 37 | 0.22 | 0.95 | 0.23 | 0.82 | 1 | Plk5 | polo like kinase 5 | 9 | 25 | 12 | 4 | 29 | 66 | 64 | 2 | 34 | 93 | 68 |
| 36 | -0.92 | 0.69 | -1.33 | 0.18 | 1 | 5330411J11Rik | | 4 | 22 | 32 | 63 | 52 | 42 | 64 | 23 | 28 | 26 | 44 |
| 2865 | -0.48 | 1.16 | -0.41 | 0.68 | 1 | Trf | transferrin | 17925 | 9059 | 4189 | 69 | 30 | 29 | 63 | 66 | 69 | 0 | 12 |
| 191 | 0.19 | 0.39 | 0.48 | 0.63 | 1 | Rhbdf2 | rhomboid 5 homolog 2 | 532 | 351 | 461 | 103 | 108 | 72 | 63 | 143 | 57 | 131 | 75 |
| 181 | 0.08 | 0.34 | 0.23 | 0.82 | 1 | Mcub | mitochondrial calcium uniporter dominant | 483 | 406 | 415 | 89 | 107 | 66 | 63 | 89 | 92 | 137 | 45 |
| 167 | 0.25 | 0.48 | 0.52 | 0.61 | 1 | Nsg1 | neuron specific gene family member 1 | 507 | 451 | 378 | 63 | 22 | 83 | 63 | 20 | 64 | 102 | 90 |
| 142 | 0.78 | 0.41 | 1.90 | 0.06 | 1 | Renbp | renin binding protein | 456 | 285 | 316 | 87 | 30 | 11 | 63 | 85 | 54 | 147 | 27 |
| 134 | -0.40 | 0.29 | -1.40 | 0.16 | 1 | Pde4c | phosphodiesterase 4C, cAMP specific | 69 | 117 | 92 | 125 | 71 | 414 | 63 | 83 | 64 | 41 | 338 |
| 117 | -0.09 | 0.43 | -0.20 | 0.84 | 1 | Krt1 | keratin 1 | 214 | 258 | 193 | 100 | 72 | 71 | 63 | 153 | 62 | 26 | 78 |
| 107 | 0.82 | 0.48 | 1.73 | 0.08 | 1 | Greb1l | growth regulation by estrogen in breast ca | 55 | 66 | 27 | 176 | 62 | 93 | 63 | 105 | 157 | 218 | 159 |
| 93 | -0.16 | 0.37 | -0.44 | 0.66 | 1 | Kiss1r | KISS1 receptor | 55 | 107 | 117 | 112 | 79 | 133 | 63 | 67 | 114 | 49 | 125 |
| 85 | -0.07 | 0.41 | -0.17 | 0.87 | 1 | Cenpt | centromere protein T | 160 | 75 | 123 | 61 | 57 | 103 | 63 | 46 | 64 | 43 | 140 |
| 80 | -0.03 | 0.40 | -0.07 | 0.95 | 1 | Gm48604 | | 63 | 133 | 111 | 56 | 90 | 81 | 63 | 67 | 64 | 61 | 94 |
| 72 | 0.33 | 0.51 | 0.64 | 0.52 | 1 | 9130604C24Rik | | 63 | 63 | 95 | 34 | 102 | 55 | 63 | 120 | 62 | 81 | 49 |
| 67 | -0.41 | 0.49 | -0.84 | 0.40 | 1 | Mpc1-ps | | 41 | 65 | 91 | 82 | 124 | 44 | 63 | 25 | 91 | 62 | 53 |
| 67 | -0.08 | 0.51 | -0.16 | 0.87 | 1 | Gm5526 | | 104 | 81 | 95 | 52 | 68 | 49 | 63 | 101 | 28 | 45 | 55 |
| 66 | -0.05 | 0.50 | -0.10 | 0.92 | 1 | Gm10399 | | 71 | 50 | 136 | 46 | 79 | 41 | 63 | 51 | 53 | 109 | 23 |
| 62 | -0.67 | 0.65 | -1.04 | 0.30 | 1 | Terb1 | telomere repeat binding bouquet formatio | 45 | 65 | 53 | 57 | 70 | 107 | 63 | 6 | 68 | 39 | 116 |
| 61 | 0.03 | 0.49 | 0.06 | 0.95 | 1 | Morf4l1-ps1 | | 114 | 126 | 48 | 42 | 40 | 38 | 63 | 36 | 29 | 102 | 38 |
| 58 | -0.05 | 0.68 | -0.08 | 0.94 | 1 | Gm50340 | | 24 | 77 | 39 | 84 | 46 | 63 | 63 | 14 | 78 | 80 | 68 |
| 56 | 0.05 | 0.42 | 0.12 | 0.90 | 1 | Gm10645 | | 51 | 28 | 57 | 50 | 67 | 55 | 63 | 50 | 58 | 85 | 54 |
| 55 | -1.23 | 0.85 | -1.44 | 0.15 | 1 | Zfp982 | zinc finger protein 982 | 76 | 78 | 68 | 79 | 53 | 63 | 63 | 39 | 59 | 0 | 32 |
| 44 | -0.54 | 0.56 | -0.97 | 0.33 | 1 | 5730414N17Rik | | 60 | 40 | 54 | 64 | 41 | 28 | 63 | 16 | 48 | 43 | 29 |
| 43 | -0.06 | 1.04 | -0.06 | 0.95 | 1 | Gm10226 | | 19 | 46 | 48 | 46 | 20 | 31 | 63 | 0 | 52 | 43 | 103 |
| 42 | -0.57 | 0.77 | -0.75 | 0.45 | 1 | Pxt1 | peroxisomal, testis specific 1 | 14 | 67 | 11 | 38 | 39 | 63 | 63 | 54 | 43 | 6 | 62 |

| | | | | | | | | | | | | | | | | | | |
|---|---|---|---|---|---|---|---|---|---|---|---|---|---|---|---|---|---|---|
| 27 | -1.64 | 0.88 | -1.85 | 0.06 | 1 | Sidt1 | SID1 transmembrane family, member 1 | 48 | 75 | 6 | 26 | 6 | 20 | 63 | 6 | 0 | 19 | 24 |
| 471 | 0.17 | 0.38 | 0.45 | 0.65 | 1 | Syk | spleen tyrosine kinase | 2901 | 975 | 632 | 98 | 88 | 46 | 62 | 105 | 63 | 166 | 48 |
| 371 | -0.02 | 0.36 | -0.04 | 0.97 | 1 | Klhl5 | kelch-like 5 | 1580 | 1151 | 974 | 40 | 31 | 52 | 62 | 86 | 34 | 53 | 21 |
| 222 | -0.30 | 0.41 | -0.72 | 0.47 | 1 | Cys1 | cystin 1 | 389 | 516 | 459 | 158 | 181 | 171 | 62 | 149 | 201 | 26 | 125 |
| 196 | -0.52 | 0.36 | -1.44 | 0.15 | 1 | Sec14l4 | SEC14-like lipid binding 4 | 313 | 685 | 428 | 122 | 128 | 117 | 62 | 88 | 105 | 71 | 39 |
| 149 | -0.36 | 0.57 | -0.63 | 0.53 | 1 | Dubr | | 399 | 433 | 244 | 126 | 38 | 87 | 62 | 105 | 73 | 8 | 66 |
| 144 | 0.56 | 0.61 | 0.92 | 0.36 | 1 | Kcnn4 | potassium intermediate/small conductanc | 816 | 179 | 292 | 23 | 26 | 19 | 62 | 96 | 33 | 21 | 23 |
| 140 | 0.18 | 0.36 | 0.50 | 0.62 | 1 | Rpain | RPA interacting protein | 130 | 181 | 196 | 70 | 108 | 230 | 62 | 149 | 92 | 43 | 274 |
| 115 | 0.28 | 0.37 | 0.77 | 0.44 | 1 | Ddb2 | damage specific DNA binding protein 2 | 55 | 95 | 98 | 38 | 63 | 323 | 62 | 66 | 88 | 62 | 317 |
| 98 | -0.81 | 0.56 | -1.46 | 0.15 | 1 | Gm1043 | predicted gene 1043 | 3 | 4 | 16 | 43 | 31 | 451 | 62 | 10 | 35 | 30 | 396 |
| 90 | -0.27 | 0.43 | -0.63 | 0.53 | 1 | Spata5l1 | | 90 | 99 | 85 | 63 | 82 | 156 | 62 | 28 | 76 | 48 | 202 |
| 84 | -0.11 | 0.42 | -0.26 | 0.80 | 1 | Mex3b | mex3 RNA binding family member B | 132 | 170 | 83 | 46 | 63 | 105 | 62 | 80 | 35 | 62 | 85 |
| 79 | 0.41 | 0.47 | 0.88 | 0.38 | 1 | Gm10069 | | 53 | 79 | 56 | 51 | 19 | 196 | 62 | 81 | 58 | 59 | 157 |
| 67 | 0.05 | 0.40 | 0.13 | 0.90 | 1 | Zfp335os | | 78 | 40 | 65 | 65 | 74 | 73 | 62 | 70 | 66 | 79 | 70 |
| 65 | -0.39 | 0.92 | -0.42 | 0.67 | 1 | Snord104 | | 96 | 38 | 64 | 69 | 59 | 75 | 62 | 68 | 55 | 0 | 128 |
| 59 | -0.19 | 0.52 | -0.36 | 0.72 | 1 | Rps12-ps4 | | 124 | 91 | 136 | 37 | 31 | 25 | 62 | 16 | 59 | 35 | 29 |
| 53 | 0.15 | 0.83 | 0.17 | 0.86 | 1 | Elavl2 | ELAV like RNA binding protein 1 | 87 | 134 | 86 | 26 | 3 | 36 | 62 | 89 | 21 | 10 | 23 |
| 52 | 0.20 | 0.44 | 0.46 | 0.65 | 1 | D930048N14Rik | | 35 | 29 | 41 | 32 | 49 | 74 | 62 | 42 | 77 | 61 | 66 |
| 49 | 0.12 | 0.56 | 0.21 | 0.83 | 1 | Gm18860 | | 102 | 69 | 35 | 49 | 25 | 25 | 62 | 57 | 60 | 48 | 13 |
| 47 | -0.77 | 0.61 | -1.27 | 0.20 | 1 | Pheta2 | PH domain containing endocytic traffickin | 146 | 123 | 75 | 13 | 15 | 21 | 62 | 6 | 10 | 27 | 18 |
| 46 | -0.20 | 0.50 | -0.39 | 0.70 | 1 | Bcl2l14 | BCL2-like 14 (apoptosis facilitator) | 58 | 27 | 34 | 72 | 36 | 38 | 62 | 64 | 30 | 49 | 37 |
| 2436 | -0.37 | 1.87 | -0.20 | 0.84 | 1 | Mpeg1 | macrophage expressed gene 1 | 11873 | 6753 | 8008 | 0 | 7 | 8 | 61 | 0 | 75 | 0 | 7 |
| 1348 | 0.35 | 0.32 | 1.08 | 0.28 | 1 | Tgm2 | transglutaminase 2, C polypeptide | 7338 | 3766 | 2977 | 103 | 100 | 63 | 61 | 110 | 157 | 76 | 77 |
| 1098 | -2.11 | 1.34 | -1.58 | 0.12 | 1 | Ets1 | E26 avian leukemia oncogene 1, 5' doma | 4637 | 4027 | 3291 | 5 | 25 | 9 | 61 | 0 | 16 | 0 | 11 |
| 215 | 0.45 | 0.30 | 1.49 | 0.14 | 1 | Poglut2 | protein O-glucosyltransferase 2 | 379 | 321 | 287 | 105 | 100 | 356 | 61 | 156 | 123 | 129 | 347 |
| 180 | -1.13 | 0.83 | -1.35 | 0.18 | 1 | Arap2 | ArfGAP with RhoGAP domain, ankyrin rep | 662 | 688 | 461 | 4 | 33 | 11 | 61 | 46 | 1 | 8 | 8 |
| 170 | -0.14 | 0.73 | -0.20 | 0.84 | 1 | A330023F24Rik | | 12 | 34 | 1 | 20 | 69 | 796 | 61 | 20 | 82 | 36 | 737 |
| 163 | -0.19 | 0.37 | -0.50 | 0.62 | 1 | Gpr137b-ps | | 426 | 410 | 375 | 94 | 68 | 85 | 61 | 35 | 84 | 73 | 82 |
| 144 | -0.49 | 0.39 | -1.25 | 0.21 | 1 | Pik3c2b | phosphatidylinositol-4-phosphate 3-kinas | 281 | 314 | 480 | 54 | 90 | 80 | 61 | 16 | 70 | 50 | 83 |
| 123 | -0.13 | 0.46 | -0.28 | 0.78 | 1 | Ephx2 | epoxide hydrolase 2, cytoplasmic | 277 | 229 | 169 | 99 | 102 | 67 | 61 | 185 | 59 | 68 | 37 |
| 122 | -0.53 | 0.36 | -1.47 | 0.14 | 1 | Msh5 | mutS homolog 5 | 40 | 45 | 46 | 46 | 70 | 513 | 61 | 31 | 64 | 26 | 399 |
| 114 | 0.05 | 0.58 | 0.08 | 0.93 | 1 | Brca1 | breast cancer 1, early onset | 283 | 103 | 109 | 107 | 82 | 110 | 61 | 190 | 69 | 32 | 106 |
| 99 | 0.41 | 0.37 | 1.11 | 0.27 | 1 | Gm11992 | predicted gene 11992 | 45 | 107 | 136 | 81 | 74 | 130 | 61 | 116 | 127 | 75 | 138 |
| 98 | 0.26 | 0.34 | 0.76 | 0.45 | 1 | Fancg | Fanconi anemia, complementation group | 144 | 136 | 176 | 46 | 58 | 129 | 61 | 70 | 78 | 70 | 115 |
| 98 | 0.74 | 0.40 | 1.85 | 0.06 | 1 | Zfp825 | zinc finger protein 825 | 162 | 83 | 65 | 73 | 62 | 85 | 61 | 75 | 111 | 135 | 162 |
| 97 | 0.34 | 0.53 | 0.65 | 0.52 | 1 | Ada | adenosine deaminase | 280 | 99 | 69 | 119 | 69 | 22 | 61 | 174 | 64 | 84 | 31 |
| 90 | -0.48 | 0.42 | -1.14 | 0.25 | 1 | Kcnab1 | potassium voltage-gated channel, shaker | 427 | 144 | 151 | 23 | 33 | 40 | 61 | 12 | 24 | 36 | 43 |
| 85 | -0.74 | 0.53 | -1.39 | 0.17 | 1 | Ttc16 | tetratricopeptide repeat domain 16 | 5 | 25 | 7 | 91 | 38 | 341 | 61 | 43 | 52 | 14 | 253 |
| 79 | 0.12 | 0.39 | 0.30 | 0.76 | 1 | Rexo5 | RNA exonuclease 5 | 63 | 77 | 34 | 53 | 58 | 165 | 61 | 61 | 58 | 68 | 177 |
| 70 | 0.63 | 0.53 | 1.20 | 0.23 | 1 | D130007C19Rik | | 33 | 31 | 35 | 54 | 60 | 66 | 61 | 103 | 51 | 218 | 55 |
| 65 | -0.33 | 0.69 | -0.48 | 0.63 | 1 | Efcab10 | EF-hand calcium binding domain 10 | 80 | 46 | 131 | 64 | 64 | 43 | 61 | 124 | 64 | 9 | 27 |
| 60 | -0.27 | 0.50 | -0.54 | 0.59 | 1 | Rpl18-ps2 | | 108 | 104 | 87 | 43 | 58 | 37 | 61 | 43 | 50 | 21 | 53 |
| 58 | 0.13 | 0.58 | 0.23 | 0.82 | 1 | Aloxe3 | arachidonate lipoxygenase 3 | 12 | 64 | 42 | 58 | 46 | 83 | 61 | 50 | 48 | 75 | 101 |
| 58 | -0.61 | 0.53 | -1.16 | 0.25 | 1 | Gm9816 | | 79 | 42 | 49 | 79 | 86 | 50 | 61 | 20 | 51 | 71 | 52 |
| 55 | 0.12 | 0.44 | 0.28 | 0.78 | 1 | Fancb | Fanconi anemia, complementation group | 72 | 44 | 81 | 25 | 37 | 80 | 61 | 32 | 60 | 49 | 69 |
| 50 | 0.02 | 0.46 | 0.04 | 0.96 | 1 | 2810032G03Rik | | 35 | 30 | 30 | 88 | 49 | 32 | 61 | 54 | 62 | 64 | 43 |
| 45 | -0.06 | 0.70 | -0.08 | 0.93 | 1 | Rab26 | RAB26, member RAS oncogene family | 21 | 101 | 63 | 24 | 26 | 38 | 61 | 41 | 53 | 18 | 45 |
| 43 | -0.11 | 0.62 | -0.17 | 0.86 | 1 | 1110013H19Rik | | 37 | 23 | 22 | 34 | 65 | 40 | 61 | 86 | 46 | 24 | 35 |
| 36 | -0.46 | 0.62 | -0.74 | 0.46 | 1 | Gm20075 | | 73 | 9 | 14 | 32 | 48 | 30 | 61 | 20 | 26 | 52 | 32 |
| 35 | 0.49 | 0.77 | 0.64 | 0.52 | 1 | Gm10366 | | 37 | 2 | 22 | 25 | 25 | 30 | 61 | 65 | 29 | 52 | 34 |
| 34 | -0.94 | 0.71 | -1.32 | 0.19 | 1 | Gm5857 | | 15 | 49 | 21 | 59 | 38 | 30 | 61 | 24 | 43 | 8 | 27 |
| 29 | -0.89 | 0.68 | -1.30 | 0.19 | 1 | Gm28455 | | 10 | 37 | 21 | 46 | 17 | 44 | 61 | 23 | 12 | 28 | 24 |
| 914 | 0.04 | 1.10 | 0.03 | 0.97 | 1 | Fzd4 | frizzled class receptor 4 | 3763 | 3291 | 2749 | 39 | 10 | 17 | 60 | 71 | 36 | 0 | 20 |
| 711 | -0.65 | 0.41 | -1.59 | 0.11 | 1 | Avpr1b | arginine vasopressin receptor 1B | 1713 | 2657 | 3032 | 45 | 65 | 71 | 60 | 89 | 48 | 33 | 11 |
| 253 | 0.52 | 0.37 | 1.42 | 0.16 | 1 | Lmo2 | LIM domain only 2 | 998 | 458 | 749 | 67 | 67 | 46 | 60 | 130 | 61 | 77 | 75 |
| 177 | -0.35 | 0.55 | -0.63 | 0.53 | 1 | Tbkbp1 | TBK1 binding protein 1 | 667 | 478 | 441 | 36 | 49 | 39 | 60 | 115 | 24 | 17 | 22 |
| 134 | -0.21 | 0.34 | -0.64 | 0.52 | 1 | Klf8 | Kruppel-like factor 8 | 195 | 216 | 293 | 121 | 78 | 153 | 60 | 83 | 97 | 44 | 136 |
| 129 | 0.11 | 0.35 | 0.31 | 0.76 | 1 | Haus3 | HAUS augmin-like complex, subunit 3 | 243 | 244 | 170 | 77 | 105 | 130 | 60 | 94 | 90 | 95 | 114 |
| 123 | 0.22 | 0.51 | 0.43 | 0.67 | 1 | Donson | downstream neighbor of SON | 221 | 127 | 130 | 82 | 38 | 231 | 60 | 149 | 59 | 26 | 235 |
| 121 | 0.41 | 0.59 | 0.69 | 0.49 | 1 | Ifit3 | interferon-induced protein with tetratricop | 316 | 202 | 454 | 20 | 43 | 14 | 60 | 3 | 90 | 103 | 24 |

| | | | | | | | | | | | | | | | | | | |
|---|---|---|---|---|---|---|---|---|---|---|---|---|---|---|---|---|---|---|
| 115 | -0.20 | 0.42 | -0.47 | 0.64 | 1 | 4930579G24Ril | RIKEN cDNA 4930579G24 gene | 137 | 81 | 219 | 134 | 164 | 90 | 60 | 105 | 107 | 100 | 66 |
| 110 | -0.47 | 0.55 | -0.86 | 0.39 | 1 | Zfp712 | zinc finger protein 712 | 195 | 245 | 83 | 46 | 143 | 154 | 60 | 31 | 85 | 70 | 94 |
| 101 | -0.22 | 0.42 | -0.51 | 0.61 | 1 | Mapk13 | mitogen-activated protein kinase 13 | 123 | 87 | 154 | 123 | 80 | 142 | 60 | 93 | 53 | 85 | 114 |
| 99 | 0.55 | 0.50 | 1.10 | 0.27 | 1 | Gucy1a2 | guanylate cyclase 1, soluble, alpha 2 | 323 | 277 | 155 | 21 | 25 | 24 | 60 | 98 | 37 | 45 | 20 |
| 92 | -0.20 | 0.45 | -0.45 | 0.65 | 1 | Rhpn1 | rhophilin, Rho GTPase binding protein 1 | 82 | 42 | 54 | 107 | 100 | 166 | 60 | 98 | 63 | 53 | 184 |
| 82 | 0.26 | 0.50 | 0.52 | 0.60 | 1 | B230377A18Rik | | 57 | 68 | 139 | 100 | 68 | 63 | 60 | 75 | 62 | 135 | 75 |
| 77 | 0.51 | 0.39 | 1.30 | 0.19 | 1 | Chrna1os | | 92 | 83 | 113 | 55 | 64 | 50 | 60 | 71 | 67 | 125 | 70 |
| 74 | -0.92 | 0.47 | -1.97 | 0.05 | 1 | Ccdc114 | coiled-coil domain containing 114 | 85 | 114 | 118 | 56 | 60 | 125 | 60 | 23 | 37 | 17 | 121 |
| 72 | 0.05 | 0.53 | 0.09 | 0.93 | 1 | Myl12b | myosin, light chain 12B, regulatory | 152 | 55 | 108 | 47 | 59 | 70 | 60 | 83 | 53 | 30 | 76 |
| 69 | -1.09 | 0.65 | -1.68 | 0.09 | 1 | Ankrd34b | ankyrin repeat domain 34B | 135 | 232 | 145 | 11 | 16 | 85 | 60 | 19 | 4 | 18 | 38 |
| 67 | -0.08 | 0.49 | -0.16 | 0.88 | 1 | Pfn4 | profilin family, member 4 | 63 | 128 | 96 | 60 | 55 | 36 | 60 | 127 | 63 | 28 | 23 |
| 64 | -0.22 | 0.44 | -0.50 | 0.62 | 1 | AI839979 | | 37 | 50 | 30 | 149 | 58 | 63 | 60 | 68 | 60 | 58 | 77 |
| 63 | -0.10 | 0.47 | -0.21 | 0.83 | 1 | Gm50455 | | 63 | 57 | 51 | 84 | 65 | 57 | 60 | 94 | 68 | 28 | 65 |
| 57 | -0.35 | 0.88 | -0.40 | 0.69 | 1 | Ifit3b | interferon-induced protein with tetratricop | 153 | 78 | 120 | 39 | 19 | 21 | 60 | 0 | 43 | 77 | 20 |
| 49 | -0.23 | 0.68 | -0.33 | 0.74 | 1 | Inca1 | inhibitor of CDK, cyclin A1 interacting prot | 23 | 64 | 43 | 57 | 46 | 44 | 60 | 16 | 23 | 75 | 83 |
| 34 | -1.05 | 0.71 | -1.49 | 0.14 | 1 | Gm3724 | | 58 | 59 | 31 | 39 | 41 | 17 | 60 | 5 | 26 | 26 | 16 |
| 33 | -0.40 | 0.63 | -0.63 | 0.53 | 1 | Catsperd | cation channel sperm associated auxiliar | 23 | 7 | 3 | 61 | 18 | 57 | 60 | 51 | 20 | 35 | 32 |
| 2259 | -0.90 | 0.55 | -1.64 | 0.10 | 1 | Mfge8 | milk fat globule-EGF factor 8 protein | 11336 | 6904 | 6292 | 61 | 50 | 38 | 59 | 41 | 35 | 6 | 29 |
| 329 | -0.58 | 0.41 | -1.42 | 0.16 | 1 | Tppp3 | tubulin polymerization-promoting protein | 1086 | 669 | 909 | 113 | 121 | 238 | 59 | 100 | 41 | 30 | 258 |
| 299 | -0.48 | 0.57 | -0.85 | 0.40 | 1 | Plekho1 | pleckstrin homology domain containing, f | 1179 | 854 | 869 | 47 | 67 | 41 | 59 | 79 | 31 | 4 | 59 |
| 247 | -0.21 | 0.37 | -0.56 | 0.58 | 1 | Pik3c2g | phosphatidylinositol-4-phosphate 3-kinas | 593 | 932 | 306 | 187 | 53 | 175 | 59 | 126 | 69 | 30 | 187 |
| 204 | 0.06 | 0.50 | 0.12 | 0.90 | 1 | Stmn1 | stathmin 1 | 407 | 215 | 256 | 232 | 277 | 110 | 59 | 291 | 207 | 77 | 118 |
| 171 | 0.59 | 0.45 | 1.32 | 0.19 | 1 | Nusap1 | nucleolar and spindle associated protein | 545 | 284 | 174 | 68 | 131 | 88 | 59 | 161 | 107 | 138 | 126 |
| 145 | 0.44 | 0.37 | 1.18 | 0.24 | 1 | Ammecr1 | Alport syndrome, mental retardation, midf | 288 | 278 | 167 | 108 | 93 | 105 | 59 | 88 | 118 | 151 | 138 |
| 137 | -1.04 | 0.53 | -1.95 | 0.05 | 1 | Atp10a | ATPase, class V, type 10A | 512 | 320 | 443 | 22 | 14 | 64 | 59 | 25 | 12 | 9 | 22 |
| 129 | -0.02 | 0.33 | -0.08 | 0.94 | 1 | Gm45805 | | 81 | 145 | 98 | 106 | 113 | 286 | 59 | 100 | 137 | 54 | 247 |
| 125 | 0.50 | 0.54 | 0.93 | 0.35 | 1 | Ncaph | non-SMC condensin I complex, subunit H | 375 | 130 | 168 | 55 | 111 | 67 | 59 | 178 | 106 | 53 | 76 |
| 122 | -0.15 | 0.41 | -0.37 | 0.71 | 1 | Smco4 | single-pass membrane protein with coilec | 476 | 289 | 289 | 32 | 57 | 10 | 59 | 37 | 26 | 48 | 18 |
| 112 | -1.07 | 0.57 | -1.89 | 0.06 | 1 | Aoc2 | amine oxidase, copper containing 2 (retin | 53 | 33 | 33 | 109 | 111 | 371 | 59 | 9 | 50 | 59 | 349 |
| 95 | -0.21 | 0.40 | -0.52 | 0.60 | 1 | H4c3 | H4 clustered histone 3 | 21 | 55 | 19 | 198 | 127 | 118 | 59 | 176 | 113 | 35 | 128 |
| 94 | -0.16 | 0.45 | -0.36 | 0.72 | 1 | Hdc | histidine decarboxylase | 255 | 185 | 140 | 35 | 64 | 79 | 59 | 24 | 76 | 68 | 47 |
| 91 | -0.22 | 0.52 | -0.43 | 0.67 | 1 | Eda2r | ectodysplasin A2 receptor | 308 | 188 | 141 | 32 | 58 | 43 | 59 | 68 | 18 | 36 | 50 |
| 84 | 0.05 | 0.39 | 0.14 | 0.89 | 1 | Fbxo47 | F-box protein 47 | 89 | 109 | 86 | 102 | 84 | 69 | 59 | 68 | 73 | 98 | 82 |
| 77 | 0.33 | 0.51 | 0.65 | 0.52 | 1 | Uba52 | ubiquitin A-52 residue ribosomal protein f | 120 | 43 | 131 | 25 | 90 | 89 | 59 | 83 | 69 | 58 | 80 |
| 71 | -1.08 | 0.68 | -1.59 | 0.11 | 1 | Chaf1b | chromatin assembly factor 1, subunit B (p | 85 | 127 | 56 | 90 | 104 | 75 | 59 | 20 | 98 | 8 | 63 |
| 65 | -0.38 | 0.49 | -0.78 | 0.44 | 1 | H3f3aos | | 73 | 110 | 30 | 116 | 59 | 52 | 59 | 54 | 28 | 67 | 71 |
| 64 | -0.47 | 0.90 | -0.52 | 0.60 | 1 | Gm38300 | | 62 | 1 | 61 | 74 | 115 | 85 | 59 | 119 | 61 | 23 | 38 |
| 61 | -0.99 | 0.79 | -1.26 | 0.21 | 1 | Gm50268 | | 16 | 12 | 4 | 73 | 38 | 200 | 59 | 12 | 56 | 5 | 198 |
| 56 | 0.28 | 0.81 | 0.35 | 0.72 | 1 | Alpk1 | alpha-kinase 1 | 172 | 116 | 125 | 6 | 9 | 16 | 59 | 5 | 31 | 10 | 62 |
| 55 | -0.41 | 0.51 | -0.80 | 0.43 | 1 | 4930502A04Rik | | 12 | 34 | 37 | 39 | 60 | 143 | 59 | 37 | 45 | 41 | 97 |
| 53 | -0.43 | 0.58 | -0.75 | 0.46 | 1 | Bmpr1b | bone morphogenetic protein receptor, typ | 45 | 76 | 82 | 88 | 19 | 53 | 59 | 78 | 21 | 28 | 39 |
| 52 | -0.78 | 0.45 | -1.73 | 0.08 | 1 | Fank1 | fibronectin type 3 and ankyrin repeat dom | 82 | 64 | 53 | 47 | 43 | 84 | 59 | 20 | 28 | 31 | 63 |
| 49 | 0.31 | 0.70 | 0.44 | 0.66 | 1 | Gm31373 | | 23 | 20 | 45 | 11 | 35 | 124 | 59 | 53 | 19 | 82 | 70 |
| 46 | -0.02 | 0.72 | -0.03 | 0.98 | 1 | Lrrc14b | leucine rich repeat containing 14B | 9 | 30 | 0 | 101 | 36 | 25 | 59 | 96 | 40 | 95 | 12 |
| 42 | -0.13 | 0.62 | -0.21 | 0.84 | 1 | B230217O12Rik | | 18 | 41 | 17 | 65 | 20 | 63 | 59 | 22 | 35 | 59 | 59 |
| 37 | 0.15 | 0.74 | 0.20 | 0.84 | 1 | Gm9333 | | 85 | 71 | 5 | 33 | 13 | 16 | 59 | 13 | 16 | 58 | 37 |
| 34 | -0.09 | 0.56 | -0.16 | 0.87 | 1 | Gm13341 | | 13 | 24 | 14 | 34 | 42 | 35 | 59 | 42 | 42 | 33 | 41 |
| 34 | -0.07 | 0.53 | -0.13 | 0.89 | 1 | 9130213A22Rik | | 26 | 25 | 9 | 43 | 28 | 30 | 59 | 34 | 28 | 62 | 29 |
| 18 | -1.61 | 0.89 | -1.80 | 0.07 | 1 | Mrap2 | melanocortin 2 receptor accessory proteir | 5 | 7 | 0 | 32 | 27 | 25 | 59 | 17 | 12 | 8 | 8 |
| 212 | -0.78 | 0.40 | -1.97 | 0.05 | 1 | Syt16 | synaptotagmin XVI | 393 | 989 | 433 | 95 | 62 | 115 | 58 | 27 | 62 | 31 | 69 |
| 121 | 0.40 | 0.50 | 0.79 | 0.43 | 1 | Zfp872 | zinc finger protein 872 | 26 | 70 | 32 | 60 | 138 | 291 | 58 | 65 | 124 | 149 | 323 |
| 97 | -1.21 | 0.75 | -1.62 | 0.11 | 1 | Kcnc2 | potassium voltage gated channel, Shaw-r | 144 | 244 | 232 | 49 | 64 | 116 | 58 | 0 | 36 | 36 | 83 |
| 91 | 0.72 | 0.50 | 1.45 | 0.15 | 1 | Rpusd3 | RNA pseudouridylate synthase domain co | 64 | 47 | 57 | 48 | 82 | 134 | 58 | 174 | 79 | 164 | 95 |
| 84 | 0.51 | 0.51 | 0.98 | 0.33 | 1 | Rnf122 | ring finger protein 122 | 254 | 128 | 116 | 19 | 51 | 34 | 58 | 24 | 100 | 120 | 22 |
| 72 | 0.22 | 0.37 | 0.60 | 0.55 | 1 | Nupr1l | nuclear protein transcriptional regulator 1 | 38 | 50 | 58 | 70 | 93 | 79 | 58 | 88 | 90 | 86 | 80 |
| 61 | 0.06 | 0.48 | 0.13 | 0.90 | 1 | Lrrc6 | leucine rich repeat containing 6 (testis) | 11 | 31 | 21 | 103 | 54 | 85 | 58 | 71 | 50 | 90 | 92 |
| 60 | -0.02 | 0.49 | -0.03 | 0.97 | 1 | Pcdhb13 | protocadherin beta 13 | 104 | 96 | 58 | 67 | 32 | 48 | 58 | 41 | 68 | 59 | 32 |
| 58 | 0.11 | 0.55 | 0.21 | 0.83 | 1 | Gm8066 | | 61 | 38 | 55 | 39 | 66 | 60 | 58 | 110 | 48 | 67 | 31 |
| 53 | -0.59 | 0.54 | -1.09 | 0.27 | 1 | 2610037D02Rik | | 43 | 87 | 46 | 38 | 25 | 106 | 58 | 22 | 21 | 21 | 117 |
| 49 | -0.32 | 0.53 | -0.60 | 0.55 | 1 | Opn3 | opsin 3 | 75 | 34 | 43 | 76 | 51 | 34 | 58 | 52 | 41 | 31 | 45 |

| | | | | | | | | | | | | | | | | | | |
|---|---|---|---|---|---|---|---|---|---|---|---|---|---|---|---|---|---|---|
| 47 | 0.02 | 0.58 | 0.03 | 0.97 | 1 | Gm37607 | | 34 | 40 | 15 | 24 | 20 | 128 | 58 | 65 | 26 | 27 | 85 |
| 42 | -0.54 | 0.85 | -0.63 | 0.53 | 1 | Gm5900 | | 78 | 36 | 55 | 29 | 42 | 26 | 58 | 0 | 40 | 67 | 30 |
| 41 | -1.00 | 0.64 | -1.57 | 0.12 | 1 | Gm10010 | | 5 | 30 | 32 | 67 | 70 | 55 | 58 | 20 | 43 | 21 | 51 |
| 37 | -1.20 | 0.81 | -1.48 | 0.14 | 1 | 2310043P16Rik | | 28 | 31 | 22 | 65 | 35 | 49 | 58 | 32 | 20 | 1 | 61 |
| 288 | -0.36 | 0.35 | -1.04 | 0.30 | 1 | Cyp2d22 | cytochrome P450, family 2, subfamily d, p | 805 | 832 | 698 | 164 | 77 | 167 | 57 | 74 | 92 | 39 | 162 |
| 123 | 0.15 | 0.42 | 0.35 | 0.72 | 1 | Aifm3 | apoptosis-inducing factor, mitochondrion- | 94 | 96 | 89 | 156 | 161 | 142 | 57 | 140 | 104 | 164 | 146 |
| 117 | -0.52 | 0.57 | -0.90 | 0.37 | 1 | Rbm11 | RNA binding motif protein 11 | 305 | 369 | 224 | 74 | 64 | 35 | 57 | 8 | 47 | 64 | 45 |
| 104 | 0.31 | 0.37 | 0.82 | 0.41 | 1 | Mrip-ps | | 185 | 133 | 180 | 74 | 68 | 88 | 57 | 61 | 79 | 129 | 94 |
| 102 | -0.28 | 0.56 | -0.50 | 0.61 | 1 | Gen1 | GEN1, Holliday junction 5' flap endonucle | 162 | 141 | 55 | 101 | 115 | 126 | 57 | 86 | 91 | 23 | 160 |
| 99 | -0.51 | 0.41 | -1.23 | 0.22 | 1 | Rasgef1a | RasGEF domain family, member 1A | 91 | 197 | 301 | 57 | 93 | 88 | 57 | 70 | 48 | 36 | 53 |
| 91 | 0.05 | 0.44 | 0.11 | 0.91 | 1 | Espnl | espin-like | 78 | 87 | 90 | 88 | 32 | 221 | 57 | 67 | 52 | 90 | 142 |
| 83 | -0.85 | 0.48 | -1.76 | 0.08 | 1 | Neu2 | neuraminidase 2 | 94 | 144 | 175 | 116 | 111 | 35 | 57 | 66 | 65 | 12 | 38 |
| 77 | 0.13 | 0.48 | 0.27 | 0.78 | 1 | Neb | nebulin | 124 | 88 | 123 | 46 | 42 | 98 | 57 | 73 | 67 | 24 | 105 |
| 75 | -0.27 | 0.43 | -0.62 | 0.54 | 1 | Pcdhgb5 | protocadherin gamma subfamily B, 5 | 116 | 130 | 133 | 44 | 93 | 49 | 57 | 30 | 78 | 64 | 34 |
| 73 | -0.03 | 0.61 | -0.05 | 0.96 | 1 | AA986860 | expressed sequence AA986860 | 214 | 113 | 131 | 39 | 38 | 37 | 57 | 60 | 66 | 12 | 32 |
| 66 | -0.66 | 0.59 | -1.12 | 0.26 | 1 | 1110006O24Rik | | 39 | 22 | 74 | 161 | 80 | 63 | 57 | 41 | 63 | 85 | 43 |
| 63 | 0.02 | 0.66 | 0.03 | 0.98 | 1 | Gm37397 | | 51 | 59 | 37 | 107 | 40 | 69 | 57 | 15 | 93 | 91 | 73 |
| 60 | -0.94 | 0.59 | -1.60 | 0.11 | 1 | Zfp786 | zinc finger protein 786 | 46 | 108 | 85 | 54 | 77 | 79 | 57 | 9 | 51 | 28 | 67 |
| 55 | 0.18 | 0.44 | 0.41 | 0.68 | 1 | Rps23-ps1 | | 94 | 89 | 90 | 30 | 32 | 39 | 57 | 38 | 42 | 55 | 40 |
| 51 | 0.54 | 0.51 | 1.05 | 0.29 | 1 | 6430503K07Rik | | 57 | 37 | 35 | 23 | 58 | 41 | 57 | 79 | 51 | 73 | 51 |
| 41 | -0.32 | 0.54 | -0.59 | 0.56 | 1 | Gm49041 | | 75 | 47 | 68 | 25 | 39 | 21 | 57 | 12 | 26 | 59 | 22 |
| 40 | -0.20 | 0.72 | -0.28 | 0.78 | 1 | D930020B18Ri | RIKEN cDNA D930020B18 gene | 13 | 11 | 0 | 50 | 43 | 64 | 57 | 92 | 33 | 39 | 40 |
| 40 | -0.07 | 1.03 | -0.07 | 0.95 | 1 | Zfp960 | zinc finger protein 960 | 75 | 56 | 10 | 34 | 22 | 42 | 57 | 59 | 34 | 0 | 48 |
| 28 | -0.39 | 0.93 | -0.42 | 0.68 | 1 | Atg4a-ps | | 21 | 17 | 27 | 37 | 10 | 30 | 57 | 42 | 49 | 3 | 19 |
| 8086 | -0.91 | 1.14 | -0.80 | 0.43 | 1 | Zg16 | zymogen granule protein 16 | 10466 | 39872 | 35802 | 1783 | 167 | 11 | 56 | 163 | 86 | 27 | 512 |
| 229 | -3.02 | 2.65 | -1.14 | 0.25 | 1 | Rac2 | Rac family small GTPase 2 | 1512 | 539 | 400 | 0 | 0 | 5 | 56 | 0 | 4 | 0 | 0 |
| 200 | -0.22 | 0.42 | -0.52 | 0.60 | 1 | Dipk1a | divergent protein kinase domain 1A | 715 | 393 | 479 | 83 | 110 | 70 | 56 | 127 | 72 | 59 | 34 |
| 119 | -0.47 | 0.52 | -0.90 | 0.37 | 1 | Gm13003 | | 23 | 63 | 17 | 77 | 41 | 498 | 56 | 20 | 57 | 32 | 421 |
| 77 | 0.15 | 0.45 | 0.33 | 0.74 | 1 | Skp2 | S-phase kinase-associated protein 2 (p45 | 170 | 87 | 67 | 33 | 35 | 142 | 56 | 39 | 63 | 49 | 108 |
| 64 | -0.13 | 0.57 | -0.23 | 0.82 | 1 | 2310068J16Rik | | 21 | 64 | 43 | 91 | 102 | 52 | 56 | 48 | 60 | 121 | 43 |
| 59 | -0.12 | 0.58 | -0.21 | 0.83 | 1 | 5730405O15Rik | | 41 | 33 | 39 | 23 | 65 | 112 | 56 | 58 | 25 | 24 | 169 |
| 58 | -0.01 | 0.50 | -0.02 | 0.98 | 1 | Gm21769 | | 54 | 57 | 61 | 25 | 34 | 134 | 56 | 46 | 24 | 62 | 89 |
| 57 | -0.29 | 0.45 | -0.64 | 0.52 | 1 | Gm6723 | | 56 | 69 | 30 | 89 | 51 | 55 | 56 | 90 | 38 | 70 | 25 |
| 55 | 0.43 | 0.48 | 0.88 | 0.38 | 1 | Gm6863 | | 94 | 104 | 77 | 20 | 39 | 27 | 56 | 46 | 39 | 63 | 38 |
| 54 | 0.28 | 0.48 | 0.58 | 0.56 | 1 | A930012O16Rik | | 28 | 25 | 24 | 55 | 62 | 55 | 56 | 110 | 63 | 41 | 72 |
| 52 | -0.14 | 0.65 | -0.22 | 0.82 | 1 | F2 | coagulation factor II | 39 | 127 | 89 | 43 | 15 | 51 | 56 | 21 | 37 | 30 | 63 |
| 48 | 0.31 | 0.54 | 0.57 | 0.57 | 1 | Zfp524 | zinc finger protein 524 | 33 | 59 | 26 | 31 | 61 | 38 | 56 | 49 | 65 | 52 | 65 |
| 48 | -0.71 | 0.49 | -1.45 | 0.15 | 1 | Gm10288 | | 77 | 77 | 80 | 69 | 34 | 25 | 56 | 37 | 29 | 22 | 19 |
| 47 | -0.45 | 0.59 | -0.76 | 0.45 | 1 | Nek11 | NIMA (never in mitosis gene a)-related ex | 21 | 40 | 65 | 34 | 50 | 72 | 56 | 41 | 61 | 12 | 65 |
| 42 | -0.25 | 0.65 | -0.39 | 0.69 | 1 | 4933428C19Rik | | 0 | 3 | 0 | 53 | 60 | 64 | 56 | 95 | 30 | 36 | 68 |
| 373 | 0.07 | 0.39 | 0.17 | 0.87 | 1 | Septin4 | septin 4 | 1370 | 1010 | 1119 | 86 | 88 | 53 | 55 | 39 | 49 | 130 | 100 |
| 185 | 0.60 | 0.34 | 1.76 | 0.08 | 1 | Pik3cg | phosphatidylinositol-4,5-bisphosphate 3-k | 817 | 332 | 426 | 44 | 43 | 43 | 55 | 73 | 77 | 67 | 59 |
| 176 | 0.20 | 0.52 | 0.38 | 0.70 | 1 | Chaf1a | chromatin assembly factor 1, subunit A (p | 420 | 199 | 229 | 146 | 121 | 163 | 55 | 301 | 56 | 102 | 150 |
| 109 | -0.19 | 0.50 | -0.37 | 0.71 | 1 | Gm10516 | | 75 | 80 | 107 | 185 | 114 | 149 | 55 | 74 | 62 | 151 | 150 |
| 70 | -0.18 | 0.58 | -0.32 | 0.75 | 1 | Zfp599 | zinc finger protein 599 | 96 | 89 | 131 | 44 | 60 | 64 | 55 | 12 | 38 | 111 | 69 |
| 67 | -0.53 | 0.53 | -1.01 | 0.31 | 1 | Gm20186 | | 47 | 79 | 78 | 68 | 101 | 76 | 55 | 97 | 76 | 12 | 53 |
| 64 | -0.69 | 0.52 | -1.33 | 0.19 | 1 | Gm5141 | predicted gene 5141 | 100 | 116 | 58 | 58 | 45 | 83 | 55 | 32 | 28 | 14 | 110 |
| 60 | -0.35 | 0.63 | -0.57 | 0.57 | 1 | 9430081H08Rik | | 50 | 36 | 83 | 54 | 72 | 81 | 55 | 89 | 46 | 10 | 86 |
| 55 | -0.38 | 0.94 | -0.41 | 0.68 | 1 | Gm10037 | predicted gene 10037 | 29 | 36 | 62 | 57 | 27 | 109 | 55 | 0 | 56 | 75 | 103 |
| 54 | -0.04 | 0.57 | -0.07 | 0.94 | 1 | Gm17484 | | 30 | 80 | 58 | 41 | 44 | 79 | 55 | 39 | 61 | 73 | 37 |
| 49 | 0.26 | 0.81 | 0.32 | 0.75 | 1 | Cdk15 | cyclin-dependent kinase 15 | 49 | 18 | 0 | 77 | 40 | 28 | 55 | 42 | 37 | 137 | 57 |
| 48 | -0.59 | 0.82 | -0.72 | 0.47 | 1 | Gm14966 | | 68 | 13 | 18 | 50 | 55 | 64 | 55 | 2 | 82 | 39 | 83 |
| 41 | 0.30 | 0.57 | 0.53 | 0.60 | 1 | 4930547M16Rik | | 37 | 40 | 38 | 33 | 26 | 31 | 55 | 80 | 25 | 64 | 23 |
| 41 | 0.20 | 0.64 | 0.31 | 0.76 | 1 | Gm3604 | predicted gene 3604 | 64 | 22 | 21 | 23 | 33 | 57 | 55 | 64 | 43 | 28 | 40 |
| 33 | -0.93 | 1.00 | -0.93 | 0.35 | 1 | Pcdha1 | protocadherin alpha 1 | 10 | 13 | 9 | 77 | 35 | 42 | 55 | 0 | 27 | 45 | 51 |
| 33 | -0.50 | 0.59 | -0.84 | 0.40 | 1 | Bod1-ps | | 27 | 32 | 49 | 47 | 35 | 14 | 55 | 23 | 35 | 24 | 18 |
| 20 | 0.43 | 1.40 | 0.31 | 0.76 | 1 | St6gal2 | beta galactoside alpha 2,6 sialyltransfera | 14 | 11 | 0 | 4 | 18 | 8 | 55 | 0 | 56 | 43 | 10 |
| 246 | -0.57 | 0.33 | -1.74 | 0.08 | 1 | Cpb2 | carboxypeptidase B2 (plasma) | 66 | 405 | 155 | 678 | 100 | 390 | 54 | 414 | 71 | 26 | 353 |
| 239 | -0.10 | 0.48 | -0.21 | 0.84 | 1 | Ston1 | stonin 1 | 1093 | 750 | 515 | 52 | 23 | 19 | 54 | 52 | 40 | 10 | 24 |
| 203 | -0.43 | 0.73 | -0.60 | 0.55 | 1 | Pcsk4 | proprotein convertase subtilisin/kexin type | 63 | 24 | 28 | 17 | 38 | 1024 | 54 | 36 | 31 | 0 | 917 |

| | | | | | | | | | | | | | | | | | | |
|---|---|---|---|---|---|---|---|---|---|---|---|---|---|---|---|---|---|---|
| 149 | 0.46 | 0.34 | 1.36 | 0.17 | 1 | Tamm41 | TAM41 mitochondrial translocator assemb | 210 | 291 | 166 | 116 | 120 | 122 | 54 | 171 | 150 | 85 | 156 |
| 141 | 0.40 | 0.52 | 0.76 | 0.45 | 1 | Ccbe1 | collagen and calcium binding EGF domai | 417 | 547 | 377 | 12 | 10 | 19 | 54 | 36 | 33 | 32 | 14 |
| 139 | -0.53 | 0.43 | -1.24 | 0.21 | 1 | Mill2 | MHC I like leukocyte 2 | 372 | 331 | 236 | 65 | 86 | 134 | 54 | 48 | 64 | 19 | 125 |
| 129 | -1.19 | 0.76 | -1.57 | 0.12 | 1 | Ppp1r17 | protein phosphatase 1, regulatory subuni | 369 | 467 | 228 | 48 | 17 | 117 | 54 | 0 | 30 | 15 | 75 |
| 118 | -1.05 | 0.92 | -1.14 | 0.25 | 1 | Bcar3 | breast cancer anti-estrogen resistance 3 | 616 | 251 | 273 | 20 | 3 | 30 | 54 | 8 | 29 | 0 | 10 |
| 101 | -0.56 | 0.50 | -1.13 | 0.26 | 1 | Pabpc1l | poly(A) binding protein, cytoplasmic 1-like | 130 | 197 | 282 | 105 | 77 | 57 | 54 | 23 | 69 | 77 | 35 |
| 90 | 0.26 | 0.46 | 0.58 | 0.56 | 1 | 5530401A14Rik | | 26 | 30 | 43 | 126 | 114 | 115 | 54 | 178 | 62 | 111 | 133 |
| 87 | 0.18 | 0.43 | 0.41 | 0.68 | 1 | Mtbp | Mdm2, transformed 3T3 cell double minut | 174 | 124 | 152 | 24 | 67 | 93 | 54 | 24 | 59 | 90 | 97 |
| 84 | -0.51 | 0.53 | -0.96 | 0.34 | 1 | Cage1 | cancer antigen 1 | 78 | 115 | 122 | 83 | 123 | 78 | 54 | 117 | 49 | 14 | 92 |
| 72 | 0.06 | 0.61 | 0.10 | 0.92 | 1 | Lrrc51 | leucine rich repeat containing 51 | 15 | 85 | 63 | 46 | 68 | 121 | 54 | 175 | 63 | 54 | 47 |
| 69 | 0.25 | 0.43 | 0.57 | 0.57 | 1 | BC055324 | cDNA sequence BC055324 | 105 | 101 | 55 | 43 | 50 | 77 | 54 | 75 | 37 | 95 | 68 |
| 69 | 0.28 | 0.53 | 0.54 | 0.59 | 1 | Gm49359 | zinc finger protein 935 | 76 | 105 | 53 | 22 | 71 | 90 | 54 | 65 | 72 | 43 | 106 |
| 60 | 0.41 | 0.62 | 0.66 | 0.51 | 1 | Gm15327 | | 18 | 36 | 50 | 46 | 66 | 62 | 54 | 17 | 98 | 121 | 90 |
| 59 | 0.46 | 0.49 | 0.94 | 0.35 | 1 | Zfp97 | zinc finger protein 97 | 66 | 90 | 36 | 39 | 29 | 67 | 54 | 98 | 48 | 36 | 82 |
| 58 | 0.20 | 0.48 | 0.43 | 0.67 | 1 | Gm2762 | | 23 | 28 | 9 | 65 | 77 | 71 | 54 | 88 | 61 | 71 | 95 |
| 54 | 0.24 | 0.60 | 0.41 | 0.68 | 1 | Zbtb8b | zinc finger and BTB domain containing 8b | 44 | 115 | 30 | 59 | 21 | 55 | 54 | 52 | 52 | 86 | 25 |
| 37 | -0.46 | 0.58 | -0.80 | 0.42 | 1 | Gm12184 | | 21 | 21 | 51 | 36 | 50 | 42 | 54 | 36 | 23 | 33 | 37 |
| 35 | -0.95 | 1.08 | -0.88 | 0.38 | 1 | Plet1os | | 1 | 5 | 18 | 44 | 60 | 32 | 54 | 108 | 36 | 0 | 30 |
| 35 | -0.57 | 0.78 | -0.73 | 0.46 | 1 | A930035D04Rik | | 12 | 23 | 69 | 60 | 20 | 36 | 54 | 25 | 16 | 37 | 36 |
| 15 | -2.21 | 1.25 | -1.77 | 0.08 | 1 | Hs3st5 | heparan sulfate (glucosamine) 3-O-sulfotr | 0 | 0 | 0 | 20 | 18 | 39 | 54 | 0 | 9 | 9 | 19 |
| 6160 | -1.84 | 1.57 | -1.17 | 0.24 | 1 | Kdr | kinase insert domain protein receptor | 22453 | 22206 | 23010 | 0 | 4 | 11 | 53 | 0 | 2 | 22 | 0 |
| 989 | 0.40 | 0.29 | 1.37 | 0.17 | 1 | Adamts2 | a disintegrin-like and metallopeptidase (re | 5443 | 2151 | 2956 | 34 | 37 | 20 | 53 | 71 | 35 | 58 | 24 |
| 266 | 0.42 | 0.37 | 1.13 | 0.26 | 1 | Entpd2 | ectonucleoside triphosphate diphosphohy | 1241 | 493 | 719 | 61 | 48 | 32 | 53 | 98 | 84 | 79 | 22 |
| 204 | 0.68 | 0.48 | 1.43 | 0.15 | 1 | Lgi4 | leucine-rich repeat LGI family, member 4 | 646 | 354 | 589 | 46 | 79 | 64 | 53 | 197 | 54 | 61 | 96 |
| 141 | -0.69 | 0.72 | -0.95 | 0.34 | 1 | Rtp4 | receptor transporter protein 4 | 359 | 345 | 542 | 32 | 57 | 27 | 53 | 22 | 71 | 0 | 38 |
| 118 | -1.25 | 0.75 | -1.67 | 0.10 | 1 | Sema4f | sema domain, immunoglobulin domain (Ig | 428 | 421 | 232 | 28 | 32 | 25 | 53 | 0 | 22 | 12 | 40 |
| 105 | -0.15 | 0.59 | -0.25 | 0.80 | 1 | Spag5 | sperm associated antigen 5 | 211 | 104 | 69 | 124 | 108 | 118 | 53 | 124 | 107 | 26 | 115 |
| 99 | 0.41 | 0.39 | 1.05 | 0.29 | 1 | Bbs12 | Bardet-Biedl syndrome 12 (human) | 94 | 125 | 85 | 89 | 41 | 168 | 53 | 110 | 103 | 50 | 172 |
| 91 | -0.02 | 0.52 | -0.04 | 0.97 | 1 | Neu3 | neuraminidase 3 | 264 | 80 | 115 | 116 | 49 | 57 | 53 | 89 | 47 | 62 | 71 |
| 74 | -0.26 | 0.40 | -0.66 | 0.51 | 1 | Ccdc96 | coiled-coil domain containing 96 | 48 | 64 | 47 | 103 | 108 | 89 | 53 | 116 | 69 | 46 | 68 |
| 65 | -0.56 | 0.44 | -1.28 | 0.20 | 1 | Wfikkn2 | WAP, follistatin/kazal, immunoglobulin, ku | 74 | 85 | 56 | 93 | 95 | 52 | 53 | 86 | 51 | 35 | 34 |
| 59 | -0.27 | 0.51 | -0.52 | 0.61 | 1 | Tas1r3 | taste receptor, type 1, member 3 | 65 | 70 | 79 | 78 | 58 | 46 | 53 | 36 | 28 | 80 | 57 |
| 59 | -0.62 | 0.53 | -1.16 | 0.24 | 1 | Carns1 | carnosine synthase 1 | 100 | 62 | 65 | 67 | 55 | 75 | 53 | 42 | 52 | 15 | 59 |
| 58 | -0.69 | 0.82 | -0.84 | 0.40 | 1 | Gins2 | GINS complex subunit 2 (Psf2 homolog) | 156 | 71 | 131 | 55 | 12 | 49 | 53 | 23 | 51 | 3 | 38 |
| 50 | -0.77 | 0.76 | -1.01 | 0.31 | 1 | Trpc7 | transient receptor potential cation channe | 37 | 28 | 67 | 77 | 81 | 41 | 53 | 81 | 46 | 3 | 43 |
| 45 | -1.96 | 1.06 | -1.85 | 0.06 | 1 | Map3k8 | mitogen-activated protein kinase kinase k | 160 | 70 | 83 | 4 | 8 | 55 | 53 | 0 | 7 | 0 | 52 |
| 42 | 1.25 | 0.73 | 1.72 | 0.09 | 1 | 4933417E11Rik | | 10 | 4 | 2 | 8 | 22 | 69 | 53 | 100 | 51 | 99 | 49 |
| 41 | 0.43 | 0.77 | 0.55 | 0.58 | 1 | Gm6916 | | 47 | 52 | 18 | 41 | 7 | 10 | 53 | 101 | 2 | 120 | 6 |
| 39 | 1.01 | 0.92 | 1.09 | 0.28 | 1 | Nos1 | nitric oxide synthase 1, neuronal | 36 | 18 | 0 | 92 | 5 | 28 | 53 | 34 | 48 | 71 | 51 |
| 28 | 0.21 | 0.74 | 0.28 | 0.78 | 1 | Gm33263 | | 0 | 0 | 0 | 31 | 33 | 25 | 53 | 17 | 60 | 52 | 41 |
| 23 | -0.23 | 1.29 | -0.18 | 0.86 | 1 | 1700030C10Rik | | 15 | 6 | 0 | 8 | 0 | 39 | 53 | 0 | 0 | 77 | 52 |
| 1119 | -0.98 | 0.82 | -1.20 | 0.23 | 1 | Ccn1 | cellular communication network factor 1 | 6882 | 2427 | 2364 | 11 | 46 | 327 | 52 | 0 | 47 | 33 | 117 |
| 386 | -0.25 | 1.09 | -0.23 | 0.82 | 1 | Jag1 | jagged 1 | 1541 | 1185 | 1129 | 53 | 36 | 45 | 52 | 0 | 124 | 52 | 35 |
| 260 | -0.48 | 0.36 | -1.33 | 0.18 | 1 | Corin | corin, serine peptidase | 726 | 590 | 1118 | 29 | 143 | 29 | 52 | 17 | 89 | 30 | 35 |
| 187 | 0.09 | 0.32 | 0.28 | 0.78 | 1 | Id4 | inhibitor of DNA binding 4 | 636 | 520 | 403 | 54 | 29 | 108 | 52 | 42 | 36 | 85 | 89 |
| 170 | 0.19 | 0.38 | 0.50 | 0.62 | 1 | Dhx58 | DEXH (Asp-Glu-X-His) box polypeptide 5 | 304 | 390 | 372 | 92 | 140 | 84 | 52 | 158 | 137 | 99 | 45 |
| 157 | 0.24 | 0.35 | 0.68 | 0.49 | 1 | Mtcp1 | mature T cell proliferation 1 | 63 | 69 | 65 | 55 | 26 | 626 | 52 | 57 | 67 | 44 | 607 |
| 99 | -0.13 | 0.47 | -0.27 | 0.79 | 1 | Mkx | mohawk homeobox | 189 | 139 | 232 | 129 | 77 | 25 | 52 | 66 | 55 | 108 | 24 |
| 85 | 0.13 | 0.41 | 0.32 | 0.75 | 1 | Rwdd2a | RWD domain containing 2A | 105 | 86 | 133 | 88 | 88 | 63 | 52 | 61 | 89 | 112 | 62 |
| 82 | 0.15 | 0.49 | 0.31 | 0.76 | 1 | 9530085L11Rik | | 52 | 134 | 63 | 70 | 49 | 111 | 52 | 185 | 59 | 22 | 107 |
| 78 | 0.23 | 0.49 | 0.48 | 0.63 | 1 | Rbm38 | RNA binding motif protein 38 | 257 | 195 | 80 | 30 | 22 | 40 | 52 | 79 | 23 | 64 | 20 |
| 75 | -0.56 | 0.46 | -1.23 | 0.22 | 1 | Syt17 | synaptotagmin XVII | 175 | 209 | 121 | 50 | 39 | 49 | 52 | 49 | 36 | 17 | 29 |
| 67 | 0.74 | 0.52 | 1.43 | 0.15 | 1 | Gm16701 | | 39 | 19 | 24 | 70 | 42 | 75 | 52 | 187 | 65 | 57 | 107 |
| 66 | 0.09 | 0.43 | 0.21 | 0.83 | 1 | Gm32856 | | 80 | 58 | 38 | 55 | 43 | 111 | 52 | 52 | 55 | 41 | 137 |
| 54 | 0.44 | 0.57 | 0.77 | 0.44 | 1 | Tspan2os | | 6 | 9 | 0 | 85 | 60 | 45 | 52 | 125 | 81 | 91 | 45 |
| 52 | 0.51 | 0.70 | 0.73 | 0.47 | 1 | Slc5a9 | solute carrier family 5 (sodium/glucose co | 14 | 60 | 75 | 46 | 26 | 39 | 52 | 119 | 32 | 84 | 26 |
| 49 | -0.39 | 0.49 | -0.79 | 0.43 | 1 | Gm6170 | | 107 | 59 | 102 | 34 | 50 | 20 | 52 | 23 | 23 | 41 | 27 |
| 47 | 0.53 | 0.58 | 0.90 | 0.37 | 1 | Uckl1os | | 0 | 2 | 0 | 40 | 60 | 56 | 52 | 90 | 94 | 76 | 49 |
| 46 | -0.03 | 0.51 | -0.06 | 0.95 | 1 | Tacr3 | tachykinin receptor 3 | 77 | 63 | 52 | 23 | 22 | 56 | 52 | 20 | 25 | 31 | 83 |
| 43 | -0.31 | 0.53 | -0.59 | 0.55 | 1 | Gm14023 | | 46 | 24 | 25 | 39 | 43 | 68 | 52 | 17 | 47 | 33 | 80 |

| | | | | | | | | | | | | | | | | | | |
|---|---|---|---|---|---|---|---|---|---|---|---|---|---|---|---|---|---|---|
| 42 | -0.34 | 0.84 | -0.40 | 0.69 | 1 | Gm50012 | | 14 | 18 | 8 | 71 | 43 | 51 | 52 | 46 | 103 | 3 | 50 |
| 38 | 0.27 | 0.67 | 0.41 | 0.68 | 1 | Plcxd1 | phosphatidylinositol-specific phospholipa | 92 | 13 | 39 | 39 | 15 | 17 | 52 | 49 | 31 | 54 | 14 |
| 26 | 0.41 | 0.97 | 0.42 | 0.67 | 1 | Gm47175 | | 18 | 14 | 26 | 4 | 43 | 8 | 52 | 24 | 3 | 81 | 15 |
| 25 | -0.54 | 0.69 | -0.77 | 0.44 | 1 | 4930405N21Rik | | 23 | 3 | 17 | 22 | 23 | 39 | 52 | 13 | 15 | 27 | 44 |
| 5946 | -0.84 | 1.25 | -0.67 | 0.50 | 1 | Ltbp4 | latent transforming growth factor beta binc | 27551 | 18820 | 18902 | 0 | 25 | 16 | 51 | 0 | 16 | 12 | 19 |
| 2544 | 0.04 | 1.05 | 0.03 | 0.97 | 1 | Amy2b | amylase 2b | 5052 | 8891 | 13385 | 285 | 39 | 3 | 51 | 63 | 32 | 46 | 138 |
| 303 | -0.51 | 0.27 | -1.87 | 0.06 | 1 | Rapgef3 | Rap guanine nucleotide exchange factor | 975 | 1024 | 1022 | 49 | 36 | 52 | 51 | 32 | 35 | 35 | 28 |
| 147 | -0.13 | 0.39 | -0.33 | 0.74 | 1 | Cercam | cerebral endothelial cell adhesion molecu | 579 | 341 | 422 | 33 | 34 | 26 | 51 | 22 | 38 | 26 | 45 |
| 93 | 0.05 | 0.41 | 0.12 | 0.90 | 1 | A230072C01Rik | | 101 | 107 | 112 | 109 | 71 | 114 | 51 | 56 | 69 | 97 | 139 |
| 77 | 0.28 | 0.41 | 0.69 | 0.49 | 1 | Gm46123 | | 8 | 32 | 7 | 109 | 47 | 168 | 51 | 131 | 68 | 67 | 161 |
| 76 | -0.27 | 0.43 | -0.63 | 0.53 | 1 | Ttll10 | tubulin tyrosine ligase-like family, membe | 31 | 83 | 80 | 50 | 49 | 207 | 51 | 59 | 50 | 30 | 151 |
| 65 | -0.09 | 0.48 | -0.18 | 0.86 | 1 | Rbpj-ps3 | | 132 | 138 | 98 | 51 | 74 | 8 | 51 | 50 | 68 | 32 | 11 |
| 60 | -0.43 | 0.69 | -0.62 | 0.53 | 1 | Gpr137c | G protein-coupled receptor 137C | 41 | 73 | 115 | 62 | 42 | 75 | 51 | 15 | 16 | 85 | 82 |
| 59 | 0.17 | 0.49 | 0.34 | 0.73 | 1 | 4930563E22Ril | RIKEN cDNA 4930563E22 gene | 14 | 27 | 41 | 36 | 60 | 123 | 51 | 54 | 45 | 58 | 146 |
| 51 | -0.36 | 0.63 | -0.57 | 0.57 | 1 | Tdg-ps2 | | 62 | 86 | 28 | 104 | 12 | 30 | 51 | 109 | 3 | 55 | 19 |
| 50 | 0.64 | 0.86 | 0.74 | 0.46 | 1 | Gm20400 | | 15 | 15 | 0 | 52 | 32 | 74 | 51 | 118 | 91 | 17 | 83 |
| 47 | -0.56 | 0.48 | -1.17 | 0.24 | 1 | Gm48069 | | 46 | 41 | 57 | 53 | 38 | 76 | 51 | 29 | 32 | 24 | 71 |
| 46 | -1.23 | 0.72 | -1.71 | 0.09 | 1 | Fhl4 | four and a half LIM domains 4 | 82 | 73 | 56 | 51 | 62 | 36 | 51 | 24 | 28 | 3 | 41 |
| 42 | -0.67 | 0.65 | -1.03 | 0.30 | 1 | Eci3 | enoyl-Coenzyme A delta isomerase 3 | 10 | 61 | 70 | 34 | 67 | 33 | 51 | 58 | 42 | 18 | 17 |
| 37 | 0.45 | 0.72 | 0.62 | 0.53 | 1 | Spsb4 | splA/ryanodine receptor domain and SOC | 19 | 13 | 47 | 32 | 26 | 31 | 51 | 68 | 18 | 43 | 63 |
| 37 | -0.20 | 0.61 | -0.34 | 0.74 | 1 | Gm16283 | | 39 | 14 | 6 | 43 | 42 | 51 | 51 | 30 | 45 | 49 | 39 |
| 33 | -0.18 | 0.58 | -0.30 | 0.76 | 1 | 6230400D17Rik | | 33 | 22 | 34 | 32 | 18 | 41 | 51 | 17 | 29 | 48 | 32 |
| 30 | 0.15 | 0.81 | 0.19 | 0.85 | 1 | Gm9768 | | 10 | 8 | 34 | 13 | 27 | 39 | 51 | 57 | 45 | 12 | 35 |
| 24 | -0.40 | 1.39 | -0.29 | 0.77 | 1 | Ppp1r14c | protein phosphatase 1, regulatory inhibito | 39 | 23 | 105 | 0 | 3 | 12 | 51 | 9 | 0 | 9 | 8 |
| 21 | -0.92 | 0.90 | -1.03 | 0.31 | 1 | Plb1 | phospholipase B1 | 45 | 9 | 10 | 14 | 10 | 23 | 51 | 10 | 0 | 41 | 16 |
| 19465 | -0.54 | 0.80 | -0.68 | 0.50 | 1 | Col3a1 | collagen, type III, alpha 1 | 102959 | 46409 | 64603 | 9 | 10 | 6 | 50 | 0 | 6 | 62 | 6 |
| 3051 | 0.18 | 1.18 | 0.15 | 0.88 | 1 | H2-Eb1 | histocompatibility 2, class II antigen E bet | 14057 | 10836 | 8421 | 27 | 6 | 23 | 50 | 0 | 82 | 33 | 25 |
| 677 | -0.01 | 0.33 | -0.04 | 0.96 | 1 | Bnip2 | BCL2/adenovirus E1B interacting protein | 2092 | 1313 | 1775 | 151 | 81 | 815 | 50 | 79 | 114 | 67 | 912 |
| 623 | -1.64 | 1.61 | -1.02 | 0.31 | 1 | Coro1a | coronin, actin binding protein 1A | 3848 | 1257 | 1646 | 19 | 1 | 6 | 50 | 0 | 19 | 0 | 6 |
| 508 | -0.45 | 0.89 | -0.50 | 0.61 | 1 | Antxr1 | anthrax toxin receptor 1 | 2730 | 1254 | 1352 | 26 | 49 | 10 | 50 | 0 | 37 | 63 | 17 |
| 194 | -0.32 | 0.38 | -0.84 | 0.40 | 1 | Phlda3 | pleckstrin homology like domain, family A | 792 | 434 | 417 | 83 | 52 | 92 | 50 | 66 | 52 | 35 | 66 |
| 167 | 0.67 | 0.78 | 0.86 | 0.39 | 1 | Cenpe | centromere protein E | 637 | 107 | 89 | 100 | 104 | 135 | 50 | 212 | 201 | 15 | 181 |
| 144 | 0.20 | 0.40 | 0.50 | 0.61 | 1 | Il18bp | interleukin 18 binding protein | 243 | 248 | 182 | 65 | 65 | 262 | 50 | 90 | 114 | 35 | 225 |
| 125 | 0.66 | 0.87 | 0.75 | 0.45 | 1 | Ppfia4 | protein tyrosine phosphatase, receptor typ | 601 | 239 | 385 | 0 | 3 | 30 | 50 | 23 | 14 | 10 | 20 |
| 107 | 0.09 | 0.42 | 0.21 | 0.84 | 1 | Zfp418 | zinc finger protein 418 | 115 | 114 | 110 | 95 | 137 | 127 | 50 | 118 | 126 | 111 | 70 |
| 102 | 0.00 | 0.36 | -0.01 | 0.99 | 1 | 2500004C02Rik | | 77 | 154 | 190 | 110 | 62 | 123 | 50 | 95 | 77 | 39 | 149 |
| 94 | -0.55 | 0.43 | -1.29 | 0.20 | 1 | Cpne5 | copine V | 120 | 263 | 141 | 112 | 93 | 43 | 50 | 57 | 76 | 58 | 17 |
| 90 | -0.63 | 0.41 | -1.53 | 0.13 | 1 | Sema4g | sema domain, immunoglobulin domain (I | 195 | 190 | 182 | 37 | 67 | 101 | 50 | 37 | 43 | 19 | 65 |
| 89 | 0.14 | 0.45 | 0.32 | 0.75 | 1 | Pinx1 | PIN2/TERF1 interacting, telomerase inhib | 124 | 135 | 99 | 62 | 72 | 107 | 50 | 80 | 105 | 28 | 119 |
| 83 | -0.07 | 0.62 | -0.11 | 0.91 | 1 | Gm26788 | | 7 | 7 | 0 | 269 | 74 | 96 | 50 | 178 | 121 | 45 | 71 |
| 73 | 0.06 | 0.46 | 0.14 | 0.89 | 1 | Zfp583 | zinc finger protein 583 | 69 | 138 | 127 | 46 | 62 | 73 | 50 | 57 | 43 | 76 | 62 |
| 72 | -0.07 | 0.52 | -0.14 | 0.89 | 1 | Pidd1 | p53 induced death domain protein 1 | 54 | 22 | 65 | 81 | 41 | 152 | 50 | 63 | 40 | 48 | 180 |
| 72 | -0.13 | 0.46 | -0.29 | 0.77 | 1 | D430040D24Rik | | 156 | 100 | 183 | 32 | 70 | 36 | 50 | 54 | 45 | 28 | 37 |
| 67 | 0.53 | 0.64 | 0.82 | 0.41 | 1 | Ttk | Ttk protein kinase | 179 | 36 | 52 | 62 | 43 | 41 | 50 | 103 | 73 | 32 | 66 |
| 60 | -2.03 | 1.07 | -1.91 | 0.06 | 1 | Gm38393 | | 11 | 18 | 11 | 25 | 22 | 256 | 50 | 0 | 25 | 0 | 241 |
| 55 | 0.57 | 0.68 | 0.84 | 0.40 | 1 | Gm15265 | | 24 | 3 | 32 | 29 | 25 | 139 | 50 | 60 | 28 | 84 | 136 |
| 55 | 0.40 | 0.48 | 0.84 | 0.40 | 1 | Snhg10 | | 38 | 34 | 71 | 50 | 33 | 66 | 50 | 45 | 68 | 70 | 85 |
| 51 | 0.21 | 0.64 | 0.34 | 0.74 | 1 | Gm48045 | | 7 | 24 | 9 | 60 | 77 | 56 | 50 | 34 | 53 | 109 | 87 |
| 51 | -0.38 | 0.53 | -0.73 | 0.47 | 1 | Ltk | leukocyte tyrosine kinase | 40 | 76 | 52 | 34 | 59 | 79 | 50 | 65 | 25 | 33 | 53 |
| 51 | 0.43 | 1.16 | 0.37 | 0.71 | 1 | Prkg2 | protein kinase, cGMP-dependent, type II | 90 | 24 | 0 | 46 | 21 | 60 | 50 | 160 | 20 | 1 | 90 |
| 44 | -0.91 | 0.94 | -0.97 | 0.33 | 1 | Gm45151 | | 27 | 46 | 22 | 40 | 87 | 68 | 50 | 0 | 57 | 37 | 56 |
| 42 | -1.11 | 0.92 | -1.21 | 0.23 | 1 | Glis1 | GLIS family zinc finger 1 | 76 | 125 | 112 | 6 | 11 | 34 | 50 | 0 | 18 | 18 | 10 |
| 39 | -0.21 | 0.99 | -0.21 | 0.83 | 1 | Gm42595 | | 27 | 15 | 35 | 25 | 54 | 48 | 50 | 66 | 46 | 0 | 62 |
| 37 | -0.01 | 0.59 | -0.01 | 0.99 | 1 | Gm41231 | | 11 | 17 | 28 | 33 | 34 | 57 | 50 | 63 | 36 | 32 | 45 |
| 34 | 0.65 | 0.73 | 0.89 | 0.37 | 1 | Dlgap2 | DLG associated protein 2 | 8 | 4 | 0 | 32 | 14 | 49 | 50 | 20 | 47 | 104 | 51 |
| 30 | -1.03 | 0.70 | -1.46 | 0.14 | 1 | Gm4895 | | 15 | 13 | 52 | 51 | 30 | 34 | 50 | 15 | 19 | 23 | 26 |
| 1174 | -1.12 | 1.73 | -0.65 | 0.52 | 1 | Anxa2 | annexin A2 | 5950 | 3203 | 3682 | 0 | 11 | 3 | 49 | 0 | 15 | 0 | 4 |
| 229 | -0.01 | 0.38 | -0.03 | 0.98 | 1 | Dock4 | dedicator of cytokinesis 4 | 709 | 405 | 736 | 100 | 67 | 117 | 49 | 57 | 115 | 84 | 78 |
| 221 | 0.35 | 0.37 | 0.95 | 0.34 | 1 | 4930523C07Ril | RIKEN cDNA 4930523C07 gene | 818 | 545 | 405 | 130 | 44 | 71 | 49 | 105 | 50 | 126 | 92 |
| 186 | -0.67 | 0.47 | -1.42 | 0.16 | 1 | Gm20554 | | 434 | 618 | 590 | 36 | 52 | 112 | 49 | 16 | 25 | 73 | 41 |

| | | | | | | | | | | | | | | | | | | |
|---|---|---|---|---|---|---|---|---|---|---|---|---|---|---|---|---|---|---|
| 177 | 0.21 | 0.36 | 0.58 | 0.56 | 1 | Gm3294 | | 26 | 17 | 45 | 109 | 95 | 651 | 49 | 122 | 142 | 54 | 638 |
| 110 | 0.08 | 0.59 | 0.13 | 0.89 | 1 | Kif20b | kinesin family member 20B | 263 | 71 | 75 | 87 | 96 | 139 | 49 | 85 | 96 | 35 | 212 |
| 107 | 0.17 | 0.78 | 0.22 | 0.82 | 1 | Ccnb1 | cyclin B1 | 408 | 83 | 27 | 81 | 94 | 89 | 49 | 130 | 108 | 24 | 84 |
| 91 | -0.56 | 0.61 | -0.91 | 0.36 | 1 | Clstn2 | calsyntenin 2 | 171 | 238 | 143 | 38 | 76 | 87 | 49 | 82 | 34 | 6 | 82 |
| 89 | -0.12 | 0.54 | -0.22 | 0.83 | 1 | Vav3 | vav 3 oncogene | 257 | 219 | 221 | 54 | 38 | 11 | 49 | 36 | 25 | 24 | 43 |
| 89 | -0.05 | 0.56 | -0.09 | 0.93 | 1 | Gm1976 | | 70 | 100 | 63 | 103 | 93 | 126 | 49 | 36 | 69 | 169 | 100 |
| 72 | 0.79 | 0.53 | 1.50 | 0.13 | 1 | 1700040D17Rik | | 35 | 93 | 132 | 46 | 41 | 56 | 49 | 102 | 95 | 102 | 47 |
| 63 | 0.20 | 0.55 | 0.38 | 0.71 | 1 | Rpl10a-ps1 | | 155 | 109 | 81 | 43 | 33 | 29 | 49 | 87 | 61 | 31 | 16 |
| 61 | 0.75 | 0.53 | 1.40 | 0.16 | 1 | Acot4 | acyl-CoA thioesterase 4 | 73 | 119 | 44 | 25 | 50 | 36 | 49 | 53 | 60 | 98 | 61 |
| 58 | -0.33 | 0.76 | -0.44 | 0.66 | 1 | Rhov | ras homolog family member V | 67 | 102 | 128 | 44 | 49 | 32 | 49 | 2 | 88 | 55 | 24 |
| 54 | 0.23 | 0.66 | 0.35 | 0.73 | 1 | Insyn1 | inhibitory synaptic factor 1 | 162 | 76 | 108 | 41 | 35 | 3 | 49 | 30 | 33 | 27 | 26 |
| 50 | 0.06 | 0.66 | 0.09 | 0.93 | 1 | Gm7984 | | 40 | 66 | 42 | 65 | 94 | 8 | 49 | 28 | 41 | 102 | 20 |
| 50 | -0.99 | 0.94 | -1.06 | 0.29 | 1 | Gm35409 | | 84 | 137 | 135 | 20 | 12 | 37 | 49 | 0 | 29 | 37 | 7 |
| 48 | 0.06 | 0.70 | 0.08 | 0.93 | 1 | A230060F14Rik | | 4 | 61 | 30 | 77 | 29 | 55 | 49 | 89 | 29 | 70 | 35 |
| 46 | 0.08 | 0.74 | 0.11 | 0.91 | 1 | 9430091E24Rik | | 5 | 44 | 51 | 69 | 29 | 52 | 49 | 37 | 42 | 76 | 48 |
| 45 | -0.82 | 0.93 | -0.88 | 0.38 | 1 | Kcnd2 | potassium voltage-gated channel, Shal-re | 84 | 195 | 74 | 21 | 9 | 13 | 49 | 0 | 6 | 35 | 14 |
| 45 | -0.88 | 0.60 | -1.48 | 0.14 | 1 | 9430065F17Rik | | 45 | 22 | 23 | 72 | 32 | 107 | 49 | 13 | 36 | 33 | 63 |
| 43 | -1.09 | 1.03 | -1.05 | 0.29 | 1 | Klhl14 | kelch-like 14 | 92 | 143 | 89 | 11 | 9 | 24 | 49 | 36 | 17 | 0 | 4 |
| 43 | 0.31 | 0.74 | 0.42 | 0.67 | 1 | Gm7435 | | 24 | 37 | 43 | 60 | 9 | 38 | 49 | 107 | 48 | 14 | 43 |
| 38 | -0.18 | 1.00 | -0.18 | 0.86 | 1 | Cfap206 | cilia and flagella associated protein 206 | 27 | 62 | 30 | 9 | 48 | 25 | 49 | 1 | 41 | 113 | 9 |
| 35 | -0.40 | 0.78 | -0.51 | 0.61 | 1 | Gm19026 | | 51 | 29 | 58 | 25 | 20 | 39 | 49 | 61 | 28 | 9 | 13 |
| 33 | -0.02 | 0.67 | -0.03 | 0.98 | 1 | Gm18227 | | 39 | 51 | 8 | 25 | 25 | 25 | 49 | 36 | 8 | 55 | 46 |
| 30 | -1.48 | 0.95 | -1.56 | 0.12 | 1 | Gm37174 | | 16 | 18 | 42 | 30 | 50 | 37 | 49 | 19 | 17 | 0 | 54 |
| 5034 | -0.80 | 1.20 | -0.67 | 0.50 | 1 | Sycn | syncollin | 7152 | 24083 | 22067 | 1080 | 129 | 4 | 48 | 219 | 42 | 19 | 530 |
| 251 | -0.74 | 0.53 | -1.39 | 0.16 | 1 | Gbp3 | guanylate binding protein 3 | 1127 | 678 | 663 | 43 | 61 | 25 | 48 | 7 | 62 | 14 | 35 |
| 187 | -0.28 | 0.50 | -0.57 | 0.57 | 1 | Mirg | | 45 | 6 | 17 | 29 | 33 | 933 | 48 | 14 | 59 | 21 | 857 |
| 134 | -0.03 | 0.53 | -0.05 | 0.96 | 1 | Cmtm7 | CKLF-like MARVEL transmembrane dom | 468 | 316 | 405 | 47 | 24 | 24 | 48 | 70 | 41 | 12 | 22 |
| 134 | -0.02 | 0.39 | -0.05 | 0.96 | 1 | Fabp5 | fatty acid binding protein 5, epidermal | 363 | 444 | 367 | 32 | 31 | 36 | 48 | 59 | 22 | 35 | 34 |
| 127 | 0.37 | 0.42 | 0.89 | 0.37 | 1 | Ccdc17 | coiled-coil domain containing 17 | 90 | 111 | 44 | 95 | 112 | 266 | 48 | 149 | 113 | 94 | 273 |
| 89 | 0.01 | 0.42 | 0.02 | 0.98 | 1 | Slc35d2 | solute carrier family 35, member D2 | 146 | 59 | 193 | 59 | 109 | 73 | 48 | 43 | 82 | 95 | 73 |
| 86 | 0.08 | 0.48 | 0.17 | 0.86 | 1 | Tmem158 | transmembrane protein 158 | 155 | 123 | 128 | 126 | 44 | 48 | 48 | 53 | 68 | 93 | 55 |
| 79 | -0.02 | 0.42 | -0.05 | 0.96 | 1 | Zfp607b | zinc finger protein 607B | 56 | 67 | 118 | 46 | 89 | 142 | 48 | 39 | 61 | 77 | 131 |
| 79 | 0.69 | 0.60 | 1.15 | 0.25 | 1 | Tmem240 | transmembrane protein 240 | 48 | 114 | 120 | 57 | 36 | 82 | 48 | 94 | 46 | 170 | 56 |
| 79 | 0.05 | 0.38 | 0.14 | 0.89 | 1 | 0610040B10Rik | | 62 | 106 | 104 | 98 | 63 | 78 | 48 | 81 | 75 | 39 | 112 |
| 73 | -0.03 | 0.56 | -0.06 | 0.96 | 1 | Bora | bora, aurora kinase A activator | 112 | 32 | 46 | 38 | 87 | 154 | 48 | 80 | 55 | 49 | 102 |
| 72 | 0.10 | 0.54 | 0.19 | 0.85 | 1 | Steap1 | six transmembrane epithelial antigen of th | 52 | 81 | 44 | 72 | 95 | 76 | 48 | 46 | 62 | 144 | 72 |
| 68 | -0.66 | 0.68 | -0.96 | 0.33 | 1 | Bard1 | BRCA1 associated RING domain 1 | 94 | 50 | 70 | 92 | 70 | 96 | 48 | 7 | 85 | 86 | 54 |
| 66 | 0.27 | 0.75 | 0.36 | 0.72 | 1 | E130102H24Rik | | 1 | 18 | 41 | 59 | 27 | 162 | 48 | 49 | 59 | 57 | 203 |
| 63 | 0.19 | 0.68 | 0.28 | 0.78 | 1 | Gm43268 | | 44 | 101 | 30 | 81 | 65 | 51 | 48 | 16 | 69 | 106 | 82 |
| 58 | -0.74 | 0.64 | -1.16 | 0.25 | 1 | Gm36371 | | 57 | 40 | 83 | 64 | 78 | 75 | 48 | 6 | 48 | 63 | 74 |
| 55 | -0.63 | 0.61 | -1.04 | 0.30 | 1 | Tmem37 | transmembrane protein 37 | 176 | 92 | 118 | 27 | 48 | 14 | 48 | 28 | 14 | 14 | 23 |
| 52 | 0.70 | 0.48 | 1.47 | 0.14 | 1 | Ppargc1b | peroxisome proliferative activated recepto | 61 | 19 | 22 | 30 | 43 | 58 | 48 | 49 | 72 | 80 | 87 |
| 52 | 0.27 | 0.59 | 0.46 | 0.65 | 1 | Gm12976 | | 5 | 56 | 38 | 67 | 58 | 36 | 48 | 92 | 73 | 57 | 39 |
| 51 | 0.28 | 0.58 | 0.49 | 0.62 | 1 | Gm15351 | | 18 | 41 | 69 | 48 | 53 | 43 | 48 | 87 | 49 | 79 | 29 |
| 49 | 0.39 | 0.53 | 0.73 | 0.46 | 1 | Ap3s1-ps1 | | 30 | 35 | 88 | 45 | 37 | 38 | 48 | 67 | 55 | 49 | 51 |
| 44 | -0.11 | 0.61 | -0.18 | 0.86 | 1 | D330050I16Rik | | 33 | 26 | 53 | 44 | 50 | 49 | 48 | 48 | 15 | 63 | 56 |
| 39 | 0.30 | 0.97 | 0.31 | 0.76 | 1 | Ryr3 | ryanodine receptor 3 | 15 | 28 | 16 | 3 | 50 | 39 | 48 | 74 | 8 | 23 | 129 |
| 37 | -0.04 | 0.62 | -0.07 | 0.94 | 1 | 2810405F17Rik | | 38 | 35 | 37 | 22 | 34 | 46 | 48 | 59 | 32 | 18 | 39 |
| 35 | -0.18 | 0.84 | -0.22 | 0.83 | 1 | Gm4875 | | 37 | 56 | 59 | 34 | 7 | 23 | 48 | 22 | 4 | 86 | 12 |
| 32 | -1.24 | 0.82 | -1.51 | 0.13 | 1 | Lhx1os | | 5 | 0 | 0 | 53 | 15 | 90 | 48 | 8 | 5 | 44 | 80 |
| 27 | -0.51 | 0.87 | -0.59 | 0.56 | 1 | Gm27030 | | 17 | 10 | 23 | 45 | 22 | 23 | 48 | 68 | 17 | 4 | 22 |
| 25 | -0.39 | 0.84 | -0.46 | 0.65 | 1 | Tekt5 | tektin 5 | 33 | 32 | 11 | 15 | 26 | 23 | 48 | 30 | 16 | 5 | 39 |
| 25 | -1.34 | 1.16 | -1.16 | 0.25 | 1 | D930007J09Rik | | 29 | 4 | 33 | 53 | 9 | 22 | 48 | 0 | 32 | 18 | 28 |
| 19 | 0.02 | 0.75 | 0.02 | 0.98 | 1 | Gm10320 | | 7 | 4 | 5 | 34 | 12 | 9 | 48 | 29 | 26 | 22 | 13 |
| 373 | 0.28 | 0.31 | 0.89 | 0.37 | 1 | Ehbp1l1 | EH domain binding protein 1-like 1 | 1607 | 914 | 1253 | 23 | 30 | 52 | 47 | 51 | 30 | 35 | 63 |
| 315 | -0.55 | 0.47 | -1.17 | 0.24 | 1 | Dtx4 | deltex 4, E3 ubiquitin ligase | 1268 | 810 | 913 | 62 | 75 | 84 | 47 | 8 | 94 | 43 | 66 |
| 170 | -0.18 | 0.46 | -0.38 | 0.70 | 1 | AK157302 | | 7 | 215 | 242 | 274 | 464 | 30 | 47 | 161 | 349 | 44 | 42 |
| 134 | 0.38 | 0.31 | 1.22 | 0.22 | 1 | Gm16973 | | 100 | 126 | 90 | 46 | 45 | 400 | 47 | 73 | 52 | 72 | 423 |
| 134 | 0.45 | 0.38 | 1.19 | 0.23 | 1 | Ptgr1 | prostaglandin reductase 1 | 373 | 193 | 379 | 42 | 60 | 77 | 47 | 81 | 78 | 82 | 59 |
| 94 | -0.08 | 0.53 | -0.14 | 0.89 | 1 | Asf1b | anti-silencing function 1B histone chapero | 242 | 150 | 54 | 81 | 97 | 72 | 47 | 94 | 71 | 76 | 53 |

| | | | | | | | | | | | | | | | | | | |
|---|---|---|---|---|---|---|---|---|---|---|---|---|---|---|---|---|---|---|
| 91 | -0.16 | 0.56 | -0.29 | 0.77 | 1 | 4930488L21Rik | | 85 | 197 | 38 | 137 | 60 | 119 | 47 | 50 | 95 | 30 | 141 |
| 85 | -0.12 | 0.41 | -0.29 | 0.77 | 1 | Dynlt1a | dynein light chain Tctex-type 1A | 136 | 179 | 135 | 59 | 72 | 74 | 47 | 41 | 45 | 73 | 76 |
| 78 | -1.16 | 1.08 | -1.08 | 0.28 | 1 | Tpbgl | trophoblast glycoprotein-like | 480 | 164 | 113 | 12 | 6 | 6 | 47 | 0 | 12 | 0 | 17 |
| 77 | -0.36 | 0.53 | -0.68 | 0.50 | 1 | Pole | polymerase (DNA directed), epsilon | 231 | 148 | 95 | 46 | 41 | 71 | 47 | 14 | 52 | 31 | 76 |
| 68 | -0.33 | 0.53 | -0.61 | 0.54 | 1 | Fam83b | family with sequence similarity 83, membe | 91 | 122 | 88 | 49 | 84 | 54 | 47 | 24 | 76 | 22 | 86 |
| 65 | -0.79 | 0.51 | -1.54 | 0.12 | 1 | Tagln3 | transgelin 3 | 88 | 199 | 136 | 38 | 40 | 62 | 47 | 25 | 39 | 26 | 18 |
| 63 | -0.30 | 0.64 | -0.48 | 0.63 | 1 | Gm46565 | | 15 | 58 | 21 | 39 | 43 | 188 | 47 | 50 | 9 | 64 | 158 |
| 63 | 0.44 | 0.47 | 0.95 | 0.34 | 1 | Gm7598 | | 104 | 74 | 67 | 41 | 25 | 81 | 47 | 59 | 61 | 35 | 97 |
| 62 | -0.49 | 0.88 | -0.55 | 0.58 | 1 | Gprc5a | G protein-coupled receptor, family C, grou | 118 | 202 | 114 | 17 | 50 | 25 | 47 | 0 | 36 | 58 | 19 |
| 62 | 0.88 | 0.45 | 1.95 | 0.05 | 1 | Alk | anaplastic lymphoma kinase | 15 | 9 | 20 | 56 | 65 | 52 | 47 | 137 | 127 | 66 | 83 |
| 60 | 0.10 | 0.58 | 0.16 | 0.87 | 1 | Gm17655 | | 64 | 36 | 55 | 99 | 41 | 51 | 47 | 34 | 86 | 48 | 95 |
| 60 | 0.61 | 0.56 | 1.09 | 0.27 | 1 | Gm20457 | | 56 | 49 | 28 | 20 | 61 | 85 | 47 | 97 | 63 | 61 | 89 |
| 56 | 0.18 | 0.63 | 0.28 | 0.78 | 1 | Gm6278 | | 124 | 57 | 102 | 75 | 40 | 2 | 47 | 93 | 41 | 31 | 4 |
| 54 | -0.12 | 0.49 | -0.24 | 0.81 | 1 | Lrrc46 | leucine rich repeat containing 46 | 40 | 60 | 74 | 67 | 46 | 51 | 47 | 90 | 46 | 44 | 27 |
| 51 | 0.39 | 0.67 | 0.58 | 0.56 | 1 | Lingo3 | leucine rich repeat and Ig domain contain | 9 | 0 | 12 | 70 | 80 | 41 | 47 | 96 | 91 | 59 | 59 |
| 49 | -0.06 | 0.56 | -0.10 | 0.92 | 1 | A930019D19Rik | | 18 | 14 | 11 | 102 | 46 | 49 | 47 | 125 | 33 | 26 | 70 |
| 46 | -1.34 | 0.79 | -1.70 | 0.09 | 1 | Gm36445 | zinc finger protein 120-like | 82 | 34 | 51 | 14 | 47 | 113 | 47 | 0 | 41 | 10 | 70 |
| 44 | 0.41 | 0.60 | 0.67 | 0.50 | 1 | Gm42778 | | 39 | 61 | 45 | 26 | 34 | 24 | 47 | 111 | 48 | 28 | 20 |
| 44 | -0.02 | 0.48 | -0.04 | 0.97 | 1 | Gpr63 | G protein-coupled receptor 63 | 32 | 55 | 18 | 60 | 40 | 44 | 47 | 54 | 43 | 45 | 43 |
| 43 | -0.63 | 0.71 | -0.89 | 0.37 | 1 | A930024E05Rik | | 72 | 7 | 47 | 23 | 21 | 83 | 47 | 2 | 22 | 43 | 108 |
| 41 | -0.94 | 0.82 | -1.14 | 0.25 | 1 | Panct2 | | 105 | 64 | 56 | 25 | 25 | 36 | 47 | 0 | 15 | 44 | 30 |
| 37 | -0.53 | 0.65 | -0.81 | 0.42 | 1 | Gm7334 | | 60 | 11 | 15 | 32 | 60 | 40 | 47 | 9 | 39 | 53 | 45 |
| 32 | -0.36 | 0.68 | -0.53 | 0.60 | 1 | Spdya | speedy/RINGO cell cycle regulator family, | 15 | 60 | 42 | 24 | 26 | 35 | 47 | 32 | 26 | 18 | 30 |
| 32 | -1.27 | 0.93 | -1.37 | 0.17 | 1 | 2700038G22Rik | | 37 | 66 | 44 | 32 | 13 | 48 | 47 | 0 | 16 | 18 | 29 |
| 29 | -0.55 | 0.84 | -0.66 | 0.51 | 1 | Gm16244 | | 14 | 35 | 41 | 26 | 24 | 25 | 47 | 38 | 17 | 4 | 46 |
| 25 | -0.73 | 0.81 | -0.90 | 0.37 | 1 | Gm37170 | | 22 | 9 | 7 | 9 | 61 | 41 | 47 | 24 | 19 | 12 | 28 |
| 25 | -0.63 | 1.09 | -0.58 | 0.56 | 1 | Gm15697 | | 12 | 54 | 12 | 15 | 9 | 41 | 47 | 22 | 17 | 0 | 49 |
| 21 | 0.04 | 0.88 | 0.04 | 0.97 | 1 | Gm37296 | | 0 | 3 | 0 | 16 | 22 | 30 | 47 | 20 | 23 | 45 | 28 |
| 14 | -0.76 | 1.10 | -0.69 | 0.49 | 1 | Gm13187 | | 4 | 0 | 2 | 2 | 16 | 18 | 47 | 0 | 15 | 10 | 35 |
| 9205 | -0.20 | 1.31 | -0.15 | 0.88 | 1 | Dmbt1 | deleted in malignant brain tumors 1 | 11608 | 54479 | 32619 | 2064 | 50 | 3 | 46 | 54 | 80 | 46 | 211 |
| 476 | 0.05 | 0.34 | 0.15 | 0.88 | 1 | Fzd1 | frizzled class receptor 1 | 2683 | 1076 | 1108 | 45 | 60 | 31 | 46 | 50 | 47 | 43 | 46 |
| 384 | -0.36 | 0.36 | -1.01 | 0.31 | 1 | Sipa1 | signal-induced proliferation associated ge | 1628 | 767 | 1095 | 99 | 103 | 163 | 46 | 78 | 58 | 48 | 142 |
| 357 | 0.19 | 0.42 | 0.44 | 0.66 | 1 | Dock2 | dedicator of cyto-kinesis 2 | 1975 | 948 | 685 | 30 | 46 | 20 | 46 | 78 | 17 | 55 | 25 |
| 249 | -0.62 | 0.67 | -0.92 | 0.36 | 1 | Tspan4 | tetraspanin 4 | 1192 | 660 | 613 | 55 | 25 | 36 | 46 | 51 | 27 | 0 | 31 |
| 154 | -0.63 | 0.72 | -0.87 | 0.38 | 1 | Mmp23 | matrix metallopeptidase 23 | 943 | 325 | 257 | 25 | 6 | 26 | 46 | 0 | 20 | 17 | 31 |
| 142 | -0.16 | 0.31 | -0.51 | 0.61 | 1 | Gm16638 | | 278 | 348 | 354 | 84 | 55 | 124 | 46 | 56 | 56 | 57 | 101 |
| 113 | 0.43 | 0.39 | 1.11 | 0.27 | 1 | Ttll11 | tubulin tyrosine ligase-like family, membe | 132 | 157 | 184 | 92 | 91 | 104 | 46 | 158 | 82 | 104 | 94 |
| 90 | -0.86 | 0.55 | -1.56 | 0.12 | 1 | Pknox2 | Pbx/knotted 1 homeobox 2 | 332 | 153 | 248 | 52 | 26 | 36 | 46 | 42 | 27 | 15 | 9 |
| 83 | 0.05 | 0.35 | 0.15 | 0.88 | 1 | Tmem147os | | 57 | 93 | 87 | 84 | 55 | 150 | 46 | 95 | 62 | 44 | 142 |
| 82 | 0.20 | 0.45 | 0.44 | 0.66 | 1 | E530011L22Rik | | 172 | 138 | 145 | 27 | 66 | 72 | 46 | 68 | 36 | 53 | 80 |
| 79 | 0.81 | 0.59 | 1.38 | 0.17 | 1 | Poc1a | POC1 centriolar protein A | 94 | 37 | 84 | 33 | 81 | 82 | 46 | 176 | 72 | 103 | 62 |
| 73 | -0.35 | 0.49 | -0.72 | 0.47 | 1 | Npas2 | neuronal PAS domain protein 2 | 154 | 84 | 116 | 69 | 82 | 46 | 46 | 68 | 72 | 50 | 17 |
| 68 | -0.27 | 0.53 | -0.51 | 0.61 | 1 | 1700030J22Rik | RIKEN cDNA 1700030J22 gene | 101 | 146 | 27 | 71 | 70 | 62 | 46 | 74 | 30 | 59 | 60 |
| 64 | 0.27 | 0.54 | 0.50 | 0.62 | 1 | Gm17733 | | 24 | 44 | 29 | 106 | 81 | 51 | 46 | 45 | 77 | 107 | 94 |
| 63 | 0.05 | 0.41 | 0.12 | 0.90 | 1 | Gm8518 | | 62 | 66 | 83 | 73 | 51 | 69 | 46 | 70 | 77 | 49 | 53 |
| 63 | -0.11 | 0.44 | -0.26 | 0.79 | 1 | Lockd | | 72 | 57 | 60 | 70 | 64 | 84 | 46 | 85 | 59 | 39 | 62 |
| 63 | -0.33 | 0.87 | -0.38 | 0.70 | 1 | Septin2 | | 32 | 14 | 34 | 5 | 51 | 216 | 46 | 45 | 16 | 3 | 233 |
| 61 | 0.32 | 0.51 | 0.62 | 0.53 | 1 | Map2k3os | | 31 | 17 | 54 | 52 | 86 | 59 | 46 | 31 | 83 | 120 | 87 |
| 57 | -0.26 | 0.44 | -0.60 | 0.55 | 1 | Tmem88b | transmembrane protein 88B | 102 | 114 | 50 | 64 | 33 | 57 | 46 | 38 | 34 | 45 | 45 |
| 56 | 0.97 | 1.01 | 0.96 | 0.34 | 1 | Gm27168 | | 0 | 0 | 28 | 56 | 65 | 45 | 46 | 173 | 64 | 85 | 60 |
| 56 | -0.17 | 0.50 | -0.34 | 0.73 | 1 | Gm26881 | | 27 | 52 | 72 | 74 | 59 | 67 | 46 | 80 | 30 | 44 | 67 |
| 51 | -1.09 | 0.83 | -1.31 | 0.19 | 1 | Togaram2 | TOG array regulator of axonemal microtul | 3 | 0 | 0 | 139 | 32 | 155 | 46 | 12 | 32 | 57 | 88 |
| 50 | -0.66 | 0.58 | -1.15 | 0.25 | 1 | Gm43597 | | 45 | 47 | 36 | 28 | 40 | 120 | 46 | 30 | 7 | 27 | 127 |
| 42 | 0.55 | 0.52 | 1.06 | 0.29 | 1 | Zfp69 | zinc finger protein 69 | 57 | 29 | 23 | 18 | 39 | 44 | 46 | 46 | 39 | 55 | 67 |
| 40 | -0.14 | 0.70 | -0.20 | 0.84 | 1 | Gm49376 | | 25 | 2 | 12 | 32 | 68 | 66 | 46 | 44 | 37 | 28 | 77 |
| 37 | 0.29 | 0.69 | 0.43 | 0.67 | 1 | 4732416N19Rik | | 35 | 16 | 35 | 44 | 9 | 46 | 46 | 24 | 43 | 50 | 58 |
| 36 | -0.07 | 0.78 | -0.09 | 0.93 | 1 | Tgm5 | transglutaminase 5 | 27 | 2 | 39 | 28 | 62 | 41 | 46 | 54 | 32 | 44 | 23 |
| 36 | -0.01 | 0.76 | -0.01 | 0.99 | 1 | 1700063J08Rik | | 5 | 16 | 34 | 71 | 24 | 33 | 46 | 29 | 32 | 67 | 40 |
| 34 | 0.48 | 0.62 | 0.78 | 0.43 | 1 | Gm14418 | | 17 | 25 | 41 | 16 | 22 | 39 | 46 | 22 | 40 | 53 | 54 |
| 31 | 0.03 | 1.14 | 0.03 | 0.98 | 1 | Rnf165 | ring finger protein 165 | 13 | 38 | 31 | 0 | 40 | 23 | 46 | 93 | 31 | 19 | 11 |

| | | | | | | | | | | | | | | | | | | |
|---|---|---|---|---|---|---|---|---|---|---|---|---|---|---|---|---|---|---|
| 17 | -0.55 | 1.11 | -0.49 | 0.62 | 1 | Gm43075 | | 18 | 12 | 4 | 25 | 10 | 17 | 46 | 29 | 11 | 0 | 20 |
| 17 | 1.01 | 1.26 | 0.80 | 0.43 | 1 | n-R5-8s1 | | 0 | 2 | 5 | 21 | 11 | 0 | 46 | 49 | 10 | 35 | 6 |
| 14 | -2.41 | 1.29 | -1.86 | 0.06 | 1 | Gm26684 | | 12 | 0 | 6 | 11 | 30 | 27 | 46 | 1 | 12 | 0 | 14 |
| 2879 | -2.23 | 1.68 | -1.33 | 0.18 | 1 | Nrp1 | neuropilin 1 | 11498 | 9489 | 10598 | 4 | 0 | 21 | 45 | 0 | 6 | 0 | 8 |
| 338 | 0.14 | 0.46 | 0.30 | 0.76 | 1 | Boc | biregional cell adhesion molecule-relatec | 1883 | 641 | 799 | 32 | 88 | 17 | 45 | 89 | 39 | 67 | 13 |
| 137 | -0.07 | 0.36 | -0.20 | 0.84 | 1 | Tax1bp3 | Tax1 (human T cell leukemia virus type I) | 333 | 294 | 404 | 88 | 60 | 52 | 45 | 49 | 68 | 48 | 67 |
| 123 | -1.37 | 0.77 | -1.78 | 0.08 | 1 | Rbm24 | RNA binding motif protein 24 | 347 | 365 | 419 | 37 | 26 | 30 | 45 | 0 | 5 | 53 | 21 |
| 100 | 0.06 | 0.56 | 0.10 | 0.92 | 1 | Dtl | denticleless E3 ubiquitin protein ligase | 253 | 113 | 63 | 78 | 120 | 83 | 45 | 68 | 135 | 66 | 74 |
| 93 | 0.58 | 0.83 | 0.70 | 0.48 | 1 | Knl1 | kinetochore scaffold 1 | 280 | 68 | 29 | 67 | 64 | 79 | 45 | 177 | 101 | 9 | 101 |
| 92 | 0.50 | 0.56 | 0.90 | 0.37 | 1 | Lmntd2 | lamin tail domain containing 2 | 9 | 44 | 31 | 18 | 38 | 320 | 45 | 63 | 73 | 37 | 331 |
| 89 | 0.65 | 0.39 | 1.66 | 0.10 | 1 | Spin2c | spindlin family, member 2C | 129 | 128 | 149 | 56 | 79 | 45 | 45 | 97 | 70 | 109 | 71 |
| 74 | -0.12 | 0.44 | -0.28 | 0.78 | 1 | 2810029C07Rik | | 60 | 60 | 60 | 55 | 74 | 146 | 45 | 50 | 88 | 23 | 153 |
| 67 | -0.83 | 0.60 | -1.37 | 0.17 | 1 | Bach2 | BTB and CNC homology, basic leucine zi | 208 | 120 | 103 | 99 | 28 | 28 | 45 | 13 | 33 | 49 | 16 |
| 59 | -0.15 | 0.49 | -0.31 | 0.76 | 1 | Gm10461 | | 40 | 105 | 87 | 63 | 58 | 57 | 45 | 43 | 48 | 66 | 41 |
| 59 | -0.48 | 0.66 | -0.72 | 0.47 | 1 | Gm13688 | | 98 | 66 | 59 | 48 | 76 | 61 | 45 | 6 | 50 | 90 | 52 |
| 57 | -1.53 | 0.94 | -1.63 | 0.10 | 1 | Kif15 | kinesin family member 15 | 225 | 80 | 24 | 57 | 52 | 46 | 45 | 0 | 20 | 23 | 60 |
| 57 | -0.69 | 0.66 | -1.05 | 0.29 | 1 | Cfap46 | cilia and flagella associated protein 46 | 7 | 5 | 32 | 21 | 38 | 221 | 45 | 21 | 45 | 5 | 187 |
| 56 | 0.23 | 0.68 | 0.34 | 0.73 | 1 | Kif18a | kinesin family member 18A | 113 | 12 | 63 | 41 | 42 | 59 | 45 | 72 | 37 | 104 | 27 |
| 56 | 0.24 | 0.91 | 0.27 | 0.79 | 1 | 1110002E22Ril | RIKEN cDNA 1110002E22 gene | 33 | 18 | 35 | 39 | 53 | 66 | 45 | 0 | 107 | 131 | 88 |
| 54 | -0.44 | 0.64 | -0.69 | 0.49 | 1 | Gm16035 | | 24 | 44 | 37 | 73 | 30 | 112 | 45 | 12 | 59 | 26 | 129 |
| 50 | -0.07 | 0.54 | -0.13 | 0.90 | 1 | Gm6712 | predicted gene 6712 | 58 | 56 | 31 | 52 | 56 | 47 | 45 | 74 | 39 | 22 | 67 |
| 47 | 0.55 | 0.54 | 1.01 | 0.31 | 1 | Gm2000 | | 111 | 104 | 44 | 35 | 17 | 14 | 45 | 36 | 49 | 43 | 21 |
| 42 | 0.45 | 0.67 | 0.67 | 0.50 | 1 | 2700054A10Rik | | 33 | 61 | 87 | 28 | 9 | 41 | 45 | 52 | 36 | 35 | 41 |
| 42 | -0.98 | 0.93 | -1.05 | 0.29 | 1 | Gm19569 | | 51 | 43 | 31 | 73 | 64 | 30 | 45 | 30 | 49 | 0 | 43 |
| 37 | -0.61 | 1.02 | -0.60 | 0.55 | 1 | A930007I19Rik | | 33 | 37 | 9 | 67 | 37 | 41 | 45 | 0 | 28 | 48 | 63 |
| 32 | 0.42 | 0.77 | 0.54 | 0.59 | 1 | Gm41760 | | 18 | 19 | 28 | 18 | 44 | 18 | 45 | 73 | 11 | 40 | 39 |
| 29 | 0.02 | 1.30 | 0.02 | 0.99 | 1 | 3110045C21Rik | | 3 | 79 | 42 | 23 | 3 | 12 | 45 | 29 | 38 | 0 | 47 |
| 28 | -0.73 | 0.72 | -1.01 | 0.31 | 1 | Mir7666 | | 32 | 13 | 14 | 54 | 22 | 26 | 45 | 39 | 26 | 33 | 4 |
| 25 | -0.14 | 0.98 | -0.15 | 0.88 | 1 | Cacna1f | calcium channel, voltage-dependent, alpl | 15 | 8 | 8 | 4 | 36 | 25 | 45 | 0 | 32 | 80 | 18 |
| 24 | 0.24 | 0.72 | 0.33 | 0.74 | 1 | Gm39271 | | 13 | 29 | 5 | 30 | 12 | 18 | 45 | 48 | 29 | 21 | 19 |
| 20 | 1.26 | 1.28 | 0.98 | 0.33 | 1 | Gm38944 | | 18 | 0 | 47 | 0 | 9 | 10 | 45 | 9 | 16 | 68 | 3 |
| 1426 | -1.81 | 1.65 | -1.10 | 0.27 | 1 | Emp1 | epithelial membrane protein 1 | 6735 | 3923 | 4902 | 14 | 0 | 27 | 44 | 9 | 0 | 0 | 27 |
| 444 | -1.30 | 0.83 | -1.56 | 0.12 | 1 | Adgrl2 | adhesion G protein-coupled receptor L2 | 1766 | 1332 | 1537 | 17 | 33 | 73 | 44 | 0 | 38 | 26 | 14 |
| 122 | -0.03 | 0.44 | -0.07 | 0.94 | 1 | Thrsp | thyroid hormone responsive | 242 | 227 | 329 | 133 | 70 | 27 | 44 | 147 | 62 | 36 | 28 |
| 114 | 0.46 | 0.44 | 1.05 | 0.29 | 1 | Lipt2 | lipoyl(octanoyl) transferase 2 (putative) | 154 | 138 | 198 | 111 | 108 | 59 | 44 | 89 | 123 | 170 | 65 |
| 81 | 0.11 | 0.46 | 0.24 | 0.81 | 1 | Pnma2 | paraneoplastic antigen MA2 | 116 | 154 | 61 | 90 | 74 | 61 | 44 | 89 | 98 | 62 | 44 |
| 80 | 0.11 | 0.48 | 0.24 | 0.81 | 1 | Tc2n | tandem C2 domains, nuclear | 145 | 237 | 112 | 66 | 31 | 50 | 44 | 30 | 61 | 55 | 50 |
| 71 | 0.11 | 0.42 | 0.27 | 0.79 | 1 | 5830448L01Rik | | 41 | 64 | 107 | 71 | 84 | 77 | 44 | 104 | 63 | 46 | 83 |
| 67 | 0.18 | 0.44 | 0.41 | 0.68 | 1 | Gm36839 | | 58 | 50 | 86 | 60 | 65 | 90 | 44 | 87 | 80 | 64 | 57 |
| 63 | -0.37 | 0.61 | -0.61 | 0.54 | 1 | Gata4 | GATA binding protein 4 | 88 | 220 | 160 | 25 | 40 | 16 | 44 | 14 | 21 | 50 | 13 |
| 60 | 0.09 | 0.57 | 0.16 | 0.87 | 1 | Fbxl12os | | 68 | 75 | 28 | 63 | 74 | 56 | 44 | 32 | 74 | 94 | 58 |
| 59 | 0.05 | 0.51 | 0.10 | 0.92 | 1 | Crybb3 | crystallin, beta B3 | 25 | 33 | 16 | 44 | 56 | 139 | 44 | 59 | 27 | 86 | 124 |
| 55 | 0.11 | 0.48 | 0.22 | 0.82 | 1 | 9330160F10Rik | | 17 | 54 | 46 | 60 | 54 | 75 | 44 | 44 | 77 | 46 | 86 |
| 51 | -0.23 | 0.55 | -0.41 | 0.68 | 1 | Mterf1a | mitochondrial transcription termination fac | 54 | 36 | 36 | 52 | 34 | 93 | 44 | 65 | 22 | 18 | 112 |
| 51 | -1.04 | 0.83 | -1.27 | 0.21 | 1 | 6530402F18Rik | | 194 | 66 | 97 | 27 | 43 | 14 | 44 | 0 | 47 | 18 | 14 |
| 48 | -0.38 | 0.61 | -0.62 | 0.53 | 1 | Gm32391 | | 22 | 75 | 56 | 69 | 20 | 70 | 44 | 80 | 37 | 12 | 47 |
| 48 | -0.57 | 0.59 | -0.97 | 0.33 | 1 | 2610528J11Rik | RIKEN cDNA 2610528J11 gene | 80 | 93 | 133 | 34 | 26 | 20 | 44 | 9 | 34 | 37 | 11 |
| 35 | 0.04 | 0.64 | 0.07 | 0.95 | 1 | Gm34654 | | 0 | 0 | 0 | 26 | 60 | 57 | 44 | 27 | 40 | 50 | 82 |
| 34 | -0.22 | 0.54 | -0.41 | 0.68 | 1 | 3010001F23Rik | | 38 | 24 | 32 | 25 | 18 | 63 | 44 | 16 | 23 | 39 | 47 |
| 33 | -1.51 | 1.13 | -1.33 | 0.18 | 1 | Gm41293 | | 3 | 0 | 21 | 50 | 44 | 52 | 44 | 0 | 77 | 49 | 26 |
| 25 | -1.31 | 0.99 | -1.32 | 0.19 | 1 | Txlnb | taxilin beta | 15 | 44 | 10 | 32 | 37 | 25 | 44 | 27 | 16 | 0 | 30 |
| 13 | 0.29 | 1.10 | 0.26 | 0.80 | 1 | Gm17324 | | 2 | 6 | 7 | 9 | 2 | 9 | 44 | 16 | 4 | 26 | 16 |
| 1787 | -1.09 | 0.87 | -1.26 | 0.21 | 1 | Itih5 | inter-alpha (globulin) inhibitor H5 | 9313 | 5309 | 4820 | 39 | 27 | 20 | 43 | 0 | 23 | 53 | 8 |
| 475 | 1.21 | 1.38 | 0.88 | 0.38 | 1 | Irf8 | interferon regulatory factor 8 | 2835 | 1253 | 1030 | 4 | 9 | 0 | 43 | 22 | 25 | 0 | 2 |
| 263 | 0.25 | 2.20 | 0.12 | 0.91 | 1 | Ikzf1 | IKAROS family zinc finger 1 | 1907 | 525 | 401 | 0 | 4 | 0 | 43 | 0 | 8 | 0 | 8 |
| 212 | 0.34 | 0.38 | 0.89 | 0.37 | 1 | Ncf1 | neutrophil cytosolic factor 1 | 1014 | 466 | 550 | 28 | 39 | 22 | 43 | 64 | 47 | 41 | 18 |
| 132 | -0.46 | 0.42 | -1.08 | 0.28 | 1 | Snhg20 | | 64 | 34 | 54 | 43 | 37 | 577 | 43 | 45 | 16 | 23 | 512 |
| 124 | 0.70 | 0.67 | 1.04 | 0.30 | 1 | A930004J17Rik | | 54 | 96 | 46 | 4 | 11 | 512 | 43 | 71 | 24 | 22 | 477 |
| 121 | 0.20 | 0.36 | 0.56 | 0.58 | 1 | Homer3 | homer scaffolding protein 3 | 383 | 257 | 315 | 48 | 44 | 36 | 43 | 67 | 37 | 61 | 37 |
| 85 | -0.28 | 0.52 | -0.53 | 0.60 | 1 | Shcbp1 | Shc SH2-domain binding protein 1 | 255 | 117 | 77 | 109 | 59 | 50 | 43 | 42 | 40 | 95 | 52 |

| | | | | | | | | | | | | | | | | | | |
|---|---|---|---|---|---|---|---|---|---|---|---|---|---|---|---|---|---|---|
| 85 | -0.80 | 0.47 | -1.68 | 0.09 | 1 | Enkd1 | enkurin domain containing 1 | 138 | 94 | 91 | 89 | 85 | 153 | 43 | 56 | 30 | 24 | 129 |
| 84 | 0.32 | 0.46 | 0.70 | 0.48 | 1 | Gm49066 | | 82 | 76 | 87 | 81 | 90 | 82 | 43 | 174 | 73 | 68 | 68 |
| 75 | 0.96 | 0.50 | 1.93 | 0.05 | 1 | Epb41l4aos | | 139 | 116 | 127 | 14 | 28 | 74 | 43 | 83 | 40 | 49 | 117 |
| 75 | -0.72 | 0.78 | -0.92 | 0.36 | 1 | H4c14 | H4 clustered histone 14 | 9 | 13 | 11 | 20 | 38 | 326 | 43 | 23 | 43 | 0 | 294 |
| 72 | 0.08 | 0.48 | 0.16 | 0.87 | 1 | Epha1 | Eph receptor A1 | 38 | 82 | 154 | 89 | 49 | 70 | 43 | 95 | 70 | 48 | 55 |
| 71 | 0.04 | 0.62 | 0.07 | 0.94 | 1 | Fbxo5 | F-box protein 5 | 132 | 26 | 61 | 38 | 79 | 121 | 43 | 81 | 28 | 64 | 105 |
| 69 | 0.75 | 0.43 | 1.75 | 0.08 | 1 | Gm3531 | | 120 | 82 | 114 | 27 | 61 | 36 | 43 | 73 | 62 | 89 | 51 |
| 65 | 0.34 | 0.51 | 0.66 | 0.51 | 1 | Gm42067 | | 41 | 52 | 72 | 54 | 58 | 83 | 43 | 140 | 57 | 59 | 53 |
| 61 | 0.03 | 0.42 | 0.08 | 0.94 | 1 | Gm47071 | | 50 | 79 | 44 | 82 | 76 | 46 | 43 | 85 | 56 | 49 | 61 |
| 52 | 0.16 | 0.54 | 0.29 | 0.77 | 1 | Ube2t | ubiquitin-conjugating enzyme E2T | 50 | 53 | 60 | 17 | 46 | 85 | 43 | 12 | 58 | 53 | 98 |
| 46 | -0.30 | 0.55 | -0.54 | 0.59 | 1 | BC049715 | cDNA sequence BC049715 | 43 | 38 | 35 | 48 | 45 | 73 | 43 | 14 | 54 | 43 | 73 |
| 45 | -0.03 | 0.57 | -0.05 | 0.96 | 1 | 4931440P22Rik | | 78 | 84 | 64 | 26 | 29 | 33 | 43 | 59 | 12 | 45 | 25 |
| 45 | -0.18 | 1.06 | -0.17 | 0.86 | 1 | 1700120C14Rik | | 17 | 45 | 57 | 52 | 29 | 51 | 43 | 0 | 54 | 117 | 26 |
| 39 | -0.73 | 0.89 | -0.82 | 0.41 | 1 | Gm31036 | | 32 | 26 | 45 | 28 | 37 | 83 | 43 | 36 | 33 | 0 | 72 |
| 35 | -1.13 | 0.58 | -1.95 | 0.05 | 1 | Rpl27a-ps4 | | 71 | 61 | 55 | 54 | 21 | 16 | 43 | 17 | 10 | 27 | 7 |
| 35 | 0.26 | 0.60 | 0.43 | 0.66 | 1 | Gm5112 | | 20 | 44 | 20 | 41 | 22 | 30 | 43 | 28 | 27 | 64 | 42 |
| 25 | -1.11 | 1.03 | -1.07 | 0.28 | 1 | 5430402O13Rik | | 52 | 17 | 7 | 33 | 21 | 31 | 43 | 0 | 25 | 17 | 26 |
| 24 | -0.57 | 0.68 | -0.84 | 0.40 | 1 | Gm44389 | | 21 | 8 | 35 | 33 | 41 | 9 | 43 | 24 | 29 | 17 | 9 |
| 5121 | -1.43 | 1.17 | -1.22 | 0.22 | 1 | Pnliprp2 | pancreatic lipase-related protein 2 | 7826 | 26908 | 20317 | 858 | 74 | 7 | 42 | 35 | 39 | 12 | 216 |
| 4042 | -1.11 | 1.86 | -0.59 | 0.55 | 1 | Col15a1 | collagen, type XV, alpha 1 | 16400 | 13703 | 14291 | 5 | 0 | 3 | 42 | 8 | 0 | 0 | 6 |
| 464 | -0.31 | 0.53 | -0.59 | 0.56 | 1 | Gas7 | growth arrest specific 7 | 2569 | 954 | 1404 | 6 | 37 | 11 | 42 | 0 | 23 | 46 | 14 |
| 364 | -0.44 | 0.98 | -0.45 | 0.66 | 1 | Igsf10 | immunoglobulin superfamily, member 10 | 2152 | 886 | 704 | 20 | 55 | 14 | 42 | 0 | 20 | 85 | 30 |
| 330 | -0.18 | 0.34 | -0.54 | 0.59 | 1 | Dmd | dystrophin, muscular dystrophy | 1179 | 1004 | 1090 | 42 | 34 | 76 | 42 | 57 | 46 | 31 | 32 |
| 105 | 0.47 | 0.48 | 0.98 | 0.33 | 1 | Fancd2 | Fanconi anemia, complementation group | 144 | 173 | 89 | 95 | 97 | 50 | 42 | 261 | 92 | 80 | 38 |
| 100 | 0.40 | 0.60 | 0.67 | 0.50 | 1 | Birc5 | baculoviral IAP repeat-containing 5 | 358 | 73 | 80 | 52 | 73 | 55 | 42 | 123 | 47 | 161 | 37 |
| 94 | -1.17 | 0.71 | -1.64 | 0.10 | 1 | Wipf3 | WAS/WASL interacting protein family, me | 347 | 272 | 164 | 66 | 33 | 28 | 42 | 2 | 43 | 14 | 19 |
| 91 | -0.16 | 0.65 | -0.24 | 0.81 | 1 | Gm10557 | | 70 | 135 | 85 | 94 | 43 | 199 | 42 | 10 | 50 | 103 | 172 |
| 88 | -0.20 | 0.55 | -0.37 | 0.71 | 1 | Ttll3 | tubulin tyrosine ligase-like family, membe | 15 | 78 | 41 | 21 | 51 | 357 | 42 | 50 | 43 | 24 | 250 |
| 87 | 0.34 | 0.47 | 0.72 | 0.47 | 1 | Shld3 | shieldin complex subunit 3 | 29 | 70 | 76 | 138 | 59 | 128 | 42 | 138 | 66 | 102 | 115 |
| 84 | -0.22 | 0.48 | -0.46 | 0.65 | 1 | Bnip5 | BCL2 interacting protein 5 | 105 | 253 | 246 | 39 | 46 | 41 | 42 | 63 | 25 | 39 | 25 |
| 74 | -0.32 | 0.41 | -0.77 | 0.44 | 1 | 9430038I01Rik | RIKEN cDNA 9430038I01 gene | 144 | 143 | 127 | 41 | 55 | 86 | 42 | 52 | 35 | 26 | 67 |
| 70 | 0.36 | 0.50 | 0.72 | 0.47 | 1 | Iqank1 | IQ motif and ankyrin repeat containing 1 | 51 | 43 | 89 | 71 | 45 | 107 | 42 | 111 | 53 | 72 | 87 |
| 70 | -0.65 | 0.55 | -1.18 | 0.24 | 1 | Ypel1 | yippee like 1 | 58 | 71 | 153 | 99 | 84 | 59 | 42 | 85 | 40 | 12 | 63 |
| 67 | 0.43 | 0.52 | 0.83 | 0.41 | 1 | Gm9843 | | 157 | 77 | 75 | 89 | 24 | 32 | 42 | 54 | 43 | 98 | 49 |
| 57 | 0.25 | 0.73 | 0.34 | 0.73 | 1 | Gm31024 | | 42 | 41 | 2 | 27 | 43 | 140 | 42 | 28 | 48 | 73 | 145 |
| 56 | -1.05 | 0.59 | -1.79 | 0.07 | 1 | Mmp16 | matrix metallopeptidase 16 | 103 | 90 | 68 | 62 | 72 | 55 | 42 | 42 | 13 | 14 | 61 |
| 52 | 0.39 | 0.53 | 0.74 | 0.46 | 1 | Myl6b | myosin, light polypeptide 6B | 51 | 95 | 124 | 35 | 41 | 19 | 42 | 42 | 35 | 55 | 38 |
| 52 | 0.10 | 0.60 | 0.17 | 0.86 | 1 | 6430562O15Rik | | 45 | 45 | 45 | 43 | 65 | 54 | 42 | 80 | 16 | 72 | 64 |
| 48 | -1.35 | 0.82 | -1.65 | 0.10 | 1 | Shroom1 | shroom family member 1 | 125 | 97 | 101 | 36 | 41 | 25 | 42 | 14 | 29 | 0 | 19 |
| 47 | 0.25 | 0.71 | 0.36 | 0.72 | 1 | Gm50394 | | 27 | 33 | 37 | 21 | 59 | 55 | 42 | 7 | 56 | 122 | 58 |
| 45 | -0.81 | 0.88 | -0.93 | 0.35 | 1 | Dnah8 | dynein, axonemal, heavy chain 8 | 94 | 54 | 83 | 29 | 62 | 31 | 42 | 25 | 40 | 0 | 38 |
| 45 | 0.24 | 0.56 | 0.42 | 0.67 | 1 | Ccdc116 | coiled-coil domain containing 116 | 9 | 24 | 36 | 38 | 37 | 74 | 42 | 43 | 35 | 46 | 112 |
| 44 | -0.33 | 0.52 | -0.64 | 0.52 | 1 | Cenpw | centromere protein W | 39 | 42 | 34 | 39 | 90 | 38 | 42 | 43 | 42 | 32 | 41 |
| 43 | -0.72 | 0.56 | -1.28 | 0.20 | 1 | Cyp46a1 | cytochrome P450, family 46, subfamily a, | 46 | 27 | 26 | 52 | 40 | 84 | 42 | 21 | 26 | 17 | 96 |
| 43 | -0.59 | 0.89 | -0.66 | 0.51 | 1 | Gm14322 | predicted gene 14322 | 28 | 22 | 49 | 29 | 32 | 90 | 42 | 0 | 23 | 68 | 90 |
| 40 | -1.42 | 1.03 | -1.38 | 0.17 | 1 | Gm26737 | | 0 | 9 | 23 | 46 | 61 | 80 | 42 | 48 | 81 | 0 | 53 |
| 39 | 1.06 | 0.64 | 1.65 | 0.10 | 1 | Gm7536 | | 70 | 55 | 64 | 9 | 15 | 16 | 42 | 64 | 30 | 41 | 24 |
| 35 | -0.08 | 0.66 | -0.12 | 0.90 | 1 | Gm6789 | | 30 | 71 | 56 | 35 | 24 | 14 | 42 | 52 | 31 | 10 | 22 |
| 33 | -0.29 | 0.57 | -0.51 | 0.61 | 1 | Rps10-ps1 | | 63 | 48 | 74 | 18 | 32 | 11 | 42 | 15 | 27 | 21 | 15 |
| 32 | 0.60 | 0.88 | 0.68 | 0.49 | 1 | Rab7-ps1 | | 22 | 17 | 18 | 12 | 40 | 20 | 42 | 73 | 51 | 4 | 49 |
| 31 | -0.23 | 1.08 | -0.22 | 0.83 | 1 | B230206L02Rik | | 8 | 43 | 52 | 24 | 24 | 32 | 42 | 0 | 34 | 53 | 32 |
| 31 | -0.27 | 0.92 | -0.29 | 0.77 | 1 | Gm45133 | | 50 | 38 | 47 | 10 | 20 | 34 | 42 | 48 | 1 | 21 | 30 |
| 29 | 0.03 | 0.76 | 0.04 | 0.97 | 1 | 4930412M03Rik | | 2 | 9 | 23 | 20 | 58 | 21 | 42 | 51 | 39 | 31 | 19 |
| 19 | -2.01 | 1.40 | -1.44 | 0.15 | 1 | Catip | ciliogenesis associated TTC17 interacting | 30 | 46 | 0 | 22 | 14 | 30 | 42 | 0 | 9 | 3 | 16 |
| 790 | -1.49 | 1.75 | -0.85 | 0.39 | 1 | Ptprc | protein tyrosine phosphatase, receptor typ | 5091 | 1635 | 1873 | 4 | 0 | 23 | 40 | 0 | 13 | 0 | 12 |
| 263 | 0.89 | 0.51 | 1.74 | 0.08 | 1 | Iigp1 | interferon inducible GTPase 1 | 1102 | 740 | 787 | 7 | 26 | 17 | 40 | 31 | 103 | 24 | 17 |
| 133 | -0.07 | 0.57 | -0.13 | 0.89 | 1 | Tex15 | testis expressed gene 15 | 11 | 12 | 6 | 388 | 34 | 274 | 40 | 341 | 54 | 18 | 282 |
| 125 | 0.51 | 0.51 | 1.01 | 0.31 | 1 | Ccnb2 | cyclin B2 | 363 | 157 | 133 | 93 | 87 | 71 | 40 | 177 | 88 | 75 | 88 |
| 116 | 0.96 | 0.49 | 1.97 | 0.05 | 1 | Coq3 | coenzyme Q3 methyltransferase | 129 | 134 | 127 | 54 | 105 | 96 | 40 | 278 | 97 | 113 | 106 |
| 102 | 0.26 | 0.47 | 0.55 | 0.58 | 1 | Gm20721 | | 154 | 224 | 191 | 51 | 84 | 75 | 40 | 49 | 77 | 121 | 55 |

| | | | | | | | | | | | | | | | | | | |
|---|---|---|---|---|---|---|---|---|---|---|---|---|---|---|---|---|---|---|
| 98 | -0.04 | 0.56 | -0.07 | 0.95 | 1 | Cdca8 | cell division cycle associated 8 | 403 | 151 | 61 | 65 | 70 | 51 | 40 | 58 | 62 | 85 | 37 |
| 92 | 0.14 | 0.43 | 0.33 | 0.74 | 1 | D330023K18Rik | | 88 | 129 | 105 | 96 | 90 | 107 | 40 | 74 | 81 | 108 | 88 |
| 90 | -0.30 | 0.73 | -0.41 | 0.68 | 1 | Neto2 | neuropilin (NRP) and tolloid (TLL)-like 2 | 150 | 296 | 170 | 65 | 31 | 57 | 40 | 5 | 38 | 103 | 29 |
| 80 | 0.05 | 0.55 | 0.10 | 0.92 | 1 | Fam78b | family with sequence similarity 78, membe | 155 | 75 | 105 | 127 | 36 | 60 | 40 | 76 | 35 | 49 | 116 |
| 78 | -0.23 | 0.37 | -0.63 | 0.53 | 1 | D330041H03Rik | | 34 | 46 | 128 | 56 | 116 | 131 | 40 | 37 | 136 | 33 | 105 |
| 78 | -0.16 | 0.68 | -0.24 | 0.81 | 1 | Cdc7 | cell division cycle 7 (S. cerevisiae) | 109 | 37 | 42 | 84 | 14 | 236 | 40 | 22 | 70 | 35 | 171 |
| 71 | -0.54 | 0.50 | -1.08 | 0.28 | 1 | Gm15927 | | 57 | 36 | 68 | 111 | 43 | 172 | 40 | 44 | 45 | 36 | 128 |
| 68 | 0.23 | 0.49 | 0.46 | 0.65 | 1 | Zfand4 | zinc finger, AN1-type domain 4 | 106 | 54 | 63 | 76 | 51 | 74 | 40 | 115 | 50 | 57 | 66 |
| 67 | 0.57 | 0.72 | 0.79 | 0.43 | 1 | Gm30239 | | 10 | 0 | 22 | 67 | 41 | 138 | 40 | 88 | 120 | 67 | 138 |
| 64 | -0.36 | 0.57 | -0.64 | 0.52 | 1 | 6820431F20Rik | | 104 | 42 | 29 | 48 | 54 | 123 | 40 | 25 | 41 | 24 | 168 |
| 63 | 0.26 | 0.48 | 0.54 | 0.59 | 1 | Slc26a11 | solute carrier family 26, member 11 | 92 | 108 | 114 | 38 | 33 | 56 | 40 | 37 | 84 | 28 | 60 |
| 62 | -0.01 | 1.00 | -0.01 | 1.00 | 1 | Nexn | nexilin | 280 | 109 | 144 | 6 | 18 | 8 | 40 | 51 | 13 | 0 | 8 |
| 61 | 0.16 | 0.50 | 0.32 | 0.75 | 1 | Gm35025 | | 78 | 92 | 47 | 52 | 54 | 56 | 40 | 110 | 39 | 31 | 67 |
| 58 | 0.01 | 0.53 | 0.03 | 0.98 | 1 | H3c4 | H3 clustered histone 4 | 4 | 17 | 8 | 43 | 71 | 155 | 40 | 44 | 55 | 58 | 140 |
| 52 | -0.74 | 0.52 | -1.43 | 0.15 | 1 | Tpm3-rs7 | | 146 | 86 | 98 | 36 | 43 | 30 | 40 | 34 | 13 | 22 | 21 |
| 50 | 0.11 | 0.50 | 0.22 | 0.83 | 1 | Ccdc89 | coiled-coil domain containing 89 | 51 | 52 | 30 | 37 | 62 | 61 | 40 | 51 | 68 | 37 | 59 |
| 46 | -0.35 | 0.54 | -0.65 | 0.52 | 1 | Ggnbp1 | gametogenetin binding protein 1 | 20 | 46 | 20 | 37 | 33 | 134 | 40 | 34 | 43 | 21 | 82 |
| 41 | -0.17 | 0.57 | -0.29 | 0.77 | 1 | Rab11b-ps2 | | 48 | 50 | 72 | 25 | 71 | 19 | 40 | 17 | 39 | 32 | 41 |
| 38 | -0.01 | 0.62 | -0.02 | 0.98 | 1 | Tsg101-ps | | 32 | 11 | 52 | 44 | 44 | 36 | 40 | 48 | 31 | 52 | 32 |
| 38 | 0.10 | 0.71 | 0.14 | 0.89 | 1 | A830035O19Rik | | 32 | 11 | 35 | 79 | 17 | 25 | 40 | 85 | 30 | 41 | 19 |
| 36 | 0.37 | 0.63 | 0.58 | 0.56 | 1 | Asb5 | ankyrin repeat and SOCs box-containing | 8 | 16 | 39 | 16 | 46 | 45 | 40 | 29 | 48 | 54 | 50 |
| 34 | -0.38 | 0.95 | -0.40 | 0.69 | 1 | Cfap45 | cilia and flagella associated protein 45 | 23 | 45 | 31 | 25 | 45 | 30 | 40 | 43 | 59 | 0 | 36 |
| 34 | 0.43 | 0.71 | 0.61 | 0.54 | 1 | Slc26a1 | solute carrier family 26 (sulfate transporte | 54 | 73 | 41 | 33 | 8 | 12 | 40 | 49 | 29 | 12 | 23 |
| 32 | -0.50 | 0.98 | -0.52 | 0.61 | 1 | Gm49785 | | 21 | 28 | 20 | 39 | 35 | 41 | 40 | 0 | 27 | 61 | 44 |
| 32 | -0.57 | 0.65 | -0.89 | 0.38 | 1 | 2210406O10Rik | | 16 | 27 | 15 | 32 | 53 | 37 | 40 | 67 | 24 | 15 | 21 |
| 31 | -0.72 | 1.02 | -0.70 | 0.48 | 1 | Gm6598 | | 14 | 5 | 34 | 33 | 26 | 57 | 40 | 31 | 29 | 0 | 67 |
| 28 | 0.03 | 0.72 | 0.05 | 0.96 | 1 | Gm16437 | | 22 | 32 | 30 | 32 | 7 | 31 | 40 | 48 | 18 | 21 | 23 |
| 27 | -0.70 | 0.85 | -0.82 | 0.41 | 1 | Gm20684 | | 9 | 40 | 9 | 16 | 61 | 30 | 40 | 14 | 31 | 36 | 12 |
| 16 | 0.29 | 1.17 | 0.25 | 0.80 | 1 | Gm5828 | | 39 | 1 | 0 | 6 | 2 | 1 | 40 | 12 | 2 | 75 | 0 |
| 14 | 0.47 | 1.40 | 0.34 | 0.74 | 1 | 1500002F19Rik | | 8 | 1 | 0 | 16 | 7 | 18 | 40 | 29 | 22 | 0 | 18 |
| 1375 | -3.60 | 2.22 | -1.62 | 0.10 | 1 | Adgrl4 | adhesion G protein-coupled receptor L4 | 4908 | 5170 | 5011 | 0 | 1 | 1 | 39 | 0 | 0 | 0 | 0 |
| 654 | -1.66 | 1.43 | -1.16 | 0.25 | 1 | Entpd1 | ectonucleoside triphosphate diphosphohy | 2852 | 2035 | 2229 | 12 | 2 | 7 | 39 | 0 | 11 | 0 | 10 |
| 645 | -0.30 | 0.34 | -0.89 | 0.38 | 1 | Cd9 | CD9 antigen | 2825 | 1617 | 2331 | 45 | 45 | 41 | 39 | 17 | 48 | 21 | 65 |
| 405 | -1.55 | 0.92 | -1.69 | 0.09 | 1 | Rnd3 | Rho family GTPase 3 | 1982 | 1002 | 1177 | 42 | 39 | 75 | 39 | 0 | 43 | 9 | 45 |
| 290 | -0.40 | 0.62 | -0.65 | 0.52 | 1 | Afap1l1 | actin filament associated protein 1-like 1 | 1102 | 1061 | 897 | 2 | 12 | 21 | 39 | 23 | 5 | 15 | 9 |
| 264 | -0.11 | 0.42 | -0.25 | 0.80 | 1 | Capg | capping protein (actin filament), gelsolin-l | 1437 | 552 | 695 | 15 | 35 | 24 | 39 | 35 | 11 | 35 | 23 |
| 219 | -0.25 | 0.44 | -0.56 | 0.57 | 1 | Septin6 | septin 6 | 1042 | 472 | 343 | 98 | 77 | 85 | 39 | 57 | 71 | 72 | 57 |
| 146 | -0.71 | 0.49 | -1.45 | 0.15 | 1 | Add2 | adducin 2 (beta) | 353 | 459 | 397 | 104 | 38 | 63 | 39 | 23 | 30 | 67 | 27 |
| 124 | 0.08 | 0.44 | 0.17 | 0.86 | 1 | Mgmt | O-6-methylguanine-DNA methyltransfera: | 235 | 333 | 194 | 79 | 98 | 74 | 39 | 109 | 37 | 88 | 81 |
| 116 | 0.84 | 0.56 | 1.49 | 0.14 | 1 | Ptp4a1 | protein tyrosine phosphatase 4a1 | 106 | 22 | 55 | 10 | 3 | 456 | 39 | 16 | 11 | 97 | 455 |
| 82 | -0.13 | 0.53 | -0.24 | 0.81 | 1 | 5031434O11Rik | | 21 | 74 | 23 | 62 | 61 | 244 | 39 | 105 | 18 | 64 | 193 |
| 82 | 0.04 | 0.41 | 0.09 | 0.93 | 1 | Mybl1 | myeloblastosis oncogene-like 1 | 128 | 140 | 83 | 72 | 64 | 99 | 39 | 80 | 72 | 48 | 74 |
| 79 | 0.44 | 0.61 | 0.73 | 0.47 | 1 | Melk | maternal embryonic leucine zipper kinase | 200 | 66 | 36 | 93 | 60 | 52 | 39 | 81 | 96 | 59 | 88 |
| 67 | 0.12 | 0.44 | 0.27 | 0.79 | 1 | Gm10602 | | 45 | 68 | 83 | 109 | 52 | 64 | 39 | 65 | 93 | 45 | 78 |
| 63 | 0.58 | 0.49 | 1.17 | 0.24 | 1 | Gm11762 | | 57 | 82 | 76 | 96 | 26 | 41 | 39 | 67 | 64 | 82 | 64 |
| 62 | -0.55 | 0.59 | -0.93 | 0.35 | 1 | Sync | syncoilin | 113 | 143 | 124 | 51 | 41 | 44 | 39 | 10 | 20 | 61 | 41 |
| 60 | 0.26 | 0.58 | 0.46 | 0.65 | 1 | A630089N07Rik | | 29 | 68 | 114 | 32 | 26 | 113 | 39 | 68 | 28 | 36 | 111 |
| 59 | 0.54 | 0.60 | 0.91 | 0.36 | 1 | 4930589O11Rik | | 42 | 69 | 82 | 23 | 63 | 64 | 39 | 78 | 34 | 98 | 56 |
| 52 | 0.35 | 0.57 | 0.61 | 0.54 | 1 | Nudt15 | nudix (nucleoside diphosphate linked mo | 26 | 21 | 27 | 45 | 76 | 49 | 39 | 53 | 46 | 148 | 45 |
| 50 | -0.94 | 0.65 | -1.45 | 0.15 | 1 | 6720489N17Ril | RIKEN cDNA 6720489N17 gene | 69 | 89 | 58 | 57 | 75 | 40 | 39 | 8 | 33 | 22 | 62 |
| 45 | -0.21 | 0.52 | -0.41 | 0.68 | 1 | Tnfrsf26 | tumor necrosis factor receptor superfamily | 102 | 122 | 79 | 41 | 15 | 10 | 39 | 32 | 24 | 28 | 6 |
| 44 | 0.08 | 0.48 | 0.16 | 0.87 | 1 | Gm5865 | | 36 | 19 | 60 | 41 | 63 | 34 | 39 | 36 | 61 | 57 | 36 |
| 42 | 0.48 | 0.63 | 0.76 | 0.45 | 1 | Psme2b | protease (prosome, macropain) activator | 101 | 54 | 107 | 15 | 14 | 19 | 39 | 44 | 28 | 21 | 22 |
| 42 | 0.17 | 0.63 | 0.27 | 0.79 | 1 | Mylpf | myosin light chain, phosphorylatable, fast | 36 | 37 | 23 | 30 | 39 | 60 | 39 | 76 | 60 | 15 | 46 |
| 39 | 0.16 | 0.81 | 0.20 | 0.84 | 1 | Cdc25c | cell division cycle 25C | 52 | 43 | 17 | 27 | 32 | 30 | 39 | 119 | 21 | 6 | 42 |
| 39 | 0.07 | 0.67 | 0.11 | 0.92 | 1 | Gm47431 | | 0 | 10 | 4 | 75 | 33 | 58 | 39 | 65 | 31 | 55 | 55 |
| 38 | 0.12 | 0.84 | 0.14 | 0.89 | 1 | Gm12715 | | 69 | 63 | 38 | 36 | 19 | 13 | 39 | 2 | 49 | 72 | 12 |
| 37 | -1.11 | 1.14 | -0.97 | 0.33 | 1 | Gm10499 | | 219 | 15 | 96 | 0 | 17 | 0 | 39 | 0 | 13 | 5 | 0 |
| 36 | 0.23 | 0.68 | 0.34 | 0.73 | 1 | Gm16253 | | 39 | 22 | 29 | 26 | 22 | 38 | 39 | 21 | 57 | 91 | 12 |
| 35 | 0.40 | 0.72 | 0.55 | 0.58 | 1 | Gm12276 | | 24 | 36 | 24 | 43 | 10 | 40 | 39 | 12 | 37 | 66 | 55 |

| | | | | | | | | | | | | | | | | | |
|---|---|---|---|---|---|---|---|---|---|---|---|---|---|---|---|---|---|
| 33 | -1.62 | 1.25 | -1.30 | 0.19 | 1 | Gm31356 | | 0 | 72 | 20 | 15 | 42 | 52 | 39 | 59 | 28 | 0 | 41 |
| 33 | -0.08 | 0.96 | -0.08 | 0.93 | 1 | Gm12299 | | 0 | 0 | 14 | 30 | 38 | 52 | 39 | 13 | 39 | 45 | 91 |
| 33 | 0.62 | 0.54 | 1.15 | 0.25 | 1 | Gm38250 | | 34 | 26 | 35 | 20 | 18 | 26 | 39 | 30 | 35 | 59 | 35 |
| 31 | -0.42 | 0.68 | -0.61 | 0.54 | 1 | Gm17530 | | 3 | 18 | 26 | 20 | 50 | 54 | 39 | 21 | 50 | 17 | 48 |
| 31 | 0.16 | 0.83 | 0.19 | 0.85 | 1 | Spcs2-ps | | 3 | 5 | 2 | 40 | 46 | 23 | 39 | 6 | 82 | 40 | 53 |
| 28 | 0.37 | 0.92 | 0.41 | 0.68 | 1 | Gm6728 | | 33 | 36 | 18 | 32 | 30 | 0 | 39 | 83 | 31 | 6 | 2 |
| 28 | 0.22 | 0.76 | 0.29 | 0.78 | 1 | Pkd1l2 | polycystic kidney disease 1 like 2 | 31 | 41 | 47 | 6 | 10 | 33 | 39 | 9 | 12 | 53 | 26 |
| 28 | 0.25 | 0.69 | 0.36 | 0.72 | 1 | Timm8a1 | translocase of inner mitochondrial membr | 54 | 34 | 25 | 18 | 17 | 19 | 39 | 32 | 30 | 13 | 25 |
| 25 | -0.08 | 1.12 | -0.07 | 0.95 | 1 | Gm7276 | | 2 | 10 | 0 | 24 | 36 | 36 | 39 | 2 | 36 | 55 | 34 |
| 22 | 0.14 | 0.78 | 0.18 | 0.86 | 1 | Gm5602 | | 16 | 9 | 26 | 4 | 25 | 21 | 39 | 6 | 38 | 14 | 40 |
| 22 | -0.52 | 1.02 | -0.51 | 0.61 | 1 | Gm26682 | | 0 | 12 | 10 | 38 | 9 | 27 | 39 | 56 | 16 | 9 | 21 |
| 21 | -0.50 | 1.00 | -0.50 | 0.62 | 1 | 1010001B22Rik | | 12 | 41 | 16 | 3 | 40 | 8 | 39 | 32 | 26 | 9 | 5 |
| 17 | -0.46 | 1.36 | -0.34 | 0.73 | 1 | Perm1 | PPARGC1 and ESRR induced regulator, | 8 | 6 | 0 | 15 | 23 | 8 | 39 | 0 | 42 | 45 | 2 |
| 17 | 0.25 | 1.19 | 0.21 | 0.83 | 1 | Gm11549 | | 22 | 0 | 2 | 31 | 2 | 14 | 39 | 27 | 22 | 18 | 8 |
| 13 | -1.49 | 1.09 | -1.37 | 0.17 | 1 | Etaa1os | | 9 | 14 | 14 | 10 | 9 | 18 | 39 | 0 | 8 | 9 | 12 |
| 508 | 0.64 | 0.41 | 1.58 | 0.11 | 1 | Adgrd1 | adhesion G protein-coupled receptor D1 | 2963 | 1346 | 696 | 107 | 37 | 46 | 38 | 74 | 86 | 128 | 64 |
| 357 | -0.42 | 0.35 | -1.23 | 0.22 | 1 | Wwc2 | WW, C2 and coiled-coil domain containin | 1434 | 1017 | 1183 | 44 | 45 | 37 | 38 | 53 | 26 | 19 | 26 |
| 125 | 0.48 | 0.43 | 1.12 | 0.26 | 1 | Tcaf2 | TRPM8 channel-associated factor 2 | 365 | 321 | 405 | 19 | 49 | 14 | 38 | 56 | 32 | 43 | 27 |
| 111 | 0.22 | 0.57 | 0.38 | 0.71 | 1 | Tubb2b | tubulin, beta 2B class IIB | 340 | 236 | 124 | 92 | 58 | 32 | 38 | 156 | 94 | 28 | 19 |
| 105 | 0.01 | 0.66 | 0.01 | 0.99 | 1 | Bub1 | BUB1, mitotic checkpoint serine/threonine | 395 | 58 | 68 | 116 | 64 | 88 | 38 | 49 | 101 | 73 | 105 |
| 80 | 0.11 | 0.63 | 0.18 | 0.86 | 1 | Phf19 | PHD finger protein 19 | 141 | 65 | 16 | 112 | 75 | 70 | 38 | 130 | 70 | 117 | 45 |
| 72 | 0.71 | 0.43 | 1.65 | 0.10 | 1 | 1810024B03Ril | RIKEN cDNA 1810024B03 gene | 57 | 114 | 59 | 53 | 60 | 59 | 38 | 98 | 117 | 73 | 61 |
| 71 | -0.33 | 0.54 | -0.60 | 0.55 | 1 | Riiad1 | regulatory subunit of type II PKA R-subun | 31 | 197 | 103 | 55 | 60 | 83 | 38 | 112 | 30 | 33 | 40 |
| 69 | -0.25 | 0.58 | -0.43 | 0.67 | 1 | Chtf18 | CTF18, chromosome transmission fidelity | 169 | 55 | 97 | 84 | 45 | 63 | 38 | 25 | 78 | 67 | 43 |
| 67 | 0.30 | 0.39 | 0.77 | 0.44 | 1 | Tcte2 | t-complex-associated testis expressed 2 | 82 | 152 | 80 | 67 | 35 | 50 | 38 | 81 | 38 | 64 | 51 |
| 65 | 0.20 | 0.76 | 0.26 | 0.79 | 1 | Stab2 | stabilin 2 | 82 | 194 | 60 | 12 | 5 | 157 | 38 | 66 | 22 | 9 | 69 |
| 60 | -0.05 | 0.48 | -0.11 | 0.91 | 1 | Gm6277 | | 98 | 92 | 105 | 46 | 49 | 46 | 38 | 38 | 23 | 46 | 77 |
| 60 | 0.81 | 0.63 | 1.29 | 0.20 | 1 | Stard6 | StAR-related lipid transfer (START) doma | 17 | 83 | 62 | 32 | 39 | 74 | 38 | 56 | 47 | 116 | 91 |
| 53 | -0.15 | 0.64 | -0.23 | 0.82 | 1 | Arhgap8 | Rho GTPase activating protein 8 | 39 | 82 | 116 | 78 | 21 | 39 | 38 | 87 | 14 | 24 | 47 |
| 50 | -0.56 | 0.91 | -0.62 | 0.54 | 1 | Spata1 | spermatogenesis associated 1 | 21 | 16 | 8 | 19 | 43 | 190 | 38 | 34 | 41 | 0 | 134 |
| 48 | -0.41 | 0.68 | -0.60 | 0.55 | 1 | Zfp773 | zinc finger protein 773 | 95 | 31 | 62 | 50 | 61 | 36 | 38 | 36 | 9 | 67 | 44 |
| 47 | 0.94 | 0.64 | 1.47 | 0.14 | 1 | Asb15 | ankyrin repeat and SOCS box-containing | 27 | 6 | 24 | 42 | 25 | 33 | 38 | 53 | 37 | 183 | 50 |
| 47 | 0.58 | 0.71 | 0.82 | 0.41 | 1 | 4921534A09Rik | | 35 | 100 | 23 | 12 | 43 | 43 | 38 | 59 | 32 | 84 | 46 |
| 47 | 0.23 | 0.58 | 0.39 | 0.70 | 1 | Cfap47 | cilia and flagella associated protein 47 | 33 | 12 | 22 | 35 | 22 | 119 | 38 | 36 | 61 | 26 | 110 |
| 46 | -0.05 | 0.53 | -0.09 | 0.93 | 1 | Gm13680 | | 117 | 115 | 53 | 20 | 33 | 20 | 38 | 34 | 22 | 33 | 20 |
| 44 | -1.46 | 0.98 | -1.49 | 0.14 | 1 | Mis18bp1 | MIS18 binding protein 1 | 130 | 40 | 23 | 53 | 41 | 58 | 38 | 0 | 34 | 12 | 53 |
| 43 | 0.26 | 0.65 | 0.41 | 0.68 | 1 | Slc22a18 | solute carrier family 22 (organic cation tra | 90 | 42 | 68 | 48 | 32 | 12 | 38 | 58 | 31 | 19 | 37 |
| 42 | 0.02 | 0.55 | 0.04 | 0.97 | 1 | Ppp1ccb | protein phosphatase 1 catalytic subunit ga | 51 | 97 | 19 | 41 | 22 | 45 | 38 | 43 | 30 | 32 | 38 |
| 40 | 0.17 | 0.54 | 0.31 | 0.76 | 1 | Gm10044 | | 9 | 12 | 24 | 42 | 33 | 67 | 38 | 35 | 48 | 41 | 87 |
| 37 | -0.48 | 0.54 | -0.88 | 0.38 | 1 | Gm7008 | | 28 | 34 | 50 | 39 | 33 | 58 | 38 | 23 | 43 | 28 | 27 |
| 36 | -0.27 | 0.67 | -0.40 | 0.69 | 1 | 9130230L23Rik | | 14 | 6 | 2 | 39 | 45 | 79 | 38 | 59 | 23 | 23 | 73 |
| 35 | 0.09 | 0.64 | 0.14 | 0.89 | 1 | A930016O22Rik | | 3 | 12 | 3 | 51 | 50 | 37 | 38 | 50 | 32 | 68 | 40 |
| 34 | -0.58 | 0.77 | -0.75 | 0.45 | 1 | Gm38391 | | 23 | 29 | 64 | 73 | 17 | 14 | 38 | 54 | 15 | 36 | 4 |
| 30 | -2.25 | 1.33 | -1.69 | 0.09 | 1 | Artn | artemin | 60 | 20 | 55 | 71 | 12 | 16 | 38 | 0 | 36 | 0 | 15 |
| 29 | -0.30 | 0.81 | -0.37 | 0.71 | 1 | Zfp541 | zinc finger protein 541 | 0 | 0 | 0 | 27 | 14 | 95 | 38 | 22 | 30 | 10 | 85 |
| 28 | 0.46 | 0.62 | 0.74 | 0.46 | 1 | Gm6767 | | 25 | 28 | 35 | 17 | 26 | 15 | 38 | 34 | 31 | 26 | 36 |
| 27 | 0.58 | 0.68 | 0.84 | 0.40 | 1 | Gm11508 | | 13 | 28 | 8 | 27 | 6 | 24 | 38 | 44 | 9 | 70 | 24 |
| 25 | -0.68 | 0.89 | -0.77 | 0.44 | 1 | Gm3807 | | 13 | 35 | 11 | 15 | 29 | 36 | 38 | 17 | 58 | 12 | 7 |
| 25 | 0.06 | 1.10 | 0.06 | 0.95 | 1 | Gm41804 | | 19 | 24 | 19 | 26 | 22 | 9 | 38 | 20 | 54 | 0 | 40 |
| 20 | -1.53 | 1.02 | -1.50 | 0.13 | 1 | 5033403F01Rik | | 16 | 12 | 36 | 7 | 10 | 45 | 38 | 0 | 18 | 4 | 29 |
| 19 | -0.28 | 0.83 | -0.34 | 0.74 | 1 | Gm18541 | | 25 | 30 | 10 | 17 | 11 | 15 | 38 | 8 | 22 | 12 | 18 |
| 16 | -0.15 | 1.57 | -0.09 | 0.93 | 1 | Cntn4 | contactin 4 | 0 | 13 | 0 | 1 | 24 | 23 | 38 | 3 | 41 | 21 | 15 |
| 13 | -0.63 | 1.12 | -0.56 | 0.57 | 1 | Rbfaos | | 0 | 0 | 3 | 24 | 11 | 8 | 38 | 6 | 29 | 18 | 10 |
| 13 | -2.05 | 1.67 | -1.23 | 0.22 | 1 | 6030442K20Rik | | 9 | 13 | 0 | 18 | 15 | 18 | 38 | 0 | 11 | 0 | 18 |
| 9271 | -2.29 | 1.80 | -1.28 | 0.20 | 1 | Igkc | | 84986 | 15100 | 1837 | 0 | 1 | 13 | 37 | 0 | 3 | 1 | 0 |
| 709 | -0.11 | 0.28 | -0.41 | 0.68 | 1 | Plk2 | polo like kinase 2 | 2711 | 2370 | 2090 | 40 | 23 | 245 | 37 | 23 | 48 | 32 | 179 |
| 427 | -0.18 | 0.39 | -0.47 | 0.64 | 1 | Steap3 | STEAP family member 3 | 2317 | 841 | 1259 | 54 | 31 | 25 | 37 | 45 | 20 | 32 | 31 |
| 133 | -1.44 | 0.75 | -1.92 | 0.05 | 1 | Gm20342 | | 18 | 79 | 50 | 60 | 32 | 566 | 37 | 8 | 46 | 0 | 569 |
| 90 | -0.62 | 0.95 | -0.65 | 0.52 | 1 | Gas2l3 | growth arrest-specific 2 like 3 | 233 | 166 | 38 | 103 | 105 | 73 | 37 | 67 | 94 | 0 | 69 |
| 78 | -0.10 | 0.40 | -0.25 | 0.80 | 1 | Ano7 | anoctamin 7 | 106 | 229 | 145 | 53 | 29 | 74 | 37 | 38 | 36 | 26 | 88 |

| | | | | | | | | | | | | | | | | | | |
|---|---|---|---|---|---|---|---|---|---|---|---|---|---|---|---|---|---|---|
| 77 | 0.01 | 0.45 | 0.02 | 0.98 | 1 | Praf2 | PRA1 domain family 2 | 130 | 94 | 161 | 32 | 106 | 53 | 37 | 25 | 100 | 71 | 38 |
| 69 | -0.46 | 0.94 | -0.49 | 0.62 | 1 | Pbld1 | phenazine biosynthesis-like protein doma | 130 | 268 | 200 | 9 | 13 | 14 | 37 | 63 | 13 | 0 | 9 |
| 62 | -0.26 | 0.52 | -0.50 | 0.61 | 1 | Gm4617 | | 112 | 56 | 124 | 77 | 46 | 55 | 37 | 65 | 36 | 33 | 45 |
| 60 | -0.56 | 0.57 | -0.98 | 0.33 | 1 | Wnt9a | wingless-type MMTV integration site famil | 238 | 111 | 130 | 16 | 27 | 19 | 37 | 14 | 24 | 41 | 3 |
| 59 | 0.94 | 0.53 | 1.77 | 0.08 | 1 | Rab17 | RAB17, member RAS oncogene family | 45 | 26 | 77 | 35 | 56 | 39 | 37 | 133 | 97 | 48 | 53 |
| 57 | 0.23 | 0.53 | 0.43 | 0.67 | 1 | Gm4032 | | 28 | 76 | 60 | 59 | 45 | 74 | 37 | 44 | 84 | 67 | 50 |
| 56 | -0.58 | 0.78 | -0.74 | 0.46 | 1 | P2ry14 | purinergic receptor P2Y, G-protein couple | 157 | 193 | 130 | 34 | 4 | 10 | 37 | 10 | 23 | 5 | 10 |
| 55 | -1.12 | 0.92 | -1.22 | 0.22 | 1 | Ntrk3 | neurotrophic tyrosine kinase, receptor, typ | 174 | 34 | 30 | 53 | 56 | 68 | 37 | 0 | 41 | 70 | 38 |
| 52 | 0.45 | 0.48 | 0.92 | 0.36 | 1 | Lix1 | limb and CNS expressed 1 | 63 | 64 | 107 | 23 | 50 | 31 | 37 | 28 | 48 | 71 | 46 |
| 47 | 0.09 | 0.45 | 0.20 | 0.84 | 1 | Apoo-ps | | 45 | 71 | 54 | 43 | 47 | 39 | 37 | 63 | 50 | 30 | 41 |
| 47 | -0.12 | 0.50 | -0.24 | 0.81 | 1 | Gm17092 | | 37 | 46 | 28 | 88 | 40 | 49 | 37 | 43 | 34 | 63 | 50 |
| 47 | -0.71 | 0.66 | -1.09 | 0.28 | 1 | Gm44349 | | 64 | 38 | 58 | 43 | 77 | 41 | 37 | 15 | 103 | 10 | 26 |
| 38 | -0.95 | 0.65 | -1.47 | 0.14 | 1 | Slc29a2 | solute carrier family 29 (nucleoside transp | 78 | 50 | 38 | 62 | 16 | 50 | 37 | 8 | 16 | 28 | 34 |
| 37 | -0.21 | 0.56 | -0.37 | 0.71 | 1 | Ebf4 | early B cell factor 4 | 33 | 56 | 41 | 39 | 34 | 36 | 37 | 44 | 28 | 48 | 15 |
| 34 | -0.66 | 0.65 | -1.02 | 0.31 | 1 | Zfp808 | zinc finger protein 808 | 14 | 24 | 50 | 37 | 51 | 40 | 37 | 32 | 20 | 12 | 53 |
| 33 | -0.30 | 0.60 | -0.50 | 0.61 | 1 | Gm17349 | | 16 | 35 | 38 | 39 | 40 | 31 | 37 | 17 | 50 | 32 | 25 |
| 31 | 0.07 | 0.59 | 0.12 | 0.91 | 1 | Gm9512 | | 54 | 30 | 46 | 17 | 24 | 27 | 37 | 32 | 25 | 22 | 25 |
| 29 | 0.58 | 0.66 | 0.89 | 0.38 | 1 | Gm4130 | | 13 | 30 | 29 | 15 | 25 | 22 | 37 | 53 | 38 | 35 | 26 |
| 29 | -0.81 | 0.69 | -1.17 | 0.24 | 1 | Gm16894 | | 20 | 21 | 50 | 61 | 33 | 18 | 37 | 20 | 15 | 24 | 20 |
| 29 | -0.01 | 0.72 | -0.01 | 0.99 | 1 | Gm6055 | | 46 | 11 | 55 | 20 | 14 | 22 | 37 | 6 | 32 | 48 | 27 |
| 28 | -0.69 | 0.72 | -0.96 | 0.34 | 1 | Crybb1 | crystallin, beta B1 | 9 | 11 | 8 | 50 | 34 | 43 | 37 | 31 | 5 | 55 | 27 |
| 28 | -0.14 | 0.96 | -0.14 | 0.89 | 1 | 4933432K03Rik | | 0 | 0 | 0 | 51 | 26 | 35 | 37 | 5 | 41 | 55 | 53 |
| 27 | -0.70 | 1.18 | -0.59 | 0.55 | 1 | Gm15841 | | 0 | 36 | 31 | 11 | 15 | 41 | 37 | 54 | 41 | 3 | 25 |
| 25 | -1.47 | 0.95 | -1.56 | 0.12 | 1 | Cfap70 | cilia and flagella associated protein 70 | 19 | 15 | 29 | 33 | 28 | 46 | 37 | 23 | 10 | 0 | 38 |
| 24 | -0.07 | 0.78 | -0.09 | 0.93 | 1 | Mtag2 | | 12 | 31 | 26 | 32 | 10 | 20 | 37 | 15 | 30 | 15 | 37 |
| 23 | 0.38 | 0.78 | 0.49 | 0.62 | 1 | Gm14703 | | 9 | 18 | 10 | 18 | 16 | 20 | 37 | 9 | 45 | 46 | 22 |
| 21 | -2.49 | 1.30 | -1.92 | 0.06 | 1 | Rdh16f2 | RDH16 family member 2 | 0 | 11 | 11 | 8 | 29 | 59 | 37 | 8 | 4 | 0 | 63 |
| 21 | -0.89 | 0.82 | -1.09 | 0.28 | 1 | Hacl1 | 2-hydroxyacyl-CoA lyase 1 | 21 | 19 | 3 | 33 | 25 | 25 | 37 | 14 | 15 | 8 | 28 |
| 19 | -0.37 | 1.05 | -0.35 | 0.72 | 1 | Gm11772 | | 20 | 15 | 0 | 18 | 17 | 18 | 37 | 3 | 14 | 52 | 13 |
| 19 | -1.30 | 0.96 | -1.36 | 0.17 | 1 | Gm35638 | | 21 | 24 | 10 | 32 | 16 | 19 | 37 | 2 | 25 | 5 | 12 |
| 15 | -0.96 | 1.04 | -0.93 | 0.35 | 1 | Gm42879 | | 4 | 22 | 17 | 27 | 9 | 8 | 37 | 3 | 7 | 12 | 19 |
| 14 | -1.99 | 1.44 | -1.38 | 0.17 | 1 | Gm17039 | | 15 | 48 | 11 | 10 | 11 | 5 | 37 | 0 | 10 | 0 | 7 |
| 1760 | -0.42 | 1.24 | -0.34 | 0.73 | 1 | Cfh | complement component factor h | 12064 | 3242 | 3996 | 0 | 2 | 6 | 36 | 1 | 10 | 1 | 3 |
| 787 | -0.16 | 0.32 | -0.49 | 0.62 | 1 | Arhgap31 | Rho GTPase activating protein 31 | 2842 | 2500 | 2572 | 182 | 92 | 94 | 36 | 118 | 78 | 59 | 86 |
| 199 | -0.36 | 0.57 | -0.63 | 0.53 | 1 | Ccdc84 | coiled-coil domain containing 84 | 66 | 120 | 70 | 53 | 84 | 862 | 36 | 7 | 115 | 32 | 743 |
| 103 | 0.04 | 0.50 | 0.08 | 0.94 | 1 | Parp3 | poly (ADP-ribose) polymerase family, mer | 377 | 232 | 268 | 12 | 38 | 42 | 36 | 27 | 54 | 36 | 15 |
| 103 | 0.29 | 0.48 | 0.61 | 0.54 | 1 | Iqgap3 | IQ motif containing GTPase activating pro | 220 | 117 | 110 | 95 | 77 | 102 | 36 | 92 | 105 | 104 | 77 |
| 91 | -0.65 | 0.82 | -0.79 | 0.43 | 1 | Abcb1b | ATP-binding cassette, sub-family B (MDR | 471 | 177 | 167 | 25 | 35 | 16 | 36 | 17 | 24 | 0 | 32 |
| 85 | -0.51 | 0.63 | -0.81 | 0.42 | 1 | Dcdc5 | doublecortin domain containing 5 | 51 | 40 | 39 | 103 | 64 | 207 | 36 | 37 | 88 | 8 | 264 |
| 78 | 0.08 | 0.40 | 0.20 | 0.84 | 1 | Cmc4 | C-x(9)-C motif containing 4 | 38 | 72 | 53 | 81 | 58 | 135 | 36 | 100 | 43 | 28 | 213 |
| 75 | -0.15 | 0.46 | -0.33 | 0.74 | 1 | Nrm | nurim (nuclear envelope membrane prote | 263 | 98 | 130 | 40 | 48 | 47 | 36 | 23 | 49 | 54 | 39 |
| 70 | -0.17 | 0.72 | -0.23 | 0.82 | 1 | Gm21811 | | 26 | 55 | 63 | 41 | 73 | 172 | 36 | 12 | 27 | 135 | 135 |
| 66 | 0.85 | 0.65 | 1.30 | 0.19 | 1 | Lonrf3 | LON peptidase N-terminal domain and rir | 82 | 34 | 7 | 68 | 63 | 34 | 36 | 159 | 85 | 100 | 53 |
| 60 | 0.76 | 0.54 | 1.42 | 0.16 | 1 | Gm49883 | | 57 | 25 | 18 | 22 | 54 | 101 | 36 | 73 | 49 | 100 | 125 |
| 57 | 0.35 | 0.48 | 0.73 | 0.47 | 1 | Plek2 | pleckstrin 2 | 67 | 68 | 27 | 58 | 56 | 52 | 36 | 67 | 57 | 73 | 63 |
| 55 | 0.72 | 0.56 | 1.29 | 0.20 | 1 | Scd3 | stearoyl-coenzyme A desaturase 3 | 28 | 26 | 73 | 46 | 70 | 31 | 36 | 123 | 50 | 76 | 45 |
| 51 | -0.61 | 0.49 | -1.24 | 0.21 | 1 | Rgs8 | regulator of G-protein signaling 8 | 84 | 124 | 47 | 91 | 24 | 34 | 36 | 54 | 13 | 18 | 36 |
| 51 | -0.72 | 0.86 | -0.84 | 0.40 | 1 | Fmo4 | flavin containing monooxygenase 4 | 54 | 62 | 65 | 45 | 29 | 102 | 36 | 0 | 30 | 39 | 95 |
| 51 | 0.22 | 0.95 | 0.23 | 0.82 | 1 | Sapcd2 | suppressor APC domain containing 2 | 89 | 8 | 3 | 54 | 72 | 48 | 36 | 110 | 55 | 17 | 63 |
| 50 | 0.29 | 0.46 | 0.62 | 0.53 | 1 | M5C1000I18Rik | | 63 | 107 | 37 | 74 | 17 | 25 | 36 | 97 | 20 | 33 | 39 |
| 50 | 0.16 | 0.67 | 0.24 | 0.81 | 1 | Zfp133-ps | | 57 | 36 | 17 | 61 | 19 | 96 | 36 | 17 | 64 | 40 | 107 |
| 50 | 0.15 | 0.80 | 0.19 | 0.85 | 1 | 9630013D21Rik | | 1 | 5 | 25 | 82 | 43 | 86 | 36 | 38 | 41 | 77 | 114 |
| 48 | 0.23 | 0.63 | 0.36 | 0.72 | 1 | Gm15853 | | 39 | 12 | 78 | 60 | 47 | 41 | 36 | 65 | 47 | 46 | 55 |
| 46 | -0.07 | 0.66 | -0.10 | 0.92 | 1 | Gm12905 | | 22 | 19 | 22 | 41 | 35 | 105 | 36 | 7 | 51 | 71 | 101 |
| 46 | 0.11 | 0.51 | 0.22 | 0.83 | 1 | 2700033N17Rik | | 51 | 64 | 90 | 32 | 35 | 41 | 36 | 58 | 30 | 37 | 32 |
| 46 | 0.42 | 0.80 | 0.53 | 0.60 | 1 | Sgo1 | shugoshin 1 | 126 | 68 | 22 | 45 | 6 | 51 | 36 | 23 | 55 | 18 | 52 |
| 43 | -0.15 | 0.64 | -0.24 | 0.81 | 1 | Fam72a | family with sequence similarity 72, memb | 62 | 80 | 77 | 12 | 54 | 28 | 36 | 17 | 33 | 15 | 59 |
| 42 | -0.07 | 0.62 | -0.11 | 0.91 | 1 | 2200002J24Rik | RIKEN cDNA 2200002J24 gene | 5 | 4 | 0 | 95 | 42 | 45 | 36 | 108 | 24 | 55 | 43 |
| 40 | -1.50 | 0.85 | -1.77 | 0.08 | 1 | Ankrd34c | ankyrin repeat domain 34C | 102 | 73 | 105 | 14 | 28 | 27 | 36 | 6 | 14 | 0 | 29 |
| 38 | -0.68 | 0.90 | -0.76 | 0.45 | 1 | Actn3 | actinin alpha 3 | 112 | 93 | 46 | 20 | 17 | 18 | 36 | 0 | 13 | 19 | 44 |

| | | | | | | | | | | | | | | | | | | |
|---|---|---|---|---|---|---|---|---|---|---|---|---|---|---|---|---|---|---|
| 37 | 0.50 | 0.58 | 0.86 | 0.39 | 1 | 4930513N10Rik | | 86 | 72 | 44 | 13 | 11 | 29 | 36 | 23 | 28 | 26 | 39 |
| 37 | -0.62 | 1.03 | -0.61 | 0.54 | 1 | 3110083C13Rik | | 64 | 30 | 19 | 70 | 13 | 48 | 36 | 0 | 31 | 63 | 26 |
| 32 | -0.75 | 0.94 | -0.79 | 0.43 | 1 | 9530018F02Rik | | 26 | 24 | 43 | 50 | 29 | 33 | 36 | 54 | 35 | 0 | 22 |
| 32 | -0.08 | 0.66 | -0.12 | 0.90 | 1 | Rab4b | RAB4B, member RAS oncogene family | 17 | 10 | 29 | 22 | 29 | 67 | 36 | 39 | 54 | 18 | 30 |
| 31 | 1.42 | 0.91 | 1.56 | 0.12 | 1 | E030030I06Rik | RIKEN cDNA E030030I06 gene | 25 | 27 | 35 | 4 | 7 | 17 | 36 | 119 | 21 | 23 | 25 |
| 31 | -0.40 | 1.02 | -0.39 | 0.70 | 1 | Ece2 | endothelin converting enzyme 2 | 3 | 10 | 11 | 18 | 8 | 106 | 36 | 0 | 17 | 36 | 92 |
| 29 | -0.66 | 0.97 | -0.67 | 0.50 | 1 | Guca1b | guanylate cyclase activator 1B | 6 | 6 | 12 | 32 | 35 | 62 | 36 | 58 | 29 | 0 | 48 |
| 29 | -1.03 | 0.67 | -1.54 | 0.12 | 1 | 1700001C19Ril | RIKEN cDNA 1700001C19 gene | 58 | 19 | 45 | 48 | 26 | 19 | 36 | 16 | 24 | 23 | 6 |
| 28 | -0.03 | 0.66 | -0.04 | 0.96 | 1 | C79130 | | 26 | 30 | 16 | 19 | 22 | 37 | 36 | 52 | 23 | 24 | 18 |
| 23 | -1.52 | 0.84 | -1.81 | 0.07 | 1 | Sms-ps | | 15 | 22 | 18 | 53 | 21 | 31 | 36 | 3 | 28 | 8 | 14 |
| 22 | -2.39 | 1.61 | -1.48 | 0.14 | 1 | Kcnj5 | potassium inwardly-rectifying channel, su | 0 | 13 | 8 | 3 | 95 | 19 | 36 | 0 | 54 | 0 | 15 |
| 22 | 0.47 | 0.81 | 0.58 | 0.56 | 1 | Gm44616 | | 5 | 17 | 4 | 26 | 4 | 24 | 36 | 52 | 16 | 44 | 11 |
| 22 | 0.58 | 1.10 | 0.53 | 0.60 | 1 | | | 0 | 12 | 12 | 7 | 16 | 23 | 36 | 9 | 54 | 45 | 22 |
| 22 | -0.46 | 1.09 | -0.42 | 0.67 | 1 | Gm3365 | | 20 | 19 | 12 | 16 | 11 | 30 | 36 | 0 | 35 | 9 | 47 |
| 18 | 0.14 | 0.91 | 0.15 | 0.88 | 1 | Gm16433 | | 9 | 7 | 27 | 6 | 7 | 25 | 36 | 14 | 17 | 10 | 35 |
| 16 | -0.99 | 1.12 | -0.89 | 0.37 | 1 | Gm19587 | | 24 | 5 | 8 | 18 | 26 | 6 | 36 | 2 | 35 | 1 | 10 |
| 14 | -0.15 | 0.80 | -0.19 | 0.85 | 1 | Gm38102 | | 3 | 2 | 4 | 9 | 20 | 16 | 36 | 14 | 17 | 15 | 19 |
| 14 | -0.52 | 1.17 | -0.45 | 0.66 | 1 | Mettl5os | | 21 | 16 | 14 | 7 | 1 | 17 | 36 | 0 | 6 | 18 | 16 |
| 1392 | -2.17 | 1.11 | -1.95 | 0.05 | 1 | Tgfbr2 | transforming growth factor, beta receptor I | 7031 | 3733 | 4430 | 7 | 22 | 21 | 35 | 0 | 12 | 0 | 17 |
| 809 | -1.09 | 1.05 | -1.03 | 0.30 | 1 | Pianp | PILR alpha associated neural protein | 265 | 2500 | 356 | 1957 | 306 | 1324 | 35 | 965 | 178 | 0 | 1015 |
| 389 | -0.60 | 0.33 | -1.82 | 0.07 | 1 | Sorbs3 | sorbin and SH3 domain containing 3 | 1571 | 1247 | 1174 | 68 | 24 | 44 | 35 | 36 | 16 | 27 | 32 |
| 237 | -0.09 | 0.43 | -0.20 | 0.84 | 1 | Slc16a2 | solute carrier family 16 (monocarboxylic a | 858 | 576 | 723 | 103 | 50 | 37 | 35 | 123 | 30 | 36 | 36 |
| 173 | -0.75 | 0.46 | -1.64 | 0.10 | 1 | D630003M21Ri | RIKEN cDNA D630003M21 gene | 738 | 374 | 449 | 110 | 38 | 32 | 35 | 45 | 23 | 30 | 23 |
| 156 | -3.29 | 2.24 | -1.47 | 0.14 | 1 | Col12a1 | collagen, type XII, alpha 1 | 619 | 491 | 528 | 0 | 4 | 5 | 35 | 0 | 30 | 0 | 0 |
| 108 | -0.44 | 0.52 | -0.84 | 0.40 | 1 | Cpne2 | copine II | 399 | 162 | 255 | 60 | 58 | 55 | 35 | 24 | 70 | 18 | 55 |
| 103 | 0.00 | 0.71 | 0.00 | 1.00 | 1 | 4632427E13Rik | | 39 | 22 | 69 | 101 | 18 | 345 | 35 | 35 | 16 | 125 | 326 |
| 85 | -0.69 | 0.90 | -0.77 | 0.44 | 1 | Cklf | chemokine-like factor | 201 | 79 | 116 | 92 | 51 | 124 | 35 | 0 | 80 | 63 | 96 |
| 79 | -0.06 | 0.58 | -0.11 | 0.91 | 1 | Ntng2 | netrin G2 | 270 | 257 | 136 | 9 | 36 | 25 | 35 | 32 | 16 | 24 | 26 |
| 74 | 0.35 | 0.60 | 0.59 | 0.56 | 1 | Kif27 | kinesin family member 27 | 51 | 27 | 125 | 88 | 54 | 83 | 35 | 156 | 94 | 32 | 69 |
| 69 | -0.67 | 0.59 | -1.13 | 0.26 | 1 | Drc3 | dynein regulatory complex subunit 3 | 73 | 125 | 72 | 37 | 103 | 112 | 35 | 17 | 25 | 44 | 120 |
| 63 | 0.87 | 0.46 | 1.88 | 0.06 | 1 | Gm48435 | | 33 | 69 | 25 | 57 | 50 | 52 | 35 | 139 | 57 | 109 | 70 |
| 61 | -0.24 | 0.99 | -0.25 | 0.81 | 1 | Kif14 | kinesin family member 14 | 181 | 50 | 44 | 71 | 59 | 44 | 35 | 64 | 71 | 0 | 49 |
| 53 | 0.33 | 0.48 | 0.68 | 0.50 | 1 | 1500004A13Rik | | 37 | 96 | 101 | 57 | 38 | 25 | 35 | 72 | 50 | 52 | 25 |
| 52 | -0.19 | 0.85 | -0.23 | 0.82 | 1 | Lrrc23 | leucine rich repeat containing 23 | 124 | 198 | 77 | 3 | 25 | 24 | 35 | 25 | 11 | 5 | 51 |
| 50 | -0.59 | 0.70 | -0.83 | 0.40 | 1 | 4732440D04Rik | | 46 | 48 | 24 | 43 | 48 | 117 | 35 | 21 | 65 | 5 | 103 |
| 49 | 0.51 | 0.63 | 0.80 | 0.42 | 1 | Cenpm | centromere protein M | 57 | 31 | 31 | 46 | 43 | 40 | 35 | 133 | 58 | 22 | 45 |
| 43 | -0.07 | 0.77 | -0.10 | 0.92 | 1 | Efcab9 | EF-hand calcium binding domain 9 | 4 | 27 | 7 | 119 | 49 | 28 | 35 | 24 | 105 | 64 | 12 |
| 42 | 0.03 | 0.63 | 0.05 | 0.96 | 1 | Mthfs | 5, 10-methenyltetrahydrofolate synthetase | 45 | 63 | 82 | 18 | 27 | 56 | 35 | 35 | 16 | 55 | 29 |
| 41 | 0.21 | 0.63 | 0.34 | 0.73 | 1 | Cd59b | CD59b antigen | 34 | 56 | 37 | 39 | 27 | 36 | 35 | 36 | 91 | 18 | 35 |
| 38 | 0.41 | 0.63 | 0.65 | 0.52 | 1 | Cnih2 | cornichon family AMPA receptor auxiliary | 51 | 42 | 47 | 25 | 15 | 41 | 35 | 12 | 41 | 73 | 39 |
| 37 | -0.49 | 0.68 | -0.73 | 0.47 | 1 | Tmem116 | transmembrane protein 116 | 24 | 46 | 64 | 53 | 42 | 29 | 35 | 8 | 41 | 31 | 38 |
| 35 | -0.43 | 0.52 | -0.84 | 0.40 | 1 | 9530080O11Rik | | 10 | 12 | 14 | 53 | 41 | 67 | 35 | 27 | 37 | 27 | 57 |
| 33 | -0.04 | 0.67 | -0.05 | 0.96 | 1 | Lysmd1 | LysM, putative peptidoglycan-binding, do | 22 | 14 | 4 | 58 | 27 | 46 | 35 | 37 | 47 | 27 | 43 |
| 31 | -1.04 | 1.02 | -1.02 | 0.31 | 1 | Gm43848 | | 64 | 14 | 43 | 46 | 15 | 25 | 35 | 0 | 31 | 62 | 9 |
| 30 | -1.03 | 0.94 | -1.09 | 0.27 | 1 | Gm31432 | | 5 | 13 | 7 | 44 | 22 | 79 | 35 | 19 | 36 | 0 | 70 |
| 30 | -0.87 | 0.63 | -1.38 | 0.17 | 1 | Gm50039 | | 25 | 15 | 25 | 69 | 33 | 30 | 35 | 38 | 19 | 30 | 10 |
| 26 | -1.20 | 1.07 | -1.13 | 0.26 | 1 | Gpr165 | G protein-coupled receptor 165 | 4 | 64 | 44 | 20 | 30 | 15 | 35 | 34 | 18 | 0 | 21 |
| 23 | -0.71 | 0.77 | -0.92 | 0.36 | 1 | Gm45629 | | 6 | 7 | 23 | 47 | 26 | 26 | 35 | 36 | 13 | 14 | 22 |
| 23 | -2.33 | 1.52 | -1.53 | 0.13 | 1 | Col25a1 | collagen, type XXV, alpha 1 | 30 | 54 | 45 | 22 | 0 | 34 | 35 | 0 | 9 | 0 | 25 |
| 23 | -0.25 | 0.88 | -0.29 | 0.77 | 1 | Amd2 | S-adenosylmethionine decarboxylase 2 | 4 | 7 | 24 | 22 | 24 | 24 | 35 | 3 | 30 | 53 | 21 |
| 20 | -1.15 | 0.93 | -1.24 | 0.21 | 1 | Nudt17 | nudix (nucleoside diphosphate linked mo | 35 | 20 | 32 | 10 | 24 | 19 | 35 | 0 | 23 | 13 | 11 |
| 18 | -1.64 | 1.12 | -1.47 | 0.14 | 1 | Gm11642 | | 4 | 19 | 25 | 25 | 22 | 25 | 35 | 0 | 11 | 23 | 7 |
| 18 | -0.64 | 1.12 | -0.57 | 0.57 | 1 | H2-Q10 | histocompatibility 2, Q region locus 10 | 75 | 1 | 47 | 1 | 14 | 4 | 35 | 5 | 5 | 5 | 2 |
| 17 | 0.19 | 1.43 | 0.13 | 0.90 | 1 | Gm16175 | | 1 | 10 | 0 | 18 | 11 | 19 | 35 | 0 | 22 | 57 | 11 |
| 11 | -0.12 | 1.44 | -0.08 | 0.93 | 1 | Gm45640 | | 16 | 1 | 4 | 6 | 0 | 8 | 35 | 0 | 13 | 17 | 15 |
| 7 | -0.49 | 2.71 | -0.18 | 0.86 | 1 | Cyp2e1 | cytochrome P450, family 2, subfamily e, p | 9 | 9 | 22 | 0 | 0 | 0 | 35 | 0 | 4 | 0 | 0 |
| 1738 | -0.99 | 1.08 | -0.92 | 0.36 | 1 | Cuzd1 | CUB and zona pellucida-like domains 1 | 2710 | 9495 | 6484 | 323 | 7 | 2 | 34 | 23 | 8 | 14 | 16 |
| 1687 | -0.96 | 1.23 | -0.78 | 0.43 | 1 | Pltp | phospholipid transfer protein | 8975 | 4422 | 5075 | 0 | 9 | 7 | 34 | 0 | 1 | 27 | 5 |
| 964 | -1.13 | 0.79 | -1.43 | 0.15 | 1 | Zfp36l1 | zinc finger protein 36, C3H type-like 1 | 5182 | 2590 | 2638 | 9 | 59 | 25 | 34 | 0 | 37 | 18 | 12 |
| 167 | -2.29 | 1.31 | -1.75 | 0.08 | 1 | Lrrn4cl | LRRN4 C-terminal like | 824 | 310 | 577 | 11 | 24 | 19 | 34 | 24 | 8 | 0 | 0 |

| | | | | | | | | | | | | | | | | | | |
|---|---|---|---|---|---|---|---|---|---|---|---|---|---|---|---|---|---|---|
| 142 | 0.26 | 0.52 | 0.50 | 0.62 | 1 | Cdk6 | cyclin-dependent kinase 6 | 571 | 239 | 457 | 56 | 9 | 36 | 34 | 34 | 26 | 43 | 56 |
| 121 | 0.28 | 0.69 | 0.40 | 0.69 | 1 | Enpp3 | ectonucleotide pyrophosphatase/phosph | 467 | 379 | 289 | 7 | 11 | 30 | 34 | 0 | 24 | 43 | 43 |
| 105 | -0.80 | 0.74 | -1.07 | 0.28 | 1 | Slc27a3 | solute carrier family 27 (fatty acid transpo | 322 | 163 | 118 | 16 | 2 | 246 | 34 | 0 | 2 | 33 | 219 |
| 87 | 0.26 | 0.49 | 0.53 | 0.60 | 1 | Mocos | molybdenum cofactor sulfurase | 228 | 115 | 121 | 104 | 63 | 19 | 34 | 87 | 68 | 90 | 22 |
| 68 | 0.31 | 0.38 | 0.84 | 0.40 | 1 | Bbof1 | basal body orientation factor 1 | 28 | 70 | 57 | 67 | 56 | 102 | 34 | 85 | 61 | 41 | 145 |
| 68 | -0.09 | 0.54 | -0.16 | 0.87 | 1 | Gm10501 | | 57 | 44 | 49 | 90 | 26 | 172 | 34 | 34 | 58 | 52 | 130 |
| 67 | -0.01 | 0.67 | -0.01 | 0.99 | 1 | B3galt1 | UDP-Gal:betaGlcNAc beta 1,3-galactosyl | 75 | 162 | 43 | 95 | 46 | 35 | 34 | 118 | 90 | 6 | 29 |
| 58 | 0.15 | 0.44 | 0.33 | 0.74 | 1 | 6330408M09Rik | | 51 | 100 | 60 | 66 | 40 | 66 | 34 | 49 | 58 | 50 | 61 |
| 56 | 1.55 | 1.31 | 1.19 | 0.24 | 1 | Gm20559 | | 135 | 85 | 112 | 2 | 0 | 74 | 34 | 72 | 23 | 0 | 74 |
| 55 | 0.59 | 0.60 | 0.99 | 0.32 | 1 | Gm26691 | | 3 | 25 | 9 | 33 | 48 | 117 | 34 | 52 | 97 | 48 | 135 |
| 55 | -0.09 | 0.45 | -0.20 | 0.84 | 1 | Eef1akmt4 | EEF1A lysine methyltransferase 4 | 68 | 84 | 50 | 65 | 53 | 54 | 34 | 54 | 59 | 37 | 42 |
| 52 | 0.14 | 0.47 | 0.31 | 0.76 | 1 | Gm32926 | | 26 | 55 | 46 | 61 | 45 | 77 | 34 | 60 | 74 | 50 | 49 |
| 52 | 0.34 | 0.53 | 0.64 | 0.52 | 1 | Zfp454 | zinc finger protein 454 | 82 | 32 | 46 | 55 | 28 | 62 | 34 | 34 | 53 | 67 | 81 |
| 51 | -0.23 | 0.57 | -0.40 | 0.69 | 1 | 1700123M08Rik | | 89 | 34 | 72 | 75 | 42 | 46 | 34 | 32 | 59 | 39 | 43 |
| 49 | 1.08 | 0.61 | 1.77 | 0.08 | 1 | Gm8186 | | 121 | 59 | 54 | 12 | 40 | 15 | 34 | 78 | 29 | 64 | 38 |
| 47 | -0.50 | 0.61 | -0.81 | 0.42 | 1 | Rybp-ps | | 55 | 114 | 60 | 32 | 47 | 36 | 34 | 83 | 19 | 18 | 16 |
| 46 | 0.31 | 0.53 | 0.59 | 0.56 | 1 | Gm19196 | | 36 | 22 | 45 | 49 | 62 | 36 | 34 | 67 | 38 | 64 | 51 |
| 46 | 0.20 | 0.55 | 0.36 | 0.72 | 1 | 1700010I14Rik | RIKEN cDNA 1700010I14 gene | 28 | 87 | 29 | 32 | 45 | 52 | 34 | 50 | 53 | 30 | 62 |
| 45 | 0.05 | 0.57 | 0.09 | 0.93 | 1 | Gm13091 | | 24 | 18 | 8 | 48 | 54 | 76 | 34 | 28 | 54 | 62 | 85 |
| 41 | -0.62 | 0.73 | -0.86 | 0.39 | 1 | Pcdhb12 | protocadherin beta 12 | 43 | 29 | 92 | 60 | 24 | 53 | 34 | 20 | 12 | 55 | 31 |
| 39 | 0.82 | 0.88 | 0.93 | 0.35 | 1 | Rnf212b | ring finger protein 212B | 54 | 59 | 133 | 9 | 7 | 14 | 34 | 53 | 4 | 37 | 19 |
| 36 | 0.74 | 0.69 | 1.06 | 0.29 | 1 | 4930515G01Rik | | 17 | 3 | 18 | 28 | 39 | 36 | 34 | 87 | 62 | 32 | 44 |
| 33 | -0.32 | 0.63 | -0.51 | 0.61 | 1 | Gm6204 | | 54 | 49 | 38 | 60 | 27 | 8 | 34 | 41 | 10 | 31 | 15 |
| 33 | -1.16 | 0.91 | -1.28 | 0.20 | 1 | Gm8034 | | 15 | 16 | 60 | 32 | 53 | 51 | 34 | 0 | 25 | 32 | 43 |
| 33 | 0.36 | 0.58 | 0.63 | 0.53 | 1 | Gm21178 | | 31 | 30 | 16 | 26 | 42 | 23 | 34 | 36 | 47 | 36 | 40 |
| 31 | -0.09 | 0.61 | -0.14 | 0.89 | 1 | Gm5814 | | 21 | 48 | 38 | 35 | 34 | 19 | 34 | 23 | 24 | 48 | 20 |
| 31 | -1.75 | 0.97 | -1.80 | 0.07 | 1 | Scml2 | Scm polycomb group protein like 2 | 29 | 14 | 49 | 61 | 26 | 52 | 34 | 0 | 14 | 21 | 40 |
| 30 | -0.73 | 0.57 | -1.28 | 0.20 | 1 | Gm5586 | | 68 | 38 | 53 | 30 | 21 | 24 | 34 | 16 | 16 | 23 | 11 |
| 27 | 0.21 | 0.87 | 0.24 | 0.81 | 1 | Gm10767 | predicted gene 10767 | 3 | 24 | 8 | 15 | 23 | 50 | 34 | 7 | 36 | 39 | 61 |
| 25 | 0.19 | 0.68 | 0.28 | 0.78 | 1 | Gm31812 | | 13 | 24 | 37 | 22 | 31 | 10 | 34 | 20 | 30 | 31 | 23 |
| 23 | 1.14 | 1.18 | 0.96 | 0.33 | 1 | Tspear | thrombospondin type laminin G domain a | 9 | 15 | 28 | 11 | 0 | 15 | 34 | 74 | 13 | 44 | 7 |
| 14 | -0.91 | 1.04 | -0.88 | 0.38 | 1 | Gm13991 | | 27 | 14 | 20 | 4 | 10 | 13 | 34 | 5 | 16 | 1 | 7 |
| 13 | 0.15 | 0.92 | 0.16 | 0.87 | 1 | Gm15590 | | 8 | 12 | 5 | 6 | 9 | 5 | 34 | 10 | 6 | 35 | 7 |
| 273 | -0.17 | 1.03 | -0.17 | 0.87 | 1 | Rab3il1 | RAB3A interacting protein (rabin3)-like 1 | 1270 | 730 | 804 | 28 | 15 | 17 | 33 | 0 | 58 | 17 | 31 |
| 180 | -0.64 | 0.60 | -1.06 | 0.29 | 1 | Atp10d | ATPase, class V, type 10D | 592 | 594 | 525 | 74 | 47 | 12 | 33 | 13 | 9 | 59 | 25 |
| 164 | -0.48 | 0.57 | -0.84 | 0.40 | 1 | Adcy1 | adenylate cyclase 1 | 595 | 473 | 608 | 4 | 15 | 24 | 33 | 2 | 18 | 21 | 11 |
| 145 | -0.34 | 0.49 | -0.71 | 0.48 | 1 | Arhgap10 | Rho GTPase activating protein 10 | 554 | 305 | 350 | 82 | 68 | 30 | 33 | 79 | 32 | 23 | 34 |
| 137 | -0.27 | 0.62 | -0.43 | 0.66 | 1 | Cldn11 | claudin 11 | 389 | 561 | 419 | 21 | 4 | 12 | 33 | 43 | 8 | 10 | 5 |
| 123 | 0.35 | 0.45 | 0.79 | 0.43 | 1 | Irak3 | interleukin-1 receptor-associated kinase 3 | 384 | 245 | 376 | 59 | 21 | 39 | 33 | 58 | 46 | 62 | 30 |
| 101 | -0.21 | 0.49 | -0.43 | 0.67 | 1 | H2-K2 | | 174 | 44 | 51 | 40 | 56 | 309 | 33 | 27 | 42 | 48 | 287 |
| 101 | -1.48 | 0.98 | -1.51 | 0.13 | 1 | Ampd3 | adenosine monophosphate deaminase 3 | 430 | 310 | 266 | 9 | 4 | 25 | 33 | 0 | 16 | 0 | 15 |
| 90 | -0.17 | 0.45 | -0.39 | 0.70 | 1 | Vnn1 | vanin 1 | 232 | 164 | 219 | 74 | 73 | 22 | 33 | 60 | 52 | 31 | 27 |
| 87 | 0.61 | 0.61 | 1.00 | 0.32 | 1 | Mms22l | MMS22-like, DNA repair protein | 244 | 78 | 95 | 54 | 65 | 61 | 33 | 134 | 98 | 35 | 64 |
| 77 | -2.28 | 1.24 | -1.85 | 0.06 | 1 | Ptpn18 | protein tyrosine phosphatase, non-recept | 274 | 172 | 224 | 49 | 1 | 49 | 33 | 0 | 31 | 0 | 17 |
| 66 | 0.44 | 0.55 | 0.80 | 0.42 | 1 | Rap1gapos | | 60 | 70 | 68 | 84 | 36 | 49 | 33 | 191 | 42 | 63 | 27 |
| 65 | 0.28 | 0.49 | 0.56 | 0.57 | 1 | Gm45051 | | 55 | 60 | 86 | 73 | 73 | 48 | 33 | 116 | 75 | 22 | 78 |
| 57 | 0.28 | 0.48 | 0.57 | 0.57 | 1 | Dnajb3 | DnaJ heat shock protein family (Hsp40) m | 69 | 56 | 76 | 58 | 60 | 38 | 33 | 82 | 42 | 71 | 38 |
| 56 | -0.57 | 0.52 | -1.08 | 0.28 | 1 | Adamtsl5 | ADAMTS-like 5 | 224 | 168 | 100 | 19 | 10 | 9 | 33 | 9 | 7 | 22 | 9 |
| 49 | 0.15 | 0.57 | 0.26 | 0.80 | 1 | 4930405A21Rik | | 9 | 21 | 23 | 69 | 68 | 61 | 33 | 32 | 89 | 80 | 51 |
| 47 | -0.18 | 0.65 | -0.28 | 0.78 | 1 | Gm9616 | | 80 | 66 | 102 | 49 | 26 | 30 | 33 | 8 | 42 | 49 | 34 |
| 42 | 0.26 | 0.60 | 0.44 | 0.66 | 1 | Zfp992 | zinc finger protein 992 | 22 | 16 | 27 | 68 | 29 | 46 | 33 | 78 | 60 | 19 | 63 |
| 40 | 0.33 | 0.61 | 0.54 | 0.59 | 1 | Gm12359 | | 31 | 26 | 7 | 46 | 55 | 30 | 33 | 49 | 55 | 67 | 39 |
| 38 | 1.21 | 0.95 | 1.27 | 0.20 | 1 | Gm17511 | | 101 | 113 | 7 | 71 | 0 | 0 | 33 | 57 | 7 | 26 | 4 |
| 38 | -0.56 | 1.03 | -0.54 | 0.59 | 1 | Gm45218 | | 24 | 51 | 16 | 81 | 26 | 61 | 33 | 0 | 53 | 31 | 42 |
| 37 | 0.05 | 0.57 | 0.09 | 0.93 | 1 | C730034F03Rik | | 33 | 47 | 28 | 43 | 13 | 60 | 33 | 43 | 29 | 17 | 63 |
| 37 | 0.42 | 0.79 | 0.53 | 0.60 | 1 | 1700012D14Rik | | 42 | 28 | 3 | 14 | 12 | 104 | 33 | 42 | 27 | 13 | 84 |
| 33 | 0.42 | 0.96 | 0.44 | 0.66 | 1 | Stpg1 | sperm tail PG rich repeat containing 1 | 29 | 56 | 1 | 13 | 49 | 20 | 33 | 53 | 50 | 22 | 39 |
| 32 | -0.23 | 0.70 | -0.33 | 0.74 | 1 | Gm16299 | | 26 | 27 | 72 | 25 | 47 | 14 | 33 | 58 | 18 | 21 | 10 |
| 28 | -0.53 | 0.71 | -0.74 | 0.46 | 1 | Gm26812 | | 2 | 0 | 0 | 35 | 52 | 61 | 33 | 34 | 27 | 15 | 54 |
| 28 | 0.25 | 0.75 | 0.33 | 0.74 | 1 | E230029C05Rik | | 23 | 16 | 47 | 31 | 13 | 20 | 33 | 58 | 21 | 18 | 24 |

| | | | | | | | | | | | | | | | | | | |
|---|---|---|---|---|---|---|---|---|---|---|---|---|---|---|---|---|---|---|
| 28 | -0.42 | 0.82 | -0.52 | 0.61 | 1 | Man2c1os | | 23 | 26 | 14 | 30 | 27 | 42 | 33 | 38 | 39 | 3 | 28 |
| 27 | 0.24 | 0.74 | 0.32 | 0.75 | 1 | Ap4b1 | adaptor-related protein complex AP-4, be | 15 | 30 | 9 | 10 | 33 | 32 | 33 | 37 | 30 | 24 | 39 |
| 25 | -1.85 | 1.04 | -1.78 | 0.08 | 1 | Gm48420 | | 10 | 30 | 39 | 46 | 44 | 27 | 33 | 0 | 8 | 28 | 11 |
| 24 | -0.34 | 1.00 | -0.34 | 0.73 | 1 | Cldn34c1 | claudin 34C1 | 39 | 14 | 0 | 31 | 27 | 27 | 33 | 15 | 34 | 32 | 14 |
| 23 | -0.24 | 1.08 | -0.23 | 0.82 | 1 | Gm14168 | | 48 | 35 | 26 | 7 | 5 | 30 | 33 | 0 | 33 | 13 | 21 |
| 22 | 0.24 | 0.68 | 0.36 | 0.72 | 1 | Gm14584 | | 13 | 14 | 14 | 22 | 10 | 26 | 33 | 20 | 24 | 23 | 40 |
| 20 | -2.25 | 1.28 | -1.76 | 0.08 | 1 | Gm45930 | | 5 | 49 | 3 | 60 | 23 | 13 | 33 | 1 | 13 | 0 | 16 |
| 18 | 0.34 | 1.10 | 0.31 | 0.76 | 1 | Klhl30 | kelch-like 30 | 37 | 9 | 4 | 11 | 16 | 4 | 33 | 0 | 30 | 43 | 14 |
| 17 | -1.89 | 1.05 | -1.81 | 0.07 | 1 | H2bu1-ps | | 15 | 31 | 27 | 16 | 18 | 18 | 33 | 0 | 11 | 13 | 3 |
| 16 | -1.44 | 1.64 | -0.88 | 0.38 | 1 | Sgca | sarcoglycan, alpha (dystrophin-associate | 46 | 9 | 18 | 26 | 0 | 7 | 33 | 14 | 0 | 0 | 17 |
| 824 | -1.49 | 0.77 | -1.93 | 0.05 | 1 | Crim1 | cysteine rich transmembrane BMP regula | 3409 | 2796 | 2656 | 42 | 39 | 33 | 32 | 0 | 25 | 19 | 15 |
| 656 | 0.19 | 0.46 | 0.41 | 0.68 | 1 | Sema3f | sema domain, immunoglobulin domain (I | 2643 | 2132 | 2261 | 10 | 28 | 14 | 32 | 36 | 23 | 5 | 29 |
| 214 | 0.09 | 0.48 | 0.18 | 0.86 | 1 | Nrros | negative regulator of reactive oxygen spe | 997 | 488 | 406 | 50 | 96 | 46 | 32 | 68 | 68 | 30 | 68 |
| 201 | -0.67 | 0.82 | -0.81 | 0.42 | 1 | Fes | feline sarcoma oncogene | 874 | 526 | 666 | 0 | 16 | 42 | 32 | 0 | 11 | 23 | 17 |
| 159 | -0.95 | 0.49 | -1.94 | 0.05 | 1 | Cacna1g | calcium channel, voltage-dependent, T ty | 631 | 483 | 326 | 48 | 43 | 68 | 32 | 30 | 7 | 54 | 23 |
| 147 | 0.20 | 0.63 | 0.31 | 0.75 | 1 | Vamp5 | vesicle-associated membrane protein 5 | 521 | 312 | 500 | 12 | 53 | 22 | 32 | 89 | 19 | 43 | 9 |
| 143 | 0.12 | 0.67 | 0.17 | 0.86 | 1 | B4galnt1 | beta-1,4-N-acetyl-galactosaminyl transfer | 659 | 373 | 434 | 4 | 6 | 15 | 32 | 20 | 20 | 8 | 5 |
| 133 | -0.27 | 0.58 | -0.48 | 0.63 | 1 | Vegfd | vascular endothelial growth factor D | 835 | 126 | 307 | 38 | 14 | 14 | 32 | 29 | 10 | 54 | 7 |
| 118 | 0.28 | 0.37 | 0.77 | 0.44 | 1 | Col28a1 | collagen, type XXVIII, alpha 1 | 324 | 349 | 309 | 17 | 22 | 70 | 32 | 24 | 24 | 35 | 90 |
| 118 | 0.81 | 0.53 | 1.51 | 0.13 | 1 | Pclaf | PCNA clamp associated factor | 606 | 148 | 127 | 47 | 30 | 38 | 32 | 107 | 53 | 66 | 44 |
| 109 | -0.56 | 0.57 | -0.98 | 0.33 | 1 | 9530082P21Rik | | 205 | 276 | 401 | 76 | 33 | 45 | 32 | 13 | 19 | 45 | 55 |
| 108 | 0.00 | 0.40 | 0.01 | 0.99 | 1 | Mycbpap | MYCBP associated protein | 171 | 246 | 238 | 45 | 48 | 154 | 32 | 64 | 30 | 53 | 104 |
| 102 | -0.39 | 0.39 | -0.99 | 0.32 | 1 | Rinl | Ras and Rab interactor-like | 226 | 263 | 265 | 50 | 76 | 54 | 32 | 54 | 32 | 30 | 44 |
| 74 | 0.35 | 0.54 | 0.64 | 0.52 | 1 | Gm16001 | | 56 | 98 | 82 | 74 | 46 | 73 | 32 | 205 | 66 | 17 | 65 |
| 74 | 1.10 | 0.97 | 1.13 | 0.26 | 1 | Cldn1 | claudin 1 | 486 | 77 | 104 | 10 | 5 | 13 | 32 | 41 | 23 | 0 | 24 |
| 66 | -0.36 | 0.46 | -0.78 | 0.44 | 1 | Dancr | | 132 | 143 | 111 | 32 | 56 | 66 | 32 | 27 | 39 | 18 | 70 |
| 59 | -0.61 | 0.68 | -0.89 | 0.37 | 1 | Gm3325 | | 146 | 83 | 28 | 94 | 60 | 47 | 32 | 17 | 35 | 43 | 68 |
| 51 | -0.05 | 0.62 | -0.08 | 0.94 | 1 | Gtse1 | G two S phase expressed protein 1 | 138 | 110 | 47 | 18 | 38 | 41 | 32 | 57 | 19 | 22 | 37 |
| 50 | 0.88 | 0.68 | 1.29 | 0.20 | 1 | Gm16675 | | 31 | 83 | 23 | 30 | 48 | 30 | 32 | 34 | 56 | 139 | 49 |
| 43 | -0.22 | 0.65 | -0.33 | 0.74 | 1 | Lrrtm4 | leucine rich repeat transmembrane neurc | 64 | 74 | 37 | 36 | 42 | 46 | 32 | 9 | 55 | 41 | 36 |
| 43 | 0.80 | 0.75 | 1.07 | 0.29 | 1 | Slc35f4 | solute carrier family 35, member F4 | 31 | 10 | 17 | 93 | 5 | 34 | 32 | 120 | 21 | 57 | 52 |
| 38 | -0.11 | 0.56 | -0.20 | 0.84 | 1 | Gm30015 | | 7 | 41 | 22 | 60 | 34 | 43 | 32 | 73 | 39 | 15 | 49 |
| 37 | -0.38 | 0.59 | -0.64 | 0.52 | 1 | 9430021M05Rik | | 36 | 64 | 64 | 42 | 27 | 32 | 32 | 32 | 12 | 36 | 24 |
| 36 | -0.54 | 1.22 | -0.44 | 0.66 | 1 | Gm14286 | | 21 | 20 | 38 | 0 | 14 | 145 | 32 | 17 | 12 | 0 | 91 |
| 35 | -0.97 | 0.92 | -1.05 | 0.29 | 1 | Camk4 | calcium/calmodulin-dependent protein kir | 94 | 66 | 82 | 18 | 4 | 34 | 32 | 0 | 4 | 28 | 27 |
| 35 | -1.02 | 0.68 | -1.51 | 0.13 | 1 | Sytl3 | synaptotagmin-like 3 | 33 | 67 | 58 | 21 | 27 | 68 | 32 | 15 | 33 | 5 | 24 |
| 35 | 0.48 | 0.76 | 0.63 | 0.53 | 1 | 3010003L21Rik | | 47 | 19 | 42 | 23 | 21 | 27 | 32 | 95 | 13 | 46 | 16 |
| 34 | -0.03 | 0.53 | -0.06 | 0.95 | 1 | Tes3-ps | | 30 | 20 | 42 | 20 | 60 | 35 | 32 | 30 | 41 | 40 | 25 |
| 34 | 0.20 | 0.74 | 0.28 | 0.78 | 1 | Dnmt3bos | | 1 | 14 | 11 | 53 | 54 | 30 | 32 | 35 | 42 | 66 | 36 |
| 34 | -0.41 | 0.65 | -0.63 | 0.53 | 1 | Sap18b | Sin3-associated polypeptide 18B | 39 | 23 | 51 | 25 | 60 | 25 | 32 | 6 | 32 | 48 | 31 |
| 34 | -0.57 | 0.62 | -0.92 | 0.36 | 1 | Gm6252 | | 63 | 69 | 62 | 16 | 38 | 19 | 32 | 8 | 23 | 18 | 21 |
| 32 | -0.73 | 0.92 | -0.79 | 0.43 | 1 | C230096K16Rik | | 21 | 33 | 25 | 15 | 41 | 63 | 32 | 25 | 27 | 0 | 73 |
| 31 | -1.35 | 1.11 | -1.22 | 0.22 | 1 | Gm13092 | | 66 | 85 | 11 | 86 | 2 | 14 | 32 | 0 | 3 | 40 | 4 |
| 30 | -0.45 | 0.71 | -0.63 | 0.53 | 1 | Gm49870 | | 7 | 10 | 28 | 46 | 20 | 61 | 32 | 43 | 38 | 14 | 29 |
| 29 | -1.11 | 0.97 | -1.14 | 0.25 | 1 | Pnma1 | paraneoplastic antigen MA1 | 19 | 41 | 57 | 47 | 43 | 14 | 32 | 1 | 8 | 36 | 23 |
| 29 | 0.23 | 0.73 | 0.31 | 0.75 | 1 | Gm4149 | | 57 | 25 | 25 | 27 | 30 | 6 | 32 | 70 | 26 | 15 | 7 |
| 28 | -0.12 | 0.76 | -0.16 | 0.87 | 1 | Gm9847 | | 15 | 22 | 25 | 51 | 17 | 16 | 32 | 46 | 10 | 64 | 7 |
| 28 | -1.48 | 0.88 | -1.69 | 0.09 | 1 | Gm15542 | | 20 | 44 | 16 | 22 | 26 | 76 | 32 | 3 | 28 | 3 | 35 |
| 25 | 0.82 | 0.88 | 0.93 | 0.35 | 1 | Rps13-ps2 | | 23 | 21 | 59 | 12 | 4 | 10 | 32 | 36 | 19 | 46 | 7 |
| 24 | -0.49 | 0.78 | -0.62 | 0.53 | 1 | Fbl-ps2 | | 15 | 29 | 56 | 15 | 18 | 26 | 32 | 19 | 22 | 5 | 28 |
| 22 | -0.55 | 1.11 | -0.50 | 0.62 | 1 | 1700102P08Ril | RIKEN cDNA 1700102P08 gene | 7 | 12 | 14 | 7 | 4 | 70 | 32 | 16 | 9 | 0 | 66 |
| 21 | -0.77 | 1.07 | -0.72 | 0.47 | 1 | BC001981 | | 20 | 62 | 20 | 18 | 21 | 8 | 32 | 15 | 26 | 0 | 10 |
| 15 | 0.40 | 0.87 | 0.46 | 0.65 | 1 | Gm16090 | | 9 | 3 | 0 | 17 | 8 | 15 | 32 | 25 | 14 | 30 | 15 |
| 14 | -1.49 | 1.24 | -1.20 | 0.23 | 1 | Gm18939 | | 0 | 8 | 10 | 47 | 7 | 6 | 32 | 25 | 10 | 1 | 6 |
| 13 | -1.64 | 1.30 | -1.26 | 0.21 | 1 | Gm20056 | | 28 | 8 | 5 | 2 | 33 | 8 | 32 | 0 | 15 | 1 | 9 |
| 11 | -0.49 | 0.86 | -0.57 | 0.57 | 1 | Galnt2 | polypeptide N-acetylgalactosaminyltransf | 6 | 3 | 6 | 11 | 10 | 9 | 32 | 10 | 10 | 15 | 6 |
| 9 | -2.17 | 2.18 | -1.00 | 0.32 | 1 | Tmem72 | transmembrane protein 72 | 0 | 17 | 0 | 3 | 20 | 8 | 32 | 0 | 12 | 0 | 10 |
| 3731 | 0.84 | 0.63 | 1.33 | 0.18 | 1 | Igfbp4 | insulin-like growth factor binding protein 4 | 24096 | 8841 | 7882 | 6 | 24 | 17 | 31 | 64 | 13 | 46 | 22 |
| 2724 | -1.83 | 1.26 | -1.45 | 0.15 | 1 | Pecam1 | platelet/endothelial cell adhesion molecu | 9960 | 10079 | 9864 | 11 | 0 | 11 | 31 | 3 | 4 | 0 | 4 |
| 474 | 0.42 | 0.39 | 1.07 | 0.28 | 1 | Gbp2 | guanylate binding protein 2 | 2207 | 1403 | 1368 | 31 | 4 | 40 | 31 | 32 | 19 | 24 | 58 |

| | | | | | | | | | | | | | | | | | | |
|---|---|---|---|---|---|---|---|---|---|---|---|---|---|---|---|---|---|---|
| 300 | 0.67 | 0.44 | 1.50 | 0.13 | 1 | Sdc1 | syndecan 1 | 1519 | 700 | 669 | 50 | 24 | 44 | 31 | 125 | 42 | 70 | 24 |
| 258 | -0.26 | 1.13 | -0.23 | 0.82 | 1 | Fat4 | FAT atypical cadherin 4 | 1023 | 779 | 875 | 10 | 5 | 25 | 31 | 0 | 36 | 4 | 52 |
| 193 | 0.58 | 0.39 | 1.49 | 0.14 | 1 | Gm12940 | | 111 | 121 | 130 | 39 | 12 | 702 | 31 | 32 | 46 | 53 | 845 |
| 164 | -0.90 | 0.46 | -1.96 | 0.05 | 1 | Col8a1 | collagen, type VIII, alpha 1 | 679 | 366 | 627 | 12 | 26 | 16 | 31 | 6 | 26 | 9 | 7 |
| 158 | -1.23 | 0.78 | -1.58 | 0.11 | 1 | Stbd1 | starch binding domain 1 | 891 | 202 | 416 | 34 | 30 | 61 | 31 | 41 | 10 | 0 | 23 |
| 86 | -0.82 | 0.88 | -0.93 | 0.35 | 1 | Plcd1 | phospholipase C, delta 1 | 342 | 229 | 175 | 4 | 47 | 39 | 31 | 46 | 12 | 1 | 19 |
| 83 | 0.69 | 0.45 | 1.53 | 0.13 | 1 | Xaf1 | XIAP associated factor 1 | 253 | 172 | 216 | 18 | 20 | 36 | 31 | 23 | 66 | 46 | 37 |
| 82 | -0.41 | 0.47 | -0.87 | 0.38 | 1 | Catsper2 | cation channel, sperm associated 2 | 39 | 52 | 34 | 45 | 46 | 318 | 31 | 39 | 50 | 10 | 238 |
| 76 | -0.03 | 0.59 | -0.06 | 0.95 | 1 | Cldn2 | claudin 2 | 71 | 119 | 181 | 54 | 82 | 67 | 31 | 21 | 48 | 99 | 62 |
| 76 | 0.08 | 0.49 | 0.17 | 0.86 | 1 | Cenpl | centromere protein L | 131 | 84 | 187 | 67 | 60 | 47 | 31 | 90 | 68 | 32 | 36 |
| 66 | -0.39 | 0.54 | -0.73 | 0.46 | 1 | Rps24-ps3 | | 153 | 142 | 121 | 79 | 30 | 39 | 31 | 23 | 31 | 59 | 23 |
| 61 | -1.54 | 0.88 | -1.75 | 0.08 | 1 | Minar2 | membrane integral NOTCH2 associated r | 95 | 213 | 164 | 71 | 27 | 17 | 31 | 0 | 13 | 12 | 27 |
| 55 | 0.55 | 0.60 | 0.92 | 0.36 | 1 | Ankrd34a | ankyrin repeat domain 34A | 100 | 101 | 80 | 80 | 12 | 14 | 31 | 50 | 46 | 82 | 13 |
| 53 | -1.19 | 0.74 | -1.61 | 0.11 | 1 | Ngb | neuroglobin | 131 | 76 | 165 | 28 | 57 | 31 | 31 | 21 | 2 | 14 | 30 |
| 49 | -0.42 | 0.55 | -0.77 | 0.44 | 1 | 1110035H17Rik | | 45 | 73 | 54 | 67 | 60 | 47 | 31 | 36 | 25 | 62 | 31 |
| 47 | 0.07 | 0.75 | 0.10 | 0.92 | 1 | Cd40 | CD40 antigen | 154 | 71 | 72 | 17 | 40 | 17 | 31 | 63 | 14 | 10 | 28 |
| 45 | 0.38 | 0.67 | 0.56 | 0.58 | 1 | Grip2 | glutamate receptor interacting protein 2 | 94 | 23 | 151 | 26 | 14 | 19 | 31 | 22 | 35 | 63 | 17 |
| 45 | 0.11 | 0.71 | 0.15 | 0.88 | 1 | 1700001G17Rik | | 12 | 29 | 62 | 88 | 47 | 30 | 31 | 67 | 17 | 57 | 49 |
| 44 | -0.50 | 0.69 | -0.72 | 0.47 | 1 | Gm12966 | | 49 | 46 | 48 | 35 | 55 | 68 | 31 | 3 | 53 | 41 | 55 |
| 42 | -0.49 | 0.79 | -0.63 | 0.53 | 1 | Hectd2os | | 21 | 46 | 27 | 67 | 49 | 45 | 31 | 44 | 85 | 1 | 44 |
| 41 | 0.16 | 1.16 | 0.14 | 0.89 | 1 | Ank1 | ankyrin 1, erythroid | 118 | 76 | 56 | 0 | 41 | 11 | 31 | 0 | 31 | 75 | 7 |
| 39 | -1.16 | 0.80 | -1.44 | 0.15 | 1 | Gm28172 | | 102 | 53 | 94 | 38 | 26 | 13 | 31 | 1 | 18 | 44 | 5 |
| 35 | -0.33 | 0.56 | -0.59 | 0.56 | 1 | Gm5835 | | 63 | 24 | 43 | 31 | 43 | 34 | 31 | 17 | 24 | 50 | 27 |
| 32 | -0.58 | 0.84 | -0.69 | 0.49 | 1 | Gm49494 | | 17 | 28 | 0 | 68 | 36 | 42 | 31 | 37 | 18 | 22 | 48 |
| 31 | -0.85 | 1.04 | -0.82 | 0.41 | 1 | Cbln4 | cerebellin 4 precursor protein | 122 | 51 | 37 | 11 | 12 | 30 | 31 | 35 | 8 | 0 | 8 |
| 31 | -0.48 | 0.63 | -0.77 | 0.44 | 1 | Gm38560 | | 16 | 41 | 9 | 40 | 43 | 36 | 31 | 60 | 18 | 26 | 21 |
| 29 | -0.23 | 0.93 | -0.24 | 0.81 | 1 | A230056P14Rik | | 36 | 36 | 53 | 10 | 19 | 37 | 31 | 60 | 2 | 14 | 25 |
| 29 | 0.72 | 0.61 | 1.18 | 0.24 | 1 | Gm12174 | | 39 | 44 | 26 | 12 | 20 | 15 | 31 | 27 | 20 | 58 | 28 |
| 29 | -2.02 | 1.20 | -1.68 | 0.09 | 1 | 1700094D03Ril | RIKEN cDNA 1700094D03 gene | 20 | 0 | 42 | 4 | 19 | 92 | 31 | 0 | 20 | 0 | 86 |
| 28 | -1.24 | 0.91 | -1.36 | 0.17 | 1 | D830025C05Rik | | 19 | 58 | 37 | 37 | 28 | 28 | 31 | 15 | 23 | 0 | 37 |
| 27 | -0.73 | 1.27 | -0.58 | 0.56 | 1 | Shisa6 | shisa family member 6 | 42 | 111 | 57 | 0 | 12 | 11 | 31 | 0 | 12 | 6 | 12 |
| 26 | -0.28 | 0.60 | -0.47 | 0.64 | 1 | Gm10608 | | 23 | 31 | 30 | 41 | 31 | 13 | 31 | 28 | 22 | 21 | 19 |
| 26 | -0.32 | 0.90 | -0.35 | 0.72 | 1 | Mir1905 | | 6 | 0 | 0 | 26 | 65 | 34 | 31 | 35 | 35 | 36 | 20 |
| 25 | -0.88 | 1.04 | -0.84 | 0.40 | 1 | Slit1 | slit guidance ligand 1 | 25 | 36 | 11 | 11 | 54 | 31 | 31 | 0 | 21 | 27 | 31 |
| 25 | -0.63 | 0.74 | -0.85 | 0.39 | 1 | 4930597O21Rik | | 9 | 6 | 9 | 44 | 30 | 36 | 31 | 10 | 11 | 50 | 35 |
| 25 | 1.70 | 1.44 | 1.18 | 0.24 | 1 | Cntd1 | cyclin N-terminal domain containing 1 | 0 | 33 | 0 | 16 | 0 | 45 | 31 | 16 | 14 | 45 | 69 |
| 24 | -0.27 | 0.63 | -0.43 | 0.67 | 1 | 1700084E18Rik | | 17 | 34 | 19 | 30 | 26 | 16 | 31 | 32 | 26 | 15 | 14 |
| 23 | 0.44 | 1.39 | 0.31 | 0.75 | 1 | 4933440M02Rik | | 6 | 0 | 0 | 26 | 20 | 36 | 31 | 67 | 25 | 0 | 45 |
| 21 | 0.71 | 0.89 | 0.80 | 0.42 | 1 | Gm10570 | | 20 | 26 | 7 | 5 | 21 | 10 | 31 | 42 | 26 | 13 | 26 |
| 21 | -1.57 | 1.31 | -1.20 | 0.23 | 1 | Kcnj4 | potassium inwardly-rectifying channel, su | 44 | 69 | 36 | 4 | 5 | 12 | 31 | 0 | 6 | 0 | 20 |
| 17 | -0.33 | 1.23 | -0.27 | 0.79 | 1 | Gm10179 | | 4 | 31 | 16 | 11 | 15 | 9 | 31 | 0 | 13 | 44 | 7 |
| 13 | -1.20 | 1.07 | -1.11 | 0.27 | 1 | Gm8399 | | 30 | 5 | 29 | 15 | 2 | 8 | 31 | 6 | 5 | 8 | 3 |
| 840 | -3.97 | 2.98 | -1.33 | 0.18 | 1 | Tek | TEK receptor tyrosine kinase | 3200 | 2951 | 3053 | 0 | 0 | 5 | 30 | 0 | 0 | 0 | 0 |
| 687 | -1.41 | 0.98 | -1.43 | 0.15 | 1 | Sema5a | sema domain, seven thrombospondin rep | 3256 | 1779 | 2198 | 81 | 53 | 50 | 30 | 71 | 12 | 0 | 27 |
| 287 | -1.09 | 0.99 | -1.10 | 0.27 | 1 | Cebpa | CCAAT/enhancer binding protein (C/EBP | 1298 | 695 | 997 | 30 | 29 | 10 | 30 | 10 | 15 | 0 | 39 |
| 247 | -0.49 | 0.40 | -1.22 | 0.22 | 1 | Kank3 | KN motif and ankyrin repeat domains 3 | 742 | 707 | 895 | 60 | 55 | 68 | 30 | 59 | 47 | 9 | 45 |
| 181 | 0.97 | 0.76 | 1.27 | 0.21 | 1 | Nsg2 | neuron specific gene family member 2 | 491 | 700 | 667 | 27 | 0 | 2 | 30 | 29 | 14 | 18 | 16 |
| 178 | -0.48 | 0.64 | -0.76 | 0.45 | 1 | Tgfb1i1 | transforming growth factor beta 1 induced | 798 | 506 | 489 | 13 | 16 | 20 | 30 | 0 | 8 | 57 | 17 |
| 172 | 0.86 | 0.45 | 1.90 | 0.06 | 1 | Csgalnact1 | chondroitin sulfate N-acetylgalactosaminy | 528 | 595 | 471 | 26 | 12 | 33 | 30 | 70 | 23 | 82 | 23 |
| 162 | 0.69 | 0.44 | 1.58 | 0.11 | 1 | Abcb1a | ATP-binding cassette, sub-family B (MDR | 479 | 580 | 404 | 15 | 38 | 36 | 30 | 74 | 42 | 52 | 31 |
| 105 | -0.15 | 0.52 | -0.30 | 0.77 | 1 | Mlkl | mixed lineage kinase domain-like | 299 | 207 | 230 | 80 | 45 | 49 | 30 | 58 | 33 | 98 | 20 |
| 99 | 0.19 | 0.46 | 0.41 | 0.68 | 1 | Sh3bp2 | SH3-domain binding protein 2 | 371 | 180 | 216 | 61 | 37 | 25 | 30 | 64 | 37 | 36 | 35 |
| 84 | 0.41 | 0.52 | 0.78 | 0.44 | 1 | Gm6136 | | 170 | 88 | 138 | 66 | 59 | 56 | 30 | 58 | 71 | 147 | 41 |
| 78 | 0.32 | 0.48 | 0.66 | 0.51 | 1 | Setd4 | SET domain containing 4 | 75 | 70 | 101 | 26 | 44 | 197 | 30 | 71 | 28 | 44 | 176 |
| 73 | 0.09 | 0.85 | 0.11 | 0.91 | 1 | Cmya5 | cardiomyopathy associated 5 | 198 | 187 | 106 | 23 | 26 | 50 | 30 | 0 | 25 | 99 | 58 |
| 72 | 0.24 | 0.49 | 0.48 | 0.63 | 1 | Gm50010 | | 82 | 45 | 120 | 33 | 55 | 140 | 30 | 52 | 31 | 55 | 146 |
| 68 | 0.78 | 0.67 | 1.16 | 0.25 | 1 | Prox2 | prospero homeobox 2 | 87 | 37 | 51 | 39 | 43 | 85 | 30 | 218 | 30 | 59 | 70 |
| 65 | 0.14 | 0.54 | 0.26 | 0.80 | 1 | Gm45902 | | 62 | 74 | 60 | 29 | 28 | 156 | 30 | 21 | 17 | 77 | 159 |
| 63 | 0.17 | 0.57 | 0.30 | 0.76 | 1 | 4921507P07Ril | RIKEN cDNA 4921507P07 gene | 83 | 49 | 49 | 111 | 56 | 45 | 30 | 46 | 78 | 91 | 53 |
| 61 | -1.46 | 0.92 | -1.58 | 0.11 | 1 | Top1mt | DNA topoisomerase 1, mitochondrial | 151 | 103 | 180 | 62 | 34 | 28 | 30 | 0 | 8 | 61 | 16 |

| | | | | | | | | | | | | | | | | | | |
|---|---|---|---|---|---|---|---|---|---|---|---|---|---|---|---|---|---|---|
| 61 | 0.15 | 0.53 | 0.28 | 0.78 | 1 | Zscan30 | | 56 | 74 | 55 | 74 | 55 | 69 | 30 | 71 | 37 | 102 | 48 |
| 57 | -1.57 | 0.84 | -1.87 | 0.06 | 1 | Xndc1 | Xrcc1 N-terminal domain containing 1 | 55 | 54 | 21 | 48 | 41 | 187 | 30 | 9 | 23 | 0 | 162 |
| 53 | -0.77 | 0.85 | -0.92 | 0.36 | 1 | Usp43 | ubiquitin specific peptidase 43 | 129 | 149 | 123 | 13 | 28 | 38 | 30 | 32 | 24 | 0 | 17 |
| 47 | 0.40 | 0.45 | 0.88 | 0.38 | 1 | D630036H23Rik | | 51 | 72 | 51 | 40 | 43 | 36 | 30 | 61 | 58 | 33 | 46 |
| 46 | -0.36 | 0.54 | -0.66 | 0.51 | 1 | Rad51ap1 | RAD51 associated protein 1 | 61 | 92 | 32 | 40 | 43 | 59 | 30 | 36 | 23 | 19 | 73 |
| 46 | 0.03 | 0.55 | 0.06 | 0.96 | 1 | Kifc5b | kinesin family member C5B | 56 | 43 | 55 | 15 | 51 | 85 | 30 | 28 | 25 | 53 | 63 |
| 43 | -0.26 | 0.72 | -0.36 | 0.72 | 1 | Gm39326 | | 16 | 21 | 50 | 18 | 19 | 129 | 30 | 10 | 6 | 43 | 135 |
| 41 | -0.21 | 0.52 | -0.41 | 0.68 | 1 | Xrra1 | X-ray radiation resistance associated 1 | 31 | 38 | 30 | 61 | 61 | 38 | 30 | 29 | 59 | 39 | 36 |
| 39 | -0.50 | 0.64 | -0.79 | 0.43 | 1 | Gm4925 | predicted gene 4925 | 27 | 49 | 44 | 25 | 70 | 54 | 30 | 64 | 23 | 15 | 32 |
| 39 | 0.47 | 0.67 | 0.70 | 0.48 | 1 | Asb9 | ankyrin repeat and SOCS box-containing | 0 | 0 | 0 | 70 | 26 | 44 | 30 | 97 | 15 | 63 | 85 |
| 35 | -0.57 | 0.85 | -0.66 | 0.51 | 1 | Dnaaf2 | dynein, axonemal assembly factor 2 | 45 | 7 | 86 | 61 | 20 | 25 | 30 | 22 | 47 | 10 | 26 |
| 34 | -0.42 | 0.79 | -0.53 | 0.60 | 1 | Znf41-ps | | 51 | 20 | 17 | 51 | 27 | 43 | 30 | 7 | 46 | 17 | 67 |
| 34 | 1.31 | 0.73 | 1.80 | 0.07 | 1 | Gm19522 | | 19 | 48 | 26 | 7 | 9 | 45 | 30 | 43 | 31 | 67 | 49 |
| 33 | -0.07 | 0.66 | -0.11 | 0.91 | 1 | Gm13594 | | 27 | 53 | 38 | 31 | 13 | 54 | 30 | 13 | 36 | 24 | 45 |
| 32 | -0.57 | 1.20 | -0.47 | 0.64 | 1 | Gm43359 | | 6 | 0 | 0 | 11 | 3 | 153 | 30 | 6 | 6 | 0 | 140 |
| 31 | -0.22 | 0.68 | -0.32 | 0.75 | 1 | Fgl1 | fibrinogen-like protein 1 | 93 | 16 | 38 | 18 | 26 | 24 | 30 | 16 | 15 | 17 | 42 |
| 29 | -0.05 | 0.60 | -0.08 | 0.94 | 1 | Gm6192 | | 45 | 25 | 41 | 29 | 40 | 13 | 30 | 32 | 26 | 22 | 20 |
| 29 | 0.06 | 0.59 | 0.09 | 0.92 | 1 | Gm5914 | | 20 | 36 | 14 | 27 | 27 | 32 | 30 | 50 | 21 | 27 | 31 |
| 27 | 0.43 | 0.72 | 0.59 | 0.56 | 1 | Gm20036 | | 17 | 28 | 44 | 10 | 26 | 19 | 30 | 8 | 49 | 45 | 21 |
| 27 | -0.64 | 0.75 | -0.86 | 0.39 | 1 | Gm15952 | | 19 | 61 | 16 | 30 | 25 | 39 | 30 | 10 | 27 | 23 | 15 |
| 26 | 0.43 | 0.61 | 0.71 | 0.48 | 1 | Gm18284 | | 35 | 9 | 19 | 11 | 38 | 17 | 30 | 14 | 50 | 39 | 25 |
| 25 | -0.37 | 0.81 | -0.45 | 0.65 | 1 | Dhrs13os | | 14 | 18 | 17 | 13 | 41 | 37 | 30 | 46 | 39 | 5 | 17 |
| 25 | -0.55 | 0.84 | -0.66 | 0.51 | 1 | Gm6444 | | 46 | 8 | 14 | 55 | 19 | 19 | 30 | 12 | 29 | 17 | 23 |
| 25 | -0.89 | 0.72 | -1.25 | 0.21 | 1 | Gm10244 | | 26 | 27 | 34 | 13 | 38 | 33 | 30 | 12 | 17 | 5 | 36 |
| 25 | -0.01 | 0.76 | -0.01 | 0.99 | 1 | Gm12854 | | 74 | 24 | 52 | 4 | 9 | 14 | 30 | 1 | 15 | 22 | 23 |
| 23 | -1.66 | 1.17 | -1.42 | 0.16 | 1 | Gm9993 | | 5 | 3 | 69 | 32 | 23 | 34 | 30 | 0 | 12 | 30 | 18 |
| 23 | -0.44 | 0.74 | -0.59 | 0.55 | 1 | Lzts1 | leucine zipper, putative tumor suppressor | 39 | 27 | 69 | 16 | 9 | 10 | 30 | 8 | 12 | 21 | 8 |
| 23 | 0.73 | 1.44 | 0.50 | 0.61 | 1 | Mxd3 | Max dimerization protein 3 | 50 | 15 | 0 | 10 | 10 | 34 | 30 | 31 | 48 | 0 | 21 |
| 22 | -0.52 | 1.06 | -0.49 | 0.62 | 1 | Slc3a1 | solute carrier family 3, member 1 | 0 | 15 | 16 | 9 | 20 | 54 | 30 | 8 | 37 | 8 | 47 |
| 20 | -1.57 | 0.94 | -1.67 | 0.10 | 1 | 1700026J14Rik | | 12 | 18 | 30 | 14 | 52 | 19 | 30 | 9 | 27 | 0 | 11 |
| 20 | -1.75 | 1.21 | -1.45 | 0.15 | 1 | Rpl21-ps8 | | 27 | 69 | 25 | 12 | 14 | 10 | 30 | 0 | 25 | 1 | 4 |
| 17 | -0.13 | 1.03 | -0.12 | 0.90 | 1 | 1700028J19Rik | | 20 | 34 | 1 | 6 | 14 | 8 | 30 | 34 | 3 | 24 | 15 |
| 16 | -0.14 | 1.01 | -0.13 | 0.89 | 1 | Gm29216 | | 11 | 34 | 7 | 8 | 7 | 25 | 30 | 3 | 19 | 21 | 12 |
| 14 | -0.54 | 1.27 | -0.42 | 0.67 | 1 | Impg2 | interphotoreceptor matrix proteoglycan 2 | 12 | 5 | 3 | 3 | 2 | 60 | 30 | 14 | 4 | 0 | 20 |
| 13 | -0.08 | 1.08 | -0.07 | 0.94 | 1 | Fbxo40 | F-box protein 40 | 0 | 0 | 0 | 38 | 7 | 8 | 30 | 19 | 22 | 14 | 10 |
| 13 | -0.59 | 1.51 | -0.39 | 0.69 | 1 | Gm42903 | | 9 | 3 | 36 | 0 | 8 | 9 | 30 | 0 | 30 | 19 | 0 |
| 12 | -0.84 | 1.30 | -0.65 | 0.51 | 1 | Rpl6l | | 13 | 29 | 8 | 15 | 1 | 14 | 30 | 0 | 8 | 6 | 12 |
| 1355 | -0.38 | 3.54 | -0.11 | 0.91 | 1 | Lama4 | laminin, alpha 4 | 5830 | 4031 | 5003 | 0 | 0 | 0 | 29 | 0 | 12 | 0 | 0 |
| 522 | -1.21 | 0.82 | -1.49 | 0.14 | 1 | Ptn | pleiotrophin | 2638 | 955 | 1945 | 31 | 32 | 32 | 29 | 0 | 14 | 50 | 17 |
| 266 | -0.23 | 0.31 | -0.76 | 0.45 | 1 | Rps7-ps3 | | 26 | 480 | 850 | 220 | 409 | 115 | 29 | 258 | 432 | 13 | 88 |
| 227 | -0.42 | 0.34 | -1.25 | 0.21 | 1 | Mfap2 | microfibrillar-associated protein 2 | 700 | 707 | 771 | 74 | 39 | 44 | 29 | 44 | 25 | 33 | 34 |
| 134 | -0.18 | 0.55 | -0.33 | 0.74 | 1 | Syne3 | spectrin repeat containing, nuclear envelo | 434 | 384 | 242 | 58 | 74 | 46 | 29 | 20 | 39 | 109 | 41 |
| 110 | 0.02 | 0.50 | 0.03 | 0.97 | 1 | Dnase1l1 | deoxyribonuclease 1-like 1 | 363 | 257 | 235 | 31 | 61 | 55 | 29 | 70 | 22 | 52 | 41 |
| 105 | -0.83 | 0.69 | -1.21 | 0.23 | 1 | R74862 | | 172 | 204 | 135 | 93 | 95 | 143 | 29 | 44 | 66 | 1 | 178 |
| 76 | -0.66 | 1.14 | -0.58 | 0.56 | 1 | Gm38020 | | 21 | 16 | 9 | 3 | 5 | 407 | 29 | 0 | 33 | 0 | 311 |
| 73 | -0.36 | 0.48 | -0.75 | 0.45 | 1 | B3gnt8 | UDP-GlcNAc:betaGal beta-1,3-N-acetylgl | 252 | 160 | 183 | 24 | 46 | 14 | 29 | 12 | 42 | 27 | 11 |
| 71 | 0.23 | 0.60 | 0.39 | 0.70 | 1 | Gna14 | guanine nucleotide binding protein, alpha | 154 | 209 | 204 | 23 | 22 | 19 | 29 | 6 | 40 | 35 | 37 |
| 56 | -0.83 | 0.86 | -0.97 | 0.33 | 1 | Ddit4l | DNA-damage-inducible transcript 4-like | 164 | 132 | 120 | 43 | 16 | 36 | 29 | 0 | 26 | 35 | 18 |
| 55 | 0.24 | 0.50 | 0.48 | 0.63 | 1 | Dtwd2 | DTW domain containing 2 | 116 | 88 | 48 | 38 | 43 | 51 | 29 | 38 | 52 | 45 | 54 |
| 53 | 0.38 | 0.75 | 0.50 | 0.62 | 1 | Med9os | | 34 | 43 | 67 | 81 | 21 | 35 | 29 | 168 | 35 | 5 | 70 |
| 53 | -0.51 | 0.86 | -0.59 | 0.55 | 1 | Arhgef38 | Rho guanine nucleotide exchange factor | 78 | 107 | 118 | 20 | 27 | 70 | 29 | 63 | 25 | 0 | 40 |
| 49 | -0.02 | 0.54 | -0.04 | 0.97 | 1 | Oasl1 | 2'-5' oligoadenylate synthetase-like 1 | 117 | 78 | 113 | 35 | 33 | 14 | 29 | 14 | 34 | 54 | 15 |
| 47 | 0.26 | 0.47 | 0.56 | 0.58 | 1 | Gm45605 | | 51 | 65 | 65 | 46 | 42 | 36 | 29 | 67 | 31 | 45 | 42 |
| 42 | 0.02 | 0.95 | 0.02 | 0.98 | 1 | 4833422M21Rik | | 66 | 131 | 68 | 10 | 27 | 25 | 29 | 0 | 27 | 49 | 30 |
| 42 | 0.16 | 0.92 | 0.17 | 0.86 | 1 | Zfp951 | zinc finger protein 951 | 60 | 81 | 32 | 9 | 17 | 76 | 29 | 74 | 26 | 1 | 52 |
| 35 | -2.58 | 1.32 | -1.95 | 0.05 | 1 | Tcf7 | transcription factor 7, T cell specific | 243 | 43 | 54 | 6 | 0 | 4 | 29 | 0 | 1 | 0 | 4 |
| 34 | -0.12 | 0.77 | -0.16 | 0.87 | 1 | Gm42690 | | 46 | 5 | 16 | 18 | 72 | 43 | 29 | 41 | 43 | 44 | 15 |
| 33 | -0.30 | 0.63 | -0.48 | 0.63 | 1 | Gm26616 | | 23 | 13 | 35 | 50 | 36 | 41 | 29 | 41 | 25 | 17 | 50 |
| 33 | -0.84 | 0.62 | -1.36 | 0.17 | 1 | Rnf26 | ring finger protein 26 | 33 | 32 | 25 | 33 | 63 | 42 | 29 | 14 | 25 | 15 | 47 |
| 32 | -0.10 | 0.66 | -0.15 | 0.88 | 1 | Gm21092 | | 34 | 26 | 54 | 18 | 17 | 61 | 29 | 35 | 17 | 13 | 48 |

| | | | | | | | | | | | | | | | | | | |
|---|---|---|---|---|---|---|---|---|---|---|---|---|---|---|---|---|---|---|
| 32 | -0.59 | 0.61 | -0.97 | 0.33 | 1 | Crry-ps | | 94 | 67 | 28 | 24 | 27 | 16 | 29 | 22 | 12 | 19 | 12 |
| 30 | -0.02 | 0.81 | -0.03 | 0.98 | 1 | 4930412C18Rik | | 21 | 59 | 57 | 4 | 33 | 30 | 29 | 36 | 17 | 21 | 21 |
| 29 | -0.70 | 0.62 | -1.13 | 0.26 | 1 | Gm10073 | | 66 | 52 | 31 | 41 | 20 | 19 | 29 | 20 | 13 | 12 | 21 |
| 28 | -0.90 | 0.76 | -1.18 | 0.24 | 1 | Psenen-ps | | 25 | 55 | 27 | 38 | 43 | 22 | 29 | 5 | 35 | 14 | 18 |
| 28 | -0.04 | 0.65 | -0.06 | 0.95 | 1 | Gm7027 | | 19 | 40 | 43 | 26 | 21 | 29 | 29 | 28 | 17 | 40 | 19 |
| 27 | -1.58 | 0.91 | -1.74 | 0.08 | 1 | Gm42902 | | 35 | 51 | 60 | 21 | 25 | 30 | 29 | 0 | 9 | 23 | 10 |
| 26 | -0.02 | 1.43 | -0.01 | 0.99 | 1 | Gm21284 | | 0 | 48 | 47 | 34 | 9 | 23 | 29 | 0 | 21 | 39 | 37 |
| 25 | -1.03 | 0.97 | -1.06 | 0.29 | 1 | A930006K02Rik | | 29 | 18 | 37 | 4 | 60 | 36 | 29 | 20 | 23 | 1 | 13 |
| 23 | -1.56 | 0.91 | -1.71 | 0.09 | 1 | Rpl9-ps6 | | 32 | 73 | 26 | 29 | 15 | 17 | 29 | 2 | 14 | 3 | 13 |
| 23 | 0.07 | 0.84 | 0.09 | 0.93 | 1 | BC030343 | | 15 | 9 | 17 | 31 | 3 | 35 | 29 | 16 | 14 | 17 | 64 |
| 17 | -0.91 | 1.07 | -0.85 | 0.39 | 1 | Ap3m1-ps | | 23 | 22 | 20 | 12 | 14 | 20 | 29 | 8 | 25 | 0 | 13 |
| 17 | 0.10 | 1.36 | 0.07 | 0.94 | 1 | 3830403N18Ril | RIKEN cDNA 3830403N18 gene | 22 | 14 | 32 | 0 | 26 | 7 | 29 | 7 | 33 | 0 | 14 |
| 17 | -0.84 | 1.12 | -0.74 | 0.46 | 1 | Gm5500 | | 2 | 6 | 21 | 13 | 24 | 19 | 29 | 20 | 31 | 0 | 20 |
| 16 | -0.54 | 0.88 | -0.62 | 0.54 | 1 | Igf2 | insulin-like growth factor 2 | 7 | 4 | 2 | 25 | 24 | 17 | 29 | 3 | 30 | 21 | 16 |
| 16 | -1.68 | 1.24 | -1.35 | 0.18 | 1 | Gm20754 | | 21 | 1 | 22 | 28 | 5 | 25 | 29 | 0 | 10 | 26 | 12 |
| 16 | -0.14 | 1.32 | -0.11 | 0.91 | 1 | Gm6724 | | 58 | 0 | 0 | 2 | 0 | 13 | 29 | 1 | 0 | 63 | 7 |
| 15 | -0.25 | 1.08 | -0.23 | 0.82 | 1 | Gm4430 | | 0 | 3 | 0 | 16 | 31 | 8 | 29 | 29 | 9 | 15 | 27 |
| 14 | 0.14 | 0.89 | 0.15 | 0.88 | 1 | Gm6988 | | 20 | 31 | 11 | 8 | 3 | 8 | 29 | 9 | 5 | 17 | 14 |
| 9 | -0.39 | 1.58 | -0.25 | 0.80 | 1 | Gm42568 | | 15 | 4 | 8 | 6 | 0 | 13 | 29 | 15 | 2 | 0 | 9 |
| 6 | -1.25 | 1.88 | -0.66 | 0.51 | 1 | Gm50015 | | 0 | 0 | 6 | 5 | 14 | 1 | 29 | 7 | 3 | 0 | 3 |
| 1916 | -1.03 | 1.89 | -0.55 | 0.59 | 1 | Ctss | cathepsin S | 9837 | 4295 | 6839 | 17 | 0 | 8 | 28 | 0 | 39 | 0 | 9 |
| 916 | -1.13 | 0.79 | -1.42 | 0.15 | 1 | Ctsc | cathepsin C | 5418 | 2255 | 2341 | 7 | 7 | 7 | 28 | 7 | 6 | 0 | 4 |
| 305 | 0.14 | 0.58 | 0.25 | 0.80 | 1 | Smo | smoothened, frizzled class receptor | 1634 | 695 | 844 | 14 | 6 | 36 | 28 | 0 | 24 | 50 | 28 |
| 236 | 0.33 | 0.54 | 0.61 | 0.54 | 1 | Rhbdf1 | rhomboid 5 homolog 1 | 964 | 552 | 759 | 21 | 32 | 54 | 28 | 107 | 21 | 28 | 29 |
| 153 | -1.43 | 1.52 | -0.94 | 0.35 | 1 | Tlr7 | toll-like receptor 7 | 824 | 386 | 440 | 0 | 3 | 1 | 28 | 2 | 3 | 0 | 0 |
| 111 | 0.36 | 0.60 | 0.60 | 0.55 | 1 | Kif4 | kinesin family member 4 | 311 | 100 | 78 | 85 | 114 | 90 | 28 | 125 | 100 | 82 | 103 |
| 110 | -0.51 | 0.70 | -0.72 | 0.47 | 1 | Ccdc141 | coiled-coil domain containing 141 | 166 | 187 | 163 | 116 | 81 | 157 | 28 | 93 | 97 | 1 | 117 |
| 94 | 0.43 | 0.49 | 0.87 | 0.38 | 1 | Ccdc15 | coiled-coil domain containing 15 | 122 | 178 | 108 | 75 | 67 | 107 | 28 | 46 | 114 | 89 | 100 |
| 89 | -0.82 | 0.90 | -0.90 | 0.37 | 1 | Gm21781 | | 3 | 74 | 38 | 12 | 13 | 379 | 28 | 43 | 14 | 0 | 376 |
| 85 | -2.06 | 1.17 | -1.76 | 0.08 | 1 | Cd22 | CD22 antigen | 563 | 204 | 41 | 35 | 12 | 21 | 28 | 0 | 28 | 0 | 6 |
| 79 | -0.07 | 0.58 | -0.13 | 0.90 | 1 | Aurkb | aurora kinase B | 343 | 86 | 104 | 46 | 29 | 67 | 28 | 39 | 31 | 21 | 73 |
| 66 | 0.55 | 0.98 | 0.56 | 0.57 | 1 | Ercc6l | excision repair cross-complementing rode | 295 | 59 | 90 | 35 | 21 | 26 | 28 | 85 | 29 | 0 | 60 |
| 66 | -0.34 | 0.63 | -0.54 | 0.59 | 1 | 1700086P04Rik | | 31 | 112 | 127 | 84 | 50 | 59 | 28 | 64 | 102 | 3 | 66 |
| 66 | -0.69 | 0.47 | -1.48 | 0.14 | 1 | Cfap300 | cilia and flagella associated protein 300 | 75 | 144 | 120 | 70 | 77 | 56 | 28 | 42 | 30 | 14 | 67 |
| 65 | 0.31 | 0.84 | 0.37 | 0.71 | 1 | E2f7 | E2F transcription factor 7 | 117 | 45 | 11 | 54 | 34 | 106 | 28 | 8 | 97 | 162 | 54 |
| 56 | -0.13 | 0.59 | -0.22 | 0.83 | 1 | Gjc2 | gap junction protein, gamma 2 | 202 | 71 | 71 | 46 | 50 | 14 | 28 | 58 | 27 | 37 | 12 |
| 52 | 0.44 | 0.55 | 0.80 | 0.42 | 1 | BC034090 | cDNA sequence BC034090 | 172 | 90 | 89 | 22 | 28 | 14 | 28 | 44 | 15 | 36 | 32 |
| 50 | -0.14 | 0.90 | -0.15 | 0.88 | 1 | A330084C13Rik | | 63 | 71 | 92 | 84 | 26 | 19 | 28 | 97 | 40 | 0 | 32 |
| 46 | -0.41 | 0.90 | -0.46 | 0.65 | 1 | Kif18b | kinesin family member 18B | 101 | 64 | 60 | 41 | 33 | 47 | 28 | 51 | 38 | 0 | 38 |
| 45 | -0.13 | 0.65 | -0.20 | 0.84 | 1 | Ube2ql1 | ubiquitin-conjugating enzyme E2Q family | 107 | 78 | 141 | 32 | 12 | 16 | 28 | 23 | 7 | 32 | 20 |
| 44 | -0.88 | 0.99 | -0.88 | 0.38 | 1 | A930004D18Rik | | 76 | 155 | 81 | 51 | 6 | 28 | 28 | 0 | 24 | 15 | 15 |
| 43 | -0.23 | 0.73 | -0.32 | 0.75 | 1 | Trpa1 | transient receptor potential cation channe | 139 | 121 | 98 | 9 | 13 | 14 | 28 | 21 | 24 | 5 | 6 |
| 42 | 0.49 | 0.62 | 0.79 | 0.43 | 1 | Kynu | kynureninase | 93 | 40 | 12 | 63 | 16 | 30 | 28 | 46 | 30 | 53 | 52 |
| 41 | -1.21 | 0.84 | -1.43 | 0.15 | 1 | Klhl38 | kelch-like 38 | 112 | 83 | 89 | 18 | 28 | 35 | 28 | 24 | 12 | 0 | 16 |
| 40 | -1.27 | 0.82 | -1.55 | 0.12 | 1 | Gm32618 | | 84 | 82 | 79 | 18 | 43 | 41 | 28 | 10 | 27 | 0 | 28 |
| 40 | 0.29 | 0.47 | 0.61 | 0.54 | 1 | Gm15918 | | 20 | 30 | 44 | 35 | 48 | 42 | 28 | 50 | 61 | 36 | 42 |
| 39 | 0.35 | 0.75 | 0.47 | 0.64 | 1 | Gm10443 | | 57 | 37 | 57 | 51 | 9 | 26 | 28 | 14 | 34 | 90 | 25 |
| 39 | -0.85 | 0.81 | -1.06 | 0.29 | 1 | Rhebl1 | Ras homolog enriched in brain like 1 | 43 | 21 | 46 | 21 | 60 | 68 | 28 | 0 | 34 | 30 | 73 |
| 38 | -0.65 | 0.68 | -0.95 | 0.34 | 1 | Gm9892 | | 33 | 73 | 33 | 34 | 43 | 48 | 28 | 17 | 38 | 5 | 61 |
| 36 | -1.77 | 1.13 | -1.56 | 0.12 | 1 | Zfp811 | zinc finger protein 811 | 79 | 138 | 57 | 11 | 20 | 25 | 28 | 0 | 19 | 0 | 23 |
| 33 | 0.54 | 0.82 | 0.66 | 0.51 | 1 | B930018H19Rik | | 12 | 21 | 0 | 33 | 34 | 37 | 28 | 54 | 46 | 44 | 53 |
| 32 | 1.06 | 0.96 | 1.11 | 0.27 | 1 | Gm28557 | | 41 | 65 | 33 | 15 | 1 | 23 | 28 | 48 | 1 | 61 | 36 |
| 31 | -0.23 | 0.70 | -0.33 | 0.74 | 1 | Bdnf | brain derived neurotrophic factor | 31 | 24 | 65 | 38 | 23 | 30 | 28 | 22 | 14 | 49 | 21 |
| 31 | -0.02 | 0.69 | -0.03 | 0.97 | 1 | Gm7895 | | 10 | 41 | 19 | 23 | 44 | 41 | 28 | 24 | 51 | 37 | 22 |
| 31 | 0.79 | 0.67 | 1.17 | 0.24 | 1 | Gm13031 | | 12 | 22 | 19 | 19 | 24 | 29 | 28 | 52 | 15 | 72 | 45 |
| 28 | -0.30 | 1.14 | -0.26 | 0.79 | 1 | Spata24 | spermatogenesis associated 24 | 21 | 29 | 2 | 34 | 34 | 37 | 28 | 0 | 28 | 43 | 51 |
| 27 | 0.72 | 1.12 | 0.64 | 0.52 | 1 | Lsmem1 | leucine-rich single-pass membrane prote | 15 | 63 | 34 | 6 | 22 | 11 | 28 | 0 | 30 | 57 | 35 |
| 27 | 0.14 | 0.65 | 0.21 | 0.83 | 1 | Ugt1a9 | UDP glucuronosyltransferase 1 family, pc | 22 | 12 | 17 | 35 | 34 | 18 | 28 | 44 | 15 | 54 | 21 |
| 27 | 0.87 | 0.84 | 1.03 | 0.30 | 1 | Gm15816 | predicted gene 15816 | 15 | 6 | 0 | 19 | 12 | 19 | 28 | 93 | 6 | 53 | 47 |
| 26 | -0.09 | 0.74 | -0.13 | 0.90 | 1 | Cbfa2t2-ps1 | | 27 | 41 | 15 | 16 | 38 | 21 | 28 | 12 | 39 | 19 | 27 |

| | | | | | | | | | | | | | | | | | | |
|---|---|---|---|---|---|---|---|---|---|---|---|---|---|---|---|---|---|---|
| 25 | -0.52 | 1.05 | -0.49 | 0.62 | 1 | Akap5 | A kinase (PRKA) anchor protein 5 | 22 | 30 | 15 | 30 | 9 | 45 | 28 | 0 | 8 | 62 | 28 |
| 19 | -0.16 | 1.12 | -0.14 | 0.89 | 1 | Zmynd10 | zinc finger, MYND domain containing 10 | 1 | 4 | 0 | 31 | 10 | 14 | 28 | 43 | 0 | 35 | 46 |
| 18 | -0.23 | 1.19 | -0.19 | 0.85 | 1 | Gm29675 | | 6 | 20 | 23 | 12 | 6 | 19 | 28 | 54 | 19 | 0 | 14 |
| 17 | -0.62 | 1.11 | -0.56 | 0.58 | 1 | Gm11914 | | 23 | 46 | 17 | 9 | 10 | 15 | 28 | 10 | 19 | 0 | 14 |
| 17 | 0.08 | 0.97 | 0.08 | 0.94 | 1 | Gm12696 | | 40 | 14 | 4 | 6 | 15 | 16 | 28 | 10 | 28 | 6 | 14 |
| 17 | 0.00 | 0.73 | 0.00 | 1.00 | 1 | Gm10702 | | 18 | 8 | 4 | 22 | 18 | 7 | 28 | 17 | 16 | 35 | 8 |
| 15 | -0.26 | 0.89 | -0.29 | 0.77 | 1 | Gm5605 | | 20 | 37 | 19 | 11 | 9 | 3 | 28 | 14 | 4 | 12 | 8 |
| 15 | -2.36 | 1.82 | -1.30 | 0.19 | 1 | Cpa5 | carboxypeptidase A5 | 33 | 40 | 50 | 0 | 1 | 3 | 28 | 0 | 0 | 0 | 5 |
| 12 | -0.62 | 1.00 | -0.62 | 0.54 | 1 | Gm5171 | | 9 | 14 | 11 | 3 | 19 | 13 | 28 | 10 | 16 | 5 | 7 |
| 11 | 0.66 | 1.83 | 0.36 | 0.72 | 1 | Gm50402 | | 18 | 0 | 8 | 7 | 0 | 11 | 28 | 25 | 16 | 0 | 6 |
| 6 | -1.87 | 2.65 | -0.71 | 0.48 | 1 | Sult6b2 | sulfotransferase family 6B, member 2 | 0 | 0 | 0 | 8 | 0 | 11 | 28 | 0 | 9 | 0 | 13 |
| 3409 | -0.53 | 0.75 | -0.71 | 0.48 | 1 | Sparcl1 | SPARC-like 1 | 16553 | 9218 | 11571 | 15 | 22 | 18 | 27 | 0 | 27 | 39 | 9 |
| 2219 | -0.65 | 1.64 | -0.40 | 0.69 | 1 | Aebp1 | AE binding protein 1 | 14091 | 5694 | 4579 | 0 | 0 | 3 | 27 | 0 | 3 | 9 | 0 |
| 179 | -1.85 | 1.77 | -1.04 | 0.30 | 1 | Rab27b | RAB27B, member RAS oncogene family | 558 | 753 | 597 | 3 | 0 | 14 | 27 | 0 | 13 | 0 | 4 |
| 145 | 0.38 | 0.77 | 0.49 | 0.63 | 1 | H2-Q7 | histocompatibility 2, Q region locus 7 | 953 | 287 | 271 | 3 | 7 | 8 | 27 | 15 | 20 | 1 | 6 |
| 126 | -0.68 | 0.46 | -1.47 | 0.14 | 1 | Ngef | neuronal guanine nucleotide exchange fa | 226 | 320 | 222 | 142 | 157 | 68 | 27 | 81 | 52 | 44 | 48 |
| 100 | -0.27 | 0.50 | -0.53 | 0.60 | 1 | Oas1a | 2'-5' oligoadenylate synthetase 1A | 241 | 163 | 218 | 103 | 73 | 57 | 27 | 95 | 46 | 53 | 28 |
| 78 | -0.30 | 0.82 | -0.37 | 0.71 | 1 | Gm43481 | | 27 | 26 | 9 | 5 | 10 | 365 | 27 | 0 | 23 | 10 | 357 |
| 48 | -0.96 | 0.82 | -1.18 | 0.24 | 1 | Shisa4 | shisa family member 4 | 195 | 69 | 91 | 25 | 36 | 16 | 27 | 0 | 34 | 21 | 13 |
| 44 | -0.56 | 0.92 | -0.61 | 0.54 | 1 | Ccdc151 | coiled-coil domain containing 151 | 45 | 50 | 16 | 50 | 43 | 93 | 27 | 59 | 39 | 0 | 66 |
| 44 | -0.43 | 0.71 | -0.60 | 0.55 | 1 | Gm8116 | | 40 | 94 | 27 | 60 | 70 | 29 | 27 | 13 | 39 | 31 | 48 |
| 42 | 0.38 | 0.70 | 0.54 | 0.59 | 1 | Gm19721 | | 3 | 5 | 12 | 34 | 40 | 96 | 27 | 32 | 23 | 104 | 87 |
| 42 | 0.25 | 0.47 | 0.53 | 0.59 | 1 | Gm12346 | | 41 | 48 | 41 | 34 | 28 | 64 | 27 | 41 | 39 | 37 | 58 |
| 40 | -0.83 | 0.56 | -1.47 | 0.14 | 1 | 4833445I07Rik | | 21 | 47 | 34 | 39 | 60 | 83 | 27 | 34 | 31 | 6 | 61 |
| 40 | 0.06 | 0.61 | 0.10 | 0.92 | 1 | Lrrc4 | leucine rich repeat containing 4 | 71 | 28 | 19 | 60 | 31 | 37 | 27 | 60 | 35 | 45 | 26 |
| 33 | -0.13 | 0.58 | -0.22 | 0.83 | 1 | Gm14303 | | 45 | 53 | 34 | 28 | 32 | 33 | 27 | 17 | 25 | 45 | 26 |
| 32 | 0.84 | 0.61 | 1.38 | 0.17 | 1 | Gm10060 | | 21 | 14 | 31 | 15 | 10 | 57 | 27 | 31 | 31 | 39 | 76 |
| 32 | -0.48 | 0.56 | -0.85 | 0.39 | 1 | Gm15972 | | 26 | 22 | 23 | 33 | 50 | 50 | 27 | 23 | 29 | 17 | 52 |
| 29 | -0.95 | 0.91 | -1.05 | 0.30 | 1 | Gm34294 | | 45 | 56 | 54 | 25 | 13 | 36 | 27 | 0 | 14 | 18 | 31 |
| 26 | -0.45 | 0.63 | -0.72 | 0.47 | 1 | C130060C02Rik | | 29 | 18 | 37 | 30 | 22 | 37 | 27 | 21 | 24 | 13 | 28 |
| 26 | 0.63 | 0.66 | 0.95 | 0.34 | 1 | Gm4353 | | 22 | 31 | 18 | 11 | 21 | 25 | 27 | 27 | 40 | 28 | 31 |
| 23 | -0.84 | 0.96 | -0.88 | 0.38 | 1 | Gm26760 | | 13 | 19 | 52 | 17 | 36 | 19 | 27 | 19 | 28 | 0 | 23 |
| 22 | -0.40 | 0.65 | -0.62 | 0.54 | 1 | Gm10250 | | 13 | 17 | 16 | 24 | 46 | 15 | 27 | 17 | 23 | 28 | 13 |
| 22 | -0.17 | 0.78 | -0.21 | 0.83 | 1 | Gm6900 | | 35 | 14 | 14 | 29 | 27 | 9 | 27 | 8 | 36 | 21 | 17 |
| 21 | 0.91 | 1.11 | 0.82 | 0.41 | 1 | Gm34121 | | 0 | 21 | 6 | 11 | 7 | 23 | 27 | 24 | 7 | 61 | 45 |
| 19 | -1.58 | 1.08 | -1.46 | 0.14 | 1 | Gm31834 | | 6 | 12 | 10 | 19 | 26 | 40 | 27 | 0 | 22 | 3 | 47 |
| 19 | -0.72 | 0.83 | -0.87 | 0.39 | 1 | Gm7327 | | 9 | 4 | 6 | 30 | 26 | 28 | 27 | 25 | 34 | 5 | 10 |
| 18 | -1.13 | 1.31 | -0.86 | 0.39 | 1 | Il20rb | interleukin 20 receptor beta | 16 | 0 | 25 | 19 | 16 | 37 | 27 | 10 | 13 | 0 | 37 |
| 14 | 1.53 | 1.53 | 1.00 | 0.32 | 1 | Lrmda | leucine rich melanocyte differentiation as | 30 | 19 | 29 | 0 | 0 | 3 | 27 | 0 | 6 | 35 | 9 |
| 13 | -2.03 | 1.33 | -1.53 | 0.13 | 1 | 1810062O18Rik | | 9 | 0 | 5 | 36 | 24 | 16 | 27 | 7 | 8 | 0 | 12 |
| 12 | -0.89 | 1.01 | -0.87 | 0.38 | 1 | Gm43775 | | 2 | 0 | 1 | 28 | 15 | 11 | 27 | 19 | 4 | 8 | 14 |
| 11 | 1.29 | 1.54 | 0.83 | 0.40 | 1 | Gm5503 | | 27 | 0 | 1 | 0 | 0 | 1 | 27 | 0 | 1 | 61 | 2 |
| 10 | -1.34 | 1.76 | -0.76 | 0.45 | 1 | Nrxn3 | neurexin III | 22 | 3 | 20 | 4 | 0 | 8 | 27 | 0 | 13 | 0 | 10 |
| 8 | -2.74 | 1.52 | -1.80 | 0.07 | 1 | Gm12743 | | 3 | 6 | 7 | 14 | 9 | 6 | 27 | 0 | 6 | 0 | 9 |
| 6 | -1.93 | 2.39 | -0.81 | 0.42 | 1 | D530033B14Rik | | 0 | 0 | 0 | 0 | 24 | 5 | 27 | 0 | 11 | 0 | 4 |
| 782 | -1.73 | 1.13 | -1.52 | 0.13 | 1 | C1qc | complement component 1, q subcompone | 4174 | 1761 | 2589 | 4 | 9 | 10 | 26 | 0 | 6 | 0 | 18 |
| 456 | -1.46 | 1.24 | -1.18 | 0.24 | 1 | Gimap6 | GTPase, IMAP family member 6 | 1766 | 1587 | 1605 | 3 | 5 | 5 | 26 | 0 | 6 | 0 | 9 |
| 313 | 0.67 | 0.51 | 1.30 | 0.19 | 1 | Tnfrsf1b | tumor necrosis factor receptor superfamily | 1577 | 949 | 801 | 5 | 4 | 9 | 26 | 31 | 11 | 23 | 4 |
| 197 | 0.16 | 0.65 | 0.24 | 0.81 | 1 | Ksr1 | kinase suppressor of ras 1 | 934 | 470 | 545 | 16 | 23 | 36 | 26 | 51 | 28 | 1 | 37 |
| 167 | -2.00 | 1.85 | -1.08 | 0.28 | 1 | Irf4 | interferon regulatory factor 4 | 1441 | 233 | 130 | 0 | 0 | 2 | 26 | 0 | 0 | 0 | 1 |
| 158 | -0.02 | 1.81 | -0.01 | 0.99 | 1 | Myo1g | myosin IG | 1021 | 365 | 286 | 2 | 0 | 7 | 26 | 30 | 0 | 0 | 6 |
| 156 | 0.61 | 0.52 | 1.19 | 0.23 | 1 | Selenbp1 | selenium binding protein 1 | 573 | 424 | 536 | 12 | 8 | 24 | 26 | 43 | 9 | 35 | 20 |
| 103 | -0.05 | 0.58 | -0.08 | 0.94 | 1 | Cdk2 | cyclin-dependent kinase 2 | 389 | 142 | 89 | 66 | 52 | 106 | 26 | 23 | 61 | 85 | 92 |
| 87 | 0.54 | 0.43 | 1.24 | 0.22 | 1 | Dnhd1 | dynein heavy chain domain 1 | 23 | 40 | 16 | 88 | 14 | 292 | 26 | 114 | 36 | 49 | 255 |
| 83 | -3.08 | 2.04 | -1.50 | 0.13 | 1 | Cxcl9 | chemokine (C-X-C motif) ligand 9 | 414 | 269 | 189 | 0 | 0 | 17 | 26 | 0 | 0 | 0 | 2 |
| 69 | 0.09 | 0.51 | 0.17 | 0.87 | 1 | Flt3l | FMS-like tyrosine kinase 3 ligand | 92 | 22 | 62 | 29 | 36 | 183 | 26 | 32 | 25 | 37 | 209 |
| 67 | -1.38 | 0.76 | -1.81 | 0.07 | 1 | Crym | crystallin, mu | 166 | 205 | 233 | 32 | 13 | 19 | 26 | 0 | 15 | 9 | 14 |
| 57 | 0.40 | 0.59 | 0.69 | 0.49 | 1 | Eaf2 | ELL associated factor 2 | 212 | 54 | 43 | 15 | 31 | 63 | 26 | 22 | 37 | 85 | 41 |
| 55 | 0.20 | 0.59 | 0.35 | 0.73 | 1 | Ribc1 | RIB43A domain with coiled-coils 1 | 39 | 103 | 42 | 51 | 59 | 59 | 26 | 46 | 37 | 88 | 51 |
| 54 | 0.02 | 0.59 | 0.04 | 0.97 | 1 | Polq | polymerase (DNA directed), theta | 156 | 81 | 29 | 38 | 29 | 67 | 26 | 39 | 29 | 68 | 31 |

| | | | | | | | | | | | | | | | | | | |
|---|---|---|---|---|---|---|---|---|---|---|---|---|---|---|---|---|---|---|
| 53 | 0.16 | 0.72 | 0.23 | 0.82 | 1 | Dbndd2 | dysbindin (dystrobrevin binding protein 1) | 118 | 63 | 154 | 32 | 7 | 52 | 26 | 54 | 9 | 18 | 49 |
| 52 | -0.25 | 0.71 | -0.36 | 0.72 | 1 | Gm43667 | | 55 | 61 | 80 | 71 | 74 | 32 | 26 | 8 | 51 | 70 | 45 |
| 52 | -1.47 | 0.88 | -1.68 | 0.09 | 1 | Gm26660 | | 105 | 49 | 141 | 67 | 40 | 53 | 26 | 0 | 33 | 19 | 38 |
| 52 | -1.18 | 0.90 | -1.32 | 0.19 | 1 | Zkscan7 | zinc finger protein 660 | 166 | 168 | 77 | 24 | 9 | 36 | 26 | 45 | 2 | 0 | 18 |
| 49 | 0.23 | 0.51 | 0.45 | 0.65 | 1 | Gm15860 | | 16 | 84 | 42 | 70 | 64 | 22 | 26 | 95 | 40 | 31 | 43 |
| 48 | 0.50 | 0.63 | 0.79 | 0.43 | 1 | Fam83d | family with sequence similarity 83, membe | 47 | 52 | 53 | 77 | 19 | 42 | 26 | 28 | 62 | 68 | 58 |
| 45 | -0.47 | 0.82 | -0.57 | 0.57 | 1 | Gm7665 | | 89 | 42 | 75 | 46 | 43 | 30 | 26 | 2 | 19 | 66 | 61 |
| 43 | 0.43 | 0.67 | 0.64 | 0.52 | 1 | Gm43668 | | 42 | 93 | 19 | 43 | 20 | 54 | 26 | 21 | 57 | 37 | 58 |
| 40 | 0.92 | 0.67 | 1.36 | 0.17 | 1 | Abcb11 | ATP-binding cassette, sub-family B (MDR | 3 | 8 | 9 | 41 | 29 | 39 | 26 | 92 | 42 | 119 | 26 |
| 39 | -0.48 | 0.89 | -0.55 | 0.59 | 1 | Nlrc4 | NLR family, CARD domain containing 4 | 103 | 57 | 105 | 18 | 10 | 38 | 26 | 20 | 18 | 0 | 35 |
| 37 | -0.12 | 0.97 | -0.12 | 0.90 | 1 | Lgi3 | leucine-rich repeat LGI family, member 3 | 186 | 79 | 37 | 1 | 9 | 15 | 26 | 0 | 9 | 36 | 8 |
| 35 | -0.23 | 0.71 | -0.33 | 0.74 | 1 | Cenph | centromere protein H | 86 | 30 | 64 | 43 | 16 | 24 | 26 | 9 | 32 | 23 | 36 |
| 32 | 0.24 | 0.70 | 0.34 | 0.74 | 1 | Gm5521 | | 10 | 48 | 28 | 36 | 38 | 25 | 26 | 16 | 43 | 44 | 37 |
| 31 | 0.12 | 0.76 | 0.16 | 0.87 | 1 | Slc12a1 | solute carrier family 12, member 1 | 3 | 22 | 35 | 36 | 18 | 52 | 26 | 59 | 26 | 19 | 50 |
| 31 | -0.39 | 0.78 | -0.50 | 0.61 | 1 | Clgn | calmegin | 36 | 14 | 52 | 53 | 25 | 30 | 26 | 25 | 7 | 40 | 36 |
| 31 | -0.87 | 0.90 | -0.96 | 0.34 | 1 | Gm18943 | | 27 | 42 | 16 | 47 | 45 | 42 | 26 | 1 | 32 | 33 | 25 |
| 31 | 0.57 | 0.71 | 0.81 | 0.42 | 1 | Gm47155 | | 27 | 23 | 16 | 11 | 26 | 42 | 26 | 9 | 52 | 67 | 37 |
| 30 | -0.04 | 0.65 | -0.06 | 0.95 | 1 | Gm47232 | | 40 | 9 | 17 | 11 | 44 | 63 | 26 | 20 | 32 | 32 | 39 |
| 30 | -0.38 | 0.93 | -0.40 | 0.69 | 1 | 6430573P05Rik | | 9 | 30 | 37 | 30 | 29 | 30 | 26 | 31 | 65 | 0 | 42 |
| 28 | -0.23 | 0.69 | -0.33 | 0.74 | 1 | Gm6493 | | 33 | 45 | 24 | 37 | 26 | 21 | 26 | 10 | 16 | 33 | 41 |
| 28 | 0.03 | 0.81 | 0.04 | 0.97 | 1 | Gm14379 | | 18 | 42 | 9 | 49 | 14 | 36 | 26 | 7 | 21 | 58 | 32 |
| 28 | -0.37 | 0.63 | -0.58 | 0.56 | 1 | Gm6210 | | 45 | 25 | 36 | 21 | 30 | 30 | 26 | 9 | 14 | 36 | 34 |
| 28 | -0.10 | 0.95 | -0.11 | 0.91 | 1 | Pcdhac1 | protocadherin alpha subfamily C, 1 | 8 | 0 | 0 | 74 | 22 | 24 | 26 | 67 | 12 | 37 | 34 |
| 28 | -0.25 | 0.76 | -0.32 | 0.75 | 1 | Gm10221 | | 52 | 17 | 16 | 39 | 17 | 37 | 26 | 30 | 37 | 15 | 15 |
| 28 | 1.40 | 1.35 | 1.04 | 0.30 | 1 | Slc22a21 | solute carrier family 22 (organic cation tra | 57 | 70 | 57 | 0 | 0 | 18 | 26 | 0 | 6 | 35 | 34 |
| 28 | 0.24 | 0.75 | 0.32 | 0.75 | 1 | 6330549D23Rik | | 8 | 72 | 21 | 29 | 21 | 18 | 26 | 25 | 21 | 28 | 34 |
| 27 | 0.79 | 1.01 | 0.78 | 0.43 | 1 | Gm44744 | | 11 | 0 | 0 | 38 | 7 | 38 | 26 | 31 | 43 | 64 | 34 |
| 26 | -0.06 | 0.81 | -0.08 | 0.94 | 1 | Gm16062 | | 12 | 2 | 42 | 37 | 26 | 27 | 26 | 30 | 42 | 23 | 24 |
| 24 | -0.69 | 1.07 | -0.65 | 0.52 | 1 | B130034C11Rik | | 32 | 32 | 63 | 20 | 18 | 7 | 26 | 7 | 24 | 0 | 35 |
| 23 | -0.82 | 0.98 | -0.84 | 0.40 | 1 | 1110046J04Rik | | 17 | 25 | 32 | 9 | 16 | 52 | 26 | 17 | 14 | 0 | 40 |
| 22 | -0.36 | 0.79 | -0.45 | 0.65 | 1 | Gm15418 | | 29 | 15 | 23 | 38 | 13 | 18 | 26 | 10 | 35 | 13 | 18 |
| 22 | -0.28 | 1.12 | -0.25 | 0.80 | 1 | Gm37035 | | 2 | 3 | 0 | 25 | 14 | 41 | 26 | 50 | 12 | 0 | 66 |
| 20 | -1.52 | 0.89 | -1.70 | 0.09 | 1 | Zfp457 | zinc finger protein 457 | 6 | 26 | 22 | 41 | 24 | 20 | 26 | 32 | 7 | 14 | 1 |
| 20 | -1.65 | 1.06 | -1.55 | 0.12 | 1 | Gm45718 | | 11 | 0 | 9 | 10 | 30 | 52 | 26 | 0 | 12 | 12 | 56 |
| 20 | -0.94 | 0.99 | -0.95 | 0.34 | 1 | Gm3055 | predicted gene 3055 | 10 | 5 | 24 | 18 | 17 | 35 | 26 | 0 | 28 | 13 | 40 |
| 19 | 0.21 | 0.94 | 0.22 | 0.83 | 1 | Gm15821 | | 5 | 9 | 1 | 10 | 21 | 31 | 26 | 5 | 34 | 21 | 51 |
| 18 | -1.53 | 1.10 | -1.39 | 0.16 | 1 | Rpl10-ps3 | | 17 | 11 | 22 | 47 | 10 | 23 | 26 | 0 | 20 | 9 | 15 |
| 18 | -0.04 | 1.42 | -0.03 | 0.98 | 1 | Dmrta1 | doublesex and mab-3 related transcriptio | 0 | 4 | 0 | 18 | 19 | 32 | 26 | 0 | 16 | 50 | 28 |
| 15 | -2.00 | 1.03 | -1.93 | 0.05 | 1 | Appbp2os | | 12 | 16 | 2 | 32 | 18 | 23 | 26 | 15 | 0 | 18 | 6 |
| 14 | -0.73 | 1.01 | -0.73 | 0.47 | 1 | 3632454L22Rik | | 7 | 2 | 0 | 32 | 19 | 15 | 26 | 12 | 12 | 8 | 23 |
| 14 | -0.36 | 1.47 | -0.24 | 0.81 | 1 | Gm46447 | | 1 | 2 | 10 | 7 | 5 | 22 | 26 | 43 | 2 | 0 | 32 |
| 14 | -1.97 | 1.58 | -1.25 | 0.21 | 1 | Spata17 | spermatogenesis associated 17 | 8 | 10 | 0 | 13 | 8 | 33 | 26 | 0 | 5 | 0 | 46 |
| 13 | -1.32 | 1.44 | -0.92 | 0.36 | 1 | 5430430B14Rik | | 11 | 15 | 14 | 13 | 16 | 6 | 26 | 0 | 28 | 0 | 14 |
| 12 | -0.36 | 1.50 | -0.24 | 0.81 | 1 | Gm43192 | | 0 | 0 | 0 | 20 | 17 | 8 | 26 | 23 | 16 | 0 | 25 |
| 12 | -0.09 | 1.38 | -0.06 | 0.95 | 1 | Hpca | hippocalcin | 28 | 17 | 7 | 0 | 9 | 4 | 26 | 0 | 8 | 22 | 5 |
| 6 | -0.48 | 2.05 | -0.23 | 0.82 | 1 | Catspere1 | cation channel sperm associated auxiliar | 10 | 0 | 0 | 0 | 2 | 14 | 26 | 0 | 13 | 3 | 1 |
| 5 | 0.27 | 2.36 | 0.11 | 0.91 | 1 | Scn4a | sodium channel, voltage-gated, type IV, a | 3 | 6 | 0 | 4 | 0 | 3 | 26 | 0 | 6 | 0 | 5 |
| 620 | -1.55 | 1.60 | -0.97 | 0.33 | 1 | Myl9 | myosin, light polypeptide 9, regulatory | 3889 | 1423 | 1462 | 4 | 3 | 0 | 25 | 0 | 3 | 0 | 5 |
| 391 | 0.39 | 0.58 | 0.68 | 0.50 | 1 | Smtn | smoothelin | 1707 | 1166 | 1157 | 46 | 13 | 25 | 25 | 3 | 34 | 85 | 39 |
| 283 | 0.08 | 1.31 | 0.06 | 0.95 | 1 | Sp100 | nuclear antigen Sp100 | 1349 | 828 | 880 | 0 | 0 | 9 | 25 | 0 | 3 | 13 | 6 |
| 264 | -0.66 | 0.51 | -1.28 | 0.20 | 1 | Hcar1 | hydrocarboxylic acid receptor 1 | 781 | 808 | 1031 | 69 | 72 | 13 | 25 | 7 | 47 | 27 | 22 |
| 235 | 0.18 | 0.40 | 0.46 | 0.65 | 1 | Parvb | parvin, beta | 916 | 691 | 778 | 27 | 8 | 38 | 25 | 21 | 22 | 33 | 29 |
| 217 | -0.41 | 1.24 | -0.33 | 0.74 | 1 | Abcg2 | ATP binding cassette subfamily G membe | 969 | 664 | 553 | 51 | 3 | 31 | 25 | 0 | 34 | 10 | 46 |
| 191 | -0.04 | 2.19 | -0.02 | 0.99 | 1 | Fcgr3 | Fc receptor, IgG, low affinity III | 998 | 486 | 571 | 0 | 1 | 0 | 25 | 0 | 2 | 0 | 13 |
| 190 | -1.31 | 3.61 | -0.36 | 0.72 | 1 | Lyz1 | lysozyme 1 | 1247 | 532 | 283 | 0 | 0 | 0 | 25 | 0 | 0 | 0 | 0 |
| 183 | -1.33 | 1.27 | -1.05 | 0.29 | 1 | Cytip | cytohesin 1 interacting protein | 1219 | 405 | 322 | 0 | 6 | 18 | 25 | 13 | 2 | 3 | 1 |
| 181 | -3.26 | 3.27 | -1.00 | 0.32 | 1 | Fcrls | Fc receptor-like S, scavenger receptor | 1090 | 250 | 618 | 0 | 0 | 0 | 25 | 0 | 0 | 0 | 3 |
| 121 | -0.51 | 0.61 | -0.84 | 0.40 | 1 | Gamt | guanidinoacetate methyltransferase | 247 | 409 | 452 | 67 | 35 | 11 | 25 | 8 | 21 | 24 | 31 |
| 86 | 0.35 | 0.63 | 0.56 | 0.57 | 1 | Ctdsp2-ps | | 16 | 174 | 102 | 153 | 31 | 87 | 25 | 200 | 18 | 59 | 81 |
| 77 | 0.15 | 0.60 | 0.25 | 0.81 | 1 | 1700110K17Rik | | 26 | 88 | 48 | 26 | 33 | 244 | 25 | 12 | 35 | 48 | 258 |

| | | | | | | | | | | | | | | | | | |
|---|---|---|---|---|---|---|---|---|---|---|---|---|---|---|---|---|---|
| 64 | -1.42 | 1.08 | -1.32 | 0.19 | 1 | Pcdh9 | protocadherin 9 | 295 | 205 | 110 | 18 | 4 | 19 | 25 | 0 | 16 | 0 | 14 |
| 56 | -0.52 | 0.73 | -0.71 | 0.48 | 1 | Gm39469 | | 53 | 63 | 14 | 58 | 90 | 94 | 25 | 14 | 61 | 23 | 121 |
| 56 | 0.32 | 0.51 | 0.63 | 0.53 | 1 | Gm9625 | | 82 | 66 | 108 | 60 | 42 | 26 | 25 | 81 | 70 | 31 | 22 |
| 50 | 0.06 | 0.63 | 0.09 | 0.93 | 1 | Kif24 | kinesin family member 24 | 26 | 25 | 96 | 70 | 32 | 72 | 25 | 50 | 33 | 43 | 73 |
| 46 | -0.07 | 0.88 | -0.08 | 0.94 | 1 | AA465934 | | 31 | 73 | 50 | 30 | 28 | 67 | 25 | 105 | 36 | 0 | 60 |
| 43 | -0.14 | 0.64 | -0.23 | 0.82 | 1 | Zp2 | zona pellucida glycoprotein 2 | 17 | 30 | 92 | 37 | 49 | 64 | 25 | 23 | 23 | 41 | 67 |
| 41 | 0.50 | 0.75 | 0.66 | 0.51 | 1 | Dnah7b | dynein, axonemal, heavy chain 7B | 28 | 32 | 85 | 36 | 11 | 51 | 25 | 61 | 9 | 33 | 79 |
| 40 | -0.44 | 1.02 | -0.43 | 0.67 | 1 | 1700055D18Rik | | 12 | 31 | 23 | 51 | 58 | 69 | 25 | 0 | 27 | 66 | 80 |
| 39 | 0.65 | 0.63 | 1.03 | 0.30 | 1 | Zpbp | zona pellucida binding protein | 27 | 24 | 42 | 57 | 21 | 27 | 25 | 44 | 29 | 77 | 51 |
| 37 | -0.49 | 0.77 | -0.63 | 0.53 | 1 | Rdh12 | retinol dehydrogenase 12 | 71 | 112 | 61 | 23 | 31 | 11 | 25 | 2 | 34 | 21 | 10 |
| 36 | -0.15 | 0.80 | -0.19 | 0.85 | 1 | Chkb | choline kinase beta | 7 | 25 | 38 | 9 | 18 | 133 | 25 | 46 | 26 | 8 | 59 |
| 36 | -0.66 | 0.84 | -0.78 | 0.44 | 1 | Prelid2 | PRELI domain containing 2 | 38 | 62 | 75 | 25 | 31 | 36 | 25 | 29 | 24 | 0 | 47 |
| 34 | -0.84 | 0.67 | -1.26 | 0.21 | 1 | Gm2614 | | 68 | 62 | 77 | 34 | 22 | 22 | 25 | 31 | 10 | 5 | 16 |
| 34 | 0.18 | 1.16 | 0.16 | 0.87 | 1 | Stx1b | syntaxin 1B | 159 | 66 | 65 | 9 | 0 | 10 | 25 | 15 | 6 | 0 | 14 |
| 32 | -0.53 | 0.70 | -0.76 | 0.45 | 1 | 2310040G07Rik | | 27 | 34 | 28 | 49 | 53 | 25 | 25 | 24 | 11 | 55 | 20 |
| 29 | -0.06 | 0.75 | -0.08 | 0.93 | 1 | Gm5105 | | 7 | 34 | 47 | 48 | 14 | 36 | 25 | 34 | 21 | 31 | 26 |
| 29 | 0.18 | 1.10 | 0.16 | 0.87 | 1 | Gzmm | granzyme M (lymphocyte met-ase 1) | 51 | 13 | 8 | 34 | 16 | 41 | 25 | 68 | 25 | 0 | 40 |
| 27 | -1.84 | 0.97 | -1.90 | 0.06 | 1 | Barx2 | BarH-like homeobox 2 | 26 | 34 | 54 | 41 | 16 | 50 | 25 | 32 | 3 | 0 | 20 |
| 27 | -0.41 | 0.65 | -0.62 | 0.53 | 1 | Rps25-ps1 | | 61 | 47 | 35 | 19 | 27 | 15 | 25 | 14 | 24 | 10 | 16 |
| 25 | 0.94 | 0.95 | 0.98 | 0.33 | 1 | 9330136K24Rik | | 5 | 62 | 11 | 4 | 6 | 30 | 25 | 22 | 10 | 31 | 66 |
| 23 | 0.33 | 0.93 | 0.35 | 0.73 | 1 | Mgarp | mitochondria localized glutamic acid rich | 6 | 11 | 43 | 23 | 21 | 14 | 25 | 52 | 7 | 14 | 42 |
| 23 | -0.37 | 1.15 | -0.32 | 0.75 | 1 | 4833408A19Rik | | 47 | 7 | 19 | 29 | 16 | 32 | 25 | 37 | 32 | 0 | 12 |
| 23 | -0.32 | 0.76 | -0.42 | 0.68 | 1 | Gm18537 | | 4 | 3 | 4 | 32 | 32 | 38 | 25 | 53 | 39 | 9 | 15 |
| 22 | 0.46 | 0.92 | 0.50 | 0.62 | 1 | Gm7730 | | 36 | 15 | 6 | 4 | 36 | 14 | 25 | 49 | 15 | 24 | 19 |
| 22 | -0.26 | 1.15 | -0.23 | 0.82 | 1 | Gm15559 | | 27 | 48 | 15 | 13 | 9 | 25 | 25 | 20 | 49 | 0 | 10 |
| 21 | 0.29 | 1.06 | 0.27 | 0.78 | 1 | Gm47414 | | 33 | 16 | 29 | 1 | 11 | 24 | 25 | 0 | 10 | 37 | 44 |
| 21 | 1.10 | 0.97 | 1.14 | 0.26 | 1 | Gm8228 | | 5 | 11 | 16 | 1 | 12 | 36 | 25 | 28 | 24 | 45 | 26 |
| 18 | -1.32 | 1.15 | -1.15 | 0.25 | 1 | Zfp520-ps | | 0 | 0 | 0 | 46 | 37 | 14 | 25 | 49 | 2 | 4 | 17 |
| 18 | -0.64 | 0.95 | -0.67 | 0.50 | 1 | Gm45774 | | 3 | 2 | 16 | 22 | 16 | 34 | 25 | 25 | 20 | 3 | 26 |
| 17 | 0.76 | 1.36 | 0.56 | 0.58 | 1 | Gm20692 | | 0 | 0 | 22 | 7 | 10 | 22 | 25 | 20 | 12 | 55 | 13 |
| 17 | -1.17 | 1.43 | -0.81 | 0.42 | 1 | Spag6l | sperm associated antigen 6-like | 0 | 26 | 18 | 20 | 15 | 22 | 25 | 0 | 31 | 4 | 23 |
| 17 | -0.93 | 1.25 | -0.74 | 0.46 | 1 | Gm7712 | | 0 | 1 | 1 | 12 | 21 | 27 | 25 | 70 | 12 | 0 | 15 |
| 16 | -1.31 | 1.04 | -1.26 | 0.21 | 1 | Gm20768 | | 19 | 29 | 17 | 24 | 19 | 7 | 25 | 3 | 12 | 1 | 20 |
| 16 | -1.84 | 1.23 | -1.50 | 0.13 | 1 | Irs3 | insulin receptor substrate 3 | 27 | 26 | 24 | 13 | 20 | 3 | 25 | 7 | 0 | 26 | 0 |
| 15 | -2.06 | 1.98 | -1.04 | 0.30 | 1 | Gm18194 | | 0 | 24 | 9 | 8 | 0 | 56 | 25 | 0 | 12 | 0 | 31 |
| 15 | 0.13 | 1.39 | 0.10 | 0.92 | 1 | Pcdha5 | protocadherin alpha 5 | 4 | 22 | 0 | 13 | 5 | 21 | 25 | 0 | 5 | 45 | 24 |
| 14 | -2.00 | 1.89 | -1.06 | 0.29 | 1 | Tmem40 | transmembrane protein 40 | 55 | 31 | 38 | 0 | 0 | 4 | 25 | 0 | 0 | 0 | 3 |
| 14 | -0.34 | 1.15 | -0.29 | 0.77 | 1 | Chst9 | carbohydrate (N-acetylgalactosamine 4-0 | 45 | 12 | 10 | 10 | 8 | 10 | 25 | 15 | 8 | 0 | 11 |
| 14 | -1.71 | 1.12 | -1.54 | 0.12 | 1 | Gm8666 | | 5 | 26 | 31 | 11 | 13 | 10 | 25 | 2 | 6 | 1 | 21 |
| 14 | 0.30 | 0.95 | 0.32 | 0.75 | 1 | Gm14493 | | 3 | 11 | 9 | 5 | 13 | 11 | 25 | 29 | 10 | 26 | 8 |
| 13 | -1.89 | 1.30 | -1.45 | 0.15 | 1 | Slc5a10 | solute carrier family 5 (sodium/glucose cc | 0 | 0 | 0 | 11 | 44 | 25 | 25 | 0 | 6 | 14 | 19 |
| 13 | 0.30 | 1.50 | 0.20 | 0.84 | 1 | Edn2 | endothelin 2 | 3 | 0 | 0 | 8 | 15 | 24 | 25 | 27 | 26 | 0 | 13 |
| 13 | -1.64 | 1.14 | -1.44 | 0.15 | 1 | Gm16196 | | 26 | 11 | 2 | 19 | 9 | 14 | 25 | 0 | 2 | 15 | 15 |
| 13 | -0.95 | 1.19 | -0.80 | 0.42 | 1 | 4930532G15Rik | | 19 | 25 | 5 | 4 | 9 | 19 | 25 | 10 | 6 | 0 | 15 |
| 12 | -2.80 | 1.69 | -1.65 | 0.10 | 1 | Zfp711 | zinc finger protein 711 | 21 | 31 | 37 | 11 | 0 | 3 | 25 | 0 | 2 | 0 | 4 |
| 12 | -0.82 | 1.81 | -0.45 | 0.65 | 1 | Ccdc106 | coiled-coil domain containing 106 | 40 | 11 | 18 | 0 | 6 | 4 | 25 | 27 | 0 | 0 | 3 |
| 12 | 1.15 | 1.77 | 0.65 | 0.52 | 1 | Gm6356 | | 9 | 0 | 0 | 0 | 0 | 4 | 25 | 0 | 0 | 84 | 11 |
| 9 | 2.67 | 2.07 | 1.29 | 0.20 | 1 | Ccdc155 | coiled-coil domain containing 155 | 10 | 0 | 3 | 0 | 0 | 13 | 25 | 15 | 12 | 0 | 18 |
| 9 | -0.82 | 1.57 | -0.52 | 0.60 | 1 | Gm5432 | | 5 | 0 | 12 | 11 | 5 | 7 | 25 | 16 | 8 | 0 | 4 |
| 7 | -1.17 | 1.99 | -0.59 | 0.56 | 1 | 2210417A02Rik | | 3 | 29 | 5 | 10 | 0 | 0 | 25 | 6 | 3 | 0 | 0 |
| 6 | -3.23 | 1.69 | -1.91 | 0.06 | 1 | Gm14154 | | 3 | 0 | 12 | 3 | 10 | 7 | 25 | 0 | 9 | 0 | 1 |
| 1055 | -1.48 | 1.31 | -1.13 | 0.26 | 1 | Cybb | cytochrome b-245, beta polypeptide | 5288 | 2799 | 3423 | 18 | 12 | 8 | 24 | 0 | 22 | 0 | 13 |
| 1006 | -2.12 | 1.27 | -1.68 | 0.09 | 1 | Mrc1 | mannose receptor, C type 1 | 6361 | 1941 | 2644 | 10 | 26 | 30 | 24 | 13 | 0 | 0 | 20 |
| 506 | -1.97 | 1.25 | -1.58 | 0.11 | 1 | Nckap1l | NCK associated protein 1 like | 2782 | 1408 | 1310 | 18 | 12 | 2 | 24 | 0 | 11 | 0 | 3 |
| 447 | -1.92 | 1.36 | -1.41 | 0.16 | 1 | Hcls1 | hematopoietic cell specific Lyn substrate | 2618 | 927 | 1310 | 0 | 12 | 11 | 24 | 0 | 6 | 0 | 6 |
| 318 | -0.79 | 0.59 | -1.33 | 0.18 | 1 | Samd5 | sterile alpha motif domain containing 5 | 986 | 1231 | 1066 | 12 | 38 | 59 | 24 | 51 | 6 | 19 | 10 |
| 309 | -2.29 | 1.43 | -1.60 | 0.11 | 1 | Snhg11 | | 52 | 20 | 52 | 66 | 20 | 1795 | 24 | 0 | 37 | 0 | 1328 |
| 294 | -2.16 | 1.18 | -1.82 | 0.07 | 1 | Lbp | lipopolysaccharide binding protein | 1736 | 598 | 799 | 14 | 28 | 7 | 24 | 0 | 14 | 0 | 9 |
| 266 | 0.41 | 0.43 | 0.96 | 0.33 | 1 | Dnah11 | dynein, axonemal, heavy chain 11 | 943 | 1075 | 687 | 11 | 26 | 31 | 24 | 52 | 30 | 26 | 19 |
| 227 | 0.23 | 0.62 | 0.37 | 0.71 | 1 | Serpine1 | serine (or cysteine) peptidase inhibitor, cl | 1189 | 481 | 601 | 9 | 20 | 59 | 24 | 21 | 40 | 53 | 5 |

| | | | | | | | | | | | | | | | | | | |
|---|---|---|---|---|---|---|---|---|---|---|---|---|---|---|---|---|---|---|
| 195 | -0.35 | 1.18 | -0.30 | 0.77 | 1 | Hic1 | hypermethylated in cancer 1 | 851 | 544 | 597 | 2 | 25 | 34 | 24 | 31 | 21 | 0 | 13 |
| 179 | -1.21 | 0.72 | -1.67 | 0.10 | 1 | Irf5 | interferon regulatory factor 5 | 953 | 534 | 423 | 3 | 10 | 6 | 24 | 0 | 2 | 13 | 5 |
| 99 | -0.33 | 0.50 | -0.66 | 0.51 | 1 | E230001N04Rik | | 156 | 232 | 173 | 77 | 61 | 132 | 24 | 25 | 31 | 61 | 120 |
| 95 | 0.51 | 0.42 | 1.23 | 0.22 | 1 | Tst | thiosulfate sulfurtransferase, mitochondria | 371 | 255 | 242 | 17 | 20 | 11 | 24 | 28 | 16 | 43 | 18 |
| 90 | 0.39 | 0.52 | 0.75 | 0.45 | 1 | Ccdc68 | coiled-coil domain containing 68 | 100 | 265 | 86 | 104 | 28 | 46 | 24 | 222 | 42 | 10 | 60 |
| 74 | 0.62 | 0.47 | 1.32 | 0.19 | 1 | Ttc12 | tetratricopeptide repeat domain 12 | 124 | 207 | 132 | 31 | 42 | 38 | 24 | 80 | 30 | 59 | 46 |
| 69 | 0.68 | 1.25 | 0.54 | 0.59 | 1 | Fgr | FGR proto-oncogene, Src family tyrosine | 450 | 194 | 68 | 0 | 0 | 0 | 24 | 0 | 2 | 23 | 0 |
| 65 | 0.01 | 0.65 | 0.01 | 0.99 | 1 | Tyro3 | TYRO3 protein tyrosine kinase 3 | 185 | 75 | 163 | 74 | 19 | 27 | 24 | 65 | 46 | 17 | 19 |
| 64 | -0.99 | 0.74 | -1.34 | 0.18 | 1 | Gm31518 | | 55 | 129 | 124 | 96 | 40 | 83 | 24 | 67 | 49 | 0 | 38 |
| 59 | 0.57 | 0.57 | 1.01 | 0.31 | 1 | 3110001I22Rik | bifunctional apoptosis regulator | 81 | 37 | 76 | 47 | 49 | 63 | 24 | 100 | 35 | 55 | 82 |
| 58 | -1.85 | 0.97 | -1.90 | 0.06 | 1 | Brip1 | BRCA1 interacting protein C-terminal heli | 209 | 23 | 88 | 78 | 51 | 66 | 24 | 0 | 41 | 15 | 39 |
| 53 | 0.62 | 0.62 | 0.99 | 0.32 | 1 | Gm6288 | | 12 | 104 | 25 | 39 | 45 | 68 | 24 | 49 | 67 | 44 | 103 |
| 43 | 0.37 | 0.58 | 0.64 | 0.52 | 1 | 4932422M17Rik | | 18 | 12 | 29 | 32 | 72 | 60 | 24 | 51 | 36 | 66 | 70 |
| 42 | 0.11 | 0.63 | 0.17 | 0.86 | 1 | AA388235 | | 35 | 73 | 28 | 28 | 42 | 65 | 24 | 46 | 31 | 62 | 29 |
| 42 | -0.37 | 0.74 | -0.50 | 0.62 | 1 | Ccdc65 | coiled-coil domain containing 65 | 50 | 23 | 60 | 73 | 39 | 41 | 24 | 42 | 51 | 5 | 56 |
| 38 | 0.24 | 0.59 | 0.41 | 0.68 | 1 | Gm9828 | | 24 | 50 | 77 | 39 | 24 | 36 | 24 | 25 | 33 | 36 | 51 |
| 37 | -1.13 | 0.87 | -1.30 | 0.19 | 1 | Krtap17-1 | keratin associated protein 17-1 | 43 | 35 | 80 | 34 | 23 | 76 | 24 | 0 | 32 | 18 | 40 |
| 36 | 0.33 | 0.86 | 0.38 | 0.70 | 1 | B4galnt3 | beta-1,4-N-acetyl-galactosaminyl transfer | 36 | 74 | 49 | 9 | 13 | 53 | 24 | 70 | 4 | 40 | 21 |
| 35 | 0.61 | 0.72 | 0.84 | 0.40 | 1 | B930094E09Rik | | 7 | 12 | 16 | 11 | 63 | 42 | 24 | 87 | 51 | 48 | 24 |
| 32 | 0.34 | 0.96 | 0.35 | 0.72 | 1 | Mutyh | mutY DNA glycosylase | 35 | 60 | 32 | 7 | 9 | 52 | 24 | 0 | 22 | 32 | 74 |
| 32 | 1.04 | 0.94 | 1.10 | 0.27 | 1 | Mapkapk5 | MAP kinase-activated protein kinase 5 | 3 | 2 | 19 | 4 | 11 | 93 | 24 | 54 | 16 | 23 | 96 |
| 31 | -1.11 | 0.98 | -1.14 | 0.26 | 1 | Arhgef33 | Rho guanine nucleotide exchange factor | 43 | 37 | 11 | 27 | 31 | 61 | 24 | 9 | 21 | 0 | 75 |
| 30 | 0.16 | 0.66 | 0.24 | 0.81 | 1 | Gm9385 | | 47 | 29 | 74 | 25 | 20 | 15 | 24 | 38 | 19 | 15 | 21 |
| 30 | 0.17 | 1.01 | 0.17 | 0.86 | 1 | Gm12655 | | 33 | 26 | 0 | 38 | 26 | 37 | 24 | 13 | 42 | 27 | 60 |
| 28 | 0.19 | 0.77 | 0.24 | 0.81 | 1 | Gm6645 | | 20 | 16 | 0 | 51 | 4 | 31 | 24 | 41 | 2 | 93 | 29 |
| 25 | -0.15 | 0.75 | -0.19 | 0.85 | 1 | Gm1848 | | 46 | 17 | 20 | 36 | 16 | 21 | 24 | 39 | 15 | 9 | 25 |
| 23 | 0.06 | 0.66 | 0.09 | 0.93 | 1 | Gm10638 | | 45 | 43 | 22 | 18 | 10 | 14 | 24 | 13 | 12 | 36 | 12 |
| 22 | 0.66 | 1.50 | 0.44 | 0.66 | 1 | Slc9a5 | solute carrier family 9 (sodium/hydrogen e | 13 | 34 | 44 | 0 | 0 | 60 | 24 | 0 | 10 | 18 | 36 |
| 21 | 0.07 | 0.73 | 0.09 | 0.93 | 1 | Gm49741 | | 7 | 10 | 31 | 21 | 17 | 23 | 24 | 17 | 40 | 22 | 19 |
| 20 | 0.30 | 0.77 | 0.38 | 0.70 | 1 | Slco5a1 | solute carrier organic anion transporter fa | 57 | 45 | 11 | 7 | 7 | 12 | 24 | 8 | 12 | 21 | 14 |
| 19 | -0.01 | 1.54 | -0.01 | 0.99 | 1 | D030025P21Rik | | 0 | 11 | 0 | 26 | 26 | 29 | 24 | 0 | 48 | 21 | 23 |
| 18 | -0.27 | 0.90 | -0.30 | 0.77 | 1 | Gm14022 | | 5 | 0 | 2 | 39 | 26 | 12 | 24 | 43 | 15 | 21 | 9 |
| 18 | -0.08 | 1.09 | -0.07 | 0.94 | 1 | Gm42743 | | 5 | 6 | 18 | 6 | 15 | 24 | 24 | 0 | 12 | 59 | 23 |
| 16 | -0.07 | 1.00 | -0.07 | 0.94 | 1 | 4930423D22Rik | | 9 | 0 | 4 | 17 | 37 | 13 | 24 | 19 | 27 | 8 | 23 |
| 16 | -1.20 | 1.12 | -1.08 | 0.28 | 1 | 9130015G15Rik | | 5 | 19 | 31 | 12 | 20 | 25 | 24 | 0 | 10 | 23 | 9 |
| 16 | 0.85 | 0.87 | 0.97 | 0.33 | 1 | Gm17115 | | 6 | 12 | 10 | 17 | 3 | 8 | 24 | 41 | 15 | 23 | 12 |
| 15 | -1.62 | 1.73 | -0.94 | 0.35 | 1 | Gm49092 | | 24 | 47 | 0 | 8 | 19 | 14 | 24 | 0 | 17 | 0 | 17 |
| 15 | -0.65 | 1.20 | -0.54 | 0.59 | 1 | Gm49300 | | 25 | 21 | 2 | 15 | 23 | 8 | 24 | 0 | 16 | 23 | 9 |
| 15 | -1.96 | 1.31 | -1.49 | 0.14 | 1 | Olfr1322 | olfactory receptor 1322 | 0 | 3 | 0 | 8 | 12 | 67 | 24 | 12 | 5 | 0 | 31 |
| 14 | -0.66 | 1.57 | -0.42 | 0.68 | 1 | Rps15a-ps6 | | 33 | 44 | 10 | 1 | 4 | 11 | 24 | 0 | 21 | 0 | 9 |
| 14 | -0.97 | 1.59 | -0.61 | 0.54 | 1 | Gm36315 | | 0 | 40 | 0 | 16 | 10 | 9 | 24 | 3 | 4 | 44 | 1 |
| 13 | 0.51 | 1.47 | 0.35 | 0.73 | 1 | Gm11836 | | 31 | 0 | 9 | 8 | 15 | 5 | 24 | 20 | 24 | 0 | 8 |
| 13 | -0.41 | 1.13 | -0.36 | 0.72 | 1 | Gm43398 | | 0 | 0 | 0 | 20 | 8 | 27 | 24 | 10 | 22 | 5 | 26 |
| 12 | 0.01 | 1.32 | 0.01 | 0.99 | 1 | Gm37784 | | 0 | 1 | 6 | 3 | 7 | 19 | 24 | 0 | 30 | 32 | 12 |
| 12 | 0.57 | 1.72 | 0.33 | 0.74 | 1 | Doc2a | double C2, alpha | 5 | 0 | 0 | 2 | 7 | 14 | 24 | 70 | 4 | 0 | 9 |
| 12 | 0.00 | 1.41 | 0.00 | 1.00 | 1 | Gm36638 | | 4 | 5 | 9 | 4 | 0 | 19 | 24 | 0 | 3 | 19 | 40 |
| 12 | -0.59 | 1.29 | -0.46 | 0.65 | 1 | Gm47525 | | 10 | 26 | 11 | 7 | 2 | 15 | 24 | 0 | 12 | 13 | 5 |
| 11 | -0.29 | 1.71 | -0.17 | 0.87 | 1 | Gm6345 | | 20 | 27 | 7 | 0 | 4 | 2 | 24 | 0 | 0 | 27 | 10 |
| 11 | 0.63 | 1.48 | 0.43 | 0.67 | 1 | Gm46136 | | 8 | 15 | 17 | 0 | 2 | 8 | 24 | 0 | 6 | 14 | 23 |
| 8 | 2.63 | 1.76 | 1.49 | 0.14 | 1 | Gm13544 | | 3 | 0 | 0 | 0 | 0 | 8 | 24 | 15 | 6 | 23 | 7 |
| 5 | -3.18 | 2.99 | -1.07 | 0.29 | 1 | H2ac21 | H2A clustered histone 21 | 0 | 0 | 10 | 0 | 0 | 11 | 24 | 0 | 4 | 0 | 1 |
| 4 | -2.94 | 2.05 | -1.43 | 0.15 | 1 | Gm15991 | | 2 | 7 | 0 | 5 | 4 | 2 | 24 | 0 | 1 | 0 | 4 |
| 1077 | -0.15 | 1.58 | -0.09 | 0.92 | 1 | Pla2g1b | phospholipase A2, group IB, pancreas | 1473 | 5557 | 4421 | 250 | 4 | 3 | 23 | 0 | 16 | 21 | 78 |
| 761 | -0.49 | 0.94 | -0.53 | 0.60 | 1 | Mmp14 | matrix metallopeptidase 14 (membrane-ir | 3899 | 1975 | 2399 | 7 | 7 | 21 | 23 | 16 | 13 | 0 | 8 |
| 448 | -0.37 | 2.05 | -0.18 | 0.86 | 1 | Vsir | V-set immunoregulatory receptor | 2559 | 954 | 1371 | 0 | 0 | 7 | 23 | 0 | 11 | 0 | 4 |
| 356 | -1.73 | 1.53 | -1.13 | 0.26 | 1 | Cfp | complement factor properdin | 2123 | 772 | 624 | 0 | 26 | 190 | 23 | 0 | 12 | 0 | 147 |
| 335 | 0.36 | 1.35 | 0.26 | 0.79 | 1 | Dhrs3 | dehydrogenase/reductase (SDR family) n | 1350 | 977 | 1293 | 0 | 7 | 3 | 23 | 7 | 21 | 0 | 4 |
| 270 | -0.68 | 0.63 | -1.07 | 0.28 | 1 | Mctp2 | multiple C2 domains, transmembrane 2 | 913 | 1228 | 609 | 20 | 51 | 19 | 23 | 59 | 25 | 24 | 0 |
| 264 | -0.38 | 1.54 | -0.24 | 0.81 | 1 | Gbp9 | guanylate-binding protein 9 | 960 | 837 | 979 | 13 | 3 | 11 | 23 | 0 | 38 | 0 | 36 |
| 247 | -0.80 | 1.70 | -0.47 | 0.64 | 1 | Gfpt2 | glutamine fructose-6-phosphate transami | 1663 | 487 | 505 | 0 | 7 | 4 | 23 | 0 | 25 | 0 | 2 |

| | | | | | | | | | | | | | | | | | | |
|---|---|---|---|---|---|---|---|---|---|---|---|---|---|---|---|---|---|---|
| 191 | -2.10 | 1.25 | -1.68 | 0.09 | 1 | Sorcs1 | sortilin-related VPS10 domain containing | 597 | 643 | 663 | 33 | 42 | 34 | 23 | 0 | 38 | 0 | 27 |
| 167 | -0.26 | 0.35 | -0.75 | 0.45 | 1 | Rpl3-ps1 | | 71 | 598 | 145 | 367 | 102 | 97 | 23 | 241 | 53 | 40 | 95 |
| 146 | 0.09 | 0.56 | 0.17 | 0.86 | 1 | Atp6v0c-ps2 | | 33 | 3 | 424 | 40 | 512 | 12 | 23 | 34 | 471 | 35 | 16 |
| 145 | -1.41 | 1.60 | -0.88 | 0.38 | 1 | Tent5d | terminal nucleotidyltransferase 5D | 385 | 718 | 430 | 0 | 9 | 6 | 23 | 0 | 0 | 19 | 4 |
| 117 | -0.29 | 0.74 | -0.39 | 0.69 | 1 | Sox6 | SRY (sex determining region Y)-box 6 | 444 | 477 | 247 | 10 | 12 | 19 | 23 | 0 | 24 | 9 | 27 |
| 104 | -0.01 | 0.57 | -0.03 | 0.98 | 1 | Diaph3 | diaphanous related formin 3 | 291 | 106 | 165 | 99 | 103 | 61 | 23 | 67 | 73 | 104 | 51 |
| 104 | 0.31 | 0.63 | 0.50 | 0.62 | 1 | Tppp | tubulin polymerization promoting protein | 403 | 331 | 188 | 41 | 3 | 25 | 23 | 27 | 11 | 75 | 11 |
| 67 | -0.03 | 0.61 | -0.05 | 0.96 | 1 | Pter | phosphotriesterase related | 141 | 67 | 193 | 56 | 40 | 46 | 23 | 54 | 27 | 63 | 23 |
| 66 | -0.09 | 0.57 | -0.16 | 0.87 | 1 | Ttc25 | tetratricopeptide repeat domain 25 | 51 | 130 | 107 | 61 | 56 | 101 | 23 | 51 | 31 | 71 | 49 |
| 66 | -1.76 | 0.99 | -1.78 | 0.08 | 1 | Cldn8 | claudin 8 | 130 | 207 | 281 | 19 | 14 | 14 | 23 | 0 | 18 | 0 | 21 |
| 53 | 0.63 | 0.49 | 1.28 | 0.20 | 1 | Tmsb15l | thymosin beta 15b like | 75 | 103 | 175 | 15 | 25 | 30 | 23 | 27 | 34 | 36 | 44 |
| 52 | 0.30 | 0.56 | 0.54 | 0.59 | 1 | Pimreg | PICALM interacting mitotic regulator | 54 | 53 | 33 | 69 | 58 | 39 | 23 | 104 | 65 | 24 | 47 |
| 45 | -0.60 | 0.58 | -1.04 | 0.30 | 1 | Rps3a2 | | 63 | 72 | 122 | 39 | 75 | 13 | 23 | 19 | 21 | 30 | 14 |
| 44 | 1.07 | 0.60 | 1.78 | 0.08 | 1 | Gpat3 | glycerol-3-phosphate acyltransferase 3 | 95 | 62 | 83 | 37 | 9 | 10 | 23 | 43 | 33 | 73 | 17 |
| 43 | -0.16 | 0.70 | -0.22 | 0.82 | 1 | Phf2os1 | | 40 | 33 | 6 | 23 | 55 | 107 | 23 | 34 | 31 | 24 | 101 |
| 42 | -0.21 | 0.53 | -0.39 | 0.70 | 1 | Dynlt1-ps1 | | 46 | 44 | 44 | 47 | 38 | 63 | 23 | 25 | 22 | 41 | 64 |
| 40 | 0.06 | 0.87 | 0.07 | 0.94 | 1 | Gm47694 | | 13 | 15 | 14 | 10 | 10 | 127 | 23 | 0 | 30 | 19 | 174 |
| 38 | 0.30 | 0.57 | 0.53 | 0.60 | 1 | Akt2-ps | | 27 | 21 | 33 | 35 | 55 | 31 | 23 | 17 | 76 | 52 | 44 |
| 36 | -0.05 | 1.05 | -0.05 | 0.96 | 1 | Hs6st2 | heparan sulfate 6-O-sulfotransferase 2 | 82 | 47 | 176 | 7 | 8 | 4 | 23 | 28 | 17 | 0 | 3 |
| 34 | 0.17 | 0.55 | 0.30 | 0.76 | 1 | Gm5905 | | 66 | 57 | 54 | 22 | 13 | 38 | 23 | 24 | 24 | 17 | 41 |
| 34 | -0.11 | 0.72 | -0.15 | 0.88 | 1 | H4c4 | H4 clustered histone 4 | 9 | 6 | 32 | 43 | 48 | 46 | 23 | 53 | 24 | 12 | 78 |
| 33 | 0.10 | 0.53 | 0.18 | 0.86 | 1 | 1700102H20Rik | | 27 | 45 | 30 | 36 | 32 | 31 | 23 | 58 | 24 | 23 | 32 |
| 33 | -0.27 | 0.96 | -0.28 | 0.78 | 1 | Dazl | deleted in azoospermia-like | 0 | 4 | 56 | 55 | 48 | 36 | 23 | 51 | 36 | 18 | 30 |
| 33 | -0.87 | 0.92 | -0.95 | 0.34 | 1 | Gm10086 | | 31 | 34 | 131 | 23 | 33 | 17 | 23 | 0 | 22 | 32 | 12 |
| 32 | -0.76 | 0.62 | -1.23 | 0.22 | 1 | Unc5cl | unc-5 family C-terminal like | 45 | 36 | 109 | 21 | 42 | 18 | 23 | 21 | 16 | 14 | 8 |
| 30 | -0.22 | 0.96 | -0.23 | 0.82 | 1 | Gm13115 | | 5 | 7 | 20 | 16 | 32 | 75 | 23 | 51 | 42 | 0 | 62 |
| 27 | 1.09 | 0.73 | 1.48 | 0.14 | 1 | Gm12409 | | 13 | 24 | 23 | 25 | 7 | 13 | 23 | 87 | 19 | 45 | 13 |
| 25 | 0.30 | 0.86 | 0.35 | 0.73 | 1 | Gm30074 | | 9 | 13 | 0 | 62 | 18 | 25 | 23 | 23 | 34 | 39 | 34 |
| 25 | 0.15 | 0.86 | 0.17 | 0.86 | 1 | Rpl3-ps2 | | 39 | 14 | 51 | 11 | 15 | 13 | 23 | 1 | 30 | 68 | 10 |
| 21 | -1.24 | 0.98 | -1.26 | 0.21 | 1 | Klhl10 | kelch-like 10 | 13 | 6 | 25 | 30 | 20 | 49 | 23 | 15 | 16 | 0 | 38 |
| 21 | -0.65 | 0.82 | -0.80 | 0.42 | 1 | Pcdhb15 | protocadherin beta 15 | 44 | 23 | 11 | 11 | 22 | 35 | 23 | 20 | 8 | 5 | 29 |
| 21 | -1.47 | 1.06 | -1.39 | 0.16 | 1 | Tdrd12 | tudor domain containing 12 | 12 | 25 | 10 | 46 | 25 | 33 | 23 | 0 | 6 | 23 | 24 |
| 20 | -1.36 | 1.29 | -1.06 | 0.29 | 1 | Gm13261 | | 9 | 14 | 18 | 9 | 7 | 60 | 23 | 0 | 26 | 0 | 59 |
| 20 | -1.01 | 0.99 | -1.02 | 0.31 | 1 | Chrnb4 | cholinergic receptor, nicotinic, beta polyp | 24 | 10 | 41 | 16 | 16 | 25 | 23 | 0 | 46 | 10 | 13 |
| 20 | 0.13 | 0.78 | 0.17 | 0.86 | 1 | Vdac3-ps1 | | 48 | 9 | 20 | 11 | 26 | 13 | 23 | 16 | 22 | 12 | 22 |
| 20 | -0.74 | 1.30 | -0.57 | 0.57 | 1 | H2ac19 | H2A clustered histone 19 | 16 | 1 | 0 | 19 | 34 | 14 | 23 | 0 | 53 | 53 | 7 |
| 19 | -0.25 | 0.68 | -0.37 | 0.71 | 1 | Gm9803 | | 25 | 14 | 9 | 19 | 9 | 31 | 23 | 12 | 8 | 19 | 36 |
| 19 | 0.56 | 0.85 | 0.65 | 0.51 | 1 | Gm30970 | | 5 | 7 | 6 | 4 | 24 | 25 | 23 | 45 | 23 | 19 | 23 |
| 18 | 0.18 | 1.33 | 0.13 | 0.89 | 1 | Gm28523 | | 21 | 12 | 43 | 0 | 8 | 31 | 23 | 7 | 26 | 0 | 24 |
| 18 | 0.12 | 0.87 | 0.14 | 0.89 | 1 | 4930426L09Rik | | 16 | 20 | 21 | 20 | 9 | 5 | 23 | 57 | 6 | 12 | 5 |
| 17 | -1.80 | 1.09 | -1.66 | 0.10 | 1 | Impdh2-ps | | 42 | 40 | 9 | 11 | 35 | 4 | 23 | 2 | 4 | 4 | 14 |
| 17 | -0.42 | 0.77 | -0.55 | 0.58 | 1 | Gm49774 | | 20 | 12 | 18 | 27 | 20 | 9 | 23 | 25 | 18 | 8 | 8 |
| 16 | -0.53 | 1.28 | -0.42 | 0.68 | 1 | Gm36283 | | 7 | 19 | 74 | 3 | 8 | 13 | 23 | 8 | 0 | 14 | 6 |
| 15 | -1.69 | 1.23 | -1.38 | 0.17 | 1 | Gm11295 | | 4 | 11 | 12 | 29 | 21 | 15 | 23 | 0 | 22 | 1 | 30 |
| 15 | -0.55 | 0.88 | -0.63 | 0.53 | 1 | Mrpl23-ps1 | | 12 | 12 | 9 | 28 | 12 | 14 | 23 | 25 | 10 | 3 | 18 |
| 15 | -0.05 | 0.88 | -0.06 | 0.95 | 1 | Gm15326 | | 9 | 18 | 15 | 8 | 13 | 15 | 23 | 20 | 4 | 22 | 15 |
| 15 | 1.06 | 1.34 | 0.79 | 0.43 | 1 | Gm4811 | | 23 | 36 | 0 | 9 | 0 | 6 | 23 | 52 | 4 | 3 | 4 |
| 13 | -0.02 | 1.36 | -0.02 | 0.99 | 1 | Gm7517 | | 17 | 20 | 1 | 2 | 16 | 12 | 23 | 0 | 19 | 24 | 11 |
| 13 | -1.92 | 1.34 | -1.43 | 0.15 | 1 | Gm21814 | | 23 | 19 | 21 | 11 | 5 | 16 | 23 | 0 | 9 | 0 | 14 |
| 13 | -0.78 | 1.18 | -0.67 | 0.50 | 1 | Gm12996 | | 3 | 5 | 9 | 13 | 10 | 18 | 23 | 0 | 28 | 22 | 5 |
| 12 | -1.52 | 1.73 | -0.88 | 0.38 | 1 | Grhl3 | grainyhead like transcription factor 3 | 34 | 14 | 0 | 8 | 13 | 13 | 23 | 0 | 14 | 0 | 14 |
| 11 | -2.16 | 1.34 | -1.61 | 0.11 | 1 | Gm13712 | | 16 | 0 | 16 | 11 | 15 | 10 | 23 | 0 | 11 | 19 | 2 |
| 11 | -0.92 | 1.33 | -0.70 | 0.49 | 1 | Gm5931 | | 30 | 8 | 30 | 1 | 11 | 3 | 23 | 1 | 6 | 0 | 9 |
| 10 | -0.42 | 1.35 | -0.31 | 0.76 | 1 | Schip1 | schwannomin interacting protein 1 | 11 | 1 | 11 | 15 | 0 | 11 | 23 | 9 | 12 | 9 | 5 |
| 9 | -0.70 | 1.71 | -0.41 | 0.68 | 1 | A830052D11Rik | | 19 | 8 | 8 | 0 | 7 | 10 | 23 | 0 | 14 | 0 | 12 |
| 9 | -1.87 | 2.03 | -0.92 | 0.36 | 1 | AI197445 | | 0 | 13 | 0 | 17 | 9 | 10 | 23 | 0 | 7 | 0 | 20 |
| 8 | -1.02 | 1.97 | -0.52 | 0.60 | 1 | Tafa2 | TAFA chemokine like family member 2 | 4 | 41 | 16 | 2 | 0 | 1 | 23 | 0 | 2 | 0 | 4 |
| 8 | -2.55 | 2.18 | -1.17 | 0.24 | 1 | Gm7514 | | 3 | 18 | 0 | 0 | 9 | 10 | 23 | 0 | 0 | 22 | 2 |
| 8 | -2.26 | 1.94 | -1.16 | 0.24 | 1 | 2610206C17Rik | | 0 | 0 | 0 | 7 | 15 | 12 | 23 | 0 | 23 | 0 | 5 |
| 6 | -1.69 | 1.74 | -0.97 | 0.33 | 1 | Gm49085 | | 1 | 0 | 1 | 3 | 5 | 12 | 23 | 0 | 10 | 0 | 14 |

| | | | | | | | | | | | | | | | | | | |
|---|---|---|---|---|---|---|---|---|---|---|---|---|---|---|---|---|---|---|
| 7446 | -2.03 | 1.45 | -1.40 | 0.16 | 1 | Gp2 | glycoprotein 2 (zymogen granule membra | 9875 | 40448 | 30071 | 1110 | 87 | 3 | 22 | 100 | 29 | 0 | 156 |
| 4619 | 0.05 | 1.61 | 0.03 | 0.97 | 1 | Nid1 | nidogen 1 | 20572 | 12767 | 17390 | 0 | 4 | 12 | 22 | 0 | 13 | 33 | 0 |
| 994 | -0.64 | 0.94 | -0.68 | 0.50 | 1 | Adamts1 | a disintegrin-like and metallopeptidase (r | 5192 | 3084 | 2490 | 33 | 37 | 7 | 22 | 39 | 19 | 0 | 8 |
| 922 | -2.09 | 1.56 | -1.34 | 0.18 | 1 | Tm4sf1 | transmembrane 4 superfamily member 1 | 3719 | 2823 | 3568 | 3 | 0 | 3 | 22 | 0 | 1 | 0 | 6 |
| 711 | 0.35 | 1.64 | 0.21 | 0.83 | 1 | Gpc3 | glypican 3 | 4322 | 1517 | 1916 | 1 | 9 | 0 | 22 | 30 | 6 | 0 | 0 |
| 460 | 0.44 | 0.48 | 0.93 | 0.35 | 1 | Pdgfb | platelet derived growth factor, B polypepti | 1611 | 1544 | 1719 | 8 | 18 | 25 | 22 | 15 | 10 | 72 | 17 |
| 244 | -2.41 | 1.29 | -1.87 | 0.06 | 1 | Mmp9 | matrix metallopeptidase 9 | 1713 | 439 | 478 | 0 | 10 | 6 | 22 | 0 | 1 | 10 | 0 |
| 140 | -1.58 | 1.87 | -0.84 | 0.40 | 1 | Sstr2 | somatostatin receptor 2 | 322 | 738 | 376 | 31 | 0 | 14 | 22 | 0 | 19 | 0 | 13 |
| 115 | -0.11 | 0.42 | -0.26 | 0.79 | 1 | Efcc1 | EF hand and coiled-coil domain containir | 315 | 308 | 276 | 55 | 27 | 91 | 22 | 42 | 20 | 45 | 65 |
| 108 | -0.12 | 0.51 | -0.24 | 0.81 | 1 | Smim6 | small integral membrane protein 6 | 185 | 162 | 482 | 46 | 101 | 25 | 22 | 17 | 48 | 55 | 39 |
| 100 | 0.58 | 0.69 | 0.85 | 0.40 | 1 | Espl1 | extra spindle pole bodies 1, separase | 426 | 99 | 116 | 67 | 26 | 75 | 22 | 98 | 92 | 40 | 38 |
| 97 | 0.21 | 0.43 | 0.48 | 0.63 | 1 | Stac2 | SH3 and cysteine rich domain 2 | 446 | 159 | 320 | 11 | 20 | 12 | 22 | 7 | 25 | 32 | 14 |
| 93 | -2.10 | 1.24 | -1.69 | 0.09 | 1 | Pou3f4 | POU domain, class 3, transcription factor | 316 | 332 | 333 | 0 | 1 | 10 | 22 | 0 | 5 | 0 | 1 |
| 72 | 0.06 | 0.60 | 0.10 | 0.92 | 1 | Dna2 | DNA replication helicase/nuclease 2 | 236 | 112 | 121 | 10 | 13 | 130 | 22 | 27 | 4 | 36 | 76 |
| 60 | -0.62 | 0.51 | -1.23 | 0.22 | 1 | Kndc1 | kinase non-catalytic C-lobe domain (KIND | 99 | 127 | 129 | 63 | 46 | 51 | 22 | 53 | 27 | 12 | 30 |
| 60 | -1.38 | 0.76 | -1.83 | 0.07 | 1 | Lmo3 | LIM domain only 3 | 120 | 157 | 250 | 11 | 30 | 25 | 22 | 17 | 17 | 0 | 6 |
| 58 | 0.52 | 0.49 | 1.05 | 0.29 | 1 | Gm43379 | | 9 | 50 | 22 | 65 | 45 | 105 | 22 | 61 | 72 | 41 | 145 |
| 56 | -1.27 | 0.69 | -1.83 | 0.07 | 1 | Cntnap1 | contactin associated protein-like 1 | 190 | 82 | 88 | 65 | 12 | 74 | 22 | 30 | 2 | 10 | 38 |
| 50 | -1.52 | 1.23 | -1.23 | 0.22 | 1 | Sgo2a | shugoshin 2A | 164 | 67 | 51 | 53 | 41 | 41 | 22 | 0 | 57 | 0 | 58 |
| 48 | -0.49 | 0.89 | -0.55 | 0.58 | 1 | Car13 | carbonic anhydrase 13 | 160 | 131 | 92 | 34 | 18 | 8 | 22 | 0 | 25 | 33 | 7 |
| 47 | -1.42 | 0.92 | -1.54 | 0.12 | 1 | Cep72 | centrosomal protein 72 | 103 | 33 | 101 | 57 | 38 | 76 | 22 | 46 | 11 | 0 | 31 |
| 47 | -0.64 | 0.64 | -1.00 | 0.32 | 1 | Hyal1 | hyaluronoglucosaminidase 1 | 130 | 60 | 120 | 38 | 19 | 42 | 22 | 6 | 26 | 23 | 30 |
| 44 | -1.24 | 1.20 | -1.03 | 0.30 | 1 | Insyn2b | inhibitory synaptic factor family member 2 | 124 | 111 | 157 | 0 | 10 | 25 | 22 | 0 | 10 | 0 | 27 |
| 43 | 0.00 | 0.53 | -0.01 | 1.00 | 1 | Tmem221 | transmembrane protein 221 | 85 | 53 | 109 | 27 | 43 | 16 | 22 | 28 | 45 | 39 | 8 |
| 40 | 0.44 | 0.70 | 0.63 | 0.53 | 1 | Fdps | farnesyl diphosphate synthetase | 23 | 17 | 25 | 15 | 28 | 97 | 22 | 6 | 84 | 55 | 65 |
| 39 | 0.92 | 0.67 | 1.38 | 0.17 | 1 | Gm9761 | | 58 | 54 | 53 | 16 | 8 | 52 | 22 | 67 | 27 | 14 | 61 |
| 37 | 0.50 | 0.55 | 0.91 | 0.36 | 1 | 4930466K18Rik | | 33 | 62 | 84 | 26 | 29 | 15 | 22 | 50 | 31 | 22 | 29 |
| 34 | -0.50 | 0.65 | -0.77 | 0.44 | 1 | Gm16008 | | 3 | 7 | 11 | 19 | 26 | 141 | 22 | 7 | 24 | 21 | 93 |
| 33 | -0.85 | 1.22 | -0.70 | 0.48 | 1 | | | 0 | 13 | 0 | 10 | 7 | 125 | 22 | 52 | 7 | 0 | 130 |
| 31 | -0.22 | 0.67 | -0.34 | 0.74 | 1 | Cox5b-ps | | 45 | 5 | 46 | 7 | 59 | 36 | 22 | 3 | 31 | 53 | 28 |
| 30 | -0.25 | 1.10 | -0.22 | 0.82 | 1 | Pcdha8 | protocadherin alpha 8 | 0 | 5 | 14 | 84 | 20 | 39 | 22 | 19 | 16 | 91 | 17 |
| 29 | 0.74 | 0.73 | 1.02 | 0.31 | 1 | Gm10125 | | 27 | 21 | 14 | 6 | 26 | 40 | 22 | 19 | 24 | 85 | 35 |
| 29 | -0.97 | 0.95 | -1.01 | 0.31 | 1 | Gm49201 | | 54 | 25 | 29 | 33 | 25 | 40 | 22 | 0 | 15 | 46 | 24 |
| 27 | -0.37 | 0.73 | -0.50 | 0.62 | 1 | Gm13033 | | 16 | 47 | 21 | 17 | 48 | 30 | 22 | 20 | 46 | 8 | 26 |
| 27 | 0.69 | 0.87 | 0.79 | 0.43 | 1 | Cstad | CSA-conditional, T cell activation-depend | 40 | 54 | 28 | 23 | 2 | 20 | 22 | 51 | 22 | 5 | 25 |
| 25 | 0.14 | 0.98 | 0.14 | 0.89 | 1 | Tssk1 | testis-specific serine kinase 1 | 24 | 21 | 0 | 19 | 34 | 21 | 22 | 59 | 20 | 28 | 21 |
| 24 | -0.58 | 0.86 | -0.67 | 0.50 | 1 | Gm43034 | | 19 | 2 | 28 | 20 | 56 | 27 | 22 | 20 | 51 | 4 | 16 |
| 24 | -0.81 | 1.10 | -0.73 | 0.46 | 1 | Gm42777 | | 16 | 69 | 30 | 10 | 17 | 17 | 22 | 64 | 0 | 8 | 9 |
| 22 | -0.38 | 0.66 | -0.58 | 0.56 | 1 | Pdf | peptide deformylase (mitochondrial) | 27 | 35 | 12 | 25 | 15 | 32 | 22 | 13 | 13 | 24 | 21 |
| 21 | 1.20 | 0.80 | 1.50 | 0.13 | 1 | Gm10039 | | 9 | 7 | 46 | 5 | 14 | 10 | 22 | 16 | 31 | 45 | 21 |
| 21 | -0.53 | 1.01 | -0.53 | 0.60 | 1 | Pcdhb5 | protocadherin beta 5 | 90 | 13 | 19 | 15 | 6 | 5 | 22 | 0 | 5 | 44 | 7 |
| 18 | 0.37 | 0.82 | 0.45 | 0.65 | 1 | Gm10785 | | 23 | 11 | 4 | 10 | 4 | 35 | 22 | 5 | 13 | 24 | 42 |
| 17 | -0.07 | 0.79 | -0.09 | 0.93 | 1 | Gm20337 | | 12 | 6 | 14 | 23 | 9 | 26 | 22 | 12 | 10 | 36 | 23 |
| 17 | -1.48 | 1.08 | -1.36 | 0.17 | 1 | Neurl2 | neuralized E3 ubiquitin protein ligase 2 | 39 | 32 | 24 | 14 | 9 | 14 | 22 | 0 | 15 | 10 | 2 |
| 15 | 0.19 | 1.15 | 0.16 | 0.87 | 1 | Hspe1-rs1 | | 8 | 7 | 26 | 1 | 17 | 19 | 22 | 23 | 9 | 3 | 29 |
| 15 | -0.37 | 1.41 | -0.27 | 0.79 | 1 | Gp9 | glycoprotein 9 (platelet) | 3 | 10 | 0 | 18 | 39 | 8 | 22 | 0 | 24 | 19 | 18 |
| 13 | -1.75 | 1.12 | -1.55 | 0.12 | 1 | B930036N10Rik | | 8 | 5 | 20 | 29 | 17 | 8 | 22 | 6 | 19 | 0 | 5 |
| 13 | -1.01 | 1.11 | -0.91 | 0.36 | 1 | Gm5294 | predicted gene 5294 | 10 | 6 | 11 | 13 | 15 | 20 | 22 | 17 | 8 | 0 | 14 |
| 12 | -1.08 | 1.57 | -0.69 | 0.49 | 1 | Hes7 | hes family bHLH transcription factor 7 | 18 | 34 | 16 | 0 | 11 | 6 | 22 | 17 | 1 | 0 | 10 |
| 12 | -2.11 | 1.62 | -1.30 | 0.19 | 1 | Mx1 | | 40 | 38 | 11 | 1 | 8 | 1 | 22 | 0 | 2 | 0 | 7 |
| 11 | 0.17 | 1.20 | 0.14 | 0.89 | 1 | Uchl4 | ubiquitin carboxyl-terminal esterase L4 | 15 | 1 | 1 | 11 | 5 | 6 | 22 | 1 | 28 | 18 | 8 |
| 10 | -3.43 | 1.86 | -1.84 | 0.07 | 1 | Gm9970 | | 0 | 31 | 23 | 10 | 7 | 4 | 22 | 0 | 2 | 0 | 13 |
| 10 | 0.19 | 1.52 | 0.12 | 0.90 | 1 | Apoa1 | apolipoprotein A-I | 1 | 22 | 1 | 4 | 4 | 8 | 22 | 0 | 4 | 18 | 25 |
| 9 | -0.83 | 1.84 | -0.45 | 0.65 | 1 | Gm28905 | | 6 | 0 | 0 | 10 | 2 | 16 | 22 | 15 | 0 | 0 | 28 |
| 8 | -0.43 | 1.25 | -0.34 | 0.73 | 1 | Gm15264 | | 11 | 1 | 3 | 0 | 13 | 10 | 22 | 0 | 14 | 10 | 6 |
| 5 | -0.72 | 2.58 | -0.28 | 0.78 | 1 | Gm45694 | | 0 | 9 | 0 | 10 | 0 | 5 | 22 | 0 | 2 | 0 | 9 |
| 4 | -1.61 | 3.00 | -0.54 | 0.59 | 1 | 4932415D10Ril | uncharacterized LOC102635990 | 0 | 0 | 0 | 4 | 0 | 3 | 22 | 0 | 4 | 0 | 10 |
| 3 | -2.10 | 3.54 | -0.59 | 0.55 | 1 | Adh4 | alcohol dehydrogenase 4 (class II), pi pol | 0 | 0 | 0 | 0 | 6 | 0 | 22 | 0 | 2 | 0 | 0 |
| 2 | -3.90 | 3.56 | -1.10 | 0.27 | 1 | Gm48356 | | 0 | 0 | 0 | 0 | 0 | 5 | 22 | 0 | 0 | 0 | 0 |
| 3570 | -2.51 | 1.37 | -1.83 | 0.07 | 1 | Fbn1 | fibrillin 1 | 16380 | 8603 | 14222 | 0 | 31 | 4 | 21 | 0 | 7 | 0 | 2 |

| | | | | | | | | | | | | | | | | | | |
|---|---|---|---|---|---|---|---|---|---|---|---|---|---|---|---|---|---|---|
| 2033 | 0.17 | 0.96 | 0.18 | 0.86 | 1 | Lama2 | laminin, alpha 2 | 8033 | 6203 | 7938 | 4 | 31 | 22 | 21 | 0 | 22 | 75 | 19 |
| 1441 | 0.39 | 1.24 | 0.31 | 0.75 | 1 | Tff2 | trefoil factor 2 (spasmolytic protein 1) | 1629 | 6829 | 6973 | 194 | 23 | 0 | 21 | 56 | 26 | 21 | 78 |
| 861 | -2.39 | 1.40 | -1.71 | 0.09 | 1 | Fcgrt | Fc receptor, IgG, alpha chain transporter | 4733 | 2033 | 2665 | 0 | 13 | 3 | 21 | 0 | 5 | 0 | 1 |
| 476 | 0.63 | 1.45 | 0.43 | 0.67 | 1 | Serpina3n | serine (or cysteine) peptidase inhibitor, cl | 3215 | 844 | 1095 | 0 | 5 | 12 | 21 | 9 | 26 | 0 | 4 |
| 315 | -1.15 | 2.20 | -0.52 | 0.60 | 1 | Gda | guanine deaminase | 2025 | 975 | 439 | 0 | 3 | 0 | 21 | 0 | 4 | 0 | 0 |
| 156 | -0.83 | 0.67 | -1.24 | 0.21 | 1 | Calhm2 | calcium homeostasis modulator family me | 467 | 434 | 474 | 61 | 89 | 37 | 21 | 41 | 59 | 0 | 32 |
| 151 | -0.74 | 0.56 | -1.31 | 0.19 | 1 | Cracr2b | calcium release activated channel regula | 524 | 396 | 523 | 37 | 24 | 50 | 21 | 2 | 20 | 36 | 29 |
| 134 | 0.34 | 0.51 | 0.66 | 0.51 | 1 | Tesc | tescalcin | 346 | 337 | 334 | 21 | 63 | 102 | 21 | 83 | 27 | 32 | 107 |
| 102 | 0.64 | 0.94 | 0.68 | 0.50 | 1 | Abca6 | ATP-binding cassette, sub-family A (ABC | 481 | 286 | 258 | 0 | 0 | 21 | 21 | 0 | 2 | 26 | 27 |
| 77 | 0.38 | 0.49 | 0.79 | 0.43 | 1 | Gm11131 | | 117 | 137 | 139 | 89 | 38 | 52 | 21 | 73 | 96 | 35 | 50 |
| 76 | -1.24 | 0.74 | -1.68 | 0.09 | 1 | Adamts10 | a disintegrin-like and metallopeptidase (r | 319 | 180 | 161 | 14 | 15 | 77 | 21 | 19 | 8 | 0 | 22 |
| 75 | -0.05 | 0.53 | -0.09 | 0.93 | 1 | Dnajc22 | DnaJ heat shock protein family (Hsp40) m | 106 | 276 | 222 | 28 | 38 | 25 | 21 | 34 | 14 | 36 | 23 |
| 69 | -0.69 | 0.92 | -0.75 | 0.45 | 1 | Caps2 | calcyphosphine 2 | 5 | 11 | 28 | 27 | 9 | 309 | 21 | 57 | 6 | 0 | 284 |
| 65 | 0.89 | 0.98 | 0.91 | 0.36 | 1 | Akr1b8 | aldo-keto reductase family 1, member B8 | 285 | 161 | 134 | 11 | 22 | 1 | 21 | 41 | 26 | 0 | 11 |
| 65 | 0.03 | 0.71 | 0.04 | 0.97 | 1 | Cox17 | cytochrome c oxidase assembly protein 1 | 16 | 44 | 4 | 15 | 19 | 242 | 21 | 7 | 18 | 27 | 299 |
| 64 | -0.82 | 0.59 | -1.38 | 0.17 | 1 | L3hypdh | L-3-hydroxyproline dehydratase (trans-) | 221 | 86 | 167 | 56 | 31 | 35 | 21 | 12 | 25 | 19 | 28 |
| 54 | -0.46 | 0.60 | -0.75 | 0.45 | 1 | Rac3 | Rac family small GTPase 3 | 108 | 62 | 116 | 61 | 41 | 48 | 21 | 28 | 14 | 30 | 65 |
| 49 | -1.13 | 0.97 | -1.16 | 0.25 | 1 | Zc3h12a | zinc finger CCCH type containing 12A | 273 | 147 | 60 | 4 | 0 | 23 | 21 | 0 | 4 | 8 | 4 |
| 48 | -0.53 | 0.81 | -0.66 | 0.51 | 1 | Zfp300 | zinc finger protein 300 | 154 | 90 | 122 | 25 | 20 | 22 | 21 | 0 | 24 | 26 | 24 |
| 45 | -0.38 | 0.74 | -0.52 | 0.60 | 1 | Cdca5 | cell division cycle associated 5 | 188 | 36 | 35 | 68 | 32 | 15 | 21 | 36 | 16 | 26 | 24 |
| 43 | 0.19 | 0.62 | 0.31 | 0.75 | 1 | Nhej1 | non-homologous end joining factor 1 | 41 | 30 | 68 | 64 | 21 | 49 | 21 | 60 | 27 | 22 | 71 |
| 42 | -0.57 | 0.76 | -0.75 | 0.45 | 1 | Scn2a | sodium channel, voltage-gated, type II, al | 112 | 91 | 110 | 12 | 18 | 35 | 21 | 31 | 20 | 8 | 4 |
| 39 | -0.09 | 0.67 | -0.13 | 0.89 | 1 | Zfp804b | zinc finger protein 804B | 30 | 36 | 14 | 39 | 55 | 58 | 21 | 21 | 23 | 57 | 80 |
| 38 | -0.40 | 0.58 | -0.68 | 0.50 | 1 | Gm4117 | | 59 | 66 | 80 | 33 | 31 | 36 | 21 | 20 | 13 | 36 | 25 |
| 38 | -0.24 | 0.65 | -0.37 | 0.71 | 1 | E130208F15Rik | | 4 | 79 | 37 | 39 | 45 | 50 | 21 | 44 | 42 | 14 | 42 |
| 32 | -0.52 | 1.03 | -0.51 | 0.61 | 1 | D630023F18Ril | RIKEN cDNA D630023F18 gene | 27 | 18 | 69 | 40 | 12 | 41 | 21 | 0 | 33 | 22 | 64 |
| 30 | -0.07 | 0.67 | -0.11 | 0.91 | 1 | Gm14165 | | 25 | 58 | 38 | 39 | 36 | 4 | 21 | 63 | 13 | 26 | 4 |
| 27 | -1.10 | 1.17 | -0.95 | 0.34 | 1 | Gm36266 | | 12 | 15 | 0 | 29 | 15 | 85 | 21 | 0 | 8 | 12 | 104 |
| 27 | -0.18 | 0.97 | -0.18 | 0.85 | 1 | Gm4247 | | 24 | 38 | 15 | 12 | 24 | 30 | 21 | 95 | 17 | 1 | 17 |
| 26 | -1.15 | 1.09 | -1.05 | 0.29 | 1 | A430103D13Rik | | 51 | 8 | 17 | 36 | 24 | 31 | 21 | 13 | 9 | 0 | 74 |
| 26 | 1.65 | 1.05 | 1.57 | 0.12 | 1 | Gm47824 | | 0 | 0 | 0 | 6 | 1 | 85 | 21 | 29 | 35 | 40 | 65 |
| 26 | -0.18 | 0.97 | -0.18 | 0.85 | 1 | Fbxw10 | F-box and WD-40 domain protein 10 | 26 | 48 | 51 | 16 | 18 | 16 | 21 | 44 | 26 | 0 | 15 |
| 25 | -0.88 | 1.01 | -0.86 | 0.39 | 1 | Lgr6 | leucine-rich repeat-containing G protein-c | 44 | 35 | 27 | 41 | 20 | 14 | 21 | 53 | 19 | 0 | 4 |
| 24 | 0.40 | 1.08 | 0.37 | 0.71 | 1 | Ppara | peroxisome proliferator activated receptor | 32 | 41 | 82 | 4 | 2 | 20 | 21 | 0 | 14 | 36 | 16 |
| 24 | 0.82 | 0.75 | 1.09 | 0.28 | 1 | A330041J22Rik | | 21 | 40 | 33 | 15 | 8 | 18 | 21 | 15 | 49 | 24 | 24 |
| 24 | 0.63 | 0.82 | 0.76 | 0.45 | 1 | Slc22a2 | solute carrier family 22 (organic cation tra | 14 | 29 | 18 | 8 | 36 | 10 | 21 | 68 | 15 | 26 | 18 |
| 23 | 0.34 | 0.88 | 0.38 | 0.70 | 1 | Cartpt | CART prepropeptide | 27 | 6 | 17 | 67 | 8 | 4 | 21 | 60 | 16 | 23 | 7 |
| 23 | -0.87 | 1.10 | -0.79 | 0.43 | 1 | Slc2a12 | solute carrier family 2 (facilitated glucose | 59 | 72 | 65 | 8 | 2 | 7 | 21 | 5 | 12 | 0 | 4 |
| 23 | -0.59 | 1.10 | -0.54 | 0.59 | 1 | Zfyve28 | zinc finger, FYVE domain containing 28 | 68 | 21 | 32 | 0 | 28 | 30 | 21 | 0 | 11 | 22 | 19 |
| 22 | -1.62 | 1.01 | -1.60 | 0.11 | 1 | Slc25a47 | solute carrier family 25, member 47 | 12 | 47 | 31 | 18 | 18 | 43 | 21 | 0 | 4 | 9 | 44 |
| 22 | 0.28 | 0.68 | 0.42 | 0.68 | 1 | Gm8203 | | 9 | 40 | 12 | 30 | 11 | 16 | 21 | 51 | 20 | 13 | 20 |
| 21 | 0.06 | 0.74 | 0.09 | 0.93 | 1 | Ccdc69 | coiled-coil domain containing 69 | 77 | 17 | 46 | 4 | 11 | 11 | 21 | 10 | 13 | 14 | 7 |
| 21 | -1.18 | 0.78 | -1.51 | 0.13 | 1 | Ncoa4-ps | | 16 | 16 | 23 | 43 | 33 | 20 | 21 | 8 | 28 | 4 | 18 |
| 21 | -0.80 | 1.01 | -0.79 | 0.43 | 1 | Gm17399 | | 34 | 44 | 33 | 6 | 15 | 30 | 21 | 16 | 0 | 15 | 15 |
| 20 | 0.13 | 1.01 | 0.13 | 0.90 | 1 | 4930599N23Rik | | 27 | 13 | 0 | 14 | 22 | 33 | 21 | 20 | 26 | 21 | 27 |
| 19 | 0.49 | 0.71 | 0.69 | 0.49 | 1 | Gm5844 | | 27 | 11 | 4 | 24 | 9 | 18 | 21 | 37 | 11 | 28 | 24 |
| 18 | -1.31 | 1.13 | -1.16 | 0.25 | 1 | Gm18180 | | 6 | 46 | 18 | 36 | 9 | 24 | 21 | 0 | 7 | 8 | 24 |
| 18 | -1.87 | 1.25 | -1.50 | 0.13 | 1 | Gm43054 | | 6 | 0 | 5 | 17 | 8 | 55 | 21 | 2 | 5 | 0 | 77 |
| 18 | 0.49 | 1.30 | 0.38 | 0.70 | 1 | Gm27003 | | 52 | 8 | 19 | 1 | 4 | 32 | 21 | 22 | 4 | 0 | 32 |
| 17 | 1.90 | 1.34 | 1.42 | 0.16 | 1 | Aadat | aminoadipate aminotransferase | 0 | 13 | 0 | 18 | 0 | 12 | 21 | 49 | 17 | 27 | 28 |
| 17 | -0.22 | 1.47 | -0.15 | 0.88 | 1 | Gm45203 | | 0 | 0 | 0 | 5 | 11 | 41 | 21 | 58 | 7 | 0 | 39 |
| 17 | 0.00 | 1.13 | 0.00 | 1.00 | 1 | Gm47512 | | 1 | 0 | 16 | 15 | 10 | 30 | 21 | 22 | 13 | 9 | 45 |
| 17 | -1.10 | 1.39 | -0.79 | 0.43 | 1 | Srrm4os | | 0 | 13 | 0 | 18 | 22 | 16 | 21 | 59 | 18 | 0 | 15 |
| 16 | 1.32 | 0.90 | 1.47 | 0.14 | 1 | Hcn1 | hyperpolarization activated cyclic nucleot | 26 | 8 | 23 | 6 | 7 | 5 | 21 | 22 | 24 | 9 | 26 |
| 16 | -1.60 | 1.37 | -1.16 | 0.24 | 1 | AV039307 | | 18 | 9 | 3 | 10 | 25 | 32 | 21 | 0 | 24 | 0 | 32 |
| 14 | 0.79 | 1.27 | 0.62 | 0.53 | 1 | 4930506C21Rik | | 10 | 12 | 0 | 8 | 4 | 15 | 21 | 23 | 39 | 13 | 3 |
| 13 | -0.39 | 1.34 | -0.29 | 0.77 | 1 | Gm6565 | | 14 | 43 | 1 | 11 | 12 | 5 | 21 | 13 | 21 | 0 | 6 |
| 13 | -2.55 | 1.31 | -1.95 | 0.05 | 1 | Kpna2-ps | | 12 | 18 | 17 | 18 | 23 | 13 | 21 | 0 | 21 | 0 | 4 |
| 13 | -2.63 | 1.47 | -1.79 | 0.07 | 1 | Pou3f1 | POU domain, class 3, transcription factor | 39 | 10 | 46 | 0 | 16 | 7 | 21 | 0 | 6 | 0 | 1 |
| 13 | -1.91 | 1.42 | -1.35 | 0.18 | 1 | Gm37637 | | 21 | 22 | 4 | 11 | 13 | 24 | 21 | 0 | 17 | 0 | 11 |

| | | | | | | | | | | | | | | | | | | |
|---|---|---|---|---|---|---|---|---|---|---|---|---|---|---|---|---|---|---|
| 12 | 0.67 | 1.00 | 0.67 | 0.50 | 1 | Atp5pb-ps | | 15 | 8 | 3 | 9 | 3 | 9 | 21 | 22 | 17 | 27 | 2 |
| 12 | -1.02 | 2.16 | -0.47 | 0.64 | 1 | Gm38192 | | 0 | 17 | 32 | 0 | 6 | 10 | 21 | 36 | 0 | 0 | 12 |
| 12 | -2.06 | 2.04 | -1.01 | 0.31 | 1 | Gm14963 | | 19 | 31 | 0 | 0 | 11 | 22 | 21 | 0 | 5 | 0 | 20 |
| 11 | 0.49 | 1.60 | 0.31 | 0.76 | 1 | Hist1h4n | histone cluster 1, H4n | 0 | 0 | 3 | 14 | 6 | 7 | 21 | 45 | 0 | 23 | 7 |
| 9 | -1.28 | 1.26 | -1.01 | 0.31 | 1 | Gm9001 | | 2 | 17 | 6 | 4 | 11 | 10 | 21 | 1 | 9 | 3 | 9 |
| 8 | -1.09 | 1.14 | -0.96 | 0.34 | 1 | Gm7476 | | 14 | 7 | 1 | 10 | 9 | 7 | 21 | 3 | 9 | 4 | 3 |
| 8 | -1.73 | 1.68 | -1.03 | 0.30 | 1 | Gm715 | predicted gene 715 | 10 | 0 | 10 | 12 | 16 | 1 | 21 | 6 | 11 | 0 | 0 |
| 8 | 0.31 | 1.41 | 0.22 | 0.83 | 1 | Gm15420 | | 2 | 0 | 2 | 0 | 10 | 5 | 21 | 0 | 8 | 15 | 20 |
| 7 | -4.02 | 2.12 | -1.89 | 0.06 | 1 | Gm49077 | | 0 | 0 | 0 | 9 | 9 | 20 | 21 | 0 | 1 | 0 | 13 |
| 5 | -1.30 | 2.04 | -0.64 | 0.52 | 1 | Gm35106 | | 10 | 7 | 1 | 0 | 7 | 3 | 21 | 0 | 8 | 0 | 2 |
| 4 | -2.63 | 2.34 | -1.12 | 0.26 | 1 | Gm44020 | | 0 | 0 | 0 | 8 | 4 | 3 | 21 | 10 | 0 | 0 | 1 |
| 4 | -1.53 | 2.07 | -0.74 | 0.46 | 1 | Gm5481 | | 4 | 8 | 1 | 1 | 0 | 5 | 21 | 0 | 1 | 0 | 5 |
| 4 | -0.70 | 3.12 | -0.22 | 0.82 | 1 | Slc22a29 | solute carrier family 22. member 29 | 0 | 0 | 6 | 0 | 0 | 3 | 21 | 0 | 9 | 0 | 4 |
| 560 | -0.11 | 0.75 | -0.14 | 0.89 | 1 | S100a6 | S100 calcium binding protein A6 (calcycli | 2638 | 1197 | 1647 | 129 | 148 | 37 | 20 | 219 | 77 | 27 | 25 |
| 468 | -0.94 | 0.94 | -1.00 | 0.32 | 1 | Prex2 | phosphatidylinositol-3,4,5-trisphosphate-c | 1760 | 1655 | 1613 | 2 | 22 | 29 | 20 | 0 | 6 | 24 | 12 |
| 432 | -1.69 | 1.67 | -1.01 | 0.31 | 1 | Tgfb1 | transforming growth factor, beta 1 | 2078 | 1102 | 1525 | 0 | 5 | 8 | 20 | 0 | 16 | 0 | 1 |
| 153 | -0.83 | 0.76 | -1.09 | 0.28 | 1 | Ctxn2 | cortexin 2 | 492 | 494 | 546 | 32 | 15 | 15 | 20 | 0 | 9 | 53 | 7 |
| 114 | 0.85 | 0.48 | 1.77 | 0.08 | 1 | 5430416N02Rik | | 52 | 81 | 121 | 30 | 47 | 330 | 20 | 61 | 41 | 90 | 383 |
| 100 | 0.43 | 0.53 | 0.81 | 0.42 | 1 | Snhg15 | | 109 | 48 | 26 | 32 | 36 | 380 | 20 | 36 | 47 | 86 | 282 |
| 95 | 0.03 | 0.52 | 0.05 | 0.96 | 1 | Rttn | rotatin | 250 | 112 | 137 | 64 | 50 | 148 | 20 | 61 | 54 | 30 | 124 |
| 95 | -1.33 | 0.92 | -1.45 | 0.15 | 1 | St6galnac2 | ST6 (alpha-N-acetyl-neuraminyl-2,3-beta | 239 | 340 | 342 | 6 | 6 | 47 | 20 | 0 | 0 | 21 | 24 |
| 90 | -0.52 | 0.52 | -1.00 | 0.32 | 1 | Smarcd3 | SWI/SNF related, matrix associated, actin | 376 | 236 | 207 | 35 | 16 | 25 | 20 | 30 | 11 | 26 | 7 |
| 72 | -0.98 | 0.87 | -1.13 | 0.26 | 1 | Klhl1 | kelch-like 1 | 99 | 116 | 8 | 166 | 7 | 119 | 20 | 80 | 3 | 0 | 177 |
| 63 | -0.52 | 0.87 | -0.60 | 0.55 | 1 | Gm45487 | | 15 | 0 | 0 | 9 | 20 | 384 | 20 | 9 | 10 | 21 | 210 |
| 59 | -1.68 | 1.04 | -1.62 | 0.11 | 1 | A530016L24Ril | RIKEN cDNA A530016L24 gene | 154 | 235 | 176 | 29 | 10 | 2 | 20 | 20 | 0 | 0 | 5 |
| 53 | -0.10 | 0.59 | -0.16 | 0.87 | 1 | Capn6 | calpain 6 | 160 | 110 | 161 | 24 | 15 | 17 | 20 | 24 | 33 | 9 | 10 |
| 47 | -1.35 | 1.33 | -1.01 | 0.31 | 1 | Plekhf1 | pleckstrin homology domain containing, f | 182 | 151 | 110 | 4 | 1 | 7 | 20 | 39 | 0 | 0 | 1 |
| 45 | -1.37 | 1.36 | -1.01 | 0.31 | 1 | Larp6 | La ribonucleoprotein domain family, mem | 233 | 107 | 100 | 0 | 9 | 6 | 20 | 16 | 0 | 0 | 3 |
| 43 | -0.41 | 0.77 | -0.53 | 0.59 | 1 | Rhof | ras homolog family member F (in filopodia | 88 | 99 | 21 | 24 | 60 | 47 | 20 | 10 | 49 | 28 | 26 |
| 41 | -0.63 | 0.99 | -0.64 | 0.52 | 1 | Btc | betacellulin, epidermal growth factor fami | 135 | 23 | 101 | 28 | 28 | 30 | 20 | 24 | 45 | 0 | 12 |
| 40 | -0.01 | 0.58 | -0.02 | 0.98 | 1 | Magee2 | melanoma antigen, family E, 2 | 77 | 54 | 65 | 54 | 28 | 17 | 20 | 32 | 28 | 45 | 14 |
| 38 | 0.14 | 0.86 | 0.16 | 0.87 | 1 | Nrgn | neurogranin | 73 | 126 | 80 | 5 | 19 | 8 | 20 | 13 | 13 | 62 | 1 |
| 38 | 0.94 | 0.80 | 1.17 | 0.24 | 1 | Sec14l2 | SEC14-like lipid binding 2 | 80 | 111 | 48 | 35 | 1 | 12 | 20 | 51 | 30 | 13 | 16 |
| 34 | 0.63 | 1.05 | 0.60 | 0.55 | 1 | Gm43815 | | 7 | 0 | 44 | 20 | 23 | 72 | 20 | 95 | 39 | 17 | 41 |
| 33 | 0.43 | 0.59 | 0.72 | 0.47 | 1 | Gm45844 | | 15 | 29 | 17 | 33 | 20 | 47 | 20 | 90 | 16 | 23 | 53 |
| 33 | -1.53 | 1.91 | -0.80 | 0.42 | 1 | Fgb | fibrinogen beta chain | 51 | 137 | 152 | 0 | 0 | 0 | 20 | 0 | 0 | 1 | 0 |
| 31 | 0.34 | 0.76 | 0.45 | 0.66 | 1 | Gm16386 | | 11 | 14 | 8 | 25 | 18 | 59 | 20 | 29 | 51 | 4 | 101 |
| 30 | 0.37 | 0.95 | 0.39 | 0.70 | 1 | Dusp15 | dual specificity phosphatase-like 15 | 33 | 8 | 0 | 23 | 19 | 28 | 20 | 95 | 11 | 80 | 15 |
| 30 | -0.46 | 0.75 | -0.62 | 0.54 | 1 | Prodh | proline dehydrogenase | 65 | 86 | 27 | 10 | 42 | 17 | 20 | 15 | 17 | 17 | 13 |
| 30 | 0.19 | 0.59 | 0.32 | 0.75 | 1 | Gm32123 | | 21 | 32 | 16 | 39 | 23 | 35 | 20 | 64 | 30 | 24 | 21 |
| 29 | 0.57 | 0.95 | 0.60 | 0.55 | 1 | Shisa7 | shisa family member 7 | 54 | 26 | 33 | 26 | 9 | 12 | 20 | 1 | 15 | 90 | 32 |
| 29 | -0.41 | 0.75 | -0.55 | 0.58 | 1 | Gm50462 | | 57 | 29 | 8 | 25 | 37 | 44 | 20 | 31 | 16 | 18 | 34 |
| 28 | -0.26 | 0.98 | -0.27 | 0.79 | 1 | Gm44717 | | 16 | 0 | 3 | 60 | 26 | 47 | 20 | 22 | 49 | 24 | 41 |
| 27 | -0.16 | 1.00 | -0.16 | 0.87 | 1 | Calca | calcitonin/calcitonin-related polypeptide, a | 81 | 38 | 83 | 8 | 11 | 9 | 20 | 23 | 14 | 0 | 8 |
| 26 | -0.94 | 0.72 | -1.30 | 0.19 | 1 | 1700022N22Rik | | 12 | 29 | 37 | 67 | 23 | 23 | 20 | 22 | 33 | 4 | 16 |
| 26 | -1.27 | 0.98 | -1.29 | 0.20 | 1 | Clvs1 | clavesin 1 | 24 | 81 | 53 | 19 | 16 | 29 | 20 | 0 | 5 | 17 | 20 |
| 25 | -0.08 | 0.95 | -0.09 | 0.93 | 1 | Gm15879 | | 33 | 29 | 15 | 7 | 9 | 67 | 20 | 0 | 10 | 41 | 50 |
| 25 | 0.34 | 0.76 | 0.44 | 0.66 | 1 | Rps6-ps3 | | 51 | 41 | 22 | 11 | 32 | 8 | 20 | 29 | 30 | 10 | 16 |
| 24 | 0.79 | 0.94 | 0.84 | 0.40 | 1 | Pcdhb10 | protocadherin beta 10 | 34 | 22 | 38 | 1 | 20 | 26 | 20 | 36 | 36 | 15 | 13 |
| 24 | -0.25 | 0.67 | -0.36 | 0.72 | 1 | A930002H24Rik | | 31 | 23 | 39 | 28 | 25 | 17 | 20 | 14 | 13 | 23 | 27 |
| 23 | 0.68 | 0.78 | 0.87 | 0.39 | 1 | Gm12663 | | 21 | 13 | 27 | 24 | 15 | 6 | 20 | 43 | 11 | 63 | 6 |
| 23 | 0.18 | 0.65 | 0.28 | 0.78 | 1 | Eef2-ps2 | | 31 | 18 | 36 | 19 | 15 | 21 | 20 | 23 | 20 | 26 | 17 |
| 22 | -0.04 | 1.28 | -0.03 | 0.97 | 1 | 4930519P11Rik | | 1 | 24 | 28 | 21 | 28 | 20 | 20 | 0 | 26 | 59 | 15 |
| 22 | -0.17 | 0.74 | -0.23 | 0.82 | 1 | 2310074N15Rik | | 3 | 4 | 6 | 45 | 32 | 16 | 20 | 43 | 24 | 39 | 6 |
| 22 | 0.21 | 1.11 | 0.19 | 0.85 | 1 | C230062I16Rik | | 28 | 37 | 4 | 13 | 6 | 40 | 20 | 20 | 19 | 0 | 50 |
| 21 | -0.04 | 0.88 | -0.04 | 0.96 | 1 | Gm47210 | | 12 | 21 | 5 | 13 | 34 | 21 | 20 | 29 | 30 | 45 | 4 |
| 21 | -1.40 | 1.04 | -1.35 | 0.18 | 1 | Slc23a4 | solute carrier family 23 member 4 | 56 | 9 | 10 | 32 | 7 | 46 | 20 | 0 | 8 | 17 | 26 |
| 21 | -0.85 | 1.02 | -0.84 | 0.40 | 1 | 5330439K02Rik | | 12 | 13 | 12 | 37 | 29 | 31 | 20 | 0 | 25 | 24 | 24 |
| 20 | 0.31 | 1.14 | 0.27 | 0.78 | 1 | Gm43457 | | 24 | 25 | 22 | 14 | 3 | 28 | 20 | 44 | 21 | 0 | 19 |
| 20 | -1.10 | 1.16 | -0.95 | 0.34 | 1 | Gm44878 | | 14 | 33 | 1 | 22 | 31 | 28 | 20 | 37 | 12 | 0 | 19 |

| | | | | | | | | | | | | | | | | | | |
|---|---|---|---|---|---|---|---|---|---|---|---|---|---|---|---|---|---|---|
| 19 | 0.83 | 1.41 | 0.59 | 0.55 | 1 | Afmid | arylformamidase | 35 | 42 | 26 | 1 | 0 | 31 | 20 | 10 | 12 | 0 | 35 |
| 19 | -0.26 | 0.79 | -0.33 | 0.74 | 1 | Hspd1-ps3 | | 24 | 5 | 24 | 11 | 31 | 22 | 20 | 13 | 10 | 37 | 12 |
| 19 | -0.14 | 1.02 | -0.14 | 0.89 | 1 | Gm41541 | | 0 | 0 | 8 | 30 | 21 | 30 | 20 | 30 | 31 | 17 | 21 |
| 18 | 0.72 | 1.09 | 0.66 | 0.51 | 1 | Gm18588 | | 14 | 2 | 30 | 5 | 9 | 22 | 20 | 46 | 20 | 19 | 5 |
| 17 | -0.68 | 1.42 | -0.48 | 0.63 | 1 | Insl6 | insulin-like 6 | 26 | 15 | 33 | 14 | 12 | 8 | 20 | 44 | 0 | 0 | 18 |
| 17 | -0.93 | 1.02 | -0.91 | 0.36 | 1 | Kcnab3os | | 4 | 4 | 33 | 3 | 37 | 28 | 20 | 0 | 9 | 14 | 35 |
| 17 | -0.60 | 0.81 | -0.74 | 0.46 | 1 | Gm26620 | | 9 | 3 | 7 | 22 | 28 | 21 | 20 | 13 | 19 | 39 | 5 |
| 16 | -0.33 | 0.92 | -0.36 | 0.72 | 1 | Gm6552 | | 15 | 27 | 38 | 15 | 8 | 8 | 20 | 3 | 28 | 10 | 7 |
| 16 | -0.52 | 0.75 | -0.69 | 0.49 | 1 | A830018L16Ril | RIKEN cDNA A830018L16 gene | 21 | 7 | 9 | 10 | 19 | 33 | 20 | 6 | 14 | 10 | 30 |
| 16 | -0.35 | 0.72 | -0.48 | 0.63 | 1 | Gm28437 | | 11 | 15 | 11 | 18 | 24 | 13 | 20 | 21 | 15 | 17 | 8 |
| 15 | -0.13 | 0.89 | -0.14 | 0.89 | 1 | Gm14269 | | 15 | 12 | 39 | 7 | 15 | 8 | 20 | 3 | 8 | 26 | 11 |
| 14 | -0.77 | 1.23 | -0.62 | 0.53 | 1 | Gm12263 | | 16 | 4 | 3 | 39 | 1 | 28 | 20 | 5 | 19 | 3 | 17 |
| 14 | 0.14 | 1.13 | 0.13 | 0.90 | 1 | Rpl22-ps1 | | 17 | 7 | 25 | 5 | 15 | 7 | 20 | 35 | 9 | 1 | 9 |
| 13 | -0.56 | 1.12 | -0.50 | 0.62 | 1 | Rps18-ps3 | | 40 | 10 | 20 | 4 | 9 | 14 | 20 | 17 | 6 | 1 | 4 |
| 13 | -1.45 | 1.19 | -1.21 | 0.23 | 1 | Slc10a4l | | 10 | 15 | 5 | 11 | 33 | 11 | 20 | 0 | 25 | 10 | 2 |
| 13 | -1.37 | 1.25 | -1.10 | 0.27 | 1 | 1700041G16Rik | | 12 | 18 | 11 | 3 | 16 | 20 | 20 | 1 | 11 | 0 | 24 |
| 12 | -0.25 | 1.42 | -0.18 | 0.86 | 1 | Gm14412 | | 0 | 8 | 0 | 4 | 10 | 21 | 20 | 16 | 22 | 1 | 31 |
| 12 | 0.46 | 2.46 | 0.19 | 0.85 | 1 | Slc6a19os | solute carrier family 6 (neurotransmitter tra | 0 | 16 | 0 | 0 | 7 | 13 | 20 | 0 | 18 | 55 | 0 |
| 12 | -0.46 | 1.52 | -0.31 | 0.76 | 1 | Rec114 | REC114 meiotic recombination protein | 25 | 8 | 11 | 0 | 30 | 6 | 20 | 14 | 7 | 0 | 7 |
| 12 | -0.41 | 0.91 | -0.46 | 0.65 | 1 | Nsa2-ps1 | | 5 | 12 | 20 | 12 | 9 | 10 | 20 | 8 | 10 | 13 | 8 |
| 11 | -0.41 | 1.08 | -0.38 | 0.71 | 1 | Gm42814 | | 2 | 5 | 11 | 10 | 11 | 11 | 20 | 1 | 13 | 15 | 16 |
| 10 | 0.57 | 1.28 | 0.45 | 0.66 | 1 | Gm26833 | | 9 | 4 | 10 | 0 | 9 | 6 | 20 | 0 | 12 | 33 | 9 |
| 9 | 0.93 | 1.45 | 0.64 | 0.52 | 1 | Spag6 | sperm associated antigen 6 | 13 | 11 | 0 | 0 | 4 | 4 | 20 | 17 | 8 | 23 | 3 |
| 9 | -0.83 | 2.65 | -0.31 | 0.76 | 1 | Gm17193 | | 0 | 0 | 0 | 0 | 0 | 38 | 20 | 0 | 1 | 0 | 40 |
| 8 | -0.53 | 1.54 | -0.34 | 0.73 | 1 | Scamp4 | secretory carrier membrane protein 4 | 0 | 0 | 0 | 4 | 15 | 6 | 20 | 0 | 9 | 15 | 15 |
| 8 | -2.13 | 1.77 | -1.21 | 0.23 | 1 | Gm49700 | | 19 | 21 | 0 | 3 | 1 | 10 | 20 | 0 | 1 | 0 | 8 |
| 7 | -2.70 | 2.10 | -1.28 | 0.20 | 1 | Gm39662 | | 2 | 0 | 11 | 7 | 0 | 11 | 20 | 0 | 19 | 0 | 11 |
| 7 | -2.80 | 1.92 | -1.46 | 0.15 | 1 | Frmpd1 | FERM and PDZ domain containing 1 | 1 | 0 | 0 | 17 | 11 | 6 | 20 | 0 | 22 | 0 | 1 |
| 5 | -4.57 | 2.45 | -1.87 | 0.06 | 1 | Gm11608 | | 0 | 0 | 10 | 11 | 9 | 2 | 20 | 0 | 0 | 1 | 0 |
| 1183 | -2.35 | 2.24 | -1.05 | 0.29 | 1 | Slfn5 | schlafen 5 | 4457 | 4109 | 4402 | 0 | 7 | 0 | 19 | 1 | 0 | 0 | 12 |
| 602 | -0.94 | 1.25 | -0.75 | 0.45 | 1 | Zeb2 | zinc finger E-box binding homeobox 2 | 3603 | 1521 | 1460 | 0 | 5 | 4 | 19 | 1 | 2 | 0 | 11 |
| 308 | 1.44 | 1.36 | 1.06 | 0.29 | 1 | Fcna | ficolin A | 2223 | 613 | 346 | 2 | 18 | 21 | 19 | 68 | 51 | 0 | 22 |
| 290 | -2.17 | 1.25 | -1.74 | 0.08 | 1 | Pla1a | phospholipase A1 member A | 935 | 892 | 1286 | 24 | 4 | 9 | 19 | 0 | 9 | 0 | 9 |
| 175 | -0.03 | 0.52 | -0.06 | 0.95 | 1 | Icosl | icos ligand | 847 | 461 | 422 | 19 | 40 | 19 | 19 | 38 | 20 | 10 | 25 |
| 126 | -1.03 | 0.66 | -1.56 | 0.12 | 1 | Sox9 | SRY (sex determining region Y)-box 9 | 364 | 351 | 560 | 12 | 23 | 14 | 19 | 21 | 13 | 0 | 5 |
| 118 | -0.60 | 0.82 | -0.73 | 0.46 | 1 | Rgs10 | regulator of G-protein signalling 10 | 666 | 200 | 308 | 15 | 21 | 16 | 19 | 30 | 20 | 0 | 4 |
| 110 | -0.59 | 0.72 | -0.82 | 0.41 | 1 | Plcl1 | phospholipase C-like 1 | 385 | 361 | 328 | 10 | 21 | 25 | 19 | 32 | 21 | 8 | 1 |
| 107 | -1.08 | 0.86 | -1.25 | 0.21 | 1 | Egr3 | early growth response 3 | 612 | 191 | 129 | 0 | 5 | 134 | 19 | 0 | 1 | 10 | 75 |
| 100 | -0.26 | 0.65 | -0.41 | 0.68 | 1 | Pqlc3 | PQ loop repeat containing | 421 | 183 | 176 | 37 | 87 | 22 | 19 | 22 | 100 | 13 | 18 |
| 98 | -1.36 | 0.96 | -1.42 | 0.16 | 1 | Pcdh8 | protocadherin 8 | 290 | 577 | 158 | 8 | 3 | 10 | 19 | 0 | 8 | 9 | 0 |
| 96 | 0.12 | 0.53 | 0.23 | 0.82 | 1 | Cyp2s1 | cytochrome P450, family 2, subfamily s, p | 275 | 219 | 137 | 71 | 50 | 61 | 19 | 38 | 53 | 79 | 50 |
| 84 | 0.27 | 0.59 | 0.46 | 0.65 | 1 | Fam227a | family with sequence similarity 227, meml | 50 | 26 | 80 | 30 | 20 | 323 | 19 | 43 | 11 | 43 | 281 |
| 73 | -0.19 | 0.70 | -0.27 | 0.79 | 1 | Wdr62 | WD repeat domain 62 | 144 | 70 | 56 | 58 | 103 | 96 | 19 | 54 | 88 | 10 | 98 |
| 67 | -0.89 | 0.65 | -1.37 | 0.17 | 1 | Cdca2 | cell division cycle associated 2 | 173 | 76 | 84 | 34 | 66 | 103 | 19 | 6 | 32 | 15 | 122 |
| 66 | -3.53 | 2.39 | -1.47 | 0.14 | 1 | Prr5 | proline rich 5 (renal) | 330 | 164 | 182 | 0 | 1 | 9 | 19 | 24 | 0 | 0 | 0 |
| 57 | 0.15 | 0.45 | 0.33 | 0.74 | 1 | Spaca1 | sperm acrosome associated 1 | 53 | 129 | 136 | 32 | 53 | 38 | 19 | 45 | 65 | 13 | 45 |
| 56 | -0.84 | 1.26 | -0.66 | 0.51 | 1 | Serpina1b | serine (or cysteine) peptidase inhibitor, c | 332 | 173 | 79 | 0 | 2 | 2 | 19 | 0 | 2 | 0 | 7 |
| 51 | -0.16 | 0.65 | -0.24 | 0.81 | 1 | Cenpn | centromere protein N | 113 | 88 | 37 | 58 | 46 | 46 | 19 | 23 | 48 | 22 | 63 |
| 48 | -0.64 | 0.91 | -0.70 | 0.48 | 1 | Prx | periaxin | 127 | 56 | 177 | 32 | 22 | 14 | 19 | 0 | 15 | 50 | 14 |
| 40 | -0.96 | 0.77 | -1.25 | 0.21 | 1 | Ankef1 | ankyrin repeat and EF-hand domain conta | 34 | 74 | 45 | 66 | 32 | 56 | 19 | 48 | 19 | 0 | 50 |
| 37 | 0.62 | 0.57 | 1.09 | 0.27 | 1 | 4930539J05Rik | | 24 | 38 | 47 | 22 | 18 | 71 | 19 | 37 | 48 | 33 | 54 |
| 37 | -0.49 | 0.74 | -0.66 | 0.51 | 1 | Cfap58 | cilia and flagella associated protein 58 | 6 | 17 | 25 | 63 | 67 | 67 | 19 | 38 | 10 | 40 | 51 |
| 35 | 0.94 | 0.67 | 1.39 | 0.16 | 1 | 3830408C21Rik | | 15 | 41 | 73 | 10 | 14 | 52 | 19 | 34 | 32 | 31 | 65 |
| 33 | -0.15 | 0.94 | -0.16 | 0.87 | 1 | Creb3l3 | cAMP responsive element binding proteir | 57 | 104 | 103 | 9 | 20 | 3 | 19 | 0 | 17 | 24 | 6 |
| 31 | 1.08 | 0.67 | 1.60 | 0.11 | 1 | 4933430I17Rik | RIKEN cDNA 4933430I17 gene | 9 | 38 | 14 | 15 | 31 | 23 | 19 | 53 | 33 | 45 | 61 |
| 29 | 0.09 | 0.99 | 0.09 | 0.93 | 1 | Akr1d1 | aldo-keto reductase family 1, member D1 | 3 | 9 | 10 | 44 | 44 | 4 | 19 | 136 | 38 | 0 | 10 |
| 28 | 0.98 | 0.77 | 1.26 | 0.21 | 1 | Gm47976 | | 20 | 3 | 6 | 14 | 3 | 90 | 19 | 28 | 19 | 44 | 64 |
| 28 | 0.30 | 0.73 | 0.42 | 0.68 | 1 | Gm47572 | | 12 | 46 | 4 | 18 | 8 | 57 | 19 | 54 | 3 | 32 | 56 |
| 28 | 0.18 | 0.74 | 0.24 | 0.81 | 1 | Jph1 | junctophilin 1 | 51 | 30 | 10 | 63 | 9 | 14 | 19 | 31 | 26 | 43 | 9 |
| 27 | -0.61 | 0.68 | -0.90 | 0.37 | 1 | Mnd1 | meiotic nuclear divisions 1 | 75 | 89 | 34 | 14 | 19 | 9 | 19 | 12 | 12 | 10 | 6 |

| | | | | | | | | | | | | | | | | | | |
|---|---|---|---|---|---|---|---|---|---|---|---|---|---|---|---|---|---|---|
| 27 | 1.07 | 0.89 | 1.20 | 0.23 | 1 | Gm37499 | | 38 | 46 | 33 | 4 | 5 | 32 | 19 | 54 | 19 | 5 | 41 |
| 27 | 0.10 | 0.60 | 0.16 | 0.87 | 1 | Gm26670 | | 25 | 18 | 42 | 28 | 29 | 25 | 19 | 28 | 24 | 24 | 31 |
| 26 | -0.39 | 0.79 | -0.50 | 0.62 | 1 | Gm8960 | | 50 | 20 | 70 | 28 | 14 | 13 | 19 | 8 | 16 | 44 | 6 |
| 26 | 0.18 | 0.95 | 0.19 | 0.85 | 1 | Gm26843 | | 7 | 56 | 7 | 4 | 15 | 73 | 19 | 17 | 11 | 32 | 46 |
| 26 | -0.48 | 0.85 | -0.56 | 0.57 | 1 | 9130230N09Rik | | 33 | 26 | 52 | 24 | 21 | 28 | 19 | 10 | 2 | 31 | 38 |
| 25 | 0.04 | 0.72 | 0.06 | 0.96 | 1 | Gm9887 | | 30 | 40 | 17 | 22 | 40 | 10 | 19 | 49 | 17 | 15 | 16 |
| 24 | 0.38 | 0.89 | 0.42 | 0.67 | 1 | Gm50149 | | 12 | 15 | 0 | 35 | 23 | 25 | 19 | 22 | 28 | 58 | 28 |
| 23 | 0.55 | 0.89 | 0.61 | 0.54 | 1 | Gm43173 | | 2 | 18 | 4 | 28 | 12 | 40 | 19 | 29 | 29 | 61 | 14 |
| 22 | -1.56 | 1.01 | -1.55 | 0.12 | 1 | Naip5 | NLR family, apoptosis inhibitory protein 5 | 83 | 24 | 42 | 18 | 6 | 27 | 19 | 3 | 8 | 0 | 15 |
| 22 | -0.87 | 0.89 | -0.98 | 0.33 | 1 | Gm6421 | | 40 | 31 | 50 | 26 | 25 | 7 | 19 | 5 | 23 | 3 | 17 |
| 22 | 0.69 | 0.80 | 0.86 | 0.39 | 1 | Gm12693 | | 5 | 10 | 8 | 17 | 16 | 18 | 19 | 76 | 39 | 6 | 22 |
| 21 | -0.08 | 0.74 | -0.11 | 0.91 | 1 | Gm49894 | | 3 | 4 | 12 | 32 | 29 | 27 | 19 | 25 | 21 | 35 | 18 |
| 20 | -0.77 | 0.84 | -0.92 | 0.36 | 1 | Gm34961 | | 29 | 19 | 19 | 22 | 28 | 19 | 19 | 6 | 23 | 4 | 31 |
| 19 | 0.06 | 0.68 | 0.09 | 0.93 | 1 | Krt8-ps | | 23 | 15 | 31 | 12 | 17 | 20 | 19 | 14 | 12 | 22 | 24 |
| 19 | -0.81 | 0.79 | -1.03 | 0.30 | 1 | Gm9517 | | 16 | 49 | 8 | 18 | 9 | 35 | 19 | 13 | 2 | 15 | 22 |
| 18 | -0.16 | 0.81 | -0.19 | 0.85 | 1 | E430024P14Rik | | 21 | 20 | 33 | 23 | 6 | 19 | 19 | 9 | 22 | 14 | 15 |
| 18 | 0.07 | 0.90 | 0.08 | 0.94 | 1 | Rps10-ps2 | | 27 | 30 | 30 | 23 | 12 | 3 | 19 | 3 | 17 | 19 | 14 |
| 18 | 0.23 | 0.73 | 0.31 | 0.75 | 1 | Gm10517 | | 8 | 18 | 22 | 29 | 15 | 8 | 19 | 22 | 24 | 21 | 13 |
| 18 | -0.72 | 0.82 | -0.89 | 0.38 | 1 | Gm44647 | | 17 | 32 | 21 | 24 | 16 | 14 | 19 | 12 | 4 | 9 | 30 |
| 18 | -0.05 | 0.96 | -0.05 | 0.96 | 1 | Mir703 | | 23 | 6 | 28 | 17 | 13 | 19 | 19 | 32 | 21 | 3 | 14 |
| 16 | 0.70 | 0.95 | 0.74 | 0.46 | 1 | Gm14292 | | 9 | 4 | 29 | 0 | 20 | 16 | 19 | 5 | 41 | 17 | 15 |
| 16 | -0.74 | 1.47 | -0.50 | 0.62 | 1 | Gm44799 | | 1 | 0 | 8 | 31 | 27 | 18 | 19 | 46 | 0 | 18 | 3 |
| 15 | -0.51 | 1.33 | -0.39 | 0.70 | 1 | Gm15634 | | 7 | 0 | 0 | 18 | 29 | 31 | 19 | 28 | 14 | 1 | 20 |
| 15 | -0.30 | 1.01 | -0.30 | 0.77 | 1 | Rbx1-ps | | 17 | 7 | 14 | 4 | 21 | 28 | 19 | 30 | 11 | 6 | 8 |
| 15 | 0.18 | 1.49 | 0.12 | 0.90 | 1 | Xlr4b | X-linked lymphocyte-regulated 4B | 9 | 6 | 4 | 1 | 0 | 41 | 19 | 0 | 0 | 41 | 39 |
| 14 | -1.00 | 0.99 | -1.01 | 0.31 | 1 | Gm45592 | | 2 | 10 | 27 | 26 | 16 | 15 | 19 | 20 | 6 | 10 | 5 |
| 14 | -0.21 | 1.21 | -0.17 | 0.86 | 1 | Vsig8 | V-set and immunoglobulin domain contai | 15 | 3 | 4 | 29 | 12 | 13 | 19 | 20 | 20 | 0 | 21 |
| 14 | -1.38 | 1.08 | -1.28 | 0.20 | 1 | Gm8494 | | 21 | 28 | 12 | 13 | 17 | 8 | 19 | 17 | 1 | 1 | 12 |
| 13 | -0.82 | 1.70 | -0.48 | 0.63 | 1 | Gm38340 | | 0 | 2 | 0 | 0 | 13 | 39 | 19 | 0 | 1 | 27 | 41 |
| 13 | 0.64 | 0.95 | 0.68 | 0.50 | 1 | Gm12989 | | 23 | 25 | 6 | 6 | 7 | 4 | 19 | 7 | 20 | 15 | 7 |
| 12 | 0.23 | 1.74 | 0.13 | 0.90 | 1 | Itk | IL2 inducible T cell kinase | 51 | 44 | 2 | 0 | 1 | 1 | 19 | 0 | 5 | 14 | 0 |
| 12 | -1.28 | 1.58 | -0.81 | 0.42 | 1 | Gm31793 | | 0 | 26 | 19 | 9 | 5 | 11 | 19 | 21 | 1 | 0 | 24 |
| 12 | -0.20 | 1.00 | -0.19 | 0.85 | 1 | Gm8712 | | 20 | 7 | 1 | 11 | 19 | 7 | 19 | 17 | 7 | 13 | 11 |
| 12 | 0.09 | 1.43 | 0.06 | 0.95 | 1 | Gm50105 | | 19 | 21 | 0 | 9 | 3 | 4 | 19 | 49 | 3 | 0 | 4 |
| 11 | -0.99 | 0.92 | -1.07 | 0.28 | 1 | Gm10819 | | 8 | 3 | 22 | 7 | 22 | 10 | 19 | 2 | 15 | 5 | 10 |
| 11 | -1.42 | 1.11 | -1.27 | 0.20 | 1 | Gm10463 | | 27 | 3 | 4 | 9 | 11 | 16 | 19 | 0 | 6 | 8 | 16 |
| 8 | -1.64 | 1.85 | -0.89 | 0.38 | 1 | 4732471J01Rik | | 0 | 13 | 0 | 4 | 4 | 25 | 19 | 1 | 5 | 0 | 20 |
| 8 | -0.23 | 1.79 | -0.13 | 0.90 | 1 | Gm5790 | | 2 | 24 | 0 | 0 | 7 | 8 | 19 | 1 | 9 | 5 | 13 |
| 8 | -0.55 | 1.22 | -0.45 | 0.65 | 1 | H3f3a-ps1 | | 9 | 3 | 1 | 1 | 18 | 8 | 19 | 7 | 10 | 10 | 2 |
| 7 | -1.47 | 1.54 | -0.96 | 0.34 | 1 | Gm48249 | | 0 | 5 | 10 | 4 | 9 | 14 | 19 | 0 | 3 | 9 | 8 |
| 7 | -1.52 | 1.22 | -1.24 | 0.21 | 1 | Gm1818 | | 5 | 1 | 6 | 4 | 10 | 12 | 19 | 0 | 4 | 17 | 3 |
| 7 | -1.16 | 1.17 | -0.99 | 0.32 | 1 | Gm8806 | | 4 | 7 | 5 | 3 | 9 | 11 | 19 | 5 | 2 | 3 | 12 |
| 7 | 0.09 | 2.21 | 0.04 | 0.97 | 1 | C230035I16Rik | | 16 | 0 | 0 | 0 | 2 | 19 | 19 | 0 | 7 | 0 | 13 |
| 4 | -2.90 | 2.17 | -1.34 | 0.18 | 1 | Gm10059 | | 1 | 5 | 10 | 1 | 2 | 1 | 19 | 0 | 3 | 0 | 0 |
| 4 | 4.87 | 3.22 | 1.51 | 0.13 | 1 | Gm22567 | | 0 | 0 | 1 | 0 | 0 | 0 | 19 | 10 | 6 | 0 | 4 |
| 2259 | -3.47 | 2.15 | -1.61 | 0.11 | 1 | Igfbp3 | insulin-like growth factor binding protein 3 | 8885 | 7024 | 8910 | 0 | 0 | 8 | 18 | 1 | 0 | 0 | 0 |
| 1332 | -1.77 | 1.18 | -1.49 | 0.14 | 1 | Plxnd1 | plexin D1 | 5799 | 4234 | 4575 | 10 | 4 | 3 | 18 | 0 | 9 | 0 | 2 |
| 892 | -2.20 | 1.45 | -1.51 | 0.13 | 1 | Exoc3l2 | exocyst complex component 3-like 2 | 3153 | 2948 | 3585 | 70 | 6 | 7 | 18 | 0 | 14 | 0 | 10 |
| 753 | -0.27 | 1.41 | -0.19 | 0.85 | 1 | Scn7a | sodium channel, voltage-gated, type VII, a | 3494 | 2208 | 2460 | 4 | 49 | 0 | 18 | 30 | 24 | 0 | 0 |
| 535 | -1.73 | 1.30 | -1.33 | 0.18 | 1 | Scara3 | scavenger receptor class A, member 3 | 2289 | 1605 | 1858 | 11 | 18 | 24 | 18 | 0 | 51 | 0 | 8 |
| 327 | 0.18 | 1.33 | 0.13 | 0.89 | 1 | Chst2 | carbohydrate sulfotransferase 2 | 974 | 1208 | 1326 | 0 | 13 | 9 | 18 | 16 | 13 | 0 | 16 |
| 278 | -0.77 | 0.56 | -1.39 | 0.16 | 1 | Igsf21 | immunoglobulin superfamily, member 21 | 900 | 1044 | 932 | 21 | 33 | 32 | 18 | 0 | 24 | 12 | 40 |
| 177 | -1.11 | 1.29 | -0.87 | 0.39 | 1 | Pou6f2 | POU domain, class 6, transcription factor | 466 | 911 | 476 | 0 | 10 | 32 | 18 | 2 | 10 | 0 | 19 |
| 173 | 0.16 | 0.49 | 0.32 | 0.75 | 1 | Lynx1 | Ly6/neurotoxin 1 | 823 | 426 | 475 | 22 | 30 | 10 | 18 | 49 | 13 | 24 | 10 |
| 145 | 0.22 | 2.10 | 0.11 | 0.92 | 1 | Slc38a1 | solute carrier family 38, member 1 | 888 | 423 | 235 | 0 | 0 | 9 | 18 | 15 | 0 | 0 | 10 |
| 126 | -0.89 | 0.85 | -1.05 | 0.29 | 1 | Fkbp7 | FK506 binding protein 7 | 576 | 256 | 339 | 0 | 19 | 95 | 18 | 7 | 5 | 0 | 76 |
| 120 | -0.53 | 1.03 | -0.52 | 0.61 | 1 | Mecom | MDS1 and EVI1 complex locus | 242 | 656 | 304 | 14 | 14 | 7 | 18 | 0 | 0 | 40 | 23 |
| 106 | -0.98 | 0.62 | -1.58 | 0.12 | 1 | Myom1 | myomesin 1 | 330 | 171 | 291 | 21 | 21 | 173 | 18 | 0 | 12 | 26 | 106 |
| 99 | -1.14 | 1.25 | -0.92 | 0.36 | 1 | Rassf9 | Ras association (RalGDS/AF-6) domain f | 370 | 312 | 345 | 3 | 1 | 4 | 18 | 0 | 0 | 39 | 0 |
| 86 | 0.62 | 0.51 | 1.21 | 0.23 | 1 | C2 | complement component 2 (within H-2S) | 361 | 150 | 150 | 32 | 15 | 46 | 18 | 42 | 33 | 59 | 34 |

| | | | | | | | | | | | | | | | | | |
|---|---|---|---|---|---|---|---|---|---|---|---|---|---|---|---|---|---|
| 75 | 1.17 | 0.69 | 1.68 | 0.09 | 1 | Tlcd2 | TLC domain containing 2 | 122 | 198 | 113 | 18 | 37 | 25 | 18 | 37 | 52 | 193 | 17 |
| 73 | -1.53 | 0.96 | -1.59 | 0.11 | 1 | Avpr1a | arginine vasopressin receptor 1A | 266 | 222 | 259 | 16 | 0 | 6 | 18 | 0 | 3 | 5 | 4 |
| 69 | -0.77 | 1.33 | -0.58 | 0.56 | 1 | Casp1 | caspase 1 | 353 | 230 | 148 | 0 | 0 | 3 | 18 | 0 | 4 | 0 | 2 |
| 68 | 0.81 | 0.72 | 1.13 | 0.26 | 1 | Lpxn | leupaxin | 331 | 83 | 115 | 8 | 19 | 41 | 18 | 61 | 26 | 17 | 27 |
| 66 | -0.24 | 1.31 | -0.18 | 0.86 | 1 | Adam8 | a disintegrin and metallopeptidase domai | 299 | 181 | 133 | 0 | 0 | 46 | 18 | 0 | 6 | 0 | 45 |
| 62 | -0.20 | 0.68 | -0.29 | 0.77 | 1 | Slc12a8 | solute carrier family 12 (potassium/chloric | 99 | 214 | 229 | 21 | 26 | 3 | 18 | 14 | 22 | 33 | 0 |
| 61 | 0.31 | 0.51 | 0.60 | 0.55 | 1 | BC065397 | | 37 | 55 | 41 | 11 | 19 | 224 | 18 | 14 | 33 | 22 | 202 |
| 51 | -0.10 | 0.49 | -0.20 | 0.84 | 1 | Tnfaip8l1 | tumor necrosis factor, alpha-induced prot | 130 | 128 | 107 | 34 | 23 | 27 | 18 | 25 | 27 | 18 | 23 |
| 50 | -1.76 | 0.89 | -1.97 | 0.05 | 1 | Svopl | SV2 related protein homolog (rat)-like | 54 | 11 | 90 | 52 | 131 | 76 | 18 | 9 | 69 | 0 | 39 |
| 45 | 0.26 | 0.65 | 0.40 | 0.69 | 1 | Gpr75 | G protein-coupled receptor 75 | 29 | 92 | 22 | 62 | 16 | 77 | 18 | 27 | 18 | 68 | 67 |
| 41 | -0.47 | 0.59 | -0.79 | 0.43 | 1 | Gm17354 | | 20 | 41 | 62 | 67 | 60 | 46 | 18 | 78 | 28 | 21 | 17 |
| 41 | -0.06 | 0.66 | -0.09 | 0.93 | 1 | Snai2 | snail family zinc finger 2 | 89 | 94 | 140 | 11 | 11 | 21 | 18 | 7 | 5 | 23 | 32 |
| 34 | 0.76 | 1.15 | 0.67 | 0.51 | 1 | Fanci | Fanconi anemia, complementation group | 102 | 55 | 5 | 18 | 18 | 25 | 18 | 37 | 51 | 0 | 49 |
| 33 | -0.02 | 0.67 | -0.03 | 0.98 | 1 | Gm42064 | | 42 | 19 | 39 | 40 | 33 | 38 | 18 | 46 | 14 | 30 | 41 |
| 32 | 0.34 | 0.78 | 0.43 | 0.67 | 1 | Ddias | DNA damage-induced apoptosis suppres | 46 | 36 | 8 | 15 | 41 | 36 | 18 | 27 | 36 | 72 | 20 |
| 31 | -1.72 | 0.89 | -1.94 | 0.05 | 1 | Zan | zonadhesin | 28 | 15 | 39 | 48 | 34 | 83 | 18 | 23 | 9 | 0 | 39 |
| 30 | -0.47 | 0.66 | -0.71 | 0.48 | 1 | Tmem95 | transmembrane protein 95 | 27 | 11 | 10 | 39 | 43 | 62 | 18 | 24 | 19 | 32 | 46 |
| 30 | -0.20 | 0.73 | -0.27 | 0.78 | 1 | Gm12216 | | 61 | 40 | 59 | 27 | 12 | 28 | 18 | 46 | 8 | 12 | 16 |
| 29 | 0.32 | 0.61 | 0.52 | 0.60 | 1 | Zfp296 | zinc finger protein 296 | 68 | 31 | 58 | 11 | 11 | 32 | 18 | 10 | 28 | 17 | 34 |
| 29 | 0.59 | 0.89 | 0.66 | 0.51 | 1 | Oas1g | 2'-5' oligoadenylate synthetase 1G | 45 | 34 | 77 | 0 | 19 | 36 | 18 | 6 | 38 | 21 | 21 |
| 29 | 0.19 | 0.82 | 0.23 | 0.82 | 1 | Gm6083 | | 3 | 0 | 0 | 33 | 50 | 40 | 18 | 59 | 23 | 46 | 43 |
| 28 | -0.77 | 0.65 | -1.19 | 0.23 | 1 | Wdr93 | WD repeat domain 93 | 33 | 34 | 8 | 50 | 25 | 55 | 18 | 24 | 12 | 17 | 34 |
| 25 | 0.06 | 0.96 | 0.06 | 0.95 | 1 | Gm11249 | | 25 | 46 | 24 | 20 | 9 | 17 | 18 | 5 | 4 | 91 | 13 |
| 24 | 0.51 | 0.66 | 0.77 | 0.44 | 1 | 4732414G09Rik | | 8 | 24 | 8 | 25 | 30 | 20 | 18 | 44 | 29 | 36 | 25 |
| 23 | 0.07 | 0.81 | 0.09 | 0.93 | 1 | Gm7862 | | 7 | 57 | 9 | 53 | 13 | 14 | 18 | 15 | 12 | 24 | 30 |
| 23 | -0.98 | 1.29 | -0.76 | 0.45 | 1 | Oas2 | 2'-5' oligoadenylate synthetase 2 | 66 | 5 | 77 | 0 | 9 | 39 | 18 | 0 | 28 | 0 | 11 |
| 23 | -0.05 | 1.00 | -0.05 | 0.96 | 1 | Mterf1b | mitochondrial transcription termination fa | 27 | 32 | 16 | 9 | 22 | 33 | 18 | 0 | 22 | 30 | 42 |
| 23 | -1.59 | 1.45 | -1.10 | 0.27 | 1 | Gm49024 | | 4 | 0 | 35 | 54 | 41 | 17 | 18 | 0 | 6 | 61 | 13 |
| 23 | -0.42 | 0.69 | -0.61 | 0.54 | 1 | Gm16580 | | 33 | 44 | 25 | 17 | 34 | 16 | 18 | 12 | 22 | 17 | 12 |
| 23 | 0.75 | 0.93 | 0.81 | 0.42 | 1 | Gm5547 | | 68 | 20 | 11 | 20 | 0 | 25 | 18 | 41 | 5 | 21 | 18 |
| 22 | -0.33 | 0.73 | -0.45 | 0.65 | 1 | Fgf20 | fibroblast growth factor 20 | 15 | 8 | 14 | 29 | 36 | 26 | 18 | 7 | 33 | 35 | 22 |
| 22 | -2.63 | 1.34 | -1.96 | 0.05 | 1 | Gm31718 | | 1 | 46 | 30 | 24 | 16 | 57 | 18 | 0 | 18 | 0 | 26 |
| 21 | -0.49 | 1.12 | -0.43 | 0.66 | 1 | Gm35696 | | 27 | 7 | 32 | 41 | 25 | 9 | 18 | 44 | 14 | 0 | 16 |
| 21 | 0.26 | 0.77 | 0.33 | 0.74 | 1 | Gm13127 | | 25 | 23 | 10 | 27 | 16 | 8 | 18 | 37 | 32 | 33 | 2 |
| 20 | 1.37 | 0.99 | 1.39 | 0.16 | 1 | 4930590J08Rik | RIKEN cDNA 4930590J08 gene | 27 | 16 | 45 | 0 | 9 | 18 | 18 | 22 | 17 | 21 | 27 |
| 19 | -0.14 | 0.83 | -0.16 | 0.87 | 1 | 1300014J16Rik | | 32 | 17 | 37 | 4 | 18 | 29 | 18 | 10 | 25 | 4 | 18 |
| 19 | -0.74 | 0.85 | -0.87 | 0.38 | 1 | Klhl35 | kelch-like 35 | 9 | 35 | 1 | 37 | 15 | 26 | 18 | 38 | 10 | 5 | 15 |
| 18 | 0.11 | 0.79 | 0.14 | 0.89 | 1 | Tceal7 | transcription elongation factor A (SII)-like | 15 | 15 | 15 | 28 | 16 | 8 | 18 | 12 | 18 | 53 | 6 |
| 18 | -1.62 | 0.94 | -1.73 | 0.08 | 1 | Gm7285 | | 25 | 15 | 18 | 12 | 11 | 46 | 18 | 1 | 5 | 3 | 48 |
| 18 | 0.19 | 0.95 | 0.20 | 0.84 | 1 | Rimbp3 | RIMS binding protein 3 | 0 | 0 | 0 | 22 | 25 | 20 | 18 | 27 | 29 | 53 | 7 |
| 17 | -1.20 | 0.98 | -1.23 | 0.22 | 1 | 1700101I11Rik | | 24 | 23 | 20 | 13 | 25 | 25 | 18 | 14 | 11 | 0 | 15 |
| 17 | -1.23 | 1.26 | -0.97 | 0.33 | 1 | Fndc11 | fibronectin type III domain containing 11 | 30 | 63 | 25 | 11 | 8 | 6 | 18 | 0 | 12 | 15 | 0 |
| 17 | 0.68 | 0.97 | 0.70 | 0.48 | 1 | Radx | RPA1 related single stranded DNA bindir | 12 | 19 | 6 | 13 | 5 | 16 | 18 | 28 | 50 | 12 | 6 |
| 17 | 1.20 | 0.92 | 1.29 | 0.20 | 1 | Gm10617 | | 2 | 12 | 12 | 6 | 9 | 12 | 18 | 35 | 22 | 39 | 14 |
| 16 | -1.36 | 1.40 | -0.97 | 0.33 | 1 | Gm45546 | | 17 | 10 | 11 | 29 | 12 | 19 | 18 | 0 | 34 | 0 | 25 |
| 16 | -1.42 | 1.56 | -0.91 | 0.36 | 1 | Gm12925 | | 45 | 26 | 5 | 0 | 23 | 19 | 18 | 27 | 2 | 0 | 8 |
| 15 | -1.08 | 1.14 | -0.94 | 0.35 | 1 | Gm10101 | | 11 | 37 | 38 | 12 | 12 | 9 | 18 | 0 | 4 | 21 | 6 |
| 15 | -1.37 | 1.15 | -1.19 | 0.23 | 1 | Gm42688 | | 15 | 19 | 23 | 1 | 11 | 37 | 18 | 0 | 4 | 4 | 34 |
| 15 | -0.97 | 0.90 | -1.08 | 0.28 | 1 | Gm6542 | | 28 | 5 | 41 | 10 | 10 | 19 | 18 | 3 | 10 | 4 | 17 |
| 15 | 0.43 | 1.22 | 0.35 | 0.73 | 1 | Stk33 | serine/threonine kinase 33 | 18 | 7 | 24 | 5 | 2 | 25 | 18 | 16 | 13 | 0 | 36 |
| 15 | -0.74 | 1.07 | -0.69 | 0.49 | 1 | Gm12669 | | 33 | 30 | 8 | 8 | 12 | 17 | 18 | 0 | 10 | 17 | 10 |
| 15 | -1.00 | 0.81 | -1.24 | 0.21 | 1 | Gm14270 | | 10 | 4 | 16 | 23 | 25 | 20 | 18 | 9 | 22 | 8 | 8 |
| 15 | -0.42 | 1.19 | -0.35 | 0.73 | 1 | Gm16364 | | 6 | 31 | 26 | 7 | 11 | 15 | 18 | 0 | 7 | 28 | 12 |
| 14 | 0.12 | 0.92 | 0.13 | 0.90 | 1 | Eif1-ps1 | | 3 | 2 | 10 | 17 | 17 | 8 | 18 | 30 | 28 | 5 | 11 |
| 14 | -0.57 | 1.45 | -0.40 | 0.69 | 1 | Gm48701 | | 0 | 6 | 11 | 11 | 12 | 28 | 18 | 0 | 22 | 35 | 5 |
| 13 | -1.84 | 1.45 | -1.27 | 0.20 | 1 | Gm15473 | | 0 | 10 | 0 | 14 | 23 | 12 | 18 | 46 | 4 | 0 | 19 |
| 12 | -0.98 | 1.65 | -0.60 | 0.55 | 1 | Gm17202 | | 11 | 19 | 16 | 25 | 0 | 1 | 18 | 36 | 4 | 0 | 0 |
| 10 | -0.80 | 0.96 | -0.82 | 0.41 | 1 | Gm4949 | | 4 | 6 | 15 | 4 | 34 | 5 | 18 | 5 | 12 | 8 | 4 |
| 10 | -0.50 | 1.19 | -0.41 | 0.68 | 1 | Gm6402 | | 20 | 23 | 7 | 4 | 9 | 7 | 18 | 0 | 9 | 10 | 7 |
| 10 | -1.40 | 1.20 | -1.17 | 0.24 | 1 | Gm4900 | | 4 | 8 | 21 | 10 | 15 | 8 | 18 | 0 | 7 | 5 | 15 |

| | | | | | | | | | | | | | | | | | | |
|---|---|---|---|---|---|---|---|---|---|---|---|---|---|---|---|---|---|---|
| 10 | 0.01 | 1.76 | 0.00 | 1.00 | 1 | Gm20667 | | 15 | 14 | 0 | 0 | 10 | 7 | 18 | 0 | 5 | 28 | 10 |
| 9 | -1.41 | 1.33 | -1.06 | 0.29 | 1 | Gm11592 | | 0 | 0 | 3 | 4 | 23 | 19 | 18 | 0 | 10 | 10 | 10 |
| 9 | -0.89 | 0.97 | -0.91 | 0.36 | 1 | 4930534H18Rik | | 9 | 5 | 1 | 16 | 7 | 12 | 18 | 6 | 5 | 10 | 6 |
| 8 | -1.56 | 1.31 | -1.19 | 0.23 | 1 | B3gat2 | beta-1,3-glucuronyltransferase 2 (glucurc | 0 | 0 | 0 | 15 | 19 | 13 | 18 | 8 | 2 | 4 | 11 |
| 8 | -1.62 | 1.40 | -1.16 | 0.24 | 1 | Tmem74b | transmembrane protein 74B | 4 | 7 | 14 | 16 | 6 | 4 | 18 | 0 | 1 | 15 | 3 |
| 7 | -0.94 | 1.64 | -0.57 | 0.57 | 1 | Gstm6 | glutathione S-transferase, mu 6 | 26 | 1 | 9 | 0 | 3 | 8 | 18 | 0 | 3 | 0 | 9 |
| 7 | -3.24 | 2.34 | -1.39 | 0.17 | 1 | Gm25604 | | 19 | 8 | 17 | 0 | 0 | 12 | 18 | 0 | 2 | 0 | 0 |
| 6 | 0.41 | 1.84 | 0.22 | 0.83 | 1 | Gm9514 | | 0 | 3 | 0 | 2 | 1 | 14 | 18 | 0 | 8 | 12 | 12 |
| 6 | -1.15 | 1.54 | -0.75 | 0.46 | 1 | Gm4735 | | 11 | 0 | 12 | 1 | 9 | 2 | 18 | 0 | 6 | 1 | 4 |
| 6 | -0.71 | 2.18 | -0.33 | 0.74 | 1 | Etv4 | ets variant 4 | 9 | 7 | 12 | 0 | 0 | 5 | 18 | 0 | 1 | 0 | 8 |
| 4 | -3.71 | 2.67 | -1.39 | 0.17 | 1 | Snx31 | sorting nexin 31 | 10 | 0 | 7 | 0 | 8 | 2 | 18 | 0 | 0 | 0 | 0 |
| 3 | -3.70 | 3.39 | -1.09 | 0.28 | 1 | Gm26612 | | 0 | 0 | 0 | 11 | 7 | 0 | 18 | 0 | 2 | 0 | 0 |
| 3 | 1.80 | 3.49 | 0.52 | 0.61 | 1 | Gm17772 | | 0 | 6 | 0 | 2 | 0 | 0 | 18 | 0 | 1 | 0 | 5 |
| 2631 | -1.14 | 1.33 | -0.85 | 0.39 | 1 | Klk1 | kallikrein 1 | 3202 | 11740 | 13277 | 333 | 35 | 7 | 17 | 49 | 35 | 0 | 250 |
| 973 | -0.75 | 1.93 | -0.39 | 0.70 | 1 | Ifitm3 | interferon induced transmembrane protei | 4553 | 2873 | 3242 | 0 | 9 | 0 | 17 | 0 | 7 | 0 | 4 |
| 756 | 0.04 | 0.46 | 0.09 | 0.93 | 1 | Olfml3 | olfactomedin-like 3 | 4983 | 1360 | 1833 | 28 | 11 | 9 | 17 | 29 | 22 | 23 | 2 |
| 339 | -0.23 | 0.47 | -0.50 | 0.62 | 1 | Yap1 | yes-associated protein 1 | 1664 | 958 | 997 | 21 | 6 | 13 | 17 | 16 | 3 | 17 | 13 |
| 324 | 0.06 | 0.47 | 0.13 | 0.89 | 1 | Nr2f2 | nuclear receptor subfamily 2, group F, me | 1671 | 804 | 890 | 37 | 32 | 12 | 17 | 39 | 11 | 24 | 27 |
| 229 | -0.14 | 0.54 | -0.27 | 0.79 | 1 | Nfam1 | Nfat activating molecule with ITAM motif 1 | 1338 | 550 | 439 | 28 | 30 | 22 | 17 | 46 | 11 | 12 | 27 |
| 224 | -1.31 | 1.34 | -0.97 | 0.33 | 1 | Neurl3 | neuralized E3 ubiquitin protein ligase 3 | 945 | 569 | 912 | 0 | 7 | 6 | 17 | 0 | 8 | 0 | 4 |
| 168 | 0.54 | 0.45 | 1.20 | 0.23 | 1 | Gm17491 | | 63 | 44 | 32 | 50 | 29 | 679 | 17 | 81 | 52 | 22 | 779 |
| 157 | 0.51 | 0.44 | 1.16 | 0.24 | 1 | Slc5a1 | solute carrier family 5 (sodium/glucose cc | 515 | 300 | 607 | 26 | 40 | 43 | 17 | 63 | 52 | 28 | 34 |
| 119 | 1.28 | 1.81 | 0.70 | 0.48 | 1 | Klhl6 | kelch-like 6 | 693 | 198 | 377 | 0 | 5 | 0 | 17 | 14 | 1 | 0 | 3 |
| 107 | -0.59 | 1.13 | -0.52 | 0.60 | 1 | Jdp2 | Jun dimerization protein 2 | 455 | 360 | 323 | 0 | 6 | 3 | 17 | 0 | 11 | 0 | 4 |
| 86 | -1.50 | 0.97 | -1.54 | 0.12 | 1 | Fam13a | family with sequence similarity 13, memb | 303 | 308 | 286 | 14 | 5 | 3 | 17 | 0 | 7 | 0 | 7 |
| 73 | -1.47 | 0.99 | -1.48 | 0.14 | 1 | Il4i1 | interleukin 4 induced 1 | 319 | 239 | 72 | 45 | 17 | 33 | 17 | 1 | 19 | 0 | 41 |
| 71 | 0.14 | 0.69 | 0.20 | 0.84 | 1 | Nhs | NHS actin remodeling regulator | 147 | 234 | 215 | 11 | 9 | 63 | 17 | 13 | 9 | 48 | 19 |
| 62 | 0.84 | 0.47 | 1.79 | 0.07 | 1 | Syn3 | synapsin III | 50 | 91 | 56 | 59 | 54 | 48 | 17 | 96 | 107 | 54 | 53 |
| 57 | 0.31 | 0.60 | 0.52 | 0.61 | 1 | Parpbp | PARP1 binding protein | 106 | 97 | 106 | 30 | 57 | 34 | 17 | 92 | 27 | 22 | 39 |
| 57 | -0.93 | 1.04 | -0.90 | 0.37 | 1 | Scube1 | signal peptide, CUB domain, EGF-like 1 | 173 | 193 | 193 | 0 | 9 | 14 | 17 | 0 | 10 | 1 | 13 |
| 56 | -2.81 | 2.69 | -1.04 | 0.30 | 1 | Siglece | sialic acid binding Ig-like lectin E | 275 | 114 | 213 | 0 | 0 | 2 | 17 | 0 | 0 | 0 | 0 |
| 55 | 0.95 | 0.93 | 1.02 | 0.31 | 1 | Gm38190 | | 21 | 43 | 4 | 0 | 9 | 232 | 17 | 34 | 14 | 30 | 203 |
| 54 | -2.47 | 2.86 | -0.86 | 0.39 | 1 | Galnt6 | polypeptide N-acetylgalactosaminyltransf | 155 | 241 | 184 | 0 | 0 | 1 | 17 | 0 | 0 | 0 | 0 |
| 54 | -1.10 | 0.57 | -1.92 | 0.05 | 1 | Gpr182 | G protein-coupled receptor 182 | 142 | 155 | 147 | 24 | 34 | 25 | 17 | 7 | 14 | 6 | 22 |
| 51 | -1.35 | 1.55 | -0.87 | 0.39 | 1 | Cdh9 | cadherin 9 | 140 | 170 | 211 | 0 | 0 | 6 | 17 | 0 | 12 | 0 | 1 |
| 49 | -0.96 | 0.79 | -1.21 | 0.23 | 1 | Gm47260 | | 26 | 4 | 15 | 12 | 7 | 240 | 17 | 0 | 4 | 22 | 197 |
| 46 | 0.12 | 0.75 | 0.16 | 0.88 | 1 | Gm50387 | | 53 | 43 | 94 | 14 | 36 | 72 | 17 | 3 | 21 | 81 | 68 |
| 45 | 1.22 | 0.68 | 1.81 | 0.07 | 1 | Esco2 | establishment of sister chromatid cohesio | 117 | 40 | 39 | 21 | 9 | 59 | 17 | 81 | 34 | 35 | 48 |
| 44 | -0.80 | 0.87 | -0.92 | 0.36 | 1 | Lst1 | leukocyte specific transcript 1 | 169 | 66 | 129 | 36 | 14 | 8 | 17 | 8 | 15 | 1 | 22 |
| 43 | 0.93 | 0.75 | 1.24 | 0.21 | 1 | Gm6883 | | 49 | 48 | 79 | 16 | 32 | 24 | 17 | 122 | 35 | 46 | 9 |
| 41 | 0.27 | 0.55 | 0.49 | 0.62 | 1 | Phtf1os | | 54 | 73 | 65 | 25 | 37 | 37 | 17 | 58 | 24 | 30 | 30 |
| 39 | -2.93 | 3.34 | -0.88 | 0.38 | 1 | Hhipl1 | hedgehog interacting protein-like 1 | 225 | 93 | 85 | 0 | 0 | 0 | 17 | 0 | 0 | 0 | 5 |
| 39 | 0.55 | 0.98 | 0.56 | 0.58 | 1 | Gm11808 | | 122 | 92 | 53 | 23 | 16 | 5 | 17 | 0 | 45 | 41 | 11 |
| 35 | 0.28 | 0.58 | 0.49 | 0.63 | 1 | Cenpp | centromere protein P | 35 | 36 | 50 | 31 | 36 | 33 | 17 | 22 | 37 | 49 | 34 |
| 33 | -1.90 | 1.06 | -1.79 | 0.07 | 1 | Gm5148 | predicted gene 5148 | 60 | 110 | 94 | 14 | 16 | 16 | 17 | 0 | 12 | 0 | 19 |
| 31 | -0.41 | 1.02 | -0.40 | 0.69 | 1 | Lca5l | Leber congenital amaurosis 5-like | 48 | 23 | 34 | 31 | 17 | 50 | 17 | 71 | 8 | 0 | 44 |
| 28 | 1.34 | 0.77 | 1.74 | 0.08 | 1 | Gm5578 | | 29 | 4 | 17 | 25 | 15 | 14 | 17 | 83 | 30 | 45 | 27 |
| 28 | 1.23 | 0.94 | 1.30 | 0.19 | 1 | Matk | megakaryocyte-associated tyrosine kinas | 99 | 18 | 7 | 0 | 5 | 46 | 17 | 20 | 5 | 24 | 62 |
| 26 | 0.07 | 1.10 | 0.07 | 0.95 | 1 | Gm2093 | | 43 | 0 | 31 | 40 | 26 | 8 | 17 | 22 | 11 | 77 | 13 |
| 26 | -0.39 | 1.68 | -0.23 | 0.82 | 1 | Gm37303 | | 7 | 31 | 22 | 0 | 0 | 91 | 17 | 0 | 5 | 0 | 110 |
| 25 | 0.10 | 1.12 | 0.09 | 0.93 | 1 | Lrrc39 | leucine rich repeat containing 39 | 42 | 54 | 16 | 29 | 19 | 9 | 17 | 0 | 62 | 24 | 8 |
| 25 | 0.02 | 0.72 | 0.03 | 0.98 | 1 | Eif3s6-ps2 | | 8 | 52 | 24 | 33 | 17 | 13 | 17 | 72 | 27 | 5 | 11 |
| 24 | 0.46 | 1.05 | 0.44 | 0.66 | 1 | Gm14486 | | 1 | 13 | 22 | 13 | 49 | 22 | 25 | 17 | 36 | 7 | 68 | 16 |
| 24 | -1.02 | 1.14 | -0.89 | 0.37 | 1 | Gm26549 | | 32 | 10 | 4 | 32 | 22 | 57 | 17 | 0 | 37 | 8 | 40 |
| 23 | -0.36 | 1.15 | -0.31 | 0.75 | 1 | Ceacam10 | carcinoembryonic antigen-related cell adl | 0 | 2 | 31 | 40 | 35 | 27 | 17 | 7 | 27 | 53 | 16 |
| 23 | 0.40 | 0.97 | 0.41 | 0.68 | 1 | Cfap77 | cilia and flagella associated protein 77 | 39 | 43 | 74 | 0 | 10 | 19 | 17 | 10 | 11 | 15 | 11 |
| 22 | -1.33 | 1.19 | -1.12 | 0.26 | 1 | Foxn1 | forkhead box N1 | 0 | 6 | 39 | 71 | 32 | 24 | 17 | 3 | 18 | 26 | 8 |
| 22 | -0.63 | 1.64 | -0.38 | 0.70 | 1 | Gm26532 | | 21 | 35 | 10 | 0 | 0 | 85 | 17 | 0 | 3 | 0 | 70 |
| 22 | 1.09 | 0.75 | 1.46 | 0.14 | 1 | Rps3a3 | | 32 | 55 | 25 | 13 | 8 | 4 | 17 | 42 | 20 | 10 | 16 |
| 21 | -2.00 | 1.23 | -1.63 | 0.10 | 1 | Fam228b | family with sequence similarity 228, meml | 13 | 14 | 0 | 39 | 29 | 53 | 17 | 20 | 3 | 0 | 47 |

| | | | | | | | | | | | | | | | | | | |
|---|---|---|---|---|---|---|---|---|---|---|---|---|---|---|---|---|---|---|
| 20 | -2.03 | 2.04 | -0.99 | 0.32 | 1 | Prss45 | protease, serine 45 | 194 | 2 | 0 | 0 | 0 | 3 | 17 | 0 | 0 | 0 | 0 |
| 19 | -0.64 | 0.85 | -0.75 | 0.45 | 1 | Gm9903 | | 19 | 17 | 19 | 32 | 19 | 22 | 17 | 5 | 40 | 8 | 16 |
| 19 | 1.24 | 1.09 | 1.13 | 0.26 | 1 | Gm11940 | | 7 | 53 | 6 | 24 | 0 | 15 | 17 | 8 | 9 | 40 | 31 |
| 19 | -1.90 | 1.28 | -1.48 | 0.14 | 1 | 4930471D02Rik | | 26 | 52 | 25 | 23 | 7 | 15 | 17 | 0 | 7 | 0 | 34 |
| 19 | 0.45 | 0.87 | 0.52 | 0.61 | 1 | Dnaaf4 | dynein axonemal assembly factor 4 | 29 | 4 | 21 | 11 | 7 | 23 | 17 | 6 | 39 | 13 | 35 |
| 18 | -0.11 | 1.10 | -0.10 | 0.92 | 1 | Gm37200 | | 11 | 0 | 0 | 11 | 26 | 18 | 17 | 2 | 22 | 57 | 35 |
| 17 | 2.27 | 1.50 | 1.51 | 0.13 | 1 | Gm35853 | | 82 | 11 | 37 | 0 | 1 | 3 | 17 | 24 | 8 | 0 | 8 |
| 17 | 0.86 | 1.15 | 0.75 | 0.46 | 1 | Gm6475 | | 27 | 17 | 1 | 2 | 20 | 8 | 17 | 2 | 24 | 40 | 23 |
| 17 | -1.19 | 0.92 | -1.29 | 0.20 | 1 | Gm7389 | | 1 | 2 | 10 | 40 | 36 | 21 | 17 | 6 | 17 | 9 | 22 |
| 17 | -0.98 | 1.20 | -0.82 | 0.41 | 1 | A230103J11Rik | | 13 | 23 | 19 | 19 | 18 | 16 | 17 | 0 | 2 | 36 | 18 |
| 16 | -0.35 | 1.07 | -0.32 | 0.75 | 1 | Gm49064 | | 18 | 27 | 18 | 9 | 12 | 19 | 17 | 0 | 22 | 17 | 12 |
| 15 | 0.05 | 0.99 | 0.05 | 0.96 | 1 | Pnpla3 | patatin-like phospholipase domain contai | 2 | 12 | 6 | 16 | 7 | 32 | 17 | 3 | 19 | 17 | 34 |
| 14 | -0.46 | 1.18 | -0.39 | 0.69 | 1 | Ccdc74a | coiled-coil domain containing 74A | 51 | 15 | 25 | 7 | 1 | 10 | 17 | 0 | 13 | 12 | 5 |
| 13 | -0.85 | 1.14 | -0.75 | 0.45 | 1 | Gm12279 | | 7 | 21 | 7 | 15 | 11 | 26 | 17 | 0 | 15 | 8 | 19 |
| 13 | -2.35 | 1.70 | -1.38 | 0.17 | 1 | Gm13032 | | 27 | 25 | 35 | 12 | 0 | 4 | 17 | 2 | 17 | 0 | 0 |
| 13 | -2.32 | 1.40 | -1.66 | 0.10 | 1 | Slc6a20a | solute carrier family 6 (neurotransmitter tr | 22 | 28 | 20 | 7 | 10 | 13 | 17 | 0 | 20 | 0 | 2 |
| 13 | -1.21 | 1.13 | -1.07 | 0.28 | 1 | Rpsa-ps2 | | 30 | 23 | 29 | 3 | 11 | 8 | 17 | 3 | 5 | 0 | 8 |
| 12 | -0.72 | 0.93 | -0.77 | 0.44 | 1 | Gm5776 | | 1 | 6 | 9 | 18 | 16 | 17 | 17 | 13 | 24 | 5 | 9 |
| 12 | -0.45 | 1.24 | -0.37 | 0.71 | 1 | Myo18b | myosin XVIIIb | 6 | 3 | 12 | 12 | 11 | 22 | 17 | 29 | 0 | 12 | 11 |
| 11 | 0.80 | 1.48 | 0.54 | 0.59 | 1 | Gm20707 | | 32 | 1 | 4 | 13 | 1 | 7 | 17 | 27 | 10 | 0 | 8 |
| 11 | -0.21 | 1.20 | -0.17 | 0.86 | 1 | Gm47079 | | 21 | 0 | 24 | 0 | 11 | 9 | 17 | 0 | 7 | 21 | 6 |
| 9 | 0.18 | 2.30 | 0.08 | 0.94 | 1 | D930036K23Rik | | 0 | 19 | 19 | 0 | 0 | 14 | 17 | 0 | 0 | 21 | 13 |
| 9 | -1.47 | 1.27 | -1.16 | 0.25 | 1 | Gm6374 | | 9 | 11 | 9 | 20 | 2 | 14 | 17 | 6 | 9 | 0 | 5 |
| 9 | -1.61 | 1.94 | -0.83 | 0.41 | 1 | Fscn2 | fascin actin-bundling protein 2 | 0 | 1 | 19 | 25 | 0 | 4 | 17 | 24 | 6 | 0 | 3 |
| 8 | -0.16 | 1.42 | -0.11 | 0.91 | 1 | Gm8623 | | 5 | 2 | 0 | 2 | 15 | 11 | 17 | 21 | 14 | 1 | 4 |
| 8 | -0.09 | 1.47 | -0.06 | 0.95 | 1 | Gm14048 | | 21 | 2 | 4 | 4 | 9 | 4 | 17 | 20 | 2 | 0 | 7 |
| 8 | -1.27 | 1.46 | -0.87 | 0.38 | 1 | Gm50163 | | 0 | 7 | 3 | 11 | 9 | 8 | 17 | 13 | 6 | 0 | 12 |
| 8 | -0.96 | 1.67 | -0.58 | 0.56 | 1 | Gm12312 | | 0 | 9 | 14 | 4 | 3 | 8 | 17 | 21 | 4 | 0 | 6 |
| 7 | -4.19 | 3.04 | -1.38 | 0.17 | 1 | Gm28941 | | 0 | 0 | 8 | 0 | 0 | 32 | 17 | 0 | 0 | 0 | 23 |
| 7 | -1.94 | 1.43 | -1.36 | 0.17 | 1 | Gm37264 | | 7 | 14 | 7 | 6 | 7 | 5 | 17 | 0 | 1 | 13 | 1 |
| 7 | -1.82 | 2.44 | -0.75 | 0.45 | 1 | Gm34376 | | 0 | 7 | 0 | 0 | 5 | 16 | 17 | 0 | 0 | 9 | 21 |
| 6 | 0.48 | 1.50 | 0.32 | 0.75 | 1 | Gm9349 | | 2 | 0 | 5 | 5 | 9 | 1 | 17 | 15 | 6 | 1 | 7 |
| 6 | -0.45 | 2.03 | -0.22 | 0.83 | 1 | Serpinc1 | serine (or cysteine) peptidase inhibitor, cl | 1 | 3 | 2 | 1 | 0 | 12 | 17 | 0 | 7 | 0 | 21 |
| 5 | 1.02 | 3.08 | 0.33 | 0.74 | 1 | Gm26461 | | 0 | 0 | 7 | 0 | 0 | 7 | 17 | 2 | 0 | 0 | 24 |
| 5 | -3.49 | 2.10 | -1.67 | 0.10 | 1 | Gm17450 | | 1 | 1 | 0 | 11 | 3 | 8 | 17 | 0 | 0 | 14 | 0 |
| 4 | 1.23 | 3.46 | 0.36 | 0.72 | 1 | BC039966 | | 0 | 0 | 0 | 0 | 0 | 3 | 17 | 21 | 0 | 0 | 6 |
| 4 | -5.83 | 3.44 | -1.70 | 0.09 | 1 | Gm43136 | | 0 | 0 | 0 | 0 | 4 | 5 | 17 | 0 | 20 | 0 | 0 |
| 4 | -0.68 | 3.37 | -0.20 | 0.84 | 1 | Gm42447 | | 0 | 0 | 0 | 0 | 0 | 10 | 17 | 0 | 0 | 5 | 6 |
| 3 | -4.27 | 2.89 | -1.48 | 0.14 | 1 | 2310016D23Rik | | 0 | 0 | 0 | 8 | 3 | 4 | 17 | 0 | 0 | 0 | 5 |
| 3 | -4.10 | 2.74 | -1.49 | 0.14 | 1 | Pde6a | phosphodiesterase 6A, cGMP-specific, ro | 2 | 4 | 0 | 0 | 6 | 5 | 17 | 0 | 0 | 0 | 3 |
| 746 | -2.93 | 1.94 | -1.51 | 0.13 | 1 | Cavin2 | caveolae associated 2 | 2995 | 2581 | 2576 | 0 | 15 | 6 | 16 | 0 | 14 | 0 | 0 |
| 386 | -1.96 | 1.60 | -1.22 | 0.22 | 1 | St8sia4 | ST8 alpha-N-acetyl-neuraminide alpha-2 | 1947 | 1138 | 1124 | 0 | 4 | 3 | 16 | 0 | 0 | 12 | 0 |
| 227 | -1.69 | 1.43 | -1.18 | 0.24 | 1 | Itga7 | integrin alpha 7 | 1058 | 658 | 748 | 0 | 5 | 2 | 16 | 0 | 2 | 0 | 3 |
| 189 | 0.29 | 0.92 | 0.31 | 0.76 | 1 | Hpgd | hydroxyprostaglandin dehydrogenase 15 | 960 | 476 | 583 | 4 | 0 | 8 | 16 | 16 | 0 | 14 | 3 |
| 182 | -0.32 | 0.48 | -0.67 | 0.50 | 1 | Sema6a | sema domain, transmembrane domain (T | 622 | 488 | 527 | 56 | 67 | 64 | 16 | 37 | 42 | 59 | 24 |
| 175 | -1.96 | 1.52 | -1.29 | 0.20 | 1 | Rcsd1 | RCSD domain containing 1 | 841 | 480 | 571 | 8 | 0 | 3 | 16 | 0 | 2 | 0 | 5 |
| 150 | 0.66 | 0.50 | 1.32 | 0.19 | 1 | Ptger4 | prostaglandin E receptor 4 (subtype EP4) | 745 | 354 | 273 | 18 | 19 | 53 | 16 | 44 | 29 | 13 | 81 |
| 107 | 0.59 | 0.74 | 0.79 | 0.43 | 1 | Slc7a7 | solute carrier family 7 (cationic amino acid | 547 | 273 | 245 | 1 | 23 | 7 | 16 | 27 | 17 | 8 | 9 |
| 76 | -1.21 | 0.70 | -1.74 | 0.08 | 1 | Hs3st1 | heparan sulfate (glucosamine) 3-O-sulfot | 341 | 228 | 206 | 11 | 5 | 9 | 16 | 0 | 6 | 8 | 4 |
| 70 | 0.18 | 0.72 | 0.25 | 0.80 | 1 | Gm49331 | | 88 | 122 | 42 | 50 | 81 | 102 | 16 | 14 | 104 | 64 | 87 |
| 69 | 0.97 | 0.60 | 1.62 | 0.10 | 1 | Ndc80 | NDC80 kinetochore complex component | 173 | 103 | 57 | 35 | 33 | 63 | 16 | 94 | 67 | 24 | 97 |
| 67 | 0.13 | 0.72 | 0.18 | 0.86 | 1 | Kif2c | kinesin family member 2C | 204 | 72 | 25 | 81 | 71 | 35 | 16 | 76 | 52 | 70 | 40 |
| 67 | -2.28 | 1.36 | -1.68 | 0.09 | 1 | Clec4a3 | C-type lectin domain family 4, member a3 | 436 | 138 | 143 | 0 | 0 | 1 | 16 | 1 | 0 | 0 | 0 |
| 64 | -1.04 | 1.29 | -0.81 | 0.42 | 1 | Dok2 | docking protein 2 | 392 | 93 | 154 | 0 | 0 | 36 | 16 | 0 | 6 | 0 | 8 |
| 57 | 0.50 | 0.61 | 0.82 | 0.41 | 1 | 2310058D17Rik | | 76 | 43 | 50 | 43 | 65 | 62 | 16 | 42 | 51 | 112 | 71 |
| 57 | -0.45 | 0.83 | -0.55 | 0.58 | 1 | Gm7694 | predicted gene 7694 | 214 | 184 | 125 | 0 | 12 | 32 | 16 | 8 | 4 | 5 | 25 |
| 56 | 0.83 | 0.52 | 1.60 | 0.11 | 1 | Neil3 | nei like 3 (E. coli) | 51 | 70 | 32 | 45 | 31 | 74 | 16 | 76 | 69 | 22 | 130 |
| 53 | 0.33 | 0.89 | 0.37 | 0.71 | 1 | E2f8 | E2F transcription factor 8 | 211 | 100 | 7 | 21 | 44 | 24 | 16 | 72 | 42 | 30 | 13 |
| 51 | 0.12 | 0.56 | 0.21 | 0.83 | 1 | Ebi3 | Epstein-Barr virus induced gene 3 | 95 | 49 | 114 | 30 | 47 | 46 | 16 | 14 | 77 | 32 | 44 |
| 46 | 0.97 | 0.59 | 1.66 | 0.10 | 1 | Tnnc1 | troponin C, cardiac/slow skeletal | 46 | 62 | 30 | 18 | 20 | 70 | 16 | 102 | 26 | 22 | 98 |

| | | | | | | | | | | | | | | | | | | |
|---|---|---|---|---|---|---|---|---|---|---|---|---|---|---|---|---|---|---|
| 44 | 0.57 | 0.82 | 0.69 | 0.49 | 1 | Gm37090 | | 22 | 40 | 24 | 1 | 22 | 161 | 16 | 10 | 16 | 40 | 136 |
| 40 | 1.19 | 0.75 | 1.58 | 0.11 | 1 | Gm960 | predicted gene 960 | 22 | 17 | 39 | 29 | 8 | 73 | 16 | 52 | 20 | 99 | 68 |
| 39 | 0.53 | 0.56 | 0.96 | 0.34 | 1 | Jhy | junctional cadherin complex regulator | 20 | 53 | 23 | 64 | 26 | 41 | 16 | 49 | 37 | 52 | 52 |
| 38 | -0.36 | 0.74 | -0.48 | 0.63 | 1 | Hypk | huntingtin interacting protein K | 37 | 42 | 12 | 16 | 25 | 111 | 16 | 7 | 7 | 44 | 100 |
| 35 | -0.46 | 0.79 | -0.59 | 0.56 | 1 | A330009N23Rik | | 104 | 59 | 73 | 16 | 29 | 14 | 16 | 21 | 8 | 49 | 2 |
| 35 | -0.51 | 0.95 | -0.54 | 0.59 | 1 | BC022960 | | 12 | 33 | 46 | 28 | 7 | 98 | 16 | 54 | 8 | 0 | 85 |
| 35 | -0.70 | 1.00 | -0.69 | 0.49 | 1 | Cfap73 | cilia and flagella associated protein 73 | 94 | 35 | 74 | 46 | 10 | 29 | 16 | 0 | 16 | 31 | 29 |
| 34 | -0.24 | 1.03 | -0.23 | 0.82 | 1 | Gm44777 | | 43 | 40 | 35 | 43 | 9 | 65 | 16 | 0 | 13 | 54 | 61 |
| 34 | -1.28 | 1.54 | -0.83 | 0.40 | 1 | Calml4 | calmodulin-like 4 | 189 | 94 | 58 | 0 | 0 | 5 | 16 | 0 | 0 | 0 | 12 |
| 32 | -0.70 | 1.07 | -0.65 | 0.52 | 1 | Stil | Scl/Tal1 interrupting locus | 136 | 10 | 56 | 3 | 47 | 38 | 16 | 9 | 21 | 0 | 15 |
| 30 | 0.00 | 0.62 | 0.01 | 1.00 | 1 | Cenpu | centromere protein U | 50 | 26 | 51 | 20 | 30 | 35 | 16 | 23 | 15 | 30 | 32 |
| 30 | -2.00 | 1.55 | -1.29 | 0.20 | 1 | Fcmr | Fc fragment of IgM receptor | 240 | 30 | 28 | 0 | 3 | 1 | 16 | 0 | 0 | 0 | 10 |
| 29 | 0.76 | 0.65 | 1.16 | 0.24 | 1 | Oip5 | Opa interacting protein 5 | 70 | 35 | 43 | 23 | 7 | 18 | 16 | 37 | 17 | 24 | 25 |
| 29 | 0.44 | 0.93 | 0.48 | 0.63 | 1 | Gm14253 | | 27 | 46 | 35 | 17 | 18 | 21 | 16 | 2 | 29 | 90 | 13 |
| 28 | -2.66 | 1.46 | -1.82 | 0.07 | 1 | H2ac7 | H2A clustered histone 7 | 0 | 0 | 0 | 75 | 20 | 99 | 16 | 0 | 19 | 0 | 81 |
| 25 | 0.61 | 1.15 | 0.53 | 0.60 | 1 | Vash2 | vasohibin 2 | 75 | 79 | 48 | 7 | 1 | 3 | 16 | 39 | 6 | 0 | 4 |
| 24 | 0.37 | 1.02 | 0.36 | 0.72 | 1 | Zfp473 | zinc finger protein 473 | 60 | 0 | 27 | 9 | 22 | 36 | 16 | 17 | 35 | 21 | 20 |
| 23 | -0.66 | 0.80 | -0.83 | 0.40 | 1 | Gm44066 | | 35 | 63 | 46 | 9 | 35 | 6 | 16 | 20 | 11 | 5 | 5 |
| 23 | -0.08 | 1.05 | -0.07 | 0.94 | 1 | Gm12992 | | 15 | 36 | 24 | 10 | 12 | 45 | 16 | 0 | 10 | 41 | 40 |
| 22 | -1.04 | 1.02 | -1.01 | 0.31 | 1 | Gm49694 | | 28 | 13 | 50 | 36 | 11 | 35 | 16 | 25 | 15 | 0 | 15 |
| 22 | -1.12 | 1.07 | -1.04 | 0.30 | 1 | Cdkl4 | cyclin-dependent kinase-like 4 | 10 | 0 | 23 | 49 | 31 | 38 | 16 | 12 | 13 | 40 | 11 |
| 21 | 0.04 | 1.05 | 0.04 | 0.97 | 1 | Hcst | hematopoietic cell signal transducer | 22 | 40 | 39 | 5 | 8 | 30 | 16 | 27 | 15 | 0 | 28 |
| 20 | 0.18 | 0.77 | 0.23 | 0.82 | 1 | Gm5786 | | 48 | 20 | 33 | 25 | 10 | 8 | 16 | 21 | 20 | 10 | 12 |
| 20 | -1.02 | 1.39 | -0.73 | 0.47 | 1 | Prr22 | proline rich 22 | 0 | 6 | 0 | 4 | 15 | 79 | 16 | 7 | 18 | 0 | 77 |
| 20 | 0.17 | 1.09 | 0.16 | 0.88 | 1 | Gm4814 | | 9 | 16 | 2 | 35 | 10 | 19 | 16 | 73 | 23 | 0 | 16 |
| 20 | -0.39 | 0.85 | -0.46 | 0.65 | 1 | Gm5628 | | 24 | 15 | 7 | 28 | 16 | 25 | 16 | 41 | 4 | 6 | 36 |
| 19 | 0.60 | 0.92 | 0.65 | 0.52 | 1 | Nkain1 | Na+/K+ transporting ATPase interacting 1 | 20 | 17 | 0 | 31 | 8 | 19 | 16 | 27 | 17 | 24 | 28 |
| 18 | -0.88 | 0.87 | -1.01 | 0.31 | 1 | Gm11767 | | 13 | 13 | 1 | 27 | 20 | 47 | 16 | 13 | 13 | 6 | 27 |
| 18 | 0.13 | 0.82 | 0.16 | 0.88 | 1 | Gm7461 | | 21 | 5 | 18 | 22 | 17 | 13 | 16 | 13 | 10 | 44 | 19 |
| 18 | 0.67 | 0.83 | 0.80 | 0.42 | 1 | Gm9392 | | 18 | 16 | 5 | 26 | 7 | 14 | 16 | 9 | 23 | 40 | 21 |
| 17 | -0.62 | 1.05 | -0.59 | 0.56 | 1 | AA474408 | | 30 | 10 | 4 | 10 | 38 | 26 | 16 | 12 | 25 | 1 | 19 |
| 17 | -1.38 | 1.69 | -0.82 | 0.41 | 1 | Slitrk1 | SLIT and NTRK-like family, member 1 | 44 | 42 | 6 | 0 | 29 | 11 | 16 | 0 | 0 | 35 | 8 |
| 17 | -0.92 | 0.98 | -0.94 | 0.35 | 1 | Gm26759 | | 9 | 5 | 5 | 17 | 15 | 52 | 16 | 0 | 11 | 18 | 41 |
| 17 | -0.81 | 1.55 | -0.52 | 0.60 | 1 | 2700099C18Rik | | 17 | 0 | 0 | 17 | 26 | 46 | 16 | 7 | 19 | 0 | 40 |
| 17 | -0.51 | 0.93 | -0.55 | 0.58 | 1 | Zcwpw2 | | 4 | 0 | 14 | 2 | 21 | 54 | 16 | 0 | 21 | 15 | 40 |
| 17 | 0.68 | 0.78 | 0.87 | 0.39 | 1 | Srsf3-ps | | 22 | 22 | 12 | 23 | 4 | 9 | 16 | 25 | 11 | 13 | 27 |
| 16 | -0.29 | 1.16 | -0.25 | 0.80 | 1 | Gm9812 | | 15 | 62 | 8 | 20 | 9 | 10 | 16 | 0 | 13 | 13 | 12 |
| 16 | -1.06 | 1.61 | -0.66 | 0.51 | 1 | Col6a4 | collagen, type VI, alpha 4 | 51 | 29 | 8 | 0 | 20 | 14 | 16 | 0 | 18 | 0 | 18 |
| 16 | -1.87 | 1.31 | -1.42 | 0.15 | 1 | Gm20033 | | 5 | 25 | 6 | 11 | 9 | 47 | 16 | 0 | 9 | 0 | 44 |
| 16 | 1.81 | 1.19 | 1.52 | 0.13 | 1 | Gm32151 | | 11 | 16 | 18 | 4 | 0 | 8 | 16 | 64 | 5 | 14 | 16 |
| 14 | -2.48 | 1.84 | -1.35 | 0.18 | 1 | Prrt4 | proline-rich transmembrane protein 4 | 71 | 10 | 50 | 0 | 0 | 4 | 16 | 0 | 0 | 0 | 1 |
| 14 | 0.48 | 1.37 | 0.35 | 0.73 | 1 | Gm5421 | | 15 | 29 | 0 | 4 | 10 | 12 | 16 | 2 | 37 | 8 | 16 |
| 14 | -0.09 | 0.99 | -0.10 | 0.92 | 1 | Gm15519 | | 20 | 19 | 9 | 14 | 10 | 10 | 16 | 13 | 16 | 1 | 20 |
| 14 | 0.66 | 1.48 | 0.45 | 0.65 | 1 | Gm15743 | | 1 | 14 | 0 | 18 | 6 | 15 | 16 | 0 | 9 | 52 | 18 |
| 13 | 0.30 | 0.97 | 0.31 | 0.76 | 1 | | | 20 | 6 | 7 | 20 | 9 | 8 | 16 | 24 | 17 | 4 | 14 |
| 13 | 0.06 | 1.29 | 0.04 | 0.97 | 1 | Slc6a15 | solute carrier family 6 (neurotransmitter tr | 6 | 11 | 15 | 32 | 5 | 1 | 16 | 42 | 7 | 0 | 8 |
| 12 | -1.05 | 1.19 | -0.88 | 0.38 | 1 | D730044K07Rik | | 8 | 36 | 11 | 11 | 12 | 8 | 16 | 0 | 9 | 9 | 5 |
| 10 | -1.79 | 1.35 | -1.33 | 0.18 | 1 | Gm16364 | | 3 | 4 | 0 | 25 | 32 | 11 | 16 | 0 | 11 | 8 | 5 |
| 10 | -1.44 | 1.51 | -0.95 | 0.34 | 1 | Gm49326 | | 18 | 28 | 0 | 4 | 12 | 10 | 16 | 10 | 4 | 0 | 5 |
| 9 | 0.84 | 1.74 | 0.48 | 0.63 | 1 | Sec14l5 | SEC14-like lipid binding 5 | 10 | 7 | 0 | 0 | 5 | 9 | 16 | 0 | 16 | 27 | 7 |
| 8 | -2.16 | 1.47 | -1.47 | 0.14 | 1 | Gm1826 | | 7 | 4 | 15 | 29 | 2 | 4 | 16 | 0 | 9 | 1 | 4 |
| 8 | -0.58 | 1.32 | -0.44 | 0.66 | 1 | Gm12799 | | 3 | 2 | 1 | 2 | 10 | 14 | 16 | 31 | 3 | 1 | 7 |
| 7 | 0.14 | 2.17 | 0.06 | 0.95 | 1 | Gm15423 | | 0 | 0 | 0 | 0 | 4 | 21 | 16 | 19 | 9 | 0 | 12 |
| 6 | 2.10 | 2.24 | 0.94 | 0.35 | 1 | Gm11520 | | 20 | 0 | 0 | 0 | 8 | 1 | 16 | 16 | 6 | 0 | 4 |
| 6 | -1.62 | 2.09 | -0.77 | 0.44 | 1 | Gm43137 | | 0 | 7 | 0 | 7 | 4 | 8 | 16 | 0 | 0 | 22 | 4 |
| 6 | -2.55 | 1.63 | -1.56 | 0.12 | 1 | Gm44545 | | 1 | 2 | 3 | 7 | 6 | 12 | 16 | 0 | 13 | 0 | 3 |
| 5 | -1.15 | 2.42 | -0.47 | 0.64 | 1 | Gm45469 | | 0 | 0 | 1 | 0 | 8 | 11 | 16 | 6 | 0 | 0 | 18 |
| 5 | -4.42 | 2.32 | -1.90 | 0.06 | 1 | Gm31513 | | 1 | 0 | 0 | 12 | 4 | 8 | 16 | 0 | 0 | 0 | 11 |
| 5 | 0.29 | 1.69 | 0.17 | 0.86 | 1 | Gm44771 | | 4 | 1 | 1 | 1 | 7 | 0 | 16 | 0 | 8 | 6 | 6 |
| 4 | -0.34 | 2.67 | -0.13 | 0.90 | 1 | Gm18280 | | 0 | 0 | 5 | 0 | 0 | 5 | 16 | 0 | 2 | 12 | 1 |

| | | | | | | | | | | | | | | | | | | |
|---|---|---|---|---|---|---|---|---|---|---|---|---|---|---|---|---|---|---|
| 4 | -3.01 | 3.05 | -0.99 | 0.32 | 1 | Csmd3 | CUB and Sushi multiple domains 3 | 3 | 5 | 15 | 0 | 0 | 2 | 16 | 0 | 0 | 0 | 0 |
| 3 | -0.39 | 2.66 | -0.15 | 0.88 | 1 | Gm5131 | | 1 | 0 | 1 | 2 | 0 | 0 | 16 | 0 | 0 | 13 | 1 |
| 3 | -2.67 | 2.86 | -0.93 | 0.35 | 1 | Serpina1c | serine (or cysteine) peptidase inhibitor, cl | 2 | 1 | 11 | 0 | 0 | 2 | 16 | 0 | 1 | 0 | 0 |
| 3 | -3.61 | 3.41 | -1.06 | 0.29 | 1 | Mir7655 | | 0 | 0 | 3 | 7 | 0 | 0 | 16 | 0 | 3 | 0 | 1 |
| 3 | -4.87 | 3.55 | -1.37 | 0.17 | 1 | Kif16bos | | 0 | 0 | 0 | 4 | 2 | 6 | 16 | 0 | 0 | 0 | 0 |
| 2894 | -1.22 | 1.50 | -0.81 | 0.42 | 1 | Ptprb | protein tyrosine phosphatase, receptor typ | 10303 | 10257 | 11219 | 0 | 3 | 19 | 15 | 0 | 18 | 1 | 2 |
| 797 | -1.32 | 3.62 | -0.37 | 0.71 | 1 | Clec14a | C-type lectin domain family 14, member a | 3132 | 2706 | 2914 | 0 | 0 | 0 | 15 | 0 | 0 | 0 | 0 |
| 599 | -0.72 | 1.68 | -0.43 | 0.67 | 1 | Chst15 | carbohydrate sulfotransferase 15 | 2209 | 2402 | 1902 | 17 | 0 | 3 | 15 | 43 | 0 | 0 | 3 |
| 508 | -2.34 | 1.71 | -1.37 | 0.17 | 1 | Kitl | kit ligand | 1964 | 1721 | 1865 | 0 | 9 | 2 | 15 | 0 | 7 | 0 | 0 |
| 385 | -0.90 | 1.20 | -0.75 | 0.45 | 1 | Cyba | cytochrome b-245, alpha polypeptide | 2242 | 836 | 1120 | 2 | 5 | 3 | 15 | 0 | 12 | 0 | 3 |
| 314 | -2.52 | 1.59 | -1.59 | 0.11 | 1 | Rgcc | regulator of cell cycle | 1274 | 1067 | 1076 | 2 | 0 | 8 | 15 | 0 | 0 | 0 | 8 |
| 244 | 0.71 | 0.46 | 1.55 | 0.12 | 1 | Csrp2 | cysteine and glycine-rich protein 2 | 1370 | 523 | 605 | 9 | 34 | 14 | 15 | 22 | 18 | 43 | 36 |
| 213 | 0.25 | 0.75 | 0.33 | 0.74 | 1 | Egr2 | early growth response 2 | 1150 | 372 | 457 | 1 | 2 | 174 | 15 | 0 | 16 | 5 | 150 |
| 203 | 1.85 | 1.39 | 1.33 | 0.18 | 1 | Arhgap45 | Rho GTPase activating protein 45 | 1289 | 441 | 416 | 0 | 0 | 14 | 15 | 0 | 13 | 33 | 16 |
| 98 | -0.02 | 0.59 | -0.03 | 0.97 | 1 | Rasgrp2 | RAS, guanyl releasing protein 2 | 516 | 241 | 157 | 13 | 16 | 38 | 15 | 17 | 20 | 5 | 37 |
| 91 | 0.01 | 0.53 | 0.02 | 0.98 | 1 | Gm14681 | | 27 | 186 | 300 | 83 | 83 | 14 | 15 | 159 | 121 | 12 | 4 |
| 73 | -1.95 | 2.10 | -0.93 | 0.35 | 1 | Kank4 | KN motif and ankyrin repeat domains 4 | 250 | 166 | 329 | 10 | 0 | 2 | 15 | 31 | 0 | 0 | 0 |
| 57 | -0.74 | 0.87 | -0.85 | 0.40 | 1 | Serpinb8 | serine (or cysteine) peptidase inhibitor, cl | 183 | 102 | 124 | 43 | 32 | 34 | 15 | 52 | 25 | 0 | 13 |
| 52 | -1.47 | 1.27 | -1.16 | 0.25 | 1 | Lhpp | phospholysine phosphohistidine inorgani | 198 | 219 | 96 | 0 | 18 | 7 | 15 | 0 | 11 | 0 | 6 |
| 50 | 0.46 | 0.47 | 0.98 | 0.33 | 1 | Zfp493 | zinc finger protein 493 | 73 | 117 | 80 | 41 | 29 | 34 | 15 | 42 | 30 | 31 | 60 |
| 50 | 0.27 | 0.56 | 0.48 | 0.63 | 1 | Car7 | carbonic anhydrase 7 | 135 | 66 | 133 | 13 | 36 | 27 | 15 | 8 | 47 | 17 | 53 |
| 49 | 0.12 | 0.84 | 0.15 | 0.88 | 1 | Cracr2a | calcium release activated channel regula | 7 | 61 | 37 | 60 | 37 | 117 | 15 | 23 | 19 | 95 | 69 |
| 43 | -0.48 | 1.34 | -0.36 | 0.72 | 1 | Gper1 | G protein-coupled estrogen receptor 1 | 129 | 207 | 102 | 0 | 1 | 6 | 15 | 0 | 4 | 0 | 10 |
| 41 | -0.99 | 1.16 | -0.86 | 0.39 | 1 | Rnf125 | ring finger protein 125 | 133 | 107 | 137 | 27 | 0 | 8 | 15 | 0 | 7 | 17 | 3 |
| 40 | 0.30 | 0.53 | 0.56 | 0.57 | 1 | C920021L13Rik | | 13 | 45 | 42 | 73 | 33 | 38 | 15 | 85 | 31 | 27 | 44 |
| 39 | -0.80 | 0.57 | -1.38 | 0.17 | 1 | Ifnlr1 | interferon lambda receptor 1 | 108 | 88 | 74 | 25 | 14 | 46 | 15 | 15 | 5 | 14 | 24 |
| 38 | -0.81 | 1.13 | -0.72 | 0.47 | 1 | Gm38372 | | 22 | 0 | 12 | 17 | 3 | 167 | 15 | 0 | 12 | 28 | 136 |
| 36 | -1.91 | 1.12 | -1.71 | 0.09 | 1 | Mesp1 | mesoderm posterior 1 | 0 | 5 | 32 | 50 | 13 | 117 | 15 | 15 | 10 | 0 | 140 |
| 35 | 0.08 | 0.63 | 0.13 | 0.90 | 1 | Rps15a-ps7 | | 54 | 22 | 94 | 36 | 35 | 19 | 15 | 25 | 36 | 30 | 20 |
| 33 | 0.64 | 0.76 | 0.83 | 0.40 | 1 | Gm10036 | | 88 | 84 | 55 | 21 | 9 | 8 | 15 | 5 | 31 | 24 | 20 |
| 33 | 0.20 | 0.65 | 0.31 | 0.75 | 1 | Slc26a8 | solute carrier family 26, member 8 | 30 | 86 | 24 | 23 | 6 | 59 | 15 | 13 | 14 | 14 | 75 |
| 32 | 0.25 | 1.00 | 0.25 | 0.80 | 1 | Sntg2 | syntrophin, gamma 2 | 43 | 79 | 79 | 20 | 6 | 23 | 15 | 30 | 0 | 44 | 15 |
| 31 | -0.65 | 1.04 | -0.63 | 0.53 | 1 | Srd5a1 | steroid 5 alpha-reductase 1 | 83 | 24 | 31 | 31 | 37 | 12 | 15 | 0 | 13 | 73 | 17 |
| 28 | 0.68 | 0.81 | 0.84 | 0.40 | 1 | Lingo4 | leucine rich repeat and Ig domain contain | 27 | 15 | 26 | 16 | 21 | 32 | 15 | 98 | 25 | 18 | 16 |
| 28 | -0.11 | 0.96 | -0.11 | 0.91 | 1 | Fibin | fin bud initiation factor homolog (zebrafish | 133 | 15 | 59 | 11 | 20 | 5 | 15 | 14 | 24 | 1 | 5 |
| 26 | 0.17 | 0.75 | 0.22 | 0.82 | 1 | Gm34574 | | 6 | 9 | 24 | 14 | 37 | 56 | 15 | 10 | 19 | 48 | 52 |
| 26 | -1.28 | 1.10 | -1.16 | 0.25 | 1 | Gm12844 | | 101 | 26 | 45 | 26 | 15 | 11 | 15 | 2 | 14 | 0 | 26 |
| 25 | -0.20 | 1.05 | -0.19 | 0.85 | 1 | Fam221b | family with sequence similarity 221, meml | 44 | 42 | 57 | 35 | 0 | 27 | 15 | 14 | 16 | 18 | 9 |
| 22 | -0.12 | 0.74 | -0.17 | 0.87 | 1 | Gm13528 | | 23 | 33 | 59 | 23 | 11 | 14 | 15 | 21 | 15 | 19 | 6 |
| 22 | 0.19 | 0.78 | 0.25 | 0.80 | 1 | Gm19531 | | 9 | 37 | 52 | 17 | 12 | 20 | 15 | 12 | 22 | 19 | 21 |
| 21 | 1.03 | 1.03 | 1.00 | 0.32 | 1 | B3gat1 | beta-1,3-glucuronyltransferase 1 (glucurc | 0 | 7 | 30 | 4 | 25 | 22 | 15 | 29 | 33 | 14 | 57 |
| 21 | 1.38 | 1.15 | 1.21 | 0.23 | 1 | Grm1 | glutamate receptor, metabotropic 1 | 0 | 11 | 0 | 6 | 15 | 19 | 15 | 36 | 20 | 62 | 47 |
| 21 | 0.92 | 0.86 | 1.06 | 0.29 | 1 | 5830408C22Rik | | 31 | 22 | 22 | 11 | 5 | 15 | 15 | 65 | 4 | 22 | 16 |
| 19 | 0.03 | 0.85 | 0.04 | 0.97 | 1 | Gm15536 | | 46 | 31 | 31 | 5 | 13 | 18 | 15 | 10 | 19 | 3 | 21 |
| 19 | -0.52 | 1.01 | -0.52 | 0.61 | 1 | Gm49745 | | 0 | 19 | 22 | 15 | 26 | 27 | 15 | 21 | 32 | 3 | 30 |
| 18 | -0.22 | 1.34 | -0.17 | 0.87 | 1 | A330102I10Rik | | 0 | 10 | 0 | 11 | 16 | 21 | 15 | 44 | 1 | 71 | 10 |
| 17 | 0.51 | 1.16 | 0.44 | 0.66 | 1 | Rpl7-ps7 | | 19 | 21 | 20 | 3 | 19 | 8 | 15 | 57 | 2 | 4 | 25 |
| 17 | 0.22 | 1.36 | 0.16 | 0.87 | 1 | H3c6 | H3 clustered histone 6 | 5 | 2 | 0 | 4 | 16 | 46 | 15 | 49 | 13 | 0 | 35 |
| 16 | -1.06 | 1.25 | -0.84 | 0.40 | 1 | Pfkfb1 | 6-phosphofructo-2-kinase/fructose-2,6-bip | 25 | 23 | 42 | 0 | 14 | 25 | 15 | 2 | 6 | 0 | 28 |
| 16 | 0.92 | 0.86 | 1.07 | 0.28 | 1 | Gm16418 | | 9 | 29 | 11 | 15 | 7 | 4 | 15 | 19 | 15 | 50 | 4 |
| 16 | -0.77 | 1.39 | -0.55 | 0.58 | 1 | Rps10-ps4 | | 34 | 13 | 0 | 17 | 28 | 11 | 15 | 0 | 16 | 32 | 9 |
| 16 | -1.75 | 1.36 | -1.29 | 0.20 | 1 | Gm45428 | | 45 | 10 | 17 | 21 | 15 | 16 | 15 | 0 | 22 | 0 | 14 |
| 16 | -0.70 | 1.57 | -0.44 | 0.66 | 1 | Cacnb4 | calcium channel, voltage-dependent, beta | 53 | 43 | 10 | 0 | 5 | 19 | 15 | 0 | 9 | 0 | 21 |
| 16 | 0.34 | 0.77 | 0.45 | 0.66 | 1 | Gm4577 | | 11 | 8 | 26 | 16 | 11 | 13 | 15 | 14 | 21 | 15 | 22 |
| 16 | 0.26 | 1.14 | 0.23 | 0.82 | 1 | Slc25a5-ps | | 0 | 19 | 2 | 21 | 4 | 23 | 15 | 31 | 1 | 31 | 24 |
| 15 | 0.34 | 1.05 | 0.33 | 0.74 | 1 | Gm50357 | | 26 | 44 | 7 | 8 | 5 | 11 | 15 | 28 | 16 | 1 | 8 |
| 15 | 1.09 | 0.99 | 1.11 | 0.27 | 1 | BC049762 | cDNA sequence BC049762 | 8 | 7 | 19 | 3 | 15 | 7 | 15 | 35 | 13 | 39 | 4 |
| 14 | -0.35 | 1.08 | -0.32 | 0.75 | 1 | Gm41031 | | 4 | 26 | 11 | 14 | 11 | 14 | 15 | 22 | 19 | 0 | 19 |
| 14 | -0.82 | 0.95 | -0.87 | 0.39 | 1 | Smarce1-ps1 | | 3 | 7 | 3 | 35 | 22 | 8 | 15 | 32 | 8 | 1 | 14 |
| 13 | -0.26 | 1.11 | -0.24 | 0.81 | 1 | Dgkeos | | 6 | 0 | 3 | 31 | 10 | 10 | 15 | 41 | 4 | 23 | 4 |

| | | | | | | | | | | | | | | | | | | |
|---|---|---|---|---|---|---|---|---|---|---|---|---|---|---|---|---|---|---|
| 13 | -0.93 | 1.28 | -0.73 | 0.47 | 1 | Gm11100 | predicted gene 11100 | 4 | 0 | 17 | 11 | 21 | 12 | 15 | 0 | 14 | 36 | 10 |
| 13 | -0.99 | 1.51 | -0.66 | 0.51 | 1 | Gm14121 | | 10 | 38 | 22 | 1 | 11 | 8 | 15 | 0 | 17 | 0 | 14 |
| 12 | 0.70 | 1.31 | 0.54 | 0.59 | 1 | Gm7785 | | 4 | 1 | 12 | 2 | 6 | 25 | 15 | 15 | 26 | 0 | 28 |
| 12 | -0.75 | 1.49 | -0.50 | 0.61 | 1 | 1700054A03Rik | | 16 | 0 | 11 | 13 | 23 | 10 | 15 | 28 | 12 | 0 | 3 |
| 12 | -1.18 | 1.58 | -0.75 | 0.45 | 1 | Tmem52 | transmembrane protein 52 | 16 | 0 | 0 | 25 | 12 | 9 | 15 | 0 | 15 | 26 | 9 |
| 12 | 0.04 | 1.17 | 0.04 | 0.97 | 1 | Tsacc | TSSK6 activating co-chaperone | 4 | 0 | 12 | 0 | 10 | 30 | 15 | 0 | 20 | 12 | 23 |
| 12 | 0.52 | 1.02 | 0.51 | 0.61 | 1 | Gm5446 | | 5 | 11 | 16 | 2 | 9 | 13 | 15 | 15 | 16 | 5 | 20 |
| 11 | -1.18 | 1.91 | -0.61 | 0.54 | 1 | Gm4787 | predicted gene 4787 | 12 | 0 | 0 | 17 | 9 | 19 | 15 | 38 | 0 | 0 | 17 |
| 11 | 0.89 | 1.55 | 0.57 | 0.57 | 1 | Gm37226 | | 15 | 34 | 15 | 0 | 1 | 9 | 15 | 13 | 11 | 0 | 12 |
| 11 | -0.26 | 0.98 | -0.26 | 0.79 | 1 | Gm6981 | | 24 | 7 | 19 | 9 | 5 | 8 | 15 | 9 | 2 | 14 | 6 |
| 11 | -1.96 | 2.04 | -0.96 | 0.34 | 1 | Mir99ahg | | 16 | 41 | 31 | 0 | 0 | 8 | 15 | 0 | 0 | 0 | 5 |
| 11 | 0.55 | 1.15 | 0.48 | 0.63 | 1 | Gm26558 | | 8 | 1 | 18 | 6 | 6 | 8 | 15 | 29 | 13 | 6 | 6 |
| 11 | -0.23 | 1.09 | -0.21 | 0.83 | 1 | Gm49335 | | 0 | 0 | 0 | 13 | 9 | 26 | 15 | 7 | 11 | 19 | 15 |
| 10 | -1.50 | 1.57 | -0.96 | 0.34 | 1 | Pdxk-ps | | 23 | 11 | 15 | 7 | 0 | 14 | 15 | 0 | 0 | 6 | 20 |
| 10 | -1.53 | 1.41 | -1.08 | 0.28 | 1 | 1810012K16Rik | | 0 | 2 | 7 | 24 | 15 | 14 | 15 | 0 | 6 | 8 | 17 |
| 9 | -1.10 | 1.22 | -0.90 | 0.37 | 1 | Gm45795 | | 4 | 7 | 4 | 11 | 11 | 16 | 15 | 0 | 15 | 10 | 4 |
| 8 | 0.88 | 1.76 | 0.50 | 0.62 | 1 | Fgf22 | fibroblast growth factor 22 | 9 | 0 | 0 | 11 | 9 | 2 | 15 | 29 | 13 | 0 | 4 |
| 8 | 1.54 | 1.59 | 0.97 | 0.33 | 1 | Ttc30a2 | tetratricopeptide repeat domain 30A2 | 9 | 13 | 5 | 0 | 4 | 1 | 15 | 0 | 12 | 22 | 8 |
| 8 | -0.42 | 1.25 | -0.33 | 0.74 | 1 | Rpl28-ps1 | | 5 | 19 | 4 | 3 | 11 | 4 | 15 | 1 | 10 | 6 | 6 |
| 8 | -1.30 | 1.56 | -0.83 | 0.41 | 1 | Nutf2-ps1 | | 5 | 22 | 11 | 7 | 2 | 3 | 15 | 13 | 0 | 0 | 6 |
| 7 | 0.95 | 1.43 | 0.67 | 0.50 | 1 | Gm48496 | | 21 | 0 | 4 | 0 | 5 | 3 | 15 | 0 | 17 | 12 | 6 |
| 7 | -1.91 | 2.25 | -0.85 | 0.40 | 1 | Gm22107 | | 18 | 9 | 0 | 0 | 8 | 10 | 15 | 0 | 14 | 0 | 1 |
| 7 | 0.48 | 1.67 | 0.28 | 0.78 | 1 | B230119M05Rik | | 12 | 6 | 0 | 0 | 10 | 2 | 15 | 1 | 12 | 8 | 6 |
| 6 | -1.41 | 1.60 | -0.89 | 0.38 | 1 | Gm20498 | predicted gene 20498 | 13 | 5 | 2 | 7 | 4 | 7 | 15 | 0 | 5 | 0 | 13 |
| 6 | 0.43 | 1.69 | 0.25 | 0.80 | 1 | Gm47735 | | 3 | 12 | 10 | 4 | 0 | 2 | 15 | 0 | 7 | 10 | 5 |
| 6 | -1.72 | 2.06 | -0.84 | 0.40 | 1 | Syna | syncytin a | 4 | 0 | 0 | 10 | 0 | 14 | 15 | 0 | 0 | 14 | 7 |
| 6 | 0.99 | 1.57 | 0.63 | 0.53 | 1 | Gm11189 | | 4 | 1 | 1 | 3 | 0 | 3 | 15 | 0 | 7 | 23 | 6 |
| 5 | -0.96 | 1.38 | -0.70 | 0.49 | 1 | Gm9755 | | 5 | 7 | 9 | 1 | 4 | 6 | 15 | 1 | 1 | 4 | 7 |
| 5 | -0.90 | 1.48 | -0.61 | 0.54 | 1 | Gm6430 | | 3 | 16 | 1 | 2 | 2 | 6 | 15 | 1 | 2 | 1 | 9 |
| 5 | -1.63 | 2.14 | -0.76 | 0.45 | 1 | Gm14335 | | 0 | 15 | 1 | 3 | 6 | 3 | 15 | 0 | 8 | 0 | 3 |
| 4 | -2.05 | 2.44 | -0.84 | 0.40 | 1 | Gm4129 | | 2 | 2 | 0 | 7 | 0 | 6 | 15 | 0 | 13 | 0 | 1 |
| 4 | -1.11 | 1.80 | -0.61 | 0.54 | 1 | Gm10029 | | 6 | 2 | 9 | 0 | 6 | 1 | 15 | 3 | 2 | 1 | 0 |
| 4 | -1.40 | 1.57 | -0.89 | 0.37 | 1 | Gm50045 | | 5 | 1 | 1 | 1 | 10 | 3 | 15 | 0 | 5 | 1 | 3 |
| 3 | -4.26 | 2.65 | -1.61 | 0.11 | 1 | Gm29629 | | 3 | 0 | 0 | 8 | 6 | 1 | 15 | 0 | 0 | 0 | 6 |
| 3 | -1.84 | 1.97 | -0.93 | 0.35 | 1 | Rpl7-ps9 | | 6 | 4 | 1 | 0 | 7 | 1 | 15 | 0 | 1 | 1 | 2 |
| 3 | -2.20 | 2.98 | -0.74 | 0.46 | 1 | A730043L09Rik | | 0 | 0 | 3 | 2 | 4 | 0 | 15 | 0 | 14 | 0 | 0 |
| 3 | -1.03 | 2.96 | -0.35 | 0.73 | 1 | Gm43889 | | 0 | 0 | 0 | 0 | 2 | 7 | 15 | 0 | 3 | 0 | 9 |
| 3 | -2.43 | 3.17 | -0.77 | 0.44 | 1 | Gm30373 | | 0 | 0 | 0 | 5 | 0 | 4 | 15 | 0 | 2 | 0 | 6 |
| 3 | -3.90 | 3.53 | -1.10 | 0.27 | 1 | Gm5819 | | 0 | 0 | 0 | 0 | 9 | 3 | 15 | 0 | 0 | 0 | 2 |
| 2 | -4.12 | 3.59 | -1.15 | 0.25 | 1 | Gm14280 | | 0 | 6 | 0 | 0 | 0 | 2 | 15 | 0 | 0 | 0 | 2 |
| 2 | -0.67 | 3.54 | -0.19 | 0.85 | 1 | Gm44013 | | 0 | 0 | 2 | 0 | 0 | 0 | 15 | 0 | 7 | 0 | 0 |
| 2 | -2.71 | 3.59 | -0.75 | 0.45 | 1 | 3110053B16Rik | | 3 | 0 | 0 | 0 | 0 | 5 | 15 | 0 | 0 | 0 | 0 |
| 2343 | -0.41 | 1.29 | -0.32 | 0.75 | 1 | Axl | AXL receptor tyrosine kinase | 12050 | 5952 | 7700 | 2 | 14 | 3 | 14 | 0 | 28 | 0 | 9 |
| 1075 | -1.21 | 1.28 | -0.94 | 0.35 | 1 | C1qb | complement component 1, q subcompone | 5293 | 2594 | 3845 | 16 | 13 | 8 | 14 | 0 | 22 | 0 | 18 |
| 283 | -0.70 | 0.61 | -1.15 | 0.25 | 1 | Egfl7 | EGF-like domain 7 | 1088 | 837 | 1040 | 15 | 42 | 19 | 14 | 0 | 11 | 28 | 24 |
| 268 | -1.04 | 1.40 | -0.74 | 0.46 | 1 | Cxcr4 | chemokine (C-X-C motif) receptor 4 | 1362 | 815 | 748 | 0 | 1 | 3 | 14 | 0 | 2 | 0 | 4 |
| 218 | -0.01 | 1.97 | 0.00 | 1.00 | 1 | Fermt3 | fermitin family member 3 | 1234 | 531 | 532 | 0 | 0 | 46 | 14 | 0 | 11 | 0 | 34 |
| 196 | -1.08 | 1.72 | -0.63 | 0.53 | 1 | Slc16a1 | solute carrier family 16 (monocarboxylic a | 879 | 668 | 566 | 11 | 0 | 3 | 14 | 0 | 12 | 0 | 3 |
| 186 | -0.80 | 0.52 | -1.55 | 0.12 | 1 | Mical1 | microtubule associated monooxygenase, | 957 | 547 | 355 | 48 | 24 | 27 | 14 | 30 | 16 | 17 | 6 |
| 148 | -0.81 | 0.56 | -1.44 | 0.15 | 1 | Fhod1 | formin homology 2 domain containing 1 | 544 | 239 | 470 | 67 | 60 | 81 | 14 | 24 | 25 | 10 | 93 |
| 132 | -0.72 | 0.86 | -0.84 | 0.40 | 1 | Plxnc1 | plexin C1 | 879 | 225 | 291 | 0 | 11 | 10 | 14 | 5 | 9 | 0 | 3 |
| 107 | 1.25 | 0.66 | 1.89 | 0.06 | 1 | 4933439C10Rik | | 31 | 75 | 20 | 5 | 1 | 500 | 14 | 16 | 17 | 24 | 471 |
| 102 | -0.21 | 0.85 | -0.25 | 0.80 | 1 | Map4k1 | mitogen-activated protein kinase kinase k | 694 | 200 | 90 | 18 | 7 | 39 | 14 | 10 | 18 | 0 | 32 |
| 92 | -0.62 | 2.05 | -0.30 | 0.76 | 1 | Tmem45a | transmembrane protein 45a | 611 | 126 | 236 | 0 | 0 | 9 | 14 | 0 | 0 | 21 | 0 |
| 89 | -0.98 | 2.30 | -0.43 | 0.67 | 1 | Gimap1 | GTPase, IMAP family member 1 | 399 | 283 | 253 | 0 | 0 | 18 | 14 | 14 | 2 | 0 | 0 |
| 83 | -0.37 | 0.56 | -0.66 | 0.51 | 1 | Trim12a | tripartite motif-containing 12A | 335 | 212 | 165 | 9 | 30 | 61 | 14 | 5 | 15 | 22 | 51 |
| 72 | -0.92 | 0.90 | -1.02 | 0.31 | 1 | Fam171a2 | family with sequence similarity 171, meml | 287 | 133 | 253 | 19 | 13 | 11 | 14 | 0 | 22 | 37 | 1 |
| 69 | 1.01 | 1.01 | 1.00 | 0.32 | 1 | Septin1 | septin 1 | 399 | 119 | 101 | 0 | 11 | 16 | 14 | 65 | 3 | 21 | 15 |
| 61 | 0.84 | 0.71 | 1.18 | 0.24 | 1 | Zcwpw1 | zinc finger, CW type with PWWP domain ' | 114 | 96 | 91 | 39 | 8 | 88 | 14 | 125 | 34 | 12 | 54 |
| 59 | -0.32 | 0.93 | -0.34 | 0.73 | 1 | Pkd1l1 | polycystic kidney disease 1 like 1 | 53 | 249 | 175 | 4 | 11 | 51 | 14 | 48 | 9 | 0 | 36 |

| | | | | | | | | | | | | | | | | | | |
|---|---|---|---|---|---|---|---|---|---|---|---|---|---|---|---|---|---|---|
| 58 | 0.44 | 0.58 | 0.76 | 0.45 | 1 | Tmem125 | transmembrane protein 125 | 40 | 58 | 139 | 49 | 77 | 41 | 14 | 75 | 32 | 35 | 72 |
| 49 | 0.22 | 0.60 | 0.36 | 0.72 | 1 | Baiap2l2 | BAI1-associated protein 2-like 2 | 57 | 35 | 33 | 34 | 60 | 84 | 14 | 24 | 76 | 31 | 90 |
| 49 | -1.62 | 1.06 | -1.53 | 0.13 | 1 | Metrn | meteorin, glial cell differentiation regulato | 247 | 81 | 155 | 7 | 2 | 14 | 14 | 0 | 9 | 0 | 6 |
| 48 | -1.47 | 0.78 | -1.89 | 0.06 | 1 | Micall2 | MICAL-like 2 | 144 | 141 | 149 | 37 | 14 | 6 | 14 | 16 | 10 | 0 | 2 |
| 48 | -0.85 | 0.88 | -0.97 | 0.33 | 1 | Cacna2d3 | calcium channel, voltage-dependent, alpl | 88 | 187 | 97 | 16 | 30 | 37 | 14 | 0 | 15 | 24 | 17 |
| 44 | -0.17 | 1.15 | -0.15 | 0.88 | 1 | Gm36684 | | 0 | 0 | 61 | 11 | 215 | 1 | 14 | 0 | 157 | 27 | 4 |
| 43 | -1.31 | 3.62 | -0.36 | 0.72 | 1 | Scimp | SLP adaptor and CSK interacting membra | 174 | 91 | 198 | 0 | 0 | 0 | 14 | 0 | 0 | 0 | 0 |
| 43 | -1.55 | 1.05 | -1.48 | 0.14 | 1 | Kcnj2 | potassium inwardly-rectifying channel, su | 108 | 173 | 102 | 10 | 2 | 37 | 14 | 0 | 7 | 0 | 24 |
| 43 | 0.36 | 0.66 | 0.54 | 0.59 | 1 | Kcnk9 | potassium channel, subfamily K, member | 7 | 0 | 10 | 9 | 17 | 137 | 14 | 10 | 19 | 15 | 238 |
| 43 | 0.38 | 0.61 | 0.63 | 0.53 | 1 | Cenpk | centromere protein K | 85 | 33 | 50 | 35 | 38 | 43 | 14 | 39 | 28 | 57 | 51 |
| 40 | 0.03 | 0.70 | 0.04 | 0.97 | 1 | Rasal1 | RAS protein activator like 1 (GAP1 like) | 70 | 13 | 80 | 32 | 29 | 59 | 14 | 30 | 30 | 15 | 62 |
| 38 | -2.44 | 1.30 | -1.87 | 0.06 | 1 | Lrrc4b | leucine rich repeat containing 4B | 156 | 71 | 166 | 0 | 5 | 2 | 14 | 0 | 3 | 0 | 0 |
| 35 | -0.23 | 0.70 | -0.33 | 0.74 | 1 | Eno4 | enolase 4 | 49 | 89 | 54 | 25 | 27 | 38 | 14 | 8 | 37 | 26 | 17 |
| 33 | -0.23 | 0.72 | -0.32 | 0.75 | 1 | Gm21399 | | 49 | 24 | 49 | 57 | 31 | 27 | 14 | 13 | 39 | 40 | 23 |
| 33 | -0.39 | 1.07 | -0.36 | 0.72 | 1 | 9330159M07Rik | | 27 | 0 | 22 | 61 | 58 | 42 | 14 | 51 | 26 | 8 | 56 |
| 32 | -1.19 | 1.64 | -0.73 | 0.47 | 1 | Cdcp3 | CUB domain containing protein 3 | 36 | 210 | 79 | 0 | 0 | 3 | 14 | 0 | 1 | 0 | 4 |
| 31 | -1.47 | 1.11 | -1.33 | 0.18 | 1 | Abcg4 | ATP binding cassette subfamily G membe | 79 | 53 | 57 | 24 | 24 | 24 | 14 | 0 | 16 | 52 | 1 |
| 31 | 1.05 | 1.29 | 0.82 | 0.41 | 1 | Cdh17 | cadherin 17 | 3 | 113 | 81 | 7 | 8 | 7 | 14 | 0 | 31 | 66 | 5 |
| 29 | 0.32 | 0.91 | 0.35 | 0.73 | 1 | Kirrel3os | | 0 | 0 | 0 | 23 | 9 | 78 | 14 | 7 | 8 | 63 | 119 |
| 27 | 0.00 | 0.79 | 0.00 | 1.00 | 1 | Gchfr | GTP cyclohydrolase I feedback regulator | 70 | 43 | 29 | 18 | 17 | 17 | 14 | 45 | 3 | 19 | 16 |
| 27 | -0.59 | 0.61 | -0.97 | 0.33 | 1 | Gm7908 | | 14 | 38 | 21 | 53 | 32 | 32 | 14 | 25 | 25 | 8 | 28 |
| 26 | -0.11 | 0.79 | -0.14 | 0.89 | 1 | Gm2999 | | 18 | 9 | 48 | 22 | 39 | 25 | 14 | 2 | 48 | 31 | 36 |
| 25 | -0.75 | 0.84 | -0.89 | 0.37 | 1 | Gm7868 | | 39 | 11 | 97 | 2 | 25 | 32 | 14 | 1 | 20 | 1 | 38 |
| 25 | 0.16 | 1.09 | 0.15 | 0.88 | 1 | Rad54b | RAD54 homolog B (S. cerevisiae) | 42 | 24 | 7 | 15 | 13 | 49 | 14 | 59 | 14 | 0 | 39 |
| 25 | -0.85 | 1.00 | -0.85 | 0.40 | 1 | Mdga2 | MAM domain containing glycosylphospha | 71 | 52 | 70 | 18 | 7 | 7 | 14 | 0 | 5 | 27 | 4 |
| 25 | 0.33 | 1.02 | 0.32 | 0.75 | 1 | Sucnr1 | succinate receptor 1 | 69 | 49 | 68 | 14 | 0 | 7 | 14 | 7 | 8 | 32 | 3 |
| 24 | 1.45 | 0.75 | 1.94 | 0.05 | 1 | Gm15429 | | 10 | 30 | 8 | 11 | 16 | 10 | 14 | 75 | 15 | 53 | 25 |
| 24 | -0.89 | 0.93 | -0.96 | 0.34 | 1 | Gm34655 | | 15 | 34 | 9 | 46 | 18 | 32 | 14 | 52 | 6 | 0 | 36 |
| 24 | -0.75 | 1.00 | -0.76 | 0.45 | 1 | Pld6 | phospholipase D family, member 6 | 43 | 46 | 29 | 28 | 31 | 15 | 14 | 0 | 21 | 18 | 17 |
| 23 | 0.06 | 1.00 | 0.06 | 0.95 | 1 | Gm42940 | | 11 | 21 | 29 | 30 | 2 | 45 | 14 | 52 | 20 | 1 | 32 |
| 23 | 0.52 | 0.66 | 0.80 | 0.42 | 1 | Gm5869 | | 27 | 30 | 32 | 15 | 23 | 12 | 14 | 16 | 49 | 17 | 16 |
| 22 | 0.24 | 0.89 | 0.28 | 0.78 | 1 | Fsbp | fibrinogen silencer binding protein | 27 | 18 | 29 | 11 | 15 | 27 | 14 | 56 | 11 | 4 | 25 |
| 21 | -2.07 | 1.49 | -1.39 | 0.16 | 1 | Gm26621 | | 0 | 0 | 3 | 4 | 2 | 96 | 14 | 0 | 11 | 0 | 96 |
| 20 | -1.92 | 1.25 | -1.53 | 0.13 | 1 | Gm45412 | | 15 | 12 | 31 | 44 | 39 | 7 | 14 | 0 | 31 | 31 | 0 |
| 20 | 1.06 | 0.83 | 1.28 | 0.20 | 1 | B3gnt2 | UDP-GlcNAc:betaGal beta-1,3-N-acetylgl | 15 | 8 | 25 | 15 | 14 | 9 | 14 | 67 | 15 | 17 | 20 |
| 20 | 0.67 | 0.98 | 0.68 | 0.49 | 1 | Eme1 | essential meiotic structure-specific endon | 19 | 20 | 0 | 11 | 15 | 15 | 14 | 51 | 9 | 36 | 25 |
| 20 | 0.54 | 1.49 | 0.36 | 0.72 | 1 | Adh7 | alcohol dehydrogenase 7 (class IV), mu c | 41 | 0 | 6 | 29 | 6 | 23 | 14 | 21 | 37 | 0 | 38 |
| 19 | -0.05 | 1.09 | -0.05 | 0.96 | 1 | Gm14443 | | 6 | 5 | 31 | 4 | 10 | 41 | 14 | 0 | 5 | 37 | 58 |
| 18 | 0.22 | 0.85 | 0.26 | 0.79 | 1 | Gm3650 | predicted gene 3650 | 13 | 11 | 7 | 16 | 17 | 19 | 14 | 70 | 8 | 10 | 17 |
| 18 | -0.90 | 1.05 | -0.85 | 0.39 | 1 | Rps19-ps3 | | 11 | 7 | 2 | 60 | 21 | 20 | 14 | 1 | 14 | 37 | 14 |
| 18 | -0.09 | 1.23 | -0.07 | 0.94 | 1 | Ptger1 | prostaglandin E receptor 1 (subtype EP1) | 46 | 63 | 15 | 0 | 2 | 30 | 14 | 10 | 5 | 1 | 11 |
| 18 | -0.58 | 0.99 | -0.59 | 0.56 | 1 | Gm35215 | | 6 | 15 | 34 | 23 | 18 | 14 | 14 | 30 | 30 | 0 | 10 |
| 17 | -0.27 | 1.08 | -0.25 | 0.80 | 1 | 1700003M07Rik | | 12 | 12 | 29 | 13 | 14 | 22 | 14 | 0 | 13 | 43 | 17 |
| 17 | 0.55 | 0.79 | 0.69 | 0.49 | 1 | Pcdha11 | protocadherin alpha 11 | 13 | 21 | 8 | 28 | 14 | 6 | 14 | 21 | 12 | 23 | 28 |
| 17 | -0.31 | 1.44 | -0.21 | 0.83 | 1 | 6720468P15Rik | | 14 | 2 | 30 | 36 | 2 | 8 | 14 | 0 | 5 | 62 | 12 |
| 17 | -0.36 | 1.12 | -0.32 | 0.75 | 1 | Zmat4 | zinc finger, matrin type 4 | 9 | 11 | 47 | 9 | 15 | 18 | 14 | 31 | 11 | 0 | 16 |
| 16 | -0.78 | 1.09 | -0.71 | 0.48 | 1 | Gabrq | gamma-aminobutyric acid (GABA) A rece | 46 | 10 | 4 | 4 | 17 | 21 | 14 | 0 | 6 | 43 | 11 |
| 15 | -0.45 | 0.92 | -0.49 | 0.63 | 1 | Tmem41b-ps | | 11 | 14 | 14 | 27 | 26 | 5 | 14 | 22 | 23 | 1 | 8 |
| 15 | -1.61 | 2.28 | -0.71 | 0.48 | 1 | Ifi206 | interferon activated gene 206 | 75 | 47 | 29 | 0 | 0 | 0 | 14 | 0 | 0 | 0 | 0 |
| 15 | -0.10 | 0.97 | -0.10 | 0.92 | 1 | Gm14567 | | 15 | 14 | 9 | 11 | 25 | 12 | 14 | 20 | 26 | 1 | 15 |
| 15 | -0.16 | 0.94 | -0.17 | 0.87 | 1 | Gm5422 | | 9 | 21 | 20 | 10 | 25 | 11 | 14 | 7 | 5 | 27 | 15 |
| 14 | -1.08 | 1.10 | -0.98 | 0.33 | 1 | Rab42 | RAB42, member RAS oncogene family | 38 | 28 | 5 | 21 | 7 | 15 | 14 | 12 | 6 | 0 | 8 |
| 14 | -0.56 | 0.87 | -0.64 | 0.52 | 1 | Gm16425 | | 9 | 10 | 22 | 22 | 24 | 5 | 14 | 20 | 17 | 3 | 6 |
| 13 | 0.94 | 1.50 | 0.62 | 0.53 | 1 | Gm15672 | | 0 | 5 | 2 | 5 | 2 | 37 | 14 | 0 | 29 | 33 | 19 |
| 13 | -0.97 | 1.48 | -0.65 | 0.51 | 1 | C530044C16Rik | | 15 | 28 | 8 | 3 | 20 | 9 | 14 | 0 | 25 | 0 | 18 |
| 13 | -1.22 | 1.52 | -0.81 | 0.42 | 1 | Gm50047 | | 1 | 0 | 0 | 41 | 22 | 4 | 14 | 38 | 1 | 0 | 18 |
| 13 | -0.09 | 1.98 | -0.05 | 0.96 | 1 | Gm5150 | predicted gene 5150 | 58 | 0 | 9 | 10 | 0 | 1 | 14 | 0 | 0 | 36 | 10 |
| 13 | -3.53 | 1.80 | -1.96 | 0.05 | 1 | H4f16 | H4 histone 16 | 13 | 23 | 25 | 9 | 0 | 20 | 14 | 0 | 0 | 0 | 34 |
| 12 | -0.33 | 0.82 | -0.40 | 0.69 | 1 | Gm10925 | | 7 | 10 | 9 | 14 | 15 | 17 | 14 | 9 | 18 | 15 | 7 |
| 12 | -2.93 | 1.92 | -1.53 | 0.13 | 1 | Tcam1 | testicular cell adhesion molecule 1 | 0 | 40 | 0 | 25 | 26 | 15 | 14 | 0 | 8 | 0 | 8 |

| | | | | | | | | | | | | | | | | | | |
|---|---|---|---|---|---|---|---|---|---|---|---|---|---|---|---|---|---|---|
| 12 | -2.96 | 1.93 | -1.53 | 0.13 | 1 | Gm28535 | | 14 | 1 | 18 | 25 | 0 | 4 | 14 | 0 | 1 | 55 | 0 |
| 12 | 0.39 | 1.14 | 0.34 | 0.73 | 1 | Gm4598 | | 3 | 0 | 0 | 29 | 6 | 3 | 14 | 50 | 4 | 19 | 4 |
| 12 | -3.01 | 1.53 | -1.96 | 0.05 | 1 | 4930512B01Rik | | 3 | 0 | 15 | 15 | 14 | 31 | 14 | 0 | 14 | 0 | 23 |
| 12 | -1.68 | 1.50 | -1.13 | 0.26 | 1 | Gm12586 | | 4 | 30 | 0 | 13 | 8 | 30 | 14 | 0 | 5 | 1 | 21 |
| 11 | -0.18 | 1.75 | -0.10 | 0.92 | 1 | 4930517E14Rik | | 0 | 17 | 7 | 0 | 11 | 10 | 14 | 20 | 18 | 0 | 28 |
| 11 | -2.57 | 1.71 | -1.50 | 0.13 | 1 | Hydin | HYDIN, axonemal central pair apparatus | 0 | 24 | 17 | 11 | 18 | 8 | 14 | 0 | 8 | 0 | 24 |
| 11 | -0.62 | 2.03 | -0.30 | 0.76 | 1 | E230020A03Rik | | 19 | 25 | 0 | 0 | 4 | 20 | 14 | 0 | 7 | 0 | 35 |
| 11 | 1.45 | 1.37 | 1.06 | 0.29 | 1 | Gm43437 | | 27 | 1 | 0 | 0 | 7 | 10 | 14 | 8 | 15 | 35 | 6 |
| 11 | 0.58 | 1.37 | 0.43 | 0.67 | 1 | Ap3s1-ps2 | | 24 | 27 | 1 | 6 | 2 | 11 | 14 | 13 | 12 | 0 | 12 |
| 11 | 1.06 | 1.13 | 0.94 | 0.35 | 1 | Oaz2-ps | | 16 | 37 | 5 | 4 | 1 | 4 | 14 | 2 | 6 | 14 | 17 |
| 11 | 1.04 | 1.22 | 0.85 | 0.39 | 1 | Aldob | aldolase B, fructose-bisphosphate | 12 | 0 | 24 | 0 | 5 | 10 | 14 | 1 | 6 | 32 | 13 |
| 10 | -0.23 | 2.41 | -0.10 | 0.92 | 1 | G630016G05Rik | | 4 | 27 | 0 | 0 | 0 | 35 | 14 | 0 | 7 | 0 | 24 |
| 9 | 0.39 | 1.42 | 0.28 | 0.78 | 1 | Gm36640 | | 3 | 0 | 0 | 24 | 0 | 9 | 14 | 9 | 5 | 31 | 7 |
| 9 | -0.34 | 1.24 | -0.27 | 0.78 | 1 | Mdh1b | malate dehydrogenase 1B, NAD (soluble | 7 | 6 | 9 | 3 | 9 | 18 | 14 | 13 | 10 | 0 | 13 |
| 9 | -2.47 | 1.50 | -1.65 | 0.10 | 1 | Gm29417 | | 15 | 6 | 3 | 21 | 13 | 8 | 14 | 0 | 5 | 0 | 16 |
| 9 | 0.22 | 1.14 | 0.19 | 0.85 | 1 | Gm5904 | | 4 | 1 | 9 | 4 | 5 | 10 | 14 | 1 | 4 | 36 | 13 |
| 9 | -1.63 | 2.00 | -0.81 | 0.42 | 1 | Gm43197 | | 42 | 11 | 11 | 0 | 0 | 12 | 14 | 0 | 0 | 0 | 8 |
| 9 | 0.09 | 1.20 | 0.08 | 0.94 | 1 | 1810049J17Rik | | 3 | 1 | 6 | 12 | 1 | 13 | 14 | 6 | 10 | 26 | 7 |
| 9 | 0.61 | 1.67 | 0.36 | 0.72 | 1 | Eif3s6-ps1 | | 1 | 7 | 7 | 0 | 15 | 5 | 14 | 16 | 0 | 23 | 9 |
| 9 | -1.76 | 1.36 | -1.29 | 0.20 | 1 | 4930563E18Rik | | 15 | 5 | 14 | 2 | 8 | 22 | 14 | 0 | 8 | 0 | 11 |
| 9 | -0.61 | 1.37 | -0.45 | 0.66 | 1 | Gm7488 | | 19 | 10 | 0 | 4 | 13 | 8 | 14 | 2 | 8 | 3 | 15 |
| 7 | 0.10 | 1.28 | 0.07 | 0.94 | 1 | Gm11889 | | 10 | 0 | 5 | 2 | 8 | 7 | 14 | 2 | 6 | 3 | 25 |
| 7 | -1.42 | 1.60 | -0.89 | 0.37 | 1 | 4930439D14Rik | | 0 | 2 | 10 | 11 | 4 | 8 | 14 | 20 | 8 | 0 | 3 |
| 7 | -0.88 | 1.24 | -0.71 | 0.48 | 1 | Gm33666 | | 2 | 1 | 0 | 13 | 15 | 3 | 14 | 3 | 2 | 18 | 7 |
| 7 | 0.06 | 1.31 | 0.05 | 0.96 | 1 | Eif1-ps2 | | 5 | 0 | 6 | 1 | 13 | 8 | 14 | 3 | 12 | 1 | 12 |
| 7 | 0.71 | 1.65 | 0.43 | 0.67 | 1 | Mir8091 | | 9 | 0 | 0 | 0 | 5 | 8 | 14 | 0 | 10 | 19 | 11 |
| 7 | -2.06 | 2.07 | -1.00 | 0.32 | 1 | Gm45437 | | 10 | 8 | 26 | 0 | 2 | 3 | 14 | 0 | 12 | 0 | 0 |
| 7 | -0.40 | 1.19 | -0.34 | 0.74 | 1 | Gm9493 | | 14 | 6 | 7 | 6 | 2 | 3 | 14 | 1 | 3 | 15 | 2 |
| 7 | -1.37 | 1.78 | -0.77 | 0.44 | 1 | Gm13066 | | 7 | 9 | 18 | 0 | 6 | 7 | 14 | 0 | 9 | 0 | 4 |
| 7 | -0.35 | 2.52 | -0.14 | 0.89 | 1 | Zfp972 | zinc finger protein 972 | 3 | 0 | 0 | 30 | 0 | 3 | 14 | 0 | 16 | 0 | 5 |
| 6 | -1.41 | 3.05 | -0.46 | 0.64 | 1 | Treml2 | triggering receptor expressed on myeloid | 19 | 20 | 11 | 0 | 0 | 0 | 14 | 0 | 0 | 0 | 0 |
| 6 | -3.21 | 2.93 | -1.10 | 0.27 | 1 | Gm42446 | | 0 | 0 | 0 | 4 | 0 | 15 | 14 | 0 | 0 | 0 | 30 |
| 5 | 0.15 | 2.13 | 0.07 | 0.95 | 1 | Gm11702 | | 1 | 9 | 0 | 7 | 0 | 2 | 14 | 0 | 0 | 15 | 8 |
| 5 | -0.58 | 3.05 | -0.19 | 0.85 | 1 | Gm37718 | | 0 | 3 | 0 | 0 | 0 | 19 | 14 | 0 | 6 | 0 | 12 |
| 5 | -2.34 | 2.75 | -0.85 | 0.39 | 1 | G630030J09Rik | | 0 | 0 | 9 | 11 | 0 | 6 | 14 | 6 | 0 | 0 | 5 |
| 4 | -0.12 | 2.95 | -0.04 | 0.97 | 1 | Gm44008 | | 0 | 11 | 0 | 0 | 1 | 3 | 14 | 0 | 16 | 0 | 3 |
| 4 | -1.59 | 1.96 | -0.81 | 0.42 | 1 | Gm16415 | | 22 | 5 | 1 | 1 | 0 | 3 | 14 | 0 | 1 | 0 | 1 |
| 4 | -2.43 | 3.08 | -0.79 | 0.43 | 1 | Gm11413 | | 4 | 9 | 11 | 0 | 0 | 1 | 14 | 0 | 0 | 0 | 0 |
| 3 | -3.66 | 3.18 | -1.15 | 0.25 | 1 | Gm29264 | | 4 | 11 | 0 | 0 | 0 | 3 | 14 | 0 | 0 | 0 | 4 |
| 2 | -2.25 | 3.55 | -0.64 | 0.53 | 1 | Ccdc36 | coiled-coil domain containing 36 | 0 | 0 | 0 | 0 | 0 | 2 | 14 | 0 | 0 | 0 | 10 |
| 2 | 0.23 | 2.86 | 0.08 | 0.94 | 1 | Gm14008 | | 7 | 0 | 0 | 0 | 0 | 1 | 14 | 1 | 0 | 1 | 1 |
| 2 | 0.52 | 3.60 | 0.14 | 0.88 | 1 | Serpina3k | serine (or cysteine) peptidase inhibitor, cl | 2 | 0 | 0 | 0 | 0 | 0 | 14 | 0 | 0 | 8 | 1 |
| 1861 | -2.06 | 1.36 | -1.52 | 0.13 | 1 | Lamb2 | laminin, beta 2 | 9727 | 4680 | 6019 | 2 | 18 | 0 | 13 | 0 | 8 | 0 | 0 |
| 1405 | -2.37 | 2.71 | -0.87 | 0.38 | 1 | Tie1 | tyrosine kinase with immunoglobulin-like | 4803 | 5014 | 5624 | 0 | 0 | 1 | 13 | 0 | 0 | 0 | 0 |
| 1010 | -1.33 | 1.27 | -1.05 | 0.30 | 1 | Pcolce | procollagen C-endopeptidase enhancer p | 5660 | 2417 | 2981 | 0 | 4 | 14 | 13 | 0 | 2 | 0 | 21 |
| 640 | -3.49 | 2.27 | -1.53 | 0.12 | 1 | Wdfy4 | WD repeat and FYVE domain containing | 3757 | 1882 | 1377 | 8 | 0 | 0 | 13 | 0 | 0 | 0 | 0 |
| 568 | 0.60 | 1.75 | 0.34 | 0.73 | 1 | S1pr1 | sphingosine-1-phosphate receptor 1 | 2177 | 2032 | 2004 | 0 | 0 | 2 | 13 | 0 | 5 | 15 | 0 |
| 539 | -0.71 | 0.84 | -0.85 | 0.40 | 1 | Cav2 | caveolin 2 | 2043 | 1758 | 2044 | 19 | 1 | 20 | 13 | 3 | 14 | 8 | 6 |
| 503 | -0.73 | 1.05 | -0.69 | 0.49 | 1 | Cgnl1 | cingulin-like 1 | 2405 | 1218 | 1636 | 61 | 54 | 26 | 13 | 48 | 64 | 0 | 9 |
| 423 | -2.73 | 1.82 | -1.50 | 0.13 | 1 | Ccr2 | chemokine (C-C motif) receptor 2 | 2563 | 1038 | 1030 | 7 | 0 | 0 | 13 | 0 | 1 | 0 | 0 |
| 395 | 0.00 | 1.29 | 0.00 | 1.00 | 1 | Tnfaip2 | tumor necrosis factor, alpha-induced prot | 1875 | 1133 | 1289 | 13 | 0 | 2 | 13 | 0 | 6 | 8 | 8 |
| 370 | -0.32 | 0.92 | -0.35 | 0.73 | 1 | Tgfb2 | transforming growth factor, beta 2 | 1393 | 1417 | 1171 | 6 | 16 | 10 | 13 | 20 | 0 | 22 | 5 |
| 370 | 0.81 | 2.27 | 0.36 | 0.72 | 1 | Cpxm2 | carboxypeptidase X 2 (M14 family) | 2246 | 584 | 1217 | 0 | 0 | 0 | 13 | 0 | 2 | 0 | 2 |
| 195 | -0.06 | 1.00 | -0.06 | 0.95 | 1 | Pdpn | podoplanin | 1172 | 508 | 433 | 0 | 0 | 2 | 13 | 0 | 0 | 8 | 4 |
| 167 | -1.20 | 3.62 | -0.33 | 0.74 | 1 | Clec10a | C-type lectin domain family 10, member A | 1167 | 258 | 396 | 0 | 0 | 0 | 13 | 0 | 0 | 0 | 0 |
| 129 | 0.09 | 3.29 | 0.03 | 0.98 | 1 | Ms4a7 | membrane-spanning 4-domains, subfami | 480 | 307 | 611 | 0 | 0 | 0 | 13 | 0 | 8 | 0 | 0 |
| 123 | -1.09 | 0.67 | -1.62 | 0.11 | 1 | Lrtm2 | leucine-rich repeats and transmembrane | 444 | 410 | 373 | 21 | 21 | 20 | 13 | 37 | 4 | 8 | 2 |
| 69 | 0.36 | 0.63 | 0.58 | 0.56 | 1 | Fignl1 | fidgetin-like 1 | 240 | 54 | 105 | 40 | 43 | 60 | 13 | 44 | 57 | 40 | 58 |
| 64 | -2.56 | 1.42 | -1.81 | 0.07 | 1 | Smad9 | SMAD family member 9 | 228 | 228 | 183 | 0 | 15 | 14 | 13 | 0 | 19 | 0 | 0 |
| 61 | 0.00 | 0.80 | 0.00 | 1.00 | 1 | Adam11 | a disintegrin and metallopeptidase domai | 199 | 176 | 167 | 12 | 14 | 16 | 13 | 0 | 20 | 37 | 12 |

| | | | | | | | | | | | | | | | | | | |
|---|---|---|---|---|---|---|---|---|---|---|---|---|---|---|---|---|---|---|
| 59 | -1.55 | 0.93 | -1.67 | 0.10 | 1 | Mycl | v-myc avian myelocytomatosis viral oncog | 298 | 159 | 123 | 23 | 9 | 3 | 13 | 0 | 7 | 17 | 0 |
| 58 | 0.73 | 0.77 | 0.95 | 0.34 | 1 | Mcm10 | minichromosome maintenance 10 replica | 251 | 107 | 25 | 18 | 19 | 48 | 13 | 14 | 63 | 31 | 49 |
| 58 | 1.95 | 1.03 | 1.89 | 0.06 | 1 | Stx11 | syntaxin 11 | 305 | 157 | 100 | 0 | 0 | 9 | 13 | 15 | 12 | 22 | 3 |
| 55 | 0.65 | 0.57 | 1.15 | 0.25 | 1 | D130040H23Ri | RIKEN cDNA D130040H23 gene | 110 | 102 | 75 | 31 | 20 | 64 | 13 | 28 | 31 | 70 | 58 |
| 54 | -0.78 | 0.78 | -1.00 | 0.32 | 1 | Slc37a2 | solute carrier family 37 (glycerol-3-phospl | 250 | 105 | 159 | 12 | 7 | 11 | 13 | 0 | 9 | 21 | 4 |
| 54 | -0.21 | 0.67 | -0.31 | 0.76 | 1 | Gm45623 | predicted gene 45623 | 95 | 117 | 131 | 17 | 81 | 25 | 13 | 50 | 23 | 24 | 11 |
| 50 | 0.14 | 0.55 | 0.26 | 0.79 | 1 | Gm43860 | | 60 | 45 | 48 | 48 | 35 | 90 | 13 | 49 | 28 | 23 | 109 |
| 50 | 0.09 | 1.01 | 0.09 | 0.93 | 1 | Wscd2 | WSC domain containing 2 | 121 | 178 | 193 | 1 | 3 | 8 | 13 | 7 | 17 | 0 | 5 |
| 49 | -0.21 | 0.83 | -0.25 | 0.80 | 1 | Rnps1-ps | | 5 | 27 | 179 | 14 | 116 | 18 | 13 | 1 | 125 | 26 | 11 |
| 39 | 0.61 | 0.67 | 0.90 | 0.37 | 1 | Gm10549 | | 15 | 17 | 63 | 49 | 29 | 39 | 13 | 93 | 31 | 18 | 63 |
| 39 | -0.69 | 0.91 | -0.76 | 0.45 | 1 | Gm36948 | | 1 | 0 | 0 | 34 | 17 | 160 | 13 | 15 | 27 | 0 | 159 |
| 38 | -1.61 | 1.17 | -1.38 | 0.17 | 1 | Gldc | glycine decarboxylase | 39 | 86 | 62 | 49 | 12 | 68 | 13 | 0 | 35 | 0 | 52 |
| 36 | -1.40 | 0.90 | -1.56 | 0.12 | 1 | Smim5 | small integral membrane protein 5 | 81 | 47 | 137 | 23 | 26 | 19 | 13 | 0 | 36 | 4 | 10 |
| 34 | -0.48 | 1.54 | -0.31 | 0.75 | 1 | Gm49204 | | 5 | 45 | 0 | 0 | 13 | 157 | 13 | 0 | 9 | 9 | 122 |
| 33 | 0.47 | 1.13 | 0.42 | 0.68 | 1 | Card9 | caspase recruitment domain family, meml | 107 | 41 | 25 | 11 | 13 | 23 | 13 | 103 | 12 | 0 | 16 |
| 32 | -2.00 | 1.17 | -1.71 | 0.09 | 1 | Gm37893 | | 5 | 0 | 10 | 2 | 2 | 182 | 13 | 0 | 2 | 0 | 141 |
| 32 | -0.25 | 0.78 | -0.32 | 0.75 | 1 | Car11 | carbonic anhydrase 11 | 33 | 84 | 70 | 13 | 29 | 36 | 13 | 2 | 22 | 23 | 30 |
| 32 | 0.06 | 0.81 | 0.08 | 0.94 | 1 | Gm8276 | | 58 | 51 | 35 | 36 | 36 | 14 | 13 | 5 | 41 | 26 | 35 |
| 31 | 0.30 | 0.64 | 0.47 | 0.64 | 1 | Cfap43 | cilia and flagella associated protein 43 | 24 | 20 | 11 | 25 | 26 | 66 | 13 | 42 | 14 | 40 | 64 |
| 31 | 0.72 | 0.64 | 1.13 | 0.26 | 1 | Gm43857 | | 27 | 40 | 14 | 21 | 22 | 45 | 13 | 37 | 31 | 48 | 40 |
| 30 | 0.81 | 0.64 | 1.26 | 0.21 | 1 | Pbx4 | pre B cell leukemia homeobox 4 | 43 | 48 | 49 | 7 | 13 | 46 | 13 | 16 | 33 | 27 | 39 |
| 29 | -0.43 | 0.84 | -0.51 | 0.61 | 1 | Gm47439 | | 11 | 20 | 63 | 56 | 49 | 16 | 13 | 6 | 31 | 33 | 18 |
| 28 | 0.81 | 0.98 | 0.83 | 0.41 | 1 | Gm7107 | | 17 | 37 | 0 | 39 | 21 | 20 | 13 | 22 | 27 | 67 | 42 |
| 27 | 1.04 | 0.69 | 1.51 | 0.13 | 1 | Gm42929 | | 27 | 24 | 11 | 7 | 7 | 57 | 13 | 9 | 19 | 39 | 79 |
| 26 | 0.50 | 0.82 | 0.61 | 0.54 | 1 | Gm49864 | | 5 | 57 | 30 | 16 | 15 | 46 | 13 | 23 | 20 | 35 | 29 |
| 25 | 0.61 | 1.11 | 0.55 | 0.58 | 1 | 9230117E06Rik | | 0 | 0 | 18 | 47 | 57 | 13 | 13 | 53 | 21 | 32 | 25 |
| 25 | 0.32 | 0.88 | 0.37 | 0.71 | 1 | Gm5913 | | 48 | 20 | 22 | 29 | 23 | 10 | 13 | 67 | 25 | 4 | 12 |
| 22 | 0.27 | 1.03 | 0.26 | 0.79 | 1 | Hrob | homologous recombination factor with OE | 36 | 48 | 26 | 14 | 16 | 14 | 13 | 0 | 36 | 24 | 18 |
| 22 | -0.54 | 2.11 | -0.26 | 0.80 | 1 | Mzb1 | marginal zone B and B1 cell-specific prot | 204 | 10 | 0 | 0 | 0 | 8 | 13 | 0 | 7 | 0 | 1 |
| 22 | -0.15 | 0.83 | -0.18 | 0.86 | 1 | Gm2223 | | 19 | 31 | 11 | 25 | 30 | 21 | 13 | 8 | 41 | 8 | 32 |
| 21 | -0.07 | 0.88 | -0.08 | 0.93 | 1 | Pdcd5-ps | | 48 | 3 | 38 | 15 | 18 | 18 | 13 | 8 | 11 | 44 | 16 |
| 21 | -0.41 | 0.83 | -0.49 | 0.62 | 1 | Gm18095 | | 34 | 26 | 18 | 6 | 49 | 22 | 13 | 12 | 19 | 21 | 10 |
| 21 | 0.16 | 0.89 | 0.19 | 0.85 | 1 | Gm9774 | | 39 | 4 | 11 | 28 | 20 | 19 | 13 | 39 | 24 | 21 | 11 |
| 21 | -0.74 | 1.04 | -0.71 | 0.48 | 1 | Haus5 | HAUS augmin-like complex, subunit 5 | 10 | 17 | 30 | 4 | 9 | 65 | 13 | 13 | 4 | 0 | 60 |
| 20 | -1.01 | 1.31 | -0.77 | 0.44 | 1 | 4933406I18Rik | | 14 | 0 | 24 | 21 | 6 | 57 | 13 | 0 | 17 | 27 | 46 |
| 20 | 0.93 | 0.84 | 1.11 | 0.27 | 1 | Gucy2e | guanylate cyclase 2e | 34 | 41 | 2 | 15 | 8 | 15 | 13 | 31 | 19 | 28 | 16 |
| 19 | 0.29 | 1.16 | 0.25 | 0.80 | 1 | Slc7a3 | solute carrier family 7 (cationic amino acid | 16 | 46 | 0 | 5 | 22 | 24 | 13 | 30 | 22 | 13 | 20 |
| 19 | -0.26 | 0.91 | -0.28 | 0.78 | 1 | Gm4189 | | 27 | 2 | 30 | 5 | 12 | 36 | 13 | 0 | 21 | 14 | 51 |
| 19 | -0.14 | 0.92 | -0.15 | 0.88 | 1 | Gm15734 | | 3 | 16 | 6 | 6 | 43 | 30 | 13 | 28 | 9 | 14 | 36 |
| 19 | -1.67 | 1.28 | -1.31 | 0.19 | 1 | 9330179D12Rik | | 32 | 22 | 61 | 21 | 9 | 11 | 13 | 0 | 21 | 0 | 14 |
| 18 | 0.58 | 1.08 | 0.53 | 0.59 | 1 | Rncr4 | | 0 | 0 | 0 | 32 | 14 | 10 | 13 | 8 | 26 | 86 | 12 |
| 18 | -0.73 | 1.08 | -0.68 | 0.50 | 1 | Gm7335 | | 7 | 1 | 28 | 32 | 17 | 17 | 13 | 1 | 50 | 14 | 19 |
| 17 | -0.43 | 0.87 | -0.49 | 0.62 | 1 | Gm20544 | | 44 | 18 | 17 | 37 | 9 | 8 | 13 | 14 | 6 | 8 | 19 |
| 17 | -0.67 | 0.84 | -0.80 | 0.43 | 1 | C2cd4d | C2 calcium-dependent domain containing | 6 | 45 | 16 | 34 | 11 | 7 | 13 | 37 | 14 | 3 | 3 |
| 17 | 1.11 | 0.99 | 1.12 | 0.26 | 1 | Gm16230 | | 15 | 18 | 44 | 4 | 13 | 4 | 13 | 25 | 5 | 10 | 36 |
| 17 | -0.23 | 0.93 | -0.24 | 0.81 | 1 | Gm12166 | | 29 | 17 | 10 | 24 | 12 | 17 | 13 | 3 | 28 | 9 | 20 |
| 16 | -1.41 | 1.04 | -1.36 | 0.18 | 1 | Gm11915 | | 6 | 45 | 17 | 38 | 15 | 19 | 13 | 1 | 7 | 14 | 5 |
| 16 | -1.31 | 1.04 | -1.26 | 0.21 | 1 | Ttc39d | tetratricopeptide repeat domain 39D | 9 | 18 | 36 | 13 | 31 | 14 | 13 | 0 | 9 | 6 | 26 |
| 15 | -1.59 | 1.84 | -0.86 | 0.39 | 1 | 2900041M22Rik | | 48 | 36 | 62 | 5 | 0 | 0 | 13 | 5 | 0 | 0 | 0 |
| 15 | -0.57 | 1.25 | -0.45 | 0.65 | 1 | Efhb | EF hand domain family, member B | 45 | 25 | 14 | 4 | 5 | 9 | 13 | 0 | 0 | 22 | 31 |
| 15 | 1.75 | 0.94 | 1.85 | 0.06 | 1 | Gpr25 | G protein-coupled receptor 25 | 0 | 0 | 0 | 8 | 13 | 4 | 13 | 36 | 25 | 53 | 15 |
| 15 | -2.11 | 1.17 | -1.80 | 0.07 | 1 | Gm18706 | | 28 | 17 | 23 | 5 | 26 | 15 | 13 | 2 | 2 | 0 | 29 |
| 15 | 0.92 | 0.91 | 1.01 | 0.31 | 1 | Gm6652 | | 41 | 15 | 18 | 2 | 10 | 7 | 13 | 20 | 15 | 13 | 7 |
| 15 | 0.25 | 1.40 | 0.18 | 0.86 | 1 | Gm36210 | predicted gene, 36210 | 29 | 24 | 58 | 0 | 6 | 2 | 13 | 0 | 1 | 17 | 10 |
| 14 | -1.29 | 1.25 | -1.04 | 0.30 | 1 | Gm2447 | | 9 | 0 | 19 | 12 | 11 | 30 | 13 | 0 | 18 | 6 | 39 |
| 14 | 0.44 | 1.18 | 0.38 | 0.71 | 1 | Gm44646 | | 14 | 22 | 19 | 5 | 5 | 12 | 13 | 31 | 17 | 0 | 11 |
| 14 | -1.73 | 1.40 | -1.23 | 0.22 | 1 | Cacng5 | calcium channel, voltage-dependent, gan | 8 | 48 | 30 | 5 | 2 | 17 | 13 | 0 | 4 | 0 | 22 |
| 13 | -2.00 | 1.29 | -1.55 | 0.12 | 1 | Pkdrej | polycystin (PKD) family receptor for egg je | 27 | 37 | 25 | 4 | 14 | 10 | 13 | 0 | 12 | 0 | 4 |
| 13 | -1.20 | 1.29 | -0.93 | 0.35 | 1 | Gm44710 | | 30 | 18 | 35 | 3 | 13 | 8 | 13 | 0 | 15 | 0 | 11 |
| 13 | -3.13 | 1.63 | -1.92 | 0.05 | 1 | A3galt2 | alpha 1,3-galactosyltransferase 2 (isoglob | 0 | 11 | 10 | 19 | 2 | 48 | 13 | 0 | 3 | 0 | 38 |

| | | | | | | | | | | | | | | | | | | |
|---|---|---|---|---|---|---|---|---|---|---|---|---|---|---|---|---|---|---|
| 13 | -0.40 | 1.37 | -0.29 | 0.77 | 1 | Gm29246 | | 13 | 4 | 0 | 20 | 17 | 14 | 13 | 0 | 18 | 21 | 22 |
| 13 | -0.23 | 1.55 | -0.15 | 0.88 | 1 | Gm37985 | | 9 | 35 | 2 | 2 | 0 | 41 | 13 | 0 | 5 | 1 | 32 |
| 13 | 0.53 | 0.97 | 0.55 | 0.58 | 1 | Fa2h | fatty acid 2-hydroxylase | 9 | 25 | 7 | 3 | 9 | 14 | 13 | 20 | 6 | 19 | 15 |
| 12 | 1.43 | 1.07 | 1.33 | 0.18 | 1 | 4931415C17Rik | | 6 | 13 | 5 | 12 | 0 | 8 | 13 | 19 | 13 | 17 | 23 |
| 12 | -0.78 | 1.23 | -0.63 | 0.53 | 1 | Glud-ps | | 20 | 23 | 30 | 2 | 9 | 8 | 13 | 1 | 14 | 0 | 9 |
| 11 | 0.08 | 1.04 | 0.07 | 0.94 | 1 | Tpt1-ps6 | | 21 | 29 | 7 | 4 | 9 | 6 | 13 | 7 | 12 | 3 | 13 |
| 11 | -1.12 | 1.44 | -0.78 | 0.44 | 1 | Gm9727 | | 17 | 43 | 5 | 1 | 9 | 11 | 13 | 1 | 11 | 0 | 7 |
| 11 | 1.69 | 1.74 | 0.97 | 0.33 | 1 | Slc6a11 | solute carrier family 6 (neurotransmitter tr | 4 | 31 | 0 | 19 | 0 | 5 | 13 | 0 | 10 | 26 | 7 |
| 11 | -0.17 | 1.34 | -0.13 | 0.90 | 1 | Rps12-ps24 | | 37 | 10 | 20 | 1 | 9 | 2 | 13 | 0 | 15 | 1 | 7 |
| 10 | -0.71 | 1.08 | -0.66 | 0.51 | 1 | Wdr63 | WD repeat domain 63 | 3 | 6 | 7 | 11 | 24 | 8 | 13 | 1 | 6 | 13 | 21 |
| 10 | -1.35 | 1.28 | -1.06 | 0.29 | 1 | Gm14057 | | 10 | 0 | 16 | 23 | 11 | 10 | 13 | 3 | 8 | 3 | 16 |
| 10 | -0.99 | 0.99 | -1.00 | 0.32 | 1 | Rpl36-ps12 | | 14 | 7 | 22 | 6 | 22 | 4 | 13 | 2 | 10 | 3 | 7 |
| 10 | 0.04 | 1.27 | 0.03 | 0.98 | 1 | Rpl29-ps2 | | 9 | 43 | 4 | 1 | 6 | 8 | 13 | 2 | 10 | 8 | 6 |
| 9 | 1.39 | 1.46 | 0.95 | 0.34 | 1 | Gm49207 | | 19 | 0 | 0 | 4 | 4 | 11 | 13 | 20 | 16 | 8 | 7 |
| 9 | -1.79 | 1.06 | -1.69 | 0.09 | 1 | Gm7899 | | 15 | 8 | 10 | 15 | 7 | 15 | 13 | 2 | 3 | 1 | 11 |
| 9 | -1.42 | 1.39 | -1.02 | 0.31 | 1 | Gm18911 | | 0 | 8 | 22 | 2 | 19 | 12 | 13 | 1 | 12 | 1 | 9 |
| 9 | 0.16 | 1.37 | 0.12 | 0.90 | 1 | Tmem150cos | | 6 | 4 | 2 | 14 | 0 | 17 | 13 | 1 | 16 | 12 | 13 |
| 9 | -1.33 | 1.65 | -0.81 | 0.42 | 1 | Gm9694 | | 3 | 9 | 8 | 0 | 8 | 19 | 13 | 29 | 3 | 0 | 4 |
| 8 | -3.20 | 1.81 | -1.76 | 0.08 | 1 | Gm48882 | | 0 | 13 | 15 | 8 | 14 | 6 | 13 | 0 | 3 | 0 | 17 |
| 8 | -0.36 | 1.64 | -0.22 | 0.83 | 1 | E130018N17Rik | | 0 | 0 | 6 | 9 | 11 | 3 | 13 | 22 | 17 | 0 | 5 |
| 8 | -1.53 | 1.46 | -1.05 | 0.30 | 1 | | | 1 | 0 | 27 | 3 | 16 | 9 | 13 | 0 | 4 | 8 | 6 |
| 8 | -0.34 | 1.51 | -0.23 | 0.82 | 1 | A230005M16Rik | | 28 | 1 | 3 | 9 | 8 | 3 | 13 | 14 | 2 | 0 | 6 |
| 8 | -2.26 | 1.90 | -1.19 | 0.23 | 1 | Gm7099 | | 1 | 38 | 6 | 5 | 0 | 14 | 13 | 0 | 2 | 0 | 6 |
| 7 | -3.25 | 1.72 | -1.89 | 0.06 | 1 | Gm50464 | | 2 | 0 | 16 | 7 | 4 | 23 | 13 | 0 | 5 | 0 | 13 |
| 7 | -0.52 | 1.89 | -0.28 | 0.78 | 1 | Gm37935 | | 12 | 7 | 4 | 0 | 5 | 8 | 13 | 22 | 0 | 0 | 10 |
| 7 | -2.00 | 1.84 | -1.09 | 0.28 | 1 | Gm37563 | | 13 | 0 | 3 | 16 | 10 | 4 | 13 | 0 | 16 | 0 | 4 |
| 7 | -2.37 | 1.49 | -1.60 | 0.11 | 1 | Gm7542 | | 11 | 23 | 5 | 11 | 2 | 6 | 13 | 5 | 3 | 0 | 0 |
| 7 | -1.25 | 1.17 | -1.07 | 0.29 | 1 | Rps6-ps1 | | 18 | 2 | 2 | 7 | 6 | 8 | 13 | 1 | 2 | 17 | 3 |
| 7 | -0.21 | 1.71 | -0.12 | 0.90 | 1 | Gm26647 | | 0 | 0 | 0 | 7 | 12 | 3 | 13 | 20 | 18 | 0 | 3 |
| 7 | -1.23 | 1.24 | -1.00 | 0.32 | 1 | Gm2225 | | 16 | 10 | 17 | 3 | 3 | 4 | 13 | 2 | 1 | 5 | 1 |
| 6 | -4.50 | 2.67 | -1.69 | 0.09 | 1 | Odaph | odontogenesis associated phosphoprotei | 0 | 19 | 22 | 2 | 15 | 0 | 13 | 0 | 0 | 0 | 0 |
| 6 | -4.51 | 2.77 | -1.63 | 0.10 | 1 | Gm37277 | | 21 | 0 | 24 | 11 | 0 | 1 | 13 | 0 | 0 | 0 | 0 |
| 6 | -0.16 | 1.35 | -0.12 | 0.91 | 1 | Gm10297 | | 7 | 0 | 5 | 1 | 8 | 14 | 13 | 3 | 8 | 1 | 9 |
| 6 | -1.63 | 2.09 | -0.78 | 0.44 | 1 | Dmrta2 | doublesex and mab-3 related transcriptio | 16 | 0 | 0 | 8 | 3 | 14 | 13 | 0 | 5 | 0 | 10 |
| 6 | -1.73 | 2.72 | -0.64 | 0.52 | 1 | Gm43747 | | 0 | 0 | 6 | 0 | 0 | 13 | 13 | 0 | 4 | 0 | 32 |
| 6 | -2.88 | 2.30 | -1.25 | 0.21 | 1 | Gm43500 | | 0 | 0 | 19 | 0 | 4 | 14 | 13 | 0 | 3 | 0 | 15 |
| 6 | -2.38 | 2.19 | -1.09 | 0.28 | 1 | Gm44729 | | 0 | 7 | 5 | 0 | 15 | 10 | 13 | 0 | 6 | 0 | 9 |
| 6 | -1.10 | 1.83 | -0.60 | 0.55 | 1 | Gm4889 | | 3 | 0 | 0 | 4 | 5 | 2 | 13 | 0 | 6 | 32 | 0 |
| 6 | -0.50 | 1.90 | -0.26 | 0.79 | 1 | Platr4 | | 5 | 11 | 0 | 5 | 1 | 7 | 13 | 0 | 3 | 19 | 0 |
| 5 | -3.43 | 2.04 | -1.68 | 0.09 | 1 | Pdzd3 | PDZ domain containing 3 | 3 | 17 | 5 | 4 | 0 | 13 | 13 | 0 | 0 | 0 | 3 |
| 5 | -2.75 | 1.52 | -1.80 | 0.07 | 1 | Gm5093 | | 6 | 18 | 3 | 8 | 3 | 2 | 13 | 1 | 0 | 1 | 1 |
| 5 | -4.07 | 2.59 | -1.57 | 0.12 | 1 | Gm17322 | | 0 | 0 | 0 | 10 | 13 | 7 | 13 | 0 | 12 | 0 | 0 |
| 5 | -2.30 | 2.31 | -1.00 | 0.32 | 1 | Gm43592 | | 0 | 8 | 9 | 0 | 6 | 6 | 13 | 0 | 2 | 0 | 10 |
| 5 | 1.74 | 2.93 | 0.59 | 0.55 | 1 | Gm6140 | | 0 | 19 | 1 | 0 | 0 | 2 | 13 | 0 | 8 | 0 | 9 |
| 5 | 0.74 | 2.61 | 0.28 | 0.78 | 1 | Gm42782 | | 0 | 0 | 0 | 0 | 0 | 11 | 13 | 0 | 6 | 5 | 16 |
| 5 | -0.62 | 2.07 | -0.30 | 0.77 | 1 | Gm6161 | | 1 | 0 | 21 | 0 | 0 | 6 | 13 | 1 | 6 | 1 | 2 |
| 5 | -3.79 | 2.74 | -1.38 | 0.17 | 1 | Gm42413 | | 0 | 0 | 8 | 8 | 0 | 5 | 13 | 0 | 1 | 15 | 0 |
| 4 | -4.06 | 2.39 | -1.70 | 0.09 | 1 | Gm4969 | predicted gene 4969 | 0 | 5 | 0 | 11 | 2 | 13 | 13 | 0 | 0 | 0 | 6 |
| 4 | -0.43 | 2.32 | -0.18 | 0.85 | 1 | Nme9 | | 8 | 2 | 0 | 10 | 0 | 4 | 13 | 0 | 4 | 0 | 5 |
| 4 | -1.51 | 1.63 | -0.93 | 0.35 | 1 | Gm7962 | | 2 | 5 | 1 | 9 | 6 | 1 | 13 | 0 | 2 | 3 | 4 |
| 4 | -3.57 | 2.97 | -1.20 | 0.23 | 1 | Zfp389 | | 2 | 0 | 9 | 0 | 0 | 19 | 13 | 0 | 2 | 0 | 0 |
| 3 | -0.26 | 1.74 | -0.15 | 0.88 | 1 | Supt4b | | 9 | 2 | 2 | 2 | 1 | 1 | 13 | 3 | 2 | 0 | 1 |
| 3 | 1.38 | 3.48 | 0.40 | 0.69 | 1 | Gm43810 | | 0 | 0 | 0 | 0 | 0 | 3 | 13 | 0 | 10 | 0 | 7 |
| 3 | -3.33 | 2.43 | -1.37 | 0.17 | 1 | Gm29793 | | 0 | 0 | 1 | 6 | 1 | 3 | 13 | 0 | 1 | 0 | 7 |
| 3 | -0.27 | 3.54 | -0.08 | 0.94 | 1 | Gm15169 | | 0 | 0 | 0 | 0 | 0 | 7 | 13 | 0 | 0 | 8 | 5 |
| 3 | -2.28 | 2.84 | -0.80 | 0.42 | 1 | Gm3888 | | 0 | 0 | 1 | 0 | 1 | 8 | 13 | 0 | 5 | 0 | 1 |
| 2 | -0.96 | 2.30 | -0.42 | 0.68 | 1 | Rpl19-ps1 | | 1 | 1 | 2 | 2 | 1 | 0 | 13 | 1 | 5 | 0 | 0 |
| 2 | -4.48 | 3.55 | -1.26 | 0.21 | 1 | Btn1a1 | butyrophilin, subfamily 1, member A1 | 0 | 0 | 0 | 6 | 4 | 1 | 13 | 0 | 0 | 0 | 0 |
| 2 | -3.40 | 3.53 | -0.96 | 0.34 | 1 | Gm23628 | | 0 | 0 | 0 | 0 | 7 | 0 | 13 | 0 | 0 | 0 | 2 |
| 2 | -0.93 | 3.61 | -0.26 | 0.80 | 1 | Gm47340 | | 3 | 0 | 0 | 0 | 0 | 0 | 13 | 0 | 0 | 0 | 0 |

| | | | | | | | | | | | | | | | | | |
|---|---|---|---|---|---|---|---|---|---|---|---|---|---|---|---|---|---|
| 1071 | -1.11 | 0.85 | -1.31 | 0.19 | 1 | Nfib | nuclear factor I/B | 4833 | 3298 | 3529 | 14 | 49 | 9 | 12 | 17 | 19 | 0 | 4 |
| 1063 | -2.26 | 2.76 | -0.82 | 0.41 | 1 | Ly6a | lymphocyte antigen 6 complex, locus A | 5306 | 2811 | 3538 | 0 | 0 | 2 | 12 | 21 | 0 | 0 | 0 |
| 1023 | -0.67 | 0.89 | -0.75 | 0.46 | 1 | Mcam | melanoma cell adhesion molecule | 4308 | 3122 | 3680 | 10 | 40 | 15 | 12 | 0 | 13 | 39 | 10 |
| 1015 | -3.66 | 2.63 | -1.39 | 0.16 | 1 | Nid2 | nidogen 2 | 3679 | 3962 | 3504 | 0 | 9 | 0 | 12 | 0 | 0 | 0 | 0 |
| 959 | -1.48 | 1.81 | -0.82 | 0.41 | 1 | Ctrc | chymotrypsin C (caldecrin) | 8709 | 877 | 617 | 24 | 2 | 1 | 12 | 0 | 0 | 18 | 291 |
| 471 | -0.60 | 1.14 | -0.53 | 0.60 | 1 | Slco2b1 | solute carrier organic anion transporter fa | 3022 | 909 | 1229 | 0 | 0 | 2 | 12 | 0 | 2 | 3 | 1 |
| 300 | -0.57 | 0.66 | -0.86 | 0.39 | 1 | Inhbb | inhibin beta-B | 986 | 1033 | 1217 | 4 | 12 | 5 | 12 | 7 | 2 | 19 | 0 |
| 292 | 0.45 | 0.42 | 1.09 | 0.28 | 1 | Adcy5 | adenylate cyclase 5 | 1489 | 730 | 790 | 11 | 26 | 36 | 12 | 29 | 19 | 18 | 48 |
| 248 | -0.96 | 0.71 | -1.35 | 0.18 | 1 | Chl1 | cell adhesion molecule L1-like | 709 | 1002 | 611 | 110 | 30 | 92 | 12 | 15 | 43 | 0 | 108 |
| 227 | 0.52 | 0.74 | 0.70 | 0.48 | 1 | Casq2 | calsequestrin 2 | 896 | 680 | 824 | 4 | 7 | 12 | 12 | 39 | 10 | 0 | 10 |
| 226 | 0.19 | 0.36 | 0.54 | 0.59 | 1 | Kcnj3 | potassium inwardly-rectifying channel, su | 741 | 789 | 749 | 22 | 30 | 30 | 12 | 20 | 35 | 23 | 28 |
| 220 | -1.72 | 1.55 | -1.11 | 0.27 | 1 | Fgfr3 | fibroblast growth factor receptor 3 | 915 | 560 | 920 | 0 | 0 | 11 | 12 | 0 | 1 | 0 | 4 |
| 213 | 0.70 | 0.41 | 1.70 | 0.09 | 1 | Nrarp | Notch-regulated ankyrin repeat protein | 837 | 675 | 616 | 27 | 26 | 13 | 12 | 31 | 38 | 57 | 11 |
| 191 | -0.19 | 0.44 | -0.44 | 0.66 | 1 | Zfp503 | zinc finger protein 503 | 935 | 473 | 537 | 19 | 27 | 22 | 12 | 15 | 13 | 17 | 28 |
| 170 | -0.01 | 0.66 | -0.02 | 0.98 | 1 | Uhrf1 | ubiquitin-like, containing PHD and RING f | 779 | 213 | 230 | 126 | 81 | 96 | 12 | 61 | 100 | 112 | 60 |
| 123 | 0.79 | 0.77 | 1.02 | 0.31 | 1 | Bche | butyrylcholinesterase | 401 | 219 | 555 | 33 | 1 | 18 | 12 | 13 | 25 | 57 | 19 |
| 118 | -1.03 | 0.72 | -1.43 | 0.15 | 1 | Hpn | hepsin | 279 | 455 | 432 | 40 | 21 | 11 | 12 | 6 | 32 | 0 | 11 |
| 111 | 0.30 | 0.59 | 0.51 | 0.61 | 1 | Ccdc62 | coiled-coil domain containing 62 | 21 | 10 | 28 | 83 | 47 | 445 | 12 | 80 | 37 | 63 | 392 |
| 74 | -3.24 | 2.33 | -1.39 | 0.16 | 1 | Rrad | Ras-related associated with diabetes | 522 | 163 | 114 | 0 | 0 | 6 | 12 | 0 | 0 | 0 | 0 |
| 63 | 0.69 | 0.58 | 1.19 | 0.23 | 1 | Cplane2 | ciliogenesis and planar polarity effector 2 | 74 | 94 | 49 | 40 | 42 | 95 | 12 | 58 | 29 | 84 | 122 |
| 63 | 0.06 | 0.79 | 0.08 | 0.94 | 1 | Cap2 | CAP, adenylate cyclase-associated prote | 278 | 169 | 138 | 19 | 1 | 21 | 12 | 5 | 14 | 26 | 8 |
| 61 | -1.13 | 1.28 | -0.89 | 0.38 | 1 | Tmem44 | transmembrane protein 44 | 173 | 231 | 209 | 16 | 0 | 8 | 12 | 0 | 20 | 0 | 6 |
| 58 | 0.24 | 0.74 | 0.33 | 0.74 | 1 | Gm15612 | | 0 | 18 | 22 | 31 | 75 | 188 | 12 | 28 | 39 | 33 | 196 |
| 53 | -1.64 | 0.95 | -1.74 | 0.08 | 1 | Ddx11 | DEAD/H (Asp-Glu-Ala-Asp/His) box helica | 135 | 36 | 65 | 39 | 57 | 121 | 12 | 0 | 47 | 5 | 66 |
| 53 | -0.14 | 0.90 | -0.16 | 0.87 | 1 | Kcp | kielin/chordin-like protein | 160 | 81 | 156 | 20 | 30 | 12 | 12 | 0 | 39 | 62 | 7 |
| 45 | -0.56 | 0.74 | -0.75 | 0.45 | 1 | Myo7b | myosin VIIB | 44 | 191 | 113 | 28 | 14 | 20 | 12 | 43 | 18 | 0 | 11 |
| 44 | -0.82 | 0.99 | -0.83 | 0.41 | 1 | Tmprss6 | transmembrane serine protease 6 | 58 | 42 | 174 | 24 | 28 | 49 | 12 | 71 | 17 | 0 | 10 |
| 42 | -1.18 | 1.09 | -1.09 | 0.28 | 1 | Gna15 | guanine nucleotide binding protein, alpha | 161 | 172 | 59 | 1 | 11 | 16 | 12 | 0 | 1 | 19 | 6 |
| 41 | 0.50 | 1.67 | 0.30 | 0.76 | 1 | Hoxb2 | homeobox B2 | 165 | 103 | 96 | 0 | 0 | 3 | 12 | 0 | 0 | 71 | 0 |
| 32 | -0.20 | 0.61 | -0.32 | 0.75 | 1 | Gm35558 | | 36 | 36 | 52 | 32 | 46 | 29 | 12 | 21 | 28 | 13 | 44 |
| 31 | 0.64 | 0.63 | 1.03 | 0.30 | 1 | Vmn1r4 | vomeronasal 1 receptor 4 | 27 | 64 | 43 | 10 | 16 | 49 | 12 | 21 | 21 | 26 | 52 |
| 31 | 0.44 | 0.58 | 0.76 | 0.45 | 1 | Proscos | | 17 | 37 | 31 | 28 | 33 | 32 | 12 | 56 | 22 | 21 | 47 |
| 29 | -0.63 | 1.06 | -0.59 | 0.55 | 1 | Recql4 | RecQ protein-like 4 | 100 | 7 | 54 | 10 | 26 | 41 | 12 | 12 | 9 | 0 | 53 |
| 28 | 0.24 | 0.72 | 0.33 | 0.74 | 1 | Troap | trophinin associated protein | 33 | 19 | 12 | 46 | 22 | 36 | 12 | 41 | 43 | 21 | 25 |
| 27 | -0.66 | 0.86 | -0.77 | 0.44 | 1 | Gm17249 | | 13 | 42 | 49 | 32 | 24 | 41 | 12 | 2 | 10 | 17 | 59 |
| 27 | 1.16 | 0.74 | 1.56 | 0.12 | 1 | Palb2 | partner and localizer of BRCA2 | 24 | 47 | 35 | 17 | 8 | 22 | 12 | 16 | 17 | 64 | 36 |
| 24 | 1.60 | 0.96 | 1.67 | 0.09 | 1 | Gm18736 | | 16 | 13 | 42 | 1 | 61 | 0 | 12 | 13 | 78 | 31 | 2 |
| 24 | 0.06 | 1.16 | 0.05 | 0.96 | 1 | 1700001G11Rik | | 15 | 16 | 23 | 53 | 6 | 33 | 12 | 0 | 15 | 41 | 54 |
| 24 | 0.01 | 1.40 | 0.01 | 0.99 | 1 | Gm26947 | | 0 | 9 | 0 | 11 | 14 | 98 | 12 | 0 | 14 | 19 | 86 |
| 24 | -0.24 | 1.15 | -0.20 | 0.84 | 1 | Ccdc24 | coiled-coil domain containing 24 | 19 | 57 | 37 | 13 | 7 | 36 | 12 | 0 | 4 | 54 | 21 |
| 23 | 0.80 | 0.69 | 1.15 | 0.25 | 1 | Gm13461 | | 37 | 30 | 33 | 7 | 20 | 19 | 12 | 30 | 17 | 17 | 35 |
| 23 | 0.90 | 0.85 | 1.06 | 0.29 | 1 | Gm12524 | | 5 | 9 | 5 | 11 | 11 | 42 | 12 | 78 | 31 | 4 | 47 |
| 22 | -0.48 | 0.93 | -0.51 | 0.61 | 1 | Gm9870 | | 14 | 18 | 25 | 64 | 17 | 19 | 12 | 6 | 10 | 41 | 19 |
| 22 | 1.15 | 1.17 | 0.98 | 0.33 | 1 | Frmd8os | | 0 | 4 | 18 | 26 | 9 | 19 | 12 | 60 | 15 | 71 | 9 |
| 22 | 0.24 | 0.84 | 0.28 | 0.78 | 1 | Phpt1 | phosphohistidine phosphatase 1 | 4 | 3 | 21 | 23 | 23 | 35 | 12 | 8 | 28 | 44 | 39 |
| 22 | 0.18 | 0.87 | 0.21 | 0.83 | 1 | Gm13436 | | 54 | 41 | 21 | 8 | 15 | 19 | 12 | 36 | 19 | 6 | 7 |
| 22 | 0.79 | 1.11 | 0.71 | 0.48 | 1 | Gm8189 | | 14 | 0 | 0 | 10 | 16 | 51 | 12 | 39 | 16 | 27 | 53 |
| 21 | -2.69 | 1.40 | -1.93 | 0.05 | 1 | Cacna1e | calcium channel, voltage-dependent, R ty | 108 | 60 | 23 | 5 | 0 | 14 | 12 | 0 | 0 | 0 | 12 |
| 21 | -0.30 | 1.05 | -0.28 | 0.78 | 1 | 8030456M14Rik | | 9 | 2 | 24 | 19 | 27 | 40 | 12 | 24 | 37 | 0 | 35 |
| 20 | -0.16 | 2.10 | -0.08 | 0.94 | 1 | Gm45838 | | 15 | 11 | 0 | 0 | 0 | 115 | 12 | 0 | 17 | 0 | 51 |
| 20 | -1.30 | 1.24 | -1.04 | 0.30 | 1 | 1700109H08Ril | RIKEN cDNA 1700109H08 gene | 0 | 5 | 5 | 4 | 6 | 89 | 12 | 1 | 12 | 0 | 83 |
| 20 | 0.16 | 1.09 | 0.15 | 0.88 | 1 | 4930413F20Rik | | 20 | 7 | 0 | 25 | 20 | 22 | 12 | 66 | 11 | 8 | 27 |
| 20 | -1.69 | 2.12 | -0.80 | 0.42 | 1 | Skint3 | selection and upkeep of intraepithelial T c | 80 | 46 | 80 | 0 | 0 | 0 | 12 | 0 | 0 | 0 | 0 |
| 20 | -1.39 | 0.84 | -1.65 | 0.10 | 1 | Slc38a8 | solute carrier family 38, member 8 | 47 | 14 | 15 | 36 | 26 | 23 | 12 | 10 | 8 | 21 | 5 |
| 19 | -1.74 | 1.26 | -1.38 | 0.17 | 1 | Gm49322 | | 3 | 0 | 5 | 1 | 23 | 83 | 12 | 0 | 0 | 6 | 75 |
| 18 | -2.82 | 1.99 | -1.42 | 0.16 | 1 | Cd226 | CD226 antigen | 119 | 66 | 3 | 0 | 2 | 0 | 12 | 0 | 0 | 0 | 0 |
| 18 | -0.11 | 0.81 | -0.14 | 0.89 | 1 | Hoga1 | 4-hydroxy-2-oxoglutarate aldolase 1 | 40 | 34 | 27 | 7 | 9 | 21 | 12 | 3 | 18 | 6 | 20 |
| 18 | -1.06 | 1.47 | -0.72 | 0.47 | 1 | Gm43789 | | 0 | 0 | 0 | 1 | 26 | 82 | 12 | 15 | 6 | 0 | 53 |
| 18 | -0.41 | 1.01 | -0.41 | 0.69 | 1 | Gm10052 | | 5 | 17 | 36 | 9 | 14 | 30 | 12 | 38 | 6 | 1 | 24 |
| 17 | 0.26 | 1.40 | 0.19 | 0.85 | 1 | Gm4118 | | 5 | 21 | 14 | 0 | 14 | 40 | 12 | 0 | 6 | 39 | 42 |

| | | | | | | | | | | | | | | | | | | |
|---|---|---|---|---|---|---|---|---|---|---|---|---|---|---|---|---|---|---|
| 17 | 1.01 | 0.95 | 1.06 | 0.29 | 1 | 9530062K07Rik | | 3 | 44 | 12 | 8 | 6 | 18 | 12 | 24 | 13 | 28 | 19 |
| 17 | 0.42 | 1.27 | 0.33 | 0.74 | 1 | Zpld1 | zona pellucida like domain containing 1 | 78 | 14 | 46 | 0 | 4 | 8 | 12 | 5 | 4 | 0 | 14 |
| 17 | -0.73 | 0.86 | -0.84 | 0.40 | 1 | Gm48933 | | 22 | 12 | 12 | 37 | 15 | 14 | 12 | 37 | 6 | 9 | 5 |
| 16 | 0.58 | 1.12 | 0.52 | 0.60 | 1 | Clec12b | C-type lectin domain family 12, member E | 0 | 4 | 0 | 4 | 23 | 14 | 12 | 71 | 12 | 18 | 22 |
| 16 | 1.39 | 1.53 | 0.91 | 0.36 | 1 | Atg9b | autophagy related 9B | 30 | 50 | 46 | 0 | 0 | 6 | 12 | 8 | 0 | 23 | 4 |
| 16 | -1.13 | 1.11 | -1.02 | 0.31 | 1 | Gm17494 | | 14 | 21 | 45 | 19 | 21 | 11 | 12 | 0 | 6 | 22 | 7 |
| 16 | -0.59 | 1.32 | -0.45 | 0.65 | 1 | Itpa-ps1 | | 47 | 13 | 0 | 13 | 25 | 14 | 12 | 1 | 26 | 8 | 17 |
| 16 | 0.16 | 1.45 | 0.11 | 0.91 | 1 | Gnasas1 | | 0 | 26 | 10 | 4 | 6 | 45 | 12 | 27 | 0 | 19 | 26 |
| 15 | -1.88 | 1.76 | -1.07 | 0.29 | 1 | Cd6 | CD6 antigen | 81 | 57 | 9 | 0 | 0 | 3 | 12 | 0 | 0 | 0 | 2 |
| 14 | 0.22 | 0.83 | 0.26 | 0.79 | 1 | Gm21596 | | 15 | 25 | 10 | 9 | 13 | 14 | 12 | 7 | 11 | 17 | 25 |
| 14 | -1.93 | 1.81 | -1.07 | 0.28 | 1 | Lypd6 | LY6/PLAUR domain containing 6 | 49 | 10 | 75 | 0 | 0 | 7 | 12 | 0 | 0 | 0 | 4 |
| 14 | -1.48 | 1.82 | -0.81 | 0.42 | 1 | B830012L14Rik | | 0 | 0 | 0 | 9 | 1 | 67 | 12 | 0 | 15 | 0 | 53 |
| 14 | -1.62 | 1.07 | -1.51 | 0.13 | 1 | Apobec4 | apolipoprotein B mRNA editing enzyme, c | 14 | 17 | 22 | 18 | 24 | 15 | 12 | 2 | 22 | 0 | 9 |
| 14 | 0.57 | 1.11 | 0.51 | 0.61 | 1 | Gm5855 | | 11 | 1 | 33 | 2 | 12 | 4 | 12 | 0 | 17 | 55 | 5 |
| 13 | 0.60 | 1.38 | 0.43 | 0.67 | 1 | Rnf17 | ring finger protein 17 | 25 | 21 | 18 | 11 | 0 | 8 | 12 | 28 | 10 | 0 | 9 |
| 13 | -0.07 | 1.61 | -0.04 | 0.97 | 1 | Scarna2 | | 15 | 0 | 0 | 15 | 9 | 32 | 12 | 12 | 12 | 0 | 36 |
| 12 | 2.31 | 1.34 | 1.73 | 0.08 | 1 | 9530052E02Rik | | 8 | 1 | 12 | 0 | 1 | 19 | 12 | 23 | 16 | 15 | 19 |
| 11 | -0.06 | 1.06 | -0.05 | 0.96 | 1 | Gm7045 | | 15 | 3 | 14 | 18 | 2 | 17 | 12 | 13 | 3 | 13 | 17 |
| 11 | -0.73 | 1.52 | -0.48 | 0.63 | 1 | Aldoa | | 12 | 10 | 3 | 11 | 3 | 25 | 12 | 0 | 21 | 0 | 26 |
| 11 | -1.59 | 1.44 | -1.10 | 0.27 | 1 | Gm45231 | | 9 | 16 | 0 | 19 | 25 | 15 | 12 | 0 | 4 | 14 | 10 |
| 11 | 0.66 | 1.80 | 0.37 | 0.71 | 1 | Gm7125 | | 70 | 0 | 0 | 0 | 0 | 6 | 12 | 0 | 0 | 15 | 15 |
| 11 | 0.20 | 1.40 | 0.14 | 0.89 | 1 | Gm807 | | 0 | 1 | 0 | 14 | 5 | 23 | 12 | 0 | 10 | 21 | 34 |
| 10 | -0.60 | 1.93 | -0.31 | 0.75 | 1 | Gm22753 | | 12 | 21 | 0 | 0 | 1 | 11 | 12 | 29 | 0 | 27 | 0 |
| 10 | -3.30 | 1.76 | -1.87 | 0.06 | 1 | Ppm1j | protein phosphatase 1J | 36 | 30 | 21 | 5 | 0 | 4 | 12 | 0 | 0 | 0 | 2 |
| 10 | -0.55 | 1.42 | -0.39 | 0.70 | 1 | Cyp17a1 | cytochrome P450, family 17, subfamily a, | 7 | 0 | 0 | 9 | 9 | 13 | 12 | 0 | 12 | 36 | 12 |
| 10 | -1.67 | 1.43 | -1.17 | 0.24 | 1 | Prlhr | prolactin releasing hormone receptor | 24 | 11 | 11 | 11 | 9 | 8 | 12 | 17 | 0 | 0 | 5 |
| 9 | 1.17 | 1.61 | 0.73 | 0.47 | 1 | Gm43813 | | 3 | 14 | 0 | 9 | 1 | 14 | 12 | 0 | 14 | 28 | 8 |
| 9 | 1.04 | 1.53 | 0.68 | 0.50 | 1 | 4930426I24Rik | | 13 | 5 | 8 | 4 | 0 | 10 | 12 | 27 | 5 | 0 | 15 |
| 9 | -0.83 | 1.98 | -0.42 | 0.68 | 1 | 1700066B19Ril | RIKEN cDNA 1700066B19 gene | 3 | 33 | 0 | 4 | 2 | 15 | 12 | 0 | 19 | 0 | 9 |
| 9 | 0.72 | 1.13 | 0.64 | 0.52 | 1 | Gm10184 | | 1 | 1 | 0 | 9 | 3 | 8 | 12 | 3 | 4 | 36 | 20 |
| 9 | -1.58 | 1.23 | -1.29 | 0.20 | 1 | Gm13651 | | 9 | 15 | 9 | 7 | 17 | 5 | 12 | 2 | 15 | 4 | 0 |
| 9 | -2.99 | 1.57 | -1.91 | 0.06 | 1 | D530018E20Rik | | 8 | 0 | 17 | 4 | 9 | 25 | 12 | 0 | 2 | 0 | 17 |
| 9 | -2.00 | 1.44 | -1.39 | 0.16 | 1 | Gm17552 | | 8 | 7 | 5 | 3 | 9 | 21 | 12 | 0 | 4 | 0 | 24 |
| 8 | -0.21 | 1.10 | -0.19 | 0.85 | 1 | Gm15013 | | 18 | 15 | 17 | 5 | 2 | 4 | 12 | 5 | 7 | 3 | 4 |
| 8 | 1.31 | 1.33 | 0.98 | 0.33 | 1 | Gm6793 | | 11 | 5 | 19 | 0 | 2 | 6 | 12 | 13 | 8 | 13 | 3 |
| 8 | -0.50 | 1.26 | -0.40 | 0.69 | 1 | Tpo | thyroid peroxidase | 0 | 7 | 6 | 5 | 17 | 11 | 12 | 12 | 3 | 10 | 7 |
| 8 | -0.92 | 1.76 | -0.52 | 0.60 | 1 | Dnali1 | dynein, axonemal, light intermediate poly | 9 | 27 | 24 | 0 | 3 | 3 | 12 | 0 | 6 | 5 | 0 |
| 7 | -1.54 | 1.32 | -1.17 | 0.24 | 1 | Gm17823 | | 13 | 2 | 20 | 5 | 9 | 3 | 12 | 0 | 12 | 1 | 3 |
| 7 | -0.84 | 1.66 | -0.51 | 0.61 | 1 | Gm6505 | | 3 | 12 | 0 | 4 | 11 | 10 | 12 | 0 | 6 | 19 | 2 |
| 7 | 1.70 | 2.10 | 0.81 | 0.42 | 1 | Tal2 | T cell acute lymphocytic leukemia 2 | 11 | 0 | 0 | 0 | 0 | 8 | 12 | 0 | 12 | 32 | 2 |
| 7 | 0.61 | 2.15 | 0.28 | 0.78 | 1 | Xkrx | X-linked Kx blood group related, X-linked | 37 | 0 | 7 | 0 | 0 | 6 | 12 | 0 | 4 | 0 | 11 |
| 7 | -0.94 | 1.16 | -0.81 | 0.42 | 1 | Gm5425 | | 2 | 3 | 3 | 4 | 9 | 17 | 12 | 5 | 15 | 1 | 5 |
| 7 | -3.41 | 2.65 | -1.29 | 0.20 | 1 | Gm40614 | | 0 | 20 | 0 | 0 | 17 | 4 | 12 | 0 | 2 | 0 | 18 |
| 6 | -2.92 | 2.46 | -1.19 | 0.24 | 1 | Gm48027 | | 0 | 6 | 0 | 0 | 14 | 23 | 12 | 0 | 4 | 0 | 11 |
| 6 | -0.20 | 1.91 | -0.10 | 0.92 | 1 | Gm5841 | | 18 | 0 | 0 | 6 | 2 | 13 | 12 | 2 | 11 | 0 | 6 |
| 6 | -0.50 | 1.15 | -0.43 | 0.67 | 1 | Gm7232 | | 3 | 4 | 4 | 3 | 6 | 12 | 12 | 5 | 6 | 3 | 9 |
| 6 | -2.04 | 1.88 | -1.09 | 0.28 | 1 | Gm13142 | | 0 | 8 | 7 | 4 | 6 | 9 | 12 | 0 | 11 | 0 | 7 |
| 6 | -2.83 | 2.77 | -1.02 | 0.31 | 1 | Gm24060 | | 9 | 0 | 11 | 0 | 0 | 3 | 12 | 0 | 0 | 0 | 27 |
| 6 | -3.34 | 2.53 | -1.32 | 0.19 | 1 | Plppr5 | phospholipid phosphatase related 5 | 32 | 0 | 0 | 4 | 4 | 2 | 12 | 0 | 7 | 0 | 0 |
| 5 | -1.08 | 2.17 | -0.50 | 0.62 | 1 | 9030624G23Ril | RIKEN cDNA 9030624G23 gene | 10 | 3 | 0 | 0 | 5 | 16 | 12 | 0 | 8 | 0 | 6 |
| 5 | -0.45 | 2.14 | -0.21 | 0.83 | 1 | Gm45185 | | 0 | 16 | 1 | 1 | 1 | 3 | 12 | 0 | 0 | 23 | 2 |
| 5 | -1.29 | 1.99 | -0.65 | 0.52 | 1 | Gm5831 | | 16 | 1 | 0 | 8 | 3 | 4 | 12 | 0 | 5 | 0 | 5 |
| 5 | -0.73 | 1.95 | -0.38 | 0.71 | 1 | Gm11953 | | 2 | 1 | 1 | 11 | 0 | 1 | 12 | 0 | 0 | 23 | 1 |
| 4 | -1.17 | 2.88 | -0.41 | 0.68 | 1 | Mbl2 | mannose-binding lectin (protein C) 2 | 0 | 0 | 0 | 7 | 0 | 6 | 12 | 0 | 7 | 0 | 17 |
| 4 | 1.26 | 2.67 | 0.47 | 0.64 | 1 | Gm13684 | | 7 | 2 | 0 | 0 | 0 | 6 | 12 | 0 | 11 | 0 | 11 |
| 4 | -0.93 | 2.07 | -0.45 | 0.65 | 1 | D430041D05Ri | RIKEN cDNA D430041D05 gene | 1 | 1 | 15 | 0 | 9 | 1 | 12 | 0 | 0 | 5 | 2 |
| 4 | -0.90 | 3.31 | -0.27 | 0.79 | 1 | Gm36888 | | 7 | 0 | 0 | 0 | 0 | 12 | 12 | 0 | 0 | 12 | 0 |
| 4 | -0.33 | 1.54 | -0.22 | 0.83 | 1 | Gstm2-ps1 | | 10 | 2 | 1 | 4 | 0 | 2 | 12 | 1 | 1 | 6 | 2 |
| 4 | -2.26 | 2.01 | -1.13 | 0.26 | 1 | Tssk3 | testis-specific serine kinase 3 | 7 | 0 | 6 | 5 | 3 | 2 | 12 | 0 | 4 | 0 | 3 |
| 4 | -1.67 | 3.46 | -0.48 | 0.63 | 1 | Gm8184 | | 0 | 1 | 0 | 0 | 0 | 15 | 12 | 0 | 0 | 0 | 13 |

| | | | | | | | | | | | | | | | | | | |
|---|---|---|---|---|---|---|---|---|---|---|---|---|---|---|---|---|---|---|
| 4 | -0.02 | 1.79 | -0.01 | 0.99 | 1 | Gm10801 | | 3 | 3 | 0 | 4 | 2 | 2 | 12 | 0 | 5 | 3 | 6 |
| 3 | -3.21 | 3.23 | -0.99 | 0.32 | 1 | Gm10322 | | 0 | 0 | 20 | 0 | 3 | 0 | 12 | 0 | 1 | 0 | 1 |
| 3 | -3.27 | 3.12 | -1.05 | 0.29 | 1 | Snord8 | | 0 | 4 | 0 | 8 | 0 | 6 | 12 | 0 | 0 | 0 | 3 |
| 3 | -3.87 | 3.19 | -1.21 | 0.23 | 1 | Igf2bp1 | insulin-like growth factor 2 mRNA binding | 4 | 0 | 7 | 0 | 0 | 5 | 12 | 0 | 0 | 0 | 4 |
| 3 | -3.07 | 3.52 | -0.87 | 0.38 | 1 | Gm14685 | predicted gene 14685 | 0 | 0 | 0 | 0 | 3 | 9 | 12 | 0 | 0 | 0 | 8 |
| 3 | -2.29 | 2.72 | -0.84 | 0.40 | 1 | mt-Ti | | 2 | 0 | 1 | 2 | 0 | 3 | 12 | 0 | 0 | 0 | 12 |
| 3 | -0.66 | 3.53 | -0.19 | 0.85 | 1 | C130051F05Rik | | 0 | 0 | 0 | 0 | 0 | 8 | 12 | 0 | 1 | 0 | 9 |
| 3 | -3.73 | 2.79 | -1.34 | 0.18 | 1 | Gm14066 | | 5 | 0 | 0 | 5 | 4 | 1 | 12 | 0 | 0 | 0 | 2 |
| 3 | -2.78 | 3.48 | -0.80 | 0.42 | 1 | Gm40040 | | 0 | 0 | 7 | 0 | 0 | 3 | 12 | 0 | 1 | 0 | 5 |
| 2 | -1.31 | 3.29 | -0.40 | 0.69 | 1 | Gm45949 | | 0 | 0 | 0 | 0 | 7 | 1 | 12 | 0 | 3 | 0 | 4 |
| 2 | -1.60 | 3.60 | -0.44 | 0.66 | 1 | Lrfn5 | leucine rich repeat and fibronectin type III | 3 | 0 | 0 | 0 | 1 | 0 | 12 | 0 | 0 | 0 | 0 |
| 1 | -3.33 | 3.57 | -0.93 | 0.35 | 1 | Gm3696 | predicted gene 3696 | 0 | 0 | 0 | 0 | 0 | 4 | 12 | 0 | 0 | 0 | 0 |
| 2502 | -1.93 | 1.15 | -1.67 | 0.09 | 1 | C4b | complement component 4B (Chido blood | 16769 | 5027 | 5700 | 4 | 7 | 0 | 11 | 0 | 3 | 0 | 1 |
| 1250 | -1.60 | 1.28 | -1.25 | 0.21 | 1 | Spon1 | spondin 1, (f-spondin) extracellular matrix | 7122 | 3312 | 3254 | 12 | 21 | 2 | 11 | 0 | 13 | 0 | 6 |
| 497 | -0.37 | 1.70 | -0.22 | 0.83 | 1 | Itgb2 | integrin beta 2 | 3218 | 1351 | 843 | 11 | 0 | 7 | 11 | 0 | 9 | 0 | 16 |
| 496 | -0.25 | 0.47 | -0.54 | 0.59 | 1 | Lama3 | laminin, alpha 3 | 1523 | 2273 | 1505 | 26 | 9 | 34 | 11 | 7 | 3 | 23 | 43 |
| 397 | 0.57 | 2.24 | 0.25 | 0.80 | 1 | Hmgcs2 | 3-hydroxy-3-methylglutaryl-Coenzyme A s | 2195 | 1064 | 1095 | 0 | 0 | 0 | 11 | 0 | 2 | 0 | 1 |
| 333 | 0.19 | 0.59 | 0.33 | 0.74 | 1 | Colgalt2 | collagen beta(1-O)galactosyltransferase 2 | 1107 | 1280 | 1191 | 3 | 10 | 14 | 11 | 29 | 4 | 4 | 12 |
| 288 | -0.79 | 0.65 | -1.21 | 0.23 | 1 | Ces1d | carboxylesterase 1D | 1073 | 691 | 1346 | 9 | 10 | 4 | 11 | 1 | 11 | 0 | 10 |
| 273 | -2.06 | 1.29 | -1.60 | 0.11 | 1 | Trim30a | tripartite motif-containing 30A | 1235 | 855 | 877 | 6 | 5 | 1 | 11 | 0 | 8 | 0 | 0 |
| 268 | -0.11 | 1.51 | -0.07 | 0.94 | 1 | Gpm6b | glycoprotein m6b | 969 | 828 | 1111 | 0 | 6 | 2 | 11 | 0 | 13 | 0 | 6 |
| 222 | 0.39 | 3.25 | 0.12 | 0.91 | 1 | Tyrobp | TYRO protein tyrosine kinase binding pro | 1120 | 555 | 747 | 0 | 0 | 0 | 11 | 0 | 12 | 0 | 0 |
| 203 | -2.46 | 3.02 | -0.82 | 0.41 | 1 | Cd52 | CD52 antigen | 1120 | 598 | 504 | 0 | 0 | 0 | 11 | 0 | 0 | 0 | 2 |
| 128 | 0.23 | 0.75 | 0.31 | 0.76 | 1 | Ncapg | non-SMC condensin I complex, subunit G | 389 | 117 | 85 | 123 | 144 | 102 | 11 | 136 | 135 | 40 | 122 |
| 109 | -1.72 | 1.02 | -1.69 | 0.09 | 1 | Pdgfc | platelet-derived growth factor, C polypept | 502 | 154 | 518 | 0 | 9 | 1 | 11 | 0 | 6 | 0 | 0 |
| 105 | -1.22 | 3.62 | -0.34 | 0.74 | 1 | Cd86 | CD86 antigen | 617 | 214 | 310 | 0 | 0 | 0 | 11 | 0 | 0 | 0 | 0 |
| 103 | -0.65 | 0.76 | -0.85 | 0.40 | 1 | Adrb2 | adrenergic receptor, beta 2 | 395 | 314 | 316 | 4 | 4 | 40 | 11 | 0 | 10 | 1 | 36 |
| 102 | -0.92 | 0.64 | -1.45 | 0.15 | 1 | Dtna | dystrobrevin alpha | 256 | 371 | 266 | 61 | 34 | 33 | 11 | 39 | 33 | 12 | 2 |
| 71 | 2.44 | 2.50 | 0.98 | 0.33 | 1 | Jaml | junction adhesion molecule like | 433 | 247 | 69 | 0 | 0 | 0 | 11 | 15 | 0 | 0 | 4 |
| 70 | -2.00 | 1.21 | -1.65 | 0.10 | 1 | Grap2 | GRB2-related adaptor protein 2 | 414 | 164 | 164 | 0 | 3 | 3 | 11 | 0 | 0 | 0 | 7 |
| 65 | -2.48 | 3.03 | -0.82 | 0.41 | 1 | Asb2 | ankyrin repeat and SOCS box-containing | 316 | 242 | 149 | 0 | 0 | 0 | 11 | 0 | 0 | 0 | 2 |
| 48 | -0.13 | 0.72 | -0.18 | 0.86 | 1 | Dynlrb2 | dynein light chain roadblock-type 2 | 103 | 125 | 150 | 17 | 21 | 22 | 11 | 51 | 6 | 5 | 19 |
| 48 | 0.68 | 0.62 | 1.11 | 0.27 | 1 | Prss41 | protease, serine 41 | 8 | 54 | 47 | 27 | 89 | 55 | 11 | 44 | 83 | 40 | 69 |
| 47 | 0.86 | 0.68 | 1.27 | 0.20 | 1 | Fanca | Fanconi anemia, complementation group | 97 | 27 | 54 | 29 | 21 | 72 | 11 | 54 | 48 | 61 | 45 |
| 46 | -1.49 | 1.24 | -1.19 | 0.23 | 1 | Mir5125 | | 0 | 0 | 10 | 1 | 5 | 222 | 11 | 0 | 10 | 0 | 243 |
| 41 | -2.58 | 1.31 | -1.97 | 0.05 | 1 | Slc27a2 | solute carrier family 27 (fatty acid transpor | 167 | 158 | 104 | 0 | 7 | 3 | 11 | 0 | 1 | 0 | 2 |
| 40 | -0.01 | 0.61 | -0.01 | 0.99 | 1 | E230016M11Rik | | 25 | 31 | 12 | 12 | 15 | 151 | 11 | 12 | 19 | 9 | 137 |
| 38 | -1.31 | 1.08 | -1.21 | 0.23 | 1 | Gm35339 | predicted gene, 35339 | 9 | 18 | 21 | 1 | 4 | 175 | 11 | 0 | 4 | 0 | 175 |
| 37 | 0.60 | 0.89 | 0.68 | 0.50 | 1 | Cenpi | centromere protein I | 82 | 11 | 5 | 16 | 38 | 43 | 11 | 16 | 82 | 85 | 23 |
| 36 | 0.47 | 0.64 | 0.74 | 0.46 | 1 | Pcdhb8 | protocadherin beta 8 | 71 | 57 | 51 | 28 | 13 | 40 | 11 | 16 | 25 | 44 | 38 |
| 35 | -2.30 | 1.57 | -1.46 | 0.14 | 1 | Il2rb | interleukin 2 receptor, beta chain | 280 | 92 | 6 | 0 | 0 | 1 | 11 | 0 | 0 | 0 | 0 |
| 34 | 1.33 | 1.41 | 0.94 | 0.35 | 1 | Gm45137 | | 0 | 0 | 0 | 0 | 0 | 198 | 11 | 0 | 14 | 19 | 135 |
| 34 | -0.56 | 1.43 | -0.39 | 0.70 | 1 | Gm38253 | | 15 | 0 | 9 | 2 | 0 | 150 | 11 | 0 | 12 | 0 | 171 |
| 33 | 0.21 | 0.64 | 0.32 | 0.75 | 1 | Mybl2 | myeloblastosis oncogene-like 2 | 101 | 42 | 31 | 7 | 9 | 76 | 11 | 14 | 8 | 19 | 48 |
| 32 | -0.01 | 0.76 | -0.01 | 0.99 | 1 | Gm37123 | | 21 | 7 | 15 | 9 | 26 | 99 | 11 | 23 | 9 | 6 | 124 |
| 30 | 0.05 | 0.78 | 0.06 | 0.95 | 1 | Myl4 | myosin, light polypeptide 4 | 123 | 26 | 63 | 18 | 12 | 19 | 11 | 23 | 14 | 6 | 17 |
| 29 | 0.27 | 0.71 | 0.38 | 0.70 | 1 | Gm43566 | | 38 | 12 | 38 | 34 | 31 | 31 | 11 | 43 | 32 | 23 | 28 |
| 27 | -0.73 | 0.96 | -0.76 | 0.45 | 1 | Car5b | carbonic anhydrase 5b, mitochondrial | 108 | 71 | 59 | 11 | 9 | 3 | 11 | 0 | 11 | 15 | 1 |
| 27 | 1.51 | 1.22 | 1.24 | 0.21 | 1 | Morrbid | | 67 | 74 | 61 | 5 | 0 | 6 | 11 | 0 | 18 | 46 | 8 |
| 26 | 0.57 | 1.03 | 0.55 | 0.58 | 1 | Ska3 | spindle and kinetochore associated comp | 56 | 12 | 0 | 29 | 19 | 19 | 11 | 63 | 18 | 41 | 22 |
| 26 | -0.19 | 0.72 | -0.26 | 0.79 | 1 | Gm16279 | | 22 | 50 | 16 | 18 | 26 | 48 | 11 | 49 | 11 | 10 | 29 |
| 26 | 0.09 | 0.88 | 0.10 | 0.92 | 1 | Gm11963 | | 35 | 36 | 26 | 30 | 19 | 26 | 11 | 3 | 33 | 53 | 15 |
| 25 | -0.79 | 0.70 | -1.13 | 0.26 | 1 | Rims1 | regulating synaptic membrane exocytosis | 21 | 51 | 29 | 53 | 29 | 18 | 11 | 25 | 7 | 8 | 24 |
| 25 | 0.03 | 0.92 | 0.03 | 0.97 | 1 | Oprm1 | opioid receptor, mu 1 | 0 | 4 | 0 | 34 | 55 | 29 | 11 | 16 | 88 | 10 | 27 |
| 24 | 0.08 | 1.23 | 0.07 | 0.95 | 1 | Tigd3 | tigger transposable element derived 3 | 65 | 8 | 48 | 6 | 25 | 24 | 11 | 46 | 6 | 0 | 27 |
| 24 | -2.13 | 1.42 | -1.50 | 0.13 | 1 | Mir344-2 | | 7 | 21 | 0 | 19 | 16 | 109 | 11 | 0 | 9 | 0 | 73 |
| 24 | 2.69 | 1.37 | 1.97 | 0.05 | 1 | Bend6 | BEN domain containing 6 | 86 | 34 | 29 | 0 | 0 | 2 | 11 | 0 | 7 | 86 | 3 |
| 23 | -1.27 | 1.47 | -0.87 | 0.39 | 1 | Gm13648 | | 9 | 6 | 0 | 2 | 16 | 113 | 11 | 0 | 17 | 0 | 83 |
| 23 | 0.95 | 1.24 | 0.76 | 0.45 | 1 | Rubcnl | RUN and cysteine rich domain containing | 174 | 21 | 31 | 1 | 2 | 1 | 11 | 2 | 12 | 0 | 2 |
| 23 | -2.17 | 1.66 | -1.30 | 0.19 | 1 | Gm46560 | | 59 | 86 | 91 | 0 | 2 | 1 | 11 | 0 | 0 | 0 | 4 |

| | | | | | | | | | | | | | | | | | | |
|---|---|---|---|---|---|---|---|---|---|---|---|---|---|---|---|---|---|---|
| 23 | -0.29 | 1.00 | -0.29 | 0.77 | 1 | 1700034H15Rik | | 11 | 24 | 21 | 15 | 9 | 68 | 11 | 0 | 11 | 22 | 57 |
| 22 | 1.13 | 0.84 | 1.34 | 0.18 | 1 | BC043934 | | 54 | 48 | 59 | 3 | 9 | 2 | 11 | 15 | 13 | 30 | 1 |
| 22 | -2.82 | 1.57 | -1.80 | 0.07 | 1 | Trac | | 111 | 70 | 36 | 7 | 0 | 1 | 11 | 0 | 0 | 0 | 2 |
| 22 | -0.29 | 0.72 | -0.41 | 0.68 | 1 | Zfp750 | zinc finger protein 750 | 49 | 48 | 37 | 23 | 19 | 5 | 11 | 17 | 7 | 15 | 6 |
| 21 | 0.10 | 0.80 | 0.12 | 0.90 | 1 | Gm20522 | | 12 | 30 | 15 | 15 | 41 | 22 | 11 | 16 | 19 | 35 | 19 |
| 21 | -1.60 | 1.98 | -0.81 | 0.42 | 1 | Ryr1 | ryanodine receptor 1, skeletal muscle | 142 | 47 | 32 | 0 | 0 | 0 | 11 | 0 | 0 | 0 | 0 |
| 19 | -1.76 | 1.50 | -1.17 | 0.24 | 1 | Gm44168 | | 0 | 2 | 23 | 1 | 3 | 85 | 11 | 0 | 6 | 0 | 83 |
| 19 | -1.76 | 1.35 | -1.30 | 0.19 | 1 | Apol10b | apolipoprotein L 10B | 52 | 65 | 69 | 7 | 3 | 0 | 11 | 0 | 4 | 0 | 1 |
| 18 | 0.81 | 0.83 | 0.97 | 0.33 | 1 | Syt12 | synaptotagmin XII | 26 | 5 | 4 | 24 | 12 | 12 | 11 | 32 | 26 | 35 | 15 |
| 18 | -0.45 | 1.13 | -0.39 | 0.69 | 1 | Gm6226 | | 17 | 15 | 15 | 8 | 41 | 13 | 11 | 0 | 26 | 49 | 4 |
| 17 | -0.09 | 1.02 | -0.09 | 0.93 | 1 | Nupl1 | | 15 | 21 | 33 | 11 | 19 | 15 | 11 | 0 | 20 | 19 | 24 |
| 17 | -1.40 | 1.32 | -1.06 | 0.29 | 1 | A930033H14Ri | RIKEN cDNA A930033H14 gene | 28 | 67 | 36 | 0 | 22 | 9 | 11 | 0 | 6 | 3 | 8 |
| 16 | -0.03 | 1.09 | -0.03 | 0.98 | 1 | Gm17538 | | 15 | 47 | 11 | 13 | 14 | 8 | 11 | 1 | 5 | 41 | 12 |
| 16 | -0.28 | 1.19 | -0.24 | 0.81 | 1 | Gm10145 | | 21 | 42 | 8 | 30 | 12 | 5 | 11 | 0 | 8 | 33 | 6 |
| 16 | 0.97 | 0.96 | 1.01 | 0.31 | 1 | Gm47438 | | 6 | 15 | 0 | 9 | 9 | 14 | 11 | 43 | 10 | 31 | 22 |
| 15 | -1.72 | 0.96 | -1.79 | 0.07 | 1 | Prss16 | protease, serine 16 (thymus) | 6 | 14 | 45 | 20 | 36 | 8 | 11 | 6 | 14 | 0 | 7 |
| 15 | -1.20 | 1.29 | -0.93 | 0.35 | 1 | Gm47140 | | 1 | 11 | 0 | 19 | 32 | 23 | 11 | 13 | 16 | 0 | 34 |
| 14 | -0.70 | 1.11 | -0.63 | 0.53 | 1 | Lipo4 | lipase, member O4 | 15 | 9 | 7 | 30 | 20 | 12 | 11 | 17 | 13 | 0 | 19 |
| 14 | -1.10 | 1.16 | -0.95 | 0.34 | 1 | Ak7 | adenylate kinase 7 | 5 | 19 | 43 | 4 | 23 | 19 | 11 | 0 | 4 | 12 | 14 |
| 14 | 0.12 | 1.29 | 0.10 | 0.92 | 1 | Gm5129 | | 3 | 17 | 12 | 10 | 4 | 36 | 11 | 0 | 30 | 17 | 13 |
| 14 | -1.41 | 1.35 | -1.05 | 0.29 | 1 | Rtn4r | reticulon 4 receptor | 19 | 1 | 0 | 49 | 26 | 8 | 11 | 2 | 21 | 9 | 6 |
| 14 | -0.64 | 1.52 | -0.42 | 0.67 | 1 | Mpp4 | membrane protein, palmitoylated 4 (MAG | 0 | 3 | 10 | 0 | 8 | 68 | 11 | 0 | 3 | 9 | 40 |
| 14 | -0.77 | 1.41 | -0.55 | 0.58 | 1 | Gm19705 | | 44 | 18 | 4 | 0 | 19 | 18 | 11 | 0 | 3 | 15 | 16 |
| 13 | -0.46 | 1.15 | -0.40 | 0.69 | 1 | Gm29585 | | 7 | 4 | 3 | 18 | 30 | 11 | 11 | 0 | 25 | 18 | 17 |
| 13 | -0.75 | 1.34 | -0.56 | 0.57 | 1 | Gm18562 | | 0 | 0 | 15 | 16 | 19 | 29 | 11 | 2 | 7 | 18 | 24 |
| 13 | -0.67 | 1.41 | -0.48 | 0.63 | 1 | Gm42966 | | 3 | 0 | 0 | 18 | 10 | 16 | 11 | 0 | 16 | 59 | 7 |
| 13 | -0.37 | 1.62 | -0.23 | 0.82 | 1 | Gm49760 | | 32 | 30 | 9 | 4 | 0 | 12 | 11 | 37 | 0 | 0 | 5 |
| 13 | 0.57 | 1.32 | 0.43 | 0.67 | 1 | D730045B01Rik | | 4 | 29 | 12 | 4 | 3 | 22 | 11 | 0 | 10 | 32 | 11 |
| 12 | -0.16 | 1.28 | -0.12 | 0.90 | 1 | Gm43372 | | 1 | 2 | 0 | 10 | 29 | 8 | 11 | 35 | 31 | 0 | 8 |
| 12 | 0.41 | 1.36 | 0.30 | 0.77 | 1 | Rn7sk | | 1 | 9 | 12 | 9 | 9 | 10 | 11 | 0 | 19 | 46 | 6 |
| 12 | 0.50 | 2.37 | 0.21 | 0.83 | 1 | Tbx6 | T-box 6 | 0 | 0 | 9 | 0 | 0 | 35 | 11 | 37 | 3 | 0 | 37 |
| 12 | -0.06 | 0.94 | -0.06 | 0.95 | 1 | Gm6180 | | 23 | 24 | 21 | 11 | 4 | 7 | 11 | 5 | 14 | 5 | 7 |
| 12 | 0.12 | 1.35 | 0.09 | 0.93 | 1 | Gm5867 | | 8 | 7 | 2 | 4 | 11 | 9 | 11 | 63 | 9 | 0 | 5 |
| 11 | -3.02 | 1.99 | -1.52 | 0.13 | 1 | Fcer2a | Fc receptor, IgE, low affinity II, alpha polyp | 83 | 2 | 10 | 12 | 0 | 1 | 11 | 0 | 2 | 0 | 0 |
| 11 | -1.42 | 1.45 | -0.97 | 0.33 | 1 | Gm671 | | 1 | 13 | 16 | 15 | 1 | 32 | 11 | 0 | 18 | 3 | 10 |
| 11 | -2.02 | 1.85 | -1.09 | 0.27 | 1 | Gm24632 | | 0 | 0 | 0 | 24 | 13 | 17 | 11 | 0 | 13 | 0 | 41 |
| 11 | -2.17 | 1.65 | -1.31 | 0.19 | 1 | Tmem45b | transmembrane protein 45b | 24 | 33 | 33 | 4 | 0 | 4 | 11 | 0 | 1 | 0 | 6 |
| 10 | -1.36 | 1.14 | -1.19 | 0.23 | 1 | Gm25262 | | 21 | 15 | 11 | 24 | 7 | 6 | 11 | 13 | 3 | 0 | 4 |
| 10 | -1.31 | 2.43 | -0.54 | 0.59 | 1 | Ptger2 | prostaglandin E receptor 2 (subtype EP2) | 75 | 12 | 16 | 0 | 0 | 0 | 11 | 0 | 0 | 0 | 0 |
| 10 | 0.06 | 1.70 | 0.04 | 0.97 | 1 | Gm19265 | | 27 | 0 | 31 | 0 | 12 | 13 | 11 | 5 | 3 | 0 | 11 |
| 10 | -0.16 | 1.25 | -0.13 | 0.90 | 1 | Smarca5-ps | | 2 | 2 | 15 | 19 | 5 | 11 | 11 | 21 | 1 | 8 | 15 |
| 10 | -1.45 | 1.75 | -0.83 | 0.41 | 1 | Gm47015 | | 21 | 15 | 0 | 1 | 3 | 32 | 11 | 0 | 3 | 0 | 21 |
| 9 | -1.86 | 1.48 | -1.26 | 0.21 | 1 | Gm13071 | | 8 | 20 | 7 | 4 | 10 | 11 | 11 | 0 | 0 | 5 | 25 |
| 9 | 0.94 | 1.56 | 0.60 | 0.55 | 1 | Ascl4 | achaete-scute family bHLH transcription f | 5 | 6 | 15 | 1 | 1 | 8 | 11 | 39 | 3 | 0 | 12 |
| 9 | -1.66 | 1.40 | -1.19 | 0.24 | 1 | H2ac8 | H2A clustered histone 8 | 14 | 9 | 20 | 7 | 5 | 10 | 11 | 0 | 7 | 0 | 15 |
| 9 | -0.60 | 1.61 | -0.37 | 0.71 | 1 | Gm10766 | | 4 | 0 | 0 | 16 | 7 | 19 | 11 | 29 | 3 | 0 | 9 |
| 9 | -0.41 | 1.39 | -0.29 | 0.77 | 1 | Gm29642 | | 0 | 5 | 7 | 7 | 11 | 7 | 11 | 16 | 16 | 0 | 15 |
| 9 | -0.81 | 1.29 | -0.63 | 0.53 | 1 | Gm45889 | | 3 | 9 | 23 | 8 | 9 | 5 | 11 | 0 | 10 | 4 | 13 |
| 8 | -1.24 | 1.83 | -0.68 | 0.50 | 1 | Gm5087 | | 0 | 0 | 24 | 9 | 5 | 12 | 11 | 20 | 6 | 0 | 6 |
| 8 | -0.29 | 1.27 | -0.23 | 0.82 | 1 | Gm47392 | | 0 | 0 | 0 | 19 | 5 | 14 | 11 | 9 | 9 | 19 | 4 |
| 8 | -0.14 | 1.14 | -0.12 | 0.91 | 1 | Gm5963 | | 11 | 13 | 8 | 2 | 10 | 6 | 11 | 2 | 17 | 4 | 5 |
| 8 | -0.16 | 1.58 | -0.10 | 0.92 | 1 | Gm38160 | | 17 | 6 | 0 | 6 | 12 | 4 | 11 | 21 | 5 | 0 | 7 |
| 8 | -0.07 | 1.70 | -0.04 | 0.97 | 1 | 4930592C13Rik | | 2 | 8 | 1 | 10 | 0 | 9 | 11 | 0 | 0 | 15 | 30 |
| 8 | -0.58 | 2.16 | -0.27 | 0.79 | 1 | Gm31532 | | 0 | 12 | 0 | 23 | 0 | 12 | 11 | 0 | 0 | 9 | 17 |
| 7 | -2.36 | 2.05 | -1.15 | 0.25 | 1 | Gm5824 | | 55 | 0 | 0 | 4 | 0 | 5 | 11 | 0 | 0 | 5 | 1 |
| 7 | -0.35 | 1.88 | -0.19 | 0.85 | 1 | Gm14122 | | 8 | 16 | 0 | 0 | 9 | 8 | 11 | 13 | 5 | 0 | 11 |
| 7 | -0.02 | 1.36 | -0.01 | 0.99 | 1 | Gm19582 | | 0 | 9 | 9 | 4 | 8 | 7 | 11 | 15 | 3 | 9 | 5 |
| 7 | 0.42 | 1.77 | 0.24 | 0.81 | 1 | Meiob | meiosis specific with OB domains | 14 | 30 | 4 | 3 | 1 | 1 | 11 | 0 | 7 | 0 | 8 |
| 7 | -1.94 | 2.21 | -0.88 | 0.38 | 1 | A130051J06Rik | | 3 | 12 | 14 | 0 | 4 | 2 | 11 | 0 | 33 | 0 | 0 |
| 7 | -2.38 | 1.80 | -1.32 | 0.19 | 1 | Gpr3 | G-protein coupled receptor 3 | 12 | 8 | 20 | 10 | 0 | 8 | 11 | 0 | 4 | 0 | 5 |

| | | | | | | | | | | | | | | | | | | |
|---|---|---|---|---|---|---|---|---|---|---|---|---|---|---|---|---|---|---|
| 7 | -0.91 | 1.45 | -0.63 | 0.53 | 1 | H2-Q1 | histocompatibility 2, Q region locus 1 | 17 | 3 | 28 | 2 | 7 | 0 | 11 | 1 | 7 | 1 | 0 |
| 7 | -0.32 | 1.55 | -0.20 | 0.84 | 1 | Cby3 | | 0 | 0 | 2 | 6 | 11 | 7 | 11 | 25 | 1 | 1 | 10 |
| 7 | -1.40 | 1.68 | -0.83 | 0.41 | 1 | Gm6520 | | 17 | 6 | 18 | 11 | 2 | 0 | 11 | 0 | 1 | 4 | 5 |
| 6 | -2.57 | 1.87 | -1.37 | 0.17 | 1 | Gm7443 | | 0 | 14 | 17 | 6 | 4 | 8 | 11 | 0 | 7 | 0 | 4 |
| 6 | -0.94 | 1.76 | -0.53 | 0.59 | 1 | Gm4219 | | 0 | 0 | 4 | 3 | 5 | 16 | 11 | 0 | 1 | 19 | 10 |
| 6 | -2.83 | 2.41 | -1.18 | 0.24 | 1 | Gm37941 | | 3 | 0 | 0 | 0 | 5 | 33 | 11 | 0 | 0 | 0 | 16 |
| 6 | -1.77 | 2.18 | -0.81 | 0.42 | 1 | Gm15644 | | 0 | 0 | 6 | 0 | 3 | 28 | 11 | 0 | 3 | 0 | 17 |
| 6 | -0.87 | 2.81 | -0.31 | 0.76 | 1 | Gm44851 | | 0 | 0 | 31 | 0 | 4 | 3 | 11 | 0 | 0 | 18 | 0 |
| 6 | -3.02 | 1.97 | -1.53 | 0.13 | 1 | Rps13-ps4 | | 0 | 10 | 3 | 2 | 7 | 17 | 11 | 0 | 0 | 1 | 15 |
| 6 | -2.02 | 2.12 | -0.95 | 0.34 | 1 | Gm50315 | | 12 | 0 | 16 | 3 | 0 | 8 | 11 | 0 | 2 | 0 | 14 |
| 6 | -2.21 | 2.71 | -0.82 | 0.41 | 1 | Cbln2 | cerebellin 2 precursor protein | 31 | 7 | 12 | 0 | 0 | 0 | 11 | 0 | 0 | 0 | 4 |
| 6 | -3.36 | 2.90 | -1.16 | 0.25 | 1 | Gm36527 | | 0 | 0 | 0 | 0 | 7 | 21 | 11 | 0 | 0 | 0 | 26 |
| 6 | -1.71 | 2.22 | -0.77 | 0.44 | 1 | Gm12249 | | 0 | 0 | 0 | 19 | 5 | 6 | 11 | 0 | 14 | 0 | 9 |
| 6 | 0.52 | 2.29 | 0.23 | 0.82 | 1 | 4833417C18Rik | | 18 | 26 | 1 | 0 | 0 | 2 | 11 | 0 | 3 | 0 | 3 |
| 6 | -2.50 | 2.18 | -1.15 | 0.25 | 1 | A930038B10Rik | | 14 | 0 | 30 | 0 | 3 | 4 | 11 | 0 | 0 | 0 | 2 |
| 5 | -2.24 | 1.83 | -1.22 | 0.22 | 1 | Glod5 | glyoxalase domain containing 5 | 0 | 0 | 0 | 18 | 11 | 5 | 11 | 0 | 3 | 5 | 2 |
| 5 | -3.83 | 2.14 | -1.79 | 0.07 | 1 | Gm42031 | | 3 | 8 | 6 | 0 | 21 | 3 | 11 | 0 | 2 | 0 | 0 |
| 5 | -1.61 | 2.54 | -0.63 | 0.53 | 1 | 1700112D23Rik | | 0 | 7 | 0 | 13 | 0 | 13 | 11 | 0 | 4 | 0 | 6 |
| 5 | -1.31 | 3.23 | -0.41 | 0.68 | 1 | 6430584L05Rik | | 17 | 15 | 8 | 0 | 0 | 0 | 11 | 0 | 0 | 0 | 0 |
| 4 | 0.16 | 2.70 | 0.06 | 0.95 | 1 | Gm16099 | | 0 | 0 | 4 | 5 | 0 | 0 | 11 | 7 | 9 | 0 | 12 |
| 4 | -2.60 | 2.22 | -1.17 | 0.24 | 1 | Gast | gastrin | 4 | 5 | 8 | 2 | 0 | 7 | 11 | 0 | 0 | 0 | 9 |
| 4 | -4.56 | 2.58 | -1.77 | 0.08 | 1 | Cacng3 | calcium channel, voltage-dependent, gan | 0 | 10 | 0 | 11 | 10 | 1 | 11 | 0 | 3 | 0 | 0 |
| 4 | -3.56 | 3.14 | -1.13 | 0.26 | 1 | Six1 | sine oculis-related homeobox 1 | 0 | 3 | 22 | 0 | 0 | 7 | 11 | 0 | 2 | 0 | 0 |
| 4 | -1.03 | 2.20 | -0.47 | 0.64 | 1 | Gm16284 | | 10 | 0 | 10 | 0 | 1 | 4 | 11 | 0 | 6 | 0 | 1 |
| 4 | -1.10 | 2.36 | -0.47 | 0.64 | 1 | Gm7854 | | 9 | 18 | 1 | 0 | 0 | 2 | 11 | 0 | 1 | 0 | 1 |
| 4 | -4.35 | 2.96 | -1.47 | 0.14 | 1 | Gm15489 | | 0 | 0 | 0 | 8 | 8 | 1 | 11 | 0 | 0 | 0 | 14 |
| 4 | -3.35 | 2.87 | -1.17 | 0.24 | 1 | Smim18 | small integral membrane protein 18 | 0 | 0 | 0 | 15 | 4 | 2 | 11 | 0 | 10 | 0 | 0 |
| 4 | -1.90 | 2.12 | -0.90 | 0.37 | 1 | Gm47205 | | 3 | 12 | 0 | 9 | 0 | 2 | 11 | 3 | 0 | 0 | 1 |
| 3 | -0.19 | 2.30 | -0.08 | 0.93 | 1 | Gm12057 | | 6 | 5 | 0 | 1 | 0 | 3 | 11 | 10 | 0 | 0 | 2 |
| 3 | 0.39 | 1.98 | 0.19 | 0.85 | 1 | Gm6397 | | 0 | 2 | 7 | 1 | 0 | 3 | 11 | 3 | 2 | 3 | 5 |
| 3 | -1.18 | 3.49 | -0.34 | 0.73 | 1 | Gm37914 | | 1 | 0 | 0 | 0 | 0 | 13 | 11 | 0 | 0 | 0 | 12 |
| 3 | -3.47 | 3.21 | -1.08 | 0.28 | 1 | Nags | N-acetylglutamate synthase | 1 | 2 | 18 | 0 | 0 | 5 | 11 | 0 | 0 | 0 | 0 |
| 3 | -0.33 | 3.55 | -0.09 | 0.92 | 1 | Gm10727 | | 0 | 0 | 0 | 0 | 0 | 3 | 11 | 0 | 0 | 21 | 0 |
| 3 | -3.47 | 2.67 | -1.30 | 0.19 | 1 | 1700022I11Rik | RIKEN cDNA 1700022I11 gene | 0 | 1 | 8 | 4 | 0 | 2 | 11 | 0 | 1 | 0 | 1 |
| 3 | -3.37 | 3.31 | -1.02 | 0.31 | 1 | Gm6659 | | 6 | 7 | 0 | 1 | 3 | 0 | 11 | 0 | 0 | 0 | 0 |
| 3 | -1.65 | 2.25 | -0.73 | 0.46 | 1 | Rpl30-ps10 | | 0 | 1 | 1 | 4 | 0 | 2 | 11 | 2 | 5 | 1 | 0 |
| 2 | -2.80 | 3.58 | -0.78 | 0.43 | 1 | Pzp | PZP, alpha-2-macroglobulin like | 6 | 0 | 0 | 0 | 9 | 0 | 11 | 0 | 0 | 0 | 0 |
| 2 | -0.25 | 3.20 | -0.08 | 0.94 | 1 | Gm42970 | | 5 | 0 | 0 | 0 | 0 | 3 | 11 | 0 | 0 | 3 | 4 |
| 2 | -1.12 | 2.43 | -0.46 | 0.64 | 1 | Uqcrh-ps2 | | 1 | 1 | 2 | 1 | 3 | 0 | 11 | 2 | 0 | 0 | 4 |
| 2 | -3.00 | 3.52 | -0.85 | 0.39 | 1 | Vmn1r53 | vomeronasal 1 receptor 53 | 0 | 0 | 3 | 2 | 0 | 1 | 11 | 0 | 7 | 0 | 0 |
| 2 | -2.99 | 3.57 | -0.84 | 0.40 | 1 | Gm44746 | | 0 | 0 | 0 | 0 | 0 | 3 | 11 | 0 | 0 | 0 | 6 |
| 2 | -1.85 | 2.75 | -0.67 | 0.50 | 1 | Gm4828 | | 0 | 2 | 1 | 1 | 3 | 0 | 11 | 0 | 1 | 1 | 0 |
| 2 | -0.78 | 3.56 | -0.22 | 0.83 | 1 | Rpl31-ps12 | ribosomal protein L31, pseudogene 1 2 | 1 | 0 | 3 | 0 | 0 | 0 | 11 | 0 | 2 | 0 | 0 |
| 1 | -2.31 | 3.54 | -0.65 | 0.51 | 1 | 4933436E23Rik | | 0 | 0 | 0 | 2 | 0 | 0 | 11 | 0 | 0 | 0 | 3 |
| 1 | -0.01 | 3.57 | 0.00 | 1.00 | 1 | Gm15151 | | 0 | 0 | 0 | 0 | 0 | 0 | 11 | 0 | 2 | 1 | 0 |
| 965 | 0.14 | 1.38 | 0.10 | 0.92 | 1 | Stab1 | stabilin 1 | 4851 | 2396 | 3280 | 0 | 5 | 15 | 10 | 0 | 0 | 26 | 32 |
| 842 | -3.38 | 2.49 | -1.36 | 0.17 | 1 | Dysf | dysferlin | 3293 | 2933 | 3024 | 7 | 0 | 0 | 10 | 0 | 0 | 0 | 0 |
| 679 | -1.65 | 1.16 | -1.42 | 0.16 | 1 | 1810028F09Rik | | 724 | 3925 | 2580 | 169 | 9 | 1 | 10 | 34 | 3 | 0 | 9 |
| 626 | 0.25 | 1.66 | 0.15 | 0.88 | 1 | Plau | plasminogen activator, urokinase | 2412 | 1878 | 2569 | 0 | 3 | 0 | 10 | 0 | 0 | 14 | 2 |
| 479 | -1.29 | 3.62 | -0.36 | 0.72 | 1 | Col13a1 | collagen, type XIII, alpha 1 | 1560 | 1864 | 1832 | 0 | 0 | 0 | 10 | 0 | 0 | 0 | 0 |
| 454 | -1.82 | 1.41 | -1.30 | 0.19 | 1 | Maf | avian musculoaponeurotic fibrosarcoma ( | 2759 | 1104 | 1109 | 0 | 5 | 1 | 10 | 0 | 1 | 0 | 2 |
| 410 | -0.18 | 0.97 | -0.19 | 0.85 | 1 | Adcyap1r1 | adenylate cyclase activating polypeptide | 1769 | 1327 | 1310 | 12 | 10 | 8 | 10 | 0 | 8 | 54 | 5 |
| 397 | -1.01 | 1.32 | -0.77 | 0.44 | 1 | Lgals9 | lectin, galactose binding, soluble 9 | 1784 | 1034 | 1498 | 0 | 14 | 5 | 10 | 0 | 13 | 0 | 5 |
| 349 | 2.52 | 2.22 | 1.13 | 0.26 | 1 | Arhgdib | Rho, GDP dissociation inhibitor (GDI) bet: | 1993 | 961 | 832 | 0 | 0 | 1 | 10 | 30 | 0 | 0 | 10 |
| 318 | -1.22 | 3.62 | -0.34 | 0.74 | 1 | Lepr | leptin receptor | 1784 | 904 | 796 | 0 | 0 | 0 | 10 | 0 | 0 | 0 | 0 |
| 283 | -1.26 | 3.62 | -0.35 | 0.73 | 1 | Cd38 | CD38 antigen | 1278 | 957 | 863 | 0 | 0 | 0 | 10 | 0 | 0 | 0 | 0 |
| 269 | -0.81 | 1.12 | -0.72 | 0.47 | 1 | Ppp1r18 | protein phosphatase 1, regulatory subuni | 1386 | 902 | 631 | 4 | 4 | 4 | 10 | 13 | 0 | 0 | 5 |
| 256 | -1.23 | 3.62 | -0.34 | 0.73 | 1 | Rftn1 | raftlin lipid raft linker 1 | 1374 | 663 | 768 | 0 | 0 | 0 | 10 | 0 | 0 | 0 | 0 |
| 224 | -3.23 | 2.47 | -1.31 | 0.19 | 1 | Ccl11 | chemokine (C-C motif) ligand 11 | 860 | 723 | 865 | 5 | 0 | 0 | 10 | 0 | 0 | 0 | 0 |
| 211 | -1.31 | 0.73 | -1.78 | 0.07 | 1 | Hhatl | hedgehog acyltransferase-like | 850 | 595 | 812 | 8 | 9 | 17 | 10 | 0 | 10 | 0 | 12 |

| | | | | | | | | | | | | | | | | | | |
|---|---|---|---|---|---|---|---|---|---|---|---|---|---|---|---|---|---|---|
| 168 | -1.01 | 2.12 | -0.48 | 0.63 | 1 | Icam2 | intercellular adhesion molecule 2 | 666 | 497 | 673 | 0 | 0 | 1 | 10 | 0 | 0 | 0 | 3 |
| 150 | -2.21 | 2.42 | -0.91 | 0.36 | 1 | Afap1l2 | actin filament associated protein 1-like 2 | 637 | 436 | 566 | 0 | 1 | 0 | 10 | 0 | 0 | 0 | 0 |
| 146 | 1.51 | 1.94 | 0.78 | 0.44 | 1 | Tmem200b | transmembrane protein 200B | 484 | 501 | 539 | 0 | 0 | 3 | 10 | 0 | 2 | 68 | 0 |
| 144 | -0.56 | 0.63 | -0.88 | 0.38 | 1 | Blnk | B cell linker | 854 | 270 | 332 | 18 | 22 | 12 | 10 | 6 | 15 | 44 | 2 |
| 142 | -1.06 | 1.89 | -0.56 | 0.57 | 1 | Fgf2 | fibroblast growth factor 2 | 649 | 441 | 453 | 0 | 3 | 0 | 10 | 0 | 4 | 0 | 0 |
| 141 | -1.08 | 1.12 | -0.97 | 0.33 | 1 | Cd209f | CD209f antigen | 1062 | 205 | 268 | 0 | 1 | 1 | 10 | 0 | 0 | 0 | 5 |
| 119 | -1.17 | 3.62 | -0.32 | 0.75 | 1 | Timd4 | T cell immunoglobulin and mucin domain | 800 | 289 | 213 | 0 | 0 | 0 | 10 | 0 | 0 | 0 | 0 |
| 109 | 1.16 | 1.65 | 0.70 | 0.48 | 1 | Smagp | small cell adhesion glycoprotein | 412 | 382 | 379 | 0 | 0 | 0 | 10 | 0 | 0 | 14 | 3 |
| 95 | -0.17 | 0.69 | -0.24 | 0.81 | 1 | Rnf152 | ring finger protein 152 | 337 | 227 | 220 | 75 | 19 | 36 | 10 | 20 | 64 | 14 | 22 |
| 82 | -0.55 | 0.78 | -0.71 | 0.48 | 1 | Prom2 | prominin 2 | 276 | 342 | 235 | 4 | 5 | 9 | 10 | 13 | 2 | 0 | 7 |
| 64 | -0.52 | 0.68 | -0.76 | 0.45 | 1 | Zwilch | zwilch kinetochore protein | 102 | 70 | 74 | 83 | 70 | 103 | 10 | 16 | 41 | 43 | 91 |
| 63 | -1.52 | 1.41 | -1.08 | 0.28 | 1 | Pik3r6 | phosphoinositide-3-kinase regulatory sub | 245 | 195 | 243 | 0 | 0 | 1 | 10 | 0 | 0 | 3 | 0 |
| 60 | 0.64 | 0.68 | 0.94 | 0.35 | 1 | Gsta4 | glutathione S-transferase, alpha 4 | 248 | 175 | 104 | 18 | 11 | 8 | 10 | 59 | 11 | 6 | 13 |
| 53 | -0.26 | 0.80 | -0.33 | 0.74 | 1 | Snhg16 | | 63 | 79 | 98 | 3 | 26 | 123 | 10 | 0 | 24 | 8 | 146 |
| 51 | -0.53 | 2.34 | -0.23 | 0.82 | 1 | F2rl3 | coagulation factor II (thrombin) receptor-li | 195 | 129 | 187 | 0 | 0 | 6 | 10 | 37 | 0 | 0 | 2 |
| 51 | -0.79 | 0.83 | -0.95 | 0.34 | 1 | Ulbp1 | UL16 binding protein 1 | 185 | 80 | 90 | 104 | 18 | 12 | 10 | 10 | 13 | 14 | 23 |
| 42 | 0.79 | 0.60 | 1.32 | 0.19 | 1 | 1700028E10Rik | | 43 | 65 | 47 | 21 | 35 | 51 | 10 | 63 | 68 | 21 | 40 |
| 38 | -0.49 | 0.78 | -0.63 | 0.53 | 1 | Fam78a | family with sequence similarity 78, membe | 172 | 49 | 89 | 29 | 13 | 9 | 10 | 20 | 3 | 12 | 11 |
| 35 | 0.77 | 0.96 | 0.81 | 0.42 | 1 | Lag3 | lymphocyte-activation gene 3 | 187 | 73 | 31 | 7 | 3 | 16 | 10 | 44 | 11 | 4 | 4 |
| 35 | 0.26 | 1.11 | 0.23 | 0.81 | 1 | Dynlt1c | dynein light chain Tctex-type 1C | 15 | 112 | 21 | 33 | 6 | 88 | 10 | 0 | 32 | 18 | 50 |
| 34 | 0.54 | 0.66 | 0.82 | 0.41 | 1 | 4930505A04Ril | RIKEN cDNA 4930505A04 gene | 22 | 82 | 22 | 39 | 32 | 27 | 10 | 37 | 33 | 41 | 30 |
| 34 | -1.21 | 3.62 | -0.34 | 0.74 | 1 | Phf11b | PHD finger protein 11B | 190 | 72 | 100 | 0 | 0 | 0 | 10 | 0 | 0 | 0 | 0 |
| 33 | -1.08 | 0.90 | -1.20 | 0.23 | 1 | Gm15441 | | 37 | 49 | 82 | 34 | 30 | 53 | 10 | 0 | 17 | 14 | 42 |
| 31 | 0.16 | 1.17 | 0.13 | 0.89 | 1 | Nos1ap | nitric oxide synthase 1 (neuronal) adaptor | 55 | 119 | 98 | 0 | 7 | 11 | 10 | 17 | 6 | 0 | 15 |
| 30 | -1.14 | 1.05 | -1.08 | 0.28 | 1 | Rmi2 | RecQ mediated genome instability 2 | 69 | 43 | 14 | 42 | 55 | 28 | 10 | 24 | 16 | 0 | 30 |
| 30 | -2.35 | 1.25 | -1.88 | 0.06 | 1 | Csrnp3 | cysteine-serine-rich nuclear protein 3 | 100 | 64 | 56 | 61 | 4 | 14 | 10 | 0 | 6 | 0 | 12 |
| 30 | 0.24 | 1.43 | 0.17 | 0.87 | 1 | Rgs14 | regulator of G-protein signaling 14 | 172 | 48 | 76 | 0 | 0 | 7 | 10 | 0 | 5 | 0 | 10 |
| 30 | 1.10 | 0.75 | 1.47 | 0.14 | 1 | Sptbn5 | spectrin beta, non-erythrocytic 5 | 18 | 14 | 9 | 26 | 9 | 41 | 10 | 78 | 5 | 54 | 61 |
| 29 | -0.28 | 1.06 | -0.27 | 0.79 | 1 | Dnd1 | DND microRNA-mediated repression inhi | 9 | 75 | 43 | 8 | 10 | 53 | 10 | 5 | 57 | 0 | 48 |
| 28 | -0.37 | 3.48 | -0.11 | 0.91 | 1 | Gm43843 | | 0 | 0 | 0 | 0 | 0 | 168 | 10 | 0 | 5 | 0 | 129 |
| 28 | 0.35 | 0.82 | 0.43 | 0.67 | 1 | Zfp385b | zinc finger protein 385B | 89 | 95 | 64 | 4 | 8 | 3 | 10 | 19 | 3 | 5 | 8 |
| 27 | 0.75 | 0.80 | 0.95 | 0.34 | 1 | Gm13620 | | 14 | 22 | 63 | 42 | 9 | 15 | 10 | 38 | 37 | 32 | 15 |
| 26 | 0.25 | 1.31 | 0.19 | 0.85 | 1 | Ticrr | TOPBP1-interacting checkpoint and replic | 153 | 9 | 5 | 30 | 7 | 15 | 10 | 16 | 28 | 0 | 16 |
| 26 | -2.05 | 1.07 | -1.92 | 0.06 | 1 | Rims4 | regulating synaptic membrane exocytosis | 38 | 141 | 32 | 36 | 14 | 5 | 10 | 0 | 5 | 1 | 7 |
| 26 | -3.17 | 1.78 | -1.78 | 0.08 | 1 | 6430511E19Rik | | 9 | 16 | 0 | 0 | 4 | 119 | 10 | 0 | 0 | 0 | 128 |
| 25 | 0.63 | 0.64 | 0.99 | 0.32 | 1 | Tmem139 | transmembrane protein 139 | 9 | 30 | 24 | 42 | 12 | 19 | 10 | 52 | 20 | 19 | 30 |
| 24 | -1.22 | 1.21 | -1.01 | 0.31 | 1 | Gm43579 | | 19 | 94 | 47 | 15 | 16 | 18 | 10 | 0 | 29 | 0 | 20 |
| 24 | 0.35 | 0.82 | 0.43 | 0.67 | 1 | Psrc1 | proline/serine-rich coiled-coil 1 | 39 | 40 | 11 | 5 | 26 | 39 | 10 | 12 | 20 | 23 | 41 |
| 23 | 0.40 | 1.07 | 0.38 | 0.71 | 1 | Pcdhgc5 | protocadherin gamma subfamily C, 5 | 76 | 34 | 34 | 6 | 5 | 35 | 10 | 24 | 16 | 0 | 17 |
| 23 | -0.43 | 1.39 | -0.31 | 0.76 | 1 | Fhl3 | four and a half LIM domains 3 | 82 | 78 | 57 | 0 | 6 | 3 | 10 | 0 | 0 | 17 | 4 |
| 21 | -0.33 | 0.88 | -0.38 | 0.71 | 1 | Ankrd9 | ankyrin repeat domain 9 | 65 | 47 | 18 | 12 | 14 | 15 | 10 | 36 | 7 | 5 | 6 |
| 20 | 0.74 | 0.93 | 0.80 | 0.42 | 1 | Gm10232 | | 28 | 16 | 26 | 33 | 2 | 15 | 10 | 21 | 31 | 27 | 13 |
| 20 | 1.69 | 1.18 | 1.44 | 0.15 | 1 | Avil | advillin | 11 | 12 | 25 | 0 | 9 | 39 | 10 | 29 | 3 | 50 | 35 |
| 20 | 0.56 | 0.71 | 0.80 | 0.42 | 1 | Ankmy1 | ankyrin repeat and MYND domain contair | 22 | 14 | 17 | 11 | 23 | 26 | 10 | 27 | 17 | 19 | 37 |
| 20 | -1.05 | 1.41 | -0.74 | 0.46 | 1 | Haspin | histone H3 associated protein kinase | 63 | 25 | 15 | 21 | 7 | 25 | 10 | 0 | 36 | 0 | 18 |
| 20 | -0.47 | 1.00 | -0.47 | 0.64 | 1 | Rpl5-ps2 | | 45 | 44 | 36 | 31 | 2 | 14 | 10 | 3 | 17 | 5 | 11 |
| 19 | 1.10 | 1.08 | 1.01 | 0.31 | 1 | Gm12843 | | 6 | 2 | 30 | 7 | 16 | 16 | 10 | 80 | 10 | 6 | 26 |
| 19 | -1.62 | 1.16 | -1.40 | 0.16 | 1 | Sec31b | Sec31 homolog B (S. cerevisiae) | 23 | 36 | 19 | 9 | 10 | 50 | 10 | 0 | 10 | 0 | 41 |
| 18 | 0.21 | 1.29 | 0.16 | 0.87 | 1 | Srcin1 | SRC kinase signaling inhibitor 1 | 42 | 14 | 53 | 0 | 17 | 19 | 10 | 9 | 30 | 0 | 7 |
| 18 | -0.93 | 1.81 | -0.51 | 0.61 | 1 | Gm47863 | | 84 | 41 | 61 | 0 | 0 | 1 | 10 | 0 | 0 | 0 | 4 |
| 17 | -0.13 | 1.26 | -0.10 | 0.92 | 1 | Gm47817 | | 8 | 13 | 18 | 0 | 8 | 59 | 10 | 10 | 7 | 0 | 59 |
| 17 | -2.83 | 1.98 | -1.43 | 0.15 | 1 | Prdm5 | PR domain containing 5 | 45 | 72 | 62 | 0 | 0 | 2 | 10 | 0 | 0 | 0 | 0 |
| 17 | -0.15 | 1.60 | -0.10 | 0.92 | 1 | Zfp433 | zinc finger protein 433 | 23 | 0 | 33 | 25 | 7 | 8 | 10 | 57 | 1 | 0 | 26 |
| 17 | 0.64 | 0.92 | 0.69 | 0.49 | 1 | Lncppara | | 2 | 3 | 22 | 9 | 40 | 19 | 10 | 22 | 13 | 27 | 23 |
| 17 | 0.09 | 0.83 | 0.11 | 0.91 | 1 | Gm42918 | | 13 | 13 | 15 | 14 | 29 | 16 | 10 | 5 | 28 | 31 | 15 |
| 17 | 0.25 | 1.13 | 0.22 | 0.83 | 1 | Gstp-ps | | 38 | 75 | 16 | 5 | 8 | 3 | 10 | 13 | 8 | 0 | 13 |
| 17 | -1.74 | 1.30 | -1.34 | 0.18 | 1 | 4932438H23Ril | RIKEN cDNA 4932438H23 gene | 11 | 67 | 0 | 51 | 7 | 17 | 10 | 5 | 4 | 0 | 15 |
| 16 | 0.80 | 1.07 | 0.75 | 0.45 | 1 | Nrg2 | neuregulin 2 | 67 | 14 | 53 | 1 | 2 | 7 | 10 | 9 | 2 | 4 | 12 |
| 16 | 0.01 | 1.24 | 0.01 | 0.99 | 1 | Gm6545 | | 13 | 43 | 9 | 33 | 15 | 3 | 10 | 0 | 8 | 39 | 7 |
| 16 | -1.69 | 1.34 | -1.27 | 0.21 | 1 | Gm42829 | | 32 | 42 | 17 | 11 | 25 | 8 | 10 | 0 | 31 | 0 | 4 |

| | | | | | | | | | | | | | | | | | | |
|---|---|---|---|---|---|---|---|---|---|---|---|---|---|---|---|---|---|---|
| 16 | -0.03 | 1.11 | -0.03 | 0.98 | 1 | Gm9949 | predicted gene 9949 | 22 | 67 | 12 | 5 | 5 | 10 | 10 | 32 | 7 | 0 | 7 |
| 16 | -0.26 | 1.14 | -0.23 | 0.82 | 1 | Aipl1 | aryl hydrocarbon receptor-interacting prot | 30 | 20 | 23 | 18 | 4 | 15 | 10 | 0 | 14 | 6 | 32 |
| 16 | -0.30 | 0.95 | -0.31 | 0.76 | 1 | Gm14539 | | 21 | 4 | 25 | 26 | 20 | 8 | 10 | 7 | 12 | 14 | 24 |
| 15 | 0.18 | 1.33 | 0.14 | 0.89 | 1 | 9430069I07Rik | RIKEN cDNA 9430069I07 gene | 0 | 0 | 0 | 4 | 21 | 32 | 10 | 15 | 28 | 0 | 59 |
| 15 | 0.11 | 1.30 | 0.08 | 0.93 | 1 | Lncpint | | 6 | 12 | 14 | 0 | 6 | 68 | 10 | 0 | 9 | 15 | 26 |
| 14 | -1.47 | 1.56 | -0.94 | 0.35 | 1 | 4930509G22Rik | | 2 | 0 | 0 | 7 | 16 | 55 | 10 | 0 | 23 | 0 | 42 |
| 13 | -1.05 | 1.25 | -0.84 | 0.40 | 1 | Gm26876 | | 20 | 13 | 2 | 18 | 11 | 19 | 10 | 43 | 2 | 0 | 9 |
| 13 | -1.42 | 1.38 | -1.03 | 0.30 | 1 | 2610020C07Rik | | 3 | 18 | 17 | 15 | 15 | 15 | 10 | 37 | 0 | 0 | 16 |
| 13 | -1.88 | 1.81 | -1.04 | 0.30 | 1 | Gm13613 | | 6 | 0 | 0 | 17 | 31 | 23 | 10 | 0 | 21 | 0 | 36 |
| 13 | -0.48 | 1.52 | -0.31 | 0.75 | 1 | Krt77 | keratin 77 | 30 | 46 | 5 | 0 | 3 | 12 | 10 | 27 | 1 | 0 | 7 |
| 13 | 0.38 | 0.90 | 0.42 | 0.68 | 1 | Gm8566 | | 27 | 8 | 9 | 9 | 16 | 7 | 10 | 10 | 20 | 8 | 14 |
| 12 | -0.95 | 1.37 | -0.70 | 0.49 | 1 | Gm14634 | | 13 | 3 | 31 | 21 | 5 | 9 | 10 | 0 | 2 | 14 | 28 |
| 12 | -1.18 | 1.46 | -0.81 | 0.42 | 1 | Ube2v1 | ubiquitin-conjugating enzyme E2 variant | 31 | 16 | 3 | 6 | 15 | 15 | 10 | 0 | 16 | 0 | 23 |
| 12 | -0.25 | 1.19 | -0.21 | 0.83 | 1 | Rnasek | ribonuclease, RNase K | 10 | 9 | 6 | 19 | 2 | 25 | 10 | 0 | 6 | 14 | 34 |
| 12 | -0.88 | 1.50 | -0.59 | 0.56 | 1 | Gm17276 | | 0 | 0 | 11 | 10 | 6 | 30 | 10 | 0 | 20 | 10 | 31 |
| 12 | 1.19 | 1.31 | 0.91 | 0.36 | 1 | Gm38137 | | 3 | 6 | 12 | 0 | 9 | 11 | 10 | 1 | 6 | 52 | 18 |
| 11 | -1.94 | 1.57 | -1.24 | 0.22 | 1 | Gm48796 | | 2 | 0 | 0 | 3 | 15 | 36 | 10 | 0 | 5 | 0 | 54 |
| 11 | -1.50 | 2.06 | -0.73 | 0.47 | 1 | Gm42467 | | 4 | 3 | 0 | 0 | 0 | 58 | 10 | 0 | 0 | 0 | 47 |
| 11 | -1.62 | 1.87 | -0.87 | 0.39 | 1 | Gm37844 | | 50 | 6 | 44 | 2 | 0 | 1 | 10 | 0 | 9 | 0 | 0 |
| 11 | -1.62 | 1.57 | -1.03 | 0.30 | 1 | Kcnt1 | potassium channel, subfamily T, member | 15 | 24 | 23 | 4 | 0 | 21 | 10 | 0 | 3 | 0 | 19 |
| 11 | -1.35 | 1.94 | -0.69 | 0.49 | 1 | Gm47585 | | 21 | 6 | 32 | 0 | 0 | 20 | 10 | 0 | 0 | 0 | 29 |
| 11 | -0.32 | 1.04 | -0.31 | 0.75 | 1 | Gm6316 | | 15 | 17 | 16 | 4 | 10 | 10 | 10 | 1 | 19 | 6 | 6 |
| 11 | -1.21 | 1.34 | -0.90 | 0.37 | 1 | Gm43823 | | 2 | 30 | 7 | 15 | 10 | 12 | 10 | 5 | 0 | 21 | 4 |
| 10 | -0.39 | 1.63 | -0.24 | 0.81 | 1 | Gm31749 | | 3 | 10 | 0 | 6 | 0 | 35 | 10 | 0 | 0 | 12 | 38 |
| 10 | -2.28 | 2.00 | -1.14 | 0.25 | 1 | Slc25a31 | solute carrier family 25 (mitochondrial car | 16 | 0 | 0 | 0 | 26 | 39 | 10 | 0 | 2 | 0 | 18 |
| 10 | -3.31 | 2.17 | -1.52 | 0.13 | 1 | Gm44645 | | 36 | 18 | 34 | 9 | 0 | 0 | 10 | 0 | 0 | 0 | 2 |
| 10 | -0.55 | 0.93 | -0.59 | 0.56 | 1 | Gm7816 | | 8 | 5 | 2 | 11 | 5 | 29 | 10 | 5 | 5 | 6 | 20 |
| 10 | -0.08 | 0.98 | -0.08 | 0.94 | 1 | Gm10157 | | 9 | 2 | 3 | 17 | 10 | 10 | 10 | 12 | 13 | 6 | 13 |
| 10 | 1.09 | 1.55 | 0.70 | 0.48 | 1 | 1700016K19Ril | RIKEN cDNA 1700016K19 gene | 0 | 3 | 11 | 0 | 17 | 8 | 10 | 24 | 4 | 18 | 9 |
| 9 | -0.65 | 1.23 | -0.53 | 0.60 | 1 | Gm42517 | predicted gene 42517 | 34 | 11 | 4 | 7 | 8 | 4 | 10 | 0 | 9 | 14 | 2 |
| 9 | -1.01 | 1.54 | -0.66 | 0.51 | 1 | Gm10752 | | 18 | 0 | 6 | 1 | 13 | 12 | 10 | 0 | 21 | 21 | 0 |
| 9 | -0.27 | 1.59 | -0.17 | 0.87 | 1 | Gm30698 | | 24 | 0 | 6 | 22 | 0 | 11 | 10 | 12 | 3 | 3 | 10 |
| 9 | -0.58 | 1.20 | -0.48 | 0.63 | 1 | Ccdc22 | coiled-coil domain containing 22 | 5 | 13 | 5 | 9 | 10 | 14 | 10 | 0 | 9 | 9 | 14 |
| 9 | 1.16 | 1.21 | 0.96 | 0.34 | 1 | Gm5874 | | 22 | 1 | 25 | 0 | 5 | 2 | 10 | 1 | 5 | 22 | 4 |
| 9 | 0.75 | 1.39 | 0.54 | 0.59 | 1 | Gm13140 | | 0 | 0 | 3 | 4 | 9 | 7 | 10 | 14 | 10 | 39 | 1 |
| 9 | -2.49 | 1.77 | -1.41 | 0.16 | 1 | Gm37307 | | 6 | 13 | 42 | 9 | 0 | 6 | 10 | 0 | 4 | 0 | 6 |
| 9 | -0.79 | 1.54 | -0.51 | 0.61 | 1 | Kctd19 | potassium channel tetramerisation domai | 3 | 0 | 0 | 16 | 19 | 14 | 10 | 17 | 9 | 0 | 8 |
| 8 | 1.22 | 1.07 | 1.14 | 0.25 | 1 | Psma5-ps | | 6 | 1 | 16 | 2 | 5 | 3 | 10 | 6 | 19 | 21 | 4 |
| 8 | 1.04 | 1.42 | 0.74 | 0.46 | 1 | Gm2810 | | 22 | 21 | 1 | 0 | 1 | 7 | 10 | 1 | 3 | 14 | 10 |
| 8 | -1.55 | 1.55 | -1.01 | 0.31 | 1 | Gm8188 | | 13 | 1 | 18 | 6 | 5 | 8 | 10 | 0 | 11 | 0 | 15 |
| 8 | -0.64 | 2.48 | -0.26 | 0.79 | 1 | Gm47258 | | 0 | 2 | 0 | 0 | 0 | 31 | 10 | 12 | 0 | 0 | 29 |
| 8 | 0.19 | 1.46 | 0.13 | 0.90 | 1 | Gm23119 | | 12 | 9 | 6 | 0 | 3 | 13 | 10 | 0 | 3 | 10 | 17 |
| 8 | -1.40 | 1.89 | -0.74 | 0.46 | 1 | Gm45307 | | 0 | 2 | 0 | 6 | 16 | 12 | 10 | 0 | 25 | 0 | 11 |
| 7 | -0.02 | 1.43 | -0.02 | 0.99 | 1 | Gm49651 | | 0 | 0 | 0 | 11 | 7 | 6 | 10 | 2 | 2 | 28 | 15 |
| 7 | -1.49 | 1.92 | -0.77 | 0.44 | 1 | Slc15a1 | solute carrier family 15 (oligopeptide trans | 14 | 26 | 3 | 0 | 12 | 3 | 10 | 0 | 10 | 0 | 2 |
| 7 | -4.00 | 2.18 | -1.83 | 0.07 | 1 | Gm47113 | | 0 | 31 | 18 | 1 | 7 | 6 | 10 | 0 | 0 | 0 | 2 |
| 7 | -1.33 | 1.91 | -0.70 | 0.49 | 1 | Gm6254 | | 3 | 0 | 0 | 4 | 11 | 11 | 10 | 0 | 9 | 0 | 25 |
| 7 | -0.43 | 1.93 | -0.22 | 0.82 | 1 | 4933425D22Rik | | 0 | 9 | 4 | 0 | 9 | 14 | 10 | 0 | 3 | 14 | 10 |
| 7 | 0.44 | 1.84 | 0.24 | 0.81 | 1 | 4930548J01Rik | | 0 | 0 | 0 | 12 | 1 | 7 | 10 | 0 | 9 | 24 | 9 |
| 7 | 1.31 | 1.71 | 0.77 | 0.44 | 1 | A930006L05Rik | | 0 | 0 | 3 | 0 | 10 | 2 | 10 | 25 | 12 | 8 | 2 |
| 6 | 1.47 | 2.37 | 0.62 | 0.54 | 1 | Gm15681 | | 0 | 0 | 0 | 0 | 0 | 16 | 10 | 0 | 5 | 30 | 9 |
| 6 | -0.54 | 1.87 | -0.29 | 0.77 | 1 | Alox12b | arachidonate 12-lipoxygenase, 12R type | 0 | 0 | 0 | 0 | 23 | 8 | 10 | 0 | 17 | 10 | 2 |
| 6 | 1.26 | 1.71 | 0.74 | 0.46 | 1 | Gm34307 | | 0 | 0 | 0 | 1 | 1 | 5 | 10 | 32 | 4 | 14 | 1 |
| 6 | -0.84 | 1.13 | -0.74 | 0.46 | 1 | Gm12009 | | 5 | 3 | 14 | 4 | 8 | 7 | 10 | 5 | 6 | 3 | 3 |
| 6 | -3.39 | 2.03 | -1.67 | 0.10 | 1 | Ly6g6d | lymphocyte antigen 6 complex, locus G6D | 7 | 18 | 18 | 0 | 8 | 3 | 10 | 0 | 2 | 0 | 0 |
| 6 | -2.37 | 1.82 | -1.30 | 0.19 | 1 | Gm38408 | | 0 | 8 | 1 | 6 | 5 | 9 | 10 | 17 | 0 | 0 | 7 |
| 5 | -1.33 | 1.51 | -0.88 | 0.38 | 1 | Gm6457 | | 8 | 9 | 4 | 1 | 10 | 3 | 10 | 5 | 3 | 0 | 2 |
| 5 | -4.34 | 3.36 | -1.29 | 0.20 | 1 | Nup210l | nucleoporin 210-like | 0 | 0 | 15 | 0 | 0 | 22 | 10 | 0 | 0 | 0 | 8 |
| 5 | -1.14 | 1.90 | -0.60 | 0.55 | 1 | Gm13605 | | 0 | 0 | 0 | 12 | 9 | 3 | 10 | 2 | 11 | 0 | 6 |
| 5 | -3.21 | 1.99 | -1.61 | 0.11 | 1 | Gm18134 | | 0 | 0 | 5 | 7 | 14 | 3 | 10 | 0 | 8 | 0 | 3 |

| | | | | | | | | | | | | | | | | | | |
|---|---|---|---|---|---|---|---|---|---|---|---|---|---|---|---|---|---|---|
| 4 | -1.21 | 2.28 | -0.53 | 0.59 | 1 | 9530056K15Rik | | 12 | 5 | 0 | 0 | 3 | 10 | 10 | 0 | 4 | 0 | 4 |
| 4 | -1.04 | 2.23 | -0.47 | 0.64 | 1 | Gm5637 | | 9 | 2 | 17 | 0 | 1 | 1 | 10 | 0 | 8 | 0 | 0 |
| 4 | -0.51 | 2.07 | -0.25 | 0.81 | 1 | Gm40709 | | 0 | 0 | 3 | 4 | 1 | 4 | 10 | 21 | 2 | 0 | 3 |
| 4 | -0.81 | 2.72 | -0.30 | 0.77 | 1 | Gm14705 | | 0 | 9 | 0 | 0 | 2 | 4 | 10 | 0 | 0 | 15 | 6 |
| 4 | -0.54 | 2.01 | -0.27 | 0.79 | 1 | Mmel1 | membrane metallo-endopeptidase-like 1 | 6 | 8 | 2 | 0 | 6 | 1 | 10 | 0 | 6 | 8 | 0 |
| 4 | -4.66 | 3.07 | -1.52 | 0.13 | 1 | 9930004E17Rik | | 3 | 0 | 20 | 12 | 0 | 1 | 10 | 0 | 0 | 0 | 0 |
| 4 | -0.55 | 2.57 | -0.21 | 0.83 | 1 | Gm3435 | predicted gene 3435 | 1 | 10 | 0 | 0 | 7 | 1 | 10 | 0 | 15 | 0 | 2 |
| 4 | -0.49 | 3.39 | -0.15 | 0.88 | 1 | Gm47728 | | 0 | 0 | 0 | 0 | 0 | 14 | 10 | 13 | 0 | 0 | 9 |
| 4 | -5.27 | 3.01 | -1.75 | 0.08 | 1 | Gm34248 | | 0 | 8 | 0 | 5 | 0 | 4 | 10 | 0 | 0 | 0 | 17 |
| 4 | 1.43 | 3.37 | 0.43 | 0.67 | 1 | Gm12676 | | 0 | 17 | 0 | 0 | 0 | 3 | 10 | 0 | 5 | 0 | 9 |
| 4 | -1.99 | 2.06 | -0.96 | 0.33 | 1 | Gm12168 | | 1 | 8 | 0 | 11 | 3 | 3 | 10 | 0 | 3 | 0 | 2 |
| 4 | -4.05 | 3.37 | -1.20 | 0.23 | 1 | Gm47141 | | 0 | 0 | 0 | 0 | 14 | 12 | 10 | 0 | 0 | 0 | 4 |
| 4 | -1.37 | 1.79 | -0.76 | 0.45 | 1 | Gm18859 | | 8 | 8 | 1 | 8 | 0 | 1 | 10 | 1 | 0 | 1 | 2 |
| 3 | 0.53 | 3.39 | 0.16 | 0.88 | 1 | Gm47405 | | 0 | 0 | 0 | 0 | 0 | 3 | 10 | 0 | 0 | 21 | 4 |
| 3 | -2.57 | 3.59 | -0.72 | 0.47 | 1 | Fzd9 | frizzled class receptor 9 | 12 | 0 | 12 | 0 | 0 | 2 | 10 | 0 | 0 | 0 | 0 |
| 3 | -4.30 | 2.88 | -1.49 | 0.14 | 1 | Gm15762 | | 0 | 0 | 0 | 11 | 11 | 1 | 10 | 1 | 0 | 0 | 0 |
| 3 | -1.52 | 3.55 | -0.43 | 0.67 | 1 | Gm48639 | | 0 | 0 | 0 | 0 | 0 | 14 | 10 | 0 | 0 | 0 | 10 |
| 3 | -1.55 | 2.71 | -0.57 | 0.57 | 1 | Tcfl5 | transcription factor-like 5 (basic helix-loop | 10 | 5 | 3 | 0 | 0 | 3 | 10 | 0 | 0 | 0 | 2 |
| 3 | -4.52 | 3.28 | -1.38 | 0.17 | 1 | Mir8116 | | 0 | 12 | 3 | 0 | 5 | 3 | 10 | 0 | 0 | 0 | 0 |
| 3 | -1.50 | 2.07 | -0.72 | 0.47 | 1 | Gm4824 | | 1 | 0 | 4 | 7 | 1 | 1 | 10 | 2 | 1 | 0 | 6 |
| 3 | 3.38 | 3.02 | 1.12 | 0.26 | 1 | BC048644 | cDNA sequence BC048644 | 0 | 0 | 8 | 0 | 2 | 0 | 10 | 5 | 2 | 0 | 6 |
| 3 | -1.68 | 2.56 | -0.66 | 0.51 | 1 | Gm7079 | | 0 | 4 | 0 | 4 | 0 | 5 | 10 | 1 | 7 | 0 | 1 |
| 3 | -0.68 | 3.54 | -0.19 | 0.85 | 1 | Gm45797 | | 3 | 0 | 0 | 0 | 0 | 7 | 10 | 0 | 0 | 0 | 13 |
| 3 | -2.67 | 3.53 | -0.76 | 0.45 | 1 | Gm45464 | | 0 | 0 | 0 | 0 | 2 | 9 | 10 | 0 | 0 | 0 | 11 |
| 3 | -0.94 | 2.03 | -0.46 | 0.64 | 1 | Gm5550 | | 3 | 1 | 2 | 7 | 1 | 0 | 10 | 1 | 2 | 0 | 4 |
| 3 | -2.89 | 3.58 | -0.81 | 0.42 | 1 | Gm31621 | | 9 | 7 | 0 | 0 | 0 | 0 | 10 | 0 | 0 | 0 | 5 |
| 3 | -2.46 | 3.14 | -0.78 | 0.43 | 1 | Cd27 | CD27 antigen | 14 | 3 | 2 | 0 | 0 | 3 | 10 | 0 | 0 | 0 | 0 |
| 3 | -0.04 | 3.15 | -0.01 | 0.99 | 1 | 1700018M17Rik | | 0 | 5 | 0 | 0 | 1 | 2 | 10 | 0 | 8 | 0 | 3 |
| 3 | -0.12 | 3.32 | -0.03 | 0.97 | 1 | Slc24a5 | solute carrier family 24, member 5 | 0 | 1 | 0 | 0 | 6 | 0 | 10 | 0 | 10 | 0 | 2 |
| 3 | -1.54 | 2.31 | -0.67 | 0.50 | 1 | Gm47273 | | 6 | 4 | 0 | 0 | 4 | 1 | 10 | 0 | 1 | 1 | 1 |
| 3 | -1.29 | 2.34 | -0.55 | 0.58 | 1 | Crabp1 | cellular retinoic acid binding protein I | 0 | 0 | 0 | 4 | 3 | 2 | 10 | 0 | 3 | 1 | 5 |
| 2 | -1.91 | 2.56 | -0.75 | 0.45 | 1 | Slc6a20b | solute carrier family 6 (neurotransmitter tra | 0 | 1 | 0 | 1 | 1 | 5 | 10 | 0 | 8 | 0 | 1 |
| 2 | -1.40 | 2.35 | -0.59 | 0.55 | 1 | Gm48100 | | 0 | 0 | 0 | 3 | 5 | 2 | 10 | 1 | 3 | 0 | 3 |
| 2 | -3.57 | 3.57 | -1.00 | 0.32 | 1 | BC065403 | | 3 | 0 | 9 | 0 | 0 | 1 | 10 | 0 | 0 | 0 | 2 |
| 2 | -6.28 | 3.55 | -1.77 | 0.08 | 1 | Fam163b | family with sequence similarity 163, meml | 0 | 0 | 11 | 1 | 0 | 0 | 10 | 0 | 0 | 0 | 2 |
| 2 | -0.74 | 3.54 | -0.21 | 0.83 | 1 | Il20ra | interleukin 20 receptor, alpha | 0 | 0 | 0 | 0 | 4 | 0 | 10 | 0 | 7 | 0 | 0 |
| 2 | -1.65 | 2.58 | -0.64 | 0.52 | 1 | Gm6594 | | 3 | 1 | 0 | 1 | 0 | 3 | 10 | 0 | 0 | 1 | 1 |
| 2 | -2.89 | 3.55 | -0.82 | 0.41 | 1 | Gm11457 | | 0 | 0 | 0 | 0 | 5 | 2 | 10 | 1 | 0 | 0 | 1 |
| 2 | -2.82 | 3.57 | -0.79 | 0.43 | 1 | Barhl2 | BarH like homeobox 2 | 0 | 0 | 0 | 7 | 0 | 0 | 10 | 0 | 1 | 0 | 0 |
| 2 | -0.77 | 3.58 | -0.21 | 0.83 | 1 | Gm37551 | | 5 | 0 | 0 | 0 | 0 | 1 | 10 | 0 | 0 | 0 | 1 |
| 1 | -1.15 | 3.61 | -0.32 | 0.75 | 1 | Tas1r2 | taste receptor, type 1, member 2 | 4 | 2 | 0 | 0 | 0 | 0 | 10 | 0 | 0 | 0 | 0 |
| 1 | 0.20 | 3.61 | 0.06 | 0.96 | 1 | Hpd | 4-hydroxyphenylpyruvic acid dioxygenas | 5 | 0 | 0 | 0 | 0 | 0 | 10 | 0 | 1 | 0 | 0 |
| 1 | -0.35 | 3.54 | -0.10 | 0.92 | 1 | 9530046B11Rik | | 0 | 0 | 0 | 0 | 0 | 2 | 10 | 0 | 1 | 0 | 3 |
| 1 | -3.21 | 3.56 | -0.90 | 0.37 | 1 | Gm43124 | | 0 | 0 | 0 | 1 | 4 | 0 | 10 | 0 | 0 | 0 | 1 |
| 1 | -3.27 | 3.58 | -0.92 | 0.36 | 1 | Gm45601 | | 0 | 1 | 0 | 0 | 0 | 4 | 10 | 0 | 0 | 0 | 0 |
| 1 | -3.31 | 3.57 | -0.92 | 0.35 | 1 | Gm47541 | | 0 | 0 | 0 | 0 | 0 | 5 | 10 | 0 | 0 | 0 | 0 |
| 1 | -0.90 | 3.60 | -0.25 | 0.80 | 1 | Gm29019 | | 0 | 0 | 0 | 0 | 0 | 1 | 10 | 0 | 0 | 4 | 0 |
| 1 | -2.11 | 3.57 | -0.59 | 0.56 | 1 | Dnaaf1 | dynein, axonemal assembly factor 1 | 0 | 1 | 0 | 0 | 0 | 3 | 10 | 0 | 0 | 0 | 1 |
| 1 | -2.62 | 3.60 | -0.73 | 0.47 | 1 | Gm5997 | | 1 | 0 | 0 | 0 | 0 | 3 | 10 | 0 | 0 | 0 | 0 |
| 1 | -2.98 | 3.59 | -0.83 | 0.41 | 1 | Gm48883 | | 0 | 1 | 0 | 0 | 0 | 3 | 10 | 0 | 0 | 0 | 0 |
| 1 | -3.07 | 3.58 | -0.86 | 0.39 | 1 | Mir7116 | | 0 | 0 | 0 | 0 | 0 | 3 | 10 | 0 | 0 | 0 | 0 |
| 1 | 0.13 | 3.58 | 0.04 | 0.97 | 1 | Cps1 | carbamoyl-phosphate synthetase 1 | 1 | 0 | 0 | 0 | 0 | 0 | 10 | 0 | 2 | 0 | 0 |
| 1 | -2.24 | 3.61 | -0.62 | 0.53 | 1 | Gm37203 | | 0 | 0 | 0 | 0 | 0 | 1 | 10 | 0 | 0 | 0 | 0 |
| 9910 | -2.41 | 1.80 | -1.34 | 0.18 | 1 | Dcn | decorin | 62276 | 23829 | 22880 | 14 | 0 | 0 | 9 | 0 | 0 | 3 | 0 |
| 5620 | -0.01 | 1.86 | -0.01 | 0.99 | 1 | Rgs5 | regulator of G-protein signaling 5 | 23964 | 18999 | 18647 | 0 | 31 | 25 | 9 | 0 | 76 | 0 | 65 |
| 2311 | -1.40 | 1.01 | -1.39 | 0.17 | 1 | Pdia2 | protein disulfide isomerase associated 2 | 3034 | 12153 | 9474 | 450 | 40 | 48 | 9 | 66 | 23 | 0 | 125 |
| 1579 | -1.28 | 3.63 | -0.35 | 0.73 | 1 | Cdh5 | cadherin 5 | 5365 | 5962 | 6033 | 0 | 0 | 0 | 9 | 0 | 0 | 0 | 0 |
| 950 | -0.64 | 1.56 | -0.41 | 0.68 | 1 | Acvrl1 | activin A receptor, type II-like 1 | 3779 | 3039 | 3570 | 31 | 0 | 0 | 9 | 19 | 1 | 0 | 1 |
| 815 | -1.67 | 1.30 | -1.28 | 0.20 | 1 | Cxcl13 | chemokine (C-X-C motif) ligand 13 | 5874 | 2935 | 100 | 13 | 3 | 19 | 9 | 0 | 5 | 6 | 1 |
| 554 | -0.19 | 1.45 | -0.13 | 0.90 | 1 | Tinagl1 | tubulointerstitial nephritis antigen-like 1 | 2296 | 1774 | 2003 | 0 | 2 | 2 | 9 | 0 | 4 | 0 | 7 |

| | | | | | | | | | | | | | | | | | | |
|---|---|---|---|---|---|---|---|---|---|---|---|---|---|---|---|---|---|---|
| 383 | -0.62 | 2.73 | -0.23 | 0.82 | 1 | Flrt2 | fibronectin leucine rich transmembrane pı | 2201 | 1222 | 784 | 0 | 0 | 0 | 9 | 0 | 0 | 0 | 1 |
| 313 | 0.19 | 3.60 | 0.05 | 0.96 | 1 | Igkv9-129 | | 3432 | 0 | 0 | 0 | 0 | 0 | 9 | 0 | 0 | 0 | 0 |
| 313 | -1.36 | 1.79 | -0.76 | 0.45 | 1 | Ptgs1 | prostaglandin-endoperoxide synthase 1 | 1643 | 674 | 1106 | 0 | 0 | 3 | 9 | 0 | 0 | 0 | 4 |
| 271 | -3.42 | 2.03 | -1.68 | 0.09 | 1 | Hecw2 | HECT, C2 and WW domain containing E3 | 941 | 925 | 1081 | 8 | 0 | 7 | 9 | 0 | 11 | 0 | 0 |
| 263 | 0.50 | 0.65 | 0.77 | 0.44 | 1 | Mdfic | MyoD family inhibitor domain containing | 1138 | 656 | 1022 | 9 | 12 | 3 | 9 | 29 | 13 | 0 | 6 |
| 250 | -0.56 | 2.42 | -0.23 | 0.82 | 1 | Myct1 | myc target 1 | 952 | 792 | 996 | 0 | 0 | 0 | 9 | 0 | 0 | 3 | 0 |
| 189 | -0.51 | 1.43 | -0.35 | 0.72 | 1 | Adamts9 | a disintegrin-like and metallopeptidase (r | 625 | 816 | 516 | 27 | 0 | 37 | 9 | 23 | 22 | 0 | 5 |
| 175 | -0.36 | 0.47 | -0.78 | 0.44 | 1 | Nos3 | nitric oxide synthase 3, endothelial cell | 572 | 502 | 714 | 25 | 30 | 11 | 9 | 24 | 12 | 10 | 10 |
| 164 | 0.65 | 0.99 | 0.66 | 0.51 | 1 | Spn | sialophorin | 1116 | 408 | 257 | 0 | 0 | 2 | 9 | 0 | 3 | 14 | 0 |
| 158 | 0.32 | 0.41 | 0.77 | 0.44 | 1 | Gm43305 | predicted gene 43305 | 51 | 321 | 258 | 214 | 218 | 71 | 9 | 283 | 196 | 15 | 104 |
| 147 | -2.74 | 2.08 | -1.32 | 0.19 | 1 | Gimap8 | GTPase, IMAP family member 8 | 447 | 617 | 541 | 0 | 0 | 4 | 9 | 0 | 1 | 0 | 0 |
| 129 | 0.70 | 0.90 | 0.78 | 0.44 | 1 | Loxl3 | lysyl oxidase-like 3 | 576 | 349 | 399 | 0 | 26 | 4 | 9 | 0 | 33 | 13 | 12 |
| 128 | -0.88 | 2.06 | -0.43 | 0.67 | 1 | Spats2l | spermatogenesis associated, serine-rich | 533 | 455 | 392 | 0 | 6 | 0 | 9 | 0 | 9 | 0 | 0 |
| 127 | 0.25 | 0.73 | 0.34 | 0.73 | 1 | Cdc42ep2 | CDC42 effector protein (Rho GTPase bin | 329 | 486 | 409 | 17 | 22 | 13 | 9 | 74 | 23 | 0 | 10 |
| 114 | -0.64 | 2.88 | -0.22 | 0.82 | 1 | Gjc1 | gap junction protein, gamma 1 | 520 | 337 | 385 | 0 | 0 | 0 | 9 | 0 | 0 | 0 | 1 |
| 92 | -0.33 | 0.70 | -0.47 | 0.64 | 1 | Clspn | claspin | 333 | 102 | 145 | 96 | 78 | 61 | 9 | 51 | 56 | 24 | 56 |
| 89 | 0.57 | 0.92 | 0.62 | 0.54 | 1 | Vstm4 | V-set and transmembrane domain contair | 339 | 327 | 261 | 0 | 8 | 4 | 9 | 12 | 14 | 0 | 5 |
| 84 | -3.00 | 2.41 | -1.24 | 0.21 | 1 | Rem1 | rad and gem related GTP binding protein | 332 | 326 | 249 | 0 | 4 | 0 | 9 | 0 | 0 | 0 | 0 |
| 84 | -0.95 | 1.29 | -0.74 | 0.46 | 1 | Fas | Fas (TNF receptor superfamily member 6) | 531 | 197 | 169 | 0 | 3 | 0 | 9 | 0 | 0 | 1 | 9 |
| 83 | -1.56 | 0.87 | -1.79 | 0.07 | 1 | Cacna2d4 | calcium channel, voltage-dependent, alpl | 268 | 254 | 290 | 6 | 27 | 23 | 9 | 0 | 15 | 0 | 18 |
| 82 | -0.13 | 0.73 | -0.17 | 0.86 | 1 | Slc2a4rg-ps | | 16 | 49 | 4 | 15 | 30 | 403 | 9 | 8 | 30 | 12 | 328 |
| 78 | 0.24 | 0.66 | 0.36 | 0.72 | 1 | Cdt1 | chromatin licensing and DNA replication f | 232 | 109 | 162 | 62 | 40 | 49 | 9 | 23 | 67 | 62 | 43 |
| 78 | -2.24 | 3.02 | -0.74 | 0.46 | 1 | Cd300ld | CD300 molecule like family member d | 465 | 200 | 179 | 0 | 0 | 0 | 9 | 0 | 0 | 0 | 4 |
| 68 | -2.92 | 2.30 | -1.27 | 0.21 | 1 | Cd69 | CD69 antigen | 444 | 150 | 145 | 0 | 0 | 4 | 9 | 0 | 0 | 0 | 0 |
| 68 | -0.02 | 0.89 | -0.02 | 0.98 | 1 | Fgf11 | fibroblast growth factor 11 | 190 | 119 | 267 | 8 | 33 | 31 | 9 | 48 | 12 | 0 | 32 |
| 58 | -0.60 | 0.83 | -0.71 | 0.47 | 1 | Aldh1l1 | aldehyde dehydrogenase 1 family, memb | 132 | 229 | 214 | 11 | 14 | 3 | 9 | 0 | 7 | 6 | 13 |
| 54 | -0.88 | 0.75 | -1.18 | 0.24 | 1 | Gm44210 | | 136 | 153 | 235 | 14 | 9 | 11 | 9 | 15 | 6 | 0 | 6 |
| 50 | 1.04 | 0.77 | 1.35 | 0.18 | 1 | Itgae | integrin alpha E, epithelial-associated | 233 | 129 | 120 | 1 | 3 | 8 | 9 | 14 | 2 | 17 | 9 |
| 44 | -2.79 | 1.53 | -1.83 | 0.07 | 1 | Cd4 | CD4 antigen | 258 | 135 | 79 | 0 | 0 | 2 | 9 | 0 | 0 | 0 | 0 |
| 43 | -0.28 | 1.39 | -0.20 | 0.84 | 1 | Flrt3 | fibronectin leucine rich transmembrane pı | 229 | 86 | 136 | 0 | 0 | 7 | 9 | 0 | 8 | 0 | 3 |
| 43 | -0.36 | 1.45 | -0.25 | 0.80 | 1 | 2310001H17Rik | | 104 | 156 | 175 | 0 | 0 | 14 | 9 | 9 | 0 | 0 | 8 |
| 43 | -0.15 | 0.60 | -0.24 | 0.81 | 1 | 1700029I15Rik | RIKEN cDNA 1700029I15 gene | 49 | 42 | 39 | 32 | 40 | 94 | 9 | 24 | 19 | 21 | 101 |
| 42 | -0.89 | 1.11 | -0.80 | 0.42 | 1 | Itga10 | integrin, alpha 10 | 44 | 49 | 31 | 3 | 0 | 155 | 9 | 0 | 3 | 0 | 165 |
| 41 | 0.63 | 0.69 | 0.91 | 0.36 | 1 | Rhod | ras homolog family member D | 117 | 59 | 128 | 22 | 18 | 8 | 9 | 43 | 26 | 8 | 13 |
| 40 | 0.53 | 1.08 | 0.49 | 0.63 | 1 | Pole2 | polymerase (DNA directed), epsilon 2 (p5 | 86 | 61 | 56 | 41 | 8 | 32 | 9 | 0 | 33 | 79 | 30 |
| 37 | 0.71 | 0.89 | 0.80 | 0.43 | 1 | Cdca7 | cell division cycle associated 7 | 207 | 87 | 37 | 7 | 3 | 8 | 9 | 0 | 11 | 22 | 15 |
| 35 | -1.07 | 0.95 | -1.13 | 0.26 | 1 | Nradd | neurotrophin receptor associated death d | 98 | 125 | 39 | 12 | 20 | 40 | 9 | 0 | 5 | 22 | 20 |
| 35 | 0.14 | 0.98 | 0.14 | 0.89 | 1 | Gm8325 | | 67 | 53 | 30 | 46 | 25 | 24 | 9 | 1 | 41 | 70 | 20 |
| 34 | -2.62 | 1.54 | -1.70 | 0.09 | 1 | Pstpip1 | proline-serine-threonine phosphatase-int | 260 | 42 | 67 | 0 | 0 | 2 | 9 | 0 | 0 | 0 | 0 |
| 34 | 0.05 | 1.54 | 0.03 | 0.98 | 1 | Sema3a | sema domain, immunoglobulin domain (Ig | 202 | 125 | 11 | 0 | 1 | 6 | 9 | 0 | 11 | 0 | 4 |
| 30 | 0.64 | 0.70 | 0.92 | 0.36 | 1 | Cks2 | CDC28 protein kinase regulatory subunit | 70 | 37 | 23 | 23 | 18 | 26 | 9 | 23 | 31 | 45 | 23 |
| 30 | -1.80 | 1.33 | -1.36 | 0.17 | 1 | Gm36989 | | 5 | 0 | 11 | 11 | 3 | 130 | 9 | 0 | 13 | 0 | 146 |
| 29 | -0.07 | 1.02 | -0.07 | 0.95 | 1 | Car15 | carbonic anhydrase 15 | 38 | 0 | 6 | 19 | 43 | 68 | 9 | 23 | 21 | 37 | 57 |
| 29 | -0.19 | 0.70 | -0.26 | 0.79 | 1 | Cdh24 | cadherin-like 24 | 7 | 16 | 22 | 25 | 15 | 96 | 9 | 37 | 6 | 8 | 80 |
| 28 | 1.59 | 0.98 | 1.62 | 0.11 | 1 | C4bp | complement component 4 binding proteir | 48 | 61 | 75 | 2 | 4 | 13 | 9 | 67 | 8 | 21 | 6 |
| 28 | -0.61 | 1.07 | -0.57 | 0.57 | 1 | C230037L18Rik | | 49 | 29 | 24 | 12 | 11 | 62 | 9 | 0 | 27 | 1 | 85 |
| 28 | 0.24 | 1.27 | 0.19 | 0.85 | 1 | Gm45110 | | 3 | 13 | 9 | 0 | 18 | 111 | 9 | 0 | 9 | 26 | 112 |
| 27 | -1.17 | 1.29 | -0.90 | 0.37 | 1 | Pde9a | phosphodiesterase 9A | 88 | 108 | 72 | 6 | 0 | 4 | 9 | 1 | 9 | 0 | 1 |
| 27 | -0.91 | 0.80 | -1.14 | 0.25 | 1 | Gm7628 | | 21 | 35 | 53 | 47 | 44 | 15 | 9 | 38 | 8 | 3 | 17 |
| 26 | 0.78 | 0.61 | 1.27 | 0.20 | 1 | Rpl39-ps | | 33 | 36 | 53 | 15 | 26 | 12 | 9 | 32 | 28 | 22 | 19 |
| 26 | -1.81 | 1.12 | -1.62 | 0.11 | 1 | Pcdhb7 | protocadherin beta 7 | 79 | 85 | 48 | 8 | 23 | 13 | 9 | 0 | 14 | 0 | 8 |
| 25 | -0.86 | 3.54 | -0.24 | 0.81 | 1 | Tgfbr3l | transforming growth factor, beta receptor I | 3 | 0 | 0 | 0 | 0 | 128 | 9 | 0 | 0 | 0 | 135 |
| 23 | 0.49 | 1.48 | 0.33 | 0.74 | 1 | BB557941 | | 0 | 34 | 56 | 48 | 13 | 12 | 9 | 0 | 17 | 44 | 23 |
| 23 | -0.85 | 1.08 | -0.79 | 0.43 | 1 | Bcas1os1 | | 55 | 96 | 55 | 4 | 7 | 6 | 9 | 17 | 5 | 0 | 1 |
| 23 | -1.24 | 0.99 | -1.26 | 0.21 | 1 | Gm9530 | | 15 | 12 | 16 | 32 | 22 | 62 | 9 | 0 | 10 | 10 | 65 |
| 23 | 0.08 | 1.16 | 0.07 | 0.94 | 1 | Gm47798 | | 1 | 0 | 17 | 2 | 5 | 107 | 9 | 0 | 14 | 22 | 76 |
| 23 | -1.33 | 1.49 | -0.89 | 0.37 | 1 | Tspan1 | tetraspanin 1 | 12 | 104 | 91 | 0 | 7 | 8 | 9 | 0 | 13 | 0 | 4 |
| 21 | 0.00 | 0.88 | 0.00 | 1.00 | 1 | Agbl2 | ATP/GTP binding protein-like 2 | 39 | 18 | 3 | 26 | 11 | 42 | 9 | 29 | 15 | 10 | 24 |
| 21 | -0.68 | 1.48 | -0.46 | 0.65 | 1 | Gm44167 | | 3 | 14 | 0 | 0 | 6 | 94 | 9 | 6 | 5 | 0 | 89 |
| 20 | -1.61 | 1.57 | -1.02 | 0.31 | 1 | Fbp2 | fructose bisphosphatase 2 | 0 | 18 | 35 | 32 | 17 | 8 | 9 | 100 | 0 | 0 | 4 |

| | | | | | | | | | | | | | | | | | |
|---|---|---|---|---|---|---|---|---|---|---|---|---|---|---|---|---|---|
| 20 | 0.25 | 1.27 | 0.20 | 0.84 | 1 | Nuggc | nuclear GTPase, germinal center associa | 92 | 55 | 32 | 4 | 0 | 10 | 9 | 7 | 9 | 0 | 4 |
| 20 | 1.05 | 1.14 | 0.92 | 0.36 | 1 | Fetub | fetuin beta | 61 | 12 | 46 | 7 | 9 | 6 | 9 | 34 | 16 | 0 | 16 |
| 19 | 0.56 | 0.86 | 0.65 | 0.51 | 1 | 1700108N11Rik | | 0 | 0 | 0 | 41 | 26 | 14 | 9 | 31 | 42 | 19 | 25 |
| 19 | -1.06 | 1.05 | -1.01 | 0.31 | 1 | Gm15408 | | 18 | 26 | 6 | 32 | 38 | 21 | 9 | 15 | 20 | 0 | 19 |
| 18 | 0.67 | 0.73 | 0.93 | 0.35 | 1 | Rps6-ps4 | | 27 | 30 | 32 | 18 | 9 | 6 | 9 | 17 | 19 | 24 | 7 |
| 18 | 0.91 | 1.14 | 0.80 | 0.43 | 1 | Gm17344 | | 3 | 24 | 0 | 9 | 11 | 14 | 9 | 27 | 5 | 75 | 21 |
| 18 | 0.38 | 0.81 | 0.47 | 0.64 | 1 | Gm10254 | | 17 | 10 | 11 | 20 | 14 | 20 | 9 | 42 | 7 | 12 | 34 |
| 17 | 0.38 | 1.72 | 0.22 | 0.83 | 1 | Cngb1 | cyclic nucleotide gated channel beta 1 | 7 | 42 | 56 | 10 | 12 | 0 | 9 | 0 | 0 | 55 | 1 |
| 17 | -1.32 | 1.09 | -1.21 | 0.23 | 1 | Wfikkn1 | WAP, FS, Ig, KU, and NTR-containing pro | 0 | 18 | 10 | 17 | 37 | 30 | 9 | 29 | 20 | 0 | 17 |
| 17 | 0.20 | 1.15 | 0.17 | 0.86 | 1 | Gm10800 | | 18 | 2 | 22 | 10 | 8 | 18 | 9 | 0 | 12 | 63 | 25 |
| 17 | 1.25 | 0.85 | 1.46 | 0.14 | 1 | Tpd52-ps | | 3 | 16 | 14 | 25 | 10 | 5 | 9 | 43 | 12 | 33 | 16 |
| 17 | 0.17 | 1.04 | 0.16 | 0.87 | 1 | 4930520O04Rik | | 15 | 0 | 21 | 18 | 22 | 24 | 9 | 21 | 21 | 17 | 19 |
| 17 | -3.33 | 2.34 | -1.42 | 0.15 | 1 | Gm49396 | | 0 | 5 | 0 | 0 | 0 | 102 | 9 | 0 | 0 | 0 | 67 |
| 16 | 0.57 | 0.76 | 0.75 | 0.45 | 1 | Gm42884 | | 20 | 28 | 8 | 11 | 9 | 16 | 9 | 20 | 7 | 22 | 25 |
| 16 | 0.96 | 0.91 | 1.05 | 0.29 | 1 | Gm32687 | predicted gene, 32687 | 3 | 10 | 5 | 11 | 9 | 10 | 9 | 68 | 19 | 3 | 24 |
| 16 | -0.08 | 1.09 | -0.07 | 0.94 | 1 | Gm43570 | | 7 | 0 | 4 | 30 | 34 | 11 | 9 | 35 | 20 | 5 | 16 |
| 15 | 0.95 | 0.90 | 1.05 | 0.29 | 1 | Gm45925 | | 3 | 11 | 12 | 18 | 6 | 16 | 9 | 19 | 8 | 35 | 31 |
| 15 | -1.25 | 1.00 | -1.25 | 0.21 | 1 | Gm5555 | | 27 | 11 | 22 | 29 | 11 | 24 | 9 | 6 | 18 | 1 | 9 |
| 15 | -0.10 | 1.94 | -0.05 | 0.96 | 1 | Cerox1 | | 58 | 31 | 22 | 0 | 0 | 11 | 9 | 0 | 0 | 30 | 0 |
| 14 | 0.02 | 1.06 | 0.02 | 0.98 | 1 | Gm7600 | | 31 | 21 | 17 | 4 | 33 | 3 | 9 | 2 | 7 | 22 | 9 |
| 14 | 0.02 | 1.19 | 0.02 | 0.99 | 1 | Gm20655 | | 7 | 0 | 0 | 22 | 20 | 11 | 9 | 27 | 5 | 33 | 22 |
| 14 | -2.68 | 1.59 | -1.69 | 0.09 | 1 | Gm43080 | | 1 | 5 | 8 | 0 | 8 | 79 | 9 | 0 | 1 | 0 | 44 |
| 14 | -1.57 | 1.41 | -1.11 | 0.27 | 1 | Gm37352 | | 25 | 19 | 23 | 0 | 4 | 39 | 9 | 0 | 2 | 0 | 31 |
| 13 | -0.50 | 1.81 | -0.28 | 0.78 | 1 | Gm20518 | | 0 | 0 | 0 | 0 | 26 | 42 | 9 | 17 | 8 | 0 | 44 |
| 13 | -1.14 | 1.30 | -0.88 | 0.38 | 1 | Rxfp2 | relaxin/insulin-like family peptide recepto | 9 | 0 | 44 | 8 | 22 | 14 | 9 | 0 | 10 | 23 | 9 |
| 13 | -0.41 | 1.53 | -0.27 | 0.79 | 1 | Gm6705 | | 21 | 2 | 0 | 17 | 16 | 19 | 9 | 45 | 5 | 0 | 13 |
| 13 | -0.05 | 1.31 | -0.04 | 0.97 | 1 | Nnt | nicotinamide nucleotide transhydrogenas | 22 | 1 | 3 | 4 | 13 | 45 | 9 | 7 | 21 | 0 | 19 |
| 13 | -0.47 | 1.29 | -0.37 | 0.71 | 1 | Ccdc87 | coiled-coil domain containing 87 | 21 | 45 | 4 | 3 | 12 | 14 | 9 | 13 | 16 | 0 | 5 |
| 13 | 1.22 | 1.62 | 0.75 | 0.45 | 1 | | | 2 | 0 | 0 | 0 | 0 | 57 | 9 | 0 | 8 | 8 | 59 |
| 12 | 1.64 | 1.65 | 0.99 | 0.32 | 1 | Gm34590 | | 11 | 0 | 0 | 5 | 5 | 21 | 9 | 34 | 18 | 0 | 30 |
| 12 | 0.12 | 0.91 | 0.14 | 0.89 | 1 | BC037039 | | 33 | 25 | 16 | 4 | 9 | 7 | 9 | 6 | 11 | 8 | 6 |
| 12 | 1.10 | 1.16 | 0.94 | 0.35 | 1 | 9930120I10Rik | | 7 | 18 | 24 | 7 | 0 | 10 | 9 | 25 | 8 | 8 | 14 |
| 12 | -2.22 | 1.27 | -1.75 | 0.08 | 1 | Gm26885 | | 12 | 24 | 36 | 4 | 13 | 15 | 9 | 0 | 11 | 0 | 4 |
| 12 | -1.01 | 1.26 | -0.80 | 0.42 | 1 | Gm17334 | | 9 | 24 | 11 | 13 | 17 | 11 | 9 | 0 | 3 | 26 | 6 |
| 12 | 0.54 | 0.87 | 0.62 | 0.53 | 1 | Bnip3l-ps | | 17 | 8 | 12 | 12 | 6 | 9 | 9 | 7 | 16 | 17 | 15 |
| 12 | 0.28 | 0.83 | 0.34 | 0.73 | 1 | Gm28661 | | 6 | 12 | 5 | 13 | 12 | 13 | 9 | 13 | 18 | 12 | 14 |
| 11 | 2.01 | 1.37 | 1.47 | 0.14 | 1 | Cdsn | corneodesmosin | 2 | 0 | 0 | 11 | 0 | 10 | 9 | 27 | 29 | 28 | 10 |
| 11 | -0.48 | 1.01 | -0.48 | 0.63 | 1 | Selenok-ps8 | | 8 | 3 | 8 | 17 | 19 | 13 | 9 | 8 | 23 | 1 | 16 |
| 11 | -2.56 | 1.86 | -1.38 | 0.17 | 1 | Gm11529 | | 7 | 0 | 0 | 24 | 27 | 25 | 9 | 0 | 12 | 0 | 20 |
| 11 | -1.50 | 1.29 | -1.17 | 0.24 | 1 | Lexm | lymphocyte expansion molecule | 9 | 0 | 8 | 11 | 17 | 22 | 9 | 0 | 19 | 19 | 6 |
| 11 | -1.04 | 1.41 | -0.74 | 0.46 | 1 | Depdc1a | DEP domain containing 1a | 0 | 0 | 0 | 12 | 24 | 30 | 9 | 0 | 13 | 13 | 19 |
| 11 | -0.47 | 1.53 | -0.30 | 0.76 | 1 | Gm47936 | | 0 | 0 | 0 | 12 | 22 | 4 | 9 | 28 | 42 | 0 | 1 |
| 11 | -0.50 | 1.28 | -0.40 | 0.69 | 1 | Gm12791 | | 9 | 16 | 20 | 1 | 20 | 12 | 9 | 0 | 6 | 18 | 7 |
| 10 | -0.82 | 1.20 | -0.68 | 0.50 | 1 | Gm37027 | | 13 | 11 | 20 | 8 | 7 | 17 | 9 | 0 | 10 | 18 | 3 |
| 10 | -0.24 | 1.40 | -0.17 | 0.86 | 1 | Gm10175 | | 6 | 20 | 10 | 26 | 0 | 7 | 9 | 16 | 6 | 0 | 13 |
| 10 | 0.60 | 1.58 | 0.38 | 0.70 | 1 | D7Ertd128e | | 25 | 38 | 0 | 2 | 5 | 3 | 9 | 0 | 7 | 18 | 6 |
| 10 | 1.47 | 1.70 | 0.87 | 0.39 | 1 | 4930470P17Rik | | 0 | 7 | 0 | 10 | 7 | 6 | 9 | 0 | 33 | 33 | 8 |
| 10 | -0.99 | 1.35 | -0.73 | 0.46 | 1 | Ccdc190 | coiled-coil domain containing 190 | 32 | 0 | 5 | 8 | 10 | 6 | 9 | 0 | 7 | 28 | 7 |
| 10 | -0.17 | 1.76 | -0.10 | 0.92 | 1 | Gm11934 | | 0 | 18 | 17 | 20 | 0 | 3 | 9 | 27 | 6 | 0 | 13 |
| 10 | -1.24 | 1.15 | -1.08 | 0.28 | 1 | Gm10575 | | 22 | 11 | 26 | 8 | 12 | 4 | 9 | 0 | 3 | 13 | 3 |
| 10 | 0.81 | 1.73 | 0.47 | 0.64 | 1 | Mak | male germ cell-associated kinase | 33 | 14 | 23 | 0 | 0 | 8 | 9 | 0 | 1 | 14 | 9 |
| 10 | -0.97 | 2.37 | -0.41 | 0.68 | 1 | Gm38104 | | 4 | 0 | 0 | 0 | 0 | 50 | 9 | 0 | 0 | 0 | 45 |
| 10 | -0.94 | 1.97 | -0.48 | 0.63 | 1 | Frmpd3 | FERM and PDZ domain containing 3 | 19 | 32 | 27 | 12 | 0 | 0 | 9 | 0 | 5 | 0 | 3 |
| 9 | -2.85 | 1.91 | -1.49 | 0.14 | 1 | Gm10478 | | 18 | 5 | 0 | 18 | 0 | 30 | 9 | 0 | 0 | 0 | 23 |
| 9 | -0.14 | 1.29 | -0.11 | 0.91 | 1 | Gm5242 | | 12 | 5 | 21 | 0 | 12 | 14 | 9 | 0 | 9 | 14 | 7 |
| 9 | 1.34 | 1.85 | 0.73 | 0.47 | 1 | H4c1 | H4 clustered histone 1 | 3 | 0 | 19 | 0 | 2 | 19 | 9 | 17 | 10 | 0 | 23 |
| 9 | 1.11 | 1.57 | 0.70 | 0.48 | 1 | Gm2991 | | 18 | 23 | 5 | 0 | 2 | 5 | 9 | 24 | 7 | 0 | 6 |
| 9 | -0.77 | 1.54 | -0.50 | 0.62 | 1 | Gm45112 | | 0 | 12 | 12 | 30 | 8 | 4 | 9 | 0 | 9 | 5 | 10 |
| 9 | -0.43 | 1.38 | -0.31 | 0.75 | 1 | Adam4 | a disintegrin and metallopeptidase domai | 7 | 0 | 0 | 26 | 1 | 24 | 9 | 10 | 3 | 5 | 14 |
| 9 | -1.92 | 1.57 | -1.22 | 0.22 | 1 | Trim17 | tripartite motif-containing 17 | 0 | 2 | 0 | 12 | 24 | 8 | 9 | 22 | 1 | 0 | 20 |

| | | | | | | | | | | | | | | | | | | |
|---|---|---|---|---|---|---|---|---|---|---|---|---|---|---|---|---|---|---|
| 9 | -0.46 | 1.39 | -0.33 | 0.74 | 1 | Gm36236 | | 2 | 0 | 6 | 16 | 9 | 11 | 9 | 15 | 16 | 0 | 12 |
| 9 | -0.50 | 1.38 | -0.36 | 0.72 | 1 | Gm17661 | | 4 | 7 | 32 | 2 | 4 | 12 | 9 | 0 | 5 | 19 | 2 |
| 9 | -0.97 | 1.68 | -0.57 | 0.57 | 1 | Hif3a | hypoxia inducible factor 3, alpha subunit | 5 | 53 | 0 | 4 | 3 | 8 | 9 | 1 | 6 | 0 | 6 |
| 8 | -0.11 | 1.12 | -0.09 | 0.93 | 1 | Rps15-ps2 | | 9 | 17 | 15 | 1 | 10 | 7 | 9 | 3 | 8 | 4 | 9 |
| 8 | -2.35 | 2.18 | -1.08 | 0.28 | 1 | Bcan | brevican | 31 | 0 | 32 | 7 | 4 | 0 | 9 | 0 | 8 | 0 | 0 |
| 8 | -0.91 | 1.29 | -0.71 | 0.48 | 1 | Sbk3 | SH3 domain binding kinase family, memb | 7 | 19 | 5 | 4 | 8 | 17 | 9 | 0 | 5 | 10 | 5 |
| 8 | -1.48 | 2.01 | -0.74 | 0.46 | 1 | 2810455O05Rik | | 0 | 13 | 0 | 22 | 6 | 14 | 9 | 0 | 11 | 0 | 11 |
| 8 | -0.63 | 1.65 | -0.38 | 0.70 | 1 | 2810442N19Rik | | 4 | 1 | 15 | 7 | 0 | 15 | 9 | 0 | 3 | 22 | 9 |
| 8 | -0.08 | 1.33 | -0.06 | 0.95 | 1 | Gm5903 | | 3 | 0 | 11 | 7 | 3 | 10 | 9 | 1 | 8 | 17 | 15 |
| 8 | -1.73 | 1.52 | -1.14 | 0.25 | 1 | Gabrd | gamma-aminobutyric acid (GABA) A rece | 2 | 4 | 5 | 10 | 9 | 15 | 9 | 0 | 0 | 8 | 21 |
| 7 | 1.64 | 1.45 | 1.13 | 0.26 | 1 | Snhg9 | | 4 | 0 | 2 | 3 | 0 | 14 | 9 | 15 | 9 | 5 | 20 |
| 7 | -1.21 | 2.22 | -0.54 | 0.59 | 1 | C130080G10Rik | | 0 | 0 | 0 | 4 | 0 | 29 | 9 | 0 | 0 | 5 | 28 |
| 7 | -2.66 | 2.48 | -1.07 | 0.28 | 1 | Hsd3b7 | hydroxy-delta-5-steroid dehydrogenase, 3 | 0 | 0 | 5 | 0 | 7 | 27 | 9 | 0 | 0 | 0 | 27 |
| 7 | -0.90 | 1.61 | -0.56 | 0.58 | 1 | Gemin4 | gem nuclear organelle associated proteir | 9 | 0 | 7 | 4 | 14 | 12 | 9 | 10 | 7 | 0 | 2 |
| 7 | -0.46 | 1.31 | -0.35 | 0.73 | 1 | Gm9294 | | 15 | 24 | 2 | 4 | 5 | 3 | 9 | 1 | 6 | 3 | 3 |
| 7 | -1.20 | 1.24 | -0.97 | 0.33 | 1 | Gm5620 | | 12 | 13 | 7 | 12 | 3 | 4 | 9 | 1 | 3 | 1 | 7 |
| 7 | -0.44 | 1.20 | -0.37 | 0.71 | 1 | Rps12-ps10 | | 9 | 6 | 20 | 3 | 12 | 1 | 9 | 2 | 5 | 3 | 4 |
| 6 | -3.08 | 2.49 | -1.24 | 0.22 | 1 | Tsx | testis specific X-linked gene | 0 | 0 | 2 | 0 | 9 | 29 | 9 | 0 | 0 | 0 | 20 |
| 6 | -1.09 | 1.98 | -0.55 | 0.58 | 1 | Gm43800 | | 1 | 8 | 7 | 10 | 0 | 5 | 9 | 0 | 0 | 5 | 23 |
| 6 | 1.29 | 1.88 | 0.69 | 0.49 | 1 | Wee2 | WEE1 homolog 2 (S. pombe) | 21 | 1 | 0 | 0 | 0 | 9 | 9 | 0 | 3 | 19 | 4 |
| 6 | -3.27 | 2.18 | -1.50 | 0.13 | 1 | Samd11 | sterile alpha motif domain containing 11 | 21 | 5 | 28 | 0 | 1 | 3 | 9 | 0 | 0 | 0 | 0 |
| 6 | -0.59 | 1.52 | -0.39 | 0.70 | 1 | Gm49896 | | 1 | 2 | 3 | 0 | 9 | 17 | 9 | 0 | 3 | 8 | 13 |
| 6 | 2.82 | 1.87 | 1.51 | 0.13 | 1 | Gm49708 | | 9 | 0 | 0 | 0 | 0 | 7 | 9 | 5 | 4 | 23 | 7 |
| 6 | -0.44 | 1.89 | -0.23 | 0.82 | 1 | Gm10120 | | 3 | 21 | 2 | 1 | 1 | 8 | 9 | 0 | 10 | 0 | 8 |
| 6 | 2.03 | 2.75 | 0.74 | 0.46 | 1 | Gm49486 | | 0 | 0 | 0 | 0 | 0 | 5 | 9 | 34 | 10 | 0 | 5 |
| 6 | 0.86 | 1.72 | 0.50 | 0.62 | 1 | Gm3571 | | 18 | 0 | 0 | 5 | 4 | 2 | 9 | 10 | 6 | 1 | 5 |
| 6 | 0.33 | 2.22 | 0.15 | 0.88 | 1 | 1700104B16Rik | | 7 | 0 | 0 | 0 | 8 | 5 | 9 | 22 | 6 | 4 | 0 |
| 5 | -2.04 | 2.48 | -0.82 | 0.41 | 1 | 5330439B14Rik | | 0 | 0 | 0 | 16 | 0 | 14 | 9 | 0 | 0 | 3 | 18 |
| 5 | -1.34 | 1.98 | -0.68 | 0.50 | 1 | Gm49329 | | 0 | 1 | 7 | 0 | 15 | 6 | 9 | 0 | 14 | 0 | 7 |
| 5 | -0.61 | 1.30 | -0.47 | 0.64 | 1 | Gm7459 | | 3 | 1 | 0 | 4 | 9 | 6 | 9 | 6 | 2 | 15 | 3 |
| 5 | 0.29 | 2.30 | 0.13 | 0.90 | 1 | Gm47547 | | 11 | 4 | 0 | 0 | 3 | 5 | 9 | 0 | 18 | 0 | 7 |
| 5 | -4.06 | 2.25 | -1.80 | 0.07 | 1 | Gm3336 | predicted gene 3336 | 28 | 12 | 0 | 4 | 2 | 2 | 9 | 0 | 0 | 0 | 0 |
| 5 | -0.38 | 2.77 | -0.14 | 0.89 | 1 | Card14 | caspase recruitment domain family, meml | 0 | 0 | 0 | 7 | 0 | 3 | 9 | 19 | 20 | 0 | 0 |
| 5 | -2.34 | 2.48 | -0.94 | 0.35 | 1 | 5033406O09Rik | | 0 | 18 | 2 | 0 | 6 | 5 | 9 | 0 | 10 | 0 | 1 |
| 5 | -0.03 | 1.91 | -0.01 | 0.99 | 1 | Tmem217 | transmembrane protein 217 | 5 | 0 | 0 | 15 | 2 | 2 | 9 | 12 | 4 | 0 | 2 |
| 4 | -3.97 | 3.47 | -1.14 | 0.25 | 1 | Gm47856 | | 0 | 13 | 0 | 0 | 0 | 16 | 9 | 0 | 0 | 0 | 10 |
| 4 | 0.71 | 1.45 | 0.49 | 0.62 | 1 | Gm6155 | | 4 | 5 | 7 | 2 | 1 | 1 | 9 | 2 | 2 | 5 | 8 |
| 4 | 2.46 | 1.66 | 1.48 | 0.14 | 1 | Gm5216 | | 3 | 1 | 0 | 0 | 1 | 1 | 9 | 2 | 8 | 15 | 6 |
| 4 | -1.32 | 2.34 | -0.57 | 0.57 | 1 | H2bc18 | H2B clustered histone 18 | 0 | 0 | 4 | 0 | 10 | 5 | 9 | 0 | 13 | 0 | 4 |
| 4 | -1.05 | 2.65 | -0.40 | 0.69 | 1 | 2310047D07Rik | | 0 | 0 | 0 | 0 | 8 | 11 | 9 | 0 | 8 | 0 | 10 |
| 4 | -3.04 | 2.73 | -1.11 | 0.27 | 1 | 4933433G15Rik | | 3 | 0 | 0 | 5 | 0 | 21 | 9 | 0 | 0 | 0 | 7 |
| 4 | -1.31 | 2.25 | -0.58 | 0.56 | 1 | Gm48655 | | 22 | 3 | 0 | 0 | 2 | 4 | 9 | 0 | 1 | 0 | 3 |
| 4 | -2.81 | 2.38 | -1.18 | 0.24 | 1 | Gm3739 | predicted gene 3739 | 4 | 0 | 7 | 0 | 9 | 9 | 9 | 0 | 0 | 0 | 5 |
| 4 | -3.30 | 2.46 | -1.34 | 0.18 | 1 | Gm42460 | | 0 | 1 | 14 | 5 | 0 | 5 | 9 | 0 | 1 | 0 | 9 |
| 4 | -1.95 | 2.40 | -0.81 | 0.42 | 1 | Gm44560 | | 0 | 0 | 0 | 9 | 4 | 7 | 9 | 0 | 7 | 0 | 8 |
| 4 | -0.21 | 1.91 | -0.11 | 0.91 | 1 | Gm5265 | | 1 | 7 | 4 | 0 | 2 | 5 | 9 | 0 | 1 | 9 | 5 |
| 4 | -1.27 | 3.46 | -0.37 | 0.71 | 1 | Armc3 | armadillo repeat containing 3 | 12 | 10 | 10 | 0 | 0 | 0 | 9 | 0 | 0 | 0 | 0 |
| 4 | -0.28 | 1.69 | -0.16 | 0.87 | 1 | Gm13408 | | 5 | 4 | 2 | 1 | 2 | 3 | 9 | 0 | 12 | 3 | 1 |
| 4 | -5.44 | 3.22 | -1.69 | 0.09 | 1 | Gm17382 | | 0 | 0 | 0 | 5 | 21 | 5 | 9 | 0 | 0 | 0 | 0 |
| 4 | -2.46 | 2.63 | -0.94 | 0.35 | 1 | Gm4316 | | 17 | 5 | 0 | 0 | 2 | 3 | 9 | 0 | 0 | 0 | 3 |
| 4 | -2.10 | 3.47 | -0.60 | 0.55 | 1 | Gm43625 | | 0 | 1 | 0 | 0 | 0 | 21 | 9 | 0 | 0 | 0 | 8 |
| 4 | -0.86 | 1.74 | -0.50 | 0.62 | 1 | Gm50390 | | 0 | 0 | 7 | 1 | 8 | 3 | 9 | 1 | 5 | 3 | 2 |
| 4 | 1.35 | 2.07 | 0.65 | 0.51 | 1 | mt-Tt | | 1 | 2 | 0 | 7 | 0 | 1 | 9 | 0 | 4 | 9 | 6 |
| 3 | -2.41 | 2.92 | -0.83 | 0.41 | 1 | Gm15742 | | 19 | 0 | 0 | 0 | 6 | 1 | 9 | 0 | 0 | 4 | 0 |
| 3 | -2.13 | 2.92 | -0.73 | 0.47 | 1 | Cpa6 | carboxypeptidase A6 | 0 | 0 | 0 | 0 | 7 | 6 | 9 | 0 | 1 | 0 | 15 |
| 3 | -4.52 | 3.40 | -1.33 | 0.18 | 1 | Gm37786 | | 0 | 4 | 3 | 0 | 0 | 4 | 9 | 0 | 17 | 0 | 0 |
| 3 | -1.47 | 2.76 | -0.53 | 0.60 | 1 | Gm44421 | | 0 | 5 | 0 | 0 | 2 | 8 | 9 | 0 | 0 | 8 | 6 |
| 3 | -0.79 | 2.19 | -0.36 | 0.72 | 1 | Lta | lymphotoxin A | 3 | 6 | 3 | 4 | 0 | 2 | 9 | 0 | 9 | 0 | 2 |
| 3 | -0.24 | 3.54 | -0.07 | 0.95 | 1 | | | 0 | 0 | 0 | 0 | 0 | 6 | 9 | 0 | 0 | 21 | 0 |
| 3 | -3.20 | 2.84 | -1.13 | 0.26 | 1 | Hmgb1-ps3 | | 0 | 2 | 6 | 0 | 8 | 3 | 9 | 0 | 0 | 0 | 7 |

| | | | | | | | | | | | | | | | | | | |
|---|---|---|---|---|---|---|---|---|---|---|---|---|---|---|---|---|---|---|
| 3 | -2.02 | 2.95 | -0.68 | 0.49 | 1 | Gpc5 | glypican 5 | 7 | 0 | 5 | 6 | 0 | 0 | 9 | 0 | 3 | 0 | 4 |
| 3 | -2.57 | 2.55 | -1.01 | 0.31 | 1 | Gm11185 | | 0 | 8 | 1 | 0 | 7 | 1 | 9 | 7 | 0 | 1 | 0 |
| 3 | -1.97 | 2.38 | -0.83 | 0.41 | 1 | Gm12525 | | 0 | 2 | 0 | 3 | 9 | 2 | 9 | 0 | 0 | 6 | 2 |
| 3 | -1.63 | 2.93 | -0.56 | 0.58 | 1 | Ahsg | alpha-2-HS-glycoprotein | 1 | 4 | 16 | 0 | 1 | 0 | 9 | 1 | 0 | 0 | 0 |
| 3 | 0.92 | 3.54 | 0.26 | 0.79 | 1 | Rspo4os | | 0 | 0 | 0 | 0 | 0 | 0 | 9 | 22 | 0 | 0 | 0 |
| 3 | -6.11 | 3.57 | -1.71 | 0.09 | 1 | Lrrc63 | leucine rich repeat containing 63 | 0 | 0 | 16 | 0 | 0 | 0 | 9 | 0 | 6 | 0 | 0 |
| 3 | -6.28 | 3.31 | -1.90 | 0.06 | 1 | Dusp13 | dual specificity phosphatase 13 | 0 | 4 | 0 | 0 | 2 | 12 | 9 | 0 | 0 | 0 | 3 |
| 3 | -1.61 | 3.56 | -0.45 | 0.65 | 1 | Mir374b | | 0 | 0 | 0 | 0 | 0 | 13 | 9 | 0 | 0 | 0 | 7 |
| 3 | -3.80 | 2.75 | -1.38 | 0.17 | 1 | Fam71b | family with sequence similarity 71, memb | 1 | 0 | 0 | 11 | 2 | 3 | 9 | 0 | 2 | 0 | 0 |
| 2 | -0.25 | 3.54 | -0.07 | 0.94 | 1 | Rnf183 | ring finger protein 183 | 0 | 0 | 0 | 0 | 0 | 7 | 9 | 0 | 0 | 6 | 4 |
| 2 | -4.37 | 3.55 | -1.23 | 0.22 | 1 | Slc17a6 | solute carrier family 17 (sodium-depende | 0 | 0 | 0 | 1 | 14 | 1 | 9 | 0 | 0 | 0 | 0 |
| 2 | -3.59 | 3.49 | -1.03 | 0.30 | 1 | Gm8319 | | 0 | 9 | 1 | 3 | 0 | 3 | 9 | 0 | 0 | 0 | 0 |
| 2 | -3.34 | 3.34 | -1.00 | 0.32 | 1 | Hmgb1-ps2 | | 4 | 0 | 0 | 0 | 6 | 3 | 9 | 0 | 0 | 1 | 0 |
| 2 | -4.00 | 3.26 | -1.23 | 0.22 | 1 | Gm33936 | | 0 | 0 | 0 | 4 | 5 | 5 | 9 | 0 | 0 | 1 | 0 |
| 2 | -4.13 | 3.30 | -1.25 | 0.21 | 1 | Gm10728 | | 0 | 0 | 0 | 9 | 4 | 1 | 9 | 0 | 0 | 0 | 1 |
| 2 | -0.03 | 3.51 | -0.01 | 0.99 | 1 | Gm13489 | | 0 | 0 | 6 | 0 | 0 | 0 | 9 | 0 | 6 | 0 | 2 |
| 2 | 0.58 | 2.39 | 0.24 | 0.81 | 1 | Hmgb1-ps5 | | 3 | 1 | 2 | 2 | 0 | 0 | 9 | 2 | 2 | 0 | 1 |
| 2 | -1.28 | 3.56 | -0.36 | 0.72 | 1 | G630018N14Rik | | 0 | 0 | 0 | 0 | 0 | 5 | 9 | 0 | 0 | 0 | 6 |
| 2 | -0.21 | 3.60 | -0.06 | 0.95 | 1 | Car6 | carbonic anhydrase 6 | 1 | 5 | 0 | 0 | 0 | 0 | 9 | 0 | 0 | 5 | 0 |
| 2 | -3.28 | 3.57 | -0.92 | 0.36 | 1 | Gm20512 | | 0 | 0 | 2 | 0 | 0 | 1 | 9 | 0 | 0 | 0 | 8 |
| 2 | -1.98 | 3.59 | -0.55 | 0.58 | 1 | Adam21 | a disintegrin and metallopeptidase domai | 8 | 0 | 0 | 0 | 0 | 3 | 9 | 0 | 0 | 0 | 0 |
| 2 | 0.37 | 3.54 | 0.10 | 0.92 | 1 | Gm21269 | | 0 | 0 | 0 | 0 | 0 | 0 | 9 | 0 | 0 | 0 | 10 |
| 2 | -0.31 | 3.53 | -0.09 | 0.93 | 1 | Lhfpl3 | lipoma HMGIC fusion partner-like 3 | 4 | 0 | 0 | 0 | 0 | 2 | 9 | 0 | 4 | 0 | 0 |
| 2 | -2.85 | 3.57 | -0.80 | 0.42 | 1 | Gm37738 | | 0 | 0 | 0 | 0 | 0 | 3 | 9 | 0 | 0 | 0 | 6 |
| 2 | 0.18 | 3.55 | 0.05 | 0.96 | 1 | 4933421A08Rik | | 4 | 0 | 0 | 0 | 0 | 0 | 9 | 0 | 0 | 0 | 5 |
| 2 | 0.23 | 3.54 | 0.07 | 0.95 | 1 | 4921507G05Rik | | 0 | 0 | 0 | 0 | 0 | 0 | 9 | 0 | 9 | 0 | 0 |
| 2 | -1.66 | 3.57 | -0.46 | 0.64 | 1 | Gm14810 | | 0 | 0 | 0 | 0 | 0 | 6 | 9 | 0 | 0 | 0 | 3 |
| 2 | -2.99 | 3.57 | -0.84 | 0.40 | 1 | Gm37106 | | 0 | 0 | 0 | 0 | 0 | 3 | 9 | 0 | 0 | 0 | 5 |
| 2 | -0.69 | 3.55 | -0.20 | 0.85 | 1 | Gm16549 | | 3 | 0 | 0 | 0 | 0 | 3 | 9 | 0 | 0 | 1 | 2 |
| 2 | -2.03 | 2.50 | -0.81 | 0.42 | 1 | Gm8864 | | 1 | 2 | 0 | 1 | 1 | 1 | 9 | 1 | 0 | 1 | 0 |
| 1 | -2.59 | 3.46 | -0.75 | 0.45 | 1 | Gm14277 | | 0 | 2 | 0 | 0 | 1 | 2 | 9 | 1 | 0 | 0 | 1 |
| 1 | -2.45 | 3.54 | -0.69 | 0.49 | 1 | Rpl10-ps1 | | 0 | 0 | 1 | 1 | 0 | 1 | 9 | 0 | 4 | 0 | 0 |
| 1 | -1.55 | 3.53 | -0.44 | 0.66 | 1 | H2bc14 | H2B clustered histone 14 | 0 | 0 | 0 | 3 | 0 | 0 | 9 | 0 | 4 | 0 | 0 |
| 1 | -0.72 | 2.93 | -0.25 | 0.81 | 1 | Gm6065 | | 1 | 1 | 1 | 0 | 1 | 0 | 9 | 0 | 1 | 1 | 0 |
| 1 | -4.08 | 3.56 | -1.15 | 0.25 | 1 | 1810014P07Rik | | 0 | 0 | 0 | 1 | 0 | 0 | 9 | 0 | 0 | 0 | 5 |
| 1 | -1.28 | 3.56 | -0.36 | 0.72 | 1 | Gm5173 | | 0 | 0 | 0 | 0 | 0 | 3 | 9 | 0 | 0 | 0 | 3 |
| 1 | -1.49 | 3.58 | -0.42 | 0.68 | 1 | Lenep | lens epithelial protein | 0 | 4 | 0 | 0 | 0 | 0 | 9 | 0 | 1 | 0 | 0 |
| 1 | -2.21 | 3.61 | -0.61 | 0.54 | 1 | Gm15467 | | 0 | 0 | 0 | 0 | 0 | 1 | 9 | 0 | 0 | 0 | 0 |
| 1 | -1.31 | 3.63 | -0.36 | 0.72 | 1 | Olfr1252 | olfactory receptor 1252 | 0 | 0 | 0 | 0 | 0 | 0 | 9 | 0 | 0 | 0 | 0 |
| 1 | -1.31 | 3.63 | -0.36 | 0.72 | 1 | Tbx20 | T-box 20 | 0 | 0 | 0 | 0 | 0 | 0 | 9 | 0 | 0 | 0 | 0 |
| 1 | -1.31 | 3.63 | -0.36 | 0.72 | 1 | Gm26071 | | 0 | 0 | 0 | 0 | 0 | 0 | 9 | 0 | 0 | 0 | 0 |
| 1718 | -2.12 | 2.39 | -0.89 | 0.37 | 1 | Cd34 | CD34 antigen | 8013 | 5254 | 5617 | 0 | 1 | 0 | 8 | 0 | 0 | 0 | 0 |
| 1044 | -2.09 | 1.55 | -1.35 | 0.18 | 1 | Serpini2 | serine (or cysteine) peptidase inhibitor, cl | 1313 | 6574 | 3355 | 179 | 21 | 3 | 8 | 0 | 18 | 0 | 13 |
| 1025 | -1.78 | 1.25 | -1.43 | 0.15 | 1 | Col18a1 | collagen, type XVIII, alpha 1 | 5612 | 2677 | 2893 | 18 | 33 | 6 | 8 | 0 | 24 | 0 | 6 |
| 1021 | -1.25 | 3.63 | -0.35 | 0.73 | 1 | Cd300lg | CD300 molecule like family member G | 3593 | 3653 | 3977 | 0 | 0 | 0 | 8 | 0 | 0 | 0 | 0 |
| 556 | -1.10 | 1.41 | -0.78 | 0.43 | 1 | Piezo1 | piezo-type mechanosensitive ion channe | 2424 | 1848 | 1789 | 0 | 12 | 13 | 8 | 0 | 14 | 0 | 7 |
| 504 | -0.83 | 1.09 | -0.77 | 0.44 | 1 | Inmt | indolethylamine N-methyltransferase | 3787 | 542 | 1192 | 3 | 3 | 2 | 8 | 0 | 5 | 0 | 3 |
| 487 | -1.24 | 3.63 | -0.34 | 0.73 | 1 | Rasgrp3 | RAS, guanyl releasing protein 3 | 1876 | 1729 | 1748 | 0 | 0 | 0 | 8 | 0 | 0 | 0 | 0 |
| 463 | -2.11 | 1.58 | -1.33 | 0.18 | 1 | Zcchc24 | zinc finger, CCHC domain containing 24 | 2690 | 1134 | 1248 | 0 | 9 | 3 | 8 | 0 | 6 | 0 | 0 |
| 435 | -0.77 | 1.22 | -0.64 | 0.52 | 1 | Lrrc8c | leucine rich repeat containing 8 family, m | 1727 | 1446 | 1586 | 1 | 1 | 6 | 8 | 0 | 7 | 0 | 3 |
| 398 | -2.83 | 2.60 | -1.09 | 0.28 | 1 | Gask1b | golgi associated kinase 1B | 1659 | 1323 | 1379 | 0 | 0 | 3 | 8 | 0 | 0 | 0 | 0 |
| 378 | 1.20 | 1.59 | 0.75 | 0.45 | 1 | Plce1 | phospholipase C, epsilon 1 | 1703 | 1036 | 1265 | 0 | 33 | 7 | 8 | 60 | 11 | 0 | 31 |
| 356 | -1.03 | 1.58 | -0.65 | 0.52 | 1 | Cspg4 | chondroitin sulfate proteoglycan 4 | 1664 | 1005 | 1230 | 0 | 0 | 1 | 8 | 0 | 0 | 1 | 1 |
| 333 | -1.31 | 1.68 | -0.78 | 0.44 | 1 | Sulf1 | sulfatase 1 | 1889 | 810 | 942 | 0 | 0 | 4 | 8 | 0 | 0 | 0 | 5 |
| 266 | -1.00 | 1.11 | -0.90 | 0.37 | 1 | Rgma | repulsive guidance molecule family meml | 1068 | 972 | 706 | 42 | 23 | 30 | 8 | 0 | 10 | 52 | 12 |
| 235 | -1.24 | 3.63 | -0.34 | 0.73 | 1 | Hlx | H2.0-like homeobox | 921 | 646 | 1009 | 0 | 0 | 0 | 8 | 0 | 0 | 0 | 0 |
| 194 | -0.31 | 0.80 | -0.39 | 0.70 | 1 | Fgfr2 | fibroblast growth factor receptor 2 | 1010 | 477 | 560 | 11 | 7 | 9 | 8 | 0 | 39 | 5 | 4 |
| 193 | -1.65 | 1.01 | -1.63 | 0.10 | 1 | Rin3 | Ras and Rab interactor 3 | 841 | 595 | 666 | 6 | 1 | 1 | 8 | 0 | 5 | 0 | 0 |
| 158 | -4.55 | 3.07 | -1.48 | 0.14 | 1 | Slc39a5 | solute carrier family 39 (metal ion transpo | 231 | 739 | 703 | 59 | 0 | 0 | 8 | 0 | 0 | 0 | 0 |
| 149 | -0.32 | 1.99 | -0.16 | 0.87 | 1 | Gfra2 | glial cell line derived neurotrophic factor f | 1035 | 255 | 332 | 0 | 4 | 0 | 8 | 0 | 9 | 0 | 0 |

| | | | | | | | | | | | | | | | | | | |
|---|---|---|---|---|---|---|---|---|---|---|---|---|---|---|---|---|---|---|
| 111 | -2.17 | 3.12 | -0.70 | 0.49 | 1 | Cd180 | CD180 antigen | 637 | 344 | 226 | 0 | 0 | 0 | 8 | 0 | 0 | 0 | 8 |
| 97 | -1.19 | 3.63 | -0.33 | 0.74 | 1 | Gpr65 | G-protein coupled receptor 65 | 516 | 264 | 275 | 0 | 0 | 0 | 8 | 0 | 0 | 0 | 0 |
| 91 | 0.77 | 1.40 | 0.55 | 0.58 | 1 | Ms4a6b | membrane-spanning 4-domains, subfami | 404 | 281 | 253 | 0 | 0 | 6 | 8 | 0 | 0 | 53 | 1 |
| 90 | 1.66 | 1.92 | 0.87 | 0.39 | 1 | Tnfrsf13b | tumor necrosis factor receptor superfamily | 664 | 161 | 145 | 0 | 1 | 0 | 8 | 0 | 10 | 0 | 3 |
| 86 | -0.62 | 0.77 | -0.80 | 0.42 | 1 | Reep1 | receptor accessory protein 1 | 211 | 209 | 293 | 71 | 32 | 22 | 8 | 32 | 21 | 0 | 48 |
| 82 | 0.24 | 0.72 | 0.33 | 0.74 | 1 | Prr11 | proline rich 11 | 305 | 123 | 69 | 62 | 81 | 35 | 8 | 61 | 52 | 49 | 54 |
| 79 | 3.06 | 2.63 | 1.16 | 0.24 | 1 | Arhgap42 | Rho GTPase activating protein 42 | 353 | 288 | 203 | 0 | 0 | 0 | 8 | 0 | 18 | 0 | 4 |
| 79 | 0.72 | 0.66 | 1.09 | 0.28 | 1 | Cdc6 | cell division cycle 6 | 273 | 98 | 82 | 60 | 39 | 46 | 8 | 53 | 49 | 88 | 76 |
| 78 | 2.08 | 2.34 | 0.89 | 0.37 | 1 | Serpinb6b | serine (or cysteine) peptidase inhibitor, cl | 458 | 155 | 222 | 0 | 0 | 0 | 8 | 0 | 7 | 0 | 3 |
| 72 | -0.30 | 0.63 | -0.47 | 0.64 | 1 | Aff3 | AF4/FMR2 family, member 3 | 442 | 105 | 122 | 10 | 20 | 27 | 8 | 9 | 10 | 10 | 24 |
| 64 | 0.49 | 0.93 | 0.53 | 0.60 | 1 | Adamts4 | a disintegrin-like and metallopeptidase (r | 368 | 124 | 131 | 17 | 0 | 11 | 8 | 2 | 20 | 14 | 11 |
| 62 | 1.79 | 1.07 | 1.67 | 0.09 | 1 | Isg15 | ISG15 ubiquitin-like modifier | 233 | 119 | 251 | 0 | 27 | 0 | 8 | 20 | 5 | 17 | 6 |
| 60 | -1.18 | 3.63 | -0.32 | 0.75 | 1 | Kmo | kynurenine 3-monooxygenase (kynurenir | 341 | 182 | 133 | 0 | 0 | 0 | 8 | 0 | 0 | 0 | 0 |
| 51 | -2.07 | 1.34 | -1.54 | 0.12 | 1 | Kcnj10 | potassium inwardly-rectifying channel, su | 232 | 86 | 203 | 24 | 2 | 0 | 8 | 0 | 9 | 0 | 0 |
| 47 | -0.46 | 1.50 | -0.31 | 0.76 | 1 | Ccl2 | chemokine (C-C motif) ligand 2 | 280 | 133 | 83 | 0 | 0 | 1 | 8 | 0 | 0 | 0 | 10 |
| 44 | -1.97 | 1.04 | -1.89 | 0.06 | 1 | Crhr2 | corticotropin releasing hormone receptor | 131 | 127 | 81 | 27 | 24 | 42 | 8 | 29 | 0 | 0 | 18 |
| 44 | -0.50 | 1.45 | -0.34 | 0.73 | 1 | Gm10134 | | 175 | 145 | 123 | 4 | 0 | 1 | 8 | 0 | 0 | 28 | 0 |
| 44 | 0.64 | 0.60 | 1.08 | 0.28 | 1 | Sfn | stratifin | 104 | 99 | 83 | 32 | 20 | 15 | 8 | 37 | 39 | 32 | 12 |
| 44 | -1.26 | 1.15 | -1.09 | 0.28 | 1 | Slc52a3 | solute carrier protein family 52, member 3 | 179 | 90 | 178 | 10 | 7 | 0 | 8 | 0 | 8 | 0 | 2 |
| 42 | -0.21 | 0.67 | -0.32 | 0.75 | 1 | Gm7993 | | 1 | 40 | 77 | 57 | 96 | 19 | 8 | 71 | 61 | 3 | 32 |
| 41 | -0.13 | 0.83 | -0.16 | 0.87 | 1 | Gm15608 | | 20 | 24 | 47 | 64 | 67 | 50 | 8 | 13 | 22 | 67 | 65 |
| 40 | -1.16 | 1.10 | -1.05 | 0.29 | 1 | 4933407K13Rik | | 45 | 24 | 60 | 14 | 3 | 117 | 8 | 0 | 13 | 0 | 156 |
| 39 | 1.04 | 1.21 | 0.86 | 0.39 | 1 | S1pr4 | sphingosine-1-phosphate receptor 4 | 173 | 95 | 118 | 0 | 0 | 5 | 8 | 0 | 2 | 15 | 6 |
| 38 | -0.48 | 1.11 | -0.43 | 0.67 | 1 | Lax1 | lymphocyte transmembrane adaptor 1 | 269 | 29 | 15 | 0 | 4 | 35 | 8 | 0 | 0 | 14 | 40 |
| 37 | 1.02 | 0.84 | 1.22 | 0.22 | 1 | Exo1 | exonuclease 1 | 113 | 25 | 33 | 31 | 6 | 34 | 8 | 90 | 12 | 41 | 18 |
| 36 | -1.92 | 1.21 | -1.58 | 0.11 | 1 | 3930402G23Rik | | 78 | 120 | 85 | 5 | 28 | 29 | 8 | 32 | 0 | 0 | 8 |
| 35 | 0.89 | 0.68 | 1.31 | 0.19 | 1 | Tfpi2 | tissue factor pathway inhibitor 2 | 109 | 44 | 87 | 18 | 14 | 11 | 8 | 36 | 28 | 13 | 18 |
| 35 | 0.11 | 0.86 | 0.13 | 0.90 | 1 | Anxa9 | annexin A9 | 111 | 43 | 136 | 5 | 3 | 20 | 8 | 0 | 5 | 15 | 34 |
| 34 | -1.60 | 1.47 | -1.08 | 0.28 | 1 | Nrip2 | nuclear receptor interacting protein 2 | 159 | 110 | 84 | 0 | 3 | 3 | 8 | 0 | 6 | 0 | 0 |
| 34 | 0.25 | 1.35 | 0.19 | 0.85 | 1 | Gimap3 | GTPase, IMAP family member 3 | 185 | 103 | 51 | 0 | 1 | 3 | 8 | 0 | 0 | 13 | 6 |
| 33 | -0.36 | 1.56 | -0.23 | 0.82 | 1 | Muc13 | mucin 13, epithelial transmembrane | 0 | 119 | 152 | 12 | 0 | 4 | 8 | 57 | 10 | 0 | 4 |
| 33 | -0.92 | 3.62 | -0.26 | 0.80 | 1 | Ighv2-9-1 | | 293 | 13 | 50 | 0 | 0 | 0 | 8 | 0 | 0 | 0 | 0 |
| 33 | 0.73 | 1.48 | 0.50 | 0.62 | 1 | Gm45221 | | 25 | 0 | 5 | 0 | 0 | 151 | 8 | 0 | 9 | 0 | 159 |
| 29 | -0.11 | 1.08 | -0.10 | 0.92 | 1 | Efcab12 | EF-hand calcium binding domain 12 | 37 | 26 | 16 | 39 | 26 | 47 | 8 | 0 | 38 | 43 | 41 |
| 28 | -0.96 | 1.24 | -0.77 | 0.44 | 1 | Ptprcap | protein tyrosine phosphatase, receptor typ | 190 | 62 | 24 | 0 | 4 | 8 | 8 | 0 | 4 | 4 | 1 |
| 26 | 0.69 | 0.87 | 0.79 | 0.43 | 1 | Gm19426 | | 20 | 7 | 78 | 31 | 24 | 11 | 8 | 53 | 19 | 14 | 23 |
| 25 | 0.58 | 0.82 | 0.71 | 0.48 | 1 | Gm9800 | | 64 | 16 | 16 | 38 | 11 | 15 | 8 | 24 | 29 | 39 | 17 |
| 23 | 0.57 | 0.79 | 0.73 | 0.47 | 1 | Gm12338 | | 15 | 56 | 38 | 29 | 18 | 4 | 8 | 35 | 12 | 36 | 4 |
| 23 | -1.61 | 1.64 | -0.98 | 0.33 | 1 | Casc1 | cancer susceptibility candidate 1 | 14 | 84 | 0 | 11 | 20 | 45 | 8 | 0 | 0 | 19 | 53 |
| 23 | -0.50 | 1.06 | -0.47 | 0.64 | 1 | Slc28a2b | solute carrier family 28 member 2b | 7 | 9 | 25 | 25 | 2 | 60 | 8 | 13 | 10 | 0 | 91 |
| 22 | -2.44 | 1.90 | -1.29 | 0.20 | 1 | Tns4 | tensin 4 | 55 | 52 | 128 | 0 | 0 | 1 | 8 | 0 | 0 | 0 | 0 |
| 22 | 1.19 | 1.38 | 0.87 | 0.39 | 1 | Gm12962 | | 0 | 0 | 0 | 0 | 10 | 60 | 8 | 63 | 12 | 5 | 81 |
| 21 | -0.31 | 1.12 | -0.28 | 0.78 | 1 | Armc2 | armadillo repeat containing 2 | 30 | 48 | 12 | 14 | 26 | 19 | 8 | 23 | 0 | 41 | 14 |
| 21 | -0.30 | 0.92 | -0.33 | 0.74 | 1 | Gm39526 | | 0 | 18 | 7 | 23 | 9 | 60 | 8 | 48 | 17 | 1 | 42 |
| 21 | -0.14 | 1.48 | -0.10 | 0.92 | 1 | Naip6 | NLR family, apoptosis inhibitory protein 6 | 73 | 99 | 28 | 0 | 0 | 9 | 8 | 0 | 0 | 10 | 4 |
| 21 | 1.44 | 1.57 | 0.92 | 0.36 | 1 | Lpar5 | lysophosphatidic acid receptor 5 | 61 | 44 | 75 | 0 | 0 | 2 | 8 | 28 | 1 | 9 | 0 |
| 21 | -0.63 | 1.06 | -0.60 | 0.55 | 1 | Cyp4x1 | cytochrome P450, family 4, subfamily x, p | 42 | 46 | 16 | 25 | 14 | 22 | 8 | 0 | 8 | 19 | 25 |
| 20 | 0.23 | 1.82 | 0.12 | 0.90 | 1 | Cd28 | CD28 antigen | 87 | 42 | 46 | 0 | 0 | 4 | 8 | 0 | 0 | 36 | 0 |
| 19 | 0.25 | 1.04 | 0.24 | 0.81 | 1 | Misp3 | MISP family member 3 | 18 | 0 | 7 | 25 | 32 | 14 | 8 | 19 | 48 | 22 | 17 |
| 18 | 0.00 | 0.83 | 0.00 | 1.00 | 1 | Gm12059 | | 16 | 4 | 19 | 25 | 25 | 21 | 8 | 12 | 18 | 23 | 31 |
| 18 | 0.12 | 0.98 | 0.12 | 0.91 | 1 | Gm11847 | | 9 | 10 | 4 | 25 | 18 | 24 | 8 | 64 | 25 | 1 | 13 |
| 17 | -2.82 | 1.50 | -1.88 | 0.06 | 1 | Sult1b1 | sulfotransferase family 1B, member 1 | 0 | 66 | 23 | 22 | 40 | 13 | 8 | 0 | 10 | 0 | 11 |
| 17 | 0.26 | 2.19 | 0.12 | 0.91 | 1 | Tmprss5 | transmembrane protease, serine 5 (spine | 33 | 57 | 78 | 0 | 0 | 0 | 8 | 0 | 15 | 0 | 0 |
| 17 | 0.01 | 1.05 | 0.01 | 0.99 | 1 | Gm30238 | | 8 | 20 | 3 | 9 | 8 | 44 | 8 | 12 | 14 | 0 | 66 |
| 17 | 0.55 | 0.89 | 0.62 | 0.54 | 1 | Gm33195 | | 15 | 23 | 41 | 4 | 7 | 33 | 8 | 10 | 4 | 18 | 28 |
| 17 | 0.53 | 1.23 | 0.43 | 0.67 | 1 | Camk2a | calcium/calmodulin-dependent protein kir | 77 | 47 | 15 | 5 | 8 | 0 | 8 | 0 | 7 | 22 | 2 |
| 17 | 1.12 | 1.07 | 1.04 | 0.30 | 1 | Gm49874 | | 7 | 10 | 0 | 47 | 1 | 17 | 8 | 43 | 20 | 15 | 13 |
| 16 | 0.60 | 0.79 | 0.76 | 0.44 | 1 | Gm5436 | | 9 | 23 | 10 | 22 | 10 | 16 | 8 | 19 | 21 | 27 | 13 |
| 16 | 1.23 | 0.99 | 1.24 | 0.21 | 1 | Gm50203 | | 9 | 6 | 22 | 5 | 9 | 19 | 8 | 56 | 18 | 17 | 8 |
| 16 | 0.51 | 1.00 | 0.51 | 0.61 | 1 | Nme2 | NME/NM23 nucleoside diphosphate kina | 6 | 1 | 21 | 20 | 11 | 20 | 8 | 43 | 16 | 9 | 20 |

| | | | | | | | | | | | | | | | | | |
|---|---|---|---|---|---|---|---|---|---|---|---|---|---|---|---|---|---|
| 15 | 1.14 | 0.95 | 1.20 | 0.23 | 1 | Rps11-ps1 | | 24 | 6 | 4 | 5 | 15 | 12 | 8 | 41 | 12 | 22 | 19 |
| 15 | -0.58 | 1.70 | -0.34 | 0.73 | 1 | Gm29609 | | 0 | 16 | 9 | 8 | 3 | 48 | 8 | 0 | 0 | 33 | 42 |
| 15 | -1.90 | 1.24 | -1.52 | 0.13 | 1 | Gm49417 | | 19 | 61 | 47 | 5 | 4 | 13 | 8 | 0 | 6 | 0 | 5 |
| 15 | -0.93 | 1.55 | -0.60 | 0.55 | 1 | Gm37045 | | 3 | 0 | 25 | 30 | 2 | 19 | 8 | 0 | 46 | 22 | 12 |
| 14 | -0.41 | 1.48 | -0.28 | 0.78 | 1 | Gm45696 | | 0 | 13 | 0 | 13 | 14 | 25 | 8 | 17 | 50 | 0 | 18 |
| 14 | 1.22 | 1.19 | 1.02 | 0.31 | 1 | Rbm3os | | 15 | 14 | 12 | 1 | 5 | 26 | 8 | 0 | 22 | 33 | 20 |
| 14 | -0.32 | 1.20 | -0.27 | 0.79 | 1 | Gm42815 | | 33 | 8 | 14 | 26 | 13 | 8 | 8 | 12 | 23 | 0 | 11 |
| 14 | -1.73 | 1.26 | -1.38 | 0.17 | 1 | Gm37285 | | 3 | 0 | 10 | 36 | 20 | 32 | 8 | 7 | 12 | 0 | 23 |
| 14 | -2.25 | 1.59 | -1.42 | 0.16 | 1 | 2810408A11Ril | RIKEN cDNA 2810408A11 gene | 31 | 22 | 28 | 25 | 0 | 17 | 8 | 0 | 3 | 0 | 16 |
| 13 | 0.43 | 2.27 | 0.19 | 0.85 | 1 | Cd3g | CD3 antigen, gamma polypeptide | 69 | 49 | 14 | 0 | 0 | 0 | 8 | 0 | 5 | 0 | 0 |
| 13 | -1.87 | 1.76 | -1.06 | 0.29 | 1 | H2-Q5 | | 48 | 22 | 34 | 0 | 7 | 4 | 8 | 0 | 0 | 17 | 0 |
| 12 | -0.68 | 1.68 | -0.41 | 0.69 | 1 | Vstm5 | V-set and transmembrane domain contail | 20 | 51 | 10 | 10 | 0 | 10 | 8 | 13 | 15 | 0 | 0 |
| 12 | -0.21 | 1.01 | -0.20 | 0.84 | 1 | Gm8430 | | 16 | 19 | 17 | 4 | 25 | 6 | 8 | 7 | 10 | 3 | 15 |
| 12 | -2.54 | 1.37 | -1.85 | 0.06 | 1 | Gm12254 | | 21 | 5 | 4 | 41 | 11 | 16 | 8 | 0 | 15 | 5 | 1 |
| 11 | 1.73 | 1.89 | 0.92 | 0.36 | 1 | Sycp2 | synaptonemal complex protein 2 | 0 | 0 | 0 | 0 | 0 | 36 | 8 | 0 | 7 | 19 | 54 |
| 11 | 0.15 | 1.28 | 0.12 | 0.91 | 1 | Gm10190 | | 0 | 17 | 0 | 17 | 11 | 8 | 8 | 35 | 8 | 12 | 8 |
| 11 | 0.44 | 1.18 | 0.38 | 0.71 | 1 | Gpr1 | G protein-coupled receptor 1 | 25 | 12 | 16 | 11 | 5 | 5 | 8 | 17 | 9 | 0 | 15 |
| 11 | 0.13 | 1.31 | 0.10 | 0.92 | 1 | Xkr5 | X-linked Kx blood group related 5 | 27 | 12 | 19 | 4 | 5 | 7 | 8 | 31 | 5 | 0 | 3 |
| 11 | 0.13 | 1.08 | 0.12 | 0.91 | 1 | Gm6807 | | 16 | 26 | 8 | 3 | 9 | 8 | 8 | 23 | 3 | 3 | 13 |
| 11 | -0.24 | 1.33 | -0.18 | 0.86 | 1 | Cyp4f41-ps | | 3 | 0 | 19 | 13 | 7 | 27 | 8 | 20 | 2 | 6 | 15 |
| 11 | -0.93 | 1.72 | -0.54 | 0.59 | 1 | Tsga10ip | testis specific 10 interacting protein | 10 | 41 | 24 | 0 | 11 | 3 | 8 | 0 | 8 | 0 | 14 |
| 11 | -1.69 | 1.31 | -1.29 | 0.20 | 1 | Gm4221 | | 13 | 11 | 19 | 8 | 13 | 19 | 8 | 0 | 14 | 0 | 14 |
| 11 | -0.74 | 1.61 | -0.46 | 0.64 | 1 | Kcnk10 | potassium channel, subfamily K, member | 0 | 0 | 0 | 23 | 5 | 25 | 8 | 0 | 18 | 3 | 37 |
| 10 | 0.49 | 2.23 | 0.22 | 0.83 | 1 | Gm42549 | | 1 | 0 | 0 | 0 | 0 | 54 | 8 | 0 | 10 | 0 | 42 |
| 10 | 0.99 | 1.06 | 0.94 | 0.35 | 1 | | | 0 | 4 | 4 | 12 | 6 | 8 | 8 | 34 | 10 | 19 | 9 |
| 10 | 0.21 | 1.07 | 0.20 | 0.84 | 1 | Tespa1 | thymocyte expressed, positive selection a | 25 | 16 | 3 | 8 | 9 | 5 | 8 | 5 | 19 | 12 | 3 |
| 10 | -0.41 | 1.31 | -0.31 | 0.75 | 1 | Gm10658 | | 11 | 34 | 6 | 11 | 6 | 8 | 8 | 0 | 10 | 14 | 2 |
| 10 | 1.32 | 1.57 | 0.84 | 0.40 | 1 | Gm15703 | | 0 | 9 | 6 | 0 | 6 | 14 | 8 | 22 | 9 | 31 | 4 |
| 10 | -0.83 | 1.60 | -0.52 | 0.60 | 1 | Rfx4 | regulatory factor X, 4 (influences HLA clas | 5 | 0 | 0 | 20 | 16 | 15 | 8 | 8 | 8 | 0 | 27 |
| 10 | 0.77 | 2.46 | 0.31 | 0.75 | 1 | Gm31166 | | 15 | 13 | 0 | 0 | 0 | 19 | 8 | 0 | 30 | 0 | 21 |
| 10 | 0.73 | 1.42 | 0.52 | 0.61 | 1 | Gm19409 | | 4 | 0 | 0 | 0 | 3 | 25 | 8 | 0 | 3 | 30 | 34 |
| 10 | -0.24 | 1.00 | -0.24 | 0.81 | 1 | Gm12350 | | 3 | 7 | 20 | 13 | 7 | 9 | 8 | 7 | 19 | 4 | 9 |
| 10 | 0.27 | 1.28 | 0.21 | 0.83 | 1 | Gm5801 | | 18 | 3 | 21 | 2 | 25 | 0 | 8 | 1 | 16 | 4 | 8 |
| 10 | -0.65 | 1.79 | -0.36 | 0.72 | 1 | Gm44901 | | 2 | 0 | 0 | 0 | 7 | 36 | 8 | 0 | 0 | 21 | 31 |
| 9 | -0.52 | 1.83 | -0.28 | 0.78 | 1 | AA386476 | translation initiation factor IF-2-like | 0 | 1 | 6 | 4 | 2 | 25 | 8 | 28 | 0 | 0 | 29 |
| 9 | -0.62 | 1.43 | -0.43 | 0.67 | 1 | Zfp781 | zinc finger protein 781 | 12 | 0 | 0 | 3 | 7 | 21 | 8 | 0 | 6 | 33 | 12 |
| 9 | -3.62 | 1.98 | -1.83 | 0.07 | 1 | Lrrc70 | | 27 | 35 | 1 | 0 | 6 | 17 | 8 | 0 | 0 | 0 | 5 |
| 9 | 0.17 | 1.10 | 0.15 | 0.88 | 1 | Ybx1-ps2 | | 21 | 22 | 6 | 4 | 5 | 4 | 8 | 1 | 4 | 15 | 6 |
| 9 | -0.44 | 1.54 | -0.29 | 0.78 | 1 | Gm36278 | | 0 | 10 | 0 | 16 | 10 | 4 | 8 | 24 | 12 | 0 | 11 |
| 9 | 0.17 | 1.26 | 0.14 | 0.89 | 1 | 2210411M09Rik | | 8 | 4 | 6 | 12 | 4 | 8 | 8 | 0 | 11 | 22 | 11 |
| 9 | -3.36 | 1.99 | -1.68 | 0.09 | 1 | Gm5781 | | 40 | 1 | 5 | 20 | 0 | 13 | 8 | 0 | 5 | 0 | 1 |
| 8 | -0.45 | 1.16 | -0.39 | 0.70 | 1 | Ptges3l | prostaglandin E synthase 3 like | 9 | 5 | 10 | 3 | 4 | 24 | 8 | 0 | 4 | 12 | 14 |
| 8 | 1.64 | 1.98 | 0.83 | 0.41 | 1 | Brinp1 | bone morphogenic protein/retinoic acid ir | 16 | 4 | 9 | 0 | 0 | 8 | 8 | 0 | 10 | 36 | 0 |
| 8 | 1.10 | 2.03 | 0.54 | 0.59 | 1 | Gm40438 | | 0 | 0 | 4 | 0 | 0 | 20 | 8 | 0 | 15 | 9 | 35 |
| 8 | -1.72 | 1.49 | -1.16 | 0.25 | 1 | A230083N12Rik | | 3 | 5 | 7 | 21 | 9 | 9 | 8 | 0 | 18 | 0 | 9 |
| 8 | -3.30 | 1.80 | -1.84 | 0.07 | 1 | A330074K22Rik | | 41 | 12 | 11 | 7 | 0 | 6 | 8 | 0 | 0 | 0 | 2 |
| 8 | -2.26 | 1.49 | -1.51 | 0.13 | 1 | Gm50322 | | 2 | 3 | 6 | 26 | 13 | 7 | 8 | 0 | 11 | 0 | 11 |
| 8 | -1.48 | 2.79 | -0.53 | 0.60 | 1 | Serpina1a | serine (or cysteine) peptidase inhibitor, cl | 12 | 48 | 20 | 0 | 0 | 0 | 8 | 0 | 0 | 0 | 0 |
| 8 | 1.04 | 1.51 | 0.69 | 0.49 | 1 | Gm5921 | | 7 | 2 | 1 | 2 | 1 | 5 | 8 | 49 | 2 | 10 | 0 |
| 8 | 1.92 | 2.40 | 0.80 | 0.42 | 1 | Elobl | elongin B-like | 3 | 0 | 10 | 4 | 0 | 0 | 8 | 42 | 0 | 19 | 0 |
| 7 | -1.92 | 1.59 | -1.21 | 0.23 | 1 | 8030423F21Rik | | 8 | 15 | 18 | 19 | 3 | 2 | 8 | 0 | 0 | 5 | 3 |
| 7 | -3.55 | 2.10 | -1.70 | 0.09 | 1 | Albfm1 | albumin superfamily member 1 | 0 | 0 | 0 | 28 | 4 | 14 | 8 | 0 | 1 | 0 | 23 |
| 7 | 1.20 | 1.33 | 0.91 | 0.37 | 1 | Thap8 | | 3 | 0 | 0 | 7 | 8 | 1 | 8 | 20 | 6 | 18 | 6 |
| 7 | -1.18 | 1.74 | -0.68 | 0.50 | 1 | Gm4754 | | 9 | 12 | 24 | 0 | 8 | 3 | 8 | 0 | 9 | 0 | 2 |
| 7 | -1.92 | 1.99 | -0.97 | 0.33 | 1 | 4833411C07Rik | | 19 | 17 | 11 | 2 | 0 | 3 | 8 | 0 | 0 | 0 | 12 |
| 7 | 0.36 | 1.48 | 0.24 | 0.81 | 1 | Gm38098 | | 5 | 5 | 10 | 0 | 6 | 9 | 8 | 0 | 11 | 13 | 5 |
| 6 | -2.79 | 2.09 | -1.33 | 0.18 | 1 | Grpr | gastrin releasing peptide receptor | 2 | 45 | 5 | 4 | 0 | 3 | 8 | 0 | 0 | 0 | 2 |
| 6 | -2.23 | 1.71 | -1.30 | 0.19 | 1 | 1700016J18Rik | | 14 | 1 | 0 | 10 | 12 | 6 | 8 | 0 | 3 | 1 | 11 |
| 6 | -1.78 | 1.36 | -1.31 | 0.19 | 1 | Gm19221 | | 6 | 5 | 11 | 12 | 3 | 8 | 8 | 2 | 4 | 0 | 4 |
| 6 | -1.10 | 1.81 | -0.61 | 0.54 | 1 | Gm47513 | | 1 | 0 | 3 | 5 | 9 | 5 | 8 | 17 | 13 | 0 | 0 |

| | | | | | | | | | | | | | | | | | | |
|---|---|---|---|---|---|---|---|---|---|---|---|---|---|---|---|---|---|---|
| 6 | 0.10 | 1.19 | 0.08 | 0.93 | 1 | Rpl36a-ps3 | | 1 | 5 | 7 | 6 | 6 | 3 | 8 | 8 | 5 | 6 | 5 |
| 6 | -1.97 | 1.47 | -1.34 | 0.18 | 1 | Gm17709 | | 5 | 11 | 20 | 3 | 4 | 4 | 8 | 1 | 4 | 0 | 1 |
| 5 | -0.46 | 1.79 | -0.26 | 0.80 | 1 | Gm43361 | | 0 | 0 | 0 | 4 | 8 | 14 | 8 | 8 | 0 | 10 | 9 |
| 5 | -1.99 | 1.97 | -1.01 | 0.31 | 1 | D630033A02Rik | | 3 | 0 | 0 | 4 | 7 | 18 | 8 | 0 | 4 | 0 | 15 |
| 5 | -2.93 | 1.85 | -1.58 | 0.11 | 1 | Cfap57 | cilia and flagella associated protein 57 | 1 | 3 | 8 | 0 | 27 | 5 | 8 | 0 | 5 | 0 | 1 |
| 5 | -1.34 | 2.61 | -0.51 | 0.61 | 1 | Gm50373 | | 0 | 17 | 21 | 0 | 0 | 8 | 8 | 0 | 1 | 0 | 4 |
| 5 | -1.18 | 1.44 | -0.82 | 0.41 | 1 | Gm11521 | | 1 | 1 | 0 | 6 | 8 | 7 | 8 | 22 | 1 | 4 | 1 |
| 5 | -2.68 | 2.69 | -1.00 | 0.32 | 1 | Gm50323 | | 0 | 13 | 0 | 19 | 0 | 8 | 8 | 0 | 0 | 0 | 10 |
| 5 | -2.56 | 3.34 | -0.77 | 0.44 | 1 | C430042M11Rik | | 21 | 22 | 0 | 0 | 0 | 0 | 8 | 0 | 0 | 0 | 6 |
| 5 | -2.34 | 1.96 | -1.19 | 0.23 | 1 | 4930568A12Rik | | 0 | 2 | 0 | 8 | 10 | 6 | 8 | 16 | 7 | 0 | 0 |
| 5 | -0.11 | 1.94 | -0.05 | 0.96 | 1 | Gm15829 | | 0 | 10 | 0 | 0 | 7 | 7 | 8 | 7 | 6 | 5 | 7 |
| 5 | -0.04 | 1.21 | -0.04 | 0.97 | 1 | Gm4977 | | 4 | 9 | 6 | 4 | 4 | 3 | 8 | 2 | 5 | 4 | 7 |
| 5 | 0.09 | 1.37 | 0.07 | 0.95 | 1 | Gm16470 | | 8 | 13 | 4 | 5 | 0 | 3 | 8 | 5 | 1 | 5 | 4 |
| 5 | -2.58 | 3.03 | -0.85 | 0.40 | 1 | Gm12089 | | 0 | 0 | 0 | 0 | 3 | 14 | 8 | 0 | 0 | 0 | 30 |
| 5 | 0.58 | 1.77 | 0.33 | 0.74 | 1 | Gm13292 | | 11 | 1 | 1 | 11 | 1 | 0 | 8 | 0 | 4 | 15 | 2 |
| 5 | 0.11 | 1.49 | 0.07 | 0.94 | 1 | Gm4691 | | 12 | 6 | 7 | 2 | 1 | 2 | 8 | 0 | 2 | 5 | 8 |
| 5 | 0.37 | 1.82 | 0.20 | 0.84 | 1 | Gm12960 | | 13 | 0 | 7 | 0 | 15 | 0 | 8 | 1 | 6 | 1 | 1 |
| 5 | -1.45 | 2.02 | -0.72 | 0.47 | 1 | H2-Ke6 | H2-K region expressed gene 6 | 3 | 11 | 2 | 0 | 1 | 13 | 8 | 0 | 1 | 0 | 13 |
| 5 | -0.56 | 1.77 | -0.32 | 0.75 | 1 | Gm17807 | | 1 | 5 | 1 | 3 | 0 | 14 | 8 | 1 | 3 | 0 | 16 |
| 5 | -1.43 | 2.29 | -0.63 | 0.53 | 1 | BC024139 | cDNA sequence BC024139 | 9 | 0 | 0 | 0 | 9 | 7 | 8 | 0 | 0 | 15 | 4 |
| 5 | -1.50 | 1.65 | -0.91 | 0.36 | 1 | Gm4887 | | 2 | 0 | 8 | 11 | 7 | 2 | 8 | 1 | 1 | 3 | 10 |
| 5 | -1.66 | 2.09 | -0.79 | 0.43 | 1 | Gbp10 | guanylate-binding protein 10 | 20 | 4 | 15 | 0 | 2 | 1 | 8 | 0 | 2 | 0 | 0 |
| 5 | -0.07 | 2.31 | -0.03 | 0.97 | 1 | Gm7054 | | 0 | 11 | 0 | 6 | 0 | 8 | 8 | 1 | 5 | 0 | 11 |
| 4 | -3.96 | 2.05 | -1.93 | 0.05 | 1 | Rpl7a-ps5 | | 2 | 7 | 12 | 7 | 7 | 1 | 8 | 0 | 0 | 0 | 4 |
| 4 | 0.94 | 1.61 | 0.59 | 0.56 | 1 | Gm4707 | | 1 | 1 | 1 | 1 | 1 | 2 | 8 | 8 | 0 | 23 | 1 |
| 4 | -2.17 | 1.37 | -1.59 | 0.11 | 1 | Gm46209 | | 5 | 4 | 4 | 10 | 9 | 2 | 8 | 1 | 2 | 1 | 1 |
| 4 | -1.64 | 2.02 | -0.81 | 0.42 | 1 | Gm28982 | | 11 | 0 | 4 | 0 | 11 | 5 | 8 | 0 | 2 | 0 | 4 |
| 4 | 1.29 | 2.00 | 0.65 | 0.52 | 1 | Gm42893 | | 15 | 0 | 4 | 0 | 0 | 2 | 8 | 0 | 5 | 9 | 3 |
| 4 | -0.60 | 1.87 | -0.32 | 0.75 | 1 | Gm26771 | | 9 | 0 | 3 | 1 | 4 | 5 | 8 | 0 | 7 | 0 | 8 |
| 4 | -0.49 | 2.71 | -0.18 | 0.85 | 1 | Gm39327 | | 0 | 0 | 0 | 0 | 6 | 8 | 8 | 0 | 0 | 14 | 9 |
| 4 | -2.44 | 3.41 | -0.72 | 0.47 | 1 | Csn3 | casein kappa | 0 | 0 | 0 | 0 | 6 | 12 | 8 | 0 | 0 | 19 | 0 |
| 4 | 0.56 | 3.29 | 0.17 | 0.86 | 1 | H3c2 | H3 clustered histone 2 | 0 | 0 | 0 | 0 | 0 | 7 | 8 | 0 | 0 | 19 | 7 |
| 4 | 1.83 | 2.34 | 0.78 | 0.43 | 1 | Gm8538 | | 2 | 0 | 0 | 0 | 1 | 4 | 8 | 17 | 5 | 0 | 3 |
| 4 | -2.92 | 3.40 | -0.86 | 0.39 | 1 | Gm42448 | | 0 | 0 | 0 | 4 | 0 | 12 | 8 | 0 | 0 | 0 | 15 |
| 3 | 0.10 | 1.80 | 0.06 | 0.95 | 1 | Gm12497 | | 3 | 0 | 2 | 3 | 4 | 2 | 8 | 5 | 3 | 0 | 7 |
| 3 | -1.82 | 3.15 | -0.58 | 0.56 | 1 | Gm42503 | | 0 | 0 | 0 | 0 | 6 | 6 | 8 | 14 | 0 | 0 | 2 |
| 3 | -2.93 | 2.17 | -1.35 | 0.18 | 1 | Gm15491 | | 3 | 0 | 5 | 10 | 2 | 2 | 8 | 0 | 5 | 0 | 1 |
| 3 | -0.50 | 2.38 | -0.21 | 0.83 | 1 | Gm26995 | | 0 | 0 | 0 | 0 | 6 | 7 | 8 | 0 | 5 | 6 | 2 |
| 3 | -0.73 | 3.53 | -0.21 | 0.84 | 1 | Gm25128 | | 0 | 0 | 0 | 0 | 0 | 13 | 8 | 0 | 1 | 0 | 11 |
| 3 | -1.28 | 1.69 | -0.76 | 0.45 | 1 | Gm37070 | | 4 | 6 | 3 | 2 | 3 | 1 | 8 | 1 | 3 | 1 | 0 |
| 3 | -3.52 | 2.29 | -1.54 | 0.12 | 1 | Gm13413 | | 0 | 1 | 0 | 4 | 12 | 4 | 8 | 0 | 0 | 1 | 2 |
| 3 | -3.28 | 3.52 | -0.93 | 0.35 | 1 | Fam228a | family with sequence similarity 228, meml | 0 | 0 | 0 | 9 | 0 | 4 | 8 | 0 | 0 | 0 | 11 |
| 3 | -3.05 | 2.90 | -1.05 | 0.29 | 1 | B230398E01Rik | | 0 | 12 | 0 | 0 | 3 | 3 | 8 | 2 | 0 | 0 | 4 |
| 3 | -4.31 | 3.04 | -1.42 | 0.16 | 1 | Rpl30-ps9 | | 1 | 2 | 0 | 18 | 0 | 3 | 8 | 0 | 0 | 0 | 0 |
| 3 | -2.75 | 2.73 | -1.01 | 0.31 | 1 | Gm11532 | | 0 | 0 | 0 | 5 | 2 | 4 | 8 | 0 | 1 | 0 | 11 |
| 2 | 0.09 | 1.94 | 0.05 | 0.96 | 1 | Gm29667 | | 3 | 2 | 0 | 3 | 2 | 0 | 8 | 3 | 1 | 1 | 3 |
| 2 | -1.42 | 3.61 | -0.39 | 0.69 | 1 | Il27 | interleukin 27 | 6 | 0 | 11 | 0 | 0 | 0 | 8 | 0 | 0 | 0 | 0 |
| 2 | -1.89 | 2.88 | -0.66 | 0.51 | 1 | Gm13162 | | 0 | 0 | 3 | 0 | 4 | 4 | 8 | 0 | 3 | 0 | 2 |
| 2 | -3.01 | 3.32 | -0.91 | 0.36 | 1 | Gm12367 | | 1 | 0 | 0 | 0 | 4 | 8 | 8 | 0 | 0 | 0 | 3 |
| 2 | -1.84 | 3.52 | -0.52 | 0.60 | 1 | Dusp9 | dual specificity phosphatase 9 | 0 | 10 | 0 | 0 | 1 | 0 | 8 | 0 | 5 | 0 | 0 |
| 2 | -4.94 | 3.50 | -1.41 | 0.16 | 1 | Gm14296 | predicted gene 14296 | 0 | 0 | 4 | 0 | 1 | 7 | 8 | 0 | 0 | 0 | 4 |
| 2 | -1.73 | 2.22 | -0.78 | 0.44 | 1 | Gm26648 | | 3 | 0 | 0 | 4 | 3 | 1 | 8 | 0 | 2 | 1 | 1 |
| 2 | -0.07 | 3.52 | -0.02 | 0.98 | 1 | Rpl19-ps6 | | 2 | 6 | 0 | 2 | 0 | 0 | 8 | 0 | 3 | 0 | 0 |
| 2 | -3.08 | 3.58 | -0.86 | 0.39 | 1 | Gm47132 | | 0 | 0 | 8 | 0 | 0 | 0 | 8 | 0 | 0 | 0 | 3 |
| 2 | 1.43 | 3.53 | 0.41 | 0.68 | 1 | Hopxos | | 3 | 0 | 0 | 0 | 0 | 1 | 8 | 3 | 0 | 0 | 4 |
| 2 | -3.16 | 3.59 | -0.88 | 0.38 | 1 | Gm35172 | | 0 | 6 | 0 | 0 | 0 | 3 | 8 | 0 | 0 | 0 | 0 |
| 2 | -3.69 | 3.56 | -1.03 | 0.30 | 1 | Gm3081 | | 2 | 0 | 0 | 0 | 5 | 3 | 8 | 0 | 0 | 0 | 0 |
| 2 | -0.94 | 3.60 | -0.26 | 0.80 | 1 | Gm8423 | | 4 | 1 | 0 | 0 | 0 | 1 | 8 | 0 | 0 | 3 | 0 |
| 1 | -1.57 | 3.57 | -0.44 | 0.66 | 1 | Mir1192 | | 0 | 0 | 0 | 0 | 0 | 5 | 8 | 0 | 0 | 0 | 3 |
| 1 | -0.71 | 3.58 | -0.20 | 0.84 | 1 | Nanp | N-acetylneuraminic acid phosphatase | 5 | 0 | 0 | 0 | 0 | 1 | 8 | 0 | 0 | 0 | 1 |

| | | | | | | | | | | | | | | | | | | |
|---|---|---|---|---|---|---|---|---|---|---|---|---|---|---|---|---|---|---|
| 1 | -3.16 | 3.58 | -0.88 | 0.38 | 1 | Gm19637 | | 0 | 0 | 0 | 7 | 0 | 0 | 8 | 0 | 0 | 0 | 0 |
| 1 | 0.02 | 3.51 | 0.01 | 1.00 | 1 | Gm5576 | | 2 | 2 | 0 | 0 | 0 | 1 | 8 | 0 | 1 | 0 | 1 |
| 1 | -0.88 | 3.56 | -0.25 | 0.81 | 1 | Mup7 | major urinary protein 7 | 0 | 0 | 0 | 0 | 2 | 0 | 8 | 0 | 3 | 0 | 0 |
| 1 | -2.94 | 3.59 | -0.82 | 0.41 | 1 | Gm37490 | | 0 | 0 | 0 | 0 | 5 | 0 | 8 | 0 | 0 | 0 | 0 |
| 1 | -1.29 | 3.62 | -0.36 | 0.72 | 1 | Gm5780 | | 2 | 2 | 0 | 0 | 0 | 0 | 8 | 0 | 0 | 0 | 0 |
| 1 | -2.74 | 3.59 | -0.76 | 0.44 | 1 | n-R5s192 | | 0 | 0 | 0 | 0 | 0 | 3 | 8 | 0 | 0 | 0 | 0 |
| 1 | -1.49 | 3.59 | -0.42 | 0.68 | 1 | Gm11658 | | 0 | 0 | 0 | 1 | 0 | 0 | 8 | 0 | 1 | 0 | 0 |
| 1 | -1.29 | 3.59 | -0.36 | 0.72 | 1 | C230053D17Rik | | 0 | 0 | 0 | 0 | 0 | 1 | 8 | 0 | 0 | 0 | 1 |
| 1 | -0.60 | 3.61 | -0.17 | 0.87 | 1 | Gm4853 | | 0 | 0 | 0 | 0 | 0 | 0 | 8 | 0 | 0 | 0 | 1 |
| 1 | -2.17 | 3.62 | -0.60 | 0.55 | 1 | Ftcd | formiminotransferase cyclodeaminase | 0 | 0 | 0 | 0 | 0 | 1 | 8 | 0 | 0 | 0 | 0 |
| 1 | -1.29 | 3.63 | -0.36 | 0.72 | 1 | Gm42853 | | 0 | 0 | 0 | 0 | 0 | 0 | 8 | 0 | 0 | 0 | 0 |
| 1 | -1.29 | 3.63 | -0.36 | 0.72 | 1 | Slc28a1 | solute carrier family 28 (sodium-coupled r | 0 | 0 | 0 | 0 | 0 | 0 | 8 | 0 | 0 | 0 | 0 |
| 1 | -1.29 | 3.63 | -0.36 | 0.72 | 1 | Tex36 | testis expressed 36 | 0 | 0 | 0 | 0 | 0 | 0 | 8 | 0 | 0 | 0 | 0 |
| 1 | -1.29 | 3.63 | -0.36 | 0.72 | 1 | Rpl13-ps2 | | 0 | 0 | 0 | 0 | 0 | 0 | 8 | 0 | 0 | 0 | 0 |
| 1 | -1.29 | 3.63 | -0.36 | 0.72 | 1 | Gm16758 | | 0 | 0 | 0 | 0 | 0 | 0 | 8 | 0 | 0 | 0 | 0 |
| 1 | -1.29 | 3.63 | -0.36 | 0.72 | 1 | Tuba-rs1 | | 0 | 0 | 0 | 0 | 0 | 0 | 8 | 0 | 0 | 0 | 0 |
| 1 | -1.29 | 3.63 | -0.36 | 0.72 | 1 | Xlr5b | X-linked lymphocyte-regulated 5B | 0 | 0 | 0 | 0 | 0 | 0 | 8 | 0 | 0 | 0 | 0 |
| 2351 | -2.01 | 1.07 | -1.89 | 0.06 | 1 | Lamb1 | laminin B1 | 9144 | 7737 | 8943 | 4 | 9 | 5 | 7 | 0 | 2 | 0 | 9 |
| 1927 | -2.12 | 1.24 | -1.71 | 0.09 | 1 | Col5a1 | collagen, type V, alpha 1 | 9238 | 4617 | 7319 | 0 | 9 | 2 | 7 | 0 | 4 | 0 | 0 |
| 724 | -0.36 | 2.06 | -0.17 | 0.86 | 1 | Fabp4 | fatty acid binding protein 4, adipocyte | 2273 | 2579 | 3089 | 0 | 0 | 3 | 7 | 0 | 0 | 12 | 0 |
| 531 | 0.82 | 2.07 | 0.39 | 0.69 | 1 | Rarres2 | retinoic acid receptor responder (tazarote | 2898 | 1387 | 1551 | 0 | 0 | 0 | 7 | 0 | 3 | 0 | 1 |
| 473 | -1.04 | 3.62 | -0.29 | 0.77 | 1 | Fmod | fibromodulin | 3626 | 729 | 840 | 0 | 0 | 0 | 7 | 0 | 0 | 0 | 0 |
| 471 | 0.68 | 1.23 | 0.56 | 0.58 | 1 | Ptpn14 | protein tyrosine phosphatase, non-recept | 2125 | 1607 | 1280 | 0 | 15 | 61 | 7 | 0 | 24 | 26 | 37 |
| 449 | -1.15 | 3.63 | -0.32 | 0.75 | 1 | Emilin2 | elastin microfibril interfacer 2 | 2562 | 1049 | 1326 | 0 | 0 | 0 | 7 | 0 | 0 | 0 | 0 |
| 307 | 2.47 | 1.53 | 1.61 | 0.11 | 1 | Filip1l | filamin A interacting protein 1-like | 1865 | 846 | 613 | 0 | 0 | 5 | 7 | 16 | 4 | 0 | 19 |
| 282 | 0.54 | 2.86 | 0.19 | 0.85 | 1 | She | src homology 2 domain-containing transfo | 1123 | 962 | 994 | 0 | 0 | 0 | 7 | 0 | 0 | 14 | 0 |
| 233 | -0.40 | 1.62 | -0.24 | 0.81 | 1 | Ciita | class II transactivator | 1375 | 658 | 515 | 0 | 3 | 0 | 7 | 0 | 0 | 4 | 5 |
| 183 | -0.11 | 1.56 | -0.07 | 0.95 | 1 | Hacd4 | 3-hydroxyacyl-CoA dehydratase 4 | 853 | 539 | 586 | 0 | 9 | 1 | 7 | 0 | 10 | 0 | 8 |
| 178 | -2.92 | 2.22 | -1.32 | 0.19 | 1 | Clec7a | C-type lectin domain family 7, member a | 1058 | 416 | 462 | 9 | 0 | 0 | 7 | 0 | 0 | 0 | 2 |
| 167 | -2.07 | 3.09 | -0.67 | 0.50 | 1 | Mmp19 | matrix metallopeptidase 19 | 803 | 523 | 496 | 0 | 0 | 0 | 7 | 0 | 0 | 0 | 7 |
| 146 | 0.75 | 0.57 | 1.32 | 0.19 | 1 | Pdzrn3 | PDZ domain containing RING finger 3 | 598 | 334 | 388 | 48 | 22 | 27 | 7 | 96 | 32 | 21 | 30 |
| 105 | -0.56 | 2.53 | -0.22 | 0.83 | 1 | Clec12a | C-type lectin domain family 12, member a | 674 | 207 | 261 | 0 | 0 | 0 | 7 | 0 | 0 | 0 | 1 |
| 100 | 0.06 | 0.80 | 0.08 | 0.94 | 1 | Nek8 | NIMA (never in mitosis gene a)-related ex | 275 | 231 | 230 | 65 | 36 | 60 | 7 | 70 | 49 | 0 | 79 |
| 98 | -1.83 | 1.13 | -1.63 | 0.10 | 1 | Cd84 | CD84 antigen | 468 | 218 | 343 | 10 | 7 | 2 | 7 | 0 | 0 | 22 | 0 |
| 91 | -2.02 | 1.20 | -1.68 | 0.09 | 1 | Sostdc1 | sclerostin domain containing 1 | 255 | 378 | 346 | 13 | 0 | 0 | 7 | 0 | 2 | 0 | 1 |
| 88 | -2.31 | 1.30 | -1.77 | 0.08 | 1 | Lpcat2 | lysophosphatidylcholine acyltransferase 2 | 458 | 278 | 218 | 7 | 0 | 1 | 7 | 0 | 0 | 0 | 3 |
| 88 | -0.46 | 0.71 | -0.64 | 0.52 | 1 | Tnfrsf10b | tumor necrosis factor receptor superfamily | 243 | 264 | 339 | 25 | 11 | 20 | 7 | 15 | 8 | 28 | 2 |
| 86 | -0.83 | 0.92 | -0.91 | 0.37 | 1 | Fhl2 | four and a half LIM domains 2 | 481 | 112 | 207 | 33 | 9 | 37 | 7 | 39 | 7 | 0 | 9 |
| 75 | 0.03 | 2.46 | 0.01 | 0.99 | 1 | Runx1 | runt related transcription factor 1 | 353 | 227 | 226 | 0 | 0 | 1 | 7 | 0 | 0 | 15 | 0 |
| 62 | 0.75 | 1.31 | 0.57 | 0.57 | 1 | Arhgap20 | Rho GTPase activating protein 20 | 299 | 197 | 150 | 0 | 1 | 4 | 7 | 0 | 0 | 19 | 9 |
| 60 | 0.82 | 0.72 | 1.14 | 0.25 | 1 | Tk1 | thymidine kinase 1 | 261 | 87 | 85 | 36 | 9 | 29 | 7 | 16 | 35 | 61 | 36 |
| 56 | -1.17 | 3.63 | -0.32 | 0.75 | 1 | Rspo3 | R-spondin 3 | 282 | 230 | 95 | 0 | 0 | 0 | 7 | 0 | 0 | 0 | 0 |
| 55 | 2.11 | 1.12 | 1.89 | 0.06 | 1 | Rab30 | RAB30, member RAS oncogene family | 274 | 162 | 86 | 4 | 0 | 7 | 7 | 25 | 27 | 0 | 10 |
| 54 | -0.61 | 0.87 | -0.70 | 0.48 | 1 | ccdc198 | coiled-coil domain containing 198 | 165 | 96 | 236 | 26 | 18 | 6 | 7 | 7 | 20 | 0 | 16 |
| 53 | 0.26 | 3.00 | 0.09 | 0.93 | 1 | Slc25a45 | solute carrier family 25, member 45 | 235 | 175 | 157 | 0 | 0 | 0 | 7 | 0 | 5 | 0 | 0 |
| 52 | 0.42 | 0.70 | 0.59 | 0.55 | 1 | Socs1 | suppressor of cytokine signaling 1 | 178 | 44 | 156 | 26 | 19 | 24 | 7 | 13 | 26 | 48 | 29 |
| 50 | -1.61 | 1.06 | -1.51 | 0.13 | 1 | Myoc | myocilin | 438 | 41 | 30 | 2 | 9 | 8 | 7 | 0 | 0 | 12 | 6 |
| 50 | -0.27 | 0.86 | -0.32 | 0.75 | 1 | Eppk1 | epiplakin 1 | 82 | 127 | 126 | 31 | 19 | 62 | 7 | 16 | 19 | 52 | 5 |
| 48 | -0.20 | 1.30 | -0.15 | 0.88 | 1 | Acss3 | acyl-CoA synthetase short-chain family m | 257 | 140 | 113 | 0 | 0 | 4 | 7 | 0 | 4 | 0 | 3 |
| 46 | -2.34 | 1.52 | -1.54 | 0.12 | 1 | Ccn4 | cellular communication network factor 4 | 287 | 61 | 151 | 0 | 0 | 1 | 7 | 0 | 0 | 0 | 0 |
| 43 | -0.35 | 1.24 | -0.29 | 0.77 | 1 | Epn3 | epsin 3 | 166 | 98 | 171 | 6 | 0 | 3 | 7 | 0 | 0 | 13 | 6 |
| 43 | 0.42 | 0.66 | 0.64 | 0.52 | 1 | Gm26826 | | 34 | 26 | 85 | 39 | 42 | 63 | 7 | 42 | 39 | 40 | 51 |
| 41 | 0.34 | 0.85 | 0.39 | 0.69 | 1 | Cgas | cyclic GMP-AMP synthase | 116 | 57 | 131 | 9 | 10 | 41 | 7 | 44 | 3 | 10 | 21 |
| 40 | -1.93 | 1.36 | -1.42 | 0.16 | 1 | Cd1d1 | CD1d1 antigen | 172 | 108 | 137 | 0 | 4 | 3 | 7 | 0 | 0 | 0 | 12 |
| 38 | 1.16 | 0.91 | 1.28 | 0.20 | 1 | Foxj1 | forkhead box J1 | 108 | 71 | 96 | 19 | 0 | 29 | 7 | 32 | 22 | 17 | 19 |
| 37 | 0.26 | 0.69 | 0.38 | 0.70 | 1 | A230065N10Rik | | 33 | 30 | 50 | 57 | 39 | 32 | 7 | 88 | 36 | 18 | 21 |
| 36 | -0.27 | 0.79 | -0.34 | 0.73 | 1 | 5930403N24Rik | | 50 | 73 | 94 | 24 | 30 | 38 | 7 | 12 | 13 | 45 | 13 |
| 34 | -1.97 | 1.69 | -1.17 | 0.24 | 1 | Nt5e | 5' nucleotidase, ecto | 199 | 74 | 57 | 0 | 0 | 36 | 7 | 0 | 3 | 0 | 0 |
| 33 | -2.27 | 1.31 | -1.74 | 0.08 | 1 | Ticam2 | toll-like receptor adaptor molecule 2 | 139 | 122 | 85 | 0 | 2 | 8 | 7 | 0 | 0 | 0 | 5 |
| 33 | -1.29 | 1.10 | -1.18 | 0.24 | 1 | Trip13 | thyroid hormone receptor interactor 13 | 116 | 83 | 14 | 31 | 40 | 25 | 7 | 0 | 22 | 15 | 9 |

| | | | | | | | | | | | | | | | | | | |
|---|---|---|---|---|---|---|---|---|---|---|---|---|---|---|---|---|---|---|
| 33 | -1.01 | 0.74 | -1.36 | 0.17 | 1 | Deup1 | deuterosome assembly protein 1 | 53 | 132 | 75 | 25 | 19 | 12 | 7 | 24 | 2 | 4 | 7 |
| 33 | 0.99 | 0.74 | 1.34 | 0.18 | 1 | Junos | | 41 | 55 | 77 | 10 | 13 | 40 | 7 | 43 | 10 | 31 | 31 |
| 30 | -2.26 | 1.22 | -1.86 | 0.06 | 1 | Agxt2 | alanine-glyoxylate aminotransferase 2 | 1 | 0 | 8 | 4 | 4 | 150 | 7 | 0 | 2 | 0 | 154 |
| 30 | 0.04 | 0.74 | 0.05 | 0.96 | 1 | Disc1 | disrupted in schizophrenia 1 | 83 | 43 | 91 | 18 | 10 | 14 | 7 | 5 | 12 | 13 | 29 |
| 29 | -1.85 | 1.44 | -1.29 | 0.20 | 1 | Ccdc18 | coiled-coil domain containing 18 | 84 | 69 | 3 | 53 | 30 | 22 | 7 | 0 | 19 | 0 | 34 |
| 28 | -0.12 | 1.65 | -0.07 | 0.94 | 1 | Hoxa3 | homeobox A3 | 130 | 108 | 61 | 0 | 0 | 0 | 7 | 0 | 0 | 5 | 0 |
| 27 | 0.15 | 0.89 | 0.17 | 0.86 | 1 | Gm44430 | | 4 | 42 | 24 | 35 | 34 | 28 | 7 | 73 | 3 | 19 | 26 |
| 27 | -0.35 | 0.83 | -0.42 | 0.67 | 1 | Gm16712 | | 19 | 5 | 24 | 25 | 60 | 36 | 7 | 8 | 46 | 49 | 16 |
| 27 | 0.91 | 1.08 | 0.85 | 0.40 | 1 | 3300002I08Rik | RIKEN cDNA 3300002I08 gene | 30 | 24 | 46 | 6 | 9 | 46 | 7 | 64 | 29 | 0 | 29 |
| 26 | 0.28 | 1.15 | 0.24 | 0.81 | 1 | Slc7a11 | solute carrier family 7 (cationic amino acid | 58 | 37 | 18 | 4 | 43 | 9 | 7 | 0 | 28 | 70 | 8 |
| 25 | -0.24 | 1.17 | -0.21 | 0.84 | 1 | Noxred1 | NADP+ dependent oxidoreductase doma | 63 | 16 | 2 | 25 | 5 | 71 | 7 | 0 | 12 | 22 | 56 |
| 24 | 0.66 | 0.77 | 0.85 | 0.40 | 1 | 1600023N17Rik | | 45 | 51 | 70 | 4 | 7 | 29 | 7 | 7 | 11 | 22 | 17 |
| 24 | -0.27 | 0.86 | -0.32 | 0.75 | 1 | Snrpert | | 73 | 9 | 56 | 10 | 34 | 11 | 7 | 1 | 28 | 23 | 13 |
| 24 | -1.11 | 3.63 | -0.31 | 0.76 | 1 | Susd3 | sushi domain containing 3 | 154 | 57 | 48 | 0 | 0 | 0 | 7 | 0 | 0 | 0 | 0 |
| 23 | -0.85 | 1.77 | -0.48 | 0.63 | 1 | Nat8l | N-acetyltransferase 8-like | 131 | 21 | 91 | 0 | 0 | 0 | 7 | 0 | 1 | 0 | 0 |
| 22 | -1.13 | 1.51 | -0.75 | 0.45 | 1 | Cd209g | CD209g antigen | 181 | 25 | 26 | 0 | 0 | 3 | 7 | 0 | 0 | 0 | 3 |
| 21 | 0.13 | 1.47 | 0.09 | 0.93 | 1 | Rgs9bp | regulator of G-protein signalling 9 binding | 35 | 0 | 78 | 20 | 23 | 9 | 7 | 15 | 0 | 35 | 14 |
| 21 | -0.35 | 1.06 | -0.33 | 0.74 | 1 | Gm45456 | | 0 | 16 | 9 | 20 | 40 | 16 | 7 | 20 | 78 | 0 | 23 |
| 21 | -0.04 | 0.97 | -0.04 | 0.97 | 1 | Smim17 | small integral membrane protein 17 | 16 | 25 | 18 | 29 | 15 | 19 | 7 | 56 | 11 | 0 | 29 |
| 20 | 0.62 | 1.09 | 0.57 | 0.57 | 1 | Traip | TRAF-interacting protein | 30 | 17 | 20 | 0 | 38 | 33 | 7 | 21 | 22 | 17 | 19 |
| 20 | -1.01 | 1.08 | -0.93 | 0.35 | 1 | Gm32699 | | 28 | 42 | 10 | 10 | 20 | 60 | 7 | 0 | 13 | 12 | 16 |
| 19 | 1.00 | 0.97 | 1.04 | 0.30 | 1 | Gm7332 | | 36 | 1 | 19 | 7 | 27 | 14 | 7 | 22 | 23 | 19 | 32 |
| 19 | -1.72 | 1.08 | -1.59 | 0.11 | 1 | Gm15760 | | 21 | 30 | 10 | 39 | 27 | 36 | 7 | 0 | 12 | 8 | 12 |
| 17 | -0.55 | 1.27 | -0.44 | 0.66 | 1 | Fxyd2 | FXYD domain-containing ion transport reg | 65 | 54 | 42 | 0 | 3 | 6 | 7 | 5 | 1 | 0 | 5 |
| 17 | -0.06 | 1.14 | -0.05 | 0.96 | 1 | Gm5909 | | 3 | 44 | 2 | 25 | 8 | 37 | 7 | 1 | 7 | 15 | 34 |
| 16 | -0.67 | 1.16 | -0.58 | 0.56 | 1 | 8430422M14Rik | | 4 | 18 | 0 | 28 | 8 | 55 | 7 | 25 | 10 | 0 | 25 |
| 16 | -2.81 | 1.68 | -1.67 | 0.10 | 1 | Gm32063 | | 3 | 2 | 0 | 14 | 0 | 86 | 7 | 0 | 0 | 0 | 64 |
| 16 | -1.33 | 0.96 | -1.38 | 0.17 | 1 | Mfsd4b5 | major facilitator superfamily domain conta | 11 | 28 | 24 | 18 | 26 | 17 | 7 | 29 | 5 | 1 | 4 |
| 15 | -2.72 | 1.69 | -1.62 | 0.11 | 1 | Comp | cartilage oligomeric matrix protein | 98 | 3 | 21 | 7 | 10 | 3 | 7 | 0 | 20 | 0 | 0 |
| 15 | -0.26 | 1.05 | -0.24 | 0.81 | 1 | Gm43205 | | 11 | 24 | 9 | 35 | 3 | 14 | 7 | 44 | 4 | 0 | 15 |
| 15 | -1.59 | 1.79 | -0.89 | 0.37 | 1 | Gm17396 | | 39 | 42 | 64 | 0 | 0 | 7 | 7 | 0 | 0 | 0 | 5 |
| 14 | -0.44 | 1.16 | -0.38 | 0.70 | 1 | Gm10353 | | 34 | 4 | 21 | 15 | 6 | 18 | 7 | 0 | 10 | 19 | 22 |
| 14 | 0.60 | 1.28 | 0.47 | 0.64 | 1 | Fbxo24 | F-box protein 24 | 27 | 9 | 5 | 0 | 8 | 25 | 7 | 0 | 4 | 37 | 32 |
| 14 | -1.67 | 1.36 | -1.23 | 0.22 | 1 | Creb3l4 | cAMP responsive element binding protein | 35 | 61 | 22 | 9 | 2 | 3 | 7 | 0 | 0 | 3 | 10 |
| 13 | -0.57 | 1.78 | -0.32 | 0.75 | 1 | F10 | coagulation factor X | 75 | 37 | 11 | 0 | 0 | 6 | 7 | 8 | 0 | 0 | 1 |
| 13 | -0.50 | 1.79 | -0.28 | 0.78 | 1 | Gm47814 | | 16 | 14 | 0 | 0 | 13 | 33 | 7 | 44 | 4 | 0 | 14 |
| 13 | -0.13 | 0.83 | -0.15 | 0.88 | 1 | Gm2962 | | 3 | 15 | 19 | 20 | 16 | 9 | 7 | 16 | 27 | 6 | 5 |
| 13 | -3.63 | 1.86 | -1.95 | 0.05 | 1 | Gdf15 | growth differentiation factor 15 | 60 | 32 | 25 | 4 | 9 | 0 | 7 | 0 | 0 | 0 | 7 |
| 13 | -0.47 | 1.45 | -0.32 | 0.75 | 1 | Gm16984 | | 15 | 4 | 28 | 2 | 14 | 13 | 7 | 0 | 0 | 49 | 10 |
| 13 | -0.61 | 1.18 | -0.51 | 0.61 | 1 | Gm47508 | | 20 | 10 | 10 | 1 | 7 | 38 | 7 | 0 | 1 | 12 | 36 |
| 13 | -2.47 | 1.45 | -1.70 | 0.09 | 1 | Gm21992 | predicted gene 21992 | 2 | 0 | 11 | 11 | 17 | 31 | 7 | 0 | 11 | 0 | 50 |
| 13 | -0.54 | 1.17 | -0.46 | 0.65 | 1 | Anp32b-ps1 | | 17 | 5 | 16 | 21 | 17 | 12 | 7 | 25 | 11 | 0 | 9 |
| 13 | -1.14 | 2.02 | -0.56 | 0.57 | 1 | Gm12406 | | 0 | 0 | 0 | 3 | 0 | 63 | 7 | 0 | 5 | 0 | 62 |
| 13 | -0.43 | 1.32 | -0.32 | 0.75 | 1 | Slc5a5 | solute carrier family 5 (sodium iodide sym | 36 | 16 | 19 | 10 | 8 | 3 | 7 | 0 | 11 | 1 | 28 |
| 12 | 1.07 | 1.39 | 0.77 | 0.44 | 1 | Iqsec3 | IQ motif and Sec7 domain 3 | 35 | 51 | 8 | 4 | 0 | 3 | 7 | 16 | 8 | 0 | 3 |
| 12 | 0.15 | 1.20 | 0.13 | 0.90 | 1 | Gm5530 | | 41 | 1 | 5 | 11 | 1 | 7 | 7 | 2 | 3 | 50 | 6 |
| 12 | 0.44 | 1.09 | 0.40 | 0.69 | 1 | Gm6627 | | 11 | 18 | 3 | 4 | 6 | 29 | 7 | 1 | 10 | 21 | 22 |
| 12 | 0.08 | 1.23 | 0.06 | 0.95 | 1 | Coch | cochlin | 13 | 5 | 0 | 18 | 18 | 10 | 7 | 5 | 29 | 9 | 17 |
| 12 | -1.62 | 1.88 | -0.86 | 0.39 | 1 | Gm15234 | | 10 | 0 | 0 | 15 | 11 | 33 | 7 | 0 | 12 | 0 | 42 |
| 12 | -1.46 | 1.25 | -1.17 | 0.24 | 1 | 1810041H14Rik | | 9 | 4 | 22 | 26 | 11 | 13 | 7 | 0 | 11 | 3 | 23 |
| 11 | 0.03 | 1.48 | 0.02 | 0.99 | 1 | Gm5915 | | 39 | 20 | 0 | 19 | 3 | 8 | 7 | 12 | 7 | 0 | 10 |
| 11 | -0.32 | 1.78 | -0.18 | 0.86 | 1 | Gm43182 | | 21 | 2 | 50 | 8 | 5 | 0 | 7 | 31 | 1 | 0 | 0 |
| 11 | -0.73 | 1.19 | -0.62 | 0.54 | 1 | Tnfsfm13 | tumor necrosis factor (ligand) superfamily | 16 | 2 | 41 | 7 | 13 | 6 | 7 | 0 | 7 | 18 | 6 |
| 11 | 0.64 | 1.49 | 0.43 | 0.67 | 1 | Gm33051 | | 8 | 11 | 27 | 0 | 10 | 6 | 7 | 34 | 13 | 0 | 6 |
| 11 | -0.41 | 1.61 | -0.25 | 0.80 | 1 | Cend1 | cell cycle exit and neuronal differentiation | 19 | 41 | 36 | 2 | 4 | 0 | 7 | 0 | 8 | 0 | 2 |
| 11 | -1.26 | 1.67 | -0.75 | 0.45 | 1 | 2900064F13Rik | | 0 | 11 | 11 | 0 | 26 | 9 | 7 | 25 | 24 | 0 | 3 |
| 11 | 0.15 | 1.25 | 0.12 | 0.90 | 1 | Gm3436 | | 4 | 37 | 3 | 15 | 11 | 3 | 7 | 1 | 5 | 19 | 9 |
| 11 | -0.14 | 1.14 | -0.12 | 0.90 | 1 | Prkcz2 | | 11 | 24 | 4 | 30 | 6 | 3 | 7 | 6 | 14 | 1 | 9 |
| 10 | 0.72 | 1.17 | 0.61 | 0.54 | 1 | Acoxl | acyl-Coenzyme A oxidase-like | 0 | 0 | 3 | 18 | 11 | 7 | 7 | 30 | 10 | 23 | 6 |
| 10 | -2.14 | 1.70 | -1.26 | 0.21 | 1 | Popdc2 | popeye domain containing 2 | 74 | 5 | 24 | 0 | 1 | 3 | 7 | 0 | 0 | 0 | 1 |

| | | | | | | | | | | | | | | | | | | |
|---|---|---|---|---|---|---|---|---|---|---|---|---|---|---|---|---|---|---|
| 10 | 1.00 | 1.39 | 0.72 | 0.47 | 1 | 4921525O09Rik | | 9 | 4 | 7 | 0 | 10 | 14 | 7 | 23 | 21 | 10 | 1 |
| 10 | 0.14 | 1.64 | 0.09 | 0.93 | 1 | Hsf2bp | heat shock transcription factor 2 binding p | 3 | 11 | 7 | 10 | 0 | 16 | 7 | 0 | 2 | 45 | 5 |
| 9 | 0.40 | 1.09 | 0.37 | 0.71 | 1 | Gm3379 | | 9 | 25 | 8 | 6 | 12 | 2 | 7 | 3 | 14 | 6 | 8 |
| 9 | 0.90 | 1.29 | 0.69 | 0.49 | 1 | Gjd3 | gap junction protein, delta 3 | 4 | 0 | 0 | 6 | 16 | 7 | 7 | 22 | 16 | 6 | 16 |
| 9 | 0.09 | 1.25 | 0.07 | 0.94 | 1 | Gm7278 | | 11 | 1 | 14 | 3 | 4 | 12 | 7 | 0 | 4 | 33 | 11 |
| 9 | 2.56 | 1.58 | 1.62 | 0.11 | 1 | Alpk3 | alpha-kinase 3 | 20 | 14 | 0 | 0 | 0 | 2 | 7 | 13 | 0 | 31 | 11 |
| 9 | -3.06 | 1.68 | -1.82 | 0.07 | 1 | Ube2u | ubiquitin-conjugating enzyme E2U (putati | 3 | 18 | 0 | 18 | 15 | 22 | 7 | 0 | 3 | 0 | 9 |
| 8 | -2.39 | 1.57 | -1.52 | 0.13 | 1 | A530088E08Rik | | 10 | 25 | 4 | 4 | 17 | 7 | 7 | 0 | 3 | 0 | 14 |
| 8 | 1.74 | 1.67 | 1.04 | 0.30 | 1 | D930030I03Rik | | 19 | 17 | 0 | 4 | 0 | 7 | 7 | 0 | 4 | 18 | 14 |
| 8 | -0.22 | 2.24 | -0.10 | 0.92 | 1 | Scx | scleraxis | 44 | 3 | 28 | 0 | 3 | 0 | 7 | 5 | 0 | 0 | 0 |
| 8 | 0.45 | 1.61 | 0.28 | 0.78 | 1 | Gm15541 | | 9 | 13 | 0 | 4 | 4 | 13 | 7 | 8 | 21 | 0 | 8 |
| 8 | -0.85 | 1.88 | -0.45 | 0.65 | 1 | Satb2 | special AT-rich sequence binding protein | 18 | 30 | 8 | 12 | 0 | 1 | 7 | 0 | 8 | 0 | 1 |
| 8 | -1.02 | 2.03 | -0.51 | 0.61 | 1 | Gm17040 | | 0 | 7 | 12 | 0 | 10 | 13 | 7 | 0 | 17 | 0 | 17 |
| 8 | -0.51 | 2.43 | -0.21 | 0.84 | 1 | Gm43290 | | 3 | 0 | 8 | 0 | 0 | 41 | 7 | 7 | 0 | 0 | 16 |
| 7 | -4.11 | 2.10 | -1.96 | 0.05 | 1 | C030017B01Rik | | 15 | 23 | 7 | 0 | 27 | 1 | 7 | 0 | 0 | 0 | 1 |
| 7 | -0.98 | 1.35 | -0.72 | 0.47 | 1 | Gm12286 | | 4 | 13 | 5 | 4 | 31 | 2 | 7 | 1 | 2 | 6 | 6 |
| 7 | 0.46 | 1.65 | 0.28 | 0.78 | 1 | Gm8546 | | 15 | 21 | 2 | 0 | 3 | 5 | 7 | 0 | 4 | 21 | 2 |
| 7 | 1.13 | 1.80 | 0.63 | 0.53 | 1 | Gm14321 | | 12 | 9 | 0 | 12 | 0 | 8 | 7 | 0 | 7 | 8 | 14 |
| 7 | 0.45 | 2.70 | 0.17 | 0.87 | 1 | Gm37780 | | 3 | 0 | 19 | 0 | 0 | 8 | 7 | 31 | 0 | 0 | 8 |
| 7 | -0.36 | 1.61 | -0.22 | 0.82 | 1 | Gm6685 | | 35 | 5 | 0 | 0 | 5 | 4 | 7 | 2 | 1 | 13 | 2 |
| 7 | -0.93 | 1.34 | -0.70 | 0.49 | 1 | Ccdc73 | coiled-coil domain containing 73 | 3 | 7 | 3 | 3 | 12 | 14 | 7 | 8 | 12 | 0 | 4 |
| 7 | -4.17 | 2.55 | -1.64 | 0.10 | 1 | Gm50253 | | 0 | 0 | 6 | 0 | 1 | 39 | 7 | 0 | 0 | 0 | 19 |
| 7 | -0.78 | 1.59 | -0.49 | 0.62 | 1 | Gm16066 | | 1 | 0 | 0 | 11 | 8 | 8 | 7 | 0 | 4 | 28 | 6 |
| 6 | -2.27 | 1.77 | -1.28 | 0.20 | 1 | A130006I12Rik | | 11 | 0 | 12 | 6 | 11 | 5 | 7 | 0 | 16 | 0 | 2 |
| 6 | -0.61 | 1.62 | -0.38 | 0.70 | 1 | Gm29395 | | 5 | 0 | 0 | 9 | 3 | 6 | 7 | 0 | 3 | 32 | 6 |
| 6 | -3.17 | 1.85 | -1.71 | 0.09 | 1 | Gm5493 | | 0 | 0 | 3 | 11 | 20 | 10 | 7 | 0 | 6 | 0 | 13 |
| 6 | -1.80 | 1.72 | -1.05 | 0.29 | 1 | Pde11a | phosphodiesterase 11A | 7 | 7 | 19 | 0 | 9 | 8 | 7 | 0 | 2 | 0 | 11 |
| 6 | 0.22 | 2.24 | 0.10 | 0.92 | 1 | Gm43403 | | 12 | 17 | 0 | 7 | 0 | 7 | 7 | 0 | 10 | 0 | 8 |
| 6 | -0.89 | 1.86 | -0.48 | 0.63 | 1 | Omp | olfactory marker protein | 7 | 1 | 0 | 3 | 11 | 8 | 7 | 0 | 12 | 0 | 19 |
| 6 | -1.74 | 2.54 | -0.68 | 0.49 | 1 | Gabrr2 | gamma-aminobutyric acid (GABA) C rece | 0 | 0 | 0 | 7 | 0 | 24 | 7 | 0 | 4 | 0 | 25 |
| 6 | -0.36 | 1.33 | -0.27 | 0.79 | 1 | Gm10282 | | 3 | 3 | 3 | 1 | 13 | 10 | 7 | 13 | 3 | 1 | 9 |
| 6 | -1.09 | 2.54 | -0.43 | 0.67 | 1 | Themis | thymocyte selection associated | 35 | 19 | 0 | 0 | 0 | 3 | 7 | 0 | 0 | 0 | 3 |
| 6 | -0.07 | 1.87 | -0.04 | 0.97 | 1 | Gm12430 | | 2 | 0 | 0 | 21 | 0 | 8 | 7 | 2 | 17 | 1 | 8 |
| 6 | -0.86 | 1.48 | -0.58 | 0.56 | 1 | Gm5896 | | 1 | 8 | 1 | 25 | 0 | 4 | 7 | 13 | 0 | 1 | 6 |
| 6 | -2.46 | 1.49 | -1.64 | 0.10 | 1 | Gm37820 | | 7 | 14 | 5 | 11 | 4 | 10 | 7 | 0 | 0 | 4 | 4 |
| 6 | -0.39 | 2.14 | -0.18 | 0.85 | 1 | Tnni3k | TNNI3 interacting kinase | 0 | 0 | 0 | 0 | 13 | 16 | 7 | 6 | 15 | 0 | 7 |
| 6 | 0.15 | 1.55 | 0.10 | 0.92 | 1 | Gm3362 | | 13 | 6 | 4 | 6 | 0 | 6 | 7 | 0 | 4 | 5 | 12 |
| 6 | -0.61 | 1.52 | -0.40 | 0.69 | 1 | 4933413C19Rik | | 0 | 0 | 0 | 4 | 6 | 19 | 7 | 3 | 2 | 19 | 3 |
| 6 | -0.94 | 2.16 | -0.44 | 0.66 | 1 | Hist1h2bp | histone cluster 1, H2bp | 7 | 0 | 0 | 9 | 8 | 6 | 7 | 0 | 19 | 0 | 8 |
| 6 | -0.85 | 2.26 | -0.38 | 0.71 | 1 | 4933417C20Rik | | 4 | 0 | 8 | 10 | 0 | 5 | 7 | 0 | 0 | 10 | 18 |
| 6 | -0.52 | 1.84 | -0.28 | 0.78 | 1 | Gm4950 | | 3 | 0 | 4 | 7 | 0 | 10 | 7 | 0 | 12 | 9 | 10 |
| 6 | -2.51 | 1.46 | -1.72 | 0.09 | 1 | Gm6054 | | 1 | 9 | 12 | 8 | 9 | 7 | 7 | 2 | 1 | 0 | 5 |
| 6 | -2.92 | 2.43 | -1.20 | 0.23 | 1 | 1700001L19Ri | RIKEN cDNA 1700001L19 gene | 3 | 0 | 17 | 12 | 0 | 1 | 7 | 0 | 6 | 0 | 16 |
| 6 | -0.57 | 2.34 | -0.24 | 0.81 | 1 | Gm11451 | | 6 | 2 | 0 | 18 | 0 | 6 | 7 | 0 | 18 | 0 | 4 |
| 6 | -1.04 | 2.13 | -0.49 | 0.63 | 1 | Gm27017 | | 0 | 5 | 0 | 3 | 3 | 18 | 7 | 0 | 8 | 0 | 17 |
| 6 | -5.52 | 2.89 | -1.91 | 0.06 | 1 | Gm43696 | | 0 | 8 | 0 | 0 | 2 | 11 | 7 | 0 | 0 | 0 | 32 |
| 5 | 0.82 | 3.47 | 0.24 | 0.81 | 1 | Gm26569 | | 0 | 0 | 12 | 0 | 0 | 0 | 7 | 30 | 10 | 0 | 0 |
| 5 | 0.50 | 1.94 | 0.26 | 0.80 | 1 | Mdm4-ps | | 15 | 2 | 0 | 0 | 1 | 14 | 7 | 0 | 9 | 1 | 10 |
| 5 | 0.35 | 1.95 | 0.18 | 0.86 | 1 | Cryba4 | crystallin, beta A4 | 5 | 23 | 6 | 0 | 0 | 3 | 7 | 8 | 3 | 0 | 2 |
| 5 | -2.48 | 2.20 | -1.13 | 0.26 | 1 | Slc34a1 | solute carrier family 34 (sodium phosphat | 0 | 12 | 0 | 10 | 14 | 3 | 7 | 0 | 4 | 0 | 7 |
| 5 | -3.96 | 2.11 | -1.88 | 0.06 | 1 | Htr1d | 5-hydroxytryptamine (serotonin) receptor | 1 | 0 | 22 | 4 | 16 | 4 | 7 | 0 | 0 | 0 | 3 |
| 5 | -3.52 | 2.72 | -1.29 | 0.20 | 1 | 0610025J13Rik | | 0 | 5 | 8 | 0 | 3 | 2 | 7 | 32 | 0 | 0 | 0 |
| 5 | -0.87 | 2.31 | -0.38 | 0.71 | 1 | Gm20616 | | 0 | 0 | 17 | 3 | 5 | 3 | 7 | 0 | 0 | 21 | 2 |
| 5 | -2.81 | 2.07 | -1.35 | 0.18 | 1 | Cacng8 | calcium channel, voltage-dependent, gan | 9 | 3 | 15 | 0 | 9 | 5 | 7 | 0 | 0 | 0 | 7 |
| 5 | 1.36 | 2.83 | 0.48 | 0.63 | 1 | Gm16104 | | 10 | 15 | 0 | 0 | 0 | 0 | 7 | 22 | 0 | 0 | 1 |
| 5 | -0.78 | 1.73 | -0.45 | 0.65 | 1 | Gm39318 | | 11 | 4 | 2 | 4 | 0 | 9 | 7 | 0 | 0 | 10 | 7 |
| 5 | 0.00 | 1.94 | 0.00 | 1.00 | 1 | Igf2os | | 8 | 0 | 4 | 6 | 0 | 8 | 7 | 7 | 2 | 0 | 12 |
| 5 | -0.67 | 1.37 | -0.49 | 0.62 | 1 | Aldoart1 | aldolase 1 A, retrogene 1 | 9 | 4 | 1 | 5 | 4 | 5 | 7 | 5 | 0 | 6 | 8 |
| 5 | -3.78 | 2.31 | -1.64 | 0.10 | 1 | Atcayos | | 10 | 9 | 19 | 0 | 2 | 7 | 7 | 0 | 0 | 0 | 0 |
| 5 | 0.29 | 1.57 | 0.18 | 0.86 | 1 | Gm4654 | | 2 | 19 | 0 | 2 | 1 | 5 | 7 | 5 | 3 | 3 | 6 |

| | | | | | | | | | | | | | | | | | | |
|---|---|---|---|---|---|---|---|---|---|---|---|---|---|---|---|---|---|---|
| 5 | -0.82 | 1.58 | -0.52 | 0.60 | 1 | Gm4734 | | 2 | 2 | 17 | 0 | 6 | 9 | 7 | 2 | 1 | 1 | 5 |
| 5 | -2.46 | 2.38 | -1.04 | 0.30 | 1 | Gm18190 | | 1 | 4 | 0 | 4 | 0 | 20 | 7 | 0 | 0 | 0 | 16 |
| 5 | 0.71 | 3.20 | 0.22 | 0.83 | 1 | Cyp2f2 | cytochrome P450, family 2, subfamily f, pc | 15 | 0 | 12 | 0 | 0 | 0 | 7 | 0 | 0 | 17 | 0 |
| 5 | -2.51 | 2.51 | -1.00 | 0.32 | 1 | 2610306O10Rik | | 2 | 18 | 7 | 0 | 0 | 15 | 7 | 0 | 0 | 0 | 2 |
| 5 | -4.50 | 2.92 | -1.54 | 0.12 | 1 | Cavin4 | caveolae associated 4 | 0 | 14 | 17 | 0 | 9 | 4 | 7 | 0 | 0 | 0 | 0 |
| 5 | -1.42 | 2.14 | -0.67 | 0.51 | 1 | Gm12766 | | 16 | 2 | 18 | 0 | 1 | 2 | 7 | 0 | 4 | 0 | 0 |
| 4 | 0.02 | 2.38 | 0.01 | 0.99 | 1 | Gm45890 | | 2 | 7 | 7 | 0 | 0 | 7 | 7 | 0 | 12 | 0 | 7 |
| 4 | -0.67 | 3.12 | -0.21 | 0.83 | 1 | Mir1968 | | 9 | 0 | 0 | 0 | 0 | 17 | 7 | 0 | 0 | 0 | 16 |
| 4 | -0.67 | 1.35 | -0.49 | 0.62 | 1 | Gm9824 | | 3 | 0 | 2 | 6 | 8 | 6 | 7 | 5 | 2 | 5 | 5 |
| 4 | -2.51 | 2.30 | -1.09 | 0.28 | 1 | Eddm3b | epididymal protein 3B | 0 | 0 | 0 | 8 | 9 | 8 | 7 | 0 | 4 | 0 | 12 |
| 4 | 0.84 | 2.23 | 0.38 | 0.71 | 1 | Gm37748 | | 9 | 0 | 0 | 0 | 4 | 11 | 7 | 5 | 4 | 0 | 8 |
| 4 | -1.97 | 1.94 | -1.02 | 0.31 | 1 | Gm7324 | | 2 | 5 | 0 | 4 | 3 | 16 | 7 | 0 | 4 | 0 | 5 |
| 4 | -0.54 | 2.03 | -0.27 | 0.79 | 1 | Gm43143 | | 3 | 6 | 0 | 3 | 2 | 8 | 7 | 0 | 7 | 0 | 10 |
| 4 | -2.44 | 2.16 | -1.13 | 0.26 | 1 | Gm45022 | | 0 | 4 | 0 | 7 | 7 | 2 | 7 | 16 | 0 | 0 | 3 |
| 4 | -1.66 | 1.53 | -1.08 | 0.28 | 1 | Rps15a-ps5 | | 9 | 7 | 6 | 3 | 7 | 1 | 7 | 3 | 1 | 1 | 0 |
| 4 | -2.94 | 2.85 | -1.03 | 0.30 | 1 | Mir3058 | | 0 | 13 | 0 | 11 | 0 | 9 | 7 | 0 | 0 | 0 | 3 |
| 4 | -1.28 | 2.27 | -0.56 | 0.57 | 1 | Gm13423 | | 8 | 0 | 0 | 7 | 0 | 12 | 7 | 0 | 3 | 1 | 5 |
| 4 | 0.20 | 1.42 | 0.14 | 0.89 | 1 | Gm12912 | | 4 | 3 | 6 | 6 | 1 | 1 | 7 | 3 | 4 | 3 | 4 |
| 4 | 0.01 | 3.37 | 0.00 | 1.00 | 1 | D230049E03Rik | | 0 | 0 | 0 | 0 | 0 | 11 | 7 | 0 | 2 | 0 | 22 |
| 4 | 0.07 | 1.60 | 0.04 | 0.97 | 1 | Gm4335 | | 3 | 0 | 10 | 1 | 9 | 1 | 7 | 2 | 4 | 1 | 3 |
| 4 | -6.02 | 3.29 | -1.83 | 0.07 | 1 | Grid2 | glutamate receptor, ionotropic, delta 2 | 6 | 0 | 15 | 11 | 0 | 0 | 7 | 0 | 0 | 0 | 2 |
| 4 | -1.42 | 1.98 | -0.72 | 0.47 | 1 | Gm6776 | | 0 | 13 | 0 | 2 | 5 | 3 | 7 | 1 | 1 | 1 | 7 |
| 4 | 0.58 | 3.44 | 0.17 | 0.87 | 1 | Gm44613 | | 0 | 0 | 0 | 0 | 0 | 3 | 7 | 25 | 4 | 0 | 0 |
| 4 | 0.61 | 2.44 | 0.25 | 0.80 | 1 | Gm8495 | | 10 | 6 | 2 | 6 | 0 | 0 | 7 | 0 | 4 | 0 | 3 |
| 3 | -1.24 | 1.69 | -0.73 | 0.46 | 1 | Gm10074 | | 3 | 10 | 4 | 1 | 4 | 3 | 7 | 0 | 2 | 1 | 3 |
| 3 | -1.53 | 2.80 | -0.55 | 0.59 | 1 | Gm47428 | | 0 | 0 | 0 | 0 | 10 | 8 | 7 | 0 | 5 | 0 | 7 |
| 3 | 1.38 | 3.06 | 0.45 | 0.65 | 1 | Platr27 | | 6 | 0 | 0 | 0 | 0 | 6 | 7 | 0 | 3 | 0 | 14 |
| 3 | -3.36 | 2.94 | -1.14 | 0.25 | 1 | Gm47701 | | 0 | 0 | 0 | 4 | 2 | 14 | 7 | 0 | 0 | 0 | 10 |
| 3 | -1.29 | 1.95 | -0.66 | 0.51 | 1 | Gm8717 | | 3 | 0 | 16 | 0 | 4 | 3 | 7 | 0 | 2 | 1 | 1 |
| 3 | -0.99 | 1.76 | -0.56 | 0.58 | 1 | Rpl9-ps7 | | 3 | 8 | 5 | 1 | 3 | 2 | 7 | 0 | 1 | 5 | 1 |
| 3 | -1.24 | 1.87 | -0.66 | 0.51 | 1 | Gm5827 | | 12 | 3 | 0 | 2 | 1 | 6 | 7 | 2 | 1 | 0 | 1 |
| 3 | -3.80 | 2.55 | -1.49 | 0.14 | 1 | Gm12580 | | 3 | 7 | 7 | 3 | 8 | 0 | 7 | 0 | 0 | 0 | 0 |
| 3 | -2.24 | 1.79 | -1.25 | 0.21 | 1 | Gm6433 | | 7 | 2 | 8 | 2 | 3 | 2 | 7 | 0 | 2 | 1 | 0 |
| 3 | -0.32 | 3.07 | -0.10 | 0.92 | 1 | Gm48940 | | 0 | 0 | 0 | 0 | 3 | 3 | 7 | 0 | 17 | 0 | 2 |
| 3 | 0.73 | 2.07 | 0.35 | 0.72 | 1 | Rpsa-ps5 | | 2 | 1 | 8 | 0 | 3 | 0 | 7 | 3 | 5 | 3 | 0 |
| 3 | -1.18 | 2.30 | -0.52 | 0.61 | 1 | Gm5510 | | 0 | 1 | 0 | 10 | 2 | 2 | 7 | 0 | 4 | 6 | 0 |
| 3 | -0.35 | 3.44 | -0.10 | 0.92 | 1 | Slc6a5 | solute carrier family 6 (neurotransmitter tr | 0 | 10 | 0 | 0 | 0 | 4 | 7 | 0 | 7 | 0 | 3 |
| 3 | -1.71 | 2.05 | -0.83 | 0.40 | 1 | Gm14200 | | 5 | 4 | 0 | 3 | 2 | 1 | 7 | 1 | 0 | 6 | 0 |
| 3 | -4.22 | 3.51 | -1.20 | 0.23 | 1 | B430218F22Rik | | 0 | 3 | 0 | 0 | 0 | 4 | 7 | 0 | 14 | 0 | 0 |
| 3 | -1.68 | 3.17 | -0.53 | 0.60 | 1 | Gm49586 | | 9 | 0 | 3 | 0 | 0 | 8 | 7 | 0 | 0 | 0 | 2 |
| 3 | 1.37 | 3.07 | 0.45 | 0.65 | 1 | Gm9316 | | 4 | 6 | 0 | 0 | 0 | 2 | 7 | 0 | 4 | 0 | 5 |
| 3 | -3.39 | 3.43 | -0.99 | 0.32 | 1 | Gm48161 | | 0 | 8 | 9 | 0 | 2 | 2 | 7 | 0 | 0 | 0 | 0 |
| 3 | -1.88 | 2.20 | -0.85 | 0.39 | 1 | Ccdc50-ps | | 2 | 2 | 3 | 0 | 4 | 4 | 7 | 0 | 1 | 0 | 5 |
| 3 | 1.09 | 2.87 | 0.38 | 0.70 | 1 | 1700037C06Rik | | 9 | 0 | 3 | 0 | 0 | 1 | 7 | 0 | 4 | 0 | 4 |
| 2 | -3.38 | 3.48 | -0.97 | 0.33 | 1 | Gm31819 | | 0 | 0 | 6 | 0 | 0 | 4 | 7 | 6 | 4 | 0 | 0 |
| 2 | -0.99 | 2.44 | -0.41 | 0.68 | 1 | Gm12226 | | 0 | 1 | 10 | 1 | 0 | 1 | 7 | 0 | 3 | 1 | 1 |
| 2 | 1.00 | 2.09 | 0.48 | 0.63 | 1 | Gm6368 | | 1 | 1 | 2 | 0 | 0 | 3 | 7 | 1 | 1 | 3 | 7 |
| 2 | -1.41 | 3.55 | -0.40 | 0.69 | 1 | Gm42769 | | 0 | 0 | 0 | 0 | 0 | 2 | 7 | 0 | 0 | 0 | 16 |
| 2 | -1.13 | 2.98 | -0.38 | 0.70 | 1 | Gm46593 | | 0 | 5 | 0 | 1 | 0 | 3 | 7 | 2 | 0 | 0 | 6 |
| 2 | -1.81 | 2.06 | -0.88 | 0.38 | 1 | Gm15796 | | 9 | 1 | 1 | 1 | 1 | 3 | 7 | 1 | 0 | 0 | 1 |
| 2 | -2.66 | 3.46 | -0.77 | 0.44 | 1 | Gm43343 | | 3 | 0 | 0 | 0 | 7 | 3 | 7 | 0 | 0 | 0 | 5 |
| 2 | -6.92 | 3.52 | -1.97 | 0.05 | 1 | 1810007C17Rik | | 0 | 3 | 0 | 0 | 8 | 3 | 7 | 0 | 0 | 0 | 3 |
| 2 | -0.85 | 2.09 | -0.41 | 0.68 | 1 | Rpl15-ps6 | | 3 | 2 | 3 | 0 | 2 | 2 | 7 | 1 | 1 | 0 | 2 |
| 2 | -0.60 | 3.58 | -0.17 | 0.87 | 1 | Gm10046 | | 7 | 0 | 7 | 0 | 0 | 0 | 7 | 0 | 1 | 0 | 0 |
| 2 | -2.16 | 3.10 | -0.70 | 0.49 | 1 | Gm5471 | | 3 | 0 | 1 | 8 | 0 | 0 | 7 | 0 | 0 | 1 | 1 |
| 2 | -2.61 | 3.56 | -0.73 | 0.46 | 1 | Gm7972 | predicted gene 7972 | 0 | 0 | 0 | 0 | 0 | 14 | 7 | 0 | 0 | 0 | 1 |
| 2 | 1.92 | 3.52 | 0.55 | 0.58 | 1 | Gm44760 | | 3 | 0 | 0 | 0 | 0 | 1 | 7 | 0 | 6 | 0 | 4 |
| 2 | -3.26 | 2.94 | -1.11 | 0.27 | 1 | Gm38317 | | 1 | 1 | 0 | 5 | 0 | 6 | 7 | 1 | 0 | 0 | 0 |
| 2 | -4.28 | 3.52 | -1.22 | 0.22 | 1 | Gm49172 | | 0 | 0 | 3 | 0 | 1 | 3 | 7 | 5 | 0 | 0 | 0 |
| 2 | -5.54 | 3.52 | -1.57 | 0.12 | 1 | Gm17853 | | 0 | 0 | 0 | 2 | 0 | 3 | 7 | 0 | 7 | 0 | 0 |

| | | | | | | | | | | | | | | | | | | |
|---|---|---|---|---|---|---|---|---|---|---|---|---|---|---|---|---|---|---|
| 2 | -1.67 | 2.94 | -0.57 | 0.57 | 1 | Gm6682 | | 0 | 2 | 4 | 1 | 0 | 2 | 7 | 1 | 0 | 1 | 0 |
| 2 | -1.88 | 3.55 | -0.53 | 0.60 | 1 | Gm43231 | | 0 | 1 | 5 | 0 | 0 | 1 | 7 | 0 | 4 | 0 | 0 |
| 2 | -3.41 | 3.57 | -0.96 | 0.34 | 1 | Gm50019 | | 4 | 0 | 0 | 0 | 4 | 3 | 7 | 0 | 0 | 0 | 0 |
| 2 | -0.29 | 3.54 | -0.08 | 0.93 | 1 | Gm43873 | | 0 | 0 | 0 | 0 | 0 | 3 | 7 | 0 | 1 | 0 | 6 |
| 2 | -2.82 | 3.58 | -0.79 | 0.43 | 1 | Gm35755 | | 0 | 2 | 0 | 0 | 0 | 0 | 7 | 0 | 0 | 0 | 8 |
| 2 | -1.54 | 3.51 | -0.44 | 0.66 | 1 | 2010110E17Rik | | 0 | 0 | 0 | 3 | 0 | 0 | 7 | 0 | 1 | 0 | 6 |
| 2 | -4.03 | 3.56 | -1.13 | 0.26 | 1 | Xkr9 | X-linked Kx blood group related 9 | 0 | 0 | 0 | 4 | 0 | 5 | 7 | 0 | 0 | 0 | 0 |
| 1 | 0.70 | 3.54 | 0.20 | 0.84 | 1 | Gm13844 | | 3 | 0 | 0 | 0 | 0 | 2 | 7 | 0 | 2 | 0 | 2 |
| 1 | -0.71 | 3.60 | -0.20 | 0.84 | 1 | Mtarc1 | mitochondrial amidoxime reducing compo | 1 | 7 | 0 | 0 | 0 | 0 | 7 | 0 | 1 | 0 | 0 |
| 1 | -1.31 | 3.58 | -0.37 | 0.71 | 1 | Rpl35a-ps5 | | 6 | 0 | 0 | 0 | 2 | 0 | 7 | 0 | 0 | 0 | 1 |
| 1 | -2.46 | 3.57 | -0.69 | 0.49 | 1 | Gm43760 | | 0 | 0 | 0 | 0 | 0 | 3 | 7 | 0 | 0 | 0 | 6 |
| 1 | -1.98 | 3.55 | -0.56 | 0.58 | 1 | Gm44434 | | 0 | 0 | 0 | 0 | 1 | 3 | 7 | 0 | 0 | 0 | 5 |
| 1 | -0.38 | 3.61 | -0.11 | 0.92 | 1 | Rpl31-ps14 | | 4 | 1 | 2 | 0 | 0 | 0 | 7 | 0 | 0 | 0 | 1 |
| 1 | -0.76 | 3.57 | -0.21 | 0.83 | 1 | Dpys | dihydropyrimidinase | 5 | 0 | 0 | 1 | 0 | 0 | 7 | 0 | 0 | 0 | 2 |
| 1 | -0.97 | 3.62 | -0.27 | 0.79 | 1 | Cyp8b1 | cytochrome P450, family 8, subfamily b, p | 5 | 0 | 2 | 0 | 0 | 0 | 7 | 0 | 0 | 0 | 0 |
| 1 | -3.07 | 3.58 | -0.86 | 0.39 | 1 | Gm50311 | | 0 | 0 | 0 | 7 | 0 | 0 | 7 | 0 | 0 | 0 | 0 |
| 1 | -1.84 | 3.54 | -0.52 | 0.60 | 1 | Gm50263 | | 0 | 0 | 0 | 0 | 0 | 3 | 7 | 0 | 4 | 0 | 0 |
| 1 | -5.25 | 3.55 | -1.48 | 0.14 | 1 | Gm8616 | | 0 | 0 | 2 | 1 | 0 | 0 | 7 | 0 | 0 | 0 | 3 |
| 1 | -1.80 | 3.56 | -0.51 | 0.61 | 1 | Gm8146 | | 0 | 0 | 0 | 0 | 1 | 1 | 7 | 0 | 0 | 0 | 4 |
| 1 | -1.53 | 3.54 | -0.43 | 0.67 | 1 | St8sia3os | | 0 | 0 | 0 | 0 | 0 | 2 | 7 | 0 | 4 | 0 | 0 |
| 1 | -0.73 | 3.62 | -0.20 | 0.84 | 1 | Gm29439 | | 4 | 0 | 0 | 0 | 0 | 0 | 7 | 0 | 0 | 0 | 0 |
| 1 | -1.00 | 3.56 | -0.28 | 0.78 | 1 | Gm20379 | | 0 | 0 | 0 | 0 | 0 | 1 | 7 | 1 | 1 | 0 | 0 |
| 1 | -2.07 | 3.59 | -0.58 | 0.56 | 1 | Gm3150 | | 0 | 0 | 0 | 0 | 0 | 2 | 7 | 1 | 0 | 0 | 0 |
| 1 | -1.56 | 3.60 | -0.43 | 0.66 | 1 | Gm14202 | | 0 | 0 | 0 | 0 | 0 | 2 | 7 | 0 | 0 | 0 | 1 |
| 1 | -1.56 | 3.60 | -0.43 | 0.66 | 1 | Gm37240 | ADP-ribosylation factor interacting protein | 0 | 0 | 0 | 0 | 0 | 2 | 7 | 0 | 0 | 0 | 1 |
| 1 | -2.64 | 3.59 | -0.74 | 0.46 | 1 | Gm45837 | phosphodiesterase 2A, cGMP-stimulated | 0 | 0 | 0 | 0 | 0 | 3 | 7 | 0 | 0 | 0 | 0 |
| 1 | -1.27 | 3.60 | -0.35 | 0.72 | 1 | Gm12973 | | 0 | 0 | 0 | 0 | 0 | 1 | 7 | 0 | 0 | 0 | 1 |
| 1 | -1.71 | 3.62 | -0.47 | 0.64 | 1 | Gm47553 | | 1 | 0 | 0 | 0 | 1 | 0 | 7 | 0 | 0 | 0 | 0 |
| 1 | -2.42 | 3.60 | -0.67 | 0.50 | 1 | Gm47212 | | 0 | 0 | 0 | 0 | 0 | 2 | 7 | 0 | 0 | 0 | 0 |
| 1 | -2.05 | 3.62 | -0.57 | 0.57 | 1 | Gm7690 | | 0 | 0 | 0 | 1 | 0 | 0 | 7 | 0 | 0 | 0 | 0 |
| 1 | -1.03 | 3.62 | -0.28 | 0.78 | 1 | Enthd1 | ENTH domain containing 1 | 1 | 0 | 0 | 0 | 0 | 0 | 7 | 0 | 0 | 0 | 0 |
| 1 | -1.03 | 3.62 | -0.28 | 0.78 | 1 | Cyp2d9 | cytochrome P450, family 2, subfamily d, p | 1 | 0 | 0 | 0 | 0 | 0 | 7 | 0 | 0 | 0 | 0 |
| 1 | -2.07 | 3.62 | -0.57 | 0.57 | 1 | Gm20110 | | 0 | 0 | 0 | 0 | 0 | 1 | 7 | 0 | 0 | 0 | 0 |
| 1 | -1.28 | 3.63 | -0.35 | 0.73 | 1 | n-R5s164 | | 0 | 0 | 0 | 0 | 0 | 0 | 7 | 0 | 0 | 0 | 0 |
| 1 | -1.28 | 3.63 | -0.35 | 0.73 | 1 | Nlrp12 | NLR family, pyrin domain containing 12 | 0 | 0 | 0 | 0 | 0 | 0 | 7 | 0 | 0 | 0 | 0 |
| 1 | -1.28 | 3.63 | -0.35 | 0.73 | 1 | Gm44684 | | 0 | 0 | 0 | 0 | 0 | 0 | 7 | 0 | 0 | 0 | 0 |
| 1 | -1.28 | 3.63 | -0.35 | 0.73 | 1 | Gm16177 | | 0 | 0 | 0 | 0 | 0 | 0 | 7 | 0 | 0 | 0 | 0 |
| 1 | -1.28 | 3.63 | -0.35 | 0.73 | 1 | 1700034J04Rik | | 0 | 0 | 0 | 0 | 0 | 0 | 7 | 0 | 0 | 0 | 0 |
| 1 | -1.28 | 3.63 | -0.35 | 0.73 | 1 | Mir154 | | 0 | 0 | 0 | 0 | 0 | 0 | 7 | 0 | 0 | 0 | 0 |
| 1 | -1.28 | 3.63 | -0.35 | 0.73 | 1 | Rpl7a-ps2 | | 0 | 0 | 0 | 0 | 0 | 0 | 7 | 0 | 0 | 0 | 0 |
| 1 | -1.28 | 3.63 | -0.35 | 0.73 | 1 | Gm47658 | | 0 | 0 | 0 | 0 | 0 | 0 | 7 | 0 | 0 | 0 | 0 |
| 1 | -1.28 | 3.63 | -0.35 | 0.73 | 1 | Trav1 | | 0 | 0 | 0 | 0 | 0 | 0 | 7 | 0 | 0 | 0 | 0 |
| 1 | -1.28 | 3.63 | -0.35 | 0.73 | 1 | Lrrc15 | leucine rich repeat containing 15 | 0 | 0 | 0 | 0 | 0 | 0 | 7 | 0 | 0 | 0 | 0 |
| 2449 | -2.77 | 1.58 | -1.76 | 0.08 | 1 | Tnxb | tenascin XB | 13072 | 6367 | 7478 | 7 | 0 | 3 | 6 | 0 | 0 | 0 | 3 |
| 1261 | -1.24 | 1.25 | -0.99 | 0.32 | 1 | Pdgfrb | platelet derived growth factor receptor, be | 5869 | 3856 | 4111 | 0 | 9 | 7 | 6 | 0 | 5 | 0 | 6 |
| 1105 | -0.31 | 1.83 | -0.17 | 0.87 | 1 | Aqp1 | aquaporin 1 | 4894 | 3418 | 3826 | 0 | 0 | 3 | 6 | 0 | 6 | 0 | 1 |
| 964 | -0.82 | 1.79 | -0.46 | 0.65 | 1 | Slc43a3 | solute carrier family 43, member 3 | 4658 | 3278 | 2657 | 0 | 1 | 0 | 6 | 0 | 2 | 0 | 0 |
| 937 | 0.45 | 1.06 | 0.42 | 0.67 | 1 | Itga1 | integrin alpha 1 | 3481 | 3792 | 2940 | 26 | 6 | 6 | 6 | 0 | 27 | 17 | 10 |
| 536 | 0.11 | 1.48 | 0.07 | 0.94 | 1 | Sfrp1 | secreted frizzled-related protein 1 | 3484 | 959 | 1397 | 0 | 13 | 8 | 6 | 14 | 15 | 0 | 1 |
| 519 | -2.91 | 1.70 | -1.71 | 0.09 | 1 | Calcrl | calcitonin receptor-like | 2090 | 1720 | 1879 | 9 | 0 | 2 | 6 | 0 | 0 | 0 | 3 |
| 399 | -0.25 | 0.56 | -0.45 | 0.65 | 1 | Aldh1a1 | aldehyde dehydrogenase family 1, subfar | 2090 | 934 | 1308 | 8 | 8 | 5 | 6 | 0 | 4 | 18 | 5 |
| 340 | 0.02 | 1.53 | 0.01 | 0.99 | 1 | Icam1 | intercellular adhesion molecule 1 | 2169 | 626 | 897 | 0 | 8 | 2 | 6 | 0 | 7 | 0 | 22 |
| 223 | 1.02 | 2.68 | 0.38 | 0.71 | 1 | Erg | ETS transcription factor | 758 | 938 | 719 | 0 | 0 | 0 | 6 | 0 | 0 | 28 | 0 |
| 207 | -0.57 | 2.62 | -0.22 | 0.83 | 1 | Pla2r1 | phospholipase A2 receptor 1 | 1034 | 559 | 672 | 0 | 0 | 0 | 6 | 0 | 0 | 0 | 1 |
| 202 | -2.34 | 2.14 | -1.09 | 0.27 | 1 | Myo1f | myosin IF | 1042 | 577 | 580 | 19 | 0 | 0 | 6 | 0 | 2 | 0 | 0 |
| 194 | -0.07 | 1.05 | -0.07 | 0.94 | 1 | Selplg | selectin, platelet (p-selectin) ligand | 1171 | 404 | 541 | 0 | 0 | 1 | 6 | 0 | 0 | 4 | 2 |
| 182 | -1.35 | 1.05 | -1.28 | 0.20 | 1 | Siglec1 | sialic acid binding Ig-like lectin 1, sialoadl | 975 | 527 | 485 | 0 | 0 | 4 | 6 | 0 | 0 | 0 | 4 |
| 178 | -1.03 | 0.85 | -1.21 | 0.23 | 1 | Pcolce2 | procollagen C-endopeptidase enhancer 2 | 1160 | 330 | 406 | 7 | 9 | 13 | 6 | 0 | 7 | 0 | 21 |
| 152 | -0.37 | 0.69 | -0.54 | 0.59 | 1 | Rab32 | RAB32, member RAS oncogene family | 817 | 328 | 349 | 46 | 25 | 13 | 6 | 59 | 6 | 14 | 7 |
| 148 | -0.42 | 1.57 | -0.27 | 0.79 | 1 | Ncf2 | neutrophil cytosolic factor 2 | 882 | 270 | 431 | 5 | 0 | 5 | 6 | 0 | 13 | 13 | 0 |

| | | | | | | | | | | | | | | | | | | |
|---|---|---|---|---|---|---|---|---|---|---|---|---|---|---|---|---|---|---|
| 128 | -1.06 | 3.63 | -0.29 | 0.77 | 1 | Lmod1 | leiomodin 1 (smooth muscle) | 874 | 274 | 254 | 0 | 0 | 0 | 6 | 0 | 0 | 0 | 0 |
| 122 | -1.97 | 2.41 | -0.82 | 0.41 | 1 | Gypc | glycophorin C | 654 | 370 | 314 | 0 | 0 | 1 | 6 | 0 | 0 | 0 | 0 |
| 122 | -3.04 | 2.20 | -1.38 | 0.17 | 1 | Grap | GRB2-related adaptor protein | 645 | 283 | 397 | 0 | 8 | 0 | 6 | 0 | 0 | 0 | 0 |
| 119 | -0.84 | 1.05 | -0.80 | 0.42 | 1 | Ifi47 | interferon gamma inducible protein 47 | 546 | 342 | 381 | 6 | 7 | 0 | 6 | 0 | 0 | 22 | 0 |
| 112 | -1.13 | 3.63 | -0.31 | 0.76 | 1 | Pkhd1l1 | polycystic kidney and hepatic disease 1-li | 579 | 478 | 174 | 0 | 0 | 0 | 6 | 0 | 0 | 0 | 0 |
| 109 | -0.29 | 1.64 | -0.18 | 0.86 | 1 | Lgals4 | lectin, galactose binding, soluble 4 | 3 | 255 | 255 | 0 | 17 | 357 | 6 | 0 | 4 | 13 | 288 |
| 107 | 0.37 | 0.59 | 0.64 | 0.52 | 1 | Cdca7l | cell division cycle associated 7 like | 481 | 246 | 322 | 11 | 17 | 22 | 6 | 28 | 22 | 12 | 10 |
| 92 | 0.32 | 0.59 | 0.54 | 0.59 | 1 | A630072M18Rik | | 192 | 192 | 180 | 81 | 39 | 82 | 6 | 86 | 24 | 35 | 95 |
| 92 | -0.30 | 2.82 | -0.10 | 0.92 | 1 | Gm1966 | | 548 | 305 | 150 | 0 | 0 | 0 | 6 | 0 | 0 | 0 | 3 |
| 91 | 0.30 | 1.20 | 0.25 | 0.80 | 1 | Glipr2 | GLI pathogenesis-related 2 | 494 | 172 | 263 | 10 | 0 | 4 | 6 | 0 | 0 | 49 | 7 |
| 87 | 0.02 | 0.70 | 0.03 | 0.98 | 1 | Mapk12 | mitogen-activated protein kinase 12 | 298 | 221 | 353 | 9 | 7 | 19 | 6 | 25 | 3 | 6 | 8 |
| 85 | -0.88 | 0.61 | -1.45 | 0.15 | 1 | Eya2 | EYA transcriptional coactivator and phosp | 238 | 255 | 271 | 43 | 53 | 14 | 6 | 23 | 11 | 10 | 10 |
| 84 | -2.20 | 3.44 | -0.64 | 0.52 | 1 | 1810018F18Rik | | 88 | 439 | 388 | 0 | 0 | 0 | 6 | 0 | 0 | 0 | 6 |
| 82 | -0.58 | 2.71 | -0.21 | 0.83 | 1 | Cysltr1 | cysteinyl leukotriene receptor 1 | 374 | 268 | 253 | 0 | 0 | 0 | 6 | 0 | 0 | 0 | 1 |
| 78 | -1.18 | 3.63 | -0.32 | 0.75 | 1 | Trim16 | tripartite motif-containing 16 | 339 | 219 | 298 | 0 | 0 | 0 | 6 | 0 | 0 | 0 | 0 |
| 76 | -1.07 | 0.95 | -1.12 | 0.26 | 1 | Rest | RE1-silencing transcription factor | 320 | 212 | 276 | 2 | 0 | 10 | 6 | 0 | 3 | 0 | 5 |
| 76 | -0.53 | 0.99 | -0.54 | 0.59 | 1 | Pkn3 | protein kinase N3 | 286 | 184 | 296 | 10 | 9 | 6 | 6 | 17 | 0 | 0 | 16 |
| 72 | 0.77 | 1.35 | 0.57 | 0.57 | 1 | Hk2 | hexokinase 2 | 432 | 182 | 171 | 0 | 0 | 0 | 6 | 0 | 3 | 0 | 2 |
| 69 | 0.49 | 0.67 | 0.74 | 0.46 | 1 | Pde1b | phosphodiesterase 1B, Ca2+-calmodulin | 262 | 190 | 191 | 6 | 8 | 30 | 6 | 29 | 9 | 5 | 18 |
| 61 | -1.27 | 1.11 | -1.15 | 0.25 | 1 | Hebp2 | heme binding protein 2 | 165 | 171 | 212 | 47 | 8 | 8 | 6 | 0 | 3 | 46 | 2 |
| 61 | 1.19 | 0.92 | 1.29 | 0.20 | 1 | Gmfg | glia maturation factor, gamma | 305 | 136 | 177 | 6 | 3 | 0 | 6 | 9 | 0 | 21 | 5 |
| 57 | -0.13 | 0.58 | -0.23 | 0.82 | 1 | Acot1 | acyl-CoA thioesterase 1 | 100 | 230 | 200 | 20 | 24 | 4 | 6 | 16 | 11 | 8 | 8 |
| 55 | -1.27 | 1.11 | -1.14 | 0.25 | 1 | Gm42664 | | 5 | 8 | 14 | 6 | 0 | 315 | 6 | 0 | 5 | 0 | 244 |
| 54 | 0.39 | 0.68 | 0.58 | 0.57 | 1 | Tlr12 | toll-like receptor 12 | 208 | 82 | 208 | 8 | 21 | 8 | 6 | 20 | 8 | 13 | 10 |
| 53 | -0.99 | 3.63 | -0.27 | 0.79 | 1 | Traf1 | TNF receptor-associated factor 1 | 411 | 89 | 74 | 0 | 0 | 0 | 6 | 0 | 0 | 0 | 0 |
| 52 | -2.15 | 1.27 | -1.69 | 0.09 | 1 | Pou2f2 | POU domain, class 2, transcription factor | 255 | 177 | 126 | 3 | 1 | 1 | 6 | 0 | 0 | 0 | 2 |
| 49 | 0.74 | 0.75 | 0.99 | 0.32 | 1 | Spdl1 | spindle apparatus coiled-coil protein 1 | 142 | 63 | 32 | 24 | 33 | 50 | 6 | 20 | 58 | 55 | 56 |
| 47 | -0.57 | 1.00 | -0.57 | 0.57 | 1 | Mir100hg | | 101 | 154 | 97 | 4 | 15 | 63 | 6 | 42 | 3 | 0 | 29 |
| 46 | -0.53 | 0.75 | -0.71 | 0.48 | 1 | Nsl1 | NSL1, MIS12 kinetochore complex compo | 132 | 51 | 94 | 52 | 42 | 38 | 6 | 32 | 17 | 9 | 36 |
| 46 | -1.43 | 1.11 | -1.29 | 0.20 | 1 | Gm13563 | | 23 | 16 | 19 | 0 | 13 | 264 | 6 | 0 | 6 | 0 | 159 |
| 44 | 0.43 | 1.13 | 0.38 | 0.71 | 1 | Sbsn | suprabasin | 107 | 98 | 152 | 0 | 5 | 50 | 6 | 25 | 4 | 0 | 37 |
| 43 | -0.26 | 0.77 | -0.34 | 0.73 | 1 | Exoc3l4 | exocyst complex component 3-like 4 | 192 | 72 | 112 | 25 | 15 | 3 | 6 | 13 | 11 | 23 | 1 |
| 42 | -1.57 | 1.38 | -1.14 | 0.26 | 1 | Gimap9 | GTPase, IMAP family member 9 | 208 | 126 | 115 | 0 | 8 | 0 | 6 | 0 | 2 | 0 | 1 |
| 42 | -2.44 | 1.35 | -1.81 | 0.07 | 1 | Hk3 | hexokinase 3 | 164 | 153 | 126 | 0 | 4 | 3 | 6 | 0 | 0 | 0 | 3 |
| 36 | -0.46 | 1.22 | -0.38 | 0.70 | 1 | Gm38077 | | 0 | 6 | 0 | 0 | 12 | 190 | 6 | 12 | 4 | 4 | 162 |
| 31 | -2.38 | 1.59 | -1.49 | 0.14 | 1 | Ighd | | 218 | 89 | 22 | 0 | 0 | 2 | 6 | 0 | 0 | 0 | 0 |
| 30 | -0.75 | 0.91 | -0.82 | 0.41 | 1 | Rpgrip1 | retinitis pigmentosa GTPase regulator inte | 44 | 34 | 14 | 44 | 23 | 74 | 6 | 6 | 31 | 4 | 54 |
| 30 | 0.38 | 0.99 | 0.38 | 0.70 | 1 | Msantd1 | Myb/SANT-like DNA-binding domain cont | 35 | 40 | 56 | 0 | 8 | 84 | 6 | 0 | 13 | 8 | 79 |
| 30 | -0.75 | 1.05 | -0.72 | 0.47 | 1 | Fam110c | family with sequence similarity 110, meml | 68 | 122 | 82 | 19 | 4 | 4 | 6 | 0 | 8 | 12 | 1 |
| 29 | -2.77 | 1.70 | -1.63 | 0.10 | 1 | Gcsam | germinal center associated, signaling anc | 154 | 124 | 32 | 0 | 3 | 0 | 6 | 0 | 0 | 0 | 0 |
| 27 | 2.07 | 1.43 | 1.44 | 0.15 | 1 | Serpina3c | serine (or cysteine) peptidase inhibitor, cl | 202 | 30 | 32 | 6 | 0 | 0 | 6 | 10 | 12 | 0 | 2 |
| 27 | 0.86 | 1.04 | 0.83 | 0.40 | 1 | Mx2 | MX dynamin-like GTPase 2 | 18 | 29 | 34 | 0 | 24 | 66 | 6 | 27 | 8 | 17 | 63 |
| 26 | -0.40 | 1.26 | -0.32 | 0.75 | 1 | 6720427I07Rik | | 17 | 13 | 0 | 9 | 25 | 105 | 6 | 0 | 17 | 19 | 79 |
| 25 | -0.91 | 1.78 | -0.51 | 0.61 | 1 | Ms4a4a | membrane-spanning 4-domains, subfami | 124 | 50 | 98 | 0 | 0 | 0 | 6 | 0 | 0 | 0 | 1 |
| 25 | 0.03 | 1.19 | 0.02 | 0.98 | 1 | Cct6b | chaperonin containing Tcp1, subunit 6b (: | 17 | 0 | 0 | 42 | 20 | 46 | 6 | 27 | 19 | 32 | 65 |
| 25 | 0.94 | 0.87 | 1.08 | 0.28 | 1 | 4930500F10Rik | | 44 | 57 | 10 | 26 | 6 | 25 | 6 | 31 | 34 | 24 | 10 |
| 25 | -0.11 | 0.78 | -0.14 | 0.89 | 1 | Tmc8 | transmembrane channel-like gene family | 103 | 45 | 44 | 9 | 7 | 19 | 6 | 13 | 3 | 5 | 19 |
| 25 | -2.06 | 1.38 | -1.49 | 0.14 | 1 | Tpbg | trophoblast glycoprotein | 188 | 38 | 11 | 15 | 6 | 2 | 6 | 0 | 5 | 0 | 1 |
| 25 | 0.40 | 0.74 | 0.55 | 0.58 | 1 | Abhd11os | | 54 | 42 | 94 | 11 | 6 | 13 | 6 | 14 | 12 | 12 | 9 |
| 25 | -0.66 | 1.11 | -0.60 | 0.55 | 1 | Gm16576 | | 89 | 47 | 21 | 6 | 45 | 12 | 6 | 6 | 20 | 0 | 18 |
| 24 | 1.24 | 0.93 | 1.34 | 0.18 | 1 | Cdkn2a | cyclin dependent kinase inhibitor 2A | 50 | 8 | 14 | 10 | 26 | 19 | 6 | 39 | 55 | 9 | 32 |
| 24 | -0.58 | 1.33 | -0.44 | 0.66 | 1 | Enkur | enkurin, TRPC channel interacting proteir | 77 | 51 | 46 | 8 | 1 | 32 | 6 | 0 | 23 | 0 | 23 |
| 24 | -0.04 | 1.19 | -0.03 | 0.97 | 1 | Gm37902 | | 10 | 21 | 0 | 17 | 8 | 88 | 6 | 15 | 16 | 0 | 83 |
| 24 | 0.81 | 0.97 | 0.83 | 0.41 | 1 | Bfsp1 | beaded filament structural protein 1, in ler | 125 | 46 | 52 | 0 | 1 | 12 | 6 | 2 | 4 | 8 | 8 |
| 24 | -2.85 | 1.76 | -1.62 | 0.11 | 1 | Gm45457 | | 0 | 0 | 15 | 4 | 0 | 136 | 6 | 0 | 4 | 0 | 98 |
| 24 | -3.29 | 1.85 | -1.78 | 0.07 | 1 | Rasef | RAS and EF hand domain containing | 27 | 126 | 98 | 0 | 7 | 0 | 6 | 0 | 0 | 0 | 0 |
| 23 | 0.14 | 1.08 | 0.13 | 0.90 | 1 | Gm19391 | | 34 | 71 | 46 | 3 | 9 | 23 | 6 | 6 | 39 | 0 | 12 |
| 22 | -0.90 | 0.89 | -1.00 | 0.32 | 1 | Ak8 | adenylate kinase 8 | 66 | 57 | 29 | 22 | 16 | 14 | 6 | 2 | 5 | 19 | 9 |
| 22 | 0.29 | 1.19 | 0.25 | 0.81 | 1 | Rep15 | RAB15 effector protein | 39 | 32 | 82 | 0 | 19 | 14 | 6 | 0 | 8 | 28 | 11 |
| 21 | -1.60 | 1.33 | -1.21 | 0.23 | 1 | 1700020D05Ril | RIKEN cDNA 1700020D05 gene | 0 | 0 | 0 | 6 | 12 | 117 | 6 | 0 | 16 | 0 | 73 |

| | | | | | | | | | | | | | | | | | | |
|---|---|---|---|---|---|---|---|---|---|---|---|---|---|---|---|---|---|---|
| 20 | 0.53 | 1.24 | 0.43 | 0.67 | 1 | P2rx6 | purinergic receptor P2X, ligand-gated ion | 66 | 40 | 46 | 13 | 0 | 8 | 6 | 27 | 3 | 0 | 13 |
| 20 | 0.69 | 1.18 | 0.58 | 0.56 | 1 | Pif1 | PIF1 5'-to-3' DNA helicase | 45 | 9 | 8 | 16 | 12 | 25 | 6 | 28 | 29 | 0 | 40 |
| 20 | 2.20 | 1.20 | 1.83 | 0.07 | 1 | Cd46 | CD46 antigen, complement regulatory pro | 9 | 0 | 0 | 3 | 0 | 69 | 6 | 7 | 14 | 31 | 76 |
| 19 | 0.91 | 1.39 | 0.65 | 0.51 | 1 | Arhgef39 | Rho guanine nucleotide exchange factor | 24 | 11 | 28 | 0 | 26 | 25 | 6 | 44 | 25 | 0 | 24 |
| 19 | -0.40 | 1.11 | -0.36 | 0.72 | 1 | Gm49747 | | 19 | 25 | 11 | 10 | 27 | 33 | 6 | 0 | 8 | 22 | 50 |
| 19 | -1.39 | 2.03 | -0.69 | 0.49 | 1 | Hrc | histidine rich calcium binding protein | 104 | 34 | 67 | 0 | 0 | 0 | 6 | 0 | 0 | 0 | 0 |
| 19 | -0.25 | 1.28 | -0.19 | 0.85 | 1 | Cyp27b1 | cytochrome P450, family 27, subfamily b, | 13 | 0 | 29 | 6 | 10 | 70 | 6 | 13 | 9 | 0 | 52 |
| 19 | 2.04 | 1.43 | 1.43 | 0.15 | 1 | Tm6sf2 | transmembrane 6 superfamily member 2 | 5 | 0 | 0 | 0 | 0 | 83 | 6 | 0 | 6 | 40 | 65 |
| 18 | 0.64 | 1.07 | 0.60 | 0.55 | 1 | Dynlt1b | dynein light chain Tctex-type 1B | 33 | 2 | 0 | 11 | 7 | 35 | 6 | 19 | 8 | 43 | 38 |
| 18 | -1.42 | 1.05 | -1.36 | 0.17 | 1 | Gm5611 | | 21 | 67 | 28 | 26 | 15 | 11 | 6 | 1 | 18 | 5 | 1 |
| 18 | -0.63 | 0.85 | -0.74 | 0.46 | 1 | 1700013F07Rik | RIKEN cDNA 1700013F07 gene | 27 | 31 | 15 | 18 | 22 | 30 | 6 | 6 | 11 | 18 | 13 |
| 17 | 0.03 | 1.04 | 0.03 | 0.98 | 1 | Mtfr2 | mitochondrial fission regulator 2 | 33 | 27 | 15 | 7 | 9 | 25 | 6 | 20 | 6 | 0 | 35 |
| 16 | -0.08 | 0.93 | -0.09 | 0.93 | 1 | 5730522E02Rik | | 20 | 1 | 30 | 2 | 23 | 33 | 6 | 0 | 20 | 17 | 29 |
| 16 | -0.61 | 0.97 | -0.63 | 0.53 | 1 | Gm43814 | | 48 | 11 | 30 | 29 | 6 | 14 | 6 | 7 | 7 | 6 | 13 |
| 16 | 0.24 | 1.52 | 0.16 | 0.88 | 1 | Gm5464 | | 0 | 30 | 18 | 5 | 21 | 31 | 6 | 0 | 30 | 24 | 8 |
| 15 | -0.93 | 1.64 | -0.57 | 0.57 | 1 | Chn1os3 | | 15 | 3 | 0 | 0 | 2 | 104 | 6 | 0 | 3 | 0 | 37 |
| 15 | 0.26 | 1.51 | 0.17 | 0.87 | 1 | 1700056N10Rik | | 5 | 28 | 0 | 0 | 29 | 31 | 6 | 7 | 21 | 10 | 28 |
| 14 | 0.56 | 0.91 | 0.61 | 0.54 | 1 | Gm5822 | | 22 | 19 | 23 | 6 | 11 | 8 | 6 | 10 | 8 | 33 | 5 |
| 14 | -1.68 | 1.15 | -1.46 | 0.15 | 1 | Tceal5 | transcription elongation factor A (SII)-like | 41 | 58 | 20 | 7 | 8 | 3 | 6 | 0 | 4 | 3 | 1 |
| 14 | -1.13 | 2.17 | -0.52 | 0.60 | 1 | 5430427O19Rik | RIKEN cDNA 5430427O19 gene | 111 | 12 | 20 | 0 | 0 | 0 | 6 | 0 | 0 | 0 | 0 |
| 14 | 0.47 | 0.91 | 0.52 | 0.61 | 1 | Gm43966 | | 10 | 6 | 29 | 12 | 17 | 9 | 6 | 21 | 10 | 19 | 8 |
| 13 | -0.10 | 1.03 | -0.10 | 0.92 | 1 | B230319C09Rik | | 0 | 9 | 11 | 46 | 20 | 4 | 6 | 20 | 11 | 10 | 6 |
| 13 | -0.15 | 0.96 | -0.15 | 0.88 | 1 | Gm9531 | | 25 | 20 | 9 | 12 | 11 | 15 | 6 | 3 | 21 | 8 | 10 |
| 13 | -1.24 | 1.51 | -0.82 | 0.41 | 1 | Gm49890 | | 0 | 2 | 6 | 2 | 2 | 52 | 6 | 0 | 7 | 0 | 64 |
| 13 | -0.09 | 0.89 | -0.10 | 0.92 | 1 | Gm7666 | | 18 | 30 | 12 | 16 | 11 | 8 | 6 | 6 | 14 | 6 | 11 |
| 13 | 0.87 | 0.89 | 0.98 | 0.33 | 1 | 4930413G21Rik | | 2 | 10 | 9 | 16 | 9 | 12 | 6 | 32 | 12 | 19 | 12 |
| 12 | -1.55 | 1.74 | -0.89 | 0.37 | 1 | Sspo | SCO-spondin | 15 | 5 | 10 | 0 | 0 | 63 | 6 | 0 | 0 | 0 | 36 |
| 12 | -0.29 | 0.94 | -0.31 | 0.76 | 1 | Gm5560 | | 3 | 11 | 37 | 12 | 13 | 14 | 6 | 17 | 8 | 4 | 9 |
| 12 | -2.27 | 2.00 | -1.14 | 0.26 | 1 | Klra17 | killer cell lectin-like receptor, subfamily A, | 75 | 23 | 26 | 0 | 2 | 0 | 6 | 0 | 0 | 0 | 0 |
| 12 | -1.40 | 1.40 | -1.00 | 0.32 | 1 | Gm31816 | | 0 | 3 | 7 | 51 | 23 | 6 | 6 | 0 | 14 | 3 | 15 |
| 12 | -0.13 | 1.15 | -0.11 | 0.91 | 1 | Gm8539 | | 8 | 5 | 6 | 24 | 11 | 10 | 6 | 32 | 16 | 0 | 8 |
| 12 | -1.13 | 1.15 | -0.98 | 0.33 | 1 | Gm6467 | | 28 | 9 | 2 | 10 | 24 | 9 | 6 | 2 | 3 | 27 | 6 |
| 11 | 1.68 | 1.27 | 1.32 | 0.19 | 1 | Gm12251 | | 6 | 0 | 12 | 5 | 4 | 8 | 6 | 46 | 15 | 4 | 18 |
| 11 | 1.06 | 0.99 | 1.06 | 0.29 | 1 | Gm4833 | | 14 | 2 | 12 | 8 | 11 | 3 | 6 | 9 | 10 | 43 | 7 |
| 11 | -1.91 | 1.94 | -0.98 | 0.32 | 1 | Hfm1 | HFM1, ATP-dependent DNA helicase hor | 0 | 30 | 0 | 23 | 23 | 14 | 6 | 0 | 19 | 0 | 8 |
| 11 | 0.10 | 0.90 | 0.11 | 0.91 | 1 | Gm12999 | | 10 | 17 | 9 | 15 | 9 | 9 | 6 | 12 | 12 | 17 | 4 |
| 11 | 0.60 | 1.08 | 0.55 | 0.58 | 1 | Gm5525 | | 29 | 4 | 2 | 13 | 2 | 8 | 6 | 8 | 8 | 32 | 5 |
| 10 | -1.56 | 1.63 | -0.95 | 0.34 | 1 | Plcd4 | phospholipase C, delta 4 | 0 | 8 | 0 | 11 | 30 | 18 | 6 | 1 | 29 | 0 | 11 |
| 10 | -0.08 | 1.42 | -0.06 | 0.95 | 1 | Gm9449 | | 33 | 8 | 11 | 17 | 2 | 4 | 6 | 2 | 17 | 0 | 9 |
| 10 | -0.04 | 1.47 | -0.03 | 0.98 | 1 | Gm16170 | | 0 | 7 | 11 | 9 | 18 | 5 | 6 | 34 | 4 | 15 | 0 |
| 10 | -0.71 | 1.77 | -0.40 | 0.69 | 1 | Vstm2a | V-set and transmembrane domain contair | 0 | 2 | 0 | 13 | 9 | 18 | 6 | 0 | 20 | 0 | 42 |
| 10 | -1.50 | 1.42 | -1.06 | 0.29 | 1 | Cyp2r1 | cytochrome P450, family 2, subfamily r, pc | 19 | 15 | 9 | 8 | 17 | 8 | 6 | 0 | 11 | 0 | 14 |
| 10 | -1.06 | 1.12 | -0.94 | 0.34 | 1 | Gm26714 | | 23 | 2 | 18 | 4 | 18 | 10 | 6 | 0 | 8 | 8 | 9 |
| 10 | -0.23 | 1.53 | -0.15 | 0.88 | 1 | Gm12932 | | 0 | 3 | 4 | 13 | 7 | 22 | 6 | 0 | 5 | 33 | 11 |
| 10 | -1.52 | 1.63 | -0.93 | 0.35 | 1 | Gm15747 | | 0 | 2 | 1 | 14 | 11 | 32 | 6 | 0 | 0 | 12 | 26 |
| 9 | -1.49 | 2.72 | -0.55 | 0.58 | 1 | Gm16136 | | 7 | 17 | 73 | 0 | 0 | 0 | 6 | 0 | 0 | 0 | 0 |
| 9 | -1.48 | 1.45 | -1.02 | 0.31 | 1 | Gm11973 | | 3 | 10 | 7 | 18 | 12 | 9 | 6 | 0 | 13 | 0 | 23 |
| 9 | 1.37 | 2.13 | 0.64 | 0.52 | 1 | Gm37274 | | 0 | 0 | 17 | 0 | 0 | 34 | 6 | 23 | 7 | 5 | 10 |
| 9 | -0.29 | 1.42 | -0.21 | 0.84 | 1 | Gm26566 | | 2 | 16 | 5 | 3 | 29 | 8 | 6 | 0 | 9 | 14 | 9 |
| 9 | 3.30 | 1.86 | 1.77 | 0.08 | 1 | Gm16380 | | 18 | 4 | 0 | 4 | 0 | 1 | 6 | 58 | 10 | 0 | 1 |
| 9 | 0.28 | 1.63 | 0.17 | 0.86 | 1 | Gm32461 | | 10 | 9 | 38 | 0 | 2 | 8 | 6 | 20 | 6 | 0 | 2 |
| 9 | -1.24 | 1.73 | -0.72 | 0.47 | 1 | Tmem212 | transmembrane protein 212 | 9 | 31 | 0 | 4 | 2 | 25 | 6 | 0 | 3 | 0 | 21 |
| 9 | -0.77 | 1.48 | -0.52 | 0.60 | 1 | Gm16006 | | 1 | 0 | 0 | 18 | 21 | 9 | 6 | 0 | 16 | 18 | 9 |
| 9 | 0.42 | 2.02 | 0.21 | 0.83 | 1 | Gm40477 | | 0 | 18 | 7 | 0 | 0 | 24 | 6 | 7 | 5 | 0 | 31 |
| 9 | 0.09 | 0.96 | 0.09 | 0.93 | 1 | Gm19810 | | 13 | 10 | 14 | 7 | 15 | 3 | 6 | 10 | 6 | 9 | 5 |
| 9 | -0.75 | 1.62 | -0.46 | 0.64 | 1 | Gm31597 | | 51 | 14 | 0 | 1 | 5 | 3 | 6 | 0 | 1 | 14 | 2 |
| 9 | 1.14 | 1.58 | 0.72 | 0.47 | 1 | Gm42928 | | 19 | 19 | 0 | 1 | 1 | 14 | 6 | 0 | 4 | 19 | 13 |
| 9 | 0.24 | 1.21 | 0.20 | 0.84 | 1 | Chchd2-ps | | 9 | 2 | 12 | 6 | 8 | 10 | 6 | 22 | 14 | 1 | 4 |
| 9 | -0.18 | 1.95 | -0.09 | 0.93 | 1 | 4930432K21Rik | RIKEN cDNA 4930432K21 gene | 21 | 34 | 12 | 0 | 0 | 9 | 6 | 0 | 5 | 0 | 6 |
| 8 | -2.37 | 1.36 | -1.75 | 0.08 | 1 | Gm6813 | | 18 | 15 | 10 | 6 | 13 | 8 | 6 | 0 | 1 | 1 | 13 |

| | | | | | | | | | | | | | | | | | |
|---|---|---|---|---|---|---|---|---|---|---|---|---|---|---|---|---|---|
| 8 | -0.99 | 1.92 | -0.52 | 0.61 | 1 | Gm8098 | | 6 | 0 | 16 | 2 | 6 | 26 | 6 | 21 | 0 | 0 | 10 |
| 8 | -0.46 | 1.05 | -0.43 | 0.67 | 1 | Pcna-ps2 | | 21 | 4 | 8 | 7 | 7 | 13 | 6 | 2 | 8 | 4 | 12 |
| 8 | 0.17 | 1.66 | 0.10 | 0.92 | 1 | Gm16565 | | 0 | 0 | 0 | 6 | 1 | 31 | 6 | 0 | 6 | 14 | 25 |
| 8 | -2.12 | 1.98 | -1.07 | 0.28 | 1 | 1700095J03Rik | | 21 | 13 | 0 | 11 | 0 | 14 | 6 | 0 | 0 | 0 | 24 |
| 8 | -2.28 | 2.53 | -0.90 | 0.37 | 1 | Ros1 | Ros1 proto-oncogene | 0 | 59 | 0 | 0 | 5 | 12 | 6 | 0 | 4 | 0 | 4 |
| 8 | -0.35 | 1.72 | -0.20 | 0.84 | 1 | Gm5820 | predicted gene 5820 | 17 | 12 | 15 | 6 | 0 | 6 | 6 | 0 | 0 | 21 | 6 |
| 8 | 0.65 | 1.87 | 0.35 | 0.73 | 1 | Gm3636 | predicted gene 3636 | 0 | 0 | 6 | 0 | 12 | 14 | 6 | 15 | 11 | 0 | 23 |
| 8 | -0.07 | 1.42 | -0.05 | 0.96 | 1 | Rps18-ps1 | | 25 | 23 | 5 | 12 | 1 | 2 | 6 | 0 | 4 | 3 | 7 |
| 8 | -1.34 | 3.08 | -0.44 | 0.66 | 1 | Gm37776 | | 0 | 0 | 0 | 0 | 0 | 43 | 6 | 0 | 0 | 0 | 39 |
| 8 | -0.52 | 1.79 | -0.29 | 0.77 | 1 | Trim38 | tripartite motif-containing 38 | 0 | 0 | 3 | 25 | 1 | 13 | 6 | 0 | 9 | 13 | 17 |
| 8 | 0.98 | 1.45 | 0.67 | 0.50 | 1 | Gm1943 | | 9 | 11 | 4 | 4 | 1 | 2 | 6 | 42 | 0 | 1 | 6 |
| 8 | 0.15 | 1.39 | 0.11 | 0.91 | 1 | Gm44702 | | 5 | 1 | 5 | 4 | 4 | 26 | 6 | 13 | 14 | 0 | 7 |
| 8 | -0.26 | 1.46 | -0.18 | 0.86 | 1 | Tldc2 | TBC/LysM associated domain containing | 0 | 0 | 1 | 4 | 16 | 12 | 6 | 10 | 9 | 0 | 25 |
| 8 | -0.34 | 2.17 | -0.15 | 0.88 | 1 | Gabra5 | gamma-aminobutyric acid (GABA) A rece | 0 | 25 | 23 | 0 | 5 | 3 | 6 | 0 | 14 | 0 | 7 |
| 7 | -1.31 | 1.91 | -0.69 | 0.49 | 1 | 0610043K17Rik | | 0 | 20 | 5 | 5 | 16 | 7 | 6 | 0 | 0 | 6 | 16 |
| 7 | -0.64 | 1.56 | -0.41 | 0.68 | 1 | Gm20511 | | 31 | 19 | 0 | 1 | 0 | 16 | 6 | 3 | 0 | 3 | 3 |
| 7 | -1.95 | 1.77 | -1.10 | 0.27 | 1 | Lrrc74b | leucine rich repeat containing 74B | 25 | 0 | 2 | 4 | 20 | 9 | 6 | 0 | 9 | 0 | 7 |
| 7 | 0.11 | 1.57 | 0.07 | 0.95 | 1 | Slc8a3 | solute carrier family 8 (sodium/calcium ex | 28 | 14 | 7 | 8 | 0 | 3 | 6 | 0 | 6 | 3 | 5 |
| 7 | 0.75 | 1.62 | 0.46 | 0.64 | 1 | Gm37201 | | 6 | 30 | 9 | 0 | 4 | 3 | 6 | 0 | 8 | 10 | 3 |
| 7 | 0.62 | 1.34 | 0.46 | 0.64 | 1 | Gm8181 | | 14 | 2 | 0 | 2 | 9 | 7 | 6 | 10 | 3 | 10 | 16 |
| 7 | 1.01 | 2.23 | 0.45 | 0.65 | 1 | Gm12802 | | 5 | 25 | 0 | 0 | 0 | 7 | 6 | 0 | 0 | 27 | 7 |
| 7 | -0.80 | 1.81 | -0.44 | 0.66 | 1 | Gm10286 | | 8 | 17 | 14 | 0 | 4 | 7 | 6 | 8 | 0 | 0 | 10 |
| 6 | 0.62 | 1.10 | 0.57 | 0.57 | 1 | Gm13268 | | 9 | 4 | 10 | 2 | 8 | 3 | 6 | 5 | 8 | 14 | 2 |
| 6 | -0.24 | 2.68 | -0.09 | 0.93 | 1 | Gm826 | predicted gene 826 | 26 | 12 | 11 | 0 | 0 | 0 | 6 | 0 | 0 | 0 | 14 |
| 6 | -1.45 | 2.46 | -0.59 | 0.56 | 1 | Gm6566 | | 3 | 0 | 4 | 0 | 0 | 32 | 6 | 0 | 0 | 0 | 24 |
| 6 | 0.09 | 1.68 | 0.05 | 0.96 | 1 | 6330562C20Rik | | 3 | 2 | 0 | 20 | 0 | 9 | 6 | 10 | 2 | 0 | 16 |
| 6 | 0.73 | 1.12 | 0.65 | 0.52 | 1 | Gm15421 | | 4 | 10 | 3 | 5 | 3 | 3 | 6 | 9 | 3 | 5 | 16 |
| 6 | -0.61 | 1.74 | -0.35 | 0.72 | 1 | A830009L08Rik | | 0 | 0 | 2 | 23 | 3 | 6 | 6 | 0 | 9 | 8 | 12 |
| 6 | -1.98 | 1.90 | -1.04 | 0.30 | 1 | Mfsd2a | major facilitator superfamily domain conta | 13 | 2 | 0 | 7 | 18 | 6 | 6 | 0 | 8 | 0 | 8 |
| 6 | 1.00 | 2.06 | 0.49 | 0.63 | 1 | Slc39a1-ps | | 1 | 13 | 1 | 4 | 4 | 0 | 6 | 0 | 0 | 35 | 3 |
| 6 | -1.30 | 2.35 | -0.55 | 0.58 | 1 | Gm21451 | predicted gene, 21451 | 0 | 0 | 0 | 0 | 26 | 11 | 6 | 0 | 16 | 0 | 8 |
| 6 | -2.91 | 1.70 | -1.71 | 0.09 | 1 | D030047H15Rik | | 2 | 23 | 14 | 4 | 2 | 13 | 6 | 0 | 3 | 0 | 1 |
| 6 | -1.07 | 1.61 | -0.66 | 0.51 | 1 | Clec2l | C-type lectin domain family 2, member L | 3 | 0 | 0 | 11 | 5 | 13 | 6 | 12 | 0 | 13 | 4 |
| 6 | 1.62 | 2.36 | 0.69 | 0.49 | 1 | Gm42596 | | 0 | 0 | 14 | 2 | 0 | 4 | 6 | 25 | 0 | 9 | 5 |
| 6 | -1.63 | 1.50 | -1.09 | 0.28 | 1 | Rpsa-ps9 | | 6 | 5 | 19 | 1 | 6 | 13 | 6 | 1 | 7 | 1 | 0 |
| 6 | -2.89 | 2.00 | -1.44 | 0.15 | 1 | 1700084C06Rik | | 6 | 16 | 0 | 6 | 9 | 3 | 6 | 0 | 1 | 0 | 16 |
| 6 | -1.89 | 2.44 | -0.77 | 0.44 | 1 | Gm26935 | | 0 | 0 | 0 | 0 | 6 | 19 | 6 | 0 | 1 | 0 | 30 |
| 6 | 0.74 | 1.24 | 0.60 | 0.55 | 1 | Gm13410 | | 16 | 2 | 1 | 2 | 5 | 2 | 6 | 2 | 5 | 17 | 4 |
| 5 | -0.77 | 1.92 | -0.40 | 0.69 | 1 | Iqch | IQ motif containing H | 4 | 0 | 4 | 0 | 13 | 9 | 6 | 0 | 0 | 5 | 17 |
| 5 | -2.24 | 2.14 | -1.05 | 0.30 | 1 | Dnase2b | deoxyribonuclease II beta | 0 | 7 | 0 | 9 | 25 | 2 | 6 | 0 | 8 | 0 | 3 |
| 5 | -1.56 | 2.65 | -0.59 | 0.56 | 1 | Ccdc33 | coiled-coil domain containing 33 | 0 | 13 | 7 | 0 | 0 | 20 | 6 | 0 | 7 | 0 | 5 |
| 5 | 0.65 | 1.52 | 0.43 | 0.67 | 1 | Gm10817 | | 3 | 0 | 4 | 2 | 2 | 13 | 6 | 16 | 5 | 3 | 4 |
| 5 | 0.24 | 1.98 | 0.12 | 0.91 | 1 | Gm19345 | predicted gene, 19345 | 3 | 0 | 15 | 0 | 7 | 8 | 6 | 9 | 2 | 0 | 7 |
| 5 | -4.57 | 3.34 | -1.37 | 0.17 | 1 | Ms4a4b | membrane-spanning 4-domains, subfami | 45 | 0 | 0 | 3 | 0 | 0 | 6 | 0 | 0 | 0 | 3 |
| 5 | -2.07 | 2.15 | -0.96 | 0.34 | 1 | Gm26901 | | 0 | 3 | 0 | 5 | 6 | 16 | 6 | 0 | 0 | 18 | 3 |
| 5 | -0.60 | 2.32 | -0.26 | 0.80 | 1 | Gm25047 | | 0 | 26 | 0 | 8 | 3 | 2 | 6 | 0 | 6 | 0 | 6 |
| 5 | -0.72 | 2.56 | -0.28 | 0.78 | 1 | Pkd1l3 | polycystic kidney disease 1 like 3 | 0 | 4 | 0 | 11 | 0 | 11 | 6 | 0 | 18 | 0 | 7 |
| 5 | -0.93 | 1.39 | -0.67 | 0.50 | 1 | Hspa9-ps1 | | 3 | 8 | 9 | 14 | 2 | 2 | 6 | 2 | 2 | 1 | 7 |
| 5 | -1.21 | 2.58 | -0.47 | 0.64 | 1 | Egfl8 | EGF-like domain 8 | 17 | 0 | 7 | 0 | 0 | 15 | 6 | 0 | 0 | 0 | 10 |
| 5 | -3.02 | 2.52 | -1.20 | 0.23 | 1 | Gm48372 | | 8 | 11 | 0 | 0 | 5 | 9 | 6 | 0 | 0 | 0 | 16 |
| 5 | -1.13 | 2.74 | -0.41 | 0.68 | 1 | 5430400D12Rik | | 0 | 0 | 4 | 0 | 0 | 25 | 6 | 0 | 4 | 0 | 16 |
| 5 | -0.90 | 2.76 | -0.33 | 0.74 | 1 | Gm44863 | | 0 | 3 | 0 | 0 | 0 | 21 | 6 | 12 | 0 | 0 | 13 |
| 5 | -1.95 | 1.82 | -1.07 | 0.28 | 1 | 4930417H01Rik | | 16 | 0 | 0 | 4 | 6 | 5 | 6 | 0 | 2 | 12 | 2 |
| 5 | -1.31 | 2.51 | -0.52 | 0.60 | 1 | Ctcfl | CCCTC-binding factor (zinc finger protein | 2 | 16 | 20 | 0 | 0 | 4 | 6 | 0 | 0 | 0 | 5 |
| 5 | -0.92 | 2.04 | -0.45 | 0.65 | 1 | Slitrk3 | SLIT and NTRK-like family, member 3 | 21 | 1 | 6 | 0 | 4 | 1 | 6 | 0 | 0 | 14 | 0 |
| 5 | 0.82 | 3.01 | 0.27 | 0.79 | 1 | | | 0 | 0 | 0 | 0 | 0 | 11 | 6 | 0 | 0 | 18 | 17 |
| 5 | 1.30 | 1.77 | 0.73 | 0.46 | 1 | Gm10054 | | 0 | 1 | 7 | 0 | 2 | 5 | 6 | 1 | 7 | 21 | 2 |
| 5 | 0.04 | 1.90 | 0.02 | 0.98 | 1 | Gm10563 | | 6 | 3 | 11 | 0 | 1 | 7 | 6 | 0 | 0 | 10 | 7 |
| 5 | 0.29 | 1.28 | 0.23 | 0.82 | 1 | Gm13578 | | 3 | 6 | 10 | 3 | 2 | 3 | 6 | 7 | 4 | 3 | 4 |
| 5 | -5.02 | 2.85 | -1.76 | 0.08 | 1 | 1700061A03Rik | | 10 | 0 | 0 | 8 | 0 | 8 | 6 | 0 | 0 | 0 | 19 |

| | | | | | | | | | | | | | | | | | |
|---|---|---|---|---|---|---|---|---|---|---|---|---|---|---|---|---|---|
| 5 | 0.58 | 2.14 | 0.27 | 0.79 | 1 | Gm44793 | | 0 | 3 | 0 | 11 | 0 | 9 | 6 | 0 | 4 | 9 | 7 |
| 5 | -4.21 | 2.24 | -1.89 | 0.06 | 1 | Meikin | meiotic kinetochore factor | 3 | 0 | 7 | 12 | 6 | 5 | 6 | 0 | 0 | 0 | 10 |
| 5 | -0.86 | 2.52 | -0.34 | 0.73 | 1 | Ifitm10 | interferon induced transmembrane protei | 16 | 12 | 11 | 0 | 0 | 1 | 6 | 0 | 3 | 0 | 0 |
| 4 | -1.23 | 2.07 | -0.60 | 0.55 | 1 | Gm42998 | | 3 | 0 | 0 | 3 | 5 | 11 | 6 | 0 | 5 | 0 | 17 |
| 4 | -1.30 | 2.02 | -0.64 | 0.52 | 1 | Nudt8 | nudix (nucleoside diphosphate linked mo | 12 | 0 | 16 | 0 | 7 | 3 | 6 | 0 | 1 | 0 | 5 |
| 4 | -3.13 | 2.20 | -1.42 | 0.15 | 1 | Moxd1 | monooxygenase, DBH-like 1 | 9 | 0 | 0 | 9 | 5 | 14 | 6 | 0 | 3 | 0 | 2 |
| 4 | -3.55 | 3.15 | -1.13 | 0.26 | 1 | Mir700 | | 0 | 0 | 3 | 4 | 0 | 5 | 6 | 0 | 0 | 30 | 0 |
| 4 | -0.33 | 1.94 | -0.17 | 0.87 | 1 | Gm37783 | | 0 | 0 | 0 | 8 | 2 | 6 | 6 | 7 | 3 | 15 | 0 |
| 4 | -0.88 | 3.56 | -0.25 | 0.80 | 1 | Gm18857 | | 0 | 0 | 0 | 0 | 0 | 10 | 6 | 0 | 0 | 0 | 30 |
| 4 | 0.06 | 2.49 | 0.02 | 0.98 | 1 | Gm49173 | | 0 | 0 | 0 | 3 | 10 | 0 | 6 | 0 | 1 | 14 | 12 |
| 4 | -5.11 | 2.85 | -1.79 | 0.07 | 1 | 4930444P10Ril | RIKEN cDNA 4930444P10 gene | 0 | 0 | 19 | 6 | 8 | 7 | 6 | 0 | 0 | 0 | 0 |
| 4 | -3.95 | 3.55 | -1.11 | 0.27 | 1 | Gm12781 | | 0 | 0 | 16 | 0 | 0 | 15 | 6 | 0 | 0 | 0 | 7 |
| 4 | -2.38 | 3.29 | -0.72 | 0.47 | 1 | Ccdc178 | coiled coil domain containing 178 | 0 | 0 | 0 | 0 | 2 | 19 | 6 | 0 | 0 | 0 | 16 |
| 4 | -0.50 | 2.31 | -0.22 | 0.83 | 1 | 4933437G19Rik | | 8 | 0 | 0 | 2 | 3 | 5 | 6 | 0 | 15 | 0 | 4 |
| 4 | 1.49 | 2.70 | 0.55 | 0.58 | 1 | 2410004P03Ril | RIKEN cDNA 2410004P03 gene | 14 | 15 | 0 | 4 | 0 | 0 | 6 | 0 | 2 | 0 | 2 |
| 4 | -4.01 | 2.32 | -1.73 | 0.08 | 1 | Gm3222 | | 4 | 5 | 7 | 7 | 9 | 0 | 6 | 0 | 0 | 0 | 3 |
| 4 | -2.10 | 2.71 | -0.77 | 0.44 | 1 | Gm44386 | | 0 | 12 | 0 | 0 | 6 | 2 | 6 | 10 | 6 | 0 | 0 |
| 4 | -2.74 | 2.42 | -1.13 | 0.26 | 1 | Gm48633 | | 4 | 13 | 0 | 0 | 4 | 8 | 6 | 0 | 0 | 5 | 1 |
| 4 | -2.96 | 1.82 | -1.62 | 0.10 | 1 | Gad1-ps | | 5 | 5 | 2 | 8 | 6 | 3 | 6 | 1 | 0 | 0 | 6 |
| 4 | -1.85 | 2.17 | -0.85 | 0.39 | 1 | Rslcan18 | regulator of sex-limitation candidate 18 | 3 | 1 | 4 | 13 | 0 | 3 | 6 | 0 | 4 | 0 | 6 |
| 4 | -0.69 | 2.35 | -0.29 | 0.77 | 1 | Gm43714 | | 13 | 3 | 6 | 0 | 3 | 0 | 6 | 0 | 1 | 0 | 8 |
| 4 | -1.67 | 2.55 | -0.66 | 0.51 | 1 | Dydc2 | DPY30 domain containing 2 | 9 | 15 | 0 | 4 | 0 | 1 | 6 | 0 | 0 | 0 | 4 |
| 4 | -0.82 | 2.24 | -0.37 | 0.71 | 1 | Gm4462 | | 1 | 2 | 10 | 4 | 0 | 3 | 6 | 10 | 0 | 0 | 3 |
| 4 | -2.85 | 2.98 | -0.96 | 0.34 | 1 | 4930432B10Rik | | 0 | 0 | 0 | 5 | 0 | 20 | 6 | 0 | 5 | 0 | 2 |
| 3 | -0.93 | 3.01 | -0.31 | 0.76 | 1 | 4930519F16Rik | RIKEN cDNA 4930519F16 gene | 0 | 0 | 0 | 8 | 0 | 3 | 6 | 0 | 3 | 18 | 0 |
| 3 | -1.34 | 3.56 | -0.38 | 0.71 | 1 | Gm43200 | | 0 | 0 | 0 | 0 | 0 | 17 | 6 | 0 | 0 | 0 | 15 |
| 3 | -3.35 | 2.84 | -1.18 | 0.24 | 1 | Gm12522 | | 0 | 7 | 5 | 11 | 0 | 2 | 6 | 0 | 0 | 0 | 6 |
| 3 | -0.66 | 2.32 | -0.28 | 0.78 | 1 | Gm20796 | | 19 | 1 | 7 | 0 | 0 | 1 | 6 | 0 | 1 | 1 | 0 |
| 3 | -2.12 | 1.88 | -1.13 | 0.26 | 1 | Gm29254 | | 0 | 5 | 6 | 4 | 7 | 3 | 6 | 0 | 1 | 1 | 3 |
| 3 | -0.21 | 3.56 | -0.06 | 0.95 | 1 | Apoc3 | apolipoprotein C-III | 15 | 0 | 14 | 0 | 0 | 0 | 6 | 0 | 2 | 0 | 0 |
| 3 | -3.82 | 2.46 | -1.55 | 0.12 | 1 | 1700010G06Rik | | 3 | 0 | 14 | 6 | 5 | 1 | 6 | 0 | 0 | 0 | 2 |
| 3 | -1.19 | 1.54 | -0.78 | 0.44 | 1 | Gm2546 | | 8 | 2 | 2 | 2 | 3 | 5 | 6 | 0 | 2 | 4 | 2 |
| 3 | -4.54 | 3.07 | -1.48 | 0.14 | 1 | Gm35611 | | 0 | 0 | 16 | 4 | 7 | 3 | 6 | 0 | 0 | 0 | 0 |
| 3 | -0.74 | 2.06 | -0.36 | 0.72 | 1 | Gm48583 | | 10 | 0 | 1 | 9 | 1 | 1 | 6 | 1 | 5 | 0 | 1 |
| 3 | 1.57 | 2.62 | 0.60 | 0.55 | 1 | Gm17139 | | 0 | 0 | 0 | 0 | 0 | 6 | 6 | 10 | 1 | 10 | 1 |
| 3 | 0.24 | 2.79 | 0.09 | 0.93 | 1 | Gm43868 | | 3 | 0 | 12 | 0 | 0 | 4 | 6 | 1 | 0 | 0 | 7 |
| 3 | -3.50 | 2.85 | -1.23 | 0.22 | 1 | Gm12953 | | 0 | 2 | 4 | 10 | 0 | 3 | 6 | 0 | 0 | 0 | 9 |
| 3 | -3.84 | 3.25 | -1.18 | 0.24 | 1 | Tmprss7 | transmembrane serine protease 7 | 0 | 12 | 9 | 0 | 3 | 3 | 6 | 0 | 0 | 0 | 0 |
| 3 | -0.89 | 3.54 | -0.25 | 0.80 | 1 | D630014O11Rik | | 8 | 0 | 0 | 0 | 0 | 13 | 6 | 0 | 0 | 0 | 7 |
| 3 | -4.24 | 3.01 | -1.41 | 0.16 | 1 | D6Ertd527e | DNA segment, Chr 6, ERATO Doi 527, ex | 0 | 0 | 0 | 8 | 8 | 8 | 6 | 0 | 0 | 0 | 4 |
| 3 | -3.71 | 2.28 | -1.63 | 0.10 | 1 | Gm20508 | | 0 | 4 | 3 | 4 | 11 | 2 | 6 | 0 | 2 | 0 | 0 |
| 3 | -3.73 | 2.93 | -1.27 | 0.20 | 1 | Rpl35a-ps4 | | 7 | 6 | 0 | 6 | 0 | 4 | 6 | 0 | 3 | 0 | 0 |
| 3 | -2.31 | 2.87 | -0.80 | 0.42 | 1 | Ace3 | angiotensin I converting enzyme (peptidy | 11 | 3 | 0 | 0 | 9 | 1 | 6 | 0 | 2 | 0 | 0 |
| 3 | 0.39 | 1.98 | 0.20 | 0.84 | 1 | Gm6576 | | 6 | 0 | 5 | 4 | 0 | 1 | 6 | 5 | 2 | 1 | 2 |
| 3 | -5.24 | 3.30 | -1.59 | 0.11 | 1 | Gm35404 | | 0 | 0 | 16 | 0 | 5 | 2 | 6 | 0 | 0 | 0 | 3 |
| 3 | -5.71 | 3.32 | -1.72 | 0.09 | 1 | Gm5784 | | 10 | 0 | 0 | 10 | 0 | 2 | 6 | 0 | 0 | 0 | 2 |
| 3 | 0.69 | 2.15 | 0.32 | 0.75 | 1 | Gm6543 | | 7 | 0 | 4 | 0 | 2 | 2 | 6 | 2 | 0 | 3 | 3 |
| 3 | -2.28 | 3.15 | -0.72 | 0.47 | 1 | Gm48780 | | 0 | 0 | 0 | 0 | 8 | 7 | 6 | 0 | 1 | 0 | 7 |
| 3 | -0.08 | 3.55 | -0.02 | 0.98 | 1 | Gm24497 | | 0 | 0 | 0 | 0 | 0 | 4 | 6 | 0 | 0 | 18 | 0 |
| 3 | -1.67 | 2.72 | -0.61 | 0.54 | 1 | Gm6375 | | 9 | 5 | 1 | 5 | 0 | 0 | 6 | 1 | 0 | 0 | 0 |
| 2 | -0.62 | 3.28 | -0.19 | 0.85 | 1 | Gm35248 | | 11 | 2 | 0 | 0 | 0 | 3 | 6 | 0 | 0 | 0 | 4 |
| 2 | -1.03 | 2.47 | -0.42 | 0.68 | 1 | Gm47441 | | 0 | 0 | 3 | 0 | 6 | 3 | 6 | 1 | 2 | 0 | 4 |
| 2 | -2.49 | 3.60 | -0.69 | 0.49 | 1 | Grm8 | glutamate receptor, metabotropic 8 | 9 | 7 | 0 | 0 | 0 | 3 | 6 | 0 | 0 | 0 | 0 |
| 2 | -0.01 | 1.85 | -0.01 | 0.99 | 1 | Gm6085 | | 5 | 2 | 1 | 1 | 1 | 2 | 6 | 1 | 2 | 0 | 4 |
| 2 | -0.86 | 3.55 | -0.24 | 0.81 | 1 | Poln | DNA polymerase N | 8 | 0 | 0 | 0 | 0 | 7 | 6 | 0 | 0 | 0 | 4 |
| 2 | -0.02 | 2.94 | -0.01 | 0.99 | 1 | Gm5528 | | 3 | 1 | 2 | 0 | 1 | 0 | 6 | 0 | 0 | 12 | 0 |
| 2 | -2.19 | 3.54 | -0.62 | 0.54 | 1 | 2900072N19Rik | | 0 | 6 | 7 | 0 | 0 | 2 | 6 | 0 | 3 | 0 | 0 |
| 2 | -3.42 | 3.58 | -0.96 | 0.34 | 1 | Hils1 | histone H1-like protein in spermatids 1 | 0 | 10 | 0 | 0 | 0 | 3 | 6 | 0 | 0 | 0 | 4 |
| 2 | -3.67 | 3.57 | -1.03 | 0.30 | 1 | Rpsa-ps12 | | 2 | 7 | 0 | 0 | 9 | 0 | 6 | 0 | 0 | 0 | 0 |
| 2 | 0.09 | 2.71 | 0.03 | 0.97 | 1 | Gm10155 | | 3 | 0 | 0 | 1 | 1 | 1 | 6 | 0 | 10 | 0 | 1 |

| | | | | | | | | | | | | | | | | | | |
|---|---|---|---|---|---|---|---|---|---|---|---|---|---|---|---|---|---|---|
| 2 | -4.35 | 3.32 | -1.31 | 0.19 | 1 | Krt87 | keratin 87 | 0 | 2 | 0 | 11 | 4 | 1 | 6 | 0 | 0 | 0 | 0 |
| 2 | -3.91 | 3.37 | -1.16 | 0.25 | 1 | Gm26861 | | 0 | 0 | 0 | 10 | 3 | 3 | 6 | 0 | 0 | 0 | 2 |
| 2 | -0.61 | 3.54 | -0.17 | 0.86 | 1 | 2900027M19Rik | | 0 | 0 | 0 | 0 | 0 | 9 | 6 | 0 | 0 | 8 | 0 |
| 2 | 2.86 | 3.55 | 0.81 | 0.42 | 1 | Gm8705 | | 8 | 0 | 0 | 0 | 0 | 0 | 6 | 0 | 6 | 0 | 3 |
| 2 | -0.36 | 1.75 | -0.20 | 0.84 | 1 | Rpl31-ps15 | | 1 | 1 | 0 | 3 | 2 | 2 | 6 | 2 | 4 | 1 | 1 |
| 2 | -0.43 | 2.32 | -0.18 | 0.85 | 1 | Gm9795 | | 1 | 2 | 0 | 3 | 0 | 4 | 6 | 0 | 3 | 1 | 2 |
| 2 | -1.37 | 3.57 | -0.38 | 0.70 | 1 | Gm18811 | | 0 | 0 | 0 | 0 | 0 | 8 | 6 | 0 | 0 | 0 | 7 |
| 2 | -1.07 | 3.56 | -0.30 | 0.76 | 1 | Gm47097 | | 0 | 1 | 0 | 0 | 0 | 4 | 6 | 0 | 0 | 0 | 10 |
| 2 | -1.10 | 2.33 | -0.47 | 0.64 | 1 | Gm4929 | | 3 | 2 | 2 | 1 | 4 | 0 | 6 | 0 | 3 | 0 | 1 |
| 2 | -5.92 | 3.56 | -1.66 | 0.10 | 1 | 4930513O06Ril | RIKEN cDNA 4930513O06 gene | 0 | 9 | 0 | 0 | 1 | 3 | 6 | 0 | 0 | 0 | 2 |
| 2 | -2.71 | 3.56 | -0.76 | 0.45 | 1 | Gm37531 | | 4 | 0 | 0 | 0 | 6 | 3 | 6 | 0 | 1 | 0 | 0 |
| 2 | 0.08 | 2.40 | 0.03 | 0.97 | 1 | Gm9308 | | 5 | 0 | 1 | 0 | 2 | 2 | 6 | 2 | 1 | 0 | 1 |
| 2 | -2.66 | 2.42 | -1.10 | 0.27 | 1 | Gm7803 | | 0 | 1 | 3 | 2 | 1 | 4 | 6 | 0 | 2 | 0 | 1 |
| 2 | -6.35 | 3.53 | -1.80 | 0.07 | 1 | Gm17029 | | 0 | 0 | 6 | 3 | 0 | 2 | 6 | 0 | 0 | 0 | 3 |
| 2 | -3.63 | 3.53 | -1.03 | 0.30 | 1 | Gm15201 | | 0 | 0 | 0 | 7 | 2 | 3 | 6 | 0 | 0 | 0 | 2 |
| 2 | 0.00 | 3.51 | 0.00 | 1.00 | 1 | Gm45733 | | 0 | 0 | 7 | 0 | 0 | 0 | 6 | 0 | 2 | 0 | 4 |
| 2 | -2.33 | 3.55 | -0.66 | 0.51 | 1 | Gm5936 | predicted gene 5936 | 0 | 0 | 5 | 0 | 2 | 2 | 6 | 0 | 4 | 0 | 0 |
| 2 | -1.06 | 2.72 | -0.39 | 0.70 | 1 | Gm46312 | | 1 | 0 | 0 | 2 | 1 | 1 | 6 | 0 | 7 | 1 | 0 |
| 2 | -1.63 | 3.58 | -0.46 | 0.65 | 1 | Gm37482 | | 0 | 0 | 0 | 0 | 0 | 8 | 6 | 0 | 0 | 0 | 4 |
| 2 | -1.39 | 2.91 | -0.48 | 0.63 | 1 | Gad1os | | 0 | 2 | 4 | 0 | 1 | 2 | 6 | 0 | 1 | 0 | 2 |
| 2 | 0.99 | 2.73 | 0.36 | 0.72 | 1 | Gm14650 | | 2 | 0 | 0 | 4 | 0 | 0 | 6 | 2 | 2 | 1 | 1 |
| 2 | -3.76 | 3.56 | -1.06 | 0.29 | 1 | Gm47693 | | 0 | 0 | 0 | 11 | 0 | 1 | 6 | 0 | 0 | 0 | 0 |
| 2 | -1.09 | 3.56 | -0.31 | 0.76 | 1 | Gm45483 | | 0 | 0 | 0 | 0 | 0 | 4 | 6 | 0 | 0 | 0 | 7 |
| 2 | -2.68 | 3.54 | -0.76 | 0.45 | 1 | Nr1i3 | nuclear receptor subfamily 1, group I, mer | 1 | 0 | 0 | 0 | 5 | 3 | 6 | 0 | 0 | 0 | 3 |
| 2 | -0.04 | 2.19 | -0.02 | 0.99 | 1 | Gm14480 | | 2 | 2 | 0 | 2 | 1 | 0 | 6 | 1 | 1 | 1 | 1 |
| 2 | -1.80 | 3.57 | -0.50 | 0.61 | 1 | Gm10241 | | 5 | 0 | 0 | 4 | 0 | 1 | 6 | 1 | 0 | 0 | 0 |
| 1 | -1.47 | 3.57 | -0.41 | 0.68 | 1 | Gm20535 | | 0 | 0 | 0 | 0 | 0 | 6 | 6 | 0 | 0 | 0 | 4 |
| 1 | 0.89 | 3.55 | 0.25 | 0.80 | 1 | Gm11281 | | 3 | 0 | 0 | 0 | 0 | 1 | 6 | 1 | 0 | 0 | 5 |
| 1 | -5.49 | 3.56 | -1.54 | 0.12 | 1 | 1700064H15Rik | | 1 | 0 | 4 | 3 | 0 | 0 | 6 | 0 | 0 | 0 | 1 |
| 1 | -0.27 | 3.08 | -0.09 | 0.93 | 1 | Gm5457 | | 1 | 2 | 1 | 0 | 0 | 1 | 6 | 0 | 1 | 3 | 0 |
| 1 | 0.77 | 3.51 | 0.22 | 0.83 | 1 | Gm19726 | | 0 | 0 | 0 | 1 | 0 | 0 | 6 | 0 | 3 | 0 | 4 |
| 1 | -2.06 | 3.57 | -0.58 | 0.56 | 1 | Gm37465 | | 0 | 0 | 0 | 0 | 0 | 2 | 6 | 0 | 0 | 0 | 6 |
| 1 | -3.25 | 3.59 | -0.91 | 0.36 | 1 | Gm16275 | | 0 | 3 | 0 | 0 | 0 | 2 | 6 | 0 | 0 | 0 | 3 |
| 1 | -0.09 | 3.52 | -0.03 | 0.98 | 1 | Gm8010 | | 1 | 0 | 0 | 0 | 0 | 3 | 6 | 0 | 3 | 0 | 1 |
| 1 | -0.61 | 3.56 | -0.17 | 0.86 | 1 | Gm6689 | | 0 | 3 | 1 | 1 | 0 | 0 | 6 | 1 | 0 | 1 | 0 |
| 1 | -5.00 | 3.55 | -1.41 | 0.16 | 1 | Gm46344 | | 1 | 0 | 2 | 2 | 0 | 0 | 6 | 0 | 0 | 0 | 2 |
| 1 | -1.11 | 3.57 | -0.31 | 0.76 | 1 | | | 0 | 0 | 0 | 0 | 0 | 3 | 6 | 0 | 0 | 0 | 4 |
| 1 | -0.64 | 3.56 | -0.18 | 0.86 | 1 | Gm10193 | | 1 | 0 | 2 | 1 | 0 | 0 | 6 | 1 | 1 | 0 | 0 |
| 1 | -0.75 | 3.57 | -0.21 | 0.83 | 1 | Gm20545 | | 2 | 1 | 0 | 0 | 0 | 1 | 6 | 0 | 0 | 0 | 2 |
| 1 | -1.23 | 3.57 | -0.34 | 0.73 | 1 | Gm19410 | predicted gene, 19410 | 0 | 0 | 0 | 0 | 0 | 3 | 6 | 0 | 0 | 0 | 3 |
| 1 | -1.23 | 3.57 | -0.34 | 0.73 | 1 | Fam71d | family with sequence similarity 71, membe | 0 | 0 | 0 | 0 | 0 | 3 | 6 | 0 | 0 | 0 | 3 |
| 1 | -1.86 | 3.57 | -0.52 | 0.60 | 1 | Gm13231 | | 1 | 0 | 0 | 1 | 3 | 0 | 6 | 0 | 1 | 0 | 0 |
| 1 | -1.84 | 3.55 | -0.52 | 0.60 | 1 | Gm34882 | | 0 | 0 | 0 | 0 | 0 | 3 | 6 | 0 | 3 | 0 | 0 |
| 1 | 0.15 | 3.61 | 0.04 | 0.97 | 1 | Gm49959 | | 4 | 0 | 0 | 0 | 0 | 0 | 6 | 0 | 0 | 0 | 1 |
| 1 | 0.13 | 3.55 | 0.04 | 0.97 | 1 | Gm48771 | | 0 | 0 | 0 | 0 | 0 | 0 | 6 | 0 | 0 | 0 | 5 |
| 1 | -0.84 | 3.53 | -0.24 | 0.81 | 1 | 4930544F09Rik | | 0 | 0 | 0 | 0 | 2 | 0 | 6 | 0 | 1 | 0 | 2 |
| 1 | -2.20 | 3.56 | -0.62 | 0.54 | 1 | Gm10655 | | 0 | 0 | 0 | 0 | 2 | 1 | 6 | 0 | 0 | 0 | 2 |
| 1 | -0.05 | 3.61 | -0.01 | 0.99 | 1 | Gm50098 | | 1 | 0 | 0 | 0 | 0 | 0 | 6 | 0 | 0 | 4 | 0 |
| 1 | -2.67 | 3.59 | -0.74 | 0.46 | 1 | Gm15892 | | 0 | 1 | 0 | 0 | 0 | 0 | 6 | 0 | 0 | 0 | 3 |
| 1 | -1.68 | 3.56 | -0.47 | 0.64 | 1 | Gm42669 | | 0 | 0 | 0 | 0 | 0 | 2 | 6 | 0 | 2 | 0 | 0 |
| 1 | -2.53 | 3.60 | -0.70 | 0.48 | 1 | Gm36388 | | 0 | 0 | 0 | 0 | 0 | 3 | 6 | 0 | 0 | 0 | 0 |
| 1 | -2.16 | 3.61 | -0.60 | 0.55 | 1 | Gm13310 | | 0 | 1 | 0 | 0 | 0 | 1 | 6 | 0 | 0 | 0 | 0 |
| 1 | -0.54 | 3.61 | -0.15 | 0.88 | 1 | Gm12316 | | 0 | 0 | 0 | 0 | 0 | 0 | 6 | 0 | 1 | 0 | 0 |
| 1 | -1.96 | 3.63 | -0.54 | 0.59 | 1 | Mup20 | major urinary protein 20 | 0 | 0 | 0 | 0 | 0 | 1 | 6 | 0 | 0 | 0 | 0 |
| 1 | -1.96 | 3.63 | -0.54 | 0.59 | 1 | Gm31897 | | 0 | 0 | 0 | 0 | 0 | 1 | 6 | 0 | 0 | 0 | 0 |
| 1 | -1.96 | 3.63 | -0.54 | 0.59 | 1 | Gm40394 | | 0 | 0 | 0 | 0 | 0 | 1 | 6 | 0 | 0 | 0 | 0 |
| 1 | -1.96 | 3.63 | -0.54 | 0.59 | 1 | Gm11349 | | 0 | 0 | 0 | 0 | 0 | 1 | 6 | 0 | 0 | 0 | 0 |
| 1 | -1.26 | 3.64 | -0.35 | 0.73 | 1 | Gm28382 | | 0 | 0 | 0 | 0 | 0 | 0 | 6 | 0 | 0 | 0 | 0 |
| 1 | -1.26 | 3.64 | -0.35 | 0.73 | 1 | Gdf5 | growth differentiation factor 5 | 0 | 0 | 0 | 0 | 0 | 0 | 6 | 0 | 0 | 0 | 0 |
| 1 | -1.26 | 3.64 | -0.35 | 0.73 | 1 | Gm30292 | | 0 | 0 | 0 | 0 | 0 | 0 | 6 | 0 | 0 | 0 | 0 |

| | | | | | | | | | | | | | | | | | | |
|---|---|---|---|---|---|---|---|---|---|---|---|---|---|---|---|---|---|---|
| 1 | -1.26 | 3.64 | -0.35 | 0.73 | 1 | Fezf1 | Fez family zinc finger 1 | 0 | 0 | 0 | 0 | 0 | 0 | 6 | 0 | 0 | 0 | 0 |
| 1 | -1.26 | 3.64 | -0.35 | 0.73 | 1 | Gm28053 | | 0 | 0 | 0 | 0 | 0 | 0 | 6 | 0 | 0 | 0 | 0 |
| 1 | -1.26 | 3.64 | -0.35 | 0.73 | 1 | 5830468F06Rik | | 0 | 0 | 0 | 0 | 0 | 0 | 6 | 0 | 0 | 0 | 0 |
| 1 | -1.26 | 3.64 | -0.35 | 0.73 | 1 | Gm16204 | | 0 | 0 | 0 | 0 | 0 | 0 | 6 | 0 | 0 | 0 | 0 |
| 1 | -1.26 | 3.64 | -0.35 | 0.73 | 1 | Gm45895 | | 0 | 0 | 0 | 0 | 0 | 0 | 6 | 0 | 0 | 0 | 0 |
| 1 | -1.26 | 3.64 | -0.35 | 0.73 | 1 | Popdc3 | popeye domain containing 3 | 0 | 0 | 0 | 0 | 0 | 0 | 6 | 0 | 0 | 0 | 0 |
| 1 | -1.26 | 3.64 | -0.35 | 0.73 | 1 | Odf3l2 | outer dense fiber of sperm tails 3-like 2 | 0 | 0 | 0 | 0 | 0 | 0 | 6 | 0 | 0 | 0 | 0 |
| 1 | -1.26 | 3.64 | -0.35 | 0.73 | 1 | H3c3 | H3 clustered histone 3 | 0 | 0 | 0 | 0 | 0 | 0 | 6 | 0 | 0 | 0 | 0 |
| 1 | -1.26 | 3.64 | -0.35 | 0.73 | 1 | Gm18367 | | 0 | 0 | 0 | 0 | 0 | 0 | 6 | 0 | 0 | 0 | 0 |
| 3807 | -1.64 | 1.08 | -1.52 | 0.13 | 1 | Jchain | immunoglobulin joining chain | 33332 | 8102 | 421 | 6 | 1 | 4 | 5 | 0 | 0 | 5 | 2 |
| 721 | -2.79 | 1.81 | -1.54 | 0.12 | 1 | Lgals1 | lectin, galactose binding, soluble 1 | 3865 | 2498 | 1541 | 17 | 0 | 2 | 5 | 0 | 3 | 0 | 0 |
| 704 | 0.65 | 3.03 | 0.21 | 0.83 | 1 | Efemp1 | epidermal growth factor-containing fibulin | 4151 | 1594 | 1989 | 0 | 0 | 0 | 5 | 0 | 9 | 0 | 0 |
| 640 | -1.49 | 1.40 | -1.07 | 0.29 | 1 | Ephb4 | Eph receptor B4 | 2423 | 2155 | 2420 | 0 | 16 | 5 | 5 | 0 | 14 | 0 | 1 |
| 614 | -2.03 | 1.49 | -1.36 | 0.17 | 1 | Anxa3 | annexin A3 | 2922 | 1442 | 2381 | 0 | 0 | 3 | 5 | 0 | 0 | 0 | 1 |
| 572 | -0.24 | 2.76 | -0.09 | 0.93 | 1 | Des | desmin | 3493 | 1659 | 1128 | 0 | 0 | 0 | 5 | 0 | 0 | 0 | 3 |
| 533 | -3.08 | 1.71 | -1.80 | 0.07 | 1 | Abca9 | ATP-binding cassette, sub-family A (ABC1 | 2901 | 1241 | 1693 | 13 | 0 | 3 | 5 | 0 | 0 | 0 | 3 |
| 496 | -0.95 | 1.49 | -0.64 | 0.52 | 1 | Arap3 | ArfGAP with RhoGAP domain, ankyrin rep | 1782 | 1692 | 1971 | 0 | 0 | 3 | 5 | 0 | 1 | 0 | 2 |
| 476 | -0.87 | 1.97 | -0.44 | 0.66 | 1 | Gimap4 | GTPase, IMAP family member 4 | 2219 | 1534 | 1455 | 0 | 0 | 9 | 5 | 8 | 0 | 0 | 1 |
| 466 | 1.32 | 2.88 | 0.46 | 0.65 | 1 | Tmem204 | transmembrane protein 204 | 1879 | 1556 | 1640 | 0 | 0 | 0 | 5 | 0 | 0 | 48 | 0 |
| 431 | -2.30 | 1.25 | -1.84 | 0.07 | 1 | Tpm2 | tropomyosin 2, beta | 2933 | 835 | 965 | 1 | 0 | 0 | 5 | 0 | 0 | 0 | 0 |
| 269 | 0.03 | 0.71 | 0.04 | 0.97 | 1 | Fxyd5 | FXYD domain-containing ion transport reg | 1325 | 719 | 860 | 4 | 16 | 2 | 5 | 0 | 4 | 21 | 5 |
| 260 | -1.37 | 0.74 | -1.85 | 0.06 | 1 | Cd68 | CD68 antigen | 1360 | 496 | 923 | 0 | 31 | 28 | 5 | 0 | 5 | 5 | 12 |
| 232 | -0.12 | 1.60 | -0.08 | 0.94 | 1 | Pah | phenylalanine hydroxylase | 596 | 1018 | 871 | 34 | 0 | 2 | 5 | 0 | 10 | 1 | 12 |
| 222 | -0.42 | 2.77 | -0.15 | 0.88 | 1 | Enpp6 | ectonucleotide pyrophosphatase/phospho | 824 | 963 | 644 | 0 | 0 | 0 | 5 | 0 | 0 | 0 | 2 |
| 216 | -1.61 | 1.22 | -1.32 | 0.19 | 1 | Cd55 | CD55 molecule, decay accelerating facto | 1444 | 370 | 403 | 62 | 29 | 13 | 5 | 17 | 4 | 0 | 28 |
| 210 | 0.33 | 0.71 | 0.46 | 0.64 | 1 | Fam49a | family with sequence similarity 49, membe | 1137 | 626 | 461 | 10 | 14 | 8 | 5 | 25 | 11 | 0 | 13 |
| 191 | 0.29 | 0.83 | 0.35 | 0.73 | 1 | Alpl | alkaline phosphatase, liver/bone/kidney | 985 | 584 | 469 | 3 | 4 | 8 | 5 | 0 | 6 | 36 | 1 |
| 181 | -2.73 | 2.29 | -1.19 | 0.23 | 1 | Fads3 | fatty acid desaturase 3 | 877 | 552 | 506 | 0 | 9 | 2 | 5 | 38 | 0 | 0 | 0 |
| 126 | -2.63 | 2.32 | -1.13 | 0.26 | 1 | P2rx7 | purinergic receptor P2X, ligand-gated ion | 600 | 331 | 450 | 0 | 0 | 4 | 5 | 0 | 0 | 0 | 0 |
| 111 | 0.29 | 0.77 | 0.38 | 0.70 | 1 | Itpripl1 | inositol 1,4,5-triphosphate receptor intera | 548 | 312 | 209 | 54 | 6 | 2 | 5 | 36 | 9 | 40 | 0 |
| 109 | 0.74 | 0.88 | 0.84 | 0.40 | 1 | Fbn2 | fibrillin 2 | 513 | 204 | 395 | 0 | 21 | 10 | 5 | 10 | 17 | 0 | 22 |
| 106 | -0.11 | 2.07 | -0.05 | 0.96 | 1 | Reg3d | regenerating islet-derived 3 delta | 154 | 557 | 434 | 5 | 0 | 0 | 5 | 0 | 5 | 0 | 9 |
| 103 | 0.03 | 0.89 | 0.04 | 0.97 | 1 | Napsa | napsin A aspartic peptidase | 701 | 200 | 214 | 0 | 1 | 4 | 5 | 0 | 0 | 5 | 6 |
| 102 | -1.10 | 3.64 | -0.30 | 0.76 | 1 | Wfdc17 | WAP four-disulfide core domain 17 | 530 | 327 | 265 | 0 | 0 | 0 | 5 | 0 | 0 | 0 | 0 |
| 96 | -1.07 | 3.64 | -0.29 | 0.77 | 1 | Adamts7 | a disintegrin-like and metallopeptidase (r | 558 | 243 | 255 | 0 | 0 | 0 | 5 | 0 | 0 | 0 | 0 |
| 93 | 0.66 | 1.05 | 0.63 | 0.53 | 1 | Thsd1 | thrombospondin, type I, domain 1 | 352 | 361 | 274 | 0 | 0 | 7 | 5 | 6 | 0 | 0 | 15 |
| 89 | -3.61 | 2.68 | -1.34 | 0.18 | 1 | Prss3 | protease, serine 3 | 242 | 345 | 347 | 35 | 0 | 0 | 5 | 0 | 0 | 0 | 7 |
| 69 | -0.48 | 0.66 | -0.72 | 0.47 | 1 | Mboat1 | membrane bound O-acyltransferase dom | 222 | 231 | 200 | 16 | 20 | 21 | 5 | 21 | 15 | 8 | 3 |
| 60 | -2.01 | 1.08 | -1.86 | 0.06 | 1 | Clec2d | C-type lectin domain family 2, member d | 236 | 157 | 220 | 0 | 3 | 26 | 5 | 0 | 0 | 0 | 16 |
| 58 | -1.02 | 3.63 | -0.28 | 0.78 | 1 | Ptpn7 | protein tyrosine phosphatase, non-recept | 393 | 127 | 116 | 0 | 0 | 0 | 5 | 0 | 0 | 0 | 0 |
| 51 | 0.57 | 1.34 | 0.42 | 0.67 | 1 | C1qtnf9 | C1q and tumor necrosis factor related pro | 152 | 205 | 168 | 0 | 0 | 8 | 5 | 0 | 0 | 12 | 7 |
| 47 | -1.09 | 3.64 | -0.30 | 0.76 | 1 | Slamf8 | SLAM family member 8 | 255 | 168 | 94 | 0 | 0 | 0 | 5 | 0 | 0 | 0 | 0 |
| 40 | -1.07 | 3.64 | -0.29 | 0.77 | 1 | Aoah | acyloxyacyl hydrolase | 227 | 96 | 106 | 0 | 0 | 0 | 5 | 0 | 0 | 0 | 0 |
| 39 | -1.68 | 1.26 | -1.34 | 0.18 | 1 | Egln3 | egl-9 family hypoxia-inducible factor 3 | 173 | 75 | 56 | 40 | 18 | 12 | 5 | 0 | 0 | 37 | 16 |
| 38 | -0.26 | 1.16 | -0.22 | 0.82 | 1 | Nfkbid | nuclear factor of kappa light polypeptide g | 212 | 37 | 33 | 0 | 0 | 57 | 5 | 0 | 1 | 0 | 77 |
| 37 | -1.51 | 1.35 | -1.11 | 0.27 | 1 | Rad54l | RAD54 like (S. cerevisiae) | 202 | 12 | 32 | 29 | 17 | 36 | 5 | 0 | 21 | 0 | 50 |
| 37 | -0.68 | 1.10 | -0.62 | 0.54 | 1 | Iyd | iodotyrosine deiodinase | 241 | 55 | 36 | 6 | 22 | 8 | 5 | 10 | 3 | 0 | 16 |
| 34 | -1.43 | 0.94 | -1.53 | 0.13 | 1 | Acrbp | proacrosin binding protein | 17 | 3 | 7 | 6 | 14 | 167 | 5 | 0 | 5 | 3 | 151 |
| 34 | -0.29 | 1.15 | -0.25 | 0.80 | 1 | Gm28875 | | 85 | 83 | 73 | 17 | 10 | 11 | 5 | 0 | 11 | 73 | 2 |
| 32 | -0.47 | 0.73 | -0.64 | 0.52 | 1 | Sez6 | seizure related gene 6 | 37 | 35 | 44 | 50 | 68 | 24 | 5 | 31 | 34 | 9 | 19 |
| 28 | 1.35 | 1.28 | 1.05 | 0.29 | 1 | Acsm3 | acyl-CoA synthetase medium-chain family | 45 | 109 | 117 | 8 | 0 | 0 | 5 | 16 | 7 | 0 | 5 |
| 28 | -0.26 | 0.94 | -0.27 | 0.79 | 1 | Prdx2-ps1 | | 0 | 83 | 29 | 35 | 8 | 36 | 5 | 78 | 10 | 1 | 25 |
| 27 | -1.19 | 3.64 | -0.33 | 0.74 | 1 | Rtp3 | receptor transporter protein 3 | 84 | 97 | 114 | 0 | 0 | 0 | 5 | 0 | 0 | 0 | 0 |
| 26 | -2.66 | 1.80 | -1.48 | 0.14 | 1 | Rasgrp4 | RAS guanyl releasing protein 4 | 180 | 76 | 18 | 0 | 7 | 0 | 5 | 0 | 0 | 0 | 1 |
| 25 | -2.48 | 1.39 | -1.79 | 0.07 | 1 | Zfp286 | zinc finger protein 286 | 94 | 77 | 44 | 14 | 17 | 2 | 5 | 21 | 0 | 0 | 0 |
| 24 | -0.41 | 1.79 | -0.23 | 0.82 | 1 | A730011C13Rik | | 0 | 4 | 0 | 0 | 3 | 132 | 5 | 0 | 14 | 0 | 110 |
| 24 | -0.41 | 1.49 | -0.27 | 0.78 | 1 | Adra1a | adrenergic receptor, alpha 1a | 105 | 104 | 45 | 0 | 0 | 2 | 5 | 0 | 1 | 0 | 3 |
| 24 | -0.04 | 1.00 | -0.04 | 0.97 | 1 | Gm43682 | | 64 | 30 | 32 | 19 | 16 | 25 | 5 | 20 | 19 | 0 | 32 |
| 24 | 0.01 | 1.04 | 0.01 | 0.99 | 1 | Gm48427 | | 82 | 34 | 37 | 14 | 3 | 29 | 5 | 0 | 7 | 26 | 25 |
| 24 | -1.46 | 1.14 | -1.28 | 0.20 | 1 | Nlgn3 | neuroligin 3 | 50 | 60 | 32 | 25 | 22 | 25 | 5 | 0 | 3 | 31 | 9 |

| | | | | | | | | | | | | | | | | | | |
|---|---|---|---|---|---|---|---|---|---|---|---|---|---|---|---|---|---|---|
| 23 | -1.64 | 1.40 | -1.17 | 0.24 | 1 | Gm45250 | | 23 | 0 | 50 | 16 | 21 | 60 | 5 | 8 | 1 | 0 | 74 |
| 23 | -0.34 | 1.08 | -0.31 | 0.75 | 1 | Gm20045 | | 10 | 29 | 19 | 23 | 16 | 68 | 5 | 0 | 10 | 24 | 51 |
| 23 | -1.10 | 3.64 | -0.30 | 0.76 | 1 | Sbspon | somatomedin B and thrombospondin, typ | 106 | 14 | 127 | 0 | 0 | 0 | 5 | 0 | 0 | 0 | 0 |
| 23 | -0.13 | 1.22 | -0.10 | 0.92 | 1 | C78859 | | 0 | 0 | 0 | 11 | 6 | 121 | 5 | 0 | 13 | 15 | 80 |
| 22 | 0.39 | 0.86 | 0.45 | 0.65 | 1 | Elavl3 | ELAV like RNA binding protein 3 | 84 | 52 | 46 | 13 | 2 | 7 | 5 | 9 | 9 | 14 | 3 |
| 22 | 1.21 | 0.81 | 1.48 | 0.14 | 1 | 2610318N02Ril | RIKEN cDNA 2610318N02 gene | 39 | 13 | 42 | 4 | 17 | 23 | 5 | 23 | 10 | 22 | 44 |
| 20 | 0.94 | 1.30 | 0.72 | 0.47 | 1 | Gm38140 | | 24 | 0 | 0 | 3 | 22 | 25 | 5 | 25 | 16 | 82 | 15 |
| 19 | 0.50 | 1.25 | 0.40 | 0.69 | 1 | Gm44148 | | 15 | 31 | 26 | 15 | 9 | 12 | 5 | 0 | 9 | 84 | 9 |
| 19 | 1.42 | 1.28 | 1.11 | 0.27 | 1 | Tex38 | testis expressed 38 | 17 | 0 | 0 | 8 | 8 | 23 | 5 | 80 | 7 | 37 | 27 |
| 19 | -0.21 | 0.95 | -0.22 | 0.83 | 1 | Gm5870 | | 27 | 7 | 11 | 25 | 26 | 23 | 5 | 3 | 25 | 28 | 26 |
| 18 | 0.16 | 0.97 | 0.16 | 0.87 | 1 | Gm49544 | | 3 | 7 | 9 | 18 | 36 | 29 | 5 | 5 | 20 | 41 | 25 |
| 18 | -1.14 | 2.00 | -0.57 | 0.57 | 1 | Siglecg | sialic acid binding Ig-like lectin G | 142 | 21 | 25 | 0 | 0 | 0 | 5 | 0 | 0 | 0 | 0 |
| 17 | 1.93 | 1.66 | 1.17 | 0.24 | 1 | Pnoc | prepronociceptin | 122 | 31 | 17 | 0 | 0 | 0 | 5 | 0 | 5 | 10 | 0 |
| 17 | -1.39 | 1.87 | -0.74 | 0.46 | 1 | Gm14616 | | 45 | 52 | 85 | 0 | 0 | 1 | 5 | 0 | 0 | 0 | 1 |
| 16 | -0.31 | 0.87 | -0.35 | 0.72 | 1 | Atad3aos | | 15 | 13 | 6 | 31 | 17 | 30 | 5 | 21 | 18 | 5 | 18 |
| 16 | -1.43 | 2.20 | -0.65 | 0.51 | 1 | Nlrp1b | NLR family, pyrin domain containing 1B | 51 | 97 | 26 | 0 | 0 | 0 | 5 | 0 | 0 | 0 | 0 |
| 16 | -0.61 | 1.01 | -0.60 | 0.55 | 1 | Gm8129 | | 30 | 20 | 26 | 36 | 16 | 5 | 5 | 5 | 9 | 5 | 21 |
| 16 | -0.15 | 1.77 | -0.08 | 0.93 | 1 | Spag8 | sperm associated antigen 8 | 26 | 47 | 33 | 0 | 0 | 36 | 5 | 0 | 14 | 0 | 16 |
| 16 | 0.60 | 1.02 | 0.59 | 0.55 | 1 | Gm10614 | | 9 | 15 | 0 | 18 | 15 | 21 | 5 | 31 | 16 | 24 | 20 |
| 16 | -0.09 | 1.50 | -0.06 | 0.95 | 1 | Gm10814 | | 6 | 22 | 0 | 14 | 28 | 31 | 5 | 0 | 22 | 32 | 14 |
| 16 | -1.09 | 2.07 | -0.53 | 0.60 | 1 | Ltb4r1 | leukotriene B4 receptor 1 | 128 | 27 | 12 | 0 | 0 | 0 | 5 | 0 | 0 | 0 | 0 |
| 15 | -0.59 | 1.01 | -0.58 | 0.56 | 1 | Awat2 | acyl-CoA wax alcohol acyltransferase 2 | 4 | 11 | 27 | 24 | 21 | 14 | 5 | 43 | 14 | 0 | 7 |
| 15 | 0.17 | 1.07 | 0.15 | 0.88 | 1 | 9330161L09Rik | | 12 | 8 | 27 | 9 | 15 | 19 | 5 | 57 | 10 | 3 | 6 |
| 15 | 1.64 | 1.06 | 1.55 | 0.12 | 1 | Gm24157 | | 6 | 0 | 16 | 12 | 10 | 8 | 5 | 37 | 23 | 26 | 22 |
| 15 | -0.35 | 0.91 | -0.39 | 0.70 | 1 | Gm41077 | | 12 | 19 | 10 | 13 | 9 | 37 | 5 | 20 | 1 | 5 | 35 |
| 15 | 0.41 | 1.47 | 0.28 | 0.78 | 1 | Slc17a4 | solute carrier family 17 (sodium phosphat | 23 | 17 | 69 | 6 | 4 | 2 | 5 | 23 | 0 | 0 | 14 |
| 14 | -1.23 | 1.33 | -0.92 | 0.36 | 1 | Gm26779 | | 14 | 0 | 5 | 18 | 9 | 39 | 5 | 0 | 7 | 12 | 50 |
| 13 | -1.44 | 1.19 | -1.22 | 0.22 | 1 | Zfp575 | zinc finger protein 575 | 46 | 46 | 16 | 11 | 7 | 7 | 5 | 0 | 7 | 1 | 3 |
| 13 | -1.68 | 1.49 | -1.13 | 0.26 | 1 | Gm42748 | | 3 | 5 | 0 | 2 | 5 | 60 | 5 | 0 | 2 | 0 | 66 |
| 13 | -3.35 | 1.81 | -1.85 | 0.07 | 1 | Wif1 | Wnt inhibitory factor 1 | 81 | 20 | 11 | 0 | 23 | 3 | 5 | 0 | 0 | 0 | 3 |
| 13 | -0.34 | 1.16 | -0.29 | 0.77 | 1 | Mdga1 | MAM domain containing glycosylphospha | 35 | 27 | 30 | 8 | 9 | 3 | 5 | 0 | 11 | 17 | 1 |
| 13 | -2.39 | 1.47 | -1.63 | 0.10 | 1 | Zbtb32 | zinc finger and BTB domain containing 32 | 88 | 17 | 5 | 8 | 9 | 5 | 5 | 0 | 2 | 0 | 7 |
| 13 | 0.27 | 1.57 | 0.17 | 0.86 | 1 | Zmynd12 | zinc finger, MYND domain containing 12 | 0 | 28 | 5 | 32 | 7 | 15 | 5 | 0 | 5 | 35 | 12 |
| 13 | 0.80 | 1.08 | 0.74 | 0.46 | 1 | Gm13205 | | 17 | 30 | 1 | 8 | 10 | 9 | 5 | 8 | 16 | 12 | 21 |
| 12 | -1.11 | 1.47 | -0.75 | 0.45 | 1 | Gm17971 | | 33 | 13 | 14 | 0 | 10 | 37 | 5 | 0 | 10 | 0 | 15 |
| 12 | 2.77 | 1.61 | 1.72 | 0.09 | 1 | AW046200 | | 17 | 35 | 0 | 0 | 0 | 26 | 5 | 14 | 7 | 17 | 14 |
| 12 | 2.15 | 1.19 | 1.81 | 0.07 | 1 | Gm2453 | | 0 | 0 | 0 | 10 | 1 | 8 | 5 | 32 | 9 | 57 | 11 |
| 12 | 0.64 | 1.37 | 0.47 | 0.64 | 1 | Gm5621 | | 22 | 3 | 27 | 6 | 29 | 1 | 5 | 28 | 2 | 1 | 8 |
| 12 | -0.80 | 1.27 | -0.63 | 0.53 | 1 | Gm49086 | | 32 | 12 | 0 | 11 | 25 | 11 | 5 | 12 | 5 | 13 | 7 |
| 12 | 0.73 | 1.39 | 0.53 | 0.60 | 1 | Calhm6 | calcium homeostasis modulator family me | 65 | 11 | 22 | 0 | 2 | 2 | 5 | 0 | 2 | 22 | 1 |
| 12 | 0.53 | 1.18 | 0.45 | 0.65 | 1 | Gm43808 | | 5 | 20 | 15 | 2 | 10 | 12 | 5 | 23 | 10 | 0 | 28 |
| 12 | -0.23 | 1.31 | -0.18 | 0.86 | 1 | H3c11 | H3 clustered histone 11 | 9 | 0 | 5 | 5 | 8 | 19 | 5 | 0 | 4 | 43 | 29 |
| 11 | -0.11 | 1.30 | -0.09 | 0.93 | 1 | Kcnc1 | potassium voltage gated channel, Shaw-r | 5 | 15 | 0 | 23 | 6 | 5 | 5 | 56 | 3 | 0 | 8 |
| 11 | 0.80 | 1.53 | 0.52 | 0.60 | 1 | Kcnj13 | potassium inwardly-rectifying channel, su | 23 | 22 | 10 | 0 | 6 | 13 | 5 | 2 | 9 | 33 | 0 |
| 11 | -0.96 | 1.56 | -0.61 | 0.54 | 1 | Scube2 | signal peptide, CUB domain, EGF-like 2 | 28 | 27 | 14 | 0 | 9 | 15 | 5 | 0 | 8 | 0 | 14 |
| 11 | -0.03 | 1.31 | -0.03 | 0.98 | 1 | Rbm3-ps | | 11 | 2 | 27 | 4 | 22 | 3 | 5 | 0 | 3 | 30 | 12 |
| 11 | -0.62 | 0.97 | -0.64 | 0.52 | 1 | Gm6142 | | 10 | 14 | 16 | 6 | 17 | 19 | 5 | 3 | 3 | 8 | 18 |
| 11 | -0.58 | 1.26 | -0.46 | 0.65 | 1 | Gm10053 | | 11 | 13 | 0 | 27 | 8 | 19 | 5 | 2 | 8 | 18 | 7 |
| 11 | -1.38 | 1.84 | -0.75 | 0.45 | 1 | Gm43863 | | 36 | 23 | 0 | 14 | 5 | 12 | 5 | 0 | 16 | 0 | 5 |
| 10 | -3.14 | 1.72 | -1.82 | 0.07 | 1 | 6330576A10Rik | | 17 | 38 | 37 | 0 | 4 | 9 | 5 | 0 | 0 | 0 | 2 |
| 10 | 0.87 | 1.02 | 0.85 | 0.39 | 1 | Gm18101 | | 21 | 3 | 7 | 10 | 4 | 5 | 5 | 7 | 12 | 32 | 6 |
| 10 | 1.27 | 1.22 | 1.04 | 0.30 | 1 | Myh7b | myosin, heavy chain 7B, cardiac muscle, | 16 | 8 | 4 | 3 | 8 | 6 | 5 | 0 | 26 | 19 | 17 |
| 10 | -0.67 | 1.53 | -0.44 | 0.66 | 1 | Phf11c | PHD finger protein 11C | 15 | 0 | 11 | 0 | 12 | 34 | 5 | 0 | 11 | 0 | 23 |
| 10 | 0.87 | 1.11 | 0.78 | 0.44 | 1 | Arhgap27os2 | | 40 | 6 | 8 | 1 | 10 | 4 | 5 | 9 | 10 | 13 | 3 |
| 10 | -1.62 | 1.58 | -1.02 | 0.31 | 1 | Dmkn | dermokine | 71 | 10 | 7 | 0 | 4 | 8 | 5 | 0 | 2 | 0 | 2 |
| 10 | -0.71 | 1.70 | -0.42 | 0.68 | 1 | Insm2 | insulinoma-associated 2 | 8 | 48 | 20 | 0 | 7 | 5 | 5 | 0 | 11 | 0 | 6 |
| 10 | -2.43 | 1.84 | -1.32 | 0.19 | 1 | Gm44037 | | 28 | 7 | 0 | 34 | 7 | 15 | 5 | 0 | 8 | 0 | 4 |
| 10 | -1.17 | 1.56 | -0.75 | 0.45 | 1 | Pdzd7 | PDZ domain containing 7 | 14 | 18 | 18 | 7 | 0 | 23 | 5 | 0 | 0 | 5 | 18 |
| 10 | -3.48 | 1.82 | -1.92 | 0.06 | 1 | C130050O18Ri | RIKEN cDNA C130050O18 gene | 36 | 27 | 22 | 13 | 0 | 3 | 5 | 0 | 0 | 0 | 1 |
| 10 | 0.54 | 0.99 | 0.55 | 0.58 | 1 | Gm11560 | | 9 | 12 | 15 | 6 | 12 | 5 | 5 | 5 | 8 | 22 | 8 |

| | | | | | | | | | | | | | | | | | | |
|---|---|---|---|---|---|---|---|---|---|---|---|---|---|---|---|---|---|---|
| 10 | -3.68 | 1.87 | -1.97 | 0.05 | 1 | Slitrk5 | SLIT and NTRK-like family, member 5 | 8 | 45 | 33 | 12 | 0 | 3 | 5 | 0 | 1 | 0 | 0 |
| 10 | 1.57 | 1.29 | 1.22 | 0.22 | 1 | 1810059C17Rik | | 3 | 3 | 18 | 5 | 0 | 14 | 5 | 16 | 8 | 15 | 19 |
| 9 | 1.00 | 1.42 | 0.70 | 0.48 | 1 | Gm16938 | | 17 | 2 | 2 | 12 | 0 | 10 | 5 | 1 | 16 | 22 | 14 |
| 9 | -2.67 | 1.44 | -1.86 | 0.06 | 1 | 2900045O20Rik | | 3 | 4 | 25 | 25 | 15 | 6 | 5 | 0 | 8 | 0 | 7 |
| 8 | -1.33 | 1.87 | -0.71 | 0.48 | 1 | Gm10344 | | 6 | 15 | 0 | 7 | 13 | 7 | 5 | 0 | 0 | 33 | 6 |
| 8 | -2.64 | 2.62 | -1.01 | 0.31 | 1 | Gm47963 | | 0 | 2 | 33 | 0 | 0 | 22 | 5 | 0 | 0 | 0 | 28 |
| 8 | 1.24 | 1.12 | 1.11 | 0.27 | 1 | Rps15-ps3 | | 14 | 4 | 22 | 2 | 3 | 6 | 5 | 9 | 10 | 3 | 13 |
| 8 | -0.01 | 1.50 | -0.01 | 0.99 | 1 | Gm14038 | | 1 | 9 | 17 | 12 | 2 | 1 | 5 | 39 | 2 | 0 | 2 |
| 8 | -1.57 | 1.50 | -1.05 | 0.30 | 1 | Gm20426 | | 3 | 0 | 5 | 16 | 13 | 13 | 5 | 0 | 3 | 24 | 7 |
| 8 | -1.93 | 2.40 | -0.80 | 0.42 | 1 | Mag | myelin-associated glycoprotein | 27 | 20 | 35 | 0 | 0 | 1 | 5 | 0 | 0 | 0 | 0 |
| 8 | 0.61 | 1.33 | 0.46 | 0.65 | 1 | Llph-ps2 | | 7 | 0 | 11 | 9 | 12 | 2 | 5 | 6 | 4 | 22 | 9 |
| 8 | -3.00 | 1.88 | -1.60 | 0.11 | 1 | Gm37339 | | 0 | 0 | 12 | 16 | 5 | 16 | 5 | 0 | 20 | 0 | 12 |
| 8 | -3.30 | 1.68 | -1.97 | 0.05 | 1 | Atp10b | ATPase, class V, type 10B | 1 | 1 | 28 | 12 | 21 | 1 | 5 | 0 | 17 | 0 | 0 |
| 8 | -1.66 | 1.98 | -0.84 | 0.40 | 1 | Mir344i | | 0 | 0 | 0 | 8 | 12 | 13 | 5 | 8 | 0 | 0 | 39 |
| 8 | 1.59 | 1.88 | 0.84 | 0.40 | 1 | Rnaset2b | ribonuclease T2B | 9 | 0 | 14 | 0 | 8 | 6 | 5 | 25 | 9 | 0 | 8 |
| 8 | 1.27 | 1.65 | 0.77 | 0.44 | 1 | Il11 | interleukin 11 | 16 | 8 | 10 | 0 | 0 | 12 | 5 | 0 | 2 | 9 | 20 |
| 8 | -2.97 | 2.11 | -1.41 | 0.16 | 1 | Tiam2 | T cell lymphoma invasion and metastasis | 29 | 14 | 31 | 1 | 0 | 3 | 5 | 0 | 0 | 0 | 0 |
| 7 | -0.74 | 1.38 | -0.54 | 0.59 | 1 | Gm11335 | | 0 | 8 | 20 | 18 | 7 | 5 | 5 | 1 | 9 | 4 | 6 |
| 7 | -1.73 | 1.43 | -1.21 | 0.23 | 1 | Gm14398 | | 8 | 10 | 3 | 1 | 18 | 18 | 5 | 2 | 2 | 0 | 14 |
| 7 | -2.50 | 1.54 | -1.62 | 0.11 | 1 | Efcab5 | EF-hand calcium binding domain 5 | 19 | 1 | 17 | 7 | 8 | 12 | 5 | 0 | 3 | 0 | 10 |
| 7 | -2.13 | 1.91 | -1.12 | 0.26 | 1 | Gm31105 | | 26 | 0 | 28 | 2 | 9 | 1 | 5 | 0 | 8 | 0 | 0 |
| 7 | -0.29 | 1.66 | -0.18 | 0.86 | 1 | Gm28800 | | 15 | 30 | 5 | 0 | 6 | 3 | 5 | 0 | 4 | 1 | 9 |
| 7 | -0.54 | 2.00 | -0.27 | 0.79 | 1 | Capn8 | calpain 8 | 13 | 1 | 0 | 0 | 10 | 22 | 5 | 0 | 12 | 0 | 15 |
| 7 | -0.23 | 1.88 | -0.12 | 0.90 | 1 | 4930431P22Rik | | 15 | 6 | 0 | 0 | 25 | 4 | 5 | 0 | 6 | 9 | 8 |
| 7 | -0.75 | 1.31 | -0.57 | 0.57 | 1 | Gm31784 | | 0 | 0 | 0 | 8 | 9 | 25 | 5 | 2 | 6 | 5 | 15 |
| 7 | -1.96 | 1.93 | -1.02 | 0.31 | 1 | Plppr3 | phospholipid phosphatase related 3 | 24 | 9 | 25 | 1 | 0 | 7 | 5 | 0 | 0 | 0 | 5 |
| 7 | 0.87 | 1.49 | 0.58 | 0.56 | 1 | Gm44705 | | 3 | 9 | 3 | 0 | 5 | 2 | 5 | 37 | 7 | 3 | 1 |
| 7 | -3.93 | 2.01 | -1.95 | 0.05 | 1 | Gm44957 | | 0 | 0 | 22 | 19 | 4 | 13 | 5 | 0 | 7 | 0 | 5 |
| 7 | 1.55 | 1.83 | 0.84 | 0.40 | 1 | Ern2 | endoplasmic reticulum (ER) to nucleus si | 8 | 0 | 20 | 0 | 0 | 5 | 5 | 0 | 5 | 26 | 5 |
| 7 | 1.02 | 1.35 | 0.76 | 0.45 | 1 | Gpr150 | G protein-coupled receptor 150 | 12 | 1 | 15 | 0 | 2 | 8 | 5 | 0 | 6 | 13 | 11 |
| 7 | -2.01 | 2.09 | -0.96 | 0.34 | 1 | Gm43679 | | 0 | 5 | 18 | 3 | 0 | 20 | 5 | 0 | 3 | 0 | 19 |
| 7 | -2.12 | 2.41 | -0.88 | 0.38 | 1 | Aicda | activation-induced cytidine deaminase | 49 | 7 | 2 | 0 | 5 | 0 | 5 | 0 | 0 | 0 | 5 |
| 7 | 0.52 | 1.46 | 0.35 | 0.72 | 1 | Scrt2 | scratch family zinc finger 2 | 2 | 0 | 3 | 3 | 8 | 13 | 5 | 13 | 11 | 0 | 16 |
| 7 | -1.11 | 1.76 | -0.63 | 0.53 | 1 | Gm36279 | | 1 | 9 | 0 | 4 | 3 | 2 | 5 | 46 | 2 | 0 | 0 |
| 6 | -2.68 | 1.84 | -1.46 | 0.15 | 1 | Gm43690 | | 10 | 29 | 0 | 9 | 4 | 7 | 5 | 0 | 1 | 0 | 5 |
| 6 | -0.63 | 1.82 | -0.34 | 0.73 | 1 | 4930461C15Rik | | 0 | 0 | 7 | 3 | 7 | 12 | 5 | 0 | 3 | 28 | 4 |
| 6 | -1.61 | 2.04 | -0.79 | 0.43 | 1 | Arl9 | ADP-ribosylation factor-like 9 | 0 | 0 | 5 | 0 | 5 | 34 | 5 | 0 | 5 | 0 | 14 |
| 6 | 0.01 | 1.17 | 0.01 | 1.00 | 1 | Gm5644 | | 6 | 6 | 16 | 4 | 11 | 2 | 5 | 5 | 3 | 9 | 2 |
| 6 | -1.14 | 2.06 | -0.55 | 0.58 | 1 | 4930592I03Rik | | 0 | 0 | 0 | 23 | 15 | 1 | 5 | 0 | 1 | 10 | 12 |
| 6 | -2.56 | 1.98 | -1.29 | 0.20 | 1 | 2610017A05Rik | | 2 | 0 | 17 | 14 | 7 | 9 | 5 | 13 | 0 | 0 | 1 |
| 6 | 0.83 | 1.70 | 0.49 | 0.63 | 1 | Gm4835 | | 3 | 1 | 5 | 0 | 14 | 3 | 5 | 21 | 8 | 0 | 6 |
| 6 | -2.63 | 2.55 | -1.03 | 0.30 | 1 | Nxpe1-ps | | 34 | 18 | 4 | 0 | 4 | 0 | 5 | 0 | 0 | 0 | 0 |
| 6 | -1.31 | 2.23 | -0.59 | 0.56 | 1 | Gm8177 | | 0 | 9 | 22 | 3 | 0 | 5 | 5 | 0 | 6 | 0 | 14 |
| 6 | -0.99 | 1.15 | -0.86 | 0.39 | 1 | Gm6352 | | 5 | 3 | 4 | 11 | 9 | 8 | 5 | 8 | 2 | 3 | 5 |
| 6 | -0.77 | 1.92 | -0.40 | 0.69 | 1 | Vwc2 | von Willebrand factor C domain containin | 2 | 4 | 22 | 7 | 7 | 0 | 5 | 0 | 1 | 15 | 0 |
| 6 | -2.39 | 1.80 | -1.33 | 0.18 | 1 | Gm28651 | | 4 | 7 | 0 | 13 | 7 | 4 | 5 | 21 | 1 | 0 | 0 |
| 6 | -1.64 | 3.44 | -0.48 | 0.63 | 1 | 1520401A03Ril | RIKEN cDNA 1520401A03 gene | 0 | 0 | 0 | 0 | 0 | 41 | 5 | 0 | 0 | 0 | 16 |
| 6 | 0.14 | 2.02 | 0.07 | 0.95 | 1 | Prox2os | | 4 | 1 | 0 | 0 | 1 | 22 | 5 | 0 | 4 | 0 | 24 |
| 5 | -2.91 | 2.59 | -1.12 | 0.26 | 1 | Art5 | ADP-ribosyltransferase 5 | 0 | 0 | 5 | 13 | 0 | 6 | 5 | 28 | 2 | 0 | 0 |
| 5 | -2.54 | 1.87 | -1.36 | 0.17 | 1 | Aunip | aurora kinase A and ninein interacting pr | 11 | 7 | 6 | 0 | 16 | 8 | 5 | 0 | 4 | 0 | 1 |
| 5 | 0.11 | 3.05 | 0.04 | 0.97 | 1 | Itgb1bp2 | integrin beta 1 binding protein 2 | 0 | 0 | 0 | 0 | 0 | 25 | 5 | 0 | 7 | 0 | 21 |
| 5 | -2.09 | 1.80 | -1.16 | 0.25 | 1 | Cdv3-ps | | 2 | 0 | 20 | 11 | 9 | 1 | 5 | 0 | 4 | 1 | 6 |
| 5 | -2.45 | 1.77 | -1.39 | 0.17 | 1 | Gm34510 | | 0 | 6 | 4 | 18 | 9 | 6 | 5 | 0 | 5 | 0 | 6 |
| 5 | -2.73 | 2.31 | -1.18 | 0.24 | 1 | Gm37317 | | 0 | 0 | 3 | 5 | 0 | 25 | 5 | 0 | 3 | 0 | 16 |
| 5 | -0.44 | 2.52 | -0.17 | 0.86 | 1 | Gm33050 | | 0 | 0 | 0 | 0 | 20 | 3 | 5 | 0 | 20 | 0 | 9 |
| 5 | -0.47 | 1.21 | -0.39 | 0.70 | 1 | Gm28439 | | 3 | 9 | 2 | 7 | 7 | 7 | 5 | 2 | 6 | 5 | 4 |
| 5 | -0.33 | 2.53 | -0.13 | 0.89 | 1 | Gm43665 | | 0 | 4 | 0 | 0 | 1 | 16 | 5 | 0 | 4 | 0 | 26 |
| 5 | 1.59 | 2.30 | 0.69 | 0.49 | 1 | Rtl9 | retrotransposon Gag like 9 | 0 | 14 | 4 | 15 | 0 | 0 | 5 | 0 | 7 | 9 | 2 |
| 5 | 1.50 | 2.61 | 0.57 | 0.57 | 1 | Gm2654 | | 0 | 0 | 0 | 0 | 0 | 8 | 5 | 0 | 1 | 39 | 4 |
| 5 | -0.97 | 2.26 | -0.43 | 0.67 | 1 | Gm15411 | | 0 | 0 | 0 | 5 | 10 | 6 | 5 | 19 | 0 | 0 | 11 |

| | | | | | | | | | | | | | | | | | | |
|---|---|---|---|---|---|---|---|---|---|---|---|---|---|---|---|---|---|---|
| 5 | -2.12 | 2.15 | -0.98 | 0.32 | 1 | Gm11752 | | 0 | 0 | 0 | 5 | 18 | 7 | 5 | 0 | 7 | 0 | 14 |
| 5 | -0.42 | 1.99 | -0.21 | 0.83 | 1 | Gm5176 | | 1 | 0 | 16 | 0 | 3 | 8 | 5 | 0 | 8 | 0 | 16 |
| 5 | 0.76 | 1.62 | 0.47 | 0.64 | 1 | Gm6581 | | 6 | 0 | 14 | 0 | 3 | 3 | 5 | 0 | 10 | 13 | 2 |
| 5 | 1.44 | 1.99 | 0.72 | 0.47 | 1 | Gm32849 | | 18 | 3 | 0 | 3 | 0 | 8 | 5 | 7 | 6 | 0 | 4 |
| 5 | -3.76 | 2.01 | -1.87 | 0.06 | 1 | DXBay18 | DNA segment, Chr X, Baylor 18 | 1 | 2 | 0 | 29 | 1 | 11 | 5 | 0 | 0 | 0 | 4 |
| 5 | -3.31 | 3.36 | -0.99 | 0.32 | 1 | Gm15445 | | 0 | 0 | 11 | 0 | 0 | 18 | 5 | 0 | 0 | 0 | 18 |
| 5 | -0.47 | 1.97 | -0.24 | 0.81 | 1 | Gm9996 | | 0 | 0 | 0 | 4 | 8 | 5 | 5 | 14 | 1 | 0 | 15 |
| 5 | -0.87 | 3.17 | -0.27 | 0.78 | 1 | Lrrc2 | leucine rich repeat containing 2 | 35 | 1 | 9 | 0 | 0 | 0 | 5 | 0 | 0 | 0 | 0 |
| 5 | 2.34 | 2.69 | 0.87 | 0.38 | 1 | BC028777 | | 0 | 0 | 0 | 0 | 0 | 3 | 5 | 16 | 0 | 24 | 2 |
| 5 | -2.23 | 2.11 | -1.06 | 0.29 | 1 | Scd4 | stearoyl-coenzyme A desaturase 4 | 3 | 0 | 0 | 6 | 13 | 9 | 5 | 0 | 9 | 0 | 4 |
| 5 | -0.04 | 1.28 | -0.03 | 0.98 | 1 | Gm5539 | | 6 | 1 | 5 | 1 | 9 | 5 | 5 | 2 | 3 | 6 | 5 |
| 4 | -1.41 | 3.57 | -0.39 | 0.69 | 1 | Gm37062 | | 0 | 0 | 0 | 0 | 0 | 25 | 5 | 0 | 0 | 0 | 18 |
| 4 | 0.67 | 2.63 | 0.25 | 0.80 | 1 | Gm27320 | | 0 | 8 | 0 | 10 | 0 | 3 | 5 | 0 | 6 | 0 | 16 |
| 4 | -0.42 | 2.84 | -0.15 | 0.88 | 1 | Gm28536 | | 0 | 16 | 0 | 0 | 7 | 2 | 5 | 0 | 5 | 13 | 0 |
| 4 | -1.54 | 2.85 | -0.54 | 0.59 | 1 | Gm42444 | | 0 | 0 | 0 | 5 | 0 | 18 | 5 | 0 | 9 | 0 | 9 |
| 4 | 0.44 | 2.46 | 0.18 | 0.86 | 1 | Gm18646 | | 2 | 0 | 10 | 3 | 0 | 1 | 5 | 0 | 0 | 17 | 8 |
| 4 | -1.28 | 2.82 | -0.45 | 0.65 | 1 | Gm44187 | | 2 | 5 | 0 | 0 | 0 | 14 | 5 | 0 | 0 | 0 | 19 |
| 4 | -0.95 | 2.28 | -0.42 | 0.68 | 1 | Gm43689 | | 0 | 5 | 3 | 9 | 0 | 3 | 5 | 12 | 0 | 0 | 7 |
| 4 | 1.45 | 2.33 | 0.62 | 0.53 | 1 | Gm11620 | | 3 | 0 | 0 | 0 | 3 | 1 | 5 | 24 | 0 | 5 | 2 |
| 4 | -2.77 | 2.54 | -1.09 | 0.28 | 1 | Crhbp | corticotropin releasing hormone binding p | 6 | 0 | 1 | 5 | 0 | 16 | 5 | 0 | 0 | 0 | 10 |
| 4 | 3.01 | 3.24 | 0.93 | 0.35 | 1 | 0610038B21Rik | | 27 | 0 | 0 | 0 | 5 | 0 | 5 | 5 | 0 | 0 | 2 |
| 4 | -1.38 | 2.72 | -0.51 | 0.61 | 1 | Snora17 | | 7 | 1 | 0 | 0 | 0 | 23 | 5 | 0 | 0 | 0 | 7 |
| 4 | -1.04 | 3.41 | -0.31 | 0.76 | 1 | Fth-ps2 | | 0 | 1 | 0 | 0 | 0 | 11 | 5 | 0 | 0 | 0 | 25 |
| 4 | -0.15 | 1.87 | -0.08 | 0.94 | 1 | Gm12444 | | 15 | 8 | 2 | 1 | 1 | 2 | 5 | 0 | 3 | 0 | 5 |
| 4 | -3.72 | 2.74 | -1.36 | 0.17 | 1 | Gm8682 | | 3 | 0 | 18 | 0 | 9 | 7 | 5 | 0 | 0 | 0 | 0 |
| 4 | -0.80 | 2.39 | -0.33 | 0.74 | 1 | Kng2 | kininogen 2 | 3 | 3 | 0 | 0 | 9 | 3 | 5 | 0 | 5 | 0 | 14 |
| 4 | -3.93 | 2.50 | -1.57 | 0.12 | 1 | Gm11431 | | 0 | 0 | 6 | 14 | 5 | 2 | 5 | 0 | 9 | 0 | 0 |
| 4 | -5.89 | 3.07 | -1.91 | 0.06 | 1 | 9430034N14Rik | | 0 | 5 | 0 | 0 | 4 | 13 | 5 | 0 | 0 | 0 | 14 |
| 4 | 0.04 | 1.60 | 0.02 | 0.98 | 1 | Gm5845 | | 2 | 7 | 7 | 2 | 2 | 3 | 5 | 3 | 0 | 4 | 6 |
| 4 | 1.04 | 3.61 | 0.29 | 0.77 | 1 | C130071C03Rik | | 0 | 0 | 24 | 0 | 0 | 0 | 5 | 0 | 0 | 12 | 0 |
| 4 | -1.10 | 2.14 | -0.51 | 0.61 | 1 | 1700042O10Rik | | 10 | 5 | 0 | 5 | 5 | 1 | 5 | 0 | 4 | 0 | 5 |
| 4 | -0.16 | 1.64 | -0.10 | 0.92 | 1 | Gm4939 | | 2 | 2 | 3 | 5 | 4 | 1 | 5 | 1 | 11 | 0 | 6 |
| 4 | 1.41 | 3.43 | 0.41 | 0.68 | 1 | Slc25a18 | solute carrier family 25 (mitochondrial car | 0 | 0 | 0 | 0 | 0 | 3 | 5 | 16 | 16 | 0 | 0 |
| 4 | -0.48 | 2.89 | -0.17 | 0.87 | 1 | Slco4c1 | solute carrier organic anion transporter fa | 18 | 0 | 4 | 0 | 0 | 3 | 5 | 0 | 0 | 9 | 0 |
| 4 | 1.10 | 1.42 | 0.78 | 0.44 | 1 | Rps8-ps5 | | 3 | 1 | 1 | 2 | 1 | 2 | 5 | 7 | 1 | 14 | 2 |
| 4 | -0.80 | 2.11 | -0.38 | 0.71 | 1 | Gm10434 | | 5 | 8 | 2 | 0 | 5 | 3 | 5 | 0 | 6 | 0 | 4 |
| 3 | 0.23 | 1.65 | 0.14 | 0.89 | 1 | Gm7618 | | 4 | 0 | 8 | 2 | 3 | 3 | 5 | 7 | 3 | 1 | 2 |
| 3 | -2.58 | 2.91 | -0.89 | 0.37 | 1 | 1700064M15Rik | | 0 | 0 | 19 | 0 | 7 | 3 | 5 | 0 | 5 | 0 | 0 |
| 3 | -0.21 | 1.72 | -0.12 | 0.90 | 1 | Gm15793 | | 3 | 1 | 8 | 4 | 0 | 3 | 5 | 1 | 2 | 9 | 1 |
| 3 | -0.51 | 2.08 | -0.24 | 0.81 | 1 | Gm12841 | | 2 | 3 | 0 | 6 | 1 | 6 | 5 | 0 | 5 | 0 | 9 |
| 3 | -2.24 | 2.09 | -1.07 | 0.28 | 1 | Lypd6b | LY6/PLAUR domain containing 6B | 0 | 4 | 7 | 8 | 2 | 3 | 5 | 0 | 6 | 0 | 2 |
| 3 | -2.99 | 3.51 | -0.85 | 0.39 | 1 | A530001N23Rik | | 0 | 0 | 0 | 6 | 0 | 15 | 5 | 0 | 0 | 0 | 9 |
| 3 | -2.29 | 2.73 | -0.84 | 0.40 | 1 | Taar4 | trace amine-associated receptor 4 | 0 | 0 | 1 | 7 | 0 | 8 | 5 | 0 | 5 | 0 | 9 |
| 3 | 1.28 | 3.52 | 0.36 | 0.72 | 1 | Gm44366 | | 0 | 0 | 0 | 0 | 0 | 2 | 5 | 0 | 0 | 22 | 6 |
| 3 | -4.66 | 2.87 | -1.62 | 0.10 | 1 | Gm42872 | | 0 | 0 | 0 | 13 | 3 | 13 | 5 | 0 | 1 | 0 | 0 |
| 3 | -3.95 | 3.20 | -1.24 | 0.22 | 1 | mt-Nd3 | NADH dehydrogenase subunit 3 | 0 | 0 | 2 | 17 | 0 | 3 | 5 | 0 | 8 | 0 | 0 |
| 3 | -3.70 | 3.52 | -1.05 | 0.29 | 1 | 4930553P18Rik | | 0 | 5 | 0 | 0 | 0 | 3 | 5 | 0 | 21 | 0 | 0 |
| 3 | -2.04 | 3.37 | -0.61 | 0.54 | 1 | Armh1 | armadillo-like helical domain containing 1 | 0 | 5 | 17 | 0 | 0 | 2 | 5 | 0 | 5 | 0 | 0 |
| 3 | 0.28 | 1.80 | 0.15 | 0.88 | 1 | Myo3b | myosin IIIB | 11 | 2 | 2 | 0 | 3 | 1 | 5 | 0 | 3 | 4 | 2 |
| 3 | -0.97 | 1.89 | -0.51 | 0.61 | 1 | Gm13270 | | 0 | 3 | 4 | 3 | 1 | 5 | 5 | 7 | 2 | 0 | 3 |
| 3 | -1.85 | 3.55 | -0.52 | 0.60 | 1 | Gm831 | | 0 | 0 | 0 | 2 | 0 | 3 | 5 | 0 | 0 | 0 | 23 |
| 3 | -2.51 | 2.50 | -1.00 | 0.32 | 1 | Gm35867 | | 0 | 0 | 0 | 3 | 15 | 3 | 5 | 0 | 3 | 0 | 4 |
| 3 | -0.74 | 2.19 | -0.34 | 0.74 | 1 | Rbm6-ps1 | | 7 | 2 | 0 | 2 | 0 | 6 | 5 | 0 | 0 | 4 | 6 |
| 3 | 0.88 | 3.54 | 0.25 | 0.80 | 1 | Gm37238 | | 0 | 0 | 0 | 0 | 0 | 3 | 5 | 0 | 0 | 18 | 5 |
| 3 | -1.74 | 3.61 | -0.48 | 0.63 | 1 | Ckm | creatine kinase, muscle | 17 | 8 | 0 | 0 | 0 | 1 | 5 | 0 | 0 | 0 | 0 |
| 3 | -1.24 | 3.64 | -0.34 | 0.73 | 1 | D7Ertd443e | DNA segment, Chr 7, ERATO Doi 443, ex | 4 | 15 | 6 | 0 | 0 | 0 | 5 | 0 | 0 | 0 | 0 |
| 3 | 0.14 | 2.51 | 0.06 | 0.95 | 1 | Gm8097 | | 3 | 5 | 6 | 0 | 0 | 3 | 5 | 0 | 5 | 0 | 3 |
| 3 | -4.24 | 2.93 | -1.45 | 0.15 | 1 | Gm9115 | | 1 | 15 | 0 | 1 | 8 | 1 | 5 | 0 | 0 | 0 | 0 |
| 3 | 0.80 | 2.04 | 0.39 | 0.70 | 1 | Gm16559 | | 11 | 0 | 1 | 0 | 1 | 1 | 5 | 0 | 1 | 5 | 5 |
| 3 | -3.13 | 2.41 | -1.30 | 0.19 | 1 | Gm50383 | | 3 | 0 | 9 | 2 | 3 | 5 | 5 | 0 | 0 | 0 | 3 |

| | | | | | | | | | | | | | | | | | | |
|---|---|---|---|---|---|---|---|---|---|---|---|---|---|---|---|---|---|---|
| 3 | 0.33 | 1.62 | 0.21 | 0.84 | 1 | Ftl1-ps2 | | 0 | 2 | 1 | 4 | 1 | 3 | 5 | 3 | 4 | 4 | 3 |
| 3 | -0.51 | 3.28 | -0.15 | 0.88 | 1 | Rpl27a-ps2 | | 12 | 0 | 8 | 0 | 0 | 1 | 5 | 0 | 0 | 0 | 3 |
| 3 | -3.72 | 2.81 | -1.33 | 0.19 | 1 | Gm39302 | | 4 | 0 | 0 | 13 | 1 | 3 | 5 | 0 | 0 | 0 | 2 |
| 3 | -0.93 | 1.72 | -0.54 | 0.59 | 1 | Ndufs6b | | 8 | 0 | 1 | 1 | 3 | 4 | 5 | 1 | 1 | 4 | 1 |
| 3 | -1.21 | 3.64 | -0.33 | 0.74 | 1 | Gal3st3 | galactose-3-O-sulfotransferase 3 | 5 | 5 | 14 | 0 | 0 | 0 | 5 | 0 | 0 | 0 | 0 |
| 3 | 0.53 | 2.83 | 0.19 | 0.85 | 1 | 4833412C05Rik | | 3 | 0 | 3 | 0 | 0 | 3 | 5 | 0 | 9 | 0 | 5 |
| 3 | -6.54 | 3.51 | -1.86 | 0.06 | 1 | A430028G04Rik | | 0 | 0 | 0 | 8 | 0 | 7 | 5 | 0 | 8 | 0 | 0 |
| 2 | 0.44 | 2.30 | 0.19 | 0.85 | 1 | Gm5469 | | 0 | 4 | 0 | 1 | 0 | 6 | 5 | 1 | 5 | 1 | 4 |
| 2 | 0.01 | 3.54 | 0.00 | 1.00 | 1 | Gm44623 | | 11 | 0 | 0 | 0 | 0 | 3 | 5 | 0 | 0 | 0 | 8 |
| 2 | -1.83 | 2.07 | -0.88 | 0.38 | 1 | Gm14448 | | 1 | 1 | 11 | 1 | 1 | 3 | 5 | 0 | 2 | 0 | 2 |
| 2 | -3.67 | 3.59 | -1.02 | 0.31 | 1 | C230057M02Rik | | 0 | 16 | 0 | 0 | 0 | 3 | 5 | 0 | 0 | 0 | 2 |
| 2 | 1.66 | 3.59 | 0.46 | 0.64 | 1 | Cyp4a14 | cytochrome P450, family 4, subfamily a, p | 0 | 0 | 0 | 0 | 0 | 0 | 5 | 0 | 2 | 19 | 0 |
| 2 | -0.53 | 2.89 | -0.18 | 0.86 | 1 | Gm8899 | | 0 | 0 | 5 | 0 | 2 | 6 | 5 | 0 | 0 | 5 | 3 |
| 2 | -3.44 | 3.26 | -1.06 | 0.29 | 1 | Gm32220 | | 0 | 0 | 0 | 4 | 3 | 8 | 5 | 0 | 0 | 0 | 6 |
| 2 | -1.05 | 3.57 | -0.29 | 0.77 | 1 | Gm45191 | | 0 | 0 | 0 | 0 | 0 | 8 | 5 | 0 | 0 | 0 | 13 |
| 2 | -4.14 | 3.26 | -1.27 | 0.20 | 1 | Gm14302 | | 0 | 0 | 0 | 11 | 6 | 2 | 5 | 0 | 0 | 0 | 2 |
| 2 | -2.15 | 3.01 | -0.72 | 0.47 | 1 | Sbpl | spermine binding protein-like | 0 | 2 | 11 | 0 | 1 | 4 | 5 | 0 | 0 | 0 | 2 |
| 2 | -1.85 | 3.20 | -0.58 | 0.56 | 1 | Gm28449 | | 0 | 0 | 0 | 0 | 9 | 5 | 5 | 0 | 4 | 0 | 2 |
| 2 | -1.34 | 2.25 | -0.60 | 0.55 | 1 | Gm32256 | | 10 | 0 | 4 | 1 | 1 | 1 | 5 | 0 | 0 | 1 | 2 |
| 2 | 0.67 | 2.20 | 0.30 | 0.76 | 1 | Wsb2-ps | | 0 | 0 | 4 | 0 | 4 | 2 | 5 | 2 | 2 | 3 | 3 |
| 2 | -1.20 | 2.53 | -0.47 | 0.64 | 1 | Gm20570 | | 2 | 1 | 4 | 1 | 4 | 0 | 5 | 0 | 7 | 0 | 0 |
| 2 | 1.65 | 3.57 | 0.46 | 0.64 | 1 | Gm12945 | | 0 | 0 | 0 | 0 | 0 | 0 | 5 | 0 | 0 | 14 | 4 |
| 2 | -0.34 | 3.61 | -0.09 | 0.93 | 1 | Slc10a4 | solute carrier family 10 (sodium/bile acid ( | 18 | 0 | 0 | 0 | 0 | 0 | 5 | 0 | 0 | 0 | 0 |
| 2 | 0.14 | 3.60 | 0.04 | 0.97 | 1 | Gm16268 | | 0 | 0 | 0 | 0 | 0 | 1 | 5 | 0 | 0 | 17 | 0 |
| 2 | 0.21 | 3.04 | 0.07 | 0.94 | 1 | Gm46142 | | 12 | 0 | 1 | 1 | 0 | 0 | 5 | 2 | 0 | 1 | 0 |
| 2 | -2.36 | 2.92 | -0.81 | 0.42 | 1 | Ubap1l | ubiquitin-associated protein 1-like | 3 | 0 | 0 | 1 | 2 | 5 | 5 | 0 | 0 | 0 | 7 |
| 2 | 2.58 | 3.57 | 0.72 | 0.47 | 1 | Gm15321 | | 0 | 9 | 3 | 0 | 0 | 0 | 5 | 0 | 4 | 0 | 1 |
| 2 | -1.52 | 3.58 | -0.43 | 0.67 | 1 | Gm26403 | | 0 | 0 | 0 | 0 | 0 | 11 | 5 | 0 | 0 | 0 | 6 |
| 2 | -1.85 | 3.58 | -0.52 | 0.60 | 1 | Gm15538 | | 0 | 0 | 0 | 0 | 0 | 14 | 5 | 0 | 0 | 0 | 3 |
| 2 | 0.13 | 3.49 | 0.04 | 0.97 | 1 | Gm2106 | | 0 | 2 | 0 | 0 | 0 | 5 | 5 | 0 | 3 | 0 | 6 |
| 2 | -3.02 | 3.53 | -0.86 | 0.39 | 1 | Gm42685 | | 0 | 0 | 0 | 0 | 5 | 8 | 5 | 0 | 0 | 0 | 3 |
| 2 | -0.75 | 3.50 | -0.21 | 0.83 | 1 | 6430710M23Rik | | 0 | 0 | 0 | 0 | 1 | 5 | 5 | 0 | 9 | 0 | 1 |
| 2 | -1.99 | 3.55 | -0.56 | 0.57 | 1 | Gm27004 | | 0 | 0 | 0 | 0 | 9 | 1 | 5 | 0 | 5 | 0 | 0 |
| 2 | -1.03 | 3.57 | -0.29 | 0.77 | 1 | Gm46350 | | 0 | 0 | 0 | 0 | 0 | 5 | 5 | 0 | 0 | 0 | 9 |
| 2 | -2.68 | 3.58 | -0.75 | 0.45 | 1 | Gm5134 | predicted gene 5134 | 0 | 6 | 0 | 0 | 0 | 1 | 5 | 0 | 0 | 0 | 7 |
| 2 | -0.34 | 3.55 | -0.09 | 0.92 | 1 | Gm36401 | | 1 | 0 | 0 | 0 | 0 | 2 | 5 | 0 | 0 | 0 | 11 |
| 2 | -1.82 | 2.83 | -0.64 | 0.52 | 1 | Gm14091 | | 3 | 5 | 2 | 0 | 1 | 1 | 5 | 0 | 0 | 0 | 2 |
| 2 | -3.27 | 3.07 | -1.06 | 0.29 | 1 | Gm14019 | | 2 | 0 | 0 | 1 | 3 | 7 | 5 | 0 | 1 | 0 | 0 |
| 2 | -3.07 | 3.54 | -0.87 | 0.38 | 1 | Gm33016 | | 0 | 0 | 0 | 5 | 0 | 6 | 5 | 0 | 0 | 0 | 2 |
| 2 | -4.27 | 3.36 | -1.27 | 0.20 | 1 | Gm12411 | | 1 | 1 | 9 | 1 | 0 | 0 | 5 | 0 | 0 | 0 | 1 |
| 2 | -3.34 | 2.66 | -1.25 | 0.21 | 1 | Gm6651 | | 2 | 1 | 0 | 4 | 5 | 1 | 5 | 0 | 0 | 1 | 0 |
| 2 | 1.45 | 3.54 | 0.41 | 0.68 | 1 | Gm49025 | | 7 | 0 | 0 | 0 | 0 | 1 | 5 | 2 | 0 | 0 | 3 |
| 2 | 1.99 | 3.57 | 0.56 | 0.58 | 1 | Gm40493 | | 0 | 0 | 0 | 0 | 0 | 0 | 5 | 0 | 3 | 9 | 1 |
| 2 | -1.36 | 2.52 | -0.54 | 0.59 | 1 | Gm11604 | | 1 | 2 | 3 | 0 | 1 | 3 | 5 | 0 | 2 | 1 | 0 |
| 2 | 0.74 | 3.56 | 0.21 | 0.84 | 1 | Gm50217 | | 0 | 0 | 0 | 0 | 0 | 1 | 5 | 0 | 0 | 8 | 3 |
| 2 | -4.05 | 3.56 | -1.14 | 0.25 | 1 | 3110021N24Rik | | 0 | 0 | 0 | 0 | 5 | 7 | 5 | 0 | 0 | 0 | 0 |
| 2 | -0.75 | 3.62 | -0.21 | 0.83 | 1 | Gm49124 | | 9 | 2 | 0 | 0 | 0 | 0 | 5 | 0 | 0 | 0 | 0 |
| 1 | -3.93 | 3.56 | -1.11 | 0.27 | 1 | Gm13792 | | 0 | 0 | 0 | 9 | 0 | 3 | 5 | 0 | 0 | 0 | 0 |
| 1 | -0.55 | 3.55 | -0.15 | 0.88 | 1 | Gm23037 | | 0 | 0 | 0 | 0 | 0 | 5 | 5 | 0 | 1 | 0 | 5 |
| 1 | -0.62 | 2.88 | -0.22 | 0.83 | 1 | Gm9703 | | 1 | 3 | 2 | 2 | 0 | 0 | 5 | 2 | 1 | 0 | 0 |
| 1 | -0.74 | 2.26 | -0.33 | 0.74 | 1 | Gm6263 | | 3 | 1 | 1 | 2 | 1 | 0 | 5 | 0 | 1 | 1 | 1 |
| 1 | 1.16 | 3.56 | 0.32 | 0.75 | 1 | 2900092N22Rik | | 0 | 0 | 0 | 0 | 0 | 0 | 5 | 0 | 0 | 6 | 4 |
| 1 | -0.46 | 3.61 | -0.13 | 0.90 | 1 | Aldh3a1 | aldehyde dehydrogenase family 3, subfar | 10 | 0 | 0 | 0 | 0 | 0 | 5 | 0 | 0 | 0 | 0 |
| 1 | -1.36 | 3.54 | -0.38 | 0.70 | 1 | Gm11510 | | 0 | 0 | 0 | 0 | 1 | 5 | 5 | 0 | 1 | 0 | 3 |
| 1 | -0.38 | 3.56 | -0.11 | 0.91 | 1 | Slc25a2 | solute carrier family 25 (mitochondrial car | 2 | 1 | 0 | 0 | 0 | 1 | 5 | 0 | 0 | 0 | 6 |
| 1 | 2.76 | 3.56 | 0.78 | 0.44 | 1 | Gm25732 | | 0 | 6 | 0 | 0 | 0 | 0 | 5 | 0 | 2 | 0 | 1 |
| 1 | -1.98 | 3.62 | -0.55 | 0.58 | 1 | Gm6749 | | 5 | 1 | 1 | 0 | 0 | 2 | 5 | 0 | 0 | 0 | 0 |
| 1 | -4.88 | 3.55 | -1.37 | 0.17 | 1 | Gm13146 | | 0 | 0 | 0 | 0 | 0 | 4 | 5 | 5 | 0 | 0 | 0 |
| 1 | -3.49 | 3.57 | -0.98 | 0.33 | 1 | 1110028F18Rik | | 0 | 0 | 0 | 8 | 0 | 1 | 5 | 0 | 0 | 0 | 0 |
| 1 | -0.41 | 3.56 | -0.12 | 0.91 | 1 | Gm12070 | | 1 | 0 | 2 | 0 | 1 | 0 | 5 | 0 | 4 | 0 | 0 |

| | | | | | | | | | | | | | | | | | | |
|---|---|---|---|---|---|---|---|---|---|---|---|---|---|---|---|---|---|---|
| 1 | -1.22 | 2.94 | -0.42 | 0.68 | 1 | Gm7299 | | 2 | 1 | 0 | 2 | 1 | 0 | 5 | 1 | 0 | 1 | 0 |
| 1 | -2.31 | 3.54 | -0.65 | 0.51 | 1 | Gm12955 | | 0 | 0 | 0 | 0 | 4 | 0 | 5 | 0 | 0 | 0 | 4 |
| 1 | -1.20 | 3.58 | -0.33 | 0.74 | 1 | Grm4 | glutamate receptor, metabotropic 4 | 0 | 0 | 0 | 0 | 0 | 3 | 5 | 0 | 0 | 0 | 4 |
| 1 | 2.03 | 3.53 | 0.57 | 0.57 | 1 | Gm6641 | | 0 | 0 | 1 | 0 | 1 | 0 | 5 | 1 | 2 | 0 | 2 |
| 1 | 0.31 | 3.56 | 0.09 | 0.93 | 1 | Gm34967 | | 0 | 0 | 0 | 0 | 0 | 0 | 5 | 0 | 7 | 0 | 0 |
| 1 | -0.53 | 3.62 | -0.15 | 0.88 | 1 | Gm7926 | | 1 | 4 | 1 | 0 | 0 | 0 | 5 | 0 | 0 | 0 | 1 |
| 1 | -3.47 | 3.56 | -0.97 | 0.33 | 1 | 4933405D12Rik | | 0 | 0 | 0 | 0 | 5 | 2 | 5 | 0 | 0 | 0 | 0 |
| 1 | -2.77 | 3.60 | -0.77 | 0.44 | 1 | Nwd2 | NACHT and WD repeat domain containin | 0 | 0 | 0 | 6 | 0 | 0 | 5 | 0 | 0 | 0 | 0 |
| 1 | -0.57 | 3.62 | -0.16 | 0.87 | 1 | Spink2 | serine peptidase inhibitor, Kazal type 2 | 6 | 0 | 0 | 0 | 0 | 0 | 5 | 0 | 0 | 0 | 0 |
| 1 | -0.94 | 3.57 | -0.26 | 0.79 | 1 | Amd-ps7 | | 0 | 0 | 0 | 0 | 0 | 2 | 5 | 0 | 0 | 0 | 4 |
| 1 | -0.61 | 3.57 | -0.17 | 0.86 | 1 | Gm11450 | | 2 | 0 | 0 | 1 | 1 | 0 | 5 | 1 | 0 | 0 | 1 |
| 1 | -1.30 | 3.57 | -0.36 | 0.72 | 1 | Gm7867 | | 2 | 0 | 0 | 0 | 1 | 1 | 5 | 0 | 0 | 0 | 2 |
| 1 | -2.79 | 3.56 | -0.78 | 0.43 | 1 | Gm15569 | | 0 | 0 | 0 | 2 | 0 | 3 | 5 | 0 | 1 | 0 | 0 |
| 1 | 0.30 | 3.56 | 0.08 | 0.93 | 1 | Gm11745 | | 0 | 0 | 0 | 0 | 0 | 0 | 5 | 0 | 0 | 0 | 5 |
| 1 | -2.77 | 3.59 | -0.77 | 0.44 | 1 | Tafa4 | TAFA chemokine like family member 4 | 0 | 0 | 0 | 0 | 0 | 5 | 5 | 0 | 0 | 0 | 0 |
| 1 | 4.33 | 3.56 | 1.22 | 0.22 | 1 | Gm47003 | | 0 | 0 | 0 | 0 | 0 | 0 | 5 | 0 | 2 | 0 | 3 |
| 1 | -2.66 | 3.60 | -0.74 | 0.46 | 1 | Gm26766 | | 0 | 1 | 0 | 0 | 0 | 3 | 5 | 0 | 0 | 0 | 0 |
| 1 | 0.13 | 3.56 | 0.04 | 0.97 | 1 | Slc17a2 | solute carrier family 17 (sodium phosphat | 0 | 0 | 0 | 0 | 0 | 0 | 5 | 0 | 0 | 0 | 4 |
| 1 | -0.66 | 3.60 | -0.18 | 0.85 | 1 | Gm15267 | | 1 | 0 | 2 | 0 | 0 | 0 | 5 | 0 | 1 | 0 | 0 |
| 1 | -0.82 | 3.58 | -0.23 | 0.82 | 1 | Gm37160 | | 0 | 0 | 0 | 0 | 0 | 1 | 5 | 0 | 0 | 0 | 3 |
| 1 | -1.23 | 3.59 | -0.34 | 0.73 | 1 | Gm11284 | | 1 | 0 | 0 | 0 | 0 | 2 | 5 | 0 | 0 | 0 | 1 |
| 1 | -2.54 | 3.60 | -0.71 | 0.48 | 1 | Gm43020 | | 0 | 0 | 0 | 0 | 0 | 3 | 5 | 0 | 0 | 0 | 0 |
| 1 | -2.54 | 3.60 | -0.71 | 0.48 | 1 | Pax2 | paired box 2 | 0 | 0 | 0 | 0 | 0 | 3 | 5 | 0 | 0 | 0 | 0 |
| 1 | -0.68 | 3.58 | -0.19 | 0.85 | 1 | Nat8 | N-acetyltransferase 8 (GCN5-related) | 0 | 0 | 1 | 0 | 0 | 0 | 5 | 0 | 2 | 0 | 0 |
| 1 | 0.37 | 3.59 | 0.10 | 0.92 | 1 | Sult2a8 | sulfotransferase family 2A, dehydroepian | 1 | 0 | 0 | 0 | 0 | 0 | 5 | 0 | 2 | 0 | 0 |
| 1 | -0.95 | 3.59 | -0.27 | 0.79 | 1 | Pde6b | phosphodiesterase 6B, cGMP, rod recept | 0 | 0 | 0 | 0 | 0 | 1 | 5 | 0 | 0 | 0 | 2 |
| 1 | -1.00 | 3.63 | -0.27 | 0.78 | 1 | Dcc | deleted in colorectal carcinoma | 2 | 1 | 0 | 0 | 0 | 0 | 5 | 0 | 0 | 0 | 0 |
| 1 | -1.50 | 3.61 | -0.42 | 0.68 | 1 | Gm28818 | | 0 | 0 | 0 | 0 | 0 | 2 | 5 | 0 | 0 | 0 | 1 |
| 1 | -2.73 | 3.59 | -0.76 | 0.45 | 1 | Gm20531 | | 0 | 0 | 0 | 2 | 0 | 1 | 5 | 0 | 0 | 0 | 0 |
| 1 | -0.41 | 3.62 | -0.11 | 0.91 | 1 | Rpl31-ps9 | | 1 | 0 | 0 | 0 | 0 | 0 | 5 | 0 | 0 | 1 | 0 |
| 1 | -0.19 | 3.61 | -0.05 | 0.96 | 1 | Gm5335 | | 1 | 0 | 0 | 0 | 0 | 0 | 5 | 0 | 0 | 0 | 1 |
| 1 | -1.63 | 3.63 | -0.45 | 0.65 | 1 | Gm15906 | | 1 | 0 | 0 | 0 | 1 | 0 | 5 | 0 | 0 | 0 | 0 |
| 1 | -0.53 | 3.62 | -0.15 | 0.88 | 1 | Slc27a5 | solute carrier family 27 (fatty acid transpor | 0 | 0 | 0 | 0 | 0 | 0 | 5 | 0 | 0 | 0 | 1 |
| 1 | -1.85 | 3.63 | -0.51 | 0.61 | 1 | Gm38235 | | 0 | 0 | 0 | 0 | 0 | 1 | 5 | 0 | 0 | 0 | 0 |
| 1 | -1.85 | 3.63 | -0.51 | 0.61 | 1 | Gm16546 | | 0 | 0 | 0 | 0 | 0 | 1 | 5 | 0 | 0 | 0 | 0 |
| 0 | -1.23 | 3.65 | -0.34 | 0.74 | 1 | Gm18300 | | 0 | 0 | 0 | 0 | 0 | 0 | 5 | 0 | 0 | 0 | 0 |
| 0 | -1.23 | 3.65 | -0.34 | 0.74 | 1 | Gm23169 | | 0 | 0 | 0 | 0 | 0 | 0 | 5 | 0 | 0 | 0 | 0 |
| 0 | -1.23 | 3.65 | -0.34 | 0.74 | 1 | Gm24976 | | 0 | 0 | 0 | 0 | 0 | 0 | 5 | 0 | 0 | 0 | 0 |
| 0 | -1.23 | 3.65 | -0.34 | 0.74 | 1 | Gm43522 | | 0 | 0 | 0 | 0 | 0 | 0 | 5 | 0 | 0 | 0 | 0 |
| 0 | -1.23 | 3.65 | -0.34 | 0.74 | 1 | E130116L18Rik | | 0 | 0 | 0 | 0 | 0 | 0 | 5 | 0 | 0 | 0 | 0 |
| 0 | -1.23 | 3.65 | -0.34 | 0.74 | 1 | Gm16055 | | 0 | 0 | 0 | 0 | 0 | 0 | 5 | 0 | 0 | 0 | 0 |
| 0 | -1.23 | 3.65 | -0.34 | 0.74 | 1 | Gm44153 | | 0 | 0 | 0 | 0 | 0 | 0 | 5 | 0 | 0 | 0 | 0 |
| 0 | -1.23 | 3.65 | -0.34 | 0.74 | 1 | Gm31848 | | 0 | 0 | 0 | 0 | 0 | 0 | 5 | 0 | 0 | 0 | 0 |
| 0 | -1.23 | 3.65 | -0.34 | 0.74 | 1 | Lrrc72 | leucine rich repeat containing 72 | 0 | 0 | 0 | 0 | 0 | 0 | 5 | 0 | 0 | 0 | 0 |
| 0 | -1.23 | 3.65 | -0.34 | 0.74 | 1 | Nkx2-1 | NK2 homeobox 1 | 0 | 0 | 0 | 0 | 0 | 0 | 5 | 0 | 0 | 0 | 0 |
| 0 | -1.23 | 3.65 | -0.34 | 0.74 | 1 | Gm47659 | | 0 | 0 | 0 | 0 | 0 | 0 | 5 | 0 | 0 | 0 | 0 |
| 0 | -1.23 | 3.65 | -0.34 | 0.74 | 1 | Olfr720 | olfactory receptor 720 | 0 | 0 | 0 | 0 | 0 | 0 | 5 | 0 | 0 | 0 | 0 |
| 0 | -1.23 | 3.65 | -0.34 | 0.74 | 1 | Trav8d-1 | | 0 | 0 | 0 | 0 | 0 | 0 | 5 | 0 | 0 | 0 | 0 |
| 0 | -1.23 | 3.65 | -0.34 | 0.74 | 1 | Gm49786 | | 0 | 0 | 0 | 0 | 0 | 0 | 5 | 0 | 0 | 0 | 0 |
| 4104 | -3.46 | 1.81 | -1.92 | 0.06 | 1 | Bgn | biglycan | 24838 | 8548 | 11741 | 9 | 5 | 0 | 4 | 0 | 0 | 0 | 2 |
| 2109 | -0.32 | 0.80 | -0.39 | 0.69 | 1 | Fstl1 | follistatin-like 1 | 11121 | 5014 | 7022 | 3 | 16 | 3 | 4 | 0 | 7 | 6 | 7 |
| 1789 | -0.38 | 1.80 | -0.21 | 0.83 | 1 | C1s1 | complement component 1, s subcompone | 10745 | 4459 | 4468 | 0 | 0 | 1 | 4 | 0 | 0 | 0 | 5 |
| 1199 | -2.45 | 2.21 | -1.11 | 0.27 | 1 | Mmrn2 | multimerin 2 | 4316 | 4126 | 4745 | 0 | 0 | 3 | 4 | 0 | 0 | 0 | 0 |
| 960 | -1.65 | 1.36 | -1.22 | 0.22 | 1 | Ednrb | endothelin receptor type B | 3517 | 3623 | 3381 | 0 | 8 | 15 | 4 | 0 | 7 | 0 | 2 |
| 590 | -1.00 | 1.15 | -0.87 | 0.38 | 1 | Jcad | junctional cadherin 5 associated | 2435 | 1882 | 2143 | 5 | 6 | 1 | 4 | 0 | 4 | 0 | 6 |
| 570 | -2.22 | 1.36 | -1.63 | 0.10 | 1 | Atp1a2 | ATPase, Na+/K+ transporting, alpha 2 pol | 2744 | 1665 | 1806 | 8 | 26 | 1 | 4 | 0 | 16 | 0 | 0 |
| 497 | -3.01 | 2.56 | -1.18 | 0.24 | 1 | Mrc2 | mannose receptor, C type 2 | 2374 | 1425 | 1634 | 18 | 0 | 0 | 4 | 0 | 0 | 0 | 7 |
| 423 | 0.12 | 2.14 | 0.05 | 0.96 | 1 | Fgd5 | FYVE, RhoGEF and PH domain containin | 1615 | 1586 | 1442 | 0 | 0 | 0 | 4 | 0 | 0 | 4 | 0 |
| 421 | -2.08 | 1.15 | -1.82 | 0.07 | 1 | Pi16 | peptidase inhibitor 16 | 2532 | 504 | 1551 | 11 | 9 | 9 | 4 | 0 | 4 | 0 | 9 |
| 400 | -1.66 | 1.63 | -1.02 | 0.31 | 1 | Cpxm1 | carboxypeptidase X 1 (M14 family) | 2599 | 838 | 925 | 11 | 0 | 8 | 4 | 0 | 0 | 19 | 1 |

| | | | | | | | | | | | | | | | | | | |
|---|---|---|---|---|---|---|---|---|---|---|---|---|---|---|---|---|---|---|
| 364 | -0.37 | 2.76 | -0.14 | 0.89 | 1 | Pcdh17 | protocadherin 17 | 1203 | 1127 | 1668 | 0 | 0 | 0 | 4 | 0 | 0 | 0 | 2 |
| 361 | 0.65 | 0.44 | 1.48 | 0.14 | 1 | Gdpd3 | glycerophosphodiester phosphodiesteras | 42 | 528 | 673 | 572 | 525 | 82 | 4 | 664 | 761 | 45 | 71 |
| 295 | -1.13 | 3.65 | -0.31 | 0.76 | 1 | Car4 | carbonic anhydrase 4 | 1006 | 1083 | 1147 | 0 | 0 | 0 | 4 | 0 | 0 | 0 | 0 |
| 253 | -1.84 | 1.42 | -1.30 | 0.19 | 1 | Id1 | inhibitor of DNA binding 1, HLH protein | 969 | 820 | 944 | 10 | 1 | 19 | 4 | 2 | 11 | 0 | 0 |
| 249 | 1.64 | 2.28 | 0.72 | 0.47 | 1 | Epha2 | Eph receptor A2 | 1168 | 886 | 677 | 0 | 0 | 0 | 4 | 0 | 8 | 0 | 1 |
| 214 | -1.02 | 1.24 | -0.82 | 0.41 | 1 | Cd302 | CD302 antigen | 1209 | 534 | 576 | 11 | 2 | 4 | 4 | 0 | 8 | 0 | 4 |
| 200 | 0.30 | 1.76 | 0.17 | 0.86 | 1 | Sla | src-like adaptor | 1143 | 530 | 508 | 0 | 0 | 3 | 4 | 0 | 7 | 0 | 2 |
| 175 | 0.29 | 0.53 | 0.54 | 0.59 | 1 | Igsf3 | immunoglobulin superfamily, member 3 | 742 | 580 | 410 | 41 | 22 | 21 | 4 | 31 | 22 | 28 | 20 |
| 171 | 0.74 | 0.83 | 0.90 | 0.37 | 1 | Emp3 | epithelial membrane protein 3 | 884 | 442 | 486 | 0 | 3 | 24 | 4 | 0 | 4 | 15 | 20 |
| 165 | -2.69 | 2.42 | -1.11 | 0.27 | 1 | Pcsk5 | proprotein convertase subtilisin/kexin type | 1110 | 233 | 450 | 10 | 0 | 0 | 4 | 0 | 0 | 0 | 4 |
| 163 | 1.63 | 0.93 | 1.75 | 0.08 | 1 | Nr4a3 | nuclear receptor subfamily 4, group A, me | 1195 | 281 | 157 | 12 | 9 | 8 | 4 | 85 | 19 | 0 | 20 |
| 160 | 0.18 | 0.77 | 0.24 | 0.81 | 1 | Tmem47 | transmembrane protein 47 | 815 | 493 | 346 | 21 | 9 | 13 | 4 | 15 | 27 | 0 | 12 |
| 150 | -0.47 | 1.19 | -0.40 | 0.69 | 1 | H2-DMb1 | histocompatibility 2, class II, locus Mb1 | 753 | 438 | 447 | 0 | 4 | 1 | 4 | 0 | 8 | 0 | 0 |
| 146 | -2.74 | 2.05 | -1.34 | 0.18 | 1 | Lgals3 | lectin, galactose binding, soluble 3 | 749 | 323 | 525 | 0 | 7 | 0 | 4 | 0 | 0 | 0 | 0 |
| 142 | -0.62 | 1.88 | -0.33 | 0.74 | 1 | Pmaip1 | phorbol-12-myristate-13-acetate-induced | 892 | 347 | 285 | 4 | 0 | 2 | 4 | 25 | 0 | 0 | 0 |
| 139 | -1.04 | 3.65 | -0.28 | 0.78 | 1 | Ccr5 | chemokine (C-C motif) receptor 5 | 745 | 284 | 494 | 0 | 0 | 0 | 4 | 0 | 0 | 0 | 0 |
| 132 | -2.03 | 1.21 | -1.68 | 0.09 | 1 | Fxyd1 | FXYD domain-containing ion transport reg | 565 | 267 | 606 | 4 | 0 | 1 | 4 | 0 | 2 | 0 | 0 |
| 116 | -3.41 | 1.81 | -1.88 | 0.06 | 1 | Gpx8 | glutathione peroxidase 8 (putative) | 611 | 307 | 343 | 5 | 8 | 0 | 4 | 0 | 0 | 0 | 3 |
| 116 | -1.14 | 3.65 | -0.31 | 0.76 | 1 | Gata2 | GATA binding protein 2 | 371 | 373 | 525 | 0 | 0 | 0 | 4 | 0 | 0 | 0 | 0 |
| 102 | -1.04 | 3.65 | -0.28 | 0.78 | 1 | Ifi205 | interferon activated gene 205 | 554 | 316 | 253 | 0 | 0 | 0 | 4 | 0 | 0 | 0 | 0 |
| 99 | -2.81 | 1.93 | -1.46 | 0.15 | 1 | Ecscr | endothelial cell surface expressed chemo | 375 | 314 | 376 | 7 | 0 | 5 | 4 | 0 | 8 | 0 | 0 |
| 97 | -2.89 | 2.37 | -1.22 | 0.22 | 1 | Sash3 | SAM and SH3 domain containing 3 | 619 | 221 | 210 | 0 | 9 | 0 | 4 | 0 | 0 | 0 | 0 |
| 90 | -0.40 | 1.26 | -0.32 | 0.75 | 1 | Trarg1 | trafficking regulator of GLUT4 (SLC2A4) 1 | 389 | 372 | 215 | 0 | 0 | 2 | 4 | 0 | 0 | 0 | 6 |
| 88 | -0.61 | 1.10 | -0.56 | 0.58 | 1 | Ablim3 | actin binding LIM protein family, member | 353 | 289 | 307 | 0 | 3 | 6 | 4 | 0 | 7 | 0 | 2 |
| 88 | 1.60 | 1.08 | 1.48 | 0.14 | 1 | Gm43360 | | 0 | 2 | 6 | 1 | 5 | 466 | 4 | 0 | 12 | 72 | 400 |
| 87 | 0.14 | 1.09 | 0.13 | 0.90 | 1 | Kcnk6 | potassium inwardly-rectifying channel, su | 467 | 252 | 201 | 8 | 0 | 5 | 4 | 0 | 9 | 0 | 13 |
| 82 | -1.57 | 2.04 | -0.77 | 0.44 | 1 | Chrm2 | cholinergic receptor, muscarinic 2, cardia | 262 | 302 | 297 | 3 | 0 | 8 | 4 | 0 | 0 | 26 | 0 |
| 78 | -0.58 | 1.30 | -0.44 | 0.66 | 1 | A4galt | alpha 1,4-galactosyltransferase | 402 | 269 | 181 | 0 | 0 | 1 | 4 | 0 | 0 | 0 | 3 |
| 75 | -1.12 | 3.65 | -0.31 | 0.76 | 1 | Ccdc102a | coiled-coil domain containing 102A | 276 | 226 | 321 | 0 | 0 | 0 | 4 | 0 | 0 | 0 | 0 |
| 69 | 0.15 | 0.90 | 0.17 | 0.87 | 1 | Fgfrl1 | fibroblast growth factor receptor-like 1 | 305 | 203 | 164 | 25 | 3 | 10 | 4 | 1 | 17 | 26 | 5 |
| 68 | -0.52 | 1.30 | -0.40 | 0.69 | 1 | Flt3 | FMS-like tyrosine kinase 3 | 413 | 250 | 79 | 0 | 0 | 1 | 4 | 0 | 0 | 0 | 3 |
| 67 | -1.78 | 1.05 | -1.69 | 0.09 | 1 | Ccdc85a | coiled-coil domain containing 85A | 251 | 219 | 228 | 19 | 6 | 3 | 4 | 0 | 10 | 0 | 1 |
| 63 | -2.31 | 1.42 | -1.63 | 0.10 | 1 | Bank1 | B cell scaffold protein with ankyrin repeats | 388 | 211 | 90 | 0 | 0 | 2 | 4 | 0 | 0 | 0 | 0 |
| 62 | 1.13 | 2.19 | 0.52 | 0.61 | 1 | Galnt17 | polypeptide N-acetylgalactosaminyltransf | 281 | 124 | 238 | 9 | 0 | 0 | 4 | 0 | 27 | 0 | 2 |
| 61 | 1.37 | 1.34 | 1.02 | 0.31 | 1 | Ugt1a7c | UDP glucuronosyltransferase 1 family, po | 344 | 238 | 58 | 0 | 0 | 4 | 4 | 0 | 6 | 0 | 13 |
| 60 | -1.77 | 0.93 | -1.90 | 0.06 | 1 | Stap2 | signal transducing adaptor family membe | 234 | 156 | 161 | 10 | 18 | 42 | 4 | 0 | 8 | 0 | 29 |
| 56 | 1.99 | 1.19 | 1.67 | 0.10 | 1 | Fbxl7 | F-box and leucine-rich repeat protein 7 | 243 | 127 | 219 | 0 | 0 | 1 | 4 | 0 | 6 | 18 | 0 |
| 56 | -2.58 | 1.45 | -1.78 | 0.08 | 1 | Ngf | nerve growth factor | 248 | 158 | 198 | 0 | 0 | 3 | 4 | 0 | 0 | 0 | 0 |
| 55 | -1.25 | 1.38 | -0.91 | 0.36 | 1 | Ntf3 | neurotrophin 3 | 198 | 181 | 212 | 0 | 0 | 4 | 4 | 0 | 3 | 0 | 0 |
| 54 | -0.29 | 1.17 | -0.25 | 0.80 | 1 | Ccr7 | chemokine (C-C motif) receptor 7 | 350 | 161 | 69 | 0 | 0 | 3 | 4 | 0 | 0 | 1 | 4 |
| 53 | -2.34 | 1.46 | -1.61 | 0.11 | 1 | Dusp2 | dual specificity phosphatase 2 | 293 | 150 | 138 | 0 | 0 | 2 | 4 | 0 | 0 | 0 | 0 |
| 51 | -0.62 | 1.06 | -0.58 | 0.56 | 1 | Kcnc4 | potassium voltage gated channel, Shaw-r | 201 | 190 | 114 | 9 | 0 | 19 | 4 | 0 | 8 | 1 | 9 |
| 49 | -0.28 | 1.10 | -0.25 | 0.80 | 1 | Exoc3l | exocyst complex component 3-like | 223 | 127 | 168 | 1 | 0 | 7 | 4 | 0 | 4 | 0 | 5 |
| 49 | -0.94 | 3.64 | -0.26 | 0.80 | 1 | Sfrp4 | secreted frizzled-related protein 4 | 341 | 57 | 134 | 0 | 0 | 0 | 4 | 0 | 0 | 0 | 0 |
| 49 | 0.65 | 0.67 | 0.98 | 0.33 | 1 | Cep55 | centrosomal protein 55 | 171 | 105 | 61 | 13 | 9 | 54 | 4 | 14 | 9 | 39 | 55 |
| 48 | -1.43 | 0.92 | -1.55 | 0.12 | 1 | Itgb6 | integrin beta 6 | 136 | 134 | 201 | 4 | 12 | 22 | 4 | 0 | 9 | 0 | 10 |
| 46 | -0.42 | 0.71 | -0.59 | 0.55 | 1 | Fndc4 | fibronectin type III domain containing 4 | 175 | 138 | 117 | 9 | 18 | 10 | 4 | 10 | 4 | 10 | 6 |
| 45 | -0.40 | 1.10 | -0.37 | 0.71 | 1 | Bcat1 | branched chain aminotransferase 1, cytos | 283 | 76 | 101 | 14 | 0 | 3 | 4 | 7 | 2 | 0 | 4 |
| 45 | -2.02 | 1.30 | -1.55 | 0.12 | 1 | Adrb1 | adrenergic receptor, beta 1 | 215 | 128 | 121 | 9 | 0 | 9 | 4 | 0 | 5 | 0 | 1 |
| 43 | 0.50 | 0.81 | 0.62 | 0.54 | 1 | Gpi-ps | | 45 | 19 | 64 | 23 | 112 | 17 | 4 | 61 | 98 | 10 | 15 |
| 41 | 0.54 | 0.77 | 0.70 | 0.48 | 1 | Gm17753 | | 60 | 57 | 129 | 29 | 21 | 34 | 4 | 24 | 32 | 49 | 15 |
| 41 | -1.31 | 1.45 | -0.90 | 0.37 | 1 | Gm44220 | | 129 | 209 | 99 | 0 | 0 | 3 | 4 | 0 | 0 | 0 | 3 |
| 38 | 0.25 | 1.59 | 0.16 | 0.88 | 1 | Hoxb4 | homeobox B4 | 154 | 132 | 110 | 0 | 0 | 3 | 4 | 0 | 0 | 17 | 0 |
| 38 | 0.21 | 1.25 | 0.17 | 0.87 | 1 | Zbp1 | Z-DNA binding protein 1 | 154 | 62 | 184 | 0 | 4 | 0 | 4 | 0 | 6 | 0 | 2 |
| 35 | 0.86 | 0.77 | 1.11 | 0.27 | 1 | 9330162G02Rik | | 16 | 19 | 62 | 36 | 27 | 48 | 4 | 36 | 22 | 44 | 74 |
| 35 | 0.65 | 0.74 | 0.88 | 0.38 | 1 | Oas1b | 2'-5' oligoadenylate synthetase 1B | 55 | 35 | 80 | 16 | 13 | 69 | 4 | 34 | 14 | 22 | 41 |
| 34 | 0.53 | 1.48 | 0.35 | 0.72 | 1 | Rps2-ps10 | | 0 | 12 | 186 | 0 | 77 | 0 | 4 | 30 | 61 | 1 | 0 |
| 34 | 0.63 | 0.75 | 0.83 | 0.41 | 1 | Got2-ps1 | | 42 | 35 | 39 | 37 | 30 | 30 | 4 | 67 | 31 | 36 | 18 |
| 31 | -1.06 | 1.02 | -1.04 | 0.30 | 1 | Gm43389 | | 16 | 12 | 31 | 0 | 8 | 149 | 4 | 1 | 2 | 0 | 123 |
| 30 | -2.15 | 1.18 | -1.83 | 0.07 | 1 | Fign | fidgetin | 123 | 77 | 97 | 15 | 8 | 2 | 4 | 0 | 3 | 0 | 3 |

| | | | | | | | | | | | | | | | | | | |
|---|---|---|---|---|---|---|---|---|---|---|---|---|---|---|---|---|---|---|
| 30 | 0.64 | 0.99 | 0.65 | 0.52 | 1 | Zfp456 | zinc finger protein 456 | 79 | 107 | 47 | 2 | 16 | 17 | 4 | 1 | 22 | 22 | 11 |
| 29 | -1.41 | 1.15 | -1.23 | 0.22 | 1 | Mmd2 | monocyte to macrophage differentiation-a | 86 | 58 | 142 | 0 | 14 | 7 | 4 | 0 | 5 | 0 | 5 |
| 28 | -1.06 | 1.46 | -0.73 | 0.47 | 1 | BC064078 | | 162 | 57 | 73 | 0 | 9 | 0 | 4 | 0 | 3 | 0 | 1 |
| 28 | -1.68 | 1.45 | -1.16 | 0.24 | 1 | Dennd1c | DENN/MADD domain containing 1C | 150 | 45 | 82 | 5 | 0 | 8 | 4 | 0 | 8 | 0 | 1 |
| 25 | -1.58 | 1.01 | -1.56 | 0.12 | 1 | Gm47283 | | 10 | 14 | 22 | 9 | 16 | 94 | 4 | 0 | 5 | 1 | 104 |
| 25 | -1.00 | 0.95 | -1.06 | 0.29 | 1 | Lrrc9 | leucine rich repeat containing 9 | 38 | 65 | 35 | 32 | 36 | 18 | 4 | 13 | 28 | 0 | 9 |
| 25 | -0.22 | 1.76 | -0.13 | 0.90 | 1 | Sema5b | sema domain, seven thrombospondin rep | 75 | 83 | 95 | 0 | 0 | 1 | 4 | 0 | 0 | 0 | 14 |
| 24 | -2.39 | 1.72 | -1.39 | 0.17 | 1 | Il7r | interleukin 7 receptor | 118 | 55 | 83 | 0 | 0 | 3 | 4 | 0 | 0 | 0 | 0 |
| 23 | 0.40 | 0.95 | 0.43 | 0.67 | 1 | Gm14546 | | 2 | 43 | 1 | 23 | 31 | 34 | 4 | 37 | 26 | 10 | 41 |
| 23 | -1.70 | 1.46 | -1.17 | 0.24 | 1 | Gm7061 | | 15 | 14 | 12 | 33 | 0 | 94 | 4 | 0 | 5 | 0 | 72 |
| 22 | -1.71 | 1.95 | -0.88 | 0.38 | 1 | 9630001P10Rik | | 67 | 119 | 34 | 0 | 0 | 10 | 4 | 0 | 6 | 0 | 0 |
| 21 | -0.90 | 0.93 | -0.97 | 0.33 | 1 | Gm5815 | | 28 | 43 | 38 | 18 | 29 | 23 | 4 | 3 | 21 | 1 | 22 |
| 21 | -1.47 | 1.24 | -1.18 | 0.24 | 1 | D7Bwg0826e | | 10 | 4 | 20 | 18 | 8 | 60 | 4 | 0 | 12 | 0 | 94 |
| 21 | 0.82 | 1.23 | 0.66 | 0.51 | 1 | Tlr5 | toll-like receptor 5 | 105 | 28 | 42 | 0 | 2 | 15 | 4 | 0 | 4 | 24 | 4 |
| 21 | 1.36 | 0.95 | 1.44 | 0.15 | 1 | Pxmp2 | peroxisomal membrane protein 2 | 43 | 31 | 16 | 8 | 15 | 3 | 4 | 39 | 5 | 54 | 7 |
| 20 | 0.28 | 1.36 | 0.21 | 0.84 | 1 | Vsig2 | V-set and immunoglobulin domain contai | 58 | 52 | 46 | 25 | 0 | 7 | 4 | 0 | 13 | 13 | 6 |
| 20 | -1.75 | 1.19 | -1.47 | 0.14 | 1 | Acaa1b | acetyl-Coenzyme A acyltransferase 1B | 11 | 23 | 16 | 47 | 14 | 31 | 4 | 21 | 0 | 0 | 53 |
| 19 | -1.23 | 2.00 | -0.61 | 0.54 | 1 | Ear2 | eosinophil-associated, ribonuclease A fa | 105 | 41 | 62 | 0 | 0 | 0 | 4 | 0 | 0 | 0 | 0 |
| 18 | -2.86 | 1.58 | -1.81 | 0.07 | 1 | Srpk3 | serine/arginine-rich protein specific kinas | 50 | 32 | 69 | 9 | 0 | 21 | 4 | 0 | 0 | 0 | 18 |
| 18 | 0.97 | 1.51 | 0.64 | 0.52 | 1 | Pld5 | phospholipase D family, member 5 | 57 | 38 | 63 | 0 | 6 | 0 | 4 | 0 | 4 | 24 | 0 |
| 18 | 0.36 | 1.39 | 0.26 | 0.80 | 1 | Ing4 | inhibitor of growth family, member 4 | 30 | 16 | 7 | 0 | 0 | 64 | 4 | 0 | 0 | 6 | 68 |
| 18 | -2.11 | 1.53 | -1.38 | 0.17 | 1 | Hspa1l | heat shock protein 1-like | 49 | 36 | 57 | 18 | 0 | 11 | 4 | 0 | 0 | 1 | 18 |
| 17 | 1.56 | 1.10 | 1.42 | 0.15 | 1 | Lgr5 | leucine rich repeat containing G protein c | 26 | 28 | 20 | 9 | 4 | 8 | 4 | 54 | 20 | 0 | 16 |
| 17 | -1.24 | 2.10 | -0.59 | 0.55 | 1 | Tlr11 | toll-like receptor 11 | 78 | 71 | 35 | 0 | 0 | 0 | 4 | 0 | 0 | 0 | 0 |
| 17 | 0.42 | 0.94 | 0.45 | 0.65 | 1 | Gm10293 | | 8 | 2 | 2 | 19 | 28 | 25 | 4 | 36 | 20 | 14 | 29 |
| 17 | 0.11 | 0.88 | 0.12 | 0.90 | 1 | Bloc1s6os | | 4 | 29 | 16 | 10 | 39 | 18 | 4 | 25 | 11 | 8 | 22 |
| 17 | -0.18 | 1.20 | -0.15 | 0.88 | 1 | Ybx2 | Y box protein 2 | 18 | 4 | 2 | 14 | 9 | 53 | 4 | 23 | 6 | 0 | 50 |
| 17 | -0.72 | 1.36 | -0.53 | 0.60 | 1 | Wdr66 | WD repeat domain 66 | 100 | 21 | 25 | 0 | 5 | 11 | 4 | 0 | 3 | 0 | 14 |
| 17 | -1.19 | 0.88 | -1.36 | 0.18 | 1 | Gm6477 | | 19 | 29 | 54 | 20 | 23 | 9 | 4 | 2 | 13 | 3 | 7 |
| 16 | 2.07 | 1.14 | 1.82 | 0.07 | 1 | Rps26-ps1 | | 29 | 9 | 24 | 1 | 9 | 9 | 4 | 29 | 2 | 52 | 12 |
| 16 | 1.71 | 1.03 | 1.65 | 0.10 | 1 | Krtap16-1 | keratin associated protein 16-1 | 10 | 26 | 3 | 12 | 1 | 32 | 4 | 22 | 20 | 24 | 15 |
| 15 | -1.32 | 1.53 | -0.86 | 0.39 | 1 | Gm9008 | | 53 | 22 | 22 | 24 | 10 | 1 | 4 | 0 | 0 | 15 | 11 |
| 15 | 1.09 | 1.09 | 1.00 | 0.32 | 1 | Gm45053 | | 0 | 5 | 7 | 4 | 10 | 14 | 4 | 61 | 33 | 1 | 20 |
| 15 | -1.19 | 2.20 | -0.54 | 0.59 | 1 | Kcna3 | potassium voltage-gated channel, shaker | 75 | 75 | 7 | 0 | 0 | 0 | 4 | 0 | 0 | 0 | 0 |
| 15 | -2.77 | 1.99 | -1.39 | 0.16 | 1 | Gm43032 | | 52 | 36 | 26 | 0 | 7 | 6 | 4 | 29 | 0 | 0 | 0 |
| 14 | 0.63 | 1.52 | 0.41 | 0.68 | 1 | Slc35f3 | solute carrier family 35, member F3 | 27 | 43 | 9 | 12 | 0 | 9 | 4 | 35 | 0 | 19 | 0 |
| 14 | -1.08 | 1.31 | -0.83 | 0.41 | 1 | Erdr1 | | 6 | 5 | 12 | 9 | 11 | 48 | 4 | 16 | 0 | 0 | 46 |
| 14 | -2.44 | 1.58 | -1.55 | 0.12 | 1 | Nup62cl | nucleoporin 62 C-terminal like | 18 | 41 | 57 | 4 | 9 | 4 | 4 | 16 | 0 | 0 | 0 |
| 14 | -0.96 | 2.25 | -0.43 | 0.67 | 1 | Ifitm6 | interferon induced transmembrane protei | 102 | 44 | 2 | 0 | 0 | 0 | 4 | 0 | 0 | 0 | 0 |
| 13 | -0.40 | 1.42 | -0.28 | 0.78 | 1 | Gm28229 | | 12 | 0 | 9 | 24 | 8 | 36 | 4 | 14 | 17 | 0 | 24 |
| 13 | -1.02 | 2.21 | -0.46 | 0.64 | 1 | Clec4b1 | C-type lectin domain family 4, member b1 | 97 | 8 | 37 | 0 | 0 | 0 | 4 | 0 | 0 | 0 | 0 |
| 13 | -3.29 | 2.09 | -1.57 | 0.12 | 1 | Gm43362 | | 13 | 12 | 0 | 0 | 12 | 51 | 4 | 0 | 0 | 0 | 54 |
| 13 | 1.99 | 2.27 | 0.88 | 0.38 | 1 | Gm22748 | | 11 | 0 | 0 | 0 | 0 | 37 | 4 | 22 | 0 | 0 | 69 |
| 13 | 2.28 | 1.62 | 1.40 | 0.16 | 1 | Slco1a4 | solute carrier organic anion transporter fa | 18 | 27 | 43 | 0 | 0 | 7 | 4 | 0 | 10 | 27 | 3 |
| 13 | -0.82 | 1.54 | -0.53 | 0.60 | 1 | Gm29666 | | 5 | 17 | 10 | 0 | 13 | 36 | 4 | 0 | 14 | 0 | 38 |
| 12 | 1.47 | 1.24 | 1.18 | 0.24 | 1 | Ttll9 | tubulin tyrosine ligase-like family, membe | 9 | 0 | 10 | 12 | 3 | 12 | 4 | 48 | 7 | 17 | 15 |
| 12 | 0.56 | 1.84 | 0.30 | 0.76 | 1 | Gm15889 | | 20 | 76 | 21 | 0 | 0 | 3 | 4 | 0 | 6 | 0 | 5 |
| 12 | 1.34 | 1.50 | 0.90 | 0.37 | 1 | Mettl24 | methyltransferase like 24 | 71 | 9 | 16 | 0 | 6 | 0 | 4 | 0 | 3 | 14 | 7 |
| 12 | -1.92 | 1.80 | -1.07 | 0.29 | 1 | Gm39460 | | 35 | 35 | 47 | 3 | 0 | 1 | 4 | 0 | 0 | 0 | 3 |
| 12 | -0.47 | 1.18 | -0.40 | 0.69 | 1 | Smc2os | | 27 | 21 | 8 | 10 | 14 | 11 | 4 | 17 | 9 | 0 | 6 |
| 11 | 1.73 | 1.10 | 1.58 | 0.12 | 1 | Tpt1-ps5 | | 31 | 19 | 6 | 2 | 5 | 3 | 4 | 3 | 20 | 30 | 3 |
| 11 | -1.07 | 1.19 | -0.90 | 0.37 | 1 | Gm29773 | | 0 | 13 | 3 | 18 | 16 | 22 | 4 | 10 | 9 | 0 | 29 |
| 11 | -1.05 | 1.46 | -0.72 | 0.47 | 1 | Parvaos | | 3 | 2 | 0 | 5 | 14 | 31 | 4 | 0 | 1 | 37 | 24 |
| 11 | 1.92 | 1.06 | 1.81 | 0.07 | 1 | Gm15772 | | 17 | 25 | 32 | 2 | 3 | 1 | 4 | 20 | 11 | 4 | 4 |
| 11 | -0.90 | 1.11 | -0.81 | 0.42 | 1 | Gm20762 | | 15 | 26 | 46 | 11 | 4 | 3 | 4 | 2 | 1 | 4 | 4 |
| 11 | -2.48 | 1.74 | -1.43 | 0.15 | 1 | Cd247 | CD247 antigen | 21 | 46 | 0 | 11 | 15 | 11 | 4 | 0 | 4 | 0 | 8 |
| 11 | -0.74 | 1.63 | -0.46 | 0.65 | 1 | Ttc21a | tetratricopeptide repeat domain 21A | 27 | 25 | 3 | 25 | 1 | 14 | 4 | 0 | 11 | 0 | 11 |
| 11 | 0.56 | 1.61 | 0.35 | 0.73 | 1 | Gm48949 | | 2 | 4 | 0 | 0 | 8 | 26 | 4 | 12 | 9 | 0 | 52 |
| 10 | -2.05 | 1.41 | -1.45 | 0.15 | 1 | Gm50433 | | 2 | 17 | 14 | 35 | 6 | 15 | 4 | 0 | 8 | 0 | 14 |
| 10 | 0.45 | 2.18 | 0.21 | 0.84 | 1 | Gm44200 | | 7 | 11 | 0 | 0 | 0 | 46 | 4 | 0 | 5 | 0 | 41 |

| | | | | | | | | | | | | | | | | | |
|---|---|---|---|---|---|---|---|---|---|---|---|---|---|---|---|---|---|
| 10 | -0.74 | 1.27 | -0.58 | 0.56 | 1 | Gm10849 | | 11 | 13 | 16 | 18 | 10 | 10 | 4 | 0 | 3 | 18 | 10 |
| 10 | -0.41 | 1.48 | -0.28 | 0.78 | 1 | Prdm14 | PR domain containing 14 | 3 | 4 | 0 | 26 | 25 | 7 | 4 | 0 | 11 | 24 | 9 |
| 10 | -2.72 | 2.00 | -1.36 | 0.17 | 1 | Gm15261 | | 2 | 53 | 45 | 0 | 5 | 2 | 4 | 0 | 0 | 0 | 2 |
| 10 | 0.73 | 1.69 | 0.43 | 0.66 | 1 | Liph | lipase, member H | 4 | 45 | 23 | 0 | 8 | 2 | 4 | 0 | 1 | 17 | 8 |
| 10 | 0.88 | 1.53 | 0.58 | 0.56 | 1 | Gm31728 | | 13 | 9 | 8 | 21 | 0 | 8 | 4 | 0 | 9 | 17 | 20 |
| 10 | -1.89 | 2.29 | -0.83 | 0.41 | 1 | Gm34934 | | 15 | 43 | 47 | 0 | 0 | 1 | 4 | 0 | 0 | 0 | 0 |
| 10 | -3.15 | 2.66 | -1.18 | 0.24 | 1 | Gm42463 | | 0 | 7 | 0 | 0 | 0 | 62 | 4 | 0 | 0 | 0 | 34 |
| 10 | -1.64 | 1.42 | -1.16 | 0.25 | 1 | Tescl | tescalcin-like | 7 | 0 | 3 | 12 | 16 | 20 | 4 | 0 | 15 | 24 | 4 |
| 9 | -0.27 | 1.15 | -0.24 | 0.81 | 1 | Fcnaos | | 5 | 2 | 11 | 12 | 22 | 9 | 4 | 21 | 3 | 12 | 3 |
| 9 | 1.34 | 1.42 | 0.94 | 0.35 | 1 | 1700008O03Ril | RIKEN cDNA 1700008O03 gene | 9 | 40 | 16 | 4 | 0 | 2 | 4 | 13 | 0 | 13 | 3 |
| 9 | -0.08 | 1.64 | -0.05 | 0.96 | 1 | Gm12296 | | 31 | 10 | 18 | 0 | 9 | 5 | 4 | 0 | 0 | 19 | 7 |
| 9 | 0.11 | 1.62 | 0.07 | 0.95 | 1 | Ccnb2-ps | | 3 | 15 | 0 | 4 | 1 | 37 | 4 | 0 | 9 | 1 | 27 |
| 9 | -1.15 | 1.40 | -0.82 | 0.41 | 1 | Gm21960 | | 14 | 19 | 22 | 5 | 13 | 4 | 4 | 0 | 12 | 0 | 8 |
| 9 | 2.28 | 1.73 | 1.32 | 0.19 | 1 | Gm7180 | | 3 | 19 | 11 | 0 | 6 | 0 | 4 | 38 | 13 | 0 | 5 |
| 9 | -1.21 | 1.59 | -0.76 | 0.45 | 1 | Gm26850 | | 18 | 7 | 0 | 19 | 11 | 14 | 4 | 0 | 11 | 3 | 11 |
| 9 | -0.01 | 1.54 | -0.01 | 1.00 | 1 | Strc | stereocilin | 1 | 0 | 0 | 13 | 12 | 7 | 4 | 28 | 0 | 28 | 5 |
| 9 | 2.22 | 1.55 | 1.43 | 0.15 | 1 | Gm20628 | | 13 | 5 | 6 | 1 | 0 | 14 | 4 | 8 | 21 | 0 | 24 |
| 9 | -0.63 | 1.62 | -0.39 | 0.70 | 1 | Ccdc158 | coiled-coil domain containing 158 | 7 | 17 | 25 | 6 | 2 | 7 | 4 | 0 | 0 | 24 | 4 |
| 9 | 1.46 | 1.74 | 0.84 | 0.40 | 1 | Wdr38 | WD repeat domain 38 | 9 | 18 | 18 | 0 | 0 | 12 | 4 | 21 | 5 | 0 | 9 |
| 9 | -1.13 | 1.79 | -0.63 | 0.53 | 1 | Gm38387 | | 0 | 0 | 2 | 0 | 8 | 39 | 4 | 0 | 5 | 0 | 39 |
| 9 | 0.99 | 1.60 | 0.62 | 0.54 | 1 | Klhl41 | kelch-like 41 | 23 | 13 | 0 | 2 | 7 | 5 | 4 | 0 | 19 | 9 | 13 |
| 9 | 1.54 | 1.50 | 1.03 | 0.30 | 1 | Gm12164 | | 3 | 18 | 0 | 6 | 3 | 2 | 4 | 1 | 5 | 49 | 3 |
| 8 | 0.89 | 1.54 | 0.58 | 0.56 | 1 | Gm42997 | | 29 | 9 | 29 | 0 | 0 | 7 | 4 | 0 | 4 | 5 | 5 |
| 8 | -2.21 | 1.77 | -1.25 | 0.21 | 1 | Gm43797 | | 0 | 0 | 35 | 15 | 10 | 9 | 4 | 0 | 9 | 5 | 2 |
| 8 | -3.03 | 1.95 | -1.55 | 0.12 | 1 | Gm15983 | | 6 | 0 | 0 | 15 | 29 | 19 | 4 | 0 | 7 | 0 | 10 |
| 8 | 0.55 | 1.39 | 0.40 | 0.69 | 1 | Gm45418 | | 2 | 6 | 10 | 18 | 0 | 12 | 4 | 5 | 8 | 15 | 10 |
| 8 | 0.91 | 1.06 | 0.86 | 0.39 | 1 | Gm6123 | | 20 | 18 | 5 | 3 | 2 | 8 | 4 | 5 | 6 | 5 | 13 |
| 8 | 1.88 | 1.45 | 1.30 | 0.19 | 1 | Gm6992 | | 3 | 1 | 18 | 11 | 1 | 1 | 4 | 19 | 2 | 24 | 6 |
| 8 | -1.78 | 2.05 | -0.87 | 0.39 | 1 | Tenm1 | teneurin transmembrane protein 1 | 48 | 5 | 25 | 0 | 0 | 4 | 4 | 1 | 0 | 0 | 0 |
| 8 | -0.08 | 1.96 | -0.04 | 0.97 | 1 | Zfp114 | zinc finger protein 114 | 1 | 50 | 9 | 0 | 0 | 8 | 4 | 2 | 9 | 0 | 4 |
| 8 | 0.63 | 1.28 | 0.49 | 0.62 | 1 | Gm40190 | | 11 | 1 | 0 | 8 | 9 | 8 | 4 | 10 | 12 | 12 | 12 |
| 8 | 1.76 | 1.58 | 1.12 | 0.26 | 1 | Gm15753 | | 8 | 15 | 11 | 0 | 6 | 1 | 4 | 24 | 11 | 0 | 6 |
| 8 | -0.07 | 1.35 | -0.05 | 0.96 | 1 | Gm48857 | | 4 | 1 | 4 | 12 | 8 | 6 | 4 | 0 | 22 | 18 | 5 |
| 8 | -1.74 | 1.41 | -1.23 | 0.22 | 1 | Gm12115 | | 1 | 17 | 2 | 12 | 17 | 15 | 4 | 0 | 6 | 1 | 8 |
| 7 | -2.25 | 2.63 | -0.85 | 0.39 | 1 | B230303O12Rik | | 0 | 0 | 0 | 0 | 4 | 24 | 4 | 0 | 0 | 0 | 51 |
| 7 | -1.07 | 1.47 | -0.73 | 0.47 | 1 | 6330537M06Rik | | 14 | 1 | 2 | 1 | 5 | 25 | 4 | 0 | 0 | 10 | 19 |
| 7 | 0.71 | 1.16 | 0.61 | 0.54 | 1 | Spata48 | spermatogenesis associated 48 | 9 | 3 | 5 | 13 | 3 | 2 | 4 | 17 | 7 | 14 | 1 |
| 7 | -3.26 | 2.23 | -1.46 | 0.14 | 1 | Fgf13 | fibroblast growth factor 13 | 51 | 1 | 16 | 3 | 0 | 3 | 4 | 0 | 0 | 0 | 0 |
| 7 | -0.59 | 1.48 | -0.40 | 0.69 | 1 | A430073D23Rik | | 11 | 0 | 11 | 6 | 12 | 3 | 4 | 0 | 14 | 9 | 5 |
| 7 | -0.34 | 1.78 | -0.19 | 0.85 | 1 | Gm7507 | | 3 | 15 | 4 | 1 | 13 | 3 | 4 | 0 | 8 | 23 | 0 |
| 7 | 0.12 | 1.58 | 0.08 | 0.94 | 1 | Tmem202 | transmembrane protein 202 | 1 | 8 | 15 | 0 | 10 | 6 | 4 | 10 | 8 | 0 | 12 |
| 7 | -0.10 | 1.63 | -0.06 | 0.95 | 1 | 9030404E10Rik | | 4 | 18 | 8 | 0 | 5 | 5 | 4 | 22 | 2 | 0 | 5 |
| 7 | 0.81 | 1.88 | 0.43 | 0.67 | 1 | Gm15832 | | 11 | 6 | 0 | 10 | 0 | 13 | 4 | 0 | 9 | 3 | 18 |
| 7 | -3.46 | 1.76 | -1.96 | 0.05 | 1 | 1500012K07Rik | | 25 | 3 | 8 | 10 | 15 | 3 | 4 | 0 | 5 | 0 | 0 |
| 6 | -3.88 | 2.62 | -1.48 | 0.14 | 1 | Gata3 | GATA binding protein 3 | 32 | 0 | 23 | 8 | 5 | 0 | 4 | 0 | 0 | 0 | 0 |
| 6 | 0.10 | 1.95 | 0.05 | 0.96 | 1 | Gm48062 | | 5 | 4 | 15 | 0 | 0 | 18 | 4 | 0 | 6 | 0 | 19 |
| 6 | 0.84 | 1.52 | 0.55 | 0.58 | 1 | Gm6394 | | 23 | 15 | 3 | 1 | 2 | 3 | 4 | 0 | 13 | 3 | 4 |
| 6 | -0.08 | 1.14 | -0.07 | 0.94 | 1 | Rps19-ps6 | | 17 | 12 | 8 | 3 | 5 | 4 | 4 | 3 | 3 | 3 | 6 |
| 6 | -1.67 | 1.54 | -1.08 | 0.28 | 1 | B4gat1 | beta-1,4-glucuronyltransferase 1 | 6 | 0 | 17 | 5 | 13 | 4 | 4 | 0 | 8 | 9 | 1 |
| 6 | 0.03 | 1.51 | 0.02 | 0.98 | 1 | Gm5424 | | 24 | 1 | 19 | 3 | 5 | 1 | 4 | 2 | 5 | 0 | 4 |
| 6 | 1.98 | 1.96 | 1.01 | 0.31 | 1 | Crisp2 | cysteine-rich secretory protein 2 | 14 | 0 | 0 | 2 | 0 | 3 | 4 | 0 | 12 | 22 | 10 |
| 6 | 0.11 | 1.40 | 0.08 | 0.93 | 1 | Gm10131 | | 7 | 12 | 2 | 5 | 6 | 6 | 4 | 0 | 9 | 5 | 10 |
| 6 | -3.25 | 2.25 | -1.45 | 0.15 | 1 | Ccr10 | chemokine (C-C motif) receptor 10 | 33 | 2 | 7 | 11 | 0 | 3 | 4 | 0 | 4 | 0 | 0 |
| 6 | -1.06 | 1.97 | -0.54 | 0.59 | 1 | Gm37370 | | 9 | 0 | 11 | 1 | 0 | 20 | 4 | 0 | 2 | 0 | 17 |
| 6 | -2.77 | 1.64 | -1.69 | 0.09 | 1 | Gm12176 | | 19 | 4 | 2 | 11 | 7 | 11 | 4 | 0 | 2 | 0 | 5 |
| 6 | 0.74 | 2.56 | 0.29 | 0.77 | 1 | Gm6787 | | 0 | 12 | 0 | 11 | 0 | 6 | 4 | 0 | 16 | 0 | 13 |
| 6 | -0.86 | 3.49 | -0.25 | 0.81 | 1 | Ankrd23 | ankyrin repeat domain 23 | 0 | 0 | 0 | 0 | 0 | 16 | 4 | 0 | 0 | 0 | 42 |
| 6 | 0.87 | 1.75 | 0.49 | 0.62 | 1 | B230317F23Rik | | 0 | 0 | 1 | 0 | 4 | 17 | 4 | 0 | 14 | 10 | 11 |
| 6 | -1.89 | 2.61 | -0.72 | 0.47 | 1 | Gm30978 | | 19 | 34 | 3 | 0 | 0 | 1 | 4 | 0 | 0 | 0 | 0 |
| 5 | 0.54 | 2.37 | 0.23 | 0.82 | 1 | Gm45222 | | 0 | 14 | 0 | 0 | 0 | 19 | 4 | 7 | 0 | 8 | 8 |

| | | | | | | | | | | | | | | | | | | |
|---|---|---|---|---|---|---|---|---|---|---|---|---|---|---|---|---|---|---|
| 5 | 0.48 | 1.36 | 0.35 | 0.72 | 1 | Gm6266 | | 3 | 3 | 2 | 5 | 3 | 3 | 4 | 1 | 1 | 30 | 5 |
| 5 | -1.47 | 1.86 | -0.79 | 0.43 | 1 | Gm48840 | | 0 | 5 | 23 | 1 | 6 | 6 | 4 | 0 | 5 | 0 | 10 |
| 5 | 0.67 | 1.79 | 0.37 | 0.71 | 1 | Gm15900 | | 1 | 21 | 5 | 2 | 0 | 8 | 4 | 0 | 7 | 3 | 9 |
| 5 | -0.56 | 1.63 | -0.34 | 0.73 | 1 | Gm16998 | | 3 | 2 | 0 | 19 | 3 | 8 | 4 | 3 | 12 | 0 | 5 |
| 5 | 2.36 | 1.90 | 1.24 | 0.21 | 1 | Serpina1e | serine (or cysteine) peptidase inhibitor, cl | 15 | 9 | 1 | 0 | 0 | 1 | 4 | 0 | 2 | 24 | 2 |
| 5 | -1.67 | 2.38 | -0.70 | 0.48 | 1 | Gm9988 | | 3 | 2 | 0 | 0 | 10 | 3 | 4 | 35 | 0 | 0 | 1 |
| 5 | -1.63 | 1.86 | -0.87 | 0.38 | 1 | Gm8172 | | 14 | 16 | 15 | 5 | 0 | 1 | 4 | 1 | 2 | 0 | 0 |
| 5 | -0.09 | 1.77 | -0.05 | 0.96 | 1 | Gm16439 | | 8 | 0 | 7 | 11 | 0 | 5 | 4 | 3 | 4 | 1 | 12 |
| 5 | -1.06 | 3.02 | -0.35 | 0.72 | 1 | Anxa10 | annexin A10 | 31 | 15 | 0 | 0 | 0 | 1 | 4 | 0 | 5 | 0 | 0 |
| 5 | -1.74 | 2.26 | -0.77 | 0.44 | 1 | Gys2 | glycogen synthase 2 | 0 | 25 | 7 | 0 | 3 | 4 | 4 | 9 | 0 | 0 | 3 |
| 5 | 0.49 | 1.82 | 0.27 | 0.79 | 1 | Rpl17-ps5 | | 29 | 5 | 1 | 0 | 4 | 1 | 4 | 0 | 5 | 1 | 5 |
| 5 | -4.80 | 2.71 | -1.77 | 0.08 | 1 | Gm42725 | | 15 | 0 | 0 | 6 | 0 | 19 | 4 | 0 | 0 | 0 | 10 |
| 5 | -0.64 | 2.30 | -0.28 | 0.78 | 1 | Gm13169 | | 7 | 18 | 0 | 6 | 0 | 8 | 4 | 0 | 7 | 0 | 4 |
| 5 | -2.90 | 3.03 | -0.96 | 0.34 | 1 | Gm38319 | | 0 | 0 | 0 | 9 | 0 | 19 | 4 | 0 | 0 | 0 | 23 |
| 5 | -1.16 | 3.12 | -0.37 | 0.71 | 1 | Gm11650 | | 15 | 13 | 22 | 0 | 0 | 0 | 4 | 0 | 0 | 0 | 0 |
| 5 | -0.99 | 3.05 | -0.32 | 0.75 | 1 | Izumo1r | IZUMO1 receptor, JUNO | 31 | 13 | 6 | 0 | 0 | 0 | 4 | 0 | 0 | 0 | 0 |
| 5 | -1.99 | 1.99 | -1.00 | 0.32 | 1 | Gm43692 | | 0 | 0 | 7 | 1 | 11 | 11 | 4 | 0 | 3 | 0 | 16 |
| 5 | -0.71 | 2.99 | -0.24 | 0.81 | 1 | Gm26513 | | 7 | 0 | 0 | 0 | 0 | 25 | 4 | 0 | 0 | 0 | 18 |
| 5 | 0.62 | 1.29 | 0.48 | 0.63 | 1 | Rbm8a2 | | 3 | 5 | 8 | 1 | 2 | 9 | 4 | 3 | 8 | 4 | 5 |
| 5 | -1.63 | 2.40 | -0.68 | 0.50 | 1 | | | 0 | 2 | 0 | 0 | 2 | 17 | 4 | 0 | 1 | 0 | 27 |
| 5 | -3.74 | 2.50 | -1.50 | 0.13 | 1 | Gm11452 | | 4 | 0 | 0 | 5 | 16 | 4 | 4 | 0 | 19 | 0 | 0 |
| 5 | -1.28 | 1.87 | -0.68 | 0.49 | 1 | Commd5 | COMM domain containing 5 | 0 | 6 | 8 | 4 | 6 | 4 | 4 | 0 | 13 | 0 | 7 |
| 5 | 0.38 | 1.48 | 0.25 | 0.80 | 1 | Gm14429 | | 10 | 0 | 6 | 1 | 2 | 6 | 4 | 0 | 5 | 4 | 14 |
| 5 | 1.90 | 1.57 | 1.21 | 0.23 | 1 | Prkag2os2 | | 0 | 2 | 0 | 1 | 3 | 1 | 4 | 16 | 4 | 18 | 3 |
| 5 | -0.59 | 2.89 | -0.21 | 0.84 | 1 | Gm9922 | | 6 | 0 | 0 | 13 | 0 | 1 | 4 | 28 | 0 | 0 | 0 |
| 5 | -0.75 | 2.37 | -0.32 | 0.75 | 1 | Pcdh10 | protocadherin 10 | 19 | 6 | 17 | 0 | 0 | 2 | 4 | 0 | 0 | 0 | 4 |
| 5 | -0.13 | 2.13 | -0.06 | 0.95 | 1 | 4930404N11Ril | RIKEN cDNA 4930404N11 gene | 11 | 10 | 6 | 0 | 0 | 10 | 4 | 0 | 3 | 0 | 6 |
| 5 | -3.87 | 2.71 | -1.43 | 0.15 | 1 | 4930558J18Rik | | 15 | 0 | 0 | 7 | 9 | 2 | 4 | 14 | 0 | 0 | 0 |
| 5 | 1.01 | 2.41 | 0.42 | 0.68 | 1 | 1700026N04Rik | | 0 | 8 | 4 | 0 | 3 | 1 | 4 | 0 | 18 | 0 | 12 |
| 5 | 0.07 | 2.31 | 0.03 | 0.97 | 1 | Gm26852 | | 0 | 2 | 4 | 0 | 7 | 7 | 4 | 0 | 0 | 18 | 8 |
| 4 | -5.40 | 2.79 | -1.94 | 0.05 | 1 | Drd3 | dopamine receptor D3 | 0 | 0 | 12 | 22 | 7 | 4 | 4 | 0 | 0 | 0 | 0 |
| 4 | -0.70 | 1.85 | -0.38 | 0.70 | 1 | Macroh2a3 | | 11 | 5 | 12 | 0 | 5 | 3 | 4 | 0 | 3 | 0 | 6 |
| 4 | 1.86 | 2.00 | 0.93 | 0.35 | 1 | Gm37120 | | 15 | 0 | 11 | 0 | 0 | 1 | 4 | 0 | 3 | 9 | 5 |
| 4 | 2.05 | 2.03 | 1.01 | 0.31 | 1 | Gm6051 | | 9 | 0 | 1 | 0 | 5 | 3 | 4 | 8 | 8 | 0 | 9 |
| 4 | -2.33 | 2.04 | -1.14 | 0.25 | 1 | Nkx2-2os | | 2 | 6 | 18 | 10 | 3 | 0 | 4 | 0 | 2 | 0 | 3 |
| 4 | -2.26 | 1.72 | -1.32 | 0.19 | 1 | BC021767 | | 13 | 0 | 1 | 13 | 9 | 2 | 4 | 1 | 2 | 1 | 1 |
| 4 | 0.72 | 2.38 | 0.30 | 0.76 | 1 | 4933408N05Rik | | 0 | 0 | 4 | 1 | 0 | 3 | 4 | 25 | 5 | 0 | 3 |
| 4 | -2.61 | 2.13 | -1.22 | 0.22 | 1 | Gm45311 | | 2 | 10 | 0 | 5 | 2 | 3 | 4 | 19 | 0 | 0 | 0 |
| 4 | -0.94 | 2.05 | -0.46 | 0.65 | 1 | Gm47230 | | 2 | 0 | 1 | 0 | 14 | 8 | 4 | 0 | 6 | 0 | 10 |
| 4 | 1.81 | 1.94 | 0.93 | 0.35 | 1 | Gm42971 | | 3 | 0 | 0 | 0 | 1 | 3 | 4 | 0 | 7 | 21 | 6 |
| 4 | 0.34 | 3.33 | 0.10 | 0.92 | 1 | H3c7 | H3 clustered histone 7 | 0 | 0 | 0 | 0 | 0 | 16 | 4 | 0 | 7 | 0 | 17 |
| 4 | 1.22 | 2.59 | 0.47 | 0.64 | 1 | Gm9974 | | 9 | 0 | 0 | 1 | 0 | 7 | 4 | 14 | 0 | 0 | 10 |
| 4 | -1.62 | 1.45 | -1.12 | 0.26 | 1 | Gm8130 | | 3 | 7 | 5 | 8 | 5 | 4 | 4 | 3 | 4 | 0 | 1 |
| 4 | -2.92 | 2.26 | -1.29 | 0.20 | 1 | 2900089D17Rik | | 1 | 10 | 3 | 0 | 2 | 19 | 4 | 0 | 0 | 0 | 4 |
| 4 | -0.28 | 1.81 | -0.16 | 0.88 | 1 | Mir8102 | | 2 | 0 | 7 | 8 | 2 | 2 | 4 | 12 | 2 | 5 | 0 |
| 4 | -4.28 | 3.04 | -1.40 | 0.16 | 1 | Cplx1 | complexin 1 | 23 | 0 | 0 | 12 | 0 | 2 | 4 | 0 | 1 | 0 | 0 |
| 4 | -1.80 | 3.31 | -0.55 | 0.59 | 1 | Gm45828 | | 0 | 0 | 0 | 0 | 1 | 19 | 4 | 0 | 0 | 0 | 17 |
| 4 | 2.07 | 2.76 | 0.75 | 0.45 | 1 | Gm9108 | | 1 | 1 | 17 | 0 | 0 | 0 | 4 | 0 | 0 | 15 | 3 |
| 4 | -2.19 | 3.52 | -0.62 | 0.53 | 1 | Gm10741 | | 0 | 5 | 0 | 29 | 0 | 0 | 4 | 0 | 0 | 0 | 3 |
| 4 | -0.09 | 1.28 | -0.07 | 0.94 | 1 | Gm8055 | | 3 | 2 | 2 | 3 | 5 | 6 | 4 | 3 | 2 | 5 | 6 |
| 4 | -2.57 | 2.08 | -1.24 | 0.22 | 1 | Gm10043 | | 1 | 4 | 2 | 18 | 0 | 6 | 4 | 0 | 1 | 0 | 4 |
| 4 | 1.31 | 2.49 | 0.53 | 0.60 | 1 | Hpx | hemopexin | 8 | 7 | 0 | 0 | 0 | 2 | 4 | 0 | 0 | 12 | 8 |
| 4 | -4.40 | 2.51 | -1.75 | 0.08 | 1 | 8030443G20Rik | | 1 | 0 | 0 | 9 | 14 | 4 | 4 | 0 | 0 | 0 | 8 |
| 4 | 1.16 | 2.41 | 0.48 | 0.63 | 1 | 1700025J12Rik | | 0 | 0 | 0 | 0 | 2 | 3 | 4 | 0 | 8 | 21 | 1 |
| 4 | -2.32 | 2.90 | -0.80 | 0.42 | 1 | Trbc1 | | 17 | 11 | 3 | 0 | 0 | 3 | 4 | 0 | 0 | 0 | 0 |
| 3 | 1.61 | 2.07 | 0.78 | 0.44 | 1 | 4930483J18Rik | | 7 | 0 | 4 | 0 | 3 | 0 | 4 | 0 | 3 | 15 | 2 |
| 3 | -2.22 | 2.96 | -0.75 | 0.45 | 1 | Gm47350 | | 0 | 2 | 0 | 4 | 10 | 0 | 4 | 17 | 0 | 0 | 0 |
| 3 | -0.74 | 3.06 | -0.24 | 0.81 | 1 | Gm11827 | | 9 | 0 | 18 | 0 | 0 | 2 | 4 | 0 | 0 | 0 | 5 |
| 3 | -0.32 | 2.66 | -0.12 | 0.90 | 1 | Snora30 | | 0 | 4 | 0 | 7 | 0 | 4 | 4 | 0 | 2 | 0 | 16 |
| 3 | 0.10 | 3.63 | 0.03 | 0.98 | 1 | Gm4804 | | 9 | 23 | 0 | 0 | 0 | 0 | 4 | 0 | 1 | 0 | 0 |

| | | | | | | | | | | | | | | | | | | |
|---|---|---|---|---|---|---|---|---|---|---|---|---|---|---|---|---|---|---|
| 3 | 0.09 | 2.90 | 0.03 | 0.98 | 1 | Myh7 | myosin, heavy polypeptide 7, cardiac mus | 14 | 0 | 0 | 4 | 0 | 3 | 4 | 0 | 9 | 0 | 3 |
| 3 | -0.21 | 3.54 | -0.06 | 0.95 | 1 | Gm4779 | predicted gene 4779 | 1 | 0 | 0 | 0 | 0 | 3 | 4 | 0 | 0 | 0 | 28 |
| 3 | 1.40 | 2.61 | 0.54 | 0.59 | 1 | Gm12097 | | 0 | 0 | 0 | 0 | 0 | 11 | 4 | 13 | 2 | 1 | 5 |
| 3 | -3.01 | 2.05 | -1.47 | 0.14 | 1 | Gm42727 | | 3 | 0 | 14 | 4 | 3 | 6 | 4 | 0 | 1 | 0 | 2 |
| 3 | 1.62 | 2.56 | 0.63 | 0.53 | 1 | Gm4800 | | 3 | 0 | 0 | 0 | 0 | 7 | 4 | 0 | 5 | 12 | 3 |
| 3 | 0.28 | 3.54 | 0.08 | 0.94 | 1 | Gm15477 | | 0 | 0 | 0 | 0 | 0 | 4 | 4 | 0 | 0 | 24 | 0 |
| 3 | -2.07 | 2.54 | -0.82 | 0.41 | 1 | Gm26631 | | 0 | 7 | 4 | 0 | 3 | 2 | 4 | 10 | 1 | 0 | 0 |
| 3 | -1.11 | 3.65 | -0.30 | 0.76 | 1 | Amelx | amelogenin, X-linked | 9 | 10 | 7 | 0 | 0 | 0 | 4 | 0 | 0 | 0 | 0 |
| 3 | -0.80 | 3.05 | -0.26 | 0.79 | 1 | Gm5475 | | 0 | 0 | 0 | 11 | 0 | 3 | 4 | 10 | 0 | 0 | 3 |
| 3 | 0.33 | 3.51 | 0.09 | 0.93 | 1 | Maskbp3 | | 0 | 0 | 0 | 0 | 0 | 8 | 4 | 0 | 4 | 0 | 13 |
| 3 | -3.18 | 3.58 | -0.89 | 0.37 | 1 | 3110080O07Rik | | 0 | 0 | 5 | 0 | 0 | 9 | 4 | 0 | 0 | 0 | 11 |
| 3 | -1.86 | 2.60 | -0.71 | 0.48 | 1 | H1f5 | H1.5 linker histone, cluster member | 0 | 14 | 0 | 3 | 3 | 2 | 4 | 0 | 3 | 0 | 1 |
| 3 | -3.69 | 3.23 | -1.14 | 0.25 | 1 | Gm3227 | | 0 | 0 | 0 | 5 | 7 | 4 | 4 | 0 | 0 | 0 | 9 |
| 3 | -2.29 | 2.99 | -0.77 | 0.44 | 1 | Gm29292 | | 7 | 0 | 10 | 3 | 0 | 1 | 4 | 0 | 0 | 0 | 4 |
| 3 | -1.31 | 3.14 | -0.42 | 0.68 | 1 | Gm12247 | | 0 | 0 | 0 | 11 | 4 | 0 | 4 | 1 | 8 | 0 | 0 |
| 3 | 0.47 | 2.89 | 0.16 | 0.87 | 1 | Gm11583 | | 15 | 0 | 0 | 0 | 1 | 3 | 4 | 0 | 4 | 0 | 2 |
| 3 | 1.62 | 2.08 | 0.78 | 0.44 | 1 | Rps2-ps11 | | 1 | 6 | 4 | 0 | 1 | 0 | 4 | 1 | 7 | 3 | 1 |
| 3 | -0.83 | 2.72 | -0.30 | 0.76 | 1 | Gm11966 | | 4 | 0 | 0 | 3 | 0 | 2 | 4 | 0 | 0 | 14 | 1 |
| 2 | -0.62 | 3.56 | -0.17 | 0.86 | 1 | 6720482G16Rik | | 2 | 0 | 0 | 0 | 0 | 8 | 4 | 0 | 0 | 0 | 13 |
| 2 | -3.18 | 3.53 | -0.90 | 0.37 | 1 | E230015B07Rik | | 2 | 0 | 5 | 0 | 0 | 6 | 4 | 10 | 0 | 0 | 0 |
| 2 | -1.77 | 3.33 | -0.53 | 0.60 | 1 | Gm24507 | | 0 | 5 | 0 | 2 | 0 | 5 | 4 | 0 | 0 | 0 | 11 |
| 2 | -3.26 | 3.49 | -0.93 | 0.35 | 1 | 4930568G15Rik | | 0 | 6 | 11 | 0 | 0 | 3 | 4 | 0 | 0 | 0 | 2 |
| 2 | -0.08 | 3.50 | -0.02 | 0.98 | 1 | C530043K16Rik | | 0 | 0 | 0 | 0 | 2 | 0 | 4 | 21 | 0 | 0 | 0 |
| 2 | 2.85 | 2.59 | 1.10 | 0.27 | 1 | Gm10291 | | 1 | 0 | 1 | 0 | 1 | 0 | 4 | 10 | 8 | 0 | 1 |
| 2 | 0.62 | 2.82 | 0.22 | 0.82 | 1 | Clic3 | chloride intracellular channel 3 | 7 | 1 | 0 | 0 | 0 | 2 | 4 | 0 | 1 | 12 | 0 |
| 2 | -0.33 | 3.55 | -0.09 | 0.93 | 1 | Pmepa1os | | 0 | 0 | 0 | 0 | 0 | 1 | 4 | 0 | 0 | 0 | 21 |
| 2 | -1.27 | 3.06 | -0.42 | 0.68 | 1 | Gm18584 | | 0 | 0 | 5 | 0 | 0 | 9 | 4 | 2 | 4 | 0 | 1 |
| 2 | -3.24 | 2.91 | -1.12 | 0.26 | 1 | 9630014M24Rik | | 0 | 7 | 0 | 4 | 3 | 3 | 4 | 0 | 0 | 0 | 4 |
| 2 | -1.26 | 2.23 | -0.57 | 0.57 | 1 | Ppp1r2-ps2 | | 3 | 0 | 1 | 2 | 1 | 2 | 4 | 0 | 1 | 12 | 0 |
| 2 | -5.70 | 3.10 | -1.84 | 0.07 | 1 | Gm39307 | | 0 | 1 | 12 | 4 | 0 | 3 | 4 | 0 | 0 | 0 | 1 |
| 2 | -1.68 | 2.23 | -0.76 | 0.45 | 1 | Gm13880 | | 0 | 0 | 1 | 12 | 0 | 3 | 4 | 1 | 2 | 1 | 1 |
| 2 | 1.05 | 1.92 | 0.55 | 0.58 | 1 | Gm9234 | | 3 | 0 | 5 | 0 | 2 | 1 | 4 | 1 | 4 | 3 | 2 |
| 2 | 0.15 | 3.35 | 0.05 | 0.96 | 1 | Tgm4 | transglutaminase 4 (prostate) | 6 | 0 | 0 | 0 | 0 | 8 | 4 | 0 | 1 | 0 | 5 |
| 2 | 0.01 | 2.06 | 0.00 | 1.00 | 1 | Gm14130 | | 5 | 0 | 1 | 3 | 0 | 2 | 4 | 1 | 3 | 3 | 1 |
| 2 | -2.72 | 3.58 | -0.76 | 0.45 | 1 | 2610318M16Rik | | 0 | 0 | 8 | 0 | 0 | 2 | 4 | 0 | 0 | 0 | 7 |
| 2 | -2.01 | 3.60 | -0.56 | 0.58 | 1 | Gm16892 | | 3 | 4 | 8 | 0 | 0 | 0 | 4 | 0 | 0 | 0 | 2 |
| 2 | 0.64 | 3.48 | 0.18 | 0.85 | 1 | 1700012B07Ril | RIKEN cDNA 1700012B07 gene | 0 | 0 | 0 | 1 | 0 | 1 | 4 | 0 | 5 | 0 | 10 |
| 2 | -3.41 | 3.59 | -0.95 | 0.34 | 1 | 1300017J02Rik | RIKEN cDNA 1300017J02 gene | 0 | 9 | 0 | 0 | 0 | 4 | 4 | 0 | 0 | 0 | 3 |
| 2 | -0.46 | 3.57 | -0.13 | 0.90 | 1 | Gm15950 | | 0 | 3 | 0 | 0 | 0 | 0 | 4 | 13 | 0 | 0 | 0 |
| 2 | -3.69 | 3.57 | -1.03 | 0.30 | 1 | Syce1l | synaptonemal complex central element p | 0 | 4 | 0 | 5 | 0 | 6 | 4 | 0 | 0 | 0 | 0 |
| 2 | 1.19 | 3.55 | 0.34 | 0.74 | 1 | Gm45212 | | 0 | 0 | 0 | 0 | 0 | 0 | 4 | 0 | 15 | 0 | 0 |
| 2 | -1.30 | 3.57 | -0.37 | 0.71 | 1 | Gm7495 | | 0 | 0 | 7 | 0 | 7 | 0 | 4 | 1 | 0 | 0 | 0 |
| 2 | -0.97 | 3.63 | -0.27 | 0.79 | 1 | Gm30108 | | 8 | 0 | 7 | 0 | 0 | 0 | 4 | 0 | 0 | 0 | 0 |
| 2 | -1.15 | 3.58 | -0.32 | 0.75 | 1 | Gm12527 | | 0 | 0 | 0 | 0 | 0 | 7 | 4 | 0 | 0 | 0 | 8 |
| 2 | -0.54 | 3.50 | -0.16 | 0.88 | 1 | Gm50354 | | 0 | 0 | 0 | 0 | 0 | 3 | 4 | 0 | 9 | 0 | 2 |
| 2 | -2.71 | 3.56 | -0.76 | 0.45 | 1 | Gm44130 | | 0 | 0 | 8 | 0 | 0 | 5 | 4 | 0 | 1 | 0 | 0 |
| 2 | 0.50 | 3.18 | 0.16 | 0.87 | 1 | Gm12161 | | 0 | 0 | 0 | 0 | 1 | 1 | 4 | 5 | 6 | 1 | 0 |
| 2 | -0.69 | 2.22 | -0.31 | 0.76 | 1 | Gm44609 | | 3 | 3 | 2 | 0 | 2 | 1 | 4 | 0 | 1 | 1 | 1 |
| 2 | -0.71 | 3.57 | -0.20 | 0.84 | 1 | Hsf5 | heat shock transcription factor family mem | 0 | 0 | 0 | 0 | 0 | 3 | 4 | 0 | 0 | 0 | 11 |
| 2 | -0.88 | 3.26 | -0.27 | 0.79 | 1 | Gm49970 | | 3 | 6 | 2 | 0 | 0 | 1 | 4 | 0 | 0 | 0 | 2 |
| 2 | 0.20 | 3.55 | 0.06 | 0.96 | 1 | Gm22953 | | 0 | 0 | 3 | 0 | 3 | 0 | 4 | 0 | 2 | 5 | 0 |
| 2 | 0.76 | 3.61 | 0.21 | 0.83 | 1 | Hrg | histidine-rich glycoprotein | 0 | 0 | 0 | 0 | 0 | 0 | 4 | 0 | 0 | 13 | 0 |
| 1 | -0.20 | 3.29 | -0.06 | 0.95 | 1 | Gm8775 | | 1 | 0 | 1 | 0 | 0 | 4 | 4 | 0 | 3 | 0 | 3 |
| 1 | -2.38 | 3.54 | -0.67 | 0.50 | 1 | Gm43790 | | 0 | 0 | 0 | 0 | 3 | 5 | 4 | 0 | 0 | 0 | 4 |
| 1 | -1.63 | 2.87 | -0.57 | 0.57 | 1 | Gm10051 | | 3 | 1 | 3 | 0 | 1 | 2 | 4 | 0 | 2 | 0 | 0 |
| 1 | -0.37 | 3.56 | -0.10 | 0.92 | 1 | Ankrd61 | ankyrin repeat domain 61 | 3 | 0 | 0 | 0 | 0 | 3 | 4 | 0 | 0 | 0 | 6 |
| 1 | -1.81 | 3.52 | -0.51 | 0.61 | 1 | Gm4294 | | 0 | 0 | 0 | 6 | 0 | 1 | 4 | 1 | 3 | 0 | 0 |
| 1 | -2.65 | 3.30 | -0.80 | 0.42 | 1 | Gm44170 | | 0 | 0 | 0 | 6 | 1 | 2 | 4 | 0 | 1 | 0 | 1 |
| 1 | -3.77 | 3.56 | -1.06 | 0.29 | 1 | Gm16208 | | 0 | 0 | 0 | 8 | 0 | 3 | 4 | 0 | 0 | 0 | 0 |
| 1 | -1.19 | 3.61 | -0.33 | 0.74 | 1 | Gm18774 | | 9 | 0 | 0 | 0 | 0 | 1 | 4 | 0 | 0 | 0 | 0 |

| | | | | | | | | | | | | | | | | | |
|---|---|---|---|---|---|---|---|---|---|---|---|---|---|---|---|---|---|
| 1 | -0.13 | 3.57 | -0.04 | 0.97 | 1 | Gm8152 | | 0 | 1 | 2 | 0 | 0 | 0 | 4 | 0 | 7 | 0 | 0 |
| 1 | 0.96 | 3.55 | 0.27 | 0.79 | 1 | Gm33843 | | 0 | 0 | 0 | 0 | 0 | 0 | 4 | 0 | 0 | 0 | 10 |
| 1 | 0.46 | 3.53 | 0.13 | 0.90 | 1 | Gm43126 | | 0 | 0 | 0 | 0 | 0 | 2 | 4 | 0 | 2 | 0 | 6 |
| 1 | -3.70 | 3.56 | -1.04 | 0.30 | 1 | Gm50334 | | 0 | 0 | 0 | 0 | 3 | 7 | 4 | 0 | 0 | 0 | 0 |
| 1 | -1.47 | 3.08 | -0.48 | 0.63 | 1 | Gm4849 | | 0 | 1 | 1 | 0 | 3 | 1 | 4 | 0 | 1 | 1 | 0 |
| 1 | 0.03 | 3.56 | 0.01 | 0.99 | 1 | Gm49490 | | 1 | 0 | 0 | 0 | 0 | 0 | 4 | 0 | 0 | 0 | 7 |
| 1 | 0.96 | 3.55 | 0.27 | 0.79 | 1 | Mrpl27-ps | | 0 | 3 | 0 | 0 | 0 | 0 | 4 | 1 | 2 | 1 | 0 |
| 1 | -1.53 | 3.59 | -0.43 | 0.67 | 1 | Gm48870 | | 0 | 0 | 0 | 0 | 0 | 5 | 4 | 0 | 0 | 0 | 2 |
| 1 | -1.83 | 3.58 | -0.51 | 0.61 | 1 | Gm37756 | | 0 | 0 | 5 | 0 | 0 | 1 | 4 | 0 | 1 | 0 | 0 |
| 1 | -0.32 | 3.55 | -0.09 | 0.93 | 1 | Gm11449 | | 0 | 0 | 0 | 1 | 0 | 0 | 4 | 0 | 1 | 0 | 5 |
| 1 | -0.83 | 3.58 | -0.23 | 0.82 | 1 | Gm33940 | | 0 | 0 | 0 | 0 | 0 | 2 | 4 | 0 | 0 | 0 | 5 |
| 1 | -0.32 | 3.56 | -0.09 | 0.93 | 1 | Gm49727 | | 0 | 0 | 0 | 0 | 0 | 3 | 4 | 0 | 1 | 1 | 1 |
| 1 | -1.25 | 3.58 | -0.35 | 0.73 | 1 | Gm16053 | | 0 | 1 | 0 | 0 | 0 | 3 | 4 | 0 | 0 | 0 | 3 |
| 1 | 4.20 | 3.57 | 1.18 | 0.24 | 1 | 4930543K20Rik | | 0 | 0 | 0 | 0 | 0 | 0 | 4 | 0 | 5 | 0 | 1 |
| 1 | -3.63 | 3.57 | -1.02 | 0.31 | 1 | Gm37566 | | 0 | 0 | 0 | 0 | 1 | 0 | 4 | 0 | 0 | 0 | 5 |
| 1 | -0.52 | 3.62 | -0.14 | 0.89 | 1 | Gm44956 | | 6 | 0 | 0 | 0 | 0 | 0 | 4 | 0 | 0 | 0 | 0 |
| 1 | -1.30 | 3.62 | -0.36 | 0.72 | 1 | Grin3b | glutamate receptor, ionotropic, NMDA3B | 5 | 0 | 0 | 0 | 0 | 1 | 4 | 0 | 0 | 0 | 0 |
| 1 | -1.58 | 3.58 | -0.44 | 0.66 | 1 | Gm1833 | | 0 | 0 | 0 | 1 | 0 | 3 | 4 | 0 | 0 | 3 | 0 |
| 1 | -1.68 | 3.61 | -0.46 | 0.64 | 1 | Glra3 | glycine receptor, alpha 3 subunit | 4 | 0 | 0 | 0 | 0 | 2 | 4 | 0 | 0 | 0 | 0 |
| 1 | -0.29 | 3.54 | -0.08 | 0.93 | 1 | Gm19445 | | 0 | 0 | 0 | 0 | 0 | 2 | 4 | 0 | 3 | 0 | 1 |
| 1 | -3.21 | 3.57 | -0.90 | 0.37 | 1 | Gm17122 | | 0 | 0 | 0 | 0 | 2 | 3 | 4 | 0 | 0 | 0 | 0 |
| 1 | -2.14 | 3.58 | -0.60 | 0.55 | 1 | Gucy1b2 | guanylate cyclase 1, soluble, beta 2 | 0 | 0 | 0 | 0 | 0 | 4 | 4 | 0 | 1 | 0 | 0 |
| 1 | -2.32 | 3.60 | -0.64 | 0.52 | 1 | C330018A13Rik | | 0 | 0 | 3 | 0 | 0 | 2 | 4 | 0 | 0 | 0 | 0 |
| 1 | -1.01 | 3.59 | -0.28 | 0.78 | 1 | Gm47615 | | 0 | 0 | 0 | 0 | 0 | 2 | 4 | 0 | 0 | 0 | 3 |
| 1 | -1.40 | 3.59 | -0.39 | 0.70 | 1 | Gm13641 | | 2 | 0 | 0 | 0 | 2 | 0 | 4 | 0 | 0 | 0 | 1 |
| 1 | -2.39 | 3.61 | -0.66 | 0.51 | 1 | Dll3 | delta like canonical Notch ligand 3 | 0 | 0 | 0 | 0 | 4 | 0 | 4 | 0 | 0 | 0 | 0 |
| 1 | -2.39 | 3.61 | -0.66 | 0.51 | 1 | Gm15645 | | 0 | 0 | 0 | 0 | 4 | 0 | 4 | 0 | 0 | 0 | 0 |
| 1 | -2.08 | 3.58 | -0.58 | 0.56 | 1 | Gm50218 | | 0 | 0 | 0 | 0 | 2 | 1 | 4 | 0 | 0 | 0 | 1 |
| 1 | -2.37 | 3.61 | -0.66 | 0.51 | 1 | Gm37329 | | 0 | 0 | 0 | 0 | 0 | 3 | 4 | 0 | 0 | 0 | 0 |
| 1 | -0.31 | 3.62 | -0.09 | 0.93 | 1 | Mup3 | major urinary protein 3 | 1 | 0 | 1 | 0 | 0 | 0 | 4 | 0 | 0 | 0 | 1 |
| 1 | -2.64 | 3.59 | -0.74 | 0.46 | 1 | Rps2-ps13 | | 0 | 1 | 0 | 0 | 1 | 1 | 4 | 0 | 0 | 0 | 0 |
| 1 | -1.50 | 3.63 | -0.41 | 0.68 | 1 | Gm6579 | | 2 | 0 | 0 | 0 | 0 | 1 | 4 | 0 | 0 | 0 | 0 |
| 1 | -0.21 | 3.58 | -0.06 | 0.95 | 1 | Gm12736 | | 0 | 0 | 0 | 0 | 0 | 0 | 4 | 0 | 0 | 0 | 2 |
| 1 | -0.11 | 3.60 | -0.03 | 0.98 | 1 | Nphs2 | nephrosis 2, podocin | 0 | 0 | 0 | 0 | 0 | 0 | 4 | 0 | 2 | 0 | 0 |
| 1 | -0.11 | 3.60 | -0.03 | 0.98 | 1 | E030018B13Ril | RIKEN cDNA E030018B13 gene | 0 | 0 | 0 | 0 | 0 | 0 | 4 | 0 | 2 | 0 | 0 |
| 1 | -1.92 | 3.62 | -0.53 | 0.60 | 1 | Gm37392 | | 0 | 1 | 0 | 0 | 0 | 1 | 4 | 0 | 0 | 0 | 0 |
| 1 | -2.01 | 3.63 | -0.55 | 0.58 | 1 | Gm13570 | | 0 | 0 | 0 | 0 | 0 | 2 | 4 | 0 | 0 | 0 | 0 |
| 1 | -2.01 | 3.63 | -0.55 | 0.58 | 1 | Rtl1 | retrotransposon Gaglike 1 | 0 | 0 | 0 | 0 | 0 | 2 | 4 | 0 | 0 | 0 | 0 |
| 0 | -1.70 | 3.64 | -0.47 | 0.64 | 1 | Gm5943 | | 0 | 0 | 0 | 1 | 0 | 0 | 4 | 0 | 0 | 0 | 0 |
| 0 | -0.89 | 3.64 | -0.25 | 0.81 | 1 | Gm6702 | | 1 | 0 | 0 | 0 | 0 | 0 | 4 | 0 | 0 | 0 | 0 |
| 0 | -1.73 | 3.64 | -0.48 | 0.63 | 1 | Gm17099 | | 0 | 0 | 0 | 0 | 0 | 1 | 4 | 0 | 0 | 0 | 0 |
| 0 | -1.18 | 3.65 | -0.32 | 0.75 | 1 | Olfr362 | olfactory receptor 362 | 0 | 0 | 0 | 0 | 0 | 0 | 4 | 0 | 0 | 0 | 0 |
| 0 | -1.18 | 3.65 | -0.32 | 0.75 | 1 | Olfr1168 | olfactory receptor 1168 | 0 | 0 | 0 | 0 | 0 | 0 | 4 | 0 | 0 | 0 | 0 |
| 0 | -1.18 | 3.65 | -0.32 | 0.75 | 1 | F830045P16Ril | RIKEN cDNA F830045P16 gene | 0 | 0 | 0 | 0 | 0 | 0 | 4 | 0 | 0 | 0 | 0 |
| 0 | -1.18 | 3.65 | -0.32 | 0.75 | 1 | Gm12540 | | 0 | 0 | 0 | 0 | 0 | 0 | 4 | 0 | 0 | 0 | 0 |
| 0 | -1.18 | 3.65 | -0.32 | 0.75 | 1 | Gm18432 | | 0 | 0 | 0 | 0 | 0 | 0 | 4 | 0 | 0 | 0 | 0 |
| 0 | -1.18 | 3.65 | -0.32 | 0.75 | 1 | Gm43329 | | 0 | 0 | 0 | 0 | 0 | 0 | 4 | 0 | 0 | 0 | 0 |
| 0 | -1.18 | 3.65 | -0.32 | 0.75 | 1 | Etnppl | ethanolamine phosphate phospholyase | 0 | 0 | 0 | 0 | 0 | 0 | 4 | 0 | 0 | 0 | 0 |
| 0 | -1.18 | 3.65 | -0.32 | 0.75 | 1 | Gm36520 | | 0 | 0 | 0 | 0 | 0 | 0 | 4 | 0 | 0 | 0 | 0 |
| 0 | -1.18 | 3.65 | -0.32 | 0.75 | 1 | Gm43229 | | 0 | 0 | 0 | 0 | 0 | 0 | 4 | 0 | 0 | 0 | 0 |
| 0 | -1.18 | 3.65 | -0.32 | 0.75 | 1 | Trbv13-2 | | 0 | 0 | 0 | 0 | 0 | 0 | 4 | 0 | 0 | 0 | 0 |
| 0 | -1.18 | 3.65 | -0.32 | 0.75 | 1 | Neurod6 | neurogenic differentiation 6 | 0 | 0 | 0 | 0 | 0 | 0 | 4 | 0 | 0 | 0 | 0 |
| 0 | -1.18 | 3.65 | -0.32 | 0.75 | 1 | Gm6786 | | 0 | 0 | 0 | 0 | 0 | 0 | 4 | 0 | 0 | 0 | 0 |
| 0 | -1.18 | 3.65 | -0.32 | 0.75 | 1 | Gm8862 | | 0 | 0 | 0 | 0 | 0 | 0 | 4 | 0 | 0 | 0 | 0 |
| 0 | -1.18 | 3.65 | -0.32 | 0.75 | 1 | Gm26604 | | 0 | 0 | 0 | 0 | 0 | 0 | 4 | 0 | 0 | 0 | 0 |
| 0 | -1.18 | 3.65 | -0.32 | 0.75 | 1 | BC030500 | cDNA sequence BC030500 | 0 | 0 | 0 | 0 | 0 | 0 | 4 | 0 | 0 | 0 | 0 |
| 0 | -1.18 | 3.65 | -0.32 | 0.75 | 1 | H2ac5-ps | | 0 | 0 | 0 | 0 | 0 | 0 | 4 | 0 | 0 | 0 | 0 |
| 0 | -1.18 | 3.65 | -0.32 | 0.75 | 1 | Gm49668 | | 0 | 0 | 0 | 0 | 0 | 0 | 4 | 0 | 0 | 0 | 0 |
| 0 | -1.18 | 3.65 | -0.32 | 0.75 | 1 | Gm47242 | | 0 | 0 | 0 | 0 | 0 | 0 | 4 | 0 | 0 | 0 | 0 |
| 0 | -1.18 | 3.65 | -0.32 | 0.75 | 1 | Gm24772 | | 0 | 0 | 0 | 0 | 0 | 0 | 4 | 0 | 0 | 0 | 0 |

| | | | | | | | | | | | | | | | | | | |
|---|---|---|---|---|---|---|---|---|---|---|---|---|---|---|---|---|---|---|
| 2358 | -0.21 | 1.39 | -0.15 | 0.88 | 1 | Myh11 | myosin, heavy polypeptide 11, smooth mu | 15547 | 5798 | 4496 | 10 | 5 | 10 | 3 | 0 | 0 | 53 | 20 |
| 1588 | -1.05 | 1.38 | -0.77 | 0.44 | 1 | Cp | ceruloplasmin | 7757 | 4697 | 4918 | 5 | 31 | 2 | 3 | 0 | 51 | 0 | 1 |
| 1573 | -0.87 | 3.65 | -0.24 | 0.81 | 1 | Acta2 | actin, alpha 2, smooth muscle, aorta | 11094 | 3369 | 2842 | 0 | 0 | 0 | 3 | 0 | 0 | 0 | 0 |
| 1063 | -0.05 | 1.33 | -0.03 | 0.97 | 1 | Cxcl12 | chemokine (C-X-C motif) ligand 12 | 4996 | 4210 | 2388 | 16 | 35 | 1 | 3 | 0 | 6 | 35 | 7 |
| 1003 | -2.59 | 1.59 | -1.63 | 0.10 | 1 | Tagln | transgelin | 7169 | 2027 | 1798 | 0 | 10 | 11 | 3 | 0 | 0 | 0 | 10 |
| 996 | -2.40 | 1.47 | -1.63 | 0.10 | 1 | Ddr2 | discoidin domain receptor family, membe | 5458 | 2548 | 2936 | 6 | 0 | 3 | 3 | 0 | 0 | 0 | 3 |
| 708 | 0.09 | 1.14 | 0.08 | 0.93 | 1 | Postn | periostin, osteoblast specific factor | 3662 | 1875 | 2220 | 3 | 11 | 0 | 3 | 0 | 12 | 1 | 4 |
| 394 | 0.95 | 2.17 | 0.44 | 0.66 | 1 | Prrx1 | paired related homeobox 1 | 2266 | 990 | 1066 | 0 | 0 | 0 | 3 | 0 | 0 | 12 | 0 |
| 372 | -0.92 | 1.06 | -0.87 | 0.39 | 1 | Adgra2 | adhesion G protein-coupled receptor A2 | 2147 | 905 | 987 | 10 | 9 | 9 | 3 | 19 | 3 | 0 | 2 |
| 362 | 1.02 | 3.18 | 0.32 | 0.75 | 1 | Vcam1 | vascular cell adhesion molecule 1 | 2305 | 928 | 725 | 0 | 0 | 0 | 3 | 0 | 18 | 0 | 0 |
| 345 | -0.12 | 1.39 | -0.08 | 0.93 | 1 | Olfml2a | olfactomedin-like 2A | 1263 | 1171 | 1320 | 0 | 11 | 5 | 3 | 19 | 7 | 0 | 1 |
| 304 | -2.42 | 2.38 | -1.02 | 0.31 | 1 | Cd53 | CD53 antigen | 1591 | 807 | 931 | 0 | 12 | 0 | 3 | 0 | 0 | 0 | 4 |
| 286 | -1.31 | 2.13 | -0.61 | 0.54 | 1 | Itgb4 | integrin beta 4 | 1131 | 801 | 1199 | 8 | 0 | 0 | 3 | 0 | 5 | 0 | 0 |
| 267 | 0.74 | 1.22 | 0.61 | 0.54 | 1 | Ndrg2 | N-myc downstream regulated gene 2 | 1455 | 527 | 842 | 5 | 26 | 7 | 3 | 45 | 8 | 0 | 19 |
| 243 | -2.25 | 1.87 | -1.20 | 0.23 | 1 | Erp27 | endoplasmic reticulum protein 27 | 269 | 1160 | 1163 | 35 | 6 | 0 | 3 | 0 | 3 | 0 | 37 |
| 195 | -0.38 | 1.43 | -0.27 | 0.79 | 1 | H2-Q6 | histocompatibility 2, Q region locus 6 | 1256 | 437 | 347 | 0 | 53 | 6 | 3 | 0 | 24 | 1 | 13 |
| 194 | -0.06 | 2.84 | -0.02 | 0.98 | 1 | Pi15 | peptidase inhibitor 15 | 999 | 500 | 624 | 0 | 0 | 0 | 3 | 0 | 0 | 0 | 7 |
| 177 | -0.13 | 1.01 | -0.13 | 0.90 | 1 | Adam19 | a disintegrin and metallopeptidase domai | 1048 | 450 | 441 | 0 | 0 | 3 | 3 | 0 | 0 | 0 | 7 |
| 175 | -0.03 | 0.76 | -0.04 | 0.97 | 1 | Cyp4b1 | cytochrome P450, family 4, subfamily b, p | 534 | 518 | 559 | 63 | 36 | 52 | 3 | 79 | 42 | 0 | 39 |
| 169 | 1.89 | 2.30 | 0.82 | 0.41 | 1 | Cd33 | CD33 antigen | 746 | 580 | 519 | 0 | 0 | 0 | 3 | 0 | 12 | 0 | 1 |
| 165 | -2.50 | 1.73 | -1.45 | 0.15 | 1 | Lfng | LFNG O-fucosylpeptide 3-beta-N-acetylgl | 710 | 476 | 611 | 9 | 0 | 1 | 3 | 0 | 0 | 0 | 2 |
| 150 | -1.67 | 2.29 | -0.73 | 0.47 | 1 | Unc45b | unc-45 myosin chaperone B | 404 | 632 | 607 | 0 | 0 | 1 | 3 | 0 | 0 | 0 | 0 |
| 147 | -2.49 | 2.10 | -1.19 | 0.24 | 1 | Themis2 | thymocyte selection associated family me | 797 | 383 | 417 | 0 | 11 | 0 | 3 | 0 | 0 | 0 | 1 |
| 142 | -1.25 | 1.04 | -1.20 | 0.23 | 1 | Sytl2 | synaptotagmin-like 2 | 626 | 412 | 496 | 0 | 4 | 9 | 3 | 0 | 0 | 0 | 15 |
| 142 | -0.39 | 0.72 | -0.54 | 0.59 | 1 | Alox12 | arachidonate 12-lipoxygenase | 402 | 544 | 557 | 14 | 8 | 8 | 3 | 0 | 5 | 14 | 7 |
| 124 | 0.91 | 0.78 | 1.16 | 0.24 | 1 | Gulp1 | GULP, engulfment adaptor PTB domain c | 631 | 270 | 432 | 3 | 3 | 3 | 3 | 12 | 6 | 0 | 5 |
| 111 | -2.47 | 2.04 | -1.21 | 0.23 | 1 | Kcnk5 | potassium channel, subfamily K, member | 523 | 325 | 366 | 0 | 5 | 0 | 3 | 0 | 0 | 0 | 0 |
| 102 | -0.41 | 0.97 | -0.42 | 0.68 | 1 | Tgif1 | TGFB-induced factor homeobox 1 | 579 | 231 | 293 | 3 | 0 | 4 | 3 | 0 | 1 | 0 | 10 |
| 99 | -1.01 | 0.89 | -1.14 | 0.26 | 1 | Dok3 | docking protein 3 | 188 | 79 | 159 | 7 | 26 | 281 | 3 | 0 | 20 | 0 | 331 |
| 99 | 1.44 | 1.23 | 1.17 | 0.24 | 1 | Glipr1 | GLI pathogenesis-related 1 (glioma) | 652 | 210 | 200 | 0 | 0 | 1 | 3 | 0 | 0 | 19 | 0 |
| 91 | -0.77 | 1.14 | -0.68 | 0.50 | 1 | Nfatc4 | nuclear factor of activated T cells, cytopla: | 353 | 315 | 297 | 19 | 0 | 3 | 3 | 0 | 5 | 0 | 6 |
| 87 | -3.53 | 2.02 | -1.75 | 0.08 | 1 | Trpv2 | transient receptor potential cation channe | 390 | 272 | 261 | 8 | 7 | 0 | 3 | 0 | 0 | 0 | 12 |
| 79 | -1.51 | 0.77 | -1.96 | 0.05 | 1 | Prag1 | PEAK1 related kinase activating pseudok | 296 | 229 | 279 | 18 | 15 | 12 | 3 | 0 | 5 | 4 | 9 |
| 76 | -2.63 | 1.34 | -1.96 | 0.05 | 1 | Atoh8 | atonal bHLH transcription factor 8 | 299 | 198 | 328 | 0 | 2 | 1 | 3 | 0 | 0 | 0 | 0 |
| 74 | 0.38 | 0.97 | 0.39 | 0.70 | 1 | Adap1 | ArfGAP with dual PH domains 1 | 504 | 135 | 141 | 0 | 8 | 8 | 3 | 5 | 9 | 0 | 6 |
| 72 | -0.16 | 1.32 | -0.12 | 0.91 | 1 | Serpind1 | serine (or cysteine) peptidase inhibitor, cl | 207 | 383 | 141 | 0 | 14 | 6 | 3 | 0 | 7 | 32 | 0 |
| 70 | -1.94 | 1.32 | -1.47 | 0.14 | 1 | Hal | histidine ammonia lyase | 471 | 227 | 51 | 0 | 4 | 4 | 3 | 0 | 0 | 0 | 5 |
| 67 | -2.80 | 2.41 | -1.16 | 0.25 | 1 | Ifi209 | interferon activated gene 209 | 340 | 135 | 250 | 0 | 13 | 0 | 3 | 0 | 0 | 0 | 0 |
| 65 | -1.87 | 1.28 | -1.46 | 0.14 | 1 | Lrmp | lymphoid-restricted membrane protein | 364 | 231 | 107 | 0 | 0 | 4 | 3 | 0 | 0 | 0 | 1 |
| 63 | 0.63 | 0.69 | 0.92 | 0.36 | 1 | Gm5577 | | 0 | 65 | 27 | 23 | 13 | 266 | 3 | 51 | 28 | 5 | 215 |
| 63 | -1.84 | 1.47 | -1.25 | 0.21 | 1 | Sp110 | Sp110 nuclear body protein | 377 | 157 | 157 | 0 | 0 | 1 | 3 | 0 | 0 | 0 | 0 |
| 59 | -1.33 | 1.47 | -0.91 | 0.37 | 1 | Sphk1 | sphingosine kinase 1 | 221 | 206 | 215 | 4 | 0 | 0 | 3 | 0 | 0 | 0 | 5 |
| 59 | -0.15 | 3.61 | -0.04 | 0.97 | 1 | Igkv3-2 | | 638 | 2 | 5 | 0 | 0 | 0 | 3 | 0 | 0 | 0 | 0 |
| 58 | 0.08 | 1.03 | 0.08 | 0.94 | 1 | Rnd1 | Rho family GTPase 1 | 226 | 104 | 204 | 25 | 7 | 8 | 3 | 0 | 5 | 37 | 16 |
| 49 | 0.02 | 0.93 | 0.02 | 0.98 | 1 | Amz1 | archaelysin family metallopeptidase 1 | 209 | 69 | 223 | 4 | 9 | 1 | 3 | 0 | 5 | 5 | 11 |
| 49 | -0.88 | 0.88 | -1.00 | 0.32 | 1 | Chn2 | chimerin 2 | 220 | 153 | 106 | 15 | 6 | 13 | 3 | 13 | 4 | 0 | 4 |
| 49 | -1.83 | 1.53 | -1.20 | 0.23 | 1 | Gpr183 | G protein-coupled receptor 183 | 263 | 103 | 165 | 0 | 0 | 1 | 3 | 0 | 0 | 0 | 0 |
| 46 | -0.96 | 3.65 | -0.26 | 0.79 | 1 | Gpr141b | G protein-coupled receptor 141B | 235 | 213 | 53 | 0 | 0 | 0 | 3 | 0 | 0 | 0 | 0 |
| 44 | -0.96 | 3.66 | -0.26 | 0.79 | 1 | Hp | haptoglobin | 231 | 168 | 77 | 0 | 0 | 0 | 3 | 0 | 0 | 0 | 0 |
| 43 | -0.56 | 1.06 | -0.53 | 0.60 | 1 | Pcdhb4 | protocadherin beta 4 | 135 | 153 | 58 | 36 | 25 | 10 | 3 | 0 | 28 | 5 | 15 |
| 42 | 0.18 | 1.48 | 0.12 | 0.90 | 1 | Fstl4 | follistatin-like 4 | 212 | 159 | 81 | 3 | 0 | 0 | 3 | 0 | 8 | 0 | 0 |
| 42 | -0.93 | 3.65 | -0.25 | 0.80 | 1 | Gpr34 | G protein-coupled receptor 34 | 249 | 75 | 131 | 0 | 0 | 0 | 3 | 0 | 0 | 0 | 0 |
| 40 | -1.86 | 1.04 | -1.79 | 0.07 | 1 | Gm9917 | | 128 | 150 | 133 | 11 | 6 | 4 | 3 | 2 | 0 | 0 | 7 |
| 40 | -1.11 | 0.76 | -1.46 | 0.15 | 1 | 4930581F22Rik | | 21 | 62 | 47 | 12 | 13 | 137 | 3 | 1 | 4 | 3 | 140 |
| 40 | -2.06 | 1.53 | -1.35 | 0.18 | 1 | Meox2 | mesenchyme homeobox 2 | 231 | 150 | 51 | 0 | 0 | 2 | 3 | 0 | 0 | 0 | 0 |
| 38 | 1.47 | 0.83 | 1.77 | 0.08 | 1 | Has3 | hyaluronan synthase 3 | 65 | 68 | 100 | 22 | 5 | 25 | 3 | 37 | 9 | 57 | 31 |
| 33 | -2.00 | 1.58 | -1.27 | 0.21 | 1 | Ltb | lymphotoxin B | 220 | 64 | 77 | 0 | 0 | 2 | 3 | 0 | 0 | 0 | 0 |
| 31 | 0.18 | 1.48 | 0.13 | 0.90 | 1 | Cth | cystathionase (cystathionine gamma-lyas | 26 | 165 | 109 | 0 | 0 | 14 | 3 | 0 | 7 | 0 | 12 |
| 30 | 0.32 | 0.81 | 0.39 | 0.70 | 1 | Gm12474 | | 23 | 27 | 29 | 38 | 15 | 65 | 3 | 30 | 8 | 32 | 57 |
| 30 | 1.58 | 1.02 | 1.55 | 0.12 | 1 | Gm37859 | | 9 | 21 | 10 | 0 | 18 | 97 | 3 | 38 | 25 | 17 | 87 |

| | | | | | | | | | | | | | | | | | | |
|---|---|---|---|---|---|---|---|---|---|---|---|---|---|---|---|---|---|---|
| 29 | -0.88 | 0.92 | -0.95 | 0.34 | 1 | Gins1 | GINS complex subunit 1 (Psf1 homolog) | 70 | 88 | 52 | 21 | 25 | 19 | 3 | 22 | 8 | 0 | 12 |
| 28 | -1.18 | 1.70 | -0.70 | 0.49 | 1 | Gm44243 | | 23 | 36 | 0 | 0 | 10 | 132 | 3 | 0 | 6 | 0 | 102 |
| 28 | 1.11 | 1.54 | 0.72 | 0.47 | 1 | Dqx1 | DEAQ RNA-dependent ATPase | 0 | 16 | 18 | 4 | 0 | 114 | 3 | 0 | 3 | 24 | 121 |
| 27 | -0.18 | 1.34 | -0.14 | 0.89 | 1 | Nrg1 | neuregulin 1 | 77 | 86 | 95 | 0 | 0 | 22 | 3 | 0 | 3 | 0 | 14 |
| 27 | -0.18 | 1.26 | -0.15 | 0.88 | 1 | Cidea | cell death-inducing DNA fragmentation fa | 65 | 70 | 72 | 12 | 11 | 3 | 3 | 16 | 0 | 41 | 0 |
| 26 | 1.88 | 1.37 | 1.38 | 0.17 | 1 | Chsy3 | chondroitin sulfate synthase 3 | 45 | 52 | 37 | 0 | 16 | 19 | 3 | 32 | 0 | 49 | 38 |
| 25 | -0.20 | 1.76 | -0.11 | 0.91 | 1 | I830077J02Rik | RIKEN cDNA I830077J02 gene | 116 | 42 | 117 | 0 | 0 | 0 | 3 | 0 | 0 | 0 | 2 |
| 22 | 0.73 | 1.09 | 0.67 | 0.50 | 1 | Gm49975 | | 12 | 3 | 8 | 52 | 2 | 39 | 3 | 45 | 5 | 50 | 23 |
| 22 | -0.51 | 1.55 | -0.33 | 0.74 | 1 | 1810017P11Rik | | 1 | 5 | 8 | 0 | 16 | 109 | 3 | 0 | 0 | 12 | 88 |
| 22 | 0.02 | 1.29 | 0.01 | 0.99 | 1 | Gm43300 | | 3 | 1 | 1 | 0 | 5 | 117 | 3 | 0 | 13 | 0 | 94 |
| 20 | -2.32 | 1.22 | -1.91 | 0.06 | 1 | Fsd1 | fibronectin type 3 and SPRY domain-cont | 56 | 82 | 61 | 8 | 3 | 4 | 3 | 0 | 1 | 0 | 3 |
| 20 | 0.70 | 1.28 | 0.55 | 0.58 | 1 | Crybg2 | crystallin beta-gamma domain containing | 36 | 45 | 3 | 6 | 22 | 19 | 3 | 0 | 40 | 18 | 26 |
| 20 | 2.09 | 1.09 | 1.91 | 0.06 | 1 | AW822252 | | 0 | 2 | 11 | 4 | 7 | 40 | 3 | 71 | 12 | 13 | 54 |
| 19 | 0.28 | 0.96 | 0.30 | 0.77 | 1 | Gm37570 | | 17 | 27 | 1 | 29 | 11 | 37 | 3 | 16 | 12 | 18 | 41 |
| 18 | -2.74 | 1.53 | -1.80 | 0.07 | 1 | Gal | galanin and GMAP prepropeptide | 27 | 95 | 65 | 3 | 7 | 1 | 3 | 0 | 0 | 0 | 3 |
| 17 | -1.78 | 1.28 | -1.39 | 0.16 | 1 | Lrrc8e | leucine rich repeat containing 8 family, m | 49 | 49 | 50 | 14 | 7 | 6 | 3 | 0 | 11 | 0 | 2 |
| 16 | -0.20 | 1.20 | -0.16 | 0.87 | 1 | Gm6983 | | 15 | 5 | 16 | 29 | 26 | 12 | 3 | 38 | 10 | 0 | 24 |
| 16 | -2.81 | 1.80 | -1.56 | 0.12 | 1 | Ntng1 | netrin G1 | 45 | 44 | 61 | 18 | 0 | 1 | 3 | 0 | 0 | 0 | 4 |
| 16 | 0.12 | 1.01 | 0.12 | 0.90 | 1 | Gm10180 | | 16 | 38 | 33 | 9 | 14 | 4 | 3 | 44 | 6 | 1 | 3 |
| 15 | 1.31 | 1.73 | 0.75 | 0.45 | 1 | Cers1 | growth differentiation factor 1 | 50 | 69 | 24 | 0 | 0 | 4 | 3 | 0 | 10 | 0 | 9 |
| 15 | 1.58 | 1.03 | 1.53 | 0.13 | 1 | 1700001O22Ril | RIKEN cDNA 1700001O22 gene | 6 | 2 | 1 | 1 | 13 | 39 | 3 | 8 | 36 | 19 | 39 |
| 15 | 0.78 | 0.98 | 0.80 | 0.42 | 1 | Gm15787 | | 11 | 16 | 35 | 1 | 21 | 20 | 3 | 16 | 22 | 5 | 13 |
| 15 | 0.48 | 1.14 | 0.42 | 0.67 | 1 | Kbtbd6 | kelch repeat and BTB (POZ) domain cont | 10 | 1 | 3 | 12 | 18 | 24 | 3 | 57 | 5 | 13 | 18 |
| 15 | -2.70 | 1.44 | -1.87 | 0.06 | 1 | Gm37233 | | 48 | 19 | 65 | 0 | 6 | 18 | 3 | 0 | 0 | 0 | 5 |
| 15 | -2.10 | 1.40 | -1.50 | 0.13 | 1 | Gm11821 | | 0 | 11 | 35 | 13 | 7 | 48 | 3 | 0 | 9 | 0 | 35 |
| 14 | 0.45 | 1.22 | 0.37 | 0.71 | 1 | 4930426D05Ril | RIKEN cDNA 4930426D05 gene | 17 | 23 | 19 | 11 | 0 | 43 | 3 | 2 | 7 | 17 | 13 |
| 14 | 0.97 | 1.55 | 0.63 | 0.53 | 1 | Gm9908 | | 9 | 18 | 18 | 0 | 13 | 20 | 3 | 23 | 1 | 45 | 1 |
| 14 | -0.15 | 1.15 | -0.13 | 0.90 | 1 | Gm4459 | | 24 | 3 | 44 | 21 | 7 | 11 | 3 | 16 | 9 | 1 | 12 |
| 13 | 1.63 | 1.43 | 1.14 | 0.25 | 1 | Gm13327 | | 0 | 0 | 0 | 32 | 26 | 0 | 3 | 19 | 25 | 36 | 7 |
| 13 | -0.38 | 2.17 | -0.18 | 0.86 | 1 | Gm37309 | | 0 | 8 | 0 | 0 | 0 | 66 | 3 | 0 | 1 | 0 | 68 |
| 13 | 2.19 | 1.87 | 1.17 | 0.24 | 1 | Gm47218 | | 0 | 0 | 0 | 0 | 0 | 68 | 3 | 0 | 13 | 15 | 47 |
| 13 | 0.57 | 1.14 | 0.50 | 0.62 | 1 | Gm17705 | | 39 | 20 | 14 | 5 | 6 | 10 | 3 | 0 | 8 | 24 | 15 |
| 13 | 1.41 | 0.91 | 1.55 | 0.12 | 1 | Gm2387 | | 11 | 29 | 15 | 13 | 5 | 5 | 3 | 9 | 18 | 14 | 17 |
| 13 | -1.08 | 2.31 | -0.47 | 0.64 | 1 | Bcl2a1a | B cell leukemia/lymphoma 2 related prote | 47 | 32 | 57 | 0 | 0 | 0 | 3 | 0 | 0 | 0 | 0 |
| 13 | -2.69 | 1.55 | -1.73 | 0.08 | 1 | D930019O06Rik | | 17 | 28 | 55 | 11 | 0 | 18 | 3 | 0 | 1 | 0 | 6 |
| 13 | 2.57 | 1.39 | 1.85 | 0.06 | 1 | Gm19710 | | 9 | 0 | 10 | 2 | 2 | 22 | 3 | 54 | 11 | 17 | 8 |
| 12 | -1.57 | 1.84 | -0.85 | 0.39 | 1 | Gm37891 | | 1 | 0 | 20 | 0 | 2 | 58 | 3 | 0 | 0 | 0 | 53 |
| 12 | -3.39 | 1.77 | -1.91 | 0.06 | 1 | Fbxo48 | F-box protein 48 | 24 | 25 | 0 | 11 | 17 | 23 | 3 | 0 | 1 | 0 | 31 |
| 12 | -0.58 | 2.14 | -0.27 | 0.78 | 1 | Gm43890 | | 13 | 0 | 0 | 0 | 0 | 68 | 3 | 0 | 0 | 0 | 51 |
| 12 | -0.54 | 1.40 | -0.39 | 0.70 | 1 | Gm6225 | | 0 | 8 | 9 | 5 | 2 | 49 | 3 | 23 | 1 | 0 | 31 |
| 12 | -1.79 | 1.77 | -1.01 | 0.31 | 1 | Gdf3 | growth differentiation factor 3 | 29 | 3 | 77 | 5 | 2 | 1 | 3 | 0 | 0 | 10 | 0 |
| 12 | -0.48 | 1.87 | -0.26 | 0.80 | 1 | Thpo | thrombopoietin | 8 | 0 | 0 | 0 | 9 | 56 | 3 | 0 | 7 | 0 | 46 |
| 12 | -0.10 | 1.23 | -0.08 | 0.93 | 1 | Gm44550 | | 0 | 9 | 4 | 24 | 6 | 15 | 3 | 8 | 27 | 0 | 30 |
| 11 | 1.14 | 0.92 | 1.23 | 0.22 | 1 | Gm7105 | | 33 | 25 | 17 | 5 | 4 | 3 | 3 | 13 | 9 | 6 | 6 |
| 11 | -1.05 | 1.10 | -0.96 | 0.34 | 1 | Wfdc3 | WAP four-disulfide core domain 3 | 11 | 17 | 32 | 16 | 8 | 14 | 3 | 10 | 8 | 0 | 4 |
| 11 | -0.08 | 1.09 | -0.07 | 0.94 | 1 | A330040F15Rik | | 2 | 6 | 10 | 18 | 8 | 17 | 3 | 36 | 13 | 0 | 10 |
| 11 | -2.01 | 1.78 | -1.13 | 0.26 | 1 | Galnt5 | polypeptide N-acetylgalactosaminyltransf | 18 | 43 | 49 | 4 | 0 | 2 | 3 | 0 | 0 | 0 | 4 |
| 11 | 1.18 | 1.16 | 1.02 | 0.31 | 1 | Gm6392 | | 20 | 11 | 12 | 11 | 9 | 1 | 3 | 32 | 3 | 5 | 13 |
| 11 | -0.76 | 1.39 | -0.54 | 0.59 | 1 | Bloc1s1 | biogenesis of lysosomal organelles comp | 19 | 14 | 14 | 17 | 2 | 12 | 3 | 14 | 0 | 0 | 26 |
| 11 | 3.02 | 1.83 | 1.65 | 0.10 | 1 | 9930012K11Ril | RIKEN cDNA 9930012K11 gene | 40 | 2 | 16 | 0 | 0 | 8 | 3 | 29 | 3 | 0 | 16 |
| 11 | -0.29 | 1.44 | -0.20 | 0.84 | 1 | Gm14176 | | 17 | 6 | 0 | 30 | 7 | 13 | 3 | 2 | 27 | 6 | 6 |
| 11 | 0.23 | 1.35 | 0.17 | 0.86 | 1 | Fbxw24 | F-box and WD-40 domain protein 24 | 10 | 23 | 28 | 14 | 0 | 7 | 3 | 14 | 6 | 0 | 10 |
| 10 | -1.67 | 1.91 | -0.88 | 0.38 | 1 | Gm45426 | | 0 | 0 | 0 | 19 | 1 | 45 | 3 | 0 | 6 | 0 | 39 |
| 10 | 0.74 | 1.03 | 0.72 | 0.47 | 1 | Rpl34-ps2 | | 15 | 5 | 27 | 7 | 7 | 7 | 3 | 13 | 8 | 14 | 5 |
| 10 | -0.60 | 1.27 | -0.47 | 0.64 | 1 | 2810429I04Rik | | 4 | 29 | 28 | 6 | 15 | 5 | 3 | 0 | 4 | 6 | 6 |
| 10 | -2.84 | 1.66 | -1.71 | 0.09 | 1 | Gm10544 | | 5 | 35 | 39 | 7 | 2 | 5 | 3 | 0 | 0 | 0 | 10 |
| 10 | 1.45 | 1.18 | 1.23 | 0.22 | 1 | Tubb4b-ps1 | | 37 | 2 | 19 | 1 | 5 | 2 | 3 | 3 | 3 | 28 | 3 |
| 9 | -2.86 | 1.48 | -1.93 | 0.05 | 1 | 5430414B19Rik | | 33 | 20 | 7 | 5 | 10 | 14 | 3 | 0 | 1 | 0 | 10 |
| 9 | 1.16 | 1.48 | 0.78 | 0.43 | 1 | D930015M05Rik | | 15 | 0 | 4 | 10 | 0 | 14 | 3 | 6 | 9 | 22 | 18 |
| 9 | -1.54 | 1.43 | -1.08 | 0.28 | 1 | Il23a | interleukin 23, alpha subunit p19 | 21 | 5 | 8 | 2 | 17 | 18 | 3 | 0 | 6 | 0 | 19 |

| | | | | | | | | | | | | | | | | | | |
|---|---|---|---|---|---|---|---|---|---|---|---|---|---|---|---|---|---|---|
| 9 | -0.90 | 1.42 | -0.63 | 0.53 | 1 | Gm9929 | | 24 | 2 | 7 | 19 | 5 | 10 | 3 | 0 | 13 | 6 | 7 |
| 9 | 1.87 | 1.17 | 1.60 | 0.11 | 1 | 6530409C15Rik | | 3 | 8 | 14 | 9 | 9 | 0 | 3 | 22 | 10 | 15 | 3 |
| 9 | 0.34 | 1.19 | 0.29 | 0.77 | 1 | Gm10269 | | 41 | 13 | 9 | 5 | 1 | 4 | 3 | 7 | 2 | 1 | 6 |
| 8 | -1.04 | 1.56 | -0.66 | 0.51 | 1 | Zfp82 | zinc finger protein 82 | 0 | 11 | 4 | 19 | 0 | 24 | 3 | 6 | 3 | 0 | 23 |
| 8 | -0.28 | 1.74 | -0.16 | 0.87 | 1 | Ifit1bl2 | interferon induced protein with tetratricop | 29 | 9 | 22 | 0 | 3 | 6 | 3 | 0 | 20 | 0 | 1 |
| 8 | 0.29 | 1.73 | 0.17 | 0.87 | 1 | Gm17597 | | 0 | 12 | 8 | 19 | 3 | 7 | 3 | 0 | 3 | 32 | 5 |
| 8 | -0.26 | 1.67 | -0.16 | 0.87 | 1 | Gm36406 | | 0 | 1 | 0 | 39 | 0 | 12 | 3 | 8 | 23 | 1 | 4 |
| 8 | -1.42 | 1.53 | -0.93 | 0.35 | 1 | Gm13524 | | 9 | 5 | 0 | 13 | 12 | 18 | 3 | 0 | 2 | 17 | 12 |
| 8 | 0.83 | 2.43 | 0.34 | 0.73 | 1 | Slitrk2 | SLIT and NTRK-like family, member 2 | 38 | 16 | 24 | 0 | 0 | 0 | 3 | 0 | 9 | 0 | 0 |
| 8 | -2.63 | 2.26 | -1.17 | 0.24 | 1 | Faim2 | Fas apoptotic inhibitory molecule 2 | 20 | 41 | 18 | 8 | 0 | 0 | 3 | 0 | 0 | 0 | 0 |
| 8 | -0.85 | 1.60 | -0.53 | 0.59 | 1 | Gm34388 | | 0 | 19 | 0 | 7 | 13 | 8 | 3 | 16 | 7 | 0 | 14 |
| 8 | 0.49 | 1.82 | 0.27 | 0.79 | 1 | Tekt4 | tektin 4 | 17 | 13 | 39 | 0 | 0 | 3 | 3 | 0 | 3 | 0 | 9 |
| 8 | -1.17 | 1.33 | -0.87 | 0.38 | 1 | Prr29 | proline rich 29 | 1 | 2 | 0 | 24 | 23 | 8 | 3 | 5 | 14 | 0 | 9 |
| 8 | 0.32 | 1.32 | 0.25 | 0.81 | 1 | Gm8515 | | 13 | 3 | 2 | 15 | 4 | 3 | 3 | 22 | 9 | 12 | 0 |
| 8 | 3.09 | 2.06 | 1.50 | 0.13 | 1 | 9130002K18Rik | | 11 | 11 | 0 | 0 | 0 | 15 | 3 | 19 | 8 | 0 | 17 |
| 8 | -1.98 | 1.80 | -1.10 | 0.27 | 1 | Gm9895 | | 9 | 23 | 0 | 10 | 6 | 11 | 3 | 0 | 2 | 0 | 19 |
| 7 | -2.65 | 2.42 | -1.09 | 0.28 | 1 | Fgf21 | fibroblast growth factor 21 | 7 | 43 | 25 | 0 | 5 | 0 | 3 | 0 | 0 | 0 | 0 |
| 7 | -0.13 | 1.15 | -0.11 | 0.91 | 1 | Gm10080 | | 18 | 13 | 19 | 1 | 3 | 12 | 3 | 2 | 3 | 3 | 5 |
| 7 | -2.46 | 1.79 | -1.38 | 0.17 | 1 | Mlf1 | myeloid leukemia factor 1 | 15 | 26 | 5 | 8 | 0 | 14 | 3 | 0 | 0 | 0 | 9 |
| 7 | -0.73 | 1.88 | -0.39 | 0.70 | 1 | Gimap1os | | 1 | 38 | 15 | 6 | 0 | 4 | 3 | 0 | 7 | 0 | 6 |
| 7 | -0.45 | 1.36 | -0.33 | 0.74 | 1 | Bloc1s2-ps | | 0 | 0 | 15 | 5 | 20 | 7 | 3 | 2 | 17 | 3 | 7 |
| 7 | -0.62 | 3.37 | -0.18 | 0.85 | 1 | Trp73 | transformation related protein 73 | 57 | 0 | 14 | 0 | 0 | 0 | 3 | 0 | 0 | 0 | 0 |
| 7 | -0.22 | 2.27 | -0.10 | 0.92 | 1 | Gm13657 | | 9 | 40 | 0 | 0 | 4 | 5 | 3 | 0 | 4 | 9 | 0 |
| 7 | 0.92 | 1.25 | 0.74 | 0.46 | 1 | Gm14373 | | 8 | 6 | 1 | 5 | 6 | 4 | 3 | 20 | 7 | 1 | 12 |
| 7 | -0.90 | 2.77 | -0.32 | 0.75 | 1 | Gm41396 | | 0 | 23 | 0 | 0 | 0 | 3 | 3 | 39 | 4 | 0 | 0 |
| 7 | 0.13 | 2.22 | 0.06 | 0.95 | 1 | Gm29093 | | 11 | 11 | 0 | 3 | 0 | 17 | 3 | 0 | 14 | 0 | 13 |
| 6 | -0.39 | 1.90 | -0.21 | 0.84 | 1 | B230112G18Rik | | 4 | 13 | 9 | 11 | 0 | 5 | 3 | 0 | 15 | 0 | 8 |
| 6 | 1.16 | 1.49 | 0.78 | 0.44 | 1 | Gm6039 | | 3 | 3 | 7 | 3 | 1 | 6 | 3 | 31 | 5 | 0 | 6 |
| 6 | -1.08 | 2.29 | -0.47 | 0.64 | 1 | Gm10382 | | 2 | 0 | 3 | 0 | 8 | 18 | 3 | 30 | 0 | 0 | 5 |
| 6 | -0.20 | 1.92 | -0.10 | 0.92 | 1 | Gm13791 | | 0 | 0 | 0 | 0 | 18 | 19 | 3 | 0 | 5 | 12 | 11 |
| 6 | -1.35 | 1.30 | -1.04 | 0.30 | 1 | Gm45855 | | 17 | 11 | 9 | 5 | 5 | 8 | 3 | 2 | 0 | 3 | 5 |
| 6 | -1.88 | 2.02 | -0.93 | 0.35 | 1 | Gm13830 | | 0 | 17 | 0 | 12 | 7 | 14 | 3 | 0 | 3 | 0 | 12 |
| 6 | -0.36 | 1.19 | -0.30 | 0.76 | 1 | Gm6540 | | 4 | 13 | 4 | 9 | 10 | 4 | 3 | 3 | 3 | 9 | 4 |
| 6 | -0.19 | 2.88 | -0.07 | 0.95 | 1 | A730091E23Rik | | 0 | 14 | 0 | 0 | 0 | 27 | 3 | 0 | 4 | 0 | 19 |
| 6 | -1.06 | 2.07 | -0.51 | 0.61 | 1 | Gm28496 | | 0 | 0 | 30 | 0 | 9 | 8 | 3 | 0 | 11 | 0 | 7 |
| 6 | -3.15 | 1.98 | -1.59 | 0.11 | 1 | E030037K01Rik | | 0 | 0 | 20 | 15 | 10 | 5 | 3 | 0 | 9 | 0 | 5 |
| 6 | 1.07 | 1.81 | 0.59 | 0.55 | 1 | Gm7589 | | 9 | 11 | 5 | 0 | 9 | 1 | 3 | 1 | 0 | 22 | 5 |
| 6 | 3.25 | 2.51 | 1.29 | 0.20 | 1 | 9530036O11Rik | | 0 | 8 | 0 | 0 | 0 | 4 | 3 | 0 | 1 | 44 | 6 |
| 6 | -2.85 | 1.82 | -1.56 | 0.12 | 1 | Gm18225 | | 4 | 6 | 0 | 4 | 7 | 30 | 3 | 0 | 1 | 0 | 10 |
| 6 | -2.23 | 2.36 | -0.94 | 0.34 | 1 | Rpl31-ps11 | | 15 | 2 | 23 | 11 | 0 | 0 | 3 | 0 | 1 | 0 | 9 |
| 6 | -2.01 | 2.51 | -0.80 | 0.42 | 1 | Ugt1a6b | UDP glucuronosyltransferase 1 family, po | 29 | 11 | 18 | 0 | 0 | 3 | 3 | 0 | 0 | 0 | 0 |
| 6 | -1.35 | 2.15 | -0.63 | 0.53 | 1 | Gm13565 | | 0 | 10 | 0 | 4 | 20 | 4 | 3 | 0 | 13 | 0 | 7 |
| 5 | 1.70 | 1.77 | 0.96 | 0.34 | 1 | Gm13855 | | 3 | 0 | 0 | 0 | 8 | 7 | 3 | 16 | 11 | 4 | 8 |
| 5 | 0.64 | 1.14 | 0.56 | 0.57 | 1 | Trmt112-ps2 | | 4 | 11 | 5 | 4 | 3 | 4 | 3 | 8 | 5 | 3 | 8 |
| 5 | 0.82 | 2.12 | 0.39 | 0.70 | 1 | Gm2155 | | 0 | 0 | 0 | 0 | 7 | 5 | 3 | 0 | 5 | 36 | 3 |
| 5 | -1.94 | 2.30 | -0.84 | 0.40 | 1 | Dbndd1 | dysbindin (dystrobrevin binding protein 1) | 13 | 0 | 0 | 11 | 4 | 4 | 3 | 21 | 0 | 0 | 1 |
| 5 | 1.54 | 1.41 | 1.09 | 0.27 | 1 | Gm13803 | | 0 | 8 | 1 | 3 | 2 | 3 | 3 | 15 | 12 | 3 | 8 |
| 5 | -3.57 | 2.57 | -1.39 | 0.16 | 1 | Gm31019 | | 0 | 33 | 0 | 11 | 2 | 4 | 3 | 0 | 4 | 0 | 0 |
| 5 | -2.00 | 1.78 | -1.12 | 0.26 | 1 | Gm47870 | | 0 | 1 | 8 | 6 | 5 | 11 | 3 | 0 | 6 | 0 | 16 |
| 5 | 1.11 | 1.76 | 0.63 | 0.53 | 1 | Gm48432 | | 11 | 2 | 2 | 9 | 1 | 1 | 3 | 0 | 14 | 1 | 11 |
| 5 | -2.14 | 2.14 | -1.00 | 0.32 | 1 | 9330198N18Rik | | 8 | 6 | 6 | 0 | 17 | 4 | 3 | 0 | 11 | 0 | 0 |
| 5 | -1.85 | 1.90 | -0.97 | 0.33 | 1 | Gm44418 | | 0 | 9 | 4 | 18 | 1 | 9 | 3 | 0 | 6 | 0 | 5 |
| 5 | -2.55 | 2.09 | -1.22 | 0.22 | 1 | Gm17546 | | 3 | 0 | 0 | 12 | 10 | 6 | 3 | 0 | 8 | 13 | 0 |
| 5 | -0.99 | 3.58 | -0.28 | 0.78 | 1 | Gm42576 | | 0 | 0 | 0 | 0 | 0 | 23 | 3 | 0 | 0 | 0 | 28 |
| 5 | 0.63 | 1.24 | 0.51 | 0.61 | 1 | Hmgb1-ps7 | | 14 | 8 | 6 | 2 | 2 | 3 | 3 | 3 | 4 | 6 | 2 |
| 5 | -0.26 | 2.86 | -0.09 | 0.93 | 1 | Apcs | serum amyloid P-component | 2 | 14 | 31 | 0 | 0 | 0 | 3 | 0 | 3 | 0 | 0 |
| 5 | -0.58 | 2.47 | -0.23 | 0.82 | 1 | Gm36677 | | 2 | 0 | 0 | 10 | 0 | 14 | 3 | 0 | 5 | 0 | 19 |
| 5 | -1.76 | 1.36 | -1.30 | 0.19 | 1 | Gm5470 | | 3 | 3 | 2 | 12 | 15 | 3 | 3 | 1 | 7 | 1 | 1 |
| 5 | 0.16 | 1.66 | 0.09 | 0.93 | 1 | Gm14857 | | 8 | 1 | 5 | 3 | 9 | 3 | 3 | 13 | 0 | 1 | 6 |
| 5 | 0.80 | 2.00 | 0.40 | 0.69 | 1 | Gm21988 | | 2 | 0 | 0 | 3 | 0 | 11 | 3 | 0 | 3 | 22 | 8 |

| | | | | | | | | | | | | | | | | | | |
|---|---|---|---|---|---|---|---|---|---|---|---|---|---|---|---|---|---|---|
| 5 | 0.22 | 3.16 | 0.07 | 0.95 | 1 | Hist1h4m | histone cluster 1, H4m | 0 | 0 | 0 | 0 | 0 | 24 | 3 | 0 | 6 | 0 | 18 |
| 5 | -3.58 | 2.15 | -1.67 | 0.10 | 1 | Gm4291 | | 0 | 0 | 7 | 25 | 9 | 2 | 3 | 0 | 3 | 0 | 2 |
| 5 | -3.94 | 2.16 | -1.82 | 0.07 | 1 | Rad21l | RAD21-like (S. pombe) | 0 | 4 | 7 | 15 | 5 | 9 | 3 | 0 | 0 | 0 | 7 |
| 5 | -3.63 | 2.36 | -1.54 | 0.12 | 1 | C630004L07Rik | | 9 | 7 | 0 | 1 | 18 | 5 | 3 | 0 | 0 | 0 | 6 |
| 5 | 0.36 | 3.21 | 0.11 | 0.91 | 1 | Gm42462 | | 0 | 0 | 0 | 0 | 0 | 21 | 3 | 0 | 8 | 0 | 17 |
| 4 | -1.15 | 2.29 | -0.50 | 0.61 | 1 | Gm42840 | | 0 | 0 | 0 | 8 | 4 | 10 | 3 | 0 | 11 | 0 | 13 |
| 4 | 0.80 | 2.97 | 0.27 | 0.79 | 1 | Gm10640 | | 1 | 0 | 0 | 0 | 0 | 8 | 3 | 31 | 0 | 0 | 5 |
| 4 | -3.13 | 3.41 | -0.92 | 0.36 | 1 | Gm45286 | | 0 | 11 | 0 | 0 | 0 | 19 | 3 | 0 | 0 | 0 | 14 |
| 4 | 0.49 | 2.96 | 0.17 | 0.87 | 1 | Mir205hg | | 9 | 18 | 10 | 0 | 0 | 0 | 3 | 0 | 7 | 0 | 0 |
| 4 | -0.11 | 1.33 | -0.08 | 0.93 | 1 | Gm5312 | | 1 | 4 | 5 | 2 | 9 | 5 | 3 | 6 | 3 | 1 | 8 |
| 4 | -1.53 | 2.83 | -0.54 | 0.59 | 1 | Gm12902 | | 0 | 0 | 6 | 0 | 4 | 8 | 3 | 0 | 0 | 0 | 24 |
| 4 | -0.43 | 2.25 | -0.19 | 0.85 | 1 | Gm15283 | | 12 | 2 | 3 | 1 | 0 | 3 | 3 | 0 | 0 | 22 | 0 |
| 4 | 0.17 | 2.12 | 0.08 | 0.94 | 1 | Dcaf12l2 | DDB1 and CUL4 associated factor 12-like | 0 | 5 | 10 | 6 | 0 | 4 | 3 | 0 | 3 | 10 | 3 |
| 4 | 2.25 | 2.28 | 0.99 | 0.32 | 1 | Gm28040 | KiSS-1 metastasis-suppressor | 0 | 6 | 0 | 5 | 0 | 3 | 3 | 0 | 7 | 9 | 10 |
| 4 | -0.96 | 1.59 | -0.60 | 0.55 | 1 | Gm4217 | | 20 | 2 | 1 | 5 | 1 | 5 | 3 | 1 | 3 | 1 | 1 |
| 4 | 1.98 | 2.82 | 0.70 | 0.48 | 1 | Gm38973 | | 0 | 0 | 0 | 0 | 8 | 0 | 3 | 0 | 2 | 24 | 5 |
| 4 | -1.43 | 1.93 | -0.74 | 0.46 | 1 | Gm8483 | | 1 | 12 | 6 | 0 | 8 | 6 | 3 | 1 | 0 | 3 | 2 |
| 4 | 5.29 | 2.73 | 1.94 | 0.05 | 1 | Gm15848 | | 1 | 6 | 0 | 0 | 0 | 0 | 3 | 0 | 8 | 18 | 5 |
| 4 | -3.82 | 3.59 | -1.06 | 0.29 | 1 | Gm13483 | | 0 | 24 | 0 | 0 | 0 | 10 | 3 | 0 | 0 | 0 | 3 |
| 4 | -2.31 | 2.10 | -1.10 | 0.27 | 1 | Lct | lactase | 0 | 0 | 1 | 21 | 5 | 3 | 3 | 6 | 0 | 0 | 2 |
| 4 | 0.91 | 2.16 | 0.42 | 0.67 | 1 | B230206H07Rik | | 10 | 3 | 2 | 0 | 2 | 3 | 3 | 9 | 7 | 0 | 0 |
| 4 | -0.05 | 1.66 | -0.03 | 0.98 | 1 | Gm14388 | | 0 | 3 | 2 | 2 | 5 | 3 | 3 | 10 | 6 | 0 | 5 |
| 4 | 3.19 | 2.37 | 1.35 | 0.18 | 1 | Rpl31-ps16 | | 13 | 3 | 7 | 0 | 0 | 0 | 3 | 1 | 6 | 0 | 6 |
| 3 | -1.27 | 2.61 | -0.49 | 0.63 | 1 | Gm43513 | | 2 | 0 | 8 | 0 | 0 | 18 | 3 | 0 | 5 | 0 | 2 |
| 3 | -4.28 | 2.57 | -1.67 | 0.10 | 1 | Gm26744 | | 5 | 5 | 7 | 11 | 0 | 6 | 3 | 0 | 0 | 0 | 0 |
| 3 | -6.05 | 3.28 | -1.84 | 0.07 | 1 | Gm3248 | | 0 | 17 | 0 | 0 | 3 | 5 | 3 | 0 | 0 | 0 | 9 |
| 3 | -1.46 | 2.33 | -0.63 | 0.53 | 1 | Gm38285 | | 3 | 0 | 0 | 5 | 11 | 2 | 3 | 0 | 8 | 0 | 5 |
| 3 | 1.14 | 3.60 | 0.32 | 0.75 | 1 | Mab21l3 | mab-21-like 3 | 31 | 1 | 0 | 0 | 0 | 0 | 3 | 0 | 2 | 0 | 0 |
| 3 | -0.42 | 1.70 | -0.25 | 0.81 | 1 | Gm5879 | | 12 | 3 | 2 | 0 | 5 | 4 | 3 | 3 | 1 | 1 | 2 |
| 3 | -1.55 | 2.10 | -0.73 | 0.46 | 1 | Gm14633 | | 11 | 4 | 9 | 5 | 0 | 1 | 3 | 1 | 2 | 0 | 0 |
| 3 | 0.03 | 3.47 | 0.01 | 0.99 | 1 | Gm13400 | | 0 | 0 | 0 | 0 | 0 | 14 | 3 | 0 | 2 | 0 | 17 |
| 3 | -4.22 | 2.59 | -1.63 | 0.10 | 1 | 4930447K03Rik | | 9 | 0 | 9 | 4 | 10 | 1 | 3 | 0 | 0 | 0 | 0 |
| 3 | 0.28 | 2.02 | 0.14 | 0.89 | 1 | Gm8514 | | 2 | 1 | 7 | 2 | 0 | 2 | 3 | 0 | 1 | 18 | 1 |
| 3 | -0.23 | 1.95 | -0.12 | 0.91 | 1 | Gm4609 | | 7 | 15 | 1 | 0 | 2 | 2 | 3 | 1 | 3 | 3 | 0 |
| 3 | -1.32 | 2.40 | -0.55 | 0.58 | 1 | Rps18-ps4 | | 5 | 0 | 0 | 6 | 7 | 4 | 3 | 7 | 4 | 0 | 0 |
| 3 | 1.28 | 2.26 | 0.57 | 0.57 | 1 | Gm43766 | | 2 | 1 | 7 | 0 | 0 | 3 | 3 | 12 | 7 | 0 | 1 |
| 3 | -2.81 | 3.49 | -0.81 | 0.42 | 1 | Gm13029 | | 0 | 0 | 0 | 9 | 0 | 12 | 3 | 0 | 0 | 0 | 12 |
| 3 | -1.24 | 1.67 | -0.74 | 0.46 | 1 | Gm29257 | | 2 | 12 | 2 | 4 | 1 | 4 | 3 | 1 | 1 | 0 | 5 |
| 3 | 0.22 | 1.59 | 0.14 | 0.89 | 1 | Gm3671 | | 2 | 3 | 2 | 2 | 1 | 3 | 3 | 16 | 1 | 1 | 1 |
| 3 | -0.30 | 1.91 | -0.16 | 0.88 | 1 | Gm6828 | | 3 | 13 | 0 | 6 | 1 | 3 | 3 | 0 | 2 | 4 | 1 |
| 3 | -1.14 | 2.23 | -0.51 | 0.61 | 1 | Gm45328 | | 0 | 1 | 9 | 0 | 5 | 6 | 3 | 0 | 8 | 0 | 3 |
| 3 | -0.40 | 3.11 | -0.13 | 0.90 | 1 | Gm17077 | | 0 | 0 | 0 | 1 | 0 | 9 | 3 | 20 | 0 | 0 | 2 |
| 3 | -1.36 | 3.25 | -0.42 | 0.68 | 1 | Ect2l | epithelial cell transforming sequence 2 or | 0 | 18 | 5 | 0 | 0 | 5 | 3 | 0 | 0 | 0 | 3 |
| 3 | 0.52 | 2.55 | 0.21 | 0.84 | 1 | Dnah12 | dynein, axonemal, heavy chain 12 | 12 | 0 | 0 | 2 | 0 | 8 | 3 | 1 | 6 | 0 | 2 |
| 3 | -3.33 | 3.50 | -0.95 | 0.34 | 1 | Gm27045 | | 0 | 0 | 0 | 8 | 0 | 19 | 3 | 0 | 0 | 0 | 3 |
| 3 | -1.18 | 3.32 | -0.36 | 0.72 | 1 | Gm15285 | | 0 | 11 | 15 | 0 | 0 | 3 | 3 | 0 | 0 | 0 | 2 |
| 3 | -4.30 | 2.81 | -1.53 | 0.13 | 1 | Gm4798 | | 15 | 0 | 0 | 2 | 8 | 3 | 3 | 0 | 2 | 0 | 0 |
| 3 | 0.24 | 1.58 | 0.15 | 0.88 | 1 | Gm4076 | | 5 | 1 | 0 | 4 | 2 | 3 | 3 | 2 | 4 | 4 | 4 |
| 3 | 0.28 | 1.92 | 0.15 | 0.88 | 1 | Gm8991 | | 2 | 0 | 1 | 11 | 0 | 1 | 3 | 10 | 2 | 1 | 1 |
| 3 | 0.18 | 1.49 | 0.12 | 0.90 | 1 | Gm9276 | | 2 | 1 | 7 | 1 | 4 | 3 | 3 | 3 | 3 | 3 | 2 |
| 3 | -3.07 | 3.50 | -0.88 | 0.38 | 1 | A230009B12Rik | | 0 | 0 | 0 | 19 | 0 | 3 | 3 | 0 | 7 | 0 | 0 |
| 3 | -0.06 | 1.60 | -0.03 | 0.97 | 1 | Gm6382 | | 9 | 1 | 3 | 3 | 3 | 1 | 3 | 2 | 1 | 1 | 5 |
| 3 | -2.48 | 2.19 | -1.13 | 0.26 | 1 | Gm17638 | | 1 | 0 | 15 | 4 | 4 | 1 | 3 | 0 | 2 | 0 | 2 |
| 3 | 1.55 | 3.00 | 0.52 | 0.61 | 1 | Gm46210 | | 0 | 11 | 0 | 2 | 0 | 3 | 3 | 0 | 3 | 9 | 0 |
| 3 | -0.31 | 3.32 | -0.09 | 0.93 | 1 | Gm7378 | | 0 | 0 | 7 | 0 | 0 | 6 | 3 | 0 | 12 | 3 | 0 |
| 3 | -3.57 | 2.43 | -1.47 | 0.14 | 1 | B3gnt4 | UDP-GlcNAc:betaGal beta-1,3-N-acetylgl | 0 | 0 | 7 | 11 | 3 | 4 | 3 | 0 | 1 | 0 | 2 |
| 3 | 0.10 | 3.14 | 0.03 | 0.98 | 1 | | | 0 | 0 | 0 | 0 | 1 | 8 | 3 | 0 | 13 | 0 | 5 |
| 3 | -2.59 | 3.53 | -0.73 | 0.46 | 1 | Gm2213 | | 0 | 0 | 5 | 12 | 0 | 0 | 3 | 0 | 10 | 0 | 0 |
| 3 | 0.84 | 3.55 | 0.24 | 0.81 | 1 | Gm20732 | | 0 | 0 | 0 | 4 | 0 | 0 | 3 | 22 | 0 | 0 | 1 |
| 3 | 0.34 | 1.89 | 0.18 | 0.86 | 1 | Gm6974 | | 1 | 0 | 10 | 0 | 2 | 4 | 3 | 1 | 2 | 1 | 5 |

| | | | | | | | | | | | | | | | | | | |
|---|---|---|---|---|---|---|---|---|---|---|---|---|---|---|---|---|---|---|
| 3 | -0.74 | 2.52 | -0.29 | 0.77 | 1 | Gm10392 | | 3 | 0 | 0 | 11 | 2 | 2 | 3 | 0 | 6 | 0 | 4 |
| 3 | -2.18 | 3.54 | -0.62 | 0.54 | 1 | 6720483E21Rik | | 0 | 0 | 0 | 0 | 3 | 13 | 3 | 0 | 0 | 0 | 11 |
| 3 | -2.06 | 3.63 | -0.57 | 0.57 | 1 | Plg | plasminogen | 4 | 21 | 0 | 0 | 0 | 0 | 3 | 0 | 0 | 0 | 1 |
| 3 | -1.87 | 3.25 | -0.58 | 0.56 | 1 | 4930542M03Rik | | 0 | 0 | 0 | 10 | 0 | 8 | 3 | 0 | 5 | 0 | 4 |
| 3 | 0.74 | 1.83 | 0.40 | 0.69 | 1 | Gm8213 | | 11 | 2 | 0 | 1 | 2 | 1 | 3 | 1 | 3 | 1 | 4 |
| 3 | -1.55 | 3.39 | -0.46 | 0.65 | 1 | Gm13427 | | 0 | 7 | 12 | 3 | 0 | 0 | 3 | 0 | 4 | 0 | 0 |
| 3 | 0.02 | 2.29 | 0.01 | 0.99 | 1 | Gm15191 | | 9 | 0 | 1 | 2 | 0 | 3 | 3 | 1 | 8 | 3 | 0 |
| 3 | -1.06 | 3.66 | -0.29 | 0.77 | 1 | Azgp1 | alpha-2-glycoprotein 1, zinc | 0 | 12 | 14 | 0 | 0 | 0 | 3 | 0 | 0 | 0 | 0 |
| 3 | 1.24 | 3.49 | 0.36 | 0.72 | 1 | Gm9902 | | 0 | 0 | 0 | 4 | 0 | 0 | 3 | 0 | 0 | 14 | 7 |
| 2 | -0.04 | 2.00 | -0.02 | 0.98 | 1 | Gm10156 | | 3 | 0 | 3 | 4 | 0 | 3 | 3 | 3 | 5 | 1 | 1 |
| 2 | -0.41 | 1.89 | -0.21 | 0.83 | 1 | Cox6c2 | | 1 | 0 | 1 | 3 | 3 | 3 | 3 | 3 | 8 | 0 | 1 |
| 2 | 0.59 | 3.53 | 0.17 | 0.87 | 1 | Gm14232 | | 0 | 18 | 0 | 0 | 0 | 0 | 3 | 0 | 5 | 0 | 0 |
| 2 | 0.23 | 3.33 | 0.07 | 0.94 | 1 | Platr26 | | 0 | 0 | 0 | 0 | 9 | 1 | 3 | 0 | 0 | 8 | 6 |
| 2 | -2.85 | 2.24 | -1.27 | 0.20 | 1 | Gm13339 | | 9 | 4 | 1 | 6 | 1 | 1 | 3 | 0 | 0 | 0 | 1 |
| 2 | -0.76 | 2.55 | -0.30 | 0.76 | 1 | Gm11263 | | 3 | 3 | 4 | 1 | 3 | 0 | 3 | 0 | 9 | 0 | 0 |
| 2 | -1.73 | 3.35 | -0.52 | 0.61 | 1 | Gm46409 | | 4 | 0 | 0 | 0 | 3 | 5 | 3 | 0 | 0 | 0 | 10 |
| 2 | 2.30 | 3.55 | 0.65 | 0.52 | 1 | 4930509H03Rik | | 0 | 0 | 2 | 0 | 0 | 0 | 3 | 0 | 0 | 14 | 6 |
| 2 | -0.63 | 2.58 | -0.24 | 0.81 | 1 | Gm6257 | | 3 | 0 | 9 | 0 | 0 | 6 | 3 | 1 | 1 | 0 | 1 |
| 2 | -1.29 | 2.00 | -0.64 | 0.52 | 1 | Gm5529 | | 3 | 2 | 7 | 2 | 2 | 2 | 3 | 0 | 0 | 3 | 2 |
| 2 | -1.50 | 3.63 | -0.41 | 0.68 | 1 | Kcne1l | potassium voltage-gated channel, Isk-rela | 5 | 0 | 16 | 0 | 1 | 0 | 3 | 0 | 0 | 0 | 0 |
| 2 | -6.06 | 3.58 | -1.69 | 0.09 | 1 | Gm13001 | | 0 | 0 | 15 | 1 | 0 | 5 | 3 | 0 | 0 | 0 | 1 |
| 2 | -2.24 | 2.79 | -0.80 | 0.42 | 1 | Gm27321 | | 0 | 0 | 0 | 7 | 7 | 2 | 3 | 0 | 3 | 0 | 3 |
| 2 | -2.49 | 2.01 | -1.24 | 0.22 | 1 | Rps8-ps1 | | 1 | 5 | 2 | 4 | 4 | 2 | 3 | 1 | 0 | 0 | 2 |
| 2 | 2.58 | 3.54 | 0.73 | 0.47 | 1 | Gm12352 | | 4 | 0 | 4 | 0 | 0 | 0 | 3 | 9 | 3 | 0 | 0 |
| 2 | -1.11 | 3.64 | -0.31 | 0.76 | 1 | Spata45 | spermatogenesis associated 45 | 5 | 0 | 16 | 0 | 0 | 0 | 3 | 0 | 0 | 0 | 0 |
| 2 | -2.46 | 2.64 | -0.93 | 0.35 | 1 | Pgam1-ps2 | | 1 | 5 | 0 | 1 | 2 | 4 | 3 | 0 | 0 | 0 | 7 |
| 2 | 0.75 | 1.84 | 0.41 | 0.68 | 1 | Rpl31-ps17 | | 3 | 1 | 2 | 4 | 1 | 0 | 3 | 1 | 3 | 3 | 3 |
| 2 | 1.02 | 3.52 | 0.29 | 0.77 | 1 | Gm28402 | | 0 | 0 | 1 | 0 | 0 | 1 | 3 | 0 | 0 | 17 | 1 |
| 2 | -1.34 | 3.58 | -0.37 | 0.71 | 1 | Gm45737 | | 0 | 0 | 0 | 0 | 0 | 14 | 3 | 0 | 0 | 0 | 6 |
| 2 | 0.36 | 3.54 | 0.10 | 0.92 | 1 | Gm19552 | | 0 | 0 | 0 | 0 | 0 | 8 | 3 | 0 | 0 | 8 | 4 |
| 2 | -1.77 | 2.05 | -0.86 | 0.39 | 1 | Gm13882 | | 4 | 3 | 2 | 1 | 4 | 2 | 3 | 0 | 0 | 3 | 1 |
| 2 | -0.14 | 3.05 | -0.05 | 0.96 | 1 | Gm16464 | | 6 | 0 | 4 | 0 | 0 | 2 | 3 | 0 | 5 | 1 | 0 |
| 2 | 0.42 | 2.03 | 0.21 | 0.83 | 1 | Gm6977 | | 3 | 0 | 2 | 3 | 3 | 0 | 3 | 3 | 1 | 3 | 1 |
| 2 | -2.36 | 3.54 | -0.67 | 0.51 | 1 | Mir8094 | | 0 | 0 | 0 | 0 | 3 | 10 | 3 | 0 | 0 | 0 | 5 |
| 2 | -1.31 | 2.10 | -0.62 | 0.53 | 1 | Gm18078 | | 4 | 4 | 3 | 1 | 1 | 2 | 3 | 0 | 1 | 0 | 2 |
| 2 | 0.81 | 3.41 | 0.24 | 0.81 | 1 | 7530414M10Rik | | 1 | 0 | 0 | 0 | 3 | 0 | 3 | 0 | 5 | 9 | 0 |
| 2 | -0.69 | 3.55 | -0.19 | 0.85 | 1 | Gm37568 | | 7 | 0 | 0 | 0 | 1 | 2 | 3 | 0 | 0 | 0 | 8 |
| 2 | -0.61 | 2.67 | -0.23 | 0.82 | 1 | Gm12112 | | 0 | 0 | 1 | 4 | 2 | 0 | 3 | 1 | 4 | 0 | 5 |
| 2 | -1.47 | 2.11 | -0.70 | 0.48 | 1 | Gm9169 | | 0 | 2 | 4 | 2 | 2 | 3 | 3 | 0 | 2 | 1 | 1 |
| 2 | -0.10 | 3.59 | -0.03 | 0.98 | 1 | A730060N03Rik | | 0 | 6 | 7 | 0 | 0 | 0 | 3 | 0 | 4 | 0 | 0 |
| 2 | -2.46 | 3.59 | -0.69 | 0.49 | 1 | Gm4681 | | 0 | 9 | 0 | 0 | 0 | 2 | 3 | 0 | 0 | 0 | 6 |
| 2 | -0.46 | 3.57 | -0.13 | 0.90 | 1 | Gm9939 | | 2 | 0 | 2 | 0 | 0 | 3 | 3 | 0 | 0 | 0 | 10 |
| 2 | -2.68 | 3.11 | -0.86 | 0.39 | 1 | Gm17819 | | 3 | 0 | 1 | 9 | 3 | 0 | 3 | 1 | 0 | 0 | 0 |
| 2 | -2.59 | 3.53 | -0.73 | 0.46 | 1 | Gm42944 | | 0 | 0 | 0 | 0 | 8 | 7 | 3 | 0 | 1 | 0 | 1 |
| 2 | -0.82 | 3.56 | -0.23 | 0.82 | 1 | Gapdh-ps14 | | 9 | 0 | 0 | 4 | 0 | 0 | 3 | 1 | 0 | 1 | 0 |
| 2 | 5.46 | 3.56 | 1.54 | 0.12 | 1 | Gm47904 | | 0 | 0 | 0 | 0 | 0 | 0 | 3 | 0 | 2 | 0 | 14 |
| 2 | -2.18 | 2.98 | -0.73 | 0.46 | 1 | Gm6532 | | 0 | 0 | 0 | 2 | 4 | 6 | 3 | 0 | 0 | 3 | 2 |
| 2 | 0.38 | 3.50 | 0.11 | 0.91 | 1 | Gm48374 | | 0 | 0 | 0 | 0 | 0 | 2 | 3 | 0 | 12 | 0 | 2 |
| 2 | -0.84 | 3.65 | -0.23 | 0.82 | 1 | Gm13307 | | 9 | 0 | 7 | 0 | 0 | 0 | 3 | 0 | 0 | 0 | 0 |
| 2 | -1.78 | 3.55 | -0.50 | 0.62 | 1 | Gm7252 | | 0 | 0 | 3 | 0 | 2 | 3 | 3 | 0 | 8 | 0 | 0 |
| 2 | -2.65 | 3.59 | -0.74 | 0.46 | 1 | Gm15893 | | 6 | 0 | 5 | 0 | 2 | 3 | 3 | 0 | 0 | 0 | 0 |
| 2 | -2.60 | 3.60 | -0.72 | 0.47 | 1 | Zyg11a | zyg-11 family member A, cell cycle regula | 0 | 10 | 1 | 0 | 0 | 4 | 3 | 0 | 0 | 0 | 0 |
| 2 | 0.12 | 2.17 | 0.06 | 0.96 | 1 | Gm17669 | | 4 | 2 | 2 | 1 | 2 | 0 | 3 | 1 | 2 | 0 | 1 |
| 2 | -1.28 | 3.57 | -0.36 | 0.72 | 1 | Gm9613 | | 3 | 0 | 0 | 3 | 0 | 1 | 3 | 0 | 0 | 9 | 0 |
| 2 | -0.66 | 3.51 | -0.19 | 0.85 | 1 | Mxra8os | | 3 | 0 | 0 | 4 | 0 | 0 | 3 | 0 | 8 | 0 | 0 |
| 2 | -0.72 | 1.82 | -0.40 | 0.69 | 1 | Gm8927 | | 1 | 2 | 3 | 2 | 1 | 2 | 3 | 1 | 1 | 1 | 1 |
| 2 | -0.84 | 2.24 | -0.38 | 0.71 | 1 | Gm6758 | | 2 | 3 | 3 | 0 | 1 | 3 | 3 | 1 | 1 | 1 | 0 |
| 2 | 0.50 | 2.81 | 0.18 | 0.86 | 1 | Gm12760 | | 3 | 1 | 1 | 0 | 3 | 0 | 3 | 0 | 6 | 0 | 1 |
| 2 | -2.28 | 2.82 | -0.81 | 0.42 | 1 | H3f3c | | 2 | 4 | 0 | 2 | 4 | 1 | 3 | 0 | 2 | 0 | 0 |
| 2 | -0.26 | 2.81 | -0.09 | 0.92 | 1 | Rpl23a-ps5 | | 8 | 2 | 1 | 0 | 1 | 0 | 3 | 0 | 0 | 1 | 1 |

| | | | | | | | | | | | | | | | | | | |
|---|---|---|---|---|---|---|---|---|---|---|---|---|---|---|---|---|---|---|
| 2 | -3.41 | 2.73 | -1.25 | 0.21 | 1 | Gm5946 | | 1 | 0 | 1 | 6 | 3 | 2 | 3 | 0 | 0 | 1 | 0 |
| 2 | -1.09 | 3.57 | -0.31 | 0.76 | 1 | Myo15 | myosin XV | 9 | 0 | 0 | 0 | 0 | 4 | 3 | 0 | 0 | 0 | 1 |
| 2 | 0.95 | 2.34 | 0.41 | 0.68 | 1 | Gm5830 | | 1 | 0 | 0 | 1 | 0 | 3 | 3 | 2 | 1 | 4 | 2 |
| 1 | -4.18 | 3.61 | -1.16 | 0.25 | 1 | Gm24727 | | 0 | 0 | 7 | 0 | 0 | 5 | 3 | 0 | 0 | 0 | 1 |
| 1 | -0.29 | 3.62 | -0.08 | 0.94 | 1 | Gm16027 | | 13 | 0 | 0 | 0 | 0 | 0 | 3 | 0 | 0 | 0 | 0 |
| 1 | -3.53 | 3.57 | -0.99 | 0.32 | 1 | Gm47692 | | 1 | 0 | 1 | 0 | 7 | 4 | 3 | 0 | 0 | 0 | 0 |
| 1 | -0.88 | 3.41 | -0.26 | 0.80 | 1 | Gm24270 | | 0 | 0 | 2 | 7 | 0 | 0 | 3 | 1 | 0 | 1 | 1 |
| 1 | 0.18 | 2.91 | 0.06 | 0.95 | 1 | Gm12182 | | 1 | 2 | 0 | 1 | 0 | 3 | 3 | 0 | 2 | 0 | 4 |
| 1 | 0.17 | 2.84 | 0.06 | 0.95 | 1 | Gm3355 | | 0 | 0 | 1 | 1 | 3 | 0 | 3 | 1 | 5 | 0 | 1 |
| 1 | -0.41 | 2.40 | -0.17 | 0.87 | 1 | Gm5064 | | 0 | 0 | 0 | 1 | 2 | 3 | 3 | 1 | 1 | 1 | 3 |
| 1 | -4.94 | 3.55 | -1.39 | 0.16 | 1 | Gm43167 | | 0 | 0 | 0 | 0 | 0 | 8 | 3 | 0 | 4 | 0 | 0 |
| 1 | -2.47 | 3.56 | -0.69 | 0.49 | 1 | Gm16103 | | 0 | 0 | 0 | 0 | 9 | 1 | 3 | 0 | 0 | 1 | 0 |
| 1 | 0.04 | 2.95 | 0.01 | 0.99 | 1 | Gm14052 | | 1 | 2 | 0 | 2 | 0 | 2 | 3 | 0 | 2 | 0 | 3 |
| 1 | -1.11 | 2.56 | -0.43 | 0.66 | 1 | Rpl13-ps3 | | 3 | 1 | 3 | 0 | 1 | 2 | 3 | 0 | 1 | 0 | 1 |
| 1 | -0.49 | 3.57 | -0.14 | 0.89 | 1 | Gm9945 | | 2 | 0 | 0 | 0 | 0 | 3 | 3 | 0 | 0 | 0 | 6 |
| 1 | -0.49 | 3.57 | -0.14 | 0.89 | 1 | Crip3 | cysteine-rich protein 3 | 2 | 0 | 0 | 0 | 0 | 3 | 3 | 0 | 0 | 0 | 6 |
| 1 | -0.68 | 3.57 | -0.19 | 0.85 | 1 | Gm19202 | | 0 | 0 | 0 | 0 | 0 | 1 | 3 | 0 | 0 | 0 | 10 |
| 1 | -0.42 | 3.53 | -0.12 | 0.91 | 1 | Gm20109 | | 0 | 0 | 2 | 0 | 0 | 2 | 3 | 0 | 5 | 0 | 2 |
| 1 | -0.57 | 3.56 | -0.16 | 0.87 | 1 | Epha10 | Eph receptor A10 | 3 | 0 | 0 | 1 | 0 | 0 | 3 | 0 | 0 | 0 | 6 |
| 1 | -3.64 | 3.56 | -1.02 | 0.31 | 1 | Csmd2os | CUB and Sushi multiple domains 2, oppo | 0 | 0 | 0 | 0 | 8 | 3 | 3 | 0 | 0 | 0 | 0 |
| 1 | 0.09 | 3.58 | 0.03 | 0.98 | 1 | Gm14049 | | 0 | 1 | 3 | 0 | 0 | 0 | 3 | 6 | 0 | 0 | 0 |
| 1 | 0.48 | 3.61 | 0.13 | 0.89 | 1 | mt-Tq | | 9 | 0 | 0 | 0 | 0 | 0 | 3 | 0 | 0 | 0 | 1 |
| 1 | -2.25 | 3.63 | -0.62 | 0.53 | 1 | Gm11511 | | 0 | 0 | 8 | 0 | 0 | 0 | 3 | 0 | 0 | 0 | 1 |
| 1 | -2.25 | 3.63 | -0.62 | 0.53 | 1 | A930002C04Rik | | 0 | 0 | 8 | 0 | 0 | 0 | 3 | 0 | 0 | 0 | 1 |
| 1 | -2.91 | 3.48 | -0.84 | 0.40 | 1 | Gm7468 | | 1 | 0 | 0 | 1 | 3 | 3 | 3 | 0 | 0 | 0 | 1 |
| 1 | -1.31 | 3.11 | -0.42 | 0.67 | 1 | Gm17828 | | 1 | 1 | 3 | 0 | 1 | 1 | 3 | 0 | 2 | 0 | 0 |
| 1 | -1.48 | 3.59 | -0.41 | 0.68 | 1 | Bhlha9 | basic helix-loop-helix family, member a9 | 0 | 0 | 0 | 0 | 0 | 7 | 3 | 0 | 0 | 0 | 2 |
| 1 | -0.33 | 3.58 | -0.09 | 0.93 | 1 | Gm9081 | | 0 | 1 | 3 | 0 | 0 | 1 | 3 | 0 | 1 | 0 | 2 |
| 1 | -1.43 | 3.60 | -0.40 | 0.69 | 1 | Rps8-ps4 | | 2 | 0 | 4 | 1 | 0 | 0 | 3 | 0 | 1 | 0 | 0 |
| 1 | -2.44 | 3.59 | -0.68 | 0.50 | 1 | Gm43497 | | 0 | 0 | 0 | 0 | 0 | 8 | 3 | 0 | 0 | 0 | 0 |
| 1 | -2.44 | 3.59 | -0.68 | 0.50 | 1 | Gm43880 | | 0 | 0 | 0 | 0 | 0 | 8 | 3 | 0 | 0 | 0 | 0 |
| 1 | -2.07 | 3.56 | -0.58 | 0.56 | 1 | Gm10961 | | 0 | 0 | 0 | 0 | 3 | 3 | 3 | 0 | 2 | 0 | 0 |
| 1 | -0.92 | 3.59 | -0.26 | 0.80 | 1 | 4933411G06Rik | | 0 | 0 | 0 | 0 | 0 | 3 | 3 | 0 | 0 | 0 | 4 |
| 1 | -2.05 | 3.56 | -0.57 | 0.57 | 1 | Gm12204 | | 0 | 0 | 0 | 4 | 0 | 0 | 3 | 0 | 0 | 0 | 2 |
| 1 | -0.22 | 3.57 | -0.06 | 0.95 | 1 | 1600017P15Rik | | 3 | 0 | 0 | 0 | 0 | 2 | 3 | 0 | 2 | 0 | 0 |
| 1 | -2.20 | 3.58 | -0.61 | 0.54 | 1 | Gm37711 | | 0 | 0 | 0 | 5 | 0 | 0 | 3 | 0 | 0 | 0 | 1 |
| 1 | -1.80 | 3.57 | -0.51 | 0.61 | 1 | Gm4852 | | 0 | 0 | 0 | 2 | 1 | 1 | 3 | 0 | 0 | 3 | 0 |
| 1 | -1.37 | 3.33 | -0.41 | 0.68 | 1 | Gm27018 | | 0 | 1 | 0 | 1 | 1 | 1 | 3 | 0 | 0 | 1 | 1 |
| 1 | -1.01 | 3.59 | -0.28 | 0.78 | 1 | Gm48880 | | 0 | 0 | 0 | 0 | 0 | 3 | 3 | 0 | 0 | 0 | 3 |
| 1 | -1.97 | 3.62 | -0.54 | 0.59 | 1 | Gm5455 | | 3 | 0 | 0 | 0 | 1 | 1 | 3 | 0 | 0 | 0 | 0 |
| 1 | -3.03 | 3.58 | -0.85 | 0.40 | 1 | Myrfl | myelin regulatory factor-like | 0 | 0 | 0 | 4 | 0 | 2 | 3 | 0 | 0 | 0 | 0 |
| 1 | 0.67 | 3.57 | 0.19 | 0.85 | 1 | Gm48408 | | 0 | 0 | 0 | 0 | 0 | 0 | 3 | 0 | 0 | 0 | 5 |
| 1 | 0.39 | 3.58 | 0.11 | 0.91 | 1 | Lrrc75aos2 | | 0 | 0 | 0 | 0 | 0 | 0 | 3 | 0 | 5 | 0 | 0 |
| 1 | -2.29 | 3.61 | -0.63 | 0.53 | 1 | Fer1l5 | fer-1-like 5 (C. elegans) | 0 | 0 | 0 | 4 | 0 | 0 | 3 | 0 | 0 | 0 | 0 |
| 1 | -0.51 | 3.63 | -0.14 | 0.89 | 1 | Gm42847 | | 4 | 0 | 0 | 0 | 0 | 0 | 3 | 0 | 0 | 0 | 0 |
| 1 | 0.10 | 3.60 | 0.03 | 0.98 | 1 | Ppp1r2-ps3 | | 0 | 1 | 1 | 0 | 0 | 0 | 3 | 1 | 1 | 0 | 0 |
| 1 | 0.52 | 3.57 | 0.15 | 0.88 | 1 | A030012G06Rik | | 0 | 0 | 0 | 0 | 0 | 0 | 3 | 0 | 0 | 0 | 4 |
| 1 | 0.30 | 3.58 | 0.08 | 0.93 | 1 | Gm26783 | | 0 | 0 | 0 | 0 | 0 | 0 | 3 | 0 | 4 | 0 | 0 |
| 1 | -1.95 | 3.62 | -0.54 | 0.59 | 1 | Gm18734 | | 0 | 0 | 1 | 0 | 3 | 0 | 3 | 0 | 0 | 0 | 0 |
| 1 | -2.16 | 3.62 | -0.60 | 0.55 | 1 | Gm38116 | | 0 | 0 | 0 | 0 | 0 | 3 | 3 | 0 | 0 | 0 | 0 |
| 1 | 0.39 | 3.63 | 0.11 | 0.92 | 1 | Rps4l-ps | | 0 | 0 | 1 | 0 | 0 | 0 | 3 | 0 | 0 | 1 | 1 |
| 1 | -1.95 | 3.60 | -0.54 | 0.59 | 1 | Gm38804 | | 0 | 0 | 0 | 0 | 0 | 1 | 3 | 0 | 0 | 0 | 2 |
| 1 | -0.86 | 3.65 | -0.24 | 0.81 | 1 | Gm46432 | | 2 | 0 | 1 | 0 | 0 | 0 | 3 | 0 | 0 | 0 | 0 |
| 1 | -2.06 | 3.61 | -0.57 | 0.57 | 1 | Gm37368 | | 0 | 1 | 0 | 0 | 0 | 2 | 3 | 0 | 0 | 0 | 0 |
| 1 | -1.75 | 3.63 | -0.48 | 0.63 | 1 | Isl2 | insulin related protein 2 (islet 2) | 1 | 0 | 0 | 0 | 0 | 2 | 3 | 0 | 0 | 0 | 0 |
| 1 | -2.02 | 3.63 | -0.56 | 0.58 | 1 | Gm15385 | | 0 | 0 | 0 | 0 | 0 | 3 | 3 | 0 | 0 | 0 | 0 |
| 0 | 0.00 | 3.59 | 0.00 | 1.00 | 1 | Gm29811 | | 0 | 0 | 0 | 0 | 0 | 0 | 3 | 0 | 0 | 0 | 2 |
| 0 | -1.14 | 3.62 | -0.31 | 0.75 | 1 | Gm7478 | | 0 | 0 | 0 | 0 | 0 | 1 | 3 | 1 | 0 | 0 | 0 |
| 0 | -2.17 | 3.61 | -0.60 | 0.55 | 1 | Gm31651 | | 0 | 0 | 0 | 0 | 1 | 1 | 3 | 0 | 0 | 0 | 0 |
| 0 | -1.47 | 3.64 | -0.40 | 0.69 | 1 | Gm5597 | | 1 | 0 | 0 | 1 | 0 | 0 | 3 | 0 | 0 | 0 | 0 |

| | | | | | | | | | | | | | | | | | | |
|---|---|---|---|---|---|---|---|---|---|---|---|---|---|---|---|---|---|---|
| 0 | -0.69 | 3.64 | -0.19 | 0.85 | 1 | Gpx6 | glutathione peroxidase 6 | 2 | 0 | 0 | 0 | 0 | 0 | 3 | 0 | 0 | 0 | 0 |
| 0 | -1.84 | 3.63 | -0.51 | 0.61 | 1 | Gm45857 | | 0 | 0 | 0 | 0 | 0 | 2 | 3 | 0 | 0 | 0 | 0 |
| 0 | -1.84 | 3.63 | -0.51 | 0.61 | 1 | Gm49660 | | 0 | 0 | 0 | 0 | 0 | 2 | 3 | 0 | 0 | 0 | 0 |
| 0 | -0.43 | 3.64 | -0.12 | 0.91 | 1 | Gm47599 | | 0 | 0 | 0 | 0 | 0 | 0 | 3 | 0 | 0 | 0 | 1 |
| 0 | -0.27 | 3.64 | -0.08 | 0.94 | 1 | Gm5217 | | 0 | 0 | 0 | 0 | 0 | 0 | 3 | 0 | 1 | 0 | 0 |
| 0 | -1.55 | 3.64 | -0.43 | 0.67 | 1 | Ccnb1ip1 | cyclin B1 interacting protein 1 | 0 | 0 | 0 | 1 | 0 | 0 | 3 | 0 | 0 | 0 | 0 |
| 0 | -0.82 | 3.65 | -0.22 | 0.82 | 1 | Gm14140 | | 1 | 0 | 0 | 0 | 0 | 0 | 3 | 0 | 0 | 0 | 0 |
| 0 | -0.82 | 3.65 | -0.22 | 0.82 | 1 | Gm15751 | | 1 | 0 | 0 | 0 | 0 | 0 | 3 | 0 | 0 | 0 | 0 |
| 0 | -0.82 | 3.65 | -0.22 | 0.82 | 1 | Gm30584 | | 1 | 0 | 0 | 0 | 0 | 0 | 3 | 0 | 0 | 0 | 0 |
| 0 | -1.58 | 3.64 | -0.43 | 0.66 | 1 | 0610005C13Rik | | 0 | 0 | 0 | 0 | 0 | 1 | 3 | 0 | 0 | 0 | 0 |
| 0 | -1.58 | 3.64 | -0.43 | 0.66 | 1 | Gm11989 | | 0 | 0 | 0 | 0 | 0 | 1 | 3 | 0 | 0 | 0 | 0 |
| 0 | -1.58 | 3.64 | -0.43 | 0.66 | 1 | Gm8401 | | 0 | 0 | 0 | 0 | 0 | 1 | 3 | 0 | 0 | 0 | 0 |
| 0 | -1.03 | 3.66 | -0.28 | 0.78 | 1 | Gm5106 | | 0 | 0 | 0 | 0 | 0 | 0 | 3 | 0 | 0 | 0 | 0 |
| 0 | -1.03 | 3.66 | -0.28 | 0.78 | 1 | A930003O13Rik | | 0 | 0 | 0 | 0 | 0 | 0 | 3 | 0 | 0 | 0 | 0 |
| 0 | -1.03 | 3.66 | -0.28 | 0.78 | 1 | Gm42830 | | 0 | 0 | 0 | 0 | 0 | 0 | 3 | 0 | 0 | 0 | 0 |
| 0 | -1.03 | 3.66 | -0.28 | 0.78 | 1 | Gm43661 | | 0 | 0 | 0 | 0 | 0 | 0 | 3 | 0 | 0 | 0 | 0 |
| 0 | -1.03 | 3.66 | -0.28 | 0.78 | 1 | Defb14 | defensin beta 14 | 0 | 0 | 0 | 0 | 0 | 0 | 3 | 0 | 0 | 0 | 0 |
| 0 | -1.03 | 3.66 | -0.28 | 0.78 | 1 | Mir5615-1 | | 0 | 0 | 0 | 0 | 0 | 0 | 3 | 0 | 0 | 0 | 0 |
| 0 | -1.03 | 3.66 | -0.28 | 0.78 | 1 | Gm47200 | | 0 | 0 | 0 | 0 | 0 | 0 | 3 | 0 | 0 | 0 | 0 |
| 0 | -1.03 | 3.66 | -0.28 | 0.78 | 1 | Gm12224 | | 0 | 0 | 0 | 0 | 0 | 0 | 3 | 0 | 0 | 0 | 0 |
| 0 | -1.03 | 3.66 | -0.28 | 0.78 | 1 | Slc47a2 | solute carrier family 47, member 2 | 0 | 0 | 0 | 0 | 0 | 0 | 3 | 0 | 0 | 0 | 0 |
| 0 | -1.03 | 3.66 | -0.28 | 0.78 | 1 | Gm25448 | | 0 | 0 | 0 | 0 | 0 | 0 | 3 | 0 | 0 | 0 | 0 |
| 0 | -1.03 | 3.66 | -0.28 | 0.78 | 1 | 4930543I03Rik | | 0 | 0 | 0 | 0 | 0 | 0 | 3 | 0 | 0 | 0 | 0 |
| 0 | -1.03 | 3.66 | -0.28 | 0.78 | 1 | Gm20443 | | 0 | 0 | 0 | 0 | 0 | 0 | 3 | 0 | 0 | 0 | 0 |
| 0 | -1.03 | 3.66 | -0.28 | 0.78 | 1 | Gm49834 | | 0 | 0 | 0 | 0 | 0 | 0 | 3 | 0 | 0 | 0 | 0 |
| 0 | -1.03 | 3.66 | -0.28 | 0.78 | 1 | Gm23512 | | 0 | 0 | 0 | 0 | 0 | 0 | 3 | 0 | 0 | 0 | 0 |
| 0 | -1.03 | 3.66 | -0.28 | 0.78 | 1 | Amd-ps1 | | 0 | 0 | 0 | 0 | 0 | 0 | 3 | 0 | 0 | 0 | 0 |
| 1153 | 0.11 | 1.00 | 0.11 | 0.91 | 1 | Ehd2 | EH-domain containing 2 | 5049 | 3614 | 3950 | 14 | 6 | 10 | 2 | 0 | 6 | 19 | 15 |
| 752 | -0.47 | 1.09 | -0.43 | 0.67 | 1 | Steap4 | STEAP family member 4 | 3783 | 2304 | 2133 | 0 | 3 | 15 | 2 | 0 | 1 | 0 | 29 |
| 708 | -0.22 | 1.20 | -0.19 | 0.85 | 1 | Fndc1 | fibronectin type III domain containing 1 | 4094 | 1788 | 1905 | 0 | 0 | 1 | 2 | 0 | 0 | 0 | 2 |
| 700 | 0.59 | 1.24 | 0.48 | 0.63 | 1 | Plxdc2 | plexin domain containing 2 | 3233 | 1984 | 2362 | 19 | 5 | 14 | 2 | 0 | 24 | 46 | 8 |
| 408 | -0.37 | 1.41 | -0.26 | 0.79 | 1 | Ackr3 | atypical chemokine receptor 3 | 1985 | 1031 | 1467 | 0 | 0 | 0 | 2 | 0 | 1 | 0 | 0 |
| 371 | -0.74 | 3.66 | -0.20 | 0.84 | 1 | Abi3bp | ABI gene family, member 3 (NESH) bindi | 2381 | 1001 | 698 | 0 | 0 | 0 | 2 | 0 | 0 | 0 | 0 |
| 331 | -0.76 | 3.66 | -0.21 | 0.84 | 1 | Aspn | asporin | 1722 | 1072 | 843 | 0 | 0 | 0 | 2 | 0 | 0 | 0 | 0 |
| 311 | -3.19 | 1.80 | -1.77 | 0.08 | 1 | Pear1 | platelet endothelial aggregation receptor | 1639 | 716 | 1048 | 7 | 0 | 12 | 2 | 0 | 2 | 0 | 0 |
| 219 | 1.59 | 1.18 | 1.35 | 0.18 | 1 | Astn2 | astrotactin 2 | 742 | 781 | 774 | 9 | 5 | 5 | 2 | 0 | 19 | 68 | 8 |
| 214 | -2.94 | 1.69 | -1.74 | 0.08 | 1 | Inf2 | inverted formin, FH2 and WH2 domain co | 1094 | 700 | 538 | 0 | 12 | 2 | 2 | 0 | 0 | 0 | 1 |
| 184 | -2.21 | 1.32 | -1.67 | 0.09 | 1 | Ggt5 | gamma-glutamyltransferase 5 | 839 | 557 | 625 | 0 | 0 | 3 | 2 | 0 | 0 | 0 | 0 |
| 164 | -0.46 | 1.50 | -0.31 | 0.76 | 1 | Stc1 | stanniocalcin 1 | 400 | 773 | 598 | 6 | 0 | 5 | 2 | 5 | 12 | 0 | 0 |
| 138 | 0.03 | 0.90 | 0.03 | 0.97 | 1 | S1pr2 | sphingosine-1-phosphate receptor 2 | 709 | 365 | 328 | 4 | 23 | 16 | 2 | 50 | 3 | 9 | 4 |
| 136 | -0.42 | 0.81 | -0.52 | 0.60 | 1 | Serpinb1a | serine (or cysteine) peptidase inhibitor, cl | 380 | 630 | 404 | 18 | 15 | 9 | 2 | 0 | 4 | 13 | 19 |
| 120 | -0.24 | 2.12 | -0.11 | 0.91 | 1 | Sh3tc1 | SH3 domain and tetratricopeptide repeats | 524 | 421 | 374 | 0 | 0 | 0 | 2 | 0 | 0 | 0 | 1 |
| 108 | 0.09 | 1.75 | 0.05 | 0.96 | 1 | Sntb1 | syntrophin, basic 1 | 251 | 583 | 325 | 0 | 11 | 2 | 2 | 0 | 0 | 12 | 4 |
| 102 | -2.30 | 1.30 | -1.77 | 0.08 | 1 | Pf4 | platelet factor 4 | 589 | 213 | 311 | 0 | 4 | 2 | 2 | 0 | 0 | 0 | 1 |
| 100 | -0.70 | 3.66 | -0.19 | 0.85 | 1 | Gpnmb | glycoprotein (transmembrane) nmb | 772 | 141 | 184 | 0 | 0 | 0 | 2 | 0 | 0 | 0 | 0 |
| 96 | -1.51 | 1.42 | -1.06 | 0.29 | 1 | Nkd1 | naked cuticle 1 | 623 | 313 | 118 | 0 | 0 | 1 | 2 | 0 | 0 | 0 | 0 |
| 91 | -1.89 | 1.41 | -1.34 | 0.18 | 1 | Bmper | BMP-binding endothelial regulator | 522 | 205 | 266 | 0 | 0 | 2 | 2 | 0 | 0 | 0 | 0 |
| 90 | -2.39 | 2.42 | -0.99 | 0.32 | 1 | Ccdc3 | coiled-coil domain containing 3 | 537 | 304 | 139 | 0 | 6 | 0 | 2 | 0 | 0 | 0 | 0 |
| 79 | 0.39 | 2.54 | 0.15 | 0.88 | 1 | Cd79b | CD79B antigen | 656 | 130 | 80 | 0 | 0 | 0 | 2 | 0 | 0 | 0 | 4 |
| 63 | -0.73 | 3.66 | -0.20 | 0.84 | 1 | Xcr1 | chemokine (C motif) receptor 1 | 420 | 201 | 71 | 0 | 0 | 0 | 2 | 0 | 0 | 0 | 0 |
| 47 | -1.58 | 1.57 | -1.01 | 0.31 | 1 | Tspan11 | tetraspanin 11 | 195 | 108 | 215 | 0 | 0 | 1 | 2 | 0 | 0 | 0 | 0 |
| 46 | -0.29 | 1.06 | -0.28 | 0.78 | 1 | Mmp11 | matrix metallopeptidase 11 | 255 | 102 | 111 | 4 | 16 | 3 | 2 | 13 | 3 | 0 | 3 |
| 45 | -0.78 | 3.66 | -0.21 | 0.83 | 1 | Hoxd8 | homeobox D8 | 194 | 139 | 155 | 0 | 0 | 0 | 2 | 0 | 0 | 0 | 0 |
| 41 | -0.11 | 0.79 | -0.14 | 0.89 | 1 | Gm15506 | | 28 | 49 | 44 | 67 | 59 | 57 | 2 | 60 | 52 | 21 | 16 |
| 37 | -0.76 | 3.66 | -0.21 | 0.84 | 1 | Art4 | ADP-ribosyltransferase 4 | 183 | 58 | 162 | 0 | 0 | 0 | 2 | 0 | 0 | 0 | 0 |
| 33 | 0.63 | 0.85 | 0.74 | 0.46 | 1 | Arntl2 | aryl hydrocarbon receptor nuclear translo | 47 | 55 | 65 | 4 | 36 | 50 | 2 | 8 | 24 | 30 | 37 |
| 32 | 0.59 | 0.92 | 0.64 | 0.52 | 1 | Prdm9 | PR domain containing 9 | 28 | 28 | 10 | 29 | 29 | 53 | 2 | 12 | 31 | 64 | 62 |
| 31 | -1.75 | 1.61 | -1.09 | 0.28 | 1 | Ccl24 | chemokine (C-C motif) ligand 24 | 200 | 82 | 55 | 0 | 0 | 2 | 2 | 0 | 0 | 0 | 0 |
| 31 | -0.35 | 1.07 | -0.32 | 0.75 | 1 | Snora31 | | 4 | 58 | 30 | 4 | 5 | 96 | 2 | 0 | 11 | 0 | 125 |
| 28 | -1.26 | 1.06 | -1.19 | 0.23 | 1 | Dscc1 | DNA replication and sister chromatid coh | 77 | 44 | 30 | 46 | 22 | 39 | 2 | 10 | 11 | 0 | 30 |

| | | | | | | | | | | | | | | | | | | |
|---|---|---|---|---|---|---|---|---|---|---|---|---|---|---|---|---|---|---|
| 28 | -2.69 | 1.49 | -1.81 | 0.07 | 1 | Mslnl | mesothelin-like | 6 | 20 | 0 | 33 | 65 | 110 | 2 | 0 | 14 | 0 | 56 |
| 27 | 0.21 | 1.48 | 0.14 | 0.89 | 1 | Olfr1033 | olfactory receptor 1033 | 111 | 59 | 72 | 17 | 0 | 5 | 2 | 0 | 0 | 18 | 13 |
| 26 | 1.61 | 0.91 | 1.77 | 0.08 | 1 | AV099323 | | 34 | 9 | 47 | 21 | 8 | 21 | 2 | 57 | 39 | 21 | 24 |
| 25 | 2.03 | 1.13 | 1.80 | 0.07 | 1 | Lif | leukemia inhibitory factor | 97 | 12 | 73 | 8 | 0 | 12 | 2 | 24 | 17 | 19 | 9 |
| 24 | -0.76 | 3.66 | -0.21 | 0.83 | 1 | Atp8b4 | ATPase, class I, type 8B, member 4 | 118 | 42 | 106 | 0 | 0 | 0 | 2 | 0 | 0 | 0 | 0 |
| 24 | 0.74 | 0.89 | 0.83 | 0.41 | 1 | Gm2981 | | 34 | 15 | 32 | 8 | 39 | 30 | 2 | 28 | 30 | 26 | 17 |
| 23 | -2.03 | 1.77 | -1.15 | 0.25 | 1 | Smim38 | small integral membrane protein 38 | 44 | 114 | 75 | 11 | 0 | 0 | 2 | 0 | 0 | 0 | 3 |
| 22 | -0.79 | 3.66 | -0.22 | 0.83 | 1 | 9930111J21Rik | RIKEN cDNA 9930111J21 gene 1 | 73 | 33 | 134 | 0 | 0 | 0 | 2 | 0 | 0 | 0 | 0 |
| 20 | -0.17 | 0.92 | -0.18 | 0.85 | 1 | Gm9118 | | 9 | 77 | 55 | 18 | 26 | 4 | 2 | 3 | 15 | 4 | 11 |
| 20 | -0.71 | 1.14 | -0.62 | 0.53 | 1 | Frat2 | frequently rearranged in advanced T cell | 76 | 27 | 55 | 4 | 12 | 14 | 2 | 0 | 2 | 23 | 7 |
| 20 | 0.00 | 1.09 | 0.00 | 1.00 | 1 | Teddm2 | transmembrane epididymal family membe | 18 | 61 | 35 | 16 | 7 | 36 | 2 | 0 | 16 | 8 | 23 |
| 17 | 2.41 | 1.42 | 1.69 | 0.09 | 1 | Adam1a | a disintegrin and metallopeptidase domai | 28 | 27 | 15 | 0 | 0 | 42 | 2 | 0 | 9 | 10 | 53 |
| 17 | 0.95 | 1.00 | 0.95 | 0.34 | 1 | Gm11400 | | 8 | 8 | 5 | 8 | 6 | 58 | 2 | 43 | 14 | 3 | 28 |
| 16 | -2.46 | 1.42 | -1.73 | 0.08 | 1 | F630042J09Rik | | 21 | 66 | 35 | 16 | 24 | 1 | 2 | 0 | 11 | 0 | 0 |
| 15 | 2.28 | 1.80 | 1.26 | 0.21 | 1 | Gm42528 | | 44 | 34 | 76 | 0 | 0 | 0 | 2 | 0 | 6 | 0 | 7 |
| 15 | -0.93 | 1.17 | -0.80 | 0.42 | 1 | B930059L03Rik | | 23 | 10 | 10 | 18 | 14 | 31 | 2 | 1 | 3 | 39 | 15 |
| 15 | 1.73 | 1.75 | 0.98 | 0.32 | 1 | Gm30122 | | 9 | 0 | 0 | 0 | 6 | 74 | 2 | 5 | 18 | 0 | 53 |
| 14 | 0.12 | 1.07 | 0.11 | 0.91 | 1 | Gm46336 | | 0 | 10 | 1 | 20 | 45 | 8 | 2 | 21 | 40 | 5 | 5 |
| 14 | -0.93 | 1.29 | -0.72 | 0.47 | 1 | Gm49032 | | 32 | 32 | 12 | 8 | 4 | 25 | 2 | 0 | 5 | 0 | 37 |
| 14 | -2.26 | 1.52 | -1.49 | 0.14 | 1 | Kcnmb4 | potassium large conductance calcium-act | 15 | 63 | 47 | 0 | 12 | 7 | 2 | 0 | 1 | 0 | 4 |
| 13 | -1.33 | 1.34 | -0.99 | 0.32 | 1 | Cfap52 | cilia and flagella associated protein 52 | 5 | 42 | 60 | 9 | 2 | 8 | 2 | 0 | 3 | 0 | 16 |
| 13 | 1.47 | 0.90 | 1.63 | 0.10 | 1 | Gm50341 | | 4 | 4 | 9 | 7 | 9 | 24 | 2 | 27 | 22 | 15 | 22 |
| 13 | -2.40 | 2.15 | -1.12 | 0.26 | 1 | Gm28043 | | 0 | 0 | 0 | 0 | 8 | 74 | 2 | 0 | 0 | 0 | 57 |
| 13 | -0.73 | 1.87 | -0.39 | 0.70 | 1 | Gnat3 | guanine nucleotide binding protein, alpha | 42 | 11 | 22 | 11 | 3 | 0 | 2 | 0 | 0 | 46 | 0 |
| 12 | -0.67 | 1.48 | -0.45 | 0.65 | 1 | Gm38228 | | 7 | 0 | 5 | 0 | 15 | 51 | 2 | 0 | 8 | 0 | 48 |
| 12 | -0.20 | 1.32 | -0.16 | 0.88 | 1 | Gm42715 | | 5 | 16 | 0 | 17 | 11 | 22 | 2 | 19 | 8 | 0 | 35 |
| 12 | -0.03 | 1.27 | -0.02 | 0.98 | 1 | Apobec2 | apolipoprotein B mRNA editing enzyme, c | 15 | 7 | 20 | 6 | 14 | 22 | 2 | 31 | 11 | 0 | 6 |
| 12 | -4.34 | 2.21 | -1.97 | 0.05 | 1 | Gm14817 | | 8 | 0 | 0 | 12 | 0 | 67 | 2 | 0 | 0 | 0 | 40 |
| 11 | 1.12 | 1.75 | 0.64 | 0.52 | 1 | Gm21986 | | 0 | 44 | 0 | 11 | 4 | 20 | 2 | 0 | 10 | 6 | 29 |
| 11 | -1.24 | 1.81 | -0.68 | 0.49 | 1 | Gm20324 | | 24 | 0 | 44 | 13 | 9 | 9 | 2 | 17 | 0 | 0 | 7 |
| 11 | -2.95 | 1.83 | -1.61 | 0.11 | 1 | Mfsd7a | major facilitator superfamily domain conta | 58 | 2 | 50 | 0 | 4 | 3 | 2 | 0 | 0 | 0 | 0 |
| 10 | -0.74 | 1.93 | -0.38 | 0.70 | 1 | Rph3a | rabphilin 3A | 0 | 0 | 10 | 24 | 2 | 12 | 2 | 0 | 13 | 48 | 4 |
| 10 | -3.82 | 2.21 | -1.73 | 0.08 | 1 | Gm26542 | | 13 | 7 | 46 | 16 | 0 | 13 | 2 | 0 | 0 | 17 | 0 |
| 10 | -1.44 | 1.81 | -0.79 | 0.43 | 1 | 2810408I11Rik | | 11 | 8 | 29 | 32 | 0 | 9 | 2 | 0 | 9 | 0 | 12 |
| 10 | -0.12 | 1.58 | -0.07 | 0.94 | 1 | Pla2g4c | phospholipase A2, group IVC (cytosolic, c | 27 | 27 | 17 | 0 | 3 | 16 | 2 | 0 | 8 | 0 | 11 |
| 10 | 0.55 | 1.22 | 0.45 | 0.65 | 1 | Gm38293 | | 11 | 9 | 0 | 30 | 6 | 4 | 2 | 14 | 8 | 18 | 8 |
| 10 | 2.14 | 1.33 | 1.61 | 0.11 | 1 | Plch2 | phospholipase C, eta 2 | 13 | 3 | 15 | 5 | 0 | 9 | 2 | 36 | 9 | 8 | 7 |
| 10 | 2.23 | 1.34 | 1.67 | 0.10 | 1 | Pmfbp1 | polyamine modulated factor 1 binding pro | 0 | 3 | 4 | 3 | 4 | 13 | 2 | 37 | 12 | 26 | 3 |
| 10 | -0.75 | 1.92 | -0.39 | 0.70 | 1 | Cited1 | Cbp/p300-interacting transactivator with C | 31 | 37 | 29 | 0 | 0 | 3 | 2 | 0 | 0 | 0 | 3 |
| 10 | -0.50 | 1.45 | -0.35 | 0.73 | 1 | Gm14276 | | 19 | 15 | 4 | 5 | 11 | 14 | 2 | 25 | 8 | 0 | 1 |
| 9 | -0.68 | 1.81 | -0.38 | 0.71 | 1 | E030022I16Rik | | 2 | 9 | 4 | 0 | 0 | 57 | 2 | 0 | 1 | 0 | 29 |
| 9 | 0.50 | 1.64 | 0.31 | 0.76 | 1 | Gm9725 | | 2 | 1 | 17 | 0 | 11 | 21 | 2 | 29 | 4 | 0 | 15 |
| 9 | -0.48 | 1.84 | -0.26 | 0.79 | 1 | Gm48627 | | 4 | 0 | 5 | 0 | 0 | 55 | 2 | 0 | 1 | 0 | 34 |
| 9 | 1.99 | 1.74 | 1.14 | 0.25 | 1 | 1700087I21Rik | | 1 | 0 | 8 | 0 | 13 | 8 | 2 | 28 | 6 | 0 | 35 |
| 9 | -0.20 | 1.71 | -0.12 | 0.91 | 1 | Slfn10-ps | | 15 | 12 | 25 | 11 | 0 | 7 | 2 | 19 | 0 | 0 | 9 |
| 9 | 2.56 | 1.55 | 1.65 | 0.10 | 1 | Gm22965 | | 26 | 2 | 10 | 0 | 3 | 6 | 2 | 14 | 0 | 22 | 15 |
| 9 | -0.04 | 1.32 | -0.03 | 0.98 | 1 | Synpo2l | synaptopodin 2-like | 24 | 1 | 14 | 17 | 5 | 5 | 2 | 3 | 16 | 4 | 7 |
| 9 | 1.67 | 1.13 | 1.47 | 0.14 | 1 | Gm8337 | | 7 | 2 | 7 | 1 | 6 | 10 | 2 | 2 | 11 | 36 | 12 |
| 9 | 1.98 | 1.56 | 1.27 | 0.20 | 1 | Gm43568 | | 3 | 25 | 18 | 0 | 3 | 3 | 2 | 16 | 12 | 13 | 0 |
| 9 | 0.97 | 1.31 | 0.74 | 0.46 | 1 | Gm5791 | | 12 | 0 | 17 | 2 | 6 | 6 | 2 | 1 | 7 | 36 | 7 |
| 9 | 0.51 | 1.39 | 0.37 | 0.71 | 1 | Rpl32l | ribosomal protein L32-like | 9 | 9 | 4 | 10 | 14 | 1 | 2 | 2 | 1 | 37 | 6 |
| 9 | 0.02 | 1.65 | 0.01 | 0.99 | 1 | Mccc1os | | 10 | 0 | 0 | 2 | 10 | 9 | 2 | 0 | 22 | 35 | 4 |
| 9 | 0.17 | 2.03 | 0.09 | 0.93 | 1 | Lef1 | lymphoid enhancer binding factor 1 | 59 | 13 | 5 | 0 | 0 | 3 | 2 | 0 | 0 | 12 | 0 |
| 8 | 1.93 | 1.24 | 1.55 | 0.12 | 1 | Gm12504 | | 24 | 10 | 22 | 2 | 2 | 1 | 2 | 3 | 12 | 1 | 12 |
| 8 | -2.43 | 2.81 | -0.87 | 0.39 | 1 | Gm45591 | | 0 | 0 | 10 | 0 | 0 | 39 | 2 | 0 | 0 | 0 | 40 |
| 8 | 0.71 | 2.12 | 0.33 | 0.74 | 1 | Tnfsf13os | | 0 | 8 | 19 | 0 | 2 | 7 | 2 | 37 | 0 | 0 | 16 |
| 8 | -0.72 | 1.37 | -0.53 | 0.60 | 1 | Gm7390 | | 9 | 2 | 0 | 4 | 17 | 20 | 2 | 1 | 5 | 15 | 14 |
| 8 | -1.27 | 1.56 | -0.82 | 0.41 | 1 | Gm7846 | | 18 | 16 | 9 | 3 | 9 | 13 | 2 | 15 | 0 | 0 | 5 |
| 8 | -0.46 | 1.54 | -0.30 | 0.77 | 1 | Gm44036 | | 17 | 12 | 8 | 2 | 13 | 6 | 2 | 0 | 19 | 0 | 9 |
| 8 | 0.76 | 1.63 | 0.47 | 0.64 | 1 | Gm47905 | | 0 | 0 | 0 | 19 | 1 | 10 | 2 | 27 | 13 | 0 | 16 |

| | | | | | | | | | | | | | | | | | | |
|---|---|---|---|---|---|---|---|---|---|---|---|---|---|---|---|---|---|---|
| 8 | -0.91 | 1.56 | -0.58 | 0.56 | 1 | Gm37660 | | 5 | 27 | 14 | 1 | 15 | 5 | 2 | 0 | 15 | 0 | 4 |
| 8 | -0.18 | 1.49 | -0.12 | 0.90 | 1 | Gm47270 | | 6 | 4 | 20 | 0 | 5 | 30 | 2 | 1 | 13 | 0 | 7 |
| 8 | 1.23 | 1.43 | 0.86 | 0.39 | 1 | Gm28592 | | 7 | 0 | 1 | 4 | 5 | 9 | 2 | 38 | 7 | 10 | 3 |
| 8 | -1.61 | 1.66 | -0.97 | 0.33 | 1 | Gm10548 | | 12 | 6 | 29 | 18 | 1 | 5 | 2 | 0 | 4 | 0 | 10 |
| 8 | 1.06 | 2.01 | 0.53 | 0.60 | 1 | Gm26533 | | 0 | 13 | 19 | 7 | 10 | 2 | 2 | 0 | 0 | 21 | 12 |
| 8 | 1.17 | 1.29 | 0.91 | 0.36 | 1 | Rps4x-ps | | 17 | 42 | 5 | 1 | 2 | 2 | 2 | 2 | 4 | 3 | 6 |
| 8 | 0.19 | 1.57 | 0.12 | 0.90 | 1 | Gm36355 | | 0 | 13 | 3 | 15 | 2 | 24 | 2 | 0 | 7 | 5 | 14 |
| 8 | -0.11 | 1.73 | -0.06 | 0.95 | 1 | Gm2308 | | 0 | 1 | 20 | 9 | 7 | 8 | 2 | 0 | 8 | 27 | 4 |
| 7 | 0.54 | 1.26 | 0.43 | 0.67 | 1 | Rps27a-ps2 | | 16 | 9 | 10 | 9 | 1 | 7 | 2 | 1 | 5 | 14 | 7 |
| 7 | -3.17 | 1.93 | -1.64 | 0.10 | 1 | Vmn2r57 | vomeronasal 2, receptor 57 | 0 | 0 | 17 | 28 | 8 | 10 | 2 | 0 | 10 | 0 | 6 |
| 7 | 0.31 | 1.22 | 0.25 | 0.80 | 1 | Gm7808 | | 8 | 19 | 11 | 3 | 9 | 3 | 2 | 6 | 3 | 1 | 15 |
| 7 | 2.85 | 2.20 | 1.29 | 0.20 | 1 | Mroh6 | maestro heat-like repeat family member 6 | 0 | 17 | 0 | 10 | 0 | 14 | 2 | 0 | 11 | 22 | 4 |
| 7 | -2.49 | 1.87 | -1.34 | 0.18 | 1 | Gm45552 | | 26 | 18 | 21 | 0 | 5 | 5 | 2 | 0 | 0 | 0 | 2 |
| 7 | 1.36 | 1.41 | 0.97 | 0.33 | 1 | Dnajc19-ps | | 15 | 10 | 2 | 4 | 3 | 3 | 2 | 0 | 7 | 24 | 8 |
| 7 | 1.71 | 2.74 | 0.62 | 0.53 | 1 | Chia1 | chitinase, acidic 1 | 0 | 0 | 0 | 0 | 0 | 13 | 2 | 0 | 0 | 36 | 25 |
| 7 | 0.84 | 1.18 | 0.71 | 0.48 | 1 | Gm5881 | | 4 | 9 | 6 | 2 | 6 | 11 | 2 | 1 | 13 | 5 | 16 |
| 7 | 0.83 | 2.22 | 0.37 | 0.71 | 1 | Gm26797 | | 0 | 0 | 0 | 0 | 0 | 41 | 2 | 0 | 2 | 3 | 28 |
| 7 | -1.07 | 3.15 | -0.34 | 0.73 | 1 | Pla2g4b | phospholipase A2, group IVB (cytosolic) | 0 | 0 | 0 | 0 | 0 | 47 | 2 | 0 | 0 | 0 | 25 |
| 7 | 3.59 | 2.50 | 1.44 | 0.15 | 1 | Tssk6 | testis-specific serine kinase 6 | 0 | 0 | 8 | 0 | 6 | 1 | 2 | 16 | 0 | 41 | 0 |
| 7 | -0.70 | 1.58 | -0.44 | 0.66 | 1 | Mboat4 | membrane bound O-acyltransferase dom | 15 | 15 | 11 | 9 | 4 | 3 | 2 | 0 | 7 | 0 | 8 |
| 7 | 0.65 | 1.78 | 0.37 | 0.71 | 1 | Gm1866 | | 22 | 0 | 20 | 0 | 12 | 1 | 2 | 1 | 11 | 0 | 2 |
| 6 | -0.44 | 1.31 | -0.34 | 0.74 | 1 | Gm8934 | | 11 | 18 | 7 | 2 | 8 | 6 | 2 | 12 | 1 | 3 | 2 |
| 6 | -0.40 | 1.22 | -0.33 | 0.74 | 1 | Gm6035 | | 3 | 6 | 19 | 14 | 5 | 3 | 2 | 2 | 11 | 3 | 3 |
| 6 | -2.75 | 1.69 | -1.63 | 0.10 | 1 | H4c2 | H4 clustered histone 2 | 2 | 0 | 6 | 10 | 15 | 20 | 2 | 0 | 1 | 12 | 2 |
| 6 | 0.41 | 1.83 | 0.22 | 0.82 | 1 | Gm50321 | | 7 | 0 | 0 | 0 | 8 | 17 | 2 | 0 | 7 | 17 | 12 |
| 6 | 0.36 | 1.66 | 0.21 | 0.83 | 1 | Taco1os | | 1 | 0 | 27 | 2 | 8 | 6 | 2 | 12 | 5 | 0 | 7 |
| 6 | 2.66 | 1.92 | 1.39 | 0.17 | 1 | Gm2199 | | 3 | 0 | 0 | 2 | 3 | 0 | 2 | 0 | 17 | 33 | 8 |
| 6 | -0.73 | 2.49 | -0.29 | 0.77 | 1 | Rsph14 | radial spoke head homolog 14 (Chlamyd | 26 | 17 | 8 | 0 | 0 | 3 | 2 | 0 | 12 | 0 | 0 |
| 6 | -1.34 | 2.14 | -0.63 | 0.53 | 1 | Gm13080 | | 9 | 0 | 0 | 1 | 15 | 3 | 2 | 0 | 0 | 14 | 22 |
| 6 | -4.79 | 2.62 | -1.83 | 0.07 | 1 | Gm44270 | | 0 | 6 | 0 | 0 | 3 | 38 | 2 | 0 | 0 | 0 | 18 |
| 6 | 2.52 | 2.23 | 1.13 | 0.26 | 1 | Gm49503 | | 8 | 0 | 0 | 0 | 8 | 3 | 2 | 21 | 20 | 0 | 5 |
| 6 | 3.44 | 3.07 | 1.12 | 0.26 | 1 | Frmd3 | FERM domain containing 3 | 13 | 0 | 20 | 0 | 0 | 0 | 2 | 0 | 0 | 31 | 0 |
| 6 | 1.77 | 2.52 | 0.70 | 0.48 | 1 | Gm15409 | | 0 | 0 | 5 | 7 | 0 | 0 | 2 | 34 | 2 | 0 | 15 |
| 6 | 1.09 | 1.24 | 0.88 | 0.38 | 1 | Gm24951 | | 6 | 4 | 2 | 7 | 2 | 3 | 2 | 28 | 4 | 3 | 4 |
| 6 | -1.41 | 1.89 | -0.75 | 0.45 | 1 | Gm38227 | | 0 | 7 | 14 | 9 | 10 | 10 | 2 | 0 | 0 | 4 | 9 |
| 6 | -0.53 | 2.51 | -0.21 | 0.83 | 1 | Gm37785 | | 19 | 15 | 0 | 0 | 0 | 11 | 2 | 0 | 0 | 0 | 16 |
| 6 | -1.12 | 2.11 | -0.53 | 0.59 | 1 | 1700105P06Rik | | 0 | 0 | 0 | 8 | 11 | 8 | 2 | 0 | 28 | 0 | 6 |
| 6 | 0.16 | 1.81 | 0.09 | 0.93 | 1 | Nkpd1 | NTPase, KAP family P-loop domain conta | 15 | 10 | 4 | 0 | 4 | 6 | 2 | 0 | 6 | 0 | 15 |
| 6 | 0.10 | 1.51 | 0.07 | 0.95 | 1 | Rpl23a-ps3 | | 6 | 5 | 26 | 3 | 1 | 3 | 2 | 1 | 11 | 5 | 0 |
| 6 | -0.62 | 1.62 | -0.39 | 0.70 | 1 | Rpl17-ps3 | | 13 | 15 | 15 | 6 | 0 | 4 | 2 | 0 | 3 | 1 | 3 |
| 6 | -1.78 | 2.13 | -0.84 | 0.40 | 1 | Gm8894 | | 15 | 0 | 4 | 27 | 5 | 1 | 2 | 0 | 5 | 0 | 3 |
| 6 | -3.35 | 1.95 | -1.71 | 0.09 | 1 | Ptp4a1 | protein tyrosine phosphatase 4a1 | 1 | 1 | 4 | 8 | 6 | 15 | 2 | 0 | 0 | 0 | 24 |
| 6 | 0.70 | 1.53 | 0.46 | 0.65 | 1 | Gm6030 | | 11 | 2 | 2 | 2 | 1 | 16 | 2 | 2 | 8 | 0 | 14 |
| 6 | 1.24 | 1.75 | 0.71 | 0.48 | 1 | Gm3788 | | 15 | 16 | 0 | 0 | 3 | 3 | 2 | 9 | 5 | 1 | 5 |
| 5 | 1.17 | 2.71 | 0.43 | 0.67 | 1 | Gm49600 | | 4 | 0 | 0 | 0 | 0 | 30 | 2 | 0 | 8 | 0 | 16 |
| 5 | 2.10 | 2.21 | 0.95 | 0.34 | 1 | Gm21691 | | 0 | 8 | 10 | 0 | 0 | 8 | 2 | 0 | 2 | 12 | 18 |
| 5 | 1.37 | 1.55 | 0.88 | 0.38 | 1 | 4930449I04Rik | | 3 | 14 | 0 | 9 | 0 | 8 | 2 | 3 | 2 | 13 | 6 |
| 5 | 0.35 | 2.02 | 0.17 | 0.86 | 1 | Pitpnm2os1 | | 4 | 9 | 0 | 0 | 9 | 10 | 2 | 0 | 5 | 12 | 7 |
| 5 | -0.94 | 1.50 | -0.62 | 0.53 | 1 | 4930583K01Rik | | 4 | 17 | 5 | 11 | 5 | 1 | 2 | 5 | 1 | 0 | 7 |
| 5 | 1.38 | 2.02 | 0.68 | 0.49 | 1 | Tfr2 | transferrin receptor 2 | 5 | 0 | 16 | 0 | 5 | 3 | 2 | 19 | 4 | 0 | 4 |
| 5 | -0.73 | 2.23 | -0.33 | 0.74 | 1 | Far2 | fatty acyl CoA reductase 2 | 16 | 21 | 2 | 0 | 8 | 0 | 2 | 7 | 1 | 0 | 0 |
| 5 | 0.14 | 1.46 | 0.10 | 0.92 | 1 | Rpsa-ps7 | | 9 | 5 | 16 | 4 | 0 | 8 | 2 | 1 | 5 | 3 | 5 |
| 5 | 0.84 | 1.52 | 0.55 | 0.58 | 1 | Gm8258 | | 3 | 0 | 1 | 9 | 1 | 2 | 2 | 2 | 2 | 28 | 5 |
| 5 | -0.37 | 1.34 | -0.28 | 0.78 | 1 | Gm8797 | | 7 | 4 | 8 | 11 | 2 | 5 | 2 | 1 | 4 | 3 | 8 |
| 5 | 1.47 | 2.23 | 0.66 | 0.51 | 1 | Gm15801 | | 0 | 0 | 1 | 1 | 9 | 1 | 2 | 0 | 0 | 35 | 7 |
| 5 | 1.36 | 1.51 | 0.91 | 0.37 | 1 | Gm12091 | | 3 | 4 | 2 | 1 | 1 | 3 | 2 | 32 | 2 | 1 | 2 |
| 5 | 2.53 | 2.29 | 1.10 | 0.27 | 1 | Gm15899 | | 20 | 13 | 0 | 9 | 0 | 0 | 2 | 2 | 4 | 0 | 4 |
| 5 | -0.91 | 1.97 | -0.46 | 0.65 | 1 | A930005G22Rik | | 5 | 6 | 8 | 4 | 9 | 0 | 2 | 0 | 6 | 0 | 14 |
| 5 | -0.33 | 1.59 | -0.21 | 0.83 | 1 | Gm6851 | | 9 | 14 | 5 | 0 | 10 | 3 | 2 | 3 | 5 | 1 | 1 |
| 5 | -1.96 | 1.85 | -1.06 | 0.29 | 1 | Gm20753 | | 0 | 12 | 2 | 4 | 13 | 9 | 2 | 0 | 7 | 0 | 3 |

| | | | | | | | | | | | | | | | | | | |
|---|---|---|---|---|---|---|---|---|---|---|---|---|---|---|---|---|---|---|
| 5 | -0.17 | 1.74 | -0.10 | 0.92 | 1 | Gm26810 | | 3 | 2 | 5 | 0 | 9 | 11 | 2 | 13 | 3 | 0 | 5 |
| 5 | -1.51 | 2.08 | -0.73 | 0.47 | 1 | Ppp1r14d | protein phosphatase 1, regulatory inhibitc | 12 | 8 | 0 | 0 | 15 | 8 | 2 | 2 | 2 | 0 | 3 |
| 5 | 2.13 | 2.11 | 1.01 | 0.31 | 1 | Gm5507 | | 2 | 1 | 4 | 0 | 11 | 0 | 2 | 28 | 0 | 4 | 1 |
| 5 | -0.44 | 1.62 | -0.27 | 0.79 | 1 | Gm11993 | | 1 | 5 | 17 | 4 | 0 | 11 | 2 | 1 | 7 | 1 | 4 |
| 5 | -2.54 | 2.40 | -1.06 | 0.29 | 1 | Gm12816 | | 27 | 19 | 1 | 0 | 1 | 3 | 2 | 0 | 0 | 0 | 0 |
| 5 | -1.03 | 2.98 | -0.35 | 0.73 | 1 | A530083I20Rik | | 0 | 33 | 0 | 0 | 0 | 12 | 2 | 0 | 1 | 0 | 4 |
| 5 | -0.03 | 1.48 | -0.02 | 0.99 | 1 | Txn-ps1 | | 11 | 3 | 1 | 15 | 1 | 4 | 2 | 2 | 6 | 3 | 4 |
| 5 | -0.39 | 2.76 | -0.14 | 0.89 | 1 | Gm37795 | | 0 | 0 | 10 | 0 | 0 | 14 | 2 | 0 | 9 | 0 | 16 |
| 5 | -0.30 | 2.09 | -0.14 | 0.89 | 1 | 4930422I22Rik | | 0 | 0 | 0 | 9 | 6 | 3 | 2 | 0 | 1 | 28 | 3 |
| 5 | -0.58 | 1.43 | -0.41 | 0.68 | 1 | Gm44126 | | 5 | 1 | 14 | 4 | 2 | 10 | 2 | 1 | 1 | 3 | 9 |
| 5 | 1.22 | 2.26 | 0.54 | 0.59 | 1 | Gm42735 | | 7 | 0 | 3 | 0 | 0 | 10 | 2 | 0 | 0 | 17 | 12 |
| 5 | -0.08 | 1.62 | -0.05 | 0.96 | 1 | Gm5047 | | 8 | 7 | 4 | 0 | 5 | 8 | 2 | 1 | 1 | 12 | 2 |
| 5 | -0.81 | 1.60 | -0.51 | 0.61 | 1 | Rpl7a-ps10 | | 15 | 2 | 16 | 3 | 2 | 4 | 2 | 3 | 2 | 1 | 0 |
| 4 | -0.33 | 2.57 | -0.13 | 0.90 | 1 | Gm44639 | | 0 | 0 | 0 | 0 | 2 | 21 | 2 | 0 | 0 | 4 | 20 |
| 4 | 1.04 | 1.32 | 0.79 | 0.43 | 1 | Gm12933 | | 2 | 3 | 2 | 3 | 3 | 3 | 2 | 21 | 4 | 1 | 5 |
| 4 | -1.14 | 1.60 | -0.71 | 0.48 | 1 | Gm13192 | | 7 | 8 | 16 | 1 | 4 | 6 | 2 | 3 | 1 | 1 | 0 |
| 4 | -5.21 | 2.86 | -1.82 | 0.07 | 1 | Gm44597 | | 0 | 2 | 0 | 0 | 8 | 19 | 2 | 0 | 0 | 0 | 18 |
| 4 | -0.45 | 1.52 | -0.29 | 0.77 | 1 | Gm8784 | | 1 | 2 | 23 | 2 | 7 | 3 | 2 | 1 | 1 | 1 | 6 |
| 4 | -1.31 | 1.61 | -0.81 | 0.42 | 1 | Gm45201 | | 0 | 1 | 0 | 14 | 8 | 8 | 2 | 6 | 5 | 0 | 4 |
| 4 | -0.93 | 1.85 | -0.50 | 0.62 | 1 | Gm44696 | | 12 | 5 | 1 | 6 | 6 | 3 | 2 | 0 | 10 | 0 | 3 |
| 4 | -1.19 | 2.37 | -0.50 | 0.62 | 1 | Gm50209 | | 0 | 3 | 10 | 0 | 2 | 8 | 2 | 17 | 5 | 0 | 0 |
| 4 | -0.67 | 1.48 | -0.46 | 0.65 | 1 | Gm43712 | | 10 | 3 | 10 | 7 | 1 | 4 | 2 | 3 | 3 | 1 | 1 |
| 4 | -2.06 | 1.68 | -1.23 | 0.22 | 1 | Gm49218 | | 0 | 0 | 1 | 16 | 6 | 13 | 2 | 3 | 2 | 0 | 4 |
| 4 | 2.59 | 2.81 | 0.92 | 0.36 | 1 | Gm38563 | | 0 | 0 | 0 | 0 | 0 | 9 | 2 | 15 | 4 | 0 | 16 |
| 4 | 1.65 | 1.61 | 1.03 | 0.30 | 1 | Gm37108 | | 20 | 5 | 4 | 0 | 2 | 1 | 2 | 3 | 2 | 1 | 6 |
| 4 | -0.01 | 1.97 | -0.01 | 1.00 | 1 | Atp5l2-ps | | 8 | 1 | 1 | 8 | 5 | 1 | 2 | 0 | 14 | 0 | 7 |
| 4 | -1.14 | 1.81 | -0.63 | 0.53 | 1 | Gm37025 | | 0 | 6 | 26 | 1 | 4 | 3 | 2 | 1 | 1 | 0 | 3 |
| 4 | 0.40 | 1.26 | 0.32 | 0.75 | 1 | Gm8880 | | 8 | 3 | 8 | 2 | 4 | 4 | 2 | 2 | 4 | 4 | 5 |
| 4 | 1.22 | 3.21 | 0.38 | 0.70 | 1 | Hmgb1-rs16 | | 0 | 0 | 0 | 0 | 0 | 11 | 2 | 0 | 0 | 22 | 11 |
| 4 | 0.08 | 1.71 | 0.05 | 0.96 | 1 | Gm2694 | | 1 | 1 | 0 | 9 | 4 | 4 | 2 | 0 | 3 | 15 | 6 |
| 4 | -0.94 | 3.59 | -0.26 | 0.79 | 1 | Mir7-1 | | 0 | 0 | 0 | 0 | 0 | 25 | 2 | 0 | 0 | 0 | 18 |
| 4 | -2.20 | 1.69 | -1.30 | 0.19 | 1 | Gm9134 | | 1 | 5 | 0 | 13 | 8 | 7 | 2 | 3 | 0 | 1 | 4 |
| 4 | 0.71 | 2.70 | 0.26 | 0.79 | 1 | 9430002A10Rik | | 0 | 16 | 0 | 3 | 0 | 6 | 2 | 0 | 5 | 0 | 12 |
| 4 | -1.20 | 2.27 | -0.53 | 0.60 | 1 | Gm9429 | | 0 | 0 | 0 | 1 | 12 | 10 | 2 | 0 | 10 | 0 | 8 |
| 4 | -3.38 | 2.83 | -1.19 | 0.23 | 1 | H2bc15 | H2B clustered histone 15 | 0 | 0 | 0 | 8 | 4 | 14 | 2 | 0 | 0 | 0 | 16 |
| 4 | -0.01 | 1.77 | -0.01 | 1.00 | 1 | Gm10129 | | 4 | 0 | 5 | 5 | 6 | 2 | 2 | 7 | 11 | 0 | 1 |
| 4 | 0.49 | 3.20 | 0.15 | 0.88 | 1 | Gm39041 | | 0 | 0 | 0 | 0 | 0 | 17 | 2 | 0 | 0 | 4 | 20 |
| 4 | -3.67 | 2.04 | -1.80 | 0.07 | 1 | Gm9923 | | 5 | 0 | 1 | 19 | 6 | 4 | 2 | 1 | 0 | 4 | 0 |
| 4 | -0.84 | 3.05 | -0.27 | 0.78 | 1 | Gm15688 | | 0 | 17 | 16 | 0 | 0 | 3 | 2 | 0 | 0 | 0 | 4 |
| 4 | 0.26 | 1.70 | 0.15 | 0.88 | 1 | Gm20568 | | 3 | 2 | 11 | 2 | 1 | 3 | 2 | 1 | 3 | 13 | 0 |
| 4 | -0.33 | 1.67 | -0.20 | 0.84 | 1 | M6pr-ps | | 6 | 2 | 7 | 9 | 5 | 0 | 2 | 1 | 2 | 3 | 5 |
| 4 | 1.50 | 1.75 | 0.86 | 0.39 | 1 | Gm10356 | | 6 | 1 | 4 | 1 | 0 | 3 | 2 | 0 | 4 | 15 | 4 |
| 4 | -2.25 | 2.28 | -0.99 | 0.32 | 1 | 4930473A02Rik | | 8 | 0 | 0 | 4 | 6 | 2 | 2 | 0 | 0 | 15 | 4 |
| 4 | 0.33 | 1.72 | 0.19 | 0.85 | 1 | Gm4596 | | 9 | 1 | 8 | 1 | 1 | 7 | 2 | 2 | 6 | 3 | 0 |
| 4 | 0.45 | 2.00 | 0.22 | 0.82 | 1 | 4933428P19Rik | | 0 | 0 | 0 | 10 | 2 | 2 | 2 | 15 | 4 | 0 | 5 |
| 4 | 0.41 | 2.05 | 0.20 | 0.84 | 1 | Gm8741 | | 11 | 5 | 6 | 2 | 0 | 2 | 2 | 5 | 0 | 0 | 6 |
| 3 | -1.04 | 3.59 | -0.29 | 0.77 | 1 | Mir7b | | 0 | 0 | 0 | 0 | 0 | 23 | 2 | 0 | 0 | 0 | 13 |
| 3 | -0.35 | 1.42 | -0.24 | 0.81 | 1 | Gm2451 | | 7 | 3 | 3 | 4 | 5 | 3 | 2 | 1 | 2 | 3 | 6 |
| 3 | -0.17 | 2.09 | -0.08 | 0.94 | 1 | Gm11599 | | 3 | 21 | 3 | 0 | 3 | 0 | 2 | 3 | 1 | 1 | 0 |
| 3 | -1.51 | 1.85 | -0.82 | 0.41 | 1 | Gm6440 | | 10 | 5 | 2 | 4 | 2 | 2 | 2 | 1 | 0 | 9 | 0 |
| 3 | 1.24 | 3.03 | 0.41 | 0.68 | 1 | Gm15232 | | 9 | 3 | 4 | 0 | 0 | 0 | 2 | 0 | 0 | 18 | 0 |
| 3 | -3.59 | 2.66 | -1.35 | 0.18 | 1 | Gm26805 | | 18 | 3 | 1 | 0 | 4 | 8 | 2 | 0 | 0 | 0 | 0 |
| 3 | 0.79 | 3.48 | 0.23 | 0.82 | 1 | Gm12958 | | 0 | 0 | 0 | 0 | 9 | 0 | 2 | 0 | 4 | 21 | 0 |
| 3 | -0.26 | 1.80 | -0.14 | 0.89 | 1 | Gm5910 | | 12 | 3 | 0 | 2 | 7 | 1 | 2 | 1 | 3 | 1 | 3 |
| 3 | -0.76 | 3.57 | -0.21 | 0.83 | 1 | Gm45159 | | 15 | 13 | 4 | 0 | 0 | 0 | 2 | 0 | 0 | 0 | 0 |
| 3 | 1.10 | 3.05 | 0.36 | 0.72 | 1 | Gm17111 | | 0 | 0 | 0 | 0 | 13 | 0 | 2 | 10 | 8 | 0 | 1 |
| 3 | -4.10 | 3.51 | -1.17 | 0.24 | 1 | Spata9 | spermatogenesis associated 9 | 0 | 10 | 0 | 0 | 6 | 3 | 2 | 0 | 0 | 14 | 0 |
| 3 | -1.78 | 2.80 | -0.63 | 0.53 | 1 | Gm4974 | | 20 | 0 | 0 | 8 | 0 | 3 | 2 | 1 | 1 | 0 | 0 |
| 3 | -0.39 | 2.01 | -0.20 | 0.84 | 1 | Rpl7-ps8 | | 11 | 2 | 2 | 0 | 10 | 1 | 2 | 2 | 2 | 0 | 1 |
| 3 | -3.57 | 2.66 | -1.34 | 0.18 | 1 | 2010016I18Rik | | 0 | 0 | 8 | 6 | 5 | 9 | 2 | 0 | 0 | 0 | 3 |

| | | | | | | | | | | | | | | | | | | |
|---|---|---|---|---|---|---|---|---|---|---|---|---|---|---|---|---|---|---|
| 3 | -0.22 | 1.69 | -0.13 | 0.90 | 1 | Gm6159 | | 11 | 4 | 1 | 3 | 3 | 1 | 2 | 2 | 4 | 3 | 0 |
| 3 | -1.84 | 2.53 | -0.73 | 0.47 | 1 | Gm15890 | | 2 | 6 | 0 | 0 | 16 | 3 | 2 | 0 | 3 | 0 | 2 |
| 3 | -2.62 | 3.03 | -0.86 | 0.39 | 1 | Gm11575 | | 0 | 0 | 17 | 7 | 0 | 2 | 2 | 1 | 0 | 0 | 4 |
| 3 | -2.96 | 3.06 | -0.97 | 0.33 | 1 | Gm44361 | | 0 | 0 | 0 | 10 | 1 | 7 | 2 | 0 | 0 | 0 | 13 |
| 3 | 1.34 | 1.62 | 0.83 | 0.41 | 1 | Gm9013 | | 8 | 5 | 1 | 2 | 0 | 3 | 2 | 5 | 2 | 3 | 3 |
| 3 | -3.28 | 2.16 | -1.52 | 0.13 | 1 | Gm7986 | | 9 | 0 | 2 | 4 | 8 | 4 | 2 | 0 | 0 | 1 | 2 |
| 3 | -0.96 | 2.67 | -0.36 | 0.72 | 1 | Gm42991 | | 8 | 11 | 1 | 0 | 0 | 6 | 2 | 0 | 0 | 0 | 4 |
| 3 | -0.45 | 3.64 | -0.12 | 0.90 | 1 | Gbx1 | gastrulation brain homeobox 1 | 26 | 4 | 0 | 0 | 0 | 0 | 2 | 0 | 0 | 0 | 0 |
| 3 | -0.93 | 3.21 | -0.29 | 0.77 | 1 | Gm15739 | | 0 | 10 | 0 | 0 | 0 | 8 | 2 | 7 | 0 | 0 | 4 |
| 3 | -2.41 | 1.99 | -1.21 | 0.23 | 1 | Gm6428 | | 7 | 0 | 3 | 9 | 4 | 3 | 2 | 0 | 2 | 1 | 1 |
| 3 | 1.18 | 1.71 | 0.69 | 0.49 | 1 | Gm37486 | | 3 | 1 | 9 | 1 | 3 | 0 | 2 | 2 | 7 | 1 | 1 |
| 3 | -0.28 | 1.93 | -0.14 | 0.89 | 1 | Gm10290 | | 15 | 1 | 3 | 2 | 0 | 3 | 2 | 1 | 0 | 4 | 1 |
| 3 | -1.52 | 2.02 | -0.75 | 0.45 | 1 | Gm7446 | | 2 | 13 | 1 | 3 | 3 | 3 | 2 | 0 | 1 | 4 | 0 |
| 3 | 1.51 | 2.39 | 0.63 | 0.53 | 1 | | | 6 | 3 | 0 | 0 | 0 | 7 | 2 | 0 | 2 | 6 | 4 |
| 3 | 0.63 | 2.25 | 0.28 | 0.78 | 1 | Gm6877 | | 3 | 0 | 0 | 0 | 4 | 3 | 2 | 0 | 6 | 10 | 2 |
| 3 | -3.84 | 2.29 | -1.68 | 0.09 | 1 | Gm7094 | | 0 | 1 | 5 | 9 | 10 | 2 | 2 | 0 | 1 | 0 | 0 |
| 3 | -0.17 | 1.82 | -0.10 | 0.92 | 1 | Gm38574 | | 3 | 7 | 1 | 1 | 3 | 3 | 2 | 0 | 6 | 1 | 2 |
| 3 | -2.03 | 2.25 | -0.90 | 0.37 | 1 | Gm4865 | | 3 | 2 | 10 | 5 | 0 | 4 | 2 | 0 | 0 | 1 | 2 |
| 3 | -0.93 | 2.02 | -0.46 | 0.64 | 1 | Gm4374 | | 3 | 0 | 6 | 7 | 1 | 2 | 2 | 1 | 6 | 0 | 2 |
| 3 | -0.43 | 2.10 | -0.21 | 0.84 | 1 | Gm18234 | | 0 | 1 | 0 | 2 | 4 | 5 | 2 | 0 | 1 | 13 | 2 |
| 3 | -0.14 | 3.55 | -0.04 | 0.97 | 1 | Gpr21 | G protein-coupled receptor 21 | 16 | 0 | 0 | 0 | 0 | 5 | 2 | 0 | 0 | 0 | 6 |
| 3 | -1.61 | 3.16 | -0.51 | 0.61 | 1 | Kcnmb4os2 | | 13 | 5 | 7 | 0 | 0 | 2 | 2 | 0 | 0 | 0 | 0 |
| 3 | -2.64 | 1.95 | -1.35 | 0.18 | 1 | Gm7935 | | 5 | 3 | 1 | 3 | 8 | 4 | 2 | 0 | 1 | 0 | 2 |
| 3 | 0.62 | 2.40 | 0.26 | 0.80 | 1 | Gm41041 | | 3 | 1 | 18 | 0 | 0 | 1 | 2 | 1 | 0 | 1 | 1 |
| 3 | 0.36 | 1.79 | 0.20 | 0.84 | 1 | Gm9027 | | 1 | 5 | 1 | 7 | 3 | 0 | 2 | 1 | 4 | 1 | 3 |
| 3 | -0.81 | 3.66 | -0.22 | 0.82 | 1 | Mpig6b | megakaryocyte and platelet inhibitory rec | 5 | 15 | 5 | 0 | 0 | 0 | 2 | 0 | 0 | 0 | 0 |
| 2 | 0.38 | 3.59 | 0.11 | 0.92 | 1 | Gm2109 | | 15 | 6 | 0 | 0 | 0 | 0 | 2 | 0 | 0 | 0 | 3 |
| 2 | -1.30 | 2.02 | -0.64 | 0.52 | 1 | Gm5451 | | 3 | 4 | 1 | 2 | 2 | 6 | 2 | 0 | 4 | 0 | 2 |
| 2 | -1.55 | 2.65 | -0.59 | 0.56 | 1 | Gm12571 | | 1 | 13 | 0 | 0 | 2 | 2 | 2 | 2 | 0 | 0 | 4 |
| 2 | 1.31 | 3.49 | 0.38 | 0.71 | 1 | Gm4869 | predicted gene 4869 | 0 | 9 | 0 | 0 | 0 | 3 | 2 | 0 | 5 | 0 | 7 |
| 2 | -0.82 | 3.30 | -0.25 | 0.80 | 1 | Gm16192 | | 6 | 0 | 6 | 0 | 0 | 6 | 2 | 0 | 0 | 0 | 5 |
| 2 | 0.66 | 1.77 | 0.37 | 0.71 | 1 | Gm4482 | | 3 | 1 | 0 | 3 | 3 | 1 | 2 | 1 | 6 | 3 | 3 |
| 2 | -0.30 | 3.36 | -0.09 | 0.93 | 1 | Gm43193 | | 0 | 0 | 0 | 1 | 0 | 8 | 2 | 0 | 4 | 0 | 10 |
| 2 | -2.23 | 3.52 | -0.63 | 0.53 | 1 | Gm25360 | | 0 | 0 | 0 | 0 | 9 | 2 | 2 | 0 | 0 | 0 | 11 |
| 2 | -2.68 | 3.11 | -0.86 | 0.39 | 1 | Gm12345 | | 6 | 0 | 6 | 8 | 0 | 1 | 2 | 0 | 0 | 1 | 0 |
| 2 | -2.85 | 2.63 | -1.08 | 0.28 | 1 | Gm6283 | | 10 | 0 | 1 | 4 | 1 | 3 | 2 | 0 | 0 | 0 | 2 |
| 2 | -0.51 | 1.77 | -0.29 | 0.77 | 1 | Rpl10-ps2 | | 1 | 10 | 2 | 2 | 2 | 1 | 2 | 1 | 1 | 1 | 1 |
| 2 | -3.23 | 3.53 | -0.91 | 0.36 | 1 | Gm48914 | | 0 | 0 | 0 | 0 | 10 | 9 | 2 | 0 | 0 | 0 | 2 |
| 2 | -0.07 | 2.31 | -0.03 | 0.98 | 1 | Gm6644 | | 4 | 1 | 10 | 0 | 0 | 3 | 2 | 0 | 1 | 1 | 1 |
| 2 | -0.31 | 2.39 | -0.13 | 0.90 | 1 | Rpl13-ps6 | | 2 | 6 | 2 | 0 | 2 | 2 | 2 | 0 | 0 | 3 | 5 |
| 2 | -0.58 | 3.34 | -0.17 | 0.86 | 1 | Gm11537 | | 0 | 0 | 0 | 0 | 5 | 5 | 2 | 0 | 8 | 0 | 3 |
| 2 | -1.90 | 2.04 | -0.93 | 0.35 | 1 | Gm8557 | | 4 | 0 | 1 | 3 | 2 | 7 | 2 | 0 | 2 | 1 | 1 |
| 2 | -0.86 | 3.67 | -0.23 | 0.81 | 1 | Art2a | | 0 | 9 | 11 | 0 | 0 | 0 | 2 | 0 | 0 | 0 | 0 |
| 2 | -0.63 | 3.58 | -0.18 | 0.86 | 1 | Gm42722 | | 1 | 0 | 0 | 0 | 0 | 8 | 2 | 0 | 0 | 0 | 11 |
| 2 | 0.36 | 3.23 | 0.11 | 0.91 | 1 | Gm12568 | | 3 | 0 | 3 | 0 | 0 | 3 | 2 | 9 | 0 | 0 | 2 |
| 2 | -1.84 | 2.09 | -0.88 | 0.38 | 1 | Gm15773 | | 7 | 4 | 1 | 2 | 1 | 3 | 2 | 1 | 0 | 1 | 0 |
| 2 | -1.46 | 3.44 | -0.42 | 0.67 | 1 | Gm12974 | | 4 | 0 | 0 | 0 | 1 | 9 | 2 | 0 | 0 | 0 | 5 |
| 2 | 0.16 | 2.16 | 0.07 | 0.94 | 1 | Gm8606 | | 1 | 3 | 2 | 1 | 6 | 0 | 2 | 0 | 1 | 4 | 2 |
| 2 | -1.56 | 3.27 | -0.48 | 0.63 | 1 | Gm13611 | | 0 | 2 | 4 | 0 | 1 | 2 | 2 | 10 | 0 | 0 | 0 |
| 2 | 2.27 | 3.12 | 0.73 | 0.47 | 1 | 4930451E10Rik | | 2 | 3 | 1 | 0 | 0 | 0 | 2 | 9 | 4 | 0 | 0 |
| 2 | -2.44 | 3.56 | -0.69 | 0.49 | 1 | Gm48772 | | 0 | 0 | 8 | 0 | 3 | 6 | 2 | 0 | 2 | 0 | 0 |
| 2 | -0.44 | 2.62 | -0.17 | 0.87 | 1 | Gm14017 | | 3 | 1 | 0 | 2 | 0 | 3 | 2 | 1 | 0 | 9 | 0 |
| 2 | 0.04 | 1.83 | 0.02 | 0.98 | 1 | Gm4928 | | 1 | 0 | 2 | 4 | 2 | 1 | 2 | 3 | 2 | 1 | 2 |
| 2 | -4.63 | 3.60 | -1.29 | 0.20 | 1 | 4930500M09Rik | | 0 | 0 | 17 | 0 | 0 | 0 | 2 | 0 | 2 | 0 | 0 |
| 2 | -0.95 | 3.65 | -0.26 | 0.80 | 1 | 1700060C16Rik | | 3 | 15 | 0 | 0 | 0 | 0 | 2 | 0 | 0 | 0 | 0 |
| 2 | 1.43 | 3.21 | 0.45 | 0.66 | 1 | Gm28530 | | 0 | 2 | 1 | 0 | 0 | 1 | 2 | 0 | 0 | 12 | 3 |
| 2 | -3.51 | 3.50 | -1.00 | 0.32 | 1 | Gm17207 | | 0 | 0 | 0 | 4 | 9 | 3 | 2 | 0 | 0 | 0 | 3 |
| 2 | -0.43 | 3.64 | -0.12 | 0.91 | 1 | Myh13 | myosin, heavy polypeptide 13, skeletal m | 16 | 0 | 2 | 0 | 0 | 0 | 2 | 0 | 0 | 0 | 0 |
| 2 | 3.86 | 3.55 | 1.09 | 0.28 | 1 | Gm16059 | | 0 | 10 | 0 | 0 | 0 | 0 | 2 | 0 | 7 | 0 | 1 |
| 2 | -1.52 | 3.62 | -0.42 | 0.67 | 1 | Gm6745 | | 1 | 14 | 1 | 0 | 0 | 0 | 2 | 0 | 0 | 0 | 2 |

| | | | | | | | | | | | | | | | | | | |
|---|---|---|---|---|---|---|---|---|---|---|---|---|---|---|---|---|---|---|
| 2 | -1.59 | 2.76 | -0.57 | 0.57 | 1 | Rpl17-ps4 | | 9 | 0 | 1 | 4 | 1 | 0 | 2 | 0 | 1 | 1 | 0 |
| 2 | -1.06 | 3.54 | -0.30 | 0.76 | 1 | Gm50351 | | 0 | 0 | 8 | 0 | 2 | 2 | 2 | 0 | 0 | 0 | 6 |
| 2 | -1.26 | 3.64 | -0.35 | 0.73 | 1 | Gm7527 | | 13 | 0 | 4 | 0 | 0 | 1 | 2 | 0 | 0 | 0 | 0 |
| 2 | -2.56 | 3.60 | -0.71 | 0.48 | 1 | Gm11425 | | 9 | 0 | 4 | 1 | 0 | 3 | 2 | 0 | 0 | 0 | 0 |
| 2 | -2.06 | 2.35 | -0.88 | 0.38 | 1 | Ftl2-ps | | 0 | 0 | 1 | 8 | 1 | 3 | 2 | 1 | 2 | 1 | 0 |
| 2 | -1.62 | 3.57 | -0.45 | 0.65 | 1 | Gm15066 | | 0 | 2 | 2 | 12 | 0 | 0 | 2 | 0 | 0 | 1 | 0 |
| 2 | -0.92 | 2.18 | -0.42 | 0.67 | 1 | Gm5518 | | 2 | 5 | 3 | 1 | 2 | 1 | 2 | 0 | 3 | 0 | 1 |
| 2 | 1.17 | 1.94 | 0.60 | 0.55 | 1 | Rpl36-ps2 | | 1 | 1 | 4 | 2 | 1 | 0 | 2 | 3 | 2 | 1 | 2 |
| 2 | -1.40 | 2.26 | -0.62 | 0.54 | 1 | Gm15190 | | 1 | 0 | 0 | 3 | 5 | 3 | 2 | 1 | 4 | 1 | 0 |
| 2 | 0.98 | 3.05 | 0.32 | 0.75 | 1 | Gm11865 | | 0 | 0 | 1 | 0 | 0 | 5 | 2 | 0 | 1 | 4 | 6 |
| 2 | -4.08 | 3.62 | -1.13 | 0.26 | 1 | Gm45719 | | 0 | 10 | 0 | 0 | 0 | 6 | 2 | 0 | 0 | 0 | 1 |
| 2 | -0.98 | 2.76 | -0.35 | 0.72 | 1 | Gm6517 | | 6 | 1 | 4 | 1 | 0 | 1 | 2 | 0 | 0 | 0 | 4 |
| 2 | 0.31 | 1.97 | 0.16 | 0.87 | 1 | Gm6023 | | 3 | 1 | 2 | 1 | 1 | 2 | 2 | 5 | 1 | 0 | 2 |
| 2 | 0.32 | 2.82 | 0.11 | 0.91 | 1 | Gm8805 | | 0 | 9 | 0 | 1 | 1 | 0 | 2 | 2 | 1 | 3 | 0 |
| 2 | -2.32 | 2.43 | -0.96 | 0.34 | 1 | Rpl21-ps12 | | 3 | 3 | 0 | 2 | 3 | 3 | 2 | 0 | 1 | 1 | 0 |
| 2 | 0.36 | 3.56 | 0.10 | 0.92 | 1 | Gm5575 | | 0 | 0 | 0 | 0 | 0 | 3 | 2 | 0 | 0 | 14 | 0 |
| 2 | -0.86 | 3.67 | -0.23 | 0.81 | 1 | Tmprss13 | transmembrane protease, serine 13 | 0 | 5 | 11 | 0 | 0 | 0 | 2 | 0 | 0 | 0 | 0 |
| 2 | -2.06 | 2.64 | -0.78 | 0.43 | 1 | Gm34432 | | 3 | 0 | 6 | 1 | 4 | 1 | 2 | 0 | 2 | 0 | 0 |
| 2 | -0.68 | 3.35 | -0.20 | 0.84 | 1 | Gm17802 | | 0 | 10 | 0 | 0 | 0 | 3 | 2 | 1 | 1 | 0 | 1 |
| 2 | 1.71 | 3.53 | 0.49 | 0.63 | 1 | Gm44180 | | 0 | 2 | 0 | 1 | 0 | 0 | 2 | 0 | 0 | 8 | 5 |
| 2 | -4.22 | 3.55 | -1.19 | 0.23 | 1 | Adtrp | androgen dependent TFPI regulating prot | 0 | 1 | 0 | 0 | 9 | 5 | 2 | 0 | 0 | 0 | 0 |
| 2 | -3.04 | 3.57 | -0.85 | 0.39 | 1 | Gm40910 | | 0 | 4 | 0 | 9 | 0 | 3 | 2 | 0 | 0 | 0 | 0 |
| 1 | -2.36 | 3.54 | -0.67 | 0.51 | 1 | Gm26781 | | 0 | 0 | 0 | 12 | 0 | 0 | 2 | 0 | 0 | 0 | 2 |
| 1 | -1.16 | 2.95 | -0.39 | 0.69 | 1 | Gm48338 | | 0 | 2 | 4 | 0 | 4 | 1 | 2 | 2 | 0 | 0 | 1 |
| 1 | 1.99 | 3.27 | 0.61 | 0.54 | 1 | Gm8318 | | 2 | 0 | 2 | 0 | 0 | 0 | 2 | 1 | 1 | 8 | 0 |
| 1 | 0.51 | 2.61 | 0.19 | 0.85 | 1 | Gm9416 | | 1 | 0 | 1 | 4 | 0 | 0 | 2 | 1 | 1 | 1 | 3 |
| 1 | 0.88 | 3.52 | 0.25 | 0.80 | 1 | Gm43748 | | 0 | 0 | 2 | 0 | 0 | 2 | 2 | 6 | 0 | 0 | 3 |
| 1 | -1.45 | 3.64 | -0.40 | 0.69 | 1 | Gm44732 | | 0 | 5 | 6 | 0 | 1 | 0 | 2 | 0 | 0 | 0 | 0 |
| 1 | -1.61 | 3.60 | -0.45 | 0.65 | 1 | Gm44021 | | 0 | 6 | 0 | 0 | 0 | 0 | 2 | 0 | 0 | 0 | 6 |
| 1 | -3.23 | 3.60 | -0.90 | 0.37 | 1 | Gm19792 | | 0 | 0 | 3 | 0 | 0 | 6 | 2 | 0 | 0 | 0 | 3 |
| 1 | 0.00 | 1.90 | 0.00 | 1.00 | 1 | Gm12017 | | 1 | 1 | 1 | 1 | 1 | 2 | 2 | 1 | 2 | 1 | 1 |
| 1 | -2.75 | 2.89 | -0.95 | 0.34 | 1 | 4930579G18Rik | | 2 | 1 | 0 | 2 | 3 | 3 | 2 | 0 | 0 | 0 | 2 |
| 1 | -0.04 | 2.11 | -0.02 | 0.98 | 1 | Gm8825 | | 2 | 1 | 1 | 1 | 1 | 1 | 2 | 0 | 3 | 1 | 1 |
| 1 | 1.22 | 3.47 | 0.35 | 0.73 | 1 | Gm12804 | | 2 | 2 | 0 | 0 | 0 | 1 | 2 | 0 | 2 | 5 | 0 |
| 1 | -2.57 | 3.59 | -0.72 | 0.47 | 1 | Gm10787 | | 0 | 0 | 0 | 11 | 0 | 0 | 2 | 0 | 0 | 0 | 0 |
| 1 | -2.15 | 3.00 | -0.72 | 0.47 | 1 | Gm14639 | | 4 | 2 | 0 | 2 | 1 | 1 | 2 | 0 | 0 | 0 | 1 |
| 1 | 1.76 | 3.52 | 0.50 | 0.62 | 1 | Gm17108 | | 0 | 0 | 0 | 0 | 0 | 1 | 2 | 7 | 1 | 0 | 2 |
| 1 | -2.24 | 3.55 | -0.63 | 0.53 | 1 | 2400006E01Rik | | 0 | 0 | 0 | 5 | 0 | 3 | 2 | 0 | 0 | 0 | 3 |
| 1 | -3.29 | 3.58 | -0.92 | 0.36 | 1 | Mnd1-ps | | 0 | 1 | 0 | 4 | 0 | 5 | 2 | 0 | 0 | 0 | 0 |
| 1 | 0.86 | 3.03 | 0.28 | 0.78 | 1 | Selenok-ps5 | | 0 | 0 | 1 | 1 | 1 | 0 | 2 | 2 | 4 | 0 | 1 |
| 1 | -3.47 | 3.57 | -0.97 | 0.33 | 1 | 1700025C18Rik | | 0 | 0 | 0 | 0 | 8 | 3 | 2 | 0 | 0 | 0 | 0 |
| 1 | -0.41 | 2.74 | -0.15 | 0.88 | 1 | Gm3617 | | 1 | 2 | 2 | 0 | 1 | 1 | 2 | 0 | 0 | 1 | 2 |
| 1 | -1.65 | 3.55 | -0.47 | 0.64 | 1 | Fut4-ps1 | | 0 | 0 | 0 | 0 | 2 | 5 | 2 | 0 | 3 | 0 | 0 |
| 1 | -2.22 | 3.08 | -0.72 | 0.47 | 1 | Gm5055 | | 3 | 1 | 0 | 1 | 2 | 2 | 2 | 0 | 1 | 0 | 0 |
| 1 | -1.42 | 3.64 | -0.39 | 0.70 | 1 | Gm5239 | | 2 | 1 | 6 | 0 | 0 | 1 | 2 | 0 | 0 | 0 | 0 |
| 1 | -1.44 | 3.13 | -0.46 | 0.64 | 1 | Gm14000 | | 0 | 1 | 1 | 1 | 1 | 1 | 2 | 0 | 5 | 0 | 0 |
| 1 | -1.58 | 3.60 | -0.44 | 0.66 | 1 | Gm14502 | | 0 | 0 | 0 | 0 | 0 | 8 | 2 | 0 | 0 | 0 | 1 |
| 1 | -2.62 | 3.60 | -0.73 | 0.47 | 1 | Gm44587 | | 0 | 5 | 0 | 2 | 0 | 3 | 2 | 0 | 0 | 0 | 0 |
| 1 | -1.28 | 3.53 | -0.36 | 0.72 | 1 | Gm15720 | | 0 | 0 | 0 | 1 | 0 | 5 | 2 | 1 | 2 | 0 | 0 |
| 1 | -0.89 | 3.14 | -0.28 | 0.78 | 1 | Gm5276 | | 0 | 1 | 1 | 2 | 0 | 2 | 2 | 2 | 0 | 1 | 0 |
| 1 | 0.63 | 3.55 | 0.18 | 0.86 | 1 | Gm9857 | | 0 | 0 | 0 | 0 | 0 | 2 | 2 | 2 | 0 | 0 | 5 |
| 1 | -0.76 | 2.82 | -0.27 | 0.79 | 1 | Rpl21-ps13 | | 1 | 2 | 1 | 0 | 1 | 1 | 2 | 2 | 0 | 0 | 1 |
| 1 | 0.83 | 2.75 | 0.30 | 0.76 | 1 | Gm13750 | | 1 | 1 | 0 | 1 | 1 | 0 | 2 | 0 | 2 | 1 | 2 |
| 1 | -0.30 | 3.60 | -0.08 | 0.93 | 1 | 4930516B21Rik | | 0 | 0 | 0 | 0 | 0 | 1 | 2 | 0 | 0 | 0 | 8 |
| 1 | -1.34 | 3.59 | -0.37 | 0.71 | 1 | Gm50087 | | 0 | 1 | 0 | 0 | 0 | 6 | 2 | 0 | 0 | 0 | 2 |
| 1 | -0.48 | 3.13 | -0.15 | 0.88 | 1 | Gm29739 | | 1 | 0 | 0 | 1 | 3 | 0 | 2 | 0 | 1 | 1 | 2 |
| 1 | -1.77 | 3.62 | -0.49 | 0.63 | 1 | Acat3 | acetyl-Coenzyme A acetyltransferase 3 | 6 | 0 | 0 | 0 | 3 | 0 | 2 | 0 | 0 | 0 | 0 |
| 1 | -1.85 | 3.26 | -0.57 | 0.57 | 1 | Gm15946 | | 1 | 1 | 1 | 0 | 1 | 3 | 2 | 2 | 0 | 0 | 0 |
| 1 | -1.78 | 3.57 | -0.50 | 0.62 | 1 | Gm5684 | | 0 | 0 | 1 | 2 | 5 | 0 | 2 | 1 | 0 | 0 | 0 |
| 1 | -1.20 | 3.58 | -0.34 | 0.74 | 1 | Gm44291 | | 1 | 0 | 0 | 0 | 1 | 3 | 2 | 0 | 0 | 0 | 4 |

| | | | | | | | | | | | | | | | | | | |
|---|---|---|---|---|---|---|---|---|---|---|---|---|---|---|---|---|---|---|
| 1 | -1.43 | 3.60 | -0.40 | 0.69 | 1 | Gm14138 | | 3 | 2 | 0 | 0 | 0 | 3 | 2 | 1 | 0 | 0 | 0 |
| 1 | 0.24 | 3.64 | 0.07 | 0.95 | 1 | Gm48088 | | 5 | 1 | 1 | 0 | 0 | 0 | 2 | 0 | 1 | 0 | 0 |
| 1 | 0.57 | 3.59 | 0.16 | 0.87 | 1 | Mug1 | murinoglobulin 1 | 0 | 0 | 1 | 0 | 1 | 0 | 2 | 0 | 5 | 0 | 1 |
| 1 | -1.19 | 3.57 | -0.33 | 0.74 | 1 | Gm15886 | | 0 | 0 | 4 | 0 | 0 | 1 | 2 | 0 | 1 | 0 | 2 |
| 1 | -2.40 | 3.57 | -0.67 | 0.50 | 1 | Gm26749 | | 0 | 0 | 0 | 0 | 5 | 2 | 2 | 0 | 0 | 0 | 1 |
| 1 | 0.00 | 3.55 | 0.00 | 1.00 | 1 | Gm2076 | | 0 | 0 | 0 | 0 | 1 | 2 | 2 | 1 | 0 | 3 | 1 |
| 1 | -0.87 | 3.59 | -0.24 | 0.81 | 1 | Gm36043 | | 2 | 0 | 0 | 0 | 0 | 3 | 2 | 0 | 0 | 0 | 2 |
| 1 | 0.81 | 3.60 | 0.22 | 0.82 | 1 | Rpl21-ps3 | | 2 | 2 | 0 | 0 | 0 | 0 | 2 | 0 | 0 | 1 | 2 |
| 1 | -0.47 | 3.60 | -0.13 | 0.90 | 1 | 4930549G23Rik | | 0 | 1 | 1 | 0 | 0 | 1 | 2 | 0 | 0 | 0 | 4 |
| 1 | -3.54 | 3.56 | -0.99 | 0.32 | 1 | Gm20659 | | 0 | 0 | 0 | 0 | 0 | 2 | 2 | 0 | 5 | 0 | 0 |
| 1 | -0.92 | 3.57 | -0.26 | 0.80 | 1 | Gm11598 | | 1 | 0 | 0 | 1 | 0 | 3 | 2 | 1 | 0 | 1 | 0 |
| 1 | -0.42 | 3.56 | -0.12 | 0.91 | 1 | Gm8524 | | 1 | 0 | 0 | 2 | 1 | 0 | 2 | 0 | 2 | 0 | 1 |
| 1 | -0.85 | 3.58 | -0.24 | 0.81 | 1 | Gm5040 | | 0 | 0 | 1 | 0 | 2 | 1 | 2 | 0 | 1 | 1 | 0 |
| 1 | 0.85 | 3.60 | 0.24 | 0.81 | 1 | Gm232 | | 0 | 1 | 0 | 0 | 0 | 0 | 2 | 0 | 5 | 0 | 0 |
| 1 | -0.34 | 3.63 | -0.09 | 0.92 | 1 | Gm42839 | | 6 | 0 | 0 | 0 | 0 | 0 | 2 | 0 | 0 | 0 | 0 |
| 1 | 1.27 | 3.60 | 0.35 | 0.72 | 1 | 4931431C16Rik | | 3 | 0 | 0 | 0 | 0 | 0 | 2 | 0 | 3 | 0 | 0 |
| 1 | -0.78 | 3.60 | -0.22 | 0.83 | 1 | Cyp4a32 | cytochrome P450, family 4, subfamily a, p | 0 | 0 | 0 | 0 | 0 | 3 | 2 | 0 | 0 | 0 | 3 |
| 1 | -1.59 | 3.63 | -0.44 | 0.66 | 1 | Gm8801 | | 3 | 0 | 0 | 0 | 2 | 0 | 2 | 0 | 0 | 0 | 0 |
| 1 | -0.80 | 3.60 | -0.22 | 0.82 | 1 | Gm15956 | | 3 | 0 | 0 | 0 | 0 | 2 | 2 | 0 | 0 | 0 | 1 |
| 1 | -0.09 | 3.63 | -0.02 | 0.98 | 1 | Rpl17-ps1 | | 1 | 1 | 0 | 0 | 1 | 0 | 2 | 0 | 0 | 1 | 1 |
| 1 | -2.01 | 3.62 | -0.56 | 0.58 | 1 | Gm45033 | | 0 | 0 | 0 | 0 | 0 | 5 | 2 | 0 | 0 | 0 | 0 |
| 1 | 2.57 | 3.60 | 0.71 | 0.48 | 1 | Gm11633 | | 0 | 1 | 0 | 0 | 0 | 0 | 2 | 0 | 1 | 0 | 3 |
| 1 | -0.01 | 3.63 | 0.00 | 1.00 | 1 | Gm10222 | | 1 | 1 | 0 | 0 | 0 | 1 | 2 | 0 | 1 | 1 | 0 |
| 1 | 0.49 | 3.60 | 0.14 | 0.89 | 1 | Gm43442 | | 1 | 0 | 0 | 0 | 1 | 0 | 2 | 1 | 0 | 0 | 2 |
| 1 | -1.72 | 3.60 | -0.48 | 0.63 | 1 | Gm7085 | | 0 | 0 | 0 | 0 | 3 | 1 | 2 | 0 | 0 | 1 | 0 |
| 1 | 0.47 | 3.58 | 0.13 | 0.90 | 1 | Gm8577 | | 0 | 0 | 0 | 2 | 0 | 0 | 2 | 1 | 1 | 0 | 1 |
| 1 | -0.01 | 3.63 | 0.00 | 1.00 | 1 | Gm7002 | | 3 | 0 | 1 | 0 | 0 | 0 | 2 | 0 | 0 | 1 | 0 |
| 1 | -2.84 | 3.58 | -0.79 | 0.43 | 1 | Gm44101 | | 0 | 1 | 0 | 0 | 2 | 2 | 2 | 0 | 0 | 0 | 0 |
| 1 | -1.99 | 3.62 | -0.55 | 0.58 | 1 | Gm37645 | | 0 | 0 | 0 | 0 | 0 | 4 | 2 | 0 | 0 | 0 | 0 |
| 1 | 0.76 | 3.62 | 0.21 | 0.83 | 1 | Gm8757 | | 2 | 0 | 0 | 0 | 0 | 0 | 2 | 0 | 1 | 1 | 0 |
| 1 | -0.07 | 3.66 | -0.02 | 0.98 | 1 | Gm8624 | | 1 | 1 | 1 | 0 | 0 | 0 | 2 | 0 | 1 | 0 | 0 |
| 1 | -2.18 | 3.61 | -0.61 | 0.55 | 1 | 1700094J05Rik | | 0 | 1 | 0 | 0 | 3 | 0 | 2 | 0 | 0 | 0 | 0 |
| 1 | 0.88 | 3.62 | 0.24 | 0.81 | 1 | Gm14056 | | 0 | 0 | 0 | 0 | 0 | 0 | 2 | 0 | 0 | 3 | 1 |
| 0 | -0.11 | 3.60 | -0.03 | 0.98 | 1 | Gm8814 | | 0 | 0 | 0 | 0 | 1 | 0 | 2 | 1 | 1 | 0 | 0 |
| 0 | -1.18 | 3.62 | -0.33 | 0.74 | 1 | Gm9387 | | 0 | 1 | 0 | 0 | 0 | 1 | 2 | 1 | 0 | 0 | 0 |
| 0 | 0.19 | 3.61 | 0.05 | 0.96 | 1 | Gm42858 | | 1 | 0 | 0 | 0 | 0 | 0 | 2 | 0 | 0 | 0 | 2 |
| 0 | -0.59 | 3.61 | -0.16 | 0.87 | 1 | Gm44335 | | 0 | 0 | 0 | 0 | 0 | 1 | 2 | 0 | 0 | 0 | 2 |
| 0 | -0.64 | 3.62 | -0.18 | 0.86 | 1 | Rps19-ps12 | | 0 | 0 | 0 | 0 | 0 | 1 | 2 | 0 | 2 | 0 | 0 |
| 0 | -0.51 | 3.64 | -0.14 | 0.89 | 1 | Gm12196 | | 3 | 0 | 0 | 0 | 0 | 0 | 2 | 0 | 0 | 0 | 0 |
| 0 | -1.81 | 3.63 | -0.50 | 0.62 | 1 | Gm48936 | | 0 | 0 | 0 | 0 | 0 | 3 | 2 | 0 | 0 | 0 | 0 |
| 0 | -0.89 | 3.63 | -0.24 | 0.81 | 1 | Gm13450 | | 0 | 0 | 0 | 0 | 1 | 0 | 2 | 1 | 0 | 0 | 0 |
| 0 | 0.30 | 3.60 | 0.08 | 0.93 | 1 | Gm11748 | | 0 | 0 | 0 | 0 | 0 | 0 | 2 | 0 | 0 | 0 | 2 |
| 0 | -0.76 | 3.66 | -0.21 | 0.84 | 1 | Gm5043 | | 1 | 0 | 1 | 0 | 0 | 0 | 2 | 0 | 0 | 0 | 0 |
| 0 | -0.80 | 3.63 | -0.22 | 0.83 | 1 | Gm28892 | | 0 | 0 | 0 | 0 | 0 | 1 | 2 | 0 | 0 | 0 | 1 |
| 0 | -1.52 | 3.64 | -0.42 | 0.68 | 1 | Gm13679 | | 0 | 1 | 0 | 0 | 0 | 1 | 2 | 0 | 0 | 0 | 0 |
| 0 | -1.62 | 3.64 | -0.45 | 0.66 | 1 | Gm24368 | | 0 | 0 | 0 | 2 | 0 | 0 | 2 | 0 | 0 | 0 | 0 |
| 0 | -0.06 | 3.64 | -0.02 | 0.99 | 1 | 4930592A05Rik | | 0 | 0 | 0 | 0 | 0 | 0 | 2 | 0 | 0 | 1 | 0 |
| 0 | -0.16 | 3.65 | -0.04 | 0.97 | 1 | Fgg | fibrinogen gamma chain | 0 | 0 | 0 | 0 | 0 | 0 | 2 | 1 | 0 | 0 | 0 |
| 0 | -0.16 | 3.65 | -0.04 | 0.97 | 1 | Gm4978 | | 0 | 0 | 0 | 0 | 0 | 0 | 2 | 1 | 0 | 0 | 0 |
| 0 | -0.11 | 3.66 | -0.03 | 0.98 | 1 | Rn7s1 | | 0 | 0 | 0 | 0 | 0 | 0 | 2 | 0 | 1 | 0 | 0 |
| 0 | -1.36 | 3.65 | -0.37 | 0.71 | 1 | Gm18737 | | 0 | 0 | 0 | 0 | 1 | 0 | 2 | 0 | 0 | 0 | 0 |
| 0 | -0.71 | 3.66 | -0.19 | 0.85 | 1 | Gm16379 | | 1 | 0 | 0 | 0 | 0 | 0 | 2 | 0 | 0 | 0 | 0 |
| 0 | -0.71 | 3.66 | -0.19 | 0.85 | 1 | Gm35229 | | 1 | 0 | 0 | 0 | 0 | 0 | 2 | 0 | 0 | 0 | 0 |
| 0 | -1.38 | 3.65 | -0.38 | 0.71 | 1 | Gm38241 | | 0 | 0 | 0 | 0 | 0 | 1 | 2 | 0 | 0 | 0 | 0 |
| 0 | -1.38 | 3.65 | -0.38 | 0.71 | 1 | Gm42972 | | 0 | 0 | 0 | 0 | 0 | 1 | 2 | 0 | 0 | 0 | 0 |
| 0 | -1.38 | 3.65 | -0.38 | 0.71 | 1 | Gm47162 | | 0 | 0 | 0 | 0 | 0 | 1 | 2 | 0 | 0 | 0 | 0 |
| 0 | -1.38 | 3.65 | -0.38 | 0.71 | 1 | Gm34609 | | 0 | 0 | 0 | 0 | 0 | 1 | 2 | 0 | 0 | 0 | 0 |
| 0 | -0.81 | 3.66 | -0.22 | 0.83 | 1 | Hc | hemolytic complement | 0 | 0 | 0 | 0 | 0 | 0 | 2 | 0 | 0 | 0 | 0 |
| 0 | -0.81 | 3.66 | -0.22 | 0.83 | 1 | Fau-ps1 | | 0 | 0 | 0 | 0 | 0 | 0 | 2 | 0 | 0 | 0 | 0 |
| 0 | -0.81 | 3.66 | -0.22 | 0.83 | 1 | Gm42514 | | 0 | 0 | 0 | 0 | 0 | 0 | 2 | 0 | 0 | 0 | 0 |

| | | | | | | | | | | | | | | | | | |
|---|---|---|---|---|---|---|---|---|---|---|---|---|---|---|---|---|---|
| 0 | -0.81 | 3.66 | -0.22 | 0.83 | 1 | Cyp4a10 | cytochrome P450, family 4, subfamily a, p | 0 | 0 | 0 | 0 | 0 | 0 | 2 | 0 | 0 | 0 | 0 |
| 0 | -0.81 | 3.66 | -0.22 | 0.83 | 1 | Drd5 | dopamine receptor D5 | 0 | 0 | 0 | 0 | 0 | 0 | 2 | 0 | 0 | 0 | 0 |
| 0 | -0.81 | 3.66 | -0.22 | 0.83 | 1 | Trmt112-ps1 | | 0 | 0 | 0 | 0 | 0 | 0 | 2 | 0 | 0 | 0 | 0 |
| 0 | -0.81 | 3.66 | -0.22 | 0.83 | 1 | Gm42620 | | 0 | 0 | 0 | 0 | 0 | 0 | 2 | 0 | 0 | 0 | 0 |
| 0 | -0.81 | 3.66 | -0.22 | 0.83 | 1 | Gm44442 | | 0 | 0 | 0 | 0 | 0 | 0 | 2 | 0 | 0 | 0 | 0 |
| 0 | -0.81 | 3.66 | -0.22 | 0.83 | 1 | Gm5596 | | 0 | 0 | 0 | 0 | 0 | 0 | 2 | 0 | 0 | 0 | 0 |
| 0 | -0.81 | 3.66 | -0.22 | 0.83 | 1 | 1700042D02Rik | | 0 | 0 | 0 | 0 | 0 | 0 | 2 | 0 | 0 | 0 | 0 |
| 0 | -0.81 | 3.66 | -0.22 | 0.83 | 1 | Gm18996 | | 0 | 0 | 0 | 0 | 0 | 0 | 2 | 0 | 0 | 0 | 0 |
| 0 | -0.81 | 3.66 | -0.22 | 0.83 | 1 | Atoh7 | atonal bHLH transcription factor 7 | 0 | 0 | 0 | 0 | 0 | 0 | 2 | 0 | 0 | 0 | 0 |
| 0 | -0.81 | 3.66 | -0.22 | 0.83 | 1 | Gm48605 | | 0 | 0 | 0 | 0 | 0 | 0 | 2 | 0 | 0 | 0 | 0 |
| 0 | -0.81 | 3.66 | -0.22 | 0.83 | 1 | Gm48366 | | 0 | 0 | 0 | 0 | 0 | 0 | 2 | 0 | 0 | 0 | 0 |
| 0 | -0.81 | 3.66 | -0.22 | 0.83 | 1 | Gm29587 | | 0 | 0 | 0 | 0 | 0 | 0 | 2 | 0 | 0 | 0 | 0 |
| 0 | -0.81 | 3.66 | -0.22 | 0.83 | 1 | Vmn1r202 | vomeronasal 1 receptor 202 | 0 | 0 | 0 | 0 | 0 | 0 | 2 | 0 | 0 | 0 | 0 |
| 0 | -0.81 | 3.66 | -0.22 | 0.83 | 1 | Mir802 | | 0 | 0 | 0 | 0 | 0 | 0 | 2 | 0 | 0 | 0 | 0 |
| 0 | -0.81 | 3.66 | -0.22 | 0.83 | 1 | Mir692-1 | | 0 | 0 | 0 | 0 | 0 | 0 | 2 | 0 | 0 | 0 | 0 |
| 0 | -0.81 | 3.66 | -0.22 | 0.83 | 1 | Gm15238 | | 0 | 0 | 0 | 0 | 0 | 0 | 2 | 0 | 0 | 0 | 0 |
| 2140 | -0.15 | 3.64 | -0.04 | 0.97 | 1 | Ighg2c | | 22505 | 457 | 577 | 0 | 0 | 0 | 1 | 0 | 0 | 0 | 0 |
| 1077 | 2.03 | 1.33 | 1.52 | 0.13 | 1 | Mmp2 | matrix metallopeptidase 2 | 5686 | 2680 | 3471 | 0 | 0 | 0 | 1 | 0 | 1 | 6 | 0 |
| 987 | -2.19 | 1.41 | -1.55 | 0.12 | 1 | Mylk | myosin, light polypeptide kinase | 5649 | 2491 | 2683 | 10 | 4 | 12 | 1 | 0 | 7 | 0 | 1 |
| 964 | 0.28 | 1.45 | 0.19 | 0.85 | 1 | Hmcn2 | hemicentin 2 | 6084 | 2278 | 2235 | 0 | 0 | 0 | 1 | 0 | 0 | 0 | 1 |
| 602 | -0.41 | 3.67 | -0.11 | 0.91 | 1 | Islr | immunoglobulin superfamily containing le | 3244 | 1495 | 1878 | 0 | 0 | 0 | 1 | 0 | 0 | 0 | 0 |
| 589 | -1.77 | 1.32 | -1.34 | 0.18 | 1 | C130074G19Ri | RIKEN cDNA C130074G19 gene | 2611 | 1910 | 1950 | 0 | 0 | 3 | 1 | 0 | 0 | 0 | 0 |
| 588 | -1.12 | 1.43 | -0.79 | 0.43 | 1 | Robo4 | roundabout guidance receptor 4 | 2239 | 2151 | 2075 | 0 | 0 | 1 | 1 | 0 | 0 | 0 | 0 |
| 443 | -1.01 | 2.17 | -0.47 | 0.64 | 1 | Galnt15 | polypeptide N-acetylgalactosaminyltransf | 2680 | 949 | 1239 | 0 | 4 | 0 | 1 | 0 | 0 | 0 | 5 |
| 382 | -0.44 | 3.68 | -0.12 | 0.90 | 1 | Pcdh12 | protocadherin 12 | 1316 | 1372 | 1516 | 0 | 0 | 0 | 1 | 0 | 0 | 0 | 0 |
| 309 | 0.13 | 1.42 | 0.09 | 0.93 | 1 | Pdgfd | platelet-derived growth factor, D polypept | 1263 | 924 | 1189 | 0 | 5 | 1 | 1 | 0 | 2 | 0 | 11 |
| 299 | 0.04 | 3.62 | 0.01 | 0.99 | 1 | Ighv14-4 | | 3251 | 32 | 0 | 0 | 0 | 0 | 1 | 0 | 0 | 0 | 0 |
| 215 | -1.33 | 1.04 | -1.28 | 0.20 | 1 | Cd14 | CD14 antigen | 1303 | 376 | 671 | 0 | 0 | 7 | 1 | 0 | 1 | 0 | 1 |
| 173 | 0.26 | 1.80 | 0.14 | 0.89 | 1 | Spi1 | spleen focus forming virus (SFFV) provira | 1013 | 430 | 454 | 0 | 3 | 0 | 1 | 0 | 6 | 0 | 0 |
| 169 | -2.41 | 1.29 | -1.86 | 0.06 | 1 | C3ar1 | complement component 3a receptor 1 | 812 | 415 | 628 | 0 | 3 | 1 | 1 | 0 | 0 | 0 | 0 |
| 147 | -0.01 | 0.95 | -0.01 | 0.99 | 1 | Stard9 | START domain containing 9 | 650 | 464 | 431 | 18 | 0 | 25 | 1 | 5 | 10 | 0 | 18 |
| 147 | -0.43 | 0.74 | -0.59 | 0.56 | 1 | Nr5a2 | nuclear receptor subfamily 5, group A, me | 356 | 675 | 511 | 20 | 9 | 12 | 1 | 20 | 9 | 0 | 4 |
| 140 | 1.06 | 0.85 | 1.25 | 0.21 | 1 | Smad6 | SMAD family member 6 | 588 | 285 | 636 | 4 | 1 | 2 | 1 | 0 | 3 | 13 | 5 |
| 124 | -0.33 | 0.96 | -0.35 | 0.73 | 1 | Rbp1 | retinol binding protein 1, cellular | 750 | 216 | 229 | 11 | 43 | 37 | 1 | 14 | 17 | 0 | 49 |
| 121 | 0.28 | 1.53 | 0.18 | 0.86 | 1 | Kif26a | kinesin family member 26A | 470 | 429 | 431 | 0 | 0 | 0 | 1 | 0 | 1 | 0 | 0 |
| 100 | 0.61 | 0.91 | 0.67 | 0.50 | 1 | Col11a2 | collagen, type XI, alpha 2 | 17 | 26 | 11 | 0 | 4 | 494 | 1 | 0 | 5 | 4 | 537 |
| 96 | -0.83 | 1.05 | -0.79 | 0.43 | 1 | Ifi44 | interferon-induced protein 44 | 338 | 284 | 375 | 13 | 20 | 2 | 1 | 0 | 16 | 0 | 7 |
| 91 | -0.98 | 2.38 | -0.41 | 0.68 | 1 | Hhip | Hedgehog-interacting protein | 926 | 33 | 36 | 0 | 1 | 0 | 1 | 0 | 0 | 0 | 0 |
| 90 | 1.68 | 1.10 | 1.53 | 0.13 | 1 | Cyp26b1 | cytochrome P450, family 26, subfamily b, | 256 | 608 | 95 | 0 | 0 | 5 | 1 | 0 | 2 | 0 | 20 |
| 75 | 0.59 | 2.00 | 0.30 | 0.77 | 1 | Card11 | caspase recruitment domain family, meml | 387 | 222 | 204 | 9 | 0 | 0 | 1 | 0 | 3 | 0 | 5 |
| 61 | -0.21 | 0.91 | -0.23 | 0.82 | 1 | Dlk1 | delta like non-canonical Notch ligand 1 | 85 | 179 | 174 | 27 | 26 | 64 | 1 | 14 | 42 | 0 | 54 |
| 57 | -0.75 | 1.02 | -0.74 | 0.46 | 1 | Tcea3 | transcription elongation factor A (SII), 3 | 155 | 192 | 212 | 17 | 18 | 6 | 1 | 21 | 3 | 0 | 5 |
| 56 | -0.69 | 1.22 | -0.57 | 0.57 | 1 | Naaladl2 | N-acetylated alpha-linked acidic dipeptid | 255 | 181 | 164 | 0 | 6 | 3 | 1 | 0 | 2 | 0 | 5 |
| 56 | 0.77 | 1.23 | 0.63 | 0.53 | 1 | Rnase6 | ribonuclease, RNase A family, 6 | 287 | 186 | 128 | 0 | 0 | 3 | 1 | 0 | 0 | 0 | 10 |
| 55 | -0.42 | 1.21 | -0.35 | 0.73 | 1 | Gm15344 | | 3 | 1 | 0 | 9 | 2 | 298 | 1 | 0 | 9 | 0 | 281 |
| 55 | -1.09 | 1.57 | -0.69 | 0.49 | 1 | Cdh6 | cadherin 6 | 290 | 101 | 207 | 0 | 0 | 1 | 1 | 0 | 0 | 0 | 0 |
| 49 | 0.14 | 0.99 | 0.14 | 0.89 | 1 | P2rx3 | purinergic receptor P2X, ligand-gated ion | 48 | 77 | 24 | 43 | 36 | 106 | 1 | 56 | 38 | 0 | 116 |
| 48 | -0.31 | 3.66 | -0.09 | 0.93 | 1 | Actg2 | actin, gamma 2, smooth muscle, enteric | 450 | 61 | 11 | 0 | 0 | 0 | 1 | 0 | 0 | 0 | 0 |
| 45 | 0.11 | 1.07 | 0.10 | 0.92 | 1 | Lrrtm1 | leucine rich repeat transmembrane neuro | 181 | 168 | 132 | 3 | 1 | 3 | 1 | 0 | 3 | 0 | 6 |
| 45 | 2.03 | 1.47 | 1.38 | 0.17 | 1 | Gli2 | GLI-Kruppel family member GLI2 | 232 | 126 | 130 | 0 | 0 | 0 | 1 | 0 | 2 | 0 | 3 |
| 45 | -2.88 | 1.50 | -1.92 | 0.05 | 1 | Il21r | interleukin 21 receptor | 323 | 120 | 37 | 9 | 0 | 3 | 1 | 0 | 1 | 0 | 0 |
| 43 | -0.65 | 1.01 | -0.64 | 0.52 | 1 | Mycn | v-myc avian myelocytomatosis viral relate | 132 | 128 | 157 | 6 | 10 | 17 | 1 | 0 | 4 | 17 | 5 |
| 43 | 0.13 | 1.05 | 0.12 | 0.90 | 1 | Rimklb | ribosomal modification protein rimK-like f | 124 | 137 | 140 | 3 | 6 | 31 | 1 | 0 | 5 | 22 | 8 |
| 42 | -1.66 | 1.56 | -1.06 | 0.29 | 1 | Ms4a6d | membrane-spanning 4-domains, subfami | 172 | 113 | 176 | 0 | 0 | 3 | 1 | 0 | 0 | 0 | 0 |
| 41 | 0.34 | 0.75 | 0.45 | 0.65 | 1 | Gm7809 | | 17 | 142 | 10 | 76 | 14 | 54 | 1 | 61 | 23 | 5 | 48 |
| 41 | -0.38 | 3.67 | -0.10 | 0.92 | 1 | Trim30d | tripartite motif-containing 30D | 295 | 62 | 89 | 0 | 0 | 0 | 1 | 0 | 0 | 0 | 0 |
| 39 | -1.54 | 1.60 | -0.96 | 0.34 | 1 | AI504432 | | 163 | 124 | 137 | 2 | 0 | 0 | 1 | 0 | 0 | 0 | 0 |
| 39 | 1.55 | 1.35 | 1.15 | 0.25 | 1 | Unc13d | unc-13 homolog D | 161 | 96 | 57 | 0 | 0 | 50 | 1 | 0 | 0 | 21 | 39 |
| 38 | -0.44 | 3.68 | -0.12 | 0.90 | 1 | Cysltr2 | cysteinyl leukotriene receptor 2 | 95 | 130 | 187 | 0 | 0 | 0 | 1 | 0 | 0 | 0 | 0 |
| 34 | -1.03 | 1.65 | -0.62 | 0.53 | 1 | Aim2 | absent in melanoma 2 | 245 | 79 | 44 | 0 | 0 | 1 | 1 | 0 | 0 | 0 | 0 |

| | | | | | | | | | | | | | | | | | | |
|---|---|---|---|---|---|---|---|---|---|---|---|---|---|---|---|---|---|---|
| 32 | 0.85 | 1.62 | 0.52 | 0.60 | 1 | Foxc1 | forkhead box C1 | 206 | 52 | 90 | 0 | 0 | 0 | 1 | 0 | 0 | 0 | 3 |
| 30 | -1.67 | 1.44 | -1.16 | 0.25 | 1 | Rab38 | RAB38, member RAS oncogene family | 124 | 119 | 75 | 0 | 3 | 3 | 1 | 0 | 0 | 0 | 2 |
| 28 | -2.10 | 1.29 | -1.63 | 0.10 | 1 | Mbnl3 | muscleblind like splicing factor 3 | 106 | 37 | 62 | 17 | 41 | 9 | 1 | 0 | 19 | 14 | 0 |
| 28 | -0.44 | 3.68 | -0.12 | 0.90 | 1 | Ackr4 | atypical chemokine receptor 4 | 94 | 70 | 140 | 0 | 0 | 0 | 1 | 0 | 0 | 0 | 0 |
| 26 | 0.82 | 1.02 | 0.80 | 0.42 | 1 | 4930526A20Rik | | 38 | 54 | 42 | 2 | 29 | 40 | 1 | 17 | 8 | 21 | 34 |
| 24 | -0.19 | 1.11 | -0.17 | 0.86 | 1 | Rgs20 | regulator of G-protein signaling 20 | 62 | 73 | 65 | 9 | 20 | 5 | 1 | 0 | 19 | 3 | 12 |
| 24 | 0.54 | 0.98 | 0.55 | 0.58 | 1 | Sco2 | SCO2 cytochrome c oxidase assembly pr | 35 | 31 | 30 | 13 | 29 | 28 | 1 | 52 | 25 | 3 | 20 |
| 24 | -2.41 | 1.82 | -1.32 | 0.19 | 1 | Fcgr4 | Fc receptor, IgG, low affinity IV | 79 | 114 | 64 | 0 | 8 | 0 | 1 | 0 | 0 | 0 | 0 |
| 22 | -1.44 | 1.83 | -0.79 | 0.43 | 1 | Erbb4 | erb-b2 receptor tyrosine kinase 4 | 45 | 43 | 132 | 4 | 0 | 5 | 1 | 0 | 0 | 14 | 0 |
| 22 | -0.03 | 1.24 | -0.03 | 0.98 | 1 | Snx22 | sorting nexin 22 | 93 | 38 | 73 | 2 | 8 | 3 | 1 | 0 | 15 | 0 | 5 |
| 20 | -0.69 | 1.48 | -0.47 | 0.64 | 1 | Sh2b2 | SH2B adaptor protein 2 | 111 | 75 | 27 | 0 | 0 | 8 | 1 | 0 | 1 | 0 | 2 |
| 19 | -1.34 | 1.26 | -1.06 | 0.29 | 1 | Gm11944 | | 21 | 7 | 23 | 44 | 15 | 54 | 1 | 0 | 16 | 12 | 21 |
| 19 | -0.94 | 1.15 | -0.82 | 0.41 | 1 | 9430060I03Rik | | 20 | 2 | 23 | 62 | 26 | 25 | 1 | 15 | 7 | 8 | 23 |
| 18 | -1.38 | 1.84 | -0.75 | 0.45 | 1 | Pdcd1 | programmed cell death 1 | 96 | 68 | 30 | 0 | 0 | 2 | 1 | 0 | 0 | 0 | 0 |
| 17 | -0.13 | 1.46 | -0.09 | 0.93 | 1 | Mamstr | MEF2 activating motif and SAP domain cc | 63 | 21 | 60 | 13 | 0 | 3 | 1 | 3 | 3 | 21 | 0 |
| 17 | 0.72 | 1.37 | 0.52 | 0.60 | 1 | Hsd17b1 | hydroxysteroid (17-beta) dehydrogenase | 3 | 3 | 6 | 31 | 2 | 46 | 1 | 0 | 19 | 21 | 56 |
| 16 | -1.77 | 1.89 | -0.94 | 0.35 | 1 | Pipox | pipecolic acid oxidase | 47 | 50 | 76 | 0 | 0 | 4 | 1 | 0 | 0 | 0 | 0 |
| 16 | -1.34 | 1.44 | -0.93 | 0.35 | 1 | Gm44698 | | 20 | 20 | 7 | 16 | 23 | 38 | 1 | 0 | 29 | 0 | 22 |
| 16 | -0.16 | 1.45 | -0.11 | 0.91 | 1 | Gm12265 | | 20 | 33 | 17 | 14 | 9 | 14 | 1 | 0 | 11 | 54 | 1 |
| 15 | -0.02 | 1.73 | -0.01 | 0.99 | 1 | Gm6166 | | 63 | 19 | 31 | 0 | 9 | 1 | 1 | 0 | 1 | 45 | 0 |
| 15 | -0.13 | 1.35 | -0.09 | 0.92 | 1 | Rpl5-ps1 | | 15 | 51 | 12 | 21 | 9 | 19 | 1 | 0 | 21 | 12 | 3 |
| 15 | 0.59 | 1.18 | 0.50 | 0.62 | 1 | Phf20-ps | | 2 | 9 | 1 | 53 | 4 | 7 | 1 | 36 | 7 | 40 | 1 |
| 14 | 0.74 | 1.19 | 0.62 | 0.53 | 1 | Lrfn2 | leucine rich repeat and fibronectin type III | 23 | 30 | 22 | 15 | 5 | 7 | 1 | 31 | 11 | 0 | 9 |
| 13 | 0.09 | 1.96 | 0.05 | 0.96 | 1 | Gm12511 | | 38 | 32 | 69 | 0 | 0 | 2 | 1 | 0 | 6 | 0 | 0 |
| 13 | -0.93 | 1.50 | -0.62 | 0.54 | 1 | 2200002D01Ril | RIKEN cDNA 2200002D01 gene | 42 | 26 | 62 | 0 | 3 | 5 | 1 | 0 | 3 | 0 | 1 |
| 13 | -2.29 | 1.37 | -1.67 | 0.09 | 1 | Cpne7 | copine VII | 40 | 8 | 35 | 6 | 15 | 20 | 1 | 0 | 7 | 0 | 5 |
| 12 | 0.80 | 1.67 | 0.48 | 0.63 | 1 | Gm17936 | | 3 | 54 | 3 | 32 | 1 | 3 | 1 | 0 | 0 | 19 | 11 |
| 11 | 0.92 | 1.26 | 0.73 | 0.47 | 1 | Gm8242 | | 1 | 32 | 18 | 2 | 23 | 8 | 1 | 8 | 16 | 12 | 5 |
| 11 | -1.26 | 1.47 | -0.86 | 0.39 | 1 | Cox7a1 | cytochrome c oxidase subunit 7A1 | 14 | 6 | 15 | 9 | 34 | 7 | 1 | 0 | 21 | 0 | 15 |
| 11 | 1.87 | 1.43 | 1.31 | 0.19 | 1 | Gm24563 | | 1 | 0 | 14 | 11 | 3 | 9 | 1 | 16 | 11 | 43 | 12 |
| 11 | -0.35 | 2.41 | -0.14 | 0.89 | 1 | Slc14a1 | solute carrier family 14 (urea transporter), | 94 | 20 | 3 | 0 | 0 | 0 | 1 | 0 | 0 | 0 | 0 |
| 11 | -1.75 | 2.12 | -0.83 | 0.41 | 1 | Gm15413 | | 21 | 59 | 32 | 4 | 0 | 0 | 1 | 0 | 0 | 0 | 0 |
| 10 | -1.79 | 1.37 | -1.31 | 0.19 | 1 | Gm13217 | | 16 | 9 | 12 | 20 | 24 | 9 | 1 | 0 | 8 | 1 | 10 |
| 10 | -0.36 | 2.44 | -0.15 | 0.88 | 1 | Cd207 | CD207 antigen | 82 | 6 | 20 | 0 | 0 | 0 | 1 | 0 | 0 | 0 | 0 |
| 10 | 0.98 | 1.34 | 0.73 | 0.47 | 1 | Gm32633 | | 5 | 8 | 11 | 9 | 9 | 8 | 1 | 0 | 10 | 23 | 23 |
| 9 | -1.70 | 2.16 | -0.79 | 0.43 | 1 | 9130008F23Ril | RIKEN cDNA 9130008F23 gene | 52 | 8 | 38 | 0 | 4 | 0 | 1 | 0 | 0 | 0 | 0 |
| 9 | -3.18 | 1.90 | -1.67 | 0.09 | 1 | Rprm | reprimo, TP53 dependent G2 arrest media | 28 | 48 | 16 | 4 | 0 | 8 | 1 | 0 | 0 | 0 | 0 |
| 9 | 0.61 | 1.85 | 0.33 | 0.74 | 1 | Nkain2 | Na+/K+ transporting ATPase interacting 2 | 15 | 52 | 27 | 4 | 0 | 0 | 1 | 0 | 3 | 0 | 2 |
| 9 | -0.14 | 1.10 | -0.13 | 0.90 | 1 | Gm7143 | | 14 | 6 | 16 | 11 | 20 | 4 | 1 | 7 | 13 | 5 | 4 |
| 9 | 0.04 | 1.34 | 0.03 | 0.97 | 1 | Gm17102 | | 7 | 13 | 4 | 8 | 9 | 8 | 1 | 41 | 2 | 0 | 8 |
| 9 | 2.17 | 1.61 | 1.35 | 0.18 | 1 | Gm48151 | | 4 | 0 | 4 | 11 | 0 | 2 | 1 | 20 | 7 | 46 | 2 |
| 9 | -0.94 | 1.89 | -0.50 | 0.62 | 1 | Gm42933 | | 0 | 0 | 0 | 48 | 0 | 21 | 1 | 9 | 2 | 0 | 14 |
| 9 | 1.74 | 1.26 | 1.39 | 0.17 | 1 | Gm6158 | | 15 | 4 | 1 | 2 | 3 | 8 | 1 | 12 | 6 | 40 | 4 |
| 8 | 0.74 | 1.24 | 0.59 | 0.55 | 1 | Gm11539 | | 22 | 10 | 26 | 1 | 4 | 8 | 1 | 7 | 1 | 3 | 9 |
| 8 | 1.50 | 1.75 | 0.86 | 0.39 | 1 | Gm15484 | | 4 | 30 | 20 | 0 | 10 | 0 | 1 | 2 | 17 | 0 | 6 |
| 8 | -0.10 | 2.20 | -0.04 | 0.96 | 1 | 1700049E15Rik | | 9 | 31 | 0 | 0 | 6 | 17 | 1 | 0 | 20 | 0 | 7 |
| 8 | 1.48 | 2.50 | 0.59 | 0.55 | 1 | Ccr9 | chemokine (C-C motif) receptor 9 | 32 | 22 | 12 | 0 | 0 | 0 | 1 | 23 | 0 | 0 | 0 |
| 8 | 0.99 | 1.57 | 0.63 | 0.53 | 1 | Amdhd1 | amidohydrolase domain containing 1 | 31 | 15 | 9 | 0 | 7 | 4 | 1 | 7 | 3 | 0 | 13 |
| 8 | -2.40 | 1.95 | -1.23 | 0.22 | 1 | Gm43088 | | 3 | 14 | 2 | 0 | 7 | 36 | 1 | 0 | 0 | 0 | 26 |
| 8 | -2.03 | 1.83 | -1.11 | 0.27 | 1 | Henmt1 | HEN1 methyltransferase homolog 1 (Arak | 0 | 0 | 24 | 10 | 29 | 3 | 1 | 0 | 13 | 0 | 10 |
| 8 | 2.51 | 1.58 | 1.58 | 0.11 | 1 | Mab21l1 | mab-21-like 1 | 27 | 14 | 11 | 0 | 2 | 1 | 1 | 0 | 3 | 22 | 7 |
| 8 | 0.25 | 1.29 | 0.19 | 0.85 | 1 | Gm7451 | | 1 | 15 | 1 | 5 | 7 | 16 | 1 | 16 | 10 | 0 | 14 |
| 8 | -1.83 | 1.54 | -1.19 | 0.23 | 1 | C230012O17Rik | | 26 | 22 | 11 | 11 | 6 | 2 | 1 | 0 | 2 | 5 | 0 |
| 8 | -1.43 | 1.51 | -0.94 | 0.35 | 1 | Gm14148 | | 6 | 26 | 2 | 17 | 10 | 6 | 1 | 0 | 1 | 3 | 13 |
| 8 | -0.09 | 1.74 | -0.05 | 0.96 | 1 | Gm14830 | | 1 | 2 | 0 | 1 | 0 | 39 | 1 | 2 | 0 | 0 | 37 |
| 7 | 0.83 | 2.27 | 0.36 | 0.72 | 1 | Gm5540 | | 11 | 30 | 1 | 0 | 8 | 0 | 1 | 0 | 2 | 28 | 0 |
| 7 | -0.55 | 1.70 | -0.33 | 0.74 | 1 | A430108G06Rik | | 15 | 10 | 9 | 13 | 2 | 7 | 1 | 0 | 17 | 0 | 6 |
| 7 | 0.14 | 3.61 | 0.04 | 0.97 | 1 | Igkv3-7 | | 80 | 0 | 0 | 0 | 0 | 0 | 1 | 0 | 0 | 0 | 0 |
| 7 | -3.48 | 2.26 | -1.54 | 0.12 | 1 | Gm16201 | | 38 | 20 | 6 | 0 | 12 | 2 | 1 | 0 | 0 | 0 | 0 |
| 7 | -1.01 | 2.51 | -0.40 | 0.69 | 1 | Cd8b1 | CD8 antigen, beta chain 1 | 37 | 0 | 37 | 0 | 0 | 3 | 1 | 0 | 1 | 0 | 0 |

| | | | | | | | | | | | | | | | | | | |
|---|---|---|---|---|---|---|---|---|---|---|---|---|---|---|---|---|---|---|
| 7 | -1.61 | 1.95 | -0.82 | 0.41 | 1 | Gm49272 | | 0 | 0 | 0 | 7 | 2 | 36 | 1 | 0 | 2 | 0 | 30 |
| 7 | -0.94 | 1.67 | -0.56 | 0.57 | 1 | Gm2573 | | 3 | 13 | 18 | 10 | 0 | 17 | 1 | 0 | 1 | 1 | 15 |
| 7 | 2.45 | 2.80 | 0.88 | 0.38 | 1 | Gm7420 | | 0 | 11 | 0 | 0 | 0 | 22 | 1 | 0 | 0 | 31 | 10 |
| 7 | -0.18 | 1.27 | -0.14 | 0.89 | 1 | Gm3815 | | 7 | 20 | 14 | 11 | 3 | 3 | 1 | 2 | 10 | 1 | 2 |
| 7 | 2.97 | 1.59 | 1.87 | 0.06 | 1 | Gm11223 | | 9 | 13 | 1 | 0 | 1 | 4 | 1 | 1 | 6 | 30 | 8 |
| 7 | -1.01 | 2.32 | -0.44 | 0.66 | 1 | Areg | amphiregulin | 43 | 0 | 7 | 9 | 0 | 4 | 1 | 0 | 6 | 0 | 3 |
| 7 | -0.41 | 1.35 | -0.30 | 0.76 | 1 | Gm11977 | | 1 | 3 | 25 | 5 | 7 | 6 | 1 | 1 | 11 | 0 | 13 |
| 7 | 1.65 | 2.06 | 0.80 | 0.42 | 1 | Rasl2-9 | RAS-like, family 2, locus 9 | 0 | 11 | 9 | 0 | 13 | 3 | 1 | 0 | 3 | 21 | 11 |
| 6 | -2.03 | 2.06 | -0.98 | 0.32 | 1 | Gm38342 | | 6 | 1 | 0 | 18 | 6 | 8 | 1 | 0 | 24 | 5 | 0 |
| 6 | 2.47 | 2.12 | 1.16 | 0.24 | 1 | Fcor | Foxo1 corepressor | 13 | 7 | 31 | 0 | 0 | 1 | 1 | 12 | 0 | 0 | 5 |
| 6 | 0.74 | 1.98 | 0.37 | 0.71 | 1 | Gm42600 | | 24 | 6 | 3 | 8 | 0 | 4 | 1 | 0 | 6 | 0 | 16 |
| 6 | 2.33 | 1.87 | 1.25 | 0.21 | 1 | AW495222 | | 13 | 19 | 1 | 5 | 0 | 0 | 1 | 15 | 1 | 0 | 11 |
| 6 | -3.06 | 1.71 | -1.79 | 0.07 | 1 | Gm15501 | predicted pseudogene 15501 | 18 | 13 | 9 | 2 | 16 | 5 | 1 | 0 | 1 | 0 | 1 |
| 6 | 0.02 | 1.39 | 0.01 | 0.99 | 1 | Rpsa-ps4 | | 3 | 40 | 1 | 6 | 2 | 3 | 1 | 3 | 2 | 1 | 3 |
| 6 | -0.79 | 1.58 | -0.50 | 0.62 | 1 | Rps15a-ps4 | | 26 | 7 | 5 | 5 | 5 | 2 | 1 | 2 | 1 | 0 | 9 |
| 6 | -0.21 | 1.57 | -0.13 | 0.89 | 1 | Gm5687 | | 11 | 1 | 14 | 0 | 7 | 14 | 1 | 0 | 9 | 1 | 4 |
| 6 | 0.45 | 2.24 | 0.20 | 0.84 | 1 | Gm7004 | | 0 | 5 | 9 | 1 | 0 | 4 | 1 | 32 | 8 | 0 | 0 |
| 6 | 0.26 | 1.88 | 0.14 | 0.89 | 1 | Gm12482 | | 6 | 7 | 0 | 1 | 14 | 2 | 1 | 21 | 8 | 0 | 1 |
| 5 | 0.39 | 2.18 | 0.18 | 0.86 | 1 | Gm5466 | | 3 | 0 | 1 | 0 | 0 | 25 | 1 | 0 | 1 | 0 | 28 |
| 5 | -0.28 | 1.56 | -0.18 | 0.86 | 1 | Gm8925 | | 7 | 4 | 5 | 1 | 19 | 2 | 1 | 1 | 8 | 12 | 0 |
| 5 | 2.08 | 1.65 | 1.26 | 0.21 | 1 | Gm43653 | | 19 | 11 | 6 | 1 | 0 | 3 | 1 | 5 | 4 | 0 | 9 |
| 5 | -1.69 | 2.32 | -0.73 | 0.47 | 1 | Hsp25-ps1 | | 20 | 23 | 12 | 0 | 1 | 1 | 1 | 0 | 0 | 0 | 0 |
| 5 | 2.41 | 2.00 | 1.20 | 0.23 | 1 | Gm10146 | | 12 | 12 | 1 | 0 | 0 | 5 | 1 | 20 | 2 | 0 | 3 |
| 5 | -1.58 | 1.86 | -0.85 | 0.40 | 1 | Gm11464 | | 4 | 16 | 9 | 9 | 4 | 1 | 1 | 12 | 0 | 0 | 0 |
| 5 | -1.06 | 1.94 | -0.55 | 0.58 | 1 | Ccdc146 | coiled-coil domain containing 146 | 3 | 0 | 5 | 9 | 6 | 6 | 1 | 0 | 12 | 0 | 13 |
| 5 | 1.47 | 1.80 | 0.82 | 0.41 | 1 | Gm6139 | | 0 | 10 | 18 | 7 | 2 | 0 | 1 | 1 | 6 | 3 | 7 |
| 5 | -1.48 | 2.04 | -0.72 | 0.47 | 1 | Gm10472 | | 8 | 13 | 0 | 12 | 3 | 8 | 1 | 0 | 6 | 0 | 3 |
| 5 | -0.62 | 2.42 | -0.26 | 0.80 | 1 | Gm44694 | | 0 | 0 | 17 | 4 | 0 | 10 | 1 | 0 | 4 | 15 | 3 |
| 5 | -1.11 | 1.80 | -0.62 | 0.54 | 1 | Gm5619 | | 13 | 2 | 0 | 10 | 7 | 3 | 1 | 0 | 2 | 9 | 7 |
| 5 | 0.64 | 1.63 | 0.39 | 0.70 | 1 | Gm6781 | | 0 | 8 | 6 | 1 | 3 | 6 | 1 | 23 | 1 | 1 | 3 |
| 5 | -2.66 | 1.87 | -1.42 | 0.16 | 1 | Gm8902 | | 4 | 19 | 5 | 10 | 5 | 4 | 1 | 0 | 4 | 0 | 0 |
| 5 | -1.39 | 1.93 | -0.72 | 0.47 | 1 | Gm18336 | | 2 | 9 | 10 | 11 | 0 | 10 | 1 | 0 | 4 | 0 | 5 |
| 5 | 0.06 | 1.76 | 0.03 | 0.97 | 1 | Ccdc152 | coiled-coil domain containing 152 | 7 | 2 | 3 | 4 | 1 | 8 | 1 | 0 | 6 | 0 | 20 |
| 5 | -1.70 | 1.76 | -0.96 | 0.33 | 1 | Gm8326 | | 3 | 0 | 3 | 23 | 4 | 7 | 1 | 2 | 2 | 0 | 7 |
| 5 | 1.38 | 2.06 | 0.67 | 0.50 | 1 | Gm5120 | | 2 | 2 | 1 | 0 | 2 | 3 | 1 | 0 | 0 | 31 | 9 |
| 5 | 0.12 | 2.82 | 0.04 | 0.97 | 1 | 2010008C14Rik | | 25 | 16 | 0 | 0 | 0 | 3 | 1 | 0 | 0 | 6 | 0 |
| 5 | -1.27 | 1.84 | -0.69 | 0.49 | 1 | Gm12492 | | 8 | 7 | 1 | 4 | 11 | 5 | 1 | 0 | 11 | 0 | 3 |
| 5 | -0.24 | 3.65 | -0.06 | 0.95 | 1 | Ighv1-2 | | 45 | 5 | 0 | 0 | 0 | 0 | 1 | 0 | 0 | 0 | 0 |
| 4 | -0.08 | 1.54 | -0.05 | 0.96 | 1 | Gm8253 | | 10 | 1 | 22 | 3 | 2 | 3 | 1 | 3 | 2 | 1 | 1 |
| 4 | 1.96 | 1.66 | 1.18 | 0.24 | 1 | Gm5566 | | 1 | 1 | 10 | 4 | 0 | 3 | 1 | 6 | 4 | 12 | 7 |
| 4 | 2.34 | 1.96 | 1.19 | 0.23 | 1 | Rpl10-ps6 | | 6 | 6 | 1 | 0 | 0 | 12 | 1 | 1 | 15 | 1 | 5 |
| 4 | -0.90 | 2.78 | -0.32 | 0.75 | 1 | Gm14612 | | 1 | 0 | 0 | 0 | 25 | 1 | 1 | 0 | 21 | 0 | 0 |
| 4 | 1.33 | 1.59 | 0.84 | 0.40 | 1 | Gm7363 | | 3 | 8 | 0 | 1 | 1 | 4 | 1 | 22 | 1 | 1 | 6 |
| 4 | -0.53 | 2.34 | -0.23 | 0.82 | 1 | Gm5864 | | 21 | 14 | 6 | 0 | 0 | 3 | 1 | 0 | 0 | 0 | 2 |
| 4 | -0.55 | 2.47 | -0.22 | 0.83 | 1 | Myo1a | myosin IA | 2 | 29 | 1 | 0 | 0 | 7 | 1 | 0 | 0 | 0 | 7 |
| 4 | -2.56 | 2.31 | -1.11 | 0.27 | 1 | Gm5873 | | 0 | 0 | 16 | 13 | 1 | 5 | 1 | 0 | 8 | 0 | 3 |
| 4 | -0.11 | 2.34 | -0.05 | 0.96 | 1 | Gm14411 | | 1 | 0 | 0 | 15 | 0 | 4 | 1 | 0 | 8 | 17 | 1 |
| 4 | 0.18 | 1.56 | 0.12 | 0.91 | 1 | Rpl10-ps5 | | 2 | 2 | 4 | 11 | 2 | 2 | 1 | 3 | 1 | 17 | 1 |
| 4 | 2.05 | 1.95 | 1.05 | 0.29 | 1 | S100a4 | S100 calcium binding protein A4 | 9 | 1 | 4 | 0 | 8 | 1 | 1 | 13 | 4 | 0 | 5 |
| 4 | -1.34 | 1.72 | -0.78 | 0.43 | 1 | Rps24-ps2 | | 15 | 0 | 1 | 6 | 4 | 9 | 1 | 1 | 2 | 1 | 5 |
| 4 | 1.48 | 2.65 | 0.56 | 0.58 | 1 | Gm49730 | | 0 | 4 | 5 | 0 | 0 | 8 | 1 | 0 | 0 | 10 | 16 |
| 4 | 3.35 | 2.19 | 1.53 | 0.13 | 1 | Atf1-ps | | 9 | 0 | 8 | 0 | 0 | 2 | 1 | 16 | 1 | 1 | 7 |
| 4 | 2.81 | 1.68 | 1.67 | 0.10 | 1 | Gm3511 | | 9 | 1 | 5 | 0 | 1 | 3 | 1 | 7 | 9 | 8 | 1 |
| 4 | -1.82 | 1.82 | -1.00 | 0.32 | 1 | Actg-ps1 | | 13 | 6 | 5 | 6 | 5 | 3 | 1 | 1 | 5 | 0 | 0 |
| 4 | -0.49 | 1.43 | -0.34 | 0.73 | 1 | Gm5846 | | 10 | 2 | 5 | 4 | 3 | 7 | 1 | 1 | 1 | 4 | 6 |
| 4 | -0.55 | 2.29 | -0.24 | 0.81 | 1 | Gm31544 | | 3 | 18 | 11 | 7 | 0 | 0 | 1 | 2 | 0 | 0 | 1 |
| 4 | -1.73 | 1.78 | -0.97 | 0.33 | 1 | Gm31266 | | 9 | 4 | 6 | 7 | 2 | 7 | 1 | 0 | 2 | 0 | 5 |
| 4 | 0.67 | 2.92 | 0.23 | 0.82 | 1 | Gm14863 | | 6 | 0 | 7 | 16 | 0 | 0 | 1 | 0 | 9 | 0 | 4 |
| 4 | -0.27 | 2.11 | -0.13 | 0.90 | 1 | 9330182O14Ril | RIKEN cDNA 9330182O14 gene | 0 | 0 | 0 | 1 | 10 | 8 | 1 | 0 | 2 | 13 | 6 |
| 4 | 0.50 | 1.71 | 0.29 | 0.77 | 1 | Selenok-ps1 | | 2 | 0 | 1 | 3 | 1 | 9 | 1 | 2 | 0 | 9 | 13 |

| | | | | | | | | | | | | | | | | | | |
|---|---|---|---|---|---|---|---|---|---|---|---|---|---|---|---|---|---|---|
| 4 | 3.34 | 2.71 | 1.24 | 0.22 | 1 | Olfr920 | olfactory receptor 920 | 2 | 0 | 0 | 0 | 0 | 19 | 1 | 12 | 4 | 0 | 4 |
| 4 | -0.09 | 2.04 | -0.04 | 0.97 | 1 | Gm8231 | | 2 | 30 | 2 | 1 | 0 | 1 | 1 | 1 | 0 | 0 | 2 |
| 4 | 0.94 | 2.40 | 0.39 | 0.69 | 1 | Rnf170-ps | | 9 | 0 | 4 | 0 | 1 | 4 | 1 | 1 | 0 | 18 | 0 |
| 4 | 0.43 | 2.52 | 0.17 | 0.86 | 1 | C130089K02Rik | | 2 | 0 | 0 | 0 | 4 | 6 | 1 | 0 | 5 | 0 | 21 |
| 3 | -1.89 | 2.29 | -0.83 | 0.41 | 1 | Gm9645 | | 0 | 0 | 10 | 8 | 5 | 2 | 1 | 0 | 9 | 0 | 3 |
| 3 | -0.66 | 2.48 | -0.27 | 0.79 | 1 | Gm17110 | | 12 | 7 | 0 | 0 | 2 | 3 | 1 | 10 | 0 | 0 | 2 |
| 3 | 2.11 | 2.56 | 0.82 | 0.41 | 1 | Gm7780 | | 15 | 0 | 0 | 0 | 0 | 3 | 1 | 3 | 0 | 13 | 2 |
| 3 | 2.08 | 1.98 | 1.05 | 0.29 | 1 | Gm13827 | | 1 | 21 | 4 | 0 | 1 | 0 | 1 | 3 | 1 | 3 | 2 |
| 3 | -4.13 | 2.35 | -1.76 | 0.08 | 1 | Gm13778 | | 2 | 0 | 5 | 13 | 5 | 8 | 1 | 0 | 3 | 0 | 0 |
| 3 | -1.87 | 2.26 | -0.83 | 0.41 | 1 | Gm5487 | | 16 | 3 | 3 | 0 | 4 | 6 | 1 | 0 | 0 | 0 | 4 |
| 3 | -1.33 | 2.91 | -0.46 | 0.65 | 1 | Gm49838 | | 0 | 0 | 2 | 1 | 0 | 17 | 1 | 0 | 0 | 0 | 16 |
| 3 | -0.39 | 1.64 | -0.24 | 0.81 | 1 | Gm14519 | | 1 | 10 | 3 | 1 | 5 | 4 | 1 | 7 | 2 | 0 | 3 |
| 3 | -0.35 | 3.06 | -0.11 | 0.91 | 1 | Gm49413 | | 0 | 0 | 0 | 0 | 7 | 9 | 1 | 0 | 0 | 17 | 3 |
| 3 | -1.03 | 1.61 | -0.64 | 0.52 | 1 | Gm5837 | | 3 | 1 | 0 | 4 | 6 | 12 | 1 | 2 | 3 | 3 | 2 |
| 3 | 1.35 | 1.60 | 0.85 | 0.40 | 1 | Gm8942 | | 8 | 4 | 6 | 3 | 2 | 0 | 1 | 5 | 2 | 5 | 1 |
| 3 | 2.28 | 2.16 | 1.06 | 0.29 | 1 | Gm16409 | | 7 | 1 | 3 | 2 | 0 | 0 | 1 | 1 | 0 | 18 | 3 |
| 3 | -1.59 | 2.07 | -0.77 | 0.44 | 1 | Rps12-ps1 | | 5 | 0 | 7 | 12 | 3 | 1 | 1 | 1 | 1 | 0 | 4 |
| 3 | -2.49 | 3.12 | -0.80 | 0.43 | 1 | Gm16072 | | 12 | 18 | 0 | 0 | 1 | 3 | 1 | 0 | 0 | 0 | 0 |
| 3 | -2.32 | 3.10 | -0.75 | 0.45 | 1 | Epha5 | Eph receptor A5 | 0 | 0 | 0 | 24 | 0 | 7 | 1 | 0 | 2 | 0 | 2 |
| 3 | -0.91 | 2.24 | -0.41 | 0.69 | 1 | Gm3219 | | 15 | 13 | 1 | 0 | 2 | 2 | 1 | 0 | 1 | 0 | 1 |
| 3 | 1.44 | 3.28 | 0.44 | 0.66 | 1 | Rab5a-ps | | 23 | 0 | 0 | 0 | 3 | 0 | 1 | 1 | 7 | 0 | 0 |
| 3 | -1.44 | 2.46 | -0.59 | 0.56 | 1 | Atp5l-ps1 | | 1 | 0 | 0 | 18 | 2 | 1 | 1 | 0 | 2 | 0 | 10 |
| 3 | -1.09 | 2.10 | -0.52 | 0.60 | 1 | Gm13653 | | 9 | 5 | 14 | 0 | 3 | 2 | 1 | 1 | 0 | 1 | 0 |
| 3 | -1.45 | 2.15 | -0.67 | 0.50 | 1 | Gm9207 | | 5 | 1 | 0 | 3 | 5 | 2 | 1 | 0 | 0 | 17 | 2 |
| 3 | -1.40 | 2.01 | -0.70 | 0.49 | 1 | Gm5787 | | 4 | 0 | 7 | 10 | 1 | 4 | 1 | 0 | 2 | 1 | 4 |
| 3 | -1.95 | 2.57 | -0.76 | 0.45 | 1 | Gm26407 | | 3 | 0 | 6 | 0 | 9 | 9 | 1 | 0 | 0 | 0 | 5 |
| 3 | -0.30 | 2.12 | -0.14 | 0.89 | 1 | Rps2-ps7 | | 14 | 7 | 4 | 2 | 0 | 2 | 1 | 0 | 1 | 0 | 4 |
| 3 | 0.79 | 1.85 | 0.43 | 0.67 | 1 | Mif-ps | | 1 | 0 | 1 | 4 | 4 | 1 | 1 | 0 | 6 | 14 | 3 |
| 3 | -0.44 | 3.58 | -0.12 | 0.90 | 1 | Gm13717 | | 3 | 14 | 16 | 0 | 0 | 0 | 1 | 0 | 0 | 0 | 0 |
| 3 | 0.45 | 2.58 | 0.17 | 0.86 | 1 | Gm46416 | | 0 | 8 | 5 | 3 | 11 | 0 | 1 | 0 | 0 | 4 | 2 |
| 3 | -1.73 | 3.34 | -0.52 | 0.60 | 1 | mt-Tp | | 0 | 3 | 2 | 11 | 0 | 0 | 1 | 0 | 0 | 0 | 16 |
| 3 | 0.43 | 1.48 | 0.29 | 0.77 | 1 | Gm7331 | | 4 | 2 | 3 | 3 | 3 | 3 | 1 | 5 | 6 | 3 | 1 |
| 3 | 0.35 | 1.69 | 0.21 | 0.84 | 1 | Gm3534 | | 3 | 5 | 6 | 2 | 1 | 3 | 1 | 0 | 2 | 1 | 9 |
| 3 | -0.08 | 2.07 | -0.04 | 0.97 | 1 | Gm7424 | | 4 | 2 | 3 | 7 | 3 | 0 | 1 | 10 | 0 | 3 | 0 |
| 3 | 1.35 | 1.93 | 0.70 | 0.49 | 1 | Gm48067 | | 0 | 0 | 0 | 2 | 3 | 3 | 1 | 5 | 1 | 14 | 5 |
| 3 | -0.15 | 2.64 | -0.06 | 0.95 | 1 | Gm10343 | | 15 | 0 | 1 | 10 | 0 | 0 | 1 | 1 | 0 | 1 | 4 |
| 3 | 0.54 | 2.48 | 0.22 | 0.83 | 1 | Gm5937 | | 1 | 18 | 8 | 0 | 0 | 1 | 1 | 0 | 0 | 1 | 2 |
| 3 | -1.00 | 1.86 | -0.54 | 0.59 | 1 | Gm6913 | | 14 | 5 | 0 | 4 | 3 | 1 | 1 | 1 | 1 | 1 | 1 |
| 3 | -0.80 | 3.05 | -0.26 | 0.79 | 1 | | | 12 | 4 | 0 | 0 | 0 | 11 | 1 | 0 | 0 | 0 | 4 |
| 3 | 1.50 | 2.01 | 0.75 | 0.46 | 1 | Gm26636 | | 3 | 0 | 0 | 1 | 1 | 3 | 1 | 0 | 6 | 13 | 5 |
| 3 | -4.43 | 3.55 | -1.25 | 0.21 | 1 | Gm15597 | | 0 | 0 | 0 | 25 | 5 | 1 | 1 | 0 | 0 | 0 | 0 |
| 3 | -0.11 | 2.75 | -0.04 | 0.97 | 1 | Gm7623 | | 1 | 16 | 0 | 0 | 1 | 8 | 1 | 0 | 3 | 1 | 0 |
| 3 | -1.16 | 1.96 | -0.59 | 0.56 | 1 | Gm5616 | | 3 | 10 | 12 | 1 | 1 | 2 | 1 | 0 | 1 | 0 | 1 |
| 3 | 3.95 | 3.29 | 1.20 | 0.23 | 1 | Spr-ps1 | | 20 | 0 | 0 | 0 | 0 | 0 | 1 | 3 | 1 | 0 | 6 |
| 3 | 0.42 | 1.90 | 0.22 | 0.83 | 1 | Rps12-ps9 | | 12 | 8 | 2 | 2 | 0 | 1 | 1 | 1 | 0 | 1 | 3 |
| 3 | -5.69 | 2.92 | -1.95 | 0.05 | 1 | Gm13034 | | 0 | 17 | 0 | 1 | 5 | 5 | 1 | 0 | 0 | 0 | 2 |
| 3 | 0.81 | 1.69 | 0.48 | 0.63 | 1 | Gm5778 | | 4 | 1 | 0 | 2 | 2 | 4 | 1 | 1 | 5 | 4 | 6 |
| 3 | 0.95 | 2.46 | 0.39 | 0.70 | 1 | Gm5643 | | 3 | 0 | 17 | 0 | 0 | 3 | 1 | 1 | 1 | 0 | 5 |
| 3 | -2.64 | 2.28 | -1.16 | 0.25 | 1 | Gm11652 | | 12 | 1 | 0 | 1 | 7 | 5 | 1 | 0 | 0 | 1 | 2 |
| 3 | -1.69 | 2.03 | -0.83 | 0.41 | 1 | Gm12922 | | 4 | 0 | 1 | 11 | 4 | 2 | 1 | 2 | 0 | 1 | 3 |
| 3 | 1.79 | 3.31 | 0.54 | 0.59 | 1 | Gm3145 | | 0 | 1 | 0 | 0 | 0 | 1 | 1 | 1 | 0 | 26 | 0 |
| 3 | -3.56 | 2.56 | -1.39 | 0.16 | 1 | Gm8069 | | 9 | 2 | 1 | 12 | 4 | 0 | 1 | 0 | 0 | 0 | 1 |
| 3 | -1.24 | 3.00 | -0.41 | 0.68 | 1 | Gm13181 | | 0 | 0 | 1 | 0 | 2 | 3 | 1 | 22 | 1 | 0 | 0 |
| 3 | 0.12 | 2.58 | 0.05 | 0.96 | 1 | H4c6 | H4 clustered histone 6 | 0 | 0 | 5 | 0 | 2 | 5 | 1 | 0 | 4 | 0 | 12 |
| 3 | 0.37 | 1.77 | 0.21 | 0.84 | 1 | Rplp0-ps1 | | 4 | 3 | 5 | 3 | 1 | 2 | 1 | 1 | 0 | 5 | 4 |
| 3 | 1.02 | 1.73 | 0.59 | 0.56 | 1 | Gm7367 | | 2 | 9 | 0 | 2 | 1 | 2 | 1 | 5 | 1 | 5 | 2 |
| 3 | -1.89 | 2.13 | -0.89 | 0.38 | 1 | Gm13161 | | 0 | 6 | 2 | 3 | 11 | 2 | 1 | 0 | 3 | 0 | 1 |
| 3 | 0.38 | 1.99 | 0.19 | 0.85 | 1 | Gm12906 | | 1 | 1 | 0 | 1 | 2 | 3 | 1 | 0 | 2 | 17 | 1 |
| 3 | -1.05 | 2.92 | -0.36 | 0.72 | 1 | Gm8371 | | 15 | 1 | 1 | 0 | 9 | 0 | 1 | 0 | 2 | 0 | 0 |
| 3 | 1.95 | 2.93 | 0.67 | 0.50 | 1 | Gm6568 | | 0 | 0 | 1 | 0 | 0 | 6 | 1 | 0 | 2 | 17 | 1 |

| | | | | | | | | | | | | | | | | | | |
|---|---|---|---|---|---|---|---|---|---|---|---|---|---|---|---|---|---|---|
| 3 | -0.05 | 2.78 | -0.02 | 0.99 | 1 | Gm11221 | | 14 | 2 | 0 | 0 | 0 | 8 | 1 | 1 | 0 | 0 | 2 |
| 2 | 1.62 | 1.67 | 0.97 | 0.33 | 1 | Gm16089 | | 7 | 2 | 1 | 1 | 1 | 2 | 1 | 5 | 4 | 1 | 3 |
| 2 | 0.82 | 2.34 | 0.35 | 0.73 | 1 | Gm8141 | | 4 | 2 | 1 | 0 | 2 | 2 | 1 | 10 | 0 | 0 | 5 |
| 2 | -0.28 | 2.34 | -0.12 | 0.90 | 1 | Gm12583 | | 3 | 1 | 16 | 2 | 2 | 0 | 1 | 0 | 0 | 3 | 1 |
| 2 | -0.41 | 3.67 | -0.11 | 0.91 | 1 | F2rl2 | coagulation factor II (thrombin) receptor-li | 12 | 11 | 3 | 0 | 0 | 0 | 1 | 0 | 0 | 0 | 0 |
| 2 | 2.73 | 3.52 | 0.77 | 0.44 | 1 | Gm24245 | | 0 | 0 | 0 | 2 | 0 | 0 | 1 | 0 | 0 | 19 | 5 |
| 2 | -0.70 | 2.43 | -0.29 | 0.77 | 1 | Gm14698 | | 7 | 1 | 3 | 6 | 0 | 1 | 1 | 0 | 1 | 0 | 7 |
| 2 | -1.37 | 3.58 | -0.38 | 0.70 | 1 | Gm8532 | | 0 | 0 | 16 | 0 | 0 | 8 | 1 | 0 | 0 | 0 | 2 |
| 2 | -2.35 | 3.33 | -0.71 | 0.48 | 1 | Lrrd1 | leucine rich repeats and death domain cc | 8 | 0 | 0 | 0 | 4 | 4 | 1 | 0 | 0 | 10 | 0 |
| 2 | 1.07 | 2.85 | 0.37 | 0.71 | 1 | 3110006O06Rik | | 0 | 0 | 2 | 0 | 0 | 3 | 1 | 0 | 17 | 1 | 2 |
| 2 | 2.31 | 3.35 | 0.69 | 0.49 | 1 | Gabra1 | gamma-aminobutyric acid (GABA) A rece | 3 | 0 | 7 | 0 | 0 | 0 | 1 | 0 | 0 | 14 | 1 |
| 2 | -2.08 | 2.15 | -0.97 | 0.33 | 1 | Gm6245 | | 0 | 1 | 0 | 6 | 12 | 2 | 1 | 1 | 1 | 3 | 0 |
| 2 | -1.39 | 3.39 | -0.41 | 0.68 | 1 | Kcna4 | potassium voltage-gated channel, shaker | 5 | 0 | 4 | 0 | 15 | 0 | 1 | 0 | 0 | 0 | 1 |
| 2 | 1.78 | 2.38 | 0.75 | 0.46 | 1 | Gm47655 | | 0 | 1 | 10 | 0 | 3 | 1 | 1 | 1 | 0 | 4 | 5 |
| 2 | -6.31 | 3.54 | -1.78 | 0.07 | 1 | Gm41505 | | 0 | 8 | 0 | 0 | 9 | 4 | 1 | 0 | 0 | 0 | 4 |
| 2 | -1.02 | 2.33 | -0.44 | 0.66 | 1 | Gm6190 | | 2 | 0 | 7 | 2 | 3 | 3 | 1 | 0 | 0 | 8 | 1 |
| 2 | -0.05 | 3.60 | -0.01 | 0.99 | 1 | Gm45756 | | 0 | 15 | 0 | 0 | 0 | 0 | 1 | 9 | 0 | 0 | 0 |
| 2 | -0.47 | 1.82 | -0.26 | 0.80 | 1 | Gm17415 | | 2 | 2 | 1 | 4 | 1 | 6 | 1 | 0 | 2 | 4 | 2 |
| 2 | -2.17 | 2.92 | -0.74 | 0.46 | 1 | Gm20492 | | 4 | 0 | 2 | 3 | 0 | 12 | 1 | 0 | 0 | 0 | 3 |
| 2 | -0.11 | 2.95 | -0.04 | 0.97 | 1 | 1700040L08Rik | | 0 | 0 | 6 | 0 | 2 | 9 | 1 | 2 | 0 | 0 | 4 |
| 2 | -0.61 | 1.87 | -0.33 | 0.74 | 1 | Gm13577 | | 3 | 1 | 2 | 4 | 4 | 2 | 1 | 7 | 1 | 0 | 1 |
| 2 | -1.67 | 2.03 | -0.82 | 0.41 | 1 | Stxbp3-ps | | 3 | 1 | 2 | 1 | 8 | 3 | 1 | 0 | 1 | 0 | 5 |
| 2 | 3.45 | 3.07 | 1.12 | 0.26 | 1 | Gm16416 | | 6 | 7 | 0 | 0 | 0 | 0 | 1 | 1 | 5 | 0 | 4 |
| 2 | -1.57 | 3.32 | -0.47 | 0.64 | 1 | Gm37376 | | 0 | 1 | 0 | 0 | 2 | 9 | 1 | 0 | 0 | 0 | 11 |
| 2 | 0.72 | 2.17 | 0.33 | 0.74 | 1 | Gm11960 | | 1 | 1 | 0 | 1 | 3 | 0 | 1 | 1 | 1 | 1 | 14 |
| 2 | -2.64 | 2.85 | -0.93 | 0.35 | 1 | Gm28294 | | 0 | 1 | 7 | 0 | 10 | 3 | 1 | 0 | 1 | 0 | 0 |
| 2 | -0.24 | 2.06 | -0.12 | 0.91 | 1 | Gm7293 | | 3 | 4 | 8 | 2 | 2 | 0 | 1 | 0 | 1 | 1 | 1 |
| 2 | 0.90 | 2.52 | 0.36 | 0.72 | 1 | Gm10651 | | 4 | 0 | 0 | 1 | 0 | 5 | 1 | 0 | 2 | 1 | 9 |
| 2 | -0.02 | 2.34 | -0.01 | 0.99 | 1 | Gm19353 | | 3 | 6 | 6 | 1 | 0 | 1 | 1 | 1 | 4 | 0 | 0 |
| 2 | 1.85 | 2.90 | 0.64 | 0.52 | 1 | Gm6296 | | 4 | 0 | 4 | 0 | 0 | 1 | 1 | 0 | 3 | 0 | 10 |
| 2 | 0.70 | 1.76 | 0.39 | 0.69 | 1 | Gm3839 | | 9 | 1 | 1 | 1 | 2 | 1 | 1 | 2 | 1 | 3 | 2 |
| 2 | 0.38 | 1.88 | 0.20 | 0.84 | 1 | Gm6485 | | 3 | 0 | 6 | 1 | 4 | 1 | 1 | 1 | 1 | 3 | 2 |
| 2 | -0.06 | 2.10 | -0.03 | 0.98 | 1 | Gm4705 | | 5 | 2 | 4 | 1 | 1 | 2 | 1 | 0 | 5 | 0 | 2 |
| 2 | -1.18 | 3.57 | -0.33 | 0.74 | 1 | Speer2 | spermatogenesis associated glutamate (E | 0 | 0 | 0 | 2 | 0 | 7 | 1 | 0 | 0 | 0 | 13 |
| 2 | -0.48 | 2.05 | -0.24 | 0.81 | 1 | Selenok-ps4 | | 1 | 3 | 1 | 6 | 0 | 3 | 1 | 1 | 1 | 0 | 5 |
| 2 | 0.92 | 3.27 | 0.28 | 0.78 | 1 | Gm16234 | | 5 | 0 | 0 | 0 | 1 | 2 | 1 | 13 | 1 | 0 | 0 |
| 2 | 1.08 | 1.98 | 0.55 | 0.58 | 1 | Gm10689 | | 0 | 1 | 2 | 0 | 3 | 3 | 1 | 5 | 5 | 1 | 2 |
| 2 | -1.10 | 2.01 | -0.55 | 0.58 | 1 | Gm13493 | | 2 | 3 | 0 | 3 | 2 | 6 | 1 | 1 | 1 | 0 | 4 |
| 2 | -0.04 | 2.69 | -0.01 | 0.99 | 1 | Gm8756 | | 8 | 0 | 2 | 2 | 0 | 2 | 1 | 0 | 2 | 0 | 6 |
| 2 | -0.96 | 2.43 | -0.39 | 0.69 | 1 | Gm16020 | | 1 | 1 | 1 | 5 | 0 | 3 | 1 | 2 | 7 | 0 | 0 |
| 2 | -1.16 | 2.79 | -0.42 | 0.68 | 1 | Gm5829 | | 0 | 1 | 6 | 5 | 0 | 1 | 1 | 0 | 6 | 1 | 0 |
| 2 | -0.35 | 3.08 | -0.11 | 0.91 | 1 | Gm10335 | | 9 | 8 | 1 | 0 | 0 | 1 | 1 | 0 | 1 | 0 | 0 |
| 2 | -0.49 | 3.60 | -0.14 | 0.89 | 1 | 4930572G02Rik | | 0 | 0 | 0 | 0 | 0 | 10 | 1 | 0 | 0 | 0 | 10 |
| 2 | 2.59 | 2.40 | 1.08 | 0.28 | 1 | Gm9575 | | 2 | 1 | 0 | 0 | 1 | 1 | 1 | 3 | 7 | 0 | 5 |
| 2 | 0.70 | 1.91 | 0.36 | 0.72 | 1 | Gm13328 | | 0 | 6 | 1 | 1 | 1 | 3 | 1 | 2 | 3 | 1 | 2 |
| 2 | -0.23 | 3.60 | -0.07 | 0.95 | 1 | Gm37879 | | 0 | 0 | 0 | 0 | 0 | 7 | 1 | 0 | 0 | 0 | 13 |
| 2 | -0.64 | 3.57 | -0.18 | 0.86 | 1 | Gm11453 | | 1 | 0 | 7 | 0 | 0 | 2 | 1 | 0 | 10 | 0 | 0 |
| 2 | -0.28 | 1.86 | -0.15 | 0.88 | 1 | Gm3756 | | 1 | 3 | 4 | 1 | 3 | 2 | 1 | 2 | 3 | 0 | 1 |
| 2 | 1.00 | 2.45 | 0.41 | 0.68 | 1 | Gm25432 | | 1 | 0 | 1 | 1 | 0 | 3 | 1 | 1 | 1 | 12 | 0 |
| 2 | 0.08 | 2.19 | 0.04 | 0.97 | 1 | Rpl31-ps13 | | 2 | 6 | 1 | 3 | 3 | 0 | 1 | 0 | 1 | 3 | 1 |
| 2 | -2.23 | 3.59 | -0.62 | 0.53 | 1 | Gm10362 | | 0 | 0 | 16 | 2 | 0 | 0 | 1 | 0 | 0 | 1 | 0 |
| 2 | 1.03 | 1.75 | 0.59 | 0.55 | 1 | Gm11512 | | 1 | 2 | 1 | 3 | 1 | 1 | 1 | 1 | 2 | 4 | 3 |
| 2 | 0.52 | 1.99 | 0.26 | 0.79 | 1 | Gm43028 | | 3 | 1 | 3 | 4 | 0 | 1 | 1 | 1 | 2 | 1 | 2 |
| 2 | 0.69 | 2.27 | 0.30 | 0.76 | 1 | Gm9845 | | 3 | 0 | 3 | 0 | 1 | 3 | 1 | 0 | 4 | 1 | 3 |
| 2 | 0.40 | 2.19 | 0.18 | 0.85 | 1 | Gm8652 | | 9 | 1 | 0 | 1 | 0 | 3 | 1 | 1 | 1 | 1 | 1 |
| 2 | 0.59 | 2.12 | 0.28 | 0.78 | 1 | Gm13123 | | 3 | 4 | 0 | 1 | 0 | 4 | 1 | 2 | 1 | 1 | 1 |
| 2 | -0.15 | 2.11 | -0.07 | 0.94 | 1 | Gm6665 | | 1 | 1 | 1 | 4 | 0 | 3 | 1 | 3 | 1 | 0 | 3 |
| 2 | 0.51 | 2.89 | 0.18 | 0.86 | 1 | Gm16209 | | 0 | 9 | 0 | 0 | 1 | 2 | 1 | 0 | 1 | 1 | 4 |
| 2 | -0.22 | 3.60 | -0.06 | 0.95 | 1 | Gm45297 | | 0 | 0 | 0 | 0 | 0 | 6 | 1 | 0 | 0 | 0 | 12 |
| 2 | -0.22 | 3.60 | -0.06 | 0.95 | 1 | Gm12294 | | 0 | 0 | 0 | 0 | 0 | 6 | 1 | 0 | 0 | 0 | 12 |

| | | | | | | | | | | | | | | | | | | |
|---|---|---|---|---|---|---|---|---|---|---|---|---|---|---|---|---|---|---|
| 2 | 0.35 | 3.06 | 0.11 | 0.91 | 1 | 9130409I23Rik | RIKEN cDNA 9130409I23 gene | 0 | 0 | 0 | 0 | 6 | 2 | 1 | 5 | 2 | 0 | 4 |
| 2 | -1.82 | 3.23 | -0.56 | 0.57 | 1 | BC085271 | | 0 | 1 | 0 | 0 | 6 | 3 | 1 | 7 | 0 | 0 | 1 |
| 2 | 0.14 | 1.89 | 0.07 | 0.94 | 1 | Rpl9-ps4 | | 1 | 0 | 3 | 2 | 3 | 1 | 1 | 1 | 5 | 1 | 1 |
| 2 | -0.50 | 2.79 | -0.18 | 0.86 | 1 | Gm11793 | | 2 | 1 | 3 | 1 | 0 | 1 | 1 | 0 | 0 | 0 | 10 |
| 2 | -1.63 | 3.53 | -0.46 | 0.64 | 1 | Gm16308 | | 0 | 0 | 0 | 0 | 11 | 0 | 1 | 0 | 0 | 0 | 6 |
| 2 | 1.21 | 1.94 | 0.63 | 0.53 | 1 | Rpl21-ps10 | | 3 | 2 | 2 | 0 | 2 | 1 | 1 | 2 | 2 | 3 | 1 |
| 2 | -1.09 | 2.56 | -0.42 | 0.67 | 1 | Gm29787 | | 1 | 2 | 0 | 1 | 10 | 0 | 1 | 1 | 1 | 0 | 1 |
| 2 | -0.06 | 2.23 | -0.03 | 0.98 | 1 | Gm10481 | | 3 | 1 | 2 | 2 | 0 | 3 | 1 | 0 | 5 | 1 | 1 |
| 2 | -0.65 | 2.24 | -0.29 | 0.77 | 1 | Gm18726 | | 4 | 0 | 0 | 2 | 5 | 1 | 1 | 1 | 1 | 1 | 2 |
| 2 | -1.53 | 3.23 | -0.47 | 0.63 | 1 | Gm12005 | | 1 | 0 | 0 | 1 | 3 | 1 | 1 | 0 | 0 | 12 | 0 |
| 2 | 1.63 | 2.22 | 0.73 | 0.46 | 1 | Tpi-rs11 | | 3 | 4 | 0 | 1 | 1 | 0 | 1 | 2 | 3 | 1 | 1 |
| 2 | 0.91 | 2.42 | 0.38 | 0.71 | 1 | Gm7551 | | 1 | 0 | 2 | 1 | 3 | 0 | 1 | 1 | 4 | 0 | 5 |
| 2 | -0.99 | 3.12 | -0.32 | 0.75 | 1 | Gm6197 | | 0 | 0 | 3 | 0 | 9 | 1 | 1 | 0 | 2 | 0 | 1 |
| 2 | -1.19 | 3.60 | -0.33 | 0.74 | 1 | Gm44542 | | 0 | 0 | 0 | 0 | 0 | 14 | 1 | 0 | 0 | 0 | 2 |
| 2 | -0.72 | 2.93 | -0.25 | 0.81 | 1 | Gm6919 | | 0 | 0 | 7 | 2 | 3 | 1 | 1 | 0 | 0 | 3 | 1 |
| 2 | -0.73 | 2.95 | -0.25 | 0.80 | 1 | Gm7464 | | 3 | 4 | 1 | 0 | 6 | 0 | 1 | 0 | 1 | 0 | 1 |
| 2 | -1.00 | 3.58 | -0.28 | 0.78 | 1 | 4930449E18Rik | | 0 | 0 | 7 | 0 | 0 | 2 | 1 | 0 | 7 | 0 | 0 |
| 2 | 0.07 | 2.19 | 0.03 | 0.98 | 1 | Gm11964 | | 0 | 1 | 3 | 1 | 1 | 3 | 1 | 0 | 4 | 1 | 2 |
| 2 | -1.35 | 3.57 | -0.38 | 0.70 | 1 | Gm49282 | | 3 | 0 | 0 | 12 | 0 | 0 | 1 | 0 | 1 | 0 | 0 |
| 2 | -1.10 | 2.94 | -0.37 | 0.71 | 1 | Gm5871 | | 1 | 1 | 2 | 0 | 3 | 3 | 1 | 0 | 0 | 6 | 0 |
| 2 | 0.46 | 2.80 | 0.16 | 0.87 | 1 | Gm2445 | | 2 | 2 | 0 | 1 | 0 | 3 | 1 | 5 | 0 | 4 | 0 |
| 1 | -3.16 | 3.58 | -0.88 | 0.38 | 1 | Gm5353 | | 9 | 0 | 0 | 1 | 1 | 5 | 1 | 0 | 0 | 0 | 0 |
| 1 | -0.13 | 3.60 | -0.04 | 0.97 | 1 | Pip5kl1 | phosphatidylinositol-4-phosphate 5-kinas | 0 | 0 | 0 | 0 | 0 | 4 | 1 | 0 | 0 | 0 | 11 |
| 1 | 3.24 | 2.48 | 1.31 | 0.19 | 1 | Gm48974 | | 3 | 2 | 1 | 0 | 0 | 0 | 1 | 2 | 4 | 1 | 2 |
| 1 | 2.15 | 2.87 | 0.75 | 0.45 | 1 | Gm7432 | | 3 | 1 | 0 | 0 | 1 | 0 | 1 | 0 | 5 | 3 | 2 |
| 1 | -2.54 | 3.55 | -0.71 | 0.47 | 1 | Gm45184 | | 0 | 0 | 0 | 8 | 0 | 5 | 1 | 0 | 0 | 0 | 2 |
| 1 | 0.56 | 3.00 | 0.19 | 0.85 | 1 | Gm3809 | | 2 | 0 | 1 | 1 | 6 | 0 | 1 | 5 | 0 | 0 | 1 |
| 1 | 1.78 | 2.94 | 0.60 | 0.55 | 1 | E030024N20Rik | | 2 | 1 | 0 | 0 | 2 | 0 | 1 | 6 | 0 | 1 | 3 |
| 1 | -0.60 | 2.89 | -0.21 | 0.83 | 1 | Rpl7a-ps7 | | 2 | 3 | 3 | 0 | 2 | 1 | 1 | 0 | 4 | 0 | 0 |
| 1 | -0.70 | 3.35 | -0.21 | 0.84 | 1 | Gm8290 | | 1 | 7 | 1 | 0 | 0 | 3 | 1 | 0 | 0 | 0 | 2 |
| 1 | -0.66 | 2.37 | -0.28 | 0.78 | 1 | Gm7498 | | 2 | 1 | 3 | 4 | 0 | 1 | 1 | 0 | 1 | 1 | 1 |
| 1 | -0.49 | 3.60 | -0.13 | 0.89 | 1 | Gm27029 | | 8 | 2 | 0 | 0 | 0 | 3 | 1 | 0 | 0 | 0 | 2 |
| 1 | 0.74 | 3.50 | 0.21 | 0.83 | 1 | Rpl15-ps2 | | 0 | 7 | 2 | 0 | 0 | 1 | 1 | 0 | 3 | 0 | 1 |
| 1 | -0.10 | 3.62 | -0.03 | 0.98 | 1 | Gm11759 | | 12 | 0 | 0 | 0 | 1 | 0 | 1 | 0 | 0 | 0 | 1 |
| 1 | 2.32 | 3.59 | 0.65 | 0.52 | 1 | Gm13624 | | 8 | 1 | 0 | 0 | 0 | 0 | 1 | 1 | 4 | 0 | 0 |
| 1 | 1.00 | 2.30 | 0.44 | 0.66 | 1 | Gm15466 | | 1 | 1 | 1 | 0 | 2 | 1 | 1 | 1 | 5 | 0 | 2 |
| 1 | -3.59 | 3.56 | -1.01 | 0.31 | 1 | C630028M04Rik | | 0 | 0 | 0 | 10 | 0 | 4 | 1 | 0 | 0 | 0 | 0 |
| 1 | -0.88 | 2.63 | -0.33 | 0.74 | 1 | Gm9525 | | 2 | 2 | 0 | 1 | 1 | 4 | 1 | 0 | 3 | 0 | 1 |
| 1 | -3.10 | 2.89 | -1.07 | 0.28 | 1 | Gm14111 | | 1 | 2 | 2 | 7 | 1 | 1 | 1 | 0 | 0 | 0 | 0 |
| 1 | 0.46 | 3.41 | 0.14 | 0.89 | 1 | Gm6722 | | 0 | 0 | 0 | 1 | 0 | 3 | 1 | 1 | 8 | 0 | 1 |
| 1 | -2.82 | 2.85 | -0.99 | 0.32 | 1 | Gm13961 | | 0 | 1 | 1 | 4 | 1 | 6 | 1 | 0 | 0 | 0 | 1 |
| 1 | -2.77 | 3.56 | -0.78 | 0.44 | 1 | Gm5641 | | 0 | 0 | 1 | 7 | 0 | 4 | 1 | 1 | 0 | 0 | 0 |
| 1 | -1.21 | 3.43 | -0.35 | 0.72 | 1 | Kat2b-ps | | 0 | 0 | 0 | 4 | 1 | 1 | 1 | 0 | 1 | 0 | 6 |
| 1 | 0.16 | 3.61 | 0.04 | 0.97 | 1 | Gm5558 | | 1 | 0 | 5 | 0 | 1 | 0 | 1 | 0 | 6 | 0 | 0 |
| 1 | -0.54 | 3.61 | -0.15 | 0.88 | 1 | Gm5912 | | 0 | 1 | 5 | 0 | 4 | 0 | 1 | 0 | 3 | 0 | 0 |
| 1 | -1.41 | 3.60 | -0.39 | 0.70 | 1 | Gm8250 | | 0 | 2 | 1 | 9 | 0 | 0 | 1 | 1 | 0 | 0 | 0 |
| 1 | 0.00 | 2.39 | 0.00 | 1.00 | 1 | Gm5939 | | 1 | 3 | 3 | 2 | 0 | 1 | 1 | 0 | 1 | 1 | 1 |
| 1 | -2.07 | 3.62 | -0.57 | 0.57 | 1 | Gm12194 | | 1 | 9 | 0 | 1 | 2 | 0 | 1 | 0 | 0 | 0 | 0 |
| 1 | -0.21 | 3.49 | -0.06 | 0.95 | 1 | Gm4045 | | 1 | 0 | 0 | 0 | 2 | 2 | 1 | 1 | 0 | 0 | 7 |
| 1 | -0.33 | 2.24 | -0.15 | 0.88 | 1 | Gm5863 | | 0 | 0 | 1 | 4 | 1 | 2 | 1 | 1 | 1 | 1 | 2 |
| 1 | 1.57 | 2.62 | 0.60 | 0.55 | 1 | Gm5944 | | 1 | 2 | 0 | 1 | 0 | 1 | 1 | 0 | 1 | 4 | 3 |
| 1 | -0.42 | 2.62 | -0.16 | 0.87 | 1 | Gm13422 | | 1 | 1 | 1 | 0 | 1 | 3 | 1 | 1 | 0 | 0 | 4 |
| 1 | -0.85 | 3.00 | -0.28 | 0.78 | 1 | Gm5160 | | 3 | 4 | 0 | 2 | 2 | 0 | 1 | 1 | 1 | 0 | 0 |
| 1 | -1.82 | 3.57 | -0.51 | 0.61 | 1 | Gm15666 | | 0 | 0 | 0 | 3 | 0 | 7 | 1 | 0 | 0 | 0 | 3 |
| 1 | -2.51 | 3.59 | -0.70 | 0.48 | 1 | Rapgef4os2 | | 0 | 0 | 0 | 0 | 0 | 1 | 1 | 12 | 0 | 0 | 0 |
| 1 | -0.47 | 2.32 | -0.20 | 0.84 | 1 | Gm11557 | | 1 | 0 | 1 | 3 | 1 | 1 | 1 | 1 | 1 | 4 | 0 |
| 1 | 2.84 | 2.65 | 1.07 | 0.28 | 1 | Gm7418 | | 1 | 1 | 1 | 0 | 0 | 0 | 1 | 1 | 4 | 3 | 1 |
| 1 | 0.59 | 2.35 | 0.25 | 0.80 | 1 | Gm11410 | | 2 | 1 | 0 | 0 | 1 | 3 | 1 | 1 | 1 | 1 | 2 |
| 1 | -1.07 | 3.06 | -0.35 | 0.73 | 1 | Gm12328 | | 2 | 2 | 2 | 0 | 2 | 1 | 1 | 0 | 0 | 0 | 3 |
| 1 | 0.97 | 3.08 | 0.32 | 0.75 | 1 | Gm5866 | | 3 | 1 | 1 | 0 | 0 | 1 | 1 | 1 | 0 | 0 | 4 |

| | | | | | | | | | | | | | | | | | | |
|---|---|---|---|---|---|---|---|---|---|---|---|---|---|---|---|---|---|---|
| 1 | 1.18 | 2.18 | 0.54 | 0.59 | 1 | Gm5883 | | 1 | 1 | 2 | 1 | 1 | 0 | 1 | 1 | 1 | 3 | 1 |
| 1 | 0.58 | 3.52 | 0.16 | 0.87 | 1 | Gm23493 | | 1 | 1 | 0 | 0 | 0 | 3 | 1 | 0 | 1 | 0 | 6 |
| 1 | -0.87 | 3.60 | -0.24 | 0.81 | 1 | Gm47075 | | 0 | 0 | 0 | 0 | 0 | 8 | 1 | 0 | 0 | 0 | 3 |
| 1 | -0.55 | 2.31 | -0.24 | 0.81 | 1 | Gm49933 | | 1 | 0 | 0 | 1 | 2 | 3 | 1 | 1 | 1 | 1 | 1 |
| 1 | -2.21 | 2.66 | -0.83 | 0.41 | 1 | Gm5070 | | 1 | 1 | 3 | 1 | 2 | 3 | 1 | 1 | 0 | 0 | 0 |
| 1 | -1.56 | 3.56 | -0.44 | 0.66 | 1 | Gm12362 | | 0 | 1 | 0 | 0 | 8 | 1 | 1 | 0 | 1 | 0 | 1 |
| 1 | 1.89 | 3.14 | 0.60 | 0.55 | 1 | Gm4784 | | 1 | 5 | 1 | 0 | 0 | 0 | 1 | 0 | 1 | 1 | 2 |
| 1 | -1.17 | 3.60 | -0.32 | 0.75 | 1 | Gm10136 | | 0 | 0 | 1 | 8 | 0 | 0 | 1 | 2 | 0 | 0 | 0 |
| 1 | -0.23 | 3.63 | -0.06 | 0.95 | 1 | Gm5256 | | 1 | 9 | 0 | 0 | 0 | 0 | 1 | 1 | 0 | 0 | 0 |
| 1 | 0.33 | 3.56 | 0.09 | 0.93 | 1 | Gm5389 | | 0 | 0 | 1 | 0 | 2 | 1 | 1 | 1 | 0 | 0 | 6 |
| 1 | -2.84 | 3.56 | -0.80 | 0.43 | 1 | Gm8663 | | 3 | 0 | 0 | 5 | 2 | 0 | 1 | 0 | 0 | 0 | 1 |
| 1 | -2.06 | 3.56 | -0.58 | 0.56 | 1 | Gm567 | | 0 | 0 | 0 | 7 | 0 | 2 | 1 | 0 | 0 | 0 | 2 |
| 1 | 0.55 | 3.63 | 0.15 | 0.88 | 1 | Tpi-rs10 | | 6 | 2 | 0 | 0 | 0 | 0 | 1 | 0 | 0 | 3 | 0 |
| 1 | 1.31 | 3.60 | 0.37 | 0.71 | 1 | Gm4972 | | 5 | 0 | 1 | 0 | 0 | 0 | 1 | 0 | 0 | 1 | 3 |
| 1 | -0.72 | 2.79 | -0.26 | 0.80 | 1 | Gm5826 | | 3 | 0 | 1 | 2 | 1 | 1 | 1 | 2 | 1 | 0 | 0 |
| 1 | 1.51 | 3.19 | 0.47 | 0.64 | 1 | Gm9658 | | 0 | 0 | 3 | 0 | 0 | 2 | 1 | 2 | 1 | 1 | 1 |
| 1 | 1.03 | 3.60 | 0.29 | 0.78 | 1 | Gm27972 | | 0 | 0 | 0 | 0 | 0 | 0 | 1 | 10 | 0 | 0 | 0 |
| 1 | -2.74 | 3.54 | -0.77 | 0.44 | 1 | Gm12473 | | 1 | 0 | 0 | 3 | 4 | 1 | 1 | 0 | 0 | 0 | 2 |
| 1 | 0.04 | 2.43 | 0.02 | 0.99 | 1 | Gm7688 | | 1 | 1 | 0 | 1 | 1 | 1 | 1 | 1 | 0 | 1 | 3 |
| 1 | 0.53 | 2.63 | 0.20 | 0.84 | 1 | Gm9242 | | 0 | 0 | 0 | 1 | 1 | 2 | 1 | 1 | 2 | 1 | 2 |
| 1 | -0.89 | 3.61 | -0.25 | 0.81 | 1 | Gm7684 | | 0 | 3 | 3 | 2 | 0 | 0 | 1 | 0 | 0 | 0 | 2 |
| 1 | 1.42 | 3.62 | 0.39 | 0.70 | 1 | Gm13216 | | 1 | 3 | 1 | 0 | 0 | 0 | 1 | 2 | 0 | 3 | 0 |
| 1 | -1.70 | 3.59 | -0.47 | 0.64 | 1 | Gm3086 | | 2 | 0 | 0 | 7 | 0 | 0 | 1 | 0 | 0 | 0 | 1 |
| 1 | -0.60 | 3.57 | -0.17 | 0.87 | 1 | Col2a1 | collagen, type II, alpha 1 | 0 | 0 | 0 | 0 | 0 | 8 | 1 | 0 | 2 | 0 | 0 |
| 1 | -0.36 | 3.25 | -0.11 | 0.91 | 1 | Gm12013 | | 2 | 1 | 1 | 0 | 1 | 1 | 1 | 0 | 4 | 0 | 0 |
| 1 | -0.13 | 3.60 | -0.04 | 0.97 | 1 | 2310016G11Rik | | 8 | 0 | 0 | 0 | 0 | 1 | 1 | 0 | 0 | 0 | 1 |
| 1 | -0.45 | 3.23 | -0.14 | 0.89 | 1 | Gm49017 | | 2 | 1 | 1 | 0 | 4 | 0 | 1 | 0 | 1 | 0 | 1 |
| 1 | -1.13 | 3.59 | -0.31 | 0.75 | 1 | Gm2467 | | 2 | 0 | 1 | 4 | 1 | 0 | 1 | 2 | 0 | 0 | 0 |
| 1 | -0.34 | 3.26 | -0.11 | 0.92 | 1 | Gm7290 | | 0 | 4 | 1 | 1 | 1 | 0 | 1 | 1 | 0 | 1 | 0 |
| 1 | -1.76 | 3.58 | -0.49 | 0.62 | 1 | Gm11222 | | 1 | 1 | 0 | 0 | 1 | 5 | 1 | 0 | 0 | 1 | 0 |
| 1 | -0.41 | 3.63 | -0.11 | 0.91 | 1 | Gm5917 | | 6 | 1 | 0 | 0 | 0 | 1 | 1 | 0 | 0 | 1 | 0 |
| 1 | -0.41 | 3.61 | -0.11 | 0.91 | 1 | H3c13 | H3 clustered histone 13 | 0 | 4 | 1 | 1 | 0 | 0 | 1 | 0 | 0 | 0 | 3 |
| 1 | 0.27 | 3.62 | 0.08 | 0.94 | 1 | Gm5356 | | 8 | 0 | 0 | 0 | 0 | 0 | 1 | 0 | 0 | 1 | 0 |
| 1 | 1.49 | 3.58 | 0.42 | 0.68 | 1 | Gm12470 | | 3 | 0 | 0 | 0 | 0 | 1 | 1 | 2 | 0 | 1 | 1 |
| 1 | -0.64 | 2.55 | -0.25 | 0.80 | 1 | Gm11605 | | 1 | 1 | 1 | 1 | 1 | 1 | 1 | 0 | 0 | 1 | 2 |
| 1 | -0.19 | 3.59 | -0.05 | 0.96 | 1 | Cfap65 | cilia and flagella associated protein 65 | 5 | 0 | 0 | 0 | 0 | 2 | 1 | 0 | 0 | 0 | 2 |
| 1 | 2.97 | 3.57 | 0.83 | 0.41 | 1 | Gm6663 | | 0 | 0 | 0 | 0 | 0 | 0 | 1 | 3 | 3 | 1 | 1 |
| 1 | -1.86 | 3.62 | -0.51 | 0.61 | 1 | Gm13349 | | 0 | 0 | 6 | 0 | 0 | 3 | 1 | 0 | 0 | 0 | 0 |
| 1 | -1.91 | 3.25 | -0.59 | 0.56 | 1 | Gm8618 | | 0 | 2 | 2 | 2 | 1 | 1 | 1 | 0 | 1 | 0 | 0 |
| 1 | -2.02 | 3.61 | -0.56 | 0.58 | 1 | Gm6625 | | 6 | 0 | 0 | 2 | 0 | 1 | 1 | 0 | 0 | 0 | 0 |
| 1 | -2.06 | 3.60 | -0.57 | 0.57 | 1 | Gm12778 | | 0 | 0 | 0 | 0 | 9 | 0 | 1 | 0 | 0 | 0 | 0 |
| 1 | -2.06 | 3.60 | -0.57 | 0.57 | 1 | Gm15124 | | 0 | 0 | 0 | 0 | 9 | 0 | 1 | 0 | 0 | 0 | 0 |
| 1 | -0.40 | 3.67 | -0.11 | 0.91 | 1 | 2010001K21Rik | | 3 | 5 | 0 | 0 | 0 | 0 | 1 | 0 | 0 | 0 | 0 |
| 1 | -1.39 | 3.57 | -0.39 | 0.70 | 1 | Gm9105 | | 1 | 0 | 0 | 1 | 3 | 1 | 1 | 0 | 3 | 0 | 0 |
| 1 | -0.23 | 3.33 | -0.07 | 0.94 | 1 | Gm6222 | | 2 | 0 | 0 | 1 | 1 | 1 | 1 | 0 | 3 | 0 | 1 |
| 1 | -3.06 | 3.58 | -0.86 | 0.39 | 1 | Gm6736 | | 0 | 3 | 0 | 0 | 2 | 3 | 1 | 0 | 0 | 0 | 0 |
| 1 | 1.18 | 3.27 | 0.36 | 0.72 | 1 | Gm18904 | | 2 | 0 | 1 | 0 | 1 | 0 | 1 | 0 | 1 | 3 | 1 |
| 1 | -0.51 | 3.61 | -0.14 | 0.89 | 1 | Gm29459 | | 0 | 0 | 0 | 0 | 0 | 4 | 1 | 0 | 0 | 0 | 4 |
| 1 | 0.72 | 3.57 | 0.20 | 0.84 | 1 | Gm2383 | | 1 | 0 | 0 | 0 | 0 | 2 | 1 | 0 | 0 | 3 | 3 |
| 1 | 1.00 | 3.35 | 0.30 | 0.76 | 1 | Gm7206 | | 1 | 0 | 2 | 1 | 0 | 0 | 1 | 0 | 1 | 1 | 2 |
| 1 | 0.21 | 3.63 | 0.06 | 0.95 | 1 | Gm6681 | | 2 | 4 | 0 | 0 | 0 | 0 | 1 | 2 | 0 | 0 | 0 |
| 1 | -1.15 | 3.60 | -0.32 | 0.75 | 1 | Gm7368 | | 4 | 0 | 0 | 1 | 2 | 0 | 1 | 0 | 1 | 0 | 0 |
| 1 | -0.42 | 2.95 | -0.14 | 0.89 | 1 | Gm7638 | | 2 | 1 | 1 | 1 | 1 | 0 | 1 | 0 | 0 | 1 | 1 |
| 1 | -0.57 | 3.61 | -0.16 | 0.87 | 1 | Gm9521 | | 1 | 3 | 0 | 0 | 1 | 0 | 1 | 0 | 0 | 0 | 3 |
| 1 | -3.20 | 3.57 | -0.90 | 0.37 | 1 | Usp50 | ubiquitin specific peptidase 50 | 0 | 0 | 0 | 5 | 1 | 2 | 1 | 0 | 0 | 0 | 0 |
| 1 | 0.82 | 3.40 | 0.24 | 0.81 | 1 | Gm28438 | | 1 | 0 | 1 | 2 | 0 | 0 | 1 | 1 | 2 | 0 | 1 |
| 1 | -0.46 | 3.67 | -0.13 | 0.90 | 1 | Gm5459 | | 2 | 6 | 0 | 0 | 0 | 0 | 1 | 0 | 0 | 0 | 0 |
| 1 | 0.14 | 3.62 | 0.04 | 0.97 | 1 | Gm4575 | | 1 | 1 | 1 | 0 | 1 | 0 | 1 | 0 | 0 | 4 | 0 |
| 1 | -1.01 | 3.59 | -0.28 | 0.78 | 1 | Gm15454 | | 1 | 0 | 0 | 2 | 1 | 0 | 1 | 0 | 0 | 4 | 0 |
| 1 | 1.47 | 3.62 | 0.41 | 0.68 | 1 | Gm9436 | | 1 | 1 | 1 | 0 | 0 | 0 | 1 | 3 | 1 | 0 | 0 |

| | | | | | | | | | | | | | | | | | | |
|---|---|---|---|---|---|---|---|---|---|---|---|---|---|---|---|---|---|---|
| 1 | 1.02 | 3.57 | 0.29 | 0.78 | 1 | Gm9644 | | 0 | 1 | 0 | 2 | 0 | 0 | 1 | 0 | 0 | 3 | 2 |
| 1 | 0.48 | 3.35 | 0.14 | 0.89 | 1 | Gm6181 | | 0 | 0 | 1 | 1 | 1 | 0 | 1 | 1 | 2 | 1 | 0 |
| 1 | 0.39 | 3.62 | 0.11 | 0.91 | 1 | Gm43330 | | 0 | 0 | 0 | 0 | 0 | 1 | 1 | 0 | 0 | 6 | 0 |
| 1 | -0.59 | 3.66 | -0.16 | 0.87 | 1 | Gm14706 | | 0 | 0 | 7 | 0 | 0 | 0 | 1 | 0 | 0 | 0 | 0 |
| 1 | -0.10 | 3.58 | -0.03 | 0.98 | 1 | Gm15599 | | 0 | 0 | 0 | 0 | 0 | 4 | 1 | 0 | 2 | 0 | 1 |
| 1 | -0.98 | 3.58 | -0.27 | 0.78 | 1 | Gm5981 | | 0 | 1 | 0 | 2 | 2 | 0 | 1 | 0 | 0 | 3 | 0 |
| 1 | 0.88 | 3.61 | 0.24 | 0.81 | 1 | Gm49588 | | 5 | 0 | 0 | 0 | 0 | 0 | 1 | 0 | 0 | 0 | 2 |
| 1 | -0.21 | 3.59 | -0.06 | 0.95 | 1 | Gm11407 | | 2 | 0 | 0 | 0 | 1 | 1 | 1 | 0 | 0 | 3 | 1 |
| 1 | 0.95 | 3.63 | 0.26 | 0.79 | 1 | Gm37421 | | 0 | 1 | 0 | 0 | 0 | 0 | 1 | 0 | 6 | 0 | 0 |
| 1 | 0.93 | 3.65 | 0.26 | 0.80 | 1 | Gm4968 | | 2 | 2 | 1 | 0 | 0 | 0 | 1 | 0 | 1 | 1 | 0 |
| 1 | -0.56 | 3.57 | -0.16 | 0.87 | 1 | Gm15432 | | 0 | 0 | 1 | 3 | 1 | 0 | 1 | 1 | 0 | 1 | 0 |
| 1 | -0.72 | 3.46 | -0.21 | 0.84 | 1 | Gm7940 | | 0 | 1 | 2 | 0 | 1 | 1 | 1 | 1 | 1 | 0 | 0 |
| 1 | -0.93 | 3.52 | -0.27 | 0.79 | 1 | Gm12186 | | 1 | 2 | 1 | 0 | 1 | 1 | 1 | 0 | 0 | 1 | 0 |
| 1 | -2.52 | 3.59 | -0.70 | 0.48 | 1 | Cidec | cell death-inducing DFFA-like effector c | 0 | 0 | 0 | 0 | 0 | 1 | 1 | 0 | 6 | 0 | 0 |
| 1 | 0.28 | 3.58 | 0.08 | 0.94 | 1 | Gm2986 | | 0 | 0 | 0 | 1 | 0 | 2 | 1 | 0 | 1 | 1 | 2 |
| 1 | -0.34 | 3.48 | -0.10 | 0.92 | 1 | Gm4936 | | 1 | 2 | 0 | 1 | 1 | 0 | 1 | 1 | 1 | 0 | 0 |
| 1 | -0.94 | 3.62 | -0.26 | 0.80 | 1 | Hmgb1-ps6 | | 2 | 0 | 2 | 0 | 1 | 1 | 1 | 0 | 0 | 1 | 0 |
| 1 | -1.27 | 3.59 | -0.35 | 0.72 | 1 | Gm12181 | | 0 | 0 | 0 | 0 | 6 | 0 | 1 | 1 | 0 | 0 | 0 |
| 1 | 1.71 | 3.62 | 0.47 | 0.64 | 1 | Rpl21-ps4 | | 2 | 0 | 1 | 0 | 0 | 0 | 1 | 0 | 3 | 0 | 1 |
| 1 | -0.47 | 3.59 | -0.13 | 0.90 | 1 | Gm44911 | | 0 | 0 | 0 | 0 | 1 | 3 | 1 | 0 | 0 | 1 | 2 |
| 1 | -0.49 | 3.65 | -0.13 | 0.89 | 1 | Gm11970 | | 2 | 2 | 1 | 0 | 0 | 1 | 1 | 1 | 0 | 0 | 0 |
| 1 | -0.58 | 3.48 | -0.17 | 0.87 | 1 | Gm12726 | | 1 | 1 | 0 | 2 | 0 | 1 | 1 | 0 | 0 | 1 | 1 |
| 1 | -0.10 | 3.61 | -0.03 | 0.98 | 1 | 2210039B01Rik | | 0 | 0 | 0 | 0 | 0 | 2 | 1 | 0 | 0 | 0 | 5 |
| 1 | 0.77 | 3.60 | 0.21 | 0.83 | 1 | Anp32-ps | | 0 | 0 | 0 | 0 | 0 | 3 | 1 | 1 | 1 | 0 | 2 |
| 1 | -1.48 | 3.60 | -0.41 | 0.68 | 1 | Gm7847 | | 0 | 0 | 1 | 1 | 4 | 0 | 1 | 0 | 1 | 0 | 0 |
| 1 | -1.02 | 3.47 | -0.30 | 0.77 | 1 | Gm5802 | | 2 | 0 | 0 | 1 | 1 | 1 | 1 | 0 | 1 | 1 | 0 |
| 1 | 0.89 | 3.57 | 0.25 | 0.80 | 1 | Gm7436 | | 0 | 0 | 0 | 0 | 2 | 0 | 1 | 1 | 1 | 3 | 0 |
| 1 | -2.45 | 3.50 | -0.70 | 0.48 | 1 | Gm19774 | | 1 | 1 | 1 | 1 | 2 | 1 | 1 | 0 | 0 | 0 | 0 |
| 1 | -2.29 | 3.61 | -0.63 | 0.53 | 1 | Gm7985 | | 1 | 0 | 1 | 0 | 2 | 3 | 1 | 0 | 0 | 0 | 0 |
| 1 | 0.68 | 3.60 | 0.19 | 0.85 | 1 | Gm7776 | | 0 | 1 | 0 | 1 | 0 | 0 | 1 | 2 | 0 | 0 | 2 |
| 1 | 1.69 | 3.62 | 0.47 | 0.64 | 1 | Gm14828 | | 0 | 0 | 0 | 0 | 0 | 0 | 1 | 2 | 0 | 4 | 0 |
| 1 | -1.28 | 3.64 | -0.35 | 0.72 | 1 | Gm43471 | | 3 | 0 | 1 | 0 | 0 | 2 | 1 | 0 | 0 | 0 | 0 |
| 1 | 0.42 | 3.61 | 0.12 | 0.91 | 1 | Smbd1 | somatomedin B domain containing 1 | 0 | 0 | 0 | 0 | 0 | 0 | 1 | 0 | 0 | 0 | 6 |
| 1 | 1.39 | 3.61 | 0.39 | 0.70 | 1 | Gm16198 | | 1 | 0 | 0 | 0 | 0 | 0 | 1 | 0 | 5 | 0 | 0 |
| 1 | 0.28 | 3.68 | 0.08 | 0.94 | 1 | Gm17541 | | 2 | 2 | 1 | 0 | 0 | 0 | 1 | 0 | 0 | 0 | 1 |
| 1 | -1.31 | 3.60 | -0.36 | 0.72 | 1 | Gm18669 | | 0 | 0 | 0 | 0 | 5 | 0 | 1 | 0 | 0 | 0 | 1 |
| 1 | -0.42 | 3.61 | -0.12 | 0.91 | 1 | Gm2065 | | 0 | 0 | 0 | 0 | 0 | 3 | 1 | 0 | 0 | 0 | 3 |
| 1 | -0.42 | 3.61 | -0.12 | 0.91 | 1 | 4933433G19Rik | | 0 | 0 | 0 | 0 | 0 | 3 | 1 | 0 | 0 | 0 | 3 |
| 1 | 2.33 | 3.59 | 0.65 | 0.52 | 1 | Gm7129 | | 0 | 0 | 0 | 0 | 0 | 0 | 1 | 1 | 2 | 1 | 1 |
| 1 | -2.63 | 3.59 | -0.73 | 0.46 | 1 | Gm42427 | | 0 | 0 | 0 | 0 | 3 | 3 | 1 | 0 | 0 | 0 | 0 |
| 1 | -1.43 | 3.64 | -0.39 | 0.69 | 1 | 1110032F04Rik | RIKEN cDNA 1110032F04 gene | 3 | 0 | 0 | 0 | 2 | 0 | 1 | 0 | 0 | 0 | 0 |
| 1 | -0.92 | 3.61 | -0.26 | 0.80 | 1 | Gm12178 | | 0 | 0 | 0 | 0 | 2 | 1 | 1 | 0 | 0 | 3 | 0 |
| 1 | -1.56 | 3.63 | -0.43 | 0.67 | 1 | Gm11531 | | 1 | 1 | 0 | 0 | 0 | 3 | 1 | 0 | 0 | 0 | 0 |
| 1 | -1.90 | 3.63 | -0.52 | 0.60 | 1 | Gm7192 | | 0 | 0 | 0 | 5 | 0 | 0 | 1 | 0 | 0 | 0 | 0 |
| 1 | -1.01 | 3.61 | -0.28 | 0.78 | 1 | Xlr3c | X-linked lymphocyte-regulated 3C | 0 | 0 | 0 | 0 | 0 | 4 | 1 | 0 | 0 | 0 | 1 |
| 1 | -0.02 | 3.63 | -0.01 | 1.00 | 1 | Gm11443 | | 0 | 0 | 3 | 0 | 1 | 0 | 1 | 1 | 0 | 0 | 0 |
| 1 | 1.31 | 3.60 | 0.36 | 0.72 | 1 | Gm49877 | | 0 | 0 | 0 | 1 | 0 | 0 | 1 | 0 | 1 | 1 | 2 |
| 1 | -0.55 | 3.67 | -0.15 | 0.88 | 1 | Gm4468 | | 0 | 0 | 5 | 0 | 0 | 0 | 1 | 0 | 0 | 0 | 0 |
| 1 | 0.29 | 3.60 | 0.08 | 0.94 | 1 | Gm8100 | | 1 | 0 | 0 | 0 | 0 | 1 | 1 | 3 | 0 | 0 | 0 |
| 1 | -0.36 | 3.65 | -0.10 | 0.92 | 1 | Gm7456 | | 1 | 1 | 1 | 0 | 1 | 0 | 1 | 0 | 0 | 1 | 0 |
| 1 | 0.28 | 3.68 | 0.08 | 0.94 | 1 | Gm10257 | | 0 | 3 | 1 | 0 | 0 | 0 | 1 | 0 | 1 | 0 | 0 |
| 1 | -1.61 | 3.62 | -0.44 | 0.66 | 1 | Gm43571 | | 0 | 0 | 0 | 0 | 0 | 5 | 1 | 0 | 0 | 0 | 0 |
| 1 | 0.89 | 3.62 | 0.25 | 0.81 | 1 | Gm13932 | | 0 | 0 | 0 | 0 | 0 | 0 | 1 | 0 | 5 | 0 | 0 |
| 1 | -0.16 | 3.61 | -0.05 | 0.96 | 1 | Fabp3-ps1 | | 0 | 0 | 0 | 1 | 0 | 1 | 1 | 2 | 1 | 0 | 0 |
| 1 | -1.27 | 3.63 | -0.35 | 0.73 | 1 | Gm15516 | | 0 | 0 | 2 | 0 | 3 | 0 | 1 | 0 | 0 | 0 | 0 |
| 1 | -0.06 | 3.60 | -0.02 | 0.99 | 1 | Gm7977 | | 0 | 0 | 0 | 0 | 1 | 1 | 1 | 0 | 2 | 0 | 1 |
| 1 | -0.72 | 3.63 | -0.20 | 0.84 | 1 | Gm8722 | | 0 | 0 | 2 | 0 | 0 | 2 | 1 | 0 | 0 | 0 | 1 |
| 1 | -1.32 | 3.61 | -0.37 | 0.71 | 1 | Eif4a3l2 | | 1 | 0 | 0 | 0 | 1 | 2 | 1 | 0 | 0 | 1 | 0 |
| 1 | 0.00 | 3.61 | 0.00 | 1.00 | 1 | Oprd1 | opioid receptor, delta 1 | 1 | 0 | 0 | 0 | 0 | 1 | 1 | 0 | 0 | 0 | 3 |
| 1 | -1.83 | 3.63 | -0.50 | 0.61 | 1 | Tubb4b-ps2 | | 0 | 0 | 0 | 0 | 5 | 0 | 1 | 0 | 0 | 0 | 0 |

| | | | | | | | | | | | | | | | | | | |
|---|---|---|---|---|---|---|---|---|---|---|---|---|---|---|---|---|---|---|
| 1 | -0.82 | 3.61 | -0.23 | 0.82 | 1 | Gm39147 | predicted gene, 39147 | 1 | 0 | 0 | 0 | 1 | 1 | 1 | 0 | 0 | 0 | 2 |
| 1 | -0.87 | 3.63 | -0.24 | 0.81 | 1 | Pafah1b1-ps1 | | 2 | 0 | 0 | 1 | 0 | 1 | 1 | 1 | 0 | 0 | 0 |
| 1 | -1.49 | 3.61 | -0.41 | 0.68 | 1 | Gm8990 | | 0 | 0 | 0 | 0 | 2 | 2 | 1 | 0 | 1 | 0 | 0 |
| 1 | -0.08 | 3.61 | -0.02 | 0.98 | 1 | Gm45405 | | 2 | 0 | 0 | 0 | 0 | 1 | 1 | 0 | 0 | 0 | 2 |
| 1 | -1.93 | 3.62 | -0.53 | 0.59 | 1 | Rps11-ps3 | | 0 | 2 | 0 | 1 | 0 | 2 | 1 | 0 | 0 | 0 | 0 |
| 1 | -1.43 | 3.60 | -0.40 | 0.69 | 1 | Gm13592 | | 0 | 0 | 0 | 3 | 1 | 0 | 1 | 0 | 1 | 0 | 0 |
| 0 | 2.04 | 3.61 | 0.57 | 0.57 | 1 | Gm43432 | | 0 | 0 | 0 | 0 | 0 | 0 | 1 | 1 | 1 | 1 | 1 |
| 0 | 0.56 | 3.64 | 0.15 | 0.88 | 1 | Gm38186 | | 3 | 0 | 0 | 0 | 0 | 0 | 1 | 0 | 0 | 0 | 1 |
| 0 | 0.27 | 3.64 | 0.07 | 0.94 | 1 | Gm4525 | | 0 | 1 | 0 | 0 | 1 | 0 | 1 | 0 | 1 | 1 | 0 |
| 0 | 0.28 | 3.68 | 0.08 | 0.94 | 1 | Gm8317 | | 0 | 1 | 2 | 0 | 0 | 0 | 1 | 1 | 0 | 0 | 0 |
| 0 | 0.97 | 3.65 | 0.26 | 0.79 | 1 | Rpsa-ps8 | | 1 | 1 | 0 | 0 | 0 | 0 | 1 | 0 | 1 | 1 | 0 |
| 0 | 0.36 | 3.64 | 0.10 | 0.92 | 1 | Gm11877 | | 0 | 0 | 2 | 0 | 0 | 0 | 1 | 0 | 0 | 0 | 2 |
| 0 | 0.40 | 3.62 | 0.11 | 0.91 | 1 | Gm44686 | | 0 | 0 | 0 | 0 | 0 | 0 | 1 | 0 | 0 | 0 | 4 |
| 0 | 0.85 | 3.65 | 0.23 | 0.82 | 1 | Gm6975 | | 0 | 1 | 0 | 0 | 0 | 0 | 1 | 0 | 3 | 0 | 0 |
| 0 | 0.80 | 3.63 | 0.22 | 0.82 | 1 | Gm24105 | | 0 | 0 | 0 | 0 | 0 | 0 | 1 | 0 | 4 | 0 | 0 |
| 0 | -0.54 | 3.64 | -0.15 | 0.88 | 1 | Gm5626 | | 0 | 2 | 0 | 0 | 1 | 0 | 1 | 0 | 1 | 0 | 0 |
| 0 | -0.48 | 3.62 | -0.13 | 0.89 | 1 | Gm8508 | | 0 | 0 | 0 | 0 | 1 | 1 | 1 | 1 | 0 | 0 | 1 |
| 0 | 0.48 | 3.62 | 0.13 | 0.89 | 1 | Gm8580 | | 1 | 0 | 0 | 0 | 0 | 0 | 1 | 0 | 0 | 0 | 3 |
| 0 | -0.05 | 3.62 | -0.01 | 0.99 | 1 | Gm13437 | | 0 | 0 | 0 | 0 | 0 | 1 | 1 | 0 | 0 | 0 | 3 |
| 0 | -0.47 | 3.65 | -0.13 | 0.90 | 1 | Gm13196 | | 1 | 0 | 1 | 0 | 1 | 0 | 1 | 0 | 0 | 0 | 1 |
| 0 | -0.45 | 3.62 | -0.12 | 0.90 | 1 | Gm5779 | | 1 | 0 | 0 | 0 | 1 | 0 | 1 | 0 | 0 | 0 | 2 |
| 0 | -0.34 | 3.65 | -0.09 | 0.93 | 1 | Gm37009 | | 1 | 1 | 0 | 0 | 1 | 0 | 1 | 0 | 1 | 0 | 0 |
| 0 | -1.14 | 3.63 | -0.32 | 0.75 | 1 | Gm9523 | | 0 | 0 | 1 | 1 | 0 | 1 | 1 | 0 | 0 | 0 | 1 |
| 0 | -0.42 | 3.67 | -0.11 | 0.91 | 1 | Hmgb1-ps9 | | 2 | 1 | 1 | 0 | 0 | 0 | 1 | 0 | 0 | 0 | 0 |
| 0 | -0.60 | 3.63 | -0.17 | 0.87 | 1 | Gm8526 | | 1 | 0 | 0 | 2 | 0 | 0 | 1 | 1 | 0 | 0 | 0 |
| 0 | 0.61 | 3.62 | 0.17 | 0.87 | 1 | Umodl1 | uromodulin-like 1 | 2 | 0 | 0 | 0 | 0 | 0 | 1 | 0 | 0 | 0 | 2 |
| 0 | -0.69 | 3.63 | -0.19 | 0.85 | 1 | Gm14251 | | 0 | 0 | 1 | 0 | 0 | 2 | 1 | 0 | 0 | 0 | 1 |
| 0 | 1.30 | 3.63 | 0.36 | 0.72 | 1 | Rpl31-ps21 | | 2 | 0 | 0 | 0 | 0 | 0 | 1 | 0 | 1 | 0 | 1 |
| 0 | -1.23 | 3.62 | -0.34 | 0.73 | 1 | Gm12749 | | 1 | 0 | 0 | 1 | 1 | 0 | 1 | 0 | 0 | 0 | 1 |
| 0 | -2.17 | 3.61 | -0.60 | 0.55 | 1 | Gm7833 | | 0 | 0 | 0 | 0 | 3 | 1 | 1 | 0 | 0 | 0 | 0 |
| 0 | -2.17 | 3.61 | -0.60 | 0.55 | 1 | Slc22a28 | solute carrier family 22, member 28 | 0 | 0 | 0 | 0 | 3 | 1 | 1 | 0 | 0 | 0 | 0 |
| 0 | -1.06 | 3.66 | -0.29 | 0.77 | 1 | Gm4960 | | 2 | 0 | 1 | 1 | 0 | 0 | 1 | 0 | 0 | 0 | 0 |
| 0 | -0.22 | 3.65 | -0.06 | 0.95 | 1 | Gm19366 | | 2 | 0 | 0 | 1 | 0 | 0 | 1 | 0 | 1 | 0 | 0 |
| 0 | 1.20 | 3.63 | 0.33 | 0.74 | 1 | Gm8976 | | 0 | 0 | 0 | 0 | 0 | 0 | 1 | 0 | 1 | 3 | 0 |
| 0 | -1.65 | 3.64 | -0.45 | 0.65 | 1 | Gm6043 | | 2 | 0 | 0 | 1 | 1 | 0 | 1 | 0 | 0 | 0 | 0 |
| 0 | 0.65 | 3.64 | 0.18 | 0.86 | 1 | Gm9873 | | 0 | 0 | 0 | 0 | 0 | 0 | 1 | 3 | 0 | 0 | 0 |
| 0 | 0.00 | 3.63 | 0.00 | 1.00 | 1 | Gm5403 | | 0 | 0 | 0 | 1 | 0 | 0 | 1 | 0 | 0 | 3 | 0 |
| 0 | -1.57 | 3.63 | -0.43 | 0.67 | 1 | Gm18916 | | 0 | 0 | 0 | 0 | 0 | 3 | 1 | 0 | 0 | 0 | 0 |
| 0 | 0.70 | 3.66 | 0.19 | 0.85 | 1 | Gm5809 | | 0 | 0 | 1 | 0 | 0 | 0 | 1 | 2 | 0 | 0 | 0 |
| 0 | 0.92 | 3.66 | 0.25 | 0.80 | 1 | H3f3a-ps2 | | 0 | 1 | 0 | 0 | 0 | 0 | 1 | 0 | 1 | 1 | 0 |
| 0 | 0.96 | 3.67 | 0.26 | 0.79 | 1 | Gm46379 | | 0 | 0 | 1 | 0 | 0 | 0 | 1 | 1 | 0 | 0 | 1 |
| 0 | 0.28 | 3.65 | 0.08 | 0.94 | 1 | Gm12834 | | 1 | 0 | 1 | 0 | 0 | 0 | 1 | 0 | 0 | 1 | 0 |
| 0 | -0.06 | 3.64 | -0.02 | 0.99 | 1 | Rpl26-ps2 | | 0 | 0 | 2 | 0 | 0 | 0 | 1 | 0 | 1 | 0 | 0 |
| 0 | 0.36 | 3.64 | 0.10 | 0.92 | 1 | Rpl21-ps14 | | 0 | 0 | 1 | 0 | 0 | 0 | 1 | 0 | 0 | 0 | 2 |
| 0 | 0.86 | 3.66 | 0.23 | 0.81 | 1 | Gm6407 | | 0 | 0 | 1 | 0 | 0 | 0 | 1 | 0 | 1 | 0 | 1 |
| 0 | 0.39 | 3.63 | 0.11 | 0.91 | 1 | Gm15806 | | 0 | 0 | 0 | 0 | 0 | 0 | 1 | 0 | 0 | 0 | 3 |
| 0 | -0.40 | 3.67 | -0.11 | 0.91 | 1 | Gm43247 | | 0 | 0 | 1 | 0 | 0 | 1 | 1 | 1 | 0 | 0 | 0 |
| 0 | -0.53 | 3.67 | -0.14 | 0.89 | 1 | Gm10231 | | 0 | 3 | 0 | 0 | 0 | 0 | 1 | 0 | 0 | 0 | 0 |
| 0 | -0.38 | 3.66 | -0.10 | 0.92 | 1 | Gm11363 | | 1 | 0 | 0 | 0 | 1 | 0 | 1 | 1 | 0 | 0 | 0 |
| 0 | -0.66 | 3.63 | -0.18 | 0.86 | 1 | Gm25920 | | 0 | 0 | 0 | 0 | 2 | 0 | 1 | 0 | 1 | 0 | 0 |
| 0 | -1.08 | 3.67 | -0.30 | 0.77 | 1 | Gm9434 | | 0 | 1 | 1 | 0 | 0 | 1 | 1 | 0 | 0 | 0 | 0 |
| 0 | 0.55 | 3.65 | 0.15 | 0.88 | 1 | Gm30747 | | 2 | 0 | 0 | 0 | 0 | 0 | 1 | 1 | 0 | 0 | 0 |
| 0 | -0.50 | 3.65 | -0.14 | 0.89 | 1 | Gm9568 | | 1 | 0 | 0 | 0 | 1 | 0 | 1 | 0 | 0 | 0 | 1 |
| 0 | -1.53 | 3.64 | -0.42 | 0.67 | 1 | Gm12380 | | 0 | 0 | 1 | 2 | 0 | 0 | 1 | 0 | 0 | 0 | 0 |
| 0 | -1.08 | 3.67 | -0.29 | 0.77 | 1 | Gm27219 | | 1 | 0 | 1 | 1 | 0 | 0 | 1 | 0 | 0 | 0 | 0 |
| 0 | -1.02 | 3.62 | -0.28 | 0.78 | 1 | Gm7075 | | 0 | 0 | 0 | 1 | 0 | 1 | 1 | 0 | 0 | 0 | 1 |
| 0 | -1.62 | 3.63 | -0.45 | 0.66 | 1 | Gm5269 | | 0 | 1 | 0 | 1 | 0 | 1 | 1 | 0 | 0 | 0 | 0 |
| 0 | -1.61 | 3.64 | -0.44 | 0.66 | 1 | Gm49867 | | 1 | 0 | 0 | 0 | 1 | 1 | 1 | 0 | 0 | 0 | 0 |
| 0 | -1.60 | 3.63 | -0.44 | 0.66 | 1 | Gm11224 | | 1 | 0 | 0 | 1 | 0 | 1 | 1 | 0 | 0 | 0 | 0 |
| 0 | 0.63 | 3.64 | 0.17 | 0.86 | 1 | Gm7369 | | 0 | 0 | 0 | 0 | 0 | 0 | 1 | 0 | 0 | 3 | 0 |

| | | | | | | | | | | | | | | | | | | |
|---|---|---|---|---|---|---|---|---|---|---|---|---|---|---|---|---|---|---|
| 0 | -0.32 | 3.66 | -0.09 | 0.93 | 1 | Gm20513 | | 3 | 0 | 0 | 0 | 0 | 0 | 1 | 0 | 0 | 0 | 0 |
| 0 | 0.52 | 3.65 | 0.14 | 0.89 | 1 | Gm6059 | | 0 | 0 | 0 | 0 | 0 | 0 | 1 | 2 | 0 | 0 | 0 |
| 0 | 0.90 | 3.65 | 0.25 | 0.81 | 1 | Gm6386 | | 0 | 0 | 0 | 0 | 0 | 0 | 1 | 0 | 0 | 1 | 1 |
| 0 | 0.90 | 3.65 | 0.25 | 0.81 | 1 | Gm15204 | | 0 | 0 | 0 | 0 | 0 | 0 | 1 | 0 | 0 | 1 | 1 |
| 0 | 0.29 | 3.66 | 0.08 | 0.94 | 1 | Gm13831 | | 0 | 1 | 0 | 0 | 0 | 0 | 1 | 0 | 0 | 1 | 0 |
| 0 | 0.89 | 3.65 | 0.24 | 0.81 | 1 | Gm14006 | | 0 | 0 | 0 | 0 | 0 | 0 | 1 | 0 | 1 | 1 | 0 |
| 0 | 0.95 | 3.66 | 0.26 | 0.80 | 1 | Gm1969 | | 0 | 0 | 0 | 0 | 0 | 0 | 1 | 1 | 1 | 0 | 0 |
| 0 | -0.33 | 3.65 | -0.09 | 0.93 | 1 | Gm8940 | | 0 | 0 | 0 | 1 | 0 | 0 | 1 | 0 | 0 | 1 | 0 |
| 0 | 0.28 | 3.64 | 0.08 | 0.94 | 1 | Gm17199 | | 1 | 0 | 0 | 0 | 0 | 0 | 1 | 0 | 0 | 1 | 0 |
| 0 | -0.33 | 3.65 | -0.09 | 0.93 | 1 | Gm5638 | | 0 | 0 | 0 | 0 | 0 | 1 | 1 | 0 | 0 | 1 | 0 |
| 0 | 0.07 | 3.65 | 0.02 | 0.98 | 1 | Gm15526 | | 0 | 0 | 1 | 0 | 0 | 0 | 1 | 0 | 1 | 0 | 0 |
| 0 | -0.36 | 3.65 | -0.10 | 0.92 | 1 | Gm17068 | | 0 | 0 | 0 | 1 | 0 | 0 | 1 | 1 | 0 | 0 | 0 |
| 0 | 0.36 | 3.64 | 0.10 | 0.92 | 1 | Gm10550 | | 0 | 0 | 0 | 0 | 0 | 0 | 1 | 0 | 0 | 0 | 2 |
| 0 | 0.28 | 3.68 | 0.08 | 0.94 | 1 | Gm2367 | | 0 | 1 | 0 | 0 | 0 | 0 | 1 | 0 | 1 | 0 | 0 |
| 0 | 0.28 | 3.68 | 0.08 | 0.94 | 1 | Gm17795 | | 0 | 1 | 0 | 0 | 0 | 0 | 1 | 0 | 1 | 0 | 0 |
| 0 | 0.56 | 3.66 | 0.15 | 0.88 | 1 | Gm8171 | | 0 | 0 | 0 | 0 | 0 | 0 | 1 | 0 | 2 | 0 | 0 |
| 0 | 0.56 | 3.66 | 0.15 | 0.88 | 1 | 4930483K19Rik | | 0 | 0 | 0 | 0 | 0 | 0 | 1 | 0 | 2 | 0 | 0 |
| 0 | -0.47 | 3.66 | -0.13 | 0.90 | 1 | Pin1rt1 | protein (peptidyl-prolyl cis/trans isomeras | 0 | 0 | 0 | 0 | 0 | 1 | 1 | 1 | 0 | 0 | 0 |
| 0 | -1.04 | 3.66 | -0.28 | 0.78 | 1 | Gm42694 | | 0 | 0 | 1 | 0 | 1 | 0 | 1 | 0 | 0 | 0 | 0 |
| 0 | -1.04 | 3.66 | -0.28 | 0.78 | 1 | Gm4987 | | 0 | 0 | 1 | 0 | 1 | 0 | 1 | 0 | 0 | 0 | 0 |
| 0 | -1.21 | 3.66 | -0.33 | 0.74 | 1 | Gm4034 | | 0 | 1 | 0 | 0 | 1 | 0 | 1 | 0 | 0 | 0 | 0 |
| 0 | -0.38 | 3.65 | -0.10 | 0.92 | 1 | Gm9034 | | 0 | 0 | 0 | 0 | 1 | 0 | 1 | 0 | 1 | 0 | 0 |
| 0 | -0.41 | 3.67 | -0.11 | 0.91 | 1 | Gm8101 | | 1 | 0 | 1 | 0 | 0 | 0 | 1 | 0 | 0 | 0 | 0 |
| 0 | -0.41 | 3.67 | -0.11 | 0.91 | 1 | Gm5876 | | 1 | 0 | 1 | 0 | 0 | 0 | 1 | 0 | 0 | 0 | 0 |
| 0 | -0.41 | 3.67 | -0.11 | 0.91 | 1 | Gm17786 | | 1 | 0 | 1 | 0 | 0 | 0 | 1 | 0 | 0 | 0 | 0 |
| 0 | -0.52 | 3.66 | -0.14 | 0.89 | 1 | Gm7611 | | 0 | 0 | 0 | 1 | 0 | 0 | 1 | 0 | 0 | 0 | 1 |
| 0 | -1.35 | 3.65 | -0.37 | 0.71 | 1 | Gm43636 | | 0 | 0 | 0 | 0 | 2 | 0 | 1 | 0 | 0 | 0 | 0 |
| 0 | -1.04 | 3.66 | -0.28 | 0.78 | 1 | Gm8731 | | 0 | 1 | 0 | 1 | 0 | 0 | 1 | 0 | 0 | 0 | 0 |
| 0 | 0.42 | 3.66 | 0.11 | 0.91 | 1 | Ube2nl | | 1 | 0 | 0 | 0 | 0 | 0 | 1 | 0 | 1 | 0 | 0 |
| 0 | -0.43 | 3.64 | -0.12 | 0.91 | 1 | Gm26716 | | 0 | 0 | 0 | 0 | 0 | 1 | 1 | 0 | 0 | 0 | 1 |
| 0 | -0.43 | 3.64 | -0.12 | 0.91 | 1 | Gm6640 | | 0 | 0 | 0 | 0 | 0 | 1 | 1 | 0 | 0 | 0 | 1 |
| 0 | -1.13 | 3.66 | -0.31 | 0.76 | 1 | Gm9109 | | 0 | 1 | 0 | 0 | 0 | 1 | 1 | 0 | 0 | 0 | 0 |
| 0 | -0.43 | 3.66 | -0.12 | 0.91 | 1 | Gm13858 | | 0 | 0 | 0 | 0 | 0 | 1 | 1 | 0 | 1 | 0 | 0 |
| 0 | -0.43 | 3.66 | -0.12 | 0.91 | 1 | Gm2791 | | 0 | 0 | 0 | 0 | 0 | 1 | 1 | 0 | 1 | 0 | 0 |
| 0 | -1.61 | 3.64 | -0.44 | 0.66 | 1 | Gm38139 | | 0 | 0 | 0 | 0 | 1 | 1 | 1 | 0 | 0 | 0 | 0 |
| 0 | -1.33 | 3.65 | -0.37 | 0.71 | 1 | Gm6794 | | 0 | 0 | 0 | 2 | 0 | 0 | 1 | 0 | 0 | 0 | 0 |
| 0 | -1.02 | 3.66 | -0.28 | 0.78 | 1 | Gm11663 | | 1 | 0 | 0 | 1 | 0 | 0 | 1 | 0 | 0 | 0 | 0 |
| 0 | -1.02 | 3.66 | -0.28 | 0.78 | 1 | Gm50041 | | 1 | 0 | 0 | 1 | 0 | 0 | 1 | 0 | 0 | 0 | 0 |
| 0 | -1.60 | 3.64 | -0.44 | 0.66 | 1 | Gm23061 | | 0 | 0 | 0 | 1 | 0 | 1 | 1 | 0 | 0 | 0 | 0 |
| 0 | -0.35 | 3.67 | -0.10 | 0.92 | 1 | Rpl19-ps10 | | 2 | 0 | 0 | 0 | 0 | 0 | 1 | 0 | 0 | 0 | 0 |
| 0 | -0.35 | 3.67 | -0.10 | 0.92 | 1 | Gm8586 | | 2 | 0 | 0 | 0 | 0 | 0 | 1 | 0 | 0 | 0 | 0 |
| 0 | -0.99 | 3.65 | -0.27 | 0.79 | 1 | Gm12838 | | 1 | 0 | 0 | 0 | 0 | 1 | 1 | 0 | 0 | 0 | 0 |
| 0 | 0.29 | 3.66 | 0.08 | 0.94 | 1 | Rps2-ps2 | | 0 | 0 | 0 | 0 | 0 | 0 | 1 | 0 | 0 | 1 | 0 |
| 0 | 0.29 | 3.66 | 0.08 | 0.94 | 1 | Gm7561 | | 0 | 0 | 0 | 0 | 0 | 0 | 1 | 0 | 0 | 1 | 0 |
| 0 | 0.28 | 3.68 | 0.08 | 0.94 | 1 | Gm13566 | | 0 | 0 | 0 | 0 | 0 | 0 | 1 | 1 | 0 | 0 | 0 |
| 0 | 0.28 | 3.68 | 0.08 | 0.94 | 1 | Gm43726 | | 0 | 0 | 0 | 0 | 0 | 0 | 1 | 1 | 0 | 0 | 0 |
| 0 | 0.28 | 3.68 | 0.08 | 0.94 | 1 | Prdx2-ps2 | | 0 | 0 | 0 | 0 | 0 | 0 | 1 | 1 | 0 | 0 | 0 |
| 0 | 0.28 | 3.68 | 0.08 | 0.94 | 1 | Gm43368 | | 0 | 0 | 0 | 0 | 0 | 0 | 1 | 1 | 0 | 0 | 0 |
| 0 | -0.44 | 3.68 | -0.12 | 0.90 | 1 | Gm12464 | | 0 | 0 | 1 | 0 | 0 | 0 | 1 | 0 | 0 | 0 | 0 |
| 0 | -0.44 | 3.68 | -0.12 | 0.90 | 1 | Gm5051 | | 0 | 0 | 1 | 0 | 0 | 0 | 1 | 0 | 0 | 0 | 0 |
| 0 | -0.44 | 3.68 | -0.12 | 0.90 | 1 | Gm10015 | | 0 | 0 | 1 | 0 | 0 | 0 | 1 | 0 | 0 | 0 | 0 |
| 0 | -0.44 | 3.68 | -0.12 | 0.90 | 1 | Rpl19-ps2 | | 0 | 0 | 1 | 0 | 0 | 0 | 1 | 0 | 0 | 0 | 0 |
| 0 | -0.44 | 3.68 | -0.12 | 0.90 | 1 | Gm46496 | | 0 | 0 | 1 | 0 | 0 | 0 | 1 | 0 | 0 | 0 | 0 |
| 0 | -0.44 | 3.68 | -0.12 | 0.90 | 1 | Gm50307 | | 0 | 0 | 1 | 0 | 0 | 0 | 1 | 0 | 0 | 0 | 0 |
| 0 | 0.28 | 3.68 | 0.08 | 0.94 | 1 | Gm13009 | | 0 | 0 | 0 | 0 | 0 | 0 | 1 | 0 | 0 | 0 | 1 |
| 0 | 0.28 | 3.68 | 0.08 | 0.94 | 1 | Ovol3 | ovo like zinc finger 3 | 0 | 0 | 0 | 0 | 0 | 0 | 1 | 0 | 0 | 0 | 1 |
| 0 | 0.28 | 3.68 | 0.08 | 0.94 | 1 | Gm47681 | | 0 | 0 | 0 | 0 | 0 | 0 | 1 | 0 | 0 | 0 | 1 |
| 0 | -0.44 | 3.68 | -0.12 | 0.90 | 1 | Gm12421 | | 0 | 1 | 0 | 0 | 0 | 0 | 1 | 0 | 0 | 0 | 0 |
| 0 | -0.44 | 3.68 | -0.12 | 0.90 | 1 | Gm5292 | | 0 | 1 | 0 | 0 | 0 | 0 | 1 | 0 | 0 | 0 | 0 |
| 0 | -0.44 | 3.68 | -0.12 | 0.90 | 1 | Gm8767 | | 0 | 1 | 0 | 0 | 0 | 0 | 1 | 0 | 0 | 0 | 0 |

| | | | | | | | | | | | | | | | | | | |
|---|---|---|---|---|---|---|---|---|---|---|---|---|---|---|---|---|---|---|
| 0 | -0.44 | 3.68 | -0.12 | 0.90 | 1 | Gm8185 | | 0 | 1 | 0 | 0 | 0 | 0 | 1 | 0 | 0 | 0 | 0 |
| 0 | -0.44 | 3.68 | -0.12 | 0.90 | 1 | Gm18335 | | 0 | 1 | 0 | 0 | 0 | 0 | 1 | 0 | 0 | 0 | 0 |
| 0 | 0.28 | 3.68 | 0.08 | 0.94 | 1 | Gm13368 | | 0 | 0 | 0 | 0 | 0 | 0 | 1 | 0 | 1 | 0 | 0 |
| 0 | 0.28 | 3.68 | 0.08 | 0.94 | 1 | Gm5738 | | 0 | 0 | 0 | 0 | 0 | 0 | 1 | 0 | 1 | 0 | 0 |
| 0 | 0.28 | 3.68 | 0.08 | 0.94 | 1 | Gm18079 | | 0 | 0 | 0 | 0 | 0 | 0 | 1 | 0 | 1 | 0 | 0 |
| 0 | 0.28 | 3.68 | 0.08 | 0.94 | 1 | Gm4821 | | 0 | 0 | 0 | 0 | 0 | 0 | 1 | 0 | 1 | 0 | 0 |
| 0 | 0.28 | 3.68 | 0.08 | 0.94 | 1 | Gm8702 | | 0 | 0 | 0 | 0 | 0 | 0 | 1 | 0 | 1 | 0 | 0 |
| 0 | 0.28 | 3.68 | 0.08 | 0.94 | 1 | Rps12-ps3 | | 0 | 0 | 0 | 0 | 0 | 0 | 1 | 0 | 1 | 0 | 0 |
| 0 | -1.04 | 3.66 | -0.28 | 0.78 | 1 | Gm7329 | | 0 | 0 | 0 | 0 | 1 | 0 | 1 | 0 | 0 | 0 | 0 |
| 0 | -1.04 | 3.66 | -0.28 | 0.78 | 1 | Gm12579 | | 0 | 0 | 0 | 0 | 1 | 0 | 1 | 0 | 0 | 0 | 0 |
| 0 | -1.04 | 3.66 | -0.28 | 0.78 | 1 | Gm6434 | | 0 | 0 | 0 | 0 | 1 | 0 | 1 | 0 | 0 | 0 | 0 |
| 0 | -1.04 | 3.66 | -0.28 | 0.78 | 1 | Gm17134 | | 0 | 0 | 0 | 0 | 1 | 0 | 1 | 0 | 0 | 0 | 0 |
| 0 | -1.03 | 3.66 | -0.28 | 0.78 | 1 | Gm2623 | | 0 | 0 | 0 | 1 | 0 | 0 | 1 | 0 | 0 | 0 | 0 |
| 0 | -1.03 | 3.66 | -0.28 | 0.78 | 1 | Gm8918 | | 0 | 0 | 0 | 1 | 0 | 0 | 1 | 0 | 0 | 0 | 0 |
| 0 | -1.03 | 3.66 | -0.28 | 0.78 | 1 | Gm6128 | | 0 | 0 | 0 | 1 | 0 | 0 | 1 | 0 | 0 | 0 | 0 |
| 0 | -1.03 | 3.66 | -0.28 | 0.78 | 1 | Gm5213 | | 0 | 0 | 0 | 1 | 0 | 0 | 1 | 0 | 0 | 0 | 0 |
| 0 | -1.03 | 3.66 | -0.28 | 0.78 | 1 | Gm36804 | | 0 | 0 | 0 | 1 | 0 | 0 | 1 | 0 | 0 | 0 | 0 |
| 0 | -0.38 | 3.67 | -0.10 | 0.92 | 1 | Gm26983 | | 1 | 0 | 0 | 0 | 0 | 0 | 1 | 0 | 0 | 0 | 0 |
| 0 | -0.38 | 3.67 | -0.10 | 0.92 | 1 | Gm12943 | | 1 | 0 | 0 | 0 | 0 | 0 | 1 | 0 | 0 | 0 | 0 |
| 0 | -0.38 | 3.67 | -0.10 | 0.92 | 1 | Gm10845 | | 1 | 0 | 0 | 0 | 0 | 0 | 1 | 0 | 0 | 0 | 0 |
| 0 | -0.38 | 3.67 | -0.10 | 0.92 | 1 | Gm26977 | | 1 | 0 | 0 | 0 | 0 | 0 | 1 | 0 | 0 | 0 | 0 |
| 0 | -1.05 | 3.66 | -0.29 | 0.78 | 1 | Shisal2a | shisa like 2A | 0 | 0 | 0 | 0 | 0 | 1 | 1 | 0 | 0 | 0 | 0 |
| 0 | -1.05 | 3.66 | -0.29 | 0.78 | 1 | Gm47808 | | 0 | 0 | 0 | 0 | 0 | 1 | 1 | 0 | 0 | 0 | 0 |
| 0 | -1.05 | 3.66 | -0.29 | 0.78 | 1 | Gm46453 | | 0 | 0 | 0 | 0 | 0 | 1 | 1 | 0 | 0 | 0 | 0 |
| 0 | -1.05 | 3.66 | -0.29 | 0.78 | 1 | Gm47505 | | 0 | 0 | 0 | 0 | 0 | 1 | 1 | 0 | 0 | 0 | 0 |
| 0 | -1.05 | 3.66 | -0.29 | 0.78 | 1 | Gm5688 | | 0 | 0 | 0 | 0 | 0 | 1 | 1 | 0 | 0 | 0 | 0 |
| 3888 | -0.78 | 3.62 | -0.21 | 0.83 | 1 | Col6a1 | collagen, type VI, alpha 1 | 19462 | 10536 | 12771 | 0 | 0 | 4 | 0 | 0 | 0 | 0 | 0 |
| 3794 | 1.34 | 3.60 | 0.37 | 0.71 | 1 | Igha | | 40560 | 824 | 306 | 0 | 0 | 0 | 0 | 0 | 0 | 39 | 0 |
| 2023 | 1.51 | 3.65 | 0.41 | 0.68 | 1 | Svep1 | sushi, von Willebrand factor type A, EGF a | 10386 | 5239 | 6625 | 0 | 0 | 0 | 0 | 0 | 0 | 4 | 0 |
| 1951 | -0.11 | 2.18 | -0.05 | 0.96 | 1 | Fbln2 | fibulin 2 | 9388 | 5712 | 6353 | 0 | 7 | 0 | 0 | 0 | 4 | 0 | 0 |
| 1779 | -1.46 | 1.84 | -0.79 | 0.43 | 1 | C7 | complement component 7 | 13656 | 2340 | 3554 | 0 | 5 | 6 | 0 | 0 | 0 | 0 | 5 |
| 1634 | 1.00 | 3.68 | 0.27 | 0.79 | 1 | Col5a2 | collagen, type V, alpha 2 | 7647 | 4261 | 6066 | 0 | 0 | 0 | 0 | 1 | 0 | 0 | 0 |
| 1604 | 3.95 | 2.56 | 1.54 | 0.12 | 1 | Dpt | dermatopontin | 10772 | 3121 | 3727 | 0 | 0 | 0 | 0 | 0 | 0 | 22 | 6 |
| 1539 | -1.37 | 1.74 | -0.79 | 0.43 | 1 | Notch3 | notch 3 | 7556 | 4639 | 4720 | 0 | 5 | 3 | 0 | 0 | 0 | 0 | 3 |
| 1525 | -0.69 | 1.72 | -0.40 | 0.69 | 1 | Slit3 | slit guidance ligand 3 | 9164 | 3527 | 4080 | 3 | 0 | 2 | 0 | 0 | 0 | 0 | 4 |
| 1498 | -2.97 | 2.42 | -1.23 | 0.22 | 1 | Crispld2 | cysteine-rich secretory protein LCCL dom | 7979 | 3609 | 4868 | 11 | 4 | 0 | 0 | 0 | 0 | 0 | 3 |
| 1440 | 1.75 | 3.64 | 0.48 | 0.63 | 1 | Gdf10 | growth differentiation factor 10 | 8975 | 2789 | 4067 | 0 | 0 | 0 | 0 | 0 | 5 | 0 | 0 |
| 1321 | -2.83 | 1.73 | -1.64 | 0.10 | 1 | Fbln1 | fibulin 1 | 9485 | 2107 | 2918 | 6 | 4 | 7 | 0 | 0 | 0 | 0 | 7 |
| 1265 | 0.00 | 3.68 | 0.00 | 1.00 | 1 | Lum | lumican | 8151 | 2813 | 2948 | 0 | 0 | 0 | 0 | 0 | 0 | 0 | 0 |
| 1234 | 0.00 | 3.68 | 0.00 | 1.00 | 1 | Pdgfra | platelet derived growth factor receptor, alp | 7042 | 2811 | 3720 | 0 | 0 | 0 | 0 | 0 | 0 | 0 | 0 |
| 1195 | 0.45 | 2.76 | 0.16 | 0.87 | 1 | Eln | elastin | 6285 | 2288 | 4547 | 9 | 0 | 0 | 0 | 15 | 0 | 0 | 0 |
| 1131 | 2.91 | 3.13 | 0.93 | 0.35 | 1 | Prelp | proline arginine-rich end leucine-rich repe | 5365 | 3373 | 3696 | 0 | 0 | 2 | 0 | 0 | 9 | 0 | 0 |
| 1064 | -0.34 | 3.12 | -0.11 | 0.91 | 1 | Smoc2 | SPARC related modular calcium binding | 3907 | 3364 | 4420 | 13 | 0 | 0 | 0 | 0 | 0 | 3 | 0 |
| 1056 | 0.25 | 2.05 | 0.12 | 0.90 | 1 | Vwf | Von Willebrand factor | 4984 | 1971 | 4641 | 0 | 5 | 3 | 0 | 14 | 0 | 0 | 2 |
| 1024 | 1.32 | 3.63 | 0.36 | 0.72 | 1 | Ogn | osteoglycin | 6280 | 2242 | 2734 | 0 | 0 | 0 | 0 | 0 | 0 | 0 | 4 |
| 984 | 0.82 | 3.65 | 0.22 | 0.82 | 1 | Igkv5-39 | | 487 | 9352 | 983 | 0 | 0 | 0 | 0 | 1 | 0 | 0 | 0 |
| 922 | -2.01 | 2.34 | -0.86 | 0.39 | 1 | Dpep1 | dipeptidase 1 | 4253 | 2383 | 3484 | 7 | 0 | 3 | 0 | 0 | 0 | 0 | 14 |
| 853 | 1.37 | 1.53 | 0.90 | 0.37 | 1 | Col5a3 | collagen, type V, alpha 3 | 2472 | 2756 | 4152 | 2 | 0 | 0 | 0 | 0 | 3 | 0 | 2 |
| 810 | 3.04 | 1.65 | 1.84 | 0.07 | 1 | Cygb | cytoglobin | 5255 | 1562 | 2082 | 0 | 0 | 0 | 0 | 0 | 0 | 5 | 4 |
| 805 | -1.29 | 3.63 | -0.36 | 0.72 | 1 | Htra1 | HtrA serine peptidase 1 | 4186 | 2173 | 2489 | 8 | 0 | 0 | 0 | 0 | 0 | 0 | 0 |
| 784 | -0.06 | 2.05 | -0.03 | 0.97 | 1 | Mgp | matrix Gla protein | 3635 | 2223 | 2765 | 0 | 0 | 3 | 0 | 0 | 1 | 0 | 0 |
| 784 | 0.97 | 3.67 | 0.26 | 0.79 | 1 | Igfbp6 | insulin-like growth factor binding protein 6 | 4695 | 1895 | 2030 | 0 | 0 | 0 | 0 | 0 | 0 | 1 | 0 |
| 776 | -0.40 | 3.67 | -0.11 | 0.91 | 1 | Cyyr1 | cysteine and tyrosine-rich protein 1 | 2488 | 3147 | 2905 | 0 | 0 | 1 | 0 | 0 | 0 | 0 | 0 |
| 774 | 0.00 | 3.68 | 0.00 | 1.00 | 1 | Scara5 | scavenger receptor class A, member 5 | 4803 | 1594 | 2114 | 0 | 0 | 0 | 0 | 0 | 0 | 0 | 0 |
| 768 | 0.00 | 3.68 | 0.00 | 1.00 | 1 | Ighg2b | | 6265 | 1065 | 1113 | 0 | 0 | 0 | 0 | 0 | 0 | 0 | 0 |
| 764 | 0.39 | 1.58 | 0.25 | 0.81 | 1 | Clec3b | C-type lectin domain family 3, member b | 4973 | 1282 | 2133 | 0 | 5 | 2 | 0 | 0 | 5 | 0 | 3 |
| 744 | 0.98 | 3.68 | 0.27 | 0.79 | 1 | Htra3 | HtrA serine peptidase 3 | 3576 | 2120 | 2484 | 0 | 0 | 0 | 0 | 0 | 1 | 0 | 0 |
| 711 | -0.73 | 3.64 | -0.20 | 0.84 | 1 | Ly6c1 | lymphocyte antigen 6 complex, locus C1 | 2710 | 2354 | 2753 | 0 | 0 | 3 | 0 | 0 | 0 | 0 | 0 |
| 695 | -1.68 | 1.52 | -1.11 | 0.27 | 1 | C1ra | complement component 1, r subcompone | 4316 | 1748 | 1491 | 14 | 45 | 4 | 0 | 3 | 4 | 0 | 13 |
| 684 | -0.59 | 2.94 | -0.20 | 0.84 | 1 | Serpinf1 | serine (or cysteine) peptidase inhibitor, cl | 4117 | 1364 | 2028 | 9 | 0 | 0 | 0 | 0 | 1 | 0 | 0 |

| | | | | | | | | | | | | | | | | | |
|---|---|---|---|---|---|---|---|---|---|---|---|---|---|---|---|---|---|
| 602 | 2.23 | 1.86 | 1.20 | 0.23 | 1 | Xdh | xanthine dehydrogenase | 3065 | 1779 | 1702 | 0 | 14 | 5 | 0 | 28 | 0 | 22 | 11 |
| 602 | -2.32 | 3.27 | -0.71 | 0.48 | 1 | Anxa1 | annexin A1 | 3493 | 1534 | 1586 | 7 | 0 | 0 | 0 | 0 | 0 | 0 | 4 |
| 602 | 0.00 | 3.68 | 0.00 | 1.00 | 1 | Flt4 | FMS-like tyrosine kinase 4 | 2744 | 1802 | 2072 | 0 | 0 | 0 | 0 | 0 | 0 | 0 | 0 |
| 599 | -2.03 | 1.49 | -1.36 | 0.17 | 1 | Wwtr1 | WW domain containing transcription regu | 2740 | 1894 | 1943 | 3 | 6 | 3 | 0 | 0 | 3 | 0 | 0 |
| 591 | -2.33 | 1.57 | -1.48 | 0.14 | 1 | Nes | nestin | 2277 | 1996 | 2189 | 10 | 19 | 2 | 0 | 0 | 2 | 0 | 6 |
| 576 | -1.75 | 2.85 | -0.62 | 0.54 | 1 | Colec12 | collectin sub-family member 12 | 3355 | 1568 | 1401 | 4 | 0 | 0 | 0 | 0 | 0 | 0 | 2 |
| 571 | -2.02 | 1.61 | -1.25 | 0.21 | 1 | Aoc3 | amine oxidase, copper containing 3 | 2786 | 1668 | 1794 | 5 | 0 | 22 | 0 | 0 | 0 | 0 | 7 |
| 561 | -0.09 | 1.17 | -0.07 | 0.94 | 1 | Gas1 | growth arrest specific 1 | 3070 | 1653 | 1420 | 0 | 2 | 14 | 0 | 0 | 2 | 0 | 10 |
| 548 | 0.00 | 3.68 | 0.00 | 1.00 | 1 | Adamts5 | a disintegrin-like and metallopeptidase (r | 2452 | 1659 | 1916 | 0 | 0 | 0 | 0 | 0 | 0 | 0 | 0 |
| 529 | -0.44 | 3.68 | -0.12 | 0.90 | 1 | Igf1 | insulin-like growth factor 1 | 3525 | 1147 | 1147 | 0 | 1 | 0 | 0 | 0 | 0 | 0 | 0 |
| 514 | -2.62 | 1.62 | -1.62 | 0.11 | 1 | Ano1 | anoctamin 1, calcium activated chloride c | 2630 | 1640 | 1312 | 35 | 15 | 14 | 0 | 0 | 5 | 0 | 7 |
| 509 | 2.03 | 2.54 | 0.80 | 0.42 | 1 | Dchs1 | dachsous cadherin related 1 | 2203 | 1583 | 1792 | 11 | 0 | 0 | 0 | 0 | 4 | 0 | 3 |
| 503 | 3.19 | 2.53 | 1.26 | 0.21 | 1 | Lhfp | lipoma HMGIC fusion partner | 2477 | 1505 | 1531 | 0 | 4 | 0 | 0 | 13 | 0 | 0 | 6 |
| 497 | -1.16 | 3.65 | -0.32 | 0.75 | 1 | Lamc3 | laminin gamma 3 | 2226 | 1163 | 2070 | 4 | 0 | 0 | 0 | 0 | 0 | 0 | 0 |
| 470 | -0.26 | 2.97 | -0.09 | 0.93 | 1 | Abcc9 | ATP-binding cassette, sub-family C (CFTI | 2479 | 1593 | 1082 | 4 | 0 | 0 | 0 | 0 | 0 | 0 | 7 |
| 444 | -0.41 | 1.58 | -0.26 | 0.80 | 1 | Pld4 | phospholipase D family, member 4 | 2435 | 981 | 1441 | 0 | 11 | 7 | 0 | 0 | 6 | 0 | 5 |
| 438 | 0.98 | 1.05 | 0.94 | 0.35 | 1 | Sod3 | superoxide dismutase 3, extracellular | 2427 | 1121 | 1199 | 11 | 7 | 6 | 0 | 22 | 1 | 17 | 12 |
| 423 | -1.36 | 1.99 | -0.68 | 0.49 | 1 | Sema7a | sema domain, immunoglobulin domain (I | 1458 | 1450 | 1731 | 0 | 9 | 3 | 0 | 0 | 0 | 0 | 4 |
| 419 | 0.00 | 3.68 | 0.00 | 1.00 | 1 | Cyp1b1 | cytochrome P450, family 1, subfamily b, p | 2328 | 1429 | 853 | 0 | 0 | 0 | 0 | 0 | 0 | 0 | 0 |
| 418 | 0.00 | 3.68 | 0.00 | 1.00 | 1 | Igkv8-24 | | 4313 | 290 | 0 | 0 | 0 | 0 | 0 | 0 | 0 | 0 | 0 |
| 418 | 0.05 | 1.63 | 0.03 | 0.97 | 1 | Cdh11 | cadherin 11 | 2758 | 1171 | 646 | 0 | 4 | 8 | 0 | 0 | 4 | 0 | 6 |
| 415 | -3.09 | 2.50 | -1.24 | 0.22 | 1 | Itgax | integrin alpha X | 2128 | 1199 | 1226 | 7 | 5 | 0 | 0 | 0 | 0 | 0 | 0 |
| 415 | -1.88 | 1.64 | -1.15 | 0.25 | 1 | Dpysl3 | dihydropyrimidinase-like 3 | 2410 | 966 | 1178 | 0 | 3 | 3 | 0 | 0 | 0 | 1 | 0 |
| 412 | 0.00 | 3.68 | 0.00 | 1.00 | 1 | Upk3b | uroplakin 3B | 2147 | 1554 | 836 | 0 | 0 | 0 | 0 | 0 | 0 | 0 | 0 |
| 397 | -0.44 | 3.68 | -0.12 | 0.90 | 1 | Lox | lysyl oxidase | 2323 | 956 | 1087 | 0 | 1 | 0 | 0 | 0 | 0 | 0 | 0 |
| 394 | -1.18 | 3.65 | -0.32 | 0.75 | 1 | Itgbl1 | integrin, beta-like 1 | 2737 | 551 | 1045 | 4 | 0 | 0 | 0 | 0 | 0 | 0 | 0 |
| 388 | 1.62 | 1.63 | 0.99 | 0.32 | 1 | S1pr3 | sphingosine-1-phosphate receptor 3 | 1897 | 1112 | 1248 | 0 | 3 | 0 | 0 | 0 | 5 | 5 | 0 |
| 387 | -0.76 | 1.76 | -0.43 | 0.67 | 1 | Ecm2 | extracellular matrix protein 2, female orga | 1854 | 1237 | 1155 | 0 | 0 | 7 | 0 | 0 | 0 | 0 | 1 |
| 380 | -1.80 | 1.91 | -0.94 | 0.35 | 1 | Ebf1 | early B cell factor 1 | 1424 | 1540 | 1215 | 0 | 2 | 2 | 0 | 0 | 0 | 0 | 0 |
| 379 | 0.99 | 3.68 | 0.27 | 0.79 | 1 | Ighv3-1 | | 284 | 3059 | 828 | 0 | 0 | 0 | 0 | 0 | 1 | 0 | 0 |
| 367 | -0.52 | 1.56 | -0.33 | 0.74 | 1 | Arhgef15 | Rho guanine nucleotide exchange factor | 1098 | 1410 | 1527 | 0 | 3 | 2 | 0 | 0 | 3 | 0 | 0 |
| 361 | -0.80 | 3.66 | -0.22 | 0.83 | 1 | Sox18 | SRY (sex determining region Y)-box 18 | 1153 | 1353 | 1467 | 0 | 2 | 0 | 0 | 0 | 0 | 0 | 0 |
| 359 | 0.48 | 0.96 | 0.50 | 0.62 | 1 | Itga5 | integrin alpha 5 (fibronectin receptor alph | 1269 | 1303 | 1359 | 0 | 6 | 1 | 0 | 0 | 8 | 1 | 0 |
| 353 | 3.88 | 3.34 | 1.16 | 0.25 | 1 | Cx3cr1 | chemokine (C-X3-C motif) receptor 1 | 1519 | 931 | 1406 | 0 | 0 | 3 | 0 | 0 | 28 | 0 | 0 |
| 341 | 0.00 | 3.68 | 0.00 | 1.00 | 1 | Hsd11b1 | hydroxysteroid 11-beta dehydrogenase 1 | 1510 | 962 | 1280 | 0 | 0 | 0 | 0 | 0 | 0 | 0 | 0 |
| 336 | -1.75 | 1.78 | -0.98 | 0.33 | 1 | Heyl | hairy/enhancer-of-split related with YRPW | 1381 | 1009 | 1286 | 7 | 0 | 8 | 0 | 0 | 0 | 0 | 5 |
| 336 | -0.73 | 1.60 | -0.46 | 0.65 | 1 | Cdc42ep1 | CDC42 effector protein (Rho GTPase bin | 1277 | 1121 | 1280 | 6 | 6 | 0 | 0 | 0 | 5 | 0 | 1 |
| 333 | -2.61 | 1.71 | -1.53 | 0.13 | 1 | Plscr2 | phospholipid scramblase 2 | 1126 | 1241 | 1260 | 13 | 9 | 4 | 0 | 9 | 0 | 0 | 0 |
| 319 | 0.00 | 3.68 | 0.00 | 1.00 | 1 | Piezo2 | piezo-type mechanosensitive ion channe | 1306 | 942 | 1261 | 0 | 0 | 0 | 0 | 0 | 0 | 0 | 0 |
| 312 | 0.95 | 3.67 | 0.26 | 0.80 | 1 | Osmr | oncostatin M receptor | 1359 | 1197 | 878 | 0 | 0 | 0 | 0 | 0 | 0 | 0 | 1 |
| 312 | 1.02 | 1.11 | 0.91 | 0.36 | 1 | Ptgis | prostaglandin I2 (prostacyclin) synthase | 1939 | 710 | 777 | 0 | 0 | 2 | 0 | 1 | 0 | 0 | 3 |
| 312 | 0.00 | 3.68 | 0.00 | 1.00 | 1 | Ces2e | carboxylesterase 2E | 912 | 994 | 1526 | 0 | 0 | 0 | 0 | 0 | 0 | 0 | 0 |
| 305 | 1.00 | 3.68 | 0.27 | 0.79 | 1 | Bdkrb2 | bradykinin receptor, beta 2 | 1179 | 821 | 1352 | 0 | 0 | 0 | 0 | 0 | 0 | 1 | 0 |
| 295 | 0.00 | 3.68 | 0.00 | 1.00 | 1 | Igkv12-46 | | 3242 | 0 | 0 | 0 | 0 | 0 | 0 | 0 | 0 | 0 | 0 |
| 289 | 0.00 | 3.68 | 0.00 | 1.00 | 1 | Aplnr | apelin receptor | 1053 | 1057 | 1069 | 0 | 0 | 0 | 0 | 0 | 0 | 0 | 0 |
| 288 | 0.90 | 2.43 | 0.37 | 0.71 | 1 | Ppp1r16b | protein phosphatase 1, regulatory subuni | 1238 | 732 | 1182 | 0 | 12 | 0 | 0 | 3 | 0 | 0 | 1 |
| 286 | -2.14 | 1.55 | -1.38 | 0.17 | 1 | Col4a5 | collagen, type IV, alpha 5 | 1740 | 630 | 654 | 12 | 18 | 54 | 0 | 0 | 7 | 0 | 29 |
| 281 | 0.00 | 3.68 | 0.00 | 1.00 | 1 | Ar | androgen receptor | 1480 | 806 | 804 | 0 | 0 | 0 | 0 | 0 | 0 | 0 | 0 |
| 275 | 0.00 | 3.68 | 0.00 | 1.00 | 1 | Lrrc32 | leucine rich repeat containing 32 | 1213 | 751 | 1062 | 0 | 0 | 0 | 0 | 0 | 0 | 0 | 0 |
| 272 | -2.00 | 1.86 | -1.07 | 0.28 | 1 | Ctsk | cathepsin K | 1307 | 617 | 987 | 7 | 8 | 20 | 0 | 0 | 0 | 46 | 3 |
| 270 | 0.57 | 1.85 | 0.31 | 0.76 | 1 | Itm2a | integral membrane protein 2A | 1260 | 691 | 986 | 0 | 12 | 1 | 0 | 0 | 8 | 13 | 0 |
| 270 | 1.30 | 3.64 | 0.36 | 0.72 | 1 | Pou2af1 | POU domain, class 2, associating factor 1 | 2362 | 518 | 88 | 0 | 0 | 0 | 0 | 0 | 0 | 0 | 3 |
| 266 | 0.21 | 1.30 | 0.16 | 0.87 | 1 | Cckar | cholecystokinin A receptor | 279 | 1611 | 997 | 12 | 0 | 6 | 0 | 12 | 0 | 0 | 9 |
| 263 | -0.77 | 1.36 | -0.57 | 0.57 | 1 | Trim47 | tripartite motif-containing 47 | 1118 | 725 | 1029 | 4 | 5 | 1 | 0 | 0 | 4 | 0 | 2 |
| 262 | 0.01 | 1.03 | 0.01 | 0.99 | 1 | Adamts12 | a disintegrin-like and metallopeptidase (r | 806 | 915 | 1159 | 0 | 0 | 3 | 0 | 0 | 0 | 0 | 3 |
| 260 | -0.73 | 3.64 | -0.20 | 0.84 | 1 | Cmklr1 | chemokine-like receptor 1 | 1121 | 834 | 902 | 0 | 0 | 3 | 0 | 0 | 0 | 0 | 0 |
| 258 | -0.83 | 2.52 | -0.33 | 0.74 | 1 | Aldh1a2 | aldehyde dehydrogenase family 1, subfar | 1224 | 824 | 767 | 7 | 0 | 1 | 0 | 0 | 10 | 0 | 0 |
| 252 | 0.00 | 3.68 | 0.00 | 1.00 | 1 | Mfap4 | microfibrillar-associated protein 4 | 1794 | 407 | 575 | 0 | 0 | 0 | 0 | 0 | 0 | 0 | 0 |
| 249 | -2.67 | 2.53 | -1.06 | 0.29 | 1 | Lims2 | LIM and senescent cell antigen like doma | 1006 | 783 | 936 | 9 | 0 | 2 | 0 | 0 | 0 | 0 | 0 |
| 247 | 0.00 | 3.68 | 0.00 | 1.00 | 1 | Cxcl14 | chemokine (C-X-C motif) ligand 14 | 1147 | 721 | 852 | 0 | 0 | 0 | 0 | 0 | 0 | 0 | 0 |

| | | | | | | | | | | | | | | | | | | |
|---|---|---|---|---|---|---|---|---|---|---|---|---|---|---|---|---|---|---|
| 246 | 0.00 | 3.68 | 0.00 | 1.00 | 1 | Ighv14-2 | | 2708 | 3 | 0 | 0 | 0 | 0 | 0 | 0 | 0 | 0 | 0 |
| 242 | 0.00 | 3.68 | 0.00 | 1.00 | 1 | Iglc2 | | 894 | 1748 | 20 | 0 | 0 | 0 | 0 | 0 | 0 | 0 | 0 |
| 242 | 0.41 | 1.97 | 0.21 | 0.84 | 1 | Il10ra | interleukin 10 receptor, alpha | 1278 | 526 | 796 | 15 | 4 | 0 | 0 | 0 | 7 | 32 | 0 |
| 241 | 0.00 | 3.68 | 0.00 | 1.00 | 1 | Slpi | secretory leukocyte peptidase inhibitor | 1311 | 750 | 585 | 0 | 0 | 0 | 0 | 0 | 0 | 0 | 0 |
| 240 | -0.51 | 1.20 | -0.42 | 0.67 | 1 | Adamtsl1 | ADAMTS-like 1 | 1530 | 540 | 567 | 0 | 0 | 3 | 0 | 0 | 0 | 0 | 1 |
| 238 | -2.22 | 1.84 | -1.20 | 0.23 | 1 | Synm | synemin, intermediate filament protein | 1421 | 681 | 472 | 2 | 26 | 4 | 0 | 0 | 11 | 0 | 0 |
| 237 | 1.00 | 3.68 | 0.27 | 0.79 | 1 | Tmem119 | transmembrane protein 119 | 1176 | 899 | 529 | 0 | 0 | 0 | 0 | 0 | 1 | 0 | 0 |
| 237 | -0.52 | 1.65 | -0.31 | 0.75 | 1 | Itgal | integrin alpha L | 1525 | 555 | 479 | 4 | 3 | 6 | 0 | 0 | 0 | 28 | 4 |
| 235 | -3.10 | 2.44 | -1.27 | 0.20 | 1 | Naalad2 | N-acetylated alpha-linked acidic dipeptid | 1205 | 520 | 845 | 11 | 5 | 0 | 0 | 0 | 0 | 0 | 3 |
| 227 | 0.28 | 1.35 | 0.21 | 0.84 | 1 | Tril | TLR4 interactor with leucine-rich repeats | 879 | 871 | 743 | 0 | 0 | 1 | 0 | 0 | 0 | 0 | 1 |
| 225 | -1.11 | 1.82 | -0.61 | 0.54 | 1 | Adgrg6 | adhesion G protein-coupled receptor G6 | 1128 | 592 | 739 | 0 | 7 | 2 | 0 | 0 | 3 | 0 | 0 |
| 224 | -0.16 | 2.42 | -0.07 | 0.95 | 1 | Ces2g | carboxylesterase 2G | 912 | 757 | 789 | 0 | 8 | 0 | 0 | 0 | 4 | 0 | 0 |
| 224 | -0.69 | 3.66 | -0.19 | 0.85 | 1 | Ifi203 | interferon activated gene 203 | 831 | 864 | 771 | 0 | 0 | 2 | 0 | 0 | 0 | 0 | 0 |
| 223 | 0.00 | 3.68 | 0.00 | 1.00 | 1 | Mrvi1 | MRV integration site 1 | 1156 | 828 | 473 | 0 | 0 | 0 | 0 | 0 | 0 | 0 | 0 |
| 223 | -1.36 | 2.11 | -0.64 | 0.52 | 1 | Mmrn1 | multimerin 1 | 2023 | 102 | 314 | 0 | 9 | 1 | 0 | 0 | 2 | 0 | 0 |
| 221 | 0.57 | 1.87 | 0.30 | 0.76 | 1 | Cadm3 | cell adhesion molecule 3 | 1280 | 545 | 595 | 0 | 4 | 0 | 0 | 0 | 3 | 0 | 1 |
| 219 | 0.00 | 3.68 | 0.00 | 1.00 | 1 | Ighv4-2 | | 2412 | 0 | 0 | 0 | 0 | 0 | 0 | 0 | 0 | 0 | 0 |
| 215 | -0.05 | 2.20 | -0.02 | 0.98 | 1 | Apln | apelin | 1170 | 650 | 494 | 0 | 20 | 5 | 0 | 0 | 0 | 15 | 6 |
| 212 | -1.11 | 3.63 | -0.31 | 0.76 | 1 | Mgl2 | macrophage galactose N-acetyl-galactos | 1081 | 413 | 829 | 0 | 6 | 0 | 0 | 0 | 0 | 0 | 0 |
| 211 | 0.00 | 3.68 | 0.00 | 1.00 | 1 | Mfap5 | microfibrillar associated protein 5 | 1101 | 396 | 823 | 0 | 0 | 0 | 0 | 0 | 0 | 0 | 0 |
| 210 | 1.00 | 3.61 | 0.28 | 0.78 | 1 | Ighv14-1 | | 2226 | 75 | 0 | 0 | 0 | 0 | 0 | 0 | 0 | 10 | 0 |
| 208 | 1.30 | 3.64 | 0.36 | 0.72 | 1 | Vgll3 | vestigial like family member 3 | 1056 | 515 | 719 | 0 | 0 | 0 | 0 | 0 | 0 | 0 | 3 |
| 208 | 0.00 | 3.68 | 0.00 | 1.00 | 1 | Iglv2 | | 315 | 1968 | 0 | 0 | 0 | 0 | 0 | 0 | 0 | 0 | 0 |
| 207 | 1.53 | 0.94 | 1.63 | 0.10 | 1 | Tbx2 | T-box 2 | 980 | 508 | 774 | 0 | 0 | 3 | 0 | 0 | 0 | 0 | 14 |
| 206 | -0.46 | 1.73 | -0.27 | 0.79 | 1 | Rnf144a | ring finger protein 144A | 785 | 833 | 633 | 8 | 0 | 5 | 0 | 0 | 4 | 0 | 3 |
| 204 | -1.15 | 2.03 | -0.57 | 0.57 | 1 | Daam2 | dishevelled associated activator of morph | 888 | 580 | 748 | 12 | 0 | 8 | 0 | 0 | 7 | 0 | 2 |
| 204 | 0.00 | 3.68 | 0.00 | 1.00 | 1 | Adcy4 | adenylate cyclase 4 | 643 | 889 | 707 | 0 | 0 | 0 | 0 | 0 | 0 | 0 | 0 |
| 203 | -0.81 | 3.61 | -0.22 | 0.82 | 1 | Ushbp1 | USH1 protein network component harmo | 737 | 640 | 852 | 0 | 0 | 8 | 0 | 0 | 0 | 0 | 0 |
| 201 | 0.00 | 3.68 | 0.00 | 1.00 | 1 | Col4a4 | collagen, type IV, alpha 4 | 1061 | 481 | 670 | 0 | 0 | 0 | 0 | 0 | 0 | 0 | 0 |
| 201 | 0.00 | 3.68 | 0.00 | 1.00 | 1 | Itga8 | integrin alpha 8 | 1246 | 558 | 402 | 0 | 0 | 0 | 0 | 0 | 0 | 0 | 0 |
| 199 | -1.27 | 3.62 | -0.35 | 0.73 | 1 | Csf2rb | colony stimulating factor 2 receptor, beta, | 1322 | 485 | 367 | 0 | 9 | 0 | 0 | 0 | 0 | 0 | 0 |
| 197 | -0.58 | 1.36 | -0.42 | 0.67 | 1 | Ifit2 | interferon-induced protein with tetratricop | 705 | 561 | 787 | 12 | 37 | 8 | 0 | 1 | 43 | 0 | 12 |
| 197 | -2.80 | 2.51 | -1.12 | 0.26 | 1 | Gng11 | guanine nucleotide binding protein (G pro | 956 | 587 | 610 | 0 | 8 | 3 | 0 | 0 | 0 | 0 | 0 |
| 196 | 1.09 | 1.33 | 0.82 | 0.41 | 1 | Gpc6 | glypican 6 | 945 | 588 | 538 | 6 | 5 | 8 | 0 | 49 | 3 | 0 | 11 |
| 193 | 1.53 | 1.24 | 1.23 | 0.22 | 1 | Col4a3 | collagen, type IV, alpha 3 | 837 | 563 | 720 | 0 | 0 | 1 | 0 | 0 | 1 | 0 | 3 |
| 192 | 0.13 | 1.50 | 0.09 | 0.93 | 1 | Acacb | acetyl-Coenzyme A carboxylase beta | 607 | 773 | 605 | 17 | 26 | 14 | 0 | 0 | 17 | 48 | 11 |
| 192 | 0.00 | 3.68 | 0.00 | 1.00 | 1 | Hmcn1 | hemicentin 1 | 1114 | 582 | 414 | 0 | 0 | 0 | 0 | 0 | 0 | 0 | 0 |
| 191 | 2.71 | 2.29 | 1.18 | 0.24 | 1 | St8sia6 | ST8 alpha-N-acetyl-neuraminide alpha-2 | 1134 | 573 | 389 | 0 | 0 | 0 | 0 | 0 | 3 | 0 | 3 |
| 191 | 1.30 | 3.64 | 0.36 | 0.72 | 1 | Nkx2-3 | NK2 homeobox 3 | 774 | 621 | 701 | 0 | 0 | 0 | 0 | 0 | 0 | 0 | 3 |
| 188 | 0.00 | 3.68 | 0.00 | 1.00 | 1 | Gpihbp1 | GPI-anchored HDL-binding protein 1 | 713 | 687 | 670 | 0 | 0 | 0 | 0 | 0 | 0 | 0 | 0 |
| 187 | -2.18 | 2.27 | -0.96 | 0.34 | 1 | Rgs1 | regulator of G-protein signaling 1 | 1124 | 381 | 545 | 0 | 4 | 2 | 0 | 0 | 0 | 0 | 0 |
| 185 | 0.00 | 3.68 | 0.00 | 1.00 | 1 | Ighv4-1 | | 2035 | 0 | 0 | 0 | 0 | 0 | 0 | 0 | 0 | 0 | 0 |
| 185 | -0.20 | 1.59 | -0.12 | 0.90 | 1 | Pla2g7 | phospholipase A2, group VII (platelet-acti | 1162 | 370 | 480 | 0 | 7 | 4 | 0 | 0 | 5 | 0 | 3 |
| 181 | 0.00 | 3.68 | 0.00 | 1.00 | 1 | Iglv1 | | 1603 | 358 | 33 | 0 | 0 | 0 | 0 | 0 | 0 | 0 | 0 |
| 180 | -3.11 | 2.03 | -1.53 | 0.13 | 1 | Plxna4 | plexin A4 | 1017 | 483 | 434 | 10 | 12 | 8 | 0 | 14 | 0 | 0 | 0 |
| 177 | -1.93 | 2.06 | -0.94 | 0.35 | 1 | Ptgs2 | prostaglandin-endoperoxide synthase 2 | 1481 | 197 | 243 | 0 | 9 | 13 | 0 | 0 | 0 | 0 | 7 |
| 176 | -3.39 | 2.66 | -1.27 | 0.20 | 1 | Slfn2 | schlafen 2 | 886 | 445 | 590 | 0 | 12 | 4 | 0 | 0 | 0 | 0 | 0 |
| 174 | 2.02 | 1.56 | 1.30 | 0.20 | 1 | AW112010 | | 767 | 559 | 516 | 0 | 9 | 12 | 0 | 22 | 14 | 0 | 18 |
| 174 | 0.00 | 3.68 | 0.00 | 1.00 | 1 | Ighv9-3 | | 1332 | 583 | 0 | 0 | 0 | 0 | 0 | 0 | 0 | 0 | 0 |
| 173 | -1.89 | 2.63 | -0.72 | 0.47 | 1 | Lrrn4 | leucine rich repeat neuronal 4 | 961 | 726 | 214 | 0 | 4 | 1 | 0 | 0 | 0 | 0 | 0 |
| 172 | 1.70 | 2.01 | 0.85 | 0.40 | 1 | Fli1 | Friend leukemia integration 1 | 756 | 523 | 604 | 0 | 0 | 2 | 0 | 5 | 0 | 1 | 0 |
| 171 | -3.17 | 2.75 | -1.15 | 0.25 | 1 | Apbb1ip | amyloid beta (A4) precursor protein-bindi | 962 | 404 | 499 | 0 | 14 | 3 | 0 | 0 | 0 | 0 | 0 |
| 170 | 1.21 | 3.07 | 0.39 | 0.69 | 1 | Vav1 | vav 1 oncogene | 888 | 438 | 527 | 0 | 0 | 1 | 0 | 0 | 0 | 22 | 0 |
| 169 | -1.37 | 1.53 | -0.90 | 0.37 | 1 | Enpp1 | ectonucleotide pyrophosphatase/phosph | 842 | 463 | 479 | 11 | 12 | 16 | 0 | 0 | 7 | 0 | 28 |
| 168 | 1.30 | 3.66 | 0.36 | 0.72 | 1 | Hbb-bs | hemoglobin, beta adult major chain | 618 | 428 | 802 | 0 | 0 | 0 | 0 | 0 | 2 | 0 | 0 |
| 167 | 0.00 | 3.68 | 0.00 | 1.00 | 1 | Igkv1-110 | | 1628 | 204 | 0 | 0 | 0 | 0 | 0 | 0 | 0 | 0 | 0 |
| 166 | 0.00 | 3.68 | 0.00 | 1.00 | 1 | Ighg3 | | 1705 | 115 | 3 | 0 | 0 | 0 | 0 | 0 | 0 | 0 | 0 |
| 164 | 0.00 | 3.68 | 0.00 | 1.00 | 1 | Rbp7 | retinol binding protein 7, cellular | 575 | 335 | 898 | 0 | 0 | 0 | 0 | 0 | 0 | 0 | 0 |
| 164 | -0.78 | 3.62 | -0.21 | 0.83 | 1 | Epha4 | Eph receptor A4 | 643 | 515 | 638 | 0 | 0 | 4 | 0 | 0 | 0 | 0 | 0 |
| 160 | 0.00 | 3.68 | 0.00 | 1.00 | 1 | Ifi207 | interferon activated gene 207 | 728 | 512 | 518 | 0 | 0 | 0 | 0 | 0 | 0 | 0 | 0 |

| | | | | | | | | | | | | | | | | | | |
|---|---|---|---|---|---|---|---|---|---|---|---|---|---|---|---|---|---|---|
| 160 | -1.29 | 3.61 | -0.36 | 0.72 | 1 | Thbs2 | thrombospondin 2 | 1034 | 328 | 382 | 0 | 11 | 0 | 0 | 0 | 0 | 0 | 0 |
| 156 | 0.00 | 3.64 | 0.00 | 1.00 | 1 | Igkv6-17 | | 1700 | 0 | 11 | 0 | 0 | 0 | 0 | 0 | 0 | 0 | 0 |
| 154 | 0.99 | 1.76 | 0.56 | 0.57 | 1 | Syde1 | synapse defective 1, Rho GTPase, homol | 658 | 451 | 505 | 0 | 13 | 17 | 0 | 0 | 13 | 31 | 4 |
| 153 | 0.73 | 1.08 | 0.67 | 0.50 | 1 | Slc11a1 | solute carrier family 11 (proton-coupled d | 894 | 478 | 298 | 0 | 0 | 3 | 0 | 0 | 0 | 0 | 5 |
| 152 | 1.66 | 3.62 | 0.46 | 0.65 | 1 | Hvcn1 | hydrogen voltage-gated channel 1 | 1014 | 320 | 329 | 0 | 0 | 0 | 0 | 0 | 0 | 10 | 0 |
| 149 | -1.91 | 2.22 | -0.86 | 0.39 | 1 | Ldb2 | LIM domain binding 2 | 565 | 526 | 509 | 26 | 0 | 4 | 0 | 0 | 0 | 0 | 3 |
| 148 | -1.08 | 1.79 | -0.60 | 0.55 | 1 | Gja4 | gap junction protein, alpha 4 | 595 | 459 | 560 | 4 | 0 | 3 | 0 | 0 | 0 | 0 | 4 |
| 147 | -0.03 | 2.45 | -0.01 | 0.99 | 1 | Casp12 | caspase 12 | 815 | 321 | 474 | 0 | 2 | 0 | 0 | 0 | 0 | 0 | 2 |
| 145 | -1.37 | 2.31 | -0.59 | 0.55 | 1 | Ly86 | lymphocyte antigen 86 | 687 | 374 | 527 | 5 | 0 | 2 | 0 | 0 | 5 | 0 | 0 |
| 145 | -2.80 | 2.32 | -1.21 | 0.23 | 1 | Aqp12 | aquaporin 12 | 153 | 817 | 600 | 13 | 8 | 0 | 0 | 0 | 0 | 0 | 5 |
| 145 | 0.95 | 3.67 | 0.26 | 0.80 | 1 | Tbxa2r | thromboxane A2 receptor | 599 | 426 | 563 | 0 | 0 | 0 | 0 | 0 | 0 | 0 | 1 |
| 143 | -2.52 | 1.80 | -1.40 | 0.16 | 1 | Sardh | sarcosine dehydrogenase | 579 | 581 | 398 | 10 | 0 | 5 | 0 | 0 | 0 | 0 | 1 |
| 143 | -1.23 | 1.20 | -1.03 | 0.30 | 1 | Sult1a1 | sulfotransferase family 1A, phenol-preferr | 577 | 408 | 577 | 0 | 5 | 3 | 0 | 0 | 3 | 0 | 0 |
| 143 | -0.51 | 1.19 | -0.43 | 0.67 | 1 | Sting1 | stimulator of interferon response cGAMP i | 738 | 410 | 414 | 0 | 3 | 3 | 0 | 0 | 0 | 0 | 5 |
| 142 | 2.05 | 3.41 | 0.60 | 0.55 | 1 | Abi3 | ABI gene family, member 3 | 590 | 422 | 498 | 3 | 0 | 0 | 0 | 0 | 0 | 44 | 0 |
| 141 | -1.70 | 2.28 | -0.74 | 0.46 | 1 | Lrrc17 | leucine rich repeat containing 17 | 750 | 402 | 395 | 0 | 3 | 1 | 0 | 0 | 0 | 0 | 0 |
| 141 | 0.00 | 3.68 | 0.00 | 1.00 | 1 | Kcne4 | potassium voltage-gated channel, Isk-rela | 995 | 226 | 326 | 0 | 0 | 0 | 0 | 0 | 0 | 0 | 0 |
| 140 | 1.04 | 1.11 | 0.94 | 0.35 | 1 | Ssc5d | scavenger receptor cysteine rich family, 5 | 811 | 343 | 276 | 0 | 0 | 56 | 0 | 0 | 3 | 0 | 53 |
| 138 | 1.19 | 3.00 | 0.40 | 0.69 | 1 | Flnc | filamin C, gamma | 688 | 419 | 391 | 0 | 0 | 1 | 0 | 0 | 0 | 17 | 0 |
| 138 | 3.07 | 2.03 | 1.51 | 0.13 | 1 | Pcdh19 | protocadherin 19 | 600 | 479 | 410 | 0 | 0 | 1 | 0 | 17 | 0 | 0 | 5 |
| 137 | 0.00 | 3.68 | 0.00 | 1.00 | 1 | Igkv1-117 | | 1448 | 58 | 0 | 0 | 0 | 0 | 0 | 0 | 0 | 0 | 0 |
| 135 | 0.00 | 3.68 | 0.00 | 1.00 | 1 | Ighv5-17 | | 1484 | 0 | 0 | 0 | 0 | 0 | 0 | 0 | 0 | 0 | 0 |
| 133 | -1.13 | 3.65 | -0.31 | 0.76 | 1 | Rab7b | RAB7B, member RAS oncogene family | 780 | 446 | 237 | 0 | 4 | 0 | 0 | 0 | 0 | 0 | 0 |
| 131 | 0.55 | 1.89 | 0.29 | 0.77 | 1 | Fblim1 | filamin binding LIM protein 1 | 602 | 344 | 484 | 3 | 0 | 0 | 0 | 1 | 0 | 0 | 3 |
| 130 | 1.25 | 3.66 | 0.34 | 0.73 | 1 | Syt15 | synaptotagmin XV | 582 | 373 | 477 | 0 | 0 | 0 | 0 | 0 | 0 | 0 | 2 |
| 130 | -2.19 | 1.27 | -1.73 | 0.08 | 1 | Itgb7 | integrin beta 7 | 812 | 421 | 182 | 6 | 8 | 2 | 0 | 0 | 2 | 0 | 1 |
| 130 | 0.97 | 3.67 | 0.26 | 0.79 | 1 | Hey2 | hairy/enhancer-of-split related with YRPW | 583 | 336 | 512 | 0 | 0 | 0 | 0 | 0 | 1 | 0 | 0 |
| 130 | 3.23 | 2.57 | 1.26 | 0.21 | 1 | Pcdh18 | protocadherin 18 | 692 | 373 | 353 | 0 | 0 | 0 | 0 | 10 | 0 | 0 | 2 |
| 130 | 0.95 | 3.67 | 0.26 | 0.80 | 1 | Hck | hemopoietic cell kinase | 566 | 349 | 509 | 0 | 0 | 0 | 0 | 0 | 0 | 0 | 1 |
| 129 | 0.00 | 3.62 | 0.00 | 1.00 | 1 | Igkv9-124 | | 1417 | 3 | 0 | 0 | 0 | 0 | 0 | 0 | 0 | 0 | 0 |
| 128 | 1.79 | 1.00 | 1.79 | 0.07 | 1 | Angpt1 | angiopoietin 1 | 456 | 358 | 488 | 3 | 20 | 3 | 0 | 43 | 27 | 0 | 9 |
| 128 | 1.16 | 2.62 | 0.44 | 0.66 | 1 | Pcdh7 | protocadherin 7 | 646 | 342 | 409 | 0 | 0 | 1 | 0 | 0 | 5 | 0 | 0 |
| 126 | 0.00 | 3.68 | 0.00 | 1.00 | 1 | Clec1a | C-type lectin domain family 1, member a | 581 | 404 | 401 | 0 | 0 | 0 | 0 | 0 | 0 | 0 | 0 |
| 126 | 2.15 | 1.62 | 1.33 | 0.18 | 1 | Gm13889 | predicted gene 13889 | 595 | 354 | 357 | 7 | 0 | 13 | 0 | 28 | 0 | 24 | 6 |
| 126 | 0.00 | 3.68 | 0.00 | 1.00 | 1 | Itga11 | integrin alpha 11 | 640 | 251 | 492 | 0 | 0 | 0 | 0 | 0 | 0 | 0 | 0 |
| 126 | 0.00 | 3.68 | 0.00 | 1.00 | 1 | Cdh13 | cadherin 13 | 807 | 251 | 323 | 0 | 0 | 0 | 0 | 0 | 0 | 0 | 0 |
| 125 | 0.00 | 3.68 | 0.00 | 1.00 | 1 | Slamf7 | SLAM family member 7 | 773 | 332 | 269 | 0 | 0 | 0 | 0 | 0 | 0 | 0 | 0 |
| 125 | -0.85 | 1.91 | -0.45 | 0.66 | 1 | Mustn1 | musculoskeletal, embryonic nuclear prote | 672 | 340 | 348 | 0 | 0 | 10 | 0 | 0 | 0 | 0 | 1 |
| 124 | 0.00 | 3.68 | 0.00 | 1.00 | 1 | Ighv7-1 | | 1269 | 95 | 0 | 0 | 0 | 0 | 0 | 0 | 0 | 0 | 0 |
| 124 | 0.00 | 3.68 | 0.00 | 1.00 | 1 | Ccl21a | chemokine (C-C motif) ligand 21A (serine | 1046 | 26 | 289 | 0 | 0 | 0 | 0 | 0 | 0 | 0 | 0 |
| 123 | 0.00 | 3.68 | 0.00 | 1.00 | 1 | Igkv14-111 | | 1247 | 102 | 0 | 0 | 0 | 0 | 0 | 0 | 0 | 0 | 0 |
| 121 | -0.84 | 2.05 | -0.41 | 0.68 | 1 | Pld2 | phospholipase D2 | 599 | 403 | 318 | 0 | 3 | 10 | 0 | 0 | 3 | 0 | 0 |
| 121 | 0.61 | 2.55 | 0.24 | 0.81 | 1 | Aif1 | allograft inflammatory factor 1 | 631 | 346 | 353 | 0 | 0 | 3 | 0 | 0 | 2 | 0 | 0 |
| 121 | 1.43 | 1.06 | 1.35 | 0.18 | 1 | Lair1 | leukocyte-associated Ig-like receptor 1 | 448 | 317 | 550 | 0 | 0 | 7 | 0 | 0 | 4 | 0 | 9 |
| 121 | 1.25 | 3.66 | 0.34 | 0.73 | 1 | Srpx2 | sushi-repeat-containing protein, X-linked | 516 | 375 | 432 | 0 | 0 | 0 | 0 | 0 | 0 | 0 | 2 |
| 120 | 1.37 | 3.61 | 0.38 | 0.70 | 1 | Ifi204 | interferon activated gene 204 | 510 | 453 | 346 | 0 | 0 | 0 | 0 | 0 | 0 | 0 | 9 |
| 120 | -2.18 | 2.36 | -0.92 | 0.36 | 1 | Marchf1 | membrane associated ring-CH-type finge | 744 | 277 | 289 | 0 | 4 | 2 | 0 | 0 | 0 | 0 | 0 |
| 119 | -0.40 | 3.67 | -0.11 | 0.91 | 1 | Slamf9 | SLAM family member 9 | 573 | 251 | 483 | 0 | 0 | 1 | 0 | 0 | 0 | 0 | 0 |
| 119 | 0.84 | 1.23 | 0.68 | 0.49 | 1 | AU020206 | | 540 | 405 | 340 | 0 | 5 | 2 | 0 | 0 | 6 | 0 | 6 |
| 118 | 0.00 | 3.68 | 0.00 | 1.00 | 1 | C1qtnf7 | C1q and tumor necrosis factor related pro | 625 | 346 | 328 | 0 | 0 | 0 | 0 | 0 | 0 | 0 | 0 |
| 117 | 1.30 | 3.64 | 0.36 | 0.72 | 1 | Marco | macrophage receptor with collagenous st | 968 | 262 | 56 | 0 | 0 | 0 | 0 | 0 | 0 | 0 | 3 |
| 117 | 0.00 | 3.68 | 0.00 | 1.00 | 1 | Ighv6-7 | | 1285 | 0 | 0 | 0 | 0 | 0 | 0 | 0 | 0 | 0 | 0 |
| 116 | 0.00 | 3.68 | 0.00 | 1.00 | 1 | Art3 | ADP-ribosyltransferase 3 | 502 | 343 | 435 | 0 | 0 | 0 | 0 | 0 | 0 | 0 | 0 |
| 115 | 0.00 | 3.68 | 0.00 | 1.00 | 1 | Iglc1 | | 1195 | 56 | 18 | 0 | 0 | 0 | 0 | 0 | 0 | 0 | 0 |
| 115 | -0.40 | 3.67 | -0.11 | 0.91 | 1 | Dnase1l3 | deoxyribonuclease 1-like 3 | 638 | 343 | 280 | 0 | 0 | 1 | 0 | 0 | 0 | 0 | 0 |
| 115 | 0.00 | 3.68 | 0.00 | 1.00 | 1 | Adamtsl3 | ADAMTS-like 3 | 358 | 360 | 545 | 0 | 0 | 0 | 0 | 0 | 0 | 0 | 0 |
| 114 | -2.07 | 1.29 | -1.61 | 0.11 | 1 | P2rx1 | purinergic receptor P2X, ligand-gated ion | 464 | 481 | 304 | 4 | 0 | 3 | 0 | 0 | 0 | 0 | 1 |
| 114 | 2.29 | 2.00 | 1.15 | 0.25 | 1 | Erbb2 | erb-b2 receptor tyrosine kinase 2 | 508 | 322 | 401 | 4 | 0 | 0 | 0 | 9 | 12 | 0 | 0 |
| 114 | 0.95 | 3.67 | 0.26 | 0.80 | 1 | Wfdc1 | WAP four-disulfide core domain 1 | 698 | 285 | 269 | 0 | 0 | 0 | 0 | 0 | 0 | 0 | 1 |
| 114 | 0.00 | 3.68 | 0.00 | 1.00 | 1 | Vit | vitrin | 999 | 156 | 95 | 0 | 0 | 0 | 0 | 0 | 0 | 0 | 0 |

| | | | | | | | | | | | | | | | | | | |
|---|---|---|---|---|---|---|---|---|---|---|---|---|---|---|---|---|---|---|
| 112 | -0.68 | 2.48 | -0.27 | 0.79 | 1 | Slc15a3 | solute carrier family 15, member 3 | 449 | 333 | 433 | 0 | 10 | 0 | 0 | 0 | 2 | 0 | 0 |
| 112 | -2.86 | 2.67 | -1.07 | 0.29 | 1 | Rasl12 | RAS-like, family 12 | 554 | 323 | 336 | 12 | 0 | 2 | 0 | 0 | 0 | 0 | 0 |
| 112 | 2.06 | 1.26 | 1.64 | 0.10 | 1 | Ajuba | ajuba LIM protein | 427 | 389 | 398 | 0 | 0 | 3 | 0 | 0 | 6 | 0 | 4 |
| 111 | -1.48 | 1.29 | -1.14 | 0.25 | 1 | Cyp27a1 | cytochrome P450, family 27, subfamily a, | 698 | 223 | 277 | 1 | 11 | 5 | 0 | 0 | 8 | 0 | 0 |
| 110 | -0.08 | 2.47 | -0.03 | 0.97 | 1 | Evi2a | ecotropic viral integration site 2a | 619 | 272 | 313 | 0 | 7 | 0 | 0 | 0 | 4 | 0 | 0 |
| 109 | 0.00 | 3.68 | 0.00 | 1.00 | 1 | Cd300a | CD300A molecule | 496 | 295 | 409 | 0 | 0 | 0 | 0 | 0 | 0 | 0 | 0 |
| 109 | 0.12 | 0.97 | 0.12 | 0.91 | 1 | Scarf1 | scavenger receptor class F, member 1 | 483 | 320 | 372 | 0 | 0 | 12 | 0 | 0 | 0 | 0 | 12 |
| 108 | 0.00 | 3.68 | 0.00 | 1.00 | 1 | Lcp2 | lymphocyte cytosolic protein 2 | 642 | 294 | 258 | 0 | 0 | 0 | 0 | 0 | 0 | 0 | 0 |
| 108 | 0.00 | 3.68 | 0.00 | 1.00 | 1 | Agtr1a | angiotensin II receptor, type 1a | 399 | 404 | 383 | 0 | 0 | 0 | 0 | 0 | 0 | 0 | 0 |
| 107 | -1.43 | 1.25 | -1.14 | 0.25 | 1 | Ptpre | protein tyrosine phosphatase, receptor typ | 614 | 254 | 294 | 8 | 0 | 7 | 0 | 1 | 0 | 0 | 4 |
| 107 | -1.18 | 3.63 | -0.32 | 0.75 | 1 | Il34 | interleukin 34 | 572 | 372 | 232 | 0 | 6 | 0 | 0 | 0 | 0 | 0 | 0 |
| 107 | -0.18 | 1.20 | -0.15 | 0.88 | 1 | Hand2 | heart and neural crest derivatives express | 621 | 223 | 310 | 3 | 9 | 0 | 0 | 6 | 3 | 0 | 0 |
| 106 | -0.41 | 3.53 | -0.11 | 0.91 | 1 | Cd48 | CD48 antigen | 642 | 304 | 185 | 0 | 16 | 0 | 0 | 0 | 0 | 22 | 0 |
| 106 | -1.47 | 1.27 | -1.16 | 0.25 | 1 | Cd37 | CD37 antigen | 623 | 335 | 196 | 1 | 3 | 2 | 0 | 0 | 0 | 0 | 2 |
| 105 | 0.00 | 3.68 | 0.00 | 1.00 | 1 | Parvg | parvin, gamma | 531 | 352 | 276 | 0 | 0 | 0 | 0 | 0 | 0 | 0 | 0 |
| 105 | 2.37 | 1.24 | 1.91 | 0.06 | 1 | Efhd1 | EF hand domain containing 1 | 382 | 319 | 417 | 10 | 0 | 0 | 0 | 15 | 13 | 0 | 2 |
| 104 | 0.35 | 3.60 | 0.10 | 0.92 | 1 | Igkv6-32 | | 1135 | 0 | 0 | 0 | 0 | 0 | 0 | 0 | 0 | 8 | 0 |
| 103 | -0.46 | 1.12 | -0.40 | 0.69 | 1 | Was | Wiskott-Aldrich syndrome | 696 | 232 | 201 | 0 | 0 | 6 | 0 | 0 | 0 | 0 | 3 |
| 103 | 0.00 | 3.68 | 0.00 | 1.00 | 1 | Gxylt2 | glucoside xylosyltransferase 2 | 795 | 160 | 176 | 0 | 0 | 0 | 0 | 0 | 0 | 0 | 0 |
| 103 | 0.00 | 3.68 | 0.00 | 1.00 | 1 | Ighv9-1 | | 839 | 289 | 0 | 0 | 0 | 0 | 0 | 0 | 0 | 0 | 0 |
| 102 | -2.44 | 2.82 | -0.87 | 0.39 | 1 | Stum | mechanosensory transduction mediator | 589 | 212 | 311 | 12 | 0 | 1 | 0 | 0 | 0 | 0 | 0 |
| 102 | 0.00 | 3.68 | 0.00 | 1.00 | 1 | Jph2 | junctophilin 2 | 619 | 188 | 315 | 0 | 0 | 0 | 0 | 0 | 0 | 0 | 0 |
| 102 | -0.63 | 2.25 | -0.28 | 0.78 | 1 | Tagap | T cell activation Rho GTPase activating pr | 505 | 265 | 312 | 0 | 13 | 8 | 0 | 0 | 16 | 0 | 0 |
| 101 | 0.00 | 3.68 | 0.00 | 1.00 | 1 | Igkv11-125 | | 1018 | 96 | 0 | 0 | 0 | 0 | 0 | 0 | 0 | 0 | 0 |
| 101 | -3.19 | 2.61 | -1.22 | 0.22 | 1 | Chrdl1 | chordin-like 1 | 878 | 131 | 83 | 16 | 0 | 1 | 0 | 0 | 0 | 0 | 2 |
| 100 | 0.00 | 3.68 | 0.00 | 1.00 | 1 | Igkv14-126 | | 200 | 902 | 0 | 0 | 0 | 0 | 0 | 0 | 0 | 0 | 0 |
| 100 | 0.00 | 3.68 | 0.00 | 1.00 | 1 | Gngt2 | guanine nucleotide binding protein (G pro | 589 | 190 | 322 | 0 | 0 | 0 | 0 | 0 | 0 | 0 | 0 |
| 100 | 0.00 | 3.68 | 0.00 | 1.00 | 1 | AU021092 | expressed sequence AU021092 | 353 | 277 | 469 | 0 | 0 | 0 | 0 | 0 | 0 | 0 | 0 |
| 99 | 0.00 | 3.68 | 0.00 | 1.00 | 1 | Adm | adrenomedullin | 555 | 223 | 315 | 0 | 0 | 0 | 0 | 0 | 0 | 0 | 0 |
| 99 | -0.69 | 3.66 | -0.19 | 0.85 | 1 | Dkk2 | dickkopf WNT signaling pathway inhibitor | 524 | 176 | 388 | 0 | 0 | 2 | 0 | 0 | 0 | 0 | 0 |
| 99 | -0.69 | 3.66 | -0.19 | 0.85 | 1 | Arhgap25 | Rho GTPase activating protein 25 | 574 | 209 | 303 | 0 | 0 | 2 | 0 | 0 | 0 | 0 | 0 |
| 98 | -0.40 | 3.67 | -0.11 | 0.91 | 1 | Msr1 | macrophage scavenger receptor 1 | 619 | 270 | 188 | 0 | 0 | 1 | 0 | 0 | 0 | 0 | 0 |
| 98 | 0.48 | 2.32 | 0.21 | 0.83 | 1 | H2-T24 | histocompatibility 2, T region locus 24 | 343 | 309 | 415 | 0 | 4 | 0 | 0 | 0 | 6 | 0 | 0 |
| 97 | 0.00 | 3.68 | 0.00 | 1.00 | 1 | Igkv4-55 | | 865 | 206 | 0 | 0 | 0 | 0 | 0 | 0 | 0 | 0 | 0 |
| 97 | -2.18 | 2.56 | -0.85 | 0.39 | 1 | Acss1 | acyl-CoA synthetase short-chain family m | 558 | 256 | 245 | 0 | 7 | 1 | 0 | 0 | 0 | 0 | 0 |
| 97 | 0.00 | 3.68 | 0.00 | 1.00 | 1 | Olfr558 | olfactory receptor 558 | 498 | 376 | 191 | 0 | 0 | 0 | 0 | 0 | 0 | 0 | 0 |
| 96 | -2.94 | 2.52 | -1.17 | 0.24 | 1 | Fap | fibroblast activation protein | 309 | 292 | 443 | 8 | 0 | 3 | 0 | 0 | 0 | 0 | 0 |
| 96 | 1.74 | 1.32 | 1.32 | 0.19 | 1 | P2ry6 | pyrimidinergic receptor P2Y, G-protein co | 554 | 264 | 220 | 0 | 2 | 1 | 0 | 0 | 6 | 0 | 5 |
| 95 | 1.34 | 1.00 | 1.34 | 0.18 | 1 | Cachd1 | cache domain containing 1 | 428 | 276 | 290 | 1 | 6 | 8 | 0 | 0 | 28 | 5 | 5 |
| 94 | 0.88 | 1.48 | 0.60 | 0.55 | 1 | Tnc | tenascin C | 561 | 209 | 259 | 1 | 0 | 1 | 0 | 0 | 4 | 0 | 0 |
| 94 | 0.00 | 3.68 | 0.00 | 1.00 | 1 | Ighv1-64 | | 405 | 623 | 5 | 0 | 0 | 0 | 0 | 0 | 0 | 0 | 0 |
| 94 | -2.16 | 1.33 | -1.63 | 0.10 | 1 | Fzd2 | frizzled class receptor 2 | 613 | 170 | 230 | 7 | 6 | 2 | 0 | 0 | 4 | 0 | 0 |
| 93 | 0.78 | 0.98 | 0.80 | 0.43 | 1 | Lrrc55 | leucine rich repeat containing 55 | 256 | 220 | 308 | 51 | 13 | 25 | 0 | 89 | 1 | 23 | 36 |
| 93 | -1.16 | 3.63 | -0.32 | 0.75 | 1 | Rnf144b | ring finger protein 144B | 362 | 330 | 322 | 0 | 7 | 0 | 0 | 0 | 0 | 0 | 0 |
| 93 | 0.00 | 3.68 | 0.00 | 1.00 | 1 | Igkv19-93 | | 1018 | 0 | 0 | 0 | 0 | 0 | 0 | 0 | 0 | 0 | 0 |
| 92 | 0.57 | 1.05 | 0.55 | 0.58 | 1 | Cldn15 | claudin 15 | 353 | 328 | 313 | 0 | 0 | 7 | 0 | 0 | 0 | 0 | 11 |
| 92 | -2.83 | 3.54 | -0.80 | 0.42 | 1 | Ccn5 | cellular communication network factor 5 | 734 | 73 | 190 | 11 | 0 | 0 | 0 | 0 | 0 | 0 | 3 |
| 91 | 0.00 | 3.67 | 0.00 | 1.00 | 1 | Ighv14-3 | | 977 | 20 | 0 | 0 | 0 | 0 | 0 | 0 | 0 | 0 | 0 |
| 90 | -3.02 | 3.31 | -0.91 | 0.36 | 1 | Amy2a5 | amylase 2a5 | 88 | 532 | 352 | 16 | 0 | 0 | 0 | 0 | 0 | 0 | 7 |
| 90 | 0.00 | 3.68 | 0.00 | 1.00 | 1 | Mmp12 | matrix metallopeptidase 12 | 444 | 281 | 265 | 0 | 0 | 0 | 0 | 0 | 0 | 0 | 0 |
| 90 | 0.00 | 3.68 | 0.00 | 1.00 | 1 | Rftn2 | raftlin family member 2 | 451 | 268 | 269 | 0 | 0 | 0 | 0 | 0 | 0 | 0 | 0 |
| 90 | 0.00 | 3.68 | 0.00 | 1.00 | 1 | Ptgir | prostaglandin I receptor (IP) | 537 | 287 | 162 | 0 | 0 | 0 | 0 | 0 | 0 | 0 | 0 |
| 90 | -1.44 | 1.18 | -1.22 | 0.22 | 1 | Ppp1r1b | protein phosphatase 1, regulatory inhibitc | 233 | 269 | 460 | 7 | 9 | 0 | 0 | 0 | 6 | 0 | 0 |
| 90 | -2.95 | 2.74 | -1.08 | 0.28 | 1 | Ahnak2 | AHNAK nucleoprotein 2 | 551 | 270 | 153 | 0 | 9 | 3 | 0 | 0 | 0 | 0 | 0 |
| 89 | 0.00 | 3.68 | 0.00 | 1.00 | 1 | Pirb | paired Ig-like receptor B | 493 | 243 | 243 | 0 | 0 | 0 | 0 | 0 | 0 | 0 | 0 |
| 88 | -1.79 | 2.19 | -0.82 | 0.41 | 1 | Foxf1 | forkhead box F1 | 359 | 360 | 248 | 4 | 0 | 1 | 0 | 0 | 0 | 0 | 0 |
| 88 | 1.88 | 3.63 | 0.52 | 0.60 | 1 | Csf2rb2 | colony stimulating factor 2 receptor, beta : | 525 | 222 | 211 | 0 | 0 | 0 | 0 | 0 | 10 | 0 | 0 |
| 88 | 0.00 | 3.68 | 0.00 | 1.00 | 1 | Igkv3-10 | | 966 | 0 | 0 | 0 | 0 | 0 | 0 | 0 | 0 | 0 | 0 |
| 87 | 3.07 | 2.30 | 1.34 | 0.18 | 1 | Clec4a1 | C-type lectin domain family 4, member a1 | 542 | 158 | 229 | 0 | 0 | 1 | 0 | 28 | 0 | 0 | 4 |
| 87 | 0.00 | 3.68 | 0.00 | 1.00 | 1 | Heph | hephaestin | 438 | 247 | 267 | 0 | 0 | 0 | 0 | 0 | 0 | 0 | 0 |

| | | | | | | | | | | | | | | | | | | |
|---|---|---|---|---|---|---|---|---|---|---|---|---|---|---|---|---|---|---|
| 86 | 4.89 | 2.97 | 1.64 | 0.10 | 1 | Ms4a1 | membrane-spanning 4-domains, subfami | 480 | 356 | 47 | 0 | 0 | 0 | 0 | 58 | 0 | 0 | 9 |
| 86 | -1.10 | 3.64 | -0.30 | 0.76 | 1 | Cd209a | CD209a antigen | 444 | 197 | 295 | 0 | 5 | 0 | 0 | 0 | 0 | 0 | 0 |
| 86 | -0.44 | 1.31 | -0.34 | 0.74 | 1 | Plaur | plasminogen activator, urokinase recepto | 412 | 304 | 221 | 0 | 0 | 3 | 0 | 0 | 0 | 0 | 1 |
| 85 | -1.58 | 1.33 | -1.18 | 0.24 | 1 | Sh2d4a | SH2 domain containing 4A | 421 | 215 | 293 | 0 | 5 | 3 | 0 | 0 | 0 | 0 | 2 |
| 85 | 0.00 | 3.68 | 0.00 | 1.00 | 1 | Mrgprf | MAS-related GPR, member F | 436 | 237 | 262 | 0 | 0 | 0 | 0 | 0 | 0 | 0 | 0 |
| 84 | 0.00 | 3.68 | 0.00 | 1.00 | 1 | Olfml1 | olfactomedin-like 1 | 540 | 199 | 189 | 0 | 0 | 0 | 0 | 0 | 0 | 0 | 0 |
| 84 | 0.94 | 1.53 | 0.62 | 0.54 | 1 | Mndal | myeloid nuclear differentiation antigen lik | 291 | 364 | 268 | 0 | 0 | 1 | 0 | 2 | 0 | 0 | 0 |
| 84 | 1.25 | 3.66 | 0.34 | 0.73 | 1 | Lrg1 | leucine-rich alpha-2-glycoprotein 1 | 476 | 135 | 307 | 0 | 0 | 0 | 0 | 0 | 0 | 0 | 2 |
| 83 | -0.40 | 3.67 | -0.11 | 0.91 | 1 | H2-Ob | histocompatibility 2, O region beta locus | 561 | 248 | 104 | 0 | 0 | 1 | 0 | 0 | 0 | 0 | 0 |
| 83 | 0.00 | 3.68 | 0.00 | 1.00 | 1 | Ctla2a | cytotoxic T lymphocyte-associated proteir | 228 | 317 | 368 | 0 | 0 | 0 | 0 | 0 | 0 | 0 | 0 |
| 82 | 1.37 | 3.61 | 0.38 | 0.70 | 1 | Wt1 | Wilms tumor 1 homolog | 487 | 355 | 56 | 0 | 0 | 0 | 0 | 0 | 0 | 0 | 9 |
| 82 | -3.83 | 2.84 | -1.35 | 0.18 | 1 | Tlr9 | toll-like receptor 9 | 522 | 144 | 215 | 5 | 16 | 0 | 0 | 0 | 0 | 0 | 0 |
| 81 | -0.25 | 1.24 | -0.20 | 0.84 | 1 | Layn | layilin | 308 | 240 | 343 | 0 | 0 | 3 | 0 | 0 | 0 | 0 | 2 |
| 81 | 0.00 | 3.68 | 0.00 | 1.00 | 1 | Kcnj8 | potassium inwardly-rectifying channel, su | 356 | 293 | 247 | 0 | 0 | 0 | 0 | 0 | 0 | 0 | 0 |
| 81 | -0.71 | 1.04 | -0.68 | 0.49 | 1 | Arhgap28 | Rho GTPase activating protein 28 | 430 | 160 | 191 | 28 | 15 | 25 | 0 | 10 | 11 | 0 | 20 |
| 81 | -0.84 | 1.24 | -0.68 | 0.50 | 1 | Tead2 | TEA domain family member 2 | 205 | 350 | 307 | 8 | 6 | 0 | 0 | 0 | 2 | 0 | 8 |
| 80 | 1.91 | 1.37 | 1.39 | 0.17 | 1 | Pappa | pregnancy-associated plasma protein A | 362 | 355 | 144 | 0 | 0 | 5 | 0 | 0 | 6 | 0 | 6 |
| 80 | 0.00 | 3.68 | 0.00 | 1.00 | 1 | Mn1 | meningioma 1 | 257 | 347 | 273 | 0 | 0 | 0 | 0 | 0 | 0 | 0 | 0 |
| 80 | 0.00 | 3.68 | 0.00 | 1.00 | 1 | Nkain4 | Na+/K+ transporting ATPase interacting 4 | 492 | 279 | 107 | 0 | 0 | 0 | 0 | 0 | 0 | 0 | 0 |
| 80 | -1.10 | 3.65 | -0.30 | 0.76 | 1 | Arhgap15 | Rho GTPase activating protein 15 | 530 | 157 | 185 | 0 | 4 | 0 | 0 | 0 | 0 | 0 | 0 |
| 80 | 1.81 | 3.65 | 0.50 | 0.62 | 1 | Cxcl1 | chemokine (C-X-C motif) ligand 1 | 517 | 233 | 120 | 0 | 0 | 0 | 0 | 0 | 5 | 0 | 0 |
| 80 | -3.79 | 3.35 | -1.13 | 0.26 | 1 | Gm5409 | | 1 | 590 | 259 | 25 | 0 | 0 | 0 | 0 | 0 | 0 | 1 |
| 79 | -2.47 | 1.39 | -1.78 | 0.07 | 1 | Gpr4 | G protein-coupled receptor 4 | 322 | 250 | 296 | 0 | 3 | 3 | 0 | 0 | 0 | 0 | 0 |
| 79 | 1.55 | 3.66 | 0.42 | 0.67 | 1 | Ebf3 | early B cell factor 3 | 348 | 290 | 232 | 0 | 0 | 0 | 0 | 0 | 3 | 0 | 0 |
| 79 | -2.48 | 2.90 | -0.86 | 0.39 | 1 | Pdlim4 | PDZ and LIM domain 4 | 484 | 166 | 181 | 0 | 6 | 3 | 0 | 32 | 0 | 0 | 0 |
| 79 | 0.00 | 2.70 | 0.00 | 1.00 | 1 | Galk1 | galactokinase 1 | 480 | 133 | 250 | 3 | 0 | 0 | 0 | 0 | 2 | 0 | 0 |
| 79 | -0.69 | 3.66 | -0.19 | 0.85 | 1 | Lvrn | laeverin | 388 | 147 | 329 | 0 | 0 | 2 | 0 | 0 | 0 | 0 | 0 |
| 78 | 0.00 | 3.68 | 0.00 | 1.00 | 1 | Ebf2 | early B cell factor 2 | 416 | 229 | 217 | 0 | 0 | 0 | 0 | 0 | 0 | 0 | 0 |
| 78 | -0.02 | 1.53 | -0.01 | 0.99 | 1 | Pla2g4a | phospholipase A2, group IVA (cytosolic, c | 451 | 214 | 176 | 0 | 7 | 1 | 0 | 0 | 9 | 0 | 0 |
| 78 | 1.99 | 1.42 | 1.40 | 0.16 | 1 | Tbx3 | T-box 3 | 338 | 215 | 298 | 0 | 0 | 1 | 0 | 0 | 4 | 0 | 1 |
| 78 | 1.25 | 3.66 | 0.34 | 0.73 | 1 | BC028528 | cDNA sequence BC028528 | 376 | 247 | 228 | 0 | 0 | 0 | 0 | 0 | 0 | 0 | 2 |
| 77 | 0.99 | 2.30 | 0.43 | 0.67 | 1 | Runx3 | runt related transcription factor 3 | 462 | 163 | 221 | 0 | 0 | 1 | 0 | 0 | 0 | 5 | 0 |
| 77 | -0.40 | 3.67 | -0.11 | 0.91 | 1 | Fam167b | family with sequence similarity 167, meml | 268 | 242 | 333 | 0 | 0 | 1 | 0 | 0 | 0 | 0 | 0 |
| 76 | -0.14 | 1.41 | -0.10 | 0.92 | 1 | Spib | Spi-B transcription factor (Spi-1/PU.1 rela | 560 | 260 | 17 | 0 | 0 | 2 | 0 | 0 | 0 | 0 | 1 |
| 76 | 0.00 | 3.68 | 0.00 | 1.00 | 1 | Igkv4-70 | | 728 | 1 | 110 | 0 | 0 | 0 | 0 | 0 | 0 | 0 | 0 |
| 76 | 0.00 | 3.68 | 0.00 | 1.00 | 1 | Npy1r | neuropeptide Y receptor Y1 | 504 | 218 | 117 | 0 | 0 | 0 | 0 | 0 | 0 | 0 | 0 |
| 76 | 0.00 | 3.68 | 0.00 | 1.00 | 1 | Sema3d | sema domain, immunoglobulin domain (Ig | 476 | 185 | 176 | 0 | 0 | 0 | 0 | 0 | 0 | 0 | 0 |
| 76 | 0.00 | 3.68 | 0.00 | 1.00 | 1 | Selp | selectin, platelet | 392 | 202 | 242 | 0 | 0 | 0 | 0 | 0 | 0 | 0 | 0 |
| 75 | 1.70 | 1.30 | 1.30 | 0.19 | 1 | Havcr2 | hepatitis A virus cellular receptor 2 | 291 | 230 | 296 | 0 | 0 | 3 | 0 | 0 | 6 | 0 | 2 |
| 75 | 0.57 | 3.46 | 0.16 | 0.87 | 1 | Ngfr | nerve growth factor receptor (TNFR super | 462 | 175 | 166 | 0 | 4 | 0 | 0 | 20 | 0 | 0 | 0 |
| 75 | 0.00 | 3.68 | 0.00 | 1.00 | 1 | Bmp4 | bone morphogenetic protein 4 | 439 | 209 | 177 | 0 | 0 | 0 | 0 | 0 | 0 | 0 | 0 |
| 75 | -0.02 | 3.33 | -0.01 | 1.00 | 1 | Kcna6 | potassium voltage-gated channel, shaker | 266 | 189 | 344 | 15 | 0 | 0 | 0 | 0 | 7 | 0 | 0 |
| 74 | 0.94 | 2.42 | 0.39 | 0.70 | 1 | Habp2 | hyaluronic acid binding protein 2 | 226 | 248 | 325 | 13 | 0 | 0 | 0 | 0 | 3 | 0 | 1 |
| 74 | -0.76 | 2.07 | -0.37 | 0.72 | 1 | Ppfibp2 | PTPRF interacting protein, binding proteir | 363 | 250 | 182 | 8 | 0 | 3 | 0 | 0 | 9 | 1 | 0 |
| 74 | -3.64 | 2.74 | -1.33 | 0.18 | 1 | Cdh19 | cadherin 19, type 2 | 305 | 136 | 347 | 0 | 20 | 5 | 0 | 0 | 0 | 0 | 0 |
| 74 | 0.60 | 2.07 | 0.29 | 0.77 | 1 | Cacng7 | calcium channel, voltage-dependent, gan | 322 | 233 | 229 | 0 | 6 | 3 | 0 | 6 | 0 | 15 | 0 |
| 74 | -0.73 | 3.64 | -0.20 | 0.84 | 1 | C5ar1 | complement component 5a receptor 1 | 498 | 144 | 165 | 0 | 0 | 3 | 0 | 0 | 0 | 0 | 0 |
| 74 | 0.00 | 3.68 | 0.00 | 1.00 | 1 | Aox3 | aldehyde oxidase 3 | 606 | 150 | 54 | 0 | 0 | 0 | 0 | 0 | 0 | 0 | 0 |
| 72 | 0.00 | 3.68 | 0.00 | 1.00 | 1 | Ncf4 | neutrophil cytosolic factor 4 | 374 | 229 | 194 | 0 | 0 | 0 | 0 | 0 | 0 | 0 | 0 |
| 72 | 4.01 | 2.50 | 1.61 | 0.11 | 1 | Rspo1 | R-spondin 1 | 470 | 201 | 82 | 0 | 0 | 0 | 0 | 8 | 0 | 32 | 0 |
| 72 | 0.54 | 1.08 | 0.50 | 0.62 | 1 | Lat2 | linker for activation of T cells family, memb | 428 | 157 | 164 | 0 | 0 | 15 | 0 | 0 | 0 | 0 | 24 |
| 71 | 0.00 | 3.68 | 0.00 | 1.00 | 1 | Mal | myelin and lymphocyte protein, T cell diffe | 277 | 159 | 350 | 0 | 0 | 0 | 0 | 0 | 0 | 0 | 0 |
| 71 | 0.27 | 1.33 | 0.21 | 0.84 | 1 | Arhgap9 | Rho GTPase activating protein 9 | 460 | 133 | 183 | 0 | 0 | 2 | 0 | 0 | 0 | 0 | 2 |
| 71 | -0.03 | 1.24 | -0.03 | 0.98 | 1 | Atcay | ataxia, cerebellar, Cayman type | 258 | 314 | 165 | 0 | 12 | 8 | 0 | 14 | 3 | 0 | 3 |
| 70 | -3.36 | 2.70 | -1.25 | 0.21 | 1 | Tlr8 | toll-like receptor 8 | 396 | 170 | 193 | 0 | 11 | 4 | 0 | 0 | 0 | 0 | 0 |
| 70 | 1.68 | 3.65 | 0.46 | 0.65 | 1 | Megf6 | multiple EGF-like-domains 6 | 420 | 127 | 224 | 0 | 0 | 0 | 0 | 0 | 4 | 0 | 0 |
| 70 | -1.17 | 3.63 | -0.32 | 0.75 | 1 | Sox7 | SRY (sex determining region Y)-box 7 | 262 | 278 | 228 | 0 | 7 | 0 | 0 | 0 | 0 | 0 | 0 |
| 70 | 0.00 | 3.68 | 0.00 | 1.00 | 1 | Adgre4 | adhesion G protein-coupled receptor E4 | 467 | 226 | 80 | 0 | 0 | 0 | 0 | 0 | 0 | 0 | 0 |
| 70 | 2.36 | 1.30 | 1.82 | 0.07 | 1 | Adam12 | a disintegrin and metallopeptidase domai | 302 | 204 | 251 | 0 | 0 | 2 | 0 | 0 | 6 | 0 | 4 |
| 69 | 0.00 | 3.68 | 0.00 | 1.00 | 1 | Wnt2b | wingless-type MMTV integration site famil | 456 | 139 | 169 | 0 | 0 | 0 | 0 | 0 | 0 | 0 | 0 |

| | | | | | | | | | | | | | | | | | | |
|---|---|---|---|---|---|---|---|---|---|---|---|---|---|---|---|---|---|---|
| 69 | 0.00 | 3.68 | 0.00 | 1.00 | 1 | G0s2 | G0/G1 switch gene 2 | 256 | 261 | 245 | 0 | 0 | 0 | 0 | 0 | 0 | 0 | 0 |
| 69 | 0.00 | 3.68 | 0.00 | 1.00 | 1 | Btk | Bruton agammaglobulinemia tyrosine kin | 432 | 171 | 157 | 0 | 0 | 0 | 0 | 0 | 0 | 0 | 0 |
| 69 | -0.42 | 1.35 | -0.31 | 0.76 | 1 | Plxdc1 | plexin domain containing 1 | 354 | 205 | 193 | 0 | 0 | 3 | 0 | 0 | 0 | 0 | 1 |
| 68 | -0.73 | 3.64 | -0.20 | 0.84 | 1 | Fgd3 | FYVE, RhoGEF and PH domain containin | 342 | 225 | 183 | 0 | 0 | 3 | 0 | 0 | 0 | 0 | 0 |
| 68 | -0.84 | 1.20 | -0.70 | 0.49 | 1 | Prr5l | proline rich 5 like | 329 | 155 | 237 | 7 | 4 | 5 | 0 | 0 | 11 | 0 | 2 |
| 68 | 0.28 | 1.58 | 0.18 | 0.86 | 1 | C1qtnf2 | C1q and tumor necrosis factor related pro | 362 | 167 | 219 | 1 | 0 | 0 | 0 | 0 | 1 | 0 | 0 |
| 68 | -1.16 | 3.63 | -0.32 | 0.75 | 1 | Nkd2 | naked cuticle 2 | 334 | 170 | 237 | 0 | 7 | 0 | 0 | 0 | 0 | 0 | 0 |
| 68 | 0.47 | 1.31 | 0.36 | 0.72 | 1 | Ccm2l | cerebral cavernous malformation 2-like | 251 | 179 | 308 | 0 | 0 | 5 | 0 | 0 | 3 | 0 | 1 |
| 68 | 0.00 | 3.68 | 0.00 | 1.00 | 1 | Igkv8-30 | | 715 | 28 | 3 | 0 | 0 | 0 | 0 | 0 | 0 | 0 | 0 |
| 67 | -2.82 | 2.27 | -1.24 | 0.21 | 1 | Cabp2 | calcium binding protein 2 | 109 | 307 | 302 | 13 | 6 | 0 | 0 | 0 | 0 | 0 | 3 |
| 67 | -0.74 | 1.14 | -0.65 | 0.52 | 1 | Oprl1 | opioid receptor-like 1 | 190 | 189 | 286 | 17 | 9 | 12 | 0 | 0 | 14 | 0 | 18 |
| 67 | 0.00 | 3.68 | 0.00 | 1.00 | 1 | Upk1b | uroplakin 1B | 334 | 255 | 145 | 0 | 0 | 0 | 0 | 0 | 0 | 0 | 0 |
| 67 | 0.00 | 3.68 | 0.00 | 1.00 | 1 | Mmp28 | matrix metallopeptidase 28 (epilysin) | 268 | 216 | 249 | 0 | 0 | 0 | 0 | 0 | 0 | 0 | 0 |
| 66 | -2.60 | 1.44 | -1.81 | 0.07 | 1 | Igsf6 | immunoglobulin superfamily, member 6 | 380 | 160 | 184 | 0 | 3 | 3 | 0 | 0 | 0 | 0 | 0 |
| 66 | 1.83 | 3.65 | 0.50 | 0.62 | 1 | Col4a6 | collagen, type IV, alpha 6 | 494 | 137 | 85 | 0 | 0 | 0 | 0 | 0 | 5 | 0 | 0 |
| 65 | -2.53 | 3.18 | -0.80 | 0.43 | 1 | Lrrc25 | leucine rich repeat containing 25 | 308 | 136 | 262 | 0 | 9 | 0 | 0 | 0 | 0 | 0 | 4 |
| 65 | 0.00 | 3.68 | 0.00 | 1.00 | 1 | Has1 | hyaluronan synthase 1 | 507 | 125 | 87 | 0 | 0 | 0 | 0 | 0 | 0 | 0 | 0 |
| 65 | 0.00 | 3.68 | 0.00 | 1.00 | 1 | Ighg1 | | 691 | 27 | 0 | 0 | 0 | 0 | 0 | 0 | 0 | 0 | 0 |
| 65 | 0.00 | 3.68 | 0.00 | 1.00 | 1 | Tmem26 | transmembrane protein 26 | 286 | 148 | 282 | 0 | 0 | 0 | 0 | 0 | 0 | 0 | 0 |
| 65 | 0.95 | 1.27 | 0.75 | 0.45 | 1 | Bmp2 | bone morphogenetic protein 2 | 324 | 201 | 182 | 0 | 0 | 2 | 0 | 0 | 0 | 0 | 5 |
| 64 | 0.00 | 3.68 | 0.00 | 1.00 | 1 | Iglc3 | | 420 | 285 | 5 | 0 | 0 | 0 | 0 | 0 | 0 | 0 | 0 |
| 64 | 1.32 | 3.63 | 0.36 | 0.72 | 1 | Ctse | cathepsin E | 362 | 161 | 182 | 0 | 0 | 0 | 0 | 0 | 0 | 0 | 4 |
| 64 | -1.67 | 1.51 | -1.11 | 0.27 | 1 | Cldn10 | claudin 10 | 174 | 350 | 174 | 0 | 2 | 1 | 0 | 0 | 0 | 0 | 0 |
| 64 | 0.41 | 1.27 | 0.32 | 0.75 | 1 | Mdk | midkine | 291 | 195 | 183 | 0 | 6 | 6 | 0 | 0 | 0 | 8 | 10 |
| 63 | -2.00 | 2.57 | -0.78 | 0.44 | 1 | P2ry10 | purinergic receptor P2Y, G-protein couple | 389 | 248 | 55 | 0 | 2 | 3 | 0 | 0 | 0 | 0 | 0 |
| 63 | -0.40 | 3.67 | -0.11 | 0.91 | 1 | Prrg1 | proline rich Gla (G-carboxyglutamic acid) | 286 | 197 | 211 | 0 | 0 | 1 | 0 | 0 | 0 | 0 | 0 |
| 63 | -0.44 | 3.68 | -0.12 | 0.90 | 1 | Pln | phospholamban | 433 | 127 | 133 | 1 | 0 | 0 | 0 | 0 | 0 | 0 | 0 |
| 63 | -1.08 | 3.62 | -0.30 | 0.77 | 1 | Lypd8 | LY6/PLAUR domain containing 8 | 0 | 417 | 272 | 0 | 4 | 0 | 0 | 0 | 0 | 0 | 0 |
| 63 | -1.97 | 1.50 | -1.32 | 0.19 | 1 | St6galnac3 | ST6 (alpha-N-acetyl-neuraminyl-2,3-beta | 224 | 245 | 218 | 0 | 3 | 1 | 0 | 0 | 0 | 0 | 0 |
| 63 | -2.00 | 1.50 | -1.33 | 0.18 | 1 | Cd209b | CD209b antigen | 501 | 81 | 104 | 0 | 2 | 2 | 0 | 0 | 0 | 0 | 0 |
| 63 | -2.58 | 1.33 | -1.95 | 0.05 | 1 | Gabrg1 | gamma-aminobutyric acid (GABA) A rece | 202 | 294 | 163 | 4 | 12 | 5 | 0 | 0 | 0 | 0 | 9 |
| 62 | 0.00 | 3.68 | 0.00 | 1.00 | 1 | Angpt4 | angiopoietin 4 | 536 | 69 | 82 | 0 | 0 | 0 | 0 | 0 | 0 | 0 | 0 |
| 62 | 0.00 | 3.68 | 0.00 | 1.00 | 1 | Osr1 | odd-skipped related transcription factor 1 | 438 | 110 | 136 | 0 | 0 | 0 | 0 | 0 | 0 | 0 | 0 |
| 62 | -1.28 | 3.62 | -0.35 | 0.72 | 1 | Sh3tc2 | SH3 domain and tetratricopeptide repeats | 300 | 224 | 151 | 9 | 0 | 0 | 0 | 0 | 0 | 0 | 0 |
| 62 | -0.69 | 3.66 | -0.19 | 0.85 | 1 | Ephx3 | epoxide hydrolase 3 | 304 | 149 | 228 | 0 | 0 | 2 | 0 | 0 | 0 | 0 | 0 |
| 62 | -0.50 | 1.19 | -0.42 | 0.67 | 1 | Ankrd29 | ankyrin repeat domain 29 | 266 | 216 | 113 | 3 | 14 | 24 | 0 | 0 | 11 | 0 | 34 |
| 61 | -0.14 | 1.01 | -0.14 | 0.89 | 1 | Esr1 | estrogen receptor 1 (alpha) | 371 | 111 | 148 | 9 | 5 | 6 | 0 | 0 | 8 | 10 | 4 |
| 61 | -2.73 | 2.61 | -1.04 | 0.30 | 1 | Klhdc7a | kelch domain containing 7A | 233 | 124 | 293 | 2 | 17 | 0 | 0 | 0 | 0 | 0 | 0 |
| 61 | -0.40 | 3.67 | -0.11 | 0.91 | 1 | Dchs2 | dachsous cadherin related 2 | 170 | 207 | 291 | 0 | 0 | 1 | 0 | 0 | 0 | 0 | 0 |
| 61 | -0.26 | 1.25 | -0.21 | 0.83 | 1 | Ppp1r14a | protein phosphatase 1, regulatory inhibito | 312 | 135 | 213 | 0 | 0 | 5 | 0 | 0 | 0 | 0 | 3 |
| 61 | -2.05 | 2.52 | -0.81 | 0.42 | 1 | H2-DMb2 | histocompatibility 2, class II, locus Mb2 | 416 | 122 | 124 | 0 | 5 | 1 | 0 | 0 | 0 | 0 | 0 |
| 60 | 3.53 | 2.42 | 1.46 | 0.15 | 1 | Trpv4 | transient receptor potential cation channe | 267 | 207 | 176 | 0 | 0 | 0 | 0 | 0 | 5 | 0 | 11 |
| 60 | 0.97 | 1.08 | 0.90 | 0.37 | 1 | Gli3 | GLI-Kruppel family member GLI3 | 264 | 113 | 220 | 3 | 11 | 3 | 0 | 7 | 1 | 40 | 2 |
| 60 | -0.42 | 1.45 | -0.29 | 0.77 | 1 | Plcb2 | phospholipase C, beta 2 | 297 | 150 | 214 | 0 | 0 | 3 | 0 | 0 | 1 | 0 | 0 |
| 60 | -0.64 | 1.29 | -0.49 | 0.62 | 1 | Frem1 | Fras1 related extracellular matrix protein | 275 | 88 | 292 | 0 | 0 | 6 | 0 | 0 | 0 | 0 | 2 |
| 60 | -0.69 | 3.66 | -0.19 | 0.85 | 1 | Lrrc3b | leucine rich repeat containing 3B | 226 | 251 | 184 | 0 | 0 | 2 | 0 | 0 | 0 | 0 | 0 |
| 60 | 1.09 | 2.38 | 0.46 | 0.65 | 1 | Klk1b5 | kallikrein 1-related peptidase b5 | 89 | 299 | 251 | 10 | 0 | 0 | 0 | 0 | 2 | 0 | 7 |
| 60 | 0.67 | 1.28 | 0.53 | 0.60 | 1 | Slc39a4 | solute carrier family 39 (zinc transporter), | 230 | 131 | 273 | 5 | 2 | 0 | 0 | 1 | 3 | 0 | 12 |
| 59 | 0.00 | 3.68 | 0.00 | 1.00 | 1 | Tcf21 | transcription factor 21 | 388 | 100 | 163 | 0 | 0 | 0 | 0 | 0 | 0 | 0 | 0 |
| 59 | 0.00 | 3.68 | 0.00 | 1.00 | 1 | Ifitm1 | interferon induced transmembrane protei | 305 | 179 | 165 | 0 | 0 | 0 | 0 | 0 | 0 | 0 | 0 |
| 59 | 0.00 | 3.68 | 0.00 | 1.00 | 1 | Gimap5 | GTPase, IMAP family member 5 | 184 | 278 | 188 | 0 | 0 | 0 | 0 | 0 | 0 | 0 | 0 |
| 59 | 0.95 | 1.39 | 0.68 | 0.49 | 1 | Fam124a | family with sequence similarity 124, meml | 273 | 109 | 263 | 0 | 0 | 1 | 0 | 0 | 0 | 0 | 3 |
| 59 | 2.32 | 1.51 | 1.54 | 0.12 | 1 | Btla | B and T lymphocyte associated | 359 | 217 | 60 | 0 | 0 | 2 | 0 | 0 | 5 | 0 | 4 |
| 59 | 1.67 | 1.57 | 1.06 | 0.29 | 1 | Pth1r | parathyroid hormone 1 receptor | 344 | 115 | 185 | 0 | 0 | 0 | 0 | 1 | 0 | 0 | 1 |
| 59 | 0.00 | 3.68 | 0.00 | 1.00 | 1 | Crispld1 | cysteine-rich secretory protein LCCL dom | 248 | 218 | 179 | 0 | 0 | 0 | 0 | 0 | 0 | 0 | 0 |
| 59 | -0.83 | 0.96 | -0.87 | 0.39 | 1 | Acsbg1 | acyl-CoA synthetase bubblegum family m | 186 | 209 | 219 | 5 | 8 | 6 | 0 | 0 | 6 | 4 | 1 |
| 58 | -1.14 | 1.59 | -0.72 | 0.47 | 1 | Ndnf | neuron-derived neurotrophic factor | 492 | 134 | 14 | 0 | 1 | 1 | 0 | 0 | 0 | 0 | 0 |
| 58 | 0.00 | 3.68 | 0.00 | 1.00 | 1 | Map3k7cl | Map3k7 C-terminal like | 391 | 94 | 155 | 0 | 0 | 0 | 0 | 0 | 0 | 0 | 0 |
| 58 | -0.79 | 1.47 | -0.54 | 0.59 | 1 | Gjb1 | gap junction protein, beta 1 | 118 | 253 | 251 | 1 | 6 | 0 | 0 | 0 | 0 | 0 | 7 |
| 58 | 0.00 | 3.68 | 0.00 | 1.00 | 1 | Igkv4-50 | | 586 | 50 | 0 | 0 | 0 | 0 | 0 | 0 | 0 | 0 | 0 |

| | | | | | | | | | | | | | | | | | | |
|---|---|---|---|---|---|---|---|---|---|---|---|---|---|---|---|---|---|---|
| 58 | 0.00 | 3.68 | 0.00 | 1.00 | 1 | Itgam | integrin alpha M | 415 | 98 | 122 | 0 | 0 | 0 | 0 | 0 | 0 | 0 | 0 |
| 58 | -0.71 | 1.14 | -0.62 | 0.54 | 1 | Nlrc5 | NLR family, CARD domain containing 5 | 309 | 96 | 132 | 8 | 32 | 19 | 0 | 23 | 2 | 1 | 10 |
| 57 | -2.16 | 3.22 | -0.67 | 0.50 | 1 | Il2rg | interleukin 2 receptor, gamma chain | 310 | 154 | 156 | 0 | 6 | 0 | 0 | 0 | 0 | 0 | 4 |
| 57 | 0.00 | 3.68 | 0.00 | 1.00 | 1 | Mmp3 | matrix metallopeptidase 3 | 517 | 37 | 74 | 0 | 0 | 0 | 0 | 0 | 0 | 0 | 0 |
| 57 | 1.30 | 3.64 | 0.36 | 0.72 | 1 | 9930111J21Rik | RIKEN cDNA 9930111J21 gene 2 | 311 | 153 | 156 | 0 | 0 | 0 | 0 | 0 | 0 | 0 | 3 |
| 56 | 0.00 | 3.68 | 0.00 | 1.00 | 1 | Mall | mal, T cell differentiation protein-like | 182 | 161 | 275 | 0 | 0 | 0 | 0 | 0 | 0 | 0 | 0 |
| 56 | 1.65 | 3.64 | 0.45 | 0.65 | 1 | Krt23 | keratin 23 | 127 | 202 | 283 | 0 | 0 | 0 | 0 | 0 | 5 | 0 | 0 |
| 56 | 1.32 | 3.66 | 0.36 | 0.72 | 1 | Cd300c2 | CD300C molecule 2 | 280 | 116 | 217 | 0 | 0 | 0 | 0 | 0 | 2 | 0 | 0 |
| 56 | 0.65 | 1.37 | 0.47 | 0.64 | 1 | Kcnd3 | potassium voltage-gated channel, Shal-re | 259 | 194 | 138 | 0 | 3 | 6 | 0 | 0 | 10 | 0 | 3 |
| 55 | -0.40 | 3.67 | -0.11 | 0.91 | 1 | Lyl1 | lymphoblastomic leukemia 1 | 371 | 93 | 142 | 0 | 0 | 1 | 0 | 0 | 0 | 0 | 0 |
| 55 | 1.71 | 3.65 | 0.47 | 0.64 | 1 | Trem2 | triggering receptor expressed on myeloid | 347 | 139 | 116 | 0 | 0 | 0 | 0 | 0 | 4 | 0 | 0 |
| 55 | 0.00 | 3.68 | 0.00 | 1.00 | 1 | Npr3 | natriuretic peptide receptor 3 | 266 | 179 | 161 | 0 | 0 | 0 | 0 | 0 | 0 | 0 | 0 |
| 55 | -2.03 | 1.43 | -1.42 | 0.16 | 1 | Tll1 | tolloid-like | 292 | 91 | 213 | 0 | 1 | 5 | 0 | 0 | 0 | 0 | 0 |
| 55 | 0.00 | 3.68 | 0.00 | 1.00 | 1 | Hba-a1 | hemoglobin alpha, adult chain 1 | 219 | 176 | 206 | 0 | 0 | 0 | 0 | 0 | 0 | 0 | 0 |
| 55 | 0.08 | 1.21 | 0.06 | 0.95 | 1 | Arhgap4 | Rho GTPase activating protein 4 | 281 | 176 | 107 | 2 | 0 | 19 | 0 | 3 | 0 | 0 | 11 |
| 54 | -1.58 | 1.25 | -1.26 | 0.21 | 1 | Col7a1 | collagen, type VII, alpha 1 | 62 | 237 | 114 | 12 | 0 | 85 | 0 | 0 | 0 | 0 | 85 |
| 53 | 0.67 | 1.63 | 0.41 | 0.68 | 1 | Tlr1 | toll-like receptor 1 | 285 | 192 | 90 | 0 | 8 | 0 | 0 | 0 | 8 | 0 | 1 |
| 53 | 0.00 | 3.68 | 0.00 | 1.00 | 1 | Muc16 | mucin 16 | 256 | 194 | 134 | 0 | 0 | 0 | 0 | 0 | 0 | 0 | 0 |
| 53 | 0.00 | 3.68 | 0.00 | 1.00 | 1 | Zfpm2 | zinc finger protein, multitype 2 | 263 | 180 | 136 | 0 | 0 | 0 | 0 | 0 | 0 | 0 | 0 |
| 53 | 0.00 | 3.68 | 0.00 | 1.00 | 1 | Vsig4 | V-set and immunoglobulin domain contai | 395 | 161 | 23 | 0 | 0 | 0 | 0 | 0 | 0 | 0 | 0 |
| 53 | -1.48 | 1.56 | -0.95 | 0.34 | 1 | Susd5 | sushi domain containing 5 | 256 | 135 | 185 | 1 | 0 | 2 | 0 | 0 | 0 | 0 | 0 |
| 52 | 0.00 | 3.68 | 0.00 | 1.00 | 1 | Cd5l | CD5 antigen-like | 452 | 81 | 44 | 0 | 0 | 0 | 0 | 0 | 0 | 0 | 0 |
| 52 | 1.71 | 3.63 | 0.47 | 0.64 | 1 | Creb5 | cAMP responsive element binding proteir | 183 | 134 | 254 | 0 | 0 | 0 | 0 | 0 | 6 | 0 | 0 |
| 52 | 0.00 | 3.68 | 0.00 | 1.00 | 1 | Slfn3 | schlafen 3 | 240 | 158 | 177 | 0 | 0 | 0 | 0 | 0 | 0 | 0 | 0 |
| 52 | -0.25 | 1.27 | -0.20 | 0.84 | 1 | Agmo | alkylglycerol monooxygenase | 220 | 97 | 220 | 0 | 16 | 4 | 0 | 0 | 5 | 0 | 11 |
| 52 | 0.32 | 0.95 | 0.34 | 0.73 | 1 | Kcnmb1 | potassium large conductance calcium-act | 252 | 133 | 79 | 13 | 13 | 14 | 0 | 15 | 4 | 39 | 11 |
| 52 | 3.62 | 2.51 | 1.44 | 0.15 | 1 | Xpnpep2 | X-prolyl aminopeptidase (aminopeptidase | 277 | 107 | 160 | 0 | 0 | 5 | 0 | 6 | 11 | 0 | 0 |
| 51 | 1.45 | 1.23 | 1.18 | 0.24 | 1 | AI662270 | | 344 | 88 | 57 | 0 | 0 | 13 | 0 | 0 | 0 | 0 | 64 |
| 51 | 0.24 | 1.34 | 0.18 | 0.86 | 1 | Fgf7 | fibroblast growth factor 7 | 334 | 97 | 111 | 0 | 0 | 14 | 0 | 0 | 2 | 0 | 4 |
| 51 | -0.52 | 0.82 | -0.64 | 0.52 | 1 | Gm35082 | | 0 | 222 | 57 | 145 | 29 | 0 | 0 | 89 | 19 | 0 | 0 |
| 51 | -0.80 | 3.66 | -0.22 | 0.83 | 1 | Ripk3 | receptor-interacting serine-threonine kina | 311 | 116 | 133 | 2 | 0 | 0 | 0 | 0 | 0 | 0 | 0 |
| 51 | 1.32 | 3.63 | 0.36 | 0.72 | 1 | Sp140 | Sp140 nuclear body protein | 257 | 157 | 137 | 0 | 0 | 0 | 0 | 0 | 0 | 0 | 4 |
| 50 | -2.06 | 1.33 | -1.55 | 0.12 | 1 | Papln | papilin, proteoglycan-like sulfated glycopr | 180 | 153 | 209 | 1 | 7 | 3 | 0 | 0 | 0 | 0 | 2 |
| 50 | -0.96 | 1.37 | -0.70 | 0.49 | 1 | Zbtb16 | zinc finger and BTB domain containing 16 | 248 | 140 | 156 | 0 | 0 | 6 | 0 | 0 | 0 | 0 | 1 |
| 50 | -0.77 | 3.66 | -0.21 | 0.83 | 1 | Scn4b | sodium channel, type IV, beta | 296 | 170 | 81 | 2 | 0 | 0 | 0 | 0 | 0 | 0 | 0 |
| 50 | 0.00 | 3.68 | 0.00 | 1.00 | 1 | Klhl4 | kelch-like 4 | 232 | 189 | 123 | 0 | 0 | 0 | 0 | 0 | 0 | 0 | 0 |
| 49 | -1.13 | 3.63 | -0.31 | 0.75 | 1 | Arhgef26 | Rho guanine nucleotide exchange factor | 172 | 195 | 169 | 0 | 6 | 0 | 0 | 0 | 0 | 0 | 0 |
| 49 | 0.00 | 3.68 | 0.00 | 1.00 | 1 | Cnn1 | calponin 1 | 412 | 90 | 39 | 0 | 0 | 0 | 0 | 0 | 0 | 0 | 0 |
| 49 | 1.34 | 3.62 | 0.37 | 0.71 | 1 | Cd19 | CD19 antigen | 347 | 162 | 26 | 0 | 0 | 0 | 0 | 0 | 0 | 0 | 5 |
| 49 | 0.00 | 3.61 | 0.00 | 1.00 | 1 | Igkv17-121 | | 538 | 0 | 0 | 0 | 0 | 0 | 0 | 0 | 0 | 0 | 0 |
| 49 | 0.04 | 2.65 | 0.02 | 0.99 | 1 | Ghrl | ghrelin | 33 | 75 | 422 | 0 | 0 | 5 | 0 | 0 | 2 | 0 | 0 |
| 49 | -1.55 | 1.27 | -1.23 | 0.22 | 1 | Slfn8 | schlafen 8 | 226 | 131 | 157 | 0 | 3 | 12 | 0 | 0 | 0 | 0 | 5 |
| 48 | 0.95 | 1.17 | 0.81 | 0.42 | 1 | Gm43858 | | 13 | 8 | 10 | 0 | 2 | 245 | 0 | 0 | 8 | 0 | 246 |
| 48 | -1.09 | 3.64 | -0.30 | 0.77 | 1 | Pxdc1 | PX domain containing 1 | 289 | 72 | 166 | 0 | 5 | 0 | 0 | 0 | 0 | 0 | 0 |
| 48 | 0.00 | 3.68 | 0.00 | 1.00 | 1 | Fgf10 | fibroblast growth factor 10 | 236 | 167 | 128 | 0 | 0 | 0 | 0 | 0 | 0 | 0 | 0 |
| 48 | 0.00 | 3.68 | 0.00 | 1.00 | 1 | Adamts8 | a disintegrin-like and metallopeptidase (r | 357 | 66 | 106 | 0 | 0 | 0 | 0 | 0 | 0 | 0 | 0 |
| 48 | 1.00 | 3.68 | 0.27 | 0.79 | 1 | Tnfaip6 | tumor necrosis factor alpha induced prote | 421 | 58 | 48 | 0 | 0 | 0 | 0 | 0 | 1 | 0 | 0 |
| 48 | -0.99 | 1.27 | -0.78 | 0.44 | 1 | Glt8d2 | glycosyltransferase 8 domain containing 2 | 300 | 108 | 82 | 7 | 5 | 11 | 0 | 0 | 7 | 0 | 7 |
| 47 | -0.69 | 3.66 | -0.19 | 0.85 | 1 | Adamts16 | a disintegrin-like and metallopeptidase (r | 136 | 71 | 313 | 0 | 0 | 2 | 0 | 0 | 0 | 0 | 0 |
| 47 | 1.34 | 3.62 | 0.37 | 0.71 | 1 | Tmed11 | transmembrane p24 trafficking protein 11 | 70 | 239 | 205 | 0 | 0 | 0 | 0 | 0 | 0 | 0 | 5 |
| 47 | 0.00 | 3.68 | 0.00 | 1.00 | 1 | Mptx1 | mucosal pentraxin 1 | 0 | 245 | 272 | 0 | 0 | 0 | 0 | 0 | 0 | 0 | 0 |
| 47 | 0.00 | 3.68 | 0.00 | 1.00 | 1 | Mcpt4 | mast cell protease 4 | 375 | 127 | 16 | 0 | 0 | 0 | 0 | 0 | 0 | 0 | 0 |
| 47 | 0.00 | 3.68 | 0.00 | 1.00 | 1 | Clec4a2 | C-type lectin domain family 4, member a2 | 269 | 102 | 145 | 0 | 0 | 0 | 0 | 0 | 0 | 0 | 0 |
| 46 | 0.00 | 3.68 | 0.00 | 1.00 | 1 | Ccr1 | chemokine (C-C motif) receptor 1 | 352 | 103 | 54 | 0 | 0 | 0 | 0 | 0 | 0 | 0 | 0 |
| 46 | 0.00 | 3.68 | 0.00 | 1.00 | 1 | Ly6d | lymphocyte antigen 6 complex, locus D | 432 | 74 | 0 | 0 | 0 | 0 | 0 | 0 | 0 | 0 | 0 |
| 46 | -1.23 | 3.64 | -0.34 | 0.73 | 1 | Tmem100 | transmembrane protein 100 | 187 | 124 | 190 | 5 | 0 | 0 | 0 | 0 | 0 | 0 | 0 |
| 46 | 0.00 | 3.68 | 0.00 | 1.00 | 1 | Gask1a | golgi associated kinase 1A | 176 | 174 | 154 | 0 | 0 | 0 | 0 | 0 | 0 | 0 | 0 |
| 46 | 0.95 | 3.67 | 0.26 | 0.80 | 1 | Gdpd2 | glycerophosphodiester phosphodiesteras | 290 | 202 | 9 | 0 | 0 | 0 | 0 | 0 | 0 | 0 | 1 |
| 46 | -1.98 | 2.79 | -0.71 | 0.48 | 1 | 2010007H06Rik | | 94 | 194 | 182 | 0 | 3 | 7 | 0 | 0 | 0 | 22 | 0 |
| 45 | 0.00 | 3.68 | 0.00 | 1.00 | 1 | Fhl5 | four and a half LIM domains 5 | 280 | 163 | 57 | 0 | 0 | 0 | 0 | 0 | 0 | 0 | 0 |

| | | | | | | | | | | | | | | | | | | |
|---|---|---|---|---|---|---|---|---|---|---|---|---|---|---|---|---|---|---|
| 45 | 1.89 | 3.62 | 0.52 | 0.60 | 1 | Edn1 | endothelin 1 | 210 | 93 | 180 | 0 | 0 | 0 | 0 | 0 | 15 | 0 | 0 |
| 45 | 0.00 | 3.68 | 0.00 | 1.00 | 1 | P2ry2 | purinergic receptor P2Y, G-protein couple | 226 | 142 | 125 | 0 | 0 | 0 | 0 | 0 | 0 | 0 | 0 |
| 45 | -1.08 | 1.51 | -0.71 | 0.48 | 1 | Ddah1 | dimethylarginine dimethylaminohydrolase | 150 | 193 | 126 | 3 | 4 | 2 | 0 | 0 | 0 | 15 | 0 |
| 45 | 0.00 | 3.68 | 0.00 | 1.00 | 1 | Slc6a2 | solute carrier family 6 (neurotransmitter tr | 405 | 55 | 33 | 0 | 0 | 0 | 0 | 0 | 0 | 0 | 0 |
| 45 | 0.69 | 1.13 | 0.62 | 0.54 | 1 | Gpr160 | G protein-coupled receptor 160 | 193 | 101 | 118 | 4 | 13 | 13 | 0 | 25 | 8 | 0 | 15 |
| 45 | 0.00 | 3.68 | 0.00 | 1.00 | 1 | Lamb3 | laminin, beta 3 | 169 | 145 | 178 | 0 | 0 | 0 | 0 | 0 | 0 | 0 | 0 |
| 44 | 0.95 | 3.67 | 0.26 | 0.80 | 1 | Myzap | myocardial zonula adherens protein | 255 | 141 | 92 | 0 | 0 | 0 | 0 | 0 | 0 | 0 | 1 |
| 44 | 0.28 | 1.46 | 0.19 | 0.85 | 1 | P2ry13 | purinergic receptor P2Y, G-protein couple | 244 | 40 | 202 | 0 | 0 | 2 | 0 | 0 | 0 | 0 | 2 |
| 44 | -0.76 | 3.63 | -0.21 | 0.83 | 1 | Rhoh | ras homolog family member H | 353 | 69 | 62 | 0 | 0 | 3 | 0 | 0 | 0 | 0 | 0 |
| 44 | 1.21 | 1.28 | 0.94 | 0.35 | 1 | Lhx6 | LIM homeobox protein 6 | 147 | 110 | 139 | 1 | 18 | 6 | 0 | 49 | 10 | 0 | 6 |
| 44 | 1.41 | 1.61 | 0.87 | 0.38 | 1 | Gjb2 | gap junction protein, beta 2 | 55 | 267 | 144 | 0 | 0 | 7 | 0 | 0 | 0 | 6 | 3 |
| 44 | 0.00 | 3.68 | 0.00 | 1.00 | 1 | Cpa3 | carboxypeptidase A3, mast cell | 257 | 207 | 19 | 0 | 0 | 0 | 0 | 0 | 0 | 0 | 0 |
| 44 | 0.00 | 3.61 | 0.00 | 1.00 | 1 | Igkv3-9 | | 480 | 0 | 0 | 0 | 0 | 0 | 0 | 0 | 0 | 0 | 0 |
| 44 | 0.00 | 3.68 | 0.00 | 1.00 | 1 | Cxcl2 | chemokine (C-X-C motif) ligand 2 | 353 | 73 | 54 | 0 | 0 | 0 | 0 | 0 | 0 | 0 | 0 |
| 44 | 0.00 | 3.68 | 0.00 | 1.00 | 1 | Slc45a3 | solute carrier family 45, member 3 | 256 | 141 | 82 | 0 | 0 | 0 | 0 | 0 | 0 | 0 | 0 |
| 43 | -0.15 | 3.09 | -0.05 | 0.96 | 1 | Clca3a1 | chloride channel accessory 3A1 | 112 | 155 | 195 | 11 | 0 | 0 | 0 | 0 | 4 | 0 | 0 |
| 43 | -1.41 | 1.64 | -0.86 | 0.39 | 1 | Gpr37l1 | G protein-coupled receptor 37-like 1 | 148 | 161 | 147 | 0 | 18 | 0 | 0 | 0 | 2 | 0 | 0 |
| 43 | 1.84 | 3.63 | 0.51 | 0.61 | 1 | Csmd1 | CUB and Sushi multiple domains 1 | 214 | 114 | 136 | 0 | 0 | 0 | 0 | 0 | 9 | 0 | 0 |
| 43 | 0.00 | 3.67 | 0.00 | 1.00 | 1 | Ighv3-8 | | 465 | 8 | 0 | 0 | 0 | 0 | 0 | 0 | 0 | 0 | 0 |
| 43 | 0.00 | 3.63 | 0.00 | 1.00 | 1 | Igkv17-127 | | 2 | 469 | 0 | 0 | 0 | 0 | 0 | 0 | 0 | 0 | 0 |
| 43 | 1.32 | 3.63 | 0.36 | 0.72 | 1 | Lrig3 | leucine-rich repeats and immunoglobulin | 265 | 122 | 78 | 0 | 0 | 0 | 0 | 0 | 0 | 0 | 4 |
| 43 | -1.74 | 1.59 | -1.09 | 0.28 | 1 | Cd109 | CD109 antigen | 245 | 139 | 73 | 0 | 8 | 3 | 0 | 0 | 2 | 0 | 0 |
| 43 | -0.44 | 3.68 | -0.12 | 0.90 | 1 | Ephb1 | Eph receptor B1 | 166 | 157 | 144 | 0 | 1 | 0 | 0 | 0 | 0 | 0 | 0 |
| 42 | 0.00 | 3.68 | 0.00 | 1.00 | 1 | Higd1b | HIG1 domain family, member 1B | 164 | 130 | 174 | 0 | 0 | 0 | 0 | 0 | 0 | 0 | 0 |
| 42 | 0.00 | 3.68 | 0.00 | 1.00 | 1 | Col23a1 | collagen, type XXIII, alpha 1 | 365 | 33 | 68 | 0 | 0 | 0 | 0 | 0 | 0 | 0 | 0 |
| 42 | 0.00 | 3.68 | 0.00 | 1.00 | 1 | Sox17 | SRY (sex determining region Y)-box 17 | 140 | 189 | 136 | 0 | 0 | 0 | 0 | 0 | 0 | 0 | 0 |
| 42 | 0.00 | 3.68 | 0.00 | 1.00 | 1 | Hoxa5 | homeobox A5 | 275 | 138 | 51 | 0 | 0 | 0 | 0 | 0 | 0 | 0 | 0 |
| 42 | -0.40 | 3.67 | -0.11 | 0.91 | 1 | Gm6377 | predicted gene 6377 | 344 | 64 | 54 | 0 | 0 | 1 | 0 | 0 | 0 | 0 | 0 |
| 42 | 0.00 | 3.64 | 0.00 | 1.00 | 1 | Ighv8-11 | | 458 | 3 | 0 | 0 | 0 | 0 | 0 | 0 | 0 | 0 | 0 |
| 42 | -1.13 | 3.62 | -0.31 | 0.75 | 1 | Cldn5 | claudin 5 | 256 | 34 | 165 | 0 | 6 | 0 | 0 | 0 | 0 | 0 | 0 |
| 41 | 0.97 | 3.67 | 0.26 | 0.79 | 1 | Prg4 | proteoglycan 4 (megakaryocyte stimulatir | 268 | 45 | 140 | 0 | 0 | 0 | 0 | 0 | 1 | 0 | 0 |
| 41 | 0.00 | 3.68 | 0.00 | 1.00 | 1 | Nnmt | nicotinamide N-methyltransferase | 305 | 58 | 88 | 0 | 0 | 0 | 0 | 0 | 0 | 0 | 0 |
| 41 | 0.00 | 3.68 | 0.00 | 1.00 | 1 | Slc10a6 | solute carrier family 10 (sodium/bile acid | 257 | 96 | 94 | 0 | 0 | 0 | 0 | 0 | 0 | 0 | 0 |
| 40 | -0.98 | 1.19 | -0.82 | 0.41 | 1 | Pm20d1 | peptidase M20 domain containing 1 | 68 | 147 | 169 | 4 | 34 | 3 | 0 | 0 | 14 | 0 | 5 |
| 40 | 0.00 | 3.68 | 0.00 | 1.00 | 1 | Sult5a1 | sulfotransferase family 5A, member 1 | 266 | 72 | 105 | 0 | 0 | 0 | 0 | 0 | 0 | 0 | 0 |
| 40 | 1.73 | 3.64 | 0.48 | 0.63 | 1 | Smim3 | small integral membrane protein 3 | 142 | 122 | 174 | 0 | 0 | 0 | 0 | 0 | 6 | 0 | 0 |
| 40 | 0.00 | 3.68 | 0.00 | 1.00 | 1 | Fst | follistatin | 191 | 86 | 165 | 0 | 0 | 0 | 0 | 0 | 0 | 0 | 0 |
| 40 | 0.00 | 3.68 | 0.00 | 1.00 | 1 | Grrp1 | glycine/arginine rich protein 1 | 139 | 146 | 153 | 0 | 0 | 0 | 0 | 0 | 0 | 0 | 0 |
| 40 | 0.00 | 3.68 | 0.00 | 1.00 | 1 | Cyp4f18 | cytochrome P450, family 4, subfamily f, pc | 212 | 70 | 155 | 0 | 0 | 0 | 0 | 0 | 0 | 0 | 0 |
| 39 | -0.18 | 1.03 | -0.18 | 0.86 | 1 | B3gnt7 | UDP-GlcNAc:betaGal beta-1,3-N-acetylgl | 25 | 168 | 198 | 7 | 7 | 7 | 0 | 0 | 12 | 0 | 10 |
| 39 | 0.28 | 1.46 | 0.19 | 0.85 | 1 | Slc2a4 | solute carrier family 2 (facilitated glucose | 182 | 169 | 78 | 0 | 0 | 2 | 0 | 0 | 0 | 0 | 2 |
| 39 | -1.86 | 1.39 | -1.34 | 0.18 | 1 | Tlr6 | toll-like receptor 6 | 138 | 75 | 107 | 50 | 17 | 18 | 0 | 0 | 14 | 0 | 13 |
| 39 | -0.09 | 1.37 | -0.06 | 0.95 | 1 | Cftr | cystic fibrosis transmembrane conductanc | 115 | 78 | 228 | 0 | 3 | 2 | 0 | 0 | 5 | 0 | 0 |
| 39 | -1.28 | 3.63 | -0.35 | 0.72 | 1 | Slc43a1 | solute carrier family 43, member 1 | 251 | 71 | 100 | 6 | 0 | 0 | 0 | 0 | 0 | 0 | 0 |
| 39 | -0.69 | 3.66 | -0.19 | 0.85 | 1 | Crb2 | crumbs family member 2 | 247 | 130 | 49 | 0 | 0 | 2 | 0 | 0 | 0 | 0 | 0 |
| 39 | -1.04 | 3.64 | -0.29 | 0.78 | 1 | Cox4i2 | cytochrome c oxidase subunit 4I2 | 142 | 134 | 147 | 0 | 4 | 0 | 0 | 0 | 0 | 0 | 0 |
| 39 | 0.00 | 3.63 | 0.00 | 1.00 | 1 | Ighv8-8 | | 425 | 1 | 0 | 0 | 0 | 0 | 0 | 0 | 0 | 0 | 0 |
| 39 | 0.69 | 3.34 | 0.21 | 0.84 | 1 | Snai1 | snail family zinc finger 1 | 127 | 108 | 163 | 0 | 5 | 0 | 0 | 22 | 0 | 0 | 0 |
| 39 | -1.19 | 1.32 | -0.90 | 0.37 | 1 | Col26a1 | collagen, type XXVI, alpha 1 | 100 | 136 | 177 | 0 | 3 | 6 | 0 | 0 | 0 | 0 | 4 |
| 38 | 2.32 | 1.70 | 1.36 | 0.17 | 1 | Grik3 | glutamate receptor, ionotropic, kainate 3 | 240 | 98 | 70 | 0 | 0 | 6 | 0 | 3 | 4 | 0 | 0 |
| 38 | -0.40 | 3.67 | -0.11 | 0.91 | 1 | Nod2 | nucleotide-binding oligomerization doma | 136 | 113 | 170 | 0 | 0 | 1 | 0 | 0 | 0 | 0 | 0 |
| 38 | -1.16 | 3.62 | -0.32 | 0.75 | 1 | Dpysl4 | dihydropyrimidinase-like 4 | 132 | 95 | 184 | 0 | 9 | 0 | 0 | 0 | 0 | 0 | 0 |
| 38 | 0.98 | 3.68 | 0.27 | 0.79 | 1 | Mpzl2 | myelin protein zero-like 2 | 221 | 75 | 121 | 0 | 0 | 0 | 0 | 0 | 1 | 0 | 0 |
| 38 | 1.55 | 3.66 | 0.43 | 0.67 | 1 | Samsn1 | SAM domain, SH3 domain and nuclear lo | 227 | 89 | 98 | 0 | 0 | 0 | 0 | 0 | 3 | 0 | 0 |
| 38 | -1.36 | 1.64 | -0.83 | 0.41 | 1 | Siglech | sialic acid binding Ig-like lectin H | 199 | 74 | 133 | 0 | 9 | 0 | 0 | 0 | 1 | 0 | 0 |
| 38 | 0.00 | 3.68 | 0.00 | 1.00 | 1 | Ighv8-5 | | 386 | 30 | 0 | 0 | 0 | 0 | 0 | 0 | 0 | 0 | 0 |
| 38 | 0.18 | 1.33 | 0.14 | 0.89 | 1 | Gpsm3 | G-protein signalling modulator 3 (AGS3-li | 197 | 94 | 108 | 0 | 0 | 8 | 0 | 0 | 0 | 0 | 8 |
| 38 | 1.34 | 1.24 | 1.08 | 0.28 | 1 | Cybrd1 | cytochrome b reductase 1 | 196 | 107 | 94 | 0 | 2 | 3 | 0 | 0 | 4 | 6 | 2 |
| 38 | 0.00 | 3.68 | 0.00 | 1.00 | 1 | Arsi | arylsulfatase i | 191 | 81 | 140 | 0 | 0 | 0 | 0 | 0 | 0 | 0 | 0 |
| 37 | 0.47 | 1.41 | 0.33 | 0.74 | 1 | Casp4 | caspase 4, apoptosis-related cysteine pe | 243 | 89 | 73 | 0 | 0 | 3 | 0 | 0 | 0 | 0 | 4 |

| | | | | | | | | | | | | | | | | | | |
|---|---|---|---|---|---|---|---|---|---|---|---|---|---|---|---|---|---|---|
| 37 | 0.00 | 3.68 | 0.00 | 1.00 | 1 | Ackr1 | atypical chemokine receptor 1 (Duffy bloc | 208 | 112 | 88 | 0 | 0 | 0 | 0 | 0 | 0 | 0 | 0 |
| 37 | 1.10 | 1.60 | 0.69 | 0.49 | 1 | Abcc12 | ATP-binding cassette, sub-family C (CFTF | 102 | 113 | 155 | 3 | 0 | 11 | 0 | 23 | 1 | 0 | 0 |
| 37 | 0.12 | 1.45 | 0.08 | 0.93 | 1 | Vill | villin-like | 70 | 73 | 249 | 0 | 2 | 4 | 0 | 0 | 8 | 0 | 0 |
| 37 | 1.77 | 1.47 | 1.20 | 0.23 | 1 | Gm47483 | | 4 | 10 | 0 | 0 | 1 | 203 | 0 | 35 | 1 | 0 | 152 |
| 37 | -1.59 | 1.51 | -1.05 | 0.29 | 1 | Colec11 | collectin sub-family member 11 | 214 | 101 | 76 | 7 | 0 | 4 | 0 | 0 | 0 | 0 | 4 |
| 37 | -2.35 | 1.60 | -1.47 | 0.14 | 1 | Gipc3 | GIPC PDZ domain containing family, men | 138 | 75 | 185 | 0 | 7 | 1 | 0 | 0 | 0 | 0 | 0 |
| 37 | 0.00 | 3.68 | 0.00 | 1.00 | 1 | Scube3 | signal peptide, CUB domain, EGF-like 3 | 154 | 167 | 80 | 0 | 0 | 0 | 0 | 0 | 0 | 0 | 0 |
| 36 | 0.00 | 3.68 | 0.00 | 1.00 | 1 | Cnr2 | cannabinoid receptor 2 (macrophage) | 186 | 117 | 98 | 0 | 0 | 0 | 0 | 0 | 0 | 0 | 0 |
| 36 | -0.43 | 3.68 | -0.12 | 0.91 | 1 | Olfr1396 | olfactory receptor 1396 | 102 | 122 | 177 | 1 | 0 | 0 | 0 | 0 | 0 | 0 | 0 |
| 36 | 0.00 | 3.61 | 0.00 | 1.00 | 1 | Ighv1-18 | | 401 | 0 | 0 | 0 | 0 | 0 | 0 | 0 | 0 | 0 | 0 |
| 36 | 0.00 | 3.68 | 0.00 | 1.00 | 1 | H2-M2 | histocompatibility 2, M region locus 2 | 131 | 177 | 91 | 0 | 0 | 0 | 0 | 0 | 0 | 0 | 0 |
| 36 | 0.00 | 3.68 | 0.00 | 1.00 | 1 | Sult1e1 | sulfotransferase family 1E, member 1 | 219 | 121 | 60 | 0 | 0 | 0 | 0 | 0 | 0 | 0 | 0 |
| 36 | -0.37 | 3.33 | -0.11 | 0.91 | 1 | Aif1l | allograft inflammatory factor 1-like | 200 | 56 | 128 | 8 | 0 | 0 | 0 | 0 | 7 | 0 | 0 |
| 36 | 0.00 | 3.68 | 0.00 | 1.00 | 1 | Bst1 | bone marrow stromal cell antigen 1 | 248 | 97 | 52 | 0 | 0 | 0 | 0 | 0 | 0 | 0 | 0 |
| 36 | 0.08 | 1.64 | 0.05 | 0.96 | 1 | Kcna1 | potassium voltage-gated channel, shaker | 69 | 155 | 165 | 0 | 3 | 0 | 0 | 0 | 0 | 0 | 4 |
| 36 | 0.00 | 3.68 | 0.00 | 1.00 | 1 | Cxcr3 | chemokine (C-X-C motif) receptor 3 | 305 | 58 | 32 | 0 | 0 | 0 | 0 | 0 | 0 | 0 | 0 |
| 36 | 0.00 | 3.68 | 0.00 | 1.00 | 1 | Barx1 | BarH-like homeobox 1 | 239 | 73 | 82 | 0 | 0 | 0 | 0 | 0 | 0 | 0 | 0 |
| 36 | 0.00 | 3.68 | 0.00 | 1.00 | 1 | Ssu2 | ssu-2 homolog (C. elegans) | 90 | 159 | 144 | 0 | 0 | 0 | 0 | 0 | 0 | 0 | 0 |
| 36 | -0.89 | 1.51 | -0.59 | 0.56 | 1 | Sox10 | SRY (sex determining region Y)-box 10 | 157 | 95 | 124 | 6 | 4 | 0 | 0 | 0 | 6 | 0 | 0 |
| 36 | -0.22 | 1.35 | -0.16 | 0.87 | 1 | Chek2 | checkpoint kinase 2 | 134 | 51 | 85 | 8 | 17 | 21 | 0 | 0 | 42 | 0 | 34 |
| 35 | 1.00 | 3.68 | 0.27 | 0.79 | 1 | Car2 | carbonic anhydrase 2 | 115 | 133 | 140 | 0 | 0 | 0 | 0 | 1 | 0 | 0 | 0 |
| 35 | -0.17 | 1.54 | -0.11 | 0.91 | 1 | Gpr35 | G protein-coupled receptor 35 | 111 | 111 | 157 | 3 | 0 | 1 | 0 | 0 | 0 | 0 | 6 |
| 35 | -0.32 | 2.96 | -0.11 | 0.91 | 1 | Ms4a4c | membrane-spanning 4-domains, subfami | 146 | 26 | 205 | 0 | 9 | 0 | 0 | 0 | 0 | 0 | 2 |
| 35 | 0.00 | 3.68 | 0.00 | 1.00 | 1 | Veph1 | ventricular zone expressed PH domain-co | 175 | 22 | 189 | 0 | 0 | 0 | 0 | 0 | 0 | 0 | 0 |
| 35 | 0.00 | 3.68 | 0.00 | 1.00 | 1 | Igkv1-135 | | 308 | 78 | 0 | 0 | 0 | 0 | 0 | 0 | 0 | 0 | 0 |
| 35 | 0.00 | 3.68 | 0.00 | 1.00 | 1 | Bmp5 | bone morphogenetic protein 5 | 154 | 106 | 125 | 0 | 0 | 0 | 0 | 0 | 0 | 0 | 0 |
| 35 | 0.00 | 3.68 | 0.00 | 1.00 | 1 | Arhgap22 | Rho GTPase activating protein 22 | 136 | 85 | 164 | 0 | 0 | 0 | 0 | 0 | 0 | 0 | 0 |
| 35 | 0.18 | 1.37 | 0.13 | 0.90 | 1 | AB124611 | cDNA sequence AB124611 | 124 | 159 | 88 | 0 | 0 | 7 | 0 | 0 | 0 | 0 | 7 |
| 35 | 1.20 | 1.13 | 1.06 | 0.29 | 1 | C6 | complement component 6 | 238 | 43 | 49 | 2 | 9 | 3 | 0 | 12 | 1 | 22 | 6 |
| 35 | -0.40 | 3.67 | -0.11 | 0.91 | 1 | Abcg3 | ATP binding cassette subfamily G membe | 231 | 107 | 43 | 0 | 0 | 1 | 0 | 0 | 0 | 0 | 0 |
| 35 | 3.83 | 3.40 | 1.13 | 0.26 | 1 | Dapk2 | death-associated protein kinase 2 | 185 | 82 | 87 | 0 | 0 | 2 | 0 | 0 | 24 | 0 | 0 |
| 35 | 0.00 | 3.68 | 0.00 | 1.00 | 1 | Tmem132c | transmembrane protein 132C | 251 | 69 | 60 | 0 | 0 | 0 | 0 | 0 | 0 | 0 | 0 |
| 35 | -1.44 | 1.67 | -0.86 | 0.39 | 1 | Ism1 | isthmin 1, angiogenesis inhibitor | 201 | 89 | 78 | 10 | 0 | 1 | 0 | 0 | 0 | 0 | 2 |
| 34 | -0.73 | 3.64 | -0.20 | 0.84 | 1 | Lrp2 | low density lipoprotein receptor-related pı | 184 | 145 | 48 | 0 | 0 | 3 | 0 | 0 | 0 | 0 | 0 |
| 34 | -0.40 | 3.67 | -0.11 | 0.91 | 1 | Lilra5 | leukocyte immunoglobulin-like receptor, s | 165 | 48 | 165 | 0 | 0 | 1 | 0 | 0 | 0 | 0 | 0 |
| 34 | 0.00 | 3.68 | 0.00 | 1.00 | 1 | Tpsb2 | tryptase beta 2 | 214 | 144 | 20 | 0 | 0 | 0 | 0 | 0 | 0 | 0 | 0 |
| 34 | 1.45 | 3.15 | 0.46 | 0.64 | 1 | Gm36936 | | 38 | 2 | 0 | 0 | 0 | 163 | 0 | 0 | 2 | 0 | 173 |
| 34 | -0.82 | 1.57 | -0.52 | 0.60 | 1 | Cd300e | CD300E molecule | 155 | 135 | 76 | 6 | 0 | 1 | 0 | 0 | 0 | 0 | 4 |
| 34 | 0.00 | 3.68 | 0.00 | 1.00 | 1 | Igkv4-68 | | 150 | 0 | 227 | 0 | 0 | 0 | 0 | 0 | 0 | 0 | 0 |
| 34 | -0.55 | 1.39 | -0.40 | 0.69 | 1 | Bvht | | 106 | 93 | 164 | 3 | 0 | 5 | 0 | 0 | 4 | 0 | 1 |
| 34 | 0.00 | 3.68 | 0.00 | 1.00 | 1 | Tgtp2 | T cell specific GTPase 2 | 108 | 94 | 173 | 0 | 0 | 0 | 0 | 0 | 0 | 0 | 0 |
| 34 | 0.00 | 3.61 | 0.00 | 1.00 | 1 | Gm20730 | | 372 | 0 | 0 | 0 | 0 | 0 | 0 | 0 | 0 | 0 | 0 |
| 34 | 2.14 | 1.29 | 1.66 | 0.10 | 1 | Ankrd22 | ankyrin repeat domain 22 | 6 | 0 | 15 | 30 | 21 | 10 | 0 | 141 | 28 | 111 | 10 |
| 34 | 1.58 | 3.65 | 0.43 | 0.67 | 1 | Lipg | lipase, endothelial | 153 | 43 | 173 | 0 | 0 | 0 | 0 | 0 | 4 | 0 | 0 |
| 34 | 1.71 | 3.65 | 0.47 | 0.64 | 1 | Chst3 | carbohydrate sulfotransferase 3 | 219 | 80 | 69 | 0 | 0 | 0 | 0 | 0 | 4 | 0 | 0 |
| 34 | 1.61 | 1.63 | 0.99 | 0.32 | 1 | Gpr68 | G protein-coupled receptor 68 | 109 | 133 | 120 | 0 | 0 | 2 | 0 | 0 | 0 | 6 | 1 |
| 34 | -1.20 | 3.63 | -0.33 | 0.74 | 1 | Ugt2b35 | UDP glucuronosyltransferase 2 family, pc | 63 | 140 | 161 | 6 | 0 | 0 | 0 | 0 | 0 | 0 | 0 |
| 34 | 0.00 | 3.67 | 0.00 | 1.00 | 1 | Igkv4-91 | | 9 | 360 | 0 | 0 | 0 | 0 | 0 | 0 | 0 | 0 | 0 |
| 34 | 1.66 | 1.66 | 1.00 | 0.32 | 1 | Rbpms2 | RNA binding protein with multiple splicing | 190 | 97 | 69 | 0 | 1 | 0 | 0 | 0 | 0 | 12 | 0 |
| 33 | 1.73 | 3.63 | 0.48 | 0.63 | 1 | Rasl11a | RAS-like, family 11, member A | 199 | 32 | 131 | 0 | 0 | 0 | 0 | 0 | 6 | 0 | 0 |
| 33 | 0.00 | 3.68 | 0.00 | 1.00 | 1 | Magix | MAGI family member, X-linked | 91 | 139 | 138 | 0 | 0 | 0 | 0 | 0 | 0 | 0 | 0 |
| 33 | -0.79 | 1.48 | -0.53 | 0.59 | 1 | Tnfaip8l2 | tumor necrosis factor, alpha-induced prot | 129 | 80 | 151 | 4 | 2 | 0 | 0 | 0 | 3 | 0 | 0 |
| 33 | 0.00 | 3.68 | 0.00 | 1.00 | 1 | Ifi202b | interferon activated gene 202B | 213 | 1 | 154 | 0 | 0 | 0 | 0 | 0 | 0 | 0 | 0 |
| 33 | 0.00 | 3.68 | 0.00 | 1.00 | 1 | Fcrla | Fc receptor-like A | 198 | 135 | 33 | 0 | 0 | 0 | 0 | 0 | 0 | 0 | 0 |
| 33 | 0.00 | 3.68 | 0.00 | 1.00 | 1 | Alox15 | arachidonate 15-lipoxygenase | 143 | 55 | 167 | 0 | 0 | 0 | 0 | 0 | 0 | 0 | 0 |
| 33 | 0.00 | 3.68 | 0.00 | 1.00 | 1 | Gbp2b | guanylate binding protein 2b | 251 | 72 | 41 | 0 | 0 | 0 | 0 | 0 | 0 | 0 | 0 |
| 33 | 0.95 | 3.67 | 0.26 | 0.80 | 1 | C1s2 | complement component 1, s subcompone | 243 | 85 | 34 | 0 | 0 | 0 | 0 | 0 | 0 | 0 | 1 |
| 33 | 0.61 | 1.18 | 0.52 | 0.60 | 1 | Zmynd15 | zinc finger, MYND-type containing 15 | 71 | 73 | 103 | 6 | 12 | 23 | 0 | 0 | 7 | 28 | 38 |
| 33 | -0.37 | 1.51 | -0.24 | 0.81 | 1 | Inka1 | inka box actin regulator 1 | 88 | 101 | 106 | 11 | 3 | 6 | 0 | 0 | 0 | 41 | 4 |
| 33 | 0.00 | 3.68 | 0.00 | 1.00 | 1 | Dipk2b | divergent protein kinase domain 2B | 154 | 69 | 136 | 0 | 0 | 0 | 0 | 0 | 0 | 0 | 0 |

| | | | | | | | | | | | | | | | | | | |
|---|---|---|---|---|---|---|---|---|---|---|---|---|---|---|---|---|---|---|
| 33 | 0.00 | 3.68 | 0.00 | 1.00 | 1 | Ackr2 | atypical chemokine receptor 2 | 219 | 69 | 71 | 0 | 0 | 0 | 0 | 0 | 0 | 0 | 0 |
| 33 | -0.47 | 1.14 | -0.42 | 0.68 | 1 | Gm43609 | | 15 | 18 | 8 | 4 | 2 | 158 | 0 | 0 | 3 | 0 | 149 |
| 32 | -0.10 | 3.00 | -0.03 | 0.97 | 1 | Rbpjl | recombination signal binding protein for i | 60 | 124 | 158 | 3 | 0 | 0 | 0 | 0 | 0 | 0 | 13 |
| 32 | 1.89 | 3.63 | 0.52 | 0.60 | 1 | Il1rn | interleukin 1 receptor antagonist | 201 | 93 | 54 | 0 | 0 | 0 | 0 | 0 | 9 | 0 | 0 |
| 32 | 0.00 | 3.61 | 0.00 | 1.00 | 1 | Igkv4-81 | | 357 | 0 | 0 | 0 | 0 | 0 | 0 | 0 | 0 | 0 | 0 |
| 32 | -0.38 | 1.74 | -0.22 | 0.82 | 1 | Ltc4s | leukotriene C4 synthase | 162 | 109 | 81 | 0 | 2 | 0 | 0 | 0 | 0 | 0 | 1 |
| 32 | 0.00 | 3.65 | 0.00 | 1.00 | 1 | Igkv5-48 | | 352 | 3 | 0 | 0 | 0 | 0 | 0 | 0 | 0 | 0 | 0 |
| 32 | 0.00 | 3.68 | 0.00 | 1.00 | 1 | Smtnl2 | smoothelin-like 2 | 125 | 112 | 116 | 0 | 0 | 0 | 0 | 0 | 0 | 0 | 0 |
| 32 | -0.80 | 3.61 | -0.22 | 0.82 | 1 | Msx1 | msh homeobox 1 | 112 | 128 | 108 | 0 | 0 | 7 | 0 | 0 | 0 | 0 | 0 |
| 32 | 0.00 | 3.68 | 0.00 | 1.00 | 1 | Muc3 | mucin 3, intestinal | 0 | 135 | 219 | 0 | 0 | 0 | 0 | 0 | 0 | 0 | 0 |
| 32 | 1.57 | 1.12 | 1.40 | 0.16 | 1 | Rasl11b | RAS-like, family 11, member B | 100 | 33 | 83 | 9 | 5 | 19 | 0 | 25 | 8 | 66 | 6 |
| 32 | 0.00 | 3.68 | 0.00 | 1.00 | 1 | Trpc3 | transient receptor potential cation channe | 159 | 123 | 72 | 0 | 0 | 0 | 0 | 0 | 0 | 0 | 0 |
| 32 | 0.00 | 3.68 | 0.00 | 1.00 | 1 | Tmem252 | transmembrane protein 252 | 83 | 130 | 139 | 0 | 0 | 0 | 0 | 0 | 0 | 0 | 0 |
| 32 | 3.35 | 3.23 | 1.04 | 0.30 | 1 | Il2ra | interleukin 2 receptor, alpha chain | 130 | 100 | 104 | 0 | 0 | 2 | 0 | 15 | 0 | 0 | 0 |
| 32 | 0.00 | 3.68 | 0.00 | 1.00 | 1 | Tal1 | T cell acute lymphocytic leukemia 1 | 105 | 75 | 171 | 0 | 0 | 0 | 0 | 0 | 0 | 0 | 0 |
| 32 | 0.00 | 3.68 | 0.00 | 1.00 | 1 | Lpar4 | lysophosphatidic acid receptor 4 | 177 | 98 | 76 | 0 | 0 | 0 | 0 | 0 | 0 | 0 | 0 |
| 32 | 0.00 | 3.68 | 0.00 | 1.00 | 1 | Clec9a | C-type lectin domain family 9, member a | 206 | 70 | 74 | 0 | 0 | 0 | 0 | 0 | 0 | 0 | 0 |
| 31 | 1.61 | 1.81 | 0.89 | 0.38 | 1 | Stap1 | signal transducing adaptor family membe | 167 | 78 | 92 | 0 | 0 | 1 | 0 | 0 | 7 | 0 | 0 |
| 31 | 1.78 | 1.55 | 1.15 | 0.25 | 1 | Endou | endonuclease, polyU-specific | 179 | 75 | 69 | 0 | 2 | 2 | 0 | 12 | 0 | 0 | 6 |
| 31 | 0.00 | 3.68 | 0.00 | 1.00 | 1 | Kcnt2 | potassium channel, subfamily T, member | 208 | 64 | 72 | 0 | 0 | 0 | 0 | 0 | 0 | 0 | 0 |
| 31 | 0.79 | 1.54 | 0.51 | 0.61 | 1 | Prrg4 | proline rich Gla (G-carboxyglutamic acid) | 96 | 136 | 51 | 0 | 9 | 13 | 0 | 0 | 27 | 0 | 11 |
| 31 | -1.99 | 1.70 | -1.17 | 0.24 | 1 | Arl5c | ADP-ribosylation factor-like 5C | 180 | 86 | 65 | 4 | 3 | 0 | 0 | 0 | 0 | 0 | 2 |
| 31 | 0.00 | 3.68 | 0.00 | 1.00 | 1 | Ighv1-26 | | 331 | 8 | 0 | 0 | 0 | 0 | 0 | 0 | 0 | 0 | 0 |
| 31 | 0.00 | 3.68 | 0.00 | 1.00 | 1 | Pilra | paired immunoglobin-like type 2 receptor | 131 | 100 | 107 | 0 | 0 | 0 | 0 | 0 | 0 | 0 | 0 |
| 31 | 0.00 | 3.68 | 0.00 | 1.00 | 1 | H2-Eb2 | histocompatibility 2, class II antigen E bet | 263 | 50 | 26 | 0 | 0 | 0 | 0 | 0 | 0 | 0 | 0 |
| 31 | -0.69 | 3.66 | -0.19 | 0.85 | 1 | Ctnna3 | catenin (cadherin associated protein), alp | 167 | 60 | 109 | 0 | 0 | 2 | 0 | 0 | 0 | 0 | 0 |
| 31 | 0.00 | 3.61 | 0.00 | 1.00 | 1 | Ighv3-5 | | 337 | 0 | 0 | 0 | 0 | 0 | 0 | 0 | 0 | 0 | 0 |
| 31 | 3.04 | 1.63 | 1.87 | 0.06 | 1 | Zc3h12d | zinc finger CCCH type containing 12D | 217 | 14 | 72 | 0 | 0 | 3 | 0 | 0 | 9 | 22 | 0 |
| 30 | -1.06 | 3.62 | -0.29 | 0.77 | 1 | Ccl4 | chemokine (C-C motif) ligand 4 | 125 | 37 | 163 | 0 | 7 | 0 | 0 | 0 | 0 | 0 | 0 |
| 30 | 0.00 | 3.68 | 0.00 | 1.00 | 1 | Kcng4 | potassium voltage-gated channel, subfan | 119 | 135 | 78 | 0 | 0 | 0 | 0 | 0 | 0 | 0 | 0 |
| 30 | -0.12 | 1.34 | -0.09 | 0.93 | 1 | Adcy2 | adenylate cyclase 2 | 118 | 65 | 75 | 11 | 5 | 14 | 0 | 27 | 0 | 0 | 18 |
| 30 | 0.00 | 3.68 | 0.00 | 1.00 | 1 | Il6 | interleukin 6 | 264 | 44 | 24 | 0 | 0 | 0 | 0 | 0 | 0 | 0 | 0 |
| 30 | 0.53 | 0.97 | 0.55 | 0.58 | 1 | Chst8 | carbohydrate sulfotransferase 8 | 82 | 92 | 53 | 22 | 9 | 14 | 0 | 16 | 4 | 23 | 17 |
| 30 | 0.00 | 3.68 | 0.00 | 1.00 | 1 | Bnc1 | basonuclin 1 | 141 | 125 | 65 | 0 | 0 | 0 | 0 | 0 | 0 | 0 | 0 |
| 30 | 0.00 | 3.65 | 0.00 | 1.00 | 1 | Ighv1-81 | | 3 | 327 | 0 | 0 | 0 | 0 | 0 | 0 | 0 | 0 | 0 |
| 30 | -2.33 | 1.21 | -1.93 | 0.05 | 1 | Itga2b | integrin alpha 2b | 51 | 109 | 71 | 6 | 21 | 42 | 0 | 0 | 2 | 0 | 28 |
| 30 | 0.00 | 3.68 | 0.00 | 1.00 | 1 | H2-Oa | histocompatibility 2, O region alpha locus | 189 | 97 | 42 | 0 | 0 | 0 | 0 | 0 | 0 | 0 | 0 |
| 29 | -2.10 | 1.49 | -1.41 | 0.16 | 1 | Stk32c | serine/threonine kinase 32C | 104 | 51 | 151 | 7 | 1 | 6 | 0 | 0 | 0 | 0 | 5 |
| 29 | 1.73 | 1.55 | 1.12 | 0.26 | 1 | Reg2 | regenerating islet-derived 2 | 25 | 28 | 224 | 0 | 0 | 3 | 0 | 0 | 0 | 0 | 43 |
| 29 | -3.42 | 1.77 | -1.94 | 0.05 | 1 | Cyp2j9 | cytochrome P450, family 2, subfamily j, pc | 150 | 90 | 69 | 0 | 10 | 4 | 0 | 0 | 0 | 0 | 0 |
| 29 | -1.41 | 1.44 | -0.98 | 0.33 | 1 | Gm20627 | | 4 | 0 | 7 | 26 | 9 | 123 | 0 | 0 | 3 | 6 | 143 |
| 29 | -0.73 | 3.64 | -0.20 | 0.84 | 1 | Clec4n | C-type lectin domain family 4, member n | 156 | 56 | 108 | 0 | 0 | 3 | 0 | 0 | 0 | 0 | 0 |
| 29 | -1.34 | 3.61 | -0.37 | 0.71 | 1 | Hamp2 | hepcidin antimicrobial peptide 2 | 51 | 131 | 123 | 18 | 0 | 0 | 0 | 0 | 0 | 0 | 0 |
| 29 | 0.00 | 3.68 | 0.00 | 1.00 | 1 | Rarb | retinoic acid receptor, beta | 154 | 78 | 88 | 0 | 0 | 0 | 0 | 0 | 0 | 0 | 0 |
| 29 | 1.82 | 3.63 | 0.50 | 0.62 | 1 | Ankrd1 | ankyrin repeat domain 1 (cardiac muscle) | 123 | 94 | 96 | 0 | 0 | 0 | 0 | 0 | 8 | 0 | 0 |
| 29 | 0.00 | 3.68 | 0.00 | 1.00 | 1 | Gli1 | GLI-Kruppel family member GLI1 | 232 | 22 | 65 | 0 | 0 | 0 | 0 | 0 | 0 | 0 | 0 |
| 29 | 0.00 | 3.68 | 0.00 | 1.00 | 1 | Msc | musculin | 189 | 50 | 81 | 0 | 0 | 0 | 0 | 0 | 0 | 0 | 0 |
| 29 | 0.00 | 3.68 | 0.00 | 1.00 | 1 | Cdh3 | cadherin 3 | 149 | 59 | 110 | 0 | 0 | 0 | 0 | 0 | 0 | 0 | 0 |
| 29 | 0.00 | 3.68 | 0.00 | 1.00 | 1 | Sfrp2 | secreted frizzled-related protein 2 | 112 | 69 | 136 | 0 | 0 | 0 | 0 | 0 | 0 | 0 | 0 |
| 29 | 0.00 | 3.68 | 0.00 | 1.00 | 1 | Milr1 | mast cell immunoglobulin like receptor 1 | 106 | 80 | 131 | 0 | 0 | 0 | 0 | 0 | 0 | 0 | 0 |
| 29 | 0.00 | 3.68 | 0.00 | 1.00 | 1 | Ighv1-34 | | 307 | 9 | 0 | 0 | 0 | 0 | 0 | 0 | 0 | 0 | 0 |
| 29 | -0.86 | 3.66 | -0.23 | 0.81 | 1 | Hspb7 | heat shock protein family, member 7 (carc | 240 | 30 | 44 | 0 | 2 | 0 | 0 | 0 | 0 | 0 | 0 |
| 29 | 0.21 | 1.33 | 0.15 | 0.88 | 1 | Pcdhb9 | protocadherin beta 9 | 136 | 14 | 82 | 24 | 9 | 6 | 0 | 22 | 17 | 0 | 5 |
| 29 | 0.87 | 1.80 | 0.48 | 0.63 | 1 | Grhl2 | grainyhead like transcription factor 2 | 147 | 79 | 82 | 0 | 0 | 3 | 0 | 0 | 4 | 0 | 0 |
| 29 | 0.00 | 3.68 | 0.00 | 1.00 | 1 | Clec5a | C-type lectin domain family 5, member a | 142 | 89 | 82 | 0 | 0 | 0 | 0 | 0 | 0 | 0 | 0 |
| 28 | 0.00 | 3.68 | 0.00 | 1.00 | 1 | Ptgfr | prostaglandin F receptor | 166 | 69 | 78 | 0 | 0 | 0 | 0 | 0 | 0 | 0 | 0 |
| 28 | -0.49 | 1.59 | -0.31 | 0.76 | 1 | Dctd | dCMP deaminase | 75 | 15 | 71 | 16 | 22 | 27 | 0 | 61 | 0 | 0 | 24 |
| 28 | 0.00 | 3.68 | 0.00 | 1.00 | 1 | Slc16a3 | solute carrier family 16 (monocarboxylic a | 155 | 96 | 58 | 0 | 0 | 0 | 0 | 0 | 0 | 0 | 0 |
| 28 | 1.21 | 1.46 | 0.83 | 0.41 | 1 | Plxnb3 | plexin B3 | 57 | 50 | 127 | 0 | 8 | 14 | 0 | 0 | 4 | 43 | 8 |
| 28 | 1.25 | 3.66 | 0.34 | 0.73 | 1 | Osr2 | odd-skipped related 2 | 161 | 52 | 94 | 0 | 0 | 0 | 0 | 0 | 0 | 0 | 2 |

| | | | | | | | | | | | | | | | | | | |
|---|---|---|---|---|---|---|---|---|---|---|---|---|---|---|---|---|---|---|
| 28 | 0.00 | 3.68 | 0.00 | 1.00 | 1 | Has2 | hyaluronan synthase 2 | 193 | 80 | 34 | 0 | 0 | 0 | 0 | 0 | 0 | 0 | 0 |
| 28 | -0.21 | 1.41 | -0.15 | 0.88 | 1 | Trim5 | tripartite motif-containing 5 | 193 | 68 | 32 | 1 | 0 | 6 | 0 | 0 | 0 | 0 | 7 |
| 28 | 0.00 | 3.68 | 0.00 | 1.00 | 1 | Cass4 | Cas scaffolding protein family member 4 | 210 | 57 | 38 | 0 | 0 | 0 | 0 | 0 | 0 | 0 | 0 |
| 28 | 0.00 | 3.68 | 0.00 | 1.00 | 1 | Sncaip | synuclein, alpha interacting protein (synp | 124 | 55 | 126 | 0 | 0 | 0 | 0 | 0 | 0 | 0 | 0 |
| 28 | 0.00 | 3.61 | 0.00 | 1.00 | 1 | Igkv3-1 | | 304 | 0 | 0 | 0 | 0 | 0 | 0 | 0 | 0 | 0 | 0 |
| 27 | -0.26 | 1.63 | -0.16 | 0.87 | 1 | Mc4r | melanocortin 4 receptor | 70 | 46 | 140 | 0 | 13 | 9 | 0 | 0 | 0 | 15 | 4 |
| 27 | 0.00 | 3.61 | 0.00 | 1.00 | 1 | Igkv6-25 | | 298 | 0 | 0 | 0 | 0 | 0 | 0 | 0 | 0 | 0 | 0 |
| 27 | -0.43 | 3.68 | -0.12 | 0.91 | 1 | Tgif2 | TGFB-induced factor homeobox 2 | 126 | 54 | 116 | 0 | 1 | 0 | 0 | 0 | 0 | 0 | 0 |
| 27 | -0.76 | 3.66 | -0.21 | 0.84 | 1 | Clic6 | chloride intracellular channel 6 | 61 | 114 | 117 | 2 | 0 | 0 | 0 | 0 | 0 | 0 | 0 |
| 27 | 0.00 | 3.68 | 0.00 | 1.00 | 1 | Igkv4-57-1 | | 45 | 248 | 0 | 0 | 0 | 0 | 0 | 0 | 0 | 0 | 0 |
| 27 | -1.27 | 3.60 | -0.35 | 0.72 | 1 | Ccl5 | chemokine (C-C motif) ligand 5 | 137 | 10 | 134 | 0 | 11 | 0 | 0 | 0 | 0 | 0 | 0 |
| 27 | 0.00 | 3.68 | 0.00 | 1.00 | 1 | Cma1 | chymase 1, mast cell | 131 | 145 | 17 | 0 | 0 | 0 | 0 | 0 | 0 | 0 | 0 |
| 27 | 0.00 | 3.68 | 0.00 | 1.00 | 1 | Adra1b | adrenergic receptor, alpha 1b | 178 | 42 | 73 | 0 | 0 | 0 | 0 | 0 | 0 | 0 | 0 |
| 27 | 0.00 | 3.68 | 0.00 | 1.00 | 1 | Slc6a4 | solute carrier family 6 (neurotransmitter tra | 200 | 80 | 12 | 0 | 0 | 0 | 0 | 0 | 0 | 0 | 0 |
| 27 | 0.00 | 3.68 | 0.00 | 1.00 | 1 | Loxl4 | lysyl oxidase-like 4 | 130 | 70 | 92 | 0 | 0 | 0 | 0 | 0 | 0 | 0 | 0 |
| 26 | 0.00 | 3.68 | 0.00 | 1.00 | 1 | Foxs1 | forkhead box S1 | 112 | 50 | 129 | 0 | 0 | 0 | 0 | 0 | 0 | 0 | 0 |
| 26 | -1.69 | 1.77 | -0.96 | 0.34 | 1 | Espn | espin | 159 | 103 | 22 | 0 | 4 | 2 | 0 | 0 | 0 | 0 | 1 |
| 26 | -1.41 | 3.61 | -0.39 | 0.70 | 1 | Ccl7 | chemokine (C-C motif) ligand 7 | 208 | 19 | 53 | 0 | 9 | 0 | 0 | 0 | 0 | 0 | 0 |
| 26 | -0.40 | 3.67 | -0.11 | 0.91 | 1 | Ptf1a | pancreas specific transcription factor, 1a | 15 | 120 | 153 | 1 | 0 | 0 | 0 | 0 | 0 | 0 | 0 |
| 26 | 2.59 | 1.69 | 1.53 | 0.13 | 1 | Nr2f1 | nuclear receptor subfamily 2, group F, me | 101 | 77 | 104 | 0 | 0 | 0 | 0 | 0 | 3 | 0 | 2 |
| 26 | 3.21 | 1.76 | 1.82 | 0.07 | 1 | Hand2os1 | | 70 | 68 | 126 | 0 | 2 | 0 | 0 | 16 | 0 | 0 | 5 |
| 26 | -2.65 | 1.75 | -1.52 | 0.13 | 1 | Gm830 | | 152 | 37 | 88 | 0 | 7 | 2 | 0 | 0 | 0 | 0 | 0 |
| 26 | -1.49 | 1.44 | -1.03 | 0.30 | 1 | Sh3gl3 | SH3-domain GRB2-like 3 | 71 | 32 | 99 | 7 | 42 | 3 | 0 | 0 | 11 | 0 | 20 |
| 26 | 1.04 | 1.20 | 0.86 | 0.39 | 1 | Bcl2l12 | BCL2-like 12 (proline rich) | 63 | 89 | 78 | 1 | 5 | 17 | 0 | 0 | 5 | 14 | 14 |
| 26 | 0.00 | 3.68 | 0.00 | 1.00 | 1 | Treml4 | triggering receptor expressed on myeloid | 158 | 99 | 28 | 0 | 0 | 0 | 0 | 0 | 0 | 0 | 0 |
| 26 | 0.00 | 3.68 | 0.00 | 1.00 | 1 | Hoxb5 | homeobox B5 | 153 | 63 | 70 | 0 | 0 | 0 | 0 | 0 | 0 | 0 | 0 |
| 26 | -0.40 | 3.67 | -0.11 | 0.91 | 1 | Tspan32 | tetraspanin 32 | 187 | 54 | 43 | 0 | 0 | 1 | 0 | 0 | 0 | 0 | 0 |
| 26 | 0.00 | 3.68 | 0.00 | 1.00 | 1 | Matn4 | matrilin 4 | 69 | 117 | 97 | 0 | 0 | 0 | 0 | 0 | 0 | 0 | 0 |
| 26 | 2.96 | 3.33 | 0.89 | 0.37 | 1 | 9530026P05Rik | | 95 | 95 | 76 | 0 | 0 | 7 | 0 | 0 | 9 | 0 | 0 |
| 26 | -1.42 | 1.39 | -1.02 | 0.31 | 1 | Sell | selectin, lymphocyte | 182 | 28 | 41 | 5 | 4 | 7 | 0 | 0 | 0 | 10 | 4 |
| 25 | -2.60 | 1.43 | -1.82 | 0.07 | 1 | Angptl1 | angiopoietin-like 1 | 92 | 69 | 105 | 3 | 7 | 3 | 0 | 0 | 0 | 0 | 2 |
| 25 | -0.40 | 3.67 | -0.11 | 0.91 | 1 | Csf3r | colony stimulating factor 3 receptor (granu | 123 | 93 | 63 | 0 | 0 | 1 | 0 | 0 | 0 | 0 | 0 |
| 25 | 0.00 | 3.61 | 0.00 | 1.00 | 1 | Ighv1-50 | | 278 | 0 | 0 | 0 | 0 | 0 | 0 | 0 | 0 | 0 | 0 |
| 25 | 0.00 | 3.68 | 0.00 | 1.00 | 1 | Igkv3-4 | | 148 | 130 | 0 | 0 | 0 | 0 | 0 | 0 | 0 | 0 | 0 |
| 25 | 0.36 | 1.26 | 0.28 | 0.78 | 1 | Extl1 | exostosin-like glycosyltransferase 1 | 122 | 70 | 47 | 10 | 3 | 2 | 0 | 17 | 1 | 0 | 4 |
| 25 | -1.79 | 1.78 | -1.01 | 0.31 | 1 | Osm | oncostatin M | 113 | 69 | 89 | 3 | 0 | 1 | 0 | 0 | 0 | 0 | 0 |
| 25 | 0.78 | 1.49 | 0.52 | 0.60 | 1 | Lin7a | lin-7 homolog A (C. elegans) | 57 | 95 | 92 | 0 | 0 | 8 | 0 | 0 | 0 | 0 | 21 |
| 25 | 1.76 | 1.57 | 1.12 | 0.26 | 1 | Nos2 | nitric oxide synthase 2, inducible | 85 | 69 | 79 | 0 | 0 | 3 | 0 | 0 | 0 | 0 | 39 |
| 25 | 0.00 | 3.68 | 0.00 | 1.00 | 1 | Gpr88 | G-protein coupled receptor 88 | 205 | 56 | 12 | 0 | 0 | 0 | 0 | 0 | 0 | 0 | 0 |
| 25 | -2.17 | 1.33 | -1.63 | 0.10 | 1 | Pde6c | phosphodiesterase 6C, cGMP specific, cc | 47 | 105 | 73 | 9 | 10 | 19 | 0 | 0 | 0 | 4 | 5 |
| 25 | 0.00 | 3.68 | 0.00 | 1.00 | 1 | C1qtnf3 | C1q and tumor necrosis factor related pro | 202 | 21 | 49 | 0 | 0 | 0 | 0 | 0 | 0 | 0 | 0 |
| 25 | 0.00 | 3.61 | 0.00 | 1.00 | 1 | Igkv8-16 | | 0 | 272 | 0 | 0 | 0 | 0 | 0 | 0 | 0 | 0 | 0 |
| 25 | 0.00 | 3.68 | 0.00 | 1.00 | 1 | Gm4951 | predicted gene 4951 | 139 | 22 | 110 | 0 | 0 | 0 | 0 | 0 | 0 | 0 | 0 |
| 25 | 0.00 | 3.68 | 0.00 | 1.00 | 1 | Cpz | carboxypeptidase Z | 166 | 41 | 64 | 0 | 0 | 0 | 0 | 0 | 0 | 0 | 0 |
| 25 | 0.00 | 3.68 | 0.00 | 1.00 | 1 | Gm45705 | | 85 | 91 | 94 | 0 | 0 | 0 | 0 | 0 | 0 | 0 | 0 |
| 24 | 1.33 | 3.66 | 0.36 | 0.72 | 1 | P2ry10b | purinergic receptor P2Y, G-protein couple | 133 | 72 | 60 | 0 | 0 | 0 | 0 | 0 | 0 | 3 | 0 |
| 24 | 1.30 | 1.00 | 1.31 | 0.19 | 1 | Gm26590 | | 19 | 44 | 11 | 18 | 29 | 10 | 0 | 71 | 37 | 6 | 20 |
| 24 | -0.19 | 1.16 | -0.17 | 0.87 | 1 | Gm36862 | | 0 | 7 | 6 | 56 | 81 | 21 | 0 | 2 | 72 | 4 | 16 |
| 24 | 0.00 | 3.68 | 0.00 | 1.00 | 1 | Shisa3 | shisa family member 3 | 257 | 9 | 0 | 0 | 0 | 0 | 0 | 0 | 0 | 0 | 0 |
| 24 | 0.53 | 1.39 | 0.38 | 0.70 | 1 | Gm43759 | | 0 | 3 | 0 | 3 | 0 | 142 | 0 | 9 | 0 | 0 | 109 |
| 24 | 0.87 | 1.69 | 0.52 | 0.61 | 1 | Gm42979 | | 7 | 17 | 0 | 0 | 0 | 105 | 0 | 0 | 1 | 0 | 135 |
| 24 | 0.95 | 3.67 | 0.26 | 0.80 | 1 | Blk | B lymphoid kinase | 196 | 59 | 8 | 0 | 0 | 0 | 0 | 0 | 0 | 0 | 1 |
| 24 | 2.27 | 1.17 | 1.94 | 0.05 | 1 | Gm15834 | | 7 | 12 | 11 | 18 | 0 | 74 | 0 | 36 | 11 | 23 | 71 |
| 24 | 0.00 | 3.64 | 0.00 | 1.00 | 1 | Ighv3-2 | | 263 | 1 | 0 | 0 | 0 | 0 | 0 | 0 | 0 | 0 | 0 |
| 24 | -1.17 | 1.22 | -0.95 | 0.34 | 1 | Rdh5 | retinol dehydrogenase 5 | 16 | 92 | 8 | 18 | 7 | 61 | 0 | 20 | 0 | 0 | 42 |
| 24 | -1.56 | 1.79 | -0.87 | 0.38 | 1 | Traf3ip3 | TRAF3 interacting protein 3 | 168 | 56 | 37 | 2 | 0 | 1 | 0 | 0 | 0 | 0 | 0 |
| 24 | -0.41 | 1.75 | -0.23 | 0.82 | 1 | Gm44164 | | 15 | 0 | 18 | 8 | 5 | 116 | 0 | 22 | 0 | 0 | 80 |
| 24 | -0.44 | 3.68 | -0.12 | 0.91 | 1 | Cyp3a13 | cytochrome P450, family 3, subfamily a, p | 74 | 88 | 101 | 0 | 1 | 0 | 0 | 0 | 0 | 0 | 0 |
| 24 | -2.15 | 1.83 | -1.18 | 0.24 | 1 | A630001G21Ri | RIKEN cDNA A630001G21 gene | 166 | 31 | 62 | 3 | 0 | 2 | 0 | 0 | 0 | 0 | 0 |
| 24 | 0.00 | 3.68 | 0.00 | 1.00 | 1 | Htr1b | 5-hydroxytryptamine (serotonin) receptor | 103 | 110 | 50 | 0 | 0 | 0 | 0 | 0 | 0 | 0 | 0 |

| | | | | | | | | | | | | | | | | | | |
|---|---|---|---|---|---|---|---|---|---|---|---|---|---|---|---|---|---|---|
| 24 | -1.27 | 3.63 | -0.35 | 0.73 | 1 | Prkcq | protein kinase C, theta | 130 | 69 | 56 | 7 | 0 | 0 | 0 | 0 | 0 | 0 | 0 |
| 24 | -0.42 | 1.06 | -0.40 | 0.69 | 1 | Rnf39 | ring finger protein 39 | 57 | 54 | 83 | 10 | 9 | 21 | 0 | 9 | 4 | 0 | 16 |
| 24 | -0.77 | 3.66 | -0.21 | 0.83 | 1 | Stra6l | STRA6-like | 75 | 60 | 126 | 0 | 2 | 0 | 0 | 0 | 0 | 0 | 0 |
| 24 | 1.20 | 1.50 | 0.80 | 0.42 | 1 | Gm44949 | | 0 | 6 | 0 | 4 | 2 | 98 | 0 | 7 | 0 | 23 | 120 |
| 24 | -3.14 | 1.80 | -1.75 | 0.08 | 1 | Il1rl1 | interleukin 1 receptor-like 1 | 93 | 72 | 83 | 9 | 0 | 3 | 0 | 0 | 0 | 0 | 0 |
| 23 | -1.02 | 3.62 | -0.28 | 0.78 | 1 | Slc5a8 | solute carrier family 5 (iodide transporter) | 31 | 71 | 150 | 0 | 7 | 0 | 0 | 0 | 0 | 0 | 0 |
| 23 | 0.99 | 0.99 | 0.99 | 0.32 | 1 | Gm13093 | | 6 | 34 | 36 | 16 | 32 | 29 | 0 | 56 | 11 | 14 | 23 |
| 23 | 1.70 | 1.50 | 1.13 | 0.26 | 1 | Timp1 | tissue inhibitor of metalloproteinase 1 | 116 | 29 | 87 | 0 | 0 | 8 | 0 | 0 | 5 | 0 | 11 |
| 23 | 0.73 | 1.71 | 0.43 | 0.67 | 1 | Nmb | neuromedin B | 33 | 79 | 85 | 6 | 0 | 14 | 0 | 0 | 0 | 22 | 16 |
| 23 | 1.96 | 1.21 | 1.62 | 0.10 | 1 | Gm14004 | | 88 | 58 | 47 | 17 | 0 | 3 | 0 | 14 | 3 | 19 | 8 |
| 23 | 0.61 | 1.65 | 0.37 | 0.71 | 1 | Gpr171 | G protein-coupled receptor 171 | 192 | 48 | 5 | 0 | 0 | 3 | 0 | 0 | 0 | 0 | 7 |
| 23 | 0.00 | 3.68 | 0.00 | 1.00 | 1 | Colq | collagen-like tail subunit (single strand of | 187 | 51 | 18 | 0 | 0 | 0 | 0 | 0 | 0 | 0 | 0 |
| 23 | 0.00 | 3.68 | 0.00 | 1.00 | 1 | Slamf6 | SLAM family member 6 | 154 | 69 | 31 | 0 | 0 | 0 | 0 | 0 | 0 | 0 | 0 |
| 23 | 1.33 | 1.23 | 1.08 | 0.28 | 1 | Efcab11 | EF-hand calcium binding domain 11 | 22 | 14 | 43 | 19 | 6 | 31 | 0 | 58 | 27 | 0 | 32 |
| 23 | -1.06 | 3.63 | -0.29 | 0.77 | 1 | Ifi208 | interferon activated gene 208 | 41 | 112 | 95 | 0 | 5 | 0 | 0 | 0 | 0 | 0 | 0 |
| 23 | -0.78 | 3.62 | -0.21 | 0.83 | 1 | Ccdc170 | coiled-coil domain containing 170 | 145 | 67 | 36 | 0 | 0 | 4 | 0 | 0 | 0 | 0 | 0 |
| 23 | 0.00 | 3.68 | 0.00 | 1.00 | 1 | Igkv10-96 | | 124 | 128 | 0 | 0 | 0 | 0 | 0 | 0 | 0 | 0 | 0 |
| 23 | 0.00 | 3.68 | 0.00 | 1.00 | 1 | Nxpe2 | neurexophilin and PC-esterase domain fa | 48 | 76 | 128 | 0 | 0 | 0 | 0 | 0 | 0 | 0 | 0 |
| 23 | -0.78 | 3.62 | -0.21 | 0.83 | 1 | Gpr141 | G protein-coupled receptor 141 | 116 | 75 | 56 | 0 | 0 | 4 | 0 | 0 | 0 | 0 | 0 |
| 23 | 0.65 | 1.85 | 0.35 | 0.73 | 1 | S100b | S100 protein, beta polypeptide, neural | 69 | 56 | 109 | 0 | 6 | 0 | 0 | 0 | 11 | 0 | 0 |
| 23 | 0.95 | 3.67 | 0.26 | 0.80 | 1 | Haao | 3-hydroxyanthranilate 3,4-dioxygenase | 94 | 124 | 31 | 0 | 0 | 0 | 0 | 0 | 0 | 0 | 1 |
| 23 | -1.26 | 1.93 | -0.65 | 0.51 | 1 | Gm42868 | | 0 | 0 | 17 | 7 | 0 | 101 | 0 | 0 | 2 | 0 | 123 |
| 23 | 0.59 | 1.19 | 0.49 | 0.62 | 1 | Gm27042 | | 21 | 2 | 31 | 15 | 43 | 33 | 0 | 42 | 23 | 5 | 35 |
| 23 | 1.35 | 3.62 | 0.37 | 0.71 | 1 | Tnf | tumor necrosis factor | 87 | 56 | 101 | 0 | 0 | 0 | 0 | 0 | 0 | 0 | 6 |
| 23 | 1.25 | 3.66 | 0.34 | 0.73 | 1 | Slc1a3 | solute carrier family 1 (glial high affinity gl | 147 | 91 | 9 | 0 | 0 | 0 | 0 | 0 | 0 | 0 | 2 |
| 23 | 0.00 | 3.68 | 0.00 | 1.00 | 1 | Irx3 | Iroquois related homeobox 3 | 87 | 75 | 87 | 0 | 0 | 0 | 0 | 0 | 0 | 0 | 0 |
| 23 | 0.00 | 3.68 | 0.00 | 1.00 | 1 | Ccl8 | chemokine (C-C motif) ligand 8 | 181 | 40 | 28 | 0 | 0 | 0 | 0 | 0 | 0 | 0 | 0 |
| 23 | 1.96 | 1.72 | 1.14 | 0.25 | 1 | 9930017N22Rik | | 2 | 24 | 4 | 0 | 0 | 106 | 0 | 0 | 8 | 0 | 105 |
| 22 | 0.00 | 3.68 | 0.00 | 1.00 | 1 | Thbs4 | thrombospondin 4 | 38 | 86 | 123 | 0 | 0 | 0 | 0 | 0 | 0 | 0 | 0 |
| 22 | 1.15 | 1.21 | 0.95 | 0.34 | 1 | 1110017D15Ril | RIKEN cDNA 1110017D15 gene | 22 | 25 | 52 | 6 | 11 | 29 | 0 | 0 | 18 | 23 | 59 |
| 22 | 1.32 | 3.63 | 0.36 | 0.72 | 1 | C1rb | complement component 1, r subcompone | 135 | 89 | 18 | 0 | 0 | 0 | 0 | 0 | 0 | 0 | 4 |
| 22 | 0.00 | 3.68 | 0.00 | 1.00 | 1 | Slc13a3 | solute carrier family 13 (sodium-depende | 117 | 44 | 84 | 0 | 0 | 0 | 0 | 0 | 0 | 0 | 0 |
| 22 | -1.26 | 1.60 | -0.78 | 0.43 | 1 | Gm17024 | | 36 | 17 | 11 | 0 | 4 | 91 | 0 | 0 | 0 | 0 | 84 |
| 22 | 1.23 | 1.03 | 1.19 | 0.23 | 1 | A430078I02Rik | | 8 | 20 | 30 | 25 | 26 | 15 | 0 | 24 | 39 | 43 | 14 |
| 22 | 0.00 | 3.68 | 0.00 | 1.00 | 1 | Kcne3 | potassium voltage-gated channel, Isk-rela | 106 | 41 | 97 | 0 | 0 | 0 | 0 | 0 | 0 | 0 | 0 |
| 22 | 0.00 | 3.68 | 0.00 | 1.00 | 1 | Gpr18 | G protein-coupled receptor 18 | 164 | 69 | 10 | 0 | 0 | 0 | 0 | 0 | 0 | 0 | 0 |
| 22 | 0.00 | 3.68 | 0.00 | 1.00 | 1 | Foxd3 | forkhead box D3 | 89 | 51 | 103 | 0 | 0 | 0 | 0 | 0 | 0 | 0 | 0 |
| 22 | -0.40 | 3.67 | -0.11 | 0.91 | 1 | Nlrp3 | NLR family, pyrin domain containing 3 | 144 | 29 | 68 | 0 | 0 | 1 | 0 | 0 | 0 | 0 | 0 |
| 22 | -1.16 | 3.63 | -0.32 | 0.75 | 1 | Aldh1a7 | aldehyde dehydrogenase family 1, subfar | 33 | 92 | 111 | 5 | 0 | 0 | 0 | 0 | 0 | 0 | 0 |
| 22 | 1.57 | 1.21 | 1.30 | 0.19 | 1 | Adgrg2 | adhesion G protein-coupled receptor G2 | 113 | 28 | 48 | 10 | 7 | 0 | 0 | 16 | 7 | 8 | 4 |
| 22 | 0.28 | 1.75 | 0.16 | 0.87 | 1 | Tgtp1 | T cell specific GTPase 1 | 61 | 71 | 107 | 0 | 0 | 1 | 0 | 0 | 0 | 0 | 1 |
| 22 | 0.00 | 3.68 | 0.00 | 1.00 | 1 | Cnmd | chondromodulin | 174 | 43 | 24 | 0 | 0 | 0 | 0 | 0 | 0 | 0 | 0 |
| 22 | -3.00 | 1.90 | -1.58 | 0.11 | 1 | Il7 | interleukin 7 | 116 | 63 | 49 | 0 | 11 | 2 | 0 | 0 | 0 | 0 | 0 |
| 22 | 2.36 | 1.37 | 1.73 | 0.08 | 1 | Mst1r | macrophage stimulating 1 receptor (c-me | 149 | 31 | 26 | 0 | 0 | 8 | 0 | 0 | 1 | 10 | 14 |
| 22 | 0.00 | 2.14 | 0.00 | 1.00 | 1 | Alkal2 | ALK and LTK ligand 2 | 93 | 103 | 44 | 0 | 0 | 0 | 0 | 0 | 0 | 0 | 0 |
| 22 | 0.62 | 2.12 | 0.29 | 0.77 | 1 | Tgm3 | transglutaminase 3, E polypeptide | 0 | 108 | 105 | 0 | 0 | 8 | 0 | 0 | 0 | 0 | 18 |
| 22 | 0.00 | 2.14 | 0.00 | 1.00 | 1 | Trpc6 | transient receptor potential cation channe | 130 | 70 | 39 | 0 | 0 | 0 | 0 | 0 | 0 | 0 | 0 |
| 22 | 1.73 | 1.60 | 1.08 | 0.28 | 1 | Sftpa1 | surfactant associated protein A1 | 4 | 9 | 9 | 0 | 0 | 110 | 0 | 0 | 9 | 0 | 96 |
| 22 | -2.05 | 1.49 | -1.37 | 0.17 | 1 | Prdm16os | | 74 | 58 | 95 | 2 | 5 | 3 | 0 | 0 | 2 | 0 | 0 |
| 22 | -1.60 | 2.30 | -0.70 | 0.49 | 1 | Aqp5 | aquaporin 5 | 157 | 66 | 7 | 0 | 7 | 0 | 0 | 0 | 0 | 0 | 0 |
| 21 | 0.00 | 2.15 | 0.00 | 1.00 | 1 | Batf3 | basic leucine zipper transcription factor, A | 111 | 51 | 75 | 0 | 0 | 0 | 0 | 0 | 0 | 0 | 0 |
| 21 | 0.89 | 1.11 | 0.80 | 0.43 | 1 | Cdkn3 | cyclin-dependent kinase inhibitor 3 | 54 | 24 | 10 | 23 | 17 | 14 | 0 | 28 | 29 | 9 | 26 |
| 21 | 0.00 | 2.15 | 0.00 | 1.00 | 1 | Anxa8 | annexin A8 | 145 | 41 | 50 | 0 | 0 | 0 | 0 | 0 | 0 | 0 | 0 |
| 21 | 0.28 | 1.68 | 0.16 | 0.87 | 1 | Il18r1 | interleukin 18 receptor 1 | 71 | 54 | 105 | 0 | 0 | 2 | 0 | 0 | 0 | 0 | 2 |
| 21 | 0.00 | 2.16 | 0.00 | 1.00 | 1 | Arl11 | ADP-ribosylation factor-like 11 | 70 | 80 | 82 | 0 | 0 | 0 | 0 | 0 | 0 | 0 | 0 |
| 21 | -0.20 | 1.60 | -0.12 | 0.90 | 1 | Dnase1l2 | deoxyribonuclease 1-like 2 | 2 | 0 | 8 | 0 | 1 | 104 | 0 | 0 | 0 | 0 | 117 |
| 21 | 1.56 | 2.04 | 0.77 | 0.44 | 1 | Slc6a14 | solute carrier family 6 (neurotransmitter tra | 4 | 140 | 81 | 0 | 0 | 0 | 0 | 0 | 0 | 0 | 5 |
| 21 | 0.77 | 1.71 | 0.45 | 0.65 | 1 | Batf2 | basic leucine zipper transcription factor, A | 63 | 81 | 50 | 0 | 10 | 3 | 0 | 13 | 0 | 0 | 9 |
| 21 | 0.00 | 2.17 | 0.00 | 1.00 | 1 | Lcn2 | lipocalin 2 | 101 | 59 | 70 | 0 | 0 | 0 | 0 | 0 | 0 | 0 | 0 |
| 21 | -0.51 | 1.67 | -0.31 | 0.76 | 1 | Kif7 | kinesin family member 7 | 81 | 51 | 79 | 12 | 0 | 2 | 0 | 0 | 2 | 0 | 3 |

| | | | | | | | | | | | | | | | | | | |
|---|---|---|---|---|---|---|---|---|---|---|---|---|---|---|---|---|---|---|
| 21 | 0.00 | 2.85 | 0.00 | 1.00 | 1 | Ighv9-2 | | 172 | 57 | 0 | 0 | 0 | 0 | 0 | 0 | 0 | 0 | 0 |
| 21 | 0.00 | 2.18 | 0.00 | 1.00 | 1 | Adgrg5 | adhesion G protein-coupled receptor G5 | 36 | 54 | 139 | 0 | 0 | 0 | 0 | 0 | 0 | 0 | 0 |
| 21 | 0.36 | 1.46 | 0.25 | 0.80 | 1 | Trim71 | tripartite motif-containing 71 | 31 | 69 | 96 | 6 | 3 | 4 | 0 | 0 | 0 | 9 | 11 |
| 21 | 1.58 | 1.16 | 1.36 | 0.17 | 1 | K230015D01Rik | | 11 | 68 | 14 | 6 | 26 | 10 | 0 | 19 | 21 | 39 | 15 |
| 21 | -2.19 | 1.85 | -1.18 | 0.24 | 1 | BC106179 | | 78 | 48 | 53 | 6 | 2 | 5 | 0 | 0 | 0 | 36 | 0 |
| 21 | 0.00 | 2.17 | 0.00 | 1.00 | 1 | Alx4 | aristaless-like homeobox 4 | 75 | 77 | 75 | 0 | 0 | 0 | 0 | 0 | 0 | 0 | 0 |
| 21 | 0.00 | 2.19 | 0.00 | 1.00 | 1 | Folh1 | folate hydrolase 1 | 108 | 96 | 23 | 0 | 0 | 0 | 0 | 0 | 0 | 0 | 0 |
| 21 | 0.00 | 2.18 | 0.00 | 1.00 | 1 | Tead4 | TEA domain family member 4 | 75 | 71 | 81 | 0 | 0 | 0 | 0 | 0 | 0 | 0 | 0 |
| 21 | 0.00 | 2.19 | 0.00 | 1.00 | 1 | Dcstamp | dendrocyte expressed seven transmembr | 46 | 23 | 157 | 0 | 0 | 0 | 0 | 0 | 0 | 0 | 0 |
| 21 | 0.00 | 2.18 | 0.00 | 1.00 | 1 | Gpr20 | G protein-coupled receptor 20 | 113 | 43 | 71 | 0 | 0 | 0 | 0 | 0 | 0 | 0 | 0 |
| 21 | 0.00 | 2.20 | 0.00 | 1.00 | 1 | Rarres1 | retinoic acid receptor responder (tazarote | 172 | 19 | 35 | 0 | 0 | 0 | 0 | 0 | 0 | 0 | 0 |
| 21 | 0.00 | 3.61 | 0.00 | 1.00 | 1 | Igkv10-94 | | 226 | 0 | 0 | 0 | 0 | 0 | 0 | 0 | 0 | 0 | 0 |
| 21 | 1.00 | 2.05 | 0.49 | 0.62 | 1 | Cacna1s | calcium channel, voltage-dependent, L ty | 178 | 34 | 12 | 0 | 0 | 0 | 0 | 0 | 1 | 0 | 0 |
| 20 | 0.16 | 1.87 | 0.09 | 0.93 | 1 | Aspa | aspartoacylase | 34 | 67 | 116 | 0 | 3 | 0 | 0 | 0 | 0 | 0 | 5 |
| 20 | -1.80 | 1.77 | -1.02 | 0.31 | 1 | Wfdc2 | WAP four-disulfide core domain 2 | 49 | 64 | 106 | 1 | 0 | 4 | 0 | 0 | 0 | 0 | 0 |
| 20 | -0.07 | 1.58 | -0.05 | 0.96 | 1 | Kcns3 | potassium voltage-gated channel, delaye | 72 | 31 | 35 | 8 | 6 | 8 | 0 | 57 | 0 | 0 | 6 |
| 20 | -0.37 | 1.41 | -0.27 | 0.79 | 1 | Kctd14 | potassium channel tetramerisation domai | 39 | 63 | 83 | 11 | 7 | 2 | 0 | 0 | 13 | 0 | 4 |
| 20 | 0.00 | 2.20 | 0.00 | 1.00 | 1 | Omd | osteomodulin | 105 | 77 | 39 | 0 | 0 | 0 | 0 | 0 | 0 | 0 | 0 |
| 20 | 0.00 | 2.25 | 0.00 | 1.00 | 1 | Igkv2-109 | | 80 | 8 | 133 | 0 | 0 | 0 | 0 | 0 | 0 | 0 | 0 |
| 20 | 3.18 | 1.99 | 1.60 | 0.11 | 1 | Qprt | quinolinate phosphoribosyltransferase | 122 | 16 | 74 | 0 | 0 | 0 | 0 | 7 | 0 | 0 | 2 |
| 20 | -0.85 | 1.97 | -0.43 | 0.67 | 1 | Cdhr5 | cadherin-related family member 5 | 31 | 83 | 105 | 0 | 2 | 0 | 0 | 0 | 0 | 0 | 0 |
| 20 | 0.00 | 2.21 | 0.00 | 1.00 | 1 | Ccl3 | chemokine (C-C motif) ligand 3 | 125 | 21 | 74 | 0 | 0 | 0 | 0 | 0 | 0 | 0 | 0 |
| 20 | 0.00 | 2.20 | 0.00 | 1.00 | 1 | Sh2d1b1 | SH2 domain containing 1B1 | 78 | 61 | 81 | 0 | 0 | 0 | 0 | 0 | 0 | 0 | 0 |
| 20 | 1.28 | 2.14 | 0.60 | 0.55 | 1 | Gm25483 | | 0 | 0 | 0 | 0 | 0 | 132 | 0 | 0 | 4 | 0 | 83 |
| 20 | 0.00 | 2.20 | 0.00 | 1.00 | 1 | Hoxb7 | homeobox B7 | 76 | 70 | 73 | 0 | 0 | 0 | 0 | 0 | 0 | 0 | 0 |
| 20 | 1.68 | 2.03 | 0.83 | 0.41 | 1 | Trim36 | tripartite motif-containing 36 | 114 | 41 | 61 | 0 | 0 | 0 | 0 | 0 | 3 | 0 | 0 |
| 20 | -1.12 | 1.94 | -0.58 | 0.56 | 1 | Il27ra | interleukin 27 receptor, alpha | 95 | 66 | 52 | 0 | 0 | 6 | 0 | 0 | 0 | 0 | 0 |
| 20 | 0.00 | 3.61 | 0.00 | 1.00 | 1 | Igkv8-27 | | 219 | 0 | 0 | 0 | 0 | 0 | 0 | 0 | 0 | 0 | 0 |
| 20 | 0.00 | 2.20 | 0.00 | 1.00 | 1 | Rgs18 | regulator of G-protein signaling 18 | 81 | 62 | 76 | 0 | 0 | 0 | 0 | 0 | 0 | 0 | 0 |
| 20 | 1.58 | 0.94 | 1.69 | 0.09 | 1 | Gm42670 | | 6 | 15 | 45 | 10 | 14 | 30 | 0 | 31 | 26 | 13 | 28 |
| 20 | 0.00 | 2.23 | 0.00 | 1.00 | 1 | Il1a | interleukin 1 alpha | 172 | 18 | 27 | 0 | 0 | 0 | 0 | 0 | 0 | 0 | 0 |
| 20 | 0.00 | 2.21 | 0.00 | 1.00 | 1 | Dclk3 | doublecortin-like kinase 3 | 73 | 71 | 74 | 0 | 0 | 0 | 0 | 0 | 0 | 0 | 0 |
| 20 | 0.00 | 2.29 | 0.00 | 1.00 | 1 | Tnfrsf13c | tumor necrosis factor receptor superfamily | 177 | 34 | 5 | 0 | 0 | 0 | 0 | 0 | 0 | 0 | 0 |
| 20 | -2.76 | 1.91 | -1.44 | 0.15 | 1 | Flvcr2 | feline leukemia virus subgroup C cellular | 75 | 72 | 46 | 13 | 4 | 0 | 0 | 0 | 0 | 0 | 5 |
| 19 | 2.03 | 1.14 | 1.78 | 0.08 | 1 | C530008M17Ri | RIKEN cDNA C530008M17 gene | 39 | 41 | 44 | 8 | 1 | 17 | 0 | 9 | 17 | 28 | 11 |
| 19 | 0.00 | 2.22 | 0.00 | 1.00 | 1 | Cd300lb | CD300 molecule like family member B | 124 | 61 | 30 | 0 | 0 | 0 | 0 | 0 | 0 | 0 | 0 |
| 19 | -1.22 | 2.00 | -0.61 | 0.54 | 1 | Gfap | glial fibrillary acidic protein | 43 | 72 | 96 | 0 | 4 | 0 | 0 | 0 | 0 | 0 | 0 |
| 19 | 0.95 | 1.77 | 0.54 | 0.59 | 1 | Pcdh20 | protocadherin 20 | 89 | 46 | 51 | 9 | 0 | 1 | 0 | 0 | 0 | 15 | 3 |
| 19 | 0.74 | 1.80 | 0.41 | 0.68 | 1 | Rsph10b | PMS1 homolog2, mismatch repair system | 3 | 7 | 6 | 1 | 14 | 36 | 0 | 0 | 0 | 90 | 56 |
| 19 | 0.00 | 2.28 | 0.00 | 1.00 | 1 | Tnfrsf8 | tumor necrosis factor receptor superfamily | 158 | 49 | 6 | 0 | 0 | 0 | 0 | 0 | 0 | 0 | 0 |
| 19 | -1.32 | 2.12 | -0.63 | 0.53 | 1 | Islr2 | immunoglobulin superfamily containing le | 94 | 37 | 78 | 4 | 0 | 0 | 0 | 0 | 0 | 0 | 0 |
| 19 | 0.00 | 2.23 | 0.00 | 1.00 | 1 | Gm10687 | | 57 | 54 | 102 | 0 | 0 | 0 | 0 | 0 | 0 | 0 | 0 |
| 19 | -0.24 | 1.64 | -0.15 | 0.88 | 1 | C4a | | 86 | 9 | 14 | 0 | 0 | 68 | 0 | 0 | 0 | 0 | 36 |
| 19 | 0.00 | 2.24 | 0.00 | 1.00 | 1 | Mdfi | MyoD family inhibitor | 80 | 42 | 88 | 0 | 0 | 0 | 0 | 0 | 0 | 0 | 0 |
| 19 | 0.00 | 2.28 | 0.00 | 1.00 | 1 | Ighv2-9 | | 134 | 68 | 7 | 0 | 0 | 0 | 0 | 0 | 0 | 0 | 0 |
| 19 | 1.59 | 1.58 | 1.01 | 0.31 | 1 | Gm45104 | | 0 | 0 | 0 | 5 | 0 | 72 | 0 | 30 | 1 | 0 | 101 |
| 19 | 0.28 | 1.94 | 0.14 | 0.89 | 1 | Nrn1 | neuritin 1 | 63 | 72 | 55 | 0 | 2 | 1 | 0 | 0 | 0 | 15 | 0 |
| 19 | 2.34 | 2.13 | 1.10 | 0.27 | 1 | Il9r | interleukin 9 receptor | 124 | 38 | 6 | 0 | 0 | 0 | 0 | 0 | 0 | 39 | 0 |
| 19 | -0.41 | 2.01 | -0.20 | 0.84 | 1 | Hrh2 | histamine receptor H2 | 113 | 42 | 52 | 0 | 0 | 1 | 0 | 0 | 0 | 0 | 0 |
| 19 | 0.00 | 2.24 | 0.00 | 1.00 | 1 | Tcf15 | transcription factor 15 | 88 | 65 | 55 | 0 | 0 | 0 | 0 | 0 | 0 | 0 | 0 |
| 19 | 0.07 | 1.94 | 0.04 | 0.97 | 1 | Gm42572 | | 0 | 0 | 0 | 0 | 4 | 85 | 0 | 2 | 0 | 0 | 116 |
| 19 | -0.94 | 1.51 | -0.62 | 0.53 | 1 | Grik4 | glutamate receptor, ionotropic, kainate 4 | 45 | 49 | 82 | 6 | 0 | 15 | 0 | 0 | 2 | 0 | 8 |
| 19 | -0.76 | 2.04 | -0.37 | 0.71 | 1 | Dhrs9 | dehydrogenase/reductase (SDR family) n | 130 | 50 | 16 | 0 | 6 | 1 | 0 | 0 | 0 | 0 | 5 |
| 19 | -0.41 | 2.02 | -0.20 | 0.84 | 1 | BC035044 | cDNA sequence BC035044 | 110 | 65 | 30 | 0 | 0 | 1 | 0 | 0 | 0 | 0 | 0 |
| 19 | 0.00 | 2.99 | 0.00 | 1.00 | 1 | Ighv1-74 | | 35 | 169 | 0 | 0 | 0 | 0 | 0 | 0 | 0 | 0 | 0 |
| 19 | 0.97 | 2.03 | 0.48 | 0.63 | 1 | Gm36161 | | 133 | 33 | 36 | 0 | 0 | 0 | 0 | 0 | 0 | 0 | 1 |
| 19 | -0.41 | 2.02 | -0.20 | 0.84 | 1 | Colca2 | COLCA2 homolog | 60 | 63 | 80 | 0 | 0 | 1 | 0 | 0 | 0 | 0 | 0 |
| 19 | -2.08 | 1.78 | -1.17 | 0.24 | 1 | Aqp8 | aquaporin 8 | 14 | 117 | 67 | 2 | 0 | 4 | 0 | 0 | 0 | 0 | 0 |
| 18 | 1.29 | 2.08 | 0.62 | 0.53 | 1 | Car3 | carbonic anhydrase 3 | 155 | 41 | 5 | 0 | 0 | 0 | 0 | 0 | 0 | 0 | 2 |
| 18 | 0.00 | 2.27 | 0.00 | 1.00 | 1 | Chst4 | carbohydrate sulfotransferase 4 | 100 | 76 | 27 | 0 | 0 | 0 | 0 | 0 | 0 | 0 | 0 |

| | | | | | | | | | | | | | | | | | | |
|---|---|---|---|---|---|---|---|---|---|---|---|---|---|---|---|---|---|---|
| 18 | 0.00 | 2.27 | 0.00 | 1.00 | 1 | Cyp4f14 | cytochrome P450, family 4, subfamily f, po | 38 | 74 | 90 | 0 | 0 | 0 | 0 | 0 | 0 | 0 | 0 |
| 18 | 1.86 | 1.82 | 1.02 | 0.31 | 1 | Gm33378 | | 0 | 66 | 9 | 2 | 12 | 30 | 0 | 0 | 13 | 37 | 32 |
| 18 | 0.00 | 2.27 | 0.00 | 1.00 | 1 | Spaar | small regulatory polypeptide of amino aci | 41 | 111 | 49 | 0 | 0 | 0 | 0 | 0 | 0 | 0 | 0 |
| 18 | -0.70 | 1.47 | -0.48 | 0.63 | 1 | Adora3 | adenosine A3 receptor | 11 | 19 | 21 | 12 | 18 | 44 | 0 | 0 | 51 | 0 | 25 |
| 18 | 0.00 | 3.62 | 0.00 | 1.00 | 1 | Igkv3-3 | | 201 | 0 | 0 | 0 | 0 | 0 | 0 | 0 | 0 | 0 | 0 |
| 18 | -1.39 | 1.41 | -0.99 | 0.32 | 1 | Depdc1b | DEP domain containing 1B | 104 | 10 | 43 | 12 | 13 | 7 | 0 | 6 | 3 | 0 | 3 |
| 18 | 3.09 | 2.01 | 1.54 | 0.12 | 1 | Gm33869 | | 0 | 0 | 0 | 0 | 0 | 110 | 0 | 0 | 7 | 6 | 77 |
| 18 | 0.00 | 2.27 | 0.00 | 1.00 | 1 | Alas2 | aminolevulinic acid synthase 2, erythroid | 51 | 75 | 74 | 0 | 0 | 0 | 0 | 0 | 0 | 0 | 0 |
| 18 | 1.05 | 1.58 | 0.67 | 0.50 | 1 | Gabrg2 | gamma-aminobutyric acid (GABA) A rece | 57 | 62 | 37 | 22 | 0 | 1 | 0 | 5 | 0 | 12 | 3 |
| 18 | 0.00 | 2.30 | 0.00 | 1.00 | 1 | H2-Ea | histocompatibility 2, class II antigen E alpl | 141 | 13 | 45 | 0 | 0 | 0 | 0 | 0 | 0 | 0 | 0 |
| 18 | 0.00 | 2.30 | 0.00 | 1.00 | 1 | Mybpc2 | myosin binding protein C, fast-type | 112 | 14 | 73 | 0 | 0 | 0 | 0 | 0 | 0 | 0 | 0 |
| 18 | 0.00 | 2.99 | 0.00 | 1.00 | 1 | Ighv1-53 | | 160 | 37 | 0 | 0 | 0 | 0 | 0 | 0 | 0 | 0 | 0 |
| 18 | -0.08 | 1.95 | -0.04 | 0.97 | 1 | Gm44321 | | 19 | 7 | 0 | 0 | 3 | 79 | 0 | 0 | 0 | 24 | 65 |
| 18 | 1.08 | 1.78 | 0.61 | 0.54 | 1 | Lilr4b | leukocyte immunoglobulin-like receptor, s | 137 | 24 | 30 | 0 | 0 | 1 | 0 | 0 | 0 | 0 | 5 |
| 18 | 0.00 | 3.05 | 0.00 | 1.00 | 1 | Igkv4-78 | | 184 | 13 | 0 | 0 | 0 | 0 | 0 | 0 | 0 | 0 | 0 |
| 18 | 1.86 | 1.92 | 0.97 | 0.33 | 1 | Ctla4 | cytotoxic T-lymphocyte-associated proteir | 132 | 44 | 0 | 0 | 0 | 3 | 0 | 0 | 0 | 14 | 4 |
| 18 | 0.00 | 2.29 | 0.00 | 1.00 | 1 | Gm12002 | | 68 | 46 | 82 | 0 | 0 | 0 | 0 | 0 | 0 | 0 | 0 |
| 18 | 0.64 | 1.70 | 0.38 | 0.71 | 1 | Lrrc4c | leucine rich repeat containing 4C | 40 | 72 | 47 | 4 | 11 | 0 | 0 | 17 | 0 | 0 | 3 |
| 18 | 0.00 | 2.29 | 0.00 | 1.00 | 1 | Rtn4rl2 | reticulon 4 receptor-like 2 | 50 | 48 | 98 | 0 | 0 | 0 | 0 | 0 | 0 | 0 | 0 |
| 18 | 0.00 | 2.29 | 0.00 | 1.00 | 1 | Xirp1 | xin actin-binding repeat containing 1 | 104 | 52 | 39 | 0 | 0 | 0 | 0 | 0 | 0 | 0 | 0 |
| 18 | 0.00 | 3.15 | 0.00 | 1.00 | 1 | Ighv8-12 | | 194 | 1 | 0 | 0 | 0 | 0 | 0 | 0 | 0 | 0 | 0 |
| 18 | -1.14 | 2.08 | -0.55 | 0.58 | 1 | Onecut1 | one cut domain, family member 1 | 76 | 54 | 61 | 0 | 3 | 0 | 0 | 0 | 0 | 0 | 0 |
| 18 | 0.00 | 2.30 | 0.00 | 1.00 | 1 | Adamts14 | a disintegrin-like and metallopeptidase (r | 83 | 53 | 58 | 0 | 0 | 0 | 0 | 0 | 0 | 0 | 0 |
| 18 | -0.61 | 1.36 | -0.45 | 0.65 | 1 | Afm | afamin | 16 | 24 | 36 | 0 | 4 | 73 | 0 | 0 | 2 | 0 | 38 |
| 18 | 1.34 | 2.09 | 0.64 | 0.52 | 1 | Gm42659 | | 20 | 0 | 0 | 12 | 0 | 87 | 0 | 14 | 5 | 0 | 55 |
| 17 | 0.00 | 2.31 | 0.00 | 1.00 | 1 | Wnt11 | wingless-type MMTV integration site famil | 114 | 27 | 51 | 0 | 0 | 0 | 0 | 0 | 0 | 0 | 0 |
| 17 | -1.45 | 1.87 | -0.78 | 0.44 | 1 | Paqr6 | progestin and adipoQ receptor family mer | 90 | 14 | 85 | 0 | 2 | 1 | 0 | 0 | 0 | 0 | 0 |
| 17 | 0.00 | 2.31 | 0.00 | 1.00 | 1 | Eva1a | eva-1 homolog A (C. elegans) | 131 | 28 | 32 | 0 | 0 | 0 | 0 | 0 | 0 | 0 | 0 |
| 17 | 1.95 | 1.02 | 1.92 | 0.05 | 1 | Gm14264 | | 4 | 24 | 10 | 12 | 2 | 36 | 0 | 60 | 12 | 6 | 23 |
| 17 | -0.73 | 2.05 | -0.36 | 0.72 | 1 | Hoxb3 | homeobox B3 | 130 | 36 | 23 | 0 | 0 | 2 | 0 | 0 | 0 | 0 | 0 |
| 17 | 0.11 | 1.90 | 0.06 | 0.96 | 1 | Gm43039 | | 3 | 6 | 0 | 0 | 4 | 73 | 0 | 0 | 0 | 6 | 97 |
| 17 | 0.82 | 1.95 | 0.42 | 0.67 | 1 | Gm48878 | | 57 | 69 | 60 | 0 | 0 | 1 | 0 | 0 | 2 | 0 | 0 |
| 17 | 0.00 | 2.32 | 0.00 | 1.00 | 1 | Hsd11b2 | hydroxysteroid 11-beta dehydrogenase 2 | 56 | 63 | 71 | 0 | 0 | 0 | 0 | 0 | 0 | 0 | 0 |
| 17 | -1.54 | 2.19 | -0.70 | 0.48 | 1 | Wdr86 | WD repeat domain 86 | 51 | 69 | 60 | 0 | 9 | 0 | 0 | 0 | 0 | 0 | 0 |
| 17 | 0.00 | 3.05 | 0.00 | 1.00 | 1 | Ighv11-2 | | 38 | 151 | 0 | 0 | 0 | 0 | 0 | 0 | 0 | 0 | 0 |
| 17 | 0.00 | 3.61 | 0.00 | 1.00 | 1 | Ighv12-3 | | 0 | 188 | 0 | 0 | 0 | 0 | 0 | 0 | 0 | 0 | 0 |
| 17 | -1.46 | 1.61 | -0.90 | 0.37 | 1 | Gm43328 | | 2 | 31 | 18 | 8 | 0 | 74 | 0 | 0 | 0 | 0 | 56 |
| 17 | -2.17 | 1.49 | -1.45 | 0.15 | 1 | Pcdhb3 | protocadherin beta 3 | 103 | 24 | 34 | 4 | 8 | 10 | 0 | 0 | 3 | 0 | 2 |
| 17 | -0.41 | 2.09 | -0.20 | 0.84 | 1 | Glp2r | glucagon-like peptide 2 receptor | 118 | 12 | 56 | 0 | 0 | 1 | 0 | 0 | 0 | 0 | 0 |
| 17 | 0.00 | 2.33 | 0.00 | 1.00 | 1 | Slc25a48 | solute carrier family 25, member 48 | 57 | 51 | 80 | 0 | 0 | 0 | 0 | 0 | 0 | 0 | 0 |
| 17 | 0.00 | 2.33 | 0.00 | 1.00 | 1 | Hrct1 | histidine rich carboxyl terminus 1 | 77 | 46 | 64 | 0 | 0 | 0 | 0 | 0 | 0 | 0 | 0 |
| 17 | 0.80 | 1.46 | 0.55 | 0.58 | 1 | 9030025P20Ril | RIKEN cDNA 9030025P20 gene | 17 | 16 | 12 | 20 | 5 | 26 | 0 | 0 | 12 | 52 | 26 |
| 17 | 0.67 | 1.64 | 0.41 | 0.68 | 1 | Pvt1 | | 52 | 59 | 34 | 0 | 9 | 4 | 0 | 0 | 9 | 0 | 18 |
| 17 | -0.18 | 1.54 | -0.12 | 0.91 | 1 | Ager | advanced glycosylation end product-spec | 21 | 18 | 18 | 0 | 1 | 56 | 0 | 0 | 0 | 0 | 73 |
| 17 | 0.00 | 3.08 | 0.00 | 1.00 | 1 | Clca1 | chloride channel accessory 1 | 0 | 92 | 94 | 0 | 0 | 0 | 0 | 0 | 0 | 0 | 0 |
| 17 | 0.00 | 2.34 | 0.00 | 1.00 | 1 | Ly6c2 | lymphocyte antigen 6 complex, locus C2 | 118 | 50 | 18 | 0 | 0 | 0 | 0 | 0 | 0 | 0 | 0 |
| 17 | 3.07 | 2.06 | 1.49 | 0.14 | 1 | Gm37963 | | 21 | 0 | 5 | 0 | 0 | 89 | 0 | 0 | 11 | 0 | 60 |
| 17 | 1.90 | 2.18 | 0.87 | 0.38 | 1 | Runx2 | runt related transcription factor 2 | 58 | 61 | 61 | 0 | 0 | 0 | 0 | 0 | 5 | 0 | 0 |
| 17 | 0.00 | 2.34 | 0.00 | 1.00 | 1 | Cd209d | CD209d antigen | 99 | 35 | 51 | 0 | 0 | 0 | 0 | 0 | 0 | 0 | 0 |
| 17 | 0.00 | 2.34 | 0.00 | 1.00 | 1 | Pof1b | premature ovarian failure 1B | 76 | 33 | 75 | 0 | 0 | 0 | 0 | 0 | 0 | 0 | 0 |
| 17 | 0.00 | 2.34 | 0.00 | 1.00 | 1 | Padi4 | peptidyl arginine deiminase, type IV | 63 | 66 | 56 | 0 | 0 | 0 | 0 | 0 | 0 | 0 | 0 |
| 17 | 0.00 | 3.62 | 0.00 | 1.00 | 1 | Ighv1-14 | | 184 | 0 | 0 | 0 | 0 | 0 | 0 | 0 | 0 | 0 | 0 |
| 17 | 0.67 | 1.15 | 0.59 | 0.56 | 1 | Gm28424 | | 28 | 19 | 8 | 18 | 11 | 14 | 0 | 15 | 7 | 48 | 16 |
| 17 | 3.00 | 1.93 | 1.55 | 0.12 | 1 | Gata6os | | 58 | 27 | 60 | 0 | 0 | 11 | 0 | 0 | 0 | 23 | 4 |
| 17 | 0.00 | 2.34 | 0.00 | 1.00 | 1 | Batf | basic leucine zipper transcription factor, A | 100 | 55 | 29 | 0 | 0 | 0 | 0 | 0 | 0 | 0 | 0 |
| 17 | 0.00 | 2.34 | 0.00 | 1.00 | 1 | Arsj | arylsulfatase J | 89 | 30 | 64 | 0 | 0 | 0 | 0 | 0 | 0 | 0 | 0 |
| 17 | -0.96 | 1.32 | -0.72 | 0.47 | 1 | Syt6 | synaptotagmin VI | 17 | 64 | 59 | 15 | 3 | 9 | 0 | 13 | 0 | 0 | 4 |
| 17 | -0.72 | 1.79 | -0.40 | 0.69 | 1 | Gm42937 | | 16 | 1 | 3 | 6 | 0 | 64 | 0 | 0 | 1 | 0 | 91 |
| 17 | 0.00 | 2.35 | 0.00 | 1.00 | 1 | Smpdl3b | sphingomyelin phosphodiesterase, acid-l | 73 | 58 | 52 | 0 | 0 | 0 | 0 | 0 | 0 | 0 | 0 |
| 17 | 0.00 | 2.36 | 0.00 | 1.00 | 1 | Gjb5 | gap junction protein, beta 5 | 132 | 27 | 23 | 0 | 0 | 0 | 0 | 0 | 0 | 0 | 0 |

| | | | | | | | | | | | | | | | | | | |
|---|---|---|---|---|---|---|---|---|---|---|---|---|---|---|---|---|---|---|
| 17 | 1.00 | 2.11 | 0.47 | 0.64 | 1 | Foxc2 | forkhead box C2 | 78 | 33 | 70 | 0 | 0 | 0 | 0 | 1 | 0 | 0 | 0 |
| 17 | 0.09 | 1.67 | 0.05 | 0.96 | 1 | Exd1 | exonuclease 3'-5' domain containing 1 | 43 | 33 | 14 | 0 | 33 | 20 | 0 | 0 | 11 | 21 | 7 |
| 17 | 0.00 | 3.09 | 0.00 | 1.00 | 1 | Igkv4-56 | | 154 | 27 | 0 | 0 | 0 | 0 | 0 | 0 | 0 | 0 | 0 |
| 16 | 0.96 | 1.84 | 0.52 | 0.60 | 1 | Bnc2 | basonuclin 2 | 127 | 10 | 39 | 0 | 0 | 1 | 0 | 0 | 0 | 0 | 4 |
| 16 | 0.00 | 2.35 | 0.00 | 1.00 | 1 | Sost | sclerostin | 59 | 54 | 69 | 0 | 0 | 0 | 0 | 0 | 0 | 0 | 0 |
| 16 | 0.00 | 2.35 | 0.00 | 1.00 | 1 | Klrb1b | killer cell lectin-like receptor subfamily B r | 79 | 62 | 41 | 0 | 0 | 0 | 0 | 0 | 0 | 0 | 0 |
| 16 | -1.32 | 1.86 | -0.71 | 0.48 | 1 | Gm43961 | | 15 | 0 | 3 | 0 | 3 | 95 | 0 | 0 | 0 | 0 | 66 |
| 16 | 0.54 | 1.94 | 0.28 | 0.78 | 1 | Nell2 | NEL-like 2 | 42 | 37 | 95 | 2 | 0 | 0 | 0 | 0 | 5 | 0 | 0 |
| 16 | 0.00 | 2.48 | 0.00 | 1.00 | 1 | Igkv12-44 | | 162 | 4 | 15 | 0 | 0 | 0 | 0 | 0 | 0 | 0 | 0 |
| 16 | -1.23 | 2.13 | -0.58 | 0.56 | 1 | Vtcn1 | V-set domain containing T cell activation i | 36 | 74 | 67 | 0 | 4 | 0 | 0 | 0 | 0 | 0 | 0 |
| 16 | 1.41 | 2.08 | 0.68 | 0.50 | 1 | Igfbp2 | insulin-like growth factor binding protein 2 | 129 | 26 | 22 | 0 | 0 | 0 | 0 | 0 | 0 | 0 | 3 |
| 16 | 2.75 | 1.58 | 1.74 | 0.08 | 1 | Btnl7-ps | | 26 | 70 | 21 | 0 | 8 | 3 | 0 | 0 | 23 | 18 | 11 |
| 16 | 0.15 | 1.98 | 0.07 | 0.94 | 1 | Mamdc4 | MAM domain containing 4 | 4 | 0 | 0 | 0 | 8 | 79 | 0 | 0 | 8 | 0 | 80 |
| 16 | -3.36 | 1.79 | -1.87 | 0.06 | 1 | 2610028H24Ril | RIKEN cDNA 2610028H24 gene | 46 | 50 | 70 | 11 | 1 | 2 | 0 | 0 | 0 | 0 | 0 |
| 16 | -1.86 | 2.01 | -0.93 | 0.35 | 1 | Itih4 | inter alpha-trypsin inhibitor, heavy chain 4 | 76 | 40 | 50 | 8 | 0 | 1 | 0 | 0 | 0 | 0 | 5 |
| 16 | 0.00 | 2.38 | 0.00 | 1.00 | 1 | Plin4 | perilipin 4 | 136 | 28 | 16 | 0 | 0 | 0 | 0 | 0 | 0 | 0 | 0 |
| 16 | -1.77 | 2.27 | -0.78 | 0.44 | 1 | Cd80 | CD80 antigen | 109 | 58 | 2 | 11 | 0 | 0 | 0 | 0 | 0 | 0 | 0 |
| 16 | 1.42 | 2.06 | 0.69 | 0.49 | 1 | Dio2 | deiodinase, iodothyronine, type II | 70 | 58 | 48 | 0 | 0 | 0 | 0 | 0 | 0 | 0 | 3 |
| 16 | 0.00 | 3.19 | 0.00 | 1.00 | 1 | Igkv4-79 | | 177 | 2 | 0 | 0 | 0 | 0 | 0 | 0 | 0 | 0 | 0 |
| 16 | 0.00 | 2.38 | 0.00 | 1.00 | 1 | Smim41 | small integral membrane protein 41 | 78 | 24 | 76 | 0 | 0 | 0 | 0 | 0 | 0 | 0 | 0 |
| 16 | 0.00 | 2.37 | 0.00 | 1.00 | 1 | B3gnt5 | UDP-GlcNAc:betaGal beta-1,3-N-acetylgl | 61 | 86 | 31 | 0 | 0 | 0 | 0 | 0 | 0 | 0 | 0 |
| 16 | 0.00 | 2.37 | 0.00 | 1.00 | 1 | Gm11816 | | 99 | 28 | 51 | 0 | 0 | 0 | 0 | 0 | 0 | 0 | 0 |
| 16 | -1.38 | 2.25 | -0.61 | 0.54 | 1 | Mcemp1 | mast cell expressed membrane protein 1 | 72 | 71 | 30 | 0 | 5 | 0 | 0 | 0 | 0 | 0 | 0 |
| 16 | -2.00 | 2.32 | -0.86 | 0.39 | 1 | BC055308 | | 17 | 9 | 0 | 0 | 23 | 81 | 0 | 0 | 0 | 0 | 48 |
| 16 | -1.06 | 2.07 | -0.51 | 0.61 | 1 | Cd3e | CD3 antigen, epsilon polypeptide | 88 | 47 | 37 | 0 | 0 | 6 | 0 | 0 | 0 | 0 | 0 |
| 16 | -2.91 | 1.91 | -1.53 | 0.13 | 1 | Bcl11b | B cell leukemia/lymphoma 11B | 49 | 30 | 83 | 10 | 0 | 4 | 0 | 0 | 1 | 0 | 0 |
| 16 | -0.29 | 2.02 | -0.14 | 0.88 | 1 | Megf10 | multiple EGF-like-domains 10 | 45 | 81 | 38 | 0 | 9 | 0 | 0 | 0 | 4 | 0 | 0 |
| 16 | 0.00 | 2.39 | 0.00 | 1.00 | 1 | Fam180a | family with sequence similarity 180, meml | 118 | 41 | 18 | 0 | 0 | 0 | 0 | 0 | 0 | 0 | 0 |
| 16 | -2.46 | 2.02 | -1.22 | 0.22 | 1 | Nfe2l3 | nuclear factor, erythroid derived 2, like 3 | 63 | 45 | 57 | 0 | 10 | 1 | 0 | 0 | 0 | 0 | 0 |
| 16 | 4.14 | 2.49 | 1.66 | 0.10 | 1 | Rpl24 | ribosomal protein L24 | 0 | 15 | 0 | 0 | 0 | 51 | 0 | 0 | 0 | 14 | 95 |
| 16 | 0.00 | 2.40 | 0.00 | 1.00 | 1 | Ppbp | pro-platelet basic protein | 29 | 129 | 18 | 0 | 0 | 0 | 0 | 0 | 0 | 0 | 0 |
| 16 | -0.98 | 1.82 | -0.54 | 0.59 | 1 | Acap1 | ArfGAP with coiled-coil, ankyrin repeat an | 124 | 23 | 18 | 0 | 3 | 3 | 0 | 0 | 0 | 0 | 4 |
| 16 | 2.13 | 1.88 | 1.13 | 0.26 | 1 | Gja5 | gap junction protein, alpha 5 | 68 | 38 | 64 | 0 | 0 | 0 | 0 | 0 | 1 | 0 | 3 |
| 16 | 0.00 | 3.25 | 0.00 | 1.00 | 1 | Ighv1-22 | | 173 | 1 | 0 | 0 | 0 | 0 | 0 | 0 | 0 | 0 | 0 |
| 16 | -0.21 | 1.47 | -0.14 | 0.89 | 1 | Urah | urate (5-hydroxyiso-) hydrolase | 17 | 40 | 48 | 11 | 6 | 18 | 0 | 0 | 3 | 26 | 7 |
| 16 | 0.00 | 3.09 | 0.00 | 1.00 | 1 | Igkv12-38 | | 129 | 46 | 0 | 0 | 0 | 0 | 0 | 0 | 0 | 0 | 0 |
| 16 | -1.14 | 2.23 | -0.51 | 0.61 | 1 | Phf24 | PHD finger protein 24 | 94 | 26 | 51 | 3 | 0 | 0 | 0 | 0 | 0 | 0 | 0 |
| 16 | 0.00 | 2.39 | 0.00 | 1.00 | 1 | Wnt6 | wingless-type MMTV integration site famil | 106 | 31 | 36 | 0 | 0 | 0 | 0 | 0 | 0 | 0 | 0 |
| 16 | 0.00 | 2.39 | 0.00 | 1.00 | 1 | Cd244a | CD244 molecule A | 80 | 43 | 51 | 0 | 0 | 0 | 0 | 0 | 0 | 0 | 0 |
| 16 | 2.05 | 1.43 | 1.43 | 0.15 | 1 | Gm43951 | RIKEN cDNA 4933427D14 gene | 29 | 11 | 16 | 0 | 2 | 45 | 0 | 0 | 5 | 22 | 44 |
| 16 | 0.65 | 1.04 | 0.63 | 0.53 | 1 | Sdr9c7 | 4short chain dehydrogenase/reductase fa | 7 | 71 | 41 | 12 | 3 | 9 | 0 | 19 | 3 | 4 | 5 |
| 16 | -0.44 | 2.14 | -0.21 | 0.84 | 1 | Aadac | arylacetamide deacetylase | 89 | 25 | 57 | 0 | 1 | 0 | 0 | 0 | 0 | 0 | 0 |
| 16 | 1.97 | 2.19 | 0.90 | 0.37 | 1 | Wfdc18 | WAP four-disulfide core domain 18 | 52 | 61 | 53 | 0 | 0 | 0 | 0 | 0 | 6 | 0 | 0 |
| 16 | 0.00 | 2.40 | 0.00 | 1.00 | 1 | Fcrl1 | Fc receptor-like 1 | 109 | 23 | 39 | 0 | 0 | 0 | 0 | 0 | 0 | 0 | 0 |
| 16 | 0.00 | 3.17 | 0.00 | 1.00 | 1 | Iglv3 | | 156 | 15 | 0 | 0 | 0 | 0 | 0 | 0 | 0 | 0 | 0 |
| 16 | 0.00 | 2.42 | 0.00 | 1.00 | 1 | Bdh2 | 3-hydroxybutyrate dehydrogenase, type 2 | 83 | 14 | 74 | 0 | 0 | 0 | 0 | 0 | 0 | 0 | 0 |
| 16 | -0.41 | 2.13 | -0.19 | 0.85 | 1 | Gm11346 | | 23 | 64 | 83 | 0 | 0 | 1 | 0 | 0 | 0 | 0 | 0 |
| 16 | 0.13 | 1.37 | 0.09 | 0.93 | 1 | B230216N24Rik | | 20 | 14 | 54 | 0 | 2 | 46 | 0 | 0 | 3 | 0 | 31 |
| 15 | 1.44 | 2.20 | 0.66 | 0.51 | 1 | Tgm1 | transglutaminase 1, K polypeptide | 92 | 49 | 28 | 0 | 0 | 0 | 0 | 0 | 2 | 0 | 0 |
| 15 | 0.00 | 2.48 | 0.00 | 1.00 | 1 | Gpr55 | G protein-coupled receptor 55 | 123 | 45 | 3 | 0 | 0 | 0 | 0 | 0 | 0 | 0 | 0 |
| 15 | 2.10 | 1.99 | 1.05 | 0.29 | 1 | Gm37968 | | 21 | 14 | 0 | 0 | 0 | 48 | 0 | 0 | 0 | 24 | 62 |
| 15 | -0.56 | 1.66 | -0.34 | 0.73 | 1 | Gm40557 | | 18 | 48 | 63 | 16 | 0 | 10 | 0 | 0 | 3 | 0 | 12 |
| 15 | -0.26 | 1.87 | -0.14 | 0.89 | 1 | Rbfox1 | RNA binding protein, fox-1 homolog (C. e | 74 | 78 | 15 | 0 | 0 | 3 | 0 | 0 | 0 | 0 | 1 |
| 15 | 0.00 | 2.41 | 0.00 | 1.00 | 1 | Eya4 | EYA transcriptional coactivator and phosp | 54 | 68 | 48 | 0 | 0 | 0 | 0 | 0 | 0 | 0 | 0 |
| 15 | -0.41 | 2.14 | -0.19 | 0.85 | 1 | Nr0b2 | nuclear receptor subfamily 0, group B, me | 24 | 75 | 70 | 0 | 0 | 1 | 0 | 0 | 0 | 0 | 0 |
| 15 | -0.30 | 1.60 | -0.19 | 0.85 | 1 | Clcf1 | cardiotrophin-like cytokine factor 1 | 55 | 44 | 53 | 0 | 3 | 8 | 0 | 0 | 3 | 0 | 3 |
| 15 | 1.83 | 1.57 | 1.17 | 0.24 | 1 | 1810053B23Rik | | 6 | 77 | 74 | 0 | 1 | 1 | 0 | 0 | 4 | 0 | 6 |
| 15 | 0.00 | 2.43 | 0.00 | 1.00 | 1 | Fndc7 | fibronectin type III domain containing 7 | 117 | 38 | 14 | 0 | 0 | 0 | 0 | 0 | 0 | 0 | 0 |
| 15 | 0.10 | 2.09 | 0.05 | 0.96 | 1 | Kcnq5 | potassium voltage-gated channel, subfam | 78 | 49 | 34 | 4 | 0 | 0 | 0 | 0 | 3 | 0 | 0 |
| 15 | -2.29 | 1.35 | -1.69 | 0.09 | 1 | Trpm6 | transient receptor potential cation channe | 8 | 19 | 42 | 15 | 41 | 23 | 0 | 0 | 16 | 0 | 5 |

| | | | | | | | | | | | | | | | | | | |
|---|---|---|---|---|---|---|---|---|---|---|---|---|---|---|---|---|---|---|
| 15 | 0.80 | 2.12 | 0.38 | 0.71 | 1 | Drp2 | dystrophin related protein 2 | 74 | 45 | 41 | 0 | 0 | 5 | 0 | 0 | 4 | 0 | 0 |
| 15 | 0.00 | 2.43 | 0.00 | 1.00 | 1 | Gm13861 | | 79 | 24 | 64 | 0 | 0 | 0 | 0 | 0 | 0 | 0 | 0 |
| 15 | -2.55 | 2.09 | -1.22 | 0.22 | 1 | Cxcr5 | chemokine (C-X-C motif) receptor 5 | 124 | 14 | 22 | 5 | 0 | 2 | 0 | 0 | 0 | 0 | 0 |
| 15 | 1.38 | 1.66 | 0.83 | 0.40 | 1 | Cox6b2 | cytochrome c oxidase subunit 6B2 | 28 | 51 | 30 | 0 | 9 | 10 | 0 | 16 | 0 | 18 | 5 |
| 15 | 1.04 | 1.06 | 0.98 | 0.33 | 1 | Kansl2-ps | | 14 | 42 | 11 | 6 | 14 | 14 | 0 | 32 | 13 | 4 | 16 |
| 15 | 1.93 | 2.24 | 0.86 | 0.39 | 1 | Cd300lf | CD300 molecule like family member F | 74 | 47 | 36 | 0 | 0 | 4 | 0 | 0 | 6 | 0 | 0 |
| 15 | -0.68 | 2.10 | -0.32 | 0.75 | 1 | 1700048O20Rik | | 66 | 57 | 34 | 8 | 0 | 0 | 0 | 0 | 0 | 0 | 2 |
| 15 | -0.46 | 1.81 | -0.25 | 0.80 | 1 | Lcat | lecithin cholesterol acyltransferase | 33 | 48 | 45 | 24 | 0 | 6 | 0 | 0 | 7 | 0 | 5 |
| 15 | -0.31 | 1.37 | -0.23 | 0.82 | 1 | Fbxl22 | F-box and leucine-rich repeat protein 22 | 55 | 28 | 57 | 1 | 3 | 10 | 0 | 0 | 3 | 0 | 9 |
| 15 | -2.21 | 2.09 | -1.06 | 0.29 | 1 | Sfmbt2 | Scm-like with four mbt domains 2 | 90 | 31 | 23 | 0 | 10 | 2 | 0 | 0 | 0 | 0 | 9 |
| 15 | 0.00 | 2.43 | 0.00 | 1.00 | 1 | Tdh | L-threonine dehydrogenase | 59 | 72 | 34 | 0 | 0 | 0 | 0 | 0 | 0 | 0 | 0 |
| 15 | 2.06 | 2.24 | 0.92 | 0.36 | 1 | Fam181b | family with sequence similarity 181, meml | 39 | 66 | 52 | 0 | 0 | 0 | 0 | 0 | 8 | 0 | 0 |
| 15 | 0.00 | 2.43 | 0.00 | 1.00 | 1 | Ch25h | cholesterol 25-hydroxylase | 66 | 62 | 37 | 0 | 0 | 0 | 0 | 0 | 0 | 0 | 0 |
| 15 | 0.13 | 1.93 | 0.07 | 0.95 | 1 | Gm36120 | | 17 | 85 | 49 | 0 | 7 | 0 | 0 | 0 | 1 | 0 | 5 |
| 15 | -0.37 | 1.83 | -0.20 | 0.84 | 1 | Il22ra1 | interleukin 22 receptor, alpha 1 | 22 | 100 | 26 | 6 | 0 | 1 | 0 | 0 | 0 | 0 | 8 |
| 15 | 1.54 | 1.86 | 0.83 | 0.41 | 1 | Gm15246 | predicted gene 15246 | 15 | 0 | 18 | 6 | 0 | 53 | 0 | 21 | 3 | 0 | 48 |
| 15 | 0.00 | 3.20 | 0.00 | 1.00 | 1 | Lpo | lactoperoxidase | 0 | 85 | 78 | 0 | 0 | 0 | 0 | 0 | 0 | 0 | 0 |
| 15 | 0.00 | 2.45 | 0.00 | 1.00 | 1 | Stac | src homology three (SH3) and cysteine ric | 97 | 36 | 29 | 0 | 0 | 0 | 0 | 0 | 0 | 0 | 0 |
| 15 | 1.61 | 2.66 | 0.60 | 0.55 | 1 | Gpr174 | G protein-coupled receptor 174 | 122 | 12 | 0 | 0 | 0 | 0 | 0 | 0 | 0 | 28 | 0 |
| 15 | 0.00 | 2.46 | 0.00 | 1.00 | 1 | Adam33 | a disintegrin and metallopeptidase domai | 97 | 14 | 51 | 0 | 0 | 0 | 0 | 0 | 0 | 0 | 0 |
| 15 | -0.26 | 1.82 | -0.14 | 0.89 | 1 | Gm20394 | | 9 | 5 | 1 | 0 | 0 | 100 | 0 | 0 | 0 | 0 | 47 |
| 15 | 0.00 | 2.45 | 0.00 | 1.00 | 1 | Bhlhe22 | basic helix-loop-helix family, member e22 | 73 | 62 | 27 | 0 | 0 | 0 | 0 | 0 | 0 | 0 | 0 |
| 15 | -1.46 | 1.96 | -0.74 | 0.46 | 1 | Mas1 | MAS1 oncogene | 27 | 68 | 64 | 0 | 2 | 1 | 0 | 0 | 0 | 0 | 0 |
| 15 | 0.00 | 2.51 | 0.00 | 1.00 | 1 | Ptx3 | pentraxin related gene | 136 | 6 | 20 | 0 | 0 | 0 | 0 | 0 | 0 | 0 | 0 |
| 15 | -0.16 | 1.69 | -0.09 | 0.92 | 1 | 9430037G07Rik | | 13 | 17 | 0 | 17 | 29 | 19 | 0 | 7 | 36 | 0 | 23 |
| 15 | 0.00 | 2.46 | 0.00 | 1.00 | 1 | Serpina3g | serine (or cysteine) peptidase inhibitor, cl | 108 | 33 | 20 | 0 | 0 | 0 | 0 | 0 | 0 | 0 | 0 |
| 15 | -0.26 | 1.91 | -0.14 | 0.89 | 1 | Gm12250 | | 66 | 21 | 60 | 0 | 2 | 3 | 0 | 0 | 0 | 9 | 0 |
| 15 | 0.86 | 2.18 | 0.40 | 0.69 | 1 | Cd8a | CD8 antigen, alpha chain | 130 | 1 | 26 | 0 | 0 | 1 | 0 | 0 | 2 | 0 | 0 |
| 15 | 1.15 | 1.14 | 1.00 | 0.32 | 1 | Gm47920 | | 17 | 4 | 19 | 7 | 9 | 32 | 0 | 12 | 24 | 21 | 16 |
| 15 | -1.46 | 1.62 | -0.90 | 0.37 | 1 | Rps10 | ribosomal protein S10 | 22 | 7 | 0 | 4 | 7 | 60 | 0 | 0 | 1 | 4 | 55 |
| 15 | 0.00 | 2.46 | 0.00 | 1.00 | 1 | Hoxa4 | homeobox A4 | 62 | 59 | 39 | 0 | 0 | 0 | 0 | 0 | 0 | 0 | 0 |
| 14 | 0.96 | 2.18 | 0.44 | 0.66 | 1 | Nxpe5 | neurexophilin and PC-esterase domain fa | 100 | 11 | 48 | 0 | 0 | 0 | 0 | 0 | 0 | 0 | 1 |
| 14 | 3.33 | 1.74 | 1.91 | 0.06 | 1 | Fam129c | family with sequence similarity 129, meml | 15 | 6 | 9 | 0 | 0 | 66 | 0 | 13 | 4 | 0 | 46 |
| 14 | 0.00 | 2.47 | 0.00 | 1.00 | 1 | Hoxb8 | homeobox B8 | 36 | 101 | 22 | 0 | 0 | 0 | 0 | 0 | 0 | 0 | 0 |
| 14 | -1.69 | 1.75 | -0.96 | 0.33 | 1 | Gm43059 | | 12 | 16 | 0 | 11 | 2 | 63 | 0 | 0 | 1 | 0 | 55 |
| 14 | -1.30 | 2.34 | -0.56 | 0.58 | 1 | Prokr1 | prokineticin receptor 1 | 47 | 16 | 91 | 4 | 0 | 0 | 0 | 0 | 0 | 0 | 0 |
| 14 | -0.69 | 1.79 | -0.39 | 0.70 | 1 | 6430590A07Rik | | 9 | 14 | 0 | 8 | 5 | 45 | 0 | 0 | 5 | 0 | 73 |
| 14 | 0.00 | 2.49 | 0.00 | 1.00 | 1 | Klrk1 | killer cell lectin-like receptor subfamily K, | 112 | 35 | 10 | 0 | 0 | 0 | 0 | 0 | 0 | 0 | 0 |
| 14 | 3.14 | 1.95 | 1.61 | 0.11 | 1 | Zfp977 | zinc finger protein 977 | 3 | 6 | 0 | 21 | 0 | 20 | 0 | 0 | 17 | 76 | 14 |
| 14 | 2.00 | 2.28 | 0.88 | 0.38 | 1 | Best1 | bestrophin 1 | 83 | 31 | 37 | 0 | 0 | 0 | 0 | 0 | 6 | 0 | 0 |
| 14 | 0.12 | 1.79 | 0.07 | 0.95 | 1 | Gm38057 | | 28 | 14 | 3 | 0 | 0 | 58 | 0 | 0 | 0 | 0 | 54 |
| 14 | 0.00 | 2.49 | 0.00 | 1.00 | 1 | Gata5 | GATA binding protein 5 | 78 | 66 | 14 | 0 | 0 | 0 | 0 | 0 | 0 | 0 | 0 |
| 14 | 0.00 | 3.19 | 0.00 | 1.00 | 1 | Ighv1-19 | | 51 | 106 | 0 | 0 | 0 | 0 | 0 | 0 | 0 | 0 | 0 |
| 14 | -0.70 | 2.79 | -0.25 | 0.80 | 1 | Muc2 | | 0 | 26 | 129 | 0 | 0 | 2 | 0 | 0 | 0 | 0 | 0 |
| 14 | -0.75 | 1.57 | -0.48 | 0.63 | 1 | Gm48267 | | 0 | 14 | 0 | 7 | 2 | 77 | 0 | 29 | 0 | 0 | 27 |
| 14 | 2.01 | 1.24 | 1.63 | 0.10 | 1 | Gm9844 | | 33 | 22 | 22 | 6 | 2 | 4 | 0 | 52 | 7 | 5 | 2 |
| 14 | 1.40 | 2.15 | 0.65 | 0.52 | 1 | Il15 | interleukin 15 | 74 | 25 | 54 | 0 | 0 | 0 | 0 | 0 | 0 | 0 | 3 |
| 14 | 0.00 | 3.62 | 0.00 | 1.00 | 1 | Igkv2-112 | | 0 | 156 | 0 | 0 | 0 | 0 | 0 | 0 | 0 | 0 | 0 |
| 14 | 3.78 | 2.00 | 1.89 | 0.06 | 1 | Naip1 | NLR family, apoptosis inhibitory protein 1 | 47 | 21 | 30 | 1 | 0 | 0 | 0 | 52 | 0 | 0 | 4 |
| 14 | 1.60 | 1.82 | 0.88 | 0.38 | 1 | Ttpa | tocopherol (alpha) transfer protein | 43 | 57 | 47 | 0 | 2 | 0 | 0 | 0 | 4 | 0 | 3 |
| 14 | 0.00 | 2.49 | 0.00 | 1.00 | 1 | Ifi211 | interferon activated gene 211 | 76 | 58 | 21 | 0 | 0 | 0 | 0 | 0 | 0 | 0 | 0 |
| 14 | 0.00 | 2.49 | 0.00 | 1.00 | 1 | Gm41381 | | 54 | 47 | 54 | 0 | 0 | 0 | 0 | 0 | 0 | 0 | 0 |
| 14 | 1.61 | 1.72 | 0.94 | 0.35 | 1 | Gm48063 | | 1 | 0 | 10 | 4 | 0 | 64 | 0 | 20 | 9 | 0 | 46 |
| 14 | 1.20 | 1.88 | 0.64 | 0.52 | 1 | Sirpb1a | signal-regulatory protein beta 1A | 46 | 42 | 58 | 0 | 0 | 1 | 0 | 0 | 0 | 0 | 7 |
| 14 | 2.44 | 1.58 | 1.55 | 0.12 | 1 | Gm30531 | | 10 | 0 | 2 | 7 | 0 | 30 | 0 | 22 | 3 | 52 | 28 |
| 14 | 0.40 | 1.55 | 0.26 | 0.80 | 1 | Vnn3 | vanin 3 | 58 | 41 | 25 | 0 | 9 | 6 | 0 | 0 | 9 | 3 | 3 |
| 14 | 0.00 | 2.50 | 0.00 | 1.00 | 1 | Dhh | desert hedgehog | 44 | 56 | 54 | 0 | 0 | 0 | 0 | 0 | 0 | 0 | 0 |
| 14 | -1.58 | 2.43 | -0.65 | 0.51 | 1 | Gm5431 | predicted gene 5431 | 73 | 17 | 54 | 9 | 0 | 0 | 0 | 0 | 0 | 0 | 0 |
| 14 | -1.46 | 2.31 | -0.63 | 0.53 | 1 | Prss35 | protease, serine 35 | 25 | 24 | 88 | 0 | 15 | 0 | 0 | 0 | 0 | 0 | 0 |
| 14 | -0.32 | 1.92 | -0.17 | 0.87 | 1 | Gm37581 | | 2 | 0 | 1 | 5 | 0 | 75 | 0 | 0 | 5 | 0 | 64 |

| | | | | | | | | | | | | | | | | | | |
|---|---|---|---|---|---|---|---|---|---|---|---|---|---|---|---|---|---|---|
| 14 | 2.14 | 1.34 | 1.60 | 0.11 | 1 | Spaca9 | sperm acrosome associated 9 | 39 | 50 | 18 | 0 | 6 | 3 | 0 | 7 | 6 | 18 | 5 |
| 14 | 1.74 | 2.15 | 0.81 | 0.42 | 1 | Ces1e | carboxylesterase 1E | 15 | 20 | 109 | 0 | 0 | 0 | 0 | 0 | 7 | 0 | 0 |
| 14 | -1.97 | 1.40 | -1.40 | 0.16 | 1 | Gm42636 | | 9 | 11 | 10 | 32 | 47 | 19 | 0 | 0 | 10 | 5 | 7 |
| 14 | 1.81 | 2.27 | 0.80 | 0.42 | 1 | Hkdc1 | hexokinase domain containing 1 | 53 | 56 | 38 | 0 | 0 | 0 | 0 | 0 | 4 | 0 | 0 |
| 14 | -0.44 | 2.26 | -0.20 | 0.84 | 1 | Gm21188 | predicted gene, 21188 | 106 | 33 | 11 | 0 | 1 | 0 | 0 | 0 | 0 | 0 | 0 |
| 14 | 0.00 | 2.57 | 0.00 | 1.00 | 1 | Gria4 | glutamate receptor, ionotropic, AMPA4 (a | 128 | 18 | 5 | 0 | 0 | 0 | 0 | 0 | 0 | 0 | 0 |
| 14 | 0.00 | 2.52 | 0.00 | 1.00 | 1 | 5033428I22Rik | | 97 | 42 | 12 | 0 | 0 | 0 | 0 | 0 | 0 | 0 | 0 |
| 14 | 0.00 | 3.31 | 0.00 | 1.00 | 1 | Dsc3 | desmocollin 3 | 142 | 9 | 0 | 0 | 0 | 0 | 0 | 0 | 0 | 0 | 0 |
| 14 | 4.64 | 2.64 | 1.76 | 0.08 | 1 | Slc39a2 | solute carrier family 39 (zinc transporter), | 0 | 0 | 4 | 0 | 0 | 57 | 0 | 34 | 0 | 0 | 55 |
| 14 | -0.72 | 2.22 | -0.32 | 0.75 | 1 | 2610203C22Rik | | 105 | 10 | 33 | 0 | 0 | 2 | 0 | 0 | 0 | 0 | 0 |
| 14 | 0.00 | 2.53 | 0.00 | 1.00 | 1 | Twist1 | twist basic helix-loop-helix transcription fa | 64 | 21 | 63 | 0 | 0 | 0 | 0 | 0 | 0 | 0 | 0 |
| 14 | 1.42 | 2.82 | 0.50 | 0.61 | 1 | Grem1 | gremlin 1, DAN family BMP antagonist | 118 | 26 | 0 | 0 | 0 | 0 | 0 | 0 | 0 | 0 | 5 |
| 14 | 0.95 | 1.86 | 0.51 | 0.61 | 1 | Ceacam2 | carcinoembryonic antigen-related cell adl | 0 | 5 | 95 | 0 | 20 | 0 | 0 | 0 | 28 | 0 | 1 |
| 13 | 0.00 | 2.53 | 0.00 | 1.00 | 1 | 4833403J16Rik | | 28 | 74 | 46 | 0 | 0 | 0 | 0 | 0 | 0 | 0 | 0 |
| 13 | 1.28 | 2.20 | 0.58 | 0.56 | 1 | Sh3rf3 | SH3 domain containing ring finger 3 | 57 | 20 | 69 | 0 | 0 | 0 | 0 | 0 | 0 | 0 | 2 |
| 13 | 0.00 | 2.54 | 0.00 | 1.00 | 1 | Slc30a2 | solute carrier family 30 (zinc transporter), | 29 | 59 | 59 | 0 | 0 | 0 | 0 | 0 | 0 | 0 | 0 |
| 13 | 0.00 | 2.54 | 0.00 | 1.00 | 1 | Dio3 | deiodinase, iodothyronine type III | 35 | 29 | 82 | 0 | 0 | 0 | 0 | 0 | 0 | 0 | 0 |
| 13 | 1.53 | 2.30 | 0.67 | 0.51 | 1 | Sox8 | SRY (sex determining region Y)-box 8 | 51 | 86 | 4 | 0 | 0 | 3 | 0 | 0 | 3 | 0 | 0 |
| 13 | 0.00 | 2.56 | 0.00 | 1.00 | 1 | Adamts3 | a disintegrin-like and metallopeptidase (r | 112 | 12 | 22 | 0 | 0 | 0 | 0 | 0 | 0 | 0 | 0 |
| 13 | 0.28 | 2.41 | 0.12 | 0.91 | 1 | Agr2 | anterior gradient 2 | 0 | 34 | 108 | 0 | 0 | 2 | 0 | 0 | 0 | 0 | 2 |
| 13 | 0.00 | 2.63 | 0.00 | 1.00 | 1 | Twist2 | twist basic helix-loop-helix transcription fa | 106 | 38 | 2 | 0 | 0 | 0 | 0 | 0 | 0 | 0 | 0 |
| 13 | -0.21 | 2.12 | -0.10 | 0.92 | 1 | Foxd2 | forkhead box D2 | 38 | 33 | 69 | 3 | 0 | 0 | 0 | 0 | 0 | 0 | 3 |
| 13 | 0.00 | 2.55 | 0.00 | 1.00 | 1 | Bcl2a1d | B cell leukemia/lymphoma 2 related prote | 69 | 45 | 31 | 0 | 0 | 0 | 0 | 0 | 0 | 0 | 0 |
| 13 | -0.50 | 1.47 | -0.34 | 0.73 | 1 | 4933406C10Rik | | 8 | 6 | 18 | 8 | 5 | 42 | 0 | 0 | 14 | 0 | 45 |
| 13 | -0.40 | 2.82 | -0.14 | 0.89 | 1 | Gsdmc4 | gasdermin C4 | 0 | 47 | 98 | 0 | 0 | 1 | 0 | 0 | 0 | 0 | 0 |
| 13 | 0.00 | 3.31 | 0.00 | 1.00 | 1 | Ighv1-69 | | 123 | 22 | 0 | 0 | 0 | 0 | 0 | 0 | 0 | 0 | 0 |
| 13 | 3.14 | 2.16 | 1.46 | 0.15 | 1 | Epop | elongin BC and polycomb repressive com | 105 | 28 | 0 | 0 | 0 | 1 | 0 | 0 | 5 | 4 | 2 |
| 13 | -1.78 | 1.67 | -1.06 | 0.29 | 1 | Chil1 | chitinase-like 1 | 56 | 0 | 15 | 6 | 27 | 14 | 0 | 0 | 11 | 5 | 11 |
| 13 | 0.26 | 1.83 | 0.14 | 0.89 | 1 | Lat | linker for activation of T cells | 34 | 12 | 38 | 7 | 0 | 17 | 0 | 0 | 0 | 9 | 26 |
| 13 | -0.49 | 1.68 | -0.29 | 0.77 | 1 | Brd8dc | BRD8 domain containing | 9 | 0 | 11 | 0 | 5 | 52 | 0 | 0 | 1 | 0 | 65 |
| 13 | 0.00 | 2.56 | 0.00 | 1.00 | 1 | Hspb2 | heat shock protein 2 | 92 | 33 | 19 | 0 | 0 | 0 | 0 | 0 | 0 | 0 | 0 |
| 13 | 0.00 | 2.56 | 0.00 | 1.00 | 1 | Pglyrp1 | peptidoglycan recognition protein 1 | 74 | 22 | 48 | 0 | 0 | 0 | 0 | 0 | 0 | 0 | 0 |
| 13 | 1.82 | 1.43 | 1.28 | 0.20 | 1 | Catsper4 | cation channel, sperm associated 4 | 10 | 24 | 11 | 0 | 7 | 21 | 0 | 7 | 16 | 0 | 47 |
| 13 | 1.41 | 2.16 | 0.65 | 0.52 | 1 | Ccdc13 | coiled-coil domain containing 13 | 41 | 68 | 20 | 0 | 0 | 1 | 0 | 14 | 0 | 0 | 0 |
| 13 | -1.57 | 1.45 | -1.08 | 0.28 | 1 | Gm45786 | | 4 | 18 | 36 | 26 | 6 | 25 | 0 | 0 | 5 | 0 | 21 |
| 13 | 0.18 | 1.48 | 0.12 | 0.91 | 1 | A430105J06Rik | | 0 | 23 | 0 | 19 | 9 | 36 | 0 | 14 | 15 | 0 | 26 |
| 13 | 0.97 | 1.13 | 0.86 | 0.39 | 1 | Gm16574 | | 15 | 10 | 10 | 5 | 8 | 36 | 0 | 22 | 6 | 9 | 21 |
| 13 | -0.34 | 1.40 | -0.24 | 0.81 | 1 | Gbp6 | guanylate binding protein 6 | 51 | 40 | 8 | 8 | 4 | 10 | 0 | 0 | 2 | 10 | 9 |
| 13 | 0.00 | 3.31 | 0.00 | 1.00 | 1 | Ighv9-4 | | 106 | 35 | 0 | 0 | 0 | 0 | 0 | 0 | 0 | 0 | 0 |
| 13 | -0.40 | 2.88 | -0.14 | 0.89 | 1 | Sval1 | seminal vesicle antigen-like 1 | 0 | 29 | 110 | 0 | 0 | 1 | 0 | 0 | 0 | 0 | 0 |
| 13 | 0.00 | 2.60 | 0.00 | 1.00 | 1 | Saa3 | serum amyloid A 3 | 82 | 9 | 49 | 0 | 0 | 0 | 0 | 0 | 0 | 0 | 0 |
| 13 | 0.00 | 2.58 | 0.00 | 1.00 | 1 | Gm16551 | | 45 | 42 | 54 | 0 | 0 | 0 | 0 | 0 | 0 | 0 | 0 |
| 13 | 0.00 | 3.34 | 0.00 | 1.00 | 1 | Reg4 | regenerating islet-derived family, member | 0 | 45 | 96 | 0 | 0 | 0 | 0 | 0 | 0 | 0 | 0 |
| 13 | 0.00 | 2.59 | 0.00 | 1.00 | 1 | H19 | | 26 | 32 | 81 | 0 | 0 | 0 | 0 | 0 | 0 | 0 | 0 |
| 13 | -1.69 | 1.76 | -0.96 | 0.34 | 1 | Macc1 | metastasis associated in colon cancer 1 | 56 | 11 | 62 | 0 | 2 | 7 | 0 | 0 | 0 | 0 | 1 |
| 13 | 0.00 | 2.59 | 0.00 | 1.00 | 1 | 5830432E09Rik | | 65 | 48 | 26 | 0 | 0 | 0 | 0 | 0 | 0 | 0 | 0 |
| 13 | -1.52 | 1.98 | -0.77 | 0.44 | 1 | Abca14 | ATP-binding cassette, sub-family A (ABC1 | 3 | 0 | 9 | 0 | 13 | 68 | 0 | 0 | 0 | 0 | 45 |
| 13 | 1.22 | 1.19 | 1.03 | 0.30 | 1 | Gm3510 | | 4 | 17 | 6 | 7 | 6 | 24 | 0 | 32 | 13 | 0 | 28 |
| 13 | 1.44 | 1.61 | 0.90 | 0.37 | 1 | Zpbp2 | zona pellucida binding protein 2 | 0 | 0 | 4 | 3 | 17 | 14 | 0 | 73 | 17 | 0 | 10 |
| 12 | 1.92 | 1.39 | 1.39 | 0.17 | 1 | Gm7832 | | 30 | 27 | 5 | 1 | 11 | 4 | 0 | 34 | 9 | 3 | 13 |
| 12 | 0.00 | 3.62 | 0.00 | 1.00 | 1 | Ighv2-7 | | 137 | 0 | 0 | 0 | 0 | 0 | 0 | 0 | 0 | 0 | 0 |
| 12 | 0.30 | 1.78 | 0.17 | 0.86 | 1 | Mfsd2b | major facilitator superfamily domain conta | 7 | 2 | 0 | 0 | 8 | 52 | 0 | 0 | 2 | 9 | 57 |
| 12 | -0.41 | 2.28 | -0.18 | 0.86 | 1 | Phf11a | PHD finger protein 11A | 64 | 55 | 17 | 0 | 0 | 1 | 0 | 0 | 0 | 0 | 0 |
| 12 | 0.66 | 2.27 | 0.29 | 0.77 | 1 | Mmp25 | matrix metallopeptidase 25 | 75 | 25 | 26 | 3 | 0 | 0 | 0 | 0 | 7 | 0 | 0 |
| 12 | 0.25 | 3.03 | 0.08 | 0.93 | 1 | Gm27000 | | 0 | 0 | 0 | 0 | 0 | 64 | 0 | 0 | 0 | 0 | 72 |
| 12 | 1.42 | 2.59 | 0.55 | 0.58 | 1 | Gm31557 | | 0 | 0 | 0 | 0 | 0 | 66 | 0 | 0 | 3 | 0 | 67 |
| 12 | 0.00 | 2.61 | 0.00 | 1.00 | 1 | Stat4 | signal transducer and activator of transcri | 66 | 43 | 27 | 0 | 0 | 0 | 0 | 0 | 0 | 0 | 0 |
| 12 | 1.77 | 1.85 | 0.96 | 0.34 | 1 | Atp2c2 | ATPase, Ca++ transporting, type 2C, men | 5 | 79 | 41 | 0 | 0 | 3 | 0 | 0 | 5 | 0 | 3 |
| 12 | -1.00 | 2.26 | -0.44 | 0.66 | 1 | D030034A15Rik | | 40 | 54 | 35 | 0 | 0 | 6 | 0 | 0 | 0 | 0 | 0 |
| 12 | 0.00 | 2.62 | 0.00 | 1.00 | 1 | Ripor3 | RIPOR family member 3 | 46 | 17 | 72 | 0 | 0 | 0 | 0 | 0 | 0 | 0 | 0 |

| | | | | | | | | | | | | | | | | | | |
|---|---|---|---|---|---|---|---|---|---|---|---|---|---|---|---|---|---|---|
| 12 | -0.76 | 1.81 | -0.42 | 0.68 | 1 | Gm43000 | | 31 | 42 | 8 | 7 | 21 | 3 | 0 | 0 | 0 | 13 | 11 |
| 12 | -0.09 | 1.67 | -0.05 | 0.96 | 1 | Stac3 | SH3 and cysteine rich domain 3 | 0 | 9 | 14 | 0 | 3 | 56 | 0 | 0 | 2 | 0 | 52 |
| 12 | 0.38 | 2.35 | 0.16 | 0.87 | 1 | Gm46658 | | 27 | 44 | 46 | 8 | 0 | 0 | 0 | 0 | 0 | 10 | 0 |
| 12 | 0.39 | 1.70 | 0.23 | 0.82 | 1 | Tnnt1 | troponin T1, skeletal, slow | 33 | 14 | 31 | 0 | 11 | 8 | 0 | 0 | 2 | 31 | 4 |
| 12 | 0.00 | 2.63 | 0.00 | 1.00 | 1 | A530064D06Ri | RIKEN cDNA A530064D06 gene | 100 | 13 | 22 | 0 | 0 | 0 | 0 | 0 | 0 | 0 | 0 |
| 12 | -0.41 | 2.29 | -0.18 | 0.86 | 1 | Duox2 | dual oxidase 2 | 69 | 18 | 46 | 0 | 0 | 1 | 0 | 0 | 0 | 0 | 0 |
| 12 | 2.24 | 2.44 | 0.92 | 0.36 | 1 | Apoc1 | apolipoprotein C-I | 38 | 14 | 65 | 0 | 0 | 0 | 0 | 0 | 0 | 17 | 0 |
| 12 | -0.41 | 2.29 | -0.18 | 0.86 | 1 | Lsmem2 | leucine-rich single-pass membrane prote | 49 | 47 | 37 | 0 | 0 | 1 | 0 | 0 | 0 | 0 | 0 |
| 12 | 1.34 | 2.26 | 0.59 | 0.55 | 1 | Zeb2os | | 43 | 24 | 64 | 0 | 0 | 0 | 0 | 0 | 2 | 0 | 0 |
| 12 | 0.00 | 3.52 | 0.00 | 1.00 | 1 | Igkv6-23 | | 132 | 1 | 0 | 0 | 0 | 0 | 0 | 0 | 0 | 0 | 0 |
| 12 | 1.43 | 2.12 | 0.67 | 0.50 | 1 | Pla2g5 | phospholipase A2, group V | 84 | 15 | 2 | 0 | 2 | 6 | 0 | 21 | 3 | 0 | 0 |
| 12 | 1.93 | 2.46 | 0.78 | 0.43 | 1 | Gm30211 | | 103 | 4 | 4 | 17 | 0 | 0 | 0 | 0 | 4 | 0 | 1 |
| 12 | -0.15 | 1.87 | -0.08 | 0.94 | 1 | Gm37818 | | 21 | 12 | 15 | 0 | 0 | 54 | 0 | 0 | 0 | 0 | 30 |
| 12 | 0.00 | 2.64 | 0.00 | 1.00 | 1 | Aoc1 | amine oxidase, copper-containing 1 | 16 | 43 | 74 | 0 | 0 | 0 | 0 | 0 | 0 | 0 | 0 |
| 12 | 0.00 | 2.63 | 0.00 | 1.00 | 1 | Dio3os | | 55 | 50 | 28 | 0 | 0 | 0 | 0 | 0 | 0 | 0 | 0 |
| 12 | 0.00 | 2.64 | 0.00 | 1.00 | 1 | Tmigd1 | transmembrane and immunoglobulin dom | 17 | 84 | 31 | 0 | 0 | 0 | 0 | 0 | 0 | 0 | 0 |
| 12 | -1.05 | 2.06 | -0.51 | 0.61 | 1 | Gm3716 | | 32 | 36 | 62 | 1 | 1 | 0 | 0 | 0 | 0 | 0 | 0 |
| 12 | -0.42 | 1.70 | -0.25 | 0.81 | 1 | 4930518I15Rik | | 9 | 7 | 14 | 0 | 1 | 52 | 0 | 0 | 0 | 0 | 50 |
| 12 | 0.29 | 2.54 | 0.11 | 0.91 | 1 | Doc2g | double C2, gamma | 1 | 0 | 0 | 0 | 0 | 58 | 0 | 0 | 0 | 0 | 72 |
| 12 | -0.96 | 2.31 | -0.41 | 0.68 | 1 | Zfp536 | zinc finger protein 536 | 54 | 63 | 9 | 0 | 0 | 5 | 0 | 0 | 0 | 0 | 0 |
| 12 | -0.18 | 1.39 | -0.13 | 0.90 | 1 | Gm36608 | | 16 | 8 | 32 | 10 | 21 | 7 | 0 | 0 | 10 | 8 | 19 |
| 12 | -0.25 | 1.61 | -0.16 | 0.87 | 1 | Slc9b2 | solute carrier family 9, subfamily B (NHA2 | 28 | 17 | 69 | 0 | 9 | 2 | 0 | 0 | 3 | 0 | 3 |
| 12 | -2.97 | 1.85 | -1.61 | 0.11 | 1 | Gm11454 | | 3 | 70 | 6 | 9 | 23 | 5 | 0 | 0 | 0 | 0 | 14 |
| 12 | 1.27 | 1.98 | 0.64 | 0.52 | 1 | Gm49601 | | 0 | 6 | 21 | 0 | 0 | 46 | 0 | 0 | 5 | 0 | 52 |
| 12 | 0.00 | 2.65 | 0.00 | 1.00 | 1 | Adra1d | adrenergic receptor, alpha 1d | 69 | 24 | 36 | 0 | 0 | 0 | 0 | 0 | 0 | 0 | 0 |
| 12 | 0.00 | 2.68 | 0.00 | 1.00 | 1 | Hao2 | hydroxyacid oxidase 2 | 9 | 59 | 61 | 0 | 0 | 0 | 0 | 0 | 0 | 0 | 0 |
| 12 | 0.71 | 1.51 | 0.47 | 0.64 | 1 | Prmt8 | protein arginine N-methyltransferase 8 | 1 | 37 | 17 | 37 | 0 | 5 | 0 | 7 | 7 | 0 | 18 |
| 12 | -0.25 | 1.78 | -0.14 | 0.89 | 1 | Ccnjl | cyclin J-like | 37 | 27 | 29 | 17 | 1 | 2 | 0 | 0 | 0 | 13 | 3 |
| 12 | 0.00 | 3.41 | 0.00 | 1.00 | 1 | Prg2 | proteoglycan 2, bone marrow | 96 | 32 | 0 | 0 | 0 | 0 | 0 | 0 | 0 | 0 | 0 |
| 12 | 0.00 | 2.66 | 0.00 | 1.00 | 1 | Tbx3os1 | | 43 | 38 | 47 | 0 | 0 | 0 | 0 | 0 | 0 | 0 | 0 |
| 12 | -2.59 | 2.10 | -1.23 | 0.22 | 1 | 4732463B04Rik | | 8 | 41 | 72 | 5 | 0 | 3 | 0 | 0 | 0 | 0 | 0 |
| 12 | -1.57 | 1.50 | -1.04 | 0.30 | 1 | Acpp | acid phosphatase, prostate | 1 | 72 | 37 | 6 | 7 | 2 | 0 | 0 | 1 | 0 | 2 |
| 12 | 0.00 | 2.67 | 0.00 | 1.00 | 1 | Fgf18 | fibroblast growth factor 18 | 67 | 25 | 35 | 0 | 0 | 0 | 0 | 0 | 0 | 0 | 0 |
| 12 | 1.45 | 2.33 | 0.62 | 0.53 | 1 | Dcdc2b | doublecortin domain containing 2b | 2 | 0 | 0 | 0 | 0 | 67 | 0 | 0 | 3 | 0 | 56 |
| 12 | -0.71 | 2.35 | -0.30 | 0.76 | 1 | Dnaaf3 | dynein, axonemal assembly factor 3 | 96 | 9 | 20 | 0 | 0 | 2 | 0 | 0 | 0 | 0 | 0 |
| 12 | 0.00 | 3.62 | 0.00 | 1.00 | 1 | Ighv10-1 | | 0 | 127 | 0 | 0 | 0 | 0 | 0 | 0 | 0 | 0 | 0 |
| 11 | 0.00 | 2.68 | 0.00 | 1.00 | 1 | Klk8 | kallikrein related-peptidase 8 | 40 | 45 | 42 | 0 | 0 | 0 | 0 | 0 | 0 | 0 | 0 |
| 11 | 0.00 | 2.68 | 0.00 | 1.00 | 1 | Cd200r1 | CD200 receptor 1 | 67 | 35 | 24 | 0 | 0 | 0 | 0 | 0 | 0 | 0 | 0 |
| 11 | 1.62 | 2.11 | 0.77 | 0.44 | 1 | Slc34a2 | solute carrier family 34 (sodium phosphat | 69 | 20 | 35 | 0 | 0 | 0 | 0 | 0 | 1 | 0 | 1 |
| 11 | 0.00 | 2.68 | 0.00 | 1.00 | 1 | Itih3 | inter-alpha trypsin inhibitor, heavy chain 3 | 69 | 23 | 34 | 0 | 0 | 0 | 0 | 0 | 0 | 0 | 0 |
| 11 | -1.30 | 2.36 | -0.55 | 0.58 | 1 | Slc16a5 | solute carrier family 16 (monocarboxylic a | 25 | 66 | 29 | 6 | 0 | 0 | 0 | 0 | 0 | 0 | 0 |
| 11 | 1.27 | 1.90 | 0.67 | 0.50 | 1 | Gm28729 | predicted gene 28729 | 37 | 7 | 42 | 4 | 0 | 5 | 0 | 25 | 0 | 0 | 6 |
| 11 | -0.71 | 2.33 | -0.31 | 0.76 | 1 | Fam169b | family with sequence similarity 169, meml | 86 | 13 | 25 | 0 | 0 | 2 | 0 | 0 | 0 | 0 | 0 |
| 11 | 3.82 | 2.39 | 1.60 | 0.11 | 1 | Gm48562 | | 0 | 0 | 0 | 0 | 0 | 36 | 0 | 39 | 4 | 0 | 46 |
| 11 | 0.00 | 2.69 | 0.00 | 1.00 | 1 | Ccl19 | chemokine (C-C motif) ligand 19 | 80 | 18 | 27 | 0 | 0 | 0 | 0 | 0 | 0 | 0 | 0 |
| 11 | 0.00 | 2.73 | 0.00 | 1.00 | 1 | Apol7c | apolipoprotein L 7c | 106 | 7 | 11 | 0 | 0 | 0 | 0 | 0 | 0 | 0 | 0 |
| 11 | 0.15 | 1.94 | 0.08 | 0.94 | 1 | Rilp | Rab interacting lysosomal protein | 59 | 37 | 5 | 0 | 0 | 12 | 0 | 0 | 0 | 0 | 11 |
| 11 | 0.00 | 2.70 | 0.00 | 1.00 | 1 | Clec2i | C-type lectin domain family 2, member i | 35 | 75 | 15 | 0 | 0 | 0 | 0 | 0 | 0 | 0 | 0 |
| 11 | -0.88 | 1.66 | -0.53 | 0.60 | 1 | Cdr1os | | 3 | 64 | 9 | 9 | 1 | 26 | 0 | 0 | 0 | 4 | 8 |
| 11 | 2.50 | 1.37 | 1.82 | 0.07 | 1 | Nat1 | N-acetyl transferase 1 | 16 | 13 | 15 | 3 | 4 | 5 | 0 | 16 | 35 | 0 | 17 |
| 11 | 0.49 | 1.54 | 0.32 | 0.75 | 1 | Ska1 | spindle and kinetochore associated comp | 28 | 9 | 11 | 8 | 7 | 13 | 0 | 37 | 8 | 0 | 3 |
| 11 | 3.10 | 2.15 | 1.44 | 0.15 | 1 | Krt17 | keratin 17 | 55 | 23 | 35 | 0 | 0 | 0 | 0 | 0 | 8 | 0 | 2 |
| 11 | -0.07 | 1.56 | -0.04 | 0.96 | 1 | Tmem82 | transmembrane protein 82 | 15 | 19 | 7 | 8 | 8 | 7 | 0 | 12 | 2 | 46 | 0 |
| 11 | 0.36 | 1.59 | 0.22 | 0.82 | 1 | Gm20501 | | 0 | 20 | 14 | 1 | 6 | 33 | 0 | 0 | 11 | 0 | 39 |
| 11 | 0.00 | 2.71 | 0.00 | 1.00 | 1 | Tmc5 | transmembrane channel-like gene family | 40 | 29 | 53 | 0 | 0 | 0 | 0 | 0 | 0 | 0 | 0 |
| 11 | 0.89 | 1.40 | 0.63 | 0.53 | 1 | Gm32200 | | 1 | 13 | 0 | 10 | 17 | 16 | 0 | 25 | 8 | 13 | 19 |
| 11 | 0.38 | 1.40 | 0.27 | 0.79 | 1 | Gm8615 | | 2 | 1 | 0 | 3 | 7 | 44 | 0 | 5 | 7 | 0 | 55 |
| 11 | 0.00 | 2.71 | 0.00 | 1.00 | 1 | Hoxb6 | homeobox B6 | 46 | 36 | 39 | 0 | 0 | 0 | 0 | 0 | 0 | 0 | 0 |
| 11 | 1.69 | 2.38 | 0.71 | 0.48 | 1 | Wnt10b | wingless-type MMTV integration site famil | 60 | 8 | 50 | 0 | 0 | 0 | 0 | 0 | 4 | 0 | 0 |
| 11 | 1.55 | 3.01 | 0.51 | 0.61 | 1 | Tg | thyroglobulin | 115 | 4 | 0 | 0 | 0 | 0 | 0 | 0 | 0 | 0 | 3 |

| | | | | | | | | | | | | | | | | | | |
|---|---|---|---|---|---|---|---|---|---|---|---|---|---|---|---|---|---|---|
| 11 | 0.00 | 3.47 | 0.00 | 1.00 | 1 | Mcoln2 | mucolipin 2 | 70 | 52 | 0 | 0 | 0 | 0 | 0 | 0 | 0 | 0 | 0 |
| 11 | 0.08 | 1.99 | 0.04 | 0.97 | 1 | Gm15915 | | 29 | 54 | 34 | 0 | 0 | 3 | 0 | 0 | 0 | 0 | 2 |
| 11 | 0.50 | 1.62 | 0.31 | 0.76 | 1 | Jakmip2 | janus kinase and microtubule interacting | 21 | 37 | 36 | 0 | 2 | 12 | 0 | 0 | 5 | 0 | 8 |
| 11 | -4.40 | 2.23 | -1.97 | 0.05 | 1 | Gm48822 | | 5 | 0 | 0 | 5 | 2 | 67 | 0 | 0 | 0 | 0 | 42 |
| 11 | -2.53 | 1.48 | -1.71 | 0.09 | 1 | Gm10649 | | 21 | 27 | 38 | 9 | 8 | 13 | 0 | 0 | 2 | 0 | 3 |
| 11 | 0.91 | 1.23 | 0.74 | 0.46 | 1 | Yjefn3 | YjeF N-terminal domain containing 3 | 9 | 12 | 19 | 7 | 9 | 17 | 0 | 17 | 5 | 18 | 7 |
| 11 | 0.48 | 2.27 | 0.21 | 0.83 | 1 | Gm42941 | | 0 | 22 | 0 | 0 | 1 | 32 | 0 | 0 | 1 | 0 | 64 |
| 11 | 0.00 | 2.73 | 0.00 | 1.00 | 1 | Hoxa7 | homeobox A7 | 61 | 36 | 23 | 0 | 0 | 0 | 0 | 0 | 0 | 0 | 0 |
| 11 | 0.00 | 2.73 | 0.00 | 1.00 | 1 | Srms | src-related kinase lacking C-terminal regu | 57 | 33 | 30 | 0 | 0 | 0 | 0 | 0 | 0 | 0 | 0 |
| 11 | 0.00 | 3.62 | 0.00 | 1.00 | 1 | Ighv6-3 | | 120 | 0 | 0 | 0 | 0 | 0 | 0 | 0 | 0 | 0 | 0 |
| 11 | 0.00 | 2.74 | 0.00 | 1.00 | 1 | Cytl1 | cytokine-like 1 | 60 | 13 | 47 | 0 | 0 | 0 | 0 | 0 | 0 | 0 | 0 |
| 11 | 1.01 | 1.87 | 0.54 | 0.59 | 1 | Nwd1 | NACHT and WD repeat domain containin | 33 | 60 | 9 | 3 | 0 | 3 | 0 | 0 | 8 | 0 | 4 |
| 11 | 0.62 | 1.99 | 0.31 | 0.76 | 1 | 4930556M19Rik | | 57 | 22 | 34 | 0 | 0 | 2 | 0 | 0 | 0 | 0 | 4 |
| 11 | -0.08 | 1.58 | -0.05 | 0.96 | 1 | 1700011D18Rik | | 9 | 14 | 9 | 16 | 14 | 8 | 0 | 0 | 4 | 36 | 9 |
| 11 | 0.00 | 2.73 | 0.00 | 1.00 | 1 | Slc26a10 | solute carrier family 26, member 10 | 23 | 46 | 51 | 0 | 0 | 0 | 0 | 0 | 0 | 0 | 0 |
| 11 | -1.70 | 1.97 | -0.86 | 0.39 | 1 | Gm45743 | | 3 | 0 | 0 | 2 | 15 | 44 | 0 | 0 | 3 | 0 | 53 |
| 11 | 3.02 | 1.85 | 1.63 | 0.10 | 1 | Atp6v0a4 | ATPase, H+ transporting, lysosomal V0 su | 38 | 12 | 26 | 1 | 0 | 9 | 0 | 24 | 0 | 8 | 1 |
| 11 | 0.00 | 2.74 | 0.00 | 1.00 | 1 | Ifi203-ps | | 39 | 26 | 54 | 0 | 0 | 0 | 0 | 0 | 0 | 0 | 0 |
| 11 | 2.15 | 1.68 | 1.28 | 0.20 | 1 | Dpf1 | D4, zinc and double PHD fingers family 1 | 33 | 18 | 44 | 0 | 0 | 9 | 0 | 0 | 2 | 5 | 7 |
| 11 | -0.44 | 2.43 | -0.18 | 0.86 | 1 | Reg3b | regenerating islet-derived 3 beta | 21 | 13 | 84 | 1 | 0 | 0 | 0 | 0 | 0 | 0 | 0 |
| 11 | 1.84 | 1.43 | 1.29 | 0.20 | 1 | Pcdhgb8 | | 7 | 14 | 2 | 22 | 0 | 19 | 0 | 15 | 8 | 22 | 10 |
| 11 | 0.00 | 2.75 | 0.00 | 1.00 | 1 | Mmp13 | matrix metallopeptidase 13 | 66 | 12 | 41 | 0 | 0 | 0 | 0 | 0 | 0 | 0 | 0 |
| 11 | 1.34 | 1.75 | 0.77 | 0.44 | 1 | Proca1 | protein interacting with cyclin A1 | 20 | 45 | 37 | 0 | 2 | 2 | 0 | 0 | 10 | 0 | 3 |
| 11 | -0.51 | 1.65 | -0.31 | 0.76 | 1 | Krt86 | keratin 86 | 14 | 44 | 12 | 6 | 16 | 3 | 0 | 14 | 0 | 0 | 9 |
| 11 | -1.53 | 1.91 | -0.80 | 0.42 | 1 | Ano3 | anoctamin 3 | 31 | 31 | 39 | 0 | 5 | 8 | 0 | 0 | 0 | 0 | 4 |
| 11 | 0.00 | 2.75 | 0.00 | 1.00 | 1 | Il10 | interleukin 10 | 81 | 19 | 18 | 0 | 0 | 0 | 0 | 0 | 0 | 0 | 0 |
| 11 | 1.83 | 2.34 | 0.78 | 0.44 | 1 | D430013B06Rik | | 6 | 0 | 0 | 0 | 4 | 31 | 0 | 51 | 4 | 0 | 21 |
| 11 | -0.96 | 2.38 | -0.40 | 0.69 | 1 | Raver2 | ribonucleoprotein, PTB-binding 2 | 58 | 42 | 11 | 0 | 0 | 6 | 0 | 0 | 0 | 0 | 0 |
| 11 | 0.00 | 2.75 | 0.00 | 1.00 | 1 | Tlx1 | T cell leukemia, homeobox 1 | 62 | 29 | 26 | 0 | 0 | 0 | 0 | 0 | 0 | 0 | 0 |
| 11 | -0.71 | 2.36 | -0.30 | 0.76 | 1 | Cd5 | CD5 antigen | 44 | 43 | 29 | 0 | 0 | 2 | 0 | 0 | 0 | 0 | 0 |
| 11 | -1.11 | 2.08 | -0.53 | 0.59 | 1 | Gm44423 | | 9 | 0 | 6 | 3 | 0 | 69 | 0 | 0 | 1 | 0 | 28 |
| 11 | 0.00 | 2.75 | 0.00 | 1.00 | 1 | Tmem182 | transmembrane protein 182 | 31 | 48 | 38 | 0 | 0 | 0 | 0 | 0 | 0 | 0 | 0 |
| 11 | -3.36 | 2.09 | -1.61 | 0.11 | 1 | Wdr72 | WD repeat domain 72 | 15 | 68 | 15 | 11 | 0 | 8 | 0 | 0 | 0 | 0 | 0 |
| 11 | -3.09 | 2.08 | -1.49 | 0.14 | 1 | Gm49125 | | 18 | 22 | 0 | 7 | 4 | 42 | 0 | 0 | 0 | 0 | 23 |
| 11 | 0.29 | 2.81 | 0.10 | 0.92 | 1 | Gm43511 | | 0 | 10 | 0 | 0 | 0 | 46 | 0 | 0 | 0 | 0 | 60 |
| 11 | 1.27 | 1.43 | 0.88 | 0.38 | 1 | Gm12191 | | 3 | 52 | 12 | 7 | 4 | 3 | 0 | 1 | 22 | 0 | 12 |
| 11 | -0.54 | 2.01 | -0.27 | 0.79 | 1 | Pcdhb2 | protocadherin beta 2 | 33 | 30 | 14 | 11 | 0 | 3 | 0 | 0 | 0 | 23 | 1 |
| 11 | 0.37 | 1.89 | 0.20 | 0.84 | 1 | 9130017K11Rik | | 13 | 27 | 17 | 0 | 15 | 7 | 0 | 2 | 0 | 32 | 3 |
| 11 | -2.27 | 2.19 | -1.03 | 0.30 | 1 | Gucy2g | guanylate cyclase 2g | 66 | 18 | 14 | 10 | 0 | 2 | 0 | 0 | 0 | 0 | 7 |
| 11 | 0.00 | 3.62 | 0.00 | 1.00 | 1 | Ighv5-1 | | 116 | 0 | 0 | 0 | 0 | 0 | 0 | 0 | 0 | 0 | 0 |
| 11 | -1.79 | 2.69 | -0.66 | 0.51 | 1 | Gm48945 | | 0 | 0 | 0 | 5 | 0 | 83 | 0 | 0 | 0 | 0 | 27 |
| 10 | 0.00 | 3.59 | 0.00 | 1.00 | 1 | Ighv1-55 | | 106 | 9 | 0 | 0 | 0 | 0 | 0 | 0 | 0 | 0 | 0 |
| 10 | -2.53 | 1.61 | -1.58 | 0.11 | 1 | Fyb2 | FYN binding protein 2 | 24 | 32 | 38 | 3 | 3 | 12 | 0 | 0 | 0 | 0 | 3 |
| 10 | -0.85 | 3.02 | -0.28 | 0.78 | 1 | Ryr2 | ryanodine receptor 2, cardiac | 100 | 13 | 0 | 0 | 0 | 3 | 0 | 0 | 0 | 0 | 0 |
| 10 | -0.53 | 2.32 | -0.23 | 0.82 | 1 | Tacr1 | tachykinin receptor 1 | 62 | 28 | 11 | 0 | 10 | 0 | 0 | 0 | 3 | 0 | 0 |
| 10 | 0.83 | 1.47 | 0.56 | 0.57 | 1 | Prkcg | protein kinase C, gamma | 0 | 5 | 37 | 0 | 6 | 28 | 0 | 2 | 6 | 0 | 30 |
| 10 | 0.54 | 1.60 | 0.34 | 0.73 | 1 | Gm43375 | | 3 | 2 | 9 | 0 | 2 | 58 | 0 | 0 | 7 | 0 | 32 |
| 10 | 3.36 | 3.16 | 1.07 | 0.29 | 1 | Nmbr | neuromedin B receptor | 14 | 74 | 0 | 0 | 0 | 13 | 0 | 0 | 14 | 0 | 0 |
| 10 | 0.61 | 1.82 | 0.33 | 0.74 | 1 | AI854703 | | 19 | 29 | 12 | 0 | 15 | 3 | 0 | 29 | 2 | 0 | 4 |
| 10 | 2.18 | 2.00 | 1.09 | 0.28 | 1 | Tnfrsf4 | tumor necrosis factor receptor superfamily | 7 | 18 | 10 | 0 | 0 | 27 | 0 | 0 | 15 | 0 | 37 |
| 10 | 0.00 | 2.78 | 0.00 | 1.00 | 1 | Gm20467 | | 45 | 42 | 28 | 0 | 0 | 0 | 0 | 0 | 0 | 0 | 0 |
| 10 | 0.59 | 2.09 | 0.28 | 0.78 | 1 | A730046J19Rik | RIKEN cDNA A730046J19 gene | 52 | 10 | 49 | 0 | 0 | 1 | 0 | 0 | 0 | 0 | 2 |
| 10 | 0.00 | 2.78 | 0.00 | 1.00 | 1 | Gpr31c | | 53 | 14 | 47 | 0 | 0 | 0 | 0 | 0 | 0 | 0 | 0 |
| 10 | 2.91 | 2.06 | 1.41 | 0.16 | 1 | 6430550D23Ril | RIKEN cDNA 6430550D23 gene | 2 | 11 | 0 | 0 | 3 | 29 | 0 | 0 | 12 | 26 | 31 |
| 10 | -0.26 | 2.07 | -0.13 | 0.90 | 1 | Nlrc3 | NLR family, CARD domain containing 3 | 30 | 0 | 3 | 0 | 8 | 22 | 0 | 0 | 0 | 33 | 17 |
| 10 | 0.00 | 2.78 | 0.00 | 1.00 | 1 | Kcnj15 | potassium inwardly-rectifying channel, su | 36 | 26 | 51 | 0 | 0 | 0 | 0 | 0 | 0 | 0 | 0 |
| 10 | -1.29 | 2.01 | -0.64 | 0.52 | 1 | Olfr1372-ps1 | | 44 | 18 | 34 | 0 | 10 | 3 | 0 | 0 | 0 | 0 | 4 |
| 10 | 2.43 | 2.28 | 1.06 | 0.29 | 1 | Gm13212 | predicted gene 13212 | 48 | 50 | 10 | 0 | 0 | 0 | 0 | 0 | 4 | 0 | 1 |
| 10 | 0.00 | 3.55 | 0.00 | 1.00 | 1 | Btnl2 | butyrophilin-like 2 | 80 | 0 | 33 | 0 | 0 | 0 | 0 | 0 | 0 | 0 | 0 |
| 10 | 0.00 | 2.79 | 0.00 | 1.00 | 1 | Myom3 | myomesin family, member 3 | 73 | 17 | 23 | 0 | 0 | 0 | 0 | 0 | 0 | 0 | 0 |

| | | | | | | | | | | | | | | | | | | |
|---|---|---|---|---|---|---|---|---|---|---|---|---|---|---|---|---|---|---|
| 10 | 0.00 | 2.79 | 0.00 | 1.00 | 1 | Adhfe1 | alcohol dehydrogenase, iron containing, | 35 | 47 | 31 | 0 | 0 | 0 | 0 | 0 | 0 | 0 | 0 |
| 10 | 0.00 | 2.81 | 0.00 | 1.00 | 1 | Myocd | myocardin | 83 | 23 | 6 | 0 | 0 | 0 | 0 | 0 | 0 | 0 | 0 |
| 10 | 1.12 | 2.30 | 0.49 | 0.63 | 1 | Gm48838 | | 0 | 4 | 0 | 8 | 0 | 45 | 0 | 0 | 11 | 0 | 45 |
| 10 | -2.00 | 2.11 | -0.95 | 0.34 | 1 | Begain | brain-enriched guanylate kinase-associat | 39 | 44 | 25 | 2 | 0 | 3 | 0 | 0 | 0 | 0 | 0 |
| 10 | 0.00 | 2.94 | 0.00 | 1.00 | 1 | Gm4956 | | 1 | 35 | 76 | 0 | 0 | 0 | 0 | 0 | 0 | 0 | 0 |
| 10 | 0.10 | 1.99 | 0.05 | 0.96 | 1 | Gm2788 | | 27 | 44 | 30 | 0 | 0 | 6 | 0 | 0 | 0 | 0 | 5 |
| 10 | 0.73 | 2.41 | 0.30 | 0.76 | 1 | Prima1 | proline rich membrane anchor 1 | 17 | 43 | 24 | 0 | 4 | 0 | 0 | 24 | 0 | 0 | 0 |
| 10 | -0.78 | 1.83 | -0.43 | 0.67 | 1 | Col9a3 | collagen, type IX, alpha 3 | 11 | 18 | 43 | 0 | 4 | 14 | 0 | 0 | 0 | 0 | 21 |
| 10 | 2.59 | 1.95 | 1.32 | 0.19 | 1 | Zap70 | zeta-chain (TCR) associated protein kina: | 62 | 10 | 20 | 0 | 0 | 3 | 0 | 0 | 8 | 0 | 9 |
| 10 | 3.58 | 2.09 | 1.71 | 0.09 | 1 | Apol9b | apolipoprotein L 9b | 32 | 3 | 19 | 0 | 1 | 0 | 0 | 0 | 0 | 53 | 4 |
| 10 | 0.27 | 1.83 | 0.15 | 0.88 | 1 | Padi1 | peptidyl arginine deiminase, type I | 34 | 16 | 45 | 0 | 0 | 11 | 0 | 0 | 1 | 0 | 4 |
| 10 | -1.09 | 2.24 | -0.49 | 0.63 | 1 | Sox11 | SRY (sex determining region Y)-box 11 | 47 | 48 | 0 | 6 | 0 | 4 | 0 | 0 | 0 | 0 | 6 |
| 10 | -0.59 | 1.64 | -0.36 | 0.72 | 1 | Pla2g4e | phospholipase A2, group IVE | 40 | 13 | 21 | 3 | 5 | 11 | 0 | 0 | 13 | 0 | 5 |
| 10 | 1.59 | 1.63 | 0.98 | 0.33 | 1 | Spef2 | sperm flagellar 2 | 11 | 49 | 19 | 0 | 2 | 6 | 0 | 17 | 5 | 0 | 2 |
| 10 | 0.80 | 2.54 | 0.31 | 0.75 | 1 | Gm43662 | | 0 | 0 | 0 | 4 | 0 | 33 | 0 | 0 | 18 | 0 | 55 |
| 10 | 0.85 | 2.08 | 0.41 | 0.68 | 1 | 0610010K14Ril | RIKEN cDNA 0610010K14 gene | 6 | 0 | 9 | 0 | 0 | 44 | 0 | 0 | 0 | 1 | 50 |
| 10 | -1.26 | 1.88 | -0.67 | 0.50 | 1 | Tmc3 | transmembrane channel-like gene family | 27 | 53 | 15 | 6 | 0 | 5 | 0 | 0 | 0 | 0 | 5 |
| 10 | -0.51 | 2.38 | -0.22 | 0.83 | 1 | Gm48086 | | 0 | 0 | 3 | 1 | 0 | 53 | 0 | 0 | 0 | 0 | 53 |
| 10 | 0.00 | 2.82 | 0.00 | 1.00 | 1 | Col11a1 | collagen, type XI, alpha 1 | 62 | 25 | 23 | 0 | 0 | 0 | 0 | 0 | 0 | 0 | 0 |
| 10 | 0.00 | 2.81 | 0.00 | 1.00 | 1 | Slc27a6 | solute carrier family 27 (fatty acid transpo | 37 | 29 | 44 | 0 | 0 | 0 | 0 | 0 | 0 | 0 | 0 |
| 10 | 0.00 | 2.82 | 0.00 | 1.00 | 1 | Klri1 | killer cell lectin-like receptor family I meml | 39 | 59 | 12 | 0 | 0 | 0 | 0 | 0 | 0 | 0 | 0 |
| 10 | 0.00 | 3.62 | 0.00 | 1.00 | 1 | Igkv1-132 | | 110 | 0 | 0 | 0 | 0 | 0 | 0 | 0 | 0 | 0 | 0 |
| 10 | 1.49 | 2.08 | 0.72 | 0.47 | 1 | Gm7889 | | 15 | 6 | 0 | 0 | 13 | 11 | 0 | 0 | 18 | 39 | 7 |
| 10 | 0.00 | 2.82 | 0.00 | 1.00 | 1 | B4galnt2 | beta-1,4-N-acetyl-galactosaminyl transfer | 35 | 12 | 62 | 0 | 0 | 0 | 0 | 0 | 0 | 0 | 0 |
| 10 | 0.03 | 3.32 | 0.01 | 0.99 | 1 | Hnf1aos2 | | 0 | 0 | 0 | 0 | 0 | 66 | 0 | 0 | 0 | 0 | 44 |
| 10 | 0.30 | 1.61 | 0.19 | 0.85 | 1 | Gm16793 | | 18 | 27 | 46 | 0 | 3 | 4 | 0 | 0 | 2 | 0 | 9 |
| 10 | 4.33 | 2.21 | 1.95 | 0.05 | 1 | Gm46378 | | 0 | 3 | 0 | 4 | 0 | 38 | 0 | 0 | 11 | 32 | 21 |
| 10 | -0.67 | 2.56 | -0.26 | 0.79 | 1 | Lama1 | laminin, alpha 1 | 0 | 74 | 8 | 0 | 20 | 0 | 0 | 0 | 2 | 0 | 5 |
| 10 | -0.86 | 1.55 | -0.55 | 0.58 | 1 | Asprv1 | aspartic peptidase, retroviral-like 1 | 7 | 13 | 4 | 17 | 17 | 20 | 0 | 0 | 5 | 17 | 9 |
| 10 | 3.17 | 2.17 | 1.46 | 0.14 | 1 | 5330431K02Rik | | 27 | 20 | 7 | 0 | 0 | 25 | 0 | 0 | 15 | 0 | 15 |
| 10 | 2.60 | 2.07 | 1.26 | 0.21 | 1 | Egfl6 | EGF-like-domain, multiple 6 | 40 | 9 | 32 | 0 | 0 | 1 | 0 | 0 | 0 | 24 | 2 |
| 10 | 0.00 | 2.99 | 0.00 | 1.00 | 1 | 9830107B12Ril | RIKEN cDNA 9830107B12 gene | 94 | 1 | 14 | 0 | 0 | 0 | 0 | 0 | 0 | 0 | 0 |
| 10 | 0.00 | 2.83 | 0.00 | 1.00 | 1 | Calr4 | calreticulin 4 | 27 | 35 | 46 | 0 | 0 | 0 | 0 | 0 | 0 | 0 | 0 |
| 10 | -3.35 | 2.40 | -1.39 | 0.16 | 1 | Tchh | trichohyalin | 75 | 19 | 2 | 10 | 3 | 0 | 0 | 0 | 0 | 0 | 0 |
| 10 | 0.57 | 1.72 | 0.33 | 0.74 | 1 | Gm26912 | | 0 | 24 | 10 | 11 | 9 | 5 | 0 | 0 | 17 | 0 | 30 |
| 10 | 0.00 | 3.62 | 0.00 | 1.00 | 1 | Igkv14-100 | | 108 | 0 | 0 | 0 | 0 | 0 | 0 | 0 | 0 | 0 | 0 |
| 10 | 0.20 | 2.28 | 0.09 | 0.93 | 1 | Gm43029 | | 0 | 0 | 0 | 0 | 8 | 51 | 0 | 0 | 10 | 0 | 40 |
| 10 | 0.20 | 2.03 | 0.10 | 0.92 | 1 | F730016J06Rik | | 23 | 54 | 5 | 0 | 0 | 13 | 0 | 0 | 0 | 0 | 13 |
| 10 | 0.00 | 3.62 | 0.00 | 1.00 | 1 | Htr2a | 5-hydroxytryptamine (serotonin) receptor | 33 | 74 | 0 | 0 | 0 | 0 | 0 | 0 | 0 | 0 | 0 |
| 10 | 0.16 | 1.93 | 0.08 | 0.93 | 1 | Gm33732 | | 0 | 3 | 0 | 11 | 2 | 43 | 0 | 0 | 9 | 0 | 40 |
| 10 | 1.86 | 2.86 | 0.65 | 0.51 | 1 | Gm14824 | | 0 | 0 | 0 | 0 | 0 | 42 | 0 | 0 | 8 | 0 | 57 |
| 10 | -1.59 | 1.72 | -0.92 | 0.36 | 1 | Gm9884 | | 15 | 41 | 15 | 5 | 13 | 8 | 0 | 0 | 0 | 6 | 4 |
| 10 | 0.07 | 1.93 | 0.04 | 0.97 | 1 | Gm31227 | | 4 | 48 | 15 | 0 | 9 | 12 | 0 | 0 | 1 | 14 | 5 |
| 10 | -1.01 | 2.81 | -0.36 | 0.72 | 1 | Gm49405 | | 0 | 0 | 0 | 3 | 0 | 52 | 0 | 0 | 0 | 0 | 52 |
| 10 | 0.17 | 1.79 | 0.10 | 0.92 | 1 | Gm20634 | | 0 | 9 | 2 | 25 | 0 | 16 | 0 | 16 | 0 | 0 | 38 |
| 10 | 0.00 | 2.86 | 0.00 | 1.00 | 1 | Spic | Spi-C transcription factor (Spi-1/PU.1 rela | 76 | 16 | 14 | 0 | 0 | 0 | 0 | 0 | 0 | 0 | 0 |
| 10 | 0.27 | 1.89 | 0.14 | 0.89 | 1 | Gm15535 | | 0 | 24 | 7 | 5 | 3 | 21 | 0 | 0 | 0 | 3 | 43 |
| 10 | -0.66 | 1.81 | -0.37 | 0.71 | 1 | Lrriq1 | leucine-rich repeats and IQ motif containi | 15 | 0 | 12 | 9 | 18 | 24 | 0 | 15 | 6 | 0 | 7 |
| 10 | -1.66 | 1.75 | -0.95 | 0.34 | 1 | Col19a1 | collagen, type XIX, alpha 1 | 15 | 8 | 36 | 4 | 10 | 14 | 0 | 0 | 4 | 13 | 0 |
| 10 | 0.00 | 2.89 | 0.00 | 1.00 | 1 | Tnfrsf17 | tumor necrosis factor receptor superfamily | 53 | 46 | 6 | 0 | 0 | 0 | 0 | 0 | 0 | 0 | 0 |
| 10 | 0.28 | 1.67 | 0.17 | 0.87 | 1 | Gm3555 | | 0 | 13 | 4 | 29 | 3 | 15 | 0 | 2 | 0 | 4 | 35 |
| 10 | -0.92 | 1.95 | -0.47 | 0.64 | 1 | Ak4 | adenylate kinase 4 | 36 | 3 | 5 | 18 | 9 | 3 | 0 | 0 | 9 | 0 | 21 |
| 10 | 0.00 | 2.86 | 0.00 | 1.00 | 1 | Rtl3 | retrotransposon Gag like 3 | 47 | 23 | 34 | 0 | 0 | 0 | 0 | 0 | 0 | 0 | 0 |
| 10 | -2.57 | 2.31 | -1.11 | 0.27 | 1 | Ankle1 | ankyrin repeat and LEM domain containir | 16 | 0 | 0 | 11 | 5 | 45 | 0 | 0 | 2 | 0 | 26 |
| 10 | -0.57 | 1.52 | -0.37 | 0.71 | 1 | Gm34552 | | 3 | 1 | 17 | 3 | 9 | 18 | 0 | 0 | 7 | 0 | 47 |
| 10 | 1.48 | 2.49 | 0.59 | 0.55 | 1 | Lilrb4a | leukocyte immunoglobulin-like receptor, s | 69 | 8 | 22 | 0 | 0 | 0 | 0 | 0 | 0 | 0 | 6 |
| 10 | 0.00 | 3.68 | 0.00 | 1.00 | 1 | Ly6a2 | | 100 | 5 | 0 | 0 | 0 | 0 | 0 | 0 | 0 | 0 | 0 |
| 10 | -0.99 | 1.91 | -0.52 | 0.60 | 1 | Gm34680 | | 3 | 0 | 0 | 22 | 11 | 20 | 0 | 0 | 10 | 14 | 23 |
| 9 | 0.00 | 2.87 | 0.00 | 1.00 | 1 | Il12b | interleukin 12b | 64 | 15 | 25 | 0 | 0 | 0 | 0 | 0 | 0 | 0 | 0 |
| 9 | 0.96 | 2.50 | 0.38 | 0.70 | 1 | Gm5771 | predicted gene 5771 | 5 | 58 | 41 | 0 | 0 | 0 | 0 | 0 | 0 | 0 | 1 |

| | | | | | | | | | | | | | | | | | | |
|---|---|---|---|---|---|---|---|---|---|---|---|---|---|---|---|---|---|---|
| 9 | 1.57 | 1.48 | 1.06 | 0.29 | 1 | Sox30 | SRY (sex determining region Y)-box 30 | 4 | 16 | 0 | 4 | 8 | 11 | 0 | 10 | 9 | 23 | 18 |
| 9 | -0.80 | 2.15 | -0.37 | 0.71 | 1 | H2bc3 | H2B clustered histone 3 | 0 | 0 | 0 | 23 | 6 | 16 | 0 | 0 | 2 | 53 | 5 |
| 9 | 0.00 | 2.88 | 0.00 | 1.00 | 1 | Prss12 | protease, serine 12 neurotrypsin (motops | 63 | 33 | 8 | 0 | 0 | 0 | 0 | 0 | 0 | 0 | 0 |
| 9 | -0.40 | 2.48 | -0.16 | 0.87 | 1 | Htra4 | HtrA serine peptidase 4 | 70 | 9 | 24 | 0 | 0 | 1 | 0 | 0 | 0 | 0 | 0 |
| 9 | 0.67 | 2.23 | 0.30 | 0.76 | 1 | Gm48194 | | 14 | 6 | 0 | 5 | 0 | 30 | 0 | 0 | 4 | 0 | 45 |
| 9 | -0.80 | 2.47 | -0.32 | 0.75 | 1 | Plekhs1 | pleckstrin homology domain containing, fa | 45 | 8 | 49 | 0 | 0 | 3 | 0 | 0 | 0 | 0 | 0 |
| 9 | 0.00 | 2.93 | 0.00 | 1.00 | 1 | Drd1 | dopamine receptor D1 | 49 | 52 | 3 | 0 | 0 | 0 | 0 | 0 | 0 | 0 | 0 |
| 9 | 0.00 | 2.88 | 0.00 | 1.00 | 1 | Pigz | phosphatidylinositol glycan anchor biosyn | 40 | 43 | 21 | 0 | 0 | 0 | 0 | 0 | 0 | 0 | 0 |
| 9 | 0.00 | 2.88 | 0.00 | 1.00 | 1 | Madcam1 | mucosal vascular addressin cell adhesion | 44 | 15 | 45 | 0 | 0 | 0 | 0 | 0 | 0 | 0 | 0 |
| 9 | 2.28 | 1.91 | 1.19 | 0.23 | 1 | Ifit1bl1 | interferon induced protein with tetratricpe | 16 | 3 | 16 | 0 | 9 | 8 | 0 | 16 | 0 | 33 | 2 |
| 9 | -0.39 | 2.91 | -0.13 | 0.89 | 1 | Mir186 | | 0 | 8 | 0 | 0 | 0 | 71 | 0 | 0 | 0 | 0 | 24 |
| 9 | 0.00 | 3.68 | 0.00 | 1.00 | 1 | Timd2 | T cell immunoglobulin and mucin domain | 85 | 18 | 0 | 0 | 0 | 0 | 0 | 0 | 0 | 0 | 0 |
| 9 | 0.47 | 2.09 | 0.23 | 0.82 | 1 | Icos | inducible T cell co-stimulator | 52 | 33 | 12 | 0 | 0 | 2 | 0 | 0 | 0 | 0 | 3 |
| 9 | -2.24 | 1.74 | -1.29 | 0.20 | 1 | Fam83e | family with sequence similarity 83, memb | 3 | 50 | 30 | 4 | 9 | 3 | 0 | 0 | 0 | 0 | 4 |
| 9 | 0.00 | 3.68 | 0.00 | 1.00 | 1 | Ighv2-4 | | 15 | 87 | 0 | 0 | 0 | 0 | 0 | 0 | 0 | 0 | 0 |
| 9 | 2.61 | 2.08 | 1.26 | 0.21 | 1 | Ptchd1 | patched domain containing 1 | 30 | 21 | 21 | 3 | 0 | 0 | 0 | 0 | 7 | 21 | 0 |
| 9 | -2.29 | 1.97 | -1.16 | 0.24 | 1 | Vwa7 | von Willebrand factor A domain containin | 42 | 19 | 26 | 0 | 4 | 10 | 0 | 0 | 0 | 0 | 1 |
| 9 | -0.60 | 2.41 | -0.25 | 0.80 | 1 | Gm48714 | | 0 | 11 | 0 | 6 | 0 | 42 | 0 | 0 | 0 | 0 | 43 |
| 9 | 0.50 | 1.34 | 0.37 | 0.71 | 1 | 4833419F23Rik | | 13 | 2 | 5 | 18 | 5 | 9 | 0 | 13 | 5 | 24 | 8 |
| 9 | -0.20 | 1.74 | -0.11 | 0.91 | 1 | 4930412L05Rik | | 8 | 2 | 0 | 8 | 5 | 41 | 0 | 3 | 9 | 0 | 26 |
| 9 | 2.81 | 2.19 | 1.29 | 0.20 | 1 | Megf11 | multiple EGF-like-domains 11 | 3 | 35 | 16 | 11 | 7 | 0 | 0 | 0 | 0 | 27 | 4 |
| 9 | -0.39 | 1.83 | -0.21 | 0.83 | 1 | Gm49502 | | 11 | 0 | 18 | 0 | 9 | 38 | 0 | 0 | 6 | 0 | 20 |
| 9 | 1.34 | 1.59 | 0.84 | 0.40 | 1 | Tekt1 | tektin 1 | 12 | 7 | 18 | 5 | 2 | 9 | 0 | 39 | 7 | 0 | 2 |
| 9 | 0.32 | 1.32 | 0.24 | 0.81 | 1 | Fsip1 | fibrous sheath-interacting protein 1 | 3 | 8 | 8 | 7 | 22 | 7 | 0 | 29 | 13 | 0 | 5 |
| 9 | -2.16 | 1.70 | -1.27 | 0.20 | 1 | Gm30082 | | 8 | 5 | 32 | 13 | 6 | 19 | 0 | 0 | 16 | 0 | 2 |
| 9 | 1.96 | 1.77 | 1.11 | 0.27 | 1 | Gm16958 | | 21 | 30 | 9 | 0 | 9 | 1 | 0 | 19 | 11 | 0 | 2 |
| 9 | -0.78 | 2.41 | -0.32 | 0.75 | 1 | Slfn1 | schlafen 1 | 15 | 19 | 59 | 6 | 0 | 0 | 0 | 0 | 0 | 0 | 1 |
| 9 | 0.00 | 2.92 | 0.00 | 1.00 | 1 | Htr7 | 5-hydroxytryptamine (serotonin) receptor | 77 | 9 | 15 | 0 | 0 | 0 | 0 | 0 | 0 | 0 | 0 |
| 9 | -0.87 | 1.46 | -0.60 | 0.55 | 1 | Pmel | premelanosome protein | 16 | 20 | 11 | 12 | 8 | 14 | 0 | 0 | 3 | 6 | 9 |
| 9 | 0.00 | 3.68 | 0.00 | 1.00 | 1 | Gpr176 | G protein-coupled receptor 176 | 74 | 26 | 0 | 0 | 0 | 0 | 0 | 0 | 0 | 0 | 0 |
| 9 | 0.00 | 2.92 | 0.00 | 1.00 | 1 | Pax1 | paired box 1 | 41 | 12 | 47 | 0 | 0 | 0 | 0 | 0 | 0 | 0 | 0 |
| 9 | 0.96 | 3.11 | 0.31 | 0.76 | 1 | Fzd10 | frizzled class receptor 10 | 58 | 41 | 0 | 0 | 0 | 0 | 0 | 0 | 0 | 0 | 1 |
| 9 | 0.00 | 3.68 | 0.00 | 1.00 | 1 | Pilrb1 | paired immunoglobin-like type 2 receptor | 83 | 0 | 17 | 0 | 0 | 0 | 0 | 0 | 0 | 0 | 0 |
| 9 | 0.18 | 2.59 | 0.07 | 0.95 | 1 | Phex | phosphate regulating endopeptidase hom | 64 | 0 | 27 | 0 | 0 | 4 | 0 | 0 | 0 | 0 | 4 |
| 9 | 2.78 | 2.49 | 1.12 | 0.26 | 1 | Gm15903 | | 0 | 0 | 0 | 0 | 0 | 46 | 0 | 0 | 3 | 6 | 44 |
| 9 | -2.28 | 2.35 | -0.97 | 0.33 | 1 | Zfp831 | zinc finger protein 831 | 47 | 38 | 8 | 0 | 4 | 2 | 0 | 0 | 0 | 0 | 0 |
| 9 | 0.79 | 2.08 | 0.38 | 0.70 | 1 | Gm12592 | | 21 | 27 | 26 | 0 | 7 | 2 | 0 | 0 | 14 | 0 | 3 |
| 9 | 4.43 | 2.87 | 1.54 | 0.12 | 1 | Gm26944 | | 4 | 0 | 0 | 0 | 0 | 40 | 0 | 21 | 0 | 0 | 35 |
| 9 | 0.00 | 2.93 | 0.00 | 1.00 | 1 | Tmem273 | transmembrane protein 273 | 60 | 8 | 31 | 0 | 0 | 0 | 0 | 0 | 0 | 0 | 0 |
| 9 | 0.05 | 1.90 | 0.03 | 0.98 | 1 | Gm45501 | | 0 | 3 | 4 | 4 | 0 | 39 | 0 | 0 | 3 | 0 | 47 |
| 9 | -1.98 | 2.45 | -0.81 | 0.42 | 1 | Asxl3 | additional sex combs like 3, transcriptiona | 29 | 29 | 31 | 0 | 8 | 0 | 0 | 0 | 0 | 0 | 2 |
| 9 | 0.69 | 1.84 | 0.37 | 0.71 | 1 | Plekhd1 | pleckstrin homology domain containing, fa | 29 | 24 | 23 | 0 | 1 | 12 | 0 | 0 | 6 | 0 | 4 |
| 9 | -0.40 | 2.52 | -0.16 | 0.87 | 1 | Xlr | X-linked lymphocyte-regulated | 45 | 44 | 9 | 0 | 0 | 1 | 0 | 0 | 0 | 0 | 0 |
| 9 | 0.00 | 2.93 | 0.00 | 1.00 | 1 | Hrh1 | histamine receptor H1 | 31 | 26 | 42 | 0 | 0 | 0 | 0 | 0 | 0 | 0 | 0 |
| 9 | 0.03 | 1.70 | 0.02 | 0.99 | 1 | Gm15638 | | 0 | 10 | 4 | 22 | 4 | 8 | 0 | 30 | 20 | 0 | 0 |
| 9 | -0.01 | 2.08 | -0.01 | 0.99 | 1 | 4930534D22Rik | | 0 | 7 | 0 | 25 | 10 | 15 | 0 | 0 | 23 | 0 | 18 |
| 9 | 0.00 | 2.99 | 0.00 | 1.00 | 1 | Fcrl5 | Fc receptor-like 5 | 71 | 25 | 2 | 0 | 0 | 0 | 0 | 0 | 0 | 0 | 0 |
| 9 | 0.00 | 2.95 | 0.00 | 1.00 | 1 | Ifi213 | interferon activated gene 213 | 45 | 8 | 46 | 0 | 0 | 0 | 0 | 0 | 0 | 0 | 0 |
| 9 | 0.04 | 1.71 | 0.02 | 0.98 | 1 | Gm26614 | | 0 | 38 | 0 | 4 | 4 | 13 | 0 | 28 | 2 | 0 | 10 |
| 9 | -3.65 | 1.90 | -1.92 | 0.06 | 1 | Spef1l | sperm flagellar 1 like | 13 | 11 | 21 | 11 | 14 | 12 | 0 | 0 | 0 | 0 | 16 |
| 9 | 0.80 | 2.47 | 0.33 | 0.75 | 1 | Gm43031 | | 0 | 0 | 22 | 0 | 0 | 34 | 0 | 0 | 5 | 0 | 38 |
| 9 | 2.55 | 2.82 | 0.90 | 0.37 | 1 | Gm36070 | | 8 | 64 | 0 | 0 | 0 | 19 | 0 | 0 | 6 | 0 | 1 |
| 9 | 0.00 | 2.94 | 0.00 | 1.00 | 1 | Hoxa2 | homeobox A2 | 31 | 58 | 9 | 0 | 0 | 0 | 0 | 0 | 0 | 0 | 0 |
| 9 | 0.76 | 1.91 | 0.40 | 0.69 | 1 | Gm9057 | | 14 | 5 | 0 | 4 | 7 | 19 | 0 | 38 | 8 | 0 | 4 |
| 9 | 1.12 | 2.04 | 0.55 | 0.58 | 1 | Crtac1 | cartilage acidic protein 1 | 12 | 35 | 37 | 0 | 5 | 0 | 0 | 0 | 6 | 0 | 2 |
| 9 | 0.00 | 2.94 | 0.00 | 1.00 | 1 | Gvin1 | GTPase, very large interferon inducible 1 | 35 | 38 | 24 | 0 | 0 | 0 | 0 | 0 | 0 | 0 | 0 |
| 9 | -0.07 | 2.05 | -0.03 | 0.97 | 1 | Gm43339 | | 0 | 24 | 0 | 4 | 1 | 42 | 0 | 0 | 2 | 0 | 24 |
| 9 | -0.56 | 1.84 | -0.31 | 0.76 | 1 | 8430426J06Rik | | 3 | 0 | 17 | 5 | 6 | 11 | 0 | 0 | 13 | 40 | 3 |
| 9 | 0.85 | 1.51 | 0.57 | 0.57 | 1 | Fbxl13 | F-box and leucine-rich repeat protein 13 | 33 | 9 | 6 | 2 | 4 | 11 | 0 | 2 | 8 | 0 | 22 |
| 9 | 0.00 | 2.97 | 0.00 | 1.00 | 1 | Prf1 | perforin 1 (pore forming protein) | 75 | 17 | 5 | 0 | 0 | 0 | 0 | 0 | 0 | 0 | 0 |

| | | | | | | | | | | | | | | | | | |
|---|---|---|---|---|---|---|---|---|---|---|---|---|---|---|---|---|---|
| 9 | 0.04 | 2.83 | 0.01 | 0.99 | 1 | Gm35279 | | 0 | 0 | 29 | 0 | 42 | 0 | 0 | 0 | 26 | 0 | 0 |
| 9 | 0.00 | 2.95 | 0.00 | 1.00 | 1 | Timp4 | tissue inhibitor of metalloproteinase 4 | 52 | 11 | 33 | 0 | 0 | 0 | 0 | 0 | 0 | 0 | 0 |
| 9 | 0.96 | 1.93 | 0.50 | 0.62 | 1 | Gm37116 | | 1 | 10 | 28 | 0 | 0 | 31 | 0 | 0 | 3 | 0 | 23 |
| 9 | -0.11 | 1.64 | -0.07 | 0.95 | 1 | 1700028I16Rik | | 24 | 10 | 21 | 6 | 7 | 2 | 0 | 0 | 1 | 23 | 3 |
| 9 | -2.06 | 1.76 | -1.17 | 0.24 | 1 | Slc25a43 | solute carrier family 25, member 43 | 9 | 8 | 65 | 3 | 2 | 7 | 0 | 0 | 0 | 0 | 3 |
| 9 | 0.23 | 1.57 | 0.15 | 0.88 | 1 | Ldhb-ps | | 0 | 25 | 0 | 11 | 1 | 18 | 0 | 28 | 1 | 0 | 12 |
| 9 | 0.00 | 2.96 | 0.00 | 1.00 | 1 | Tnfsf15 | tumor necrosis factor (ligand) superfamily | 64 | 12 | 20 | 0 | 0 | 0 | 0 | 0 | 0 | 0 | 0 |
| 9 | 0.00 | 2.96 | 0.00 | 1.00 | 1 | Tll2 | tolloid-like 2 | 15 | 20 | 60 | 0 | 0 | 0 | 0 | 0 | 0 | 0 | 0 |
| 9 | -0.85 | 2.54 | -0.33 | 0.74 | 1 | F830016B08Ril | RIKEN cDNA F830016B08 gene | 43 | 8 | 42 | 0 | 0 | 3 | 0 | 0 | 0 | 0 | 0 |
| 9 | 0.53 | 1.76 | 0.30 | 0.76 | 1 | Gm33543 | | 12 | 13 | 0 | 4 | 7 | 8 | 0 | 17 | 0 | 15 | 19 |
| 9 | -2.02 | 1.76 | -1.15 | 0.25 | 1 | Gm45767 | | 11 | 10 | 4 | 21 | 10 | 23 | 0 | 0 | 10 | 0 | 6 |
| 9 | 2.84 | 2.33 | 1.22 | 0.22 | 1 | Itgad | integrin, alpha D | 23 | 33 | 31 | 0 | 0 | 0 | 0 | 0 | 6 | 0 | 2 |
| 9 | 0.00 | 2.96 | 0.00 | 1.00 | 1 | Hoxc5 | homeobox C5 | 35 | 26 | 34 | 0 | 0 | 0 | 0 | 0 | 0 | 0 | 0 |
| 9 | 1.27 | 2.54 | 0.50 | 0.62 | 1 | Wfdc15b | WAP four-disulfide core domain 15B | 16 | 18 | 59 | 0 | 0 | 0 | 0 | 0 | 0 | 0 | 2 |
| 9 | 0.00 | 2.98 | 0.00 | 1.00 | 1 | Bmx | BMX non-receptor tyrosine kinase | 43 | 7 | 46 | 0 | 0 | 0 | 0 | 0 | 0 | 0 | 0 |
| 9 | 0.00 | 3.00 | 0.00 | 1.00 | 1 | Ccn6 | cellular communication network factor 6 | 4 | 63 | 28 | 0 | 0 | 0 | 0 | 0 | 0 | 0 | 0 |
| 9 | 1.98 | 2.70 | 0.73 | 0.46 | 1 | Apol9a | apolipoprotein L 9a | 37 | 16 | 24 | 0 | 0 | 0 | 0 | 0 | 0 | 18 | 0 |
| 9 | -0.40 | 2.55 | -0.16 | 0.87 | 1 | Apba2 | amyloid beta (A4) precursor protein-bindi | 53 | 28 | 12 | 0 | 0 | 1 | 0 | 0 | 0 | 0 | 0 |
| 9 | -1.49 | 2.68 | -0.56 | 0.58 | 1 | Gm15138 | | 16 | 27 | 41 | 11 | 0 | 0 | 0 | 0 | 0 | 0 | 0 |
| 9 | 0.00 | 3.13 | 0.00 | 1.00 | 1 | Dok6 | docking protein 6 | 15 | 1 | 78 | 0 | 0 | 0 | 0 | 0 | 0 | 0 | 0 |
| 9 | 1.60 | 1.57 | 1.02 | 0.31 | 1 | Gm42604 | | 30 | 4 | 30 | 0 | 4 | 3 | 0 | 0 | 3 | 15 | 5 |
| 9 | -0.85 | 1.96 | -0.43 | 0.66 | 1 | Mep1a | meprin 1 alpha | 8 | 31 | 32 | 12 | 0 | 3 | 0 | 0 | 6 | 0 | 1 |
| 9 | 0.00 | 2.98 | 0.00 | 1.00 | 1 | Hbb-bt | hemoglobin, beta adult t chain | 55 | 14 | 25 | 0 | 0 | 0 | 0 | 0 | 0 | 0 | 0 |
| 9 | 0.00 | 2.98 | 0.00 | 1.00 | 1 | Klrg2 | killer cell lectin-like receptor subfamily G, | 35 | 15 | 44 | 0 | 0 | 0 | 0 | 0 | 0 | 0 | 0 |
| 9 | 0.00 | 3.00 | 0.00 | 1.00 | 1 | Gm34084 | | 75 | 14 | 5 | 0 | 0 | 0 | 0 | 0 | 0 | 0 | 0 |
| 9 | -1.50 | 2.54 | -0.59 | 0.55 | 1 | Cr2 | complement receptor 2 | 36 | 21 | 21 | 0 | 5 | 0 | 0 | 0 | 0 | 0 | 11 |
| 9 | -1.42 | 2.15 | -0.66 | 0.51 | 1 | 4930448E22Rik | | 33 | 34 | 19 | 0 | 4 | 2 | 0 | 0 | 0 | 3 | 0 |
| 9 | -1.13 | 1.45 | -0.78 | 0.44 | 1 | Rpsa-ps11 | | 20 | 13 | 3 | 16 | 19 | 6 | 0 | 1 | 6 | 4 | 6 |
| 9 | 0.00 | 3.68 | 0.00 | 1.00 | 1 | Clec2g | C-type lectin domain family 2, member g | 88 | 0 | 6 | 0 | 0 | 0 | 0 | 0 | 0 | 0 | 0 |
| 9 | -2.68 | 2.11 | -1.27 | 0.20 | 1 | Fbxo43 | F-box protein 43 | 0 | 0 | 21 | 19 | 10 | 12 | 0 | 0 | 7 | 0 | 24 |
| 9 | 0.00 | 3.00 | 0.00 | 1.00 | 1 | Slc22a3 | solute carrier family 22 (organic cation tra | 49 | 37 | 7 | 0 | 0 | 0 | 0 | 0 | 0 | 0 | 0 |
| 9 | 1.67 | 1.49 | 1.12 | 0.26 | 1 | Gm16098 | | 0 | 4 | 0 | 7 | 11 | 11 | 0 | 13 | 28 | 9 | 10 |
| 9 | -0.56 | 1.87 | -0.30 | 0.76 | 1 | Gm15884 | | 0 | 4 | 9 | 0 | 5 | 39 | 0 | 0 | 1 | 0 | 36 |
| 9 | -1.43 | 2.69 | -0.53 | 0.60 | 1 | Eprn | | 45 | 19 | 22 | 7 | 0 | 0 | 0 | 0 | 0 | 0 | 0 |
| 9 | 0.00 | 2.99 | 0.00 | 1.00 | 1 | 2810430I11Rik | | 62 | 17 | 15 | 0 | 0 | 0 | 0 | 0 | 0 | 0 | 0 |
| 8 | -0.71 | 2.21 | -0.32 | 0.75 | 1 | Bcas1 | breast carcinoma amplified sequence 1 | 27 | 26 | 30 | 0 | 6 | 1 | 0 | 0 | 3 | 0 | 0 |
| 8 | 0.12 | 2.06 | 0.06 | 0.95 | 1 | Orc1 | origin recognition complex, subunit 1 | 37 | 11 | 0 | 12 | 0 | 9 | 0 | 12 | 0 | 0 | 12 |
| 8 | 0.00 | 3.02 | 0.00 | 1.00 | 1 | Fpr2 | formyl peptide receptor 2 | 79 | 5 | 9 | 0 | 0 | 0 | 0 | 0 | 0 | 0 | 0 |
| 8 | 1.87 | 2.48 | 0.75 | 0.45 | 1 | Gm25596 | | 15 | 27 | 0 | 24 | 0 | 3 | 0 | 0 | 10 | 0 | 14 |
| 8 | 1.46 | 2.63 | 0.56 | 0.58 | 1 | Gm37084 | | 9 | 0 | 0 | 0 | 0 | 16 | 0 | 0 | 0 | 52 | 16 |
| 8 | -2.97 | 1.92 | -1.55 | 0.12 | 1 | Gm13652 | | 0 | 23 | 31 | 16 | 13 | 7 | 0 | 0 | 0 | 0 | 3 |
| 8 | 0.28 | 2.57 | 0.11 | 0.91 | 1 | Gm24119 | | 27 | 5 | 0 | 0 | 0 | 28 | 0 | 0 | 0 | 0 | 34 |
| 8 | 0.00 | 3.01 | 0.00 | 1.00 | 1 | Ighv1-82 | | 51 | 7 | 34 | 0 | 0 | 0 | 0 | 0 | 0 | 0 | 0 |
| 8 | 0.78 | 2.48 | 0.32 | 0.75 | 1 | Gm42798 | | 0 | 0 | 11 | 0 | 0 | 35 | 0 | 0 | 2 | 0 | 45 |
| 8 | -0.25 | 2.01 | -0.12 | 0.90 | 1 | Taar2 | trace amine-associated receptor 2 | 0 | 7 | 0 | 42 | 6 | 6 | 0 | 3 | 4 | 24 | 0 |
| 8 | 2.18 | 1.47 | 1.48 | 0.14 | 1 | H2ac13 | H2A clustered histone 13 | 4 | 11 | 2 | 15 | 0 | 13 | 0 | 12 | 12 | 12 | 12 |
| 8 | -0.17 | 2.62 | -0.06 | 0.95 | 1 | Gm20219 | | 18 | 34 | 0 | 0 | 0 | 28 | 0 | 0 | 0 | 0 | 12 |
| 8 | -0.80 | 1.78 | -0.45 | 0.65 | 1 | Adgrb1 | adhesion G protein-coupled receptor B1 | 35 | 14 | 8 | 4 | 6 | 7 | 0 | 14 | 0 | 0 | 4 |
| 8 | 1.23 | 2.21 | 0.56 | 0.58 | 1 | Gm37827 | | 0 | 0 | 3 | 18 | 0 | 14 | 0 | 36 | 0 | 1 | 19 |
| 8 | 0.17 | 1.99 | 0.08 | 0.93 | 1 | Gm45220 | | 0 | 0 | 11 | 0 | 5 | 30 | 0 | 0 | 3 | 0 | 43 |
| 8 | 0.00 | 3.05 | 0.00 | 1.00 | 1 | 1810010K12Rik | | 3 | 31 | 58 | 0 | 0 | 0 | 0 | 0 | 0 | 0 | 0 |
| 8 | 0.00 | 3.00 | 0.00 | 1.00 | 1 | Gm12551 | | 18 | 26 | 48 | 0 | 0 | 0 | 0 | 0 | 0 | 0 | 0 |
| 8 | 0.00 | 3.68 | 0.00 | 1.00 | 1 | Pilrb2 | paired immunoglobin-like type 2 receptor | 75 | 0 | 17 | 0 | 0 | 0 | 0 | 0 | 0 | 0 | 0 |
| 8 | -0.01 | 2.99 | 0.00 | 1.00 | 1 | Gm25082 | | 7 | 0 | 0 | 0 | 0 | 51 | 0 | 0 | 0 | 0 | 35 |
| 8 | 0.00 | 3.68 | 0.00 | 1.00 | 1 | Ighv2-3 | | 6 | 86 | 0 | 0 | 0 | 0 | 0 | 0 | 0 | 0 | 0 |
| 8 | 0.00 | 3.01 | 0.00 | 1.00 | 1 | Gdf6 | growth differentiation factor 6 | 45 | 12 | 35 | 0 | 0 | 0 | 0 | 0 | 0 | 0 | 0 |
| 8 | 1.66 | 1.96 | 0.85 | 0.40 | 1 | 4933424M12Rik | | 8 | 4 | 14 | 0 | 4 | 3 | 0 | 54 | 4 | 0 | 1 |
| 8 | 0.58 | 2.12 | 0.27 | 0.78 | 1 | Mroh2a | maestro heat-like repeat family member 2 | 21 | 38 | 17 | 0 | 0 | 5 | 0 | 0 | 0 | 0 | 11 |
| 8 | -0.01 | 2.96 | 0.00 | 1.00 | 1 | Gm44667 | | 5 | 0 | 0 | 0 | 0 | 53 | 0 | 0 | 0 | 0 | 34 |
| 8 | 0.00 | 3.01 | 0.00 | 1.00 | 1 | Rxrg | retinoid X receptor gamma | 57 | 12 | 23 | 0 | 0 | 0 | 0 | 0 | 0 | 0 | 0 |

| | | | | | | | | | | | | | | | | | | |
|---|---|---|---|---|---|---|---|---|---|---|---|---|---|---|---|---|---|---|
| 8 | 0.70 | 1.71 | 0.41 | 0.68 | 1 | Dusp27 | dual specificity phosphatase 27 (putative) | 39 | 12 | 6 | 4 | 2 | 2 | 0 | 25 | 1 | 0 | 1 |
| 8 | 0.00 | 3.68 | 0.00 | 1.00 | 1 | Gm5086 | | 55 | 0 | 36 | 0 | 0 | 0 | 0 | 0 | 0 | 0 | 0 |
| 8 | 2.44 | 2.25 | 1.08 | 0.28 | 1 | Gm12207 | | 0 | 4 | 0 | 9 | 0 | 19 | 0 | 0 | 1 | 28 | 29 |
| 8 | 1.22 | 1.96 | 0.62 | 0.53 | 1 | Gm49493 | | 4 | 3 | 7 | 0 | 0 | 44 | 0 | 0 | 4 | 0 | 28 |
| 8 | 2.81 | 1.92 | 1.46 | 0.14 | 1 | Serinc4 | serine incorporator 4 | 9 | 0 | 7 | 0 | 7 | 13 | 0 | 9 | 3 | 43 | 1 |
| 8 | -1.07 | 2.17 | -0.49 | 0.62 | 1 | D430018E03Rik | | 6 | 0 | 0 | 2 | 5 | 40 | 0 | 0 | 5 | 0 | 34 |
| 8 | -0.90 | 2.18 | -0.41 | 0.68 | 1 | | | 0 | 1 | 8 | 0 | 6 | 42 | 0 | 0 | 0 | 0 | 34 |
| 8 | -0.87 | 1.74 | -0.50 | 0.62 | 1 | Gm9937 | | 0 | 6 | 0 | 10 | 32 | 15 | 0 | 2 | 15 | 0 | 10 |
| 8 | 3.09 | 2.21 | 1.40 | 0.16 | 1 | Gm45148 | | 0 | 0 | 1 | 0 | 1 | 26 | 0 | 45 | 1 | 0 | 16 |
| 8 | 1.28 | 2.51 | 0.51 | 0.61 | 1 | Adgrf1 | adhesion G protein-coupled receptor F1 | 11 | 35 | 35 | 0 | 0 | 1 | 0 | 0 | 0 | 8 | 0 |
| 8 | 0.00 | 3.68 | 0.00 | 1.00 | 1 | Tnni2 | troponin I, skeletal, fast 2 | 59 | 0 | 31 | 0 | 0 | 0 | 0 | 0 | 0 | 0 | 0 |
| 8 | 1.39 | 2.59 | 0.54 | 0.59 | 1 | Gm16044 | | 0 | 16 | 0 | 0 | 0 | 18 | 0 | 25 | 0 | 0 | 30 |
| 8 | -1.45 | 2.68 | -0.54 | 0.59 | 1 | Klk1b4 | kallikrein 1-related pepidase b4 | 6 | 43 | 29 | 12 | 0 | 0 | 0 | 0 | 0 | 0 | 0 |
| 8 | 0.90 | 2.55 | 0.35 | 0.72 | 1 | Fam71f2 | family with sequence similarity 71, membe | 11 | 0 | 0 | 14 | 0 | 10 | 0 | 30 | 0 | 0 | 24 |
| 8 | -2.81 | 2.84 | -0.99 | 0.32 | 1 | Mss51 | MSS51 mitochondrial translational activat | 0 | 0 | 6 | 4 | 0 | 31 | 0 | 0 | 0 | 0 | 48 |
| 8 | 2.33 | 2.60 | 0.90 | 0.37 | 1 | Prrx2 | paired related homeobox 2 | 48 | 24 | 9 | 0 | 0 | 2 | 0 | 0 | 6 | 0 | 0 |
| 8 | -0.84 | 2.99 | -0.28 | 0.78 | 1 | Rhbdl1 | rhomboid like 1 | 0 | 0 | 0 | 0 | 2 | 45 | 0 | 0 | 0 | 0 | 43 |
| 8 | 1.17 | 2.11 | 0.56 | 0.58 | 1 | B230303A05Rik | | 0 | 9 | 0 | 32 | 4 | 4 | 0 | 0 | 4 | 33 | 3 |
| 8 | 3.42 | 1.92 | 1.78 | 0.07 | 1 | Gm13027 | | 0 | 0 | 0 | 0 | 2 | 21 | 0 | 28 | 10 | 4 | 24 |
| 8 | 4.51 | 2.92 | 1.55 | 0.12 | 1 | Ovgp1 | oviductal glycoprotein 1 | 0 | 0 | 3 | 0 | 0 | 29 | 0 | 20 | 0 | 0 | 38 |
| 8 | -1.04 | 2.65 | -0.39 | 0.69 | 1 | Slc18a2 | solute carrier family 18 (vesicular monoar | 15 | 25 | 46 | 3 | 0 | 0 | 0 | 0 | 0 | 0 | 0 |
| 8 | 0.00 | 3.04 | 0.00 | 1.00 | 1 | BC147527 | cDNA sequence BC147527 | 32 | 33 | 24 | 0 | 0 | 0 | 0 | 0 | 0 | 0 | 0 |
| 8 | 0.00 | 3.04 | 0.00 | 1.00 | 1 | Gjb3 | gap junction protein, beta 3 | 32 | 36 | 21 | 0 | 0 | 0 | 0 | 0 | 0 | 0 | 0 |
| 8 | 0.41 | 2.22 | 0.19 | 0.85 | 1 | Trp53-ps | | 20 | 0 | 26 | 10 | 0 | 14 | 0 | 8 | 9 | 0 | 2 |
| 8 | 0.67 | 1.92 | 0.35 | 0.73 | 1 | Gm26787 | | 0 | 11 | 0 | 11 | 4 | 7 | 0 | 32 | 0 | 19 | 4 |
| 8 | 0.94 | 2.60 | 0.36 | 0.72 | 1 | Cyp1a1 | cytochrome P450, family 1, subfamily a, p | 9 | 31 | 48 | 0 | 0 | 0 | 0 | 0 | 1 | 0 | 0 |
| 8 | 0.00 | 3.05 | 0.00 | 1.00 | 1 | Ip6k3 | inositol hexaphosphate kinase 3 | 56 | 15 | 18 | 0 | 0 | 0 | 0 | 0 | 0 | 0 | 0 |
| 8 | 1.42 | 2.02 | 0.70 | 0.48 | 1 | Gm28707 | | 0 | 26 | 7 | 23 | 1 | 1 | 0 | 0 | 9 | 0 | 21 |
| 8 | -0.41 | 2.63 | -0.15 | 0.88 | 1 | Cela3a | chymotrypsin-like elastase family, membe | 3 | 41 | 44 | 0 | 1 | 0 | 0 | 0 | 0 | 0 | 0 |
| 8 | 0.28 | 2.27 | 0.12 | 0.90 | 1 | Adarb2 | adenosine deaminase, RNA-specific, B2 | 21 | 58 | 8 | 0 | 0 | 1 | 0 | 0 | 0 | 0 | 1 |
| 8 | 0.00 | 3.68 | 0.00 | 1.00 | 1 | Prss32 | protease, serine 32 | 0 | 34 | 54 | 0 | 0 | 0 | 0 | 0 | 0 | 0 | 0 |
| 8 | -1.55 | 2.10 | -0.74 | 0.46 | 1 | Gabre | gamma-aminobutyric acid (GABA) A rece | 33 | 3 | 5 | 0 | 5 | 24 | 0 | 0 | 0 | 0 | 18 |
| 8 | -1.76 | 2.16 | -0.82 | 0.41 | 1 | 1810064F22Rik | | 14 | 33 | 26 | 13 | 0 | 1 | 0 | 1 | 0 | 0 | 0 |
| 8 | 1.26 | 2.09 | 0.60 | 0.55 | 1 | Gm10676 | | 9 | 4 | 0 | 3 | 0 | 27 | 0 | 27 | 0 | 0 | 19 |
| 8 | -2.31 | 1.68 | -1.38 | 0.17 | 1 | Odf3b | outer dense fiber of sperm tails 3B | 13 | 17 | 35 | 11 | 8 | 2 | 0 | 0 | 2 | 0 | 1 |
| 8 | 3.29 | 2.33 | 1.41 | 0.16 | 1 | Gm42559 | | 0 | 17 | 9 | 0 | 0 | 26 | 0 | 0 | 3 | 9 | 23 |
| 8 | 0.00 | 3.09 | 0.00 | 1.00 | 1 | Tnip3 | TNFAIP3 interacting protein 3 | 60 | 3 | 25 | 0 | 0 | 0 | 0 | 0 | 0 | 0 | 0 |
| 8 | -0.11 | 2.74 | -0.04 | 0.97 | 1 | Kcnh7 | potassium voltage-gated channel, subfam | 0 | 63 | 12 | 0 | 0 | 8 | 0 | 0 | 0 | 0 | 4 |
| 8 | -3.35 | 2.95 | -1.13 | 0.26 | 1 | Gm7441 | | 19 | 0 | 0 | 0 | 7 | 29 | 0 | 0 | 0 | 0 | 34 |
| 8 | 0.08 | 2.90 | 0.03 | 0.98 | 1 | Gm20775 | | 0 | 10 | 0 | 21 | 17 | 0 | 0 | 0 | 39 | 0 | 0 |
| 8 | 0.00 | 3.62 | 0.00 | 1.00 | 1 | Ighv1-47 | | 0 | 87 | 0 | 0 | 0 | 0 | 0 | 0 | 0 | 0 | 0 |
| 8 | -0.09 | 3.00 | -0.03 | 0.98 | 1 | Gm43272 | | 0 | 2 | 0 | 0 | 0 | 57 | 0 | 0 | 0 | 0 | 28 |
| 8 | 0.71 | 2.21 | 0.32 | 0.75 | 1 | Gm11427 | | 31 | 3 | 39 | 0 | 0 | 3 | 0 | 0 | 0 | 0 | 10 |
| 8 | 0.66 | 1.89 | 0.35 | 0.73 | 1 | Lrr1 | leucine rich repeat protein 1 | 23 | 13 | 0 | 5 | 10 | 4 | 0 | 10 | 13 | 0 | 7 |
| 8 | -3.16 | 2.36 | -1.34 | 0.18 | 1 | Drd2 | dopamine receptor D2 | 27 | 15 | 31 | 5 | 0 | 8 | 0 | 0 | 0 | 0 | 0 |
| 8 | 0.96 | 2.63 | 0.36 | 0.72 | 1 | Serpina3f | serine (or cysteine) peptidase inhibitor, cl | 60 | 17 | 8 | 0 | 0 | 0 | 0 | 0 | 0 | 0 | 1 |
| 8 | -1.25 | 2.02 | -0.62 | 0.54 | 1 | Vps25 | vacuolar protein sorting 25 | 9 | 1 | 0 | 8 | 3 | 28 | 0 | 7 | 0 | 0 | 30 |
| 8 | 3.28 | 2.62 | 1.25 | 0.21 | 1 | Gm37517 | | 0 | 0 | 0 | 0 | 0 | 44 | 0 | 7 | 7 | 0 | 28 |
| 8 | -0.66 | 1.98 | -0.33 | 0.74 | 1 | A830005F24Ril | RIKEN cDNA A830005F24 gene | 8 | 0 | 0 | 10 | 6 | 16 | 0 | 0 | 9 | 27 | 11 |
| 8 | 0.32 | 1.84 | 0.17 | 0.86 | 1 | Spag4 | sperm associated antigen 4 | 17 | 28 | 16 | 0 | 6 | 4 | 0 | 0 | 4 | 0 | 11 |
| 8 | 0.07 | 3.12 | 0.02 | 0.98 | 1 | Gm45527 | | 0 | 9 | 0 | 0 | 0 | 41 | 0 | 0 | 0 | 0 | 36 |
| 8 | 1.68 | 2.74 | 0.61 | 0.54 | 1 | Wnt7b | wingless-type MMTV integration site famil | 65 | 7 | 10 | 0 | 0 | 0 | 0 | 0 | 3 | 0 | 0 |
| 8 | 1.79 | 1.32 | 1.35 | 0.18 | 1 | Gm6501 | | 10 | 8 | 2 | 4 | 5 | 4 | 0 | 21 | 3 | 5 | 23 |
| 8 | 2.38 | 1.62 | 1.47 | 0.14 | 1 | Cd96 | CD96 antigen | 22 | 5 | 11 | 0 | 2 | 8 | 0 | 0 | 9 | 17 | 11 |
| 8 | -1.78 | 1.72 | -1.03 | 0.30 | 1 | Gm20541 | | 5 | 14 | 7 | 17 | 26 | 3 | 0 | 0 | 7 | 0 | 6 |
| 8 | -1.22 | 2.72 | -0.45 | 0.65 | 1 | Cmtm5 | CKLF-like MARVEL transmembrane dom | 34 | 16 | 29 | 0 | 6 | 0 | 0 | 0 | 0 | 0 | 0 |
| 8 | -2.74 | 2.24 | -1.23 | 0.22 | 1 | Mfsd13b | major facilitator superfamily domain conta | 0 | 0 | 0 | 11 | 14 | 29 | 0 | 0 | 1 | 0 | 30 |
| 8 | 0.00 | 3.10 | 0.00 | 1.00 | 1 | Spns3 | spinster homolog 3 | 50 | 8 | 27 | 0 | 0 | 0 | 0 | 0 | 0 | 0 | 0 |
| 8 | 0.00 | 3.68 | 0.00 | 1.00 | 1 | Ighv1-71 | | 24 | 61 | 0 | 0 | 0 | 0 | 0 | 0 | 0 | 0 | 0 |
| 8 | 0.00 | 3.10 | 0.00 | 1.00 | 1 | Tacstd2 | tumor-associated calcium signal transduc | 59 | 17 | 8 | 0 | 0 | 0 | 0 | 0 | 0 | 0 | 0 |

| | | | | | | | | | | | | | | | | | |
|---|---|---|---|---|---|---|---|---|---|---|---|---|---|---|---|---|---|
| 8 | 0.88 | 2.46 | 0.36 | 0.72 | 1 | Gm16487 | | 21 | 8 | 44 | 0 | 0 | 3 | 0 | 0 | 8 | 0 | 0 |
| 8 | -0.10 | 2.36 | -0.04 | 0.97 | 1 | Ncan | neurocan | 23 | 0 | 5 | 9 | 3 | 0 | 0 | 3 | 0 | 41 | 0 |
| 8 | 0.91 | 1.23 | 0.74 | 0.46 | 1 | Gm7901 | | 3 | 13 | 2 | 9 | 7 | 5 | 0 | 32 | 4 | 3 | 7 |
| 8 | 0.00 | 3.10 | 0.00 | 1.00 | 1 | Gm44751 | | 33 | 14 | 37 | 0 | 0 | 0 | 0 | 0 | 0 | 0 | 0 |
| 8 | 1.34 | 2.08 | 0.64 | 0.52 | 1 | Catsper3 | cation channel, sperm associated 3 | 7 | 0 | 0 | 0 | 23 | 8 | 0 | 14 | 6 | 17 | 10 |
| 8 | -1.13 | 2.10 | -0.54 | 0.59 | 1 | Gm19938 | | 13 | 21 | 15 | 7 | 0 | 11 | 0 | 0 | 0 | 0 | 17 |
| 8 | -0.47 | 2.19 | -0.21 | 0.83 | 1 | Map7d3 | MAP7 domain containing 3 | 30 | 17 | 24 | 0 | 5 | 1 | 0 | 0 | 0 | 0 | 7 |
| 8 | 0.28 | 2.21 | 0.13 | 0.90 | 1 | Ihh | Indian hedgehog | 21 | 19 | 37 | 0 | 0 | 3 | 0 | 0 | 0 | 0 | 3 |
| 8 | 3.32 | 2.00 | 1.66 | 0.10 | 1 | Gm20635 | | 2 | 19 | 23 | 0 | 1 | 13 | 0 | 9 | 0 | 13 | 4 |
| 8 | -0.70 | 2.68 | -0.26 | 0.79 | 1 | Trank1 | tetratricopeptide repeat and ankyrin repea | 4 | 57 | 21 | 0 | 0 | 2 | 0 | 0 | 0 | 0 | 0 |
| 8 | 0.00 | 3.68 | 0.00 | 1.00 | 1 | Tigit | T cell immunoreceptor with Ig and ITIM dc | 71 | 12 | 0 | 0 | 0 | 0 | 0 | 0 | 0 | 0 | 0 |
| 8 | -1.16 | 2.27 | -0.51 | 0.61 | 1 | Gm12405 | | 5 | 0 | 0 | 3 | 3 | 47 | 0 | 0 | 8 | 0 | 17 |
| 8 | -1.02 | 2.01 | -0.51 | 0.61 | 1 | Gm4287 | | 4 | 19 | 0 | 4 | 5 | 9 | 0 | 37 | 0 | 0 | 4 |
| 8 | 0.95 | 3.04 | 0.31 | 0.75 | 1 | Gm27028 | | 0 | 0 | 0 | 0 | 0 | 36 | 0 | 0 | 1 | 0 | 46 |
| 8 | 0.87 | 2.36 | 0.37 | 0.71 | 1 | Gm32688 | | 10 | 31 | 37 | 0 | 0 | 1 | 0 | 0 | 3 | 0 | 0 |
| 8 | 2.94 | 2.16 | 1.36 | 0.17 | 1 | Gm23027 | | 0 | 21 | 8 | 0 | 6 | 5 | 0 | 0 | 15 | 14 | 13 |
| 8 | 0.00 | 3.68 | 0.00 | 1.00 | 1 | Clec4d | C-type lectin domain family 4, member d | 65 | 0 | 18 | 0 | 0 | 0 | 0 | 0 | 0 | 0 | 0 |
| 8 | 0.00 | 3.22 | 0.00 | 1.00 | 1 | Gm5084 | | 65 | 1 | 17 | 0 | 0 | 0 | 0 | 0 | 0 | 0 | 0 |
| 8 | -0.92 | 1.99 | -0.46 | 0.64 | 1 | E330034L11Rik | | 6 | 11 | 0 | 5 | 5 | 26 | 0 | 0 | 4 | 0 | 25 |
| 8 | -0.15 | 2.64 | -0.06 | 0.95 | 1 | Gm38413 | | 14 | 0 | 0 | 0 | 10 | 31 | 0 | 0 | 15 | 0 | 12 |
| 8 | 2.50 | 2.71 | 0.92 | 0.36 | 1 | Slc2a7 | solute carrier family 2 (facilitated glucose | 0 | 45 | 19 | 0 | 0 | 0 | 0 | 17 | 2 | 0 | 0 |
| 7 | -5.11 | 2.60 | -1.97 | 0.05 | 1 | Fam220a | family with sequence similarity 220, meml | 15 | 0 | 0 | 3 | 3 | 38 | 0 | 0 | 0 | 0 | 24 |
| 7 | 1.90 | 2.73 | 0.69 | 0.49 | 1 | P4ha3 | procollagen-proline, 2-oxoglutarate 4-dio | 15 | 7 | 45 | 0 | 0 | 0 | 0 | 0 | 15 | 0 | 0 |
| 7 | 0.00 | 3.13 | 0.00 | 1.00 | 1 | Nkx3-2 | NK3 homeobox 2 | 34 | 30 | 18 | 0 | 0 | 0 | 0 | 0 | 0 | 0 | 0 |
| 7 | -3.70 | 3.02 | -1.22 | 0.22 | 1 | Prph | peripherin | 46 | 12 | 0 | 23 | 0 | 0 | 0 | 0 | 0 | 0 | 1 |
| 7 | 2.32 | 1.37 | 1.70 | 0.09 | 1 | Gm12397 | | 1 | 0 | 1 | 2 | 10 | 6 | 0 | 24 | 16 | 5 | 16 |
| 7 | 0.00 | 3.23 | 0.00 | 1.00 | 1 | Sele | selectin, endothelial cell | 53 | 27 | 1 | 0 | 0 | 0 | 0 | 0 | 0 | 0 | 0 |
| 7 | -4.08 | 2.43 | -1.68 | 0.09 | 1 | Gm31218 | | 15 | 0 | 16 | 10 | 17 | 8 | 0 | 0 | 0 | 0 | 16 |
| 7 | 0.00 | 3.63 | 0.00 | 1.00 | 1 | Ighv1-62 | | 82 | 0 | 0 | 0 | 0 | 0 | 0 | 0 | 0 | 0 | 0 |
| 7 | -2.41 | 2.20 | -1.10 | 0.27 | 1 | Gm14027 | | 16 | 0 | 8 | 23 | 5 | 12 | 0 | 0 | 2 | 0 | 15 |
| 7 | 0.02 | 1.64 | 0.01 | 0.99 | 1 | Gm10300 | | 11 | 7 | 0 | 7 | 18 | 5 | 0 | 3 | 6 | 14 | 9 |
| 7 | -0.17 | 2.02 | -0.09 | 0.93 | 1 | 4930455G09Rik | | 12 | 4 | 24 | 11 | 0 | 13 | 0 | 14 | 0 | 0 | 4 |
| 7 | 0.00 | 3.15 | 0.00 | 1.00 | 1 | C5ar2 | complement component 5a receptor 2 | 52 | 13 | 16 | 0 | 0 | 0 | 0 | 0 | 0 | 0 | 0 |
| 7 | -3.49 | 3.03 | -1.15 | 0.25 | 1 | Gm29008 | | 0 | 9 | 0 | 0 | 9 | 23 | 0 | 0 | 0 | 0 | 40 |
| 7 | -2.28 | 2.31 | -0.99 | 0.32 | 1 | Mirt1 | | 10 | 22 | 0 | 3 | 5 | 14 | 0 | 0 | 0 | 0 | 27 |
| 7 | 0.00 | 3.68 | 0.00 | 1.00 | 1 | Vxn | vexin | 40 | 41 | 0 | 0 | 0 | 0 | 0 | 0 | 0 | 0 | 0 |
| 7 | 1.80 | 2.72 | 0.66 | 0.51 | 1 | Car12 | carbonic anhydrase 12 | 33 | 11 | 30 | 0 | 0 | 0 | 0 | 0 | 6 | 0 | 0 |
| 7 | 2.40 | 1.91 | 1.26 | 0.21 | 1 | C030005K06Rik | | 6 | 5 | 0 | 4 | 0 | 14 | 0 | 32 | 2 | 0 | 17 |
| 7 | -0.37 | 1.82 | -0.20 | 0.84 | 1 | Fam131c | family with sequence similarity 131, meml | 10 | 16 | 24 | 11 | 2 | 2 | 0 | 14 | 2 | 0 | 0 |
| 7 | -0.91 | 3.09 | -0.30 | 0.77 | 1 | Gm45804 | | 0 | 0 | 0 | 0 | 2 | 44 | 0 | 0 | 0 | 0 | 35 |
| 7 | 0.02 | 1.98 | 0.01 | 0.99 | 1 | C030010L15Rik | | 3 | 13 | 2 | 23 | 5 | 0 | 0 | 30 | 0 | 0 | 4 |
| 7 | 0.00 | 3.15 | 0.00 | 1.00 | 1 | 2810459M11Ri | RIKEN cDNA 2810459M11 gene | 23 | 40 | 18 | 0 | 0 | 0 | 0 | 0 | 0 | 0 | 0 |
| 7 | 0.05 | 1.94 | 0.03 | 0.98 | 1 | Hepacam | hepatocyte cell adhesion molecule | 9 | 8 | 33 | 10 | 0 | 4 | 0 | 0 | 2 | 12 | 2 |
| 7 | 2.24 | 2.04 | 1.10 | 0.27 | 1 | A230083G16Rik | | 11 | 1 | 5 | 0 | 4 | 11 | 0 | 36 | 9 | 0 | 3 |
| 7 | 2.12 | 1.87 | 1.13 | 0.26 | 1 | Gm4651 | | 12 | 15 | 6 | 0 | 3 | 14 | 0 | 9 | 18 | 0 | 2 |
| 7 | 2.43 | 2.27 | 1.07 | 0.28 | 1 | Gm49893 | | 6 | 0 | 3 | 0 | 11 | 3 | 0 | 37 | 0 | 18 | 2 |
| 7 | 1.06 | 2.84 | 0.37 | 0.71 | 1 | Piwil2 | piwi-like RNA-mediated gene silencing 2 | 27 | 45 | 0 | 0 | 0 | 1 | 0 | 0 | 0 | 0 | 8 |
| 7 | 1.60 | 2.73 | 0.59 | 0.56 | 1 | Hoxd9 | homeobox D9 | 30 | 29 | 18 | 0 | 0 | 0 | 0 | 0 | 3 | 0 | 0 |
| 7 | 0.75 | 2.30 | 0.33 | 0.74 | 1 | Pcnx2 | pecanex homolog 2 | 45 | 25 | 6 | 0 | 0 | 1 | 0 | 0 | 0 | 0 | 3 |
| 7 | -0.06 | 2.32 | -0.03 | 0.98 | 1 | Gm27252 | | 4 | 0 | 25 | 10 | 0 | 8 | 0 | 0 | 2 | 23 | 7 |
| 7 | -1.19 | 2.12 | -0.56 | 0.57 | 1 | Gm12915 | | 0 | 22 | 0 | 10 | 20 | 4 | 0 | 12 | 3 | 9 | 0 |
| 7 | 0.00 | 3.17 | 0.00 | 1.00 | 1 | Sh2d2a | SH2 domain containing 2A | 33 | 34 | 12 | 0 | 0 | 0 | 0 | 0 | 0 | 0 | 0 |
| 7 | 0.20 | 3.60 | 0.06 | 0.96 | 1 | Gm20751 | | 0 | 0 | 0 | 0 | 0 | 40 | 0 | 0 | 0 | 0 | 40 |
| 7 | -1.00 | 2.07 | -0.48 | 0.63 | 1 | 1810020O05Rik | | 10 | 42 | 14 | 0 | 2 | 7 | 0 | 0 | 0 | 0 | 5 |
| 7 | -3.22 | 1.87 | -1.72 | 0.08 | 1 | 1810006J02Rik | | 11 | 33 | 7 | 8 | 7 | 8 | 0 | 0 | 0 | 0 | 4 |
| 7 | 2.14 | 1.72 | 1.24 | 0.21 | 1 | Mrps36-ps1 | | 7 | 18 | 8 | 0 | 4 | 6 | 0 | 10 | 21 | 0 | 4 |
| 7 | 0.00 | 3.18 | 0.00 | 1.00 | 1 | Dgki | diacylglycerol kinase, iota | 30 | 9 | 39 | 0 | 0 | 0 | 0 | 0 | 0 | 0 | 0 |
| 7 | 0.40 | 1.83 | 0.22 | 0.83 | 1 | Spx | spexin hormone | 0 | 9 | 6 | 2 | 5 | 18 | 0 | 20 | 0 | 0 | 19 |
| 7 | -2.05 | 1.89 | -1.08 | 0.28 | 1 | Gm15975 | | 15 | 2 | 22 | 13 | 15 | 2 | 0 | 0 | 5 | 0 | 5 |
| 7 | 0.00 | 3.19 | 0.00 | 1.00 | 1 | Fat2 | FAT atypical cadherin 2 | 8 | 41 | 30 | 0 | 0 | 0 | 0 | 0 | 0 | 0 | 0 |

| | | | | | | | | | | | | | | | | | |
|---|---|---|---|---|---|---|---|---|---|---|---|---|---|---|---|---|---|
| 7 | 2.68 | 2.15 | 1.25 | 0.21 | 1 | Cxcl11 | | 9 | 1 | 28 | 0 | 5 | 0 | 0 | 5 | 31 | 0 | 0 |
| 7 | 1.53 | 2.04 | 0.75 | 0.45 | 1 | Gm44325 | | 2 | 9 | 16 | 2 | 8 | 2 | 0 | 0 | 0 | 35 | 6 |
| 7 | 0.00 | 3.18 | 0.00 | 1.00 | 1 | Nlrp1a | NLR family, pyrin domain containing 1A | 36 | 16 | 26 | 0 | 0 | 0 | 0 | 0 | 0 | 0 | 0 |
| 7 | -0.34 | 2.11 | -0.16 | 0.87 | 1 | Gm38375 | | 0 | 0 | 9 | 0 | 8 | 34 | 0 | 0 | 3 | 0 | 24 |
| 7 | 1.38 | 2.25 | 0.61 | 0.54 | 1 | Gm48582 | | 0 | 4 | 0 | 0 | 4 | 25 | 0 | 25 | 17 | 0 | 3 |
| 7 | 0.00 | 3.68 | 0.00 | 1.00 | 1 | Gm15987 | | 51 | 27 | 0 | 0 | 0 | 0 | 0 | 0 | 0 | 0 | 0 |
| 7 | -1.10 | 2.79 | -0.39 | 0.69 | 1 | Bmerb1 | bMERB domain containing 1 | 9 | 47 | 20 | 0 | 3 | 0 | 0 | 0 | 0 | 0 | 0 |
| 7 | 0.00 | 3.19 | 0.00 | 1.00 | 1 | Prdm8 | PR domain containing 8 | 33 | 15 | 29 | 0 | 0 | 0 | 0 | 0 | 0 | 0 | 0 |
| 7 | 1.21 | 2.43 | 0.50 | 0.62 | 1 | D230022J07Rik | | 0 | 3 | 11 | 0 | 6 | 17 | 0 | 0 | 0 | 14 | 26 |
| 7 | 0.00 | 3.68 | 0.00 | 1.00 | 1 | Tbx21 | T-box 21 | 53 | 24 | 0 | 0 | 0 | 0 | 0 | 0 | 0 | 0 | 0 |
| 7 | 0.00 | 3.20 | 0.00 | 1.00 | 1 | Gm43820 | | 45 | 16 | 17 | 0 | 0 | 0 | 0 | 0 | 0 | 0 | 0 |
| 7 | 0.00 | 3.20 | 0.00 | 1.00 | 1 | Rapsn | receptor-associated protein of the synaps | 26 | 19 | 32 | 0 | 0 | 0 | 0 | 0 | 0 | 0 | 0 |
| 7 | 0.00 | 3.20 | 0.00 | 1.00 | 1 | Cd3d | CD3 antigen, delta polypeptide | 31 | 21 | 25 | 0 | 0 | 0 | 0 | 0 | 0 | 0 | 0 |
| 7 | 0.00 | 3.21 | 0.00 | 1.00 | 1 | 2610307P16Rik | | 52 | 11 | 14 | 0 | 0 | 0 | 0 | 0 | 0 | 0 | 0 |
| 7 | 0.00 | 3.21 | 0.00 | 1.00 | 1 | Ptgds | prostaglandin D2 synthase (brain) | 23 | 9 | 45 | 0 | 0 | 0 | 0 | 0 | 0 | 0 | 0 |
| 7 | 1.97 | 1.78 | 1.11 | 0.27 | 1 | Disp3 | dispatched RND transporter family memb | 3 | 32 | 4 | 0 | 2 | 4 | 0 | 19 | 11 | 0 | 2 |
| 7 | 1.75 | 3.18 | 0.55 | 0.58 | 1 | Gnrh1 | gonadotropin releasing hormone 1 | 0 | 0 | 0 | 0 | 0 | 28 | 0 | 0 | 4 | 0 | 45 |
| 7 | -1.25 | 2.60 | -0.48 | 0.63 | 1 | Gm46218 | | 1 | 0 | 0 | 0 | 12 | 21 | 0 | 0 | 0 | 27 | 15 |
| 7 | 0.28 | 2.45 | 0.11 | 0.91 | 1 | Lipc | lipase, hepatic | 36 | 37 | 1 | 0 | 0 | 1 | 0 | 0 | 0 | 0 | 1 |
| 7 | -0.40 | 2.73 | -0.15 | 0.88 | 1 | Ism2 | isthmin 2 | 28 | 27 | 20 | 0 | 0 | 1 | 0 | 0 | 0 | 0 | 0 |
| 7 | 0.00 | 3.22 | 0.00 | 1.00 | 1 | Peg12 | paternally expressed 12 | 33 | 33 | 9 | 0 | 0 | 0 | 0 | 0 | 0 | 0 | 0 |
| 7 | 0.00 | 3.28 | 0.00 | 1.00 | 1 | C130026I21Rik | RIKEN cDNA C130026I21 gene | 67 | 6 | 3 | 0 | 0 | 0 | 0 | 0 | 0 | 0 | 0 |
| 7 | 0.04 | 3.60 | 0.01 | 0.99 | 1 | Slco1a1 | solute carrier organic anion transporter fa | 0 | 0 | 0 | 0 | 0 | 46 | 0 | 0 | 0 | 0 | 30 |
| 7 | -1.42 | 2.86 | -0.50 | 0.62 | 1 | Prtg | protogenin | 45 | 2 | 22 | 0 | 8 | 0 | 0 | 0 | 0 | 0 | 0 |
| 7 | 0.80 | 1.62 | 0.49 | 0.62 | 1 | Gm8451 | | 5 | 13 | 6 | 13 | 12 | 0 | 0 | 17 | 1 | 3 | 5 |
| 7 | 0.23 | 2.64 | 0.09 | 0.93 | 1 | Gm15979 | | 1 | 0 | 1 | 0 | 0 | 36 | 0 | 0 | 0 | 0 | 39 |
| 7 | 0.96 | 2.74 | 0.35 | 0.73 | 1 | Gm22270 | | 9 | 0 | 0 | 0 | 0 | 32 | 0 | 0 | 0 | 13 | 22 |
| 7 | 0.00 | 3.23 | 0.00 | 1.00 | 1 | Tfec | transcription factor EC | 46 | 21 | 8 | 0 | 0 | 0 | 0 | 0 | 0 | 0 | 0 |
| 7 | 3.25 | 2.24 | 1.45 | 0.15 | 1 | Myl7 | myosin, light polypeptide 7, regulatory | 5 | 5 | 0 | 0 | 0 | 22 | 0 | 0 | 5 | 5 | 34 |
| 7 | -0.12 | 2.26 | -0.05 | 0.96 | 1 | A230107N01Rik | | 3 | 6 | 10 | 32 | 0 | 8 | 0 | 0 | 0 | 8 | 9 |
| 7 | 0.00 | 3.22 | 0.00 | 1.00 | 1 | Clca3a2 | chloride channel accessory 3A2 | 48 | 11 | 17 | 0 | 0 | 0 | 0 | 0 | 0 | 0 | 0 |
| 7 | -0.34 | 2.28 | -0.15 | 0.88 | 1 | Grid1 | glutamate receptor, ionotropic, delta 1 | 30 | 19 | 18 | 0 | 0 | 7 | 0 | 0 | 0 | 0 | 2 |
| 7 | 2.67 | 2.45 | 1.09 | 0.28 | 1 | C130040N14Rik | | 0 | 9 | 0 | 0 | 0 | 6 | 0 | 52 | 3 | 0 | 5 |
| 7 | -4.64 | 2.42 | -1.92 | 0.05 | 1 | Gm49719 | | 9 | 10 | 0 | 14 | 24 | 12 | 0 | 0 | 0 | 0 | 7 |
| 7 | -0.33 | 2.08 | -0.16 | 0.88 | 1 | Gm16278 | | 3 | 44 | 0 | 4 | 2 | 13 | 0 | 0 | 4 | 0 | 6 |
| 7 | -2.32 | 2.20 | -1.05 | 0.29 | 1 | 4930469K13Rik | | 0 | 0 | 16 | 16 | 23 | 6 | 0 | 0 | 2 | 0 | 13 |
| 7 | 0.00 | 3.24 | 0.00 | 1.00 | 1 | Trem1 | triggering receptor expressed on myeloid | 51 | 16 | 8 | 0 | 0 | 0 | 0 | 0 | 0 | 0 | 0 |
| 7 | 0.09 | 2.30 | 0.04 | 0.97 | 1 | Epor | erythropoietin receptor | 17 | 30 | 23 | 0 | 0 | 3 | 0 | 0 | 0 | 0 | 2 |
| 7 | 2.06 | 2.32 | 0.89 | 0.37 | 1 | Apoh | apolipoprotein H | 10 | 15 | 17 | 8 | 0 | 0 | 0 | 0 | 17 | 8 | 0 |
| 7 | 0.00 | 3.24 | 0.00 | 1.00 | 1 | A830082K12Rik | | 11 | 28 | 35 | 0 | 0 | 0 | 0 | 0 | 0 | 0 | 0 |
| 7 | 0.00 | 3.68 | 0.00 | 1.00 | 1 | Olr1 | oxidized low density lipoprotein (lectin-lik | 69 | 6 | 0 | 0 | 0 | 0 | 0 | 0 | 0 | 0 | 0 |
| 7 | -1.17 | 2.35 | -0.50 | 0.62 | 1 | Piwil4 | piwi-like RNA-mediated gene silencing 4 | 28 | 19 | 5 | 0 | 17 | 1 | 0 | 0 | 4 | 0 | 0 |
| 7 | 0.19 | 3.60 | 0.05 | 0.96 | 1 | Gm43138 | | 0 | 0 | 0 | 0 | 0 | 38 | 0 | 0 | 0 | 0 | 37 |
| 7 | 0.00 | 3.68 | 0.00 | 1.00 | 1 | Cdh16 | cadherin 16 | 46 | 0 | 28 | 0 | 0 | 0 | 0 | 0 | 0 | 0 | 0 |
| 7 | 1.88 | 3.04 | 0.62 | 0.54 | 1 | Tafa3 | TAFA chemokine like family member 3 | 45 | 2 | 20 | 0 | 0 | 0 | 0 | 7 | 0 | 0 | 0 |
| 7 | -1.61 | 1.82 | -0.89 | 0.38 | 1 | Gm18890 | | 9 | 12 | 16 | 21 | 2 | 7 | 0 | 0 | 6 | 0 | 2 |
| 7 | -0.40 | 3.44 | -0.12 | 0.91 | 1 | B430306N03Ri | RIKEN cDNA B430306N03 gene | 60 | 13 | 0 | 0 | 0 | 1 | 0 | 0 | 0 | 0 | 0 |
| 7 | -1.99 | 1.98 | -1.00 | 0.31 | 1 | Tslp | thymic stromal lymphopoietin | 42 | 5 | 18 | 1 | 4 | 3 | 0 | 0 | 0 | 0 | 2 |
| 7 | 0.20 | 2.29 | 0.09 | 0.93 | 1 | Scn1a | sodium channel, voltage-gated, type I, alp | 26 | 5 | 31 | 0 | 0 | 6 | 0 | 0 | 0 | 0 | 6 |
| 7 | 0.00 | 3.63 | 0.00 | 1.00 | 1 | Ighv5-4 | | 74 | 0 | 0 | 0 | 0 | 0 | 0 | 0 | 0 | 0 | 0 |
| 7 | -1.14 | 2.40 | -0.47 | 0.64 | 1 | Gm28578 | | 33 | 12 | 9 | 0 | 9 | 1 | 0 | 0 | 0 | 10 | 0 |
| 7 | 0.00 | 3.27 | 0.00 | 1.00 | 1 | Gm5535 | | 58 | 7 | 8 | 0 | 0 | 0 | 0 | 0 | 0 | 0 | 0 |
| 7 | 3.11 | 2.20 | 1.42 | 0.16 | 1 | Gm18113 | | 11 | 0 | 4 | 5 | 0 | 2 | 0 | 38 | 8 | 0 | 5 |
| 7 | 0.00 | 3.26 | 0.00 | 1.00 | 1 | Car9 | carbonic anhydrase 9 | 19 | 12 | 43 | 0 | 0 | 0 | 0 | 0 | 0 | 0 | 0 |
| 7 | -0.75 | 2.13 | -0.35 | 0.72 | 1 | Zfp385c | zinc finger protein 385C | 14 | 26 | 26 | 0 | 3 | 2 | 0 | 0 | 0 | 0 | 3 |
| 7 | 0.00 | 3.27 | 0.00 | 1.00 | 1 | Mrgprh | MAS-related GPR, member H | 42 | 26 | 5 | 0 | 0 | 0 | 0 | 0 | 0 | 0 | 0 |
| 7 | -1.28 | 3.64 | -0.35 | 0.73 | 1 | Havcr1 | hepatitis A virus cellular receptor 1 | 51 | 17 | 0 | 0 | 5 | 0 | 0 | 0 | 0 | 0 | 0 |
| 7 | 0.00 | 3.26 | 0.00 | 1.00 | 1 | Adamts13 | a disintegrin-like and metallopeptidase (r | 45 | 14 | 15 | 0 | 0 | 0 | 0 | 0 | 0 | 0 | 0 |
| 7 | -0.40 | 2.78 | -0.14 | 0.89 | 1 | Gabrb1 | gamma-aminobutyric acid (GABA) A rece | 21 | 5 | 47 | 0 | 0 | 1 | 0 | 0 | 0 | 0 | 0 |
| 7 | 0.36 | 2.55 | 0.14 | 0.89 | 1 | Gm44731 | | 0 | 0 | 6 | 10 | 0 | 14 | 0 | 0 | 6 | 26 | 11 |

| | | | | | | | | | | | | | | | | | | |
|---|---|---|---|---|---|---|---|---|---|---|---|---|---|---|---|---|---|---|
| 7 | -0.74 | 3.45 | -0.21 | 0.83 | 1 | Notum | notum palmitoleoyl-protein carboxylestera | 45 | 0 | 26 | 0 | 0 | 3 | 0 | 0 | 0 | 0 | 0 |
| 7 | 0.00 | 3.68 | 0.00 | 1.00 | 1 | Apol7a | apolipoprotein L 7a | 0 | 25 | 48 | 0 | 0 | 0 | 0 | 0 | 0 | 0 | 0 |
| 7 | -2.17 | 2.41 | -0.90 | 0.37 | 1 | 4930404H11Rik | | 0 | 4 | 0 | 4 | 5 | 25 | 0 | 0 | 0 | 0 | 35 |
| 7 | 1.03 | 2.35 | 0.44 | 0.66 | 1 | Gbp11 | guanylate binding protein 11 | 23 | 28 | 15 | 0 | 0 | 1 | 0 | 0 | 0 | 0 | 6 |
| 7 | -1.57 | 1.75 | -0.89 | 0.37 | 1 | Gm5640 | predicted gene 5640 | 13 | 15 | 22 | 10 | 6 | 2 | 0 | 0 | 2 | 0 | 4 |
| 7 | 0.00 | 3.28 | 0.00 | 1.00 | 1 | Mefv | Mediterranean fever | 52 | 12 | 8 | 0 | 0 | 0 | 0 | 0 | 0 | 0 | 0 |
| 7 | 0.00 | 3.30 | 0.00 | 1.00 | 1 | Mgat4c | MGAT4 family, member C | 3 | 53 | 17 | 0 | 0 | 0 | 0 | 0 | 0 | 0 | 0 |
| 7 | 0.20 | 2.62 | 0.08 | 0.94 | 1 | Tff3 | trefoil factor 3, intestinal | 3 | 40 | 8 | 0 | 5 | 0 | 0 | 17 | 0 | 0 | 0 |
| 7 | -1.30 | 2.06 | -0.63 | 0.53 | 1 | Npb | neuropeptide B | 15 | 14 | 23 | 0 | 3 | 12 | 0 | 0 | 0 | 0 | 6 |
| 7 | 1.10 | 2.26 | 0.49 | 0.63 | 1 | Gm37033 | | 0 | 0 | 0 | 0 | 5 | 35 | 0 | 0 | 10 | 3 | 20 |
| 7 | 0.00 | 3.28 | 0.00 | 1.00 | 1 | Kcng2 | potassium voltage-gated channel, subfam | 22 | 33 | 17 | 0 | 0 | 0 | 0 | 0 | 0 | 0 | 0 |
| 7 | 1.20 | 1.91 | 0.63 | 0.53 | 1 | Gm4017 | | 9 | 0 | 6 | 2 | 0 | 23 | 0 | 1 | 19 | 3 | 9 |
| 7 | 2.32 | 2.02 | 1.15 | 0.25 | 1 | Tesmin | testis expressed metallothionein like | 4 | 0 | 12 | 3 | 3 | 1 | 0 | 37 | 1 | 0 | 11 |
| 7 | 0.90 | 3.00 | 0.30 | 0.76 | 1 | Slc26a3 | solute carrier family 26, member 3 | 0 | 21 | 31 | 0 | 5 | 0 | 0 | 15 | 0 | 0 | 0 |
| 7 | 0.28 | 1.89 | 0.15 | 0.88 | 1 | C430014B12Rik | | 8 | 20 | 20 | 0 | 4 | 8 | 0 | 0 | 7 | 0 | 5 |
| 7 | 0.16 | 2.73 | 0.06 | 0.95 | 1 | Snora41 | | 3 | 7 | 0 | 0 | 0 | 32 | 0 | 0 | 0 | 0 | 29 |
| 7 | 1.76 | 2.85 | 0.62 | 0.54 | 1 | Lrrc26 | leucine rich repeat containing 26 | 26 | 16 | 25 | 0 | 0 | 0 | 0 | 0 | 5 | 0 | 0 |
| 7 | 0.00 | 3.28 | 0.00 | 1.00 | 1 | Rergl | RERG/RAS-like | 37 | 12 | 23 | 0 | 0 | 0 | 0 | 0 | 0 | 0 | 0 |
| 7 | 1.60 | 2.57 | 0.62 | 0.53 | 1 | Gm48137 | | 0 | 3 | 0 | 1 | 0 | 31 | 0 | 0 | 9 | 0 | 27 |
| 7 | 0.00 | 3.29 | 0.00 | 1.00 | 1 | Slc16a4 | solute carrier family 16 (monocarboxylic a | 37 | 22 | 12 | 0 | 0 | 0 | 0 | 0 | 0 | 0 | 0 |
| 7 | 0.60 | 2.77 | 0.22 | 0.83 | 1 | Gm45342 | | 0 | 0 | 19 | 0 | 0 | 25 | 0 | 0 | 2 | 0 | 26 |
| 7 | -0.40 | 2.81 | -0.14 | 0.89 | 1 | Hapln3 | hyaluronan and proteoglycan link protein | 9 | 9 | 53 | 0 | 0 | 1 | 0 | 0 | 0 | 0 | 0 |
| 7 | -2.16 | 2.48 | -0.87 | 0.38 | 1 | A530095I07Rik | | 5 | 27 | 31 | 7 | 0 | 1 | 0 | 0 | 0 | 0 | 0 |
| 6 | 0.11 | 2.59 | 0.04 | 0.97 | 1 | Aarsd1 | alanyl-tRNA synthetase domain containin | 1 | 0 | 0 | 8 | 0 | 26 | 0 | 0 | 11 | 0 | 25 |
| 6 | 0.09 | 2.98 | 0.03 | 0.98 | 1 | Gm47583 | | 4 | 0 | 0 | 0 | 8 | 15 | 0 | 30 | 0 | 0 | 14 |
| 6 | 1.38 | 2.83 | 0.49 | 0.63 | 1 | C1ql3 | C1q-like 3 | 37 | 28 | 2 | 0 | 0 | 0 | 0 | 0 | 0 | 0 | 4 |
| 6 | -1.31 | 2.46 | -0.53 | 0.59 | 1 | mt-Tw | | 42 | 6 | 0 | 0 | 3 | 5 | 0 | 0 | 0 | 14 | 1 |
| 6 | -0.17 | 3.35 | -0.05 | 0.96 | 1 | Nlrp5-ps | | 12 | 0 | 0 | 0 | 0 | 39 | 0 | 0 | 0 | 0 | 20 |
| 6 | 1.06 | 2.30 | 0.46 | 0.64 | 1 | Gm36634 | | 0 | 3 | 11 | 0 | 7 | 21 | 0 | 0 | 6 | 22 | 1 |
| 6 | 1.82 | 2.86 | 0.64 | 0.52 | 1 | Nkx6-3 | NK6 homeobox 3 | 23 | 27 | 16 | 0 | 0 | 0 | 0 | 0 | 5 | 0 | 0 |
| 6 | -1.09 | 1.97 | -0.55 | 0.58 | 1 | 1810019N24Rik | | 6 | 0 | 9 | 2 | 12 | 14 | 0 | 0 | 23 | 0 | 4 |
| 6 | -3.62 | 1.89 | -1.92 | 0.05 | 1 | Ccdc60 | coiled-coil domain containing 60 | 7 | 27 | 14 | 4 | 9 | 8 | 0 | 0 | 0 | 0 | 1 |
| 6 | 0.00 | 3.68 | 0.00 | 1.00 | 1 | Igkv4-73 | | 67 | 4 | 0 | 0 | 0 | 0 | 0 | 0 | 0 | 0 | 0 |
| 6 | 1.66 | 1.48 | 1.12 | 0.26 | 1 | Gm44837 | | 2 | 3 | 5 | 3 | 5 | 13 | 0 | 22 | 3 | 13 | 3 |
| 6 | -0.04 | 1.81 | -0.02 | 0.98 | 1 | Nlrp4f | NLR family, pyrin domain containing 4F | 8 | 20 | 0 | 9 | 7 | 6 | 0 | 10 | 5 | 0 | 6 |
| 6 | 0.00 | 3.68 | 0.00 | 1.00 | 1 | Oosp1 | oocyte secreted protein 1 | 51 | 19 | 0 | 0 | 0 | 0 | 0 | 0 | 0 | 0 | 0 |
| 6 | 0.00 | 3.31 | 0.00 | 1.00 | 1 | Prss30 | protease, serine 30 | 22 | 27 | 21 | 0 | 0 | 0 | 0 | 0 | 0 | 0 | 0 |
| 6 | 1.79 | 2.62 | 0.68 | 0.49 | 1 | Gm12941 | | 1 | 4 | 0 | 0 | 0 | 33 | 0 | 0 | 6 | 0 | 26 |
| 6 | -0.40 | 2.63 | -0.15 | 0.88 | 1 | Sstr4 | somatostatin receptor 4 | 24 | 31 | 3 | 9 | 0 | 0 | 0 | 0 | 0 | 0 | 3 |
| 6 | 1.53 | 3.31 | 0.46 | 0.64 | 1 | Gm47580 | | 0 | 0 | 0 | 0 | 0 | 35 | 0 | 0 | 4 | 0 | 31 |
| 6 | 0.06 | 2.21 | 0.03 | 0.98 | 1 | C330002G04Rik | | 0 | 0 | 3 | 22 | 6 | 9 | 0 | 0 | 1 | 13 | 16 |
| 6 | 0.62 | 2.89 | 0.22 | 0.83 | 1 | Gm45632 | | 0 | 0 | 0 | 2 | 0 | 31 | 0 | 0 | 0 | 12 | 25 |
| 6 | 1.40 | 2.24 | 0.63 | 0.53 | 1 | Acbd7 | acyl-Coenzyme A binding domain contair | 13 | 8 | 0 | 1 | 0 | 10 | 0 | 0 | 0 | 26 | 12 |
| 6 | -2.13 | 1.84 | -1.16 | 0.25 | 1 | Adgra1 | adhesion G protein-coupled receptor A1 | 11 | 10 | 27 | 5 | 11 | 1 | 0 | 0 | 1 | 0 | 3 |
| 6 | -0.86 | 1.96 | -0.44 | 0.66 | 1 | Gm30124 | | 0 | 7 | 32 | 4 | 2 | 8 | 0 | 1 | 16 | 0 | 0 |
| 6 | 0.85 | 1.74 | 0.49 | 0.63 | 1 | Gm47257 | | 0 | 4 | 10 | 6 | 4 | 16 | 0 | 0 | 11 | 4 | 14 |
| 6 | -0.92 | 1.74 | -0.53 | 0.60 | 1 | Gm16853 | | 0 | 1 | 25 | 2 | 8 | 16 | 0 | 0 | 5 | 0 | 13 |
| 6 | 0.00 | 3.68 | 0.00 | 1.00 | 1 | Sirpb1c | signal-regulatory protein beta 1C | 47 | 22 | 0 | 0 | 0 | 0 | 0 | 0 | 0 | 0 | 0 |
| 6 | 1.29 | 2.36 | 0.55 | 0.58 | 1 | Gm15690 | | 2 | 16 | 0 | 1 | 0 | 26 | 0 | 0 | 4 | 0 | 20 |
| 6 | 3.24 | 2.85 | 1.14 | 0.26 | 1 | Nrap | nebulin-related anchoring protein | 24 | 11 | 8 | 0 | 0 | 8 | 0 | 17 | 0 | 0 | 0 |
| 6 | -0.09 | 2.66 | -0.03 | 0.97 | 1 | Pck1 | phosphoenolpyruvate carboxykinase 1, c | 1 | 28 | 28 | 0 | 0 | 11 | 0 | 0 | 1 | 0 | 0 |
| 6 | 3.60 | 2.48 | 1.45 | 0.15 | 1 | Bpi | bactericidal permeablility increasing prote | 19 | 5 | 28 | 0 | 0 | 0 | 0 | 0 | 12 | 0 | 5 |
| 6 | 3.12 | 2.09 | 1.50 | 0.13 | 1 | Gm37940 | | 3 | 2 | 0 | 0 | 1 | 19 | 0 | 12 | 11 | 0 | 21 |
| 6 | 1.06 | 2.11 | 0.50 | 0.62 | 1 | Zfp57 | zinc finger protein 57 | 9 | 26 | 17 | 0 | 0 | 8 | 0 | 0 | 2 | 0 | 6 |
| 6 | 2.79 | 2.83 | 0.98 | 0.33 | 1 | Gabrb2 | gamma-aminobutyric acid (GABA) A rece | 5 | 4 | 52 | 0 | 0 | 1 | 0 | 7 | 0 | 0 | 0 |
| 6 | 0.70 | 2.76 | 0.25 | 0.80 | 1 | | | 0 | 0 | 0 | 10 | 0 | 14 | 0 | 34 | 0 | 0 | 12 |
| 6 | 0.00 | 3.33 | 0.00 | 1.00 | 1 | Asb10 | ankyrin repeat and SOCS box-containing | 20 | 27 | 22 | 0 | 0 | 0 | 0 | 0 | 0 | 0 | 0 |
| 6 | 1.15 | 2.35 | 0.49 | 0.62 | 1 | Gm49334 | | 0 | 0 | 0 | 8 | 2 | 8 | 0 | 0 | 17 | 32 | 2 |
| 6 | 0.54 | 1.94 | 0.28 | 0.78 | 1 | Gm42477 | | 1 | 25 | 11 | 0 | 2 | 10 | 0 | 7 | 0 | 0 | 12 |
| 6 | -0.40 | 2.82 | -0.14 | 0.89 | 1 | Ly6k | lymphocyte antigen 6 complex, locus K | 36 | 18 | 14 | 0 | 0 | 1 | 0 | 0 | 0 | 0 | 0 |

| | | | | | | | | | | | | | | | | | | |
|---|---|---|---|---|---|---|---|---|---|---|---|---|---|---|---|---|---|---|
| 6 | -0.15 | 2.29 | -0.07 | 0.95 | 1 | 9230116L04Rik | | 0 | 7 | 10 | 12 | 0 | 14 | 0 | 0 | 16 | 0 | 9 |
| 6 | -0.63 | 2.15 | -0.29 | 0.77 | 1 | E330037G11Rik | | 0 | 9 | 0 | 4 | 3 | 29 | 0 | 0 | 2 | 0 | 22 |
| 6 | 0.00 | 3.34 | 0.00 | 1.00 | 1 | Hk1os | | 18 | 15 | 35 | 0 | 0 | 0 | 0 | 0 | 0 | 0 | 0 |
| 6 | 3.29 | 2.90 | 1.14 | 0.26 | 1 | Lrp8os2 | | 30 | 21 | 0 | 0 | 0 | 5 | 0 | 0 | 9 | 0 | 3 |
| 6 | 0.38 | 3.42 | 0.11 | 0.91 | 1 | Amh | anti-Mullerian hormone | 0 | 0 | 11 | 0 | 0 | 25 | 0 | 0 | 0 | 0 | 32 |
| 6 | 0.00 | 3.68 | 0.00 | 1.00 | 1 | Sh3rf2 | SH3 domain containing ring finger 2 | 25 | 44 | 0 | 0 | 0 | 0 | 0 | 0 | 0 | 0 | 0 |
| 6 | -2.05 | 2.42 | -0.85 | 0.40 | 1 | Gulo | gulonolactone (L-) oxidase | 10 | 2 | 50 | 0 | 3 | 3 | 0 | 0 | 0 | 0 | 0 |
| 6 | -0.84 | 2.86 | -0.29 | 0.77 | 1 | Mucl3 | mucin like 3 | 10 | 46 | 8 | 0 | 0 | 4 | 0 | 0 | 0 | 0 | 0 |
| 6 | 1.93 | 3.01 | 0.64 | 0.52 | 1 | Pllp | plasma membrane proteolipid | 16 | 15 | 22 | 0 | 0 | 0 | 0 | 15 | 0 | 0 | 0 |
| 6 | 1.24 | 2.69 | 0.46 | 0.65 | 1 | Il12rb1 | interleukin 12 receptor, beta 1 | 26 | 31 | 0 | 0 | 0 | 5 | 0 | 0 | 2 | 0 | 4 |
| 6 | 2.61 | 2.24 | 1.16 | 0.24 | 1 | Gm47690 | | 0 | 22 | 9 | 0 | 2 | 6 | 0 | 0 | 3 | 12 | 14 |
| 6 | -2.21 | 2.02 | -1.10 | 0.27 | 1 | Hpcal4 | hippocalcin-like 4 | 19 | 19 | 10 | 5 | 3 | 3 | 0 | 0 | 0 | 0 | 9 |
| 6 | -3.32 | 1.94 | -1.71 | 0.09 | 1 | Gm37415 | | 0 | 16 | 10 | 4 | 26 | 8 | 0 | 0 | 1 | 1 | 0 |
| 6 | 1.38 | 2.82 | 0.49 | 0.63 | 1 | Smpd5 | sphingomyelin phosphodiesterase 5 | 29 | 17 | 18 | 0 | 0 | 0 | 0 | 0 | 0 | 0 | 4 |
| 6 | 1.97 | 1.66 | 1.19 | 0.23 | 1 | Gm9442 | | 2 | 7 | 7 | 8 | 0 | 10 | 0 | 13 | 3 | 14 | 4 |
| 6 | 2.94 | 2.65 | 1.11 | 0.27 | 1 | H3c8 | H3 clustered histone 8 | 0 | 0 | 0 | 3 | 0 | 5 | 0 | 0 | 6 | 50 | 4 |
| 6 | 0.00 | 3.35 | 0.00 | 1.00 | 1 | Myb | myeloblastosis oncogene | 36 | 8 | 24 | 0 | 0 | 0 | 0 | 0 | 0 | 0 | 0 |
| 6 | 0.21 | 1.99 | 0.11 | 0.92 | 1 | Gm33782 | | 1 | 26 | 9 | 0 | 7 | 8 | 0 | 0 | 6 | 0 | 11 |
| 6 | -0.15 | 2.52 | -0.06 | 0.95 | 1 | Acot3 | acyl-CoA thioesterase 3 | 33 | 13 | 14 | 0 | 6 | 0 | 0 | 0 | 3 | 0 | 0 |
| 6 | 0.85 | 1.93 | 0.44 | 0.66 | 1 | Gm26672 | | 6 | 4 | 10 | 0 | 6 | 16 | 0 | 19 | 4 | 0 | 3 |
| 6 | -0.74 | 3.53 | -0.21 | 0.83 | 1 | Lgals7 | lectin, galactose binding, soluble 7 | 26 | 40 | 0 | 0 | 0 | 3 | 0 | 0 | 0 | 0 | 0 |
| 6 | 0.00 | 3.36 | 0.00 | 1.00 | 1 | Tcf23 | transcription factor 23 | 42 | 11 | 15 | 0 | 0 | 0 | 0 | 0 | 0 | 0 | 0 |
| 6 | -4.00 | 3.02 | -1.33 | 0.18 | 1 | Gm24494 | | 2 | 0 | 0 | 10 | 0 | 38 | 0 | 0 | 0 | 0 | 18 |
| 6 | 0.12 | 3.60 | 0.03 | 0.97 | 1 | 5830487J09Rik | | 0 | 0 | 0 | 0 | 0 | 37 | 0 | 0 | 0 | 0 | 30 |
| 6 | 0.00 | 3.39 | 0.00 | 1.00 | 1 | Gm26588 | | 8 | 56 | 4 | 0 | 0 | 0 | 0 | 0 | 0 | 0 | 0 |
| 6 | 0.00 | 3.36 | 0.00 | 1.00 | 1 | Ltf | lactotransferrin | 18 | 13 | 36 | 0 | 0 | 0 | 0 | 0 | 0 | 0 | 0 |
| 6 | -2.05 | 2.76 | -0.75 | 0.46 | 1 | 4933432I09Rik | | 0 | 0 | 0 | 1 | 5 | 41 | 0 | 0 | 0 | 0 | 21 |
| 6 | 0.60 | 2.54 | 0.24 | 0.81 | 1 | Gm47826 | | 0 | 3 | 0 | 0 | 2 | 32 | 0 | 0 | 5 | 0 | 25 |
| 6 | 0.00 | 3.45 | 0.00 | 1.00 | 1 | Il12rb2 | interleukin 12 receptor, beta 2 | 43 | 23 | 1 | 0 | 0 | 0 | 0 | 0 | 0 | 0 | 0 |
| 6 | 0.14 | 2.32 | 0.06 | 0.95 | 1 | 2010308F09Rik | | 6 | 12 | 1 | 8 | 0 | 4 | 0 | 36 | 0 | 0 | 0 |
| 6 | -1.70 | 2.29 | -0.74 | 0.46 | 1 | Trim34b | tripartite motif-containing 34B | 0 | 14 | 24 | 18 | 1 | 2 | 0 | 0 | 8 | 0 | 0 |
| 6 | 0.00 | 3.38 | 0.00 | 1.00 | 1 | Gm26645 | | 51 | 5 | 11 | 0 | 0 | 0 | 0 | 0 | 0 | 0 | 0 |
| 6 | -0.29 | 3.37 | -0.09 | 0.93 | 1 | Gm45853 | | 0 | 5 | 0 | 0 | 0 | 46 | 0 | 0 | 0 | 0 | 16 |
| 6 | 2.77 | 2.27 | 1.22 | 0.22 | 1 | 2900009J06Rik | | 0 | 0 | 10 | 10 | 1 | 1 | 0 | 36 | 0 | 5 | 4 |
| 6 | 2.56 | 2.64 | 0.97 | 0.33 | 1 | Gm23849 | | 18 | 20 | 0 | 0 | 0 | 1 | 0 | 0 | 0 | 26 | 2 |
| 6 | 1.43 | 2.92 | 0.49 | 0.62 | 1 | Vwa2 | von Willebrand factor A domain containin | 3 | 40 | 18 | 0 | 0 | 0 | 0 | 0 | 0 | 0 | 7 |
| 6 | -3.24 | 2.10 | -1.54 | 0.12 | 1 | Shd | src homology 2 domain-containing transfo | 12 | 7 | 5 | 23 | 3 | 7 | 0 | 0 | 0 | 0 | 10 |
| 6 | -0.98 | 2.61 | -0.38 | 0.71 | 1 | Gm35037 | | 8 | 0 | 0 | 11 | 9 | 5 | 0 | 29 | 0 | 0 | 5 |
| 6 | 0.00 | 3.68 | 0.00 | 1.00 | 1 | Cdx1 | caudal type homeobox 1 | 0 | 33 | 33 | 0 | 0 | 0 | 0 | 0 | 0 | 0 | 0 |
| 6 | -1.35 | 2.67 | -0.50 | 0.61 | 1 | Il1bos | | 26 | 7 | 0 | 0 | 2 | 23 | 0 | 0 | 0 | 0 | 9 |
| 6 | 3.05 | 3.10 | 0.98 | 0.33 | 1 | Nxnl2 | nucleoredoxin-like 2 | 27 | 19 | 0 | 0 | 0 | 0 | 0 | 20 | 1 | 0 | 0 |
| 6 | 1.37 | 2.87 | 0.48 | 0.63 | 1 | Cfb | complement factor B | 43 | 6 | 14 | 0 | 0 | 0 | 0 | 0 | 0 | 0 | 4 |
| 6 | 2.09 | 2.46 | 0.85 | 0.40 | 1 | Gm37019 | | 2 | 0 | 22 | 0 | 0 | 3 | 0 | 0 | 28 | 0 | 11 |
| 6 | 0.00 | 3.40 | 0.00 | 1.00 | 1 | Pgam2 | phosphoglycerate mutase 2 | 49 | 12 | 5 | 0 | 0 | 0 | 0 | 0 | 0 | 0 | 0 |
| 6 | 0.96 | 2.86 | 0.33 | 0.74 | 1 | Ranbp3l | RAN binding protein 3-like | 15 | 12 | 38 | 0 | 0 | 0 | 0 | 0 | 0 | 0 | 1 |
| 6 | 0.96 | 2.90 | 0.33 | 0.74 | 1 | Lbx2 | ladybird homeobox 2 | 55 | 6 | 4 | 0 | 0 | 0 | 0 | 0 | 0 | 0 | 1 |
| 6 | 0.93 | 2.98 | 0.31 | 0.76 | 1 | Gm34237 | | 0 | 0 | 0 | 2 | 0 | 23 | 0 | 0 | 19 | 0 | 22 |
| 6 | 1.81 | 2.66 | 0.68 | 0.50 | 1 | Gm23134 | | 0 | 2 | 0 | 1 | 0 | 39 | 0 | 0 | 10 | 0 | 14 |
| 6 | 0.00 | 3.39 | 0.00 | 1.00 | 1 | Sp7 | Sp7 transcription factor 7 | 14 | 27 | 25 | 0 | 0 | 0 | 0 | 0 | 0 | 0 | 0 |
| 6 | -3.84 | 3.12 | -1.23 | 0.22 | 1 | Gm42986 | | 12 | 0 | 0 | 0 | 6 | 33 | 0 | 0 | 0 | 0 | 15 |
| 6 | -0.16 | 2.11 | -0.08 | 0.94 | 1 | Tex14 | testis expressed gene 14 | 15 | 27 | 14 | 0 | 1 | 3 | 0 | 0 | 0 | 0 | 6 |
| 6 | -3.54 | 2.46 | -1.44 | 0.15 | 1 | Gm10030 | | 4 | 0 | 10 | 27 | 2 | 17 | 0 | 0 | 0 | 0 | 5 |
| 6 | 0.00 | 3.68 | 0.00 | 1.00 | 1 | Musk | muscle, skeletal, receptor tyrosine kinase | 53 | 0 | 12 | 0 | 0 | 0 | 0 | 0 | 0 | 0 | 0 |
| 6 | -0.46 | 2.06 | -0.22 | 0.82 | 1 | BC060293 | | 0 | 3 | 15 | 0 | 5 | 22 | 0 | 0 | 1 | 0 | 20 |
| 6 | 0.09 | 1.88 | 0.05 | 0.96 | 1 | Gm9522 | | 2 | 0 | 0 | 1 | 6 | 22 | 0 | 0 | 5 | 4 | 26 |
| 6 | 0.00 | 3.68 | 0.00 | 1.00 | 1 | Fut2 | fucosyltransferase 2 | 34 | 0 | 31 | 0 | 0 | 0 | 0 | 0 | 0 | 0 | 0 |
| 6 | 3.07 | 3.08 | 1.00 | 0.32 | 1 | Gm32815 | | 0 | 20 | 19 | 0 | 0 | 0 | 0 | 25 | 0 | 0 | 1 |
| 6 | 1.33 | 2.89 | 0.46 | 0.64 | 1 | Tcap | titin-cap | 24 | 4 | 34 | 0 | 0 | 0 | 0 | 0 | 0 | 0 | 3 |
| 6 | 2.07 | 2.34 | 0.88 | 0.38 | 1 | 1700125H03Rik | | 3 | 36 | 15 | 0 | 0 | 2 | 0 | 0 | 0 | 5 | 4 |
| 6 | 0.00 | 3.42 | 0.00 | 1.00 | 1 | Fpr1 | formyl peptide receptor 1 | 33 | 27 | 5 | 0 | 0 | 0 | 0 | 0 | 0 | 0 | 0 |

| | | | | | | | | | | | | | | | | | | |
|---|---|---|---|---|---|---|---|---|---|---|---|---|---|---|---|---|---|---|
| 6 | 0.00 | 3.40 | 0.00 | 1.00 | 1 | Gm266 | predicted gene 266 | 18 | 29 | 18 | 0 | 0 | 0 | 0 | 0 | 0 | 0 | 0 |
| 6 | 3.38 | 2.24 | 1.51 | 0.13 | 1 | 1700026F02Rik | | 0 | 0 | 0 | 0 | 4 | 3 | 0 | 35 | 5 | 17 | 2 |
| 6 | 0.16 | 3.38 | 0.05 | 0.96 | 1 | Gm44982 | | 5 | 0 | 0 | 0 | 0 | 31 | 0 | 0 | 0 | 0 | 28 |
| 6 | -0.20 | 3.47 | -0.06 | 0.95 | 1 | 2900018N21Rik | | 0 | 10 | 0 | 0 | 0 | 36 | 0 | 0 | 0 | 0 | 18 |
| 6 | 0.20 | 2.69 | 0.07 | 0.94 | 1 | Gm45246 | | 5 | 12 | 25 | 0 | 11 | 0 | 0 | 0 | 11 | 0 | 0 |
| 6 | 4.47 | 2.34 | 1.91 | 0.06 | 1 | Gm12185 | predicted gene 12185 | 8 | 26 | 6 | 0 | 0 | 0 | 0 | 0 | 1 | 19 | 4 |
| 6 | 0.00 | 3.41 | 0.00 | 1.00 | 1 | Ms4a8a | membrane-spanning 4-domains, subfami | 37 | 11 | 17 | 0 | 0 | 0 | 0 | 0 | 0 | 0 | 0 |
| 6 | 0.00 | 3.41 | 0.00 | 1.00 | 1 | Nccrp1 | non-specific cytotoxic cell receptor proteir | 20 | 31 | 14 | 0 | 0 | 0 | 0 | 0 | 0 | 0 | 0 |
| 6 | 1.25 | 2.30 | 0.54 | 0.59 | 1 | 4833422C13Rik | | 11 | 27 | 17 | 0 | 0 | 3 | 0 | 0 | 5 | 0 | 1 |
| 6 | -0.32 | 2.48 | -0.13 | 0.90 | 1 | Gm47424 | | 0 | 14 | 10 | 1 | 0 | 18 | 0 | 0 | 0 | 0 | 21 |
| 6 | 0.00 | 3.43 | 0.00 | 1.00 | 1 | Gm13387 | | 6 | 32 | 26 | 0 | 0 | 0 | 0 | 0 | 0 | 0 | 0 |
| 6 | 0.00 | 3.68 | 0.00 | 1.00 | 1 | A530032D15Ri | RIKEN cDNA A530032D15Rik gene | 31 | 33 | 0 | 0 | 0 | 0 | 0 | 0 | 0 | 0 | 0 |
| 6 | 0.00 | 3.42 | 0.00 | 1.00 | 1 | Duox1 | dual oxidase 1 | 39 | 15 | 10 | 0 | 0 | 0 | 0 | 0 | 0 | 0 | 0 |
| 6 | 0.09 | 3.46 | 0.03 | 0.98 | 1 | Mir1898 | | 0 | 8 | 0 | 0 | 0 | 30 | 0 | 0 | 0 | 0 | 26 |
| 6 | 0.05 | 2.53 | 0.02 | 0.99 | 1 | Gm9955 | | 0 | 0 | 8 | 1 | 3 | 7 | 0 | 0 | 18 | 27 | 0 |
| 6 | 0.13 | 1.92 | 0.07 | 0.95 | 1 | Gm37292 | | 2 | 8 | 12 | 0 | 6 | 13 | 0 | 0 | 4 | 0 | 19 |
| 6 | -1.24 | 3.01 | -0.41 | 0.68 | 1 | Ccl12 | chemokine (C-C motif) ligand 12 | 15 | 7 | 29 | 0 | 12 | 0 | 0 | 0 | 0 | 0 | 0 |
| 6 | -3.43 | 3.12 | -1.10 | 0.27 | 1 | Gm43654 | | 0 | 11 | 0 | 0 | 3 | 35 | 0 | 0 | 0 | 0 | 15 |
| 6 | 0.32 | 2.39 | 0.13 | 0.89 | 1 | Slc4a11 | solute carrier family 4, sodium bicarbonat | 9 | 29 | 6 | 0 | 0 | 8 | 0 | 0 | 0 | 0 | 11 |
| 6 | 0.63 | 2.70 | 0.23 | 0.82 | 1 | Gm19439 | | 0 | 10 | 11 | 7 | 0 | 8 | 0 | 0 | 0 | 19 | 8 |
| 6 | 0.57 | 1.85 | 0.31 | 0.76 | 1 | Gm23833 | | 0 | 24 | 0 | 17 | 2 | 3 | 0 | 7 | 3 | 0 | 8 |
| 6 | 1.62 | 2.98 | 0.54 | 0.59 | 1 | Gm34030 | | 44 | 3 | 14 | 0 | 0 | 0 | 0 | 0 | 3 | 0 | 0 |
| 6 | 2.24 | 3.53 | 0.63 | 0.53 | 1 | Gm18073 | | 0 | 0 | 0 | 0 | 0 | 4 | 0 | 0 | 0 | 49 | 10 |
| 6 | 0.00 | 3.46 | 0.00 | 1.00 | 1 | Klk1b3 | kallikrein 1-related peptidase b3 | 4 | 17 | 42 | 0 | 0 | 0 | 0 | 0 | 0 | 0 | 0 |
| 6 | 1.93 | 3.05 | 0.63 | 0.53 | 1 | Slc1a2 | solute carrier family 1 (glial high affinity gl | 39 | 12 | 6 | 0 | 0 | 0 | 0 | 0 | 6 | 0 | 0 |
| 6 | 0.19 | 2.70 | 0.07 | 0.95 | 1 | Pate2 | prostate and testis expressed 2 | 8 | 0 | 0 | 18 | 1 | 8 | 0 | 0 | 9 | 0 | 19 |
| 6 | -0.03 | 1.94 | -0.02 | 0.99 | 1 | Gm38037 | | 0 | 11 | 3 | 14 | 6 | 7 | 0 | 0 | 12 | 0 | 10 |
| 6 | 0.62 | 2.16 | 0.29 | 0.78 | 1 | Gm37834 | | 2 | 2 | 8 | 0 | 8 | 15 | 0 | 14 | 0 | 0 | 14 |
| 6 | -0.70 | 2.90 | -0.24 | 0.81 | 1 | Gm10584 | | 16 | 30 | 15 | 0 | 0 | 2 | 0 | 0 | 0 | 0 | 0 |
| 6 | 0.00 | 3.68 | 0.00 | 1.00 | 1 | Eps8l3 | EPS8-like 3 | 0 | 21 | 42 | 0 | 0 | 0 | 0 | 0 | 0 | 0 | 0 |
| 6 | 0.00 | 3.45 | 0.00 | 1.00 | 1 | A830012C17Rik | | 29 | 24 | 9 | 0 | 0 | 0 | 0 | 0 | 0 | 0 | 0 |
| 6 | -1.50 | 3.49 | -0.43 | 0.67 | 1 | Gm20274 | | 0 | 0 | 0 | 0 | 10 | 32 | 0 | 0 | 0 | 0 | 20 |
| 6 | 4.06 | 2.37 | 1.72 | 0.09 | 1 | Sag | S-antigen, retina and pineal gland (arrest | 4 | 5 | 0 | 0 | 0 | 12 | 0 | 16 | 6 | 0 | 19 |
| 6 | 0.75 | 1.52 | 0.49 | 0.62 | 1 | Gm10327 | | 11 | 7 | 9 | 9 | 2 | 3 | 0 | 1 | 2 | 12 | 7 |
| 6 | 0.00 | 3.47 | 0.00 | 1.00 | 1 | Hs3st3a1 | heparan sulfate (glucosamine) 3-O-sulfot | 52 | 5 | 5 | 0 | 0 | 0 | 0 | 0 | 0 | 0 | 0 |
| 6 | 0.00 | 3.46 | 0.00 | 1.00 | 1 | Six2 | sine oculis-related homeobox 2 | 36 | 19 | 7 | 0 | 0 | 0 | 0 | 0 | 0 | 0 | 0 |
| 6 | 0.74 | 1.99 | 0.37 | 0.71 | 1 | 1700047K16Rik | | 5 | 11 | 15 | 0 | 4 | 8 | 0 | 0 | 8 | 0 | 12 |
| 6 | 0.00 | 3.46 | 0.00 | 1.00 | 1 | 1810007D17Rik | | 12 | 18 | 32 | 0 | 0 | 0 | 0 | 0 | 0 | 0 | 0 |
| 6 | 4.17 | 3.27 | 1.28 | 0.20 | 1 | Gm37144 | | 0 | 0 | 16 | 0 | 0 | 14 | 0 | 8 | 0 | 0 | 24 |
| 6 | 1.60 | 3.41 | 0.47 | 0.64 | 1 | Gm17195 | | 0 | 0 | 0 | 0 | 0 | 23 | 0 | 0 | 3 | 0 | 37 |
| 6 | 0.00 | 3.46 | 0.00 | 1.00 | 1 | Foxf2 | forkhead box F2 | 32 | 20 | 10 | 0 | 0 | 0 | 0 | 0 | 0 | 0 | 0 |
| 6 | -0.40 | 3.63 | -0.11 | 0.91 | 1 | C920009B18Rik | | 47 | 14 | 0 | 0 | 0 | 1 | 0 | 0 | 0 | 0 | 0 |
| 6 | 0.00 | 3.47 | 0.00 | 1.00 | 1 | Slc36a2 | solute carrier family 36 (proton/amino acid | 44 | 12 | 6 | 0 | 0 | 0 | 0 | 0 | 0 | 0 | 0 |
| 6 | -0.60 | 2.35 | -0.26 | 0.80 | 1 | Gm40828 | | 0 | 28 | 6 | 0 | 14 | 4 | 0 | 0 | 4 | 0 | 5 |
| 6 | 2.88 | 3.15 | 0.92 | 0.36 | 1 | Mir421 | | 0 | 11 | 0 | 0 | 0 | 25 | 0 | 0 | 2 | 0 | 24 |
| 6 | 1.82 | 2.19 | 0.83 | 0.41 | 1 | A030005L19Rik | | 1 | 11 | 12 | 0 | 0 | 10 | 0 | 0 | 4 | 0 | 23 |
| 6 | 1.85 | 2.25 | 0.82 | 0.41 | 1 | 4930507D05Rik | | 0 | 0 | 8 | 6 | 10 | 0 | 0 | 29 | 5 | 0 | 3 |
| 6 | 1.04 | 2.68 | 0.39 | 0.70 | 1 | Slc23a3 | solute carrier family 23 (nucleobase trans | 12 | 13 | 0 | 0 | 0 | 25 | 0 | 0 | 2 | 0 | 10 |
| 6 | 0.00 | 3.68 | 0.00 | 1.00 | 1 | Ighv1-62-2 | | 11 | 51 | 0 | 0 | 0 | 0 | 0 | 0 | 0 | 0 | 0 |
| 6 | 0.19 | 3.60 | 0.05 | 0.96 | 1 | Gm24564 | | 0 | 0 | 0 | 0 | 0 | 31 | 0 | 0 | 0 | 0 | 30 |
| 6 | -2.13 | 3.30 | -0.65 | 0.52 | 1 | Hoxa11 | homeobox A11 | 48 | 8 | 0 | 0 | 5 | 1 | 0 | 0 | 0 | 0 | 0 |
| 6 | 0.99 | 2.93 | 0.34 | 0.74 | 1 | Tmem171 | transmembrane protein 171 | 31 | 13 | 17 | 0 | 0 | 0 | 0 | 0 | 1 | 0 | 0 |
| 6 | 2.95 | 2.42 | 1.22 | 0.22 | 1 | Gm16596 | | 15 | 0 | 6 | 0 | 2 | 6 | 0 | 22 | 9 | 0 | 1 |
| 6 | 0.00 | 3.52 | 0.00 | 1.00 | 1 | Tmem156 | transmembrane protein 156 | 53 | 3 | 5 | 0 | 0 | 0 | 0 | 0 | 0 | 0 | 0 |
| 6 | 0.00 | 3.49 | 0.00 | 1.00 | 1 | Cav3 | caveolin 3 | 34 | 7 | 20 | 0 | 0 | 0 | 0 | 0 | 0 | 0 | 0 |
| 6 | 1.45 | 3.14 | 0.46 | 0.65 | 1 | Dnah6 | dynein, axonemal, heavy chain 6 | 0 | 12 | 34 | 0 | 2 | 0 | 0 | 13 | 0 | 0 | 0 |
| 6 | -1.42 | 2.46 | -0.58 | 0.56 | 1 | Gm43604 | | 3 | 0 | 0 | 2 | 14 | 4 | 0 | 0 | 0 | 32 | 6 |
| 6 | 0.00 | 3.58 | 0.00 | 1.00 | 1 | Gapt | Grb2-binding adaptor, transmembrane | 51 | 1 | 9 | 0 | 0 | 0 | 0 | 0 | 0 | 0 | 0 |
| 6 | -1.43 | 2.58 | -0.56 | 0.58 | 1 | Npc1l1 | NPC1 like intracellular cholesterol transp | 9 | 38 | 10 | 0 | 1 | 2 | 0 | 0 | 0 | 0 | 0 |
| 6 | 0.05 | 2.86 | 0.02 | 0.99 | 1 | Gm37979 | | 3 | 2 | 0 | 0 | 0 | 33 | 0 | 0 | 0 | 0 | 23 |

| | | | | | | | | | | | | | | | | | | |
|---|---|---|---|---|---|---|---|---|---|---|---|---|---|---|---|---|---|---|
| 6 | -0.40 | 3.67 | -0.11 | 0.91 | 1 | Cd101 | CD101 antigen | 0 | 10 | 50 | 0 | 0 | 1 | 0 | 0 | 0 | 0 | 0 |
| 6 | 3.02 | 2.49 | 1.21 | 0.22 | 1 | Gm48653 | | 0 | 9 | 0 | 12 | 0 | 12 | 0 | 0 | 3 | 15 | 9 |
| 6 | 0.25 | 3.46 | 0.07 | 0.94 | 1 | Gm38248 | | 5 | 0 | 0 | 0 | 0 | 26 | 0 | 0 | 0 | 0 | 29 |
| 6 | 0.00 | 3.68 | 0.00 | 1.00 | 1 | Adgrg7 | adhesion G protein-coupled receptor G7 | 0 | 21 | 39 | 0 | 0 | 0 | 0 | 0 | 0 | 0 | 0 |
| 6 | 1.74 | 2.18 | 0.80 | 0.42 | 1 | Gm18867 | | 18 | 0 | 0 | 3 | 1 | 0 | 0 | 1 | 2 | 31 | 5 |
| 6 | -2.32 | 3.50 | -0.66 | 0.51 | 1 | Snai3 | snail family zinc finger 3 | 4 | 44 | 0 | 0 | 5 | 0 | 0 | 0 | 0 | 0 | 8 |
| 6 | -0.16 | 2.44 | -0.07 | 0.95 | 1 | Tigd4 | tigger transposable element derived 4 | 20 | 8 | 23 | 0 | 0 | 7 | 0 | 0 | 0 | 0 | 3 |
| 5 | 0.30 | 3.43 | 0.09 | 0.93 | 1 | Snord72 | | 3 | 0 | 0 | 0 | 0 | 25 | 0 | 0 | 0 | 0 | 32 |
| 5 | 2.11 | 1.58 | 1.34 | 0.18 | 1 | Gm9320 | | 3 | 1 | 16 | 5 | 1 | 2 | 0 | 21 | 6 | 1 | 5 |
| 5 | 0.00 | 3.54 | 0.00 | 1.00 | 1 | Cyp2ab1 | cytochrome P450, family 2, subfamily ab, | 42 | 16 | 2 | 0 | 0 | 0 | 0 | 0 | 0 | 0 | 0 |
| 5 | 0.28 | 2.53 | 0.11 | 0.91 | 1 | Gm44067 | | 21 | 23 | 15 | 0 | 0 | 1 | 0 | 0 | 0 | 0 | 1 |
| 5 | 1.33 | 2.94 | 0.45 | 0.65 | 1 | Gfi1 | growth factor independent 1 transcription | 10 | 34 | 12 | 0 | 0 | 0 | 0 | 0 | 0 | 0 | 3 |
| 5 | -1.00 | 3.50 | -0.29 | 0.78 | 1 | Gm37559 | | 0 | 0 | 0 | 0 | 5 | 22 | 0 | 0 | 0 | 0 | 34 |
| 5 | -0.35 | 2.79 | -0.13 | 0.90 | 1 | Tub | tubby bipartite transcription factor | 13 | 16 | 15 | 14 | 0 | 0 | 0 | 0 | 0 | 3 | 0 |
| 5 | 3.68 | 3.54 | 1.04 | 0.30 | 1 | Gm37653 | | 0 | 0 | 0 | 0 | 0 | 22 | 0 | 0 | 0 | 27 | 11 |
| 5 | 3.56 | 2.86 | 1.24 | 0.21 | 1 | Kremen2 | kringle containing transmembrane proteir | 24 | 7 | 7 | 0 | 0 | 1 | 0 | 21 | 0 | 0 | 0 |
| 5 | -2.68 | 2.98 | -0.90 | 0.37 | 1 | Foxp3 | forkhead box P3 | 7 | 15 | 0 | 0 | 2 | 3 | 0 | 34 | 0 | 0 | 0 |
| 5 | 0.00 | 3.63 | 0.00 | 1.00 | 1 | Igkv3-5 | | 60 | 0 | 0 | 0 | 0 | 0 | 0 | 0 | 0 | 0 | 0 |
| 5 | 0.28 | 2.54 | 0.11 | 0.91 | 1 | Gm45187 | | 34 | 3 | 21 | 0 | 0 | 1 | 0 | 0 | 0 | 0 | 1 |
| 5 | -0.83 | 2.95 | -0.28 | 0.78 | 1 | Gm10109 | | 15 | 9 | 31 | 0 | 0 | 4 | 0 | 0 | 0 | 0 | 0 |
| 5 | -0.40 | 3.00 | -0.13 | 0.89 | 1 | Tnfrsf18 | tumor necrosis factor receptor superfamily | 45 | 2 | 12 | 0 | 0 | 1 | 0 | 0 | 0 | 0 | 0 |
| 5 | 0.49 | 1.55 | 0.32 | 0.75 | 1 | Gm27217 | | 6 | 12 | 12 | 4 | 5 | 4 | 0 | 0 | 8 | 3 | 6 |
| 5 | 3.68 | 2.88 | 1.28 | 0.20 | 1 | 4632418H02Rik | | 11 | 5 | 0 | 0 | 0 | 15 | 0 | 0 | 11 | 0 | 17 |
| 5 | 0.00 | 3.51 | 0.00 | 1.00 | 1 | Gm15270 | | 24 | 23 | 12 | 0 | 0 | 0 | 0 | 0 | 0 | 0 | 0 |
| 5 | 1.02 | 2.91 | 0.35 | 0.73 | 1 | C4bp-ps1 | | 0 | 0 | 0 | 0 | 1 | 30 | 0 | 0 | 8 | 0 | 20 |
| 5 | 0.00 | 3.52 | 0.00 | 1.00 | 1 | Mmp7 | matrix metallopeptidase 7 | 34 | 20 | 5 | 0 | 0 | 0 | 0 | 0 | 0 | 0 | 0 |
| 5 | -2.28 | 2.29 | -1.00 | 0.32 | 1 | Gm13534 | | 0 | 5 | 1 | 11 | 8 | 14 | 0 | 22 | 0 | 0 | 0 |
| 5 | 0.00 | 3.66 | 0.00 | 1.00 | 1 | Ifng | interferon gamma | 1 | 59 | 0 | 0 | 0 | 0 | 0 | 0 | 0 | 0 | 0 |
| 5 | 0.03 | 2.36 | 0.01 | 0.99 | 1 | Nkain3 | Na+/K+ transporting ATPase interacting 3 | 22 | 14 | 8 | 0 | 5 | 1 | 0 | 0 | 9 | 0 | 0 |
| 5 | 2.85 | 2.73 | 1.04 | 0.30 | 1 | 2600014E21Rik | | 9 | 13 | 1 | 0 | 0 | 25 | 0 | 0 | 11 | 0 | 1 |
| 5 | -3.29 | 2.67 | -1.23 | 0.22 | 1 | Gm16548 | | 22 | 14 | 9 | 0 | 9 | 5 | 0 | 0 | 0 | 0 | 0 |
| 5 | 0.53 | 2.89 | 0.19 | 0.85 | 1 | Actn2 | actinin alpha 2 | 0 | 0 | 0 | 4 | 0 | 28 | 0 | 13 | 0 | 0 | 15 |
| 5 | 0.34 | 3.44 | 0.10 | 0.92 | 1 | Gm9278 | | 10 | 0 | 0 | 0 | 24 | 0 | 0 | 0 | 6 | 0 | 19 |
| 5 | -2.71 | 3.08 | -0.88 | 0.38 | 1 | Gm19784 | | 27 | 22 | 0 | 7 | 0 | 3 | 0 | 0 | 0 | 0 | 0 |
| 5 | 0.00 | 3.53 | 0.00 | 1.00 | 1 | Crocc2 | ciliary rootlet coiled-coil, rootletin family m | 21 | 27 | 10 | 0 | 0 | 0 | 0 | 0 | 0 | 0 | 0 |
| 5 | 0.00 | 3.68 | 0.00 | 1.00 | 1 | Cubn | cubilin (intrinsic factor-cobalamin recepto | 25 | 0 | 34 | 0 | 0 | 0 | 0 | 0 | 0 | 0 | 0 |
| 5 | 2.17 | 2.25 | 0.97 | 0.33 | 1 | Foxq1 | forkhead box Q1 | 12 | 13 | 25 | 0 | 0 | 1 | 0 | 0 | 4 | 0 | 4 |
| 5 | -0.76 | 2.96 | -0.26 | 0.80 | 1 | Hoxb3os | | 15 | 19 | 22 | 0 | 0 | 3 | 0 | 0 | 0 | 0 | 0 |
| 5 | 0.00 | 3.68 | 0.00 | 1.00 | 1 | Chrm1 | cholinergic receptor, muscarinic 1, CNS | 40 | 0 | 19 | 0 | 0 | 0 | 0 | 0 | 0 | 0 | 0 |
| 5 | -1.25 | 3.63 | -0.34 | 0.73 | 1 | Olfr78 | olfactory receptor 78 | 40 | 15 | 0 | 4 | 0 | 0 | 0 | 0 | 0 | 0 | 0 |
| 5 | 2.41 | 3.33 | 0.72 | 0.47 | 1 | Gm48702 | | 0 | 0 | 0 | 7 | 1 | 0 | 0 | 0 | 0 | 49 | 2 |
| 5 | -0.42 | 2.53 | -0.17 | 0.87 | 1 | Snora73b | | 12 | 1 | 3 | 0 | 0 | 35 | 0 | 0 | 0 | 0 | 8 |
| 5 | -0.83 | 3.06 | -0.27 | 0.78 | 1 | Adgrf4 | adhesion G protein-coupled receptor F4 | 46 | 2 | 6 | 0 | 0 | 4 | 0 | 0 | 0 | 0 | 0 |
| 5 | 3.53 | 3.21 | 1.10 | 0.27 | 1 | Grk1 | G protein-coupled receptor kinase 1 | 0 | 0 | 1 | 0 | 0 | 1 | 0 | 0 | 4 | 53 | 0 |
| 5 | 0.55 | 3.02 | 0.18 | 0.86 | 1 | Gm49027 | | 7 | 0 | 0 | 7 | 0 | 14 | 0 | 0 | 8 | 0 | 22 |
| 5 | 0.52 | 3.58 | 0.15 | 0.88 | 1 | Gm29107 | | 13 | 0 | 0 | 0 | 0 | 15 | 0 | 0 | 0 | 0 | 30 |
| 5 | -0.46 | 2.22 | -0.21 | 0.84 | 1 | Trim6 | tripartite motif-containing 6 | 2 | 19 | 30 | 0 | 1 | 3 | 0 | 0 | 0 | 0 | 3 |
| 5 | 0.00 | 3.55 | 0.00 | 1.00 | 1 | Klrd1 | killer cell lectin-like receptor, subfamily D, | 36 | 17 | 5 | 0 | 0 | 0 | 0 | 0 | 0 | 0 | 0 |
| 5 | -3.20 | 2.29 | -1.40 | 0.16 | 1 | Rfx8 | regulatory factor X 8 | 22 | 23 | 3 | 5 | 1 | 3 | 0 | 0 | 0 | 0 | 0 |
| 5 | -0.16 | 1.93 | -0.08 | 0.93 | 1 | Gm37194 | | 10 | 0 | 12 | 7 | 4 | 1 | 0 | 1 | 1 | 21 | 1 |
| 5 | 0.08 | 2.98 | 0.03 | 0.98 | 1 | Dmc1 | DNA meiotic recombinase 1 | 6 | 0 | 17 | 0 | 0 | 20 | 0 | 0 | 0 | 0 | 15 |
| 5 | 1.76 | 3.54 | 0.50 | 0.62 | 1 | Gm29427 | | 0 | 0 | 0 | 0 | 0 | 24 | 0 | 0 | 14 | 0 | 20 |
| 5 | -3.91 | 3.33 | -1.17 | 0.24 | 1 | Rbp2 | retinol binding protein 2, cellular | 0 | 0 | 0 | 9 | 15 | 6 | 0 | 0 | 0 | 28 | 0 |
| 5 | 0.28 | 3.06 | 0.09 | 0.93 | 1 | Gm42856 | | 7 | 0 | 36 | 0 | 0 | 7 | 0 | 0 | 0 | 0 | 8 |
| 5 | 0.00 | 3.56 | 0.00 | 1.00 | 1 | Tm4sf5 | transmembrane 4 superfamily member 5 | 34 | 5 | 19 | 0 | 0 | 0 | 0 | 0 | 0 | 0 | 0 |
| 5 | -0.39 | 2.14 | -0.18 | 0.85 | 1 | Gm32950 | | 12 | 3 | 20 | 5 | 0 | 7 | 0 | 0 | 2 | 0 | 9 |
| 5 | -1.21 | 3.09 | -0.39 | 0.70 | 1 | Ctsw | cathepsin W | 21 | 23 | 9 | 0 | 5 | 0 | 0 | 0 | 0 | 0 | 0 |
| 5 | 0.01 | 3.10 | 0.00 | 1.00 | 1 | Gm45809 | | 0 | 0 | 0 | 12 | 0 | 15 | 0 | 0 | 4 | 0 | 26 |
| 5 | 0.00 | 3.56 | 0.00 | 1.00 | 1 | Hoxc4 | homeobox C4 | 30 | 7 | 21 | 0 | 0 | 0 | 0 | 0 | 0 | 0 | 0 |
| 5 | 0.65 | 1.39 | 0.47 | 0.64 | 1 | Gm9260 | | 3 | 7 | 5 | 2 | 10 | 8 | 0 | 2 | 9 | 3 | 9 |

| | | | | | | | | | | | | | | | | | |
|---|---|---|---|---|---|---|---|---|---|---|---|---|---|---|---|---|---|
| 5 | 0.00 | 3.68 | 0.00 | 1.00 | 1 | Gm28809 | | 0 | 18 | 39 | 0 | 0 | 0 | 0 | 0 | 0 | 0 | 0 |
| 5 | -0.85 | 2.36 | -0.36 | 0.72 | 1 | Shc3 | src homology 2 domain-containing transfo | 25 | 14 | 6 | 0 | 4 | 3 | 0 | 0 | 0 | 0 | 6 |
| 5 | -0.41 | 1.75 | -0.23 | 0.82 | 1 | Gm14117 | | 9 | 1 | 3 | 7 | 6 | 8 | 0 | 0 | 14 | 3 | 7 |
| 5 | -4.21 | 2.65 | -1.59 | 0.11 | 1 | Gm16191 | | 7 | 7 | 0 | 18 | 10 | 2 | 0 | 0 | 0 | 0 | 13 |
| 5 | -3.29 | 3.25 | -1.01 | 0.31 | 1 | Gm49783 | | 0 | 0 | 6 | 6 | 0 | 18 | 0 | 0 | 0 | 0 | 27 |
| 5 | -0.92 | 1.93 | -0.48 | 0.63 | 1 | Rpsa-ps1 | | 3 | 1 | 16 | 8 | 9 | 3 | 0 | 1 | 0 | 15 | 1 |
| 5 | 0.53 | 3.24 | 0.16 | 0.87 | 1 | Gm9077 | | 0 | 0 | 0 | 0 | 6 | 10 | 0 | 34 | 8 | 0 | 0 |
| 5 | 3.61 | 3.32 | 1.09 | 0.28 | 1 | 2810428J06Rik | | 17 | 0 | 0 | 0 | 0 | 22 | 0 | 0 | 7 | 0 | 11 |
| 5 | 1.20 | 1.76 | 0.68 | 0.50 | 1 | Gm5763 | | 5 | 2 | 1 | 7 | 5 | 2 | 0 | 0 | 15 | 15 | 5 |
| 5 | 1.57 | 3.03 | 0.52 | 0.60 | 1 | Sorcs3 | sortilin-related VPS10 domain containing | 15 | 24 | 15 | 0 | 0 | 0 | 0 | 0 | 3 | 0 | 0 |
| 5 | 2.20 | 1.79 | 1.23 | 0.22 | 1 | Gm7102 | predicted gene 7102 | 9 | 8 | 10 | 4 | 1 | 0 | 0 | 20 | 1 | 1 | 2 |
| 5 | -0.90 | 2.41 | -0.37 | 0.71 | 1 | Draxin | dorsal inhibitory axon guidance protein | 26 | 12 | 9 | 6 | 0 | 1 | 0 | 0 | 0 | 0 | 3 |
| 5 | 1.06 | 2.97 | 0.36 | 0.72 | 1 | Gm27010 | | 9 | 0 | 0 | 0 | 0 | 19 | 0 | 0 | 0 | 8 | 22 |
| 5 | 0.00 | 3.58 | 0.00 | 1.00 | 1 | Sapcd1 | suppressor APC domain containing 1 | 16 | 0 | 0 | 0 | 0 | 24 | 0 | 0 | 0 | 0 | 17 |
| 5 | -2.25 | 2.97 | -0.76 | 0.45 | 1 | 4930483O08Rik | | 17 | 0 | 28 | 4 | 0 | 3 | 0 | 0 | 5 | 0 | 0 |
| 5 | 1.37 | 2.25 | 0.61 | 0.54 | 1 | Gm49327 | | 3 | 6 | 12 | 0 | 0 | 17 | 0 | 0 | 5 | 0 | 13 |
| 5 | 0.00 | 3.57 | 0.00 | 1.00 | 1 | Gm38410 | | 21 | 26 | 9 | 0 | 0 | 0 | 0 | 0 | 0 | 0 | 0 |
| 5 | 0.73 | 2.01 | 0.36 | 0.72 | 1 | Gm43062 | | 9 | 4 | 9 | 0 | 3 | 11 | 0 | 0 | 7 | 0 | 14 |
| 5 | 0.60 | 3.03 | 0.20 | 0.84 | 1 | Fosl1 | fos-like antigen 1 | 20 | 16 | 0 | 0 | 0 | 6 | 0 | 0 | 0 | 0 | 15 |
| 5 | 0.57 | 2.55 | 0.22 | 0.82 | 1 | Gm16046 | | 13 | 13 | 28 | 0 | 0 | 1 | 0 | 0 | 0 | 0 | 2 |
| 5 | -1.54 | 3.52 | -0.44 | 0.66 | 1 | H2bc1 | H2B clustered histone 1 | 0 | 0 | 0 | 39 | 9 | 0 | 0 | 0 | 8 | 0 | 0 |
| 5 | -1.17 | 2.25 | -0.52 | 0.60 | 1 | Dnah7a | dynein, axonemal, heavy chain 7A | 13 | 25 | 5 | 5 | 0 | 4 | 0 | 0 | 0 | 0 | 4 |
| 5 | -1.48 | 3.18 | -0.47 | 0.64 | 1 | Tmem196 | transmembrane protein 196 | 7 | 12 | 23 | 15 | 0 | 0 | 0 | 0 | 0 | 0 | 0 |
| 5 | 0.00 | 3.63 | 0.00 | 1.00 | 1 | Ighv6-6 | | 57 | 0 | 0 | 0 | 0 | 0 | 0 | 0 | 0 | 0 | 0 |
| 5 | -1.07 | 2.17 | -0.49 | 0.62 | 1 | Kcnrg | potassium channel regulator | 1 | 6 | 7 | 23 | 0 | 8 | 0 | 0 | 1 | 0 | 10 |
| 5 | -0.05 | 3.04 | -0.02 | 0.99 | 1 | Tnfrsf25 | tumor necrosis factor receptor superfamily | 6 | 0 | 22 | 0 | 0 | 19 | 0 | 0 | 0 | 0 | 10 |
| 5 | 2.69 | 2.14 | 1.26 | 0.21 | 1 | A730082K24Rik | | 8 | 1 | 2 | 0 | 11 | 0 | 0 | 0 | 14 | 14 | 6 |
| 5 | 0.27 | 1.80 | 0.15 | 0.88 | 1 | Gm5512 | | 0 | 1 | 0 | 10 | 10 | 8 | 0 | 1 | 18 | 6 | 2 |
| 5 | 0.17 | 2.86 | 0.06 | 0.95 | 1 | Vinac1 | vinculin/alpha-catenin family member 1 | 7 | 0 | 0 | 0 | 8 | 18 | 0 | 0 | 19 | 0 | 5 |
| 5 | -0.76 | 1.87 | -0.41 | 0.68 | 1 | Gm14648 | | 9 | 0 | 9 | 5 | 18 | 2 | 0 | 2 | 1 | 1 | 8 |
| 5 | 1.81 | 1.96 | 0.92 | 0.36 | 1 | Rpl19-ps7 | | 3 | 23 | 0 | 1 | 2 | 4 | 0 | 1 | 5 | 15 | 1 |
| 5 | 0.00 | 3.61 | 0.00 | 1.00 | 1 | Tbx4 | T-box 4 | 36 | 17 | 3 | 0 | 0 | 0 | 0 | 0 | 0 | 0 | 0 |
| 5 | -0.16 | 3.59 | -0.05 | 0.96 | 1 | Gm37899 | | 0 | 0 | 7 | 0 | 0 | 34 | 0 | 0 | 0 | 0 | 15 |
| 5 | 0.00 | 3.68 | 0.00 | 1.00 | 1 | Galnt6os | | 0 | 37 | 19 | 0 | 0 | 0 | 0 | 0 | 0 | 0 | 0 |
| 5 | 0.00 | 3.61 | 0.00 | 1.00 | 1 | Tnfsf8 | tumor necrosis factor (ligand) superfamily | 44 | 5 | 7 | 0 | 0 | 0 | 0 | 0 | 0 | 0 | 0 |
| 5 | 2.39 | 2.33 | 1.03 | 0.31 | 1 | Gm14207 | | 15 | 16 | 10 | 0 | 0 | 2 | 0 | 0 | 6 | 0 | 7 |
| 5 | 0.56 | 2.90 | 0.19 | 0.85 | 1 | Gm48349 | | 0 | 0 | 0 | 0 | 6 | 17 | 0 | 0 | 10 | 0 | 23 |
| 5 | 0.41 | 2.49 | 0.17 | 0.87 | 1 | Snord49a | | 3 | 7 | 12 | 0 | 0 | 13 | 0 | 0 | 0 | 0 | 20 |
| 5 | 2.46 | 1.62 | 1.52 | 0.13 | 1 | Gm11675 | | 3 | 1 | 19 | 1 | 2 | 4 | 0 | 5 | 1 | 9 | 11 |
| 5 | 0.44 | 2.02 | 0.22 | 0.83 | 1 | Gdf9 | growth differentiation factor 9 | 11 | 2 | 16 | 0 | 8 | 3 | 0 | 0 | 6 | 0 | 10 |
| 5 | 0.00 | 3.68 | 0.00 | 1.00 | 1 | Has2os | | 40 | 0 | 16 | 0 | 0 | 0 | 0 | 0 | 0 | 0 | 0 |
| 5 | 0.00 | 3.62 | 0.00 | 1.00 | 1 | Cd7 | CD7 antigen | 27 | 25 | 3 | 0 | 0 | 0 | 0 | 0 | 0 | 0 | 0 |
| 5 | 0.13 | 3.03 | 0.04 | 0.96 | 1 | Gm45094 | | 0 | 0 | 0 | 3 | 0 | 28 | 0 | 0 | 4 | 0 | 21 |
| 5 | 1.07 | 2.27 | 0.47 | 0.64 | 1 | Serpinf2 | serine (or cysteine) peptidase inhibitor, cl | 7 | 7 | 22 | 0 | 0 | 11 | 0 | 0 | 4 | 0 | 5 |
| 5 | -1.56 | 2.28 | -0.68 | 0.49 | 1 | Gm12762 | | 8 | 3 | 0 | 5 | 10 | 5 | 0 | 0 | 5 | 19 | 0 |
| 5 | -0.61 | 3.62 | -0.17 | 0.87 | 1 | Nlrp10 | NLR family, pyrin domain containing 10 | 55 | 0 | 0 | 0 | 0 | 1 | 0 | 0 | 0 | 0 | 0 |
| 5 | 0.00 | 3.60 | 0.00 | 1.00 | 1 | Ces2b | carboxyesterase 2B | 25 | 13 | 18 | 0 | 0 | 0 | 0 | 0 | 0 | 0 | 0 |
| 5 | 2.15 | 2.29 | 0.94 | 0.35 | 1 | Gm37475 | | 9 | 1 | 0 | 3 | 0 | 7 | 0 | 0 | 16 | 15 | 5 |
| 5 | 0.00 | 3.62 | 0.00 | 1.00 | 1 | Hoxd3os1 | | 36 | 3 | 17 | 0 | 0 | 0 | 0 | 0 | 0 | 0 | 0 |
| 5 | 1.00 | 3.47 | 0.29 | 0.77 | 1 | Gm47769 | | 0 | 14 | 0 | 6 | 0 | 4 | 0 | 0 | 0 | 31 | 0 |
| 5 | -0.40 | 3.67 | -0.11 | 0.91 | 1 | Gldn | gliomedin | 17 | 0 | 37 | 0 | 0 | 1 | 0 | 0 | 0 | 0 | 0 |
| 5 | -2.14 | 2.28 | -0.93 | 0.35 | 1 | Nr1i2 | nuclear receptor subfamily 1, group I, mer | 0 | 8 | 14 | 4 | 5 | 19 | 0 | 0 | 6 | 0 | 0 |
| 5 | 0.00 | 3.68 | 0.00 | 1.00 | 1 | Ermap | erythroblast membrane-associated protei | 30 | 24 | 1 | 0 | 0 | 0 | 0 | 0 | 0 | 0 | 0 |
| 5 | -0.69 | 3.66 | -0.19 | 0.85 | 1 | Gm17501 | | 18 | 35 | 0 | 0 | 0 | 2 | 0 | 0 | 0 | 0 | 0 |
| 5 | 0.00 | 3.63 | 0.00 | 1.00 | 1 | Ccr6 | chemokine (C-C motif) receptor 6 | 38 | 12 | 5 | 0 | 0 | 0 | 0 | 0 | 0 | 0 | 0 |
| 5 | -1.73 | 2.27 | -0.76 | 0.45 | 1 | Gm26608 | | 0 | 6 | 0 | 11 | 9 | 17 | 0 | 0 | 2 | 0 | 9 |
| 5 | 0.00 | 3.62 | 0.00 | 1.00 | 1 | Tmem150b | transmembrane protein 150B | 24 | 11 | 20 | 0 | 0 | 0 | 0 | 0 | 0 | 0 | 0 |
| 5 | 0.48 | 2.42 | 0.20 | 0.84 | 1 | Gm4275 | predicted gene 4275 | 8 | 3 | 0 | 4 | 0 | 6 | 0 | 0 | 0 | 30 | 5 |
| 5 | -0.30 | 2.10 | -0.14 | 0.89 | 1 | Gm41442 | | 9 | 19 | 11 | 5 | 0 | 3 | 0 | 0 | 1 | 0 | 5 |
| 5 | 0.20 | 2.54 | 0.08 | 0.94 | 1 | Kif6 | kinesin family member 6 | 1 | 0 | 2 | 18 | 0 | 6 | 0 | 16 | 12 | 0 | 0 |

| | | | | | | | | | | | | | | | | | | |
|---|---|---|---|---|---|---|---|---|---|---|---|---|---|---|---|---|---|---|
| 5 | 0.41 | 2.46 | 0.17 | 0.87 | 1 | Rsf1os2 | | 0 | 18 | 0 | 5 | 8 | 3 | 0 | 0 | 11 | 0 | 9 |
| 5 | -1.11 | 2.13 | -0.52 | 0.60 | 1 | Rorb | RAR-related orphan receptor beta | 1 | 0 | 0 | 4 | 7 | 15 | 0 | 0 | 1 | 19 | 8 |
| 5 | -0.62 | 2.30 | -0.27 | 0.79 | 1 | Mgat5b | mannoside acetylglucosaminyltransferase | 0 | 16 | 20 | 0 | 7 | 6 | 0 | 0 | 4 | 0 | 2 |
| 5 | 0.00 | 3.66 | 0.00 | 1.00 | 1 | Tmem200c | transmembrane protein 200C | 27 | 3 | 24 | 0 | 0 | 0 | 0 | 0 | 0 | 0 | 0 |
| 5 | 0.13 | 3.58 | 0.04 | 0.97 | 1 | 4930503B20Ril | RIKEN cDNA 4930503B20 gene | 0 | 0 | 11 | 0 | 0 | 24 | 0 | 0 | 0 | 0 | 19 |
| 5 | 2.46 | 1.66 | 1.49 | 0.14 | 1 | | | 1 | 3 | 20 | 1 | 4 | 2 | 0 | 1 | 2 | 9 | 12 |
| 5 | 2.12 | 1.83 | 1.16 | 0.25 | 1 | Zfp993 | | 0 | 4 | 0 | 11 | 7 | 1 | 0 | 7 | 10 | 9 | 6 |
| 5 | -1.31 | 2.22 | -0.59 | 0.55 | 1 | Trpm1 | transient receptor potential cation channe | 5 | 0 | 21 | 11 | 4 | 3 | 0 | 3 | 7 | 0 | 0 |
| 5 | 0.96 | 3.05 | 0.31 | 0.75 | 1 | Rasl10a | RAS-like, family 10, member A | 12 | 17 | 24 | 0 | 0 | 0 | 0 | 0 | 0 | 0 | 1 |
| 5 | 0.00 | 3.68 | 0.00 | 1.00 | 1 | Fam83a | family with sequence similarity 83, memb | 37 | 17 | 0 | 0 | 0 | 0 | 0 | 0 | 0 | 0 | 0 |
| 5 | 0.00 | 3.65 | 0.00 | 1.00 | 1 | Gm35154 | | 27 | 19 | 7 | 0 | 0 | 0 | 0 | 0 | 0 | 0 | 0 |
| 5 | 2.47 | 3.50 | 0.71 | 0.48 | 1 | Clec1b | C-type lectin domain family 1, member b | 1 | 47 | 0 | 0 | 0 | 3 | 0 | 0 | 3 | 0 | 0 |
| 5 | 1.64 | 2.18 | 0.75 | 0.45 | 1 | D9Wsu149 | | 0 | 8 | 0 | 1 | 12 | 2 | 0 | 1 | 6 | 21 | 3 |
| 5 | -4.19 | 2.81 | -1.49 | 0.14 | 1 | Gm30091 | | 0 | 21 | 0 | 2 | 5 | 12 | 0 | 0 | 0 | 0 | 14 |
| 5 | 2.27 | 2.85 | 0.79 | 0.43 | 1 | Gm47096 | | 34 | 8 | 4 | 0 | 0 | 3 | 0 | 5 | 0 | 0 | 0 |
| 5 | -2.04 | 2.84 | -0.72 | 0.47 | 1 | Gm15708 | | 0 | 0 | 15 | 8 | 0 | 21 | 0 | 0 | 3 | 0 | 7 |
| 5 | 0.00 | 3.65 | 0.00 | 1.00 | 1 | Cd9-ps | | 29 | 16 | 8 | 0 | 0 | 0 | 0 | 0 | 0 | 0 | 0 |
| 5 | 1.44 | 2.04 | 0.70 | 0.48 | 1 | Gm2541 | | 6 | 16 | 19 | 0 | 1 | 2 | 0 | 0 | 5 | 0 | 5 |
| 5 | -1.08 | 3.55 | -0.31 | 0.76 | 1 | Gm49706 | | 0 | 0 | 0 | 0 | 9 | 18 | 0 | 0 | 0 | 0 | 27 |
| 5 | -0.93 | 2.60 | -0.36 | 0.72 | 1 | 9530052C20Rik | | 3 | 20 | 0 | 11 | 0 | 10 | 0 | 0 | 0 | 0 | 10 |
| 5 | 0.22 | 2.17 | 0.10 | 0.92 | 1 | A730071L15Rik | | 0 | 5 | 11 | 0 | 16 | 9 | 0 | 0 | 2 | 3 | 7 |
| 5 | 0.93 | 3.14 | 0.30 | 0.77 | 1 | Sycp1 | synaptonemal complex protein 1 | 19 | 0 | 22 | 0 | 0 | 2 | 0 | 0 | 0 | 0 | 11 |
| 5 | -1.29 | 3.30 | -0.39 | 0.70 | 1 | Gm47862 | | 0 | 4 | 34 | 13 | 0 | 0 | 0 | 0 | 2 | 0 | 0 |
| 5 | 1.10 | 2.33 | 0.47 | 0.64 | 1 | Kcnj9 | potassium inwardly-rectifying channel, su | 17 | 0 | 12 | 5 | 0 | 3 | 0 | 0 | 4 | 4 | 8 |
| 5 | -5.02 | 2.79 | -1.80 | 0.07 | 1 | Gm15627 | | 0 | 19 | 0 | 3 | 8 | 17 | 0 | 0 | 0 | 0 | 7 |
| 5 | -1.12 | 2.19 | -0.51 | 0.61 | 1 | Gm39197 | | 5 | 11 | 20 | 11 | 3 | 0 | 0 | 0 | 1 | 0 | 3 |
| 5 | 0.13 | 2.63 | 0.05 | 0.96 | 1 | Gm47797 | | 5 | 25 | 0 | 0 | 3 | 11 | 0 | 0 | 7 | 0 | 2 |
| 5 | 3.71 | 2.71 | 1.37 | 0.17 | 1 | Itpr3os | | 0 | 4 | 4 | 0 | 0 | 1 | 0 | 0 | 2 | 41 | 1 |
| 5 | -1.00 | 2.59 | -0.39 | 0.70 | 1 | Coro6 | coronin 6 | 3 | 0 | 26 | 0 | 2 | 18 | 0 | 0 | 0 | 0 | 5 |
| 5 | 0.89 | 3.15 | 0.28 | 0.78 | 1 | Gm42913 | | 0 | 0 | 0 | 0 | 1 | 24 | 0 | 0 | 0 | 10 | 18 |
| 5 | 3.80 | 2.64 | 1.44 | 0.15 | 1 | Gm44573 | | 0 | 2 | 0 | 0 | 0 | 3 | 0 | 34 | 8 | 0 | 6 |
| 5 | 0.00 | 3.68 | 0.00 | 1.00 | 1 | Fgf16 | fibroblast growth factor 16 | 48 | 5 | 0 | 0 | 0 | 0 | 0 | 0 | 0 | 0 | 0 |
| 5 | -1.45 | 2.72 | -0.53 | 0.59 | 1 | 4930556J24Rik | | 15 | 0 | 14 | 0 | 9 | 8 | 0 | 0 | 0 | 0 | 7 |
| 5 | 0.15 | 3.30 | 0.04 | 0.96 | 1 | 1810008B01Rik | | 0 | 21 | 27 | 2 | 0 | 0 | 0 | 0 | 0 | 0 | 3 |
| 5 | 0.00 | 3.68 | 0.00 | 1.00 | 1 | Gm43102 | | 38 | 10 | 5 | 0 | 0 | 0 | 0 | 0 | 0 | 0 | 0 |
| 5 | -0.69 | 3.07 | -0.23 | 0.82 | 1 | Foxd2os | | 21 | 15 | 16 | 0 | 0 | 2 | 0 | 0 | 0 | 0 | 0 |
| 5 | 1.26 | 3.07 | 0.41 | 0.68 | 1 | Wnt2 | wingless-type MMTV integration site famil | 28 | 8 | 15 | 0 | 0 | 0 | 0 | 0 | 0 | 0 | 2 |
| 5 | -0.05 | 3.60 | -0.01 | 0.99 | 1 | Slc12a3 | solute carrier family 12, member 3 | 0 | 0 | 0 | 0 | 0 | 35 | 0 | 0 | 0 | 0 | 18 |
| 5 | 2.67 | 3.44 | 0.78 | 0.44 | 1 | Pnp2 | purine-nucleoside phosphorylase 2 | 27 | 18 | 0 | 0 | 0 | 0 | 0 | 0 | 7 | 0 | 1 |
| 5 | 1.92 | 3.13 | 0.61 | 0.54 | 1 | Gm42658 | | 1 | 0 | 0 | 0 | 0 | 19 | 0 | 0 | 6 | 0 | 26 |
| 5 | 3.24 | 3.55 | 0.91 | 0.36 | 1 | Gm25835 | | 0 | 0 | 0 | 0 | 0 | 25 | 0 | 0 | 0 | 18 | 10 |
| 5 | 0.88 | 1.99 | 0.44 | 0.66 | 1 | Ccdc78 | coiled-coil domain containing 78 | 3 | 9 | 0 | 6 | 0 | 13 | 0 | 6 | 0 | 10 | 6 |
| 5 | 1.19 | 2.59 | 0.46 | 0.65 | 1 | Gm25777 | | 3 | 0 | 7 | 0 | 0 | 19 | 0 | 0 | 4 | 0 | 19 |
| 5 | 1.38 | 2.31 | 0.59 | 0.55 | 1 | Gm1141 | | 10 | 6 | 0 | 1 | 0 | 10 | 0 | 7 | 0 | 0 | 18 |
| 5 | 0.96 | 3.11 | 0.31 | 0.76 | 1 | 4933416E03Rik | | 7 | 40 | 5 | 0 | 0 | 0 | 0 | 0 | 0 | 0 | 1 |
| 5 | 0.00 | 3.68 | 0.00 | 1.00 | 1 | Psma8 | proteasome subunit alpha 8 | 21 | 4 | 27 | 0 | 0 | 0 | 0 | 0 | 0 | 0 | 0 |
| 5 | 0.86 | 2.61 | 0.33 | 0.74 | 1 | Mypopos | | 4 | 0 | 7 | 0 | 0 | 24 | 0 | 0 | 2 | 0 | 15 |
| 5 | 0.37 | 2.53 | 0.15 | 0.88 | 1 | Gm27002 | | 5 | 0 | 0 | 4 | 0 | 6 | 0 | 0 | 4 | 31 | 2 |
| 5 | 4.46 | 2.57 | 1.74 | 0.08 | 1 | Pcdha7 | protocadherin alpha 7 | 0 | 3 | 0 | 1 | 0 | 3 | 0 | 0 | 4 | 35 | 7 |
| 5 | 0.00 | 3.68 | 0.00 | 1.00 | 1 | Il17rb | interleukin 17 receptor B | 13 | 0 | 39 | 0 | 0 | 0 | 0 | 0 | 0 | 0 | 0 |
| 5 | 2.11 | 2.80 | 0.75 | 0.45 | 1 | Mroh8 | maestro heat-like repeat family member 8 | 6 | 0 | 0 | 0 | 9 | 3 | 0 | 16 | 1 | 0 | 16 |
| 5 | 4.06 | 2.78 | 1.46 | 0.14 | 1 | B230208H11Rik | | 3 | 1 | 0 | 0 | 0 | 1 | 0 | 0 | 7 | 40 | 0 |
| 5 | 3.33 | 2.69 | 1.24 | 0.22 | 1 | Hhipl2 | hedgehog interacting protein-like 2 | 0 | 0 | 0 | 0 | 1 | 9 | 0 | 6 | 0 | 13 | 23 |
| 5 | 1.63 | 3.58 | 0.46 | 0.65 | 1 | Eqtn | equatorin, sperm acrosome associated | 0 | 0 | 0 | 0 | 0 | 18 | 0 | 0 | 3 | 0 | 31 |
| 5 | -1.31 | 1.94 | -0.68 | 0.50 | 1 | Gm26725 | | 6 | 15 | 3 | 18 | 3 | 3 | 0 | 0 | 3 | 0 | 2 |
| 5 | 0.00 | 3.68 | 0.00 | 1.00 | 1 | Asgr2 | asialoglycoprotein receptor 2 | 43 | 1 | 8 | 0 | 0 | 0 | 0 | 0 | 0 | 0 | 0 |
| 5 | 0.07 | 2.31 | 0.03 | 0.98 | 1 | Gm2449 | | 1 | 15 | 11 | 4 | 0 | 11 | 0 | 0 | 0 | 5 | 5 |
| 5 | 0.00 | 3.68 | 0.00 | 1.00 | 1 | Gm16118 | | 27 | 8 | 17 | 0 | 0 | 0 | 0 | 0 | 0 | 0 | 0 |
| 5 | 1.90 | 3.21 | 0.59 | 0.55 | 1 | Lrrc31 | leucine rich repeat containing 31 | 21 | 11 | 10 | 0 | 0 | 0 | 0 | 0 | 9 | 0 | 0 |
| 5 | 1.88 | 3.26 | 0.58 | 0.56 | 1 | Gm30054 | | 20 | 0 | 0 | 11 | 0 | 3 | 0 | 0 | 5 | 0 | 14 |

| | | | | | | | | | | | | | | | | | | |
|---|---|---|---|---|---|---|---|---|---|---|---|---|---|---|---|---|---|---|
| 5 | 0.00 | 3.68 | 0.00 | 1.00 | 1 | Mef2b | myocyte enhancer factor 2B | 34 | 0 | 18 | 0 | 0 | 0 | 0 | 0 | 0 | 0 | 0 |
| 5 | -1.58 | 3.56 | -0.44 | 0.66 | 1 | Gm23458 | | 0 | 0 | 0 | 6 | 0 | 30 | 0 | 0 | 0 | 0 | 16 |
| 5 | -1.11 | 3.58 | -0.31 | 0.76 | 1 | Gm38366 | | 0 | 0 | 0 | 3 | 0 | 28 | 0 | 0 | 0 | 0 | 21 |
| 5 | 0.36 | 2.34 | 0.15 | 0.88 | 1 | Stpg3 | sperm tail PG rich repeat containing 3 | 7 | 18 | 4 | 0 | 5 | 3 | 0 | 0 | 0 | 10 | 4 |
| 5 | 0.00 | 3.68 | 0.00 | 1.00 | 1 | Hoxb9 | homeobox B9 | 18 | 25 | 8 | 0 | 0 | 0 | 0 | 0 | 0 | 0 | 0 |
| 5 | -0.60 | 2.80 | -0.21 | 0.83 | 1 | Mcmdc2 | minichromosome maintenance domain co | 16 | 25 | 0 | 6 | 0 | 1 | 0 | 0 | 0 | 0 | 3 |
| 5 | 1.50 | 1.95 | 0.77 | 0.44 | 1 | Chtf8 | CTF8, chromosome transmission fidelity f | 6 | 5 | 2 | 0 | 2 | 18 | 0 | 0 | 9 | 3 | 7 |
| 5 | 0.00 | 3.68 | 0.00 | 1.00 | 1 | Klk1b11 | kallikrein 1-related peptidase b11 | 5 | 14 | 32 | 0 | 0 | 0 | 0 | 0 | 0 | 0 | 0 |
| 5 | 1.38 | 2.72 | 0.51 | 0.61 | 1 | Gm45059 | | 5 | 0 | 5 | 0 | 0 | 21 | 0 | 0 | 0 | 8 | 12 |
| 5 | -0.83 | 2.44 | -0.34 | 0.73 | 1 | 2410080I02Rik | | 0 | 0 | 4 | 19 | 0 | 15 | 0 | 5 | 3 | 0 | 5 |
| 5 | -2.21 | 1.99 | -1.11 | 0.27 | 1 | Gm16759 | | 6 | 2 | 1 | 12 | 4 | 16 | 0 | 8 | 0 | 0 | 2 |
| 5 | 0.00 | 3.68 | 0.00 | 1.00 | 1 | H2-M9 | histocompatibility 2, M region locus 9 | 19 | 32 | 0 | 0 | 0 | 0 | 0 | 0 | 0 | 0 | 0 |
| 5 | 0.00 | 3.68 | 0.00 | 1.00 | 1 | Cyp2j5 | cytochrome P450, family 2, subfamily j, po | 13 | 3 | 35 | 0 | 0 | 0 | 0 | 0 | 0 | 0 | 0 |
| 5 | 0.43 | 3.60 | 0.12 | 0.90 | 1 | Gm37677 | | 0 | 0 | 0 | 0 | 0 | 19 | 0 | 0 | 0 | 0 | 32 |
| 5 | 0.00 | 3.68 | 0.00 | 1.00 | 1 | A230001M10Rik | | 6 | 26 | 19 | 0 | 0 | 0 | 0 | 0 | 0 | 0 | 0 |
| 5 | -2.75 | 2.72 | -1.01 | 0.31 | 1 | Nog | noggin | 15 | 9 | 17 | 8 | 3 | 0 | 0 | 0 | 0 | 0 | 0 |
| 5 | -0.69 | 3.11 | -0.22 | 0.82 | 1 | 8430419K02Rik | | 9 | 24 | 16 | 0 | 0 | 2 | 0 | 0 | 0 | 0 | 0 |
| 5 | 0.80 | 2.83 | 0.28 | 0.78 | 1 | Mir212 | | 26 | 0 | 11 | 0 | 0 | 5 | 0 | 0 | 0 | 8 | 1 |
| 5 | -0.40 | 3.67 | -0.11 | 0.91 | 1 | Stmn4 | stathmin-like 4 | 28 | 0 | 22 | 0 | 0 | 1 | 0 | 0 | 0 | 0 | 0 |
| 5 | 1.23 | 2.12 | 0.58 | 0.56 | 1 | Gm11588 | | 0 | 0 | 0 | 9 | 1 | 7 | 0 | 16 | 7 | 0 | 11 |
| 5 | 1.23 | 3.58 | 0.34 | 0.73 | 1 | Gm48478 | | 0 | 0 | 0 | 0 | 0 | 25 | 0 | 0 | 2 | 0 | 24 |
| 5 | -1.43 | 3.55 | -0.40 | 0.69 | 1 | Gm50481 | | 0 | 0 | 0 | 10 | 0 | 22 | 0 | 0 | 0 | 0 | 19 |
| 5 | -0.06 | 3.58 | -0.02 | 0.99 | 1 | Gm37154 | | 0 | 10 | 0 | 0 | 0 | 25 | 0 | 0 | 0 | 0 | 16 |
| 5 | -0.03 | 2.48 | -0.01 | 0.99 | 1 | 1700069B07Rik | | 0 | 15 | 0 | 11 | 7 | 3 | 0 | 0 | 12 | 0 | 3 |
| 5 | 3.42 | 3.00 | 1.14 | 0.25 | 1 | Klkb1 | kallikrein B, plasma 1 | 11 | 14 | 0 | 0 | 0 | 9 | 0 | 0 | 8 | 0 | 8 |
| 5 | -0.62 | 3.09 | -0.20 | 0.84 | 1 | Rnu73b | | 0 | 0 | 0 | 4 | 0 | 28 | 0 | 0 | 1 | 0 | 18 |
| 5 | -1.53 | 3.22 | -0.47 | 0.64 | 1 | Gm2237 | predicted gene 2237 | 1 | 0 | 0 | 0 | 10 | 21 | 0 | 0 | 0 | 0 | 18 |
| 5 | -1.52 | 1.90 | -0.80 | 0.42 | 1 | Gm42470 | | 3 | 11 | 2 | 6 | 9 | 9 | 0 | 0 | 6 | 0 | 3 |
| 5 | 0.94 | 2.17 | 0.43 | 0.66 | 1 | Gm44116 | | 7 | 9 | 2 | 0 | 3 | 8 | 0 | 0 | 10 | 0 | 11 |
| 5 | -0.86 | 3.18 | -0.27 | 0.79 | 1 | 2900052N01Rik | | 25 | 12 | 4 | 0 | 0 | 9 | 0 | 0 | 0 | 0 | 0 |
| 5 | 3.06 | 2.94 | 1.04 | 0.30 | 1 | Gm43412 | | 9 | 21 | 0 | 0 | 0 | 8 | 0 | 0 | 4 | 0 | 8 |
| 5 | -0.87 | 3.03 | -0.29 | 0.77 | 1 | Capn11 | calpain 11 | 0 | 0 | 0 | 0 | 10 | 28 | 0 | 0 | 3 | 0 | 9 |
| 5 | 0.00 | 3.68 | 0.00 | 1.00 | 1 | 4930511M06Rik | | 15 | 15 | 20 | 0 | 0 | 0 | 0 | 0 | 0 | 0 | 0 |
| 5 | -1.54 | 2.32 | -0.66 | 0.51 | 1 | Klf1 | Kruppel-like factor 1 (erythroid) | 3 | 2 | 0 | 20 | 8 | 5 | 0 | 0 | 6 | 0 | 6 |
| 5 | 0.00 | 3.68 | 0.00 | 1.00 | 1 | Spta1 | spectrin alpha, erythrocytic 1 | 13 | 0 | 37 | 0 | 0 | 0 | 0 | 0 | 0 | 0 | 0 |
| 5 | 1.32 | 3.12 | 0.42 | 0.67 | 1 | Ccno | cyclin O | 10 | 19 | 18 | 0 | 0 | 0 | 0 | 0 | 0 | 0 | 3 |
| 5 | -0.30 | 2.57 | -0.12 | 0.91 | 1 | Gm10357 | | 0 | 0 | 15 | 16 | 0 | 3 | 0 | 10 | 5 | 0 | 1 |
| 5 | 1.26 | 3.13 | 0.40 | 0.69 | 1 | Ankrd2 | ankyrin repeat domain 2 (stretch responsi | 11 | 18 | 19 | 0 | 0 | 0 | 0 | 0 | 0 | 0 | 2 |
| 5 | 0.00 | 3.68 | 0.00 | 1.00 | 1 | 2610528A11Ril | RIKEN cDNA 2610528A11 gene | 12 | 13 | 25 | 0 | 0 | 0 | 0 | 0 | 0 | 0 | 0 |
| 5 | 1.26 | 3.13 | 0.40 | 0.69 | 1 | Serpina3h | | 27 | 6 | 15 | 0 | 0 | 0 | 0 | 0 | 0 | 0 | 2 |
| 5 | 3.91 | 3.38 | 1.16 | 0.25 | 1 | AF357399 | | 0 | 2 | 0 | 0 | 0 | 25 | 0 | 0 | 0 | 10 | 13 |
| 5 | -1.68 | 3.22 | -0.52 | 0.60 | 1 | Kcnq3 | potassium voltage-gated channel, subfan | 8 | 0 | 0 | 13 | 17 | 0 | 0 | 0 | 9 | 0 | 3 |
| 5 | -2.81 | 3.37 | -0.83 | 0.40 | 1 | Pla2g3 | phospholipase A2, group III | 3 | 0 | 0 | 5 | 0 | 8 | 0 | 0 | 0 | 0 | 35 |
| 5 | 1.86 | 3.24 | 0.57 | 0.57 | 1 | Hpse2 | heparanase 2 | 27 | 8 | 8 | 0 | 0 | 0 | 0 | 0 | 6 | 0 | 0 |
| 5 | 0.00 | 3.68 | 0.00 | 1.00 | 1 | 9130208D14Rik | | 8 | 15 | 27 | 0 | 0 | 0 | 0 | 0 | 0 | 0 | 0 |
| 5 | 1.64 | 3.11 | 0.53 | 0.60 | 1 | Masp2 | mannan-binding lectin serine peptidase 2 | 3 | 0 | 0 | 0 | 0 | 12 | 0 | 0 | 0 | 14 | 21 |
| 5 | 0.00 | 3.68 | 0.00 | 1.00 | 1 | Abcg8 | ATP binding cassette subfamily G membe | 0 | 41 | 9 | 0 | 0 | 0 | 0 | 0 | 0 | 0 | 0 |
| 5 | 1.33 | 3.53 | 0.38 | 0.71 | 1 | Gm23338 | | 0 | 0 | 0 | 0 | 0 | 9 | 0 | 41 | 0 | 0 | 0 |
| 5 | 0.00 | 3.68 | 0.00 | 1.00 | 1 | Tnfsf11 | tumor necrosis factor (ligand) superfamily | 21 | 28 | 0 | 0 | 0 | 0 | 0 | 0 | 0 | 0 | 0 |
| 5 | 0.36 | 3.60 | 0.10 | 0.92 | 1 | 4633401B06Rik | | 0 | 0 | 0 | 0 | 0 | 20 | 0 | 0 | 0 | 0 | 29 |
| 5 | 0.00 | 3.68 | 0.00 | 1.00 | 1 | Gm42922 | | 18 | 19 | 12 | 0 | 0 | 0 | 0 | 0 | 0 | 0 | 0 |
| 5 | -2.16 | 2.35 | -0.92 | 0.36 | 1 | Gm41664 | | 0 | 0 | 12 | 11 | 7 | 10 | 0 | 0 | 5 | 0 | 4 |
| 5 | 0.02 | 3.60 | 0.01 | 1.00 | 1 | 1110003F10Rik | | 0 | 0 | 0 | 0 | 0 | 30 | 0 | 0 | 0 | 0 | 19 |
| 5 | -0.40 | 3.15 | -0.13 | 0.90 | 1 | Zfp804a | zinc finger protein 804A | 33 | 10 | 5 | 0 | 0 | 1 | 0 | 0 | 0 | 0 | 0 |
| 5 | 1.66 | 3.58 | 0.46 | 0.64 | 1 | 4833412K13Rik | | 0 | 0 | 0 | 0 | 0 | 16 | 0 | 0 | 3 | 0 | 30 |
| 4 | -3.10 | 2.27 | -1.37 | 0.17 | 1 | Mir7688 | | 26 | 5 | 3 | 5 | 5 | 3 | 0 | 0 | 0 | 0 | 3 |
| 4 | 0.96 | 2.36 | 0.41 | 0.68 | 1 | 5430425K12Rik | | 0 | 0 | 0 | 9 | 5 | 4 | 0 | 0 | 19 | 6 | 6 |
| 4 | 2.45 | 2.33 | 1.05 | 0.29 | 1 | 7630403G23Rik | | 0 | 0 | 0 | 1 | 2 | 5 | 0 | 0 | 15 | 19 | 7 |
| 4 | -3.89 | 3.51 | -1.11 | 0.27 | 1 | Gm43011 | | 0 | 0 | 14 | 11 | 0 | 9 | 0 | 0 | 0 | 0 | 15 |
| 4 | -0.04 | 3.60 | -0.01 | 0.99 | 1 | Gm6614 | predicted gene 6614 | 0 | 0 | 0 | 0 | 0 | 32 | 0 | 0 | 0 | 0 | 17 |

| | | | | | | | | | | | | | | | | | | |
|---|---|---|---|---|---|---|---|---|---|---|---|---|---|---|---|---|---|---|
| 4 | 1.71 | 3.19 | 0.54 | 0.59 | 1 | S100a14 | S100 calcium binding protein A14 | 5 | 11 | 25 | 0 | 0 | 0 | 0 | 0 | 8 | 0 | 0 |
| 4 | 1.04 | 2.60 | 0.40 | 0.69 | 1 | Tceal6 | transcription elongation factor A (SII)-like | 1 | 0 | 0 | 1 | 1 | 4 | 0 | 0 | 5 | 37 | 0 |
| 4 | -0.01 | 3.17 | 0.00 | 1.00 | 1 | Gm44104 | | 0 | 13 | 0 | 7 | 0 | 7 | 0 | 0 | 0 | 0 | 22 |
| 4 | -1.13 | 2.07 | -0.55 | 0.58 | 1 | Gm14150 | | 9 | 2 | 18 | 1 | 1 | 11 | 0 | 0 | 0 | 0 | 8 |
| 4 | -1.02 | 3.55 | -0.29 | 0.77 | 1 | Gm43413 | | 0 | 0 | 0 | 0 | 8 | 15 | 0 | 0 | 0 | 0 | 26 |
| 4 | 0.00 | 3.68 | 0.00 | 1.00 | 1 | Gm32219 | | 7 | 12 | 30 | 0 | 0 | 0 | 0 | 0 | 0 | 0 | 0 |
| 4 | 0.99 | 3.19 | 0.31 | 0.76 | 1 | Adamts20 | a disintegrin-like and metallopeptidase (r | 4 | 32 | 11 | 0 | 0 | 0 | 0 | 0 | 1 | 0 | 0 |
| 4 | -3.76 | 3.38 | -1.11 | 0.27 | 1 | Cfap157 | cilia and flagella associated protein 157 | 3 | 0 | 0 | 0 | 9 | 18 | 0 | 0 | 0 | 0 | 20 |
| 4 | 0.00 | 3.68 | 0.00 | 1.00 | 1 | 2610027K06Rik | | 39 | 1 | 9 | 0 | 0 | 0 | 0 | 0 | 0 | 0 | 0 |
| 4 | -2.71 | 3.32 | -0.82 | 0.41 | 1 | Gm45083 | | 0 | 24 | 14 | 0 | 9 | 2 | 0 | 0 | 0 | 0 | 0 |
| 4 | -0.44 | 2.03 | -0.22 | 0.83 | 1 | Gm6451 | | 9 | 2 | 27 | 0 | 5 | 3 | 0 | 0 | 1 | 0 | 3 |
| 4 | 0.40 | 2.48 | 0.16 | 0.87 | 1 | Emx1 | empty spiracles homeobox 1 | 12 | 7 | 10 | 0 | 12 | 0 | 0 | 0 | 4 | 0 | 3 |
| 4 | 0.00 | 3.68 | 0.00 | 1.00 | 1 | Duoxa1 | dual oxidase maturation factor 1 | 34 | 0 | 15 | 0 | 0 | 0 | 0 | 0 | 0 | 0 | 0 |
| 4 | 2.16 | 2.34 | 0.92 | 0.36 | 1 | Gm48146 | | 0 | 0 | 6 | 0 | 4 | 8 | 0 | 0 | 6 | 9 | 15 |
| 4 | 1.23 | 2.55 | 0.48 | 0.63 | 1 | Gm19680 | | 0 | 10 | 0 | 1 | 5 | 5 | 0 | 0 | 5 | 22 | 1 |
| 4 | 4.32 | 3.56 | 1.21 | 0.23 | 1 | Gm23346 | | 0 | 7 | 5 | 0 | 0 | 11 | 0 | 25 | 0 | 0 | 0 |
| 4 | 0.26 | 3.13 | 0.08 | 0.93 | 1 | Gm43462 | | 3 | 9 | 0 | 0 | 0 | 17 | 0 | 0 | 0 | 0 | 19 |
| 4 | 0.00 | 3.68 | 0.00 | 1.00 | 1 | Serpinb10 | serine (or cysteine) peptidase inhibitor, cl | 23 | 17 | 8 | 0 | 0 | 0 | 0 | 0 | 0 | 0 | 0 |
| 4 | -2.65 | 3.30 | -0.80 | 0.42 | 1 | Gm37020 | | 0 | 34 | 0 | 10 | 4 | 1 | 0 | 0 | 0 | 0 | 0 |
| 4 | -0.40 | 1.97 | -0.20 | 0.84 | 1 | Gm6063 | | 9 | 0 | 8 | 3 | 3 | 6 | 0 | 0 | 3 | 1 | 15 |
| 4 | 0.00 | 3.68 | 0.00 | 1.00 | 1 | Ocstamp | osteoclast stimulatory transmembrane prc | 9 | 17 | 23 | 0 | 0 | 0 | 0 | 0 | 0 | 0 | 0 |
| 4 | 0.00 | 3.68 | 0.00 | 1.00 | 1 | Fut7 | fucosyltransferase 7 | 15 | 9 | 25 | 0 | 0 | 0 | 0 | 0 | 0 | 0 | 0 |
| 4 | 0.66 | 2.59 | 0.25 | 0.80 | 1 | Gm26696 | | 6 | 0 | 11 | 3 | 0 | 4 | 0 | 0 | 5 | 0 | 19 |
| 4 | 0.00 | 3.68 | 0.00 | 1.00 | 1 | Ahrr | aryl-hydrocarbon receptor repressor | 8 | 28 | 12 | 0 | 0 | 0 | 0 | 0 | 0 | 0 | 0 |
| 4 | 0.05 | 3.60 | 0.01 | 0.99 | 1 | Rho | rhodopsin | 0 | 4 | 0 | 0 | 0 | 26 | 0 | 0 | 0 | 0 | 18 |
| 4 | -0.99 | 2.60 | -0.38 | 0.70 | 1 | Gm44997 | | 0 | 0 | 0 | 2 | 24 | 6 | 0 | 0 | 12 | 0 | 5 |
| 4 | 2.15 | 1.44 | 1.49 | 0.14 | 1 | Gm7308 | | 5 | 10 | 7 | 3 | 1 | 2 | 0 | 7 | 4 | 3 | 7 |
| 4 | 0.00 | 3.68 | 0.00 | 1.00 | 1 | Gm9913 | | 8 | 34 | 6 | 0 | 0 | 0 | 0 | 0 | 0 | 0 | 0 |
| 4 | -1.32 | 3.58 | -0.37 | 0.71 | 1 | 2610028D06Rik | | 0 | 0 | 0 | 0 | 3 | 31 | 0 | 0 | 0 | 0 | 14 |
| 4 | 0.18 | 3.60 | 0.05 | 0.96 | 1 | Gm6939 | | 0 | 0 | 4 | 0 | 0 | 23 | 0 | 0 | 0 | 0 | 21 |
| 4 | 0.62 | 3.60 | 0.17 | 0.86 | 1 | Gm47573 | | 0 | 0 | 0 | 0 | 0 | 13 | 0 | 0 | 0 | 0 | 36 |
| 4 | -2.46 | 2.79 | -0.88 | 0.38 | 1 | Gm37125 | | 16 | 14 | 10 | 0 | 5 | 3 | 0 | 0 | 0 | 0 | 0 |
| 4 | 1.35 | 3.17 | 0.43 | 0.67 | 1 | Gm10561 | | 21 | 10 | 12 | 0 | 0 | 0 | 0 | 0 | 0 | 0 | 4 |
| 4 | 0.00 | 3.68 | 0.00 | 1.00 | 1 | Rasgef1c | RasGEF domain family, member 1C | 15 | 24 | 8 | 0 | 0 | 0 | 0 | 0 | 0 | 0 | 0 |
| 4 | 0.00 | 3.68 | 0.00 | 1.00 | 1 | A130077B15Rik | | 27 | 7 | 14 | 0 | 0 | 0 | 0 | 0 | 0 | 0 | 0 |
| 4 | -0.34 | 3.60 | -0.09 | 0.93 | 1 | Psg29 | pregnancy-specific glycoprotein 29 | 0 | 0 | 0 | 0 | 36 | 0 | 0 | 0 | 12 | 0 | 0 |
| 4 | 4.35 | 3.48 | 1.25 | 0.21 | 1 | Gm15667 | | 8 | 0 | 0 | 0 | 0 | 14 | 0 | 10 | 0 | 0 | 16 |
| 4 | 3.75 | 2.98 | 1.26 | 0.21 | 1 | Tmem74 | transmembrane protein 74 | 19 | 0 | 5 | 0 | 0 | 8 | 0 | 0 | 14 | 0 | 2 |
| 4 | -0.40 | 3.67 | -0.11 | 0.91 | 1 | Gm13010 | | 0 | 30 | 17 | 0 | 0 | 1 | 0 | 0 | 0 | 0 | 0 |
| 4 | 3.46 | 3.36 | 1.03 | 0.30 | 1 | Gm27616 | | 0 | 5 | 0 | 0 | 0 | 19 | 0 | 0 | 4 | 0 | 19 |
| 4 | -0.87 | 2.26 | -0.39 | 0.70 | 1 | Gm12454 | | 9 | 8 | 10 | 11 | 0 | 4 | 0 | 0 | 3 | 0 | 2 |
| 4 | -0.29 | 2.46 | -0.12 | 0.91 | 1 | Rpl17-ps10 | | 3 | 15 | 5 | 0 | 6 | 7 | 0 | 0 | 2 | 10 | 0 |
| 4 | 0.00 | 3.68 | 0.00 | 1.00 | 1 | Podnl1 | podocan-like 1 | 39 | 9 | 0 | 0 | 0 | 0 | 0 | 0 | 0 | 0 | 0 |
| 4 | 1.37 | 3.24 | 0.42 | 0.67 | 1 | Cdh18 | cadherin 18 | 2 | 25 | 16 | 0 | 0 | 0 | 0 | 0 | 0 | 0 | 5 |
| 4 | 0.69 | 3.27 | 0.21 | 0.83 | 1 | Gm4134 | | 18 | 0 | 20 | 0 | 3 | 0 | 0 | 0 | 7 | 0 | 0 |
| 4 | 0.00 | 3.68 | 0.00 | 1.00 | 1 | Gm9968 | | 25 | 10 | 12 | 0 | 0 | 0 | 0 | 0 | 0 | 0 | 0 |
| 4 | 0.00 | 3.68 | 0.00 | 1.00 | 1 | Serpinb2 | serine (or cysteine) peptidase inhibitor, cl | 36 | 0 | 11 | 0 | 0 | 0 | 0 | 0 | 0 | 0 | 0 |
| 4 | 2.80 | 2.80 | 1.00 | 0.32 | 1 | Gm43852 | | 0 | 0 | 8 | 0 | 0 | 11 | 0 | 6 | 5 | 0 | 17 |
| 4 | 1.79 | 2.74 | 0.65 | 0.51 | 1 | Gm10419 | | 0 | 17 | 3 | 6 | 0 | 2 | 0 | 0 | 0 | 14 | 5 |
| 4 | 0.44 | 3.60 | 0.12 | 0.90 | 1 | Gm44136 | | 0 | 0 | 0 | 0 | 0 | 17 | 0 | 0 | 0 | 0 | 30 |
| 4 | 1.76 | 3.33 | 0.53 | 0.60 | 1 | Ppp1r18os | | 4 | 21 | 17 | 0 | 0 | 0 | 0 | 0 | 0 | 5 | 0 |
| 4 | -0.52 | 2.55 | -0.20 | 0.84 | 1 | Gm15710 | | 15 | 5 | 5 | 0 | 7 | 6 | 0 | 0 | 9 | 0 | 0 |
| 4 | 0.61 | 2.52 | 0.24 | 0.81 | 1 | Gm32358 | | 0 | 0 | 6 | 0 | 5 | 11 | 0 | 0 | 10 | 0 | 15 |
| 4 | -0.69 | 3.19 | -0.22 | 0.83 | 1 | Efcab6 | EF-hand calcium binding domain 6 | 13 | 22 | 10 | 0 | 0 | 2 | 0 | 0 | 0 | 0 | 0 |
| 4 | 3.61 | 3.28 | 1.10 | 0.27 | 1 | Krt27 | keratin 27 | 0 | 0 | 0 | 0 | 0 | 7 | 0 | 31 | 3 | 0 | 6 |
| 4 | 0.00 | 3.68 | 0.00 | 1.00 | 1 | Elfn1 | leucine rich repeat and fibronectin type III | 42 | 3 | 2 | 0 | 0 | 0 | 0 | 0 | 0 | 0 | 0 |
| 4 | -0.26 | 2.94 | -0.09 | 0.93 | 1 | Gm19696 | | 0 | 7 | 20 | 0 | 11 | 1 | 0 | 0 | 0 | 0 | 8 |
| 4 | 0.41 | 2.40 | 0.17 | 0.86 | 1 | Gm44065 | | 16 | 21 | 4 | 0 | 0 | 3 | 0 | 0 | 1 | 0 | 1 |
| 4 | 0.11 | 3.21 | 0.03 | 0.97 | 1 | Gckr | glucokinase regulatory protein | 15 | 14 | 0 | 0 | 0 | 10 | 0 | 0 | 0 | 0 | 8 |
| 4 | 0.38 | 3.60 | 0.10 | 0.92 | 1 | Gm15539 | | 0 | 0 | 0 | 0 | 0 | 19 | 0 | 0 | 0 | 0 | 28 |

| | | | | | | | | | | | | | | | | | | |
|---|---|---|---|---|---|---|---|---|---|---|---|---|---|---|---|---|---|---|
| 4 | 0.79 | 3.19 | 0.25 | 0.80 | 1 | Gm13373 | | 0 | 10 | 0 | 0 | 4 | 9 | 0 | 0 | 0 | 6 | 17 |
| 4 | 2.52 | 3.55 | 0.71 | 0.48 | 1 | C330022C24Rik | | 0 | 0 | 0 | 0 | 0 | 2 | 0 | 45 | 0 | 0 | 0 |
| 4 | 1.33 | 2.27 | 0.59 | 0.56 | 1 | Gm22540 | | 3 | 0 | 7 | 8 | 0 | 4 | 0 | 13 | 3 | 0 | 9 |
| 4 | 0.74 | 3.29 | 0.23 | 0.82 | 1 | Gm42611 | | 0 | 8 | 0 | 0 | 0 | 25 | 0 | 0 | 1 | 0 | 13 |
| 4 | -0.83 | 3.22 | -0.26 | 0.80 | 1 | 6030466F02Rik | | 22 | 7 | 11 | 0 | 0 | 6 | 0 | 0 | 0 | 0 | 0 |
| 4 | 2.70 | 2.47 | 1.09 | 0.27 | 1 | Gm8349 | | 3 | 20 | 0 | 1 | 6 | 0 | 0 | 0 | 5 | 5 | 7 |
| 4 | 0.09 | 3.60 | 0.03 | 0.98 | 1 | Gm47370 | | 0 | 2 | 0 | 0 | 0 | 25 | 0 | 0 | 0 | 0 | 19 |
| 4 | 0.00 | 3.68 | 0.00 | 1.00 | 1 | Unc45bos | | 3 | 16 | 27 | 0 | 0 | 0 | 0 | 0 | 0 | 0 | 0 |
| 4 | -0.18 | 2.65 | -0.07 | 0.95 | 1 | Catsperg1 | cation channel sperm associated auxiliar | 4 | 14 | 4 | 0 | 0 | 17 | 0 | 0 | 0 | 0 | 7 |
| 4 | -2.68 | 3.52 | -0.76 | 0.45 | 1 | Gm45353 | | 0 | 18 | 19 | 0 | 7 | 0 | 0 | 0 | 0 | 0 | 3 |
| 4 | -1.28 | 2.76 | -0.46 | 0.64 | 1 | Ppp1r32 | protein phosphatase 1, regulatory subuni | 0 | 4 | 9 | 0 | 9 | 14 | 0 | 0 | 0 | 0 | 10 |
| 4 | 0.69 | 2.23 | 0.31 | 0.76 | 1 | Gm26887 | | 0 | 4 | 10 | 2 | 2 | 3 | 0 | 0 | 1 | 22 | 2 |
| 4 | 0.00 | 3.68 | 0.00 | 1.00 | 1 | D630033O11Ri | RIKEN cDNA D630033O11 gene | 35 | 8 | 3 | 0 | 0 | 0 | 0 | 0 | 0 | 0 | 0 |
| 4 | 0.25 | 3.60 | 0.07 | 0.95 | 1 | Gm45823 | | 0 | 0 | 0 | 0 | 0 | 22 | 0 | 0 | 0 | 0 | 24 |
| 4 | 0.00 | 3.63 | 0.00 | 1.00 | 1 | Trp63 | transformation related protein 63 | 46 | 0 | 0 | 0 | 0 | 0 | 0 | 0 | 0 | 0 | 0 |
| 4 | 0.00 | 3.68 | 0.00 | 1.00 | 1 | Krt76 | keratin 76 | 35 | 11 | 0 | 0 | 0 | 0 | 0 | 0 | 0 | 0 | 0 |
| 4 | -5.76 | 3.43 | -1.68 | 0.09 | 1 | Gm16092 | | 0 | 0 | 0 | 22 | 3 | 3 | 0 | 0 | 18 | 0 | 0 |
| 4 | -0.40 | 3.67 | -0.11 | 0.91 | 1 | Gm20485 | | 12 | 33 | 0 | 0 | 0 | 1 | 0 | 0 | 0 | 0 | 0 |
| 4 | -0.38 | 1.99 | -0.19 | 0.85 | 1 | Gm45885 | | 6 | 1 | 8 | 2 | 8 | 3 | 0 | 0 | 7 | 0 | 11 |
| 4 | 0.23 | 2.34 | 0.10 | 0.92 | 1 | Gm10224 | | 0 | 12 | 1 | 0 | 13 | 2 | 0 | 7 | 1 | 0 | 10 |
| 4 | 1.79 | 2.79 | 0.64 | 0.52 | 1 | Gm44954 | | 9 | 6 | 0 | 0 | 0 | 8 | 0 | 0 | 0 | 13 | 10 |
| 4 | 1.27 | 2.06 | 0.61 | 0.54 | 1 | Gtf2a1l | general transcription factor IIA, 1-like | 7 | 0 | 9 | 4 | 2 | 3 | 0 | 14 | 4 | 0 | 4 |
| 4 | -1.36 | 2.94 | -0.46 | 0.64 | 1 | Pnldc1 | poly(A)-specific ribonuclease (PARN)-like | 6 | 0 | 6 | 3 | 0 | 21 | 0 | 0 | 0 | 0 | 10 |
| 4 | -1.01 | 2.89 | -0.35 | 0.73 | 1 | Scarna13 | | 10 | 0 | 3 | 2 | 0 | 18 | 0 | 0 | 0 | 0 | 13 |
| 4 | 0.00 | 3.16 | 0.00 | 1.00 | 1 | 5730521K06Rik | | 3 | 1 | 0 | 0 | 0 | 26 | 0 | 0 | 0 | 0 | 16 |
| 4 | 0.21 | 3.37 | 0.06 | 0.95 | 1 | Gm45449 | | 15 | 0 | 0 | 14 | 0 | 3 | 0 | 0 | 9 | 0 | 4 |
| 4 | -1.08 | 3.56 | -0.30 | 0.76 | 1 | Gm50186 | | 0 | 0 | 0 | 0 | 6 | 17 | 0 | 0 | 0 | 0 | 23 |
| 4 | -5.43 | 3.54 | -1.53 | 0.13 | 1 | Gm47359 | | 0 | 0 | 0 | 25 | 10 | 10 | 0 | 0 | 0 | 0 | 0 |
| 4 | -1.13 | 2.90 | -0.39 | 0.70 | 1 | E030044B06Rik | | 0 | 27 | 6 | 4 | 0 | 6 | 0 | 0 | 2 | 0 | 0 |
| 4 | -0.75 | 3.24 | -0.23 | 0.82 | 1 | Tvp23bos | | 5 | 9 | 29 | 0 | 0 | 3 | 0 | 0 | 0 | 0 | 0 |
| 4 | -0.56 | 2.83 | -0.20 | 0.84 | 1 | Gm9267 | | 0 | 2 | 1 | 0 | 16 | 3 | 0 | 0 | 4 | 19 | 0 |
| 4 | 1.33 | 3.58 | 0.37 | 0.71 | 1 | Gm21297 | | 0 | 0 | 0 | 15 | 0 | 0 | 0 | 29 | 0 | 0 | 2 |
| 4 | 4.89 | 3.31 | 1.48 | 0.14 | 1 | Gm47772 | | 31 | 1 | 0 | 2 | 0 | 0 | 0 | 0 | 7 | 0 | 5 |
| 4 | 0.40 | 2.69 | 0.15 | 0.88 | 1 | Ppp1r36 | protein phosphatase 1, regulatory subuni | 8 | 2 | 23 | 0 | 0 | 5 | 0 | 0 | 0 | 0 | 8 |
| 4 | 0.51 | 2.05 | 0.25 | 0.80 | 1 | Kdm4d | lysine (K)-specific demethylase 4D | 0 | 10 | 7 | 7 | 4 | 5 | 0 | 0 | 3 | 6 | 3 |
| 4 | 1.57 | 3.60 | 0.44 | 0.66 | 1 | Gm9626 | | 0 | 4 | 0 | 0 | 0 | 0 | 0 | 42 | 0 | 0 | 0 |
| 4 | 4.15 | 3.27 | 1.27 | 0.20 | 1 | Kng1 | kininogen 1 | 3 | 0 | 1 | 0 | 0 | 0 | 0 | 0 | 20 | 22 | 0 |
| 4 | 1.16 | 3.65 | 0.32 | 0.75 | 1 | Clrn3 | clarin 3 | 15 | 0 | 29 | 0 | 0 | 0 | 0 | 0 | 2 | 0 | 0 |
| 4 | 1.68 | 3.56 | 0.47 | 0.64 | 1 | Gm16540 | | 0 | 0 | 0 | 0 | 0 | 20 | 0 | 0 | 6 | 0 | 19 |
| 4 | 1.70 | 3.59 | 0.47 | 0.64 | 1 | Mybpc1 | myosin binding protein C, slow-type | 0 | 42 | 0 | 0 | 0 | 0 | 0 | 0 | 0 | 0 | 4 |
| 4 | 0.56 | 2.31 | 0.24 | 0.81 | 1 | Psd2 | pleckstrin and Sec7 domain containing 2 | 0 | 4 | 0 | 11 | 5 | 8 | 0 | 0 | 5 | 12 | 2 |
| 4 | 2.35 | 1.86 | 1.26 | 0.21 | 1 | Gm44419 | | 4 | 3 | 12 | 0 | 3 | 2 | 0 | 0 | 6 | 9 | 6 |
| 4 | -2.25 | 3.07 | -0.73 | 0.46 | 1 | Gm9888 | | 23 | 7 | 4 | 7 | 0 | 0 | 0 | 0 | 0 | 0 | 4 |
| 4 | 0.00 | 3.68 | 0.00 | 1.00 | 1 | Art2b | ADP-ribosyltransferase 2b | 34 | 11 | 0 | 0 | 0 | 0 | 0 | 0 | 0 | 0 | 0 |
| 4 | 0.00 | 3.68 | 0.00 | 1.00 | 1 | Aqp9 | aquaporin 9 | 18 | 24 | 3 | 0 | 0 | 0 | 0 | 0 | 0 | 0 | 0 |
| 4 | -0.14 | 3.60 | -0.04 | 0.97 | 1 | Gm24644 | | 0 | 0 | 0 | 0 | 0 | 32 | 0 | 0 | 0 | 0 | 13 |
| 4 | 1.43 | 3.58 | 0.40 | 0.69 | 1 | Gm37447 | | 0 | 0 | 0 | 0 | 0 | 22 | 0 | 0 | 3 | 0 | 20 |
| 4 | 2.96 | 2.63 | 1.13 | 0.26 | 1 | Gm9256 | | 1 | 0 | 18 | 0 | 3 | 0 | 0 | 0 | 1 | 22 | 1 |
| 4 | 0.00 | 2.15 | 0.00 | 1.00 | 1 | Gm9805 | | 1 | 5 | 0 | 10 | 4 | 3 | 0 | 10 | 12 | 0 | 0 |
| 4 | -0.44 | 3.05 | -0.14 | 0.89 | 1 | Gm9722 | | 5 | 0 | 6 | 0 | 5 | 3 | 0 | 0 | 0 | 26 | 0 |
| 4 | 0.76 | 3.39 | 0.22 | 0.82 | 1 | | | 0 | 0 | 0 | 11 | 0 | 4 | 0 | 0 | 4 | 0 | 26 |
| 4 | 4.46 | 3.56 | 1.25 | 0.21 | 1 | Gm23442 | | 0 | 0 | 0 | 0 | 0 | 8 | 0 | 25 | 0 | 0 | 11 |
| 4 | -2.37 | 2.84 | -0.83 | 0.40 | 1 | Gm16485 | | 2 | 35 | 0 | 5 | 1 | 2 | 0 | 0 | 0 | 0 | 0 |
| 4 | -1.26 | 2.47 | -0.51 | 0.61 | 1 | Obscn | obscurin, cytoskeletal calmodulin and titir | 0 | 19 | 14 | 4 | 0 | 7 | 0 | 0 | 1 | 0 | 1 |
| 4 | 0.00 | 3.68 | 0.00 | 1.00 | 1 | Spdef | SAM pointed domain containing ets trans | 15 | 21 | 8 | 0 | 0 | 0 | 0 | 0 | 0 | 0 | 0 |
| 4 | -0.60 | 3.17 | -0.19 | 0.85 | 1 | Gm14001 | | 0 | 0 | 7 | 0 | 2 | 24 | 0 | 0 | 0 | 0 | 12 |
| 4 | -0.13 | 2.59 | -0.05 | 0.96 | 1 | Rpl27-ps1 | | 0 | 15 | 5 | 0 | 6 | 2 | 0 | 16 | 0 | 0 | 1 |
| 4 | -1.89 | 2.61 | -0.72 | 0.47 | 1 | Trpv6 | transient receptor potential cation channe | 11 | 9 | 12 | 5 | 2 | 0 | 0 | 0 | 0 | 0 | 5 |
| 4 | -0.01 | 3.25 | 0.00 | 1.00 | 1 | Gm47813 | | 11 | 12 | 0 | 0 | 0 | 14 | 0 | 0 | 0 | 0 | 8 |
| 4 | 4.19 | 3.55 | 1.18 | 0.24 | 1 | Gm43872 | | 0 | 0 | 0 | 0 | 0 | 8 | 0 | 0 | 0 | 28 | 8 |

| | | | | | | | | | | | | | | | | | |
|---|---|---|---|---|---|---|---|---|---|---|---|---|---|---|---|---|---|
| 4 | -2.85 | 3.55 | -0.80 | 0.42 | 1 | Gm49616 | | 0 | 0 | 0 | 12 | 0 | 9 | 0 | 0 | 0 | 0 | 23 |
| 4 | -1.84 | 2.21 | -0.83 | 0.40 | 1 | Syngr4 | synaptogyrin 4 | 10 | 2 | 0 | 4 | 16 | 2 | 0 | 0 | 2 | 1 | 7 |
| 4 | 1.13 | 2.67 | 0.42 | 0.67 | 1 | Tssk4 | testis-specific serine kinase 4 | 2 | 8 | 0 | 6 | 0 | 8 | 0 | 0 | 10 | 0 | 11 |
| 4 | 0.00 | 3.68 | 0.00 | 1.00 | 1 | Nodal | nodal | 22 | 13 | 9 | 0 | 0 | 0 | 0 | 0 | 0 | 0 | 0 |
| 4 | -0.72 | 2.09 | -0.34 | 0.73 | 1 | Gm8565 | | 5 | 0 | 1 | 3 | 9 | 2 | 0 | 0 | 1 | 17 | 7 |
| 4 | 0.00 | 3.68 | 0.00 | 1.00 | 1 | Gdf2 | growth differentiation factor 2 | 9 | 0 | 35 | 0 | 0 | 0 | 0 | 0 | 0 | 0 | 0 |
| 4 | 1.31 | 2.50 | 0.52 | 0.60 | 1 | Lyzl4 | lysozyme-like 4 | 11 | 6 | 12 | 0 | 0 | 6 | 0 | 0 | 7 | 0 | 2 |
| 4 | 0.00 | 3.68 | 0.00 | 1.00 | 1 | Pdcd1lg2 | programmed cell death 1 ligand 2 | 39 | 6 | 0 | 0 | 0 | 0 | 0 | 0 | 0 | 0 | 0 |
| 4 | -1.80 | 2.84 | -0.64 | 0.53 | 1 | Gm20597 | | 5 | 13 | 21 | 0 | 5 | 1 | 0 | 0 | 0 | 0 | 0 |
| 4 | 0.00 | 3.63 | 0.00 | 1.00 | 1 | Ighv1-7 | | 0 | 45 | 0 | 0 | 0 | 0 | 0 | 0 | 0 | 0 | 0 |
| 4 | -1.08 | 2.43 | -0.44 | 0.66 | 1 | Gm33037 | | 0 | 9 | 8 | 11 | 5 | 1 | 0 | 0 | 10 | 0 | 0 |
| 4 | -0.54 | 3.62 | -0.15 | 0.88 | 1 | 5430437J10Rik | | 0 | 0 | 44 | 0 | 0 | 1 | 0 | 0 | 0 | 0 | 0 |
| 4 | 0.00 | 3.68 | 0.00 | 1.00 | 1 | Clca3b | chloride channel accessory 3B | 0 | 9 | 35 | 0 | 0 | 0 | 0 | 0 | 0 | 0 | 0 |
| 4 | 0.89 | 2.62 | 0.34 | 0.73 | 1 | Gm11843 | | 16 | 8 | 6 | 0 | 6 | 0 | 0 | 0 | 7 | 0 | 1 |
| 4 | -0.95 | 2.76 | -0.34 | 0.73 | 1 | Gm18665 | | 0 | 0 | 0 | 5 | 1 | 27 | 0 | 0 | 5 | 0 | 6 |
| 4 | 0.16 | 3.28 | 0.05 | 0.96 | 1 | Myo1h | myosin 1H | 22 | 9 | 0 | 0 | 0 | 7 | 0 | 0 | 0 | 0 | 6 |
| 4 | 0.00 | 3.68 | 0.00 | 1.00 | 1 | E030013I19Rik | | 31 | 9 | 4 | 0 | 0 | 0 | 0 | 0 | 0 | 0 | 0 |
| 4 | 0.00 | 3.68 | 0.00 | 1.00 | 1 | Ugt2b36 | UDP glucuronosyltransferase 2 family, pc | 2 | 7 | 35 | 0 | 0 | 0 | 0 | 0 | 0 | 0 | 0 |
| 4 | -1.44 | 3.57 | -0.40 | 0.69 | 1 | Tmod4 | tropomodulin 4 | 0 | 0 | 0 | 4 | 0 | 25 | 0 | 0 | 0 | 0 | 15 |
| 4 | 3.55 | 3.03 | 1.17 | 0.24 | 1 | Gm38376 | | 0 | 11 | 4 | 0 | 0 | 14 | 0 | 0 | 10 | 0 | 5 |
| 4 | 0.56 | 3.29 | 0.17 | 0.86 | 1 | Sh2d4b | SH2 domain containing 4B | 11 | 0 | 16 | 0 | 0 | 5 | 0 | 0 | 0 | 0 | 12 |
| 4 | -0.80 | 2.89 | -0.28 | 0.78 | 1 | Gm4356 | | 7 | 7 | 0 | 0 | 2 | 12 | 0 | 0 | 0 | 0 | 16 |
| 4 | 3.43 | 3.54 | 0.97 | 0.33 | 1 | Gm18917 | | 0 | 0 | 0 | 0 | 0 | 4 | 0 | 0 | 0 | 39 | 1 |
| 4 | -0.08 | 3.38 | -0.02 | 0.98 | 1 | Gm15483 | | 2 | 30 | 0 | 0 | 0 | 8 | 0 | 0 | 0 | 0 | 4 |
| 4 | 0.13 | 3.60 | 0.04 | 0.97 | 1 | Eif4ebp3 | eukaryotic translation initiation factor 4E b | 3 | 0 | 0 | 0 | 0 | 22 | 0 | 0 | 0 | 0 | 18 |
| 4 | 0.00 | 3.68 | 0.00 | 1.00 | 1 | Tinag | tubulointerstitial nephritis antigen | 5 | 37 | 1 | 0 | 0 | 0 | 0 | 0 | 0 | 0 | 0 |
| 4 | 3.76 | 3.06 | 1.23 | 0.22 | 1 | Gm8036 | | 0 | 0 | 1 | 0 | 1 | 2 | 0 | 36 | 4 | 0 | 0 |
| 4 | -0.11 | 3.59 | -0.03 | 0.98 | 1 | Xntrpc | Xndc1-transient receptor potential cation | 0 | 5 | 0 | 0 | 0 | 25 | 0 | 0 | 0 | 0 | 13 |
| 4 | -0.69 | 3.31 | -0.21 | 0.83 | 1 | Tmem132e | transmembrane protein 132E | 9 | 29 | 3 | 0 | 0 | 2 | 0 | 0 | 0 | 0 | 0 |
| 4 | 3.45 | 3.58 | 0.96 | 0.34 | 1 | Gm47389 | | 0 | 15 | 10 | 0 | 0 | 3 | 0 | 0 | 0 | 15 | 0 |
| 4 | -3.86 | 2.97 | -1.30 | 0.19 | 1 | Unc5d | unc-5 netrin receptor D | 15 | 0 | 16 | 3 | 6 | 4 | 0 | 0 | 0 | 0 | 0 |
| 4 | -1.73 | 3.48 | -0.50 | 0.62 | 1 | Wnt3 | wingless-type MMTV integration site famil | 0 | 0 | 0 | 0 | 5 | 8 | 0 | 0 | 0 | 28 | 2 |
| 4 | 0.00 | 3.68 | 0.00 | 1.00 | 1 | Gm10710 | predicted gene 10710 | 3 | 16 | 24 | 0 | 0 | 0 | 0 | 0 | 0 | 0 | 0 |
| 4 | 0.00 | 3.68 | 0.00 | 1.00 | 1 | Gm11695 | | 6 | 35 | 2 | 0 | 0 | 0 | 0 | 0 | 0 | 0 | 0 |
| 4 | 0.14 | 3.48 | 0.04 | 0.97 | 1 | Gm40770 | | 0 | 0 | 15 | 10 | 0 | 0 | 0 | 0 | 3 | 0 | 16 |
| 4 | 0.00 | 3.68 | 0.00 | 1.00 | 1 | Mgst2 | microsomal glutathione S-transferase 2 | 30 | 3 | 10 | 0 | 0 | 0 | 0 | 0 | 0 | 0 | 0 |
| 4 | -4.92 | 2.69 | -1.83 | 0.07 | 1 | Gm13568 | | 13 | 0 | 0 | 6 | 5 | 9 | 0 | 0 | 0 | 1 | 9 |
| 4 | -2.21 | 2.65 | -0.83 | 0.40 | 1 | Pinlyp | phospholipase A2 inhibitor and LY6/PLAU | 4 | 13 | 0 | 12 | 2 | 2 | 0 | 0 | 0 | 0 | 10 |
| 4 | -0.34 | 2.96 | -0.11 | 0.91 | 1 | A730061H03Rik | | 0 | 24 | 0 | 6 | 1 | 1 | 0 | 0 | 0 | 0 | 11 |
| 4 | 0.30 | 3.60 | 0.08 | 0.93 | 1 | Gm47234 | | 5 | 0 | 0 | 0 | 0 | 17 | 0 | 0 | 0 | 0 | 21 |
| 4 | -0.50 | 2.83 | -0.18 | 0.86 | 1 | Gm45223 | | 10 | 9 | 0 | 0 | 1 | 12 | 0 | 0 | 0 | 0 | 11 |
| 4 | 0.00 | 3.68 | 0.00 | 1.00 | 1 | Evi2 | ecotropic viral integration site 2 | 4 | 16 | 23 | 0 | 0 | 0 | 0 | 0 | 0 | 0 | 0 |
| 4 | 0.80 | 3.58 | 0.22 | 0.82 | 1 | Pglyrp2 | peptidoglycan recognition protein 2 | 39 | 0 | 0 | 0 | 0 | 1 | 0 | 0 | 0 | 0 | 3 |
| 4 | 0.38 | 3.14 | 0.12 | 0.90 | 1 | Gm37580 | | 0 | 7 | 0 | 11 | 0 | 13 | 0 | 0 | 11 | 0 | 1 |
| 4 | 0.00 | 3.68 | 0.00 | 1.00 | 1 | Gm11707 | | 31 | 5 | 7 | 0 | 0 | 0 | 0 | 0 | 0 | 0 | 0 |
| 4 | 3.55 | 3.29 | 1.08 | 0.28 | 1 | Gm20517 | predicted gene 20517 | 0 | 0 | 0 | 0 | 0 | 11 | 0 | 0 | 4 | 13 | 15 |
| 4 | -1.76 | 2.74 | -0.64 | 0.52 | 1 | Gm15696 | | 6 | 0 | 5 | 4 | 0 | 20 | 0 | 0 | 6 | 0 | 2 |
| 4 | -0.69 | 3.66 | -0.19 | 0.85 | 1 | Trbc2 | | 23 | 18 | 0 | 0 | 0 | 2 | 0 | 0 | 0 | 0 | 0 |
| 4 | 0.00 | 3.68 | 0.00 | 1.00 | 1 | Mettl7b | methyltransferase like 7B | 9 | 25 | 8 | 0 | 0 | 0 | 0 | 0 | 0 | 0 | 0 |
| 4 | 0.00 | 3.68 | 0.00 | 1.00 | 1 | Ly6i | lymphocyte antigen 6 complex, locus I | 27 | 0 | 16 | 0 | 0 | 0 | 0 | 0 | 0 | 0 | 0 |
| 4 | 0.00 | 3.68 | 0.00 | 1.00 | 1 | Reg3g | regenerating islet-derived 3 gamma | 13 | 0 | 30 | 0 | 0 | 0 | 0 | 0 | 0 | 0 | 0 |
| 4 | 0.13 | 2.80 | 0.05 | 0.96 | 1 | Gm45081 | | 0 | 0 | 0 | 0 | 15 | 11 | 0 | 0 | 4 | 8 | 5 |
| 4 | 0.00 | 3.68 | 0.00 | 1.00 | 1 | Klrb1a | killer cell lectin-like receptor subfamily B r | 33 | 2 | 8 | 0 | 0 | 0 | 0 | 0 | 0 | 0 | 0 |
| 4 | -0.06 | 2.48 | -0.03 | 0.98 | 1 | Gm36486 | | 9 | 0 | 7 | 0 | 13 | 3 | 0 | 0 | 6 | 0 | 5 |
| 4 | -0.32 | 3.22 | -0.10 | 0.92 | 1 | Prok1 | prokineticin 1 | 0 | 15 | 0 | 11 | 0 | 1 | 0 | 16 | 0 | 0 | 0 |
| 4 | -0.08 | 3.07 | -0.03 | 0.98 | 1 | Gm15401 | | 0 | 18 | 9 | 0 | 1 | 10 | 0 | 0 | 4 | 0 | 0 |
| 4 | 0.00 | 3.68 | 0.00 | 1.00 | 1 | Gm4841 | predicted gene 4841 | 34 | 0 | 8 | 0 | 0 | 0 | 0 | 0 | 0 | 0 | 0 |
| 4 | 0.28 | 1.67 | 0.17 | 0.87 | 1 | Cdiptos | CDIP transferase, opposite strand | 3 | 2 | 9 | 2 | 7 | 4 | 0 | 0 | 9 | 4 | 3 |
| 4 | 0.00 | 3.68 | 0.00 | 1.00 | 1 | Gm19935 | | 20 | 3 | 20 | 0 | 0 | 0 | 0 | 0 | 0 | 0 | 0 |
| 4 | -0.19 | 3.60 | -0.05 | 0.96 | 1 | Gm38268 | | 0 | 0 | 0 | 0 | 0 | 31 | 0 | 0 | 0 | 0 | 11 |

| | | | | | | | | | | | | | | | | | | |
|---|---|---|---|---|---|---|---|---|---|---|---|---|---|---|---|---|---|---|
| 4 | -0.40 | 3.31 | -0.12 | 0.90 | 1 | Oit3 | oncoprotein induced transcript 3 | 19 | 7 | 16 | 0 | 0 | 1 | 0 | 0 | 0 | 0 | 0 |
| 4 | 1.92 | 3.14 | 0.61 | 0.54 | 1 | Gm26571 | | 0 | 1 | 0 | 2 | 0 | 5 | 0 | 0 | 0 | 33 | 1 |
| 4 | 0.99 | 3.59 | 0.28 | 0.78 | 1 | C1ql1 | complement component 1, q subcompon | 9 | 0 | 19 | 1 | 0 | 0 | 0 | 0 | 0 | 14 | 0 |
| 4 | 0.63 | 1.97 | 0.32 | 0.75 | 1 | Gm16238 | | 2 | 16 | 8 | 1 | 8 | 0 | 0 | 2 | 4 | 1 | 0 |
| 4 | 0.61 | 3.60 | 0.17 | 0.87 | 1 | Ldha-ps2 | | 1 | 0 | 0 | 0 | 0 | 11 | 0 | 0 | 0 | 0 | 30 |
| 4 | 1.58 | 3.55 | 0.45 | 0.66 | 1 | Gm31727 | | 0 | 0 | 0 | 0 | 2 | 0 | 0 | 39 | 1 | 0 | 0 |
| 4 | 0.00 | 3.68 | 0.00 | 1.00 | 1 | 2810457G06Rik | | 3 | 24 | 15 | 0 | 0 | 0 | 0 | 0 | 0 | 0 | 0 |
| 4 | -0.70 | 2.86 | -0.24 | 0.81 | 1 | Gm15964 | | 0 | 7 | 21 | 0 | 9 | 1 | 0 | 0 | 5 | 0 | 0 |
| 4 | 0.00 | 3.68 | 0.00 | 1.00 | 1 | Gm765 | predicted gene 765 | 13 | 29 | 0 | 0 | 0 | 0 | 0 | 0 | 0 | 0 | 0 |
| 4 | -4.06 | 3.30 | -1.23 | 0.22 | 1 | Gm38329 | | 0 | 0 | 0 | 10 | 14 | 14 | 0 | 0 | 0 | 0 | 4 |
| 4 | -3.53 | 2.74 | -1.29 | 0.20 | 1 | Siah3 | siah E3 ubiquitin protein ligase family me | 0 | 13 | 10 | 11 | 5 | 3 | 0 | 0 | 0 | 0 | 0 |
| 4 | 0.00 | 3.68 | 0.00 | 1.00 | 1 | Ighv11-1 | | 19 | 23 | 0 | 0 | 0 | 0 | 0 | 0 | 0 | 0 | 0 |
| 4 | 0.00 | 3.68 | 0.00 | 1.00 | 1 | Cd209c | CD209c antigen | 21 | 3 | 18 | 0 | 0 | 0 | 0 | 0 | 0 | 0 | 0 |
| 4 | 0.61 | 2.76 | 0.22 | 0.82 | 1 | 1190001M18Rik | | 3 | 3 | 0 | 6 | 0 | 10 | 0 | 0 | 10 | 0 | 10 |
| 4 | 2.91 | 2.74 | 1.06 | 0.29 | 1 | Gm47956 | | 0 | 0 | 0 | 0 | 1 | 3 | 0 | 1 | 16 | 0 | 21 |
| 4 | -2.31 | 3.04 | -0.76 | 0.45 | 1 | Gm37986 | | 0 | 0 | 6 | 12 | 0 | 14 | 0 | 0 | 8 | 0 | 2 |
| 4 | 2.05 | 2.47 | 0.83 | 0.41 | 1 | Tnfsf13 | tumor necrosis factor (ligand) superfamily | 3 | 7 | 6 | 0 | 0 | 6 | 0 | 0 | 6 | 0 | 14 |
| 4 | 0.00 | 3.68 | 0.00 | 1.00 | 1 | Ighv1-72 | | 3 | 40 | 0 | 0 | 0 | 0 | 0 | 0 | 0 | 0 | 0 |
| 4 | -0.33 | 2.85 | -0.11 | 0.91 | 1 | H1f1 | H1.1 linker histone, cluster member | 0 | 0 | 5 | 20 | 3 | 3 | 0 | 0 | 0 | 6 | 4 |
| 4 | 0.00 | 3.64 | 0.00 | 1.00 | 1 | Ighv2-2 | | 42 | 0 | 0 | 0 | 0 | 0 | 0 | 0 | 0 | 0 | 0 |
| 4 | -3.20 | 3.30 | -0.97 | 0.33 | 1 | Gm20539 | | 0 | 0 | 0 | 13 | 3 | 17 | 0 | 0 | 0 | 0 | 9 |
| 4 | -0.90 | 2.41 | -0.37 | 0.71 | 1 | Gm16741 | | 5 | 5 | 0 | 8 | 6 | 5 | 0 | 0 | 9 | 0 | 4 |
| 4 | -5.39 | 3.46 | -1.56 | 0.12 | 1 | Gm15694 | | 0 | 0 | 0 | 2 | 6 | 7 | 0 | 28 | 0 | 0 | 0 |
| 4 | 2.04 | 1.94 | 1.05 | 0.29 | 1 | Tbx15 | T-box 15 | 9 | 3 | 8 | 0 | 1 | 5 | 0 | 0 | 7 | 4 | 5 |
| 4 | 0.00 | 3.68 | 0.00 | 1.00 | 1 | Acnat1 | acyl-coenzyme A amino acid N-acyltransf | 7 | 16 | 19 | 0 | 0 | 0 | 0 | 0 | 0 | 0 | 0 |
| 4 | 0.00 | 3.68 | 0.00 | 1.00 | 1 | Mrgprb1 | MAS-related GPR, member B1 | 28 | 2 | 11 | 0 | 0 | 0 | 0 | 0 | 0 | 0 | 0 |
| 4 | 0.03 | 3.60 | 0.01 | 0.99 | 1 | Gm12784 | | 0 | 0 | 0 | 0 | 25 | 0 | 0 | 0 | 17 | 0 | 0 |
| 4 | 0.00 | 3.68 | 0.00 | 1.00 | 1 | Ifi214 | interferon activated gene 214 | 19 | 0 | 23 | 0 | 0 | 0 | 0 | 0 | 0 | 0 | 0 |
| 4 | -1.74 | 3.64 | -0.48 | 0.63 | 1 | Gm4876 | | 19 | 19 | 0 | 0 | 2 | 2 | 0 | 0 | 0 | 0 | 0 |
| 4 | 0.22 | 2.57 | 0.08 | 0.93 | 1 | Tymp | thymidine phosphorylase | 0 | 0 | 0 | 4 | 5 | 6 | 0 | 13 | 1 | 13 | 0 |
| 4 | -1.09 | 3.62 | -0.30 | 0.76 | 1 | Serpina9 | serine (or cysteine) peptidase inhibitor, cl | 17 | 19 | 0 | 5 | 0 | 0 | 0 | 0 | 0 | 0 | 0 |
| 4 | 0.00 | 3.68 | 0.00 | 1.00 | 1 | Gm15931 | | 13 | 13 | 16 | 0 | 0 | 0 | 0 | 0 | 0 | 0 | 0 |
| 4 | 0.15 | 2.77 | 0.06 | 0.96 | 1 | Itih5l-ps | inter-alpha (globulin) inhibitor H5-like, ps | 3 | 24 | 8 | 0 | 0 | 3 | 0 | 0 | 0 | 0 | 3 |
| 4 | 1.98 | 3.59 | 0.55 | 0.58 | 1 | D030045P18Rik | | 7 | 0 | 11 | 0 | 0 | 0 | 0 | 0 | 23 | 0 | 0 |
| 4 | 0.00 | 3.68 | 0.00 | 1.00 | 1 | Ms4a14 | membrane-spanning 4-domains, subfami | 24 | 10 | 7 | 0 | 0 | 0 | 0 | 0 | 0 | 0 | 0 |
| 4 | 0.00 | 3.68 | 0.00 | 1.00 | 1 | Adra2b | adrenergic receptor, alpha 2b | 30 | 3 | 8 | 0 | 0 | 0 | 0 | 0 | 0 | 0 | 0 |
| 4 | -2.53 | 3.27 | -0.77 | 0.44 | 1 | Gm16199 | | 22 | 9 | 0 | 4 | 0 | 3 | 0 | 0 | 3 | 0 | 0 |
| 4 | 0.00 | 3.60 | 0.00 | 1.00 | 1 | Gm8113 | | 3 | 0 | 0 | 0 | 0 | 24 | 0 | 0 | 0 | 0 | 14 |
| 4 | -0.41 | 2.80 | -0.15 | 0.88 | 1 | Gm12108 | | 15 | 6 | 15 | 0 | 0 | 5 | 0 | 0 | 0 | 0 | 1 |
| 4 | -0.04 | 3.05 | -0.01 | 0.99 | 1 | Gm37488 | | 0 | 14 | 0 | 0 | 4 | 10 | 0 | 0 | 3 | 0 | 10 |
| 4 | 2.92 | 3.47 | 0.84 | 0.40 | 1 | Rpl35a-ps3 | | 1 | 0 | 1 | 0 | 0 | 0 | 0 | 0 | 2 | 37 | 0 |
| 4 | 0.00 | 3.68 | 0.00 | 1.00 | 1 | Trim30c | tripartite motif-containing 30C | 31 | 3 | 7 | 0 | 0 | 0 | 0 | 0 | 0 | 0 | 0 |
| 4 | 0.00 | 3.64 | 0.00 | 1.00 | 1 | Igkv4-77 | | 41 | 0 | 0 | 0 | 0 | 0 | 0 | 0 | 0 | 0 | 0 |
| 4 | -3.23 | 3.49 | -0.92 | 0.35 | 1 | Pirt | phosphoinositide-interacting regulator of | 24 | 5 | 0 | 7 | 0 | 5 | 0 | 0 | 0 | 0 | 0 |
| 4 | -1.72 | 2.56 | -0.67 | 0.50 | 1 | Nsun7 | NOL1/NOP2/Sun domain family, member | 3 | 8 | 3 | 0 | 8 | 13 | 0 | 0 | 0 | 0 | 7 |
| 4 | 0.00 | 3.68 | 0.00 | 1.00 | 1 | Wfdc6a | WAP four-disulfide core domain 6A | 9 | 20 | 11 | 0 | 0 | 0 | 0 | 0 | 0 | 0 | 0 |
| 4 | 0.00 | 3.68 | 0.00 | 1.00 | 1 | Prss1 | protease, serine 1 (trypsin 1) | 0 | 17 | 24 | 0 | 0 | 0 | 0 | 0 | 0 | 0 | 0 |
| 4 | 0.00 | 3.68 | 0.00 | 1.00 | 1 | Slamf1 | signaling lymphocytic activation molecule | 21 | 10 | 10 | 0 | 0 | 0 | 0 | 0 | 0 | 0 | 0 |
| 4 | 4.77 | 3.57 | 1.34 | 0.18 | 1 | Gm44432 | | 0 | 0 | 2 | 0 | 0 | 9 | 0 | 27 | 0 | 0 | 3 |
| 4 | -3.08 | 3.22 | -0.96 | 0.34 | 1 | Gm27343 | | 7 | 15 | 0 | 0 | 9 | 3 | 0 | 0 | 0 | 0 | 7 |
| 4 | -0.59 | 3.60 | -0.16 | 0.87 | 1 | Gm37140 | | 0 | 0 | 0 | 0 | 1 | 23 | 0 | 0 | 0 | 0 | 17 |
| 4 | 0.00 | 3.68 | 0.00 | 1.00 | 1 | Lgals2 | lectin, galactose-binding, soluble 2 | 20 | 21 | 0 | 0 | 0 | 0 | 0 | 0 | 0 | 0 | 0 |
| 4 | 0.80 | 3.43 | 0.23 | 0.82 | 1 | 9330175E14Rik | | 19 | 17 | 0 | 0 | 0 | 1 | 0 | 0 | 0 | 0 | 4 |
| 4 | 0.03 | 2.92 | 0.01 | 0.99 | 1 | Gm42706 | | 7 | 18 | 0 | 3 | 0 | 2 | 0 | 12 | 0 | 0 | 0 |
| 4 | 2.49 | 3.56 | 0.70 | 0.48 | 1 | D030055H07Rik | | 0 | 0 | 0 | 0 | 0 | 1 | 0 | 37 | 0 | 0 | 3 |
| 4 | -1.55 | 2.16 | -0.72 | 0.47 | 1 | Gm6218 | | 15 | 5 | 2 | 8 | 1 | 6 | 0 | 3 | 1 | 0 | 0 |
| 4 | -0.82 | 1.90 | -0.43 | 0.67 | 1 | Gm12502 | | 7 | 0 | 7 | 6 | 8 | 3 | 0 | 1 | 2 | 1 | 6 |
| 4 | -0.95 | 3.61 | -0.26 | 0.79 | 1 | Malrd1 | MAM and LDL receptor class A domain c | 3 | 32 | 0 | 0 | 0 | 5 | 0 | 0 | 0 | 0 | 0 |
| 4 | -3.09 | 2.72 | -1.14 | 0.26 | 1 | Gm43935 | | 3 | 4 | 0 | 6 | 7 | 3 | 0 | 0 | 0 | 0 | 17 |
| 4 | -0.24 | 2.96 | -0.08 | 0.93 | 1 | Gm43877 | | 0 | 0 | 0 | 3 | 3 | 4 | 0 | 30 | 0 | 0 | 1 |

| | | | | | | | | | | | | | | | | | | |
|---|---|---|---|---|---|---|---|---|---|---|---|---|---|---|---|---|---|---|
| 4 | 3.53 | 3.56 | 0.99 | 0.32 | 1 | Gm14400 | | 0 | 0 | 0 | 0 | 0 | 0 | 0 | 38 | 0 | 3 | 0 |
| 4 | 4.45 | 3.51 | 1.27 | 0.21 | 1 | Gm48908 | | 0 | 0 | 0 | 0 | 0 | 7 | 0 | 0 | 12 | 22 | 0 |
| 4 | 0.14 | 2.34 | 0.06 | 0.95 | 1 | Gm44075 | | 0 | 1 | 6 | 9 | 8 | 0 | 0 | 8 | 9 | 0 | 0 |
| 4 | -1.97 | 2.85 | -0.69 | 0.49 | 1 | Gm38257 | | 3 | 3 | 0 | 9 | 0 | 20 | 0 | 0 | 0 | 0 | 6 |
| 4 | 0.14 | 3.59 | 0.04 | 0.97 | 1 | Gm26202 | | 9 | 0 | 0 | 0 | 0 | 16 | 0 | 0 | 0 | 0 | 15 |
| 4 | -6.04 | 3.08 | -1.96 | 0.05 | 1 | Gm17613 | | 0 | 13 | 0 | 11 | 10 | 2 | 0 | 0 | 0 | 0 | 4 |
| 4 | -2.06 | 2.89 | -0.71 | 0.47 | 1 | Gm26868 | | 8 | 0 | 0 | 18 | 2 | 3 | 0 | 0 | 1 | 0 | 9 |
| 4 | 1.86 | 3.04 | 0.61 | 0.54 | 1 | Gm26796 | | 0 | 15 | 9 | 7 | 0 | 0 | 0 | 0 | 6 | 0 | 3 |
| 4 | 4.86 | 3.06 | 1.59 | 0.11 | 1 | Iqca1l | IQ motif containing with AAA domain 1 like | 11 | 4 | 0 | 0 | 0 | 2 | 0 | 20 | 0 | 0 | 4 |
| 4 | 0.22 | 3.13 | 0.07 | 0.94 | 1 | 9530056E24Rik | | 10 | 0 | 0 | 4 | 0 | 14 | 0 | 9 | 0 | 0 | 4 |
| 4 | 0.00 | 3.68 | 0.00 | 1.00 | 1 | Dmrt3 | doublesex and mab-3 related transcriptio | 17 | 16 | 7 | 0 | 0 | 0 | 0 | 0 | 0 | 0 | 0 |
| 4 | 0.27 | 3.45 | 0.08 | 0.94 | 1 | Dlx1 | distal-less homeobox 1 | 0 | 31 | 0 | 5 | 0 | 1 | 0 | 0 | 0 | 0 | 3 |
| 4 | -2.60 | 2.87 | -0.91 | 0.36 | 1 | Eif3s6-ps4 | | 1 | 0 | 23 | 12 | 0 | 3 | 0 | 0 | 1 | 0 | 1 |
| 4 | 0.00 | 3.68 | 0.00 | 1.00 | 1 | Elfn2 | leucine rich repeat and fibronectin type III | 9 | 10 | 22 | 0 | 0 | 0 | 0 | 0 | 0 | 0 | 0 |
| 4 | 0.00 | 3.63 | 0.00 | 1.00 | 1 | Gm39464 | | 0 | 0 | 41 | 0 | 0 | 0 | 0 | 0 | 0 | 0 | 0 |
| 4 | -0.76 | 2.08 | -0.36 | 0.72 | 1 | 9230111E07Rik | | 0 | 3 | 6 | 4 | 7 | 6 | 0 | 0 | 3 | 0 | 12 |
| 4 | 0.20 | 3.27 | 0.06 | 0.95 | 1 | Stpg2 | sperm tail PG rich repeat containing 2 | 0 | 0 | 0 | 0 | 8 | 13 | 0 | 0 | 9 | 0 | 11 |
| 4 | -0.52 | 2.27 | -0.23 | 0.82 | 1 | AY512931 | | 6 | 20 | 3 | 0 | 3 | 4 | 0 | 0 | 1 | 0 | 3 |
| 4 | -0.54 | 3.35 | -0.16 | 0.87 | 1 | Cdk3 | | 0 | 0 | 3 | 0 | 5 | 14 | 0 | 0 | 0 | 0 | 18 |
| 4 | 3.05 | 2.60 | 1.18 | 0.24 | 1 | Gm13523 | | 2 | 0 | 11 | 0 | 9 | 0 | 0 | 5 | 9 | 0 | 4 |
| 4 | 0.00 | 3.68 | 0.00 | 1.00 | 1 | Gm39822 | | 15 | 8 | 18 | 0 | 0 | 0 | 0 | 0 | 0 | 0 | 0 |
| 4 | -3.71 | 3.51 | -1.06 | 0.29 | 1 | 4930556H04Rik | | 12 | 0 | 0 | 0 | 4 | 3 | 0 | 22 | 0 | 0 | 0 |
| 4 | 0.00 | 3.64 | 0.00 | 1.00 | 1 | Igkv1-133 | | 40 | 0 | 0 | 0 | 0 | 0 | 0 | 0 | 0 | 0 | 0 |
| 4 | -0.70 | 2.95 | -0.24 | 0.81 | 1 | Gm40309 | | 16 | 0 | 4 | 1 | 0 | 12 | 0 | 0 | 0 | 0 | 7 |
| 4 | 1.87 | 3.01 | 0.62 | 0.53 | 1 | Ccdc177 | coiled-coil domain containing 177 | 13 | 15 | 0 | 0 | 0 | 1 | 0 | 0 | 0 | 10 | 1 |
| 4 | 3.84 | 3.58 | 1.07 | 0.28 | 1 | Gm40617 | | 0 | 21 | 0 | 0 | 0 | 7 | 0 | 0 | 6 | 0 | 6 |
| 4 | 0.00 | 3.68 | 0.00 | 1.00 | 1 | Gm5485 | | 0 | 13 | 27 | 0 | 0 | 0 | 0 | 0 | 0 | 0 | 0 |
| 4 | 0.00 | 3.68 | 0.00 | 1.00 | 1 | Gm34829 | | 31 | 2 | 7 | 0 | 0 | 0 | 0 | 0 | 0 | 0 | 0 |
| 4 | 0.00 | 3.68 | 0.00 | 1.00 | 1 | Gm10334 | predicted gene 10334 | 1 | 8 | 31 | 0 | 0 | 0 | 0 | 0 | 0 | 0 | 0 |
| 4 | 0.28 | 2.81 | 0.10 | 0.92 | 1 | 2600006K01Rik | | 13 | 15 | 8 | 0 | 0 | 2 | 0 | 0 | 0 | 0 | 2 |
| 4 | -1.68 | 2.61 | -0.64 | 0.52 | 1 | Gm10840 | | 5 | 0 | 0 | 3 | 2 | 7 | 0 | 0 | 0 | 21 | 3 |
| 4 | 0.33 | 3.59 | 0.09 | 0.93 | 1 | Gm43294 | | 0 | 0 | 8 | 0 | 0 | 14 | 0 | 0 | 0 | 0 | 17 |
| 4 | 1.98 | 3.55 | 0.56 | 0.58 | 1 | Gm37673 | | 0 | 0 | 0 | 0 | 0 | 3 | 0 | 0 | 0 | 37 | 0 |
| 4 | -0.28 | 2.89 | -0.10 | 0.92 | 1 | Gm27353 | | 0 | 1 | 0 | 4 | 0 | 26 | 0 | 0 | 2 | 0 | 7 |
| 4 | 0.97 | 3.50 | 0.28 | 0.78 | 1 | Lncenc1 | | 0 | 13 | 19 | 0 | 0 | 1 | 0 | 0 | 0 | 0 | 7 |
| 4 | -0.86 | 2.66 | -0.32 | 0.75 | 1 | Gm50338 | | 11 | 5 | 0 | 3 | 0 | 17 | 0 | 0 | 2 | 0 | 2 |
| 4 | 0.39 | 3.34 | 0.12 | 0.91 | 1 | 9430041J12Rik | | 1 | 4 | 0 | 0 | 0 | 14 | 0 | 0 | 0 | 0 | 21 |
| 4 | 0.62 | 3.59 | 0.17 | 0.86 | 1 | Gm45698 | | 0 | 7 | 0 | 0 | 0 | 9 | 0 | 0 | 0 | 0 | 23 |
| 4 | -1.98 | 3.14 | -0.63 | 0.53 | 1 | Robo3 | roundabout guidance receptor 3 | 11 | 6 | 0 | 0 | 7 | 12 | 0 | 0 | 0 | 0 | 4 |
| 4 | 0.95 | 3.39 | 0.28 | 0.78 | 1 | Ceacam20 | carcinoembryonic antigen-related cell adh | 8 | 7 | 24 | 0 | 0 | 0 | 0 | 0 | 0 | 0 | 1 |
| 4 | -2.17 | 3.46 | -0.63 | 0.53 | 1 | Gm10747 | | 0 | 0 | 0 | 2 | 10 | 4 | 0 | 0 | 0 | 0 | 23 |
| 4 | -0.64 | 2.83 | -0.23 | 0.82 | 1 | 1700063D05Rik | | 0 | 0 | 0 | 11 | 3 | 10 | 0 | 0 | 7 | 0 | 9 |
| 4 | 0.22 | 1.75 | 0.12 | 0.90 | 1 | Gm13136 | | 1 | 1 | 1 | 2 | 8 | 8 | 0 | 14 | 1 | 4 | 1 |
| 4 | 3.77 | 3.55 | 1.06 | 0.29 | 1 | Tmem253 | transmembrane protein 253 | 0 | 0 | 0 | 0 | 0 | 3 | 0 | 0 | 0 | 36 | 1 |
| 4 | 2.08 | 3.55 | 0.59 | 0.56 | 1 | Gm25323 | | 0 | 0 | 0 | 0 | 0 | 3 | 0 | 37 | 0 | 0 | 0 |
| 4 | -0.03 | 3.60 | -0.01 | 0.99 | 1 | Gm45493 | | 0 | 0 | 0 | 0 | 0 | 25 | 0 | 0 | 0 | 0 | 14 |
| 4 | 0.00 | 3.63 | 0.00 | 1.00 | 1 | Ighv1-43 | | 0 | 0 | 39 | 0 | 0 | 0 | 0 | 0 | 0 | 0 | 0 |
| 4 | 3.71 | 3.57 | 1.04 | 0.30 | 1 | Gm13707 | | 8 | 0 | 0 | 0 | 0 | 19 | 0 | 0 | 8 | 0 | 5 |
| 4 | 0.00 | 3.64 | 0.00 | 1.00 | 1 | Slc30a10 | solute carrier family 30, member 10 | 39 | 0 | 0 | 0 | 0 | 0 | 0 | 0 | 0 | 0 | 0 |
| 4 | -0.40 | 2.82 | -0.14 | 0.89 | 1 | Gm13710 | | 8 | 15 | 5 | 0 | 0 | 9 | 0 | 0 | 0 | 0 | 2 |
| 4 | -1.68 | 2.27 | -0.74 | 0.46 | 1 | 4933425M03Rik | | 2 | 0 | 5 | 3 | 5 | 11 | 0 | 0 | 13 | 0 | 1 |
| 4 | 1.31 | 1.97 | 0.66 | 0.51 | 1 | Gm47766 | | 2 | 1 | 8 | 0 | 4 | 8 | 0 | 6 | 3 | 0 | 8 |
| 4 | 0.18 | 3.60 | 0.05 | 0.96 | 1 | Gm14320 | | 7 | 0 | 0 | 0 | 0 | 15 | 0 | 0 | 0 | 0 | 17 |
| 4 | 3.98 | 3.55 | 1.12 | 0.26 | 1 | Gm44913 | | 0 | 0 | 0 | 0 | 0 | 8 | 0 | 27 | 0 | 0 | 5 |
| 4 | -3.26 | 3.59 | -0.91 | 0.36 | 1 | Papolb | poly (A) polymerase beta (testis specific) | 0 | 27 | 0 | 0 | 3 | 5 | 0 | 0 | 0 | 0 | 4 |
| 4 | 0.11 | 3.42 | 0.03 | 0.97 | 1 | Gm36899 | | 0 | 15 | 10 | 0 | 0 | 8 | 0 | 0 | 0 | 0 | 6 |
| 4 | 0.36 | 2.57 | 0.14 | 0.89 | 1 | Acot5 | acyl-CoA thioesterase 5 | 0 | 4 | 18 | 0 | 5 | 2 | 0 | 8 | 3 | 0 | 0 |
| 4 | 0.00 | 3.68 | 0.00 | 1.00 | 1 | Dhx58os | | 25 | 6 | 8 | 0 | 0 | 0 | 0 | 0 | 0 | 0 | 0 |
| 4 | 0.48 | 3.60 | 0.13 | 0.89 | 1 | Gm43336 | | 0 | 4 | 0 | 0 | 0 | 12 | 0 | 0 | 0 | 0 | 23 |
| 4 | 0.00 | 3.68 | 0.00 | 1.00 | 1 | Gm4070 | predicted gene 4070 | 9 | 21 | 9 | 0 | 0 | 0 | 0 | 0 | 0 | 0 | 0 |

| | | | | | | | | | | | | | | | | | | |
|---|---|---|---|---|---|---|---|---|---|---|---|---|---|---|---|---|---|---|
| 4 | -2.61 | 3.19 | -0.82 | 0.41 | 1 | Gm15722 | | 8 | 0 | 5 | 14 | 0 | 10 | 0 | 0 | 0 | 0 | 2 |
| 4 | 0.00 | 3.68 | 0.00 | 1.00 | 1 | Trim50 | tripartite motif-containing 50 | 0 | 16 | 23 | 0 | 0 | 0 | 0 | 0 | 0 | 0 | 0 |
| 4 | -3.68 | 3.57 | -1.03 | 0.30 | 1 | Gm23301 | | 11 | 0 | 0 | 5 | 0 | 14 | 0 | 0 | 0 | 0 | 9 |
| 4 | -2.94 | 3.57 | -0.82 | 0.41 | 1 | Gm48552 | deoxyhypusine hydroxylase/monooxyger | 10 | 0 | 0 | 5 | 0 | 4 | 0 | 0 | 0 | 0 | 19 |
| 4 | 3.27 | 3.58 | 0.91 | 0.36 | 1 | Slc22a13b | | 0 | 13 | 0 | 0 | 0 | 13 | 0 | 0 | 3 | 0 | 10 |
| 4 | 0.00 | 3.68 | 0.00 | 1.00 | 1 | Gm5532 | | 11 | 7 | 21 | 0 | 0 | 0 | 0 | 0 | 0 | 0 | 0 |
| 4 | 0.00 | 3.68 | 0.00 | 1.00 | 1 | Hoxb13 | homeobox B13 | 0 | 19 | 20 | 0 | 0 | 0 | 0 | 0 | 0 | 0 | 0 |
| 4 | 0.03 | 3.41 | 0.01 | 0.99 | 1 | Tdrd5 | tudor domain containing 5 | 8 | 13 | 0 | 0 | 0 | 11 | 0 | 0 | 0 | 0 | 7 |
| 4 | -1.31 | 3.07 | -0.43 | 0.67 | 1 | 3300002A11Rik | | 0 | 0 | 5 | 9 | 0 | 12 | 0 | 0 | 6 | 0 | 7 |
| 4 | 1.30 | 3.44 | 0.38 | 0.70 | 1 | Dppa3 | developmental pluripotency-associated 3 | 7 | 4 | 25 | 0 | 0 | 0 | 0 | 0 | 0 | 0 | 3 |
| 4 | -0.09 | 3.43 | -0.03 | 0.98 | 1 | Dnah7c | dynein, axonemal, heavy chain 7C | 13 | 0 | 14 | 0 | 0 | 8 | 0 | 0 | 0 | 0 | 4 |
| 4 | -0.40 | 3.41 | -0.12 | 0.91 | 1 | Efna4 | ephrin A4 | 10 | 12 | 16 | 0 | 0 | 1 | 0 | 0 | 0 | 0 | 0 |
| 4 | 0.00 | 3.68 | 0.00 | 1.00 | 1 | Atp6v0d2 | ATPase, H+ transporting, lysosomal V0 su | 4 | 29 | 5 | 0 | 0 | 0 | 0 | 0 | 0 | 0 | 0 |
| 4 | 0.66 | 2.65 | 0.25 | 0.80 | 1 | C230038L03Rik | | 0 | 0 | 0 | 5 | 2 | 5 | 0 | 0 | 10 | 15 | 1 |
| 4 | -0.88 | 3.56 | -0.25 | 0.81 | 1 | Gm35256 | | 0 | 0 | 0 | 0 | 4 | 13 | 0 | 0 | 0 | 0 | 22 |
| 4 | 0.54 | 3.60 | 0.15 | 0.88 | 1 | Gm6047 | | 0 | 0 | 5 | 0 | 11 | 0 | 0 | 0 | 22 | 0 | 0 |
| 4 | 0.00 | 3.68 | 0.00 | 1.00 | 1 | Trhde | TRH-degrading enzyme | 17 | 8 | 14 | 0 | 0 | 0 | 0 | 0 | 0 | 0 | 0 |
| 4 | -0.13 | 2.70 | -0.05 | 0.96 | 1 | Slc4a5 | solute carrier family 4, sodium bicarbonat | 0 | 20 | 0 | 0 | 6 | 3 | 0 | 2 | 2 | 0 | 5 |
| 4 | 0.51 | 2.84 | 0.18 | 0.86 | 1 | Necab3 | N-terminal EF-hand calcium binding prote | 0 | 0 | 4 | 0 | 3 | 8 | 0 | 0 | 21 | 0 | 2 |
| 4 | 0.09 | 3.56 | 0.03 | 0.98 | 1 | Amy2a1 | amylase 2a1 | 0 | 19 | 17 | 2 | 0 | 0 | 0 | 0 | 0 | 0 | 1 |
| 4 | -2.74 | 2.68 | -1.02 | 0.31 | 1 | Pramef12 | PRAME family member 12 | 3 | 6 | 12 | 8 | 0 | 8 | 0 | 0 | 2 | 0 | 0 |
| 4 | 0.00 | 3.64 | 0.00 | 1.00 | 1 | Zfp853 | zinc finger protein 853 | 39 | 0 | 0 | 0 | 0 | 0 | 0 | 0 | 0 | 0 | 0 |
| 4 | 0.00 | 3.68 | 0.00 | 1.00 | 1 | Gm7538 | | 7 | 18 | 14 | 0 | 0 | 0 | 0 | 0 | 0 | 0 | 0 |
| 4 | -1.27 | 3.16 | -0.40 | 0.69 | 1 | Gpat2 | glycerol-3-phosphate acyltransferase 2, n | 11 | 0 | 0 | 0 | 9 | 12 | 0 | 0 | 4 | 0 | 3 |
| 4 | 0.16 | 3.60 | 0.04 | 0.96 | 1 | Gm37008 | | 0 | 0 | 0 | 0 | 0 | 20 | 0 | 0 | 0 | 0 | 18 |
| 4 | 0.00 | 3.68 | 0.00 | 1.00 | 1 | Dmp1 | dentin matrix protein 1 | 17 | 16 | 5 | 0 | 0 | 0 | 0 | 0 | 0 | 0 | 0 |
| 4 | 1.74 | 3.59 | 0.48 | 0.63 | 1 | Mucl1 | mucin-like 1 | 33 | 0 | 0 | 0 | 0 | 0 | 0 | 0 | 0 | 0 | 5 |
| 3 | 0.00 | 3.68 | 0.00 | 1.00 | 1 | Mrap | melanocortin 2 receptor accessory proteir | 28 | 9 | 1 | 0 | 0 | 0 | 0 | 0 | 0 | 0 | 0 |
| 3 | 0.00 | 3.68 | 0.00 | 1.00 | 1 | Igfbpl1 | insulin-like growth factor binding protein-l | 13 | 15 | 10 | 0 | 0 | 0 | 0 | 0 | 0 | 0 | 0 |
| 3 | -0.40 | 3.48 | -0.11 | 0.91 | 1 | Gm6212 | | 14 | 1 | 23 | 0 | 0 | 1 | 0 | 0 | 0 | 0 | 0 |
| 3 | 4.17 | 3.42 | 1.22 | 0.22 | 1 | Ano2 | anoctamin 2 | 0 | 0 | 0 | 0 | 0 | 3 | 0 | 21 | 6 | 0 | 8 |
| 3 | -1.07 | 3.48 | -0.31 | 0.76 | 1 | G530011O06Rik | | 17 | 7 | 10 | 0 | 4 | 0 | 0 | 0 | 0 | 0 | 0 |
| 3 | 2.61 | 3.43 | 0.76 | 0.45 | 1 | Gm8719 | | 19 | 0 | 1 | 0 | 0 | 0 | 0 | 0 | 0 | 15 | 3 |
| 3 | 0.57 | 3.60 | 0.16 | 0.87 | 1 | Gm15328 | | 0 | 0 | 0 | 0 | 0 | 11 | 0 | 0 | 0 | 0 | 27 |
| 3 | 1.57 | 2.37 | 0.66 | 0.51 | 1 | Rpl18a-ps1 | | 8 | 2 | 21 | 1 | 1 | 0 | 0 | 3 | 0 | 3 | 0 |
| 3 | 1.10 | 3.60 | 0.31 | 0.76 | 1 | Gm10605 | | 18 | 0 | 11 | 0 | 0 | 1 | 0 | 0 | 8 | 0 | 0 |
| 3 | -3.22 | 2.79 | -1.15 | 0.25 | 1 | Gm13194 | predicted gene 13194 | 10 | 4 | 0 | 11 | 3 | 3 | 0 | 0 | 0 | 0 | 8 |
| 3 | -1.85 | 2.08 | -0.89 | 0.37 | 1 | Gm4600 | | 2 | 8 | 1 | 11 | 9 | 3 | 0 | 0 | 4 | 0 | 1 |
| 3 | 0.87 | 3.50 | 0.25 | 0.80 | 1 | AI838599 | | 19 | 0 | 14 | 0 | 0 | 1 | 0 | 0 | 0 | 0 | 5 |
| 3 | 0.00 | 3.68 | 0.00 | 1.00 | 1 | Ccl17 | chemokine (C-C motif) ligand 17 | 10 | 2 | 26 | 0 | 0 | 0 | 0 | 0 | 0 | 0 | 0 |
| 3 | 0.00 | 3.68 | 0.00 | 1.00 | 1 | B130024G19Rik | | 18 | 20 | 0 | 0 | 0 | 0 | 0 | 0 | 0 | 0 | 0 |
| 3 | 3.73 | 2.97 | 1.25 | 0.21 | 1 | Gm12129 | | 1 | 0 | 0 | 0 | 0 | 1 | 0 | 2 | 2 | 32 | 0 |
| 3 | 1.37 | 3.57 | 0.38 | 0.70 | 1 | Gm2824 | | 0 | 0 | 0 | 0 | 0 | 22 | 0 | 0 | 4 | 0 | 12 |
| 3 | -0.98 | 3.50 | -0.28 | 0.78 | 1 | Tph1 | tryptophan hydroxylase 1 | 2 | 10 | 20 | 0 | 7 | 0 | 0 | 0 | 0 | 0 | 0 |
| 3 | 0.00 | 2.88 | 0.00 | 1.00 | 1 | Cebpe | CCAAT/enhancer binding protein (C/EBP | 7 | 18 | 10 | 0 | 0 | 2 | 0 | 0 | 0 | 0 | 1 |
| 3 | 0.92 | 3.57 | 0.26 | 0.80 | 1 | Ly6g5b | lymphocyte antigen 6 complex, locus G5E | 0 | 28 | 0 | 0 | 0 | 2 | 0 | 0 | 0 | 0 | 8 |
| 3 | 3.46 | 3.56 | 0.97 | 0.33 | 1 | Snord53 | | 0 | 0 | 0 | 0 | 0 | 15 | 0 | 13 | 0 | 0 | 10 |
| 3 | 0.84 | 3.45 | 0.24 | 0.81 | 1 | Gm45309 | | 4 | 0 | 0 | 0 | 0 | 19 | 0 | 0 | 1 | 0 | 14 |
| 3 | 2.45 | 3.57 | 0.68 | 0.49 | 1 | A430010J10Rik | | 0 | 0 | 0 | 3 | 0 | 0 | 0 | 32 | 0 | 0 | 3 |
| 3 | 0.26 | 3.60 | 0.07 | 0.94 | 1 | Gm44041 | | 0 | 0 | 0 | 0 | 0 | 18 | 0 | 0 | 0 | 0 | 20 |
| 3 | 4.38 | 3.51 | 1.25 | 0.21 | 1 | 4930505N22Rik | | 0 | 0 | 0 | 0 | 0 | 4 | 0 | 10 | 0 | 19 | 4 |
| 3 | -0.76 | 3.63 | -0.21 | 0.83 | 1 | Gng8 | guanine nucleotide binding protein (G prc | 23 | 0 | 11 | 0 | 0 | 3 | 0 | 0 | 0 | 0 | 0 |
| 3 | 0.71 | 3.03 | 0.23 | 0.81 | 1 | Gm48242 | | 9 | 2 | 20 | 0 | 0 | 2 | 0 | 0 | 5 | 0 | 0 |
| 3 | -0.74 | 2.32 | -0.32 | 0.75 | 1 | Gm14221 | | 1 | 10 | 12 | 8 | 0 | 3 | 0 | 0 | 1 | 0 | 3 |
| 3 | -0.74 | 3.53 | -0.21 | 0.83 | 1 | Phkg1 | phosphorylase kinase gamma 1 | 29 | 1 | 5 | 0 | 0 | 3 | 0 | 0 | 0 | 0 | 0 |
| 3 | 4.08 | 3.59 | 1.13 | 0.26 | 1 | Gm29707 | | 0 | 17 | 0 | 0 | 0 | 3 | 0 | 0 | 2 | 0 | 15 |
| 3 | -0.75 | 3.43 | -0.22 | 0.83 | 1 | Gm48357 | | 3 | 0 | 0 | 0 | 1 | 21 | 0 | 0 | 0 | 0 | 13 |
| 3 | -2.01 | 2.91 | -0.69 | 0.49 | 1 | Gm43660 | | 0 | 2 | 0 | 11 | 3 | 5 | 0 | 17 | 0 | 0 | 0 |
| 3 | 1.31 | 3.66 | 0.36 | 0.72 | 1 | Gm12688 | | 0 | 23 | 12 | 0 | 0 | 0 | 0 | 0 | 2 | 0 | 0 |
| 3 | 0.18 | 3.60 | 0.05 | 0.96 | 1 | Gm42548 | | 0 | 0 | 0 | 0 | 0 | 19 | 0 | 0 | 0 | 0 | 18 |

| | | | | | | | | | | | | | | | | | | |
|---|---|---|---|---|---|---|---|---|---|---|---|---|---|---|---|---|---|---|
| 3 | -1.26 | 3.10 | -0.41 | 0.68 | 1 | | | 6 | 3 | 0 | 0 | 4 | 13 | 0 | 0 | 0 | 0 | 12 |
| 3 | 1.56 | 3.02 | 0.52 | 0.60 | 1 | Icam5 | intercellular adhesion molecule 5, telence | 19 | 0 | 8 | 0 | 0 | 3 | 0 | 0 | 4 | 0 | 3 |
| 3 | 0.00 | 3.68 | 0.00 | 1.00 | 1 | Gm50405 | | 21 | 16 | 0 | 0 | 0 | 0 | 0 | 0 | 0 | 0 | 0 |
| 3 | 0.72 | 2.84 | 0.25 | 0.80 | 1 | Fem1al | RIKEN cDNA 4931440F15 gene | 13 | 1 | 0 | 0 | 1 | 13 | 0 | 0 | 7 | 0 | 3 |
| 3 | 0.36 | 3.60 | 0.10 | 0.92 | 1 | Gm37689 | | 0 | 0 | 0 | 0 | 0 | 15 | 0 | 0 | 0 | 0 | 22 |
| 3 | 4.31 | 3.56 | 1.21 | 0.23 | 1 | Gm43018 | | 0 | 0 | 0 | 0 | 0 | 0 | 0 | 0 | 0 | 33 | 4 |
| 3 | -1.25 | 3.60 | -0.35 | 0.73 | 1 | Casq1 | calsequestrin 1 | 24 | 0 | 0 | 0 | 7 | 1 | 0 | 0 | 6 | 0 | 0 |
| 3 | 0.00 | 3.68 | 0.00 | 1.00 | 1 | Gm18345 | | 11 | 24 | 2 | 0 | 0 | 0 | 0 | 0 | 0 | 0 | 0 |
| 3 | 0.43 | 3.57 | 0.12 | 0.90 | 1 | Gm44237 | | 0 | 0 | 19 | 0 | 0 | 8 | 0 | 0 | 0 | 0 | 11 |
| 3 | 0.00 | 3.68 | 0.00 | 1.00 | 1 | Gm13056 | | 29 | 0 | 8 | 0 | 0 | 0 | 0 | 0 | 0 | 0 | 0 |
| 3 | -0.42 | 3.53 | -0.12 | 0.90 | 1 | AA914427 | | 0 | 11 | 12 | 0 | 0 | 12 | 0 | 0 | 0 | 0 | 2 |
| 3 | 0.00 | 3.68 | 0.00 | 1.00 | 1 | Gm21013 | | 8 | 15 | 15 | 0 | 0 | 0 | 0 | 0 | 0 | 0 | 0 |
| 3 | -0.14 | 3.48 | -0.04 | 0.97 | 1 | Gm5103 | | 0 | 0 | 0 | 12 | 0 | 10 | 0 | 7 | 8 | 0 | 0 |
| 3 | 3.93 | 2.98 | 1.32 | 0.19 | 1 | Gm38004 | | 0 | 2 | 0 | 0 | 0 | 2 | 0 | 19 | 12 | 0 | 3 |
| 3 | -3.75 | 3.12 | -1.20 | 0.23 | 1 | B9d1os | | 7 | 0 | 0 | 4 | 6 | 5 | 0 | 0 | 0 | 15 | 0 |
| 3 | 0.00 | 3.68 | 0.00 | 1.00 | 1 | Sectm1a | secreted and transmembrane 1A | 34 | 0 | 3 | 0 | 0 | 0 | 0 | 0 | 0 | 0 | 0 |
| 3 | -2.59 | 3.50 | -0.74 | 0.46 | 1 | 1700020L24Rik | RIKEN cDNA 1700020L24 gene | 0 | 0 | 0 | 9 | 4 | 2 | 0 | 0 | 0 | 23 | 0 |
| 3 | 0.95 | 2.22 | 0.43 | 0.67 | 1 | Gm15455 | predicted gene 15455 | 2 | 0 | 4 | 0 | 4 | 9 | 0 | 0 | 2 | 10 | 6 |
| 3 | 1.47 | 2.71 | 0.54 | 0.59 | 1 | Gm47980 | | 3 | 0 | 0 | 0 | 5 | 8 | 0 | 3 | 3 | 0 | 14 |
| 3 | 3.61 | 2.73 | 1.32 | 0.19 | 1 | Gm37452 | | 0 | 1 | 0 | 3 | 0 | 5 | 0 | 0 | 9 | 15 | 4 |
| 3 | 0.11 | 3.58 | 0.03 | 0.98 | 1 | Gm37900 | | 18 | 0 | 0 | 0 | 0 | 10 | 0 | 0 | 0 | 0 | 9 |
| 3 | 3.81 | 3.51 | 1.09 | 0.28 | 1 | Gm5075 | | 0 | 17 | 0 | 0 | 14 | 0 | 0 | 0 | 0 | 4 | 2 |
| 3 | 0.00 | 3.68 | 0.00 | 1.00 | 1 | Igsf23 | immunoglobulin superfamily, member 23 | 5 | 3 | 29 | 0 | 0 | 0 | 0 | 0 | 0 | 0 | 0 |
| 3 | 0.00 | 3.68 | 0.00 | 1.00 | 1 | Fendrr | | 19 | 14 | 4 | 0 | 0 | 0 | 0 | 0 | 0 | 0 | 0 |
| 3 | -3.55 | 3.56 | -1.00 | 0.32 | 1 | E430014B02Rik | | 0 | 0 | 0 | 15 | 0 | 10 | 0 | 0 | 0 | 0 | 12 |
| 3 | 0.00 | 3.68 | 0.00 | 1.00 | 1 | Gm14548 | paired-Ig-like receptor A11 | 24 | 13 | 0 | 0 | 0 | 0 | 0 | 0 | 0 | 0 | 0 |
| 3 | 1.21 | 2.12 | 0.57 | 0.57 | 1 | Abcb6 | ATP-binding cassette, sub-family B (MDR | 1 | 3 | 5 | 4 | 0 | 9 | 0 | 0 | 3 | 5 | 7 |
| 3 | 1.68 | 2.32 | 0.73 | 0.47 | 1 | Gm15784 | | 1 | 13 | 0 | 4 | 0 | 3 | 0 | 1 | 0 | 9 | 6 |
| 3 | 2.88 | 2.39 | 1.20 | 0.23 | 1 | Cpne9 | copine family member IX | 0 | 2 | 9 | 0 | 1 | 3 | 0 | 0 | 5 | 10 | 6 |
| 3 | -2.37 | 2.42 | -0.98 | 0.33 | 1 | Dnajb6-ps | | 11 | 7 | 0 | 4 | 5 | 8 | 0 | 0 | 1 | 0 | 2 |
| 3 | 0.00 | 3.68 | 0.00 | 1.00 | 1 | A530040E14Rik | | 19 | 18 | 0 | 0 | 0 | 0 | 0 | 0 | 0 | 0 | 0 |
| 3 | 2.99 | 2.98 | 1.00 | 0.32 | 1 | Vpreb1 | pre-B lymphocyte gene 1 | 5 | 0 | 0 | 0 | 0 | 5 | 0 | 0 | 6 | 17 | 4 |
| 3 | 0.00 | 3.68 | 0.00 | 1.00 | 1 | Spink4 | serine peptidase inhibitor, Kazal type 4 | 8 | 29 | 0 | 0 | 0 | 0 | 0 | 0 | 0 | 0 | 0 |
| 3 | 0.00 | 3.68 | 0.00 | 1.00 | 1 | Hba-a2 | hemoglobin alpha, adult chain 2 | 11 | 2 | 24 | 0 | 0 | 0 | 0 | 0 | 0 | 0 | 0 |
| 3 | 0.43 | 2.38 | 0.18 | 0.86 | 1 | A430027C01Rik | | 1 | 0 | 7 | 0 | 7 | 5 | 0 | 0 | 6 | 0 | 11 |
| 3 | 0.01 | 3.60 | 0.00 | 1.00 | 1 | Gm43588 | | 0 | 0 | 0 | 0 | 0 | 23 | 0 | 0 | 0 | 0 | 14 |
| 3 | 0.04 | 3.49 | 0.01 | 0.99 | 1 | Tac1 | tachykinin 1 | 9 | 0 | 20 | 0 | 5 | 0 | 0 | 0 | 3 | 0 | 0 |
| 3 | 2.48 | 3.58 | 0.69 | 0.49 | 1 | Gm12026 | | 0 | 0 | 0 | 1 | 0 | 0 | 0 | 35 | 0 | 1 | 0 |
| 3 | 0.00 | 3.68 | 0.00 | 1.00 | 1 | Chrdl2 | chordin-like 2 | 0 | 22 | 15 | 0 | 0 | 0 | 0 | 0 | 0 | 0 | 0 |
| 3 | 0.02 | 3.59 | 0.00 | 1.00 | 1 | Gm37166 | | 8 | 0 | 0 | 0 | 0 | 17 | 0 | 0 | 0 | 0 | 12 |
| 3 | -1.32 | 2.36 | -0.56 | 0.57 | 1 | Gm11523 | | 2 | 11 | 0 | 6 | 7 | 5 | 0 | 0 | 3 | 0 | 3 |
| 3 | -1.09 | 3.15 | -0.34 | 0.73 | 1 | Gm16025 | | 0 | 0 | 0 | 1 | 5 | 8 | 0 | 0 | 0 | 21 | 2 |
| 3 | -1.23 | 3.00 | -0.41 | 0.68 | 1 | Rnf225 | ring finger protein 225 | 8 | 4 | 0 | 6 | 0 | 8 | 0 | 0 | 0 | 0 | 11 |
| 3 | 0.00 | 3.68 | 0.00 | 1.00 | 1 | Gm16263 | | 21 | 15 | 0 | 0 | 0 | 0 | 0 | 0 | 0 | 0 | 0 |
| 3 | 0.85 | 3.58 | 0.24 | 0.81 | 1 | Znrd2 | zinc ribbon domain containing 2 | 0 | 0 | 10 | 0 | 0 | 6 | 0 | 0 | 0 | 0 | 20 |
| 3 | 0.91 | 2.21 | 0.41 | 0.68 | 1 | Cntf | ciliary neurotrophic factor | 8 | 0 | 3 | 4 | 2 | 2 | 0 | 13 | 3 | 0 | 2 |
| 3 | -1.14 | 3.10 | -0.37 | 0.71 | 1 | Gm45091 | | 0 | 12 | 15 | 2 | 0 | 5 | 0 | 0 | 3 | 0 | 0 |
| 3 | 0.00 | 3.68 | 0.00 | 1.00 | 1 | a | nonagouti | 12 | 15 | 9 | 0 | 0 | 0 | 0 | 0 | 0 | 0 | 0 |
| 3 | -0.87 | 3.22 | -0.27 | 0.79 | 1 | A930001C03Rik | | 0 | 7 | 0 | 0 | 12 | 3 | 0 | 12 | 0 | 0 | 3 |
| 3 | -2.38 | 3.55 | -0.67 | 0.50 | 1 | Gm25259 | | 0 | 0 | 0 | 11 | 0 | 22 | 0 | 0 | 0 | 0 | 4 |
| 3 | 0.11 | 3.60 | 0.03 | 0.97 | 1 | Gm37983 | | 0 | 0 | 0 | 0 | 0 | 20 | 0 | 0 | 0 | 0 | 16 |
| 3 | 0.36 | 3.61 | 0.10 | 0.92 | 1 | Gm1965 | | 26 | 0 | 0 | 0 | 5 | 0 | 0 | 0 | 6 | 0 | 0 |
| 3 | 4.40 | 3.57 | 1.23 | 0.22 | 1 | Gm42841 | | 0 | 0 | 11 | 0 | 0 | 4 | 0 | 21 | 0 | 0 | 0 |
| 3 | -2.51 | 3.59 | -0.70 | 0.48 | 1 | Gria1 | glutamate receptor, ionotropic, AMPA1 (a | 16 | 0 | 6 | 13 | 0 | 1 | 0 | 0 | 0 | 0 | 0 |
| 3 | 0.00 | 3.63 | 0.00 | 1.00 | 1 | Gm15056 | predicted gene 15056 | 0 | 36 | 0 | 0 | 0 | 0 | 0 | 0 | 0 | 0 | 0 |
| 3 | -0.58 | 3.43 | -0.17 | 0.87 | 1 | 4930404H24Rik | | 11 | 0 | 0 | 9 | 0 | 3 | 0 | 0 | 11 | 0 | 2 |
| 3 | 1.50 | 2.39 | 0.63 | 0.53 | 1 | Gm11405 | | 0 | 3 | 2 | 0 | 1 | 2 | 0 | 27 | 1 | 0 | 1 |
| 3 | 1.36 | 3.55 | 0.38 | 0.70 | 1 | Gm15063 | | 0 | 0 | 10 | 0 | 17 | 0 | 0 | 0 | 0 | 9 | 0 |
| 3 | 0.00 | 3.68 | 0.00 | 1.00 | 1 | Cst7 | cystatin F (leukocystatin) | 28 | 8 | 0 | 0 | 0 | 0 | 0 | 0 | 0 | 0 | 0 |
| 3 | 1.61 | 3.64 | 0.44 | 0.66 | 1 | Gsg1l | GSG1-like | 24 | 0 | 9 | 0 | 0 | 0 | 0 | 0 | 3 | 0 | 0 |

| | | | | | | | | | | | | | | | | | | |
|---|---|---|---|---|---|---|---|---|---|---|---|---|---|---|---|---|---|---|
| 3 | 0.05 | 2.36 | 0.02 | 0.98 | 1 | Gm4518 | | 2 | 0 | 0 | 7 | 2 | 1 | 0 | 6 | 0 | 18 | 1 |
| 3 | -0.32 | 1.67 | -0.19 | 0.85 | 1 | Rpl13a-ps1 | | 3 | 6 | 3 | 11 | 4 | 1 | 0 | 2 | 2 | 1 | 2 |
| 3 | -2.19 | 2.40 | -0.91 | 0.36 | 1 | Gm2693 | | 8 | 2 | 11 | 2 | 9 | 1 | 0 | 0 | 0 | 0 | 3 |
| 3 | 1.21 | 2.93 | 0.41 | 0.68 | 1 | Gm43267 | | 8 | 0 | 2 | 0 | 0 | 10 | 0 | 0 | 0 | 5 | 11 |
| 3 | -1.28 | 3.58 | -0.36 | 0.72 | 1 | Foxa1 | forkhead box A1 | 20 | 7 | 4 | 5 | 0 | 0 | 0 | 0 | 0 | 0 | 0 |
| 3 | -3.13 | 3.44 | -0.91 | 0.36 | 1 | Gm12764 | | 13 | 1 | 0 | 0 | 11 | 1 | 0 | 0 | 0 | 0 | 10 |
| 3 | -2.40 | 3.30 | -0.73 | 0.47 | 1 | Mchr1 | melanin-concentrating hormone receptor | 1 | 1 | 0 | 0 | 4 | 1 | 0 | 0 | 0 | 30 | 0 |
| 3 | 0.00 | 3.68 | 0.00 | 1.00 | 1 | Msmp | microseminoprotein, prostate associated | 10 | 9 | 17 | 0 | 0 | 0 | 0 | 0 | 0 | 0 | 0 |
| 3 | 0.00 | 3.64 | 0.00 | 1.00 | 1 | Ighv5-15 | | 36 | 0 | 0 | 0 | 0 | 0 | 0 | 0 | 0 | 0 | 0 |
| 3 | 2.28 | 2.68 | 0.85 | 0.40 | 1 | Gm48048 | | 0 | 0 | 2 | 9 | 0 | 0 | 0 | 21 | 2 | 1 | 1 |
| 3 | -2.40 | 3.59 | -0.67 | 0.50 | 1 | Gm48163 | | 0 | 14 | 14 | 0 | 7 | 2 | 0 | 0 | 0 | 0 | 0 |
| 3 | 0.75 | 2.24 | 0.33 | 0.74 | 1 | Gm13572 | | 0 | 2 | 9 | 3 | 3 | 1 | 0 | 0 | 11 | 0 | 7 |
| 3 | 1.25 | 3.50 | 0.36 | 0.72 | 1 | Obp2b | odorant binding protein 2B | 5 | 15 | 14 | 0 | 0 | 0 | 0 | 0 | 0 | 0 | 2 |
| 3 | 2.52 | 2.34 | 1.08 | 0.28 | 1 | Rnaset2a | ribonuclease T2A | 9 | 2 | 2 | 0 | 0 | 9 | 0 | 0 | 5 | 4 | 4 |
| 3 | 0.44 | 2.62 | 0.17 | 0.87 | 1 | Amd-ps4 | | 0 | 2 | 26 | 1 | 0 | 1 | 0 | 0 | 4 | 0 | 2 |
| 3 | 2.66 | 3.54 | 0.75 | 0.45 | 1 | Gm37420 | | 0 | 0 | 0 | 0 | 0 | 9 | 0 | 25 | 0 | 0 | 1 |
| 3 | -3.07 | 3.58 | -0.86 | 0.39 | 1 | Snord73a | | 6 | 0 | 0 | 0 | 3 | 13 | 0 | 0 | 0 | 0 | 14 |
| 3 | 0.00 | 3.68 | 0.00 | 1.00 | 1 | Gm29491 | | 13 | 0 | 23 | 0 | 0 | 0 | 0 | 0 | 0 | 0 | 0 |
| 3 | 0.48 | 3.52 | 0.14 | 0.89 | 1 | Eid3 | EP300 interacting inhibitor of differentiatio | 1 | 0 | 7 | 0 | 0 | 9 | 0 | 0 | 0 | 0 | 18 |
| 3 | 1.33 | 3.58 | 0.37 | 0.71 | 1 | Gm11870 | | 0 | 0 | 0 | 0 | 0 | 14 | 0 | 0 | 2 | 0 | 19 |
| 3 | -1.62 | 3.61 | -0.45 | 0.65 | 1 | Kcnj14 | potassium inwardly-rectifying channel, su | 18 | 14 | 0 | 3 | 0 | 1 | 0 | 0 | 0 | 0 | 0 |
| 3 | 1.56 | 3.56 | 0.44 | 0.66 | 1 | Igdcc3 | immunoglobulin superfamily, DCC subcla | 11 | 14 | 7 | 0 | 0 | 0 | 0 | 0 | 3 | 0 | 0 |
| 3 | -3.31 | 3.56 | -0.93 | 0.35 | 1 | Gm26756 | | 0 | 21 | 0 | 6 | 6 | 3 | 0 | 0 | 0 | 0 | 0 |
| 3 | -4.15 | 3.01 | -1.38 | 0.17 | 1 | Prr18 | proline rich 18 | 0 | 2 | 0 | 10 | 17 | 6 | 0 | 0 | 0 | 0 | 1 |
| 3 | -3.48 | 3.29 | -1.06 | 0.29 | 1 | Gm49519 | | 5 | 0 | 4 | 11 | 0 | 7 | 0 | 0 | 0 | 0 | 8 |
| 3 | 0.95 | 3.67 | 0.26 | 0.80 | 1 | Greb1 | gene regulated by estrogen in breast can | 27 | 7 | 0 | 0 | 0 | 0 | 0 | 0 | 0 | 0 | 1 |
| 3 | 0.05 | 3.59 | 0.01 | 0.99 | 1 | Gm22637 | | 0 | 0 | 8 | 0 | 0 | 16 | 0 | 0 | 0 | 0 | 11 |
| 3 | -2.37 | 3.05 | -0.78 | 0.44 | 1 | Rapgef3os2 | | 8 | 5 | 15 | 0 | 6 | 3 | 0 | 0 | 0 | 0 | 0 |
| 3 | 0.32 | 3.60 | 0.09 | 0.93 | 1 | Gm43931 | | 0 | 0 | 0 | 0 | 0 | 15 | 0 | 0 | 0 | 0 | 20 |
| 3 | -2.04 | 2.58 | -0.79 | 0.43 | 1 | Gm48961 | | 0 | 0 | 8 | 8 | 4 | 8 | 0 | 0 | 4 | 0 | 3 |
| 3 | 0.00 | 3.68 | 0.00 | 1.00 | 1 | Cnksr2 | connector enhancer of kinase suppressor | 23 | 5 | 7 | 0 | 0 | 0 | 0 | 0 | 0 | 0 | 0 |
| 3 | 4.05 | 3.57 | 1.13 | 0.26 | 1 | Gm17096 | | 0 | 4 | 0 | 0 | 0 | 10 | 0 | 0 | 6 | 0 | 15 |
| 3 | 0.00 | 3.68 | 0.00 | 1.00 | 1 | Prcd | photoreceptor disc component | 2 | 21 | 12 | 0 | 0 | 0 | 0 | 0 | 0 | 0 | 0 |
| 3 | 0.00 | 3.68 | 0.00 | 1.00 | 1 | Palm3 | paralemmin 3 | 18 | 9 | 8 | 0 | 0 | 0 | 0 | 0 | 0 | 0 | 0 |
| 3 | 6.09 | 3.54 | 1.72 | 0.09 | 1 | 4930447F24Rik | | 0 | 0 | 0 | 0 | 15 | 0 | 0 | 16 | 0 | 0 | 4 |
| 3 | 0.00 | 3.68 | 0.00 | 1.00 | 1 | Igkj3 | | 17 | 18 | 0 | 0 | 0 | 0 | 0 | 0 | 0 | 0 | 0 |
| 3 | 3.39 | 3.59 | 0.95 | 0.34 | 1 | Cck | cholecystokinin | 0 | 13 | 7 | 0 | 0 | 0 | 0 | 0 | 0 | 13 | 2 |
| 3 | 0.28 | 3.60 | 0.08 | 0.94 | 1 | C230085N15Rik | | 0 | 0 | 0 | 0 | 0 | 16 | 0 | 0 | 0 | 0 | 19 |
| 3 | -0.39 | 2.56 | -0.15 | 0.88 | 1 | Gm10863 | | 15 | 12 | 2 | 0 | 1 | 3 | 0 | 0 | 0 | 0 | 3 |
| 3 | -0.40 | 3.53 | -0.11 | 0.91 | 1 | Bmp8b | bone morphogenetic protein 8b | 18 | 6 | 10 | 0 | 0 | 1 | 0 | 0 | 0 | 0 | 0 |
| 3 | -1.05 | 3.62 | -0.29 | 0.77 | 1 | Chad | chondroadherin | 12 | 9 | 10 | 0 | 4 | 0 | 0 | 0 | 0 | 0 | 0 |
| 3 | 4.03 | 3.56 | 1.13 | 0.26 | 1 | Gm28055 | | 0 | 0 | 0 | 0 | 0 | 0 | 0 | 0 | 0 | 32 | 3 |
| 3 | -0.04 | 3.54 | -0.01 | 0.99 | 1 | Gm47819 | | 0 | 0 | 0 | 7 | 0 | 5 | 0 | 23 | 0 | 0 | 0 |
| 3 | 1.62 | 2.67 | 0.61 | 0.54 | 1 | Gm1070 | predicted gene 1070 | 0 | 1 | 0 | 0 | 2 | 8 | 0 | 0 | 2 | 17 | 5 |
| 3 | 0.00 | 3.64 | 0.00 | 1.00 | 1 | Ptprv | protein tyrosine phosphatase, receptor ty | 35 | 0 | 0 | 0 | 0 | 0 | 0 | 0 | 0 | 0 | 0 |
| 3 | 1.92 | 3.59 | 0.53 | 0.59 | 1 | Gm45237 | | 0 | 9 | 5 | 0 | 0 | 0 | 0 | 21 | 0 | 0 | 0 |
| 3 | 0.00 | 3.68 | 0.00 | 1.00 | 1 | Zfp652os | | 13 | 22 | 0 | 0 | 0 | 0 | 0 | 0 | 0 | 0 | 0 |
| 3 | -0.01 | 3.30 | 0.00 | 1.00 | 1 | Gm6334 | | 21 | 5 | 0 | 0 | 2 | 2 | 0 | 0 | 5 | 0 | 0 |
| 3 | -0.40 | 3.59 | -0.11 | 0.91 | 1 | Gm2814 | | 2 | 27 | 5 | 0 | 0 | 1 | 0 | 0 | 0 | 0 | 0 |
| 3 | 1.60 | 2.19 | 0.73 | 0.47 | 1 | Rps2-ps6 | | 3 | 2 | 2 | 3 | 0 | 3 | 0 | 2 | 2 | 18 | 0 |
| 3 | -0.55 | 3.19 | -0.17 | 0.86 | 1 | Gm30889 | | 0 | 0 | 8 | 6 | 0 | 3 | 0 | 0 | 11 | 0 | 6 |
| 3 | 1.46 | 2.62 | 0.56 | 0.58 | 1 | Gm11205 | | 2 | 9 | 0 | 5 | 0 | 1 | 0 | 15 | 3 | 0 | 0 |
| 3 | 0.43 | 3.60 | 0.12 | 0.90 | 1 | Gm42672 | | 0 | 0 | 0 | 0 | 0 | 13 | 0 | 0 | 0 | 0 | 22 |
| 3 | 0.43 | 3.60 | 0.12 | 0.90 | 1 | AA413626 | | 0 | 0 | 0 | 0 | 0 | 13 | 0 | 0 | 0 | 0 | 22 |
| 3 | 0.00 | 3.68 | 0.00 | 1.00 | 1 | Hoxaas2 | | 21 | 5 | 9 | 0 | 0 | 0 | 0 | 0 | 0 | 0 | 0 |
| 3 | 1.91 | 3.04 | 0.63 | 0.53 | 1 | E230020D15Rik | | 1 | 1 | 27 | 0 | 0 | 0 | 0 | 0 | 5 | 0 | 1 |
| 3 | 0.00 | 3.68 | 0.00 | 1.00 | 1 | Ggt6 | gamma-glutamyltransferase 6 | 0 | 19 | 16 | 0 | 0 | 0 | 0 | 0 | 0 | 0 | 0 |
| 3 | -2.54 | 2.93 | -0.87 | 0.39 | 1 | Gm9067 | | 18 | 0 | 0 | 3 | 8 | 3 | 0 | 0 | 2 | 0 | 2 |
| 3 | 0.52 | 1.88 | 0.27 | 0.78 | 1 | Pabpn1l | poly(A)binding protein nuclear 1-like | 7 | 5 | 0 | 2 | 5 | 3 | 0 | 2 | 4 | 3 | 4 |
| 3 | -1.23 | 3.60 | -0.34 | 0.73 | 1 | Olig1 | oligodendrocyte transcription factor 1 | 0 | 31 | 0 | 0 | 0 | 3 | 0 | 0 | 0 | 0 | 0 |

| | | | | | | | | | | | | | | | | | | |
|---|---|---|---|---|---|---|---|---|---|---|---|---|---|---|---|---|---|---|
| 3 | -4.00 | 3.56 | -1.13 | 0.26 | 1 | Gm47509 | | 0 | 0 | 10 | 4 | 6 | 15 | 0 | 0 | 0 | 0 | 0 |
| 3 | 0.00 | 3.68 | 0.00 | 1.00 | 1 | C730027H18Rik | | 0 | 22 | 12 | 0 | 0 | 0 | 0 | 0 | 0 | 0 | 0 |
| 3 | 0.00 | 3.68 | 0.00 | 1.00 | 1 | Sectm1b | secreted and transmembrane 1B | 0 | 23 | 11 | 0 | 0 | 0 | 0 | 0 | 0 | 0 | 0 |
| 3 | 0.00 | 3.68 | 0.00 | 1.00 | 1 | Serpina3m | serine (or cysteine) peptidase inhibitor, cl | 27 | 8 | 0 | 0 | 0 | 0 | 0 | 0 | 0 | 0 | 0 |
| 3 | 0.03 | 2.93 | 0.01 | 0.99 | 1 | Erfe | erythroferrone | 6 | 2 | 0 | 8 | 0 | 8 | 0 | 0 | 5 | 0 | 6 |
| 3 | -3.42 | 2.42 | -1.41 | 0.16 | 1 | Gm6169 | | 3 | 2 | 7 | 9 | 9 | 2 | 0 | 0 | 0 | 0 | 2 |
| 3 | 0.75 | 2.82 | 0.26 | 0.79 | 1 | Gm42655 | | 0 | 0 | 5 | 3 | 3 | 5 | 0 | 0 | 0 | 8 | 11 |
| 3 | 1.95 | 3.06 | 0.64 | 0.52 | 1 | Defa39 | defensin, alpha, 39 | 0 | 0 | 8 | 1 | 2 | 0 | 0 | 17 | 6 | 0 | 0 |
| 3 | -1.54 | 3.61 | -0.43 | 0.67 | 1 | Sct | secretin | 0 | 9 | 21 | 0 | 4 | 1 | 0 | 0 | 0 | 0 | 0 |
| 3 | -2.50 | 2.93 | -0.85 | 0.39 | 1 | Gm8054 | | 3 | 4 | 0 | 16 | 5 | 0 | 0 | 0 | 0 | 1 | 6 |
| 3 | 0.09 | 3.55 | 0.03 | 0.98 | 1 | Gm20605 | | 0 | 1 | 5 | 0 | 0 | 16 | 0 | 0 | 0 | 0 | 12 |
| 3 | -1.38 | 2.57 | -0.54 | 0.59 | 1 | Gm10373 | | 0 | 0 | 7 | 5 | 6 | 5 | 0 | 0 | 7 | 0 | 4 |
| 3 | -2.35 | 3.60 | -0.65 | 0.51 | 1 | Adamts19 | a disintegrin-like and metallopeptidase (re | 26 | 0 | 0 | 0 | 2 | 2 | 0 | 0 | 0 | 0 | 5 |
| 3 | -0.59 | 3.49 | -0.17 | 0.86 | 1 | Gm49223 | | 0 | 0 | 0 | 0 | 8 | 16 | 0 | 5 | 0 | 0 | 6 |
| 3 | -0.40 | 3.56 | -0.11 | 0.91 | 1 | Gm11730 | | 12 | 14 | 7 | 0 | 0 | 1 | 0 | 0 | 0 | 0 | 0 |
| 3 | 0.00 | 3.68 | 0.00 | 1.00 | 1 | Trdc | | 23 | 11 | 0 | 0 | 0 | 0 | 0 | 0 | 0 | 0 | 0 |
| 3 | 1.00 | 3.59 | 0.28 | 0.78 | 1 | Gm13777 | | 0 | 1 | 0 | 0 | 0 | 1 | 0 | 32 | 0 | 0 | 0 |
| 3 | 0.00 | 3.63 | 0.00 | 1.00 | 1 | Gsdmc2 | gasdermin C2 | 0 | 0 | 34 | 0 | 0 | 0 | 0 | 0 | 0 | 0 | 0 |
| 3 | -1.03 | 2.78 | -0.37 | 0.71 | 1 | Gm33035 | | 3 | 18 | 2 | 0 | 5 | 2 | 0 | 0 | 0 | 0 | 4 |
| 3 | -0.03 | 3.60 | -0.01 | 0.99 | 1 | Gm37886 | | 0 | 0 | 0 | 0 | 0 | 22 | 0 | 0 | 0 | 0 | 12 |
| 3 | 0.00 | 3.68 | 0.00 | 1.00 | 1 | Col17a1 | collagen, type XVII, alpha 1 | 24 | 10 | 0 | 0 | 0 | 0 | 0 | 0 | 0 | 0 | 0 |
| 3 | 0.00 | 3.68 | 0.00 | 1.00 | 1 | Ptgdr2 | prostaglandin D2 receptor 2 | 3 | 25 | 6 | 0 | 0 | 0 | 0 | 0 | 0 | 0 | 0 |
| 3 | -0.24 | 2.33 | -0.11 | 0.92 | 1 | Gm5737 | predicted gene 5737 | 1 | 11 | 9 | 1 | 2 | 3 | 0 | 0 | 0 | 6 | 1 |
| 3 | 0.00 | 3.68 | 0.00 | 1.00 | 1 | | | 15 | 3 | 16 | 0 | 0 | 0 | 0 | 0 | 0 | 0 | 0 |
| 3 | 0.00 | 3.68 | 0.00 | 1.00 | 1 | Krtdap | keratinocyte differentiation associated prc | 26 | 0 | 8 | 0 | 0 | 0 | 0 | 0 | 0 | 0 | 0 |
| 3 | 0.17 | 3.60 | 0.05 | 0.96 | 1 | Cpne6 | copine VI | 0 | 0 | 0 | 0 | 0 | 18 | 0 | 0 | 0 | 0 | 16 |
| 3 | 1.34 | 3.58 | 0.38 | 0.71 | 1 | Fgfbp1 | fibroblast growth factor binding protein 1 | 15 | 5 | 8 | 0 | 0 | 0 | 0 | 0 | 0 | 0 | 5 |
| 3 | 0.00 | 3.68 | 0.00 | 1.00 | 1 | Gm29340 | | 11 | 5 | 18 | 0 | 0 | 0 | 0 | 0 | 0 | 0 | 0 |
| 3 | -4.53 | 3.61 | -1.25 | 0.21 | 1 | Gm20468 | | 9 | 0 | 0 | 0 | 22 | 2 | 0 | 0 | 1 | 0 | 0 |
| 3 | 0.39 | 3.60 | 0.11 | 0.91 | 1 | Clhc1 | clathrin heavy chain linker domain contai | 1 | 0 | 0 | 0 | 0 | 13 | 0 | 0 | 0 | 0 | 20 |
| 3 | 0.38 | 3.59 | 0.11 | 0.92 | 1 | Gm37509 | | 10 | 0 | 0 | 0 | 0 | 9 | 0 | 0 | 0 | 0 | 14 |
| 3 | 3.84 | 3.56 | 1.08 | 0.28 | 1 | Gm15704 | | 0 | 0 | 0 | 0 | 0 | 7 | 0 | 0 | 0 | 22 | 5 |
| 3 | 4.68 | 3.61 | 1.30 | 0.19 | 1 | Gm41030 | | 0 | 0 | 6 | 0 | 0 | 1 | 0 | 27 | 0 | 0 | 0 |
| 3 | -2.30 | 2.99 | -0.77 | 0.44 | 1 | 1500032F14Rik | | 2 | 20 | 3 | 5 | 0 | 3 | 0 | 0 | 0 | 0 | 0 |
| 3 | 1.32 | 3.62 | 0.37 | 0.71 | 1 | 2900060B14Rik | | 7 | 3 | 20 | 0 | 0 | 0 | 0 | 0 | 0 | 0 | 4 |
| 3 | 0.00 | 3.68 | 0.00 | 1.00 | 1 | Clec4e | C-type lectin domain family 4, member e | 15 | 18 | 0 | 0 | 0 | 0 | 0 | 0 | 0 | 0 | 0 |
| 3 | 0.57 | 3.49 | 0.16 | 0.87 | 1 | Pde6g | phosphodiesterase 6G, cGMP-specific, rc | 0 | 0 | 0 | 0 | 4 | 9 | 0 | 10 | 0 | 0 | 10 |
| 3 | 4.25 | 3.57 | 1.19 | 0.23 | 1 | Gm11945 | | 0 | 0 | 6 | 0 | 0 | 7 | 0 | 0 | 0 | 21 | 0 |
| 3 | 0.00 | 3.68 | 0.00 | 1.00 | 1 | Krt20 | keratin 20 | 6 | 19 | 8 | 0 | 0 | 0 | 0 | 0 | 0 | 0 | 0 |
| 3 | -3.80 | 3.52 | -1.08 | 0.28 | 1 | Gm48558 | | 0 | 0 | 0 | 18 | 8 | 3 | 0 | 0 | 0 | 0 | 5 |
| 3 | 0.03 | 3.60 | 0.01 | 0.99 | 1 | Gm44836 | | 0 | 0 | 0 | 0 | 0 | 20 | 0 | 0 | 0 | 0 | 13 |
| 3 | 1.36 | 3.56 | 0.38 | 0.70 | 1 | Gm48139 | | 0 | 0 | 0 | 0 | 0 | 20 | 0 | 0 | 6 | 0 | 7 |
| 3 | 0.00 | 3.64 | 0.00 | 1.00 | 1 | Htr2b | 5-hydroxytryptamine (serotonin) receptor | 33 | 0 | 0 | 0 | 0 | 0 | 0 | 0 | 0 | 0 | 0 |
| 3 | 0.00 | 3.64 | 0.00 | 1.00 | 1 | Igkv2-116 | | 33 | 0 | 0 | 0 | 0 | 0 | 0 | 0 | 0 | 0 | 0 |
| 3 | 0.07 | 2.01 | 0.03 | 0.97 | 1 | Gm7204 | | 1 | 1 | 7 | 2 | 6 | 2 | 0 | 0 | 8 | 0 | 7 |
| 3 | -0.31 | 3.57 | -0.09 | 0.93 | 1 | 4933407L21Rik | | 0 | 26 | 0 | 0 | 0 | 5 | 0 | 0 | 0 | 0 | 2 |
| 3 | 0.00 | 3.68 | 0.00 | 1.00 | 1 | Gm44089 | | 3 | 6 | 24 | 0 | 0 | 0 | 0 | 0 | 0 | 0 | 0 |
| 3 | 0.46 | 1.68 | 0.27 | 0.78 | 1 | Gm3200 | | 4 | 2 | 4 | 4 | 2 | 3 | 0 | 1 | 1 | 4 | 8 |
| 3 | 0.65 | 3.60 | 0.18 | 0.86 | 1 | Gm49754 | | 0 | 7 | 10 | 0 | 5 | 0 | 0 | 0 | 11 | 0 | 0 |
| 3 | 1.38 | 3.61 | 0.38 | 0.70 | 1 | Acod1 | aconitate decarboxylase 1 | 12 | 10 | 0 | 0 | 0 | 0 | 0 | 0 | 0 | 0 | 11 |
| 3 | 0.34 | 2.96 | 0.12 | 0.91 | 1 | Ppef2 | protein phosphatase, EF hand calcium-bi | 8 | 5 | 8 | 0 | 0 | 5 | 0 | 0 | 0 | 0 | 7 |
| 3 | 1.02 | 1.91 | 0.53 | 0.59 | 1 | Gm9791 | | 8 | 4 | 12 | 1 | 1 | 1 | 0 | 0 | 3 | 1 | 2 |
| 3 | -2.18 | 3.59 | -0.61 | 0.54 | 1 | Morn5 | MORN repeat containing 5 | 0 | 25 | 0 | 0 | 3 | 0 | 0 | 0 | 0 | 0 | 5 |
| 3 | 2.57 | 2.90 | 0.88 | 0.38 | 1 | 1700020N18Rik | | 7 | 14 | 1 | 0 | 0 | 5 | 0 | 0 | 5 | 0 | 1 |
| 3 | 0.00 | 3.68 | 0.00 | 1.00 | 1 | 1700009J07Rik | | 25 | 0 | 8 | 0 | 0 | 0 | 0 | 0 | 0 | 0 | 0 |
| 3 | 0.44 | 3.60 | 0.12 | 0.90 | 1 | Gm45343 | | 0 | 0 | 0 | 0 | 0 | 12 | 0 | 0 | 0 | 0 | 21 |
| 3 | -0.61 | 2.25 | -0.27 | 0.79 | 1 | Gm10420 | | 11 | 6 | 2 | 3 | 1 | 5 | 0 | 1 | 4 | 0 | 0 |
| 3 | 1.36 | 1.96 | 0.70 | 0.49 | 1 | Gm5523 | | 0 | 3 | 1 | 2 | 1 | 4 | 0 | 15 | 4 | 0 | 3 |
| 3 | 0.00 | 3.68 | 0.00 | 1.00 | 1 | Ccdc180 | coiled-coil domain containing 180 | 9 | 14 | 10 | 0 | 0 | 0 | 0 | 0 | 0 | 0 | 0 |
| 3 | 4.55 | 3.56 | 1.28 | 0.20 | 1 | 4833405L11Rik | | 0 | 0 | 0 | 0 | 0 | 0 | 0 | 15 | 0 | 18 | 0 |

| | | | | | | | | | | | | | | | | | | |
|---|---|---|---|---|---|---|---|---|---|---|---|---|---|---|---|---|---|---|
| 3 | -2.20 | 2.36 | -0.93 | 0.35 | 1 | Gm7114 | | 3 | 0 | 2 | 4 | 9 | 8 | 0 | 0 | 5 | 0 | 2 |
| 3 | -5.83 | 3.29 | -1.77 | 0.08 | 1 | Gm9967 | | 0 | 7 | 0 | 11 | 7 | 1 | 0 | 0 | 0 | 0 | 7 |
| 3 | -1.89 | 3.02 | -0.63 | 0.53 | 1 | Gm37320 | | 6 | 6 | 0 | 4 | 0 | 14 | 0 | 0 | 0 | 0 | 3 |
| 3 | 0.00 | 3.68 | 0.00 | 1.00 | 1 | Nfe2 | nuclear factor, erythroid derived 2 | 14 | 19 | 0 | 0 | 0 | 0 | 0 | 0 | 0 | 0 | 0 |
| 3 | -1.26 | 3.55 | -0.36 | 0.72 | 1 | Efcab8 | EF-hand calcium binding domain 8 | 0 | 0 | 0 | 10 | 0 | 10 | 0 | 0 | 0 | 0 | 13 |
| 3 | -3.00 | 3.40 | -0.88 | 0.38 | 1 | Gm14488 | | 0 | 0 | 10 | 17 | 0 | 1 | 0 | 0 | 4 | 0 | 1 |
| 3 | -0.87 | 3.60 | -0.24 | 0.81 | 1 | Gm38377 | | 19 | 3 | 0 | 0 | 0 | 11 | 0 | 0 | 0 | 0 | 0 |
| 3 | 0.05 | 2.09 | 0.02 | 0.98 | 1 | Gm18949 | | 0 | 0 | 0 | 4 | 10 | 4 | 0 | 2 | 7 | 1 | 4 |
| 3 | 0.00 | 3.68 | 0.00 | 1.00 | 1 | Gm10384 | | 0 | 15 | 18 | 0 | 0 | 0 | 0 | 0 | 0 | 0 | 0 |
| 3 | 0.64 | 2.61 | 0.24 | 0.81 | 1 | Platr31 | | 11 | 6 | 0 | 2 | 2 | 1 | 0 | 0 | 7 | 0 | 4 |
| 3 | 2.30 | 3.55 | 0.65 | 0.52 | 1 | A430110L20Rik | | 0 | 0 | 0 | 0 | 0 | 3 | 0 | 0 | 0 | 19 | 10 |
| 3 | 4.48 | 3.38 | 1.33 | 0.18 | 1 | Gm3507 | | 0 | 1 | 1 | 0 | 0 | 2 | 0 | 0 | 0 | 27 | 2 |
| 3 | 1.10 | 3.59 | 0.31 | 0.76 | 1 | Gm38245 | | 0 | 0 | 0 | 0 | 0 | 19 | 0 | 0 | 2 | 0 | 12 |
| 3 | -0.25 | 3.60 | -0.07 | 0.94 | 1 | Gm25271 | | 0 | 2 | 0 | 0 | 0 | 24 | 0 | 0 | 0 | 0 | 7 |
| 3 | 1.20 | 3.49 | 0.34 | 0.73 | 1 | Rps23rg1 | ribosomal protein S23, retrogene 1 | 0 | 0 | 0 | 16 | 0 | 1 | 0 | 0 | 14 | 0 | 2 |
| 3 | 0.00 | 3.68 | 0.00 | 1.00 | 1 | Ear-ps2 | | 21 | 12 | 0 | 0 | 0 | 0 | 0 | 0 | 0 | 0 | 0 |
| 3 | 1.14 | 3.54 | 0.32 | 0.75 | 1 | Ascl3 | achaete-scute family bHLH transcription f | 0 | 0 | 0 | 0 | 6 | 0 | 0 | 0 | 0 | 27 | 0 |
| 3 | 0.17 | 3.60 | 0.05 | 0.96 | 1 | Gm48804 | | 5 | 0 | 0 | 0 | 0 | 14 | 0 | 0 | 0 | 0 | 13 |
| 3 | 0.00 | 3.68 | 0.00 | 1.00 | 1 | Gm17745 | | 16 | 8 | 8 | 0 | 0 | 0 | 0 | 0 | 0 | 0 | 0 |
| 3 | 0.15 | 3.60 | 0.04 | 0.97 | 1 | Gm12981 | | 4 | 0 | 0 | 0 | 0 | 15 | 0 | 0 | 0 | 0 | 13 |
| 3 | -0.04 | 2.59 | -0.01 | 0.99 | 1 | A530058N18Rik | | 0 | 0 | 0 | 1 | 4 | 11 | 0 | 0 | 1 | 4 | 12 |
| 3 | 0.00 | 3.68 | 0.00 | 1.00 | 1 | Gm9947 | | 21 | 7 | 4 | 0 | 0 | 0 | 0 | 0 | 0 | 0 | 0 |
| 3 | -0.54 | 3.60 | -0.15 | 0.88 | 1 | Gm49658 | | 1 | 0 | 0 | 0 | 0 | 29 | 0 | 0 | 0 | 0 | 3 |
| 3 | 3.68 | 3.52 | 1.04 | 0.30 | 1 | Gm50222 | | 0 | 0 | 0 | 4 | 0 | 0 | 0 | 7 | 0 | 14 | 7 |
| 3 | 0.00 | 3.64 | 0.00 | 1.00 | 1 | A2m | alpha-2-macroglobulin | 33 | 0 | 0 | 0 | 0 | 0 | 0 | 0 | 0 | 0 | 0 |
| 3 | 0.00 | 3.64 | 0.00 | 1.00 | 1 | Ighv1-5 | | 33 | 0 | 0 | 0 | 0 | 0 | 0 | 0 | 0 | 0 | 0 |
| 3 | 4.59 | 3.59 | 1.28 | 0.20 | 1 | Gm13814 | | 8 | 0 | 0 | 0 | 0 | 2 | 0 | 23 | 0 | 0 | 0 |
| 3 | 0.28 | 3.60 | 0.08 | 0.94 | 1 | B130021K23Rik | | 1 | 0 | 0 | 0 | 0 | 14 | 0 | 0 | 0 | 0 | 17 |
| 3 | 1.62 | 3.55 | 0.46 | 0.65 | 1 | AA387883 | | 0 | 0 | 16 | 0 | 0 | 2 | 0 | 0 | 6 | 0 | 9 |
| 3 | -2.85 | 3.18 | -0.90 | 0.37 | 1 | Ooep | oocyte expressed protein | 6 | 0 | 12 | 1 | 12 | 1 | 0 | 0 | 0 | 0 | 0 |
| 3 | -1.81 | 2.76 | -0.65 | 0.51 | 1 | Rpl28-ps3 | | 17 | 5 | 2 | 0 | 4 | 3 | 0 | 0 | 0 | 0 | 1 |
| 3 | 0.25 | 3.60 | 0.07 | 0.94 | 1 | Gm42436 | | 0 | 0 | 0 | 0 | 0 | 15 | 0 | 0 | 0 | 0 | 17 |
| 3 | 0.83 | 3.59 | 0.23 | 0.82 | 1 | Gm25112 | | 0 | 7 | 0 | 0 | 0 | 5 | 0 | 0 | 0 | 0 | 20 |
| 3 | 0.15 | 3.58 | 0.04 | 0.97 | 1 | Gm23136 | | 0 | 12 | 0 | 0 | 0 | 10 | 0 | 0 | 0 | 0 | 10 |
| 3 | 0.00 | 3.68 | 0.00 | 1.00 | 1 | Klhl40 | kelch-like 40 | 22 | 10 | 0 | 0 | 0 | 0 | 0 | 0 | 0 | 0 | 0 |
| 3 | 3.92 | 3.54 | 1.11 | 0.27 | 1 | Nxf3 | nuclear RNA export factor 3 | 0 | 0 | 0 | 0 | 0 | 1 | 0 | 25 | 2 | 0 | 4 |
| 3 | -0.50 | 3.18 | -0.16 | 0.88 | 1 | Gm28417 | | 13 | 12 | 0 | 4 | 0 | 1 | 0 | 0 | 0 | 0 | 3 |
| 3 | -4.28 | 3.61 | -1.19 | 0.24 | 1 | Gm34069 | | 0 | 20 | 0 | 0 | 9 | 2 | 0 | 0 | 0 | 0 | 1 |
| 3 | -1.86 | 2.80 | -0.66 | 0.51 | 1 | Gm16731 | | 9 | 7 | 2 | 0 | 8 | 5 | 0 | 0 | 0 | 0 | 2 |
| 3 | 0.00 | 3.68 | 0.00 | 1.00 | 1 | Gm6556 | | 7 | 17 | 8 | 0 | 0 | 0 | 0 | 0 | 0 | 0 | 0 |
| 3 | -3.51 | 3.44 | -1.02 | 0.31 | 1 | Ak9 | adenylate kinase 9 | 3 | 8 | 0 | 0 | 13 | 3 | 0 | 0 | 0 | 0 | 5 |
| 3 | -1.07 | 2.57 | -0.41 | 0.68 | 1 | Rpl36a-ps1 | | 6 | 0 | 4 | 3 | 1 | 1 | 0 | 0 | 1 | 17 | 0 |
| 3 | 0.00 | 3.68 | 0.00 | 1.00 | 1 | Nat8f3 | N-acetyltransferase 8 (GCN5-related) fam | 9 | 0 | 23 | 0 | 0 | 0 | 0 | 0 | 0 | 0 | 0 |
| 3 | 1.25 | 3.66 | 0.34 | 0.73 | 1 | Gm12840 | | 12 | 17 | 1 | 0 | 0 | 0 | 0 | 0 | 0 | 0 | 2 |
| 3 | -0.26 | 3.27 | -0.08 | 0.94 | 1 | Ndufb4c | NADH:ubiquinone oxidoreductase subun | 11 | 0 | 0 | 1 | 4 | 0 | 0 | 2 | 0 | 14 | 0 |
| 3 | 0.92 | 1.68 | 0.55 | 0.58 | 1 | Gm6533 | | 6 | 7 | 5 | 1 | 2 | 3 | 0 | 2 | 2 | 1 | 3 |
| 3 | -1.23 | 2.87 | -0.43 | 0.67 | 1 | Gm28959 | | 8 | 0 | 0 | 1 | 11 | 4 | 0 | 0 | 5 | 0 | 3 |
| 3 | -0.22 | 3.00 | -0.07 | 0.94 | 1 | Mir5710 | | 3 | 7 | 3 | 0 | 0 | 14 | 0 | 0 | 0 | 0 | 5 |
| 3 | 0.86 | 3.54 | 0.24 | 0.81 | 1 | Gm31706 | | 0 | 0 | 11 | 0 | 0 | 8 | 0 | 0 | 5 | 0 | 8 |
| 3 | -5.63 | 3.26 | -1.73 | 0.08 | 1 | Gm14197 | | 0 | 0 | 7 | 9 | 6 | 3 | 0 | 0 | 0 | 0 | 7 |
| 3 | 0.39 | 3.60 | 0.11 | 0.91 | 1 | 4930593A02Rik | | 0 | 3 | 9 | 0 | 0 | 8 | 0 | 0 | 0 | 0 | 12 |
| 3 | 0.00 | 3.68 | 0.00 | 1.00 | 1 | A730020M07Rik | | 30 | 0 | 2 | 0 | 0 | 0 | 0 | 0 | 0 | 0 | 0 |
| 3 | 0.00 | 3.68 | 0.00 | 1.00 | 1 | Gm15050 | | 25 | 2 | 5 | 0 | 0 | 0 | 0 | 0 | 0 | 0 | 0 |
| 3 | -0.74 | 3.03 | -0.24 | 0.81 | 1 | Gm47950 | | 1 | 0 | 0 | 0 | 3 | 20 | 0 | 0 | 1 | 0 | 7 |
| 3 | -0.57 | 3.04 | -0.19 | 0.85 | 1 | Gm4759 | | 4 | 7 | 10 | 0 | 0 | 9 | 0 | 0 | 0 | 0 | 1 |
| 3 | 0.00 | 3.68 | 0.00 | 1.00 | 1 | Hoxa13 | homeobox A13 | 4 | 11 | 17 | 0 | 0 | 0 | 0 | 0 | 0 | 0 | 0 |
| 3 | -2.74 | 3.57 | -0.77 | 0.44 | 1 | E430021H15Rik | | 6 | 0 | 0 | 4 | 0 | 3 | 0 | 0 | 0 | 0 | 18 |
| 3 | 1.49 | 3.54 | 0.42 | 0.67 | 1 | Gm43183 | | 0 | 0 | 0 | 0 | 0 | 4 | 0 | 28 | 0 | 0 | 0 |
| 3 | 3.32 | 3.59 | 0.92 | 0.36 | 1 | Gm31342 | | 0 | 8 | 9 | 0 | 0 | 2 | 0 | 0 | 0 | 13 | 0 |
| 3 | 0.20 | 2.75 | 0.07 | 0.94 | 1 | Gm11733 | | 1 | 10 | 10 | 0 | 0 | 8 | 0 | 1 | 0 | 0 | 1 |

| | | | | | | | | | | | | | | | | | |
|---|---|---|---|---|---|---|---|---|---|---|---|---|---|---|---|---|---|
| 3 | 2.03 | 3.58 | 0.57 | 0.57 | 1 | Snord65 | | 0 | 0 | 0 | 0 | 0 | 8 | 0 | 0 | 5 | 0 | 19 |
| 3 | -2.15 | 3.61 | -0.59 | 0.55 | 1 | Stra6 | stimulated by retinoic acid gene 6 | 16 | 9 | 0 | 0 | 6 | 1 | 0 | 0 | 0 | 0 | 0 |
| 3 | 0.00 | 3.68 | 0.00 | 1.00 | 1 | Gm43489 | | 9 | 14 | 8 | 0 | 0 | 0 | 0 | 0 | 0 | 0 | 0 |
| 3 | 3.02 | 3.57 | 0.85 | 0.40 | 1 | Gm37682 | | 0 | 0 | 0 | 0 | 0 | 0 | 0 | 0 | 0 | 31 | 1 |
| 3 | 4.22 | 3.58 | 1.18 | 0.24 | 1 | Gm11743 | | 9 | 0 | 0 | 0 | 0 | 5 | 0 | 0 | 5 | 0 | 13 |
| 3 | 0.00 | 3.68 | 0.00 | 1.00 | 1 | Arhgap15os | | 16 | 0 | 16 | 0 | 0 | 0 | 0 | 0 | 0 | 0 | 0 |
| 3 | 0.00 | 3.68 | 0.00 | 1.00 | 1 | Frmd7 | FERM domain containing 7 | 26 | 4 | 2 | 0 | 0 | 0 | 0 | 0 | 0 | 0 | 0 |
| 3 | 0.05 | 3.60 | 0.01 | 0.99 | 1 | Gm11948 | | 1 | 1 | 0 | 0 | 0 | 18 | 0 | 0 | 0 | 0 | 12 |
| 3 | 0.00 | 3.68 | 0.00 | 1.00 | 1 | 5031425F14Rik | | 4 | 21 | 6 | 0 | 0 | 0 | 0 | 0 | 0 | 0 | 0 |
| 3 | 0.00 | 3.68 | 0.00 | 1.00 | 1 | Gm44763 | | 0 | 16 | 16 | 0 | 0 | 0 | 0 | 0 | 0 | 0 | 0 |
| 3 | -0.11 | 3.59 | -0.03 | 0.98 | 1 | Snord71 | | 0 | 10 | 0 | 0 | 0 | 14 | 0 | 0 | 0 | 0 | 8 |
| 3 | 0.00 | 3.64 | 0.00 | 1.00 | 1 | Ighv1-20 | | 32 | 0 | 0 | 0 | 0 | 0 | 0 | 0 | 0 | 0 | 0 |
| 3 | 0.00 | 3.68 | 0.00 | 1.00 | 1 | Ptgs2os2 | | 0 | 18 | 14 | 0 | 0 | 0 | 0 | 0 | 0 | 0 | 0 |
| 3 | -2.33 | 3.60 | -0.65 | 0.52 | 1 | Gm36757 | | 1 | 26 | 0 | 0 | 3 | 2 | 0 | 0 | 0 | 0 | 0 |
| 3 | -0.12 | 2.44 | -0.05 | 0.96 | 1 | Gm2048 | | 1 | 2 | 0 | 11 | 2 | 3 | 0 | 0 | 9 | 0 | 3 |
| 3 | 0.52 | 3.60 | 0.14 | 0.89 | 1 | Gm3052 | | 4 | 11 | 0 | 0 | 0 | 5 | 0 | 0 | 0 | 0 | 11 |
| 3 | -0.92 | 3.54 | -0.26 | 0.79 | 1 | Gm7318 | | 0 | 1 | 1 | 1 | 0 | 2 | 0 | 0 | 0 | 27 | 0 |
| 3 | 0.36 | 3.01 | 0.12 | 0.90 | 1 | Mir6976 | | 3 | 3 | 7 | 0 | 0 | 8 | 0 | 0 | 0 | 0 | 11 |
| 3 | 0.00 | 3.68 | 0.00 | 1.00 | 1 | 1600010M07Rik | | 15 | 16 | 0 | 0 | 0 | 0 | 0 | 0 | 0 | 0 | 0 |
| 3 | -2.93 | 3.39 | -0.86 | 0.39 | 1 | Spag16 | sperm associated antigen 16 | 0 | 0 | 15 | 11 | 0 | 3 | 0 | 0 | 1 | 0 | 3 |
| 3 | 1.28 | 3.57 | 0.36 | 0.72 | 1 | Gm49839 | | 0 | 0 | 0 | 0 | 0 | 19 | 0 | 0 | 4 | 0 | 8 |
| 3 | -2.31 | 3.60 | -0.64 | 0.52 | 1 | Tunar | | 15 | 10 | 0 | 4 | 0 | 3 | 0 | 0 | 0 | 0 | 0 |
| 3 | 0.00 | 3.68 | 0.00 | 1.00 | 1 | Gm26862 | | 0 | 24 | 7 | 0 | 0 | 0 | 0 | 0 | 0 | 0 | 0 |
| 3 | 0.00 | 3.68 | 0.00 | 1.00 | 1 | Gm11266 | | 21 | 10 | 0 | 0 | 0 | 0 | 0 | 0 | 0 | 0 | 0 |
| 3 | -3.16 | 3.58 | -0.88 | 0.38 | 1 | Mir17hg | | 3 | 0 | 0 | 0 | 4 | 10 | 0 | 0 | 0 | 0 | 14 |
| 3 | 1.32 | 3.63 | 0.36 | 0.72 | 1 | Rgs22 | regulator of G-protein signalling 22 | 22 | 2 | 3 | 0 | 0 | 0 | 0 | 0 | 0 | 0 | 4 |
| 3 | -2.84 | 3.15 | -0.90 | 0.37 | 1 | Scn11a | sodium channel, voltage-gated, type XI, a | 3 | 6 | 8 | 4 | 0 | 10 | 0 | 0 | 0 | 0 | 0 |
| 3 | -1.46 | 3.60 | -0.41 | 0.68 | 1 | Gm41414 | | 13 | 0 | 7 | 0 | 11 | 0 | 0 | 0 | 0 | 0 | 0 |
| 3 | -0.70 | 3.66 | -0.19 | 0.85 | 1 | Gm48073 | | 0 | 16 | 14 | 2 | 0 | 0 | 0 | 0 | 0 | 0 | 0 |
| 3 | 1.76 | 3.17 | 0.56 | 0.58 | 1 | Icam4 | intercellular adhesion molecule 4, Landst | 11 | 0 | 8 | 0 | 3 | 0 | 0 | 0 | 6 | 0 | 3 |
| 3 | 0.00 | 3.64 | 0.00 | 1.00 | 1 | Gm15444 | | 0 | 31 | 0 | 0 | 0 | 0 | 0 | 0 | 0 | 0 | 0 |
| 3 | -0.92 | 3.55 | -0.26 | 0.79 | 1 | Gm44154 | | 0 | 0 | 0 | 0 | 6 | 8 | 0 | 0 | 0 | 0 | 17 |
| 3 | 0.00 | 3.68 | 0.00 | 1.00 | 1 | Sctr | secretin receptor | 3 | 5 | 23 | 0 | 0 | 0 | 0 | 0 | 0 | 0 | 0 |
| 3 | 0.00 | 3.68 | 0.00 | 1.00 | 1 | Apol6 | apolipoprotein L 6 | 28 | 3 | 0 | 0 | 0 | 0 | 0 | 0 | 0 | 0 | 0 |
| 3 | 0.80 | 3.60 | 0.22 | 0.82 | 1 | A930014E10Rik | | 0 | 0 | 0 | 7 | 0 | 0 | 0 | 24 | 0 | 0 | 0 |
| 3 | 0.00 | 3.68 | 0.00 | 1.00 | 1 | Myh1 | myosin, heavy polypeptide 1, skeletal mu | 6 | 25 | 0 | 0 | 0 | 0 | 0 | 0 | 0 | 0 | 0 |
| 3 | -4.79 | 3.55 | -1.35 | 0.18 | 1 | Gm37247 | | 0 | 0 | 0 | 8 | 6 | 18 | 0 | 0 | 0 | 0 | 0 |
| 3 | 0.28 | 3.05 | 0.09 | 0.93 | 1 | A930015P04Rik | | 5 | 17 | 5 | 0 | 0 | 2 | 0 | 0 | 0 | 0 | 2 |
| 3 | 0.00 | 3.68 | 0.00 | 1.00 | 1 | Clca2 | chloride channel accessory 2 | 23 | 8 | 0 | 0 | 0 | 0 | 0 | 0 | 0 | 0 | 0 |
| 3 | -0.13 | 3.25 | -0.04 | 0.97 | 1 | Gng3 | guanine nucleotide binding protein (G prc | 4 | 11 | 6 | 4 | 0 | 0 | 0 | 0 | 0 | 0 | 6 |
| 3 | -0.49 | 3.64 | -0.14 | 0.89 | 1 | Gm48801 | | 3 | 2 | 24 | 0 | 2 | 0 | 0 | 0 | 0 | 0 | 0 |
| 3 | 0.83 | 3.34 | 0.25 | 0.80 | 1 | Gm12104 | | 12 | 0 | 0 | 0 | 3 | 4 | 0 | 0 | 6 | 0 | 6 |
| 3 | 4.48 | 3.57 | 1.25 | 0.21 | 1 | Mybph | myosin binding protein H | 0 | 6 | 0 | 0 | 0 | 6 | 0 | 0 | 14 | 0 | 5 |
| 3 | 0.11 | 3.53 | 0.03 | 0.98 | 1 | Gm17949 | | 0 | 0 | 0 | 0 | 8 | 8 | 0 | 0 | 7 | 0 | 8 |
| 3 | 5.84 | 3.56 | 1.64 | 0.10 | 1 | Rps6-ps2 | | 0 | 8 | 1 | 0 | 0 | 2 | 0 | 0 | 1 | 19 | 0 |
| 3 | 2.83 | 3.61 | 0.78 | 0.43 | 1 | Aldh3b3 | aldehyde dehydrogenase 3 family, memb | 11 | 2 | 0 | 2 | 0 | 0 | 0 | 16 | 0 | 0 | 0 |
| 3 | -0.50 | 2.60 | -0.19 | 0.85 | 1 | Gng10 | guanine nucleotide binding protein (G prc | 9 | 2 | 0 | 2 | 2 | 7 | 0 | 0 | 7 | 0 | 3 |
| 3 | 2.12 | 2.31 | 0.92 | 0.36 | 1 | Rpl36-ps3 | | 2 | 1 | 9 | 1 | 0 | 6 | 0 | 2 | 6 | 4 | 0 |
| 3 | -5.68 | 3.54 | -1.61 | 0.11 | 1 | Tdrd6 | tudor domain containing 6 | 0 | 0 | 0 | 4 | 0 | 14 | 0 | 0 | 13 | 0 | 0 |
| 3 | 0.00 | 3.68 | 0.00 | 1.00 | 1 | Tnni1 | troponin I, skeletal, slow 1 | 8 | 23 | 0 | 0 | 0 | 0 | 0 | 0 | 0 | 0 | 0 |
| 3 | -4.36 | 3.52 | -1.24 | 0.22 | 1 | Gm11688 | | 8 | 1 | 0 | 14 | 0 | 4 | 0 | 0 | 4 | 0 | 0 |
| 3 | -2.33 | 2.72 | -0.85 | 0.39 | 1 | Gm44658 | | 0 | 3 | 2 | 4 | 7 | 7 | 0 | 0 | 0 | 0 | 8 |
| 3 | 0.56 | 3.44 | 0.16 | 0.87 | 1 | 9330111N05Rik | | 4 | 0 | 0 | 5 | 0 | 4 | 0 | 0 | 11 | 0 | 6 |
| 3 | -0.71 | 3.59 | -0.20 | 0.84 | 1 | Npm2 | nucleophosmin/nucleoplasmin 2 | 0 | 0 | 0 | 0 | 2 | 12 | 0 | 0 | 0 | 0 | 17 |
| 3 | -0.66 | 2.79 | -0.24 | 0.81 | 1 | Gm2238 | | 8 | 5 | 0 | 4 | 0 | 6 | 0 | 0 | 0 | 8 | 1 |
| 3 | -0.40 | 3.67 | -0.11 | 0.91 | 1 | Igf2bp3 | insulin-like growth factor 2 mRNA binding | 24 | 6 | 0 | 0 | 0 | 1 | 0 | 0 | 0 | 0 | 0 |
| 3 | 5.11 | 3.31 | 1.54 | 0.12 | 1 | Gm45444 | | 2 | 10 | 0 | 0 | 0 | 1 | 0 | 0 | 0 | 14 | 4 |
| 3 | -1.11 | 3.60 | -0.31 | 0.76 | 1 | 5730507C01Ril | RIKEN cDNA 5730507C01 gene | 1 | 23 | 0 | 0 | 0 | 7 | 0 | 0 | 0 | 0 | 0 |
| 3 | 1.36 | 3.60 | 0.38 | 0.71 | 1 | Hs3st2 | heparan sulfate (glucosamine) 3-O-sulfot | 8 | 0 | 0 | 0 | 0 | 0 | 0 | 0 | 0 | 23 | 0 |
| 3 | 0.00 | 3.64 | 0.00 | 1.00 | 1 | Sh2d1b2 | SH2 domain containing 1B2 | 31 | 0 | 0 | 0 | 0 | 0 | 0 | 0 | 0 | 0 | 0 |

| | | | | | | | | | | | | | | | | | |
|---|---|---|---|---|---|---|---|---|---|---|---|---|---|---|---|---|---|
| 3 | 0.00 | 3.64 | 0.00 | 1.00 | 1 | Ighv1-80 | | 31 | 0 | 0 | 0 | 0 | 0 | 0 | 0 | 0 | 0 | 0 |
| 3 | 0.61 | 3.33 | 0.18 | 0.86 | 1 | Akap3 | A kinase (PRKA) anchor protein 3 | 9 | 0 | 0 | 0 | 5 | 3 | 0 | 0 | 5 | 0 | 9 |
| 3 | 2.56 | 2.50 | 1.02 | 0.31 | 1 | Gm5587 | | 15 | 1 | 6 | 1 | 0 | 0 | 0 | 1 | 2 | 0 | 4 |
| 3 | 0.00 | 3.68 | 0.00 | 1.00 | 1 | Slc6a13 | solute carrier family 6 (neurotransmitter tra | 10 | 12 | 8 | 0 | 0 | 0 | 0 | 0 | 0 | 0 | 0 |
| 3 | -0.95 | 3.58 | -0.26 | 0.79 | 1 | Rtbdn | retbindin | 0 | 0 | 0 | 0 | 3 | 13 | 0 | 0 | 0 | 0 | 15 |
| 3 | 1.64 | 3.22 | 0.51 | 0.61 | 1 | Tmem269 | transmembrane protein 269 | 5 | 0 | 4 | 0 | 0 | 4 | 0 | 0 | 0 | 14 | 3 |
| 3 | -0.71 | 3.28 | -0.22 | 0.83 | 1 | 4930404I05Rik | | 5 | 0 | 0 | 0 | 6 | 4 | 0 | 0 | 0 | 12 | 4 |
| 3 | -0.58 | 2.40 | -0.24 | 0.81 | 1 | Gm6363 | | 2 | 8 | 0 | 0 | 7 | 6 | 0 | 1 | 1 | 5 | 1 |
| 3 | -2.10 | 3.55 | -0.59 | 0.55 | 1 | Gm37052 | | 0 | 0 | 0 | 11 | 0 | 15 | 0 | 0 | 0 | 0 | 4 |
| 3 | -2.78 | 2.94 | -0.94 | 0.35 | 1 | Gm19241 | | 1 | 5 | 7 | 11 | 0 | 1 | 0 | 0 | 0 | 0 | 6 |
| 3 | 0.58 | 3.60 | 0.16 | 0.87 | 1 | Gm27326 | | 4 | 0 | 5 | 0 | 0 | 6 | 0 | 0 | 0 | 0 | 15 |
| 3 | 0.00 | 3.11 | 0.00 | 1.00 | 1 | Gm30270 | | 9 | 17 | 2 | 0 | 0 | 2 | 0 | 0 | 0 | 0 | 1 |
| 3 | 0.00 | 3.68 | 0.00 | 1.00 | 1 | Tbx18 | T-box18 | 5 | 23 | 2 | 0 | 0 | 0 | 0 | 0 | 0 | 0 | 0 |
| 3 | -1.26 | 2.72 | -0.46 | 0.64 | 1 | Dbil5 | diazepam binding inhibitor-like 5 | 6 | 15 | 3 | 4 | 0 | 2 | 0 | 0 | 1 | 0 | 0 |
| 3 | 1.58 | 3.39 | 0.46 | 0.64 | 1 | Rpl17-ps7 | | 10 | 13 | 0 | 0 | 1 | 0 | 0 | 0 | 2 | 0 | 4 |
| 3 | 3.91 | 3.52 | 1.11 | 0.27 | 1 | Gm33677 | | 0 | 0 | 0 | 0 | 0 | 5 | 0 | 0 | 6 | 19 | 0 |
| 3 | -1.22 | 2.11 | -0.58 | 0.56 | 1 | Gm36964 | | 13 | 2 | 1 | 1 | 7 | 3 | 0 | 1 | 0 | 1 | 2 |
| 3 | 1.14 | 2.33 | 0.49 | 0.62 | 1 | Gm10923 | | 8 | 0 | 7 | 0 | 1 | 4 | 0 | 0 | 1 | 5 | 4 |
| 3 | -0.40 | 3.67 | -0.11 | 0.91 | 1 | Ndst3 | N-deacetylase/N-sulfotransferase (hepara | 8 | 4 | 18 | 0 | 1 | 0 | 0 | 0 | 0 | 0 | 0 |
| 3 | -0.40 | 3.67 | -0.11 | 0.91 | 1 | Gm45894 | | 0 | 24 | 5 | 0 | 0 | 1 | 0 | 0 | 0 | 0 | 0 |
| 3 | 0.00 | 3.68 | 0.00 | 1.00 | 1 | 9330158H04Rik | | 8 | 10 | 12 | 0 | 0 | 0 | 0 | 0 | 0 | 0 | 0 |
| 3 | -1.77 | 2.72 | -0.65 | 0.51 | 1 | 1700016A09Rik | | 1 | 5 | 0 | 4 | 2 | 4 | 0 | 14 | 0 | 0 | 0 |
| 3 | -2.36 | 3.15 | -0.75 | 0.45 | 1 | Gm7645 | | 0 | 4 | 0 | 10 | 4 | 7 | 0 | 0 | 0 | 0 | 6 |
| 3 | -0.69 | 3.66 | -0.19 | 0.85 | 1 | Gm36953 | | 15 | 13 | 0 | 0 | 0 | 2 | 0 | 0 | 0 | 0 | 0 |
| 3 | 0.18 | 3.60 | 0.05 | 0.96 | 1 | Gm37621 | | 1 | 0 | 0 | 0 | 0 | 15 | 0 | 0 | 0 | 0 | 14 |
| 3 | 0.00 | 3.68 | 0.00 | 1.00 | 1 | Ces1c | carboxylesterase 1C | 9 | 0 | 21 | 0 | 0 | 0 | 0 | 0 | 0 | 0 | 0 |
| 3 | -0.96 | 3.25 | -0.30 | 0.77 | 1 | Snora57 | | 0 | 6 | 4 | 0 | 9 | 3 | 0 | 0 | 8 | 0 | 0 |
| 3 | -1.71 | 3.19 | -0.54 | 0.59 | 1 | 1810044K17Rik | | 0 | 13 | 0 | 11 | 2 | 2 | 0 | 0 | 0 | 0 | 2 |
| 3 | -4.85 | 3.29 | -1.47 | 0.14 | 1 | Gm20383 | | 3 | 1 | 0 | 18 | 6 | 3 | 0 | 0 | 0 | 0 | 0 |
| 3 | 1.12 | 3.55 | 0.32 | 0.75 | 1 | Gm37962 | | 0 | 0 | 6 | 0 | 0 | 8 | 0 | 0 | 4 | 0 | 12 |
| 3 | -1.07 | 3.21 | -0.33 | 0.74 | 1 | Rps19-ps4 | | 2 | 18 | 8 | 0 | 1 | 1 | 0 | 0 | 0 | 0 | 0 |
| 3 | -6.11 | 3.25 | -1.88 | 0.06 | 1 | Gm48734 | | 0 | 9 | 0 | 6 | 10 | 3 | 0 | 0 | 0 | 0 | 1 |
| 3 | 0.00 | 3.64 | 0.00 | 1.00 | 1 | Ighv1-9 | | 30 | 0 | 0 | 0 | 0 | 0 | 0 | 0 | 0 | 0 | 0 |
| 3 | -1.06 | 3.56 | -0.30 | 0.76 | 1 | Gm8850 | | 0 | 0 | 1 | 0 | 4 | 16 | 0 | 0 | 0 | 0 | 9 |
| 3 | -0.60 | 3.64 | -0.17 | 0.87 | 1 | Gm7879 | | 29 | 0 | 0 | 0 | 0 | 1 | 0 | 0 | 0 | 0 | 0 |
| 3 | -5.76 | 3.52 | -1.64 | 0.10 | 1 | Gm48913 | | 0 | 0 | 0 | 4 | 4 | 3 | 0 | 0 | 0 | 19 | 0 |
| 3 | -2.24 | 3.58 | -0.62 | 0.53 | 1 | Foxr2 | forkhead box R2 | 0 | 9 | 0 | 18 | 0 | 3 | 0 | 0 | 0 | 0 | 0 |
| 3 | -0.20 | 3.60 | -0.06 | 0.96 | 1 | Gm13054 | | 3 | 0 | 0 | 0 | 0 | 20 | 0 | 0 | 0 | 0 | 7 |
| 3 | 0.00 | 3.68 | 0.00 | 1.00 | 1 | Smyd1 | SET and MYND domain containing 1 | 9 | 17 | 4 | 0 | 0 | 0 | 0 | 0 | 0 | 0 | 0 |
| 3 | 0.30 | 3.60 | 0.08 | 0.93 | 1 | Gm48865 | | 0 | 1 | 0 | 0 | 0 | 13 | 0 | 0 | 0 | 0 | 16 |
| 3 | 1.57 | 2.71 | 0.58 | 0.56 | 1 | Acot12 | acyl-CoA thioesterase 12 | 8 | 4 | 10 | 0 | 0 | 2 | 0 | 0 | 4 | 0 | 2 |
| 3 | -1.14 | 3.58 | -0.32 | 0.75 | 1 | Krt88 | keratin 88 | 0 | 0 | 0 | 20 | 0 | 2 | 0 | 8 | 0 | 0 | 0 |
| 3 | -1.50 | 3.22 | -0.47 | 0.64 | 1 | BC016579 | cDNA sequence, BC016579 | 14 | 9 | 0 | 4 | 0 | 2 | 0 | 0 | 0 | 0 | 1 |
| 3 | 0.00 | 3.68 | 0.00 | 1.00 | 1 | Ido1 | indoleamine 2,3-dioxygenase 1 | 3 | 27 | 0 | 0 | 0 | 0 | 0 | 0 | 0 | 0 | 0 |
| 3 | 6.89 | 3.54 | 1.95 | 0.05 | 1 | Gm40332 | | 0 | 7 | 0 | 0 | 0 | 5 | 0 | 0 | 1 | 17 | 0 |
| 3 | -2.00 | 3.56 | -0.56 | 0.57 | 1 | Gm31805 | | 0 | 16 | 0 | 5 | 0 | 8 | 0 | 0 | 0 | 0 | 0 |
| 3 | 0.38 | 2.43 | 0.16 | 0.87 | 1 | Gm16372 | | 6 | 4 | 3 | 6 | 1 | 0 | 0 | 6 | 0 | 4 | 0 |
| 3 | -1.22 | 3.58 | -0.34 | 0.73 | 1 | Gm17893 | | 0 | 0 | 0 | 0 | 3 | 17 | 0 | 0 | 0 | 0 | 10 |
| 3 | -0.78 | 3.62 | -0.21 | 0.83 | 1 | G930009F23Rik | | 16 | 0 | 9 | 0 | 0 | 4 | 0 | 0 | 0 | 0 | 0 |
| 3 | -0.24 | 3.62 | -0.07 | 0.95 | 1 | Apom | apolipoprotein M | 0 | 19 | 6 | 0 | 0 | 3 | 0 | 0 | 0 | 0 | 1 |
| 3 | 3.44 | 3.57 | 0.97 | 0.33 | 1 | Gm13199 | | 0 | 0 | 0 | 0 | 0 | 10 | 0 | 0 | 0 | 12 | 8 |
| 3 | 0.17 | 3.49 | 0.05 | 0.96 | 1 | Gm37519 | | 0 | 0 | 0 | 8 | 0 | 7 | 0 | 0 | 5 | 0 | 10 |
| 3 | -2.36 | 3.60 | -0.65 | 0.51 | 1 | 2810405F15Rik | | 10 | 11 | 0 | 0 | 8 | 1 | 0 | 0 | 0 | 0 | 0 |
| 3 | 0.00 | 3.68 | 0.00 | 1.00 | 1 | Hsd17b2 | hydroxysteroid (17-beta) dehydrogenase | 0 | 12 | 18 | 0 | 0 | 0 | 0 | 0 | 0 | 0 | 0 |
| 3 | 0.28 | 3.61 | 0.08 | 0.94 | 1 | Cpa4 | carboxypeptidase A4 | 5 | 17 | 0 | 0 | 0 | 3 | 0 | 0 | 0 | 0 | 4 |
| 3 | 0.61 | 2.27 | 0.27 | 0.79 | 1 | Gm50432 | | 1 | 0 | 2 | 4 | 2 | 6 | 0 | 8 | 1 | 6 | 0 |
| 3 | 4.77 | 3.25 | 1.47 | 0.14 | 1 | Gm15247 | | 4 | 0 | 0 | 0 | 0 | 1 | 0 | 8 | 10 | 6 | 0 |
| 3 | -2.11 | 2.83 | -0.75 | 0.46 | 1 | Gm34808 | | 0 | 0 | 14 | 4 | 3 | 7 | 0 | 0 | 0 | 1 | 1 |
| 3 | -0.96 | 2.66 | -0.36 | 0.72 | 1 | Gm26520 | | 0 | 1 | 0 | 8 | 6 | 5 | 0 | 0 | 3 | 0 | 7 |
| 3 | -3.52 | 3.57 | -0.99 | 0.32 | 1 | Dyrk4 | dual-specificity tyrosine-(Y)-phosphorylati | 0 | 0 | 11 | 7 | 0 | 4 | 0 | 0 | 0 | 0 | 7 |

| | | | | | | | | | | | | | | | | | | |
|---|---|---|---|---|---|---|---|---|---|---|---|---|---|---|---|---|---|---|
| 3 | -0.11 | 3.59 | -0.03 | 0.98 | 1 | Gm48624 | | 0 | 8 | 0 | 0 | 0 | 14 | 0 | 0 | 0 | 0 | 8 |
| 3 | 0.00 | 3.68 | 0.00 | 1.00 | 1 | Mylk4 | myosin light chain kinase family, member | 6 | 6 | 18 | 0 | 0 | 0 | 0 | 0 | 0 | 0 | 0 |
| 3 | 0.25 | 3.60 | 0.07 | 0.95 | 1 | Stag3 | stromal antigen 3 | 3 | 0 | 3 | 0 | 0 | 11 | 0 | 0 | 0 | 0 | 12 |
| 3 | 3.47 | 3.58 | 0.97 | 0.33 | 1 | Gm45873 | | 9 | 0 | 0 | 0 | 0 | 10 | 0 | 0 | 5 | 0 | 5 |
| 3 | 0.00 | 3.68 | 0.00 | 1.00 | 1 | Gm21955 | | 11 | 6 | 12 | 0 | 0 | 0 | 0 | 0 | 0 | 0 | 0 |
| 3 | 2.55 | 3.35 | 0.76 | 0.45 | 1 | Trat1 | T cell receptor associated transmembrane | 13 | 5 | 4 | 0 | 0 | 3 | 0 | 0 | 5 | 0 | 0 |
| 3 | 4.14 | 3.55 | 1.17 | 0.24 | 1 | Gm45024 | | 0 | 0 | 0 | 0 | 1 | 0 | 0 | 0 | 0 | 21 | 8 |
| 3 | -4.20 | 3.53 | -1.19 | 0.23 | 1 | Hmgb1-rs17 | | 13 | 1 | 0 | 7 | 0 | 7 | 0 | 0 | 2 | 0 | 0 |
| 3 | -0.76 | 3.63 | -0.21 | 0.83 | 1 | Gm11961 | | 6 | 20 | 0 | 0 | 0 | 3 | 0 | 0 | 0 | 0 | 0 |
| 3 | 2.90 | 3.51 | 0.82 | 0.41 | 1 | 1700006F04Rik | | 3 | 14 | 0 | 0 | 0 | 6 | 0 | 0 | 5 | 0 | 1 |
| 3 | -0.05 | 3.60 | -0.01 | 0.99 | 1 | Gm43313 | | 0 | 0 | 0 | 0 | 0 | 19 | 0 | 0 | 0 | 0 | 10 |
| 3 | -0.37 | 3.58 | -0.10 | 0.92 | 1 | Gm38147 | | 0 | 0 | 0 | 11 | 0 | 2 | 0 | 17 | 0 | 0 | 0 |
| 3 | 0.24 | 3.57 | 0.07 | 0.95 | 1 | L3mbtl1 | L3MBTL1 histone methyl-lysine binding p | 0 | 20 | 0 | 0 | 0 | 4 | 0 | 0 | 0 | 0 | 5 |
| 3 | 0.86 | 3.28 | 0.26 | 0.79 | 1 | Esrrb | estrogen related receptor, beta | 0 | 23 | 2 | 0 | 1 | 0 | 0 | 1 | 0 | 0 | 2 |
| 3 | 0.63 | 3.56 | 0.18 | 0.86 | 1 | Grm2 | glutamate receptor, metabotropic 2 | 0 | 0 | 19 | 0 | 0 | 2 | 0 | 0 | 8 | 0 | 1 |
| 3 | 4.38 | 3.55 | 1.23 | 0.22 | 1 | Tas1r1 | taste receptor, type 1, member 1 | 1 | 0 | 0 | 0 | 1 | 0 | 0 | 19 | 0 | 0 | 9 |
| 3 | 0.10 | 3.25 | 0.03 | 0.98 | 1 | Gm34418 | | 3 | 15 | 4 | 3 | 0 | 0 | 0 | 0 | 0 | 0 | 4 |
| 3 | -0.43 | 3.67 | -0.12 | 0.91 | 1 | E230016K23Rik | | 27 | 2 | 0 | 0 | 0 | 1 | 0 | 0 | 0 | 0 | 0 |
| 3 | 0.63 | 2.22 | 0.28 | 0.78 | 1 | Gm7384 | | 3 | 6 | 9 | 2 | 0 | 4 | 0 | 1 | 3 | 1 | 0 |
| 3 | -0.69 | 3.66 | -0.19 | 0.85 | 1 | Snord123 | | 0 | 11 | 17 | 0 | 0 | 2 | 0 | 0 | 0 | 0 | 0 |
| 3 | 0.00 | 3.68 | 0.00 | 1.00 | 1 | Gm37824 | | 6 | 17 | 6 | 0 | 0 | 0 | 0 | 0 | 0 | 0 | 0 |
| 3 | -0.07 | 3.06 | -0.02 | 0.98 | 1 | Gm46440 | | 0 | 0 | 4 | 0 | 1 | 15 | 0 | 0 | 1 | 0 | 8 |
| 3 | 0.00 | 3.68 | 0.00 | 1.00 | 1 | 9430062P05Rik | | 18 | 0 | 11 | 0 | 0 | 0 | 0 | 0 | 0 | 0 | 0 |
| 3 | -0.57 | 3.01 | -0.19 | 0.85 | 1 | Gm42501 | | 3 | 0 | 0 | 13 | 2 | 2 | 0 | 0 | 4 | 0 | 6 |
| 3 | 0.00 | 3.68 | 0.00 | 1.00 | 1 | Hotairm1 | | 3 | 0 | 26 | 0 | 0 | 0 | 0 | 0 | 0 | 0 | 0 |
| 3 | -3.46 | 2.39 | -1.45 | 0.15 | 1 | Gm28902 | | 1 | 3 | 5 | 5 | 10 | 3 | 0 | 0 | 0 | 1 | 0 |
| 3 | 1.04 | 2.38 | 0.44 | 0.66 | 1 | Gm12418 | | 1 | 13 | 1 | 0 | 1 | 7 | 0 | 0 | 1 | 3 | 3 |
| 3 | 1.99 | 3.32 | 0.60 | 0.55 | 1 | Gm9165 | | 0 | 1 | 1 | 0 | 0 | 1 | 0 | 25 | 0 | 0 | 1 |
| 3 | 1.07 | 2.56 | 0.42 | 0.68 | 1 | Gm43353 | | 2 | 0 | 6 | 2 | 1 | 3 | 0 | 8 | 8 | 0 | 0 |
| 3 | -3.66 | 3.56 | -1.03 | 0.30 | 1 | Gm37790 | | 0 | 0 | 0 | 0 | 2 | 2 | 0 | 0 | 0 | 26 | 0 |
| 3 | -0.56 | 2.99 | -0.19 | 0.85 | 1 | Gm47761 | | 2 | 0 | 0 | 1 | 5 | 5 | 0 | 14 | 0 | 0 | 3 |
| 3 | -1.29 | 2.84 | -0.45 | 0.65 | 1 | Gm14113 | | 10 | 9 | 1 | 7 | 0 | 1 | 0 | 0 | 0 | 0 | 1 |
| 3 | 0.84 | 3.60 | 0.23 | 0.82 | 1 | Gm9866 | | 0 | 0 | 0 | 6 | 0 | 0 | 0 | 23 | 0 | 0 | 0 |
| 3 | -1.45 | 2.76 | -0.53 | 0.60 | 1 | Wbscr25 | | 4 | 0 | 10 | 1 | 2 | 7 | 0 | 0 | 0 | 0 | 5 |
| 3 | 0.49 | 3.54 | 0.14 | 0.89 | 1 | Gm49067 | | 0 | 0 | 0 | 0 | 7 | 3 | 0 | 0 | 5 | 0 | 14 |
| 3 | 0.00 | 3.68 | 0.00 | 1.00 | 1 | Ecrg4 | ECRG4 augurin precursor | 19 | 0 | 10 | 0 | 0 | 0 | 0 | 0 | 0 | 0 | 0 |
| 3 | 0.00 | 3.68 | 0.00 | 1.00 | 1 | Tmem178 | transmembrane protein 178 | 19 | 0 | 10 | 0 | 0 | 0 | 0 | 0 | 0 | 0 | 0 |
| 3 | 0.78 | 2.62 | 0.30 | 0.77 | 1 | Amd-ps3 | | 0 | 0 | 1 | 0 | 9 | 4 | 0 | 0 | 4 | 1 | 10 |
| 3 | -2.27 | 3.58 | -0.64 | 0.53 | 1 | Zfp980 | zinc finger protein 980 | 9 | 0 | 7 | 11 | 0 | 0 | 0 | 0 | 2 | 0 | 0 |
| 3 | 2.14 | 3.54 | 0.61 | 0.54 | 1 | Btnl1 | butyrophilin-like 1 | 0 | 0 | 0 | 4 | 0 | 0 | 0 | 12 | 14 | 0 | 0 |
| 3 | -3.25 | 3.17 | -1.03 | 0.30 | 1 | Gm44697 | | 0 | 0 | 3 | 8 | 9 | 7 | 0 | 0 | 0 | 0 | 2 |
| 3 | 1.25 | 3.66 | 0.34 | 0.73 | 1 | Erich2 | glutamate rich 2 | 7 | 17 | 3 | 0 | 0 | 0 | 0 | 0 | 0 | 0 | 2 |
| 3 | 0.00 | 3.68 | 0.00 | 1.00 | 1 | Slc6a12 | solute carrier family 6 (neurotransmitter tr | 24 | 0 | 5 | 0 | 0 | 0 | 0 | 0 | 0 | 0 | 0 |
| 3 | -1.29 | 3.59 | -0.36 | 0.72 | 1 | 9630050E16Rik | | 0 | 23 | 0 | 0 | 0 | 6 | 0 | 0 | 0 | 0 | 0 |
| 3 | 2.19 | 3.60 | 0.61 | 0.54 | 1 | Axdnd1 | axonemal dynein light chain domain cont | 0 | 21 | 0 | 0 | 0 | 6 | 0 | 0 | 2 | 0 | 0 |
| 3 | 0.00 | 3.68 | 0.00 | 1.00 | 1 | Gm36377 | | 8 | 13 | 8 | 0 | 0 | 0 | 0 | 0 | 0 | 0 | 0 |
| 3 | -0.98 | 3.62 | -0.27 | 0.79 | 1 | 2210407C18Ril | RIKEN cDNA 2210407C18 gene | 6 | 2 | 15 | 0 | 7 | 0 | 0 | 0 | 0 | 0 | 0 |
| 3 | 1.83 | 3.55 | 0.52 | 0.61 | 1 | Gm10571 | | 0 | 0 | 0 | 0 | 0 | 3 | 0 | 27 | 0 | 0 | 0 |
| 3 | 1.83 | 3.55 | 0.52 | 0.61 | 1 | Ces4a | carboxylesterase 4A | 0 | 0 | 0 | 0 | 0 | 3 | 0 | 27 | 0 | 0 | 0 |
| 3 | -1.13 | 3.59 | -0.31 | 0.75 | 1 | A930012L18Rik | | 0 | 0 | 25 | 0 | 0 | 4 | 0 | 0 | 0 | 0 | 0 |
| 3 | 2.40 | 3.55 | 0.68 | 0.50 | 1 | Gm17021 | | 0 | 0 | 0 | 0 | 0 | 16 | 0 | 0 | 0 | 9 | 4 |
| 3 | -3.67 | 3.57 | -1.03 | 0.30 | 1 | Gm29462 | | 2 | 0 | 8 | 15 | 0 | 4 | 0 | 0 | 0 | 0 | 0 |
| 3 | -1.39 | 3.61 | -0.39 | 0.70 | 1 | Ntf5 | neurotrophin 5 | 0 | 17 | 6 | 0 | 6 | 0 | 0 | 0 | 0 | 0 | 0 |
| 3 | 3.22 | 2.32 | 1.39 | 0.17 | 1 | Gm5564 | | 3 | 1 | 2 | 1 | 0 | 3 | 0 | 7 | 7 | 5 | 0 |
| 3 | -0.18 | 2.77 | -0.07 | 0.95 | 1 | A230052G05Rik | | 4 | 0 | 10 | 0 | 5 | 3 | 0 | 0 | 2 | 5 | 0 |
| 3 | -1.28 | 3.59 | -0.36 | 0.72 | 1 | Gm48314 | | 0 | 0 | 0 | 0 | 2 | 21 | 0 | 0 | 0 | 0 | 6 |
| 3 | 0.31 | 3.60 | 0.09 | 0.93 | 1 | Gm49659 | | 2 | 4 | 0 | 0 | 0 | 10 | 0 | 0 | 0 | 0 | 13 |
| 3 | 0.07 | 3.60 | 0.02 | 0.98 | 1 | Gm25945 | | 0 | 0 | 0 | 0 | 0 | 17 | 0 | 0 | 0 | 0 | 12 |
| 3 | 0.07 | 3.60 | 0.02 | 0.98 | 1 | Mir3060 | | 0 | 0 | 0 | 0 | 0 | 17 | 0 | 0 | 0 | 0 | 12 |
| 3 | 1.77 | 2.67 | 0.66 | 0.51 | 1 | Mm2pr | | 9 | 1 | 0 | 0 | 2 | 3 | 0 | 8 | 2 | 0 | 4 |

| | | | | | | | | | | | | | | | | | | |
|---|---|---|---|---|---|---|---|---|---|---|---|---|---|---|---|---|---|---|
| 3 | 0.00 | 3.68 | 0.00 | 1.00 | 1 | | predicted gene, 21104 | 5 | 0 | 24 | 0 | 0 | 0 | 0 | 0 | 0 | 0 | 0 |
| 3 | -0.64 | 3.54 | -0.18 | 0.86 | 1 | H2ac4 | H2A clustered histone 4 | 0 | 0 | 0 | 0 | 14 | 7 | 0 | 0 | 5 | 0 | 3 |
| 3 | -0.15 | 3.60 | -0.04 | 0.97 | 1 | Gm19325 | | 0 | 0 | 4 | 0 | 0 | 18 | 0 | 0 | 0 | 0 | 7 |
| 3 | -3.56 | 3.58 | -0.99 | 0.32 | 1 | Gm44053 | | 2 | 0 | 0 | 0 | 4 | 14 | 0 | 0 | 0 | 0 | 9 |
| 3 | -1.96 | 3.39 | -0.58 | 0.56 | 1 | Gm28874 | | 10 | 0 | 0 | 0 | 4 | 12 | 0 | 0 | 1 | 0 | 2 |
| 3 | -2.49 | 3.60 | -0.69 | 0.49 | 1 | Gm12098 | | 10 | 0 | 11 | 4 | 3 | 0 | 0 | 0 | 0 | 0 | 0 |
| 3 | 0.02 | 3.60 | 0.01 | 1.00 | 1 | Gm48798 | | 0 | 0 | 0 | 0 | 0 | 18 | 0 | 0 | 0 | 0 | 11 |
| 3 | 0.00 | 3.68 | 0.00 | 1.00 | 1 | Lpar3 | lysophosphatidic acid receptor 3 | 21 | 0 | 8 | 0 | 0 | 0 | 0 | 0 | 0 | 0 | 0 |
| 3 | 2.78 | 2.06 | 1.35 | 0.18 | 1 | Gm4342 | | 3 | 2 | 3 | 0 | 1 | 1 | 0 | 15 | 2 | 1 | 1 |
| 3 | 0.00 | 3.68 | 0.00 | 1.00 | 1 | Pira2 | paired-Ig-like receptor A2 | 26 | 0 | 3 | 0 | 0 | 0 | 0 | 0 | 0 | 0 | 0 |
| 3 | -2.78 | 3.58 | -0.78 | 0.44 | 1 | Gad2 | glutamic acid decarboxylase 2 | 5 | 10 | 0 | 0 | 9 | 0 | 0 | 0 | 0 | 0 | 4 |
| 3 | 0.00 | 3.68 | 0.00 | 1.00 | 1 | Phgr1 | proline/histidine/glycine-rich 1 | 0 | 13 | 16 | 0 | 0 | 0 | 0 | 0 | 0 | 0 | 0 |
| 3 | 0.11 | 3.59 | 0.03 | 0.97 | 1 | 6030400A10Rik | | 9 | 0 | 0 | 0 | 0 | 10 | 0 | 0 | 0 | 0 | 9 |
| 3 | -0.19 | 3.54 | -0.05 | 0.96 | 1 | Gm44117 | | 0 | 12 | 0 | 0 | 0 | 8 | 0 | 8 | 0 | 0 | 0 |
| 3 | -0.11 | 3.58 | -0.03 | 0.98 | 1 | Tsix | | 0 | 23 | 0 | 0 | 0 | 3 | 0 | 0 | 0 | 0 | 2 |
| 3 | 0.00 | 3.68 | 0.00 | 1.00 | 1 | Rbm46 | RNA binding motif protein 46 | 12 | 0 | 17 | 0 | 0 | 0 | 0 | 0 | 0 | 0 | 0 |
| 3 | 0.82 | 2.90 | 0.28 | 0.78 | 1 | Tcte1 | t-complex-associated testis expressed 1 | 4 | 11 | 0 | 4 | 1 | 0 | 0 | 0 | 0 | 8 | 1 |
| 3 | -2.87 | 3.38 | -0.85 | 0.40 | 1 | Majin | membrane anchored junction protein | 4 | 14 | 1 | 0 | 6 | 3 | 0 | 0 | 0 | 0 | 0 |
| 3 | 1.10 | 3.31 | 0.33 | 0.74 | 1 | Gm26873 | | 4 | 13 | 5 | 0 | 0 | 1 | 0 | 0 | 5 | 0 | 0 |
| 3 | 0.44 | 3.54 | 0.12 | 0.90 | 1 | Bpifc | BPI fold containing family C | 0 | 0 | 14 | 0 | 0 | 6 | 0 | 0 | 7 | 0 | 2 |
| 3 | 0.00 | 3.68 | 0.00 | 1.00 | 1 | Slc26a7 | solute carrier family 26, member 7 | 21 | 7 | 0 | 0 | 0 | 0 | 0 | 0 | 0 | 0 | 0 |
| 3 | 0.56 | 3.58 | 0.16 | 0.88 | 1 | Gm48405 | | 0 | 0 | 11 | 0 | 0 | 6 | 0 | 0 | 0 | 0 | 11 |
| 3 | -0.69 | 3.66 | -0.19 | 0.85 | 1 | Slc25a34 | solute carrier family 25, member 34 | 0 | 8 | 19 | 0 | 0 | 2 | 0 | 0 | 0 | 0 | 0 |
| 3 | -3.80 | 3.61 | -1.05 | 0.29 | 1 | Necab1 | N-terminal EF-hand calcium binding prote | 21 | 0 | 0 | 4 | 0 | 3 | 0 | 0 | 0 | 0 | 1 |
| 3 | -0.86 | 3.26 | -0.26 | 0.79 | 1 | 4930417O13Rik | | 12 | 0 | 4 | 0 | 3 | 3 | 0 | 0 | 0 | 0 | 6 |
| 3 | -0.74 | 3.55 | -0.21 | 0.83 | 1 | 4930535L15Rik | | 0 | 0 | 16 | 0 | 7 | 4 | 0 | 0 | 0 | 0 | 2 |
| 3 | -4.43 | 3.55 | -1.25 | 0.21 | 1 | Gm13061 | | 0 | 0 | 0 | 6 | 0 | 2 | 0 | 0 | 0 | 21 | 0 |
| 3 | 0.15 | 3.60 | 0.04 | 0.97 | 1 | Gm43482 | | 0 | 0 | 0 | 0 | 0 | 15 | 0 | 0 | 0 | 0 | 13 |
| 3 | 0.65 | 3.60 | 0.18 | 0.86 | 1 | Gm37558 | | 1 | 10 | 0 | 0 | 0 | 4 | 0 | 0 | 0 | 0 | 13 |
| 3 | -0.30 | 3.12 | -0.10 | 0.92 | 1 | Gm15912 | | 5 | 5 | 4 | 0 | 0 | 11 | 0 | 0 | 0 | 0 | 3 |
| 3 | -1.32 | 2.76 | -0.48 | 0.63 | 1 | Gm50336 | | 0 | 0 | 5 | 4 | 2 | 7 | 0 | 0 | 2 | 0 | 8 |
| 3 | 0.00 | 3.64 | 0.00 | 1.00 | 1 | Gm43525 | | 0 | 28 | 0 | 0 | 0 | 0 | 0 | 0 | 0 | 0 | 0 |
| 3 | 0.00 | 3.64 | 0.00 | 1.00 | 1 | Gm4764 | | 0 | 28 | 0 | 0 | 0 | 0 | 0 | 0 | 0 | 0 | 0 |
| 3 | 0.00 | 3.64 | 0.00 | 1.00 | 1 | 6030498E09Ril | RIKEN cDNA 6030498E09 gene | 0 | 28 | 0 | 0 | 0 | 0 | 0 | 0 | 0 | 0 | 0 |
| 3 | 0.73 | 3.60 | 0.20 | 0.84 | 1 | Gm27626 | | 0 | 0 | 19 | 0 | 7 | 0 | 0 | 0 | 2 | 0 | 1 |
| 3 | 0.40 | 3.51 | 0.12 | 0.91 | 1 | Foxh1 | forkhead box H1 | 0 | 0 | 15 | 4 | 0 | 0 | 0 | 0 | 0 | 10 | 0 |
| 3 | 1.24 | 3.61 | 0.34 | 0.73 | 1 | Gal3st2b | galactose-3-O-sulfotransferase 2B | 10 | 0 | 0 | 0 | 0 | 0 | 0 | 0 | 0 | 18 | 0 |
| 3 | 0.00 | 3.64 | 0.00 | 1.00 | 1 | Igkv4-62 | | 28 | 0 | 0 | 0 | 0 | 0 | 0 | 0 | 0 | 0 | 0 |
| 3 | 0.00 | 3.64 | 0.00 | 1.00 | 1 | Raet1e | retinoic acid early transcript 1E | 28 | 0 | 0 | 0 | 0 | 0 | 0 | 0 | 0 | 0 | 0 |
| 3 | 0.00 | 3.64 | 0.00 | 1.00 | 1 | Ighj1 | | 28 | 0 | 0 | 0 | 0 | 0 | 0 | 0 | 0 | 0 | 0 |
| 3 | 0.00 | 3.68 | 0.00 | 1.00 | 1 | Vstm2b | V-set and transmembrane domain contai | 12 | 16 | 0 | 0 | 0 | 0 | 0 | 0 | 0 | 0 | 0 |
| 3 | 1.20 | 3.58 | 0.33 | 0.74 | 1 | Gm31333 | | 0 | 19 | 0 | 0 | 0 | 1 | 0 | 0 | 0 | 0 | 8 |
| 3 | -1.00 | 2.79 | -0.36 | 0.72 | 1 | Gm13245 | | 0 | 0 | 3 | 11 | 4 | 2 | 0 | 0 | 5 | 0 | 4 |
| 3 | 0.00 | 3.68 | 0.00 | 1.00 | 1 | Smc1b | structural maintenance of chromosomes ' | 1 | 27 | 0 | 0 | 0 | 0 | 0 | 0 | 0 | 0 | 0 |
| 3 | 3.18 | 3.62 | 0.88 | 0.38 | 1 | Gm13522 | | 0 | 22 | 0 | 0 | 0 | 1 | 0 | 0 | 5 | 0 | 0 |
| 3 | -0.56 | 1.88 | -0.30 | 0.76 | 1 | Ndufab1-ps | | 0 | 1 | 4 | 6 | 8 | 3 | 0 | 3 | 1 | 1 | 1 |
| 3 | 0.01 | 3.61 | 0.00 | 1.00 | 1 | Slc19a3 | solute carrier family 19, member 3 | 20 | 3 | 0 | 0 | 0 | 3 | 0 | 0 | 0 | 0 | 2 |
| 3 | 0.05 | 2.46 | 0.02 | 0.98 | 1 | Gm12416 | | 1 | 3 | 15 | 0 | 5 | 2 | 0 | 0 | 0 | 1 | 2 |
| 3 | 0.62 | 2.83 | 0.22 | 0.83 | 1 | Gpbar1 | G protein-coupled bile acid receptor 1 | 8 | 0 | 1 | 0 | 3 | 3 | 0 | 0 | 3 | 0 | 10 |
| 3 | 0.37 | 3.60 | 0.10 | 0.92 | 1 | Insl3 | insulin-like 3 | 0 | 8 | 3 | 0 | 0 | 7 | 0 | 0 | 0 | 0 | 10 |
| 3 | 0.00 | 3.68 | 0.00 | 1.00 | 1 | Lilra6 | leukocyte immunoglobulin-like receptor, s | 19 | 5 | 4 | 0 | 0 | 0 | 0 | 0 | 0 | 0 | 0 |
| 3 | 1.22 | 2.36 | 0.52 | 0.61 | 1 | Gm5461 | | 0 | 1 | 7 | 0 | 6 | 2 | 0 | 0 | 3 | 1 | 8 |
| 3 | 2.54 | 3.60 | 0.71 | 0.48 | 1 | Ly6g6e | lymphocyte antigen 6 complex, locus G6E | 9 | 3 | 0 | 0 | 0 | 0 | 0 | 0 | 1 | 15 | 0 |
| 3 | -1.82 | 3.58 | -0.51 | 0.61 | 1 | Rskr | ribosomal protein S6 kinase related | 0 | 0 | 0 | 5 | 0 | 18 | 0 | 0 | 0 | 0 | 5 |
| 3 | 2.95 | 3.54 | 0.83 | 0.40 | 1 | 4933406B17Rik | | 0 | 0 | 0 | 0 | 0 | 7 | 0 | 8 | 2 | 0 | 11 |
| 3 | 0.46 | 3.60 | 0.13 | 0.90 | 1 | Npff | neuropeptide FF-amide peptide precurso | 0 | 1 | 5 | 0 | 0 | 8 | 0 | 0 | 0 | 0 | 14 |
| 3 | 1.25 | 3.66 | 0.34 | 0.73 | 1 | Frmpd2 | FERM and PDZ domain containing 2 | 8 | 18 | 0 | 0 | 0 | 0 | 0 | 0 | 0 | 0 | 2 |
| 3 | -2.59 | 3.54 | -0.73 | 0.46 | 1 | Gm12348 | | 0 | 0 | 9 | 6 | 0 | 3 | 0 | 0 | 9 | 0 | 0 |
| 3 | 4.47 | 3.56 | 1.25 | 0.21 | 1 | 9330159N22Rik | | 0 | 0 | 0 | 0 | 0 | 0 | 0 | 0 | 0 | 17 | 11 |
| 3 | -0.02 | 3.60 | 0.00 | 1.00 | 1 | Xist | | 0 | 0 | 0 | 0 | 0 | 18 | 0 | 0 | 0 | 0 | 10 |

| | | | | | | | | | | | | | | | | | | |
|---|---|---|---|---|---|---|---|---|---|---|---|---|---|---|---|---|---|---|
| 3 | -0.22 | 3.59 | -0.06 | 0.95 | 1 | Gm13110 | | 0 | 9 | 0 | 0 | 0 | 13 | 0 | 0 | 0 | 0 | 6 |
| 3 | 0.00 | 3.68 | 0.00 | 1.00 | 1 | Gm6133 | | 1 | 0 | 27 | 0 | 0 | 0 | 0 | 0 | 0 | 0 | 0 |
| 3 | -0.98 | 3.58 | -0.27 | 0.78 | 1 | Mir7-2 | | 0 | 0 | 0 | 0 | 3 | 12 | 0 | 0 | 0 | 0 | 13 |
| 3 | 0.00 | 3.68 | 0.00 | 1.00 | 1 | Pvrig | poliovirus receptor related immunoglobul | 21 | 0 | 7 | 0 | 0 | 0 | 0 | 0 | 0 | 0 | 0 |
| 3 | -0.86 | 2.33 | -0.37 | 0.71 | 1 | Hpdl | 4-hydroxyphenylpyruvate dioxygenase-lil | 0 | 7 | 2 | 11 | 1 | 2 | 0 | 3 | 0 | 0 | 2 |
| 3 | 0.15 | 3.49 | 0.04 | 0.97 | 1 | Gm48853 | | 5 | 0 | 0 | 0 | 5 | 1 | 0 | 15 | 2 | 0 | 0 |
| 3 | 0.00 | 3.68 | 0.00 | 1.00 | 1 | Ighv1-59 | | 26 | 2 | 0 | 0 | 0 | 0 | 0 | 0 | 0 | 0 | 0 |
| 3 | 0.12 | 3.62 | 0.03 | 0.97 | 1 | Gm13532 | | 23 | 0 | 1 | 0 | 0 | 3 | 0 | 0 | 1 | 0 | 0 |
| 3 | 0.54 | 3.60 | 0.15 | 0.88 | 1 | 1700001J03Rik | RIKEN cDNA 1700001J03 gene | 0 | 0 | 0 | 0 | 0 | 8 | 0 | 0 | 0 | 0 | 19 |
| 3 | 0.00 | 3.68 | 0.00 | 1.00 | 1 | 1810073O08Rik | | 11 | 2 | 15 | 0 | 0 | 0 | 0 | 0 | 0 | 0 | 0 |
| 3 | 2.17 | 2.94 | 0.74 | 0.46 | 1 | Rpl35a-ps2 | | 3 | 3 | 5 | 0 | 0 | 1 | 0 | 15 | 1 | 0 | 0 |
| 3 | -4.47 | 3.48 | -1.29 | 0.20 | 1 | Uba52-ps | | 9 | 0 | 1 | 9 | 9 | 1 | 0 | 0 | 0 | 0 | 0 |
| 3 | 0.00 | 3.68 | 0.00 | 1.00 | 1 | Gm26902 | | 16 | 2 | 9 | 0 | 0 | 0 | 0 | 0 | 0 | 0 | 0 |
| 3 | -0.17 | 3.53 | -0.05 | 0.96 | 1 | Gm42716 | | 0 | 0 | 0 | 0 | 0 | 16 | 0 | 0 | 0 | 12 | 0 |
| 3 | 0.00 | 3.68 | 0.00 | 1.00 | 1 | Igkv4-86 | | 27 | 1 | 0 | 0 | 0 | 0 | 0 | 0 | 0 | 0 | 0 |
| 3 | 4.22 | 3.58 | 1.18 | 0.24 | 1 | F630040K05Rik | | 0 | 0 | 10 | 0 | 0 | 3 | 0 | 0 | 0 | 8 | 6 |
| 3 | -1.15 | 3.46 | -0.33 | 0.74 | 1 | Pip5k1bos | | 0 | 0 | 8 | 10 | 0 | 3 | 0 | 0 | 4 | 0 | 3 |
| 3 | 0.45 | 1.74 | 0.26 | 0.80 | 1 | Gm7429 | predicted pseudogene 7429 | 3 | 3 | 5 | 2 | 4 | 2 | 0 | 2 | 1 | 1 | 4 |
| 3 | 1.73 | 3.30 | 0.52 | 0.60 | 1 | Gm14820 | | 0 | 0 | 0 | 4 | 0 | 6 | 0 | 0 | 2 | 9 | 7 |
| 3 | 0.00 | 3.68 | 0.00 | 1.00 | 1 | Cd40lg | CD40 ligand | 1 | 12 | 15 | 0 | 0 | 0 | 0 | 0 | 0 | 0 | 0 |
| 3 | -2.61 | 2.76 | -0.95 | 0.34 | 1 | Gm7535 | | 1 | 9 | 10 | 1 | 4 | 3 | 0 | 0 | 0 | 0 | 0 |
| 3 | 0.00 | 3.68 | 0.00 | 1.00 | 1 | Gm26510 | | 17 | 0 | 10 | 0 | 0 | 0 | 0 | 0 | 0 | 0 | 0 |
| 2 | 1.17 | 3.59 | 0.33 | 0.74 | 1 | Marchf10 | membrane associated ring-CH-type finge | 0 | 0 | 0 | 0 | 0 | 1 | 0 | 27 | 0 | 0 | 0 |
| 2 | -0.73 | 3.64 | -0.20 | 0.84 | 1 | Mro | maestro | 14 | 8 | 3 | 0 | 0 | 3 | 0 | 0 | 0 | 0 | 0 |
| 2 | 0.00 | 3.68 | 0.00 | 1.00 | 1 | Gm9796 | | 8 | 0 | 20 | 0 | 0 | 0 | 0 | 0 | 0 | 0 | 0 |
| 2 | -1.12 | 3.41 | -0.33 | 0.74 | 1 | Gm16751 | | 0 | 5 | 0 | 4 | 5 | 5 | 0 | 0 | 9 | 0 | 0 |
| 2 | -0.87 | 3.57 | -0.24 | 0.81 | 1 | Dnah10 | dynein, axonemal, heavy chain 10 | 0 | 0 | 0 | 0 | 4 | 8 | 0 | 0 | 0 | 0 | 15 |
| 2 | 0.00 | 3.64 | 0.00 | 1.00 | 1 | Cntn6 | contactin 6 | 27 | 0 | 0 | 0 | 0 | 0 | 0 | 0 | 0 | 0 | 0 |
| 2 | 0.00 | 3.64 | 0.00 | 1.00 | 1 | Tbc1d10c | TBC1 domain family, member 10c | 27 | 0 | 0 | 0 | 0 | 0 | 0 | 0 | 0 | 0 | 0 |
| 2 | 0.00 | 3.68 | 0.00 | 1.00 | 1 | Ttc6 | tetratricopeptide repeat domain 6 | 16 | 11 | 0 | 0 | 0 | 0 | 0 | 0 | 0 | 0 | 0 |
| 2 | 0.00 | 3.68 | 0.00 | 1.00 | 1 | Dnajb13 | DnaJ heat shock protein family (Hsp40) m | 11 | 16 | 0 | 0 | 0 | 0 | 0 | 0 | 0 | 0 | 0 |
| 2 | 0.00 | 3.19 | 0.00 | 1.00 | 1 | Prickle4 | prickle planar cell polarity protein 4 | 3 | 2 | 15 | 0 | 0 | 5 | 0 | 0 | 0 | 0 | 3 |
| 2 | 2.87 | 2.49 | 1.15 | 0.25 | 1 | Gm8692 | | 0 | 1 | 3 | 1 | 1 | 0 | 0 | 16 | 1 | 0 | 4 |
| 2 | 2.66 | 3.45 | 0.77 | 0.44 | 1 | Gm44567 | | 3 | 10 | 0 | 0 | 0 | 8 | 0 | 0 | 3 | 0 | 3 |
| 2 | -0.24 | 2.51 | -0.10 | 0.92 | 1 | Tmsb15b2 | thymosin beta 15b2 | 4 | 1 | 3 | 0 | 4 | 7 | 0 | 1 | 0 | 0 | 7 |
| 2 | 0.44 | 3.59 | 0.12 | 0.90 | 1 | Gm37612 | | 11 | 0 | 0 | 0 | 0 | 6 | 0 | 0 | 0 | 0 | 10 |
| 2 | -0.58 | 2.53 | -0.23 | 0.82 | 1 | Gm48133 | | 3 | 6 | 3 | 0 | 4 | 7 | 0 | 0 | 3 | 0 | 1 |
| 2 | 1.37 | 3.38 | 0.41 | 0.68 | 1 | Gm12735 | | 3 | 0 | 0 | 3 | 0 | 2 | 0 | 16 | 0 | 0 | 4 |
| 2 | -2.20 | 3.30 | -0.67 | 0.51 | 1 | Gm17146 | | 0 | 9 | 0 | 2 | 4 | 8 | 0 | 0 | 0 | 0 | 4 |
| 2 | 1.34 | 1.96 | 0.68 | 0.49 | 1 | Rps2-ps8 | | 3 | 2 | 7 | 2 | 5 | 0 | 0 | 1 | 3 | 3 | 2 |
| 2 | 5.53 | 3.56 | 1.56 | 0.12 | 1 | Gm36500 | | 0 | 11 | 0 | 0 | 0 | 2 | 0 | 1 | 12 | 0 | 1 |
| 2 | -1.04 | 2.48 | -0.42 | 0.68 | 1 | 4930429F24Rik | | 6 | 0 | 3 | 1 | 2 | 10 | 0 | 0 | 2 | 0 | 3 |
| 2 | -0.34 | 3.60 | -0.09 | 0.93 | 1 | Mir341 | | 0 | 0 | 0 | 0 | 0 | 22 | 0 | 0 | 0 | 0 | 5 |
| 2 | 0.00 | 3.68 | 0.00 | 1.00 | 1 | Gm14812 | | 14 | 8 | 5 | 0 | 0 | 0 | 0 | 0 | 0 | 0 | 0 |
| 2 | -2.02 | 2.76 | -0.73 | 0.46 | 1 | A730081D07Rik | | 0 | 7 | 5 | 2 | 6 | 4 | 0 | 0 | 0 | 0 | 3 |
| 2 | 0.95 | 3.67 | 0.26 | 0.80 | 1 | 1200007C13Rik | | 9 | 11 | 6 | 0 | 0 | 0 | 0 | 0 | 0 | 0 | 1 |
| 2 | 4.03 | 3.57 | 1.13 | 0.26 | 1 | A930027P06Rik | | 4 | 0 | 0 | 0 | 0 | 8 | 0 | 0 | 11 | 0 | 3 |
| 2 | 0.42 | 3.60 | 0.12 | 0.91 | 1 | Slc17a8 | solute carrier family 17 (sodium-depende | 2 | 9 | 0 | 0 | 0 | 6 | 0 | 0 | 0 | 0 | 10 |
| 2 | 0.48 | 3.60 | 0.13 | 0.89 | 1 | Gm28187 | | 0 | 4 | 0 | 0 | 0 | 8 | 0 | 0 | 0 | 0 | 15 |
| 2 | 1.87 | 3.55 | 0.53 | 0.60 | 1 | Gm37672 | | 0 | 0 | 0 | 0 | 0 | 18 | 0 | 0 | 6 | 0 | 3 |
| 2 | -1.04 | 3.53 | -0.30 | 0.77 | 1 | Snord70 | | 0 | 0 | 0 | 9 | 0 | 11 | 0 | 0 | 1 | 0 | 6 |
| 2 | 2.96 | 3.44 | 0.86 | 0.39 | 1 | Pla2g4f | phospholipase A2, group IVF | 8 | 1 | 0 | 0 | 0 | 1 | 0 | 0 | 5 | 0 | 12 |
| 2 | 1.25 | 3.66 | 0.34 | 0.73 | 1 | Gm16499 | | 9 | 16 | 0 | 0 | 0 | 0 | 0 | 0 | 0 | 0 | 2 |
| 2 | 3.09 | 3.56 | 0.87 | 0.39 | 1 | Gm45738 | | 0 | 0 | 0 | 0 | 0 | 10 | 0 | 0 | 0 | 12 | 5 |
| 2 | -1.16 | 3.62 | -0.32 | 0.75 | 1 | Gm42793 | | 0 | 10 | 11 | 5 | 0 | 0 | 0 | 0 | 0 | 0 | 0 |
| 2 | 0.95 | 3.61 | 0.26 | 0.79 | 1 | Gm17783 | predicted gene, 17783 | 0 | 0 | 19 | 0 | 0 | 0 | 0 | 0 | 8 | 0 | 0 |
| 2 | -0.66 | 3.58 | -0.18 | 0.85 | 1 | Pga5 | pepsinogen 5, group I | 0 | 0 | 6 | 6 | 0 | 2 | 0 | 13 | 0 | 0 | 0 |
| 2 | 0.00 | 3.68 | 0.00 | 1.00 | 1 | Klk13 | kallikrein related-peptidase 13 | 6 | 0 | 21 | 0 | 0 | 0 | 0 | 0 | 0 | 0 | 0 |
| 2 | -0.50 | 3.59 | -0.14 | 0.89 | 1 | Gm44816 | | 0 | 0 | 9 | 0 | 0 | 14 | 0 | 0 | 0 | 0 | 3 |
| 2 | -4.84 | 3.54 | -1.37 | 0.17 | 1 | Gm26596 | | 0 | 0 | 0 | 4 | 1 | 4 | 0 | 0 | 17 | 0 | 0 |

| | | | | | | | | | | | | | | | | | |
|---|---|---|---|---|---|---|---|---|---|---|---|---|---|---|---|---|---|
| 2 | 0.00 | 3.68 | 0.00 | 1.00 | 1 | Gm16685 | | 15 | 12 | 0 | 0 | 0 | 0 | 0 | 0 | 0 | 0 | 0 |
| 2 | 0.00 | 3.68 | 0.00 | 1.00 | 1 | Ms4a2 | membrane-spanning 4-domains, subfami | 15 | 12 | 0 | 0 | 0 | 0 | 0 | 0 | 0 | 0 | 0 |
| 2 | -0.90 | 3.62 | -0.25 | 0.80 | 1 | Gm15596 | | 0 | 21 | 2 | 0 | 0 | 3 | 0 | 0 | 0 | 0 | 0 |
| 2 | -0.38 | 3.50 | -0.11 | 0.91 | 1 | Gm41307 | | 11 | 0 | 0 | 11 | 0 | 0 | 0 | 0 | 5 | 0 | 0 |
| 2 | 0.39 | 1.71 | 0.22 | 0.82 | 1 | Gm4799 | | 2 | 3 | 2 | 3 | 5 | 2 | 0 | 3 | 5 | 1 | 1 |
| 2 | 0.00 | 3.68 | 0.00 | 1.00 | 1 | Gm49767 | | 16 | 0 | 10 | 0 | 0 | 0 | 0 | 0 | 0 | 0 | 0 |
| 2 | 0.00 | 3.68 | 0.00 | 1.00 | 1 | Clec2h | C-type lectin domain family 2, member h | 11 | 2 | 14 | 0 | 0 | 0 | 0 | 0 | 0 | 0 | 0 |
| 2 | -0.11 | 3.60 | -0.03 | 0.97 | 1 | Gm17999 | | 3 | 0 | 0 | 0 | 0 | 16 | 0 | 0 | 0 | 0 | 7 |
| 2 | 0.00 | 3.64 | 0.00 | 1.00 | 1 | Igkv13-84 | | 27 | 0 | 0 | 0 | 0 | 0 | 0 | 0 | 0 | 0 | 0 |
| 2 | 0.00 | 3.64 | 0.00 | 1.00 | 1 | P2rx5 | purinergic receptor P2X, ligand-gated ion | 27 | 0 | 0 | 0 | 0 | 0 | 0 | 0 | 0 | 0 | 0 |
| 2 | 0.32 | 3.12 | 0.10 | 0.92 | 1 | 1700037F03Rik | | 0 | 0 | 6 | 0 | 6 | 3 | 0 | 0 | 10 | 0 | 2 |
| 2 | -4.16 | 3.59 | -1.16 | 0.25 | 1 | Gm48296 | | 0 | 13 | 0 | 0 | 4 | 8 | 0 | 0 | 0 | 0 | 2 |
| 2 | 0.84 | 3.60 | 0.23 | 0.82 | 1 | Gm48735 | | 0 | 0 | 0 | 0 | 0 | 4 | 0 | 0 | 0 | 0 | 22 |
| 2 | 3.59 | 3.52 | 1.02 | 0.31 | 1 | Gm11839 | | 0 | 0 | 0 | 0 | 0 | 8 | 0 | 0 | 6 | 13 | 0 |
| 2 | -2.86 | 3.59 | -0.80 | 0.43 | 1 | Mkrn3 | makorin, ring finger protein, 3 | 3 | 0 | 15 | 4 | 0 | 6 | 0 | 0 | 0 | 0 | 0 |
| 2 | -3.45 | 2.92 | -1.18 | 0.24 | 1 | Gm49328 | | 3 | 0 | 6 | 7 | 4 | 6 | 0 | 0 | 0 | 0 | 1 |
| 2 | 0.93 | 2.03 | 0.46 | 0.65 | 1 | Gm6997 | | 3 | 1 | 0 | 3 | 1 | 2 | 0 | 1 | 3 | 12 | 1 |
| 2 | 2.16 | 2.66 | 0.81 | 0.42 | 1 | Gm47942 | | 0 | 14 | 0 | 3 | 2 | 0 | 0 | 1 | 1 | 3 | 3 |
| 2 | 5.76 | 3.54 | 1.63 | 0.10 | 1 | Mir328 | | 0 | 0 | 0 | 8 | 0 | 0 | 0 | 0 | 0 | 15 | 3 |
| 2 | -1.81 | 3.29 | -0.55 | 0.58 | 1 | Nrg4 | neuregulin 4 | 0 | 1 | 1 | 0 | 8 | 13 | 0 | 0 | 0 | 0 | 4 |
| 2 | 1.61 | 3.63 | 0.44 | 0.66 | 1 | Arg2 | arginase type II | 2 | 5 | 12 | 0 | 0 | 0 | 0 | 0 | 7 | 0 | 0 |
| 2 | 0.00 | 3.68 | 0.00 | 1.00 | 1 | Adad1 | adenosine deaminase domain containinç | 13 | 2 | 11 | 0 | 0 | 0 | 0 | 0 | 0 | 0 | 0 |
| 2 | -3.27 | 3.49 | -0.94 | 0.35 | 1 | Gm7466 | | 4 | 0 | 0 | 0 | 7 | 3 | 0 | 12 | 0 | 1 | 0 |
| 2 | 0.00 | 3.68 | 0.00 | 1.00 | 1 | Lrat | lecithin-retinol acyltransferase (phosphati | 17 | 9 | 0 | 0 | 0 | 0 | 0 | 0 | 0 | 0 | 0 |
| 2 | -3.62 | 3.58 | -1.01 | 0.31 | 1 | Gm36638 | | 9 | 0 | 0 | 0 | 6 | 6 | 0 | 0 | 0 | 0 | 6 |
| 2 | 0.00 | 3.68 | 0.00 | 1.00 | 1 | Fzd10os | | 18 | 3 | 5 | 0 | 0 | 0 | 0 | 0 | 0 | 0 | 0 |
| 2 | 2.26 | 3.52 | 0.64 | 0.52 | 1 | Gm10284 | | 0 | 2 | 1 | 0 | 3 | 0 | 0 | 2 | 0 | 18 | 0 |
| 2 | -2.11 | 3.59 | -0.59 | 0.56 | 1 | 1700108F19Rik | | 1 | 0 | 14 | 1 | 0 | 11 | 0 | 0 | 0 | 0 | 0 |
| 2 | 0.47 | 3.59 | 0.13 | 0.90 | 1 | Gm20100 | | 12 | 0 | 0 | 0 | 0 | 5 | 0 | 0 | 0 | 0 | 9 |
| 2 | 0.34 | 3.60 | 0.09 | 0.93 | 1 | Gm23084 | | 0 | 0 | 0 | 0 | 0 | 11 | 0 | 0 | 0 | 0 | 15 |
| 2 | -0.36 | 2.41 | -0.15 | 0.88 | 1 | Gm7729 | | 17 | 2 | 1 | 1 | 1 | 1 | 0 | 0 | 2 | 1 | 0 |
| 2 | 0.00 | 3.68 | 0.00 | 1.00 | 1 | Alox12e | arachidonate lipoxygenase, epidermal | 12 | 14 | 0 | 0 | 0 | 0 | 0 | 0 | 0 | 0 | 0 |
| 2 | 0.00 | 3.68 | 0.00 | 1.00 | 1 | Cdh12 | cadherin 12 | 8 | 9 | 9 | 0 | 0 | 0 | 0 | 0 | 0 | 0 | 0 |
| 2 | 1.10 | 3.60 | 0.31 | 0.76 | 1 | Gm4535 | | 0 | 1 | 0 | 8 | 0 | 0 | 0 | 16 | 0 | 0 | 1 |
| 2 | 1.23 | 3.54 | 0.35 | 0.73 | 1 | Gm44802 | | 0 | 0 | 0 | 0 | 0 | 4 | 0 | 0 | 0 | 22 | 0 |
| 2 | -2.75 | 3.56 | -0.77 | 0.44 | 1 | Rps12-ps5 | | 1 | 0 | 2 | 20 | 0 | 0 | 0 | 0 | 3 | 0 | 0 |
| 2 | 0.28 | 3.60 | 0.08 | 0.94 | 1 | Ccdc169 | coiled-coil domain containing 169 | 0 | 0 | 0 | 0 | 0 | 12 | 0 | 0 | 0 | 0 | 14 |
| 2 | -0.04 | 3.60 | -0.01 | 0.99 | 1 | Gm14827 | | 0 | 0 | 0 | 0 | 0 | 17 | 0 | 0 | 0 | 0 | 9 |
| 2 | 1.90 | 3.59 | 0.53 | 0.60 | 1 | Hsd3b2 | hydroxy-delta-5-steroid dehydrogenase, 3 | 0 | 0 | 0 | 0 | 4 | 0 | 0 | 0 | 20 | 0 | 2 |
| 2 | -0.40 | 3.60 | -0.11 | 0.91 | 1 | Gm42648 | | 0 | 0 | 0 | 0 | 0 | 22 | 0 | 0 | 0 | 0 | 4 |
| 2 | -0.62 | 2.00 | -0.31 | 0.76 | 1 | Fkbp1a-ps1 | | 1 | 2 | 2 | 7 | 1 | 7 | 0 | 0 | 2 | 1 | 3 |
| 2 | -0.69 | 3.66 | -0.19 | 0.85 | 1 | Gpr17 | G protein-coupled receptor 17 | 11 | 13 | 0 | 0 | 0 | 2 | 0 | 0 | 0 | 0 | 0 |
| 2 | 2.40 | 3.04 | 0.79 | 0.43 | 1 | Gm12928 | | 2 | 3 | 0 | 3 | 0 | 2 | 0 | 0 | 4 | 13 | 0 |
| 2 | -0.08 | 3.55 | -0.02 | 0.98 | 1 | Gm37885 | | 0 | 0 | 16 | 0 | 6 | 2 | 0 | 0 | 0 | 0 | 3 |
| 2 | 3.50 | 3.53 | 0.99 | 0.32 | 1 | Cabyr | calcium-binding tyrosine-(Y)-phosphoryla | 0 | 0 | 0 | 0 | 0 | 8 | 0 | 9 | 7 | 0 | 2 |
| 2 | -4.94 | 3.61 | -1.37 | 0.17 | 1 | Gm15528 | | 10 | 0 | 0 | 0 | 3 | 12 | 0 | 0 | 0 | 0 | 1 |
| 2 | 0.00 | 3.68 | 0.00 | 1.00 | 1 | Rln3 | relaxin 3 | 15 | 2 | 9 | 0 | 0 | 0 | 0 | 0 | 0 | 0 | 0 |
| 2 | 0.00 | 3.68 | 0.00 | 1.00 | 1 | Gstp3 | glutathione S-transferase pi 3 | 5 | 0 | 21 | 0 | 0 | 0 | 0 | 0 | 0 | 0 | 0 |
| 2 | 1.45 | 3.57 | 0.41 | 0.68 | 1 | Gm19327 | | 0 | 0 | 0 | 0 | 0 | 13 | 0 | 0 | 4 | 0 | 9 |
| 2 | -1.53 | 3.58 | -0.43 | 0.67 | 1 | Gm12353 | | 0 | 0 | 0 | 4 | 0 | 14 | 0 | 0 | 0 | 0 | 7 |
| 2 | -0.06 | 3.60 | -0.02 | 0.99 | 1 | Gm44033 | | 7 | 0 | 1 | 0 | 0 | 12 | 0 | 0 | 0 | 0 | 6 |
| 2 | 3.62 | 3.54 | 1.02 | 0.31 | 1 | Gm16343 | | 8 | 0 | 0 | 0 | 4 | 0 | 0 | 9 | 5 | 0 | 0 |
| 2 | 1.53 | 3.25 | 0.47 | 0.64 | 1 | Nanos3 | nanos C2HC-type zinc finger 3 | 0 | 0 | 0 | 4 | 2 | 0 | 0 | 10 | 2 | 0 | 7 |
| 2 | 0.26 | 3.34 | 0.08 | 0.94 | 1 | Gm15616 | | 3 | 14 | 6 | 0 | 1 | 0 | 0 | 0 | 1 | 0 | 0 |
| 2 | 1.35 | 3.62 | 0.37 | 0.71 | 1 | Cfap299 | cilia and flagella associated protein 299 | 5 | 0 | 15 | 0 | 0 | 0 | 0 | 0 | 0 | 0 | 6 |
| 2 | -0.21 | 3.61 | -0.06 | 0.95 | 1 | 4930415O20Ril | RIKEN cDNA 4930415O20 gene | 0 | 22 | 0 | 4 | 0 | 0 | 0 | 0 | 0 | 0 | 0 |
| 2 | 0.00 | 3.68 | 0.00 | 1.00 | 1 | Gm46212 | | 9 | 11 | 5 | 0 | 0 | 0 | 0 | 0 | 0 | 0 | 0 |
| 2 | -0.55 | 3.45 | -0.16 | 0.87 | 1 | Zfp978 | zinc finger protein 978 | 0 | 7 | 7 | 2 | 0 | 3 | 0 | 0 | 7 | 0 | 0 |
| 2 | 0.16 | 3.60 | 0.04 | 0.96 | 1 | Gm37003 | | 0 | 0 | 0 | 0 | 0 | 14 | 0 | 0 | 0 | 0 | 12 |
| 2 | -3.44 | 3.56 | -0.97 | 0.33 | 1 | Gm27011 | | 0 | 0 | 0 | 0 | 3 | 1 | 0 | 22 | 0 | 0 | 0 |

| | | | | | | | | | | | | | | | | | | |
|---|---|---|---|---|---|---|---|---|---|---|---|---|---|---|---|---|---|---|
| 2 | 0.09 | 3.54 | 0.03 | 0.98 | 1 | Gm5655 | | 0 | 0 | 0 | 4 | 0 | 5 | 0 | 0 | 1 | 0 | 15 |
| 2 | 0.49 | 3.60 | 0.14 | 0.89 | 1 | Gm43751 | | 9 | 0 | 0 | 0 | 0 | 6 | 0 | 0 | 0 | 0 | 11 |
| 2 | -0.76 | 3.63 | -0.21 | 0.83 | 1 | 4930458D05Rik | | 12 | 6 | 4 | 0 | 0 | 3 | 0 | 0 | 0 | 0 | 0 |
| 2 | -4.55 | 3.59 | -1.27 | 0.20 | 1 | Gm47652 | | 0 | 0 | 6 | 13 | 0 | 4 | 0 | 0 | 0 | 0 | 2 |
| 2 | 0.66 | 3.50 | 0.19 | 0.85 | 1 | C130083M11Rik | | 0 | 0 | 0 | 3 | 0 | 7 | 0 | 0 | 7 | 0 | 9 |
| 2 | -0.79 | 3.62 | -0.22 | 0.83 | 1 | Mylk3 | myosin light chain kinase 3 | 11 | 0 | 9 | 0 | 0 | 5 | 0 | 0 | 0 | 0 | 0 |
| 2 | -2.66 | 3.57 | -0.75 | 0.46 | 1 | Gm38009 | | 0 | 0 | 3 | 0 | 12 | 10 | 0 | 0 | 0 | 0 | 0 |
| 2 | 0.43 | 3.60 | 0.12 | 0.91 | 1 | Gm42869 | | 0 | 0 | 0 | 0 | 0 | 9 | 0 | 0 | 0 | 0 | 16 |
| 2 | 0.00 | 3.68 | 0.00 | 1.00 | 1 | Capn13 | calpain 13 | 2 | 4 | 20 | 0 | 0 | 0 | 0 | 0 | 0 | 0 | 0 |
| 2 | 1.63 | 3.33 | 0.49 | 0.62 | 1 | Angptl8 | angiopoietin-like 8 | 0 | 3 | 14 | 0 | 0 | 1 | 0 | 0 | 6 | 0 | 2 |
| 2 | 3.86 | 3.55 | 1.09 | 0.28 | 1 | Gm36742 | | 0 | 0 | 0 | 0 | 0 | 1 | 0 | 19 | 5 | 0 | 1 |
| 2 | 2.04 | 3.08 | 0.66 | 0.51 | 1 | Mlana | melan-A | 1 | 4 | 1 | 11 | 0 | 0 | 0 | 0 | 5 | 0 | 3 |
| 2 | 0.00 | 3.68 | 0.00 | 1.00 | 1 | Gsdmc3 | gasdermin C3 | 0 | 21 | 4 | 0 | 0 | 0 | 0 | 0 | 0 | 0 | 0 |
| 2 | 0.56 | 3.59 | 0.16 | 0.88 | 1 | Gm43263 | | 12 | 0 | 0 | 0 | 0 | 4 | 0 | 0 | 0 | 0 | 9 |
| 2 | 0.04 | 3.60 | 0.01 | 0.99 | 1 | Gm43133 | | 0 | 0 | 0 | 0 | 0 | 15 | 0 | 0 | 0 | 0 | 10 |
| 2 | 0.04 | 3.60 | 0.01 | 0.99 | 1 | Gm34081 | | 0 | 0 | 0 | 0 | 0 | 15 | 0 | 0 | 0 | 0 | 10 |
| 2 | 0.71 | 3.60 | 0.20 | 0.84 | 1 | Gm9860 | | 0 | 0 | 0 | 0 | 0 | 15 | 0 | 0 | 1 | 0 | 9 |
| 2 | -0.70 | 3.54 | -0.20 | 0.84 | 1 | Gm5447 | | 0 | 10 | 0 | 5 | 0 | 6 | 0 | 0 | 0 | 0 | 4 |
| 2 | -2.64 | 3.38 | -0.78 | 0.43 | 1 | Mir1938 | | 6 | 2 | 0 | 7 | 0 | 9 | 0 | 0 | 0 | 0 | 1 |
| 2 | 2.04 | 3.57 | 0.57 | 0.57 | 1 | Gm26509 | | 9 | 0 | 0 | 0 | 2 | 1 | 0 | 0 | 10 | 0 | 3 |
| 2 | 0.00 | 3.64 | 0.00 | 1.00 | 1 | Lrrc66 | leucine rich repeat containing 66 | 0 | 25 | 0 | 0 | 0 | 0 | 0 | 0 | 0 | 0 | 0 |
| 2 | 0.00 | 3.68 | 0.00 | 1.00 | 1 | Ifi27l2b | interferon, alpha-inducible protein 27 like | 13 | 1 | 11 | 0 | 0 | 0 | 0 | 0 | 0 | 0 | 0 |
| 2 | -2.87 | 2.60 | -1.10 | 0.27 | 1 | Ywhaq-ps3 | | 1 | 5 | 0 | 6 | 8 | 3 | 0 | 0 | 0 | 1 | 1 |
| 2 | 3.33 | 3.53 | 0.94 | 0.35 | 1 | Gm7580 | | 0 | 0 | 0 | 0 | 2 | 0 | 0 | 0 | 0 | 19 | 4 |
| 2 | 0.00 | 3.68 | 0.00 | 1.00 | 1 | Sox2 | SRY (sex determining region Y)-box 2 | 3 | 1 | 21 | 0 | 0 | 0 | 0 | 0 | 0 | 0 | 0 |
| 2 | -2.41 | 3.16 | -0.76 | 0.45 | 1 | Gm43110 | | 2 | 1 | 0 | 8 | 9 | 0 | 0 | 0 | 1 | 0 | 5 |
| 2 | -0.80 | 3.61 | -0.22 | 0.82 | 1 | Gm39121 | | 11 | 0 | 7 | 0 | 0 | 7 | 0 | 0 | 0 | 0 | 0 |
| 2 | 3.82 | 3.58 | 1.07 | 0.29 | 1 | 1700082M22Rik | | 0 | 7 | 0 | 0 | 0 | 6 | 0 | 0 | 5 | 0 | 7 |
| 2 | -1.61 | 3.32 | -0.48 | 0.63 | 1 | Gm26618 | | 4 | 9 | 8 | 2 | 0 | 2 | 0 | 0 | 0 | 0 | 0 |
| 2 | 1.16 | 3.38 | 0.34 | 0.73 | 1 | 9530053A07Ril | RIKEN cDNA 9530053A07 gene | 2 | 1 | 7 | 0 | 0 | 1 | 0 | 0 | 0 | 0 | 14 |
| 2 | 0.00 | 3.68 | 0.00 | 1.00 | 1 | Rufy4 | RUN and FYVE domain containing 4 | 4 | 1 | 20 | 0 | 0 | 0 | 0 | 0 | 0 | 0 | 0 |
| 2 | 2.35 | 2.98 | 0.79 | 0.43 | 1 | Gm13213 | | 0 | 13 | 1 | 1 | 0 | 1 | 0 | 0 | 5 | 0 | 4 |
| 2 | -0.22 | 3.55 | -0.06 | 0.95 | 1 | Gpr6 | G protein-coupled receptor 6 | 0 | 0 | 2 | 0 | 3 | 6 | 0 | 0 | 0 | 0 | 14 |
| 2 | 0.50 | 3.54 | 0.14 | 0.89 | 1 | Gm13783 | | 0 | 0 | 0 | 0 | 2 | 11 | 0 | 0 | 5 | 0 | 7 |
| 2 | 0.25 | 3.60 | 0.07 | 0.95 | 1 | Gm37960 | | 0 | 0 | 0 | 0 | 0 | 12 | 0 | 0 | 0 | 0 | 13 |
| 2 | -0.40 | 3.67 | -0.11 | 0.91 | 1 | Gm34611 | | 3 | 8 | 14 | 0 | 0 | 1 | 0 | 0 | 0 | 0 | 0 |
| 2 | 0.00 | 3.68 | 0.00 | 1.00 | 1 | Igkv5-43 | | 2 | 23 | 0 | 0 | 0 | 0 | 0 | 0 | 0 | 0 | 0 |
| 2 | 2.61 | 3.54 | 0.74 | 0.46 | 1 | Gm5340 | | 0 | 0 | 0 | 0 | 7 | 0 | 0 | 17 | 1 | 0 | 0 |
| 2 | -5.97 | 3.54 | -1.69 | 0.09 | 1 | 4933424G06Ril | RIKEN cDNA 4933424G06 gene | 0 | 0 | 0 | 9 | 0 | 10 | 0 | 0 | 6 | 0 | 0 |
| 2 | 0.87 | 3.61 | 0.24 | 0.81 | 1 | Ces5a | carboxylesterase 5A | 3 | 0 | 16 | 0 | 0 | 1 | 0 | 0 | 0 | 0 | 5 |
| 2 | 0.00 | 3.64 | 0.00 | 1.00 | 1 | 1700029F12Ril | RIKEN cDNA 1700029F12 gene | 0 | 0 | 25 | 0 | 0 | 0 | 0 | 0 | 0 | 0 | 0 |
| 2 | 0.26 | 3.44 | 0.08 | 0.94 | 1 | Gm19815 | | 0 | 4 | 8 | 0 | 9 | 0 | 0 | 0 | 1 | 0 | 3 |
| 2 | 0.00 | 3.68 | 0.00 | 1.00 | 1 | Ctrcos | | 15 | 3 | 7 | 0 | 0 | 0 | 0 | 0 | 0 | 0 | 0 |
| 2 | -0.33 | 2.56 | -0.13 | 0.90 | 1 | Rps2-ps5 | | 13 | 2 | 2 | 0 | 1 | 3 | 0 | 0 | 0 | 3 | 1 |
| 2 | 0.00 | 3.68 | 0.00 | 1.00 | 1 | Gpr84 | G protein-coupled receptor 84 | 4 | 7 | 14 | 0 | 0 | 0 | 0 | 0 | 0 | 0 | 0 |
| 2 | -1.46 | 2.27 | -0.65 | 0.52 | 1 | Gm11353 | | 2 | 2 | 1 | 8 | 4 | 3 | 0 | 0 | 3 | 0 | 2 |
| 2 | 0.00 | 3.65 | 0.00 | 1.00 | 1 | Hoxa10 | homeobox A10 | 25 | 0 | 0 | 0 | 0 | 0 | 0 | 0 | 0 | 0 | 0 |
| 2 | 0.00 | 3.65 | 0.00 | 1.00 | 1 | Cryaa | crystallin, alpha A | 25 | 0 | 0 | 0 | 0 | 0 | 0 | 0 | 0 | 0 | 0 |
| 2 | -0.76 | 3.63 | -0.21 | 0.83 | 1 | Chodl | chondrolectin | 16 | 0 | 5 | 0 | 0 | 3 | 0 | 0 | 0 | 0 | 0 |
| 2 | 0.92 | 3.61 | 0.26 | 0.80 | 1 | Vwa3a | von Willebrand factor A domain containin | 13 | 5 | 0 | 0 | 0 | 1 | 0 | 0 | 0 | 0 | 6 |
| 2 | 0.40 | 3.54 | 0.11 | 0.91 | 1 | 6030460B20Rik | | 0 | 0 | 8 | 0 | 0 | 8 | 0 | 0 | 6 | 0 | 2 |
| 2 | -0.15 | 3.60 | -0.04 | 0.97 | 1 | Gm37553 | | 0 | 0 | 0 | 0 | 0 | 18 | 0 | 0 | 0 | 0 | 7 |
| 2 | 0.00 | 3.68 | 0.00 | 1.00 | 1 | Gm36569 | | 3 | 22 | 0 | 0 | 0 | 0 | 0 | 0 | 0 | 0 | 0 |
| 2 | 0.80 | 3.04 | 0.26 | 0.79 | 1 | Gm13414 | | 0 | 1 | 1 | 0 | 7 | 3 | 0 | 0 | 0 | 8 | 5 |
| 2 | -0.96 | 3.53 | -0.27 | 0.79 | 1 | Gm50103 | | 0 | 0 | 0 | 19 | 0 | 3 | 0 | 0 | 1 | 0 | 2 |
| 2 | -0.44 | 3.24 | -0.13 | 0.89 | 1 | Gm11264 | | 0 | 0 | 0 | 2 | 6 | 4 | 0 | 0 | 5 | 0 | 8 |
| 2 | -0.16 | 3.39 | -0.05 | 0.96 | 1 | Gm37422 | | 13 | 0 | 5 | 0 | 1 | 2 | 0 | 0 | 0 | 0 | 4 |
| 2 | 1.82 | 3.60 | 0.50 | 0.61 | 1 | Lrrc7 | leucine rich repeat containing 7 | 0 | 12 | 0 | 0 | 0 | 0 | 0 | 12 | 0 | 0 | 1 |
| 2 | -0.53 | 3.63 | -0.15 | 0.88 | 1 | Iqcm | IQ motif containing M | 0 | 0 | 24 | 0 | 0 | 1 | 0 | 0 | 0 | 0 | 0 |
| 2 | 0.00 | 3.68 | 0.00 | 1.00 | 1 | Gm43101 | | 21 | 0 | 4 | 0 | 0 | 0 | 0 | 0 | 0 | 0 | 0 |

| | | | | | | | | | | | | | | | | | | |
|---|---|---|---|---|---|---|---|---|---|---|---|---|---|---|---|---|---|---|
| 2 | -0.51 | 3.49 | -0.15 | 0.88 | 1 | Gm14227 | | 0 | 1 | 0 | 0 | 8 | 1 | 0 | 1 | 0 | 14 | 0 |
| 2 | 1.45 | 3.01 | 0.48 | 0.63 | 1 | Gm17190 | | 0 | 0 | 18 | 1 | 1 | 0 | 0 | 1 | 1 | 0 | 3 |
| 2 | 0.00 | 3.68 | 0.00 | 1.00 | 1 | C330024D21Rik | | 3 | 21 | 0 | 0 | 0 | 0 | 0 | 0 | 0 | 0 | 0 |
| 2 | 0.13 | 3.60 | 0.04 | 0.97 | 1 | Gm38067 | | 0 | 0 | 0 | 0 | 0 | 14 | 0 | 0 | 0 | 0 | 11 |
| 2 | -0.24 | 2.42 | -0.10 | 0.92 | 1 | Gm9619 | | 1 | 2 | 10 | 0 | 6 | 2 | 0 | 0 | 2 | 0 | 2 |
| 2 | 0.00 | 3.68 | 0.00 | 1.00 | 1 | 4930430E12Rik | | 5 | 8 | 11 | 0 | 0 | 0 | 0 | 0 | 0 | 0 | 0 |
| 2 | -0.78 | 3.62 | -0.21 | 0.83 | 1 | Gm13330 | | 0 | 13 | 7 | 0 | 0 | 4 | 0 | 0 | 0 | 0 | 0 |
| 2 | 3.32 | 3.56 | 0.93 | 0.35 | 1 | Gm49168 | | 9 | 1 | 0 | 1 | 0 | 0 | 0 | 0 | 4 | 0 | 10 |
| 2 | -1.37 | 3.53 | -0.39 | 0.70 | 1 | Cntnap5a | contactin associated protein-like 5A | 7 | 0 | 0 | 16 | 0 | 0 | 0 | 0 | 2 | 0 | 0 |
| 2 | -3.52 | 3.07 | -1.15 | 0.25 | 1 | Gm15387 | | 10 | 1 | 0 | 5 | 3 | 4 | 0 | 0 | 0 | 0 | 1 |
| 2 | -2.83 | 3.59 | -0.79 | 0.43 | 1 | Gm29724 | | 9 | 0 | 5 | 0 | 7 | 3 | 0 | 0 | 0 | 0 | 0 |
| 2 | 0.00 | 3.68 | 0.00 | 1.00 | 1 | Spata20 | spermatogenesis associated 20 | 5 | 10 | 9 | 0 | 0 | 0 | 0 | 0 | 0 | 0 | 0 |
| 2 | 0.00 | 3.21 | 0.00 | 1.00 | 1 | Izumo1 | izumo sperm-egg fusion 1 | 0 | 0 | 9 | 0 | 7 | 3 | 0 | 0 | 3 | 0 | 3 |
| 2 | 1.32 | 2.93 | 0.45 | 0.65 | 1 | Gm7286 | | 1 | 1 | 19 | 0 | 1 | 0 | 0 | 0 | 1 | 0 | 2 |
| 2 | 0.00 | 3.68 | 0.00 | 1.00 | 1 | Cyp4x1os | | 6 | 7 | 11 | 0 | 0 | 0 | 0 | 0 | 0 | 0 | 0 |
| 2 | 0.06 | 3.60 | 0.02 | 0.99 | 1 | Gm23205 | | 0 | 0 | 0 | 0 | 0 | 14 | 0 | 0 | 0 | 0 | 10 |
| 2 | -0.30 | 3.60 | -0.08 | 0.93 | 1 | Gm43879 | | 0 | 0 | 0 | 0 | 0 | 19 | 0 | 0 | 0 | 0 | 5 |
| 2 | 0.00 | 3.68 | 0.00 | 1.00 | 1 | Gm36535 | | 6 | 8 | 10 | 0 | 0 | 0 | 0 | 0 | 0 | 0 | 0 |
| 2 | 2.14 | 3.55 | 0.60 | 0.55 | 1 | Gm42717 | | 0 | 0 | 0 | 0 | 0 | 14 | 0 | 0 | 7 | 0 | 3 |
| 2 | 0.00 | 3.68 | 0.00 | 1.00 | 1 | Gm43618 | | 1 | 10 | 14 | 0 | 0 | 0 | 0 | 0 | 0 | 0 | 0 |
| 2 | -1.31 | 3.59 | -0.36 | 0.72 | 1 | Gm48375 | | 0 | 15 | 0 | 0 | 0 | 9 | 0 | 0 | 0 | 0 | 0 |
| 2 | -2.89 | 3.40 | -0.85 | 0.40 | 1 | C230034O21Rik | | 4 | 5 | 3 | 4 | 9 | 0 | 0 | 0 | 0 | 0 | 0 |
| 2 | 0.00 | 3.68 | 0.00 | 1.00 | 1 | BC048559 | | 0 | 18 | 6 | 0 | 0 | 0 | 0 | 0 | 0 | 0 | 0 |
| 2 | -3.40 | 3.56 | -0.95 | 0.34 | 1 | Gm45413 | | 0 | 0 | 0 | 0 | 12 | 5 | 0 | 0 | 0 | 0 | 7 |
| 2 | -0.40 | 3.67 | -0.11 | 0.91 | 1 | Rgs13 | regulator of G-protein signaling 13 | 9 | 14 | 0 | 0 | 0 | 1 | 0 | 0 | 0 | 0 | 0 |
| 2 | 0.28 | 3.56 | 0.08 | 0.94 | 1 | Gng13 | guanine nucleotide binding protein (G prc | 0 | 0 | 0 | 0 | 1 | 14 | 0 | 0 | 2 | 0 | 7 |
| 2 | 0.00 | 3.68 | 0.00 | 1.00 | 1 | D630011A20Rik | | 0 | 20 | 4 | 0 | 0 | 0 | 0 | 0 | 0 | 0 | 0 |
| 2 | 0.00 | 3.68 | 0.00 | 1.00 | 1 | Akr1c20 | aldo-keto reductase family 1, member C2 | 3 | 1 | 21 | 0 | 0 | 0 | 0 | 0 | 0 | 0 | 0 |
| 2 | 0.34 | 3.60 | 0.09 | 0.92 | 1 | Gm44735 | | 0 | 0 | 0 | 0 | 0 | 10 | 0 | 0 | 0 | 0 | 14 |
| 2 | 3.68 | 3.59 | 1.03 | 0.31 | 1 | Gm17873 | | 0 | 1 | 1 | 0 | 0 | 2 | 0 | 0 | 0 | 21 | 0 |
| 2 | -1.95 | 3.06 | -0.64 | 0.52 | 1 | Gm42917 | | 4 | 0 | 4 | 1 | 5 | 3 | 0 | 0 | 0 | 8 | 0 |
| 2 | -2.89 | 3.40 | -0.85 | 0.39 | 1 | Tomt | transmembrane O-methyltransferase | 3 | 3 | 6 | 0 | 6 | 7 | 0 | 0 | 0 | 0 | 0 |
| 2 | -2.23 | 3.37 | -0.66 | 0.51 | 1 | Gm22877 | | 0 | 0 | 9 | 4 | 4 | 2 | 0 | 0 | 5 | 0 | 0 |
| 2 | 4.78 | 3.59 | 1.33 | 0.18 | 1 | Gm43637 | | 0 | 4 | 0 | 0 | 0 | 2 | 0 | 0 | 0 | 15 | 3 |
| 2 | -0.62 | 2.96 | -0.21 | 0.83 | 1 | Cfap53 | cilia and flagella associated protein 53 | 3 | 3 | 4 | 0 | 6 | 2 | 0 | 0 | 0 | 0 | 7 |
| 2 | 3.08 | 3.53 | 0.87 | 0.38 | 1 | Gm6611 | | 0 | 10 | 2 | 0 | 0 | 6 | 0 | 0 | 4 | 0 | 2 |
| 2 | 0.00 | 3.68 | 0.00 | 1.00 | 1 | 4732419C18Rik | | 12 | 11 | 1 | 0 | 0 | 0 | 0 | 0 | 0 | 0 | 0 |
| 2 | 1.96 | 3.60 | 0.55 | 0.59 | 1 | Hgd | homogentisate 1, 2-dioxygenase | 0 | 8 | 0 | 0 | 0 | 0 | 0 | 15 | 0 | 0 | 1 |
| 2 | 3.05 | 3.55 | 0.86 | 0.39 | 1 | Gm5117 | | 0 | 0 | 0 | 0 | 0 | 1 | 0 | 16 | 0 | 0 | 7 |
| 2 | 0.00 | 3.68 | 0.00 | 1.00 | 1 | Sec1 | secretory blood group 1 | 9 | 0 | 16 | 0 | 0 | 0 | 0 | 0 | 0 | 0 | 0 |
| 2 | 0.00 | 3.68 | 0.00 | 1.00 | 1 | Crtam | cytotoxic and regulatory T cell molecule | 9 | 0 | 16 | 0 | 0 | 0 | 0 | 0 | 0 | 0 | 0 |
| 2 | 3.15 | 3.52 | 0.89 | 0.37 | 1 | Fezf2 | Fez family zinc finger 2 | 0 | 0 | 0 | 0 | 0 | 3 | 0 | 19 | 3 | 0 | 0 |
| 2 | -0.51 | 3.05 | -0.17 | 0.87 | 1 | Gm2805 | | 12 | 5 | 2 | 0 | 1 | 2 | 0 | 2 | 0 | 0 | 0 |
| 2 | 0.00 | 3.68 | 0.00 | 1.00 | 1 | Trim30b | tripartite motif-containing 30B | 2 | 18 | 4 | 0 | 0 | 0 | 0 | 0 | 0 | 0 | 0 |
| 2 | -0.02 | 3.35 | 0.00 | 1.00 | 1 | Snord55 | | 2 | 0 | 4 | 0 | 1 | 5 | 0 | 0 | 0 | 0 | 12 |
| 2 | -3.94 | 3.52 | -1.12 | 0.26 | 1 | Spaca4 | sperm acrosome associated 4 | 9 | 0 | 0 | 8 | 0 | 2 | 0 | 0 | 5 | 0 | 0 |
| 2 | 0.21 | 3.60 | 0.06 | 0.95 | 1 | Gm17082 | | 1 | 0 | 1 | 0 | 0 | 11 | 0 | 0 | 0 | 0 | 11 |
| 2 | 0.00 | 3.65 | 0.00 | 1.00 | 1 | Myh3 | myosin, heavy polypeptide 3, skeletal mu | 24 | 0 | 0 | 0 | 0 | 0 | 0 | 0 | 0 | 0 | 0 |
| 2 | 0.00 | 3.68 | 0.00 | 1.00 | 1 | Asic4 | acid-sensing (proton-gated) ion channel f | 13 | 11 | 0 | 0 | 0 | 0 | 0 | 0 | 0 | 0 | 0 |
| 2 | -0.11 | 3.58 | -0.03 | 0.98 | 1 | 9930024M15Rik | | 0 | 13 | 0 | 0 | 0 | 7 | 0 | 0 | 0 | 0 | 4 |
| 2 | 4.16 | 3.57 | 1.16 | 0.24 | 1 | Gm33373 | | 0 | 1 | 0 | 0 | 0 | 7 | 0 | 0 | 0 | 14 | 2 |
| 2 | 0.50 | 3.61 | 0.14 | 0.89 | 1 | Gm47122 | | 5 | 0 | 8 | 0 | 0 | 3 | 0 | 0 | 0 | 0 | 7 |
| 2 | -0.64 | 3.52 | -0.18 | 0.86 | 1 | Gm12917 | | 0 | 0 | 0 | 3 | 0 | 15 | 0 | 0 | 3 | 0 | 3 |
| 2 | -4.61 | 3.54 | -1.30 | 0.19 | 1 | Gm37229 | | 0 | 0 | 0 | 5 | 1 | 3 | 0 | 0 | 15 | 0 | 0 |
| 2 | 0.00 | 3.64 | 0.00 | 1.00 | 1 | Gm11732 | | 0 | 0 | 24 | 0 | 0 | 0 | 0 | 0 | 0 | 0 | 0 |
| 2 | 3.86 | 3.57 | 1.08 | 0.28 | 1 | Thegl | theg spermatid protein like | 0 | 0 | 0 | 0 | 0 | 0 | 0 | 21 | 0 | 0 | 3 |
| 2 | -1.53 | 3.55 | -0.43 | 0.67 | 1 | H2bc11 | H2B clustered histone 11 | 0 | 0 | 3 | 0 | 9 | 9 | 0 | 0 | 0 | 0 | 2 |
| 2 | -0.83 | 2.58 | -0.32 | 0.75 | 1 | Rps23-ps2 | | 3 | 1 | 16 | 1 | 1 | 1 | 0 | 0 | 2 | 0 | 0 |
| 2 | -0.56 | 3.52 | -0.16 | 0.87 | 1 | Gm38336 | | 0 | 0 | 0 | 0 | 4 | 13 | 0 | 2 | 0 | 0 | 5 |
| 2 | 1.13 | 3.59 | 0.32 | 0.75 | 1 | B230362B09Rik | | 0 | 0 | 0 | 0 | 0 | 13 | 0 | 0 | 2 | 0 | 9 |

| | | | | | | | | | | | | | | | | | | |
|---|---|---|---|---|---|---|---|---|---|---|---|---|---|---|---|---|---|---|
| 2 | 1.77 | 3.56 | 0.50 | 0.62 | 1 | Gm37268 | | 0 | 0 | 0 | 0 | 0 | 8 | 0 | 0 | 8 | 0 | 8 |
| 2 | 1.34 | 3.02 | 0.44 | 0.66 | 1 | Gm4795 | | 0 | 16 | 0 | 2 | 0 | 3 | 0 | 0 | 1 | 1 | 1 |
| 2 | 1.81 | 3.61 | 0.50 | 0.62 | 1 | Klk1b24 | kallikrein 1-related peptidase b24 | 0 | 0 | 20 | 0 | 0 | 0 | 0 | 0 | 0 | 0 | 4 |
| 2 | 0.00 | 3.68 | 0.00 | 1.00 | 1 | Bdkrb1 | bradykinin receptor, beta 1 | 10 | 0 | 14 | 0 | 0 | 0 | 0 | 0 | 0 | 0 | 0 |
| 2 | 0.98 | 3.54 | 0.28 | 0.78 | 1 | AU017674 | | 0 | 0 | 0 | 0 | 8 | 0 | 0 | 0 | 6 | 0 | 10 |
| 2 | 2.30 | 3.57 | 0.65 | 0.52 | 1 | Gm12040 | | 0 | 0 | 0 | 1 | 0 | 0 | 0 | 0 | 1 | 22 | 0 |
| 2 | 0.00 | 3.68 | 0.00 | 1.00 | 1 | Arl14 | ADP-ribosylation factor-like 14 | 9 | 15 | 0 | 0 | 0 | 0 | 0 | 0 | 0 | 0 | 0 |
| 2 | 1.48 | 2.94 | 0.50 | 0.61 | 1 | Gm6336 | | 3 | 1 | 9 | 0 | 0 | 2 | 0 | 1 | 0 | 0 | 7 |
| 2 | 0.00 | 3.68 | 0.00 | 1.00 | 1 | Slc30a3 | solute carrier family 30 (zinc transporter), | 10 | 2 | 11 | 0 | 0 | 0 | 0 | 0 | 0 | 0 | 0 |
| 2 | -2.01 | 3.55 | -0.57 | 0.57 | 1 | A530064N14Rik | | 0 | 0 | 0 | 5 | 2 | 3 | 0 | 0 | 0 | 0 | 13 |
| 2 | 0.00 | 3.68 | 0.00 | 1.00 | 1 | Arhgap27os3 | | 1 | 0 | 23 | 0 | 0 | 0 | 0 | 0 | 0 | 0 | 0 |
| 2 | -0.42 | 3.55 | -0.12 | 0.91 | 1 | Gm43627 | | 0 | 0 | 7 | 0 | 2 | 8 | 0 | 0 | 0 | 0 | 6 |
| 2 | 0.00 | 3.68 | 0.00 | 1.00 | 1 | Ren1 | renin 1 structural | 21 | 0 | 3 | 0 | 0 | 0 | 0 | 0 | 0 | 0 | 0 |
| 2 | 0.44 | 3.60 | 0.12 | 0.90 | 1 | Gm22728 | | 0 | 0 | 0 | 0 | 0 | 8 | 0 | 0 | 0 | 0 | 15 |
| 2 | -1.52 | 3.52 | -0.43 | 0.67 | 1 | Gm37933 | | 0 | 0 | 0 | 15 | 2 | 1 | 0 | 0 | 6 | 0 | 0 |
| 2 | 0.09 | 3.60 | 0.02 | 0.98 | 1 | Khdc3 | KH domain containing 3, subcortical mate | 0 | 0 | 0 | 0 | 0 | 14 | 0 | 0 | 0 | 0 | 10 |
| 2 | 0.09 | 3.60 | 0.02 | 0.98 | 1 | Gm19046 | | 0 | 0 | 0 | 0 | 0 | 14 | 0 | 0 | 0 | 0 | 10 |
| 2 | -0.82 | 3.45 | -0.24 | 0.81 | 1 | Gm13496 | | 13 | 0 | 5 | 0 | 2 | 2 | 0 | 0 | 0 | 0 | 2 |
| 2 | 2.09 | 3.62 | 0.58 | 0.56 | 1 | Spata25 | spermatogenesis associated 25 | 0 | 4 | 15 | 0 | 0 | 0 | 0 | 0 | 1 | 0 | 4 |
| 2 | 5.92 | 3.54 | 1.67 | 0.09 | 1 | Uroc1 | urocanase domain containing 1 | 0 | 0 | 0 | 7 | 0 | 0 | 0 | 0 | 0 | 12 | 5 |
| 2 | 1.88 | 3.59 | 0.52 | 0.60 | 1 | Pou5f2 | POU domain class 5, transcription factor 2 | 0 | 0 | 14 | 0 | 0 | 0 | 0 | 0 | 0 | 0 | 10 |
| 2 | 0.55 | 3.36 | 0.16 | 0.87 | 1 | Gm47467 | | 4 | 5 | 11 | 0 | 0 | 1 | 0 | 0 | 0 | 0 | 2 |
| 2 | -0.25 | 3.60 | -0.07 | 0.94 | 1 | Gm37639 | | 1 | 0 | 5 | 0 | 0 | 14 | 0 | 0 | 0 | 0 | 4 |
| 2 | -1.07 | 3.00 | -0.36 | 0.72 | 1 | Gm48557 | | 1 | 15 | 1 | 2 | 4 | 0 | 0 | 0 | 1 | 0 | 0 |
| 2 | 0.00 | 3.68 | 0.00 | 1.00 | 1 | Slc13a5 | solute carrier family 13 (sodium-depende | 11 | 0 | 12 | 0 | 0 | 0 | 0 | 0 | 0 | 0 | 0 |
| 2 | -0.61 | 2.93 | -0.21 | 0.84 | 1 | Gm10371 | | 1 | 1 | 10 | 0 | 7 | 2 | 0 | 0 | 0 | 0 | 3 |
| 2 | -4.90 | 3.60 | -1.36 | 0.17 | 1 | 0610040F04Rik | | 0 | 3 | 0 | 0 | 17 | 3 | 0 | 0 | 0 | 0 | 1 |
| 2 | 0.95 | 3.67 | 0.26 | 0.80 | 1 | Tfap2a | transcription factor AP-2, alpha | 11 | 0 | 11 | 0 | 0 | 0 | 0 | 0 | 0 | 0 | 1 |
| 2 | 4.47 | 3.58 | 1.25 | 0.21 | 1 | Rdh7 | retinol dehydrogenase 7 | 7 | 0 | 0 | 0 | 0 | 3 | 0 | 0 | 6 | 0 | 8 |
| 2 | -0.15 | 3.33 | -0.05 | 0.96 | 1 | Phyhd1 | phytanoyl-CoA dioxygenase domain cont | 5 | 4 | 7 | 0 | 0 | 5 | 0 | 0 | 0 | 0 | 2 |
| 2 | 0.00 | 3.68 | 0.00 | 1.00 | 1 | Slc22a14 | solute carrier family 22 (organic cation tra | 16 | 0 | 7 | 0 | 0 | 0 | 0 | 0 | 0 | 0 | 0 |
| 2 | 1.30 | 3.64 | 0.36 | 0.72 | 1 | Gm44860 | | 0 | 10 | 10 | 0 | 0 | 0 | 0 | 0 | 0 | 0 | 3 |
| 2 | 1.34 | 3.62 | 0.37 | 0.71 | 1 | Nxph2 | neurexophilin 2 | 0 | 9 | 9 | 0 | 0 | 0 | 0 | 0 | 0 | 0 | 5 |
| 2 | -0.40 | 3.67 | -0.11 | 0.91 | 1 | Vpreb3 | pre-B lymphocyte gene 3 | 15 | 8 | 0 | 0 | 0 | 1 | 0 | 0 | 0 | 0 | 0 |
| 2 | -0.56 | 3.65 | -0.15 | 0.88 | 1 | Bfsp2 | beaded filament structural protein 2, phak | 0 | 7 | 15 | 0 | 2 | 0 | 0 | 0 | 0 | 0 | 0 |
| 2 | 0.28 | 3.64 | 0.08 | 0.94 | 1 | Gm15396 | | 3 | 18 | 0 | 0 | 0 | 1 | 0 | 0 | 0 | 0 | 1 |
| 2 | 0.02 | 3.60 | 0.01 | 0.99 | 1 | Gm37105 | | 0 | 0 | 0 | 0 | 0 | 14 | 0 | 0 | 0 | 0 | 9 |
| 2 | -3.71 | 3.62 | -1.02 | 0.31 | 1 | Gm32624 | | 0 | 17 | 0 | 0 | 2 | 3 | 0 | 0 | 0 | 0 | 1 |
| 2 | -0.37 | 3.60 | -0.10 | 0.92 | 1 | Gm43061 | | 0 | 0 | 0 | 0 | 0 | 19 | 0 | 0 | 0 | 0 | 4 |
| 2 | 0.34 | 3.58 | 0.09 | 0.93 | 1 | Snord7 | | 0 | 13 | 0 | 0 | 0 | 4 | 0 | 0 | 0 | 0 | 6 |
| 2 | 1.34 | 3.62 | 0.37 | 0.71 | 1 | Gm48449 | | 0 | 12 | 6 | 0 | 0 | 0 | 0 | 0 | 0 | 0 | 5 |
| 2 | 0.00 | 3.68 | 0.00 | 1.00 | 1 | Paqr5 | progestin and adipoQ receptor family mer | 12 | 0 | 11 | 0 | 0 | 0 | 0 | 0 | 0 | 0 | 0 |
| 2 | -4.26 | 3.63 | -1.17 | 0.24 | 1 | Mst1 | macrophage stimulating 1 (hepatocyte gr | 0 | 12 | 0 | 0 | 1 | 9 | 0 | 0 | 0 | 0 | 1 |
| 2 | -1.72 | 2.91 | -0.59 | 0.55 | 1 | Gm10566 | | 0 | 0 | 12 | 2 | 4 | 3 | 0 | 0 | 1 | 0 | 1 |
| 2 | 0.00 | 3.68 | 0.00 | 1.00 | 1 | Slitrk4 | SLIT and NTRK-like family, member 4 | 2 | 5 | 17 | 0 | 0 | 0 | 0 | 0 | 0 | 0 | 0 |
| 2 | -2.45 | 3.00 | -0.82 | 0.41 | 1 | Gm12033 | | 1 | 16 | 1 | 1 | 2 | 3 | 0 | 0 | 0 | 0 | 0 |
| 2 | 1.30 | 3.59 | 0.36 | 0.72 | 1 | Gm44091 | | 4 | 0 | 0 | 0 | 0 | 6 | 0 | 0 | 1 | 0 | 12 |
| 2 | -4.82 | 3.59 | -1.34 | 0.18 | 1 | Gm7622 | | 3 | 0 | 8 | 9 | 2 | 0 | 0 | 0 | 0 | 0 | 1 |
| 2 | 0.74 | 3.52 | 0.21 | 0.83 | 1 | Gm30934 | | 0 | 0 | 0 | 5 | 0 | 3 | 0 | 13 | 2 | 0 | 0 |
| 2 | -0.76 | 3.63 | -0.21 | 0.83 | 1 | Cep295nl | CEP295 N-terminal like | 9 | 0 | 11 | 0 | 0 | 3 | 0 | 0 | 0 | 0 | 0 |
| 2 | 1.41 | 3.67 | 0.38 | 0.70 | 1 | | | 16 | 5 | 0 | 0 | 0 | 0 | 0 | 0 | 2 | 0 | 0 |
| 2 | 0.28 | 3.64 | 0.08 | 0.94 | 1 | Shisa8 | shisa family member 8 | 4 | 17 | 0 | 0 | 0 | 1 | 0 | 0 | 0 | 0 | 1 |
| 2 | -0.36 | 3.61 | -0.10 | 0.92 | 1 | Gm49195 | | 0 | 15 | 2 | 0 | 0 | 5 | 0 | 0 | 0 | 0 | 1 |
| 2 | 1.30 | 3.58 | 0.36 | 0.72 | 1 | Tpt1-ps7 | | 1 | 1 | 0 | 1 | 0 | 0 | 0 | 0 | 0 | 21 | 0 |
| 2 | 3.69 | 3.58 | 1.03 | 0.30 | 1 | Gm38162 | | 2 | 0 | 0 | 0 | 0 | 8 | 0 | 0 | 5 | 0 | 8 |
| 2 | -1.28 | 3.40 | -0.38 | 0.71 | 1 | Gm4322 | | 6 | 2 | 0 | 3 | 0 | 8 | 0 | 0 | 0 | 0 | 5 |
| 2 | 0.16 | 3.60 | 0.04 | 0.96 | 1 | Gm44510 | | 0 | 4 | 0 | 0 | 0 | 10 | 0 | 0 | 0 | 0 | 9 |
| 2 | -0.11 | 3.58 | -0.03 | 0.98 | 1 | 4930447M23Rik | | 0 | 15 | 0 | 0 | 0 | 5 | 0 | 0 | 0 | 0 | 3 |
| 2 | 1.71 | 3.60 | 0.48 | 0.63 | 1 | Gm38299 | | 0 | 18 | 0 | 0 | 0 | 0 | 0 | 0 | 0 | 0 | 5 |
| 2 | 0.00 | 3.64 | 0.00 | 1.00 | 1 | Grin3a | glutamate receptor ionotropic, NMDA3A | 0 | 23 | 0 | 0 | 0 | 0 | 0 | 0 | 0 | 0 | 0 |

| | | | | | | | | | | | | | | | | | |
|---|---|---|---|---|---|---|---|---|---|---|---|---|---|---|---|---|---|
| 2 | 0.00 | 3.64 | 0.00 | 1.00 | 1 | Ighv1-4 | | 0 | 23 | 0 | 0 | 0 | 0 | 0 | 0 | 0 | 0 | 0 |
| 2 | 2.54 | 3.24 | 0.78 | 0.43 | 1 | Gm6157 | | 0 | 10 | 0 | 1 | 5 | 0 | 0 | 1 | 0 | 1 | 5 |
| 2 | -0.50 | 3.54 | -0.14 | 0.89 | 1 | Gm42783 | | 3 | 0 | 0 | 3 | 0 | 14 | 0 | 3 | 0 | 0 | 1 |
| 2 | -4.52 | 3.55 | -1.27 | 0.20 | 1 | Gm11629 | | 3 | 0 | 1 | 11 | 6 | 3 | 0 | 0 | 0 | 0 | 0 |
| 2 | 0.00 | 3.68 | 0.00 | 1.00 | 1 | 1700012C14Rik | | 1 | 20 | 2 | 0 | 0 | 0 | 0 | 0 | 0 | 0 | 0 |
| 2 | -3.60 | 3.59 | -1.00 | 0.32 | 1 | Gm8738 | | 4 | 0 | 0 | 0 | 3 | 11 | 0 | 0 | 0 | 0 | 5 |
| 2 | 0.63 | 3.60 | 0.17 | 0.86 | 1 | Gm43633 | | 0 | 0 | 0 | 0 | 0 | 6 | 0 | 0 | 0 | 0 | 17 |
| 2 | 0.00 | 3.65 | 0.00 | 1.00 | 1 | Gm8818 | | 23 | 0 | 0 | 0 | 0 | 0 | 0 | 0 | 0 | 0 | 0 |
| 2 | 0.00 | 3.65 | 0.00 | 1.00 | 1 | Smim40 | small integral membrane protein 40 | 23 | 0 | 0 | 0 | 0 | 0 | 0 | 0 | 0 | 0 | 0 |
| 2 | 0.00 | 3.65 | 0.00 | 1.00 | 1 | Ndp | Norrie disease (pseudoglioma) (human) | 23 | 0 | 0 | 0 | 0 | 0 | 0 | 0 | 0 | 0 | 0 |
| 2 | 0.00 | 3.68 | 0.00 | 1.00 | 1 | Cideb | cell death-inducing DNA fragmentation fa | 8 | 6 | 9 | 0 | 0 | 0 | 0 | 0 | 0 | 0 | 0 |
| 2 | -0.62 | 3.64 | -0.17 | 0.87 | 1 | Gm36447 | | 0 | 22 | 0 | 0 | 0 | 1 | 0 | 0 | 0 | 0 | 0 |
| 2 | -0.62 | 3.64 | -0.17 | 0.87 | 1 | Gm39059 | | 0 | 22 | 0 | 0 | 0 | 1 | 0 | 0 | 0 | 0 | 0 |
| 2 | 3.36 | 3.53 | 0.95 | 0.34 | 1 | 1700122E12Rik | | 0 | 0 | 0 | 0 | 0 | 4 | 0 | 0 | 5 | 8 | 6 |
| 2 | 0.80 | 3.54 | 0.23 | 0.82 | 1 | Gm29791 | | 0 | 0 | 0 | 0 | 3 | 5 | 0 | 0 | 9 | 0 | 6 |
| 2 | -2.25 | 3.61 | -0.62 | 0.53 | 1 | Cdh15 | cadherin 15 | 10 | 0 | 7 | 2 | 4 | 0 | 0 | 0 | 0 | 0 | 0 |
| 2 | -0.23 | 2.52 | -0.09 | 0.93 | 1 | Gm6341 | | 3 | 0 | 6 | 7 | 1 | 1 | 0 | 1 | 0 | 1 | 3 |
| 2 | 0.79 | 3.62 | 0.22 | 0.83 | 1 | Gm5920 | | 17 | 1 | 0 | 0 | 0 | 1 | 0 | 0 | 0 | 0 | 4 |
| 2 | 4.31 | 3.56 | 1.21 | 0.23 | 1 | Gm42567 | | 0 | 2 | 0 | 0 | 0 | 0 | 0 | 9 | 4 | 8 | 0 |
| 2 | -0.69 | 3.66 | -0.19 | 0.85 | 1 | Gm47108 | | 0 | 17 | 4 | 0 | 0 | 2 | 0 | 0 | 0 | 0 | 0 |
| 2 | 0.00 | 3.68 | 0.00 | 1.00 | 1 | Gpr156 | G protein-coupled receptor 156 | 9 | 3 | 11 | 0 | 0 | 0 | 0 | 0 | 0 | 0 | 0 |
| 2 | 0.56 | 2.39 | 0.23 | 0.82 | 1 | Gm4859 | | 9 | 2 | 2 | 2 | 3 | 0 | 0 | 3 | 0 | 1 | 1 |
| 2 | 0.56 | 3.60 | 0.15 | 0.88 | 1 | Snord98 | | 0 | 0 | 0 | 0 | 0 | 7 | 0 | 0 | 0 | 0 | 16 |
| 2 | 3.57 | 3.16 | 1.13 | 0.26 | 1 | Gm16726 | | 3 | 1 | 0 | 0 | 0 | 5 | 0 | 9 | 4 | 0 | 1 |
| 2 | -0.07 | 3.58 | -0.02 | 0.99 | 1 | Gstp2 | glutathione S-transferase, pi 2 | 17 | 0 | 0 | 1 | 0 | 1 | 0 | 0 | 0 | 0 | 4 |
| 2 | -0.66 | 1.94 | -0.34 | 0.73 | 1 | Gm6311 | | 3 | 2 | 1 | 1 | 6 | 4 | 0 | 2 | 1 | 1 | 1 |
| 2 | 1.34 | 3.63 | 0.37 | 0.71 | 1 | 2900076G11Rik | | 4 | 0 | 15 | 0 | 0 | 0 | 0 | 0 | 0 | 0 | 4 |
| 2 | 0.00 | 3.68 | 0.00 | 1.00 | 1 | Glb1l3 | galactosidase, beta 1 like 3 | 15 | 0 | 8 | 0 | 0 | 0 | 0 | 0 | 0 | 0 | 0 |
| 2 | 3.00 | 3.55 | 0.84 | 0.40 | 1 | Gm26736 | | 3 | 3 | 6 | 0 | 0 | 0 | 0 | 0 | 0 | 9 | 2 |
| 2 | 0.00 | 3.68 | 0.00 | 1.00 | 1 | Gm46224 | | 3 | 11 | 8 | 0 | 0 | 0 | 0 | 0 | 0 | 0 | 0 |
| 2 | -1.94 | 3.52 | -0.55 | 0.58 | 1 | Mettl4-ps1 | | 4 | 1 | 7 | 0 | 9 | 1 | 0 | 0 | 0 | 0 | 0 |
| 2 | 1.22 | 3.58 | 0.34 | 0.73 | 1 | Gm14327 | | 3 | 10 | 0 | 0 | 0 | 4 | 0 | 0 | 2 | 0 | 3 |
| 2 | 0.97 | 3.60 | 0.27 | 0.79 | 1 | A930028O11Rik | | 0 | 0 | 0 | 0 | 0 | 3 | 0 | 0 | 0 | 0 | 20 |
| 2 | 0.11 | 3.60 | 0.03 | 0.97 | 1 | Gm22063 | | 0 | 0 | 0 | 0 | 0 | 13 | 0 | 0 | 0 | 0 | 10 |
| 2 | -0.27 | 3.60 | -0.07 | 0.94 | 1 | Mir1188 | | 0 | 0 | 0 | 0 | 0 | 18 | 0 | 0 | 0 | 0 | 5 |
| 2 | 0.51 | 3.01 | 0.17 | 0.86 | 1 | | | 6 | 0 | 4 | 0 | 5 | 1 | 0 | 0 | 4 | 0 | 3 |
| 2 | 0.88 | 2.63 | 0.33 | 0.74 | 1 | Gm6649 | | 7 | 1 | 0 | 0 | 2 | 3 | 0 | 0 | 2 | 3 | 5 |
| 2 | 0.00 | 3.68 | 0.00 | 1.00 | 1 | Gm10824 | | 20 | 1 | 2 | 0 | 0 | 0 | 0 | 0 | 0 | 0 | 0 |
| 2 | -0.05 | 3.59 | -0.01 | 0.99 | 1 | Gm45877 | | 0 | 10 | 0 | 0 | 0 | 8 | 0 | 0 | 0 | 0 | 5 |
| 2 | -0.76 | 3.63 | -0.21 | 0.83 | 1 | I830134H01Rik | | 0 | 12 | 7 | 0 | 0 | 3 | 0 | 0 | 0 | 0 | 0 |
| 2 | -1.18 | 3.62 | -0.32 | 0.75 | 1 | E330020D12Rik | | 0 | 20 | 0 | 0 | 0 | 3 | 0 | 0 | 0 | 0 | 0 |
| 2 | 1.34 | 3.66 | 0.37 | 0.71 | 1 | Cyp4f15 | cytochrome P450, family 4, subfamily f, po | 15 | 2 | 4 | 0 | 0 | 0 | 0 | 0 | 2 | 0 | 0 |
| 2 | 4.41 | 3.59 | 1.23 | 0.22 | 1 | 4930470H14Rik | | 0 | 0 | 0 | 0 | 0 | 2 | 0 | 0 | 0 | 18 | 3 |
| 2 | 0.00 | 3.68 | 0.00 | 1.00 | 1 | Ces2a | carboxylesterase 2A | 0 | 4 | 19 | 0 | 0 | 0 | 0 | 0 | 0 | 0 | 0 |
| 2 | 0.00 | 3.68 | 0.00 | 1.00 | 1 | Acer1 | alkaline ceramidase 1 | 0 | 4 | 19 | 0 | 0 | 0 | 0 | 0 | 0 | 0 | 0 |
| 2 | 1.94 | 3.53 | 0.55 | 0.58 | 1 | Cntnap4 | contactin associated protein-like 4 | 0 | 3 | 14 | 0 | 0 | 1 | 0 | 2 | 0 | 0 | 3 |
| 2 | 0.00 | 3.68 | 0.00 | 1.00 | 1 | Gm4610 | | 4 | 10 | 8 | 0 | 0 | 0 | 0 | 0 | 0 | 0 | 0 |
| 2 | 0.00 | 3.68 | 0.00 | 1.00 | 1 | Gm44005 | | 0 | 5 | 18 | 0 | 0 | 0 | 0 | 0 | 0 | 0 | 0 |
| 2 | 3.58 | 3.56 | 1.00 | 0.32 | 1 | Gm37074 | | 0 | 0 | 0 | 0 | 0 | 4 | 0 | 0 | 0 | 15 | 3 |
| 2 | -0.67 | 3.60 | -0.19 | 0.85 | 1 | Gm16570 | | 0 | 0 | 1 | 1 | 0 | 13 | 0 | 0 | 0 | 0 | 8 |
| 2 | 0.00 | 3.68 | 0.00 | 1.00 | 1 | 2310057J18Rik | RIKEN cDNA 2310057J18 gene | 15 | 0 | 7 | 0 | 0 | 0 | 0 | 0 | 0 | 0 | 0 |
| 2 | -0.57 | 3.60 | -0.16 | 0.88 | 1 | 1700015F17Rik | RIKEN cDNA 1700015F17 gene | 0 | 0 | 10 | 0 | 12 | 0 | 0 | 0 | 0 | 0 | 0 |
| 2 | 0.05 | 3.60 | 0.01 | 0.99 | 1 | Gm22596 | | 0 | 0 | 0 | 0 | 0 | 14 | 0 | 0 | 0 | 0 | 9 |
| 2 | -1.90 | 3.57 | -0.53 | 0.59 | 1 | Gm6580 | | 2 | 0 | 2 | 17 | 1 | 0 | 0 | 0 | 0 | 1 | 0 |
| 2 | 4.10 | 3.57 | 1.15 | 0.25 | 1 | Gm12020 | | 0 | 0 | 0 | 0 | 0 | 0 | 0 | 2 | 0 | 19 | 1 |
| 2 | -0.40 | 3.67 | -0.11 | 0.91 | 1 | Gm45867 | | 9 | 13 | 0 | 0 | 0 | 1 | 0 | 0 | 0 | 0 | 0 |
| 2 | 2.77 | 3.54 | 0.78 | 0.43 | 1 | Amn | amnionless | 0 | 0 | 0 | 0 | 0 | 9 | 0 | 0 | 4 | 5 | 4 |
| 2 | -1.08 | 3.67 | -0.30 | 0.77 | 1 | Slc1a6 | solute carrier family 1 (high affinity asparta | 9 | 12 | 0 | 0 | 1 | 1 | 0 | 0 | 0 | 0 | 0 |
| 2 | 0.00 | 3.68 | 0.00 | 1.00 | 1 | Gm37121 | | 4 | 18 | 0 | 0 | 0 | 0 | 0 | 0 | 0 | 0 | 0 |
| 2 | -0.12 | 2.16 | -0.06 | 0.95 | 1 | Gm10252 | | 1 | 10 | 1 | 1 | 2 | 3 | 0 | 0 | 1 | 1 | 2 |

| | | | | | | | | | | | | | | | | | | |
|---|---|---|---|---|---|---|---|---|---|---|---|---|---|---|---|---|---|---|
| 2 | 4.92 | 3.41 | 1.44 | 0.15 | 1 | Mir5620 | | 3 | 0 | 0 | 0 | 0 | 0 | 0 | 6 | 3 | 5 | 5 |
| 2 | -0.45 | 3.60 | -0.12 | 0.90 | 1 | Jmjd7 | jumonji domain containing 7 | 0 | 0 | 0 | 0 | 0 | 19 | 0 | 0 | 0 | 0 | 3 |
| 2 | -1.29 | 3.59 | -0.36 | 0.72 | 1 | Tmem30c | transmembrane protein 30C | 0 | 13 | 0 | 0 | 0 | 9 | 0 | 0 | 0 | 0 | 0 |
| 2 | -3.44 | 3.62 | -0.95 | 0.34 | 1 | Gm5445 | | 0 | 12 | 0 | 0 | 8 | 2 | 0 | 0 | 1 | 0 | 0 |
| 2 | 0.53 | 3.50 | 0.15 | 0.88 | 1 | Gm45069 | | 1 | 0 | 0 | 3 | 0 | 3 | 0 | 0 | 0 | 10 | 6 |
| 2 | -1.52 | 3.56 | -0.43 | 0.67 | 1 | Tiaf2 | | 0 | 0 | 0 | 7 | 0 | 9 | 0 | 0 | 0 | 0 | 6 |
| 2 | -0.51 | 3.61 | -0.14 | 0.89 | 1 | Gm10916 | | 9 | 6 | 1 | 0 | 6 | 0 | 0 | 0 | 0 | 0 | 1 |
| 2 | -5.32 | 3.54 | -1.50 | 0.13 | 1 | Gm43593 | | 0 | 0 | 0 | 7 | 0 | 5 | 0 | 0 | 0 | 10 | 0 |
| 2 | -0.55 | 3.59 | -0.15 | 0.88 | 1 | Gm49216 | | 0 | 0 | 9 | 0 | 0 | 11 | 0 | 0 | 0 | 0 | 2 |
| 2 | -1.81 | 2.32 | -0.78 | 0.43 | 1 | Gm3695 | | 2 | 3 | 1 | 2 | 8 | 4 | 0 | 0 | 2 | 0 | 1 |
| 2 | -1.68 | 3.60 | -0.47 | 0.64 | 1 | Gm29397 | | 0 | 0 | 14 | 2 | 0 | 0 | 0 | 0 | 0 | 0 | 7 |
| 2 | 0.00 | 3.68 | 0.00 | 1.00 | 1 | Gm44929 | | 17 | 0 | 5 | 0 | 0 | 0 | 0 | 0 | 0 | 0 | 0 |
| 2 | 0.00 | 3.68 | 0.00 | 1.00 | 1 | Tnfsf14 | tumor necrosis factor (ligand) superfamily | 17 | 0 | 5 | 0 | 0 | 0 | 0 | 0 | 0 | 0 | 0 |
| 2 | -0.46 | 3.68 | -0.12 | 0.90 | 1 | Gm6445 | | 0 | 16 | 5 | 0 | 1 | 0 | 0 | 0 | 0 | 0 | 0 |
| 2 | -1.56 | 3.60 | -0.43 | 0.66 | 1 | Gm46313 | | 0 | 15 | 0 | 5 | 2 | 0 | 0 | 0 | 0 | 0 | 0 |
| 2 | 2.02 | 3.57 | 0.57 | 0.57 | 1 | Gm44492 | | 0 | 0 | 0 | 0 | 0 | 5 | 0 | 0 | 6 | 0 | 11 |
| 2 | 0.00 | 3.65 | 0.00 | 1.00 | 1 | Adamdec1 | ADAM-like, decysin 1 | 22 | 0 | 0 | 0 | 0 | 0 | 0 | 0 | 0 | 0 | 0 |
| 2 | 0.00 | 3.65 | 0.00 | 1.00 | 1 | Cd200r4 | CD200 receptor 4 | 22 | 0 | 0 | 0 | 0 | 0 | 0 | 0 | 0 | 0 | 0 |
| 2 | 0.00 | 3.68 | 0.00 | 1.00 | 1 | Gm12426 | | 11 | 11 | 0 | 0 | 0 | 0 | 0 | 0 | 0 | 0 | 0 |
| 2 | 0.00 | 3.68 | 0.00 | 1.00 | 1 | Ribc2 | RIB43A domain with coiled-coils 2 | 6 | 16 | 0 | 0 | 0 | 0 | 0 | 0 | 0 | 0 | 0 |
| 2 | 0.21 | 3.60 | 0.06 | 0.95 | 1 | Cpne1 | copine I | 0 | 0 | 0 | 0 | 0 | 11 | 0 | 0 | 0 | 0 | 11 |
| 2 | 0.37 | 3.60 | 0.10 | 0.92 | 1 | Gm50023 | | 0 | 7 | 0 | 0 | 0 | 6 | 0 | 0 | 0 | 0 | 9 |
| 2 | 0.07 | 3.59 | 0.02 | 0.98 | 1 | Gm20405 | | 0 | 11 | 0 | 0 | 0 | 6 | 0 | 0 | 0 | 0 | 5 |
| 2 | 4.44 | 3.56 | 1.25 | 0.21 | 1 | Gm43948 | | 0 | 0 | 0 | 14 | 0 | 0 | 0 | 0 | 1 | 0 | 7 |
| 2 | 0.81 | 3.58 | 0.23 | 0.82 | 1 | Uchl1os | | 6 | 1 | 10 | 0 | 0 | 1 | 0 | 0 | 0 | 4 | 0 |
| 2 | -4.26 | 3.61 | -1.18 | 0.24 | 1 | Gm11724 | | 0 | 0 | 12 | 5 | 0 | 3 | 0 | 0 | 0 | 0 | 1 |
| 2 | -0.09 | 3.48 | -0.03 | 0.98 | 1 | Krtap29-1 | | 3 | 9 | 0 | 2 | 0 | 2 | 0 | 0 | 0 | 0 | 6 |
| 2 | -1.19 | 3.39 | -0.35 | 0.73 | 1 | Gm14401 | | 0 | 0 | 0 | 6 | 4 | 5 | 0 | 0 | 3 | 0 | 4 |
| 2 | 2.65 | 3.55 | 0.75 | 0.46 | 1 | Gm37423 | | 0 | 0 | 0 | 0 | 0 | 8 | 0 | 12 | 0 | 0 | 2 |
| 2 | 5.70 | 3.55 | 1.60 | 0.11 | 1 | Gm9214 | | 1 | 0 | 0 | 0 | 0 | 3 | 0 | 17 | 0 | 1 | 0 |
| 2 | 0.32 | 3.60 | 0.09 | 0.93 | 1 | Gm43466 | | 4 | 0 | 0 | 0 | 0 | 8 | 0 | 0 | 0 | 0 | 10 |
| 2 | -0.81 | 3.59 | -0.23 | 0.82 | 1 | Gm25193 | | 0 | 0 | 0 | 2 | 0 | 10 | 0 | 0 | 0 | 0 | 10 |
| 2 | 1.46 | 3.60 | 0.41 | 0.68 | 1 | Gm8162 | | 0 | 0 | 2 | 0 | 4 | 0 | 0 | 0 | 15 | 0 | 1 |
| 2 | 0.00 | 3.68 | 0.00 | 1.00 | 1 | Gm9465 | | 3 | 5 | 14 | 0 | 0 | 0 | 0 | 0 | 0 | 0 | 0 |
| 2 | -1.74 | 3.23 | -0.54 | 0.59 | 1 | Gm7511 | | 9 | 1 | 0 | 1 | 2 | 7 | 0 | 0 | 2 | 0 | 0 |
| 2 | 0.53 | 3.60 | 0.15 | 0.88 | 1 | Gm37314 | | 0 | 0 | 0 | 0 | 0 | 7 | 0 | 0 | 0 | 0 | 15 |
| 2 | 0.14 | 3.60 | 0.04 | 0.97 | 1 | Gm26109 | | 0 | 0 | 0 | 0 | 0 | 12 | 0 | 0 | 0 | 0 | 10 |
| 2 | 2.54 | 3.56 | 0.71 | 0.48 | 1 | Capn3 | calpain 3 | 0 | 0 | 0 | 0 | 0 | 2 | 0 | 0 | 12 | 0 | 8 |
| 2 | 2.98 | 3.54 | 0.84 | 0.40 | 1 | Arf4os | | 0 | 0 | 0 | 0 | 0 | 6 | 0 | 0 | 6 | 4 | 6 |
| 2 | 0.00 | 3.68 | 0.00 | 1.00 | 1 | A730065G17Rik | | 8 | 14 | 0 | 0 | 0 | 0 | 0 | 0 | 0 | 0 | 0 |
| 2 | -0.56 | 3.54 | -0.16 | 0.87 | 1 | Gm14541 | | 0 | 0 | 0 | 0 | 6 | 10 | 0 | 0 | 3 | 0 | 3 |
| 2 | 0.45 | 3.51 | 0.13 | 0.90 | 1 | Gm12689 | | 0 | 0 | 0 | 0 | 3 | 7 | 0 | 0 | 0 | 5 | 7 |
| 2 | 0.16 | 3.60 | 0.04 | 0.96 | 1 | Snord83b | | 3 | 0 | 0 | 0 | 0 | 10 | 0 | 0 | 0 | 0 | 9 |
| 2 | 0.11 | 3.60 | 0.03 | 0.97 | 1 | Gm32914 | | 0 | 0 | 7 | 0 | 9 | 0 | 0 | 0 | 6 | 0 | 0 |
| 2 | 0.95 | 3.60 | 0.26 | 0.79 | 1 | Gm22574 | | 0 | 0 | 0 | 0 | 0 | 3 | 0 | 0 | 0 | 0 | 19 |
| 2 | 0.46 | 3.60 | 0.13 | 0.90 | 1 | Mir7027 | | 0 | 0 | 0 | 0 | 0 | 8 | 0 | 0 | 0 | 0 | 14 |
| 2 | 0.07 | 3.60 | 0.02 | 0.98 | 1 | 1700028K03Ril | RIKEN cDNA 1700028K03 gene | 0 | 0 | 0 | 0 | 0 | 13 | 0 | 0 | 0 | 0 | 9 |
| 2 | 0.88 | 3.54 | 0.25 | 0.80 | 1 | Zfp967 | zinc finger protein 968 | 0 | 0 | 0 | 0 | 0 | 5 | 0 | 0 | 0 | 17 | 0 |
| 2 | 0.83 | 3.60 | 0.23 | 0.82 | 1 | Gm6426 | | 3 | 1 | 1 | 1 | 0 | 0 | 0 | 0 | 0 | 15 | 0 |
| 2 | 1.24 | 2.82 | 0.44 | 0.66 | 1 | Rpl35a-ps7 | | 4 | 3 | 1 | 0 | 1 | 4 | 0 | 1 | 7 | 0 | 0 |
| 2 | -0.34 | 2.34 | -0.14 | 0.89 | 1 | Gm8543 | | 5 | 5 | 2 | 2 | 1 | 2 | 0 | 0 | 1 | 0 | 4 |
| 2 | 0.00 | 3.68 | 0.00 | 1.00 | 1 | Unc13c | unc-13 homolog C | 9 | 13 | 0 | 0 | 0 | 0 | 0 | 0 | 0 | 0 | 0 |
| 2 | 4.66 | 3.56 | 1.31 | 0.19 | 1 | Gm6153 | | 12 | 0 | 0 | 0 | 0 | 3 | 0 | 0 | 5 | 1 | 0 |
| 2 | 0.41 | 3.53 | 0.12 | 0.91 | 1 | 1700099I09Rik | | 0 | 10 | 0 | 0 | 4 | 0 | 0 | 6 | 0 | 0 | 2 |
| 2 | 0.00 | 3.68 | 0.00 | 1.00 | 1 | Ralyl | RALY RNA binding protein-like | 4 | 10 | 7 | 0 | 0 | 0 | 0 | 0 | 0 | 0 | 0 |
| 2 | 0.38 | 3.60 | 0.11 | 0.92 | 1 | Gm30717 | | 0 | 0 | 0 | 0 | 0 | 8 | 0 | 0 | 0 | 0 | 13 |
| 2 | 0.00 | 3.60 | 0.00 | 1.00 | 1 | Gm37860 | | 0 | 0 | 0 | 0 | 0 | 14 | 0 | 0 | 0 | 0 | 8 |
| 2 | 0.00 | 3.60 | 0.00 | 1.00 | 1 | 3110062G12Rik | | 0 | 0 | 0 | 0 | 0 | 14 | 0 | 0 | 0 | 0 | 8 |
| 2 | -4.37 | 3.52 | -1.24 | 0.21 | 1 | Gm44077 | | 3 | 0 | 0 | 5 | 3 | 5 | 0 | 0 | 5 | 0 | 0 |
| 2 | -1.22 | 3.61 | -0.34 | 0.74 | 1 | Nppc | natriuretic peptide type C | 0 | 18 | 0 | 0 | 0 | 3 | 0 | 0 | 0 | 0 | 0 |

| | | | | | | | | | | | | | | | | | | |
|---|---|---|---|---|---|---|---|---|---|---|---|---|---|---|---|---|---|---|
| 2 | 0.00 | 3.68 | 0.00 | 1.00 | 1 | Gm1968 | | 0 | 8 | 14 | 0 | 0 | 0 | 0 | 0 | 0 | 0 | 0 |
| 2 | 0.63 | 3.55 | 0.18 | 0.86 | 1 | 1810012K08Rik | | 2 | 0 | 0 | 2 | 0 | 4 | 0 | 14 | 0 | 0 | 0 |
| 2 | -0.42 | 3.54 | -0.12 | 0.90 | 1 | Gm48236 | | 7 | 0 | 0 | 0 | 6 | 1 | 0 | 0 | 3 | 5 | 0 |
| 2 | 3.12 | 3.53 | 0.88 | 0.38 | 1 | Gm42992 | | 0 | 0 | 0 | 0 | 0 | 6 | 0 | 0 | 3 | 12 | 1 |
| 2 | 0.00 | 3.68 | 0.00 | 1.00 | 1 | Nlrp1c-ps | | 15 | 6 | 0 | 0 | 0 | 0 | 0 | 0 | 0 | 0 | 0 |
| 2 | 0.00 | 3.68 | 0.00 | 1.00 | 1 | Gm15318 | | 11 | 1 | 9 | 0 | 0 | 0 | 0 | 0 | 0 | 0 | 0 |
| 2 | 3.13 | 3.58 | 0.87 | 0.38 | 1 | 5330406M23Rik | | 0 | 0 | 8 | 0 | 0 | 7 | 0 | 0 | 0 | 6 | 0 |
| 2 | -1.79 | 3.58 | -0.50 | 0.62 | 1 | Gm6365 | | 2 | 10 | 0 | 0 | 2 | 7 | 0 | 0 | 0 | 0 | 1 |
| 2 | -0.73 | 3.64 | -0.20 | 0.84 | 1 | Ccdc150 | coiled-coil domain containing 150 | 9 | 0 | 10 | 0 | 0 | 3 | 0 | 0 | 0 | 0 | 0 |
| 2 | 1.18 | 3.55 | 0.33 | 0.74 | 1 | Gm26658 | | 0 | 0 | 0 | 0 | 2 | 3 | 0 | 0 | 8 | 0 | 8 |
| 2 | -3.35 | 3.57 | -0.94 | 0.35 | 1 | Gm35405 | | 0 | 0 | 0 | 15 | 0 | 3 | 0 | 0 | 0 | 0 | 4 |
| 2 | 0.00 | 3.65 | 0.00 | 1.00 | 1 | Mylk2 | myosin, light polypeptide kinase 2, skeleta | 21 | 0 | 0 | 0 | 0 | 0 | 0 | 0 | 0 | 0 | 0 |
| 2 | 0.00 | 3.65 | 0.00 | 1.00 | 1 | 4932441J04Rik | | 21 | 0 | 0 | 0 | 0 | 0 | 0 | 0 | 0 | 0 | 0 |
| 2 | -0.35 | 3.58 | -0.10 | 0.92 | 1 | Gm29502 | | 6 | 6 | 2 | 5 | 0 | 0 | 0 | 0 | 0 | 0 | 2 |
| 2 | 0.48 | 3.54 | 0.14 | 0.89 | 1 | Lypd10 | Ly6/PLAUR domain containing 10 | 0 | 0 | 0 | 0 | 8 | 2 | 0 | 0 | 8 | 0 | 4 |
| 2 | -2.68 | 2.81 | -0.95 | 0.34 | 1 | Gm46522 | | 2 | 1 | 2 | 11 | 1 | 2 | 0 | 0 | 0 | 3 | 0 |
| 2 | -1.18 | 3.59 | -0.33 | 0.74 | 1 | Gm37248 | | 0 | 0 | 10 | 0 | 0 | 11 | 0 | 0 | 0 | 0 | 0 |
| 2 | 0.00 | 3.68 | 0.00 | 1.00 | 1 | Clrn1 | clarin 1 | 12 | 0 | 9 | 0 | 0 | 0 | 0 | 0 | 0 | 0 | 0 |
| 2 | 0.00 | 3.68 | 0.00 | 1.00 | 1 | 4833407H14Rik | | 12 | 0 | 9 | 0 | 0 | 0 | 0 | 0 | 0 | 0 | 0 |
| 2 | 0.00 | 3.68 | 0.00 | 1.00 | 1 | Il5ra | interleukin 5 receptor, alpha | 0 | 18 | 3 | 0 | 0 | 0 | 0 | 0 | 0 | 0 | 0 |
| 2 | -1.21 | 3.59 | -0.34 | 0.74 | 1 | Gm38043 | | 0 | 0 | 0 | 0 | 2 | 14 | 0 | 0 | 0 | 0 | 5 |
| 2 | -2.15 | 3.62 | -0.60 | 0.55 | 1 | E330011O21Rik | | 12 | 4 | 0 | 0 | 2 | 3 | 0 | 0 | 0 | 0 | 0 |
| 2 | 0.00 | 3.68 | 0.00 | 1.00 | 1 | Rps11-ps5 | | 0 | 20 | 1 | 0 | 0 | 0 | 0 | 0 | 0 | 0 | 0 |
| 2 | 1.46 | 3.56 | 0.41 | 0.68 | 1 | Gm36933 | | 0 | 0 | 0 | 0 | 0 | 10 | 0 | 0 | 6 | 0 | 5 |
| 2 | 0.00 | 3.64 | 0.00 | 1.00 | 1 | F630028O10Rik | | 0 | 21 | 0 | 0 | 0 | 0 | 0 | 0 | 0 | 0 | 0 |
| 2 | -0.14 | 3.61 | -0.04 | 0.97 | 1 | Ugt1a6a | UDP glucuronosyltransferase 1 family, pc | 7 | 0 | 7 | 0 | 0 | 5 | 0 | 0 | 0 | 0 | 2 |
| 2 | 0.85 | 3.52 | 0.24 | 0.81 | 1 | Gm50242 | | 0 | 0 | 0 | 3 | 0 | 3 | 0 | 0 | 3 | 0 | 12 |
| 2 | 0.28 | 3.60 | 0.08 | 0.94 | 1 | Gm48632 | | 3 | 1 | 0 | 0 | 0 | 8 | 0 | 0 | 0 | 0 | 9 |
| 2 | 0.55 | 3.61 | 0.15 | 0.88 | 1 | 1700016C15Ril | RIKEN cDNA 1700016C15 gene | 4 | 11 | 0 | 0 | 0 | 2 | 0 | 0 | 0 | 0 | 4 |
| 2 | -1.77 | 3.57 | -0.50 | 0.62 | 1 | Gm45344 | | 0 | 0 | 0 | 6 | 0 | 11 | 0 | 0 | 0 | 0 | 4 |
| 2 | -1.39 | 3.56 | -0.39 | 0.70 | 1 | Gm17203 | | 0 | 0 | 0 | 0 | 7 | 8 | 0 | 0 | 0 | 0 | 6 |
| 2 | 1.21 | 3.60 | 0.33 | 0.74 | 1 | Vmn1r90 | vomeronasal 1 receptor 90 | 0 | 0 | 0 | 0 | 0 | 1 | 0 | 0 | 0 | 0 | 20 |
| 2 | -3.39 | 3.60 | -0.94 | 0.35 | 1 | Gm37969 | | 9 | 0 | 0 | 0 | 2 | 7 | 0 | 0 | 0 | 0 | 3 |
| 2 | 0.57 | 3.55 | 0.16 | 0.87 | 1 | Gm45708 | | 1 | 0 | 7 | 0 | 0 | 6 | 0 | 0 | 6 | 0 | 1 |
| 2 | 0.00 | 3.68 | 0.00 | 1.00 | 1 | Tulp2 | tubby-like protein 2 | 18 | 0 | 3 | 0 | 0 | 0 | 0 | 0 | 0 | 0 | 0 |
| 2 | 0.00 | 3.68 | 0.00 | 1.00 | 1 | Mir142hg | | 12 | 9 | 0 | 0 | 0 | 0 | 0 | 0 | 0 | 0 | 0 |
| 2 | 0.82 | 2.13 | 0.39 | 0.70 | 1 | Gm6305 | | 0 | 1 | 1 | 9 | 1 | 1 | 0 | 1 | 4 | 1 | 2 |
| 2 | 2.35 | 3.56 | 0.66 | 0.51 | 1 | Gm12990 | | 0 | 0 | 0 | 0 | 0 | 11 | 0 | 0 | 7 | 0 | 3 |
| 2 | 1.15 | 3.01 | 0.38 | 0.70 | 1 | Gm10913 | | 1 | 1 | 12 | 0 | 0 | 2 | 0 | 0 | 1 | 0 | 4 |
| 2 | 2.30 | 3.47 | 0.66 | 0.51 | 1 | Gm11527 | | 4 | 0 | 0 | 0 | 0 | 2 | 0 | 0 | 1 | 9 | 5 |
| 2 | -2.59 | 3.55 | -0.73 | 0.47 | 1 | Gm50136 | | 0 | 0 | 0 | 0 | 12 | 2 | 0 | 0 | 0 | 0 | 7 |
| 2 | -0.27 | 3.49 | -0.08 | 0.94 | 1 | Rpl7a-ps8 | | 1 | 0 | 11 | 1 | 3 | 0 | 0 | 0 | 5 | 0 | 0 |
| 2 | 0.00 | 3.68 | 0.00 | 1.00 | 1 | E130215H24Rik | | 18 | 3 | 0 | 0 | 0 | 0 | 0 | 0 | 0 | 0 | 0 |
| 2 | 1.71 | 3.56 | 0.48 | 0.63 | 1 | 4932443L11Rik | | 0 | 9 | 3 | 0 | 0 | 2 | 0 | 0 | 2 | 0 | 5 |
| 2 | 0.00 | 3.68 | 0.00 | 1.00 | 1 | Gm29686 | | 14 | 0 | 7 | 0 | 0 | 0 | 0 | 0 | 0 | 0 | 0 |
| 2 | 2.88 | 3.53 | 0.81 | 0.42 | 1 | Gm6044 | | 0 | 0 | 0 | 0 | 0 | 5 | 0 | 0 | 2 | 13 | 1 |
| 2 | 0.05 | 3.60 | 0.01 | 0.99 | 1 | Gm25117 | | 0 | 1 | 0 | 0 | 0 | 12 | 0 | 0 | 0 | 0 | 8 |
| 2 | 0.00 | 3.68 | 0.00 | 1.00 | 1 | Calcb | calcitonin-related polypeptide, beta | 4 | 0 | 17 | 0 | 0 | 0 | 0 | 0 | 0 | 0 | 0 |
| 2 | 2.98 | 3.55 | 0.84 | 0.40 | 1 | Gm34354 | | 0 | 0 | 0 | 0 | 0 | 3 | 0 | 1 | 11 | 0 | 6 |
| 2 | 0.00 | 3.68 | 0.00 | 1.00 | 1 | Kcnv1 | potassium channel, subfamily V, member | 8 | 13 | 0 | 0 | 0 | 0 | 0 | 0 | 0 | 0 | 0 |
| 2 | -3.54 | 3.57 | -0.99 | 0.32 | 1 | Gm43048 | | 0 | 0 | 0 | 18 | 0 | 3 | 0 | 0 | 0 | 0 | 0 |
| 2 | -0.52 | 3.57 | -0.15 | 0.88 | 1 | Rpl30-ps6 | | 0 | 11 | 0 | 8 | 0 | 1 | 0 | 0 | 0 | 0 | 1 |
| 2 | 0.00 | 3.68 | 0.00 | 1.00 | 1 | B430305J03Rik | | 4 | 3 | 14 | 0 | 0 | 0 | 0 | 0 | 0 | 0 | 0 |
| 2 | 0.43 | 3.60 | 0.12 | 0.91 | 1 | Apc-ps1 | | 0 | 0 | 0 | 0 | 0 | 8 | 0 | 0 | 0 | 0 | 13 |
| 2 | 0.43 | 3.60 | 0.12 | 0.91 | 1 | Gm16133 | | 0 | 0 | 0 | 0 | 0 | 8 | 0 | 0 | 0 | 0 | 13 |
| 2 | 0.70 | 3.50 | 0.20 | 0.84 | 1 | Gm47322 | | 3 | 8 | 6 | 0 | 0 | 1 | 0 | 0 | 0 | 0 | 3 |
| 2 | 0.00 | 3.64 | 0.00 | 1.00 | 1 | Gm19434 | | 0 | 0 | 21 | 0 | 0 | 0 | 0 | 0 | 0 | 0 | 0 |
| 2 | 0.00 | 3.64 | 0.00 | 1.00 | 1 | Btn2a2 | butyrophilin, subfamily 2, member A2 | 0 | 0 | 21 | 0 | 0 | 0 | 0 | 0 | 0 | 0 | 0 |
| 2 | 3.42 | 3.58 | 0.96 | 0.34 | 1 | Gm44780 | | 0 | 5 | 0 | 0 | 0 | 8 | 0 | 0 | 5 | 0 | 3 |
| 2 | 4.37 | 3.56 | 1.23 | 0.22 | 1 | Iqub | IQ motif and ubiquitin domain containing | 6 | 2 | 0 | 0 | 0 | 0 | 0 | 5 | 4 | 0 | 4 |

| | | | | | | | | | | | | | | | | | |
|---|---|---|---|---|---|---|---|---|---|---|---|---|---|---|---|---|---|
| 2 | -0.76 | 3.63 | -0.21 | 0.83 | 1 | Gm38832 | | 10 | 7 | 0 | 0 | 0 | 3 | 0 | 0 | 0 | 0 | 0 |
| 2 | -0.82 | 2.41 | -0.34 | 0.73 | 1 | Gm10313 | | 2 | 0 | 10 | 2 | 2 | 2 | 0 | 0 | 1 | 1 | 1 |
| 2 | 0.00 | 3.68 | 0.00 | 1.00 | 1 | Eomes | eomesodermin | 5 | 0 | 16 | 0 | 0 | 0 | 0 | 0 | 0 | 0 | 0 |
| 2 | 0.92 | 3.61 | 0.26 | 0.80 | 1 | Gm15843 | | 9 | 0 | 5 | 0 | 0 | 1 | 0 | 0 | 0 | 0 | 6 |
| 2 | 0.03 | 2.94 | 0.01 | 0.99 | 1 | Gm8696 | | 1 | 0 | 5 | 0 | 1 | 8 | 0 | 0 | 1 | 0 | 5 |
| 2 | 1.80 | 3.61 | 0.50 | 0.62 | 1 | Gm13977 | | 0 | 0 | 17 | 0 | 0 | 0 | 0 | 0 | 0 | 0 | 4 |
| 2 | 0.00 | 3.68 | 0.00 | 1.00 | 1 | Gm43573 | | 9 | 8 | 3 | 0 | 0 | 0 | 0 | 0 | 0 | 0 | 0 |
| 2 | -0.05 | 3.60 | -0.01 | 0.99 | 1 | Gm43203 | | 0 | 0 | 0 | 0 | 0 | 14 | 0 | 0 | 0 | 0 | 7 |
| 2 | -0.05 | 3.60 | -0.01 | 0.99 | 1 | Mir7669 | | 0 | 0 | 0 | 0 | 0 | 14 | 0 | 0 | 0 | 0 | 7 |
| 2 | -0.05 | 3.60 | -0.01 | 0.99 | 1 | Gm44974 | | 0 | 0 | 0 | 0 | 0 | 14 | 0 | 0 | 0 | 0 | 7 |
| 2 | 0.00 | 3.68 | 0.00 | 1.00 | 1 | Gm15958 | | 0 | 6 | 15 | 0 | 0 | 0 | 0 | 0 | 0 | 0 | 0 |
| 2 | -0.69 | 3.59 | -0.19 | 0.85 | 1 | Gm12403 | | 5 | 0 | 1 | 1 | 0 | 8 | 0 | 0 | 0 | 0 | 5 |
| 2 | 0.00 | 3.65 | 0.00 | 1.00 | 1 | Syndig1 | synapse differentiation inducing 1 | 21 | 0 | 0 | 0 | 0 | 0 | 0 | 0 | 0 | 0 | 0 |
| 2 | 0.00 | 3.65 | 0.00 | 1.00 | 1 | Gkn3 | gastrokine 3 | 21 | 0 | 0 | 0 | 0 | 0 | 0 | 0 | 0 | 0 | 0 |
| 2 | 0.28 | 3.61 | 0.08 | 0.94 | 1 | Gm26448 | | 7 | 0 | 6 | 0 | 0 | 3 | 0 | 0 | 0 | 0 | 4 |
| 2 | -1.76 | 3.64 | -0.48 | 0.63 | 1 | Art1 | ADP-ribosyltransferase 1 | 8 | 5 | 3 | 0 | 4 | 1 | 0 | 0 | 0 | 0 | 0 |
| 2 | 0.42 | 2.03 | 0.21 | 0.84 | 1 | Gm6929 | | 0 | 2 | 1 | 4 | 2 | 3 | 0 | 1 | 2 | 3 | 2 |
| 2 | 1.26 | 3.06 | 0.41 | 0.68 | 1 | Gm6978 | | 3 | 0 | 9 | 0 | 0 | 3 | 0 | 0 | 2 | 1 | 1 |
| 2 | 1.61 | 3.66 | 0.44 | 0.66 | 1 | Gm4793 | | 5 | 13 | 0 | 0 | 0 | 0 | 0 | 1 | 1 | 0 | 0 |
| 2 | -0.84 | 3.62 | -0.23 | 0.82 | 1 | Gm13039 | | 1 | 15 | 1 | 0 | 0 | 3 | 0 | 0 | 0 | 0 | 0 |
| 2 | 0.28 | 3.60 | 0.08 | 0.94 | 1 | 6820402A03Rik | | 0 | 0 | 0 | 0 | 0 | 9 | 0 | 0 | 0 | 0 | 11 |
| 2 | 0.28 | 3.60 | 0.08 | 0.94 | 1 | Gm24608 | | 0 | 0 | 0 | 0 | 0 | 9 | 0 | 0 | 0 | 0 | 11 |
| 2 | -0.82 | 3.59 | -0.23 | 0.82 | 1 | Gm37584 | | 0 | 5 | 0 | 0 | 0 | 14 | 0 | 0 | 0 | 0 | 1 |
| 2 | 0.73 | 3.58 | 0.20 | 0.84 | 1 | Gm11799 | | 0 | 13 | 0 | 0 | 0 | 4 | 0 | 0 | 1 | 0 | 2 |
| 2 | 1.26 | 3.66 | 0.35 | 0.73 | 1 | Kbtbd12 | kelch repeat and BTB (POZ) domain conta | 16 | 0 | 2 | 0 | 0 | 0 | 0 | 0 | 0 | 0 | 2 |
| 2 | 0.00 | 3.68 | 0.00 | 1.00 | 1 | Gm8598 | | 11 | 4 | 5 | 0 | 0 | 0 | 0 | 0 | 0 | 0 | 0 |
| 2 | 0.00 | 3.68 | 0.00 | 1.00 | 1 | Dlk2 | delta like non-canonical Notch ligand 2 | 0 | 15 | 5 | 0 | 0 | 0 | 0 | 0 | 0 | 0 | 0 |
| 2 | -1.25 | 3.56 | -0.35 | 0.73 | 1 | Gm23969 | | 0 | 0 | 0 | 0 | 6 | 8 | 0 | 0 | 0 | 0 | 7 |
| 2 | -1.03 | 3.59 | -0.29 | 0.77 | 1 | 4933412A08Rik | | 0 | 0 | 5 | 0 | 9 | 2 | 0 | 0 | 4 | 0 | 0 |
| 2 | -1.00 | 3.62 | -0.28 | 0.78 | 1 | Gm19684 | | 6 | 9 | 0 | 5 | 0 | 0 | 0 | 0 | 0 | 0 | 0 |
| 2 | 2.10 | 3.60 | 0.58 | 0.56 | 1 | Gm13586 | | 0 | 0 | 1 | 0 | 0 | 0 | 0 | 0 | 0 | 19 | 0 |
| 2 | 0.00 | 3.68 | 0.00 | 1.00 | 1 | Phf11 | | 16 | 4 | 0 | 0 | 0 | 0 | 0 | 0 | 0 | 0 | 0 |
| 2 | 0.00 | 3.68 | 0.00 | 1.00 | 1 | Otogl | otogelin-like | 7 | 1 | 12 | 0 | 0 | 0 | 0 | 0 | 0 | 0 | 0 |
| 2 | -2.10 | 3.52 | -0.60 | 0.55 | 1 | Gm32816 | | 0 | 0 | 0 | 7 | 3 | 3 | 0 | 0 | 7 | 0 | 0 |
| 2 | 1.16 | 3.59 | 0.32 | 0.75 | 1 | Ankrd31 | ankyrin repeat domain 31 | 0 | 0 | 0 | 0 | 0 | 10 | 0 | 0 | 2 | 0 | 8 |
| 2 | 3.24 | 3.55 | 0.91 | 0.36 | 1 | Gm5374 | | 0 | 0 | 4 | 9 | 0 | 0 | 0 | 0 | 0 | 1 | 6 |
| 2 | 3.69 | 3.58 | 1.03 | 0.30 | 1 | Gm8388 | | 3 | 0 | 2 | 0 | 0 | 0 | 0 | 12 | 4 | 0 | 0 |
| 2 | 0.00 | 3.68 | 0.00 | 1.00 | 1 | Dmrt2 | doublesex and mab-3 related transcriptio | 17 | 0 | 3 | 0 | 0 | 0 | 0 | 0 | 0 | 0 | 0 |
| 2 | 0.04 | 3.59 | 0.01 | 0.99 | 1 | Gm23908 | | 0 | 11 | 0 | 0 | 0 | 5 | 0 | 0 | 0 | 0 | 4 |
| 2 | -1.09 | 3.53 | -0.31 | 0.76 | 1 | 4930527F14Rik | | 7 | 0 | 0 | 0 | 3 | 3 | 0 | 0 | 0 | 5 | 2 |
| 2 | 0.00 | 3.64 | 0.00 | 1.00 | 1 | Htr6 | 5-hydroxytryptamine (serotonin) receptor | 0 | 20 | 0 | 0 | 0 | 0 | 0 | 0 | 0 | 0 | 0 |
| 2 | 0.00 | 3.64 | 0.00 | 1.00 | 1 | Oosp2 | oocyte secreted protein 2 | 0 | 20 | 0 | 0 | 0 | 0 | 0 | 0 | 0 | 0 | 0 |
| 2 | 1.79 | 3.58 | 0.50 | 0.62 | 1 | Gm19801 | | 0 | 0 | 0 | 0 | 0 | 6 | 0 | 0 | 0 | 5 | 9 |
| 2 | -1.06 | 3.62 | -0.29 | 0.77 | 1 | Gm833 | | 0 | 0 | 18 | 0 | 0 | 3 | 0 | 0 | 0 | 0 | 0 |
| 2 | 0.00 | 3.68 | 0.00 | 1.00 | 1 | Ighv5-12 | | 17 | 3 | 0 | 0 | 0 | 0 | 0 | 0 | 0 | 0 | 0 |
| 2 | 0.64 | 3.18 | 0.20 | 0.84 | 1 | Gm14730 | | 9 | 1 | 0 | 0 | 1 | 2 | 0 | 0 | 1 | 0 | 6 |
| 2 | -3.26 | 3.58 | -0.91 | 0.36 | 1 | Gm5590 | | 0 | 1 | 7 | 6 | 0 | 2 | 0 | 0 | 0 | 0 | 4 |
| 2 | 0.56 | 3.60 | 0.15 | 0.88 | 1 | Gm42930 | | 0 | 0 | 0 | 0 | 0 | 6 | 0 | 0 | 0 | 0 | 14 |
| 2 | 0.00 | 3.68 | 0.00 | 1.00 | 1 | Snorc | secondary ossification center associated | 2 | 11 | 7 | 0 | 0 | 0 | 0 | 0 | 0 | 0 | 0 |
| 2 | 0.13 | 3.60 | 0.04 | 0.97 | 1 | Gm38220 | | 0 | 0 | 0 | 0 | 0 | 11 | 0 | 0 | 0 | 0 | 9 |
| 2 | 1.18 | 3.55 | 0.33 | 0.74 | 1 | Tsks | testis-specific serine kinase substrate | 0 | 0 | 0 | 0 | 0 | 3 | 0 | 0 | 0 | 17 | 0 |
| 2 | 1.48 | 3.59 | 0.41 | 0.68 | 1 | Gm49932 | | 0 | 0 | 0 | 0 | 0 | 6 | 0 | 0 | 2 | 0 | 12 |
| 2 | -1.31 | 3.54 | -0.37 | 0.71 | 1 | Gm14455 | | 3 | 0 | 0 | 0 | 8 | 5 | 0 | 0 | 4 | 0 | 0 |
| 2 | 1.76 | 3.56 | 0.50 | 0.62 | 1 | Gm23183 | | 0 | 0 | 0 | 0 | 0 | 6 | 0 | 0 | 8 | 0 | 6 |
| 2 | 4.13 | 3.04 | 1.36 | 0.18 | 1 | Gm29668 | | 1 | 1 | 0 | 0 | 0 | 0 | 0 | 14 | 2 | 1 | 1 |
| 2 | 0.78 | 3.59 | 0.22 | 0.83 | 1 | Gm5724 | predicted gene 5724 | 0 | 16 | 0 | 0 | 0 | 1 | 0 | 0 | 0 | 0 | 3 |
| 2 | -0.62 | 3.64 | -0.17 | 0.87 | 1 | Klrg1 | killer cell lectin-like receptor subfamily G, | 0 | 19 | 0 | 0 | 0 | 1 | 0 | 0 | 0 | 0 | 0 |
| 2 | -1.32 | 3.64 | -0.36 | 0.72 | 1 | Gm13771 | | 0 | 7 | 10 | 1 | 0 | 2 | 0 | 0 | 0 | 0 | 0 |
| 2 | -0.24 | 3.61 | -0.07 | 0.95 | 1 | Gm48369 | | 1 | 0 | 10 | 0 | 0 | 7 | 0 | 0 | 0 | 0 | 2 |
| 2 | 1.95 | 3.56 | 0.55 | 0.58 | 1 | | predicted gene, 31160 | 0 | 2 | 4 | 0 | 0 | 2 | 0 | 0 | 7 | 0 | 5 |

| | | | | | | | | | | | | | | | | | | |
|---|---|---|---|---|---|---|---|---|---|---|---|---|---|---|---|---|---|---|
| 2 | 3.75 | 3.58 | 1.05 | 0.29 | 1 | Gm42457 | | 0 | 0 | 4 | 0 | 0 | 5 | 0 | 0 | 0 | 8 | 3 |
| 2 | 3.23 | 3.58 | 0.90 | 0.37 | 1 | Gm48551 | | 3 | 0 | 0 | 0 | 0 | 9 | 0 | 0 | 4 | 0 | 4 |
| 2 | 1.05 | 3.60 | 0.29 | 0.77 | 1 | Gm26493 | | 0 | 0 | 0 | 0 | 0 | 2 | 0 | 0 | 0 | 0 | 18 |
| 2 | -4.08 | 3.52 | -1.16 | 0.25 | 1 | Gm9419 | | 3 | 0 | 0 | 9 | 0 | 3 | 0 | 0 | 6 | 0 | 0 |
| 2 | -3.21 | 3.58 | -0.90 | 0.37 | 1 | Omg | oligodendrocyte myelin glycoprotein | 9 | 0 | 0 | 9 | 0 | 3 | 0 | 0 | 0 | 0 | 0 |
| 2 | 1.68 | 3.57 | 0.47 | 0.64 | 1 | Gm10369 | | 0 | 5 | 4 | 0 | 0 | 3 | 0 | 0 | 4 | 0 | 4 |
| 2 | -1.03 | 3.18 | -0.32 | 0.75 | 1 | Gm10132 | | 10 | 3 | 1 | 0 | 1 | 3 | 0 | 1 | 0 | 0 | 0 |
| 2 | -0.65 | 3.49 | -0.18 | 0.85 | 1 | Vmn2r1 | vomeronasal 2, receptor 1 | 0 | 0 | 0 | 3 | 5 | 3 | 0 | 0 | 4 | 0 | 5 |
| 2 | 0.38 | 3.55 | 0.11 | 0.91 | 1 | Gm45441 | | 0 | 0 | 10 | 0 | 0 | 3 | 0 | 0 | 5 | 0 | 1 |
| 2 | -0.94 | 3.56 | -0.26 | 0.79 | 1 | 4933427D14Ril | RIKEN cDNA 4933427D14 gene | 0 | 0 | 5 | 0 | 1 | 13 | 0 | 0 | 1 | 0 | 0 |
| 2 | 0.00 | 3.68 | 0.00 | 1.00 | 1 | Prkag3 | protein kinase, AMP-activated, gamma 3 r | 9 | 0 | 10 | 0 | 0 | 0 | 0 | 0 | 0 | 0 | 0 |
| 2 | 0.78 | 3.61 | 0.22 | 0.83 | 1 | Gm11802 | | 0 | 15 | 0 | 0 | 0 | 0 | 0 | 5 | 0 | 0 | 0 |
| 2 | 0.02 | 2.12 | 0.01 | 0.99 | 1 | Gm16385 | | 3 | 1 | 2 | 2 | 1 | 5 | 0 | 2 | 1 | 0 | 3 |
| 2 | 0.00 | 3.68 | 0.00 | 1.00 | 1 | Fcer1a | Fc receptor, IgE, high affinity I, alpha poly | 14 | 6 | 0 | 0 | 0 | 0 | 0 | 0 | 0 | 0 | 0 |
| 2 | 0.00 | 3.68 | 0.00 | 1.00 | 1 | Serpina3i | serine (or cysteine) peptidase inhibitor, cl | 4 | 5 | 10 | 0 | 0 | 0 | 0 | 0 | 0 | 0 | 0 |
| 2 | 0.00 | 3.64 | 0.00 | 1.00 | 1 | Igfbp1 | insulin-like growth factor binding protein 1 | 0 | 0 | 20 | 0 | 0 | 0 | 0 | 0 | 0 | 0 | 0 |
| 2 | 0.00 | 3.65 | 0.00 | 1.00 | 1 | Baalc | brain and acute leukemia, cytoplasmic | 20 | 0 | 0 | 0 | 0 | 0 | 0 | 0 | 0 | 0 | 0 |
| 2 | -5.05 | 3.54 | -1.42 | 0.15 | 1 | Gm6906 | | 0 | 0 | 0 | 0 | 3 | 8 | 0 | 9 | 0 | 0 | 0 |
| 2 | 0.00 | 3.68 | 0.00 | 1.00 | 1 | Ptk6 | PTK6 protein tyrosine kinase 6 | 0 | 2 | 18 | 0 | 0 | 0 | 0 | 0 | 0 | 0 | 0 |
| 2 | -0.74 | 3.59 | -0.21 | 0.84 | 1 | Gm36745 | | 0 | 0 | 0 | 0 | 2 | 8 | 0 | 0 | 0 | 0 | 10 |
| 2 | 0.00 | 3.68 | 0.00 | 1.00 | 1 | Kcnd3os | | 15 | 5 | 0 | 0 | 0 | 0 | 0 | 0 | 0 | 0 | 0 |
| 2 | 3.58 | 3.61 | 0.99 | 0.32 | 1 | Bhmt | betaine-homocysteine methyltransferase | 0 | 1 | 1 | 0 | 0 | 1 | 0 | 0 | 0 | 17 | 0 |
| 2 | 0.00 | 3.68 | 0.00 | 1.00 | 1 | Guca2a | guanylate cyclase activator 2a (guanylin) | 0 | 4 | 16 | 0 | 0 | 0 | 0 | 0 | 0 | 0 | 0 |
| 2 | 0.00 | 3.68 | 0.00 | 1.00 | 1 | Gpr83 | G protein-coupled receptor 83 | 0 | 4 | 16 | 0 | 0 | 0 | 0 | 0 | 0 | 0 | 0 |
| 2 | 1.74 | 3.59 | 0.49 | 0.63 | 1 | Platr14 | | 0 | 0 | 0 | 0 | 0 | 3 | 0 | 0 | 2 | 0 | 14 |
| 2 | -0.11 | 3.60 | -0.03 | 0.98 | 1 | Gm22571 | | 0 | 0 | 0 | 0 | 0 | 14 | 0 | 0 | 0 | 0 | 6 |
| 2 | 1.08 | 3.63 | 0.30 | 0.77 | 1 | Phospho1 | phosphatase, orphan 1 | 14 | 2 | 0 | 0 | 0 | 0 | 0 | 0 | 0 | 4 | 0 |
| 2 | 0.89 | 3.55 | 0.25 | 0.80 | 1 | Fam209 | family with sequence similarity 209 | 0 | 0 | 7 | 0 | 0 | 4 | 0 | 0 | 3 | 0 | 5 |
| 2 | 3.50 | 3.58 | 0.98 | 0.33 | 1 | Gm49330 | | 0 | 0 | 7 | 0 | 0 | 4 | 0 | 8 | 0 | 0 | 0 |
| 2 | -0.76 | 3.63 | -0.21 | 0.83 | 1 | Gm20689 | | 11 | 5 | 0 | 0 | 0 | 3 | 0 | 0 | 0 | 0 | 0 |
| 2 | -0.43 | 3.59 | -0.12 | 0.91 | 1 | Gm43301 | | 0 | 15 | 0 | 0 | 0 | 3 | 0 | 0 | 0 | 0 | 1 |
| 2 | 3.86 | 3.58 | 1.08 | 0.28 | 1 | Rps12-ps2 | | 0 | 6 | 0 | 0 | 0 | 3 | 0 | 0 | 9 | 0 | 1 |
| 2 | 0.94 | 3.67 | 0.25 | 0.80 | 1 | Saa1 | serum amyloid A 1 | 0 | 11 | 7 | 0 | 0 | 0 | 0 | 1 | 0 | 0 | 0 |
| 2 | 1.43 | 3.63 | 0.39 | 0.69 | 1 | Nipal4 | NIPA-like domain containing 4 | 5 | 0 | 9 | 0 | 0 | 0 | 0 | 0 | 5 | 0 | 0 |
| 2 | -1.89 | 3.59 | -0.52 | 0.60 | 1 | Gm42425 | | 0 | 6 | 1 | 11 | 2 | 0 | 0 | 0 | 0 | 0 | 0 |
| 2 | 0.00 | 3.68 | 0.00 | 1.00 | 1 | Ugt2a3 | UDP glucuronosyltransferase 2 family, pc | 4 | 15 | 0 | 0 | 0 | 0 | 0 | 0 | 0 | 0 | 0 |
| 2 | 1.37 | 3.58 | 0.38 | 0.70 | 1 | Gm37342 | | 0 | 0 | 0 | 0 | 0 | 9 | 0 | 0 | 3 | 0 | 7 |
| 2 | 0.00 | 3.68 | 0.00 | 1.00 | 1 | Htr4 | 5 hydroxytryptamine (serotonin) receptor | 0 | 12 | 7 | 0 | 0 | 0 | 0 | 0 | 0 | 0 | 0 |
| 2 | -0.21 | 3.59 | -0.06 | 0.95 | 1 | Mir6937 | | 0 | 13 | 0 | 0 | 0 | 4 | 0 | 0 | 0 | 0 | 2 |
| 2 | 2.28 | 3.55 | 0.64 | 0.52 | 1 | Gm43425 | | 0 | 0 | 0 | 0 | 0 | 9 | 0 | 0 | 8 | 0 | 2 |
| 2 | -1.06 | 3.53 | -0.30 | 0.76 | 1 | Gm2518 | | 0 | 0 | 0 | 4 | 0 | 11 | 0 | 0 | 2 | 0 | 2 |
| 2 | -0.73 | 3.54 | -0.21 | 0.84 | 1 | Gm36584 | | 3 | 0 | 0 | 0 | 6 | 5 | 0 | 0 | 1 | 0 | 5 |
| 2 | -0.40 | 3.67 | -0.11 | 0.91 | 1 | Gm43719 | | 9 | 9 | 0 | 0 | 0 | 1 | 0 | 0 | 0 | 0 | 0 |
| 2 | 3.66 | 3.58 | 1.02 | 0.31 | 1 | Spem1 | sperm maturation 1 | 0 | 0 | 5 | 0 | 0 | 5 | 0 | 8 | 0 | 0 | 1 |
| 2 | 1.11 | 3.60 | 0.31 | 0.76 | 1 | C630004M23Rik | | 6 | 0 | 0 | 0 | 0 | 4 | 0 | 0 | 0 | 0 | 9 |
| 2 | -0.17 | 3.57 | -0.05 | 0.96 | 1 | Gm16288 | | 0 | 4 | 9 | 2 | 2 | 0 | 0 | 2 | 0 | 0 | 0 |
| 2 | 1.30 | 3.64 | 0.36 | 0.72 | 1 | Slc17a7 | solute carrier family 17 (sodium-depende | 5 | 11 | 0 | 0 | 0 | 0 | 0 | 0 | 0 | 0 | 3 |
| 2 | 0.61 | 3.60 | 0.17 | 0.86 | 1 | Mir5100 | | 0 | 0 | 0 | 0 | 0 | 5 | 0 | 0 | 0 | 0 | 14 |
| 2 | 0.00 | 3.68 | 0.00 | 1.00 | 1 | Gpr31b | G protein-coupled receptor 31, D17Leh66 | 12 | 0 | 7 | 0 | 0 | 0 | 0 | 0 | 0 | 0 | 0 |
| 2 | 0.16 | 3.60 | 0.04 | 0.96 | 1 | Gm50141 | | 0 | 0 | 0 | 0 | 0 | 10 | 0 | 0 | 0 | 0 | 9 |
| 2 | 1.75 | 3.56 | 0.49 | 0.62 | 1 | Cdc34b | | 9 | 0 | 0 | 2 | 0 | 0 | 0 | 6 | 1 | 1 | 0 |
| 2 | 0.00 | 3.68 | 0.00 | 1.00 | 1 | Nlrp6 | NLR family, pyrin domain containing 6 | 0 | 18 | 1 | 0 | 0 | 0 | 0 | 0 | 0 | 0 | 0 |
| 2 | 3.27 | 3.59 | 0.91 | 0.36 | 1 | Nyx | nyctalopin | 0 | 7 | 0 | 0 | 0 | 5 | 0 | 0 | 3 | 0 | 4 |
| 2 | 1.64 | 3.62 | 0.45 | 0.65 | 1 | D030018L15Rik | | 0 | 16 | 0 | 0 | 0 | 0 | 0 | 0 | 0 | 0 | 3 |
| 2 | 1.50 | 3.63 | 0.41 | 0.68 | 1 | Gm15025 | | 0 | 17 | 0 | 0 | 0 | 0 | 0 | 0 | 0 | 0 | 2 |
| 2 | 0.00 | 3.65 | 0.00 | 1.00 | 1 | Platr16 | | 0 | 19 | 0 | 0 | 0 | 0 | 0 | 0 | 0 | 0 | 0 |
| 2 | 0.00 | 3.65 | 0.00 | 1.00 | 1 | Hsd17b3 | hydroxysteroid (17-beta) dehydrogenase | 0 | 19 | 0 | 0 | 0 | 0 | 0 | 0 | 0 | 0 | 0 |
| 2 | 3.78 | 3.61 | 1.05 | 0.30 | 1 | Gm12034 | | 10 | 1 | 0 | 0 | 0 | 1 | 0 | 0 | 6 | 0 | 1 |
| 2 | 0.35 | 3.58 | 0.10 | 0.92 | 1 | 4930445N18Rik | | 6 | 0 | 7 | 0 | 2 | 0 | 0 | 0 | 0 | 0 | 4 |
| 2 | 0.00 | 3.68 | 0.00 | 1.00 | 1 | 2410017I17Rik | | 3 | 0 | 17 | 0 | 0 | 0 | 0 | 0 | 0 | 0 | 0 |

| | | | | | | | | | | | | | | | | | |
|---|---|---|---|---|---|---|---|---|---|---|---|---|---|---|---|---|---|
| 2 | 0.00 | 3.68 | 0.00 | 1.00 | 1 | Ubash3a | ubiquitin associated and SH3 domain cor | 12 | 4 | 3 | 0 | 0 | 0 | 0 | 0 | 0 | 0 | 0 |
| 2 | -0.14 | 3.60 | -0.04 | 0.97 | 1 | Gm44093 | | 0 | 0 | 0 | 1 | 0 | 5 | 0 | 0 | 0 | 0 | 13 |
| 2 | 0.29 | 3.56 | 0.08 | 0.94 | 1 | A330015K06Rik | | 0 | 8 | 0 | 6 | 0 | 1 | 0 | 0 | 4 | 0 | 0 |
| 2 | -3.02 | 3.58 | -0.84 | 0.40 | 1 | Gm49590 | | 1 | 0 | 0 | 0 | 5 | 3 | 0 | 0 | 0 | 0 | 10 |
| 2 | 0.08 | 3.60 | 0.02 | 0.98 | 1 | Gm25492 | | 0 | 0 | 0 | 0 | 0 | 11 | 0 | 0 | 0 | 0 | 8 |
| 2 | 0.22 | 3.50 | 0.06 | 0.95 | 1 | D5Ertd615e | | 0 | 0 | 0 | 4 | 0 | 4 | 0 | 0 | 0 | 6 | 4 |
| 2 | 0.00 | 3.68 | 0.00 | 1.00 | 1 | Gm45912 | | 13 | 0 | 6 | 0 | 0 | 0 | 0 | 0 | 0 | 0 | 0 |
| 2 | 0.00 | 3.68 | 0.00 | 1.00 | 1 | Igkj2 | | 12 | 7 | 0 | 0 | 0 | 0 | 0 | 0 | 0 | 0 | 0 |
| 2 | 2.09 | 3.59 | 0.58 | 0.56 | 1 | Gm37760 | | 0 | 1 | 0 | 0 | 0 | 16 | 0 | 0 | 2 | 0 | 0 |
| 2 | 3.67 | 3.57 | 1.03 | 0.30 | 1 | Cyp2d26 | cytochrome P450, family 2, subfamily d, p | 0 | 2 | 0 | 0 | 0 | 7 | 0 | 0 | 0 | 10 | 0 |
| 2 | 0.18 | 3.60 | 0.05 | 0.96 | 1 | Gm15937 | | 0 | 0 | 6 | 0 | 0 | 7 | 0 | 0 | 0 | 0 | 6 |
| 2 | -3.31 | 3.58 | -0.92 | 0.36 | 1 | Gm14002 | | 9 | 0 | 0 | 0 | 7 | 1 | 0 | 0 | 0 | 0 | 3 |
| 2 | 0.00 | 3.68 | 0.00 | 1.00 | 1 | Retnlg | resistin like gamma | 2 | 12 | 5 | 0 | 0 | 0 | 0 | 0 | 0 | 0 | 0 |
| 2 | -1.01 | 3.61 | -0.28 | 0.78 | 1 | Plin5 | perilipin 5 | 5 | 0 | 7 | 0 | 7 | 0 | 0 | 0 | 0 | 0 | 0 |
| 2 | -0.33 | 3.60 | -0.09 | 0.93 | 1 | Gm25291 | | 0 | 0 | 4 | 0 | 0 | 12 | 0 | 0 | 0 | 0 | 3 |
| 2 | 0.00 | 3.68 | 0.00 | 1.00 | 1 | G6pc | glucose-6-phosphatase, catalytic | 3 | 0 | 16 | 0 | 0 | 0 | 0 | 0 | 0 | 0 | 0 |
| 2 | 0.00 | 3.68 | 0.00 | 1.00 | 1 | Krt14 | keratin 14 | 7 | 12 | 0 | 0 | 0 | 0 | 0 | 0 | 0 | 0 | 0 |
| 2 | 0.88 | 3.66 | 0.24 | 0.81 | 1 | Slc9a4 | solute carrier family 9 (sodium/hydrogen e | 3 | 0 | 15 | 0 | 0 | 0 | 0 | 0 | 1 | 0 | 0 |
| 2 | 1.30 | 3.64 | 0.36 | 0.72 | 1 | Gm9767 | | 2 | 13 | 1 | 0 | 0 | 0 | 0 | 0 | 0 | 0 | 3 |
| 2 | 0.44 | 3.60 | 0.12 | 0.90 | 1 | | | 0 | 0 | 0 | 0 | 0 | 7 | 0 | 0 | 0 | 0 | 12 |
| 2 | 0.44 | 3.60 | 0.12 | 0.90 | 1 | Gm37183 | | 0 | 0 | 0 | 0 | 0 | 7 | 0 | 0 | 0 | 0 | 12 |
| 2 | 0.00 | 3.68 | 0.00 | 1.00 | 1 | Banf2os | | 13 | 6 | 0 | 0 | 0 | 0 | 0 | 0 | 0 | 0 | 0 |
| 2 | 0.41 | 3.60 | 0.11 | 0.91 | 1 | 4930533D04Rik | | 0 | 1 | 0 | 0 | 0 | 7 | 0 | 0 | 0 | 0 | 11 |
| 2 | 4.65 | 3.05 | 1.52 | 0.13 | 1 | Gm12818 | | 1 | 1 | 0 | 0 | 0 | 0 | 0 | 5 | 5 | 1 | 6 |
| 2 | 0.00 | 3.60 | 0.00 | 1.00 | 1 | Gm29036 | | 0 | 0 | 0 | 0 | 0 | 12 | 0 | 0 | 0 | 0 | 7 |
| 2 | 1.33 | 3.63 | 0.37 | 0.71 | 1 | Apol8 | apolipoprotein L 8 | 3 | 0 | 11 | 0 | 0 | 0 | 0 | 0 | 0 | 0 | 4 |
| 2 | 0.00 | 3.65 | 0.00 | 1.00 | 1 | Gnrhr | gonadotropin releasing hormone recepto | 19 | 0 | 0 | 0 | 0 | 0 | 0 | 0 | 0 | 0 | 0 |
| 2 | 0.00 | 3.65 | 0.00 | 1.00 | 1 | Igkv4-63 | | 19 | 0 | 0 | 0 | 0 | 0 | 0 | 0 | 0 | 0 | 0 |
| 2 | 0.00 | 3.65 | 0.00 | 1.00 | 1 | Colec10 | collectin sub-family member 10 | 19 | 0 | 0 | 0 | 0 | 0 | 0 | 0 | 0 | 0 | 0 |
| 2 | 0.00 | 3.68 | 0.00 | 1.00 | 1 | Cers3 | ceramide synthase 3 | 8 | 11 | 0 | 0 | 0 | 0 | 0 | 0 | 0 | 0 | 0 |
| 2 | 0.00 | 3.68 | 0.00 | 1.00 | 1 | Trim29 | tripartite motif-containing 29 | 8 | 11 | 0 | 0 | 0 | 0 | 0 | 0 | 0 | 0 | 0 |
| 2 | 0.00 | 3.68 | 0.00 | 1.00 | 1 | Rbbp8nl | RBBP8 N-terminal like | 3 | 6 | 9 | 0 | 0 | 0 | 0 | 0 | 0 | 0 | 0 |
| 2 | 0.23 | 3.60 | 0.06 | 0.95 | 1 | Snord49b | | 0 | 0 | 3 | 0 | 0 | 8 | 0 | 0 | 0 | 0 | 8 |
| 2 | 0.07 | 3.60 | 0.02 | 0.98 | 1 | D830026I12Rik | | 4 | 0 | 0 | 0 | 0 | 8 | 0 | 0 | 0 | 0 | 6 |
| 2 | 4.36 | 3.59 | 1.21 | 0.22 | 1 | Spin3-ps | | 6 | 0 | 0 | 0 | 0 | 2 | 0 | 0 | 8 | 0 | 3 |
| 2 | -0.73 | 3.64 | -0.20 | 0.84 | 1 | Ddn | dendrin | 5 | 11 | 0 | 0 | 0 | 3 | 0 | 0 | 0 | 0 | 0 |
| 2 | -1.14 | 3.59 | -0.32 | 0.75 | 1 | Gm48334 | | 0 | 0 | 0 | 0 | 2 | 12 | 0 | 0 | 0 | 0 | 5 |
| 2 | -0.08 | 3.60 | -0.02 | 0.98 | 1 | Gm15381 | | 0 | 0 | 0 | 0 | 0 | 13 | 0 | 0 | 0 | 0 | 6 |
| 2 | 0.37 | 3.60 | 0.10 | 0.92 | 1 | Polr2k-ps | | 10 | 0 | 0 | 0 | 0 | 3 | 0 | 0 | 0 | 0 | 5 |
| 2 | 0.00 | 3.68 | 0.00 | 1.00 | 1 | Gm26577 | | 3 | 11 | 4 | 0 | 0 | 0 | 0 | 0 | 0 | 0 | 0 |
| 2 | 0.00 | 3.68 | 0.00 | 1.00 | 1 | Optc | opticin | 9 | 10 | 0 | 0 | 0 | 0 | 0 | 0 | 0 | 0 | 0 |
| 2 | 0.00 | 3.64 | 0.00 | 1.00 | 1 | Mab21l4 | mab-21-like 4 | 0 | 0 | 19 | 0 | 0 | 0 | 0 | 0 | 0 | 0 | 0 |
| 2 | 0.00 | 3.64 | 0.00 | 1.00 | 1 | Gm14133 | | 0 | 0 | 19 | 0 | 0 | 0 | 0 | 0 | 0 | 0 | 0 |
| 2 | 0.00 | 3.64 | 0.00 | 1.00 | 1 | Chil3 | chitinase-like 3 | 0 | 0 | 19 | 0 | 0 | 0 | 0 | 0 | 0 | 0 | 0 |
| 2 | 0.00 | 3.64 | 0.00 | 1.00 | 1 | Gm19557 | | 0 | 0 | 19 | 0 | 0 | 0 | 0 | 0 | 0 | 0 | 0 |
| 2 | 0.39 | 3.61 | 0.11 | 0.91 | 1 | Gm37242 | | 2 | 0 | 10 | 0 | 0 | 3 | 0 | 0 | 0 | 0 | 4 |
| 2 | -2.14 | 3.55 | -0.60 | 0.55 | 1 | Stx19 | syntaxin 19 | 0 | 0 | 5 | 10 | 0 | 1 | 0 | 0 | 3 | 0 | 0 |
| 2 | 0.02 | 3.60 | 0.01 | 1.00 | 1 | Gm26397 | | 6 | 0 | 0 | 0 | 0 | 8 | 0 | 0 | 0 | 0 | 5 |
| 2 | -0.03 | 3.60 | -0.01 | 0.99 | 1 | 2210412B16Rik | | 4 | 0 | 0 | 0 | 0 | 9 | 0 | 0 | 0 | 0 | 5 |
| 2 | -0.03 | 3.60 | -0.01 | 0.99 | 1 | Gm48673 | | 4 | 0 | 0 | 0 | 0 | 9 | 0 | 0 | 0 | 0 | 5 |
| 2 | -0.04 | 3.54 | -0.01 | 0.99 | 1 | Gm37521 | | 0 | 0 | 5 | 0 | 6 | 2 | 0 | 0 | 0 | 0 | 6 |
| 2 | -0.36 | 3.60 | -0.10 | 0.92 | 1 | Gm3468 | | 0 | 0 | 0 | 1 | 0 | 8 | 0 | 0 | 0 | 0 | 10 |
| 2 | 1.57 | 3.61 | 0.43 | 0.66 | 1 | Gm22980 | | 2 | 5 | 0 | 0 | 0 | 2 | 0 | 0 | 1 | 0 | 9 |
| 2 | 0.76 | 3.60 | 0.21 | 0.83 | 1 | Gm15513 | | 0 | 1 | 0 | 0 | 0 | 3 | 0 | 0 | 0 | 0 | 14 |
| 2 | 0.28 | 3.60 | 0.08 | 0.94 | 1 | Gm37357 | | 0 | 0 | 0 | 0 | 0 | 8 | 0 | 0 | 0 | 0 | 10 |
| 2 | 0.28 | 3.60 | 0.08 | 0.94 | 1 | Gm49286 | | 0 | 0 | 0 | 0 | 0 | 8 | 0 | 0 | 0 | 0 | 10 |
| 2 | 0.94 | 3.56 | 0.26 | 0.79 | 1 | Gm13874 | | 0 | 4 | 0 | 11 | 0 | 0 | 0 | 0 | 4 | 0 | 0 |
| 2 | -0.76 | 3.63 | -0.21 | 0.83 | 1 | Gm32552 | | 11 | 4 | 0 | 0 | 0 | 3 | 0 | 0 | 0 | 0 | 0 |
| 2 | 4.17 | 3.56 | 1.17 | 0.24 | 1 | Gm30881 | | 0 | 0 | 0 | 0 | 0 | 0 | 0 | 0 | 11 | 6 | 1 |
| 2 | 2.55 | 3.56 | 0.72 | 0.47 | 1 | Gm43541 | | 0 | 0 | 0 | 0 | 0 | 8 | 0 | 0 | 7 | 0 | 3 |

| | | | | | | | | | | | | | | | | | |
|---|---|---|---|---|---|---|---|---|---|---|---|---|---|---|---|---|---|
| 2 | 1.95 | 3.56 | 0.55 | 0.58 | 1 | Gm43274 | | 0 | 0 | 0 | 0 | 0 | 3 | 0 | 0 | 10 | 0 | 5 |
| 2 | 2.57 | 3.57 | 0.72 | 0.47 | 1 | Gm11979 | | 0 | 0 | 0 | 0 | 1 | 0 | 0 | 1 | 1 | 15 | 0 |
| 2 | 0.89 | 3.55 | 0.25 | 0.80 | 1 | Rgsl1 | regulator of G-protein signaling like 1 | 1 | 0 | 0 | 0 | 3 | 2 | 0 | 0 | 4 | 0 | 9 |
| 2 | 1.06 | 3.50 | 0.30 | 0.76 | 1 | H2ac15 | H2A clustered histone 15 | 0 | 0 | 0 | 9 | 0 | 2 | 0 | 0 | 4 | 0 | 4 |
| 2 | 0.42 | 2.99 | 0.14 | 0.89 | 1 | D830039M14Rik | | 0 | 0 | 0 | 5 | 2 | 1 | 0 | 3 | 6 | 0 | 1 |
| 2 | 0.00 | 3.68 | 0.00 | 1.00 | 1 | Raet1d | retinoic acid early transcript 1, alpha | 0 | 7 | 11 | 0 | 0 | 0 | 0 | 0 | 0 | 0 | 0 |
| 2 | 3.50 | 3.59 | 0.98 | 0.33 | 1 | Gm43097 | | 0 | 1 | 0 | 0 | 0 | 0 | 0 | 0 | 0 | 15 | 2 |
| 2 | 0.00 | 3.68 | 0.00 | 1.00 | 1 | Gm12022 | | 0 | 8 | 10 | 0 | 0 | 0 | 0 | 0 | 0 | 0 | 0 |
| 2 | 0.00 | 3.68 | 0.00 | 1.00 | 1 | Gm38198 | | 4 | 14 | 0 | 0 | 0 | 0 | 0 | 0 | 0 | 0 | 0 |
| 2 | 0.19 | 3.57 | 0.05 | 0.96 | 1 | Zfa-ps | | 0 | 0 | 0 | 0 | 9 | 1 | 0 | 0 | 8 | 0 | 1 |
| 2 | -3.07 | 3.59 | -0.85 | 0.39 | 1 | Gm6058 | | 3 | 2 | 0 | 10 | 0 | 4 | 0 | 0 | 0 | 0 | 0 |
| 2 | 2.54 | 3.59 | 0.71 | 0.48 | 1 | Mansc4 | MANSC domain containing 4 | 0 | 0 | 8 | 0 | 0 | 0 | 0 | 0 | 4 | 0 | 6 |
| 2 | 0.20 | 3.60 | 0.05 | 0.96 | 1 | Mir6955 | | 0 | 0 | 0 | 0 | 0 | 9 | 0 | 0 | 0 | 0 | 9 |
| 2 | 0.95 | 3.67 | 0.26 | 0.80 | 1 | Il22ra2 | interleukin 22 receptor, alpha 2 | 0 | 10 | 7 | 0 | 0 | 0 | 0 | 0 | 0 | 0 | 1 |
| 2 | 0.95 | 3.67 | 0.26 | 0.80 | 1 | Dnah5 | dynein, axonemal, heavy chain 5 | 0 | 10 | 7 | 0 | 0 | 0 | 0 | 0 | 0 | 0 | 1 |
| 2 | 0.00 | 3.68 | 0.00 | 1.00 | 1 | Hoxd13 | homeobox D13 | 0 | 11 | 7 | 0 | 0 | 0 | 0 | 0 | 0 | 0 | 0 |
| 2 | 2.08 | 3.59 | 0.58 | 0.56 | 1 | Gm9877 | | 0 | 10 | 0 | 0 | 0 | 0 | 0 | 1 | 0 | 0 | 7 |
| 2 | 0.39 | 3.63 | 0.11 | 0.91 | 1 | Rps12-ps19 | | 4 | 8 | 2 | 1 | 0 | 0 | 0 | 0 | 0 | 0 | 3 |
| 2 | -0.37 | 3.67 | -0.10 | 0.92 | 1 | Crnde | | 6 | 0 | 11 | 0 | 1 | 0 | 0 | 0 | 0 | 0 | 0 |
| 2 | 0.00 | 3.68 | 0.00 | 1.00 | 1 | Gm10684 | | 0 | 13 | 5 | 0 | 0 | 0 | 0 | 0 | 0 | 0 | 0 |
| 2 | 0.00 | 3.68 | 0.00 | 1.00 | 1 | Gm1123 | predicted gene 1123 | 0 | 13 | 5 | 0 | 0 | 0 | 0 | 0 | 0 | 0 | 0 |
| 2 | 0.03 | 3.35 | 0.01 | 0.99 | 1 | Gm9354 | | 8 | 0 | 1 | 4 | 0 | 1 | 0 | 1 | 0 | 0 | 4 |
| 2 | -5.18 | 3.54 | -1.46 | 0.14 | 1 | Gm4319 | | 0 | 0 | 0 | 7 | 0 | 4 | 0 | 0 | 7 | 0 | 0 |
| 2 | 0.00 | 3.68 | 0.00 | 1.00 | 1 | 1810009J06Rik | RIKEN cDNA 1810009J06 gene | 7 | 0 | 11 | 0 | 0 | 0 | 0 | 0 | 0 | 0 | 0 |
| 2 | -4.25 | 3.59 | -1.19 | 0.24 | 1 | Lrrc10 | leucine rich repeat containing 10 | 3 | 0 | 0 | 0 | 9 | 3 | 0 | 0 | 0 | 0 | 2 |
| 2 | 1.99 | 2.69 | 0.74 | 0.46 | 1 | Gm35013 | | 0 | 1 | 2 | 0 | 6 | 0 | 0 | 1 | 6 | 1 | 1 |
| 2 | -0.38 | 3.60 | -0.10 | 0.92 | 1 | Gm33707 | | 0 | 0 | 0 | 0 | 0 | 15 | 0 | 0 | 0 | 0 | 3 |
| 2 | 0.00 | 3.68 | 0.00 | 1.00 | 1 | Slc22a8 | solute carrier family 22 (organic anion tra | 12 | 0 | 6 | 0 | 0 | 0 | 0 | 0 | 0 | 0 | 0 |
| 2 | 3.44 | 3.15 | 1.09 | 0.27 | 1 | Gm8643 | | 2 | 0 | 1 | 0 | 0 | 1 | 0 | 2 | 2 | 10 | 0 |
| 2 | 0.00 | 3.68 | 0.00 | 1.00 | 1 | Gm10552 | | 11 | 7 | 0 | 0 | 0 | 0 | 0 | 0 | 0 | 0 | 0 |
| 2 | 0.00 | 3.65 | 0.00 | 1.00 | 1 | Hsd3b3 | hydroxy-delta-5-steroid dehydrogenase, 3 | 0 | 18 | 0 | 0 | 0 | 0 | 0 | 0 | 0 | 0 | 0 |
| 2 | 0.00 | 3.65 | 0.00 | 1.00 | 1 | Gm19385 | | 0 | 18 | 0 | 0 | 0 | 0 | 0 | 0 | 0 | 0 | 0 |
| 2 | 0.00 | 3.65 | 0.00 | 1.00 | 1 | Casp14 | caspase 14 | 0 | 18 | 0 | 0 | 0 | 0 | 0 | 0 | 0 | 0 | 0 |
| 2 | 0.00 | 3.65 | 0.00 | 1.00 | 1 | Gm34983 | | 0 | 18 | 0 | 0 | 0 | 0 | 0 | 0 | 0 | 0 | 0 |
| 2 | 0.00 | 3.65 | 0.00 | 1.00 | 1 | Gm38407 | | 0 | 18 | 0 | 0 | 0 | 0 | 0 | 0 | 0 | 0 | 0 |
| 2 | 0.00 | 3.65 | 0.00 | 1.00 | 1 | Fbxo15 | F-box protein 15 | 0 | 18 | 0 | 0 | 0 | 0 | 0 | 0 | 0 | 0 | 0 |
| 2 | 3.75 | 3.58 | 1.05 | 0.29 | 1 | Gm22422 | | 0 | 0 | 0 | 0 | 0 | 0 | 0 | 0 | 0 | 14 | 4 |
| 2 | -0.28 | 2.48 | -0.11 | 0.91 | 1 | Gm11362 | | 2 | 3 | 1 | 2 | 1 | 3 | 0 | 0 | 0 | 5 | 2 |
| 2 | 2.22 | 3.55 | 0.63 | 0.53 | 1 | Gm26903 | | 0 | 2 | 0 | 0 | 0 | 2 | 0 | 10 | 1 | 0 | 3 |
| 2 | -1.20 | 3.56 | -0.34 | 0.74 | 1 | Gm30541 | | 0 | 0 | 0 | 8 | 0 | 4 | 0 | 0 | 0 | 0 | 6 |
| 2 | -0.47 | 3.60 | -0.13 | 0.90 | 1 | Rnf223 | ring finger 223 | 7 | 0 | 0 | 0 | 0 | 9 | 0 | 0 | 0 | 0 | 2 |
| 2 | 0.87 | 3.59 | 0.24 | 0.81 | 1 | Gm37204 | | 0 | 0 | 10 | 0 | 0 | 2 | 0 | 0 | 0 | 0 | 6 |
| 2 | 1.62 | 3.59 | 0.45 | 0.65 | 1 | Gm5517 | | 0 | 13 | 0 | 0 | 1 | 0 | 0 | 0 | 3 | 0 | 1 |
| 2 | 0.00 | 3.68 | 0.00 | 1.00 | 1 | Gm15156 | | 2 | 7 | 9 | 0 | 0 | 0 | 0 | 0 | 0 | 0 | 0 |
| 2 | 1.72 | 2.75 | 0.63 | 0.53 | 1 | Rpl18-ps1 | | 2 | 3 | 8 | 1 | 0 | 0 | 0 | 1 | 2 | 0 | 1 |
| 2 | 2.59 | 3.13 | 0.83 | 0.41 | 1 | Gm6474 | | 3 | 2 | 8 | 0 | 0 | 0 | 0 | 1 | 3 | 0 | 1 |
| 2 | 0.95 | 3.67 | 0.26 | 0.80 | 1 | Gm14230 | | 7 | 6 | 4 | 0 | 0 | 0 | 0 | 0 | 0 | 0 | 1 |
| 2 | 1.03 | 3.57 | 0.29 | 0.77 | 1 | Gm8802 | | 0 | 0 | 8 | 0 | 5 | 0 | 0 | 0 | 2 | 0 | 3 |
| 2 | -2.79 | 3.56 | -0.78 | 0.43 | 1 | Snora34 | | 0 | 0 | 0 | 0 | 11 | 2 | 0 | 0 | 0 | 0 | 5 |
| 2 | 0.14 | 3.64 | 0.04 | 0.97 | 1 | Gm7588 | | 0 | 7 | 8 | 0 | 0 | 2 | 0 | 0 | 1 | 0 | 0 |
| 2 | 1.02 | 3.59 | 0.29 | 0.78 | 1 | Dnah17 | dynein, axonemal, heavy chain 17 | 0 | 0 | 7 | 0 | 0 | 2 | 0 | 0 | 0 | 0 | 9 |
| 2 | 0.49 | 3.60 | 0.14 | 0.89 | 1 | Gm12259 | | 0 | 0 | 0 | 0 | 0 | 6 | 0 | 0 | 0 | 0 | 12 |
| 2 | 0.42 | 3.60 | 0.12 | 0.91 | 1 | Gm13689 | | 0 | 2 | 0 | 0 | 0 | 6 | 0 | 0 | 0 | 0 | 10 |
| 2 | 3.27 | 3.59 | 0.91 | 0.36 | 1 | Gnat1 | guanine nucleotide binding protein, alpha | 0 | 0 | 0 | 0 | 0 | 0 | 0 | 15 | 3 | 0 | 0 |
| 2 | -0.45 | 3.54 | -0.13 | 0.90 | 1 | Gm7457 | | 0 | 0 | 0 | 0 | 12 | 0 | 0 | 6 | 0 | 0 | 0 |
| 2 | -1.64 | 3.25 | -0.50 | 0.61 | 1 | Rpl31-ps1 | | 1 | 10 | 0 | 2 | 1 | 3 | 0 | 0 | 1 | 0 | 0 |
| 2 | -0.22 | 3.55 | -0.06 | 0.95 | 1 | Gm37402 | | 0 | 0 | 0 | 0 | 8 | 3 | 0 | 0 | 5 | 0 | 2 |
| 2 | 0.10 | 3.58 | 0.03 | 0.98 | 1 | Gm10654 | predicted gene 10654 | 1 | 0 | 11 | 0 | 0 | 2 | 0 | 0 | 4 | 0 | 0 |
| 2 | 1.83 | 2.48 | 0.74 | 0.46 | 1 | Rps2-ps9 | | 1 | 2 | 1 | 0 | 1 | 1 | 0 | 9 | 1 | 0 | 2 |
| 2 | 0.00 | 3.65 | 0.00 | 1.00 | 1 | Wnt10a | wingless-type MMTV integration site famil | 18 | 0 | 0 | 0 | 0 | 0 | 0 | 0 | 0 | 0 | 0 |

| | | | | | | | | | | | | | | | | | | |
|---|---|---|---|---|---|---|---|---|---|---|---|---|---|---|---|---|---|---|
| 2 | 0.00 | 3.65 | 0.00 | 1.00 | 1 | Gm14406 | | 18 | 0 | 0 | 0 | 0 | 0 | 0 | 0 | 0 | 0 | 0 |
| 2 | 0.00 | 3.65 | 0.00 | 1.00 | 1 | Angptl3 | angiopoietin-like 3 | 18 | 0 | 0 | 0 | 0 | 0 | 0 | 0 | 0 | 0 | 0 |
| 2 | 0.00 | 3.65 | 0.00 | 1.00 | 1 | Gm16350 | | 18 | 0 | 0 | 0 | 0 | 0 | 0 | 0 | 0 | 0 | 0 |
| 2 | 0.39 | 2.86 | 0.14 | 0.89 | 1 | Gm10653 | | 3 | 0 | 6 | 3 | 1 | 0 | 0 | 1 | 1 | 3 | 0 |
| 2 | -3.17 | 3.57 | -0.89 | 0.37 | 1 | Gm44714 | | 0 | 0 | 0 | 11 | 0 | 3 | 0 | 0 | 0 | 0 | 4 |
| 2 | -2.61 | 3.53 | -0.74 | 0.46 | 1 | Gm25238 | | 0 | 0 | 0 | 4 | 4 | 7 | 0 | 0 | 3 | 0 | 0 |
| 2 | 0.41 | 3.60 | 0.11 | 0.91 | 1 | Snora23 | | 0 | 0 | 0 | 0 | 0 | 7 | 0 | 0 | 0 | 0 | 11 |
| 2 | 2.21 | 3.49 | 0.63 | 0.53 | 1 | Gm8225 | | 3 | 0 | 4 | 8 | 0 | 0 | 0 | 0 | 1 | 1 | 1 |
| 2 | 0.00 | 3.68 | 0.00 | 1.00 | 1 | Sirpb1b | signal-regulatory protein beta 1B | 2 | 16 | 0 | 0 | 0 | 0 | 0 | 0 | 0 | 0 | 0 |
| 2 | 1.78 | 3.56 | 0.50 | 0.62 | 1 | Gm16351 | | 0 | 0 | 0 | 0 | 0 | 2 | 0 | 16 | 0 | 0 | 0 |
| 2 | -4.67 | 3.61 | -1.29 | 0.20 | 1 | Hnrnpa1l2-ps2 | heterogeneous nuclear ribonucleoprotein | 0 | 4 | 0 | 0 | 9 | 3 | 0 | 0 | 0 | 0 | 1 |
| 2 | 0.23 | 2.78 | 0.08 | 0.94 | 1 | Gm14131 | | 0 | 1 | 1 | 2 | 3 | 1 | 0 | 0 | 9 | 1 | 0 |
| 2 | -1.42 | 3.60 | -0.39 | 0.69 | 1 | Mei4 | meiotic double-stranded break formation | 0 | 14 | 0 | 2 | 2 | 0 | 0 | 0 | 0 | 0 | 0 |
| 2 | -0.98 | 3.57 | -0.27 | 0.78 | 1 | Gm42895 | | 0 | 0 | 0 | 0 | 4 | 6 | 0 | 0 | 0 | 0 | 8 |
| 2 | 2.52 | 3.58 | 0.70 | 0.48 | 1 | Gm37500 | | 0 | 0 | 0 | 1 | 0 | 0 | 0 | 14 | 3 | 0 | 0 |
| 2 | 2.70 | 3.59 | 0.75 | 0.45 | 1 | Fam186a | family with sequence similarity 186, meml | 0 | 0 | 0 | 0 | 0 | 0 | 0 | 0 | 0 | 3 | 15 |
| 2 | -1.44 | 3.56 | -0.41 | 0.68 | 1 | Gm6020 | | 0 | 0 | 0 | 11 | 0 | 3 | 0 | 0 | 0 | 0 | 3 |
| 2 | -4.16 | 3.55 | -1.17 | 0.24 | 1 | Gm13257 | | 0 | 0 | 0 | 5 | 7 | 6 | 0 | 0 | 0 | 0 | 0 |
| 2 | 0.00 | 3.68 | 0.00 | 1.00 | 1 | Fam71a | family with sequence similarity 71, memb | 14 | 4 | 0 | 0 | 0 | 0 | 0 | 0 | 0 | 0 | 0 |
| 2 | 0.00 | 3.68 | 0.00 | 1.00 | 1 | Gm11545 | | 4 | 1 | 12 | 0 | 0 | 0 | 0 | 0 | 0 | 0 | 0 |
| 2 | 1.57 | 3.54 | 0.44 | 0.66 | 1 | Sycp3 | synaptonemal complex protein 3 | 4 | 0 | 0 | 4 | 0 | 1 | 0 | 0 | 1 | 0 | 8 |
| 2 | 0.88 | 3.60 | 0.24 | 0.81 | 1 | Prrt2 | proline-rich transmembrane protein 2 | 0 | 0 | 0 | 0 | 0 | 3 | 0 | 0 | 0 | 0 | 15 |
| 2 | 0.00 | 3.56 | 0.00 | 1.00 | 1 | Skor1 | SKI family transcriptional corepressor 1 | 0 | 0 | 6 | 0 | 0 | 8 | 0 | 0 | 1 | 0 | 2 |
| 2 | 2.45 | 3.59 | 0.68 | 0.49 | 1 | Gm6419 | | 0 | 0 | 11 | 0 | 1 | 0 | 0 | 0 | 0 | 1 | 4 |
| 2 | -0.31 | 3.26 | -0.10 | 0.92 | 1 | Rps12-ps26 | | 5 | 0 | 1 | 1 | 0 | 8 | 0 | 0 | 1 | 0 | 2 |
| 2 | -1.32 | 3.63 | -0.36 | 0.72 | 1 | Cox7c-ps1 | | 11 | 3 | 0 | 4 | 0 | 0 | 0 | 0 | 0 | 0 | 0 |
| 2 | 0.00 | 3.65 | 0.00 | 1.00 | 1 | Gm5420 | | 0 | 0 | 18 | 0 | 0 | 0 | 0 | 0 | 0 | 0 | 0 |
| 2 | 0.00 | 3.65 | 0.00 | 1.00 | 1 | Cyp4f40 | cytochrome P450, family 4, subfamily f, pc | 0 | 0 | 18 | 0 | 0 | 0 | 0 | 0 | 0 | 0 | 0 |
| 2 | -1.44 | 3.57 | -0.40 | 0.69 | 1 | Gm43882 | | 0 | 0 | 0 | 0 | 10 | 4 | 0 | 0 | 3 | 0 | 0 |
| 2 | -3.39 | 3.56 | -0.95 | 0.34 | 1 | Acsbg3 | acyl-CoA synthetase bubblegum family m | 0 | 0 | 1 | 3 | 12 | 2 | 0 | 0 | 0 | 0 | 0 |
| 2 | 1.34 | 3.65 | 0.37 | 0.71 | 1 | Rrh | retinal pigment epithelium derived rhodop | 15 | 1 | 0 | 0 | 0 | 0 | 0 | 0 | 0 | 0 | 2 |
| 2 | 0.00 | 3.68 | 0.00 | 1.00 | 1 | Gm13012 | | 15 | 3 | 0 | 0 | 0 | 0 | 0 | 0 | 0 | 0 | 0 |
| 2 | 0.00 | 3.68 | 0.00 | 1.00 | 1 | Siglecf | sialic acid binding Ig-like lectin F | 5 | 0 | 12 | 0 | 0 | 0 | 0 | 0 | 0 | 0 | 0 |
| 2 | 0.00 | 3.68 | 0.00 | 1.00 | 1 | Gzmb | granzyme B | 0 | 2 | 16 | 0 | 0 | 0 | 0 | 0 | 0 | 0 | 0 |
| 2 | 0.00 | 3.68 | 0.00 | 1.00 | 1 | Gm336 | | 3 | 14 | 0 | 0 | 0 | 0 | 0 | 0 | 0 | 0 | 0 |
| 2 | 0.76 | 3.60 | 0.21 | 0.83 | 1 | Mir697 | | 0 | 0 | 0 | 0 | 0 | 3 | 0 | 0 | 0 | 0 | 14 |
| 2 | 0.26 | 3.61 | 0.07 | 0.94 | 1 | Gm9918 | | 8 | 0 | 0 | 0 | 0 | 1 | 0 | 0 | 0 | 9 | 0 |
| 2 | 1.40 | 3.57 | 0.39 | 0.70 | 1 | A430057M04Ri | RIKEN cDNA A430057M04 gene | 0 | 0 | 0 | 0 | 0 | 8 | 0 | 0 | 5 | 0 | 4 |
| 2 | 1.79 | 3.61 | 0.49 | 0.62 | 1 | Best2 | bestrophin 2 | 0 | 0 | 14 | 0 | 0 | 0 | 0 | 0 | 0 | 0 | 4 |
| 2 | 0.48 | 3.60 | 0.13 | 0.89 | 1 | Fam166b | family with sequence similarity 166, meml | 0 | 0 | 6 | 0 | 0 | 4 | 0 | 0 | 0 | 0 | 7 |
| 2 | -2.79 | 3.58 | -0.78 | 0.44 | 1 | 4930509E16Rik | | 0 | 0 | 7 | 4 | 0 | 2 | 0 | 0 | 0 | 0 | 5 |
| 2 | -2.85 | 3.53 | -0.81 | 0.42 | 1 | Hapln2 | hyaluronan and proteoglycan link protein | 0 | 0 | 0 | 9 | 5 | 0 | 0 | 0 | 0 | 0 | 4 |
| 2 | 1.68 | 3.61 | 0.47 | 0.64 | 1 | Iqcn | IQ motif containing N | 7 | 0 | 0 | 0 | 0 | 3 | 0 | 0 | 0 | 0 | 8 |
| 2 | -1.04 | 3.56 | -0.29 | 0.77 | 1 | 1700067K01Ril | RIKEN cDNA 1700067K01 gene | 0 | 0 | 0 | 7 | 0 | 3 | 0 | 0 | 0 | 0 | 7 |
| 2 | 0.29 | 2.80 | 0.10 | 0.92 | 1 | Gm10247 | | 4 | 6 | 3 | 1 | 1 | 0 | 0 | 1 | 0 | 0 | 1 |
| 2 | 0.94 | 3.55 | 0.27 | 0.79 | 1 | Foxo6os | | 0 | 0 | 3 | 0 | 0 | 4 | 0 | 0 | 8 | 0 | 2 |
| 2 | -0.01 | 3.60 | 0.00 | 1.00 | 1 | Casp16 | | 0 | 0 | 0 | 1 | 0 | 3 | 0 | 0 | 0 | 0 | 13 |
| 2 | 0.00 | 3.68 | 0.00 | 1.00 | 1 | Gm28177 | | 0 | 8 | 9 | 0 | 0 | 0 | 0 | 0 | 0 | 0 | 0 |
| 2 | 1.09 | 3.52 | 0.31 | 0.76 | 1 | Ddx4 | DEAD (Asp-Glu-Ala-Asp) box polypeptide | 0 | 0 | 0 | 2 | 0 | 3 | 0 | 0 | 8 | 0 | 5 |
| 2 | -0.43 | 3.68 | -0.12 | 0.91 | 1 | Amy2-ps1 | | 0 | 3 | 14 | 1 | 0 | 0 | 0 | 0 | 0 | 0 | 0 |
| 2 | 0.15 | 3.60 | 0.04 | 0.97 | 1 | Gm23318 | | 0 | 0 | 0 | 0 | 0 | 9 | 0 | 0 | 0 | 0 | 8 |
| 2 | 0.00 | 3.68 | 0.00 | 1.00 | 1 | Gm32401 | | 11 | 0 | 6 | 0 | 0 | 0 | 0 | 0 | 0 | 0 | 0 |
| 2 | 0.00 | 3.68 | 0.00 | 1.00 | 1 | Igkj5 | | 10 | 7 | 0 | 0 | 0 | 0 | 0 | 0 | 0 | 0 | 0 |
| 2 | 1.50 | 2.82 | 0.53 | 0.60 | 1 | Gm3226 | | 4 | 4 | 1 | 0 | 0 | 3 | 0 | 2 | 0 | 1 | 1 |
| 2 | 1.57 | 3.57 | 0.44 | 0.66 | 1 | Gm20402 | | 3 | 0 | 4 | 0 | 0 | 2 | 0 | 0 | 6 | 0 | 2 |
| 2 | 0.00 | 3.68 | 0.00 | 1.00 | 1 | Gm8013 | | 0 | 11 | 6 | 0 | 0 | 0 | 0 | 0 | 0 | 0 | 0 |
| 2 | 1.66 | 3.59 | 0.46 | 0.64 | 1 | Gm47918 | | 0 | 0 | 4 | 0 | 0 | 0 | 0 | 0 | 0 | 0 | 13 |
| 2 | -1.75 | 3.62 | -0.48 | 0.63 | 1 | Prss50 | protease, serine 50 | 9 | 0 | 5 | 0 | 2 | 2 | 0 | 0 | 0 | 0 | 0 |
| 2 | -0.66 | 3.60 | -0.18 | 0.85 | 1 | Gm6829 | | 0 | 1 | 0 | 0 | 1 | 9 | 0 | 0 | 0 | 0 | 6 |
| 2 | 0.75 | 3.61 | 0.21 | 0.84 | 1 | Gm15943 | | 6 | 0 | 0 | 0 | 0 | 4 | 0 | 0 | 0 | 0 | 7 |

| | | | | | | | | | | | | | | | | | | |
|---|---|---|---|---|---|---|---|---|---|---|---|---|---|---|---|---|---|---|
| 2 | -1.39 | 2.59 | -0.54 | 0.59 | 1 | Gm8302 | | 1 | 1 | 1 | 1 | 10 | 1 | 0 | 0 | 0 | 1 | 1 |
| 2 | 0.00 | 3.68 | 0.00 | 1.00 | 1 | Gm12407 | | 5 | 11 | 1 | 0 | 0 | 0 | 0 | 0 | 0 | 0 | 0 |
| 2 | 5.16 | 3.61 | 1.43 | 0.15 | 1 | Tmem238l | | 3 | 0 | 0 | 0 | 0 | 1 | 0 | 7 | 0 | 0 | 6 |
| 2 | 1.39 | 3.57 | 0.39 | 0.70 | 1 | U2af1l4 | U2 small nuclear RNA auxiliary factor 1-li | 0 | 0 | 0 | 0 | 0 | 8 | 0 | 5 | 0 | 0 | 5 |
| 2 | -0.07 | 3.61 | -0.02 | 0.98 | 1 | B130046B21Rik | | 0 | 0 | 14 | 1 | 2 | 0 | 0 | 0 | 0 | 0 | 1 |
| 2 | -3.50 | 3.57 | -0.98 | 0.33 | 1 | Tmprss9 | transmembrane protease, serine 9 | 2 | 0 | 0 | 11 | 0 | 5 | 0 | 0 | 0 | 0 | 0 |
| 2 | 2.15 | 2.84 | 0.76 | 0.45 | 1 | Gm13135 | | 6 | 4 | 1 | 0 | 0 | 1 | 0 | 0 | 2 | 1 | 2 |
| 2 | 4.54 | 3.59 | 1.26 | 0.21 | 1 | Gm36955 | | 3 | 0 | 0 | 0 | 0 | 2 | 0 | 0 | 7 | 0 | 5 |
| 2 | 0.28 | 3.61 | 0.08 | 0.94 | 1 | Gm48231 | | 3 | 9 | 0 | 0 | 0 | 3 | 0 | 0 | 0 | 0 | 3 |
| 2 | 1.39 | 3.61 | 0.39 | 0.70 | 1 | Gm26640 | | 0 | 0 | 0 | 0 | 0 | 0 | 0 | 0 | 0 | 0 | 17 |
| 2 | 0.27 | 2.81 | 0.09 | 0.92 | 1 | Gm12712 | | 2 | 6 | 2 | 4 | 0 | 1 | 0 | 0 | 1 | 0 | 2 |
| 2 | 0.55 | 3.60 | 0.15 | 0.88 | 1 | Acr | acrosin prepropeptide | 0 | 0 | 0 | 0 | 0 | 5 | 0 | 0 | 0 | 0 | 12 |
| 2 | 0.55 | 3.60 | 0.15 | 0.88 | 1 | Snora78 | | 0 | 0 | 0 | 0 | 0 | 5 | 0 | 0 | 0 | 0 | 12 |
| 2 | 2.80 | 3.58 | 0.78 | 0.43 | 1 | Gm29229 | | 0 | 0 | 0 | 0 | 0 | 1 | 0 | 0 | 0 | 10 | 6 |
| 2 | 3.56 | 3.58 | 0.99 | 0.32 | 1 | Gm4065 | | 0 | 0 | 0 | 0 | 0 | 0 | 0 | 0 | 6 | 0 | 11 |
| 2 | 0.04 | 3.60 | 0.01 | 0.99 | 1 | Gm25630 | | 0 | 8 | 0 | 0 | 0 | 5 | 0 | 0 | 0 | 0 | 4 |
| 2 | 2.84 | 3.59 | 0.79 | 0.43 | 1 | Nxnl1 | nucleoredoxin-like 1 | 0 | 0 | 0 | 0 | 0 | 0 | 0 | 16 | 0 | 0 | 1 |
| 2 | 1.64 | 3.62 | 0.45 | 0.65 | 1 | Gm43072 | | 0 | 14 | 0 | 0 | 0 | 0 | 0 | 0 | 0 | 0 | 3 |
| 2 | -0.69 | 3.66 | -0.19 | 0.85 | 1 | Srarp | steroid receptor associated and regulatec | 0 | 2 | 14 | 0 | 0 | 2 | 0 | 0 | 0 | 0 | 0 |
| 2 | 1.96 | 3.63 | 0.54 | 0.59 | 1 | Gm11972 | | 0 | 15 | 0 | 0 | 0 | 0 | 0 | 0 | 1 | 0 | 1 |
| 2 | -0.76 | 3.63 | -0.21 | 0.83 | 1 | Dleu7 | deleted in lymphocytic leukemia, 7 | 8 | 6 | 0 | 0 | 0 | 3 | 0 | 0 | 0 | 0 | 0 |
| 2 | 0.00 | 3.65 | 0.00 | 1.00 | 1 | Gm6264 | | 17 | 0 | 0 | 0 | 0 | 0 | 0 | 0 | 0 | 0 | 0 |
| 2 | 0.00 | 3.65 | 0.00 | 1.00 | 1 | Lcn11 | lipocalin 11 | 17 | 0 | 0 | 0 | 0 | 0 | 0 | 0 | 0 | 0 | 0 |
| 2 | 0.00 | 3.65 | 0.00 | 1.00 | 1 | Iglj1 | | 17 | 0 | 0 | 0 | 0 | 0 | 0 | 0 | 0 | 0 | 0 |
| 2 | -0.40 | 3.67 | -0.11 | 0.91 | 1 | Gm2808 | predicted gene 2808 | 0 | 12 | 4 | 0 | 0 | 1 | 0 | 0 | 0 | 0 | 0 |
| 2 | 1.25 | 3.66 | 0.34 | 0.73 | 1 | Gm49482 | | 6 | 9 | 0 | 0 | 0 | 0 | 0 | 0 | 0 | 0 | 2 |
| 2 | 1.63 | 3.56 | 0.46 | 0.65 | 1 | C430039J01Rik | | 0 | 0 | 0 | 0 | 0 | 2 | 0 | 0 | 0 | 15 | 0 |
| 2 | 0.00 | 3.68 | 0.00 | 1.00 | 1 | Gm9748 | | 2 | 6 | 9 | 0 | 0 | 0 | 0 | 0 | 0 | 0 | 0 |
| 2 | -0.59 | 3.65 | -0.16 | 0.87 | 1 | Tnni3 | troponin I, cardiac 3 | 16 | 0 | 0 | 0 | 0 | 1 | 0 | 0 | 0 | 0 | 0 |
| 2 | 0.28 | 3.68 | 0.08 | 0.94 | 1 | | | 11 | 1 | 3 | 0 | 0 | 1 | 0 | 0 | 1 | 0 | 0 |
| 2 | 0.25 | 3.61 | 0.07 | 0.94 | 1 | Gm8219 | | 3 | 0 | 0 | 0 | 7 | 0 | 0 | 0 | 7 | 0 | 0 |
| 2 | 2.42 | 3.55 | 0.68 | 0.49 | 1 | Gm12366 | | 2 | 2 | 0 | 0 | 0 | 7 | 0 | 0 | 4 | 3 | 0 |
| 2 | 0.24 | 3.58 | 0.07 | 0.95 | 1 | 0610033M10Rik | | 0 | 1 | 9 | 0 | 0 | 2 | 0 | 0 | 5 | 0 | 0 |
| 2 | -1.86 | 3.61 | -0.52 | 0.61 | 1 | | | 5 | 7 | 0 | 0 | 3 | 0 | 0 | 0 | 0 | 0 | 2 |
| 2 | -0.03 | 3.60 | -0.01 | 0.99 | 1 | Fgf17 | fibroblast growth factor 17 | 0 | 0 | 0 | 0 | 0 | 11 | 0 | 0 | 0 | 0 | 6 |
| 2 | 0.00 | 3.68 | 0.00 | 1.00 | 1 | Hoxa6 | homeobox A6 | 12 | 5 | 0 | 0 | 0 | 0 | 0 | 0 | 0 | 0 | 0 |
| 2 | -0.23 | 3.55 | -0.06 | 0.95 | 1 | Gm43686 | | 0 | 0 | 7 | 0 | 0 | 7 | 0 | 0 | 3 | 0 | 0 |
| 2 | 1.14 | 3.60 | 0.32 | 0.75 | 1 | 4930555B11Rik | | 0 | 0 | 0 | 0 | 0 | 1 | 0 | 16 | 0 | 0 | 0 |
| 2 | 1.64 | 3.57 | 0.46 | 0.64 | 1 | Mir670 | | 0 | 0 | 0 | 0 | 0 | 6 | 0 | 0 | 6 | 0 | 5 |
| 2 | -1.41 | 3.58 | -0.39 | 0.69 | 1 | Gm47073 | | 0 | 0 | 0 | 4 | 0 | 8 | 0 | 0 | 0 | 0 | 5 |
| 2 | 0.58 | 3.60 | 0.16 | 0.87 | 1 | Tex22 | testis expressed gene 22 | 0 | 14 | 0 | 0 | 0 | 1 | 0 | 0 | 0 | 0 | 2 |
| 2 | 2.79 | 3.58 | 0.78 | 0.44 | 1 | Gm49308 | | 0 | 0 | 0 | 0 | 0 | 1 | 0 | 0 | 10 | 0 | 6 |
| 2 | -0.30 | 3.27 | -0.09 | 0.93 | 1 | Gm5678 | | 1 | 4 | 5 | 3 | 0 | 1 | 0 | 3 | 0 | 0 | 0 |
| 2 | -0.61 | 3.64 | -0.17 | 0.87 | 1 | Gm49313 | | 0 | 16 | 0 | 0 | 0 | 1 | 0 | 0 | 0 | 0 | 0 |
| 2 | 0.00 | 3.68 | 0.00 | 1.00 | 1 | Gm15675 | | 3 | 3 | 11 | 0 | 0 | 0 | 0 | 0 | 0 | 0 | 0 |
| 2 | 2.39 | 3.58 | 0.67 | 0.50 | 1 | Gm42651 | | 0 | 0 | 0 | 0 | 0 | 1 | 0 | 0 | 13 | 0 | 3 |
| 2 | -0.73 | 3.64 | -0.20 | 0.84 | 1 | Sptssb | serine palmitoyltransferase, small subunit | 0 | 2 | 12 | 0 | 0 | 3 | 0 | 0 | 0 | 0 | 0 |
| 2 | 0.90 | 3.33 | 0.27 | 0.79 | 1 | Gm9083 | | 3 | 1 | 10 | 1 | 0 | 0 | 0 | 1 | 0 | 0 | 1 |
| 2 | 0.00 | 3.68 | 0.00 | 1.00 | 1 | Fam131b | family with sequence similarity 131, meml | 13 | 0 | 4 | 0 | 0 | 0 | 0 | 0 | 0 | 0 | 0 |
| 2 | 0.28 | 2.48 | 0.11 | 0.91 | 1 | Gm7287 | | 3 | 2 | 2 | 2 | 0 | 3 | 0 | 0 | 1 | 1 | 2 |
| 2 | 2.26 | 3.63 | 0.62 | 0.53 | 1 | Aprt-ps | | 3 | 0 | 9 | 0 | 0 | 0 | 0 | 0 | 2 | 0 | 3 |
| 2 | 0.00 | 3.68 | 0.00 | 1.00 | 1 | Nkg7 | natural killer cell group 7 sequence | 7 | 10 | 0 | 0 | 0 | 0 | 0 | 0 | 0 | 0 | 0 |
| 2 | 0.00 | 3.68 | 0.00 | 1.00 | 1 | Gm18852 | | 7 | 10 | 0 | 0 | 0 | 0 | 0 | 0 | 0 | 0 | 0 |
| 2 | 1.52 | 3.56 | 0.43 | 0.67 | 1 | S100a3 | S100 calcium binding protein A3 | 0 | 0 | 9 | 2 | 0 | 0 | 0 | 0 | 2 | 4 | 0 |
| 2 | -1.20 | 3.63 | -0.33 | 0.74 | 1 | Myl1 | myosin, light polypeptide 1 | 9 | 4 | 0 | 4 | 0 | 0 | 0 | 0 | 0 | 0 | 0 |
| 2 | -2.07 | 3.62 | -0.57 | 0.57 | 1 | Gm14719 | | 10 | 2 | 0 | 1 | 4 | 0 | 0 | 0 | 0 | 0 | 0 |
| 2 | 1.38 | 3.63 | 0.38 | 0.71 | 1 | Lhcgr | luteinizing hormone/choriogonadotropin r | 13 | 0 | 1 | 0 | 0 | 0 | 0 | 0 | 0 | 0 | 3 |
| 2 | -2.82 | 3.59 | -0.79 | 0.43 | 1 | Gm46233 | | 0 | 3 | 0 | 0 | 2 | 6 | 0 | 0 | 0 | 0 | 6 |
| 2 | 1.36 | 3.30 | 0.41 | 0.68 | 1 | Gm9480 | | 3 | 2 | 1 | 0 | 0 | 2 | 0 | 0 | 0 | 8 | 1 |
| 2 | 0.37 | 3.60 | 0.10 | 0.92 | 1 | Gpc2 | glypican 2 (cerebroglycan) | 0 | 0 | 0 | 0 | 0 | 7 | 0 | 0 | 0 | 0 | 10 |

| | | | | | | | | | | | | | | | | | | |
|---|---|---|---|---|---|---|---|---|---|---|---|---|---|---|---|---|---|---|
| 2 | 0.00 | 3.68 | 0.00 | 1.00 | 1 | Afp | alpha fetoprotein | 9 | 0 | 8 | 0 | 0 | 0 | 0 | 0 | 0 | 0 | 0 |
| 2 | 0.00 | 3.68 | 0.00 | 1.00 | 1 | Gm16200 | | 9 | 0 | 8 | 0 | 0 | 0 | 0 | 0 | 0 | 0 | 0 |
| 2 | -1.09 | 3.61 | -0.30 | 0.76 | 1 | Gm22806 | | 0 | 0 | 14 | 0 | 0 | 3 | 0 | 0 | 0 | 0 | 0 |
| 2 | 2.52 | 3.56 | 0.71 | 0.48 | 1 | BC026513 | | 0 | 0 | 0 | 0 | 0 | 7 | 0 | 0 | 8 | 0 | 2 |
| 2 | -1.01 | 3.63 | -0.28 | 0.78 | 1 | Gm19248 | | 0 | 15 | 0 | 0 | 0 | 2 | 0 | 0 | 0 | 0 | 0 |
| 2 | -2.17 | 3.63 | -0.60 | 0.55 | 1 | Akap14 | A kinase (PRKA) anchor protein 14 | 6 | 6 | 0 | 0 | 4 | 0 | 0 | 0 | 0 | 0 | 1 |
| 2 | -1.50 | 3.58 | -0.42 | 0.68 | 1 | Gm50352 | | 0 | 0 | 0 | 0 | 3 | 11 | 0 | 0 | 0 | 0 | 3 |
| 2 | -3.52 | 3.58 | -0.98 | 0.33 | 1 | Gm44822 | | 5 | 0 | 0 | 6 | 0 | 3 | 0 | 0 | 0 | 0 | 3 |
| 2 | 0.00 | 3.68 | 0.00 | 1.00 | 1 | S100a9 | S100 calcium binding protein A9 (calgran | 8 | 9 | 0 | 0 | 0 | 0 | 0 | 0 | 0 | 0 | 0 |
| 2 | 2.78 | 3.59 | 0.78 | 0.44 | 1 | Gm16010 | | 7 | 0 | 0 | 0 | 0 | 6 | 0 | 0 | 3 | 0 | 1 |
| 2 | 0.57 | 3.61 | 0.16 | 0.87 | 1 | Gm41253 | | 0 | 0 | 0 | 0 | 6 | 0 | 0 | 0 | 11 | 0 | 0 |
| 2 | 0.51 | 3.60 | 0.14 | 0.89 | 1 | Gm42969 | | 3 | 0 | 0 | 0 | 0 | 4 | 0 | 0 | 0 | 0 | 9 |
| 2 | -0.15 | 3.58 | -0.04 | 0.97 | 1 | Gm7407 | | 0 | 13 | 0 | 2 | 0 | 1 | 0 | 0 | 0 | 0 | 1 |
| 2 | -3.44 | 3.58 | -0.96 | 0.34 | 1 | Gm26511 | | 0 | 0 | 3 | 9 | 0 | 1 | 0 | 0 | 0 | 0 | 4 |
| 2 | 0.00 | 3.68 | 0.00 | 1.00 | 1 | Gm11627 | | 14 | 3 | 0 | 0 | 0 | 0 | 0 | 0 | 0 | 0 | 0 |
| 2 | 2.26 | 3.42 | 0.66 | 0.51 | 1 | Gm4740 | | 0 | 0 | 7 | 1 | 3 | 0 | 0 | 1 | 0 | 3 | 2 |
| 2 | 0.28 | 3.60 | 0.08 | 0.94 | 1 | Gm42851 | | 0 | 0 | 0 | 0 | 0 | 8 | 0 | 0 | 0 | 0 | 9 |
| 2 | 0.28 | 3.60 | 0.08 | 0.94 | 1 | 1700023H06Rik | | 0 | 0 | 0 | 0 | 0 | 8 | 0 | 0 | 0 | 0 | 9 |
| 2 | -0.23 | 3.60 | -0.06 | 0.95 | 1 | Snora47 | | 0 | 0 | 0 | 0 | 0 | 13 | 0 | 0 | 0 | 0 | 4 |
| 2 | -1.16 | 3.62 | -0.32 | 0.75 | 1 | Mbl1 | mannose-binding lectin (protein A) 1 | 0 | 14 | 0 | 0 | 0 | 3 | 0 | 0 | 0 | 0 | 0 |
| 2 | 0.92 | 3.55 | 0.26 | 0.80 | 1 | Rnu11 | | 0 | 0 | 6 | 0 | 0 | 3 | 0 | 0 | 3 | 0 | 4 |
| 2 | -2.50 | 3.54 | -0.71 | 0.48 | 1 | Zxda | | 4 | 0 | 2 | 4 | 4 | 0 | 0 | 0 | 0 | 0 | 3 |
| 2 | 0.00 | 3.68 | 0.00 | 1.00 | 1 | 9430078K24Rik | | 3 | 10 | 3 | 0 | 0 | 0 | 0 | 0 | 0 | 0 | 0 |
| 2 | -0.89 | 3.58 | -0.25 | 0.80 | 1 | Gm5191 | | 0 | 3 | 1 | 2 | 0 | 7 | 0 | 0 | 0 | 0 | 4 |
| 2 | 0.00 | 3.68 | 0.00 | 1.00 | 1 | Gm15998 | | 3 | 11 | 2 | 0 | 0 | 0 | 0 | 0 | 0 | 0 | 0 |
| 2 | -0.63 | 3.55 | -0.18 | 0.86 | 1 | Gm48071 | | 0 | 3 | 0 | 7 | 0 | 3 | 0 | 0 | 0 | 0 | 4 |
| 2 | 0.00 | 3.65 | 0.00 | 1.00 | 1 | Oprk1 | opioid receptor, kappa 1 | 0 | 0 | 17 | 0 | 0 | 0 | 0 | 0 | 0 | 0 | 0 |
| 2 | 0.00 | 3.65 | 0.00 | 1.00 | 1 | 6330415B21Rik | | 0 | 0 | 17 | 0 | 0 | 0 | 0 | 0 | 0 | 0 | 0 |
| 2 | 0.00 | 3.65 | 0.00 | 1.00 | 1 | Krt15 | keratin 15 | 0 | 0 | 17 | 0 | 0 | 0 | 0 | 0 | 0 | 0 | 0 |
| 2 | 0.00 | 3.65 | 0.00 | 1.00 | 1 | Gm31517 | | 0 | 0 | 17 | 0 | 0 | 0 | 0 | 0 | 0 | 0 | 0 |
| 2 | 0.00 | 3.65 | 0.00 | 1.00 | 1 | Apol10c-ps | | 0 | 0 | 17 | 0 | 0 | 0 | 0 | 0 | 0 | 0 | 0 |
| 2 | 0.00 | 3.65 | 0.00 | 1.00 | 1 | I730030J21Rik | | 0 | 0 | 17 | 0 | 0 | 0 | 0 | 0 | 0 | 0 | 0 |
| 2 | 0.00 | 3.65 | 0.00 | 1.00 | 1 | Gm6553 | | 0 | 0 | 17 | 0 | 0 | 0 | 0 | 0 | 0 | 0 | 0 |
| 2 | 2.99 | 3.56 | 0.84 | 0.40 | 1 | 4930465K10Rik | | 0 | 0 | 0 | 0 | 0 | 1 | 0 | 13 | 3 | 0 | 0 |
| 2 | -1.01 | 3.37 | -0.30 | 0.76 | 1 | Gm6335 | | 0 | 1 | 1 | 3 | 1 | 1 | 0 | 0 | 0 | 0 | 10 |
| 2 | -1.84 | 3.43 | -0.54 | 0.59 | 1 | Gm10171 | | 4 | 0 | 0 | 5 | 2 | 1 | 0 | 0 | 3 | 1 | 0 |
| 2 | 2.45 | 3.51 | 0.70 | 0.49 | 1 | Cox7b2 | cytochrome c oxidase subunit 7B2 | 0 | 0 | 0 | 4 | 0 | 1 | 0 | 0 | 3 | 5 | 4 |
| 2 | -0.70 | 3.66 | -0.19 | 0.85 | 1 | Shisa9 | shisa family member 9 | 13 | 2 | 0 | 0 | 0 | 2 | 0 | 0 | 0 | 0 | 0 |
| 2 | -3.96 | 3.59 | -1.10 | 0.27 | 1 | Gm44775 | | 4 | 0 | 0 | 4 | 0 | 7 | 0 | 0 | 0 | 0 | 2 |
| 2 | 0.74 | 3.60 | 0.20 | 0.84 | 1 | Snord58b | | 0 | 0 | 0 | 0 | 0 | 3 | 0 | 0 | 0 | 0 | 13 |
| 2 | 0.45 | 3.60 | 0.12 | 0.90 | 1 | Mir6942 | | 0 | 7 | 0 | 0 | 0 | 3 | 0 | 0 | 0 | 0 | 6 |
| 2 | 2.01 | 3.58 | 0.56 | 0.58 | 1 | Gm44718 | | 0 | 0 | 0 | 0 | 0 | 3 | 0 | 0 | 5 | 0 | 8 |
| 2 | -3.45 | 3.61 | -0.96 | 0.34 | 1 | Gm5406 | | 9 | 0 | 0 | 5 | 0 | 1 | 0 | 0 | 0 | 0 | 1 |
| 2 | -0.79 | 3.62 | -0.22 | 0.83 | 1 | Gm41639 | | 9 | 2 | 0 | 0 | 0 | 5 | 0 | 0 | 0 | 0 | 0 |
| 2 | 2.04 | 2.26 | 0.90 | 0.37 | 1 | Gm15148 | | 2 | 2 | 3 | 1 | 0 | 1 | 0 | 5 | 1 | 1 | 1 |
| 2 | 4.33 | 3.62 | 1.20 | 0.23 | 1 | Gm43099 | | 9 | 0 | 0 | 0 | 0 | 1 | 0 | 0 | 2 | 0 | 5 |
| 2 | 0.80 | 3.50 | 0.23 | 0.82 | 1 | Gm16537 | | 0 | 0 | 0 | 0 | 3 | 3 | 0 | 8 | 2 | 0 | 1 |
| 2 | 0.88 | 3.61 | 0.24 | 0.81 | 1 | Gm28181 | | 0 | 0 | 12 | 0 | 0 | 0 | 0 | 0 | 4 | 0 | 0 |
| 1 | 1.02 | 2.56 | 0.40 | 0.69 | 1 | Gm11824 | | 2 | 5 | 0 | 1 | 1 | 1 | 0 | 1 | 2 | 4 | 0 |
| 1 | 0.27 | 2.55 | 0.11 | 0.91 | 1 | Rpl31-ps10 | | 3 | 5 | 1 | 4 | 0 | 1 | 0 | 1 | 1 | 1 | 0 |
| 1 | -0.21 | 3.61 | -0.06 | 0.95 | 1 | Nrg3 | neuregulin 3 | 3 | 0 | 7 | 0 | 2 | 1 | 0 | 0 | 3 | 0 | 0 |
| 1 | -0.83 | 3.59 | -0.23 | 0.82 | 1 | Gm49512 | | 0 | 0 | 0 | 2 | 0 | 8 | 0 | 0 | 0 | 0 | 7 |
| 1 | -0.47 | 3.55 | -0.13 | 0.90 | 1 | A630081D01Rik | | 0 | 0 | 0 | 0 | 4 | 8 | 0 | 0 | 2 | 0 | 3 |
| 1 | -2.09 | 3.29 | -0.63 | 0.53 | 1 | Gm5270 | | 1 | 0 | 1 | 9 | 1 | 3 | 0 | 2 | 0 | 0 | 0 |
| 1 | 0.00 | 3.68 | 0.00 | 1.00 | 1 | C030034L19Rik | | 1 | 0 | 16 | 0 | 0 | 0 | 0 | 0 | 0 | 0 | 0 |
| 1 | -0.71 | 3.66 | -0.19 | 0.85 | 1 | Ptprh | protein tyrosine phosphatase, receptor typ | 3 | 12 | 0 | 0 | 0 | 2 | 0 | 0 | 0 | 0 | 0 |
| 1 | 0.00 | 3.68 | 0.00 | 1.00 | 1 | Akr1c18 | aldo-keto reductase family 1, member C1 | 0 | 8 | 8 | 0 | 0 | 0 | 0 | 0 | 0 | 0 | 0 |
| 1 | 0.62 | 3.60 | 0.17 | 0.86 | 1 | Fdx2 | ferredoxin 2 | 0 | 0 | 0 | 0 | 0 | 4 | 0 | 0 | 0 | 0 | 12 |
| 1 | 0.10 | 3.60 | 0.03 | 0.98 | 1 | Mir384 | | 0 | 0 | 0 | 0 | 0 | 9 | 0 | 0 | 0 | 0 | 7 |
| 1 | -0.71 | 3.56 | -0.20 | 0.84 | 1 | T | brachyury, T-box transcription factor T | 0 | 0 | 0 | 0 | 5 | 3 | 0 | 0 | 0 | 0 | 9 |

| | | | | | | | | | | | | | | | | | | |
|---|---|---|---|---|---|---|---|---|---|---|---|---|---|---|---|---|---|---|
| 1 | 0.75 | 3.61 | 0.21 | 0.84 | 1 | Gm43445 | | 0 | 5 | 0 | 0 | 0 | 4 | 0 | 0 | 0 | 0 | 7 |
| 1 | -2.59 | 3.56 | -0.73 | 0.47 | 1 | Gm33467 | | 0 | 0 | 0 | 10 | 0 | 2 | 0 | 0 | 0 | 0 | 5 |
| 1 | 1.44 | 3.64 | 0.40 | 0.69 | 1 | Gm12335 | | 2 | 10 | 0 | 0 | 1 | 0 | 0 | 0 | 0 | 3 | 1 |
| 1 | 3.62 | 3.58 | 1.01 | 0.31 | 1 | Gm16156 | | 0 | 4 | 0 | 0 | 0 | 4 | 0 | 0 | 6 | 0 | 2 |
| 1 | -0.58 | 3.60 | -0.16 | 0.87 | 1 | Gm31406 | | 0 | 0 | 0 | 0 | 9 | 1 | 0 | 0 | 6 | 0 | 0 |
| 1 | 0.00 | 3.68 | 0.00 | 1.00 | 1 | Gm45238 | | 11 | 0 | 5 | 0 | 0 | 0 | 0 | 0 | 0 | 0 | 0 |
| 1 | 0.00 | 3.68 | 0.00 | 1.00 | 1 | BC049352 | cDNA sequence BC049352 | 11 | 0 | 5 | 0 | 0 | 0 | 0 | 0 | 0 | 0 | 0 |
| 1 | -1.18 | 3.56 | -0.33 | 0.74 | 1 | Gm46563 | | 0 | 0 | 0 | 8 | 0 | 3 | 0 | 0 | 0 | 0 | 5 |
| 1 | -0.67 | 3.60 | -0.19 | 0.85 | 1 | Gm29361 | | 7 | 0 | 0 | 0 | 0 | 8 | 0 | 0 | 0 | 0 | 1 |
| 1 | 2.08 | 3.55 | 0.59 | 0.56 | 1 | Gm33201 | | 0 | 3 | 0 | 0 | 0 | 5 | 0 | 2 | 0 | 4 | 2 |
| 1 | 0.00 | 3.65 | 0.00 | 1.00 | 1 | Slc7a13 | solute carrier family 7, (cationic amino aci | 16 | 0 | 0 | 0 | 0 | 0 | 0 | 0 | 0 | 0 | 0 |
| 1 | 0.00 | 3.65 | 0.00 | 1.00 | 1 | Tecrl | trans-2,3-enoyl-CoA reductase-like | 16 | 0 | 0 | 0 | 0 | 0 | 0 | 0 | 0 | 0 | 0 |
| 1 | 0.00 | 3.65 | 0.00 | 1.00 | 1 | Igkv3-12 | | 16 | 0 | 0 | 0 | 0 | 0 | 0 | 0 | 0 | 0 | 0 |
| 1 | 1.14 | 3.45 | 0.33 | 0.74 | 1 | Gm7172 | | 2 | 1 | 0 | 2 | 0 | 2 | 0 | 0 | 7 | 0 | 3 |
| 1 | -0.98 | 3.64 | -0.27 | 0.79 | 1 | Gabrr1 | gamma-aminobutyric acid (GABA) C rece | 15 | 0 | 0 | 0 | 0 | 2 | 0 | 0 | 0 | 0 | 0 |
| 1 | 0.00 | 3.68 | 0.00 | 1.00 | 1 | Gm10791 | | 1 | 7 | 8 | 0 | 0 | 0 | 0 | 0 | 0 | 0 | 0 |
| 1 | -1.15 | 3.61 | -0.32 | 0.75 | 1 | Tgm7 | transglutaminase 7 | 13 | 0 | 0 | 0 | 0 | 3 | 0 | 0 | 0 | 0 | 0 |
| 1 | 0.00 | 3.68 | 0.00 | 1.00 | 1 | Gm5452 | | 11 | 4 | 1 | 0 | 0 | 0 | 0 | 0 | 0 | 0 | 0 |
| 1 | 0.52 | 3.60 | 0.14 | 0.89 | 1 | Gm37906 | | 0 | 0 | 0 | 0 | 0 | 5 | 0 | 0 | 0 | 0 | 11 |
| 1 | 0.52 | 3.60 | 0.14 | 0.89 | 1 | Gm37411 | | 0 | 0 | 0 | 0 | 0 | 5 | 0 | 0 | 0 | 0 | 11 |
| 1 | 0.52 | 3.60 | 0.14 | 0.89 | 1 | Gm29438 | | 0 | 0 | 0 | 0 | 0 | 5 | 0 | 0 | 0 | 0 | 11 |
| 1 | -0.93 | 3.63 | -0.25 | 0.80 | 1 | Gm30055 | | 0 | 0 | 15 | 0 | 0 | 2 | 0 | 0 | 0 | 0 | 0 |
| 1 | 0.00 | 3.60 | 0.00 | 1.00 | 1 | Gm16093 | | 0 | 0 | 0 | 0 | 0 | 10 | 0 | 0 | 0 | 0 | 6 |
| 1 | -0.41 | 3.56 | -0.12 | 0.91 | 1 | Amhr2 | anti-Mullerian hormone type 2 receptor | 0 | 4 | 1 | 4 | 0 | 2 | 0 | 0 | 0 | 0 | 5 |
| 1 | -0.07 | 3.60 | -0.02 | 0.99 | 1 | Gm14537 | | 0 | 1 | 0 | 0 | 0 | 10 | 0 | 0 | 0 | 0 | 5 |
| 1 | 0.34 | 3.60 | 0.09 | 0.92 | 1 | Mir1198 | | 0 | 4 | 0 | 0 | 0 | 5 | 0 | 0 | 0 | 0 | 7 |
| 1 | 0.52 | 3.61 | 0.15 | 0.88 | 1 | Gm29458 | | 0 | 0 | 8 | 0 | 7 | 0 | 0 | 0 | 0 | 1 | 0 |
| 1 | 1.67 | 3.61 | 0.46 | 0.64 | 1 | Gm50128 | | 0 | 12 | 0 | 0 | 0 | 0 | 0 | 0 | 0 | 0 | 4 |
| 1 | -0.76 | 3.65 | -0.21 | 0.83 | 1 | 1700020G03Rik | | 0 | 1 | 14 | 0 | 0 | 2 | 0 | 0 | 0 | 0 | 0 |
| 1 | 0.00 | 3.65 | 0.00 | 1.00 | 1 | Ccr3 | chemokine (C-C motif) receptor 3 | 0 | 16 | 0 | 0 | 0 | 0 | 0 | 0 | 0 | 0 | 0 |
| 1 | 0.00 | 3.65 | 0.00 | 1.00 | 1 | Prtn3 | proteinase 3 | 0 | 16 | 0 | 0 | 0 | 0 | 0 | 0 | 0 | 0 | 0 |
| 1 | 0.00 | 3.65 | 0.00 | 1.00 | 1 | Gm10735 | | 0 | 16 | 0 | 0 | 0 | 0 | 0 | 0 | 0 | 0 | 0 |
| 1 | 0.00 | 3.65 | 0.00 | 1.00 | 1 | Traj7 | | 0 | 16 | 0 | 0 | 0 | 0 | 0 | 0 | 0 | 0 | 0 |
| 1 | 0.52 | 3.60 | 0.14 | 0.89 | 1 | Gm32089 | | 9 | 0 | 0 | 0 | 0 | 3 | 0 | 0 | 0 | 0 | 5 |
| 1 | 0.37 | 3.61 | 0.10 | 0.92 | 1 | Rbm42 | RNA binding motif protein 42 | 8 | 0 | 0 | 0 | 0 | 3 | 0 | 0 | 0 | 0 | 5 |
| 1 | -3.23 | 3.58 | -0.90 | 0.37 | 1 | Gm12356 | | 0 | 0 | 0 | 11 | 6 | 0 | 0 | 0 | 0 | 0 | 0 |
| 1 | 0.88 | 3.56 | 0.25 | 0.81 | 1 | Gm22009 | | 9 | 0 | 0 | 0 | 0 | 2 | 0 | 0 | 0 | 4 | 2 |
| 1 | 1.46 | 3.59 | 0.41 | 0.68 | 1 | Gm15549 | | 0 | 0 | 0 | 0 | 0 | 8 | 0 | 1 | 0 | 1 | 6 |
| 1 | 1.30 | 3.64 | 0.36 | 0.72 | 1 | Grm3 | glutamate receptor, metabotropic 3 | 7 | 0 | 6 | 0 | 0 | 0 | 0 | 0 | 0 | 0 | 3 |
| 1 | 0.00 | 3.68 | 0.00 | 1.00 | 1 | Daw1 | dynein assembly factor with WDR repeat | 6 | 10 | 0 | 0 | 0 | 0 | 0 | 0 | 0 | 0 | 0 |
| 1 | 0.00 | 3.68 | 0.00 | 1.00 | 1 | Gm12796 | | 6 | 10 | 0 | 0 | 0 | 0 | 0 | 0 | 0 | 0 | 0 |
| 1 | -1.11 | 3.23 | -0.34 | 0.73 | 1 | Gm13864 | | 3 | 1 | 6 | 0 | 2 | 3 | 0 | 0 | 0 | 0 | 1 |
| 1 | 0.78 | 3.60 | 0.22 | 0.83 | 1 | Gm17891 | | 0 | 0 | 9 | 0 | 0 | 2 | 0 | 0 | 0 | 0 | 5 |
| 1 | 0.08 | 3.58 | 0.02 | 0.98 | 1 | Gm36501 | | 0 | 11 | 0 | 0 | 1 | 1 | 0 | 1 | 2 | 0 | 0 |
| 1 | -2.57 | 3.60 | -0.71 | 0.47 | 1 | Gm12657 | | 0 | 2 | 6 | 1 | 2 | 5 | 0 | 0 | 0 | 0 | 0 |
| 1 | 0.54 | 3.54 | 0.15 | 0.88 | 1 | Gm50068 | | 0 | 0 | 0 | 0 | 6 | 0 | 0 | 10 | 0 | 0 | 0 |
| 1 | -0.07 | 3.60 | -0.02 | 0.99 | 1 | Gm26347 | | 1 | 0 | 0 | 0 | 0 | 10 | 0 | 0 | 0 | 0 | 5 |
| 1 | 0.42 | 3.60 | 0.12 | 0.91 | 1 | Gm26457 | | 0 | 0 | 0 | 0 | 0 | 6 | 0 | 0 | 0 | 0 | 10 |
| 1 | 0.42 | 3.60 | 0.12 | 0.91 | 1 | Gm11969 | | 0 | 0 | 0 | 0 | 0 | 6 | 0 | 0 | 0 | 0 | 10 |
| 1 | -0.01 | 3.62 | 0.00 | 1.00 | 1 | Spp2 | secreted phosphoprotein 2 | 0 | 15 | 0 | 1 | 0 | 0 | 0 | 0 | 0 | 0 | 0 |
| 1 | -1.89 | 3.27 | -0.58 | 0.56 | 1 | Rpl27a-ps1 | | 4 | 2 | 6 | 1 | 1 | 2 | 0 | 0 | 0 | 0 | 0 |
| 1 | 1.00 | 3.58 | 0.28 | 0.78 | 1 | Mir1943 | | 0 | 0 | 0 | 0 | 0 | 11 | 0 | 0 | 3 | 0 | 2 |
| 1 | 0.78 | 3.60 | 0.22 | 0.83 | 1 | Gm27457 | | 0 | 12 | 0 | 0 | 0 | 1 | 0 | 0 | 0 | 0 | 3 |
| 1 | 1.27 | 3.60 | 0.35 | 0.72 | 1 | 2310015A16Rik | | 5 | 0 | 3 | 0 | 0 | 2 | 0 | 0 | 1 | 0 | 5 |
| 1 | -1.51 | 3.56 | -0.42 | 0.67 | 1 | Gm44214 | | 0 | 0 | 0 | 9 | 0 | 4 | 0 | 0 | 0 | 0 | 3 |
| 1 | -1.43 | 3.58 | -0.40 | 0.69 | 1 | Gm20548 | | 0 | 0 | 0 | 4 | 0 | 8 | 0 | 0 | 0 | 0 | 4 |
| 1 | 1.26 | 3.59 | 0.35 | 0.73 | 1 | Nr5a1 | nuclear receptor subfamily 5, group A, me | 0 | 0 | 0 | 0 | 1 | 0 | 0 | 15 | 0 | 0 | 0 |
| 1 | -3.01 | 3.58 | -0.84 | 0.40 | 1 | Gm47664 | | 0 | 0 | 0 | 13 | 0 | 1 | 0 | 0 | 0 | 0 | 2 |
| 1 | -5.28 | 3.55 | -1.49 | 0.14 | 1 | Gm15530 | | 0 | 0 | 0 | 8 | 0 | 4 | 0 | 0 | 0 | 4 | 0 |
| 1 | 2.90 | 3.55 | 0.82 | 0.41 | 1 | 4930406D18Rik | | 0 | 0 | 0 | 2 | 0 | 0 | 0 | 0 | 7 | 0 | 7 |

| | | | | | | | | | | | | | | | | | | |
|---|---|---|---|---|---|---|---|---|---|---|---|---|---|---|---|---|---|---|
| 1 | -1.97 | 3.58 | -0.55 | 0.58 | 1 | Gm12264 | | 0 | 0 | 6 | 0 | 4 | 6 | 0 | 0 | 0 | 0 | 0 |
| 1 | 0.00 | 3.68 | 0.00 | 1.00 | 1 | 6030445D17Rik | | 8 | 4 | 4 | 0 | 0 | 0 | 0 | 0 | 0 | 0 | 0 |
| 1 | -1.12 | 3.60 | -0.31 | 0.75 | 1 | Gm10827 | | 0 | 0 | 8 | 0 | 0 | 8 | 0 | 0 | 0 | 0 | 0 |
| 1 | 0.98 | 3.60 | 0.27 | 0.79 | 1 | Gm37678 | | 0 | 0 | 0 | 0 | 0 | 2 | 0 | 0 | 0 | 0 | 14 |
| 1 | 1.96 | 3.61 | 0.54 | 0.59 | 1 | Gm11517 | | 2 | 0 | 1 | 0 | 0 | 0 | 0 | 0 | 13 | 0 | 0 |
| 1 | 0.33 | 3.60 | 0.09 | 0.93 | 1 | 1700093K21Ril | RIKEN cDNA 1700093K21 gene | 0 | 0 | 0 | 0 | 0 | 7 | 0 | 0 | 0 | 0 | 9 |
| 1 | 0.00 | 3.68 | 0.00 | 1.00 | 1 | Gm21816 | | 2 | 14 | 0 | 0 | 0 | 0 | 0 | 0 | 0 | 0 | 0 |
| 1 | -0.20 | 3.60 | -0.06 | 0.95 | 1 | Gm24616 | | 0 | 0 | 0 | 0 | 0 | 12 | 0 | 0 | 0 | 0 | 4 |
| 1 | 1.42 | 3.58 | 0.40 | 0.69 | 1 | Gm23864 | | 0 | 0 | 0 | 0 | 0 | 7 | 0 | 0 | 3 | 0 | 6 |
| 1 | -1.71 | 3.57 | -0.48 | 0.63 | 1 | Gm37973 | | 0 | 0 | 0 | 10 | 0 | 4 | 0 | 0 | 0 | 0 | 2 |
| 1 | -1.21 | 3.60 | -0.34 | 0.74 | 1 | Abo | ABO blood group (transferase A, alpha 1- | 0 | 9 | 0 | 0 | 0 | 7 | 0 | 0 | 0 | 0 | 0 |
| 1 | 0.00 | 3.68 | 0.00 | 1.00 | 1 | 4930443O20Rik | | 3 | 1 | 11 | 0 | 0 | 0 | 0 | 0 | 0 | 0 | 0 |
| 1 | 1.25 | 3.66 | 0.34 | 0.73 | 1 | A330032B11Rik | | 3 | 0 | 10 | 0 | 0 | 0 | 0 | 0 | 0 | 0 | 2 |
| 1 | 0.00 | 3.68 | 0.00 | 1.00 | 1 | Olfr56 | olfactory receptor 56 | 3 | 2 | 10 | 0 | 0 | 0 | 0 | 0 | 0 | 0 | 0 |
| 1 | -0.99 | 3.60 | -0.27 | 0.78 | 1 | Gm42445 | | 0 | 0 | 0 | 0 | 1 | 12 | 0 | 0 | 0 | 0 | 3 |
| 1 | 0.28 | 3.61 | 0.08 | 0.94 | 1 | Gm43430 | | 1 | 0 | 9 | 0 | 0 | 3 | 0 | 0 | 0 | 0 | 3 |
| 1 | 0.52 | 3.61 | 0.14 | 0.89 | 1 | Gm35585 | | 0 | 0 | 0 | 0 | 6 | 0 | 0 | 0 | 10 | 0 | 0 |
| 1 | -1.60 | 3.58 | -0.45 | 0.65 | 1 | Gm16272 | | 0 | 0 | 3 | 10 | 0 | 1 | 0 | 1 | 0 | 0 | 1 |
| 1 | 0.00 | 3.68 | 0.00 | 1.00 | 1 | Gm37412 | | 14 | 2 | 0 | 0 | 0 | 0 | 0 | 0 | 0 | 0 | 0 |
| 1 | -1.18 | 3.58 | -0.33 | 0.74 | 1 | Grin1os | glutamate receptor, ionotropic, NMDA1 (z | 0 | 0 | 6 | 1 | 2 | 2 | 0 | 0 | 5 | 0 | 0 |
| 1 | 0.00 | 3.68 | 0.00 | 1.00 | 1 | Pla2g12b | phospholipase A2, group XIIB | 3 | 6 | 6 | 0 | 0 | 0 | 0 | 0 | 0 | 0 | 0 |
| 1 | -0.28 | 3.61 | -0.08 | 0.94 | 1 | Gm36423 | | 0 | 5 | 1 | 0 | 0 | 8 | 0 | 0 | 0 | 0 | 2 |
| 1 | 0.83 | 3.60 | 0.23 | 0.82 | 1 | Gm25970 | | 0 | 0 | 0 | 0 | 0 | 3 | 0 | 0 | 0 | 0 | 13 |
| 1 | -0.32 | 3.60 | -0.09 | 0.93 | 1 | Gm43024 | | 0 | 0 | 0 | 0 | 0 | 13 | 0 | 0 | 0 | 0 | 3 |
| 1 | 0.00 | 3.68 | 0.00 | 1.00 | 1 | Dipk1c | divergent protein kinase domain 1C | 4 | 0 | 11 | 0 | 0 | 0 | 0 | 0 | 0 | 0 | 0 |
| 1 | -0.76 | 3.63 | -0.21 | 0.83 | 1 | Tm4sf20 | transmembrane 4 L six family member 20 | 0 | 5 | 7 | 0 | 0 | 3 | 0 | 0 | 0 | 0 | 0 |
| 1 | 2.11 | 3.58 | 0.59 | 0.56 | 1 | Gm14421 | | 0 | 0 | 0 | 0 | 0 | 3 | 0 | 0 | 5 | 0 | 8 |
| 1 | 0.37 | 3.61 | 0.10 | 0.92 | 1 | Mir6907 | | 0 | 1 | 6 | 0 | 0 | 3 | 0 | 0 | 0 | 0 | 5 |
| 1 | -3.59 | 3.57 | -1.01 | 0.31 | 1 | Gm50439 | | 0 | 0 | 0 | 11 | 1 | 3 | 0 | 0 | 0 | 0 | 0 |
| 1 | -1.15 | 3.62 | -0.32 | 0.75 | 1 | Rex2 | reduced expression 2 | 0 | 13 | 0 | 0 | 0 | 3 | 0 | 0 | 0 | 0 | 0 |
| 1 | -1.15 | 3.62 | -0.32 | 0.75 | 1 | Gm15938 | | 0 | 13 | 0 | 0 | 0 | 3 | 0 | 0 | 0 | 0 | 0 |
| 1 | -0.76 | 3.63 | -0.21 | 0.83 | 1 | Gm15537 | | 10 | 2 | 0 | 0 | 0 | 3 | 0 | 0 | 0 | 0 | 0 |
| 1 | 0.95 | 3.67 | 0.26 | 0.80 | 1 | Aspg | asparaginase | 9 | 6 | 0 | 0 | 0 | 0 | 0 | 0 | 0 | 0 | 1 |
| 1 | 0.00 | 3.68 | 0.00 | 1.00 | 1 | Gm43923 | | 9 | 7 | 0 | 0 | 0 | 0 | 0 | 0 | 0 | 0 | 0 |
| 1 | 0.02 | 3.60 | 0.01 | 0.99 | 1 | Gm9042 | | 0 | 1 | 0 | 0 | 1 | 3 | 0 | 0 | 0 | 0 | 11 |
| 1 | -0.19 | 3.60 | -0.05 | 0.96 | 1 | Zfp264 | | 0 | 0 | 4 | 0 | 0 | 8 | 0 | 0 | 0 | 0 | 3 |
| 1 | 3.62 | 3.58 | 1.01 | 0.31 | 1 | Gm19918 | | 0 | 0 | 0 | 0 | 0 | 0 | 0 | 12 | 0 | 0 | 4 |
| 1 | -3.54 | 3.62 | -0.98 | 0.33 | 1 | Alms1-ps1 | | 9 | 0 | 0 | 4 | 0 | 2 | 0 | 0 | 0 | 0 | 1 |
| 1 | 0.66 | 3.61 | 0.18 | 0.86 | 1 | Fam229a | family with sequence similarity 229, meml | 1 | 0 | 5 | 3 | 0 | 0 | 0 | 7 | 0 | 0 | 0 |
| 1 | 1.11 | 3.65 | 0.30 | 0.76 | 1 | Gm41392 | | 15 | 0 | 0 | 0 | 0 | 0 | 0 | 0 | 0 | 0 | 1 |
| 1 | -0.78 | 3.61 | -0.22 | 0.83 | 1 | Npffr1 | neuropeptide FF receptor 1 | 1 | 0 | 6 | 0 | 9 | 0 | 0 | 0 | 0 | 0 | 0 |
| 1 | 0.00 | 3.68 | 0.00 | 1.00 | 1 | Gm14108 | | 15 | 1 | 0 | 0 | 0 | 0 | 0 | 0 | 0 | 0 | 0 |
| 1 | -1.07 | 3.59 | -0.30 | 0.77 | 1 | Snord42b | | 0 | 5 | 0 | 0 | 4 | 2 | 0 | 0 | 5 | 0 | 0 |
| 1 | 0.00 | 3.65 | 0.00 | 1.00 | 1 | Igf1os | | 0 | 0 | 16 | 0 | 0 | 0 | 0 | 0 | 0 | 0 | 0 |
| 1 | 0.00 | 3.65 | 0.00 | 1.00 | 1 | Gm18169 | | 0 | 0 | 16 | 0 | 0 | 0 | 0 | 0 | 0 | 0 | 0 |
| 1 | -4.46 | 3.55 | -1.26 | 0.21 | 1 | Gm50048 | | 0 | 0 | 0 | 0 | 4 | 3 | 0 | 9 | 0 | 0 | 0 |
| 1 | 0.71 | 3.60 | 0.20 | 0.84 | 1 | Gm37436 | | 0 | 0 | 0 | 0 | 0 | 3 | 0 | 0 | 0 | 0 | 12 |
| 1 | 0.00 | 3.68 | 0.00 | 1.00 | 1 | Slc16a14 | solute carrier family 16 (monocarboxylic a | 4 | 6 | 5 | 0 | 0 | 0 | 0 | 0 | 0 | 0 | 0 |
| 1 | 0.13 | 3.60 | 0.04 | 0.97 | 1 | Gm37864 | | 0 | 0 | 0 | 0 | 0 | 8 | 0 | 0 | 0 | 0 | 7 |
| 1 | 0.59 | 3.55 | 0.17 | 0.87 | 1 | Ift88os | | 0 | 0 | 0 | 5 | 0 | 0 | 0 | 0 | 0 | 10 | 0 |
| 1 | 0.25 | 3.60 | 0.07 | 0.95 | 1 | Snord23 | | 0 | 8 | 0 | 0 | 0 | 3 | 0 | 0 | 0 | 0 | 4 |
| 1 | -1.16 | 3.57 | -0.33 | 0.74 | 1 | Gm45047 | | 0 | 0 | 0 | 4 | 0 | 5 | 0 | 0 | 0 | 0 | 6 |
| 1 | -0.48 | 3.60 | -0.13 | 0.89 | 1 | Miat | | 10 | 0 | 0 | 0 | 0 | 4 | 0 | 0 | 0 | 0 | 1 |
| 1 | -0.14 | 3.47 | -0.04 | 0.97 | 1 | Ccnd3-ps | | 1 | 6 | 0 | 0 | 3 | 2 | 0 | 0 | 1 | 0 | 3 |
| 1 | -0.69 | 3.66 | -0.19 | 0.85 | 1 | Gm15984 | | 3 | 0 | 10 | 0 | 0 | 2 | 0 | 0 | 0 | 0 | 0 |
| 1 | 0.00 | 3.68 | 0.00 | 1.00 | 1 | Cldn22 | claudin 22 | 9 | 6 | 0 | 0 | 0 | 0 | 0 | 0 | 0 | 0 | 0 |
| 1 | 0.00 | 3.68 | 0.00 | 1.00 | 1 | Gm17025 | | 9 | 6 | 0 | 0 | 0 | 0 | 0 | 0 | 0 | 0 | 0 |
| 1 | 0.00 | 3.68 | 0.00 | 1.00 | 1 | Gm2629 | | 9 | 6 | 0 | 0 | 0 | 0 | 0 | 0 | 0 | 0 | 0 |
| 1 | 0.00 | 3.68 | 0.00 | 1.00 | 1 | 1500005C15Rik | | 10 | 0 | 5 | 0 | 0 | 0 | 0 | 0 | 0 | 0 | 0 |
| 1 | 0.00 | 3.68 | 0.00 | 1.00 | 1 | C430049B03Rik | | 5 | 2 | 8 | 0 | 0 | 0 | 0 | 0 | 0 | 0 | 0 |

| | | | | | | | | | | | | | | | | | | |
|---|---|---|---|---|---|---|---|---|---|---|---|---|---|---|---|---|---|---|
| 1 | -0.36 | 3.58 | -0.10 | 0.92 | 1 | Gm13047 | | 8 | 0 | 0 | 1 | 3 | 0 | 0 | 0 | 4 | 0 | 0 |
| 1 | -0.78 | 3.62 | -0.21 | 0.83 | 1 | Gm44985 | | 0 | 7 | 4 | 0 | 0 | 4 | 0 | 0 | 0 | 0 | 0 |
| 1 | 0.46 | 3.34 | 0.14 | 0.89 | 1 | Rps15a-ps3 | | 9 | 1 | 1 | 0 | 0 | 3 | 0 | 0 | 0 | 1 | 1 |
| 1 | 0.00 | 3.66 | 0.00 | 1.00 | 1 | Igfn1 | immunoglobulin-like and fibronectin type | 15 | 0 | 0 | 0 | 0 | 0 | 0 | 0 | 0 | 0 | 0 |
| 1 | 0.00 | 3.66 | 0.00 | 1.00 | 1 | Igkv9-123 | | 15 | 0 | 0 | 0 | 0 | 0 | 0 | 0 | 0 | 0 | 0 |
| 1 | 0.00 | 3.66 | 0.00 | 1.00 | 1 | Igkj4 | | 15 | 0 | 0 | 0 | 0 | 0 | 0 | 0 | 0 | 0 | 0 |
| 1 | 0.00 | 3.66 | 0.00 | 1.00 | 1 | Gm48276 | | 15 | 0 | 0 | 0 | 0 | 0 | 0 | 0 | 0 | 0 | 0 |
| 1 | 0.00 | 3.66 | 0.00 | 1.00 | 1 | Ighv1-39 | | 15 | 0 | 0 | 0 | 0 | 0 | 0 | 0 | 0 | 0 | 0 |
| 1 | 0.00 | 3.66 | 0.00 | 1.00 | 1 | 4930448D08Rik | | 15 | 0 | 0 | 0 | 0 | 0 | 0 | 0 | 0 | 0 | 0 |
| 1 | -5.03 | 3.55 | -1.42 | 0.16 | 1 | Gm47558 | | 0 | 0 | 0 | 7 | 0 | 3 | 0 | 0 | 5 | 0 | 0 |
| 1 | 0.00 | 3.68 | 0.00 | 1.00 | 1 | Ighj4 | | 4 | 11 | 0 | 0 | 0 | 0 | 0 | 0 | 0 | 0 | 0 |
| 1 | 0.00 | 3.68 | 0.00 | 1.00 | 1 | Gpr82 | G protein-coupled receptor 82 | 4 | 11 | 0 | 0 | 0 | 0 | 0 | 0 | 0 | 0 | 0 |
| 1 | 0.67 | 3.60 | 0.19 | 0.85 | 1 | Gm22581 | | 1 | 0 | 0 | 0 | 0 | 3 | 0 | 0 | 0 | 0 | 11 |
| 1 | 0.00 | 3.68 | 0.00 | 1.00 | 1 | Lmx1a | LIM homeobox transcription factor 1 alpha | 1 | 0 | 15 | 0 | 0 | 0 | 0 | 0 | 0 | 0 | 0 |
| 1 | 0.00 | 3.68 | 0.00 | 1.00 | 1 | Sohlh2 | spermatogenesis and oogenesis specific | 0 | 7 | 8 | 0 | 0 | 0 | 0 | 0 | 0 | 0 | 0 |
| 1 | 0.07 | 3.60 | 0.02 | 0.98 | 1 | Gm38150 | | 1 | 0 | 0 | 0 | 0 | 8 | 0 | 0 | 0 | 0 | 6 |
| 1 | -0.81 | 3.63 | -0.22 | 0.82 | 1 | Gm37694 | | 2 | 11 | 0 | 0 | 0 | 3 | 0 | 0 | 0 | 0 | 0 |
| 1 | 2.37 | 3.56 | 0.67 | 0.51 | 1 | Gm9202 | | 2 | 0 | 0 | 0 | 0 | 3 | 0 | 1 | 2 | 0 | 7 |
| 1 | 0.53 | 3.61 | 0.15 | 0.88 | 1 | Gm44084 | | 0 | 5 | 0 | 0 | 0 | 4 | 0 | 0 | 0 | 0 | 6 |
| 1 | -2.14 | 3.54 | -0.61 | 0.54 | 1 | Gm13610 | | 0 | 0 | 0 | 8 | 4 | 2 | 0 | 0 | 1 | 0 | 1 |
| 1 | 0.92 | 3.30 | 0.28 | 0.78 | 1 | Gm5527 | | 3 | 1 | 0 | 1 | 0 | 2 | 0 | 0 | 0 | 5 | 4 |
| 1 | 0.00 | 3.68 | 0.00 | 1.00 | 1 | Igkv4-71 | | 10 | 5 | 0 | 0 | 0 | 0 | 0 | 0 | 0 | 0 | 0 |
| 1 | 0.00 | 3.68 | 0.00 | 1.00 | 1 | Apon | apolipoprotein N | 6 | 0 | 9 | 0 | 0 | 0 | 0 | 0 | 0 | 0 | 0 |
| 1 | 0.52 | 3.62 | 0.14 | 0.89 | 1 | Dcst2 | DC-STAMP domain containing 2 | 0 | 14 | 0 | 0 | 0 | 0 | 0 | 1 | 0 | 0 | 0 |
| 1 | 0.00 | 3.68 | 0.00 | 1.00 | 1 | Gm4632 | | 0 | 10 | 5 | 0 | 0 | 0 | 0 | 0 | 0 | 0 | 0 |
| 1 | -0.71 | 3.60 | -0.20 | 0.84 | 1 | Gm22661 | | 0 | 0 | 0 | 0 | 1 | 9 | 0 | 0 | 0 | 0 | 5 |
| 1 | -3.21 | 3.55 | -0.90 | 0.37 | 1 | Gm43684 | | 0 | 0 | 0 | 5 | 6 | 3 | 0 | 0 | 0 | 0 | 1 |
| 1 | 1.14 | 3.54 | 0.32 | 0.75 | 1 | Gm48342 | | 0 | 3 | 0 | 3 | 0 | 3 | 0 | 0 | 2 | 0 | 5 |
| 1 | -2.99 | 3.58 | -0.84 | 0.40 | 1 | Dscaml1 | DS cell adhesion molecule like 1 | 3 | 0 | 0 | 0 | 7 | 0 | 0 | 0 | 0 | 0 | 6 |
| 1 | 1.41 | 3.66 | 0.38 | 0.70 | 1 | Rpl30-ps11 | | 11 | 1 | 1 | 0 | 0 | 0 | 0 | 0 | 2 | 0 | 0 |
| 1 | 1.19 | 3.60 | 0.33 | 0.74 | 1 | Mir148a | | 0 | 0 | 7 | 0 | 0 | 1 | 0 | 0 | 0 | 0 | 7 |
| 1 | 0.48 | 3.60 | 0.13 | 0.89 | 1 | Gm10912 | | 0 | 0 | 0 | 0 | 0 | 5 | 0 | 0 | 0 | 0 | 10 |
| 1 | 1.55 | 3.60 | 0.43 | 0.67 | 1 | Cyp2d37-ps | | 0 | 4 | 0 | 0 | 0 | 0 | 0 | 0 | 0 | 0 | 11 |
| 1 | 0.39 | 3.60 | 0.11 | 0.91 | 1 | Ldhal6b | lactate dehydrogenase A-like 6B | 0 | 2 | 0 | 0 | 0 | 5 | 0 | 0 | 0 | 0 | 8 |
| 1 | -0.07 | 3.60 | -0.02 | 0.99 | 1 | Gm26287 | | 0 | 0 | 0 | 0 | 0 | 10 | 0 | 0 | 0 | 0 | 5 |
| 1 | 2.59 | 3.45 | 0.75 | 0.45 | 1 | Gm11703 | | 0 | 1 | 1 | 2 | 0 | 0 | 0 | 0 | 9 | 1 | 1 |
| 1 | 1.10 | 3.65 | 0.30 | 0.76 | 1 | Ccdc105 | coiled-coil domain containing 105 | 0 | 14 | 0 | 0 | 0 | 0 | 0 | 0 | 0 | 0 | 1 |
| 1 | 0.00 | 3.65 | 0.00 | 1.00 | 1 | Gm32460 | | 0 | 15 | 0 | 0 | 0 | 0 | 0 | 0 | 0 | 0 | 0 |
| 1 | 0.00 | 3.65 | 0.00 | 1.00 | 1 | Gm13183 | | 0 | 15 | 0 | 0 | 0 | 0 | 0 | 0 | 0 | 0 | 0 |
| 1 | 0.00 | 3.65 | 0.00 | 1.00 | 1 | Gm33370 | | 0 | 15 | 0 | 0 | 0 | 0 | 0 | 0 | 0 | 0 | 0 |
| 1 | 0.00 | 3.65 | 0.00 | 1.00 | 1 | Gm11338 | | 0 | 15 | 0 | 0 | 0 | 0 | 0 | 0 | 0 | 0 | 0 |
| 1 | 2.82 | 3.60 | 0.78 | 0.43 | 1 | Gm34150 | | 0 | 0 | 0 | 0 | 0 | 0 | 0 | 0 | 0 | 14 | 1 |
| 1 | 2.17 | 3.61 | 0.60 | 0.55 | 1 | Gm49271 | | 3 | 0 | 0 | 0 | 0 | 0 | 0 | 0 | 0 | 12 | 1 |
| 1 | 3.79 | 3.58 | 1.06 | 0.29 | 1 | Gm36448 | | 1 | 1 | 0 | 0 | 0 | 0 | 0 | 2 | 7 | 4 | 0 |
| 1 | -2.97 | 3.58 | -0.83 | 0.41 | 1 | Gm45234 | | 0 | 0 | 0 | 12 | 0 | 1 | 0 | 0 | 0 | 0 | 2 |
| 1 | 0.81 | 3.55 | 0.23 | 0.82 | 1 | Nkx6-2 | NK6 homeobox 2 | 0 | 0 | 0 | 0 | 1 | 5 | 0 | 0 | 7 | 0 | 2 |
| 1 | -3.29 | 3.58 | -0.92 | 0.36 | 1 | Gm44144 | | 0 | 0 | 0 | 0 | 7 | 8 | 0 | 0 | 0 | 0 | 0 |
| 1 | 1.14 | 3.61 | 0.32 | 0.75 | 1 | Gm44699 | | 0 | 0 | 0 | 0 | 0 | 1 | 0 | 0 | 0 | 0 | 14 |
| 1 | 0.85 | 3.60 | 0.24 | 0.81 | 1 | Gm7444 | | 0 | 0 | 7 | 0 | 0 | 2 | 0 | 0 | 0 | 0 | 6 |
| 1 | 0.38 | 3.60 | 0.11 | 0.92 | 1 | 5530400K19Rik | | 0 | 0 | 0 | 0 | 0 | 6 | 0 | 0 | 0 | 0 | 9 |
| 1 | 0.38 | 3.60 | 0.11 | 0.92 | 1 | Gm43693 | | 0 | 0 | 0 | 0 | 0 | 6 | 0 | 0 | 0 | 0 | 9 |
| 1 | 0.38 | 3.60 | 0.11 | 0.92 | 1 | Gm38344 | | 0 | 0 | 0 | 0 | 0 | 6 | 0 | 0 | 0 | 0 | 9 |
| 1 | 0.95 | 3.67 | 0.26 | 0.80 | 1 | D130058E05Rik | | 12 | 2 | 0 | 0 | 0 | 0 | 0 | 0 | 0 | 0 | 1 |
| 1 | -0.18 | 3.60 | -0.05 | 0.96 | 1 | Gm45442 | | 0 | 0 | 0 | 0 | 0 | 11 | 0 | 0 | 0 | 0 | 4 |
| 1 | -0.18 | 3.60 | -0.05 | 0.96 | 1 | Gm44830 | | 0 | 0 | 0 | 0 | 0 | 11 | 0 | 0 | 0 | 0 | 4 |
| 1 | -0.18 | 3.60 | -0.05 | 0.96 | 1 | Gm45289 | | 0 | 0 | 0 | 0 | 0 | 11 | 0 | 0 | 0 | 0 | 4 |
| 1 | -0.18 | 3.60 | -0.05 | 0.96 | 1 | Gm47588 | | 0 | 0 | 0 | 0 | 0 | 11 | 0 | 0 | 0 | 0 | 4 |
| 1 | -2.35 | 3.59 | -0.65 | 0.51 | 1 | Gm6087 | | 5 | 0 | 1 | 0 | 4 | 0 | 0 | 0 | 0 | 0 | 5 |
| 1 | 0.00 | 3.68 | 0.00 | 1.00 | 1 | Gm34256 | | 1 | 14 | 0 | 0 | 0 | 0 | 0 | 0 | 0 | 0 | 0 |
| 1 | 0.03 | 3.59 | 0.01 | 0.99 | 1 | Gm35455 | | 0 | 8 | 0 | 0 | 0 | 6 | 0 | 0 | 1 | 0 | 0 |

| | | | | | | | | | | | | | | | | | | |
|---|---|---|---|---|---|---|---|---|---|---|---|---|---|---|---|---|---|---|
| 1 | -0.88 | 3.58 | -0.25 | 0.80 | 1 | Pcdha6 | protocadherin alpha 6 | 0 | 0 | 0 | 0 | 3 | 5 | 0 | 0 | 0 | 0 | 7 |
| 1 | -0.81 | 3.61 | -0.22 | 0.82 | 1 | Upk1a | uroplakin 1A | 4 | 0 | 3 | 0 | 0 | 8 | 0 | 0 | 0 | 0 | 0 |
| 1 | 0.57 | 3.63 | 0.16 | 0.87 | 1 | Gm11581 | | 14 | 0 | 0 | 0 | 0 | 0 | 0 | 0 | 0 | 1 | 0 |
| 1 | 1.15 | 3.60 | 0.32 | 0.75 | 1 | Cysrt1 | cysteine rich tail 1 | 0 | 0 | 0 | 0 | 0 | 1 | 0 | 0 | 0 | 14 | 0 |
| 1 | 0.00 | 3.68 | 0.00 | 1.00 | 1 | BB123696 | | 3 | 2 | 10 | 0 | 0 | 0 | 0 | 0 | 0 | 0 | 0 |
| 1 | -1.11 | 3.06 | -0.36 | 0.72 | 1 | Rpl17-ps9 | | 3 | 1 | 2 | 6 | 0 | 1 | 0 | 0 | 1 | 1 | 0 |
| 1 | -1.03 | 3.59 | -0.29 | 0.77 | 1 | D930007P13Rik | | 8 | 0 | 0 | 0 | 1 | 4 | 0 | 0 | 0 | 0 | 2 |
| 1 | 1.33 | 3.58 | 0.37 | 0.71 | 1 | Gm13981 | | 2 | 0 | 1 | 0 | 0 | 5 | 0 | 0 | 3 | 0 | 4 |
| 1 | 0.87 | 3.61 | 0.24 | 0.81 | 1 | Mogat2 | monoacylglycerol O-acyltransferase 2 | 2 | 0 | 7 | 0 | 0 | 1 | 0 | 0 | 0 | 0 | 5 |
| 1 | 1.49 | 3.64 | 0.41 | 0.68 | 1 | D630029K05Rik | | 13 | 0 | 0 | 0 | 0 | 0 | 0 | 0 | 0 | 0 | 2 |
| 1 | 0.91 | 3.61 | 0.25 | 0.80 | 1 | Gm37078 | | 0 | 2 | 0 | 0 | 0 | 2 | 0 | 0 | 0 | 0 | 11 |
| 1 | 0.00 | 3.68 | 0.00 | 1.00 | 1 | Kcns1 | K+ voltage-gated channel, subfamily S, 1 | 2 | 13 | 0 | 0 | 0 | 0 | 0 | 0 | 0 | 0 | 0 |
| 1 | -0.47 | 3.66 | -0.13 | 0.90 | 1 | Gm45113 | | 1 | 13 | 0 | 0 | 0 | 1 | 0 | 0 | 0 | 0 | 0 |
| 1 | 0.00 | 3.68 | 0.00 | 1.00 | 1 | Gabra2 | gamma-aminobutyric acid (GABA) A rece | 8 | 5 | 2 | 0 | 0 | 0 | 0 | 0 | 0 | 0 | 0 |
| 1 | 1.24 | 3.36 | 0.37 | 0.71 | 1 | Gm5297 | | 0 | 0 | 1 | 1 | 0 | 3 | 0 | 2 | 7 | 0 | 1 |
| 1 | 4.04 | 3.59 | 1.12 | 0.26 | 1 | Gm37012 | | 0 | 5 | 0 | 0 | 0 | 2 | 0 | 0 | 5 | 0 | 3 |
| 1 | 3.38 | 3.58 | 0.94 | 0.35 | 1 | Olfr316 | olfactory receptor 316 | 0 | 0 | 0 | 0 | 0 | 2 | 0 | 0 | 12 | 0 | 1 |
| 1 | -0.76 | 3.63 | -0.21 | 0.83 | 1 | Cyp2j8 | cytochrome P450, family 2, subfamily j, pc | 9 | 2 | 0 | 0 | 0 | 3 | 0 | 0 | 0 | 0 | 0 |
| 1 | 0.00 | 3.68 | 0.00 | 1.00 | 1 | Gm39556 | | 9 | 0 | 6 | 0 | 0 | 0 | 0 | 0 | 0 | 0 | 0 |
| 1 | 0.23 | 3.61 | 0.06 | 0.95 | 1 | Gm7452 | | 0 | 1 | 0 | 0 | 3 | 1 | 0 | 0 | 10 | 0 | 0 |
| 1 | -0.69 | 3.66 | -0.19 | 0.85 | 1 | Dgkk | diacylglycerol kinase kappa | 7 | 0 | 6 | 0 | 0 | 2 | 0 | 0 | 0 | 0 | 0 |
| 1 | 3.31 | 3.59 | 0.92 | 0.36 | 1 | Zic4 | zinc finger protein of the cerebellum 4 | 0 | 0 | 0 | 0 | 0 | 0 | 0 | 13 | 0 | 0 | 2 |
| 1 | -0.96 | 3.61 | -0.27 | 0.79 | 1 | Gm26642 | | 0 | 12 | 0 | 2 | 0 | 1 | 0 | 0 | 0 | 0 | 0 |
| 1 | 1.12 | 3.60 | 0.31 | 0.76 | 1 | 1190005I06Rik | RIKEN cDNA 1190005I06 gene | 0 | 0 | 0 | 0 | 0 | 1 | 0 | 14 | 0 | 0 | 0 |
| 1 | 1.45 | 3.56 | 0.41 | 0.68 | 1 | Gm25394 | | 0 | 0 | 2 | 0 | 0 | 3 | 0 | 0 | 6 | 0 | 4 |
| 1 | 0.26 | 3.62 | 0.07 | 0.94 | 1 | Gm17430 | | 13 | 0 | 0 | 0 | 0 | 1 | 0 | 0 | 0 | 0 | 1 |
| 1 | 1.62 | 3.65 | 0.44 | 0.66 | 1 | Gm2607 | | 5 | 0 | 7 | 0 | 0 | 0 | 0 | 0 | 0 | 1 | 1 |
| 1 | -1.84 | 3.62 | -0.51 | 0.61 | 1 | Gm37419 | | 1 | 0 | 10 | 3 | 0 | 1 | 0 | 0 | 0 | 0 | 0 |
| 1 | 0.28 | 3.58 | 0.08 | 0.94 | 1 | Gm7859 | | 7 | 1 | 0 | 0 | 0 | 2 | 0 | 0 | 0 | 5 | 0 |
| 1 | 2.06 | 3.58 | 0.58 | 0.56 | 1 | Gm30929 | | 0 | 0 | 0 | 0 | 0 | 3 | 0 | 0 | 5 | 0 | 7 |
| 1 | -1.16 | 3.44 | -0.34 | 0.74 | 1 | Gm5430 | | 2 | 1 | 1 | 4 | 0 | 4 | 0 | 2 | 0 | 0 | 0 |
| 1 | 0.33 | 3.55 | 0.09 | 0.93 | 1 | Gm11750 | | 0 | 0 | 0 | 0 | 6 | 0 | 0 | 0 | 0 | 9 | 0 |
| 1 | 0.00 | 3.68 | 0.00 | 1.00 | 1 | Gm27477 | | 4 | 0 | 10 | 0 | 0 | 0 | 0 | 0 | 0 | 0 | 0 |
| 1 | 0.00 | 3.68 | 0.00 | 1.00 | 1 | Gm15523 | | 4 | 0 | 10 | 0 | 0 | 0 | 0 | 0 | 0 | 0 | 0 |
| 1 | 1.25 | 3.40 | 0.37 | 0.71 | 1 | Gm48690 | | 8 | 2 | 1 | 0 | 0 | 1 | 0 | 0 | 1 | 0 | 2 |
| 1 | -0.69 | 3.66 | -0.19 | 0.85 | 1 | Gm50100 | | 3 | 0 | 10 | 0 | 0 | 2 | 0 | 0 | 0 | 0 | 0 |
| 1 | -1.02 | 3.61 | -0.28 | 0.78 | 1 | Gm7909 | | 0 | 9 | 1 | 0 | 1 | 3 | 0 | 0 | 0 | 0 | 1 |
| 1 | 0.00 | 3.68 | 0.00 | 1.00 | 1 | Gm30329 | | 9 | 0 | 5 | 0 | 0 | 0 | 0 | 0 | 0 | 0 | 0 |
| 1 | -3.26 | 3.57 | -0.91 | 0.36 | 1 | Fmo6 | flavin containing monooxygenase 6 | 0 | 0 | 0 | 0 | 10 | 4 | 0 | 0 | 0 | 0 | 0 |
| 1 | 1.17 | 2.86 | 0.41 | 0.68 | 1 | Csl | citrate synthase like | 3 | 0 | 2 | 0 | 2 | 1 | 0 | 0 | 2 | 1 | 3 |
| 1 | 0.28 | 3.60 | 0.08 | 0.94 | 1 | Gm13398 | | 3 | 0 | 0 | 0 | 0 | 5 | 0 | 0 | 0 | 0 | 6 |
| 1 | 1.64 | 2.95 | 0.56 | 0.58 | 1 | Gm7059 | | 3 | 1 | 0 | 3 | 0 | 1 | 0 | 3 | 2 | 0 | 2 |
| 1 | 0.00 | 3.66 | 0.00 | 1.00 | 1 | A530013C23Rik | | 15 | 0 | 0 | 0 | 0 | 0 | 0 | 0 | 0 | 0 | 0 |
| 1 | 0.00 | 3.66 | 0.00 | 1.00 | 1 | Gm43327 | | 15 | 0 | 0 | 0 | 0 | 0 | 0 | 0 | 0 | 0 | 0 |
| 1 | 0.00 | 3.66 | 0.00 | 1.00 | 1 | Gm15448 | paired-Ig-like receptor A6 | 15 | 0 | 0 | 0 | 0 | 0 | 0 | 0 | 0 | 0 | 0 |
| 1 | 0.00 | 3.66 | 0.00 | 1.00 | 1 | Apol7e | apolipoprotein L 7e | 15 | 0 | 0 | 0 | 0 | 0 | 0 | 0 | 0 | 0 | 0 |
| 1 | 0.00 | 3.66 | 0.00 | 1.00 | 1 | Tulp1 | tubby like protein 1 | 15 | 0 | 0 | 0 | 0 | 0 | 0 | 0 | 0 | 0 | 0 |
| 1 | 0.00 | 3.66 | 0.00 | 1.00 | 1 | Cyp4f37 | cytochrome P450, family 4, subfamily f, pc | 15 | 0 | 0 | 0 | 0 | 0 | 0 | 0 | 0 | 0 | 0 |
| 1 | 0.00 | 3.66 | 0.00 | 1.00 | 1 | Pcdh11x | protocadherin 11 X-linked | 15 | 0 | 0 | 0 | 0 | 0 | 0 | 0 | 0 | 0 | 0 |
| 1 | -0.09 | 3.62 | -0.03 | 0.98 | 1 | Gm15790 | | 11 | 0 | 0 | 0 | 0 | 1 | 0 | 0 | 0 | 3 | 0 |
| 1 | 0.00 | 3.65 | 0.00 | 1.00 | 1 | 9130019P16Rik | | 0 | 0 | 15 | 0 | 0 | 0 | 0 | 0 | 0 | 0 | 0 |
| 1 | 0.00 | 3.65 | 0.00 | 1.00 | 1 | 5430421F17Rik | | 0 | 0 | 15 | 0 | 0 | 0 | 0 | 0 | 0 | 0 | 0 |
| 1 | 0.00 | 3.65 | 0.00 | 1.00 | 1 | Prss57 | protease, serine 57 | 0 | 0 | 15 | 0 | 0 | 0 | 0 | 0 | 0 | 0 | 0 |
| 1 | 0.00 | 3.65 | 0.00 | 1.00 | 1 | 1700054O19Rik | | 0 | 0 | 15 | 0 | 0 | 0 | 0 | 0 | 0 | 0 | 0 |
| 1 | 0.00 | 3.65 | 0.00 | 1.00 | 1 | Gm50269 | | 0 | 0 | 15 | 0 | 0 | 0 | 0 | 0 | 0 | 0 | 0 |
| 1 | 0.07 | 3.60 | 0.02 | 0.98 | 1 | Scarna6 | | 0 | 0 | 0 | 0 | 0 | 8 | 0 | 0 | 0 | 0 | 6 |
| 1 | 0.07 | 3.60 | 0.02 | 0.98 | 1 | Gm37289 | | 0 | 0 | 0 | 0 | 0 | 8 | 0 | 0 | 0 | 0 | 6 |
| 1 | 0.07 | 3.60 | 0.02 | 0.98 | 1 | Gm42876 | | 0 | 0 | 0 | 0 | 0 | 8 | 0 | 0 | 0 | 0 | 6 |
| 1 | 0.07 | 3.60 | 0.02 | 0.98 | 1 | Gm16558 | | 0 | 0 | 0 | 0 | 0 | 8 | 0 | 0 | 0 | 0 | 6 |
| 1 | 0.07 | 3.60 | 0.02 | 0.98 | 1 | Brms1 | breast cancer metastasis-suppressor 1 | 0 | 0 | 0 | 0 | 0 | 8 | 0 | 0 | 0 | 0 | 6 |

| | | | | | | | | | | | | | | | | | | |
|---|---|---|---|---|---|---|---|---|---|---|---|---|---|---|---|---|---|---|
| 1 | 1.87 | 3.57 | 0.52 | 0.60 | 1 | Gm9143 | | 0 | 0 | 1 | 0 | 0 | 5 | 0 | 2 | 1 | 0 | 5 |
| 1 | 0.00 | 3.68 | 0.00 | 1.00 | 1 | Fabp2 | fatty acid binding protein 2, intestinal | 0 | 2 | 12 | 0 | 0 | 0 | 0 | 0 | 0 | 0 | 0 |
| 1 | 3.37 | 2.96 | 1.14 | 0.25 | 1 | Gm13082 | | 0 | 1 | 2 | 1 | 0 | 0 | 0 | 2 | 1 | 5 | 2 |
| 1 | -4.93 | 3.55 | -1.39 | 0.16 | 1 | Gm24610 | | 0 | 0 | 0 | 4 | 0 | 6 | 0 | 0 | 5 | 0 | 0 |
| 1 | 1.35 | 3.53 | 0.38 | 0.70 | 1 | H2ac20 | H2A clustered histone 20 | 0 | 0 | 0 | 0 | 3 | 1 | 0 | 0 | 1 | 8 | 2 |
| 1 | 1.51 | 3.60 | 0.42 | 0.67 | 1 | Gm44260 | | 4 | 0 | 0 | 0 | 0 | 0 | 0 | 0 | 0 | 0 | 10 |
| 1 | -0.36 | 3.61 | -0.10 | 0.92 | 1 | Nxf2 | nuclear RNA export factor 2 | 4 | 4 | 0 | 0 | 0 | 5 | 0 | 0 | 0 | 0 | 1 |
| 1 | 0.51 | 3.60 | 0.14 | 0.89 | 1 | Bricd5 | BRICHOS domain containing 5 | 0 | 0 | 1 | 0 | 0 | 4 | 0 | 0 | 0 | 0 | 9 |
| 1 | 0.55 | 3.60 | 0.15 | 0.88 | 1 | Gm24139 | | 0 | 0 | 0 | 0 | 0 | 4 | 0 | 0 | 0 | 0 | 10 |
| 1 | 0.55 | 3.60 | 0.15 | 0.88 | 1 | Gm20427 | | 0 | 0 | 0 | 0 | 0 | 4 | 0 | 0 | 0 | 0 | 10 |
| 1 | 0.28 | 3.61 | 0.08 | 0.94 | 1 | 1700019A02Ril | RIKEN cDNA 1700019A02 gene | 1 | 6 | 0 | 0 | 0 | 3 | 0 | 0 | 0 | 0 | 4 |
| 1 | -0.03 | 3.60 | -0.01 | 0.99 | 1 | Gm22990 | | 0 | 0 | 0 | 0 | 0 | 9 | 0 | 0 | 0 | 0 | 5 |
| 1 | -2.70 | 3.55 | -0.76 | 0.45 | 1 | Gm47887 | | 0 | 0 | 0 | 6 | 3 | 3 | 0 | 0 | 0 | 0 | 2 |
| 1 | 2.40 | 3.61 | 0.67 | 0.51 | 1 | Fam90a1b | family with sequence similarity 90, membe | 0 | 0 | 7 | 0 | 0 | 0 | 0 | 0 | 3 | 0 | 4 |
| 1 | -2.23 | 3.57 | -0.62 | 0.53 | 1 | 1700001D01Rik | | 0 | 0 | 0 | 0 | 5 | 3 | 0 | 0 | 0 | 0 | 7 |
| 1 | 0.00 | 3.68 | 0.00 | 1.00 | 1 | Gm14617 | | 5 | 5 | 4 | 0 | 0 | 0 | 0 | 0 | 0 | 0 | 0 |
| 1 | 0.00 | 3.68 | 0.00 | 1.00 | 1 | Gm4961 | | 5 | 5 | 4 | 0 | 0 | 0 | 0 | 0 | 0 | 0 | 0 |
| 1 | 2.31 | 3.62 | 0.64 | 0.52 | 1 | Lmntd1 | lamin tail domain containing 1 | 5 | 0 | 4 | 0 | 0 | 0 | 0 | 0 | 4 | 0 | 1 |
| 1 | 2.88 | 3.57 | 0.81 | 0.42 | 1 | Gm31274 | | 0 | 0 | 0 | 0 | 0 | 4 | 0 | 0 | 8 | 0 | 2 |
| 1 | -0.07 | 3.00 | -0.02 | 0.98 | 1 | Gm6450 | | 1 | 3 | 2 | 4 | 1 | 0 | 0 | 0 | 1 | 0 | 2 |
| 1 | 0.20 | 3.59 | 0.05 | 0.96 | 1 | Gm44759 | | 0 | 7 | 0 | 4 | 0 | 1 | 0 | 0 | 2 | 0 | 0 |
| 1 | -1.41 | 3.59 | -0.39 | 0.69 | 1 | Gm43799 | | 0 | 0 | 0 | 0 | 3 | 8 | 0 | 0 | 0 | 0 | 3 |
| 1 | -0.41 | 3.64 | -0.11 | 0.91 | 1 | S1pr5 | sphingosine-1-phosphate receptor 5 | 0 | 2 | 10 | 0 | 2 | 0 | 0 | 0 | 0 | 0 | 0 |
| 1 | 1.89 | 3.38 | 0.56 | 0.58 | 1 | Gm5511 | | 0 | 1 | 4 | 0 | 3 | 0 | 0 | 1 | 2 | 0 | 3 |
| 1 | -0.55 | 3.61 | -0.15 | 0.88 | 1 | Pnpla5 | patatin-like phospholipase domain contai | 0 | 8 | 0 | 6 | 0 | 0 | 0 | 0 | 0 | 0 | 0 |
| 1 | 4.82 | 3.55 | 1.36 | 0.17 | 1 | Gm36975 | | 0 | 0 | 0 | 6 | 0 | 0 | 0 | 0 | 6 | 0 | 2 |
| 1 | 0.51 | 3.60 | 0.14 | 0.89 | 1 | Gm23887 | | 1 | 0 | 0 | 0 | 0 | 4 | 0 | 0 | 0 | 0 | 9 |
| 1 | -0.31 | 3.55 | -0.09 | 0.93 | 1 | Gm13168 | | 0 | 0 | 0 | 0 | 7 | 3 | 0 | 0 | 3 | 0 | 2 |
| 1 | 1.39 | 3.61 | 0.38 | 0.70 | 1 | 8030474K03Ril | RIKEN cDNA 8030474K03 gene | 0 | 0 | 0 | 0 | 0 | 0 | 0 | 0 | 0 | 0 | 14 |
| 1 | -0.15 | 3.60 | -0.04 | 0.97 | 1 | Mir7085 | | 0 | 0 | 0 | 0 | 0 | 10 | 0 | 0 | 0 | 0 | 4 |
| 1 | 1.59 | 3.60 | 0.44 | 0.66 | 1 | Gm45014 | | 0 | 5 | 0 | 0 | 0 | 0 | 0 | 0 | 0 | 0 | 9 |
| 1 | -2.04 | 3.53 | -0.58 | 0.56 | 1 | Gm31323 | | 0 | 0 | 0 | 5 | 0 | 6 | 0 | 0 | 3 | 0 | 0 |
| 1 | -3.91 | 3.61 | -1.08 | 0.28 | 1 | Gm13166 | | 7 | 0 | 0 | 0 | 4 | 3 | 0 | 0 | 0 | 0 | 1 |
| 1 | 1.62 | 3.63 | 0.45 | 0.65 | 1 | 9230104M06Rik | | 0 | 11 | 0 | 0 | 0 | 0 | 0 | 0 | 0 | 0 | 3 |
| 1 | 0.00 | 3.65 | 0.00 | 1.00 | 1 | Gm37168 | | 0 | 14 | 0 | 0 | 0 | 0 | 0 | 0 | 0 | 0 | 0 |
| 1 | 0.00 | 3.65 | 0.00 | 1.00 | 1 | Rpl13-ps1 | | 0 | 14 | 0 | 0 | 0 | 0 | 0 | 0 | 0 | 0 | 0 |
| 1 | 0.00 | 3.65 | 0.00 | 1.00 | 1 | Gm33497 | | 0 | 14 | 0 | 0 | 0 | 0 | 0 | 0 | 0 | 0 | 0 |
| 1 | 1.35 | 3.57 | 0.38 | 0.70 | 1 | Gm13552 | | 0 | 0 | 5 | 1 | 0 | 0 | 0 | 0 | 4 | 0 | 4 |
| 1 | 1.85 | 3.62 | 0.51 | 0.61 | 1 | Gm5544 | predicted gene 5544 | 0 | 0 | 0 | 0 | 0 | 0 | 0 | 0 | 0 | 14 | 0 |
| 1 | 1.85 | 3.62 | 0.51 | 0.61 | 1 | Gm16054 | | 0 | 0 | 0 | 0 | 0 | 0 | 0 | 0 | 0 | 14 | 0 |
| 1 | 1.85 | 3.62 | 0.51 | 0.61 | 1 | Gm20715 | | 0 | 0 | 0 | 0 | 0 | 0 | 0 | 0 | 0 | 14 | 0 |
| 1 | 0.12 | 3.61 | 0.03 | 0.97 | 1 | Snord43 | | 8 | 0 | 0 | 0 | 0 | 3 | 0 | 0 | 0 | 0 | 3 |
| 1 | 0.00 | 3.68 | 0.00 | 1.00 | 1 | Cd300c | CD300C molecule | 7 | 0 | 7 | 0 | 0 | 0 | 0 | 0 | 0 | 0 | 0 |
| 1 | 1.12 | 3.56 | 0.32 | 0.75 | 1 | Gm16796 | | 0 | 0 | 0 | 0 | 0 | 3 | 0 | 0 | 0 | 12 | 0 |
| 1 | -0.40 | 3.67 | -0.11 | 0.91 | 1 | Gm45734 | | 5 | 8 | 0 | 0 | 0 | 1 | 0 | 0 | 0 | 0 | 0 |
| 1 | -0.39 | 3.56 | -0.11 | 0.91 | 1 | Sun3 | Sad1 and UNC84 domain containing 3 | 0 | 0 | 0 | 0 | 4 | 4 | 0 | 0 | 1 | 0 | 5 |
| 1 | 0.22 | 3.60 | 0.06 | 0.95 | 1 | Gm21057 | | 0 | 0 | 2 | 0 | 0 | 6 | 0 | 0 | 0 | 0 | 6 |
| 1 | 0.79 | 3.62 | 0.22 | 0.83 | 1 | Gm2253 | | 0 | 7 | 2 | 0 | 0 | 1 | 0 | 0 | 0 | 0 | 4 |
| 1 | 0.67 | 3.60 | 0.19 | 0.85 | 1 | Gm22486 | | 0 | 0 | 8 | 0 | 0 | 2 | 0 | 0 | 0 | 0 | 4 |
| 1 | 0.28 | 3.60 | 0.08 | 0.94 | 1 | Gm25395 | | 0 | 0 | 1 | 0 | 0 | 6 | 0 | 0 | 0 | 0 | 7 |
| 1 | -0.79 | 3.56 | -0.22 | 0.82 | 1 | Gzmk | granzyme K | 0 | 0 | 0 | 5 | 0 | 2 | 0 | 0 | 0 | 0 | 7 |
| 1 | -1.55 | 3.58 | -0.43 | 0.66 | 1 | Gm38252 | | 0 | 0 | 0 | 0 | 11 | 2 | 0 | 0 | 0 | 0 | 1 |
| 1 | -0.27 | 3.60 | -0.08 | 0.94 | 1 | Gm37334 | | 0 | 0 | 0 | 0 | 0 | 11 | 0 | 0 | 0 | 0 | 3 |
| 1 | -0.69 | 3.66 | -0.19 | 0.85 | 1 | Gm16251 | | 0 | 5 | 7 | 0 | 0 | 2 | 0 | 0 | 0 | 0 | 0 |
| 1 | 0.60 | 3.62 | 0.17 | 0.87 | 1 | Gm37407 | | 11 | 0 | 0 | 0 | 0 | 1 | 0 | 0 | 0 | 0 | 2 |
| 1 | 0.00 | 3.68 | 0.00 | 1.00 | 1 | Gm17877 | | 1 | 13 | 0 | 0 | 0 | 0 | 0 | 0 | 0 | 0 | 0 |
| 1 | 0.00 | 3.68 | 0.00 | 1.00 | 1 | Gm16166 | | 1 | 13 | 0 | 0 | 0 | 0 | 0 | 0 | 0 | 0 | 0 |
| 1 | 0.00 | 3.68 | 0.00 | 1.00 | 1 | Gm28932 | | 1 | 13 | 0 | 0 | 0 | 0 | 0 | 0 | 0 | 0 | 0 |
| 1 | 0.58 | 3.61 | 0.16 | 0.87 | 1 | Akr1c21 | aldo-keto reductase family 1, member C2 | 0 | 11 | 0 | 0 | 0 | 1 | 0 | 0 | 0 | 0 | 2 |
| 1 | -0.61 | 3.65 | -0.17 | 0.87 | 1 | Pram1 | PML-RAR alpha-regulated adaptor molec | 0 | 13 | 0 | 0 | 0 | 1 | 0 | 0 | 0 | 0 | 0 |

| | | | | | | | | | | | | | | | | | | |
|---|---|---|---|---|---|---|---|---|---|---|---|---|---|---|---|---|---|---|
| 1 | 1.57 | 3.57 | 0.44 | 0.66 | 1 | Adgrf3 | adhesion G protein-coupled receptor F3 | 0 | 0 | 0 | 0 | 3 | 0 | 0 | 0 | 6 | 0 | 5 |
| 1 | -2.31 | 3.60 | -0.64 | 0.52 | 1 | 4930593C16Rik | | 0 | 0 | 0 | 13 | 0 | 1 | 0 | 0 | 0 | 0 | 0 |
| 1 | -1.05 | 3.63 | -0.29 | 0.77 | 1 | Galr2 | galanin receptor 2 | 0 | 0 | 11 | 0 | 0 | 3 | 0 | 0 | 0 | 0 | 0 |
| 1 | -2.91 | 3.59 | -0.81 | 0.42 | 1 | Ccer2 | coiled-coil glutamate-rich protein 2 | 1 | 4 | 0 | 0 | 6 | 3 | 0 | 0 | 0 | 0 | 0 |
| 1 | 0.00 | 3.68 | 0.00 | 1.00 | 1 | Gm13749 | | 8 | 0 | 6 | 0 | 0 | 0 | 0 | 0 | 0 | 0 | 0 |
| 1 | 0.00 | 3.68 | 0.00 | 1.00 | 1 | Fam3b | family with sequence similarity 3, member | 8 | 0 | 6 | 0 | 0 | 0 | 0 | 0 | 0 | 0 | 0 |
| 1 | 0.00 | 3.68 | 0.00 | 1.00 | 1 | Gm38079 | | 7 | 7 | 0 | 0 | 0 | 0 | 0 | 0 | 0 | 0 | 0 |
| 1 | -0.40 | 3.67 | -0.11 | 0.91 | 1 | Gm5320 | | 7 | 0 | 6 | 0 | 0 | 1 | 0 | 0 | 0 | 0 | 0 |
| 1 | 1.68 | 3.57 | 0.47 | 0.64 | 1 | Gm26724 | | 0 | 0 | 0 | 0 | 1 | 1 | 0 | 0 | 9 | 0 | 3 |
| 1 | -0.27 | 3.66 | -0.07 | 0.94 | 1 | Wnt9b | wingless-type MMTV integration site famil | 3 | 0 | 10 | 0 | 1 | 0 | 0 | 0 | 0 | 0 | 0 |
| 1 | 0.00 | 3.68 | 0.00 | 1.00 | 1 | Fam124b | family with sequence similarity 124, meml | 13 | 0 | 1 | 0 | 0 | 0 | 0 | 0 | 0 | 0 | 0 |
| 1 | 1.89 | 2.60 | 0.73 | 0.47 | 1 | Gm7251 | | 2 | 1 | 2 | 1 | 0 | 1 | 0 | 0 | 3 | 1 | 3 |
| 1 | 1.67 | 2.33 | 0.71 | 0.48 | 1 | Gm8959 | | 3 | 2 | 2 | 1 | 0 | 1 | 0 | 1 | 1 | 1 | 2 |
| 1 | -3.12 | 3.58 | -0.87 | 0.38 | 1 | Gm44234 | | 0 | 0 | 0 | 11 | 0 | 3 | 0 | 0 | 0 | 0 | 0 |
| 1 | 0.00 | 3.68 | 0.00 | 1.00 | 1 | Rpl19-ps12 | | 8 | 3 | 3 | 0 | 0 | 0 | 0 | 0 | 0 | 0 | 0 |
| 1 | 0.47 | 3.60 | 0.13 | 0.90 | 1 | Gm10616 | | 0 | 0 | 7 | 0 | 0 | 3 | 0 | 0 | 0 | 0 | 4 |
| 1 | 0.22 | 3.60 | 0.06 | 0.95 | 1 | Gm46436 | | 0 | 0 | 0 | 0 | 0 | 7 | 0 | 0 | 0 | 0 | 7 |
| 1 | -0.42 | 3.60 | -0.12 | 0.91 | 1 | Gm45206 | | 0 | 0 | 0 | 0 | 0 | 12 | 0 | 0 | 0 | 0 | 2 |
| 1 | -0.46 | 3.66 | -0.13 | 0.90 | 1 | Gm12704 | | 1 | 12 | 0 | 0 | 0 | 1 | 0 | 0 | 0 | 0 | 0 |
| 1 | 2.19 | 3.57 | 0.61 | 0.54 | 1 | Gm18605 | | 0 | 0 | 0 | 0 | 0 | 7 | 0 | 0 | 5 | 0 | 2 |
| 1 | -0.10 | 3.61 | -0.03 | 0.98 | 1 | Gm10634 | | 0 | 11 | 0 | 0 | 0 | 2 | 0 | 0 | 0 | 0 | 1 |
| 1 | -0.32 | 3.58 | -0.09 | 0.93 | 1 | Gm48667 | | 0 | 0 | 3 | 3 | 0 | 0 | 0 | 0 | 8 | 0 | 0 |
| 1 | -0.20 | 3.57 | -0.06 | 0.95 | 1 | Gm9752 | | 0 | 0 | 8 | 0 | 1 | 3 | 0 | 0 | 0 | 0 | 2 |
| 1 | 0.00 | 3.68 | 0.00 | 1.00 | 1 | Gm12159 | | 8 | 6 | 0 | 0 | 0 | 0 | 0 | 0 | 0 | 0 | 0 |
| 1 | -0.73 | 3.58 | -0.20 | 0.84 | 1 | Gm37716 | | 0 | 0 | 0 | 0 | 1 | 12 | 0 | 0 | 1 | 0 | 0 |
| 1 | -1.08 | 3.56 | -0.30 | 0.76 | 1 | Gm18087 | | 0 | 0 | 0 | 0 | 4 | 1 | 0 | 0 | 0 | 0 | 9 |
| 1 | 1.63 | 3.62 | 0.45 | 0.65 | 1 | Gm5130 | | 1 | 0 | 0 | 0 | 0 | 0 | 0 | 0 | 0 | 13 | 0 |
| 1 | 0.75 | 3.40 | 0.22 | 0.83 | 1 | Gm35857 | | 1 | 1 | 2 | 0 | 5 | 0 | 0 | 0 | 4 | 0 | 1 |
| 1 | 0.00 | 3.66 | 0.00 | 1.00 | 1 | Brinp3 | bone morphogenetic protein/retinoic acid | 14 | 0 | 0 | 0 | 0 | 0 | 0 | 0 | 0 | 0 | 0 |
| 1 | 0.00 | 3.66 | 0.00 | 1.00 | 1 | Elf5 | E74-like factor 5 | 14 | 0 | 0 | 0 | 0 | 0 | 0 | 0 | 0 | 0 | 0 |
| 1 | 0.00 | 3.66 | 0.00 | 1.00 | 1 | Gm43514 | | 14 | 0 | 0 | 0 | 0 | 0 | 0 | 0 | 0 | 0 | 0 |
| 1 | 0.00 | 3.66 | 0.00 | 1.00 | 1 | Igkv4-80 | | 14 | 0 | 0 | 0 | 0 | 0 | 0 | 0 | 0 | 0 | 0 |
| 1 | 0.00 | 3.66 | 0.00 | 1.00 | 1 | Gm44812 | | 14 | 0 | 0 | 0 | 0 | 0 | 0 | 0 | 0 | 0 | 0 |
| 1 | 0.00 | 3.66 | 0.00 | 1.00 | 1 | Gm44930 | | 14 | 0 | 0 | 0 | 0 | 0 | 0 | 0 | 0 | 0 | 0 |
| 1 | 0.00 | 3.66 | 0.00 | 1.00 | 1 | 2410004I01Rik | | 14 | 0 | 0 | 0 | 0 | 0 | 0 | 0 | 0 | 0 | 0 |
| 1 | 0.00 | 3.66 | 0.00 | 1.00 | 1 | Dsg1b | desmoglein 1 beta | 14 | 0 | 0 | 0 | 0 | 0 | 0 | 0 | 0 | 0 | 0 |
| 1 | 0.00 | 3.68 | 0.00 | 1.00 | 1 | Hoxaas3 | | 3 | 11 | 0 | 0 | 0 | 0 | 0 | 0 | 0 | 0 | 0 |
| 1 | 0.00 | 3.68 | 0.00 | 1.00 | 1 | Klri2 | killer cell lectin-like receptor family I meml | 3 | 11 | 0 | 0 | 0 | 0 | 0 | 0 | 0 | 0 | 0 |
| 1 | 0.78 | 3.60 | 0.22 | 0.83 | 1 | Gm22952 | | 0 | 0 | 0 | 0 | 0 | 3 | 0 | 0 | 0 | 0 | 11 |
| 1 | 0.78 | 3.60 | 0.22 | 0.83 | 1 | Gm45579 | | 0 | 0 | 0 | 0 | 0 | 3 | 0 | 0 | 0 | 0 | 11 |
| 1 | 0.12 | 3.60 | 0.03 | 0.97 | 1 | Xlr4d-ps | | 0 | 0 | 0 | 0 | 0 | 8 | 0 | 0 | 0 | 0 | 6 |
| 1 | 1.69 | 3.59 | 0.47 | 0.64 | 1 | Snord17 | | 0 | 0 | 0 | 0 | 0 | 3 | 0 | 0 | 2 | 0 | 9 |
| 1 | -1.33 | 3.56 | -0.37 | 0.71 | 1 | Gm37593 | | 0 | 0 | 0 | 0 | 5 | 1 | 0 | 0 | 0 | 0 | 8 |
| 1 | 1.05 | 3.59 | 0.29 | 0.77 | 1 | Gm48382 | | 0 | 0 | 0 | 0 | 0 | 8 | 0 | 0 | 2 | 0 | 4 |
| 1 | 2.72 | 3.61 | 0.75 | 0.45 | 1 | Gm18119 | | 1 | 8 | 0 | 0 | 0 | 2 | 0 | 0 | 2 | 0 | 1 |
| 1 | 0.39 | 3.60 | 0.11 | 0.91 | 1 | Gm10933 | | 0 | 7 | 0 | 0 | 0 | 3 | 0 | 0 | 0 | 0 | 4 |
| 1 | 1.62 | 3.62 | 0.45 | 0.65 | 1 | Atp5l-ps2 | | 0 | 0 | 10 | 0 | 0 | 1 | 0 | 1 | 0 | 1 | 0 |
| 1 | -1.14 | 3.62 | -0.31 | 0.75 | 1 | Gm28373 | | 0 | 11 | 0 | 0 | 0 | 3 | 0 | 0 | 0 | 0 | 0 |
| 1 | -1.14 | 3.62 | -0.31 | 0.75 | 1 | Gm3558 | predicted gene 3558 | 0 | 11 | 0 | 0 | 0 | 3 | 0 | 0 | 0 | 0 | 0 |
| 1 | 0.09 | 3.60 | 0.02 | 0.98 | 1 | Gm49625 | | 0 | 10 | 0 | 0 | 0 | 3 | 0 | 0 | 1 | 0 | 0 |
| 1 | 1.25 | 3.60 | 0.35 | 0.73 | 1 | Gm14016 | | 2 | 1 | 0 | 0 | 0 | 3 | 0 | 0 | 0 | 1 | 7 |
| 1 | -0.72 | 2.97 | -0.24 | 0.81 | 1 | Gm12727 | | 2 | 0 | 0 | 2 | 2 | 3 | 0 | 1 | 2 | 3 | 0 |
| 1 | 0.00 | 3.68 | 0.00 | 1.00 | 1 | A730056A06Rik | | 9 | 5 | 0 | 0 | 0 | 0 | 0 | 0 | 0 | 0 | 0 |
| 1 | -0.23 | 3.58 | -0.06 | 0.95 | 1 | Zfp600 | zinc finger protein 600 | 0 | 3 | 0 | 0 | 3 | 2 | 0 | 0 | 6 | 0 | 0 |
| 1 | 1.04 | 3.54 | 0.29 | 0.77 | 1 | Rpl30-ps1 | | 6 | 1 | 1 | 0 | 0 | 3 | 0 | 0 | 2 | 0 | 1 |
| 1 | -1.56 | 3.56 | -0.44 | 0.66 | 1 | 1700124L16Rik | | 0 | 0 | 0 | 0 | 6 | 1 | 0 | 0 | 0 | 0 | 7 |
| 1 | 1.37 | 3.61 | 0.38 | 0.70 | 1 | Gm4860 | | 0 | 0 | 0 | 1 | 0 | 0 | 0 | 13 | 0 | 0 | 0 |
| 1 | -1.43 | 3.61 | -0.40 | 0.69 | 1 | Gm29125 | | 1 | 0 | 6 | 0 | 6 | 1 | 0 | 0 | 0 | 0 | 0 |
| 1 | -1.02 | 3.60 | -0.28 | 0.78 | 1 | Pjvk | pejvakin | 5 | 0 | 0 | 0 | 0 | 8 | 0 | 0 | 0 | 0 | 0 |
| 1 | -1.60 | 3.60 | -0.45 | 0.66 | 1 | Ankfn1 | ankyrin-repeat and fibronectin type III don | 0 | 0 | 6 | 0 | 6 | 2 | 0 | 0 | 0 | 0 | 0 |

| | | | | | | | | | | | | | | | | | | |
|---|---|---|---|---|---|---|---|---|---|---|---|---|---|---|---|---|---|---|
| 1 | 0.00 | 3.68 | 0.00 | 1.00 | 1 | S100a8 | S100 calcium binding protein A8 (calgran | 9 | 0 | 4 | 0 | 0 | 0 | 0 | 0 | 0 | 0 | 0 |
| 1 | 0.00 | 3.68 | 0.00 | 1.00 | 1 | Bean1 | brain expressed, associated with Nedd4, | 9 | 0 | 4 | 0 | 0 | 0 | 0 | 0 | 0 | 0 | 0 |
| 1 | 1.11 | 3.60 | 0.31 | 0.76 | 1 | Hnf4aos | | 0 | 0 | 0 | 0 | 0 | 1 | 0 | 13 | 0 | 0 | 0 |
| 1 | 0.00 | 3.68 | 0.00 | 1.00 | 1 | Gm30984 | | 3 | 10 | 0 | 0 | 0 | 0 | 0 | 0 | 0 | 0 | 0 |
| 1 | -2.99 | 3.56 | -0.84 | 0.40 | 1 | Gm16274 | | 0 | 0 | 0 | 6 | 5 | 2 | 0 | 0 | 0 | 0 | 1 |
| 1 | -0.40 | 3.67 | -0.11 | 0.91 | 1 | Fmr1nb | Fmr1 neighbor | 3 | 3 | 6 | 0 | 0 | 1 | 0 | 0 | 0 | 0 | 0 |
| 1 | 0.74 | 3.55 | 0.21 | 0.84 | 1 | Gm19209 | | 0 | 0 | 0 | 0 | 2 | 3 | 0 | 0 | 7 | 0 | 2 |
| 1 | 0.64 | 3.60 | 0.18 | 0.86 | 1 | Gm48805 | | 0 | 0 | 0 | 0 | 0 | 3 | 0 | 0 | 0 | 0 | 10 |
| 1 | 0.00 | 3.60 | 0.00 | 1.00 | 1 | Gm38431 | | 0 | 0 | 0 | 0 | 0 | 8 | 0 | 0 | 0 | 0 | 5 |
| 1 | 1.80 | 3.58 | 0.50 | 0.62 | 1 | Gm43266 | | 0 | 0 | 0 | 0 | 0 | 3 | 0 | 0 | 4 | 0 | 6 |
| 1 | 0.00 | 3.68 | 0.00 | 1.00 | 1 | Tmem132d | transmembrane protein 132D | 9 | 3 | 1 | 0 | 0 | 0 | 0 | 0 | 0 | 0 | 0 |
| 1 | 1.37 | 3.56 | 0.39 | 0.70 | 1 | Lhx3 | LIM homeobox protein 3 | 0 | 0 | 0 | 0 | 0 | 8 | 0 | 0 | 4 | 0 | 1 |
| 1 | -0.50 | 3.55 | -0.14 | 0.89 | 1 | Trpv1 | transient receptor potential cation channe | 0 | 0 | 0 | 0 | 9 | 0 | 0 | 0 | 0 | 0 | 4 |
| 1 | 0.00 | 3.65 | 0.00 | 1.00 | 1 | Gm42533 | | 0 | 0 | 14 | 0 | 0 | 0 | 0 | 0 | 0 | 0 | 0 |
| 1 | 0.00 | 3.65 | 0.00 | 1.00 | 1 | Nanog | Nanog homeobox | 0 | 0 | 14 | 0 | 0 | 0 | 0 | 0 | 0 | 0 | 0 |
| 1 | 0.00 | 3.65 | 0.00 | 1.00 | 1 | n-R5s151 | | 0 | 0 | 14 | 0 | 0 | 0 | 0 | 0 | 0 | 0 | 0 |
| 1 | 0.00 | 3.65 | 0.00 | 1.00 | 1 | Gm15348 | | 0 | 0 | 14 | 0 | 0 | 0 | 0 | 0 | 0 | 0 | 0 |
| 1 | 0.00 | 3.65 | 0.00 | 1.00 | 1 | Gm32352 | | 0 | 0 | 14 | 0 | 0 | 0 | 0 | 0 | 0 | 0 | 0 |
| 1 | 0.00 | 3.65 | 0.00 | 1.00 | 1 | H60b | histocompatibility 60b | 0 | 0 | 14 | 0 | 0 | 0 | 0 | 0 | 0 | 0 | 0 |
| 1 | 0.00 | 3.65 | 0.00 | 1.00 | 1 | Gm48703 | | 0 | 0 | 14 | 0 | 0 | 0 | 0 | 0 | 0 | 0 | 0 |
| 1 | -0.06 | 3.55 | -0.02 | 0.99 | 1 | Gm22935 | | 0 | 0 | 0 | 0 | 8 | 1 | 0 | 0 | 2 | 0 | 3 |
| 1 | 0.00 | 3.68 | 0.00 | 1.00 | 1 | Gm47754 | | 5 | 0 | 8 | 0 | 0 | 0 | 0 | 0 | 0 | 0 | 0 |
| 1 | -0.78 | 3.55 | -0.22 | 0.83 | 1 | Vmn2r-ps19 | | 0 | 0 | 0 | 0 | 10 | 0 | 0 | 0 | 0 | 0 | 3 |
| 1 | 1.60 | 3.57 | 0.45 | 0.65 | 1 | Gm5652 | | 1 | 0 | 2 | 2 | 0 | 0 | 0 | 0 | 0 | 8 | 1 |
| 1 | 0.00 | 3.68 | 0.00 | 1.00 | 1 | Serpinb5 | serine (or cysteine) peptidase inhibitor, cl | 10 | 0 | 3 | 0 | 0 | 0 | 0 | 0 | 0 | 0 | 0 |
| 1 | 0.00 | 3.68 | 0.00 | 1.00 | 1 | Dlx5 | distal-less homeobox 5 | 10 | 0 | 3 | 0 | 0 | 0 | 0 | 0 | 0 | 0 | 0 |
| 1 | 0.51 | 3.60 | 0.14 | 0.89 | 1 | Gm12610 | | 0 | 0 | 0 | 0 | 0 | 4 | 0 | 0 | 0 | 0 | 9 |
| 1 | 0.51 | 3.60 | 0.14 | 0.89 | 1 | 1700003G18Rik | | 0 | 0 | 0 | 0 | 0 | 4 | 0 | 0 | 0 | 0 | 9 |
| 1 | 0.51 | 3.60 | 0.14 | 0.89 | 1 | Gm11753 | | 0 | 0 | 0 | 0 | 0 | 4 | 0 | 0 | 0 | 0 | 9 |
| 1 | -0.12 | 3.60 | -0.03 | 0.97 | 1 | Gm33533 | | 0 | 0 | 0 | 0 | 0 | 9 | 0 | 0 | 0 | 0 | 4 |
| 1 | -3.76 | 3.56 | -1.05 | 0.29 | 1 | Gm6961 | | 0 | 0 | 0 | 5 | 5 | 3 | 0 | 0 | 0 | 0 | 0 |
| 1 | 0.00 | 3.68 | 0.00 | 1.00 | 1 | Il31ra | interleukin 31 receptor A | 5 | 4 | 4 | 0 | 0 | 0 | 0 | 0 | 0 | 0 | 0 |
| 1 | 1.72 | 3.60 | 0.48 | 0.63 | 1 | Gm19744 | | 0 | 0 | 6 | 0 | 0 | 0 | 0 | 0 | 0 | 0 | 7 |
| 1 | 1.59 | 3.57 | 0.44 | 0.66 | 1 | 4930556G22Rik | | 0 | 0 | 0 | 0 | 0 | 4 | 0 | 0 | 6 | 0 | 3 |
| 1 | 0.00 | 3.68 | 0.00 | 1.00 | 1 | Gm11476 | | 10 | 3 | 0 | 0 | 0 | 0 | 0 | 0 | 0 | 0 | 0 |
| 1 | 0.00 | 3.68 | 0.00 | 1.00 | 1 | Ubd | ubiquitin D | 10 | 3 | 0 | 0 | 0 | 0 | 0 | 0 | 0 | 0 | 0 |
| 1 | 0.00 | 3.68 | 0.00 | 1.00 | 1 | Rd3 | retinal degeneration 3 | 1 | 0 | 12 | 0 | 0 | 0 | 0 | 0 | 0 | 0 | 0 |
| 1 | 1.42 | 3.57 | 0.40 | 0.69 | 1 | Rnu2-10 | | 0 | 0 | 0 | 0 | 0 | 2 | 0 | 0 | 0 | 12 | 0 |
| 1 | 0.00 | 3.68 | 0.00 | 1.00 | 1 | Fam25c | family with sequence similarity 25, memb | 6 | 0 | 7 | 0 | 0 | 0 | 0 | 0 | 0 | 0 | 0 |
| 1 | 0.00 | 3.68 | 0.00 | 1.00 | 1 | Gm26717 | | 6 | 0 | 7 | 0 | 0 | 0 | 0 | 0 | 0 | 0 | 0 |
| 1 | -1.15 | 3.66 | -0.31 | 0.75 | 1 | Rpl29-ps5 | | 9 | 1 | 1 | 0 | 1 | 1 | 0 | 0 | 0 | 0 | 0 |
| 1 | 1.53 | 3.57 | 0.43 | 0.67 | 1 | Gm8288 | | 0 | 0 | 0 | 0 | 0 | 2 | 0 | 12 | 0 | 0 | 0 |
| 1 | -0.67 | 3.60 | -0.19 | 0.85 | 1 | Gm43698 | | 0 | 0 | 0 | 12 | 0 | 0 | 0 | 0 | 0 | 0 | 1 |
| 1 | 0.99 | 3.63 | 0.27 | 0.79 | 1 | Rps12l1 | | 10 | 0 | 0 | 0 | 1 | 0 | 0 | 0 | 1 | 0 | 1 |
| 1 | -2.38 | 3.52 | -0.68 | 0.50 | 1 | Baat | bile acid-Coenzyme A: amino acid N-acyl | 2 | 0 | 0 | 4 | 5 | 0 | 0 | 0 | 0 | 1 | 2 |
| 1 | -1.67 | 3.58 | -0.47 | 0.64 | 1 | Ttn | titin | 0 | 0 | 0 | 7 | 0 | 4 | 0 | 0 | 0 | 0 | 2 |
| 1 | -1.67 | 3.58 | -0.47 | 0.64 | 1 | Hspe1-ps3 | | 0 | 0 | 0 | 7 | 0 | 4 | 0 | 0 | 0 | 0 | 2 |
| 1 | 0.00 | 3.68 | 0.00 | 1.00 | 1 | Bex6 | brain expressed family member 6 | 5 | 8 | 0 | 0 | 0 | 0 | 0 | 0 | 0 | 0 | 0 |
| 1 | 0.28 | 3.64 | 0.08 | 0.94 | 1 | Gm47400 | | 0 | 3 | 8 | 0 | 0 | 1 | 0 | 0 | 0 | 0 | 1 |
| 1 | 1.45 | 3.58 | 0.41 | 0.69 | 1 | Gm22767 | | 0 | 0 | 0 | 0 | 0 | 5 | 0 | 0 | 3 | 0 | 5 |
| 1 | 1.64 | 3.61 | 0.45 | 0.65 | 1 | Gm49058 | | 0 | 8 | 0 | 0 | 0 | 5 | 0 | 0 | 0 | 0 | 5 |
| 1 | -0.25 | 3.60 | -0.07 | 0.95 | 1 | Gm43702 | | 0 | 6 | 0 | 0 | 0 | 5 | 0 | 0 | 0 | 0 | 2 |
| 1 | -0.71 | 3.57 | -0.20 | 0.84 | 1 | Gm9701 | | 3 | 1 | 0 | 1 | 6 | 0 | 0 | 0 | 1 | 0 | 2 |
| 1 | 0.00 | 3.65 | 0.00 | 1.00 | 1 | Fgf5 | fibroblast growth factor 5 | 0 | 13 | 0 | 0 | 0 | 0 | 0 | 0 | 0 | 0 | 0 |
| 1 | 0.00 | 3.65 | 0.00 | 1.00 | 1 | Mrgprx2 | MAS-related GPR, member X2 | 0 | 13 | 0 | 0 | 0 | 0 | 0 | 0 | 0 | 0 | 0 |
| 1 | 0.00 | 3.65 | 0.00 | 1.00 | 1 | Gstt4 | glutathione S-transferase, theta 4 | 0 | 13 | 0 | 0 | 0 | 0 | 0 | 0 | 0 | 0 | 0 |
| 1 | 0.00 | 3.65 | 0.00 | 1.00 | 1 | Gm27544 | | 0 | 13 | 0 | 0 | 0 | 0 | 0 | 0 | 0 | 0 | 0 |
| 1 | 0.00 | 3.65 | 0.00 | 1.00 | 1 | Gm7727 | | 0 | 13 | 0 | 0 | 0 | 0 | 0 | 0 | 0 | 0 | 0 |
| 1 | 0.00 | 3.65 | 0.00 | 1.00 | 1 | 3300005D01Rik | | 0 | 13 | 0 | 0 | 0 | 0 | 0 | 0 | 0 | 0 | 0 |
| 1 | 0.00 | 3.65 | 0.00 | 1.00 | 1 | Gm41545 | | 0 | 13 | 0 | 0 | 0 | 0 | 0 | 0 | 0 | 0 | 0 |

| | | | | | | | | | | | | | | | | | | |
|---|---|---|---|---|---|---|---|---|---|---|---|---|---|---|---|---|---|---|
| 1 | 0.00 | 3.65 | 0.00 | 1.00 | 1 | Olfr111 | olfactory receptor 111 | 0 | 13 | 0 | 0 | 0 | 0 | 0 | 0 | 0 | 0 | 0 |
| 1 | -1.38 | 3.59 | -0.38 | 0.70 | 1 | Plcz1 | phospholipase C, zeta 1 | 0 | 0 | 0 | 0 | 9 | 2 | 0 | 0 | 2 | 0 | 0 |
| 1 | -0.69 | 3.66 | -0.19 | 0.85 | 1 | Sit1 | suppression inducing transmembrane ad | 9 | 2 | 0 | 0 | 0 | 2 | 0 | 0 | 0 | 0 | 0 |
| 1 | -0.21 | 3.56 | -0.06 | 0.95 | 1 | Boll | boule homolog, RNA binding protein | 0 | 0 | 0 | 0 | 3 | 4 | 0 | 0 | 6 | 0 | 0 |
| 1 | -2.26 | 3.59 | -0.63 | 0.53 | 1 | Gm16577 | | 0 | 0 | 0 | 0 | 12 | 1 | 0 | 0 | 0 | 0 | 0 |
| 1 | -1.36 | 3.62 | -0.38 | 0.71 | 1 | Gm45702 | | 0 | 0 | 0 | 13 | 0 | 0 | 0 | 0 | 0 | 0 | 0 |
| 1 | -1.36 | 3.62 | -0.38 | 0.71 | 1 | Mettl7a3 | methyltransferase like 7A3 | 0 | 0 | 0 | 13 | 0 | 0 | 0 | 0 | 0 | 0 | 0 |
| 1 | -1.18 | 3.61 | -0.33 | 0.74 | 1 | Morc1 | microrchidia 1 | 1 | 6 | 0 | 0 | 1 | 4 | 0 | 0 | 0 | 0 | 1 |
| 1 | -0.69 | 3.65 | -0.19 | 0.85 | 1 | Gm16295 | | 0 | 12 | 0 | 0 | 1 | 0 | 0 | 0 | 0 | 0 | 0 |
| 1 | 0.00 | 3.68 | 0.00 | 1.00 | 1 | Fam47e | family with sequence similarity 47, membe | 7 | 0 | 6 | 0 | 0 | 0 | 0 | 0 | 0 | 0 | 0 |
| 1 | 0.00 | 3.68 | 0.00 | 1.00 | 1 | AU022754 | | 6 | 7 | 0 | 0 | 0 | 0 | 0 | 0 | 0 | 0 | 0 |
| 1 | -4.98 | 3.55 | -1.40 | 0.16 | 1 | Ppp1r2-ps4 | | 2 | 0 | 0 | 1 | 5 | 2 | 0 | 0 | 0 | 0 | 4 |
| 1 | 0.00 | 3.68 | 0.00 | 1.00 | 1 | 4930555A03Rik | | 12 | 0 | 1 | 0 | 0 | 0 | 0 | 0 | 0 | 0 | 0 |
| 1 | -0.76 | 3.63 | -0.21 | 0.83 | 1 | Gm37510 | | 3 | 7 | 0 | 0 | 0 | 3 | 0 | 0 | 0 | 0 | 0 |
| 1 | -2.38 | 3.56 | -0.67 | 0.50 | 1 | Gja6 | gap junction protein, alpha 6 | 3 | 0 | 0 | 2 | 1 | 1 | 0 | 0 | 6 | 0 | 0 |
| 1 | 0.28 | 3.64 | 0.08 | 0.94 | 1 | Gm11975 | | 0 | 10 | 1 | 0 | 0 | 1 | 0 | 0 | 0 | 0 | 1 |
| 1 | 1.11 | 3.66 | 0.30 | 0.76 | 1 | Smkr-ps | | 12 | 0 | 0 | 0 | 0 | 0 | 0 | 0 | 0 | 0 | 1 |
| 1 | 0.28 | 3.60 | 0.08 | 0.94 | 1 | Aanat | arylalkylamine N-acetyltransferase | 0 | 0 | 0 | 0 | 0 | 6 | 0 | 0 | 0 | 0 | 7 |
| 1 | 0.00 | 3.68 | 0.00 | 1.00 | 1 | Agtr1b | angiotensin II receptor, type 1b | 12 | 1 | 0 | 0 | 0 | 0 | 0 | 0 | 0 | 0 | 0 |
| 1 | 0.00 | 3.68 | 0.00 | 1.00 | 1 | Krt35 | keratin 35 | 12 | 1 | 0 | 0 | 0 | 0 | 0 | 0 | 0 | 0 | 0 |
| 1 | 1.63 | 3.60 | 0.45 | 0.65 | 1 | Gm34106 | | 0 | 0 | 0 | 0 | 2 | 0 | 0 | 0 | 10 | 0 | 1 |
| 1 | 1.35 | 3.57 | 0.38 | 0.71 | 1 | Gm37006 | | 0 | 0 | 0 | 0 | 0 | 6 | 0 | 0 | 4 | 0 | 3 |
| 1 | -1.75 | 3.58 | -0.49 | 0.62 | 1 | Gm12408 | | 0 | 0 | 0 | 0 | 9 | 3 | 0 | 0 | 0 | 0 | 1 |
| 1 | -0.61 | 3.65 | -0.17 | 0.87 | 1 | Gm16130 | predicted gene 16130 | 0 | 12 | 0 | 0 | 0 | 1 | 0 | 0 | 0 | 0 | 0 |
| 1 | -0.61 | 3.65 | -0.17 | 0.87 | 1 | Gm2379 | | 0 | 12 | 0 | 0 | 0 | 1 | 0 | 0 | 0 | 0 | 0 |
| 1 | 0.28 | 2.41 | 0.12 | 0.91 | 1 | Gm4973 | | 1 | 1 | 2 | 3 | 1 | 1 | 0 | 2 | 0 | 1 | 1 |
| 1 | -0.37 | 3.61 | -0.10 | 0.92 | 1 | Gm16141 | | 9 | 0 | 0 | 0 | 0 | 3 | 0 | 0 | 0 | 0 | 1 |
| 1 | -0.31 | 3.61 | -0.09 | 0.93 | 1 | Gm20481 | | 3 | 2 | 3 | 0 | 0 | 4 | 0 | 0 | 0 | 0 | 1 |
| 1 | 1.00 | 3.61 | 0.28 | 0.78 | 1 | Gm48986 | | 6 | 0 | 0 | 0 | 0 | 1 | 0 | 0 | 0 | 0 | 6 |
| 1 | -0.86 | 2.70 | -0.32 | 0.75 | 1 | Gm5559 | | 1 | 2 | 1 | 2 | 3 | 1 | 0 | 0 | 1 | 3 | 0 |
| 1 | 0.64 | 3.61 | 0.18 | 0.86 | 1 | Cilp2 | cartilage intermediate layer protein 2 | 0 | 3 | 3 | 0 | 0 | 2 | 0 | 0 | 0 | 0 | 5 |
| 1 | -2.91 | 3.59 | -0.81 | 0.42 | 1 | Gm38021 | | 4 | 0 | 0 | 5 | 0 | 3 | 0 | 0 | 0 | 0 | 0 |
| 1 | 0.42 | 3.06 | 0.14 | 0.89 | 1 | Gm24276 | | 3 | 2 | 1 | 1 | 2 | 0 | 0 | 3 | 1 | 0 | 0 |
| 1 | 0.00 | 3.68 | 0.00 | 1.00 | 1 | Gm7667 | | 2 | 10 | 1 | 0 | 0 | 0 | 0 | 0 | 0 | 0 | 0 |
| 1 | 1.84 | 3.62 | 0.51 | 0.61 | 1 | Nr6a1os | | 0 | 0 | 0 | 0 | 0 | 0 | 0 | 0 | 0 | 13 | 0 |
| 1 | 1.84 | 3.62 | 0.51 | 0.61 | 1 | Gm5551 | | 0 | 0 | 0 | 0 | 0 | 0 | 0 | 0 | 0 | 13 | 0 |
| 1 | 0.00 | 3.66 | 0.00 | 1.00 | 1 | Gm37858 | | 13 | 0 | 0 | 0 | 0 | 0 | 0 | 0 | 0 | 0 | 0 |
| 1 | 0.00 | 3.66 | 0.00 | 1.00 | 1 | Hemgn | hemogen | 13 | 0 | 0 | 0 | 0 | 0 | 0 | 0 | 0 | 0 | 0 |
| 1 | 0.00 | 3.66 | 0.00 | 1.00 | 1 | Igkj1 | | 13 | 0 | 0 | 0 | 0 | 0 | 0 | 0 | 0 | 0 | 0 |
| 1 | 0.00 | 3.66 | 0.00 | 1.00 | 1 | Gm43914 | | 13 | 0 | 0 | 0 | 0 | 0 | 0 | 0 | 0 | 0 | 0 |
| 1 | 0.00 | 3.66 | 0.00 | 1.00 | 1 | A630009H07Rik | | 13 | 0 | 0 | 0 | 0 | 0 | 0 | 0 | 0 | 0 | 0 |
| 1 | 0.00 | 3.66 | 0.00 | 1.00 | 1 | Gm45540 | | 13 | 0 | 0 | 0 | 0 | 0 | 0 | 0 | 0 | 0 | 0 |
| 1 | 0.00 | 3.66 | 0.00 | 1.00 | 1 | Cd300ld2 | CD300 molecule like family member D2 | 13 | 0 | 0 | 0 | 0 | 0 | 0 | 0 | 0 | 0 | 0 |
| 1 | 0.00 | 3.66 | 0.00 | 1.00 | 1 | Gm30042 | | 13 | 0 | 0 | 0 | 0 | 0 | 0 | 0 | 0 | 0 | 0 |
| 1 | 0.00 | 3.66 | 0.00 | 1.00 | 1 | Dntt | deoxynucleotidyltransferase, terminal | 13 | 0 | 0 | 0 | 0 | 0 | 0 | 0 | 0 | 0 | 0 |
| 1 | 1.00 | 3.58 | 0.28 | 0.78 | 1 | Gm45575 | | 0 | 0 | 0 | 0 | 2 | 1 | 0 | 0 | 9 | 0 | 1 |
| 1 | 0.16 | 3.60 | 0.05 | 0.96 | 1 | Gm42770 | | 0 | 0 | 0 | 0 | 0 | 7 | 0 | 0 | 0 | 0 | 6 |
| 1 | 0.16 | 3.60 | 0.05 | 0.96 | 1 | Snord91a | | 0 | 0 | 0 | 0 | 0 | 7 | 0 | 0 | 0 | 0 | 6 |
| 1 | 0.16 | 3.60 | 0.05 | 0.96 | 1 | Gm22200 | | 0 | 0 | 0 | 0 | 0 | 7 | 0 | 0 | 0 | 0 | 6 |
| 1 | -0.58 | 3.61 | -0.16 | 0.87 | 1 | Gm44086 | | 0 | 0 | 0 | 0 | 0 | 12 | 0 | 0 | 0 | 0 | 1 |
| 1 | 2.29 | 2.98 | 0.77 | 0.44 | 1 | Rpl15-ps5 | | 3 | 0 | 2 | 0 | 0 | 2 | 0 | 1 | 1 | 1 | 3 |
| 1 | 1.66 | 3.61 | 0.46 | 0.65 | 1 | Gm11687 | | 1 | 2 | 1 | 0 | 7 | 0 | 0 | 0 | 0 | 1 | 1 |
| 1 | 1.03 | 3.56 | 0.29 | 0.77 | 1 | Gm37266 | | 0 | 0 | 0 | 0 | 0 | 3 | 0 | 0 | 0 | 10 | 0 |
| 1 | -1.08 | 3.62 | -0.30 | 0.77 | 1 | Gm47657 | | 10 | 0 | 0 | 0 | 0 | 3 | 0 | 0 | 0 | 0 | 0 |
| 1 | -0.85 | 3.55 | -0.24 | 0.81 | 1 | Chn1os1 | | 0 | 0 | 0 | 6 | 0 | 3 | 0 | 0 | 0 | 1 | 2 |
| 1 | 0.00 | 3.68 | 0.00 | 1.00 | 1 | Gm36787 | | 8 | 5 | 0 | 0 | 0 | 0 | 0 | 0 | 0 | 0 | 0 |
| 1 | 0.00 | 3.68 | 0.00 | 1.00 | 1 | Gm30571 | | 3 | 0 | 9 | 0 | 0 | 0 | 0 | 0 | 0 | 0 | 0 |
| 1 | -0.21 | 3.58 | -0.06 | 0.95 | 1 | Gm48894 | | 0 | 0 | 0 | 3 | 0 | 0 | 0 | 0 | 0 | 0 | 10 |
| 1 | 0.24 | 3.55 | 0.07 | 0.95 | 1 | Gm49704 | | 0 | 0 | 0 | 1 | 4 | 0 | 0 | 0 | 3 | 0 | 5 |
| 1 | 3.29 | 3.57 | 0.92 | 0.36 | 1 | Gm17036 | | 0 | 0 | 0 | 0 | 0 | 1 | 0 | 1 | 3 | 8 | 0 |

| | | | | | | | | | | | | | | | | | | |
|---|---|---|---|---|---|---|---|---|---|---|---|---|---|---|---|---|---|---|
| 1 | 5.52 | 3.54 | 1.56 | 0.12 | 1 | Gm15165 | | 0 | 0 | 0 | 4 | 0 | 0 | 0 | 0 | 4 | 1 | 3 |
| 1 | -0.98 | 3.60 | -0.27 | 0.79 | 1 | A230059L01Rik | | 4 | 0 | 0 | 0 | 0 | 8 | 0 | 0 | 0 | 0 | 0 |
| 1 | -0.44 | 3.68 | -0.12 | 0.90 | 1 | Slc9a2 | solute carrier family 9 (sodium/hydrogen | 3 | 5 | 4 | 0 | 1 | 0 | 0 | 0 | 0 | 0 | 0 |
| 1 | 0.00 | 3.68 | 0.00 | 1.00 | 1 | Selenbp2 | selenium binding protein 2 | 9 | 0 | 4 | 0 | 0 | 0 | 0 | 0 | 0 | 0 | 0 |
| 1 | 0.00 | 3.68 | 0.00 | 1.00 | 1 | Prss23os | | 9 | 0 | 4 | 0 | 0 | 0 | 0 | 0 | 0 | 0 | 0 |
| 1 | 0.28 | 3.61 | 0.08 | 0.94 | 1 | Gm11628 | | 3 | 0 | 0 | 0 | 0 | 4 | 0 | 0 | 0 | 0 | 5 |
| 1 | -0.60 | 3.56 | -0.17 | 0.87 | 1 | Gm13077 | | 0 | 0 | 0 | 9 | 1 | 0 | 0 | 0 | 3 | 0 | 0 |
| 1 | 2.16 | 3.56 | 0.61 | 0.54 | 1 | Gm13252 | | 0 | 0 | 0 | 0 | 0 | 3 | 0 | 1 | 0 | 9 | 0 |
| 1 | 3.04 | 3.58 | 0.85 | 0.40 | 1 | Gm49169 | | 0 | 0 | 0 | 0 | 0 | 1 | 0 | 6 | 6 | 0 | 0 |
| 1 | 0.00 | 3.68 | 0.00 | 1.00 | 1 | Gm13604 | | 3 | 10 | 0 | 0 | 0 | 0 | 0 | 0 | 0 | 0 | 0 |
| 1 | 0.74 | 3.60 | 0.21 | 0.84 | 1 | Gm6238 | | 0 | 0 | 0 | 0 | 0 | 3 | 0 | 0 | 0 | 0 | 10 |
| 1 | -0.02 | 3.60 | -0.01 | 1.00 | 1 | Gm43196 | | 0 | 0 | 7 | 0 | 0 | 3 | 0 | 0 | 0 | 0 | 2 |
| 1 | 0.04 | 3.60 | 0.01 | 0.99 | 1 | Gm43172 | | 0 | 0 | 0 | 0 | 0 | 8 | 0 | 0 | 0 | 0 | 5 |
| 1 | 1.64 | 3.59 | 0.46 | 0.65 | 1 | Gm24144 | | 0 | 0 | 0 | 0 | 0 | 3 | 0 | 0 | 2 | 0 | 8 |
| 1 | -0.82 | 3.61 | -0.23 | 0.82 | 1 | Mir344 | | 0 | 0 | 0 | 0 | 0 | 13 | 0 | 0 | 0 | 0 | 0 |
| 1 | 1.80 | 3.59 | 0.50 | 0.62 | 1 | Glipr1l3 | GLI pathogenesis-related 1 like 3 | 0 | 0 | 0 | 0 | 0 | 3 | 0 | 0 | 3 | 0 | 7 |
| 1 | -1.01 | 3.58 | -0.28 | 0.78 | 1 | Gm29808 | | 1 | 0 | 0 | 11 | 0 | 0 | 0 | 0 | 0 | 1 | 0 |
| 1 | -0.75 | 3.64 | -0.21 | 0.84 | 1 | Gm13498 | | 3 | 7 | 0 | 0 | 1 | 1 | 0 | 1 | 0 | 0 | 0 |
| 1 | 0.00 | 3.68 | 0.00 | 1.00 | 1 | Gm29376 | | 4 | 0 | 8 | 0 | 0 | 0 | 0 | 0 | 0 | 0 | 0 |
| 1 | 0.00 | 3.68 | 0.00 | 1.00 | 1 | AI463229 | | 4 | 0 | 8 | 0 | 0 | 0 | 0 | 0 | 0 | 0 | 0 |
| 1 | -0.44 | 3.55 | -0.12 | 0.90 | 1 | Gm829 | predicted gene 829 | 0 | 0 | 0 | 0 | 9 | 0 | 0 | 0 | 0 | 0 | 4 |
| 1 | 0.00 | 3.68 | 0.00 | 1.00 | 1 | Gm38843 | | 9 | 0 | 3 | 0 | 0 | 0 | 0 | 0 | 0 | 0 | 0 |
| 1 | -0.16 | 3.61 | -0.05 | 0.96 | 1 | Gm45474 | | 0 | 0 | 0 | 0 | 9 | 0 | 0 | 0 | 4 | 0 | 0 |
| 1 | -1.96 | 3.54 | -0.55 | 0.58 | 1 | Gm7902 | | 0 | 0 | 0 | 6 | 0 | 3 | 0 | 0 | 3 | 0 | 0 |
| 1 | -1.34 | 3.56 | -0.38 | 0.71 | 1 | Gm8261 | | 2 | 0 | 0 | 2 | 0 | 7 | 0 | 0 | 0 | 1 | 1 |
| 1 | 0.00 | 3.68 | 0.00 | 1.00 | 1 | Gjc3 | gap junction protein, gamma 3 | 4 | 3 | 5 | 0 | 0 | 0 | 0 | 0 | 0 | 0 | 0 |
| 1 | 0.60 | 3.60 | 0.17 | 0.87 | 1 | Gm43158 | | 0 | 0 | 0 | 0 | 0 | 3 | 0 | 0 | 0 | 0 | 9 |
| 1 | 0.60 | 3.60 | 0.17 | 0.87 | 1 | Mir7651 | | 0 | 0 | 0 | 0 | 0 | 3 | 0 | 0 | 0 | 0 | 9 |
| 1 | 0.60 | 3.60 | 0.17 | 0.87 | 1 | Sstr5 | somatostatin receptor 5 | 0 | 0 | 0 | 0 | 0 | 3 | 0 | 0 | 0 | 0 | 9 |
| 1 | -0.08 | 3.60 | -0.02 | 0.98 | 1 | Gm37745 | | 0 | 0 | 0 | 0 | 0 | 8 | 0 | 0 | 0 | 0 | 4 |
| 1 | -0.08 | 3.60 | -0.02 | 0.98 | 1 | Gm45807 | | 0 | 0 | 0 | 0 | 0 | 8 | 0 | 0 | 0 | 0 | 4 |
| 1 | -0.08 | 3.60 | -0.02 | 0.98 | 1 | Gm50468 | | 0 | 0 | 0 | 0 | 0 | 8 | 0 | 0 | 0 | 0 | 4 |
| 1 | 1.17 | 3.60 | 0.33 | 0.75 | 1 | 1700081N11Rik | | 0 | 2 | 0 | 0 | 0 | 3 | 0 | 0 | 1 | 0 | 6 |
| 1 | -2.80 | 3.58 | -0.78 | 0.43 | 1 | Gm43180 | | 0 | 0 | 0 | 10 | 0 | 1 | 0 | 0 | 0 | 0 | 2 |
| 1 | 1.72 | 3.58 | 0.48 | 0.63 | 1 | Mir434 | | 0 | 0 | 0 | 0 | 0 | 3 | 0 | 0 | 5 | 0 | 4 |
| 1 | -0.11 | 3.57 | -0.03 | 0.97 | 1 | Gm9104 | | 1 | 0 | 3 | 3 | 0 | 2 | 0 | 1 | 0 | 0 | 3 |
| 1 | 0.20 | 3.58 | 0.06 | 0.96 | 1 | Gm47136 | | 0 | 0 | 7 | 3 | 0 | 0 | 0 | 0 | 0 | 3 | 0 |
| 1 | 0.00 | 3.68 | 0.00 | 1.00 | 1 | Gm50130 | | 9 | 3 | 0 | 0 | 0 | 0 | 0 | 0 | 0 | 0 | 0 |
| 1 | 2.64 | 3.60 | 0.73 | 0.46 | 1 | 4930511A02Rik | | 0 | 0 | 0 | 0 | 0 | 0 | 0 | 2 | 0 | 0 | 10 |
| 1 | 0.00 | 3.65 | 0.00 | 1.00 | 1 | Gm13985 | | 0 | 0 | 12 | 0 | 0 | 0 | 0 | 0 | 0 | 0 | 0 |
| 1 | 0.00 | 3.65 | 0.00 | 1.00 | 1 | Gm36548 | | 0 | 0 | 12 | 0 | 0 | 0 | 0 | 0 | 0 | 0 | 0 |
| 1 | 0.00 | 3.65 | 0.00 | 1.00 | 1 | Gm4744 | | 0 | 0 | 12 | 0 | 0 | 0 | 0 | 0 | 0 | 0 | 0 |
| 1 | 0.00 | 3.65 | 0.00 | 1.00 | 1 | Gm8209 | | 0 | 0 | 12 | 0 | 0 | 0 | 0 | 0 | 0 | 0 | 0 |
| 1 | 0.00 | 3.65 | 0.00 | 1.00 | 1 | Gm33586 | | 0 | 0 | 12 | 0 | 0 | 0 | 0 | 0 | 0 | 0 | 0 |
| 1 | 0.00 | 3.65 | 0.00 | 1.00 | 1 | Slc10a2 | solute carrier family 10, member 2 | 0 | 0 | 12 | 0 | 0 | 0 | 0 | 0 | 0 | 0 | 0 |
| 1 | 0.00 | 3.65 | 0.00 | 1.00 | 1 | Gm10825 | | 0 | 0 | 12 | 0 | 0 | 0 | 0 | 0 | 0 | 0 | 0 |
| 1 | 0.00 | 3.65 | 0.00 | 1.00 | 1 | Gm53 | | 0 | 0 | 12 | 0 | 0 | 0 | 0 | 0 | 0 | 0 | 0 |
| 1 | 0.00 | 3.65 | 0.00 | 1.00 | 1 | Cyp2d34 | cytochrome P450, family 2, subfamily d, p | 0 | 0 | 12 | 0 | 0 | 0 | 0 | 0 | 0 | 0 | 0 |
| 1 | 0.00 | 3.65 | 0.00 | 1.00 | 1 | Gm9028 | | 0 | 0 | 12 | 0 | 0 | 0 | 0 | 0 | 0 | 0 | 0 |
| 1 | 0.26 | 3.63 | 0.07 | 0.94 | 1 | Gm2027 | | 3 | 6 | 0 | 0 | 1 | 1 | 0 | 0 | 1 | 0 | 1 |
| 1 | 1.34 | 3.61 | 0.37 | 0.71 | 1 | EU599041 | expressed sequence EU599041 | 0 | 0 | 0 | 1 | 0 | 0 | 0 | 12 | 0 | 0 | 0 |
| 1 | 3.18 | 3.60 | 0.88 | 0.38 | 1 | Gm31138 | | 0 | 0 | 0 | 0 | 0 | 0 | 0 | 10 | 0 | 0 | 2 |
| 1 | 0.26 | 3.56 | 0.07 | 0.94 | 1 | Adprhl1 | ADP-ribosylhydrolase like 1 | 0 | 0 | 0 | 0 | 3 | 3 | 0 | 0 | 6 | 0 | 1 |
| 1 | 1.61 | 3.62 | 0.45 | 0.66 | 1 | Gm12099 | | 1 | 0 | 0 | 0 | 0 | 0 | 0 | 0 | 0 | 12 | 0 |
| 1 | 1.18 | 3.65 | 0.32 | 0.75 | 1 | Rnf222 | ring finger protein 222 | 0 | 0 | 11 | 0 | 0 | 0 | 0 | 0 | 0 | 0 | 1 |
| 1 | 1.18 | 3.65 | 0.32 | 0.75 | 1 | Gm48500 | | 0 | 0 | 11 | 0 | 0 | 0 | 0 | 0 | 0 | 0 | 1 |
| 1 | 0.27 | 3.61 | 0.07 | 0.94 | 1 | Gm44364 | | 7 | 0 | 0 | 0 | 0 | 3 | 0 | 0 | 0 | 0 | 3 |
| 1 | 1.13 | 3.60 | 0.31 | 0.75 | 1 | Gm49646 | | 0 | 0 | 0 | 0 | 0 | 1 | 0 | 0 | 0 | 12 | 0 |
| 1 | 2.66 | 3.61 | 0.74 | 0.46 | 1 | Gm3160 | | 1 | 0 | 6 | 0 | 0 | 0 | 0 | 0 | 2 | 1 | 2 |
| 1 | 1.55 | 3.64 | 0.43 | 0.67 | 1 | Plekhd1os | | 0 | 0 | 10 | 0 | 0 | 0 | 0 | 0 | 0 | 0 | 2 |

| | | | | | | | | | | | | | | | | | | |
|---|---|---|---|---|---|---|---|---|---|---|---|---|---|---|---|---|---|---|
| 1 | 3.18 | 3.57 | 0.89 | 0.37 | 1 | Gm14472 | | 0 | 0 | 0 | 0 | 0 | 1 | 0 | 1 | 8 | 1 | 1 |
| 1 | -3.44 | 3.55 | -0.97 | 0.33 | 1 | Rpl7a-ps3 | | 3 | 0 | 0 | 1 | 4 | 2 | 0 | 3 | 0 | 0 | 0 |
| 1 | 0.00 | 3.68 | 0.00 | 1.00 | 1 | 5033403H07Rik | | 4 | 8 | 0 | 0 | 0 | 0 | 0 | 0 | 0 | 0 | 0 |
| 1 | 2.28 | 3.53 | 0.64 | 0.52 | 1 | Gm8885 | | 0 | 2 | 0 | 1 | 5 | 0 | 0 | 1 | 0 | 3 | 1 |
| 1 | 0.12 | 3.62 | 0.03 | 0.97 | 1 | Gm29797 | predicted gene, 29797 | 2 | 6 | 0 | 0 | 0 | 3 | 0 | 0 | 0 | 0 | 2 |
| 1 | 0.47 | 3.60 | 0.13 | 0.90 | 1 | Gm42671 | | 0 | 0 | 0 | 0 | 0 | 4 | 0 | 0 | 0 | 0 | 8 |
| 1 | 0.47 | 3.60 | 0.13 | 0.90 | 1 | Gm48670 | | 0 | 0 | 0 | 0 | 0 | 4 | 0 | 0 | 0 | 0 | 8 |
| 1 | 0.47 | 3.60 | 0.13 | 0.90 | 1 | Gm10113 | | 0 | 0 | 0 | 0 | 0 | 4 | 0 | 0 | 0 | 0 | 8 |
| 1 | -0.22 | 3.60 | -0.06 | 0.95 | 1 | Gm42890 | | 0 | 0 | 0 | 0 | 0 | 9 | 0 | 0 | 0 | 0 | 3 |
| 1 | -0.22 | 3.60 | -0.06 | 0.95 | 1 | Gm45084 | | 0 | 0 | 0 | 0 | 0 | 9 | 0 | 0 | 0 | 0 | 3 |
| 1 | -0.22 | 3.60 | -0.06 | 0.95 | 1 | Gm45205 | | 0 | 0 | 0 | 0 | 0 | 9 | 0 | 0 | 0 | 0 | 3 |
| 1 | -0.22 | 3.60 | -0.06 | 0.95 | 1 | Gm24366 | | 0 | 0 | 0 | 0 | 0 | 9 | 0 | 0 | 0 | 0 | 3 |
| 1 | -0.22 | 3.60 | -0.06 | 0.95 | 1 | Gm48978 | | 0 | 0 | 0 | 0 | 0 | 9 | 0 | 0 | 0 | 0 | 3 |
| 1 | 1.36 | 3.59 | 0.38 | 0.71 | 1 | Gm46339 | | 0 | 0 | 0 | 0 | 0 | 4 | 0 | 0 | 2 | 0 | 6 |
| 1 | -1.81 | 3.63 | -0.50 | 0.62 | 1 | Gm8425 | | 1 | 3 | 4 | 2 | 0 | 3 | 0 | 0 | 0 | 0 | 0 |
| 1 | -0.36 | 2.71 | -0.13 | 0.89 | 1 | Gm12728 | | 1 | 3 | 1 | 1 | 1 | 3 | 0 | 0 | 2 | 0 | 1 |
| 1 | 1.71 | 3.60 | 0.47 | 0.64 | 1 | Zfp683 | | 0 | 0 | 6 | 0 | 0 | 0 | 0 | 0 | 0 | 0 | 6 |
| 1 | 0.64 | 3.63 | 0.18 | 0.86 | 1 | Gm14173 | | 4 | 0 | 4 | 0 | 0 | 1 | 0 | 0 | 3 | 0 | 0 |
| 1 | 0.00 | 3.68 | 0.00 | 1.00 | 1 | Ighv5-6 | | 10 | 2 | 0 | 0 | 0 | 0 | 0 | 0 | 0 | 0 | 0 |
| 1 | -1.11 | 3.60 | -0.31 | 0.76 | 1 | Gm44541 | | 0 | 0 | 0 | 0 | 9 | 1 | 0 | 0 | 2 | 0 | 0 |
| 1 | 0.00 | 3.68 | 0.00 | 1.00 | 1 | Gm10603 | | 0 | 6 | 6 | 0 | 0 | 0 | 0 | 0 | 0 | 0 | 0 |
| 1 | -2.85 | 3.58 | -0.80 | 0.43 | 1 | Gm15410 | | 3 | 0 | 0 | 0 | 6 | 0 | 0 | 0 | 0 | 0 | 4 |
| 1 | 1.68 | 3.60 | 0.47 | 0.64 | 1 | Gm5449 | | 0 | 0 | 5 | 0 | 0 | 0 | 0 | 0 | 0 | 0 | 7 |
| 1 | -0.53 | 3.65 | -0.14 | 0.88 | 1 | Ces1b | carboxylesterase 1B | 0 | 0 | 11 | 0 | 0 | 1 | 0 | 0 | 0 | 0 | 0 |
| 1 | 0.93 | 3.53 | 0.26 | 0.79 | 1 | Gm4349 | | 0 | 0 | 0 | 4 | 0 | 1 | 0 | 0 | 0 | 4 | 4 |
| 1 | -1.43 | 3.58 | -0.40 | 0.69 | 1 | Gm26583 | | 3 | 0 | 0 | 2 | 0 | 6 | 0 | 0 | 0 | 0 | 2 |
| 1 | 1.02 | 3.67 | 0.28 | 0.78 | 1 | Gm14267 | | 1 | 0 | 10 | 0 | 0 | 0 | 0 | 0 | 0 | 0 | 1 |
| 1 | 0.00 | 3.68 | 0.00 | 1.00 | 1 | Gm12631 | | 0 | 8 | 4 | 0 | 0 | 0 | 0 | 0 | 0 | 0 | 0 |
| 1 | -0.18 | 3.61 | -0.05 | 0.96 | 1 | Gm37953 | | 5 | 0 | 0 | 0 | 0 | 5 | 0 | 0 | 0 | 0 | 2 |
| 1 | 0.00 | 3.68 | 0.00 | 1.00 | 1 | Nr0b1 | nuclear receptor subfamily 0, group B, me | 5 | 7 | 0 | 0 | 0 | 0 | 0 | 0 | 0 | 0 | 0 |
| 1 | 0.00 | 3.68 | 0.00 | 1.00 | 1 | 4930519A11Rik | | 0 | 9 | 3 | 0 | 0 | 0 | 0 | 0 | 0 | 0 | 0 |
| 1 | 0.00 | 3.68 | 0.00 | 1.00 | 1 | Pla2g10 | phospholipase A2, group X | 0 | 9 | 3 | 0 | 0 | 0 | 0 | 0 | 0 | 0 | 0 |
| 1 | -2.27 | 3.60 | -0.63 | 0.53 | 1 | Nudt12os | | 0 | 0 | 0 | 11 | 0 | 1 | 0 | 0 | 0 | 0 | 0 |
| 1 | -1.03 | 3.60 | -0.29 | 0.78 | 1 | Tnn | tenascin N | 0 | 0 | 0 | 1 | 0 | 9 | 0 | 0 | 0 | 0 | 2 |
| 1 | -0.73 | 3.64 | -0.20 | 0.84 | 1 | 1700023B13Rik | | 3 | 0 | 6 | 0 | 0 | 3 | 0 | 0 | 0 | 0 | 0 |
| 1 | 0.00 | 3.68 | 0.00 | 1.00 | 1 | Ighv1-56 | | 11 | 0 | 1 | 0 | 0 | 0 | 0 | 0 | 0 | 0 | 0 |
| 1 | 0.97 | 3.60 | 0.27 | 0.79 | 1 | Gm37674 | | 0 | 0 | 7 | 0 | 0 | 1 | 0 | 0 | 0 | 0 | 4 |
| 1 | 0.97 | 3.60 | 0.27 | 0.79 | 1 | Gm42133 | | 0 | 0 | 7 | 0 | 0 | 1 | 0 | 0 | 0 | 0 | 4 |
| 1 | -2.90 | 3.59 | -0.81 | 0.42 | 1 | Mybphl | myosin binding protein H-like | 0 | 0 | 0 | 10 | 0 | 3 | 0 | 0 | 0 | 0 | 0 |
| 1 | -2.90 | 3.59 | -0.81 | 0.42 | 1 | Gm26652 | | 0 | 0 | 0 | 10 | 0 | 3 | 0 | 0 | 0 | 0 | 0 |
| 1 | 1.20 | 3.61 | 0.33 | 0.74 | 1 | Gm18283 | | 0 | 4 | 0 | 0 | 0 | 0 | 0 | 8 | 0 | 0 | 0 |
| 1 | 1.10 | 3.65 | 0.30 | 0.76 | 1 | Gm47061 | | 0 | 11 | 0 | 0 | 0 | 0 | 0 | 0 | 0 | 0 | 1 |
| 1 | -0.24 | 3.62 | -0.07 | 0.95 | 1 | Gm9130 | | 1 | 0 | 7 | 0 | 2 | 1 | 0 | 0 | 0 | 1 | 0 |
| 1 | 0.00 | 3.66 | 0.00 | 1.00 | 1 | Gm29453 | | 0 | 12 | 0 | 0 | 0 | 0 | 0 | 0 | 0 | 0 | 0 |
| 1 | 0.00 | 3.66 | 0.00 | 1.00 | 1 | Mindy4b-ps | | 0 | 12 | 0 | 0 | 0 | 0 | 0 | 0 | 0 | 0 | 0 |
| 1 | 0.00 | 3.66 | 0.00 | 1.00 | 1 | Gm11902 | | 0 | 12 | 0 | 0 | 0 | 0 | 0 | 0 | 0 | 0 | 0 |
| 1 | 0.00 | 3.66 | 0.00 | 1.00 | 1 | Gm42543 | | 0 | 12 | 0 | 0 | 0 | 0 | 0 | 0 | 0 | 0 | 0 |
| 1 | 0.00 | 3.66 | 0.00 | 1.00 | 1 | Gm38708 | | 0 | 12 | 0 | 0 | 0 | 0 | 0 | 0 | 0 | 0 | 0 |
| 1 | 0.00 | 3.66 | 0.00 | 1.00 | 1 | Peak1os | | 0 | 12 | 0 | 0 | 0 | 0 | 0 | 0 | 0 | 0 | 0 |
| 1 | 0.00 | 3.66 | 0.00 | 1.00 | 1 | Prss44 | protease, serine 44 | 0 | 12 | 0 | 0 | 0 | 0 | 0 | 0 | 0 | 0 | 0 |
| 1 | 0.00 | 3.66 | 0.00 | 1.00 | 1 | Treml1 | triggering receptor expressed on myeloid | 0 | 12 | 0 | 0 | 0 | 0 | 0 | 0 | 0 | 0 | 0 |
| 1 | -1.95 | 3.59 | -0.54 | 0.59 | 1 | 4932702P03Rik | | 0 | 0 | 0 | 4 | 0 | 7 | 0 | 0 | 0 | 0 | 1 |
| 1 | 1.17 | 3.65 | 0.32 | 0.75 | 1 | Gm4258 | | 0 | 11 | 0 | 0 | 0 | 0 | 0 | 0 | 1 | 0 | 0 |
| 1 | 1.17 | 3.65 | 0.32 | 0.75 | 1 | Gm20147 | | 0 | 11 | 0 | 0 | 0 | 0 | 0 | 0 | 1 | 0 | 0 |
| 1 | 1.91 | 3.62 | 0.53 | 0.60 | 1 | Arhgap27os1 | | 0 | 0 | 0 | 0 | 0 | 0 | 0 | 0 | 12 | 0 | 0 |
| 1 | -2.97 | 3.58 | -0.83 | 0.41 | 1 | 4930567H12Rik | | 0 | 0 | 0 | 9 | 0 | 3 | 0 | 0 | 0 | 0 | 0 |
| 1 | 1.54 | 3.61 | 0.43 | 0.67 | 1 | Mir3091 | | 0 | 0 | 0 | 0 | 0 | 2 | 0 | 0 | 0 | 1 | 9 |
| 1 | 0.30 | 3.61 | 0.08 | 0.93 | 1 | Gm37613 | | 0 | 0 | 5 | 0 | 0 | 1 | 0 | 0 | 6 | 0 | 0 |
| 1 | 1.56 | 3.61 | 0.43 | 0.67 | 1 | Gm16048 | | 6 | 0 | 0 | 0 | 0 | 0 | 0 | 0 | 0 | 0 | 6 |
| 1 | -0.67 | 3.66 | -0.18 | 0.86 | 1 | Hjv | hemojuvelin BMP co-receptor | 11 | 0 | 0 | 0 | 1 | 0 | 0 | 0 | 0 | 0 | 0 |

| | | | | | | | | | | | | | | | | | | |
|---|---|---|---|---|---|---|---|---|---|---|---|---|---|---|---|---|---|---|
| 1 | -1.27 | 3.57 | -0.36 | 0.72 | 1 | Snord89 | | 0 | 0 | 0 | 0 | 6 | 3 | 0 | 0 | 0 | 0 | 3 |
| 1 | -0.19 | 3.56 | -0.05 | 0.96 | 1 | Gm42800 | | 0 | 0 | 4 | 0 | 0 | 6 | 0 | 0 | 2 | 0 | 0 |
| 1 | -0.40 | 3.67 | -0.11 | 0.91 | 1 | Klrb1f | killer cell lectin-like receptor subfamily B r | 1 | 0 | 10 | 0 | 0 | 1 | 0 | 0 | 0 | 0 | 0 |
| 1 | -0.92 | 3.64 | -0.25 | 0.80 | 1 | Gm45346 | | 0 | 0 | 10 | 0 | 0 | 2 | 0 | 0 | 0 | 0 | 0 |
| 1 | 1.94 | 3.61 | 0.54 | 0.59 | 1 | Gm8375 | | 0 | 0 | 0 | 0 | 1 | 0 | 0 | 0 | 10 | 0 | 1 |
| 1 | -0.68 | 3.61 | -0.19 | 0.85 | 1 | Tmem262 | transmembrane protein 262 | 0 | 5 | 0 | 7 | 0 | 0 | 0 | 0 | 0 | 0 | 0 |
| 1 | 0.55 | 3.61 | 0.15 | 0.88 | 1 | Gm128 | predicted gene 128 | 3 | 0 | 0 | 0 | 0 | 3 | 0 | 0 | 0 | 0 | 6 |
| 1 | -1.94 | 3.62 | -0.53 | 0.59 | 1 | Gm49521 | | 9 | 0 | 0 | 0 | 1 | 3 | 0 | 0 | 0 | 0 | 0 |
| 1 | 0.08 | 3.56 | 0.02 | 0.98 | 1 | Tmem91 | transmembrane protein 91 | 0 | 0 | 0 | 7 | 0 | 0 | 0 | 0 | 5 | 0 | 0 |
| 1 | 0.00 | 3.68 | 0.00 | 1.00 | 1 | Gm26685 | | 7 | 0 | 5 | 0 | 0 | 0 | 0 | 0 | 0 | 0 | 0 |
| 1 | 2.57 | 3.61 | 0.71 | 0.48 | 1 | Gm17794 | | 8 | 0 | 0 | 0 | 0 | 0 | 0 | 0 | 2 | 1 | 1 |
| 1 | 1.40 | 3.56 | 0.39 | 0.69 | 1 | Rpl21-ps7 | | 2 | 2 | 3 | 0 | 0 | 1 | 0 | 0 | 0 | 1 | 3 |
| 1 | 0.72 | 3.61 | 0.20 | 0.84 | 1 | Gm15921 | predicted gene 15921 | 3 | 2 | 0 | 0 | 0 | 3 | 0 | 1 | 0 | 0 | 2 |
| 1 | 1.02 | 3.54 | 0.29 | 0.77 | 1 | Gm9299 | | 1 | 0 | 0 | 2 | 1 | 0 | 0 | 0 | 1 | 6 | 1 |
| 1 | -0.05 | 3.63 | -0.01 | 0.99 | 1 | Gm11534 | | 0 | 11 | 0 | 1 | 0 | 0 | 0 | 0 | 0 | 0 | 0 |
| 1 | 1.09 | 3.61 | 0.30 | 0.76 | 1 | Gm5066 | | 0 | 0 | 0 | 0 | 0 | 1 | 0 | 0 | 0 | 0 | 11 |
| 1 | -2.79 | 3.60 | -0.77 | 0.44 | 1 | Fsd2 | fibronectin type III and SPRY domain cont | 3 | 2 | 0 | 0 | 3 | 2 | 0 | 0 | 0 | 0 | 2 |
| 1 | 1.51 | 3.15 | 0.48 | 0.63 | 1 | Gm16061 | | 2 | 0 | 3 | 1 | 1 | 0 | 0 | 2 | 1 | 0 | 2 |
| 1 | 1.77 | 3.53 | 0.50 | 0.62 | 1 | Gm7614 | | 1 | 1 | 3 | 0 | 1 | 0 | 0 | 0 | 4 | 0 | 2 |
| 1 | 0.00 | 3.66 | 0.00 | 1.00 | 1 | Padi3 | peptidyl arginine deiminase, type III | 12 | 0 | 0 | 0 | 0 | 0 | 0 | 0 | 0 | 0 | 0 |
| 1 | 0.00 | 3.66 | 0.00 | 1.00 | 1 | Ereg | epiregulin | 12 | 0 | 0 | 0 | 0 | 0 | 0 | 0 | 0 | 0 | 0 |
| 1 | 0.00 | 3.66 | 0.00 | 1.00 | 1 | A630023P12Rik | | 12 | 0 | 0 | 0 | 0 | 0 | 0 | 0 | 0 | 0 | 0 |
| 1 | 0.00 | 3.66 | 0.00 | 1.00 | 1 | Igkv4-61 | | 12 | 0 | 0 | 0 | 0 | 0 | 0 | 0 | 0 | 0 | 0 |
| 1 | 0.00 | 3.66 | 0.00 | 1.00 | 1 | Gm15353 | | 12 | 0 | 0 | 0 | 0 | 0 | 0 | 0 | 0 | 0 | 0 |
| 1 | 0.00 | 3.66 | 0.00 | 1.00 | 1 | Gm8093 | | 12 | 0 | 0 | 0 | 0 | 0 | 0 | 0 | 0 | 0 | 0 |
| 1 | 0.00 | 3.66 | 0.00 | 1.00 | 1 | Gm8873 | | 12 | 0 | 0 | 0 | 0 | 0 | 0 | 0 | 0 | 0 | 0 |
| 1 | 0.00 | 3.66 | 0.00 | 1.00 | 1 | Gm36660 | | 12 | 0 | 0 | 0 | 0 | 0 | 0 | 0 | 0 | 0 | 0 |
| 1 | 0.00 | 3.66 | 0.00 | 1.00 | 1 | Grp | gastrin releasing peptide | 12 | 0 | 0 | 0 | 0 | 0 | 0 | 0 | 0 | 0 | 0 |
| 1 | 0.00 | 3.66 | 0.00 | 1.00 | 1 | Irs4 | insulin receptor substrate 4 | 12 | 0 | 0 | 0 | 0 | 0 | 0 | 0 | 0 | 0 | 0 |
| 1 | 0.22 | 3.60 | 0.06 | 0.95 | 1 | Gm12737 | | 0 | 0 | 0 | 0 | 0 | 6 | 0 | 0 | 0 | 0 | 6 |
| 1 | -0.82 | 3.61 | -0.23 | 0.82 | 1 | Gm13511 | | 0 | 1 | 0 | 0 | 0 | 11 | 0 | 0 | 0 | 0 | 0 |
| 1 | 1.26 | 3.58 | 0.35 | 0.73 | 1 | Gm37165 | | 0 | 0 | 0 | 0 | 0 | 6 | 0 | 0 | 3 | 0 | 3 |
| 1 | 1.34 | 3.57 | 0.38 | 0.71 | 1 | Muc3a | mucin 3A, cell surface associated | 0 | 0 | 0 | 0 | 0 | 2 | 0 | 0 | 0 | 10 | 0 |
| 1 | 0.24 | 3.62 | 0.07 | 0.95 | 1 | Gm14139 | predicted gene 14139 | 0 | 10 | 0 | 0 | 0 | 1 | 0 | 0 | 0 | 0 | 1 |
| 1 | -0.61 | 3.65 | -0.17 | 0.87 | 1 | Gm15443 | | 0 | 11 | 0 | 0 | 0 | 1 | 0 | 0 | 0 | 0 | 0 |
| 1 | -0.95 | 3.57 | -0.27 | 0.79 | 1 | Stmnd1 | stathmin domain containing 1 | 0 | 0 | 0 | 0 | 5 | 4 | 0 | 0 | 3 | 0 | 0 |
| 1 | 0.24 | 3.65 | 0.07 | 0.95 | 1 | 1700018A23Rik | | 0 | 10 | 0 | 0 | 0 | 1 | 0 | 0 | 1 | 0 | 0 |
| 1 | -2.14 | 3.61 | -0.59 | 0.55 | 1 | Gm9595 | | 0 | 1 | 2 | 7 | 2 | 0 | 0 | 0 | 0 | 0 | 0 |
| 1 | 3.18 | 3.62 | 0.88 | 0.38 | 1 | Esr2 | estrogen receptor 2 (beta) | 0 | 6 | 0 | 0 | 0 | 1 | 0 | 0 | 5 | 0 | 0 |
| 1 | 0.46 | 3.55 | 0.13 | 0.90 | 1 | Gm21998 | | 0 | 0 | 0 | 0 | 0 | 4 | 0 | 0 | 0 | 8 | 0 |
| 1 | -1.07 | 3.61 | -0.30 | 0.77 | 1 | Cyb5r2 | cytochrome b5 reductase 2 | 8 | 0 | 0 | 0 | 0 | 4 | 0 | 0 | 0 | 0 | 0 |
| 1 | 0.00 | 3.68 | 0.00 | 1.00 | 1 | Gm45512 | | 3 | 0 | 9 | 0 | 0 | 0 | 0 | 0 | 0 | 0 | 0 |
| 1 | 0.00 | 3.68 | 0.00 | 1.00 | 1 | Tm4sf19 | transmembrane 4 L six family member 19 | 3 | 0 | 9 | 0 | 0 | 0 | 0 | 0 | 0 | 0 | 0 |
| 1 | -0.19 | 3.58 | -0.05 | 0.96 | 1 | Gm45363 | | 3 | 0 | 0 | 0 | 2 | 3 | 0 | 0 | 1 | 0 | 3 |
| 1 | -0.10 | 3.61 | -0.03 | 0.98 | 1 | 1810009A15Ril | RIKEN cDNA 1810009A15 gene | 0 | 0 | 2 | 0 | 0 | 7 | 0 | 0 | 0 | 0 | 3 |
| 1 | 0.00 | 3.68 | 0.00 | 1.00 | 1 | Gm49990 | | 8 | 0 | 4 | 0 | 0 | 0 | 0 | 0 | 0 | 0 | 0 |
| 1 | -0.20 | 3.61 | -0.06 | 0.96 | 1 | Gm27019 | | 0 | 0 | 8 | 0 | 0 | 3 | 0 | 0 | 0 | 0 | 1 |
| 1 | 0.00 | 3.61 | 0.00 | 1.00 | 1 | Snora69 | | 2 | 2 | 0 | 0 | 0 | 5 | 0 | 0 | 0 | 0 | 3 |
| 1 | 0.09 | 3.60 | 0.02 | 0.98 | 1 | AU022133 | | 0 | 0 | 0 | 0 | 0 | 7 | 0 | 0 | 0 | 0 | 5 |
| 1 | 0.09 | 3.60 | 0.02 | 0.98 | 1 | A930030B08Rik | | 0 | 0 | 0 | 0 | 0 | 7 | 0 | 0 | 0 | 0 | 5 |
| 1 | 0.09 | 3.60 | 0.02 | 0.98 | 1 | Gm18042 | | 0 | 0 | 0 | 0 | 0 | 7 | 0 | 0 | 0 | 0 | 5 |
| 1 | -0.82 | 3.61 | -0.23 | 0.82 | 1 | Gm44524 | | 0 | 0 | 0 | 0 | 0 | 12 | 0 | 0 | 0 | 0 | 0 |
| 1 | -0.82 | 3.61 | -0.23 | 0.82 | 1 | Gm37589 | | 0 | 0 | 0 | 0 | 0 | 12 | 0 | 0 | 0 | 0 | 0 |
| 1 | -0.26 | 3.61 | -0.07 | 0.94 | 1 | Gm19500 | | 0 | 3 | 0 | 0 | 0 | 7 | 0 | 0 | 0 | 0 | 2 |
| 1 | 1.01 | 3.59 | 0.28 | 0.78 | 1 | Gm43658 | | 0 | 0 | 0 | 0 | 0 | 7 | 0 | 0 | 2 | 0 | 3 |
| 1 | 0.03 | 3.56 | 0.01 | 0.99 | 1 | E530001F21Rik | | 0 | 0 | 4 | 0 | 2 | 2 | 0 | 0 | 0 | 0 | 4 |
| 1 | -0.55 | 3.56 | -0.16 | 0.88 | 1 | A230028O05Rik | | 0 | 0 | 0 | 9 | 0 | 0 | 0 | 0 | 0 | 0 | 3 |
| 1 | -0.22 | 3.56 | -0.06 | 0.95 | 1 | Gm48035 | | 0 | 0 | 0 | 9 | 0 | 0 | 0 | 0 | 3 | 0 | 0 |
| 1 | -1.19 | 3.62 | -0.33 | 0.74 | 1 | Gm7392 | | 1 | 7 | 0 | 1 | 1 | 1 | 0 | 1 | 0 | 0 | 0 |
| 1 | 0.17 | 3.55 | 0.05 | 0.96 | 1 | 4930598N05Rik | | 0 | 0 | 0 | 0 | 6 | 0 | 0 | 0 | 0 | 0 | 6 |

| | | | | | | | | | | | | | | | | | | |
|---|---|---|---|---|---|---|---|---|---|---|---|---|---|---|---|---|---|---|
| 1 | 0.00 | 3.68 | 0.00 | 1.00 | 1 | Clec4g | C-type lectin domain family 4, member g | 3 | 0 | 8 | 0 | 0 | 0 | 0 | 0 | 0 | 0 | 0 |
| 1 | -5.95 | 3.55 | -1.67 | 0.09 | 1 | Hrk | harakiri, BCL2 interacting protein (contair | 3 | 0 | 0 | 4 | 4 | 0 | 0 | 0 | 0 | 0 | 1 |
| 1 | -0.32 | 3.62 | -0.09 | 0.93 | 1 | Gm45114 | | 0 | 9 | 0 | 3 | 0 | 0 | 0 | 0 | 0 | 0 | 0 |
| 1 | -0.28 | 3.61 | -0.08 | 0.94 | 1 | Gm5754 | | 0 | 0 | 2 | 0 | 0 | 8 | 0 | 0 | 0 | 0 | 2 |
| 1 | -0.66 | 3.58 | -0.19 | 0.85 | 1 | Gm4606 | predicted gene 4606 | 5 | 0 | 0 | 0 | 3 | 2 | 0 | 0 | 1 | 0 | 1 |
| 1 | 2.60 | 3.61 | 0.72 | 0.47 | 1 | Gm34091 | | 3 | 0 | 0 | 0 | 0 | 0 | 0 | 1 | 0 | 0 | 7 |
| 1 | 0.00 | 3.68 | 0.00 | 1.00 | 1 | Gm15133 | | 9 | 0 | 3 | 0 | 0 | 0 | 0 | 0 | 0 | 0 | 0 |
| 1 | 0.00 | 3.68 | 0.00 | 1.00 | 1 | Igkv8-19 | | 3 | 9 | 0 | 0 | 0 | 0 | 0 | 0 | 0 | 0 | 0 |
| 1 | -0.11 | 3.58 | -0.03 | 0.98 | 1 | Gm43767 | | 0 | 7 | 0 | 1 | 0 | 2 | 0 | 0 | 0 | 0 | 2 |
| 1 | -0.04 | 3.60 | -0.01 | 0.99 | 1 | Gm44253 | | 0 | 0 | 0 | 0 | 0 | 8 | 0 | 0 | 0 | 0 | 4 |
| 1 | -0.04 | 3.60 | -0.01 | 0.99 | 1 | Gm45041 | | 0 | 0 | 0 | 0 | 0 | 8 | 0 | 0 | 0 | 0 | 4 |
| 1 | -0.04 | 3.60 | -0.01 | 0.99 | 1 | Gm45893 | | 0 | 0 | 0 | 0 | 0 | 8 | 0 | 0 | 0 | 0 | 4 |
| 1 | -0.04 | 3.60 | -0.01 | 0.99 | 1 | 2610001A08Rik | | 0 | 0 | 0 | 0 | 0 | 8 | 0 | 0 | 0 | 0 | 4 |
| 1 | -3.55 | 3.56 | -0.99 | 0.32 | 1 | Gm48890 | | 0 | 0 | 0 | 4 | 5 | 3 | 0 | 0 | 0 | 0 | 0 |
| 1 | 0.41 | 3.56 | 0.12 | 0.91 | 1 | 2600002D14Rik | | 0 | 3 | 0 | 4 | 0 | 0 | 0 | 0 | 0 | 0 | 5 |
| 1 | 0.77 | 3.61 | 0.21 | 0.83 | 1 | Gm13487 | | 8 | 0 | 0 | 0 | 0 | 1 | 0 | 0 | 0 | 0 | 3 |
| 1 | -3.27 | 3.61 | -0.90 | 0.37 | 1 | B430219N15Rik | | 3 | 0 | 1 | 6 | 0 | 1 | 0 | 0 | 0 | 0 | 1 |
| 1 | 1.83 | 3.62 | 0.51 | 0.61 | 1 | Gm29783 | | 0 | 0 | 0 | 0 | 0 | 0 | 0 | 0 | 0 | 12 | 0 |
| 1 | 1.83 | 3.62 | 0.51 | 0.61 | 1 | C130012C08Rik | | 0 | 0 | 0 | 0 | 0 | 0 | 0 | 0 | 0 | 12 | 0 |
| 1 | 1.81 | 3.62 | 0.50 | 0.62 | 1 | 4933430L12Rik | | 0 | 0 | 0 | 0 | 0 | 0 | 0 | 12 | 0 | 0 | 0 |
| 1 | 1.81 | 3.62 | 0.50 | 0.62 | 1 | Gm11714 | | 0 | 0 | 0 | 0 | 0 | 0 | 0 | 12 | 0 | 0 | 0 |
| 1 | 1.49 | 3.57 | 0.42 | 0.68 | 1 | Gm44996 | | 3 | 0 | 0 | 0 | 2 | 0 | 0 | 0 | 3 | 0 | 4 |
| 1 | 1.08 | 3.55 | 0.30 | 0.76 | 1 | Gm8330 | | 1 | 1 | 0 | 0 | 4 | 0 | 0 | 2 | 1 | 3 | 0 |
| 1 | 0.55 | 3.61 | 0.15 | 0.88 | 1 | Gm23201 | | 0 | 0 | 0 | 0 | 0 | 3 | 0 | 0 | 0 | 0 | 8 |
| 1 | 0.55 | 3.61 | 0.15 | 0.88 | 1 | Gm13238 | | 0 | 0 | 0 | 0 | 0 | 3 | 0 | 0 | 0 | 0 | 8 |
| 1 | -1.05 | 3.61 | -0.29 | 0.77 | 1 | Gm24091 | | 0 | 0 | 7 | 0 | 0 | 4 | 0 | 0 | 0 | 0 | 0 |
| 1 | -0.18 | 3.60 | -0.05 | 0.96 | 1 | Lrch4 | leucine-rich repeats and calponin homolc | 0 | 0 | 0 | 0 | 0 | 8 | 0 | 0 | 0 | 0 | 3 |
| 1 | 0.81 | 3.60 | 0.23 | 0.82 | 1 | Gm45817 | | 0 | 5 | 0 | 0 | 0 | 3 | 0 | 0 | 1 | 0 | 2 |
| 1 | -0.40 | 3.61 | -0.11 | 0.91 | 1 | Clcn1 | chloride channel, voltage-sensitive 1 | 0 | 0 | 6 | 0 | 0 | 4 | 0 | 0 | 0 | 0 | 1 |
| 1 | 1.47 | 3.64 | 0.40 | 0.69 | 1 | Mab21l2 | mab-21-like 2 | 9 | 0 | 0 | 0 | 0 | 0 | 0 | 0 | 0 | 0 | 2 |
| 1 | 1.47 | 3.64 | 0.40 | 0.69 | 1 | Gm43152 | | 9 | 0 | 0 | 0 | 0 | 0 | 0 | 0 | 0 | 0 | 2 |
| 1 | 0.00 | 3.68 | 0.00 | 1.00 | 1 | Duoxa2 | dual oxidase maturation factor 2 | 9 | 2 | 0 | 0 | 0 | 0 | 0 | 0 | 0 | 0 | 0 |
| 1 | 0.00 | 3.68 | 0.00 | 1.00 | 1 | Bcl2a1b | B cell leukemia/lymphoma 2 related prote | 9 | 2 | 0 | 0 | 0 | 0 | 0 | 0 | 0 | 0 | 0 |
| 1 | 0.00 | 3.68 | 0.00 | 1.00 | 1 | Gm47567 | | 9 | 2 | 0 | 0 | 0 | 0 | 0 | 0 | 0 | 0 | 0 |
| 1 | -2.08 | 3.57 | -0.58 | 0.56 | 1 | Npffr2 | neuropeptide FF receptor 2 | 0 | 0 | 0 | 9 | 0 | 2 | 0 | 0 | 1 | 0 | 0 |
| 1 | -1.63 | 3.64 | -0.45 | 0.65 | 1 | Gm5121 | | 2 | 3 | 3 | 0 | 2 | 2 | 0 | 0 | 0 | 0 | 0 |
| 1 | 0.67 | 3.56 | 0.19 | 0.85 | 1 | Tubb2a-ps2 | | 1 | 0 | 0 | 4 | 0 | 0 | 0 | 0 | 7 | 0 | 0 |
| 1 | 0.00 | 3.66 | 0.00 | 1.00 | 1 | Entpd8 | ectonucleoside triphosphate diphosphohy | 0 | 0 | 11 | 0 | 0 | 0 | 0 | 0 | 0 | 0 | 0 |
| 1 | 0.00 | 3.66 | 0.00 | 1.00 | 1 | Sh2d6 | SH2 domain containing 6 | 0 | 0 | 11 | 0 | 0 | 0 | 0 | 0 | 0 | 0 | 0 |
| 1 | 0.00 | 3.66 | 0.00 | 1.00 | 1 | Capn12 | calpain 12 | 0 | 0 | 11 | 0 | 0 | 0 | 0 | 0 | 0 | 0 | 0 |
| 1 | 0.00 | 3.66 | 0.00 | 1.00 | 1 | Gm2996 | | 0 | 0 | 11 | 0 | 0 | 0 | 0 | 0 | 0 | 0 | 0 |
| 1 | 0.00 | 3.66 | 0.00 | 1.00 | 1 | Gm15503 | | 0 | 0 | 11 | 0 | 0 | 0 | 0 | 0 | 0 | 0 | 0 |
| 1 | 0.00 | 3.66 | 0.00 | 1.00 | 1 | Gm45643 | | 0 | 0 | 11 | 0 | 0 | 0 | 0 | 0 | 0 | 0 | 0 |
| 1 | 0.00 | 3.66 | 0.00 | 1.00 | 1 | Calb2 | calbindin 2 | 0 | 0 | 11 | 0 | 0 | 0 | 0 | 0 | 0 | 0 | 0 |
| 1 | 0.00 | 3.66 | 0.00 | 1.00 | 1 | Mmp8 | matrix metallopeptidase 8 | 0 | 0 | 11 | 0 | 0 | 0 | 0 | 0 | 0 | 0 | 0 |
| 1 | 0.00 | 3.66 | 0.00 | 1.00 | 1 | Gm39323 | | 0 | 0 | 11 | 0 | 0 | 0 | 0 | 0 | 0 | 0 | 0 |
| 1 | 0.00 | 3.66 | 0.00 | 1.00 | 1 | Gm18032 | | 0 | 0 | 11 | 0 | 0 | 0 | 0 | 0 | 0 | 0 | 0 |
| 1 | 0.00 | 3.66 | 0.00 | 1.00 | 1 | Slc9c1 | solute carrier family 9, subfamily C (Na+-t | 0 | 0 | 11 | 0 | 0 | 0 | 0 | 0 | 0 | 0 | 0 |
| 1 | -1.78 | 3.59 | -0.50 | 0.62 | 1 | Gm20644 | | 0 | 0 | 0 | 8 | 0 | 3 | 0 | 0 | 0 | 0 | 1 |
| 1 | 1.18 | 3.65 | 0.32 | 0.75 | 1 | Gm16096 | | 0 | 0 | 10 | 0 | 0 | 0 | 0 | 0 | 0 | 0 | 1 |
| 1 | 0.62 | 3.63 | 0.17 | 0.86 | 1 | Gm17890 | | 0 | 0 | 10 | 0 | 0 | 0 | 0 | 0 | 1 | 0 | 0 |
| 1 | 2.16 | 3.60 | 0.60 | 0.55 | 1 | 5430420F09Rik | | 5 | 0 | 0 | 0 | 0 | 4 | 0 | 0 | 2 | 0 | 0 |
| 1 | -0.49 | 3.59 | -0.14 | 0.89 | 1 | Gm44571 | | 0 | 0 | 0 | 2 | 0 | 3 | 0 | 0 | 0 | 0 | 7 |
| 1 | 0.00 | 3.68 | 0.00 | 1.00 | 1 | Gm32282 | | 5 | 0 | 6 | 0 | 0 | 0 | 0 | 0 | 0 | 0 | 0 |
| 1 | 0.00 | 3.68 | 0.00 | 1.00 | 1 | Tmem236 | transmembrane protein 236 | 0 | 2 | 9 | 0 | 0 | 0 | 0 | 0 | 0 | 0 | 0 |
| 1 | 1.99 | 3.60 | 0.55 | 0.58 | 1 | Gm11542 | | 1 | 0 | 4 | 0 | 0 | 1 | 0 | 3 | 0 | 0 | 2 |
| 1 | -2.24 | 3.60 | -0.62 | 0.53 | 1 | Gm37017 | | 0 | 0 | 0 | 11 | 0 | 1 | 0 | 0 | 0 | 0 | 0 |
| 1 | 0.41 | 3.60 | 0.11 | 0.91 | 1 | Gm9353 | | 0 | 0 | 0 | 0 | 0 | 4 | 0 | 0 | 0 | 0 | 7 |
| 1 | 0.41 | 3.60 | 0.11 | 0.91 | 1 | Gm47687 | | 0 | 0 | 0 | 0 | 0 | 4 | 0 | 0 | 0 | 0 | 7 |
| 1 | 0.41 | 3.60 | 0.11 | 0.91 | 1 | Krt74 | | 0 | 0 | 0 | 0 | 0 | 4 | 0 | 0 | 0 | 0 | 7 |

| | | | | | | | | | | | | | | | | | | |
|---|---|---|---|---|---|---|---|---|---|---|---|---|---|---|---|---|---|---|
| 1 | 1.40 | 3.57 | 0.39 | 0.69 | 1 | Gm45779 | | 0 | 0 | 0 | 0 | 0 | 4 | 0 | 0 | 5 | 0 | 2 |
| 1 | -1.40 | 3.57 | -0.39 | 0.69 | 1 | Gm16618 | | 0 | 0 | 0 | 4 | 0 | 1 | 0 | 0 | 0 | 0 | 6 |
| 1 | 0.00 | 3.68 | 0.00 | 1.00 | 1 | Ighv10-3 | | 10 | 1 | 0 | 0 | 0 | 0 | 0 | 0 | 0 | 0 | 0 |
| 1 | -2.13 | 3.58 | -0.60 | 0.55 | 1 | Gm9509 | | 1 | 0 | 3 | 4 | 0 | 3 | 0 | 0 | 0 | 1 | 0 |
| 1 | 0.00 | 3.68 | 0.00 | 1.00 | 1 | Reg3a | regenerating islet-derived 3 alpha | 0 | 5 | 6 | 0 | 0 | 0 | 0 | 0 | 0 | 0 | 0 |
| 1 | 0.00 | 3.68 | 0.00 | 1.00 | 1 | Lncbate6 | | 0 | 5 | 6 | 0 | 0 | 0 | 0 | 0 | 0 | 0 | 0 |
| 1 | 0.48 | 3.61 | 0.13 | 0.89 | 1 | Gm7083 | | 3 | 1 | 0 | 4 | 0 | 0 | 0 | 2 | 0 | 0 | 1 |
| 1 | 0.00 | 3.68 | 0.00 | 1.00 | 1 | Mir423 | | 1 | 0 | 10 | 0 | 0 | 0 | 0 | 0 | 0 | 0 | 0 |
| 1 | 0.00 | 3.68 | 0.00 | 1.00 | 1 | Gm30938 | | 1 | 0 | 10 | 0 | 0 | 0 | 0 | 0 | 0 | 0 | 0 |
| 1 | 0.00 | 3.68 | 0.00 | 1.00 | 1 | Nmur1 | neuromedin U receptor 1 | 0 | 7 | 4 | 0 | 0 | 0 | 0 | 0 | 0 | 0 | 0 |
| 1 | 0.00 | 3.68 | 0.00 | 1.00 | 1 | Cd70 | CD70 antigen | 0 | 7 | 4 | 0 | 0 | 0 | 0 | 0 | 0 | 0 | 0 |
| 1 | 1.99 | 3.60 | 0.55 | 0.58 | 1 | Gm5054 | | 1 | 0 | 5 | 0 | 0 | 1 | 0 | 2 | 1 | 0 | 1 |
| 1 | -0.53 | 3.65 | -0.14 | 0.89 | 1 | Cfap100 | cilia and flagella associated protein 100 | 0 | 0 | 10 | 0 | 0 | 1 | 0 | 0 | 0 | 0 | 0 |
| 1 | 1.25 | 3.66 | 0.34 | 0.73 | 1 | Dydc1 | DPY30 domain containing 1 | 0 | 6 | 3 | 0 | 0 | 0 | 0 | 0 | 0 | 0 | 2 |
| 1 | 0.00 | 3.68 | 0.00 | 1.00 | 1 | Ccr4 | chemokine (C-C motif) receptor 4 | 5 | 6 | 0 | 0 | 0 | 0 | 0 | 0 | 0 | 0 | 0 |
| 1 | 0.00 | 3.68 | 0.00 | 1.00 | 1 | Pitx1 | paired-like homeodomain transcription fa | 5 | 6 | 0 | 0 | 0 | 0 | 0 | 0 | 0 | 0 | 0 |
| 1 | 2.93 | 3.60 | 0.81 | 0.42 | 1 | Gapdhs | glyceraldehyde-3-phosphate dehydroger | 5 | 0 | 0 | 0 | 0 | 0 | 0 | 0 | 4 | 0 | 2 |
| 1 | -2.90 | 3.59 | -0.81 | 0.42 | 1 | Gm7364 | | 0 | 0 | 0 | 6 | 0 | 5 | 0 | 0 | 0 | 0 | 0 |
| 1 | -0.77 | 3.59 | -0.22 | 0.83 | 1 | Gm22027 | | 0 | 0 | 0 | 0 | 2 | 4 | 0 | 0 | 0 | 0 | 5 |
| 1 | -2.05 | 3.62 | -0.57 | 0.57 | 1 | A730013G03Rik | | 7 | 0 | 0 | 0 | 1 | 3 | 0 | 0 | 0 | 0 | 0 |
| 1 | -0.40 | 3.67 | -0.11 | 0.91 | 1 | 4933415A04Rik | | 5 | 0 | 5 | 0 | 0 | 1 | 0 | 0 | 0 | 0 | 0 |
| 1 | -2.06 | 3.60 | -0.57 | 0.57 | 1 | A430110C17Rik | | 0 | 0 | 0 | 0 | 2 | 9 | 0 | 0 | 0 | 0 | 0 |
| 1 | 1.30 | 3.64 | 0.36 | 0.72 | 1 | Gm12536 | | 0 | 7 | 1 | 0 | 0 | 0 | 0 | 0 | 0 | 0 | 3 |
| 1 | 1.38 | 3.61 | 0.38 | 0.70 | 1 | Gm43414 | | 0 | 0 | 0 | 0 | 0 | 0 | 0 | 0 | 0 | 0 | 11 |
| 1 | 1.38 | 3.61 | 0.38 | 0.70 | 1 | Gm43340 | | 0 | 0 | 0 | 0 | 0 | 0 | 0 | 0 | 0 | 0 | 11 |
| 1 | 0.28 | 3.60 | 0.08 | 0.94 | 1 | Gm37522 | | 0 | 0 | 0 | 0 | 0 | 5 | 0 | 0 | 0 | 0 | 6 |
| 1 | 0.28 | 3.60 | 0.08 | 0.94 | 1 | Gm12850 | | 0 | 0 | 0 | 0 | 0 | 5 | 0 | 0 | 0 | 0 | 6 |
| 1 | 0.28 | 3.60 | 0.08 | 0.94 | 1 | Gm22362 | | 0 | 0 | 0 | 0 | 0 | 5 | 0 | 0 | 0 | 0 | 6 |
| 1 | 0.28 | 3.60 | 0.08 | 0.94 | 1 | Gm27186 | | 0 | 0 | 0 | 0 | 0 | 5 | 0 | 0 | 0 | 0 | 6 |
| 1 | 0.28 | 3.60 | 0.08 | 0.94 | 1 | Gm11274 | | 0 | 0 | 0 | 0 | 0 | 5 | 0 | 0 | 0 | 0 | 6 |
| 1 | 0.28 | 3.60 | 0.08 | 0.94 | 1 | Gm27271 | | 0 | 0 | 0 | 0 | 0 | 5 | 0 | 0 | 0 | 0 | 6 |
| 1 | -1.46 | 3.58 | -0.41 | 0.68 | 1 | 1700113H08Ril | RIKEN cDNA 1700113H08 gene | 2 | 0 | 0 | 2 | 6 | 0 | 0 | 0 | 2 | 0 | 0 |
| 1 | 0.12 | 3.61 | 0.03 | 0.97 | 1 | Gm45120 | | 0 | 2 | 0 | 0 | 0 | 5 | 0 | 0 | 0 | 0 | 4 |
| 1 | -0.55 | 3.61 | -0.15 | 0.88 | 1 | Fbxw15 | F-box and WD-40 domain protein 15 | 0 | 0 | 0 | 0 | 0 | 10 | 0 | 0 | 0 | 0 | 1 |
| 1 | 1.17 | 3.59 | 0.32 | 0.75 | 1 | Gm45124 | | 0 | 0 | 0 | 0 | 0 | 5 | 0 | 0 | 2 | 0 | 4 |
| 1 | 0.00 | 3.66 | 0.00 | 1.00 | 1 | Gm2619 | | 11 | 0 | 0 | 0 | 0 | 0 | 0 | 0 | 0 | 0 | 0 |
| 1 | 0.00 | 3.66 | 0.00 | 1.00 | 1 | Fmo3 | flavin containing monooxygenase 3 | 11 | 0 | 0 | 0 | 0 | 0 | 0 | 0 | 0 | 0 | 0 |
| 1 | 0.00 | 3.66 | 0.00 | 1.00 | 1 | Sp5 | trans-acting transcription factor 5 | 11 | 0 | 0 | 0 | 0 | 0 | 0 | 0 | 0 | 0 | 0 |
| 1 | 0.00 | 3.66 | 0.00 | 1.00 | 1 | Gm27715 | | 11 | 0 | 0 | 0 | 0 | 0 | 0 | 0 | 0 | 0 | 0 |
| 1 | 0.00 | 3.66 | 0.00 | 1.00 | 1 | Tmem54 | transmembrane protein 54 | 11 | 0 | 0 | 0 | 0 | 0 | 0 | 0 | 0 | 0 | 0 |
| 1 | 0.00 | 3.66 | 0.00 | 1.00 | 1 | Gm5309 | | 11 | 0 | 0 | 0 | 0 | 0 | 0 | 0 | 0 | 0 | 0 |
| 1 | 0.00 | 3.66 | 0.00 | 1.00 | 1 | Igkv6-15 | | 11 | 0 | 0 | 0 | 0 | 0 | 0 | 0 | 0 | 0 | 0 |
| 1 | 0.00 | 3.66 | 0.00 | 1.00 | 1 | Clec4a4 | C-type lectin domain family 4, member a4 | 11 | 0 | 0 | 0 | 0 | 0 | 0 | 0 | 0 | 0 | 0 |
| 1 | 0.00 | 3.66 | 0.00 | 1.00 | 1 | Smco3 | single-pass membrane protein with coilec | 11 | 0 | 0 | 0 | 0 | 0 | 0 | 0 | 0 | 0 | 0 |
| 1 | 0.00 | 3.66 | 0.00 | 1.00 | 1 | Gm44861 | | 11 | 0 | 0 | 0 | 0 | 0 | 0 | 0 | 0 | 0 | 0 |
| 1 | 0.00 | 3.66 | 0.00 | 1.00 | 1 | Gm39090 | | 11 | 0 | 0 | 0 | 0 | 0 | 0 | 0 | 0 | 0 | 0 |
| 1 | 0.00 | 3.66 | 0.00 | 1.00 | 1 | Cd209e | CD209e antigen | 11 | 0 | 0 | 0 | 0 | 0 | 0 | 0 | 0 | 0 | 0 |
| 1 | 0.00 | 3.66 | 0.00 | 1.00 | 1 | Gm35572 | | 11 | 0 | 0 | 0 | 0 | 0 | 0 | 0 | 0 | 0 | 0 |
| 1 | 0.00 | 3.66 | 0.00 | 1.00 | 1 | Ighv15-2 | | 11 | 0 | 0 | 0 | 0 | 0 | 0 | 0 | 0 | 0 | 0 |
| 1 | 0.00 | 3.66 | 0.00 | 1.00 | 1 | Ctsg | cathepsin G | 11 | 0 | 0 | 0 | 0 | 0 | 0 | 0 | 0 | 0 | 0 |
| 1 | 0.00 | 3.66 | 0.00 | 1.00 | 1 | Gm5970 | | 11 | 0 | 0 | 0 | 0 | 0 | 0 | 0 | 0 | 0 | 0 |
| 1 | 0.00 | 3.66 | 0.00 | 1.00 | 1 | Gm43581 | | 0 | 11 | 0 | 0 | 0 | 0 | 0 | 0 | 0 | 0 | 0 |
| 1 | 0.00 | 3.66 | 0.00 | 1.00 | 1 | Gm43142 | | 0 | 11 | 0 | 0 | 0 | 0 | 0 | 0 | 0 | 0 | 0 |
| 1 | 0.00 | 3.66 | 0.00 | 1.00 | 1 | Gm42456 | | 0 | 11 | 0 | 0 | 0 | 0 | 0 | 0 | 0 | 0 | 0 |
| 1 | 0.00 | 3.66 | 0.00 | 1.00 | 1 | Sec14l3 | SEC14-like lipid binding 3 | 0 | 11 | 0 | 0 | 0 | 0 | 0 | 0 | 0 | 0 | 0 |
| 1 | 0.00 | 3.66 | 0.00 | 1.00 | 1 | CJ186046Rik | | 0 | 11 | 0 | 0 | 0 | 0 | 0 | 0 | 0 | 0 | 0 |
| 1 | 0.19 | 3.55 | 0.05 | 0.96 | 1 | 2310020H05Rik | | 0 | 0 | 0 | 0 | 0 | 5 | 0 | 0 | 6 | 0 | 0 |
| 1 | -2.68 | 3.59 | -0.75 | 0.46 | 1 | Gm33474 | | 0 | 0 | 0 | 0 | 9 | 2 | 0 | 0 | 0 | 0 | 0 |
| 1 | -2.80 | 3.59 | -0.78 | 0.44 | 1 | F7 | coagulation factor VII | 0 | 0 | 0 | 4 | 0 | 7 | 0 | 0 | 0 | 0 | 0 |
| 1 | 0.00 | 3.68 | 0.00 | 1.00 | 1 | Gm2399 | | 6 | 3 | 2 | 0 | 0 | 0 | 0 | 0 | 0 | 0 | 0 |

| | | | | | | | | | | | | | | | | | | |
|---|---|---|---|---|---|---|---|---|---|---|---|---|---|---|---|---|---|---|
| 1 | 2.00 | 3.09 | 0.65 | 0.52 | 1 | Gm4948 | | 1 | 1 | 0 | 1 | 0 | 1 | 0 | 3 | 2 | 0 | 2 |
| 1 | 0.62 | 3.56 | 0.17 | 0.86 | 1 | Gm37829 | | 0 | 0 | 0 | 0 | 0 | 3 | 0 | 0 | 0 | 8 | 0 |
| 1 | -1.05 | 3.62 | -0.29 | 0.77 | 1 | Gm45216 | | 8 | 0 | 0 | 0 | 0 | 3 | 0 | 0 | 0 | 0 | 0 |
| 1 | -0.98 | 3.62 | -0.27 | 0.79 | 1 | Gm9431 | | 2 | 4 | 0 | 3 | 0 | 2 | 0 | 0 | 1 | 0 | 0 |
| 1 | 1.48 | 3.59 | 0.41 | 0.68 | 1 | Aox2 | aldehyde oxidase 2 | 0 | 0 | 4 | 0 | 0 | 1 | 0 | 0 | 1 | 0 | 5 |
| 1 | -1.04 | 3.61 | -0.29 | 0.77 | 1 | Gm23722 | | 7 | 0 | 0 | 0 | 0 | 4 | 0 | 0 | 0 | 0 | 0 |
| 1 | 0.83 | 3.60 | 0.23 | 0.82 | 1 | Gm7701 | | 1 | 0 | 6 | 0 | 0 | 2 | 0 | 0 | 1 | 1 | 0 |
| 1 | -0.26 | 3.24 | -0.08 | 0.94 | 1 | Gm7413 | | 2 | 1 | 1 | 0 | 2 | 2 | 0 | 1 | 0 | 3 | 0 |
| 1 | -1.03 | 3.61 | -0.28 | 0.78 | 1 | Gm15764 | | 6 | 0 | 0 | 0 | 0 | 5 | 0 | 0 | 0 | 0 | 0 |
| 1 | 2.75 | 3.61 | 0.76 | 0.45 | 1 | Gm13785 | | 6 | 0 | 0 | 0 | 0 | 0 | 0 | 0 | 2 | 0 | 3 |
| 1 | 0.95 | 3.67 | 0.26 | 0.80 | 1 | Gm6756 | | 6 | 4 | 0 | 0 | 0 | 0 | 0 | 0 | 0 | 0 | 1 |
| 1 | 0.00 | 3.68 | 0.00 | 1.00 | 1 | Ctxn3 | cortexin 3 | 6 | 5 | 0 | 0 | 0 | 0 | 0 | 0 | 0 | 0 | 0 |
| 1 | 0.00 | 3.68 | 0.00 | 1.00 | 1 | Gm5127 | predicted gene 5127 | 6 | 5 | 0 | 0 | 0 | 0 | 0 | 0 | 0 | 0 | 0 |
| 1 | 0.00 | 3.68 | 0.00 | 1.00 | 1 | Abcg5 | ATP binding cassette subfamily G membe | 2 | 0 | 9 | 0 | 0 | 0 | 0 | 0 | 0 | 0 | 0 |
| 1 | -1.00 | 3.60 | -0.28 | 0.78 | 1 | A230087F16Rik | | 5 | 0 | 0 | 0 | 0 | 6 | 0 | 0 | 0 | 0 | 0 |
| 1 | -0.40 | 3.67 | -0.11 | 0.91 | 1 | Gm14393 | predicted gene 14393 | 1 | 0 | 9 | 0 | 0 | 1 | 0 | 0 | 0 | 0 | 0 |
| 1 | -0.75 | 3.59 | -0.21 | 0.84 | 1 | Gm41836 | | 0 | 0 | 0 | 2 | 0 | 4 | 0 | 0 | 0 | 0 | 5 |
| 1 | -1.25 | 3.58 | -0.35 | 0.73 | 1 | Naaladl1 | N-acetylated alpha-linked acidic dipeptid | 0 | 0 | 0 | 0 | 4 | 4 | 0 | 0 | 0 | 0 | 3 |
| 1 | -0.91 | 3.64 | -0.25 | 0.80 | 1 | Mir207 | | 0 | 0 | 9 | 0 | 0 | 2 | 0 | 0 | 0 | 0 | 0 |
| 1 | -0.91 | 3.64 | -0.25 | 0.80 | 1 | Gm45016 | | 0 | 0 | 9 | 0 | 0 | 2 | 0 | 0 | 0 | 0 | 0 |
| 1 | 4.48 | 3.57 | 1.26 | 0.21 | 1 | Gm22875 | | 0 | 0 | 0 | 0 | 4 | 0 | 0 | 1 | 0 | 0 | 6 |
| 1 | 0.00 | 3.68 | 0.00 | 1.00 | 1 | Gm43324 | | 7 | 0 | 4 | 0 | 0 | 0 | 0 | 0 | 0 | 0 | 0 |
| 1 | 0.00 | 3.68 | 0.00 | 1.00 | 1 | Gm45866 | | 7 | 0 | 4 | 0 | 0 | 0 | 0 | 0 | 0 | 0 | 0 |
| 1 | 0.00 | 3.68 | 0.00 | 1.00 | 1 | Gm45494 | | 7 | 0 | 4 | 0 | 0 | 0 | 0 | 0 | 0 | 0 | 0 |
| 1 | 1.06 | 3.61 | 0.29 | 0.77 | 1 | Gm25134 | | 0 | 0 | 0 | 0 | 0 | 1 | 0 | 0 | 0 | 0 | 10 |
| 1 | 1.73 | 3.60 | 0.48 | 0.63 | 1 | Lhfpl1 | lipoma HMGIC fusion partner-like 1 | 0 | 0 | 0 | 0 | 0 | 1 | 0 | 0 | 1 | 0 | 9 |
| 1 | 0.14 | 3.60 | 0.04 | 0.97 | 1 | A730090N16Rik | | 0 | 0 | 0 | 0 | 0 | 6 | 0 | 0 | 0 | 0 | 5 |
| 1 | 0.14 | 3.60 | 0.04 | 0.97 | 1 | 2310026L22Rik | | 0 | 0 | 0 | 0 | 0 | 6 | 0 | 0 | 0 | 0 | 5 |
| 1 | 0.14 | 3.60 | 0.04 | 0.97 | 1 | Aurkc | aurora kinase C | 0 | 0 | 0 | 0 | 0 | 6 | 0 | 0 | 0 | 0 | 5 |
| 1 | 0.14 | 3.60 | 0.04 | 0.97 | 1 | Gm17981 | | 0 | 0 | 0 | 0 | 0 | 6 | 0 | 0 | 0 | 0 | 5 |
| 1 | -3.51 | 3.61 | -0.97 | 0.33 | 1 | Gm17149 | | 3 | 0 | 0 | 1 | 6 | 0 | 0 | 0 | 1 | 0 | 0 |
| 1 | -0.82 | 3.61 | -0.23 | 0.82 | 1 | Arl4aos | | 0 | 0 | 0 | 0 | 0 | 11 | 0 | 0 | 0 | 0 | 0 |
| 1 | 2.18 | 3.59 | 0.61 | 0.54 | 1 | Gm8237 | | 0 | 0 | 0 | 0 | 0 | 1 | 0 | 0 | 3 | 0 | 7 |
| 1 | 0.00 | 3.68 | 0.00 | 1.00 | 1 | 4930595D18Rik | | 1 | 10 | 0 | 0 | 0 | 0 | 0 | 0 | 0 | 0 | 0 |
| 1 | 0.77 | 3.61 | 0.21 | 0.83 | 1 | Gm11722 | | 0 | 7 | 0 | 0 | 0 | 1 | 0 | 0 | 0 | 0 | 3 |
| 1 | 1.09 | 3.57 | 0.31 | 0.76 | 1 | Gm44967 | | 0 | 0 | 0 | 0 | 0 | 6 | 0 | 0 | 4 | 0 | 1 |
| 1 | 2.35 | 3.60 | 0.65 | 0.51 | 1 | Slc26a5 | solute carrier family 26, member 5 | 0 | 0 | 0 | 0 | 0 | 1 | 0 | 0 | 6 | 0 | 4 |
| 1 | 2.14 | 3.59 | 0.60 | 0.55 | 1 | Gm13380 | | 0 | 0 | 0 | 0 | 0 | 1 | 0 | 0 | 8 | 0 | 2 |
| 1 | -0.47 | 3.62 | -0.13 | 0.90 | 1 | B230110C06Rik | | 1 | 0 | 7 | 0 | 3 | 0 | 0 | 0 | 0 | 0 | 0 |
| 1 | -1.16 | 3.55 | -0.33 | 0.74 | 1 | Gm9396 | | 0 | 3 | 0 | 1 | 3 | 0 | 0 | 1 | 0 | 0 | 3 |
| 1 | -2.32 | 3.56 | -0.65 | 0.51 | 1 | Gstm3 | glutathione S-transferase, mu 3 | 0 | 0 | 0 | 4 | 3 | 2 | 0 | 0 | 0 | 0 | 2 |
| 1 | -0.40 | 3.67 | -0.11 | 0.91 | 1 | Gm6201 | | 6 | 4 | 0 | 0 | 0 | 1 | 0 | 0 | 0 | 0 | 0 |
| 1 | -0.05 | 3.58 | -0.02 | 0.99 | 1 | Gm13184 | | 0 | 0 | 6 | 0 | 1 | 2 | 0 | 0 | 0 | 0 | 2 |
| 1 | 0.95 | 3.67 | 0.26 | 0.80 | 1 | Gm7357 | | 2 | 6 | 2 | 0 | 0 | 0 | 0 | 0 | 0 | 0 | 1 |
| 1 | 1.55 | 3.66 | 0.42 | 0.67 | 1 | Gm45667 | | 2 | 4 | 2 | 0 | 0 | 0 | 0 | 0 | 3 | 0 | 0 |
| 1 | 0.96 | 3.57 | 0.27 | 0.79 | 1 | Gm15522 | | 0 | 0 | 0 | 0 | 4 | 0 | 0 | 0 | 3 | 0 | 4 |
| 1 | 1.13 | 3.61 | 0.31 | 0.75 | 1 | Gm14807 | | 0 | 1 | 0 | 0 | 0 | 1 | 0 | 0 | 0 | 9 | 0 |
| 1 | -0.22 | 3.65 | -0.06 | 0.95 | 1 | Gm30262 | | 1 | 9 | 0 | 1 | 0 | 0 | 0 | 0 | 0 | 0 | 0 |
| 1 | 1.67 | 3.53 | 0.47 | 0.64 | 1 | Gm21738 | | 0 | 0 | 0 | 3 | 0 | 1 | 0 | 0 | 3 | 1 | 3 |
| 1 | 0.00 | 3.68 | 0.00 | 1.00 | 1 | Gm35161 | | 8 | 0 | 3 | 0 | 0 | 0 | 0 | 0 | 0 | 0 | 0 |
| 1 | 0.00 | 3.68 | 0.00 | 1.00 | 1 | Armc12 | armadillo repeat containing 12 | 2 | 9 | 0 | 0 | 0 | 0 | 0 | 0 | 0 | 0 | 0 |
| 1 | 0.80 | 3.61 | 0.22 | 0.82 | 1 | Gm38304 | | 0 | 1 | 0 | 0 | 0 | 2 | 0 | 0 | 0 | 0 | 8 |
| 1 | 0.00 | 3.61 | 0.00 | 1.00 | 1 | 9430031K09Rik | | 0 | 0 | 0 | 0 | 0 | 7 | 0 | 0 | 0 | 0 | 4 |
| 1 | 0.64 | 3.61 | 0.18 | 0.86 | 1 | 4930502E09Rik | | 0 | 4 | 0 | 0 | 0 | 2 | 0 | 0 | 0 | 0 | 5 |
| 1 | 2.01 | 3.59 | 0.56 | 0.58 | 1 | Gm42791 | | 0 | 0 | 0 | 0 | 0 | 2 | 0 | 0 | 5 | 0 | 4 |
| 1 | -0.48 | 3.65 | -0.13 | 0.89 | 1 | E130201H02Rik | | 1 | 6 | 1 | 1 | 1 | 0 | 0 | 0 | 0 | 0 | 1 |
| 1 | 2.25 | 3.62 | 0.62 | 0.53 | 1 | Gm23421 | | 0 | 7 | 0 | 0 | 0 | 2 | 0 | 0 | 2 | 0 | 0 |
| 1 | 2.63 | 3.61 | 0.73 | 0.46 | 1 | Gm15567 | | 0 | 6 | 0 | 0 | 0 | 2 | 0 | 0 | 3 | 0 | 0 |
| 1 | 2.63 | 3.61 | 0.73 | 0.46 | 1 | Gm11437 | predicted gene 11437 | 0 | 6 | 0 | 0 | 0 | 2 | 0 | 0 | 3 | 0 | 0 |
| 1 | -2.36 | 3.59 | -0.66 | 0.51 | 1 | Gm37503 | | 0 | 1 | 0 | 0 | 3 | 1 | 0 | 0 | 0 | 0 | 6 |

| | | | | | | | | | | | | | | | | | | |
|---|---|---|---|---|---|---|---|---|---|---|---|---|---|---|---|---|---|---|
| 1 | -1.01 | 3.58 | -0.28 | 0.78 | 1 | Gm12961 | | 0 | 1 | 0 | 8 | 0 | 1 | 0 | 0 | 0 | 0 | 1 |
| 1 | -0.15 | 3.23 | -0.05 | 0.96 | 1 | Gm44768 | | 1 | 3 | 1 | 0 | 1 | 3 | 0 | 0 | 0 | 1 | 1 |
| 1 | 2.55 | 3.59 | 0.71 | 0.48 | 1 | Chrna10 | cholinergic receptor, nicotinic, alpha poly | 3 | 0 | 0 | 0 | 0 | 4 | 0 | 0 | 3 | 0 | 0 |
| 1 | -2.06 | 3.63 | -0.57 | 0.57 | 1 | Gm10268 | | 0 | 6 | 0 | 0 | 2 | 2 | 0 | 0 | 1 | 0 | 0 |
| 1 | -0.76 | 3.64 | -0.21 | 0.83 | 1 | Gm26586 | | 1 | 0 | 7 | 0 | 0 | 3 | 0 | 0 | 0 | 0 | 0 |
| 1 | -0.40 | 3.67 | -0.11 | 0.91 | 1 | Cyp2c70 | cytochrome P450, family 2, subfamily c, p | 3 | 1 | 6 | 0 | 0 | 1 | 0 | 0 | 0 | 0 | 0 |
| 1 | -0.10 | 3.61 | -0.03 | 0.98 | 1 | Cpt1b | carnitine palmitoyltransferase 1b, muscle | 1 | 0 | 0 | 0 | 0 | 7 | 0 | 0 | 0 | 0 | 3 |
| 1 | 0.66 | 3.61 | 0.18 | 0.85 | 1 | Gm29848 | | 0 | 0 | 0 | 0 | 0 | 3 | 0 | 0 | 0 | 0 | 8 |
| 1 | 0.00 | 3.68 | 0.00 | 1.00 | 1 | Ddx43 | DEAD (Asp-Glu-Ala-Asp) box polypeptide | 9 | 0 | 2 | 0 | 0 | 0 | 0 | 0 | 0 | 0 | 0 |
| 1 | -1.02 | 3.62 | -0.28 | 0.78 | 1 | Gm42731 | | 0 | 0 | 7 | 0 | 0 | 3 | 0 | 0 | 0 | 0 | 0 |
| 1 | -0.15 | 3.61 | -0.04 | 0.97 | 1 | Gm25411 | | 0 | 0 | 0 | 0 | 0 | 8 | 0 | 0 | 0 | 0 | 3 |
| 1 | -0.15 | 3.61 | -0.04 | 0.97 | 1 | Gm37478 | | 0 | 0 | 0 | 0 | 0 | 8 | 0 | 0 | 0 | 0 | 3 |
| 1 | -0.15 | 3.61 | -0.04 | 0.97 | 1 | Gm25219 | | 0 | 0 | 0 | 0 | 0 | 8 | 0 | 0 | 0 | 0 | 3 |
| 1 | -0.15 | 3.61 | -0.04 | 0.97 | 1 | Gm35876 | | 0 | 0 | 0 | 0 | 0 | 8 | 0 | 0 | 0 | 0 | 3 |
| 1 | -0.15 | 3.61 | -0.04 | 0.97 | 1 | Mir1929 | | 0 | 0 | 0 | 0 | 0 | 8 | 0 | 0 | 0 | 0 | 3 |
| 1 | -0.15 | 3.61 | -0.04 | 0.97 | 1 | Taf9 | TATA-box binding protein associated fact | 0 | 0 | 0 | 0 | 0 | 8 | 0 | 0 | 0 | 0 | 3 |
| 1 | -0.48 | 3.61 | -0.13 | 0.89 | 1 | Gm38163 | | 0 | 2 | 0 | 0 | 0 | 8 | 0 | 0 | 0 | 0 | 1 |
| 1 | 1.68 | 3.59 | 0.47 | 0.64 | 1 | Gm13357 | | 0 | 0 | 0 | 0 | 0 | 3 | 0 | 0 | 3 | 0 | 5 |
| 1 | 0.96 | 3.56 | 0.27 | 0.79 | 1 | 4930543N07Rik | | 0 | 0 | 0 | 0 | 0 | 3 | 0 | 8 | 0 | 0 | 0 |
| 1 | 0.77 | 3.58 | 0.22 | 0.83 | 1 | A630001O12Rik | | 0 | 0 | 0 | 0 | 0 | 8 | 0 | 0 | 3 | 0 | 0 |
| 1 | -3.55 | 3.56 | -1.00 | 0.32 | 1 | 1700022H01Rik | | 1 | 0 | 0 | 5 | 3 | 2 | 0 | 0 | 0 | 0 | 0 |
| 1 | 3.15 | 3.60 | 0.88 | 0.38 | 1 | Atp5c1-ps | | 0 | 1 | 3 | 0 | 0 | 0 | 0 | 1 | 3 | 1 | 1 |
| 1 | 1.37 | 3.65 | 0.38 | 0.71 | 1 | Gm37696 | | 3 | 0 | 4 | 0 | 0 | 0 | 0 | 0 | 3 | 0 | 0 |
| 1 | 2.79 | 3.61 | 0.77 | 0.44 | 1 | Gm48596 | | 0 | 5 | 0 | 0 | 0 | 2 | 0 | 0 | 0 | 4 | 0 |
| 1 | -1.32 | 3.59 | -0.37 | 0.71 | 1 | Cnga1 | cyclic nucleotide gated channel alpha 1 | 0 | 0 | 0 | 9 | 0 | 1 | 0 | 0 | 0 | 0 | 1 |
| 1 | -0.44 | 3.62 | -0.12 | 0.90 | 1 | Hipk4 | homeodomain interacting protein kinase | 0 | 7 | 0 | 4 | 0 | 0 | 0 | 0 | 0 | 0 | 0 |
| 1 | 2.50 | 3.62 | 0.69 | 0.49 | 1 | Slfnl1 | schlafen like 1 | 0 | 3 | 0 | 0 | 0 | 0 | 0 | 3 | 0 | 0 | 4 |
| 1 | 0.96 | 3.56 | 0.27 | 0.79 | 1 | Gm8444 | | 0 | 0 | 1 | 0 | 1 | 3 | 0 | 1 | 3 | 0 | 1 |
| 1 | -1.11 | 3.59 | -0.31 | 0.76 | 1 | Gm16374 | | 3 | 1 | 0 | 1 | 1 | 2 | 0 | 3 | 0 | 0 | 0 |
| 1 | -2.48 | 3.61 | -0.69 | 0.49 | 1 | Gm2099 | | 1 | 0 | 0 | 9 | 1 | 0 | 0 | 0 | 0 | 0 | 0 |
| 1 | -0.01 | 3.61 | 0.00 | 1.00 | 1 | Gm2531 | | 1 | 1 | 0 | 1 | 0 | 2 | 0 | 0 | 0 | 0 | 6 |
| 1 | 0.00 | 3.68 | 0.00 | 1.00 | 1 | 2310015D24Rik | | 4 | 0 | 6 | 0 | 0 | 0 | 0 | 0 | 0 | 0 | 0 |
| 1 | -2.13 | 3.63 | -0.59 | 0.56 | 1 | Zfp985 | zinc finger protein 985 | 7 | 0 | 0 | 3 | 0 | 0 | 0 | 0 | 0 | 0 | 1 |
| 1 | -1.07 | 3.67 | -0.29 | 0.77 | 1 | Gm20528 | | 3 | 1 | 5 | 1 | 0 | 1 | 0 | 0 | 0 | 0 | 0 |
| 1 | 0.50 | 3.61 | 0.14 | 0.89 | 1 | Gm28784 | | 0 | 0 | 0 | 0 | 0 | 3 | 0 | 0 | 0 | 0 | 7 |
| 1 | 0.50 | 3.61 | 0.14 | 0.89 | 1 | Gm17224 | | 0 | 0 | 0 | 0 | 0 | 3 | 0 | 0 | 0 | 0 | 7 |
| 1 | 0.50 | 3.61 | 0.14 | 0.89 | 1 | Mir453 | | 0 | 0 | 0 | 0 | 0 | 3 | 0 | 0 | 0 | 0 | 7 |
| 1 | -0.32 | 3.61 | -0.09 | 0.93 | 1 | Mir30c-1 | | 0 | 0 | 0 | 0 | 0 | 8 | 0 | 0 | 0 | 0 | 2 |
| 1 | -0.32 | 3.61 | -0.09 | 0.93 | 1 | Gm48437 | | 0 | 0 | 0 | 0 | 0 | 8 | 0 | 0 | 0 | 0 | 2 |
| 1 | 0.28 | 3.61 | 0.08 | 0.94 | 1 | A930041C12Rik | | 0 | 3 | 0 | 0 | 0 | 3 | 0 | 0 | 0 | 0 | 4 |
| 1 | 0.16 | 3.61 | 0.05 | 0.96 | 1 | 1810013D15Rik | | 0 | 4 | 0 | 0 | 0 | 3 | 0 | 0 | 0 | 0 | 3 |
| 1 | -2.44 | 3.59 | -0.68 | 0.50 | 1 | Gm43067 | | 2 | 0 | 0 | 0 | 3 | 1 | 0 | 0 | 0 | 0 | 5 |
| 1 | 1.34 | 3.57 | 0.37 | 0.71 | 1 | Mageb3 | melanoma antigen, family B, 3 | 0 | 0 | 0 | 0 | 0 | 3 | 0 | 0 | 6 | 0 | 1 |
| 1 | -0.51 | 3.60 | -0.14 | 0.89 | 1 | Gm26705 | | 0 | 0 | 0 | 0 | 9 | 0 | 0 | 0 | 0 | 0 | 1 |
| 1 | -0.53 | 3.59 | -0.15 | 0.88 | 1 | Rpl30-ps8 | | 4 | 1 | 0 | 2 | 1 | 0 | 0 | 1 | 0 | 1 | 0 |
| 1 | -2.62 | 3.60 | -0.73 | 0.47 | 1 | Actl10 | actin-like 10 | 0 | 0 | 0 | 9 | 0 | 2 | 0 | 0 | 0 | 0 | 0 |
| 1 | -2.62 | 3.60 | -0.73 | 0.47 | 1 | Gm50009 | | 0 | 0 | 0 | 9 | 0 | 2 | 0 | 0 | 0 | 0 | 0 |
| 1 | -0.41 | 3.67 | -0.11 | 0.91 | 1 | Gm13597 | | 3 | 4 | 2 | 1 | 0 | 0 | 0 | 0 | 0 | 0 | 0 |
| 1 | 1.84 | 3.64 | 0.51 | 0.61 | 1 | Gm8115 | | 1 | 0 | 7 | 0 | 0 | 0 | 0 | 0 | 0 | 1 | 1 |
| 1 | 3.02 | 3.59 | 0.84 | 0.40 | 1 | Snora21 | | 0 | 0 | 0 | 0 | 0 | 2 | 0 | 0 | 0 | 8 | 1 |
| 1 | 1.80 | 3.62 | 0.50 | 0.62 | 1 | Ube4bos3 | | 0 | 0 | 0 | 0 | 0 | 0 | 0 | 10 | 0 | 0 | 0 |
| 1 | 1.80 | 3.62 | 0.50 | 0.62 | 1 | 4930560O18Rik | | 0 | 0 | 0 | 0 | 0 | 0 | 0 | 10 | 0 | 0 | 0 |
| 1 | 1.80 | 3.62 | 0.50 | 0.62 | 1 | Gm29154 | | 0 | 0 | 0 | 0 | 0 | 0 | 0 | 10 | 0 | 0 | 0 |
| 1 | 1.80 | 3.62 | 0.50 | 0.62 | 1 | 5730435O14Rik | | 0 | 0 | 0 | 0 | 0 | 0 | 0 | 10 | 0 | 0 | 0 |
| 1 | 0.30 | 3.20 | 0.09 | 0.93 | 1 | Rpl21-ps6 | | 0 | 1 | 1 | 0 | 1 | 4 | 0 | 1 | 1 | 0 | 1 |
| 1 | -1.68 | 3.59 | -0.47 | 0.64 | 1 | Gm47726 | | 0 | 0 | 0 | 3 | 0 | 7 | 0 | 0 | 0 | 0 | 1 |
| 1 | -1.25 | 3.62 | -0.34 | 0.73 | 1 | Gm42962 | | 0 | 0 | 0 | 0 | 10 | 0 | 0 | 0 | 0 | 0 | 0 |
| 1 | 0.00 | 3.66 | 0.00 | 1.00 | 1 | 9430097D07Rik | | 0 | 0 | 10 | 0 | 0 | 0 | 0 | 0 | 0 | 0 | 0 |
| 1 | 0.00 | 3.66 | 0.00 | 1.00 | 1 | Gm14102 | | 0 | 0 | 10 | 0 | 0 | 0 | 0 | 0 | 0 | 0 | 0 |
| 1 | 0.00 | 3.66 | 0.00 | 1.00 | 1 | 4921539H07Rik | | 0 | 0 | 10 | 0 | 0 | 0 | 0 | 0 | 0 | 0 | 0 |

| | | | | | | | | | | | | | | | | | | |
|---|---|---|---|---|---|---|---|---|---|---|---|---|---|---|---|---|---|---|
| 1 | 0.00 | 3.66 | 0.00 | 1.00 | 1 | 1700009C05Rik | | 0 | 0 | 10 | 0 | 0 | 0 | 0 | 0 | 0 | 0 | 0 |
| 1 | 0.00 | 3.66 | 0.00 | 1.00 | 1 | Gm9375 | | 0 | 0 | 10 | 0 | 0 | 0 | 0 | 0 | 0 | 0 | 0 |
| 1 | 0.00 | 3.66 | 0.00 | 1.00 | 1 | Gm10161 | | 0 | 0 | 10 | 0 | 0 | 0 | 0 | 0 | 0 | 0 | 0 |
| 1 | 0.00 | 3.66 | 0.00 | 1.00 | 1 | Gm29794 | | 0 | 0 | 10 | 0 | 0 | 0 | 0 | 0 | 0 | 0 | 0 |
| 1 | 0.00 | 3.66 | 0.00 | 1.00 | 1 | A530084C06Rik | | 0 | 0 | 10 | 0 | 0 | 0 | 0 | 0 | 0 | 0 | 0 |
| 1 | 0.00 | 3.66 | 0.00 | 1.00 | 1 | Gm32067 | | 0 | 0 | 10 | 0 | 0 | 0 | 0 | 0 | 0 | 0 | 0 |
| 1 | 0.00 | 3.66 | 0.00 | 1.00 | 1 | Gm4813 | | 0 | 0 | 10 | 0 | 0 | 0 | 0 | 0 | 0 | 0 | 0 |
| 1 | 0.00 | 3.66 | 0.00 | 1.00 | 1 | Gzma | granzyme A | 0 | 0 | 10 | 0 | 0 | 0 | 0 | 0 | 0 | 0 | 0 |
| 1 | 0.00 | 3.66 | 0.00 | 1.00 | 1 | Gm49256 | | 0 | 0 | 10 | 0 | 0 | 0 | 0 | 0 | 0 | 0 | 0 |
| 1 | 0.00 | 3.66 | 0.00 | 1.00 | 1 | Ugt3a2 | UDP glycosyltransferases 3 family, polype | 0 | 0 | 10 | 0 | 0 | 0 | 0 | 0 | 0 | 0 | 0 |
| 1 | 0.00 | 3.66 | 0.00 | 1.00 | 1 | B630019A10Rik | | 0 | 0 | 10 | 0 | 0 | 0 | 0 | 0 | 0 | 0 | 0 |
| 1 | 0.00 | 3.66 | 0.00 | 1.00 | 1 | Gm15665 | | 0 | 0 | 10 | 0 | 0 | 0 | 0 | 0 | 0 | 0 | 0 |
| 1 | 0.00 | 3.66 | 0.00 | 1.00 | 1 | Gm6425 | | 0 | 0 | 10 | 0 | 0 | 0 | 0 | 0 | 0 | 0 | 0 |
| 1 | 0.00 | 3.68 | 0.00 | 1.00 | 1 | Pkp1 | plakophilin 1 | 4 | 6 | 0 | 0 | 0 | 0 | 0 | 0 | 0 | 0 | 0 |
| 1 | 0.00 | 3.68 | 0.00 | 1.00 | 1 | Gm7298 | | 0 | 1 | 9 | 0 | 0 | 0 | 0 | 0 | 0 | 0 | 0 |
| 1 | 3.49 | 3.57 | 0.98 | 0.33 | 1 | Gm42683 | | 0 | 1 | 0 | 0 | 2 | 0 | 0 | 0 | 0 | 6 | 1 |
| 1 | -0.15 | 3.58 | -0.04 | 0.97 | 1 | Gm47201 | | 0 | 2 | 1 | 5 | 0 | 0 | 0 | 0 | 0 | 0 | 2 |
| 1 | 1.53 | 3.64 | 0.42 | 0.67 | 1 | Gm41556 | | 0 | 0 | 8 | 0 | 0 | 0 | 0 | 0 | 0 | 0 | 2 |
| 1 | 0.00 | 3.68 | 0.00 | 1.00 | 1 | Gm44764 | | 5 | 0 | 5 | 0 | 0 | 0 | 0 | 0 | 0 | 0 | 0 |
| 1 | 0.00 | 3.68 | 0.00 | 1.00 | 1 | Gm41611 | | 5 | 0 | 5 | 0 | 0 | 0 | 0 | 0 | 0 | 0 | 0 |
| 1 | 0.00 | 3.68 | 0.00 | 1.00 | 1 | Gsc | goosecoid homeobox | 0 | 2 | 8 | 0 | 0 | 0 | 0 | 0 | 0 | 0 | 0 |
| 1 | -2.03 | 3.62 | -0.56 | 0.57 | 1 | Edar | ectodysplasin-A receptor | 6 | 0 | 0 | 0 | 1 | 3 | 0 | 0 | 0 | 0 | 0 |
| 1 | 1.65 | 3.63 | 0.46 | 0.65 | 1 | 4930428N03Rik | | 0 | 0 | 7 | 0 | 0 | 0 | 0 | 0 | 0 | 0 | 3 |
| 1 | 0.35 | 3.61 | 0.10 | 0.92 | 1 | Gm43793 | | 0 | 0 | 0 | 0 | 0 | 4 | 0 | 0 | 0 | 0 | 6 |
| 1 | 0.35 | 3.61 | 0.10 | 0.92 | 1 | Gm43844 | | 0 | 0 | 0 | 0 | 0 | 4 | 0 | 0 | 0 | 0 | 6 |
| 1 | 0.35 | 3.61 | 0.10 | 0.92 | 1 | Gm20519 | | 0 | 0 | 0 | 0 | 0 | 4 | 0 | 0 | 0 | 0 | 6 |
| 1 | -0.82 | 3.61 | -0.23 | 0.82 | 1 | Gm38214 | | 0 | 1 | 0 | 0 | 0 | 9 | 0 | 0 | 0 | 0 | 0 |
| 1 | -0.42 | 3.60 | -0.12 | 0.91 | 1 | Gm20150 | | 0 | 5 | 0 | 0 | 0 | 4 | 0 | 0 | 0 | 0 | 1 |
| 1 | 1.82 | 3.63 | 0.50 | 0.62 | 1 | Tbx3os2 | | 0 | 0 | 0 | 0 | 0 | 0 | 0 | 0 | 0 | 10 | 0 |
| 1 | 1.82 | 3.63 | 0.50 | 0.62 | 1 | Eif4e1b | eukaryotic translation initiation factor 4E fa | 0 | 0 | 0 | 0 | 0 | 0 | 0 | 0 | 0 | 10 | 0 |
| 1 | -2.18 | 3.60 | -0.61 | 0.55 | 1 | Gm38114 | | 0 | 0 | 0 | 0 | 9 | 1 | 0 | 0 | 0 | 0 | 0 |
| 1 | 0.00 | 3.67 | 0.00 | 1.00 | 1 | D630023O14Rik | | 10 | 0 | 0 | 0 | 0 | 0 | 0 | 0 | 0 | 0 | 0 |
| 1 | 0.00 | 3.67 | 0.00 | 1.00 | 1 | Banf2 | barrier to autointegration factor 2 | 10 | 0 | 0 | 0 | 0 | 0 | 0 | 0 | 0 | 0 | 0 |
| 1 | 0.00 | 3.67 | 0.00 | 1.00 | 1 | AI646519 | | 10 | 0 | 0 | 0 | 0 | 0 | 0 | 0 | 0 | 0 | 0 |
| 1 | 0.00 | 3.67 | 0.00 | 1.00 | 1 | Zfp663 | zinc finger protein 663 | 10 | 0 | 0 | 0 | 0 | 0 | 0 | 0 | 0 | 0 | 0 |
| 1 | 0.00 | 3.67 | 0.00 | 1.00 | 1 | Bcl2l15 | BCLl2-like 15 | 10 | 0 | 0 | 0 | 0 | 0 | 0 | 0 | 0 | 0 | 0 |
| 1 | 0.00 | 3.67 | 0.00 | 1.00 | 1 | Spocd1 | SPOC domain containing 1 | 10 | 0 | 0 | 0 | 0 | 0 | 0 | 0 | 0 | 0 | 0 |
| 1 | 0.00 | 3.67 | 0.00 | 1.00 | 1 | Gm8773 | | 10 | 0 | 0 | 0 | 0 | 0 | 0 | 0 | 0 | 0 | 0 |
| 1 | 0.00 | 3.67 | 0.00 | 1.00 | 1 | Trbv16 | | 10 | 0 | 0 | 0 | 0 | 0 | 0 | 0 | 0 | 0 | 0 |
| 1 | 0.00 | 3.67 | 0.00 | 1.00 | 1 | Igkv13-85 | | 10 | 0 | 0 | 0 | 0 | 0 | 0 | 0 | 0 | 0 | 0 |
| 1 | 0.00 | 3.67 | 0.00 | 1.00 | 1 | Lypd3 | Ly6/Plaur domain containing 3 | 10 | 0 | 0 | 0 | 0 | 0 | 0 | 0 | 0 | 0 | 0 |
| 1 | 0.00 | 3.67 | 0.00 | 1.00 | 1 | Gm18853 | | 10 | 0 | 0 | 0 | 0 | 0 | 0 | 0 | 0 | 0 | 0 |
| 1 | 0.00 | 3.67 | 0.00 | 1.00 | 1 | Cd163l1 | CD163 molecule-like 1 | 10 | 0 | 0 | 0 | 0 | 0 | 0 | 0 | 0 | 0 | 0 |
| 1 | 0.00 | 3.67 | 0.00 | 1.00 | 1 | Gm25661 | | 10 | 0 | 0 | 0 | 0 | 0 | 0 | 0 | 0 | 0 | 0 |
| 1 | 0.00 | 3.67 | 0.00 | 1.00 | 1 | Gm17169 | | 10 | 0 | 0 | 0 | 0 | 0 | 0 | 0 | 0 | 0 | 0 |
| 1 | 0.00 | 3.67 | 0.00 | 1.00 | 1 | Tdgf1 | teratocarcinoma-derived growth factor 1 | 10 | 0 | 0 | 0 | 0 | 0 | 0 | 0 | 0 | 0 | 0 |
| 1 | 0.00 | 3.67 | 0.00 | 1.00 | 1 | Gm12221 | | 10 | 0 | 0 | 0 | 0 | 0 | 0 | 0 | 0 | 0 | 0 |
| 1 | 0.00 | 3.67 | 0.00 | 1.00 | 1 | Ighv3-6 | | 10 | 0 | 0 | 0 | 0 | 0 | 0 | 0 | 0 | 0 | 0 |
| 1 | 0.00 | 3.67 | 0.00 | 1.00 | 1 | Ankdd1b | ankyrin repeat and death domain contain | 10 | 0 | 0 | 0 | 0 | 0 | 0 | 0 | 0 | 0 | 0 |
| 1 | 0.00 | 3.67 | 0.00 | 1.00 | 1 | Gm49123 | | 10 | 0 | 0 | 0 | 0 | 0 | 0 | 0 | 0 | 0 | 0 |
| 1 | 0.00 | 3.67 | 0.00 | 1.00 | 1 | Trav6-5 | | 10 | 0 | 0 | 0 | 0 | 0 | 0 | 0 | 0 | 0 | 0 |
| 1 | 0.00 | 3.67 | 0.00 | 1.00 | 1 | Gm34643 | | 10 | 0 | 0 | 0 | 0 | 0 | 0 | 0 | 0 | 0 | 0 |
| 1 | 0.00 | 3.67 | 0.00 | 1.00 | 1 | Cyp2c23 | cytochrome P450, family 2, subfamily c, p | 10 | 0 | 0 | 0 | 0 | 0 | 0 | 0 | 0 | 0 | 0 |
| 1 | -2.74 | 3.60 | -0.76 | 0.45 | 1 | 4930596I21Rik | | 0 | 0 | 0 | 4 | 0 | 6 | 0 | 0 | 0 | 0 | 0 |
| 1 | -2.74 | 3.60 | -0.76 | 0.45 | 1 | Gm45358 | | 0 | 0 | 0 | 4 | 0 | 6 | 0 | 0 | 0 | 0 | 0 |
| 1 | 2.66 | 3.61 | 0.74 | 0.46 | 1 | Gm42974 | | 0 | 0 | 0 | 0 | 0 | 0 | 0 | 9 | 0 | 0 | 1 |
| 1 | -0.57 | 3.66 | -0.15 | 0.88 | 1 | Gm13486 | | 9 | 0 | 0 | 0 | 0 | 1 | 0 | 0 | 0 | 0 | 0 |
| 1 | -0.65 | 3.56 | -0.18 | 0.86 | 1 | Gm26703 | | 0 | 0 | 0 | 4 | 0 | 1 | 0 | 0 | 5 | 0 | 0 |
| 1 | 0.00 | 3.68 | 0.00 | 1.00 | 1 | 1810065E05Ril | RIKEN cDNA 1810065E05 gene | 0 | 5 | 5 | 0 | 0 | 0 | 0 | 0 | 0 | 0 | 0 |
| 1 | -1.01 | 3.63 | -0.28 | 0.78 | 1 | Gm14637 | | 8 | 0 | 0 | 0 | 0 | 3 | 0 | 0 | 0 | 0 | 0 |

| | | | | | | | | | | | | | | | | | | |
|---|---|---|---|---|---|---|---|---|---|---|---|---|---|---|---|---|---|---|
| 1 | -1.01 | 3.63 | -0.28 | 0.78 | 1 | Gpr37 | G protein-coupled receptor 37 | 8 | 0 | 0 | 0 | 0 | 3 | 0 | 0 | 0 | 0 | 0 |
| 1 | 1.61 | 3.61 | 0.45 | 0.66 | 1 | Zbtb9 | zinc finger and BTB domain containing 9 | 0 | 0 | 4 | 0 | 0 | 0 | 0 | 0 | 0 | 0 | 6 |
| 1 | -1.02 | 3.62 | -0.28 | 0.78 | 1 | Gm12963 | | 7 | 0 | 0 | 0 | 0 | 3 | 0 | 0 | 0 | 0 | 0 |
| 1 | 5.52 | 3.59 | 1.54 | 0.12 | 1 | Gm13215 | | 0 | 3 | 0 | 0 | 0 | 1 | 0 | 0 | 2 | 1 | 3 |
| 1 | -2.28 | 3.60 | -0.63 | 0.53 | 1 | Gm43544 | | 0 | 0 | 0 | 3 | 0 | 8 | 0 | 0 | 0 | 0 | 0 |
| 1 | -1.01 | 3.61 | -0.28 | 0.78 | 1 | Gm26674 | | 6 | 0 | 0 | 0 | 0 | 4 | 0 | 0 | 0 | 0 | 0 |
| 1 | 0.25 | 3.56 | 0.07 | 0.94 | 1 | Gm23966 | | 0 | 0 | 0 | 0 | 1 | 4 | 0 | 0 | 5 | 0 | 0 |
| 1 | 2.41 | 3.61 | 0.67 | 0.50 | 1 | Ankrd65 | ankyrin repeat domain 65 | 0 | 0 | 3 | 0 | 0 | 0 | 0 | 0 | 2 | 0 | 5 |
| 1 | 0.00 | 3.68 | 0.00 | 1.00 | 1 | Irgc1 | immunity-related GTPase family, cinema | 5 | 5 | 0 | 0 | 0 | 0 | 0 | 0 | 0 | 0 | 0 |
| 1 | 0.00 | 3.68 | 0.00 | 1.00 | 1 | E230025N22Ri | Riken cDNA E230025N22 gene | 5 | 5 | 0 | 0 | 0 | 0 | 0 | 0 | 0 | 0 | 0 |
| 1 | 0.10 | 3.60 | 0.03 | 0.98 | 1 | Gm16074 | | 1 | 0 | 1 | 1 | 0 | 3 | 0 | 0 | 1 | 0 | 3 |
| 1 | -0.53 | 3.66 | -0.14 | 0.89 | 1 | Gm11855 | | 0 | 0 | 9 | 0 | 0 | 1 | 0 | 0 | 0 | 0 | 0 |
| 1 | -0.53 | 3.66 | -0.14 | 0.89 | 1 | 4930433N12Rik | | 0 | 0 | 9 | 0 | 0 | 1 | 0 | 0 | 0 | 0 | 0 |
| 1 | -0.53 | 3.66 | -0.14 | 0.89 | 1 | Platr7 | | 0 | 0 | 9 | 0 | 0 | 1 | 0 | 0 | 0 | 0 | 0 |
| 1 | 0.58 | 3.61 | 0.16 | 0.87 | 1 | Gm6623 | | 7 | 0 | 0 | 1 | 0 | 0 | 0 | 1 | 0 | 1 | 0 |
| 1 | -0.81 | 3.61 | -0.22 | 0.82 | 1 | Gm37928 | | 2 | 0 | 0 | 0 | 0 | 8 | 0 | 0 | 0 | 0 | 0 |
| 1 | -0.84 | 3.63 | -0.23 | 0.82 | 1 | Gm14279 | | 0 | 1 | 5 | 1 | 1 | 1 | 0 | 0 | 0 | 1 | 0 |
| 1 | 1.38 | 3.61 | 0.38 | 0.70 | 1 | Gm43886 | | 0 | 0 | 0 | 0 | 0 | 0 | 0 | 0 | 0 | 0 | 10 |
| 1 | 1.38 | 3.61 | 0.38 | 0.70 | 1 | Gm34093 | | 0 | 0 | 0 | 0 | 0 | 0 | 0 | 0 | 0 | 0 | 10 |
| 1 | 1.38 | 3.61 | 0.38 | 0.70 | 1 | Gm50044 | | 0 | 0 | 0 | 0 | 0 | 0 | 0 | 0 | 0 | 0 | 10 |
| 1 | 0.20 | 3.61 | 0.06 | 0.95 | 1 | Gm28512 | | 0 | 0 | 0 | 0 | 0 | 5 | 0 | 0 | 0 | 0 | 5 |
| 1 | 0.20 | 3.61 | 0.06 | 0.95 | 1 | Vmn2r29 | vomeronasal 2, receptor 29 | 0 | 0 | 0 | 0 | 0 | 5 | 0 | 0 | 0 | 0 | 5 |
| 1 | 0.20 | 3.61 | 0.06 | 0.95 | 1 | Gm38355 | | 0 | 0 | 0 | 0 | 0 | 5 | 0 | 0 | 0 | 0 | 5 |
| 1 | 0.20 | 3.61 | 0.06 | 0.95 | 1 | Gm24400 | | 0 | 0 | 0 | 0 | 0 | 5 | 0 | 0 | 0 | 0 | 5 |
| 1 | 1.51 | 3.61 | 0.42 | 0.68 | 1 | Gm8540 | | 0 | 4 | 0 | 0 | 0 | 0 | 0 | 0 | 0 | 0 | 6 |
| 1 | 2.79 | 3.61 | 0.77 | 0.44 | 1 | Gm18126 | | 0 | 0 | 0 | 0 | 0 | 0 | 0 | 0 | 3 | 0 | 7 |
| 1 | -2.43 | 3.60 | -0.68 | 0.50 | 1 | 1700001P01Ril | RIKEN cDNA 1700001P01 gene | 0 | 0 | 4 | 2 | 0 | 4 | 0 | 0 | 0 | 0 | 0 |
| 1 | 1.10 | 3.66 | 0.30 | 0.76 | 1 | Plin1 | perilipin 1 | 0 | 9 | 0 | 0 | 0 | 0 | 0 | 0 | 0 | 0 | 1 |
| 1 | 4.73 | 3.58 | 1.32 | 0.19 | 1 | Gm12599 | | 0 | 0 | 3 | 0 | 0 | 3 | 0 | 1 | 0 | 1 | 2 |
| 1 | 0.00 | 3.66 | 0.00 | 1.00 | 1 | Gm37573 | | 0 | 10 | 0 | 0 | 0 | 0 | 0 | 0 | 0 | 0 | 0 |
| 1 | 0.00 | 3.66 | 0.00 | 1.00 | 1 | Lrrc19 | leucine rich repeat containing 19 | 0 | 10 | 0 | 0 | 0 | 0 | 0 | 0 | 0 | 0 | 0 |
| 1 | 0.00 | 3.66 | 0.00 | 1.00 | 1 | 4933427I22Rik | | 0 | 10 | 0 | 0 | 0 | 0 | 0 | 0 | 0 | 0 | 0 |
| 1 | 0.00 | 3.66 | 0.00 | 1.00 | 1 | Dnah3 | dynein, axonemal, heavy chain 3 | 0 | 10 | 0 | 0 | 0 | 0 | 0 | 0 | 0 | 0 | 0 |
| 1 | 0.00 | 3.66 | 0.00 | 1.00 | 1 | Atp2a1 | ATPase, Ca++ transporting, cardiac musc | 0 | 10 | 0 | 0 | 0 | 0 | 0 | 0 | 0 | 0 | 0 |
| 1 | 0.00 | 3.66 | 0.00 | 1.00 | 1 | Gm44891 | | 0 | 10 | 0 | 0 | 0 | 0 | 0 | 0 | 0 | 0 | 0 |
| 1 | 0.00 | 3.66 | 0.00 | 1.00 | 1 | Ccdc194 | coiled-coil domain containing 194 | 0 | 10 | 0 | 0 | 0 | 0 | 0 | 0 | 0 | 0 | 0 |
| 1 | 0.00 | 3.66 | 0.00 | 1.00 | 1 | Ces2c | carboxylesterase 2C | 0 | 10 | 0 | 0 | 0 | 0 | 0 | 0 | 0 | 0 | 0 |
| 1 | 0.00 | 3.66 | 0.00 | 1.00 | 1 | Gm32515 | | 0 | 10 | 0 | 0 | 0 | 0 | 0 | 0 | 0 | 0 | 0 |
| 1 | 0.00 | 3.66 | 0.00 | 1.00 | 1 | 4732465J04Rik | | 0 | 10 | 0 | 0 | 0 | 0 | 0 | 0 | 0 | 0 | 0 |
| 1 | 0.00 | 3.66 | 0.00 | 1.00 | 1 | Gm15642 | | 0 | 10 | 0 | 0 | 0 | 0 | 0 | 0 | 0 | 0 | 0 |
| 1 | 0.00 | 3.66 | 0.00 | 1.00 | 1 | Tshr | thyroid stimulating hormone receptor | 0 | 10 | 0 | 0 | 0 | 0 | 0 | 0 | 0 | 0 | 0 |
| 1 | 0.00 | 3.66 | 0.00 | 1.00 | 1 | Gm48733 | | 0 | 10 | 0 | 0 | 0 | 0 | 0 | 0 | 0 | 0 | 0 |
| 1 | 0.00 | 3.66 | 0.00 | 1.00 | 1 | Irx4 | Iroquois homeobox 4 | 0 | 10 | 0 | 0 | 0 | 0 | 0 | 0 | 0 | 0 | 0 |
| 1 | 0.00 | 3.66 | 0.00 | 1.00 | 1 | Trav14d-3-dv8 | | 0 | 10 | 0 | 0 | 0 | 0 | 0 | 0 | 0 | 0 | 0 |
| 1 | 0.00 | 3.66 | 0.00 | 1.00 | 1 | Ppnr | | 0 | 10 | 0 | 0 | 0 | 0 | 0 | 0 | 0 | 0 | 0 |
| 1 | 0.00 | 3.66 | 0.00 | 1.00 | 1 | Gm14699 | | 0 | 10 | 0 | 0 | 0 | 0 | 0 | 0 | 0 | 0 | 0 |
| 1 | 1.17 | 3.65 | 0.32 | 0.75 | 1 | Gm37092 | | 0 | 9 | 0 | 0 | 0 | 0 | 0 | 0 | 1 | 0 | 0 |
| 1 | 3.05 | 3.61 | 0.84 | 0.40 | 1 | H1f6 | H1.6 linker histone, cluster member | 0 | 0 | 0 | 0 | 0 | 0 | 0 | 0 | 7 | 0 | 3 |
| 1 | 0.82 | 3.61 | 0.23 | 0.82 | 1 | Gm43788 | | 0 | 0 | 6 | 0 | 0 | 1 | 0 | 0 | 0 | 0 | 3 |
| 1 | 1.89 | 3.63 | 0.52 | 0.60 | 1 | Gm45174 | | 0 | 0 | 0 | 0 | 0 | 0 | 0 | 0 | 10 | 0 | 0 |
| 1 | -2.73 | 3.59 | -0.76 | 0.45 | 1 | Gm42766 | | 0 | 0 | 0 | 0 | 8 | 3 | 0 | 0 | 0 | 0 | 0 |
| 1 | -0.54 | 3.67 | -0.15 | 0.88 | 1 | Gm14414 | | 0 | 8 | 1 | 0 | 1 | 0 | 0 | 0 | 0 | 0 | 0 |
| 1 | 0.24 | 3.65 | 0.06 | 0.95 | 1 | Gm18722 | | 0 | 8 | 0 | 0 | 1 | 0 | 0 | 0 | 0 | 0 | 1 |
| 1 | 0.31 | 3.62 | 0.09 | 0.93 | 1 | Gm5513 | | 0 | 1 | 5 | 0 | 3 | 0 | 0 | 0 | 0 | 0 | 1 |
| 1 | 0.00 | 3.68 | 0.00 | 1.00 | 1 | Gm13815 | | 6 | 4 | 0 | 0 | 0 | 0 | 0 | 0 | 0 | 0 | 0 |
| 1 | 0.00 | 3.68 | 0.00 | 1.00 | 1 | 5330411O13Rik | | 6 | 4 | 0 | 0 | 0 | 0 | 0 | 0 | 0 | 0 | 0 |
| 1 | 0.00 | 3.68 | 0.00 | 1.00 | 1 | Gm30873 | | 6 | 4 | 0 | 0 | 0 | 0 | 0 | 0 | 0 | 0 | 0 |
| 1 | 0.00 | 3.68 | 0.00 | 1.00 | 1 | Gm11682 | | 6 | 4 | 0 | 0 | 0 | 0 | 0 | 0 | 0 | 0 | 0 |
| 1 | 0.82 | 3.62 | 0.23 | 0.82 | 1 | Gm42432 | | 5 | 0 | 0 | 0 | 0 | 1 | 0 | 0 | 0 | 0 | 4 |
| 1 | -0.39 | 3.56 | -0.11 | 0.91 | 1 | Ifi44l | interferon-induced protein 44 like | 0 | 0 | 0 | 7 | 0 | 0 | 0 | 0 | 0 | 0 | 3 |

| | | | | | | | | | | | | | | | | | | |
|---|---|---|---|---|---|---|---|---|---|---|---|---|---|---|---|---|---|---|
| 1 | -0.40 | 3.67 | -0.11 | 0.91 | 1 | Gm38312 | | 0 | 6 | 3 | 0 | 0 | 1 | 0 | 0 | 0 | 0 | 0 |
| 1 | -0.40 | 3.67 | -0.11 | 0.91 | 1 | Gm48295 | | 0 | 6 | 3 | 0 | 0 | 1 | 0 | 0 | 0 | 0 | 0 |
| 1 | 3.51 | 3.57 | 0.98 | 0.33 | 1 | Myoz1 | myozenin 1 | 0 | 0 | 0 | 7 | 0 | 0 | 0 | 0 | 2 | 0 | 1 |
| 1 | 0.00 | 3.68 | 0.00 | 1.00 | 1 | Gm13919 | | 2 | 0 | 8 | 0 | 0 | 0 | 0 | 0 | 0 | 0 | 0 |
| 1 | 0.81 | 3.62 | 0.22 | 0.82 | 1 | Smt3h2-ps | | 1 | 0 | 4 | 0 | 1 | 0 | 0 | 0 | 4 | 0 | 0 |
| 1 | -2.76 | 3.56 | -0.78 | 0.44 | 1 | Gm15795 | | 1 | 0 | 0 | 4 | 1 | 3 | 0 | 0 | 0 | 1 | 0 |
| 1 | -1.80 | 3.61 | -0.50 | 0.62 | 1 | Gm37706 | | 0 | 5 | 0 | 0 | 2 | 0 | 0 | 0 | 0 | 0 | 3 |
| 1 | 1.04 | 3.61 | 0.29 | 0.77 | 1 | Gm25821 | | 0 | 0 | 0 | 0 | 0 | 1 | 0 | 0 | 0 | 0 | 9 |
| 1 | 1.04 | 3.61 | 0.29 | 0.77 | 1 | Tomm6 | translocase of outer mitochondrial membr | 0 | 0 | 0 | 0 | 0 | 1 | 0 | 0 | 0 | 0 | 9 |
| 1 | 0.00 | 3.68 | 0.00 | 1.00 | 1 | Il18rap | interleukin 18 receptor accessory protein | 7 | 0 | 3 | 0 | 0 | 0 | 0 | 0 | 0 | 0 | 0 |
| 1 | 0.00 | 3.68 | 0.00 | 1.00 | 1 | 9530034E10Rik | | 7 | 0 | 3 | 0 | 0 | 0 | 0 | 0 | 0 | 0 | 0 |
| 1 | 1.01 | 3.61 | 0.28 | 0.78 | 1 | 3110067C02Rik | | 0 | 2 | 1 | 0 | 0 | 1 | 0 | 0 | 6 | 0 | 0 |
| 1 | 1.17 | 3.60 | 0.33 | 0.74 | 1 | Olfr330 | olfactory receptor 330 | 0 | 0 | 0 | 1 | 0 | 0 | 0 | 0 | 9 | 0 | 0 |
| 1 | 0.06 | 3.61 | 0.02 | 0.99 | 1 | Gm43480 | | 0 | 0 | 0 | 0 | 0 | 6 | 0 | 0 | 0 | 0 | 4 |
| 1 | 0.06 | 3.61 | 0.02 | 0.99 | 1 | Gm7265 | | 0 | 0 | 0 | 0 | 0 | 6 | 0 | 0 | 0 | 0 | 4 |
| 1 | 0.06 | 3.61 | 0.02 | 0.99 | 1 | Hus1b | HUS1 checkpoint clamp component B | 0 | 0 | 0 | 0 | 0 | 6 | 0 | 0 | 0 | 0 | 4 |
| 1 | 1.98 | 3.60 | 0.55 | 0.58 | 1 | Gm12339 | | 0 | 0 | 0 | 0 | 0 | 1 | 0 | 0 | 2 | 0 | 7 |
| 1 | 2.13 | 3.60 | 0.59 | 0.55 | 1 | Gm9670 | | 0 | 0 | 0 | 0 | 0 | 1 | 0 | 0 | 3 | 0 | 6 |
| 1 | 0.99 | 3.58 | 0.28 | 0.78 | 1 | Gm47767 | | 0 | 0 | 0 | 0 | 0 | 6 | 0 | 0 | 3 | 0 | 1 |
| 1 | -0.59 | 3.65 | -0.16 | 0.87 | 1 | Gm45140 | | 0 | 9 | 0 | 0 | 0 | 1 | 0 | 0 | 0 | 0 | 0 |
| 1 | 3.26 | 3.63 | 0.90 | 0.37 | 1 | 4931406G06Rik | | 0 | 6 | 0 | 0 | 0 | 1 | 0 | 0 | 2 | 0 | 1 |
| 1 | -1.17 | 3.58 | -0.33 | 0.74 | 1 | 2510017J16Rik | | 0 | 0 | 0 | 4 | 0 | 3 | 0 | 0 | 0 | 0 | 3 |
| 1 | 1.20 | 3.60 | 0.33 | 0.74 | 1 | Gm37443 | | 0 | 0 | 0 | 0 | 0 | 1 | 0 | 0 | 9 | 0 | 0 |
| 1 | -1.24 | 3.59 | -0.35 | 0.73 | 1 | Gm27184 | | 0 | 0 | 0 | 1 | 1 | 5 | 0 | 0 | 0 | 0 | 3 |
| 1 | -0.90 | 3.57 | -0.25 | 0.80 | 1 | Xlr4c | X-linked lymphocyte-regulated 4C | 0 | 0 | 0 | 0 | 3 | 5 | 0 | 0 | 2 | 0 | 0 |
| 1 | 1.07 | 3.25 | 0.33 | 0.74 | 1 | Gm14794 | | 1 | 0 | 0 | 3 | 1 | 0 | 0 | 1 | 1 | 1 | 2 |
| 1 | -0.73 | 3.61 | -0.20 | 0.84 | 1 | Gm7616 | predicted gene 7616 | 0 | 0 | 0 | 0 | 1 | 6 | 0 | 0 | 0 | 0 | 3 |
| 1 | -2.61 | 3.60 | -0.72 | 0.47 | 1 | Gm16982 | | 0 | 0 | 0 | 6 | 4 | 0 | 0 | 0 | 0 | 0 | 0 |
| 1 | 0.00 | 3.68 | 0.00 | 1.00 | 1 | Hes2 | hes family bHLH transcription factor 2 | 7 | 3 | 0 | 0 | 0 | 0 | 0 | 0 | 0 | 0 | 0 |
| 1 | -2.93 | 3.59 | -0.82 | 0.41 | 1 | Smim9 | small integral membrane protein 9 | 1 | 0 | 0 | 7 | 0 | 0 | 0 | 0 | 0 | 0 | 2 |
| 1 | 0.95 | 3.61 | 0.26 | 0.79 | 1 | Gm11551 | | 0 | 0 | 0 | 0 | 4 | 0 | 0 | 0 | 5 | 0 | 1 |
| 1 | -0.58 | 3.61 | -0.16 | 0.87 | 1 | Gm37364 | | 0 | 0 | 4 | 0 | 6 | 0 | 0 | 0 | 0 | 0 | 0 |
| 1 | -0.02 | 3.62 | -0.01 | 0.99 | 1 | Hmgb1-ps1 | | 0 | 7 | 0 | 1 | 0 | 1 | 0 | 0 | 1 | 0 | 0 |
| 1 | -0.06 | 3.64 | -0.02 | 0.99 | 1 | Gm5388 | | 2 | 0 | 5 | 2 | 0 | 0 | 0 | 1 | 0 | 0 | 0 |
| 1 | 0.00 | 3.68 | 0.00 | 1.00 | 1 | Igkv12-41 | | 2 | 8 | 0 | 0 | 0 | 0 | 0 | 0 | 0 | 0 | 0 |
| 1 | -3.32 | 3.57 | -0.93 | 0.35 | 1 | Gm12248 | predicted gene 12248 | 0 | 0 | 0 | 4 | 4 | 3 | 0 | 0 | 0 | 0 | 0 |
| 1 | 1.19 | 3.57 | 0.33 | 0.74 | 1 | Gm16754 | | 0 | 0 | 1 | 0 | 0 | 2 | 0 | 0 | 6 | 0 | 1 |
| 1 | 0.80 | 3.61 | 0.22 | 0.82 | 1 | Gm44625 | | 0 | 0 | 0 | 0 | 0 | 2 | 0 | 0 | 0 | 0 | 8 |
| 1 | 0.80 | 3.61 | 0.22 | 0.82 | 1 | Gm23645 | | 0 | 0 | 0 | 0 | 0 | 2 | 0 | 0 | 0 | 0 | 8 |
| 1 | -2.48 | 3.59 | -0.69 | 0.49 | 1 | Gm15583 | | 0 | 0 | 0 | 1 | 0 | 1 | 0 | 0 | 8 | 0 | 0 |
| 1 | 0.17 | 3.55 | 0.05 | 0.96 | 1 | Gm27206 | | 0 | 0 | 0 | 0 | 5 | 0 | 0 | 0 | 0 | 0 | 5 |
| 1 | 0.85 | 3.62 | 0.23 | 0.81 | 1 | Gm23094 | | 0 | 5 | 0 | 0 | 0 | 2 | 0 | 0 | 0 | 0 | 3 |
| 1 | 0.00 | 3.68 | 0.00 | 1.00 | 1 | 5830428M24Rik | | 8 | 0 | 2 | 0 | 0 | 0 | 0 | 0 | 0 | 0 | 0 |
| 1 | -0.95 | 3.64 | -0.26 | 0.79 | 1 | Gm4755 | | 0 | 8 | 0 | 0 | 0 | 2 | 0 | 0 | 0 | 0 | 0 |
| 1 | -0.95 | 3.64 | -0.26 | 0.79 | 1 | Gm40918 | | 0 | 8 | 0 | 0 | 0 | 2 | 0 | 0 | 0 | 0 | 0 |
| 1 | 0.59 | 3.57 | 0.17 | 0.87 | 1 | Gm16505 | | 0 | 0 | 0 | 0 | 3 | 0 | 0 | 7 | 0 | 0 | 0 |
| 1 | -0.80 | 3.57 | -0.22 | 0.82 | 1 | Gm5928 | | 0 | 1 | 2 | 1 | 4 | 0 | 0 | 0 | 0 | 0 | 2 |
| 1 | 1.12 | 3.65 | 0.31 | 0.76 | 1 | Gm48909 | | 7 | 0 | 0 | 1 | 0 | 0 | 0 | 0 | 1 | 0 | 1 |
| 1 | -0.65 | 3.60 | -0.18 | 0.86 | 1 | Gm7229 | | 0 | 0 | 0 | 9 | 0 | 0 | 0 | 0 | 1 | 0 | 0 |
| 1 | 2.91 | 3.61 | 0.81 | 0.42 | 1 | Gm15802 | | 0 | 0 | 0 | 0 | 0 | 0 | 0 | 0 | 0 | 8 | 2 |
| 1 | 0.00 | 3.61 | 0.00 | 1.00 | 1 | Gm49098 | | 0 | 0 | 0 | 0 | 3 | 1 | 0 | 0 | 6 | 0 | 0 |
| 1 | -1.30 | 3.59 | -0.36 | 0.72 | 1 | Gm40264 | | 0 | 0 | 0 | 8 | 0 | 1 | 0 | 0 | 0 | 0 | 1 |
| 1 | -0.12 | 3.60 | -0.03 | 0.97 | 1 | Gm9570 | | 0 | 0 | 0 | 0 | 8 | 0 | 0 | 1 | 1 | 0 | 0 |
| 1 | 1.91 | 3.57 | 0.53 | 0.59 | 1 | Gm16523 | | 0 | 0 | 0 | 0 | 0 | 3 | 0 | 0 | 5 | 1 | 0 |
| 1 | -0.86 | 3.62 | -0.24 | 0.81 | 1 | Ccnb1-ps | | 0 | 6 | 0 | 0 | 1 | 2 | 0 | 0 | 1 | 0 | 0 |
| 1 | -1.39 | 3.62 | -0.38 | 0.70 | 1 | Gm4402 | | 3 | 1 | 0 | 5 | 0 | 0 | 0 | 0 | 0 | 0 | 0 |
| 1 | -1.74 | 3.60 | -0.48 | 0.63 | 1 | Gm2735 | | 0 | 0 | 0 | 6 | 0 | 3 | 0 | 0 | 0 | 0 | 1 |
| 1 | -0.79 | 3.62 | -0.22 | 0.83 | 1 | St6galnac2 | | 3 | 2 | 0 | 0 | 0 | 5 | 0 | 0 | 0 | 0 | 0 |
| 1 | 1.25 | 3.66 | 0.34 | 0.73 | 1 | Aknad1 | AKNA domain containing 1 | 3 | 5 | 0 | 0 | 0 | 0 | 0 | 0 | 0 | 0 | 2 |
| 1 | 0.55 | 3.61 | 0.15 | 0.88 | 1 | Gm50456 | | 0 | 0 | 1 | 0 | 0 | 3 | 0 | 0 | 0 | 0 | 6 |

| | | | | | | | | | | | | | | | | | |
|---|---|---|---|---|---|---|---|---|---|---|---|---|---|---|---|---|---|
| 1 | 0.00 | 3.68 | 0.00 | 1.00 | 1 | Utf1 | undifferentiated embryonic cell transcripti | 3 | 0 | 6 | 0 | 0 | 0 | 0 | 0 | 0 | 0 | 0 |
| 1 | 1.40 | 3.67 | 0.38 | 0.70 | 1 | Lmod3 | leiomodin 3 (fetal) | 3 | 5 | 0 | 0 | 0 | 0 | 0 | 0 | 2 | 0 | 0 |
| 1 | -1.81 | 3.60 | -0.50 | 0.62 | 1 | Gm12868 | | 0 | 0 | 0 | 5 | 0 | 3 | 0 | 0 | 0 | 0 | 1 |
| 1 | 0.61 | 3.61 | 0.17 | 0.87 | 1 | Trim43a | tripartite motif-containing 43A | 0 | 0 | 0 | 0 | 0 | 3 | 0 | 0 | 0 | 0 | 7 |
| 1 | 0.61 | 3.61 | 0.17 | 0.87 | 1 | Gm19056 | | 0 | 0 | 0 | 0 | 0 | 3 | 0 | 0 | 0 | 0 | 7 |
| 1 | 0.61 | 3.61 | 0.17 | 0.87 | 1 | Gm20683 | | 0 | 0 | 0 | 0 | 0 | 3 | 0 | 0 | 0 | 0 | 7 |
| 1 | -0.28 | 3.61 | -0.08 | 0.94 | 1 | Gm26049 | | 0 | 0 | 0 | 0 | 0 | 8 | 0 | 0 | 0 | 0 | 2 |
| 1 | -0.64 | 3.57 | -0.18 | 0.86 | 1 | Gm43965 | | 0 | 0 | 0 | 0 | 5 | 1 | 0 | 0 | 0 | 0 | 4 |
| 1 | 1.65 | 3.59 | 0.46 | 0.64 | 1 | Gm15941 | | 0 | 0 | 0 | 0 | 0 | 3 | 0 | 0 | 4 | 0 | 3 |
| 1 | -1.33 | 3.62 | -0.37 | 0.71 | 1 | Ugt1a2 | UDP glucuronosyltransferase 1 family, pc | 0 | 0 | 0 | 10 | 0 | 0 | 0 | 0 | 0 | 0 | 0 |
| 1 | -1.33 | 3.62 | -0.37 | 0.71 | 1 | Gm20652 | | 0 | 0 | 0 | 10 | 0 | 0 | 0 | 0 | 0 | 0 | 0 |
| 1 | -1.33 | 3.62 | -0.37 | 0.71 | 1 | Mir6400 | | 0 | 0 | 0 | 10 | 0 | 0 | 0 | 0 | 0 | 0 | 0 |
| 1 | -1.33 | 3.62 | -0.37 | 0.71 | 1 | Gm44605 | | 0 | 0 | 0 | 10 | 0 | 0 | 0 | 0 | 0 | 0 | 0 |
| 1 | -1.33 | 3.62 | -0.37 | 0.71 | 1 | Gm8281 | | 0 | 0 | 0 | 10 | 0 | 0 | 0 | 0 | 0 | 0 | 0 |
| 1 | -1.05 | 3.63 | -0.29 | 0.77 | 1 | Gm29488 | | 0 | 7 | 0 | 0 | 0 | 3 | 0 | 0 | 0 | 0 | 0 |
| 1 | -0.99 | 3.62 | -0.27 | 0.78 | 1 | Gm48872 | | 0 | 0 | 6 | 0 | 0 | 3 | 0 | 0 | 0 | 0 | 0 |
| 1 | -0.51 | 3.67 | -0.14 | 0.89 | 1 | F9 | coagulation factor IX | 8 | 1 | 0 | 1 | 0 | 0 | 0 | 0 | 0 | 0 | 0 |
| 1 | 0.00 | 3.68 | 0.00 | 1.00 | 1 | Gm49288 | | 9 | 1 | 0 | 0 | 0 | 0 | 0 | 0 | 0 | 0 | 0 |
| 1 | -0.76 | 3.57 | -0.21 | 0.83 | 1 | Gm26165 | | 0 | 0 | 0 | 0 | 6 | 1 | 0 | 0 | 0 | 0 | 3 |
| 1 | 0.36 | 3.57 | 0.10 | 0.92 | 1 | Gm5147 | | 0 | 0 | 0 | 0 | 1 | 3 | 0 | 0 | 6 | 0 | 0 |
| 1 | 1.32 | 3.60 | 0.37 | 0.71 | 1 | Gm48849 | | 0 | 0 | 5 | 1 | 0 | 0 | 0 | 1 | 1 | 1 | 0 |
| 1 | 1.43 | 3.61 | 0.40 | 0.69 | 1 | Gm17191 | | 3 | 0 | 0 | 0 | 0 | 0 | 0 | 0 | 0 | 0 | 6 |
| 1 | -0.64 | 3.66 | -0.18 | 0.86 | 1 | Gm14044 | | 9 | 0 | 0 | 0 | 1 | 0 | 0 | 0 | 0 | 0 | 0 |
| 1 | -3.51 | 3.56 | -0.99 | 0.32 | 1 | Gm10253 | | 0 | 0 | 0 | 2 | 0 | 2 | 0 | 0 | 6 | 0 | 0 |
| 1 | 0.55 | 3.61 | 0.15 | 0.88 | 1 | Snord37 | | 1 | 0 | 0 | 0 | 0 | 3 | 0 | 0 | 0 | 0 | 6 |
| 1 | 0.00 | 3.68 | 0.00 | 1.00 | 1 | Gm44000 | | 4 | 0 | 5 | 0 | 0 | 0 | 0 | 0 | 0 | 0 | 0 |
| 1 | -0.69 | 3.66 | -0.19 | 0.85 | 1 | Hesx1 | homeobox gene expressed in ES cells | 2 | 6 | 0 | 0 | 0 | 2 | 0 | 0 | 0 | 0 | 0 |
| 1 | 0.31 | 3.55 | 0.09 | 0.93 | 1 | Aldoart2 | aldolase 1 A, retrogene 2 | 0 | 2 | 0 | 1 | 0 | 3 | 0 | 0 | 4 | 0 | 0 |
| 1 | 0.44 | 3.61 | 0.12 | 0.90 | 1 | Gm42907 | | 0 | 0 | 0 | 0 | 0 | 3 | 0 | 0 | 0 | 0 | 6 |
| 1 | 0.44 | 3.61 | 0.12 | 0.90 | 1 | Gm24888 | | 0 | 0 | 0 | 0 | 0 | 3 | 0 | 0 | 0 | 0 | 6 |
| 1 | -0.50 | 3.61 | -0.14 | 0.89 | 1 | Mir343 | | 0 | 0 | 0 | 0 | 0 | 8 | 0 | 0 | 0 | 0 | 1 |
| 1 | 1.42 | 3.59 | 0.40 | 0.69 | 1 | Gm47857 | | 0 | 0 | 0 | 0 | 0 | 3 | 0 | 0 | 3 | 0 | 3 |
| 1 | 1.42 | 3.59 | 0.40 | 0.69 | 1 | Gm48302 | | 0 | 0 | 0 | 0 | 0 | 3 | 0 | 0 | 3 | 0 | 3 |
| 1 | 0.33 | 3.56 | 0.09 | 0.93 | 1 | Gm48856 | | 0 | 0 | 0 | 4 | 0 | 0 | 0 | 0 | 0 | 0 | 5 |
| 1 | -1.24 | 3.62 | -0.34 | 0.73 | 1 | B3gnt6 | UDP-GlcNAc:betaGal beta-1,3-N-acetylgl | 0 | 0 | 0 | 0 | 9 | 0 | 0 | 0 | 0 | 0 | 0 |
| 1 | -1.24 | 3.62 | -0.34 | 0.73 | 1 | Gm47169 | | 0 | 0 | 0 | 0 | 9 | 0 | 0 | 0 | 0 | 0 | 0 |
| 1 | -1.24 | 3.62 | -0.34 | 0.73 | 1 | D030024E09Rik | | 0 | 0 | 0 | 0 | 9 | 0 | 0 | 0 | 0 | 0 | 0 |
| 1 | -1.24 | 3.62 | -0.34 | 0.73 | 1 | Opn1mw | opsin 1 (cone pigments), medium-wave-s | 0 | 0 | 0 | 0 | 9 | 0 | 0 | 0 | 0 | 0 | 0 |
| 1 | -1.81 | 3.61 | -0.50 | 0.62 | 1 | Cdh10 | cadherin 10 | 0 | 0 | 2 | 0 | 6 | 2 | 0 | 0 | 0 | 0 | 0 |
| 1 | 0.00 | 3.67 | 0.00 | 1.00 | 1 | Gm28884 | | 9 | 0 | 0 | 0 | 0 | 0 | 0 | 0 | 0 | 0 | 0 |
| 1 | 0.00 | 3.67 | 0.00 | 1.00 | 1 | Vis1 | | 9 | 0 | 0 | 0 | 0 | 0 | 0 | 0 | 0 | 0 | 0 |
| 1 | 0.00 | 3.67 | 0.00 | 1.00 | 1 | Ankub1 | ankrin repeat and ubiquitin domain conta | 9 | 0 | 0 | 0 | 0 | 0 | 0 | 0 | 0 | 0 | 0 |
| 1 | 0.00 | 3.67 | 0.00 | 1.00 | 1 | Bach2os | | 9 | 0 | 0 | 0 | 0 | 0 | 0 | 0 | 0 | 0 | 0 |
| 1 | 0.00 | 3.67 | 0.00 | 1.00 | 1 | Mir5120 | | 9 | 0 | 0 | 0 | 0 | 0 | 0 | 0 | 0 | 0 | 0 |
| 1 | 0.00 | 3.67 | 0.00 | 1.00 | 1 | Gm10582 | | 9 | 0 | 0 | 0 | 0 | 0 | 0 | 0 | 0 | 0 | 0 |
| 1 | 0.00 | 3.67 | 0.00 | 1.00 | 1 | 4930407G08Rik | | 9 | 0 | 0 | 0 | 0 | 0 | 0 | 0 | 0 | 0 | 0 |
| 1 | 0.00 | 3.67 | 0.00 | 1.00 | 1 | Gm8879 | | 9 | 0 | 0 | 0 | 0 | 0 | 0 | 0 | 0 | 0 | 0 |
| 1 | 0.00 | 3.67 | 0.00 | 1.00 | 1 | Gm26953 | | 9 | 0 | 0 | 0 | 0 | 0 | 0 | 0 | 0 | 0 | 0 |
| 1 | 0.00 | 3.67 | 0.00 | 1.00 | 1 | Doxl1 | diamine oxidase-like protein 1 | 9 | 0 | 0 | 0 | 0 | 0 | 0 | 0 | 0 | 0 | 0 |
| 1 | 0.00 | 3.67 | 0.00 | 1.00 | 1 | Igkv9-119 | | 9 | 0 | 0 | 0 | 0 | 0 | 0 | 0 | 0 | 0 | 0 |
| 1 | 0.00 | 3.67 | 0.00 | 1.00 | 1 | M1ap | meiosis 1 associated protein | 9 | 0 | 0 | 0 | 0 | 0 | 0 | 0 | 0 | 0 | 0 |
| 1 | 0.00 | 3.67 | 0.00 | 1.00 | 1 | Vmn2r26 | vomeronasal 2, receptor 26 | 9 | 0 | 0 | 0 | 0 | 0 | 0 | 0 | 0 | 0 | 0 |
| 1 | 0.00 | 3.67 | 0.00 | 1.00 | 1 | Gm44135 | | 9 | 0 | 0 | 0 | 0 | 0 | 0 | 0 | 0 | 0 | 0 |
| 1 | 0.00 | 3.67 | 0.00 | 1.00 | 1 | Gm45778 | | 9 | 0 | 0 | 0 | 0 | 0 | 0 | 0 | 0 | 0 | 0 |
| 1 | 0.00 | 3.67 | 0.00 | 1.00 | 1 | Mir7073 | | 9 | 0 | 0 | 0 | 0 | 0 | 0 | 0 | 0 | 0 | 0 |
| 1 | 0.00 | 3.67 | 0.00 | 1.00 | 1 | Gm48814 | | 9 | 0 | 0 | 0 | 0 | 0 | 0 | 0 | 0 | 0 | 0 |
| 1 | 0.00 | 3.67 | 0.00 | 1.00 | 1 | Gm23323 | | 9 | 0 | 0 | 0 | 0 | 0 | 0 | 0 | 0 | 0 | 0 |
| 1 | 0.00 | 3.67 | 0.00 | 1.00 | 1 | 1700120B22Rik | | 9 | 0 | 0 | 0 | 0 | 0 | 0 | 0 | 0 | 0 | 0 |
| 1 | 0.00 | 3.67 | 0.00 | 1.00 | 1 | Ttll6 | tubulin tyrosine ligase-like family, membe | 9 | 0 | 0 | 0 | 0 | 0 | 0 | 0 | 0 | 0 | 0 |
| 1 | 0.00 | 3.67 | 0.00 | 1.00 | 1 | Gm33680 | | 9 | 0 | 0 | 0 | 0 | 0 | 0 | 0 | 0 | 0 | 0 |

| | | | | | | | | | | | | | | | | | |
|---|---|---|---|---|---|---|---|---|---|---|---|---|---|---|---|---|---|
| 1 | 0.00 | 3.67 | 0.00 | 1.00 | 1 | Gm48746 | | 9 | 0 | 0 | 0 | 0 | 0 | 0 | 0 | 0 | 0 | 0 |
| 1 | 0.00 | 3.67 | 0.00 | 1.00 | 1 | Gm47709 | | 9 | 0 | 0 | 0 | 0 | 0 | 0 | 0 | 0 | 0 | 0 |
| 1 | 0.00 | 3.67 | 0.00 | 1.00 | 1 | Gm8221 | | 9 | 0 | 0 | 0 | 0 | 0 | 0 | 0 | 0 | 0 | 0 |
| 1 | 0.00 | 3.67 | 0.00 | 1.00 | 1 | Pvalb | parvalbumin | 9 | 0 | 0 | 0 | 0 | 0 | 0 | 0 | 0 | 0 | 0 |
| 1 | 0.00 | 3.67 | 0.00 | 1.00 | 1 | 1700097N02Rik | | 9 | 0 | 0 | 0 | 0 | 0 | 0 | 0 | 0 | 0 | 0 |
| 1 | 0.00 | 3.67 | 0.00 | 1.00 | 1 | H2-M11 | histocompatibility 2, M region locus 11 | 9 | 0 | 0 | 0 | 0 | 0 | 0 | 0 | 0 | 0 | 0 |
| 1 | 0.00 | 3.67 | 0.00 | 1.00 | 1 | Themis3 | thymocyte selection associated family me | 9 | 0 | 0 | 0 | 0 | 0 | 0 | 0 | 0 | 0 | 0 |
| 1 | 0.00 | 3.67 | 0.00 | 1.00 | 1 | 1700065O20Rik | | 9 | 0 | 0 | 0 | 0 | 0 | 0 | 0 | 0 | 0 | 0 |
| 1 | -2.02 | 3.61 | -0.56 | 0.57 | 1 | Slc47a1 | solute carrier family 47, member 1 | 3 | 0 | 0 | 0 | 1 | 6 | 0 | 0 | 0 | 0 | 0 |
| 1 | -0.90 | 3.64 | -0.25 | 0.80 | 1 | Gm12709 | | 8 | 0 | 0 | 0 | 0 | 2 | 0 | 0 | 0 | 0 | 0 |
| 1 | -0.90 | 3.64 | -0.25 | 0.80 | 1 | Gm41517 | | 8 | 0 | 0 | 0 | 0 | 2 | 0 | 0 | 0 | 0 | 0 |
| 1 | -0.98 | 3.63 | -0.27 | 0.79 | 1 | Gm43230 | | 7 | 0 | 0 | 0 | 0 | 3 | 0 | 0 | 0 | 0 | 0 |
| 1 | 1.02 | 3.66 | 0.28 | 0.78 | 1 | Gm8174 | | 0 | 1 | 5 | 1 | 0 | 0 | 0 | 0 | 0 | 1 | 1 |
| 1 | 0.00 | 3.68 | 0.00 | 1.00 | 1 | Gdnf | glial cell line derived neurotrophic factor | 4 | 4 | 1 | 0 | 0 | 0 | 0 | 0 | 0 | 0 | 0 |
| 1 | -0.08 | 3.61 | -0.02 | 0.98 | 1 | Gm43795 | | 0 | 0 | 3 | 0 | 0 | 4 | 0 | 0 | 0 | 0 | 2 |
| 1 | 1.00 | 3.68 | 0.27 | 0.79 | 1 | Gm5321 | | 1 | 2 | 5 | 0 | 0 | 0 | 0 | 0 | 0 | 1 | 0 |
| 1 | 0.06 | 3.58 | 0.02 | 0.99 | 1 | Gm45162 | | 2 | 0 | 0 | 0 | 2 | 2 | 0 | 0 | 1 | 0 | 3 |
| 1 | 0.00 | 3.66 | 0.00 | 1.00 | 1 | Gm29100 | | 0 | 0 | 9 | 0 | 0 | 0 | 0 | 0 | 0 | 0 | 0 |
| 1 | 0.00 | 3.66 | 0.00 | 1.00 | 1 | Car1 | carbonic anhydrase 1 | 0 | 0 | 9 | 0 | 0 | 0 | 0 | 0 | 0 | 0 | 0 |
| 1 | 0.00 | 3.66 | 0.00 | 1.00 | 1 | Gm9273 | | 0 | 0 | 9 | 0 | 0 | 0 | 0 | 0 | 0 | 0 | 0 |
| 1 | 0.00 | 3.66 | 0.00 | 1.00 | 1 | Gm42776 | | 0 | 0 | 9 | 0 | 0 | 0 | 0 | 0 | 0 | 0 | 0 |
| 1 | 0.00 | 3.66 | 0.00 | 1.00 | 1 | Lao1 | L-amino acid oxidase 1 | 0 | 0 | 9 | 0 | 0 | 0 | 0 | 0 | 0 | 0 | 0 |
| 1 | 0.00 | 3.66 | 0.00 | 1.00 | 1 | Gm9954 | | 0 | 0 | 9 | 0 | 0 | 0 | 0 | 0 | 0 | 0 | 0 |
| 1 | 0.00 | 3.66 | 0.00 | 1.00 | 1 | Zbed4-ps3 | | 0 | 0 | 9 | 0 | 0 | 0 | 0 | 0 | 0 | 0 | 0 |
| 1 | 0.00 | 3.66 | 0.00 | 1.00 | 1 | Olfr539 | olfactory receptor 539 | 0 | 0 | 9 | 0 | 0 | 0 | 0 | 0 | 0 | 0 | 0 |
| 1 | 0.00 | 3.66 | 0.00 | 1.00 | 1 | Gm39128 | | 0 | 0 | 9 | 0 | 0 | 0 | 0 | 0 | 0 | 0 | 0 |
| 1 | 0.00 | 3.66 | 0.00 | 1.00 | 1 | Gm47697 | | 0 | 0 | 9 | 0 | 0 | 0 | 0 | 0 | 0 | 0 | 0 |
| 1 | 0.00 | 3.66 | 0.00 | 1.00 | 1 | Gm48610 | | 0 | 0 | 9 | 0 | 0 | 0 | 0 | 0 | 0 | 0 | 0 |
| 1 | 0.00 | 3.66 | 0.00 | 1.00 | 1 | Gm46323 | | 0 | 0 | 9 | 0 | 0 | 0 | 0 | 0 | 0 | 0 | 0 |
| 1 | 0.00 | 3.66 | 0.00 | 1.00 | 1 | Gm35240 | | 0 | 0 | 9 | 0 | 0 | 0 | 0 | 0 | 0 | 0 | 0 |
| 1 | 0.00 | 3.66 | 0.00 | 1.00 | 1 | Gm32834 | | 0 | 0 | 9 | 0 | 0 | 0 | 0 | 0 | 0 | 0 | 0 |
| 1 | 0.00 | 3.66 | 0.00 | 1.00 | 1 | Gm49663 | | 0 | 0 | 9 | 0 | 0 | 0 | 0 | 0 | 0 | 0 | 0 |
| 1 | 0.00 | 3.66 | 0.00 | 1.00 | 1 | Gm5145 | | 0 | 0 | 9 | 0 | 0 | 0 | 0 | 0 | 0 | 0 | 0 |
| 1 | 0.00 | 3.66 | 0.00 | 1.00 | 1 | Glyat | glycine-N-acyltransferase | 0 | 0 | 9 | 0 | 0 | 0 | 0 | 0 | 0 | 0 | 0 |
| 1 | 0.00 | 3.66 | 0.00 | 1.00 | 1 | A630012P03Rik | | 0 | 0 | 9 | 0 | 0 | 0 | 0 | 0 | 0 | 0 | 0 |
| 1 | 0.00 | 3.66 | 0.00 | 1.00 | 1 | Gm2174 | | 0 | 0 | 9 | 0 | 0 | 0 | 0 | 0 | 0 | 0 | 0 |
| 1 | -2.14 | 3.61 | -0.59 | 0.55 | 1 | Gm48996 | | 0 | 0 | 0 | 0 | 9 | 1 | 0 | 0 | 0 | 0 | 0 |
| 1 | -1.97 | 3.61 | -0.55 | 0.58 | 1 | Il2 | interleukin 2 | 0 | 0 | 1 | 0 | 7 | 2 | 0 | 0 | 0 | 0 | 0 |
| 1 | 0.28 | 3.61 | 0.08 | 0.94 | 1 | Gm23690 | | 0 | 0 | 0 | 0 | 0 | 4 | 0 | 0 | 0 | 0 | 5 |
| 1 | 0.28 | 3.61 | 0.08 | 0.94 | 1 | Gm4714 | | 0 | 0 | 0 | 0 | 0 | 4 | 0 | 0 | 0 | 0 | 5 |
| 1 | 0.28 | 3.61 | 0.08 | 0.94 | 1 | Gm12208 | | 0 | 0 | 0 | 0 | 0 | 4 | 0 | 0 | 0 | 0 | 5 |
| 1 | 0.28 | 3.61 | 0.08 | 0.94 | 1 | Gm15186 | | 0 | 0 | 0 | 0 | 0 | 4 | 0 | 0 | 0 | 0 | 5 |
| 1 | 0.00 | 3.68 | 0.00 | 1.00 | 1 | Gm44855 | | 0 | 2 | 7 | 0 | 0 | 0 | 0 | 0 | 0 | 0 | 0 |
| 1 | -0.82 | 3.61 | -0.23 | 0.82 | 1 | Cntn2 | contactin 2 | 0 | 0 | 0 | 0 | 0 | 9 | 0 | 0 | 0 | 0 | 0 |
| 1 | -0.82 | 3.61 | -0.23 | 0.82 | 1 | Gm47087 | | 0 | 0 | 0 | 0 | 0 | 9 | 0 | 0 | 0 | 0 | 0 |
| 1 | -1.05 | 3.61 | -0.29 | 0.77 | 1 | Gm42797 | | 0 | 5 | 0 | 0 | 0 | 4 | 0 | 0 | 0 | 0 | 0 |
| 1 | -1.24 | 3.62 | -0.34 | 0.73 | 1 | Gm12929 | | 3 | 0 | 1 | 0 | 6 | 0 | 0 | 0 | 0 | 0 | 0 |
| 1 | 1.79 | 3.63 | 0.49 | 0.62 | 1 | Gm4810 | | 0 | 0 | 0 | 0 | 0 | 0 | 0 | 9 | 0 | 0 | 0 |
| 1 | -2.05 | 3.61 | -0.57 | 0.57 | 1 | Cntnap3 | contactin associated protein-like 3 | 0 | 0 | 4 | 1 | 0 | 4 | 0 | 0 | 0 | 0 | 0 |
| 1 | 1.62 | 3.62 | 0.45 | 0.65 | 1 | 4931403E22Rik | | 0 | 0 | 5 | 0 | 0 | 0 | 0 | 0 | 0 | 0 | 4 |
| 1 | -0.71 | 3.58 | -0.20 | 0.84 | 1 | Gm8648 | | 0 | 2 | 0 | 1 | 1 | 3 | 0 | 0 | 1 | 0 | 1 |
| 1 | 0.00 | 3.68 | 0.00 | 1.00 | 1 | Hoxa9 | homeobox A9 | 5 | 2 | 2 | 0 | 0 | 0 | 0 | 0 | 0 | 0 | 0 |
| 1 | -2.53 | 3.60 | -0.70 | 0.48 | 1 | Gm9712 | | 0 | 0 | 0 | 0 | 8 | 2 | 0 | 0 | 0 | 0 | 0 |
| 1 | 0.00 | 3.68 | 0.00 | 1.00 | 1 | Gm13344 | | 0 | 4 | 5 | 0 | 0 | 0 | 0 | 0 | 0 | 0 | 0 |
| 1 | 0.00 | 3.68 | 0.00 | 1.00 | 1 | D330050G23Rik | | 0 | 4 | 5 | 0 | 0 | 0 | 0 | 0 | 0 | 0 | 0 |
| 1 | 0.00 | 3.68 | 0.00 | 1.00 | 1 | C730027H18Rik | | 0 | 4 | 5 | 0 | 0 | 0 | 0 | 0 | 0 | 0 | 0 |
| 1 | 1.65 | 3.55 | 0.46 | 0.64 | 1 | Cbx3-ps7 | | 0 | 0 | 0 | 2 | 0 | 1 | 0 | 2 | 1 | 1 | 2 |
| 1 | 2.04 | 3.60 | 0.57 | 0.57 | 1 | Gm7565 | | 1 | 3 | 0 | 1 | 0 | 0 | 0 | 1 | 2 | 1 | 0 |
| 1 | -0.95 | 3.61 | -0.26 | 0.79 | 1 | Gm37432 | | 0 | 0 | 4 | 0 | 0 | 5 | 0 | 0 | 0 | 0 | 0 |
| 1 | 1.92 | 3.62 | 0.53 | 0.60 | 1 | Gm47885 | | 3 | 0 | 1 | 0 | 0 | 1 | 0 | 1 | 0 | 3 | 1 |

| | | | | | | | | | | | | | | | | | | |
|---|---|---|---|---|---|---|---|---|---|---|---|---|---|---|---|---|---|---|
| 1 | 1.71 | 3.58 | 0.48 | 0.63 | 1 | Gm6937 | | 0 | 4 | 0 | 0 | 0 | 2 | 0 | 1 | 1 | 1 | 0 |
| 1 | 0.00 | 3.68 | 0.00 | 1.00 | 1 | Gm37712 | | 5 | 4 | 0 | 0 | 0 | 0 | 0 | 0 | 0 | 0 | 0 |
| 1 | 0.00 | 3.68 | 0.00 | 1.00 | 1 | Rhox5 | reproductive homeobox 5 | 1 | 0 | 8 | 0 | 0 | 0 | 0 | 0 | 0 | 0 | 0 |
| 1 | 2.88 | 3.60 | 0.80 | 0.42 | 1 | Gm26446 | | 3 | 0 | 0 | 0 | 0 | 2 | 0 | 0 | 4 | 0 | 0 |
| 1 | -0.19 | 3.62 | -0.05 | 0.96 | 1 | Gm12732 | | 0 | 0 | 7 | 0 | 2 | 0 | 0 | 0 | 0 | 0 | 0 |
| 1 | -0.53 | 3.66 | -0.14 | 0.89 | 1 | Gm5977 | | 0 | 0 | 8 | 0 | 0 | 1 | 0 | 0 | 0 | 0 | 0 |
| 1 | 2.85 | 3.59 | 0.79 | 0.43 | 1 | Gm5234 | | 3 | 0 | 0 | 0 | 0 | 3 | 0 | 0 | 4 | 0 | 0 |
| 1 | 0.35 | 3.57 | 0.10 | 0.92 | 1 | Gm13363 | | 0 | 0 | 1 | 0 | 0 | 5 | 0 | 0 | 3 | 0 | 0 |
| 1 | 2.67 | 3.62 | 0.74 | 0.46 | 1 | 6720475M21Rik | | 0 | 0 | 1 | 0 | 0 | 0 | 0 | 0 | 5 | 0 | 3 |
| 1 | 1.37 | 3.61 | 0.38 | 0.70 | 1 | Gm20005 | | 0 | 0 | 0 | 0 | 0 | 0 | 0 | 0 | 0 | 0 | 9 |
| 1 | 1.37 | 3.61 | 0.38 | 0.70 | 1 | Gm24412 | | 0 | 0 | 0 | 0 | 0 | 0 | 0 | 0 | 0 | 0 | 9 |
| 1 | 1.37 | 3.61 | 0.38 | 0.70 | 1 | Gm44549 | | 0 | 0 | 0 | 0 | 0 | 0 | 0 | 0 | 0 | 0 | 9 |
| 1 | 1.37 | 3.61 | 0.38 | 0.70 | 1 | Gm36235 | | 0 | 0 | 0 | 0 | 0 | 0 | 0 | 0 | 0 | 0 | 9 |
| 1 | 1.37 | 3.61 | 0.38 | 0.70 | 1 | Gm47254 | | 0 | 0 | 0 | 0 | 0 | 0 | 0 | 0 | 0 | 0 | 9 |
| 1 | 0.12 | 3.61 | 0.03 | 0.97 | 1 | Proz | protein Z, vitamin K-dependent plasma gl | 0 | 0 | 0 | 0 | 0 | 5 | 0 | 0 | 0 | 0 | 4 |
| 1 | 0.12 | 3.61 | 0.03 | 0.97 | 1 | Gm20694 | | 0 | 0 | 0 | 0 | 0 | 5 | 0 | 0 | 0 | 0 | 4 |
| 1 | 0.12 | 3.61 | 0.03 | 0.97 | 1 | Gm19967 | | 0 | 0 | 0 | 0 | 0 | 5 | 0 | 0 | 0 | 0 | 4 |
| 1 | 0.12 | 3.61 | 0.03 | 0.97 | 1 | 4930447C04Ril | RIKEN cDNA 4930447C04 gene | 0 | 0 | 0 | 0 | 0 | 5 | 0 | 0 | 0 | 0 | 4 |
| 1 | 0.12 | 3.61 | 0.03 | 0.97 | 1 | Gm15712 | | 0 | 0 | 0 | 0 | 0 | 5 | 0 | 0 | 0 | 0 | 4 |
| 1 | 1.52 | 3.63 | 0.42 | 0.68 | 1 | 6330410L21Rik | | 0 | 6 | 0 | 0 | 0 | 0 | 0 | 0 | 0 | 0 | 3 |
| 1 | -1.01 | 3.61 | -0.28 | 0.78 | 1 | Eif2s3x-ps1 | | 0 | 4 | 0 | 0 | 0 | 5 | 0 | 0 | 0 | 0 | 0 |
| 1 | 0.00 | 3.66 | 0.00 | 1.00 | 1 | Gm15851 | | 0 | 9 | 0 | 0 | 0 | 0 | 0 | 0 | 0 | 0 | 0 |
| 1 | 0.00 | 3.66 | 0.00 | 1.00 | 1 | 3830422I06Rik | | 0 | 9 | 0 | 0 | 0 | 0 | 0 | 0 | 0 | 0 | 0 |
| 1 | 0.00 | 3.66 | 0.00 | 1.00 | 1 | Gm43351 | | 0 | 9 | 0 | 0 | 0 | 0 | 0 | 0 | 0 | 0 | 0 |
| 1 | 0.00 | 3.66 | 0.00 | 1.00 | 1 | G630022F23Rik | | 0 | 9 | 0 | 0 | 0 | 0 | 0 | 0 | 0 | 0 | 0 |
| 1 | 0.00 | 3.66 | 0.00 | 1.00 | 1 | Trbv19 | | 0 | 9 | 0 | 0 | 0 | 0 | 0 | 0 | 0 | 0 | 0 |
| 1 | 0.00 | 3.66 | 0.00 | 1.00 | 1 | Gm32389 | | 0 | 9 | 0 | 0 | 0 | 0 | 0 | 0 | 0 | 0 | 0 |
| 1 | 0.00 | 3.66 | 0.00 | 1.00 | 1 | Il5 | interleukin 5 | 0 | 9 | 0 | 0 | 0 | 0 | 0 | 0 | 0 | 0 | 0 |
| 1 | 0.00 | 3.66 | 0.00 | 1.00 | 1 | Cacng1 | calcium channel, voltage-dependent, gan | 0 | 9 | 0 | 0 | 0 | 0 | 0 | 0 | 0 | 0 | 0 |
| 1 | 0.00 | 3.66 | 0.00 | 1.00 | 1 | Trav4-2 | | 0 | 9 | 0 | 0 | 0 | 0 | 0 | 0 | 0 | 0 | 0 |
| 1 | 0.00 | 3.66 | 0.00 | 1.00 | 1 | 4930556J02Rik | | 0 | 9 | 0 | 0 | 0 | 0 | 0 | 0 | 0 | 0 | 0 |
| 1 | 0.00 | 3.66 | 0.00 | 1.00 | 1 | Gm10865 | | 0 | 9 | 0 | 0 | 0 | 0 | 0 | 0 | 0 | 0 | 0 |
| 1 | 0.00 | 3.66 | 0.00 | 1.00 | 1 | Gm50037 | | 0 | 9 | 0 | 0 | 0 | 0 | 0 | 0 | 0 | 0 | 0 |
| 1 | 2.77 | 3.61 | 0.77 | 0.44 | 1 | Gm12811 | | 0 | 1 | 0 | 0 | 0 | 0 | 0 | 0 | 7 | 0 | 1 |
| 1 | 1.87 | 3.63 | 0.52 | 0.61 | 1 | Gm10444 | | 0 | 0 | 0 | 0 | 0 | 0 | 0 | 0 | 9 | 0 | 0 |
| 1 | -0.46 | 3.56 | -0.13 | 0.90 | 1 | Gm8492 | | 2 | 0 | 0 | 1 | 2 | 0 | 0 | 0 | 0 | 3 | 2 |
| 1 | -0.73 | 3.64 | -0.20 | 0.84 | 1 | F730035M05Rik | | 3 | 0 | 3 | 0 | 0 | 3 | 0 | 0 | 0 | 0 | 0 |
| 1 | -2.33 | 3.57 | -0.65 | 0.51 | 1 | Gm47586 | | 0 | 0 | 0 | 2 | 4 | 3 | 0 | 0 | 0 | 0 | 1 |
| 1 | 1.58 | 3.58 | 0.44 | 0.66 | 1 | Gpx2-ps1 | | 0 | 0 | 1 | 0 | 0 | 2 | 0 | 0 | 5 | 1 | 0 |
| 1 | 0.44 | 3.62 | 0.12 | 0.90 | 1 | Gm2895 | | 4 | 0 | 0 | 0 | 0 | 2 | 0 | 0 | 0 | 0 | 3 |
| 1 | -1.50 | 3.59 | -0.42 | 0.68 | 1 | Gm11491 | | 0 | 0 | 0 | 0 | 4 | 4 | 0 | 0 | 1 | 0 | 0 |
| 1 | 1.80 | 3.63 | 0.50 | 0.62 | 1 | Gm11099 | | 0 | 0 | 0 | 0 | 0 | 0 | 0 | 0 | 0 | 9 | 0 |
| 1 | 1.80 | 3.63 | 0.50 | 0.62 | 1 | Gm15462 | | 0 | 0 | 0 | 0 | 0 | 0 | 0 | 0 | 0 | 9 | 0 |
| 1 | 1.80 | 3.63 | 0.50 | 0.62 | 1 | Gja10 | gap junction protein, alpha 10 | 0 | 0 | 0 | 0 | 0 | 0 | 0 | 0 | 0 | 9 | 0 |
| 1 | 1.80 | 3.63 | 0.50 | 0.62 | 1 | Cyp3a11 | cytochrome P450, family 3, subfamily a, p | 0 | 0 | 0 | 0 | 0 | 0 | 0 | 0 | 0 | 9 | 0 |
| 1 | 1.80 | 3.63 | 0.50 | 0.62 | 1 | Gm6852 | | 0 | 0 | 0 | 0 | 0 | 0 | 0 | 0 | 0 | 9 | 0 |
| 1 | 1.80 | 3.63 | 0.50 | 0.62 | 1 | Gm50316 | | 0 | 0 | 0 | 0 | 0 | 0 | 0 | 0 | 0 | 9 | 0 |
| 1 | 2.41 | 3.63 | 0.67 | 0.51 | 1 | Gm7591 | | 1 | 0 | 2 | 0 | 0 | 0 | 0 | 0 | 4 | 0 | 2 |
| 1 | 2.08 | 3.64 | 0.57 | 0.57 | 1 | Gm20900 | | 1 | 2 | 2 | 0 | 0 | 0 | 0 | 0 | 3 | 0 | 1 |
| 1 | -2.60 | 3.60 | -0.72 | 0.47 | 1 | Gm23639 | | 2 | 0 | 0 | 0 | 5 | 3 | 0 | 0 | 0 | 0 | 0 |
| 1 | -0.43 | 3.62 | -0.12 | 0.91 | 1 | Gm18076 | | 0 | 0 | 5 | 0 | 4 | 0 | 0 | 0 | 0 | 0 | 0 |
| 1 | 0.89 | 3.61 | 0.25 | 0.81 | 1 | H2-Q3 | | 0 | 0 | 0 | 0 | 2 | 0 | 0 | 0 | 7 | 0 | 0 |
| 1 | 1.01 | 3.61 | 0.28 | 0.78 | 1 | Gm36546 | | 0 | 0 | 0 | 0 | 0 | 1 | 0 | 0 | 0 | 0 | 8 |
| 1 | 1.01 | 3.61 | 0.28 | 0.78 | 1 | Gm47833 | | 0 | 0 | 0 | 0 | 0 | 1 | 0 | 0 | 0 | 0 | 8 |
| 1 | -1.16 | 3.65 | -0.32 | 0.75 | 1 | Gm7543 | | 0 | 1 | 6 | 1 | 0 | 1 | 0 | 0 | 0 | 0 | 0 |
| 1 | -0.05 | 3.61 | -0.02 | 0.99 | 1 | Gm44437 | | 0 | 0 | 0 | 0 | 0 | 6 | 0 | 0 | 0 | 0 | 3 |
| 1 | -0.05 | 3.61 | -0.02 | 0.99 | 1 | Gm49972 | | 0 | 0 | 0 | 0 | 0 | 6 | 0 | 0 | 0 | 0 | 3 |
| 1 | 0.00 | 3.68 | 0.00 | 1.00 | 1 | 6430710C18Rik | | 1 | 8 | 0 | 0 | 0 | 0 | 0 | 0 | 0 | 0 | 0 |
| 1 | 0.00 | 3.68 | 0.00 | 1.00 | 1 | Ighv1-85 | | 1 | 8 | 0 | 0 | 0 | 0 | 0 | 0 | 0 | 0 | 0 |
| 1 | 2.14 | 3.60 | 0.59 | 0.55 | 1 | 1700034P13Rik | | 0 | 0 | 0 | 0 | 0 | 1 | 0 | 0 | 4 | 0 | 4 |

| | | | | | | | | | | | | | | | | | | |
|---|---|---|---|---|---|---|---|---|---|---|---|---|---|---|---|---|---|---|
| 1 | -0.58 | 3.65 | -0.16 | 0.87 | 1 | Tubal3 | tubulin, alpha-like 3 | 0 | 8 | 0 | 0 | 0 | 1 | 0 | 0 | 0 | 0 | 0 |
| 1 | -0.58 | 3.65 | -0.16 | 0.87 | 1 | Tmprss3 | transmembrane protease, serine 3 | 0 | 8 | 0 | 0 | 0 | 1 | 0 | 0 | 0 | 0 | 0 |
| 1 | -1.34 | 3.59 | -0.37 | 0.71 | 1 | Gm50375 | | 0 | 0 | 0 | 4 | 0 | 3 | 0 | 0 | 0 | 0 | 2 |
| 1 | 1.18 | 3.61 | 0.33 | 0.74 | 1 | Gm28551 | | 0 | 0 | 0 | 0 | 0 | 1 | 0 | 0 | 8 | 0 | 0 |
| 1 | -2.03 | 3.61 | -0.56 | 0.57 | 1 | Gm44085 | | 0 | 2 | 0 | 4 | 0 | 3 | 0 | 0 | 0 | 0 | 0 |
| 1 | -0.19 | 3.56 | -0.05 | 0.96 | 1 | Gm38120 | | 0 | 0 | 0 | 0 | 0 | 5 | 0 | 0 | 0 | 4 | 0 |
| 1 | 1.18 | 3.56 | 0.33 | 0.74 | 1 | Rpl27-ps3 | | 0 | 1 | 0 | 1 | 0 | 2 | 0 | 2 | 3 | 0 | 0 |
| 1 | -1.35 | 3.59 | -0.38 | 0.71 | 1 | Gm17833 | | 0 | 0 | 0 | 4 | 0 | 3 | 0 | 0 | 0 | 0 | 2 |
| 1 | 1.41 | 3.64 | 0.39 | 0.70 | 1 | Gm6569 | predicted gene 6569 | 7 | 0 | 0 | 0 | 0 | 0 | 0 | 0 | 0 | 0 | 2 |
| 1 | -0.47 | 3.67 | -0.13 | 0.90 | 1 | Gm7289 | | 1 | 7 | 0 | 0 | 1 | 0 | 0 | 0 | 0 | 0 | 0 |
| 1 | 0.00 | 3.68 | 0.00 | 1.00 | 1 | Gm10309 | | 7 | 2 | 0 | 0 | 0 | 0 | 0 | 0 | 0 | 0 | 0 |
| 1 | -1.13 | 3.61 | -0.31 | 0.76 | 1 | Gm12630 | | 4 | 0 | 0 | 0 | 2 | 2 | 0 | 0 | 1 | 0 | 0 |
| 1 | 0.55 | 3.63 | 0.15 | 0.88 | 1 | Gm46329 | | 6 | 0 | 0 | 0 | 0 | 1 | 0 | 0 | 0 | 0 | 2 |
| 1 | -0.07 | 3.63 | -0.02 | 0.99 | 1 | 4930405O22Rik | | 0 | 4 | 0 | 0 | 1 | 1 | 0 | 0 | 3 | 0 | 0 |
| 1 | 0.55 | 3.61 | 0.15 | 0.88 | 1 | Gm42957 | | 0 | 0 | 3 | 0 | 0 | 2 | 0 | 0 | 0 | 0 | 4 |
| 1 | -0.79 | 3.62 | -0.22 | 0.83 | 1 | Foxd1 | forkhead box D1 | 2 | 0 | 2 | 0 | 0 | 5 | 0 | 0 | 0 | 0 | 0 |
| 1 | 0.56 | 3.59 | 0.16 | 0.88 | 1 | Gm47021 | | 0 | 2 | 2 | 1 | 0 | 1 | 0 | 0 | 2 | 0 | 1 |
| 1 | 0.64 | 3.61 | 0.18 | 0.86 | 1 | Gm44664 | | 0 | 0 | 2 | 0 | 0 | 2 | 0 | 0 | 0 | 0 | 5 |
| 1 | -0.08 | 3.61 | -0.02 | 0.98 | 1 | Gm37666 | | 3 | 0 | 0 | 0 | 0 | 4 | 0 | 0 | 0 | 0 | 2 |
| 1 | 0.01 | 3.63 | 0.00 | 1.00 | 1 | Gm13935 | | 0 | 5 | 1 | 0 | 0 | 2 | 0 | 0 | 0 | 0 | 1 |
| 1 | 0.00 | 3.68 | 0.00 | 1.00 | 1 | Igkv16-104 | | 2 | 7 | 0 | 0 | 0 | 0 | 0 | 0 | 0 | 0 | 0 |
| 1 | 0.76 | 3.61 | 0.21 | 0.83 | 1 | D630030B08Rik | | 0 | 0 | 0 | 0 | 0 | 2 | 0 | 0 | 0 | 0 | 7 |
| 1 | 0.76 | 3.61 | 0.21 | 0.83 | 1 | Gm45570 | | 0 | 0 | 0 | 0 | 0 | 2 | 0 | 0 | 0 | 0 | 7 |
| 1 | 0.76 | 3.61 | 0.21 | 0.83 | 1 | Gm47004 | | 0 | 0 | 0 | 0 | 0 | 2 | 0 | 0 | 0 | 0 | 7 |
| 1 | 1.91 | 3.60 | 0.53 | 0.60 | 1 | Gm10540 | | 1 | 0 | 0 | 0 | 0 | 1 | 0 | 0 | 6 | 0 | 1 |
| 1 | -0.93 | 3.64 | -0.26 | 0.80 | 1 | Dsg3 | desmoglein 3 | 0 | 7 | 0 | 0 | 0 | 2 | 0 | 0 | 0 | 0 | 0 |
| 1 | 1.54 | 3.58 | 0.43 | 0.67 | 1 | Gm19195 | | 0 | 0 | 0 | 0 | 0 | 2 | 0 | 0 | 6 | 0 | 1 |
| 1 | 1.12 | 3.57 | 0.31 | 0.75 | 1 | Gm24430 | | 0 | 0 | 0 | 0 | 0 | 2 | 0 | 0 | 7 | 0 | 0 |
| 1 | -0.96 | 3.63 | -0.26 | 0.79 | 1 | Gm42908 | | 0 | 0 | 6 | 0 | 0 | 3 | 0 | 0 | 0 | 0 | 0 |
| 1 | -0.96 | 3.63 | -0.26 | 0.79 | 1 | Ssmem1 | serine-rich single-pass membrane proteir | 0 | 0 | 6 | 0 | 0 | 3 | 0 | 0 | 0 | 0 | 0 |
| 1 | -0.96 | 3.63 | -0.26 | 0.79 | 1 | Gm50348 | | 0 | 0 | 6 | 0 | 0 | 3 | 0 | 0 | 0 | 0 | 0 |
| 1 | 2.48 | 3.59 | 0.69 | 0.49 | 1 | Gm28913 | | 0 | 0 | 0 | 0 | 0 | 1 | 0 | 0 | 4 | 4 | 0 |
| 1 | -1.32 | 3.62 | -0.36 | 0.72 | 1 | Gm7721 | | 0 | 0 | 0 | 9 | 0 | 0 | 0 | 0 | 0 | 0 | 0 |
| 1 | -1.32 | 3.62 | -0.36 | 0.72 | 1 | Rps8-ps3 | | 0 | 0 | 0 | 9 | 0 | 0 | 0 | 0 | 0 | 0 | 0 |
| 1 | -1.32 | 3.62 | -0.36 | 0.72 | 1 | Gm16348 | | 0 | 0 | 0 | 9 | 0 | 0 | 0 | 0 | 0 | 0 | 0 |
| 1 | -1.32 | 3.62 | -0.36 | 0.72 | 1 | Gm10744 | | 0 | 0 | 0 | 9 | 0 | 0 | 0 | 0 | 0 | 0 | 0 |
| 1 | -1.32 | 3.62 | -0.36 | 0.72 | 1 | Mir320 | | 0 | 0 | 0 | 9 | 0 | 0 | 0 | 0 | 0 | 0 | 0 |
| 1 | -0.42 | 3.59 | -0.12 | 0.91 | 1 | Gm32899 | | 0 | 0 | 5 | 0 | 0 | 3 | 0 | 0 | 1 | 0 | 0 |
| 1 | -0.65 | 3.58 | -0.18 | 0.85 | 1 | Gm44140 | | 0 | 0 | 0 | 0 | 2 | 2 | 0 | 0 | 0 | 5 | 0 |
| 1 | 0.84 | 3.38 | 0.25 | 0.80 | 1 | Gm12538 | | 1 | 1 | 0 | 0 | 1 | 2 | 0 | 1 | 1 | 0 | 2 |
| 1 | -0.18 | 3.58 | -0.05 | 0.96 | 1 | Gm9369 | | 0 | 0 | 0 | 6 | 0 | 0 | 0 | 0 | 0 | 3 | 0 |
| 1 | -2.44 | 3.61 | -0.68 | 0.50 | 1 | G6pd2 | glucose-6-phosphate dehydrogenase 2 | 0 | 0 | 0 | 7 | 0 | 2 | 0 | 0 | 0 | 0 | 0 |
| 1 | -0.31 | 3.61 | -0.09 | 0.93 | 1 | Gm47644 | | 3 | 0 | 0 | 0 | 0 | 4 | 0 | 0 | 0 | 0 | 1 |
| 1 | 2.61 | 3.62 | 0.72 | 0.47 | 1 | Gm37409 | | 0 | 0 | 0 | 0 | 0 | 0 | 0 | 0 | 0 | 3 | 6 |
| 1 | -0.49 | 3.59 | -0.14 | 0.89 | 1 | 4931429P17Rik | | 0 | 0 | 0 | 0 | 2 | 2 | 0 | 0 | 0 | 0 | 5 |
| 1 | 0.95 | 3.67 | 0.26 | 0.80 | 1 | Shh | sonic hedgehog | 3 | 5 | 0 | 0 | 0 | 0 | 0 | 0 | 0 | 0 | 1 |
| 1 | -0.40 | 3.64 | -0.11 | 0.91 | 1 | Gm6443 | | 2 | 1 | 3 | 0 | 1 | 1 | 0 | 0 | 0 | 0 | 1 |
| 1 | 0.05 | 3.61 | 0.01 | 0.99 | 1 | Gm15083 | | 0 | 0 | 0 | 0 | 7 | 0 | 0 | 0 | 1 | 0 | 1 |
| 1 | -2.57 | 3.60 | -0.71 | 0.48 | 1 | Gm15823 | | 0 | 0 | 0 | 5 | 0 | 3 | 0 | 0 | 0 | 0 | 0 |
| 1 | 0.55 | 3.61 | 0.15 | 0.88 | 1 | n-R5s197 | | 0 | 0 | 0 | 0 | 0 | 3 | 0 | 0 | 0 | 0 | 6 |
| 1 | 0.55 | 3.61 | 0.15 | 0.88 | 1 | B230208B08Rik | | 0 | 0 | 0 | 0 | 0 | 3 | 0 | 0 | 0 | 0 | 6 |
| 1 | 0.55 | 3.61 | 0.15 | 0.88 | 1 | Gm33609 | | 0 | 0 | 0 | 0 | 0 | 3 | 0 | 0 | 0 | 0 | 6 |
| 1 | 0.55 | 3.61 | 0.15 | 0.88 | 1 | Mir7647 | | 0 | 0 | 0 | 0 | 0 | 3 | 0 | 0 | 0 | 0 | 6 |
| 1 | 0.55 | 3.61 | 0.15 | 0.88 | 1 | Gm36262 | | 0 | 0 | 0 | 0 | 0 | 3 | 0 | 0 | 0 | 0 | 6 |
| 1 | 0.55 | 3.61 | 0.15 | 0.88 | 1 | Gm16310 | | 0 | 0 | 0 | 0 | 0 | 3 | 0 | 0 | 0 | 0 | 6 |
| 1 | 0.55 | 3.61 | 0.15 | 0.88 | 1 | Mir6986 | | 0 | 0 | 0 | 0 | 0 | 3 | 0 | 0 | 0 | 0 | 6 |
| 1 | 0.00 | 3.68 | 0.00 | 1.00 | 1 | Hoxd10 | homeobox D10 | 3 | 0 | 5 | 0 | 0 | 0 | 0 | 0 | 0 | 0 | 0 |
| 1 | 0.00 | 3.68 | 0.00 | 1.00 | 1 | Gm34425 | | 3 | 0 | 5 | 0 | 0 | 0 | 0 | 0 | 0 | 0 | 0 |
| 1 | -0.48 | 3.61 | -0.13 | 0.89 | 1 | Gm26708 | | 0 | 0 | 0 | 0 | 0 | 8 | 0 | 0 | 0 | 0 | 1 |
| 1 | -0.48 | 3.61 | -0.13 | 0.89 | 1 | Gm23482 | | 0 | 0 | 0 | 0 | 0 | 8 | 0 | 0 | 0 | 0 | 1 |

| | | | | | | | | | | | | | | | | | | |
|---|---|---|---|---|---|---|---|---|---|---|---|---|---|---|---|---|---|---|
| 1 | -0.48 | 3.61 | -0.13 | 0.89 | 1 | Gm31499 | | 0 | 0 | 0 | 0 | 0 | 8 | 0 | 0 | 0 | 0 | 1 |
| 1 | 0.12 | 3.62 | 0.03 | 0.97 | 1 | Gm25076 | | 0 | 4 | 0 | 0 | 0 | 3 | 0 | 0 | 0 | 0 | 2 |
| 1 | -2.69 | 3.61 | -0.75 | 0.46 | 1 | Gm12523 | | 0 | 0 | 3 | 2 | 0 | 2 | 0 | 0 | 0 | 0 | 2 |
| 1 | -0.69 | 3.58 | -0.19 | 0.85 | 1 | Gm43874 | | 0 | 0 | 0 | 0 | 3 | 2 | 0 | 0 | 0 | 0 | 4 |
| 1 | 0.00 | 3.67 | 0.00 | 1.00 | 1 | Gm29040 | | 9 | 0 | 0 | 0 | 0 | 0 | 0 | 0 | 0 | 0 | 0 |
| 1 | 0.00 | 3.67 | 0.00 | 1.00 | 1 | Gm3364 | | 9 | 0 | 0 | 0 | 0 | 0 | 0 | 0 | 0 | 0 | 0 |
| 1 | 0.00 | 3.67 | 0.00 | 1.00 | 1 | Gm6213 | | 9 | 0 | 0 | 0 | 0 | 0 | 0 | 0 | 0 | 0 | 0 |
| 1 | 0.00 | 3.67 | 0.00 | 1.00 | 1 | Spata4 | spermatogenesis associated 4 | 9 | 0 | 0 | 0 | 0 | 0 | 0 | 0 | 0 | 0 | 0 |
| 1 | 0.00 | 3.67 | 0.00 | 1.00 | 1 | Gm7850 | | 9 | 0 | 0 | 0 | 0 | 0 | 0 | 0 | 0 | 0 | 0 |
| 1 | 0.00 | 3.67 | 0.00 | 1.00 | 1 | Mir24-2 | | 9 | 0 | 0 | 0 | 0 | 0 | 0 | 0 | 0 | 0 | 0 |
| 1 | 0.00 | 3.67 | 0.00 | 1.00 | 1 | Cntn5 | contactin 5 | 9 | 0 | 0 | 0 | 0 | 0 | 0 | 0 | 0 | 0 | 0 |
| 1 | 0.00 | 3.67 | 0.00 | 1.00 | 1 | F930017D23Rik | | 9 | 0 | 0 | 0 | 0 | 0 | 0 | 0 | 0 | 0 | 0 |
| 1 | 0.00 | 3.67 | 0.00 | 1.00 | 1 | 4933404K13Rik | | 9 | 0 | 0 | 0 | 0 | 0 | 0 | 0 | 0 | 0 | 0 |
| 1 | 0.00 | 3.67 | 0.00 | 1.00 | 1 | Slc36a3os | | 9 | 0 | 0 | 0 | 0 | 0 | 0 | 0 | 0 | 0 | 0 |
| 1 | 0.00 | 3.67 | 0.00 | 1.00 | 1 | Gm16342 | | 9 | 0 | 0 | 0 | 0 | 0 | 0 | 0 | 0 | 0 | 0 |
| 1 | 0.00 | 3.67 | 0.00 | 1.00 | 1 | 9930038B18Rik | | 9 | 0 | 0 | 0 | 0 | 0 | 0 | 0 | 0 | 0 | 0 |
| 1 | 0.00 | 3.67 | 0.00 | 1.00 | 1 | Nkx2-9 | NK2 homeobox 9 | 9 | 0 | 0 | 0 | 0 | 0 | 0 | 0 | 0 | 0 | 0 |
| 1 | 0.00 | 3.67 | 0.00 | 1.00 | 1 | 4930474H06Rik | | 9 | 0 | 0 | 0 | 0 | 0 | 0 | 0 | 0 | 0 | 0 |
| 1 | 0.00 | 3.67 | 0.00 | 1.00 | 1 | Gm21939 | | 9 | 0 | 0 | 0 | 0 | 0 | 0 | 0 | 0 | 0 | 0 |
| 1 | 0.00 | 3.67 | 0.00 | 1.00 | 1 | Ighv1-61 | | 9 | 0 | 0 | 0 | 0 | 0 | 0 | 0 | 0 | 0 | 0 |
| 1 | 0.00 | 3.67 | 0.00 | 1.00 | 1 | Gm2670 | | 9 | 0 | 0 | 0 | 0 | 0 | 0 | 0 | 0 | 0 | 0 |
| 1 | 0.00 | 3.67 | 0.00 | 1.00 | 1 | Gm49236 | | 9 | 0 | 0 | 0 | 0 | 0 | 0 | 0 | 0 | 0 | 0 |
| 1 | 0.00 | 3.67 | 0.00 | 1.00 | 1 | Apol11a | apolipoprotein L 11a | 9 | 0 | 0 | 0 | 0 | 0 | 0 | 0 | 0 | 0 | 0 |
| 1 | 0.00 | 3.67 | 0.00 | 1.00 | 1 | Gm26634 | | 9 | 0 | 0 | 0 | 0 | 0 | 0 | 0 | 0 | 0 | 0 |
| 1 | 0.00 | 3.67 | 0.00 | 1.00 | 1 | Tmdd1 | | 9 | 0 | 0 | 0 | 0 | 0 | 0 | 0 | 0 | 0 | 0 |
| 1 | 0.00 | 3.67 | 0.00 | 1.00 | 1 | Ifitm7 | interferon induced transmembrane protei | 9 | 0 | 0 | 0 | 0 | 0 | 0 | 0 | 0 | 0 | 0 |
| 1 | 0.00 | 3.67 | 0.00 | 1.00 | 1 | A530030E21Rik | | 9 | 0 | 0 | 0 | 0 | 0 | 0 | 0 | 0 | 0 | 0 |
| 1 | 0.00 | 3.67 | 0.00 | 1.00 | 1 | Gm15157 | | 9 | 0 | 0 | 0 | 0 | 0 | 0 | 0 | 0 | 0 | 0 |
| 1 | -0.49 | 3.60 | -0.13 | 0.89 | 1 | Gm41693 | | 0 | 0 | 0 | 0 | 8 | 0 | 0 | 0 | 0 | 0 | 1 |
| 1 | 0.79 | 3.61 | 0.22 | 0.83 | 1 | Gm6564 | | 0 | 0 | 0 | 0 | 0 | 4 | 0 | 0 | 0 | 1 | 3 |
| 1 | -0.88 | 3.64 | -0.24 | 0.81 | 1 | Serpinb9b | serine (or cysteine) peptidase inhibitor, cl | 7 | 0 | 0 | 0 | 0 | 2 | 0 | 0 | 0 | 0 | 0 |
| 1 | -0.95 | 3.63 | -0.26 | 0.79 | 1 | Gm44017 | | 6 | 0 | 0 | 0 | 0 | 3 | 0 | 0 | 0 | 0 | 0 |
| 1 | -0.95 | 3.63 | -0.26 | 0.79 | 1 | Gm15556 | | 6 | 0 | 0 | 0 | 0 | 3 | 0 | 0 | 0 | 0 | 0 |
| 1 | 0.53 | 3.62 | 0.15 | 0.88 | 1 | Gm8844 | | 2 | 0 | 0 | 0 | 4 | 0 | 0 | 0 | 2 | 0 | 1 |
| 1 | -1.63 | 3.56 | -0.46 | 0.65 | 1 | Gm45080 | | 0 | 0 | 0 | 3 | 2 | 2 | 0 | 0 | 1 | 1 | 0 |
| 1 | -1.23 | 3.62 | -0.34 | 0.73 | 1 | Gm22286 | | 0 | 0 | 0 | 0 | 9 | 0 | 0 | 0 | 0 | 0 | 0 |
| 1 | -1.23 | 3.62 | -0.34 | 0.73 | 1 | Rps17-ps1 | | 0 | 0 | 0 | 0 | 9 | 0 | 0 | 0 | 0 | 0 | 0 |
| 1 | 0.00 | 3.68 | 0.00 | 1.00 | 1 | A830008E24Rik | | 3 | 5 | 0 | 0 | 0 | 0 | 0 | 0 | 0 | 0 | 0 |
| 1 | 2.60 | 3.58 | 0.72 | 0.47 | 1 | H2bc7 | H2B clustered histone 7 | 0 | 0 | 0 | 0 | 0 | 1 | 0 | 0 | 3 | 3 | 2 |
| 1 | 0.44 | 3.62 | 0.12 | 0.90 | 1 | Gm37728 | | 2 | 2 | 0 | 0 | 0 | 2 | 0 | 0 | 0 | 0 | 3 |
| 1 | 3.07 | 3.60 | 0.85 | 0.39 | 1 | Gm15726 | | 2 | 0 | 0 | 0 | 0 | 2 | 0 | 0 | 5 | 0 | 0 |
| 1 | 0.37 | 3.61 | 0.10 | 0.92 | 1 | Gm16024 | | 0 | 0 | 0 | 0 | 0 | 3 | 0 | 0 | 0 | 0 | 5 |
| 1 | 0.37 | 3.61 | 0.10 | 0.92 | 1 | Gm43371 | | 0 | 0 | 0 | 0 | 0 | 3 | 0 | 0 | 0 | 0 | 5 |
| 1 | 0.37 | 3.61 | 0.10 | 0.92 | 1 | Gm15809 | | 0 | 0 | 0 | 0 | 0 | 3 | 0 | 0 | 0 | 0 | 5 |
| 1 | -0.81 | 3.61 | -0.22 | 0.82 | 1 | Gm24379 | | 0 | 0 | 0 | 0 | 0 | 8 | 0 | 0 | 0 | 0 | 0 |
| 1 | -0.81 | 3.61 | -0.22 | 0.82 | 1 | 5430434F05Rik | | 0 | 0 | 0 | 0 | 0 | 8 | 0 | 0 | 0 | 0 | 0 |
| 1 | -0.81 | 3.61 | -0.22 | 0.82 | 1 | Gm44589 | | 0 | 0 | 0 | 0 | 0 | 8 | 0 | 0 | 0 | 0 | 0 |
| 1 | -0.81 | 3.61 | -0.22 | 0.82 | 1 | Gm26971 | | 0 | 0 | 0 | 0 | 0 | 8 | 0 | 0 | 0 | 0 | 0 |
| 1 | -0.81 | 3.61 | -0.22 | 0.82 | 1 | Gm48147 | | 0 | 0 | 0 | 0 | 0 | 8 | 0 | 0 | 0 | 0 | 0 |
| 1 | -0.81 | 3.61 | -0.22 | 0.82 | 1 | Gm36101 | | 0 | 0 | 0 | 0 | 0 | 8 | 0 | 0 | 0 | 0 | 0 |
| 1 | -0.81 | 3.61 | -0.22 | 0.82 | 1 | Gm29394 | | 0 | 0 | 0 | 0 | 0 | 8 | 0 | 0 | 0 | 0 | 0 |
| 1 | -0.81 | 3.61 | -0.22 | 0.82 | 1 | Gm49802 | | 0 | 0 | 0 | 0 | 0 | 8 | 0 | 0 | 0 | 0 | 0 |
| 1 | -0.81 | 3.61 | -0.22 | 0.82 | 1 | AU022751 | expressed sequence AU022751 | 0 | 0 | 0 | 0 | 0 | 8 | 0 | 0 | 0 | 0 | 0 |
| 1 | 1.20 | 3.59 | 0.33 | 0.74 | 1 | Mir26a-1 | | 0 | 0 | 0 | 0 | 0 | 3 | 0 | 0 | 2 | 0 | 3 |
| 1 | 0.18 | 3.57 | 0.05 | 0.96 | 1 | Mid1-ps1 | | 0 | 0 | 0 | 4 | 0 | 0 | 0 | 0 | 4 | 0 | 0 |
| 1 | -0.95 | 3.59 | -0.27 | 0.79 | 1 | Mir153 | | 0 | 0 | 0 | 0 | 3 | 3 | 0 | 0 | 0 | 0 | 3 |
| 1 | 1.52 | 3.63 | 0.42 | 0.68 | 1 | Rps27a-ps1 | | 0 | 0 | 6 | 0 | 0 | 0 | 0 | 1 | 1 | 0 | 0 |
| 1 | -0.31 | 3.61 | -0.09 | 0.93 | 1 | Gm18147 | | 1 | 0 | 0 | 0 | 1 | 3 | 0 | 0 | 0 | 0 | 4 |
| 1 | -0.63 | 3.58 | -0.17 | 0.86 | 1 | Gm7117 | | 0 | 0 | 1 | 3 | 0 | 3 | 0 | 1 | 0 | 0 | 1 |
| 1 | 1.25 | 3.66 | 0.34 | 0.73 | 1 | Gm6069 | | 4 | 1 | 1 | 0 | 0 | 0 | 0 | 0 | 0 | 0 | 2 |

| | | | | | | | | | | | | | | | | | | |
|---|---|---|---|---|---|---|---|---|---|---|---|---|---|---|---|---|---|---|
| 1 | 2.70 | 3.58 | 0.76 | 0.45 | 1 | Gm5138 | | 0 | 1 | 0 | 0 | 0 | 1 | 0 | 1 | 2 | 1 | 2 |
| 1 | 2.44 | 3.39 | 0.72 | 0.47 | 1 | Gm8641 | | 1 | 1 | 0 | 0 | 0 | 1 | 0 | 2 | 1 | 1 | 1 |
| 1 | -1.07 | 3.67 | -0.29 | 0.77 | 1 | Gm5257 | | 3 | 2 | 1 | 1 | 1 | 0 | 0 | 0 | 0 | 0 | 0 |
| 1 | 0.16 | 3.62 | 0.04 | 0.96 | 1 | Cbx3-ps6 | | 1 | 2 | 1 | 0 | 0 | 3 | 0 | 0 | 1 | 0 | 0 |
| 1 | -2.50 | 3.60 | -0.69 | 0.49 | 1 | Gm5750 | | 0 | 0 | 2 | 4 | 2 | 1 | 0 | 0 | 0 | 0 | 0 |
| 1 | 0.00 | 3.66 | 0.00 | 1.00 | 1 | Trim55 | tripartite motif-containing 55 | 0 | 0 | 8 | 0 | 0 | 0 | 0 | 0 | 0 | 0 | 0 |
| 1 | 0.00 | 3.66 | 0.00 | 1.00 | 1 | Nppa | natriuretic peptide type A | 0 | 0 | 8 | 0 | 0 | 0 | 0 | 0 | 0 | 0 | 0 |
| 1 | 0.00 | 3.66 | 0.00 | 1.00 | 1 | A430072P03Rik | | 0 | 0 | 8 | 0 | 0 | 0 | 0 | 0 | 0 | 0 | 0 |
| 1 | 0.00 | 3.66 | 0.00 | 1.00 | 1 | Ncr1 | natural cytotoxicity triggering receptor 1 | 0 | 0 | 8 | 0 | 0 | 0 | 0 | 0 | 0 | 0 | 0 |
| 1 | 0.00 | 3.66 | 0.00 | 1.00 | 1 | Gm45264 | | 0 | 0 | 8 | 0 | 0 | 0 | 0 | 0 | 0 | 0 | 0 |
| 1 | 0.00 | 3.66 | 0.00 | 1.00 | 1 | Gm29521 | | 0 | 0 | 8 | 0 | 0 | 0 | 0 | 0 | 0 | 0 | 0 |
| 1 | 0.00 | 3.66 | 0.00 | 1.00 | 1 | Gm33104 | | 0 | 0 | 8 | 0 | 0 | 0 | 0 | 0 | 0 | 0 | 0 |
| 1 | 0.00 | 3.66 | 0.00 | 1.00 | 1 | Gm4898 | | 0 | 0 | 8 | 0 | 0 | 0 | 0 | 0 | 0 | 0 | 0 |
| 1 | 0.00 | 3.66 | 0.00 | 1.00 | 1 | Fndc8 | fibronectin type III domain containing 8 | 0 | 0 | 8 | 0 | 0 | 0 | 0 | 0 | 0 | 0 | 0 |
| 1 | 0.00 | 3.66 | 0.00 | 1.00 | 1 | Ighv1-15 | | 0 | 0 | 8 | 0 | 0 | 0 | 0 | 0 | 0 | 0 | 0 |
| 1 | 0.00 | 3.66 | 0.00 | 1.00 | 1 | Gm29590 | | 0 | 0 | 8 | 0 | 0 | 0 | 0 | 0 | 0 | 0 | 0 |
| 1 | 0.00 | 3.66 | 0.00 | 1.00 | 1 | Slc6a18 | solute carrier family 6 (neurotransmitter tr | 0 | 0 | 8 | 0 | 0 | 0 | 0 | 0 | 0 | 0 | 0 |
| 1 | 0.00 | 3.66 | 0.00 | 1.00 | 1 | Trav3-3 | | 0 | 0 | 8 | 0 | 0 | 0 | 0 | 0 | 0 | 0 | 0 |
| 1 | 0.00 | 3.66 | 0.00 | 1.00 | 1 | Gm31107 | | 0 | 0 | 8 | 0 | 0 | 0 | 0 | 0 | 0 | 0 | 0 |
| 1 | 0.00 | 3.66 | 0.00 | 1.00 | 1 | Gm49518 | | 0 | 0 | 8 | 0 | 0 | 0 | 0 | 0 | 0 | 0 | 0 |
| 1 | 0.00 | 3.66 | 0.00 | 1.00 | 1 | Dppa2 | developmental pluripotency associated 2 | 0 | 0 | 8 | 0 | 0 | 0 | 0 | 0 | 0 | 0 | 0 |
| 1 | 0.00 | 3.66 | 0.00 | 1.00 | 1 | Gm8974 | | 0 | 0 | 8 | 0 | 0 | 0 | 0 | 0 | 0 | 0 | 0 |
| 1 | 0.85 | 3.66 | 0.23 | 0.82 | 1 | Gm12322 | | 6 | 1 | 0 | 0 | 0 | 0 | 0 | 0 | 0 | 1 | 0 |
| 1 | -0.18 | 3.56 | -0.05 | 0.96 | 1 | Gm20172 | predicted gene, 20172 | 0 | 0 | 0 | 5 | 0 | 0 | 0 | 0 | 0 | 0 | 3 |
| 1 | 1.14 | 3.65 | 0.31 | 0.75 | 1 | 4933431K23Rik | | 0 | 0 | 7 | 0 | 0 | 0 | 0 | 0 | 0 | 0 | 1 |
| 1 | 1.14 | 3.65 | 0.31 | 0.75 | 1 | Tecta | tectorin alpha | 0 | 0 | 7 | 0 | 0 | 0 | 0 | 0 | 0 | 0 | 1 |
| 1 | 1.14 | 3.65 | 0.31 | 0.75 | 1 | Gm18270 | | 0 | 0 | 7 | 0 | 0 | 0 | 0 | 0 | 0 | 0 | 1 |
| 1 | 0.19 | 3.61 | 0.05 | 0.96 | 1 | Gm22888 | | 0 | 0 | 0 | 0 | 0 | 4 | 0 | 0 | 0 | 0 | 4 |
| 1 | 0.19 | 3.61 | 0.05 | 0.96 | 1 | Gm5953 | | 0 | 0 | 0 | 0 | 0 | 4 | 0 | 0 | 0 | 0 | 4 |
| 1 | 1.21 | 3.61 | 0.34 | 0.74 | 1 | Gm29570 | | 0 | 0 | 3 | 0 | 0 | 1 | 0 | 0 | 3 | 1 | 0 |
| 1 | 0.77 | 3.61 | 0.21 | 0.83 | 1 | Gm43077 | | 0 | 0 | 0 | 0 | 0 | 4 | 0 | 0 | 1 | 0 | 3 |
| 1 | 0.10 | 3.61 | 0.03 | 0.98 | 1 | Hnrnpa1l2-ps | | 0 | 1 | 1 | 0 | 0 | 5 | 0 | 1 | 0 | 0 | 0 |
| 1 | -1.58 | 3.60 | -0.44 | 0.66 | 1 | Glra4 | glycine receptor, alpha 4 subunit | 0 | 0 | 0 | 0 | 5 | 3 | 0 | 0 | 0 | 0 | 1 |
| 1 | 1.49 | 3.64 | 0.41 | 0.68 | 1 | Gm37336 | | 0 | 0 | 6 | 0 | 0 | 0 | 0 | 0 | 0 | 0 | 2 |
| 1 | -0.24 | 3.58 | -0.07 | 0.95 | 1 | Gm10243 | | 0 | 0 | 0 | 1 | 0 | 5 | 0 | 2 | 0 | 0 | 0 |
| 1 | 3.24 | 3.59 | 0.90 | 0.37 | 1 | Gm7741 | | 0 | 0 | 0 | 0 | 0 | 0 | 0 | 1 | 4 | 0 | 3 |
| 1 | 0.62 | 3.63 | 0.17 | 0.86 | 1 | Gm44002 | | 0 | 7 | 0 | 0 | 0 | 0 | 0 | 1 | 0 | 0 | 0 |
| 1 | -1.67 | 3.60 | -0.46 | 0.64 | 1 | Gm5732 | | 1 | 1 | 0 | 1 | 1 | 3 | 0 | 1 | 0 | 0 | 0 |
| 1 | 0.00 | 3.68 | 0.00 | 1.00 | 1 | 1110002J07Rik | | 5 | 2 | 1 | 0 | 0 | 0 | 0 | 0 | 0 | 0 | 0 |
| 1 | 0.39 | 3.63 | 0.11 | 0.91 | 1 | Gm49761 | | 4 | 0 | 0 | 1 | 0 | 0 | 0 | 0 | 0 | 0 | 3 |
| 1 | 2.82 | 3.61 | 0.78 | 0.44 | 1 | Gm47675 | | 0 | 0 | 0 | 0 | 0 | 0 | 0 | 0 | 0 | 5 | 3 |
| 1 | -0.33 | 3.58 | -0.09 | 0.93 | 1 | Gm4777 | | 2 | 0 | 0 | 2 | 0 | 2 | 0 | 0 | 2 | 0 | 1 |
| 1 | 0.00 | 3.68 | 0.00 | 1.00 | 1 | Gm16556 | | 0 | 5 | 3 | 0 | 0 | 0 | 0 | 0 | 0 | 0 | 0 |
| 1 | 0.69 | 3.62 | 0.19 | 0.85 | 1 | Lrrc34 | leucine rich repeat containing 34 | 4 | 0 | 0 | 0 | 0 | 1 | 0 | 0 | 0 | 0 | 3 |
| 1 | -0.84 | 3.62 | -0.23 | 0.82 | 1 | Gm12860 | | 0 | 2 | 0 | 6 | 0 | 0 | 0 | 0 | 0 | 0 | 0 |
| 1 | -0.84 | 3.62 | -0.23 | 0.82 | 1 | Gm13841 | | 0 | 2 | 0 | 6 | 0 | 0 | 0 | 0 | 0 | 0 | 0 |
| 1 | 2.57 | 3.60 | 0.72 | 0.47 | 1 | 4921516A02Rik | | 3 | 0 | 0 | 0 | 0 | 3 | 0 | 0 | 3 | 0 | 0 |
| 1 | 1.85 | 3.64 | 0.51 | 0.61 | 1 | Gm7198 | | 1 | 3 | 1 | 0 | 0 | 0 | 0 | 1 | 0 | 0 | 2 |
| 1 | 1.37 | 3.61 | 0.38 | 0.71 | 1 | 1700072G22Rik | | 0 | 0 | 0 | 0 | 0 | 0 | 0 | 0 | 0 | 0 | 8 |
| 1 | 1.37 | 3.61 | 0.38 | 0.71 | 1 | Gm45630 | | 0 | 0 | 0 | 0 | 0 | 0 | 0 | 0 | 0 | 0 | 8 |
| 1 | 1.37 | 3.61 | 0.38 | 0.71 | 1 | B930025P03Rik | | 0 | 0 | 0 | 0 | 0 | 0 | 0 | 0 | 0 | 0 | 8 |
| 1 | 1.37 | 3.61 | 0.38 | 0.71 | 1 | Mir6905 | | 0 | 0 | 0 | 0 | 0 | 0 | 0 | 0 | 0 | 0 | 8 |
| 1 | 1.37 | 3.61 | 0.38 | 0.71 | 1 | Pnma5 | paraneoplastic antigen family 5 | 0 | 0 | 0 | 0 | 0 | 0 | 0 | 0 | 0 | 0 | 8 |
| 1 | 2.16 | 3.61 | 0.60 | 0.55 | 1 | 1700034I23Rik | | 0 | 0 | 0 | 0 | 0 | 0 | 0 | 0 | 1 | 0 | 7 |
| 1 | 0.00 | 3.61 | 0.00 | 1.00 | 1 | Gm24837 | | 0 | 0 | 0 | 0 | 0 | 5 | 0 | 0 | 0 | 0 | 3 |
| 1 | 0.00 | 3.61 | 0.00 | 1.00 | 1 | Spatc1l | spermatogenesis and centriole associate | 0 | 0 | 0 | 0 | 0 | 5 | 0 | 0 | 0 | 0 | 3 |
| 1 | 0.00 | 3.61 | 0.00 | 1.00 | 1 | Gm11755 | | 0 | 0 | 0 | 0 | 0 | 5 | 0 | 0 | 0 | 0 | 3 |
| 1 | 0.00 | 3.61 | 0.00 | 1.00 | 1 | Gm49968 | | 0 | 0 | 0 | 0 | 0 | 5 | 0 | 0 | 0 | 0 | 3 |
| 1 | 1.43 | 3.62 | 0.40 | 0.69 | 1 | Prodh2 | proline dehydrogenase (oxidase) 2 | 0 | 3 | 0 | 0 | 0 | 0 | 0 | 0 | 0 | 0 | 5 |
| 1 | 2.73 | 3.62 | 0.75 | 0.45 | 1 | Gm3307 | | 0 | 0 | 0 | 0 | 0 | 0 | 0 | 0 | 3 | 0 | 5 |

| | | | | | | | | | | | | | | | | | | |
|---|---|---|---|---|---|---|---|---|---|---|---|---|---|---|---|---|---|---|
| 1 | 1.49 | 3.63 | 0.41 | 0.68 | 1 | Gm44953 | | 0 | 5 | 0 | 0 | 0 | 0 | 0 | 0 | 0 | 0 | 3 |
| 1 | 1.77 | 3.63 | 0.49 | 0.63 | 1 | A630076J17Rik | RIKEN cDNA A630076J17 gene | 0 | 0 | 0 | 0 | 0 | 0 | 0 | 8 | 0 | 0 | 0 |
| 1 | 1.77 | 3.63 | 0.49 | 0.63 | 1 | 1700047A11Rik | | 0 | 0 | 0 | 0 | 0 | 0 | 0 | 8 | 0 | 0 | 0 |
| 1 | 1.42 | 3.64 | 0.39 | 0.70 | 1 | Gm49924 | | 0 | 6 | 0 | 0 | 0 | 0 | 0 | 0 | 0 | 0 | 2 |
| 1 | 1.07 | 3.66 | 0.29 | 0.77 | 1 | Gm37811 | | 0 | 7 | 0 | 0 | 0 | 0 | 0 | 0 | 0 | 0 | 1 |
| 1 | 2.54 | 3.61 | 0.70 | 0.48 | 1 | Spink10 | serine peptidase inhibitor, Kazal type 10 | 0 | 4 | 0 | 0 | 0 | 0 | 0 | 0 | 2 | 0 | 2 |
| 1 | 2.79 | 3.62 | 0.77 | 0.44 | 1 | Gm14376 | | 0 | 0 | 0 | 0 | 0 | 0 | 0 | 0 | 5 | 0 | 3 |
| 1 | 2.79 | 3.62 | 0.77 | 0.44 | 1 | Gm32184 | | 0 | 0 | 0 | 0 | 0 | 0 | 0 | 0 | 5 | 0 | 3 |
| 1 | 0.00 | 3.67 | 0.00 | 1.00 | 1 | Gm28342 | | 0 | 8 | 0 | 0 | 0 | 0 | 0 | 0 | 0 | 0 | 0 |
| 1 | 0.00 | 3.67 | 0.00 | 1.00 | 1 | Gm37516 | | 0 | 8 | 0 | 0 | 0 | 0 | 0 | 0 | 0 | 0 | 0 |
| 1 | 0.00 | 3.67 | 0.00 | 1.00 | 1 | Gm19817 | | 0 | 8 | 0 | 0 | 0 | 0 | 0 | 0 | 0 | 0 | 0 |
| 1 | 0.00 | 3.67 | 0.00 | 1.00 | 1 | Pakap | paralemmin A kinase anchor protein | 0 | 8 | 0 | 0 | 0 | 0 | 0 | 0 | 0 | 0 | 0 |
| 1 | 0.00 | 3.67 | 0.00 | 1.00 | 1 | Gm42981 | | 0 | 8 | 0 | 0 | 0 | 0 | 0 | 0 | 0 | 0 | 0 |
| 1 | 0.00 | 3.67 | 0.00 | 1.00 | 1 | Fkbp6 | FK506 binding protein 6 | 0 | 8 | 0 | 0 | 0 | 0 | 0 | 0 | 0 | 0 | 0 |
| 1 | 0.00 | 3.67 | 0.00 | 1.00 | 1 | Gm9951 | | 0 | 8 | 0 | 0 | 0 | 0 | 0 | 0 | 0 | 0 | 0 |
| 1 | 0.00 | 3.67 | 0.00 | 1.00 | 1 | Gm32743 | | 0 | 8 | 0 | 0 | 0 | 0 | 0 | 0 | 0 | 0 | 0 |
| 1 | 0.00 | 3.67 | 0.00 | 1.00 | 1 | Gm44787 | | 0 | 8 | 0 | 0 | 0 | 0 | 0 | 0 | 0 | 0 | 0 |
| 1 | 0.00 | 3.67 | 0.00 | 1.00 | 1 | Gm20556 | | 0 | 8 | 0 | 0 | 0 | 0 | 0 | 0 | 0 | 0 | 0 |
| 1 | 0.00 | 3.67 | 0.00 | 1.00 | 1 | Gm49595 | | 0 | 8 | 0 | 0 | 0 | 0 | 0 | 0 | 0 | 0 | 0 |
| 1 | 0.00 | 3.67 | 0.00 | 1.00 | 1 | Gm15336 | | 0 | 8 | 0 | 0 | 0 | 0 | 0 | 0 | 0 | 0 | 0 |
| 1 | 0.00 | 3.67 | 0.00 | 1.00 | 1 | Nxf7 | nuclear RNA export factor 7 | 0 | 8 | 0 | 0 | 0 | 0 | 0 | 0 | 0 | 0 | 0 |
| 1 | 1.86 | 3.63 | 0.51 | 0.61 | 1 | Gm29460 | | 0 | 0 | 0 | 0 | 0 | 0 | 0 | 0 | 8 | 0 | 0 |
| 1 | 1.86 | 3.63 | 0.51 | 0.61 | 1 | Serpinb13 | serine (or cysteine) peptidase inhibitor, cl | 0 | 0 | 0 | 0 | 0 | 0 | 0 | 0 | 8 | 0 | 0 |
| 1 | 1.86 | 3.63 | 0.51 | 0.61 | 1 | 4930540M05Rik | | 0 | 0 | 0 | 0 | 0 | 0 | 0 | 0 | 8 | 0 | 0 |
| 1 | 1.86 | 3.63 | 0.51 | 0.61 | 1 | Gm48362 | | 0 | 0 | 0 | 0 | 0 | 0 | 0 | 0 | 8 | 0 | 0 |
| 1 | 1.86 | 3.63 | 0.51 | 0.61 | 1 | Gm48455 | | 0 | 0 | 0 | 0 | 0 | 0 | 0 | 0 | 8 | 0 | 0 |
| 1 | -2.81 | 3.60 | -0.78 | 0.43 | 1 | Gm8807 | | 3 | 0 | 0 | 2 | 2 | 0 | 0 | 0 | 1 | 0 | 0 |
| 1 | 1.25 | 3.61 | 0.35 | 0.73 | 1 | Gm13464 | | 0 | 0 | 1 | 0 | 0 | 2 | 0 | 0 | 0 | 1 | 4 |
| 1 | 2.56 | 3.62 | 0.71 | 0.48 | 1 | Gm14336 | | 3 | 0 | 0 | 0 | 0 | 0 | 0 | 2 | 0 | 1 | 1 |
| 1 | 0.76 | 3.61 | 0.21 | 0.83 | 1 | Gm6034 | predicted gene 6034 | 0 | 0 | 4 | 0 | 0 | 1 | 0 | 0 | 0 | 0 | 3 |
| 1 | -0.83 | 3.61 | -0.23 | 0.82 | 1 | Gm26167 | | 0 | 0 | 0 | 0 | 1 | 5 | 0 | 0 | 0 | 0 | 2 |
| 1 | -0.76 | 3.63 | -0.21 | 0.83 | 1 | Gm12537 | | 4 | 0 | 0 | 2 | 0 | 1 | 0 | 1 | 0 | 0 | 0 |
| 1 | -1.45 | 3.62 | -0.40 | 0.69 | 1 | Gm5854 | | 0 | 0 | 1 | 7 | 0 | 0 | 0 | 0 | 0 | 0 | 0 |
| 1 | -0.36 | 3.62 | -0.10 | 0.92 | 1 | Gm13612 | | 0 | 0 | 5 | 0 | 3 | 0 | 0 | 0 | 0 | 0 | 0 |
| 1 | 0.00 | 3.68 | 0.00 | 1.00 | 1 | Hcrtr1 | hypocretin (orexin) receptor 1 | 6 | 2 | 0 | 0 | 0 | 0 | 0 | 0 | 0 | 0 | 0 |
| 1 | 1.34 | 3.62 | 0.37 | 0.71 | 1 | Gm15195 | | 1 | 0 | 2 | 0 | 0 | 0 | 0 | 0 | 0 | 0 | 5 |
| 1 | 0.31 | 3.62 | 0.09 | 0.93 | 1 | Gm34544 | | 0 | 0 | 3 | 0 | 0 | 1 | 0 | 0 | 4 | 0 | 0 |
| 1 | -0.69 | 3.66 | -0.19 | 0.85 | 1 | Gm29187 | | 4 | 2 | 0 | 0 | 0 | 2 | 0 | 0 | 0 | 0 | 0 |
| 1 | 0.00 | 3.68 | 0.00 | 1.00 | 1 | Gm29683 | | 1 | 5 | 2 | 0 | 0 | 0 | 0 | 0 | 0 | 0 | 0 |
| 1 | 0.12 | 3.62 | 0.03 | 0.97 | 1 | Gm3928 | | 3 | 0 | 0 | 0 | 0 | 3 | 0 | 0 | 0 | 0 | 2 |
| 1 | 1.58 | 3.66 | 0.43 | 0.67 | 1 | Gm29770 | | 2 | 2 | 2 | 0 | 0 | 0 | 0 | 1 | 1 | 0 | 0 |
| 1 | -0.96 | 3.64 | -0.26 | 0.79 | 1 | Gm10364 | | 5 | 0 | 0 | 1 | 1 | 0 | 0 | 0 | 0 | 0 | 1 |
| 1 | -0.08 | 3.61 | -0.02 | 0.98 | 1 | Gm12846 | | 2 | 0 | 0 | 0 | 0 | 4 | 0 | 0 | 0 | 0 | 2 |
| 1 | 0.14 | 3.66 | 0.04 | 0.97 | 1 | Gm15419 | | 6 | 0 | 0 | 0 | 1 | 0 | 0 | 0 | 1 | 0 | 0 |
| 1 | 0.97 | 3.61 | 0.27 | 0.79 | 1 | Gm20778 | | 0 | 0 | 0 | 0 | 0 | 1 | 0 | 0 | 0 | 0 | 7 |
| 1 | 0.00 | 3.68 | 0.00 | 1.00 | 1 | Arl14epl | ADP-ribosylation factor-like 14 effector pr | 1 | 7 | 0 | 0 | 0 | 0 | 0 | 0 | 0 | 0 | 0 |
| 1 | -2.38 | 3.62 | -0.66 | 0.51 | 1 | Gm16069 | | 3 | 0 | 0 | 4 | 0 | 0 | 0 | 0 | 0 | 0 | 1 |
| 1 | -0.20 | 3.61 | -0.06 | 0.96 | 1 | Gm13828 | | 0 | 0 | 0 | 0 | 0 | 6 | 0 | 0 | 0 | 0 | 2 |
| 1 | -0.20 | 3.61 | -0.06 | 0.96 | 1 | Gm20696 | | 0 | 0 | 0 | 0 | 0 | 6 | 0 | 0 | 0 | 0 | 2 |
| 1 | -0.20 | 3.61 | -0.06 | 0.96 | 1 | Gm44553 | | 0 | 0 | 0 | 0 | 0 | 6 | 0 | 0 | 0 | 0 | 2 |
| 1 | -0.20 | 3.61 | -0.06 | 0.96 | 1 | Gm45845 | | 0 | 0 | 0 | 0 | 0 | 6 | 0 | 0 | 0 | 0 | 2 |
| 1 | -0.20 | 3.61 | -0.06 | 0.96 | 1 | Cdhr4 | cadherin-related family member 4 | 0 | 0 | 0 | 0 | 0 | 6 | 0 | 0 | 0 | 0 | 2 |
| 1 | -0.20 | 3.61 | -0.06 | 0.96 | 1 | Gm44314 | | 0 | 0 | 0 | 0 | 0 | 6 | 0 | 0 | 0 | 0 | 2 |
| 1 | -0.20 | 3.61 | -0.06 | 0.96 | 1 | Gm36931 | | 0 | 0 | 0 | 0 | 0 | 6 | 0 | 0 | 0 | 0 | 2 |
| 1 | -2.01 | 3.62 | -0.56 | 0.58 | 1 | Gm40692 | | 0 | 0 | 0 | 7 | 1 | 0 | 0 | 0 | 0 | 0 | 0 |
| 1 | 1.87 | 3.60 | 0.52 | 0.60 | 1 | Gm13856 | | 0 | 0 | 0 | 0 | 0 | 1 | 0 | 0 | 2 | 0 | 5 |
| 1 | 1.94 | 3.63 | 0.53 | 0.59 | 1 | Gm50156 | | 1 | 0 | 0 | 0 | 0 | 0 | 0 | 0 | 7 | 0 | 0 |
| 1 | 0.94 | 3.60 | 0.26 | 0.79 | 1 | Gm11235 | | 0 | 0 | 1 | 0 | 5 | 0 | 0 | 1 | 0 | 0 | 1 |
| 1 | 0.83 | 3.47 | 0.24 | 0.81 | 1 | Gm11488 | | 0 | 1 | 1 | 1 | 0 | 2 | 0 | 0 | 1 | 1 | 1 |
| 1 | -2.99 | 3.58 | -0.84 | 0.40 | 1 | Gm3565 | | 0 | 0 | 0 | 4 | 2 | 3 | 0 | 0 | 0 | 0 | 0 |

| | | | | | | | | | | | | | | | | | | |
|---|---|---|---|---|---|---|---|---|---|---|---|---|---|---|---|---|---|---|
| 1 | 0.00 | 3.68 | 0.00 | 1.00 | 1 | Ecel1 | endothelin converting enzyme-like 1 | 7 | 0 | 1 | 0 | 0 | 0 | 0 | 0 | 0 | 0 | 0 |
| 1 | 0.00 | 3.68 | 0.00 | 1.00 | 1 | Gm45669 | | 7 | 0 | 1 | 0 | 0 | 0 | 0 | 0 | 0 | 0 | 0 |
| 1 | 1.05 | 3.66 | 0.29 | 0.77 | 1 | Gm42615 | | 7 | 0 | 0 | 0 | 0 | 0 | 0 | 0 | 0 | 0 | 1 |
| 1 | 1.05 | 3.66 | 0.29 | 0.77 | 1 | Gm48990 | | 7 | 0 | 0 | 0 | 0 | 0 | 0 | 0 | 0 | 0 | 1 |
| 1 | 0.28 | 3.62 | 0.08 | 0.94 | 1 | 1700092M07Ri | RIKEN cDNA 1700092M07 gene | 0 | 0 | 4 | 0 | 0 | 2 | 0 | 0 | 0 | 0 | 2 |
| 1 | -1.31 | 3.63 | -0.36 | 0.72 | 1 | Gm28071 | | 0 | 0 | 0 | 8 | 0 | 0 | 0 | 0 | 0 | 0 | 0 |
| 1 | -1.31 | 3.63 | -0.36 | 0.72 | 1 | Gm42705 | | 0 | 0 | 0 | 8 | 0 | 0 | 0 | 0 | 0 | 0 | 0 |
| 1 | -1.31 | 3.63 | -0.36 | 0.72 | 1 | Htr5a | 5-hydroxytryptamine (serotonin) receptor | 0 | 0 | 0 | 8 | 0 | 0 | 0 | 0 | 0 | 0 | 0 |
| 1 | -1.31 | 3.63 | -0.36 | 0.72 | 1 | Gm8930 | | 0 | 0 | 0 | 8 | 0 | 0 | 0 | 0 | 0 | 0 | 0 |
| 1 | -0.27 | 3.58 | -0.08 | 0.94 | 1 | Gm7422 | | 0 | 0 | 0 | 0 | 5 | 1 | 0 | 0 | 0 | 1 | 1 |
| 1 | -2.02 | 3.61 | -0.56 | 0.58 | 1 | 1810063I02Rik | | 4 | 0 | 0 | 0 | 2 | 2 | 0 | 0 | 0 | 0 | 0 |
| 1 | 0.26 | 3.64 | 0.07 | 0.94 | 1 | Gm44022 | | 6 | 0 | 0 | 0 | 0 | 1 | 0 | 0 | 0 | 0 | 1 |
| 1 | -2.10 | 3.62 | -0.58 | 0.56 | 1 | Gm15352 | | 0 | 0 | 0 | 7 | 0 | 1 | 0 | 0 | 0 | 0 | 0 |
| 1 | -2.10 | 3.62 | -0.58 | 0.56 | 1 | Gm8849 | | 0 | 0 | 0 | 7 | 0 | 1 | 0 | 0 | 0 | 0 | 0 |
| 1 | 1.63 | 3.67 | 0.44 | 0.66 | 1 | Rps15a-ps8 | | 3 | 1 | 2 | 0 | 0 | 0 | 0 | 1 | 0 | 0 | 1 |
| 1 | 0.40 | 3.57 | 0.11 | 0.91 | 1 | Ywhaq-ps2 | | 0 | 0 | 0 | 0 | 4 | 0 | 0 | 0 | 1 | 0 | 3 |
| 1 | 1.23 | 3.61 | 0.34 | 0.73 | 1 | Gm8670 | | 2 | 1 | 0 | 1 | 0 | 0 | 0 | 1 | 0 | 0 | 3 |
| 1 | 0.00 | 3.68 | 0.00 | 1.00 | 1 | Ighv2-6-8 | | 2 | 6 | 0 | 0 | 0 | 0 | 0 | 0 | 0 | 0 | 0 |
| 1 | 0.00 | 3.68 | 0.00 | 1.00 | 1 | Gm46526 | | 2 | 6 | 0 | 0 | 0 | 0 | 0 | 0 | 0 | 0 | 0 |
| 1 | 0.36 | 3.61 | 0.10 | 0.92 | 1 | Gm4246 | | 0 | 0 | 1 | 0 | 5 | 0 | 0 | 0 | 1 | 0 | 1 |
| 1 | 0.70 | 3.61 | 0.19 | 0.85 | 1 | Gm37602 | | 0 | 0 | 0 | 0 | 0 | 2 | 0 | 0 | 0 | 0 | 6 |
| 1 | 0.70 | 3.61 | 0.19 | 0.85 | 1 | F830115B05Rik | | 0 | 0 | 0 | 0 | 0 | 2 | 0 | 0 | 0 | 0 | 6 |
| 1 | 0.70 | 3.61 | 0.19 | 0.85 | 1 | Gm15842 | | 0 | 0 | 0 | 0 | 0 | 2 | 0 | 0 | 0 | 0 | 6 |
| 1 | 0.70 | 3.61 | 0.19 | 0.85 | 1 | Lman1l | lectin, mannose-binding 1 like | 0 | 0 | 0 | 0 | 0 | 2 | 0 | 0 | 0 | 0 | 6 |
| 1 | 0.70 | 3.61 | 0.19 | 0.85 | 1 | Gm12066 | | 0 | 0 | 0 | 0 | 0 | 2 | 0 | 0 | 0 | 0 | 6 |
| 1 | 0.70 | 3.61 | 0.19 | 0.85 | 1 | Gm22106 | | 0 | 0 | 0 | 0 | 0 | 2 | 0 | 0 | 0 | 0 | 6 |
| 1 | 0.70 | 3.61 | 0.19 | 0.85 | 1 | Gm49435 | | 0 | 0 | 0 | 0 | 0 | 2 | 0 | 0 | 0 | 0 | 6 |
| 1 | 0.70 | 3.61 | 0.19 | 0.85 | 1 | | | 0 | 0 | 0 | 0 | 0 | 2 | 0 | 0 | 0 | 0 | 6 |
| 1 | 0.64 | 3.61 | 0.18 | 0.86 | 1 | Gm16287 | | 0 | 1 | 0 | 0 | 0 | 2 | 0 | 0 | 0 | 0 | 5 |
| 1 | 0.55 | 3.61 | 0.15 | 0.88 | 1 | Gm48960 | | 0 | 2 | 0 | 0 | 0 | 2 | 0 | 0 | 0 | 0 | 4 |
| 1 | 0.55 | 3.61 | 0.15 | 0.88 | 1 | E330032C10Rik | | 0 | 2 | 0 | 0 | 0 | 2 | 0 | 0 | 0 | 0 | 4 |
| 1 | -0.45 | 3.61 | -0.12 | 0.90 | 1 | Gm43875 | | 0 | 0 | 0 | 0 | 0 | 7 | 0 | 0 | 0 | 0 | 1 |
| 1 | 1.25 | 3.61 | 0.35 | 0.73 | 1 | Gm13416 | | 0 | 1 | 0 | 0 | 0 | 2 | 0 | 0 | 1 | 0 | 4 |
| 1 | 0.14 | 3.61 | 0.04 | 0.97 | 1 | Gm22043 | | 0 | 0 | 0 | 0 | 0 | 7 | 0 | 0 | 1 | 0 | 0 |
| 1 | 0.00 | 3.68 | 0.00 | 1.00 | 1 | Foxl1 | forkhead box L1 | 3 | 0 | 5 | 0 | 0 | 0 | 0 | 0 | 0 | 0 | 0 |
| 1 | 0.00 | 3.68 | 0.00 | 1.00 | 1 | Tpsab1 | tryptase alpha/beta 1 | 3 | 0 | 5 | 0 | 0 | 0 | 0 | 0 | 0 | 0 | 0 |
| 1 | -0.91 | 3.64 | -0.25 | 0.80 | 1 | Wfdc10 | WAP four-disulfide core domain 10 | 0 | 6 | 0 | 0 | 0 | 2 | 0 | 0 | 0 | 0 | 0 |
| 1 | 2.24 | 3.62 | 0.62 | 0.54 | 1 | Gm13558 | | 0 | 4 | 0 | 0 | 0 | 2 | 0 | 0 | 2 | 0 | 0 |
| 1 | 1.49 | 3.59 | 0.41 | 0.68 | 1 | Gm16436 | | 0 | 0 | 0 | 0 | 0 | 2 | 0 | 0 | 5 | 0 | 1 |
| 1 | 3.08 | 3.60 | 0.86 | 0.39 | 1 | Rpl7a-ps1 | | 0 | 1 | 0 | 0 | 0 | 2 | 0 | 0 | 5 | 0 | 0 |
| 1 | 1.00 | 3.58 | 0.28 | 0.78 | 1 | Gm44737 | | 0 | 0 | 0 | 0 | 0 | 2 | 0 | 0 | 6 | 0 | 0 |
| 1 | 1.00 | 3.58 | 0.28 | 0.78 | 1 | Gm10545 | | 0 | 0 | 0 | 0 | 0 | 2 | 0 | 0 | 6 | 0 | 0 |
| 1 | 1.86 | 3.60 | 0.52 | 0.61 | 1 | Gm47639 | | 0 | 0 | 0 | 0 | 1 | 0 | 0 | 6 | 0 | 0 | 1 |
| 1 | -1.48 | 3.61 | -0.41 | 0.68 | 1 | Gm7558 | | 1 | 2 | 0 | 0 | 2 | 2 | 0 | 0 | 0 | 1 | 0 |
| 1 | 0.95 | 3.58 | 0.27 | 0.79 | 1 | Gm6061 | | 0 | 1 | 0 | 1 | 0 | 3 | 0 | 0 | 1 | 1 | 1 |
| 1 | -0.92 | 3.63 | -0.25 | 0.80 | 1 | Gm42589 | | 0 | 0 | 5 | 0 | 0 | 3 | 0 | 0 | 0 | 0 | 0 |
| 1 | 1.78 | 3.64 | 0.49 | 0.62 | 1 | Gm38055 | | 0 | 0 | 0 | 0 | 0 | 0 | 0 | 0 | 0 | 8 | 0 |
| 1 | 0.00 | 3.68 | 0.00 | 1.00 | 1 | Gm37618 | | 8 | 0 | 0 | 0 | 0 | 0 | 0 | 0 | 0 | 0 | 0 |
| 1 | 0.00 | 3.68 | 0.00 | 1.00 | 1 | Gm2427 | | 8 | 0 | 0 | 0 | 0 | 0 | 0 | 0 | 0 | 0 | 0 |
| 1 | 0.00 | 3.68 | 0.00 | 1.00 | 1 | Gm16587 | | 8 | 0 | 0 | 0 | 0 | 0 | 0 | 0 | 0 | 0 | 0 |
| 1 | 0.00 | 3.68 | 0.00 | 1.00 | 1 | Gm16701 | | 8 | 0 | 0 | 0 | 0 | 0 | 0 | 0 | 0 | 0 | 0 |
| 1 | 0.00 | 3.68 | 0.00 | 1.00 | 1 | Hoxd11 | homeobox D11 | 8 | 0 | 0 | 0 | 0 | 0 | 0 | 0 | 0 | 0 | 0 |
| 1 | 0.00 | 3.68 | 0.00 | 1.00 | 1 | Rpl29-ps1 | | 8 | 0 | 0 | 0 | 0 | 0 | 0 | 0 | 0 | 0 | 0 |
| 1 | 0.00 | 3.68 | 0.00 | 1.00 | 1 | 8030451A03Rik | | 8 | 0 | 0 | 0 | 0 | 0 | 0 | 0 | 0 | 0 | 0 |
| 1 | 0.00 | 3.68 | 0.00 | 1.00 | 1 | 9130410C08Rik | | 8 | 0 | 0 | 0 | 0 | 0 | 0 | 0 | 0 | 0 | 0 |
| 1 | 0.00 | 3.68 | 0.00 | 1.00 | 1 | Gm22656 | | 8 | 0 | 0 | 0 | 0 | 0 | 0 | 0 | 0 | 0 | 0 |
| 1 | 0.00 | 3.68 | 0.00 | 1.00 | 1 | Cib4 | calcium and integrin binding family memb | 8 | 0 | 0 | 0 | 0 | 0 | 0 | 0 | 0 | 0 | 0 |
| 1 | 0.00 | 3.68 | 0.00 | 1.00 | 1 | Morn3 | MORN repeat containing 3 | 8 | 0 | 0 | 0 | 0 | 0 | 0 | 0 | 0 | 0 | 0 |
| 1 | 0.00 | 3.68 | 0.00 | 1.00 | 1 | Gm42442 | | 8 | 0 | 0 | 0 | 0 | 0 | 0 | 0 | 0 | 0 | 0 |
| 1 | 0.00 | 3.68 | 0.00 | 1.00 | 1 | Gm43013 | | 8 | 0 | 0 | 0 | 0 | 0 | 0 | 0 | 0 | 0 | 0 |

| | | | | | | | | | | | | | | | | | | |
|---|---|---|---|---|---|---|---|---|---|---|---|---|---|---|---|---|---|---|
| 1 | 0.00 | 3.68 | 0.00 | 1.00 | 1 | Igkv4-75 | | 8 | 0 | 0 | 0 | 0 | 0 | 0 | 0 | 0 | 0 | 0 |
| 1 | 0.00 | 3.68 | 0.00 | 1.00 | 1 | Igkv8-28 | | 8 | 0 | 0 | 0 | 0 | 0 | 0 | 0 | 0 | 0 | 0 |
| 1 | 0.00 | 3.68 | 0.00 | 1.00 | 1 | Klrc1 | killer cell lectin-like receptor subfamily C, | 8 | 0 | 0 | 0 | 0 | 0 | 0 | 0 | 0 | 0 | 0 |
| 1 | 0.00 | 3.68 | 0.00 | 1.00 | 1 | Psg22 | pregnancy-specific glycoprotein 22 | 8 | 0 | 0 | 0 | 0 | 0 | 0 | 0 | 0 | 0 | 0 |
| 1 | 0.00 | 3.68 | 0.00 | 1.00 | 1 | Gm50483 | | 8 | 0 | 0 | 0 | 0 | 0 | 0 | 0 | 0 | 0 | 0 |
| 1 | 0.00 | 3.68 | 0.00 | 1.00 | 1 | Gm7870 | | 8 | 0 | 0 | 0 | 0 | 0 | 0 | 0 | 0 | 0 | 0 |
| 1 | 0.00 | 3.68 | 0.00 | 1.00 | 1 | Fbll1 | fibrillarin-like 1 | 8 | 0 | 0 | 0 | 0 | 0 | 0 | 0 | 0 | 0 | 0 |
| 1 | 0.00 | 3.68 | 0.00 | 1.00 | 1 | Dhrs7c | dehydrogenase/reductase (SDR family) m | 8 | 0 | 0 | 0 | 0 | 0 | 0 | 0 | 0 | 0 | 0 |
| 1 | 0.00 | 3.68 | 0.00 | 1.00 | 1 | Rab11fip4os1 | | 8 | 0 | 0 | 0 | 0 | 0 | 0 | 0 | 0 | 0 | 0 |
| 1 | 0.00 | 3.68 | 0.00 | 1.00 | 1 | Gm23444 | | 8 | 0 | 0 | 0 | 0 | 0 | 0 | 0 | 0 | 0 | 0 |
| 1 | 0.00 | 3.68 | 0.00 | 1.00 | 1 | Gm525 | predicted gene 525 | 8 | 0 | 0 | 0 | 0 | 0 | 0 | 0 | 0 | 0 | 0 |
| 1 | 0.00 | 3.68 | 0.00 | 1.00 | 1 | Gm11638 | | 8 | 0 | 0 | 0 | 0 | 0 | 0 | 0 | 0 | 0 | 0 |
| 1 | 0.00 | 3.68 | 0.00 | 1.00 | 1 | Ighe | | 8 | 0 | 0 | 0 | 0 | 0 | 0 | 0 | 0 | 0 | 0 |
| 1 | 0.00 | 3.68 | 0.00 | 1.00 | 1 | Ighv1-33 | | 8 | 0 | 0 | 0 | 0 | 0 | 0 | 0 | 0 | 0 | 0 |
| 1 | 0.00 | 3.68 | 0.00 | 1.00 | 1 | Gm4066 | | 8 | 0 | 0 | 0 | 0 | 0 | 0 | 0 | 0 | 0 | 0 |
| 1 | 0.00 | 3.68 | 0.00 | 1.00 | 1 | Gm31577 | | 8 | 0 | 0 | 0 | 0 | 0 | 0 | 0 | 0 | 0 | 0 |
| 1 | 0.00 | 3.68 | 0.00 | 1.00 | 1 | Gm10385 | | 8 | 0 | 0 | 0 | 0 | 0 | 0 | 0 | 0 | 0 | 0 |
| 1 | 0.00 | 3.68 | 0.00 | 1.00 | 1 | Gm21917 | | 8 | 0 | 0 | 0 | 0 | 0 | 0 | 0 | 0 | 0 | 0 |
| 1 | 0.00 | 3.68 | 0.00 | 1.00 | 1 | Gm28505 | | 8 | 0 | 0 | 0 | 0 | 0 | 0 | 0 | 0 | 0 | 0 |
| 1 | 0.00 | 3.68 | 0.00 | 1.00 | 1 | Gm15320 | | 8 | 0 | 0 | 0 | 0 | 0 | 0 | 0 | 0 | 0 | 0 |
| 1 | 0.00 | 3.68 | 0.00 | 1.00 | 1 | Gm25348 | | 8 | 0 | 0 | 0 | 0 | 0 | 0 | 0 | 0 | 0 | 0 |
| 1 | 0.00 | 3.68 | 0.00 | 1.00 | 1 | 2700046A07Rik | | 8 | 0 | 0 | 0 | 0 | 0 | 0 | 0 | 0 | 0 | 0 |
| 1 | 0.00 | 3.68 | 0.00 | 1.00 | 1 | Il1rapl1 | interleukin 1 receptor accessory protein-li | 8 | 0 | 0 | 0 | 0 | 0 | 0 | 0 | 0 | 0 | 0 |
| 1 | -0.52 | 3.66 | -0.14 | 0.89 | 1 | Gm29155 | | 7 | 0 | 0 | 0 | 0 | 1 | 0 | 0 | 0 | 0 | 0 |
| 1 | -0.52 | 3.66 | -0.14 | 0.89 | 1 | Gm37691 | | 7 | 0 | 0 | 0 | 0 | 1 | 0 | 0 | 0 | 0 | 0 |
| 1 | -0.52 | 3.66 | -0.14 | 0.89 | 1 | 1700019L13Rik | | 7 | 0 | 0 | 0 | 0 | 1 | 0 | 0 | 0 | 0 | 0 |
| 1 | -0.52 | 3.66 | -0.14 | 0.89 | 1 | Gm36169 | | 7 | 0 | 0 | 0 | 0 | 1 | 0 | 0 | 0 | 0 | 0 |
| 1 | -0.60 | 3.62 | -0.17 | 0.87 | 1 | Gm12447 | | 0 | 1 | 0 | 0 | 6 | 0 | 0 | 0 | 1 | 0 | 0 |
| 1 | 0.60 | 3.58 | 0.17 | 0.87 | 1 | Gm9246 | | 0 | 0 | 0 | 4 | 0 | 0 | 0 | 1 | 3 | 0 | 0 |
| 1 | -0.91 | 3.60 | -0.25 | 0.80 | 1 | Rps7-ps2 | | 0 | 1 | 1 | 0 | 4 | 1 | 0 | 0 | 0 | 0 | 1 |
| 1 | 0.00 | 3.68 | 0.00 | 1.00 | 1 | Gm9247 | | 3 | 3 | 2 | 0 | 0 | 0 | 0 | 0 | 0 | 0 | 0 |
| 1 | -0.34 | 3.62 | -0.09 | 0.92 | 1 | Gm12988 | | 1 | 0 | 0 | 0 | 6 | 0 | 0 | 1 | 0 | 0 | 0 |
| 1 | -0.91 | 3.62 | -0.25 | 0.80 | 1 | Gm3948 | | 4 | 0 | 0 | 0 | 0 | 3 | 0 | 0 | 0 | 0 | 0 |
| 1 | 1.73 | 3.59 | 0.48 | 0.63 | 1 | Gm7370 | | 2 | 0 | 0 | 2 | 0 | 0 | 0 | 1 | 2 | 0 | 1 |
| 1 | 0.28 | 3.61 | 0.08 | 0.94 | 1 | Mir375 | | 0 | 0 | 2 | 0 | 0 | 3 | 0 | 0 | 0 | 0 | 3 |
| 1 | -0.47 | 3.61 | -0.13 | 0.90 | 1 | Plpp4 | phospholipid phosphatase 4 | 0 | 0 | 0 | 0 | 7 | 0 | 0 | 0 | 0 | 0 | 1 |
| 1 | -0.47 | 3.61 | -0.13 | 0.90 | 1 | Gm46332 | | 0 | 0 | 0 | 0 | 7 | 0 | 0 | 0 | 0 | 0 | 1 |
| 1 | 1.25 | 3.61 | 0.35 | 0.73 | 1 | Gm6494 | | 1 | 0 | 0 | 0 | 0 | 2 | 0 | 0 | 1 | 0 | 4 |
| 1 | 0.48 | 3.61 | 0.13 | 0.89 | 1 | Gm3513 | | 0 | 0 | 0 | 0 | 0 | 3 | 0 | 0 | 0 | 0 | 5 |
| 1 | 0.48 | 3.61 | 0.13 | 0.89 | 1 | Mir1960 | | 0 | 0 | 0 | 0 | 0 | 3 | 0 | 0 | 0 | 0 | 5 |
| 1 | 0.48 | 3.61 | 0.13 | 0.89 | 1 | Gm42728 | | 0 | 0 | 0 | 0 | 0 | 3 | 0 | 0 | 0 | 0 | 5 |
| 1 | 0.48 | 3.61 | 0.13 | 0.89 | 1 | 2700029L08Rik | | 0 | 0 | 0 | 0 | 0 | 3 | 0 | 0 | 0 | 0 | 5 |
| 1 | 0.48 | 3.61 | 0.13 | 0.89 | 1 | Tcp10a | t-complex protein 10a | 0 | 0 | 0 | 0 | 0 | 3 | 0 | 0 | 0 | 0 | 5 |
| 1 | -0.81 | 3.61 | -0.22 | 0.82 | 1 | Gm38024 | | 0 | 0 | 0 | 0 | 0 | 8 | 0 | 0 | 0 | 0 | 0 |
| 1 | -0.81 | 3.61 | -0.22 | 0.82 | 1 | 9630010A21Rik | | 0 | 0 | 0 | 0 | 0 | 8 | 0 | 0 | 0 | 0 | 0 |
| 1 | -0.81 | 3.61 | -0.22 | 0.82 | 1 | Gm43103 | | 0 | 0 | 0 | 0 | 0 | 8 | 0 | 0 | 0 | 0 | 0 |
| 1 | -0.81 | 3.61 | -0.22 | 0.82 | 1 | Gm17875 | | 0 | 0 | 0 | 0 | 0 | 8 | 0 | 0 | 0 | 0 | 0 |
| 1 | -0.81 | 3.61 | -0.22 | 0.82 | 1 | Gm42763 | | 0 | 0 | 0 | 0 | 0 | 8 | 0 | 0 | 0 | 0 | 0 |
| 1 | -0.99 | 3.63 | -0.27 | 0.79 | 1 | Gm43474 | | 0 | 5 | 0 | 0 | 0 | 3 | 0 | 0 | 0 | 0 | 0 |
| 1 | 2.59 | 3.60 | 0.72 | 0.47 | 1 | Gm45542 | | 0 | 2 | 0 | 0 | 0 | 3 | 0 | 0 | 3 | 0 | 0 |
| 1 | 0.56 | 3.57 | 0.16 | 0.87 | 1 | Gm47503 | | 0 | 0 | 0 | 0 | 0 | 3 | 0 | 0 | 5 | 0 | 0 |
| 1 | -0.42 | 3.57 | -0.12 | 0.91 | 1 | Gm13904 | | 0 | 0 | 1 | 1 | 1 | 2 | 0 | 0 | 2 | 0 | 1 |
| 1 | -0.35 | 3.64 | -0.10 | 0.92 | 1 | Gm5777 | | 3 | 1 | 0 | 0 | 3 | 0 | 0 | 1 | 0 | 0 | 0 |
| 1 | 0.48 | 3.58 | 0.13 | 0.89 | 1 | Gm7253 | | 0 | 0 | 0 | 1 | 0 | 3 | 0 | 0 | 2 | 1 | 0 |
| 1 | -1.21 | 3.62 | -0.33 | 0.74 | 1 | Ldlrad2 | low density lipoprotein receptor class A do | 0 | 0 | 0 | 0 | 8 | 0 | 0 | 0 | 0 | 0 | 0 |
| 1 | -1.21 | 3.62 | -0.33 | 0.74 | 1 | C1ql4 | complement component 1, q subcompone | 0 | 0 | 0 | 0 | 8 | 0 | 0 | 0 | 0 | 0 | 0 |
| 1 | -1.49 | 3.60 | -0.41 | 0.68 | 1 | Gm13529 | | 0 | 0 | 1 | 1 | 4 | 1 | 0 | 0 | 1 | 0 | 0 |
| 1 | -0.73 | 3.64 | -0.20 | 0.84 | 1 | Gm48952 | | 1 | 0 | 4 | 0 | 0 | 3 | 0 | 0 | 0 | 0 | 0 |
| 1 | -1.33 | 3.57 | -0.37 | 0.71 | 1 | Gm10075 | | 0 | 0 | 0 | 2 | 1 | 3 | 0 | 0 | 1 | 1 | 0 |
| 1 | -2.01 | 3.62 | -0.56 | 0.58 | 1 | Gm5767 | predicted gene 5767 | 0 | 0 | 0 | 0 | 7 | 1 | 0 | 0 | 0 | 0 | 0 |

| | | | | | | | | | | | | | | | | | |
|---|---|---|---|---|---|---|---|---|---|---|---|---|---|---|---|---|---|
| 1 | -2.01 | 3.62 | -0.56 | 0.58 | 1 | Gm9432 | | 0 | 0 | 0 | 0 | 7 | 1 | 0 | 0 | 0 | 0 | 0 |
| 1 | -1.89 | 3.62 | -0.52 | 0.60 | 1 | Gm44987 | | 0 | 0 | 1 | 0 | 5 | 2 | 0 | 0 | 0 | 0 | 0 |
| 1 | 1.85 | 3.64 | 0.51 | 0.61 | 1 | Trim69 | tripartite motif-containing 69 | 0 | 1 | 0 | 0 | 0 | 0 | 0 | 0 | 0 | 6 | 0 |
| 1 | -1.03 | 3.57 | -0.29 | 0.77 | 1 | Gm14753 | | 0 | 0 | 1 | 2 | 1 | 2 | 0 | 0 | 1 | 0 | 1 |
| 1 | -0.85 | 3.62 | -0.24 | 0.81 | 1 | Rpl12-ps1 | | 2 | 0 | 1 | 0 | 3 | 1 | 0 | 0 | 1 | 0 | 0 |
| 1 | 0.28 | 3.61 | 0.08 | 0.94 | 1 | Gm37255 | | 0 | 0 | 0 | 0 | 0 | 3 | 0 | 0 | 0 | 0 | 4 |
| 1 | 0.28 | 3.61 | 0.08 | 0.94 | 1 | Bnipl | BCL2/adenovirus E1B 19kD interacting p | 0 | 0 | 0 | 0 | 0 | 3 | 0 | 0 | 0 | 0 | 4 |
| 1 | 0.28 | 3.61 | 0.08 | 0.94 | 1 | Gm22940 | | 0 | 0 | 0 | 0 | 0 | 3 | 0 | 0 | 0 | 0 | 4 |
| 1 | 0.28 | 3.61 | 0.08 | 0.94 | 1 | Mir7015 | | 0 | 0 | 0 | 0 | 0 | 3 | 0 | 0 | 0 | 0 | 4 |
| 1 | 0.28 | 3.61 | 0.08 | 0.94 | 1 | Gm44103 | | 0 | 0 | 0 | 0 | 0 | 3 | 0 | 0 | 0 | 0 | 4 |
| 1 | 0.28 | 3.61 | 0.08 | 0.94 | 1 | Snord32a | | 0 | 0 | 0 | 0 | 0 | 3 | 0 | 0 | 0 | 0 | 4 |
| 1 | 0.28 | 3.61 | 0.08 | 0.94 | 1 | Gm22314 | | 0 | 0 | 0 | 0 | 0 | 3 | 0 | 0 | 0 | 0 | 4 |
| 1 | 0.28 | 3.61 | 0.08 | 0.94 | 1 | Gm19337 | | 0 | 0 | 0 | 0 | 0 | 3 | 0 | 0 | 0 | 0 | 4 |
| 1 | 0.28 | 3.61 | 0.08 | 0.94 | 1 | Il3 | interleukin 3 | 0 | 0 | 0 | 0 | 0 | 3 | 0 | 0 | 0 | 0 | 4 |
| 1 | 0.28 | 3.61 | 0.08 | 0.94 | 1 | Gm48017 | | 0 | 0 | 0 | 0 | 0 | 3 | 0 | 0 | 0 | 0 | 4 |
| 1 | 0.28 | 3.61 | 0.08 | 0.94 | 1 | Gm48566 | | 0 | 0 | 0 | 0 | 0 | 3 | 0 | 0 | 0 | 0 | 4 |
| 1 | 0.28 | 3.61 | 0.08 | 0.94 | 1 | Snora19 | | 0 | 0 | 0 | 0 | 0 | 3 | 0 | 0 | 0 | 0 | 4 |
| 1 | -1.53 | 3.58 | -0.43 | 0.67 | 1 | Gm4478 | | 0 | 0 | 0 | 1 | 3 | 2 | 0 | 0 | 0 | 0 | 2 |
| 1 | 0.19 | 3.56 | 0.05 | 0.96 | 1 | Gm49394 | | 0 | 0 | 0 | 0 | 0 | 3 | 0 | 0 | 4 | 0 | 0 |
| 1 | -0.05 | 3.57 | -0.01 | 0.99 | 1 | Gm13840 | | 0 | 0 | 0 | 4 | 0 | 0 | 0 | 0 | 0 | 0 | 3 |
| 1 | -0.05 | 3.57 | -0.01 | 0.99 | 1 | Rhox3f | reproductive homeobox 3F | 0 | 0 | 0 | 4 | 0 | 0 | 0 | 0 | 0 | 0 | 3 |
| 1 | -1.08 | 3.62 | -0.30 | 0.76 | 1 | Gm8099 | | 1 | 1 | 0 | 3 | 2 | 0 | 0 | 0 | 1 | 0 | 0 |
| 1 | -2.30 | 3.61 | -0.64 | 0.52 | 1 | Ucma | upper zone of growth plate and cartilage r | 0 | 0 | 0 | 0 | 6 | 2 | 0 | 0 | 0 | 0 | 0 |
| 1 | -2.30 | 3.61 | -0.64 | 0.52 | 1 | Gm16155 | | 0 | 0 | 0 | 0 | 6 | 2 | 0 | 0 | 0 | 0 | 0 |
| 1 | -1.83 | 3.63 | -0.50 | 0.61 | 1 | Gm12031 | | 2 | 1 | 0 | 0 | 4 | 1 | 0 | 0 | 0 | 0 | 0 |
| 1 | -1.04 | 3.58 | -0.29 | 0.77 | 1 | Rdh16 | retinol dehydrogenase 16 | 0 | 0 | 0 | 3 | 0 | 2 | 0 | 0 | 3 | 0 | 0 |
| 1 | 1.41 | 3.63 | 0.39 | 0.70 | 1 | n-R5s29 | | 4 | 0 | 0 | 0 | 0 | 0 | 0 | 0 | 0 | 0 | 3 |
| 1 | 0.00 | 3.68 | 0.00 | 1.00 | 1 | Sla2 | Src-like-adaptor 2 | 4 | 3 | 0 | 0 | 0 | 0 | 0 | 0 | 0 | 0 | 0 |
| 1 | 0.00 | 3.68 | 0.00 | 1.00 | 1 | Abcc6 | ATP-binding cassette, sub-family C (CFTF | 4 | 3 | 0 | 0 | 0 | 0 | 0 | 0 | 0 | 0 | 0 |
| 1 | 0.00 | 3.68 | 0.00 | 1.00 | 1 | Gm49476 | | 4 | 3 | 0 | 0 | 0 | 0 | 0 | 0 | 0 | 0 | 0 |
| 1 | 1.52 | 3.58 | 0.42 | 0.67 | 1 | Gm6506 | | 0 | 0 | 0 | 1 | 1 | 0 | 0 | 1 | 0 | 1 | 3 |
| 1 | 2.69 | 3.63 | 0.74 | 0.46 | 1 | Sox2ot | | 3 | 0 | 0 | 0 | 0 | 1 | 0 | 0 | 3 | 0 | 0 |
| 1 | -0.51 | 3.61 | -0.14 | 0.89 | 1 | Gm4935 | | 0 | 0 | 0 | 1 | 0 | 3 | 0 | 0 | 0 | 0 | 3 |
| 1 | 0.84 | 3.63 | 0.23 | 0.82 | 1 | Gm11516 | | 0 | 0 | 4 | 0 | 0 | 1 | 0 | 0 | 1 | 1 | 0 |
| 1 | 1.14 | 3.61 | 0.32 | 0.75 | 1 | Gm8520 | | 1 | 0 | 1 | 0 | 1 | 1 | 0 | 2 | 0 | 1 | 0 |
| 1 | -1.52 | 3.58 | -0.42 | 0.67 | 1 | 1700030C12Rik | | 0 | 0 | 0 | 3 | 1 | 2 | 0 | 0 | 0 | 0 | 2 |
| 1 | -0.69 | 3.60 | -0.19 | 0.85 | 1 | Gm19503 | | 1 | 1 | 0 | 1 | 0 | 3 | 0 | 1 | 0 | 0 | 0 |
| 1 | 1.06 | 3.62 | 0.29 | 0.77 | 1 | Gm22969 | | 0 | 0 | 0 | 0 | 0 | 1 | 0 | 0 | 0 | 6 | 0 |
| 1 | 1.06 | 3.62 | 0.29 | 0.77 | 1 | Gm2862 | | 0 | 0 | 0 | 0 | 0 | 1 | 0 | 0 | 0 | 6 | 0 |
| 1 | 0.00 | 3.67 | 0.00 | 1.00 | 1 | Gm28386 | | 0 | 0 | 7 | 0 | 0 | 0 | 0 | 0 | 0 | 0 | 0 |
| 1 | 0.00 | 3.67 | 0.00 | 1.00 | 1 | Gm35048 | | 0 | 0 | 7 | 0 | 0 | 0 | 0 | 0 | 0 | 0 | 0 |
| 1 | 0.00 | 3.67 | 0.00 | 1.00 | 1 | Gal3st2c | galactose-3-O-sulfotransferase 2C | 0 | 0 | 7 | 0 | 0 | 0 | 0 | 0 | 0 | 0 | 0 |
| 1 | 0.00 | 3.67 | 0.00 | 1.00 | 1 | Mymk | myomaker, myoblast fusion factor | 0 | 0 | 7 | 0 | 0 | 0 | 0 | 0 | 0 | 0 | 0 |
| 1 | 0.00 | 3.67 | 0.00 | 1.00 | 1 | Kcng1 | potassium voltage-gated channel, subfam | 0 | 0 | 7 | 0 | 0 | 0 | 0 | 0 | 0 | 0 | 0 |
| 1 | 0.00 | 3.67 | 0.00 | 1.00 | 1 | Gm42629 | | 0 | 0 | 7 | 0 | 0 | 0 | 0 | 0 | 0 | 0 | 0 |
| 1 | 0.00 | 3.67 | 0.00 | 1.00 | 1 | Gm12828 | | 0 | 0 | 7 | 0 | 0 | 0 | 0 | 0 | 0 | 0 | 0 |
| 1 | 0.00 | 3.67 | 0.00 | 1.00 | 1 | Gm7932 | predicted gene 7932 | 0 | 0 | 7 | 0 | 0 | 0 | 0 | 0 | 0 | 0 | 0 |
| 1 | 0.00 | 3.67 | 0.00 | 1.00 | 1 | Gm44090 | | 0 | 0 | 7 | 0 | 0 | 0 | 0 | 0 | 0 | 0 | 0 |
| 1 | 0.00 | 3.67 | 0.00 | 1.00 | 1 | Ceacam9 | carcinoembryonic antigen-related cell adl | 0 | 0 | 7 | 0 | 0 | 0 | 0 | 0 | 0 | 0 | 0 |
| 1 | 0.00 | 3.67 | 0.00 | 1.00 | 1 | Gm18959 | | 0 | 0 | 7 | 0 | 0 | 0 | 0 | 0 | 0 | 0 | 0 |
| 1 | 0.00 | 3.67 | 0.00 | 1.00 | 1 | Acsm5 | acyl-CoA synthetase medium-chain famil | 0 | 0 | 7 | 0 | 0 | 0 | 0 | 0 | 0 | 0 | 0 |
| 1 | 0.00 | 3.67 | 0.00 | 1.00 | 1 | Gm48068 | | 0 | 0 | 7 | 0 | 0 | 0 | 0 | 0 | 0 | 0 | 0 |
| 1 | 0.00 | 3.67 | 0.00 | 1.00 | 1 | Gm48827 | | 0 | 0 | 7 | 0 | 0 | 0 | 0 | 0 | 0 | 0 | 0 |
| 1 | 0.00 | 3.67 | 0.00 | 1.00 | 1 | Gm48290 | | 0 | 0 | 7 | 0 | 0 | 0 | 0 | 0 | 0 | 0 | 0 |
| 1 | 0.00 | 3.67 | 0.00 | 1.00 | 1 | Wnt1 | wingless-type MMTV integration site famil | 0 | 0 | 7 | 0 | 0 | 0 | 0 | 0 | 0 | 0 | 0 |
| 1 | 0.00 | 3.67 | 0.00 | 1.00 | 1 | Pnpla1 | patatin-like phospholipase domain contai | 0 | 0 | 7 | 0 | 0 | 0 | 0 | 0 | 0 | 0 | 0 |
| 1 | 0.00 | 3.67 | 0.00 | 1.00 | 1 | Gm50288 | | 0 | 0 | 7 | 0 | 0 | 0 | 0 | 0 | 0 | 0 | 0 |
| 1 | 0.08 | 3.61 | 0.02 | 0.98 | 1 | Gm38392 | | 0 | 0 | 0 | 0 | 0 | 4 | 0 | 0 | 0 | 0 | 3 |
| 1 | 0.08 | 3.61 | 0.02 | 0.98 | 1 | Cort | cortistatin | 0 | 0 | 0 | 0 | 0 | 4 | 0 | 0 | 0 | 0 | 3 |
| 1 | 0.08 | 3.61 | 0.02 | 0.98 | 1 | Gm8493 | | 0 | 0 | 0 | 0 | 0 | 4 | 0 | 0 | 0 | 0 | 3 |

| | | | | | | | | | | | | | | | | | | |
|---|---|---|---|---|---|---|---|---|---|---|---|---|---|---|---|---|---|---|
| 1 | 0.08 | 3.61 | 0.02 | 0.98 | 1 | 4930479D17Rik | | 0 | 0 | 0 | 0 | 0 | 4 | 0 | 0 | 0 | 0 | 3 |
| 1 | 0.08 | 3.61 | 0.02 | 0.98 | 1 | Gm24189 | | 0 | 0 | 0 | 0 | 0 | 4 | 0 | 0 | 0 | 0 | 3 |
| 1 | 0.08 | 3.61 | 0.02 | 0.98 | 1 | Gm47445 | | 0 | 0 | 0 | 0 | 0 | 4 | 0 | 0 | 0 | 0 | 3 |
| 1 | 0.08 | 3.61 | 0.02 | 0.98 | 1 | Gm24241 | | 0 | 0 | 0 | 0 | 0 | 4 | 0 | 0 | 0 | 0 | 3 |
| 1 | 0.08 | 3.61 | 0.02 | 0.98 | 1 | Gm22667 | | 0 | 0 | 0 | 0 | 0 | 4 | 0 | 0 | 0 | 0 | 3 |
| 1 | -0.08 | 3.61 | -0.02 | 0.98 | 1 | Wdr20rt | WD repeat domain 20, retrogene | 0 | 1 | 0 | 0 | 0 | 4 | 0 | 0 | 0 | 0 | 2 |
| 1 | -2.36 | 3.61 | -0.65 | 0.51 | 1 | Gm50399 | | 0 | 0 | 0 | 0 | 5 | 3 | 0 | 0 | 0 | 0 | 0 |
| 1 | 0.00 | 3.68 | 0.00 | 1.00 | 1 | Kpna7 | karyopherin alpha 7 (importin alpha 8) | 0 | 1 | 6 | 0 | 0 | 0 | 0 | 0 | 0 | 0 | 0 |
| 1 | 0.65 | 3.63 | 0.18 | 0.86 | 1 | Gm50058 | | 0 | 6 | 0 | 0 | 0 | 0 | 0 | 1 | 0 | 0 | 0 |
| 1 | 1.47 | 3.64 | 0.40 | 0.69 | 1 | Cdx2 | caudal type homeobox 2 | 0 | 0 | 5 | 0 | 0 | 0 | 0 | 0 | 0 | 0 | 2 |
| 1 | 1.47 | 3.64 | 0.40 | 0.69 | 1 | Drc7 | dynein regulatory complex subunit 7 | 0 | 0 | 5 | 0 | 0 | 0 | 0 | 0 | 0 | 0 | 2 |
| 1 | 1.50 | 3.62 | 0.41 | 0.68 | 1 | Gm13431 | | 0 | 0 | 3 | 0 | 0 | 0 | 0 | 0 | 0 | 0 | 4 |
| 1 | 0.95 | 3.67 | 0.26 | 0.80 | 1 | Gm15131 | | 5 | 1 | 0 | 0 | 0 | 0 | 0 | 0 | 0 | 0 | 1 |
| 1 | 1.84 | 3.65 | 0.51 | 0.61 | 1 | Gm9836 | | 5 | 0 | 0 | 0 | 0 | 0 | 0 | 0 | 1 | 0 | 1 |
| 1 | 0.00 | 3.68 | 0.00 | 1.00 | 1 | Plxna4os1 | | 5 | 2 | 0 | 0 | 0 | 0 | 0 | 0 | 0 | 0 | 0 |
| 1 | 0.00 | 3.68 | 0.00 | 1.00 | 1 | Slc7a10 | solute carrier family 7 (cationic amino acid | 5 | 2 | 0 | 0 | 0 | 0 | 0 | 0 | 0 | 0 | 0 |
| 1 | 0.00 | 3.68 | 0.00 | 1.00 | 1 | Lbx1 | ladybird homeobox 1 | 5 | 2 | 0 | 0 | 0 | 0 | 0 | 0 | 0 | 0 | 0 |
| 1 | 0.95 | 3.67 | 0.26 | 0.80 | 1 | 9930022D16Rik | | 0 | 3 | 3 | 0 | 0 | 0 | 0 | 0 | 0 | 0 | 1 |
| 1 | 0.00 | 3.68 | 0.00 | 1.00 | 1 | Brs3 | bombesin-like receptor 3 | 0 | 4 | 3 | 0 | 0 | 0 | 0 | 0 | 0 | 0 | 0 |
| 1 | 1.12 | 3.67 | 0.30 | 0.76 | 1 | Gm8655 | | 6 | 0 | 0 | 0 | 0 | 0 | 0 | 1 | 0 | 0 | 0 |
| 1 | 1.49 | 3.64 | 0.41 | 0.68 | 1 | Hmgb1-ps4 | | 2 | 1 | 0 | 0 | 0 | 1 | 0 | 0 | 1 | 3 | 0 |
| 1 | -2.34 | 3.61 | -0.65 | 0.52 | 1 | Gm44951 | | 0 | 0 | 0 | 0 | 4 | 3 | 0 | 0 | 0 | 0 | 0 |
| 1 | -2.34 | 3.61 | -0.65 | 0.52 | 1 | Adgrg4 | adhesion G protein-coupled receptor G4 | 0 | 0 | 0 | 0 | 4 | 3 | 0 | 0 | 0 | 0 | 0 |
| 1 | -0.69 | 3.61 | -0.19 | 0.85 | 1 | Llph-ps1 | | 3 | 0 | 0 | 1 | 0 | 3 | 0 | 1 | 0 | 0 | 0 |
| 1 | 0.01 | 3.61 | 0.00 | 1.00 | 1 | Gm38365 | | 2 | 0 | 0 | 0 | 0 | 3 | 0 | 0 | 0 | 0 | 2 |
| 1 | -0.54 | 3.58 | -0.15 | 0.88 | 1 | Gm16235 | | 0 | 0 | 0 | 1 | 0 | 4 | 0 | 0 | 2 | 0 | 0 |
| 1 | -0.08 | 3.61 | -0.02 | 0.98 | 1 | Tcp10b | t-complex protein 10b | 1 | 0 | 0 | 0 | 0 | 4 | 0 | 0 | 0 | 0 | 2 |
| 1 | 1.36 | 3.62 | 0.38 | 0.71 | 1 | Gm38164 | | 0 | 0 | 0 | 0 | 0 | 0 | 0 | 0 | 0 | 0 | 7 |
| 1 | 1.36 | 3.62 | 0.38 | 0.71 | 1 | | | 0 | 0 | 0 | 0 | 0 | 0 | 0 | 0 | 0 | 0 | 7 |
| 1 | 1.36 | 3.62 | 0.38 | 0.71 | 1 | Gm48353 | | 0 | 0 | 0 | 0 | 0 | 0 | 0 | 0 | 0 | 0 | 7 |
| 1 | 1.36 | 3.62 | 0.38 | 0.71 | 1 | Gm11196 | | 0 | 0 | 0 | 0 | 0 | 0 | 0 | 0 | 0 | 0 | 7 |
| 1 | 1.36 | 3.62 | 0.38 | 0.71 | 1 | Mir494 | | 0 | 0 | 0 | 0 | 0 | 0 | 0 | 0 | 0 | 0 | 7 |
| 1 | 1.36 | 3.62 | 0.38 | 0.71 | 1 | Mir652 | | 0 | 0 | 0 | 0 | 0 | 0 | 0 | 0 | 0 | 0 | 7 |
| 1 | 1.35 | 3.62 | 0.37 | 0.71 | 1 | Pgam1-ps1 | | 0 | 1 | 0 | 0 | 0 | 0 | 0 | 0 | 0 | 0 | 6 |
| 1 | 1.34 | 3.62 | 0.37 | 0.71 | 1 | Gm49227 | | 0 | 2 | 0 | 0 | 0 | 0 | 0 | 0 | 0 | 0 | 5 |
| 1 | -0.14 | 3.61 | -0.04 | 0.97 | 1 | Gm25290 | | 0 | 0 | 0 | 0 | 0 | 5 | 0 | 0 | 0 | 0 | 2 |
| 1 | -0.14 | 3.61 | -0.04 | 0.97 | 1 | Gm43518 | | 0 | 0 | 0 | 0 | 0 | 5 | 0 | 0 | 0 | 0 | 2 |
| 1 | -0.14 | 3.61 | -0.04 | 0.97 | 1 | Gm35648 | | 0 | 0 | 0 | 0 | 0 | 5 | 0 | 0 | 0 | 0 | 2 |
| 1 | -0.14 | 3.61 | -0.04 | 0.97 | 1 | 4933411K16Ril | RIKEN cDNA 4933411K16 gene | 0 | 0 | 0 | 0 | 0 | 5 | 0 | 0 | 0 | 0 | 2 |
| 1 | 1.44 | 3.63 | 0.40 | 0.69 | 1 | F730311O21Rik | | 0 | 4 | 0 | 0 | 0 | 0 | 0 | 0 | 0 | 0 | 3 |
| 1 | 0.62 | 3.59 | 0.17 | 0.86 | 1 | Gm43792 | | 0 | 0 | 0 | 0 | 0 | 5 | 0 | 0 | 2 | 0 | 0 |
| 1 | 0.00 | 3.68 | 0.00 | 1.00 | 1 | A630031M04Rik | | 1 | 0 | 6 | 0 | 0 | 0 | 0 | 0 | 0 | 0 | 0 |
| 1 | 0.00 | 3.68 | 0.00 | 1.00 | 1 | Gm38901 | | 1 | 0 | 6 | 0 | 0 | 0 | 0 | 0 | 0 | 0 | 0 |
| 1 | 2.66 | 3.62 | 0.73 | 0.46 | 1 | 4930512H18Rik | | 0 | 0 | 0 | 0 | 0 | 0 | 0 | 0 | 4 | 0 | 3 |
| 1 | 2.66 | 3.62 | 0.73 | 0.46 | 1 | Gm10637 | | 0 | 0 | 0 | 0 | 0 | 0 | 0 | 0 | 4 | 0 | 3 |
| 1 | 0.00 | 3.67 | 0.00 | 1.00 | 1 | Fcrlb | Fc receptor-like B | 0 | 7 | 0 | 0 | 0 | 0 | 0 | 0 | 0 | 0 | 0 |
| 1 | 0.00 | 3.67 | 0.00 | 1.00 | 1 | Slc44a5 | solute carrier family 44, member 5 | 0 | 7 | 0 | 0 | 0 | 0 | 0 | 0 | 0 | 0 | 0 |
| 1 | 0.00 | 3.67 | 0.00 | 1.00 | 1 | 4930595O18Rik | | 0 | 7 | 0 | 0 | 0 | 0 | 0 | 0 | 0 | 0 | 0 |
| 1 | 0.00 | 3.67 | 0.00 | 1.00 | 1 | Gm10693 | | 0 | 7 | 0 | 0 | 0 | 0 | 0 | 0 | 0 | 0 | 0 |
| 1 | 0.00 | 3.67 | 0.00 | 1.00 | 1 | Gm45074 | | 0 | 7 | 0 | 0 | 0 | 0 | 0 | 0 | 0 | 0 | 0 |
| 1 | 0.00 | 3.67 | 0.00 | 1.00 | 1 | Gm20786 | | 0 | 7 | 0 | 0 | 0 | 0 | 0 | 0 | 0 | 0 | 0 |
| 1 | 0.00 | 3.67 | 0.00 | 1.00 | 1 | 1700011L03Rik | | 0 | 7 | 0 | 0 | 0 | 0 | 0 | 0 | 0 | 0 | 0 |
| 1 | 0.00 | 3.67 | 0.00 | 1.00 | 1 | Gm16322 | | 0 | 7 | 0 | 0 | 0 | 0 | 0 | 0 | 0 | 0 | 0 |
| 1 | 0.00 | 3.67 | 0.00 | 1.00 | 1 | Gm47770 | | 0 | 7 | 0 | 0 | 0 | 0 | 0 | 0 | 0 | 0 | 0 |
| 1 | 0.00 | 3.67 | 0.00 | 1.00 | 1 | Gm23105 | | 0 | 7 | 0 | 0 | 0 | 0 | 0 | 0 | 0 | 0 | 0 |
| 1 | 0.00 | 3.67 | 0.00 | 1.00 | 1 | Hormad2 | HORMA domain containing 2 | 0 | 7 | 0 | 0 | 0 | 0 | 0 | 0 | 0 | 0 | 0 |
| 1 | 0.00 | 3.67 | 0.00 | 1.00 | 1 | Gm5144 | | 0 | 7 | 0 | 0 | 0 | 0 | 0 | 0 | 0 | 0 | 0 |
| 1 | 0.00 | 3.67 | 0.00 | 1.00 | 1 | Gm37176 | | 0 | 7 | 0 | 0 | 0 | 0 | 0 | 0 | 0 | 0 | 0 |
| 1 | 0.00 | 3.67 | 0.00 | 1.00 | 1 | Gm35551 | | 0 | 7 | 0 | 0 | 0 | 0 | 0 | 0 | 0 | 0 | 0 |
| 1 | 0.00 | 3.67 | 0.00 | 1.00 | 1 | Gm50123 | | 0 | 7 | 0 | 0 | 0 | 0 | 0 | 0 | 0 | 0 | 0 |

| | | | | | | | | | | | | | | | | | | |
|---|---|---|---|---|---|---|---|---|---|---|---|---|---|---|---|---|---|---|
| 1 | 0.00 | 3.67 | 0.00 | 1.00 | 1 | Gm15345 | | 0 | 7 | 0 | 0 | 0 | 0 | 0 | 0 | 0 | 0 | 0 |
| 1 | 0.00 | 3.67 | 0.00 | 1.00 | 1 | Rhox3h | reproductive homeobox 3H | 0 | 7 | 0 | 0 | 0 | 0 | 0 | 0 | 0 | 0 | 0 |
| 1 | 0.00 | 3.67 | 0.00 | 1.00 | 1 | Tex16 | testis expressed gene 16 | 0 | 7 | 0 | 0 | 0 | 0 | 0 | 0 | 0 | 0 | 0 |
| 1 | 1.84 | 3.63 | 0.51 | 0.61 | 1 | Mael | maelstrom spermatogenic transposon sile | 0 | 0 | 0 | 0 | 0 | 0 | 0 | 0 | 7 | 0 | 0 |
| 1 | 1.84 | 3.63 | 0.51 | 0.61 | 1 | Gm13997 | | 0 | 0 | 0 | 0 | 0 | 0 | 0 | 0 | 7 | 0 | 0 |
| 1 | 1.84 | 3.63 | 0.51 | 0.61 | 1 | Gm21984 | | 0 | 0 | 0 | 0 | 0 | 0 | 0 | 0 | 7 | 0 | 0 |
| 1 | 1.84 | 3.63 | 0.51 | 0.61 | 1 | 5330439A09Rik | | 0 | 0 | 0 | 0 | 0 | 0 | 0 | 0 | 7 | 0 | 0 |
| 1 | 1.84 | 3.63 | 0.51 | 0.61 | 1 | Gm47725 | | 0 | 0 | 0 | 0 | 0 | 0 | 0 | 0 | 7 | 0 | 0 |
| 1 | 1.56 | 3.61 | 0.43 | 0.67 | 1 | Fscn3 | fascin actin-bundling protein 3 | 0 | 0 | 0 | 0 | 0 | 1 | 0 | 1 | 0 | 0 | 5 |
| 1 | 1.56 | 3.61 | 0.43 | 0.67 | 1 | Gm16471 | | 0 | 0 | 0 | 0 | 0 | 1 | 0 | 1 | 0 | 0 | 5 |
| 1 | -0.50 | 3.66 | -0.14 | 0.89 | 1 | Gm15689 | | 0 | 0 | 6 | 0 | 0 | 1 | 0 | 0 | 0 | 0 | 0 |
| 1 | -0.50 | 3.66 | -0.14 | 0.89 | 1 | Gm25415 | | 0 | 0 | 6 | 0 | 0 | 1 | 0 | 0 | 0 | 0 | 0 |
| 1 | -0.50 | 3.66 | -0.14 | 0.89 | 1 | Gm49552 | | 0 | 0 | 6 | 0 | 0 | 1 | 0 | 0 | 0 | 0 | 0 |
| 1 | 0.42 | 3.61 | 0.12 | 0.91 | 1 | Hao1 | hydroxyacid oxidase 1, liver | 0 | 0 | 0 | 0 | 1 | 0 | 0 | 0 | 0 | 0 | 6 |
| 1 | 1.57 | 3.61 | 0.44 | 0.66 | 1 | Gm4986 | | 0 | 1 | 0 | 0 | 1 | 0 | 0 | 0 | 2 | 0 | 3 |
| 1 | 1.03 | 3.66 | 0.28 | 0.78 | 1 | Gm36287 | | 6 | 0 | 0 | 0 | 0 | 0 | 0 | 0 | 0 | 0 | 1 |
| 1 | 0.00 | 3.68 | 0.00 | 1.00 | 1 | Gm9733 | predicted gene 9733 | 6 | 1 | 0 | 0 | 0 | 0 | 0 | 0 | 0 | 0 | 0 |
| 1 | 0.00 | 3.68 | 0.00 | 1.00 | 1 | Cd300ld3 | CD300 molecule like family member D3 | 1 | 3 | 3 | 0 | 0 | 0 | 0 | 0 | 0 | 0 | 0 |
| 1 | -1.29 | 3.63 | -0.36 | 0.72 | 1 | Iqca | IQ motif containing with AAA domain | 0 | 0 | 0 | 7 | 0 | 0 | 0 | 0 | 0 | 0 | 0 |
| 1 | -1.29 | 3.63 | -0.36 | 0.72 | 1 | 2210418O10Ril | RIKEN cDNA 2210418O10 gene | 0 | 0 | 0 | 7 | 0 | 0 | 0 | 0 | 0 | 0 | 0 |
| 1 | -1.29 | 3.63 | -0.36 | 0.72 | 1 | Gm11225 | | 0 | 0 | 0 | 7 | 0 | 0 | 0 | 0 | 0 | 0 | 0 |
| 1 | -1.29 | 3.63 | -0.36 | 0.72 | 1 | Gm23498 | | 0 | 0 | 0 | 7 | 0 | 0 | 0 | 0 | 0 | 0 | 0 |
| 1 | -1.29 | 3.63 | -0.36 | 0.72 | 1 | Gm27169 | | 0 | 0 | 0 | 7 | 0 | 0 | 0 | 0 | 0 | 0 | 0 |
| 1 | -1.29 | 3.63 | -0.36 | 0.72 | 1 | BC050972 | | 0 | 0 | 0 | 7 | 0 | 0 | 0 | 0 | 0 | 0 | 0 |
| 1 | -1.29 | 3.63 | -0.36 | 0.72 | 1 | Gm6986 | | 0 | 0 | 0 | 7 | 0 | 0 | 0 | 0 | 0 | 0 | 0 |
| 1 | -1.29 | 3.63 | -0.36 | 0.72 | 1 | Gm34743 | | 0 | 0 | 0 | 7 | 0 | 0 | 0 | 0 | 0 | 0 | 0 |
| 1 | -0.92 | 3.65 | -0.25 | 0.80 | 1 | Gm14618 | | 0 | 4 | 1 | 0 | 2 | 0 | 0 | 0 | 0 | 0 | 0 |
| 1 | -2.00 | 3.63 | -0.55 | 0.58 | 1 | Gm6420 | | 0 | 0 | 0 | 6 | 0 | 1 | 0 | 0 | 0 | 0 | 0 |
| 1 | 0.42 | 3.61 | 0.12 | 0.91 | 1 | Vmn2r79 | vomeronasal 2, receptor 79 | 0 | 0 | 0 | 1 | 0 | 0 | 0 | 0 | 0 | 0 | 6 |
| 1 | 0.87 | 3.61 | 0.24 | 0.81 | 1 | Slc45a1 | solute carrier family 45, member 1 | 0 | 0 | 1 | 0 | 0 | 1 | 0 | 0 | 0 | 0 | 5 |
| 1 | 1.60 | 3.64 | 0.44 | 0.66 | 1 | Gm37401 | | 1 | 0 | 1 | 0 | 0 | 0 | 0 | 0 | 5 | 0 | 0 |
| 1 | 1.75 | 3.63 | 0.48 | 0.63 | 1 | Gm14419 | | 0 | 0 | 0 | 0 | 0 | 0 | 0 | 7 | 0 | 0 | 0 |
| 1 | 1.35 | 3.62 | 0.37 | 0.71 | 1 | Gm42967 | | 1 | 0 | 0 | 0 | 0 | 0 | 0 | 0 | 0 | 0 | 6 |
| 1 | 1.35 | 3.62 | 0.37 | 0.71 | 1 | Slc6a1 | solute carrier family 6 (neurotransmitter tr | 1 | 0 | 0 | 0 | 0 | 0 | 0 | 0 | 0 | 0 | 6 |
| 1 | -0.07 | 3.65 | -0.02 | 0.98 | 1 | Rps13-ps5 | | 2 | 1 | 1 | 0 | 2 | 0 | 0 | 0 | 0 | 1 | 0 |
| 1 | 0.92 | 3.61 | 0.26 | 0.80 | 1 | Gm44508 | | 0 | 0 | 0 | 0 | 0 | 1 | 0 | 0 | 0 | 0 | 6 |
| 1 | 0.92 | 3.61 | 0.26 | 0.80 | 1 | Tspan10 | tetraspanin 10 | 0 | 0 | 0 | 0 | 0 | 1 | 0 | 0 | 0 | 0 | 6 |
| 1 | 0.00 | 3.68 | 0.00 | 1.00 | 1 | Lhx2 | LIM homeobox protein 2 | 1 | 6 | 0 | 0 | 0 | 0 | 0 | 0 | 0 | 0 | 0 |
| 1 | 0.00 | 3.68 | 0.00 | 1.00 | 1 | Gm8893 | | 1 | 6 | 0 | 0 | 0 | 0 | 0 | 0 | 0 | 0 | 0 |
| 1 | 0.79 | 3.62 | 0.22 | 0.83 | 1 | Tsnaxip1 | translin-associated factor X (Tsnax) intera | 0 | 2 | 0 | 0 | 0 | 1 | 0 | 0 | 0 | 0 | 4 |
| 1 | -0.41 | 3.61 | -0.11 | 0.91 | 1 | Gm25788 | | 0 | 0 | 0 | 0 | 0 | 6 | 0 | 0 | 0 | 0 | 1 |
| 1 | -0.41 | 3.61 | -0.11 | 0.91 | 1 | Gm25654 | | 0 | 0 | 0 | 0 | 0 | 6 | 0 | 0 | 0 | 0 | 1 |
| 1 | -0.41 | 3.61 | -0.11 | 0.91 | 1 | 4833412C15Rik | | 0 | 0 | 0 | 0 | 0 | 6 | 0 | 0 | 0 | 0 | 1 |
| 1 | -0.41 | 3.61 | -0.11 | 0.91 | 1 | Gm38109 | | 0 | 0 | 0 | 0 | 0 | 6 | 0 | 0 | 0 | 0 | 1 |
| 1 | -0.54 | 3.66 | -0.15 | 0.88 | 1 | Gm29358 | | 0 | 6 | 0 | 0 | 0 | 1 | 0 | 0 | 0 | 0 | 0 |
| 1 | -0.54 | 3.66 | -0.15 | 0.88 | 1 | Gm38251 | | 0 | 6 | 0 | 0 | 0 | 1 | 0 | 0 | 0 | 0 | 0 |
| 1 | -0.54 | 3.66 | -0.15 | 0.88 | 1 | Gm20673 | | 0 | 6 | 0 | 0 | 0 | 1 | 0 | 0 | 0 | 0 | 0 |
| 1 | 1.74 | 3.61 | 0.48 | 0.63 | 1 | Slc5a2 | solute carrier family 5 (sodium/glucose co | 0 | 0 | 0 | 0 | 0 | 1 | 0 | 0 | 5 | 0 | 1 |
| 1 | 1.13 | 3.61 | 0.31 | 0.75 | 1 | Gm1335 | | 0 | 0 | 0 | 0 | 0 | 1 | 0 | 0 | 6 | 0 | 0 |
| 1 | 1.13 | 3.61 | 0.31 | 0.75 | 1 | Rnu5g | | 0 | 0 | 0 | 0 | 0 | 1 | 0 | 0 | 6 | 0 | 0 |
| 1 | 1.13 | 3.61 | 0.31 | 0.75 | 1 | Gm12278 | | 0 | 0 | 0 | 0 | 0 | 1 | 0 | 0 | 6 | 0 | 0 |
| 1 | 1.13 | 3.61 | 0.31 | 0.75 | 1 | Gm26877 | | 0 | 0 | 0 | 0 | 0 | 1 | 0 | 0 | 6 | 0 | 0 |
| 1 | 0.00 | 3.68 | 0.00 | 1.00 | 1 | Fxyd7 | FXYD domain-containing ion transport re | 2 | 0 | 5 | 0 | 0 | 0 | 0 | 0 | 0 | 0 | 0 |
| 1 | 0.00 | 3.68 | 0.00 | 1.00 | 1 | Gm6052 | | 2 | 0 | 5 | 0 | 0 | 0 | 0 | 0 | 0 | 0 | 0 |
| 1 | 0.14 | 3.61 | 0.04 | 0.97 | 1 | Gm5990 | | 0 | 0 | 0 | 3 | 1 | 0 | 0 | 2 | 0 | 0 | 1 |
| 1 | 0.11 | 3.62 | 0.03 | 0.98 | 1 | Gm46546 | | 2 | 1 | 0 | 1 | 0 | 1 | 0 | 1 | 0 | 1 | 0 |
| 1 | -0.84 | 3.65 | -0.23 | 0.82 | 1 | Gm15856 | | 0 | 0 | 5 | 0 | 0 | 2 | 0 | 0 | 0 | 0 | 0 |
| 1 | -0.84 | 3.65 | -0.23 | 0.82 | 1 | Trpv3 | transient receptor potential cation channe | 0 | 0 | 5 | 0 | 0 | 2 | 0 | 0 | 0 | 0 | 0 |
| 1 | -0.84 | 3.65 | -0.23 | 0.82 | 1 | Gm11630 | | 0 | 0 | 5 | 0 | 0 | 2 | 0 | 0 | 0 | 0 | 0 |
| 1 | -0.84 | 3.65 | -0.23 | 0.82 | 1 | Gm47836 | | 0 | 0 | 5 | 0 | 0 | 2 | 0 | 0 | 0 | 0 | 0 |

| | | | | | | | | | | | | | | | | | | |
|---|---|---|---|---|---|---|---|---|---|---|---|---|---|---|---|---|---|---|
| 1 | 0.36 | 3.64 | 0.10 | 0.92 | 1 | Gm13226 | | 1 | 3 | 0 | 0 | 1 | 0 | 0 | 0 | 0 | 0 | 2 |
| 1 | 0.00 | 3.68 | 0.00 | 1.00 | 1 | Tnfsf4 | tumor necrosis factor (ligand) superfamily | 7 | 0 | 0 | 0 | 0 | 0 | 0 | 0 | 0 | 0 | 0 |
| 1 | 0.00 | 3.68 | 0.00 | 1.00 | 1 | Btbd18 | BTB (POZ) domain containing 18 | 7 | 0 | 0 | 0 | 0 | 0 | 0 | 0 | 0 | 0 | 0 |
| 1 | 0.00 | 3.68 | 0.00 | 1.00 | 1 | Egfem1 | EGF-like and EMI domain containing 1 | 7 | 0 | 0 | 0 | 0 | 0 | 0 | 0 | 0 | 0 | 0 |
| 1 | 0.00 | 3.68 | 0.00 | 1.00 | 1 | Gm42752 | | 7 | 0 | 0 | 0 | 0 | 0 | 0 | 0 | 0 | 0 | 0 |
| 1 | 0.00 | 3.68 | 0.00 | 1.00 | 1 | Gm26871 | | 7 | 0 | 0 | 0 | 0 | 0 | 0 | 0 | 0 | 0 | 0 |
| 1 | 0.00 | 3.68 | 0.00 | 1.00 | 1 | 6530403H02Rik | | 7 | 0 | 0 | 0 | 0 | 0 | 0 | 0 | 0 | 0 | 0 |
| 1 | 0.00 | 3.68 | 0.00 | 1.00 | 1 | 4930428O21Rik | | 7 | 0 | 0 | 0 | 0 | 0 | 0 | 0 | 0 | 0 | 0 |
| 1 | 0.00 | 3.68 | 0.00 | 1.00 | 1 | Ppp1r3a | protein phosphatase 1, regulatory subuni | 7 | 0 | 0 | 0 | 0 | 0 | 0 | 0 | 0 | 0 | 0 |
| 1 | 0.00 | 3.68 | 0.00 | 1.00 | 1 | 9530036M11Rik | | 7 | 0 | 0 | 0 | 0 | 0 | 0 | 0 | 0 | 0 | 0 |
| 1 | 0.00 | 3.68 | 0.00 | 1.00 | 1 | Cacng6 | calcium channel, voltage-dependent, gan | 7 | 0 | 0 | 0 | 0 | 0 | 0 | 0 | 0 | 0 | 0 |
| 1 | 0.00 | 3.68 | 0.00 | 1.00 | 1 | Pgpep1l | pyroglutamyl-peptidase I-like | 7 | 0 | 0 | 0 | 0 | 0 | 0 | 0 | 0 | 0 | 0 |
| 1 | 0.00 | 3.68 | 0.00 | 1.00 | 1 | Olfr701 | olfactory receptor 701 | 7 | 0 | 0 | 0 | 0 | 0 | 0 | 0 | 0 | 0 | 0 |
| 1 | 0.00 | 3.68 | 0.00 | 1.00 | 1 | Gm45670 | | 7 | 0 | 0 | 0 | 0 | 0 | 0 | 0 | 0 | 0 | 0 |
| 1 | 0.00 | 3.68 | 0.00 | 1.00 | 1 | Ascl2 | achaete-scute family bHLH transcription f | 7 | 0 | 0 | 0 | 0 | 0 | 0 | 0 | 0 | 0 | 0 |
| 1 | 0.00 | 3.68 | 0.00 | 1.00 | 1 | Gypa | glycophorin A | 7 | 0 | 0 | 0 | 0 | 0 | 0 | 0 | 0 | 0 | 0 |
| 1 | 0.00 | 3.68 | 0.00 | 1.00 | 1 | A530010L16Rik | | 7 | 0 | 0 | 0 | 0 | 0 | 0 | 0 | 0 | 0 | 0 |
| 1 | 0.00 | 3.68 | 0.00 | 1.00 | 1 | Slc51b | solute carrier family 51, beta subunit | 7 | 0 | 0 | 0 | 0 | 0 | 0 | 0 | 0 | 0 | 0 |
| 1 | 0.00 | 3.68 | 0.00 | 1.00 | 1 | Mir677 | | 7 | 0 | 0 | 0 | 0 | 0 | 0 | 0 | 0 | 0 | 0 |
| 1 | 0.00 | 3.68 | 0.00 | 1.00 | 1 | Gm12253 | predicted gene 12253 | 7 | 0 | 0 | 0 | 0 | 0 | 0 | 0 | 0 | 0 | 0 |
| 1 | 0.00 | 3.68 | 0.00 | 1.00 | 1 | Krt36 | keratin 36 | 7 | 0 | 0 | 0 | 0 | 0 | 0 | 0 | 0 | 0 | 0 |
| 1 | 0.00 | 3.68 | 0.00 | 1.00 | 1 | Gm48482 | | 7 | 0 | 0 | 0 | 0 | 0 | 0 | 0 | 0 | 0 | 0 |
| 1 | 0.00 | 3.68 | 0.00 | 1.00 | 1 | Gm5184 | | 7 | 0 | 0 | 0 | 0 | 0 | 0 | 0 | 0 | 0 | 0 |
| 1 | 0.00 | 3.68 | 0.00 | 1.00 | 1 | Gm47796 | | 7 | 0 | 0 | 0 | 0 | 0 | 0 | 0 | 0 | 0 | 0 |
| 1 | 0.00 | 3.68 | 0.00 | 1.00 | 1 | Gm47507 | | 7 | 0 | 0 | 0 | 0 | 0 | 0 | 0 | 0 | 0 | 0 |
| 1 | 0.00 | 3.68 | 0.00 | 1.00 | 1 | 4933404K08Rik | | 7 | 0 | 0 | 0 | 0 | 0 | 0 | 0 | 0 | 0 | 0 |
| 1 | 0.00 | 3.68 | 0.00 | 1.00 | 1 | Gm21762 | | 7 | 0 | 0 | 0 | 0 | 0 | 0 | 0 | 0 | 0 | 0 |
| 1 | 0.00 | 3.68 | 0.00 | 1.00 | 1 | 5430440P10Rik | | 7 | 0 | 0 | 0 | 0 | 0 | 0 | 0 | 0 | 0 | 0 |
| 1 | 0.00 | 3.68 | 0.00 | 1.00 | 1 | Olfr166 | olfactory receptor 166 | 7 | 0 | 0 | 0 | 0 | 0 | 0 | 0 | 0 | 0 | 0 |
| 1 | 0.00 | 3.68 | 0.00 | 1.00 | 1 | Vmn2r96 | vomeronasal 2, receptor 96 | 7 | 0 | 0 | 0 | 0 | 0 | 0 | 0 | 0 | 0 | 0 |
| 1 | 0.00 | 3.68 | 0.00 | 1.00 | 1 | Gm28347 | | 7 | 0 | 0 | 0 | 0 | 0 | 0 | 0 | 0 | 0 | 0 |
| 1 | 0.00 | 3.68 | 0.00 | 1.00 | 1 | Tslrn1 | | 7 | 0 | 0 | 0 | 0 | 0 | 0 | 0 | 0 | 0 | 0 |
| 1 | 0.00 | 3.68 | 0.00 | 1.00 | 1 | n-R5s8 | | 7 | 0 | 0 | 0 | 0 | 0 | 0 | 0 | 0 | 0 | 0 |
| 1 | 0.56 | 3.57 | 0.16 | 0.88 | 1 | Gm19028 | | 0 | 0 | 0 | 1 | 1 | 1 | 0 | 1 | 2 | 0 | 1 |
| 1 | 0.81 | 3.63 | 0.22 | 0.82 | 1 | Gm6129 | | 3 | 0 | 1 | 0 | 1 | 0 | 0 | 0 | 1 | 1 | 0 |
| 1 | -0.49 | 3.66 | -0.13 | 0.89 | 1 | Ldlrad1 | low density lipoprotein receptor class A d | 6 | 0 | 0 | 0 | 0 | 1 | 0 | 0 | 0 | 0 | 0 |
| 1 | -0.49 | 3.66 | -0.13 | 0.89 | 1 | Gm13121 | | 6 | 0 | 0 | 0 | 0 | 1 | 0 | 0 | 0 | 0 | 0 |
| 1 | -0.49 | 3.66 | -0.13 | 0.89 | 1 | Gm49373 | | 6 | 0 | 0 | 0 | 0 | 1 | 0 | 0 | 0 | 0 | 0 |
| 1 | -0.49 | 3.66 | -0.13 | 0.89 | 1 | Gm49097 | | 6 | 0 | 0 | 0 | 0 | 1 | 0 | 0 | 0 | 0 | 0 |
| 1 | 0.92 | 3.62 | 0.25 | 0.80 | 1 | Gm7593 | | 1 | 0 | 0 | 0 | 1 | 1 | 0 | 1 | 3 | 0 | 0 |
| 1 | -0.69 | 3.66 | -0.19 | 0.85 | 1 | Rpl21-ps5 | | 0 | 2 | 3 | 0 | 0 | 2 | 0 | 0 | 0 | 0 | 0 |
| 1 | -0.03 | 3.57 | -0.01 | 0.99 | 1 | Gm31462 | | 0 | 0 | 0 | 0 | 4 | 0 | 0 | 0 | 0 | 0 | 3 |
| 1 | -0.86 | 3.62 | -0.24 | 0.81 | 1 | Gm44995 | | 3 | 0 | 0 | 0 | 0 | 3 | 0 | 0 | 0 | 0 | 0 |
| 1 | -0.36 | 3.58 | -0.10 | 0.92 | 1 | Gm48078 | | 0 | 0 | 0 | 0 | 0 | 4 | 0 | 0 | 0 | 3 | 0 |
| 1 | 0.87 | 3.61 | 0.24 | 0.81 | 1 | Gm3045 | predicted gene 3045 | 1 | 0 | 0 | 0 | 0 | 1 | 0 | 0 | 0 | 0 | 5 |
| 1 | 0.00 | 3.68 | 0.00 | 1.00 | 1 | Gm48898 | | 2 | 5 | 0 | 0 | 0 | 0 | 0 | 0 | 0 | 0 | 0 |
| 1 | 0.64 | 3.61 | 0.18 | 0.86 | 1 | Gm42892 | | 0 | 0 | 0 | 0 | 0 | 2 | 0 | 0 | 0 | 0 | 5 |
| 1 | 0.64 | 3.61 | 0.18 | 0.86 | 1 | Gm23245 | | 0 | 0 | 0 | 0 | 0 | 2 | 0 | 0 | 0 | 0 | 5 |
| 1 | 0.64 | 3.61 | 0.18 | 0.86 | 1 | Gm16163 | | 0 | 0 | 0 | 0 | 0 | 2 | 0 | 0 | 0 | 0 | 5 |
| 1 | 0.64 | 3.61 | 0.18 | 0.86 | 1 | Olfr317 | olfactory receptor 317 | 0 | 0 | 0 | 0 | 0 | 2 | 0 | 0 | 0 | 0 | 5 |
| 1 | 0.64 | 3.61 | 0.18 | 0.86 | 1 | Gm29819 | | 0 | 0 | 0 | 0 | 0 | 2 | 0 | 0 | 0 | 0 | 5 |
| 1 | 1.21 | 3.61 | 0.33 | 0.74 | 1 | Ankrd60 | ankyrin repeat domain 60 | 0 | 0 | 0 | 0 | 0 | 2 | 0 | 0 | 1 | 0 | 4 |
| 1 | 0.44 | 3.62 | 0.12 | 0.90 | 1 | Gm15057 | | 0 | 2 | 0 | 0 | 0 | 2 | 0 | 0 | 0 | 0 | 3 |
| 1 | -0.80 | 3.61 | -0.22 | 0.82 | 1 | Gm37383 | | 0 | 0 | 0 | 0 | 0 | 7 | 0 | 0 | 0 | 0 | 0 |
| 1 | -0.80 | 3.61 | -0.22 | 0.82 | 1 | Gm37876 | | 0 | 0 | 0 | 0 | 0 | 7 | 0 | 0 | 0 | 0 | 0 |
| 1 | -0.80 | 3.61 | -0.22 | 0.82 | 1 | Gm12472 | | 0 | 0 | 0 | 0 | 0 | 7 | 0 | 0 | 0 | 0 | 0 |
| 1 | -0.80 | 3.61 | -0.22 | 0.82 | 1 | Gm25982 | | 0 | 0 | 0 | 0 | 0 | 7 | 0 | 0 | 0 | 0 | 0 |
| 1 | -0.80 | 3.61 | -0.22 | 0.82 | 1 | Gm42551 | | 0 | 0 | 0 | 0 | 0 | 7 | 0 | 0 | 0 | 0 | 0 |
| 1 | -0.80 | 3.61 | -0.22 | 0.82 | 1 | Gm43936 | | 0 | 0 | 0 | 0 | 0 | 7 | 0 | 0 | 0 | 0 | 0 |
| 1 | -0.80 | 3.61 | -0.22 | 0.82 | 1 | Gm10648 | | 0 | 0 | 0 | 0 | 0 | 7 | 0 | 0 | 0 | 0 | 0 |

| | | | | | | | | | | | | | | | | | |
|---|---|---|---|---|---|---|---|---|---|---|---|---|---|---|---|---|---|
| 1 | -0.80 | 3.61 | -0.22 | 0.82 | 1 | BC046251 | | 0 | 0 | 0 | 0 | 0 | 7 | 0 | 0 | 0 | 0 | 0 |
| 1 | -0.80 | 3.61 | -0.22 | 0.82 | 1 | Gm14382 | | 0 | 0 | 0 | 0 | 0 | 7 | 0 | 0 | 0 | 0 | 0 |
| 1 | -0.80 | 3.61 | -0.22 | 0.82 | 1 | Gm18600 | | 0 | 0 | 0 | 0 | 0 | 7 | 0 | 0 | 0 | 0 | 0 |
| 1 | -0.80 | 3.61 | -0.22 | 0.82 | 1 | Gm33053 | | 0 | 0 | 0 | 0 | 0 | 7 | 0 | 0 | 0 | 0 | 0 |
| 1 | -0.80 | 3.61 | -0.22 | 0.82 | 1 | Gm48215 | | 0 | 0 | 0 | 0 | 0 | 7 | 0 | 0 | 0 | 0 | 0 |
| 1 | -0.80 | 3.61 | -0.22 | 0.82 | 1 | Gm47594 | | 0 | 0 | 0 | 0 | 0 | 7 | 0 | 0 | 0 | 0 | 0 |
| 1 | -0.80 | 3.61 | -0.22 | 0.82 | 1 | Gm47794 | | 0 | 0 | 0 | 0 | 0 | 7 | 0 | 0 | 0 | 0 | 0 |
| 1 | 0.00 | 3.63 | 0.00 | 1.00 | 1 | Gm42303 | | 0 | 4 | 0 | 0 | 0 | 2 | 0 | 0 | 0 | 0 | 1 |
| 1 | -0.89 | 3.64 | -0.24 | 0.81 | 1 | Gm9962 | | 0 | 5 | 0 | 0 | 0 | 2 | 0 | 0 | 0 | 0 | 0 |
| 1 | 0.27 | 3.60 | 0.08 | 0.94 | 1 | Gm7556 | | 1 | 0 | 2 | 0 | 1 | 1 | 0 | 0 | 1 | 0 | 1 |
| 1 | 0.86 | 3.58 | 0.24 | 0.81 | 1 | Mir7090 | | 0 | 0 | 0 | 0 | 0 | 2 | 0 | 0 | 5 | 0 | 0 |
| 1 | 0.39 | 3.63 | 0.11 | 0.91 | 1 | Gm12229 | | 2 | 1 | 0 | 0 | 1 | 0 | 0 | 0 | 0 | 0 | 3 |
| 1 | 0.04 | 3.61 | 0.01 | 0.99 | 1 | Gm5218 | | 1 | 0 | 0 | 0 | 1 | 1 | 0 | 0 | 0 | 0 | 4 |
| 1 | -0.88 | 3.63 | -0.24 | 0.81 | 1 | Gm10775 | | 0 | 0 | 4 | 0 | 0 | 3 | 0 | 0 | 0 | 0 | 0 |
| 1 | -1.03 | 3.64 | -0.28 | 0.78 | 1 | Rps2-ps4 | | 1 | 1 | 1 | 0 | 4 | 0 | 0 | 0 | 0 | 0 | 0 |
| 1 | -0.23 | 3.59 | -0.07 | 0.95 | 1 | Gm44026 | | 0 | 0 | 0 | 3 | 0 | 0 | 0 | 0 | 0 | 0 | 4 |
| 1 | 0.00 | 3.68 | 0.00 | 1.00 | 1 | D730045A05Rik | | 3 | 2 | 2 | 0 | 0 | 0 | 0 | 0 | 0 | 0 | 0 |
| 1 | -0.40 | 3.59 | -0.11 | 0.91 | 1 | Gm17829 | | 0 | 0 | 0 | 0 | 4 | 1 | 0 | 0 | 1 | 0 | 1 |
| 1 | 0.33 | 3.61 | 0.09 | 0.93 | 1 | Gm9826 | | 1 | 0 | 2 | 0 | 0 | 2 | 0 | 0 | 2 | 0 | 0 |
| 1 | -0.65 | 3.60 | -0.18 | 0.86 | 1 | Gm12043 | | 0 | 0 | 0 | 0 | 2 | 2 | 0 | 0 | 0 | 0 | 3 |
| 1 | 0.00 | 3.68 | 0.00 | 1.00 | 1 | Kbtbd13 | kelch repeat and BTB (POZ) domain conta | 3 | 4 | 0 | 0 | 0 | 0 | 0 | 0 | 0 | 0 | 0 |
| 1 | -0.20 | 3.61 | -0.05 | 0.96 | 1 | Gm9182 | | 0 | 1 | 0 | 1 | 0 | 2 | 0 | 0 | 0 | 0 | 3 |
| 1 | -1.20 | 3.63 | -0.33 | 0.74 | 1 | Gm37688 | | 0 | 0 | 0 | 0 | 7 | 0 | 0 | 0 | 0 | 0 | 0 |
| 1 | -1.20 | 3.63 | -0.33 | 0.74 | 1 | Gm43106 | | 0 | 0 | 0 | 0 | 7 | 0 | 0 | 0 | 0 | 0 | 0 |
| 1 | -1.20 | 3.63 | -0.33 | 0.74 | 1 | Gm15982 | | 0 | 0 | 0 | 0 | 7 | 0 | 0 | 0 | 0 | 0 | 0 |
| 1 | -1.20 | 3.63 | -0.33 | 0.74 | 1 | Rpl35a-ps6 | | 0 | 0 | 0 | 0 | 7 | 0 | 0 | 0 | 0 | 0 | 0 |
| 1 | -1.20 | 3.63 | -0.33 | 0.74 | 1 | Gm7546 | | 0 | 0 | 0 | 0 | 7 | 0 | 0 | 0 | 0 | 0 | 0 |
| 1 | -1.20 | 3.63 | -0.33 | 0.74 | 1 | Gm47990 | | 0 | 0 | 0 | 0 | 7 | 0 | 0 | 0 | 0 | 0 | 0 |
| 1 | -1.20 | 3.63 | -0.33 | 0.74 | 1 | Gm48300 | | 0 | 0 | 0 | 0 | 7 | 0 | 0 | 0 | 0 | 0 | 0 |
| 1 | -1.20 | 3.63 | -0.33 | 0.74 | 1 | AU015836 | | 0 | 0 | 0 | 0 | 7 | 0 | 0 | 0 | 0 | 0 | 0 |
| 1 | 0.39 | 3.61 | 0.11 | 0.91 | 1 | Mir6999 | | 0 | 0 | 0 | 0 | 0 | 3 | 0 | 0 | 0 | 0 | 4 |
| 1 | 0.39 | 3.61 | 0.11 | 0.91 | 1 | Gm12609 | | 0 | 0 | 0 | 0 | 0 | 3 | 0 | 0 | 0 | 0 | 4 |
| 1 | 0.39 | 3.61 | 0.11 | 0.91 | 1 | A130012E19Rik | | 0 | 0 | 0 | 0 | 0 | 3 | 0 | 0 | 0 | 0 | 4 |
| 1 | 0.95 | 3.61 | 0.26 | 0.79 | 1 | Gm49355 | RIKEN cDNA 3830406C13 gene | 0 | 0 | 0 | 0 | 0 | 3 | 0 | 0 | 1 | 0 | 3 |
| 1 | -0.14 | 3.62 | -0.04 | 0.97 | 1 | Akp-ps1 | | 0 | 3 | 0 | 0 | 0 | 3 | 0 | 0 | 0 | 0 | 1 |
| 1 | 1.17 | 3.60 | 0.33 | 0.74 | 1 | Gm15518 | | 0 | 0 | 0 | 0 | 0 | 3 | 0 | 0 | 2 | 0 | 2 |
| 1 | 0.40 | 3.57 | 0.11 | 0.91 | 1 | BC051077 | | 0 | 0 | 0 | 0 | 0 | 3 | 0 | 0 | 4 | 0 | 0 |
| 1 | 0.15 | 3.62 | 0.04 | 0.97 | 1 | Gm14843 | | 0 | 0 | 1 | 0 | 0 | 4 | 0 | 0 | 0 | 1 | 0 |
| 1 | -1.94 | 3.63 | -0.53 | 0.59 | 1 | Spdye4b | speedy/RINGO cell cycle regulator family | 0 | 0 | 0 | 0 | 6 | 1 | 0 | 0 | 0 | 0 | 0 |
| 1 | -1.94 | 3.63 | -0.53 | 0.59 | 1 | Zcchc13 | zinc finger, CCHC domain containing 13 | 0 | 0 | 0 | 0 | 6 | 1 | 0 | 0 | 0 | 0 | 0 |
| 1 | -0.85 | 3.62 | -0.23 | 0.82 | 1 | Gm37297 | | 0 | 0 | 3 | 0 | 0 | 3 | 0 | 0 | 0 | 0 | 0 |
| 1 | -0.85 | 3.62 | -0.23 | 0.82 | 1 | Aldh3b2 | aldehyde dehydrogenase 3 family, memb | 0 | 0 | 3 | 0 | 0 | 3 | 0 | 0 | 0 | 0 | 0 |
| 1 | -2.09 | 3.62 | -0.58 | 0.56 | 1 | Gm16116 | | 0 | 0 | 0 | 2 | 5 | 0 | 0 | 0 | 0 | 0 | 0 |
| 1 | 0.00 | 3.68 | 0.00 | 1.00 | 1 | Gm8979 | | 3 | 3 | 0 | 0 | 0 | 0 | 0 | 0 | 0 | 0 | 0 |
| 1 | 0.00 | 3.68 | 0.00 | 1.00 | 1 | Gm31378 | | 3 | 3 | 0 | 0 | 0 | 0 | 0 | 0 | 0 | 0 | 0 |
| 1 | -1.16 | 3.63 | -0.32 | 0.75 | 1 | Gm6028 | | 1 | 1 | 0 | 3 | 1 | 0 | 0 | 0 | 0 | 0 | 1 |
| 1 | 1.76 | 3.64 | 0.48 | 0.63 | 1 | Gm10518 | | 0 | 0 | 0 | 0 | 0 | 0 | 0 | 0 | 0 | 6 | 0 |
| 1 | 1.76 | 3.64 | 0.48 | 0.63 | 1 | Prok2 | prokineticin 2 | 0 | 0 | 0 | 0 | 0 | 0 | 0 | 0 | 0 | 6 | 0 |
| 1 | 1.76 | 3.64 | 0.48 | 0.63 | 1 | Gm5509 | | 0 | 0 | 0 | 0 | 0 | 0 | 0 | 0 | 0 | 6 | 0 |
| 1 | 0.16 | 3.62 | 0.04 | 0.96 | 1 | Gm12969 | | 1 | 0 | 1 | 0 | 0 | 3 | 0 | 1 | 0 | 0 | 0 |
| 1 | 0.16 | 3.61 | 0.05 | 0.96 | 1 | Gm8251 | | 0 | 0 | 0 | 0 | 0 | 3 | 0 | 0 | 0 | 0 | 3 |
| 1 | 0.16 | 3.61 | 0.05 | 0.96 | 1 | Gm43628 | | 0 | 0 | 0 | 0 | 0 | 3 | 0 | 0 | 0 | 0 | 3 |
| 1 | 0.16 | 3.61 | 0.05 | 0.96 | 1 | Gm18981 | | 0 | 0 | 0 | 0 | 0 | 3 | 0 | 0 | 0 | 0 | 3 |
| 1 | 0.16 | 3.61 | 0.05 | 0.96 | 1 | Gm11723 | | 0 | 0 | 0 | 0 | 0 | 3 | 0 | 0 | 0 | 0 | 3 |
| 1 | 0.01 | 3.61 | 0.00 | 1.00 | 1 | Ckb-ps2 | | 0 | 1 | 0 | 0 | 0 | 3 | 0 | 0 | 0 | 0 | 2 |
| 1 | 0.01 | 3.61 | 0.00 | 1.00 | 1 | mt-Tm | | 0 | 1 | 0 | 0 | 0 | 3 | 0 | 0 | 0 | 0 | 2 |
| 1 | -0.07 | 3.64 | -0.02 | 0.99 | 1 | Gm45663 | | 4 | 0 | 0 | 0 | 0 | 1 | 0 | 0 | 0 | 1 | 0 |
| 1 | 0.93 | 3.60 | 0.26 | 0.80 | 1 | Gm8206 | | 0 | 0 | 0 | 0 | 0 | 3 | 0 | 0 | 2 | 0 | 1 |
| 1 | -1.32 | 3.59 | -0.37 | 0.71 | 1 | Gm9231 | | 0 | 0 | 0 | 2 | 1 | 2 | 0 | 0 | 0 | 0 | 2 |
| 1 | 0.79 | 3.59 | 0.22 | 0.82 | 1 | Gm37390 | | 0 | 0 | 0 | 0 | 0 | 3 | 0 | 0 | 3 | 0 | 0 |

| | | | | | | | | | | | | | | | | | |
|---|---|---|---|---|---|---|---|---|---|---|---|---|---|---|---|---|---|
| 1 | 0.79 | 3.59 | 0.22 | 0.82 | 1 | Ucp3 | uncoupling protein 3 (mitochondrial, protc | 0 | 0 | 0 | 0 | 0 | 3 | 0 | 0 | 3 | 0 | 0 |
| 1 | 0.79 | 3.59 | 0.22 | 0.82 | 1 | Gm32828 | | 0 | 0 | 0 | 0 | 0 | 3 | 0 | 0 | 3 | 0 | 0 |
| 1 | 0.79 | 3.59 | 0.22 | 0.82 | 1 | Gm48885 | | 0 | 0 | 0 | 0 | 0 | 3 | 0 | 0 | 3 | 0 | 0 |
| 1 | -1.34 | 3.65 | -0.37 | 0.71 | 1 | Gm17241 | | 2 | 1 | 1 | 0 | 1 | 2 | 0 | 0 | 0 | 0 | 0 |
| 1 | -0.63 | 3.63 | -0.17 | 0.86 | 1 | Gm16585 | | 2 | 0 | 1 | 0 | 1 | 2 | 0 | 0 | 1 | 0 | 0 |
| 1 | 0.00 | 3.68 | 0.00 | 1.00 | 1 | Gm9754 | | 4 | 0 | 2 | 0 | 0 | 0 | 0 | 0 | 0 | 0 | 0 |
| 1 | 1.99 | 3.65 | 0.55 | 0.58 | 1 | Gm5175 | | 2 | 0 | 1 | 0 | 0 | 0 | 0 | 2 | 0 | 1 | 0 |
| 1 | -1.69 | 3.63 | -0.47 | 0.64 | 1 | Psca | prostate stem cell antigen | 0 | 2 | 0 | 3 | 0 | 2 | 0 | 0 | 0 | 0 | 0 |
| 1 | 2.49 | 3.62 | 0.69 | 0.49 | 1 | Gm15464 | | 1 | 1 | 0 | 0 | 0 | 0 | 0 | 1 | 2 | 1 | 0 |
| 1 | -2.15 | 3.62 | -0.60 | 0.55 | 1 | Gm17035 | | 0 | 0 | 0 | 4 | 3 | 0 | 0 | 0 | 0 | 0 | 0 |
| 1 | -1.00 | 3.66 | -0.27 | 0.78 | 1 | Gm9132 | | 3 | 2 | 0 | 1 | 0 | 1 | 0 | 0 | 0 | 0 | 0 |
| 1 | 1.10 | 3.67 | 0.30 | 0.77 | 1 | Gm11127 | predicted gene 11127 | 5 | 0 | 0 | 0 | 0 | 0 | 0 | 1 | 0 | 0 | 0 |
| 1 | 0.27 | 3.62 | 0.07 | 0.94 | 1 | Gm5599 | | 0 | 2 | 0 | 1 | 1 | 0 | 0 | 1 | 0 | 1 | 0 |
| 1 | 0.01 | 3.61 | 0.00 | 1.00 | 1 | Hm629797 | | 1 | 0 | 0 | 0 | 0 | 3 | 0 | 0 | 0 | 0 | 2 |
| 1 | 0.01 | 3.61 | 0.00 | 1.00 | 1 | Asgr1 | asialoglycoprotein receptor 1 | 1 | 0 | 0 | 0 | 0 | 3 | 0 | 0 | 0 | 0 | 2 |
| 1 | 2.36 | 3.63 | 0.65 | 0.51 | 1 | Gm13690 | | 0 | 0 | 1 | 0 | 0 | 0 | 0 | 1 | 0 | 0 | 4 |
| 1 | -0.08 | 3.61 | -0.02 | 0.98 | 1 | Gm13574 | | 0 | 0 | 0 | 0 | 0 | 4 | 0 | 0 | 0 | 0 | 2 |
| 1 | -0.08 | 3.61 | -0.02 | 0.98 | 1 | Cbx3-ps3 | | 0 | 0 | 0 | 0 | 0 | 4 | 0 | 0 | 0 | 0 | 2 |
| 1 | -0.08 | 3.61 | -0.02 | 0.98 | 1 | Gm43490 | | 0 | 0 | 0 | 0 | 0 | 4 | 0 | 0 | 0 | 0 | 2 |
| 1 | -0.08 | 3.61 | -0.02 | 0.98 | 1 | 4930520P13Rik | | 0 | 0 | 0 | 0 | 0 | 4 | 0 | 0 | 0 | 0 | 2 |
| 1 | -0.08 | 3.61 | -0.02 | 0.98 | 1 | Gm49297 | | 0 | 0 | 0 | 0 | 0 | 4 | 0 | 0 | 0 | 0 | 2 |
| 1 | 0.16 | 3.62 | 0.04 | 0.96 | 1 | Gm38082 | | 2 | 0 | 0 | 0 | 0 | 3 | 0 | 1 | 0 | 0 | 0 |
| 1 | -0.83 | 3.62 | -0.23 | 0.82 | 1 | Gm3912 | | 0 | 2 | 0 | 0 | 0 | 4 | 0 | 0 | 0 | 0 | 0 |
| 1 | 1.59 | 3.66 | 0.43 | 0.66 | 1 | Gm11878 | | 1 | 1 | 2 | 0 | 0 | 0 | 0 | 0 | 1 | 1 | 0 |
| 1 | 0.00 | 3.67 | 0.00 | 1.00 | 1 | Hoxd4 | homeobox D4 | 0 | 0 | 6 | 0 | 0 | 0 | 0 | 0 | 0 | 0 | 0 |
| 1 | 0.00 | 3.67 | 0.00 | 1.00 | 1 | A730089K16Rik | | 0 | 0 | 6 | 0 | 0 | 0 | 0 | 0 | 0 | 0 | 0 |
| 1 | 0.00 | 3.67 | 0.00 | 1.00 | 1 | 1700034G24Rik | | 0 | 0 | 6 | 0 | 0 | 0 | 0 | 0 | 0 | 0 | 0 |
| 1 | 0.00 | 3.67 | 0.00 | 1.00 | 1 | 5330437M03Rik | | 0 | 0 | 6 | 0 | 0 | 0 | 0 | 0 | 0 | 0 | 0 |
| 1 | 0.00 | 3.67 | 0.00 | 1.00 | 1 | Nanos2 | nanos C2HC-type zinc finger 2 | 0 | 0 | 6 | 0 | 0 | 0 | 0 | 0 | 0 | 0 | 0 |
| 1 | 0.00 | 3.67 | 0.00 | 1.00 | 1 | Gm44689 | | 0 | 0 | 6 | 0 | 0 | 0 | 0 | 0 | 0 | 0 | 0 |
| 1 | 0.00 | 3.67 | 0.00 | 1.00 | 1 | Smim31 | small integral membrane protein 31 | 0 | 0 | 6 | 0 | 0 | 0 | 0 | 0 | 0 | 0 | 0 |
| 1 | 0.00 | 3.67 | 0.00 | 1.00 | 1 | Gm47625 | | 0 | 0 | 6 | 0 | 0 | 0 | 0 | 0 | 0 | 0 | 0 |
| 1 | 0.00 | 3.67 | 0.00 | 1.00 | 1 | Gm47093 | | 0 | 0 | 6 | 0 | 0 | 0 | 0 | 0 | 0 | 0 | 0 |
| 1 | 0.00 | 3.67 | 0.00 | 1.00 | 1 | Gm11951 | | 0 | 0 | 6 | 0 | 0 | 0 | 0 | 0 | 0 | 0 | 0 |
| 1 | 0.00 | 3.67 | 0.00 | 1.00 | 1 | Gm11615 | | 0 | 0 | 6 | 0 | 0 | 0 | 0 | 0 | 0 | 0 | 0 |
| 1 | 0.00 | 3.67 | 0.00 | 1.00 | 1 | BC006965 | | 0 | 0 | 6 | 0 | 0 | 0 | 0 | 0 | 0 | 0 | 0 |
| 1 | 0.00 | 3.67 | 0.00 | 1.00 | 1 | Gm36635 | | 0 | 0 | 6 | 0 | 0 | 0 | 0 | 0 | 0 | 0 | 0 |
| 1 | 0.00 | 3.67 | 0.00 | 1.00 | 1 | D030007L05Rik | | 0 | 0 | 6 | 0 | 0 | 0 | 0 | 0 | 0 | 0 | 0 |
| 1 | 0.00 | 3.67 | 0.00 | 1.00 | 1 | Gm5796 | predicted gene 5796 | 0 | 0 | 6 | 0 | 0 | 0 | 0 | 0 | 0 | 0 | 0 |
| 1 | 0.00 | 3.67 | 0.00 | 1.00 | 1 | Gm15222 | | 0 | 0 | 6 | 0 | 0 | 0 | 0 | 0 | 0 | 0 | 0 |
| 1 | 0.00 | 3.67 | 0.00 | 1.00 | 1 | Trdv4 | | 0 | 0 | 6 | 0 | 0 | 0 | 0 | 0 | 0 | 0 | 0 |
| 1 | 0.00 | 3.67 | 0.00 | 1.00 | 1 | Krt5 | keratin 5 | 0 | 0 | 6 | 0 | 0 | 0 | 0 | 0 | 0 | 0 | 0 |
| 1 | 0.00 | 3.67 | 0.00 | 1.00 | 1 | Gm36200 | | 0 | 0 | 6 | 0 | 0 | 0 | 0 | 0 | 0 | 0 | 0 |
| 1 | -2.14 | 3.62 | -0.59 | 0.55 | 1 | Gm37802 | | 0 | 0 | 0 | 0 | 3 | 3 | 0 | 0 | 0 | 0 | 0 |
| 1 | -2.14 | 3.62 | -0.59 | 0.55 | 1 | Gm4971 | | 0 | 0 | 0 | 0 | 3 | 3 | 0 | 0 | 0 | 0 | 0 |
| 1 | 0.31 | 3.65 | 0.08 | 0.93 | 1 | Gm7434 | | 0 | 2 | 2 | 0 | 1 | 0 | 0 | 1 | 0 | 0 | 0 |
| 1 | 0.74 | 3.63 | 0.21 | 0.84 | 1 | Gm18913 | | 0 | 0 | 2 | 1 | 1 | 0 | 0 | 0 | 0 | 1 | 1 |
| 1 | 0.00 | 3.68 | 0.00 | 1.00 | 1 | Fam43b | family with sequence similarity 43, membe | 0 | 2 | 4 | 0 | 0 | 0 | 0 | 0 | 0 | 0 | 0 |
| 1 | 1.55 | 3.62 | 0.43 | 0.67 | 1 | Gm7860 | | 1 | 1 | 0 | 0 | 0 | 1 | 0 | 1 | 0 | 1 | 1 |
| 1 | -2.39 | 3.60 | -0.66 | 0.51 | 1 | Gm46515 | | 0 | 0 | 0 | 4 | 1 | 1 | 0 | 0 | 0 | 0 | 0 |
| 1 | 0.00 | 3.68 | 0.00 | 1.00 | 1 | Fam178b | family with sequence similarity 178, meml | 0 | 4 | 2 | 0 | 0 | 0 | 0 | 0 | 0 | 0 | 0 |
| 1 | -1.27 | 3.63 | -0.35 | 0.73 | 1 | Mir8120 | | 0 | 0 | 0 | 6 | 0 | 0 | 0 | 0 | 0 | 0 | 0 |
| 1 | -1.27 | 3.63 | -0.35 | 0.73 | 1 | Gm15788 | | 0 | 0 | 0 | 6 | 0 | 0 | 0 | 0 | 0 | 0 | 0 |
| 1 | -1.27 | 3.63 | -0.35 | 0.73 | 1 | Gm44832 | | 0 | 0 | 0 | 6 | 0 | 0 | 0 | 0 | 0 | 0 | 0 |
| 1 | -1.27 | 3.63 | -0.35 | 0.73 | 1 | Olfr312 | olfactory receptor 312 | 0 | 0 | 0 | 6 | 0 | 0 | 0 | 0 | 0 | 0 | 0 |
| 1 | -1.27 | 3.63 | -0.35 | 0.73 | 1 | Gm47849 | | 0 | 0 | 0 | 6 | 0 | 0 | 0 | 0 | 0 | 0 | 0 |
| 1 | -1.27 | 3.63 | -0.35 | 0.73 | 1 | Hrh4 | histamine receptor H4 | 0 | 0 | 0 | 6 | 0 | 0 | 0 | 0 | 0 | 0 | 0 |
| 1 | -1.98 | 3.62 | -0.55 | 0.59 | 1 | H2al1m | H2A histone family member L1M | 0 | 0 | 0 | 0 | 2 | 4 | 0 | 0 | 0 | 0 | 0 |
| 1 | -1.37 | 3.64 | -0.38 | 0.71 | 1 | Rhoay-ps3 | | 1 | 0 | 0 | 5 | 0 | 0 | 0 | 0 | 0 | 0 | 0 |
| 1 | -0.31 | 3.61 | -0.09 | 0.93 | 1 | Gm17112 | | 1 | 0 | 0 | 0 | 0 | 4 | 0 | 0 | 0 | 0 | 1 |

| | | | | | | | | | | | | | | | | | |
|---|---|---|---|---|---|---|---|---|---|---|---|---|---|---|---|---|---|
| 1 | -1.91 | 3.63 | -0.53 | 0.60 | 1 | Gm16229 | | 0 | 0 | 0 | 5 | 0 | 1 | 0 | 0 | 0 | 0 | 0 |
| 1 | -1.91 | 3.63 | -0.53 | 0.60 | 1 | Gm49009 | | 0 | 0 | 0 | 5 | 0 | 1 | 0 | 0 | 0 | 0 | 0 |
| 1 | 1.35 | 3.62 | 0.37 | 0.71 | 1 | Gm37315 | | 0 | 0 | 0 | 0 | 0 | 0 | 0 | 0 | 0 | 0 | 6 |
| 1 | 1.35 | 3.62 | 0.37 | 0.71 | 1 | Il17a | interleukin 17A | 0 | 0 | 0 | 0 | 0 | 0 | 0 | 0 | 0 | 0 | 6 |
| 1 | 1.35 | 3.62 | 0.37 | 0.71 | 1 | Gm25035 | | 0 | 0 | 0 | 0 | 0 | 0 | 0 | 0 | 0 | 0 | 6 |
| 1 | 1.35 | 3.62 | 0.37 | 0.71 | 1 | Gm37101 | | 0 | 0 | 0 | 0 | 0 | 0 | 0 | 0 | 0 | 0 | 6 |
| 1 | 1.35 | 3.62 | 0.37 | 0.71 | 1 | 1700003F12Ri | RIKEN cDNA 1700003F12 gene | 0 | 0 | 0 | 0 | 0 | 0 | 0 | 0 | 0 | 0 | 6 |
| 1 | 1.35 | 3.62 | 0.37 | 0.71 | 1 | 2610316D01Rik | | 0 | 0 | 0 | 0 | 0 | 0 | 0 | 0 | 0 | 0 | 6 |
| 1 | 1.35 | 3.62 | 0.37 | 0.71 | 1 | Gm17951 | | 0 | 0 | 0 | 0 | 0 | 0 | 0 | 0 | 0 | 0 | 6 |
| 1 | 1.35 | 3.62 | 0.37 | 0.71 | 1 | Gm43560 | | 0 | 0 | 0 | 0 | 0 | 0 | 0 | 0 | 0 | 0 | 6 |
| 1 | 1.35 | 3.62 | 0.37 | 0.71 | 1 | Mos | Moloney sarcoma oncogene | 0 | 0 | 0 | 0 | 0 | 0 | 0 | 0 | 0 | 0 | 6 |
| 1 | 1.35 | 3.62 | 0.37 | 0.71 | 1 | 2010310C07Rik | | 0 | 0 | 0 | 0 | 0 | 0 | 0 | 0 | 0 | 0 | 6 |
| 1 | 1.35 | 3.62 | 0.37 | 0.71 | 1 | Mir704 | | 0 | 0 | 0 | 0 | 0 | 0 | 0 | 0 | 0 | 0 | 6 |
| 1 | 1.35 | 3.62 | 0.37 | 0.71 | 1 | Gm44433 | | 0 | 0 | 0 | 0 | 0 | 0 | 0 | 0 | 0 | 0 | 6 |
| 1 | 1.35 | 3.62 | 0.37 | 0.71 | 1 | Gm44438 | | 0 | 0 | 0 | 0 | 0 | 0 | 0 | 0 | 0 | 0 | 6 |
| 1 | 1.35 | 3.62 | 0.37 | 0.71 | 1 | Gm44045 | | 0 | 0 | 0 | 0 | 0 | 0 | 0 | 0 | 0 | 0 | 6 |
| 1 | 1.35 | 3.62 | 0.37 | 0.71 | 1 | Gm44189 | | 0 | 0 | 0 | 0 | 0 | 0 | 0 | 0 | 0 | 0 | 6 |
| 1 | 1.35 | 3.62 | 0.37 | 0.71 | 1 | Gm43635 | | 0 | 0 | 0 | 0 | 0 | 0 | 0 | 0 | 0 | 0 | 6 |
| 1 | 1.35 | 3.62 | 0.37 | 0.71 | 1 | Hamp | hepcidin antimicrobial peptide | 0 | 0 | 0 | 0 | 0 | 0 | 0 | 0 | 0 | 0 | 6 |
| 1 | 1.35 | 3.62 | 0.37 | 0.71 | 1 | Trim58 | tripartite motif-containing 58 | 0 | 0 | 0 | 0 | 0 | 0 | 0 | 0 | 0 | 0 | 6 |
| 1 | 1.35 | 3.62 | 0.37 | 0.71 | 1 | Krt28 | keratin 28 | 0 | 0 | 0 | 0 | 0 | 0 | 0 | 0 | 0 | 0 | 6 |
| 1 | 1.35 | 3.62 | 0.37 | 0.71 | 1 | 1100001I12Rik | | 0 | 0 | 0 | 0 | 0 | 0 | 0 | 0 | 0 | 0 | 6 |
| 1 | 1.35 | 3.62 | 0.37 | 0.71 | 1 | Dsg1c | desmoglein 1 gamma | 0 | 0 | 0 | 0 | 0 | 0 | 0 | 0 | 0 | 0 | 6 |
| 1 | 1.35 | 3.62 | 0.37 | 0.71 | 1 | Gm28374 | | 0 | 0 | 0 | 0 | 0 | 0 | 0 | 0 | 0 | 0 | 6 |
| 1 | 1.35 | 3.62 | 0.37 | 0.71 | 1 | Trex2 | three prime repair exonuclease 2 | 0 | 0 | 0 | 0 | 0 | 0 | 0 | 0 | 0 | 0 | 6 |
| 1 | -0.57 | 3.67 | -0.16 | 0.88 | 1 | Gm26761 | | 5 | 0 | 0 | 0 | 1 | 0 | 0 | 0 | 0 | 0 | 0 |
| 1 | -0.57 | 3.67 | -0.16 | 0.88 | 1 | Pnma3 | paraneoplastic antigen MA3 | 5 | 0 | 0 | 0 | 1 | 0 | 0 | 0 | 0 | 0 | 0 |
| 1 | 1.32 | 3.63 | 0.36 | 0.72 | 1 | Gm47480 | | 0 | 2 | 0 | 0 | 0 | 0 | 0 | 0 | 0 | 0 | 4 |
| 1 | -0.36 | 3.61 | -0.10 | 0.92 | 1 | Gm5709 | predicted gene 5709 | 0 | 0 | 0 | 0 | 0 | 5 | 0 | 0 | 0 | 0 | 1 |
| 1 | -0.36 | 3.61 | -0.10 | 0.92 | 1 | Gm22518 | | 0 | 0 | 0 | 0 | 0 | 5 | 0 | 0 | 0 | 0 | 1 |
| 1 | -0.36 | 3.61 | -0.10 | 0.92 | 1 | Gm45269 | | 0 | 0 | 0 | 0 | 0 | 5 | 0 | 0 | 0 | 0 | 1 |
| 1 | -0.36 | 3.61 | -0.10 | 0.92 | 1 | Gm45360 | | 0 | 0 | 0 | 0 | 0 | 5 | 0 | 0 | 0 | 0 | 1 |
| 1 | -0.36 | 3.61 | -0.10 | 0.92 | 1 | Gm25694 | | 0 | 0 | 0 | 0 | 0 | 5 | 0 | 0 | 0 | 0 | 1 |
| 1 | -0.36 | 3.61 | -0.10 | 0.92 | 1 | Gm22771 | | 0 | 0 | 0 | 0 | 0 | 5 | 0 | 0 | 0 | 0 | 1 |
| 1 | -0.36 | 3.61 | -0.10 | 0.92 | 1 | Mir1291 | | 0 | 0 | 0 | 0 | 0 | 5 | 0 | 0 | 0 | 0 | 1 |
| 1 | -0.36 | 3.61 | -0.10 | 0.92 | 1 | Gm22858 | | 0 | 0 | 0 | 0 | 0 | 5 | 0 | 0 | 0 | 0 | 1 |
| 1 | 2.26 | 3.63 | 0.62 | 0.53 | 1 | Gm12094 | | 0 | 1 | 0 | 0 | 0 | 0 | 0 | 0 | 1 | 0 | 4 |
| 1 | -0.79 | 3.62 | -0.22 | 0.83 | 1 | Mir541 | | 0 | 1 | 0 | 0 | 0 | 5 | 0 | 0 | 0 | 0 | 0 |
| 1 | 2.52 | 3.63 | 0.69 | 0.49 | 1 | Gm41662 | | 0 | 0 | 0 | 0 | 0 | 0 | 0 | 0 | 3 | 0 | 3 |
| 1 | 2.52 | 3.63 | 0.69 | 0.49 | 1 | Gm6938 | | 0 | 0 | 0 | 0 | 0 | 0 | 0 | 0 | 3 | 0 | 3 |
| 1 | 0.99 | 3.65 | 0.27 | 0.79 | 1 | Gm4963 | | 0 | 2 | 1 | 0 | 0 | 1 | 0 | 1 | 1 | 0 | 0 |
| 1 | 1.02 | 3.66 | 0.28 | 0.78 | 1 | B230334C09Rik | | 0 | 5 | 0 | 0 | 0 | 0 | 0 | 0 | 0 | 0 | 1 |
| 1 | 2.23 | 3.64 | 0.61 | 0.54 | 1 | Gm43096 | | 1 | 0 | 0 | 0 | 0 | 0 | 0 | 1 | 0 | 0 | 4 |
| 1 | 0.00 | 3.68 | 0.00 | 1.00 | 1 | Gm29695 | | 0 | 6 | 0 | 0 | 0 | 0 | 0 | 0 | 0 | 0 | 0 |
| 1 | 0.00 | 3.68 | 0.00 | 1.00 | 1 | Gm15850 | | 0 | 6 | 0 | 0 | 0 | 0 | 0 | 0 | 0 | 0 | 0 |
| 1 | 0.00 | 3.68 | 0.00 | 1.00 | 1 | Gm13446 | | 0 | 6 | 0 | 0 | 0 | 0 | 0 | 0 | 0 | 0 | 0 |
| 1 | 0.00 | 3.68 | 0.00 | 1.00 | 1 | Kcna10 | potassium voltage-gated channel, shaker | 0 | 6 | 0 | 0 | 0 | 0 | 0 | 0 | 0 | 0 | 0 |
| 1 | 0.00 | 3.68 | 0.00 | 1.00 | 1 | Gm12679 | | 0 | 6 | 0 | 0 | 0 | 0 | 0 | 0 | 0 | 0 | 0 |
| 1 | 0.00 | 3.68 | 0.00 | 1.00 | 1 | | | 0 | 6 | 0 | 0 | 0 | 0 | 0 | 0 | 0 | 0 | 0 |
| 1 | 0.00 | 3.68 | 0.00 | 1.00 | 1 | Rnf212 | ring finger protein 212 | 0 | 6 | 0 | 0 | 0 | 0 | 0 | 0 | 0 | 0 | 0 |
| 1 | 0.00 | 3.68 | 0.00 | 1.00 | 1 | Gm30003 | | 0 | 6 | 0 | 0 | 0 | 0 | 0 | 0 | 0 | 0 | 0 |
| 1 | 0.00 | 3.68 | 0.00 | 1.00 | 1 | Cd177 | CD177 antigen | 0 | 6 | 0 | 0 | 0 | 0 | 0 | 0 | 0 | 0 | 0 |
| 1 | 0.00 | 3.68 | 0.00 | 1.00 | 1 | Gm36864 | predicted gene, 36864 | 0 | 6 | 0 | 0 | 0 | 0 | 0 | 0 | 0 | 0 | 0 |
| 1 | 0.00 | 3.68 | 0.00 | 1.00 | 1 | Tpte | transmembrane phosphatase with tensin | 0 | 6 | 0 | 0 | 0 | 0 | 0 | 0 | 0 | 0 | 0 |
| 1 | 0.00 | 3.68 | 0.00 | 1.00 | 1 | Gm48385 | | 0 | 6 | 0 | 0 | 0 | 0 | 0 | 0 | 0 | 0 | 0 |
| 1 | 0.00 | 3.68 | 0.00 | 1.00 | 1 | Mfsd4b2 | | 0 | 6 | 0 | 0 | 0 | 0 | 0 | 0 | 0 | 0 | 0 |
| 1 | 0.00 | 3.68 | 0.00 | 1.00 | 1 | Gm47398 | | 0 | 6 | 0 | 0 | 0 | 0 | 0 | 0 | 0 | 0 | 0 |
| 1 | 0.00 | 3.68 | 0.00 | 1.00 | 1 | 4930442H23Rik | | 0 | 6 | 0 | 0 | 0 | 0 | 0 | 0 | 0 | 0 | 0 |
| 1 | 0.00 | 3.68 | 0.00 | 1.00 | 1 | Gm11204 | | 0 | 6 | 0 | 0 | 0 | 0 | 0 | 0 | 0 | 0 | 0 |
| 1 | 0.00 | 3.68 | 0.00 | 1.00 | 1 | Gm9315 | | 0 | 6 | 0 | 0 | 0 | 0 | 0 | 0 | 0 | 0 | 0 |

| | | | | | | | | | | | | | | | | | | |
|---|---|---|---|---|---|---|---|---|---|---|---|---|---|---|---|---|---|---|
| 1 | 0.00 | 3.68 | 0.00 | 1.00 | 1 | Ankrd33 | ankyrin repeat domain 33 | 0 | 6 | 0 | 0 | 0 | 0 | 0 | 0 | 0 | 0 | 0 |
| 1 | 0.00 | 3.68 | 0.00 | 1.00 | 1 | Mir5107 | | 0 | 6 | 0 | 0 | 0 | 0 | 0 | 0 | 0 | 0 | 0 |
| 1 | 0.00 | 3.68 | 0.00 | 1.00 | 1 | E330013P04Rik | | 0 | 6 | 0 | 0 | 0 | 0 | 0 | 0 | 0 | 0 | 0 |
| 1 | 1.14 | 3.67 | 0.31 | 0.76 | 1 | Gm34653 | predicted gene, 34653 | 0 | 5 | 0 | 0 | 0 | 0 | 0 | 0 | 1 | 0 | 0 |
| 1 | 2.36 | 3.63 | 0.65 | 0.52 | 1 | Gm15573 | | 0 | 0 | 0 | 0 | 0 | 0 | 0 | 0 | 5 | 0 | 1 |
| 1 | 1.80 | 3.64 | 0.49 | 0.62 | 1 | 9430024E24Rik | | 0 | 0 | 0 | 0 | 0 | 0 | 0 | 0 | 6 | 0 | 0 |
| 1 | 1.80 | 3.64 | 0.49 | 0.62 | 1 | Gm8579 | | 0 | 0 | 0 | 0 | 0 | 0 | 0 | 0 | 6 | 0 | 0 |
| 1 | 1.80 | 3.64 | 0.49 | 0.62 | 1 | Gm44288 | | 0 | 0 | 0 | 0 | 0 | 0 | 0 | 0 | 6 | 0 | 0 |
| 1 | 1.80 | 3.64 | 0.49 | 0.62 | 1 | Gm25780 | | 0 | 0 | 0 | 0 | 0 | 0 | 0 | 0 | 6 | 0 | 0 |
| 1 | 1.80 | 3.64 | 0.49 | 0.62 | 1 | Gm22513 | | 0 | 0 | 0 | 0 | 0 | 0 | 0 | 0 | 6 | 0 | 0 |
| 1 | 1.80 | 3.64 | 0.49 | 0.62 | 1 | Mir1197 | | 0 | 0 | 0 | 0 | 0 | 0 | 0 | 0 | 6 | 0 | 0 |
| 1 | 1.80 | 3.64 | 0.49 | 0.62 | 1 | Gm49743 | | 0 | 0 | 0 | 0 | 0 | 0 | 0 | 0 | 6 | 0 | 0 |
| 1 | 1.80 | 3.64 | 0.49 | 0.62 | 1 | 4833415N18Rik | | 0 | 0 | 0 | 0 | 0 | 0 | 0 | 0 | 6 | 0 | 0 |
| 1 | 1.80 | 3.64 | 0.49 | 0.62 | 1 | 2010320O07Rik | | 0 | 0 | 0 | 0 | 0 | 0 | 0 | 0 | 6 | 0 | 0 |
| 1 | -2.07 | 3.63 | -0.57 | 0.57 | 1 | Gm14123 | | 0 | 0 | 0 | 4 | 0 | 2 | 0 | 0 | 0 | 0 | 0 |
| 1 | 0.00 | 3.68 | 0.00 | 1.00 | 1 | Rpl10a-ps4 | | 1 | 0 | 5 | 0 | 0 | 0 | 0 | 0 | 0 | 0 | 0 |
| 1 | 0.00 | 3.68 | 0.00 | 1.00 | 1 | Gm19585 | | 1 | 0 | 5 | 0 | 0 | 0 | 0 | 0 | 0 | 0 | 0 |
| 1 | -0.47 | 3.66 | -0.13 | 0.90 | 1 | Gm28178 | | 0 | 0 | 5 | 0 | 0 | 1 | 0 | 0 | 0 | 0 | 0 |
| 1 | 0.41 | 3.62 | 0.11 | 0.91 | 1 | 1700021J08Rik | | 0 | 0 | 0 | 0 | 1 | 0 | 0 | 0 | 0 | 0 | 5 |
| 1 | 0.41 | 3.62 | 0.11 | 0.91 | 1 | Gm18555 | | 0 | 0 | 0 | 0 | 1 | 0 | 0 | 0 | 0 | 0 | 5 |
| 1 | 0.41 | 3.62 | 0.11 | 0.91 | 1 | Gm12230 | | 0 | 0 | 0 | 0 | 1 | 0 | 0 | 0 | 0 | 0 | 5 |
| 1 | 1.16 | 3.62 | 0.32 | 0.75 | 1 | Gm11760 | | 0 | 1 | 0 | 0 | 1 | 0 | 0 | 0 | 1 | 0 | 3 |
| 1 | 0.00 | 3.68 | 0.00 | 1.00 | 1 | 4931408C20Ril | RIKEN cDNA 4931408C20 gene | 6 | 0 | 0 | 0 | 0 | 0 | 0 | 0 | 0 | 0 | 0 |
| 1 | 0.00 | 3.68 | 0.00 | 1.00 | 1 | Gbx2 | gastrulation brain homeobox 2 | 6 | 0 | 0 | 0 | 0 | 0 | 0 | 0 | 0 | 0 | 0 |
| 1 | 0.00 | 3.68 | 0.00 | 1.00 | 1 | Olfr1034 | olfactory receptor 1034 | 6 | 0 | 0 | 0 | 0 | 0 | 0 | 0 | 0 | 0 | 0 |
| 1 | 0.00 | 3.68 | 0.00 | 1.00 | 1 | Gm42699 | | 6 | 0 | 0 | 0 | 0 | 0 | 0 | 0 | 0 | 0 | 0 |
| 1 | 0.00 | 3.68 | 0.00 | 1.00 | 1 | Gm42639 | | 6 | 0 | 0 | 0 | 0 | 0 | 0 | 0 | 0 | 0 | 0 |
| 1 | 0.00 | 3.68 | 0.00 | 1.00 | 1 | Cxcl3 | chemokine (C-X-C motif) ligand 3 | 6 | 0 | 0 | 0 | 0 | 0 | 0 | 0 | 0 | 0 | 0 |
| 1 | 0.00 | 3.68 | 0.00 | 1.00 | 1 | Gm13836 | | 6 | 0 | 0 | 0 | 0 | 0 | 0 | 0 | 0 | 0 | 0 |
| 1 | 0.00 | 3.68 | 0.00 | 1.00 | 1 | Trbv29 | | 6 | 0 | 0 | 0 | 0 | 0 | 0 | 0 | 0 | 0 | 0 |
| 1 | 0.00 | 3.68 | 0.00 | 1.00 | 1 | Igkv1-131 | | 6 | 0 | 0 | 0 | 0 | 0 | 0 | 0 | 0 | 0 | 0 |
| 1 | 0.00 | 3.68 | 0.00 | 1.00 | 1 | Nat8f7 | N-acetyltransferase 8 (GCN5-related) fam | 6 | 0 | 0 | 0 | 0 | 0 | 0 | 0 | 0 | 0 | 0 |
| 1 | 0.00 | 3.68 | 0.00 | 1.00 | 1 | Trim30e-ps1 | | 6 | 0 | 0 | 0 | 0 | 0 | 0 | 0 | 0 | 0 | 0 |
| 1 | 0.00 | 3.68 | 0.00 | 1.00 | 1 | Gm27530 | | 6 | 0 | 0 | 0 | 0 | 0 | 0 | 0 | 0 | 0 | 0 |
| 1 | 0.00 | 3.68 | 0.00 | 1.00 | 1 | Gm23811 | | 6 | 0 | 0 | 0 | 0 | 0 | 0 | 0 | 0 | 0 | 0 |
| 1 | 0.00 | 3.68 | 0.00 | 1.00 | 1 | Bcl2l10 | Bcl2-like 10 | 6 | 0 | 0 | 0 | 0 | 0 | 0 | 0 | 0 | 0 | 0 |
| 1 | 0.00 | 3.68 | 0.00 | 1.00 | 1 | Col10a1 | collagen, type X, alpha 1 | 6 | 0 | 0 | 0 | 0 | 0 | 0 | 0 | 0 | 0 | 0 |
| 1 | 0.00 | 3.68 | 0.00 | 1.00 | 1 | Gm26576 | | 6 | 0 | 0 | 0 | 0 | 0 | 0 | 0 | 0 | 0 | 0 |
| 1 | 0.00 | 3.68 | 0.00 | 1.00 | 1 | Cfd | complement factor D (adipsin) | 6 | 0 | 0 | 0 | 0 | 0 | 0 | 0 | 0 | 0 | 0 |
| 1 | 0.00 | 3.68 | 0.00 | 1.00 | 1 | Nxph4 | neurexophilin 4 | 6 | 0 | 0 | 0 | 0 | 0 | 0 | 0 | 0 | 0 | 0 |
| 1 | 0.00 | 3.68 | 0.00 | 1.00 | 1 | Tac2 | tachykinin 2 | 6 | 0 | 0 | 0 | 0 | 0 | 0 | 0 | 0 | 0 | 0 |
| 1 | 0.00 | 3.68 | 0.00 | 1.00 | 1 | Neurod4 | neurogenic differentiation 4 | 6 | 0 | 0 | 0 | 0 | 0 | 0 | 0 | 0 | 0 | 0 |
| 1 | 0.00 | 3.68 | 0.00 | 1.00 | 1 | Scgb3a1 | secretoglobin, family 3A, member 1 | 6 | 0 | 0 | 0 | 0 | 0 | 0 | 0 | 0 | 0 | 0 |
| 1 | 0.00 | 3.68 | 0.00 | 1.00 | 1 | Gltpd2 | glycolipid transfer protein domain contain | 6 | 0 | 0 | 0 | 0 | 0 | 0 | 0 | 0 | 0 | 0 |
| 1 | 0.00 | 3.68 | 0.00 | 1.00 | 1 | Gm12349 | | 6 | 0 | 0 | 0 | 0 | 0 | 0 | 0 | 0 | 0 | 0 |
| 1 | 0.00 | 3.68 | 0.00 | 1.00 | 1 | Slc4a1 | solute carrier family 4 (anion exchanger), | 6 | 0 | 0 | 0 | 0 | 0 | 0 | 0 | 0 | 0 | 0 |
| 1 | 0.00 | 3.68 | 0.00 | 1.00 | 1 | Slc26a4 | solute carrier family 26, member 4 | 6 | 0 | 0 | 0 | 0 | 0 | 0 | 0 | 0 | 0 | 0 |
| 1 | 0.00 | 3.68 | 0.00 | 1.00 | 1 | Gpr33 | G protein-coupled receptor 33 | 6 | 0 | 0 | 0 | 0 | 0 | 0 | 0 | 0 | 0 | 0 |
| 1 | 0.00 | 3.68 | 0.00 | 1.00 | 1 | Gm36756 | | 6 | 0 | 0 | 0 | 0 | 0 | 0 | 0 | 0 | 0 | 0 |
| 1 | 0.00 | 3.68 | 0.00 | 1.00 | 1 | B430119L08Rik | | 6 | 0 | 0 | 0 | 0 | 0 | 0 | 0 | 0 | 0 | 0 |
| 1 | 0.00 | 3.68 | 0.00 | 1.00 | 1 | Ighv1-25 | | 6 | 0 | 0 | 0 | 0 | 0 | 0 | 0 | 0 | 0 | 0 |
| 1 | 0.00 | 3.68 | 0.00 | 1.00 | 1 | Gm34471 | | 6 | 0 | 0 | 0 | 0 | 0 | 0 | 0 | 0 | 0 | 0 |
| 1 | 0.00 | 3.68 | 0.00 | 1.00 | 1 | Gm3715 | | 6 | 0 | 0 | 0 | 0 | 0 | 0 | 0 | 0 | 0 | 0 |
| 1 | 0.00 | 3.68 | 0.00 | 1.00 | 1 | Gm7604 | | 6 | 0 | 0 | 0 | 0 | 0 | 0 | 0 | 0 | 0 | 0 |
| 1 | 0.00 | 3.68 | 0.00 | 1.00 | 1 | Ear1 | eosinophil-associated, ribonuclease A fa | 6 | 0 | 0 | 0 | 0 | 0 | 0 | 0 | 0 | 0 | 0 |
| 1 | 0.00 | 3.68 | 0.00 | 1.00 | 1 | Abra | actin-binding Rho activating protein | 6 | 0 | 0 | 0 | 0 | 0 | 0 | 0 | 0 | 0 | 0 |
| 1 | 0.00 | 3.68 | 0.00 | 1.00 | 1 | Gm43388 | | 6 | 0 | 0 | 0 | 0 | 0 | 0 | 0 | 0 | 0 | 0 |
| 1 | 0.00 | 3.68 | 0.00 | 1.00 | 1 | Gm3320 | | 6 | 0 | 0 | 0 | 0 | 0 | 0 | 0 | 0 | 0 | 0 |
| 1 | 0.00 | 3.68 | 0.00 | 1.00 | 1 | Prss34 | protease, serine 34 | 6 | 0 | 0 | 0 | 0 | 0 | 0 | 0 | 0 | 0 | 0 |
| 1 | 0.00 | 3.68 | 0.00 | 1.00 | 1 | Gm33055 | | 6 | 0 | 0 | 0 | 0 | 0 | 0 | 0 | 0 | 0 | 0 |

| | | | | | | | | | | | | | | | | | | |
|---|---|---|---|---|---|---|---|---|---|---|---|---|---|---|---|---|---|---|
| 1 | 0.00 | 3.68 | 0.00 | 1.00 | 1 | BC051408 | | 6 | 0 | 0 | 0 | 0 | 0 | 0 | 0 | 0 | 0 | 0 |
| 1 | 0.00 | 3.68 | 0.00 | 1.00 | 1 | Gm550 | predicted gene 550 | 6 | 0 | 0 | 0 | 0 | 0 | 0 | 0 | 0 | 0 | 0 |
| 1 | 0.00 | 3.68 | 0.00 | 1.00 | 1 | Gm8369 | predicted gene 8369 | 6 | 0 | 0 | 0 | 0 | 0 | 0 | 0 | 0 | 0 | 0 |
| 1 | 0.00 | 3.68 | 0.00 | 1.00 | 1 | Glra2 | glycine receptor, alpha 2 subunit | 6 | 0 | 0 | 0 | 0 | 0 | 0 | 0 | 0 | 0 | 0 |
| 1 | 0.54 | 3.63 | 0.15 | 0.88 | 1 | Gm12266 | | 0 | 0 | 3 | 0 | 0 | 1 | 0 | 0 | 0 | 0 | 2 |
| 1 | -0.47 | 3.66 | -0.13 | 0.90 | 1 | A930018P22Ril | RIKEN cDNA A930018P22 gene | 5 | 0 | 0 | 0 | 0 | 1 | 0 | 0 | 0 | 0 | 0 |
| 1 | -0.47 | 3.66 | -0.13 | 0.90 | 1 | H2ap | H2A.P histone | 5 | 0 | 0 | 0 | 0 | 1 | 0 | 0 | 0 | 0 | 0 |
| 1 | 0.33 | 3.63 | 0.09 | 0.93 | 1 | Gm6109 | predicted gene 6109 | 1 | 0 | 2 | 0 | 2 | 0 | 0 | 1 | 0 | 0 | 0 |
| 1 | 0.37 | 3.61 | 0.10 | 0.92 | 1 | Gm12242 | | 0 | 1 | 0 | 0 | 1 | 1 | 0 | 1 | 2 | 0 | 0 |
| 1 | 0.63 | 3.63 | 0.17 | 0.86 | 1 | Gm5853 | | 0 | 1 | 1 | 2 | 0 | 0 | 0 | 1 | 0 | 0 | 1 |
| 1 | -0.83 | 3.63 | -0.23 | 0.82 | 1 | Ninj2 | ninjurin 2 | 3 | 0 | 0 | 0 | 0 | 3 | 0 | 0 | 0 | 0 | 0 |
| 1 | 0.01 | 3.63 | 0.00 | 1.00 | 1 | Gm49879 | | 0 | 1 | 0 | 0 | 1 | 2 | 0 | 0 | 0 | 1 | 1 |
| 1 | 0.02 | 3.60 | 0.00 | 1.00 | 1 | Gm44983 | | 0 | 0 | 0 | 0 | 2 | 0 | 0 | 0 | 0 | 0 | 4 |
| 1 | -0.78 | 3.62 | -0.21 | 0.83 | 1 | Gm42608 | | 2 | 0 | 0 | 0 | 0 | 4 | 0 | 0 | 0 | 0 | 0 |
| 1 | -0.78 | 3.62 | -0.21 | 0.83 | 1 | Gm42789 | | 2 | 0 | 0 | 0 | 0 | 4 | 0 | 0 | 0 | 0 | 0 |
| 1 | -0.78 | 3.62 | -0.21 | 0.83 | 1 | Gm48766 | | 2 | 0 | 0 | 0 | 0 | 4 | 0 | 0 | 0 | 0 | 0 |
| 1 | 0.46 | 3.62 | 0.13 | 0.90 | 1 | Gm9061 | | 0 | 1 | 0 | 0 | 2 | 0 | 0 | 0 | 3 | 0 | 0 |
| 1 | 0.37 | 3.60 | 0.10 | 0.92 | 1 | Gm45473 | | 0 | 0 | 0 | 0 | 2 | 1 | 0 | 1 | 0 | 0 | 2 |
| 1 | -0.37 | 3.63 | -0.10 | 0.92 | 1 | Gm18025 | | 0 | 1 | 2 | 1 | 1 | 0 | 0 | 0 | 1 | 0 | 0 |
| 1 | 0.87 | 3.61 | 0.24 | 0.81 | 1 | Gm38380 | | 0 | 0 | 0 | 0 | 0 | 1 | 0 | 0 | 0 | 0 | 5 |
| 1 | 0.87 | 3.61 | 0.24 | 0.81 | 1 | Gm9823 | | 0 | 0 | 0 | 0 | 0 | 1 | 0 | 0 | 0 | 0 | 5 |
| 1 | 0.87 | 3.61 | 0.24 | 0.81 | 1 | Gm14104 | | 0 | 0 | 0 | 0 | 0 | 1 | 0 | 0 | 0 | 0 | 5 |
| 1 | 0.87 | 3.61 | 0.24 | 0.81 | 1 | Gm37353 | | 0 | 0 | 0 | 0 | 0 | 1 | 0 | 0 | 0 | 0 | 5 |
| 1 | 0.87 | 3.61 | 0.24 | 0.81 | 1 | Gm43387 | | 0 | 0 | 0 | 0 | 0 | 1 | 0 | 0 | 0 | 0 | 5 |
| 1 | 0.87 | 3.61 | 0.24 | 0.81 | 1 | Gm13274 | | 0 | 0 | 0 | 0 | 0 | 1 | 0 | 0 | 0 | 0 | 5 |
| 1 | 0.87 | 3.61 | 0.24 | 0.81 | 1 | Yipf7 | Yip1 domain family, member 7 | 0 | 0 | 0 | 0 | 0 | 1 | 0 | 0 | 0 | 0 | 5 |
| 1 | 0.87 | 3.61 | 0.24 | 0.81 | 1 | Gm28719 | | 0 | 0 | 0 | 0 | 0 | 1 | 0 | 0 | 0 | 0 | 5 |
| 1 | 0.87 | 3.61 | 0.24 | 0.81 | 1 | Snora62 | | 0 | 0 | 0 | 0 | 0 | 1 | 0 | 0 | 0 | 0 | 5 |
| 1 | 0.87 | 3.61 | 0.24 | 0.81 | 1 | Gm49298 | | 0 | 0 | 0 | 0 | 0 | 1 | 0 | 0 | 0 | 0 | 5 |
| 1 | 0.00 | 3.68 | 0.00 | 1.00 | 1 | Gm26574 | | 1 | 5 | 0 | 0 | 0 | 0 | 0 | 0 | 0 | 0 | 0 |
| 1 | 0.00 | 3.68 | 0.00 | 1.00 | 1 | Gm11830 | | 1 | 5 | 0 | 0 | 0 | 0 | 0 | 0 | 0 | 0 | 0 |
| 1 | 0.00 | 3.68 | 0.00 | 1.00 | 1 | Gm8848 | | 1 | 5 | 0 | 0 | 0 | 0 | 0 | 0 | 0 | 0 | 0 |
| 1 | 0.79 | 3.62 | 0.22 | 0.83 | 1 | Shcbp1l | Shc SH2-domain binding protein 1-like | 0 | 1 | 0 | 0 | 0 | 1 | 0 | 0 | 0 | 0 | 4 |
| 1 | 0.79 | 3.62 | 0.22 | 0.83 | 1 | Gm12986 | | 0 | 1 | 0 | 0 | 0 | 1 | 0 | 0 | 0 | 0 | 4 |
| 1 | -0.80 | 3.62 | -0.22 | 0.83 | 1 | Gm29156 | | 0 | 0 | 0 | 0 | 0 | 6 | 0 | 0 | 0 | 0 | 0 |
| 1 | -0.80 | 3.62 | -0.22 | 0.83 | 1 | Gm29596 | | 0 | 0 | 0 | 0 | 0 | 6 | 0 | 0 | 0 | 0 | 0 |
| 1 | -0.80 | 3.62 | -0.22 | 0.83 | 1 | Gm37733 | | 0 | 0 | 0 | 0 | 0 | 6 | 0 | 0 | 0 | 0 | 0 |
| 1 | -0.80 | 3.62 | -0.22 | 0.83 | 1 | Gm37529 | | 0 | 0 | 0 | 0 | 0 | 6 | 0 | 0 | 0 | 0 | 0 |
| 1 | -0.80 | 3.62 | -0.22 | 0.83 | 1 | Gm37258 | | 0 | 0 | 0 | 0 | 0 | 6 | 0 | 0 | 0 | 0 | 0 |
| 1 | -0.80 | 3.62 | -0.22 | 0.83 | 1 | Gm43128 | | 0 | 0 | 0 | 0 | 0 | 6 | 0 | 0 | 0 | 0 | 0 |
| 1 | -0.80 | 3.62 | -0.22 | 0.83 | 1 | Mir7001 | | 0 | 0 | 0 | 0 | 0 | 6 | 0 | 0 | 0 | 0 | 0 |
| 1 | -0.80 | 3.62 | -0.22 | 0.83 | 1 | Gm24081 | | 0 | 0 | 0 | 0 | 0 | 6 | 0 | 0 | 0 | 0 | 0 |
| 1 | -0.80 | 3.62 | -0.22 | 0.83 | 1 | Gm43312 | | 0 | 0 | 0 | 0 | 0 | 6 | 0 | 0 | 0 | 0 | 0 |
| 1 | -0.80 | 3.62 | -0.22 | 0.83 | 1 | Gm43883 | | 0 | 0 | 0 | 0 | 0 | 6 | 0 | 0 | 0 | 0 | 0 |
| 1 | -0.80 | 3.62 | -0.22 | 0.83 | 1 | Gm44157 | | 0 | 0 | 0 | 0 | 0 | 6 | 0 | 0 | 0 | 0 | 0 |
| 1 | -0.80 | 3.62 | -0.22 | 0.83 | 1 | Mir7049 | | 0 | 0 | 0 | 0 | 0 | 6 | 0 | 0 | 0 | 0 | 0 |
| 1 | -0.80 | 3.62 | -0.22 | 0.83 | 1 | Gm44797 | | 0 | 0 | 0 | 0 | 0 | 6 | 0 | 0 | 0 | 0 | 0 |
| 1 | -0.80 | 3.62 | -0.22 | 0.83 | 1 | n-R5s96 | | 0 | 0 | 0 | 0 | 0 | 6 | 0 | 0 | 0 | 0 | 0 |
| 1 | -0.80 | 3.62 | -0.22 | 0.83 | 1 | Gm31614 | | 0 | 0 | 0 | 0 | 0 | 6 | 0 | 0 | 0 | 0 | 0 |
| 1 | -0.80 | 3.62 | -0.22 | 0.83 | 1 | Gm47000 | | 0 | 0 | 0 | 0 | 0 | 6 | 0 | 0 | 0 | 0 | 0 |
| 1 | -0.80 | 3.62 | -0.22 | 0.83 | 1 | Adgrf2 | adhesion G protein-coupled receptor F2 | 0 | 0 | 0 | 0 | 0 | 6 | 0 | 0 | 0 | 0 | 0 |
| 1 | 0.27 | 3.63 | 0.08 | 0.94 | 1 | Gm9358 | | 0 | 2 | 0 | 1 | 0 | 1 | 0 | 1 | 1 | 0 | 0 |
| 1 | 1.87 | 3.65 | 0.51 | 0.61 | 1 | Trbv12-2 | | 1 | 0 | 0 | 0 | 0 | 0 | 0 | 0 | 5 | 0 | 0 |
| 1 | 1.69 | 3.61 | 0.47 | 0.64 | 1 | Gm12105 | | 0 | 0 | 0 | 0 | 0 | 1 | 0 | 0 | 2 | 0 | 3 |
| 1 | 1.75 | 3.61 | 0.48 | 0.63 | 1 | 1700065J18Rik | | 0 | 0 | 0 | 0 | 0 | 1 | 0 | 0 | 3 | 0 | 2 |
| 1 | -0.52 | 3.66 | -0.14 | 0.89 | 1 | 1700039I01Rik | | 0 | 5 | 0 | 0 | 0 | 1 | 0 | 0 | 0 | 0 | 0 |
| 1 | 1.46 | 3.62 | 0.40 | 0.69 | 1 | Gm43794 | | 0 | 2 | 0 | 0 | 0 | 1 | 0 | 0 | 2 | 0 | 1 |
| 1 | 1.67 | 3.62 | 0.46 | 0.64 | 1 | Gm15343 | | 0 | 0 | 0 | 0 | 0 | 1 | 0 | 0 | 4 | 0 | 1 |
| 1 | 1.09 | 3.62 | 0.30 | 0.76 | 1 | Rnf151 | ring finger protein 151 | 0 | 0 | 0 | 0 | 0 | 1 | 0 | 0 | 5 | 0 | 0 |
| 1 | -0.44 | 3.65 | -0.12 | 0.90 | 1 | Rpl31-ps20 | | 0 | 2 | 1 | 1 | 1 | 0 | 0 | 0 | 0 | 0 | 1 |

| | | | | | | | | | | | | | | | | | | |
|---|---|---|---|---|---|---|---|---|---|---|---|---|---|---|---|---|---|---|
| 1 | 0.39 | 3.59 | 0.11 | 0.91 | 1 | Calr-ps | | 0 | 0 | 0 | 1 | 0 | 2 | 0 | 2 | 0 | 0 | 1 |
| 1 | 0.18 | 3.58 | 0.05 | 0.96 | 1 | Mir3473d | | 0 | 0 | 0 | 0 | 3 | 0 | 0 | 0 | 0 | 0 | 3 |
| 1 | 0.18 | 3.58 | 0.05 | 0.96 | 1 | Gm20082 | | 0 | 0 | 0 | 0 | 3 | 0 | 0 | 0 | 0 | 0 | 3 |
| 1 | 0.59 | 3.64 | 0.16 | 0.87 | 1 | Gm11191 | | 1 | 1 | 1 | 0 | 1 | 0 | 0 | 0 | 2 | 0 | 0 |
| 1 | 0.67 | 3.62 | 0.18 | 0.85 | 1 | Gm43435 | | 0 | 0 | 0 | 0 | 3 | 0 | 0 | 0 | 2 | 0 | 1 |
| 1 | 0.00 | 3.68 | 0.00 | 1.00 | 1 | Adra2c | adrenergic receptor, alpha 2c | 2 | 0 | 4 | 0 | 0 | 0 | 0 | 0 | 0 | 0 | 0 |
| 1 | -0.81 | 3.65 | -0.22 | 0.82 | 1 | Gm47112 | | 0 | 0 | 4 | 0 | 0 | 2 | 0 | 0 | 0 | 0 | 0 |
| 1 | 0.92 | 3.67 | 0.25 | 0.80 | 1 | Gm15032 | | 2 | 0 | 3 | 0 | 0 | 0 | 0 | 0 | 1 | 0 | 0 |
| 1 | -1.36 | 3.64 | -0.37 | 0.71 | 1 | Kdm6bos | | 0 | 4 | 0 | 0 | 1 | 1 | 0 | 0 | 0 | 0 | 0 |
| 1 | -0.07 | 3.60 | -0.02 | 0.98 | 1 | Gm12608 | | 0 | 0 | 0 | 2 | 0 | 0 | 0 | 0 | 0 | 0 | 4 |
| 1 | -0.07 | 3.60 | -0.02 | 0.98 | 1 | Gm45251 | | 0 | 0 | 0 | 2 | 0 | 0 | 0 | 0 | 0 | 0 | 4 |
| 1 | -0.91 | 3.60 | -0.25 | 0.80 | 1 | Gm4742 | | 0 | 1 | 0 | 1 | 2 | 0 | 0 | 0 | 0 | 0 | 2 |
| 1 | 0.46 | 3.60 | 0.13 | 0.90 | 1 | Cyp2c55 | cytochrome P450, family 2, subfamily c, p | 0 | 2 | 0 | 2 | 0 | 0 | 0 | 0 | 0 | 0 | 2 |
| 1 | -0.69 | 3.66 | -0.19 | 0.85 | 1 | Gm11716 | | 0 | 2 | 2 | 0 | 0 | 2 | 0 | 0 | 0 | 0 | 0 |
| 1 | 1.71 | 3.64 | 0.47 | 0.64 | 1 | Gm37832 | | 0 | 0 | 0 | 0 | 0 | 0 | 0 | 6 | 0 | 0 | 0 |
| 1 | 1.71 | 3.64 | 0.47 | 0.64 | 1 | 5730596B20Rik | | 0 | 0 | 0 | 0 | 0 | 0 | 0 | 6 | 0 | 0 | 0 |
| 1 | 1.71 | 3.64 | 0.47 | 0.64 | 1 | Gm47992 | | 0 | 0 | 0 | 0 | 0 | 0 | 0 | 6 | 0 | 0 | 0 |
| 1 | -0.44 | 3.60 | -0.12 | 0.90 | 1 | Rdh1 | retinol dehydrogenase 1 (all trans) | 0 | 0 | 0 | 0 | 2 | 1 | 0 | 0 | 0 | 0 | 3 |
| 1 | -0.14 | 3.62 | -0.04 | 0.97 | 1 | Gm35558 | | 0 | 0 | 0 | 0 | 2 | 1 | 0 | 0 | 3 | 0 | 0 |
| 1 | 0.00 | 3.68 | 0.00 | 1.00 | 1 | Gm44758 | | 2 | 4 | 0 | 0 | 0 | 0 | 0 | 0 | 0 | 0 | 0 |
| 1 | 0.69 | 3.62 | 0.19 | 0.85 | 1 | Gm17383 | | 1 | 1 | 0 | 0 | 0 | 1 | 0 | 0 | 0 | 0 | 3 |
| 1 | 0.55 | 3.61 | 0.15 | 0.88 | 1 | Gm37909 | | 0 | 0 | 0 | 0 | 0 | 2 | 0 | 0 | 0 | 0 | 4 |
| 1 | 0.55 | 3.61 | 0.15 | 0.88 | 1 | Gm25612 | | 0 | 0 | 0 | 0 | 0 | 2 | 0 | 0 | 0 | 0 | 4 |
| 1 | 0.55 | 3.61 | 0.15 | 0.88 | 1 | Gm37060 | | 0 | 0 | 0 | 0 | 0 | 2 | 0 | 0 | 0 | 0 | 4 |
| 1 | 0.55 | 3.61 | 0.15 | 0.88 | 1 | Gm13627 | | 0 | 0 | 0 | 0 | 0 | 2 | 0 | 0 | 0 | 0 | 4 |
| 1 | 0.55 | 3.61 | 0.15 | 0.88 | 1 | Gm12463 | | 0 | 0 | 0 | 0 | 0 | 2 | 0 | 0 | 0 | 0 | 4 |
| 1 | 0.55 | 3.61 | 0.15 | 0.88 | 1 | Gm2719 | | 0 | 0 | 0 | 0 | 0 | 2 | 0 | 0 | 0 | 0 | 4 |
| 1 | 0.55 | 3.61 | 0.15 | 0.88 | 1 | Gm45537 | | 0 | 0 | 0 | 0 | 0 | 2 | 0 | 0 | 0 | 0 | 4 |
| 1 | 0.55 | 3.61 | 0.15 | 0.88 | 1 | Gm45518 | | 0 | 0 | 0 | 0 | 0 | 2 | 0 | 0 | 0 | 0 | 4 |
| 1 | 0.55 | 3.61 | 0.15 | 0.88 | 1 | Gm37195 | | 0 | 0 | 0 | 0 | 0 | 2 | 0 | 0 | 0 | 0 | 4 |
| 1 | 0.55 | 3.61 | 0.15 | 0.88 | 1 | Gm23156 | | 0 | 0 | 0 | 0 | 0 | 2 | 0 | 0 | 0 | 0 | 4 |
| 1 | 0.55 | 3.61 | 0.15 | 0.88 | 1 | Gm17196 | | 0 | 0 | 0 | 0 | 0 | 2 | 0 | 0 | 0 | 0 | 4 |
| 1 | 0.55 | 3.61 | 0.15 | 0.88 | 1 | Gm47972 | | 0 | 0 | 0 | 0 | 0 | 2 | 0 | 0 | 0 | 0 | 4 |
| 1 | 0.55 | 3.61 | 0.15 | 0.88 | 1 | Gm47279 | | 0 | 0 | 0 | 0 | 0 | 2 | 0 | 0 | 0 | 0 | 4 |
| 1 | 0.55 | 3.61 | 0.15 | 0.88 | 1 | Mir6941 | | 0 | 0 | 0 | 0 | 0 | 2 | 0 | 0 | 0 | 0 | 4 |
| 1 | 0.55 | 3.61 | 0.15 | 0.88 | 1 | Gm24620 | | 0 | 0 | 0 | 0 | 0 | 2 | 0 | 0 | 0 | 0 | 4 |
| 1 | 0.55 | 3.61 | 0.15 | 0.88 | 1 | Gm48824 | | 0 | 0 | 0 | 0 | 0 | 2 | 0 | 0 | 0 | 0 | 4 |
| 1 | 0.27 | 3.64 | 0.07 | 0.94 | 1 | Gm13623 | | 2 | 0 | 2 | 0 | 1 | 0 | 0 | 0 | 1 | 0 | 0 |
| 1 | 1.12 | 3.63 | 0.31 | 0.76 | 1 | Gm14383 | | 1 | 0 | 0 | 0 | 0 | 1 | 0 | 0 | 4 | 0 | 0 |
| 1 | 1.10 | 3.61 | 0.30 | 0.76 | 1 | Gm5832 | | 0 | 0 | 0 | 0 | 0 | 2 | 0 | 0 | 1 | 0 | 3 |
| 1 | -0.85 | 3.65 | -0.23 | 0.82 | 1 | C230031I18Rik | | 0 | 4 | 0 | 0 | 0 | 2 | 0 | 0 | 0 | 0 | 0 |
| 1 | -0.85 | 3.65 | -0.23 | 0.82 | 1 | 2010106G04Rik | | 0 | 4 | 0 | 0 | 0 | 2 | 0 | 0 | 0 | 0 | 0 |
| 1 | -0.85 | 3.60 | -0.24 | 0.81 | 1 | Selenot-ps | | 0 | 0 | 0 | 2 | 1 | 0 | 0 | 0 | 0 | 0 | 3 |
| 1 | -0.85 | 3.60 | -0.24 | 0.81 | 1 | Gm16301 | | 0 | 0 | 0 | 2 | 1 | 0 | 0 | 0 | 0 | 0 | 3 |
| 1 | 0.70 | 3.58 | 0.20 | 0.85 | 1 | Gm43721 | | 0 | 0 | 0 | 0 | 0 | 2 | 0 | 0 | 4 | 0 | 0 |
| 1 | 1.53 | 3.65 | 0.42 | 0.67 | 1 | Gm5548 | | 3 | 0 | 0 | 0 | 0 | 0 | 0 | 0 | 0 | 1 | 1 |
| 1 | 0.14 | 3.64 | 0.04 | 0.97 | 1 | Rps15a-ps2 | | 2 | 0 | 1 | 0 | 0 | 2 | 0 | 0 | 0 | 1 | 0 |
| 1 | 0.36 | 3.64 | 0.10 | 0.92 | 1 | Gm14036 | | 2 | 0 | 1 | 0 | 1 | 0 | 0 | 0 | 0 | 0 | 2 |
| 1 | -0.70 | 3.60 | -0.20 | 0.84 | 1 | Gm49026 | | 0 | 0 | 0 | 0 | 3 | 1 | 0 | 0 | 0 | 0 | 2 |
| 1 | 0.29 | 3.66 | 0.08 | 0.94 | 1 | Gm6029 | | 2 | 0 | 0 | 1 | 1 | 0 | 0 | 1 | 0 | 0 | 1 |
| 1 | -1.91 | 3.62 | -0.53 | 0.60 | 1 | Gm42744 | | 0 | 2 | 0 | 0 | 3 | 1 | 0 | 0 | 0 | 0 | 0 |
| 1 | 0.00 | 3.68 | 0.00 | 1.00 | 1 | Gm9925 | | 3 | 0 | 3 | 0 | 0 | 0 | 0 | 0 | 0 | 0 | 0 |
| 1 | -0.25 | 3.59 | -0.07 | 0.94 | 1 | Gm43607 | | 0 | 0 | 0 | 3 | 0 | 0 | 0 | 0 | 0 | 0 | 3 |
| 1 | -1.18 | 3.63 | -0.32 | 0.75 | 1 | Gm23814 | | 0 | 0 | 0 | 0 | 6 | 0 | 0 | 0 | 0 | 0 | 0 |
| 1 | -1.18 | 3.63 | -0.32 | 0.75 | 1 | Chil4 | chitinase-like 4 | 0 | 0 | 0 | 0 | 6 | 0 | 0 | 0 | 0 | 0 | 0 |
| 1 | -1.18 | 3.63 | -0.32 | 0.75 | 1 | Taf9-ps | TATA-box binding protein associated fact | 0 | 0 | 0 | 0 | 6 | 0 | 0 | 0 | 0 | 0 | 0 |
| 1 | -1.18 | 3.63 | -0.32 | 0.75 | 1 | Gm7596 | | 0 | 0 | 0 | 0 | 6 | 0 | 0 | 0 | 0 | 0 | 0 |
| 1 | -1.18 | 3.63 | -0.32 | 0.75 | 1 | Csnk2a1-ps1 | | 0 | 0 | 0 | 0 | 6 | 0 | 0 | 0 | 0 | 0 | 0 |
| 1 | -1.18 | 3.63 | -0.32 | 0.75 | 1 | Gm7473 | | 0 | 0 | 0 | 0 | 6 | 0 | 0 | 0 | 0 | 0 | 0 |
| 1 | -1.18 | 3.63 | -0.32 | 0.75 | 1 | Prrxl1 | paired related homeobox protein-like 1 | 0 | 0 | 0 | 0 | 6 | 0 | 0 | 0 | 0 | 0 | 0 |

| | | | | | | | | | | | | | | | | | | |
|---|---|---|---|---|---|---|---|---|---|---|---|---|---|---|---|---|---|---|
| 1 | -1.18 | 3.63 | -0.32 | 0.75 | 1 | Gm50029 | | 0 | 0 | 0 | 0 | 6 | 0 | 0 | 0 | 0 | 0 | 0 |
| 1 | 0.00 | 3.68 | 0.00 | 1.00 | 1 | D430036J16Rik | | 3 | 1 | 2 | 0 | 0 | 0 | 0 | 0 | 0 | 0 | 0 |
| 1 | 0.27 | 3.59 | 0.08 | 0.94 | 1 | Bloodlinc | | 0 | 0 | 0 | 3 | 0 | 0 | 0 | 0 | 3 | 0 | 0 |
| 1 | -1.06 | 3.61 | -0.29 | 0.77 | 1 | Pde6h | phosphodiesterase 6H, cGMP-specific, cc | 0 | 0 | 0 | 0 | 4 | 1 | 0 | 0 | 0 | 0 | 1 |
| 1 | 0.00 | 3.68 | 0.00 | 1.00 | 1 | Cyp26a1 | cytochrome P450, family 26, subfamily a, | 3 | 2 | 1 | 0 | 0 | 0 | 0 | 0 | 0 | 0 | 0 |
| 1 | 3.00 | 3.59 | 0.84 | 0.40 | 1 | Gm12109 | | 0 | 0 | 0 | 0 | 0 | 0 | 0 | 2 | 1 | 1 | 1 |
| 1 | 0.00 | 3.68 | 0.00 | 1.00 | 1 | Mmp27 | matrix metallopeptidase 27 | 3 | 3 | 0 | 0 | 0 | 0 | 0 | 0 | 0 | 0 | 0 |
| 1 | -0.78 | 3.61 | -0.22 | 0.83 | 1 | Oaz3 | ornithine decarboxylase antizyme 3 | 0 | 0 | 0 | 0 | 2 | 2 | 0 | 0 | 0 | 0 | 2 |
| 1 | -0.14 | 3.62 | -0.04 | 0.97 | 1 | Gm4832 | predicted gene 4832 | 0 | 1 | 1 | 0 | 0 | 3 | 0 | 0 | 0 | 0 | 1 |
| 1 | 2.54 | 3.61 | 0.71 | 0.48 | 1 | Gm49096 | | 1 | 0 | 0 | 0 | 0 | 2 | 0 | 0 | 3 | 0 | 0 |
| 1 | 0.28 | 3.61 | 0.08 | 0.94 | 1 | Gm37221 | | 0 | 0 | 0 | 0 | 0 | 3 | 0 | 0 | 0 | 0 | 3 |
| 1 | 0.28 | 3.61 | 0.08 | 0.94 | 1 | Gm43496 | | 0 | 0 | 0 | 0 | 0 | 3 | 0 | 0 | 0 | 0 | 3 |
| 1 | 0.28 | 3.61 | 0.08 | 0.94 | 1 | Gm43773 | | 0 | 0 | 0 | 0 | 0 | 3 | 0 | 0 | 0 | 0 | 3 |
| 1 | 0.28 | 3.61 | 0.08 | 0.94 | 1 | Gm12678 | | 0 | 0 | 0 | 0 | 0 | 3 | 0 | 0 | 0 | 0 | 3 |
| 1 | 0.28 | 3.61 | 0.08 | 0.94 | 1 | Mir3473e | | 0 | 0 | 0 | 0 | 0 | 3 | 0 | 0 | 0 | 0 | 3 |
| 1 | 0.28 | 3.61 | 0.08 | 0.94 | 1 | Tex45 | testis expressed 45 | 0 | 0 | 0 | 0 | 0 | 3 | 0 | 0 | 0 | 0 | 3 |
| 1 | 0.28 | 3.61 | 0.08 | 0.94 | 1 | Gm47171 | | 0 | 0 | 0 | 0 | 0 | 3 | 0 | 0 | 0 | 0 | 3 |
| 1 | 0.28 | 3.61 | 0.08 | 0.94 | 1 | Mir6913 | | 0 | 0 | 0 | 0 | 0 | 3 | 0 | 0 | 0 | 0 | 3 |
| 1 | 0.28 | 3.61 | 0.08 | 0.94 | 1 | Gm12239 | | 0 | 0 | 0 | 0 | 0 | 3 | 0 | 0 | 0 | 0 | 3 |
| 1 | 0.28 | 3.61 | 0.08 | 0.94 | 1 | Gm35239 | | 0 | 0 | 0 | 0 | 0 | 3 | 0 | 0 | 0 | 0 | 3 |
| 1 | 0.28 | 3.61 | 0.08 | 0.94 | 1 | D930032P07Rik | | 0 | 0 | 0 | 0 | 0 | 3 | 0 | 0 | 0 | 0 | 3 |
| 1 | -1.06 | 3.63 | -0.29 | 0.77 | 1 | Mkrn1-ps1 | | 1 | 0 | 1 | 1 | 1 | 1 | 0 | 0 | 0 | 0 | 1 |
| 1 | -0.87 | 3.63 | -0.24 | 0.81 | 1 | Gm48565 | | 0 | 3 | 0 | 0 | 0 | 3 | 0 | 0 | 0 | 0 | 0 |
| 1 | -1.86 | 3.63 | -0.51 | 0.61 | 1 | Trh | thyrotropin releasing hormone | 0 | 0 | 0 | 0 | 5 | 1 | 0 | 0 | 0 | 0 | 0 |
| 1 | -1.86 | 3.63 | -0.51 | 0.61 | 1 | Gm35595 | | 0 | 0 | 0 | 0 | 5 | 1 | 0 | 0 | 0 | 0 | 0 |
| 1 | 0.52 | 3.61 | 0.15 | 0.88 | 1 | Gm12501 | | 0 | 0 | 0 | 0 | 0 | 3 | 0 | 1 | 0 | 0 | 1 |
| 1 | -0.79 | 3.61 | -0.22 | 0.83 | 1 | Gm14853 | | 0 | 0 | 0 | 1 | 0 | 3 | 0 | 0 | 0 | 1 | 0 |
| 1 | -0.18 | 3.58 | -0.05 | 0.96 | 1 | Gm15379 | | 0 | 0 | 0 | 4 | 0 | 0 | 0 | 0 | 0 | 0 | 2 |
| 1 | -1.48 | 3.61 | -0.41 | 0.68 | 1 | Gm12565 | | 0 | 0 | 0 | 3 | 2 | 0 | 0 | 0 | 0 | 0 | 1 |
| 1 | 0.15 | 3.62 | 0.04 | 0.97 | 1 | Gm46391 | | 0 | 0 | 0 | 0 | 0 | 4 | 0 | 0 | 0 | 1 | 0 |
| 1 | 0.00 | 3.68 | 0.00 | 1.00 | 1 | Mrgprb2 | MAS-related GPR, member B2 | 3 | 0 | 2 | 0 | 0 | 0 | 0 | 0 | 0 | 0 | 0 |
| 1 | 0.00 | 3.68 | 0.00 | 1.00 | 1 | Gm48495 | | 3 | 0 | 2 | 0 | 0 | 0 | 0 | 0 | 0 | 0 | 0 |
| 1 | -0.05 | 3.62 | -0.01 | 0.99 | 1 | Gm15511 | | 1 | 0 | 0 | 3 | 0 | 0 | 0 | 0 | 2 | 0 | 0 |
| 0 | -0.69 | 3.66 | -0.19 | 0.85 | 1 | Gm49449 | | 2 | 0 | 2 | 0 | 0 | 2 | 0 | 0 | 0 | 0 | 0 |
| 0 | 1.86 | 3.64 | 0.51 | 0.61 | 1 | Gm7091 | | 1 | 1 | 0 | 0 | 0 | 0 | 0 | 2 | 0 | 1 | 0 |
| 0 | -2.00 | 3.63 | -0.55 | 0.58 | 1 | Gm43871 | | 0 | 0 | 0 | 0 | 4 | 2 | 0 | 0 | 0 | 0 | 0 |
| 0 | 0.95 | 3.67 | 0.26 | 0.80 | 1 | Gm4739 | | 3 | 1 | 0 | 0 | 0 | 0 | 0 | 0 | 0 | 0 | 1 |
| 0 | 0.00 | 3.68 | 0.00 | 1.00 | 1 | Slfn5os | | 3 | 2 | 0 | 0 | 0 | 0 | 0 | 0 | 0 | 0 | 0 |
| 0 | 0.00 | 3.68 | 0.00 | 1.00 | 1 | Gm34215 | | 3 | 2 | 0 | 0 | 0 | 0 | 0 | 0 | 0 | 0 | 0 |
| 0 | 0.01 | 3.63 | 0.00 | 1.00 | 1 | Gm13379 | | 1 | 0 | 0 | 1 | 0 | 2 | 0 | 0 | 1 | 0 | 1 |
| 0 | -0.24 | 3.62 | -0.07 | 0.95 | 1 | Gm7787 | | 0 | 0 | 1 | 0 | 0 | 3 | 0 | 0 | 0 | 0 | 1 |
| 0 | 0.18 | 3.63 | 0.05 | 0.96 | 1 | Gm18705 | | 1 | 1 | 0 | 0 | 0 | 3 | 0 | 0 | 1 | 0 | 0 |
| 0 | 0.01 | 3.61 | 0.00 | 1.00 | 1 | Gm37250 | | 0 | 0 | 0 | 0 | 0 | 3 | 0 | 0 | 0 | 0 | 2 |
| 0 | 0.01 | 3.61 | 0.00 | 1.00 | 1 | Mir7682 | | 0 | 0 | 0 | 0 | 0 | 3 | 0 | 0 | 0 | 0 | 2 |
| 0 | 0.01 | 3.61 | 0.00 | 1.00 | 1 | Gm25632 | | 0 | 0 | 0 | 0 | 0 | 3 | 0 | 0 | 0 | 0 | 2 |
| 0 | 0.01 | 3.61 | 0.00 | 1.00 | 1 | Gm14425 | | 0 | 0 | 0 | 0 | 0 | 3 | 0 | 0 | 0 | 0 | 2 |
| 0 | 0.01 | 3.61 | 0.00 | 1.00 | 1 | n-R5s200 | | 0 | 0 | 0 | 0 | 0 | 3 | 0 | 0 | 0 | 0 | 2 |
| 0 | 0.01 | 3.61 | 0.00 | 1.00 | 1 | Gm26425 | | 0 | 0 | 0 | 0 | 0 | 3 | 0 | 0 | 0 | 0 | 2 |
| 0 | 0.01 | 3.61 | 0.00 | 1.00 | 1 | Gm24196 | | 0 | 0 | 0 | 0 | 0 | 3 | 0 | 0 | 0 | 0 | 2 |
| 0 | 0.01 | 3.61 | 0.00 | 1.00 | 1 | Gm43273 | | 0 | 0 | 0 | 0 | 0 | 3 | 0 | 0 | 0 | 0 | 2 |
| 0 | 0.01 | 3.61 | 0.00 | 1.00 | 1 | Gm25108 | | 0 | 0 | 0 | 0 | 0 | 3 | 0 | 0 | 0 | 0 | 2 |
| 0 | 0.01 | 3.61 | 0.00 | 1.00 | 1 | Gm44190 | | 0 | 0 | 0 | 0 | 0 | 3 | 0 | 0 | 0 | 0 | 2 |
| 0 | 0.01 | 3.61 | 0.00 | 1.00 | 1 | 4933405O20Ril | RIKEN cDNA 4933405O20 gene | 0 | 0 | 0 | 0 | 0 | 3 | 0 | 0 | 0 | 0 | 2 |
| 0 | 0.01 | 3.61 | 0.00 | 1.00 | 1 | Gm24923 | | 0 | 0 | 0 | 0 | 0 | 3 | 0 | 0 | 0 | 0 | 2 |
| 0 | 0.01 | 3.61 | 0.00 | 1.00 | 1 | Gm26189 | | 0 | 0 | 0 | 0 | 0 | 3 | 0 | 0 | 0 | 0 | 2 |
| 0 | 0.01 | 3.61 | 0.00 | 1.00 | 1 | Gm25545 | | 0 | 0 | 0 | 0 | 0 | 3 | 0 | 0 | 0 | 0 | 2 |
| 0 | 0.01 | 3.61 | 0.00 | 1.00 | 1 | Aire | autoimmune regulator (autoimmune polye | 0 | 0 | 0 | 0 | 0 | 3 | 0 | 0 | 0 | 0 | 2 |
| 0 | 0.01 | 3.61 | 0.00 | 1.00 | 1 | Snora5c | | 0 | 0 | 0 | 0 | 0 | 3 | 0 | 0 | 0 | 0 | 2 |
| 0 | 0.01 | 3.61 | 0.00 | 1.00 | 1 | Gm44577 | | 0 | 0 | 0 | 0 | 0 | 3 | 0 | 0 | 0 | 0 | 2 |
| 0 | -0.52 | 3.62 | -0.14 | 0.89 | 1 | Gm6009 | | 0 | 0 | 0 | 4 | 0 | 0 | 0 | 0 | 0 | 0 | 1 |

| | | | | | | | | | | | | | | | | | | |
|---|---|---|---|---|---|---|---|---|---|---|---|---|---|---|---|---|---|---|
| 0 | 0.99 | 3.62 | 0.27 | 0.79 | 1 | Gm12138 | | 1 | 0 | 0 | 0 | 1 | 0 | 0 | 0 | 1 | 3 | 0 |
| 0 | 1.84 | 3.64 | 0.51 | 0.61 | 1 | Gm4997 | | 0 | 1 | 1 | 0 | 0 | 0 | 0 | 2 | 1 | 0 | 0 |
| 0 | -1.08 | 3.67 | -0.30 | 0.77 | 1 | Gm8157 | | 3 | 1 | 0 | 0 | 1 | 1 | 0 | 0 | 0 | 0 | 0 |
| 0 | 1.65 | 3.63 | 0.46 | 0.65 | 1 | Selenok-ps7 | | 0 | 0 | 1 | 1 | 0 | 0 | 0 | 1 | 0 | 1 | 1 |
| 0 | -2.01 | 3.63 | -0.55 | 0.58 | 1 | Gm20540 | | 0 | 0 | 0 | 0 | 3 | 3 | 0 | 0 | 0 | 0 | 0 |
| 0 | 1.94 | 3.65 | 0.53 | 0.60 | 1 | Gm13007 | | 0 | 1 | 1 | 0 | 0 | 0 | 0 | 0 | 2 | 1 | 0 |
| 0 | -0.92 | 3.62 | -0.26 | 0.80 | 1 | Gm38025 | | 0 | 0 | 0 | 0 | 1 | 3 | 0 | 0 | 0 | 0 | 1 |
| 0 | -1.78 | 3.63 | -0.49 | 0.62 | 1 | Gm8343 | | 0 | 1 | 0 | 0 | 1 | 3 | 0 | 0 | 0 | 0 | 0 |
| 0 | -0.67 | 3.61 | -0.19 | 0.85 | 1 | Tmem181c-ps | | 0 | 0 | 0 | 0 | 1 | 3 | 0 | 0 | 1 | 0 | 0 |
| 0 | 2.44 | 3.63 | 0.67 | 0.50 | 1 | Gm7583 | | 1 | 0 | 0 | 0 | 0 | 0 | 0 | 1 | 0 | 1 | 2 |
| 0 | 1.56 | 3.65 | 0.43 | 0.67 | 1 | Gm7783 | | 1 | 2 | 0 | 0 | 0 | 0 | 0 | 1 | 0 | 1 | 0 |
| 0 | 0.97 | 3.67 | 0.27 | 0.79 | 1 | Txk | TXK tyrosine kinase | 4 | 0 | 0 | 0 | 0 | 0 | 0 | 0 | 0 | 0 | 1 |
| 0 | 0.00 | 3.68 | 0.00 | 1.00 | 1 | Igkv1-88 | | 4 | 1 | 0 | 0 | 0 | 0 | 0 | 0 | 0 | 0 | 0 |
| 0 | 0.00 | 3.68 | 0.00 | 1.00 | 1 | Hoxc6 | homeobox C6 | 4 | 1 | 0 | 0 | 0 | 0 | 0 | 0 | 0 | 0 | 0 |
| 0 | 1.08 | 3.67 | 0.29 | 0.77 | 1 | Mettl7a2 | methyltransferase like 7A2 | 4 | 0 | 0 | 0 | 0 | 0 | 0 | 0 | 1 | 0 | 0 |
| 0 | -2.36 | 3.60 | -0.65 | 0.51 | 1 | Gm46204 | | 0 | 0 | 0 | 1 | 2 | 3 | 0 | 0 | 0 | 0 | 0 |
| 0 | 1.77 | 3.62 | 0.49 | 0.62 | 1 | Gpx4-ps1 | | 0 | 0 | 0 | 0 | 1 | 0 | 0 | 2 | 0 | 0 | 2 |
| 0 | -1.92 | 3.63 | -0.53 | 0.60 | 1 | Potegl | fem-1 homolog A like | 0 | 0 | 0 | 0 | 2 | 3 | 0 | 0 | 0 | 0 | 0 |
| 0 | 1.01 | 3.68 | 0.27 | 0.78 | 1 | Gm47724 | | 1 | 0 | 3 | 0 | 0 | 0 | 0 | 0 | 0 | 1 | 0 |
| 0 | -0.24 | 3.62 | -0.07 | 0.95 | 1 | Gm6967 | | 1 | 0 | 0 | 0 | 0 | 3 | 0 | 0 | 0 | 0 | 1 |
| 0 | -1.25 | 3.64 | -0.34 | 0.73 | 1 | 1700121C08Rik | | 0 | 0 | 0 | 5 | 0 | 0 | 0 | 0 | 0 | 0 | 0 |
| 0 | -1.25 | 3.64 | -0.34 | 0.73 | 1 | Gm43319 | | 0 | 0 | 0 | 5 | 0 | 0 | 0 | 0 | 0 | 0 | 0 |
| 0 | -1.25 | 3.64 | -0.34 | 0.73 | 1 | Gm32158 | | 0 | 0 | 0 | 5 | 0 | 0 | 0 | 0 | 0 | 0 | 0 |
| 0 | -1.25 | 3.64 | -0.34 | 0.73 | 1 | Mir344f | | 0 | 0 | 0 | 5 | 0 | 0 | 0 | 0 | 0 | 0 | 0 |
| 0 | -1.25 | 3.64 | -0.34 | 0.73 | 1 | Gm20681 | | 0 | 0 | 0 | 5 | 0 | 0 | 0 | 0 | 0 | 0 | 0 |
| 0 | -1.25 | 3.64 | -0.34 | 0.73 | 1 | Gm49769 | | 0 | 0 | 0 | 5 | 0 | 0 | 0 | 0 | 0 | 0 | 0 |
| 0 | 0.16 | 3.62 | 0.04 | 0.96 | 1 | Hspe1-ps1 | | 1 | 0 | 0 | 0 | 0 | 3 | 0 | 0 | 1 | 0 | 0 |
| 0 | -0.31 | 3.61 | -0.09 | 0.93 | 1 | Gm37161 | | 0 | 0 | 0 | 0 | 0 | 4 | 0 | 0 | 0 | 0 | 1 |
| 0 | -0.31 | 3.61 | -0.09 | 0.93 | 1 | 2310001K24Rik | | 0 | 0 | 0 | 0 | 0 | 4 | 0 | 0 | 0 | 0 | 1 |
| 0 | -0.31 | 3.61 | -0.09 | 0.93 | 1 | Gm27899 | | 0 | 0 | 0 | 0 | 0 | 4 | 0 | 0 | 0 | 0 | 1 |
| 0 | -0.31 | 3.61 | -0.09 | 0.93 | 1 | Gm37978 | | 0 | 0 | 0 | 0 | 0 | 4 | 0 | 0 | 0 | 0 | 1 |
| 0 | -0.31 | 3.61 | -0.09 | 0.93 | 1 | Gm44907 | | 0 | 0 | 0 | 0 | 0 | 4 | 0 | 0 | 0 | 0 | 1 |
| 0 | -0.31 | 3.61 | -0.09 | 0.93 | 1 | Gm31786 | | 0 | 0 | 0 | 0 | 0 | 4 | 0 | 0 | 0 | 0 | 1 |
| 0 | -0.31 | 3.61 | -0.09 | 0.93 | 1 | Gm47160 | | 0 | 0 | 0 | 0 | 0 | 4 | 0 | 0 | 0 | 0 | 1 |
| 0 | -0.31 | 3.61 | -0.09 | 0.93 | 1 | Gm3211 | | 0 | 0 | 0 | 0 | 0 | 4 | 0 | 0 | 0 | 0 | 1 |
| 0 | -0.31 | 3.61 | -0.09 | 0.93 | 1 | Gm40884 | | 0 | 0 | 0 | 0 | 0 | 4 | 0 | 0 | 0 | 0 | 1 |
| 0 | -0.31 | 3.61 | -0.09 | 0.93 | 1 | Timm8a2 | translocase of inner mitochondrial membr | 0 | 0 | 0 | 0 | 0 | 4 | 0 | 0 | 0 | 0 | 1 |
| 0 | -0.31 | 3.61 | -0.09 | 0.93 | 1 | Gm24907 | | 0 | 0 | 0 | 0 | 0 | 4 | 0 | 0 | 0 | 0 | 1 |
| 0 | 0.15 | 3.62 | 0.04 | 0.97 | 1 | Gm25902 | | 0 | 0 | 0 | 0 | 0 | 4 | 0 | 0 | 1 | 0 | 0 |
| 0 | -0.55 | 3.67 | -0.15 | 0.88 | 1 | Gm43578 | | 4 | 0 | 0 | 0 | 1 | 0 | 0 | 0 | 0 | 0 | 0 |
| 0 | 1.34 | 3.65 | 0.37 | 0.71 | 1 | Gm6244 | | 0 | 0 | 1 | 1 | 0 | 0 | 0 | 0 | 0 | 1 | 2 |
| 0 | -1.81 | 3.63 | -0.50 | 0.62 | 1 | Gm24082 | | 0 | 0 | 0 | 4 | 0 | 1 | 0 | 0 | 0 | 0 | 0 |
| 0 | 0.72 | 3.64 | 0.20 | 0.84 | 1 | Gm9797 | | 0 | 4 | 0 | 0 | 0 | 0 | 0 | 1 | 0 | 0 | 0 |
| 0 | 1.59 | 3.66 | 0.44 | 0.66 | 1 | Rps19-ps7 | | 1 | 1 | 1 | 0 | 0 | 0 | 0 | 0 | 1 | 1 | 0 |
| 0 | 0.00 | 3.68 | 0.00 | 1.00 | 1 | Gm37469 | | 0 | 0 | 5 | 0 | 0 | 0 | 0 | 0 | 0 | 0 | 0 |
| 0 | 0.00 | 3.68 | 0.00 | 1.00 | 1 | Olfr335-ps | | 0 | 0 | 5 | 0 | 0 | 0 | 0 | 0 | 0 | 0 | 0 |
| 0 | 0.00 | 3.68 | 0.00 | 1.00 | 1 | Gm22261 | | 0 | 0 | 5 | 0 | 0 | 0 | 0 | 0 | 0 | 0 | 0 |
| 0 | 0.00 | 3.68 | 0.00 | 1.00 | 1 | Gm42772 | | 0 | 0 | 5 | 0 | 0 | 0 | 0 | 0 | 0 | 0 | 0 |
| 0 | 0.00 | 3.68 | 0.00 | 1.00 | 1 | Gm42882 | | 0 | 0 | 5 | 0 | 0 | 0 | 0 | 0 | 0 | 0 | 0 |
| 0 | 0.00 | 3.68 | 0.00 | 1.00 | 1 | Gm3054 | | 0 | 0 | 5 | 0 | 0 | 0 | 0 | 0 | 0 | 0 | 0 |
| 0 | 0.00 | 3.68 | 0.00 | 1.00 | 1 | 9530018H14Rik | | 0 | 0 | 5 | 0 | 0 | 0 | 0 | 0 | 0 | 0 | 0 |
| 0 | 0.00 | 3.68 | 0.00 | 1.00 | 1 | Gm44712 | | 0 | 0 | 5 | 0 | 0 | 0 | 0 | 0 | 0 | 0 | 0 |
| 0 | 0.00 | 3.68 | 0.00 | 1.00 | 1 | Gm44914 | | 0 | 0 | 5 | 0 | 0 | 0 | 0 | 0 | 0 | 0 | 0 |
| 0 | 0.00 | 3.68 | 0.00 | 1.00 | 1 | Spaca7 | sperm acrosome associated 7 | 0 | 0 | 5 | 0 | 0 | 0 | 0 | 0 | 0 | 0 | 0 |
| 0 | 0.00 | 3.68 | 0.00 | 1.00 | 1 | Ccdc110 | coiled-coil domain containing 110 | 0 | 0 | 5 | 0 | 0 | 0 | 0 | 0 | 0 | 0 | 0 |
| 0 | 0.00 | 3.68 | 0.00 | 1.00 | 1 | Gm48293 | | 0 | 0 | 5 | 0 | 0 | 0 | 0 | 0 | 0 | 0 | 0 |
| 0 | 0.00 | 3.68 | 0.00 | 1.00 | 1 | 4932413F04Rik | | 0 | 0 | 5 | 0 | 0 | 0 | 0 | 0 | 0 | 0 | 0 |
| 0 | 0.00 | 3.68 | 0.00 | 1.00 | 1 | Gm47094 | | 0 | 0 | 5 | 0 | 0 | 0 | 0 | 0 | 0 | 0 | 0 |
| 0 | 0.00 | 3.68 | 0.00 | 1.00 | 1 | Gm47456 | | 0 | 0 | 5 | 0 | 0 | 0 | 0 | 0 | 0 | 0 | 0 |
| 0 | 0.00 | 3.68 | 0.00 | 1.00 | 1 | Gm16086 | | 0 | 0 | 5 | 0 | 0 | 0 | 0 | 0 | 0 | 0 | 0 |

| | | | | | | | | | | | | | | | | | | |
|---|---|---|---|---|---|---|---|---|---|---|---|---|---|---|---|---|---|---|
| 0 | 0.00 | 3.68 | 0.00 | 1.00 | 1 | Gm48783 | | 0 | 0 | 5 | 0 | 0 | 0 | 0 | 0 | 0 | 0 | 0 |
| 0 | 0.00 | 3.68 | 0.00 | 1.00 | 1 | Trav14-1 | | 0 | 0 | 5 | 0 | 0 | 0 | 0 | 0 | 0 | 0 | 0 |
| 0 | 0.00 | 3.68 | 0.00 | 1.00 | 1 | Fam216b | family with sequence similarity 216, meml | 0 | 0 | 5 | 0 | 0 | 0 | 0 | 0 | 0 | 0 | 0 |
| 0 | 0.00 | 3.68 | 0.00 | 1.00 | 1 | Cldn14 | claudin 14 | 0 | 0 | 5 | 0 | 0 | 0 | 0 | 0 | 0 | 0 | 0 |
| 0 | 0.00 | 3.68 | 0.00 | 1.00 | 1 | Gm20496 | | 0 | 0 | 5 | 0 | 0 | 0 | 0 | 0 | 0 | 0 | 0 |
| 0 | 0.00 | 3.68 | 0.00 | 1.00 | 1 | Gm9598 | | 0 | 0 | 5 | 0 | 0 | 0 | 0 | 0 | 0 | 0 | 0 |
| 0 | 0.00 | 3.68 | 0.00 | 1.00 | 1 | Gm15337 | | 0 | 0 | 5 | 0 | 0 | 0 | 0 | 0 | 0 | 0 | 0 |
| 0 | -1.95 | 3.63 | -0.54 | 0.59 | 1 | Gm7781 | | 0 | 0 | 0 | 4 | 0 | 2 | 0 | 0 | 0 | 0 | 0 |
| 0 | 1.06 | 3.66 | 0.29 | 0.77 | 1 | Gm48966 | | 0 | 0 | 4 | 0 | 0 | 0 | 0 | 0 | 0 | 0 | 1 |
| 0 | 0.08 | 3.65 | 0.02 | 0.98 | 1 | Gm9224 | | 0 | 2 | 1 | 0 | 1 | 0 | 0 | 1 | 0 | 0 | 0 |
| 0 | -0.54 | 3.67 | -0.15 | 0.88 | 1 | Gm3331 | | 4 | 0 | 0 | 1 | 0 | 0 | 0 | 0 | 0 | 0 | 0 |
| 0 | -1.97 | 3.63 | -0.54 | 0.59 | 1 | D830013O20Rik | | 0 | 0 | 0 | 3 | 0 | 3 | 0 | 0 | 0 | 0 | 0 |
| 0 | -1.97 | 3.63 | -0.54 | 0.59 | 1 | H2bc13 | H2B clustered histone 13 | 0 | 0 | 0 | 3 | 0 | 3 | 0 | 0 | 0 | 0 | 0 |
| 0 | 1.72 | 3.65 | 0.47 | 0.64 | 1 | Gm49397 | | 0 | 0 | 0 | 0 | 0 | 0 | 0 | 0 | 0 | 5 | 0 |
| 0 | 0.00 | 3.68 | 0.00 | 1.00 | 1 | Gm33280 | | 5 | 0 | 0 | 0 | 0 | 0 | 0 | 0 | 0 | 0 | 0 |
| 0 | 0.00 | 3.68 | 0.00 | 1.00 | 1 | Trpm8 | transient receptor potential cation channe | 5 | 0 | 0 | 0 | 0 | 0 | 0 | 0 | 0 | 0 | 0 |
| 0 | 0.00 | 3.68 | 0.00 | 1.00 | 1 | Gm28404 | | 5 | 0 | 0 | 0 | 0 | 0 | 0 | 0 | 0 | 0 | 0 |
| 0 | 0.00 | 3.68 | 0.00 | 1.00 | 1 | Fcamr | Fc receptor, IgA, IgM, high affinity | 5 | 0 | 0 | 0 | 0 | 0 | 0 | 0 | 0 | 0 | 0 |
| 0 | 0.00 | 3.68 | 0.00 | 1.00 | 1 | Tnfsf18 | tumor necrosis factor (ligand) superfamily | 5 | 0 | 0 | 0 | 0 | 0 | 0 | 0 | 0 | 0 | 0 |
| 0 | 0.00 | 3.68 | 0.00 | 1.00 | 1 | Fasl | Fas ligand (TNF superfamily, member 6) | 5 | 0 | 0 | 0 | 0 | 0 | 0 | 0 | 0 | 0 | 0 |
| 0 | 0.00 | 3.68 | 0.00 | 1.00 | 1 | Il1f10 | interleukin 1 family, member 10 | 5 | 0 | 0 | 0 | 0 | 0 | 0 | 0 | 0 | 0 | 0 |
| 0 | 0.00 | 3.68 | 0.00 | 1.00 | 1 | Gm14029 | | 5 | 0 | 0 | 0 | 0 | 0 | 0 | 0 | 0 | 0 | 0 |
| 0 | 0.00 | 3.68 | 0.00 | 1.00 | 1 | Gm22245 | | 5 | 0 | 0 | 0 | 0 | 0 | 0 | 0 | 0 | 0 | 0 |
| 0 | 0.00 | 3.68 | 0.00 | 1.00 | 1 | AW047730 | | 5 | 0 | 0 | 0 | 0 | 0 | 0 | 0 | 0 | 0 | 0 |
| 0 | 0.00 | 3.68 | 0.00 | 1.00 | 1 | Gm38078 | | 5 | 0 | 0 | 0 | 0 | 0 | 0 | 0 | 0 | 0 | 0 |
| 0 | 0.00 | 3.68 | 0.00 | 1.00 | 1 | Pitx2 | paired-like homeodomain transcription fa | 5 | 0 | 0 | 0 | 0 | 0 | 0 | 0 | 0 | 0 | 0 |
| 0 | 0.00 | 3.68 | 0.00 | 1.00 | 1 | Gm43428 | | 5 | 0 | 0 | 0 | 0 | 0 | 0 | 0 | 0 | 0 | 0 |
| 0 | 0.00 | 3.68 | 0.00 | 1.00 | 1 | Gm13223 | | 5 | 0 | 0 | 0 | 0 | 0 | 0 | 0 | 0 | 0 | 0 |
| 0 | 0.00 | 3.68 | 0.00 | 1.00 | 1 | Gm42802 | | 5 | 0 | 0 | 0 | 0 | 0 | 0 | 0 | 0 | 0 | 0 |
| 0 | 0.00 | 3.68 | 0.00 | 1.00 | 1 | Csn1s2b | casein alpha s2-like B | 5 | 0 | 0 | 0 | 0 | 0 | 0 | 0 | 0 | 0 | 0 |
| 0 | 0.00 | 3.68 | 0.00 | 1.00 | 1 | Vmn1r30 | vomeronasal 1 receptor 30 | 5 | 0 | 0 | 0 | 0 | 0 | 0 | 0 | 0 | 0 | 0 |
| 0 | 0.00 | 3.68 | 0.00 | 1.00 | 1 | Igkv4-83 | | 5 | 0 | 0 | 0 | 0 | 0 | 0 | 0 | 0 | 0 | 0 |
| 0 | 0.00 | 3.68 | 0.00 | 1.00 | 1 | Igkv4-74 | | 5 | 0 | 0 | 0 | 0 | 0 | 0 | 0 | 0 | 0 | 0 |
| 0 | 0.00 | 3.68 | 0.00 | 1.00 | 1 | Gm43218 | | 5 | 0 | 0 | 0 | 0 | 0 | 0 | 0 | 0 | 0 | 0 |
| 0 | 0.00 | 3.68 | 0.00 | 1.00 | 1 | Igkv6-20 | | 5 | 0 | 0 | 0 | 0 | 0 | 0 | 0 | 0 | 0 | 0 |
| 0 | 0.00 | 3.68 | 0.00 | 1.00 | 1 | Gm15862 | | 5 | 0 | 0 | 0 | 0 | 0 | 0 | 0 | 0 | 0 | 0 |
| 0 | 0.00 | 3.68 | 0.00 | 1.00 | 1 | Gsg1 | germ cell associated 1 | 5 | 0 | 0 | 0 | 0 | 0 | 0 | 0 | 0 | 0 | 0 |
| 0 | 0.00 | 3.68 | 0.00 | 1.00 | 1 | Tarm1 | T cell-interacting, activating receptor on m | 5 | 0 | 0 | 0 | 0 | 0 | 0 | 0 | 0 | 0 | 0 |
| 0 | 0.00 | 3.68 | 0.00 | 1.00 | 1 | Dkkl1 | dickkopf-like 1 | 5 | 0 | 0 | 0 | 0 | 0 | 0 | 0 | 0 | 0 | 0 |
| 0 | 0.00 | 3.68 | 0.00 | 1.00 | 1 | Gm9885 | | 5 | 0 | 0 | 0 | 0 | 0 | 0 | 0 | 0 | 0 | 0 |
| 0 | 0.00 | 3.68 | 0.00 | 1.00 | 1 | Gm26747 | | 5 | 0 | 0 | 0 | 0 | 0 | 0 | 0 | 0 | 0 | 0 |
| 0 | 0.00 | 3.68 | 0.00 | 1.00 | 1 | Jsrp1 | junctional sarcoplasmic reticulum protein | 5 | 0 | 0 | 0 | 0 | 0 | 0 | 0 | 0 | 0 | 0 |
| 0 | 0.00 | 3.68 | 0.00 | 1.00 | 1 | Gm40723 | | 5 | 0 | 0 | 0 | 0 | 0 | 0 | 0 | 0 | 0 | 0 |
| 0 | 0.00 | 3.68 | 0.00 | 1.00 | 1 | Slc25a21 | solute carrier family 25 (mitochondrial oxo | 5 | 0 | 0 | 0 | 0 | 0 | 0 | 0 | 0 | 0 | 0 |
| 0 | 0.00 | 3.68 | 0.00 | 1.00 | 1 | Gm9238 | | 5 | 0 | 0 | 0 | 0 | 0 | 0 | 0 | 0 | 0 | 0 |
| 0 | 0.00 | 3.68 | 0.00 | 1.00 | 1 | Trav8n-2 | | 5 | 0 | 0 | 0 | 0 | 0 | 0 | 0 | 0 | 0 | 0 |
| 0 | 0.00 | 3.68 | 0.00 | 1.00 | 1 | Gm536 | predicted gene 536 | 5 | 0 | 0 | 0 | 0 | 0 | 0 | 0 | 0 | 0 | 0 |
| 0 | 0.00 | 3.68 | 0.00 | 1.00 | 1 | Gm22146 | | 5 | 0 | 0 | 0 | 0 | 0 | 0 | 0 | 0 | 0 | 0 |
| 0 | 0.00 | 3.68 | 0.00 | 1.00 | 1 | Gm50127 | | 5 | 0 | 0 | 0 | 0 | 0 | 0 | 0 | 0 | 0 | 0 |
| 0 | 0.00 | 3.68 | 0.00 | 1.00 | 1 | Gm46606 | | 5 | 0 | 0 | 0 | 0 | 0 | 0 | 0 | 0 | 0 | 0 |
| 0 | 0.00 | 3.68 | 0.00 | 1.00 | 1 | Gm50428 | | 5 | 0 | 0 | 0 | 0 | 0 | 0 | 0 | 0 | 0 | 0 |
| 0 | 0.18 | 3.62 | 0.05 | 0.96 | 1 | Gm12508 | | 0 | 0 | 1 | 0 | 1 | 1 | 0 | 2 | 0 | 0 | 0 |
| 0 | 1.43 | 3.63 | 0.39 | 0.69 | 1 | Hspd1-ps5 | | 0 | 1 | 0 | 0 | 1 | 0 | 0 | 1 | 1 | 0 | 1 |
| 0 | 0.00 | 3.68 | 0.00 | 1.00 | 1 | Gm14523 | | 0 | 2 | 3 | 0 | 0 | 0 | 0 | 0 | 0 | 0 | 0 |
| 0 | -0.44 | 3.67 | -0.12 | 0.90 | 1 | Gm37202 | | 4 | 0 | 0 | 0 | 0 | 1 | 0 | 0 | 0 | 0 | 0 |
| 0 | -0.44 | 3.67 | -0.12 | 0.90 | 1 | Pdzd9 | PDZ domain containing 9 | 4 | 0 | 0 | 0 | 0 | 1 | 0 | 0 | 0 | 0 | 0 |
| 0 | -0.77 | 3.65 | -0.21 | 0.83 | 1 | 2810407A14Rik | | 3 | 0 | 0 | 0 | 0 | 2 | 0 | 0 | 0 | 0 | 0 |
| 0 | 0.48 | 3.61 | 0.13 | 0.89 | 1 | Gm9487 | | 0 | 0 | 0 | 0 | 1 | 1 | 0 | 2 | 1 | 0 | 0 |
| 0 | 1.32 | 3.63 | 0.36 | 0.72 | 1 | Gm6331 | | 0 | 0 | 1 | 0 | 0 | 0 | 0 | 0 | 0 | 0 | 4 |
| 0 | 0.00 | 3.68 | 0.00 | 1.00 | 1 | Gm5969 | | 0 | 4 | 1 | 0 | 0 | 0 | 0 | 0 | 0 | 0 | 0 |

| | | | | | | | | | | | | | | | | | | |
|---|---|---|---|---|---|---|---|---|---|---|---|---|---|---|---|---|---|---|
| 0 | 1.38 | 3.63 | 0.38 | 0.70 | 1 | Gm5187 | | 0 | 0 | 1 | 0 | 0 | 0 | 0 | 0 | 4 | 0 | 0 |
| 0 | -0.78 | 3.62 | -0.21 | 0.83 | 1 | Gm26814 | | 1 | 0 | 0 | 0 | 0 | 4 | 0 | 0 | 0 | 0 | 0 |
| 0 | -0.78 | 3.62 | -0.21 | 0.83 | 1 | LOC665443 | | 1 | 0 | 0 | 0 | 0 | 4 | 0 | 0 | 0 | 0 | 0 |
| 0 | 1.34 | 3.62 | 0.37 | 0.71 | 1 | Gm16107 | | 0 | 0 | 0 | 0 | 0 | 0 | 0 | 0 | 0 | 0 | 5 |
| 0 | 1.34 | 3.62 | 0.37 | 0.71 | 1 | Gm37246 | | 0 | 0 | 0 | 0 | 0 | 0 | 0 | 0 | 0 | 0 | 5 |
| 0 | 1.34 | 3.62 | 0.37 | 0.71 | 1 | Gm28035 | | 0 | 0 | 0 | 0 | 0 | 0 | 0 | 0 | 0 | 0 | 5 |
| 0 | 1.34 | 3.62 | 0.37 | 0.71 | 1 | Gm25184 | | 0 | 0 | 0 | 0 | 0 | 0 | 0 | 0 | 0 | 0 | 5 |
| 0 | 1.34 | 3.62 | 0.37 | 0.71 | 1 | Gm38034 | | 0 | 0 | 0 | 0 | 0 | 0 | 0 | 0 | 0 | 0 | 5 |
| 0 | 1.34 | 3.62 | 0.37 | 0.71 | 1 | Gm42696 | | 0 | 0 | 0 | 0 | 0 | 0 | 0 | 0 | 0 | 0 | 5 |
| 0 | 1.34 | 3.62 | 0.37 | 0.71 | 1 | Gm43111 | | 0 | 0 | 0 | 0 | 0 | 0 | 0 | 0 | 0 | 0 | 5 |
| 0 | 1.34 | 3.62 | 0.37 | 0.71 | 1 | Gm29681 | | 0 | 0 | 0 | 0 | 0 | 0 | 0 | 0 | 0 | 0 | 5 |
| 0 | 1.34 | 3.62 | 0.37 | 0.71 | 1 | Gm5315 | | 0 | 0 | 0 | 0 | 0 | 0 | 0 | 0 | 0 | 0 | 5 |
| 0 | 1.34 | 3.62 | 0.37 | 0.71 | 1 | Gm24307 | | 0 | 0 | 0 | 0 | 0 | 0 | 0 | 0 | 0 | 0 | 5 |
| 0 | 1.34 | 3.62 | 0.37 | 0.71 | 1 | Gm6518 | | 0 | 0 | 0 | 0 | 0 | 0 | 0 | 0 | 0 | 0 | 5 |
| 0 | 1.34 | 3.62 | 0.37 | 0.71 | 1 | Snord34 | | 0 | 0 | 0 | 0 | 0 | 0 | 0 | 0 | 0 | 0 | 5 |
| 0 | 1.34 | 3.62 | 0.37 | 0.71 | 1 | Gm23917 | | 0 | 0 | 0 | 0 | 0 | 0 | 0 | 0 | 0 | 0 | 5 |
| 0 | 1.34 | 3.62 | 0.37 | 0.71 | 1 | Gm18958 | | 0 | 0 | 0 | 0 | 0 | 0 | 0 | 0 | 0 | 0 | 5 |
| 0 | 1.34 | 3.62 | 0.37 | 0.71 | 1 | B930086L07Rik | | 0 | 0 | 0 | 0 | 0 | 0 | 0 | 0 | 0 | 0 | 5 |
| 0 | 1.34 | 3.62 | 0.37 | 0.71 | 1 | Gm45785 | | 0 | 0 | 0 | 0 | 0 | 0 | 0 | 0 | 0 | 0 | 5 |
| 0 | 1.34 | 3.62 | 0.37 | 0.71 | 1 | Gm45490 | | 0 | 0 | 0 | 0 | 0 | 0 | 0 | 0 | 0 | 0 | 5 |
| 0 | 1.34 | 3.62 | 0.37 | 0.71 | 1 | 4930578M07Rik | | 0 | 0 | 0 | 0 | 0 | 0 | 0 | 0 | 0 | 0 | 5 |
| 0 | 1.34 | 3.62 | 0.37 | 0.71 | 1 | Gm47050 | | 0 | 0 | 0 | 0 | 0 | 0 | 0 | 0 | 0 | 0 | 5 |
| 0 | 1.34 | 3.62 | 0.37 | 0.71 | 1 | Gm30228 | | 0 | 0 | 0 | 0 | 0 | 0 | 0 | 0 | 0 | 0 | 5 |
| 0 | 1.34 | 3.62 | 0.37 | 0.71 | 1 | Gm8793 | | 0 | 0 | 0 | 0 | 0 | 0 | 0 | 0 | 0 | 0 | 5 |
| 0 | 1.34 | 3.62 | 0.37 | 0.71 | 1 | E230014E18Rik | | 0 | 0 | 0 | 0 | 0 | 0 | 0 | 0 | 0 | 0 | 5 |
| 0 | 1.34 | 3.62 | 0.37 | 0.71 | 1 | Gm49628 | | 0 | 0 | 0 | 0 | 0 | 0 | 0 | 0 | 0 | 0 | 5 |
| 0 | 1.34 | 3.62 | 0.37 | 0.71 | 1 | Mir5621 | | 0 | 0 | 0 | 0 | 0 | 0 | 0 | 0 | 0 | 0 | 5 |
| 0 | 1.34 | 3.62 | 0.37 | 0.71 | 1 | Gm48542 | | 0 | 0 | 0 | 0 | 0 | 0 | 0 | 0 | 0 | 0 | 5 |
| 0 | 1.34 | 3.62 | 0.37 | 0.71 | 1 | 4930474N09Rik | | 0 | 0 | 0 | 0 | 0 | 0 | 0 | 0 | 0 | 0 | 5 |
| 0 | 1.34 | 3.62 | 0.37 | 0.71 | 1 | Gm20621 | | 0 | 0 | 0 | 0 | 0 | 0 | 0 | 0 | 0 | 0 | 5 |
| 0 | 1.34 | 3.62 | 0.37 | 0.71 | 1 | Gm26562 | | 0 | 0 | 0 | 0 | 0 | 0 | 0 | 0 | 0 | 0 | 5 |
| 0 | 1.34 | 3.62 | 0.37 | 0.71 | 1 | 1700116B05Rik | | 0 | 0 | 0 | 0 | 0 | 0 | 0 | 0 | 0 | 0 | 5 |
| 0 | 1.34 | 3.62 | 0.37 | 0.71 | 1 | Gm50050 | | 0 | 0 | 0 | 0 | 0 | 0 | 0 | 0 | 0 | 0 | 5 |
| 0 | 1.34 | 3.62 | 0.37 | 0.71 | 1 | Gm17097 | | 0 | 0 | 0 | 0 | 0 | 0 | 0 | 0 | 0 | 0 | 5 |
| 0 | 1.34 | 3.62 | 0.37 | 0.71 | 1 | Gm19853 | | 0 | 0 | 0 | 0 | 0 | 0 | 0 | 0 | 0 | 0 | 5 |
| 0 | 1.34 | 3.62 | 0.37 | 0.71 | 1 | Gm19035 | | 0 | 0 | 0 | 0 | 0 | 0 | 0 | 0 | 0 | 0 | 5 |
| 0 | 1.34 | 3.62 | 0.37 | 0.71 | 1 | Gm49971 | | 0 | 0 | 0 | 0 | 0 | 0 | 0 | 0 | 0 | 0 | 5 |
| 0 | 1.43 | 3.67 | 0.39 | 0.70 | 1 | Gm19620 | | 2 | 0 | 1 | 0 | 0 | 0 | 0 | 2 | 0 | 0 | 0 |
| 0 | 1.30 | 3.64 | 0.36 | 0.72 | 1 | Rhox3b-ps | | 0 | 2 | 0 | 0 | 0 | 0 | 0 | 0 | 0 | 0 | 3 |
| 0 | -0.79 | 3.62 | -0.22 | 0.83 | 1 | Gm37686 | | 0 | 0 | 0 | 0 | 0 | 5 | 0 | 0 | 0 | 0 | 0 |
| 0 | -0.79 | 3.62 | -0.22 | 0.83 | 1 | Rbm44 | RNA binding motif protein 44 | 0 | 0 | 0 | 0 | 0 | 5 | 0 | 0 | 0 | 0 | 0 |
| 0 | -0.79 | 3.62 | -0.22 | 0.83 | 1 | Gm10537 | | 0 | 0 | 0 | 0 | 0 | 5 | 0 | 0 | 0 | 0 | 0 |
| 0 | -0.79 | 3.62 | -0.22 | 0.83 | 1 | Gm23608 | | 0 | 0 | 0 | 0 | 0 | 5 | 0 | 0 | 0 | 0 | 0 |
| 0 | -0.79 | 3.62 | -0.22 | 0.83 | 1 | Gm22192 | | 0 | 0 | 0 | 0 | 0 | 5 | 0 | 0 | 0 | 0 | 0 |
| 0 | -0.79 | 3.62 | -0.22 | 0.83 | 1 | | | 0 | 0 | 0 | 0 | 0 | 5 | 0 | 0 | 0 | 0 | 0 |
| 0 | -0.79 | 3.62 | -0.22 | 0.83 | 1 | Lime1 | Lck interacting transmembrane adaptor 1 | 0 | 0 | 0 | 0 | 0 | 5 | 0 | 0 | 0 | 0 | 0 |
| 0 | -0.79 | 3.62 | -0.22 | 0.83 | 1 | Gm38200 | | 0 | 0 | 0 | 0 | 0 | 5 | 0 | 0 | 0 | 0 | 0 |
| 0 | -0.79 | 3.62 | -0.22 | 0.83 | 1 | Gm44043 | | 0 | 0 | 0 | 0 | 0 | 5 | 0 | 0 | 0 | 0 | 0 |
| 0 | -0.79 | 3.62 | -0.22 | 0.83 | 1 | Snord15a | | 0 | 0 | 0 | 0 | 0 | 5 | 0 | 0 | 0 | 0 | 0 |
| 0 | -0.79 | 3.62 | -0.22 | 0.83 | 1 | Gm45399 | | 0 | 0 | 0 | 0 | 0 | 5 | 0 | 0 | 0 | 0 | 0 |
| 0 | -0.79 | 3.62 | -0.22 | 0.83 | 1 | Gm15718 | | 0 | 0 | 0 | 0 | 0 | 5 | 0 | 0 | 0 | 0 | 0 |
| 0 | -0.79 | 3.62 | -0.22 | 0.83 | 1 | Gm48207 | | 0 | 0 | 0 | 0 | 0 | 5 | 0 | 0 | 0 | 0 | 0 |
| 0 | -0.79 | 3.62 | -0.22 | 0.83 | 1 | Mir3971 | | 0 | 0 | 0 | 0 | 0 | 5 | 0 | 0 | 0 | 0 | 0 |
| 0 | -0.79 | 3.62 | -0.22 | 0.83 | 1 | 1700028N14Rik | | 0 | 0 | 0 | 0 | 0 | 5 | 0 | 0 | 0 | 0 | 0 |
| 0 | -0.79 | 3.62 | -0.22 | 0.83 | 1 | Gm48889 | | 0 | 0 | 0 | 0 | 0 | 5 | 0 | 0 | 0 | 0 | 0 |
| 0 | -0.79 | 3.62 | -0.22 | 0.83 | 1 | Vmn1r203 | vomeronasal 1 receptor 203 | 0 | 0 | 0 | 0 | 0 | 5 | 0 | 0 | 0 | 0 | 0 |
| 0 | -0.79 | 3.62 | -0.22 | 0.83 | 1 | Gm34939 | | 0 | 0 | 0 | 0 | 0 | 5 | 0 | 0 | 0 | 0 | 0 |
| 0 | -0.79 | 3.62 | -0.22 | 0.83 | 1 | 1700031F05Ri | RIKEN cDNA 1700031F05 gene | 0 | 0 | 0 | 0 | 0 | 5 | 0 | 0 | 0 | 0 | 0 |
| 0 | -0.79 | 3.62 | -0.22 | 0.83 | 1 | Gm14870 | | 0 | 0 | 0 | 0 | 0 | 5 | 0 | 0 | 0 | 0 | 0 |
| 0 | 1.10 | 3.62 | 0.30 | 0.76 | 1 | Gm15235 | | 0 | 1 | 0 | 2 | 0 | 0 | 0 | 0 | 0 | 1 | 1 |

| | | | | | | | | | | | | | | | | | | |
|---|---|---|---|---|---|---|---|---|---|---|---|---|---|---|---|---|---|---|
| 0 | 1.48 | 3.60 | 0.41 | 0.68 | 1 | Gm38166 | | 0 | 0 | 0 | 2 | 0 | 0 | 0 | 0 | 1 | 1 | 1 |
| 0 | 0.00 | 3.68 | 0.00 | 1.00 | 1 | Gm16070 | | 0 | 5 | 0 | 0 | 0 | 0 | 0 | 0 | 0 | 0 | 0 |
| 0 | 0.00 | 3.68 | 0.00 | 1.00 | 1 | Agxt | alanine-glyoxylate aminotransferase | 0 | 5 | 0 | 0 | 0 | 0 | 0 | 0 | 0 | 0 | 0 |
| 0 | 0.00 | 3.68 | 0.00 | 1.00 | 1 | Mybpc3 | myosin binding protein C, cardiac | 0 | 5 | 0 | 0 | 0 | 0 | 0 | 0 | 0 | 0 | 0 |
| 0 | 0.00 | 3.68 | 0.00 | 1.00 | 1 | Gm43577 | | 0 | 5 | 0 | 0 | 0 | 0 | 0 | 0 | 0 | 0 | 0 |
| 0 | 0.00 | 3.68 | 0.00 | 1.00 | 1 | 1700054K19Rik | | 0 | 5 | 0 | 0 | 0 | 0 | 0 | 0 | 0 | 0 | 0 |
| 0 | 0.00 | 3.68 | 0.00 | 1.00 | 1 | Cyp2b10 | cytochrome P450, family 2, subfamily b, p | 0 | 5 | 0 | 0 | 0 | 0 | 0 | 0 | 0 | 0 | 0 |
| 0 | 0.00 | 3.68 | 0.00 | 1.00 | 1 | Gm32255 | | 0 | 5 | 0 | 0 | 0 | 0 | 0 | 0 | 0 | 0 | 0 |
| 0 | 0.00 | 3.68 | 0.00 | 1.00 | 1 | Phxr2 | | 0 | 5 | 0 | 0 | 0 | 0 | 0 | 0 | 0 | 0 | 0 |
| 0 | 0.00 | 3.68 | 0.00 | 1.00 | 1 | Gm4425 | | 0 | 5 | 0 | 0 | 0 | 0 | 0 | 0 | 0 | 0 | 0 |
| 0 | 0.00 | 3.68 | 0.00 | 1.00 | 1 | Ighv1-42 | | 0 | 5 | 0 | 0 | 0 | 0 | 0 | 0 | 0 | 0 | 0 |
| 0 | 0.00 | 3.68 | 0.00 | 1.00 | 1 | Calm4 | calmodulin 4 | 0 | 5 | 0 | 0 | 0 | 0 | 0 | 0 | 0 | 0 | 0 |
| 0 | 0.00 | 3.68 | 0.00 | 1.00 | 1 | Ltb4r2 | leukotriene B4 receptor 2 | 0 | 5 | 0 | 0 | 0 | 0 | 0 | 0 | 0 | 0 | 0 |
| 0 | 0.00 | 3.68 | 0.00 | 1.00 | 1 | Gm15945 | | 0 | 5 | 0 | 0 | 0 | 0 | 0 | 0 | 0 | 0 | 0 |
| 0 | 0.00 | 3.68 | 0.00 | 1.00 | 1 | Gm41492 | | 0 | 5 | 0 | 0 | 0 | 0 | 0 | 0 | 0 | 0 | 0 |
| 0 | 0.00 | 3.68 | 0.00 | 1.00 | 1 | Gm16312 | | 0 | 5 | 0 | 0 | 0 | 0 | 0 | 0 | 0 | 0 | 0 |
| 0 | 0.00 | 3.68 | 0.00 | 1.00 | 1 | Gm5384 | | 0 | 5 | 0 | 0 | 0 | 0 | 0 | 0 | 0 | 0 | 0 |
| 0 | 1.75 | 3.64 | 0.48 | 0.63 | 1 | Dnah14 | dynein, axonemal, heavy chain 14 | 0 | 0 | 0 | 0 | 0 | 0 | 0 | 0 | 5 | 0 | 0 |
| 0 | 1.75 | 3.64 | 0.48 | 0.63 | 1 | Gm43064 | | 0 | 0 | 0 | 0 | 0 | 0 | 0 | 0 | 5 | 0 | 0 |
| 0 | 1.75 | 3.64 | 0.48 | 0.63 | 1 | Gm9364 | | 0 | 0 | 0 | 0 | 0 | 0 | 0 | 0 | 5 | 0 | 0 |
| 0 | 1.75 | 3.64 | 0.48 | 0.63 | 1 | Cbln1 | cerebellin 1 precursor protein | 0 | 0 | 0 | 0 | 0 | 0 | 0 | 0 | 5 | 0 | 0 |
| 0 | 1.75 | 3.64 | 0.48 | 0.63 | 1 | Gm35850 | | 0 | 0 | 0 | 0 | 0 | 0 | 0 | 0 | 5 | 0 | 0 |
| 0 | 1.75 | 3.64 | 0.48 | 0.63 | 1 | Gm45709 | | 0 | 0 | 0 | 0 | 0 | 0 | 0 | 0 | 5 | 0 | 0 |
| 0 | 1.75 | 3.64 | 0.48 | 0.63 | 1 | Gm23668 | | 0 | 0 | 0 | 0 | 0 | 0 | 0 | 0 | 5 | 0 | 0 |
| 0 | 1.75 | 3.64 | 0.48 | 0.63 | 1 | 4930432O09Rik | | 0 | 0 | 0 | 0 | 0 | 0 | 0 | 0 | 5 | 0 | 0 |
| 0 | 1.75 | 3.64 | 0.48 | 0.63 | 1 | Nt5c1b | 5'-nucleotidase, cytosolic IB | 0 | 0 | 0 | 0 | 0 | 0 | 0 | 0 | 5 | 0 | 0 |
| 0 | 1.75 | 3.64 | 0.48 | 0.63 | 1 | Gm16677 | | 0 | 0 | 0 | 0 | 0 | 0 | 0 | 0 | 5 | 0 | 0 |
| 0 | 1.75 | 3.64 | 0.48 | 0.63 | 1 | Mlc1 | megalencephalic leukoencephalopathy w | 0 | 0 | 0 | 0 | 0 | 0 | 0 | 0 | 5 | 0 | 0 |
| 0 | 1.75 | 3.64 | 0.48 | 0.63 | 1 | Gm9164 | | 0 | 0 | 0 | 0 | 0 | 0 | 0 | 0 | 5 | 0 | 0 |
| 0 | -0.55 | 3.67 | -0.15 | 0.88 | 1 | Gm42888 | | 0 | 0 | 4 | 1 | 0 | 0 | 0 | 0 | 0 | 0 | 0 |
| 0 | -0.35 | 3.63 | -0.10 | 0.92 | 1 | Gm29725 | | 0 | 1 | 1 | 1 | 1 | 0 | 0 | 1 | 0 | 0 | 0 |
| 0 | -0.31 | 3.62 | -0.08 | 0.93 | 1 | Gm28659 | | 0 | 0 | 0 | 2 | 1 | 0 | 0 | 2 | 0 | 0 | 0 |
| 0 | 1.38 | 3.61 | 0.38 | 0.70 | 1 | Gm37164 | | 0 | 0 | 0 | 0 | 0 | 2 | 0 | 0 | 1 | 1 | 1 |
| 0 | -0.44 | 3.67 | -0.12 | 0.90 | 1 | Gm8153 | | 0 | 0 | 4 | 0 | 0 | 1 | 0 | 0 | 0 | 0 | 0 |
| 0 | -0.44 | 3.67 | -0.12 | 0.90 | 1 | Gm23918 | | 0 | 0 | 4 | 0 | 0 | 1 | 0 | 0 | 0 | 0 | 0 |
| 0 | -0.60 | 3.66 | -0.16 | 0.87 | 1 | Gm13602 | | 0 | 4 | 0 | 0 | 1 | 0 | 0 | 0 | 0 | 0 | 0 |
| 0 | -0.40 | 3.67 | -0.11 | 0.91 | 1 | Serpinb6c | serine (or cysteine) peptidase inhibitor, cl | 0 | 1 | 3 | 0 | 0 | 1 | 0 | 0 | 0 | 0 | 0 |
| 0 | -0.99 | 3.62 | -0.27 | 0.78 | 1 | Gm6195 | | 0 | 0 | 0 | 0 | 3 | 1 | 0 | 0 | 0 | 1 | 0 |
| 0 | 0.95 | 3.67 | 0.26 | 0.80 | 1 | Gm13668 | | 1 | 1 | 2 | 0 | 0 | 0 | 0 | 0 | 0 | 0 | 1 |
| 0 | 0.29 | 3.65 | 0.08 | 0.94 | 1 | Gm15682 | | 1 | 1 | 1 | 1 | 0 | 0 | 0 | 1 | 0 | 0 | 0 |
| 0 | 0.28 | 3.68 | 0.08 | 0.94 | 1 | Gm7239 | | 1 | 0 | 2 | 0 | 0 | 1 | 0 | 1 | 0 | 0 | 0 |
| 0 | 0.00 | 3.60 | 0.00 | 1.00 | 1 | Mir433 | | 0 | 0 | 0 | 0 | 2 | 0 | 0 | 0 | 0 | 0 | 3 |
| 0 | 1.19 | 3.62 | 0.33 | 0.74 | 1 | Gm6621 | | 0 | 0 | 0 | 2 | 0 | 0 | 0 | 1 | 1 | 0 | 1 |
| 0 | 1.34 | 3.62 | 0.37 | 0.71 | 1 | Gm7664 | | 0 | 0 | 0 | 1 | 0 | 0 | 0 | 0 | 1 | 0 | 3 |
| 0 | 0.74 | 3.61 | 0.20 | 0.84 | 1 | Gm11826 | | 0 | 0 | 0 | 0 | 2 | 0 | 0 | 0 | 1 | 0 | 2 |
| 0 | 1.46 | 3.65 | 0.40 | 0.69 | 1 | Gm11470 | | 1 | 0 | 1 | 0 | 0 | 0 | 0 | 0 | 3 | 0 | 0 |
| 0 | 0.70 | 3.62 | 0.19 | 0.85 | 1 | Gm41780 | | 0 | 2 | 0 | 1 | 0 | 0 | 0 | 0 | 0 | 0 | 2 |
| 0 | -0.36 | 3.64 | -0.10 | 0.92 | 1 | Gm12852 | | 0 | 1 | 1 | 1 | 0 | 1 | 0 | 1 | 0 | 0 | 0 |
| 0 | 0.46 | 3.62 | 0.13 | 0.90 | 1 | Gm45760 | | 0 | 0 | 0 | 0 | 2 | 0 | 0 | 0 | 3 | 0 | 0 |
| 0 | 0.00 | 3.68 | 0.00 | 1.00 | 1 | Gm47922 | | 1 | 4 | 0 | 0 | 0 | 0 | 0 | 0 | 0 | 0 | 0 |
| 0 | 0.79 | 3.62 | 0.22 | 0.83 | 1 | Gm37588 | | 0 | 0 | 0 | 0 | 0 | 1 | 0 | 0 | 0 | 0 | 4 |
| 0 | 0.79 | 3.62 | 0.22 | 0.83 | 1 | Gm13625 | | 0 | 0 | 0 | 0 | 0 | 1 | 0 | 0 | 0 | 0 | 4 |
| 0 | 0.79 | 3.62 | 0.22 | 0.83 | 1 | Gm42842 | | 0 | 0 | 0 | 0 | 0 | 1 | 0 | 0 | 0 | 0 | 4 |
| 0 | 0.79 | 3.62 | 0.22 | 0.83 | 1 | Gm20731 | | 0 | 0 | 0 | 0 | 0 | 1 | 0 | 0 | 0 | 0 | 4 |
| 0 | 0.79 | 3.62 | 0.22 | 0.83 | 1 | Gm43320 | | 0 | 0 | 0 | 0 | 0 | 1 | 0 | 0 | 0 | 0 | 4 |
| 0 | 0.79 | 3.62 | 0.22 | 0.83 | 1 | Gm37321 | | 0 | 0 | 0 | 0 | 0 | 1 | 0 | 0 | 0 | 0 | 4 |
| 0 | 0.79 | 3.62 | 0.22 | 0.83 | 1 | Gm20705 | | 0 | 0 | 0 | 0 | 0 | 1 | 0 | 0 | 0 | 0 | 4 |
| 0 | 0.79 | 3.62 | 0.22 | 0.83 | 1 | Gm44584 | | 0 | 0 | 0 | 0 | 0 | 1 | 0 | 0 | 0 | 0 | 4 |
| 0 | 0.79 | 3.62 | 0.22 | 0.83 | 1 | Gm37257 | | 0 | 0 | 0 | 0 | 0 | 1 | 0 | 0 | 0 | 0 | 4 |
| 0 | 0.79 | 3.62 | 0.22 | 0.83 | 1 | Gm6740 | | 0 | 0 | 0 | 0 | 0 | 1 | 0 | 0 | 0 | 0 | 4 |

| | | | | | | | | | | | | | | | | | | |
|---|---|---|---|---|---|---|---|---|---|---|---|---|---|---|---|---|---|---|
| 0 | 0.69 | 3.62 | 0.19 | 0.85 | 1 | Gm3835 | | 0 | 1 | 0 | 0 | 0 | 1 | 0 | 0 | 0 | 0 | 3 |
| 0 | 1.75 | 3.65 | 0.48 | 0.63 | 1 | Gm28379 | | 1 | 0 | 0 | 0 | 0 | 0 | 0 | 0 | 4 | 0 | 0 |
| 0 | 1.36 | 3.62 | 0.38 | 0.71 | 1 | Mir7052 | | 0 | 0 | 0 | 0 | 0 | 1 | 0 | 0 | 1 | 0 | 3 |
| 0 | 0.28 | 3.64 | 0.08 | 0.94 | 1 | Gm17201 | | 0 | 3 | 0 | 0 | 0 | 1 | 0 | 0 | 0 | 0 | 1 |
| 0 | 0.28 | 3.64 | 0.08 | 0.94 | 1 | Asmt | acetylserotonin O-methyltransferase | 0 | 3 | 0 | 0 | 0 | 1 | 0 | 0 | 0 | 0 | 1 |
| 0 | -0.48 | 3.66 | -0.13 | 0.90 | 1 | 1700120K04Rik | | 0 | 4 | 0 | 0 | 0 | 1 | 0 | 0 | 0 | 0 | 0 |
| 0 | -0.44 | 3.65 | -0.12 | 0.90 | 1 | Gm5492 | | 1 | 1 | 0 | 1 | 0 | 1 | 0 | 0 | 0 | 1 | 0 |
| 0 | 1.63 | 3.66 | 0.45 | 0.66 | 1 | Cnga3 | cyclic nucleotide gated channel alpha 3 | 0 | 1 | 0 | 0 | 0 | 0 | 0 | 0 | 0 | 4 | 0 |
| 0 | 0.14 | 3.62 | 0.04 | 0.97 | 1 | Gm11434 | | 0 | 0 | 0 | 1 | 1 | 0 | 0 | 0 | 3 | 0 | 0 |
| 0 | -0.05 | 3.62 | -0.01 | 0.99 | 1 | Gm8870 | | 0 | 0 | 0 | 0 | 1 | 1 | 0 | 0 | 0 | 0 | 3 |
| 0 | -0.05 | 3.62 | -0.01 | 0.99 | 1 | Gm26175 | | 0 | 0 | 0 | 0 | 1 | 1 | 0 | 0 | 0 | 0 | 3 |
| 0 | 0.88 | 3.63 | 0.24 | 0.81 | 1 | Gm13502 | | 0 | 0 | 0 | 0 | 1 | 0 | 0 | 0 | 0 | 4 | 0 |
| 0 | -0.36 | 3.62 | -0.10 | 0.92 | 1 | Gm14822 | | 0 | 0 | 0 | 0 | 4 | 0 | 0 | 0 | 0 | 0 | 1 |
| 0 | -1.30 | 3.64 | -0.36 | 0.72 | 1 | Gm4883 | | 0 | 1 | 0 | 0 | 4 | 0 | 0 | 0 | 0 | 0 | 0 |
| 0 | -0.28 | 3.63 | -0.08 | 0.94 | 1 | Gm12919 | | 0 | 0 | 0 | 4 | 0 | 0 | 0 | 0 | 0 | 1 | 0 |
| 0 | 0.01 | 3.63 | 0.00 | 1.00 | 1 | Gm49733 | | 0 | 0 | 2 | 0 | 0 | 2 | 0 | 0 | 0 | 0 | 1 |
| 0 | 0.44 | 3.66 | 0.12 | 0.90 | 1 | Gm12460 | | 1 | 2 | 0 | 1 | 0 | 0 | 0 | 0 | 1 | 0 | 0 |
| 0 | -0.05 | 3.62 | -0.01 | 0.99 | 1 | Gm6612 | | 0 | 0 | 0 | 1 | 0 | 1 | 0 | 0 | 0 | 0 | 3 |
| 0 | -0.26 | 3.66 | -0.07 | 0.94 | 1 | Gm42573 | | 2 | 0 | 0 | 0 | 2 | 0 | 0 | 1 | 0 | 0 | 0 |
| 0 | 0.00 | 3.68 | 0.00 | 1.00 | 1 | Gm6637 | | 2 | 3 | 0 | 0 | 0 | 0 | 0 | 0 | 0 | 0 | 0 |
| 0 | 0.00 | 3.68 | 0.00 | 1.00 | 1 | Ceacam15 | carcinoembryonic antigen-related cell adl | 2 | 3 | 0 | 0 | 0 | 0 | 0 | 0 | 0 | 0 | 0 |
| 0 | 2.09 | 3.63 | 0.57 | 0.57 | 1 | Gm16581 | | 2 | 0 | 0 | 0 | 0 | 0 | 0 | 0 | 2 | 0 | 1 |
| 0 | 0.29 | 3.62 | 0.08 | 0.94 | 1 | Gm4837 | | 1 | 0 | 0 | 1 | 0 | 1 | 0 | 1 | 0 | 0 | 1 |
| 0 | 0.69 | 3.62 | 0.19 | 0.85 | 1 | Gm12987 | | 1 | 0 | 0 | 0 | 0 | 1 | 0 | 0 | 0 | 0 | 3 |
| 0 | 0.69 | 3.62 | 0.19 | 0.85 | 1 | Gm49594 | | 1 | 0 | 0 | 0 | 0 | 1 | 0 | 0 | 0 | 0 | 3 |
| 0 | 1.25 | 3.63 | 0.34 | 0.73 | 1 | Gm4880 | | 1 | 0 | 0 | 0 | 0 | 1 | 0 | 0 | 1 | 0 | 2 |
| 0 | 0.85 | 3.66 | 0.23 | 0.82 | 1 | Gm8216 | | 1 | 1 | 0 | 0 | 0 | 1 | 0 | 0 | 2 | 0 | 0 |
| 0 | 0.44 | 3.62 | 0.12 | 0.90 | 1 | Mir103-2 | | 0 | 0 | 0 | 0 | 0 | 2 | 0 | 0 | 0 | 0 | 3 |
| 0 | 0.44 | 3.62 | 0.12 | 0.90 | 1 | Mir7013 | | 0 | 0 | 0 | 0 | 0 | 2 | 0 | 0 | 0 | 0 | 3 |
| 0 | 0.44 | 3.62 | 0.12 | 0.90 | 1 | Gm26387 | | 0 | 0 | 0 | 0 | 0 | 2 | 0 | 0 | 0 | 0 | 3 |
| 0 | 0.44 | 3.62 | 0.12 | 0.90 | 1 | Sval2 | seminal vesicle antigen-like 2 | 0 | 0 | 0 | 0 | 0 | 2 | 0 | 0 | 0 | 0 | 3 |
| 0 | 0.44 | 3.62 | 0.12 | 0.90 | 1 | Mir7042 | | 0 | 0 | 0 | 0 | 0 | 2 | 0 | 0 | 0 | 0 | 3 |
| 0 | 0.44 | 3.62 | 0.12 | 0.90 | 1 | Snord88c | | 0 | 0 | 0 | 0 | 0 | 2 | 0 | 0 | 0 | 0 | 3 |
| 0 | 0.44 | 3.62 | 0.12 | 0.90 | 1 | Gm9801 | | 0 | 0 | 0 | 0 | 0 | 2 | 0 | 0 | 0 | 0 | 3 |
| 0 | 0.44 | 3.62 | 0.12 | 0.90 | 1 | Gm16202 | | 0 | 0 | 0 | 0 | 0 | 2 | 0 | 0 | 0 | 0 | 3 |
| 0 | 0.44 | 3.62 | 0.12 | 0.90 | 1 | Mir8103 | | 0 | 0 | 0 | 0 | 0 | 2 | 0 | 0 | 0 | 0 | 3 |
| 0 | 0.44 | 3.62 | 0.12 | 0.90 | 1 | Spata32 | spermatogenesis associated 32 | 0 | 0 | 0 | 0 | 0 | 2 | 0 | 0 | 0 | 0 | 3 |
| 0 | 0.44 | 3.62 | 0.12 | 0.90 | 1 | Gm11653 | | 0 | 0 | 0 | 0 | 0 | 2 | 0 | 0 | 0 | 0 | 3 |
| 0 | 0.44 | 3.62 | 0.12 | 0.90 | 1 | H3c1 | H3 clustered histone 1 | 0 | 0 | 0 | 0 | 0 | 2 | 0 | 0 | 0 | 0 | 3 |
| 0 | 0.44 | 3.62 | 0.12 | 0.90 | 1 | Gm46540 | | 0 | 0 | 0 | 0 | 0 | 2 | 0 | 0 | 0 | 0 | 3 |
| 0 | 0.44 | 3.62 | 0.12 | 0.90 | 1 | Gm4786 | | 0 | 0 | 0 | 0 | 0 | 2 | 0 | 0 | 0 | 0 | 3 |
| 0 | 0.44 | 3.62 | 0.12 | 0.90 | 1 | Gm14636 | | 0 | 0 | 0 | 0 | 0 | 2 | 0 | 0 | 0 | 0 | 3 |
| 0 | 0.95 | 3.62 | 0.26 | 0.79 | 1 | 4933417D19Rik | | 0 | 0 | 0 | 0 | 0 | 2 | 0 | 0 | 1 | 0 | 2 |
| 0 | 0.74 | 3.63 | 0.20 | 0.84 | 1 | Gm35781 | | 0 | 1 | 0 | 0 | 0 | 2 | 0 | 0 | 1 | 0 | 1 |
| 0 | -0.81 | 3.65 | -0.22 | 0.83 | 1 | Tprg | transformation related protein 63 regulate | 0 | 3 | 0 | 0 | 0 | 2 | 0 | 0 | 0 | 0 | 0 |
| 0 | 0.78 | 3.62 | 0.22 | 0.83 | 1 | Gm37676 | | 0 | 1 | 0 | 0 | 0 | 2 | 0 | 0 | 2 | 0 | 0 |
| 0 | 0.83 | 3.60 | 0.23 | 0.82 | 1 | Gm7838 | | 0 | 0 | 0 | 0 | 0 | 2 | 0 | 0 | 3 | 0 | 0 |
| 0 | 0.83 | 3.60 | 0.23 | 0.82 | 1 | Atp8b3 | ATPase, class I, type 8B, member 3 | 0 | 0 | 0 | 0 | 0 | 2 | 0 | 0 | 3 | 0 | 0 |
| 0 | -1.13 | 3.64 | -0.31 | 0.76 | 1 | Olah | oleoyl-ACP hydrolase | 0 | 0 | 0 | 0 | 5 | 0 | 0 | 0 | 0 | 0 | 0 |
| 0 | -1.13 | 3.64 | -0.31 | 0.76 | 1 | Accsl | 1-aminocyclopropane-1-carboxylate syntl | 0 | 0 | 0 | 0 | 5 | 0 | 0 | 0 | 0 | 0 | 0 |
| 0 | -1.13 | 3.64 | -0.31 | 0.76 | 1 | Zfp345 | zinc finger protein 345 | 0 | 0 | 0 | 0 | 5 | 0 | 0 | 0 | 0 | 0 | 0 |
| 0 | -1.13 | 3.64 | -0.31 | 0.76 | 1 | Gm37675 | | 0 | 0 | 0 | 0 | 5 | 0 | 0 | 0 | 0 | 0 | 0 |
| 0 | -1.13 | 3.64 | -0.31 | 0.76 | 1 | S100a2 | S100 calcium binding protein A2 | 0 | 0 | 0 | 0 | 5 | 0 | 0 | 0 | 0 | 0 | 0 |
| 0 | -1.13 | 3.64 | -0.31 | 0.76 | 1 | Trim63 | tripartite motif-containing 63 | 0 | 0 | 0 | 0 | 5 | 0 | 0 | 0 | 0 | 0 | 0 |
| 0 | -1.13 | 3.64 | -0.31 | 0.76 | 1 | Gm43870 | | 0 | 0 | 0 | 0 | 5 | 0 | 0 | 0 | 0 | 0 | 0 |
| 0 | -1.13 | 3.64 | -0.31 | 0.76 | 1 | Hoxa11os | | 0 | 0 | 0 | 0 | 5 | 0 | 0 | 0 | 0 | 0 | 0 |
| 0 | -1.13 | 3.64 | -0.31 | 0.76 | 1 | Gm44755 | | 0 | 0 | 0 | 0 | 5 | 0 | 0 | 0 | 0 | 0 | 0 |
| 0 | -1.13 | 3.64 | -0.31 | 0.76 | 1 | Sp8 | trans-acting transcription factor 8 | 0 | 0 | 0 | 0 | 5 | 0 | 0 | 0 | 0 | 0 | 0 |
| 0 | -1.13 | 3.64 | -0.31 | 0.76 | 1 | Gm49015 | | 0 | 0 | 0 | 0 | 5 | 0 | 0 | 0 | 0 | 0 | 0 |
| 0 | -1.13 | 3.64 | -0.31 | 0.76 | 1 | Gm1604a | | 0 | 0 | 0 | 0 | 5 | 0 | 0 | 0 | 0 | 0 | 0 |

| | | | | | | | | | | | | | | | | | | |
|---|---|---|---|---|---|---|---|---|---|---|---|---|---|---|---|---|---|---|
| 0 | -1.13 | 3.64 | -0.31 | 0.76 | 1 | Pnpla1os | | 0 | 0 | 0 | 0 | 5 | 0 | 0 | 0 | 0 | 0 | 0 |
| 0 | -1.13 | 3.64 | -0.31 | 0.76 | 1 | Ppef1 | protein phosphatase with EF hand calciu | 0 | 0 | 0 | 0 | 5 | 0 | 0 | 0 | 0 | 0 | 0 |
| 0 | 0.96 | 3.64 | 0.26 | 0.79 | 1 | Gm37975 | | 0 | 0 | 0 | 0 | 0 | 1 | 0 | 0 | 0 | 4 | 0 |
| 0 | 0.16 | 3.62 | 0.04 | 0.96 | 1 | Gm37967 | | 0 | 0 | 0 | 0 | 0 | 3 | 0 | 0 | 0 | 1 | 0 |
| 0 | 0.16 | 3.62 | 0.04 | 0.96 | 1 | Tdg-ps | thymine DNA glycosylase, pseudogene | 0 | 0 | 0 | 0 | 0 | 3 | 0 | 0 | 0 | 1 | 0 |
| 0 | -0.27 | 3.60 | -0.08 | 0.94 | 1 | Prdx6b | peroxiredoxin 6B | 0 | 0 | 0 | 3 | 0 | 0 | 0 | 0 | 0 | 0 | 2 |
| 0 | -0.27 | 3.60 | -0.08 | 0.94 | 1 | Mir484 | | 0 | 0 | 0 | 3 | 0 | 0 | 0 | 0 | 0 | 0 | 2 |
| 0 | -1.69 | 3.64 | -0.46 | 0.64 | 1 | Gm13378 | | 0 | 0 | 0 | 1 | 4 | 0 | 0 | 0 | 0 | 0 | 0 |
| 0 | -0.62 | 3.63 | -0.17 | 0.86 | 1 | Gm6872 | | 0 | 2 | 0 | 3 | 0 | 0 | 0 | 0 | 0 | 0 | 0 |
| 0 | 0.10 | 3.60 | 0.03 | 0.98 | 1 | Mir762 | | 0 | 0 | 0 | 3 | 0 | 0 | 0 | 0 | 2 | 0 | 0 |
| 0 | -0.22 | 3.62 | -0.06 | 0.95 | 1 | Gm11685 | | 1 | 0 | 0 | 2 | 0 | 0 | 0 | 0 | 0 | 0 | 2 |
| 0 | 1.65 | 3.65 | 0.45 | 0.65 | 1 | Gm43042 | | 0 | 0 | 0 | 0 | 0 | 0 | 0 | 5 | 0 | 0 | 0 |
| 0 | -0.67 | 3.66 | -0.18 | 0.85 | 1 | Gm17212 | | 1 | 2 | 0 | 2 | 0 | 0 | 0 | 0 | 0 | 0 | 0 |
| 0 | 0.27 | 3.64 | 0.07 | 0.94 | 1 | 1700003N22Rik | | 1 | 0 | 0 | 2 | 0 | 0 | 0 | 0 | 2 | 0 | 0 |
| 0 | -1.72 | 3.64 | -0.47 | 0.64 | 1 | Gm2287 | | 0 | 0 | 0 | 0 | 4 | 1 | 0 | 0 | 0 | 0 | 0 |
| 0 | -1.72 | 3.64 | -0.47 | 0.64 | 1 | Mir7051 | | 0 | 0 | 0 | 0 | 4 | 1 | 0 | 0 | 0 | 0 | 0 |
| 0 | -1.91 | 3.64 | -0.52 | 0.60 | 1 | Gm6378 | | 2 | 0 | 0 | 0 | 2 | 0 | 0 | 0 | 0 | 0 | 1 |
| 0 | -0.40 | 3.67 | -0.11 | 0.91 | 1 | Gm13666 | | 2 | 1 | 1 | 0 | 0 | 1 | 0 | 0 | 0 | 0 | 0 |
| 0 | -0.85 | 3.63 | -0.23 | 0.81 | 1 | Gm9009 | | 1 | 0 | 0 | 0 | 2 | 1 | 0 | 0 | 0 | 0 | 1 |
| 0 | 2.21 | 3.64 | 0.61 | 0.54 | 1 | Gm8134 | | 0 | 0 | 0 | 0 | 0 | 0 | 0 | 0 | 0 | 3 | 2 |
| 0 | 1.25 | 3.66 | 0.34 | 0.73 | 1 | Gm50230 | | 3 | 0 | 0 | 0 | 0 | 0 | 0 | 0 | 0 | 0 | 2 |
| 0 | -0.08 | 3.65 | -0.02 | 0.98 | 1 | Gm9347 | | 1 | 0 | 0 | 1 | 0 | 1 | 0 | 0 | 2 | 0 | 0 |
| 0 | 1.48 | 3.66 | 0.40 | 0.69 | 1 | Gm6822 | | 3 | 0 | 0 | 0 | 0 | 0 | 0 | 0 | 2 | 0 | 0 |
| 0 | -1.02 | 3.62 | -0.28 | 0.78 | 1 | 9430064I24Rik | | 0 | 0 | 0 | 0 | 2 | 2 | 0 | 0 | 0 | 0 | 1 |
| 0 | -1.40 | 3.65 | -0.38 | 0.70 | 1 | Rpl26-ps4 | | 1 | 0 | 1 | 2 | 1 | 0 | 0 | 0 | 0 | 0 | 0 |
| 0 | -0.14 | 3.62 | -0.04 | 0.97 | 1 | Gm43322 | | 0 | 0 | 1 | 0 | 0 | 3 | 0 | 0 | 0 | 0 | 1 |
| 0 | 0.74 | 3.63 | 0.20 | 0.84 | 1 | Ptchd4 | patched domain containing 4 | 1 | 0 | 0 | 0 | 0 | 2 | 0 | 0 | 1 | 0 | 1 |
| 0 | -1.06 | 3.63 | -0.29 | 0.77 | 1 | Gm17060 | | 1 | 0 | 0 | 2 | 1 | 0 | 0 | 0 | 0 | 0 | 1 |
| 0 | 0.12 | 3.62 | 0.03 | 0.97 | 1 | C2cd6 | C2 calcium dependent domain containing | 0 | 0 | 0 | 0 | 0 | 3 | 0 | 0 | 0 | 0 | 2 |
| 0 | 0.12 | 3.62 | 0.03 | 0.97 | 1 | Gm37349 | | 0 | 0 | 0 | 0 | 0 | 3 | 0 | 0 | 0 | 0 | 2 |
| 0 | 0.12 | 3.62 | 0.03 | 0.97 | 1 | Gm38105 | | 0 | 0 | 0 | 0 | 0 | 3 | 0 | 0 | 0 | 0 | 2 |
| 0 | 0.12 | 3.62 | 0.03 | 0.97 | 1 | Gm48183 | | 0 | 0 | 0 | 0 | 0 | 3 | 0 | 0 | 0 | 0 | 2 |
| 0 | 0.12 | 3.62 | 0.03 | 0.97 | 1 | Gm24607 | | 0 | 0 | 0 | 0 | 0 | 3 | 0 | 0 | 0 | 0 | 2 |
| 0 | 0.12 | 3.62 | 0.03 | 0.97 | 1 | Skint5 | selection and upkeep of intraepithelial T c | 0 | 0 | 0 | 0 | 0 | 3 | 0 | 0 | 0 | 0 | 2 |
| 0 | 0.12 | 3.62 | 0.03 | 0.97 | 1 | Gm42466 | | 0 | 0 | 0 | 0 | 0 | 3 | 0 | 0 | 0 | 0 | 2 |
| 0 | 0.12 | 3.62 | 0.03 | 0.97 | 1 | Gm43195 | | 0 | 0 | 0 | 0 | 0 | 3 | 0 | 0 | 0 | 0 | 2 |
| 0 | 0.12 | 3.62 | 0.03 | 0.97 | 1 | Gm45799 | translocase of inner mitochondrial membr | 0 | 0 | 0 | 0 | 0 | 3 | 0 | 0 | 0 | 0 | 2 |
| 0 | 0.12 | 3.62 | 0.03 | 0.97 | 1 | Mir7061 | | 0 | 0 | 0 | 0 | 0 | 3 | 0 | 0 | 0 | 0 | 2 |
| 0 | 0.12 | 3.62 | 0.03 | 0.97 | 1 | 4833427G06Ril | RIKEN cDNA 4833427G06 gene | 0 | 0 | 0 | 0 | 0 | 3 | 0 | 0 | 0 | 0 | 2 |
| 0 | 0.12 | 3.62 | 0.03 | 0.97 | 1 | Gm23567 | | 0 | 0 | 0 | 0 | 0 | 3 | 0 | 0 | 0 | 0 | 2 |
| 0 | 0.12 | 3.62 | 0.03 | 0.97 | 1 | Gm36995 | | 0 | 0 | 0 | 0 | 0 | 3 | 0 | 0 | 0 | 0 | 2 |
| 0 | 0.65 | 3.61 | 0.18 | 0.86 | 1 | Gm11874 | | 0 | 0 | 0 | 0 | 0 | 3 | 0 | 0 | 2 | 0 | 0 |
| 0 | 0.65 | 3.61 | 0.18 | 0.86 | 1 | Gm44862 | | 0 | 0 | 0 | 0 | 0 | 3 | 0 | 0 | 2 | 0 | 0 |
| 0 | 0.65 | 3.61 | 0.18 | 0.86 | 1 | Gm39228 | | 0 | 0 | 0 | 0 | 0 | 3 | 0 | 0 | 2 | 0 | 0 |
| 0 | 0.65 | 3.61 | 0.18 | 0.86 | 1 | Mir679 | | 0 | 0 | 0 | 0 | 0 | 3 | 0 | 0 | 2 | 0 | 0 |
| 0 | 0.65 | 3.61 | 0.18 | 0.86 | 1 | Gm29331 | | 0 | 0 | 0 | 0 | 0 | 3 | 0 | 0 | 2 | 0 | 0 |
| 0 | 0.16 | 3.62 | 0.04 | 0.96 | 1 | Gm43752 | | 0 | 0 | 0 | 0 | 0 | 3 | 0 | 1 | 0 | 0 | 0 |
| 0 | -1.12 | 3.63 | -0.31 | 0.76 | 1 | Mrps36-ps2 | | 1 | 0 | 0 | 1 | 1 | 1 | 0 | 0 | 0 | 0 | 1 |
| 0 | -1.30 | 3.65 | -0.36 | 0.72 | 1 | Rpl23a-ps14 | | 1 | 0 | 1 | 0 | 1 | 2 | 0 | 0 | 0 | 0 | 0 |
| 0 | -1.83 | 3.63 | -0.50 | 0.61 | 1 | Gm7516 | | 0 | 0 | 0 | 0 | 3 | 2 | 0 | 0 | 0 | 0 | 0 |
| 0 | -1.83 | 3.63 | -0.50 | 0.61 | 1 | Gm13608 | | 0 | 0 | 0 | 0 | 3 | 2 | 0 | 0 | 0 | 0 | 0 |
| 0 | -1.83 | 3.63 | -0.50 | 0.61 | 1 | Gssos2 | | 0 | 0 | 0 | 0 | 3 | 2 | 0 | 0 | 0 | 0 | 0 |
| 0 | -1.83 | 3.63 | -0.50 | 0.61 | 1 | Gm2420 | | 0 | 0 | 0 | 0 | 3 | 2 | 0 | 0 | 0 | 0 | 0 |
| 0 | -1.83 | 3.63 | -0.50 | 0.61 | 1 | Olfr1349 | olfactory receptor 1349 | 0 | 0 | 0 | 0 | 3 | 2 | 0 | 0 | 0 | 0 | 0 |
| 0 | -0.45 | 3.63 | -0.13 | 0.90 | 1 | Nrl | neural retina leucine zipper gene | 0 | 0 | 0 | 4 | 0 | 0 | 0 | 0 | 0 | 0 | 1 |
| 0 | -0.31 | 3.62 | -0.08 | 0.93 | 1 | C230024C17Rik | | 0 | 0 | 0 | 4 | 0 | 0 | 0 | 0 | 1 | 0 | 0 |
| 0 | 2.43 | 3.62 | 0.67 | 0.50 | 1 | Gm5497 | | 0 | 0 | 1 | 0 | 0 | 0 | 0 | 1 | 0 | 1 | 1 |
| 0 | -0.91 | 3.65 | -0.25 | 0.80 | 1 | Gm9402 | | 1 | 1 | 0 | 3 | 0 | 0 | 0 | 0 | 0 | 0 | 0 |
| 0 | -0.82 | 3.62 | -0.23 | 0.82 | 1 | Gm49974 | | 0 | 0 | 0 | 0 | 1 | 3 | 0 | 0 | 0 | 0 | 1 |
| 0 | -0.65 | 3.61 | -0.18 | 0.86 | 1 | Psmb6-ps2 | | 0 | 0 | 0 | 0 | 1 | 3 | 0 | 0 | 1 | 0 | 0 |

| | | | | | | | | | | | | | | | | | | |
|---|---|---|---|---|---|---|---|---|---|---|---|---|---|---|---|---|---|---|
| 0 | 2.10 | 3.64 | 0.58 | 0.56 | 1 | 4933431M02Rik | | 0 | 0 | 0 | 0 | 0 | 0 | 0 | 3 | 1 | 0 | 0 |
| 0 | -1.31 | 3.65 | -0.36 | 0.72 | 1 | Gm5561 | | 1 | 1 | 0 | 2 | 0 | 1 | 0 | 0 | 0 | 0 | 0 |
| 0 | 0.28 | 3.68 | 0.08 | 0.94 | 1 | Gm45371 | | 3 | 0 | 0 | 1 | 0 | 0 | 0 | 0 | 0 | 0 | 1 |
| 0 | 0.28 | 3.68 | 0.08 | 0.94 | 1 | Gm15294 | | 3 | 0 | 0 | 1 | 0 | 0 | 0 | 0 | 0 | 0 | 1 |
| 0 | -0.99 | 3.61 | -0.28 | 0.78 | 1 | Gm16550 | | 0 | 0 | 0 | 2 | 0 | 2 | 0 | 0 | 0 | 0 | 1 |
| 0 | -0.99 | 3.61 | -0.28 | 0.78 | 1 | Zfhx2os | | 0 | 0 | 0 | 2 | 0 | 2 | 0 | 0 | 0 | 0 | 1 |
| 0 | 0.95 | 3.67 | 0.26 | 0.80 | 1 | 1600014C23Ril | RIKEN cDNA 1600014C23 gene | 3 | 0 | 0 | 0 | 0 | 0 | 0 | 0 | 0 | 0 | 1 |
| 0 | -1.67 | 3.64 | -0.46 | 0.65 | 1 | Gm37915 | | 0 | 0 | 0 | 4 | 1 | 0 | 0 | 0 | 0 | 0 | 0 |
| 0 | 0.00 | 3.68 | 0.00 | 1.00 | 1 | Gm14220 | | 3 | 1 | 0 | 0 | 0 | 0 | 0 | 0 | 0 | 0 | 0 |
| 0 | -1.80 | 3.63 | -0.50 | 0.62 | 1 | Gm44798 | | 0 | 0 | 0 | 0 | 2 | 3 | 0 | 0 | 0 | 0 | 0 |
| 0 | -1.80 | 3.63 | -0.50 | 0.62 | 1 | Gm47807 | | 0 | 0 | 0 | 0 | 2 | 3 | 0 | 0 | 0 | 0 | 0 |
| 0 | -0.76 | 3.63 | -0.21 | 0.83 | 1 | Gm13370 | | 0 | 0 | 1 | 0 | 0 | 3 | 0 | 0 | 0 | 0 | 0 |
| 0 | -0.76 | 3.63 | -0.21 | 0.83 | 1 | Gm43912 | | 0 | 0 | 1 | 0 | 0 | 3 | 0 | 0 | 0 | 0 | 0 |
| 0 | -0.76 | 3.63 | -0.21 | 0.83 | 1 | Gm32842 | | 0 | 0 | 1 | 0 | 0 | 3 | 0 | 0 | 0 | 0 | 0 |
| 0 | -0.76 | 3.63 | -0.21 | 0.83 | 1 | Gm34045 | | 0 | 0 | 1 | 0 | 0 | 3 | 0 | 0 | 0 | 0 | 0 |
| 0 | -0.24 | 3.62 | -0.07 | 0.95 | 1 | Gm6162 | | 0 | 0 | 0 | 0 | 0 | 3 | 0 | 0 | 0 | 0 | 1 |
| 0 | -0.24 | 3.62 | -0.07 | 0.95 | 1 | C030017G13Rik | | 0 | 0 | 0 | 0 | 0 | 3 | 0 | 0 | 0 | 0 | 1 |
| 0 | -0.24 | 3.62 | -0.07 | 0.95 | 1 | 9330198I05Rik | | 0 | 0 | 0 | 0 | 0 | 3 | 0 | 0 | 0 | 0 | 1 |
| 0 | -0.24 | 3.62 | -0.07 | 0.95 | 1 | Gm39027 | | 0 | 0 | 0 | 0 | 0 | 3 | 0 | 0 | 0 | 0 | 1 |
| 0 | -0.24 | 3.62 | -0.07 | 0.95 | 1 | Plekhg4 | pleckstrin homology domain containing, fa | 0 | 0 | 0 | 0 | 0 | 3 | 0 | 0 | 0 | 0 | 1 |
| 0 | -0.24 | 3.62 | -0.07 | 0.95 | 1 | Gm16349 | | 0 | 0 | 0 | 0 | 0 | 3 | 0 | 0 | 0 | 0 | 1 |
| 0 | -0.24 | 3.62 | -0.07 | 0.95 | 1 | Gm20444 | | 0 | 0 | 0 | 0 | 0 | 3 | 0 | 0 | 0 | 0 | 1 |
| 0 | -0.24 | 3.62 | -0.07 | 0.95 | 1 | Gm22000 | | 0 | 0 | 0 | 0 | 0 | 3 | 0 | 0 | 0 | 0 | 1 |
| 0 | -0.24 | 3.62 | -0.07 | 0.95 | 1 | Gm47166 | | 0 | 0 | 0 | 0 | 0 | 3 | 0 | 0 | 0 | 0 | 1 |
| 0 | -0.24 | 3.62 | -0.07 | 0.95 | 1 | Gm23654 | | 0 | 0 | 0 | 0 | 0 | 3 | 0 | 0 | 0 | 0 | 1 |
| 0 | -0.76 | 3.63 | -0.21 | 0.83 | 1 | Gm6012 | | 0 | 1 | 0 | 0 | 0 | 3 | 0 | 0 | 0 | 0 | 0 |
| 0 | -0.76 | 3.63 | -0.21 | 0.83 | 1 | 2900078I11Rik | | 0 | 1 | 0 | 0 | 0 | 3 | 0 | 0 | 0 | 0 | 0 |
| 0 | 0.16 | 3.62 | 0.04 | 0.96 | 1 | Gm13886 | | 0 | 0 | 0 | 0 | 0 | 3 | 0 | 0 | 1 | 0 | 0 |
| 0 | 0.16 | 3.62 | 0.04 | 0.96 | 1 | Gm10308 | | 0 | 0 | 0 | 0 | 0 | 3 | 0 | 0 | 1 | 0 | 0 |
| 0 | 1.81 | 3.65 | 0.50 | 0.62 | 1 | Gm4953 | | 0 | 0 | 2 | 0 | 0 | 0 | 0 | 0 | 0 | 1 | 1 |
| 0 | 1.61 | 3.66 | 0.44 | 0.66 | 1 | Gm13665 | | 0 | 0 | 2 | 0 | 0 | 0 | 0 | 0 | 1 | 1 | 0 |
| 0 | -0.52 | 3.67 | -0.14 | 0.89 | 1 | Gm11906 | | 3 | 0 | 0 | 0 | 1 | 0 | 0 | 0 | 0 | 0 | 0 |
| 0 | -1.20 | 3.64 | -0.33 | 0.74 | 1 | Crabp2 | cellular retinoic acid binding protein II | 0 | 0 | 0 | 4 | 0 | 0 | 0 | 0 | 0 | 0 | 0 |
| 0 | -1.20 | 3.64 | -0.33 | 0.74 | 1 | Spag17 | sperm associated antigen 17 | 0 | 0 | 0 | 4 | 0 | 0 | 0 | 0 | 0 | 0 | 0 |
| 0 | -1.20 | 3.64 | -0.33 | 0.74 | 1 | 4930447N08Rik | | 0 | 0 | 0 | 4 | 0 | 0 | 0 | 0 | 0 | 0 | 0 |
| 0 | -1.20 | 3.64 | -0.33 | 0.74 | 1 | 4930548H24Ril | RIKEN cDNA 4930548H24 gene | 0 | 0 | 0 | 4 | 0 | 0 | 0 | 0 | 0 | 0 | 0 |
| 0 | -1.20 | 3.64 | -0.33 | 0.74 | 1 | Gm26611 | | 0 | 0 | 0 | 4 | 0 | 0 | 0 | 0 | 0 | 0 | 0 |
| 0 | -1.20 | 3.64 | -0.33 | 0.74 | 1 | Gm17229 | | 0 | 0 | 0 | 4 | 0 | 0 | 0 | 0 | 0 | 0 | 0 |
| 0 | -1.20 | 3.64 | -0.33 | 0.74 | 1 | 4933408J17Rik | | 0 | 0 | 0 | 4 | 0 | 0 | 0 | 0 | 0 | 0 | 0 |
| 0 | -1.20 | 3.64 | -0.33 | 0.74 | 1 | Gm48438 | | 0 | 0 | 0 | 4 | 0 | 0 | 0 | 0 | 0 | 0 | 0 |
| 0 | -1.20 | 3.64 | -0.33 | 0.74 | 1 | Gm34567 | | 0 | 0 | 0 | 4 | 0 | 0 | 0 | 0 | 0 | 0 | 0 |
| 0 | -1.20 | 3.64 | -0.33 | 0.74 | 1 | E130008D07Rik | | 0 | 0 | 0 | 4 | 0 | 0 | 0 | 0 | 0 | 0 | 0 |
| 0 | -1.20 | 3.64 | -0.33 | 0.74 | 1 | Gm29992 | | 0 | 0 | 0 | 4 | 0 | 0 | 0 | 0 | 0 | 0 | 0 |
| 0 | 2.04 | 3.64 | 0.56 | 0.58 | 1 | Ces3a | carboxylesterase 3A | 0 | 0 | 1 | 0 | 0 | 0 | 0 | 0 | 0 | 1 | 2 |
| 0 | -1.70 | 3.64 | -0.47 | 0.64 | 1 | Nlk-ps1 | | 0 | 0 | 0 | 4 | 0 | 1 | 0 | 0 | 0 | 0 | 0 |
| 0 | 1.01 | 3.64 | 0.28 | 0.78 | 1 | Gm15730 | | 0 | 1 | 0 | 1 | 0 | 0 | 0 | 1 | 0 | 1 | 0 |
| 0 | 1.98 | 3.65 | 0.54 | 0.59 | 1 | Gm2676 | | 0 | 1 | 0 | 0 | 0 | 0 | 0 | 0 | 0 | 1 | 2 |
| 0 | 2.44 | 3.63 | 0.67 | 0.50 | 1 | Atp6v0c-ps1 | | 0 | 0 | 0 | 0 | 0 | 0 | 0 | 0 | 2 | 1 | 1 |
| 0 | -1.56 | 3.63 | -0.43 | 0.67 | 1 | Gm20125 | | 0 | 0 | 0 | 0 | 1 | 3 | 0 | 0 | 0 | 0 | 0 |
| 0 | -1.81 | 3.63 | -0.50 | 0.62 | 1 | Gm37328 | | 0 | 0 | 0 | 3 | 0 | 2 | 0 | 0 | 0 | 0 | 0 |
| 0 | -1.81 | 3.63 | -0.50 | 0.62 | 1 | Gm11701 | | 0 | 0 | 0 | 3 | 0 | 2 | 0 | 0 | 0 | 0 | 0 |
| 0 | 2.17 | 3.64 | 0.60 | 0.55 | 1 | Gm2926 | | 1 | 0 | 0 | 0 | 0 | 0 | 0 | 1 | 1 | 1 | 0 |
| 0 | -0.15 | 3.64 | -0.04 | 0.97 | 1 | Gm3852 | | 0 | 0 | 0 | 1 | 0 | 1 | 0 | 0 | 0 | 3 | 0 |
| 0 | 0.00 | 3.68 | 0.00 | 1.00 | 1 | Khdc1a | KH domain containing 1A | 4 | 0 | 0 | 0 | 0 | 0 | 0 | 0 | 0 | 0 | 0 |
| 0 | 0.00 | 3.68 | 0.00 | 1.00 | 1 | Col9a1 | collagen, type IX, alpha 1 | 4 | 0 | 0 | 0 | 0 | 0 | 0 | 0 | 0 | 0 | 0 |
| 0 | 0.00 | 3.68 | 0.00 | 1.00 | 1 | Gm18342 | | 4 | 0 | 0 | 0 | 0 | 0 | 0 | 0 | 0 | 0 | 0 |
| 0 | 0.00 | 3.68 | 0.00 | 1.00 | 1 | Slc39a12 | solute carrier family 39 (zinc transporter), | 4 | 0 | 0 | 0 | 0 | 0 | 0 | 0 | 0 | 0 | 0 |
| 0 | 0.00 | 3.68 | 0.00 | 1.00 | 1 | Dbh | dopamine beta hydroxylase | 4 | 0 | 0 | 0 | 0 | 0 | 0 | 0 | 0 | 0 | 0 |
| 0 | 0.00 | 3.68 | 0.00 | 1.00 | 1 | Mir6997 | | 4 | 0 | 0 | 0 | 0 | 0 | 0 | 0 | 0 | 0 | 0 |
| 0 | 0.00 | 3.68 | 0.00 | 1.00 | 1 | C230014O12Rik | | 4 | 0 | 0 | 0 | 0 | 0 | 0 | 0 | 0 | 0 | 0 |

| | | | | | | | | | | | | | | | | | | |
|---|---|---|---|---|---|---|---|---|---|---|---|---|---|---|---|---|---|---|
| 0 | 0.00 | 3.68 | 0.00 | 1.00 | 1 | Gm13477 | | 4 | 0 | 0 | 0 | 0 | 0 | 0 | 0 | 0 | 0 | 0 |
| 0 | 0.00 | 3.68 | 0.00 | 1.00 | 1 | B230312C02Rik | | 4 | 0 | 0 | 0 | 0 | 0 | 0 | 0 | 0 | 0 | 0 |
| 0 | 0.00 | 3.68 | 0.00 | 1.00 | 1 | Gm42923 | | 4 | 0 | 0 | 0 | 0 | 0 | 0 | 0 | 0 | 0 | 0 |
| 0 | 0.00 | 3.68 | 0.00 | 1.00 | 1 | Gm38124 | | 4 | 0 | 0 | 0 | 0 | 0 | 0 | 0 | 0 | 0 | 0 |
| 0 | 0.00 | 3.68 | 0.00 | 1.00 | 1 | L1td1 | LINE-1 type transposase domain containi | 4 | 0 | 0 | 0 | 0 | 0 | 0 | 0 | 0 | 0 | 0 |
| 0 | 0.00 | 3.68 | 0.00 | 1.00 | 1 | Zfp990 | zinc finger protein 990 | 4 | 0 | 0 | 0 | 0 | 0 | 0 | 0 | 0 | 0 | 0 |
| 0 | 0.00 | 3.68 | 0.00 | 1.00 | 1 | Ccdc27 | coiled-coil domain containing 27 | 4 | 0 | 0 | 0 | 0 | 0 | 0 | 0 | 0 | 0 | 0 |
| 0 | 0.00 | 3.68 | 0.00 | 1.00 | 1 | Mir7031 | | 4 | 0 | 0 | 0 | 0 | 0 | 0 | 0 | 0 | 0 | 0 |
| 0 | 0.00 | 3.68 | 0.00 | 1.00 | 1 | Gm36551 | | 4 | 0 | 0 | 0 | 0 | 0 | 0 | 0 | 0 | 0 | 0 |
| 0 | 0.00 | 3.68 | 0.00 | 1.00 | 1 | Calcr | calcitonin receptor | 4 | 0 | 0 | 0 | 0 | 0 | 0 | 0 | 0 | 0 | 0 |
| 0 | 0.00 | 3.68 | 0.00 | 1.00 | 1 | Trbv2 | | 4 | 0 | 0 | 0 | 0 | 0 | 0 | 0 | 0 | 0 | 0 |
| 0 | 0.00 | 3.68 | 0.00 | 1.00 | 1 | Hoxa1 | homeobox A1 | 4 | 0 | 0 | 0 | 0 | 0 | 0 | 0 | 0 | 0 | 0 |
| 0 | 0.00 | 3.68 | 0.00 | 1.00 | 1 | Gm44050 | | 4 | 0 | 0 | 0 | 0 | 0 | 0 | 0 | 0 | 0 | 0 |
| 0 | 0.00 | 3.68 | 0.00 | 1.00 | 1 | Gm44427 | | 4 | 0 | 0 | 0 | 0 | 0 | 0 | 0 | 0 | 0 | 0 |
| 0 | 0.00 | 3.68 | 0.00 | 1.00 | 1 | Tmem52b | transmembrane protein 52B | 4 | 0 | 0 | 0 | 0 | 0 | 0 | 0 | 0 | 0 | 0 |
| 0 | 0.00 | 3.68 | 0.00 | 1.00 | 1 | Gm15922 | paired-Ig-like receptor A1 | 4 | 0 | 0 | 0 | 0 | 0 | 0 | 0 | 0 | 0 | 0 |
| 0 | 0.00 | 3.68 | 0.00 | 1.00 | 1 | Klk4 | kallikrein related-peptidase 4 (prostase, e | 4 | 0 | 0 | 0 | 0 | 0 | 0 | 0 | 0 | 0 | 0 |
| 0 | 0.00 | 3.68 | 0.00 | 1.00 | 1 | Myod1 | myogenic differentiation 1 | 4 | 0 | 0 | 0 | 0 | 0 | 0 | 0 | 0 | 0 | 0 |
| 0 | 0.00 | 3.68 | 0.00 | 1.00 | 1 | Gm42742 | PAXIP1 associated glutamate rich protein | 4 | 0 | 0 | 0 | 0 | 0 | 0 | 0 | 0 | 0 | 0 |
| 0 | 0.00 | 3.68 | 0.00 | 1.00 | 1 | Gm37082 | | 4 | 0 | 0 | 0 | 0 | 0 | 0 | 0 | 0 | 0 | 0 |
| 0 | 0.00 | 3.68 | 0.00 | 1.00 | 1 | Slc24a1 | solute carrier family 24 (sodium/potassiur | 4 | 0 | 0 | 0 | 0 | 0 | 0 | 0 | 0 | 0 | 0 |
| 0 | 0.00 | 3.68 | 0.00 | 1.00 | 1 | Mlip | muscular LMNA-interacting protein | 4 | 0 | 0 | 0 | 0 | 0 | 0 | 0 | 0 | 0 | 0 |
| 0 | 0.00 | 3.68 | 0.00 | 1.00 | 1 | Gm36993 | | 4 | 0 | 0 | 0 | 0 | 0 | 0 | 0 | 0 | 0 | 0 |
| 0 | 0.00 | 3.68 | 0.00 | 1.00 | 1 | Cxcr6 | chemokine (C-X-C motif) receptor 6 | 4 | 0 | 0 | 0 | 0 | 0 | 0 | 0 | 0 | 0 | 0 |
| 0 | 0.00 | 3.68 | 0.00 | 1.00 | 1 | Gm40634 | | 4 | 0 | 0 | 0 | 0 | 0 | 0 | 0 | 0 | 0 | 0 |
| 0 | 0.00 | 3.68 | 0.00 | 1.00 | 1 | Gm19624 | | 4 | 0 | 0 | 0 | 0 | 0 | 0 | 0 | 0 | 0 | 0 |
| 0 | 0.00 | 3.68 | 0.00 | 1.00 | 1 | Nts | neurotensin | 4 | 0 | 0 | 0 | 0 | 0 | 0 | 0 | 0 | 0 | 0 |
| 0 | 0.00 | 3.68 | 0.00 | 1.00 | 1 | Gm12065 | | 4 | 0 | 0 | 0 | 0 | 0 | 0 | 0 | 0 | 0 | 0 |
| 0 | 0.00 | 3.68 | 0.00 | 1.00 | 1 | Gm12111 | | 4 | 0 | 0 | 0 | 0 | 0 | 0 | 0 | 0 | 0 | 0 |
| 0 | 0.00 | 3.68 | 0.00 | 1.00 | 1 | Mir142 | | 4 | 0 | 0 | 0 | 0 | 0 | 0 | 0 | 0 | 0 | 0 |
| 0 | 0.00 | 3.68 | 0.00 | 1.00 | 1 | Dlx3 | distal-less homeobox 3 | 4 | 0 | 0 | 0 | 0 | 0 | 0 | 0 | 0 | 0 | 0 |
| 0 | 0.00 | 3.68 | 0.00 | 1.00 | 1 | Gm11689 | | 4 | 0 | 0 | 0 | 0 | 0 | 0 | 0 | 0 | 0 | 0 |
| 0 | 0.00 | 3.68 | 0.00 | 1.00 | 1 | Gpr22 | G protein-coupled receptor 22 | 4 | 0 | 0 | 0 | 0 | 0 | 0 | 0 | 0 | 0 | 0 |
| 0 | 0.00 | 3.68 | 0.00 | 1.00 | 1 | Ighd4-1 | | 4 | 0 | 0 | 0 | 0 | 0 | 0 | 0 | 0 | 0 | 0 |
| 0 | 0.00 | 3.68 | 0.00 | 1.00 | 1 | Ighv1-62-3 | | 4 | 0 | 0 | 0 | 0 | 0 | 0 | 0 | 0 | 0 | 0 |
| 0 | 0.00 | 3.68 | 0.00 | 1.00 | 1 | Gm48928 | | 4 | 0 | 0 | 0 | 0 | 0 | 0 | 0 | 0 | 0 | 0 |
| 0 | 0.00 | 3.68 | 0.00 | 1.00 | 1 | 4932411K12Rik | | 4 | 0 | 0 | 0 | 0 | 0 | 0 | 0 | 0 | 0 | 0 |
| 0 | 0.00 | 3.68 | 0.00 | 1.00 | 1 | 4930405A10Rik | | 4 | 0 | 0 | 0 | 0 | 0 | 0 | 0 | 0 | 0 | 0 |
| 0 | 0.00 | 3.68 | 0.00 | 1.00 | 1 | Gm4383 | | 4 | 0 | 0 | 0 | 0 | 0 | 0 | 0 | 0 | 0 | 0 |
| 0 | 0.00 | 3.68 | 0.00 | 1.00 | 1 | 4833419O12Rik | | 4 | 0 | 0 | 0 | 0 | 0 | 0 | 0 | 0 | 0 | 0 |
| 0 | 0.00 | 3.68 | 0.00 | 1.00 | 1 | Csnka2ip | casein kinase 2, alpha prime interacting p | 4 | 0 | 0 | 0 | 0 | 0 | 0 | 0 | 0 | 0 | 0 |
| 0 | 0.00 | 3.68 | 0.00 | 1.00 | 1 | Sim2 | single-minded family bHLH transcription f | 4 | 0 | 0 | 0 | 0 | 0 | 0 | 0 | 0 | 0 | 0 |
| 0 | 0.00 | 3.68 | 0.00 | 1.00 | 1 | 2300002M23Ri | RIKEN cDNA 2300002M23 gene | 4 | 0 | 0 | 0 | 0 | 0 | 0 | 0 | 0 | 0 | 0 |
| 0 | 0.00 | 3.68 | 0.00 | 1.00 | 1 | Gm9284 | | 4 | 0 | 0 | 0 | 0 | 0 | 0 | 0 | 0 | 0 | 0 |
| 0 | 0.00 | 3.68 | 0.00 | 1.00 | 1 | Gm41720 | | 4 | 0 | 0 | 0 | 0 | 0 | 0 | 0 | 0 | 0 | 0 |
| 0 | 0.00 | 3.68 | 0.00 | 1.00 | 1 | Psmb7-ps2 | | 4 | 0 | 0 | 0 | 0 | 0 | 0 | 0 | 0 | 0 | 0 |
| 0 | -1.78 | 3.63 | -0.49 | 0.62 | 1 | Dbx2 | developing brain homeobox 2 | 0 | 0 | 0 | 2 | 0 | 3 | 0 | 0 | 0 | 0 | 0 |
| 0 | -1.78 | 3.63 | -0.49 | 0.62 | 1 | Map6d1 | MAP6 domain containing 1 | 0 | 0 | 0 | 2 | 0 | 3 | 0 | 0 | 0 | 0 | 0 |
| 0 | 0.43 | 3.64 | 0.12 | 0.91 | 1 | Gm18733 | | 1 | 0 | 0 | 0 | 0 | 1 | 0 | 0 | 0 | 3 | 0 |
| 0 | -0.41 | 3.67 | -0.11 | 0.91 | 1 | Gm13675 | | 3 | 0 | 0 | 0 | 0 | 1 | 0 | 0 | 0 | 0 | 0 |
| 0 | -0.41 | 3.67 | -0.11 | 0.91 | 1 | 4930537H20Rik | | 3 | 0 | 0 | 0 | 0 | 1 | 0 | 0 | 0 | 0 | 0 |
| 0 | -0.41 | 3.67 | -0.11 | 0.91 | 1 | Gm38379 | | 3 | 0 | 0 | 0 | 0 | 1 | 0 | 0 | 0 | 0 | 0 |
| 0 | 0.75 | 3.62 | 0.21 | 0.84 | 1 | Rny1 | | 0 | 0 | 0 | 0 | 0 | 2 | 0 | 0 | 0 | 3 | 0 |
| 0 | -0.71 | 3.66 | -0.19 | 0.85 | 1 | 4930544L04Rik | | 3 | 0 | 0 | 0 | 0 | 2 | 0 | 0 | 0 | 0 | 0 |
| 0 | 1.40 | 3.64 | 0.38 | 0.70 | 1 | Anapc15-ps | | 0 | 0 | 0 | 0 | 1 | 0 | 0 | 0 | 0 | 1 | 2 |
| 0 | -1.55 | 3.63 | -0.43 | 0.67 | 1 | Gm43215 | | 0 | 0 | 0 | 1 | 0 | 3 | 0 | 0 | 0 | 0 | 0 |
| 0 | -1.55 | 3.63 | -0.43 | 0.67 | 1 | Lamp3 | lysosomal-associated membrane protein | 0 | 0 | 0 | 1 | 0 | 3 | 0 | 0 | 0 | 0 | 0 |
| 0 | -1.55 | 3.63 | -0.43 | 0.67 | 1 | Gm30961 | | 0 | 0 | 0 | 1 | 0 | 3 | 0 | 0 | 0 | 0 | 0 |
| 0 | -0.73 | 3.64 | -0.20 | 0.84 | 1 | F11 | coagulation factor XI | 2 | 0 | 0 | 0 | 0 | 3 | 0 | 0 | 0 | 0 | 0 |
| 0 | -0.76 | 3.63 | -0.21 | 0.83 | 1 | Gm43162 | | 1 | 0 | 0 | 0 | 0 | 3 | 0 | 0 | 0 | 0 | 0 |

| | | | | | | | | | | | | | | | | | |
|---|---|---|---|---|---|---|---|---|---|---|---|---|---|---|---|---|---|
| 0 | -0.76 | 3.63 | -0.21 | 0.83 | 1 | Gm17938 | | 1 | 0 | 0 | 0 | 0 | 3 | 0 | 0 | 0 | 0 | 0 |
| 0 | -0.78 | 3.62 | -0.21 | 0.83 | 1 | Gm37211 | | 0 | 0 | 0 | 0 | 0 | 4 | 0 | 0 | 0 | 0 | 0 |
| 0 | -0.78 | 3.62 | -0.21 | 0.83 | 1 | Gm37868 | | 0 | 0 | 0 | 0 | 0 | 4 | 0 | 0 | 0 | 0 | 0 |
| 0 | -0.78 | 3.62 | -0.21 | 0.83 | 1 | Gm28883 | | 0 | 0 | 0 | 0 | 0 | 4 | 0 | 0 | 0 | 0 | 0 |
| 0 | -0.78 | 3.62 | -0.21 | 0.83 | 1 | Platr23 | | 0 | 0 | 0 | 0 | 0 | 4 | 0 | 0 | 0 | 0 | 0 |
| 0 | -0.78 | 3.62 | -0.21 | 0.83 | 1 | Lcn5 | lipocalin 5 | 0 | 0 | 0 | 0 | 0 | 4 | 0 | 0 | 0 | 0 | 0 |
| 0 | -0.78 | 3.62 | -0.21 | 0.83 | 1 | Gm25966 | | 0 | 0 | 0 | 0 | 0 | 4 | 0 | 0 | 0 | 0 | 0 |
| 0 | -0.78 | 3.62 | -0.21 | 0.83 | 1 | Gm22865 | | 0 | 0 | 0 | 0 | 0 | 4 | 0 | 0 | 0 | 0 | 0 |
| 0 | -0.78 | 3.62 | -0.21 | 0.83 | 1 | Gm25516 | | 0 | 0 | 0 | 0 | 0 | 4 | 0 | 0 | 0 | 0 | 0 |
| 0 | -0.78 | 3.62 | -0.21 | 0.83 | 1 | Gm37726 | | 0 | 0 | 0 | 0 | 0 | 4 | 0 | 0 | 0 | 0 | 0 |
| 0 | -0.78 | 3.62 | -0.21 | 0.83 | 1 | Gm43768 | | 0 | 0 | 0 | 0 | 0 | 4 | 0 | 0 | 0 | 0 | 0 |
| 0 | -0.78 | 3.62 | -0.21 | 0.83 | 1 | Gm12745 | | 0 | 0 | 0 | 0 | 0 | 4 | 0 | 0 | 0 | 0 | 0 |
| 0 | -0.78 | 3.62 | -0.21 | 0.83 | 1 | Gm12904 | | 0 | 0 | 0 | 0 | 0 | 4 | 0 | 0 | 0 | 0 | 0 |
| 0 | -0.78 | 3.62 | -0.21 | 0.83 | 1 | Gm15587 | | 0 | 0 | 0 | 0 | 0 | 4 | 0 | 0 | 0 | 0 | 0 |
| 0 | -0.78 | 3.62 | -0.21 | 0.83 | 1 | Gm43016 | | 0 | 0 | 0 | 0 | 0 | 4 | 0 | 0 | 0 | 0 | 0 |
| 0 | -0.78 | 3.62 | -0.21 | 0.83 | 1 | Gm43084 | | 0 | 0 | 0 | 0 | 0 | 4 | 0 | 0 | 0 | 0 | 0 |
| 0 | -0.78 | 3.62 | -0.21 | 0.83 | 1 | Gm43547 | | 0 | 0 | 0 | 0 | 0 | 4 | 0 | 0 | 0 | 0 | 0 |
| 0 | -0.78 | 3.62 | -0.21 | 0.83 | 1 | Gm42866 | | 0 | 0 | 0 | 0 | 0 | 4 | 0 | 0 | 0 | 0 | 0 |
| 0 | -0.78 | 3.62 | -0.21 | 0.83 | 1 | Ankrd7 | ankyrin repeat domain 7 | 0 | 0 | 0 | 0 | 0 | 4 | 0 | 0 | 0 | 0 | 0 |
| 0 | -0.78 | 3.62 | -0.21 | 0.83 | 1 | Gm43864 | | 0 | 0 | 0 | 0 | 0 | 4 | 0 | 0 | 0 | 0 | 0 |
| 0 | -0.78 | 3.62 | -0.21 | 0.83 | 1 | Gm43982 | | 0 | 0 | 0 | 0 | 0 | 4 | 0 | 0 | 0 | 0 | 0 |
| 0 | -0.78 | 3.62 | -0.21 | 0.83 | 1 | Olfr1348 | olfactory receptor 1348 | 0 | 0 | 0 | 0 | 0 | 4 | 0 | 0 | 0 | 0 | 0 |
| 0 | -0.78 | 3.62 | -0.21 | 0.83 | 1 | Gm20482 | | 0 | 0 | 0 | 0 | 0 | 4 | 0 | 0 | 0 | 0 | 0 |
| 0 | -0.78 | 3.62 | -0.21 | 0.83 | 1 | Prr19 | proline rich 19 | 0 | 0 | 0 | 0 | 0 | 4 | 0 | 0 | 0 | 0 | 0 |
| 0 | -0.78 | 3.62 | -0.21 | 0.83 | 1 | Mir1964 | | 0 | 0 | 0 | 0 | 0 | 4 | 0 | 0 | 0 | 0 | 0 |
| 0 | -0.78 | 3.62 | -0.21 | 0.83 | 1 | Gfy | golgi-associated olfactory signaling regul | 0 | 0 | 0 | 0 | 0 | 4 | 0 | 0 | 0 | 0 | 0 |
| 0 | -0.78 | 3.62 | -0.21 | 0.83 | 1 | Gm23873 | | 0 | 0 | 0 | 0 | 0 | 4 | 0 | 0 | 0 | 0 | 0 |
| 0 | -0.78 | 3.62 | -0.21 | 0.83 | 1 | C130083A15Rik | | 0 | 0 | 0 | 0 | 0 | 4 | 0 | 0 | 0 | 0 | 0 |
| 0 | -0.78 | 3.62 | -0.21 | 0.83 | 1 | Mir7064 | | 0 | 0 | 0 | 0 | 0 | 4 | 0 | 0 | 0 | 0 | 0 |
| 0 | -0.78 | 3.62 | -0.21 | 0.83 | 1 | 4933422A05Rik | | 0 | 0 | 0 | 0 | 0 | 4 | 0 | 0 | 0 | 0 | 0 |
| 0 | -0.78 | 3.62 | -0.21 | 0.83 | 1 | Gm10647 | | 0 | 0 | 0 | 0 | 0 | 4 | 0 | 0 | 0 | 0 | 0 |
| 0 | -0.78 | 3.62 | -0.21 | 0.83 | 1 | Mir7089 | | 0 | 0 | 0 | 0 | 0 | 4 | 0 | 0 | 0 | 0 | 0 |
| 0 | -0.78 | 3.62 | -0.21 | 0.83 | 1 | Mir7663 | | 0 | 0 | 0 | 0 | 0 | 4 | 0 | 0 | 0 | 0 | 0 |
| 0 | -0.78 | 3.62 | -0.21 | 0.83 | 1 | Gm15197 | | 0 | 0 | 0 | 0 | 0 | 4 | 0 | 0 | 0 | 0 | 0 |
| 0 | -0.78 | 3.62 | -0.21 | 0.83 | 1 | Gm47006 | | 0 | 0 | 0 | 0 | 0 | 4 | 0 | 0 | 0 | 0 | 0 |
| 0 | -0.78 | 3.62 | -0.21 | 0.83 | 1 | Gm47673 | | 0 | 0 | 0 | 0 | 0 | 4 | 0 | 0 | 0 | 0 | 0 |
| 0 | -0.78 | 3.62 | -0.21 | 0.83 | 1 | Gm47540 | | 0 | 0 | 0 | 0 | 0 | 4 | 0 | 0 | 0 | 0 | 0 |
| 0 | -0.78 | 3.62 | -0.21 | 0.83 | 1 | Gm22611 | | 0 | 0 | 0 | 0 | 0 | 4 | 0 | 0 | 0 | 0 | 0 |
| 0 | -0.78 | 3.62 | -0.21 | 0.83 | 1 | Gm20682 | | 0 | 0 | 0 | 0 | 0 | 4 | 0 | 0 | 0 | 0 | 0 |
| 0 | -0.78 | 3.62 | -0.21 | 0.83 | 1 | Gm27596 | | 0 | 0 | 0 | 0 | 0 | 4 | 0 | 0 | 0 | 0 | 0 |
| 0 | -0.78 | 3.62 | -0.21 | 0.83 | 1 | Gm23553 | | 0 | 0 | 0 | 0 | 0 | 4 | 0 | 0 | 0 | 0 | 0 |
| 0 | -0.78 | 3.62 | -0.21 | 0.83 | 1 | Gm44484 | | 0 | 0 | 0 | 0 | 0 | 4 | 0 | 0 | 0 | 0 | 0 |
| 0 | -0.78 | 3.62 | -0.21 | 0.83 | 1 | Mir6943 | | 0 | 0 | 0 | 0 | 0 | 4 | 0 | 0 | 0 | 0 | 0 |
| 0 | -0.78 | 3.62 | -0.21 | 0.83 | 1 | Gm18398 | | 0 | 0 | 0 | 0 | 0 | 4 | 0 | 0 | 0 | 0 | 0 |
| 0 | -0.78 | 3.62 | -0.21 | 0.83 | 1 | Gm24041 | | 0 | 0 | 0 | 0 | 0 | 4 | 0 | 0 | 0 | 0 | 0 |
| 0 | -0.78 | 3.62 | -0.21 | 0.83 | 1 | Gm30159 | | 0 | 0 | 0 | 0 | 0 | 4 | 0 | 0 | 0 | 0 | 0 |
| 0 | -0.78 | 3.62 | -0.21 | 0.83 | 1 | Gm49410 | | 0 | 0 | 0 | 0 | 0 | 4 | 0 | 0 | 0 | 0 | 0 |
| 0 | -0.78 | 3.62 | -0.21 | 0.83 | 1 | Gm49500 | | 0 | 0 | 0 | 0 | 0 | 4 | 0 | 0 | 0 | 0 | 0 |
| 0 | -0.78 | 3.62 | -0.21 | 0.83 | 1 | Gm4566 | | 0 | 0 | 0 | 0 | 0 | 4 | 0 | 0 | 0 | 0 | 0 |
| 0 | -0.78 | 3.62 | -0.21 | 0.83 | 1 | Gm25406 | | 0 | 0 | 0 | 0 | 0 | 4 | 0 | 0 | 0 | 0 | 0 |
| 0 | -0.78 | 3.62 | -0.21 | 0.83 | 1 | Sult6b1 | sulfotransferase family, cytosolic, 6B, mer | 0 | 0 | 0 | 0 | 0 | 4 | 0 | 0 | 0 | 0 | 0 |
| 0 | -0.78 | 3.62 | -0.21 | 0.83 | 1 | Vaultrc5 | | 0 | 0 | 0 | 0 | 0 | 4 | 0 | 0 | 0 | 0 | 0 |
| 0 | -0.78 | 3.62 | -0.21 | 0.83 | 1 | Scarna17 | | 0 | 0 | 0 | 0 | 0 | 4 | 0 | 0 | 0 | 0 | 0 |
| 0 | -0.78 | 3.62 | -0.21 | 0.83 | 1 | Csnk2a1-ps | | 0 | 0 | 0 | 0 | 0 | 4 | 0 | 0 | 0 | 0 | 0 |
| 0 | -0.78 | 3.62 | -0.21 | 0.83 | 1 | Gm6292 | | 0 | 0 | 0 | 0 | 0 | 4 | 0 | 0 | 0 | 0 | 0 |
| 0 | 0.96 | 3.67 | 0.26 | 0.79 | 1 | Gm7399 | | 0 | 0 | 1 | 1 | 0 | 0 | 0 | 0 | 0 | 1 | 1 |
| 0 | 2.05 | 3.64 | 0.56 | 0.57 | 1 | Gm8556 | | 0 | 0 | 0 | 0 | 0 | 0 | 0 | 1 | 0 | 0 | 3 |
| 0 | 2.05 | 3.64 | 0.56 | 0.57 | 1 | Gm45454 | | 0 | 0 | 0 | 0 | 0 | 0 | 0 | 1 | 0 | 0 | 3 |
| 0 | 0.87 | 3.66 | 0.24 | 0.81 | 1 | Gm5230 | | 0 | 0 | 1 | 0 | 0 | 1 | 0 | 2 | 0 | 0 | 0 |
| 0 | 1.55 | 3.65 | 0.42 | 0.67 | 1 | Gm5050 | | 1 | 0 | 1 | 0 | 0 | 0 | 0 | 0 | 1 | 1 | 0 |

| | | | | | | | | | | | | | | | | | | |
|---|---|---|---|---|---|---|---|---|---|---|---|---|---|---|---|---|---|---|
| 0 | 1.91 | 3.64 | 0.52 | 0.60 | 1 | Gm19807 | | 0 | 1 | 0 | 0 | 0 | 0 | 0 | 1 | 2 | 0 | 0 |
| 0 | 1.26 | 3.64 | 0.35 | 0.73 | 1 | Gm12419 | | 0 | 1 | 0 | 1 | 0 | 0 | 0 | 0 | 0 | 1 | 1 |
| 0 | 2.03 | 3.64 | 0.56 | 0.58 | 1 | Gm17045 | | 1 | 0 | 0 | 0 | 0 | 0 | 0 | 2 | 0 | 0 | 1 |
| 0 | 1.58 | 3.66 | 0.43 | 0.67 | 1 | Gm7191 | | 1 | 1 | 0 | 0 | 0 | 0 | 0 | 0 | 0 | 1 | 1 |
| 0 | 0.00 | 3.68 | 0.00 | 1.00 | 1 | 7530428D23Rik | | 0 | 0 | 4 | 0 | 0 | 0 | 0 | 0 | 0 | 0 | 0 |
| 0 | 0.00 | 3.68 | 0.00 | 1.00 | 1 | Mecomos | | 0 | 0 | 4 | 0 | 0 | 0 | 0 | 0 | 0 | 0 | 0 |
| 0 | 0.00 | 3.68 | 0.00 | 1.00 | 1 | Gm42660 | | 0 | 0 | 4 | 0 | 0 | 0 | 0 | 0 | 0 | 0 | 0 |
| 0 | 0.00 | 3.68 | 0.00 | 1.00 | 1 | Garem2 | GRB2 associated regulator of MAPK1 sub | 0 | 0 | 4 | 0 | 0 | 0 | 0 | 0 | 0 | 0 | 0 |
| 0 | 0.00 | 3.68 | 0.00 | 1.00 | 1 | Gm43685 | | 0 | 0 | 4 | 0 | 0 | 0 | 0 | 0 | 0 | 0 | 0 |
| 0 | 0.00 | 3.68 | 0.00 | 1.00 | 1 | Gm6583 | predicted gene 6583 | 0 | 0 | 4 | 0 | 0 | 0 | 0 | 0 | 0 | 0 | 0 |
| 0 | 0.00 | 3.68 | 0.00 | 1.00 | 1 | Gm22740 | | 0 | 0 | 4 | 0 | 0 | 0 | 0 | 0 | 0 | 0 | 0 |
| 0 | 0.00 | 3.68 | 0.00 | 1.00 | 1 | Igkv4-69 | | 0 | 0 | 4 | 0 | 0 | 0 | 0 | 0 | 0 | 0 | 0 |
| 0 | 0.00 | 3.68 | 0.00 | 1.00 | 1 | Minpp1-ps | | 0 | 0 | 4 | 0 | 0 | 0 | 0 | 0 | 0 | 0 | 0 |
| 0 | 0.00 | 3.68 | 0.00 | 1.00 | 1 | Gm8989 | | 0 | 0 | 4 | 0 | 0 | 0 | 0 | 0 | 0 | 0 | 0 |
| 0 | 0.00 | 3.68 | 0.00 | 1.00 | 1 | BC051019 | cDNA sequence BC051019 | 0 | 0 | 4 | 0 | 0 | 0 | 0 | 0 | 0 | 0 | 0 |
| 0 | 0.00 | 3.68 | 0.00 | 1.00 | 1 | Hs3st4 | heparan sulfate (glucosamine) 3-O-sulfot | 0 | 0 | 4 | 0 | 0 | 0 | 0 | 0 | 0 | 0 | 0 |
| 0 | 0.00 | 3.68 | 0.00 | 1.00 | 1 | Gm12292 | | 0 | 0 | 4 | 0 | 0 | 0 | 0 | 0 | 0 | 0 | 0 |
| 0 | 0.00 | 3.68 | 0.00 | 1.00 | 1 | 4732490B19Rik | | 0 | 0 | 4 | 0 | 0 | 0 | 0 | 0 | 0 | 0 | 0 |
| 0 | 0.00 | 3.68 | 0.00 | 1.00 | 1 | Ppp1r27 | protein phosphatase 1, regulatory subuni | 0 | 0 | 4 | 0 | 0 | 0 | 0 | 0 | 0 | 0 | 0 |
| 0 | 0.00 | 3.68 | 0.00 | 1.00 | 1 | Gm15512 | | 0 | 0 | 4 | 0 | 0 | 0 | 0 | 0 | 0 | 0 | 0 |
| 0 | 0.00 | 3.68 | 0.00 | 1.00 | 1 | Gm24540 | | 0 | 0 | 4 | 0 | 0 | 0 | 0 | 0 | 0 | 0 | 0 |
| 0 | 0.00 | 3.68 | 0.00 | 1.00 | 1 | C130023A14Rik | | 0 | 0 | 4 | 0 | 0 | 0 | 0 | 0 | 0 | 0 | 0 |
| 0 | 0.00 | 3.68 | 0.00 | 1.00 | 1 | Gm45879 | | 0 | 0 | 4 | 0 | 0 | 0 | 0 | 0 | 0 | 0 | 0 |
| 0 | 0.00 | 3.68 | 0.00 | 1.00 | 1 | Vwce | von Willebrand factor C and EGF domains | 0 | 0 | 4 | 0 | 0 | 0 | 0 | 0 | 0 | 0 | 0 |
| 0 | 0.30 | 3.66 | 0.08 | 0.93 | 1 | Gm8277 | | 0 | 1 | 1 | 0 | 1 | 0 | 0 | 1 | 0 | 0 | 0 |
| 0 | 1.01 | 3.63 | 0.28 | 0.78 | 1 | Gm6750 | | 0 | 0 | 1 | 0 | 1 | 0 | 0 | 1 | 1 | 0 | 0 |
| 0 | 1.01 | 3.67 | 0.28 | 0.78 | 1 | Gm25588 | | 0 | 0 | 3 | 0 | 0 | 0 | 0 | 0 | 0 | 0 | 1 |
| 0 | 0.83 | 3.66 | 0.23 | 0.82 | 1 | Gm11893 | | 0 | 1 | 0 | 0 | 1 | 0 | 0 | 1 | 0 | 0 | 1 |
| 0 | 0.00 | 3.68 | 0.00 | 1.00 | 1 | Gm43399 | | 0 | 1 | 3 | 0 | 0 | 0 | 0 | 0 | 0 | 0 | 0 |
| 0 | 0.00 | 3.68 | 0.00 | 1.00 | 1 | Gm4945 | | 0 | 1 | 3 | 0 | 0 | 0 | 0 | 0 | 0 | 0 | 0 |
| 0 | 1.32 | 3.65 | 0.36 | 0.72 | 1 | 4933411E08Rik | | 0 | 0 | 2 | 0 | 0 | 0 | 0 | 0 | 0 | 0 | 2 |
| 0 | 1.00 | 3.68 | 0.27 | 0.79 | 1 | Gm12757 | | 1 | 0 | 2 | 0 | 0 | 0 | 0 | 1 | 0 | 0 | 0 |
| 0 | 1.30 | 3.64 | 0.36 | 0.72 | 1 | BC046401 | | 0 | 0 | 1 | 0 | 0 | 0 | 0 | 0 | 0 | 0 | 3 |
| 0 | 1.30 | 3.64 | 0.36 | 0.72 | 1 | Gm11841 | | 0 | 0 | 1 | 0 | 0 | 0 | 0 | 0 | 0 | 0 | 3 |
| 0 | 1.32 | 3.63 | 0.36 | 0.72 | 1 | Gm37051 | | 0 | 0 | 0 | 0 | 0 | 0 | 0 | 0 | 0 | 0 | 4 |
| 0 | 1.32 | 3.63 | 0.36 | 0.72 | 1 | Gm38260 | | 0 | 0 | 0 | 0 | 0 | 0 | 0 | 0 | 0 | 0 | 4 |
| 0 | 1.32 | 3.63 | 0.36 | 0.72 | 1 | Gm37036 | | 0 | 0 | 0 | 0 | 0 | 0 | 0 | 0 | 0 | 0 | 4 |
| 0 | 1.32 | 3.63 | 0.36 | 0.72 | 1 | Gm38026 | | 0 | 0 | 0 | 0 | 0 | 0 | 0 | 0 | 0 | 0 | 4 |
| 0 | 1.32 | 3.63 | 0.36 | 0.72 | 1 | Gm28693 | | 0 | 0 | 0 | 0 | 0 | 0 | 0 | 0 | 0 | 0 | 4 |
| 0 | 1.32 | 3.63 | 0.36 | 0.72 | 1 | Gm37107 | | 0 | 0 | 0 | 0 | 0 | 0 | 0 | 0 | 0 | 0 | 4 |
| 0 | 1.32 | 3.63 | 0.36 | 0.72 | 1 | Gm37894 | | 0 | 0 | 0 | 0 | 0 | 0 | 0 | 0 | 0 | 0 | 4 |
| 0 | 1.32 | 3.63 | 0.36 | 0.72 | 1 | Gm37139 | | 0 | 0 | 0 | 0 | 0 | 0 | 0 | 0 | 0 | 0 | 4 |
| 0 | 1.32 | 3.63 | 0.36 | 0.72 | 1 | Gm13596 | | 0 | 0 | 0 | 0 | 0 | 0 | 0 | 0 | 0 | 0 | 4 |
| 0 | 1.32 | 3.63 | 0.36 | 0.72 | 1 | Bpifb3 | BPI fold containing family B, member 3 | 0 | 0 | 0 | 0 | 0 | 0 | 0 | 0 | 0 | 0 | 4 |
| 0 | 1.32 | 3.63 | 0.36 | 0.72 | 1 | Gm14246 | | 0 | 0 | 0 | 0 | 0 | 0 | 0 | 0 | 0 | 0 | 4 |
| 0 | 1.32 | 3.63 | 0.36 | 0.72 | 1 | Gm42196 | | 0 | 0 | 0 | 0 | 0 | 0 | 0 | 0 | 0 | 0 | 4 |
| 0 | 1.32 | 3.63 | 0.36 | 0.72 | 1 | 9430025C20Rik | | 0 | 0 | 0 | 0 | 0 | 0 | 0 | 0 | 0 | 0 | 4 |
| 0 | 1.32 | 3.63 | 0.36 | 0.72 | 1 | Gm22690 | | 0 | 0 | 0 | 0 | 0 | 0 | 0 | 0 | 0 | 0 | 4 |
| 0 | 1.32 | 3.63 | 0.36 | 0.72 | 1 | Gm22335 | | 0 | 0 | 0 | 0 | 0 | 0 | 0 | 0 | 0 | 0 | 4 |
| 0 | 1.32 | 3.63 | 0.36 | 0.72 | 1 | Olfr1342 | olfactory receptor 1342 | 0 | 0 | 0 | 0 | 0 | 0 | 0 | 0 | 0 | 0 | 4 |
| 0 | 1.32 | 3.63 | 0.36 | 0.72 | 1 | Gm42490 | | 0 | 0 | 0 | 0 | 0 | 0 | 0 | 0 | 0 | 0 | 4 |
| 0 | 1.32 | 3.63 | 0.36 | 0.72 | 1 | Clnk | cytokine-dependent hematopoietic cell lir | 0 | 0 | 0 | 0 | 0 | 0 | 0 | 0 | 0 | 0 | 4 |
| 0 | 1.32 | 3.63 | 0.36 | 0.72 | 1 | Gm43321 | | 0 | 0 | 0 | 0 | 0 | 0 | 0 | 0 | 0 | 0 | 4 |
| 0 | 1.32 | 3.63 | 0.36 | 0.72 | 1 | Gm42461 | | 0 | 0 | 0 | 0 | 0 | 0 | 0 | 0 | 0 | 0 | 4 |
| 0 | 1.32 | 3.63 | 0.36 | 0.72 | 1 | D130004A15Rik | | 0 | 0 | 0 | 0 | 0 | 0 | 0 | 0 | 0 | 0 | 4 |
| 0 | 1.32 | 3.63 | 0.36 | 0.72 | 1 | Atp6v1fnb | Atp6v1f neighbor | 0 | 0 | 0 | 0 | 0 | 0 | 0 | 0 | 0 | 0 | 4 |
| 0 | 1.32 | 3.63 | 0.36 | 0.72 | 1 | Gm44105 | | 0 | 0 | 0 | 0 | 0 | 0 | 0 | 0 | 0 | 0 | 4 |
| 0 | 1.32 | 3.63 | 0.36 | 0.72 | 1 | Gm50492 | | 0 | 0 | 0 | 0 | 0 | 0 | 0 | 0 | 0 | 0 | 4 |
| 0 | 1.32 | 3.63 | 0.36 | 0.72 | 1 | Gm45415 | | 0 | 0 | 0 | 0 | 0 | 0 | 0 | 0 | 0 | 0 | 4 |
| 0 | 1.32 | 3.63 | 0.36 | 0.72 | 1 | Mir7048 | | 0 | 0 | 0 | 0 | 0 | 0 | 0 | 0 | 0 | 0 | 4 |

| | | | | | | | | | | | | | | | | | | |
|---|---|---|---|---|---|---|---|---|---|---|---|---|---|---|---|---|---|---|
| 0 | 1.32 | 3.63 | 0.36 | 0.72 | 1 | Mir344d-1 | | 0 | 0 | 0 | 0 | 0 | 0 | 0 | 0 | 0 | 0 | 4 |
| 0 | 1.32 | 3.63 | 0.36 | 0.72 | 1 | Gm33882 | | 0 | 0 | 0 | 0 | 0 | 0 | 0 | 0 | 0 | 0 | 4 |
| 0 | 1.32 | 3.63 | 0.36 | 0.72 | 1 | Gm45155 | | 0 | 0 | 0 | 0 | 0 | 0 | 0 | 0 | 0 | 0 | 4 |
| 0 | 1.32 | 3.63 | 0.36 | 0.72 | 1 | Tspan32os | | 0 | 0 | 0 | 0 | 0 | 0 | 0 | 0 | 0 | 0 | 4 |
| 0 | 1.32 | 3.63 | 0.36 | 0.72 | 1 | Gm45534 | | 0 | 0 | 0 | 0 | 0 | 0 | 0 | 0 | 0 | 0 | 4 |
| 0 | 1.32 | 3.63 | 0.36 | 0.72 | 1 | Tepp | testis, prostate and placenta expressed | 0 | 0 | 0 | 0 | 0 | 0 | 0 | 0 | 0 | 0 | 4 |
| 0 | 1.32 | 3.63 | 0.36 | 0.72 | 1 | Gm26815 | | 0 | 0 | 0 | 0 | 0 | 0 | 0 | 0 | 0 | 0 | 4 |
| 0 | 1.32 | 3.63 | 0.36 | 0.72 | 1 | Gm19324 | | 0 | 0 | 0 | 0 | 0 | 0 | 0 | 0 | 0 | 0 | 4 |
| 0 | 1.32 | 3.63 | 0.36 | 0.72 | 1 | Gm2087 | | 0 | 0 | 0 | 0 | 0 | 0 | 0 | 0 | 0 | 0 | 4 |
| 0 | 1.32 | 3.63 | 0.36 | 0.72 | 1 | Gm39388 | | 0 | 0 | 0 | 0 | 0 | 0 | 0 | 0 | 0 | 0 | 4 |
| 0 | 1.32 | 3.63 | 0.36 | 0.72 | 1 | Gm38297 | | 0 | 0 | 0 | 0 | 0 | 0 | 0 | 0 | 0 | 0 | 4 |
| 0 | 1.32 | 3.63 | 0.36 | 0.72 | 1 | Gm24860 | | 0 | 0 | 0 | 0 | 0 | 0 | 0 | 0 | 0 | 0 | 4 |
| 0 | 1.32 | 3.63 | 0.36 | 0.72 | 1 | Gm22105 | | 0 | 0 | 0 | 0 | 0 | 0 | 0 | 0 | 0 | 0 | 4 |
| 0 | 1.32 | 3.63 | 0.36 | 0.72 | 1 | Gm48065 | | 0 | 0 | 0 | 0 | 0 | 0 | 0 | 0 | 0 | 0 | 4 |
| 0 | 1.32 | 3.63 | 0.36 | 0.72 | 1 | Il22 | interleukin 22 | 0 | 0 | 0 | 0 | 0 | 0 | 0 | 0 | 0 | 0 | 4 |
| 0 | 1.32 | 3.63 | 0.36 | 0.72 | 1 | Gm25856 | | 0 | 0 | 0 | 0 | 0 | 0 | 0 | 0 | 0 | 0 | 4 |
| 0 | 1.32 | 3.63 | 0.36 | 0.72 | 1 | Mir6940 | | 0 | 0 | 0 | 0 | 0 | 0 | 0 | 0 | 0 | 0 | 4 |
| 0 | 1.32 | 3.63 | 0.36 | 0.72 | 1 | C030017D09Rik | | 0 | 0 | 0 | 0 | 0 | 0 | 0 | 0 | 0 | 0 | 4 |
| 0 | 1.32 | 3.63 | 0.36 | 0.72 | 1 | Gm20242 | | 0 | 0 | 0 | 0 | 0 | 0 | 0 | 0 | 0 | 0 | 4 |
| 0 | 1.32 | 3.63 | 0.36 | 0.72 | 1 | Gm49109 | | 0 | 0 | 0 | 0 | 0 | 0 | 0 | 0 | 0 | 0 | 4 |
| 0 | 1.32 | 3.63 | 0.36 | 0.72 | 1 | Gm17232 | | 0 | 0 | 0 | 0 | 0 | 0 | 0 | 0 | 0 | 0 | 4 |
| 0 | 1.32 | 3.63 | 0.36 | 0.72 | 1 | Gm4775 | | 0 | 0 | 0 | 0 | 0 | 0 | 0 | 0 | 0 | 0 | 4 |
| 0 | 1.32 | 3.63 | 0.36 | 0.72 | 1 | Mir6951 | | 0 | 0 | 0 | 0 | 0 | 0 | 0 | 0 | 0 | 0 | 4 |
| 0 | 1.32 | 3.63 | 0.36 | 0.72 | 1 | Gm32886 | | 0 | 0 | 0 | 0 | 0 | 0 | 0 | 0 | 0 | 0 | 4 |
| 0 | 1.32 | 3.63 | 0.36 | 0.72 | 1 | Gm50295 | | 0 | 0 | 0 | 0 | 0 | 0 | 0 | 0 | 0 | 0 | 4 |
| 0 | 1.32 | 3.63 | 0.36 | 0.72 | 1 | Gm4710 | | 0 | 0 | 0 | 0 | 0 | 0 | 0 | 0 | 0 | 0 | 4 |
| 0 | 1.32 | 3.63 | 0.36 | 0.72 | 1 | Gm4840 | | 0 | 0 | 0 | 0 | 0 | 0 | 0 | 0 | 0 | 0 | 4 |
| 0 | 1.32 | 3.63 | 0.36 | 0.72 | 1 | Gm26115 | | 0 | 0 | 0 | 0 | 0 | 0 | 0 | 0 | 0 | 0 | 4 |
| 0 | 1.32 | 3.63 | 0.36 | 0.72 | 1 | Mir101b | | 0 | 0 | 0 | 0 | 0 | 0 | 0 | 0 | 0 | 0 | 4 |
| 0 | 1.32 | 3.63 | 0.36 | 0.72 | 1 | Gm19200 | | 0 | 0 | 0 | 0 | 0 | 0 | 0 | 0 | 0 | 0 | 4 |
| 0 | 1.32 | 3.63 | 0.36 | 0.72 | 1 | Tspy-ps | | 0 | 0 | 0 | 0 | 0 | 0 | 0 | 0 | 0 | 0 | 4 |
| 0 | -0.52 | 3.63 | -0.14 | 0.89 | 1 | Hspd1-ps4 | | 0 | 1 | 0 | 0 | 2 | 0 | 0 | 1 | 0 | 0 | 0 |
| 0 | 1.49 | 3.63 | 0.41 | 0.68 | 1 | Gm46544 | | 0 | 0 | 0 | 1 | 0 | 0 | 0 | 1 | 1 | 0 | 1 |
| 0 | 1.30 | 3.64 | 0.36 | 0.72 | 1 | Gm44916 | | 0 | 1 | 0 | 0 | 0 | 0 | 0 | 0 | 0 | 0 | 3 |
| 0 | 2.04 | 3.64 | 0.56 | 0.58 | 1 | Gm12389 | | 0 | 0 | 0 | 0 | 0 | 0 | 0 | 0 | 1 | 0 | 3 |
| 0 | 1.25 | 3.66 | 0.34 | 0.73 | 1 | Slc22a1 | solute carrier family 22 (organic cation tra | 0 | 2 | 0 | 0 | 0 | 0 | 0 | 0 | 0 | 0 | 2 |
| 0 | 1.05 | 3.65 | 0.29 | 0.77 | 1 | Gm10123 | | 0 | 0 | 1 | 0 | 0 | 1 | 0 | 1 | 1 | 0 | 0 |
| 0 | 0.00 | 3.68 | 0.00 | 1.00 | 1 | Gm38039 | | 0 | 4 | 0 | 0 | 0 | 0 | 0 | 0 | 0 | 0 | 0 |
| 0 | 0.00 | 3.68 | 0.00 | 1.00 | 1 | Mannr | | 0 | 4 | 0 | 0 | 0 | 0 | 0 | 0 | 0 | 0 | 0 |
| 0 | 0.00 | 3.68 | 0.00 | 1.00 | 1 | Gm43012 | | 0 | 4 | 0 | 0 | 0 | 0 | 0 | 0 | 0 | 0 | 0 |
| 0 | 0.00 | 3.68 | 0.00 | 1.00 | 1 | Gm16043 | | 0 | 4 | 0 | 0 | 0 | 0 | 0 | 0 | 0 | 0 | 0 |
| 0 | 0.00 | 3.68 | 0.00 | 1.00 | 1 | Mgam2-ps | | 0 | 4 | 0 | 0 | 0 | 0 | 0 | 0 | 0 | 0 | 0 |
| 0 | 0.00 | 3.68 | 0.00 | 1.00 | 1 | Klk6 | kallikrein related-peptidase 6 | 0 | 4 | 0 | 0 | 0 | 0 | 0 | 0 | 0 | 0 | 0 |
| 0 | 0.00 | 3.68 | 0.00 | 1.00 | 1 | Klk1b8 | kallikrein 1-related peptidase b8 | 0 | 4 | 0 | 0 | 0 | 0 | 0 | 0 | 0 | 0 | 0 |
| 0 | 0.00 | 3.68 | 0.00 | 1.00 | 1 | Tbata | thymus, brain and testes associated | 0 | 4 | 0 | 0 | 0 | 0 | 0 | 0 | 0 | 0 | 0 |
| 0 | 0.00 | 3.68 | 0.00 | 1.00 | 1 | Gm12212 | | 0 | 4 | 0 | 0 | 0 | 0 | 0 | 0 | 0 | 0 | 0 |
| 0 | 0.00 | 3.68 | 0.00 | 1.00 | 1 | Ccl1 | chemokine (C-C motif) ligand 1 | 0 | 4 | 0 | 0 | 0 | 0 | 0 | 0 | 0 | 0 | 0 |
| 0 | 0.00 | 3.68 | 0.00 | 1.00 | 1 | Gm11713 | | 0 | 4 | 0 | 0 | 0 | 0 | 0 | 0 | 0 | 0 | 0 |
| 0 | 0.00 | 3.68 | 0.00 | 1.00 | 1 | Gm27299 | | 0 | 4 | 0 | 0 | 0 | 0 | 0 | 0 | 0 | 0 | 0 |
| 0 | 0.00 | 3.68 | 0.00 | 1.00 | 1 | Acox2 | acyl-Coenzyme A oxidase 2, branched ch | 0 | 4 | 0 | 0 | 0 | 0 | 0 | 0 | 0 | 0 | 0 |
| 0 | 0.00 | 3.68 | 0.00 | 1.00 | 1 | Gm28033 | | 0 | 4 | 0 | 0 | 0 | 0 | 0 | 0 | 0 | 0 | 0 |
| 0 | 0.00 | 3.68 | 0.00 | 1.00 | 1 | Sult1c1 | sulfotransferase family, cytosolic, 1C, mer | 0 | 4 | 0 | 0 | 0 | 0 | 0 | 0 | 0 | 0 | 0 |
| 0 | 0.00 | 3.68 | 0.00 | 1.00 | 1 | Gm50008 | | 0 | 4 | 0 | 0 | 0 | 0 | 0 | 0 | 0 | 0 | 0 |
| 0 | 0.00 | 3.68 | 0.00 | 1.00 | 1 | Gm50215 | | 0 | 4 | 0 | 0 | 0 | 0 | 0 | 0 | 0 | 0 | 0 |
| 0 | 0.00 | 3.68 | 0.00 | 1.00 | 1 | 1810058N15Rik | | 0 | 4 | 0 | 0 | 0 | 0 | 0 | 0 | 0 | 0 | 0 |
| 0 | 0.00 | 3.68 | 0.00 | 1.00 | 1 | Cyp2c69 | cytochrome P450, family 2, subfamily c, p | 0 | 4 | 0 | 0 | 0 | 0 | 0 | 0 | 0 | 0 | 0 |
| 0 | 1.07 | 3.67 | 0.29 | 0.77 | 1 | Gm8459 | | 0 | 3 | 0 | 0 | 0 | 0 | 0 | 0 | 1 | 0 | 0 |
| 0 | 1.48 | 3.65 | 0.40 | 0.69 | 1 | Gm13013 | | 0 | 2 | 0 | 0 | 0 | 0 | 0 | 0 | 2 | 0 | 0 |
| 0 | 1.68 | 3.65 | 0.46 | 0.65 | 1 | Gm14009 | | 0 | 0 | 0 | 0 | 0 | 0 | 0 | 0 | 4 | 0 | 0 |
| 0 | 1.68 | 3.65 | 0.46 | 0.65 | 1 | Zfp966 | zinc finger protein 966 | 0 | 0 | 0 | 0 | 0 | 0 | 0 | 0 | 4 | 0 | 0 |

| | | | | | | | | | | | | | | | | | | |
|---|---|---|---|---|---|---|---|---|---|---|---|---|---|---|---|---|---|---|
| 0 | 1.68 | 3.65 | 0.46 | 0.65 | 1 | Rxfp1 | relaxin/insulin-like family peptide receptor | 0 | 0 | 0 | 0 | 0 | 0 | 0 | 0 | 4 | 0 | 0 |
| 0 | 1.68 | 3.65 | 0.46 | 0.65 | 1 | 1700016H13Ri | RIKEN cDNA 1700016H13 gene | 0 | 0 | 0 | 0 | 0 | 0 | 0 | 0 | 4 | 0 | 0 |
| 0 | 1.68 | 3.65 | 0.46 | 0.65 | 1 | 2810432F15Rik | | 0 | 0 | 0 | 0 | 0 | 0 | 0 | 0 | 4 | 0 | 0 |
| 0 | 1.68 | 3.65 | 0.46 | 0.65 | 1 | Gm10619 | | 0 | 0 | 0 | 0 | 0 | 0 | 0 | 0 | 4 | 0 | 0 |
| 0 | 1.68 | 3.65 | 0.46 | 0.65 | 1 | Akirin1-ps | | 0 | 0 | 0 | 0 | 0 | 0 | 0 | 0 | 4 | 0 | 0 |
| 0 | 1.68 | 3.65 | 0.46 | 0.65 | 1 | Qrich2 | glutamine rich 2 | 0 | 0 | 0 | 0 | 0 | 0 | 0 | 0 | 4 | 0 | 0 |
| 0 | 1.68 | 3.65 | 0.46 | 0.65 | 1 | Gm47705 | | 0 | 0 | 0 | 0 | 0 | 0 | 0 | 0 | 4 | 0 | 0 |
| 0 | 1.68 | 3.65 | 0.46 | 0.65 | 1 | B230220B15Rik | | 0 | 0 | 0 | 0 | 0 | 0 | 0 | 0 | 4 | 0 | 0 |
| 0 | 1.68 | 3.65 | 0.46 | 0.65 | 1 | Gm2244 | predicted gene 2244 | 0 | 0 | 0 | 0 | 0 | 0 | 0 | 0 | 4 | 0 | 0 |
| 0 | 1.68 | 3.65 | 0.46 | 0.65 | 1 | Gm20026 | | 0 | 0 | 0 | 0 | 0 | 0 | 0 | 0 | 4 | 0 | 0 |
| 0 | 1.68 | 3.65 | 0.46 | 0.65 | 1 | Calhm3 | | 0 | 0 | 0 | 0 | 0 | 0 | 0 | 0 | 4 | 0 | 0 |
| 0 | 0.95 | 3.67 | 0.26 | 0.80 | 1 | Gm29228 | | 2 | 0 | 1 | 0 | 0 | 0 | 0 | 0 | 0 | 1 | 0 |
| 0 | -0.38 | 3.67 | -0.10 | 0.92 | 1 | Gm6867 | | 0 | 1 | 2 | 0 | 1 | 0 | 0 | 0 | 0 | 0 | 0 |
| 0 | 0.92 | 3.63 | 0.25 | 0.80 | 1 | Gm29718 | | 0 | 1 | 0 | 0 | 0 | 1 | 0 | 1 | 0 | 0 | 1 |
| 0 | 0.21 | 3.64 | 0.06 | 0.95 | 1 | Gm4761 | | 0 | 0 | 1 | 0 | 0 | 2 | 0 | 0 | 0 | 1 | 0 |
| 0 | 0.38 | 3.67 | 0.10 | 0.92 | 1 | Gm4796 | | 1 | 0 | 1 | 0 | 1 | 0 | 0 | 1 | 0 | 0 | 0 |
| 0 | 1.15 | 3.63 | 0.32 | 0.75 | 1 | Gm21963 | | 0 | 0 | 0 | 0 | 1 | 0 | 0 | 0 | 1 | 0 | 2 |
| 0 | 0.26 | 3.66 | 0.07 | 0.94 | 1 | Gm17090 | | 0 | 2 | 0 | 0 | 1 | 0 | 0 | 0 | 1 | 0 | 0 |
| 0 | 0.00 | 3.68 | 0.00 | 1.00 | 1 | Gm6285 | | 1 | 0 | 3 | 0 | 0 | 0 | 0 | 0 | 0 | 0 | 0 |
| 0 | -0.41 | 3.67 | -0.11 | 0.91 | 1 | Gm8599 | | 0 | 1 | 2 | 1 | 0 | 0 | 0 | 0 | 0 | 0 | 0 |
| 0 | -0.40 | 3.67 | -0.11 | 0.91 | 1 | Rpl27-ps2 | | 0 | 0 | 3 | 0 | 0 | 1 | 0 | 0 | 0 | 0 | 0 |
| 0 | -0.40 | 3.67 | -0.11 | 0.91 | 1 | Gm3512 | | 0 | 0 | 3 | 0 | 0 | 1 | 0 | 0 | 0 | 0 | 0 |
| 0 | 0.95 | 3.67 | 0.26 | 0.80 | 1 | Gm8062 | | 1 | 0 | 2 | 0 | 0 | 0 | 0 | 0 | 0 | 0 | 1 |
| 0 | 1.25 | 3.66 | 0.34 | 0.73 | 1 | Gm11645 | | 1 | 0 | 1 | 0 | 0 | 0 | 0 | 0 | 0 | 0 | 2 |
| 0 | 1.25 | 3.66 | 0.34 | 0.73 | 1 | Gm6643 | | 1 | 0 | 1 | 0 | 0 | 0 | 0 | 0 | 0 | 0 | 2 |
| 0 | 0.59 | 3.64 | 0.16 | 0.87 | 1 | Gm6473 | | 0 | 0 | 0 | 2 | 0 | 0 | 0 | 1 | 0 | 0 | 1 |
| 0 | 0.00 | 3.68 | 0.00 | 1.00 | 1 | Gm8019 | | 1 | 2 | 1 | 0 | 0 | 0 | 0 | 0 | 0 | 0 | 0 |
| 0 | -0.03 | 3.61 | -0.01 | 0.99 | 1 | Gm26820 | | 0 | 0 | 0 | 0 | 2 | 0 | 0 | 0 | 0 | 0 | 2 |
| 0 | -0.09 | 3.64 | -0.02 | 0.98 | 1 | Aldoa-ps4 | | 0 | 1 | 0 | 0 | 2 | 0 | 0 | 0 | 1 | 0 | 0 |
| 0 | 0.90 | 3.63 | 0.25 | 0.80 | 1 | Gm12848 | | 0 | 0 | 1 | 0 | 0 | 1 | 0 | 0 | 1 | 0 | 1 |
| 0 | 0.74 | 3.64 | 0.20 | 0.84 | 1 | Gm7575 | | 0 | 0 | 0 | 1 | 0 | 0 | 0 | 0 | 3 | 0 | 0 |
| 0 | -0.12 | 3.66 | -0.03 | 0.97 | 1 | Gm7642 | | 1 | 0 | 0 | 0 | 2 | 0 | 0 | 1 | 0 | 0 | 0 |
| 0 | 1.30 | 3.64 | 0.36 | 0.72 | 1 | Gm7200 | | 1 | 0 | 0 | 0 | 0 | 0 | 0 | 0 | 0 | 0 | 3 |
| 0 | 1.30 | 3.64 | 0.36 | 0.72 | 1 | 6330409D20Ri | RIKEN cDNA 6330409D20 gene | 1 | 0 | 0 | 0 | 0 | 0 | 0 | 0 | 0 | 0 | 3 |
| 0 | 1.30 | 3.64 | 0.36 | 0.72 | 1 | Gm14115 | | 1 | 0 | 0 | 0 | 0 | 0 | 0 | 0 | 0 | 0 | 3 |
| 0 | 1.30 | 3.64 | 0.36 | 0.72 | 1 | 8030453O22Rik | | 1 | 0 | 0 | 0 | 0 | 0 | 0 | 0 | 0 | 0 | 3 |
| 0 | 1.30 | 3.64 | 0.36 | 0.72 | 1 | Gm11949 | | 1 | 0 | 0 | 0 | 0 | 0 | 0 | 0 | 0 | 0 | 3 |
| 0 | 0.28 | 3.68 | 0.08 | 0.94 | 1 | Gm3800 | | 1 | 0 | 1 | 0 | 0 | 1 | 0 | 1 | 0 | 0 | 0 |
| 0 | 0.00 | 3.68 | 0.00 | 1.00 | 1 | Gm8182 | | 1 | 3 | 0 | 0 | 0 | 0 | 0 | 0 | 0 | 0 | 0 |
| 0 | 0.00 | 3.68 | 0.00 | 1.00 | 1 | Gm15895 | | 1 | 3 | 0 | 0 | 0 | 0 | 0 | 0 | 0 | 0 | 0 |
| 0 | 1.59 | 3.66 | 0.43 | 0.66 | 1 | Gm11578 | | 1 | 0 | 0 | 0 | 0 | 0 | 0 | 0 | 3 | 0 | 0 |
| 0 | 0.69 | 3.62 | 0.19 | 0.85 | 1 | A130048G24Rik | | 0 | 0 | 0 | 0 | 0 | 1 | 0 | 0 | 0 | 0 | 3 |
| 0 | 0.69 | 3.62 | 0.19 | 0.85 | 1 | Gm38142 | | 0 | 0 | 0 | 0 | 0 | 1 | 0 | 0 | 0 | 0 | 3 |
| 0 | 0.69 | 3.62 | 0.19 | 0.85 | 1 | Gm27027 | | 0 | 0 | 0 | 0 | 0 | 1 | 0 | 0 | 0 | 0 | 3 |
| 0 | 0.69 | 3.62 | 0.19 | 0.85 | 1 | Gm14226 | nonagouti | 0 | 0 | 0 | 0 | 0 | 1 | 0 | 0 | 0 | 0 | 3 |
| 0 | 0.69 | 3.62 | 0.19 | 0.85 | 1 | Gm11455 | | 0 | 0 | 0 | 0 | 0 | 1 | 0 | 0 | 0 | 0 | 3 |
| 0 | 0.69 | 3.62 | 0.19 | 0.85 | 1 | Gm12404 | | 0 | 0 | 0 | 0 | 0 | 1 | 0 | 0 | 0 | 0 | 3 |
| 0 | 0.69 | 3.62 | 0.19 | 0.85 | 1 | Gm11210 | | 0 | 0 | 0 | 0 | 0 | 1 | 0 | 0 | 0 | 0 | 3 |
| 0 | 0.69 | 3.62 | 0.19 | 0.85 | 1 | 2700068H02Rik | | 0 | 0 | 0 | 0 | 0 | 1 | 0 | 0 | 0 | 0 | 3 |
| 0 | 0.69 | 3.62 | 0.19 | 0.85 | 1 | Gm12934 | | 0 | 0 | 0 | 0 | 0 | 1 | 0 | 0 | 0 | 0 | 3 |
| 0 | 0.69 | 3.62 | 0.19 | 0.85 | 1 | Gm13063 | | 0 | 0 | 0 | 0 | 0 | 1 | 0 | 0 | 0 | 0 | 3 |
| 0 | 0.69 | 3.62 | 0.19 | 0.85 | 1 | Gm43727 | | 0 | 0 | 0 | 0 | 0 | 1 | 0 | 0 | 0 | 0 | 3 |
| 0 | 0.69 | 3.62 | 0.19 | 0.85 | 1 | Gm22716 | | 0 | 0 | 0 | 0 | 0 | 1 | 0 | 0 | 0 | 0 | 3 |
| 0 | 0.69 | 3.62 | 0.19 | 0.85 | 1 | Gm24414 | | 0 | 0 | 0 | 0 | 0 | 1 | 0 | 0 | 0 | 0 | 3 |
| 0 | 0.69 | 3.62 | 0.19 | 0.85 | 1 | Gm45095 | | 0 | 0 | 0 | 0 | 0 | 1 | 0 | 0 | 0 | 0 | 3 |
| 0 | 0.69 | 3.62 | 0.19 | 0.85 | 1 | Gm19656 | | 0 | 0 | 0 | 0 | 0 | 1 | 0 | 0 | 0 | 0 | 3 |
| 0 | 0.69 | 3.62 | 0.19 | 0.85 | 1 | B830042I05Rik | | 0 | 0 | 0 | 0 | 0 | 1 | 0 | 0 | 0 | 0 | 3 |
| 0 | 0.69 | 3.62 | 0.19 | 0.85 | 1 | 4930524O08Rik | | 0 | 0 | 0 | 0 | 0 | 1 | 0 | 0 | 0 | 0 | 3 |
| 0 | 0.69 | 3.62 | 0.19 | 0.85 | 1 | Gm28586 | | 0 | 0 | 0 | 0 | 0 | 1 | 0 | 0 | 0 | 0 | 3 |
| 0 | 0.69 | 3.62 | 0.19 | 0.85 | 1 | Gm48297 | | 0 | 0 | 0 | 0 | 0 | 1 | 0 | 0 | 0 | 0 | 3 |

| | | | | | | | | | | | | | | | | | | |
|---|---|---|---|---|---|---|---|---|---|---|---|---|---|---|---|---|---|---|
| 0 | 0.69 | 3.62 | 0.19 | 0.85 | 1 | A930033M14Rik | | 0 | 0 | 0 | 0 | 0 | 1 | 0 | 0 | 0 | 0 | 3 |
| 0 | 0.69 | 3.62 | 0.19 | 0.85 | 1 | Lin28b | lin-28 homolog B (C. elegans) | 0 | 0 | 0 | 0 | 0 | 1 | 0 | 0 | 0 | 0 | 3 |
| 0 | 0.69 | 3.62 | 0.19 | 0.85 | 1 | Gm32364 | | 0 | 0 | 0 | 0 | 0 | 1 | 0 | 0 | 0 | 0 | 3 |
| 0 | 0.27 | 3.65 | 0.07 | 0.94 | 1 | Gm5206 | | 0 | 0 | 0 | 1 | 0 | 1 | 0 | 1 | 1 | 0 | 0 |
| 0 | 0.28 | 3.64 | 0.08 | 0.94 | 1 | Rpl30-ps5 | | 0 | 2 | 0 | 0 | 0 | 1 | 0 | 0 | 0 | 0 | 1 |
| 0 | 0.83 | 3.66 | 0.23 | 0.82 | 1 | Gm16148 | | 0 | 1 | 0 | 0 | 0 | 1 | 0 | 0 | 2 | 0 | 0 |
| 0 | 0.90 | 3.64 | 0.25 | 0.80 | 1 | AI115009 | | 0 | 0 | 0 | 0 | 0 | 1 | 0 | 0 | 3 | 0 | 0 |
| 0 | 0.90 | 3.64 | 0.25 | 0.80 | 1 | Gm17200 | | 0 | 0 | 0 | 0 | 0 | 1 | 0 | 0 | 3 | 0 | 0 |
| 0 | 1.60 | 3.66 | 0.44 | 0.66 | 1 | Gm42435 | | 0 | 0 | 0 | 0 | 0 | 0 | 0 | 0 | 0 | 4 | 0 |
| 0 | 1.60 | 3.66 | 0.44 | 0.66 | 1 | Gm44185 | | 0 | 0 | 0 | 0 | 0 | 0 | 0 | 0 | 0 | 4 | 0 |
| 0 | 1.60 | 3.66 | 0.44 | 0.66 | 1 | Acot10 | acyl-CoA thioesterase 10 | 0 | 0 | 0 | 0 | 0 | 0 | 0 | 0 | 0 | 4 | 0 |
| 0 | 0.31 | 3.67 | 0.08 | 0.93 | 1 | Rpl31-ps18 | | 1 | 0 | 1 | 0 | 1 | 0 | 0 | 0 | 0 | 0 | 1 |
| 0 | -0.26 | 3.63 | -0.07 | 0.94 | 1 | Gm36198 | | 0 | 0 | 0 | 0 | 3 | 0 | 0 | 0 | 0 | 0 | 1 |
| 0 | -1.05 | 3.66 | -0.29 | 0.77 | 1 | Gm12035 | | 0 | 2 | 0 | 1 | 1 | 0 | 0 | 0 | 0 | 0 | 0 |
| 0 | -1.01 | 3.66 | -0.28 | 0.78 | 1 | Gm14046 | | 0 | 1 | 1 | 0 | 1 | 1 | 0 | 0 | 0 | 0 | 0 |
| 0 | -0.39 | 3.63 | -0.11 | 0.91 | 1 | Gm15470 | | 0 | 0 | 1 | 0 | 1 | 1 | 0 | 0 | 1 | 0 | 0 |
| 0 | -0.39 | 3.63 | -0.11 | 0.91 | 1 | Gm6088 | | 0 | 0 | 1 | 0 | 1 | 1 | 0 | 0 | 1 | 0 | 0 |
| 0 | -1.26 | 3.65 | -0.34 | 0.73 | 1 | Gm13042 | | 0 | 2 | 0 | 0 | 1 | 1 | 0 | 0 | 0 | 0 | 0 |
| 0 | 0.00 | 3.68 | 0.00 | 1.00 | 1 | Gadl1 | glutamate decarboxylase-like 1 | 2 | 0 | 2 | 0 | 0 | 0 | 0 | 0 | 0 | 0 | 0 |
| 0 | 0.00 | 3.68 | 0.00 | 1.00 | 1 | Ldha-ps3 | | 2 | 0 | 2 | 0 | 0 | 0 | 0 | 0 | 0 | 0 | 0 |
| 0 | 0.00 | 3.68 | 0.00 | 1.00 | 1 | B430212C06Rik | | 2 | 0 | 2 | 0 | 0 | 0 | 0 | 0 | 0 | 0 | 0 |
| 0 | -0.12 | 3.62 | -0.03 | 0.97 | 1 | Gm13822 | | 0 | 0 | 0 | 2 | 0 | 0 | 0 | 0 | 0 | 0 | 2 |
| 0 | -1.07 | 3.64 | -0.29 | 0.77 | 1 | Zswim2 | zinc finger SWIM-type containing 2 | 0 | 0 | 0 | 0 | 4 | 0 | 0 | 0 | 0 | 0 | 0 |
| 0 | -1.07 | 3.64 | -0.29 | 0.77 | 1 | S100a5 | S100 calcium binding protein A5 | 0 | 0 | 0 | 0 | 4 | 0 | 0 | 0 | 0 | 0 | 0 |
| 0 | -1.07 | 3.64 | -0.29 | 0.77 | 1 | Gm28685 | | 0 | 0 | 0 | 0 | 4 | 0 | 0 | 0 | 0 | 0 | 0 |
| 0 | -1.07 | 3.64 | -0.29 | 0.77 | 1 | Gm34086 | | 0 | 0 | 0 | 0 | 4 | 0 | 0 | 0 | 0 | 0 | 0 |
| 0 | -1.07 | 3.64 | -0.29 | 0.77 | 1 | Gm43397 | | 0 | 0 | 0 | 0 | 4 | 0 | 0 | 0 | 0 | 0 | 0 |
| 0 | -1.07 | 3.64 | -0.29 | 0.77 | 1 | Erich4 | glutamate rich 4 | 0 | 0 | 0 | 0 | 4 | 0 | 0 | 0 | 0 | 0 | 0 |
| 0 | -1.07 | 3.64 | -0.29 | 0.77 | 1 | Gm34533 | | 0 | 0 | 0 | 0 | 4 | 0 | 0 | 0 | 0 | 0 | 0 |
| 0 | -1.07 | 3.64 | -0.29 | 0.77 | 1 | Gm17231 | | 0 | 0 | 0 | 0 | 4 | 0 | 0 | 0 | 0 | 0 | 0 |
| 0 | -1.07 | 3.64 | -0.29 | 0.77 | 1 | Gm5161 | | 0 | 0 | 0 | 0 | 4 | 0 | 0 | 0 | 0 | 0 | 0 |
| 0 | -1.07 | 3.64 | -0.29 | 0.77 | 1 | 2410018L13Rik | | 0 | 0 | 0 | 0 | 4 | 0 | 0 | 0 | 0 | 0 | 0 |
| 0 | -1.07 | 3.64 | -0.29 | 0.77 | 1 | Gm33785 | | 0 | 0 | 0 | 0 | 4 | 0 | 0 | 0 | 0 | 0 | 0 |
| 0 | -1.07 | 3.64 | -0.29 | 0.77 | 1 | Gm49540 | | 0 | 0 | 0 | 0 | 4 | 0 | 0 | 0 | 0 | 0 | 0 |
| 0 | -1.07 | 3.64 | -0.29 | 0.77 | 1 | 8430430B14Rik | | 0 | 0 | 0 | 0 | 4 | 0 | 0 | 0 | 0 | 0 | 0 |
| 0 | -1.07 | 3.64 | -0.29 | 0.77 | 1 | Gm14829 | | 0 | 0 | 0 | 0 | 4 | 0 | 0 | 0 | 0 | 0 | 0 |
| 0 | -0.51 | 3.64 | -0.14 | 0.89 | 1 | Gm14409 | | 0 | 2 | 0 | 2 | 0 | 0 | 0 | 0 | 0 | 0 | 0 |
| 0 | -0.68 | 3.64 | -0.19 | 0.85 | 1 | Gm8824 | | 0 | 0 | 1 | 1 | 0 | 1 | 0 | 0 | 1 | 0 | 0 |
| 0 | -0.69 | 3.66 | -0.19 | 0.85 | 1 | Gm13369 | | 0 | 0 | 2 | 0 | 0 | 2 | 0 | 0 | 0 | 0 | 0 |
| 0 | -0.40 | 3.67 | -0.11 | 0.91 | 1 | Gm12944 | | 1 | 1 | 1 | 0 | 0 | 1 | 0 | 0 | 0 | 0 | 0 |
| 0 | 0.01 | 3.63 | 0.00 | 1.00 | 1 | Gm18889 | | 0 | 0 | 1 | 0 | 0 | 2 | 0 | 0 | 0 | 0 | 1 |
| 0 | 1.69 | 3.66 | 0.46 | 0.64 | 1 | Gm17825 | | 2 | 0 | 0 | 0 | 0 | 0 | 0 | 0 | 1 | 0 | 1 |
| 0 | -0.69 | 3.66 | -0.19 | 0.85 | 1 | Rpl19-ps9 | | 0 | 1 | 1 | 0 | 0 | 2 | 0 | 0 | 0 | 0 | 0 |
| 0 | 0.00 | 3.68 | 0.00 | 1.00 | 1 | Gm9063 | | 2 | 2 | 0 | 0 | 0 | 0 | 0 | 0 | 0 | 0 | 0 |
| 0 | 0.00 | 3.68 | 0.00 | 1.00 | 1 | 2010309G21Rik | | 2 | 2 | 0 | 0 | 0 | 0 | 0 | 0 | 0 | 0 | 0 |
| 0 | 0.00 | 3.68 | 0.00 | 1.00 | 1 | Zfp474 | zinc finger protein 474 | 2 | 2 | 0 | 0 | 0 | 0 | 0 | 0 | 0 | 0 | 0 |
| 0 | 0.03 | 3.63 | 0.01 | 0.99 | 1 | Gm4613 | | 0 | 0 | 1 | 0 | 0 | 2 | 0 | 0 | 1 | 0 | 0 |
| 0 | -0.32 | 3.66 | -0.09 | 0.93 | 1 | Ckb-ps1 | | 1 | 0 | 0 | 1 | 0 | 1 | 0 | 1 | 0 | 0 | 0 |
| 0 | 1.00 | 3.68 | 0.27 | 0.78 | 1 | Gm15989 | | 3 | 0 | 0 | 0 | 0 | 0 | 0 | 1 | 0 | 0 | 0 |
| 0 | -0.40 | 3.67 | -0.11 | 0.91 | 1 | 5430427N15Rik | | 1 | 2 | 0 | 0 | 0 | 1 | 0 | 0 | 0 | 0 | 0 |
| 0 | 0.28 | 3.62 | 0.08 | 0.94 | 1 | C030007H22Rik | | 0 | 0 | 0 | 0 | 0 | 2 | 0 | 0 | 0 | 0 | 2 |
| 0 | 0.28 | 3.62 | 0.08 | 0.94 | 1 | Gm37214 | | 0 | 0 | 0 | 0 | 0 | 2 | 0 | 0 | 0 | 0 | 2 |
| 0 | 0.28 | 3.62 | 0.08 | 0.94 | 1 | Gm44283 | | 0 | 0 | 0 | 0 | 0 | 2 | 0 | 0 | 0 | 0 | 2 |
| 0 | 0.28 | 3.62 | 0.08 | 0.94 | 1 | Gm22334 | | 0 | 0 | 0 | 0 | 0 | 2 | 0 | 0 | 0 | 0 | 2 |
| 0 | 0.28 | 3.62 | 0.08 | 0.94 | 1 | Gm45477 | | 0 | 0 | 0 | 0 | 0 | 2 | 0 | 0 | 0 | 0 | 2 |
| 0 | 0.28 | 3.62 | 0.08 | 0.94 | 1 | A930036I15Rik | | 0 | 0 | 0 | 0 | 0 | 2 | 0 | 0 | 0 | 0 | 2 |
| 0 | 0.28 | 3.62 | 0.08 | 0.94 | 1 | Gm42973 | | 0 | 0 | 0 | 0 | 0 | 2 | 0 | 0 | 0 | 0 | 2 |
| 0 | 0.28 | 3.62 | 0.08 | 0.94 | 1 | Aqp3 | aquaporin 3 | 0 | 0 | 0 | 0 | 0 | 2 | 0 | 0 | 0 | 0 | 2 |
| 0 | 0.28 | 3.62 | 0.08 | 0.94 | 1 | Gm42771 | | 0 | 0 | 0 | 0 | 0 | 2 | 0 | 0 | 0 | 0 | 2 |
| 0 | 0.28 | 3.62 | 0.08 | 0.94 | 1 | Gm13859 | | 0 | 0 | 0 | 0 | 0 | 2 | 0 | 0 | 0 | 0 | 2 |

| | | | | | | | | | | | | | | | | | | |
|---|---|---|---|---|---|---|---|---|---|---|---|---|---|---|---|---|---|---|
| 0 | 0.28 | 3.62 | 0.08 | 0.94 | 1 | Gm45292 | | 0 | 0 | 0 | 0 | 0 | 2 | 0 | 0 | 0 | 0 | 2 |
| 0 | 0.28 | 3.62 | 0.08 | 0.94 | 1 | Gm42755 | | 0 | 0 | 0 | 0 | 0 | 2 | 0 | 0 | 0 | 0 | 2 |
| 0 | 0.28 | 3.62 | 0.08 | 0.94 | 1 | Cep112it | | 0 | 0 | 0 | 0 | 0 | 2 | 0 | 0 | 0 | 0 | 2 |
| 0 | 0.28 | 3.62 | 0.08 | 0.94 | 1 | Gm24233 | | 0 | 0 | 0 | 0 | 0 | 2 | 0 | 0 | 0 | 0 | 2 |
| 0 | 0.28 | 3.62 | 0.08 | 0.94 | 1 | Gm47139 | | 0 | 0 | 0 | 0 | 0 | 2 | 0 | 0 | 0 | 0 | 2 |
| 0 | 0.28 | 3.62 | 0.08 | 0.94 | 1 | Rnu12 | | 0 | 0 | 0 | 0 | 0 | 2 | 0 | 0 | 0 | 0 | 2 |
| 0 | 0.28 | 3.62 | 0.08 | 0.94 | 1 | Krt78 | keratin 78 | 0 | 0 | 0 | 0 | 0 | 2 | 0 | 0 | 0 | 0 | 2 |
| 0 | 0.28 | 3.62 | 0.08 | 0.94 | 1 | Gm49705 | | 0 | 0 | 0 | 0 | 0 | 2 | 0 | 0 | 0 | 0 | 2 |
| 0 | -1.07 | 3.67 | -0.29 | 0.77 | 1 | Gm13392 | | 1 | 0 | 1 | 1 | 1 | 0 | 0 | 0 | 0 | 0 | 0 |
| 0 | -0.72 | 3.65 | -0.20 | 0.84 | 1 | Gm13481 | | 0 | 2 | 0 | 0 | 0 | 2 | 0 | 0 | 0 | 0 | 0 |
| 0 | -0.72 | 3.65 | -0.20 | 0.84 | 1 | Gm38327 | | 0 | 2 | 0 | 0 | 0 | 2 | 0 | 0 | 0 | 0 | 0 |
| 0 | -0.72 | 3.65 | -0.20 | 0.84 | 1 | Gm5441 | | 0 | 2 | 0 | 0 | 0 | 2 | 0 | 0 | 0 | 0 | 0 |
| 0 | 0.21 | 3.64 | 0.06 | 0.95 | 1 | Gm17103 | | 0 | 1 | 0 | 0 | 0 | 2 | 0 | 0 | 1 | 0 | 0 |
| 0 | 0.67 | 3.62 | 0.19 | 0.85 | 1 | Gm28791 | | 0 | 0 | 0 | 0 | 0 | 2 | 0 | 0 | 2 | 0 | 0 |
| 0 | 0.67 | 3.62 | 0.19 | 0.85 | 1 | Gm22503 | | 0 | 0 | 0 | 0 | 0 | 2 | 0 | 0 | 2 | 0 | 0 |
| 0 | -0.90 | 3.63 | -0.25 | 0.80 | 1 | Gm28364 | | 0 | 0 | 0 | 2 | 1 | 0 | 0 | 0 | 0 | 0 | 1 |
| 0 | -1.07 | 3.66 | -0.29 | 0.77 | 1 | Gm12267 | | 1 | 1 | 0 | 1 | 1 | 0 | 0 | 0 | 0 | 0 | 0 |
| 0 | -1.57 | 3.64 | -0.43 | 0.67 | 1 | Bbox1 | butyrobetaine (gamma), 2-oxoglutarate di | 0 | 0 | 0 | 0 | 3 | 1 | 0 | 0 | 0 | 0 | 0 |
| 0 | -0.67 | 3.63 | -0.19 | 0.85 | 1 | Gm37399 | | 0 | 0 | 0 | 0 | 1 | 2 | 0 | 0 | 0 | 0 | 1 |
| 0 | -0.67 | 3.63 | -0.19 | 0.85 | 1 | Zar1 | zygote arrest 1 | 0 | 0 | 0 | 0 | 1 | 2 | 0 | 0 | 0 | 0 | 1 |
| 0 | -0.67 | 3.63 | -0.19 | 0.85 | 1 | Prss38 | protease, serine 38 | 0 | 0 | 0 | 0 | 1 | 2 | 0 | 0 | 0 | 0 | 1 |
| 0 | -0.44 | 3.68 | -0.12 | 0.90 | 1 | Gm5764 | | 2 | 0 | 1 | 1 | 0 | 0 | 0 | 0 | 0 | 0 | 0 |
| 0 | -0.73 | 3.66 | -0.20 | 0.84 | 1 | Gm49514 | | 1 | 1 | 0 | 2 | 0 | 0 | 0 | 0 | 0 | 0 | 0 |
| 0 | -1.08 | 3.67 | -0.30 | 0.77 | 1 | Gm17826 | | 1 | 0 | 1 | 1 | 0 | 1 | 0 | 0 | 0 | 0 | 0 |
| 0 | 0.00 | 3.68 | 0.00 | 1.00 | 1 | Gm9790 | | 3 | 0 | 1 | 0 | 0 | 0 | 0 | 0 | 0 | 0 | 0 |
| 0 | -0.40 | 3.67 | -0.11 | 0.91 | 1 | Gm14263 | | 2 | 0 | 1 | 0 | 0 | 1 | 0 | 0 | 0 | 0 | 0 |
| 0 | -1.50 | 3.64 | -0.41 | 0.68 | 1 | Gm14407 | | 1 | 0 | 0 | 0 | 2 | 1 | 0 | 0 | 0 | 0 | 0 |
| 0 | 1.42 | 3.67 | 0.39 | 0.70 | 1 | Gm29164 | | 0 | 1 | 0 | 0 | 0 | 0 | 0 | 0 | 0 | 3 | 0 |
| 0 | 0.00 | 3.68 | 0.00 | 1.00 | 1 | Gm9370 | | 3 | 1 | 0 | 0 | 0 | 0 | 0 | 0 | 0 | 0 | 0 |
| 0 | 1.01 | 3.68 | 0.27 | 0.78 | 1 | Gm3508 | | 3 | 0 | 0 | 0 | 0 | 0 | 0 | 0 | 1 | 0 | 0 |
| 0 | -1.64 | 3.64 | -0.45 | 0.65 | 1 | 2010109P13Rik | | 0 | 0 | 0 | 0 | 2 | 2 | 0 | 0 | 0 | 0 | 0 |
| 0 | -1.64 | 3.64 | -0.45 | 0.65 | 1 | Gm29542 | | 0 | 0 | 0 | 0 | 2 | 2 | 0 | 0 | 0 | 0 | 0 |
| 0 | -1.64 | 3.64 | -0.45 | 0.65 | 1 | 5730416F02Rik | | 0 | 0 | 0 | 0 | 2 | 2 | 0 | 0 | 0 | 0 | 0 |
| 0 | -0.73 | 3.64 | -0.20 | 0.84 | 1 | Gm8971 | | 0 | 0 | 0 | 1 | 0 | 2 | 0 | 0 | 1 | 0 | 0 |
| 0 | -0.73 | 3.64 | -0.20 | 0.84 | 1 | Gm5673 | | 0 | 0 | 0 | 1 | 0 | 2 | 0 | 0 | 1 | 0 | 0 |
| 0 | 0.28 | 3.64 | 0.08 | 0.94 | 1 | Gm16091 | | 2 | 0 | 0 | 0 | 0 | 1 | 0 | 0 | 0 | 0 | 1 |
| 0 | -0.73 | 3.64 | -0.20 | 0.84 | 1 | Gm2622 | | 0 | 0 | 1 | 0 | 0 | 3 | 0 | 0 | 0 | 0 | 0 |
| 0 | -0.73 | 3.64 | -0.20 | 0.84 | 1 | Gm7380 | | 0 | 0 | 1 | 0 | 0 | 3 | 0 | 0 | 0 | 0 | 0 |
| 0 | 0.28 | 3.68 | 0.08 | 0.94 | 1 | Gm50174 | | 2 | 0 | 0 | 0 | 0 | 1 | 0 | 0 | 1 | 0 | 0 |
| 0 | -0.14 | 3.62 | -0.04 | 0.97 | 1 | Gm37437 | | 0 | 0 | 0 | 0 | 0 | 3 | 0 | 0 | 0 | 0 | 1 |
| 0 | -0.14 | 3.62 | -0.04 | 0.97 | 1 | Gm14124 | predicted gene 14124 | 0 | 0 | 0 | 0 | 0 | 3 | 0 | 0 | 0 | 0 | 1 |
| 0 | -0.14 | 3.62 | -0.04 | 0.97 | 1 | Gm4866 | | 0 | 0 | 0 | 0 | 0 | 3 | 0 | 0 | 0 | 0 | 1 |
| 0 | -0.14 | 3.62 | -0.04 | 0.97 | 1 | Gm15547 | | 0 | 0 | 0 | 0 | 0 | 3 | 0 | 0 | 0 | 0 | 1 |
| 0 | -0.14 | 3.62 | -0.04 | 0.97 | 1 | Gm24646 | | 0 | 0 | 0 | 0 | 0 | 3 | 0 | 0 | 0 | 0 | 1 |
| 0 | -0.14 | 3.62 | -0.04 | 0.97 | 1 | Gm11496 | | 0 | 0 | 0 | 0 | 0 | 3 | 0 | 0 | 0 | 0 | 1 |
| 0 | -0.14 | 3.62 | -0.04 | 0.97 | 1 | Mir6947 | | 0 | 0 | 0 | 0 | 0 | 3 | 0 | 0 | 0 | 0 | 1 |
| 0 | -0.14 | 3.62 | -0.04 | 0.97 | 1 | Gm50301 | | 0 | 0 | 0 | 0 | 0 | 3 | 0 | 0 | 0 | 0 | 1 |
| 0 | 0.18 | 3.63 | 0.05 | 0.96 | 1 | Gm48271 | | 0 | 0 | 0 | 0 | 0 | 3 | 0 | 0 | 1 | 0 | 0 |
| 0 | -0.48 | 3.67 | -0.13 | 0.90 | 1 | Gm12090 | | 3 | 0 | 0 | 0 | 1 | 0 | 0 | 0 | 0 | 0 | 0 |
| 0 | -0.48 | 3.67 | -0.13 | 0.90 | 1 | Gm45027 | | 3 | 0 | 0 | 0 | 1 | 0 | 0 | 0 | 0 | 0 | 0 |
| 0 | -1.13 | 3.65 | -0.31 | 0.76 | 1 | Fibcd1 | fibrinogen C domain containing 1 | 0 | 0 | 0 | 4 | 0 | 0 | 0 | 0 | 0 | 0 | 0 |
| 0 | -1.13 | 3.65 | -0.31 | 0.76 | 1 | Matn1 | matrilin 1, cartilage matrix protein | 0 | 0 | 0 | 4 | 0 | 0 | 0 | 0 | 0 | 0 | 0 |
| 0 | -1.13 | 3.65 | -0.31 | 0.76 | 1 | Gm17545 | | 0 | 0 | 0 | 4 | 0 | 0 | 0 | 0 | 0 | 0 | 0 |
| 0 | -1.13 | 3.65 | -0.31 | 0.76 | 1 | Dppa5a | developmental pluripotency associated 5 | 0 | 0 | 0 | 4 | 0 | 0 | 0 | 0 | 0 | 0 | 0 |
| 0 | -1.13 | 3.65 | -0.31 | 0.76 | 1 | Gm33301 | | 0 | 0 | 0 | 4 | 0 | 0 | 0 | 0 | 0 | 0 | 0 |
| 0 | -1.13 | 3.65 | -0.31 | 0.76 | 1 | Gm26834 | | 0 | 0 | 0 | 4 | 0 | 0 | 0 | 0 | 0 | 0 | 0 |
| 0 | -1.13 | 3.65 | -0.31 | 0.76 | 1 | Gm3621 | | 0 | 0 | 0 | 4 | 0 | 0 | 0 | 0 | 0 | 0 | 0 |
| 0 | -1.13 | 3.65 | -0.31 | 0.76 | 1 | Gm8980 | | 0 | 0 | 0 | 4 | 0 | 0 | 0 | 0 | 0 | 0 | 0 |
| 0 | -1.39 | 3.65 | -0.38 | 0.70 | 1 | Gm45507 | | 1 | 0 | 0 | 0 | 1 | 2 | 0 | 0 | 0 | 0 | 0 |
| 0 | -1.39 | 3.65 | -0.38 | 0.70 | 1 | Pgk1-rs7 | | 1 | 0 | 0 | 0 | 1 | 2 | 0 | 0 | 0 | 0 | 0 |

| | | | | | | | | | | | | | | | | | | |
|---|---|---|---|---|---|---|---|---|---|---|---|---|---|---|---|---|---|---|
| 0 | 1.79 | 3.66 | 0.49 | 0.62 | 1 | Gm2574 | | 0 | 0 | 1 | 0 | 0 | 0 | 0 | 1 | 0 | 1 | 0 |
| 0 | 1.79 | 3.66 | 0.49 | 0.62 | 1 | Gm12341 | | 0 | 0 | 1 | 0 | 0 | 0 | 0 | 1 | 0 | 1 | 0 |
| 0 | 1.79 | 3.66 | 0.49 | 0.62 | 1 | Gm9174 | | 0 | 0 | 1 | 0 | 0 | 0 | 0 | 1 | 0 | 1 | 0 |
| 0 | -1.05 | 3.66 | -0.29 | 0.77 | 1 | Atp2b3 | ATPase, Ca++ transporting, plasma mem | 1 | 0 | 0 | 3 | 0 | 0 | 0 | 0 | 0 | 0 | 0 |
| 0 | 1.56 | 3.66 | 0.43 | 0.67 | 1 | Gm6706 | | 0 | 0 | 0 | 0 | 0 | 0 | 0 | 3 | 0 | 0 | 0 |
| 0 | 1.56 | 3.66 | 0.43 | 0.67 | 1 | E130114P18Rik | | 0 | 0 | 0 | 0 | 0 | 0 | 0 | 3 | 0 | 0 | 0 |
| 0 | 1.56 | 3.66 | 0.43 | 0.67 | 1 | Gm31683 | | 0 | 0 | 0 | 0 | 0 | 0 | 0 | 3 | 0 | 0 | 0 |
| 0 | -1.55 | 3.64 | -0.43 | 0.67 | 1 | | | 0 | 0 | 0 | 3 | 0 | 1 | 0 | 0 | 0 | 0 | 0 |
| 0 | -1.55 | 3.64 | -0.43 | 0.67 | 1 | Gm17041 | | 0 | 0 | 0 | 3 | 0 | 1 | 0 | 0 | 0 | 0 | 0 |
| 0 | -1.55 | 3.64 | -0.43 | 0.67 | 1 | Gm27038 | | 0 | 0 | 0 | 3 | 0 | 1 | 0 | 0 | 0 | 0 | 0 |
| 0 | -0.47 | 3.67 | -0.13 | 0.90 | 1 | Gm46577 | | 3 | 0 | 0 | 1 | 0 | 0 | 0 | 0 | 0 | 0 | 0 |
| 0 | -1.62 | 3.64 | -0.44 | 0.66 | 1 | 4930488N24Rik | | 0 | 0 | 0 | 2 | 0 | 2 | 0 | 0 | 0 | 0 | 0 |
| 0 | 0.00 | 3.68 | 0.00 | 1.00 | 1 | Crygd | crystallin, gamma D | 3 | 0 | 0 | 0 | 0 | 0 | 0 | 0 | 0 | 0 | 0 |
| 0 | 0.00 | 3.68 | 0.00 | 1.00 | 1 | Gm7582 | | 3 | 0 | 0 | 0 | 0 | 0 | 0 | 0 | 0 | 0 | 0 |
| 0 | 0.00 | 3.68 | 0.00 | 1.00 | 1 | Efhd1os | | 3 | 0 | 0 | 0 | 0 | 0 | 0 | 0 | 0 | 0 | 0 |
| 0 | 0.00 | 3.68 | 0.00 | 1.00 | 1 | Gal3st2 | galactose-3-O-sulfotransferase 2 | 3 | 0 | 0 | 0 | 0 | 0 | 0 | 0 | 0 | 0 | 0 |
| 0 | 0.00 | 3.68 | 0.00 | 1.00 | 1 | Gm37103 | | 3 | 0 | 0 | 0 | 0 | 0 | 0 | 0 | 0 | 0 | 0 |
| 0 | 0.00 | 3.68 | 0.00 | 1.00 | 1 | Gm14040 | | 3 | 0 | 0 | 0 | 0 | 0 | 0 | 0 | 0 | 0 | 0 |
| 0 | 0.00 | 3.68 | 0.00 | 1.00 | 1 | Gm12453 | | 3 | 0 | 0 | 0 | 0 | 0 | 0 | 0 | 0 | 0 | 0 |
| 0 | 0.00 | 3.68 | 0.00 | 1.00 | 1 | Orm1 | orosomucoid 1 | 3 | 0 | 0 | 0 | 0 | 0 | 0 | 0 | 0 | 0 | 0 |
| 0 | 0.00 | 3.68 | 0.00 | 1.00 | 1 | 4930551L18Rik | | 3 | 0 | 0 | 0 | 0 | 0 | 0 | 0 | 0 | 0 | 0 |
| 0 | 0.00 | 3.68 | 0.00 | 1.00 | 1 | 4933407E24Rik | | 3 | 0 | 0 | 0 | 0 | 0 | 0 | 0 | 0 | 0 | 0 |
| 0 | 0.00 | 3.68 | 0.00 | 1.00 | 1 | Gjb4 | gap junction protein, beta 4 | 3 | 0 | 0 | 0 | 0 | 0 | 0 | 0 | 0 | 0 | 0 |
| 0 | 0.00 | 3.68 | 0.00 | 1.00 | 1 | Gm26226 | | 3 | 0 | 0 | 0 | 0 | 0 | 0 | 0 | 0 | 0 | 0 |
| 0 | 0.00 | 3.68 | 0.00 | 1.00 | 1 | Ajap1 | adherens junction associated protein 1 | 3 | 0 | 0 | 0 | 0 | 0 | 0 | 0 | 0 | 0 | 0 |
| 0 | 0.00 | 3.68 | 0.00 | 1.00 | 1 | Upk3bl | uroplakin 3B-like | 3 | 0 | 0 | 0 | 0 | 0 | 0 | 0 | 0 | 0 | 0 |
| 0 | 0.00 | 3.68 | 0.00 | 1.00 | 1 | Gm36241 | | 3 | 0 | 0 | 0 | 0 | 0 | 0 | 0 | 0 | 0 | 0 |
| 0 | 0.00 | 3.68 | 0.00 | 1.00 | 1 | Trbv3 | | 3 | 0 | 0 | 0 | 0 | 0 | 0 | 0 | 0 | 0 | 0 |
| 0 | 0.00 | 3.68 | 0.00 | 1.00 | 1 | Trbv31 | | 3 | 0 | 0 | 0 | 0 | 0 | 0 | 0 | 0 | 0 | 0 |
| 0 | 0.00 | 3.68 | 0.00 | 1.00 | 1 | G930045G22Rik | | 3 | 0 | 0 | 0 | 0 | 0 | 0 | 0 | 0 | 0 | 0 |
| 0 | 0.00 | 3.68 | 0.00 | 1.00 | 1 | Igkv14-130 | | 3 | 0 | 0 | 0 | 0 | 0 | 0 | 0 | 0 | 0 | 0 |
| 0 | 0.00 | 3.68 | 0.00 | 1.00 | 1 | Igkv11-114 | | 3 | 0 | 0 | 0 | 0 | 0 | 0 | 0 | 0 | 0 | 0 |
| 0 | 0.00 | 3.68 | 0.00 | 1.00 | 1 | Igkv4-92 | | 3 | 0 | 0 | 0 | 0 | 0 | 0 | 0 | 0 | 0 | 0 |
| 0 | 0.00 | 3.68 | 0.00 | 1.00 | 1 | Igkv5-45 | | 3 | 0 | 0 | 0 | 0 | 0 | 0 | 0 | 0 | 0 | 0 |
| 0 | 0.00 | 3.68 | 0.00 | 1.00 | 1 | Nat8f6 | N-acetyltransferase 8 (GCN5-related) far | 3 | 0 | 0 | 0 | 0 | 0 | 0 | 0 | 0 | 0 | 0 |
| 0 | 0.00 | 3.68 | 0.00 | 1.00 | 1 | Gm16567 | | 3 | 0 | 0 | 0 | 0 | 0 | 0 | 0 | 0 | 0 | 0 |
| 0 | 0.00 | 3.68 | 0.00 | 1.00 | 1 | Gm44428 | | 3 | 0 | 0 | 0 | 0 | 0 | 0 | 0 | 0 | 0 | 0 |
| 0 | 0.00 | 3.68 | 0.00 | 1.00 | 1 | Gm45282 | | 3 | 0 | 0 | 0 | 0 | 0 | 0 | 0 | 0 | 0 | 0 |
| 0 | 0.00 | 3.68 | 0.00 | 1.00 | 1 | Gm32098 | | 3 | 0 | 0 | 0 | 0 | 0 | 0 | 0 | 0 | 0 | 0 |
| 0 | 0.00 | 3.68 | 0.00 | 1.00 | 1 | Isx | intestine specific homeobox | 3 | 0 | 0 | 0 | 0 | 0 | 0 | 0 | 0 | 0 | 0 |
| 0 | 0.00 | 3.68 | 0.00 | 1.00 | 1 | Gm10181 | | 3 | 0 | 0 | 0 | 0 | 0 | 0 | 0 | 0 | 0 | 0 |
| 0 | 0.00 | 3.68 | 0.00 | 1.00 | 1 | Gm9869 | | 3 | 0 | 0 | 0 | 0 | 0 | 0 | 0 | 0 | 0 | 0 |
| 0 | 0.00 | 3.68 | 0.00 | 1.00 | 1 | Prr23a3 | proline rich 23A, member 3 | 3 | 0 | 0 | 0 | 0 | 0 | 0 | 0 | 0 | 0 | 0 |
| 0 | 0.00 | 3.68 | 0.00 | 1.00 | 1 | 5830418P13Rik | | 3 | 0 | 0 | 0 | 0 | 0 | 0 | 0 | 0 | 0 | 0 |
| 0 | 0.00 | 3.68 | 0.00 | 1.00 | 1 | Gm39465 | | 3 | 0 | 0 | 0 | 0 | 0 | 0 | 0 | 0 | 0 | 0 |
| 0 | 0.00 | 3.68 | 0.00 | 1.00 | 1 | Usp44 | ubiquitin specific peptidase 44 | 3 | 0 | 0 | 0 | 0 | 0 | 0 | 0 | 0 | 0 | 0 |
| 0 | 0.00 | 3.68 | 0.00 | 1.00 | 1 | A630014C17Rik | | 3 | 0 | 0 | 0 | 0 | 0 | 0 | 0 | 0 | 0 | 0 |
| 0 | 0.00 | 3.68 | 0.00 | 1.00 | 1 | Dnah2os | | 3 | 0 | 0 | 0 | 0 | 0 | 0 | 0 | 0 | 0 | 0 |
| 0 | 0.00 | 3.68 | 0.00 | 1.00 | 1 | Sap30bpos | | 3 | 0 | 0 | 0 | 0 | 0 | 0 | 0 | 0 | 0 | 0 |
| 0 | 0.00 | 3.68 | 0.00 | 1.00 | 1 | Ighv3-7 | | 3 | 0 | 0 | 0 | 0 | 0 | 0 | 0 | 0 | 0 | 0 |
| 0 | 0.00 | 3.68 | 0.00 | 1.00 | 1 | Ighv1-54 | | 3 | 0 | 0 | 0 | 0 | 0 | 0 | 0 | 0 | 0 | 0 |
| 0 | 0.00 | 3.68 | 0.00 | 1.00 | 1 | Gm5083 | | 3 | 0 | 0 | 0 | 0 | 0 | 0 | 0 | 0 | 0 | 0 |
| 0 | 0.00 | 3.68 | 0.00 | 1.00 | 1 | Gm47040 | | 3 | 0 | 0 | 0 | 0 | 0 | 0 | 0 | 0 | 0 | 0 |
| 0 | 0.00 | 3.68 | 0.00 | 1.00 | 1 | Npy4r | neuropeptide Y receptor Y4 | 3 | 0 | 0 | 0 | 0 | 0 | 0 | 0 | 0 | 0 | 0 |
| 0 | 0.00 | 3.68 | 0.00 | 1.00 | 1 | Gm17173 | | 3 | 0 | 0 | 0 | 0 | 0 | 0 | 0 | 0 | 0 | 0 |
| 0 | 0.00 | 3.68 | 0.00 | 1.00 | 1 | Trav13-1 | | 3 | 0 | 0 | 0 | 0 | 0 | 0 | 0 | 0 | 0 | 0 |
| 0 | 0.00 | 3.68 | 0.00 | 1.00 | 1 | 4930563I02Rik | | 3 | 0 | 0 | 0 | 0 | 0 | 0 | 0 | 0 | 0 | 0 |
| 0 | 0.00 | 3.68 | 0.00 | 1.00 | 1 | Gm36246 | | 3 | 0 | 0 | 0 | 0 | 0 | 0 | 0 | 0 | 0 | 0 |
| 0 | 0.00 | 3.68 | 0.00 | 1.00 | 1 | Gm21859 | | 3 | 0 | 0 | 0 | 0 | 0 | 0 | 0 | 0 | 0 | 0 |
| 0 | 0.00 | 3.68 | 0.00 | 1.00 | 1 | Gpr15 | G protein-coupled receptor 15 | 3 | 0 | 0 | 0 | 0 | 0 | 0 | 0 | 0 | 0 | 0 |

| | | | | | | | | | | | | | | | | | | |
|---|---|---|---|---|---|---|---|---|---|---|---|---|---|---|---|---|---|---|
| 0 | 0.00 | 3.68 | 0.00 | 1.00 | 1 | Ankrd66 | ankyrin repeat domain 66 | 3 | 0 | 0 | 0 | 0 | 0 | 0 | 0 | 0 | 0 | 0 |
| 0 | 0.00 | 3.68 | 0.00 | 1.00 | 1 | Tmem232 | transmembrane protein 232 | 3 | 0 | 0 | 0 | 0 | 0 | 0 | 0 | 0 | 0 | 0 |
| 0 | 0.00 | 3.68 | 0.00 | 1.00 | 1 | Gm50124 | | 3 | 0 | 0 | 0 | 0 | 0 | 0 | 0 | 0 | 0 | 0 |
| 0 | 0.00 | 3.68 | 0.00 | 1.00 | 1 | Ccdc192 | coiled-coil domain containing 192 | 3 | 0 | 0 | 0 | 0 | 0 | 0 | 0 | 0 | 0 | 0 |
| 0 | 0.00 | 3.68 | 0.00 | 1.00 | 1 | Bmp15 | bone morphogenetic protein 15 | 3 | 0 | 0 | 0 | 0 | 0 | 0 | 0 | 0 | 0 | 0 |
| 0 | 0.82 | 3.66 | 0.22 | 0.82 | 1 | Gm6944 | | 0 | 0 | 0 | 0 | 0 | 1 | 0 | 0 | 0 | 3 | 0 |
| 0 | -0.40 | 3.67 | -0.11 | 0.91 | 1 | Gm43484 | | 3 | 0 | 0 | 0 | 0 | 1 | 0 | 0 | 0 | 0 | 0 |
| 0 | -0.40 | 3.67 | -0.11 | 0.91 | 1 | 5830416I19Rik | | 3 | 0 | 0 | 0 | 0 | 1 | 0 | 0 | 0 | 0 | 0 |
| 0 | -0.40 | 3.67 | -0.11 | 0.91 | 1 | Tmem45a2 | transmembrane protein 45A2 | 3 | 0 | 0 | 0 | 0 | 1 | 0 | 0 | 0 | 0 | 0 |
| 0 | -0.40 | 3.67 | -0.11 | 0.91 | 1 | F830208F22Rik | | 3 | 0 | 0 | 0 | 0 | 1 | 0 | 0 | 0 | 0 | 0 |
| 0 | -0.73 | 3.64 | -0.20 | 0.84 | 1 | Gm42544 | | 1 | 0 | 0 | 0 | 0 | 3 | 0 | 0 | 0 | 0 | 0 |
| 0 | 1.08 | 3.65 | 0.30 | 0.77 | 1 | Gm26976 | | 0 | 0 | 0 | 0 | 1 | 0 | 0 | 1 | 0 | 1 | 0 |
| 0 | -0.76 | 3.63 | -0.21 | 0.83 | 1 | Gm37569 | | 0 | 0 | 0 | 0 | 0 | 3 | 0 | 0 | 0 | 0 | 0 |
| 0 | -0.76 | 3.63 | -0.21 | 0.83 | 1 | Mir6897 | | 0 | 0 | 0 | 0 | 0 | 3 | 0 | 0 | 0 | 0 | 0 |
| 0 | -0.76 | 3.63 | -0.21 | 0.83 | 1 | Gm28499 | | 0 | 0 | 0 | 0 | 0 | 3 | 0 | 0 | 0 | 0 | 0 |
| 0 | -0.76 | 3.63 | -0.21 | 0.83 | 1 | Gm28188 | | 0 | 0 | 0 | 0 | 0 | 3 | 0 | 0 | 0 | 0 | 0 |
| 0 | -0.76 | 3.63 | -0.21 | 0.83 | 1 | Gm38301 | | 0 | 0 | 0 | 0 | 0 | 3 | 0 | 0 | 0 | 0 | 0 |
| 0 | -0.76 | 3.63 | -0.21 | 0.83 | 1 | Gm1627 | | 0 | 0 | 0 | 0 | 0 | 3 | 0 | 0 | 0 | 0 | 0 |
| 0 | -0.76 | 3.63 | -0.21 | 0.83 | 1 | 2610012C04Rik | | 0 | 0 | 0 | 0 | 0 | 3 | 0 | 0 | 0 | 0 | 0 |
| 0 | -0.76 | 3.63 | -0.21 | 0.83 | 1 | Gm37873 | | 0 | 0 | 0 | 0 | 0 | 3 | 0 | 0 | 0 | 0 | 0 |
| 0 | -0.76 | 3.63 | -0.21 | 0.83 | 1 | Gm38064 | | 0 | 0 | 0 | 0 | 0 | 3 | 0 | 0 | 0 | 0 | 0 |
| 0 | -0.76 | 3.63 | -0.21 | 0.83 | 1 | Mir6998 | | 0 | 0 | 0 | 0 | 0 | 3 | 0 | 0 | 0 | 0 | 0 |
| 0 | -0.76 | 3.63 | -0.21 | 0.83 | 1 | Gm14033 | | 0 | 0 | 0 | 0 | 0 | 3 | 0 | 0 | 0 | 0 | 0 |
| 0 | -0.76 | 3.63 | -0.21 | 0.83 | 1 | Mir1955 | | 0 | 0 | 0 | 0 | 0 | 3 | 0 | 0 | 0 | 0 | 0 |
| 0 | -0.76 | 3.63 | -0.21 | 0.83 | 1 | Gm13868 | | 0 | 0 | 0 | 0 | 0 | 3 | 0 | 0 | 0 | 0 | 0 |
| 0 | -0.76 | 3.63 | -0.21 | 0.83 | 1 | Lamp5 | lysosomal-associated membrane protein | 0 | 0 | 0 | 0 | 0 | 3 | 0 | 0 | 0 | 0 | 0 |
| 0 | -0.76 | 3.63 | -0.21 | 0.83 | 1 | Zfp973 | zinc finger protein 973 | 0 | 0 | 0 | 0 | 0 | 3 | 0 | 0 | 0 | 0 | 0 |
| 0 | -0.76 | 3.63 | -0.21 | 0.83 | 1 | Gm37498 | | 0 | 0 | 0 | 0 | 0 | 3 | 0 | 0 | 0 | 0 | 0 |
| 0 | -0.76 | 3.63 | -0.21 | 0.83 | 1 | Gm42843 | | 0 | 0 | 0 | 0 | 0 | 3 | 0 | 0 | 0 | 0 | 0 |
| 0 | -0.76 | 3.63 | -0.21 | 0.83 | 1 | Gm43073 | | 0 | 0 | 0 | 0 | 0 | 3 | 0 | 0 | 0 | 0 | 0 |
| 0 | -0.76 | 3.63 | -0.21 | 0.83 | 1 | Gm43824 | | 0 | 0 | 0 | 0 | 0 | 3 | 0 | 0 | 0 | 0 | 0 |
| 0 | -0.76 | 3.63 | -0.21 | 0.83 | 1 | Gm42874 | | 0 | 0 | 0 | 0 | 0 | 3 | 0 | 0 | 0 | 0 | 0 |
| 0 | -0.76 | 3.63 | -0.21 | 0.83 | 1 | Gm11842 | | 0 | 0 | 0 | 0 | 0 | 3 | 0 | 0 | 0 | 0 | 0 |
| 0 | -0.76 | 3.63 | -0.21 | 0.83 | 1 | Gm12577 | | 0 | 0 | 0 | 0 | 0 | 3 | 0 | 0 | 0 | 0 | 0 |
| 0 | -0.76 | 3.63 | -0.21 | 0.83 | 1 | Gm50475 | | 0 | 0 | 0 | 0 | 0 | 3 | 0 | 0 | 0 | 0 | 0 |
| 0 | -0.76 | 3.63 | -0.21 | 0.83 | 1 | Gm13053 | | 0 | 0 | 0 | 0 | 0 | 3 | 0 | 0 | 0 | 0 | 0 |
| 0 | -0.76 | 3.63 | -0.21 | 0.83 | 1 | Gm23303 | | 0 | 0 | 0 | 0 | 0 | 3 | 0 | 0 | 0 | 0 | 0 |
| 0 | -0.76 | 3.63 | -0.21 | 0.83 | 1 | Wdr46-ps | | 0 | 0 | 0 | 0 | 0 | 3 | 0 | 0 | 0 | 0 | 0 |
| 0 | -0.76 | 3.63 | -0.21 | 0.83 | 1 | Gm43787 | | 0 | 0 | 0 | 0 | 0 | 3 | 0 | 0 | 0 | 0 | 0 |
| 0 | -0.76 | 3.63 | -0.21 | 0.83 | 1 | Gm43421 | | 0 | 0 | 0 | 0 | 0 | 3 | 0 | 0 | 0 | 0 | 0 |
| 0 | -0.76 | 3.63 | -0.21 | 0.83 | 1 | Gm43423 | | 0 | 0 | 0 | 0 | 0 | 3 | 0 | 0 | 0 | 0 | 0 |
| 0 | -0.76 | 3.63 | -0.21 | 0.83 | 1 | Gm26692 | | 0 | 0 | 0 | 0 | 0 | 3 | 0 | 0 | 0 | 0 | 0 |
| 0 | -0.76 | 3.63 | -0.21 | 0.83 | 1 | Gm43364 | | 0 | 0 | 0 | 0 | 0 | 3 | 0 | 0 | 0 | 0 | 0 |
| 0 | -0.76 | 3.63 | -0.21 | 0.83 | 1 | Mir7028 | | 0 | 0 | 0 | 0 | 0 | 3 | 0 | 0 | 0 | 0 | 0 |
| 0 | -0.76 | 3.63 | -0.21 | 0.83 | 1 | 2610011E03Rik | | 0 | 0 | 0 | 0 | 0 | 3 | 0 | 0 | 0 | 0 | 0 |
| 0 | -0.76 | 3.63 | -0.21 | 0.83 | 1 | Gm43378 | | 0 | 0 | 0 | 0 | 0 | 3 | 0 | 0 | 0 | 0 | 0 |
| 0 | -0.76 | 3.63 | -0.21 | 0.83 | 1 | Gm42531 | | 0 | 0 | 0 | 0 | 0 | 3 | 0 | 0 | 0 | 0 | 0 |
| 0 | -0.76 | 3.63 | -0.21 | 0.83 | 1 | Gm44280 | | 0 | 0 | 0 | 0 | 0 | 3 | 0 | 0 | 0 | 0 | 0 |
| 0 | -0.76 | 3.63 | -0.21 | 0.83 | 1 | Gm8778 | | 0 | 0 | 0 | 0 | 0 | 3 | 0 | 0 | 0 | 0 | 0 |
| 0 | -0.76 | 3.63 | -0.21 | 0.83 | 1 | D6Ertd474e | | 0 | 0 | 0 | 0 | 0 | 3 | 0 | 0 | 0 | 0 | 0 |
| 0 | -0.76 | 3.63 | -0.21 | 0.83 | 1 | Cabp5 | calcium binding protein 5 | 0 | 0 | 0 | 0 | 0 | 3 | 0 | 0 | 0 | 0 | 0 |
| 0 | -0.76 | 3.63 | -0.21 | 0.83 | 1 | Snord35a | | 0 | 0 | 0 | 0 | 0 | 3 | 0 | 0 | 0 | 0 | 0 |
| 0 | -0.76 | 3.63 | -0.21 | 0.83 | 1 | Otog | otogelin | 0 | 0 | 0 | 0 | 0 | 3 | 0 | 0 | 0 | 0 | 0 |
| 0 | -0.76 | 3.63 | -0.21 | 0.83 | 1 | Gm44708 | | 0 | 0 | 0 | 0 | 0 | 3 | 0 | 0 | 0 | 0 | 0 |
| 0 | -0.76 | 3.63 | -0.21 | 0.83 | 1 | Mir344g | | 0 | 0 | 0 | 0 | 0 | 3 | 0 | 0 | 0 | 0 | 0 |
| 0 | -0.76 | 3.63 | -0.21 | 0.83 | 1 | Gm44931 | | 0 | 0 | 0 | 0 | 0 | 3 | 0 | 0 | 0 | 0 | 0 |
| 0 | -0.76 | 3.63 | -0.21 | 0.83 | 1 | Gm45088 | | 0 | 0 | 0 | 0 | 0 | 3 | 0 | 0 | 0 | 0 | 0 |
| 0 | -0.76 | 3.63 | -0.21 | 0.83 | 1 | Gm23262 | | 0 | 0 | 0 | 0 | 0 | 3 | 0 | 0 | 0 | 0 | 0 |
| 0 | -0.76 | 3.63 | -0.21 | 0.83 | 1 | Gm44661 | | 0 | 0 | 0 | 0 | 0 | 3 | 0 | 0 | 0 | 0 | 0 |
| 0 | -0.76 | 3.63 | -0.21 | 0.83 | 1 | Gm44663 | | 0 | 0 | 0 | 0 | 0 | 3 | 0 | 0 | 0 | 0 | 0 |

| | | | | | | | | | | | | | | | | | | |
|---|---|---|---|---|---|---|---|---|---|---|---|---|---|---|---|---|---|---|
| 0 | -0.76 | 3.63 | -0.21 | 0.83 | 1 | Gm24596 | | 0 | 0 | 0 | 0 | 0 | 3 | 0 | 0 | 0 | 0 | 0 |
| 0 | -0.76 | 3.63 | -0.21 | 0.83 | 1 | Btbd16 | BTB (POZ) domain containing 16 | 0 | 0 | 0 | 0 | 0 | 3 | 0 | 0 | 0 | 0 | 0 |
| 0 | -0.76 | 3.63 | -0.21 | 0.83 | 1 | Gm45416 | | 0 | 0 | 0 | 0 | 0 | 3 | 0 | 0 | 0 | 0 | 0 |
| 0 | -0.76 | 3.63 | -0.21 | 0.83 | 1 | Gm45073 | | 0 | 0 | 0 | 0 | 0 | 3 | 0 | 0 | 0 | 0 | 0 |
| 0 | -0.76 | 3.63 | -0.21 | 0.83 | 1 | 5330413D20Rik | | 0 | 0 | 0 | 0 | 0 | 3 | 0 | 0 | 0 | 0 | 0 |
| 0 | -0.76 | 3.63 | -0.21 | 0.83 | 1 | Gm45266 | | 0 | 0 | 0 | 0 | 0 | 3 | 0 | 0 | 0 | 0 | 0 |
| 0 | -0.76 | 3.63 | -0.21 | 0.83 | 1 | Gm35021 | | 0 | 0 | 0 | 0 | 0 | 3 | 0 | 0 | 0 | 0 | 0 |
| 0 | -0.76 | 3.63 | -0.21 | 0.83 | 1 | Gm45757 | | 0 | 0 | 0 | 0 | 0 | 3 | 0 | 0 | 0 | 0 | 0 |
| 0 | -0.76 | 3.63 | -0.21 | 0.83 | 1 | Gm47347 | | 0 | 0 | 0 | 0 | 0 | 3 | 0 | 0 | 0 | 0 | 0 |
| 0 | -0.76 | 3.63 | -0.21 | 0.83 | 1 | Gm19936 | | 0 | 0 | 0 | 0 | 0 | 3 | 0 | 0 | 0 | 0 | 0 |
| 0 | -0.76 | 3.63 | -0.21 | 0.83 | 1 | Gm27231 | | 0 | 0 | 0 | 0 | 0 | 3 | 0 | 0 | 0 | 0 | 0 |
| 0 | -0.76 | 3.63 | -0.21 | 0.83 | 1 | Gm47430 | | 0 | 0 | 0 | 0 | 0 | 3 | 0 | 0 | 0 | 0 | 0 |
| 0 | -0.76 | 3.63 | -0.21 | 0.83 | 1 | Gm47427 | | 0 | 0 | 0 | 0 | 0 | 3 | 0 | 0 | 0 | 0 | 0 |
| 0 | -0.76 | 3.63 | -0.21 | 0.83 | 1 | Gm47504 | | 0 | 0 | 0 | 0 | 0 | 3 | 0 | 0 | 0 | 0 | 0 |
| 0 | -0.76 | 3.63 | -0.21 | 0.83 | 1 | Mir6906 | | 0 | 0 | 0 | 0 | 0 | 3 | 0 | 0 | 0 | 0 | 0 |
| 0 | -0.76 | 3.63 | -0.21 | 0.83 | 1 | Gm47247 | | 0 | 0 | 0 | 0 | 0 | 3 | 0 | 0 | 0 | 0 | 0 |
| 0 | -0.76 | 3.63 | -0.21 | 0.83 | 1 | 1700025F24Rik | | 0 | 0 | 0 | 0 | 0 | 3 | 0 | 0 | 0 | 0 | 0 |
| 0 | -0.76 | 3.63 | -0.21 | 0.83 | 1 | Gm18755 | | 0 | 0 | 0 | 0 | 0 | 3 | 0 | 0 | 0 | 0 | 0 |
| 0 | -0.76 | 3.63 | -0.21 | 0.83 | 1 | Gm47296 | | 0 | 0 | 0 | 0 | 0 | 3 | 0 | 0 | 0 | 0 | 0 |
| 0 | -0.76 | 3.63 | -0.21 | 0.83 | 1 | Hbq1a | hemoglobin, theta 1A | 0 | 0 | 0 | 0 | 0 | 3 | 0 | 0 | 0 | 0 | 0 |
| 0 | -0.76 | 3.63 | -0.21 | 0.83 | 1 | Mir340 | | 0 | 0 | 0 | 0 | 0 | 3 | 0 | 0 | 0 | 0 | 0 |
| 0 | -0.76 | 3.63 | -0.21 | 0.83 | 1 | Gm23564 | | 0 | 0 | 0 | 0 | 0 | 3 | 0 | 0 | 0 | 0 | 0 |
| 0 | -0.76 | 3.63 | -0.21 | 0.83 | 1 | Gm49377 | | 0 | 0 | 0 | 0 | 0 | 3 | 0 | 0 | 0 | 0 | 0 |
| 0 | -0.76 | 3.63 | -0.21 | 0.83 | 1 | Gm48536 | | 0 | 0 | 0 | 0 | 0 | 3 | 0 | 0 | 0 | 0 | 0 |
| 0 | -0.76 | 3.63 | -0.21 | 0.83 | 1 | Gm25926 | | 0 | 0 | 0 | 0 | 0 | 3 | 0 | 0 | 0 | 0 | 0 |
| 0 | -0.76 | 3.63 | -0.21 | 0.83 | 1 | Gm48225 | | 0 | 0 | 0 | 0 | 0 | 3 | 0 | 0 | 0 | 0 | 0 |
| 0 | -0.76 | 3.63 | -0.21 | 0.83 | 1 | Mir431 | | 0 | 0 | 0 | 0 | 0 | 3 | 0 | 0 | 0 | 0 | 0 |
| 0 | -0.76 | 3.63 | -0.21 | 0.83 | 1 | Gm48869 | | 0 | 0 | 0 | 0 | 0 | 3 | 0 | 0 | 0 | 0 | 0 |
| 0 | -0.76 | 3.63 | -0.21 | 0.83 | 1 | Gm44394 | | 0 | 0 | 0 | 0 | 0 | 3 | 0 | 0 | 0 | 0 | 0 |
| 0 | -0.76 | 3.63 | -0.21 | 0.83 | 1 | Gm48626 | | 0 | 0 | 0 | 0 | 0 | 3 | 0 | 0 | 0 | 0 | 0 |
| 0 | -0.76 | 3.63 | -0.21 | 0.83 | 1 | D130062J10Rik | | 0 | 0 | 0 | 0 | 0 | 3 | 0 | 0 | 0 | 0 | 0 |
| 0 | -0.76 | 3.63 | -0.21 | 0.83 | 1 | Gm3500 | predicted gene 3500 | 0 | 0 | 0 | 0 | 0 | 3 | 0 | 0 | 0 | 0 | 0 |
| 0 | -0.76 | 3.63 | -0.21 | 0.83 | 1 | Gm23593 | | 0 | 0 | 0 | 0 | 0 | 3 | 0 | 0 | 0 | 0 | 0 |
| 0 | -0.76 | 3.63 | -0.21 | 0.83 | 1 | Gm7968 | | 0 | 0 | 0 | 0 | 0 | 3 | 0 | 0 | 0 | 0 | 0 |
| 0 | -0.76 | 3.63 | -0.21 | 0.83 | 1 | Gm18994 | | 0 | 0 | 0 | 0 | 0 | 3 | 0 | 0 | 0 | 0 | 0 |
| 0 | -0.76 | 3.63 | -0.21 | 0.83 | 1 | Gm19236 | | 0 | 0 | 0 | 0 | 0 | 3 | 0 | 0 | 0 | 0 | 0 |
| 0 | -0.76 | 3.63 | -0.21 | 0.83 | 1 | Gm49520 | | 0 | 0 | 0 | 0 | 0 | 3 | 0 | 0 | 0 | 0 | 0 |
| 0 | -0.76 | 3.63 | -0.21 | 0.83 | 1 | Gm36618 | | 0 | 0 | 0 | 0 | 0 | 3 | 0 | 0 | 0 | 0 | 0 |
| 0 | -0.76 | 3.63 | -0.21 | 0.83 | 1 | Fam186b | family with sequence similarity 186, meml | 0 | 0 | 0 | 0 | 0 | 3 | 0 | 0 | 0 | 0 | 0 |
| 0 | -0.76 | 3.63 | -0.21 | 0.83 | 1 | Gm49495 | | 0 | 0 | 0 | 0 | 0 | 3 | 0 | 0 | 0 | 0 | 0 |
| 0 | -0.76 | 3.63 | -0.21 | 0.83 | 1 | Gm37186 | | 0 | 0 | 0 | 0 | 0 | 3 | 0 | 0 | 0 | 0 | 0 |
| 0 | -0.76 | 3.63 | -0.21 | 0.83 | 1 | Gm22576 | | 0 | 0 | 0 | 0 | 0 | 3 | 0 | 0 | 0 | 0 | 0 |
| 0 | -0.76 | 3.63 | -0.21 | 0.83 | 1 | Snora20 | | 0 | 0 | 0 | 0 | 0 | 3 | 0 | 0 | 0 | 0 | 0 |
| 0 | -0.76 | 3.63 | -0.21 | 0.83 | 1 | Gm38099 | | 0 | 0 | 0 | 0 | 0 | 3 | 0 | 0 | 0 | 0 | 0 |
| 0 | -0.76 | 3.63 | -0.21 | 0.83 | 1 | Gm50364 | | 0 | 0 | 0 | 0 | 0 | 3 | 0 | 0 | 0 | 0 | 0 |
| 0 | -0.76 | 3.63 | -0.21 | 0.83 | 1 | Rab33a | RAB33A, member RAS oncogene family | 0 | 0 | 0 | 0 | 0 | 3 | 0 | 0 | 0 | 0 | 0 |
| 0 | -0.76 | 3.63 | -0.21 | 0.83 | 1 | Gm15016 | | 0 | 0 | 0 | 0 | 0 | 3 | 0 | 0 | 0 | 0 | 0 |
| 0 | -0.76 | 3.63 | -0.21 | 0.83 | 1 | Pih1h3b | PIH1 domain containing 3B | 0 | 0 | 0 | 0 | 0 | 3 | 0 | 0 | 0 | 0 | 0 |
| 0 | -0.76 | 3.63 | -0.21 | 0.83 | 1 | Gm27980 | | 0 | 0 | 0 | 0 | 0 | 3 | 0 | 0 | 0 | 0 | 0 |
| 0 | 1.05 | 3.67 | 0.29 | 0.77 | 1 | Gm6075 | | 0 | 0 | 2 | 0 | 0 | 0 | 0 | 0 | 0 | 1 | 0 |
| 0 | 1.47 | 3.66 | 0.40 | 0.69 | 1 | Gm5702 | | 0 | 0 | 1 | 0 | 0 | 0 | 0 | 2 | 0 | 0 | 0 |
| 0 | 1.71 | 3.66 | 0.47 | 0.64 | 1 | Gm7481 | | 0 | 0 | 1 | 0 | 0 | 0 | 0 | 0 | 0 | 1 | 1 |
| 0 | 1.93 | 3.65 | 0.53 | 0.60 | 1 | Gm14473 | | 0 | 0 | 0 | 0 | 0 | 0 | 0 | 2 | 0 | 0 | 1 |
| 0 | 1.16 | 3.65 | 0.32 | 0.75 | 1 | Gm7541 | | 0 | 1 | 0 | 0 | 0 | 0 | 0 | 2 | 0 | 0 | 0 |
| 0 | 0.94 | 3.65 | 0.26 | 0.80 | 1 | Gm13902 | | 0 | 0 | 0 | 1 | 0 | 0 | 0 | 1 | 0 | 1 | 0 |
| 0 | 0.94 | 3.65 | 0.26 | 0.80 | 1 | Gm12245 | | 0 | 0 | 0 | 1 | 0 | 0 | 0 | 1 | 0 | 1 | 0 |
| 0 | 0.94 | 3.65 | 0.26 | 0.80 | 1 | Gm6771 | | 0 | 0 | 0 | 1 | 0 | 0 | 0 | 1 | 0 | 1 | 0 |
| 0 | 2.16 | 3.64 | 0.59 | 0.55 | 1 | Gm5267 | | 0 | 0 | 0 | 0 | 0 | 0 | 0 | 0 | 1 | 1 | 1 |
| 0 | 1.00 | 3.68 | 0.27 | 0.79 | 1 | Gm8661 | | 0 | 2 | 0 | 0 | 0 | 0 | 0 | 0 | 0 | 1 | 0 |
| 0 | 0.28 | 3.68 | 0.08 | 0.94 | 1 | Gm10240 | | 0 | 1 | 0 | 0 | 1 | 0 | 0 | 0 | 0 | 1 | 0 |

| | | | | | | | | | | | | | | | | | | |
|---|---|---|---|---|---|---|---|---|---|---|---|---|---|---|---|---|---|---|
| 0 | 1.59 | 3.66 | 0.43 | 0.66 | 1 | Gm8529 | | 0 | 0 | 1 | 0 | 0 | 0 | 0 | 1 | 1 | 0 | 0 |
| 0 | 0.28 | 3.68 | 0.08 | 0.94 | 1 | Rpl30-ps2 | | 0 | 0 | 1 | 1 | 0 | 0 | 0 | 0 | 0 | 1 | 0 |
| 0 | 1.88 | 3.65 | 0.52 | 0.61 | 1 | Gm43245 | | 0 | 0 | 0 | 0 | 0 | 0 | 0 | 1 | 0 | 0 | 2 |
| 0 | 1.88 | 3.65 | 0.52 | 0.61 | 1 | Gm8314 | | 0 | 0 | 0 | 0 | 0 | 0 | 0 | 1 | 0 | 0 | 2 |
| 0 | 1.55 | 3.65 | 0.42 | 0.67 | 1 | Gm5263 | | 0 | 1 | 0 | 0 | 0 | 0 | 0 | 1 | 0 | 0 | 1 |
| 0 | 1.55 | 3.65 | 0.42 | 0.67 | 1 | Rpl31-ps6 | | 0 | 1 | 0 | 0 | 0 | 0 | 0 | 1 | 0 | 0 | 1 |
| 0 | 1.55 | 3.65 | 0.42 | 0.67 | 1 | Gm6275 | | 0 | 1 | 0 | 0 | 0 | 0 | 0 | 1 | 0 | 0 | 1 |
| 0 | 2.15 | 3.64 | 0.59 | 0.55 | 1 | Gm7858 | | 0 | 0 | 0 | 0 | 0 | 0 | 0 | 1 | 1 | 0 | 1 |
| 0 | 1.42 | 3.67 | 0.39 | 0.70 | 1 | Gm10237 | | 1 | 0 | 0 | 0 | 0 | 0 | 0 | 2 | 0 | 0 | 0 |
| 0 | 0.71 | 3.65 | 0.19 | 0.85 | 1 | Tmem229b-ps | | 0 | 0 | 0 | 0 | 0 | 1 | 0 | 2 | 0 | 0 | 0 |
| 0 | 1.55 | 3.66 | 0.43 | 0.67 | 1 | Gm6276 | | 1 | 0 | 0 | 0 | 0 | 0 | 0 | 0 | 0 | 1 | 1 |
| 0 | 0.98 | 3.68 | 0.27 | 0.79 | 1 | Gm15602 | | 1 | 1 | 0 | 0 | 0 | 0 | 0 | 0 | 0 | 1 | 0 |
| 0 | 0.99 | 3.64 | 0.27 | 0.79 | 1 | Gm9341 | | 0 | 0 | 0 | 0 | 0 | 1 | 0 | 0 | 0 | 1 | 1 |
| 0 | 0.56 | 3.65 | 0.15 | 0.88 | 1 | Rpl21-ps11 | | 0 | 0 | 1 | 0 | 1 | 0 | 0 | 1 | 0 | 0 | 0 |
| 0 | 0.00 | 3.68 | 0.00 | 1.00 | 1 | Ccl20 | chemokine (C-C motif) ligand 20 | 0 | 0 | 3 | 0 | 0 | 0 | 0 | 0 | 0 | 0 | 0 |
| 0 | 0.00 | 3.68 | 0.00 | 1.00 | 1 | Otos | otospiralin | 0 | 0 | 3 | 0 | 0 | 0 | 0 | 0 | 0 | 0 | 0 |
| 0 | 0.00 | 3.68 | 0.00 | 1.00 | 1 | Cfap221 | cilia and flagella associated protein 221 | 0 | 0 | 3 | 0 | 0 | 0 | 0 | 0 | 0 | 0 | 0 |
| 0 | 0.00 | 3.68 | 0.00 | 1.00 | 1 | Gm10530 | | 0 | 0 | 3 | 0 | 0 | 0 | 0 | 0 | 0 | 0 | 0 |
| 0 | 0.00 | 3.68 | 0.00 | 1.00 | 1 | Gm37267 | | 0 | 0 | 3 | 0 | 0 | 0 | 0 | 0 | 0 | 0 | 0 |
| 0 | 0.00 | 3.68 | 0.00 | 1.00 | 1 | Gm17167 | | 0 | 0 | 3 | 0 | 0 | 0 | 0 | 0 | 0 | 0 | 0 |
| 0 | 0.00 | 3.68 | 0.00 | 1.00 | 1 | Gm42790 | | 0 | 0 | 3 | 0 | 0 | 0 | 0 | 0 | 0 | 0 | 0 |
| 0 | 0.00 | 3.68 | 0.00 | 1.00 | 1 | Phox2a | paired-like homeobox 2a | 0 | 0 | 3 | 0 | 0 | 0 | 0 | 0 | 0 | 0 | 0 |
| 0 | 0.00 | 3.68 | 0.00 | 1.00 | 1 | Gm19514 | | 0 | 0 | 3 | 0 | 0 | 0 | 0 | 0 | 0 | 0 | 0 |
| 0 | 0.00 | 3.68 | 0.00 | 1.00 | 1 | Gm16159 | | 0 | 0 | 3 | 0 | 0 | 0 | 0 | 0 | 0 | 0 | 0 |
| 0 | 0.00 | 3.68 | 0.00 | 1.00 | 1 | Vmo1 | vitelline membrane outer layer 1 homolog | 0 | 0 | 3 | 0 | 0 | 0 | 0 | 0 | 0 | 0 | 0 |
| 0 | 0.00 | 3.68 | 0.00 | 1.00 | 1 | Agr3 | anterior gradient 3 | 0 | 0 | 3 | 0 | 0 | 0 | 0 | 0 | 0 | 0 | 0 |
| 0 | 0.00 | 3.68 | 0.00 | 1.00 | 1 | Ighv1-52 | | 0 | 0 | 3 | 0 | 0 | 0 | 0 | 0 | 0 | 0 | 0 |
| 0 | 0.00 | 3.68 | 0.00 | 1.00 | 1 | Gm48526 | | 0 | 0 | 3 | 0 | 0 | 0 | 0 | 0 | 0 | 0 | 0 |
| 0 | 0.00 | 3.68 | 0.00 | 1.00 | 1 | Trav13-5 | | 0 | 0 | 3 | 0 | 0 | 0 | 0 | 0 | 0 | 0 | 0 |
| 0 | 0.00 | 3.68 | 0.00 | 1.00 | 1 | Ly6g | lymphocyte antigen 6 complex, locus G | 0 | 0 | 3 | 0 | 0 | 0 | 0 | 0 | 0 | 0 | 0 |
| 0 | 0.00 | 3.68 | 0.00 | 1.00 | 1 | Pla2g10os | | 0 | 0 | 3 | 0 | 0 | 0 | 0 | 0 | 0 | 0 | 0 |
| 0 | 0.00 | 3.68 | 0.00 | 1.00 | 1 | Gm38593 | | 0 | 0 | 3 | 0 | 0 | 0 | 0 | 0 | 0 | 0 | 0 |
| 0 | 0.00 | 3.68 | 0.00 | 1.00 | 1 | Gm18763 | | 0 | 0 | 3 | 0 | 0 | 0 | 0 | 0 | 0 | 0 | 0 |
| 0 | 0.00 | 3.68 | 0.00 | 1.00 | 1 | Gykl1 | glycerol kinase-like 1 | 0 | 0 | 3 | 0 | 0 | 0 | 0 | 0 | 0 | 0 | 0 |
| 0 | 0.00 | 3.68 | 0.00 | 1.00 | 1 | Gm14845 | | 0 | 0 | 3 | 0 | 0 | 0 | 0 | 0 | 0 | 0 | 0 |
| 0 | 0.03 | 3.65 | 0.01 | 0.99 | 1 | Gm12390 | | 0 | 1 | 0 | 0 | 1 | 0 | 0 | 1 | 0 | 0 | 0 |
| 0 | 0.03 | 3.65 | 0.01 | 0.99 | 1 | Gm14760 | | 0 | 1 | 0 | 0 | 1 | 0 | 0 | 1 | 0 | 0 | 0 |
| 0 | 0.91 | 3.65 | 0.25 | 0.80 | 1 | Gm14127 | | 0 | 0 | 0 | 0 | 1 | 0 | 0 | 1 | 1 | 0 | 0 |
| 0 | 0.91 | 3.65 | 0.25 | 0.80 | 1 | Gm17775 | | 0 | 0 | 0 | 0 | 1 | 0 | 0 | 1 | 1 | 0 | 0 |
| 0 | -0.42 | 3.66 | -0.11 | 0.91 | 1 | Gm8420 | | 0 | 0 | 0 | 1 | 1 | 0 | 0 | 0 | 0 | 1 | 0 |
| 0 | 0.95 | 3.67 | 0.26 | 0.80 | 1 | Gm5350 | | 0 | 0 | 2 | 0 | 0 | 0 | 0 | 0 | 0 | 0 | 1 |
| 0 | 0.00 | 3.68 | 0.00 | 1.00 | 1 | Gm5426 | | 0 | 1 | 2 | 0 | 0 | 0 | 0 | 0 | 0 | 0 | 0 |
| 0 | 0.88 | 3.66 | 0.24 | 0.81 | 1 | Gm13430 | | 0 | 0 | 2 | 0 | 0 | 0 | 0 | 0 | 1 | 0 | 0 |
| 0 | 0.88 | 3.66 | 0.24 | 0.81 | 1 | Gm7672 | | 0 | 0 | 2 | 0 | 0 | 0 | 0 | 0 | 1 | 0 | 0 |
| 0 | 1.25 | 3.66 | 0.34 | 0.73 | 1 | Gm17098 | | 0 | 0 | 1 | 0 | 0 | 0 | 0 | 0 | 0 | 0 | 2 |
| 0 | 1.25 | 3.66 | 0.34 | 0.73 | 1 | Gm16106 | | 0 | 0 | 1 | 0 | 0 | 0 | 0 | 0 | 0 | 0 | 2 |
| 0 | 0.95 | 3.67 | 0.26 | 0.80 | 1 | Gm11765 | | 0 | 1 | 1 | 0 | 0 | 0 | 0 | 0 | 0 | 0 | 1 |
| 0 | 1.54 | 3.65 | 0.42 | 0.67 | 1 | Olfr912 | olfactory receptor 912 | 0 | 0 | 1 | 0 | 0 | 0 | 0 | 0 | 1 | 0 | 1 |
| 0 | 0.00 | 3.68 | 0.00 | 1.00 | 1 | L3mbtl4 | L3MBTL4 histone methyl-lysine binding p | 0 | 2 | 1 | 0 | 0 | 0 | 0 | 0 | 0 | 0 | 0 |
| 0 | 0.94 | 3.67 | 0.26 | 0.80 | 1 | Gm12428 | | 0 | 1 | 1 | 0 | 0 | 0 | 0 | 0 | 1 | 0 | 0 |
| 0 | 0.94 | 3.67 | 0.26 | 0.80 | 1 | Gm40319 | | 0 | 1 | 1 | 0 | 0 | 0 | 0 | 0 | 1 | 0 | 0 |
| 0 | 0.94 | 3.67 | 0.26 | 0.80 | 1 | Gm17827 | | 0 | 1 | 1 | 0 | 0 | 0 | 0 | 0 | 1 | 0 | 0 |
| 0 | 0.94 | 3.67 | 0.26 | 0.80 | 1 | Gm5042 | | 0 | 1 | 1 | 0 | 0 | 0 | 0 | 0 | 1 | 0 | 0 |
| 0 | 1.00 | 3.68 | 0.27 | 0.79 | 1 | Gm42402 | | 1 | 0 | 1 | 0 | 0 | 0 | 0 | 1 | 0 | 0 | 0 |
| 0 | 1.00 | 3.68 | 0.27 | 0.79 | 1 | Gm2606 | | 1 | 0 | 1 | 0 | 0 | 0 | 0 | 1 | 0 | 0 | 0 |
| 0 | 0.00 | 3.64 | 0.00 | 1.00 | 1 | Gm43561 | | 0 | 0 | 0 | 0 | 2 | 0 | 0 | 1 | 0 | 0 | 0 |
| 0 | 1.30 | 3.64 | 0.36 | 0.72 | 1 | Gm37629 | | 0 | 0 | 0 | 0 | 0 | 0 | 0 | 0 | 0 | 0 | 3 |
| 0 | 1.30 | 3.64 | 0.36 | 0.72 | 1 | Gm28777 | | 0 | 0 | 0 | 0 | 0 | 0 | 0 | 0 | 0 | 0 | 3 |
| 0 | 1.30 | 3.64 | 0.36 | 0.72 | 1 | Abca12 | ATP-binding cassette, sub-family A (ABC | 0 | 0 | 0 | 0 | 0 | 0 | 0 | 0 | 0 | 0 | 3 |
| 0 | 1.30 | 3.64 | 0.36 | 0.72 | 1 | Gm37171 | | 0 | 0 | 0 | 0 | 0 | 0 | 0 | 0 | 0 | 0 | 3 |

| | | | | | | | | | | | | | | | | | |
|---|---|---|---|---|---|---|---|---|---|---|---|---|---|---|---|---|---|
| 0 | 1.30 | 3.64 | 0.36 | 0.72 | 1 | Gm38073 | | 0 | 0 | 0 | 0 | 0 | 0 | 0 | 0 | 0 | 0 | 3 |
| 0 | 1.30 | 3.64 | 0.36 | 0.72 | 1 | Gm37406 | | 0 | 0 | 0 | 0 | 0 | 0 | 0 | 0 | 0 | 0 | 3 |
| 0 | 1.30 | 3.64 | 0.36 | 0.72 | 1 | Gm13626 | | 0 | 0 | 0 | 0 | 0 | 0 | 0 | 0 | 0 | 0 | 3 |
| 0 | 1.30 | 3.64 | 0.36 | 0.72 | 1 | Gm13787 | | 0 | 0 | 0 | 0 | 0 | 0 | 0 | 0 | 0 | 0 | 3 |
| 0 | 1.30 | 3.64 | 0.36 | 0.72 | 1 | Gm14021 | | 0 | 0 | 0 | 0 | 0 | 0 | 0 | 0 | 0 | 0 | 3 |
| 0 | 1.30 | 3.64 | 0.36 | 0.72 | 1 | AA387200 | | 0 | 0 | 0 | 0 | 0 | 0 | 0 | 0 | 0 | 0 | 3 |
| 0 | 1.30 | 3.64 | 0.36 | 0.72 | 1 | Gm14238 | | 0 | 0 | 0 | 0 | 0 | 0 | 0 | 0 | 0 | 0 | 3 |
| 0 | 1.30 | 3.64 | 0.36 | 0.72 | 1 | Platr29 | | 0 | 0 | 0 | 0 | 0 | 0 | 0 | 0 | 0 | 0 | 3 |
| 0 | 1.30 | 3.64 | 0.36 | 0.72 | 1 | Gm22773 | | 0 | 0 | 0 | 0 | 0 | 0 | 0 | 0 | 0 | 0 | 3 |
| 0 | 1.30 | 3.64 | 0.36 | 0.72 | 1 | Gm12531 | | 0 | 0 | 0 | 0 | 0 | 0 | 0 | 0 | 0 | 0 | 3 |
| 0 | 1.30 | 3.64 | 0.36 | 0.72 | 1 | Gm16100 | | 0 | 0 | 0 | 0 | 0 | 0 | 0 | 0 | 0 | 0 | 3 |
| 0 | 1.30 | 3.64 | 0.36 | 0.72 | 1 | Gm12519 | | 0 | 0 | 0 | 0 | 0 | 0 | 0 | 0 | 0 | 0 | 3 |
| 0 | 1.30 | 3.64 | 0.36 | 0.72 | 1 | Gm12786 | | 0 | 0 | 0 | 0 | 0 | 0 | 0 | 0 | 0 | 0 | 3 |
| 0 | 1.30 | 3.64 | 0.36 | 0.72 | 1 | Kncn | kinocilin | 0 | 0 | 0 | 0 | 0 | 0 | 0 | 0 | 0 | 0 | 3 |
| 0 | 1.30 | 3.64 | 0.36 | 0.72 | 1 | Gm12863 | | 0 | 0 | 0 | 0 | 0 | 0 | 0 | 0 | 0 | 0 | 3 |
| 0 | 1.30 | 3.64 | 0.36 | 0.72 | 1 | Zfp986 | | 0 | 0 | 0 | 0 | 0 | 0 | 0 | 0 | 0 | 0 | 3 |
| 0 | 1.30 | 3.64 | 0.36 | 0.72 | 1 | Mir29a | | 0 | 0 | 0 | 0 | 0 | 0 | 0 | 0 | 0 | 0 | 3 |
| 0 | 1.30 | 3.64 | 0.36 | 0.72 | 1 | Tuba3a | tubulin, alpha 3A | 0 | 0 | 0 | 0 | 0 | 0 | 0 | 0 | 0 | 0 | 3 |
| 0 | 1.30 | 3.64 | 0.36 | 0.72 | 1 | Gm29638 | | 0 | 0 | 0 | 0 | 0 | 0 | 0 | 0 | 0 | 0 | 3 |
| 0 | 1.30 | 3.64 | 0.36 | 0.72 | 1 | Gm21028 | | 0 | 0 | 0 | 0 | 0 | 0 | 0 | 0 | 0 | 0 | 3 |
| 0 | 1.30 | 3.64 | 0.36 | 0.72 | 1 | 1700015G11Ril | RIKEN cDNA 1700015G11 gene | 0 | 0 | 0 | 0 | 0 | 0 | 0 | 0 | 0 | 0 | 3 |
| 0 | 1.30 | 3.64 | 0.36 | 0.72 | 1 | Gm45597 | | 0 | 0 | 0 | 0 | 0 | 0 | 0 | 0 | 0 | 0 | 3 |
| 0 | 1.30 | 3.64 | 0.36 | 0.72 | 1 | Gm32786 | | 0 | 0 | 0 | 0 | 0 | 0 | 0 | 0 | 0 | 0 | 3 |
| 0 | 1.30 | 3.64 | 0.36 | 0.72 | 1 | Got1l1 | glutamic-oxaloacetic transaminase 1-like | 0 | 0 | 0 | 0 | 0 | 0 | 0 | 0 | 0 | 0 | 3 |
| 0 | 1.30 | 3.64 | 0.36 | 0.72 | 1 | Mir181d | | 0 | 0 | 0 | 0 | 0 | 0 | 0 | 0 | 0 | 0 | 3 |
| 0 | 1.30 | 3.64 | 0.36 | 0.72 | 1 | Gapdh-ps16 | | 0 | 0 | 0 | 0 | 0 | 0 | 0 | 0 | 0 | 0 | 3 |
| 0 | 1.30 | 3.64 | 0.36 | 0.72 | 1 | Mir7081 | | 0 | 0 | 0 | 0 | 0 | 0 | 0 | 0 | 0 | 0 | 3 |
| 0 | 1.30 | 3.64 | 0.36 | 0.72 | 1 | 6530413G14Rik | | 0 | 0 | 0 | 0 | 0 | 0 | 0 | 0 | 0 | 0 | 3 |
| 0 | 1.30 | 3.64 | 0.36 | 0.72 | 1 | Gm49380 | | 0 | 0 | 0 | 0 | 0 | 0 | 0 | 0 | 0 | 0 | 3 |
| 0 | 1.30 | 3.64 | 0.36 | 0.72 | 1 | Gm8131 | | 0 | 0 | 0 | 0 | 0 | 0 | 0 | 0 | 0 | 0 | 3 |
| 0 | 1.30 | 3.64 | 0.36 | 0.72 | 1 | Gm17477 | | 0 | 0 | 0 | 0 | 0 | 0 | 0 | 0 | 0 | 0 | 3 |
| 0 | 1.30 | 3.64 | 0.36 | 0.72 | 1 | Trim42 | tripartite motif-containing 42 | 0 | 0 | 0 | 0 | 0 | 0 | 0 | 0 | 0 | 0 | 3 |
| 0 | 1.30 | 3.64 | 0.36 | 0.72 | 1 | Gm47704 | | 0 | 0 | 0 | 0 | 0 | 0 | 0 | 0 | 0 | 0 | 3 |
| 0 | 1.30 | 3.64 | 0.36 | 0.72 | 1 | Gm18514 | | 0 | 0 | 0 | 0 | 0 | 0 | 0 | 0 | 0 | 0 | 3 |
| 0 | 1.30 | 3.64 | 0.36 | 0.72 | 1 | Gm47107 | | 0 | 0 | 0 | 0 | 0 | 0 | 0 | 0 | 0 | 0 | 3 |
| 0 | 1.30 | 3.64 | 0.36 | 0.72 | 1 | Gm26602 | | 0 | 0 | 0 | 0 | 0 | 0 | 0 | 0 | 0 | 0 | 3 |
| 0 | 1.30 | 3.64 | 0.36 | 0.72 | 1 | Mir6918 | | 0 | 0 | 0 | 0 | 0 | 0 | 0 | 0 | 0 | 0 | 3 |
| 0 | 1.30 | 3.64 | 0.36 | 0.72 | 1 | Gm11967 | | 0 | 0 | 0 | 0 | 0 | 0 | 0 | 0 | 0 | 0 | 3 |
| 0 | 1.30 | 3.64 | 0.36 | 0.72 | 1 | Gm12246 | | 0 | 0 | 0 | 0 | 0 | 0 | 0 | 0 | 0 | 0 | 3 |
| 0 | 1.30 | 3.64 | 0.36 | 0.72 | 1 | Wnt3a | wingless-type MMTV integration site famil | 0 | 0 | 0 | 0 | 0 | 0 | 0 | 0 | 0 | 0 | 3 |
| 0 | 1.30 | 3.64 | 0.36 | 0.72 | 1 | Gm27493 | | 0 | 0 | 0 | 0 | 0 | 0 | 0 | 0 | 0 | 0 | 3 |
| 0 | 1.30 | 3.64 | 0.36 | 0.72 | 1 | Gm46317 | | 0 | 0 | 0 | 0 | 0 | 0 | 0 | 0 | 0 | 0 | 3 |
| 0 | 1.30 | 3.64 | 0.36 | 0.72 | 1 | 4732487G21Rik | | 0 | 0 | 0 | 0 | 0 | 0 | 0 | 0 | 0 | 0 | 3 |
| 0 | 1.30 | 3.64 | 0.36 | 0.72 | 1 | Mir337 | | 0 | 0 | 0 | 0 | 0 | 0 | 0 | 0 | 0 | 0 | 3 |
| 0 | 1.30 | 3.64 | 0.36 | 0.72 | 1 | Mir381 | | 0 | 0 | 0 | 0 | 0 | 0 | 0 | 0 | 0 | 0 | 3 |
| 0 | 1.30 | 3.64 | 0.36 | 0.72 | 1 | Mir382 | | 0 | 0 | 0 | 0 | 0 | 0 | 0 | 0 | 0 | 0 | 3 |
| 0 | 1.30 | 3.64 | 0.36 | 0.72 | 1 | Ofcc1 | orofacial cleft 1 candidate 1 | 0 | 0 | 0 | 0 | 0 | 0 | 0 | 0 | 0 | 0 | 3 |
| 0 | 1.30 | 3.64 | 0.36 | 0.72 | 1 | Mir5124a | | 0 | 0 | 0 | 0 | 0 | 0 | 0 | 0 | 0 | 0 | 3 |
| 0 | 1.30 | 3.64 | 0.36 | 0.72 | 1 | Msx2 | msh homeobox 2 | 0 | 0 | 0 | 0 | 0 | 0 | 0 | 0 | 0 | 0 | 3 |
| 0 | 1.30 | 3.64 | 0.36 | 0.72 | 1 | 1700014D04Ril | RIKEN cDNA 1700014D04 gene | 0 | 0 | 0 | 0 | 0 | 0 | 0 | 0 | 0 | 0 | 3 |
| 0 | 1.30 | 3.64 | 0.36 | 0.72 | 1 | 2310067P03Rik | | 0 | 0 | 0 | 0 | 0 | 0 | 0 | 0 | 0 | 0 | 3 |
| 0 | 1.30 | 3.64 | 0.36 | 0.72 | 1 | Gm10406 | predicted gene 10406 | 0 | 0 | 0 | 0 | 0 | 0 | 0 | 0 | 0 | 0 | 3 |
| 0 | 1.30 | 3.64 | 0.36 | 0.72 | 1 | Gm49134 | | 0 | 0 | 0 | 0 | 0 | 0 | 0 | 0 | 0 | 0 | 3 |
| 0 | 1.30 | 3.64 | 0.36 | 0.72 | 1 | Gm20555 | | 0 | 0 | 0 | 0 | 0 | 0 | 0 | 0 | 0 | 0 | 3 |
| 0 | 1.30 | 3.64 | 0.36 | 0.72 | 1 | Mir1907 | | 0 | 0 | 0 | 0 | 0 | 0 | 0 | 0 | 0 | 0 | 3 |
| 0 | 1.30 | 3.64 | 0.36 | 0.72 | 1 | Gm28876 | | 0 | 0 | 0 | 0 | 0 | 0 | 0 | 0 | 0 | 0 | 3 |
| 0 | 1.30 | 3.64 | 0.36 | 0.72 | 1 | Gm24927 | | 0 | 0 | 0 | 0 | 0 | 0 | 0 | 0 | 0 | 0 | 3 |
| 0 | 1.30 | 3.64 | 0.36 | 0.72 | 1 | 4930547E14Rik | | 0 | 0 | 0 | 0 | 0 | 0 | 0 | 0 | 0 | 0 | 3 |
| 0 | 1.30 | 3.64 | 0.36 | 0.72 | 1 | Mir691 | | 0 | 0 | 0 | 0 | 0 | 0 | 0 | 0 | 0 | 0 | 3 |
| 0 | 1.30 | 3.64 | 0.36 | 0.72 | 1 | Zscan10 | zinc finger and SCAN domain containing | 0 | 0 | 0 | 0 | 0 | 0 | 0 | 0 | 0 | 0 | 3 |

| | | | | | | | | | | | | | | | | | | |
|---|---|---|---|---|---|---|---|---|---|---|---|---|---|---|---|---|---|---|
| 0 | 1.30 | 3.64 | 0.36 | 0.72 | 1 | Gm29994 | | 0 | 0 | 0 | 0 | 0 | 0 | 0 | 0 | 0 | 0 | 3 |
| 0 | 1.30 | 3.64 | 0.36 | 0.72 | 1 | Mir6991 | | 0 | 0 | 0 | 0 | 0 | 0 | 0 | 0 | 0 | 0 | 3 |
| 0 | 1.30 | 3.64 | 0.36 | 0.72 | 1 | Mir6993 | | 0 | 0 | 0 | 0 | 0 | 0 | 0 | 0 | 0 | 0 | 3 |
| 0 | 1.30 | 3.64 | 0.36 | 0.72 | 1 | Gm23658 | | 0 | 0 | 0 | 0 | 0 | 0 | 0 | 0 | 0 | 0 | 3 |
| 0 | 1.30 | 3.64 | 0.36 | 0.72 | 1 | Rhox4a | reproductive homeobox 4A | 0 | 0 | 0 | 0 | 0 | 0 | 0 | 0 | 0 | 0 | 3 |
| 0 | 1.30 | 3.64 | 0.36 | 0.72 | 1 | Fsip2l | fibrous sheath-interacting protein 2-like | 0 | 0 | 0 | 0 | 0 | 0 | 0 | 0 | 0 | 0 | 3 |
| 0 | 1.30 | 3.64 | 0.36 | 0.72 | 1 | Gm14861 | | 0 | 0 | 0 | 0 | 0 | 0 | 0 | 0 | 0 | 0 | 3 |
| 0 | 1.30 | 3.64 | 0.36 | 0.72 | 1 | Gm15153 | | 0 | 0 | 0 | 0 | 0 | 0 | 0 | 0 | 0 | 0 | 3 |
| 0 | 1.25 | 3.66 | 0.34 | 0.73 | 1 | Gm7069 | | 0 | 1 | 0 | 0 | 0 | 0 | 0 | 0 | 0 | 0 | 2 |
| 0 | 1.86 | 3.65 | 0.51 | 0.61 | 1 | Gm16185 | | 0 | 0 | 0 | 0 | 0 | 0 | 0 | 0 | 1 | 0 | 2 |
| 0 | 1.86 | 3.65 | 0.51 | 0.61 | 1 | Gm49739 | | 0 | 0 | 0 | 0 | 0 | 0 | 0 | 0 | 1 | 0 | 2 |
| 0 | 1.61 | 3.66 | 0.44 | 0.66 | 1 | Gm7450 | | 0 | 1 | 0 | 0 | 0 | 0 | 0 | 0 | 1 | 0 | 1 |
| 0 | 0.00 | 3.68 | 0.00 | 1.00 | 1 | Gm38033 | | 0 | 3 | 0 | 0 | 0 | 0 | 0 | 0 | 0 | 0 | 0 |
| 0 | 0.00 | 3.68 | 0.00 | 1.00 | 1 | Gm5834 | | 0 | 3 | 0 | 0 | 0 | 0 | 0 | 0 | 0 | 0 | 0 |
| 0 | 0.00 | 3.68 | 0.00 | 1.00 | 1 | Fgfr3-ps | | 0 | 3 | 0 | 0 | 0 | 0 | 0 | 0 | 0 | 0 | 0 |
| 0 | 0.00 | 3.68 | 0.00 | 1.00 | 1 | Cstdc2 | cystatin domain containing 2 | 0 | 3 | 0 | 0 | 0 | 0 | 0 | 0 | 0 | 0 | 0 |
| 0 | 0.00 | 3.68 | 0.00 | 1.00 | 1 | Gm11814 | | 0 | 3 | 0 | 0 | 0 | 0 | 0 | 0 | 0 | 0 | 0 |
| 0 | 0.00 | 3.68 | 0.00 | 1.00 | 1 | Arid3c | AT rich interactive domain 3C (BRIGHT-lil | 0 | 3 | 0 | 0 | 0 | 0 | 0 | 0 | 0 | 0 | 0 |
| 0 | 0.00 | 3.68 | 0.00 | 1.00 | 1 | Gm12439 | | 0 | 3 | 0 | 0 | 0 | 0 | 0 | 0 | 0 | 0 | 0 |
| 0 | 0.00 | 3.68 | 0.00 | 1.00 | 1 | Crygn | crystallin, gamma N | 0 | 3 | 0 | 0 | 0 | 0 | 0 | 0 | 0 | 0 | 0 |
| 0 | 0.00 | 3.68 | 0.00 | 1.00 | 1 | Gm42141 | | 0 | 3 | 0 | 0 | 0 | 0 | 0 | 0 | 0 | 0 | 0 |
| 0 | 0.00 | 3.68 | 0.00 | 1.00 | 1 | Gm14508 | | 0 | 3 | 0 | 0 | 0 | 0 | 0 | 0 | 0 | 0 | 0 |
| 0 | 0.00 | 3.68 | 0.00 | 1.00 | 1 | Gm45060 | | 0 | 3 | 0 | 0 | 0 | 0 | 0 | 0 | 0 | 0 | 0 |
| 0 | 0.00 | 3.68 | 0.00 | 1.00 | 1 | Gm26802 | | 0 | 3 | 0 | 0 | 0 | 0 | 0 | 0 | 0 | 0 | 0 |
| 0 | 0.00 | 3.68 | 0.00 | 1.00 | 1 | Gm32647 | | 0 | 3 | 0 | 0 | 0 | 0 | 0 | 0 | 0 | 0 | 0 |
| 0 | 0.00 | 3.68 | 0.00 | 1.00 | 1 | Gm8291 | | 0 | 3 | 0 | 0 | 0 | 0 | 0 | 0 | 0 | 0 | 0 |
| 0 | 0.00 | 3.68 | 0.00 | 1.00 | 1 | Gm35657 | | 0 | 3 | 0 | 0 | 0 | 0 | 0 | 0 | 0 | 0 | 0 |
| 0 | 0.00 | 3.68 | 0.00 | 1.00 | 1 | Gm47542 | | 0 | 3 | 0 | 0 | 0 | 0 | 0 | 0 | 0 | 0 | 0 |
| 0 | 0.00 | 3.68 | 0.00 | 1.00 | 1 | Gm38063 | | 0 | 3 | 0 | 0 | 0 | 0 | 0 | 0 | 0 | 0 | 0 |
| 0 | 0.00 | 3.68 | 0.00 | 1.00 | 1 | Gm47590 | | 0 | 3 | 0 | 0 | 0 | 0 | 0 | 0 | 0 | 0 | 0 |
| 0 | 0.00 | 3.68 | 0.00 | 1.00 | 1 | Abca13 | ATP-binding cassette, sub-family A (ABC | 0 | 3 | 0 | 0 | 0 | 0 | 0 | 0 | 0 | 0 | 0 |
| 0 | 0.00 | 3.68 | 0.00 | 1.00 | 1 | Serpina6 | serine (or cysteine) peptidase inhibitor, cl | 0 | 3 | 0 | 0 | 0 | 0 | 0 | 0 | 0 | 0 | 0 |
| 0 | 0.00 | 3.68 | 0.00 | 1.00 | 1 | Gm47193 | | 0 | 3 | 0 | 0 | 0 | 0 | 0 | 0 | 0 | 0 | 0 |
| 0 | 0.00 | 3.68 | 0.00 | 1.00 | 1 | Gm33172 | | 0 | 3 | 0 | 0 | 0 | 0 | 0 | 0 | 0 | 0 | 0 |
| 0 | 0.00 | 3.68 | 0.00 | 1.00 | 1 | Gm47202 | | 0 | 3 | 0 | 0 | 0 | 0 | 0 | 0 | 0 | 0 | 0 |
| 0 | 0.00 | 3.68 | 0.00 | 1.00 | 1 | Gm36617 | | 0 | 3 | 0 | 0 | 0 | 0 | 0 | 0 | 0 | 0 | 0 |
| 0 | 0.00 | 3.68 | 0.00 | 1.00 | 1 | Trim31 | tripartite motif-containing 31 | 0 | 3 | 0 | 0 | 0 | 0 | 0 | 0 | 0 | 0 | 0 |
| 0 | 0.00 | 3.68 | 0.00 | 1.00 | 1 | Trem3 | triggering receptor expressed on myeloid | 0 | 3 | 0 | 0 | 0 | 0 | 0 | 0 | 0 | 0 | 0 |
| 0 | 0.00 | 3.68 | 0.00 | 1.00 | 1 | Gm45352 | | 0 | 3 | 0 | 0 | 0 | 0 | 0 | 0 | 0 | 0 | 0 |
| 0 | 0.00 | 3.68 | 0.00 | 1.00 | 1 | 1810035K13Rik | | 0 | 3 | 0 | 0 | 0 | 0 | 0 | 0 | 0 | 0 | 0 |
| 0 | 0.00 | 3.68 | 0.00 | 1.00 | 1 | Mycs | myc-like oncogene, s-myc protein | 0 | 3 | 0 | 0 | 0 | 0 | 0 | 0 | 0 | 0 | 0 |
| 0 | 1.90 | 3.65 | 0.52 | 0.60 | 1 | Hmgn2-ps1 | | 0 | 0 | 0 | 0 | 0 | 0 | 0 | 0 | 2 | 0 | 1 |
| 0 | 1.38 | 3.66 | 0.38 | 0.71 | 1 | Gm19457 | | 0 | 1 | 0 | 0 | 0 | 0 | 0 | 0 | 2 | 0 | 0 |
| 0 | 1.54 | 3.66 | 0.42 | 0.67 | 1 | Gm15852 | | 0 | 0 | 0 | 0 | 0 | 0 | 0 | 0 | 3 | 0 | 0 |
| 0 | 1.54 | 3.66 | 0.42 | 0.67 | 1 | Gm38158 | | 0 | 0 | 0 | 0 | 0 | 0 | 0 | 0 | 3 | 0 | 0 |
| 0 | 1.54 | 3.66 | 0.42 | 0.67 | 1 | Gm34866 | | 0 | 0 | 0 | 0 | 0 | 0 | 0 | 0 | 3 | 0 | 0 |
| 0 | 1.54 | 3.66 | 0.42 | 0.67 | 1 | 1700060J05Rik | | 0 | 0 | 0 | 0 | 0 | 0 | 0 | 0 | 3 | 0 | 0 |
| 0 | 1.54 | 3.66 | 0.42 | 0.67 | 1 | Gm42729 | | 0 | 0 | 0 | 0 | 0 | 0 | 0 | 0 | 3 | 0 | 0 |
| 0 | 1.54 | 3.66 | 0.42 | 0.67 | 1 | Gm26738 | | 0 | 0 | 0 | 0 | 0 | 0 | 0 | 0 | 3 | 0 | 0 |
| 0 | 1.54 | 3.66 | 0.42 | 0.67 | 1 | Gm10988 | | 0 | 0 | 0 | 0 | 0 | 0 | 0 | 0 | 3 | 0 | 0 |
| 0 | 1.54 | 3.66 | 0.42 | 0.67 | 1 | Gm8168 | | 0 | 0 | 0 | 0 | 0 | 0 | 0 | 0 | 3 | 0 | 0 |
| 0 | 1.54 | 3.66 | 0.42 | 0.67 | 1 | Gm15680 | | 0 | 0 | 0 | 0 | 0 | 0 | 0 | 0 | 3 | 0 | 0 |
| 0 | 1.54 | 3.66 | 0.42 | 0.67 | 1 | 5730403I07Rik | | 0 | 0 | 0 | 0 | 0 | 0 | 0 | 0 | 3 | 0 | 0 |
| 0 | 1.54 | 3.66 | 0.42 | 0.67 | 1 | Iqcf5 | IQ motif containing F5 | 0 | 0 | 0 | 0 | 0 | 0 | 0 | 0 | 3 | 0 | 0 |
| 0 | 1.54 | 3.66 | 0.42 | 0.67 | 1 | Gm48278 | | 0 | 0 | 0 | 0 | 0 | 0 | 0 | 0 | 3 | 0 | 0 |
| 0 | 1.54 | 3.66 | 0.42 | 0.67 | 1 | Gm48851 | | 0 | 0 | 0 | 0 | 0 | 0 | 0 | 0 | 3 | 0 | 0 |
| 0 | 1.54 | 3.66 | 0.42 | 0.67 | 1 | Gm3791 | | 0 | 0 | 0 | 0 | 0 | 0 | 0 | 0 | 3 | 0 | 0 |
| 0 | 1.54 | 3.66 | 0.42 | 0.67 | 1 | Gm48797 | | 0 | 0 | 0 | 0 | 0 | 0 | 0 | 0 | 3 | 0 | 0 |
| 0 | 1.54 | 3.66 | 0.42 | 0.67 | 1 | Gm48433 | | 0 | 0 | 0 | 0 | 0 | 0 | 0 | 0 | 3 | 0 | 0 |
| 0 | 1.54 | 3.66 | 0.42 | 0.67 | 1 | Gm5207 | | 0 | 0 | 0 | 0 | 0 | 0 | 0 | 0 | 3 | 0 | 0 |

| | | | | | | | | | | | | | | | | | | |
|---|---|---|---|---|---|---|---|---|---|---|---|---|---|---|---|---|---|---|
| 0 | 1.54 | 3.66 | 0.42 | 0.67 | 1 | Gm49442 | | 0 | 0 | 0 | 0 | 0 | 0 | 0 | 0 | 3 | 0 | 0 |
| 0 | 1.54 | 3.66 | 0.42 | 0.67 | 1 | Gm27023 | | 0 | 0 | 0 | 0 | 0 | 0 | 0 | 0 | 3 | 0 | 0 |
| 0 | 0.99 | 3.64 | 0.27 | 0.79 | 1 | Gm48254 | | 0 | 0 | 0 | 0 | 0 | 1 | 0 | 1 | 0 | 0 | 1 |
| 0 | 0.92 | 3.67 | 0.25 | 0.80 | 1 | Gng5-ps | | 2 | 0 | 0 | 0 | 0 | 0 | 0 | 0 | 0 | 1 | 0 |
| 0 | -0.41 | 3.67 | -0.11 | 0.91 | 1 | Gm7634 | | 0 | 1 | 1 | 0 | 1 | 0 | 0 | 0 | 0 | 0 | 0 |
| 0 | 0.28 | 3.64 | 0.08 | 0.94 | 1 | Gm13622 | | 0 | 0 | 1 | 0 | 1 | 0 | 0 | 0 | 1 | 0 | 0 |
| 0 | 0.28 | 3.64 | 0.08 | 0.94 | 1 | Gm15785 | | 0 | 0 | 1 | 0 | 1 | 0 | 0 | 0 | 1 | 0 | 0 |
| 0 | 0.21 | 3.67 | 0.06 | 0.95 | 1 | Gm8752 | | 1 | 0 | 0 | 0 | 0 | 1 | 0 | 0 | 0 | 1 | 0 |
| 0 | -0.35 | 3.65 | -0.10 | 0.92 | 1 | Gm13313 | | 0 | 0 | 0 | 1 | 1 | 0 | 0 | 1 | 0 | 0 | 0 |
| 0 | 0.36 | 3.64 | 0.10 | 0.92 | 1 | Gm43118 | | 0 | 0 | 0 | 0 | 1 | 0 | 0 | 0 | 0 | 0 | 2 |
| 0 | 0.36 | 3.64 | 0.10 | 0.92 | 1 | 1700126G02Rik | | 0 | 0 | 0 | 0 | 1 | 0 | 0 | 0 | 0 | 0 | 2 |
| 0 | 0.36 | 3.64 | 0.10 | 0.92 | 1 | Gm9502 | | 0 | 0 | 0 | 0 | 1 | 0 | 0 | 0 | 0 | 0 | 2 |
| 0 | 0.36 | 3.64 | 0.10 | 0.92 | 1 | Gm14970 | | 0 | 0 | 0 | 0 | 1 | 0 | 0 | 0 | 0 | 0 | 2 |
| 0 | 0.28 | 3.68 | 0.08 | 0.94 | 1 | mt-Co3 | cytochrome c oxidase subunit III | 0 | 1 | 0 | 0 | 1 | 0 | 0 | 0 | 0 | 0 | 1 |
| 0 | 0.26 | 3.66 | 0.07 | 0.94 | 1 | Rpl19-ps5 | | 0 | 1 | 0 | 0 | 1 | 0 | 0 | 0 | 1 | 0 | 0 |
| 0 | 0.26 | 3.66 | 0.07 | 0.94 | 1 | Gm4910 | | 0 | 1 | 0 | 0 | 1 | 0 | 0 | 0 | 1 | 0 | 0 |
| 0 | 0.60 | 3.64 | 0.16 | 0.87 | 1 | Lrriq4 | leucine-rich repeats and IQ motif containi | 0 | 0 | 0 | 0 | 1 | 0 | 0 | 0 | 2 | 0 | 0 |
| 0 | 0.60 | 3.64 | 0.16 | 0.87 | 1 | Gm38040 | | 0 | 0 | 0 | 0 | 1 | 0 | 0 | 0 | 2 | 0 | 0 |
| 0 | 0.28 | 3.68 | 0.08 | 0.94 | 1 | Gm5305 | | 1 | 0 | 0 | 0 | 1 | 0 | 0 | 1 | 0 | 0 | 0 |
| 0 | 0.00 | 3.68 | 0.00 | 1.00 | 1 | Gm11643 | | 1 | 0 | 2 | 0 | 0 | 0 | 0 | 0 | 0 | 0 | 0 |
| 0 | 0.00 | 3.68 | 0.00 | 1.00 | 1 | Rps19-ps5 | | 1 | 0 | 2 | 0 | 0 | 0 | 0 | 0 | 0 | 0 | 0 |
| 0 | -0.39 | 3.67 | -0.11 | 0.91 | 1 | Gm8662 | | 0 | 1 | 1 | 1 | 0 | 0 | 0 | 0 | 0 | 0 | 0 |
| 0 | 0.07 | 3.65 | 0.02 | 0.98 | 1 | Gm9711 | | 0 | 0 | 1 | 1 | 0 | 0 | 0 | 0 | 1 | 0 | 0 |
| 0 | 0.95 | 3.67 | 0.26 | 0.80 | 1 | Gm9173 | | 1 | 0 | 1 | 0 | 0 | 0 | 0 | 0 | 0 | 0 | 1 |
| 0 | 0.95 | 3.67 | 0.26 | 0.80 | 1 | Gnb1-ps2 | | 1 | 0 | 1 | 0 | 0 | 0 | 0 | 0 | 0 | 0 | 1 |
| 0 | 0.00 | 3.68 | 0.00 | 1.00 | 1 | Gm13418 | | 1 | 1 | 1 | 0 | 0 | 0 | 0 | 0 | 0 | 0 | 0 |
| 0 | 0.00 | 3.68 | 0.00 | 1.00 | 1 | Gm5552 | | 1 | 1 | 1 | 0 | 0 | 0 | 0 | 0 | 0 | 0 | 0 |
| 0 | 0.36 | 3.64 | 0.10 | 0.92 | 1 | Gm29084 | | 0 | 0 | 0 | 1 | 0 | 0 | 0 | 0 | 0 | 0 | 2 |
| 0 | 0.36 | 3.64 | 0.10 | 0.92 | 1 | Gm5967 | | 0 | 0 | 0 | 1 | 0 | 0 | 0 | 0 | 0 | 0 | 2 |
| 0 | -0.11 | 3.65 | -0.03 | 0.98 | 1 | Olfr1238 | olfactory receptor 1238 | 0 | 0 | 0 | 0 | 2 | 0 | 0 | 0 | 0 | 0 | 1 |
| 0 | -0.11 | 3.65 | -0.03 | 0.98 | 1 | Gm48458 | | 0 | 0 | 0 | 0 | 2 | 0 | 0 | 0 | 0 | 0 | 1 |
| 0 | 0.39 | 3.66 | 0.11 | 0.92 | 1 | Gm15740 | | 0 | 1 | 0 | 1 | 0 | 0 | 0 | 0 | 0 | 0 | 1 |
| 0 | -0.90 | 3.66 | -0.25 | 0.81 | 1 | Gm13094 | | 0 | 1 | 0 | 0 | 2 | 0 | 0 | 0 | 0 | 0 | 0 |
| 0 | -0.33 | 3.66 | -0.09 | 0.93 | 1 | Gm24959 | | 0 | 2 | 0 | 1 | 0 | 0 | 0 | 0 | 0 | 0 | 0 |
| 0 | 0.28 | 3.64 | 0.08 | 0.94 | 1 | Gm8132 | | 0 | 0 | 1 | 0 | 0 | 1 | 0 | 0 | 0 | 0 | 1 |
| 0 | 0.28 | 3.64 | 0.08 | 0.94 | 1 | Gm23388 | | 0 | 0 | 1 | 0 | 0 | 1 | 0 | 0 | 0 | 0 | 1 |
| 0 | 0.57 | 3.65 | 0.16 | 0.88 | 1 | Gm10401 | | 0 | 0 | 0 | 1 | 0 | 0 | 0 | 0 | 2 | 0 | 0 |
| 0 | -0.40 | 3.67 | -0.11 | 0.91 | 1 | Gm13537 | | 0 | 1 | 1 | 0 | 0 | 1 | 0 | 0 | 0 | 0 | 0 |
| 0 | -0.40 | 3.67 | -0.11 | 0.91 | 1 | Gm5434 | | 0 | 1 | 1 | 0 | 0 | 1 | 0 | 0 | 0 | 0 | 0 |
| 0 | 1.25 | 3.66 | 0.34 | 0.73 | 1 | Gm13320 | | 1 | 0 | 0 | 0 | 0 | 0 | 0 | 0 | 0 | 0 | 2 |
| 0 | 0.95 | 3.67 | 0.26 | 0.80 | 1 | Gm14231 | | 1 | 1 | 0 | 0 | 0 | 0 | 0 | 0 | 0 | 0 | 1 |
| 0 | 0.95 | 3.67 | 0.26 | 0.80 | 1 | Gm46311 | | 1 | 1 | 0 | 0 | 0 | 0 | 0 | 0 | 0 | 0 | 1 |
| 0 | 1.61 | 3.66 | 0.44 | 0.66 | 1 | Gm13332 | | 1 | 0 | 0 | 0 | 0 | 0 | 0 | 0 | 1 | 0 | 1 |
| 0 | 1.61 | 3.66 | 0.44 | 0.66 | 1 | Gm9078 | | 1 | 0 | 0 | 0 | 0 | 0 | 0 | 0 | 1 | 0 | 1 |
| 0 | 0.00 | 3.68 | 0.00 | 1.00 | 1 | Gm7351 | | 1 | 2 | 0 | 0 | 0 | 0 | 0 | 0 | 0 | 0 | 0 |
| 0 | 0.00 | 3.68 | 0.00 | 1.00 | 1 | Gm8858 | | 1 | 2 | 0 | 0 | 0 | 0 | 0 | 0 | 0 | 0 | 0 |
| 0 | 1.00 | 3.68 | 0.27 | 0.79 | 1 | Gm40598 | | 1 | 1 | 0 | 0 | 0 | 0 | 0 | 0 | 1 | 0 | 0 |
| 0 | 0.54 | 3.63 | 0.15 | 0.88 | 1 | Gm37444 | | 0 | 0 | 0 | 0 | 0 | 1 | 0 | 0 | 0 | 0 | 2 |
| 0 | 0.54 | 3.63 | 0.15 | 0.88 | 1 | Gm18800 | | 0 | 0 | 0 | 0 | 0 | 1 | 0 | 0 | 0 | 0 | 2 |
| 0 | 0.54 | 3.63 | 0.15 | 0.88 | 1 | Gm28403 | | 0 | 0 | 0 | 0 | 0 | 1 | 0 | 0 | 0 | 0 | 2 |
| 0 | 0.54 | 3.63 | 0.15 | 0.88 | 1 | Gm38169 | | 0 | 0 | 0 | 0 | 0 | 1 | 0 | 0 | 0 | 0 | 2 |
| 0 | 0.54 | 3.63 | 0.15 | 0.88 | 1 | Gm13401 | | 0 | 0 | 0 | 0 | 0 | 1 | 0 | 0 | 0 | 0 | 2 |
| 0 | 0.54 | 3.63 | 0.15 | 0.88 | 1 | Gm44027 | | 0 | 0 | 0 | 0 | 0 | 1 | 0 | 0 | 0 | 0 | 2 |
| 0 | 0.54 | 3.63 | 0.15 | 0.88 | 1 | Cyct | cytochrome c, testis | 0 | 0 | 0 | 0 | 0 | 1 | 0 | 0 | 0 | 0 | 2 |
| 0 | 0.54 | 3.63 | 0.15 | 0.88 | 1 | Gm21994 | | 0 | 0 | 0 | 0 | 0 | 1 | 0 | 0 | 0 | 0 | 2 |
| 0 | 0.54 | 3.63 | 0.15 | 0.88 | 1 | Gm25637 | | 0 | 0 | 0 | 0 | 0 | 1 | 0 | 0 | 0 | 0 | 2 |
| 0 | 0.54 | 3.63 | 0.15 | 0.88 | 1 | Gm24346 | | 0 | 0 | 0 | 0 | 0 | 1 | 0 | 0 | 0 | 0 | 2 |
| 0 | 0.54 | 3.63 | 0.15 | 0.88 | 1 | 4930477O15Rik | | 0 | 0 | 0 | 0 | 0 | 1 | 0 | 0 | 0 | 0 | 2 |
| 0 | 0.54 | 3.63 | 0.15 | 0.88 | 1 | Gm3793 | | 0 | 0 | 0 | 0 | 0 | 1 | 0 | 0 | 0 | 0 | 2 |
| 0 | 0.54 | 3.63 | 0.15 | 0.88 | 1 | Phlda2 | pleckstrin homology like domain, family A | 0 | 0 | 0 | 0 | 0 | 1 | 0 | 0 | 0 | 0 | 2 |

| | | | | | | | | | | | | | | | | | | |
|---|---|---|---|---|---|---|---|---|---|---|---|---|---|---|---|---|---|---|
| 0 | 0.54 | 3.63 | 0.15 | 0.88 | 1 | Mir1946b | | 0 | 0 | 0 | 0 | 0 | 1 | 0 | 0 | 0 | 0 | 2 |
| 0 | 0.54 | 3.63 | 0.15 | 0.88 | 1 | Gm48335 | | 0 | 0 | 0 | 0 | 0 | 1 | 0 | 0 | 0 | 0 | 2 |
| 0 | 0.54 | 3.63 | 0.15 | 0.88 | 1 | Gm22442 | | 0 | 0 | 0 | 0 | 0 | 1 | 0 | 0 | 0 | 0 | 2 |
| 0 | 0.54 | 3.63 | 0.15 | 0.88 | 1 | Gm8795 | | 0 | 0 | 0 | 0 | 0 | 1 | 0 | 0 | 0 | 0 | 2 |
| 0 | 0.54 | 3.63 | 0.15 | 0.88 | 1 | Miox | myo-inositol oxygenase | 0 | 0 | 0 | 0 | 0 | 1 | 0 | 0 | 0 | 0 | 2 |
| 0 | 0.54 | 3.63 | 0.15 | 0.88 | 1 | Gm26158 | | 0 | 0 | 0 | 0 | 0 | 1 | 0 | 0 | 0 | 0 | 2 |
| 0 | 0.54 | 3.63 | 0.15 | 0.88 | 1 | Gm8503 | | 0 | 0 | 0 | 0 | 0 | 1 | 0 | 0 | 0 | 0 | 2 |
| 0 | -0.40 | 3.67 | -0.11 | 0.91 | 1 | Gm42849 | | 0 | 2 | 0 | 0 | 0 | 1 | 0 | 0 | 0 | 0 | 0 |
| 0 | -0.40 | 3.67 | -0.11 | 0.91 | 1 | Tas2r108 | taste receptor, type 2, member 108 | 0 | 2 | 0 | 0 | 0 | 1 | 0 | 0 | 0 | 0 | 0 |
| 0 | 0.75 | 3.65 | 0.21 | 0.84 | 1 | Gm44673 | | 0 | 0 | 0 | 0 | 0 | 1 | 0 | 0 | 2 | 0 | 0 |
| 0 | 0.75 | 3.65 | 0.21 | 0.84 | 1 | Gm6524 | | 0 | 0 | 0 | 0 | 0 | 1 | 0 | 0 | 2 | 0 | 0 |
| 0 | 0.75 | 3.65 | 0.21 | 0.84 | 1 | Gm5612 | | 0 | 0 | 0 | 0 | 0 | 1 | 0 | 0 | 2 | 0 | 0 |
| 0 | 0.75 | 3.65 | 0.21 | 0.84 | 1 | Gm28370 | | 0 | 0 | 0 | 0 | 0 | 1 | 0 | 0 | 2 | 0 | 0 |
| 0 | 0.75 | 3.65 | 0.21 | 0.84 | 1 | Gm10143 | | 0 | 0 | 0 | 0 | 0 | 1 | 0 | 0 | 2 | 0 | 0 |
| 0 | 0.28 | 3.68 | 0.08 | 0.94 | 1 | Gm7553 | | 1 | 0 | 0 | 0 | 0 | 1 | 0 | 1 | 0 | 0 | 0 |
| 0 | 0.28 | 3.68 | 0.08 | 0.94 | 1 | Gm12516 | | 1 | 0 | 0 | 0 | 0 | 1 | 0 | 1 | 0 | 0 | 0 |
| 0 | 0.21 | 3.64 | 0.06 | 0.95 | 1 | C730045M19Rik | | 0 | 0 | 0 | 0 | 0 | 2 | 0 | 1 | 0 | 0 | 0 |
| 0 | -0.52 | 3.66 | -0.14 | 0.89 | 1 | Gm5210 | | 0 | 0 | 0 | 1 | 1 | 0 | 0 | 0 | 0 | 0 | 1 |
| 0 | -0.98 | 3.65 | -0.27 | 0.79 | 1 | Gm43438 | | 0 | 0 | 0 | 0 | 3 | 0 | 0 | 0 | 0 | 0 | 0 |
| 0 | -0.98 | 3.65 | -0.27 | 0.79 | 1 | 2610028E06Rik | | 0 | 0 | 0 | 0 | 3 | 0 | 0 | 0 | 0 | 0 | 0 |
| 0 | -0.98 | 3.65 | -0.27 | 0.79 | 1 | C130026L21Rik | | 0 | 0 | 0 | 0 | 3 | 0 | 0 | 0 | 0 | 0 | 0 |
| 0 | -0.98 | 3.65 | -0.27 | 0.79 | 1 | Tmem233 | transmembrane protein 233 | 0 | 0 | 0 | 0 | 3 | 0 | 0 | 0 | 0 | 0 | 0 |
| 0 | -0.98 | 3.65 | -0.27 | 0.79 | 1 | Gm6308 | | 0 | 0 | 0 | 0 | 3 | 0 | 0 | 0 | 0 | 0 | 0 |
| 0 | -0.98 | 3.65 | -0.27 | 0.79 | 1 | Mir7668 | | 0 | 0 | 0 | 0 | 3 | 0 | 0 | 0 | 0 | 0 | 0 |
| 0 | -0.98 | 3.65 | -0.27 | 0.79 | 1 | Gm44695 | | 0 | 0 | 0 | 0 | 3 | 0 | 0 | 0 | 0 | 0 | 0 |
| 0 | -0.98 | 3.65 | -0.27 | 0.79 | 1 | Gm5749 | | 0 | 0 | 0 | 0 | 3 | 0 | 0 | 0 | 0 | 0 | 0 |
| 0 | -0.98 | 3.65 | -0.27 | 0.79 | 1 | B430010I23Rik | | 0 | 0 | 0 | 0 | 3 | 0 | 0 | 0 | 0 | 0 | 0 |
| 0 | -0.98 | 3.65 | -0.27 | 0.79 | 1 | Gm44450 | | 0 | 0 | 0 | 0 | 3 | 0 | 0 | 0 | 0 | 0 | 0 |
| 0 | -0.98 | 3.65 | -0.27 | 0.79 | 1 | Gm45424 | | 0 | 0 | 0 | 0 | 3 | 0 | 0 | 0 | 0 | 0 | 0 |
| 0 | -0.98 | 3.65 | -0.27 | 0.79 | 1 | Iltifb | interleukin 10-related T cell-derived induc | 0 | 0 | 0 | 0 | 3 | 0 | 0 | 0 | 0 | 0 | 0 |
| 0 | -0.98 | 3.65 | -0.27 | 0.79 | 1 | 1700013N06Rik | | 0 | 0 | 0 | 0 | 3 | 0 | 0 | 0 | 0 | 0 | 0 |
| 0 | -0.98 | 3.65 | -0.27 | 0.79 | 1 | Gm48691 | | 0 | 0 | 0 | 0 | 3 | 0 | 0 | 0 | 0 | 0 | 0 |
| 0 | -0.98 | 3.65 | -0.27 | 0.79 | 1 | Gm41361 | | 0 | 0 | 0 | 0 | 3 | 0 | 0 | 0 | 0 | 0 | 0 |
| 0 | -0.98 | 3.65 | -0.27 | 0.79 | 1 | Krt71 | keratin 71 | 0 | 0 | 0 | 0 | 3 | 0 | 0 | 0 | 0 | 0 | 0 |
| 0 | -0.98 | 3.65 | -0.27 | 0.79 | 1 | Atf7ip2 | activating transcription factor 7 interacting | 0 | 0 | 0 | 0 | 3 | 0 | 0 | 0 | 0 | 0 | 0 |
| 0 | -0.98 | 3.65 | -0.27 | 0.79 | 1 | Gm50251 | | 0 | 0 | 0 | 0 | 3 | 0 | 0 | 0 | 0 | 0 | 0 |
| 0 | -0.98 | 3.65 | -0.27 | 0.79 | 1 | Pitx3 | paired-like homeodomain transcription fa | 0 | 0 | 0 | 0 | 3 | 0 | 0 | 0 | 0 | 0 | 0 |
| 0 | -0.98 | 3.65 | -0.27 | 0.79 | 1 | Atp1b4 | ATPase, (Na+)/K+ transporting, beta 4 po | 0 | 0 | 0 | 0 | 3 | 0 | 0 | 0 | 0 | 0 | 0 |
| 0 | -0.98 | 3.65 | -0.27 | 0.79 | 1 | Gm5392 | | 0 | 0 | 0 | 0 | 3 | 0 | 0 | 0 | 0 | 0 | 0 |
| 0 | -1.03 | 3.66 | -0.28 | 0.78 | 1 | Gm12870 | | 0 | 1 | 0 | 1 | 1 | 0 | 0 | 0 | 0 | 0 | 0 |
| 0 | -0.44 | 3.68 | -0.12 | 0.90 | 1 | Gm14214 | | 1 | 1 | 0 | 0 | 1 | 0 | 0 | 0 | 0 | 0 | 0 |
| 0 | -0.44 | 3.68 | -0.12 | 0.90 | 1 | Gm5472 | | 1 | 1 | 0 | 0 | 1 | 0 | 0 | 0 | 0 | 0 | 0 |
| 0 | 0.26 | 3.66 | 0.07 | 0.94 | 1 | Gm2710 | | 1 | 0 | 0 | 0 | 1 | 0 | 0 | 0 | 1 | 0 | 0 |
| 0 | -0.43 | 3.64 | -0.12 | 0.91 | 1 | Gm20536 | | 0 | 0 | 0 | 0 | 1 | 1 | 0 | 0 | 0 | 0 | 1 |
| 0 | -1.13 | 3.66 | -0.31 | 0.76 | 1 | Gm15877 | | 0 | 1 | 0 | 0 | 1 | 1 | 0 | 0 | 0 | 0 | 0 |
| 0 | -0.36 | 3.65 | -0.10 | 0.92 | 1 | Gm12256 | | 0 | 0 | 0 | 0 | 1 | 1 | 0 | 0 | 1 | 0 | 0 |
| 0 | -0.83 | 3.66 | -0.23 | 0.82 | 1 | Rpl21-ps1 | | 0 | 0 | 1 | 2 | 0 | 0 | 0 | 0 | 0 | 0 | 0 |
| 0 | 0.00 | 3.68 | 0.00 | 1.00 | 1 | Gmfg-ps | | 2 | 0 | 1 | 0 | 0 | 0 | 0 | 0 | 0 | 0 | 0 |
| 0 | 0.00 | 3.68 | 0.00 | 1.00 | 1 | Gm46369 | | 2 | 0 | 1 | 0 | 0 | 0 | 0 | 0 | 0 | 0 | 0 |
| 0 | 0.28 | 3.68 | 0.08 | 0.94 | 1 | Gm6317 | | 1 | 0 | 0 | 1 | 0 | 0 | 0 | 0 | 0 | 0 | 1 |
| 0 | -0.86 | 3.67 | -0.24 | 0.81 | 1 | Gm5274 | | 1 | 0 | 0 | 0 | 2 | 0 | 0 | 0 | 0 | 0 | 0 |
| 0 | -0.39 | 3.67 | -0.11 | 0.92 | 1 | Gm32536 | | 1 | 1 | 0 | 1 | 0 | 0 | 0 | 0 | 0 | 0 | 0 |
| 0 | -0.39 | 3.67 | -0.11 | 0.92 | 1 | Gm6743 | | 1 | 1 | 0 | 1 | 0 | 0 | 0 | 0 | 0 | 0 | 0 |
| 0 | -0.40 | 3.67 | -0.11 | 0.91 | 1 | Rpl30-ps3 | | 1 | 0 | 1 | 0 | 0 | 1 | 0 | 0 | 0 | 0 | 0 |
| 0 | -0.41 | 3.64 | -0.11 | 0.91 | 1 | Gm11380 | | 0 | 0 | 0 | 1 | 0 | 1 | 0 | 0 | 0 | 0 | 1 |
| 0 | -1.37 | 3.65 | -0.38 | 0.71 | 1 | Gm13743 | | 0 | 0 | 0 | 0 | 2 | 1 | 0 | 0 | 0 | 0 | 0 |
| 0 | -1.37 | 3.65 | -0.38 | 0.71 | 1 | Rpl39l | ribosomal protein L39-like | 0 | 0 | 0 | 0 | 2 | 1 | 0 | 0 | 0 | 0 | 0 |
| 0 | -0.98 | 3.65 | -0.27 | 0.79 | 1 | Gm9726 | | 0 | 1 | 0 | 1 | 0 | 1 | 0 | 0 | 0 | 0 | 0 |
| 0 | -0.40 | 3.66 | -0.11 | 0.91 | 1 | Gm18899 | | 0 | 0 | 0 | 1 | 0 | 1 | 0 | 0 | 1 | 0 | 0 |
| 0 | 0.95 | 3.67 | 0.26 | 0.80 | 1 | Gm17749 | | 2 | 0 | 0 | 0 | 0 | 0 | 0 | 0 | 0 | 0 | 1 |

| | | | | | | | | | | | | | | | | | | |
|---|---|---|---|---|---|---|---|---|---|---|---|---|---|---|---|---|---|---|
| 0 | 0.95 | 3.67 | 0.26 | 0.80 | 1 | Mir155hg | | 2 | 0 | 0 | 0 | 0 | 0 | 0 | 0 | 0 | 0 | 1 |
| 0 | 0.00 | 3.68 | 0.00 | 1.00 | 1 | Klf14 | Kruppel-like factor 14 | 2 | 1 | 0 | 0 | 0 | 0 | 0 | 0 | 0 | 0 | 0 |
| 0 | 0.00 | 3.68 | 0.00 | 1.00 | 1 | Gm49083 | predicted gene, 49083 | 2 | 1 | 0 | 0 | 0 | 0 | 0 | 0 | 0 | 0 | 0 |
| 0 | 0.00 | 3.68 | 0.00 | 1.00 | 1 | Ms4a10 | membrane-spanning 4-domains, subfami | 2 | 1 | 0 | 0 | 0 | 0 | 0 | 0 | 0 | 0 | 0 |
| 0 | 1.00 | 3.68 | 0.27 | 0.79 | 1 | Gm12319 | | 2 | 0 | 0 | 0 | 0 | 0 | 0 | 0 | 1 | 0 | 0 |
| 0 | 0.28 | 3.64 | 0.08 | 0.94 | 1 | Gm5978 | | 1 | 0 | 0 | 0 | 0 | 1 | 0 | 0 | 0 | 0 | 1 |
| 0 | 0.28 | 3.64 | 0.08 | 0.94 | 1 | Gm9311 | | 1 | 0 | 0 | 0 | 0 | 1 | 0 | 0 | 0 | 0 | 1 |
| 0 | 0.28 | 3.64 | 0.08 | 0.94 | 1 | Gm7722 | | 1 | 0 | 0 | 0 | 0 | 1 | 0 | 0 | 0 | 0 | 1 |
| 0 | -0.40 | 3.67 | -0.11 | 0.91 | 1 | Gm12733 | | 1 | 1 | 0 | 0 | 0 | 1 | 0 | 0 | 0 | 0 | 0 |
| 0 | -0.40 | 3.67 | -0.11 | 0.91 | 1 | Gm16335 | | 1 | 1 | 0 | 0 | 0 | 1 | 0 | 0 | 0 | 0 | 0 |
| 0 | 0.28 | 3.68 | 0.08 | 0.94 | 1 | Gm12734 | | 1 | 0 | 0 | 0 | 0 | 1 | 0 | 0 | 1 | 0 | 0 |
| 0 | 0.01 | 3.63 | 0.00 | 1.00 | 1 | Gm43116 | | 0 | 0 | 0 | 0 | 0 | 2 | 0 | 0 | 0 | 0 | 1 |
| 0 | 0.01 | 3.63 | 0.00 | 1.00 | 1 | Gm42867 | | 0 | 0 | 0 | 0 | 0 | 2 | 0 | 0 | 0 | 0 | 1 |
| 0 | 0.01 | 3.63 | 0.00 | 1.00 | 1 | Gm43316 | | 0 | 0 | 0 | 0 | 0 | 2 | 0 | 0 | 0 | 0 | 1 |
| 0 | 0.01 | 3.63 | 0.00 | 1.00 | 1 | Gm44555 | | 0 | 0 | 0 | 0 | 0 | 2 | 0 | 0 | 0 | 0 | 1 |
| 0 | 0.01 | 3.63 | 0.00 | 1.00 | 1 | Gm17816 | | 0 | 0 | 0 | 0 | 0 | 2 | 0 | 0 | 0 | 0 | 1 |
| 0 | 0.01 | 3.63 | 0.00 | 1.00 | 1 | Gm26064 | | 0 | 0 | 0 | 0 | 0 | 2 | 0 | 0 | 0 | 0 | 1 |
| 0 | 0.01 | 3.63 | 0.00 | 1.00 | 1 | Gm19077 | | 0 | 0 | 0 | 0 | 0 | 2 | 0 | 0 | 0 | 0 | 1 |
| 0 | 0.01 | 3.63 | 0.00 | 1.00 | 1 | Gm41409 | | 0 | 0 | 0 | 0 | 0 | 2 | 0 | 0 | 0 | 0 | 1 |
| 0 | 0.01 | 3.63 | 0.00 | 1.00 | 1 | Gm16298 | | 0 | 0 | 0 | 0 | 0 | 2 | 0 | 0 | 0 | 0 | 1 |
| 0 | -0.69 | 3.66 | -0.19 | 0.85 | 1 | Rnu1a1 | | 0 | 1 | 0 | 0 | 0 | 2 | 0 | 0 | 0 | 0 | 0 |
| 0 | -0.69 | 3.66 | -0.19 | 0.85 | 1 | Gm49699 | | 0 | 1 | 0 | 0 | 0 | 2 | 0 | 0 | 0 | 0 | 0 |
| 0 | 0.21 | 3.64 | 0.06 | 0.95 | 1 | Gm5533 | | 0 | 0 | 0 | 0 | 0 | 2 | 0 | 0 | 1 | 0 | 0 |
| 0 | 0.21 | 3.64 | 0.06 | 0.95 | 1 | Eif1-ps3 | | 0 | 0 | 0 | 0 | 0 | 2 | 0 | 0 | 1 | 0 | 0 |
| 0 | 0.21 | 3.64 | 0.06 | 0.95 | 1 | 6330419E04Rik | | 0 | 0 | 0 | 0 | 0 | 2 | 0 | 0 | 1 | 0 | 0 |
| 0 | 0.21 | 3.64 | 0.06 | 0.95 | 1 | Gm37871 | | 0 | 0 | 0 | 0 | 0 | 2 | 0 | 0 | 1 | 0 | 0 |
| 0 | 0.21 | 3.64 | 0.06 | 0.95 | 1 | 4930455C13Rik | | 0 | 0 | 0 | 0 | 0 | 2 | 0 | 0 | 1 | 0 | 0 |
| 0 | -1.33 | 3.65 | -0.36 | 0.72 | 1 | Gm4471 | | 0 | 0 | 0 | 2 | 1 | 0 | 0 | 0 | 0 | 0 | 0 |
| 0 | -0.44 | 3.68 | -0.12 | 0.90 | 1 | Rbmx2-ps | | 2 | 0 | 0 | 0 | 1 | 0 | 0 | 0 | 0 | 0 | 0 |
| 0 | -0.44 | 3.68 | -0.12 | 0.90 | 1 | Gm5322 | | 2 | 0 | 0 | 0 | 1 | 0 | 0 | 0 | 0 | 0 | 0 |
| 0 | -1.08 | 3.67 | -0.29 | 0.77 | 1 | Gm6905 | | 1 | 0 | 0 | 0 | 1 | 1 | 0 | 0 | 0 | 0 | 0 |
| 0 | -1.33 | 3.65 | -0.36 | 0.72 | 1 | Gm37055 | | 0 | 0 | 0 | 0 | 1 | 2 | 0 | 0 | 0 | 0 | 0 |
| 0 | -1.33 | 3.65 | -0.36 | 0.72 | 1 | Gm49548 | | 0 | 0 | 0 | 0 | 1 | 2 | 0 | 0 | 0 | 0 | 0 |
| 0 | -0.99 | 3.66 | -0.27 | 0.79 | 1 | Gm5696 | | 0 | 0 | 0 | 3 | 0 | 0 | 0 | 0 | 0 | 0 | 0 |
| 0 | -0.99 | 3.66 | -0.27 | 0.79 | 1 | Gm43666 | | 0 | 0 | 0 | 3 | 0 | 0 | 0 | 0 | 0 | 0 | 0 |
| 0 | -0.99 | 3.66 | -0.27 | 0.79 | 1 | Gm4248 | | 0 | 0 | 0 | 3 | 0 | 0 | 0 | 0 | 0 | 0 | 0 |
| 0 | -0.99 | 3.66 | -0.27 | 0.79 | 1 | Gm37270 | | 0 | 0 | 0 | 3 | 0 | 0 | 0 | 0 | 0 | 0 | 0 |
| 0 | -0.99 | 3.66 | -0.27 | 0.79 | 1 | Usp46os2 | | 0 | 0 | 0 | 3 | 0 | 0 | 0 | 0 | 0 | 0 | 0 |
| 0 | -0.99 | 3.66 | -0.27 | 0.79 | 1 | Wnt7a | wingless-type MMTV integration site famil | 0 | 0 | 0 | 3 | 0 | 0 | 0 | 0 | 0 | 0 | 0 |
| 0 | -0.99 | 3.66 | -0.27 | 0.79 | 1 | Gm48855 | | 0 | 0 | 0 | 3 | 0 | 0 | 0 | 0 | 0 | 0 | 0 |
| 0 | -0.99 | 3.66 | -0.27 | 0.79 | 1 | Rpl15-ps1 | | 0 | 0 | 0 | 3 | 0 | 0 | 0 | 0 | 0 | 0 | 0 |
| 0 | -1.35 | 3.65 | -0.37 | 0.71 | 1 | Gm43112 | | 0 | 0 | 0 | 2 | 0 | 1 | 0 | 0 | 0 | 0 | 0 |
| 0 | -1.35 | 3.65 | -0.37 | 0.71 | 1 | Rmst | | 0 | 0 | 0 | 2 | 0 | 1 | 0 | 0 | 0 | 0 | 0 |
| 0 | -0.44 | 3.68 | -0.12 | 0.90 | 1 | Gm17214 | | 2 | 0 | 0 | 1 | 0 | 0 | 0 | 0 | 0 | 0 | 0 |
| 0 | -1.06 | 3.67 | -0.29 | 0.77 | 1 | Gm12950 | | 1 | 0 | 0 | 1 | 0 | 1 | 0 | 0 | 0 | 0 | 0 |
| 0 | 1.38 | 3.66 | 0.38 | 0.71 | 1 | Gm25911 | | 0 | 0 | 0 | 0 | 0 | 0 | 0 | 0 | 0 | 3 | 0 |
| 0 | 1.38 | 3.66 | 0.38 | 0.71 | 1 | Gm37579 | | 0 | 0 | 0 | 0 | 0 | 0 | 0 | 0 | 0 | 3 | 0 |
| 0 | 1.38 | 3.66 | 0.38 | 0.71 | 1 | Gm15607 | | 0 | 0 | 0 | 0 | 0 | 0 | 0 | 0 | 0 | 3 | 0 |
| 0 | 1.38 | 3.66 | 0.38 | 0.71 | 1 | Gm15766 | | 0 | 0 | 0 | 0 | 0 | 0 | 0 | 0 | 0 | 3 | 0 |
| 0 | 1.38 | 3.66 | 0.38 | 0.71 | 1 | 1700112L15Rik | | 0 | 0 | 0 | 0 | 0 | 0 | 0 | 0 | 0 | 3 | 0 |
| 0 | 1.38 | 3.66 | 0.38 | 0.71 | 1 | Gm47620 | | 0 | 0 | 0 | 0 | 0 | 0 | 0 | 0 | 0 | 3 | 0 |
| 0 | 1.38 | 3.66 | 0.38 | 0.71 | 1 | Trav4-3 | | 0 | 0 | 0 | 0 | 0 | 0 | 0 | 0 | 0 | 3 | 0 |
| 0 | 1.38 | 3.66 | 0.38 | 0.71 | 1 | Gm6788 | | 0 | 0 | 0 | 0 | 0 | 0 | 0 | 0 | 0 | 3 | 0 |
| 0 | 0.00 | 3.68 | 0.00 | 1.00 | 1 | Gm16152 | | 3 | 0 | 0 | 0 | 0 | 0 | 0 | 0 | 0 | 0 | 0 |
| 0 | 0.00 | 3.68 | 0.00 | 1.00 | 1 | Gm16026 | | 3 | 0 | 0 | 0 | 0 | 0 | 0 | 0 | 0 | 0 | 0 |
| 0 | 0.00 | 3.68 | 0.00 | 1.00 | 1 | Gm16332 | | 3 | 0 | 0 | 0 | 0 | 0 | 0 | 0 | 0 | 0 | 0 |
| 0 | 0.00 | 3.68 | 0.00 | 1.00 | 1 | Ptgs2os | | 3 | 0 | 0 | 0 | 0 | 0 | 0 | 0 | 0 | 0 | 0 |
| 0 | 0.00 | 3.68 | 0.00 | 1.00 | 1 | Gm37273 | | 3 | 0 | 0 | 0 | 0 | 0 | 0 | 0 | 0 | 0 | 0 |
| 0 | 0.00 | 3.68 | 0.00 | 1.00 | 1 | Gm4846 | predicted gene 4846 | 3 | 0 | 0 | 0 | 0 | 0 | 0 | 0 | 0 | 0 | 0 |
| 0 | 0.00 | 3.68 | 0.00 | 1.00 | 1 | Gm13293 | | 3 | 0 | 0 | 0 | 0 | 0 | 0 | 0 | 0 | 0 | 0 |

| | | | | | | | | | | | | | | | | | | |
|---|---|---|---|---|---|---|---|---|---|---|---|---|---|---|---|---|---|---|
| 0 | 0.00 | 3.68 | 0.00 | 1.00 | 1 | Gm25857 | | 3 | 0 | 0 | 0 | 0 | 0 | 0 | 0 | 0 | 0 | 0 |
| 0 | 0.00 | 3.68 | 0.00 | 1.00 | 1 | Gm11474 | | 3 | 0 | 0 | 0 | 0 | 0 | 0 | 0 | 0 | 0 | 0 |
| 0 | 0.00 | 3.68 | 0.00 | 1.00 | 1 | Il21 | interleukin 21 | 3 | 0 | 0 | 0 | 0 | 0 | 0 | 0 | 0 | 0 | 0 |
| 0 | 0.00 | 3.68 | 0.00 | 1.00 | 1 | Gm42899 | | 3 | 0 | 0 | 0 | 0 | 0 | 0 | 0 | 0 | 0 | 0 |
| 0 | 0.00 | 3.68 | 0.00 | 1.00 | 1 | Myoz2 | myozenin 2 | 3 | 0 | 0 | 0 | 0 | 0 | 0 | 0 | 0 | 0 | 0 |
| 0 | 0.00 | 3.68 | 0.00 | 1.00 | 1 | Gm43010 | | 3 | 0 | 0 | 0 | 0 | 0 | 0 | 0 | 0 | 0 | 0 |
| 0 | 0.00 | 3.68 | 0.00 | 1.00 | 1 | Slc1a7 | solute carrier family 1 (glutamate transpor | 3 | 0 | 0 | 0 | 0 | 0 | 0 | 0 | 0 | 0 | 0 |
| 0 | 0.00 | 3.68 | 0.00 | 1.00 | 1 | Gm21411 | | 3 | 0 | 0 | 0 | 0 | 0 | 0 | 0 | 0 | 0 | 0 |
| 0 | 0.00 | 3.68 | 0.00 | 1.00 | 1 | Gm13160 | | 3 | 0 | 0 | 0 | 0 | 0 | 0 | 0 | 0 | 0 | 0 |
| 0 | 0.00 | 3.68 | 0.00 | 1.00 | 1 | Gm5861 | predicted gene 5861 | 3 | 0 | 0 | 0 | 0 | 0 | 0 | 0 | 0 | 0 | 0 |
| 0 | 0.00 | 3.68 | 0.00 | 1.00 | 1 | Cwh43 | cell wall biogenesis 43 C-terminal homolo | 3 | 0 | 0 | 0 | 0 | 0 | 0 | 0 | 0 | 0 | 0 |
| 0 | 0.00 | 3.68 | 0.00 | 1.00 | 1 | Ccdc63 | coiled-coil domain containing 63 | 3 | 0 | 0 | 0 | 0 | 0 | 0 | 0 | 0 | 0 | 0 |
| 0 | 0.00 | 3.68 | 0.00 | 1.00 | 1 | Gm43703 | | 3 | 0 | 0 | 0 | 0 | 0 | 0 | 0 | 0 | 0 | 0 |
| 0 | 0.00 | 3.68 | 0.00 | 1.00 | 1 | Gm43921 | | 3 | 0 | 0 | 0 | 0 | 0 | 0 | 0 | 0 | 0 | 0 |
| 0 | 0.00 | 3.68 | 0.00 | 1.00 | 1 | Gm29430 | | 3 | 0 | 0 | 0 | 0 | 0 | 0 | 0 | 0 | 0 | 0 |
| 0 | 0.00 | 3.68 | 0.00 | 1.00 | 1 | Igkv1-99 | | 3 | 0 | 0 | 0 | 0 | 0 | 0 | 0 | 0 | 0 | 0 |
| 0 | 0.00 | 3.68 | 0.00 | 1.00 | 1 | Igkv4-59 | | 3 | 0 | 0 | 0 | 0 | 0 | 0 | 0 | 0 | 0 | 0 |
| 0 | 0.00 | 3.68 | 0.00 | 1.00 | 1 | Chst13 | carbohydrate sulfotransferase 13 | 3 | 0 | 0 | 0 | 0 | 0 | 0 | 0 | 0 | 0 | 0 |
| 0 | 0.00 | 3.68 | 0.00 | 1.00 | 1 | Ceacam19 | carcinoembryonic antigen-related cell adl | 3 | 0 | 0 | 0 | 0 | 0 | 0 | 0 | 0 | 0 | 0 |
| 0 | 0.00 | 3.68 | 0.00 | 1.00 | 1 | Gm50477 | | 3 | 0 | 0 | 0 | 0 | 0 | 0 | 0 | 0 | 0 | 0 |
| 0 | 0.00 | 3.68 | 0.00 | 1.00 | 1 | Gm45179 | | 3 | 0 | 0 | 0 | 0 | 0 | 0 | 0 | 0 | 0 | 0 |
| 0 | 0.00 | 3.68 | 0.00 | 1.00 | 1 | Gm34280 | | 3 | 0 | 0 | 0 | 0 | 0 | 0 | 0 | 0 | 0 | 0 |
| 0 | 0.00 | 3.68 | 0.00 | 1.00 | 1 | Hmx2 | H6 homeobox 2 | 3 | 0 | 0 | 0 | 0 | 0 | 0 | 0 | 0 | 0 | 0 |
| 0 | 0.00 | 3.68 | 0.00 | 1.00 | 1 | Gm45397 | | 3 | 0 | 0 | 0 | 0 | 0 | 0 | 0 | 0 | 0 | 0 |
| 0 | 0.00 | 3.68 | 0.00 | 1.00 | 1 | Impg1 | interphotoreceptor matrix proteoglycan 1 | 3 | 0 | 0 | 0 | 0 | 0 | 0 | 0 | 0 | 0 | 0 |
| 0 | 0.00 | 3.68 | 0.00 | 1.00 | 1 | Samd3 | sterile alpha motif domain containing 3 | 3 | 0 | 0 | 0 | 0 | 0 | 0 | 0 | 0 | 0 | 0 |
| 0 | 0.00 | 3.68 | 0.00 | 1.00 | 1 | 4930551I15Rik | | 3 | 0 | 0 | 0 | 0 | 0 | 0 | 0 | 0 | 0 | 0 |
| 0 | 0.00 | 3.68 | 0.00 | 1.00 | 1 | Epyc | epiphycan | 3 | 0 | 0 | 0 | 0 | 0 | 0 | 0 | 0 | 0 | 0 |
| 0 | 0.00 | 3.68 | 0.00 | 1.00 | 1 | Hoxb5os | | 3 | 0 | 0 | 0 | 0 | 0 | 0 | 0 | 0 | 0 | 0 |
| 0 | 0.00 | 3.68 | 0.00 | 1.00 | 1 | Hspb9 | heat shock protein, alpha-crystallin-relate | 3 | 0 | 0 | 0 | 0 | 0 | 0 | 0 | 0 | 0 | 0 |
| 0 | 0.00 | 3.68 | 0.00 | 1.00 | 1 | Gm11694 | | 3 | 0 | 0 | 0 | 0 | 0 | 0 | 0 | 0 | 0 | 0 |
| 0 | 0.00 | 3.68 | 0.00 | 1.00 | 1 | Cdhr3 | cadherin-related family member 3 | 3 | 0 | 0 | 0 | 0 | 0 | 0 | 0 | 0 | 0 | 0 |
| 0 | 0.00 | 3.68 | 0.00 | 1.00 | 1 | 9130015A21Rik | | 3 | 0 | 0 | 0 | 0 | 0 | 0 | 0 | 0 | 0 | 0 |
| 0 | 0.00 | 3.68 | 0.00 | 1.00 | 1 | Gm40893 | | 3 | 0 | 0 | 0 | 0 | 0 | 0 | 0 | 0 | 0 | 0 |
| 0 | 0.00 | 3.68 | 0.00 | 1.00 | 1 | 9530050K03Rik | | 3 | 0 | 0 | 0 | 0 | 0 | 0 | 0 | 0 | 0 | 0 |
| 0 | 0.00 | 3.68 | 0.00 | 1.00 | 1 | Ighj3 | | 3 | 0 | 0 | 0 | 0 | 0 | 0 | 0 | 0 | 0 | 0 |
| 0 | 0.00 | 3.68 | 0.00 | 1.00 | 1 | Ighv2-5 | | 3 | 0 | 0 | 0 | 0 | 0 | 0 | 0 | 0 | 0 | 0 |
| 0 | 0.00 | 3.68 | 0.00 | 1.00 | 1 | Ighv1-17 | | 3 | 0 | 0 | 0 | 0 | 0 | 0 | 0 | 0 | 0 | 0 |
| 0 | 0.00 | 3.68 | 0.00 | 1.00 | 1 | Ighv1-21 | | 3 | 0 | 0 | 0 | 0 | 0 | 0 | 0 | 0 | 0 | 0 |
| 0 | 0.00 | 3.68 | 0.00 | 1.00 | 1 | Gm31946 | | 3 | 0 | 0 | 0 | 0 | 0 | 0 | 0 | 0 | 0 | 0 |
| 0 | 0.00 | 3.68 | 0.00 | 1.00 | 1 | 7330404K18Rik | | 3 | 0 | 0 | 0 | 0 | 0 | 0 | 0 | 0 | 0 | 0 |
| 0 | 0.00 | 3.68 | 0.00 | 1.00 | 1 | Gm34589 | | 3 | 0 | 0 | 0 | 0 | 0 | 0 | 0 | 0 | 0 | 0 |
| 0 | 0.00 | 3.68 | 0.00 | 1.00 | 1 | Gm15422 | | 3 | 0 | 0 | 0 | 0 | 0 | 0 | 0 | 0 | 0 | 0 |
| 0 | 0.00 | 3.68 | 0.00 | 1.00 | 1 | Gm17473 | | 3 | 0 | 0 | 0 | 0 | 0 | 0 | 0 | 0 | 0 | 0 |
| 0 | 0.00 | 3.68 | 0.00 | 1.00 | 1 | Gm49553 | | 3 | 0 | 0 | 0 | 0 | 0 | 0 | 0 | 0 | 0 | 0 |
| 0 | 0.00 | 3.68 | 0.00 | 1.00 | 1 | Soat2 | sterol O-acyltransferase 2 | 3 | 0 | 0 | 0 | 0 | 0 | 0 | 0 | 0 | 0 | 0 |
| 0 | 0.00 | 3.68 | 0.00 | 1.00 | 1 | Gm42870 | | 3 | 0 | 0 | 0 | 0 | 0 | 0 | 0 | 0 | 0 | 0 |
| 0 | 0.00 | 3.68 | 0.00 | 1.00 | 1 | Btnl4 | butyrophilin-like 4 | 3 | 0 | 0 | 0 | 0 | 0 | 0 | 0 | 0 | 0 | 0 |
| 0 | 0.00 | 3.68 | 0.00 | 1.00 | 1 | Gm36201 | | 3 | 0 | 0 | 0 | 0 | 0 | 0 | 0 | 0 | 0 | 0 |
| 0 | 0.00 | 3.68 | 0.00 | 1.00 | 1 | Gm47898 | | 3 | 0 | 0 | 0 | 0 | 0 | 0 | 0 | 0 | 0 | 0 |
| 0 | 0.00 | 3.68 | 0.00 | 1.00 | 1 | Carmn | | 3 | 0 | 0 | 0 | 0 | 0 | 0 | 0 | 0 | 0 | 0 |
| 0 | 0.00 | 3.68 | 0.00 | 1.00 | 1 | Esx1 | extraembryonic, spermatogenesis, homeo | 3 | 0 | 0 | 0 | 0 | 0 | 0 | 0 | 0 | 0 | 0 |
| 0 | -1.31 | 3.65 | -0.36 | 0.72 | 1 | Gm42768 | | 0 | 0 | 0 | 1 | 0 | 2 | 0 | 0 | 0 | 0 | 0 |
| 0 | -0.40 | 3.67 | -0.11 | 0.91 | 1 | Rpl31-ps24 | | 2 | 0 | 0 | 0 | 0 | 1 | 0 | 0 | 0 | 0 | 0 |
| 0 | -0.40 | 3.67 | -0.11 | 0.91 | 1 | Gm34821 | | 2 | 0 | 0 | 0 | 0 | 1 | 0 | 0 | 0 | 0 | 0 |
| 0 | -0.40 | 3.67 | -0.11 | 0.91 | 1 | Gm15798 | | 2 | 0 | 0 | 0 | 0 | 1 | 0 | 0 | 0 | 0 | 0 |
| 0 | -0.40 | 3.67 | -0.11 | 0.91 | 1 | Gm3946 | | 2 | 0 | 0 | 0 | 0 | 1 | 0 | 0 | 0 | 0 | 0 |
| 0 | -0.40 | 3.67 | -0.11 | 0.91 | 1 | Gm16265 | | 2 | 0 | 0 | 0 | 0 | 1 | 0 | 0 | 0 | 0 | 0 |
| 0 | -0.69 | 3.66 | -0.19 | 0.85 | 1 | Ak3l2-ps | | 1 | 0 | 0 | 0 | 0 | 2 | 0 | 0 | 0 | 0 | 0 |
| 0 | -0.69 | 3.66 | -0.19 | 0.85 | 1 | Gm6439 | | 1 | 0 | 0 | 0 | 0 | 2 | 0 | 0 | 0 | 0 | 0 |

| | | | | | | | | | | | | | | | | | | |
|---|---|---|---|---|---|---|---|---|---|---|---|---|---|---|---|---|---|---|
| 0 | -0.73 | 3.64 | -0.20 | 0.84 | 1 | Il17f | interleukin 17F | 0 | 0 | 0 | 0 | 0 | 3 | 0 | 0 | 0 | 0 | 0 |
| 0 | -0.73 | 3.64 | -0.20 | 0.84 | 1 | Gm37011 | | 0 | 0 | 0 | 0 | 0 | 3 | 0 | 0 | 0 | 0 | 0 |
| 0 | -0.73 | 3.64 | -0.20 | 0.84 | 1 | Gm38012 | | 0 | 0 | 0 | 0 | 0 | 3 | 0 | 0 | 0 | 0 | 0 |
| 0 | -0.73 | 3.64 | -0.20 | 0.84 | 1 | Gm25939 | | 0 | 0 | 0 | 0 | 0 | 3 | 0 | 0 | 0 | 0 | 0 |
| 0 | -0.73 | 3.64 | -0.20 | 0.84 | 1 | Gm10555 | | 0 | 0 | 0 | 0 | 0 | 3 | 0 | 0 | 0 | 0 | 0 |
| 0 | -0.73 | 3.64 | -0.20 | 0.84 | 1 | Gm29480 | | 0 | 0 | 0 | 0 | 0 | 3 | 0 | 0 | 0 | 0 | 0 |
| 0 | -0.73 | 3.64 | -0.20 | 0.84 | 1 | Mir6345 | | 0 | 0 | 0 | 0 | 0 | 3 | 0 | 0 | 0 | 0 | 0 |
| 0 | -0.73 | 3.64 | -0.20 | 0.84 | 1 | Gm17634 | | 0 | 0 | 0 | 0 | 0 | 3 | 0 | 0 | 0 | 0 | 0 |
| 0 | -0.73 | 3.64 | -0.20 | 0.84 | 1 | Gm18182 | | 0 | 0 | 0 | 0 | 0 | 3 | 0 | 0 | 0 | 0 | 0 |
| 0 | -0.73 | 3.64 | -0.20 | 0.84 | 1 | Gm3851 | | 0 | 0 | 0 | 0 | 0 | 3 | 0 | 0 | 0 | 0 | 0 |
| 0 | -0.73 | 3.64 | -0.20 | 0.84 | 1 | Gm28277 | | 0 | 0 | 0 | 0 | 0 | 3 | 0 | 0 | 0 | 0 | 0 |
| 0 | -0.73 | 3.64 | -0.20 | 0.84 | 1 | Mroh3 | maestro heat-like repeat family member 3 | 0 | 0 | 0 | 0 | 0 | 3 | 0 | 0 | 0 | 0 | 0 |
| 0 | -0.73 | 3.64 | -0.20 | 0.84 | 1 | 4930562F07Rik | | 0 | 0 | 0 | 0 | 0 | 3 | 0 | 0 | 0 | 0 | 0 |
| 0 | -0.73 | 3.64 | -0.20 | 0.84 | 1 | Nhlh1 | nescient helix loop helix 1 | 0 | 0 | 0 | 0 | 0 | 3 | 0 | 0 | 0 | 0 | 0 |
| 0 | -0.73 | 3.64 | -0.20 | 0.84 | 1 | Mir6904 | | 0 | 0 | 0 | 0 | 0 | 3 | 0 | 0 | 0 | 0 | 0 |
| 0 | -0.73 | 3.64 | -0.20 | 0.84 | 1 | Gm38359 | | 0 | 0 | 0 | 0 | 0 | 3 | 0 | 0 | 0 | 0 | 0 |
| 0 | -0.73 | 3.64 | -0.20 | 0.84 | 1 | Gm37132 | | 0 | 0 | 0 | 0 | 0 | 3 | 0 | 0 | 0 | 0 | 0 |
| 0 | -0.73 | 3.64 | -0.20 | 0.84 | 1 | Gm38014 | | 0 | 0 | 0 | 0 | 0 | 3 | 0 | 0 | 0 | 0 | 0 |
| 0 | -0.73 | 3.64 | -0.20 | 0.84 | 1 | Gm25220 | | 0 | 0 | 0 | 0 | 0 | 3 | 0 | 0 | 0 | 0 | 0 |
| 0 | -0.73 | 3.64 | -0.20 | 0.84 | 1 | Gm13397 | | 0 | 0 | 0 | 0 | 0 | 3 | 0 | 0 | 0 | 0 | 0 |
| 0 | -0.73 | 3.64 | -0.20 | 0.84 | 1 | Gm25541 | | 0 | 0 | 0 | 0 | 0 | 3 | 0 | 0 | 0 | 0 | 0 |
| 0 | -0.73 | 3.64 | -0.20 | 0.84 | 1 | Gm22516 | | 0 | 0 | 0 | 0 | 0 | 3 | 0 | 0 | 0 | 0 | 0 |
| 0 | -0.73 | 3.64 | -0.20 | 0.84 | 1 | Olfr361 | olfactory receptor 361 | 0 | 0 | 0 | 0 | 0 | 3 | 0 | 0 | 0 | 0 | 0 |
| 0 | -0.73 | 3.64 | -0.20 | 0.84 | 1 | Gm13786 | | 0 | 0 | 0 | 0 | 0 | 3 | 0 | 0 | 0 | 0 | 0 |
| 0 | -0.73 | 3.64 | -0.20 | 0.84 | 1 | Mir129-2 | | 0 | 0 | 0 | 0 | 0 | 3 | 0 | 0 | 0 | 0 | 0 |
| 0 | -0.73 | 3.64 | -0.20 | 0.84 | 1 | Gm13999 | | 0 | 0 | 0 | 0 | 0 | 3 | 0 | 0 | 0 | 0 | 0 |
| 0 | -0.73 | 3.64 | -0.20 | 0.84 | 1 | Gm14397 | | 0 | 0 | 0 | 0 | 0 | 3 | 0 | 0 | 0 | 0 | 0 |
| 0 | -0.73 | 3.64 | -0.20 | 0.84 | 1 | Mir7004 | | 0 | 0 | 0 | 0 | 0 | 3 | 0 | 0 | 0 | 0 | 0 |
| 0 | -0.73 | 3.64 | -0.20 | 0.84 | 1 | Gm12582 | | 0 | 0 | 0 | 0 | 0 | 3 | 0 | 0 | 0 | 0 | 0 |
| 0 | -0.73 | 3.64 | -0.20 | 0.84 | 1 | Gm37256 | | 0 | 0 | 0 | 0 | 0 | 3 | 0 | 0 | 0 | 0 | 0 |
| 0 | -0.73 | 3.64 | -0.20 | 0.84 | 1 | Gm37773 | | 0 | 0 | 0 | 0 | 0 | 3 | 0 | 0 | 0 | 0 | 0 |
| 0 | -0.73 | 3.64 | -0.20 | 0.84 | 1 | Gm43750 | | 0 | 0 | 0 | 0 | 0 | 3 | 0 | 0 | 0 | 0 | 0 |
| 0 | -0.73 | 3.64 | -0.20 | 0.84 | 1 | Gm42873 | | 0 | 0 | 0 | 0 | 0 | 3 | 0 | 0 | 0 | 0 | 0 |
| 0 | -0.73 | 3.64 | -0.20 | 0.84 | 1 | Gm568 | | 0 | 0 | 0 | 0 | 0 | 3 | 0 | 0 | 0 | 0 | 0 |
| 0 | -0.73 | 3.64 | -0.20 | 0.84 | 1 | Gm12867 | | 0 | 0 | 0 | 0 | 0 | 3 | 0 | 0 | 0 | 0 | 0 |
| 0 | -0.73 | 3.64 | -0.20 | 0.84 | 1 | Zfp988 | gastrula zinc finger protein XlCGF26.1-lik | 0 | 0 | 0 | 0 | 0 | 3 | 0 | 0 | 0 | 0 | 0 |
| 0 | -0.73 | 3.64 | -0.20 | 0.84 | 1 | Gm42975 | | 0 | 0 | 0 | 0 | 0 | 3 | 0 | 0 | 0 | 0 | 0 |
| 0 | -0.73 | 3.64 | -0.20 | 0.84 | 1 | Gm22887 | | 0 | 0 | 0 | 0 | 0 | 3 | 0 | 0 | 0 | 0 | 0 |
| 0 | -0.73 | 3.64 | -0.20 | 0.84 | 1 | Gm10471 | predicted gene 10471 | 0 | 0 | 0 | 0 | 0 | 3 | 0 | 0 | 0 | 0 | 0 |
| 0 | -0.73 | 3.64 | -0.20 | 0.84 | 1 | Gm43317 | | 0 | 0 | 0 | 0 | 0 | 3 | 0 | 0 | 0 | 0 | 0 |
| 0 | -0.73 | 3.64 | -0.20 | 0.84 | 1 | Gm42522 | | 0 | 0 | 0 | 0 | 0 | 3 | 0 | 0 | 0 | 0 | 0 |
| 0 | -0.73 | 3.64 | -0.20 | 0.84 | 1 | Mir5619 | | 0 | 0 | 0 | 0 | 0 | 3 | 0 | 0 | 0 | 0 | 0 |
| 0 | -0.73 | 3.64 | -0.20 | 0.84 | 1 | Gm43422 | | 0 | 0 | 0 | 0 | 0 | 3 | 0 | 0 | 0 | 0 | 0 |
| 0 | -0.73 | 3.64 | -0.20 | 0.84 | 1 | Gm43275 | | 0 | 0 | 0 | 0 | 0 | 3 | 0 | 0 | 0 | 0 | 0 |
| 0 | -0.73 | 3.64 | -0.20 | 0.84 | 1 | 1700008B11Rik | | 0 | 0 | 0 | 0 | 0 | 3 | 0 | 0 | 0 | 0 | 0 |
| 0 | -0.73 | 3.64 | -0.20 | 0.84 | 1 | Gm43461 | | 0 | 0 | 0 | 0 | 0 | 3 | 0 | 0 | 0 | 0 | 0 |
| 0 | -0.73 | 3.64 | -0.20 | 0.84 | 1 | Asz1 | ankyrin repeat, SAM and basic leucine zip | 0 | 0 | 0 | 0 | 0 | 3 | 0 | 0 | 0 | 0 | 0 |
| 0 | -0.73 | 3.64 | -0.20 | 0.84 | 1 | Gm43629 | | 0 | 0 | 0 | 0 | 0 | 3 | 0 | 0 | 0 | 0 | 0 |
| 0 | -0.73 | 3.64 | -0.20 | 0.84 | 1 | Gm43876 | | 0 | 0 | 0 | 0 | 0 | 3 | 0 | 0 | 0 | 0 | 0 |
| 0 | -0.73 | 3.64 | -0.20 | 0.84 | 1 | Gm25260 | | 0 | 0 | 0 | 0 | 0 | 3 | 0 | 0 | 0 | 0 | 0 |
| 0 | -0.73 | 3.64 | -0.20 | 0.84 | 1 | Gm40377 | | 0 | 0 | 0 | 0 | 0 | 3 | 0 | 0 | 0 | 0 | 0 |
| 0 | -0.73 | 3.64 | -0.20 | 0.84 | 1 | Nat8f5 | N-acetyltransferase 8 (GCN5-related) fam | 0 | 0 | 0 | 0 | 0 | 3 | 0 | 0 | 0 | 0 | 0 |
| 0 | -0.73 | 3.64 | -0.20 | 0.84 | 1 | Gm43946 | | 0 | 0 | 0 | 0 | 0 | 3 | 0 | 0 | 0 | 0 | 0 |
| 0 | -0.73 | 3.64 | -0.20 | 0.84 | 1 | Gm44193 | | 0 | 0 | 0 | 0 | 0 | 3 | 0 | 0 | 0 | 0 | 0 |
| 0 | -0.73 | 3.64 | -0.20 | 0.84 | 1 | Gm38947 | | 0 | 0 | 0 | 0 | 0 | 3 | 0 | 0 | 0 | 0 | 0 |
| 0 | -0.73 | 3.64 | -0.20 | 0.84 | 1 | Gm44873 | | 0 | 0 | 0 | 0 | 0 | 3 | 0 | 0 | 0 | 0 | 0 |
| 0 | -0.73 | 3.64 | -0.20 | 0.84 | 1 | Gm44627 | | 0 | 0 | 0 | 0 | 0 | 3 | 0 | 0 | 0 | 0 | 0 |
| 0 | -0.73 | 3.64 | -0.20 | 0.84 | 1 | Gm45169 | | 0 | 0 | 0 | 0 | 0 | 3 | 0 | 0 | 0 | 0 | 0 |
| 0 | -0.73 | 3.64 | -0.20 | 0.84 | 1 | Gm44704 | | 0 | 0 | 0 | 0 | 0 | 3 | 0 | 0 | 0 | 0 | 0 |
| 0 | -0.73 | 3.64 | -0.20 | 0.84 | 1 | Gm45793 | | 0 | 0 | 0 | 0 | 0 | 3 | 0 | 0 | 0 | 0 | 0 |

| | | | | | | | | | | | | | | | | | | |
|---|---|---|---|---|---|---|---|---|---|---|---|---|---|---|---|---|---|---|
| 0 | -0.73 | 3.64 | -0.20 | 0.84 | 1 | Gm45129 | | 0 | 0 | 0 | 0 | 0 | 3 | 0 | 0 | 0 | 0 | 0 |
| 0 | -0.73 | 3.64 | -0.20 | 0.84 | 1 | Gm45846 | | 0 | 0 | 0 | 0 | 0 | 3 | 0 | 0 | 0 | 0 | 0 |
| 0 | -0.73 | 3.64 | -0.20 | 0.84 | 1 | Mir7059 | | 0 | 0 | 0 | 0 | 0 | 3 | 0 | 0 | 0 | 0 | 0 |
| 0 | -0.73 | 3.64 | -0.20 | 0.84 | 1 | Gm28198 | | 0 | 0 | 0 | 0 | 0 | 3 | 0 | 0 | 0 | 0 | 0 |
| 0 | -0.73 | 3.64 | -0.20 | 0.84 | 1 | Gm44624 | | 0 | 0 | 0 | 0 | 0 | 3 | 0 | 0 | 0 | 0 | 0 |
| 0 | -0.73 | 3.64 | -0.20 | 0.84 | 1 | Gm39119 | | 0 | 0 | 0 | 0 | 0 | 3 | 0 | 0 | 0 | 0 | 0 |
| 0 | -0.73 | 3.64 | -0.20 | 0.84 | 1 | Gm27409 | | 0 | 0 | 0 | 0 | 0 | 3 | 0 | 0 | 0 | 0 | 0 |
| 0 | -0.73 | 3.64 | -0.20 | 0.84 | 1 | Gm45484 | | 0 | 0 | 0 | 0 | 0 | 3 | 0 | 0 | 0 | 0 | 0 |
| 0 | -0.73 | 3.64 | -0.20 | 0.84 | 1 | Olfr374 | olfactory receptor 374 | 0 | 0 | 0 | 0 | 0 | 3 | 0 | 0 | 0 | 0 | 0 |
| 0 | -0.73 | 3.64 | -0.20 | 0.84 | 1 | 1700011L22Rik | RIKEN cDNA 1700011L22 gene | 0 | 0 | 0 | 0 | 0 | 3 | 0 | 0 | 0 | 0 | 0 |
| 0 | -0.73 | 3.64 | -0.20 | 0.84 | 1 | Ucp1 | uncoupling protein 1 (mitochondrial, protc | 0 | 0 | 0 | 0 | 0 | 3 | 0 | 0 | 0 | 0 | 0 |
| 0 | -0.73 | 3.64 | -0.20 | 0.84 | 1 | D830024N08Rik | | 0 | 0 | 0 | 0 | 0 | 3 | 0 | 0 | 0 | 0 | 0 |
| 0 | -0.73 | 3.64 | -0.20 | 0.84 | 1 | Gm45608 | | 0 | 0 | 0 | 0 | 0 | 3 | 0 | 0 | 0 | 0 | 0 |
| 0 | -0.73 | 3.64 | -0.20 | 0.84 | 1 | Fth-ps3 | | 0 | 0 | 0 | 0 | 0 | 3 | 0 | 0 | 0 | 0 | 0 |
| 0 | -0.73 | 3.64 | -0.20 | 0.84 | 1 | Gm26739 | | 0 | 0 | 0 | 0 | 0 | 3 | 0 | 0 | 0 | 0 | 0 |
| 0 | -0.73 | 3.64 | -0.20 | 0.84 | 1 | Gm48646 | | 0 | 0 | 0 | 0 | 0 | 3 | 0 | 0 | 0 | 0 | 0 |
| 0 | -0.73 | 3.64 | -0.20 | 0.84 | 1 | Gm8049 | | 0 | 0 | 0 | 0 | 0 | 3 | 0 | 0 | 0 | 0 | 0 |
| 0 | -0.73 | 3.64 | -0.20 | 0.84 | 1 | Gm27161 | | 0 | 0 | 0 | 0 | 0 | 3 | 0 | 0 | 0 | 0 | 0 |
| 0 | -0.73 | 3.64 | -0.20 | 0.84 | 1 | Gm28119 | | 0 | 0 | 0 | 0 | 0 | 3 | 0 | 0 | 0 | 0 | 0 |
| 0 | -0.73 | 3.64 | -0.20 | 0.84 | 1 | Gm22141 | | 0 | 0 | 0 | 0 | 0 | 3 | 0 | 0 | 0 | 0 | 0 |
| 0 | -0.73 | 3.64 | -0.20 | 0.84 | 1 | Mir7087 | | 0 | 0 | 0 | 0 | 0 | 3 | 0 | 0 | 0 | 0 | 0 |
| 0 | -0.73 | 3.64 | -0.20 | 0.84 | 1 | Gm11149 | | 0 | 0 | 0 | 0 | 0 | 3 | 0 | 0 | 0 | 0 | 0 |
| 0 | -0.73 | 3.64 | -0.20 | 0.84 | 1 | Gm25064 | | 0 | 0 | 0 | 0 | 0 | 3 | 0 | 0 | 0 | 0 | 0 |
| 0 | -0.73 | 3.64 | -0.20 | 0.84 | 1 | Gm25640 | | 0 | 0 | 0 | 0 | 0 | 3 | 0 | 0 | 0 | 0 | 0 |
| 0 | -0.73 | 3.64 | -0.20 | 0.84 | 1 | Gm18291 | | 0 | 0 | 0 | 0 | 0 | 3 | 0 | 0 | 0 | 0 | 0 |
| 0 | -0.73 | 3.64 | -0.20 | 0.84 | 1 | Gm18655 | | 0 | 0 | 0 | 0 | 0 | 3 | 0 | 0 | 0 | 0 | 0 |
| 0 | -0.73 | 3.64 | -0.20 | 0.84 | 1 | Gm47303 | | 0 | 0 | 0 | 0 | 0 | 3 | 0 | 0 | 0 | 0 | 0 |
| 0 | -0.73 | 3.64 | -0.20 | 0.84 | 1 | Gm25341 | | 0 | 0 | 0 | 0 | 0 | 3 | 0 | 0 | 0 | 0 | 0 |
| 0 | -0.73 | 3.64 | -0.20 | 0.84 | 1 | | | 0 | 0 | 0 | 0 | 0 | 3 | 0 | 0 | 0 | 0 | 0 |
| 0 | -0.73 | 3.64 | -0.20 | 0.84 | 1 | Gm8249 | | 0 | 0 | 0 | 0 | 0 | 3 | 0 | 0 | 0 | 0 | 0 |
| 0 | -0.73 | 3.64 | -0.20 | 0.84 | 1 | Gm47082 | | 0 | 0 | 0 | 0 | 0 | 3 | 0 | 0 | 0 | 0 | 0 |
| 0 | -0.73 | 3.64 | -0.20 | 0.84 | 1 | 1700086D15Rik | | 0 | 0 | 0 | 0 | 0 | 3 | 0 | 0 | 0 | 0 | 0 |
| 0 | -0.73 | 3.64 | -0.20 | 0.84 | 1 | Gm11419 | | 0 | 0 | 0 | 0 | 0 | 3 | 0 | 0 | 0 | 0 | 0 |
| 0 | -0.73 | 3.64 | -0.20 | 0.84 | 1 | Gm11683 | | 0 | 0 | 0 | 0 | 0 | 3 | 0 | 0 | 0 | 0 | 0 |
| 0 | -0.73 | 3.64 | -0.20 | 0.84 | 1 | Trim80 | tripartite motif-containing 80 | 0 | 0 | 0 | 0 | 0 | 3 | 0 | 0 | 0 | 0 | 0 |
| 0 | -0.73 | 3.64 | -0.20 | 0.84 | 1 | Gm11737 | | 0 | 0 | 0 | 0 | 0 | 3 | 0 | 0 | 0 | 0 | 0 |
| 0 | -0.73 | 3.64 | -0.20 | 0.84 | 1 | Gm25948 | | 0 | 0 | 0 | 0 | 0 | 3 | 0 | 0 | 0 | 0 | 0 |
| 0 | -0.73 | 3.64 | -0.20 | 0.84 | 1 | Gm21941 | | 0 | 0 | 0 | 0 | 0 | 3 | 0 | 0 | 0 | 0 | 0 |
| 0 | -0.73 | 3.64 | -0.20 | 0.84 | 1 | Gm9292 | | 0 | 0 | 0 | 0 | 0 | 3 | 0 | 0 | 0 | 0 | 0 |
| 0 | -0.73 | 3.64 | -0.20 | 0.84 | 1 | Gm35135 | | 0 | 0 | 0 | 0 | 0 | 3 | 0 | 0 | 0 | 0 | 0 |
| 0 | -0.73 | 3.64 | -0.20 | 0.84 | 1 | Gm23296 | | 0 | 0 | 0 | 0 | 0 | 3 | 0 | 0 | 0 | 0 | 0 |
| 0 | -0.73 | 3.64 | -0.20 | 0.84 | 1 | Mir681 | | 0 | 0 | 0 | 0 | 0 | 3 | 0 | 0 | 0 | 0 | 0 |
| 0 | -0.73 | 3.64 | -0.20 | 0.84 | 1 | Gm22079 | | 0 | 0 | 0 | 0 | 0 | 3 | 0 | 0 | 0 | 0 | 0 |
| 0 | -0.73 | 3.64 | -0.20 | 0.84 | 1 | Mir409 | | 0 | 0 | 0 | 0 | 0 | 3 | 0 | 0 | 0 | 0 | 0 |
| 0 | -0.73 | 3.64 | -0.20 | 0.84 | 1 | Gm48887 | | 0 | 0 | 0 | 0 | 0 | 3 | 0 | 0 | 0 | 0 | 0 |
| 0 | -0.73 | 3.64 | -0.20 | 0.84 | 1 | Gm11290 | | 0 | 0 | 0 | 0 | 0 | 3 | 0 | 0 | 0 | 0 | 0 |
| 0 | -0.73 | 3.64 | -0.20 | 0.84 | 1 | 1700015C15Rik | | 0 | 0 | 0 | 0 | 0 | 3 | 0 | 0 | 0 | 0 | 0 |
| 0 | -0.73 | 3.64 | -0.20 | 0.84 | 1 | 5330416C01Rik | | 0 | 0 | 0 | 0 | 0 | 3 | 0 | 0 | 0 | 0 | 0 |
| 0 | -0.73 | 3.64 | -0.20 | 0.84 | 1 | Gm3298 | | 0 | 0 | 0 | 0 | 0 | 3 | 0 | 0 | 0 | 0 | 0 |
| 0 | -0.73 | 3.64 | -0.20 | 0.84 | 1 | Gm3629 | | 0 | 0 | 0 | 0 | 0 | 3 | 0 | 0 | 0 | 0 | 0 |
| 0 | -0.73 | 3.64 | -0.20 | 0.84 | 1 | Gm47921 | | 0 | 0 | 0 | 0 | 0 | 3 | 0 | 0 | 0 | 0 | 0 |
| 0 | -0.73 | 3.64 | -0.20 | 0.84 | 1 | Trav7-6 | | 0 | 0 | 0 | 0 | 0 | 3 | 0 | 0 | 0 | 0 | 0 |
| 0 | -0.73 | 3.64 | -0.20 | 0.84 | 1 | Gm49197 | | 0 | 0 | 0 | 0 | 0 | 3 | 0 | 0 | 0 | 0 | 0 |
| 0 | -0.73 | 3.64 | -0.20 | 0.84 | 1 | Gcat | glycine C-acetyltransferase (2-amino-3-ke | 0 | 0 | 0 | 0 | 0 | 3 | 0 | 0 | 0 | 0 | 0 |
| 0 | -0.73 | 3.64 | -0.20 | 0.84 | 1 | Mir6392 | | 0 | 0 | 0 | 0 | 0 | 3 | 0 | 0 | 0 | 0 | 0 |
| 0 | -0.73 | 3.64 | -0.20 | 0.84 | 1 | Gm24145 | | 0 | 0 | 0 | 0 | 0 | 3 | 0 | 0 | 0 | 0 | 0 |
| 0 | -0.73 | 3.64 | -0.20 | 0.84 | 1 | Gm15738 | | 0 | 0 | 0 | 0 | 0 | 3 | 0 | 0 | 0 | 0 | 0 |
| 0 | -0.73 | 3.64 | -0.20 | 0.84 | 1 | Gm49584 | | 0 | 0 | 0 | 0 | 0 | 3 | 0 | 0 | 0 | 0 | 0 |
| 0 | -0.73 | 3.64 | -0.20 | 0.84 | 1 | n-R5s33 | | 0 | 0 | 0 | 0 | 0 | 3 | 0 | 0 | 0 | 0 | 0 |
| 0 | -0.73 | 3.64 | -0.20 | 0.84 | 1 | Gm37606 | | 0 | 0 | 0 | 0 | 0 | 3 | 0 | 0 | 0 | 0 | 0 |

| | | | | | | | | | | | | | | | | | |
|---|---|---|---|---|---|---|---|---|---|---|---|---|---|---|---|---|---|
| 0 | -0.73 | 3.64 | -0.20 | 0.84 | 1 | Gm7912 | | 0 | 0 | 0 | 0 | 0 | 3 | 0 | 0 | 0 | 0 | 0 |
| 0 | -0.73 | 3.64 | -0.20 | 0.84 | 1 | Gm23111 | | 0 | 0 | 0 | 0 | 0 | 3 | 0 | 0 | 0 | 0 | 0 |
| 0 | -0.73 | 3.64 | -0.20 | 0.84 | 1 | Gm17228 | | 0 | 0 | 0 | 0 | 0 | 3 | 0 | 0 | 0 | 0 | 0 |
| 0 | -0.73 | 3.64 | -0.20 | 0.84 | 1 | Gm47471 | | 0 | 0 | 0 | 0 | 0 | 3 | 0 | 0 | 0 | 0 | 0 |
| 0 | -0.73 | 3.64 | -0.20 | 0.84 | 1 | Gm18581 | | 0 | 0 | 0 | 0 | 0 | 3 | 0 | 0 | 0 | 0 | 0 |
| 0 | -0.73 | 3.64 | -0.20 | 0.84 | 1 | Gm4834 | | 0 | 0 | 0 | 0 | 0 | 3 | 0 | 0 | 0 | 0 | 0 |
| 0 | -0.73 | 3.64 | -0.20 | 0.84 | 1 | A430019L02Rik | | 0 | 0 | 0 | 0 | 0 | 3 | 0 | 0 | 0 | 0 | 0 |
| 0 | -0.73 | 3.64 | -0.20 | 0.84 | 1 | Gm50382 | | 0 | 0 | 0 | 0 | 0 | 3 | 0 | 0 | 0 | 0 | 0 |
| 0 | -0.73 | 3.64 | -0.20 | 0.84 | 1 | Gm14529 | | 0 | 0 | 0 | 0 | 0 | 3 | 0 | 0 | 0 | 0 | 0 |
| 0 | -0.73 | 3.64 | -0.20 | 0.84 | 1 | Gm15008 | | 0 | 0 | 0 | 0 | 0 | 3 | 0 | 0 | 0 | 0 | 0 |
| 0 | -0.73 | 3.64 | -0.20 | 0.84 | 1 | Rhox3g | reproductive homeobox 3G | 0 | 0 | 0 | 0 | 0 | 3 | 0 | 0 | 0 | 0 | 0 |
| 0 | -0.73 | 3.64 | -0.20 | 0.84 | 1 | Gm9103 | | 0 | 0 | 0 | 0 | 0 | 3 | 0 | 0 | 0 | 0 | 0 |
| 0 | -0.73 | 3.64 | -0.20 | 0.84 | 1 | Gm25107 | | 0 | 0 | 0 | 0 | 0 | 3 | 0 | 0 | 0 | 0 | 0 |
| 0 | 1.61 | 3.66 | 0.44 | 0.66 | 1 | Gm7393 | | 0 | 0 | 0 | 0 | 0 | 0 | 0 | 1 | 0 | 1 | 0 |
| 0 | 1.00 | 3.68 | 0.27 | 0.79 | 1 | Gm49550 | | 0 | 0 | 1 | 0 | 0 | 0 | 0 | 0 | 0 | 1 | 0 |
| 0 | 1.00 | 3.68 | 0.27 | 0.79 | 1 | Gm14450 | | 0 | 0 | 1 | 0 | 0 | 0 | 0 | 0 | 0 | 1 | 0 |
| 0 | 1.36 | 3.66 | 0.37 | 0.71 | 1 | Mir804 | | 0 | 0 | 0 | 0 | 0 | 0 | 0 | 2 | 0 | 0 | 0 |
| 0 | 1.36 | 3.66 | 0.37 | 0.71 | 1 | Gm17805 | | 0 | 0 | 0 | 0 | 0 | 0 | 0 | 2 | 0 | 0 | 0 |
| 0 | 1.61 | 3.66 | 0.44 | 0.66 | 1 | Gm13359 | | 0 | 0 | 0 | 0 | 0 | 0 | 0 | 0 | 0 | 1 | 1 |
| 0 | 1.61 | 3.66 | 0.44 | 0.66 | 1 | Gm5303 | | 0 | 0 | 0 | 0 | 0 | 0 | 0 | 0 | 0 | 1 | 1 |
| 0 | 1.61 | 3.66 | 0.44 | 0.66 | 1 | Gm15763 | | 0 | 0 | 0 | 0 | 0 | 0 | 0 | 0 | 0 | 1 | 1 |
| 0 | 1.61 | 3.66 | 0.44 | 0.66 | 1 | Gm4995 | | 0 | 0 | 0 | 0 | 0 | 0 | 0 | 0 | 0 | 1 | 1 |
| 0 | 1.00 | 3.68 | 0.27 | 0.79 | 1 | Gm11686 | | 0 | 1 | 0 | 0 | 0 | 0 | 0 | 0 | 0 | 1 | 0 |
| 0 | 1.00 | 3.68 | 0.27 | 0.79 | 1 | Gm9830 | | 0 | 1 | 0 | 0 | 0 | 0 | 0 | 0 | 0 | 1 | 0 |
| 0 | 1.00 | 3.68 | 0.27 | 0.79 | 1 | Gm11343 | | 0 | 1 | 0 | 0 | 0 | 0 | 0 | 0 | 0 | 1 | 0 |
| 0 | 0.28 | 3.68 | 0.08 | 0.94 | 1 | Gm14094 | | 0 | 0 | 0 | 0 | 1 | 0 | 0 | 0 | 0 | 1 | 0 |
| 0 | 0.28 | 3.68 | 0.08 | 0.94 | 1 | Gm2157 | | 0 | 0 | 0 | 0 | 1 | 0 | 0 | 0 | 0 | 1 | 0 |
| 0 | 0.28 | 3.68 | 0.08 | 0.94 | 1 | Gm4708 | | 0 | 0 | 0 | 0 | 1 | 0 | 0 | 0 | 0 | 1 | 0 |
| 0 | 0.28 | 3.68 | 0.08 | 0.94 | 1 | Prkaca-ps1 | | 0 | 0 | 0 | 0 | 1 | 0 | 0 | 0 | 0 | 1 | 0 |
| 0 | 1.00 | 3.68 | 0.27 | 0.79 | 1 | Gm42666 | | 0 | 0 | 1 | 0 | 0 | 0 | 0 | 1 | 0 | 0 | 0 |
| 0 | 1.00 | 3.68 | 0.27 | 0.79 | 1 | Gm46649 | | 0 | 0 | 1 | 0 | 0 | 0 | 0 | 1 | 0 | 0 | 0 |
| 0 | 1.60 | 3.66 | 0.44 | 0.66 | 1 | Gm11929 | | 0 | 0 | 0 | 0 | 0 | 0 | 0 | 1 | 0 | 0 | 1 |
| 0 | 1.60 | 3.66 | 0.44 | 0.66 | 1 | Gm5370 | | 0 | 0 | 0 | 0 | 0 | 0 | 0 | 1 | 0 | 0 | 1 |
| 0 | 1.60 | 3.66 | 0.44 | 0.66 | 1 | Selenok-ps6 | | 0 | 0 | 0 | 0 | 0 | 0 | 0 | 1 | 0 | 0 | 1 |
| 0 | 0.93 | 3.67 | 0.25 | 0.80 | 1 | Gm5537 | | 0 | 1 | 0 | 0 | 0 | 0 | 0 | 1 | 0 | 0 | 0 |
| 0 | 0.93 | 3.67 | 0.25 | 0.80 | 1 | Gm5735 | | 0 | 1 | 0 | 0 | 0 | 0 | 0 | 1 | 0 | 0 | 0 |
| 0 | 0.93 | 3.67 | 0.25 | 0.80 | 1 | Gm16494 | | 0 | 1 | 0 | 0 | 0 | 0 | 0 | 1 | 0 | 0 | 0 |
| 0 | 0.93 | 3.67 | 0.25 | 0.80 | 1 | Gm16412 | | 0 | 1 | 0 | 0 | 0 | 0 | 0 | 1 | 0 | 0 | 0 |
| 0 | 1.58 | 3.66 | 0.43 | 0.67 | 1 | Gm12337 | | 0 | 0 | 0 | 0 | 0 | 0 | 0 | 1 | 1 | 0 | 0 |
| 0 | 0.28 | 3.68 | 0.08 | 0.94 | 1 | Gm17764 | | 0 | 0 | 0 | 1 | 0 | 0 | 0 | 0 | 0 | 1 | 0 |
| 0 | 0.28 | 3.68 | 0.08 | 0.94 | 1 | Gm38279 | | 0 | 0 | 0 | 1 | 0 | 0 | 0 | 0 | 0 | 1 | 0 |
| 0 | 0.28 | 3.68 | 0.08 | 0.94 | 1 | Gm7670 | | 0 | 0 | 0 | 1 | 0 | 0 | 0 | 0 | 0 | 1 | 0 |
| 0 | 0.94 | 3.67 | 0.26 | 0.80 | 1 | Gm14441 | | 1 | 0 | 0 | 0 | 0 | 0 | 0 | 0 | 0 | 1 | 0 |
| 0 | 0.94 | 3.67 | 0.26 | 0.80 | 1 | Gm27046 | | 1 | 0 | 0 | 0 | 0 | 0 | 0 | 0 | 0 | 1 | 0 |
| 0 | 0.94 | 3.67 | 0.26 | 0.80 | 1 | Gm21168 | | 1 | 0 | 0 | 0 | 0 | 0 | 0 | 0 | 0 | 1 | 0 |
| 0 | 0.94 | 3.67 | 0.26 | 0.80 | 1 | Gm8041 | | 1 | 0 | 0 | 0 | 0 | 0 | 0 | 0 | 0 | 1 | 0 |
| 0 | 0.94 | 3.67 | 0.26 | 0.80 | 1 | Gm45654 | | 1 | 0 | 0 | 0 | 0 | 0 | 0 | 0 | 0 | 1 | 0 |
| 0 | 0.94 | 3.67 | 0.26 | 0.80 | 1 | Gm7619 | | 1 | 0 | 0 | 0 | 0 | 0 | 0 | 0 | 0 | 1 | 0 |
| 0 | 0.94 | 3.67 | 0.26 | 0.80 | 1 | Gm12618 | | 1 | 0 | 0 | 0 | 0 | 0 | 0 | 0 | 0 | 1 | 0 |
| 0 | 0.94 | 3.67 | 0.26 | 0.80 | 1 | Gm7707 | | 1 | 0 | 0 | 0 | 0 | 0 | 0 | 0 | 0 | 1 | 0 |
| 0 | 0.94 | 3.67 | 0.26 | 0.80 | 1 | Gm17087 | | 1 | 0 | 0 | 0 | 0 | 0 | 0 | 0 | 0 | 1 | 0 |
| 0 | 0.28 | 3.68 | 0.08 | 0.94 | 1 | Gm2033 | | 0 | 0 | 0 | 0 | 1 | 0 | 0 | 1 | 0 | 0 | 0 |
| 0 | 0.00 | 3.68 | 0.00 | 1.00 | 1 | Gm3940 | | 0 | 0 | 2 | 0 | 0 | 0 | 0 | 0 | 0 | 0 | 0 |
| 0 | 0.00 | 3.68 | 0.00 | 1.00 | 1 | Gm6177 | | 0 | 0 | 2 | 0 | 0 | 0 | 0 | 0 | 0 | 0 | 0 |
| 0 | 0.00 | 3.68 | 0.00 | 1.00 | 1 | Fcrl6 | Fc receptor-like 6 | 0 | 0 | 2 | 0 | 0 | 0 | 0 | 0 | 0 | 0 | 0 |
| 0 | 0.00 | 3.68 | 0.00 | 1.00 | 1 | Gm37768 | | 0 | 0 | 2 | 0 | 0 | 0 | 0 | 0 | 0 | 0 | 0 |
| 0 | 0.00 | 3.68 | 0.00 | 1.00 | 1 | Gm37239 | | 0 | 0 | 2 | 0 | 0 | 0 | 0 | 0 | 0 | 0 | 0 |
| 0 | 0.00 | 3.68 | 0.00 | 1.00 | 1 | A330043C09Rik | | 0 | 0 | 2 | 0 | 0 | 0 | 0 | 0 | 0 | 0 | 0 |
| 0 | 0.00 | 3.68 | 0.00 | 1.00 | 1 | Gm43555 | | 0 | 0 | 2 | 0 | 0 | 0 | 0 | 0 | 0 | 0 | 0 |
| 0 | 0.00 | 3.68 | 0.00 | 1.00 | 1 | Gm37897 | | 0 | 0 | 2 | 0 | 0 | 0 | 0 | 0 | 0 | 0 | 0 |

| | | | | | | | | | | | | | | | | | | |
|---|---|---|---|---|---|---|---|---|---|---|---|---|---|---|---|---|---|---|
| 0 | 0.00 | 3.68 | 0.00 | 1.00 | 1 | Ifnb1 | interferon beta 1, fibroblast | 0 | 0 | 2 | 0 | 0 | 0 | 0 | 0 | 0 | 0 | 0 |
| 0 | 0.00 | 3.68 | 0.00 | 1.00 | 1 | Gm13067 | | 0 | 0 | 2 | 0 | 0 | 0 | 0 | 0 | 0 | 0 | 0 |
| 0 | 0.00 | 3.68 | 0.00 | 1.00 | 1 | Gm43279 | | 0 | 0 | 2 | 0 | 0 | 0 | 0 | 0 | 0 | 0 | 0 |
| 0 | 0.00 | 3.68 | 0.00 | 1.00 | 1 | Oas1e | 2'-5' oligoadenylate synthetase 1E | 0 | 0 | 2 | 0 | 0 | 0 | 0 | 0 | 0 | 0 | 0 |
| 0 | 0.00 | 3.68 | 0.00 | 1.00 | 1 | Gm6559 | | 0 | 0 | 2 | 0 | 0 | 0 | 0 | 0 | 0 | 0 | 0 |
| 0 | 0.00 | 3.68 | 0.00 | 1.00 | 1 | D830036C21Rik | | 0 | 0 | 2 | 0 | 0 | 0 | 0 | 0 | 0 | 0 | 0 |
| 0 | 0.00 | 3.68 | 0.00 | 1.00 | 1 | Cxcl17 | chemokine (C-X-C motif) ligand 17 | 0 | 0 | 2 | 0 | 0 | 0 | 0 | 0 | 0 | 0 | 0 |
| 0 | 0.00 | 3.68 | 0.00 | 1.00 | 1 | Catsperg2 | cation channel sperm associated auxiliar | 0 | 0 | 2 | 0 | 0 | 0 | 0 | 0 | 0 | 0 | 0 |
| 0 | 0.00 | 3.68 | 0.00 | 1.00 | 1 | Olfr557 | olfactory receptor 557 | 0 | 0 | 2 | 0 | 0 | 0 | 0 | 0 | 0 | 0 | 0 |
| 0 | 0.00 | 3.68 | 0.00 | 1.00 | 1 | Gm45499 | | 0 | 0 | 2 | 0 | 0 | 0 | 0 | 0 | 0 | 0 | 0 |
| 0 | 0.00 | 3.68 | 0.00 | 1.00 | 1 | Gm7607 | | 0 | 0 | 2 | 0 | 0 | 0 | 0 | 0 | 0 | 0 | 0 |
| 0 | 0.00 | 3.68 | 0.00 | 1.00 | 1 | Gm47235 | | 0 | 0 | 2 | 0 | 0 | 0 | 0 | 0 | 0 | 0 | 0 |
| 0 | 0.00 | 3.68 | 0.00 | 1.00 | 1 | Gm12056 | | 0 | 0 | 2 | 0 | 0 | 0 | 0 | 0 | 0 | 0 | 0 |
| 0 | 0.00 | 3.68 | 0.00 | 1.00 | 1 | Gm11515 | | 0 | 0 | 2 | 0 | 0 | 0 | 0 | 0 | 0 | 0 | 0 |
| 0 | 0.00 | 3.68 | 0.00 | 1.00 | 1 | Gm40847 | | 0 | 0 | 2 | 0 | 0 | 0 | 0 | 0 | 0 | 0 | 0 |
| 0 | 0.00 | 3.68 | 0.00 | 1.00 | 1 | Gm30409 | | 0 | 0 | 2 | 0 | 0 | 0 | 0 | 0 | 0 | 0 | 0 |
| 0 | 0.00 | 3.68 | 0.00 | 1.00 | 1 | Gm40999 | | 0 | 0 | 2 | 0 | 0 | 0 | 0 | 0 | 0 | 0 | 0 |
| 0 | 0.00 | 3.68 | 0.00 | 1.00 | 1 | Npw | neuropeptide W | 0 | 0 | 2 | 0 | 0 | 0 | 0 | 0 | 0 | 0 | 0 |
| 0 | 0.00 | 3.68 | 0.00 | 1.00 | 1 | Gm29168 | | 0 | 0 | 2 | 0 | 0 | 0 | 0 | 0 | 0 | 0 | 0 |
| 0 | 0.00 | 3.68 | 0.00 | 1.00 | 1 | Gm9435 | | 0 | 0 | 2 | 0 | 0 | 0 | 0 | 0 | 0 | 0 | 0 |
| 0 | 0.95 | 3.67 | 0.26 | 0.80 | 1 | Gm38358 | | 0 | 0 | 1 | 0 | 0 | 0 | 0 | 0 | 0 | 0 | 1 |
| 0 | 0.95 | 3.67 | 0.26 | 0.80 | 1 | Gm13870 | | 0 | 0 | 1 | 0 | 0 | 0 | 0 | 0 | 0 | 0 | 1 |
| 0 | 0.95 | 3.67 | 0.26 | 0.80 | 1 | Gm14013 | | 0 | 0 | 1 | 0 | 0 | 0 | 0 | 0 | 0 | 0 | 1 |
| 0 | 0.95 | 3.67 | 0.26 | 0.80 | 1 | Gm9023 | | 0 | 0 | 1 | 0 | 0 | 0 | 0 | 0 | 0 | 0 | 1 |
| 0 | 0.95 | 3.67 | 0.26 | 0.80 | 1 | Gm47941 | | 0 | 0 | 1 | 0 | 0 | 0 | 0 | 0 | 0 | 0 | 1 |
| 0 | 0.95 | 3.67 | 0.26 | 0.80 | 1 | Gm49105 | | 0 | 0 | 1 | 0 | 0 | 0 | 0 | 0 | 0 | 0 | 1 |
| 0 | 0.95 | 3.67 | 0.26 | 0.80 | 1 | Gm6599 | | 0 | 0 | 1 | 0 | 0 | 0 | 0 | 0 | 0 | 0 | 1 |
| 0 | 0.00 | 3.68 | 0.00 | 1.00 | 1 | Gm29138 | | 0 | 1 | 1 | 0 | 0 | 0 | 0 | 0 | 0 | 0 | 0 |
| 0 | 0.00 | 3.68 | 0.00 | 1.00 | 1 | Gm14274 | | 0 | 1 | 1 | 0 | 0 | 0 | 0 | 0 | 0 | 0 | 0 |
| 0 | 0.00 | 3.68 | 0.00 | 1.00 | 1 | Gm15737 | | 0 | 1 | 1 | 0 | 0 | 0 | 0 | 0 | 0 | 0 | 0 |
| 0 | 0.00 | 3.68 | 0.00 | 1.00 | 1 | Gm17136 | | 0 | 1 | 1 | 0 | 0 | 0 | 0 | 0 | 0 | 0 | 0 |
| 0 | 0.00 | 3.68 | 0.00 | 1.00 | 1 | Gm2976 | | 0 | 1 | 1 | 0 | 0 | 0 | 0 | 0 | 0 | 0 | 0 |
| 0 | 0.00 | 3.68 | 0.00 | 1.00 | 1 | Mc2r | melanocortin 2 receptor | 0 | 1 | 1 | 0 | 0 | 0 | 0 | 0 | 0 | 0 | 0 |
| 0 | 0.00 | 3.68 | 0.00 | 1.00 | 1 | Gm17150 | | 0 | 1 | 1 | 0 | 0 | 0 | 0 | 0 | 0 | 0 | 0 |
| 0 | 0.00 | 3.68 | 0.00 | 1.00 | 1 | Gm14925 | | 0 | 1 | 1 | 0 | 0 | 0 | 0 | 0 | 0 | 0 | 0 |
| 0 | 0.91 | 3.67 | 0.25 | 0.80 | 1 | Gm16210 | | 0 | 0 | 1 | 0 | 0 | 0 | 0 | 0 | 1 | 0 | 0 |
| 0 | 0.91 | 3.67 | 0.25 | 0.80 | 1 | Gm8513 | | 0 | 0 | 1 | 0 | 0 | 0 | 0 | 0 | 1 | 0 | 0 |
| 0 | 0.91 | 3.67 | 0.25 | 0.80 | 1 | Gm14506 | | 0 | 0 | 1 | 0 | 0 | 0 | 0 | 0 | 1 | 0 | 0 |
| 0 | 0.91 | 3.67 | 0.25 | 0.80 | 1 | Rplp1-ps1 | | 0 | 0 | 1 | 0 | 0 | 0 | 0 | 0 | 1 | 0 | 0 |
| 0 | 0.29 | 3.65 | 0.08 | 0.94 | 1 | Vma21-ps | | 0 | 0 | 0 | 1 | 0 | 0 | 0 | 1 | 0 | 0 | 0 |
| 0 | 0.29 | 3.65 | 0.08 | 0.94 | 1 | Gm4959 | | 0 | 0 | 0 | 1 | 0 | 0 | 0 | 1 | 0 | 0 | 0 |
| 0 | 0.29 | 3.65 | 0.08 | 0.94 | 1 | Gm15593 | | 0 | 0 | 0 | 1 | 0 | 0 | 0 | 1 | 0 | 0 | 0 |
| 0 | 0.29 | 3.65 | 0.08 | 0.94 | 1 | Gm8724 | | 0 | 0 | 0 | 1 | 0 | 0 | 0 | 1 | 0 | 0 | 0 |
| 0 | 0.29 | 3.65 | 0.08 | 0.94 | 1 | Gm47820 | | 0 | 0 | 0 | 1 | 0 | 0 | 0 | 1 | 0 | 0 | 0 |
| 0 | 0.29 | 3.65 | 0.08 | 0.94 | 1 | Gm20702 | | 0 | 0 | 0 | 1 | 0 | 0 | 0 | 1 | 0 | 0 | 0 |
| 0 | 1.25 | 3.66 | 0.34 | 0.73 | 1 | Gm37005 | | 0 | 0 | 0 | 0 | 0 | 0 | 0 | 0 | 0 | 0 | 2 |
| 0 | 1.25 | 3.66 | 0.34 | 0.73 | 1 | Gm7933 | | 0 | 0 | 0 | 0 | 0 | 0 | 0 | 0 | 0 | 0 | 2 |
| 0 | 1.25 | 3.66 | 0.34 | 0.73 | 1 | A930005N03Rik | | 0 | 0 | 0 | 0 | 0 | 0 | 0 | 0 | 0 | 0 | 2 |
| 0 | 1.25 | 3.66 | 0.34 | 0.73 | 1 | Gm29670 | | 0 | 0 | 0 | 0 | 0 | 0 | 0 | 0 | 0 | 0 | 2 |
| 0 | 1.25 | 3.66 | 0.34 | 0.73 | 1 | Gm25829 | | 0 | 0 | 0 | 0 | 0 | 0 | 0 | 0 | 0 | 0 | 2 |
| 0 | 1.25 | 3.66 | 0.34 | 0.73 | 1 | Flacc1 | flagellum associated containing coiled-cc | 0 | 0 | 0 | 0 | 0 | 0 | 0 | 0 | 0 | 0 | 2 |
| 0 | 1.25 | 3.66 | 0.34 | 0.73 | 1 | Gm4208 | | 0 | 0 | 0 | 0 | 0 | 0 | 0 | 0 | 0 | 0 | 2 |
| 0 | 1.25 | 3.66 | 0.34 | 0.73 | 1 | Gm37262 | | 0 | 0 | 0 | 0 | 0 | 0 | 0 | 0 | 0 | 0 | 2 |
| 0 | 1.25 | 3.66 | 0.34 | 0.73 | 1 | Gm36999 | | 0 | 0 | 0 | 0 | 0 | 0 | 0 | 0 | 0 | 0 | 2 |
| 0 | 1.25 | 3.66 | 0.34 | 0.73 | 1 | Gm37541 | | 0 | 0 | 0 | 0 | 0 | 0 | 0 | 0 | 0 | 0 | 2 |
| 0 | 1.25 | 3.66 | 0.34 | 0.73 | 1 | Gm28169 | | 0 | 0 | 0 | 0 | 0 | 0 | 0 | 0 | 0 | 0 | 2 |
| 0 | 1.25 | 3.66 | 0.34 | 0.73 | 1 | Gm37625 | | 0 | 0 | 0 | 0 | 0 | 0 | 0 | 0 | 0 | 0 | 2 |
| 0 | 1.25 | 3.66 | 0.34 | 0.73 | 1 | Gm37713 | | 0 | 0 | 0 | 0 | 0 | 0 | 0 | 0 | 0 | 0 | 2 |
| 0 | 1.25 | 3.66 | 0.34 | 0.73 | 1 | Gm37524 | | 0 | 0 | 0 | 0 | 0 | 0 | 0 | 0 | 0 | 0 | 2 |
| 0 | 1.25 | 3.66 | 0.34 | 0.73 | 1 | Atp1a4 | ATPase, Na+/K+ transporting, alpha 4 pol | 0 | 0 | 0 | 0 | 0 | 0 | 0 | 0 | 0 | 0 | 2 |

| | | | | | | | | | | | | | | | | | | |
|---|---|---|---|---|---|---|---|---|---|---|---|---|---|---|---|---|---|---|
| 0 | 1.25 | 3.66 | 0.34 | 0.73 | 1 | Mir1981 | | 0 | 0 | 0 | 0 | 0 | 0 | 0 | 0 | 0 | 0 | 2 |
| 0 | 1.25 | 3.66 | 0.34 | 0.73 | 1 | Gm44146 | | 0 | 0 | 0 | 0 | 0 | 0 | 0 | 0 | 0 | 0 | 2 |
| 0 | 1.25 | 3.66 | 0.34 | 0.73 | 1 | Gm37159 | | 0 | 0 | 0 | 0 | 0 | 0 | 0 | 0 | 0 | 0 | 2 |
| 0 | 1.25 | 3.66 | 0.34 | 0.73 | 1 | Gm13670 | | 0 | 0 | 0 | 0 | 0 | 0 | 0 | 0 | 0 | 0 | 2 |
| 0 | 1.25 | 3.66 | 0.34 | 0.73 | 1 | Olfr1233 | olfactory receptor 1233 | 0 | 0 | 0 | 0 | 0 | 0 | 0 | 0 | 0 | 0 | 2 |
| 0 | 1.25 | 3.66 | 0.34 | 0.73 | 1 | Gm23630 | | 0 | 0 | 0 | 0 | 0 | 0 | 0 | 0 | 0 | 0 | 2 |
| 0 | 1.25 | 3.66 | 0.34 | 0.73 | 1 | Gm10038 | | 0 | 0 | 0 | 0 | 0 | 0 | 0 | 0 | 0 | 0 | 2 |
| 0 | 1.25 | 3.66 | 0.34 | 0.73 | 1 | 4930527A07Rik | | 0 | 0 | 0 | 0 | 0 | 0 | 0 | 0 | 0 | 0 | 2 |
| 0 | 1.25 | 3.66 | 0.34 | 0.73 | 1 | Tgm6 | transglutaminase 6 | 0 | 0 | 0 | 0 | 0 | 0 | 0 | 0 | 0 | 0 | 2 |
| 0 | 1.25 | 3.66 | 0.34 | 0.73 | 1 | Gm20637 | | 0 | 0 | 0 | 0 | 0 | 0 | 0 | 0 | 0 | 0 | 2 |
| 0 | 1.25 | 3.66 | 0.34 | 0.73 | 1 | Gm14444 | predicted gene 14444 | 0 | 0 | 0 | 0 | 0 | 0 | 0 | 0 | 0 | 0 | 2 |
| 0 | 1.25 | 3.66 | 0.34 | 0.73 | 1 | Gm37685 | | 0 | 0 | 0 | 0 | 0 | 0 | 0 | 0 | 0 | 0 | 2 |
| 0 | 1.25 | 3.66 | 0.34 | 0.73 | 1 | Gm8813 | | 0 | 0 | 0 | 0 | 0 | 0 | 0 | 0 | 0 | 0 | 2 |
| 0 | 1.25 | 3.66 | 0.34 | 0.73 | 1 | Gm42812 | | 0 | 0 | 0 | 0 | 0 | 0 | 0 | 0 | 0 | 0 | 2 |
| 0 | 1.25 | 3.66 | 0.34 | 0.73 | 1 | Pdzk1 | PDZ domain containing 1 | 0 | 0 | 0 | 0 | 0 | 0 | 0 | 0 | 0 | 0 | 2 |
| 0 | 1.25 | 3.66 | 0.34 | 0.73 | 1 | 5730437C11Rik | | 0 | 0 | 0 | 0 | 0 | 0 | 0 | 0 | 0 | 0 | 2 |
| 0 | 1.25 | 3.66 | 0.34 | 0.73 | 1 | Gm43121 | | 0 | 0 | 0 | 0 | 0 | 0 | 0 | 0 | 0 | 0 | 2 |
| 0 | 1.25 | 3.66 | 0.34 | 0.73 | 1 | Gm25869 | | 0 | 0 | 0 | 0 | 0 | 0 | 0 | 0 | 0 | 0 | 2 |
| 0 | 1.25 | 3.66 | 0.34 | 0.73 | 1 | Pifo | primary cilia formation | 0 | 0 | 0 | 0 | 0 | 0 | 0 | 0 | 0 | 0 | 2 |
| 0 | 1.25 | 3.66 | 0.34 | 0.73 | 1 | Gm11864 | | 0 | 0 | 0 | 0 | 0 | 0 | 0 | 0 | 0 | 0 | 2 |
| 0 | 1.25 | 3.66 | 0.34 | 0.73 | 1 | Gm12730 | | 0 | 0 | 0 | 0 | 0 | 0 | 0 | 0 | 0 | 0 | 2 |
| 0 | 1.25 | 3.66 | 0.34 | 0.73 | 1 | Gm12998 | | 0 | 0 | 0 | 0 | 0 | 0 | 0 | 0 | 0 | 0 | 2 |
| 0 | 1.25 | 3.66 | 0.34 | 0.73 | 1 | Snora73a | | 0 | 0 | 0 | 0 | 0 | 0 | 0 | 0 | 0 | 0 | 2 |
| 0 | 1.25 | 3.66 | 0.34 | 0.73 | 1 | Lactbl1 | lactamase, beta-like 1 | 0 | 0 | 0 | 0 | 0 | 0 | 0 | 0 | 0 | 0 | 2 |
| 0 | 1.25 | 3.66 | 0.34 | 0.73 | 1 | 1700045H11Rik | | 0 | 0 | 0 | 0 | 0 | 0 | 0 | 0 | 0 | 0 | 2 |
| 0 | 1.25 | 3.66 | 0.34 | 0.73 | 1 | Gm7361 | predicted gene 7361 | 0 | 0 | 0 | 0 | 0 | 0 | 0 | 0 | 0 | 0 | 2 |
| 0 | 1.25 | 3.66 | 0.34 | 0.73 | 1 | Gm24878 | | 0 | 0 | 0 | 0 | 0 | 0 | 0 | 0 | 0 | 0 | 2 |
| 0 | 1.25 | 3.66 | 0.34 | 0.73 | 1 | Gm26560 | | 0 | 0 | 0 | 0 | 0 | 0 | 0 | 0 | 0 | 0 | 2 |
| 0 | 1.25 | 3.66 | 0.34 | 0.73 | 1 | Gm7205 | | 0 | 0 | 0 | 0 | 0 | 0 | 0 | 0 | 0 | 0 | 2 |
| 0 | 1.25 | 3.66 | 0.34 | 0.73 | 1 | Gm5108 | predicted gene 5108 | 0 | 0 | 0 | 0 | 0 | 0 | 0 | 0 | 0 | 0 | 2 |
| 0 | 1.25 | 3.66 | 0.34 | 0.73 | 1 | 4930430O22Rik | | 0 | 0 | 0 | 0 | 0 | 0 | 0 | 0 | 0 | 0 | 2 |
| 0 | 1.25 | 3.66 | 0.34 | 0.73 | 1 | Gsx1 | GS homeobox 1 | 0 | 0 | 0 | 0 | 0 | 0 | 0 | 0 | 0 | 0 | 2 |
| 0 | 1.25 | 3.66 | 0.34 | 0.73 | 1 | Gm3289 | | 0 | 0 | 0 | 0 | 0 | 0 | 0 | 0 | 0 | 0 | 2 |
| 0 | 1.25 | 3.66 | 0.34 | 0.73 | 1 | Gm16563 | | 0 | 0 | 0 | 0 | 0 | 0 | 0 | 0 | 0 | 0 | 2 |
| 0 | 1.25 | 3.66 | 0.34 | 0.73 | 1 | Vmn1r16 | vomeronasal 1 receptor 16 | 0 | 0 | 0 | 0 | 0 | 0 | 0 | 0 | 0 | 0 | 2 |
| 0 | 1.25 | 3.66 | 0.34 | 0.73 | 1 | Gm27896 | | 0 | 0 | 0 | 0 | 0 | 0 | 0 | 0 | 0 | 0 | 2 |
| 0 | 1.25 | 3.66 | 0.34 | 0.73 | 1 | Vax2os | | 0 | 0 | 0 | 0 | 0 | 0 | 0 | 0 | 0 | 0 | 2 |
| 0 | 1.25 | 3.66 | 0.34 | 0.73 | 1 | Gm44249 | | 0 | 0 | 0 | 0 | 0 | 0 | 0 | 0 | 0 | 0 | 2 |
| 0 | 1.25 | 3.66 | 0.34 | 0.73 | 1 | Gm17055 | | 0 | 0 | 0 | 0 | 0 | 0 | 0 | 0 | 0 | 0 | 2 |
| 0 | 1.25 | 3.66 | 0.34 | 0.73 | 1 | Klrb1c | killer cell lectin-like receptor subfamily B r | 0 | 0 | 0 | 0 | 0 | 0 | 0 | 0 | 0 | 0 | 2 |
| 0 | 1.25 | 3.66 | 0.34 | 0.73 | 1 | Gm3654 | | 0 | 0 | 0 | 0 | 0 | 0 | 0 | 0 | 0 | 0 | 2 |
| 0 | 1.25 | 3.66 | 0.34 | 0.73 | 1 | Gm31152 | | 0 | 0 | 0 | 0 | 0 | 0 | 0 | 0 | 0 | 0 | 2 |
| 0 | 1.25 | 3.66 | 0.34 | 0.73 | 1 | Gm20083 | | 0 | 0 | 0 | 0 | 0 | 0 | 0 | 0 | 0 | 0 | 2 |
| 0 | 1.25 | 3.66 | 0.34 | 0.73 | 1 | Gm10160 | | 0 | 0 | 0 | 0 | 0 | 0 | 0 | 0 | 0 | 0 | 2 |
| 0 | 1.25 | 3.66 | 0.34 | 0.73 | 1 | Tex29 | testis expressed 29 | 0 | 0 | 0 | 0 | 0 | 0 | 0 | 0 | 0 | 0 | 2 |
| 0 | 1.25 | 3.66 | 0.34 | 0.73 | 1 | Gm45349 | | 0 | 0 | 0 | 0 | 0 | 0 | 0 | 0 | 0 | 0 | 2 |
| 0 | 1.25 | 3.66 | 0.34 | 0.73 | 1 | Gm45693 | | 0 | 0 | 0 | 0 | 0 | 0 | 0 | 0 | 0 | 0 | 2 |
| 0 | 1.25 | 3.66 | 0.34 | 0.73 | 1 | Gm45758 | | 0 | 0 | 0 | 0 | 0 | 0 | 0 | 0 | 0 | 0 | 2 |
| 0 | 1.25 | 3.66 | 0.34 | 0.73 | 1 | Gm6013 | | 0 | 0 | 0 | 0 | 0 | 0 | 0 | 0 | 0 | 0 | 2 |
| 0 | 1.25 | 3.66 | 0.34 | 0.73 | 1 | Gm38384 | | 0 | 0 | 0 | 0 | 0 | 0 | 0 | 0 | 0 | 0 | 2 |
| 0 | 1.25 | 3.66 | 0.34 | 0.73 | 1 | Gm15898 | | 0 | 0 | 0 | 0 | 0 | 0 | 0 | 0 | 0 | 0 | 2 |
| 0 | 1.25 | 3.66 | 0.34 | 0.73 | 1 | Gm16302 | | 0 | 0 | 0 | 0 | 0 | 0 | 0 | 0 | 0 | 0 | 2 |
| 0 | 1.25 | 3.66 | 0.34 | 0.73 | 1 | Spata19 | spermatogenesis associated 19 | 0 | 0 | 0 | 0 | 0 | 0 | 0 | 0 | 0 | 0 | 2 |
| 0 | 1.25 | 3.66 | 0.34 | 0.73 | 1 | Gm47643 | | 0 | 0 | 0 | 0 | 0 | 0 | 0 | 0 | 0 | 0 | 2 |
| 0 | 1.25 | 3.66 | 0.34 | 0.73 | 1 | Gm47409 | | 0 | 0 | 0 | 0 | 0 | 0 | 0 | 0 | 0 | 0 | 2 |
| 0 | 1.25 | 3.66 | 0.34 | 0.73 | 1 | 1700008A23Rik | | 0 | 0 | 0 | 0 | 0 | 0 | 0 | 0 | 0 | 0 | 2 |
| 0 | 1.25 | 3.66 | 0.34 | 0.73 | 1 | Gm28979 | | 0 | 0 | 0 | 0 | 0 | 0 | 0 | 0 | 0 | 0 | 2 |
| 0 | 1.25 | 3.66 | 0.34 | 0.73 | 1 | Gm16252 | | 0 | 0 | 0 | 0 | 0 | 0 | 0 | 0 | 0 | 0 | 2 |
| 0 | 1.25 | 3.66 | 0.34 | 0.73 | 1 | Rp31-ps19 | | 0 | 0 | 0 | 0 | 0 | 0 | 0 | 0 | 0 | 0 | 2 |
| 0 | 1.25 | 3.66 | 0.34 | 0.73 | 1 | Gm37111 | | 0 | 0 | 0 | 0 | 0 | 0 | 0 | 0 | 0 | 0 | 2 |

| | | | | | | | | | | | | | | | | | | |
|---|---|---|---|---|---|---|---|---|---|---|---|---|---|---|---|---|---|---|
| 0 | 1.25 | 3.66 | 0.34 | 0.73 | 1 | Gm20470 | | 0 | 0 | 0 | 0 | 0 | 0 | 0 | 0 | 0 | 0 | 2 |
| 0 | 1.25 | 3.66 | 0.34 | 0.73 | 1 | Fam162b | family with sequence similarity 162, meml | 0 | 0 | 0 | 0 | 0 | 0 | 0 | 0 | 0 | 0 | 2 |
| 0 | 1.25 | 3.66 | 0.34 | 0.73 | 1 | Gm47611 | | 0 | 0 | 0 | 0 | 0 | 0 | 0 | 0 | 0 | 0 | 2 |
| 0 | 1.25 | 3.66 | 0.34 | 0.73 | 1 | Gm4767 | predicted gene 4767 | 0 | 0 | 0 | 0 | 0 | 0 | 0 | 0 | 0 | 0 | 2 |
| 0 | 1.25 | 3.66 | 0.34 | 0.73 | 1 | Snord59a | | 0 | 0 | 0 | 0 | 0 | 0 | 0 | 0 | 0 | 0 | 2 |
| 0 | 1.25 | 3.66 | 0.34 | 0.73 | 1 | Gm16518 | | 0 | 0 | 0 | 0 | 0 | 0 | 0 | 0 | 0 | 0 | 2 |
| 0 | 1.25 | 3.66 | 0.34 | 0.73 | 1 | Gm12123 | | 0 | 0 | 0 | 0 | 0 | 0 | 0 | 0 | 0 | 0 | 2 |
| 0 | 1.25 | 3.66 | 0.34 | 0.73 | 1 | Gm24582 | | 0 | 0 | 0 | 0 | 0 | 0 | 0 | 0 | 0 | 0 | 2 |
| 0 | 1.25 | 3.66 | 0.34 | 0.73 | 1 | Gp1ba | glycoprotein 1b, alpha polypeptide | 0 | 0 | 0 | 0 | 0 | 0 | 0 | 0 | 0 | 0 | 2 |
| 0 | 1.25 | 3.66 | 0.34 | 0.73 | 1 | Gm22297 | | 0 | 0 | 0 | 0 | 0 | 0 | 0 | 0 | 0 | 0 | 2 |
| 0 | 1.25 | 3.66 | 0.34 | 0.73 | 1 | Scpep1os | | 0 | 0 | 0 | 0 | 0 | 0 | 0 | 0 | 0 | 0 | 2 |
| 0 | 1.25 | 3.66 | 0.34 | 0.73 | 1 | Gm11770 | | 0 | 0 | 0 | 0 | 0 | 0 | 0 | 0 | 0 | 0 | 2 |
| 0 | 1.25 | 3.66 | 0.34 | 0.73 | 1 | Dnmt3aos | | 0 | 0 | 0 | 0 | 0 | 0 | 0 | 0 | 0 | 0 | 2 |
| 0 | 1.25 | 3.66 | 0.34 | 0.73 | 1 | Gm9921 | | 0 | 0 | 0 | 0 | 0 | 0 | 0 | 0 | 0 | 0 | 2 |
| 0 | 1.25 | 3.66 | 0.34 | 0.73 | 1 | 3110039M20Rik | | 0 | 0 | 0 | 0 | 0 | 0 | 0 | 0 | 0 | 0 | 2 |
| 0 | 1.25 | 3.66 | 0.34 | 0.73 | 1 | Gm48747 | | 0 | 0 | 0 | 0 | 0 | 0 | 0 | 0 | 0 | 0 | 2 |
| 0 | 1.25 | 3.66 | 0.34 | 0.73 | 1 | Gm48585 | | 0 | 0 | 0 | 0 | 0 | 0 | 0 | 0 | 0 | 0 | 2 |
| 0 | 1.25 | 3.66 | 0.34 | 0.73 | 1 | Mir6939 | | 0 | 0 | 0 | 0 | 0 | 0 | 0 | 0 | 0 | 0 | 2 |
| 0 | 1.25 | 3.66 | 0.34 | 0.73 | 1 | Gm24873 | | 0 | 0 | 0 | 0 | 0 | 0 | 0 | 0 | 0 | 0 | 2 |
| 0 | 1.25 | 3.66 | 0.34 | 0.73 | 1 | Gm44350 | | 0 | 0 | 0 | 0 | 0 | 0 | 0 | 0 | 0 | 0 | 2 |
| 0 | 1.25 | 3.66 | 0.34 | 0.73 | 1 | C630044B11Rik | | 0 | 0 | 0 | 0 | 0 | 0 | 0 | 0 | 0 | 0 | 2 |
| 0 | 1.25 | 3.66 | 0.34 | 0.73 | 1 | Gm18883 | | 0 | 0 | 0 | 0 | 0 | 0 | 0 | 0 | 0 | 0 | 2 |
| 0 | 1.25 | 3.66 | 0.34 | 0.73 | 1 | Gm48265 | | 0 | 0 | 0 | 0 | 0 | 0 | 0 | 0 | 0 | 0 | 2 |
| 0 | 1.25 | 3.66 | 0.34 | 0.73 | 1 | Gm3637 | | 0 | 0 | 0 | 0 | 0 | 0 | 0 | 0 | 0 | 0 | 2 |
| 0 | 1.25 | 3.66 | 0.34 | 0.73 | 1 | Olfr1507 | olfactory receptor 1507 | 0 | 0 | 0 | 0 | 0 | 0 | 0 | 0 | 0 | 0 | 2 |
| 0 | 1.25 | 3.66 | 0.34 | 0.73 | 1 | Gm48200 | | 0 | 0 | 0 | 0 | 0 | 0 | 0 | 0 | 0 | 0 | 2 |
| 0 | 1.25 | 3.66 | 0.34 | 0.73 | 1 | Gm16573 | | 0 | 0 | 0 | 0 | 0 | 0 | 0 | 0 | 0 | 0 | 2 |
| 0 | 1.25 | 3.66 | 0.34 | 0.73 | 1 | Gm29717 | | 0 | 0 | 0 | 0 | 0 | 0 | 0 | 0 | 0 | 0 | 2 |
| 0 | 1.25 | 3.66 | 0.34 | 0.73 | 1 | Gm29266 | | 0 | 0 | 0 | 0 | 0 | 0 | 0 | 0 | 0 | 0 | 2 |
| 0 | 1.25 | 3.66 | 0.34 | 0.73 | 1 | Gm34907 | | 0 | 0 | 0 | 0 | 0 | 0 | 0 | 0 | 0 | 0 | 2 |
| 0 | 1.25 | 3.66 | 0.34 | 0.73 | 1 | Gm35909 | | 0 | 0 | 0 | 0 | 0 | 0 | 0 | 0 | 0 | 0 | 2 |
| 0 | 1.25 | 3.66 | 0.34 | 0.73 | 1 | AA536875 | | 0 | 0 | 0 | 0 | 0 | 0 | 0 | 0 | 0 | 0 | 2 |
| 0 | 1.25 | 3.66 | 0.34 | 0.73 | 1 | Ttc23l | tetratricopeptide repeat domain 23-like | 0 | 0 | 0 | 0 | 0 | 0 | 0 | 0 | 0 | 0 | 2 |
| 0 | 1.25 | 3.66 | 0.34 | 0.73 | 1 | 9230109A22Rik | | 0 | 0 | 0 | 0 | 0 | 0 | 0 | 0 | 0 | 0 | 2 |
| 0 | 1.25 | 3.66 | 0.34 | 0.73 | 1 | Mroh5 | maestro heat-like repeat family member 5 | 0 | 0 | 0 | 0 | 0 | 0 | 0 | 0 | 0 | 0 | 2 |
| 0 | 1.25 | 3.66 | 0.34 | 0.73 | 1 | | | 0 | 0 | 0 | 0 | 0 | 0 | 0 | 0 | 0 | 0 | 2 |
| 0 | 1.25 | 3.66 | 0.34 | 0.73 | 1 | Gm22818 | | 0 | 0 | 0 | 0 | 0 | 0 | 0 | 0 | 0 | 0 | 2 |
| 0 | 1.25 | 3.66 | 0.34 | 0.73 | 1 | Gm24647 | | 0 | 0 | 0 | 0 | 0 | 0 | 0 | 0 | 0 | 0 | 2 |
| 0 | 1.25 | 3.66 | 0.34 | 0.73 | 1 | Krt72 | keratin 72 | 0 | 0 | 0 | 0 | 0 | 0 | 0 | 0 | 0 | 0 | 2 |
| 0 | 1.25 | 3.66 | 0.34 | 0.73 | 1 | Crygs | crystallin, gamma S | 0 | 0 | 0 | 0 | 0 | 0 | 0 | 0 | 0 | 0 | 2 |
| 0 | 1.25 | 3.66 | 0.34 | 0.73 | 1 | Gm15591 | | 0 | 0 | 0 | 0 | 0 | 0 | 0 | 0 | 0 | 0 | 2 |
| 0 | 1.25 | 3.66 | 0.34 | 0.73 | 1 | Unc93a2 | unc-93 homolog A2 | 0 | 0 | 0 | 0 | 0 | 0 | 0 | 0 | 0 | 0 | 2 |
| 0 | 1.25 | 3.66 | 0.34 | 0.73 | 1 | Gm23294 | | 0 | 0 | 0 | 0 | 0 | 0 | 0 | 0 | 0 | 0 | 2 |
| 0 | 1.25 | 3.66 | 0.34 | 0.73 | 1 | Olfr755-ps1 | | 0 | 0 | 0 | 0 | 0 | 0 | 0 | 0 | 0 | 0 | 2 |
| 0 | 1.25 | 3.66 | 0.34 | 0.73 | 1 | Gm49848 | | 0 | 0 | 0 | 0 | 0 | 0 | 0 | 0 | 0 | 0 | 2 |
| 0 | 1.25 | 3.66 | 0.34 | 0.73 | 1 | | | 0 | 0 | 0 | 0 | 0 | 0 | 0 | 0 | 0 | 0 | 2 |
| 0 | 1.25 | 3.66 | 0.34 | 0.73 | 1 | Gm7502 | | 0 | 0 | 0 | 0 | 0 | 0 | 0 | 0 | 0 | 0 | 2 |
| 0 | 1.25 | 3.66 | 0.34 | 0.73 | 1 | Gm50067 | | 0 | 0 | 0 | 0 | 0 | 0 | 0 | 0 | 0 | 0 | 2 |
| 0 | 1.25 | 3.66 | 0.34 | 0.73 | 1 | Gpr151 | G protein-coupled receptor 151 | 0 | 0 | 0 | 0 | 0 | 0 | 0 | 0 | 0 | 0 | 2 |
| 0 | 1.25 | 3.66 | 0.34 | 0.73 | 1 | Gm41750 | | 0 | 0 | 0 | 0 | 0 | 0 | 0 | 0 | 0 | 0 | 2 |
| 0 | 1.25 | 3.66 | 0.34 | 0.73 | 1 | Kcnk4 | potassium channel, subfamily K, member | 0 | 0 | 0 | 0 | 0 | 0 | 0 | 0 | 0 | 0 | 2 |
| 0 | 1.25 | 3.66 | 0.34 | 0.73 | 1 | Gm17227 | | 0 | 0 | 0 | 0 | 0 | 0 | 0 | 0 | 0 | 0 | 2 |
| 0 | 1.25 | 3.66 | 0.34 | 0.73 | 1 | Gm50191 | | 0 | 0 | 0 | 0 | 0 | 0 | 0 | 0 | 0 | 0 | 2 |
| 0 | 1.25 | 3.66 | 0.34 | 0.73 | 1 | Gm25520 | | 0 | 0 | 0 | 0 | 0 | 0 | 0 | 0 | 0 | 0 | 2 |
| 0 | 1.25 | 3.66 | 0.34 | 0.73 | 1 | Gm8522 | | 0 | 0 | 0 | 0 | 0 | 0 | 0 | 0 | 0 | 0 | 2 |
| 0 | 1.25 | 3.66 | 0.34 | 0.73 | 1 | Gm14766 | | 0 | 0 | 0 | 0 | 0 | 0 | 0 | 0 | 0 | 0 | 2 |
| 0 | 1.25 | 3.66 | 0.34 | 0.73 | 1 | Gm25793 | | 0 | 0 | 0 | 0 | 0 | 0 | 0 | 0 | 0 | 0 | 2 |
| 0 | 1.25 | 3.66 | 0.34 | 0.73 | 1 | Gm15211 | | 0 | 0 | 0 | 0 | 0 | 0 | 0 | 0 | 0 | 0 | 2 |
| 0 | 0.95 | 3.67 | 0.26 | 0.80 | 1 | Gm38353 | | 0 | 1 | 0 | 0 | 0 | 0 | 0 | 0 | 0 | 0 | 1 |
| 0 | 0.95 | 3.67 | 0.26 | 0.80 | 1 | Gm12901 | | 0 | 1 | 0 | 0 | 0 | 0 | 0 | 0 | 0 | 0 | 1 |

| | | | | | | | | | | | | | | | | | | |
|---|---|---|---|---|---|---|---|---|---|---|---|---|---|---|---|---|---|---|
| 0 | 0.95 | 3.67 | 0.26 | 0.80 | 1 | Gm42860 | | 0 | 1 | 0 | 0 | 0 | 0 | 0 | 0 | 0 | 0 | 1 |
| 0 | 0.95 | 3.67 | 0.26 | 0.80 | 1 | Gm5364 | | 0 | 1 | 0 | 0 | 0 | 0 | 0 | 0 | 0 | 0 | 1 |
| 0 | 0.95 | 3.67 | 0.26 | 0.80 | 1 | Gm37137 | | 0 | 1 | 0 | 0 | 0 | 0 | 0 | 0 | 0 | 0 | 1 |
| 0 | 0.95 | 3.67 | 0.26 | 0.80 | 1 | Gm9378 | | 0 | 1 | 0 | 0 | 0 | 0 | 0 | 0 | 0 | 0 | 1 |
| 0 | 0.95 | 3.67 | 0.26 | 0.80 | 1 | Gm21378 | | 0 | 1 | 0 | 0 | 0 | 0 | 0 | 0 | 0 | 0 | 1 |
| 0 | 0.95 | 3.67 | 0.26 | 0.80 | 1 | Gm16082 | | 0 | 1 | 0 | 0 | 0 | 0 | 0 | 0 | 0 | 0 | 1 |
| 0 | 0.95 | 3.67 | 0.26 | 0.80 | 1 | Gm16172 | | 0 | 1 | 0 | 0 | 0 | 0 | 0 | 0 | 0 | 0 | 1 |
| 0 | 1.58 | 3.66 | 0.43 | 0.67 | 1 | Gm28287 | | 0 | 0 | 0 | 0 | 0 | 0 | 0 | 0 | 1 | 0 | 1 |
| 0 | 1.58 | 3.66 | 0.43 | 0.67 | 1 | Gm18505 | | 0 | 0 | 0 | 0 | 0 | 0 | 0 | 0 | 1 | 0 | 1 |
| 0 | 1.58 | 3.66 | 0.43 | 0.67 | 1 | Gm13770 | | 0 | 0 | 0 | 0 | 0 | 0 | 0 | 0 | 1 | 0 | 1 |
| 0 | 1.58 | 3.66 | 0.43 | 0.67 | 1 | Cox20b | | 0 | 0 | 0 | 0 | 0 | 0 | 0 | 0 | 1 | 0 | 1 |
| 0 | 1.58 | 3.66 | 0.43 | 0.67 | 1 | Gm19178 | | 0 | 0 | 0 | 0 | 0 | 0 | 0 | 0 | 1 | 0 | 1 |
| 0 | 1.58 | 3.66 | 0.43 | 0.67 | 1 | Gm12096 | | 0 | 0 | 0 | 0 | 0 | 0 | 0 | 0 | 1 | 0 | 1 |
| 0 | 1.58 | 3.66 | 0.43 | 0.67 | 1 | Gm10784 | | 0 | 0 | 0 | 0 | 0 | 0 | 0 | 0 | 1 | 0 | 1 |
| 0 | 1.58 | 3.66 | 0.43 | 0.67 | 1 | Gm3785 | | 0 | 0 | 0 | 0 | 0 | 0 | 0 | 0 | 1 | 0 | 1 |
| 0 | 0.00 | 3.68 | 0.00 | 1.00 | 1 | Gm6119 | | 0 | 2 | 0 | 0 | 0 | 0 | 0 | 0 | 0 | 0 | 0 |
| 0 | 0.00 | 3.68 | 0.00 | 1.00 | 1 | 2700078F05Rik | | 0 | 2 | 0 | 0 | 0 | 0 | 0 | 0 | 0 | 0 | 0 |
| 0 | 0.00 | 3.68 | 0.00 | 1.00 | 1 | A430027H14Rik | | 0 | 2 | 0 | 0 | 0 | 0 | 0 | 0 | 0 | 0 | 0 |
| 0 | 0.00 | 3.68 | 0.00 | 1.00 | 1 | Gm38197 | | 0 | 2 | 0 | 0 | 0 | 0 | 0 | 0 | 0 | 0 | 0 |
| 0 | 0.00 | 3.68 | 0.00 | 1.00 | 1 | S100a7a | S100 calcium binding protein A7A | 0 | 2 | 0 | 0 | 0 | 0 | 0 | 0 | 0 | 0 | 0 |
| 0 | 0.00 | 3.68 | 0.00 | 1.00 | 1 | Tpt1-ps1 | | 0 | 2 | 0 | 0 | 0 | 0 | 0 | 0 | 0 | 0 | 0 |
| 0 | 0.00 | 3.68 | 0.00 | 1.00 | 1 | Gm12530 | | 0 | 2 | 0 | 0 | 0 | 0 | 0 | 0 | 0 | 0 | 0 |
| 0 | 0.00 | 3.68 | 0.00 | 1.00 | 1 | CK137956 | cDNA sequence CK137956 | 0 | 2 | 0 | 0 | 0 | 0 | 0 | 0 | 0 | 0 | 0 |
| 0 | 0.00 | 3.68 | 0.00 | 1.00 | 1 | 4933427G23Rik | | 0 | 2 | 0 | 0 | 0 | 0 | 0 | 0 | 0 | 0 | 0 |
| 0 | 0.00 | 3.68 | 0.00 | 1.00 | 1 | Gm42765 | | 0 | 2 | 0 | 0 | 0 | 0 | 0 | 0 | 0 | 0 | 0 |
| 0 | 0.00 | 3.68 | 0.00 | 1.00 | 1 | A730035I17Rik | | 0 | 2 | 0 | 0 | 0 | 0 | 0 | 0 | 0 | 0 | 0 |
| 0 | 0.00 | 3.68 | 0.00 | 1.00 | 1 | Gm17660 | predicted gene, 17660 | 0 | 2 | 0 | 0 | 0 | 0 | 0 | 0 | 0 | 0 | 0 |
| 0 | 0.00 | 3.68 | 0.00 | 1.00 | 1 | Gm42993 | | 0 | 2 | 0 | 0 | 0 | 0 | 0 | 0 | 0 | 0 | 0 |
| 0 | 0.00 | 3.68 | 0.00 | 1.00 | 1 | Gm20619 | | 0 | 2 | 0 | 0 | 0 | 0 | 0 | 0 | 0 | 0 | 0 |
| 0 | 0.00 | 3.68 | 0.00 | 1.00 | 1 | Klre1 | killer cell lectin-like receptor family E mem | 0 | 2 | 0 | 0 | 0 | 0 | 0 | 0 | 0 | 0 | 0 |
| 0 | 0.00 | 3.68 | 0.00 | 1.00 | 1 | Gm15723 | | 0 | 2 | 0 | 0 | 0 | 0 | 0 | 0 | 0 | 0 | 0 |
| 0 | 0.00 | 3.68 | 0.00 | 1.00 | 1 | Gm32365 | | 0 | 2 | 0 | 0 | 0 | 0 | 0 | 0 | 0 | 0 | 0 |
| 0 | 0.00 | 3.68 | 0.00 | 1.00 | 1 | 4930524B15Ril | RIKEN cDNA 4930524B15 gene | 0 | 2 | 0 | 0 | 0 | 0 | 0 | 0 | 0 | 0 | 0 |
| 0 | 0.00 | 3.68 | 0.00 | 1.00 | 1 | Myhas | | 0 | 2 | 0 | 0 | 0 | 0 | 0 | 0 | 0 | 0 | 0 |
| 0 | 0.00 | 3.68 | 0.00 | 1.00 | 1 | Sebox | SEBOX homeobox | 0 | 2 | 0 | 0 | 0 | 0 | 0 | 0 | 0 | 0 | 0 |
| 0 | 0.00 | 3.68 | 0.00 | 1.00 | 1 | Rps11-ps4 | | 0 | 2 | 0 | 0 | 0 | 0 | 0 | 0 | 0 | 0 | 0 |
| 0 | 0.00 | 3.68 | 0.00 | 1.00 | 1 | Gm48931 | | 0 | 2 | 0 | 0 | 0 | 0 | 0 | 0 | 0 | 0 | 0 |
| 0 | 0.00 | 3.68 | 0.00 | 1.00 | 1 | Wbp2nl | WBP2 N-terminal like | 0 | 2 | 0 | 0 | 0 | 0 | 0 | 0 | 0 | 0 | 0 |
| 0 | 0.00 | 3.68 | 0.00 | 1.00 | 1 | Gm50254 | | 0 | 2 | 0 | 0 | 0 | 0 | 0 | 0 | 0 | 0 | 0 |
| 0 | 0.00 | 3.68 | 0.00 | 1.00 | 1 | 2310026I22Rik | | 0 | 2 | 0 | 0 | 0 | 0 | 0 | 0 | 0 | 0 | 0 |
| 0 | 0.00 | 3.68 | 0.00 | 1.00 | 1 | Gm5821 | | 0 | 2 | 0 | 0 | 0 | 0 | 0 | 0 | 0 | 0 | 0 |
| 0 | 1.00 | 3.68 | 0.27 | 0.79 | 1 | Gm13545 | | 0 | 1 | 0 | 0 | 0 | 0 | 0 | 0 | 1 | 0 | 0 |
| 0 | 1.00 | 3.68 | 0.27 | 0.79 | 1 | Gm3745 | | 0 | 1 | 0 | 0 | 0 | 0 | 0 | 0 | 1 | 0 | 0 |
| 0 | 1.00 | 3.68 | 0.27 | 0.79 | 1 | Gm7336 | | 0 | 1 | 0 | 0 | 0 | 0 | 0 | 0 | 1 | 0 | 0 |
| 0 | 1.00 | 3.68 | 0.27 | 0.79 | 1 | Gm7106 | | 0 | 1 | 0 | 0 | 0 | 0 | 0 | 0 | 1 | 0 | 0 |
| 0 | 1.00 | 3.68 | 0.27 | 0.79 | 1 | Gm7748 | | 0 | 1 | 0 | 0 | 0 | 0 | 0 | 0 | 1 | 0 | 0 |
| 0 | 1.32 | 3.66 | 0.36 | 0.72 | 1 | Gm13315 | | 0 | 0 | 0 | 0 | 0 | 0 | 0 | 0 | 2 | 0 | 0 |
| 0 | 1.32 | 3.66 | 0.36 | 0.72 | 1 | Mir7000 | | 0 | 0 | 0 | 0 | 0 | 0 | 0 | 0 | 2 | 0 | 0 |
| 0 | 1.32 | 3.66 | 0.36 | 0.72 | 1 | Gm14014 | | 0 | 0 | 0 | 0 | 0 | 0 | 0 | 0 | 2 | 0 | 0 |
| 0 | 1.32 | 3.66 | 0.36 | 0.72 | 1 | Gm1647 | | 0 | 0 | 0 | 0 | 0 | 0 | 0 | 0 | 2 | 0 | 0 |
| 0 | 1.32 | 3.66 | 0.36 | 0.72 | 1 | Gm12869 | | 0 | 0 | 0 | 0 | 0 | 0 | 0 | 0 | 2 | 0 | 0 |
| 0 | 1.32 | 3.66 | 0.36 | 0.72 | 1 | Gm43772 | | 0 | 0 | 0 | 0 | 0 | 0 | 0 | 0 | 2 | 0 | 0 |
| 0 | 1.32 | 3.66 | 0.36 | 0.72 | 1 | Gm9497 | | 0 | 0 | 0 | 0 | 0 | 0 | 0 | 0 | 2 | 0 | 0 |
| 0 | 1.32 | 3.66 | 0.36 | 0.72 | 1 | Gm23103 | | 0 | 0 | 0 | 0 | 0 | 0 | 0 | 0 | 2 | 0 | 0 |
| 0 | 1.32 | 3.66 | 0.36 | 0.72 | 1 | Mir3569 | | 0 | 0 | 0 | 0 | 0 | 0 | 0 | 0 | 2 | 0 | 0 |
| 0 | 1.32 | 3.66 | 0.36 | 0.72 | 1 | Gm45278 | | 0 | 0 | 0 | 0 | 0 | 0 | 0 | 0 | 2 | 0 | 0 |
| 0 | 1.32 | 3.66 | 0.36 | 0.72 | 1 | Gm24649 | | 0 | 0 | 0 | 0 | 0 | 0 | 0 | 0 | 2 | 0 | 0 |
| 0 | 1.32 | 3.66 | 0.36 | 0.72 | 1 | 5830462I19Rik | | 0 | 0 | 0 | 0 | 0 | 0 | 0 | 0 | 2 | 0 | 0 |
| 0 | 1.32 | 3.66 | 0.36 | 0.72 | 1 | Hbq1b | hemoglobin, theta 1B | 0 | 0 | 0 | 0 | 0 | 0 | 0 | 0 | 2 | 0 | 0 |
| 0 | 1.32 | 3.66 | 0.36 | 0.72 | 1 | Slfn14 | schlafen 14 | 0 | 0 | 0 | 0 | 0 | 0 | 0 | 0 | 2 | 0 | 0 |

| | | | | | | | | | | | | | | | | | |
|---|---|---|---|---|---|---|---|---|---|---|---|---|---|---|---|---|---|
| 0 | 1.32 | 3.66 | 0.36 | 0.72 | 1 | Gm48905 | | 0 | 0 | 0 | 0 | 0 | 0 | 0 | 0 | 2 | 0 | 0 |
| 0 | 1.32 | 3.66 | 0.36 | 0.72 | 1 | Gm48559 | | 0 | 0 | 0 | 0 | 0 | 0 | 0 | 0 | 2 | 0 | 0 |
| 0 | 1.32 | 3.66 | 0.36 | 0.72 | 1 | Gm50469 | | 0 | 0 | 0 | 0 | 0 | 0 | 0 | 0 | 2 | 0 | 0 |
| 0 | 1.32 | 3.66 | 0.36 | 0.72 | 1 | Gm9195 | | 0 | 0 | 0 | 0 | 0 | 0 | 0 | 0 | 2 | 0 | 0 |
| 0 | 1.32 | 3.66 | 0.36 | 0.72 | 1 | Slain1os | | 0 | 0 | 0 | 0 | 0 | 0 | 0 | 0 | 2 | 0 | 0 |
| 0 | 1.32 | 3.66 | 0.36 | 0.72 | 1 | Gm15868 | | 0 | 0 | 0 | 0 | 0 | 0 | 0 | 0 | 2 | 0 | 0 |
| 0 | 1.32 | 3.66 | 0.36 | 0.72 | 1 | Gm33801 | | 0 | 0 | 0 | 0 | 0 | 0 | 0 | 0 | 2 | 0 | 0 |
| 0 | 1.32 | 3.66 | 0.36 | 0.72 | 1 | Nkx2-5 | NK2 homeobox 5 | 0 | 0 | 0 | 0 | 0 | 0 | 0 | 0 | 2 | 0 | 0 |
| 0 | 1.32 | 3.66 | 0.36 | 0.72 | 1 | Gm42723 | | 0 | 0 | 0 | 0 | 0 | 0 | 0 | 0 | 2 | 0 | 0 |
| 0 | 1.00 | 3.68 | 0.27 | 0.79 | 1 | Gm8860 | | 1 | 0 | 0 | 0 | 0 | 0 | 0 | 1 | 0 | 0 | 0 |
| 0 | 1.00 | 3.68 | 0.27 | 0.79 | 1 | Gm12918 | | 1 | 0 | 0 | 0 | 0 | 0 | 0 | 1 | 0 | 0 | 0 |
| 0 | 1.00 | 3.68 | 0.27 | 0.79 | 1 | Gm12807 | | 1 | 0 | 0 | 0 | 0 | 0 | 0 | 1 | 0 | 0 | 0 |
| 0 | 1.00 | 3.68 | 0.27 | 0.79 | 1 | Gm17909 | | 1 | 0 | 0 | 0 | 0 | 0 | 0 | 1 | 0 | 0 | 0 |
| 0 | 1.00 | 3.68 | 0.27 | 0.79 | 1 | Gm47435 | | 1 | 0 | 0 | 0 | 0 | 0 | 0 | 1 | 0 | 0 | 0 |
| 0 | 1.00 | 3.68 | 0.27 | 0.79 | 1 | Gm21959 | | 1 | 0 | 0 | 0 | 0 | 0 | 0 | 1 | 0 | 0 | 0 |
| 0 | 0.28 | 3.68 | 0.08 | 0.94 | 1 | Gm2792 | | 0 | 0 | 0 | 0 | 0 | 1 | 0 | 1 | 0 | 0 | 0 |
| 0 | -0.37 | 3.67 | -0.10 | 0.92 | 1 | Gm13466 | | 0 | 0 | 1 | 0 | 1 | 0 | 0 | 0 | 0 | 0 | 0 |
| 0 | -0.37 | 3.67 | -0.10 | 0.92 | 1 | Gm13132 | | 0 | 0 | 1 | 0 | 1 | 0 | 0 | 0 | 0 | 0 | 0 |
| 0 | -0.37 | 3.67 | -0.10 | 0.92 | 1 | Gm6655 | | 0 | 0 | 1 | 0 | 1 | 0 | 0 | 0 | 0 | 0 | 0 |
| 0 | -0.37 | 3.67 | -0.10 | 0.92 | 1 | Gm19246 | | 0 | 0 | 1 | 0 | 1 | 0 | 0 | 0 | 0 | 0 | 0 |
| 0 | -0.37 | 3.67 | -0.10 | 0.92 | 1 | Gm15568 | | 0 | 0 | 1 | 0 | 1 | 0 | 0 | 0 | 0 | 0 | 0 |
| 0 | -0.37 | 3.67 | -0.10 | 0.92 | 1 | Gm16276 | | 0 | 0 | 1 | 0 | 1 | 0 | 0 | 0 | 0 | 0 | 0 |
| 0 | 0.28 | 3.68 | 0.08 | 0.94 | 1 | Gm20421 | | 0 | 0 | 0 | 0 | 1 | 0 | 0 | 0 | 0 | 0 | 1 |
| 0 | 0.28 | 3.68 | 0.08 | 0.94 | 1 | Gm26243 | | 0 | 0 | 0 | 0 | 1 | 0 | 0 | 0 | 0 | 0 | 1 |
| 0 | 0.28 | 3.68 | 0.08 | 0.94 | 1 | Gm2645 | | 0 | 0 | 0 | 0 | 1 | 0 | 0 | 0 | 0 | 0 | 1 |
| 0 | 0.28 | 3.68 | 0.08 | 0.94 | 1 | Eif3s6-ps3 | | 0 | 0 | 0 | 0 | 1 | 0 | 0 | 0 | 0 | 0 | 1 |
| 0 | 0.28 | 3.68 | 0.08 | 0.94 | 1 | Gm7411 | | 0 | 0 | 0 | 0 | 1 | 0 | 0 | 0 | 0 | 0 | 1 |
| 0 | -0.44 | 3.68 | -0.12 | 0.90 | 1 | Gm8173 | | 0 | 1 | 0 | 0 | 1 | 0 | 0 | 0 | 0 | 0 | 0 |
| 0 | -0.44 | 3.68 | -0.12 | 0.90 | 1 | Gm13294 | | 0 | 1 | 0 | 0 | 1 | 0 | 0 | 0 | 0 | 0 | 0 |
| 0 | -0.44 | 3.68 | -0.12 | 0.90 | 1 | Gm13772 | | 0 | 1 | 0 | 0 | 1 | 0 | 0 | 0 | 0 | 0 | 0 |
| 0 | -0.44 | 3.68 | -0.12 | 0.90 | 1 | Gm5139 | | 0 | 1 | 0 | 0 | 1 | 0 | 0 | 0 | 0 | 0 | 0 |
| 0 | -0.44 | 3.68 | -0.12 | 0.90 | 1 | Gm13015 | | 0 | 1 | 0 | 0 | 1 | 0 | 0 | 0 | 0 | 0 | 0 |
| 0 | -0.44 | 3.68 | -0.12 | 0.90 | 1 | Gm7118 | | 0 | 1 | 0 | 0 | 1 | 0 | 0 | 0 | 0 | 0 | 0 |
| 0 | -0.44 | 3.68 | -0.12 | 0.90 | 1 | Gm10182 | | 0 | 1 | 0 | 0 | 1 | 0 | 0 | 0 | 0 | 0 | 0 |
| 0 | -0.44 | 3.68 | -0.12 | 0.90 | 1 | Gm16373 | | 0 | 1 | 0 | 0 | 1 | 0 | 0 | 0 | 0 | 0 | 0 |
| 0 | -0.44 | 3.68 | -0.12 | 0.90 | 1 | Gm8737 | | 0 | 1 | 0 | 0 | 1 | 0 | 0 | 0 | 0 | 0 | 0 |
| 0 | 0.26 | 3.65 | 0.07 | 0.94 | 1 | Gm12435 | | 0 | 0 | 0 | 0 | 1 | 0 | 0 | 0 | 1 | 0 | 0 |
| 0 | 0.26 | 3.65 | 0.07 | 0.94 | 1 | Gm20072 | | 0 | 0 | 0 | 0 | 1 | 0 | 0 | 0 | 1 | 0 | 0 |
| 0 | 0.26 | 3.65 | 0.07 | 0.94 | 1 | Gm45631 | | 0 | 0 | 0 | 0 | 1 | 0 | 0 | 0 | 1 | 0 | 0 |
| 0 | 0.26 | 3.65 | 0.07 | 0.94 | 1 | Gm39214 | | 0 | 0 | 0 | 0 | 1 | 0 | 0 | 0 | 1 | 0 | 0 |
| 0 | 0.26 | 3.65 | 0.07 | 0.94 | 1 | Kdm4dl | | 0 | 0 | 0 | 0 | 1 | 0 | 0 | 0 | 1 | 0 | 0 |
| 0 | 0.26 | 3.65 | 0.07 | 0.94 | 1 | Gm50006 | | 0 | 0 | 0 | 0 | 1 | 0 | 0 | 0 | 1 | 0 | 0 |
| 0 | 0.26 | 3.65 | 0.07 | 0.94 | 1 | Gm9200 | | 0 | 0 | 0 | 0 | 1 | 0 | 0 | 0 | 1 | 0 | 0 |
| 0 | 0.26 | 3.65 | 0.07 | 0.94 | 1 | Gm8832 | | 0 | 0 | 0 | 0 | 1 | 0 | 0 | 0 | 1 | 0 | 0 |
| 0 | -0.44 | 3.68 | -0.12 | 0.90 | 1 | Gm8048 | | 0 | 0 | 1 | 1 | 0 | 0 | 0 | 0 | 0 | 0 | 0 |
| 0 | -0.44 | 3.68 | -0.12 | 0.90 | 1 | Gm45522 | | 0 | 0 | 1 | 1 | 0 | 0 | 0 | 0 | 0 | 0 | 0 |
| 0 | -0.44 | 3.68 | -0.12 | 0.90 | 1 | Gm19843 | | 0 | 0 | 1 | 1 | 0 | 0 | 0 | 0 | 0 | 0 | 0 |
| 0 | -0.44 | 3.68 | -0.12 | 0.90 | 1 | Gm49119 | | 0 | 0 | 1 | 1 | 0 | 0 | 0 | 0 | 0 | 0 | 0 |
| 0 | -0.44 | 3.68 | -0.12 | 0.90 | 1 | Gm49713 | | 0 | 0 | 1 | 1 | 0 | 0 | 0 | 0 | 0 | 0 | 0 |
| 0 | -0.44 | 3.68 | -0.12 | 0.90 | 1 | Gm49720 | | 0 | 0 | 1 | 1 | 0 | 0 | 0 | 0 | 0 | 0 | 0 |
| 0 | 0.00 | 3.68 | 0.00 | 1.00 | 1 | Gm6104 | | 1 | 0 | 1 | 0 | 0 | 0 | 0 | 0 | 0 | 0 | 0 |
| 0 | 0.00 | 3.68 | 0.00 | 1.00 | 1 | Gm14107 | | 1 | 0 | 1 | 0 | 0 | 0 | 0 | 0 | 0 | 0 | 0 |
| 0 | 0.00 | 3.68 | 0.00 | 1.00 | 1 | Gm8109 | | 1 | 0 | 1 | 0 | 0 | 0 | 0 | 0 | 0 | 0 | 0 |
| 0 | 0.00 | 3.68 | 0.00 | 1.00 | 1 | Gm5847 | | 1 | 0 | 1 | 0 | 0 | 0 | 0 | 0 | 0 | 0 | 0 |
| 0 | 0.00 | 3.68 | 0.00 | 1.00 | 1 | Gm8667 | | 1 | 0 | 1 | 0 | 0 | 0 | 0 | 0 | 0 | 0 | 0 |
| 0 | 0.00 | 3.68 | 0.00 | 1.00 | 1 | Gm4823 | | 1 | 0 | 1 | 0 | 0 | 0 | 0 | 0 | 0 | 0 | 0 |
| 0 | 0.00 | 3.68 | 0.00 | 1.00 | 1 | Gm15807 | | 1 | 0 | 1 | 0 | 0 | 0 | 0 | 0 | 0 | 0 | 0 |
| 0 | 0.00 | 3.68 | 0.00 | 1.00 | 1 | Gm8118 | | 1 | 0 | 1 | 0 | 0 | 0 | 0 | 0 | 0 | 0 | 0 |
| 0 | 0.00 | 3.68 | 0.00 | 1.00 | 1 | Gm7162 | | 1 | 0 | 1 | 0 | 0 | 0 | 0 | 0 | 0 | 0 | 0 |
| 0 | 0.00 | 3.68 | 0.00 | 1.00 | 1 | Gm16519 | | 1 | 0 | 1 | 0 | 0 | 0 | 0 | 0 | 0 | 0 | 0 |

| | | | | | | | | | | | | | | | | | | |
|---|---|---|---|---|---|---|---|---|---|---|---|---|---|---|---|---|---|---|
| 0 | 0.28 | 3.68 | 0.08 | 0.94 | 1 | Gm12842 | | 0 | 0 | 0 | 1 | 0 | 0 | 0 | 0 | 0 | 0 | 1 |
| 0 | 0.28 | 3.68 | 0.08 | 0.94 | 1 | Gm12971 | | 0 | 0 | 0 | 1 | 0 | 0 | 0 | 0 | 0 | 0 | 1 |
| 0 | 0.28 | 3.68 | 0.08 | 0.94 | 1 | Gm42834 | | 0 | 0 | 0 | 1 | 0 | 0 | 0 | 0 | 0 | 0 | 1 |
| 0 | 0.28 | 3.68 | 0.08 | 0.94 | 1 | Gm20156 | | 0 | 0 | 0 | 1 | 0 | 0 | 0 | 0 | 0 | 0 | 1 |
| 0 | 0.28 | 3.68 | 0.08 | 0.94 | 1 | Cycs-ps2 | | 0 | 0 | 0 | 1 | 0 | 0 | 0 | 0 | 0 | 0 | 1 |
| 0 | 0.28 | 3.68 | 0.08 | 0.94 | 1 | Gm19040 | | 0 | 0 | 0 | 1 | 0 | 0 | 0 | 0 | 0 | 0 | 1 |
| 0 | 0.28 | 3.68 | 0.08 | 0.94 | 1 | Gm7183 | | 0 | 0 | 0 | 1 | 0 | 0 | 0 | 0 | 0 | 0 | 1 |
| 0 | 0.28 | 3.68 | 0.08 | 0.94 | 1 | Gm4938 | | 0 | 0 | 0 | 1 | 0 | 0 | 0 | 0 | 0 | 0 | 1 |
| 0 | 0.28 | 3.68 | 0.08 | 0.94 | 1 | Gm50059 | | 0 | 0 | 0 | 1 | 0 | 0 | 0 | 0 | 0 | 0 | 1 |
| 0 | 0.28 | 3.68 | 0.08 | 0.94 | 1 | Gm14776 | | 0 | 0 | 0 | 1 | 0 | 0 | 0 | 0 | 0 | 0 | 1 |
| 0 | -0.80 | 3.66 | -0.22 | 0.83 | 1 | Gm6060 | | 0 | 0 | 0 | 0 | 2 | 0 | 0 | 0 | 0 | 0 | 0 |
| 0 | -0.80 | 3.66 | -0.22 | 0.83 | 1 | Mir8114 | | 0 | 0 | 0 | 0 | 2 | 0 | 0 | 0 | 0 | 0 | 0 |
| 0 | -0.80 | 3.66 | -0.22 | 0.83 | 1 | Gm37313 | | 0 | 0 | 0 | 0 | 2 | 0 | 0 | 0 | 0 | 0 | 0 |
| 0 | -0.80 | 3.66 | -0.22 | 0.83 | 1 | Gm16599 | | 0 | 0 | 0 | 0 | 2 | 0 | 0 | 0 | 0 | 0 | 0 |
| 0 | -0.80 | 3.66 | -0.22 | 0.83 | 1 | Gm15624 | | 0 | 0 | 0 | 0 | 2 | 0 | 0 | 0 | 0 | 0 | 0 |
| 0 | -0.80 | 3.66 | -0.22 | 0.83 | 1 | Gm26554 | | 0 | 0 | 0 | 0 | 2 | 0 | 0 | 0 | 0 | 0 | 0 |
| 0 | -0.80 | 3.66 | -0.22 | 0.83 | 1 | Gm44917 | | 0 | 0 | 0 | 0 | 2 | 0 | 0 | 0 | 0 | 0 | 0 |
| 0 | -0.80 | 3.66 | -0.22 | 0.83 | 1 | Gm20650 | | 0 | 0 | 0 | 0 | 2 | 0 | 0 | 0 | 0 | 0 | 0 |
| 0 | -0.80 | 3.66 | -0.22 | 0.83 | 1 | Cdk19os | | 0 | 0 | 0 | 0 | 2 | 0 | 0 | 0 | 0 | 0 | 0 |
| 0 | -0.80 | 3.66 | -0.22 | 0.83 | 1 | 9230020A06Rik | | 0 | 0 | 0 | 0 | 2 | 0 | 0 | 0 | 0 | 0 | 0 |
| 0 | -0.80 | 3.66 | -0.22 | 0.83 | 1 | Gm31938 | | 0 | 0 | 0 | 0 | 2 | 0 | 0 | 0 | 0 | 0 | 0 |
| 0 | -0.80 | 3.66 | -0.22 | 0.83 | 1 | Hist1h2bq | histone cluster 1, H2bq | 0 | 0 | 0 | 0 | 2 | 0 | 0 | 0 | 0 | 0 | 0 |
| 0 | -0.80 | 3.66 | -0.22 | 0.83 | 1 | Gm30363 | | 0 | 0 | 0 | 0 | 2 | 0 | 0 | 0 | 0 | 0 | 0 |
| 0 | -0.80 | 3.66 | -0.22 | 0.83 | 1 | A430093F15Rik | | 0 | 0 | 0 | 0 | 2 | 0 | 0 | 0 | 0 | 0 | 0 |
| 0 | -0.36 | 3.67 | -0.10 | 0.92 | 1 | Gm7621 | | 0 | 1 | 0 | 1 | 0 | 0 | 0 | 0 | 0 | 0 | 0 |
| 0 | -0.36 | 3.67 | -0.10 | 0.92 | 1 | Gm7788 | | 0 | 1 | 0 | 1 | 0 | 0 | 0 | 0 | 0 | 0 | 0 |
| 0 | -0.36 | 3.67 | -0.10 | 0.92 | 1 | Gm50360 | | 0 | 1 | 0 | 1 | 0 | 0 | 0 | 0 | 0 | 0 | 0 |
| 0 | -0.36 | 3.67 | -0.10 | 0.92 | 1 | Gapdh-ps15 | | 0 | 1 | 0 | 1 | 0 | 0 | 0 | 0 | 0 | 0 | 0 |
| 0 | 0.28 | 3.68 | 0.08 | 0.94 | 1 | Gm7266 | | 0 | 0 | 0 | 1 | 0 | 0 | 0 | 0 | 1 | 0 | 0 |
| 0 | 0.28 | 3.68 | 0.08 | 0.94 | 1 | Gm9167 | | 0 | 0 | 0 | 1 | 0 | 0 | 0 | 0 | 1 | 0 | 0 |
| 0 | 0.28 | 3.68 | 0.08 | 0.94 | 1 | Gm6125 | | 0 | 0 | 0 | 1 | 0 | 0 | 0 | 0 | 1 | 0 | 0 |
| 0 | -0.40 | 3.67 | -0.11 | 0.91 | 1 | Gm12422 | | 0 | 0 | 1 | 0 | 0 | 1 | 0 | 0 | 0 | 0 | 0 |
| 0 | -0.40 | 3.67 | -0.11 | 0.91 | 1 | Gm7931 | | 0 | 0 | 1 | 0 | 0 | 1 | 0 | 0 | 0 | 0 | 0 |
| 0 | -0.40 | 3.67 | -0.11 | 0.91 | 1 | Gm45293 | | 0 | 0 | 1 | 0 | 0 | 1 | 0 | 0 | 0 | 0 | 0 |
| 0 | -0.40 | 3.67 | -0.11 | 0.91 | 1 | Gm21089 | | 0 | 0 | 1 | 0 | 0 | 1 | 0 | 0 | 0 | 0 | 0 |
| 0 | -0.40 | 3.67 | -0.11 | 0.91 | 1 | Gm9176 | | 0 | 0 | 1 | 0 | 0 | 1 | 0 | 0 | 0 | 0 | 0 |
| 0 | -0.40 | 3.67 | -0.11 | 0.91 | 1 | Gm4812 | | 0 | 0 | 1 | 0 | 0 | 1 | 0 | 0 | 0 | 0 | 0 |
| 0 | -0.40 | 3.67 | -0.11 | 0.91 | 1 | Gm15808 | | 0 | 0 | 1 | 0 | 0 | 1 | 0 | 0 | 0 | 0 | 0 |
| 0 | 0.95 | 3.67 | 0.26 | 0.80 | 1 | Gm34144 | | 1 | 0 | 0 | 0 | 0 | 0 | 0 | 0 | 0 | 0 | 1 |
| 0 | 0.95 | 3.67 | 0.26 | 0.80 | 1 | Gm15497 | | 1 | 0 | 0 | 0 | 0 | 0 | 0 | 0 | 0 | 0 | 1 |
| 0 | 0.95 | 3.67 | 0.26 | 0.80 | 1 | Gm47481 | | 1 | 0 | 0 | 0 | 0 | 0 | 0 | 0 | 0 | 0 | 1 |
| 0 | 0.95 | 3.67 | 0.26 | 0.80 | 1 | Gm11307 | | 1 | 0 | 0 | 0 | 0 | 0 | 0 | 0 | 0 | 0 | 1 |
| 0 | 0.95 | 3.67 | 0.26 | 0.80 | 1 | Gm36758 | | 1 | 0 | 0 | 0 | 0 | 0 | 0 | 0 | 0 | 0 | 1 |
| 0 | 0.95 | 3.67 | 0.26 | 0.80 | 1 | Gm4057 | | 1 | 0 | 0 | 0 | 0 | 0 | 0 | 0 | 0 | 0 | 1 |
| 0 | 0.95 | 3.67 | 0.26 | 0.80 | 1 | Gm5766 | | 1 | 0 | 0 | 0 | 0 | 0 | 0 | 0 | 0 | 0 | 1 |
| 0 | 0.95 | 3.67 | 0.26 | 0.80 | 1 | Gm46578 | | 1 | 0 | 0 | 0 | 0 | 0 | 0 | 0 | 0 | 0 | 1 |
| 0 | 0.95 | 3.67 | 0.26 | 0.80 | 1 | BE692007 | | 1 | 0 | 0 | 0 | 0 | 0 | 0 | 0 | 0 | 0 | 1 |
| 0 | 0.00 | 3.68 | 0.00 | 1.00 | 1 | Gm38247 | | 1 | 1 | 0 | 0 | 0 | 0 | 0 | 0 | 0 | 0 | 0 |
| 0 | 0.00 | 3.68 | 0.00 | 1.00 | 1 | Gm7774 | | 1 | 1 | 0 | 0 | 0 | 0 | 0 | 0 | 0 | 0 | 0 |
| 0 | 0.00 | 3.68 | 0.00 | 1.00 | 1 | Gm7545 | | 1 | 1 | 0 | 0 | 0 | 0 | 0 | 0 | 0 | 0 | 0 |
| 0 | 0.00 | 3.68 | 0.00 | 1.00 | 1 | Gm15661 | | 1 | 1 | 0 | 0 | 0 | 0 | 0 | 0 | 0 | 0 | 0 |
| 0 | 0.00 | 3.68 | 0.00 | 1.00 | 1 | Rps19-ps10 | | 1 | 1 | 0 | 0 | 0 | 0 | 0 | 0 | 0 | 0 | 0 |
| 0 | 0.00 | 3.68 | 0.00 | 1.00 | 1 | Gm12271 | | 1 | 1 | 0 | 0 | 0 | 0 | 0 | 0 | 0 | 0 | 0 |
| 0 | 0.00 | 3.68 | 0.00 | 1.00 | 1 | Ighv2-6 | | 1 | 1 | 0 | 0 | 0 | 0 | 0 | 0 | 0 | 0 | 0 |
| 0 | 0.00 | 3.68 | 0.00 | 1.00 | 1 | Gm5473 | | 1 | 1 | 0 | 0 | 0 | 0 | 0 | 0 | 0 | 0 | 0 |
| 0 | 0.00 | 3.68 | 0.00 | 1.00 | 1 | Gm45338 | | 1 | 1 | 0 | 0 | 0 | 0 | 0 | 0 | 0 | 0 | 0 |
| 0 | 1.00 | 3.68 | 0.27 | 0.79 | 1 | Gm5100 | | 1 | 0 | 0 | 0 | 0 | 0 | 0 | 0 | 1 | 0 | 0 |
| 0 | 1.00 | 3.68 | 0.27 | 0.79 | 1 | Gm33312 | | 1 | 0 | 0 | 0 | 0 | 0 | 0 | 0 | 1 | 0 | 0 |
| 0 | 1.00 | 3.68 | 0.27 | 0.79 | 1 | Gm17799 | | 1 | 0 | 0 | 0 | 0 | 0 | 0 | 0 | 1 | 0 | 0 |
| 0 | 1.00 | 3.68 | 0.27 | 0.79 | 1 | Gm6995 | | 1 | 0 | 0 | 0 | 0 | 0 | 0 | 0 | 1 | 0 | 0 |

| | | | | | | | | | | | | | | | | | | |
|---|---|---|---|---|---|---|---|---|---|---|---|---|---|---|---|---|---|---|
| 0 | 0.28 | 3.64 | 0.08 | 0.94 | 1 | Gm37949 | | 0 | 0 | 0 | 0 | 0 | 1 | 0 | 0 | 0 | 0 | 1 |
| 0 | 0.28 | 3.64 | 0.08 | 0.94 | 1 | Gm34068 | | 0 | 0 | 0 | 0 | 0 | 1 | 0 | 0 | 0 | 0 | 1 |
| 0 | 0.28 | 3.64 | 0.08 | 0.94 | 1 | Gm13352 | | 0 | 0 | 0 | 0 | 0 | 1 | 0 | 0 | 0 | 0 | 1 |
| 0 | 0.28 | 3.64 | 0.08 | 0.94 | 1 | Tbr1 | T-box brain gene 1 | 0 | 0 | 0 | 0 | 0 | 1 | 0 | 0 | 0 | 0 | 1 |
| 0 | 0.28 | 3.64 | 0.08 | 0.94 | 1 | Mir684-1 | | 0 | 0 | 0 | 0 | 0 | 1 | 0 | 0 | 0 | 0 | 1 |
| 0 | 0.28 | 3.64 | 0.08 | 0.94 | 1 | Gm25583 | | 0 | 0 | 0 | 0 | 0 | 1 | 0 | 0 | 0 | 0 | 1 |
| 0 | 0.28 | 3.64 | 0.08 | 0.94 | 1 | Gm24786 | | 0 | 0 | 0 | 0 | 0 | 1 | 0 | 0 | 0 | 0 | 1 |
| 0 | 0.28 | 3.64 | 0.08 | 0.94 | 1 | Gm8957 | | 0 | 0 | 0 | 0 | 0 | 1 | 0 | 0 | 0 | 0 | 1 |
| 0 | 0.28 | 3.64 | 0.08 | 0.94 | 1 | Gm12424 | | 0 | 0 | 0 | 0 | 0 | 1 | 0 | 0 | 0 | 0 | 1 |
| 0 | 0.28 | 3.64 | 0.08 | 0.94 | 1 | Gm42438 | | 0 | 0 | 0 | 0 | 0 | 1 | 0 | 0 | 0 | 0 | 1 |
| 0 | 0.28 | 3.64 | 0.08 | 0.94 | 1 | Gm25852 | | 0 | 0 | 0 | 0 | 0 | 1 | 0 | 0 | 0 | 0 | 1 |
| 0 | 0.28 | 3.64 | 0.08 | 0.94 | 1 | Gm37269 | | 0 | 0 | 0 | 0 | 0 | 1 | 0 | 0 | 0 | 0 | 1 |
| 0 | 0.28 | 3.64 | 0.08 | 0.94 | 1 | Gm20449 | | 0 | 0 | 0 | 0 | 0 | 1 | 0 | 0 | 0 | 0 | 1 |
| 0 | 0.28 | 3.64 | 0.08 | 0.94 | 1 | Magel2 | melanoma antigen, family L, 2 | 0 | 0 | 0 | 0 | 0 | 1 | 0 | 0 | 0 | 0 | 1 |
| 0 | 0.28 | 3.64 | 0.08 | 0.94 | 1 | Gm31510 | | 0 | 0 | 0 | 0 | 0 | 1 | 0 | 0 | 0 | 0 | 1 |
| 0 | 0.28 | 3.64 | 0.08 | 0.94 | 1 | Gm5373 | | 0 | 0 | 0 | 0 | 0 | 1 | 0 | 0 | 0 | 0 | 1 |
| 0 | 0.28 | 3.64 | 0.08 | 0.94 | 1 | Mir23a | | 0 | 0 | 0 | 0 | 0 | 1 | 0 | 0 | 0 | 0 | 1 |
| 0 | 0.28 | 3.64 | 0.08 | 0.94 | 1 | Gm48129 | | 0 | 0 | 0 | 0 | 0 | 1 | 0 | 0 | 0 | 0 | 1 |
| 0 | 0.28 | 3.64 | 0.08 | 0.94 | 1 | Gm22659 | | 0 | 0 | 0 | 0 | 0 | 1 | 0 | 0 | 0 | 0 | 1 |
| 0 | 0.28 | 3.64 | 0.08 | 0.94 | 1 | Gm18821 | | 0 | 0 | 0 | 0 | 0 | 1 | 0 | 0 | 0 | 0 | 1 |
| 0 | 0.28 | 3.64 | 0.08 | 0.94 | 1 | Gm16128 | | 0 | 0 | 0 | 0 | 0 | 1 | 0 | 0 | 0 | 0 | 1 |
| 0 | 0.28 | 3.64 | 0.08 | 0.94 | 1 | Gm48298 | | 0 | 0 | 0 | 0 | 0 | 1 | 0 | 0 | 0 | 0 | 1 |
| 0 | 0.28 | 3.64 | 0.08 | 0.94 | 1 | Gm24195 | | 0 | 0 | 0 | 0 | 0 | 1 | 0 | 0 | 0 | 0 | 1 |
| 0 | 0.28 | 3.64 | 0.08 | 0.94 | 1 | Gm48965 | | 0 | 0 | 0 | 0 | 0 | 1 | 0 | 0 | 0 | 0 | 1 |
| 0 | 0.28 | 3.64 | 0.08 | 0.94 | 1 | n-R5s40 | | 0 | 0 | 0 | 0 | 0 | 1 | 0 | 0 | 0 | 0 | 1 |
| 0 | 0.28 | 3.64 | 0.08 | 0.94 | 1 | Mir1947 | | 0 | 0 | 0 | 0 | 0 | 1 | 0 | 0 | 0 | 0 | 1 |
| 0 | 0.28 | 3.64 | 0.08 | 0.94 | 1 | Abca17 | ATP-binding cassette, sub-family A (ABC | 0 | 0 | 0 | 0 | 0 | 1 | 0 | 0 | 0 | 0 | 1 |
| 0 | 0.28 | 3.64 | 0.08 | 0.94 | 1 | Mir6975 | | 0 | 0 | 0 | 0 | 0 | 1 | 0 | 0 | 0 | 0 | 1 |
| 0 | -0.40 | 3.67 | -0.11 | 0.91 | 1 | Gm6187 | | 0 | 1 | 0 | 0 | 0 | 1 | 0 | 0 | 0 | 0 | 0 |
| 0 | -0.40 | 3.67 | -0.11 | 0.91 | 1 | Gm8762 | | 0 | 1 | 0 | 0 | 0 | 1 | 0 | 0 | 0 | 0 | 0 |
| 0 | -0.40 | 3.67 | -0.11 | 0.91 | 1 | Lhx9 | LIM homeobox protein 9 | 0 | 1 | 0 | 0 | 0 | 1 | 0 | 0 | 0 | 0 | 0 |
| 0 | -0.40 | 3.67 | -0.11 | 0.91 | 1 | Gm42566 | | 0 | 1 | 0 | 0 | 0 | 1 | 0 | 0 | 0 | 0 | 0 |
| 0 | -0.40 | 3.67 | -0.11 | 0.91 | 1 | Gm29763 | | 0 | 1 | 0 | 0 | 0 | 1 | 0 | 0 | 0 | 0 | 0 |
| 0 | -0.40 | 3.67 | -0.11 | 0.91 | 1 | Gm49506 | | 0 | 1 | 0 | 0 | 0 | 1 | 0 | 0 | 0 | 0 | 0 |
| 0 | -0.40 | 3.67 | -0.11 | 0.91 | 1 | Ppp1r2-ps1 | | 0 | 1 | 0 | 0 | 0 | 1 | 0 | 0 | 0 | 0 | 0 |
| 0 | -0.40 | 3.67 | -0.11 | 0.91 | 1 | H2-M5 | histocompatibility 2, M region locus 5 | 0 | 1 | 0 | 0 | 0 | 1 | 0 | 0 | 0 | 0 | 0 |
| 0 | 0.28 | 3.68 | 0.08 | 0.94 | 1 | Gm7775 | | 0 | 0 | 0 | 0 | 0 | 1 | 0 | 0 | 1 | 0 | 0 |
| 0 | 0.28 | 3.68 | 0.08 | 0.94 | 1 | Gm48110 | | 0 | 0 | 0 | 0 | 0 | 1 | 0 | 0 | 1 | 0 | 0 |
| 0 | 0.28 | 3.68 | 0.08 | 0.94 | 1 | Gm50198 | | 0 | 0 | 0 | 0 | 0 | 1 | 0 | 0 | 1 | 0 | 0 |
| 0 | -1.03 | 3.66 | -0.28 | 0.78 | 1 | Gm8596 | | 0 | 0 | 0 | 1 | 1 | 0 | 0 | 0 | 0 | 0 | 0 |
| 0 | -1.03 | 3.66 | -0.28 | 0.78 | 1 | Gm9423 | | 0 | 0 | 0 | 1 | 1 | 0 | 0 | 0 | 0 | 0 | 0 |
| 0 | -1.03 | 3.66 | -0.28 | 0.78 | 1 | Gm13172 | | 0 | 0 | 0 | 1 | 1 | 0 | 0 | 0 | 0 | 0 | 0 |
| 0 | -1.03 | 3.66 | -0.28 | 0.78 | 1 | Gm4751 | | 0 | 0 | 0 | 1 | 1 | 0 | 0 | 0 | 0 | 0 | 0 |
| 0 | -1.03 | 3.66 | -0.28 | 0.78 | 1 | Gm9729 | | 0 | 0 | 0 | 1 | 1 | 0 | 0 | 0 | 0 | 0 | 0 |
| 0 | -1.03 | 3.66 | -0.28 | 0.78 | 1 | Gnb3 | guanine nucleotide binding protein (G pro | 0 | 0 | 0 | 1 | 1 | 0 | 0 | 0 | 0 | 0 | 0 |
| 0 | -1.03 | 3.66 | -0.28 | 0.78 | 1 | Gm14630 | | 0 | 0 | 0 | 1 | 1 | 0 | 0 | 0 | 0 | 0 | 0 |
| 0 | -1.03 | 3.66 | -0.28 | 0.78 | 1 | Gm7108 | | 0 | 0 | 0 | 1 | 1 | 0 | 0 | 0 | 0 | 0 | 0 |
| 0 | -0.44 | 3.68 | -0.12 | 0.90 | 1 | Rpl32-ps | | 1 | 0 | 0 | 0 | 1 | 0 | 0 | 0 | 0 | 0 | 0 |
| 0 | -0.44 | 3.68 | -0.12 | 0.90 | 1 | Gm12517 | | 1 | 0 | 0 | 0 | 1 | 0 | 0 | 0 | 0 | 0 | 0 |
| 0 | -0.44 | 3.68 | -0.12 | 0.90 | 1 | Gm12458 | | 1 | 0 | 0 | 0 | 1 | 0 | 0 | 0 | 0 | 0 | 0 |
| 0 | -0.44 | 3.68 | -0.12 | 0.90 | 1 | Gm12936 | | 1 | 0 | 0 | 0 | 1 | 0 | 0 | 0 | 0 | 0 | 0 |
| 0 | -0.44 | 3.68 | -0.12 | 0.90 | 1 | Gm15779 | | 1 | 0 | 0 | 0 | 1 | 0 | 0 | 0 | 0 | 0 | 0 |
| 0 | -0.44 | 3.68 | -0.12 | 0.90 | 1 | Gm8512 | | 1 | 0 | 0 | 0 | 1 | 0 | 0 | 0 | 0 | 0 | 0 |
| 0 | -0.44 | 3.68 | -0.12 | 0.90 | 1 | Gm8697 | | 1 | 0 | 0 | 0 | 1 | 0 | 0 | 0 | 0 | 0 | 0 |
| 0 | -1.04 | 3.66 | -0.28 | 0.78 | 1 | Dnmt3c | | 0 | 0 | 0 | 0 | 1 | 1 | 0 | 0 | 0 | 0 | 0 |
| 0 | -1.04 | 3.66 | -0.28 | 0.78 | 1 | Gm11838 | | 0 | 0 | 0 | 0 | 1 | 1 | 0 | 0 | 0 | 0 | 0 |
| 0 | -1.04 | 3.66 | -0.28 | 0.78 | 1 | Gm44709 | | 0 | 0 | 0 | 0 | 1 | 1 | 0 | 0 | 0 | 0 | 0 |
| 0 | -1.04 | 3.66 | -0.28 | 0.78 | 1 | Gm10709 | | 0 | 0 | 0 | 0 | 1 | 1 | 0 | 0 | 0 | 0 | 0 |
| 0 | -1.04 | 3.66 | -0.28 | 0.78 | 1 | Gm11611 | | 0 | 0 | 0 | 0 | 1 | 1 | 0 | 0 | 0 | 0 | 0 |
| 0 | -1.04 | 3.66 | -0.28 | 0.78 | 1 | Gm8839 | | 0 | 0 | 0 | 0 | 1 | 1 | 0 | 0 | 0 | 0 | 0 |

| | | | | | | | | | | | | | | | | | | |
|---|---|---|---|---|---|---|---|---|---|---|---|---|---|---|---|---|---|---|
| 0 | -1.04 | 3.66 | -0.28 | 0.78 | 1 | Gm7222 | | 0 | 0 | 0 | 0 | 1 | 1 | 0 | 0 | 0 | 0 | 0 |
| 0 | -0.77 | 3.66 | -0.21 | 0.83 | 1 | Gm25312 | | 0 | 0 | 0 | 2 | 0 | 0 | 0 | 0 | 0 | 0 | 0 |
| 0 | -0.77 | 3.66 | -0.21 | 0.83 | 1 | Gm15557 | | 0 | 0 | 0 | 2 | 0 | 0 | 0 | 0 | 0 | 0 | 0 |
| 0 | -0.77 | 3.66 | -0.21 | 0.83 | 1 | 4921514A10Rik | | 0 | 0 | 0 | 2 | 0 | 0 | 0 | 0 | 0 | 0 | 0 |
| 0 | -0.77 | 3.66 | -0.21 | 0.83 | 1 | Gm26931 | | 0 | 0 | 0 | 2 | 0 | 0 | 0 | 0 | 0 | 0 | 0 |
| 0 | -0.77 | 3.66 | -0.21 | 0.83 | 1 | Gm26058 | | 0 | 0 | 0 | 2 | 0 | 0 | 0 | 0 | 0 | 0 | 0 |
| 0 | -0.77 | 3.66 | -0.21 | 0.83 | 1 | Gm16553 | | 0 | 0 | 0 | 2 | 0 | 0 | 0 | 0 | 0 | 0 | 0 |
| 0 | -0.77 | 3.66 | -0.21 | 0.83 | 1 | Htr3a | 5-hydroxytryptamine (serotonin) receptor | 0 | 0 | 0 | 2 | 0 | 0 | 0 | 0 | 0 | 0 | 0 |
| 0 | -0.77 | 3.66 | -0.21 | 0.83 | 1 | Gm47077 | | 0 | 0 | 0 | 2 | 0 | 0 | 0 | 0 | 0 | 0 | 0 |
| 0 | -0.77 | 3.66 | -0.21 | 0.83 | 1 | Gm9719 | | 0 | 0 | 0 | 2 | 0 | 0 | 0 | 0 | 0 | 0 | 0 |
| 0 | -0.77 | 3.66 | -0.21 | 0.83 | 1 | Gm48975 | | 0 | 0 | 0 | 2 | 0 | 0 | 0 | 0 | 0 | 0 | 0 |
| 0 | -0.77 | 3.66 | -0.21 | 0.83 | 1 | 9330117O12Rik | | 0 | 0 | 0 | 2 | 0 | 0 | 0 | 0 | 0 | 0 | 0 |
| 0 | -0.44 | 3.68 | -0.12 | 0.90 | 1 | Gm7924 | | 1 | 0 | 0 | 1 | 0 | 0 | 0 | 0 | 0 | 0 | 0 |
| 0 | -0.44 | 3.68 | -0.12 | 0.90 | 1 | Gm6654 | | 1 | 0 | 0 | 1 | 0 | 0 | 0 | 0 | 0 | 0 | 0 |
| 0 | -0.44 | 3.68 | -0.12 | 0.90 | 1 | Gm12003 | | 1 | 0 | 0 | 1 | 0 | 0 | 0 | 0 | 0 | 0 | 0 |
| 0 | -0.44 | 3.68 | -0.12 | 0.90 | 1 | Gm50429 | | 1 | 0 | 0 | 1 | 0 | 0 | 0 | 0 | 0 | 0 | 0 |
| 0 | -1.03 | 3.66 | -0.28 | 0.78 | 1 | Htr5b | 5-hydroxytryptamine (serotonin) receptor | 0 | 0 | 0 | 1 | 0 | 1 | 0 | 0 | 0 | 0 | 0 |
| 0 | -1.03 | 3.66 | -0.28 | 0.78 | 1 | Gm13912 | | 0 | 0 | 0 | 1 | 0 | 1 | 0 | 0 | 0 | 0 | 0 |
| 0 | -1.03 | 3.66 | -0.28 | 0.78 | 1 | Opn1sw | opsin 1 (cone pigments), short-wave-sens | 0 | 0 | 0 | 1 | 0 | 1 | 0 | 0 | 0 | 0 | 0 |
| 0 | -1.03 | 3.66 | -0.28 | 0.78 | 1 | Moap1 | modulator of apoptosis 1 | 0 | 0 | 0 | 1 | 0 | 1 | 0 | 0 | 0 | 0 | 0 |
| 0 | -1.03 | 3.66 | -0.28 | 0.78 | 1 | n-R5s41 | | 0 | 0 | 0 | 1 | 0 | 1 | 0 | 0 | 0 | 0 | 0 |
| 0 | -1.03 | 3.66 | -0.28 | 0.78 | 1 | Gm14965 | | 0 | 0 | 0 | 1 | 0 | 1 | 0 | 0 | 0 | 0 | 0 |
| 0 | -1.03 | 3.66 | -0.28 | 0.78 | 1 | Gm7416 | | 0 | 0 | 0 | 1 | 0 | 1 | 0 | 0 | 0 | 0 | 0 |
| 0 | 0.00 | 3.68 | 0.00 | 1.00 | 1 | Ugt1a8 | | 2 | 0 | 0 | 0 | 0 | 0 | 0 | 0 | 0 | 0 | 0 |
| 0 | 0.00 | 3.68 | 0.00 | 1.00 | 1 | Olfr1413 | olfactory receptor 1413 | 2 | 0 | 0 | 0 | 0 | 0 | 0 | 0 | 0 | 0 | 0 |
| 0 | 0.00 | 3.68 | 0.00 | 1.00 | 1 | Atp6v1g3 | ATPase, H+ transporting, lysosomal V1 su | 2 | 0 | 0 | 0 | 0 | 0 | 0 | 0 | 0 | 0 | 0 |
| 0 | 0.00 | 3.68 | 0.00 | 1.00 | 1 | Gm15867 | | 2 | 0 | 0 | 0 | 0 | 0 | 0 | 0 | 0 | 0 | 0 |
| 0 | 0.00 | 3.68 | 0.00 | 1.00 | 1 | AI847159 | | 2 | 0 | 0 | 0 | 0 | 0 | 0 | 0 | 0 | 0 | 0 |
| 0 | 0.00 | 3.68 | 0.00 | 1.00 | 1 | Gm14156 | | 2 | 0 | 0 | 0 | 0 | 0 | 0 | 0 | 0 | 0 | 0 |
| 0 | 0.00 | 3.68 | 0.00 | 1.00 | 1 | Gm14391 | predicted gene 14391 | 2 | 0 | 0 | 0 | 0 | 0 | 0 | 0 | 0 | 0 | 0 |
| 0 | 0.00 | 3.68 | 0.00 | 1.00 | 1 | Gm37131 | | 2 | 0 | 0 | 0 | 0 | 0 | 0 | 0 | 0 | 0 | 0 |
| 0 | 0.00 | 3.68 | 0.00 | 1.00 | 1 | 4632404M16Rik | | 2 | 0 | 0 | 0 | 0 | 0 | 0 | 0 | 0 | 0 | 0 |
| 0 | 0.00 | 3.68 | 0.00 | 1.00 | 1 | En2 | engrailed 2 | 2 | 0 | 0 | 0 | 0 | 0 | 0 | 0 | 0 | 0 | 0 |
| 0 | 0.00 | 3.68 | 0.00 | 1.00 | 1 | Gm10459 | | 2 | 0 | 0 | 0 | 0 | 0 | 0 | 0 | 0 | 0 | 0 |
| 0 | 0.00 | 3.68 | 0.00 | 1.00 | 1 | Caln1 | calneuron 1 | 2 | 0 | 0 | 0 | 0 | 0 | 0 | 0 | 0 | 0 | 0 |
| 0 | 0.00 | 3.68 | 0.00 | 1.00 | 1 | Gm42808 | | 2 | 0 | 0 | 0 | 0 | 0 | 0 | 0 | 0 | 0 | 0 |
| 0 | 0.00 | 3.68 | 0.00 | 1.00 | 1 | Gm43380 | | 2 | 0 | 0 | 0 | 0 | 0 | 0 | 0 | 0 | 0 | 0 |
| 0 | 0.00 | 3.68 | 0.00 | 1.00 | 1 | D5Ertd605e | | 2 | 0 | 0 | 0 | 0 | 0 | 0 | 0 | 0 | 0 | 0 |
| 0 | 0.00 | 3.68 | 0.00 | 1.00 | 1 | Igkv4-57 | | 2 | 0 | 0 | 0 | 0 | 0 | 0 | 0 | 0 | 0 | 0 |
| 0 | 0.00 | 3.68 | 0.00 | 1.00 | 1 | A730049H05Rik | | 2 | 0 | 0 | 0 | 0 | 0 | 0 | 0 | 0 | 0 | 0 |
| 0 | 0.00 | 3.68 | 0.00 | 1.00 | 1 | Umod | uromodulin | 2 | 0 | 0 | 0 | 0 | 0 | 0 | 0 | 0 | 0 | 0 |
| 0 | 0.00 | 3.68 | 0.00 | 1.00 | 1 | 2210406H18Rik | | 2 | 0 | 0 | 0 | 0 | 0 | 0 | 0 | 0 | 0 | 0 |
| 0 | 0.00 | 3.68 | 0.00 | 1.00 | 1 | Gm38414 | | 2 | 0 | 0 | 0 | 0 | 0 | 0 | 0 | 0 | 0 | 0 |
| 0 | 0.00 | 3.68 | 0.00 | 1.00 | 1 | Gm33023 | | 2 | 0 | 0 | 0 | 0 | 0 | 0 | 0 | 0 | 0 | 0 |
| 0 | 0.00 | 3.68 | 0.00 | 1.00 | 1 | Gm45865 | | 2 | 0 | 0 | 0 | 0 | 0 | 0 | 0 | 0 | 0 | 0 |
| 0 | 0.00 | 3.68 | 0.00 | 1.00 | 1 | Olfr921 | olfactory receptor 921 | 2 | 0 | 0 | 0 | 0 | 0 | 0 | 0 | 0 | 0 | 0 |
| 0 | 0.00 | 3.68 | 0.00 | 1.00 | 1 | Gm31698 | | 2 | 0 | 0 | 0 | 0 | 0 | 0 | 0 | 0 | 0 | 0 |
| 0 | 0.00 | 3.68 | 0.00 | 1.00 | 1 | 9630041A04Rik | | 2 | 0 | 0 | 0 | 0 | 0 | 0 | 0 | 0 | 0 | 0 |
| 0 | 0.00 | 3.68 | 0.00 | 1.00 | 1 | Gm26740 | | 2 | 0 | 0 | 0 | 0 | 0 | 0 | 0 | 0 | 0 | 0 |
| 0 | 0.00 | 3.68 | 0.00 | 1.00 | 1 | Gm16271 | | 2 | 0 | 0 | 0 | 0 | 0 | 0 | 0 | 0 | 0 | 0 |
| 0 | 0.00 | 3.68 | 0.00 | 1.00 | 1 | Gm22507 | | 2 | 0 | 0 | 0 | 0 | 0 | 0 | 0 | 0 | 0 | 0 |
| 0 | 0.00 | 3.68 | 0.00 | 1.00 | 1 | Gm11998 | | 2 | 0 | 0 | 0 | 0 | 0 | 0 | 0 | 0 | 0 | 0 |
| 0 | 0.00 | 3.68 | 0.00 | 1.00 | 1 | Sox15 | SRY (sex determining region Y)-box 15 | 2 | 0 | 0 | 0 | 0 | 0 | 0 | 0 | 0 | 0 | 0 |
| 0 | 0.00 | 3.68 | 0.00 | 1.00 | 1 | Psmb5-ps | | 2 | 0 | 0 | 0 | 0 | 0 | 0 | 0 | 0 | 0 | 0 |
| 0 | 0.00 | 3.68 | 0.00 | 1.00 | 1 | Gm48008 | | 2 | 0 | 0 | 0 | 0 | 0 | 0 | 0 | 0 | 0 | 0 |
| 0 | 0.00 | 3.68 | 0.00 | 1.00 | 1 | Tdrd9 | tudor domain containing 9 | 2 | 0 | 0 | 0 | 0 | 0 | 0 | 0 | 0 | 0 | 0 |
| 0 | 0.00 | 3.68 | 0.00 | 1.00 | 1 | Adam6b | a disintegrin and metallopeptidase domai | 2 | 0 | 0 | 0 | 0 | 0 | 0 | 0 | 0 | 0 | 0 |
| 0 | 0.00 | 3.68 | 0.00 | 1.00 | 1 | Ighv8-10 | | 2 | 0 | 0 | 0 | 0 | 0 | 0 | 0 | 0 | 0 | 0 |
| 0 | 0.00 | 3.68 | 0.00 | 1.00 | 1 | Gm6105 | | 2 | 0 | 0 | 0 | 0 | 0 | 0 | 0 | 0 | 0 | 0 |
| 0 | 0.00 | 3.68 | 0.00 | 1.00 | 1 | Ighv1-83 | | 2 | 0 | 0 | 0 | 0 | 0 | 0 | 0 | 0 | 0 | 0 |

| | | | | | | | | | | | | | | | | | | |
|---|---|---|---|---|---|---|---|---|---|---|---|---|---|---|---|---|---|---|
| 0 | 0.00 | 3.68 | 0.00 | 1.00 | 1 | Gm10790 | | 2 | 0 | 0 | 0 | 0 | 0 | 0 | 0 | 0 | 0 | 0 |
| 0 | 0.00 | 3.68 | 0.00 | 1.00 | 1 | Gm47533 | | 2 | 0 | 0 | 0 | 0 | 0 | 0 | 0 | 0 | 0 | 0 |
| 0 | 0.00 | 3.68 | 0.00 | 1.00 | 1 | Gm49194 | | 2 | 0 | 0 | 0 | 0 | 0 | 0 | 0 | 0 | 0 | 0 |
| 0 | 0.00 | 3.68 | 0.00 | 1.00 | 1 | A530076I17Rik | | 2 | 0 | 0 | 0 | 0 | 0 | 0 | 0 | 0 | 0 | 0 |
| 0 | 0.00 | 3.68 | 0.00 | 1.00 | 1 | Trav3-1 | | 2 | 0 | 0 | 0 | 0 | 0 | 0 | 0 | 0 | 0 | 0 |
| 0 | 0.00 | 3.68 | 0.00 | 1.00 | 1 | Traj44 | | 2 | 0 | 0 | 0 | 0 | 0 | 0 | 0 | 0 | 0 | 0 |
| 0 | 0.00 | 3.68 | 0.00 | 1.00 | 1 | Gm16029 | | 2 | 0 | 0 | 0 | 0 | 0 | 0 | 0 | 0 | 0 | 0 |
| 0 | 0.00 | 3.68 | 0.00 | 1.00 | 1 | Gm3787 | | 2 | 0 | 0 | 0 | 0 | 0 | 0 | 0 | 0 | 0 | 0 |
| 0 | 0.00 | 3.68 | 0.00 | 1.00 | 1 | Gm32885 | | 2 | 0 | 0 | 0 | 0 | 0 | 0 | 0 | 0 | 0 | 0 |
| 0 | 0.00 | 3.68 | 0.00 | 1.00 | 1 | Gm26800 | | 2 | 0 | 0 | 0 | 0 | 0 | 0 | 0 | 0 | 0 | 0 |
| 0 | 0.00 | 3.68 | 0.00 | 1.00 | 1 | Gpr31a | | 2 | 0 | 0 | 0 | 0 | 0 | 0 | 0 | 0 | 0 | 0 |
| 0 | 0.00 | 3.68 | 0.00 | 1.00 | 1 | Gm31087 | | 2 | 0 | 0 | 0 | 0 | 0 | 0 | 0 | 0 | 0 | 0 |
| 0 | 0.00 | 3.68 | 0.00 | 1.00 | 1 | Gm50457 | predicted gene, 51263 | 2 | 0 | 0 | 0 | 0 | 0 | 0 | 0 | 0 | 0 | 0 |
| 0 | 0.00 | 3.68 | 0.00 | 1.00 | 1 | Slc6a7 | solute carrier family 6 (neurotransmitter tr | 2 | 0 | 0 | 0 | 0 | 0 | 0 | 0 | 0 | 0 | 0 |
| 0 | 0.00 | 3.68 | 0.00 | 1.00 | 1 | Slc22a26 | solute carrier family 22 (organic cation tra | 2 | 0 | 0 | 0 | 0 | 0 | 0 | 0 | 0 | 0 | 0 |
| 0 | 0.00 | 3.68 | 0.00 | 1.00 | 1 | Il13ra2 | interleukin 13 receptor, alpha 2 | 2 | 0 | 0 | 0 | 0 | 0 | 0 | 0 | 0 | 0 | 0 |
| 0 | -0.40 | 3.67 | -0.11 | 0.91 | 1 | Gm26879 | | 1 | 0 | 0 | 0 | 0 | 1 | 0 | 0 | 0 | 0 | 0 |
| 0 | -0.40 | 3.67 | -0.11 | 0.91 | 1 | Gm13342 | | 1 | 0 | 0 | 0 | 0 | 1 | 0 | 0 | 0 | 0 | 0 |
| 0 | -0.40 | 3.67 | -0.11 | 0.91 | 1 | Gm13435 | | 1 | 0 | 0 | 0 | 0 | 1 | 0 | 0 | 0 | 0 | 0 |
| 0 | -0.40 | 3.67 | -0.11 | 0.91 | 1 | Gm13740 | | 1 | 0 | 0 | 0 | 0 | 1 | 0 | 0 | 0 | 0 | 0 |
| 0 | -0.40 | 3.67 | -0.11 | 0.91 | 1 | Gm14445 | | 1 | 0 | 0 | 0 | 0 | 1 | 0 | 0 | 0 | 0 | 0 |
| 0 | -0.40 | 3.67 | -0.11 | 0.91 | 1 | Gm12468 | | 1 | 0 | 0 | 0 | 0 | 1 | 0 | 0 | 0 | 0 | 0 |
| 0 | -0.40 | 3.67 | -0.11 | 0.91 | 1 | Gm36378 | | 1 | 0 | 0 | 0 | 0 | 1 | 0 | 0 | 0 | 0 | 0 |
| 0 | -0.40 | 3.67 | -0.11 | 0.91 | 1 | Gm5308 | | 1 | 0 | 0 | 0 | 0 | 1 | 0 | 0 | 0 | 0 | 0 |
| 0 | -0.40 | 3.67 | -0.11 | 0.91 | 1 | Smt3h2-ps2 | | 1 | 0 | 0 | 0 | 0 | 1 | 0 | 0 | 0 | 0 | 0 |
| 0 | -0.40 | 3.67 | -0.11 | 0.91 | 1 | Gm18441 | | 1 | 0 | 0 | 0 | 0 | 1 | 0 | 0 | 0 | 0 | 0 |
| 0 | -0.40 | 3.67 | -0.11 | 0.91 | 1 | Gm6343 | | 1 | 0 | 0 | 0 | 0 | 1 | 0 | 0 | 0 | 0 | 0 |
| 0 | -0.69 | 3.66 | -0.19 | 0.85 | 1 | Gm38259 | | 0 | 0 | 0 | 0 | 0 | 2 | 0 | 0 | 0 | 0 | 0 |
| 0 | -0.69 | 3.66 | -0.19 | 0.85 | 1 | Gm15604 | | 0 | 0 | 0 | 0 | 0 | 2 | 0 | 0 | 0 | 0 | 0 |
| 0 | -0.69 | 3.66 | -0.19 | 0.85 | 1 | D030040B21Rik | | 0 | 0 | 0 | 0 | 0 | 2 | 0 | 0 | 0 | 0 | 0 |
| 0 | -0.69 | 3.66 | -0.19 | 0.85 | 1 | Gm29506 | | 0 | 0 | 0 | 0 | 0 | 2 | 0 | 0 | 0 | 0 | 0 |
| 0 | -0.69 | 3.66 | -0.19 | 0.85 | 1 | Gm16150 | | 0 | 0 | 0 | 0 | 0 | 2 | 0 | 0 | 0 | 0 | 0 |
| 0 | -0.69 | 3.66 | -0.19 | 0.85 | 1 | 9130024F11Rik | | 0 | 0 | 0 | 0 | 0 | 2 | 0 | 0 | 0 | 0 | 0 |
| 0 | -0.69 | 3.66 | -0.19 | 0.85 | 1 | Snord11 | | 0 | 0 | 0 | 0 | 0 | 2 | 0 | 0 | 0 | 0 | 0 |
| 0 | -0.69 | 3.66 | -0.19 | 0.85 | 1 | Pth2r | parathyroid hormone 2 receptor | 0 | 0 | 0 | 0 | 0 | 2 | 0 | 0 | 0 | 0 | 0 |
| 0 | -0.69 | 3.66 | -0.19 | 0.85 | 1 | Gm24729 | | 0 | 0 | 0 | 0 | 0 | 2 | 0 | 0 | 0 | 0 | 0 |
| 0 | -0.69 | 3.66 | -0.19 | 0.85 | 1 | Gm37884 | | 0 | 0 | 0 | 0 | 0 | 2 | 0 | 0 | 0 | 0 | 0 |
| 0 | -0.69 | 3.66 | -0.19 | 0.85 | 1 | 2310015K22Rik | | 0 | 0 | 0 | 0 | 0 | 2 | 0 | 0 | 0 | 0 | 0 |
| 0 | -0.69 | 3.66 | -0.19 | 0.85 | 1 | Gm8403 | | 0 | 0 | 0 | 0 | 0 | 2 | 0 | 0 | 0 | 0 | 0 |
| 0 | -0.69 | 3.66 | -0.19 | 0.85 | 1 | Gm37669 | | 0 | 0 | 0 | 0 | 0 | 2 | 0 | 0 | 0 | 0 | 0 |
| 0 | -0.69 | 3.66 | -0.19 | 0.85 | 1 | Gm37115 | | 0 | 0 | 0 | 0 | 0 | 2 | 0 | 0 | 0 | 0 | 0 |
| 0 | -0.69 | 3.66 | -0.19 | 0.85 | 1 | 6720464F23Rik | | 0 | 0 | 0 | 0 | 0 | 2 | 0 | 0 | 0 | 0 | 0 |
| 0 | -0.69 | 3.66 | -0.19 | 0.85 | 1 | Gm32569 | | 0 | 0 | 0 | 0 | 0 | 2 | 0 | 0 | 0 | 0 | 0 |
| 0 | -0.69 | 3.66 | -0.19 | 0.85 | 1 | Gm37403 | | 0 | 0 | 0 | 0 | 0 | 2 | 0 | 0 | 0 | 0 | 0 |
| 0 | -0.69 | 3.66 | -0.19 | 0.85 | 1 | Gm37598 | | 0 | 0 | 0 | 0 | 0 | 2 | 0 | 0 | 0 | 0 | 0 |
| 0 | -0.69 | 3.66 | -0.19 | 0.85 | 1 | Snora36b | | 0 | 0 | 0 | 0 | 0 | 2 | 0 | 0 | 0 | 0 | 0 |
| 0 | -0.69 | 3.66 | -0.19 | 0.85 | 1 | Gm37095 | | 0 | 0 | 0 | 0 | 0 | 2 | 0 | 0 | 0 | 0 | 0 |
| 0 | -0.69 | 3.66 | -0.19 | 0.85 | 1 | Gm13350 | | 0 | 0 | 0 | 0 | 0 | 2 | 0 | 0 | 0 | 0 | 0 |
| 0 | -0.69 | 3.66 | -0.19 | 0.85 | 1 | Gm13362 | | 0 | 0 | 0 | 0 | 0 | 2 | 0 | 0 | 0 | 0 | 0 |
| 0 | -0.69 | 3.66 | -0.19 | 0.85 | 1 | Gm23546 | | 0 | 0 | 0 | 0 | 0 | 2 | 0 | 0 | 0 | 0 | 0 |
| 0 | -0.69 | 3.66 | -0.19 | 0.85 | 1 | Gm13550 | | 0 | 0 | 0 | 0 | 0 | 2 | 0 | 0 | 0 | 0 | 0 |
| 0 | -0.69 | 3.66 | -0.19 | 0.85 | 1 | Gm13618 | | 0 | 0 | 0 | 0 | 0 | 2 | 0 | 0 | 0 | 0 | 0 |
| 0 | -0.69 | 3.66 | -0.19 | 0.85 | 1 | Gm13713 | | 0 | 0 | 0 | 0 | 0 | 2 | 0 | 0 | 0 | 0 | 0 |
| 0 | -0.69 | 3.66 | -0.19 | 0.85 | 1 | | | 0 | 0 | 0 | 0 | 0 | 2 | 0 | 0 | 0 | 0 | 0 |
| 0 | -0.69 | 3.66 | -0.19 | 0.85 | 1 | Gm13954 | | 0 | 0 | 0 | 0 | 0 | 2 | 0 | 0 | 0 | 0 | 0 |
| 0 | -0.69 | 3.66 | -0.19 | 0.85 | 1 | n-R5s202 | | 0 | 0 | 0 | 0 | 0 | 2 | 0 | 0 | 0 | 0 | 0 |
| 0 | -0.69 | 3.66 | -0.19 | 0.85 | 1 | Gm14132 | | 0 | 0 | 0 | 0 | 0 | 2 | 0 | 0 | 0 | 0 | 0 |
| 0 | -0.69 | 3.66 | -0.19 | 0.85 | 1 | Gm14142 | | 0 | 0 | 0 | 0 | 0 | 2 | 0 | 0 | 0 | 0 | 0 |
| 0 | -0.69 | 3.66 | -0.19 | 0.85 | 1 | Gssos1 | | 0 | 0 | 0 | 0 | 0 | 2 | 0 | 0 | 0 | 0 | 0 |
| 0 | -0.69 | 3.66 | -0.19 | 0.85 | 1 | Gm14223 | | 0 | 0 | 0 | 0 | 0 | 2 | 0 | 0 | 0 | 0 | 0 |

| | | | | | | | | | | | | | | | | | |
|---|---|---|---|---|---|---|---|---|---|---|---|---|---|---|---|---|---|
| 0 | -0.69 | 3.66 | -0.19 | 0.85 | 1 | Gm14291 | | 0 | 0 | 0 | 0 | 0 | 2 | 0 | 0 | 0 | 0 | 0 |
| 0 | -0.69 | 3.66 | -0.19 | 0.85 | 1 | Gm14235 | | 0 | 0 | 0 | 0 | 0 | 2 | 0 | 0 | 0 | 0 | 0 |
| 0 | -0.69 | 3.66 | -0.19 | 0.85 | 1 | 1700017J07Rik | | 0 | 0 | 0 | 0 | 0 | 2 | 0 | 0 | 0 | 0 | 0 |
| 0 | -0.69 | 3.66 | -0.19 | 0.85 | 1 | Gm42437 | | 0 | 0 | 0 | 0 | 0 | 2 | 0 | 0 | 0 | 0 | 0 |
| 0 | -0.69 | 3.66 | -0.19 | 0.85 | 1 | Gm38047 | | 0 | 0 | 0 | 0 | 0 | 2 | 0 | 0 | 0 | 0 | 0 |
| 0 | -0.69 | 3.66 | -0.19 | 0.85 | 1 | Gm30735 | | 0 | 0 | 0 | 0 | 0 | 2 | 0 | 0 | 0 | 0 | 0 |
| 0 | -0.69 | 3.66 | -0.19 | 0.85 | 1 | Gm43589 | | 0 | 0 | 0 | 0 | 0 | 2 | 0 | 0 | 0 | 0 | 0 |
| 0 | -0.69 | 3.66 | -0.19 | 0.85 | 1 | 4921511C10Rik | | 0 | 0 | 0 | 0 | 0 | 2 | 0 | 0 | 0 | 0 | 0 |
| 0 | -0.69 | 3.66 | -0.19 | 0.85 | 1 | Bglap3 | bone gamma-carboxyglutamate protein 3 | 0 | 0 | 0 | 0 | 0 | 2 | 0 | 0 | 0 | 0 | 0 |
| 0 | -0.69 | 3.66 | -0.19 | 0.85 | 1 | Gm45354 | | 0 | 0 | 0 | 0 | 0 | 2 | 0 | 0 | 0 | 0 | 0 |
| 0 | -0.69 | 3.66 | -0.19 | 0.85 | 1 | Gm43774 | | 0 | 0 | 0 | 0 | 0 | 2 | 0 | 0 | 0 | 0 | 0 |
| 0 | -0.69 | 3.66 | -0.19 | 0.85 | 1 | Gm43463 | | 0 | 0 | 0 | 0 | 0 | 2 | 0 | 0 | 0 | 0 | 0 |
| 0 | -0.69 | 3.66 | -0.19 | 0.85 | 1 | Gm42515 | | 0 | 0 | 0 | 0 | 0 | 2 | 0 | 0 | 0 | 0 | 0 |
| 0 | -0.69 | 3.66 | -0.19 | 0.85 | 1 | Adh6a | alcohol dehydrogenase 6A (class V) | 0 | 0 | 0 | 0 | 0 | 2 | 0 | 0 | 0 | 0 | 0 |
| 0 | -0.69 | 3.66 | -0.19 | 0.85 | 1 | Pdha2 | pyruvate dehydrogenase E1 alpha 2 | 0 | 0 | 0 | 0 | 0 | 2 | 0 | 0 | 0 | 0 | 0 |
| 0 | -0.69 | 3.66 | -0.19 | 0.85 | 1 | Col24a1 | collagen, type XXIV, alpha 1 | 0 | 0 | 0 | 0 | 0 | 2 | 0 | 0 | 0 | 0 | 0 |
| 0 | -0.69 | 3.66 | -0.19 | 0.85 | 1 | Gm43400 | | 0 | 0 | 0 | 0 | 0 | 2 | 0 | 0 | 0 | 0 | 0 |
| 0 | -0.69 | 3.66 | -0.19 | 0.85 | 1 | Slc46a2 | solute carrier family 46, member 2 | 0 | 0 | 0 | 0 | 0 | 2 | 0 | 0 | 0 | 0 | 0 |
| 0 | -0.69 | 3.66 | -0.19 | 0.85 | 1 | Gm10305 | | 0 | 0 | 0 | 0 | 0 | 2 | 0 | 0 | 0 | 0 | 0 |
| 0 | -0.69 | 3.66 | -0.19 | 0.85 | 1 | Gm12798 | | 0 | 0 | 0 | 0 | 0 | 2 | 0 | 0 | 0 | 0 | 0 |
| 0 | -0.69 | 3.66 | -0.19 | 0.85 | 1 | Gm24967 | | 0 | 0 | 0 | 0 | 0 | 2 | 0 | 0 | 0 | 0 | 0 |
| 0 | -0.69 | 3.66 | -0.19 | 0.85 | 1 | Gm12865 | | 0 | 0 | 0 | 0 | 0 | 2 | 0 | 0 | 0 | 0 | 0 |
| 0 | -0.69 | 3.66 | -0.19 | 0.85 | 1 | Gm50484 | | 0 | 0 | 0 | 0 | 0 | 2 | 0 | 0 | 0 | 0 | 0 |
| 0 | -0.69 | 3.66 | -0.19 | 0.85 | 1 | Gm22154 | | 0 | 0 | 0 | 0 | 0 | 2 | 0 | 0 | 0 | 0 | 0 |
| 0 | -0.69 | 3.66 | -0.19 | 0.85 | 1 | 4933427I04Rik | | 0 | 0 | 0 | 0 | 0 | 2 | 0 | 0 | 0 | 0 | 0 |
| 0 | -0.69 | 3.66 | -0.19 | 0.85 | 1 | Gm2164 | | 0 | 0 | 0 | 0 | 0 | 2 | 0 | 0 | 0 | 0 | 0 |
| 0 | -0.69 | 3.66 | -0.19 | 0.85 | 1 | Tfap2e | transcription factor AP-2, epsilon | 0 | 0 | 0 | 0 | 0 | 2 | 0 | 0 | 0 | 0 | 0 |
| 0 | -0.69 | 3.66 | -0.19 | 0.85 | 1 | Gm23711 | | 0 | 0 | 0 | 0 | 0 | 2 | 0 | 0 | 0 | 0 | 0 |
| 0 | -0.69 | 3.66 | -0.19 | 0.85 | 1 | Gm23634 | | 0 | 0 | 0 | 0 | 0 | 2 | 0 | 0 | 0 | 0 | 0 |
| 0 | -0.69 | 3.66 | -0.19 | 0.85 | 1 | Gm24362 | | 0 | 0 | 0 | 0 | 0 | 2 | 0 | 0 | 0 | 0 | 0 |
| 0 | -0.69 | 3.66 | -0.19 | 0.85 | 1 | Gm22961 | | 0 | 0 | 0 | 0 | 0 | 2 | 0 | 0 | 0 | 0 | 0 |
| 0 | -0.69 | 3.66 | -0.19 | 0.85 | 1 | Vmn2r-ps18 | | 0 | 0 | 0 | 0 | 0 | 2 | 0 | 0 | 0 | 0 | 0 |
| 0 | -0.69 | 3.66 | -0.19 | 0.85 | 1 | Vmn2r-ps20 | | 0 | 0 | 0 | 0 | 0 | 2 | 0 | 0 | 0 | 0 | 0 |
| 0 | -0.69 | 3.66 | -0.19 | 0.85 | 1 | Gm24002 | | 0 | 0 | 0 | 0 | 0 | 2 | 0 | 0 | 0 | 0 | 0 |
| 0 | -0.69 | 3.66 | -0.19 | 0.85 | 1 | Gm13209 | | 0 | 0 | 0 | 0 | 0 | 2 | 0 | 0 | 0 | 0 | 0 |
| 0 | -0.69 | 3.66 | -0.19 | 0.85 | 1 | Gm44363 | | 0 | 0 | 0 | 0 | 0 | 2 | 0 | 0 | 0 | 0 | 0 |
| 0 | -0.69 | 3.66 | -0.19 | 0.85 | 1 | Gm17130 | | 0 | 0 | 0 | 0 | 0 | 2 | 0 | 0 | 0 | 0 | 0 |
| 0 | -0.69 | 3.66 | -0.19 | 0.85 | 1 | Gm43318 | | 0 | 0 | 0 | 0 | 0 | 2 | 0 | 0 | 0 | 0 | 0 |
| 0 | -0.69 | 3.66 | -0.19 | 0.85 | 1 | Gm24931 | | 0 | 0 | 0 | 0 | 0 | 2 | 0 | 0 | 0 | 0 | 0 |
| 0 | -0.69 | 3.66 | -0.19 | 0.85 | 1 | Mir7026 | | 0 | 0 | 0 | 0 | 0 | 2 | 0 | 0 | 0 | 0 | 0 |
| 0 | -0.69 | 3.66 | -0.19 | 0.85 | 1 | Gm13790 | | 0 | 0 | 0 | 0 | 0 | 2 | 0 | 0 | 0 | 0 | 0 |
| 0 | -0.69 | 3.66 | -0.19 | 0.85 | 1 | Gm43051 | | 0 | 0 | 0 | 0 | 0 | 2 | 0 | 0 | 0 | 0 | 0 |
| 0 | -0.69 | 3.66 | -0.19 | 0.85 | 1 | C230071H17Rik | | 0 | 0 | 0 | 0 | 0 | 2 | 0 | 0 | 0 | 0 | 0 |
| 0 | -0.69 | 3.66 | -0.19 | 0.85 | 1 | Gm43188 | | 0 | 0 | 0 | 0 | 0 | 2 | 0 | 0 | 0 | 0 | 0 |
| 0 | -0.69 | 3.66 | -0.19 | 0.85 | 1 | Mir339 | | 0 | 0 | 0 | 0 | 0 | 2 | 0 | 0 | 0 | 0 | 0 |
| 0 | -0.69 | 3.66 | -0.19 | 0.85 | 1 | Gm18753 | | 0 | 0 | 0 | 0 | 0 | 2 | 0 | 0 | 0 | 0 | 0 |
| 0 | -0.69 | 3.66 | -0.19 | 0.85 | 1 | Zar1l | zygote arrest 1-like | 0 | 0 | 0 | 0 | 0 | 2 | 0 | 0 | 0 | 0 | 0 |
| 0 | -0.69 | 3.66 | -0.19 | 0.85 | 1 | Gm8686 | | 0 | 0 | 0 | 0 | 0 | 2 | 0 | 0 | 0 | 0 | 0 |
| 0 | -0.69 | 3.66 | -0.19 | 0.85 | 1 | Trbv1 | | 0 | 0 | 0 | 0 | 0 | 2 | 0 | 0 | 0 | 0 | 0 |
| 0 | -0.69 | 3.66 | -0.19 | 0.85 | 1 | B230112I24Rik | | 0 | 0 | 0 | 0 | 0 | 2 | 0 | 0 | 0 | 0 | 0 |
| 0 | -0.69 | 3.66 | -0.19 | 0.85 | 1 | Gm43885 | | 0 | 0 | 0 | 0 | 0 | 2 | 0 | 0 | 0 | 0 | 0 |
| 0 | -0.69 | 3.66 | -0.19 | 0.85 | 1 | Gm15572 | | 0 | 0 | 0 | 0 | 0 | 2 | 0 | 0 | 0 | 0 | 0 |
| 0 | -0.69 | 3.66 | -0.19 | 0.85 | 1 | Gm43896 | | 0 | 0 | 0 | 0 | 0 | 2 | 0 | 0 | 0 | 0 | 0 |
| 0 | -0.69 | 3.66 | -0.19 | 0.85 | 1 | Gm44108 | | 0 | 0 | 0 | 0 | 0 | 2 | 0 | 0 | 0 | 0 | 0 |
| 0 | -0.69 | 3.66 | -0.19 | 0.85 | 1 | Mir7043 | | 0 | 0 | 0 | 0 | 0 | 2 | 0 | 0 | 0 | 0 | 0 |
| 0 | -0.69 | 3.66 | -0.19 | 0.85 | 1 | Stambp-ps1 | | 0 | 0 | 0 | 0 | 0 | 2 | 0 | 0 | 0 | 0 | 0 |
| 0 | -0.69 | 3.66 | -0.19 | 0.85 | 1 | Gm25430 | | 0 | 0 | 0 | 0 | 0 | 2 | 0 | 0 | 0 | 0 | 0 |
| 0 | -0.69 | 3.66 | -0.19 | 0.85 | 1 | Sspnos | | 0 | 0 | 0 | 0 | 0 | 2 | 0 | 0 | 0 | 0 | 0 |
| 0 | -0.69 | 3.66 | -0.19 | 0.85 | 1 | Oscar | osteoclast associated receptor | 0 | 0 | 0 | 0 | 0 | 2 | 0 | 0 | 0 | 0 | 0 |
| 0 | -0.69 | 3.66 | -0.19 | 0.85 | 1 | Gm29443 | | 0 | 0 | 0 | 0 | 0 | 2 | 0 | 0 | 0 | 0 | 0 |

| | | | | | | | | | | | | | | | | | | |
|---|---|---|---|---|---|---|---|---|---|---|---|---|---|---|---|---|---|---|
| 0 | -0.69 | 3.66 | -0.19 | 0.85 | 1 | Psg23 | pregnancy-specific glycoprotein 23 | 0 | 0 | 0 | 0 | 0 | 2 | 0 | 0 | 0 | 0 | 0 |
| 0 | -0.69 | 3.66 | -0.19 | 0.85 | 1 | Gm26891 | | 0 | 0 | 0 | 0 | 0 | 2 | 0 | 0 | 0 | 0 | 0 |
| 0 | -0.69 | 3.66 | -0.19 | 0.85 | 1 | E130304I02Rik | | 0 | 0 | 0 | 0 | 0 | 2 | 0 | 0 | 0 | 0 | 0 |
| 0 | -0.69 | 3.66 | -0.19 | 0.85 | 1 | Gm24938 | | 0 | 0 | 0 | 0 | 0 | 2 | 0 | 0 | 0 | 0 | 0 |
| 0 | -0.69 | 3.66 | -0.19 | 0.85 | 1 | Zfp936 | zinc finger protein 936 | 0 | 0 | 0 | 0 | 0 | 2 | 0 | 0 | 0 | 0 | 0 |
| 0 | -0.69 | 3.66 | -0.19 | 0.85 | 1 | Mir7054 | | 0 | 0 | 0 | 0 | 0 | 2 | 0 | 0 | 0 | 0 | 0 |
| 0 | -0.69 | 3.66 | -0.19 | 0.85 | 1 | Gm23962 | | 0 | 0 | 0 | 0 | 0 | 2 | 0 | 0 | 0 | 0 | 0 |
| 0 | -0.69 | 3.66 | -0.19 | 0.85 | 1 | Mir344d-2 | | 0 | 0 | 0 | 0 | 0 | 2 | 0 | 0 | 0 | 0 | 0 |
| 0 | -0.69 | 3.66 | -0.19 | 0.85 | 1 | Gm23431 | | 0 | 0 | 0 | 0 | 0 | 2 | 0 | 0 | 0 | 0 | 0 |
| 0 | -0.69 | 3.66 | -0.19 | 0.85 | 1 | Gm34350 | | 0 | 0 | 0 | 0 | 0 | 2 | 0 | 0 | 0 | 0 | 0 |
| 0 | -0.69 | 3.66 | -0.19 | 0.85 | 1 | Gm24541 | | 0 | 0 | 0 | 0 | 0 | 2 | 0 | 0 | 0 | 0 | 0 |
| 0 | -0.69 | 3.66 | -0.19 | 0.85 | 1 | Gm45378 | | 0 | 0 | 0 | 0 | 0 | 2 | 0 | 0 | 0 | 0 | 0 |
| 0 | -0.69 | 3.66 | -0.19 | 0.85 | 1 | Gm26147 | | 0 | 0 | 0 | 0 | 0 | 2 | 0 | 0 | 0 | 0 | 0 |
| 0 | -0.69 | 3.66 | -0.19 | 0.85 | 1 | Gm44986 | | 0 | 0 | 0 | 0 | 0 | 2 | 0 | 0 | 0 | 0 | 0 |
| 0 | -0.69 | 3.66 | -0.19 | 0.85 | 1 | Gm18258 | | 0 | 0 | 0 | 0 | 0 | 2 | 0 | 0 | 0 | 0 | 0 |
| 0 | -0.69 | 3.66 | -0.19 | 0.85 | 1 | Olfr525 | olfactory receptor 525 | 0 | 0 | 0 | 0 | 0 | 2 | 0 | 0 | 0 | 0 | 0 |
| 0 | -0.69 | 3.66 | -0.19 | 0.85 | 1 | Mir7063 | | 0 | 0 | 0 | 0 | 0 | 2 | 0 | 0 | 0 | 0 | 0 |
| 0 | -0.69 | 3.66 | -0.19 | 0.85 | 1 | Gm45224 | | 0 | 0 | 0 | 0 | 0 | 2 | 0 | 0 | 0 | 0 | 0 |
| 0 | -0.69 | 3.66 | -0.19 | 0.85 | 1 | Gm45555 | | 0 | 0 | 0 | 0 | 0 | 2 | 0 | 0 | 0 | 0 | 0 |
| 0 | -0.69 | 3.66 | -0.19 | 0.85 | 1 | Gm31545 | | 0 | 0 | 0 | 0 | 0 | 2 | 0 | 0 | 0 | 0 | 0 |
| 0 | -0.69 | 3.66 | -0.19 | 0.85 | 1 | Gm5741 | predicted gene 5741 | 0 | 0 | 0 | 0 | 0 | 2 | 0 | 0 | 0 | 0 | 0 |
| 0 | -0.69 | 3.66 | -0.19 | 0.85 | 1 | Gm45888 | | 0 | 0 | 0 | 0 | 0 | 2 | 0 | 0 | 0 | 0 | 0 |
| 0 | -0.69 | 3.66 | -0.19 | 0.85 | 1 | 1700030M09Rik | | 0 | 0 | 0 | 0 | 0 | 2 | 0 | 0 | 0 | 0 | 0 |
| 0 | -0.69 | 3.66 | -0.19 | 0.85 | 1 | Gm26592 | | 0 | 0 | 0 | 0 | 0 | 2 | 0 | 0 | 0 | 0 | 0 |
| 0 | -0.69 | 3.66 | -0.19 | 0.85 | 1 | Gm39317 | | 0 | 0 | 0 | 0 | 0 | 2 | 0 | 0 | 0 | 0 | 0 |
| 0 | -0.69 | 3.66 | -0.19 | 0.85 | 1 | Gm48081 | | 0 | 0 | 0 | 0 | 0 | 2 | 0 | 0 | 0 | 0 | 0 |
| 0 | -0.69 | 3.66 | -0.19 | 0.85 | 1 | Gm7439 | | 0 | 0 | 0 | 0 | 0 | 2 | 0 | 0 | 0 | 0 | 0 |
| 0 | -0.69 | 3.66 | -0.19 | 0.85 | 1 | Gm19427 | | 0 | 0 | 0 | 0 | 0 | 2 | 0 | 0 | 0 | 0 | 0 |
| 0 | -0.69 | 3.66 | -0.19 | 0.85 | 1 | Gm20537 | | 0 | 0 | 0 | 0 | 0 | 2 | 0 | 0 | 0 | 0 | 0 |
| 0 | -0.69 | 3.66 | -0.19 | 0.85 | 1 | Trim43c | tripartite motif-containing 43C | 0 | 0 | 0 | 0 | 0 | 2 | 0 | 0 | 0 | 0 | 0 |
| 0 | -0.69 | 3.66 | -0.19 | 0.85 | 1 | Gm38314 | | 0 | 0 | 0 | 0 | 0 | 2 | 0 | 0 | 0 | 0 | 0 |
| 0 | -0.69 | 3.66 | -0.19 | 0.85 | 1 | Gm37850 | | 0 | 0 | 0 | 0 | 0 | 2 | 0 | 0 | 0 | 0 | 0 |
| 0 | -0.69 | 3.66 | -0.19 | 0.85 | 1 | Slc22a13 | solute carrier family 22 (organic cation tra | 0 | 0 | 0 | 0 | 0 | 2 | 0 | 0 | 0 | 0 | 0 |
| 0 | -0.69 | 3.66 | -0.19 | 0.85 | 1 | Gm48743 | | 0 | 0 | 0 | 0 | 0 | 2 | 0 | 0 | 0 | 0 | 0 |
| 0 | -0.69 | 3.66 | -0.19 | 0.85 | 1 | Gm23044 | | 0 | 0 | 0 | 0 | 0 | 2 | 0 | 0 | 0 | 0 | 0 |
| 0 | -0.69 | 3.66 | -0.19 | 0.85 | 1 | Gm26528 | | 0 | 0 | 0 | 0 | 0 | 2 | 0 | 0 | 0 | 0 | 0 |
| 0 | -0.69 | 3.66 | -0.19 | 0.85 | 1 | Taar3 | trace amine-associated receptor 3 | 0 | 0 | 0 | 0 | 0 | 2 | 0 | 0 | 0 | 0 | 0 |
| 0 | -0.69 | 3.66 | -0.19 | 0.85 | 1 | Gm47598 | | 0 | 0 | 0 | 0 | 0 | 2 | 0 | 0 | 0 | 0 | 0 |
| 0 | -0.69 | 3.66 | -0.19 | 0.85 | 1 | Gm47388 | | 0 | 0 | 0 | 0 | 0 | 2 | 0 | 0 | 0 | 0 | 0 |
| 0 | -0.69 | 3.66 | -0.19 | 0.85 | 1 | Gm47622 | | 0 | 0 | 0 | 0 | 0 | 2 | 0 | 0 | 0 | 0 | 0 |
| 0 | -0.69 | 3.66 | -0.19 | 0.85 | 1 | Gm19459 | | 0 | 0 | 0 | 0 | 0 | 2 | 0 | 0 | 0 | 0 | 0 |
| 0 | -0.69 | 3.66 | -0.19 | 0.85 | 1 | Gm7530 | | 0 | 0 | 0 | 0 | 0 | 2 | 0 | 0 | 0 | 0 | 0 |
| 0 | -0.69 | 3.66 | -0.19 | 0.85 | 1 | Gm47459 | | 0 | 0 | 0 | 0 | 0 | 2 | 0 | 0 | 0 | 0 | 0 |
| 0 | -0.69 | 3.66 | -0.19 | 0.85 | 1 | Mir6911 | | 0 | 0 | 0 | 0 | 0 | 2 | 0 | 0 | 0 | 0 | 0 |
| 0 | -0.69 | 3.66 | -0.19 | 0.85 | 1 | Gm34184 | | 0 | 0 | 0 | 0 | 0 | 2 | 0 | 0 | 0 | 0 | 0 |
| 0 | -0.69 | 3.66 | -0.19 | 0.85 | 1 | B530045E10Rik | | 0 | 0 | 0 | 0 | 0 | 2 | 0 | 0 | 0 | 0 | 0 |
| 0 | -0.69 | 3.66 | -0.19 | 0.85 | 1 | Gm47009 | | 0 | 0 | 0 | 0 | 0 | 2 | 0 | 0 | 0 | 0 | 0 |
| 0 | -0.69 | 3.66 | -0.19 | 0.85 | 1 | Mir26a-2 | | 0 | 0 | 0 | 0 | 0 | 2 | 0 | 0 | 0 | 0 | 0 |
| 0 | -0.69 | 3.66 | -0.19 | 0.85 | 1 | Gm23182 | | 0 | 0 | 0 | 0 | 0 | 2 | 0 | 0 | 0 | 0 | 0 |
| 0 | -0.69 | 3.66 | -0.19 | 0.85 | 1 | Mir6916 | | 0 | 0 | 0 | 0 | 0 | 2 | 0 | 0 | 0 | 0 | 0 |
| 0 | -0.69 | 3.66 | -0.19 | 0.85 | 1 | Gm12664 | | 0 | 0 | 0 | 0 | 0 | 2 | 0 | 0 | 0 | 0 | 0 |
| 0 | -0.69 | 3.66 | -0.19 | 0.85 | 1 | Gm12088 | | 0 | 0 | 0 | 0 | 0 | 2 | 0 | 0 | 0 | 0 | 0 |
| 0 | -0.69 | 3.66 | -0.19 | 0.85 | 1 | Gm12100 | | 0 | 0 | 0 | 0 | 0 | 2 | 0 | 0 | 0 | 0 | 0 |
| 0 | -0.69 | 3.66 | -0.19 | 0.85 | 1 | Hba-x | hemoglobin X, alpha-like embryonic chai | 0 | 0 | 0 | 0 | 0 | 2 | 0 | 0 | 0 | 0 | 0 |
| 0 | -0.69 | 3.66 | -0.19 | 0.85 | 1 | Nmur2 | neuromedin U receptor 2 | 0 | 0 | 0 | 0 | 0 | 2 | 0 | 0 | 0 | 0 | 0 |
| 0 | -0.69 | 3.66 | -0.19 | 0.85 | 1 | H3f4 | H3.4 histone | 0 | 0 | 0 | 0 | 0 | 2 | 0 | 0 | 0 | 0 | 0 |
| 0 | -0.69 | 3.66 | -0.19 | 0.85 | 1 | Gm11424 | | 0 | 0 | 0 | 0 | 0 | 2 | 0 | 0 | 0 | 0 | 0 |
| 0 | -0.69 | 3.66 | -0.19 | 0.85 | 1 | Gm11440 | | 0 | 0 | 0 | 0 | 0 | 2 | 0 | 0 | 0 | 0 | 0 |
| 0 | -0.69 | 3.66 | -0.19 | 0.85 | 1 | Gsdma3 | gasdermin A3 | 0 | 0 | 0 | 0 | 0 | 2 | 0 | 0 | 0 | 0 | 0 |
| 0 | -0.69 | 3.66 | -0.19 | 0.85 | 1 | Gm11585 | | 0 | 0 | 0 | 0 | 0 | 2 | 0 | 0 | 0 | 0 | 0 |

| | | | | | | | | | | | | | | | | | |
|---|---|---|---|---|---|---|---|---|---|---|---|---|---|---|---|---|---|
| 0 | -0.69 | 3.66 | -0.19 | 0.85 | 1 | Gm25337 | | 0 | 0 | 0 | 0 | 0 | 2 | 0 | 0 | 0 | 0 | 0 |
| 0 | -0.69 | 3.66 | -0.19 | 0.85 | 1 | Gm39397 | | 0 | 0 | 0 | 0 | 0 | 2 | 0 | 0 | 0 | 0 | 0 |
| 0 | -0.69 | 3.66 | -0.19 | 0.85 | 1 | Gm16045 | | 0 | 0 | 0 | 0 | 0 | 2 | 0 | 0 | 0 | 0 | 0 |
| 0 | -0.69 | 3.66 | -0.19 | 0.85 | 1 | Gm48606 | | 0 | 0 | 0 | 0 | 0 | 2 | 0 | 0 | 0 | 0 | 0 |
| 0 | -0.69 | 3.66 | -0.19 | 0.85 | 1 | Gm36495 | | 0 | 0 | 0 | 0 | 0 | 2 | 0 | 0 | 0 | 0 | 0 |
| 0 | -0.69 | 3.66 | -0.19 | 0.85 | 1 | B430203G13Rik | | 0 | 0 | 0 | 0 | 0 | 2 | 0 | 0 | 0 | 0 | 0 |
| 0 | -0.69 | 3.66 | -0.19 | 0.85 | 1 | Gm48398 | | 0 | 0 | 0 | 0 | 0 | 2 | 0 | 0 | 0 | 0 | 0 |
| 0 | -0.69 | 3.66 | -0.19 | 0.85 | 1 | Gm9269 | | 0 | 0 | 0 | 0 | 0 | 2 | 0 | 0 | 0 | 0 | 0 |
| 0 | -0.69 | 3.66 | -0.19 | 0.85 | 1 | Gm47446 | | 0 | 0 | 0 | 0 | 0 | 2 | 0 | 0 | 0 | 0 | 0 |
| 0 | -0.69 | 3.66 | -0.19 | 0.85 | 1 | Gm48535 | | 0 | 0 | 0 | 0 | 0 | 2 | 0 | 0 | 0 | 0 | 0 |
| 0 | -0.69 | 3.66 | -0.19 | 0.85 | 1 | Gm11042 | | 0 | 0 | 0 | 0 | 0 | 2 | 0 | 0 | 0 | 0 | 0 |
| 0 | -0.69 | 3.66 | -0.19 | 0.85 | 1 | Gm48230 | | 0 | 0 | 0 | 0 | 0 | 2 | 0 | 0 | 0 | 0 | 0 |
| 0 | -0.69 | 3.66 | -0.19 | 0.85 | 1 | Gm25776 | | 0 | 0 | 0 | 0 | 0 | 2 | 0 | 0 | 0 | 0 | 0 |
| 0 | -0.69 | 3.66 | -0.19 | 0.85 | 1 | Gm47109 | | 0 | 0 | 0 | 0 | 0 | 2 | 0 | 0 | 0 | 0 | 0 |
| 0 | -0.69 | 3.66 | -0.19 | 0.85 | 1 | Gm47648 | | 0 | 0 | 0 | 0 | 0 | 2 | 0 | 0 | 0 | 0 | 0 |
| 0 | -0.69 | 3.66 | -0.19 | 0.85 | 1 | Mir412 | | 0 | 0 | 0 | 0 | 0 | 2 | 0 | 0 | 0 | 0 | 0 |
| 0 | -0.69 | 3.66 | -0.19 | 0.85 | 1 | Gm48248 | | 0 | 0 | 0 | 0 | 0 | 2 | 0 | 0 | 0 | 0 | 0 |
| 0 | -0.69 | 3.66 | -0.19 | 0.85 | 1 | Gm48286 | | 0 | 0 | 0 | 0 | 0 | 2 | 0 | 0 | 0 | 0 | 0 |
| 0 | -0.69 | 3.66 | -0.19 | 0.85 | 1 | Ighv6-4 | | 0 | 0 | 0 | 0 | 0 | 2 | 0 | 0 | 0 | 0 | 0 |
| 0 | -0.69 | 3.66 | -0.19 | 0.85 | 1 | Gm27649 | | 0 | 0 | 0 | 0 | 0 | 2 | 0 | 0 | 0 | 0 | 0 |
| 0 | -0.69 | 3.66 | -0.19 | 0.85 | 1 | Gm20043 | | 0 | 0 | 0 | 0 | 0 | 2 | 0 | 0 | 0 | 0 | 0 |
| 0 | -0.69 | 3.66 | -0.19 | 0.85 | 1 | Gm47665 | | 0 | 0 | 0 | 0 | 0 | 2 | 0 | 0 | 0 | 0 | 0 |
| 0 | -0.69 | 3.66 | -0.19 | 0.85 | 1 | Gm44346 | | 0 | 0 | 0 | 0 | 0 | 2 | 0 | 0 | 0 | 0 | 0 |
| 0 | -0.69 | 3.66 | -0.19 | 0.85 | 1 | Gm48188 | | 0 | 0 | 0 | 0 | 0 | 2 | 0 | 0 | 0 | 0 | 0 |
| 0 | -0.69 | 3.66 | -0.19 | 0.85 | 1 | F12 | coagulation factor XII (Hageman factor) | 0 | 0 | 0 | 0 | 0 | 2 | 0 | 0 | 0 | 0 | 0 |
| 0 | -0.69 | 3.66 | -0.19 | 0.85 | 1 | 1700066J03Rik | | 0 | 0 | 0 | 0 | 0 | 2 | 0 | 0 | 0 | 0 | 0 |
| 0 | -0.69 | 3.66 | -0.19 | 0.85 | 1 | Gm18413 | | 0 | 0 | 0 | 0 | 0 | 2 | 0 | 0 | 0 | 0 | 0 |
| 0 | -0.69 | 3.66 | -0.19 | 0.85 | 1 | 4930544M13Rik | | 0 | 0 | 0 | 0 | 0 | 2 | 0 | 0 | 0 | 0 | 0 |
| 0 | -0.69 | 3.66 | -0.19 | 0.85 | 1 | Gm47914 | | 0 | 0 | 0 | 0 | 0 | 2 | 0 | 0 | 0 | 0 | 0 |
| 0 | -0.69 | 3.66 | -0.19 | 0.85 | 1 | Gm2897 | predicted gene 2897 | 0 | 0 | 0 | 0 | 0 | 2 | 0 | 0 | 0 | 0 | 0 |
| 0 | -0.69 | 3.66 | -0.19 | 0.85 | 1 | 2610042L04Ri | RIKEN cDNA 2610042L04 gene | 0 | 0 | 0 | 0 | 0 | 2 | 0 | 0 | 0 | 0 | 0 |
| 0 | -0.69 | 3.66 | -0.19 | 0.85 | 1 | Gm48860 | | 0 | 0 | 0 | 0 | 0 | 2 | 0 | 0 | 0 | 0 | 0 |
| 0 | -0.69 | 3.66 | -0.19 | 0.85 | 1 | Dupd1 | dual specificity phosphatase and pro ison | 0 | 0 | 0 | 0 | 0 | 2 | 0 | 0 | 0 | 0 | 0 |
| 0 | -0.69 | 3.66 | -0.19 | 0.85 | 1 | Gm49270 | | 0 | 0 | 0 | 0 | 0 | 2 | 0 | 0 | 0 | 0 | 0 |
| 0 | -0.69 | 3.66 | -0.19 | 0.85 | 1 | Gm49199 | | 0 | 0 | 0 | 0 | 0 | 2 | 0 | 0 | 0 | 0 | 0 |
| 0 | -0.69 | 3.66 | -0.19 | 0.85 | 1 | Gm48985 | | 0 | 0 | 0 | 0 | 0 | 2 | 0 | 0 | 0 | 0 | 0 |
| 0 | -0.69 | 3.66 | -0.19 | 0.85 | 1 | Gm49260 | | 0 | 0 | 0 | 0 | 0 | 2 | 0 | 0 | 0 | 0 | 0 |
| 0 | -0.69 | 3.66 | -0.19 | 0.85 | 1 | Gm27760 | | 0 | 0 | 0 | 0 | 0 | 2 | 0 | 0 | 0 | 0 | 0 |
| 0 | -0.69 | 3.66 | -0.19 | 0.85 | 1 | Gm49162 | | 0 | 0 | 0 | 0 | 0 | 2 | 0 | 0 | 0 | 0 | 0 |
| 0 | -0.69 | 3.66 | -0.19 | 0.85 | 1 | Fam136b-ps | | 0 | 0 | 0 | 0 | 0 | 2 | 0 | 0 | 0 | 0 | 0 |
| 0 | -0.69 | 3.66 | -0.19 | 0.85 | 1 | Gm23754 | | 0 | 0 | 0 | 0 | 0 | 2 | 0 | 0 | 0 | 0 | 0 |
| 0 | -0.69 | 3.66 | -0.19 | 0.85 | 1 | Gm28221 | | 0 | 0 | 0 | 0 | 0 | 2 | 0 | 0 | 0 | 0 | 0 |
| 0 | -0.69 | 3.66 | -0.19 | 0.85 | 1 | Gm16575 | | 0 | 0 | 0 | 0 | 0 | 2 | 0 | 0 | 0 | 0 | 0 |
| 0 | -0.69 | 3.66 | -0.19 | 0.85 | 1 | | | 0 | 0 | 0 | 0 | 0 | 2 | 0 | 0 | 0 | 0 | 0 |
| 0 | -0.69 | 3.66 | -0.19 | 0.85 | 1 | Gm4825 | | 0 | 0 | 0 | 0 | 0 | 2 | 0 | 0 | 0 | 0 | 0 |
| 0 | -0.69 | 3.66 | -0.19 | 0.85 | 1 | B230214G05Rik | | 0 | 0 | 0 | 0 | 0 | 2 | 0 | 0 | 0 | 0 | 0 |
| 0 | -0.69 | 3.66 | -0.19 | 0.85 | 1 | Gm20695 | | 0 | 0 | 0 | 0 | 0 | 2 | 0 | 0 | 0 | 0 | 0 |
| 0 | -0.69 | 3.66 | -0.19 | 0.85 | 1 | Gm30251 | | 0 | 0 | 0 | 0 | 0 | 2 | 0 | 0 | 0 | 0 | 0 |
| 0 | -0.69 | 3.66 | -0.19 | 0.85 | 1 | Gm25848 | | 0 | 0 | 0 | 0 | 0 | 2 | 0 | 0 | 0 | 0 | 0 |
| 0 | -0.69 | 3.66 | -0.19 | 0.85 | 1 | Gm15711 | | 0 | 0 | 0 | 0 | 0 | 2 | 0 | 0 | 0 | 0 | 0 |
| 0 | -0.69 | 3.66 | -0.19 | 0.85 | 1 | Gm25908 | | 0 | 0 | 0 | 0 | 0 | 2 | 0 | 0 | 0 | 0 | 0 |
| 0 | -0.69 | 3.66 | -0.19 | 0.85 | 1 | Gm7831 | | 0 | 0 | 0 | 0 | 0 | 2 | 0 | 0 | 0 | 0 | 0 |
| 0 | -0.69 | 3.66 | -0.19 | 0.85 | 1 | D430001F17Rik | | 0 | 0 | 0 | 0 | 0 | 2 | 0 | 0 | 0 | 0 | 0 |
| 0 | -0.69 | 3.66 | -0.19 | 0.85 | 1 | Fam243 | family with sequence similarity 243 | 0 | 0 | 0 | 0 | 0 | 2 | 0 | 0 | 0 | 0 | 0 |
| 0 | -0.69 | 3.66 | -0.19 | 0.85 | 1 | Gm41504 | | 0 | 0 | 0 | 0 | 0 | 2 | 0 | 0 | 0 | 0 | 0 |
| 0 | -0.69 | 3.66 | -0.19 | 0.85 | 1 | Gm49681 | | 0 | 0 | 0 | 0 | 0 | 2 | 0 | 0 | 0 | 0 | 0 |
| 0 | -0.69 | 3.66 | -0.19 | 0.85 | 1 | Gm34883 | | 0 | 0 | 0 | 0 | 0 | 2 | 0 | 0 | 0 | 0 | 0 |
| 0 | -0.69 | 3.66 | -0.19 | 0.85 | 1 | Gm19283 | | 0 | 0 | 0 | 0 | 0 | 2 | 0 | 0 | 0 | 0 | 0 |
| 0 | -0.69 | 3.66 | -0.19 | 0.85 | 1 | Gm6811 | | 0 | 0 | 0 | 0 | 0 | 2 | 0 | 0 | 0 | 0 | 0 |
| 0 | -0.69 | 3.66 | -0.19 | 0.85 | 1 | Gm49935 | | 0 | 0 | 0 | 0 | 0 | 2 | 0 | 0 | 0 | 0 | 0 |

| | | | | | | | | | | | | | | | | | | |
|---|---|---|---|---|---|---|---|---|---|---|---|---|---|---|---|---|---|---|
| 0 | -0.69 | 3.66 | -0.19 | 0.85 | 1 | Gm50070 | | 0 | 0 | 0 | 0 | 0 | 2 | 0 | 0 | 0 | 0 | 0 |
| 0 | -0.69 | 3.66 | -0.19 | 0.85 | 1 | Gm16005 | | 0 | 0 | 0 | 0 | 0 | 2 | 0 | 0 | 0 | 0 | 0 |
| 0 | -0.69 | 3.66 | -0.19 | 0.85 | 1 | Gm25667 | | 0 | 0 | 0 | 0 | 0 | 2 | 0 | 0 | 0 | 0 | 0 |
| 0 | -0.69 | 3.66 | -0.19 | 0.85 | 1 | Gm7030 | predicted gene 7030 | 0 | 0 | 0 | 0 | 0 | 2 | 0 | 0 | 0 | 0 | 0 |
| 0 | -0.69 | 3.66 | -0.19 | 0.85 | 1 | Gm6934 | | 0 | 0 | 0 | 0 | 0 | 2 | 0 | 0 | 0 | 0 | 0 |
| 0 | -0.69 | 3.66 | -0.19 | 0.85 | 1 | Gm18417 | | 0 | 0 | 0 | 0 | 0 | 2 | 0 | 0 | 0 | 0 | 0 |
| 0 | -0.69 | 3.66 | -0.19 | 0.85 | 1 | Gm36594 | | 0 | 0 | 0 | 0 | 0 | 2 | 0 | 0 | 0 | 0 | 0 |
| 0 | -0.69 | 3.66 | -0.19 | 0.85 | 1 | Gm50055 | | 0 | 0 | 0 | 0 | 0 | 2 | 0 | 0 | 0 | 0 | 0 |
| 0 | -0.69 | 3.66 | -0.19 | 0.85 | 1 | Gm36858 | | 0 | 0 | 0 | 0 | 0 | 2 | 0 | 0 | 0 | 0 | 0 |
| 0 | -0.69 | 3.66 | -0.19 | 0.85 | 1 | Gm50223 | | 0 | 0 | 0 | 0 | 0 | 2 | 0 | 0 | 0 | 0 | 0 |
| 0 | -0.69 | 3.66 | -0.19 | 0.85 | 1 | Gm8629 | | 0 | 0 | 0 | 0 | 0 | 2 | 0 | 0 | 0 | 0 | 0 |
| 0 | -0.69 | 3.66 | -0.19 | 0.85 | 1 | Spink13 | serine peptidase inhibitor, Kazal type 13 | 0 | 0 | 0 | 0 | 0 | 2 | 0 | 0 | 0 | 0 | 0 |
| 0 | -0.69 | 3.66 | -0.19 | 0.85 | 1 | Gm37643 | | 0 | 0 | 0 | 0 | 0 | 2 | 0 | 0 | 0 | 0 | 0 |
| 0 | -0.69 | 3.66 | -0.19 | 0.85 | 1 | 1700003O11Rik | | 0 | 0 | 0 | 0 | 0 | 2 | 0 | 0 | 0 | 0 | 0 |
| 0 | -0.69 | 3.66 | -0.19 | 0.85 | 1 | Gm2176 | | 0 | 0 | 0 | 0 | 0 | 2 | 0 | 0 | 0 | 0 | 0 |
| 0 | -0.69 | 3.66 | -0.19 | 0.85 | 1 | Gm41790 | | 0 | 0 | 0 | 0 | 0 | 2 | 0 | 0 | 0 | 0 | 0 |
| 0 | -0.69 | 3.66 | -0.19 | 0.85 | 1 | Gm50438 | | 0 | 0 | 0 | 0 | 0 | 2 | 0 | 0 | 0 | 0 | 0 |
| 0 | -0.69 | 3.66 | -0.19 | 0.85 | 1 | Wnt8b | wingless-type MMTV integration site famil | 0 | 0 | 0 | 0 | 0 | 2 | 0 | 0 | 0 | 0 | 0 |
| 0 | -0.69 | 3.66 | -0.19 | 0.85 | 1 | Gm10490 | | 0 | 0 | 0 | 0 | 0 | 2 | 0 | 0 | 0 | 0 | 0 |
| 0 | -0.69 | 3.66 | -0.19 | 0.85 | 1 | Gm5380 | | 0 | 0 | 0 | 0 | 0 | 2 | 0 | 0 | 0 | 0 | 0 |
| 0 | -0.69 | 3.66 | -0.19 | 0.85 | 1 | Mir221 | | 0 | 0 | 0 | 0 | 0 | 2 | 0 | 0 | 0 | 0 | 0 |
| 0 | -0.69 | 3.66 | -0.19 | 0.85 | 1 | Rhox4b | reproductive homeobox 4B | 0 | 0 | 0 | 0 | 0 | 2 | 0 | 0 | 0 | 0 | 0 |
| 0 | -0.69 | 3.66 | -0.19 | 0.85 | 1 | Gm23322 | | 0 | 0 | 0 | 0 | 0 | 2 | 0 | 0 | 0 | 0 | 0 |
| 0 | -0.69 | 3.66 | -0.19 | 0.85 | 1 | Mir3473a | | 0 | 0 | 0 | 0 | 0 | 2 | 0 | 0 | 0 | 0 | 0 |
| 0 | -0.69 | 3.66 | -0.19 | 0.85 | 1 | mt-Tl1 | | 0 | 0 | 0 | 0 | 0 | 2 | 0 | 0 | 0 | 0 | 0 |
| 0 | -0.44 | 3.68 | -0.12 | 0.90 | NA | Gm15833 | | 0 | 0 | 0 | 0 | 0 | 0 | 1 | 0 | 0 | 0 | 0 |
| 0 | -0.44 | 3.68 | -0.12 | 0.90 | NA | Rps13-ps7 | | 0 | 0 | 0 | 0 | 0 | 0 | 1 | 0 | 0 | 0 | 0 |
| 0 | -0.44 | 3.68 | -0.12 | 0.90 | NA | Gm24513 | | 0 | 0 | 0 | 0 | 0 | 0 | 1 | 0 | 0 | 0 | 0 |
| 0 | -0.44 | 3.68 | -0.12 | 0.90 | NA | Gm28926 | | 0 | 0 | 0 | 0 | 0 | 0 | 1 | 0 | 0 | 0 | 0 |
| 0 | -0.44 | 3.68 | -0.12 | 0.90 | NA | Cdh20 | cadherin 20 | 0 | 0 | 0 | 0 | 0 | 0 | 1 | 0 | 0 | 0 | 0 |
| 0 | -0.44 | 3.68 | -0.12 | 0.90 | NA | Gm37470 | | 0 | 0 | 0 | 0 | 0 | 0 | 1 | 0 | 0 | 0 | 0 |
| 0 | -0.44 | 3.68 | -0.12 | 0.90 | NA | Rbm6-ps2 | | 0 | 0 | 0 | 0 | 0 | 0 | 1 | 0 | 0 | 0 | 0 |
| 0 | -0.44 | 3.68 | -0.12 | 0.90 | NA | Gm30725 | | 0 | 0 | 0 | 0 | 0 | 0 | 1 | 0 | 0 | 0 | 0 |
| 0 | -0.44 | 3.68 | -0.12 | 0.90 | NA | Gm13609 | | 0 | 0 | 0 | 0 | 0 | 0 | 1 | 0 | 0 | 0 | 0 |
| 0 | -0.44 | 3.68 | -0.12 | 0.90 | NA | Gm13669 | | 0 | 0 | 0 | 0 | 0 | 0 | 1 | 0 | 0 | 0 | 0 |
| 0 | -0.44 | 3.68 | -0.12 | 0.90 | NA | Gm14465 | | 0 | 0 | 0 | 0 | 0 | 0 | 1 | 0 | 0 | 0 | 0 |
| 0 | -0.44 | 3.68 | -0.12 | 0.90 | NA | Gm14287 | | 0 | 0 | 0 | 0 | 0 | 0 | 1 | 0 | 0 | 0 | 0 |
| 0 | -0.44 | 3.68 | -0.12 | 0.90 | NA | Gm17887 | | 0 | 0 | 0 | 0 | 0 | 0 | 1 | 0 | 0 | 0 | 0 |
| 0 | -0.44 | 3.68 | -0.12 | 0.90 | NA | Gm42647 | | 0 | 0 | 0 | 0 | 0 | 0 | 1 | 0 | 0 | 0 | 0 |
| 0 | -0.44 | 3.68 | -0.12 | 0.90 | NA | Gm12543 | | 0 | 0 | 0 | 0 | 0 | 0 | 1 | 0 | 0 | 0 | 0 |
| 0 | -0.44 | 3.68 | -0.12 | 0.90 | NA | Gm12716 | | 0 | 0 | 0 | 0 | 0 | 0 | 1 | 0 | 0 | 0 | 0 |
| 0 | -0.44 | 3.68 | -0.12 | 0.90 | NA | Gm12821 | | 0 | 0 | 0 | 0 | 0 | 0 | 1 | 0 | 0 | 0 | 0 |
| 0 | -0.44 | 3.68 | -0.12 | 0.90 | NA | Gm24045 | | 0 | 0 | 0 | 0 | 0 | 0 | 1 | 0 | 0 | 0 | 0 |
| 0 | -0.44 | 3.68 | -0.12 | 0.90 | NA | Gm13075 | | 0 | 0 | 0 | 0 | 0 | 0 | 1 | 0 | 0 | 0 | 0 |
| 0 | -0.44 | 3.68 | -0.12 | 0.90 | NA | Gm8602 | | 0 | 0 | 0 | 0 | 0 | 0 | 1 | 0 | 0 | 0 | 0 |
| 0 | -0.44 | 3.68 | -0.12 | 0.90 | NA | Gm5300 | | 0 | 0 | 0 | 0 | 0 | 0 | 1 | 0 | 0 | 0 | 0 |
| 0 | -0.44 | 3.68 | -0.12 | 0.90 | NA | Gm4877 | | 0 | 0 | 0 | 0 | 0 | 0 | 1 | 0 | 0 | 0 | 0 |
| 0 | -0.44 | 3.68 | -0.12 | 0.90 | NA | Gm31747 | | 0 | 0 | 0 | 0 | 0 | 0 | 1 | 0 | 0 | 0 | 0 |
| 0 | -0.44 | 3.68 | -0.12 | 0.90 | NA | Gm36189 | | 0 | 0 | 0 | 0 | 0 | 0 | 1 | 0 | 0 | 0 | 0 |
| 0 | -0.44 | 3.68 | -0.12 | 0.90 | NA | Mug-ps1 | | 0 | 0 | 0 | 0 | 0 | 0 | 1 | 0 | 0 | 0 | 0 |
| 0 | -0.44 | 3.68 | -0.12 | 0.90 | NA | Gm45173 | | 0 | 0 | 0 | 0 | 0 | 0 | 1 | 0 | 0 | 0 | 0 |
| 0 | -0.44 | 3.68 | -0.12 | 0.90 | NA | A330074H02Rik | | 0 | 0 | 0 | 0 | 0 | 0 | 1 | 0 | 0 | 0 | 0 |
| 0 | -0.44 | 3.68 | -0.12 | 0.90 | NA | A530021J07Ri | Riken cDNA A530021J07 gene | 0 | 0 | 0 | 0 | 0 | 0 | 1 | 0 | 0 | 0 | 0 |
| 0 | -0.44 | 3.68 | -0.12 | 0.90 | NA | Gm6007 | | 0 | 0 | 0 | 0 | 0 | 0 | 1 | 0 | 0 | 0 | 0 |
| 0 | -0.44 | 3.68 | -0.12 | 0.90 | NA | Acsm1 | acyl-CoA synthetase medium-chain famil | 0 | 0 | 0 | 0 | 0 | 0 | 1 | 0 | 0 | 0 | 0 |
| 0 | -0.44 | 3.68 | -0.12 | 0.90 | NA | Gm7381 | | 0 | 0 | 0 | 0 | 0 | 0 | 1 | 0 | 0 | 0 | 0 |
| 0 | -0.44 | 3.68 | -0.12 | 0.90 | NA | Gm10358 | | 0 | 0 | 0 | 0 | 0 | 0 | 1 | 0 | 0 | 0 | 0 |
| 0 | -0.44 | 3.68 | -0.12 | 0.90 | NA | Gm9655 | | 0 | 0 | 0 | 0 | 0 | 0 | 1 | 0 | 0 | 0 | 0 |
| 0 | -0.44 | 3.68 | -0.12 | 0.90 | NA | 4933405L10Ri | RIKEN cDNA 4933405L10 gene | 0 | 0 | 0 | 0 | 0 | 0 | 1 | 0 | 0 | 0 | 0 |
| 0 | -0.44 | 3.68 | -0.12 | 0.90 | NA | Gm9204 | | 0 | 0 | 0 | 0 | 0 | 0 | 1 | 0 | 0 | 0 | 0 |

| | | | | | | | | | | | | | | | | | | |
|---|---|---|---|---|---|---|---|---|---|---|---|---|---|---|---|---|---|---|
| 0 | -0.44 | 3.68 | -0.12 | 0.90 | NA | Gm37167 | | 0 | 0 | 0 | 0 | 0 | 0 | 1 | 0 | 0 | 0 | 0 |
| 0 | -0.44 | 3.68 | -0.12 | 0.90 | NA | Gm7097 | | 0 | 0 | 0 | 0 | 0 | 0 | 1 | 0 | 0 | 0 | 0 |
| 0 | -0.44 | 3.68 | -0.12 | 0.90 | NA | Gm19688 | | 0 | 0 | 0 | 0 | 0 | 0 | 1 | 0 | 0 | 0 | 0 |
| 0 | -0.44 | 3.68 | -0.12 | 0.90 | NA | Gm4556 | | 0 | 0 | 0 | 0 | 0 | 0 | 1 | 0 | 0 | 0 | 0 |
| 0 | -0.44 | 3.68 | -0.12 | 0.90 | NA | Gm37439 | | 0 | 0 | 0 | 0 | 0 | 0 | 1 | 0 | 0 | 0 | 0 |
| 0 | -0.44 | 3.68 | -0.12 | 0.90 | NA | Gm12136 | | 0 | 0 | 0 | 0 | 0 | 0 | 1 | 0 | 0 | 0 | 0 |
| 0 | -0.44 | 3.68 | -0.12 | 0.90 | NA | Gm12628 | | 0 | 0 | 0 | 0 | 0 | 0 | 1 | 0 | 0 | 0 | 0 |
| 0 | -0.44 | 3.68 | -0.12 | 0.90 | NA | Gm12611 | | 0 | 0 | 0 | 0 | 0 | 0 | 1 | 0 | 0 | 0 | 0 |
| 0 | -0.44 | 3.68 | -0.12 | 0.90 | NA | BC106175 | | 0 | 0 | 0 | 0 | 0 | 0 | 1 | 0 | 0 | 0 | 0 |
| 0 | -0.44 | 3.68 | -0.12 | 0.90 | NA | Gm18116 | | 0 | 0 | 0 | 0 | 0 | 0 | 1 | 0 | 0 | 0 | 0 |
| 0 | -0.44 | 3.68 | -0.12 | 0.90 | NA | Mir5135 | | 0 | 0 | 0 | 0 | 0 | 0 | 1 | 0 | 0 | 0 | 0 |
| 0 | -0.44 | 3.68 | -0.12 | 0.90 | NA | Gm16125 | | 0 | 0 | 0 | 0 | 0 | 0 | 1 | 0 | 0 | 0 | 0 |
| 0 | -0.44 | 3.68 | -0.12 | 0.90 | NA | Gm2534 | | 0 | 0 | 0 | 0 | 0 | 0 | 1 | 0 | 0 | 0 | 0 |
| 0 | -0.44 | 3.68 | -0.12 | 0.90 | NA | Gm31191 | | 0 | 0 | 0 | 0 | 0 | 0 | 1 | 0 | 0 | 0 | 0 |
| 0 | -0.44 | 3.68 | -0.12 | 0.90 | NA | Gm29767 | | 0 | 0 | 0 | 0 | 0 | 0 | 1 | 0 | 0 | 0 | 0 |
| 0 | -0.44 | 3.68 | -0.12 | 0.90 | NA | Gm48312 | | 0 | 0 | 0 | 0 | 0 | 0 | 1 | 0 | 0 | 0 | 0 |
| 0 | -0.44 | 3.68 | -0.12 | 0.90 | NA | Gm25353 | | 0 | 0 | 0 | 0 | 0 | 0 | 1 | 0 | 0 | 0 | 0 |
| 0 | -0.44 | 3.68 | -0.12 | 0.90 | NA | Gm35167 | | 0 | 0 | 0 | 0 | 0 | 0 | 1 | 0 | 0 | 0 | 0 |
| 0 | -0.44 | 3.68 | -0.12 | 0.90 | NA | Gm29697 | | 0 | 0 | 0 | 0 | 0 | 0 | 1 | 0 | 0 | 0 | 0 |
| 0 | -0.44 | 3.68 | -0.12 | 0.90 | NA | Gm15799 | | 0 | 0 | 0 | 0 | 0 | 0 | 1 | 0 | 0 | 0 | 0 |
| 0 | -0.44 | 3.68 | -0.12 | 0.90 | NA | Gm26980 | | 0 | 0 | 0 | 0 | 0 | 0 | 1 | 0 | 0 | 0 | 0 |
| 0 | -0.44 | 3.68 | -0.12 | 0.90 | NA | Gm15692 | | 0 | 0 | 0 | 0 | 0 | 0 | 1 | 0 | 0 | 0 | 0 |
| 0 | -0.44 | 3.68 | -0.12 | 0.90 | NA | Gm7043 | | 0 | 0 | 0 | 0 | 0 | 0 | 1 | 0 | 0 | 0 | 0 |
| 0 | -0.44 | 3.68 | -0.12 | 0.90 | NA | Zfp422-ps | | 0 | 0 | 0 | 0 | 0 | 0 | 1 | 0 | 0 | 0 | 0 |
| 0 | -0.44 | 3.68 | -0.12 | 0.90 | NA | Slc22a27 | solute carrier family 22, member 27 | 0 | 0 | 0 | 0 | 0 | 0 | 1 | 0 | 0 | 0 | 0 |
| 0 | -0.44 | 3.68 | -0.12 | 0.90 | NA | Gm14591 | | 0 | 0 | 0 | 0 | 0 | 0 | 1 | 0 | 0 | 0 | 0 |
| 0 | -0.44 | 3.68 | -0.12 | 0.90 | NA | Gm7076 | | 0 | 0 | 0 | 0 | 0 | 0 | 1 | 0 | 0 | 0 | 0 |
| 0 | -0.44 | 3.68 | -0.12 | 0.90 | NA | Rps12-ps7 | | 0 | 0 | 0 | 0 | 0 | 0 | 1 | 0 | 0 | 0 | 0 |
| 0 | -0.44 | 3.68 | -0.12 | 0.90 | NA | Gm16459 | | 0 | 0 | 0 | 0 | 0 | 0 | 1 | 0 | 0 | 0 | 0 |
| 0 | 1.00 | 3.68 | 0.27 | 0.79 | NA | Gm37670 | | 0 | 0 | 0 | 0 | 0 | 0 | 0 | 0 | 0 | 1 | 0 |
| 0 | 1.00 | 3.68 | 0.27 | 0.79 | NA | Gm13321 | | 0 | 0 | 0 | 0 | 0 | 0 | 0 | 0 | 0 | 1 | 0 |
| 0 | 1.00 | 3.68 | 0.27 | 0.79 | NA | Gm13719 | | 0 | 0 | 0 | 0 | 0 | 0 | 0 | 0 | 0 | 1 | 0 |
| 0 | 1.00 | 3.68 | 0.27 | 0.79 | NA | Olfr1175 | | 0 | 0 | 0 | 0 | 0 | 0 | 0 | 0 | 0 | 1 | 0 |
| 0 | 1.00 | 3.68 | 0.27 | 0.79 | NA | Gm2004 | predicted gene 2004 | 0 | 0 | 0 | 0 | 0 | 0 | 0 | 0 | 0 | 1 | 0 |
| 0 | 1.00 | 3.68 | 0.27 | 0.79 | NA | Gm7132 | | 0 | 0 | 0 | 0 | 0 | 0 | 0 | 0 | 0 | 1 | 0 |
| 0 | 1.00 | 3.68 | 0.27 | 0.79 | NA | Gm38305 | | 0 | 0 | 0 | 0 | 0 | 0 | 0 | 0 | 0 | 1 | 0 |
| 0 | 1.00 | 3.68 | 0.27 | 0.79 | NA | Gm9504 | | 0 | 0 | 0 | 0 | 0 | 0 | 0 | 0 | 0 | 1 | 0 |
| 0 | 1.00 | 3.68 | 0.27 | 0.79 | NA | Gm9632 | | 0 | 0 | 0 | 0 | 0 | 0 | 0 | 0 | 0 | 1 | 0 |
| 0 | 1.00 | 3.68 | 0.27 | 0.79 | NA | Gm5710 | | 0 | 0 | 0 | 0 | 0 | 0 | 0 | 0 | 0 | 1 | 0 |
| 0 | 1.00 | 3.68 | 0.27 | 0.79 | NA | Cct3-ps1 | | 0 | 0 | 0 | 0 | 0 | 0 | 0 | 0 | 0 | 1 | 0 |
| 0 | 1.00 | 3.68 | 0.27 | 0.79 | NA | Gm42959 | | 0 | 0 | 0 | 0 | 0 | 0 | 0 | 0 | 0 | 1 | 0 |
| 0 | 1.00 | 3.68 | 0.27 | 0.79 | NA | Gm15550 | | 0 | 0 | 0 | 0 | 0 | 0 | 0 | 0 | 0 | 1 | 0 |
| 0 | 1.00 | 3.68 | 0.27 | 0.79 | NA | Gm44163 | | 0 | 0 | 0 | 0 | 0 | 0 | 0 | 0 | 0 | 1 | 0 |
| 0 | 1.00 | 3.68 | 0.27 | 0.79 | NA | Gm5311 | | 0 | 0 | 0 | 0 | 0 | 0 | 0 | 0 | 0 | 1 | 0 |
| 0 | 1.00 | 3.68 | 0.27 | 0.79 | NA | Gm15782 | | 0 | 0 | 0 | 0 | 0 | 0 | 0 | 0 | 0 | 1 | 0 |
| 0 | 1.00 | 3.68 | 0.27 | 0.79 | NA | Gm3665 | | 0 | 0 | 0 | 0 | 0 | 0 | 0 | 0 | 0 | 1 | 0 |
| 0 | 1.00 | 3.68 | 0.27 | 0.79 | NA | Gm9211 | | 0 | 0 | 0 | 0 | 0 | 0 | 0 | 0 | 0 | 1 | 0 |
| 0 | 1.00 | 3.68 | 0.27 | 0.79 | NA | Gm45319 | | 0 | 0 | 0 | 0 | 0 | 0 | 0 | 0 | 0 | 1 | 0 |
| 0 | 1.00 | 3.68 | 0.27 | 0.79 | NA | Gm8268 | | 0 | 0 | 0 | 0 | 0 | 0 | 0 | 0 | 0 | 1 | 0 |
| 0 | 1.00 | 3.68 | 0.27 | 0.79 | NA | Gm45633 | | 0 | 0 | 0 | 0 | 0 | 0 | 0 | 0 | 0 | 1 | 0 |
| 0 | 1.00 | 3.68 | 0.27 | 0.79 | NA | Gm3635 | | 0 | 0 | 0 | 0 | 0 | 0 | 0 | 0 | 0 | 1 | 0 |
| 0 | 1.00 | 3.68 | 0.27 | 0.79 | NA | Gm18709 | | 0 | 0 | 0 | 0 | 0 | 0 | 0 | 0 | 0 | 1 | 0 |
| 0 | 1.00 | 3.68 | 0.27 | 0.79 | NA | Gm7680 | | 0 | 0 | 0 | 0 | 0 | 0 | 0 | 0 | 0 | 1 | 0 |
| 0 | 1.00 | 3.68 | 0.27 | 0.79 | NA | Gm8087 | | 0 | 0 | 0 | 0 | 0 | 0 | 0 | 0 | 0 | 1 | 0 |
| 0 | 1.00 | 3.68 | 0.27 | 0.79 | NA | Gm47894 | | 0 | 0 | 0 | 0 | 0 | 0 | 0 | 0 | 0 | 1 | 0 |
| 0 | 1.00 | 3.68 | 0.27 | 0.79 | NA | Gm12218 | | 0 | 0 | 0 | 0 | 0 | 0 | 0 | 0 | 0 | 1 | 0 |
| 0 | 1.00 | 3.68 | 0.27 | 0.79 | NA | Gm12344 | | 0 | 0 | 0 | 0 | 0 | 0 | 0 | 0 | 0 | 1 | 0 |
| 0 | 1.00 | 3.68 | 0.27 | 0.79 | NA | Gm11198 | | 0 | 0 | 0 | 0 | 0 | 0 | 0 | 0 | 0 | 1 | 0 |
| 0 | 1.00 | 3.68 | 0.27 | 0.79 | NA | Gm11509 | | 0 | 0 | 0 | 0 | 0 | 0 | 0 | 0 | 0 | 1 | 0 |
| 0 | 1.00 | 3.68 | 0.27 | 0.79 | NA | Neurod2 | neurogenic differentiation 2 | 0 | 0 | 0 | 0 | 0 | 0 | 0 | 0 | 0 | 1 | 0 |

| | | | | | | | | | | | | | | | | | | |
|---|---|---|---|---|---|---|---|---|---|---|---|---|---|---|---|---|---|---|
| 0 | 1.00 | 3.68 | 0.27 | 0.79 | NA | Gm11646 | | 0 | 0 | 0 | 0 | 0 | 0 | 0 | 0 | 0 | 1 | 0 |
| 0 | 1.00 | 3.68 | 0.27 | 0.79 | NA | Gm8438 | | 0 | 0 | 0 | 0 | 0 | 0 | 0 | 0 | 0 | 1 | 0 |
| 0 | 1.00 | 3.68 | 0.27 | 0.79 | NA | Gm47763 | | 0 | 0 | 0 | 0 | 0 | 0 | 0 | 0 | 0 | 1 | 0 |
| 0 | 1.00 | 3.68 | 0.27 | 0.79 | NA | Gm4817 | | 0 | 0 | 0 | 0 | 0 | 0 | 0 | 0 | 0 | 1 | 0 |
| 0 | 1.00 | 3.68 | 0.27 | 0.79 | NA | Gm16261 | | 0 | 0 | 0 | 0 | 0 | 0 | 0 | 0 | 0 | 1 | 0 |
| 0 | 1.00 | 3.68 | 0.27 | 0.79 | NA | Gm6847 | | 0 | 0 | 0 | 0 | 0 | 0 | 0 | 0 | 0 | 1 | 0 |
| 0 | 1.00 | 3.68 | 0.27 | 0.79 | NA | Syt10 | synaptotagmin X | 0 | 0 | 0 | 0 | 0 | 0 | 0 | 0 | 0 | 1 | 0 |
| 0 | 1.00 | 3.68 | 0.27 | 0.79 | NA | Gm49996 | | 0 | 0 | 0 | 0 | 0 | 0 | 0 | 0 | 0 | 1 | 0 |
| 0 | 1.00 | 3.68 | 0.27 | 0.79 | NA | | | 0 | 0 | 0 | 0 | 0 | 0 | 0 | 0 | 0 | 1 | 0 |
| 0 | 1.00 | 3.68 | 0.27 | 0.79 | NA | Gm14328 | | 0 | 0 | 0 | 0 | 0 | 0 | 0 | 0 | 0 | 1 | 0 |
| 0 | 1.00 | 3.68 | 0.27 | 0.79 | NA | Gm27037 | | 0 | 0 | 0 | 0 | 0 | 0 | 0 | 0 | 0 | 1 | 0 |
| 0 | 1.00 | 3.68 | 0.27 | 0.79 | NA | Gm5249 | | 0 | 0 | 0 | 0 | 0 | 0 | 0 | 0 | 0 | 1 | 0 |
| 0 | 1.00 | 3.68 | 0.27 | 0.79 | NA | Gm9051 | | 0 | 0 | 0 | 0 | 0 | 0 | 0 | 0 | 0 | 1 | 0 |
| 0 | 1.00 | 3.68 | 0.27 | 0.79 | NA | Gm6500 | | 0 | 0 | 0 | 0 | 0 | 0 | 0 | 0 | 0 | 1 | 0 |
| 0 | 1.00 | 3.68 | 0.27 | 0.79 | NA | Gm14928 | | 0 | 0 | 0 | 0 | 0 | 0 | 0 | 0 | 0 | 1 | 0 |
| 0 | 1.00 | 3.68 | 0.27 | 0.79 | NA | Gm8534 | | 0 | 0 | 0 | 0 | 0 | 0 | 0 | 0 | 0 | 1 | 0 |
| 0 | 1.00 | 3.68 | 0.27 | 0.79 | NA | mt-Atp6 | ATP synthase F0 subunit 6 | 0 | 0 | 0 | 0 | 0 | 0 | 0 | 0 | 0 | 1 | 0 |
| 0 | 1.00 | 3.68 | 0.27 | 0.79 | NA | Gm7449 | | 0 | 0 | 0 | 0 | 0 | 0 | 0 | 1 | 0 | 0 | 0 |
| 0 | 1.00 | 3.68 | 0.27 | 0.79 | NA | Gm15702 | | 0 | 0 | 0 | 0 | 0 | 0 | 0 | 1 | 0 | 0 | 0 |
| 0 | 1.00 | 3.68 | 0.27 | 0.79 | NA | Gm16102 | | 0 | 0 | 0 | 0 | 0 | 0 | 0 | 1 | 0 | 0 | 0 |
| 0 | 1.00 | 3.68 | 0.27 | 0.79 | NA | Gm28527 | | 0 | 0 | 0 | 0 | 0 | 0 | 0 | 1 | 0 | 0 | 0 |
| 0 | 1.00 | 3.68 | 0.27 | 0.79 | NA | Gm5974 | | 0 | 0 | 0 | 0 | 0 | 0 | 0 | 1 | 0 | 0 | 0 |
| 0 | 1.00 | 3.68 | 0.27 | 0.79 | NA | Gm28900 | | 0 | 0 | 0 | 0 | 0 | 0 | 0 | 1 | 0 | 0 | 0 |
| 0 | 1.00 | 3.68 | 0.27 | 0.79 | NA | Gm16305 | | 0 | 0 | 0 | 0 | 0 | 0 | 0 | 1 | 0 | 0 | 0 |
| 0 | 1.00 | 3.68 | 0.27 | 0.79 | NA | Gm13296 | | 0 | 0 | 0 | 0 | 0 | 0 | 0 | 1 | 0 | 0 | 0 |
| 0 | 1.00 | 3.68 | 0.27 | 0.79 | NA | Gm13326 | | 0 | 0 | 0 | 0 | 0 | 0 | 0 | 1 | 0 | 0 | 0 |
| 0 | 1.00 | 3.68 | 0.27 | 0.79 | NA | Gm13360 | | 0 | 0 | 0 | 0 | 0 | 0 | 0 | 1 | 0 | 0 | 0 |
| 0 | 1.00 | 3.68 | 0.27 | 0.79 | NA | Gm13433 | | 0 | 0 | 0 | 0 | 0 | 0 | 0 | 1 | 0 | 0 | 0 |
| 0 | 1.00 | 3.68 | 0.27 | 0.79 | NA | Gm44183 | | 0 | 0 | 0 | 0 | 0 | 0 | 0 | 1 | 0 | 0 | 0 |
| 0 | 1.00 | 3.68 | 0.27 | 0.79 | NA | Gm13517 | | 0 | 0 | 0 | 0 | 0 | 0 | 0 | 1 | 0 | 0 | 0 |
| 0 | 1.00 | 3.68 | 0.27 | 0.79 | NA | Gm23330 | | 0 | 0 | 0 | 0 | 0 | 0 | 0 | 1 | 0 | 0 | 0 |
| 0 | 1.00 | 3.68 | 0.27 | 0.79 | NA | Mif-ps3 | | 0 | 0 | 0 | 0 | 0 | 0 | 0 | 1 | 0 | 0 | 0 |
| 0 | 1.00 | 3.68 | 0.27 | 0.79 | NA | Gm13232 | | 0 | 0 | 0 | 0 | 0 | 0 | 0 | 1 | 0 | 0 | 0 |
| 0 | 1.00 | 3.68 | 0.27 | 0.79 | NA | Gm15469 | | 0 | 0 | 0 | 0 | 0 | 0 | 0 | 1 | 0 | 0 | 0 |
| 0 | 1.00 | 3.68 | 0.27 | 0.79 | NA | Gm15621 | | 0 | 0 | 0 | 0 | 0 | 0 | 0 | 1 | 0 | 0 | 0 |
| 0 | 1.00 | 3.68 | 0.27 | 0.79 | NA | Gm2796 | | 0 | 0 | 0 | 0 | 0 | 0 | 0 | 1 | 0 | 0 | 0 |
| 0 | 1.00 | 3.68 | 0.27 | 0.79 | NA | Gm40414 | | 0 | 0 | 0 | 0 | 0 | 0 | 0 | 1 | 0 | 0 | 0 |
| 0 | 1.00 | 3.68 | 0.27 | 0.79 | NA | | | 0 | 0 | 0 | 0 | 0 | 0 | 0 | 1 | 0 | 0 | 0 |
| 0 | 1.00 | 3.68 | 0.27 | 0.79 | NA | Gm45315 | | 0 | 0 | 0 | 0 | 0 | 0 | 0 | 1 | 0 | 0 | 0 |
| 0 | 1.00 | 3.68 | 0.27 | 0.79 | NA | Gm6112 | | 0 | 0 | 0 | 0 | 0 | 0 | 0 | 1 | 0 | 0 | 0 |
| 0 | 1.00 | 3.68 | 0.27 | 0.79 | NA | Gm15744 | | 0 | 0 | 0 | 0 | 0 | 0 | 0 | 1 | 0 | 0 | 0 |
| 0 | 1.00 | 3.68 | 0.27 | 0.79 | NA | Gm8436 | | 0 | 0 | 0 | 0 | 0 | 0 | 0 | 1 | 0 | 0 | 0 |
| 0 | 1.00 | 3.68 | 0.27 | 0.79 | NA | Gm8079 | | 0 | 0 | 0 | 0 | 0 | 0 | 0 | 1 | 0 | 0 | 0 |
| 0 | 1.00 | 3.68 | 0.27 | 0.79 | NA | Gm5614 | | 0 | 0 | 0 | 0 | 0 | 0 | 0 | 1 | 0 | 0 | 0 |
| 0 | 1.00 | 3.68 | 0.27 | 0.79 | NA | Gm16073 | | 0 | 0 | 0 | 0 | 0 | 0 | 0 | 1 | 0 | 0 | 0 |
| 0 | 1.00 | 3.68 | 0.27 | 0.79 | NA | Gm15896 | | 0 | 0 | 0 | 0 | 0 | 0 | 0 | 1 | 0 | 0 | 0 |
| 0 | 1.00 | 3.68 | 0.27 | 0.79 | NA | Gm3096 | | 0 | 0 | 0 | 0 | 0 | 0 | 0 | 1 | 0 | 0 | 0 |
| 0 | 1.00 | 3.68 | 0.27 | 0.79 | NA | Gm26445 | | 0 | 0 | 0 | 0 | 0 | 0 | 0 | 1 | 0 | 0 | 0 |
| 0 | 1.00 | 3.68 | 0.27 | 0.79 | NA | Gm11984 | | 0 | 0 | 0 | 0 | 0 | 0 | 0 | 1 | 0 | 0 | 0 |
| 0 | 1.00 | 3.68 | 0.27 | 0.79 | NA | Rpsa-ps3 | | 0 | 0 | 0 | 0 | 0 | 0 | 0 | 1 | 0 | 0 | 0 |
| 0 | 1.00 | 3.68 | 0.27 | 0.79 | NA | Rpl9-ps3 | | 0 | 0 | 0 | 0 | 0 | 0 | 0 | 1 | 0 | 0 | 0 |
| 0 | 1.00 | 3.68 | 0.27 | 0.79 | NA | Gm11494 | | 0 | 0 | 0 | 0 | 0 | 0 | 0 | 1 | 0 | 0 | 0 |
| 0 | 1.00 | 3.68 | 0.27 | 0.79 | NA | Gm11502 | | 0 | 0 | 0 | 0 | 0 | 0 | 0 | 1 | 0 | 0 | 0 |
| 0 | 1.00 | 3.68 | 0.27 | 0.79 | NA | Gm12587 | | 0 | 0 | 0 | 0 | 0 | 0 | 0 | 1 | 0 | 0 | 0 |
| 0 | 1.00 | 3.68 | 0.27 | 0.79 | NA | Gm21232 | | 0 | 0 | 0 | 0 | 0 | 0 | 0 | 1 | 0 | 0 | 0 |
| 0 | 1.00 | 3.68 | 0.27 | 0.79 | NA | Gm6211 | | 0 | 0 | 0 | 0 | 0 | 0 | 0 | 1 | 0 | 0 | 0 |
| 0 | 1.00 | 3.68 | 0.27 | 0.79 | NA | Gm5200 | | 0 | 0 | 0 | 0 | 0 | 0 | 0 | 1 | 0 | 0 | 0 |
| 0 | 1.00 | 3.68 | 0.27 | 0.79 | NA | Rpl19-ps3 | | 0 | 0 | 0 | 0 | 0 | 0 | 0 | 1 | 0 | 0 | 0 |
| 0 | 1.00 | 3.68 | 0.27 | 0.79 | NA | Gm9391 | | 0 | 0 | 0 | 0 | 0 | 0 | 0 | 1 | 0 | 0 | 0 |
| 0 | 1.00 | 3.68 | 0.27 | 0.79 | NA | Gm6361 | | 0 | 0 | 0 | 0 | 0 | 0 | 0 | 1 | 0 | 0 | 0 |

| | | | | | | | | | | | | | | | | | | |
|---|---|---|---|---|---|---|---|---|---|---|---|---|---|---|---|---|---|---|
| 0 | 1.00 | 3.68 | 0.27 | 0.79 | NA | Gm49657 | | 0 | 0 | 0 | 0 | 0 | 0 | 0 | 1 | 0 | 0 | 0 |
| 0 | 1.00 | 3.68 | 0.27 | 0.79 | NA | Vmn2r100 | vomeronasal 2, receptor 100 | 0 | 0 | 0 | 0 | 0 | 0 | 0 | 1 | 0 | 0 | 0 |
| 0 | 1.00 | 3.68 | 0.27 | 0.79 | NA | Gm19183 | | 0 | 0 | 0 | 0 | 0 | 0 | 0 | 1 | 0 | 0 | 0 |
| 0 | 1.00 | 3.68 | 0.27 | 0.79 | NA | Gm5504 | | 0 | 0 | 0 | 0 | 0 | 0 | 0 | 1 | 0 | 0 | 0 |
| 0 | 1.00 | 3.68 | 0.27 | 0.79 | NA | Lipo5 | | 0 | 0 | 0 | 0 | 0 | 0 | 0 | 1 | 0 | 0 | 0 |
| 0 | 1.00 | 3.68 | 0.27 | 0.79 | NA | Cyp2c50 | cytochrome P450, family 2, subfamily c, p | 0 | 0 | 0 | 0 | 0 | 0 | 0 | 1 | 0 | 0 | 0 |
| 0 | 1.00 | 3.68 | 0.27 | 0.79 | NA | Gm14859 | | 0 | 0 | 0 | 0 | 0 | 0 | 0 | 1 | 0 | 0 | 0 |
| 0 | 1.00 | 3.68 | 0.27 | 0.79 | NA | Gm4913 | | 0 | 0 | 0 | 0 | 0 | 0 | 0 | 1 | 0 | 0 | 0 |
| 0 | 0.00 | 3.68 | 0.00 | 1.00 | NA | Gm17970 | | 0 | 0 | 1 | 0 | 0 | 0 | 0 | 0 | 0 | 0 | 0 |
| 0 | 0.00 | 3.68 | 0.00 | 1.00 | NA | Gm3841 | | 0 | 0 | 1 | 0 | 0 | 0 | 0 | 0 | 0 | 0 | 0 |
| 0 | 0.00 | 3.68 | 0.00 | 1.00 | NA | Gm20302 | | 0 | 0 | 1 | 0 | 0 | 0 | 0 | 0 | 0 | 0 | 0 |
| 0 | 0.00 | 3.68 | 0.00 | 1.00 | NA | Gm9687 | | 0 | 0 | 1 | 0 | 0 | 0 | 0 | 0 | 0 | 0 | 0 |
| 0 | 0.00 | 3.68 | 0.00 | 1.00 | NA | Olfr433 | olfactory receptor 433 | 0 | 0 | 1 | 0 | 0 | 0 | 0 | 0 | 0 | 0 | 0 |
| 0 | 0.00 | 3.68 | 0.00 | 1.00 | NA | Gm5706 | | 0 | 0 | 1 | 0 | 0 | 0 | 0 | 0 | 0 | 0 | 0 |
| 0 | 0.00 | 3.68 | 0.00 | 1.00 | NA | Gm37069 | | 0 | 0 | 1 | 0 | 0 | 0 | 0 | 0 | 0 | 0 | 0 |
| 0 | 0.00 | 3.68 | 0.00 | 1.00 | NA | Gm2272 | | 0 | 0 | 1 | 0 | 0 | 0 | 0 | 0 | 0 | 0 | 0 |
| 0 | 0.00 | 3.68 | 0.00 | 1.00 | NA | Gm13311 | | 0 | 0 | 1 | 0 | 0 | 0 | 0 | 0 | 0 | 0 | 0 |
| 0 | 0.00 | 3.68 | 0.00 | 1.00 | NA | Gm13508 | | 0 | 0 | 1 | 0 | 0 | 0 | 0 | 0 | 0 | 0 | 0 |
| 0 | 0.00 | 3.68 | 0.00 | 1.00 | NA | Gm13607 | | 0 | 0 | 1 | 0 | 0 | 0 | 0 | 0 | 0 | 0 | 0 |
| 0 | 0.00 | 3.68 | 0.00 | 1.00 | NA | Gm13732 | | 0 | 0 | 1 | 0 | 0 | 0 | 0 | 0 | 0 | 0 | 0 |
| 0 | 0.00 | 3.68 | 0.00 | 1.00 | NA | Gm13811 | | 0 | 0 | 1 | 0 | 0 | 0 | 0 | 0 | 0 | 0 | 0 |
| 0 | 0.00 | 3.68 | 0.00 | 1.00 | NA | Gm10770 | | 0 | 0 | 1 | 0 | 0 | 0 | 0 | 0 | 0 | 0 | 0 |
| 0 | 0.00 | 3.68 | 0.00 | 1.00 | NA | Gm14497 | | 0 | 0 | 1 | 0 | 0 | 0 | 0 | 0 | 0 | 0 | 0 |
| 0 | 0.00 | 3.68 | 0.00 | 1.00 | NA | Rpl21-ps9 | | 0 | 0 | 1 | 0 | 0 | 0 | 0 | 0 | 0 | 0 | 0 |
| 0 | 0.00 | 3.68 | 0.00 | 1.00 | NA | Gm36985 | | 0 | 0 | 1 | 0 | 0 | 0 | 0 | 0 | 0 | 0 | 0 |
| 0 | 0.00 | 3.68 | 0.00 | 1.00 | NA | Gm6098 | | 0 | 0 | 1 | 0 | 0 | 0 | 0 | 0 | 0 | 0 | 0 |
| 0 | 0.00 | 3.68 | 0.00 | 1.00 | NA | Cd1d2 | CD1d2 antigen | 0 | 0 | 1 | 0 | 0 | 0 | 0 | 0 | 0 | 0 | 0 |
| 0 | 0.00 | 3.68 | 0.00 | 1.00 | NA | Gm43464 | | 0 | 0 | 1 | 0 | 0 | 0 | 0 | 0 | 0 | 0 | 0 |
| 0 | 0.00 | 3.68 | 0.00 | 1.00 | NA | Gm5711 | | 0 | 0 | 1 | 0 | 0 | 0 | 0 | 0 | 0 | 0 | 0 |
| 0 | 0.00 | 3.68 | 0.00 | 1.00 | NA | Gm16060 | | 0 | 0 | 1 | 0 | 0 | 0 | 0 | 0 | 0 | 0 | 0 |
| 0 | 0.00 | 3.68 | 0.00 | 1.00 | NA | Gm11888 | | 0 | 0 | 1 | 0 | 0 | 0 | 0 | 0 | 0 | 0 | 0 |
| 0 | 0.00 | 3.68 | 0.00 | 1.00 | NA | Gm11913 | | 0 | 0 | 1 | 0 | 0 | 0 | 0 | 0 | 0 | 0 | 0 |
| 0 | 0.00 | 3.68 | 0.00 | 1.00 | NA | Pakap | paralemmin A kinase anchor protein | 0 | 0 | 1 | 0 | 0 | 0 | 0 | 0 | 0 | 0 | 0 |
| 0 | 0.00 | 3.68 | 0.00 | 1.00 | NA | Gm11255 | | 0 | 0 | 1 | 0 | 0 | 0 | 0 | 0 | 0 | 0 | 0 |
| 0 | 0.00 | 3.68 | 0.00 | 1.00 | NA | Gm11412 | | 0 | 0 | 1 | 0 | 0 | 0 | 0 | 0 | 0 | 0 | 0 |
| 0 | 0.00 | 3.68 | 0.00 | 1.00 | NA | Gm13118 | | 0 | 0 | 1 | 0 | 0 | 0 | 0 | 0 | 0 | 0 | 0 |
| 0 | 0.00 | 3.68 | 0.00 | 1.00 | NA | Gm9506 | | 0 | 0 | 1 | 0 | 0 | 0 | 0 | 0 | 0 | 0 | 0 |
| 0 | 0.00 | 3.68 | 0.00 | 1.00 | NA | Gm7047 | | 0 | 0 | 1 | 0 | 0 | 0 | 0 | 0 | 0 | 0 | 0 |
| 0 | 0.00 | 3.68 | 0.00 | 1.00 | NA | Gm16111 | | 0 | 0 | 1 | 0 | 0 | 0 | 0 | 0 | 0 | 0 | 0 |
| 0 | 0.00 | 3.68 | 0.00 | 1.00 | NA | Gm15866 | | 0 | 0 | 1 | 0 | 0 | 0 | 0 | 0 | 0 | 0 | 0 |
| 0 | 0.00 | 3.68 | 0.00 | 1.00 | NA | Gm10135 | | 0 | 0 | 1 | 0 | 0 | 0 | 0 | 0 | 0 | 0 | 0 |
| 0 | 0.00 | 3.68 | 0.00 | 1.00 | NA | Gm15653 | | 0 | 0 | 1 | 0 | 0 | 0 | 0 | 0 | 0 | 0 | 0 |
| 0 | 0.00 | 3.68 | 0.00 | 1.00 | NA | Gm42601 | | 0 | 0 | 1 | 0 | 0 | 0 | 0 | 0 | 0 | 0 | 0 |
| 0 | 0.00 | 3.68 | 0.00 | 1.00 | NA | P2rx2 | purinergic receptor P2X, ligand-gated ion | 0 | 0 | 1 | 0 | 0 | 0 | 0 | 0 | 0 | 0 | 0 |
| 0 | 0.00 | 3.68 | 0.00 | 1.00 | NA | Gm21839 | | 0 | 0 | 1 | 0 | 0 | 0 | 0 | 0 | 0 | 0 | 0 |
| 0 | 0.00 | 3.68 | 0.00 | 1.00 | NA | Gm43649 | | 0 | 0 | 1 | 0 | 0 | 0 | 0 | 0 | 0 | 0 | 0 |
| 0 | 0.00 | 3.68 | 0.00 | 1.00 | NA | Wnt16 | wingless-type MMTV integration site famil | 0 | 0 | 1 | 0 | 0 | 0 | 0 | 0 | 0 | 0 | 0 |
| 0 | 0.00 | 3.68 | 0.00 | 1.00 | NA | Gm37301 | | 0 | 0 | 1 | 0 | 0 | 0 | 0 | 0 | 0 | 0 | 0 |
| 0 | 0.00 | 3.68 | 0.00 | 1.00 | NA | Rpl31-ps7 | | 0 | 0 | 1 | 0 | 0 | 0 | 0 | 0 | 0 | 0 | 0 |
| 0 | 0.00 | 3.68 | 0.00 | 1.00 | NA | Gm6134 | | 0 | 0 | 1 | 0 | 0 | 0 | 0 | 0 | 0 | 0 | 0 |
| 0 | 0.00 | 3.68 | 0.00 | 1.00 | NA | Eif4a3l1 | eukaryotic translation initiation factor 4A3 | 0 | 0 | 1 | 0 | 0 | 0 | 0 | 0 | 0 | 0 | 0 |
| 0 | 0.00 | 3.68 | 0.00 | 1.00 | NA | Vmn1r55 | vomeronasal 1 receptor 55 | 0 | 0 | 1 | 0 | 0 | 0 | 0 | 0 | 0 | 0 | 0 |
| 0 | 0.00 | 3.68 | 0.00 | 1.00 | NA | Wdr88 | G patch domain containing 1 | 0 | 0 | 1 | 0 | 0 | 0 | 0 | 0 | 0 | 0 | 0 |
| 0 | 0.00 | 3.68 | 0.00 | 1.00 | NA | Rpl7a-ps9 | | 0 | 0 | 1 | 0 | 0 | 0 | 0 | 0 | 0 | 0 | 0 |
| 0 | 0.00 | 3.68 | 0.00 | 1.00 | NA | Olfr628 | olfactory receptor 628 | 0 | 0 | 1 | 0 | 0 | 0 | 0 | 0 | 0 | 0 | 0 |
| 0 | 0.00 | 3.68 | 0.00 | 1.00 | NA | Snrp1c-ps1 | | 0 | 0 | 1 | 0 | 0 | 0 | 0 | 0 | 0 | 0 | 0 |
| 0 | 0.00 | 3.68 | 0.00 | 1.00 | NA | Gm45160 | | 0 | 0 | 1 | 0 | 0 | 0 | 0 | 0 | 0 | 0 | 0 |
| 0 | 0.00 | 3.68 | 0.00 | 1.00 | NA | Gm15302 | | 0 | 0 | 1 | 0 | 0 | 0 | 0 | 0 | 0 | 0 | 0 |
| 0 | 0.00 | 3.68 | 0.00 | 1.00 | NA | Gm6463 | | 0 | 0 | 1 | 0 | 0 | 0 | 0 | 0 | 0 | 0 | 0 |
| 0 | 0.00 | 3.68 | 0.00 | 1.00 | NA | Gm48640 | | 0 | 0 | 1 | 0 | 0 | 0 | 0 | 0 | 0 | 0 | 0 |

| | | | | | | | | | | | | | | | | | | |
|---|---|---|---|---|---|---|---|---|---|---|---|---|---|---|---|---|---|---|
| 0 | 0.00 | 3.68 | 0.00 | 1.00 | NA | 4930467K11Rik | | 0 | 0 | 1 | 0 | 0 | 0 | 0 | 0 | 0 | 0 | 0 |
| 0 | 0.00 | 3.68 | 0.00 | 1.00 | NA | Gm48145 | | 0 | 0 | 1 | 0 | 0 | 0 | 0 | 0 | 0 | 0 | 0 |
| 0 | 0.00 | 3.68 | 0.00 | 1.00 | NA | Gm20091 | | 0 | 0 | 1 | 0 | 0 | 0 | 0 | 0 | 0 | 0 | 0 |
| 0 | 0.00 | 3.68 | 0.00 | 1.00 | NA | Gm8796 | | 0 | 0 | 1 | 0 | 0 | 0 | 0 | 0 | 0 | 0 | 0 |
| 0 | 0.00 | 3.68 | 0.00 | 1.00 | NA | Rps15a-ps1 | | 0 | 0 | 1 | 0 | 0 | 0 | 0 | 0 | 0 | 0 | 0 |
| 0 | 0.00 | 3.68 | 0.00 | 1.00 | NA | Gm12045 | | 0 | 0 | 1 | 0 | 0 | 0 | 0 | 0 | 0 | 0 | 0 |
| 0 | 0.00 | 3.68 | 0.00 | 1.00 | NA | Gm12101 | | 0 | 0 | 1 | 0 | 0 | 0 | 0 | 0 | 0 | 0 | 0 |
| 0 | 0.00 | 3.68 | 0.00 | 1.00 | NA | Gm12261 | | 0 | 0 | 1 | 0 | 0 | 0 | 0 | 0 | 0 | 0 | 0 |
| 0 | 0.00 | 3.68 | 0.00 | 1.00 | NA | Gm48636 | | 0 | 0 | 1 | 0 | 0 | 0 | 0 | 0 | 0 | 0 | 0 |
| 0 | 0.00 | 3.68 | 0.00 | 1.00 | NA | Gm47399 | | 0 | 0 | 1 | 0 | 0 | 0 | 0 | 0 | 0 | 0 | 0 |
| 0 | 0.00 | 3.68 | 0.00 | 1.00 | NA | Ly6e-ps1 | | 0 | 0 | 1 | 0 | 0 | 0 | 0 | 0 | 0 | 0 | 0 |
| 0 | 0.00 | 3.68 | 0.00 | 1.00 | NA | Gm48513 | | 0 | 0 | 1 | 0 | 0 | 0 | 0 | 0 | 0 | 0 | 0 |
| 0 | 0.00 | 3.68 | 0.00 | 1.00 | NA | Gm47195 | | 0 | 0 | 1 | 0 | 0 | 0 | 0 | 0 | 0 | 0 | 0 |
| 0 | 0.00 | 3.68 | 0.00 | 1.00 | NA | Gm6436 | | 0 | 0 | 1 | 0 | 0 | 0 | 0 | 0 | 0 | 0 | 0 |
| 0 | 0.00 | 3.68 | 0.00 | 1.00 | NA | Gm21726 | | 0 | 0 | 1 | 0 | 0 | 0 | 0 | 0 | 0 | 0 | 0 |
| 0 | 0.00 | 3.68 | 0.00 | 1.00 | NA | Naip3 | | 0 | 0 | 1 | 0 | 0 | 0 | 0 | 0 | 0 | 0 | 0 |
| 0 | 0.00 | 3.68 | 0.00 | 1.00 | NA | Gm21370 | | 0 | 0 | 1 | 0 | 0 | 0 | 0 | 0 | 0 | 0 | 0 |
| 0 | 0.00 | 3.68 | 0.00 | 1.00 | NA | Gm41109 | | 0 | 0 | 1 | 0 | 0 | 0 | 0 | 0 | 0 | 0 | 0 |
| 0 | 0.00 | 3.68 | 0.00 | 1.00 | NA | Gm20430 | | 0 | 0 | 1 | 0 | 0 | 0 | 0 | 0 | 0 | 0 | 0 |
| 0 | 0.00 | 3.68 | 0.00 | 1.00 | NA | Gm5209 | | 0 | 0 | 1 | 0 | 0 | 0 | 0 | 0 | 0 | 0 | 0 |
| 0 | 0.00 | 3.68 | 0.00 | 1.00 | NA | AU022793 | | 0 | 0 | 1 | 0 | 0 | 0 | 0 | 0 | 0 | 0 | 0 |
| 0 | 0.00 | 3.68 | 0.00 | 1.00 | NA | Gm49212 | | 0 | 0 | 1 | 0 | 0 | 0 | 0 | 0 | 0 | 0 | 0 |
| 0 | 0.00 | 3.68 | 0.00 | 1.00 | NA | Gm6031 | | 0 | 0 | 1 | 0 | 0 | 0 | 0 | 0 | 0 | 0 | 0 |
| 0 | 0.00 | 3.68 | 0.00 | 1.00 | NA | Gm18334 | | 0 | 0 | 1 | 0 | 0 | 0 | 0 | 0 | 0 | 0 | 0 |
| 0 | 0.00 | 3.68 | 0.00 | 1.00 | NA | Gm17809 | | 0 | 0 | 1 | 0 | 0 | 0 | 0 | 0 | 0 | 0 | 0 |
| 0 | 0.00 | 3.68 | 0.00 | 1.00 | NA | Ly6g6f | lymphocyte antigen 6 complex, locus G6F | 0 | 0 | 1 | 0 | 0 | 0 | 0 | 0 | 0 | 0 | 0 |
| 0 | 0.00 | 3.68 | 0.00 | 1.00 | NA | Gm49949 | | 0 | 0 | 1 | 0 | 0 | 0 | 0 | 0 | 0 | 0 | 0 |
| 0 | 0.00 | 3.68 | 0.00 | 1.00 | NA | Gm50093 | | 0 | 0 | 1 | 0 | 0 | 0 | 0 | 0 | 0 | 0 | 0 |
| 0 | 0.00 | 3.68 | 0.00 | 1.00 | NA | Gm46611 | | 0 | 0 | 1 | 0 | 0 | 0 | 0 | 0 | 0 | 0 | 0 |
| 0 | 0.00 | 3.68 | 0.00 | 1.00 | NA | Gm8986 | | 0 | 0 | 1 | 0 | 0 | 0 | 0 | 0 | 0 | 0 | 0 |
| 0 | 0.00 | 3.68 | 0.00 | 1.00 | NA | Gm19437 | | 0 | 0 | 1 | 0 | 0 | 0 | 0 | 0 | 0 | 0 | 0 |
| 0 | 0.00 | 3.68 | 0.00 | 1.00 | NA | Emx2 | empty spiracles homeobox 2 | 0 | 0 | 1 | 0 | 0 | 0 | 0 | 0 | 0 | 0 | 0 |
| 0 | 0.00 | 3.68 | 0.00 | 1.00 | NA | Gm14890 | | 0 | 0 | 1 | 0 | 0 | 0 | 0 | 0 | 0 | 0 | 0 |
| 0 | 0.00 | 3.68 | 0.00 | 1.00 | NA | Gm8545 | | 0 | 0 | 1 | 0 | 0 | 0 | 0 | 0 | 0 | 0 | 0 |
| 0 | 0.00 | 3.68 | 0.00 | 1.00 | NA | Gm15365 | | 0 | 0 | 1 | 0 | 0 | 0 | 0 | 0 | 0 | 0 | 0 |
| 0 | 0.00 | 3.68 | 0.00 | 1.00 | NA | Rpl17-ps8 | | 0 | 0 | 1 | 0 | 0 | 0 | 0 | 0 | 0 | 0 | 0 |
| 0 | 0.00 | 3.68 | 0.00 | 1.00 | NA | Gm14813 | | 0 | 0 | 1 | 0 | 0 | 0 | 0 | 0 | 0 | 0 | 0 |
| 0 | 0.00 | 3.68 | 0.00 | 1.00 | NA | Gm6461 | | 0 | 0 | 1 | 0 | 0 | 0 | 0 | 0 | 0 | 0 | 0 |
| 0 | 0.00 | 3.68 | 0.00 | 1.00 | NA | Gm7823 | | 0 | 0 | 1 | 0 | 0 | 0 | 0 | 0 | 0 | 0 | 0 |
| 0 | 0.00 | 3.68 | 0.00 | 1.00 | NA | Gm16410 | | 0 | 0 | 1 | 0 | 0 | 0 | 0 | 0 | 0 | 0 | 0 |
| 0 | 0.00 | 3.68 | 0.00 | 1.00 | NA | Gm7973 | | 0 | 0 | 1 | 0 | 0 | 0 | 0 | 0 | 0 | 0 | 0 |
| 0 | 0.00 | 3.68 | 0.00 | 1.00 | NA | Smt3h1-ps | | 0 | 0 | 1 | 0 | 0 | 0 | 0 | 0 | 0 | 0 | 0 |
| 0 | 0.95 | 3.67 | 0.26 | 0.80 | NA | Gm37567 | | 0 | 0 | 0 | 0 | 0 | 0 | 0 | 0 | 0 | 0 | 1 |
| 0 | 0.95 | 3.67 | 0.26 | 0.80 | NA | Snord87 | | 0 | 0 | 0 | 0 | 0 | 0 | 0 | 0 | 0 | 0 | 1 |
| 0 | 0.95 | 3.67 | 0.26 | 0.80 | NA | Gm37138 | | 0 | 0 | 0 | 0 | 0 | 0 | 0 | 0 | 0 | 0 | 1 |
| 0 | 0.95 | 3.67 | 0.26 | 0.80 | NA | Tfap2b | transcription factor AP-2 beta | 0 | 0 | 0 | 0 | 0 | 0 | 0 | 0 | 0 | 0 | 1 |
| 0 | 0.95 | 3.67 | 0.26 | 0.80 | NA | Gm7784 | | 0 | 0 | 0 | 0 | 0 | 0 | 0 | 0 | 0 | 0 | 1 |
| 0 | 0.95 | 3.67 | 0.26 | 0.80 | NA | Gm5697 | | 0 | 0 | 0 | 0 | 0 | 0 | 0 | 0 | 0 | 0 | 1 |
| 0 | 0.95 | 3.67 | 0.26 | 0.80 | NA | Gm37506 | | 0 | 0 | 0 | 0 | 0 | 0 | 0 | 0 | 0 | 0 | 1 |
| 0 | 0.95 | 3.67 | 0.26 | 0.80 | NA | Gm9915 | | 0 | 0 | 0 | 0 | 0 | 0 | 0 | 0 | 0 | 0 | 1 |
| 0 | 0.95 | 3.67 | 0.26 | 0.80 | NA | 4930444A19Rik | | 0 | 0 | 0 | 0 | 0 | 0 | 0 | 0 | 0 | 0 | 1 |
| 0 | 0.95 | 3.67 | 0.26 | 0.80 | NA | Dytn | dystrotelin | 0 | 0 | 0 | 0 | 0 | 0 | 0 | 0 | 0 | 0 | 1 |
| 0 | 0.95 | 3.67 | 0.26 | 0.80 | NA | Gm37058 | | 0 | 0 | 0 | 0 | 0 | 0 | 0 | 0 | 0 | 0 | 1 |
| 0 | 0.95 | 3.67 | 0.26 | 0.80 | NA | 1700067G17Rik | | 0 | 0 | 0 | 0 | 0 | 0 | 0 | 0 | 0 | 0 | 1 |
| 0 | 0.95 | 3.67 | 0.26 | 0.80 | NA | Asb18 | ankyrin repeat and SOCS box-containing | 0 | 0 | 0 | 0 | 0 | 0 | 0 | 0 | 0 | 0 | 1 |
| 0 | 0.95 | 3.67 | 0.26 | 0.80 | NA | Gm17847 | | 0 | 0 | 0 | 0 | 0 | 0 | 0 | 0 | 0 | 0 | 1 |
| 0 | 0.95 | 3.67 | 0.26 | 0.80 | NA | Gm17752 | | 0 | 0 | 0 | 0 | 0 | 0 | 0 | 0 | 0 | 0 | 1 |
| 0 | 0.95 | 3.67 | 0.26 | 0.80 | NA | Gm8217 | | 0 | 0 | 0 | 0 | 0 | 0 | 0 | 0 | 0 | 0 | 1 |
| 0 | 0.95 | 3.67 | 0.26 | 0.80 | NA | Gm37333 | | 0 | 0 | 0 | 0 | 0 | 0 | 0 | 0 | 0 | 0 | 1 |
| 0 | 0.95 | 3.67 | 0.26 | 0.80 | NA | Gm5262 | | 0 | 0 | 0 | 0 | 0 | 0 | 0 | 0 | 0 | 0 | 1 |

| | | | | | | | | | | | | | | | | | | |
|---|---|---|---|---|---|---|---|---|---|---|---|---|---|---|---|---|---|---|
| 0 | 0.95 | 3.67 | 0.26 | 0.80 | NA | Gm28792 | | 0 | 0 | 0 | 0 | 0 | 0 | 0 | 0 | 0 | 0 | 1 |
| 0 | 0.95 | 3.67 | 0.26 | 0.80 | NA | Gm38302 | | 0 | 0 | 0 | 0 | 0 | 0 | 0 | 0 | 0 | 0 | 1 |
| 0 | 0.95 | 3.67 | 0.26 | 0.80 | NA | Gm26801 | | 0 | 0 | 0 | 0 | 0 | 0 | 0 | 0 | 0 | 0 | 1 |
| 0 | 0.95 | 3.67 | 0.26 | 0.80 | NA | Lefty2 | left-right determination factor 2 | 0 | 0 | 0 | 0 | 0 | 0 | 0 | 0 | 0 | 0 | 1 |
| 0 | 0.95 | 3.67 | 0.26 | 0.80 | NA | Gm37018 | | 0 | 0 | 0 | 0 | 0 | 0 | 0 | 0 | 0 | 0 | 1 |
| 0 | 0.95 | 3.67 | 0.26 | 0.80 | NA | Gm37177 | | 0 | 0 | 0 | 0 | 0 | 0 | 0 | 0 | 0 | 0 | 1 |
| 0 | 0.95 | 3.67 | 0.26 | 0.80 | NA | | | 0 | 0 | 0 | 0 | 0 | 0 | 0 | 0 | 0 | 0 | 1 |
| 0 | 0.95 | 3.67 | 0.26 | 0.80 | NA | Gm23153 | | 0 | 0 | 0 | 0 | 0 | 0 | 0 | 0 | 0 | 0 | 1 |
| 0 | 0.95 | 3.67 | 0.26 | 0.80 | NA | Mir3089 | | 0 | 0 | 0 | 0 | 0 | 0 | 0 | 0 | 0 | 0 | 1 |
| 0 | 0.95 | 3.67 | 0.26 | 0.80 | NA | Gm13429 | | 0 | 0 | 0 | 0 | 0 | 0 | 0 | 0 | 0 | 0 | 1 |
| 0 | 0.95 | 3.67 | 0.26 | 0.80 | NA | Gm13474 | | 0 | 0 | 0 | 0 | 0 | 0 | 0 | 0 | 0 | 0 | 1 |
| 0 | 0.95 | 3.67 | 0.26 | 0.80 | NA | Gm14032 | | 0 | 0 | 0 | 0 | 0 | 0 | 0 | 0 | 0 | 0 | 1 |
| 0 | 0.95 | 3.67 | 0.26 | 0.80 | NA | Olfr1018 | olfactory receptor 1018 | 0 | 0 | 0 | 0 | 0 | 0 | 0 | 0 | 0 | 0 | 1 |
| 0 | 0.95 | 3.67 | 0.26 | 0.80 | NA | Olfr1173 | olfactory receptor 1173 | 0 | 0 | 0 | 0 | 0 | 0 | 0 | 0 | 0 | 0 | 1 |
| 0 | 0.95 | 3.67 | 0.26 | 0.80 | NA | Gm14120 | | 0 | 0 | 0 | 0 | 0 | 0 | 0 | 0 | 0 | 0 | 1 |
| 0 | 0.95 | 3.67 | 0.26 | 0.80 | NA | Gm14413 | | 0 | 0 | 0 | 0 | 0 | 0 | 0 | 0 | 0 | 0 | 1 |
| 0 | 0.95 | 3.67 | 0.26 | 0.80 | NA | 1110015O18Rik | | 0 | 0 | 0 | 0 | 0 | 0 | 0 | 0 | 0 | 0 | 1 |
| 0 | 0.95 | 3.67 | 0.26 | 0.80 | NA | Gm8935 | | 0 | 0 | 0 | 0 | 0 | 0 | 0 | 0 | 0 | 0 | 1 |
| 0 | 0.95 | 3.67 | 0.26 | 0.80 | NA | Gm7442 | | 0 | 0 | 0 | 0 | 0 | 0 | 0 | 0 | 0 | 0 | 1 |
| 0 | 0.95 | 3.67 | 0.26 | 0.80 | NA | Gm36983 | | 0 | 0 | 0 | 0 | 0 | 0 | 0 | 0 | 0 | 0 | 1 |
| 0 | 0.95 | 3.67 | 0.26 | 0.80 | NA | Gm10259 | | 0 | 0 | 0 | 0 | 0 | 0 | 0 | 0 | 0 | 0 | 1 |
| 0 | 0.95 | 3.67 | 0.26 | 0.80 | NA | Gm37031 | | 0 | 0 | 0 | 0 | 0 | 0 | 0 | 0 | 0 | 0 | 1 |
| 0 | 0.95 | 3.67 | 0.26 | 0.80 | NA | Gm8684 | | 0 | 0 | 0 | 0 | 0 | 0 | 0 | 0 | 0 | 0 | 1 |
| 0 | 0.95 | 3.67 | 0.26 | 0.80 | NA | Gm8869 | | 0 | 0 | 0 | 0 | 0 | 0 | 0 | 0 | 0 | 0 | 1 |
| 0 | 0.95 | 3.67 | 0.26 | 0.80 | NA | Ttc24 | tetratricopeptide repeat domain 24 | 0 | 0 | 0 | 0 | 0 | 0 | 0 | 0 | 0 | 0 | 1 |
| 0 | 0.95 | 3.67 | 0.26 | 0.80 | NA | Gm26465 | | 0 | 0 | 0 | 0 | 0 | 0 | 0 | 0 | 0 | 0 | 1 |
| 0 | 0.95 | 3.67 | 0.26 | 0.80 | NA | Gm5284 | | 0 | 0 | 0 | 0 | 0 | 0 | 0 | 0 | 0 | 0 | 1 |
| 0 | 0.95 | 3.67 | 0.26 | 0.80 | NA | Nhlh2 | nescient helix loop helix 2 | 0 | 0 | 0 | 0 | 0 | 0 | 0 | 0 | 0 | 0 | 1 |
| 0 | 0.95 | 3.67 | 0.26 | 0.80 | NA | Gm43149 | | 0 | 0 | 0 | 0 | 0 | 0 | 0 | 0 | 0 | 0 | 1 |
| 0 | 0.95 | 3.67 | 0.26 | 0.80 | NA | Gm4861 | predicted gene 4861 | 0 | 0 | 0 | 0 | 0 | 0 | 0 | 0 | 0 | 0 | 1 |
| 0 | 0.95 | 3.67 | 0.26 | 0.80 | NA | Gm15460 | | 0 | 0 | 0 | 0 | 0 | 0 | 0 | 0 | 0 | 0 | 1 |
| 0 | 0.95 | 3.67 | 0.26 | 0.80 | NA | Gm43185 | | 0 | 0 | 0 | 0 | 0 | 0 | 0 | 0 | 0 | 0 | 1 |
| 0 | 0.95 | 3.67 | 0.26 | 0.80 | NA | Gm16232 | | 0 | 0 | 0 | 0 | 0 | 0 | 0 | 0 | 0 | 0 | 1 |
| 0 | 0.95 | 3.67 | 0.26 | 0.80 | NA | Gm11912 | | 0 | 0 | 0 | 0 | 0 | 0 | 0 | 0 | 0 | 0 | 1 |
| 0 | 0.95 | 3.67 | 0.26 | 0.80 | NA | Mir5106 | | 0 | 0 | 0 | 0 | 0 | 0 | 0 | 0 | 0 | 0 | 1 |
| 0 | 0.95 | 3.67 | 0.26 | 0.80 | NA | Gm12466 | | 0 | 0 | 0 | 0 | 0 | 0 | 0 | 0 | 0 | 0 | 1 |
| 0 | 0.95 | 3.67 | 0.26 | 0.80 | NA | Olfr273 | olfactory receptor 273 | 0 | 0 | 0 | 0 | 0 | 0 | 0 | 0 | 0 | 0 | 1 |
| 0 | 0.95 | 3.67 | 0.26 | 0.80 | NA | Gm12700 | | 0 | 0 | 0 | 0 | 0 | 0 | 0 | 0 | 0 | 0 | 1 |
| 0 | 0.95 | 3.67 | 0.26 | 0.80 | NA | Gm12792 | | 0 | 0 | 0 | 0 | 0 | 0 | 0 | 0 | 0 | 0 | 1 |
| 0 | 0.95 | 3.67 | 0.26 | 0.80 | NA | Gm29434 | | 0 | 0 | 0 | 0 | 0 | 0 | 0 | 0 | 0 | 0 | 1 |
| 0 | 0.95 | 3.67 | 0.26 | 0.80 | NA | Mir7016 | | 0 | 0 | 0 | 0 | 0 | 0 | 0 | 0 | 0 | 0 | 1 |
| 0 | 0.95 | 3.67 | 0.26 | 0.80 | NA | Gm13008 | | 0 | 0 | 0 | 0 | 0 | 0 | 0 | 0 | 0 | 0 | 1 |
| 0 | 0.95 | 3.67 | 0.26 | 0.80 | NA | Gm13052 | | 0 | 0 | 0 | 0 | 0 | 0 | 0 | 0 | 0 | 0 | 1 |
| 0 | 0.95 | 3.67 | 0.26 | 0.80 | NA | AAdacl4fm3 | AADACL4 family member 3 | 0 | 0 | 0 | 0 | 0 | 0 | 0 | 0 | 0 | 0 | 1 |
| 0 | 0.95 | 3.67 | 0.26 | 0.80 | NA | Gm13036 | | 0 | 0 | 0 | 0 | 0 | 0 | 0 | 0 | 0 | 0 | 1 |
| 0 | 0.95 | 3.67 | 0.26 | 0.80 | NA | Zfp989 | | 0 | 0 | 0 | 0 | 0 | 0 | 0 | 0 | 0 | 0 | 1 |
| 0 | 0.95 | 3.67 | 0.26 | 0.80 | NA | 4933438K21Rik | | 0 | 0 | 0 | 0 | 0 | 0 | 0 | 0 | 0 | 0 | 1 |
| 0 | 0.95 | 3.67 | 0.26 | 0.80 | NA | Dnaja1-ps | | 0 | 0 | 0 | 0 | 0 | 0 | 0 | 0 | 0 | 0 | 1 |
| 0 | 0.95 | 3.67 | 0.26 | 0.80 | NA | Gm19798 | | 0 | 0 | 0 | 0 | 0 | 0 | 0 | 0 | 0 | 0 | 1 |
| 0 | 0.95 | 3.67 | 0.26 | 0.80 | NA | Gm5291 | | 0 | 0 | 0 | 0 | 0 | 0 | 0 | 0 | 0 | 0 | 1 |
| 0 | 0.95 | 3.67 | 0.26 | 0.80 | NA | Gm43046 | | 0 | 0 | 0 | 0 | 0 | 0 | 0 | 0 | 0 | 0 | 1 |
| 0 | 0.95 | 3.67 | 0.26 | 0.80 | NA | Gm15981 | | 0 | 0 | 0 | 0 | 0 | 0 | 0 | 0 | 0 | 0 | 1 |
| 0 | 0.95 | 3.67 | 0.26 | 0.80 | NA | Gm43094 | | 0 | 0 | 0 | 0 | 0 | 0 | 0 | 0 | 0 | 0 | 1 |
| 0 | 0.95 | 3.67 | 0.26 | 0.80 | NA | Gm22450 | | 0 | 0 | 0 | 0 | 0 | 0 | 0 | 0 | 0 | 0 | 1 |
| 0 | 0.95 | 3.67 | 0.26 | 0.80 | NA | Gm15502 | | 0 | 0 | 0 | 0 | 0 | 0 | 0 | 0 | 0 | 0 | 1 |
| 0 | 0.95 | 3.67 | 0.26 | 0.80 | NA | Gm23960 | | 0 | 0 | 0 | 0 | 0 | 0 | 0 | 0 | 0 | 0 | 1 |
| 0 | 0.95 | 3.67 | 0.26 | 0.80 | NA | Gm43154 | | 0 | 0 | 0 | 0 | 0 | 0 | 0 | 0 | 0 | 0 | 1 |
| 0 | 0.95 | 3.67 | 0.26 | 0.80 | NA | Akr1b7 | aldo-keto reductase family 1, member B7 | 0 | 0 | 0 | 0 | 0 | 0 | 0 | 0 | 0 | 0 | 1 |
| 0 | 0.95 | 3.67 | 0.26 | 0.80 | NA | Gm6695 | | 0 | 0 | 0 | 0 | 0 | 0 | 0 | 0 | 0 | 0 | 1 |
| 0 | 0.95 | 3.67 | 0.26 | 0.80 | NA | Gm4873 | | 0 | 0 | 0 | 0 | 0 | 0 | 0 | 0 | 0 | 0 | 1 |

| | | | | | | | | | | | | | | | | | | |
|---|---|---|---|---|---|---|---|---|---|---|---|---|---|---|---|---|---|---|
| 0 | 0.95 | 3.67 | 0.26 | 0.80 | NA | Gm43292 | | 0 | 0 | 0 | 0 | 0 | 0 | 0 | 0 | 0 | 0 | 1 |
| 0 | 0.95 | 3.67 | 0.26 | 0.80 | NA | Kbtbd8os | | 0 | 0 | 0 | 0 | 0 | 0 | 0 | 0 | 0 | 0 | 1 |
| 0 | 0.95 | 3.67 | 0.26 | 0.80 | NA | Gm44102 | | 0 | 0 | 0 | 0 | 0 | 0 | 0 | 0 | 0 | 0 | 1 |
| 0 | 0.95 | 3.67 | 0.26 | 0.80 | NA | Gm17482 | | 0 | 0 | 0 | 0 | 0 | 0 | 0 | 0 | 0 | 0 | 1 |
| 0 | 0.95 | 3.67 | 0.26 | 0.80 | NA | Gm44215 | | 0 | 0 | 0 | 0 | 0 | 0 | 0 | 0 | 0 | 0 | 1 |
| 0 | 0.95 | 3.67 | 0.26 | 0.80 | NA | Gm44958 | | 0 | 0 | 0 | 0 | 0 | 0 | 0 | 0 | 0 | 0 | 1 |
| 0 | 0.95 | 3.67 | 0.26 | 0.80 | NA | Gm6855 | | 0 | 0 | 0 | 0 | 0 | 0 | 0 | 0 | 0 | 0 | 1 |
| 0 | 0.95 | 3.67 | 0.26 | 0.80 | NA | Olfr1347 | olfactory receptor 1347 | 0 | 0 | 0 | 0 | 0 | 0 | 0 | 0 | 0 | 0 | 1 |
| 0 | 0.95 | 3.67 | 0.26 | 0.80 | NA | Vmn1r72 | vomeronasal 1 receptor 72 | 0 | 0 | 0 | 0 | 0 | 0 | 0 | 0 | 0 | 0 | 1 |
| 0 | 0.95 | 3.67 | 0.26 | 0.80 | NA | Gm6946 | | 0 | 0 | 0 | 0 | 0 | 0 | 0 | 0 | 0 | 0 | 1 |
| 0 | 0.95 | 3.67 | 0.26 | 0.80 | NA | Gm7965 | | 0 | 0 | 0 | 0 | 0 | 0 | 0 | 0 | 0 | 0 | 1 |
| 0 | 0.95 | 3.67 | 0.26 | 0.80 | NA | Gm45034 | | 0 | 0 | 0 | 0 | 0 | 0 | 0 | 0 | 0 | 0 | 1 |
| 0 | 0.95 | 3.67 | 0.26 | 0.80 | NA | Gm44521 | | 0 | 0 | 0 | 0 | 0 | 0 | 0 | 0 | 0 | 0 | 1 |
| 0 | 0.95 | 3.67 | 0.26 | 0.80 | NA | Vmn2r59 | vomeronasal 2, receptor 59 | 0 | 0 | 0 | 0 | 0 | 0 | 0 | 0 | 0 | 0 | 1 |
| 0 | 0.95 | 3.67 | 0.26 | 0.80 | NA | Gm6290 | | 0 | 0 | 0 | 0 | 0 | 0 | 0 | 0 | 0 | 0 | 1 |
| 0 | 0.95 | 3.67 | 0.26 | 0.80 | NA | Gm44741 | | 0 | 0 | 0 | 0 | 0 | 0 | 0 | 0 | 0 | 0 | 1 |
| 0 | 0.95 | 3.67 | 0.26 | 0.80 | NA | Gm7675 | | 0 | 0 | 0 | 0 | 0 | 0 | 0 | 0 | 0 | 0 | 1 |
| 0 | 0.95 | 3.67 | 0.26 | 0.80 | NA | Olfr2 | olfactory receptor 2 | 0 | 0 | 0 | 0 | 0 | 0 | 0 | 0 | 0 | 0 | 1 |
| 0 | 0.95 | 3.67 | 0.26 | 0.80 | NA | Olfr488 | olfactory receptor 488 | 0 | 0 | 0 | 0 | 0 | 0 | 0 | 0 | 0 | 0 | 1 |
| 0 | 0.95 | 3.67 | 0.26 | 0.80 | NA | Gm25450 | | 0 | 0 | 0 | 0 | 0 | 0 | 0 | 0 | 0 | 0 | 1 |
| 0 | 0.95 | 3.67 | 0.26 | 0.80 | NA | Gm5338 | | 0 | 0 | 0 | 0 | 0 | 0 | 0 | 0 | 0 | 0 | 1 |
| 0 | 0.95 | 3.67 | 0.26 | 0.80 | NA | Gm45121 | | 0 | 0 | 0 | 0 | 0 | 0 | 0 | 0 | 0 | 0 | 1 |
| 0 | 0.95 | 3.67 | 0.26 | 0.80 | NA | Gm45532 | | 0 | 0 | 0 | 0 | 0 | 0 | 0 | 0 | 0 | 0 | 1 |
| 0 | 0.95 | 3.67 | 0.26 | 0.80 | NA | Gm44622 | | 0 | 0 | 0 | 0 | 0 | 0 | 0 | 0 | 0 | 0 | 1 |
| 0 | 0.95 | 3.67 | 0.26 | 0.80 | NA | Snord13 | | 0 | 0 | 0 | 0 | 0 | 0 | 0 | 0 | 0 | 0 | 1 |
| 0 | 0.95 | 3.67 | 0.26 | 0.80 | NA | Gm45660 | | 0 | 0 | 0 | 0 | 0 | 0 | 0 | 0 | 0 | 0 | 1 |
| 0 | 0.95 | 3.67 | 0.26 | 0.80 | NA | C130073E24Rik | | 0 | 0 | 0 | 0 | 0 | 0 | 0 | 0 | 0 | 0 | 1 |
| 0 | 0.95 | 3.67 | 0.26 | 0.80 | NA | Hspd1-ps2 | | 0 | 0 | 0 | 0 | 0 | 0 | 0 | 0 | 0 | 0 | 1 |
| 0 | 0.95 | 3.67 | 0.26 | 0.80 | NA | Gm18212 | | 0 | 0 | 0 | 0 | 0 | 0 | 0 | 0 | 0 | 0 | 1 |
| 0 | 0.95 | 3.67 | 0.26 | 0.80 | NA | Mir709 | | 0 | 0 | 0 | 0 | 0 | 0 | 0 | 0 | 0 | 0 | 1 |
| 0 | 0.95 | 3.67 | 0.26 | 0.80 | NA | Gm8688 | | 0 | 0 | 0 | 0 | 0 | 0 | 0 | 0 | 0 | 0 | 1 |
| 0 | 0.95 | 3.67 | 0.26 | 0.80 | NA | Gm38042 | | 0 | 0 | 0 | 0 | 0 | 0 | 0 | 0 | 0 | 0 | 1 |
| 0 | 0.95 | 3.67 | 0.26 | 0.80 | NA | Gm37497 | | 0 | 0 | 0 | 0 | 0 | 0 | 0 | 0 | 0 | 0 | 1 |
| 0 | 0.95 | 3.67 | 0.26 | 0.80 | NA | Gm6607 | | 0 | 0 | 0 | 0 | 0 | 0 | 0 | 0 | 0 | 0 | 1 |
| 0 | 0.95 | 3.67 | 0.26 | 0.80 | NA | 5830462O15Rik | | 0 | 0 | 0 | 0 | 0 | 0 | 0 | 0 | 0 | 0 | 1 |
| 0 | 0.95 | 3.67 | 0.26 | 0.80 | NA | Htr3b | 5-hydroxytryptamine (serotonin) receptor | 0 | 0 | 0 | 0 | 0 | 0 | 0 | 0 | 0 | 0 | 1 |
| 0 | 0.95 | 3.67 | 0.26 | 0.80 | NA | Gm49389 | | 0 | 0 | 0 | 0 | 0 | 0 | 0 | 0 | 0 | 0 | 1 |
| 0 | 0.95 | 3.67 | 0.26 | 0.80 | NA | Gm37663 | | 0 | 0 | 0 | 0 | 0 | 0 | 0 | 0 | 0 | 0 | 1 |
| 0 | 0.95 | 3.67 | 0.26 | 0.80 | NA | Gm37955 | | 0 | 0 | 0 | 0 | 0 | 0 | 0 | 0 | 0 | 0 | 1 |
| 0 | 0.95 | 3.67 | 0.26 | 0.80 | NA | Gm8226 | | 0 | 0 | 0 | 0 | 0 | 0 | 0 | 0 | 0 | 0 | 1 |
| 0 | 0.95 | 3.67 | 0.26 | 0.80 | NA | 4930422M22Rik | | 0 | 0 | 0 | 0 | 0 | 0 | 0 | 0 | 0 | 0 | 1 |
| 0 | 0.95 | 3.67 | 0.26 | 0.80 | NA | Gm48750 | | 0 | 0 | 0 | 0 | 0 | 0 | 0 | 0 | 0 | 0 | 1 |
| 0 | 0.95 | 3.67 | 0.26 | 0.80 | NA | Hmgb1-ps8 | | 0 | 0 | 0 | 0 | 0 | 0 | 0 | 0 | 0 | 0 | 1 |
| 0 | 0.95 | 3.67 | 0.26 | 0.80 | NA | Gm8911 | | 0 | 0 | 0 | 0 | 0 | 0 | 0 | 0 | 0 | 0 | 1 |
| 0 | 0.95 | 3.67 | 0.26 | 0.80 | NA | Gm23058 | | 0 | 0 | 0 | 0 | 0 | 0 | 0 | 0 | 0 | 0 | 1 |
| 0 | 0.95 | 3.67 | 0.26 | 0.80 | NA | Gm16135 | | 0 | 0 | 0 | 0 | 0 | 0 | 0 | 0 | 0 | 0 | 1 |
| 0 | 0.95 | 3.67 | 0.26 | 0.80 | NA | Gm47809 | | 0 | 0 | 0 | 0 | 0 | 0 | 0 | 0 | 0 | 0 | 1 |
| 0 | 0.95 | 3.67 | 0.26 | 0.80 | NA | Gm46192 | | 0 | 0 | 0 | 0 | 0 | 0 | 0 | 0 | 0 | 0 | 1 |
| 0 | 0.95 | 3.67 | 0.26 | 0.80 | NA | Mir1251 | | 0 | 0 | 0 | 0 | 0 | 0 | 0 | 0 | 0 | 0 | 1 |
| 0 | 0.95 | 3.67 | 0.26 | 0.80 | NA | Gm47958 | | 0 | 0 | 0 | 0 | 0 | 0 | 0 | 0 | 0 | 0 | 1 |
| 0 | 0.95 | 3.67 | 0.26 | 0.80 | NA | 1700025K04Rik | | 0 | 0 | 0 | 0 | 0 | 0 | 0 | 0 | 0 | 0 | 1 |
| 0 | 0.95 | 3.67 | 0.26 | 0.80 | NA | Gm6333 | | 0 | 0 | 0 | 0 | 0 | 0 | 0 | 0 | 0 | 0 | 1 |
| 0 | 0.95 | 3.67 | 0.26 | 0.80 | NA | Gm11965 | | 0 | 0 | 0 | 0 | 0 | 0 | 0 | 0 | 0 | 0 | 1 |
| 0 | 0.95 | 3.67 | 0.26 | 0.80 | NA | Gm12162 | | 0 | 0 | 0 | 0 | 0 | 0 | 0 | 0 | 0 | 0 | 1 |
| 0 | 0.95 | 3.67 | 0.26 | 0.80 | NA | Mir6920 | | 0 | 0 | 0 | 0 | 0 | 0 | 0 | 0 | 0 | 0 | 1 |
| 0 | 0.95 | 3.67 | 0.26 | 0.80 | NA | Gm44313 | | 0 | 0 | 0 | 0 | 0 | 0 | 0 | 0 | 0 | 0 | 1 |
| 0 | 0.95 | 3.67 | 0.26 | 0.80 | NA | Gm27711 | | 0 | 0 | 0 | 0 | 0 | 0 | 0 | 0 | 0 | 0 | 1 |
| 0 | 0.95 | 3.67 | 0.26 | 0.80 | NA | Gm12949 | | 0 | 0 | 0 | 0 | 0 | 0 | 0 | 0 | 0 | 0 | 1 |
| 0 | 0.95 | 3.67 | 0.26 | 0.80 | NA | Gm12318 | | 0 | 0 | 0 | 0 | 0 | 0 | 0 | 0 | 0 | 0 | 1 |
| 0 | 0.95 | 3.67 | 0.26 | 0.80 | NA | Gm26121 | | 0 | 0 | 0 | 0 | 0 | 0 | 0 | 0 | 0 | 0 | 1 |

| | | | | | | | | | | | | | | | | | | |
|---|---|---|---|---|---|---|---|---|---|---|---|---|---|---|---|---|---|---|
| 0 | 0.95 | 3.67 | 0.26 | 0.80 | NA | Otop3 | otopetrin 3 | 0 | 0 | 0 | 0 | 0 | 0 | 0 | 0 | 0 | 0 | 1 |
| 0 | 0.95 | 3.67 | 0.26 | 0.80 | NA | Ict1os | | 0 | 0 | 0 | 0 | 0 | 0 | 0 | 0 | 0 | 0 | 1 |
| 0 | 0.95 | 3.67 | 0.26 | 0.80 | NA | Rpl36-ps1 | | 0 | 0 | 0 | 0 | 0 | 0 | 0 | 0 | 0 | 0 | 1 |
| 0 | 0.95 | 3.67 | 0.26 | 0.80 | NA | Gm11742 | | 0 | 0 | 0 | 0 | 0 | 0 | 0 | 0 | 0 | 0 | 1 |
| 0 | 0.95 | 3.67 | 0.26 | 0.80 | NA | Gm11788 | | 0 | 0 | 0 | 0 | 0 | 0 | 0 | 0 | 0 | 0 | 1 |
| 0 | 0.95 | 3.67 | 0.26 | 0.80 | NA | Mir7675 | | 0 | 0 | 0 | 0 | 0 | 0 | 0 | 0 | 0 | 0 | 1 |
| 0 | 0.95 | 3.67 | 0.26 | 0.80 | NA | Gm44305 | | 0 | 0 | 0 | 0 | 0 | 0 | 0 | 0 | 0 | 0 | 1 |
| 0 | 0.95 | 3.67 | 0.26 | 0.80 | NA | Gm48779 | | 0 | 0 | 0 | 0 | 0 | 0 | 0 | 0 | 0 | 0 | 1 |
| 0 | 0.95 | 3.67 | 0.26 | 0.80 | NA | Gm24859 | | 0 | 0 | 0 | 0 | 0 | 0 | 0 | 0 | 0 | 0 | 1 |
| 0 | 0.95 | 3.67 | 0.26 | 0.80 | NA | Gm6068 | | 0 | 0 | 0 | 0 | 0 | 0 | 0 | 0 | 0 | 0 | 1 |
| 0 | 0.95 | 3.67 | 0.26 | 0.80 | NA | Gm48365 | | 0 | 0 | 0 | 0 | 0 | 0 | 0 | 0 | 0 | 0 | 1 |
| 0 | 0.95 | 3.67 | 0.26 | 0.80 | NA | Ccdc175 | coiled-coil domain containing 175 | 0 | 0 | 0 | 0 | 0 | 0 | 0 | 0 | 0 | 0 | 1 |
| 0 | 0.95 | 3.67 | 0.26 | 0.80 | NA | Gm47765 | | 0 | 0 | 0 | 0 | 0 | 0 | 0 | 0 | 0 | 0 | 1 |
| 0 | 0.95 | 3.67 | 0.26 | 0.80 | NA | Gm5435 | | 0 | 0 | 0 | 0 | 0 | 0 | 0 | 0 | 0 | 0 | 1 |
| 0 | 0.95 | 3.67 | 0.26 | 0.80 | NA | Heatr4 | HEAT repeat containing 4 | 0 | 0 | 0 | 0 | 0 | 0 | 0 | 0 | 0 | 0 | 1 |
| 0 | 0.95 | 3.67 | 0.26 | 0.80 | NA | Mir6938 | | 0 | 0 | 0 | 0 | 0 | 0 | 0 | 0 | 0 | 0 | 1 |
| 0 | 0.95 | 3.67 | 0.26 | 0.80 | NA | Gm5789 | | 0 | 0 | 0 | 0 | 0 | 0 | 0 | 0 | 0 | 0 | 1 |
| 0 | 0.95 | 3.67 | 0.26 | 0.80 | NA | 3110009F21Rik | | 0 | 0 | 0 | 0 | 0 | 0 | 0 | 0 | 0 | 0 | 1 |
| 0 | 0.95 | 3.67 | 0.26 | 0.80 | NA | Gm48621 | | 0 | 0 | 0 | 0 | 0 | 0 | 0 | 0 | 0 | 0 | 1 |
| 0 | 0.95 | 3.67 | 0.26 | 0.80 | NA | Gm47511 | | 0 | 0 | 0 | 0 | 0 | 0 | 0 | 0 | 0 | 0 | 1 |
| 0 | 0.95 | 3.67 | 0.26 | 0.80 | NA | Gm3338 | | 0 | 0 | 0 | 0 | 0 | 0 | 0 | 0 | 0 | 0 | 1 |
| 0 | 0.95 | 3.67 | 0.26 | 0.80 | NA | Gm7911 | | 0 | 0 | 0 | 0 | 0 | 0 | 0 | 0 | 0 | 0 | 1 |
| 0 | 0.95 | 3.67 | 0.26 | 0.80 | NA | Gm47636 | | 0 | 0 | 0 | 0 | 0 | 0 | 0 | 0 | 0 | 0 | 1 |
| 0 | 0.95 | 3.67 | 0.26 | 0.80 | NA | Gm24261 | | 0 | 0 | 0 | 0 | 0 | 0 | 0 | 0 | 0 | 0 | 1 |
| 0 | 0.95 | 3.67 | 0.26 | 0.80 | NA | Gm28989 | | 0 | 0 | 0 | 0 | 0 | 0 | 0 | 0 | 0 | 0 | 1 |
| 0 | 0.95 | 3.67 | 0.26 | 0.80 | NA | Gm48239 | | 0 | 0 | 0 | 0 | 0 | 0 | 0 | 0 | 0 | 0 | 1 |
| 0 | 0.95 | 3.67 | 0.26 | 0.80 | NA | Gm48781 | | 0 | 0 | 0 | 0 | 0 | 0 | 0 | 0 | 0 | 0 | 1 |
| 0 | 0.95 | 3.67 | 0.26 | 0.80 | NA | Gm49196 | | 0 | 0 | 0 | 0 | 0 | 0 | 0 | 0 | 0 | 0 | 1 |
| 0 | 0.95 | 3.67 | 0.26 | 0.80 | NA | Gm48939 | | 0 | 0 | 0 | 0 | 0 | 0 | 0 | 0 | 0 | 0 | 1 |
| 0 | 0.95 | 3.67 | 0.26 | 0.80 | NA | Gm9022 | | 0 | 0 | 0 | 0 | 0 | 0 | 0 | 0 | 0 | 0 | 1 |
| 0 | 0.95 | 3.67 | 0.26 | 0.80 | NA | Serpine3 | serpin peptidase inhibitor, clade E (nexin, | 0 | 0 | 0 | 0 | 0 | 0 | 0 | 0 | 0 | 0 | 1 |
| 0 | 0.95 | 3.67 | 0.26 | 0.80 | NA | 2900040C04Rik | | 0 | 0 | 0 | 0 | 0 | 0 | 0 | 0 | 0 | 0 | 1 |
| 0 | 0.95 | 3.67 | 0.26 | 0.80 | NA | Gm26679 | | 0 | 0 | 0 | 0 | 0 | 0 | 0 | 0 | 0 | 0 | 1 |
| 0 | 0.95 | 3.67 | 0.26 | 0.80 | NA | Zic5 | zinc finger protein of the cerebellum 5 | 0 | 0 | 0 | 0 | 0 | 0 | 0 | 0 | 0 | 0 | 1 |
| 0 | 0.95 | 3.67 | 0.26 | 0.80 | NA | Kcns2 | K+ voltage-gated channel, subfamily S, 2 | 0 | 0 | 0 | 0 | 0 | 0 | 0 | 0 | 0 | 0 | 1 |
| 0 | 0.95 | 3.67 | 0.26 | 0.80 | NA | Col22a1 | collagen, type XXII, alpha 1 | 0 | 0 | 0 | 0 | 0 | 0 | 0 | 0 | 0 | 0 | 1 |
| 0 | 0.95 | 3.67 | 0.26 | 0.80 | NA | Gm5808 | | 0 | 0 | 0 | 0 | 0 | 0 | 0 | 0 | 0 | 0 | 1 |
| 0 | 0.95 | 3.67 | 0.26 | 0.80 | NA | Gm9536 | | 0 | 0 | 0 | 0 | 0 | 0 | 0 | 0 | 0 | 0 | 1 |
| 0 | 0.95 | 3.67 | 0.26 | 0.80 | NA | Adipoq | adiponectin, C1Q and collagen domain c | 0 | 0 | 0 | 0 | 0 | 0 | 0 | 0 | 0 | 0 | 1 |
| 0 | 0.95 | 3.67 | 0.26 | 0.80 | NA | Gm49702 | | 0 | 0 | 0 | 0 | 0 | 0 | 0 | 0 | 0 | 0 | 1 |
| 0 | 0.95 | 3.67 | 0.26 | 0.80 | NA | Gm15575 | | 0 | 0 | 0 | 0 | 0 | 0 | 0 | 0 | 0 | 0 | 1 |
| 0 | 0.95 | 3.67 | 0.26 | 0.80 | NA | Gm6931 | | 0 | 0 | 0 | 0 | 0 | 0 | 0 | 0 | 0 | 0 | 1 |
| 0 | 0.95 | 3.67 | 0.26 | 0.80 | NA | Gm49577 | | 0 | 0 | 0 | 0 | 0 | 0 | 0 | 0 | 0 | 0 | 1 |
| 0 | 0.95 | 3.67 | 0.26 | 0.80 | NA | Olfr187 | olfactory receptor 187 | 0 | 0 | 0 | 0 | 0 | 0 | 0 | 0 | 0 | 0 | 1 |
| 0 | 0.95 | 3.67 | 0.26 | 0.80 | NA | Gm15425 | | 0 | 0 | 0 | 0 | 0 | 0 | 0 | 0 | 0 | 0 | 1 |
| 0 | 0.95 | 3.67 | 0.26 | 0.80 | NA | Gm6618 | | 0 | 0 | 0 | 0 | 0 | 0 | 0 | 0 | 0 | 0 | 1 |
| 0 | 0.95 | 3.67 | 0.26 | 0.80 | NA | Nme3 | NME/NM23 nucleoside diphosphate kina | 0 | 0 | 0 | 0 | 0 | 0 | 0 | 0 | 0 | 0 | 1 |
| 0 | 0.95 | 3.67 | 0.26 | 0.80 | NA | Gm8373 | | 0 | 0 | 0 | 0 | 0 | 0 | 0 | 0 | 0 | 0 | 1 |
| 0 | 0.95 | 3.67 | 0.26 | 0.80 | NA | Skiv2l-ps1 | | 0 | 0 | 0 | 0 | 0 | 0 | 0 | 0 | 0 | 0 | 1 |
| 0 | 0.95 | 3.67 | 0.26 | 0.80 | NA | Gm35097 | | 0 | 0 | 0 | 0 | 0 | 0 | 0 | 0 | 0 | 0 | 1 |
| 0 | 0.95 | 3.67 | 0.26 | 0.80 | NA | Gm9210 | | 0 | 0 | 0 | 0 | 0 | 0 | 0 | 0 | 0 | 0 | 1 |
| 0 | 0.95 | 3.67 | 0.26 | 0.80 | NA | Gm49939 | | 0 | 0 | 0 | 0 | 0 | 0 | 0 | 0 | 0 | 0 | 1 |
| 0 | 0.95 | 3.67 | 0.26 | 0.80 | NA | Gm50043 | | 0 | 0 | 0 | 0 | 0 | 0 | 0 | 0 | 0 | 0 | 1 |
| 0 | 0.95 | 3.67 | 0.26 | 0.80 | NA | Six3 | sine oculis-related homeobox 3 | 0 | 0 | 0 | 0 | 0 | 0 | 0 | 0 | 0 | 0 | 1 |
| 0 | 0.95 | 3.67 | 0.26 | 0.80 | NA | Gm6235 | | 0 | 0 | 0 | 0 | 0 | 0 | 0 | 0 | 0 | 0 | 1 |
| 0 | 0.95 | 3.67 | 0.26 | 0.80 | NA | Rpl7a-ps6 | | 0 | 0 | 0 | 0 | 0 | 0 | 0 | 0 | 0 | 0 | 1 |
| 0 | 0.95 | 3.67 | 0.26 | 0.80 | NA | Gm5240 | | 0 | 0 | 0 | 0 | 0 | 0 | 0 | 0 | 0 | 0 | 1 |
| 0 | 0.95 | 3.67 | 0.26 | 0.80 | NA | 2010110K18Rik | | 0 | 0 | 0 | 0 | 0 | 0 | 0 | 0 | 0 | 0 | 1 |
| 0 | 0.95 | 3.67 | 0.26 | 0.80 | NA | Gm16490 | | 0 | 0 | 0 | 0 | 0 | 0 | 0 | 0 | 0 | 0 | 1 |
| 0 | 0.95 | 3.67 | 0.26 | 0.80 | NA | Gm30093 | | 0 | 0 | 0 | 0 | 0 | 0 | 0 | 0 | 0 | 0 | 1 |

| | | | | | | | | | | | | | | | | | | |
|---|---|---|---|---|---|---|---|---|---|---|---|---|---|---|---|---|---|---|
| 0 | 0.95 | 3.67 | 0.26 | 0.80 | NA | Gm18086 | | 0 | 0 | 0 | 0 | 0 | 0 | 0 | 0 | 0 | 0 | 1 |
| 0 | 0.95 | 3.67 | 0.26 | 0.80 | NA | Gm6409 | | 0 | 0 | 0 | 0 | 0 | 0 | 0 | 0 | 0 | 0 | 1 |
| 0 | 0.95 | 3.67 | 0.26 | 0.80 | NA | Gm50204 | | 0 | 0 | 0 | 0 | 0 | 0 | 0 | 0 | 0 | 0 | 1 |
| 0 | 0.95 | 3.67 | 0.26 | 0.80 | NA | Mir192 | | 0 | 0 | 0 | 0 | 0 | 0 | 0 | 0 | 0 | 0 | 1 |
| 0 | 0.95 | 3.67 | 0.26 | 0.80 | NA | Foxd4 | forkhead box D4 | 0 | 0 | 0 | 0 | 0 | 0 | 0 | 0 | 0 | 0 | 1 |
| 0 | 0.95 | 3.67 | 0.26 | 0.80 | NA | Gm36419 | | 0 | 0 | 0 | 0 | 0 | 0 | 0 | 0 | 0 | 0 | 1 |
| 0 | 0.95 | 3.67 | 0.26 | 0.80 | NA | Gm50271 | | 0 | 0 | 0 | 0 | 0 | 0 | 0 | 0 | 0 | 0 | 1 |
| 0 | 0.95 | 3.67 | 0.26 | 0.80 | NA | Emx2os | | 0 | 0 | 0 | 0 | 0 | 0 | 0 | 0 | 0 | 0 | 1 |
| 0 | 0.95 | 3.67 | 0.26 | 0.80 | NA | Gm9439 | | 0 | 0 | 0 | 0 | 0 | 0 | 0 | 0 | 0 | 0 | 1 |
| 0 | 0.95 | 3.67 | 0.26 | 0.80 | NA | Gm14643 | | 0 | 0 | 0 | 0 | 0 | 0 | 0 | 0 | 0 | 0 | 1 |
| 0 | 0.95 | 3.67 | 0.26 | 0.80 | NA | Gm14652 | | 0 | 0 | 0 | 0 | 0 | 0 | 0 | 0 | 0 | 0 | 1 |
| 0 | 0.95 | 3.67 | 0.26 | 0.80 | NA | Gm14620 | | 0 | 0 | 0 | 0 | 0 | 0 | 0 | 0 | 0 | 0 | 1 |
| 0 | 0.95 | 3.67 | 0.26 | 0.80 | NA | Gm24598 | | 0 | 0 | 0 | 0 | 0 | 0 | 0 | 0 | 0 | 0 | 1 |
| 0 | 0.95 | 3.67 | 0.26 | 0.80 | NA | Ctag2l2 | CTAG2 like 2 | 0 | 0 | 0 | 0 | 0 | 0 | 0 | 0 | 0 | 0 | 1 |
| 0 | 0.95 | 3.67 | 0.26 | 0.80 | NA | Gm8876 | | 0 | 0 | 0 | 0 | 0 | 0 | 0 | 0 | 0 | 0 | 1 |
| 0 | 0.95 | 3.67 | 0.26 | 0.80 | NA | Mir7093 | | 0 | 0 | 0 | 0 | 0 | 0 | 0 | 0 | 0 | 0 | 1 |
| 0 | 0.95 | 3.67 | 0.26 | 0.80 | NA | Gm7123 | | 0 | 0 | 0 | 0 | 0 | 0 | 0 | 0 | 0 | 0 | 1 |
| 0 | 0.95 | 3.67 | 0.26 | 0.80 | NA | Gm8199 | | 0 | 0 | 0 | 0 | 0 | 0 | 0 | 0 | 0 | 0 | 1 |
| 0 | 0.95 | 3.67 | 0.26 | 0.80 | NA | Gm4919 | | 0 | 0 | 0 | 0 | 0 | 0 | 0 | 0 | 0 | 0 | 1 |
| 0 | 0.95 | 3.67 | 0.26 | 0.80 | NA | Gm26960 | | 0 | 0 | 0 | 0 | 0 | 0 | 0 | 0 | 0 | 0 | 1 |
| 0 | 0.95 | 3.67 | 0.26 | 0.80 | NA | Gm15243 | | 0 | 0 | 0 | 0 | 0 | 0 | 0 | 0 | 0 | 0 | 1 |
| 0 | 0.00 | 3.68 | 0.00 | 1.00 | NA | Gm7341 | | 0 | 1 | 0 | 0 | 0 | 0 | 0 | 0 | 0 | 0 | 0 |
| 0 | 0.00 | 3.68 | 0.00 | 1.00 | NA | Gm15452 | | 0 | 1 | 0 | 0 | 0 | 0 | 0 | 0 | 0 | 0 | 0 |
| 0 | 0.00 | 3.68 | 0.00 | 1.00 | NA | Gm28410 | | 0 | 1 | 0 | 0 | 0 | 0 | 0 | 0 | 0 | 0 | 0 |
| 0 | 0.00 | 3.68 | 0.00 | 1.00 | NA | Slc26a9 | solute carrier family 26, member 9 | 0 | 1 | 0 | 0 | 0 | 0 | 0 | 0 | 0 | 0 | 0 |
| 0 | 0.00 | 3.68 | 0.00 | 1.00 | NA | Gm29485 | | 0 | 1 | 0 | 0 | 0 | 0 | 0 | 0 | 0 | 0 | 0 |
| 0 | 0.00 | 3.68 | 0.00 | 1.00 | NA | Gm28286 | | 0 | 1 | 0 | 0 | 0 | 0 | 0 | 0 | 0 | 0 | 0 |
| 0 | 0.00 | 3.68 | 0.00 | 1.00 | NA | Gm18407 | | 0 | 1 | 0 | 0 | 0 | 0 | 0 | 0 | 0 | 0 | 0 |
| 0 | 0.00 | 3.68 | 0.00 | 1.00 | NA | 5033404E19Rik | | 0 | 1 | 0 | 0 | 0 | 0 | 0 | 0 | 0 | 0 | 0 |
| 0 | 0.00 | 3.68 | 0.00 | 1.00 | NA | Lcn6 | lipocalin 6 | 0 | 1 | 0 | 0 | 0 | 0 | 0 | 0 | 0 | 0 | 0 |
| 0 | 0.00 | 3.68 | 0.00 | 1.00 | NA | Gm13381 | | 0 | 1 | 0 | 0 | 0 | 0 | 0 | 0 | 0 | 0 | 0 |
| 0 | 0.00 | 3.68 | 0.00 | 1.00 | NA | Gm13875 | | 0 | 1 | 0 | 0 | 0 | 0 | 0 | 0 | 0 | 0 | 0 |
| 0 | 0.00 | 3.68 | 0.00 | 1.00 | NA | Gm13998 | | 0 | 1 | 0 | 0 | 0 | 0 | 0 | 0 | 0 | 0 | 0 |
| 0 | 0.00 | 3.68 | 0.00 | 1.00 | NA | Gm37098 | | 0 | 1 | 0 | 0 | 0 | 0 | 0 | 0 | 0 | 0 | 0 |
| 0 | 0.00 | 3.68 | 0.00 | 1.00 | NA | Gm43008 | | 0 | 1 | 0 | 0 | 0 | 0 | 0 | 0 | 0 | 0 | 0 |
| 0 | 0.00 | 3.68 | 0.00 | 1.00 | NA | Gm9723 | | 0 | 1 | 0 | 0 | 0 | 0 | 0 | 0 | 0 | 0 | 0 |
| 0 | 0.00 | 3.68 | 0.00 | 1.00 | NA | Gm36976 | | 0 | 1 | 0 | 0 | 0 | 0 | 0 | 0 | 0 | 0 | 0 |
| 0 | 0.00 | 3.68 | 0.00 | 1.00 | NA | Gm2136 | | 0 | 1 | 0 | 0 | 0 | 0 | 0 | 0 | 0 | 0 | 0 |
| 0 | 0.00 | 3.68 | 0.00 | 1.00 | NA | Gm5278 | | 0 | 1 | 0 | 0 | 0 | 0 | 0 | 0 | 0 | 0 | 0 |
| 0 | 0.00 | 3.68 | 0.00 | 1.00 | NA | Gm17954 | | 0 | 1 | 0 | 0 | 0 | 0 | 0 | 0 | 0 | 0 | 0 |
| 0 | 0.00 | 3.68 | 0.00 | 1.00 | NA | Gm9889 | | 0 | 1 | 0 | 0 | 0 | 0 | 0 | 0 | 0 | 0 | 0 |
| 0 | 0.00 | 3.68 | 0.00 | 1.00 | NA | Gm11884 | | 0 | 1 | 0 | 0 | 0 | 0 | 0 | 0 | 0 | 0 | 0 |
| 0 | 0.00 | 3.68 | 0.00 | 1.00 | NA | Gm11930 | | 0 | 1 | 0 | 0 | 0 | 0 | 0 | 0 | 0 | 0 | 0 |
| 0 | 0.00 | 3.68 | 0.00 | 1.00 | NA | Fam205c | family with sequence similarity 205, meml | 0 | 1 | 0 | 0 | 0 | 0 | 0 | 0 | 0 | 0 | 0 |
| 0 | 0.00 | 3.68 | 0.00 | 1.00 | NA | Gm12461 | | 0 | 1 | 0 | 0 | 0 | 0 | 0 | 0 | 0 | 0 | 0 |
| 0 | 0.00 | 3.68 | 0.00 | 1.00 | NA | Gm12475 | | 0 | 1 | 0 | 0 | 0 | 0 | 0 | 0 | 0 | 0 | 0 |
| 0 | 0.00 | 3.68 | 0.00 | 1.00 | NA | Gm12469 | | 0 | 1 | 0 | 0 | 0 | 0 | 0 | 0 | 0 | 0 | 0 |
| 0 | 0.00 | 3.68 | 0.00 | 1.00 | NA | Tpt1-ps2 | | 0 | 1 | 0 | 0 | 0 | 0 | 0 | 0 | 0 | 0 | 0 |
| 0 | 0.00 | 3.68 | 0.00 | 1.00 | NA | Hmgb1-rs18 | | 0 | 1 | 0 | 0 | 0 | 0 | 0 | 0 | 0 | 0 | 0 |
| 0 | 0.00 | 3.68 | 0.00 | 1.00 | NA | Gm13131 | | 0 | 1 | 0 | 0 | 0 | 0 | 0 | 0 | 0 | 0 | 0 |
| 0 | 0.00 | 3.68 | 0.00 | 1.00 | NA | Gm43825 | | 0 | 1 | 0 | 0 | 0 | 0 | 0 | 0 | 0 | 0 | 0 |
| 0 | 0.00 | 3.68 | 0.00 | 1.00 | NA | Gm5714 | | 0 | 1 | 0 | 0 | 0 | 0 | 0 | 0 | 0 | 0 | 0 |
| 0 | 0.00 | 3.68 | 0.00 | 1.00 | NA | Gm42428 | | 0 | 1 | 0 | 0 | 0 | 0 | 0 | 0 | 0 | 0 | 0 |
| 0 | 0.00 | 3.68 | 0.00 | 1.00 | NA | Gm43510 | | 0 | 1 | 0 | 0 | 0 | 0 | 0 | 0 | 0 | 0 | 0 |
| 0 | 0.00 | 3.68 | 0.00 | 1.00 | NA | Gm43704 | | 0 | 1 | 0 | 0 | 0 | 0 | 0 | 0 | 0 | 0 | 0 |
| 0 | 0.00 | 3.68 | 0.00 | 1.00 | NA | Lep | leptin | 0 | 1 | 0 | 0 | 0 | 0 | 0 | 0 | 0 | 0 | 0 |
| 0 | 0.00 | 3.68 | 0.00 | 1.00 | NA | Gm7463 | | 0 | 1 | 0 | 0 | 0 | 0 | 0 | 0 | 0 | 0 | 0 |
| 0 | 0.00 | 3.68 | 0.00 | 1.00 | NA | Tlx2 | T cell leukemia, homeobox 2 | 0 | 1 | 0 | 0 | 0 | 0 | 0 | 0 | 0 | 0 | 0 |
| 0 | 0.00 | 3.68 | 0.00 | 1.00 | NA | Gm44369 | | 0 | 1 | 0 | 0 | 0 | 0 | 0 | 0 | 0 | 0 | 0 |
| 0 | 0.00 | 3.68 | 0.00 | 1.00 | NA | Gm15495 | | 0 | 1 | 0 | 0 | 0 | 0 | 0 | 0 | 0 | 0 | 0 |

| | | | | | | | | | | | | | | | | | | |
|---|---|---|---|---|---|---|---|---|---|---|---|---|---|---|---|---|---|---|
| 0 | 0.00 | 3.68 | 0.00 | 1.00 | NA | 4932431P20Rik | | 0 | 1 | 0 | 0 | 0 | 0 | 0 | 0 | 0 | 0 | 0 |
| 0 | 0.00 | 3.68 | 0.00 | 1.00 | NA | Gm9146 | | 0 | 1 | 0 | 0 | 0 | 0 | 0 | 0 | 0 | 0 | 0 |
| 0 | 0.00 | 3.68 | 0.00 | 1.00 | NA | Gm45308 | | 0 | 1 | 0 | 0 | 0 | 0 | 0 | 0 | 0 | 0 | 0 |
| 0 | 0.00 | 3.68 | 0.00 | 1.00 | NA | Gm7482 | | 0 | 1 | 0 | 0 | 0 | 0 | 0 | 0 | 0 | 0 | 0 |
| 0 | 0.00 | 3.68 | 0.00 | 1.00 | NA | Gm18805 | | 0 | 1 | 0 | 0 | 0 | 0 | 0 | 0 | 0 | 0 | 0 |
| 0 | 0.00 | 3.68 | 0.00 | 1.00 | NA | Gm26633 | | 0 | 1 | 0 | 0 | 0 | 0 | 0 | 0 | 0 | 0 | 0 |
| 0 | 0.00 | 3.68 | 0.00 | 1.00 | NA | Gm26646 | | 0 | 1 | 0 | 0 | 0 | 0 | 0 | 0 | 0 | 0 | 0 |
| 0 | 0.00 | 3.68 | 0.00 | 1.00 | NA | Olfr472 | olfactory receptor 472 | 0 | 1 | 0 | 0 | 0 | 0 | 0 | 0 | 0 | 0 | 0 |
| 0 | 0.00 | 3.68 | 0.00 | 1.00 | NA | Gm9240 | | 0 | 1 | 0 | 0 | 0 | 0 | 0 | 0 | 0 | 0 | 0 |
| 0 | 0.00 | 3.68 | 0.00 | 1.00 | NA | 9130214F15Rik | | 0 | 1 | 0 | 0 | 0 | 0 | 0 | 0 | 0 | 0 | 0 |
| 0 | 0.00 | 3.68 | 0.00 | 1.00 | NA | Gm16933 | | 0 | 1 | 0 | 0 | 0 | 0 | 0 | 0 | 0 | 0 | 0 |
| 0 | 0.00 | 3.68 | 0.00 | 1.00 | NA | Gm15660 | | 0 | 1 | 0 | 0 | 0 | 0 | 0 | 0 | 0 | 0 | 0 |
| 0 | 0.00 | 3.68 | 0.00 | 1.00 | NA | 6030452D12Rik | | 0 | 1 | 0 | 0 | 0 | 0 | 0 | 0 | 0 | 0 | 0 |
| 0 | 0.00 | 3.68 | 0.00 | 1.00 | NA | Gm15655 | | 0 | 1 | 0 | 0 | 0 | 0 | 0 | 0 | 0 | 0 | 0 |
| 0 | 0.00 | 3.68 | 0.00 | 1.00 | NA | Gm5362 | | 0 | 1 | 0 | 0 | 0 | 0 | 0 | 0 | 0 | 0 | 0 |
| 0 | 0.00 | 3.68 | 0.00 | 1.00 | NA | Gm47986 | | 0 | 1 | 0 | 0 | 0 | 0 | 0 | 0 | 0 | 0 | 0 |
| 0 | 0.00 | 3.68 | 0.00 | 1.00 | NA | Gm8058 | | 0 | 1 | 0 | 0 | 0 | 0 | 0 | 0 | 0 | 0 | 0 |
| 0 | 0.00 | 3.68 | 0.00 | 1.00 | NA | Gm33054 | | 0 | 1 | 0 | 0 | 0 | 0 | 0 | 0 | 0 | 0 | 0 |
| 0 | 0.00 | 3.68 | 0.00 | 1.00 | NA | Gm48727 | | 0 | 1 | 0 | 0 | 0 | 0 | 0 | 0 | 0 | 0 | 0 |
| 0 | 0.00 | 3.68 | 0.00 | 1.00 | NA | Gm5177 | | 0 | 1 | 0 | 0 | 0 | 0 | 0 | 0 | 0 | 0 | 0 |
| 0 | 0.00 | 3.68 | 0.00 | 1.00 | NA | Gm40604 | | 0 | 1 | 0 | 0 | 0 | 0 | 0 | 0 | 0 | 0 | 0 |
| 0 | 0.00 | 3.68 | 0.00 | 1.00 | NA | Gm15272 | | 0 | 1 | 0 | 0 | 0 | 0 | 0 | 0 | 0 | 0 | 0 |
| 0 | 0.00 | 3.68 | 0.00 | 1.00 | NA | Gm47835 | | 0 | 1 | 0 | 0 | 0 | 0 | 0 | 0 | 0 | 0 | 0 |
| 0 | 0.00 | 3.68 | 0.00 | 1.00 | NA | Gm17779 | | 0 | 1 | 0 | 0 | 0 | 0 | 0 | 0 | 0 | 0 | 0 |
| 0 | 0.00 | 3.68 | 0.00 | 1.00 | NA | Gm50490 | | 0 | 1 | 0 | 0 | 0 | 0 | 0 | 0 | 0 | 0 | 0 |
| 0 | 0.00 | 3.68 | 0.00 | 1.00 | NA | Gm48106 | | 0 | 1 | 0 | 0 | 0 | 0 | 0 | 0 | 0 | 0 | 0 |
| 0 | 0.00 | 3.68 | 0.00 | 1.00 | NA | Gm11971 | | 0 | 1 | 0 | 0 | 0 | 0 | 0 | 0 | 0 | 0 | 0 |
| 0 | 0.00 | 3.68 | 0.00 | 1.00 | NA | Gm16140 | | 0 | 1 | 0 | 0 | 0 | 0 | 0 | 0 | 0 | 0 | 0 |
| 0 | 0.00 | 3.68 | 0.00 | 1.00 | NA | Gm12052 | | 0 | 1 | 0 | 0 | 0 | 0 | 0 | 0 | 0 | 0 | 0 |
| 0 | 0.00 | 3.68 | 0.00 | 1.00 | NA | Gm12626 | | 0 | 1 | 0 | 0 | 0 | 0 | 0 | 0 | 0 | 0 | 0 |
| 0 | 0.00 | 3.68 | 0.00 | 1.00 | NA | 1700023C21Rik | | 0 | 1 | 0 | 0 | 0 | 0 | 0 | 0 | 0 | 0 | 0 |
| 0 | 0.00 | 3.68 | 0.00 | 1.00 | NA | Gm11725 | | 0 | 1 | 0 | 0 | 0 | 0 | 0 | 0 | 0 | 0 | 0 |
| 0 | 0.00 | 3.68 | 0.00 | 1.00 | NA | Gm47556 | | 0 | 1 | 0 | 0 | 0 | 0 | 0 | 0 | 0 | 0 | 0 |
| 0 | 0.00 | 3.68 | 0.00 | 1.00 | NA | Gm6869 | | 0 | 1 | 0 | 0 | 0 | 0 | 0 | 0 | 0 | 0 | 0 |
| 0 | 0.00 | 3.68 | 0.00 | 1.00 | NA | Gm21056 | | 0 | 1 | 0 | 0 | 0 | 0 | 0 | 0 | 0 | 0 | 0 |
| 0 | 0.00 | 3.68 | 0.00 | 1.00 | NA | Slc24a4 | solute carrier family 24 (sodium/potassiur | 0 | 1 | 0 | 0 | 0 | 0 | 0 | 0 | 0 | 0 | 0 |
| 0 | 0.00 | 3.68 | 0.00 | 1.00 | NA | | | 0 | 1 | 0 | 0 | 0 | 0 | 0 | 0 | 0 | 0 | 0 |
| 0 | 0.00 | 3.68 | 0.00 | 1.00 | NA | Gm11285 | | 0 | 1 | 0 | 0 | 0 | 0 | 0 | 0 | 0 | 0 | 0 |
| 0 | 0.00 | 3.68 | 0.00 | 1.00 | NA | Gm11367 | | 0 | 1 | 0 | 0 | 0 | 0 | 0 | 0 | 0 | 0 | 0 |
| 0 | 0.00 | 3.68 | 0.00 | 1.00 | NA | Gm48234 | | 0 | 1 | 0 | 0 | 0 | 0 | 0 | 0 | 0 | 0 | 0 |
| 0 | 0.00 | 3.68 | 0.00 | 1.00 | NA | Gm2907 | | 0 | 1 | 0 | 0 | 0 | 0 | 0 | 0 | 0 | 0 | 0 |
| 0 | 0.00 | 3.68 | 0.00 | 1.00 | NA | Otx2 | orthodenticle homeobox 2 | 0 | 1 | 0 | 0 | 0 | 0 | 0 | 0 | 0 | 0 | 0 |
| 0 | 0.00 | 3.68 | 0.00 | 1.00 | NA | Rnase2a | ribonuclease, RNase A family, 2A (liver, e | 0 | 1 | 0 | 0 | 0 | 0 | 0 | 0 | 0 | 0 | 0 |
| 0 | 0.00 | 3.68 | 0.00 | 1.00 | NA | Gm5408 | | 0 | 1 | 0 | 0 | 0 | 0 | 0 | 0 | 0 | 0 | 0 |
| 0 | 0.00 | 3.68 | 0.00 | 1.00 | NA | Fitm1 | fat storage-inducing transmembrane prote | 0 | 1 | 0 | 0 | 0 | 0 | 0 | 0 | 0 | 0 | 0 |
| 0 | 0.00 | 3.68 | 0.00 | 1.00 | NA | Cma2 | chymase 2, mast cell | 0 | 1 | 0 | 0 | 0 | 0 | 0 | 0 | 0 | 0 | 0 |
| 0 | 0.00 | 3.68 | 0.00 | 1.00 | NA | Gm9012 | | 0 | 1 | 0 | 0 | 0 | 0 | 0 | 0 | 0 | 0 | 0 |
| 0 | 0.00 | 3.68 | 0.00 | 1.00 | NA | Gm49244 | | 0 | 1 | 0 | 0 | 0 | 0 | 0 | 0 | 0 | 0 | 0 |
| 0 | 0.00 | 3.68 | 0.00 | 1.00 | NA | Gm49205 | | 0 | 1 | 0 | 0 | 0 | 0 | 0 | 0 | 0 | 0 | 0 |
| 0 | 0.00 | 3.68 | 0.00 | 1.00 | NA | Gm8341 | | 0 | 1 | 0 | 0 | 0 | 0 | 0 | 0 | 0 | 0 | 0 |
| 0 | 0.00 | 3.68 | 0.00 | 1.00 | NA | Gm19840 | | 0 | 1 | 0 | 0 | 0 | 0 | 0 | 0 | 0 | 0 | 0 |
| 0 | 0.00 | 3.68 | 0.00 | 1.00 | NA | Apol11b | apolipoprotein L 11b | 0 | 1 | 0 | 0 | 0 | 0 | 0 | 0 | 0 | 0 | 0 |
| 0 | 0.00 | 3.68 | 0.00 | 1.00 | NA | Atp13a4 | ATPase type 13A4 | 0 | 1 | 0 | 0 | 0 | 0 | 0 | 0 | 0 | 0 | 0 |
| 0 | 0.00 | 3.68 | 0.00 | 1.00 | NA | Gm8387 | | 0 | 1 | 0 | 0 | 0 | 0 | 0 | 0 | 0 | 0 | 0 |
| 0 | 0.00 | 3.68 | 0.00 | 1.00 | NA | Csta2 | cystatin A family member 2 | 0 | 1 | 0 | 0 | 0 | 0 | 0 | 0 | 0 | 0 | 0 |
| 0 | 0.00 | 3.68 | 0.00 | 1.00 | NA | Gm5479 | | 0 | 1 | 0 | 0 | 0 | 0 | 0 | 0 | 0 | 0 | 0 |
| 0 | 0.00 | 3.68 | 0.00 | 1.00 | NA | Gm38655 | predicted gene, 38655 | 0 | 1 | 0 | 0 | 0 | 0 | 0 | 0 | 0 | 0 | 0 |
| 0 | 0.00 | 3.68 | 0.00 | 1.00 | NA | Gm7326 | | 0 | 1 | 0 | 0 | 0 | 0 | 0 | 0 | 0 | 0 | 0 |
| 0 | 0.00 | 3.68 | 0.00 | 1.00 | NA | Runx2os1 | | 0 | 1 | 0 | 0 | 0 | 0 | 0 | 0 | 0 | 0 | 0 |
| 0 | 0.00 | 3.68 | 0.00 | 1.00 | NA | Gm7497 | | 0 | 1 | 0 | 0 | 0 | 0 | 0 | 0 | 0 | 0 | 0 |

| | | | | | | | | | | | | | | | | | | |
|---|---|---|---|---|---|---|---|---|---|---|---|---|---|---|---|---|---|---|
| 0 | 0.00 | 3.68 | 0.00 | 1.00 | NA | Mospd4 | | 0 | 1 | 0 | 0 | 0 | 0 | 0 | 0 | 0 | 0 | 0 |
| 0 | 0.00 | 3.68 | 0.00 | 1.00 | NA | Gm16297 | | 0 | 1 | 0 | 0 | 0 | 0 | 0 | 0 | 0 | 0 | 0 |
| 0 | 0.00 | 3.68 | 0.00 | 1.00 | NA | Gm50327 | | 0 | 1 | 0 | 0 | 0 | 0 | 0 | 0 | 0 | 0 | 0 |
| 0 | 0.00 | 3.68 | 0.00 | 1.00 | NA | Gm18379 | | 0 | 1 | 0 | 0 | 0 | 0 | 0 | 0 | 0 | 0 | 0 |
| 0 | 0.00 | 3.68 | 0.00 | 1.00 | NA | Gm23140 | | 0 | 1 | 0 | 0 | 0 | 0 | 0 | 0 | 0 | 0 | 0 |
| 0 | 0.00 | 3.68 | 0.00 | 1.00 | NA | Gm50185 | | 0 | 1 | 0 | 0 | 0 | 0 | 0 | 0 | 0 | 0 | 0 |
| 0 | 0.00 | 3.68 | 0.00 | 1.00 | NA | Gm15033 | | 0 | 1 | 0 | 0 | 0 | 0 | 0 | 0 | 0 | 0 | 0 |
| 0 | 0.00 | 3.68 | 0.00 | 1.00 | NA | Gm6783 | | 0 | 1 | 0 | 0 | 0 | 0 | 0 | 0 | 0 | 0 | 0 |
| 0 | 0.00 | 3.68 | 0.00 | 1.00 | NA | Gm5391 | | 0 | 1 | 0 | 0 | 0 | 0 | 0 | 0 | 0 | 0 | 0 |
| 0 | 0.00 | 3.68 | 0.00 | 1.00 | NA | Gm6881 | | 0 | 1 | 0 | 0 | 0 | 0 | 0 | 0 | 0 | 0 | 0 |
| 0 | 0.00 | 3.68 | 0.00 | 1.00 | NA | Gm9095 | | 0 | 1 | 0 | 0 | 0 | 0 | 0 | 0 | 0 | 0 | 0 |
| 0 | 0.00 | 3.68 | 0.00 | 1.00 | NA | Nsa2-ps2 | | 0 | 1 | 0 | 0 | 0 | 0 | 0 | 0 | 0 | 0 | 0 |
| 0 | 0.00 | 3.68 | 0.00 | 1.00 | NA | Gm5761 | | 0 | 1 | 0 | 0 | 0 | 0 | 0 | 0 | 0 | 0 | 0 |
| 0 | 0.00 | 3.68 | 0.00 | 1.00 | NA | Gm15007 | | 0 | 1 | 0 | 0 | 0 | 0 | 0 | 0 | 0 | 0 | 0 |
| 0 | 0.00 | 3.68 | 0.00 | 1.00 | NA | Gm4760 | | 0 | 1 | 0 | 0 | 0 | 0 | 0 | 0 | 0 | 0 | 0 |
| 0 | 0.00 | 3.68 | 0.00 | 1.00 | NA | Rpl7a-ps12 | | 0 | 1 | 0 | 0 | 0 | 0 | 0 | 0 | 0 | 0 | 0 |
| 0 | 0.97 | 3.67 | 0.26 | 0.79 | NA | Gm7470 | | 0 | 0 | 0 | 0 | 0 | 0 | 0 | 0 | 1 | 0 | 0 |
| 0 | 0.97 | 3.67 | 0.26 | 0.79 | NA | Gm10558 | | 0 | 0 | 0 | 0 | 0 | 0 | 0 | 0 | 1 | 0 | 0 |
| 0 | 0.97 | 3.67 | 0.26 | 0.79 | NA | Gm37730 | | 0 | 0 | 0 | 0 | 0 | 0 | 0 | 0 | 1 | 0 | 0 |
| 0 | 0.97 | 3.67 | 0.26 | 0.79 | NA | Rpl35-ps1 | | 0 | 0 | 0 | 0 | 0 | 0 | 0 | 0 | 1 | 0 | 0 |
| 0 | 0.97 | 3.67 | 0.26 | 0.79 | NA | Gm13727 | | 0 | 0 | 0 | 0 | 0 | 0 | 0 | 0 | 1 | 0 | 0 |
| 0 | 0.97 | 3.67 | 0.26 | 0.79 | NA | Chrm5 | cholinergic receptor, muscarinic 5 | 0 | 0 | 0 | 0 | 0 | 0 | 0 | 0 | 1 | 0 | 0 |
| 0 | 0.97 | 3.67 | 0.26 | 0.79 | NA | Gm14240 | | 0 | 0 | 0 | 0 | 0 | 0 | 0 | 0 | 1 | 0 | 0 |
| 0 | 0.97 | 3.67 | 0.26 | 0.79 | NA | Adig | adipogenin | 0 | 0 | 0 | 0 | 0 | 0 | 0 | 0 | 1 | 0 | 0 |
| 0 | 0.97 | 3.67 | 0.26 | 0.79 | NA | Gm11472 | | 0 | 0 | 0 | 0 | 0 | 0 | 0 | 0 | 1 | 0 | 0 |
| 0 | 0.97 | 3.67 | 0.26 | 0.79 | NA | Gm14387 | | 0 | 0 | 0 | 0 | 0 | 0 | 0 | 0 | 1 | 0 | 0 |
| 0 | 0.97 | 3.67 | 0.26 | 0.79 | NA | Gm7799 | | 0 | 0 | 0 | 0 | 0 | 0 | 0 | 0 | 1 | 0 | 0 |
| 0 | 0.97 | 3.67 | 0.26 | 0.79 | NA | Sis | sucrase isomaltase (alpha-glucosidase) | 0 | 0 | 0 | 0 | 0 | 0 | 0 | 0 | 1 | 0 | 0 |
| 0 | 0.97 | 3.67 | 0.26 | 0.79 | NA | Gm25820 | | 0 | 0 | 0 | 0 | 0 | 0 | 0 | 0 | 1 | 0 | 0 |
| 0 | 0.97 | 3.67 | 0.26 | 0.79 | NA | 0610031O16Rik | | 0 | 0 | 0 | 0 | 0 | 0 | 0 | 0 | 1 | 0 | 0 |
| 0 | 0.97 | 3.67 | 0.26 | 0.79 | NA | Gm11829 | | 0 | 0 | 0 | 0 | 0 | 0 | 0 | 0 | 1 | 0 | 0 |
| 0 | 0.97 | 3.67 | 0.26 | 0.79 | NA | Gm13306 | predicted gene 13306 | 0 | 0 | 0 | 0 | 0 | 0 | 0 | 0 | 1 | 0 | 0 |
| 0 | 0.97 | 3.67 | 0.26 | 0.79 | NA | Gm12478 | | 0 | 0 | 0 | 0 | 0 | 0 | 0 | 0 | 1 | 0 | 0 |
| 0 | 0.97 | 3.67 | 0.26 | 0.79 | NA | Mir32 | | 0 | 0 | 0 | 0 | 0 | 0 | 0 | 0 | 1 | 0 | 0 |
| 0 | 0.97 | 3.67 | 0.26 | 0.79 | NA | Gm12420 | | 0 | 0 | 0 | 0 | 0 | 0 | 0 | 0 | 1 | 0 | 0 |
| 0 | 0.97 | 3.67 | 0.26 | 0.79 | NA | Ube4bos1 | | 0 | 0 | 0 | 0 | 0 | 0 | 0 | 0 | 1 | 0 | 0 |
| 0 | 0.97 | 3.67 | 0.26 | 0.79 | NA | Gm8929 | | 0 | 0 | 0 | 0 | 0 | 0 | 0 | 0 | 1 | 0 | 0 |
| 0 | 0.97 | 3.67 | 0.26 | 0.79 | NA | Nkx1-1 | NK1 homeobox 1 | 0 | 0 | 0 | 0 | 0 | 0 | 0 | 0 | 1 | 0 | 0 |
| 0 | 0.97 | 3.67 | 0.26 | 0.79 | NA | Gm43155 | | 0 | 0 | 0 | 0 | 0 | 0 | 0 | 0 | 1 | 0 | 0 |
| 0 | 0.97 | 3.67 | 0.26 | 0.79 | NA | Gm43150 | | 0 | 0 | 0 | 0 | 0 | 0 | 0 | 0 | 1 | 0 | 0 |
| 0 | 0.97 | 3.67 | 0.26 | 0.79 | NA | Gm5319 | | 0 | 0 | 0 | 0 | 0 | 0 | 0 | 0 | 1 | 0 | 0 |
| 0 | 0.97 | 3.67 | 0.26 | 0.79 | NA | Gm45351 | | 0 | 0 | 0 | 0 | 0 | 0 | 0 | 0 | 1 | 0 | 0 |
| 0 | 0.97 | 3.67 | 0.26 | 0.79 | NA | Gm6887 | | 0 | 0 | 0 | 0 | 0 | 0 | 0 | 0 | 1 | 0 | 0 |
| 0 | 0.97 | 3.67 | 0.26 | 0.79 | NA | Gm18988 | | 0 | 0 | 0 | 0 | 0 | 0 | 0 | 0 | 1 | 0 | 0 |
| 0 | 0.97 | 3.67 | 0.26 | 0.79 | NA | Gm15544 | | 0 | 0 | 0 | 0 | 0 | 0 | 0 | 0 | 1 | 0 | 0 |
| 0 | 0.97 | 3.67 | 0.26 | 0.79 | NA | Gm44578 | | 0 | 0 | 0 | 0 | 0 | 0 | 0 | 0 | 1 | 0 | 0 |
| 0 | 0.97 | 3.67 | 0.26 | 0.79 | NA | Olfr639 | olfactory receptor 639 | 0 | 0 | 0 | 0 | 0 | 0 | 0 | 0 | 1 | 0 | 0 |
| 0 | 0.97 | 3.67 | 0.26 | 0.79 | NA | Gm19269 | | 0 | 0 | 0 | 0 | 0 | 0 | 0 | 0 | 1 | 0 | 0 |
| 0 | 0.97 | 3.67 | 0.26 | 0.79 | NA | Gm20735 | | 0 | 0 | 0 | 0 | 0 | 0 | 0 | 0 | 1 | 0 | 0 |
| 0 | 0.97 | 3.67 | 0.26 | 0.79 | NA | Gm6921 | | 0 | 0 | 0 | 0 | 0 | 0 | 0 | 0 | 1 | 0 | 0 |
| 0 | 0.97 | 3.67 | 0.26 | 0.79 | NA | Mobp | myelin-associated oligodendrocytic basic | 0 | 0 | 0 | 0 | 0 | 0 | 0 | 0 | 1 | 0 | 0 |
| 0 | 0.97 | 3.67 | 0.26 | 0.79 | NA | Gm39463 | | 0 | 0 | 0 | 0 | 0 | 0 | 0 | 0 | 1 | 0 | 0 |
| 0 | 0.97 | 3.67 | 0.26 | 0.79 | NA | A730085K08Rik | | 0 | 0 | 0 | 0 | 0 | 0 | 0 | 0 | 1 | 0 | 0 |
| 0 | 0.97 | 3.67 | 0.26 | 0.79 | NA | Gje1 | gap junction protein, epsilon 1 | 0 | 0 | 0 | 0 | 0 | 0 | 0 | 0 | 1 | 0 | 0 |
| 0 | 0.97 | 3.67 | 0.26 | 0.79 | NA | Gm26741 | | 0 | 0 | 0 | 0 | 0 | 0 | 0 | 0 | 1 | 0 | 0 |
| 0 | 0.97 | 3.67 | 0.26 | 0.79 | NA | Gm47623 | | 0 | 0 | 0 | 0 | 0 | 0 | 0 | 0 | 1 | 0 | 0 |
| 0 | 0.97 | 3.67 | 0.26 | 0.79 | NA | Olfr1356 | olfactory receptor 1356 | 0 | 0 | 0 | 0 | 0 | 0 | 0 | 0 | 1 | 0 | 0 |
| 0 | 0.97 | 3.67 | 0.26 | 0.79 | NA | Gm25794 | | 0 | 0 | 0 | 0 | 0 | 0 | 0 | 0 | 1 | 0 | 0 |
| 0 | 0.97 | 3.67 | 0.26 | 0.79 | NA | 4930534H03Rik | | 0 | 0 | 0 | 0 | 0 | 0 | 0 | 0 | 1 | 0 | 0 |
| 0 | 0.97 | 3.67 | 0.26 | 0.79 | NA | Gm12042 | | 0 | 0 | 0 | 0 | 0 | 0 | 0 | 0 | 1 | 0 | 0 |

| | | | | | | | | | | | | | | | | | | |
|---|---|---|---|---|---|---|---|---|---|---|---|---|---|---|---|---|---|---|
| 0 | 0.97 | 3.67 | 0.26 | 0.79 | NA | Gm12140 | | 0 | 0 | 0 | 0 | 0 | 0 | 0 | 0 | 1 | 0 | 0 |
| 0 | 0.97 | 3.67 | 0.26 | 0.79 | NA | Leap2 | liver-expressed antimicrobial peptide 2 | 0 | 0 | 0 | 0 | 0 | 0 | 0 | 0 | 1 | 0 | 0 |
| 0 | 0.97 | 3.67 | 0.26 | 0.79 | NA | Gm11631 | | 0 | 0 | 0 | 0 | 0 | 0 | 0 | 0 | 1 | 0 | 0 |
| 0 | 0.97 | 3.67 | 0.26 | 0.79 | NA | Gm23875 | | 0 | 0 | 0 | 0 | 0 | 0 | 0 | 0 | 1 | 0 | 0 |
| 0 | 0.97 | 3.67 | 0.26 | 0.79 | NA | Gm48553 | | 0 | 0 | 0 | 0 | 0 | 0 | 0 | 0 | 1 | 0 | 0 |
| 0 | 0.97 | 3.67 | 0.26 | 0.79 | NA | 1700011B04Rik | | 0 | 0 | 0 | 0 | 0 | 0 | 0 | 0 | 1 | 0 | 0 |
| 0 | 0.97 | 3.67 | 0.26 | 0.79 | NA | Gm6344 | | 0 | 0 | 0 | 0 | 0 | 0 | 0 | 0 | 1 | 0 | 0 |
| 0 | 0.97 | 3.67 | 0.26 | 0.79 | NA | BC048507 | cDNA sequence BC048507 | 0 | 0 | 0 | 0 | 0 | 0 | 0 | 0 | 1 | 0 | 0 |
| 0 | 0.97 | 3.67 | 0.26 | 0.79 | NA | Gm7352 | | 0 | 0 | 0 | 0 | 0 | 0 | 0 | 0 | 1 | 0 | 0 |
| 0 | 0.97 | 3.67 | 0.26 | 0.79 | NA | Chat | choline acetyltransferase | 0 | 0 | 0 | 0 | 0 | 0 | 0 | 0 | 1 | 0 | 0 |
| 0 | 0.97 | 3.67 | 0.26 | 0.79 | NA | Gm5203 | | 0 | 0 | 0 | 0 | 0 | 0 | 0 | 0 | 1 | 0 | 0 |
| 0 | 0.97 | 3.67 | 0.26 | 0.79 | NA | Gm10233 | | 0 | 0 | 0 | 0 | 0 | 0 | 0 | 0 | 1 | 0 | 0 |
| 0 | 0.97 | 3.67 | 0.26 | 0.79 | NA | Gm33121 | | 0 | 0 | 0 | 0 | 0 | 0 | 0 | 0 | 1 | 0 | 0 |
| 0 | 0.97 | 3.67 | 0.26 | 0.79 | NA | Gm49301 | | 0 | 0 | 0 | 0 | 0 | 0 | 0 | 0 | 1 | 0 | 0 |
| 0 | 0.97 | 3.67 | 0.26 | 0.79 | NA | Gm5215 | | 0 | 0 | 0 | 0 | 0 | 0 | 0 | 0 | 1 | 0 | 0 |
| 0 | 0.97 | 3.67 | 0.26 | 0.79 | NA | Ly6g2 | lymphocyte antigen 6 complex, locus G2 | 0 | 0 | 0 | 0 | 0 | 0 | 0 | 0 | 1 | 0 | 0 |
| 0 | 0.97 | 3.67 | 0.26 | 0.79 | NA | Gm5768 | | 0 | 0 | 0 | 0 | 0 | 0 | 0 | 0 | 1 | 0 | 0 |
| 0 | 0.97 | 3.67 | 0.26 | 0.79 | NA | Gm46542 | | 0 | 0 | 0 | 0 | 0 | 0 | 0 | 0 | 1 | 0 | 0 |
| 0 | 0.97 | 3.67 | 0.26 | 0.79 | NA | Gm8497 | | 0 | 0 | 0 | 0 | 0 | 0 | 0 | 0 | 1 | 0 | 0 |
| 0 | 0.97 | 3.67 | 0.26 | 0.79 | NA | Lgals1-ps1 | | 0 | 0 | 0 | 0 | 0 | 0 | 0 | 0 | 1 | 0 | 0 |
| 0 | 0.97 | 3.67 | 0.26 | 0.79 | NA | Gm9386 | | 0 | 0 | 0 | 0 | 0 | 0 | 0 | 0 | 1 | 0 | 0 |
| 0 | 0.97 | 3.67 | 0.26 | 0.79 | NA | Gm15334 | | 0 | 0 | 0 | 0 | 0 | 0 | 0 | 0 | 1 | 0 | 0 |
| 0 | 0.97 | 3.67 | 0.26 | 0.79 | NA | Rps19-ps13 | | 0 | 0 | 0 | 0 | 0 | 0 | 0 | 0 | 1 | 0 | 0 |
| 0 | 0.97 | 3.67 | 0.26 | 0.79 | NA | Gm7569 | | 0 | 0 | 0 | 0 | 0 | 0 | 0 | 0 | 1 | 0 | 0 |
| 0 | 0.97 | 3.67 | 0.26 | 0.79 | NA | Pkd2l1 | polycystic kidney disease 2-like 1 | 0 | 0 | 0 | 0 | 0 | 0 | 0 | 0 | 1 | 0 | 0 |
| 0 | 0.97 | 3.67 | 0.26 | 0.79 | NA | Gm14513 | | 0 | 0 | 0 | 0 | 0 | 0 | 0 | 0 | 1 | 0 | 0 |
| 0 | 0.97 | 3.67 | 0.26 | 0.79 | NA | Rpl36-ps10 | | 0 | 0 | 0 | 0 | 0 | 0 | 0 | 0 | 1 | 0 | 0 |
| 0 | 0.97 | 3.67 | 0.26 | 0.79 | NA | 1700111N16Rik | | 0 | 0 | 0 | 0 | 0 | 0 | 0 | 0 | 1 | 0 | 0 |
| 0 | 0.97 | 3.67 | 0.26 | 0.79 | NA | Gm4989 | | 0 | 0 | 0 | 0 | 0 | 0 | 0 | 0 | 1 | 0 | 0 |
| 0 | 0.97 | 3.67 | 0.26 | 0.79 | NA | Gm2214 | | 0 | 0 | 0 | 0 | 0 | 0 | 0 | 0 | 1 | 0 | 0 |
| 0 | 0.97 | 3.67 | 0.26 | 0.79 | NA | Gm15212 | | 0 | 0 | 0 | 0 | 0 | 0 | 0 | 0 | 1 | 0 | 0 |
| 0 | -0.44 | 3.68 | -0.12 | 0.90 | NA | Gm2147 | | 0 | 0 | 0 | 0 | 1 | 0 | 0 | 0 | 0 | 0 | 0 |
| 0 | -0.44 | 3.68 | -0.12 | 0.90 | NA | Gm37110 | | 0 | 0 | 0 | 0 | 1 | 0 | 0 | 0 | 0 | 0 | 0 |
| 0 | -0.44 | 3.68 | -0.12 | 0.90 | NA | Gm28580 | | 0 | 0 | 0 | 0 | 1 | 0 | 0 | 0 | 0 | 0 | 0 |
| 0 | -0.44 | 3.68 | -0.12 | 0.90 | NA | Gm28501 | | 0 | 0 | 0 | 0 | 1 | 0 | 0 | 0 | 0 | 0 | 0 |
| 0 | -0.44 | 3.68 | -0.12 | 0.90 | NA | Gm13198 | | 0 | 0 | 0 | 0 | 1 | 0 | 0 | 0 | 0 | 0 | 0 |
| 0 | -0.44 | 3.68 | -0.12 | 0.90 | NA | Gm13478 | | 0 | 0 | 0 | 0 | 1 | 0 | 0 | 0 | 0 | 0 | 0 |
| 0 | -0.44 | 3.68 | -0.12 | 0.90 | NA | Gm13730 | | 0 | 0 | 0 | 0 | 1 | 0 | 0 | 0 | 0 | 0 | 0 |
| 0 | -0.44 | 3.68 | -0.12 | 0.90 | NA | Gm13921 | | 0 | 0 | 0 | 0 | 1 | 0 | 0 | 0 | 0 | 0 | 0 |
| 0 | -0.44 | 3.68 | -0.12 | 0.90 | NA | Gm9864 | | 0 | 0 | 0 | 0 | 1 | 0 | 0 | 0 | 0 | 0 | 0 |
| 0 | -0.44 | 3.68 | -0.12 | 0.90 | NA | 4933427J07Rik | | 0 | 0 | 0 | 0 | 1 | 0 | 0 | 0 | 0 | 0 | 0 |
| 0 | -0.44 | 3.68 | -0.12 | 0.90 | NA | Gm14278 | | 0 | 0 | 0 | 0 | 1 | 0 | 0 | 0 | 0 | 0 | 0 |
| 0 | -0.44 | 3.68 | -0.12 | 0.90 | NA | Gm14288 | predicted gene 14288 | 0 | 0 | 0 | 0 | 1 | 0 | 0 | 0 | 0 | 0 | 0 |
| 0 | -0.44 | 3.68 | -0.12 | 0.90 | NA | Gm8823 | | 0 | 0 | 0 | 0 | 1 | 0 | 0 | 0 | 0 | 0 | 0 |
| 0 | -0.44 | 3.68 | -0.12 | 0.90 | NA | 5930409G06Rik | | 0 | 0 | 0 | 0 | 1 | 0 | 0 | 0 | 0 | 0 | 0 |
| 0 | -0.44 | 3.68 | -0.12 | 0.90 | NA | Gm29741 | | 0 | 0 | 0 | 0 | 1 | 0 | 0 | 0 | 0 | 0 | 0 |
| 0 | -0.44 | 3.68 | -0.12 | 0.90 | NA | Gm18294 | | 0 | 0 | 0 | 0 | 1 | 0 | 0 | 0 | 0 | 0 | 0 |
| 0 | -0.44 | 3.68 | -0.12 | 0.90 | NA | Gm34780 | | 0 | 0 | 0 | 0 | 1 | 0 | 0 | 0 | 0 | 0 | 0 |
| 0 | -0.44 | 3.68 | -0.12 | 0.90 | NA | Gm19739 | | 0 | 0 | 0 | 0 | 1 | 0 | 0 | 0 | 0 | 0 | 0 |
| 0 | -0.44 | 3.68 | -0.12 | 0.90 | NA | Gm37361 | | 0 | 0 | 0 | 0 | 1 | 0 | 0 | 0 | 0 | 0 | 0 |
| 0 | -0.44 | 3.68 | -0.12 | 0.90 | NA | Gm43587 | | 0 | 0 | 0 | 0 | 1 | 0 | 0 | 0 | 0 | 0 | 0 |
| 0 | -0.44 | 3.68 | -0.12 | 0.90 | NA | Gm9381 | | 0 | 0 | 0 | 0 | 1 | 0 | 0 | 0 | 0 | 0 | 0 |
| 0 | -0.44 | 3.68 | -0.12 | 0.90 | NA | Gm42587 | | 0 | 0 | 0 | 0 | 1 | 0 | 0 | 0 | 0 | 0 | 0 |
| 0 | -0.44 | 3.68 | -0.12 | 0.90 | NA | Gm12412 | | 0 | 0 | 0 | 0 | 1 | 0 | 0 | 0 | 0 | 0 | 0 |
| 0 | -0.44 | 3.68 | -0.12 | 0.90 | NA | Gm12642 | | 0 | 0 | 0 | 0 | 1 | 0 | 0 | 0 | 0 | 0 | 0 |
| 0 | -0.44 | 3.68 | -0.12 | 0.90 | NA | Gm42585 | | 0 | 0 | 0 | 0 | 1 | 0 | 0 | 0 | 0 | 0 | 0 |
| 0 | -0.44 | 3.68 | -0.12 | 0.90 | NA | Gm42846 | | 0 | 0 | 0 | 0 | 1 | 0 | 0 | 0 | 0 | 0 | 0 |
| 0 | -0.44 | 3.68 | -0.12 | 0.90 | NA | Gm43105 | | 0 | 0 | 0 | 0 | 1 | 0 | 0 | 0 | 0 | 0 | 0 |
| 0 | -0.44 | 3.68 | -0.12 | 0.90 | NA | Gm7181 | | 0 | 0 | 0 | 0 | 1 | 0 | 0 | 0 | 0 | 0 | 0 |
| 0 | -0.44 | 3.68 | -0.12 | 0.90 | NA | Gm15727 | | 0 | 0 | 0 | 0 | 1 | 0 | 0 | 0 | 0 | 0 | 0 |

| | | | | | | | | | | | | | | | | | | |
|---|---|---|---|---|---|---|---|---|---|---|---|---|---|---|---|---|---|---|
| 0 | -0.44 | 3.68 | -0.12 | 0.90 | NA | Gm43295 | | 0 | 0 | 0 | 0 | 1 | 0 | 0 | 0 | 0 | 0 | 0 |
| 0 | -0.44 | 3.68 | -0.12 | 0.90 | NA | Gm8836 | | 0 | 0 | 0 | 0 | 1 | 0 | 0 | 0 | 0 | 0 | 0 |
| 0 | -0.44 | 3.68 | -0.12 | 0.90 | NA | Gm24601 | | 0 | 0 | 0 | 0 | 1 | 0 | 0 | 0 | 0 | 0 | 0 |
| 0 | -0.44 | 3.68 | -0.12 | 0.90 | NA | Gm15929 | | 0 | 0 | 0 | 0 | 1 | 0 | 0 | 0 | 0 | 0 | 0 |
| 0 | -0.44 | 3.68 | -0.12 | 0.90 | NA | Pafah1b1-ps2 | | 0 | 0 | 0 | 0 | 1 | 0 | 0 | 0 | 0 | 0 | 0 |
| 0 | -0.44 | 3.68 | -0.12 | 0.90 | NA | Ceacam12 | carcinoembryonic antigen-related cell adl | 0 | 0 | 0 | 0 | 1 | 0 | 0 | 0 | 0 | 0 | 0 |
| 0 | -0.44 | 3.68 | -0.12 | 0.90 | NA | Ttc9b | tetratricopeptide repeat domain 9B | 0 | 0 | 0 | 0 | 1 | 0 | 0 | 0 | 0 | 0 | 0 |
| 0 | -0.44 | 3.68 | -0.12 | 0.90 | NA | Gm32983 | | 0 | 0 | 0 | 0 | 1 | 0 | 0 | 0 | 0 | 0 | 0 |
| 0 | -0.44 | 3.68 | -0.12 | 0.90 | NA | Vmn2r66 | vomeronasal 2, receptor 66 | 0 | 0 | 0 | 0 | 1 | 0 | 0 | 0 | 0 | 0 | 0 |
| 0 | -0.44 | 3.68 | -0.12 | 0.90 | NA | Gm45678 | | 0 | 0 | 0 | 0 | 1 | 0 | 0 | 0 | 0 | 0 | 0 |
| 0 | -0.44 | 3.68 | -0.12 | 0.90 | NA | Gm47240 | | 0 | 0 | 0 | 0 | 1 | 0 | 0 | 0 | 0 | 0 | 0 |
| 0 | -0.44 | 3.68 | -0.12 | 0.90 | NA | Gm4797 | | 0 | 0 | 0 | 0 | 1 | 0 | 0 | 0 | 0 | 0 | 0 |
| 0 | -0.44 | 3.68 | -0.12 | 0.90 | NA | Gm2628 | | 0 | 0 | 0 | 0 | 1 | 0 | 0 | 0 | 0 | 0 | 0 |
| 0 | -0.44 | 3.68 | -0.12 | 0.90 | NA | Gm11950 | | 0 | 0 | 0 | 0 | 1 | 0 | 0 | 0 | 0 | 0 | 0 |
| 0 | -0.44 | 3.68 | -0.12 | 0.90 | NA | Gm12223 | | 0 | 0 | 0 | 0 | 1 | 0 | 0 | 0 | 0 | 0 | 0 |
| 0 | -0.44 | 3.68 | -0.12 | 0.90 | NA | Gm12612 | | 0 | 0 | 0 | 0 | 1 | 0 | 0 | 0 | 0 | 0 | 0 |
| 0 | -0.44 | 3.68 | -0.12 | 0.90 | NA | Slc35g3 | solute carrier family 35, member G3 | 0 | 0 | 0 | 0 | 1 | 0 | 0 | 0 | 0 | 0 | 0 |
| 0 | -0.44 | 3.68 | -0.12 | 0.90 | NA | Gm11518 | | 0 | 0 | 0 | 0 | 1 | 0 | 0 | 0 | 0 | 0 | 0 |
| 0 | -0.44 | 3.68 | -0.12 | 0.90 | NA | Gm11662 | | 0 | 0 | 0 | 0 | 1 | 0 | 0 | 0 | 0 | 0 | 0 |
| 0 | -0.44 | 3.68 | -0.12 | 0.90 | NA | Gm47750 | | 0 | 0 | 0 | 0 | 1 | 0 | 0 | 0 | 0 | 0 | 0 |
| 0 | -0.44 | 3.68 | -0.12 | 0.90 | NA | Gm18848 | | 0 | 0 | 0 | 0 | 1 | 0 | 0 | 0 | 0 | 0 | 0 |
| 0 | -0.44 | 3.68 | -0.12 | 0.90 | NA | Gm29689 | | 0 | 0 | 0 | 0 | 1 | 0 | 0 | 0 | 0 | 0 | 0 |
| 0 | -0.44 | 3.68 | -0.12 | 0.90 | NA | Gm4052 | | 0 | 0 | 0 | 0 | 1 | 0 | 0 | 0 | 0 | 0 | 0 |
| 0 | -0.44 | 3.68 | -0.12 | 0.90 | NA | 5930438M14Rik | | 0 | 0 | 0 | 0 | 1 | 0 | 0 | 0 | 0 | 0 | 0 |
| 0 | -0.44 | 3.68 | -0.12 | 0.90 | NA | Gm41073 | | 0 | 0 | 0 | 0 | 1 | 0 | 0 | 0 | 0 | 0 | 0 |
| 0 | -0.44 | 3.68 | -0.12 | 0.90 | NA | Gm20369 | | 0 | 0 | 0 | 0 | 1 | 0 | 0 | 0 | 0 | 0 | 0 |
| 0 | -0.44 | 3.68 | -0.12 | 0.90 | NA | Krt81 | keratin 81 | 0 | 0 | 0 | 0 | 1 | 0 | 0 | 0 | 0 | 0 | 0 |
| 0 | -0.44 | 3.68 | -0.12 | 0.90 | NA | Gm6630 | | 0 | 0 | 0 | 0 | 1 | 0 | 0 | 0 | 0 | 0 | 0 |
| 0 | -0.44 | 3.68 | -0.12 | 0.90 | NA | Slc51a | solute carrier family 51, alpha subunit | 0 | 0 | 0 | 0 | 1 | 0 | 0 | 0 | 0 | 0 | 0 |
| 0 | -0.44 | 3.68 | -0.12 | 0.90 | NA | Gm20463 | | 0 | 0 | 0 | 0 | 1 | 0 | 0 | 0 | 0 | 0 | 0 |
| 0 | -0.44 | 3.68 | -0.12 | 0.90 | NA | Rpl9-ps5 | | 0 | 0 | 0 | 0 | 1 | 0 | 0 | 0 | 0 | 0 | 0 |
| 0 | -0.44 | 3.68 | -0.12 | 0.90 | NA | | | 0 | 0 | 0 | 0 | 1 | 0 | 0 | 0 | 0 | 0 | 0 |
| 0 | -0.44 | 3.68 | -0.12 | 0.90 | NA | AY702103 | | 0 | 0 | 0 | 0 | 1 | 0 | 0 | 0 | 0 | 0 | 0 |
| 0 | -0.44 | 3.68 | -0.12 | 0.90 | NA | Gm17168 | | 0 | 0 | 0 | 0 | 1 | 0 | 0 | 0 | 0 | 0 | 0 |
| 0 | -0.44 | 3.68 | -0.12 | 0.90 | NA | Gm19399 | | 0 | 0 | 0 | 0 | 1 | 0 | 0 | 0 | 0 | 0 | 0 |
| 0 | -0.44 | 3.68 | -0.12 | 0.90 | NA | Gm6248 | | 0 | 0 | 0 | 0 | 1 | 0 | 0 | 0 | 0 | 0 | 0 |
| 0 | -0.44 | 3.68 | -0.12 | 0.90 | NA | Pabpc2 | poly(A) binding protein, cytoplasmic 2 | 0 | 0 | 0 | 0 | 1 | 0 | 0 | 0 | 0 | 0 | 0 |
| 0 | -0.44 | 3.68 | -0.12 | 0.90 | NA | Gm8390 | | 0 | 0 | 0 | 0 | 1 | 0 | 0 | 0 | 0 | 0 | 0 |
| 0 | -0.44 | 3.68 | -0.12 | 0.90 | NA | Gm2390 | | 0 | 0 | 0 | 0 | 1 | 0 | 0 | 0 | 0 | 0 | 0 |
| 0 | -0.44 | 3.68 | -0.12 | 0.90 | NA | Gm50434 | | 0 | 0 | 0 | 0 | 1 | 0 | 0 | 0 | 0 | 0 | 0 |
| 0 | -0.44 | 3.68 | -0.12 | 0.90 | NA | Gm42068 | | 0 | 0 | 0 | 0 | 1 | 0 | 0 | 0 | 0 | 0 | 0 |
| 0 | -0.44 | 3.68 | -0.12 | 0.90 | NA | Gm31300 | | 0 | 0 | 0 | 0 | 1 | 0 | 0 | 0 | 0 | 0 | 0 |
| 0 | -0.44 | 3.68 | -0.12 | 0.90 | NA | Gm14530 | | 0 | 0 | 0 | 0 | 1 | 0 | 0 | 0 | 0 | 0 | 0 |
| 0 | -0.44 | 3.68 | -0.12 | 0.90 | NA | Gm5385 | | 0 | 0 | 0 | 0 | 1 | 0 | 0 | 0 | 0 | 0 | 0 |
| 0 | -0.44 | 3.68 | -0.12 | 0.90 | NA | Gm14697 | | 0 | 0 | 0 | 0 | 1 | 0 | 0 | 0 | 0 | 0 | 0 |
| 0 | -0.44 | 3.68 | -0.12 | 0.90 | NA | Gm14780 | | 0 | 0 | 0 | 0 | 1 | 0 | 0 | 0 | 0 | 0 | 0 |
| 0 | -0.44 | 3.68 | -0.12 | 0.90 | NA | Gm6184 | | 0 | 0 | 0 | 0 | 1 | 0 | 0 | 0 | 0 | 0 | 0 |
| 0 | -0.44 | 3.68 | -0.12 | 0.90 | NA | Gm6298 | | 0 | 0 | 0 | 0 | 1 | 0 | 0 | 0 | 0 | 0 | 0 |
| 0 | -0.44 | 3.68 | -0.12 | 0.90 | NA | Gm8659 | | 0 | 0 | 0 | 0 | 1 | 0 | 0 | 0 | 0 | 0 | 0 |
| 0 | -0.42 | 3.68 | -0.11 | 0.91 | NA | Smt3h2-ps4 | | 0 | 0 | 0 | 1 | 0 | 0 | 0 | 0 | 0 | 0 | 0 |
| 0 | -0.42 | 3.68 | -0.11 | 0.91 | NA | Gm37331 | | 0 | 0 | 0 | 1 | 0 | 0 | 0 | 0 | 0 | 0 | 0 |
| 0 | -0.42 | 3.68 | -0.11 | 0.91 | NA | Gm28980 | | 0 | 0 | 0 | 1 | 0 | 0 | 0 | 0 | 0 | 0 | 0 |
| 0 | -0.42 | 3.68 | -0.11 | 0.91 | NA | Gm38122 | | 0 | 0 | 0 | 1 | 0 | 0 | 0 | 0 | 0 | 0 | 0 |
| 0 | -0.42 | 3.68 | -0.11 | 0.91 | NA | Gm13588 | | 0 | 0 | 0 | 1 | 0 | 0 | 0 | 0 | 0 | 0 | 0 |
| 0 | -0.42 | 3.68 | -0.11 | 0.91 | NA | Gm13758 | | 0 | 0 | 0 | 1 | 0 | 0 | 0 | 0 | 0 | 0 | 0 |
| 0 | -0.42 | 3.68 | -0.11 | 0.91 | NA | Gm11448 | | 0 | 0 | 0 | 1 | 0 | 0 | 0 | 0 | 0 | 0 | 0 |
| 0 | -0.42 | 3.68 | -0.11 | 0.91 | NA | Gm14439 | | 0 | 0 | 0 | 1 | 0 | 0 | 0 | 0 | 0 | 0 | 0 |
| 0 | -0.42 | 3.68 | -0.11 | 0.91 | NA | Gm37636 | | 0 | 0 | 0 | 1 | 0 | 0 | 0 | 0 | 0 | 0 | 0 |
| 0 | -0.42 | 3.68 | -0.11 | 0.91 | NA | Ube2l3-ps1 | | 0 | 0 | 0 | 1 | 0 | 0 | 0 | 0 | 0 | 0 | 0 |
| 0 | -0.42 | 3.68 | -0.11 | 0.91 | NA | Gm37424 | | 0 | 0 | 0 | 1 | 0 | 0 | 0 | 0 | 0 | 0 | 0 |

| | | | | | | | | | | | | | | | | | | |
|---|---|---|---|---|---|---|---|---|---|---|---|---|---|---|---|---|---|---|
| 0 | -0.42 | 3.68 | -0.11 | 0.91 | NA | Clca4b | chloride channel accessory 4B | 0 | 0 | 0 | 1 | 0 | 0 | 0 | 0 | 0 | 0 | 0 |
| 0 | -0.42 | 3.68 | -0.11 | 0.91 | NA | Msh4 | mutS homolog 4 | 0 | 0 | 0 | 1 | 0 | 0 | 0 | 0 | 0 | 0 | 0 |
| 0 | -0.42 | 3.68 | -0.11 | 0.91 | NA | Scp2-ps1 | | 0 | 0 | 0 | 1 | 0 | 0 | 0 | 0 | 0 | 0 | 0 |
| 0 | -0.42 | 3.68 | -0.11 | 0.91 | NA | Gm12513 | | 0 | 0 | 0 | 1 | 0 | 0 | 0 | 0 | 0 | 0 | 0 |
| 0 | -0.42 | 3.68 | -0.11 | 0.91 | NA | Mup22 | major urinary protein 22 | 0 | 0 | 0 | 1 | 0 | 0 | 0 | 0 | 0 | 0 | 0 |
| 0 | -0.42 | 3.68 | -0.11 | 0.91 | NA | Gm12671 | | 0 | 0 | 0 | 1 | 0 | 0 | 0 | 0 | 0 | 0 | 0 |
| 0 | -0.42 | 3.68 | -0.11 | 0.91 | NA | Gm12650 | | 0 | 0 | 0 | 1 | 0 | 0 | 0 | 0 | 0 | 0 | 0 |
| 0 | -0.42 | 3.68 | -0.11 | 0.91 | NA | Gm8984 | | 0 | 0 | 0 | 1 | 0 | 0 | 0 | 0 | 0 | 0 | 0 |
| 0 | -0.42 | 3.68 | -0.11 | 0.91 | NA | Gm43117 | | 0 | 0 | 0 | 1 | 0 | 0 | 0 | 0 | 0 | 0 | 0 |
| 0 | -0.42 | 3.68 | -0.11 | 0.91 | NA | Gm43694 | | 0 | 0 | 0 | 1 | 0 | 0 | 0 | 0 | 0 | 0 | 0 |
| 0 | -0.42 | 3.68 | -0.11 | 0.91 | NA | Gm10359 | | 0 | 0 | 0 | 1 | 0 | 0 | 0 | 0 | 0 | 0 | 0 |
| 0 | -0.42 | 3.68 | -0.11 | 0.91 | NA | Gm8365 | | 0 | 0 | 0 | 1 | 0 | 0 | 0 | 0 | 0 | 0 | 0 |
| 0 | -0.42 | 3.68 | -0.11 | 0.91 | NA | Gm15546 | | 0 | 0 | 0 | 1 | 0 | 0 | 0 | 0 | 0 | 0 | 0 |
| 0 | -0.42 | 3.68 | -0.11 | 0.91 | NA | Gm7890 | | 0 | 0 | 0 | 1 | 0 | 0 | 0 | 0 | 0 | 0 | 0 |
| 0 | -0.42 | 3.68 | -0.11 | 0.91 | NA | Gm15534 | | 0 | 0 | 0 | 1 | 0 | 0 | 0 | 0 | 0 | 0 | 0 |
| 0 | -0.42 | 3.68 | -0.11 | 0.91 | NA | Gm42818 | | 0 | 0 | 0 | 1 | 0 | 0 | 0 | 0 | 0 | 0 | 0 |
| 0 | -0.42 | 3.68 | -0.11 | 0.91 | NA | Gm14573 | | 0 | 0 | 0 | 1 | 0 | 0 | 0 | 0 | 0 | 0 | 0 |
| 0 | -0.42 | 3.68 | -0.11 | 0.91 | NA | Gm15531 | | 0 | 0 | 0 | 1 | 0 | 0 | 0 | 0 | 0 | 0 | 0 |
| 0 | -0.42 | 3.68 | -0.11 | 0.91 | NA | Gm44317 | | 0 | 0 | 0 | 1 | 0 | 0 | 0 | 0 | 0 | 0 | 0 |
| 0 | -0.42 | 3.68 | -0.11 | 0.91 | NA | Slco1b2 | solute carrier organic anion transporter fa | 0 | 0 | 0 | 1 | 0 | 0 | 0 | 0 | 0 | 0 | 0 |
| 0 | -0.42 | 3.68 | -0.11 | 0.91 | NA | Gm15767 | | 0 | 0 | 0 | 1 | 0 | 0 | 0 | 0 | 0 | 0 | 0 |
| 0 | -0.42 | 3.68 | -0.11 | 0.91 | NA | Gm15873 | | 0 | 0 | 0 | 1 | 0 | 0 | 0 | 0 | 0 | 0 | 0 |
| 0 | -0.42 | 3.68 | -0.11 | 0.91 | NA | Gm4607 | | 0 | 0 | 0 | 1 | 0 | 0 | 0 | 0 | 0 | 0 | 0 |
| 0 | -0.42 | 3.68 | -0.11 | 0.91 | NA | Gm44662 | | 0 | 0 | 0 | 1 | 0 | 0 | 0 | 0 | 0 | 0 | 0 |
| 0 | -0.42 | 3.68 | -0.11 | 0.91 | NA | Gm9127 | | 0 | 0 | 0 | 1 | 0 | 0 | 0 | 0 | 0 | 0 | 0 |
| 0 | -0.42 | 3.68 | -0.11 | 0.91 | NA | Gm5339 | | 0 | 0 | 0 | 1 | 0 | 0 | 0 | 0 | 0 | 0 | 0 |
| 0 | -0.42 | 3.68 | -0.11 | 0.91 | NA | Gm6108 | | 0 | 0 | 0 | 1 | 0 | 0 | 0 | 0 | 0 | 0 | 0 |
| 0 | -0.42 | 3.68 | -0.11 | 0.91 | NA | Gm15580 | | 0 | 0 | 0 | 1 | 0 | 0 | 0 | 0 | 0 | 0 | 0 |
| 0 | -0.42 | 3.68 | -0.11 | 0.91 | NA | Gm9457 | | 0 | 0 | 0 | 1 | 0 | 0 | 0 | 0 | 0 | 0 | 0 |
| 0 | -0.42 | 3.68 | -0.11 | 0.91 | NA | Gm4891 | | 0 | 0 | 0 | 1 | 0 | 0 | 0 | 0 | 0 | 0 | 0 |
| 0 | -0.42 | 3.68 | -0.11 | 0.91 | NA | Gm18000 | | 0 | 0 | 0 | 1 | 0 | 0 | 0 | 0 | 0 | 0 | 0 |
| 0 | -0.42 | 3.68 | -0.11 | 0.91 | NA | Gm36940 | | 0 | 0 | 0 | 1 | 0 | 0 | 0 | 0 | 0 | 0 | 0 |
| 0 | -0.42 | 3.68 | -0.11 | 0.91 | NA | Gm7213 | | 0 | 0 | 0 | 1 | 0 | 0 | 0 | 0 | 0 | 0 | 0 |
| 0 | -0.42 | 3.68 | -0.11 | 0.91 | NA | Gm8834 | | 0 | 0 | 0 | 1 | 0 | 0 | 0 | 0 | 0 | 0 | 0 |
| 0 | -0.42 | 3.68 | -0.11 | 0.91 | NA | Gm5950 | | 0 | 0 | 0 | 1 | 0 | 0 | 0 | 0 | 0 | 0 | 0 |
| 0 | -0.42 | 3.68 | -0.11 | 0.91 | NA | Gm3828 | | 0 | 0 | 0 | 1 | 0 | 0 | 0 | 0 | 0 | 0 | 0 |
| 0 | -0.42 | 3.68 | -0.11 | 0.91 | NA | Gm12079 | | 0 | 0 | 0 | 1 | 0 | 0 | 0 | 0 | 0 | 0 | 0 |
| 0 | -0.42 | 3.68 | -0.11 | 0.91 | NA | Rpl9-ps1 | | 0 | 0 | 0 | 1 | 0 | 0 | 0 | 0 | 0 | 0 | 0 |
| 0 | -0.42 | 3.68 | -0.11 | 0.91 | NA | Gm19084 | | 0 | 0 | 0 | 1 | 0 | 0 | 0 | 0 | 0 | 0 | 0 |
| 0 | -0.42 | 3.68 | -0.11 | 0.91 | NA | Gm29737 | | 0 | 0 | 0 | 1 | 0 | 0 | 0 | 0 | 0 | 0 | 0 |
| 0 | -0.42 | 3.68 | -0.11 | 0.91 | NA | Gm48643 | | 0 | 0 | 0 | 1 | 0 | 0 | 0 | 0 | 0 | 0 | 0 |
| 0 | -0.42 | 3.68 | -0.11 | 0.91 | NA | Hspe1-ps2 | | 0 | 0 | 0 | 1 | 0 | 0 | 0 | 0 | 0 | 0 | 0 |
| 0 | -0.42 | 3.68 | -0.11 | 0.91 | NA | Gm3191 | | 0 | 0 | 0 | 1 | 0 | 0 | 0 | 0 | 0 | 0 | 0 |
| 0 | -0.42 | 3.68 | -0.11 | 0.91 | NA | Gm7979 | | 0 | 0 | 0 | 1 | 0 | 0 | 0 | 0 | 0 | 0 | 0 |
| 0 | -0.42 | 3.68 | -0.11 | 0.91 | NA | Gm47463 | | 0 | 0 | 0 | 1 | 0 | 0 | 0 | 0 | 0 | 0 | 0 |
| 0 | -0.42 | 3.68 | -0.11 | 0.91 | NA | Gm5205 | | 0 | 0 | 0 | 1 | 0 | 0 | 0 | 0 | 0 | 0 | 0 |
| 0 | -0.42 | 3.68 | -0.11 | 0.91 | NA | Gm29753 | | 0 | 0 | 0 | 1 | 0 | 0 | 0 | 0 | 0 | 0 | 0 |
| 0 | -0.42 | 3.68 | -0.11 | 0.91 | NA | Gm6704 | | 0 | 0 | 0 | 1 | 0 | 0 | 0 | 0 | 0 | 0 | 0 |
| 0 | -0.42 | 3.68 | -0.11 | 0.91 | NA | Gm2675 | | 0 | 0 | 0 | 1 | 0 | 0 | 0 | 0 | 0 | 0 | 0 |
| 0 | -0.42 | 3.68 | -0.11 | 0.91 | NA | Uqcrh-ps1 | | 0 | 0 | 0 | 1 | 0 | 0 | 0 | 0 | 0 | 0 | 0 |
| 0 | -0.42 | 3.68 | -0.11 | 0.91 | NA | Rpl31-ps4 | | 0 | 0 | 0 | 1 | 0 | 0 | 0 | 0 | 0 | 0 | 0 |
| 0 | -0.42 | 3.68 | -0.11 | 0.91 | NA | Gm49954 | | 0 | 0 | 0 | 1 | 0 | 0 | 0 | 0 | 0 | 0 | 0 |
| 0 | -0.42 | 3.68 | -0.11 | 0.91 | NA | H2-T3 | histocompatibility 2, T region locus 18 | 0 | 0 | 0 | 1 | 0 | 0 | 0 | 0 | 0 | 0 | 0 |
| 0 | -0.42 | 3.68 | -0.11 | 0.91 | NA | Gm10350 | | 0 | 0 | 0 | 1 | 0 | 0 | 0 | 0 | 0 | 0 | 0 |
| 0 | -0.42 | 3.68 | -0.11 | 0.91 | NA | Gm18948 | | 0 | 0 | 0 | 1 | 0 | 0 | 0 | 0 | 0 | 0 | 0 |
| 0 | -0.42 | 3.68 | -0.11 | 0.91 | NA | Gm7532 | | 0 | 0 | 0 | 1 | 0 | 0 | 0 | 0 | 0 | 0 | 0 |
| 0 | -0.42 | 3.68 | -0.11 | 0.91 | NA | Gm5686 | | 0 | 0 | 0 | 1 | 0 | 0 | 0 | 0 | 0 | 0 | 0 |
| 0 | -0.42 | 3.68 | -0.11 | 0.91 | NA | Gm50313 | | 0 | 0 | 0 | 1 | 0 | 0 | 0 | 0 | 0 | 0 | 0 |
| 0 | -0.42 | 3.68 | -0.11 | 0.91 | NA | Gm8412 | | 0 | 0 | 0 | 1 | 0 | 0 | 0 | 0 | 0 | 0 | 0 |
| 0 | -0.42 | 3.68 | -0.11 | 0.91 | NA | 4930414N06Rik | | 0 | 0 | 0 | 1 | 0 | 0 | 0 | 0 | 0 | 0 | 0 |

| | | | | | | | | | | | | | | | | | | |
|---|---|---|---|---|---|---|---|---|---|---|---|---|---|---|---|---|---|---|
| 0 | -0.42 | 3.68 | -0.11 | 0.91 | NA | Gm8163 | | 0 | 0 | 0 | 1 | 0 | 0 | 0 | 0 | 0 | 0 | 0 |
| 0 | -0.42 | 3.68 | -0.11 | 0.91 | NA | Rpl7a-ps13 | | 0 | 0 | 0 | 1 | 0 | 0 | 0 | 0 | 0 | 0 | 0 |
| 0 | -0.42 | 3.68 | -0.11 | 0.91 | NA | Gm14755 | | 0 | 0 | 0 | 1 | 0 | 0 | 0 | 0 | 0 | 0 | 0 |
| 0 | -0.42 | 3.68 | -0.11 | 0.91 | NA | Gm14769 | | 0 | 0 | 0 | 1 | 0 | 0 | 0 | 0 | 0 | 0 | 0 |
| 0 | -0.42 | 3.68 | -0.11 | 0.91 | NA | Gm16463 | | 0 | 0 | 0 | 1 | 0 | 0 | 0 | 0 | 0 | 0 | 0 |
| 0 | -0.42 | 3.68 | -0.11 | 0.91 | NA | Gm7820 | | 0 | 0 | 0 | 1 | 0 | 0 | 0 | 0 | 0 | 0 | 0 |
| 0 | -0.42 | 3.68 | -0.11 | 0.91 | NA | Actb-ps1 | | 0 | 0 | 0 | 1 | 0 | 0 | 0 | 0 | 0 | 0 | 0 |
| 0 | -0.42 | 3.68 | -0.11 | 0.91 | NA | Gm15223 | | 0 | 0 | 0 | 1 | 0 | 0 | 0 | 0 | 0 | 0 | 0 |
| 0 | 0.00 | 3.68 | 0.00 | 1.00 | NA | Gm7512 | | 1 | 0 | 0 | 0 | 0 | 0 | 0 | 0 | 0 | 0 | 0 |
| 0 | 0.00 | 3.68 | 0.00 | 1.00 | NA | Gm28686 | | 1 | 0 | 0 | 0 | 0 | 0 | 0 | 0 | 0 | 0 | 0 |
| 0 | 0.00 | 3.68 | 0.00 | 1.00 | NA | Gm8009 | | 1 | 0 | 0 | 0 | 0 | 0 | 0 | 0 | 0 | 0 | 0 |
| 0 | 0.00 | 3.68 | 0.00 | 1.00 | NA | 1110002O04Rik | | 1 | 0 | 0 | 0 | 0 | 0 | 0 | 0 | 0 | 0 | 0 |
| 0 | 0.00 | 3.68 | 0.00 | 1.00 | NA | Gm29041 | | 1 | 0 | 0 | 0 | 0 | 0 | 0 | 0 | 0 | 0 | 0 |
| 0 | 0.00 | 3.68 | 0.00 | 1.00 | NA | Gm38056 | | 1 | 0 | 0 | 0 | 0 | 0 | 0 | 0 | 0 | 0 | 0 |
| 0 | 0.00 | 3.68 | 0.00 | 1.00 | NA | Gm7609 | predicted pseudogene 7609 | 1 | 0 | 0 | 0 | 0 | 0 | 0 | 0 | 0 | 0 | 0 |
| 0 | 0.00 | 3.68 | 0.00 | 1.00 | NA | Gm28721 | | 1 | 0 | 0 | 0 | 0 | 0 | 0 | 0 | 0 | 0 | 0 |
| 0 | 0.00 | 3.68 | 0.00 | 1.00 | NA | Gm28392 | | 1 | 0 | 0 | 0 | 0 | 0 | 0 | 0 | 0 | 0 | 0 |
| 0 | 0.00 | 3.68 | 0.00 | 1.00 | NA | Gm8964 | | 1 | 0 | 0 | 0 | 0 | 0 | 0 | 0 | 0 | 0 | 0 |
| 0 | 0.00 | 3.68 | 0.00 | 1.00 | NA | 9430083B18Rik | | 1 | 0 | 0 | 0 | 0 | 0 | 0 | 0 | 0 | 0 | 0 |
| 0 | 0.00 | 3.68 | 0.00 | 1.00 | NA | Gm13297 | | 1 | 0 | 0 | 0 | 0 | 0 | 0 | 0 | 0 | 0 | 0 |
| 0 | 0.00 | 3.68 | 0.00 | 1.00 | NA | Gm37126 | | 1 | 0 | 0 | 0 | 0 | 0 | 0 | 0 | 0 | 0 | 0 |
| 0 | 0.00 | 3.68 | 0.00 | 1.00 | NA | Gm13351 | | 1 | 0 | 0 | 0 | 0 | 0 | 0 | 0 | 0 | 0 | 0 |
| 0 | 0.00 | 3.68 | 0.00 | 1.00 | NA | Pax8 | paired box 8 | 1 | 0 | 0 | 0 | 0 | 0 | 0 | 0 | 0 | 0 | 0 |
| 0 | 0.00 | 3.68 | 0.00 | 1.00 | NA | Gm38287 | | 1 | 0 | 0 | 0 | 0 | 0 | 0 | 0 | 0 | 0 | 0 |
| 0 | 0.00 | 3.68 | 0.00 | 1.00 | NA | Gm13428 | | 1 | 0 | 0 | 0 | 0 | 0 | 0 | 0 | 0 | 0 | 0 |
| 0 | 0.00 | 3.68 | 0.00 | 1.00 | NA | Prg3 | proteoglycan 3 | 1 | 0 | 0 | 0 | 0 | 0 | 0 | 0 | 0 | 0 | 0 |
| 0 | 0.00 | 3.68 | 0.00 | 1.00 | NA | Gm13733 | | 1 | 0 | 0 | 0 | 0 | 0 | 0 | 0 | 0 | 0 | 0 |
| 0 | 0.00 | 3.68 | 0.00 | 1.00 | NA | Gm13816 | | 1 | 0 | 0 | 0 | 0 | 0 | 0 | 0 | 0 | 0 | 0 |
| 0 | 0.00 | 3.68 | 0.00 | 1.00 | NA | Wt1os | | 1 | 0 | 0 | 0 | 0 | 0 | 0 | 0 | 0 | 0 | 0 |
| 0 | 0.00 | 3.68 | 0.00 | 1.00 | NA | Platr8 | | 1 | 0 | 0 | 0 | 0 | 0 | 0 | 0 | 0 | 0 | 0 |
| 0 | 0.00 | 3.68 | 0.00 | 1.00 | NA | Gm11462 | | 1 | 0 | 0 | 0 | 0 | 0 | 0 | 0 | 0 | 0 | 0 |
| 0 | 0.00 | 3.68 | 0.00 | 1.00 | NA | Atp5k-ps2 | | 1 | 0 | 0 | 0 | 0 | 0 | 0 | 0 | 0 | 0 | 0 |
| 0 | 0.00 | 3.68 | 0.00 | 1.00 | NA | Ntsr1 | neurotensin receptor 1 | 1 | 0 | 0 | 0 | 0 | 0 | 0 | 0 | 0 | 0 | 0 |
| 0 | 0.00 | 3.68 | 0.00 | 1.00 | NA | Gm5272 | | 1 | 0 | 0 | 0 | 0 | 0 | 0 | 0 | 0 | 0 | 0 |
| 0 | 0.00 | 3.68 | 0.00 | 1.00 | NA | Gm37358 | | 1 | 0 | 0 | 0 | 0 | 0 | 0 | 0 | 0 | 0 | 0 |
| 0 | 0.00 | 3.68 | 0.00 | 1.00 | NA | Gm38088 | | 1 | 0 | 0 | 0 | 0 | 0 | 0 | 0 | 0 | 0 | 0 |
| 0 | 0.00 | 3.68 | 0.00 | 1.00 | NA | 4930529C04Rik | | 1 | 0 | 0 | 0 | 0 | 0 | 0 | 0 | 0 | 0 | 0 |
| 0 | 0.00 | 3.68 | 0.00 | 1.00 | NA | Gm40117 | | 1 | 0 | 0 | 0 | 0 | 0 | 0 | 0 | 0 | 0 | 0 |
| 0 | 0.00 | 3.68 | 0.00 | 1.00 | NA | Gpr61 | G protein-coupled receptor 61 | 1 | 0 | 0 | 0 | 0 | 0 | 0 | 0 | 0 | 0 | 0 |
| 0 | 0.00 | 3.68 | 0.00 | 1.00 | NA | Gm19166 | | 1 | 0 | 0 | 0 | 0 | 0 | 0 | 0 | 0 | 0 | 0 |
| 0 | 0.00 | 3.68 | 0.00 | 1.00 | NA | Rpl9-ps8 | | 1 | 0 | 0 | 0 | 0 | 0 | 0 | 0 | 0 | 0 | 0 |
| 0 | 0.00 | 3.68 | 0.00 | 1.00 | NA | Gm31881 | | 1 | 0 | 0 | 0 | 0 | 0 | 0 | 0 | 0 | 0 | 0 |
| 0 | 0.00 | 3.68 | 0.00 | 1.00 | NA | Gm3076 | | 1 | 0 | 0 | 0 | 0 | 0 | 0 | 0 | 0 | 0 | 0 |
| 0 | 0.00 | 3.68 | 0.00 | 1.00 | NA | Gm11810 | | 1 | 0 | 0 | 0 | 0 | 0 | 0 | 0 | 0 | 0 | 0 |
| 0 | 0.00 | 3.68 | 0.00 | 1.00 | NA | Mup14 | major urinary protein 14 | 1 | 0 | 0 | 0 | 0 | 0 | 0 | 0 | 0 | 0 | 0 |
| 0 | 0.00 | 3.68 | 0.00 | 1.00 | NA | Cdcp2 | CUB domain containing protein 2 | 1 | 0 | 0 | 0 | 0 | 0 | 0 | 0 | 0 | 0 | 0 |
| 0 | 0.00 | 3.68 | 0.00 | 1.00 | NA | Gm12739 | | 1 | 0 | 0 | 0 | 0 | 0 | 0 | 0 | 0 | 0 | 0 |
| 0 | 0.00 | 3.68 | 0.00 | 1.00 | NA | Gm12947 | | 1 | 0 | 0 | 0 | 0 | 0 | 0 | 0 | 0 | 0 | 0 |
| 0 | 0.00 | 3.68 | 0.00 | 1.00 | NA | Gm29367 | | 1 | 0 | 0 | 0 | 0 | 0 | 0 | 0 | 0 | 0 | 0 |
| 0 | 0.00 | 3.68 | 0.00 | 1.00 | NA | Gm16188 | | 1 | 0 | 0 | 0 | 0 | 0 | 0 | 0 | 0 | 0 | 0 |
| 0 | 0.00 | 3.68 | 0.00 | 1.00 | NA | Gm30652 | | 1 | 0 | 0 | 0 | 0 | 0 | 0 | 0 | 0 | 0 | 0 |
| 0 | 0.00 | 3.68 | 0.00 | 1.00 | NA | Gm8906 | | 1 | 0 | 0 | 0 | 0 | 0 | 0 | 0 | 0 | 0 | 0 |
| 0 | 0.00 | 3.68 | 0.00 | 1.00 | NA | Gm15732 | | 1 | 0 | 0 | 0 | 0 | 0 | 0 | 0 | 0 | 0 | 0 |
| 0 | 0.00 | 3.68 | 0.00 | 1.00 | NA | Gm5298 | | 1 | 0 | 0 | 0 | 0 | 0 | 0 | 0 | 0 | 0 | 0 |
| 0 | 0.00 | 3.68 | 0.00 | 1.00 | NA | Gm42521 | | 1 | 0 | 0 | 0 | 0 | 0 | 0 | 0 | 0 | 0 | 0 |
| 0 | 0.00 | 3.68 | 0.00 | 1.00 | NA | Gm21461 | | 1 | 0 | 0 | 0 | 0 | 0 | 0 | 0 | 0 | 0 | 0 |
| 0 | 0.00 | 3.68 | 0.00 | 1.00 | NA | Gm7048 | | 1 | 0 | 0 | 0 | 0 | 0 | 0 | 0 | 0 | 0 | 0 |
| 0 | 0.00 | 3.68 | 0.00 | 1.00 | NA | Gm42627 | | 1 | 0 | 0 | 0 | 0 | 0 | 0 | 0 | 0 | 0 | 0 |
| 0 | 0.00 | 3.68 | 0.00 | 1.00 | NA | Gm42443 | | 1 | 0 | 0 | 0 | 0 | 0 | 0 | 0 | 0 | 0 | 0 |
| 0 | 0.00 | 3.68 | 0.00 | 1.00 | NA | Gm26970 | | 1 | 0 | 0 | 0 | 0 | 0 | 0 | 0 | 0 | 0 | 0 |

| | | | | | | | | | | | | | | | | | | |
|---|---|---|---|---|---|---|---|---|---|---|---|---|---|---|---|---|---|---|
| 0 | 0.00 | 3.68 | 0.00 | 1.00 | NA | Gm8675 | | 1 | 0 | 0 | 0 | 0 | 0 | 0 | 0 | 0 | 0 | 0 |
| 0 | 0.00 | 3.68 | 0.00 | 1.00 | NA | Gm5110 | | 1 | 0 | 0 | 0 | 0 | 0 | 0 | 0 | 0 | 0 | 0 |
| 0 | 0.00 | 3.68 | 0.00 | 1.00 | NA | 4930412F09Rik | | 1 | 0 | 0 | 0 | 0 | 0 | 0 | 0 | 0 | 0 | 0 |
| 0 | 0.00 | 3.68 | 0.00 | 1.00 | NA | Trbv13-1 | | 1 | 0 | 0 | 0 | 0 | 0 | 0 | 0 | 0 | 0 | 0 |
| 0 | 0.00 | 3.68 | 0.00 | 1.00 | NA | Trbv13-3 | | 1 | 0 | 0 | 0 | 0 | 0 | 0 | 0 | 0 | 0 | 0 |
| 0 | 0.00 | 3.68 | 0.00 | 1.00 | NA | Gm32479 | | 1 | 0 | 0 | 0 | 0 | 0 | 0 | 0 | 0 | 0 | 0 |
| 0 | 0.00 | 3.68 | 0.00 | 1.00 | NA | Igkv6-13 | | 1 | 0 | 0 | 0 | 0 | 0 | 0 | 0 | 0 | 0 | 0 |
| 0 | 0.00 | 3.68 | 0.00 | 1.00 | NA | Gm29781 | | 1 | 0 | 0 | 0 | 0 | 0 | 0 | 0 | 0 | 0 | 0 |
| 0 | 0.00 | 3.68 | 0.00 | 1.00 | NA | Gm26728 | | 1 | 0 | 0 | 0 | 0 | 0 | 0 | 0 | 0 | 0 | 0 |
| 0 | 0.00 | 3.68 | 0.00 | 1.00 | NA | Zbed4-ps1 | | 1 | 0 | 0 | 0 | 0 | 0 | 0 | 0 | 0 | 0 | 0 |
| 0 | 0.00 | 3.68 | 0.00 | 1.00 | NA | Gm44971 | | 1 | 0 | 0 | 0 | 0 | 0 | 0 | 0 | 0 | 0 | 0 |
| 0 | 0.00 | 3.68 | 0.00 | 1.00 | NA | Gm44640 | | 1 | 0 | 0 | 0 | 0 | 0 | 0 | 0 | 0 | 0 | 0 |
| 0 | 0.00 | 3.68 | 0.00 | 1.00 | NA | Gm9142 | | 1 | 0 | 0 | 0 | 0 | 0 | 0 | 0 | 0 | 0 | 0 |
| 0 | 0.00 | 3.68 | 0.00 | 1.00 | NA | Gm12755 | | 1 | 0 | 0 | 0 | 0 | 0 | 0 | 0 | 0 | 0 | 0 |
| 0 | 0.00 | 3.68 | 0.00 | 1.00 | NA | Gm8463 | | 1 | 0 | 0 | 0 | 0 | 0 | 0 | 0 | 0 | 0 | 0 |
| 0 | 0.00 | 3.68 | 0.00 | 1.00 | NA | Gm18561 | | 1 | 0 | 0 | 0 | 0 | 0 | 0 | 0 | 0 | 0 | 0 |
| 0 | 0.00 | 3.68 | 0.00 | 1.00 | NA | Gm45204 | | 1 | 0 | 0 | 0 | 0 | 0 | 0 | 0 | 0 | 0 | 0 |
| 0 | 0.00 | 3.68 | 0.00 | 1.00 | NA | Gm16477 | | 1 | 0 | 0 | 0 | 0 | 0 | 0 | 0 | 0 | 0 | 0 |
| 0 | 0.00 | 3.68 | 0.00 | 1.00 | NA | Gm45280 | | 1 | 0 | 0 | 0 | 0 | 0 | 0 | 0 | 0 | 0 | 0 |
| 0 | 0.00 | 3.68 | 0.00 | 1.00 | NA | Gm45049 | | 1 | 0 | 0 | 0 | 0 | 0 | 0 | 0 | 0 | 0 | 0 |
| 0 | 0.00 | 3.68 | 0.00 | 1.00 | NA | Gm6070 | | 1 | 0 | 0 | 0 | 0 | 0 | 0 | 0 | 0 | 0 | 0 |
| 0 | 0.00 | 3.68 | 0.00 | 1.00 | NA | Gm7760 | | 1 | 0 | 0 | 0 | 0 | 0 | 0 | 0 | 0 | 0 | 0 |
| 0 | 0.00 | 3.68 | 0.00 | 1.00 | NA | Gm45470 | | 1 | 0 | 0 | 0 | 0 | 0 | 0 | 0 | 0 | 0 | 0 |
| 0 | 0.00 | 3.68 | 0.00 | 1.00 | NA | 5430403N17Rik | | 1 | 0 | 0 | 0 | 0 | 0 | 0 | 0 | 0 | 0 | 0 |
| 0 | 0.00 | 3.68 | 0.00 | 1.00 | NA | Gm7082 | | 1 | 0 | 0 | 0 | 0 | 0 | 0 | 0 | 0 | 0 | 0 |
| 0 | 0.00 | 3.68 | 0.00 | 1.00 | NA | Gm45422 | | 1 | 0 | 0 | 0 | 0 | 0 | 0 | 0 | 0 | 0 | 0 |
| 0 | 0.00 | 3.68 | 0.00 | 1.00 | NA | Gm45860 | | 1 | 0 | 0 | 0 | 0 | 0 | 0 | 0 | 0 | 0 | 0 |
| 0 | 0.00 | 3.68 | 0.00 | 1.00 | NA | Gm16071 | | 1 | 0 | 0 | 0 | 0 | 0 | 0 | 0 | 0 | 0 | 0 |
| 0 | 0.00 | 3.68 | 0.00 | 1.00 | NA | Gm48114 | | 1 | 0 | 0 | 0 | 0 | 0 | 0 | 0 | 0 | 0 | 0 |
| 0 | 0.00 | 3.68 | 0.00 | 1.00 | NA | Foxr1 | forkhead box R1 | 1 | 0 | 0 | 0 | 0 | 0 | 0 | 0 | 0 | 0 | 0 |
| 0 | 0.00 | 3.68 | 0.00 | 1.00 | NA | Sln | sarcolipin | 1 | 0 | 0 | 0 | 0 | 0 | 0 | 0 | 0 | 0 | 0 |
| 0 | 0.00 | 3.68 | 0.00 | 1.00 | NA | Gm20620 | | 1 | 0 | 0 | 0 | 0 | 0 | 0 | 0 | 0 | 0 | 0 |
| 0 | 0.00 | 3.68 | 0.00 | 1.00 | NA | Gm7613 | | 1 | 0 | 0 | 0 | 0 | 0 | 0 | 0 | 0 | 0 | 0 |
| 0 | 0.00 | 3.68 | 0.00 | 1.00 | NA | Gm33699 | | 1 | 0 | 0 | 0 | 0 | 0 | 0 | 0 | 0 | 0 | 0 |
| 0 | 0.00 | 3.68 | 0.00 | 1.00 | NA | Mir3109 | | 1 | 0 | 0 | 0 | 0 | 0 | 0 | 0 | 0 | 0 | 0 |
| 0 | 0.00 | 3.68 | 0.00 | 1.00 | NA | Gm5745 | | 1 | 0 | 0 | 0 | 0 | 0 | 0 | 0 | 0 | 0 | 0 |
| 0 | 0.00 | 3.68 | 0.00 | 1.00 | NA | Gm29562 | | 1 | 0 | 0 | 0 | 0 | 0 | 0 | 0 | 0 | 0 | 0 |
| 0 | 0.00 | 3.68 | 0.00 | 1.00 | NA | Arpp21 | cyclic AMP-regulated phosphoprotein, 21 | 1 | 0 | 0 | 0 | 0 | 0 | 0 | 0 | 0 | 0 | 0 |
| 0 | 0.00 | 3.68 | 0.00 | 1.00 | NA | Gm29825 | predicted gene, 29825 | 1 | 0 | 0 | 0 | 0 | 0 | 0 | 0 | 0 | 0 | 0 |
| 0 | 0.00 | 3.68 | 0.00 | 1.00 | NA | Gm9535 | | 1 | 0 | 0 | 0 | 0 | 0 | 0 | 0 | 0 | 0 | 0 |
| 0 | 0.00 | 3.68 | 0.00 | 1.00 | NA | Tpi-rs5 | | 1 | 0 | 0 | 0 | 0 | 0 | 0 | 0 | 0 | 0 | 0 |
| 0 | 0.00 | 3.68 | 0.00 | 1.00 | NA | Gm47940 | | 1 | 0 | 0 | 0 | 0 | 0 | 0 | 0 | 0 | 0 | 0 |
| 0 | 0.00 | 3.68 | 0.00 | 1.00 | NA | Gm7001 | | 1 | 0 | 0 | 0 | 0 | 0 | 0 | 0 | 0 | 0 | 0 |
| 0 | 0.00 | 3.68 | 0.00 | 1.00 | NA | Gm10118 | | 1 | 0 | 0 | 0 | 0 | 0 | 0 | 0 | 0 | 0 | 0 |
| 0 | 0.00 | 3.68 | 0.00 | 1.00 | NA | Dnmt3l | DNA (cytosine-5-)-methyltransferase 3-lik | 1 | 0 | 0 | 0 | 0 | 0 | 0 | 0 | 0 | 0 | 0 |
| 0 | 0.00 | 3.68 | 0.00 | 1.00 | NA | Tyms-ps | | 1 | 0 | 0 | 0 | 0 | 0 | 0 | 0 | 0 | 0 | 0 |
| 0 | 0.00 | 3.68 | 0.00 | 1.00 | NA | Gm48085 | | 1 | 0 | 0 | 0 | 0 | 0 | 0 | 0 | 0 | 0 | 0 |
| 0 | 0.00 | 3.68 | 0.00 | 1.00 | NA | Gm12198 | | 1 | 0 | 0 | 0 | 0 | 0 | 0 | 0 | 0 | 0 | 0 |
| 0 | 0.00 | 3.68 | 0.00 | 1.00 | NA | Gm12219 | | 1 | 0 | 0 | 0 | 0 | 0 | 0 | 0 | 0 | 0 | 0 |
| 0 | 0.00 | 3.68 | 0.00 | 1.00 | NA | Gm12298 | | 1 | 0 | 0 | 0 | 0 | 0 | 0 | 0 | 0 | 0 | 0 |
| 0 | 0.00 | 3.68 | 0.00 | 1.00 | NA | Gm12331 | | 1 | 0 | 0 | 0 | 0 | 0 | 0 | 0 | 0 | 0 | 0 |
| 0 | 0.00 | 3.68 | 0.00 | 1.00 | NA | Gm11533 | | 1 | 0 | 0 | 0 | 0 | 0 | 0 | 0 | 0 | 0 | 0 |
| 0 | 0.00 | 3.68 | 0.00 | 1.00 | NA | 6030469F06Rik | | 1 | 0 | 0 | 0 | 0 | 0 | 0 | 0 | 0 | 0 | 0 |
| 0 | 0.00 | 3.68 | 0.00 | 1.00 | NA | Gm17056 | | 1 | 0 | 0 | 0 | 0 | 0 | 0 | 0 | 0 | 0 | 0 |
| 0 | 0.00 | 3.68 | 0.00 | 1.00 | NA | Gm30738 | | 1 | 0 | 0 | 0 | 0 | 0 | 0 | 0 | 0 | 0 | 0 |
| 0 | 0.00 | 3.68 | 0.00 | 1.00 | NA | Gm15745 | | 1 | 0 | 0 | 0 | 0 | 0 | 0 | 0 | 0 | 0 | 0 |
| 0 | 0.00 | 3.68 | 0.00 | 1.00 | NA | Gm26512 | | 1 | 0 | 0 | 0 | 0 | 0 | 0 | 0 | 0 | 0 | 0 |
| 0 | 0.00 | 3.68 | 0.00 | 1.00 | NA | Ighv5-9-1 | | 1 | 0 | 0 | 0 | 0 | 0 | 0 | 0 | 0 | 0 | 0 |
| 0 | 0.00 | 3.68 | 0.00 | 1.00 | NA | Ighv5-18 | | 1 | 0 | 0 | 0 | 0 | 0 | 0 | 0 | 0 | 0 | 0 |
| 0 | 0.00 | 3.68 | 0.00 | 1.00 | NA | Ighv3-4 | | 1 | 0 | 0 | 0 | 0 | 0 | 0 | 0 | 0 | 0 | 0 |

| | | | | | | | | | | | | | | | | | | |
|---|---|---|---|---|---|---|---|---|---|---|---|---|---|---|---|---|---|---|
| 0 | 0.00 | 3.68 | 0.00 | 1.00 | NA | Ighv1-31 | | 1 | 0 | 0 | 0 | 0 | 0 | 0 | 0 | 0 | 0 | 0 |
| 0 | 0.00 | 3.68 | 0.00 | 1.00 | NA | Gm48620 | | 1 | 0 | 0 | 0 | 0 | 0 | 0 | 0 | 0 | 0 | 0 |
| 0 | 0.00 | 3.68 | 0.00 | 1.00 | NA | Gm48544 | | 1 | 0 | 0 | 0 | 0 | 0 | 0 | 0 | 0 | 0 | 0 |
| 0 | 0.00 | 3.68 | 0.00 | 1.00 | NA | Serpinb1c | serine (or cysteine) peptidase inhibitor, cl | 1 | 0 | 0 | 0 | 0 | 0 | 0 | 0 | 0 | 0 | 0 |
| 0 | 0.00 | 3.68 | 0.00 | 1.00 | NA | Gm47662 | | 1 | 0 | 0 | 0 | 0 | 0 | 0 | 0 | 0 | 0 | 0 |
| 0 | 0.00 | 3.68 | 0.00 | 1.00 | NA | Gm48629 | | 1 | 0 | 0 | 0 | 0 | 0 | 0 | 0 | 0 | 0 | 0 |
| 0 | 0.00 | 3.68 | 0.00 | 1.00 | NA | Gm30177 | | 1 | 0 | 0 | 0 | 0 | 0 | 0 | 0 | 0 | 0 | 0 |
| 0 | 0.00 | 3.68 | 0.00 | 1.00 | NA | Gm46415 | predicted gene, 46415 | 1 | 0 | 0 | 0 | 0 | 0 | 0 | 0 | 0 | 0 | 0 |
| 0 | 0.00 | 3.68 | 0.00 | 1.00 | NA | Gm3333 | | 1 | 0 | 0 | 0 | 0 | 0 | 0 | 0 | 0 | 0 | 0 |
| 0 | 0.00 | 3.68 | 0.00 | 1.00 | NA | Gm24588 | | 1 | 0 | 0 | 0 | 0 | 0 | 0 | 0 | 0 | 0 | 0 |
| 0 | 0.00 | 3.68 | 0.00 | 1.00 | NA | Gm47706 | | 1 | 0 | 0 | 0 | 0 | 0 | 0 | 0 | 0 | 0 | 0 |
| 0 | 0.00 | 3.68 | 0.00 | 1.00 | NA | Gm46396 | | 1 | 0 | 0 | 0 | 0 | 0 | 0 | 0 | 0 | 0 | 0 |
| 0 | 0.00 | 3.68 | 0.00 | 1.00 | NA | Gm32376 | | 1 | 0 | 0 | 0 | 0 | 0 | 0 | 0 | 0 | 0 | 0 |
| 0 | 0.00 | 3.68 | 0.00 | 1.00 | NA | Esd-ps | | 1 | 0 | 0 | 0 | 0 | 0 | 0 | 0 | 0 | 0 | 0 |
| 0 | 0.00 | 3.68 | 0.00 | 1.00 | NA | Rpl31-ps3 | | 1 | 0 | 0 | 0 | 0 | 0 | 0 | 0 | 0 | 0 | 0 |
| 0 | 0.00 | 3.68 | 0.00 | 1.00 | NA | Gm46448 | | 1 | 0 | 0 | 0 | 0 | 0 | 0 | 0 | 0 | 0 | 0 |
| 0 | 0.00 | 3.68 | 0.00 | 1.00 | NA | Gm8417 | | 1 | 0 | 0 | 0 | 0 | 0 | 0 | 0 | 0 | 0 | 0 |
| 0 | 0.00 | 3.68 | 0.00 | 1.00 | NA | Gm48934 | | 1 | 0 | 0 | 0 | 0 | 0 | 0 | 0 | 0 | 0 | 0 |
| 0 | 0.00 | 3.68 | 0.00 | 1.00 | NA | Gm2245 | | 1 | 0 | 0 | 0 | 0 | 0 | 0 | 0 | 0 | 0 | 0 |
| 0 | 0.00 | 3.68 | 0.00 | 1.00 | NA | Gm8238 | | 1 | 0 | 0 | 0 | 0 | 0 | 0 | 0 | 0 | 0 | 0 |
| 0 | 0.00 | 3.68 | 0.00 | 1.00 | NA | Gm49255 | | 1 | 0 | 0 | 0 | 0 | 0 | 0 | 0 | 0 | 0 | 0 |
| 0 | 0.00 | 3.68 | 0.00 | 1.00 | NA | Gm18152 | | 1 | 0 | 0 | 0 | 0 | 0 | 0 | 0 | 0 | 0 | 0 |
| 0 | 0.00 | 3.68 | 0.00 | 1.00 | NA | Gm19277 | | 1 | 0 | 0 | 0 | 0 | 0 | 0 | 0 | 0 | 0 | 0 |
| 0 | 0.00 | 3.68 | 0.00 | 1.00 | NA | Gm15859 | | 1 | 0 | 0 | 0 | 0 | 0 | 0 | 0 | 0 | 0 | 0 |
| 0 | 0.00 | 3.68 | 0.00 | 1.00 | NA | Gm7572 | | 1 | 0 | 0 | 0 | 0 | 0 | 0 | 0 | 0 | 0 | 0 |
| 0 | 0.00 | 3.68 | 0.00 | 1.00 | NA | Gm19571 | | 1 | 0 | 0 | 0 | 0 | 0 | 0 | 0 | 0 | 0 | 0 |
| 0 | 0.00 | 3.68 | 0.00 | 1.00 | NA | 4930542D17Rik | | 1 | 0 | 0 | 0 | 0 | 0 | 0 | 0 | 0 | 0 | 0 |
| 0 | 0.00 | 3.68 | 0.00 | 1.00 | NA | Gm49587 | | 1 | 0 | 0 | 0 | 0 | 0 | 0 | 0 | 0 | 0 | 0 |
| 0 | 0.00 | 3.68 | 0.00 | 1.00 | NA | Gm6032 | | 1 | 0 | 0 | 0 | 0 | 0 | 0 | 0 | 0 | 0 | 0 |
| 0 | 0.00 | 3.68 | 0.00 | 1.00 | NA | Gm49226 | | 1 | 0 | 0 | 0 | 0 | 0 | 0 | 0 | 0 | 0 | 0 |
| 0 | 0.00 | 3.68 | 0.00 | 1.00 | NA | Gm49689 | | 1 | 0 | 0 | 0 | 0 | 0 | 0 | 0 | 0 | 0 | 0 |
| 0 | 0.00 | 3.68 | 0.00 | 1.00 | NA | Gm7856 | | 1 | 0 | 0 | 0 | 0 | 0 | 0 | 0 | 0 | 0 | 0 |
| 0 | 0.00 | 3.68 | 0.00 | 1.00 | NA | Gm49795 | | 1 | 0 | 0 | 0 | 0 | 0 | 0 | 0 | 0 | 0 | 0 |
| 0 | 0.00 | 3.68 | 0.00 | 1.00 | NA | Gm17377 | | 1 | 0 | 0 | 0 | 0 | 0 | 0 | 0 | 0 | 0 | 0 |
| 0 | 0.00 | 3.68 | 0.00 | 1.00 | NA | Gm24813 | | 1 | 0 | 0 | 0 | 0 | 0 | 0 | 0 | 0 | 0 | 0 |
| 0 | 0.00 | 3.68 | 0.00 | 1.00 | NA | Gm49925 | | 1 | 0 | 0 | 0 | 0 | 0 | 0 | 0 | 0 | 0 | 0 |
| 0 | 0.00 | 3.68 | 0.00 | 1.00 | NA | Gm6291 | | 1 | 0 | 0 | 0 | 0 | 0 | 0 | 0 | 0 | 0 | 0 |
| 0 | 0.00 | 3.68 | 0.00 | 1.00 | NA | Gm7936 | | 1 | 0 | 0 | 0 | 0 | 0 | 0 | 0 | 0 | 0 | 0 |
| 0 | 0.00 | 3.68 | 0.00 | 1.00 | NA | Gm6293 | | 1 | 0 | 0 | 0 | 0 | 0 | 0 | 0 | 0 | 0 | 0 |
| 0 | 0.00 | 3.68 | 0.00 | 1.00 | NA | Keg1 | kidney expressed gene 1 | 1 | 0 | 0 | 0 | 0 | 0 | 0 | 0 | 0 | 0 | 0 |
| 0 | 0.00 | 3.68 | 0.00 | 1.00 | NA | Gm50264 | | 1 | 0 | 0 | 0 | 0 | 0 | 0 | 0 | 0 | 0 | 0 |
| 0 | 0.00 | 3.68 | 0.00 | 1.00 | NA | Gm32027 | | 1 | 0 | 0 | 0 | 0 | 0 | 0 | 0 | 0 | 0 | 0 |
| 0 | 0.00 | 3.68 | 0.00 | 1.00 | NA | Gm14538 | | 1 | 0 | 0 | 0 | 0 | 0 | 0 | 0 | 0 | 0 | 0 |
| 0 | 0.00 | 3.68 | 0.00 | 1.00 | NA | Gm14565 | | 1 | 0 | 0 | 0 | 0 | 0 | 0 | 0 | 0 | 0 | 0 |
| 0 | 0.00 | 3.68 | 0.00 | 1.00 | NA | Gm7189 | | 1 | 0 | 0 | 0 | 0 | 0 | 0 | 0 | 0 | 0 | 0 |
| 0 | 0.00 | 3.68 | 0.00 | 1.00 | NA | Gm7190 | | 1 | 0 | 0 | 0 | 0 | 0 | 0 | 0 | 0 | 0 | 0 |
| 0 | 0.00 | 3.68 | 0.00 | 1.00 | NA | Gm14576 | | 1 | 0 | 0 | 0 | 0 | 0 | 0 | 0 | 0 | 0 | 0 |
| 0 | 0.00 | 3.68 | 0.00 | 1.00 | NA | Gm7927 | | 1 | 0 | 0 | 0 | 0 | 0 | 0 | 0 | 0 | 0 | 0 |
| 0 | 0.00 | 3.68 | 0.00 | 1.00 | NA | Gm14745 | | 1 | 0 | 0 | 0 | 0 | 0 | 0 | 0 | 0 | 0 | 0 |
| 0 | 0.00 | 3.68 | 0.00 | 1.00 | NA | Gm16465 | | 1 | 0 | 0 | 0 | 0 | 0 | 0 | 0 | 0 | 0 | 0 |
| 0 | 0.00 | 3.68 | 0.00 | 1.00 | NA | Gm5760 | | 1 | 0 | 0 | 0 | 0 | 0 | 0 | 0 | 0 | 0 | 0 |
| 0 | 0.00 | 3.68 | 0.00 | 1.00 | NA | Gm14927 | | 1 | 0 | 0 | 0 | 0 | 0 | 0 | 0 | 0 | 0 | 0 |
| 0 | 0.00 | 3.68 | 0.00 | 1.00 | NA | Gm5648 | | 1 | 0 | 0 | 0 | 0 | 0 | 0 | 0 | 0 | 0 | 0 |
| 0 | 0.00 | 3.68 | 0.00 | 1.00 | NA | Gm23806 | | 1 | 0 | 0 | 0 | 0 | 0 | 0 | 0 | 0 | 0 | 0 |
| 0 | -0.40 | 3.67 | -0.11 | 0.91 | NA | Gm37079 | | 0 | 0 | 0 | 0 | 0 | 1 | 0 | 0 | 0 | 0 | 0 |
| 0 | -0.40 | 3.67 | -0.11 | 0.91 | NA | Gm38005 | | 0 | 0 | 0 | 0 | 0 | 1 | 0 | 0 | 0 | 0 | 0 |
| 0 | -0.40 | 3.67 | -0.11 | 0.91 | NA | Gm37702 | | 0 | 0 | 0 | 0 | 0 | 1 | 0 | 0 | 0 | 0 | 0 |
| 0 | -0.40 | 3.67 | -0.11 | 0.91 | NA | Gm36952 | | 0 | 0 | 0 | 0 | 0 | 1 | 0 | 0 | 0 | 0 | 0 |
| 0 | -0.40 | 3.67 | -0.11 | 0.91 | NA | Gm37905 | | 0 | 0 | 0 | 0 | 0 | 1 | 0 | 0 | 0 | 0 | 0 |
| 0 | -0.40 | 3.67 | -0.11 | 0.91 | NA | Gm19863 | | 0 | 0 | 0 | 0 | 0 | 1 | 0 | 0 | 0 | 0 | 0 |

| | | | | | | | | | | | | | | | | | | |
|---|---|---|---|---|---|---|---|---|---|---|---|---|---|---|---|---|---|---|
| 0 | -0.40 | 3.67 | -0.11 | 0.91 | NA | Gm18303 | | 0 | 0 | 0 | 0 | 0 | 1 | 0 | 0 | 0 | 0 | 0 |
| 0 | -0.40 | 3.67 | -0.11 | 0.91 | NA | Gm28320 | | 0 | 0 | 0 | 0 | 0 | 1 | 0 | 0 | 0 | 0 | 0 |
| 0 | -0.40 | 3.67 | -0.11 | 0.91 | NA | Gm25839 | | 0 | 0 | 0 | 0 | 0 | 1 | 0 | 0 | 0 | 0 | 0 |
| 0 | -0.40 | 3.67 | -0.11 | 0.91 | NA | Gm15789 | | 0 | 0 | 0 | 0 | 0 | 1 | 0 | 0 | 0 | 0 | 0 |
| 0 | -0.40 | 3.67 | -0.11 | 0.91 | NA | Gm8840 | | 0 | 0 | 0 | 0 | 0 | 1 | 0 | 0 | 0 | 0 | 0 |
| 0 | -0.40 | 3.67 | -0.11 | 0.91 | NA | Gm28112 | | 0 | 0 | 0 | 0 | 0 | 1 | 0 | 0 | 0 | 0 | 0 |
| 0 | -0.40 | 3.67 | -0.11 | 0.91 | NA | Apol7d | | 0 | 0 | 0 | 0 | 0 | 1 | 0 | 0 | 0 | 0 | 0 |
| 0 | -0.40 | 3.67 | -0.11 | 0.91 | NA | Mir26b | | 0 | 0 | 0 | 0 | 0 | 1 | 0 | 0 | 0 | 0 | 0 |
| 0 | -0.40 | 3.67 | -0.11 | 0.91 | NA | A630095N17Ri | RIKEN cDNA A630095N17 gene | 0 | 0 | 0 | 0 | 0 | 1 | 0 | 0 | 0 | 0 | 0 |
| 0 | -0.40 | 3.67 | -0.11 | 0.91 | NA | Gm37485 | | 0 | 0 | 0 | 0 | 0 | 1 | 0 | 0 | 0 | 0 | 0 |
| 0 | -0.40 | 3.67 | -0.11 | 0.91 | NA | Gm15375 | | 0 | 0 | 0 | 0 | 0 | 1 | 0 | 0 | 0 | 0 | 0 |
| 0 | -0.40 | 3.67 | -0.11 | 0.91 | NA | 4930440C22Rik | | 0 | 0 | 0 | 0 | 0 | 1 | 0 | 0 | 0 | 0 | 0 |
| 0 | -0.40 | 3.67 | -0.11 | 0.91 | NA | Gm19029 | | 0 | 0 | 0 | 0 | 0 | 1 | 0 | 0 | 0 | 0 | 0 |
| 0 | -0.40 | 3.67 | -0.11 | 0.91 | NA | 3830432H09Rik | | 0 | 0 | 0 | 0 | 0 | 1 | 0 | 0 | 0 | 0 | 0 |
| 0 | -0.40 | 3.67 | -0.11 | 0.91 | NA | C1ql2 | complement component 1, q subcompone | 0 | 0 | 0 | 0 | 0 | 1 | 0 | 0 | 0 | 0 | 0 |
| 0 | -0.40 | 3.67 | -0.11 | 0.91 | NA | 2610027F03Rik | | 0 | 0 | 0 | 0 | 0 | 1 | 0 | 0 | 0 | 0 | 0 |
| 0 | -0.40 | 3.67 | -0.11 | 0.91 | NA | Gm7241 | | 0 | 0 | 0 | 0 | 0 | 1 | 0 | 0 | 0 | 0 | 0 |
| 0 | -0.40 | 3.67 | -0.11 | 0.91 | NA | Gm8941 | | 0 | 0 | 0 | 0 | 0 | 1 | 0 | 0 | 0 | 0 | 0 |
| 0 | -0.40 | 3.67 | -0.11 | 0.91 | NA | Gm38256 | | 0 | 0 | 0 | 0 | 0 | 1 | 0 | 0 | 0 | 0 | 0 |
| 0 | -0.40 | 3.67 | -0.11 | 0.91 | NA | Scarna3a | | 0 | 0 | 0 | 0 | 0 | 1 | 0 | 0 | 0 | 0 | 0 |
| 0 | -0.40 | 3.67 | -0.11 | 0.91 | NA | Gm37731 | | 0 | 0 | 0 | 0 | 0 | 1 | 0 | 0 | 0 | 0 | 0 |
| 0 | -0.40 | 3.67 | -0.11 | 0.91 | NA | Gm37729 | | 0 | 0 | 0 | 0 | 0 | 1 | 0 | 0 | 0 | 0 | 0 |
| 0 | -0.40 | 3.67 | -0.11 | 0.91 | NA | Gm23208 | | 0 | 0 | 0 | 0 | 0 | 1 | 0 | 0 | 0 | 0 | 0 |
| 0 | -0.40 | 3.67 | -0.11 | 0.91 | NA | Gm36992 | | 0 | 0 | 0 | 0 | 0 | 1 | 0 | 0 | 0 | 0 | 0 |
| 0 | -0.40 | 3.67 | -0.11 | 0.91 | NA | Gm38146 | | 0 | 0 | 0 | 0 | 0 | 1 | 0 | 0 | 0 | 0 | 0 |
| 0 | -0.40 | 3.67 | -0.11 | 0.91 | NA | Gm26004 | | 0 | 0 | 0 | 0 | 0 | 1 | 0 | 0 | 0 | 0 | 0 |
| 0 | -0.40 | 3.67 | -0.11 | 0.91 | NA | Gm38315 | | 0 | 0 | 0 | 0 | 0 | 1 | 0 | 0 | 0 | 0 | 0 |
| 0 | -0.40 | 3.67 | -0.11 | 0.91 | NA | Gm8407 | | 0 | 0 | 0 | 0 | 0 | 1 | 0 | 0 | 0 | 0 | 0 |
| 0 | -0.40 | 3.67 | -0.11 | 0.91 | NA | Gm38036 | | 0 | 0 | 0 | 0 | 0 | 1 | 0 | 0 | 0 | 0 | 0 |
| 0 | -0.40 | 3.67 | -0.11 | 0.91 | NA | Gm13188 | | 0 | 0 | 0 | 0 | 0 | 1 | 0 | 0 | 0 | 0 | 0 |
| 0 | -0.40 | 3.67 | -0.11 | 0.91 | NA | Gm36988 | | 0 | 0 | 0 | 0 | 0 | 1 | 0 | 0 | 0 | 0 | 0 |
| 0 | -0.40 | 3.67 | -0.11 | 0.91 | NA | Gm13391 | | 0 | 0 | 0 | 0 | 0 | 1 | 0 | 0 | 0 | 0 | 0 |
| 0 | -0.40 | 3.67 | -0.11 | 0.91 | NA | Gm13312 | | 0 | 0 | 0 | 0 | 0 | 1 | 0 | 0 | 0 | 0 | 0 |
| 0 | -0.40 | 3.67 | -0.11 | 0.91 | NA | Gm38273 | | 0 | 0 | 0 | 0 | 0 | 1 | 0 | 0 | 0 | 0 | 0 |
| 0 | -0.40 | 3.67 | -0.11 | 0.91 | NA | Gm14471 | | 0 | 0 | 0 | 0 | 0 | 1 | 0 | 0 | 0 | 0 | 0 |
| 0 | -0.40 | 3.67 | -0.11 | 0.91 | NA | Gm37651 | | 0 | 0 | 0 | 0 | 0 | 1 | 0 | 0 | 0 | 0 | 0 |
| 0 | -0.40 | 3.67 | -0.11 | 0.91 | NA | Gm11084 | | 0 | 0 | 0 | 0 | 0 | 1 | 0 | 0 | 0 | 0 | 0 |
| 0 | -0.40 | 3.67 | -0.11 | 0.91 | NA | BC016548 | | 0 | 0 | 0 | 0 | 0 | 1 | 0 | 0 | 0 | 0 | 0 |
| 0 | -0.40 | 3.67 | -0.11 | 0.91 | NA | 4931422A03Ril | RIKEN cDNA 4931422A03 gene | 0 | 0 | 0 | 0 | 0 | 1 | 0 | 0 | 0 | 0 | 0 |
| 0 | -0.40 | 3.67 | -0.11 | 0.91 | NA | Gm13890 | | 0 | 0 | 0 | 0 | 0 | 1 | 0 | 0 | 0 | 0 | 0 |
| 0 | -0.40 | 3.67 | -0.11 | 0.91 | NA | Gm24115 | | 0 | 0 | 0 | 0 | 0 | 1 | 0 | 0 | 0 | 0 | 0 |
| 0 | -0.40 | 3.67 | -0.11 | 0.91 | NA | Snord110 | | 0 | 0 | 0 | 0 | 0 | 1 | 0 | 0 | 0 | 0 | 0 |
| 0 | -0.40 | 3.67 | -0.11 | 0.91 | NA | Gm11037 | | 0 | 0 | 0 | 0 | 0 | 1 | 0 | 0 | 0 | 0 | 0 |
| 0 | -0.40 | 3.67 | -0.11 | 0.91 | NA | Gm14037 | | 0 | 0 | 0 | 0 | 0 | 1 | 0 | 0 | 0 | 0 | 0 |
| 0 | -0.40 | 3.67 | -0.11 | 0.91 | NA | Gm14239 | | 0 | 0 | 0 | 0 | 0 | 1 | 0 | 0 | 0 | 0 | 0 |
| 0 | -0.40 | 3.67 | -0.11 | 0.91 | NA | Gm23463 | | 0 | 0 | 0 | 0 | 0 | 1 | 0 | 0 | 0 | 0 | 0 |
| 0 | -0.40 | 3.67 | -0.11 | 0.91 | NA | Wfdc6b | WAP four-disulfide core domain 6B | 0 | 0 | 0 | 0 | 0 | 1 | 0 | 0 | 0 | 0 | 0 |
| 0 | -0.40 | 3.67 | -0.11 | 0.91 | NA | Gm22355 | | 0 | 0 | 0 | 0 | 0 | 1 | 0 | 0 | 0 | 0 | 0 |
| 0 | -0.40 | 3.67 | -0.11 | 0.91 | NA | Spo11 | SPO11 meiotic protein covalently bound t | 0 | 0 | 0 | 0 | 0 | 1 | 0 | 0 | 0 | 0 | 0 |
| 0 | -0.40 | 3.67 | -0.11 | 0.91 | NA | Mir7006 | | 0 | 0 | 0 | 0 | 0 | 1 | 0 | 0 | 0 | 0 | 0 |
| 0 | -0.40 | 3.67 | -0.11 | 0.91 | NA | Gm16399 | | 0 | 0 | 0 | 0 | 0 | 1 | 0 | 0 | 0 | 0 | 0 |
| 0 | -0.40 | 3.67 | -0.11 | 0.91 | NA | Gm22282 | | 0 | 0 | 0 | 0 | 0 | 1 | 0 | 0 | 0 | 0 | 0 |
| 0 | -0.40 | 3.67 | -0.11 | 0.91 | NA | Gm37191 | | 0 | 0 | 0 | 0 | 0 | 1 | 0 | 0 | 0 | 0 | 0 |
| 0 | -0.40 | 3.67 | -0.11 | 0.91 | NA | Gm7733 | | 0 | 0 | 0 | 0 | 0 | 1 | 0 | 0 | 0 | 0 | 0 |
| 0 | -0.40 | 3.67 | -0.11 | 0.91 | NA | Gm32340 | | 0 | 0 | 0 | 0 | 0 | 1 | 0 | 0 | 0 | 0 | 0 |
| 0 | -0.40 | 3.67 | -0.11 | 0.91 | NA | Gm26671 | | 0 | 0 | 0 | 0 | 0 | 1 | 0 | 0 | 0 | 0 | 0 |
| 0 | -0.40 | 3.67 | -0.11 | 0.91 | NA | Gm20657 | | 0 | 0 | 0 | 0 | 0 | 1 | 0 | 0 | 0 | 0 | 0 |
| 0 | -0.40 | 3.67 | -0.11 | 0.91 | NA | Gm20691 | | 0 | 0 | 0 | 0 | 0 | 1 | 0 | 0 | 0 | 0 | 0 |
| 0 | -0.40 | 3.67 | -0.11 | 0.91 | NA | Zbbx | zinc finger, B-box domain containing | 0 | 0 | 0 | 0 | 0 | 1 | 0 | 0 | 0 | 0 | 0 |
| 0 | -0.40 | 3.67 | -0.11 | 0.91 | NA | Fga | fibrinogen alpha chain | 0 | 0 | 0 | 0 | 0 | 1 | 0 | 0 | 0 | 0 | 0 |

| | | | | | | | | | | | | | | | | | | |
|---|---|---|---|---|---|---|---|---|---|---|---|---|---|---|---|---|---|---|
| 0 | -0.40 | 3.67 | -0.11 | 0.91 | NA | Gm24584 | | 0 | 0 | 0 | 0 | 0 | 1 | 0 | 0 | 0 | 0 | 0 |
| 0 | -0.40 | 3.67 | -0.11 | 0.91 | NA | 4930565D16Rik | | 0 | 0 | 0 | 0 | 0 | 1 | 0 | 0 | 0 | 0 | 0 |
| 0 | -0.40 | 3.67 | -0.11 | 0.91 | NA | Gm23054 | | 0 | 0 | 0 | 0 | 0 | 1 | 0 | 0 | 0 | 0 | 0 |
| 0 | -0.40 | 3.67 | -0.11 | 0.91 | NA | Gm16561 | | 0 | 0 | 0 | 0 | 0 | 1 | 0 | 0 | 0 | 0 | 0 |
| 0 | -0.40 | 3.67 | -0.11 | 0.91 | NA | Gm26361 | | 0 | 0 | 0 | 0 | 0 | 1 | 0 | 0 | 0 | 0 | 0 |
| 0 | -0.40 | 3.67 | -0.11 | 0.91 | NA | Gm15471 | | 0 | 0 | 0 | 0 | 0 | 1 | 0 | 0 | 0 | 0 | 0 |
| 0 | -0.40 | 3.67 | -0.11 | 0.91 | NA | Gm42516 | | 0 | 0 | 0 | 0 | 0 | 1 | 0 | 0 | 0 | 0 | 0 |
| 0 | -0.40 | 3.67 | -0.11 | 0.91 | NA | Gm43651 | | 0 | 0 | 0 | 0 | 0 | 1 | 0 | 0 | 0 | 0 | 0 |
| 0 | -0.40 | 3.67 | -0.11 | 0.91 | NA | Gm9379 | | 0 | 0 | 0 | 0 | 0 | 1 | 0 | 0 | 0 | 0 | 0 |
| 0 | -0.40 | 3.67 | -0.11 | 0.91 | NA | Gm43765 | | 0 | 0 | 0 | 0 | 0 | 1 | 0 | 0 | 0 | 0 | 0 |
| 0 | -0.40 | 3.67 | -0.11 | 0.91 | NA | Gm35986 | | 0 | 0 | 0 | 0 | 0 | 1 | 0 | 0 | 0 | 0 | 0 |
| 0 | -0.40 | 3.67 | -0.11 | 0.91 | NA | 4930579F01Ri | RIKEN cDNA 4930579F01 gene | 0 | 0 | 0 | 0 | 0 | 1 | 0 | 0 | 0 | 0 | 0 |
| 0 | -0.40 | 3.67 | -0.11 | 0.91 | NA | Gm24260 | | 0 | 0 | 0 | 0 | 0 | 1 | 0 | 0 | 0 | 0 | 0 |
| 0 | -0.40 | 3.67 | -0.11 | 0.91 | NA | Gm18589 | | 0 | 0 | 0 | 0 | 0 | 1 | 0 | 0 | 0 | 0 | 0 |
| 0 | -0.40 | 3.67 | -0.11 | 0.91 | NA | Gm11787 | | 0 | 0 | 0 | 0 | 0 | 1 | 0 | 0 | 0 | 0 | 0 |
| 0 | -0.40 | 3.67 | -0.11 | 0.91 | NA | Mir3094 | | 0 | 0 | 0 | 0 | 0 | 1 | 0 | 0 | 0 | 0 | 0 |
| 0 | -0.40 | 3.67 | -0.11 | 0.91 | NA | Gm23257 | | 0 | 0 | 0 | 0 | 0 | 1 | 0 | 0 | 0 | 0 | 0 |
| 0 | -0.40 | 3.67 | -0.11 | 0.91 | NA | Mup6 | major urinary protein 6 | 0 | 0 | 0 | 0 | 0 | 1 | 0 | 0 | 0 | 0 | 0 |
| 0 | -0.40 | 3.67 | -0.11 | 0.91 | NA | Gm23148 | | 0 | 0 | 0 | 0 | 0 | 1 | 0 | 0 | 0 | 0 | 0 |
| 0 | -0.40 | 3.67 | -0.11 | 0.91 | NA | n-R5s189 | | 0 | 0 | 0 | 0 | 0 | 1 | 0 | 0 | 0 | 0 | 0 |
| 0 | -0.40 | 3.67 | -0.11 | 0.91 | NA | Gm12691 | | 0 | 0 | 0 | 0 | 0 | 1 | 0 | 0 | 0 | 0 | 0 |
| 0 | -0.40 | 3.67 | -0.11 | 0.91 | NA | Gm12706 | | 0 | 0 | 0 | 0 | 0 | 1 | 0 | 0 | 0 | 0 | 0 |
| 0 | -0.40 | 3.67 | -0.11 | 0.91 | NA | Lrp8os1 | | 0 | 0 | 0 | 0 | 0 | 1 | 0 | 0 | 0 | 0 | 0 |
| 0 | -0.40 | 3.67 | -0.11 | 0.91 | NA | Skint2 | selection and upkeep of intraepithelial T c | 0 | 0 | 0 | 0 | 0 | 1 | 0 | 0 | 0 | 0 | 0 |
| 0 | -0.40 | 3.67 | -0.11 | 0.91 | NA | Gm17114 | | 0 | 0 | 0 | 0 | 0 | 1 | 0 | 0 | 0 | 0 | 0 |
| 0 | -0.40 | 3.67 | -0.11 | 0.91 | NA | Gm12900 | | 0 | 0 | 0 | 0 | 0 | 1 | 0 | 0 | 0 | 0 | 0 |
| 0 | -0.40 | 3.67 | -0.11 | 0.91 | NA | Gm13022 | | 0 | 0 | 0 | 0 | 0 | 1 | 0 | 0 | 0 | 0 | 0 |
| 0 | -0.40 | 3.67 | -0.11 | 0.91 | NA | Tex46 | testis expressed 46 | 0 | 0 | 0 | 0 | 0 | 1 | 0 | 0 | 0 | 0 | 0 |
| 0 | -0.40 | 3.67 | -0.11 | 0.91 | NA | AI507597 | | 0 | 0 | 0 | 0 | 0 | 1 | 0 | 0 | 0 | 0 | 0 |
| 0 | -0.40 | 3.67 | -0.11 | 0.91 | NA | 6330411D24Rik | | 0 | 0 | 0 | 0 | 0 | 1 | 0 | 0 | 0 | 0 | 0 |
| 0 | -0.40 | 3.67 | -0.11 | 0.91 | NA | Zfp981 | zinc finger protein 981 | 0 | 0 | 0 | 0 | 0 | 1 | 0 | 0 | 0 | 0 | 0 |
| 0 | -0.40 | 3.67 | -0.11 | 0.91 | NA | Gm13201 | | 0 | 0 | 0 | 0 | 0 | 1 | 0 | 0 | 0 | 0 | 0 |
| 0 | -0.40 | 3.67 | -0.11 | 0.91 | NA | Rnf207 | ring finger protein 207 | 0 | 0 | 0 | 0 | 0 | 1 | 0 | 0 | 0 | 0 | 0 |
| 0 | -0.40 | 3.67 | -0.11 | 0.91 | NA | Gm43113 | | 0 | 0 | 0 | 0 | 0 | 1 | 0 | 0 | 0 | 0 | 0 |
| 0 | -0.40 | 3.67 | -0.11 | 0.91 | NA | Gm42597 | | 0 | 0 | 0 | 0 | 0 | 1 | 0 | 0 | 0 | 0 | 0 |
| 0 | -0.40 | 3.67 | -0.11 | 0.91 | NA | Gm43519 | | 0 | 0 | 0 | 0 | 0 | 1 | 0 | 0 | 0 | 0 | 0 |
| 0 | -0.40 | 3.67 | -0.11 | 0.91 | NA | Gm26954 | | 0 | 0 | 0 | 0 | 0 | 1 | 0 | 0 | 0 | 0 | 0 |
| 0 | -0.40 | 3.67 | -0.11 | 0.91 | NA | Gm10475 | | 0 | 0 | 0 | 0 | 0 | 1 | 0 | 0 | 0 | 0 | 0 |
| 0 | -0.40 | 3.67 | -0.11 | 0.91 | NA | Gm19666 | | 0 | 0 | 0 | 0 | 0 | 1 | 0 | 0 | 0 | 0 | 0 |
| 0 | -0.40 | 3.67 | -0.11 | 0.91 | NA | Gm50482 | | 0 | 0 | 0 | 0 | 0 | 1 | 0 | 0 | 0 | 0 | 0 |
| 0 | -0.40 | 3.67 | -0.11 | 0.91 | NA | Gm43850 | | 0 | 0 | 0 | 0 | 0 | 1 | 0 | 0 | 0 | 0 | 0 |
| 0 | -0.40 | 3.67 | -0.11 | 0.91 | NA | Gm10048 | | 0 | 0 | 0 | 0 | 0 | 1 | 0 | 0 | 0 | 0 | 0 |
| 0 | -0.40 | 3.67 | -0.11 | 0.91 | NA | Gm42614 | | 0 | 0 | 0 | 0 | 0 | 1 | 0 | 0 | 0 | 0 | 0 |
| 0 | -0.40 | 3.67 | -0.11 | 0.91 | NA | Gm42632 | | 0 | 0 | 0 | 0 | 0 | 1 | 0 | 0 | 0 | 0 | 0 |
| 0 | -0.40 | 3.67 | -0.11 | 0.91 | NA | 3110031N09Rik | | 0 | 0 | 0 | 0 | 0 | 1 | 0 | 0 | 0 | 0 | 0 |
| 0 | -0.40 | 3.67 | -0.11 | 0.91 | NA | 4930432L08Rik | | 0 | 0 | 0 | 0 | 0 | 1 | 0 | 0 | 0 | 0 | 0 |
| 0 | -0.40 | 3.67 | -0.11 | 0.91 | NA | Gm6366 | | 0 | 0 | 0 | 0 | 0 | 1 | 0 | 0 | 0 | 0 | 0 |
| 0 | -0.40 | 3.67 | -0.11 | 0.91 | NA | Gm43599 | | 0 | 0 | 0 | 0 | 0 | 1 | 0 | 0 | 0 | 0 | 0 |
| 0 | -0.40 | 3.67 | -0.11 | 0.91 | NA | Gm27553 | | 0 | 0 | 0 | 0 | 0 | 1 | 0 | 0 | 0 | 0 | 0 |
| 0 | -0.40 | 3.67 | -0.11 | 0.91 | NA | Gm20484 | | 0 | 0 | 0 | 0 | 0 | 1 | 0 | 0 | 0 | 0 | 0 |
| 0 | -0.40 | 3.67 | -0.11 | 0.91 | NA | Gm43365 | | 0 | 0 | 0 | 0 | 0 | 1 | 0 | 0 | 0 | 0 | 0 |
| 0 | -0.40 | 3.67 | -0.11 | 0.91 | NA | Mir701 | | 0 | 0 | 0 | 0 | 0 | 1 | 0 | 0 | 0 | 0 | 0 |
| 0 | -0.40 | 3.67 | -0.11 | 0.91 | NA | Gm42780 | | 0 | 0 | 0 | 0 | 0 | 1 | 0 | 0 | 0 | 0 | 0 |
| 0 | -0.40 | 3.67 | -0.11 | 0.91 | NA | Gm43676 | | 0 | 0 | 0 | 0 | 0 | 1 | 0 | 0 | 0 | 0 | 0 |
| 0 | -0.40 | 3.67 | -0.11 | 0.91 | NA | Gm24265 | | 0 | 0 | 0 | 0 | 0 | 1 | 0 | 0 | 0 | 0 | 0 |
| 0 | -0.40 | 3.67 | -0.11 | 0.91 | NA | Mir7030 | | 0 | 0 | 0 | 0 | 0 | 1 | 0 | 0 | 0 | 0 | 0 |
| 0 | -0.40 | 3.67 | -0.11 | 0.91 | NA | Gm24050 | | 0 | 0 | 0 | 0 | 0 | 1 | 0 | 0 | 0 | 0 | 0 |
| 0 | -0.40 | 3.67 | -0.11 | 0.91 | NA | Gm15749 | | 0 | 0 | 0 | 0 | 0 | 1 | 0 | 0 | 0 | 0 | 0 |
| 0 | -0.40 | 3.67 | -0.11 | 0.91 | NA | Gm42878 | | 0 | 0 | 0 | 0 | 0 | 1 | 0 | 0 | 0 | 0 | 0 |
| 0 | -0.40 | 3.67 | -0.11 | 0.91 | NA | Gm40323 | | 0 | 0 | 0 | 0 | 0 | 1 | 0 | 0 | 0 | 0 | 0 |

| | | | | | | | | | | | | | | | | | | |
|---|---|---|---|---|---|---|---|---|---|---|---|---|---|---|---|---|---|---|
| 0 | -0.40 | 3.67 | -0.11 | 0.91 | NA | Gm42875 | | 0 | 0 | 0 | 0 | 0 | 1 | 0 | 0 | 0 | 0 | 0 |
| 0 | -0.40 | 3.67 | -0.11 | 0.91 | NA | Gm42871 | | 0 | 0 | 0 | 0 | 0 | 1 | 0 | 0 | 0 | 0 | 0 |
| 0 | -0.40 | 3.67 | -0.11 | 0.91 | NA | Gm42989 | | 0 | 0 | 0 | 0 | 0 | 1 | 0 | 0 | 0 | 0 | 0 |
| 0 | -0.40 | 3.67 | -0.11 | 0.91 | NA | Gm23995 | | 0 | 0 | 0 | 0 | 0 | 1 | 0 | 0 | 0 | 0 | 0 |
| 0 | -0.40 | 3.67 | -0.11 | 0.91 | NA | Gm15407 | | 0 | 0 | 0 | 0 | 0 | 1 | 0 | 0 | 0 | 0 | 0 |
| 0 | -0.40 | 3.67 | -0.11 | 0.91 | NA | Gm43332 | | 0 | 0 | 0 | 0 | 0 | 1 | 0 | 0 | 0 | 0 | 0 |
| 0 | -0.40 | 3.67 | -0.11 | 0.91 | NA | BC028471 | | 0 | 0 | 0 | 0 | 0 | 1 | 0 | 0 | 0 | 0 | 0 |
| 0 | -0.40 | 3.67 | -0.11 | 0.91 | NA | Gm44411 | | 0 | 0 | 0 | 0 | 0 | 1 | 0 | 0 | 0 | 0 | 0 |
| 0 | -0.40 | 3.67 | -0.11 | 0.91 | NA | Gm28008 | | 0 | 0 | 0 | 0 | 0 | 1 | 0 | 0 | 0 | 0 | 0 |
| 0 | -0.40 | 3.67 | -0.11 | 0.91 | NA | Gm5302 | | 0 | 0 | 0 | 0 | 0 | 1 | 0 | 0 | 0 | 0 | 0 |
| 0 | -0.40 | 3.67 | -0.11 | 0.91 | NA | Gm7504 | | 0 | 0 | 0 | 0 | 0 | 1 | 0 | 0 | 0 | 0 | 0 |
| 0 | -0.40 | 3.67 | -0.11 | 0.91 | NA | Olfr455 | olfactory receptor 455 | 0 | 0 | 0 | 0 | 0 | 1 | 0 | 0 | 0 | 0 | 0 |
| 0 | -0.40 | 3.67 | -0.11 | 0.91 | NA | Gm43869 | | 0 | 0 | 0 | 0 | 0 | 1 | 0 | 0 | 0 | 0 | 0 |
| 0 | -0.40 | 3.67 | -0.11 | 0.91 | NA | Gm43887 | | 0 | 0 | 0 | 0 | 0 | 1 | 0 | 0 | 0 | 0 | 0 |
| 0 | -0.40 | 3.67 | -0.11 | 0.91 | NA | Gm43881 | | 0 | 0 | 0 | 0 | 0 | 1 | 0 | 0 | 0 | 0 | 0 |
| 0 | -0.40 | 3.67 | -0.11 | 0.91 | NA | Gm24325 | | 0 | 0 | 0 | 0 | 0 | 1 | 0 | 0 | 0 | 0 | 0 |
| 0 | -0.40 | 3.67 | -0.11 | 0.91 | NA | Gm43980 | | 0 | 0 | 0 | 0 | 0 | 1 | 0 | 0 | 0 | 0 | 0 |
| 0 | -0.40 | 3.67 | -0.11 | 0.91 | NA | Stk31 | serine threonine kinase 31 | 0 | 0 | 0 | 0 | 0 | 1 | 0 | 0 | 0 | 0 | 0 |
| 0 | -0.40 | 3.67 | -0.11 | 0.91 | NA | Vmn1r-ps12 | | 0 | 0 | 0 | 0 | 0 | 1 | 0 | 0 | 0 | 0 | 0 |
| 0 | -0.40 | 3.67 | -0.11 | 0.91 | NA | Gm43867 | | 0 | 0 | 0 | 0 | 0 | 1 | 0 | 0 | 0 | 0 | 0 |
| 0 | -0.40 | 3.67 | -0.11 | 0.91 | NA | Gm44770 | | 0 | 0 | 0 | 0 | 0 | 1 | 0 | 0 | 0 | 0 | 0 |
| 0 | -0.40 | 3.67 | -0.11 | 0.91 | NA | Ccdc142 | coiled-coil domain containing 142 | 0 | 0 | 0 | 0 | 0 | 1 | 0 | 0 | 0 | 0 | 0 |
| 0 | -0.40 | 3.67 | -0.11 | 0.91 | NA | Mrpl53 | mitochondrial ribosomal protein L53 | 0 | 0 | 0 | 0 | 0 | 1 | 0 | 0 | 0 | 0 | 0 |
| 0 | -0.40 | 3.67 | -0.11 | 0.91 | NA | Gm43920 | | 0 | 0 | 0 | 0 | 0 | 1 | 0 | 0 | 0 | 0 | 0 |
| 0 | -0.40 | 3.67 | -0.11 | 0.91 | NA | 4930471M09Rik | | 0 | 0 | 0 | 0 | 0 | 1 | 0 | 0 | 0 | 0 | 0 |
| 0 | -0.40 | 3.67 | -0.11 | 0.91 | NA | Gm4602 | | 0 | 0 | 0 | 0 | 0 | 1 | 0 | 0 | 0 | 0 | 0 |
| 0 | -0.40 | 3.67 | -0.11 | 0.91 | NA | Gm44199 | | 0 | 0 | 0 | 0 | 0 | 1 | 0 | 0 | 0 | 0 | 0 |
| 0 | -0.40 | 3.67 | -0.11 | 0.91 | NA | Vmn2r25 | vomeronasal 2, receptor 25 | 0 | 0 | 0 | 0 | 0 | 1 | 0 | 0 | 0 | 0 | 0 |
| 0 | -0.40 | 3.67 | -0.11 | 0.91 | NA | Gm6423 | | 0 | 0 | 0 | 0 | 0 | 1 | 0 | 0 | 0 | 0 | 0 |
| 0 | -0.40 | 3.67 | -0.11 | 0.91 | NA | Gprc5d | G protein-coupled receptor, family C, grou | 0 | 0 | 0 | 0 | 0 | 1 | 0 | 0 | 0 | 0 | 0 |
| 0 | -0.40 | 3.67 | -0.11 | 0.91 | NA | Grin2b | glutamate receptor, ionotropic, NMDA2B ( | 0 | 0 | 0 | 0 | 0 | 1 | 0 | 0 | 0 | 0 | 0 |
| 0 | -0.40 | 3.67 | -0.11 | 0.91 | NA | Gm17216 | | 0 | 0 | 0 | 0 | 0 | 1 | 0 | 0 | 0 | 0 | 0 |
| 0 | -0.40 | 3.67 | -0.11 | 0.91 | NA | Gm44586 | | 0 | 0 | 0 | 0 | 0 | 1 | 0 | 0 | 0 | 0 | 0 |
| 0 | -0.40 | 3.67 | -0.11 | 0.91 | NA | Tpt1-ps4 | | 0 | 0 | 0 | 0 | 0 | 1 | 0 | 0 | 0 | 0 | 0 |
| 0 | -0.40 | 3.67 | -0.11 | 0.91 | NA | Gm17850 | | 0 | 0 | 0 | 0 | 0 | 1 | 0 | 0 | 0 | 0 | 0 |
| 0 | -0.40 | 3.67 | -0.11 | 0.91 | NA | Gm26707 | | 0 | 0 | 0 | 0 | 0 | 1 | 0 | 0 | 0 | 0 | 0 |
| 0 | -0.40 | 3.67 | -0.11 | 0.91 | NA | Mia | melanoma inhibitory activity | 0 | 0 | 0 | 0 | 0 | 1 | 0 | 0 | 0 | 0 | 0 |
| 0 | -0.40 | 3.67 | -0.11 | 0.91 | NA | Gm19880 | | 0 | 0 | 0 | 0 | 0 | 1 | 0 | 0 | 0 | 0 | 0 |
| 0 | -0.40 | 3.67 | -0.11 | 0.91 | NA | Gm23368 | | 0 | 0 | 0 | 0 | 0 | 1 | 0 | 0 | 0 | 0 | 0 |
| 0 | -0.40 | 3.67 | -0.11 | 0.91 | NA | Usf2-ps1 | | 0 | 0 | 0 | 0 | 0 | 1 | 0 | 0 | 0 | 0 | 0 |
| 0 | -0.40 | 3.67 | -0.11 | 0.91 | NA | Klk10 | kallikrein related-peptidase 10 | 0 | 0 | 0 | 0 | 0 | 1 | 0 | 0 | 0 | 0 | 0 |
| 0 | -0.40 | 3.67 | -0.11 | 0.91 | NA | 6530437J22Rik | | 0 | 0 | 0 | 0 | 0 | 1 | 0 | 0 | 0 | 0 | 0 |
| 0 | -0.40 | 3.67 | -0.11 | 0.91 | NA | Gm31479 | | 0 | 0 | 0 | 0 | 0 | 1 | 0 | 0 | 0 | 0 | 0 |
| 0 | -0.40 | 3.67 | -0.11 | 0.91 | NA | Gm44894 | | 0 | 0 | 0 | 0 | 0 | 1 | 0 | 0 | 0 | 0 | 0 |
| 0 | -0.40 | 3.67 | -0.11 | 0.91 | NA | Gm44742 | | 0 | 0 | 0 | 0 | 0 | 1 | 0 | 0 | 0 | 0 | 0 |
| 0 | -0.40 | 3.67 | -0.11 | 0.91 | NA | Gm44978 | | 0 | 0 | 0 | 0 | 0 | 1 | 0 | 0 | 0 | 0 | 0 |
| 0 | -0.40 | 3.67 | -0.11 | 0.91 | NA | Gm38405 | | 0 | 0 | 0 | 0 | 0 | 1 | 0 | 0 | 0 | 0 | 0 |
| 0 | -0.40 | 3.67 | -0.11 | 0.91 | NA | Gm20476 | | 0 | 0 | 0 | 0 | 0 | 1 | 0 | 0 | 0 | 0 | 0 |
| 0 | -0.40 | 3.67 | -0.11 | 0.91 | NA | Olfr519 | olfactory receptor 519 | 0 | 0 | 0 | 0 | 0 | 1 | 0 | 0 | 0 | 0 | 0 |
| 0 | -0.40 | 3.67 | -0.11 | 0.91 | NA | Gm47284 | | 0 | 0 | 0 | 0 | 0 | 1 | 0 | 0 | 0 | 0 | 0 |
| 0 | -0.40 | 3.67 | -0.11 | 0.91 | NA | Far1os | | 0 | 0 | 0 | 0 | 0 | 1 | 0 | 0 | 0 | 0 | 0 |
| 0 | -0.40 | 3.67 | -0.11 | 0.91 | NA | Gm26414 | | 0 | 0 | 0 | 0 | 0 | 1 | 0 | 0 | 0 | 0 | 0 |
| 0 | -0.40 | 3.67 | -0.11 | 0.91 | NA | Gm23445 | | 0 | 0 | 0 | 0 | 0 | 1 | 0 | 0 | 0 | 0 | 0 |
| 0 | -0.40 | 3.67 | -0.11 | 0.91 | NA | Trim72 | tripartite motif-containing 72 | 0 | 0 | 0 | 0 | 0 | 1 | 0 | 0 | 0 | 0 | 0 |
| 0 | -0.40 | 3.67 | -0.11 | 0.91 | NA | Gm44518 | | 0 | 0 | 0 | 0 | 0 | 1 | 0 | 0 | 0 | 0 | 0 |
| 0 | -0.40 | 3.67 | -0.11 | 0.91 | NA | n-R5s95 | | 0 | 0 | 0 | 0 | 0 | 1 | 0 | 0 | 0 | 0 | 0 |
| 0 | -0.40 | 3.67 | -0.11 | 0.91 | NA | Gm44800 | | 0 | 0 | 0 | 0 | 0 | 1 | 0 | 0 | 0 | 0 | 0 |
| 0 | -0.40 | 3.67 | -0.11 | 0.91 | NA | Gm44801 | | 0 | 0 | 0 | 0 | 0 | 1 | 0 | 0 | 0 | 0 | 0 |
| 0 | -0.40 | 3.67 | -0.11 | 0.91 | NA | 1700128E19Rik | | 0 | 0 | 0 | 0 | 0 | 1 | 0 | 0 | 0 | 0 | 0 |
| 0 | -0.40 | 3.67 | -0.11 | 0.91 | NA | Gm15350 | | 0 | 0 | 0 | 0 | 0 | 1 | 0 | 0 | 0 | 0 | 0 |

| | | | | | | | | | | | | | | | | | | |
|---|---|---|---|---|---|---|---|---|---|---|---|---|---|---|---|---|---|---|
| 0 | -0.40 | 3.67 | -0.11 | 0.91 | NA | Gm25014 | | 0 | 0 | 0 | 0 | 0 | 1 | 0 | 0 | 0 | 0 | 0 |
| 0 | -0.40 | 3.67 | -0.11 | 0.91 | NA | Cfap97d2 | CFAP97 domain containing 2 | 0 | 0 | 0 | 0 | 0 | 1 | 0 | 0 | 0 | 0 | 0 |
| 0 | -0.40 | 3.67 | -0.11 | 0.91 | NA | Gm20946 | | 0 | 0 | 0 | 0 | 0 | 1 | 0 | 0 | 0 | 0 | 0 |
| 0 | -0.40 | 3.67 | -0.11 | 0.91 | NA | Gm45458 | | 0 | 0 | 0 | 0 | 0 | 1 | 0 | 0 | 0 | 0 | 0 |
| 0 | -0.40 | 3.67 | -0.11 | 0.91 | NA | Gm45321 | | 0 | 0 | 0 | 0 | 0 | 1 | 0 | 0 | 0 | 0 | 0 |
| 0 | -0.40 | 3.67 | -0.11 | 0.91 | NA | Gm6458 | | 0 | 0 | 0 | 0 | 0 | 1 | 0 | 0 | 0 | 0 | 0 |
| 0 | -0.40 | 3.67 | -0.11 | 0.91 | NA | Gm7328 | | 0 | 0 | 0 | 0 | 0 | 1 | 0 | 0 | 0 | 0 | 0 |
| 0 | -0.40 | 3.67 | -0.11 | 0.91 | NA | Gm10997 | | 0 | 0 | 0 | 0 | 0 | 1 | 0 | 0 | 0 | 0 | 0 |
| 0 | -0.40 | 3.67 | -0.11 | 0.91 | NA | Gm45706 | | 0 | 0 | 0 | 0 | 0 | 1 | 0 | 0 | 0 | 0 | 0 |
| 0 | -0.40 | 3.67 | -0.11 | 0.91 | NA | Gm16183 | | 0 | 0 | 0 | 0 | 0 | 1 | 0 | 0 | 0 | 0 | 0 |
| 0 | -0.40 | 3.67 | -0.11 | 0.91 | NA | Gm45594 | | 0 | 0 | 0 | 0 | 0 | 1 | 0 | 0 | 0 | 0 | 0 |
| 0 | -0.40 | 3.67 | -0.11 | 0.91 | NA | Gm24693 | | 0 | 0 | 0 | 0 | 0 | 1 | 0 | 0 | 0 | 0 | 0 |
| 0 | -0.40 | 3.67 | -0.11 | 0.91 | NA | Gm8748 | | 0 | 0 | 0 | 0 | 0 | 1 | 0 | 0 | 0 | 0 | 0 |
| 0 | -0.40 | 3.67 | -0.11 | 0.91 | NA | Agrp | agouti related neuropeptide | 0 | 0 | 0 | 0 | 0 | 1 | 0 | 0 | 0 | 0 | 0 |
| 0 | -0.40 | 3.67 | -0.11 | 0.91 | NA | Gm38318 | | 0 | 0 | 0 | 0 | 0 | 1 | 0 | 0 | 0 | 0 | 0 |
| 0 | -0.40 | 3.67 | -0.11 | 0.91 | NA | Mir7237 | | 0 | 0 | 0 | 0 | 0 | 1 | 0 | 0 | 0 | 0 | 0 |
| 0 | -0.40 | 3.67 | -0.11 | 0.91 | NA | Gm10717 | | 0 | 0 | 0 | 0 | 0 | 1 | 0 | 0 | 0 | 0 | 0 |
| 0 | -0.40 | 3.67 | -0.11 | 0.91 | NA | Gm6015 | | 0 | 0 | 0 | 0 | 0 | 1 | 0 | 0 | 0 | 0 | 0 |
| 0 | -0.40 | 3.67 | -0.11 | 0.91 | NA | Gm24339 | | 0 | 0 | 0 | 0 | 0 | 1 | 0 | 0 | 0 | 0 | 0 |
| 0 | -0.40 | 3.67 | -0.11 | 0.91 | NA | Gm15521 | | 0 | 0 | 0 | 0 | 0 | 1 | 0 | 0 | 0 | 0 | 0 |
| 0 | -0.40 | 3.67 | -0.11 | 0.91 | NA | Mfrp | membrane frizzled-related protein | 0 | 0 | 0 | 0 | 0 | 1 | 0 | 0 | 0 | 0 | 0 |
| 0 | -0.40 | 3.67 | -0.11 | 0.91 | NA | Gm47231 | | 0 | 0 | 0 | 0 | 0 | 1 | 0 | 0 | 0 | 0 | 0 |
| 0 | -0.40 | 3.67 | -0.11 | 0.91 | NA | Gm31374 | | 0 | 0 | 0 | 0 | 0 | 1 | 0 | 0 | 0 | 0 | 0 |
| 0 | -0.40 | 3.67 | -0.11 | 0.91 | NA | Gm47676 | | 0 | 0 | 0 | 0 | 0 | 1 | 0 | 0 | 0 | 0 | 0 |
| 0 | -0.40 | 3.67 | -0.11 | 0.91 | NA | Gm24289 | | 0 | 0 | 0 | 0 | 0 | 1 | 0 | 0 | 0 | 0 | 0 |
| 0 | -0.40 | 3.67 | -0.11 | 0.91 | NA | Gm22169 | | 0 | 0 | 0 | 0 | 0 | 1 | 0 | 0 | 0 | 0 | 0 |
| 0 | -0.40 | 3.67 | -0.11 | 0.91 | NA | Gm27253 | | 0 | 0 | 0 | 0 | 0 | 1 | 0 | 0 | 0 | 0 | 0 |
| 0 | -0.40 | 3.67 | -0.11 | 0.91 | NA | Gm37710 | | 0 | 0 | 0 | 0 | 0 | 1 | 0 | 0 | 0 | 0 | 0 |
| 0 | -0.40 | 3.67 | -0.11 | 0.91 | NA | Gm38215 | | 0 | 0 | 0 | 0 | 0 | 1 | 0 | 0 | 0 | 0 | 0 |
| 0 | -0.40 | 3.67 | -0.11 | 0.91 | NA | Gm38083 | | 0 | 0 | 0 | 0 | 0 | 1 | 0 | 0 | 0 | 0 | 0 |
| 0 | -0.40 | 3.67 | -0.11 | 0.91 | NA | Gm37484 | | 0 | 0 | 0 | 0 | 0 | 1 | 0 | 0 | 0 | 0 | 0 |
| 0 | -0.40 | 3.67 | -0.11 | 0.91 | NA | 1700065D16Ril | RIKEN cDNA 1700065D16 gene | 0 | 0 | 0 | 0 | 0 | 1 | 0 | 0 | 0 | 0 | 0 |
| 0 | -0.40 | 3.67 | -0.11 | 0.91 | NA | Gm29387 | | 0 | 0 | 0 | 0 | 0 | 1 | 0 | 0 | 0 | 0 | 0 |
| 0 | -0.40 | 3.67 | -0.11 | 0.91 | NA | Gm37649 | | 0 | 0 | 0 | 0 | 0 | 1 | 0 | 0 | 0 | 0 | 0 |
| 0 | -0.40 | 3.67 | -0.11 | 0.91 | NA | Gm37714 | | 0 | 0 | 0 | 0 | 0 | 1 | 0 | 0 | 0 | 0 | 0 |
| 0 | -0.40 | 3.67 | -0.11 | 0.91 | NA | Tmem89 | transmembrane protein 89 | 0 | 0 | 0 | 0 | 0 | 1 | 0 | 0 | 0 | 0 | 0 |
| 0 | -0.40 | 3.67 | -0.11 | 0.91 | NA | Ucn2 | urocortin 2 | 0 | 0 | 0 | 0 | 0 | 1 | 0 | 0 | 0 | 0 | 0 |
| 0 | -0.40 | 3.67 | -0.11 | 0.91 | NA | Gm47121 | | 0 | 0 | 0 | 0 | 0 | 1 | 0 | 0 | 0 | 0 | 0 |
| 0 | -0.40 | 3.67 | -0.11 | 0.91 | NA | Gm47135 | | 0 | 0 | 0 | 0 | 0 | 1 | 0 | 0 | 0 | 0 | 0 |
| 0 | -0.40 | 3.67 | -0.11 | 0.91 | NA | Gm47148 | | 0 | 0 | 0 | 0 | 0 | 1 | 0 | 0 | 0 | 0 | 0 |
| 0 | -0.40 | 3.67 | -0.11 | 0.91 | NA | Gm5178 | | 0 | 0 | 0 | 0 | 0 | 1 | 0 | 0 | 0 | 0 | 0 |
| 0 | -0.40 | 3.67 | -0.11 | 0.91 | NA | Gm48546 | | 0 | 0 | 0 | 0 | 0 | 1 | 0 | 0 | 0 | 0 | 0 |
| 0 | -0.40 | 3.67 | -0.11 | 0.91 | NA | Gm17230 | | 0 | 0 | 0 | 0 | 0 | 1 | 0 | 0 | 0 | 0 | 0 |
| 0 | -0.40 | 3.67 | -0.11 | 0.91 | NA | Gm26585 | | 0 | 0 | 0 | 0 | 0 | 1 | 0 | 0 | 0 | 0 | 0 |
| 0 | -0.40 | 3.67 | -0.11 | 0.91 | NA | Gm48092 | | 0 | 0 | 0 | 0 | 0 | 1 | 0 | 0 | 0 | 0 | 0 |
| 0 | -0.40 | 3.67 | -0.11 | 0.91 | NA | Gm47926 | | 0 | 0 | 0 | 0 | 0 | 1 | 0 | 0 | 0 | 0 | 0 |
| 0 | -0.40 | 3.67 | -0.11 | 0.91 | NA | Gm5949 | | 0 | 0 | 0 | 0 | 0 | 1 | 0 | 0 | 0 | 0 | 0 |
| 0 | -0.40 | 3.67 | -0.11 | 0.91 | NA | Gm47624 | | 0 | 0 | 0 | 0 | 0 | 1 | 0 | 0 | 0 | 0 | 0 |
| 0 | -0.40 | 3.67 | -0.11 | 0.91 | NA | Gm31763 | | 0 | 0 | 0 | 0 | 0 | 1 | 0 | 0 | 0 | 0 | 0 |
| 0 | -0.40 | 3.67 | -0.11 | 0.91 | NA | 4930533K18Rik | | 0 | 0 | 0 | 0 | 0 | 1 | 0 | 0 | 0 | 0 | 0 |
| 0 | -0.40 | 3.67 | -0.11 | 0.91 | NA | Gm867 | predicted gene 867 | 0 | 0 | 0 | 0 | 0 | 1 | 0 | 0 | 0 | 0 | 0 |
| 0 | -0.40 | 3.67 | -0.11 | 0.91 | NA | Gm16222 | | 0 | 0 | 0 | 0 | 0 | 1 | 0 | 0 | 0 | 0 | 0 |
| 0 | -0.40 | 3.67 | -0.11 | 0.91 | NA | Gm7842 | | 0 | 0 | 0 | 0 | 0 | 1 | 0 | 0 | 0 | 0 | 0 |
| 0 | -0.40 | 3.67 | -0.11 | 0.91 | NA | Onecut3 | one cut domain, family member 3 | 0 | 0 | 0 | 0 | 0 | 1 | 0 | 0 | 0 | 0 | 0 |
| 0 | -0.40 | 3.67 | -0.11 | 0.91 | NA | Mir1191b | | 0 | 0 | 0 | 0 | 0 | 1 | 0 | 0 | 0 | 0 | 0 |
| 0 | -0.40 | 3.67 | -0.11 | 0.91 | NA | Gm4792 | | 0 | 0 | 0 | 0 | 0 | 1 | 0 | 0 | 0 | 0 | 0 |
| 0 | -0.40 | 3.67 | -0.11 | 0.91 | NA | Gm3619 | | 0 | 0 | 0 | 0 | 0 | 1 | 0 | 0 | 0 | 0 | 0 |
| 0 | -0.40 | 3.67 | -0.11 | 0.91 | NA | Gm48203 | | 0 | 0 | 0 | 0 | 0 | 1 | 0 | 0 | 0 | 0 | 0 |
| 0 | -0.40 | 3.67 | -0.11 | 0.91 | NA | Gm48204 | | 0 | 0 | 0 | 0 | 0 | 1 | 0 | 0 | 0 | 0 | 0 |
| 0 | -0.40 | 3.67 | -0.11 | 0.91 | NA | 1700020G17Rik | | 0 | 0 | 0 | 0 | 0 | 1 | 0 | 0 | 0 | 0 | 0 |

| | | | | | | | | | | | | | | | | | | |
|---|---|---|---|---|---|---|---|---|---|---|---|---|---|---|---|---|---|---|
| 0 | -0.40 | 3.67 | -0.11 | 0.91 | NA | Gm47534 | | 0 | 0 | 0 | 0 | 0 | 1 | 0 | 0 | 0 | 0 | 0 |
| 0 | -0.40 | 3.67 | -0.11 | 0.91 | NA | Gm25190 | | 0 | 0 | 0 | 0 | 0 | 1 | 0 | 0 | 0 | 0 | 0 |
| 0 | -0.40 | 3.67 | -0.11 | 0.91 | NA | Gm47027 | | 0 | 0 | 0 | 0 | 0 | 1 | 0 | 0 | 0 | 0 | 0 |
| 0 | -0.40 | 3.67 | -0.11 | 0.91 | NA | Gm4489 | | 0 | 0 | 0 | 0 | 0 | 1 | 0 | 0 | 0 | 0 | 0 |
| 0 | -0.40 | 3.67 | -0.11 | 0.91 | NA | Rdh9 | retinol dehydrogenase 9 | 0 | 0 | 0 | 0 | 0 | 1 | 0 | 0 | 0 | 0 | 0 |
| 0 | -0.40 | 3.67 | -0.11 | 0.91 | NA | Olfr814 | olfactory receptor 814 | 0 | 0 | 0 | 0 | 0 | 1 | 0 | 0 | 0 | 0 | 0 |
| 0 | -0.40 | 3.67 | -0.11 | 0.91 | NA | Gm11401 | | 0 | 0 | 0 | 0 | 0 | 1 | 0 | 0 | 0 | 0 | 0 |
| 0 | -0.40 | 3.67 | -0.11 | 0.91 | NA | Gm28401 | | 0 | 0 | 0 | 0 | 0 | 1 | 0 | 0 | 0 | 0 | 0 |
| 0 | -0.40 | 3.67 | -0.11 | 0.91 | NA | Otx1 | orthodenticle homeobox 1 | 0 | 0 | 0 | 0 | 0 | 1 | 0 | 0 | 0 | 0 | 0 |
| 0 | -0.40 | 3.67 | -0.11 | 0.91 | NA | Gm12120 | | 0 | 0 | 0 | 0 | 0 | 1 | 0 | 0 | 0 | 0 | 0 |
| 0 | -0.40 | 3.67 | -0.11 | 0.91 | NA | Mir7671 | | 0 | 0 | 0 | 0 | 0 | 1 | 0 | 0 | 0 | 0 | 0 |
| 0 | -0.40 | 3.67 | -0.11 | 0.91 | NA | Gm12217 | | 0 | 0 | 0 | 0 | 0 | 1 | 0 | 0 | 0 | 0 | 0 |
| 0 | -0.40 | 3.67 | -0.11 | 0.91 | NA | Gm12285 | | 0 | 0 | 0 | 0 | 0 | 1 | 0 | 0 | 0 | 0 | 0 |
| 0 | -0.40 | 3.67 | -0.11 | 0.91 | NA | C030013C21Rik | | 0 | 0 | 0 | 0 | 0 | 1 | 0 | 0 | 0 | 0 | 0 |
| 0 | -0.40 | 3.67 | -0.11 | 0.91 | NA | Gm11417 | | 0 | 0 | 0 | 0 | 0 | 1 | 0 | 0 | 0 | 0 | 0 |
| 0 | -0.40 | 3.67 | -0.11 | 0.91 | NA | Gm11420 | | 0 | 0 | 0 | 0 | 0 | 1 | 0 | 0 | 0 | 0 | 0 |
| 0 | -0.40 | 3.67 | -0.11 | 0.91 | NA | Slfn4 | schlafen 4 | 0 | 0 | 0 | 0 | 0 | 1 | 0 | 0 | 0 | 0 | 0 |
| 0 | -0.40 | 3.67 | -0.11 | 0.91 | NA | Gm11439 | | 0 | 0 | 0 | 0 | 0 | 1 | 0 | 0 | 0 | 0 | 0 |
| 0 | -0.40 | 3.67 | -0.11 | 0.91 | NA | Mir152 | | 0 | 0 | 0 | 0 | 0 | 1 | 0 | 0 | 0 | 0 | 0 |
| 0 | -0.40 | 3.67 | -0.11 | 0.91 | NA | Mir5119 | | 0 | 0 | 0 | 0 | 0 | 1 | 0 | 0 | 0 | 0 | 0 |
| 0 | -0.40 | 3.67 | -0.11 | 0.91 | NA | Gsdma2 | gasdermin A2 | 0 | 0 | 0 | 0 | 0 | 1 | 0 | 0 | 0 | 0 | 0 |
| 0 | -0.40 | 3.67 | -0.11 | 0.91 | NA | Gm22059 | | 0 | 0 | 0 | 0 | 0 | 1 | 0 | 0 | 0 | 0 | 0 |
| 0 | -0.40 | 3.67 | -0.11 | 0.91 | NA | | | 0 | 0 | 0 | 0 | 0 | 1 | 0 | 0 | 0 | 0 | 0 |
| 0 | -0.40 | 3.67 | -0.11 | 0.91 | NA | Gm11625 | | 0 | 0 | 0 | 0 | 0 | 1 | 0 | 0 | 0 | 0 | 0 |
| 0 | -0.40 | 3.67 | -0.11 | 0.91 | NA | Gm11706 | | 0 | 0 | 0 | 0 | 0 | 1 | 0 | 0 | 0 | 0 | 0 |
| 0 | -0.40 | 3.67 | -0.11 | 0.91 | NA | | | 0 | 0 | 0 | 0 | 0 | 1 | 0 | 0 | 0 | 0 | 0 |
| 0 | -0.40 | 3.67 | -0.11 | 0.91 | NA | Gm11766 | | 0 | 0 | 0 | 0 | 0 | 1 | 0 | 0 | 0 | 0 | 0 |
| 0 | -0.40 | 3.67 | -0.11 | 0.91 | NA | Slc7a15 | solute carrier family 7 (cationic amino aci | 0 | 0 | 0 | 0 | 0 | 1 | 0 | 0 | 0 | 0 | 0 |
| 0 | -0.40 | 3.67 | -0.11 | 0.91 | NA | Mir6387 | | 0 | 0 | 0 | 0 | 0 | 1 | 0 | 0 | 0 | 0 | 0 |
| 0 | -0.40 | 3.67 | -0.11 | 0.91 | NA | Gm36752 | | 0 | 0 | 0 | 0 | 0 | 1 | 0 | 0 | 0 | 0 | 0 |
| 0 | -0.40 | 3.67 | -0.11 | 0.91 | NA | Gm9229 | | 0 | 0 | 0 | 0 | 0 | 1 | 0 | 0 | 0 | 0 | 0 |
| 0 | -0.40 | 3.67 | -0.11 | 0.91 | NA | Gm47713 | | 0 | 0 | 0 | 0 | 0 | 1 | 0 | 0 | 0 | 0 | 0 |
| 0 | -0.40 | 3.67 | -0.11 | 0.91 | NA | Gm31508 | | 0 | 0 | 0 | 0 | 0 | 1 | 0 | 0 | 0 | 0 | 0 |
| 0 | -0.40 | 3.67 | -0.11 | 0.91 | NA | Gm25913 | | 0 | 0 | 0 | 0 | 0 | 1 | 0 | 0 | 0 | 0 | 0 |
| 0 | -0.40 | 3.67 | -0.11 | 0.91 | NA | Gm40375 | | 0 | 0 | 0 | 0 | 0 | 1 | 0 | 0 | 0 | 0 | 0 |
| 0 | -0.40 | 3.67 | -0.11 | 0.91 | NA | Gm47506 | | 0 | 0 | 0 | 0 | 0 | 1 | 0 | 0 | 0 | 0 | 0 |
| 0 | -0.40 | 3.67 | -0.11 | 0.91 | NA | Gm48713 | | 0 | 0 | 0 | 0 | 0 | 1 | 0 | 0 | 0 | 0 | 0 |
| 0 | -0.40 | 3.67 | -0.11 | 0.91 | NA | Gm35188 | | 0 | 0 | 0 | 0 | 0 | 1 | 0 | 0 | 0 | 0 | 0 |
| 0 | -0.40 | 3.67 | -0.11 | 0.91 | NA | 4930458K08Rik | | 0 | 0 | 0 | 0 | 0 | 1 | 0 | 0 | 0 | 0 | 0 |
| 0 | -0.40 | 3.67 | -0.11 | 0.91 | NA | Gm3693 | | 0 | 0 | 0 | 0 | 0 | 1 | 0 | 0 | 0 | 0 | 0 |
| 0 | -0.40 | 3.67 | -0.11 | 0.91 | NA | Gm47080 | | 0 | 0 | 0 | 0 | 0 | 1 | 0 | 0 | 0 | 0 | 0 |
| 0 | -0.40 | 3.67 | -0.11 | 0.91 | NA | Vsx2 | visual system homeobox 2 | 0 | 0 | 0 | 0 | 0 | 1 | 0 | 0 | 0 | 0 | 0 |
| 0 | -0.40 | 3.67 | -0.11 | 0.91 | NA | Gm40548 | | 0 | 0 | 0 | 0 | 0 | 1 | 0 | 0 | 0 | 0 | 0 |
| 0 | -0.40 | 3.67 | -0.11 | 0.91 | NA | Gm35412 | | 0 | 0 | 0 | 0 | 0 | 1 | 0 | 0 | 0 | 0 | 0 |
| 0 | -0.40 | 3.67 | -0.11 | 0.91 | NA | Gm49703 | | 0 | 0 | 0 | 0 | 0 | 1 | 0 | 0 | 0 | 0 | 0 |
| 0 | -0.40 | 3.67 | -0.11 | 0.91 | NA | Gm48586 | | 0 | 0 | 0 | 0 | 0 | 1 | 0 | 0 | 0 | 0 | 0 |
| 0 | -0.40 | 3.67 | -0.11 | 0.91 | NA | Gm44324 | | 0 | 0 | 0 | 0 | 0 | 1 | 0 | 0 | 0 | 0 | 0 |
| 0 | -0.40 | 3.67 | -0.11 | 0.91 | NA | 5033424D13Rik | | 0 | 0 | 0 | 0 | 0 | 1 | 0 | 0 | 0 | 0 | 0 |
| 0 | -0.40 | 3.67 | -0.11 | 0.91 | NA | Mir3070b | | 0 | 0 | 0 | 0 | 0 | 1 | 0 | 0 | 0 | 0 | 0 |
| 0 | -0.40 | 3.67 | -0.11 | 0.91 | NA | Mir1193 | | 0 | 0 | 0 | 0 | 0 | 1 | 0 | 0 | 0 | 0 | 0 |
| 0 | -0.40 | 3.67 | -0.11 | 0.91 | NA | Mir496a | | 0 | 0 | 0 | 0 | 0 | 1 | 0 | 0 | 0 | 0 | 0 |
| 0 | -0.40 | 3.67 | -0.11 | 0.91 | NA | Mir369 | | 0 | 0 | 0 | 0 | 0 | 1 | 0 | 0 | 0 | 0 | 0 |
| 0 | -0.40 | 3.67 | -0.11 | 0.91 | NA | Gm9280 | | 0 | 0 | 0 | 0 | 0 | 1 | 0 | 0 | 0 | 0 | 0 |
| 0 | -0.40 | 3.67 | -0.11 | 0.91 | NA | Gm48530 | | 0 | 0 | 0 | 0 | 0 | 1 | 0 | 0 | 0 | 0 | 0 |
| 0 | -0.40 | 3.67 | -0.11 | 0.91 | NA | Gm48574 | | 0 | 0 | 0 | 0 | 0 | 1 | 0 | 0 | 0 | 0 | 0 |
| 0 | -0.40 | 3.67 | -0.11 | 0.91 | NA | Gm48871 | | 0 | 0 | 0 | 0 | 0 | 1 | 0 | 0 | 0 | 0 | 0 |
| 0 | -0.40 | 3.67 | -0.11 | 0.91 | NA | Gm47411 | | 0 | 0 | 0 | 0 | 0 | 1 | 0 | 0 | 0 | 0 | 0 |
| 0 | -0.40 | 3.67 | -0.11 | 0.91 | NA | Gm7042 | | 0 | 0 | 0 | 0 | 0 | 1 | 0 | 0 | 0 | 0 | 0 |
| 0 | -0.40 | 3.67 | -0.11 | 0.91 | NA | Gm47721 | | 0 | 0 | 0 | 0 | 0 | 1 | 0 | 0 | 0 | 0 | 0 |
| 0 | -0.40 | 3.67 | -0.11 | 0.91 | NA | H3c10 | H3 clustered histone 10 | 0 | 0 | 0 | 0 | 0 | 1 | 0 | 0 | 0 | 0 | 0 |

| | | | | | | | | | | | | | | | | | | |
|---|---|---|---|---|---|---|---|---|---|---|---|---|---|---|---|---|---|---|
| 0 | -0.40 | 3.67 | -0.11 | 0.91 | NA | Gm25084 | | 0 | 0 | 0 | 0 | 0 | 1 | 0 | 0 | 0 | 0 | 0 |
| 0 | -0.40 | 3.67 | -0.11 | 0.91 | NA | 4930586N03Rik | | 0 | 0 | 0 | 0 | 0 | 1 | 0 | 0 | 0 | 0 | 0 |
| 0 | -0.40 | 3.67 | -0.11 | 0.91 | NA | Sycp2l | synaptonemal complex protein 2-like | 0 | 0 | 0 | 0 | 0 | 1 | 0 | 0 | 0 | 0 | 0 |
| 0 | -0.40 | 3.67 | -0.11 | 0.91 | NA | Gm15813 | | 0 | 0 | 0 | 0 | 0 | 1 | 0 | 0 | 0 | 0 | 0 |
| 0 | -0.40 | 3.67 | -0.11 | 0.91 | NA | Gm44341 | | 0 | 0 | 0 | 0 | 0 | 1 | 0 | 0 | 0 | 0 | 0 |
| 0 | -0.40 | 3.67 | -0.11 | 0.91 | NA | Gm16248 | | 0 | 0 | 0 | 0 | 0 | 1 | 0 | 0 | 0 | 0 | 0 |
| 0 | -0.40 | 3.67 | -0.11 | 0.91 | NA | Pfn3 | profilin 3 | 0 | 0 | 0 | 0 | 0 | 1 | 0 | 0 | 0 | 0 | 0 |
| 0 | -0.40 | 3.67 | -0.11 | 0.91 | NA | Gm48441 | | 0 | 0 | 0 | 0 | 0 | 1 | 0 | 0 | 0 | 0 | 0 |
| 0 | -0.40 | 3.67 | -0.11 | 0.91 | NA | Gm6478 | | 0 | 0 | 0 | 0 | 0 | 1 | 0 | 0 | 0 | 0 | 0 |
| 0 | -0.40 | 3.67 | -0.11 | 0.91 | NA | Gm38307 | | 0 | 0 | 0 | 0 | 0 | 1 | 0 | 0 | 0 | 0 | 0 |
| 0 | -0.40 | 3.67 | -0.11 | 0.91 | NA | Gm48095 | | 0 | 0 | 0 | 0 | 0 | 1 | 0 | 0 | 0 | 0 | 0 |
| 0 | -0.40 | 3.67 | -0.11 | 0.91 | NA | Gm8278 | | 0 | 0 | 0 | 0 | 0 | 1 | 0 | 0 | 0 | 0 | 0 |
| 0 | -0.40 | 3.67 | -0.11 | 0.91 | NA | Gm8345 | | 0 | 0 | 0 | 0 | 0 | 1 | 0 | 0 | 0 | 0 | 0 |
| 0 | -0.40 | 3.67 | -0.11 | 0.91 | NA | Gm4211 | | 0 | 0 | 0 | 0 | 0 | 1 | 0 | 0 | 0 | 0 | 0 |
| 0 | -0.40 | 3.67 | -0.11 | 0.91 | NA | Gm48138 | | 0 | 0 | 0 | 0 | 0 | 1 | 0 | 0 | 0 | 0 | 0 |
| 0 | -0.40 | 3.67 | -0.11 | 0.91 | NA | Gm47052 | | 0 | 0 | 0 | 0 | 0 | 1 | 0 | 0 | 0 | 0 | 0 |
| 0 | -0.40 | 3.67 | -0.11 | 0.91 | NA | Naip3-ps1 | | 0 | 0 | 0 | 0 | 0 | 1 | 0 | 0 | 0 | 0 | 0 |
| 0 | -0.40 | 3.67 | -0.11 | 0.91 | NA | Gm20784 | | 0 | 0 | 0 | 0 | 0 | 1 | 0 | 0 | 0 | 0 | 0 |
| 0 | -0.40 | 3.67 | -0.11 | 0.91 | NA | Gm5456 | | 0 | 0 | 0 | 0 | 0 | 1 | 0 | 0 | 0 | 0 | 0 |
| 0 | -0.40 | 3.67 | -0.11 | 0.91 | NA | Gm48371 | | 0 | 0 | 0 | 0 | 0 | 1 | 0 | 0 | 0 | 0 | 0 |
| 0 | -0.40 | 3.67 | -0.11 | 0.91 | NA | Gm48430 | | 0 | 0 | 0 | 0 | 0 | 1 | 0 | 0 | 0 | 0 | 0 |
| 0 | -0.40 | 3.67 | -0.11 | 0.91 | NA | Gm48601 | | 0 | 0 | 0 | 0 | 0 | 1 | 0 | 0 | 0 | 0 | 0 |
| 0 | -0.40 | 3.67 | -0.11 | 0.91 | NA | n-R5s45 | | 0 | 0 | 0 | 0 | 0 | 1 | 0 | 0 | 0 | 0 | 0 |
| 0 | -0.40 | 3.67 | -0.11 | 0.91 | NA | Rpl19-ps4 | | 0 | 0 | 0 | 0 | 0 | 1 | 0 | 0 | 0 | 0 | 0 |
| 0 | -0.40 | 3.67 | -0.11 | 0.91 | NA | Gm47743 | | 0 | 0 | 0 | 0 | 0 | 1 | 0 | 0 | 0 | 0 | 0 |
| 0 | -0.40 | 3.67 | -0.11 | 0.91 | NA | Gm8517 | | 0 | 0 | 0 | 0 | 0 | 1 | 0 | 0 | 0 | 0 | 0 |
| 0 | -0.40 | 3.67 | -0.11 | 0.91 | NA | Gm49252 | | 0 | 0 | 0 | 0 | 0 | 1 | 0 | 0 | 0 | 0 | 0 |
| 0 | -0.40 | 3.67 | -0.11 | 0.91 | NA | Gm8224 | | 0 | 0 | 0 | 0 | 0 | 1 | 0 | 0 | 0 | 0 | 0 |
| 0 | -0.40 | 3.67 | -0.11 | 0.91 | NA | Gm49132 | | 0 | 0 | 0 | 0 | 0 | 1 | 0 | 0 | 0 | 0 | 0 |
| 0 | -0.40 | 3.67 | -0.11 | 0.91 | NA | Gm49189 | | 0 | 0 | 0 | 0 | 0 | 1 | 0 | 0 | 0 | 0 | 0 |
| 0 | -0.40 | 3.67 | -0.11 | 0.91 | NA | Trav12-1 | | 0 | 0 | 0 | 0 | 0 | 1 | 0 | 0 | 0 | 0 | 0 |
| 0 | -0.40 | 3.67 | -0.11 | 0.91 | NA | Myh6 | myosin, heavy polypeptide 6, cardiac mus | 0 | 0 | 0 | 0 | 0 | 1 | 0 | 0 | 0 | 0 | 0 |
| 0 | -0.40 | 3.67 | -0.11 | 0.91 | NA | Gm46455 | | 0 | 0 | 0 | 0 | 0 | 1 | 0 | 0 | 0 | 0 | 0 |
| 0 | -0.40 | 3.67 | -0.11 | 0.91 | NA | Gm15932 | | 0 | 0 | 0 | 0 | 0 | 1 | 0 | 0 | 0 | 0 | 0 |
| 0 | -0.40 | 3.67 | -0.11 | 0.91 | NA | 1700109G14Rik | | 0 | 0 | 0 | 0 | 0 | 1 | 0 | 0 | 0 | 0 | 0 |
| 0 | -0.40 | 3.67 | -0.11 | 0.91 | NA | Gm21430 | | 0 | 0 | 0 | 0 | 0 | 1 | 0 | 0 | 0 | 0 | 0 |
| 0 | -0.40 | 3.67 | -0.11 | 0.91 | NA | Gm19222 | | 0 | 0 | 0 | 0 | 0 | 1 | 0 | 0 | 0 | 0 | 0 |
| 0 | -0.40 | 3.67 | -0.11 | 0.91 | NA | Gm9290 | | 0 | 0 | 0 | 0 | 0 | 1 | 0 | 0 | 0 | 0 | 0 |
| 0 | -0.40 | 3.67 | -0.11 | 0.91 | NA | 4930594M22Rik | | 0 | 0 | 0 | 0 | 0 | 1 | 0 | 0 | 0 | 0 | 0 |
| 0 | -0.40 | 3.67 | -0.11 | 0.91 | NA | Gm49089 | | 0 | 0 | 0 | 0 | 0 | 1 | 0 | 0 | 0 | 0 | 0 |
| 0 | -0.40 | 3.67 | -0.11 | 0.91 | NA | Gm22208 | | 0 | 0 | 0 | 0 | 0 | 1 | 0 | 0 | 0 | 0 | 0 |
| 0 | -0.40 | 3.67 | -0.11 | 0.91 | NA | Gm35019 | | 0 | 0 | 0 | 0 | 0 | 1 | 0 | 0 | 0 | 0 | 0 |
| 0 | -0.40 | 3.67 | -0.11 | 0.91 | NA | Gm22488 | | 0 | 0 | 0 | 0 | 0 | 1 | 0 | 0 | 0 | 0 | 0 |
| 0 | -0.40 | 3.67 | -0.11 | 0.91 | NA | Gm49436 | | 0 | 0 | 0 | 0 | 0 | 1 | 0 | 0 | 0 | 0 | 0 |
| 0 | -0.40 | 3.67 | -0.11 | 0.91 | NA | Mir6959 | | 0 | 0 | 0 | 0 | 0 | 1 | 0 | 0 | 0 | 0 | 0 |
| 0 | -0.40 | 3.67 | -0.11 | 0.91 | NA | Gm15383 | | 0 | 0 | 0 | 0 | 0 | 1 | 0 | 0 | 0 | 0 | 0 |
| 0 | -0.40 | 3.67 | -0.11 | 0.91 | NA | Gm15382 | | 0 | 0 | 0 | 0 | 0 | 1 | 0 | 0 | 0 | 0 | 0 |
| 0 | -0.40 | 3.67 | -0.11 | 0.91 | NA | Gm25546 | | 0 | 0 | 0 | 0 | 0 | 1 | 0 | 0 | 0 | 0 | 0 |
| 0 | -0.40 | 3.67 | -0.11 | 0.91 | NA | Gm49273 | | 0 | 0 | 0 | 0 | 0 | 1 | 0 | 0 | 0 | 0 | 0 |
| 0 | -0.40 | 3.67 | -0.11 | 0.91 | NA | Dnase1 | deoxyribonuclease I | 0 | 0 | 0 | 0 | 0 | 1 | 0 | 0 | 0 | 0 | 0 |
| 0 | -0.40 | 3.67 | -0.11 | 0.91 | NA | Gm15897 | | 0 | 0 | 0 | 0 | 0 | 1 | 0 | 0 | 0 | 0 | 0 |
| 0 | -0.40 | 3.67 | -0.11 | 0.91 | NA | Gm25547 | | 0 | 0 | 0 | 0 | 0 | 1 | 0 | 0 | 0 | 0 | 0 |
| 0 | -0.40 | 3.67 | -0.11 | 0.91 | NA | Mir130b | | 0 | 0 | 0 | 0 | 0 | 1 | 0 | 0 | 0 | 0 | 0 |
| 0 | -0.40 | 3.67 | -0.11 | 0.91 | NA | Gm26170 | | 0 | 0 | 0 | 0 | 0 | 1 | 0 | 0 | 0 | 0 | 0 |
| 0 | -0.40 | 3.67 | -0.11 | 0.91 | NA | Gm15729 | | 0 | 0 | 0 | 0 | 0 | 1 | 0 | 0 | 0 | 0 | 0 |
| 0 | -0.40 | 3.67 | -0.11 | 0.91 | NA | 4932412D23Rik | | 0 | 0 | 0 | 0 | 0 | 1 | 0 | 0 | 0 | 0 | 0 |
| 0 | -0.40 | 3.67 | -0.11 | 0.91 | NA | Gm49701 | | 0 | 0 | 0 | 0 | 0 | 1 | 0 | 0 | 0 | 0 | 0 |
| 0 | -0.40 | 3.67 | -0.11 | 0.91 | NA | Olfr204 | olfactory receptor 204 | 0 | 0 | 0 | 0 | 0 | 1 | 0 | 0 | 0 | 0 | 0 |
| 0 | -0.40 | 3.67 | -0.11 | 0.91 | NA | Gm21833 | | 0 | 0 | 0 | 0 | 0 | 1 | 0 | 0 | 0 | 0 | 0 |
| 0 | -0.40 | 3.67 | -0.11 | 0.91 | NA | Cldn17 | claudin 17 | 0 | 0 | 0 | 0 | 0 | 1 | 0 | 0 | 0 | 0 | 0 |

| | | | | | | | | | | | | | | | | | | |
|---|---|---|---|---|---|---|---|---|---|---|---|---|---|---|---|---|---|---|
| 0 | -0.40 | 3.67 | -0.11 | 0.91 | NA | 2310043M15Rik | | 0 | 0 | 0 | 0 | 0 | 1 | 0 | 0 | 0 | 0 | 0 |
| 0 | -0.40 | 3.67 | -0.11 | 0.91 | NA | Gm15342 | | 0 | 0 | 0 | 0 | 0 | 1 | 0 | 0 | 0 | 0 | 0 |
| 0 | -0.40 | 3.67 | -0.11 | 0.91 | NA | Gm49673 | | 0 | 0 | 0 | 0 | 0 | 1 | 0 | 0 | 0 | 0 | 0 |
| 0 | -0.40 | 3.67 | -0.11 | 0.91 | NA | Gm34799 | | 0 | 0 | 0 | 0 | 0 | 1 | 0 | 0 | 0 | 0 | 0 |
| 0 | -0.40 | 3.67 | -0.11 | 0.91 | NA | 1700110C19Rik | | 0 | 0 | 0 | 0 | 0 | 1 | 0 | 0 | 0 | 0 | 0 |
| 0 | -0.40 | 3.67 | -0.11 | 0.91 | NA | Gm8597 | | 0 | 0 | 0 | 0 | 0 | 1 | 0 | 0 | 0 | 0 | 0 |
| 0 | -0.40 | 3.67 | -0.11 | 0.91 | NA | Rnu6 | | 0 | 0 | 0 | 0 | 0 | 1 | 0 | 0 | 0 | 0 | 0 |
| 0 | -0.40 | 3.67 | -0.11 | 0.91 | NA | Gm7168 | predicted gene 7168 | 0 | 0 | 0 | 0 | 0 | 1 | 0 | 0 | 0 | 0 | 0 |
| 0 | -0.40 | 3.67 | -0.11 | 0.91 | NA | Gm18564 | | 0 | 0 | 0 | 0 | 0 | 1 | 0 | 0 | 0 | 0 | 0 |
| 0 | -0.40 | 3.67 | -0.11 | 0.91 | NA | Gm5092 | | 0 | 0 | 0 | 0 | 0 | 1 | 0 | 0 | 0 | 0 | 0 |
| 0 | -0.40 | 3.67 | -0.11 | 0.91 | NA | Gm49691 | | 0 | 0 | 0 | 0 | 0 | 1 | 0 | 0 | 0 | 0 | 0 |
| 0 | -0.40 | 3.67 | -0.11 | 0.91 | NA | Gm34455 | | 0 | 0 | 0 | 0 | 0 | 1 | 0 | 0 | 0 | 0 | 0 |
| 0 | -0.40 | 3.67 | -0.11 | 0.91 | NA | Mir6968 | | 0 | 0 | 0 | 0 | 0 | 1 | 0 | 0 | 0 | 0 | 0 |
| 0 | -0.40 | 3.67 | -0.11 | 0.91 | NA | Gm49889 | | 0 | 0 | 0 | 0 | 0 | 1 | 0 | 0 | 0 | 0 | 0 |
| 0 | -0.40 | 3.67 | -0.11 | 0.91 | NA | Gm50226 | | 0 | 0 | 0 | 0 | 0 | 1 | 0 | 0 | 0 | 0 | 0 |
| 0 | -0.40 | 3.67 | -0.11 | 0.91 | NA | Gm50244 | | 0 | 0 | 0 | 0 | 0 | 1 | 0 | 0 | 0 | 0 | 0 |
| 0 | -0.40 | 3.67 | -0.11 | 0.91 | NA | Gm50248 | | 0 | 0 | 0 | 0 | 0 | 1 | 0 | 0 | 0 | 0 | 0 |
| 0 | -0.40 | 3.67 | -0.11 | 0.91 | NA | 4833413E03Rik | | 0 | 0 | 0 | 0 | 0 | 1 | 0 | 0 | 0 | 0 | 0 |
| 0 | -0.40 | 3.67 | -0.11 | 0.91 | NA | Gm20483 | | 0 | 0 | 0 | 0 | 0 | 1 | 0 | 0 | 0 | 0 | 0 |
| 0 | -0.40 | 3.67 | -0.11 | 0.91 | NA | Gm8878 | | 0 | 0 | 0 | 0 | 0 | 1 | 0 | 0 | 0 | 0 | 0 |
| 0 | -0.40 | 3.67 | -0.11 | 0.91 | NA | Olfr132 | olfactory receptor 132 | 0 | 0 | 0 | 0 | 0 | 1 | 0 | 0 | 0 | 0 | 0 |
| 0 | -0.40 | 3.67 | -0.11 | 0.91 | NA | Gm17080 | | 0 | 0 | 0 | 0 | 0 | 1 | 0 | 0 | 0 | 0 | 0 |
| 0 | -0.40 | 3.67 | -0.11 | 0.91 | NA | Slc22a7 | solute carrier family 22 (organic anion tra | 0 | 0 | 0 | 0 | 0 | 1 | 0 | 0 | 0 | 0 | 0 |
| 0 | -0.40 | 3.67 | -0.11 | 0.91 | NA | 1700122O11Ril | RIKEN cDNA 1700122O11 gene | 0 | 0 | 0 | 0 | 0 | 1 | 0 | 0 | 0 | 0 | 0 |
| 0 | -0.40 | 3.67 | -0.11 | 0.91 | NA | 1700067P10Rik | | 0 | 0 | 0 | 0 | 0 | 1 | 0 | 0 | 0 | 0 | 0 |
| 0 | -0.40 | 3.67 | -0.11 | 0.91 | NA | Gm49914 | | 0 | 0 | 0 | 0 | 0 | 1 | 0 | 0 | 0 | 0 | 0 |
| 0 | -0.40 | 3.67 | -0.11 | 0.91 | NA | Znrf4 | zinc and ring finger 4 | 0 | 0 | 0 | 0 | 0 | 1 | 0 | 0 | 0 | 0 | 0 |
| 0 | -0.40 | 3.67 | -0.11 | 0.91 | NA | Gm49888 | | 0 | 0 | 0 | 0 | 0 | 1 | 0 | 0 | 0 | 0 | 0 |
| 0 | -0.40 | 3.67 | -0.11 | 0.91 | NA | Gm15978 | | 0 | 0 | 0 | 0 | 0 | 1 | 0 | 0 | 0 | 0 | 0 |
| 0 | -0.40 | 3.67 | -0.11 | 0.91 | NA | Gm30794 | | 0 | 0 | 0 | 0 | 0 | 1 | 0 | 0 | 0 | 0 | 0 |
| 0 | -0.40 | 3.67 | -0.11 | 0.91 | NA | Rpl36-ps4 | | 0 | 0 | 0 | 0 | 0 | 1 | 0 | 0 | 0 | 0 | 0 |
| 0 | -0.40 | 3.67 | -0.11 | 0.91 | NA | Gm9407 | | 0 | 0 | 0 | 0 | 0 | 1 | 0 | 0 | 0 | 0 | 0 |
| 0 | -0.40 | 3.67 | -0.11 | 0.91 | NA | Gm26575 | | 0 | 0 | 0 | 0 | 0 | 1 | 0 | 0 | 0 | 0 | 0 |
| 0 | -0.40 | 3.67 | -0.11 | 0.91 | NA | 4930415O11Rik | | 0 | 0 | 0 | 0 | 0 | 1 | 0 | 0 | 0 | 0 | 0 |
| 0 | -0.40 | 3.67 | -0.11 | 0.91 | NA | Gm25845 | | 0 | 0 | 0 | 0 | 0 | 1 | 0 | 0 | 0 | 0 | 0 |
| 0 | -0.40 | 3.67 | -0.11 | 0.91 | NA | 0710001A04Rik | | 0 | 0 | 0 | 0 | 0 | 1 | 0 | 0 | 0 | 0 | 0 |
| 0 | -0.40 | 3.67 | -0.11 | 0.91 | NA | Gm50060 | | 0 | 0 | 0 | 0 | 0 | 1 | 0 | 0 | 0 | 0 | 0 |
| 0 | -0.40 | 3.67 | -0.11 | 0.91 | NA | Gm49966 | | 0 | 0 | 0 | 0 | 0 | 1 | 0 | 0 | 0 | 0 | 0 |
| 0 | -0.40 | 3.67 | -0.11 | 0.91 | NA | Gm37753 | | 0 | 0 | 0 | 0 | 0 | 1 | 0 | 0 | 0 | 0 | 0 |
| 0 | -0.40 | 3.67 | -0.11 | 0.91 | NA | Gm20162 | | 0 | 0 | 0 | 0 | 0 | 1 | 0 | 0 | 0 | 0 | 0 |
| 0 | -0.40 | 3.67 | -0.11 | 0.91 | NA | Gm31907 | | 0 | 0 | 0 | 0 | 0 | 1 | 0 | 0 | 0 | 0 | 0 |
| 0 | -0.40 | 3.67 | -0.11 | 0.91 | NA | Gm8642 | | 0 | 0 | 0 | 0 | 0 | 1 | 0 | 0 | 0 | 0 | 0 |
| 0 | -0.40 | 3.67 | -0.11 | 0.91 | NA | Cplx4 | complexin 4 | 0 | 0 | 0 | 0 | 0 | 1 | 0 | 0 | 0 | 0 | 0 |
| 0 | -0.40 | 3.67 | -0.11 | 0.91 | NA | A730085E03Rik | | 0 | 0 | 0 | 0 | 0 | 1 | 0 | 0 | 0 | 0 | 0 |
| 0 | -0.40 | 3.67 | -0.11 | 0.91 | NA | | | 0 | 0 | 0 | 0 | 0 | 1 | 0 | 0 | 0 | 0 | 0 |
| 0 | -0.40 | 3.67 | -0.11 | 0.91 | NA | Gm9750 | | 0 | 0 | 0 | 0 | 0 | 1 | 0 | 0 | 0 | 0 | 0 |
| 0 | -0.40 | 3.67 | -0.11 | 0.91 | NA | Gm6684 | | 0 | 0 | 0 | 0 | 0 | 1 | 0 | 0 | 0 | 0 | 0 |
| 0 | -0.40 | 3.67 | -0.11 | 0.91 | NA | Gm23095 | | 0 | 0 | 0 | 0 | 0 | 1 | 0 | 0 | 0 | 0 | 0 |
| 0 | -0.40 | 3.67 | -0.11 | 0.91 | NA | Gm18425 | | 0 | 0 | 0 | 0 | 0 | 1 | 0 | 0 | 0 | 0 | 0 |
| 0 | -0.40 | 3.67 | -0.11 | 0.91 | NA | Cyp2c52-ps | | 0 | 0 | 0 | 0 | 0 | 1 | 0 | 0 | 0 | 0 | 0 |
| 0 | -0.40 | 3.67 | -0.11 | 0.91 | NA | Gm47938 | | 0 | 0 | 0 | 0 | 0 | 1 | 0 | 0 | 0 | 0 | 0 |
| 0 | -0.40 | 3.67 | -0.11 | 0.91 | NA | Gm20395 | | 0 | 0 | 0 | 0 | 0 | 1 | 0 | 0 | 0 | 0 | 0 |
| 0 | -0.40 | 3.67 | -0.11 | 0.91 | NA | Gm6990 | | 0 | 0 | 0 | 0 | 0 | 1 | 0 | 0 | 0 | 0 | 0 |
| 0 | -0.40 | 3.67 | -0.11 | 0.91 | NA | Gm50279 | | 0 | 0 | 0 | 0 | 0 | 1 | 0 | 0 | 0 | 0 | 0 |
| 0 | -0.40 | 3.67 | -0.11 | 0.91 | NA | Gm14504 | | 0 | 0 | 0 | 0 | 0 | 1 | 0 | 0 | 0 | 0 | 0 |
| 0 | -0.40 | 3.67 | -0.11 | 0.91 | NA | Gm6024 | | 0 | 0 | 0 | 0 | 0 | 1 | 0 | 0 | 0 | 0 | 0 |
| 0 | -0.40 | 3.67 | -0.11 | 0.91 | NA | Gm5639 | | 0 | 0 | 0 | 0 | 0 | 1 | 0 | 0 | 0 | 0 | 0 |
| 0 | -0.40 | 3.67 | -0.11 | 0.91 | NA | Gm26726 | | 0 | 0 | 0 | 0 | 0 | 1 | 0 | 0 | 0 | 0 | 0 |
| 0 | -0.40 | 3.67 | -0.11 | 0.91 | NA | Gm7153 | | 0 | 0 | 0 | 0 | 0 | 1 | 0 | 0 | 0 | 0 | 0 |
| 0 | -0.40 | 3.67 | -0.11 | 0.91 | NA | Gm14788 | | 0 | 0 | 0 | 0 | 0 | 1 | 0 | 0 | 0 | 0 | 0 |

| | | | | | | | | | | | | | | | | | | | |
|---|---|---|---|---|---|---|---|---|---|---|---|---|---|---|---|---|---|---|---|
| 0 | | -0.40 | 3.67 | -0.11 | 0.91 | NA | Gm14858 | | 0 | 0 | 0 | 0 | 0 | 1 | 0 | 0 | 0 | 0 | 0 |
| 0 | | -0.40 | 3.67 | -0.11 | 0.91 | NA | Gm9157 | | 0 | 0 | 0 | 0 | 0 | 1 | 0 | 0 | 0 | 0 | 0 |
| 0 | | -0.40 | 3.67 | -0.11 | 0.91 | NA | Gm6491 | | 0 | 0 | 0 | 0 | 0 | 1 | 0 | 0 | 0 | 0 | 0 |
| 0 | | -0.40 | 3.67 | -0.11 | 0.91 | NA | Gucy2f | guanylate cyclase 2f | 0 | 0 | 0 | 0 | 0 | 1 | 0 | 0 | 0 | 0 | 0 |
| 0 | | -0.40 | 3.67 | -0.11 | 0.91 | NA | Mir3620 | | 0 | 0 | 0 | 0 | 0 | 1 | 0 | 0 | 0 | 0 | 0 |
| 0 | | -0.40 | 3.67 | -0.11 | 0.91 | NA | Gm15149 | | 0 | 0 | 0 | 0 | 0 | 1 | 0 | 0 | 0 | 0 | 0 |
| 0 | | -0.40 | 3.67 | -0.11 | 0.91 | NA | Gm5395 | | 0 | 0 | 0 | 0 | 0 | 1 | 0 | 0 | 0 | 0 | 0 |
| 0 | | -0.40 | 3.67 | -0.11 | 0.91 | NA | Gm5396 | | 0 | 0 | 0 | 0 | 0 | 1 | 0 | 0 | 0 | 0 | 0 |
| 0 | NA | NA | NA | NA | | NA | 4933401J01Rik | | 0 | 0 | 0 | 0 | 0 | 0 | 0 | 0 | 0 | 0 | 0 |
| 0 | NA | NA | NA | NA | | NA | Gm26206 | | 0 | 0 | 0 | 0 | 0 | 0 | 0 | 0 | 0 | 0 | 0 |
| 0 | NA | NA | NA | NA | | NA | Gm18956 | | 0 | 0 | 0 | 0 | 0 | 0 | 0 | 0 | 0 | 0 | 0 |
| 0 | NA | NA | NA | NA | | NA | Gm37180 | | 0 | 0 | 0 | 0 | 0 | 0 | 0 | 0 | 0 | 0 | 0 |
| 0 | NA | NA | NA | NA | | NA | Gm37363 | | 0 | 0 | 0 | 0 | 0 | 0 | 0 | 0 | 0 | 0 | 0 |
| 0 | NA | NA | NA | NA | | NA | Gm1992 | | 0 | 0 | 0 | 0 | 0 | 0 | 0 | 0 | 0 | 0 | 0 |
| 0 | NA | NA | NA | NA | | NA | Gm38148 | | 0 | 0 | 0 | 0 | 0 | 0 | 0 | 0 | 0 | 0 | 0 |
| 0 | NA | NA | NA | NA | | NA | Gm10568 | | 0 | 0 | 0 | 0 | 0 | 0 | 0 | 0 | 0 | 0 | 0 |
| 0 | NA | NA | NA | NA | | NA | Gm38385 | | 0 | 0 | 0 | 0 | 0 | 0 | 0 | 0 | 0 | 0 | 0 |
| 0 | NA | NA | NA | NA | | NA | Gm27396 | | 0 | 0 | 0 | 0 | 0 | 0 | 0 | 0 | 0 | 0 | 0 |
| 0 | NA | NA | NA | NA | | NA | Gm37381 | | 0 | 0 | 0 | 0 | 0 | 0 | 0 | 0 | 0 | 0 | 0 |
| 0 | NA | NA | NA | NA | | NA | Rp1 | retinitis pigmentosa 1 (human) | 0 | 0 | 0 | 0 | 0 | 0 | 0 | 0 | 0 | 0 | 0 |
| 0 | NA | NA | NA | NA | | NA | Gm6101 | | 0 | 0 | 0 | 0 | 0 | 0 | 0 | 0 | 0 | 0 | 0 |
| 0 | NA | NA | NA | NA | | NA | Gm37483 | | 0 | 0 | 0 | 0 | 0 | 0 | 0 | 0 | 0 | 0 | 0 |
| 0 | NA | NA | NA | NA | | NA | Gm37587 | | 0 | 0 | 0 | 0 | 0 | 0 | 0 | 0 | 0 | 0 | 0 |
| 0 | NA | NA | NA | NA | | NA | Gm22307 | | 0 | 0 | 0 | 0 | 0 | 0 | 0 | 0 | 0 | 0 | 0 |
| 0 | NA | NA | NA | NA | | NA | Gm38076 | | 0 | 0 | 0 | 0 | 0 | 0 | 0 | 0 | 0 | 0 | 0 |
| 0 | NA | NA | NA | NA | | NA | Gm37323 | | 0 | 0 | 0 | 0 | 0 | 0 | 0 | 0 | 0 | 0 | 0 |
| 0 | NA | NA | NA | NA | | NA | Gm25493 | | 0 | 0 | 0 | 0 | 0 | 0 | 0 | 0 | 0 | 0 | 0 |
| 0 | NA | NA | NA | NA | | NA | Gm2053 | | 0 | 0 | 0 | 0 | 0 | 0 | 0 | 0 | 0 | 0 | 0 |
| 0 | NA | NA | NA | NA | | NA | Gm37988 | | 0 | 0 | 0 | 0 | 0 | 0 | 0 | 0 | 0 | 0 | 0 |
| 0 | NA | NA | NA | NA | | NA | Gm17100 | | 0 | 0 | 0 | 0 | 0 | 0 | 0 | 0 | 0 | 0 | 0 |
| 0 | NA | NA | NA | NA | | NA | Gm16041 | | 0 | 0 | 0 | 0 | 0 | 0 | 0 | 0 | 0 | 0 | 0 |
| 0 | NA | NA | NA | NA | | NA | Gm17101 | | 0 | 0 | 0 | 0 | 0 | 0 | 0 | 0 | 0 | 0 | 0 |
| 0 | NA | NA | NA | NA | | NA | Gm7182 | | 0 | 0 | 0 | 0 | 0 | 0 | 0 | 0 | 0 | 0 | 0 |
| 0 | NA | NA | NA | NA | | NA | Gm38264 | | 0 | 0 | 0 | 0 | 0 | 0 | 0 | 0 | 0 | 0 | 0 |
| 0 | NA | NA | NA | NA | | NA | Gm36965 | | 0 | 0 | 0 | 0 | 0 | 0 | 0 | 0 | 0 | 0 | 0 |
| 0 | NA | NA | NA | NA | | NA | Gm22463 | | 0 | 0 | 0 | 0 | 0 | 0 | 0 | 0 | 0 | 0 | 0 |
| 0 | NA | NA | NA | NA | | NA | Gm37429 | | 0 | 0 | 0 | 0 | 0 | 0 | 0 | 0 | 0 | 0 | 0 |
| 0 | NA | NA | NA | NA | | NA | Npbwr1 | neuropeptides B/W receptor 1 | 0 | 0 | 0 | 0 | 0 | 0 | 0 | 0 | 0 | 0 | 0 |
| 0 | NA | NA | NA | NA | | NA | Gm19214 | | 0 | 0 | 0 | 0 | 0 | 0 | 0 | 0 | 0 | 0 | 0 |
| 0 | NA | NA | NA | NA | | NA | Alkal1 | ALK and LTK ligand 1 | 0 | 0 | 0 | 0 | 0 | 0 | 0 | 0 | 0 | 0 | 0 |
| 0 | NA | NA | NA | NA | | NA | Gm7417 | | 0 | 0 | 0 | 0 | 0 | 0 | 0 | 0 | 0 | 0 | 0 |
| 0 | NA | NA | NA | NA | | NA | Gm42492 | | 0 | 0 | 0 | 0 | 0 | 0 | 0 | 0 | 0 | 0 | 0 |
| 0 | NA | NA | NA | NA | | NA | Gm37275 | | 0 | 0 | 0 | 0 | 0 | 0 | 0 | 0 | 0 | 0 | 0 |
| 0 | NA | NA | NA | NA | | NA | Gm37225 | | 0 | 0 | 0 | 0 | 0 | 0 | 0 | 0 | 0 | 0 | 0 |
| 0 | NA | NA | NA | NA | | NA | Gm37489 | | 0 | 0 | 0 | 0 | 0 | 0 | 0 | 0 | 0 | 0 | 0 |
| 0 | NA | NA | NA | NA | | NA | Gm5694 | | 0 | 0 | 0 | 0 | 0 | 0 | 0 | 0 | 0 | 0 | 0 |
| 0 | NA | NA | NA | NA | | NA | Nras-ps2 | | 0 | 0 | 0 | 0 | 0 | 0 | 0 | 0 | 0 | 0 | 0 |
| 0 | NA | NA | NA | NA | | NA | Gm23274 | | 0 | 0 | 0 | 0 | 0 | 0 | 0 | 0 | 0 | 0 | 0 |
| 0 | NA | NA | NA | NA | | NA | Gm19002 | | 0 | 0 | 0 | 0 | 0 | 0 | 0 | 0 | 0 | 0 | 0 |
| 0 | NA | NA | NA | NA | | NA | Gm18984 | | 0 | 0 | 0 | 0 | 0 | 0 | 0 | 0 | 0 | 0 | 0 |
| 0 | NA | NA | NA | NA | | NA | Gm30414 | | 0 | 0 | 0 | 0 | 0 | 0 | 0 | 0 | 0 | 0 | 0 |
| 0 | NA | NA | NA | NA | | NA | Gm37791 | | 0 | 0 | 0 | 0 | 0 | 0 | 0 | 0 | 0 | 0 | 0 |
| 0 | NA | NA | NA | NA | | NA | Gm38216 | | 0 | 0 | 0 | 0 | 0 | 0 | 0 | 0 | 0 | 0 | 0 |
| 0 | NA | NA | NA | NA | | NA | Gm23358 | | 0 | 0 | 0 | 0 | 0 | 0 | 0 | 0 | 0 | 0 | 0 |
| 0 | NA | NA | NA | NA | | NA | Gm6152 | | 0 | 0 | 0 | 0 | 0 | 0 | 0 | 0 | 0 | 0 | 0 |
| 0 | NA | NA | NA | NA | | NA | Gm7445 | | 0 | 0 | 0 | 0 | 0 | 0 | 0 | 0 | 0 | 0 | 0 |
| 0 | NA | NA | NA | NA | | NA | Gm7493 | | 0 | 0 | 0 | 0 | 0 | 0 | 0 | 0 | 0 | 0 | 0 |
| 0 | NA | NA | NA | NA | | NA | Gm38008 | | 0 | 0 | 0 | 0 | 0 | 0 | 0 | 0 | 0 | 0 | 0 |
| 0 | NA | NA | NA | NA | | NA | Gm37143 | | 0 | 0 | 0 | 0 | 0 | 0 | 0 | 0 | 0 | 0 | 0 |
| 0 | NA | NA | NA | NA | | NA | Gm18299 | | 0 | 0 | 0 | 0 | 0 | 0 | 0 | 0 | 0 | 0 | 0 |

| | | | | | | | | | | | | | | | | | | |
|---|---|---|---|---|---|---|---|---|---|---|---|---|---|---|---|---|---|---|
| 0 | NA | NA | NA | NA | NA | Gm24765 | | 0 | 0 | 0 | 0 | 0 | 0 | 0 | 0 | 0 | 0 | 0 |
| 0 | NA | NA | NA | NA | NA | Gm29520 | | 0 | 0 | 0 | 0 | 0 | 0 | 0 | 0 | 0 | 0 | 0 |
| 0 | NA | NA | NA | NA | NA | Gm22607 | | 0 | 0 | 0 | 0 | 0 | 0 | 0 | 0 | 0 | 0 | 0 |
| 0 | NA | NA | NA | NA | NA | Gm15818 | | 0 | 0 | 0 | 0 | 0 | 0 | 0 | 0 | 0 | 0 | 0 |
| 0 | NA | NA | NA | NA | NA | Gm24674 | | 0 | 0 | 0 | 0 | 0 | 0 | 0 | 0 | 0 | 0 | 0 |
| 0 | NA | NA | NA | NA | NA | 1700047N06Rik | | 0 | 0 | 0 | 0 | 0 | 0 | 0 | 0 | 0 | 0 | 0 |
| 0 | NA | NA | NA | NA | NA | Gm26348 | | 0 | 0 | 0 | 0 | 0 | 0 | 0 | 0 | 0 | 0 | 0 |
| 0 | NA | NA | NA | NA | NA | Gm25253 | | 0 | 0 | 0 | 0 | 0 | 0 | 0 | 0 | 0 | 0 | 0 |
| 0 | NA | NA | NA | NA | NA | Gm37133 | | 0 | 0 | 0 | 0 | 0 | 0 | 0 | 0 | 0 | 0 | 0 |
| 0 | NA | NA | NA | NA | NA | Gm5522 | | 0 | 0 | 0 | 0 | 0 | 0 | 0 | 0 | 0 | 0 | 0 |
| 0 | NA | NA | NA | NA | NA | Gm22963 | | 0 | 0 | 0 | 0 | 0 | 0 | 0 | 0 | 0 | 0 | 0 |
| 0 | NA | NA | NA | NA | NA | Gm24173 | | 0 | 0 | 0 | 0 | 0 | 0 | 0 | 0 | 0 | 0 | 0 |
| 0 | NA | NA | NA | NA | NA | Gm37410 | | 0 | 0 | 0 | 0 | 0 | 0 | 0 | 0 | 0 | 0 | 0 |
| 0 | NA | NA | NA | NA | NA | Gm38178 | | 0 | 0 | 0 | 0 | 0 | 0 | 0 | 0 | 0 | 0 | 0 |
| 0 | NA | NA | NA | NA | NA | Gm29273 | | 0 | 0 | 0 | 0 | 0 | 0 | 0 | 0 | 0 | 0 | 0 |
| 0 | NA | NA | NA | NA | NA | Gm38069 | | 0 | 0 | 0 | 0 | 0 | 0 | 0 | 0 | 0 | 0 | 0 |
| 0 | NA | NA | NA | NA | NA | Gm7560 | | 0 | 0 | 0 | 0 | 0 | 0 | 0 | 0 | 0 | 0 | 0 |
| 0 | NA | NA | NA | NA | NA | Gm22633 | | 0 | 0 | 0 | 0 | 0 | 0 | 0 | 0 | 0 | 0 | 0 |
| 0 | NA | NA | NA | NA | NA | Mir6341 | | 0 | 0 | 0 | 0 | 0 | 0 | 0 | 0 | 0 | 0 | 0 |
| 0 | NA | NA | NA | NA | NA | Gm6216 | | 0 | 0 | 0 | 0 | 0 | 0 | 0 | 0 | 0 | 0 | 0 |
| 0 | NA | NA | NA | NA | NA | Gm17644 | | 0 | 0 | 0 | 0 | 0 | 0 | 0 | 0 | 0 | 0 | 0 |
| 0 | NA | NA | NA | NA | NA | Gm29663 | | 0 | 0 | 0 | 0 | 0 | 0 | 0 | 0 | 0 | 0 | 0 |
| 0 | NA | NA | NA | NA | NA | Gm37162 | | 0 | 0 | 0 | 0 | 0 | 0 | 0 | 0 | 0 | 0 | 0 |
| 0 | NA | NA | NA | NA | NA | Gm5250 | | 0 | 0 | 0 | 0 | 0 | 0 | 0 | 0 | 0 | 0 | 0 |
| 0 | NA | NA | NA | NA | NA | Gm29283 | | 0 | 0 | 0 | 0 | 0 | 0 | 0 | 0 | 0 | 0 | 0 |
| 0 | NA | NA | NA | NA | NA | Gm27881 | | 0 | 0 | 0 | 0 | 0 | 0 | 0 | 0 | 0 | 0 | 0 |
| 0 | NA | NA | NA | NA | NA | Gm38223 | | 0 | 0 | 0 | 0 | 0 | 0 | 0 | 0 | 0 | 0 | 0 |
| 0 | NA | NA | NA | NA | NA | Gm26273 | | 0 | 0 | 0 | 0 | 0 | 0 | 0 | 0 | 0 | 0 | 0 |
| 0 | NA | NA | NA | NA | NA | Gm36947 | | 0 | 0 | 0 | 0 | 0 | 0 | 0 | 0 | 0 | 0 | 0 |
| 0 | NA | NA | NA | NA | NA | Gm25491 | | 0 | 0 | 0 | 0 | 0 | 0 | 0 | 0 | 0 | 0 | 0 |
| 0 | NA | NA | NA | NA | NA | Gm37400 | | 0 | 0 | 0 | 0 | 0 | 0 | 0 | 0 | 0 | 0 | 0 |
| 0 | NA | NA | NA | NA | NA | Gm22616 | | 0 | 0 | 0 | 0 | 0 | 0 | 0 | 0 | 0 | 0 | 0 |
| 0 | NA | NA | NA | NA | NA | Gm26345 | | 0 | 0 | 0 | 0 | 0 | 0 | 0 | 0 | 0 | 0 | 0 |
| 0 | NA | NA | NA | NA | NA | Gm25227 | | 0 | 0 | 0 | 0 | 0 | 0 | 0 | 0 | 0 | 0 | 0 |
| 0 | NA | NA | NA | NA | NA | Gm25168 | | 0 | 0 | 0 | 0 | 0 | 0 | 0 | 0 | 0 | 0 | 0 |
| 0 | NA | NA | NA | NA | NA | Gm28669 | | 0 | 0 | 0 | 0 | 0 | 0 | 0 | 0 | 0 | 0 | 0 |
| 0 | NA | NA | NA | NA | NA | Gm23259 | | 0 | 0 | 0 | 0 | 0 | 0 | 0 | 0 | 0 | 0 | 0 |
| 0 | NA | NA | NA | NA | NA | Gm38249 | | 0 | 0 | 0 | 0 | 0 | 0 | 0 | 0 | 0 | 0 | 0 |
| 0 | NA | NA | NA | NA | NA | Gm28095 | | 0 | 0 | 0 | 0 | 0 | 0 | 0 | 0 | 0 | 0 | 0 |
| 0 | NA | NA | NA | NA | NA | Gm7568 | | 0 | 0 | 0 | 0 | 0 | 0 | 0 | 0 | 0 | 0 | 0 |
| 0 | NA | NA | NA | NA | NA | Gm7654 | | 0 | 0 | 0 | 0 | 0 | 0 | 0 | 0 | 0 | 0 | 0 |
| 0 | NA | NA | NA | NA | NA | Gm29649 | | 0 | 0 | 0 | 0 | 0 | 0 | 0 | 0 | 0 | 0 | 0 |
| 0 | NA | NA | NA | NA | NA | Gm28376 | | 0 | 0 | 0 | 0 | 0 | 0 | 0 | 0 | 0 | 0 | 0 |
| 0 | NA | NA | NA | NA | NA | 4921511E07Rik | | 0 | 0 | 0 | 0 | 0 | 0 | 0 | 0 | 0 | 0 | 0 |
| 0 | NA | NA | NA | NA | NA | Gm28783 | | 0 | 0 | 0 | 0 | 0 | 0 | 0 | 0 | 0 | 0 | 0 |
| 0 | NA | NA | NA | NA | NA | Gm28785 | | 0 | 0 | 0 | 0 | 0 | 0 | 0 | 0 | 0 | 0 | 0 |
| 0 | NA | NA | NA | NA | NA | Gm5251 | | 0 | 0 | 0 | 0 | 0 | 0 | 0 | 0 | 0 | 0 | 0 |
| 0 | NA | NA | NA | NA | NA | Gm25166 | | 0 | 0 | 0 | 0 | 0 | 0 | 0 | 0 | 0 | 0 | 0 |
| 0 | NA | NA | NA | NA | NA | Gm28154 | | 0 | 0 | 0 | 0 | 0 | 0 | 0 | 0 | 0 | 0 | 0 |
| 0 | NA | NA | NA | NA | NA | Gm28153 | | 0 | 0 | 0 | 0 | 0 | 0 | 0 | 0 | 0 | 0 | 0 |
| 0 | NA | NA | NA | NA | NA | Gm18775 | | 0 | 0 | 0 | 0 | 0 | 0 | 0 | 0 | 0 | 0 | 0 |
| 0 | NA | NA | NA | NA | NA | Gm17969 | | 0 | 0 | 0 | 0 | 0 | 0 | 0 | 0 | 0 | 0 | 0 |
| 0 | NA | NA | NA | NA | NA | Gm28756 | | 0 | 0 | 0 | 0 | 0 | 0 | 0 | 0 | 0 | 0 | 0 |
| 0 | NA | NA | NA | NA | NA | Gm28755 | | 0 | 0 | 0 | 0 | 0 | 0 | 0 | 0 | 0 | 0 | 0 |
| 0 | NA | NA | NA | NA | NA | Gm45784 | | 0 | 0 | 0 | 0 | 0 | 0 | 0 | 0 | 0 | 0 | 0 |
| 0 | NA | NA | NA | NA | NA | Crisp4 | cysteine-rich secretory protein 4 | 0 | 0 | 0 | 0 | 0 | 0 | 0 | 0 | 0 | 0 | 0 |
| 0 | NA | NA | NA | NA | NA | Defb44-ps | | 0 | 0 | 0 | 0 | 0 | 0 | 0 | 0 | 0 | 0 | 0 |
| 0 | NA | NA | NA | NA | NA | Defb18 | defensin beta 18 | 0 | 0 | 0 | 0 | 0 | 0 | 0 | 0 | 0 | 0 | 0 |
| 0 | NA | NA | NA | NA | NA | Defb41 | defensin beta 41 | 0 | 0 | 0 | 0 | 0 | 0 | 0 | 0 | 0 | 0 | 0 |
| 0 | NA | NA | NA | NA | NA | Gm17837 | | 0 | 0 | 0 | 0 | 0 | 0 | 0 | 0 | 0 | 0 | 0 |

| | | | | | | | | | | | | | | | | | | |
|---|---|---|---|---|---|---|---|---|---|---|---|---|---|---|---|---|---|---|
| 0 | NA | NA | NA | NA | NA | Gm5252 | | 0 | 0 | 0 | 0 | 0 | 0 | 0 | 0 | 0 | 0 | 0 |
| 0 | NA | NA | NA | NA | NA | Gm24075 | | 0 | 0 | 0 | 0 | 0 | 0 | 0 | 0 | 0 | 0 | 0 |
| 0 | NA | NA | NA | NA | NA | Gm28343 | | 0 | 0 | 0 | 0 | 0 | 0 | 0 | 0 | 0 | 0 | 0 |
| 0 | NA | NA | NA | NA | NA | Gm28344 | | 0 | 0 | 0 | 0 | 0 | 0 | 0 | 0 | 0 | 0 | 0 |
| 0 | NA | NA | NA | NA | NA | Gm15825 | | 0 | 0 | 0 | 0 | 0 | 0 | 0 | 0 | 0 | 0 | 0 |
| 0 | NA | NA | NA | NA | NA | Tfap2d | transcription factor AP-2, delta | 0 | 0 | 0 | 0 | 0 | 0 | 0 | 0 | 0 | 0 | 0 |
| 0 | NA | NA | NA | NA | NA | Gm28340 | | 0 | 0 | 0 | 0 | 0 | 0 | 0 | 0 | 0 | 0 | 0 |
| 0 | NA | NA | NA | NA | NA | Gm28341 | | 0 | 0 | 0 | 0 | 0 | 0 | 0 | 0 | 0 | 0 | 0 |
| 0 | NA | NA | NA | NA | NA | Gm24633 | | 0 | 0 | 0 | 0 | 0 | 0 | 0 | 0 | 0 | 0 | 0 |
| 0 | NA | NA | NA | NA | NA | Gm38382 | | 0 | 0 | 0 | 0 | 0 | 0 | 0 | 0 | 0 | 0 | 0 |
| 0 | NA | NA | NA | NA | NA | 4930486I03Rik | | 0 | 0 | 0 | 0 | 0 | 0 | 0 | 0 | 0 | 0 | 0 |
| 0 | NA | NA | NA | NA | NA | Gm24162 | | 0 | 0 | 0 | 0 | 0 | 0 | 0 | 0 | 0 | 0 | 0 |
| 0 | NA | NA | NA | NA | NA | Gm28653 | | 0 | 0 | 0 | 0 | 0 | 0 | 0 | 0 | 0 | 0 | 0 |
| 0 | NA | NA | NA | NA | NA | Mir206 | | 0 | 0 | 0 | 0 | 0 | 0 | 0 | 0 | 0 | 0 | 0 |
| 0 | NA | NA | NA | NA | NA | Mir133b | | 0 | 0 | 0 | 0 | 0 | 0 | 0 | 0 | 0 | 0 | 0 |
| 0 | NA | NA | NA | NA | NA | Gm20587 | | 0 | 0 | 0 | 0 | 0 | 0 | 0 | 0 | 0 | 0 | 0 |
| 0 | NA | NA | NA | NA | NA | Gm28065 | | 0 | 0 | 0 | 0 | 0 | 0 | 0 | 0 | 0 | 0 | 0 |
| 0 | NA | NA | NA | NA | NA | Gm24723 | | 0 | 0 | 0 | 0 | 0 | 0 | 0 | 0 | 0 | 0 | 0 |
| 0 | NA | NA | NA | NA | NA | Gm24171 | | 0 | 0 | 0 | 0 | 0 | 0 | 0 | 0 | 0 | 0 | 0 |
| 0 | NA | NA | NA | NA | NA | Gm28064 | | 0 | 0 | 0 | 0 | 0 | 0 | 0 | 0 | 0 | 0 | 0 |
| 0 | NA | NA | NA | NA | NA | Gm38224 | | 0 | 0 | 0 | 0 | 0 | 0 | 0 | 0 | 0 | 0 | 0 |
| 0 | NA | NA | NA | NA | NA | Gm28836 | | 0 | 0 | 0 | 0 | 0 | 0 | 0 | 0 | 0 | 0 | 0 |
| 0 | NA | NA | NA | NA | NA | Gm21909 | | 0 | 0 | 0 | 0 | 0 | 0 | 0 | 0 | 0 | 0 | 0 |
| 0 | NA | NA | NA | NA | NA | Gm16329 | | 0 | 0 | 0 | 0 | 0 | 0 | 0 | 0 | 0 | 0 | 0 |
| 0 | NA | NA | NA | NA | NA | Gm16327 | | 0 | 0 | 0 | 0 | 0 | 0 | 0 | 0 | 0 | 0 | 0 |
| 0 | NA | NA | NA | NA | NA | Khdc1c | KH domain containing 1C | 0 | 0 | 0 | 0 | 0 | 0 | 0 | 0 | 0 | 0 | 0 |
| 0 | NA | NA | NA | NA | NA | Gm2709 | | 0 | 0 | 0 | 0 | 0 | 0 | 0 | 0 | 0 | 0 | 0 |
| 0 | NA | NA | NA | NA | NA | Gm16533 | | 0 | 0 | 0 | 0 | 0 | 0 | 0 | 0 | 0 | 0 | 0 |
| 0 | NA | NA | NA | NA | NA | Khdc1b | KH domain containing 1B | 0 | 0 | 0 | 0 | 0 | 0 | 0 | 0 | 0 | 0 | 0 |
| 0 | NA | NA | NA | NA | NA | Gm16328 | | 0 | 0 | 0 | 0 | 0 | 0 | 0 | 0 | 0 | 0 | 0 |
| 0 | NA | NA | NA | NA | NA | Gm36949 | | 0 | 0 | 0 | 0 | 0 | 0 | 0 | 0 | 0 | 0 | 0 |
| 0 | NA | NA | NA | NA | NA | Gm38243 | | 0 | 0 | 0 | 0 | 0 | 0 | 0 | 0 | 0 | 0 | 0 |
| 0 | NA | NA | NA | NA | NA | Gm28239 | | 0 | 0 | 0 | 0 | 0 | 0 | 0 | 0 | 0 | 0 | 0 |
| 0 | NA | NA | NA | NA | NA | Gm26866 | | 0 | 0 | 0 | 0 | 0 | 0 | 0 | 0 | 0 | 0 | 0 |
| 0 | NA | NA | NA | NA | NA | Gm7761 | | 0 | 0 | 0 | 0 | 0 | 0 | 0 | 0 | 0 | 0 | 0 |
| 0 | NA | NA | NA | NA | NA | Ppp1r14bl | protein phosphatase 1, regulatory inhibitc | 0 | 0 | 0 | 0 | 0 | 0 | 0 | 0 | 0 | 0 | 0 |
| 0 | NA | NA | NA | NA | NA | Gm24176 | | 0 | 0 | 0 | 0 | 0 | 0 | 0 | 0 | 0 | 0 | 0 |
| 0 | NA | NA | NA | NA | NA | Mir30a | | 0 | 0 | 0 | 0 | 0 | 0 | 0 | 0 | 0 | 0 | 0 |
| 0 | NA | NA | NA | NA | NA | Mir30c-2 | | 0 | 0 | 0 | 0 | 0 | 0 | 0 | 0 | 0 | 0 | 0 |
| 0 | NA | NA | NA | NA | NA | Gm20954 | | 0 | 0 | 0 | 0 | 0 | 0 | 0 | 0 | 0 | 0 | 0 |
| 0 | NA | NA | NA | NA | NA | Gm28822 | | 0 | 0 | 0 | 0 | 0 | 0 | 0 | 0 | 0 | 0 | 0 |
| 0 | NA | NA | NA | NA | NA | Gm26524 | | 0 | 0 | 0 | 0 | 0 | 0 | 0 | 0 | 0 | 0 | 0 |
| 0 | NA | NA | NA | NA | NA | Gm26607 | | 0 | 0 | 0 | 0 | 0 | 0 | 0 | 0 | 0 | 0 | 0 |
| 0 | NA | NA | NA | NA | NA | Gm26418 | | 0 | 0 | 0 | 0 | 0 | 0 | 0 | 0 | 0 | 0 | 0 |
| 0 | NA | NA | NA | NA | NA | Gm28594 | | 0 | 0 | 0 | 0 | 0 | 0 | 0 | 0 | 0 | 0 | 0 |
| 0 | NA | NA | NA | NA | NA | Gm28237 | | 0 | 0 | 0 | 0 | 0 | 0 | 0 | 0 | 0 | 0 | 0 |
| 0 | NA | NA | NA | NA | NA | Gm9443 | | 0 | 0 | 0 | 0 | 0 | 0 | 0 | 0 | 0 | 0 | 0 |
| 0 | NA | NA | NA | NA | NA | Gm29414 | | 0 | 0 | 0 | 0 | 0 | 0 | 0 | 0 | 0 | 0 | 0 |
| 0 | NA | NA | NA | NA | NA | Gm22260 | | 0 | 0 | 0 | 0 | 0 | 0 | 0 | 0 | 0 | 0 | 0 |
| 0 | NA | NA | NA | NA | NA | Gm37543 | | 0 | 0 | 0 | 0 | 0 | 0 | 0 | 0 | 0 | 0 | 0 |
| 0 | NA | NA | NA | NA | NA | Gm37862 | | 0 | 0 | 0 | 0 | 0 | 0 | 0 | 0 | 0 | 0 | 0 |
| 0 | NA | NA | NA | NA | NA | Gm38286 | | 0 | 0 | 0 | 0 | 0 | 0 | 0 | 0 | 0 | 0 | 0 |
| 0 | NA | NA | NA | NA | NA | Gm37880 | | 0 | 0 | 0 | 0 | 0 | 0 | 0 | 0 | 0 | 0 | 0 |
| 0 | NA | NA | NA | NA | NA | Gm25294 | | 0 | 0 | 0 | 0 | 0 | 0 | 0 | 0 | 0 | 0 | 0 |
| 0 | NA | NA | NA | NA | NA | Gm29214 | | 0 | 0 | 0 | 0 | 0 | 0 | 0 | 0 | 0 | 0 | 0 |
| 0 | NA | NA | NA | NA | NA | Gm37628 | | 0 | 0 | 0 | 0 | 0 | 0 | 0 | 0 | 0 | 0 | 0 |
| 0 | NA | NA | NA | NA | NA | 8030445P17Rik | | 0 | 0 | 0 | 0 | 0 | 0 | 0 | 0 | 0 | 0 | 0 |
| 0 | NA | NA | NA | NA | NA | Gm37077 | | 0 | 0 | 0 | 0 | 0 | 0 | 0 | 0 | 0 | 0 | 0 |
| 0 | NA | NA | NA | NA | NA | Gm37895 | | 0 | 0 | 0 | 0 | 0 | 0 | 0 | 0 | 0 | 0 | 0 |
| 0 | NA | NA | NA | NA | NA | 2900002M20Rik | | 0 | 0 | 0 | 0 | 0 | 0 | 0 | 0 | 0 | 0 | 0 |

| | | | | | | | | | | | | | | | | | | |
|---|---|---|---|---|---|---|---|---|---|---|---|---|---|---|---|---|---|---|
| 0 | NA | NA | NA | NA | NA | Gm18443 | | 0 | 0 | 0 | 0 | 0 | 0 | 0 | 0 | 0 | 0 | 0 |
| 0 | NA | NA | NA | NA | NA | Gm19123 | | 0 | 0 | 0 | 0 | 0 | 0 | 0 | 0 | 0 | 0 | 0 |
| 0 | NA | NA | NA | NA | NA | Gm24064 | | 0 | 0 | 0 | 0 | 0 | 0 | 0 | 0 | 0 | 0 | 0 |
| 0 | NA | NA | NA | NA | NA | n-R5s209 | | 0 | 0 | 0 | 0 | 0 | 0 | 0 | 0 | 0 | 0 | 0 |
| 0 | NA | NA | NA | NA | NA | Gm19641 | | 0 | 0 | 0 | 0 | 0 | 0 | 0 | 0 | 0 | 0 | 0 |
| 0 | NA | NA | NA | NA | NA | Gm5524 | | 0 | 0 | 0 | 0 | 0 | 0 | 0 | 0 | 0 | 0 | 0 |
| 0 | NA | NA | NA | NA | NA | Gm18776 | | 0 | 0 | 0 | 0 | 0 | 0 | 0 | 0 | 0 | 0 | 0 |
| 0 | NA | NA | NA | NA | NA | Gm38089 | | 0 | 0 | 0 | 0 | 0 | 0 | 0 | 0 | 0 | 0 | 0 |
| 0 | NA | NA | NA | NA | NA | Gm597 | predicted gene 597 | 0 | 0 | 0 | 0 | 0 | 0 | 0 | 0 | 0 | 0 | 0 |
| 0 | NA | NA | NA | NA | NA | Gm6462 | | 0 | 0 | 0 | 0 | 0 | 0 | 0 | 0 | 0 | 0 | 0 |
| 0 | NA | NA | NA | NA | NA | Gm18033 | | 0 | 0 | 0 | 0 | 0 | 0 | 0 | 0 | 0 | 0 | 0 |
| 0 | NA | NA | NA | NA | NA | Gm25586 | | 0 | 0 | 0 | 0 | 0 | 0 | 0 | 0 | 0 | 0 | 0 |
| 0 | NA | NA | NA | NA | NA | Gm22679 | | 0 | 0 | 0 | 0 | 0 | 0 | 0 | 0 | 0 | 0 | 0 |
| 0 | NA | NA | NA | NA | NA | Gm23785 | | 0 | 0 | 0 | 0 | 0 | 0 | 0 | 0 | 0 | 0 | 0 |
| 0 | NA | NA | NA | NA | NA | Mir6342 | | 0 | 0 | 0 | 0 | 0 | 0 | 0 | 0 | 0 | 0 | 0 |
| 0 | NA | NA | NA | NA | NA | | | 0 | 0 | 0 | 0 | 0 | 0 | 0 | 0 | 0 | 0 | 0 |
| 0 | NA | NA | NA | NA | NA | Gm23771 | | 0 | 0 | 0 | 0 | 0 | 0 | 0 | 0 | 0 | 0 | 0 |
| 0 | NA | NA | NA | NA | NA | Gm27430 | | 0 | 0 | 0 | 0 | 0 | 0 | 0 | 0 | 0 | 0 | 0 |
| 0 | NA | NA | NA | NA | NA | Gm9898 | | 0 | 0 | 0 | 0 | 0 | 0 | 0 | 0 | 0 | 0 | 0 |
| 0 | NA | NA | NA | NA | NA | Gm2914 | | 0 | 0 | 0 | 0 | 0 | 0 | 0 | 0 | 0 | 0 | 0 |
| 0 | NA | NA | NA | NA | NA | Gm5698 | | 0 | 0 | 0 | 0 | 0 | 0 | 0 | 0 | 0 | 0 | 0 |
| 0 | NA | NA | NA | NA | NA | Gm9458 | | 0 | 0 | 0 | 0 | 0 | 0 | 0 | 0 | 0 | 0 | 0 |
| 0 | NA | NA | NA | NA | NA | 4931428L18Rik | | 0 | 0 | 0 | 0 | 0 | 0 | 0 | 0 | 0 | 0 | 0 |
| 0 | NA | NA | NA | NA | NA | Gm28644 | | 0 | 0 | 0 | 0 | 0 | 0 | 0 | 0 | 0 | 0 | 0 |
| 0 | NA | NA | NA | NA | NA | Gm27362 | | 0 | 0 | 0 | 0 | 0 | 0 | 0 | 0 | 0 | 0 | 0 |
| 0 | NA | NA | NA | NA | NA | Gm29128 | | 0 | 0 | 0 | 0 | 0 | 0 | 0 | 0 | 0 | 0 | 0 |
| 0 | NA | NA | NA | NA | NA | Gm7893 | | 0 | 0 | 0 | 0 | 0 | 0 | 0 | 0 | 0 | 0 | 0 |
| 0 | NA | NA | NA | NA | NA | Lgsn | lengsin, lens protein with glutamine synth | 0 | 0 | 0 | 0 | 0 | 0 | 0 | 0 | 0 | 0 | 0 |
| 0 | NA | NA | NA | NA | NA | Pih1d3 | PIH1 domain containing 3 | 0 | 0 | 0 | 0 | 0 | 0 | 0 | 0 | 0 | 0 | 0 |
| 0 | NA | NA | NA | NA | NA | Gm26145 | | 0 | 0 | 0 | 0 | 0 | 0 | 0 | 0 | 0 | 0 | 0 |
| 0 | NA | NA | NA | NA | NA | Gm4850 | | 0 | 0 | 0 | 0 | 0 | 0 | 0 | 0 | 0 | 0 | 0 |
| 0 | NA | NA | NA | NA | NA | Gm28645 | | 0 | 0 | 0 | 0 | 0 | 0 | 0 | 0 | 0 | 0 | 0 |
| 0 | NA | NA | NA | NA | NA | Gm6489 | | 0 | 0 | 0 | 0 | 0 | 0 | 0 | 0 | 0 | 0 | 0 |
| 0 | NA | NA | NA | NA | NA | Eef1a1-ps2 | | 0 | 0 | 0 | 0 | 0 | 0 | 0 | 0 | 0 | 0 | 0 |
| 0 | NA | NA | NA | NA | NA | Gm23032 | | 0 | 0 | 0 | 0 | 0 | 0 | 0 | 0 | 0 | 0 | 0 |
| 0 | NA | NA | NA | NA | NA | Gm7910 | | 0 | 0 | 0 | 0 | 0 | 0 | 0 | 0 | 0 | 0 | 0 |
| 0 | NA | NA | NA | NA | NA | Gm25814 | | 0 | 0 | 0 | 0 | 0 | 0 | 0 | 0 | 0 | 0 | 0 |
| 0 | NA | NA | NA | NA | NA | Gm37999 | | 0 | 0 | 0 | 0 | 0 | 0 | 0 | 0 | 0 | 0 | 0 |
| 0 | NA | NA | NA | NA | NA | Gm37911 | | 0 | 0 | 0 | 0 | 0 | 0 | 0 | 0 | 0 | 0 | 0 |
| 0 | NA | NA | NA | NA | NA | Gm9839 | predicted gene 9839 | 0 | 0 | 0 | 0 | 0 | 0 | 0 | 0 | 0 | 0 | 0 |
| 0 | NA | NA | NA | NA | NA | Gm37792 | | 0 | 0 | 0 | 0 | 0 | 0 | 0 | 0 | 0 | 0 | 0 |
| 0 | NA | NA | NA | NA | NA | Gm5415 | predicted gene 5415 | 0 | 0 | 0 | 0 | 0 | 0 | 0 | 0 | 0 | 0 | 0 |
| 0 | NA | NA | NA | NA | NA | Gm24732 | | 0 | 0 | 0 | 0 | 0 | 0 | 0 | 0 | 0 | 0 | 0 |
| 0 | NA | NA | NA | NA | NA | Gm37426 | | 0 | 0 | 0 | 0 | 0 | 0 | 0 | 0 | 0 | 0 | 0 |
| 0 | NA | NA | NA | NA | NA | Gm23024 | | 0 | 0 | 0 | 0 | 0 | 0 | 0 | 0 | 0 | 0 | 0 |
| 0 | NA | NA | NA | NA | NA | Gm37151 | | 0 | 0 | 0 | 0 | 0 | 0 | 0 | 0 | 0 | 0 | 0 |
| 0 | NA | NA | NA | NA | NA | Gm28624 | | 0 | 0 | 0 | 0 | 0 | 0 | 0 | 0 | 0 | 0 | 0 |
| 0 | NA | NA | NA | NA | NA | Gm37724 | | 0 | 0 | 0 | 0 | 0 | 0 | 0 | 0 | 0 | 0 | 0 |
| 0 | NA | NA | NA | NA | NA | Gm4785 | | 0 | 0 | 0 | 0 | 0 | 0 | 0 | 0 | 0 | 0 | 0 |
| 0 | NA | NA | NA | NA | NA | Gm24901 | | 0 | 0 | 0 | 0 | 0 | 0 | 0 | 0 | 0 | 0 | 0 |
| 0 | NA | NA | NA | NA | NA | Gm37591 | | 0 | 0 | 0 | 0 | 0 | 0 | 0 | 0 | 0 | 0 | 0 |
| 0 | NA | NA | NA | NA | NA | Gm2987 | | 0 | 0 | 0 | 0 | 0 | 0 | 0 | 0 | 0 | 0 | 0 |
| 0 | NA | NA | NA | NA | NA | Gm25792 | | 0 | 0 | 0 | 0 | 0 | 0 | 0 | 0 | 0 | 0 | 0 |
| 0 | NA | NA | NA | NA | NA | Gm18799 | | 0 | 0 | 0 | 0 | 0 | 0 | 0 | 0 | 0 | 0 | 0 |
| 0 | NA | NA | NA | NA | NA | Gm23453 | | 0 | 0 | 0 | 0 | 0 | 0 | 0 | 0 | 0 | 0 | 0 |
| 0 | NA | NA | NA | NA | NA | Gm17813 | | 0 | 0 | 0 | 0 | 0 | 0 | 0 | 0 | 0 | 0 | 0 |
| 0 | NA | NA | NA | NA | NA | Gm38173 | | 0 | 0 | 0 | 0 | 0 | 0 | 0 | 0 | 0 | 0 | 0 |
| 0 | NA | NA | NA | NA | NA | Gm37067 | | 0 | 0 | 0 | 0 | 0 | 0 | 0 | 0 | 0 | 0 | 0 |
| 0 | NA | NA | NA | NA | NA | Gm37354 | | 0 | 0 | 0 | 0 | 0 | 0 | 0 | 0 | 0 | 0 | 0 |
| 0 | NA | NA | NA | NA | NA | Gm28631 | | 0 | 0 | 0 | 0 | 0 | 0 | 0 | 0 | 0 | 0 | 0 |

| | | | | | | | | | | | | | | | | | | |
|---|---|---|---|---|---|---|---|---|---|---|---|---|---|---|---|---|---|---|
| 0 | NA | NA | NA | NA | NA | Gm23792 | | 0 | 0 | 0 | 0 | 0 | 0 | 0 | 0 | 0 | 0 | 0 |
| 0 | NA | NA | NA | NA | NA | Mir6896 | | 0 | 0 | 0 | 0 | 0 | 0 | 0 | 0 | 0 | 0 | 0 |
| 0 | NA | NA | NA | NA | NA | Gm37958 | | 0 | 0 | 0 | 0 | 0 | 0 | 0 | 0 | 0 | 0 | 0 |
| 0 | NA | NA | NA | NA | NA | Gm37385 | | 0 | 0 | 0 | 0 | 0 | 0 | 0 | 0 | 0 | 0 | 0 |
| 0 | NA | NA | NA | NA | NA | Gm5266 | | 0 | 0 | 0 | 0 | 0 | 0 | 0 | 0 | 0 | 0 | 0 |
| 0 | NA | NA | NA | NA | NA | Mir5103 | | 0 | 0 | 0 | 0 | 0 | 0 | 0 | 0 | 0 | 0 | 0 |
| 0 | NA | NA | NA | NA | NA | Gm28306 | | 0 | 0 | 0 | 0 | 0 | 0 | 0 | 0 | 0 | 0 | 0 |
| 0 | NA | NA | NA | NA | NA | Prss39 | protease, serine 39 | 0 | 0 | 0 | 0 | 0 | 0 | 0 | 0 | 0 | 0 | 0 |
| 0 | NA | NA | NA | NA | NA | Gm28416 | | 0 | 0 | 0 | 0 | 0 | 0 | 0 | 0 | 0 | 0 | 0 |
| 0 | NA | NA | NA | NA | NA | Gm28419 | | 0 | 0 | 0 | 0 | 0 | 0 | 0 | 0 | 0 | 0 | 0 |
| 0 | NA | NA | NA | NA | NA | Prss40 | protease, serine 40 | 0 | 0 | 0 | 0 | 0 | 0 | 0 | 0 | 0 | 0 | 0 |
| 0 | NA | NA | NA | NA | NA | 1700101I19Rik | | 0 | 0 | 0 | 0 | 0 | 0 | 0 | 0 | 0 | 0 | 0 |
| 0 | NA | NA | NA | NA | NA | Gm28418 | | 0 | 0 | 0 | 0 | 0 | 0 | 0 | 0 | 0 | 0 | 0 |
| 0 | NA | NA | NA | NA | NA | Gm36984 | | 0 | 0 | 0 | 0 | 0 | 0 | 0 | 0 | 0 | 0 | 0 |
| 0 | NA | NA | NA | NA | NA | Gm29730 | | 0 | 0 | 0 | 0 | 0 | 0 | 0 | 0 | 0 | 0 | 0 |
| 0 | NA | NA | NA | NA | NA | AA619741 | | 0 | 0 | 0 | 0 | 0 | 0 | 0 | 0 | 0 | 0 | 0 |
| 0 | NA | NA | NA | NA | NA | Gm37224 | | 0 | 0 | 0 | 0 | 0 | 0 | 0 | 0 | 0 | 0 | 0 |
| 0 | NA | NA | NA | NA | NA | Gm37845 | | 0 | 0 | 0 | 0 | 0 | 0 | 0 | 0 | 0 | 0 | 0 |
| 0 | NA | NA | NA | NA | NA | Arhgef4 | | 0 | 0 | 0 | 0 | 0 | 0 | 0 | 0 | 0 | 0 | 0 |
| 0 | NA | NA | NA | NA | NA | Gm37127 | | 0 | 0 | 0 | 0 | 0 | 0 | 0 | 0 | 0 | 0 | 0 |
| 0 | NA | NA | NA | NA | NA | Gm19430 | | 0 | 0 | 0 | 0 | 0 | 0 | 0 | 0 | 0 | 0 | 0 |
| 0 | NA | NA | NA | NA | NA | Gm38027 | | 0 | 0 | 0 | 0 | 0 | 0 | 0 | 0 | 0 | 0 | 0 |
| 0 | NA | NA | NA | NA | NA | Gm25096 | | 0 | 0 | 0 | 0 | 0 | 0 | 0 | 0 | 0 | 0 | 0 |
| 0 | NA | NA | NA | NA | NA | Gm37068 | | 0 | 0 | 0 | 0 | 0 | 0 | 0 | 0 | 0 | 0 | 0 |
| 0 | NA | NA | NA | NA | NA | Gm25634 | | 0 | 0 | 0 | 0 | 0 | 0 | 0 | 0 | 0 | 0 | 0 |
| 0 | NA | NA | NA | NA | NA | Gm28415 | | 0 | 0 | 0 | 0 | 0 | 0 | 0 | 0 | 0 | 0 | 0 |
| 0 | NA | NA | NA | NA | NA | Gm8022 | | 0 | 0 | 0 | 0 | 0 | 0 | 0 | 0 | 0 | 0 | 0 |
| 0 | NA | NA | NA | NA | NA | Gm37146 | | 0 | 0 | 0 | 0 | 0 | 0 | 0 | 0 | 0 | 0 | 0 |
| 0 | NA | NA | NA | NA | NA | Gm33222 | | 0 | 0 | 0 | 0 | 0 | 0 | 0 | 0 | 0 | 0 | 0 |
| 0 | NA | NA | NA | NA | NA | 4930535G08Rik | | 0 | 0 | 0 | 0 | 0 | 0 | 0 | 0 | 0 | 0 | 0 |
| 0 | NA | NA | NA | NA | NA | Gm38049 | | 0 | 0 | 0 | 0 | 0 | 0 | 0 | 0 | 0 | 0 | 0 |
| 0 | NA | NA | NA | NA | NA | Gm37335 | | 0 | 0 | 0 | 0 | 0 | 0 | 0 | 0 | 0 | 0 | 0 |
| 0 | NA | NA | NA | NA | NA | 4930403P22Rik | | 0 | 0 | 0 | 0 | 0 | 0 | 0 | 0 | 0 | 0 | 0 |
| 0 | NA | NA | NA | NA | NA | Mir6898 | | 0 | 0 | 0 | 0 | 0 | 0 | 0 | 0 | 0 | 0 | 0 |
| 0 | NA | NA | NA | NA | NA | Gm47280 | | 0 | 0 | 0 | 0 | 0 | 0 | 0 | 0 | 0 | 0 | 0 |
| 0 | NA | NA | NA | NA | NA | Gm42417 | | 0 | 0 | 0 | 0 | 0 | 0 | 0 | 0 | 0 | 0 | 0 |
| 0 | NA | NA | NA | NA | NA | Gm24133 | | 0 | 0 | 0 | 0 | 0 | 0 | 0 | 0 | 0 | 0 | 0 |
| 0 | NA | NA | NA | NA | NA | Gm18828 | | 0 | 0 | 0 | 0 | 0 | 0 | 0 | 0 | 0 | 0 | 0 |
| 0 | NA | NA | NA | NA | NA | Gm38115 | | 0 | 0 | 0 | 0 | 0 | 0 | 0 | 0 | 0 | 0 | 0 |
| 0 | NA | NA | NA | NA | NA | F830112A20Rik | | 0 | 0 | 0 | 0 | 0 | 0 | 0 | 0 | 0 | 0 | 0 |
| 0 | NA | NA | NA | NA | NA | n-R5s210 | | 0 | 0 | 0 | 0 | 0 | 0 | 0 | 0 | 0 | 0 | 0 |
| 0 | NA | NA | NA | NA | NA | 1700074A21Rik | | 0 | 0 | 0 | 0 | 0 | 0 | 0 | 0 | 0 | 0 | 0 |
| 0 | NA | NA | NA | NA | NA | 4930439A04Rik | | 0 | 0 | 0 | 0 | 0 | 0 | 0 | 0 | 0 | 0 | 0 |
| 0 | NA | NA | NA | NA | NA | Gm37135 | | 0 | 0 | 0 | 0 | 0 | 0 | 0 | 0 | 0 | 0 | 0 |
| 0 | NA | NA | NA | NA | NA | 4930594C11Rik | | 0 | 0 | 0 | 0 | 0 | 0 | 0 | 0 | 0 | 0 | 0 |
| 0 | NA | NA | NA | NA | NA | Gm43213 | | 0 | 0 | 0 | 0 | 0 | 0 | 0 | 0 | 0 | 0 | 0 |
| 0 | NA | NA | NA | NA | NA | 4930556I23Rik | | 0 | 0 | 0 | 0 | 0 | 0 | 0 | 0 | 0 | 0 | 0 |
| 0 | NA | NA | NA | NA | NA | 4930470B04Rik | | 0 | 0 | 0 | 0 | 0 | 0 | 0 | 0 | 0 | 0 | 0 |
| 0 | NA | NA | NA | NA | NA | Lyg2 | lysozyme G-like 2 | 0 | 0 | 0 | 0 | 0 | 0 | 0 | 0 | 0 | 0 | 0 |
| 0 | NA | NA | NA | NA | NA | Gm15457 | | 0 | 0 | 0 | 0 | 0 | 0 | 0 | 0 | 0 | 0 | 0 |
| 0 | NA | NA | NA | NA | NA | Lyg1 | lysozyme G-like 1 | 0 | 0 | 0 | 0 | 0 | 0 | 0 | 0 | 0 | 0 | 0 |
| 0 | NA | NA | NA | NA | NA | Gm23409 | | 0 | 0 | 0 | 0 | 0 | 0 | 0 | 0 | 0 | 0 | 0 |
| 0 | NA | NA | NA | NA | NA | Gm5099 | | 0 | 0 | 0 | 0 | 0 | 0 | 0 | 0 | 0 | 0 | 0 |
| 0 | NA | NA | NA | NA | NA | Gm16151 | | 0 | 0 | 0 | 0 | 0 | 0 | 0 | 0 | 0 | 0 | 0 |
| 0 | NA | NA | NA | NA | NA | Gm34727 | | 0 | 0 | 0 | 0 | 0 | 0 | 0 | 0 | 0 | 0 | 0 |
| 0 | NA | NA | NA | NA | NA | Gm38135 | | 0 | 0 | 0 | 0 | 0 | 0 | 0 | 0 | 0 | 0 | 0 |
| 0 | NA | NA | NA | NA | NA | 2610300A13Rik | | 0 | 0 | 0 | 0 | 0 | 0 | 0 | 0 | 0 | 0 | 0 |
| 0 | NA | NA | NA | NA | NA | Gm37707 | | 0 | 0 | 0 | 0 | 0 | 0 | 0 | 0 | 0 | 0 | 0 |
| 0 | NA | NA | NA | NA | NA | Nms | neuromedin S | 0 | 0 | 0 | 0 | 0 | 0 | 0 | 0 | 0 | 0 | 0 |
| 0 | NA | NA | NA | NA | NA | Gm37821 | | 0 | 0 | 0 | 0 | 0 | 0 | 0 | 0 | 0 | 0 | 0 |

| | | | | | | | | | | | | | | | | | | |
|---|---|---|---|---|---|---|---|---|---|---|---|---|---|---|---|---|---|---|
| 0 | NA | NA | NA | NA | NA | Gm37091 | | 0 | 0 | 0 | 0 | 0 | 0 | 0 | 0 | 0 | 0 | 0 |
| 0 | NA | NA | NA | NA | NA | Mir6349 | | 0 | 0 | 0 | 0 | 0 | 0 | 0 | 0 | 0 | 0 | 0 |
| 0 | NA | NA | NA | NA | NA | Gm20428 | | 0 | 0 | 0 | 0 | 0 | 0 | 0 | 0 | 0 | 0 | 0 |
| 0 | NA | NA | NA | NA | NA | Gm37265 | | 0 | 0 | 0 | 0 | 0 | 0 | 0 | 0 | 0 | 0 | 0 |
| 0 | NA | NA | NA | NA | NA | Gm24826 | | 0 | 0 | 0 | 0 | 0 | 0 | 0 | 0 | 0 | 0 | 0 |
| 0 | NA | NA | NA | NA | NA | Gm24826 | | 0 | 0 | 0 | 0 | 0 | 0 | 0 | 0 | 0 | 0 | 0 |
| 0 | NA | NA | NA | NA | NA | Gm3646 | predicted gene 3646 | 0 | 0 | 0 | 0 | 0 | 0 | 0 | 0 | 0 | 0 | 0 |
| 0 | NA | NA | NA | NA | NA | 1700066B17Rik | | 0 | 0 | 0 | 0 | 0 | 0 | 0 | 0 | 0 | 0 | 0 |
| 0 | NA | NA | NA | NA | NA | Gm37347 | | 0 | 0 | 0 | 0 | 0 | 0 | 0 | 0 | 0 | 0 | 0 |
| 0 | NA | NA | NA | NA | NA | Gm35801 | | 0 | 0 | 0 | 0 | 0 | 0 | 0 | 0 | 0 | 0 | 0 |
| 0 | NA | NA | NA | NA | NA | Gm37435 | | 0 | 0 | 0 | 0 | 0 | 0 | 0 | 0 | 0 | 0 | 0 |
| 0 | NA | NA | NA | NA | NA | Gm37623 | | 0 | 0 | 0 | 0 | 0 | 0 | 0 | 0 | 0 | 0 | 0 |
| 0 | NA | NA | NA | NA | NA | Gm5973 | | 0 | 0 | 0 | 0 | 0 | 0 | 0 | 0 | 0 | 0 | 0 |
| 0 | NA | NA | NA | NA | NA | 4930448I06Rik | | 0 | 0 | 0 | 0 | 0 | 0 | 0 | 0 | 0 | 0 | 0 |
| 0 | NA | NA | NA | NA | NA | Gm28634 | | 0 | 0 | 0 | 0 | 0 | 0 | 0 | 0 | 0 | 0 | 0 |
| 0 | NA | NA | NA | NA | NA | Gm29260 | | 0 | 0 | 0 | 0 | 0 | 0 | 0 | 0 | 0 | 0 | 0 |
| 0 | NA | NA | NA | NA | NA | Gm8176 | | 0 | 0 | 0 | 0 | 0 | 0 | 0 | 0 | 0 | 0 | 0 |
| 0 | NA | NA | NA | NA | NA | Gm29664 | | 0 | 0 | 0 | 0 | 0 | 0 | 0 | 0 | 0 | 0 | 0 |
| 0 | NA | NA | NA | NA | NA | Gm28175 | | 0 | 0 | 0 | 0 | 0 | 0 | 0 | 0 | 0 | 0 | 0 |
| 0 | NA | NA | NA | NA | NA | Gm28140 | | 0 | 0 | 0 | 0 | 0 | 0 | 0 | 0 | 0 | 0 | 0 |
| 0 | NA | NA | NA | NA | NA | Gm37047 | | 0 | 0 | 0 | 0 | 0 | 0 | 0 | 0 | 0 | 0 | 0 |
| 0 | NA | NA | NA | NA | NA | Pantr1 | | 0 | 0 | 0 | 0 | 0 | 0 | 0 | 0 | 0 | 0 | 0 |
| 0 | NA | NA | NA | NA | NA | 4930420N18Rik | | 0 | 0 | 0 | 0 | 0 | 0 | 0 | 0 | 0 | 0 | 0 |
| 0 | NA | NA | NA | NA | NA | Pou3f3 | POU domain, class 3, transcription factor | 0 | 0 | 0 | 0 | 0 | 0 | 0 | 0 | 0 | 0 | 0 |
| 0 | NA | NA | NA | NA | NA | Gm20646 | | 0 | 0 | 0 | 0 | 0 | 0 | 0 | 0 | 0 | 0 | 0 |
| 0 | NA | NA | NA | NA | NA | Pantr2 | | 0 | 0 | 0 | 0 | 0 | 0 | 0 | 0 | 0 | 0 | 0 |
| 0 | NA | NA | NA | NA | NA | Dalir | | 0 | 0 | 0 | 0 | 0 | 0 | 0 | 0 | 0 | 0 | 0 |
| 0 | NA | NA | NA | NA | NA | Gm23126 | | 0 | 0 | 0 | 0 | 0 | 0 | 0 | 0 | 0 | 0 | 0 |
| 0 | NA | NA | NA | NA | NA | Gm24001 | | 0 | 0 | 0 | 0 | 0 | 0 | 0 | 0 | 0 | 0 | 0 |
| 0 | NA | NA | NA | NA | NA | Gm28782 | | 0 | 0 | 0 | 0 | 0 | 0 | 0 | 0 | 0 | 0 | 0 |
| 0 | NA | NA | NA | NA | NA | Gm29610 | | 0 | 0 | 0 | 0 | 0 | 0 | 0 | 0 | 0 | 0 | 0 |
| 0 | NA | NA | NA | NA | NA | Gm37536 | | 0 | 0 | 0 | 0 | 0 | 0 | 0 | 0 | 0 | 0 | 0 |
| 0 | NA | NA | NA | NA | NA | Gm29157 | | 0 | 0 | 0 | 0 | 0 | 0 | 0 | 0 | 0 | 0 | 0 |
| 0 | NA | NA | NA | NA | NA | Gm8241 | | 0 | 0 | 0 | 0 | 0 | 0 | 0 | 0 | 0 | 0 | 0 |
| 0 | NA | NA | NA | NA | NA | Gm26352 | | 0 | 0 | 0 | 0 | 0 | 0 | 0 | 0 | 0 | 0 | 0 |
| 0 | NA | NA | NA | NA | NA | Gm24180 | | 0 | 0 | 0 | 0 | 0 | 0 | 0 | 0 | 0 | 0 | 0 |
| 0 | NA | NA | NA | NA | NA | Mettl21c | methyltransferase like 21C | 0 | 0 | 0 | 0 | 0 | 0 | 0 | 0 | 0 | 0 | 0 |
| 0 | NA | NA | NA | NA | NA | Gm15838 | | 0 | 0 | 0 | 0 | 0 | 0 | 0 | 0 | 0 | 0 | 0 |
| 0 | NA | NA | NA | NA | NA | Gm29247 | | 0 | 0 | 0 | 0 | 0 | 0 | 0 | 0 | 0 | 0 | 0 |
| 0 | NA | NA | NA | NA | NA | Mettl21e | methyltransferase like 21E | 0 | 0 | 0 | 0 | 0 | 0 | 0 | 0 | 0 | 0 | 0 |
| 0 | NA | NA | NA | NA | NA | Gm5700 | | 0 | 0 | 0 | 0 | 0 | 0 | 0 | 0 | 0 | 0 | 0 |
| 0 | NA | NA | NA | NA | NA | Gm28893 | | 0 | 0 | 0 | 0 | 0 | 0 | 0 | 0 | 0 | 0 | 0 |
| 0 | NA | NA | NA | NA | NA | Gm37626 | | 0 | 0 | 0 | 0 | 0 | 0 | 0 | 0 | 0 | 0 | 0 |
| 0 | NA | NA | NA | NA | NA | 4930521E06Rik | | 0 | 0 | 0 | 0 | 0 | 0 | 0 | 0 | 0 | 0 | 0 |
| 0 | NA | NA | NA | NA | NA | Gm38110 | | 0 | 0 | 0 | 0 | 0 | 0 | 0 | 0 | 0 | 0 | 0 |
| 0 | NA | NA | NA | NA | NA | Gm3489 | | 0 | 0 | 0 | 0 | 0 | 0 | 0 | 0 | 0 | 0 | 0 |
| 0 | NA | NA | NA | NA | NA | Gm19480 | | 0 | 0 | 0 | 0 | 0 | 0 | 0 | 0 | 0 | 0 | 0 |
| 0 | NA | NA | NA | NA | NA | Gm3496 | | 0 | 0 | 0 | 0 | 0 | 0 | 0 | 0 | 0 | 0 | 0 |
| 0 | NA | NA | NA | NA | NA | Gm5268 | | 0 | 0 | 0 | 0 | 0 | 0 | 0 | 0 | 0 | 0 | 0 |
| 0 | NA | NA | NA | NA | NA | Gm47302 | | 0 | 0 | 0 | 0 | 0 | 0 | 0 | 0 | 0 | 0 | 0 |
| 0 | NA | NA | NA | NA | NA | Gm38211 | | 0 | 0 | 0 | 0 | 0 | 0 | 0 | 0 | 0 | 0 | 0 |
| 0 | NA | NA | NA | NA | NA | Gm8296 | | 0 | 0 | 0 | 0 | 0 | 0 | 0 | 0 | 0 | 0 | 0 |
| 0 | NA | NA | NA | NA | NA | Gm7063 | | 0 | 0 | 0 | 0 | 0 | 0 | 0 | 0 | 0 | 0 | 0 |
| 0 | NA | NA | NA | NA | NA | Gm8304 | | 0 | 0 | 0 | 0 | 0 | 0 | 0 | 0 | 0 | 0 | 0 |
| 0 | NA | NA | NA | NA | NA | Gm23216 | | 0 | 0 | 0 | 0 | 0 | 0 | 0 | 0 | 0 | 0 | 0 |
| 0 | NA | NA | NA | NA | NA | Gm23974 | | 0 | 0 | 0 | 0 | 0 | 0 | 0 | 0 | 0 | 0 | 0 |
| 0 | NA | NA | NA | NA | NA | Gm8307 | | 0 | 0 | 0 | 0 | 0 | 0 | 0 | 0 | 0 | 0 | 0 |
| 0 | NA | NA | NA | NA | NA | Gm23984 | | 0 | 0 | 0 | 0 | 0 | 0 | 0 | 0 | 0 | 0 | 0 |
| 0 | NA | NA | NA | NA | NA | Gm28906 | | 0 | 0 | 0 | 0 | 0 | 0 | 0 | 0 | 0 | 0 | 0 |
| 0 | NA | NA | NA | NA | NA | Gm9496 | | 0 | 0 | 0 | 0 | 0 | 0 | 0 | 0 | 0 | 0 | 0 |

| | | | | | | | | | | | | | | | | | | |
|---|---|---|---|---|---|---|---|---|---|---|---|---|---|---|---|---|---|---|
| 0 | NA | NA | NA | NA | NA | Rpl23a-ps1 | | 0 | 0 | 0 | 0 | 0 | 0 | 0 | 0 | 0 | 0 | 0 |
| 0 | NA | NA | NA | NA | NA | Stk-ps2 | | 0 | 0 | 0 | 0 | 0 | 0 | 0 | 0 | 0 | 0 | 0 |
| 0 | NA | NA | NA | NA | NA | Gm18849 | | 0 | 0 | 0 | 0 | 0 | 0 | 0 | 0 | 0 | 0 | 0 |
| 0 | NA | NA | NA | NA | NA | n-R5s211 | | 0 | 0 | 0 | 0 | 0 | 0 | 0 | 0 | 0 | 0 | 0 |
| 0 | NA | NA | NA | NA | NA | Gm37393 | | 0 | 0 | 0 | 0 | 0 | 0 | 0 | 0 | 0 | 0 | 0 |
| 0 | NA | NA | NA | NA | NA | Gm28825 | | 0 | 0 | 0 | 0 | 0 | 0 | 0 | 0 | 0 | 0 | 0 |
| 0 | NA | NA | NA | NA | NA | Gm28826 | | 0 | 0 | 0 | 0 | 0 | 0 | 0 | 0 | 0 | 0 | 0 |
| 0 | NA | NA | NA | NA | NA | Gm38090 | | 0 | 0 | 0 | 0 | 0 | 0 | 0 | 0 | 0 | 0 | 0 |
| 0 | NA | NA | NA | NA | NA | Mir6350 | | 0 | 0 | 0 | 0 | 0 | 0 | 0 | 0 | 0 | 0 | 0 |
| 0 | NA | NA | NA | NA | NA | Gm18117 | | 0 | 0 | 0 | 0 | 0 | 0 | 0 | 0 | 0 | 0 | 0 |
| 0 | NA | NA | NA | NA | NA | Gm37196 | | 0 | 0 | 0 | 0 | 0 | 0 | 0 | 0 | 0 | 0 | 0 |
| 0 | NA | NA | NA | NA | NA | Gm22419 | | 0 | 0 | 0 | 0 | 0 | 0 | 0 | 0 | 0 | 0 | 0 |
| 0 | NA | NA | NA | NA | NA | Gm23829 | | 0 | 0 | 0 | 0 | 0 | 0 | 0 | 0 | 0 | 0 | 0 |
| 0 | NA | NA | NA | NA | NA | Gm19206 | | 0 | 0 | 0 | 0 | 0 | 0 | 0 | 0 | 0 | 0 | 0 |
| 0 | NA | NA | NA | NA | NA | Gm23240 | | 0 | 0 | 0 | 0 | 0 | 0 | 0 | 0 | 0 | 0 | 0 |
| 0 | NA | NA | NA | NA | NA | Gm28830 | | 0 | 0 | 0 | 0 | 0 | 0 | 0 | 0 | 0 | 0 | 0 |
| 0 | NA | NA | NA | NA | NA | Gm8354 | | 0 | 0 | 0 | 0 | 0 | 0 | 0 | 0 | 0 | 0 | 0 |
| 0 | NA | NA | NA | NA | NA | Gm19124 | | 0 | 0 | 0 | 0 | 0 | 0 | 0 | 0 | 0 | 0 | 0 |
| 0 | NA | NA | NA | NA | NA | Gm8357 | | 0 | 0 | 0 | 0 | 0 | 0 | 0 | 0 | 0 | 0 | 0 |
| 0 | NA | NA | NA | NA | NA | Gm22693 | | 0 | 0 | 0 | 0 | 0 | 0 | 0 | 0 | 0 | 0 | 0 |
| 0 | NA | NA | NA | NA | NA | Gm28740 | | 0 | 0 | 0 | 0 | 0 | 0 | 0 | 0 | 0 | 0 | 0 |
| 0 | NA | NA | NA | NA | NA | Gm8367 | | 0 | 0 | 0 | 0 | 0 | 0 | 0 | 0 | 0 | 0 | 0 |
| 0 | NA | NA | NA | NA | NA | C230029F24Rik | | 0 | 0 | 0 | 0 | 0 | 0 | 0 | 0 | 0 | 0 | 0 |
| 0 | NA | NA | NA | NA | NA | Gm6757 | | 0 | 0 | 0 | 0 | 0 | 0 | 0 | 0 | 0 | 0 | 0 |
| 0 | NA | NA | NA | NA | NA | Gm3605 | | 0 | 0 | 0 | 0 | 0 | 0 | 0 | 0 | 0 | 0 | 0 |
| 0 | NA | NA | NA | NA | NA | Gm28388 | | 0 | 0 | 0 | 0 | 0 | 0 | 0 | 0 | 0 | 0 | 0 |
| 0 | NA | NA | NA | NA | NA | Gm8384 | | 0 | 0 | 0 | 0 | 0 | 0 | 0 | 0 | 0 | 0 | 0 |
| 0 | NA | NA | NA | NA | NA | 4933411E06Rik | | 0 | 0 | 0 | 0 | 0 | 0 | 0 | 0 | 0 | 0 | 0 |
| 0 | NA | NA | NA | NA | NA | Gm8391 | | 0 | 0 | 0 | 0 | 0 | 0 | 0 | 0 | 0 | 0 | 0 |
| 0 | NA | NA | NA | NA | NA | Gm27330 | | 0 | 0 | 0 | 0 | 0 | 0 | 0 | 0 | 0 | 0 | 0 |
| 0 | NA | NA | NA | NA | NA | Gm29665 | | 0 | 0 | 0 | 0 | 0 | 0 | 0 | 0 | 0 | 0 | 0 |
| 0 | NA | NA | NA | NA | NA | Gm20118 | | 0 | 0 | 0 | 0 | 0 | 0 | 0 | 0 | 0 | 0 | 0 |
| 0 | NA | NA | NA | NA | NA | Gm25843 | | 0 | 0 | 0 | 0 | 0 | 0 | 0 | 0 | 0 | 0 | 0 |
| 0 | NA | NA | NA | NA | NA | Gm28321 | | 0 | 0 | 0 | 0 | 0 | 0 | 0 | 0 | 0 | 0 | 0 |
| 0 | NA | NA | NA | NA | NA | Gm28322 | | 0 | 0 | 0 | 0 | 0 | 0 | 0 | 0 | 0 | 0 | 0 |
| 0 | NA | NA | NA | NA | NA | Gm25736 | | 0 | 0 | 0 | 0 | 0 | 0 | 0 | 0 | 0 | 0 | 0 |
| 0 | NA | NA | NA | NA | NA | Gm28319 | | 0 | 0 | 0 | 0 | 0 | 0 | 0 | 0 | 0 | 0 | 0 |
| 0 | NA | NA | NA | NA | NA | Gm29325 | | 0 | 0 | 0 | 0 | 0 | 0 | 0 | 0 | 0 | 0 | 0 |
| 0 | NA | NA | NA | NA | NA | 9330175M20Rik | | 0 | 0 | 0 | 0 | 0 | 0 | 0 | 0 | 0 | 0 | 0 |
| 0 | NA | NA | NA | NA | NA | Gm5253 | | 0 | 0 | 0 | 0 | 0 | 0 | 0 | 0 | 0 | 0 | 0 |
| 0 | NA | NA | NA | NA | NA | Gm37535 | | 0 | 0 | 0 | 0 | 0 | 0 | 0 | 0 | 0 | 0 | 0 |
| 0 | NA | NA | NA | NA | NA | Gm28323 | | 0 | 0 | 0 | 0 | 0 | 0 | 0 | 0 | 0 | 0 | 0 |
| 0 | NA | NA | NA | NA | NA | Gm8419 | | 0 | 0 | 0 | 0 | 0 | 0 | 0 | 0 | 0 | 0 | 0 |
| 0 | NA | NA | NA | NA | NA | Gm27607 | | 0 | 0 | 0 | 0 | 0 | 0 | 0 | 0 | 0 | 0 | 0 |
| 0 | NA | NA | NA | NA | NA | Gm18301 | | 0 | 0 | 0 | 0 | 0 | 0 | 0 | 0 | 0 | 0 | 0 |
| 0 | NA | NA | NA | NA | NA | Gm5975 | | 0 | 0 | 0 | 0 | 0 | 0 | 0 | 0 | 0 | 0 | 0 |
| 0 | NA | NA | NA | NA | NA | Gm23460 | | 0 | 0 | 0 | 0 | 0 | 0 | 0 | 0 | 0 | 0 | 0 |
| 0 | NA | NA | NA | NA | NA | Gm5976 | | 0 | 0 | 0 | 0 | 0 | 0 | 0 | 0 | 0 | 0 | 0 |
| 0 | NA | NA | NA | NA | NA | Gm553 | | 0 | 0 | 0 | 0 | 0 | 0 | 0 | 0 | 0 | 0 | 0 |
| 0 | NA | NA | NA | NA | NA | Gm28650 | | 0 | 0 | 0 | 0 | 0 | 0 | 0 | 0 | 0 | 0 | 0 |
| 0 | NA | NA | NA | NA | NA | Gm24349 | | 0 | 0 | 0 | 0 | 0 | 0 | 0 | 0 | 0 | 0 | 0 |
| 0 | NA | NA | NA | NA | NA | Gm28778 | | 0 | 0 | 0 | 0 | 0 | 0 | 0 | 0 | 0 | 0 | 0 |
| 0 | NA | NA | NA | NA | NA | Gm50478 | ASNSD1 upstream reading frame | 0 | 0 | 0 | 0 | 0 | 0 | 0 | 0 | 0 | 0 | 0 |
| 0 | NA | NA | NA | NA | NA | Gm28779 | | 0 | 0 | 0 | 0 | 0 | 0 | 0 | 0 | 0 | 0 | 0 |
| 0 | NA | NA | NA | NA | NA | Gm6799 | | 0 | 0 | 0 | 0 | 0 | 0 | 0 | 0 | 0 | 0 | 0 |
| 0 | NA | NA | NA | NA | NA | Gm6801 | | 0 | 0 | 0 | 0 | 0 | 0 | 0 | 0 | 0 | 0 | 0 |
| 0 | NA | NA | NA | NA | NA | Gm25240 | | 0 | 0 | 0 | 0 | 0 | 0 | 0 | 0 | 0 | 0 | 0 |
| 0 | NA | NA | NA | NA | NA | Gm37863 | | 0 | 0 | 0 | 0 | 0 | 0 | 0 | 0 | 0 | 0 | 0 |
| 0 | NA | NA | NA | NA | NA | Mir7681 | | 0 | 0 | 0 | 0 | 0 | 0 | 0 | 0 | 0 | 0 | 0 |
| 0 | NA | NA | NA | NA | NA | Gm24251 | | 0 | 0 | 0 | 0 | 0 | 0 | 0 | 0 | 0 | 0 | 0 |

| | | | | | | | | | | | | | | | | | | |
|---|---|---|---|---|---|---|---|---|---|---|---|---|---|---|---|---|---|---|
| 0 | NA | NA | NA | NA | NA | Gm37633 | | 0 | 0 | 0 | 0 | 0 | 0 | 0 | 0 | 0 | 0 | 0 |
| 0 | NA | NA | NA | NA | NA | Gm34066 | chromosome 1 C2orf66 homolog | 0 | 0 | 0 | 0 | 0 | 0 | 0 | 0 | 0 | 0 | 0 |
| 0 | NA | NA | NA | NA | NA | Gm24053 | | 0 | 0 | 0 | 0 | 0 | 0 | 0 | 0 | 0 | 0 | 0 |
| 0 | NA | NA | NA | NA | NA | Gm8531 | | 0 | 0 | 0 | 0 | 0 | 0 | 0 | 0 | 0 | 0 | 0 |
| 0 | NA | NA | NA | NA | NA | Gm37382 | | 0 | 0 | 0 | 0 | 0 | 0 | 0 | 0 | 0 | 0 | 0 |
| 0 | NA | NA | NA | NA | NA | 9130227L01Rik | | 0 | 0 | 0 | 0 | 0 | 0 | 0 | 0 | 0 | 0 | 0 |
| 0 | NA | NA | NA | NA | NA | Gm36986 | | 0 | 0 | 0 | 0 | 0 | 0 | 0 | 0 | 0 | 0 | 0 |
| 0 | NA | NA | NA | NA | NA | 1700003I22Rik | | 0 | 0 | 0 | 0 | 0 | 0 | 0 | 0 | 0 | 0 | 0 |
| 0 | NA | NA | NA | NA | NA | Gm28240 | | 0 | 0 | 0 | 0 | 0 | 0 | 0 | 0 | 0 | 0 | 0 |
| 0 | NA | NA | NA | NA | NA | Hsfy2 | heat shock transcription factor, Y-linked 2 | 0 | 0 | 0 | 0 | 0 | 0 | 0 | 0 | 0 | 0 | 0 |
| 0 | NA | NA | NA | NA | NA | Gm4081 | | 0 | 0 | 0 | 0 | 0 | 0 | 0 | 0 | 0 | 0 | 0 |
| 0 | NA | NA | NA | NA | NA | Gm23943 | | 0 | 0 | 0 | 0 | 0 | 0 | 0 | 0 | 0 | 0 | 0 |
| 0 | NA | NA | NA | NA | NA | 1700126A01Rik | | 0 | 0 | 0 | 0 | 0 | 0 | 0 | 0 | 0 | 0 | 0 |
| 0 | NA | NA | NA | NA | NA | BC055402 | | 0 | 0 | 0 | 0 | 0 | 0 | 0 | 0 | 0 | 0 | 0 |
| 0 | NA | NA | NA | NA | NA | Gm5254 | | 0 | 0 | 0 | 0 | 0 | 0 | 0 | 0 | 0 | 0 | 0 |
| 0 | NA | NA | NA | NA | NA | Gm22371 | | 0 | 0 | 0 | 0 | 0 | 0 | 0 | 0 | 0 | 0 | 0 |
| 0 | NA | NA | NA | NA | NA | Gm24014 | | 0 | 0 | 0 | 0 | 0 | 0 | 0 | 0 | 0 | 0 | 0 |
| 0 | NA | NA | NA | NA | NA | Gm8581 | | 0 | 0 | 0 | 0 | 0 | 0 | 0 | 0 | 0 | 0 | 0 |
| 0 | NA | NA | NA | NA | NA | Gm37373 | | 0 | 0 | 0 | 0 | 0 | 0 | 0 | 0 | 0 | 0 | 0 |
| 0 | NA | NA | NA | NA | NA | Gm17929 | | 0 | 0 | 0 | 0 | 0 | 0 | 0 | 0 | 0 | 0 | 0 |
| 0 | NA | NA | NA | NA | NA | Gm17234 | | 0 | 0 | 0 | 0 | 0 | 0 | 0 | 0 | 0 | 0 | 0 |
| 0 | NA | NA | NA | NA | NA | Gm24548 | | 0 | 0 | 0 | 0 | 0 | 0 | 0 | 0 | 0 | 0 | 0 |
| 0 | NA | NA | NA | NA | NA | Aox4 | aldehyde oxidase 4 | 0 | 0 | 0 | 0 | 0 | 0 | 0 | 0 | 0 | 0 | 0 |
| 0 | NA | NA | NA | NA | NA | Gm15759 | | 0 | 0 | 0 | 0 | 0 | 0 | 0 | 0 | 0 | 0 | 0 |
| 0 | NA | NA | NA | NA | NA | Gm29389 | | 0 | 0 | 0 | 0 | 0 | 0 | 0 | 0 | 0 | 0 | 0 |
| 0 | NA | NA | NA | NA | NA | Gm18802 | | 0 | 0 | 0 | 0 | 0 | 0 | 0 | 0 | 0 | 0 | 0 |
| 0 | NA | NA | NA | NA | NA | Gm28803 | | 0 | 0 | 0 | 0 | 0 | 0 | 0 | 0 | 0 | 0 | 0 |
| 0 | NA | NA | NA | NA | NA | Gm28802 | | 0 | 0 | 0 | 0 | 0 | 0 | 0 | 0 | 0 | 0 | 0 |
| 0 | NA | NA | NA | NA | NA | | | 0 | 0 | 0 | 0 | 0 | 0 | 0 | 0 | 0 | 0 | 0 |
| 0 | NA | NA | NA | NA | NA | Gm29018 | | 0 | 0 | 0 | 0 | 0 | 0 | 0 | 0 | 0 | 0 | 0 |
| 0 | NA | NA | NA | NA | NA | Gm25502 | | 0 | 0 | 0 | 0 | 0 | 0 | 0 | 0 | 0 | 0 | 0 |
| 0 | NA | NA | NA | NA | NA | Gm29017 | | 0 | 0 | 0 | 0 | 0 | 0 | 0 | 0 | 0 | 0 | 0 |
| 0 | NA | NA | NA | NA | NA | Gm19125 | | 0 | 0 | 0 | 0 | 0 | 0 | 0 | 0 | 0 | 0 | 0 |
| 0 | NA | NA | NA | NA | NA | Gm29016 | | 0 | 0 | 0 | 0 | 0 | 0 | 0 | 0 | 0 | 0 | 0 |
| 0 | NA | NA | NA | NA | NA | Gm37737 | | 0 | 0 | 0 | 0 | 0 | 0 | 0 | 0 | 0 | 0 | 0 |
| 0 | NA | NA | NA | NA | NA | Gm26813 | | 0 | 0 | 0 | 0 | 0 | 0 | 0 | 0 | 0 | 0 | 0 |
| 0 | NA | NA | NA | NA | NA | Gm28411 | | 0 | 0 | 0 | 0 | 0 | 0 | 0 | 0 | 0 | 0 | 0 |
| 0 | NA | NA | NA | NA | NA | 1700122D07Rik | | 0 | 0 | 0 | 0 | 0 | 0 | 0 | 0 | 0 | 0 | 0 |
| 0 | NA | NA | NA | NA | NA | Gm24545 | | 0 | 0 | 0 | 0 | 0 | 0 | 0 | 0 | 0 | 0 | 0 |
| 0 | NA | NA | NA | NA | NA | Gm26293 | | 0 | 0 | 0 | 0 | 0 | 0 | 0 | 0 | 0 | 0 | 0 |
| 0 | NA | NA | NA | NA | NA | Gm38125 | | 0 | 0 | 0 | 0 | 0 | 0 | 0 | 0 | 0 | 0 | 0 |
| 0 | NA | NA | NA | NA | NA | Gm37977 | | 0 | 0 | 0 | 0 | 0 | 0 | 0 | 0 | 0 | 0 | 0 |
| 0 | NA | NA | NA | NA | NA | Gm18829 | | 0 | 0 | 0 | 0 | 0 | 0 | 0 | 0 | 0 | 0 | 0 |
| 0 | NA | NA | NA | NA | NA | Gm29332 | | 0 | 0 | 0 | 0 | 0 | 0 | 0 | 0 | 0 | 0 | 0 |
| 0 | NA | NA | NA | NA | NA | Gm11576 | | 0 | 0 | 0 | 0 | 0 | 0 | 0 | 0 | 0 | 0 | 0 |
| 0 | NA | NA | NA | NA | NA | Gm38388 | | 0 | 0 | 0 | 0 | 0 | 0 | 0 | 0 | 0 | 0 | 0 |
| 0 | NA | NA | NA | NA | NA | Gm23762 | | 0 | 0 | 0 | 0 | 0 | 0 | 0 | 0 | 0 | 0 | 0 |
| 0 | NA | NA | NA | NA | NA | Gm11577 | | 0 | 0 | 0 | 0 | 0 | 0 | 0 | 0 | 0 | 0 | 0 |
| 0 | NA | NA | NA | NA | NA | Gm11579 | | 0 | 0 | 0 | 0 | 0 | 0 | 0 | 0 | 0 | 0 | 0 |
| 0 | NA | NA | NA | NA | NA | Gm37198 | | 0 | 0 | 0 | 0 | 0 | 0 | 0 | 0 | 0 | 0 | 0 |
| 0 | NA | NA | NA | NA | NA | Gm11609 | | 0 | 0 | 0 | 0 | 0 | 0 | 0 | 0 | 0 | 0 | 0 |
| 0 | NA | NA | NA | NA | NA | Gm11582 | | 0 | 0 | 0 | 0 | 0 | 0 | 0 | 0 | 0 | 0 | 0 |
| 0 | NA | NA | NA | NA | NA | Gm37039 | | 0 | 0 | 0 | 0 | 0 | 0 | 0 | 0 | 0 | 0 | 0 |
| 0 | NA | NA | NA | NA | NA | Gm11590 | | 0 | 0 | 0 | 0 | 0 | 0 | 0 | 0 | 0 | 0 | 0 |
| 0 | NA | NA | NA | NA | NA | Gm11591 | | 0 | 0 | 0 | 0 | 0 | 0 | 0 | 0 | 0 | 0 | 0 |
| 0 | NA | NA | NA | NA | NA | Gm28083 | | 0 | 0 | 0 | 0 | 0 | 0 | 0 | 0 | 0 | 0 | 0 |
| 0 | NA | NA | NA | NA | NA | Gm11587 | | 0 | 0 | 0 | 0 | 0 | 0 | 0 | 0 | 0 | 0 | 0 |
| 0 | NA | NA | NA | NA | NA | 9530026F06Rik | | 0 | 0 | 0 | 0 | 0 | 0 | 0 | 0 | 0 | 0 | 0 |
| 0 | NA | NA | NA | NA | NA | Gm36943 | | 0 | 0 | 0 | 0 | 0 | 0 | 0 | 0 | 0 | 0 | 0 |
| 0 | NA | NA | NA | NA | NA | Gm28094 | | 0 | 0 | 0 | 0 | 0 | 0 | 0 | 0 | 0 | 0 | 0 |

| | | | | | | | | | | | | | | | | | | |
|---|---|---|---|---|---|---|---|---|---|---|---|---|---|---|---|---|---|---|
| 0 | NA | NA | NA | NA | NA | 4930587A21Rik | | 0 | 0 | 0 | 0 | 0 | 0 | 0 | 0 | 0 | 0 | 0 |
| 0 | NA | NA | NA | NA | NA | Gm37205 | | 0 | 0 | 0 | 0 | 0 | 0 | 0 | 0 | 0 | 0 | 0 |
| 0 | NA | NA | NA | NA | NA | Gm38278 | | 0 | 0 | 0 | 0 | 0 | 0 | 0 | 0 | 0 | 0 | 0 |
| 0 | NA | NA | NA | NA | NA | Pard3bos1 | | 0 | 0 | 0 | 0 | 0 | 0 | 0 | 0 | 0 | 0 | 0 |
| 0 | NA | NA | NA | NA | NA | Gm38075 | | 0 | 0 | 0 | 0 | 0 | 0 | 0 | 0 | 0 | 0 | 0 |
| 0 | NA | NA | NA | NA | NA | Gm29640 | | 0 | 0 | 0 | 0 | 0 | 0 | 0 | 0 | 0 | 0 | 0 |
| 0 | NA | NA | NA | NA | NA | Gm29641 | | 0 | 0 | 0 | 0 | 0 | 0 | 0 | 0 | 0 | 0 | 0 |
| 0 | NA | NA | NA | NA | NA | Pard3bos2 | | 0 | 0 | 0 | 0 | 0 | 0 | 0 | 0 | 0 | 0 | 0 |
| 0 | NA | NA | NA | NA | NA | Gm38351 | | 0 | 0 | 0 | 0 | 0 | 0 | 0 | 0 | 0 | 0 | 0 |
| 0 | NA | NA | NA | NA | NA | Pard3bos3 | | 0 | 0 | 0 | 0 | 0 | 0 | 0 | 0 | 0 | 0 | 0 |
| 0 | NA | NA | NA | NA | NA | Gm11600 | | 0 | 0 | 0 | 0 | 0 | 0 | 0 | 0 | 0 | 0 | 0 |
| 0 | NA | NA | NA | NA | NA | Gm29083 | | 0 | 0 | 0 | 0 | 0 | 0 | 0 | 0 | 0 | 0 | 0 |
| 0 | NA | NA | NA | NA | NA | Gm44468 | | 0 | 0 | 0 | 0 | 0 | 0 | 0 | 0 | 0 | 0 | 0 |
| 0 | NA | NA | NA | NA | NA | Gm24942 | | 0 | 0 | 0 | 0 | 0 | 0 | 0 | 0 | 0 | 0 | 0 |
| 0 | NA | NA | NA | NA | NA | Rps27-ps1 | | 0 | 0 | 0 | 0 | 0 | 0 | 0 | 0 | 0 | 0 | 0 |
| 0 | NA | NA | NA | NA | NA | Gm24557 | | 0 | 0 | 0 | 0 | 0 | 0 | 0 | 0 | 0 | 0 | 0 |
| 0 | NA | NA | NA | NA | NA | Gm27390 | | 0 | 0 | 0 | 0 | 0 | 0 | 0 | 0 | 0 | 0 | 0 |
| 0 | NA | NA | NA | NA | NA | Gm26457 | | 0 | 0 | 0 | 0 | 0 | 0 | 0 | 0 | 0 | 0 | 0 |
| 0 | NA | NA | NA | NA | NA | Gm27512 | | 0 | 0 | 0 | 0 | 0 | 0 | 0 | 0 | 0 | 0 | 0 |
| 0 | NA | NA | NA | NA | NA | Platr12 | | 0 | 0 | 0 | 0 | 0 | 0 | 0 | 0 | 0 | 0 | 0 |
| 0 | NA | NA | NA | NA | NA | Gm23448 | | 0 | 0 | 0 | 0 | 0 | 0 | 0 | 0 | 0 | 0 | 0 |
| 0 | NA | NA | NA | NA | NA | Gm11606 | | 0 | 0 | 0 | 0 | 0 | 0 | 0 | 0 | 0 | 0 | 0 |
| 0 | NA | NA | NA | NA | NA | Gm22281 | | 0 | 0 | 0 | 0 | 0 | 0 | 0 | 0 | 0 | 0 | 0 |
| 0 | NA | NA | NA | NA | NA | Gm39653 | predicted gene, 39653 | 0 | 0 | 0 | 0 | 0 | 0 | 0 | 0 | 0 | 0 | 0 |
| 0 | NA | NA | NA | NA | NA | Gm12748 | | 0 | 0 | 0 | 0 | 0 | 0 | 0 | 0 | 0 | 0 | 0 |
| 0 | NA | NA | NA | NA | NA | 4933402D24Ril | RIKEN cDNA 4933402D24 gene | 0 | 0 | 0 | 0 | 0 | 0 | 0 | 0 | 0 | 0 | 0 |
| 0 | NA | NA | NA | NA | NA | Gm13751 | | 0 | 0 | 0 | 0 | 0 | 0 | 0 | 0 | 0 | 0 | 0 |
| 0 | NA | NA | NA | NA | NA | Gm13754 | | 0 | 0 | 0 | 0 | 0 | 0 | 0 | 0 | 0 | 0 | 0 |
| 0 | NA | NA | NA | NA | NA | Mir6899 | | 0 | 0 | 0 | 0 | 0 | 0 | 0 | 0 | 0 | 0 | 0 |
| 0 | NA | NA | NA | NA | NA | Gm13748 | | 0 | 0 | 0 | 0 | 0 | 0 | 0 | 0 | 0 | 0 | 0 |
| 0 | NA | NA | NA | NA | NA | Gm28981 | | 0 | 0 | 0 | 0 | 0 | 0 | 0 | 0 | 0 | 0 | 0 |
| 0 | NA | NA | NA | NA | NA | Gm25748 | | 0 | 0 | 0 | 0 | 0 | 0 | 0 | 0 | 0 | 0 | 0 |
| 0 | NA | NA | NA | NA | NA | Gm29152 | | 0 | 0 | 0 | 0 | 0 | 0 | 0 | 0 | 0 | 0 | 0 |
| 0 | NA | NA | NA | NA | NA | Gm24788 | | 0 | 0 | 0 | 0 | 0 | 0 | 0 | 0 | 0 | 0 | 0 |
| 0 | NA | NA | NA | NA | NA | Akr1cl | aldo-keto reductase family 1, member C-l | 0 | 0 | 0 | 0 | 0 | 0 | 0 | 0 | 0 | 0 | 0 |
| 0 | NA | NA | NA | NA | NA | Gm8550 | | 0 | 0 | 0 | 0 | 0 | 0 | 0 | 0 | 0 | 0 | 0 |
| 0 | NA | NA | NA | NA | NA | Cryge | crystallin, gamma E | 0 | 0 | 0 | 0 | 0 | 0 | 0 | 0 | 0 | 0 | 0 |
| 0 | NA | NA | NA | NA | NA | Crygc | crystallin, gamma C | 0 | 0 | 0 | 0 | 0 | 0 | 0 | 0 | 0 | 0 | 0 |
| 0 | NA | NA | NA | NA | NA | Crygb | crystallin, gamma B | 0 | 0 | 0 | 0 | 0 | 0 | 0 | 0 | 0 | 0 | 0 |
| 0 | NA | NA | NA | NA | NA | Cryga | crystallin, gamma A | 0 | 0 | 0 | 0 | 0 | 0 | 0 | 0 | 0 | 0 | 0 |
| 0 | NA | NA | NA | NA | NA | Gm28845 | | 0 | 0 | 0 | 0 | 0 | 0 | 0 | 0 | 0 | 0 | 0 |
| 0 | NA | NA | NA | NA | NA | Gm28927 | | 0 | 0 | 0 | 0 | 0 | 0 | 0 | 0 | 0 | 0 | 0 |
| 0 | NA | NA | NA | NA | NA | Gm28925 | | 0 | 0 | 0 | 0 | 0 | 0 | 0 | 0 | 0 | 0 | 0 |
| 0 | NA | NA | NA | NA | NA | Gm28923 | | 0 | 0 | 0 | 0 | 0 | 0 | 0 | 0 | 0 | 0 | 0 |
| 0 | NA | NA | NA | NA | NA | Gm28922 | | 0 | 0 | 0 | 0 | 0 | 0 | 0 | 0 | 0 | 0 | 0 |
| 0 | NA | NA | NA | NA | NA | Gm28728 | | 0 | 0 | 0 | 0 | 0 | 0 | 0 | 0 | 0 | 0 | 0 |
| 0 | NA | NA | NA | NA | NA | Gm5255 | | 0 | 0 | 0 | 0 | 0 | 0 | 0 | 0 | 0 | 0 | 0 |
| 0 | NA | NA | NA | NA | NA | Crygf | crystallin, gamma F | 0 | 0 | 0 | 0 | 0 | 0 | 0 | 0 | 0 | 0 | 0 |
| 0 | NA | NA | NA | NA | NA | Gm8809 | | 0 | 0 | 0 | 0 | 0 | 0 | 0 | 0 | 0 | 0 | 0 |
| 0 | NA | NA | NA | NA | NA | Gm15659 | | 0 | 0 | 0 | 0 | 0 | 0 | 0 | 0 | 0 | 0 | 0 |
| 0 | NA | NA | NA | NA | NA | Gm8812 | | 0 | 0 | 0 | 0 | 0 | 0 | 0 | 0 | 0 | 0 | 0 |
| 0 | NA | NA | NA | NA | NA | Gm37181 | | 0 | 0 | 0 | 0 | 0 | 0 | 0 | 0 | 0 | 0 | 0 |
| 0 | NA | NA | NA | NA | NA | Gm37755 | | 0 | 0 | 0 | 0 | 0 | 0 | 0 | 0 | 0 | 0 | 0 |
| 0 | NA | NA | NA | NA | NA | Gm38218 | | 0 | 0 | 0 | 0 | 0 | 0 | 0 | 0 | 0 | 0 | 0 |
| 0 | NA | NA | NA | NA | NA | Gm27934 | | 0 | 0 | 0 | 0 | 0 | 0 | 0 | 0 | 0 | 0 | 0 |
| 0 | NA | NA | NA | NA | NA | Gm25832 | | 0 | 0 | 0 | 0 | 0 | 0 | 0 | 0 | 0 | 0 | 0 |
| 0 | NA | NA | NA | NA | NA | Gm15826 | | 0 | 0 | 0 | 0 | 0 | 0 | 0 | 0 | 0 | 0 | 0 |
| 0 | NA | NA | NA | NA | NA | Gm15668 | | 0 | 0 | 0 | 0 | 0 | 0 | 0 | 0 | 0 | 0 | 0 |
| 0 | NA | NA | NA | NA | NA | Gm26342 | | 0 | 0 | 0 | 0 | 0 | 0 | 0 | 0 | 0 | 0 | 0 |
| 0 | NA | NA | NA | NA | NA | Kif22-ps | | 0 | 0 | 0 | 0 | 0 | 0 | 0 | 0 | 0 | 0 | 0 |

| | | | | | | | | | | | | | | | | | |
|---|---|---|---|---|---|---|---|---|---|---|---|---|---|---|---|---|---|
| 0 | NA | NA | NA | NA | NA | Gm22548 | | 0 | 0 | 0 | 0 | 0 | 0 | 0 | 0 | 0 | 0 | 0 |
| 0 | NA | NA | NA | NA | NA | Gm15669 | | 0 | 0 | 0 | 0 | 0 | 0 | 0 | 0 | 0 | 0 | 0 |
| 0 | NA | NA | NA | NA | NA | Gm37957 | | 0 | 0 | 0 | 0 | 0 | 0 | 0 | 0 | 0 | 0 | 0 |
| 0 | NA | NA | NA | NA | NA | Gm23189 | | 0 | 0 | 0 | 0 | 0 | 0 | 0 | 0 | 0 | 0 | 0 |
| 0 | NA | NA | NA | NA | NA | Gm15671 | | 0 | 0 | 0 | 0 | 0 | 0 | 0 | 0 | 0 | 0 | 0 |
| 0 | NA | NA | NA | NA | NA | Gm37100 | | 0 | 0 | 0 | 0 | 0 | 0 | 0 | 0 | 0 | 0 | 0 |
| 0 | NA | NA | NA | NA | NA | Gm15670 | | 0 | 0 | 0 | 0 | 0 | 0 | 0 | 0 | 0 | 0 | 0 |
| 0 | NA | NA | NA | NA | NA | Gm25542 | | 0 | 0 | 0 | 0 | 0 | 0 | 0 | 0 | 0 | 0 | 0 |
| 0 | NA | NA | NA | NA | NA | Gm19335 | | 0 | 0 | 0 | 0 | 0 | 0 | 0 | 0 | 0 | 0 | 0 |
| 0 | NA | NA | NA | NA | NA | Gm37735 | | 0 | 0 | 0 | 0 | 0 | 0 | 0 | 0 | 0 | 0 | 0 |
| 0 | NA | NA | NA | NA | NA | Gm37769 | | 0 | 0 | 0 | 0 | 0 | 0 | 0 | 0 | 0 | 0 | 0 |
| 0 | NA | NA | NA | NA | NA | Gm37061 | | 0 | 0 | 0 | 0 | 0 | 0 | 0 | 0 | 0 | 0 | 0 |
| 0 | NA | NA | NA | NA | NA | Gm16076 | | 0 | 0 | 0 | 0 | 0 | 0 | 0 | 0 | 0 | 0 | 0 |
| 0 | NA | NA | NA | NA | NA | Pced1c-ps | | 0 | 0 | 0 | 0 | 0 | 0 | 0 | 0 | 0 | 0 | 0 |
| 0 | NA | NA | NA | NA | NA | Gm28497 | | 0 | 0 | 0 | 0 | 0 | 0 | 0 | 0 | 0 | 0 | 0 |
| 0 | NA | NA | NA | NA | NA | Gm29113 | | 0 | 0 | 0 | 0 | 0 | 0 | 0 | 0 | 0 | 0 | 0 |
| 0 | NA | NA | NA | NA | NA | Gm29114 | | 0 | 0 | 0 | 0 | 0 | 0 | 0 | 0 | 0 | 0 | 0 |
| 0 | NA | NA | NA | NA | NA | Gm29112 | | 0 | 0 | 0 | 0 | 0 | 0 | 0 | 0 | 0 | 0 | 0 |
| 0 | NA | NA | NA | NA | NA | Gm37930 | | 0 | 0 | 0 | 0 | 0 | 0 | 0 | 0 | 0 | 0 | 0 |
| 0 | NA | NA | NA | NA | NA | Gm22665 | | 0 | 0 | 0 | 0 | 0 | 0 | 0 | 0 | 0 | 0 | 0 |
| 0 | NA | NA | NA | NA | NA | Gm23879 | | 0 | 0 | 0 | 0 | 0 | 0 | 0 | 0 | 0 | 0 | 0 |
| 0 | NA | NA | NA | NA | NA | Gm38272 | | 0 | 0 | 0 | 0 | 0 | 0 | 0 | 0 | 0 | 0 | 0 |
| 0 | NA | NA | NA | NA | NA | Gm37456 | | 0 | 0 | 0 | 0 | 0 | 0 | 0 | 0 | 0 | 0 | 0 |
| 0 | NA | NA | NA | NA | NA | Gm23422 | | 0 | 0 | 0 | 0 | 0 | 0 | 0 | 0 | 0 | 0 | 0 |
| 0 | NA | NA | NA | NA | NA | Vwc2l | von Willebrand factor C domain-containin | 0 | 0 | 0 | 0 | 0 | 0 | 0 | 0 | 0 | 0 | 0 |
| 0 | NA | NA | NA | NA | NA | Gm16236 | | 0 | 0 | 0 | 0 | 0 | 0 | 0 | 0 | 0 | 0 | 0 |
| 0 | NA | NA | NA | NA | NA | Gm6947 | | 0 | 0 | 0 | 0 | 0 | 0 | 0 | 0 | 0 | 0 | 0 |
| 0 | NA | NA | NA | NA | NA | Gm37217 | | 0 | 0 | 0 | 0 | 0 | 0 | 0 | 0 | 0 | 0 | 0 |
| 0 | NA | NA | NA | NA | NA | Gm8883 | | 0 | 0 | 0 | 0 | 0 | 0 | 0 | 0 | 0 | 0 | 0 |
| 0 | NA | NA | NA | NA | NA | Gm23472 | | 0 | 0 | 0 | 0 | 0 | 0 | 0 | 0 | 0 | 0 | 0 |
| 0 | NA | NA | NA | NA | NA | Ankar | ankyrin and armadillo repeat containing | 0 | 0 | 0 | 0 | 0 | 0 | 0 | 0 | 0 | 0 | 0 |
| 0 | NA | NA | NA | NA | NA | Gm18035 | | 0 | 0 | 0 | 0 | 0 | 0 | 0 | 0 | 0 | 0 | 0 |
| 0 | NA | NA | NA | NA | NA | Gm45859 | | 0 | 0 | 0 | 0 | 0 | 0 | 0 | 0 | 0 | 0 | 0 |
| 0 | NA | NA | NA | NA | NA | 1700027A15Rik | | 0 | 0 | 0 | 0 | 0 | 0 | 0 | 0 | 0 | 0 | 0 |
| 0 | NA | NA | NA | NA | NA | Tnp1 | transition protein 1 | 0 | 0 | 0 | 0 | 0 | 0 | 0 | 0 | 0 | 0 | 0 |
| 0 | NA | NA | NA | NA | NA | Gm38106 | | 0 | 0 | 0 | 0 | 0 | 0 | 0 | 0 | 0 | 0 | 0 |
| 0 | NA | NA | NA | NA | NA | Gm29185 | | 0 | 0 | 0 | 0 | 0 | 0 | 0 | 0 | 0 | 0 | 0 |
| 0 | NA | NA | NA | NA | NA | Gm29184 | | 0 | 0 | 0 | 0 | 0 | 0 | 0 | 0 | 0 | 0 | 0 |
| 0 | NA | NA | NA | NA | NA | Pinc | | 0 | 0 | 0 | 0 | 0 | 0 | 0 | 0 | 0 | 0 | 0 |
| 0 | NA | NA | NA | NA | NA | Gm27315 | | 0 | 0 | 0 | 0 | 0 | 0 | 0 | 0 | 0 | 0 | 0 |
| 0 | NA | NA | NA | NA | NA | D530049I02Rik | | 0 | 0 | 0 | 0 | 0 | 0 | 0 | 0 | 0 | 0 | 0 |
| 0 | NA | NA | NA | NA | NA | Gm9553 | | 0 | 0 | 0 | 0 | 0 | 0 | 0 | 0 | 0 | 0 | 0 |
| 0 | NA | NA | NA | NA | NA | C530043A13Rik | | 0 | 0 | 0 | 0 | 0 | 0 | 0 | 0 | 0 | 0 | 0 |
| 0 | NA | NA | NA | NA | NA | Gm29186 | | 0 | 0 | 0 | 0 | 0 | 0 | 0 | 0 | 0 | 0 | 0 |
| 0 | NA | NA | NA | NA | NA | 6030407O03Rik | | 0 | 0 | 0 | 0 | 0 | 0 | 0 | 0 | 0 | 0 | 0 |
| 0 | NA | NA | NA | NA | NA | Gm29183 | | 0 | 0 | 0 | 0 | 0 | 0 | 0 | 0 | 0 | 0 | 0 |
| 0 | NA | NA | NA | NA | NA | Gm25329 | | 0 | 0 | 0 | 0 | 0 | 0 | 0 | 0 | 0 | 0 | 0 |
| 0 | NA | NA | NA | NA | NA | Mir6351 | | 0 | 0 | 0 | 0 | 0 | 0 | 0 | 0 | 0 | 0 | 0 |
| 0 | NA | NA | NA | NA | NA | Cxcr2 | chemokine (C-X-C motif) receptor 2 | 0 | 0 | 0 | 0 | 0 | 0 | 0 | 0 | 0 | 0 | 0 |
| 0 | NA | NA | NA | NA | NA | Cxcr1 | chemokine (C-X-C motif) receptor 1 | 0 | 0 | 0 | 0 | 0 | 0 | 0 | 0 | 0 | 0 | 0 |
| 0 | NA | NA | NA | NA | NA | Mir1928 | | 0 | 0 | 0 | 0 | 0 | 0 | 0 | 0 | 0 | 0 | 0 |
| 0 | NA | NA | NA | NA | NA | Gm25524 | | 0 | 0 | 0 | 0 | 0 | 0 | 0 | 0 | 0 | 0 | 0 |
| 0 | NA | NA | NA | NA | NA | Gm37931 | | 0 | 0 | 0 | 0 | 0 | 0 | 0 | 0 | 0 | 0 | 0 |
| 0 | NA | NA | NA | NA | NA | Gm27507 | | 0 | 0 | 0 | 0 | 0 | 0 | 0 | 0 | 0 | 0 | 0 |
| 0 | NA | NA | NA | NA | NA | Gm29539 | | 0 | 0 | 0 | 0 | 0 | 0 | 0 | 0 | 0 | 0 | 0 |
| 0 | NA | NA | NA | NA | NA | Gm16582 | | 0 | 0 | 0 | 0 | 0 | 0 | 0 | 0 | 0 | 0 | 0 |
| 0 | NA | NA | NA | NA | NA | Gm37744 | | 0 | 0 | 0 | 0 | 0 | 0 | 0 | 0 | 0 | 0 | 0 |
| 0 | NA | NA | NA | NA | NA | Gm29253 | | 0 | 0 | 0 | 0 | 0 | 0 | 0 | 0 | 0 | 0 | 0 |
| 0 | NA | NA | NA | NA | NA | Gm27943 | | 0 | 0 | 0 | 0 | 0 | 0 | 0 | 0 | 0 | 0 | 0 |
| 0 | NA | NA | NA | NA | NA | Gm50472 | | 0 | 0 | 0 | 0 | 0 | 0 | 0 | 0 | 0 | 0 | 0 |

| | | | | | | | | | | | | | | | | | | |
|---|---|---|---|---|---|---|---|---|---|---|---|---|---|---|---|---|---|---|
| 0 | NA | NA | NA | NA | NA | Gm15179 | | 0 | 0 | 0 | 0 | 0 | 0 | 0 | 0 | 0 | 0 | 0 |
| 0 | NA | NA | NA | NA | NA | Gm15178 | | 0 | 0 | 0 | 0 | 0 | 0 | 0 | 0 | 0 | 0 | 0 |
| 0 | NA | NA | NA | NA | NA | Gm15181 | | 0 | 0 | 0 | 0 | 0 | 0 | 0 | 0 | 0 | 0 | 0 |
| 0 | NA | NA | NA | NA | NA | Gm15183 | | 0 | 0 | 0 | 0 | 0 | 0 | 0 | 0 | 0 | 0 | 0 |
| 0 | NA | NA | NA | NA | NA | Gm15180 | | 0 | 0 | 0 | 0 | 0 | 0 | 0 | 0 | 0 | 0 | 0 |
| 0 | NA | NA | NA | NA | NA | Gm15182 | | 0 | 0 | 0 | 0 | 0 | 0 | 0 | 0 | 0 | 0 | 0 |
| 0 | NA | NA | NA | NA | NA | Gm29065 | | 0 | 0 | 0 | 0 | 0 | 0 | 0 | 0 | 0 | 0 | 0 |
| 0 | NA | NA | NA | NA | NA | Gm29068 | | 0 | 0 | 0 | 0 | 0 | 0 | 0 | 0 | 0 | 0 | 0 |
| 0 | NA | NA | NA | NA | NA | Gm29069 | | 0 | 0 | 0 | 0 | 0 | 0 | 0 | 0 | 0 | 0 | 0 |
| 0 | NA | NA | NA | NA | NA | Gm17751 | | 0 | 0 | 0 | 0 | 0 | 0 | 0 | 0 | 0 | 0 | 0 |
| 0 | NA | NA | NA | NA | NA | Mir6343 | | 0 | 0 | 0 | 0 | 0 | 0 | 0 | 0 | 0 | 0 | 0 |
| 0 | NA | NA | NA | NA | NA | Gm816 | | 0 | 0 | 0 | 0 | 0 | 0 | 0 | 0 | 0 | 0 | 0 |
| 0 | NA | NA | NA | NA | NA | Gm38265 | | 0 | 0 | 0 | 0 | 0 | 0 | 0 | 0 | 0 | 0 | 0 |
| 0 | NA | NA | NA | NA | NA | Gm25974 | | 0 | 0 | 0 | 0 | 0 | 0 | 0 | 0 | 0 | 0 | 0 |
| 0 | NA | NA | NA | NA | NA | Mir6352 | | 0 | 0 | 0 | 0 | 0 | 0 | 0 | 0 | 0 | 0 | 0 |
| 0 | NA | NA | NA | NA | NA | Gm28385 | | 0 | 0 | 0 | 0 | 0 | 0 | 0 | 0 | 0 | 0 | 0 |
| 0 | NA | NA | NA | NA | NA | Gm26263 | | 0 | 0 | 0 | 0 | 0 | 0 | 0 | 0 | 0 | 0 | 0 |
| 0 | NA | NA | NA | NA | NA | Gm28387 | | 0 | 0 | 0 | 0 | 0 | 0 | 0 | 0 | 0 | 0 | 0 |
| 0 | NA | NA | NA | NA | NA | Pax3 | paired box 3 | 0 | 0 | 0 | 0 | 0 | 0 | 0 | 0 | 0 | 0 | 0 |
| 0 | NA | NA | NA | NA | NA | Gm2102 | | 0 | 0 | 0 | 0 | 0 | 0 | 0 | 0 | 0 | 0 | 0 |
| 0 | NA | NA | NA | NA | NA | Gm28995 | | 0 | 0 | 0 | 0 | 0 | 0 | 0 | 0 | 0 | 0 | 0 |
| 0 | NA | NA | NA | NA | NA | BC035947 | | 0 | 0 | 0 | 0 | 0 | 0 | 0 | 0 | 0 | 0 | 0 |
| 0 | NA | NA | NA | NA | NA | 5730419F03Rik | | 0 | 0 | 0 | 0 | 0 | 0 | 0 | 0 | 0 | 0 | 0 |
| 0 | NA | NA | NA | NA | NA | Gm29611 | | 0 | 0 | 0 | 0 | 0 | 0 | 0 | 0 | 0 | 0 | 0 |
| 0 | NA | NA | NA | NA | NA | Gm29536 | | 0 | 0 | 0 | 0 | 0 | 0 | 0 | 0 | 0 | 0 | 0 |
| 0 | NA | NA | NA | NA | NA | Gm19027 | | 0 | 0 | 0 | 0 | 0 | 0 | 0 | 0 | 0 | 0 | 0 |
| 0 | NA | NA | NA | NA | NA | Gm24160 | | 0 | 0 | 0 | 0 | 0 | 0 | 0 | 0 | 0 | 0 | 0 |
| 0 | NA | NA | NA | NA | NA | Gm28058 | | 0 | 0 | 0 | 0 | 0 | 0 | 0 | 0 | 0 | 0 | 0 |
| 0 | NA | NA | NA | NA | NA | Gm23934 | | 0 | 0 | 0 | 0 | 0 | 0 | 0 | 0 | 0 | 0 | 0 |
| 0 | NA | NA | NA | NA | NA | Gm38339 | | 0 | 0 | 0 | 0 | 0 | 0 | 0 | 0 | 0 | 0 | 0 |
| 0 | NA | NA | NA | NA | NA | 9830004L10Rik | | 0 | 0 | 0 | 0 | 0 | 0 | 0 | 0 | 0 | 0 | 0 |
| 0 | NA | NA | NA | NA | NA | Gm37932 | | 0 | 0 | 0 | 0 | 0 | 0 | 0 | 0 | 0 | 0 | 0 |
| 0 | NA | NA | NA | NA | NA | Gm6189 | | 0 | 0 | 0 | 0 | 0 | 0 | 0 | 0 | 0 | 0 | 0 |
| 0 | NA | NA | NA | NA | NA | Gm45261 | | 0 | 0 | 0 | 0 | 0 | 0 | 0 | 0 | 0 | 0 | 0 |
| 0 | NA | NA | NA | NA | NA | Gm38062 | | 0 | 0 | 0 | 0 | 0 | 0 | 0 | 0 | 0 | 0 | 0 |
| 0 | NA | NA | NA | NA | NA | Gm29124 | | 0 | 0 | 0 | 0 | 0 | 0 | 0 | 0 | 0 | 0 | 0 |
| 0 | NA | NA | NA | NA | NA | Gm28066 | | 0 | 0 | 0 | 0 | 0 | 0 | 0 | 0 | 0 | 0 | 0 |
| 0 | NA | NA | NA | NA | NA | Gm37210 | | 0 | 0 | 0 | 0 | 0 | 0 | 0 | 0 | 0 | 0 | 0 |
| 0 | NA | NA | NA | NA | NA | Gm6198 | | 0 | 0 | 0 | 0 | 0 | 0 | 0 | 0 | 0 | 0 | 0 |
| 0 | NA | NA | NA | NA | NA | Gm37114 | | 0 | 0 | 0 | 0 | 0 | 0 | 0 | 0 | 0 | 0 | 0 |
| 0 | NA | NA | NA | NA | NA | Mir6344 | | 0 | 0 | 0 | 0 | 0 | 0 | 0 | 0 | 0 | 0 | 0 |
| 0 | NA | NA | NA | NA | NA | Gm9747 | | 0 | 0 | 0 | 0 | 0 | 0 | 0 | 0 | 0 | 0 | 0 |
| 0 | NA | NA | NA | NA | NA | Gm28940 | | 0 | 0 | 0 | 0 | 0 | 0 | 0 | 0 | 0 | 0 | 0 |
| 0 | NA | NA | NA | NA | NA | Gm22112 | | 0 | 0 | 0 | 0 | 0 | 0 | 0 | 0 | 0 | 0 | 0 |
| 0 | NA | NA | NA | NA | NA | n-R5s213 | | 0 | 0 | 0 | 0 | 0 | 0 | 0 | 0 | 0 | 0 | 0 |
| 0 | NA | NA | NA | NA | NA | Gm47791 | | 0 | 0 | 0 | 0 | 0 | 0 | 0 | 0 | 0 | 0 | 0 |
| 0 | NA | NA | NA | NA | NA | Gm19257 | | 0 | 0 | 0 | 0 | 0 | 0 | 0 | 0 | 0 | 0 | 0 |
| 0 | NA | NA | NA | NA | NA | Gm28942 | | 0 | 0 | 0 | 0 | 0 | 0 | 0 | 0 | 0 | 0 | 0 |
| 0 | NA | NA | NA | NA | NA | Gm22396 | | 0 | 0 | 0 | 0 | 0 | 0 | 0 | 0 | 0 | 0 | 0 |
| 0 | NA | NA | NA | NA | NA | A030014E15Rik | | 0 | 0 | 0 | 0 | 0 | 0 | 0 | 0 | 0 | 0 | 0 |
| 0 | NA | NA | NA | NA | NA | Gm6217 | | 0 | 0 | 0 | 0 | 0 | 0 | 0 | 0 | 0 | 0 | 0 |
| 0 | NA | NA | NA | NA | NA | A030003K21Rik | | 0 | 0 | 0 | 0 | 0 | 0 | 0 | 0 | 0 | 0 | 0 |
| 0 | NA | NA | NA | NA | NA | Gm7544 | | 0 | 0 | 0 | 0 | 0 | 0 | 0 | 0 | 0 | 0 | 0 |
| 0 | NA | NA | NA | NA | NA | Gm47955 | | 0 | 0 | 0 | 0 | 0 | 0 | 0 | 0 | 0 | 0 | 0 |
| 0 | NA | NA | NA | NA | NA | Gm47959 | | 0 | 0 | 0 | 0 | 0 | 0 | 0 | 0 | 0 | 0 | 0 |
| 0 | NA | NA | NA | NA | NA | Gm7539 | | 0 | 0 | 0 | 0 | 0 | 0 | 0 | 0 | 0 | 0 | 0 |
| 0 | NA | NA | NA | NA | NA | Krtap28-10 | | 0 | 0 | 0 | 0 | 0 | 0 | 0 | 0 | 0 | 0 | 0 |
| 0 | NA | NA | NA | NA | NA | Gm47969 | | 0 | 0 | 0 | 0 | 0 | 0 | 0 | 0 | 0 | 0 | 0 |
| 0 | NA | NA | NA | NA | NA | A030005K14Rik | | 0 | 0 | 0 | 0 | 0 | 0 | 0 | 0 | 0 | 0 | 0 |
| 0 | NA | NA | NA | NA | NA | Krtap28-13 | keratin associated protein 28-13 | 0 | 0 | 0 | 0 | 0 | 0 | 0 | 0 | 0 | 0 | 0 |

| | | | | | | | | | | | | | | | | | | |
|---|---|---|---|---|---|---|---|---|---|---|---|---|---|---|---|---|---|---|
| 0 | NA | NA | NA | NA | NA | Gm29518 | | 0 | 0 | 0 | 0 | 0 | 0 | 0 | 0 | 0 | 0 | 0 |
| 0 | NA | NA | NA | NA | NA | Gm29519 | | 0 | 0 | 0 | 0 | 0 | 0 | 0 | 0 | 0 | 0 | 0 |
| 0 | NA | NA | NA | NA | NA | n-R5s214 | | 0 | 0 | 0 | 0 | 0 | 0 | 0 | 0 | 0 | 0 | 0 |
| 0 | NA | NA | NA | NA | NA | Gm37989 | | 0 | 0 | 0 | 0 | 0 | 0 | 0 | 0 | 0 | 0 | 0 |
| 0 | NA | NA | NA | NA | NA | 4933436I20Rik | | 0 | 0 | 0 | 0 | 0 | 0 | 0 | 0 | 0 | 0 | 0 |
| 0 | NA | NA | NA | NA | NA | Gm25754 | | 0 | 0 | 0 | 0 | 0 | 0 | 0 | 0 | 0 | 0 | 0 |
| 0 | NA | NA | NA | NA | NA | Gm24555 | | 0 | 0 | 0 | 0 | 0 | 0 | 0 | 0 | 0 | 0 | 0 |
| 0 | NA | NA | NA | NA | NA | Gm18304 | | 0 | 0 | 0 | 0 | 0 | 0 | 0 | 0 | 0 | 0 | 0 |
| 0 | NA | NA | NA | NA | NA | Mir6353 | | 0 | 0 | 0 | 0 | 0 | 0 | 0 | 0 | 0 | 0 | 0 |
| 0 | NA | NA | NA | NA | NA | Gm37959 | | 0 | 0 | 0 | 0 | 0 | 0 | 0 | 0 | 0 | 0 | 0 |
| 0 | NA | NA | NA | NA | NA | Gm37925 | | 0 | 0 | 0 | 0 | 0 | 0 | 0 | 0 | 0 | 0 | 0 |
| 0 | NA | NA | NA | NA | NA | Mir5126 | | 0 | 0 | 0 | 0 | 0 | 0 | 0 | 0 | 0 | 0 | 0 |
| 0 | NA | NA | NA | NA | NA | Gm38374 | | 0 | 0 | 0 | 0 | 0 | 0 | 0 | 0 | 0 | 0 | 0 |
| 0 | NA | NA | NA | NA | NA | | | 0 | 0 | 0 | 0 | 0 | 0 | 0 | 0 | 0 | 0 | 0 |
| 0 | NA | NA | NA | NA | NA | Gm29284 | | 0 | 0 | 0 | 0 | 0 | 0 | 0 | 0 | 0 | 0 | 0 |
| 0 | NA | NA | NA | NA | NA | Gm27907 | | 0 | 0 | 0 | 0 | 0 | 0 | 0 | 0 | 0 | 0 | 0 |
| 0 | NA | NA | NA | NA | NA | | | 0 | 0 | 0 | 0 | 0 | 0 | 0 | 0 | 0 | 0 | 0 |
| 0 | NA | NA | NA | NA | NA | Gm2389 | | 0 | 0 | 0 | 0 | 0 | 0 | 0 | 0 | 0 | 0 | 0 |
| 0 | NA | NA | NA | NA | NA | Gm10553 | | 0 | 0 | 0 | 0 | 0 | 0 | 0 | 0 | 0 | 0 | 0 |
| 0 | NA | NA | NA | NA | NA | Gm16038 | | 0 | 0 | 0 | 0 | 0 | 0 | 0 | 0 | 0 | 0 | 0 |
| 0 | NA | NA | NA | NA | NA | Gm7592 | | 0 | 0 | 0 | 0 | 0 | 0 | 0 | 0 | 0 | 0 | 0 |
| 0 | NA | NA | NA | NA | NA | | | 0 | 0 | 0 | 0 | 0 | 0 | 0 | 0 | 0 | 0 | 0 |
| 0 | NA | NA | NA | NA | NA | Gm16028 | | 0 | 0 | 0 | 0 | 0 | 0 | 0 | 0 | 0 | 0 | 0 |
| 0 | NA | NA | NA | NA | NA | Gm7281 | | 0 | 0 | 0 | 0 | 0 | 0 | 0 | 0 | 0 | 0 | 0 |
| 0 | NA | NA | NA | NA | NA | | | 0 | 0 | 0 | 0 | 0 | 0 | 0 | 0 | 0 | 0 | 0 |
| 0 | NA | NA | NA | NA | NA | | predicted gene 7592 | 0 | 0 | 0 | 0 | 0 | 0 | 0 | 0 | 0 | 0 | 0 |
| 0 | NA | NA | NA | NA | NA | | | 0 | 0 | 0 | 0 | 0 | 0 | 0 | 0 | 0 | 0 | 0 |
| 0 | NA | NA | NA | NA | NA | G530012D18Rik | | 0 | 0 | 0 | 0 | 0 | 0 | 0 | 0 | 0 | 0 | 0 |
| 0 | NA | NA | NA | NA | NA | Gm16094 | | 0 | 0 | 0 | 0 | 0 | 0 | 0 | 0 | 0 | 0 | 0 |
| 0 | NA | NA | NA | NA | NA | n-R5s215 | | 0 | 0 | 0 | 0 | 0 | 0 | 0 | 0 | 0 | 0 | 0 |
| 0 | NA | NA | NA | NA | NA | Mir8096 | | 0 | 0 | 0 | 0 | 0 | 0 | 0 | 0 | 0 | 0 | 0 |
| 0 | NA | NA | NA | NA | NA | Gm25472 | | 0 | 0 | 0 | 0 | 0 | 0 | 0 | 0 | 0 | 0 | 0 |
| 0 | NA | NA | NA | NA | NA | 9930111H07Rik | | 0 | 0 | 0 | 0 | 0 | 0 | 0 | 0 | 0 | 0 | 0 |
| 0 | NA | NA | NA | NA | NA | Gm22906 | | 0 | 0 | 0 | 0 | 0 | 0 | 0 | 0 | 0 | 0 | 0 |
| 0 | NA | NA | NA | NA | NA | Gm28100 | | 0 | 0 | 0 | 0 | 0 | 0 | 0 | 0 | 0 | 0 | 0 |
| 0 | NA | NA | NA | NA | NA | Spata3 | spermatogenesis associated 3 | 0 | 0 | 0 | 0 | 0 | 0 | 0 | 0 | 0 | 0 | 0 |
| 0 | NA | NA | NA | NA | NA | Gm22786 | | 0 | 0 | 0 | 0 | 0 | 0 | 0 | 0 | 0 | 0 | 0 |
| 0 | NA | NA | NA | NA | NA | Gm21972 | | 0 | 0 | 0 | 0 | 0 | 0 | 0 | 0 | 0 | 0 | 0 |
| 0 | NA | NA | NA | NA | NA | Gm16341 | | 0 | 0 | 0 | 0 | 0 | 0 | 0 | 0 | 0 | 0 | 0 |
| 0 | NA | NA | NA | NA | NA | Gm28626 | | 0 | 0 | 0 | 0 | 0 | 0 | 0 | 0 | 0 | 0 | 0 |
| 0 | NA | NA | NA | NA | NA | Gm5258 | | 0 | 0 | 0 | 0 | 0 | 0 | 0 | 0 | 0 | 0 | 0 |
| 0 | NA | NA | NA | NA | NA | Gm28627 | | 0 | 0 | 0 | 0 | 0 | 0 | 0 | 0 | 0 | 0 | 0 |
| 0 | NA | NA | NA | NA | NA | Snora75 | | 0 | 0 | 0 | 0 | 0 | 0 | 0 | 0 | 0 | 0 | 0 |
| 0 | NA | NA | NA | NA | NA | Mir3535 | | 0 | 0 | 0 | 0 | 0 | 0 | 0 | 0 | 0 | 0 | 0 |
| 0 | NA | NA | NA | NA | NA | Gm24148 | | 0 | 0 | 0 | 0 | 0 | 0 | 0 | 0 | 0 | 0 | 0 |
| 0 | NA | NA | NA | NA | NA | Snord82 | | 0 | 0 | 0 | 0 | 0 | 0 | 0 | 0 | 0 | 0 | 0 |
| 0 | NA | NA | NA | NA | NA | Gm45908 | | 0 | 0 | 0 | 0 | 0 | 0 | 0 | 0 | 0 | 0 | 0 |
| 0 | NA | NA | NA | NA | NA | Tex44 | testis expressed 44 | 0 | 0 | 0 | 0 | 0 | 0 | 0 | 0 | 0 | 0 | 0 |
| 0 | NA | NA | NA | NA | NA | Gm37152 | | 0 | 0 | 0 | 0 | 0 | 0 | 0 | 0 | 0 | 0 | 0 |
| 0 | NA | NA | NA | NA | NA | Gm22955 | | 0 | 0 | 0 | 0 | 0 | 0 | 0 | 0 | 0 | 0 | 0 |
| 0 | NA | NA | NA | NA | NA | Gm37022 | | 0 | 0 | 0 | 0 | 0 | 0 | 0 | 0 | 0 | 0 | 0 |
| 0 | NA | NA | NA | NA | NA | Gm29097 | | 0 | 0 | 0 | 0 | 0 | 0 | 0 | 0 | 0 | 0 | 0 |
| 0 | NA | NA | NA | NA | NA | Gm29055 | | 0 | 0 | 0 | 0 | 0 | 0 | 0 | 0 | 0 | 0 | 0 |
| 0 | NA | NA | NA | NA | NA | Gm38010 | | 0 | 0 | 0 | 0 | 0 | 0 | 0 | 0 | 0 | 0 | 0 |
| 0 | NA | NA | NA | NA | NA | Gm28375 | | 0 | 0 | 0 | 0 | 0 | 0 | 0 | 0 | 0 | 0 | 0 |
| 0 | NA | NA | NA | NA | NA | Gm29371 | | 0 | 0 | 0 | 0 | 0 | 0 | 0 | 0 | 0 | 0 | 0 |
| 0 | NA | NA | NA | NA | NA | Alppl2 | alkaline phosphatase, placental-like 2 | 0 | 0 | 0 | 0 | 0 | 0 | 0 | 0 | 0 | 0 | 0 |
| 0 | NA | NA | NA | NA | NA | Alpi | alkaline phosphatase, intestinal | 0 | 0 | 0 | 0 | 0 | 0 | 0 | 0 | 0 | 0 | 0 |
| 0 | NA | NA | NA | NA | NA | Gm29372 | | 0 | 0 | 0 | 0 | 0 | 0 | 0 | 0 | 0 | 0 | 0 |
| 0 | NA | NA | NA | NA | NA | Akp3 | alkaline phosphatase 3, intestine, not Mn | 0 | 0 | 0 | 0 | 0 | 0 | 0 | 0 | 0 | 0 | 0 |

| | | | | | | | | | | | | | | | | | | |
|---|---|---|---|---|---|---|---|---|---|---|---|---|---|---|---|---|---|---|
| 0 | NA | NA | NA | NA | NA | Gm29374 | | 0 | 0 | 0 | 0 | 0 | 0 | 0 | 0 | 0 | 0 | 0 |
| 0 | NA | NA | NA | NA | NA | Prss56 | protease, serine 56 | 0 | 0 | 0 | 0 | 0 | 0 | 0 | 0 | 0 | 0 | 0 |
| 0 | NA | NA | NA | NA | NA | Chrnd | cholinergic receptor, nicotinic, delta polyp | 0 | 0 | 0 | 0 | 0 | 0 | 0 | 0 | 0 | 0 | 0 |
| 0 | NA | NA | NA | NA | NA | Chrng | cholinergic receptor, nicotinic, gamma po | 0 | 0 | 0 | 0 | 0 | 0 | 0 | 0 | 0 | 0 | 0 |
| 0 | NA | NA | NA | NA | NA | Gm26187 | | 0 | 0 | 0 | 0 | 0 | 0 | 0 | 0 | 0 | 0 | 0 |
| 0 | NA | NA | NA | NA | NA | Gm22549 | | 0 | 0 | 0 | 0 | 0 | 0 | 0 | 0 | 0 | 0 | 0 |
| 0 | NA | NA | NA | NA | NA | Gm23504 | | 0 | 0 | 0 | 0 | 0 | 0 | 0 | 0 | 0 | 0 | 0 |
| 0 | NA | NA | NA | NA | NA | Gm17181 | | 0 | 0 | 0 | 0 | 0 | 0 | 0 | 0 | 0 | 0 | 0 |
| 0 | NA | NA | NA | NA | NA | 4833421G17Rik | | 0 | 0 | 0 | 0 | 0 | 0 | 0 | 0 | 0 | 0 | 0 |
| 0 | NA | NA | NA | NA | NA | Gm25198 | | 0 | 0 | 0 | 0 | 0 | 0 | 0 | 0 | 0 | 0 | 0 |
| 0 | NA | NA | NA | NA | NA | | | 0 | 0 | 0 | 0 | 0 | 0 | 0 | 0 | 0 | 0 | 0 |
| 0 | NA | NA | NA | NA | NA | Ugt1a10 | UDP glycosyltransferase 1 family, polype | 0 | 0 | 0 | 0 | 0 | 0 | 0 | 0 | 0 | 0 | 0 |
| 0 | NA | NA | NA | NA | NA | | | 0 | 0 | 0 | 0 | 0 | 0 | 0 | 0 | 0 | 0 | 0 |
| 0 | NA | NA | NA | NA | NA | Gm15373 | | 0 | 0 | 0 | 0 | 0 | 0 | 0 | 0 | 0 | 0 | 0 |
| 0 | NA | NA | NA | NA | NA | Gm15372 | | 0 | 0 | 0 | 0 | 0 | 0 | 0 | 0 | 0 | 0 | 0 |
| 0 | NA | NA | NA | NA | NA | Gm15370 | | 0 | 0 | 0 | 0 | 0 | 0 | 0 | 0 | 0 | 0 | 0 |
| 0 | NA | NA | NA | NA | NA | Gm15369 | | 0 | 0 | 0 | 0 | 0 | 0 | 0 | 0 | 0 | 0 | 0 |
| 0 | NA | NA | NA | NA | NA | Gm15374 | | 0 | 0 | 0 | 0 | 0 | 0 | 0 | 0 | 0 | 0 | 0 |
| 0 | NA | NA | NA | NA | NA | Gm15371 | | 0 | 0 | 0 | 0 | 0 | 0 | 0 | 0 | 0 | 0 | 0 |
| 0 | NA | NA | NA | NA | NA | Ugt1a5 | UDP glucuronosyltransferase 1 family, pc | 0 | 0 | 0 | 0 | 0 | 0 | 0 | 0 | 0 | 0 | 0 |
| 0 | NA | NA | NA | NA | NA | | | 0 | 0 | 0 | 0 | 0 | 0 | 0 | 0 | 0 | 0 | 0 |
| 0 | NA | NA | NA | NA | NA | Gm15368 | | 0 | 0 | 0 | 0 | 0 | 0 | 0 | 0 | 0 | 0 | 0 |
| 0 | NA | NA | NA | NA | NA | Gm15376 | | 0 | 0 | 0 | 0 | 0 | 0 | 0 | 0 | 0 | 0 | 0 |
| 0 | NA | NA | NA | NA | NA | 4930453O03Rik | | 0 | 0 | 0 | 0 | 0 | 0 | 0 | 0 | 0 | 0 | 0 |
| 0 | NA | NA | NA | NA | NA | Gm29538 | | 0 | 0 | 0 | 0 | 0 | 0 | 0 | 0 | 0 | 0 | 0 |
| 0 | NA | NA | NA | NA | NA | Gm28888 | | 0 | 0 | 0 | 0 | 0 | 0 | 0 | 0 | 0 | 0 | 0 |
| 0 | NA | NA | NA | NA | NA | Glrp1 | glutamine repeat protein 1 | 0 | 0 | 0 | 0 | 0 | 0 | 0 | 0 | 0 | 0 | 0 |
| 0 | NA | NA | NA | NA | NA | Gm19589 | | 0 | 0 | 0 | 0 | 0 | 0 | 0 | 0 | 0 | 0 | 0 |
| 0 | NA | NA | NA | NA | NA | Gm38130 | | 0 | 0 | 0 | 0 | 0 | 0 | 0 | 0 | 0 | 0 | 0 |
| 0 | NA | NA | NA | NA | NA | Gm29336 | | 0 | 0 | 0 | 0 | 0 | 0 | 0 | 0 | 0 | 0 | 0 |
| 0 | NA | NA | NA | NA | NA | Gm29335 | | 0 | 0 | 0 | 0 | 0 | 0 | 0 | 0 | 0 | 0 | 0 |
| 0 | NA | NA | NA | NA | NA | Platr5 | | 0 | 0 | 0 | 0 | 0 | 0 | 0 | 0 | 0 | 0 | 0 |
| 0 | NA | NA | NA | NA | NA | Gm29337 | | 0 | 0 | 0 | 0 | 0 | 0 | 0 | 0 | 0 | 0 | 0 |
| 0 | NA | NA | NA | NA | NA | Gm4753 | | 0 | 0 | 0 | 0 | 0 | 0 | 0 | 0 | 0 | 0 | 0 |
| 0 | NA | NA | NA | NA | NA | Gm18699 | | 0 | 0 | 0 | 0 | 0 | 0 | 0 | 0 | 0 | 0 | 0 |
| 0 | NA | NA | NA | NA | NA | Gm5259 | | 0 | 0 | 0 | 0 | 0 | 0 | 0 | 0 | 0 | 0 | 0 |
| 0 | NA | NA | NA | NA | NA | Gm25180 | | 0 | 0 | 0 | 0 | 0 | 0 | 0 | 0 | 0 | 0 | 0 |
| 0 | NA | NA | NA | NA | NA | 4933400F21Rik | | 0 | 0 | 0 | 0 | 0 | 0 | 0 | 0 | 0 | 0 | 0 |
| 0 | NA | NA | NA | NA | NA | Gm26037 | | 0 | 0 | 0 | 0 | 0 | 0 | 0 | 0 | 0 | 0 | 0 |
| 0 | NA | NA | NA | NA | NA | 4930434B07Rik | | 0 | 0 | 0 | 0 | 0 | 0 | 0 | 0 | 0 | 0 | 0 |
| 0 | NA | NA | NA | NA | NA | Gm38277 | | 0 | 0 | 0 | 0 | 0 | 0 | 0 | 0 | 0 | 0 | 0 |
| 0 | NA | NA | NA | NA | NA | Gm28723 | | 0 | 0 | 0 | 0 | 0 | 0 | 0 | 0 | 0 | 0 | 0 |
| 0 | NA | NA | NA | NA | NA | Gm27648 | | 0 | 0 | 0 | 0 | 0 | 0 | 0 | 0 | 0 | 0 | 0 |
| 0 | NA | NA | NA | NA | NA | Gm28722 | | 0 | 0 | 0 | 0 | 0 | 0 | 0 | 0 | 0 | 0 | 0 |
| 0 | NA | NA | NA | NA | NA | Gm9991 | | 0 | 0 | 0 | 0 | 0 | 0 | 0 | 0 | 0 | 0 | 0 |
| 0 | NA | NA | NA | NA | NA | Gm36970 | | 0 | 0 | 0 | 0 | 0 | 0 | 0 | 0 | 0 | 0 | 0 |
| 0 | NA | NA | NA | NA | NA | 4930474B08Rik | | 0 | 0 | 0 | 0 | 0 | 0 | 0 | 0 | 0 | 0 | 0 |
| 0 | NA | NA | NA | NA | NA | Prlh | prolactin releasing hormone | 0 | 0 | 0 | 0 | 0 | 0 | 0 | 0 | 0 | 0 | 0 |
| 0 | NA | NA | NA | NA | NA | Gm28199 | | 0 | 0 | 0 | 0 | 0 | 0 | 0 | 0 | 0 | 0 | 0 |
| 0 | NA | NA | NA | NA | NA | Gm28381 | | 0 | 0 | 0 | 0 | 0 | 0 | 0 | 0 | 0 | 0 | 0 |
| 0 | NA | NA | NA | NA | NA | Gm19085 | | 0 | 0 | 0 | 0 | 0 | 0 | 0 | 0 | 0 | 0 | 0 |
| 0 | NA | NA | NA | NA | NA | Gm28380 | | 0 | 0 | 0 | 0 | 0 | 0 | 0 | 0 | 0 | 0 | 0 |
| 0 | NA | NA | NA | NA | NA | Gm29101 | | 0 | 0 | 0 | 0 | 0 | 0 | 0 | 0 | 0 | 0 | 0 |
| 0 | NA | NA | NA | NA | NA | Gm37600 | | 0 | 0 | 0 | 0 | 0 | 0 | 0 | 0 | 0 | 0 | 0 |
| 0 | NA | NA | NA | NA | NA | Gm37048 | | 0 | 0 | 0 | 0 | 0 | 0 | 0 | 0 | 0 | 0 | 0 |
| 0 | NA | NA | NA | NA | NA | Gm29099 | | 0 | 0 | 0 | 0 | 0 | 0 | 0 | 0 | 0 | 0 | 0 |
| 0 | NA | NA | NA | NA | NA | Gm7837 | | 0 | 0 | 0 | 0 | 0 | 0 | 0 | 0 | 0 | 0 | 0 |
| 0 | NA | NA | NA | NA | NA | Mir6900 | | 0 | 0 | 0 | 0 | 0 | 0 | 0 | 0 | 0 | 0 | 0 |
| 0 | NA | NA | NA | NA | NA | Olfr1416 | olfactory receptor 1416 | 0 | 0 | 0 | 0 | 0 | 0 | 0 | 0 | 0 | 0 | 0 |
| 0 | NA | NA | NA | NA | NA | Olfr1415 | olfactory receptor 1415 | 0 | 0 | 0 | 0 | 0 | 0 | 0 | 0 | 0 | 0 | 0 |

| | | | | | | | | | | | | | | | | | | |
|---|---|---|---|---|---|---|---|---|---|---|---|---|---|---|---|---|---|---|
| 0 | NA | NA | NA | NA | NA | Olfr1414 | olfactory receptor 1414 | 0 | 0 | 0 | 0 | 0 | 0 | 0 | 0 | 0 | 0 | 0 |
| 0 | NA | NA | NA | NA | NA | 1700054K02Rik | | 0 | 0 | 0 | 0 | 0 | 0 | 0 | 0 | 0 | 0 | 0 |
| 0 | NA | NA | NA | NA | NA | Gm28478 | | 0 | 0 | 0 | 0 | 0 | 0 | 0 | 0 | 0 | 0 | 0 |
| 0 | NA | NA | NA | NA | NA | Olfr1412 | olfactory receptor 1412 | 0 | 0 | 0 | 0 | 0 | 0 | 0 | 0 | 0 | 0 | 0 |
| 0 | NA | NA | NA | NA | NA | Gm44098 | | 0 | 0 | 0 | 0 | 0 | 0 | 0 | 0 | 0 | 0 | 0 |
| 0 | NA | NA | NA | NA | NA | Olfr1411 | olfactory receptor 1411 | 0 | 0 | 0 | 0 | 0 | 0 | 0 | 0 | 0 | 0 | 0 |
| 0 | NA | NA | NA | NA | NA | Gm44099 | | 0 | 0 | 0 | 0 | 0 | 0 | 0 | 0 | 0 | 0 | 0 |
| 0 | NA | NA | NA | NA | NA | Olfr1410 | olfactory receptor 1410 | 0 | 0 | 0 | 0 | 0 | 0 | 0 | 0 | 0 | 0 | 0 |
| 0 | NA | NA | NA | NA | NA | Olfr12 | olfactory receptor 12 | 0 | 0 | 0 | 0 | 0 | 0 | 0 | 0 | 0 | 0 | 0 |
| 0 | NA | NA | NA | NA | NA | Gm29483 | | 0 | 0 | 0 | 0 | 0 | 0 | 0 | 0 | 0 | 0 | 0 |
| 0 | NA | NA | NA | NA | NA | Gm29482 | | 0 | 0 | 0 | 0 | 0 | 0 | 0 | 0 | 0 | 0 | 0 |
| 0 | NA | NA | NA | NA | NA | Gm29481 | | 0 | 0 | 0 | 0 | 0 | 0 | 0 | 0 | 0 | 0 | 0 |
| 0 | NA | NA | NA | NA | NA | Mir149 | | 0 | 0 | 0 | 0 | 0 | 0 | 0 | 0 | 0 | 0 | 0 |
| 0 | NA | NA | NA | NA | NA | Gm18697 | | 0 | 0 | 0 | 0 | 0 | 0 | 0 | 0 | 0 | 0 | 0 |
| 0 | NA | NA | NA | NA | NA | Gm28086 | | 0 | 0 | 0 | 0 | 0 | 0 | 0 | 0 | 0 | 0 | 0 |
| 0 | NA | NA | NA | NA | NA | Mir6901 | | 0 | 0 | 0 | 0 | 0 | 0 | 0 | 0 | 0 | 0 | 0 |
| 0 | NA | NA | NA | NA | NA | Mir6902 | | 0 | 0 | 0 | 0 | 0 | 0 | 0 | 0 | 0 | 0 | 0 |
| 0 | NA | NA | NA | NA | NA | Septin2 | septin 2 | 0 | 0 | 0 | 0 | 0 | 0 | 0 | 0 | 0 | 0 | 0 |
| 0 | NA | NA | NA | NA | NA | Gm22089 | | 0 | 0 | 0 | 0 | 0 | 0 | 0 | 0 | 0 | 0 | 0 |
| 0 | NA | NA | NA | NA | NA | Gm18666 | | 0 | 0 | 0 | 0 | 0 | 0 | 0 | 0 | 0 | 0 | 0 |
| 0 | NA | NA | NA | NA | NA | Neu4 | sialidase 4 | 0 | 0 | 0 | 0 | 0 | 0 | 0 | 0 | 0 | 0 | 0 |
| 0 | NA | NA | NA | NA | NA | Gm7891 | | 0 | 0 | 0 | 0 | 0 | 0 | 0 | 0 | 0 | 0 | 0 |
| 0 | NA | NA | NA | NA | NA | Gm23389 | | 0 | 0 | 0 | 0 | 0 | 0 | 0 | 0 | 0 | 0 | 0 |
| 0 | NA | NA | NA | NA | NA | Gm23181 | | 0 | 0 | 0 | 0 | 0 | 0 | 0 | 0 | 0 | 0 | 0 |
| 0 | NA | NA | NA | NA | NA | Gm5264 | | 0 | 0 | 0 | 0 | 0 | 0 | 0 | 0 | 0 | 0 | 0 |
| 0 | NA | NA | NA | NA | NA | Gm37642 | | 0 | 0 | 0 | 0 | 0 | 0 | 0 | 0 | 0 | 0 | 0 |
| 0 | NA | NA | NA | NA | NA | 4930598F16Rik | | 0 | 0 | 0 | 0 | 0 | 0 | 0 | 0 | 0 | 0 | 0 |
| 0 | NA | NA | NA | NA | NA | Gm37761 | | 0 | 0 | 0 | 0 | 0 | 0 | 0 | 0 | 0 | 0 | 0 |
| 0 | NA | NA | NA | NA | NA | Gm22360 | | 0 | 0 | 0 | 0 | 0 | 0 | 0 | 0 | 0 | 0 | 0 |
| 0 | NA | NA | NA | NA | NA | 1700063A18Rik | | 0 | 0 | 0 | 0 | 0 | 0 | 0 | 0 | 0 | 0 | 0 |
| 0 | NA | NA | NA | NA | NA | Gm24586 | | 0 | 0 | 0 | 0 | 0 | 0 | 0 | 0 | 0 | 0 | 0 |
| 0 | NA | NA | NA | NA | NA | Gm22917 | | 0 | 0 | 0 | 0 | 0 | 0 | 0 | 0 | 0 | 0 | 0 |
| 0 | NA | NA | NA | NA | NA | Gm37076 | | 0 | 0 | 0 | 0 | 0 | 0 | 0 | 0 | 0 | 0 | 0 |
| 0 | NA | NA | NA | NA | NA | Gm29601 | | 0 | 0 | 0 | 0 | 0 | 0 | 0 | 0 | 0 | 0 | 0 |
| 0 | NA | NA | NA | NA | NA | 4930533P14Rik | | 0 | 0 | 0 | 0 | 0 | 0 | 0 | 0 | 0 | 0 | 0 |
| 0 | NA | NA | NA | NA | NA | Gm26245 | | 0 | 0 | 0 | 0 | 0 | 0 | 0 | 0 | 0 | 0 | 0 |
| 0 | NA | NA | NA | NA | NA | Gm26078 | | 0 | 0 | 0 | 0 | 0 | 0 | 0 | 0 | 0 | 0 | 0 |
| 0 | NA | NA | NA | NA | NA | Slco6b1 | | 0 | 0 | 0 | 0 | 0 | 0 | 0 | 0 | 0 | 0 | 0 |
| 0 | NA | NA | NA | NA | NA | Gm9499 | | 0 | 0 | 0 | 0 | 0 | 0 | 0 | 0 | 0 | 0 | 0 |
| 0 | NA | NA | NA | NA | NA | Gm28901 | | 0 | 0 | 0 | 0 | 0 | 0 | 0 | 0 | 0 | 0 | 0 |
| 0 | NA | NA | NA | NA | NA | Slco6c1 | solute carrier organic anion transporter fa | 0 | 0 | 0 | 0 | 0 | 0 | 0 | 0 | 0 | 0 | 0 |
| 0 | NA | NA | NA | NA | NA | Gm7133 | predicted gene 7133 | 0 | 0 | 0 | 0 | 0 | 0 | 0 | 0 | 0 | 0 | 0 |
| 0 | NA | NA | NA | NA | NA | Gm7135 | predicted gene 7135 | 0 | 0 | 0 | 0 | 0 | 0 | 0 | 0 | 0 | 0 | 0 |
| 0 | NA | NA | NA | NA | NA | Gm25413 | | 0 | 0 | 0 | 0 | 0 | 0 | 0 | 0 | 0 | 0 | 0 |
| 0 | NA | NA | NA | NA | NA | Gm24465 | | 0 | 0 | 0 | 0 | 0 | 0 | 0 | 0 | 0 | 0 | 0 |
| 0 | NA | NA | NA | NA | NA | Gm29461 | | 0 | 0 | 0 | 0 | 0 | 0 | 0 | 0 | 0 | 0 | 0 |
| 0 | NA | NA | NA | NA | NA | Gm7952 | | 0 | 0 | 0 | 0 | 0 | 0 | 0 | 0 | 0 | 0 | 0 |
| 0 | NA | NA | NA | NA | NA | Slco6d1 | solute carrier organic anion transporter fa | 0 | 0 | 0 | 0 | 0 | 0 | 0 | 0 | 0 | 0 | 0 |
| 0 | NA | NA | NA | NA | NA | Gm26134 | | 0 | 0 | 0 | 0 | 0 | 0 | 0 | 0 | 0 | 0 | 0 |
| 0 | NA | NA | NA | NA | NA | Gm23434 | | 0 | 0 | 0 | 0 | 0 | 0 | 0 | 0 | 0 | 0 | 0 |
| 0 | NA | NA | NA | NA | NA | Gm18248 | | 0 | 0 | 0 | 0 | 0 | 0 | 0 | 0 | 0 | 0 | 0 |
| 0 | NA | NA | NA | NA | NA | Gm37877 | | 0 | 0 | 0 | 0 | 0 | 0 | 0 | 0 | 0 | 0 | 0 |
| 0 | NA | NA | NA | NA | NA | Gm23602 | | 0 | 0 | 0 | 0 | 0 | 0 | 0 | 0 | 0 | 0 | 0 |
| 0 | NA | NA | NA | NA | NA | Gm18700 | | 0 | 0 | 0 | 0 | 0 | 0 | 0 | 0 | 0 | 0 | 0 |
| 0 | NA | NA | NA | NA | NA | Gm18185 | | 0 | 0 | 0 | 0 | 0 | 0 | 0 | 0 | 0 | 0 | 0 |
| 0 | NA | NA | NA | NA | NA | Gm29334 | | 0 | 0 | 0 | 0 | 0 | 0 | 0 | 0 | 0 | 0 | 0 |
| 0 | NA | NA | NA | NA | NA | Gm25537 | | 0 | 0 | 0 | 0 | 0 | 0 | 0 | 0 | 0 | 0 | 0 |
| 0 | NA | NA | NA | NA | NA | Gm20268 | | 0 | 0 | 0 | 0 | 0 | 0 | 0 | 0 | 0 | 0 | 0 |
| 0 | NA | NA | NA | NA | NA | Gm28801 | | 0 | 0 | 0 | 0 | 0 | 0 | 0 | 0 | 0 | 0 | 0 |
| 0 | NA | NA | NA | NA | NA | Gm18969 | | 0 | 0 | 0 | 0 | 0 | 0 | 0 | 0 | 0 | 0 | 0 |

| | | | | | | | | | | | | | | | | | | |
|---|---|---|---|---|---|---|---|---|---|---|---|---|---|---|---|---|---|---|
| 0 | NA | NA | NA | NA | NA | Gm24122 | | 0 | 0 | 0 | 0 | 0 | 0 | 0 | 0 | 0 | 0 | 0 |
| 0 | NA | NA | NA | NA | NA | Gm28623 | | 0 | 0 | 0 | 0 | 0 | 0 | 0 | 0 | 0 | 0 | 0 |
| 0 | NA | NA | NA | NA | NA | Gm23036 | | 0 | 0 | 0 | 0 | 0 | 0 | 0 | 0 | 0 | 0 | 0 |
| 0 | NA | NA | NA | NA | NA | Gm3116 | | 0 | 0 | 0 | 0 | 0 | 0 | 0 | 0 | 0 | 0 | 0 |
| 0 | NA | NA | NA | NA | NA | Gm20281 | | 0 | 0 | 0 | 0 | 0 | 0 | 0 | 0 | 0 | 0 | 0 |
| 0 | NA | NA | NA | NA | NA | Gm22034 | | 0 | 0 | 0 | 0 | 0 | 0 | 0 | 0 | 0 | 0 | 0 |
| 0 | NA | NA | NA | NA | NA | Gm18728 | | 0 | 0 | 0 | 0 | 0 | 0 | 0 | 0 | 0 | 0 | 0 |
| 0 | NA | NA | NA | NA | NA | Gm23965 | | 0 | 0 | 0 | 0 | 0 | 0 | 0 | 0 | 0 | 0 | 0 |
| 0 | NA | NA | NA | NA | NA | Gm29141 | | 0 | 0 | 0 | 0 | 0 | 0 | 0 | 0 | 0 | 0 | 0 |
| 0 | NA | NA | NA | NA | NA | Gm37789 | | 0 | 0 | 0 | 0 | 0 | 0 | 0 | 0 | 0 | 0 | 0 |
| 0 | NA | NA | NA | NA | NA | Gm18447 | | 0 | 0 | 0 | 0 | 0 | 0 | 0 | 0 | 0 | 0 | 0 |
| 0 | NA | NA | NA | NA | NA | Gm29140 | | 0 | 0 | 0 | 0 | 0 | 0 | 0 | 0 | 0 | 0 | 0 |
| 0 | NA | NA | NA | NA | NA | Gm23537 | | 0 | 0 | 0 | 0 | 0 | 0 | 0 | 0 | 0 | 0 | 0 |
| 0 | NA | NA | NA | NA | NA | Gm5260 | | 0 | 0 | 0 | 0 | 0 | 0 | 0 | 0 | 0 | 0 | 0 |
| 0 | NA | NA | NA | NA | NA | Gm29592 | | 0 | 0 | 0 | 0 | 0 | 0 | 0 | 0 | 0 | 0 | 0 |
| 0 | NA | NA | NA | NA | NA | Gm29011 | | 0 | 0 | 0 | 0 | 0 | 0 | 0 | 0 | 0 | 0 | 0 |
| 0 | NA | NA | NA | NA | NA | Gm38031 | | 0 | 0 | 0 | 0 | 0 | 0 | 0 | 0 | 0 | 0 | 0 |
| 0 | NA | NA | NA | NA | NA | n-R5s216 | | 0 | 0 | 0 | 0 | 0 | 0 | 0 | 0 | 0 | 0 | 0 |
| 0 | NA | NA | NA | NA | NA | Gm36969 | | 0 | 0 | 0 | 0 | 0 | 0 | 0 | 0 | 0 | 0 | 0 |
| 0 | NA | NA | NA | NA | NA | Gm15699 | | 0 | 0 | 0 | 0 | 0 | 0 | 0 | 0 | 0 | 0 | 0 |
| 0 | NA | NA | NA | NA | NA | Gm29012 | | 0 | 0 | 0 | 0 | 0 | 0 | 0 | 0 | 0 | 0 | 0 |
| 0 | NA | NA | NA | NA | NA | Gm29013 | | 0 | 0 | 0 | 0 | 0 | 0 | 0 | 0 | 0 | 0 | 0 |
| 0 | NA | NA | NA | NA | NA | Gm29014 | | 0 | 0 | 0 | 0 | 0 | 0 | 0 | 0 | 0 | 0 | 0 |
| 0 | NA | NA | NA | NA | NA | Gm22426 | | 0 | 0 | 0 | 0 | 0 | 0 | 0 | 0 | 0 | 0 | 0 |
| 0 | NA | NA | NA | NA | NA | Gm8004 | | 0 | 0 | 0 | 0 | 0 | 0 | 0 | 0 | 0 | 0 | 0 |
| 0 | NA | NA | NA | NA | NA | Gm37779 | | 0 | 0 | 0 | 0 | 0 | 0 | 0 | 0 | 0 | 0 | 0 |
| 0 | NA | NA | NA | NA | NA | Gm26214 | | 0 | 0 | 0 | 0 | 0 | 0 | 0 | 0 | 0 | 0 | 0 |
| 0 | NA | NA | NA | NA | NA | Gm18801 | | 0 | 0 | 0 | 0 | 0 | 0 | 0 | 0 | 0 | 0 | 0 |
| 0 | NA | NA | NA | NA | NA | Gm37372 | | 0 | 0 | 0 | 0 | 0 | 0 | 0 | 0 | 0 | 0 | 0 |
| 0 | NA | NA | NA | NA | NA | A530053M12Rik | | 0 | 0 | 0 | 0 | 0 | 0 | 0 | 0 | 0 | 0 | 0 |
| 0 | NA | NA | NA | NA | NA | D630008O14Rik | | 0 | 0 | 0 | 0 | 0 | 0 | 0 | 0 | 0 | 0 | 0 |
| 0 | NA | NA | NA | NA | NA | Mir3473f | | 0 | 0 | 0 | 0 | 0 | 0 | 0 | 0 | 0 | 0 | 0 |
| 0 | NA | NA | NA | NA | NA | Gm37053 | | 0 | 0 | 0 | 0 | 0 | 0 | 0 | 0 | 0 | 0 | 0 |
| 0 | NA | NA | NA | NA | NA | Pou2f3-rs1 | | 0 | 0 | 0 | 0 | 0 | 0 | 0 | 0 | 0 | 0 | 0 |
| 0 | NA | NA | NA | NA | NA | Serpinb12 | serine (or cysteine) peptidase inhibitor, cl | 0 | 0 | 0 | 0 | 0 | 0 | 0 | 0 | 0 | 0 | 0 |
| 0 | NA | NA | NA | NA | NA | Gm8080 | | 0 | 0 | 0 | 0 | 0 | 0 | 0 | 0 | 0 | 0 | 0 |
| 0 | NA | NA | NA | NA | NA | Serpinb3a | serine (or cysteine) peptidase inhibitor, cl | 0 | 0 | 0 | 0 | 0 | 0 | 0 | 0 | 0 | 0 | 0 |
| 0 | NA | NA | NA | NA | NA | Serpinb3d | serine (or cysteine) peptidase inhibitor, cl | 0 | 0 | 0 | 0 | 0 | 0 | 0 | 0 | 0 | 0 | 0 |
| 0 | NA | NA | NA | NA | NA | Gm15388 | | 0 | 0 | 0 | 0 | 0 | 0 | 0 | 0 | 0 | 0 | 0 |
| 0 | NA | NA | NA | NA | NA | Serpinb3b | serine (or cysteine) peptidase inhibitor, cl | 0 | 0 | 0 | 0 | 0 | 0 | 0 | 0 | 0 | 0 | 0 |
| 0 | NA | NA | NA | NA | NA | Gm8084 | | 0 | 0 | 0 | 0 | 0 | 0 | 0 | 0 | 0 | 0 | 0 |
| 0 | NA | NA | NA | NA | NA | Gm15390 | | 0 | 0 | 0 | 0 | 0 | 0 | 0 | 0 | 0 | 0 | 0 |
| 0 | NA | NA | NA | NA | NA | Serpinb3c | serine (or cysteine) peptidase inhibitor, cl | 0 | 0 | 0 | 0 | 0 | 0 | 0 | 0 | 0 | 0 | 0 |
| 0 | NA | NA | NA | NA | NA | Gm8089 | | 0 | 0 | 0 | 0 | 0 | 0 | 0 | 0 | 0 | 0 | 0 |
| 0 | NA | NA | NA | NA | NA | Serpinb11 | serine (or cysteine) peptidase inhibitor, cl | 0 | 0 | 0 | 0 | 0 | 0 | 0 | 0 | 0 | 0 | 0 |
| 0 | NA | NA | NA | NA | NA | Gm15391 | | 0 | 0 | 0 | 0 | 0 | 0 | 0 | 0 | 0 | 0 | 0 |
| 0 | NA | NA | NA | NA | NA | Serpinb7 | serine (or cysteine) peptidase inhibitor, cl | 0 | 0 | 0 | 0 | 0 | 0 | 0 | 0 | 0 | 0 | 0 |
| 0 | NA | NA | NA | NA | NA | Gm24553 | | 0 | 0 | 0 | 0 | 0 | 0 | 0 | 0 | 0 | 0 | 0 |
| 0 | NA | NA | NA | NA | NA | Gm15389 | | 0 | 0 | 0 | 0 | 0 | 0 | 0 | 0 | 0 | 0 | 0 |
| 0 | NA | NA | NA | NA | NA | Gm24267 | | 0 | 0 | 0 | 0 | 0 | 0 | 0 | 0 | 0 | 0 | 0 |
| 0 | NA | NA | NA | NA | NA | Gm22438 | | 0 | 0 | 0 | 0 | 0 | 0 | 0 | 0 | 0 | 0 | 0 |
| 0 | NA | NA | NA | NA | NA | D830032E09Rik | | 0 | 0 | 0 | 0 | 0 | 0 | 0 | 0 | 0 | 0 | 0 |
| 0 | NA | NA | NA | NA | NA | Gm18448 | | 0 | 0 | 0 | 0 | 0 | 0 | 0 | 0 | 0 | 0 | 0 |
| 0 | NA | NA | NA | NA | NA | Gm28282 | | 0 | 0 | 0 | 0 | 0 | 0 | 0 | 0 | 0 | 0 | 0 |
| 0 | NA | NA | NA | NA | NA | Gm38212 | | 0 | 0 | 0 | 0 | 0 | 0 | 0 | 0 | 0 | 0 | 0 |
| 0 | NA | NA | NA | NA | NA | Gm28283 | | 0 | 0 | 0 | 0 | 0 | 0 | 0 | 0 | 0 | 0 | 0 |
| 0 | NA | NA | NA | NA | NA | Gm6679 | | 0 | 0 | 0 | 0 | 0 | 0 | 0 | 0 | 0 | 0 | 0 |
| 0 | NA | NA | NA | NA | NA | Gm28281 | | 0 | 0 | 0 | 0 | 0 | 0 | 0 | 0 | 0 | 0 | 0 |
| 0 | NA | NA | NA | NA | NA | Gm22159 | | 0 | 0 | 0 | 0 | 0 | 0 | 0 | 0 | 0 | 0 | 0 |
| 0 | NA | NA | NA | NA | NA | Gm25293 | | 0 | 0 | 0 | 0 | 0 | 0 | 0 | 0 | 0 | 0 | 0 |

| | | | | | | | | | | | | | | | | | |
|---|---|---|---|---|---|---|---|---|---|---|---|---|---|---|---|---|---|
| 0 | NA | NA | NA | NA | NA | Gm18552 | 0 | 0 | 0 | 0 | 0 | 0 | 0 | 0 | 0 | 0 | 0 |
| 0 | NA | NA | NA | NA | NA | Gm3332 | 0 | 0 | 0 | 0 | 0 | 0 | 0 | 0 | 0 | 0 | 0 |
| 0 | NA | NA | NA | NA | NA | Gm6693 | 0 | 0 | 0 | 0 | 0 | 0 | 0 | 0 | 0 | 0 | 0 |
| 0 | NA | NA | NA | NA | NA | Gm38255 | 0 | 0 | 0 | 0 | 0 | 0 | 0 | 0 | 0 | 0 | 0 |
| 0 | NA | NA | NA | NA | NA | Gm36944 | 0 | 0 | 0 | 0 | 0 | 0 | 0 | 0 | 0 | 0 | 0 |
| 0 | NA | NA | NA | NA | NA | Gm29088 | 0 | 0 | 0 | 0 | 0 | 0 | 0 | 0 | 0 | 0 | 0 |
| 0 | NA | NA | NA | NA | NA | Gm19927 | 0 | 0 | 0 | 0 | 0 | 0 | 0 | 0 | 0 | 0 | 0 |
| 0 | NA | NA | NA | NA | NA | Gm37934 | 0 | 0 | 0 | 0 | 0 | 0 | 0 | 0 | 0 | 0 | 0 |
| 0 | NA | NA | NA | NA | NA | Gm6717 | 0 | 0 | 0 | 0 | 0 | 0 | 0 | 0 | 0 | 0 | 0 |
| 0 | NA | NA | NA | NA | NA | Gm19086 | 0 | 0 | 0 | 0 | 0 | 0 | 0 | 0 | 0 | 0 | 0 |
| 0 | NA | NA | NA | NA | NA | Gm18406 | 0 | 0 | 0 | 0 | 0 | 0 | 0 | 0 | 0 | 0 | 0 |
| 0 | NA | NA | NA | NA | NA | Gm22331 | 0 | 0 | 0 | 0 | 0 | 0 | 0 | 0 | 0 | 0 | 0 |
| 0 | NA | NA | NA | NA | NA | Gm8204 | 0 | 0 | 0 | 0 | 0 | 0 | 0 | 0 | 0 | 0 | 0 |
| 0 | NA | NA | NA | NA | NA | Gm28190 | 0 | 0 | 0 | 0 | 0 | 0 | 0 | 0 | 0 | 0 | 0 |
| 0 | NA | NA | NA | NA | NA | Gm28189 | 0 | 0 | 0 | 0 | 0 | 0 | 0 | 0 | 0 | 0 | 0 |
| 0 | NA | NA | NA | NA | NA | Gm17946 | 0 | 0 | 0 | 0 | 0 | 0 | 0 | 0 | 0 | 0 | 0 |
| 0 | NA | NA | NA | NA | NA | Gm18181 | 0 | 0 | 0 | 0 | 0 | 0 | 0 | 0 | 0 | 0 | 0 |
| 0 | NA | NA | NA | NA | NA | Gm24937 | 0 | 0 | 0 | 0 | 0 | 0 | 0 | 0 | 0 | 0 | 0 |
| 0 | NA | NA | NA | NA | NA | 9330185C12Rik | 0 | 0 | 0 | 0 | 0 | 0 | 0 | 0 | 0 | 0 | 0 |
| 0 | NA | NA | NA | NA | NA | Gm4854 | 0 | 0 | 0 | 0 | 0 | 0 | 0 | 0 | 0 | 0 | 0 |
| 0 | NA | NA | NA | NA | NA | Gm17916 | 0 | 0 | 0 | 0 | 0 | 0 | 0 | 0 | 0 | 0 | 0 |
| 0 | NA | NA | NA | NA | NA | Gm23393 | 0 | 0 | 0 | 0 | 0 | 0 | 0 | 0 | 0 | 0 | 0 |
| 0 | NA | NA | NA | NA | NA | Gm28263 | 0 | 0 | 0 | 0 | 0 | 0 | 0 | 0 | 0 | 0 | 0 |
| 0 | NA | NA | NA | NA | NA | Gm19965 | 0 | 0 | 0 | 0 | 0 | 0 | 0 | 0 | 0 | 0 | 0 |
| 0 | NA | NA | NA | NA | NA | Gm17777 | 0 | 0 | 0 | 0 | 0 | 0 | 0 | 0 | 0 | 0 | 0 |
| 0 | NA | NA | NA | NA | NA | Gm24030 | 0 | 0 | 0 | 0 | 0 | 0 | 0 | 0 | 0 | 0 | 0 |
| 0 | NA | NA | NA | NA | NA | Gm27785 | 0 | 0 | 0 | 0 | 0 | 0 | 0 | 0 | 0 | 0 | 0 |
| 0 | NA | NA | NA | NA | NA | Gm28361 | 0 | 0 | 0 | 0 | 0 | 0 | 0 | 0 | 0 | 0 | 0 |
| 0 | NA | NA | NA | NA | NA | Zfp813-ps | 0 | 0 | 0 | 0 | 0 | 0 | 0 | 0 | 0 | 0 | 0 |
| 0 | NA | NA | NA | NA | NA | Zfp813-ps | 0 | 0 | 0 | 0 | 0 | 0 | 0 | 0 | 0 | 0 | 0 |
| 0 | NA | NA | NA | NA | NA | Gm28362 | 0 | 0 | 0 | 0 | 0 | 0 | 0 | 0 | 0 | 0 | 0 |
| 0 | NA | NA | NA | NA | NA | Gm28363 | 0 | 0 | 0 | 0 | 0 | 0 | 0 | 0 | 0 | 0 | 0 |
| 0 | NA | NA | NA | NA | NA | Gm28359 | 0 | 0 | 0 | 0 | 0 | 0 | 0 | 0 | 0 | 0 | 0 |
| 0 | NA | NA | NA | NA | NA | B020011L13Rik | 0 | 0 | 0 | 0 | 0 | 0 | 0 | 0 | 0 | 0 | 0 |
| 0 | NA | NA | NA | NA | NA | Gm28360 | 0 | 0 | 0 | 0 | 0 | 0 | 0 | 0 | 0 | 0 | 0 |
| 0 | NA | NA | NA | NA | NA | Gm7195 | 0 | 0 | 0 | 0 | 0 | 0 | 0 | 0 | 0 | 0 | 0 |
| 0 | NA | NA | NA | NA | NA | Gm28168 | 0 | 0 | 0 | 0 | 0 | 0 | 0 | 0 | 0 | 0 | 0 |
| 0 | NA | NA | NA | NA | NA | Gm7145 | 0 | 0 | 0 | 0 | 0 | 0 | 0 | 0 | 0 | 0 | 0 |
| 0 | NA | NA | NA | NA | NA | Gm28358 | 0 | 0 | 0 | 0 | 0 | 0 | 0 | 0 | 0 | 0 | 0 |
| 0 | NA | NA | NA | NA | NA | Gm29489 | 0 | 0 | 0 | 0 | 0 | 0 | 0 | 0 | 0 | 0 | 0 |
| 0 | NA | NA | NA | NA | NA | Gm29490 | 0 | 0 | 0 | 0 | 0 | 0 | 0 | 0 | 0 | 0 | 0 |
| 0 | NA | NA | NA | NA | NA | Gm29104 | 0 | 0 | 0 | 0 | 0 | 0 | 0 | 0 | 0 | 0 | 0 |
| 0 | NA | NA | NA | NA | NA | Gm29106 | 0 | 0 | 0 | 0 | 0 | 0 | 0 | 0 | 0 | 0 | 0 |
| 0 | NA | NA | NA | NA | NA | Gm28867 | 0 | 0 | 0 | 0 | 0 | 0 | 0 | 0 | 0 | 0 | 0 |
| 0 | NA | NA | NA | NA | NA | Gm26080 | 0 | 0 | 0 | 0 | 0 | 0 | 0 | 0 | 0 | 0 | 0 |
| 0 | NA | NA | NA | NA | NA | Gm22710 | 0 | 0 | 0 | 0 | 0 | 0 | 0 | 0 | 0 | 0 | 0 |
| 0 | NA | NA | NA | NA | NA | Gm28466 | 0 | 0 | 0 | 0 | 0 | 0 | 0 | 0 | 0 | 0 | 0 |
| 0 | NA | NA | NA | NA | NA | Gm22094 | 0 | 0 | 0 | 0 | 0 | 0 | 0 | 0 | 0 | 0 | 0 |
| 0 | NA | NA | NA | NA | NA | Gm28882 | 0 | 0 | 0 | 0 | 0 | 0 | 0 | 0 | 0 | 0 | 0 |
| 0 | NA | NA | NA | NA | NA | Gm23497 | 0 | 0 | 0 | 0 | 0 | 0 | 0 | 0 | 0 | 0 | 0 |
| 0 | NA | NA | NA | NA | NA | Gm28467 | 0 | 0 | 0 | 0 | 0 | 0 | 0 | 0 | 0 | 0 | 0 |
| 0 | NA | NA | NA | NA | NA | Mir6346 | 0 | 0 | 0 | 0 | 0 | 0 | 0 | 0 | 0 | 0 | 0 |
| 0 | NA | NA | NA | NA | NA | Gm26831 | 0 | 0 | 0 | 0 | 0 | 0 | 0 | 0 | 0 | 0 | 0 |
| 0 | NA | NA | NA | NA | NA | Gm8321 | 0 | 0 | 0 | 0 | 0 | 0 | 0 | 0 | 0 | 0 | 0 |
| 0 | NA | NA | NA | NA | NA | Gm29456 | 0 | 0 | 0 | 0 | 0 | 0 | 0 | 0 | 0 | 0 | 0 |
| 0 | NA | NA | NA | NA | NA | Gm29455 | 0 | 0 | 0 | 0 | 0 | 0 | 0 | 0 | 0 | 0 | 0 |
| 0 | NA | NA | NA | NA | NA | Gm29454 | 0 | 0 | 0 | 0 | 0 | 0 | 0 | 0 | 0 | 0 | 0 |
| 0 | NA | NA | NA | NA | NA | Gm25322 | 0 | 0 | 0 | 0 | 0 | 0 | 0 | 0 | 0 | 0 | 0 |
| 0 | NA | NA | NA | NA | NA | Gm37219 | 0 | 0 | 0 | 0 | 0 | 0 | 0 | 0 | 0 | 0 | 0 |
| 0 | NA | NA | NA | NA | NA | Gm29657 | 0 | 0 | 0 | 0 | 0 | 0 | 0 | 0 | 0 | 0 | 0 |

| | | | | | | | | | | | | | | | | | |
|---|---|---|---|---|---|---|---|---|---|---|---|---|---|---|---|---|---|
| 0 | NA | NA | NA | NA | NA | Gm8338 | | 0 | 0 | 0 | 0 | 0 | 0 | 0 | 0 | 0 | 0 | 0 |
| 0 | NA | NA | NA | NA | NA | Gm29658 | | 0 | 0 | 0 | 0 | 0 | 0 | 0 | 0 | 0 | 0 | 0 |
| 0 | NA | NA | NA | NA | NA | Gm3551 | | 0 | 0 | 0 | 0 | 0 | 0 | 0 | 0 | 0 | 0 | 0 |
| 0 | NA | NA | NA | NA | NA | Gm28209 | | 0 | 0 | 0 | 0 | 0 | 0 | 0 | 0 | 0 | 0 | 0 |
| 0 | NA | NA | NA | NA | NA | Gm17973 | | 0 | 0 | 0 | 0 | 0 | 0 | 0 | 0 | 0 | 0 | 0 |
| 0 | NA | NA | NA | NA | NA | 1700012E03Rik | | 0 | 0 | 0 | 0 | 0 | 0 | 0 | 0 | 0 | 0 | 0 |
| 0 | NA | NA | NA | NA | NA | En1 | engrailed 1 | 0 | 0 | 0 | 0 | 0 | 0 | 0 | 0 | 0 | 0 | 0 |
| 0 | NA | NA | NA | NA | NA | Gm29346 | | 0 | 0 | 0 | 0 | 0 | 0 | 0 | 0 | 0 | 0 | 0 |
| 0 | NA | NA | NA | NA | NA | Gm29345 | | 0 | 0 | 0 | 0 | 0 | 0 | 0 | 0 | 0 | 0 | 0 |
| 0 | NA | NA | NA | NA | NA | Gm29344 | | 0 | 0 | 0 | 0 | 0 | 0 | 0 | 0 | 0 | 0 | 0 |
| 0 | NA | NA | NA | NA | NA | Gm29348 | | 0 | 0 | 0 | 0 | 0 | 0 | 0 | 0 | 0 | 0 | 0 |
| 0 | NA | NA | NA | NA | NA | Gm29347 | | 0 | 0 | 0 | 0 | 0 | 0 | 0 | 0 | 0 | 0 | 0 |
| 0 | NA | NA | NA | NA | NA | Gm37043 | | 0 | 0 | 0 | 0 | 0 | 0 | 0 | 0 | 0 | 0 | 0 |
| 0 | NA | NA | NA | NA | NA | Celrr | | 0 | 0 | 0 | 0 | 0 | 0 | 0 | 0 | 0 | 0 | 0 |
| 0 | NA | NA | NA | NA | NA | Gm29359 | | 0 | 0 | 0 | 0 | 0 | 0 | 0 | 0 | 0 | 0 | 0 |
| 0 | NA | NA | NA | NA | NA | Gm38283 | | 0 | 0 | 0 | 0 | 0 | 0 | 0 | 0 | 0 | 0 | 0 |
| 0 | NA | NA | NA | NA | NA | 2210011K15Rik | | 0 | 0 | 0 | 0 | 0 | 0 | 0 | 0 | 0 | 0 | 0 |
| 0 | NA | NA | NA | NA | NA | Gm15392 | | 0 | 0 | 0 | 0 | 0 | 0 | 0 | 0 | 0 | 0 | 0 |
| 0 | NA | NA | NA | NA | NA | Gm15393 | | 0 | 0 | 0 | 0 | 0 | 0 | 0 | 0 | 0 | 0 | 0 |
| 0 | NA | NA | NA | NA | NA | Gm28591 | | 0 | 0 | 0 | 0 | 0 | 0 | 0 | 0 | 0 | 0 | 0 |
| 0 | NA | NA | NA | NA | NA | Gm37920 | | 0 | 0 | 0 | 0 | 0 | 0 | 0 | 0 | 0 | 0 | 0 |
| 0 | NA | NA | NA | NA | NA | Gm15394 | | 0 | 0 | 0 | 0 | 0 | 0 | 0 | 0 | 0 | 0 | 0 |
| 0 | NA | NA | NA | NA | NA | Gm8392 | | 0 | 0 | 0 | 0 | 0 | 0 | 0 | 0 | 0 | 0 | 0 |
| 0 | NA | NA | NA | NA | NA | Gm26472 | | 0 | 0 | 0 | 0 | 0 | 0 | 0 | 0 | 0 | 0 | 0 |
| 0 | NA | NA | NA | NA | NA | Gm28590 | | 0 | 0 | 0 | 0 | 0 | 0 | 0 | 0 | 0 | 0 | 0 |
| 0 | NA | NA | NA | NA | NA | Gm24547 | | 0 | 0 | 0 | 0 | 0 | 0 | 0 | 0 | 0 | 0 | 0 |
| 0 | NA | NA | NA | NA | NA | Gm36982 | | 0 | 0 | 0 | 0 | 0 | 0 | 0 | 0 | 0 | 0 | 0 |
| 0 | NA | NA | NA | NA | NA | Gm25771 | | 0 | 0 | 0 | 0 | 0 | 0 | 0 | 0 | 0 | 0 | 0 |
| 0 | NA | NA | NA | NA | NA | Gm29192 | | 0 | 0 | 0 | 0 | 0 | 0 | 0 | 0 | 0 | 0 | 0 |
| 0 | NA | NA | NA | NA | NA | Gm37970 | | 0 | 0 | 0 | 0 | 0 | 0 | 0 | 0 | 0 | 0 | 0 |
| 0 | NA | NA | NA | NA | NA | Gm28911 | | 0 | 0 | 0 | 0 | 0 | 0 | 0 | 0 | 0 | 0 | 0 |
| 0 | NA | NA | NA | NA | NA | Gm20058 | | 0 | 0 | 0 | 0 | 0 | 0 | 0 | 0 | 0 | 0 | 0 |
| 0 | NA | NA | NA | NA | NA | Gm28912 | | 0 | 0 | 0 | 0 | 0 | 0 | 0 | 0 | 0 | 0 | 0 |
| 0 | NA | NA | NA | NA | NA | Gm24791 | | 0 | 0 | 0 | 0 | 0 | 0 | 0 | 0 | 0 | 0 | 0 |
| 0 | NA | NA | NA | NA | NA | Gm37066 | | 0 | 0 | 0 | 0 | 0 | 0 | 0 | 0 | 0 | 0 | 0 |
| 0 | NA | NA | NA | NA | NA | Gm37781 | | 0 | 0 | 0 | 0 | 0 | 0 | 0 | 0 | 0 | 0 | 0 |
| 0 | NA | NA | NA | NA | NA | Gm37396 | | 0 | 0 | 0 | 0 | 0 | 0 | 0 | 0 | 0 | 0 | 0 |
| 0 | NA | NA | NA | NA | NA | Gm38263 | | 0 | 0 | 0 | 0 | 0 | 0 | 0 | 0 | 0 | 0 | 0 |
| 0 | NA | NA | NA | NA | NA | Gm28929 | | 0 | 0 | 0 | 0 | 0 | 0 | 0 | 0 | 0 | 0 | 0 |
| 0 | NA | NA | NA | NA | NA | Gm18875 | | 0 | 0 | 0 | 0 | 0 | 0 | 0 | 0 | 0 | 0 | 0 |
| 0 | NA | NA | NA | NA | NA | Gm25578 | | 0 | 0 | 0 | 0 | 0 | 0 | 0 | 0 | 0 | 0 | 0 |
| 0 | NA | NA | NA | NA | NA | Gm28299 | | 0 | 0 | 0 | 0 | 0 | 0 | 0 | 0 | 0 | 0 | 0 |
| 0 | NA | NA | NA | NA | NA | Gm37717 | | 0 | 0 | 0 | 0 | 0 | 0 | 0 | 0 | 0 | 0 | 0 |
| 0 | NA | NA | NA | NA | NA | 1700121L03Rik | | 0 | 0 | 0 | 0 | 0 | 0 | 0 | 0 | 0 | 0 | 0 |
| 0 | NA | NA | NA | NA | NA | Gm28706 | | 0 | 0 | 0 | 0 | 0 | 0 | 0 | 0 | 0 | 0 | 0 |
| 0 | NA | NA | NA | NA | NA | Gm18444 | | 0 | 0 | 0 | 0 | 0 | 0 | 0 | 0 | 0 | 0 | 0 |
| 0 | NA | NA | NA | NA | NA | Gm37936 | | 0 | 0 | 0 | 0 | 0 | 0 | 0 | 0 | 0 | 0 | 0 |
| 0 | NA | NA | NA | NA | NA | Gm38177 | | 0 | 0 | 0 | 0 | 0 | 0 | 0 | 0 | 0 | 0 | 0 |
| 0 | NA | NA | NA | NA | NA | 4930599A14Rik | | 0 | 0 | 0 | 0 | 0 | 0 | 0 | 0 | 0 | 0 | 0 |
| 0 | NA | NA | NA | NA | NA | Gm5261 | | 0 | 0 | 0 | 0 | 0 | 0 | 0 | 0 | 0 | 0 | 0 |
| 0 | NA | NA | NA | NA | NA | Gm37772 | | 0 | 0 | 0 | 0 | 0 | 0 | 0 | 0 | 0 | 0 | 0 |
| 0 | NA | NA | NA | NA | NA | Gm23370 | | 0 | 0 | 0 | 0 | 0 | 0 | 0 | 0 | 0 | 0 | 0 |
| 0 | NA | NA | NA | NA | NA | Gm23370 | | 0 | 0 | 0 | 0 | 0 | 0 | 0 | 0 | 0 | 0 | 0 |
| 0 | NA | NA | NA | NA | NA | Gm23734 | | 0 | 0 | 0 | 0 | 0 | 0 | 0 | 0 | 0 | 0 | 0 |
| 0 | NA | NA | NA | NA | NA | Acmsd | amino carboxymuconate semialdehyde d | 0 | 0 | 0 | 0 | 0 | 0 | 0 | 0 | 0 | 0 | 0 |
| 0 | NA | NA | NA | NA | NA | Gm29383 | | 0 | 0 | 0 | 0 | 0 | 0 | 0 | 0 | 0 | 0 | 0 |
| 0 | NA | NA | NA | NA | NA | Mir128-1 | | 0 | 0 | 0 | 0 | 0 | 0 | 0 | 0 | 0 | 0 | 0 |
| 0 | NA | NA | NA | NA | NA | Gm23902 | | 0 | 0 | 0 | 0 | 0 | 0 | 0 | 0 | 0 | 0 | 0 |
| 0 | NA | NA | NA | NA | NA | Gm29598 | | 0 | 0 | 0 | 0 | 0 | 0 | 0 | 0 | 0 | 0 | 0 |
| 0 | NA | NA | NA | NA | NA | Gm29597 | | 0 | 0 | 0 | 0 | 0 | 0 | 0 | 0 | 0 | 0 | 0 |

| | | | | | | | | | | | | | | | | | | |
|---|---|---|---|---|---|---|---|---|---|---|---|---|---|---|---|---|---|---|
| 0 | NA | NA | NA | NA | NA | Gm25384 | | 0 | 0 | 0 | 0 | 0 | 0 | 0 | 0 | 0 | 0 | 0 |
| 0 | NA | NA | NA | NA | NA | Gm29599 | | 0 | 0 | 0 | 0 | 0 | 0 | 0 | 0 | 0 | 0 | 0 |
| 0 | NA | NA | NA | NA | NA | Gm26686 | | 0 | 0 | 0 | 0 | 0 | 0 | 0 | 0 | 0 | 0 | 0 |
| 0 | NA | NA | NA | NA | NA | Gm23056 | | 0 | 0 | 0 | 0 | 0 | 0 | 0 | 0 | 0 | 0 | 0 |
| 0 | NA | NA | NA | NA | NA | Thsd7b | thrombospondin, type I, domain containin | 0 | 0 | 0 | 0 | 0 | 0 | 0 | 0 | 0 | 0 | 0 |
| 0 | NA | NA | NA | NA | NA | Gm38240 | | 0 | 0 | 0 | 0 | 0 | 0 | 0 | 0 | 0 | 0 | 0 |
| 0 | NA | NA | NA | NA | NA | Gm38330 | | 0 | 0 | 0 | 0 | 0 | 0 | 0 | 0 | 0 | 0 | 0 |
| 0 | NA | NA | NA | NA | NA | Gm36951 | | 0 | 0 | 0 | 0 | 0 | 0 | 0 | 0 | 0 | 0 | 0 |
| 0 | NA | NA | NA | NA | NA | Gm37479 | | 0 | 0 | 0 | 0 | 0 | 0 | 0 | 0 | 0 | 0 | 0 |
| 0 | NA | NA | NA | NA | NA | Gm37064 | | 0 | 0 | 0 | 0 | 0 | 0 | 0 | 0 | 0 | 0 | 0 |
| 0 | NA | NA | NA | NA | NA | Gm37278 | | 0 | 0 | 0 | 0 | 0 | 0 | 0 | 0 | 0 | 0 | 0 |
| 0 | NA | NA | NA | NA | NA | Gm29428 | | 0 | 0 | 0 | 0 | 0 | 0 | 0 | 0 | 0 | 0 | 0 |
| 0 | NA | NA | NA | NA | NA | Gm16081 | | 0 | 0 | 0 | 0 | 0 | 0 | 0 | 0 | 0 | 0 | 0 |
| 0 | NA | NA | NA | NA | NA | Cd55b | CD55 molecule, decay accelerating facto | 0 | 0 | 0 | 0 | 0 | 0 | 0 | 0 | 0 | 0 | 0 |
| 0 | NA | NA | NA | NA | NA | Cd55os | | 0 | 0 | 0 | 0 | 0 | 0 | 0 | 0 | 0 | 0 | 0 |
| 0 | NA | NA | NA | NA | NA | Gm16083 | | 0 | 0 | 0 | 0 | 0 | 0 | 0 | 0 | 0 | 0 | 0 |
| 0 | NA | NA | NA | NA | NA | Gm37109 | | 0 | 0 | 0 | 0 | 0 | 0 | 0 | 0 | 0 | 0 | 0 |
| 0 | NA | NA | NA | NA | NA | Gm37283 | | 0 | 0 | 0 | 0 | 0 | 0 | 0 | 0 | 0 | 0 | 0 |
| 0 | NA | NA | NA | NA | NA | Zp3r | zona pellucida 3 receptor | 0 | 0 | 0 | 0 | 0 | 0 | 0 | 0 | 0 | 0 | 0 |
| 0 | NA | NA | NA | NA | NA | Gm23623 | | 0 | 0 | 0 | 0 | 0 | 0 | 0 | 0 | 0 | 0 | 0 |
| 0 | NA | NA | NA | NA | NA | Gm37163 | | 0 | 0 | 0 | 0 | 0 | 0 | 0 | 0 | 0 | 0 | 0 |
| 0 | NA | NA | NA | NA | NA | Gm28857 | | 0 | 0 | 0 | 0 | 0 | 0 | 0 | 0 | 0 | 0 | 0 |
| 0 | NA | NA | NA | NA | NA | Gm28856 | | 0 | 0 | 0 | 0 | 0 | 0 | 0 | 0 | 0 | 0 | 0 |
| 0 | NA | NA | NA | NA | NA | Il24 | interleukin 24 | 0 | 0 | 0 | 0 | 0 | 0 | 0 | 0 | 0 | 0 | 0 |
| 0 | NA | NA | NA | NA | NA | Il20 | interleukin 20 | 0 | 0 | 0 | 0 | 0 | 0 | 0 | 0 | 0 | 0 | 0 |
| 0 | NA | NA | NA | NA | NA | Il19 | interleukin 19 | 0 | 0 | 0 | 0 | 0 | 0 | 0 | 0 | 0 | 0 | 0 |
| 0 | NA | NA | NA | NA | NA | Gm25549 | | 0 | 0 | 0 | 0 | 0 | 0 | 0 | 0 | 0 | 0 | 0 |
| 0 | NA | NA | NA | NA | NA | Gm25717 | | 0 | 0 | 0 | 0 | 0 | 0 | 0 | 0 | 0 | 0 | 0 |
| 0 | NA | NA | NA | NA | NA | Gm17796 | | 0 | 0 | 0 | 0 | 0 | 0 | 0 | 0 | 0 | 0 | 0 |
| 0 | NA | NA | NA | NA | NA | Gm28914 | | 0 | 0 | 0 | 0 | 0 | 0 | 0 | 0 | 0 | 0 | 0 |
| 0 | NA | NA | NA | NA | NA | Gm38262 | | 0 | 0 | 0 | 0 | 0 | 0 | 0 | 0 | 0 | 0 | 0 |
| 0 | NA | NA | NA | NA | NA | Gm37954 | | 0 | 0 | 0 | 0 | 0 | 0 | 0 | 0 | 0 | 0 | 0 |
| 0 | NA | NA | NA | NA | NA | Gm29487 | | 0 | 0 | 0 | 0 | 0 | 0 | 0 | 0 | 0 | 0 | 0 |
| 0 | NA | NA | NA | NA | NA | Gm24273 | | 0 | 0 | 0 | 0 | 0 | 0 | 0 | 0 | 0 | 0 | 0 |
| 0 | NA | NA | NA | NA | NA | Gm29105 | | 0 | 0 | 0 | 0 | 0 | 0 | 0 | 0 | 0 | 0 | 0 |
| 0 | NA | NA | NA | NA | NA | Gm29103 | | 0 | 0 | 0 | 0 | 0 | 0 | 0 | 0 | 0 | 0 | 0 |
| 0 | NA | NA | NA | NA | NA | Gm26034 | | 0 | 0 | 0 | 0 | 0 | 0 | 0 | 0 | 0 | 0 | 0 |
| 0 | NA | NA | NA | NA | NA | Lemd1 | LEM domain containing 1 | 0 | 0 | 0 | 0 | 0 | 0 | 0 | 0 | 0 | 0 | 0 |
| 0 | NA | NA | NA | NA | NA | Mir135b | | 0 | 0 | 0 | 0 | 0 | 0 | 0 | 0 | 0 | 0 | 0 |
| 0 | NA | NA | NA | NA | NA | Gm10188 | | 0 | 0 | 0 | 0 | 0 | 0 | 0 | 0 | 0 | 0 | 0 |
| 0 | NA | NA | NA | NA | NA | Gm29630 | | 0 | 0 | 0 | 0 | 0 | 0 | 0 | 0 | 0 | 0 | 0 |
| 0 | NA | NA | NA | NA | NA | Gm15849 | | 0 | 0 | 0 | 0 | 0 | 0 | 0 | 0 | 0 | 0 | 0 |
| 0 | NA | NA | NA | NA | NA | Gm10538 | | 0 | 0 | 0 | 0 | 0 | 0 | 0 | 0 | 0 | 0 | 0 |
| 0 | NA | NA | NA | NA | NA | Gm22609 | | 0 | 0 | 0 | 0 | 0 | 0 | 0 | 0 | 0 | 0 | 0 |
| 0 | NA | NA | NA | NA | NA | Gm44300 | | 0 | 0 | 0 | 0 | 0 | 0 | 0 | 0 | 0 | 0 | 0 |
| 0 | NA | NA | NA | NA | NA | Gm38308 | | 0 | 0 | 0 | 0 | 0 | 0 | 0 | 0 | 0 | 0 | 0 |
| 0 | NA | NA | NA | NA | NA | Gm28609 | | 0 | 0 | 0 | 0 | 0 | 0 | 0 | 0 | 0 | 0 | 0 |
| 0 | NA | NA | NA | NA | NA | Gm28040 | | 0 | 0 | 0 | 0 | 0 | 0 | 0 | 0 | 0 | 0 | 0 |
| 0 | NA | NA | NA | NA | NA | Kiss1 | KiSS-1 metastasis-suppressor | 0 | 0 | 0 | 0 | 0 | 0 | 0 | 0 | 0 | 0 | 0 |
| 0 | NA | NA | NA | NA | NA | Gm23939 | | 0 | 0 | 0 | 0 | 0 | 0 | 0 | 0 | 0 | 0 | 0 |
| 0 | NA | NA | NA | NA | NA | Zc3h11a | zinc finger CCCH type containing 11A | 0 | 0 | 0 | 0 | 0 | 0 | 0 | 0 | 0 | 0 | 0 |
| 0 | NA | NA | NA | NA | NA | Mir6903 | | 0 | 0 | 0 | 0 | 0 | 0 | 0 | 0 | 0 | 0 | 0 |
| 0 | NA | NA | NA | NA | NA | Gm28441 | | 0 | 0 | 0 | 0 | 0 | 0 | 0 | 0 | 0 | 0 | 0 |
| 0 | NA | NA | NA | NA | NA | Gm18970 | | 0 | 0 | 0 | 0 | 0 | 0 | 0 | 0 | 0 | 0 | 0 |
| 0 | NA | NA | NA | NA | NA | Chit1 | chitinase 1 (chitotriosidase) | 0 | 0 | 0 | 0 | 0 | 0 | 0 | 0 | 0 | 0 | 0 |
| 0 | NA | NA | NA | NA | NA | Gm24517 | | 0 | 0 | 0 | 0 | 0 | 0 | 0 | 0 | 0 | 0 | 0 |
| 0 | NA | NA | NA | NA | NA | Myog | myogenin | 0 | 0 | 0 | 0 | 0 | 0 | 0 | 0 | 0 | 0 | 0 |
| 0 | NA | NA | NA | NA | NA | Mgat4f | MGAT4 family, member F | 0 | 0 | 0 | 0 | 0 | 0 | 0 | 0 | 0 | 0 | 0 |
| 0 | NA | NA | NA | NA | NA | Platr1 | | 0 | 0 | 0 | 0 | 0 | 0 | 0 | 0 | 0 | 0 | 0 |
| 0 | NA | NA | NA | NA | NA | Mgat4e | MGAT4 family, member E | 0 | 0 | 0 | 0 | 0 | 0 | 0 | 0 | 0 | 0 | 0 |

| | | | | | | | | | | | | | | | | | |
|---|---|---|---|---|---|---|---|---|---|---|---|---|---|---|---|---|---|
| 0 | NA | NA | NA | NA | NA | Gm22232 | | 0 | 0 | 0 | 0 | 0 | 0 | 0 | 0 | 0 | 0 | 0 |
| 0 | NA | NA | NA | NA | NA | Gm10535 | | 0 | 0 | 0 | 0 | 0 | 0 | 0 | 0 | 0 | 0 | 0 |
| 0 | NA | NA | NA | NA | NA | Gm26280 | | 0 | 0 | 0 | 0 | 0 | 0 | 0 | 0 | 0 | 0 | 0 |
| 0 | NA | NA | NA | NA | NA | Gm22796 | | 0 | 0 | 0 | 0 | 0 | 0 | 0 | 0 | 0 | 0 | 0 |
| 0 | NA | NA | NA | NA | NA | Mir1231 | | 0 | 0 | 0 | 0 | 0 | 0 | 0 | 0 | 0 | 0 | 0 |
| 0 | NA | NA | NA | NA | NA | Gm36938 | | 0 | 0 | 0 | 0 | 0 | 0 | 0 | 0 | 0 | 0 | 0 |
| 0 | NA | NA | NA | NA | NA | Ascl5 | achaete-scute family bHLH transcription f | 0 | 0 | 0 | 0 | 0 | 0 | 0 | 0 | 0 | 0 | 0 |
| 0 | NA | NA | NA | NA | NA | Gm37014 | | 0 | 0 | 0 | 0 | 0 | 0 | 0 | 0 | 0 | 0 | 0 |
| 0 | NA | NA | NA | NA | NA | Gm37759 | | 0 | 0 | 0 | 0 | 0 | 0 | 0 | 0 | 0 | 0 | 0 |
| 0 | NA | NA | NA | NA | NA | Gm26568 | | 0 | 0 | 0 | 0 | 0 | 0 | 0 | 0 | 0 | 0 | 0 |
| 0 | NA | NA | NA | NA | NA | A430034D21Rik | | 0 | 0 | 0 | 0 | 0 | 0 | 0 | 0 | 0 | 0 | 0 |
| 0 | NA | NA | NA | NA | NA | Gm37552 | | 0 | 0 | 0 | 0 | 0 | 0 | 0 | 0 | 0 | 0 | 0 |
| 0 | NA | NA | NA | NA | NA | Gm29486 | | 0 | 0 | 0 | 0 | 0 | 0 | 0 | 0 | 0 | 0 | 0 |
| 0 | NA | NA | NA | NA | NA | Gm37799 | | 0 | 0 | 0 | 0 | 0 | 0 | 0 | 0 | 0 | 0 | 0 |
| 0 | NA | NA | NA | NA | NA | Gm33994 | | 0 | 0 | 0 | 0 | 0 | 0 | 0 | 0 | 0 | 0 | 0 |
| 0 | NA | NA | NA | NA | NA | 4933409D19Rik | | 0 | 0 | 0 | 0 | 0 | 0 | 0 | 0 | 0 | 0 | 0 |
| 0 | NA | NA | NA | NA | NA | Platr22 | | 0 | 0 | 0 | 0 | 0 | 0 | 0 | 0 | 0 | 0 | 0 |
| 0 | NA | NA | NA | NA | NA | Gm24086 | | 0 | 0 | 0 | 0 | 0 | 0 | 0 | 0 | 0 | 0 | 0 |
| 0 | NA | NA | NA | NA | NA | Gm22132 | | 0 | 0 | 0 | 0 | 0 | 0 | 0 | 0 | 0 | 0 | 0 |
| 0 | NA | NA | NA | NA | NA | Gm17781 | | 0 | 0 | 0 | 0 | 0 | 0 | 0 | 0 | 0 | 0 | 0 |
| 0 | NA | NA | NA | NA | NA | Gm23763 | | 0 | 0 | 0 | 0 | 0 | 0 | 0 | 0 | 0 | 0 | 0 |
| 0 | NA | NA | NA | NA | NA | Gm25609 | | 0 | 0 | 0 | 0 | 0 | 0 | 0 | 0 | 0 | 0 | 0 |
| 0 | NA | NA | NA | NA | NA | Gm37057 | | 0 | 0 | 0 | 0 | 0 | 0 | 0 | 0 | 0 | 0 | 0 |
| 0 | NA | NA | NA | NA | NA | 1700006P03Rik | | 0 | 0 | 0 | 0 | 0 | 0 | 0 | 0 | 0 | 0 | 0 |
| 0 | NA | NA | NA | NA | NA | Gm23534 | | 0 | 0 | 0 | 0 | 0 | 0 | 0 | 0 | 0 | 0 | 0 |
| 0 | NA | NA | NA | NA | NA | Gm37903 | | 0 | 0 | 0 | 0 | 0 | 0 | 0 | 0 | 0 | 0 | 0 |
| 0 | NA | NA | NA | NA | NA | Gm22727 | | 0 | 0 | 0 | 0 | 0 | 0 | 0 | 0 | 0 | 0 | 0 |
| 0 | NA | NA | NA | NA | NA | Gm37099 | | 0 | 0 | 0 | 0 | 0 | 0 | 0 | 0 | 0 | 0 | 0 |
| 0 | NA | NA | NA | NA | NA | Gm37293 | | 0 | 0 | 0 | 0 | 0 | 0 | 0 | 0 | 0 | 0 | 0 |
| 0 | NA | NA | NA | NA | NA | Gm37298 | | 0 | 0 | 0 | 0 | 0 | 0 | 0 | 0 | 0 | 0 | 0 |
| 0 | NA | NA | NA | NA | NA | Gm26979 | | 0 | 0 | 0 | 0 | 0 | 0 | 0 | 0 | 0 | 0 | 0 |
| 0 | NA | NA | NA | NA | NA | Gm37124 | | 0 | 0 | 0 | 0 | 0 | 0 | 0 | 0 | 0 | 0 | 0 |
| 0 | NA | NA | NA | NA | NA | A130050O07Rik | | 0 | 0 | 0 | 0 | 0 | 0 | 0 | 0 | 0 | 0 | 0 |
| 0 | NA | NA | NA | NA | NA | Gm26936 | | 0 | 0 | 0 | 0 | 0 | 0 | 0 | 0 | 0 | 0 | 0 |
| 0 | NA | NA | NA | NA | NA | Gm26930 | | 0 | 0 | 0 | 0 | 0 | 0 | 0 | 0 | 0 | 0 | 0 |
| 0 | NA | NA | NA | NA | NA | D130019J16Rik | | 0 | 0 | 0 | 0 | 0 | 0 | 0 | 0 | 0 | 0 | 0 |
| 0 | NA | NA | NA | NA | NA | Gm37534 | | 0 | 0 | 0 | 0 | 0 | 0 | 0 | 0 | 0 | 0 | 0 |
| 0 | NA | NA | NA | NA | NA | Mir181a-1 | | 0 | 0 | 0 | 0 | 0 | 0 | 0 | 0 | 0 | 0 | 0 |
| 0 | NA | NA | NA | NA | NA | Mir181a-1hg | | 0 | 0 | 0 | 0 | 0 | 0 | 0 | 0 | 0 | 0 | 0 |
| 0 | NA | NA | NA | NA | NA | Mir181b-1 | | 0 | 0 | 0 | 0 | 0 | 0 | 0 | 0 | 0 | 0 | 0 |
| 0 | NA | NA | NA | NA | NA | Gm37632 | | 0 | 0 | 0 | 0 | 0 | 0 | 0 | 0 | 0 | 0 | 0 |
| 0 | NA | NA | NA | NA | NA | Gm23782 | | 0 | 0 | 0 | 0 | 0 | 0 | 0 | 0 | 0 | 0 | 0 |
| 0 | NA | NA | NA | NA | NA | A130071D04Rik | | 0 | 0 | 0 | 0 | 0 | 0 | 0 | 0 | 0 | 0 | 0 |
| 0 | NA | NA | NA | NA | NA | Gm18667 | | 0 | 0 | 0 | 0 | 0 | 0 | 0 | 0 | 0 | 0 | 0 |
| 0 | NA | NA | NA | NA | NA | Gm28556 | | 0 | 0 | 0 | 0 | 0 | 0 | 0 | 0 | 0 | 0 | 0 |
| 0 | NA | NA | NA | NA | NA | 1700113B19Rik | | 0 | 0 | 0 | 0 | 0 | 0 | 0 | 0 | 0 | 0 | 0 |
| 0 | NA | NA | NA | NA | NA | Gm28500 | | 0 | 0 | 0 | 0 | 0 | 0 | 0 | 0 | 0 | 0 | 0 |
| 0 | NA | NA | NA | NA | NA | Gm37080 | | 0 | 0 | 0 | 0 | 0 | 0 | 0 | 0 | 0 | 0 | 0 |
| 0 | NA | NA | NA | NA | NA | Gm5833 | | 0 | 0 | 0 | 0 | 0 | 0 | 0 | 0 | 0 | 0 | 0 |
| 0 | NA | NA | NA | NA | NA | Gm8790 | | 0 | 0 | 0 | 0 | 0 | 0 | 0 | 0 | 0 | 0 | 0 |
| 0 | NA | NA | NA | NA | NA | 1700019P21Rik | | 0 | 0 | 0 | 0 | 0 | 0 | 0 | 0 | 0 | 0 | 0 |
| 0 | NA | NA | NA | NA | NA | n-R5s217 | | 0 | 0 | 0 | 0 | 0 | 0 | 0 | 0 | 0 | 0 | 0 |
| 0 | NA | NA | NA | NA | NA | Gm3933 | | 0 | 0 | 0 | 0 | 0 | 0 | 0 | 0 | 0 | 0 | 0 |
| 0 | NA | NA | NA | NA | NA | Gm18553 | | 0 | 0 | 0 | 0 | 0 | 0 | 0 | 0 | 0 | 0 | 0 |
| 0 | NA | NA | NA | NA | NA | Gm37655 | | 0 | 0 | 0 | 0 | 0 | 0 | 0 | 0 | 0 | 0 | 0 |
| 0 | NA | NA | NA | NA | NA | Gm16304 | | 0 | 0 | 0 | 0 | 0 | 0 | 0 | 0 | 0 | 0 | 0 |
| 0 | NA | NA | NA | NA | NA | Gm34816 | | 0 | 0 | 0 | 0 | 0 | 0 | 0 | 0 | 0 | 0 | 0 |
| 0 | NA | NA | NA | NA | NA | F13b | coagulation factor XIII, beta subunit | 0 | 0 | 0 | 0 | 0 | 0 | 0 | 0 | 0 | 0 | 0 |
| 0 | NA | NA | NA | NA | NA | Cfhr1 | complement factor H-related 1 | 0 | 0 | 0 | 0 | 0 | 0 | 0 | 0 | 0 | 0 | 0 |
| 0 | NA | NA | NA | NA | NA | Cfhr3 | | 0 | 0 | 0 | 0 | 0 | 0 | 0 | 0 | 0 | 0 | 0 |

| | | | | | | | | | | | | | | | | | | |
|---|---|---|---|---|---|---|---|---|---|---|---|---|---|---|---|---|---|---|
| 0 | NA | NA | NA | NA | NA | Gm26271 | | 0 | 0 | 0 | 0 | 0 | 0 | 0 | 0 | 0 | 0 | 0 |
| 0 | NA | NA | NA | NA | NA | Gm4788 | predicted gene 4788 | 0 | 0 | 0 | 0 | 0 | 0 | 0 | 0 | 0 | 0 | 0 |
| 0 | NA | NA | NA | NA | NA | Gm25619 | | 0 | 0 | 0 | 0 | 0 | 0 | 0 | 0 | 0 | 0 | 0 |
| 0 | NA | NA | NA | NA | NA | Cfhr2 | complement factor H-related 2 | 0 | 0 | 0 | 0 | 0 | 0 | 0 | 0 | 0 | 0 | 0 |
| 0 | NA | NA | NA | NA | NA | Gm38266 | | 0 | 0 | 0 | 0 | 0 | 0 | 0 | 0 | 0 | 0 | 0 |
| 0 | NA | NA | NA | NA | NA | Gm26048 | | 0 | 0 | 0 | 0 | 0 | 0 | 0 | 0 | 0 | 0 | 0 |
| 0 | NA | NA | NA | NA | NA | Gm38019 | | 0 | 0 | 0 | 0 | 0 | 0 | 0 | 0 | 0 | 0 | 0 |
| 0 | NA | NA | NA | NA | NA | Gm37777 | | 0 | 0 | 0 | 0 | 0 | 0 | 0 | 0 | 0 | 0 | 0 |
| 0 | NA | NA | NA | NA | NA | 4930590L20Rik | | 0 | 0 | 0 | 0 | 0 | 0 | 0 | 0 | 0 | 0 | 0 |
| 0 | NA | NA | NA | NA | NA | Gm37371 | | 0 | 0 | 0 | 0 | 0 | 0 | 0 | 0 | 0 | 0 | 0 |
| 0 | NA | NA | NA | NA | NA | Gm4845 | | 0 | 0 | 0 | 0 | 0 | 0 | 0 | 0 | 0 | 0 | 0 |
| 0 | NA | NA | NA | NA | NA | Gm25457 | | 0 | 0 | 0 | 0 | 0 | 0 | 0 | 0 | 0 | 0 | 0 |
| 0 | NA | NA | NA | NA | NA | Gm22016 | | 0 | 0 | 0 | 0 | 0 | 0 | 0 | 0 | 0 | 0 | 0 |
| 0 | NA | NA | NA | NA | NA | Gm37599 | | 0 | 0 | 0 | 0 | 0 | 0 | 0 | 0 | 0 | 0 | 0 |
| 0 | NA | NA | NA | NA | NA | Gm37142 | | 0 | 0 | 0 | 0 | 0 | 0 | 0 | 0 | 0 | 0 | 0 |
| 0 | NA | NA | NA | NA | NA | Gm37604 | | 0 | 0 | 0 | 0 | 0 | 0 | 0 | 0 | 0 | 0 | 0 |
| 0 | NA | NA | NA | NA | NA | Gm18183 | | 0 | 0 | 0 | 0 | 0 | 0 | 0 | 0 | 0 | 0 | 0 |
| 0 | NA | NA | NA | NA | NA | Gm28724 | | 0 | 0 | 0 | 0 | 0 | 0 | 0 | 0 | 0 | 0 | 0 |
| 0 | NA | NA | NA | NA | NA | Gm28724 | | 0 | 0 | 0 | 0 | 0 | 0 | 0 | 0 | 0 | 0 | 0 |
| 0 | NA | NA | NA | NA | NA | Gm37796 | | 0 | 0 | 0 | 0 | 0 | 0 | 0 | 0 | 0 | 0 | 0 |
| 0 | NA | NA | NA | NA | NA | Gm25663 | | 0 | 0 | 0 | 0 | 0 | 0 | 0 | 0 | 0 | 0 | 0 |
| 0 | NA | NA | NA | NA | NA | Rgs21 | regulator of G-protein signalling 21 | 0 | 0 | 0 | 0 | 0 | 0 | 0 | 0 | 0 | 0 | 0 |
| 0 | NA | NA | NA | NA | NA | Gm23672 | | 0 | 0 | 0 | 0 | 0 | 0 | 0 | 0 | 0 | 0 | 0 |
| 0 | NA | NA | NA | NA | NA | Gm8856 | | 0 | 0 | 0 | 0 | 0 | 0 | 0 | 0 | 0 | 0 | 0 |
| 0 | NA | NA | NA | NA | NA | Gm22681 | | 0 | 0 | 0 | 0 | 0 | 0 | 0 | 0 | 0 | 0 | 0 |
| 0 | NA | NA | NA | NA | NA | Gm25708 | | 0 | 0 | 0 | 0 | 0 | 0 | 0 | 0 | 0 | 0 | 0 |
| 0 | NA | NA | NA | NA | NA | Gm6550 | | 0 | 0 | 0 | 0 | 0 | 0 | 0 | 0 | 0 | 0 | 0 |
| 0 | NA | NA | NA | NA | NA | Gm25460 | | 0 | 0 | 0 | 0 | 0 | 0 | 0 | 0 | 0 | 0 | 0 |
| 0 | NA | NA | NA | NA | NA | Gm38269 | | 0 | 0 | 0 | 0 | 0 | 0 | 0 | 0 | 0 | 0 | 0 |
| 0 | NA | NA | NA | NA | NA | Gm29514 | | 0 | 0 | 0 | 0 | 0 | 0 | 0 | 0 | 0 | 0 | 0 |
| 0 | NA | NA | NA | NA | NA | Gm38187 | | 0 | 0 | 0 | 0 | 0 | 0 | 0 | 0 | 0 | 0 | 0 |
| 0 | NA | NA | NA | NA | NA | Gm15584 | | 0 | 0 | 0 | 0 | 0 | 0 | 0 | 0 | 0 | 0 | 0 |
| 0 | NA | NA | NA | NA | NA | Gm37749 | | 0 | 0 | 0 | 0 | 0 | 0 | 0 | 0 | 0 | 0 | 0 |
| 0 | NA | NA | NA | NA | NA | Gm9931 | | 0 | 0 | 0 | 0 | 0 | 0 | 0 | 0 | 0 | 0 | 0 |
| 0 | NA | NA | NA | NA | NA | Gm17867 | | 0 | 0 | 0 | 0 | 0 | 0 | 0 | 0 | 0 | 0 | 0 |
| 0 | NA | NA | NA | NA | NA | Gm38284 | | 0 | 0 | 0 | 0 | 0 | 0 | 0 | 0 | 0 | 0 | 0 |
| 0 | NA | NA | NA | NA | NA | Gm29516 | | 0 | 0 | 0 | 0 | 0 | 0 | 0 | 0 | 0 | 0 | 0 |
| 0 | NA | NA | NA | NA | NA | Gm29515 | | 0 | 0 | 0 | 0 | 0 | 0 | 0 | 0 | 0 | 0 | 0 |
| 0 | NA | NA | NA | NA | NA | Gm22966 | | 0 | 0 | 0 | 0 | 0 | 0 | 0 | 0 | 0 | 0 | 0 |
| 0 | NA | NA | NA | NA | NA | Gm19891 | | 0 | 0 | 0 | 0 | 0 | 0 | 0 | 0 | 0 | 0 | 0 |
| 0 | NA | NA | NA | NA | NA | Gm29022 | | 0 | 0 | 0 | 0 | 0 | 0 | 0 | 0 | 0 | 0 | 0 |
| 0 | NA | NA | NA | NA | NA | Gm5703 | | 0 | 0 | 0 | 0 | 0 | 0 | 0 | 0 | 0 | 0 | 0 |
| 0 | NA | NA | NA | NA | NA | Gm18751 | | 0 | 0 | 0 | 0 | 0 | 0 | 0 | 0 | 0 | 0 | 0 |
| 0 | NA | NA | NA | NA | NA | Gm29020 | | 0 | 0 | 0 | 0 | 0 | 0 | 0 | 0 | 0 | 0 | 0 |
| 0 | NA | NA | NA | NA | NA | Gm18701 | | 0 | 0 | 0 | 0 | 0 | 0 | 0 | 0 | 0 | 0 | 0 |
| 0 | NA | NA | NA | NA | NA | Gm37630 | | 0 | 0 | 0 | 0 | 0 | 0 | 0 | 0 | 0 | 0 | 0 |
| 0 | NA | NA | NA | NA | NA | Gm38189 | | 0 | 0 | 0 | 0 | 0 | 0 | 0 | 0 | 0 | 0 | 0 |
| 0 | NA | NA | NA | NA | NA | Gm29398 | | 0 | 0 | 0 | 0 | 0 | 0 | 0 | 0 | 0 | 0 | 0 |
| 0 | NA | NA | NA | NA | NA | Gm37096 | | 0 | 0 | 0 | 0 | 0 | 0 | 0 | 0 | 0 | 0 | 0 |
| 0 | NA | NA | NA | NA | NA | Gm37775 | | 0 | 0 | 0 | 0 | 0 | 0 | 0 | 0 | 0 | 0 | 0 |
| 0 | NA | NA | NA | NA | NA | Gm23535 | | 0 | 0 | 0 | 0 | 0 | 0 | 0 | 0 | 0 | 0 | 0 |
| 0 | NA | NA | NA | NA | NA | n-R5s218 | | 0 | 0 | 0 | 0 | 0 | 0 | 0 | 0 | 0 | 0 | 0 |
| 0 | NA | NA | NA | NA | NA | n-R5s219 | | 0 | 0 | 0 | 0 | 0 | 0 | 0 | 0 | 0 | 0 | 0 |
| 0 | NA | NA | NA | NA | NA | Gm19136 | | 0 | 0 | 0 | 0 | 0 | 0 | 0 | 0 | 0 | 0 | 0 |
| 0 | NA | NA | NA | NA | NA | Gm44329 | | 0 | 0 | 0 | 0 | 0 | 0 | 0 | 0 | 0 | 0 | 0 |
| 0 | NA | NA | NA | NA | NA | Pdc | phosducin | 0 | 0 | 0 | 0 | 0 | 0 | 0 | 0 | 0 | 0 | 0 |
| 0 | NA | NA | NA | NA | NA | 2310030A07Rik | | 0 | 0 | 0 | 0 | 0 | 0 | 0 | 0 | 0 | 0 | 0 |
| 0 | NA | NA | NA | NA | NA | Gm37764 | | 0 | 0 | 0 | 0 | 0 | 0 | 0 | 0 | 0 | 0 | 0 |
| 0 | NA | NA | NA | NA | NA | Ska2l-ps | | 0 | 0 | 0 | 0 | 0 | 0 | 0 | 0 | 0 | 0 | 0 |
| 0 | NA | NA | NA | NA | NA | Gm20631 | | 0 | 0 | 0 | 0 | 0 | 0 | 0 | 0 | 0 | 0 | 0 |

| | | | | | | | | | | | | | | | | | | |
|---|---|---|---|---|---|---|---|---|---|---|---|---|---|---|---|---|---|---|
| 0 | NA | NA | NA | NA | NA | Gm29188 | | 0 | 0 | 0 | 0 | 0 | 0 | 0 | 0 | 0 | 0 | 0 |
| 0 | NA | NA | NA | NA | NA | Gm22756 | | 0 | 0 | 0 | 0 | 0 | 0 | 0 | 0 | 0 | 0 | 0 |
| 0 | NA | NA | NA | NA | NA | Gm37671 | | 0 | 0 | 0 | 0 | 0 | 0 | 0 | 0 | 0 | 0 | 0 |
| 0 | NA | NA | NA | NA | NA | Gm19087 | | 0 | 0 | 0 | 0 | 0 | 0 | 0 | 0 | 0 | 0 | 0 |
| 0 | NA | NA | NA | NA | NA | C730036E19Rik | | 0 | 0 | 0 | 0 | 0 | 0 | 0 | 0 | 0 | 0 | 0 |
| 0 | NA | NA | NA | NA | NA | Gm47985 | RIKEN cDNA 3110040M04 gene | 0 | 0 | 0 | 0 | 0 | 0 | 0 | 0 | 0 | 0 | 0 |
| 0 | NA | NA | NA | NA | NA | Gm8947 | | 0 | 0 | 0 | 0 | 0 | 0 | 0 | 0 | 0 | 0 | 0 |
| 0 | NA | NA | NA | NA | NA | Gm47995 | | 0 | 0 | 0 | 0 | 0 | 0 | 0 | 0 | 0 | 0 | 0 |
| 0 | NA | NA | NA | NA | NA | 3110040M04Rik | | 0 | 0 | 0 | 0 | 0 | 0 | 0 | 0 | 0 | 0 | 0 |
| 0 | NA | NA | NA | NA | NA | Gm47996 | | 0 | 0 | 0 | 0 | 0 | 0 | 0 | 0 | 0 | 0 | 0 |
| 0 | NA | NA | NA | NA | NA | Gm24402 | | 0 | 0 | 0 | 0 | 0 | 0 | 0 | 0 | 0 | 0 | 0 |
| 0 | NA | NA | NA | NA | NA | Gm29072 | | 0 | 0 | 0 | 0 | 0 | 0 | 0 | 0 | 0 | 0 | 0 |
| 0 | NA | NA | NA | NA | NA | Gm25279 | | 0 | 0 | 0 | 0 | 0 | 0 | 0 | 0 | 0 | 0 | 0 |
| 0 | NA | NA | NA | NA | NA | Gm28610 | | 0 | 0 | 0 | 0 | 0 | 0 | 0 | 0 | 0 | 0 | 0 |
| 0 | NA | NA | NA | NA | NA | Gm38238 | | 0 | 0 | 0 | 0 | 0 | 0 | 0 | 0 | 0 | 0 | 0 |
| 0 | NA | NA | NA | NA | NA | Gm15618 | | 0 | 0 | 0 | 0 | 0 | 0 | 0 | 0 | 0 | 0 | 0 |
| 0 | NA | NA | NA | NA | NA | Gm15479 | | 0 | 0 | 0 | 0 | 0 | 0 | 0 | 0 | 0 | 0 | 0 |
| 0 | NA | NA | NA | NA | NA | 1700040K01Rik | | 0 | 0 | 0 | 0 | 0 | 0 | 0 | 0 | 0 | 0 | 0 |
| 0 | NA | NA | NA | NA | NA | Gm28960 | | 0 | 0 | 0 | 0 | 0 | 0 | 0 | 0 | 0 | 0 | 0 |
| 0 | NA | NA | NA | NA | NA | Gm29529 | | 0 | 0 | 0 | 0 | 0 | 0 | 0 | 0 | 0 | 0 | 0 |
| 0 | NA | NA | NA | NA | NA | Gm28513 | | 0 | 0 | 0 | 0 | 0 | 0 | 0 | 0 | 0 | 0 | 0 |
| 0 | NA | NA | NA | NA | NA | Teddm1b | transmembrane epididymal protein 1B | 0 | 0 | 0 | 0 | 0 | 0 | 0 | 0 | 0 | 0 | 0 |
| 0 | NA | NA | NA | NA | NA | Gm38352 | | 0 | 0 | 0 | 0 | 0 | 0 | 0 | 0 | 0 | 0 | 0 |
| 0 | NA | NA | NA | NA | NA | Teddm1a | transmembrane epididymal protein 1A | 0 | 0 | 0 | 0 | 0 | 0 | 0 | 0 | 0 | 0 | 0 |
| 0 | NA | NA | NA | NA | NA | Gm29290 | | 0 | 0 | 0 | 0 | 0 | 0 | 0 | 0 | 0 | 0 | 0 |
| 0 | NA | NA | NA | NA | NA | Gm29291 | | 0 | 0 | 0 | 0 | 0 | 0 | 0 | 0 | 0 | 0 | 0 |
| 0 | NA | NA | NA | NA | NA | Gm25336 | | 0 | 0 | 0 | 0 | 0 | 0 | 0 | 0 | 0 | 0 | 0 |
| 0 | NA | NA | NA | NA | NA | Zfp648 | zinc finger protein 648 | 0 | 0 | 0 | 0 | 0 | 0 | 0 | 0 | 0 | 0 | 0 |
| 0 | NA | NA | NA | NA | NA | 4930452N14Rik | | 0 | 0 | 0 | 0 | 0 | 0 | 0 | 0 | 0 | 0 | 0 |
| 0 | NA | NA | NA | NA | NA | 4930532M18Rik | | 0 | 0 | 0 | 0 | 0 | 0 | 0 | 0 | 0 | 0 | 0 |
| 0 | NA | NA | NA | NA | NA | n-R5s220 | | 0 | 0 | 0 | 0 | 0 | 0 | 0 | 0 | 0 | 0 | 0 |
| 0 | NA | NA | NA | NA | NA | Gm18727 | | 0 | 0 | 0 | 0 | 0 | 0 | 0 | 0 | 0 | 0 | 0 |
| 0 | NA | NA | NA | NA | NA | Gm29282 | | 0 | 0 | 0 | 0 | 0 | 0 | 0 | 0 | 0 | 0 | 0 |
| 0 | NA | NA | NA | NA | NA | Gm29441 | | 0 | 0 | 0 | 0 | 0 | 0 | 0 | 0 | 0 | 0 | 0 |
| 0 | NA | NA | NA | NA | NA | Gm37622 | | 0 | 0 | 0 | 0 | 0 | 0 | 0 | 0 | 0 | 0 | 0 |
| 0 | NA | NA | NA | NA | NA | Gm22569 | | 0 | 0 | 0 | 0 | 0 | 0 | 0 | 0 | 0 | 0 | 0 |
| 0 | NA | NA | NA | NA | NA | Lhx4 | LIM homeobox protein 4 | 0 | 0 | 0 | 0 | 0 | 0 | 0 | 0 | 0 | 0 | 0 |
| 0 | NA | NA | NA | NA | NA | Gm37089 | | 0 | 0 | 0 | 0 | 0 | 0 | 0 | 0 | 0 | 0 | 0 |
| 0 | NA | NA | NA | NA | NA | Gm37539 | | 0 | 0 | 0 | 0 | 0 | 0 | 0 | 0 | 0 | 0 | 0 |
| 0 | NA | NA | NA | NA | NA | Gm37571 | | 0 | 0 | 0 | 0 | 0 | 0 | 0 | 0 | 0 | 0 | 0 |
| 0 | NA | NA | NA | NA | NA | Gm38038 | | 0 | 0 | 0 | 0 | 0 | 0 | 0 | 0 | 0 | 0 | 0 |
| 0 | NA | NA | NA | NA | NA | Gm36934 | | 0 | 0 | 0 | 0 | 0 | 0 | 0 | 0 | 0 | 0 | 0 |
| 0 | NA | NA | NA | NA | NA | Gm23744 | | 0 | 0 | 0 | 0 | 0 | 0 | 0 | 0 | 0 | 0 | 0 |
| 0 | NA | NA | NA | NA | NA | Gm38113 | | 0 | 0 | 0 | 0 | 0 | 0 | 0 | 0 | 0 | 0 | 0 |
| 0 | NA | NA | NA | NA | NA | A430050A11Rik | | 0 | 0 | 0 | 0 | 0 | 0 | 0 | 0 | 0 | 0 | 0 |
| 0 | NA | NA | NA | NA | NA | 4930518J20Rik | | 0 | 0 | 0 | 0 | 0 | 0 | 0 | 0 | 0 | 0 | 0 |
| 0 | NA | NA | NA | NA | NA | Gm37648 | | 0 | 0 | 0 | 0 | 0 | 0 | 0 | 0 | 0 | 0 | 0 |
| 0 | NA | NA | NA | NA | NA | Gm15428 | | 0 | 0 | 0 | 0 | 0 | 0 | 0 | 0 | 0 | 0 | 0 |
| 0 | NA | NA | NA | NA | NA | Gm37943 | | 0 | 0 | 0 | 0 | 0 | 0 | 0 | 0 | 0 | 0 | 0 |
| 0 | NA | NA | NA | NA | NA | Gm28694 | | 0 | 0 | 0 | 0 | 0 | 0 | 0 | 0 | 0 | 0 | 0 |
| 0 | NA | NA | NA | NA | NA | Gm10531 | | 0 | 0 | 0 | 0 | 0 | 0 | 0 | 0 | 0 | 0 | 0 |
| 0 | NA | NA | NA | NA | NA | Tex35 | testis expressed 35 | 0 | 0 | 0 | 0 | 0 | 0 | 0 | 0 | 0 | 0 | 0 |
| 0 | NA | NA | NA | NA | NA | Gm30694 | | 0 | 0 | 0 | 0 | 0 | 0 | 0 | 0 | 0 | 0 | 0 |
| 0 | NA | NA | NA | NA | NA | Gm37788 | | 0 | 0 | 0 | 0 | 0 | 0 | 0 | 0 | 0 | 0 | 0 |
| 0 | NA | NA | NA | NA | NA | Gm37578 | | 0 | 0 | 0 | 0 | 0 | 0 | 0 | 0 | 0 | 0 | 0 |
| 0 | NA | NA | NA | NA | NA | Gm7412 | | 0 | 0 | 0 | 0 | 0 | 0 | 0 | 0 | 0 | 0 | 0 |
| 0 | NA | NA | NA | NA | NA | Gm38213 | | 0 | 0 | 0 | 0 | 0 | 0 | 0 | 0 | 0 | 0 | 0 |
| 0 | NA | NA | NA | NA | NA | Gm37861 | | 0 | 0 | 0 | 0 | 0 | 0 | 0 | 0 | 0 | 0 | 0 |
| 0 | NA | NA | NA | NA | NA | Gm37679 | | 0 | 0 | 0 | 0 | 0 | 0 | 0 | 0 | 0 | 0 | 0 |
| 0 | NA | NA | NA | NA | NA | Mir488 | | 0 | 0 | 0 | 0 | 0 | 0 | 0 | 0 | 0 | 0 | 0 |

| | | | | | | | | | | | | | | | | | | |
|---|---|---|---|---|---|---|---|---|---|---|---|---|---|---|---|---|---|---|
| 0 | NA | NA | NA | NA | NA | Gm37367 | | 0 | 0 | 0 | 0 | 0 | 0 | 0 | 0 | 0 | 0 | 0 |
| 0 | NA | NA | NA | NA | NA | Gm37848 | | 0 | 0 | 0 | 0 | 0 | 0 | 0 | 0 | 0 | 0 | 0 |
| 0 | NA | NA | NA | NA | NA | 1600012P17Rik | | 0 | 0 | 0 | 0 | 0 | 0 | 0 | 0 | 0 | 0 | 0 |
| 0 | NA | NA | NA | NA | NA | 4930562F17Rik | | 0 | 0 | 0 | 0 | 0 | 0 | 0 | 0 | 0 | 0 | 0 |
| 0 | NA | NA | NA | NA | NA | Gm7430 | | 0 | 0 | 0 | 0 | 0 | 0 | 0 | 0 | 0 | 0 | 0 |
| 0 | NA | NA | NA | NA | NA | Gm37088 | | 0 | 0 | 0 | 0 | 0 | 0 | 0 | 0 | 0 | 0 | 0 |
| 0 | NA | NA | NA | NA | NA | Mir1843b | | 0 | 0 | 0 | 0 | 0 | 0 | 0 | 0 | 0 | 0 | 0 |
| 0 | NA | NA | NA | NA | NA | Gm31256 | | 0 | 0 | 0 | 0 | 0 | 0 | 0 | 0 | 0 | 0 | 0 |
| 0 | NA | NA | NA | NA | NA | Gm37757 | | 0 | 0 | 0 | 0 | 0 | 0 | 0 | 0 | 0 | 0 | 0 |
| 0 | NA | NA | NA | NA | NA | Gm24719 | | 0 | 0 | 0 | 0 | 0 | 0 | 0 | 0 | 0 | 0 | 0 |
| 0 | NA | NA | NA | NA | NA | Gm37294 | | 0 | 0 | 0 | 0 | 0 | 0 | 0 | 0 | 0 | 0 | 0 |
| 0 | NA | NA | NA | NA | NA | Mir1927 | | 0 | 0 | 0 | 0 | 0 | 0 | 0 | 0 | 0 | 0 | 0 |
| 0 | NA | NA | NA | NA | NA | 9430087J23Rik | | 0 | 0 | 0 | 0 | 0 | 0 | 0 | 0 | 0 | 0 | 0 |
| 0 | NA | NA | NA | NA | NA | Gm37212 | | 0 | 0 | 0 | 0 | 0 | 0 | 0 | 0 | 0 | 0 | 0 |
| 0 | NA | NA | NA | NA | NA | Gm37215 | | 0 | 0 | 0 | 0 | 0 | 0 | 0 | 0 | 0 | 0 | 0 |
| 0 | NA | NA | NA | NA | NA | Gm23528 | | 0 | 0 | 0 | 0 | 0 | 0 | 0 | 0 | 0 | 0 | 0 |
| 0 | NA | NA | NA | NA | NA | Gm38126 | | 0 | 0 | 0 | 0 | 0 | 0 | 0 | 0 | 0 | 0 | 0 |
| 0 | NA | NA | NA | NA | NA | Gm37083 | | 0 | 0 | 0 | 0 | 0 | 0 | 0 | 0 | 0 | 0 | 0 |
| 0 | NA | NA | NA | NA | NA | Gm38060 | | 0 | 0 | 0 | 0 | 0 | 0 | 0 | 0 | 0 | 0 | 0 |
| 0 | NA | NA | NA | NA | NA | Gm24988 | | 0 | 0 | 0 | 0 | 0 | 0 | 0 | 0 | 0 | 0 | 0 |
| 0 | NA | NA | NA | NA | NA | Gm37809 | | 0 | 0 | 0 | 0 | 0 | 0 | 0 | 0 | 0 | 0 | 0 |
| 0 | NA | NA | NA | NA | NA | Gm37072 | | 0 | 0 | 0 | 0 | 0 | 0 | 0 | 0 | 0 | 0 | 0 |
| 0 | NA | NA | NA | NA | NA | Gm37362 | | 0 | 0 | 0 | 0 | 0 | 0 | 0 | 0 | 0 | 0 | 0 |
| 0 | NA | NA | NA | NA | NA | Gm26224 | | 0 | 0 | 0 | 0 | 0 | 0 | 0 | 0 | 0 | 0 | 0 |
| 0 | NA | NA | NA | NA | NA | Gm22489 | | 0 | 0 | 0 | 0 | 0 | 0 | 0 | 0 | 0 | 0 | 0 |
| 0 | NA | NA | NA | NA | NA | Gm22357 | | 0 | 0 | 0 | 0 | 0 | 0 | 0 | 0 | 0 | 0 | 0 |
| 0 | NA | NA | NA | NA | NA | Gm23212 | | 0 | 0 | 0 | 0 | 0 | 0 | 0 | 0 | 0 | 0 | 0 |
| 0 | NA | NA | NA | NA | NA | Snord78 | | 0 | 0 | 0 | 0 | 0 | 0 | 0 | 0 | 0 | 0 | 0 |
| 0 | NA | NA | NA | NA | NA | Gm50452 | | 0 | 0 | 0 | 0 | 0 | 0 | 0 | 0 | 0 | 0 | 0 |
| 0 | NA | NA | NA | NA | NA | Snord80 | | 0 | 0 | 0 | 0 | 0 | 0 | 0 | 0 | 0 | 0 | 0 |
| 0 | NA | NA | NA | NA | NA | Snord80 | | 0 | 0 | 0 | 0 | 0 | 0 | 0 | 0 | 0 | 0 | 0 |
| 0 | NA | NA | NA | NA | NA | Snord47 | | 0 | 0 | 0 | 0 | 0 | 0 | 0 | 0 | 0 | 0 | 0 |
| 0 | NA | NA | NA | NA | NA | Gm25789 | | 0 | 0 | 0 | 0 | 0 | 0 | 0 | 0 | 0 | 0 | 0 |
| 0 | NA | NA | NA | NA | NA | Gm37634 | | 0 | 0 | 0 | 0 | 0 | 0 | 0 | 0 | 0 | 0 | 0 |
| 0 | NA | NA | NA | NA | NA | Tex50 | testis expressed 50 | 0 | 0 | 0 | 0 | 0 | 0 | 0 | 0 | 0 | 0 | 0 |
| 0 | NA | NA | NA | NA | NA | Gm6185 | | 0 | 0 | 0 | 0 | 0 | 0 | 0 | 0 | 0 | 0 | 0 |
| 0 | NA | NA | NA | NA | NA | Gm19057 | | 0 | 0 | 0 | 0 | 0 | 0 | 0 | 0 | 0 | 0 | 0 |
| 0 | NA | NA | NA | NA | NA | Gm25488 | | 0 | 0 | 0 | 0 | 0 | 0 | 0 | 0 | 0 | 0 | 0 |
| 0 | NA | NA | NA | NA | NA | Gm37638 | | 0 | 0 | 0 | 0 | 0 | 0 | 0 | 0 | 0 | 0 | 0 |
| 0 | NA | NA | NA | NA | NA | Gm5049 | | 0 | 0 | 0 | 0 | 0 | 0 | 0 | 0 | 0 | 0 | 0 |
| 0 | NA | NA | NA | NA | NA | Gm7496 | | 0 | 0 | 0 | 0 | 0 | 0 | 0 | 0 | 0 | 0 | 0 |
| 0 | NA | NA | NA | NA | NA | Gm31925 | | 0 | 0 | 0 | 0 | 0 | 0 | 0 | 0 | 0 | 0 | 0 |
| 0 | NA | NA | NA | NA | NA | 4930558K02Ril | RIKEN cDNA 4930558K02 gene | 0 | 0 | 0 | 0 | 0 | 0 | 0 | 0 | 0 | 0 | 0 |
| 0 | NA | NA | NA | NA | NA | Gm37767 | | 0 | 0 | 0 | 0 | 0 | 0 | 0 | 0 | 0 | 0 | 0 |
| 0 | NA | NA | NA | NA | NA | Mir199a-2 | | 0 | 0 | 0 | 0 | 0 | 0 | 0 | 0 | 0 | 0 | 0 |
| 0 | NA | NA | NA | NA | NA | Mir214 | | 0 | 0 | 0 | 0 | 0 | 0 | 0 | 0 | 0 | 0 | 0 |
| 0 | NA | NA | NA | NA | NA | Gm16101 | | 0 | 0 | 0 | 0 | 0 | 0 | 0 | 0 | 0 | 0 | 0 |
| 0 | NA | NA | NA | NA | NA | Gm10176 | | 0 | 0 | 0 | 0 | 0 | 0 | 0 | 0 | 0 | 0 | 0 |
| 0 | NA | NA | NA | NA | NA | Gm5705 | | 0 | 0 | 0 | 0 | 0 | 0 | 0 | 0 | 0 | 0 | 0 |
| 0 | NA | NA | NA | NA | NA | Myocos | myocilin opposite strand | 0 | 0 | 0 | 0 | 0 | 0 | 0 | 0 | 0 | 0 | 0 |
| 0 | NA | NA | NA | NA | NA | Mroh9 | maestro heat-like repeat family member 9 | 0 | 0 | 0 | 0 | 0 | 0 | 0 | 0 | 0 | 0 | 0 |
| 0 | NA | NA | NA | NA | NA | Gm22671 | | 0 | 0 | 0 | 0 | 0 | 0 | 0 | 0 | 0 | 0 | 0 |
| 0 | NA | NA | NA | NA | NA | Gm37049 | | 0 | 0 | 0 | 0 | 0 | 0 | 0 | 0 | 0 | 0 | 0 |
| 0 | NA | NA | NA | NA | NA | Gm22434 | | 0 | 0 | 0 | 0 | 0 | 0 | 0 | 0 | 0 | 0 | 0 |
| 0 | NA | NA | NA | NA | NA | Gm20471 | | 0 | 0 | 0 | 0 | 0 | 0 | 0 | 0 | 0 | 0 | 0 |
| 0 | NA | NA | NA | NA | NA | Gm37644 | | 0 | 0 | 0 | 0 | 0 | 0 | 0 | 0 | 0 | 0 | 0 |
| 0 | NA | NA | NA | NA | NA | Gm24940 | | 0 | 0 | 0 | 0 | 0 | 0 | 0 | 0 | 0 | 0 | 0 |
| 0 | NA | NA | NA | NA | NA | Gm29500 | | 0 | 0 | 0 | 0 | 0 | 0 | 0 | 0 | 0 | 0 | 0 |
| 0 | NA | NA | NA | NA | NA | Mettl11b | methyltransferase like 11B | 0 | 0 | 0 | 0 | 0 | 0 | 0 | 0 | 0 | 0 | 0 |
| 0 | NA | NA | NA | NA | NA | Gm2343 | | 0 | 0 | 0 | 0 | 0 | 0 | 0 | 0 | 0 | 0 | 0 |

| | | | | | | | | | | | | | | | | | | |
|---|---|---|---|---|---|---|---|---|---|---|---|---|---|---|---|---|---|---|
| 0 | NA | NA | NA | NA | NA | Gm37119 | | 0 | 0 | 0 | 0 | 0 | 0 | 0 | 0 | 0 | 0 | 0 |
| 0 | NA | NA | NA | NA | NA | Gm16588 | | 0 | 0 | 0 | 0 | 0 | 0 | 0 | 0 | 0 | 0 | 0 |
| 0 | NA | NA | NA | NA | NA | Gm37732 | | 0 | 0 | 0 | 0 | 0 | 0 | 0 | 0 | 0 | 0 | 0 |
| 0 | NA | NA | NA | NA | NA | Gm37754 | | 0 | 0 | 0 | 0 | 0 | 0 | 0 | 0 | 0 | 0 | 0 |
| 0 | NA | NA | NA | NA | NA | Gm37853 | | 0 | 0 | 0 | 0 | 0 | 0 | 0 | 0 | 0 | 0 | 0 |
| 0 | NA | NA | NA | NA | NA | Gm37184 | | 0 | 0 | 0 | 0 | 0 | 0 | 0 | 0 | 0 | 0 | 0 |
| 0 | NA | NA | NA | NA | NA | Gm36945 | | 0 | 0 | 0 | 0 | 0 | 0 | 0 | 0 | 0 | 0 | 0 |
| 0 | NA | NA | NA | NA | NA | Gm20743 | | 0 | 0 | 0 | 0 | 0 | 0 | 0 | 0 | 0 | 0 | 0 |
| 0 | NA | NA | NA | NA | NA | Xcl1 | chemokine (C motif) ligand 1 | 0 | 0 | 0 | 0 | 0 | 0 | 0 | 0 | 0 | 0 | 0 |
| 0 | NA | NA | NA | NA | NA | 1700029M03Rik | | 0 | 0 | 0 | 0 | 0 | 0 | 0 | 0 | 0 | 0 | 0 |
| 0 | NA | NA | NA | NA | NA | Gm26042 | | 0 | 0 | 0 | 0 | 0 | 0 | 0 | 0 | 0 | 0 | 0 |
| 0 | NA | NA | NA | NA | NA | Tbx19 | T-box 19 | 0 | 0 | 0 | 0 | 0 | 0 | 0 | 0 | 0 | 0 | 0 |
| 0 | NA | NA | NA | NA | NA | Gm38368 | | 0 | 0 | 0 | 0 | 0 | 0 | 0 | 0 | 0 | 0 | 0 |
| 0 | NA | NA | NA | NA | NA | Gm24637 | | 0 | 0 | 0 | 0 | 0 | 0 | 0 | 0 | 0 | 0 | 0 |
| 0 | NA | NA | NA | NA | NA | Adcy10 | adenylate cyclase 10 | 0 | 0 | 0 | 0 | 0 | 0 | 0 | 0 | 0 | 0 | 0 |
| 0 | NA | NA | NA | NA | NA | Gm18186 | | 0 | 0 | 0 | 0 | 0 | 0 | 0 | 0 | 0 | 0 | 0 |
| 0 | NA | NA | NA | NA | NA | Gm38325 | | 0 | 0 | 0 | 0 | 0 | 0 | 0 | 0 | 0 | 0 | 0 |
| 0 | NA | NA | NA | NA | NA | Gm23599 | | 0 | 0 | 0 | 0 | 0 | 0 | 0 | 0 | 0 | 0 | 0 |
| 0 | NA | NA | NA | NA | NA | Gm38210 | | 0 | 0 | 0 | 0 | 0 | 0 | 0 | 0 | 0 | 0 | 0 |
| 0 | NA | NA | NA | NA | NA | Gm44458 | | 0 | 0 | 0 | 0 | 0 | 0 | 0 | 0 | 0 | 0 | 0 |
| 0 | NA | NA | NA | NA | NA | Gm37073 | | 0 | 0 | 0 | 0 | 0 | 0 | 0 | 0 | 0 | 0 | 0 |
| 0 | NA | NA | NA | NA | NA | Gm34767 | | 0 | 0 | 0 | 0 | 0 | 0 | 0 | 0 | 0 | 0 | 0 |
| 0 | NA | NA | NA | NA | NA | Gm36972 | | 0 | 0 | 0 | 0 | 0 | 0 | 0 | 0 | 0 | 0 | 0 |
| 0 | NA | NA | NA | NA | NA | Gm24808 | | 0 | 0 | 0 | 0 | 0 | 0 | 0 | 0 | 0 | 0 | 0 |
| 0 | NA | NA | NA | NA | NA | Gm23402 | | 0 | 0 | 0 | 0 | 0 | 0 | 0 | 0 | 0 | 0 | 0 |
| 0 | NA | NA | NA | NA | NA | Gm32999 | | 0 | 0 | 0 | 0 | 0 | 0 | 0 | 0 | 0 | 0 | 0 |
| 0 | NA | NA | NA | NA | NA | Gm37856 | | 0 | 0 | 0 | 0 | 0 | 0 | 0 | 0 | 0 | 0 | 0 |
| 0 | NA | NA | NA | NA | NA | Gm17976 | | 0 | 0 | 0 | 0 | 0 | 0 | 0 | 0 | 0 | 0 | 0 |
| 0 | NA | NA | NA | NA | NA | Gm26665 | | 0 | 0 | 0 | 0 | 0 | 0 | 0 | 0 | 0 | 0 | 0 |
| 0 | NA | NA | NA | NA | NA | Gm7626 | | 0 | 0 | 0 | 0 | 0 | 0 | 0 | 0 | 0 | 0 | 0 |
| 0 | NA | NA | NA | NA | NA | Gm37923 | | 0 | 0 | 0 | 0 | 0 | 0 | 0 | 0 | 0 | 0 | 0 |
| 0 | NA | NA | NA | NA | NA | 5330438I03Rik | | 0 | 0 | 0 | 0 | 0 | 0 | 0 | 0 | 0 | 0 | 0 |
| 0 | NA | NA | NA | NA | NA | Gm33354 | | 0 | 0 | 0 | 0 | 0 | 0 | 0 | 0 | 0 | 0 | 0 |
| 0 | NA | NA | NA | NA | NA | Gm22992 | | 0 | 0 | 0 | 0 | 0 | 0 | 0 | 0 | 0 | 0 | 0 |
| 0 | NA | NA | NA | NA | NA | Gm4847 | predicted gene 4847 | 0 | 0 | 0 | 0 | 0 | 0 | 0 | 0 | 0 | 0 | 0 |
| 0 | NA | NA | NA | NA | NA | Fmo9 | flavin containing monooxygenase 9 | 0 | 0 | 0 | 0 | 0 | 0 | 0 | 0 | 0 | 0 | 0 |
| 0 | NA | NA | NA | NA | NA | Gm4954 | | 0 | 0 | 0 | 0 | 0 | 0 | 0 | 0 | 0 | 0 | 0 |
| 0 | NA | NA | NA | NA | NA | Gm37904 | | 0 | 0 | 0 | 0 | 0 | 0 | 0 | 0 | 0 | 0 | 0 |
| 0 | NA | NA | NA | NA | NA | Gm17868 | | 0 | 0 | 0 | 0 | 0 | 0 | 0 | 0 | 0 | 0 | 0 |
| 0 | NA | NA | NA | NA | NA | Gm37994 | | 0 | 0 | 0 | 0 | 0 | 0 | 0 | 0 | 0 | 0 | 0 |
| 0 | NA | NA | NA | NA | NA | Gm37982 | | 0 | 0 | 0 | 0 | 0 | 0 | 0 | 0 | 0 | 0 | 0 |
| 0 | NA | NA | NA | NA | NA | Mir6347 | | 0 | 0 | 0 | 0 | 0 | 0 | 0 | 0 | 0 | 0 | 0 |
| 0 | NA | NA | NA | NA | NA | Gm37232 | | 0 | 0 | 0 | 0 | 0 | 0 | 0 | 0 | 0 | 0 | 0 |
| 0 | NA | NA | NA | NA | NA | Gm24182 | | 0 | 0 | 0 | 0 | 0 | 0 | 0 | 0 | 0 | 0 | 0 |
| 0 | NA | NA | NA | NA | NA | Lrrc52 | leucine rich repeat containing 52 | 0 | 0 | 0 | 0 | 0 | 0 | 0 | 0 | 0 | 0 | 0 |
| 0 | NA | NA | NA | NA | NA | Gm20711 | | 0 | 0 | 0 | 0 | 0 | 0 | 0 | 0 | 0 | 0 | 0 |
| 0 | NA | NA | NA | NA | NA | Gm38381 | | 0 | 0 | 0 | 0 | 0 | 0 | 0 | 0 | 0 | 0 | 0 |
| 0 | NA | NA | NA | NA | NA | Mir6348 | | 0 | 0 | 0 | 0 | 0 | 0 | 0 | 0 | 0 | 0 | 0 |
| 0 | NA | NA | NA | NA | NA | 1700063I16Rik | | 0 | 0 | 0 | 0 | 0 | 0 | 0 | 0 | 0 | 0 | 0 |
| 0 | NA | NA | NA | NA | NA | Mir6354 | | 0 | 0 | 0 | 0 | 0 | 0 | 0 | 0 | 0 | 0 | 0 |
| 0 | NA | NA | NA | NA | NA | Gm37839 | | 0 | 0 | 0 | 0 | 0 | 0 | 0 | 0 | 0 | 0 | 0 |
| 0 | NA | NA | NA | NA | NA | Gm18900 | | 0 | 0 | 0 | 0 | 0 | 0 | 0 | 0 | 0 | 0 | 0 |
| 0 | NA | NA | NA | NA | NA | Gm36980 | | 0 | 0 | 0 | 0 | 0 | 0 | 0 | 0 | 0 | 0 | 0 |
| 0 | NA | NA | NA | NA | NA | Gm22114 | | 0 | 0 | 0 | 0 | 0 | 0 | 0 | 0 | 0 | 0 | 0 |
| 0 | NA | NA | NA | NA | NA | Gm37586 | | 0 | 0 | 0 | 0 | 0 | 0 | 0 | 0 | 0 | 0 | 0 |
| 0 | NA | NA | NA | NA | NA | Gm37502 | | 0 | 0 | 0 | 0 | 0 | 0 | 0 | 0 | 0 | 0 | 0 |
| 0 | NA | NA | NA | NA | NA | Spata46 | spermatogenesis associated 46 | 0 | 0 | 0 | 0 | 0 | 0 | 0 | 0 | 0 | 0 | 0 |
| 0 | NA | NA | NA | NA | NA | Gm23523 | | 0 | 0 | 0 | 0 | 0 | 0 | 0 | 0 | 0 | 0 | 0 |
| 0 | NA | NA | NA | NA | NA | Gm25235 | | 0 | 0 | 0 | 0 | 0 | 0 | 0 | 0 | 0 | 0 | 0 |
| 0 | NA | NA | NA | NA | NA | Gm37030 | | 0 | 0 | 0 | 0 | 0 | 0 | 0 | 0 | 0 | 0 | 0 |

| | | | | | | | | | | | | | | | | | | |
|---|---|---|---|---|---|---|---|---|---|---|---|---|---|---|---|---|---|---|
| 0 | NA | NA | NA | NA | NA | Gm38204 | | 0 | 0 | 0 | 0 | 0 | 0 | 0 | 0 | 0 | 0 | 0 |
| 0 | NA | NA | NA | NA | NA | Gm10522 | | 0 | 0 | 0 | 0 | 0 | 0 | 0 | 0 | 0 | 0 | 0 |
| 0 | NA | NA | NA | NA | NA | Gm26110 | | 0 | 0 | 0 | 0 | 0 | 0 | 0 | 0 | 0 | 0 | 0 |
| 0 | NA | NA | NA | NA | NA | Mir6546 | | 0 | 0 | 0 | 0 | 0 | 0 | 0 | 0 | 0 | 0 | 0 |
| 0 | NA | NA | NA | NA | NA | Gm27552 | | 0 | 0 | 0 | 0 | 0 | 0 | 0 | 0 | 0 | 0 | 0 |
| 0 | NA | NA | NA | NA | NA | Gm27844 | | 0 | 0 | 0 | 0 | 0 | 0 | 0 | 0 | 0 | 0 | 0 |
| 0 | NA | NA | NA | NA | NA | Gm25151 | | 0 | 0 | 0 | 0 | 0 | 0 | 0 | 0 | 0 | 0 | 0 |
| 0 | NA | NA | NA | NA | NA | Gm37097 | | 0 | 0 | 0 | 0 | 0 | 0 | 0 | 0 | 0 | 0 | 0 |
| 0 | NA | NA | NA | NA | NA | Gm24871 | | 0 | 0 | 0 | 0 | 0 | 0 | 0 | 0 | 0 | 0 | 0 |
| 0 | NA | NA | NA | NA | NA | Gm9765 | | 0 | 0 | 0 | 0 | 0 | 0 | 0 | 0 | 0 | 0 | 0 |
| 0 | NA | NA | NA | NA | NA | Itln1 | intelectin 1 (galactofuranose binding) | 0 | 0 | 0 | 0 | 0 | 0 | 0 | 0 | 0 | 0 | 0 |
| 0 | NA | NA | NA | NA | NA | Gm37450 | | 0 | 0 | 0 | 0 | 0 | 0 | 0 | 0 | 0 | 0 | 0 |
| 0 | NA | NA | NA | NA | NA | Gm37787 | | 0 | 0 | 0 | 0 | 0 | 0 | 0 | 0 | 0 | 0 | 0 |
| 0 | NA | NA | NA | NA | NA | Mir7683 | | 0 | 0 | 0 | 0 | 0 | 0 | 0 | 0 | 0 | 0 | 0 |
| 0 | NA | NA | NA | NA | NA | A630035G10Rik | | 0 | 0 | 0 | 0 | 0 | 0 | 0 | 0 | 0 | 0 | 0 |
| 0 | NA | NA | NA | NA | NA | Gm37010 | | 0 | 0 | 0 | 0 | 0 | 0 | 0 | 0 | 0 | 0 | 0 |
| 0 | NA | NA | NA | NA | NA | Gm37693 | | 0 | 0 | 0 | 0 | 0 | 0 | 0 | 0 | 0 | 0 | 0 |
| 0 | NA | NA | NA | NA | NA | Gm19030 | | 0 | 0 | 0 | 0 | 0 | 0 | 0 | 0 | 0 | 0 | 0 |
| 0 | NA | NA | NA | NA | NA | Gm26412 | | 0 | 0 | 0 | 0 | 0 | 0 | 0 | 0 | 0 | 0 | 0 |
| 0 | NA | NA | NA | NA | NA | Gm22950 | | 0 | 0 | 0 | 0 | 0 | 0 | 0 | 0 | 0 | 0 | 0 |
| 0 | NA | NA | NA | NA | NA | Gm10521 | | 0 | 0 | 0 | 0 | 0 | 0 | 0 | 0 | 0 | 0 | 0 |
| 0 | NA | NA | NA | NA | NA | Gm37065 | | 0 | 0 | 0 | 0 | 0 | 0 | 0 | 0 | 0 | 0 | 0 |
| 0 | NA | NA | NA | NA | NA | Gm37950 | | 0 | 0 | 0 | 0 | 0 | 0 | 0 | 0 | 0 | 0 | 0 |
| 0 | NA | NA | NA | NA | NA | Gm36937 | | 0 | 0 | 0 | 0 | 0 | 0 | 0 | 0 | 0 | 0 | 0 |
| 0 | NA | NA | NA | NA | NA | Gm37145 | | 0 | 0 | 0 | 0 | 0 | 0 | 0 | 0 | 0 | 0 | 0 |
| 0 | NA | NA | NA | NA | NA | Gm7804 | | 0 | 0 | 0 | 0 | 0 | 0 | 0 | 0 | 0 | 0 | 0 |
| 0 | NA | NA | NA | NA | NA | Gm22383 | | 0 | 0 | 0 | 0 | 0 | 0 | 0 | 0 | 0 | 0 | 0 |
| 0 | NA | NA | NA | NA | NA | Gm38051 | | 0 | 0 | 0 | 0 | 0 | 0 | 0 | 0 | 0 | 0 | 0 |
| 0 | NA | NA | NA | NA | NA | Gm24817 | | 0 | 0 | 0 | 0 | 0 | 0 | 0 | 0 | 0 | 0 | 0 |
| 0 | NA | NA | NA | NA | NA | Gm26077 | | 0 | 0 | 0 | 0 | 0 | 0 | 0 | 0 | 0 | 0 | 0 |
| 0 | NA | NA | NA | NA | NA | Olfr16 | olfactory receptor 16 | 0 | 0 | 0 | 0 | 0 | 0 | 0 | 0 | 0 | 0 | 0 |
| 0 | NA | NA | NA | NA | NA | Gm42472 | | 0 | 0 | 0 | 0 | 0 | 0 | 0 | 0 | 0 | 0 | 0 |
| 0 | NA | NA | NA | NA | NA | Gm42475 | | 0 | 0 | 0 | 0 | 0 | 0 | 0 | 0 | 0 | 0 | 0 |
| 0 | NA | NA | NA | NA | NA | Olfr1409-ps1 | | 0 | 0 | 0 | 0 | 0 | 0 | 0 | 0 | 0 | 0 | 0 |
| 0 | NA | NA | NA | NA | NA | Olfr1408 | olfactory receptor 1408 | 0 | 0 | 0 | 0 | 0 | 0 | 0 | 0 | 0 | 0 | 0 |
| 0 | NA | NA | NA | NA | NA | Olfr1407-ps1 | | 0 | 0 | 0 | 0 | 0 | 0 | 0 | 0 | 0 | 0 | 0 |
| 0 | NA | NA | NA | NA | NA | Olfr1406 | olfactory receptor 1406 | 0 | 0 | 0 | 0 | 0 | 0 | 0 | 0 | 0 | 0 | 0 |
| 0 | NA | NA | NA | NA | NA | Olfr218 | olfactory receptor 218 | 0 | 0 | 0 | 0 | 0 | 0 | 0 | 0 | 0 | 0 | 0 |
| 0 | NA | NA | NA | NA | NA | Olfr1404 | olfactory receptor 1404 | 0 | 0 | 0 | 0 | 0 | 0 | 0 | 0 | 0 | 0 | 0 |
| 0 | NA | NA | NA | NA | NA | Olfr418 | olfactory receptor 418 | 0 | 0 | 0 | 0 | 0 | 0 | 0 | 0 | 0 | 0 | 0 |
| 0 | NA | NA | NA | NA | NA | Mptx2 | mucosal pentraxin 2 | 0 | 0 | 0 | 0 | 0 | 0 | 0 | 0 | 0 | 0 | 0 |
| 0 | NA | NA | NA | NA | NA | Gm44481 | | 0 | 0 | 0 | 0 | 0 | 0 | 0 | 0 | 0 | 0 | 0 |
| 0 | NA | NA | NA | NA | NA | Gm44339 | | 0 | 0 | 0 | 0 | 0 | 0 | 0 | 0 | 0 | 0 | 0 |
| 0 | NA | NA | NA | NA | NA | Gm18752 | | 0 | 0 | 0 | 0 | 0 | 0 | 0 | 0 | 0 | 0 | 0 |
| 0 | NA | NA | NA | NA | NA | Gm7871 | | 0 | 0 | 0 | 0 | 0 | 0 | 0 | 0 | 0 | 0 | 0 |
| 0 | NA | NA | NA | NA | NA | Gm7897 | | 0 | 0 | 0 | 0 | 0 | 0 | 0 | 0 | 0 | 0 | 0 |
| 0 | NA | NA | NA | NA | NA | Gm38050 | | 0 | 0 | 0 | 0 | 0 | 0 | 0 | 0 | 0 | 0 | 0 |
| 0 | NA | NA | NA | NA | NA | Gm18445 | | 0 | 0 | 0 | 0 | 0 | 0 | 0 | 0 | 0 | 0 | 0 |
| 0 | NA | NA | NA | NA | NA | Olfr432 | olfactory receptor 432 | 0 | 0 | 0 | 0 | 0 | 0 | 0 | 0 | 0 | 0 | 0 |
| 0 | NA | NA | NA | NA | NA | Olfr431-ps1 | | 0 | 0 | 0 | 0 | 0 | 0 | 0 | 0 | 0 | 0 | 0 |
| 0 | NA | NA | NA | NA | NA | Olfr430 | olfactory receptor 430 | 0 | 0 | 0 | 0 | 0 | 0 | 0 | 0 | 0 | 0 | 0 |
| 0 | NA | NA | NA | NA | NA | Olfr429 | olfactory receptor 429 | 0 | 0 | 0 | 0 | 0 | 0 | 0 | 0 | 0 | 0 | 0 |
| 0 | NA | NA | NA | NA | NA | Olfr428-ps1 | | 0 | 0 | 0 | 0 | 0 | 0 | 0 | 0 | 0 | 0 | 0 |
| 0 | NA | NA | NA | NA | NA | Olfr427 | olfactory receptor 427 | 0 | 0 | 0 | 0 | 0 | 0 | 0 | 0 | 0 | 0 | 0 |
| 0 | NA | NA | NA | NA | NA | Gm37417 | | 0 | 0 | 0 | 0 | 0 | 0 | 0 | 0 | 0 | 0 | 0 |
| 0 | NA | NA | NA | NA | NA | Olfr231 | olfactory receptor 231 | 0 | 0 | 0 | 0 | 0 | 0 | 0 | 0 | 0 | 0 | 0 |
| 0 | NA | NA | NA | NA | NA | Olfr424 | olfactory receptor 424 | 0 | 0 | 0 | 0 | 0 | 0 | 0 | 0 | 0 | 0 | 0 |
| 0 | NA | NA | NA | NA | NA | Olfr258-ps1 | | 0 | 0 | 0 | 0 | 0 | 0 | 0 | 0 | 0 | 0 | 0 |
| 0 | NA | NA | NA | NA | NA | Olfr421-ps1 | olfactory receptor 421, pseudogene 1 | 0 | 0 | 0 | 0 | 0 | 0 | 0 | 0 | 0 | 0 | 0 |
| 0 | NA | NA | NA | NA | NA | Olfr420 | olfactory receptor 420 | 0 | 0 | 0 | 0 | 0 | 0 | 0 | 0 | 0 | 0 | 0 |

| | | | | | | | | | | | | | | | | | |
|---|---|---|---|---|---|---|---|---|---|---|---|---|---|---|---|---|---|
| 0 | NA | NA | NA | NA | NA | Gm43440 | | 0 | 0 | 0 | 0 | 0 | 0 | 0 | 0 | 0 | 0 | 0 |
| 0 | NA | NA | NA | NA | NA | Olfr419 | olfactory receptor 419 | 0 | 0 | 0 | 0 | 0 | 0 | 0 | 0 | 0 | 0 | 0 |
| 0 | NA | NA | NA | NA | NA | Olfr417 | olfactory receptor 417 | 0 | 0 | 0 | 0 | 0 | 0 | 0 | 0 | 0 | 0 | 0 |
| 0 | NA | NA | NA | NA | NA | Olfr416-ps1 | | 0 | 0 | 0 | 0 | 0 | 0 | 0 | 0 | 0 | 0 | 0 |
| 0 | NA | NA | NA | NA | NA | Olfr248 | olfactory receptor 248 | 0 | 0 | 0 | 0 | 0 | 0 | 0 | 0 | 0 | 0 | 0 |
| 0 | NA | NA | NA | NA | NA | Olfr414 | olfactory receptor 414 | 0 | 0 | 0 | 0 | 0 | 0 | 0 | 0 | 0 | 0 | 0 |
| 0 | NA | NA | NA | NA | NA | Olfr220 | olfactory receptor 220 | 0 | 0 | 0 | 0 | 0 | 0 | 0 | 0 | 0 | 0 | 0 |
| 0 | NA | NA | NA | NA | NA | 5730408A14Rik | | 0 | 0 | 0 | 0 | 0 | 0 | 0 | 0 | 0 | 0 | 0 |
| 0 | NA | NA | NA | NA | NA | Gm36307 | | 0 | 0 | 0 | 0 | 0 | 0 | 0 | 0 | 0 | 0 | 0 |
| 0 | NA | NA | NA | NA | NA | Wdr64 | WD repeat domain 64 | 0 | 0 | 0 | 0 | 0 | 0 | 0 | 0 | 0 | 0 | 0 |
| 0 | NA | NA | NA | NA | NA | Gm25560 | | 0 | 0 | 0 | 0 | 0 | 0 | 0 | 0 | 0 | 0 | 0 |
| 0 | NA | NA | NA | NA | NA | Gm23805 | | 0 | 0 | 0 | 0 | 0 | 0 | 0 | 0 | 0 | 0 | 0 |
| 0 | NA | NA | NA | NA | NA | Gm38100 | beclin 2 | 0 | 0 | 0 | 0 | 0 | 0 | 0 | 0 | 0 | 0 | 0 |
| 0 | NA | NA | NA | NA | NA | Gm37668 | | 0 | 0 | 0 | 0 | 0 | 0 | 0 | 0 | 0 | 0 | 0 |
| 0 | NA | NA | NA | NA | NA | Gm38081 | | 0 | 0 | 0 | 0 | 0 | 0 | 0 | 0 | 0 | 0 | 0 |
| 0 | NA | NA | NA | NA | NA | Gm37916 | | 0 | 0 | 0 | 0 | 0 | 0 | 0 | 0 | 0 | 0 | 0 |
| 0 | NA | NA | NA | NA | NA | Gm38101 | | 0 | 0 | 0 | 0 | 0 | 0 | 0 | 0 | 0 | 0 | 0 |
| 0 | NA | NA | NA | NA | NA | Gm17965 | | 0 | 0 | 0 | 0 | 0 | 0 | 0 | 0 | 0 | 0 | 0 |
| 0 | NA | NA | NA | NA | NA | Gm38152 | | 0 | 0 | 0 | 0 | 0 | 0 | 0 | 0 | 0 | 0 | 0 |
| 0 | NA | NA | NA | NA | NA | Gm36536 | | 0 | 0 | 0 | 0 | 0 | 0 | 0 | 0 | 0 | 0 | 0 |
| 0 | NA | NA | NA | NA | NA | Gm17966 | | 0 | 0 | 0 | 0 | 0 | 0 | 0 | 0 | 0 | 0 | 0 |
| 0 | NA | NA | NA | NA | NA | Gm26104 | | 0 | 0 | 0 | 0 | 0 | 0 | 0 | 0 | 0 | 0 | 0 |
| 0 | NA | NA | NA | NA | NA | Mir350 | | 0 | 0 | 0 | 0 | 0 | 0 | 0 | 0 | 0 | 0 | 0 |
| 0 | NA | NA | NA | NA | NA | Gm25288 | | 0 | 0 | 0 | 0 | 0 | 0 | 0 | 0 | 0 | 0 | 0 |
| 0 | NA | NA | NA | NA | NA | Gm38053 | | 0 | 0 | 0 | 0 | 0 | 0 | 0 | 0 | 0 | 0 | 0 |
| 0 | NA | NA | NA | NA | NA | Gm25993 | | 0 | 0 | 0 | 0 | 0 | 0 | 0 | 0 | 0 | 0 | 0 |
| 0 | NA | NA | NA | NA | NA | Gm37463 | | 0 | 0 | 0 | 0 | 0 | 0 | 0 | 0 | 0 | 0 | 0 |
| 0 | NA | NA | NA | NA | NA | Gm38328 | | 0 | 0 | 0 | 0 | 0 | 0 | 0 | 0 | 0 | 0 | 0 |
| 0 | NA | NA | NA | NA | NA | Gm29856 | | 0 | 0 | 0 | 0 | 0 | 0 | 0 | 0 | 0 | 0 | 0 |
| 0 | NA | NA | NA | NA | NA | Gm37306 | | 0 | 0 | 0 | 0 | 0 | 0 | 0 | 0 | 0 | 0 | 0 |
| 0 | NA | NA | NA | NA | NA | Gm37280 | | 0 | 0 | 0 | 0 | 0 | 0 | 0 | 0 | 0 | 0 | 0 |
| 0 | NA | NA | NA | NA | NA | 2310043L19Rik | | 0 | 0 | 0 | 0 | 0 | 0 | 0 | 0 | 0 | 0 | 0 |
| 0 | NA | NA | NA | NA | NA | Gm8023 | | 0 | 0 | 0 | 0 | 0 | 0 | 0 | 0 | 0 | 0 | 0 |
| 0 | NA | NA | NA | NA | NA | 4930527J03Rik | | 0 | 0 | 0 | 0 | 0 | 0 | 0 | 0 | 0 | 0 | 0 |
| 0 | NA | NA | NA | NA | NA | Gm27458 | | 0 | 0 | 0 | 0 | 0 | 0 | 0 | 0 | 0 | 0 | 0 |
| 0 | NA | NA | NA | NA | NA | Gm24919 | | 0 | 0 | 0 | 0 | 0 | 0 | 0 | 0 | 0 | 0 | 0 |
| 0 | NA | NA | NA | NA | NA | Gm24405 | | 0 | 0 | 0 | 0 | 0 | 0 | 0 | 0 | 0 | 0 | 0 |
| 0 | NA | NA | NA | NA | NA | Gm34176 | | 0 | 0 | 0 | 0 | 0 | 0 | 0 | 0 | 0 | 0 | 0 |
| 0 | NA | NA | NA | NA | NA | Gm16564 | | 0 | 0 | 0 | 0 | 0 | 0 | 0 | 0 | 0 | 0 | 0 |
| 0 | NA | NA | NA | NA | NA | Gm15480 | | 0 | 0 | 0 | 0 | 0 | 0 | 0 | 0 | 0 | 0 | 0 |
| 0 | NA | NA | NA | NA | NA | Kif28 | kinesin family member 28 | 0 | 0 | 0 | 0 | 0 | 0 | 0 | 0 | 0 | 0 | 0 |
| 0 | NA | NA | NA | NA | NA | Gm38331 | | 0 | 0 | 0 | 0 | 0 | 0 | 0 | 0 | 0 | 0 | 0 |
| 0 | NA | NA | NA | NA | NA | Gm37984 | | 0 | 0 | 0 | 0 | 0 | 0 | 0 | 0 | 0 | 0 | 0 |
| 0 | NA | NA | NA | NA | NA | Mixl1 | Mix1 homeobox-like 1 (Xenopus laevis) | 0 | 0 | 0 | 0 | 0 | 0 | 0 | 0 | 0 | 0 | 0 |
| 0 | NA | NA | NA | NA | NA | Gm24836 | | 0 | 0 | 0 | 0 | 0 | 0 | 0 | 0 | 0 | 0 | 0 |
| 0 | NA | NA | NA | NA | NA | Gm20305 | | 0 | 0 | 0 | 0 | 0 | 0 | 0 | 0 | 0 | 0 | 0 |
| 0 | NA | NA | NA | NA | NA | Gm18036 | | 0 | 0 | 0 | 0 | 0 | 0 | 0 | 0 | 0 | 0 | 0 |
| 0 | NA | NA | NA | NA | NA | Gm36961 | | 0 | 0 | 0 | 0 | 0 | 0 | 0 | 0 | 0 | 0 | 0 |
| 0 | NA | NA | NA | NA | NA | Gm37431 | | 0 | 0 | 0 | 0 | 0 | 0 | 0 | 0 | 0 | 0 | 0 |
| 0 | NA | NA | NA | NA | NA | Gm37086 | | 0 | 0 | 0 | 0 | 0 | 0 | 0 | 0 | 0 | 0 | 0 |
| 0 | NA | NA | NA | NA | NA | Gm17967 | | 0 | 0 | 0 | 0 | 0 | 0 | 0 | 0 | 0 | 0 | 0 |
| 0 | NA | NA | NA | NA | NA | Gm6606 | | 0 | 0 | 0 | 0 | 0 | 0 | 0 | 0 | 0 | 0 | 0 |
| 0 | NA | NA | NA | NA | NA | Cnih3 | cornichon family AMPA receptor auxiliary | 0 | 0 | 0 | 0 | 0 | 0 | 0 | 0 | 0 | 0 | 0 |
| 0 | NA | NA | NA | NA | NA | 2900069G24Rik | | 0 | 0 | 0 | 0 | 0 | 0 | 0 | 0 | 0 | 0 | 0 |
| 0 | NA | NA | NA | NA | NA | Gm16547 | | 0 | 0 | 0 | 0 | 0 | 0 | 0 | 0 | 0 | 0 | 0 |
| 0 | NA | NA | NA | NA | NA | Gm16539 | | 0 | 0 | 0 | 0 | 0 | 0 | 0 | 0 | 0 | 0 | 0 |
| 0 | NA | NA | NA | NA | NA | Ccdc121 | coiled-coil domain containing 121 | 0 | 0 | 0 | 0 | 0 | 0 | 0 | 0 | 0 | 0 | 0 |
| 0 | NA | NA | NA | NA | NA | Gm37664 | | 0 | 0 | 0 | 0 | 0 | 0 | 0 | 0 | 0 | 0 | 0 |
| 0 | NA | NA | NA | NA | NA | Gm17968 | | 0 | 0 | 0 | 0 | 0 | 0 | 0 | 0 | 0 | 0 | 0 |
| 0 | NA | NA | NA | NA | NA | Vmn1r1 | vomeronasal 1 receptor 1 | 0 | 0 | 0 | 0 | 0 | 0 | 0 | 0 | 0 | 0 | 0 |

| | | | | | | | | | | | | | | | | | |
|---|---|---|---|---|---|---|---|---|---|---|---|---|---|---|---|---|---|
| 0 | NA | NA | NA | NA | NA | 1700047M11Rik | | 0 | 0 | 0 | 0 | 0 | 0 | 0 | 0 | 0 | 0 | 0 |
| 0 | NA | NA | NA | NA | NA | Gm8197 | | 0 | 0 | 0 | 0 | 0 | 0 | 0 | 0 | 0 | 0 | 0 |
| 0 | NA | NA | NA | NA | NA | Gm37218 | | 0 | 0 | 0 | 0 | 0 | 0 | 0 | 0 | 0 | 0 | 0 |
| 0 | NA | NA | NA | NA | NA | Ccdc185 | coiled-coil domain containing 185 | 0 | 0 | 0 | 0 | 0 | 0 | 0 | 0 | 0 | 0 | 0 |
| 0 | NA | NA | NA | NA | NA | Gm37542 | | 0 | 0 | 0 | 0 | 0 | 0 | 0 | 0 | 0 | 0 | 0 |
| 0 | NA | NA | NA | NA | NA | Gm24975 | | 0 | 0 | 0 | 0 | 0 | 0 | 0 | 0 | 0 | 0 | 0 |
| 0 | NA | NA | NA | NA | NA | | | 0 | 0 | 0 | 0 | 0 | 0 | 0 | 0 | 0 | 0 | 0 |
| 0 | NA | NA | NA | NA | NA | Gm33973 | | 0 | 0 | 0 | 0 | 0 | 0 | 0 | 0 | 0 | 0 | 0 |
| 0 | NA | NA | NA | NA | NA | Gm8214 | | 0 | 0 | 0 | 0 | 0 | 0 | 0 | 0 | 0 | 0 | 0 |
| 0 | NA | NA | NA | NA | NA | Gm38332 | | 0 | 0 | 0 | 0 | 0 | 0 | 0 | 0 | 0 | 0 | 0 |
| 0 | NA | NA | NA | NA | NA | Gm38108 | | 0 | 0 | 0 | 0 | 0 | 0 | 0 | 0 | 0 | 0 | 0 |
| 0 | NA | NA | NA | NA | NA | Gm37223 | | 0 | 0 | 0 | 0 | 0 | 0 | 0 | 0 | 0 | 0 | 0 |
| 0 | NA | NA | NA | NA | NA | 2900092O11Rik | | 0 | 0 | 0 | 0 | 0 | 0 | 0 | 0 | 0 | 0 | 0 |
| 0 | NA | NA | NA | NA | NA | 1700112H15Rik | | 0 | 0 | 0 | 0 | 0 | 0 | 0 | 0 | 0 | 0 | 0 |
| 0 | NA | NA | NA | NA | NA | Gm23349 | | 0 | 0 | 0 | 0 | 0 | 0 | 0 | 0 | 0 | 0 | 0 |
| 0 | NA | NA | NA | NA | NA | Gm37800 | | 0 | 0 | 0 | 0 | 0 | 0 | 0 | 0 | 0 | 0 | 0 |
| 0 | NA | NA | NA | NA | NA | 4930488B22Rik | | 0 | 0 | 0 | 0 | 0 | 0 | 0 | 0 | 0 | 0 | 0 |
| 0 | NA | NA | NA | NA | NA | Gm34342 | | 0 | 0 | 0 | 0 | 0 | 0 | 0 | 0 | 0 | 0 | 0 |
| 0 | NA | NA | NA | NA | NA | Gm23780 | | 0 | 0 | 0 | 0 | 0 | 0 | 0 | 0 | 0 | 0 | 0 |
| 0 | NA | NA | NA | NA | NA | Gm34732 | | 0 | 0 | 0 | 0 | 0 | 0 | 0 | 0 | 0 | 0 | 0 |
| 0 | NA | NA | NA | NA | NA | 4930433J02Rik | | 0 | 0 | 0 | 0 | 0 | 0 | 0 | 0 | 0 | 0 | 0 |
| 0 | NA | NA | NA | NA | NA | Gm37662 | | 0 | 0 | 0 | 0 | 0 | 0 | 0 | 0 | 0 | 0 | 0 |
| 0 | NA | NA | NA | NA | NA | Mir664 | | 0 | 0 | 0 | 0 | 0 | 0 | 0 | 0 | 0 | 0 | 0 |
| 0 | NA | NA | NA | NA | NA | 2010103J01Rik | | 0 | 0 | 0 | 0 | 0 | 0 | 0 | 0 | 0 | 0 | 0 |
| 0 | NA | NA | NA | NA | NA | Mir194-1 | | 0 | 0 | 0 | 0 | 0 | 0 | 0 | 0 | 0 | 0 | 0 |
| 0 | NA | NA | NA | NA | NA | Mir215 | | 0 | 0 | 0 | 0 | 0 | 0 | 0 | 0 | 0 | 0 | 0 |
| 0 | NA | NA | NA | NA | NA | 9630028B13Rik | | 0 | 0 | 0 | 0 | 0 | 0 | 0 | 0 | 0 | 0 | 0 |
| 0 | NA | NA | NA | NA | NA | Gm2061 | | 0 | 0 | 0 | 0 | 0 | 0 | 0 | 0 | 0 | 0 | 0 |
| 0 | NA | NA | NA | NA | NA | Gm38093 | | 0 | 0 | 0 | 0 | 0 | 0 | 0 | 0 | 0 | 0 | 0 |
| 0 | NA | NA | NA | NA | NA | Gm23821 | | 0 | 0 | 0 | 0 | 0 | 0 | 0 | 0 | 0 | 0 | 0 |
| 0 | NA | NA | NA | NA | NA | Gm37134 | | 0 | 0 | 0 | 0 | 0 | 0 | 0 | 0 | 0 | 0 | 0 |
| 0 | NA | NA | NA | NA | NA | Gm37272 | | 0 | 0 | 0 | 0 | 0 | 0 | 0 | 0 | 0 | 0 | 0 |
| 0 | NA | NA | NA | NA | NA | Gm37491 | | 0 | 0 | 0 | 0 | 0 | 0 | 0 | 0 | 0 | 0 | 0 |
| 0 | NA | NA | NA | NA | NA | A730004F24Rik | | 0 | 0 | 0 | 0 | 0 | 0 | 0 | 0 | 0 | 0 | 0 |
| 0 | NA | NA | NA | NA | NA | Gm26169 | | 0 | 0 | 0 | 0 | 0 | 0 | 0 | 0 | 0 | 0 | 0 |
| 0 | NA | NA | NA | NA | NA | Gm37849 | | 0 | 0 | 0 | 0 | 0 | 0 | 0 | 0 | 0 | 0 | 0 |
| 0 | NA | NA | NA | NA | NA | 1700023G09Rik | | 0 | 0 | 0 | 0 | 0 | 0 | 0 | 0 | 0 | 0 | 0 |
| 0 | NA | NA | NA | NA | NA | Gm37912 | | 0 | 0 | 0 | 0 | 0 | 0 | 0 | 0 | 0 | 0 | 0 |
| 0 | NA | NA | NA | NA | NA | Gm19058 | | 0 | 0 | 0 | 0 | 0 | 0 | 0 | 0 | 0 | 0 | 0 |
| 0 | NA | NA | NA | NA | NA | Gm15508 | | 0 | 0 | 0 | 0 | 0 | 0 | 0 | 0 | 0 | 0 | 0 |
| 0 | NA | NA | NA | NA | NA | Gm15509 | | 0 | 0 | 0 | 0 | 0 | 0 | 0 | 0 | 0 | 0 | 0 |
| 0 | NA | NA | NA | NA | NA | 1700007P06Rik | | 0 | 0 | 0 | 0 | 0 | 0 | 0 | 0 | 0 | 0 | 0 |
| 0 | NA | NA | NA | NA | NA | Gm38155 | | 0 | 0 | 0 | 0 | 0 | 0 | 0 | 0 | 0 | 0 | 0 |
| 0 | NA | NA | NA | NA | NA | Gm37896 | | 0 | 0 | 0 | 0 | 0 | 0 | 0 | 0 | 0 | 0 | 0 |
| 0 | NA | NA | NA | NA | NA | Gm37929 | | 0 | 0 | 0 | 0 | 0 | 0 | 0 | 0 | 0 | 0 | 0 |
| 0 | NA | NA | NA | NA | NA | | | 0 | 0 | 0 | 0 | 0 | 0 | 0 | 0 | 0 | 0 | 0 |
| 0 | NA | NA | NA | NA | NA | 9330162B11Rik | | 0 | 0 | 0 | 0 | 0 | 0 | 0 | 0 | 0 | 0 | 0 |
| 0 | NA | NA | NA | NA | NA | Gm25095 | | 0 | 0 | 0 | 0 | 0 | 0 | 0 | 0 | 0 | 0 | 0 |
| 0 | NA | NA | NA | NA | NA | Gm25269 | | 0 | 0 | 0 | 0 | 0 | 0 | 0 | 0 | 0 | 0 | 0 |
| 0 | NA | NA | NA | NA | NA | Gm37178 | | 0 | 0 | 0 | 0 | 0 | 0 | 0 | 0 | 0 | 0 | 0 |
| 0 | NA | NA | NA | NA | NA | Gm18698 | | 0 | 0 | 0 | 0 | 0 | 0 | 0 | 0 | 0 | 0 | 0 |
| 0 | NA | NA | NA | NA | NA | 2900042K21Rik | | 0 | 0 | 0 | 0 | 0 | 0 | 0 | 0 | 0 | 0 | 0 |
| 0 | NA | NA | NA | NA | NA | Gm24092 | | 0 | 0 | 0 | 0 | 0 | 0 | 0 | 0 | 0 | 0 | 0 |
| 0 | NA | NA | NA | NA | NA | Gm30280 | | 0 | 0 | 0 | 0 | 0 | 0 | 0 | 0 | 0 | 0 | 0 |
| 0 | NA | NA | NA | NA | NA | Gm29678 | | 0 | 0 | 0 | 0 | 0 | 0 | 0 | 0 | 0 | 0 | 0 |
| 0 | NA | NA | NA | NA | NA | Gm47294 | | 0 | 0 | 0 | 0 | 0 | 0 | 0 | 0 | 0 | 0 | 0 |
| 0 | NA | NA | NA | NA | NA | Gm30932 | | 0 | 0 | 0 | 0 | 0 | 0 | 0 | 0 | 0 | 0 | 0 |
| 0 | NA | NA | NA | NA | NA | A230020J21Rik | | 0 | 0 | 0 | 0 | 0 | 0 | 0 | 0 | 0 | 0 | 0 |
| 0 | NA | NA | NA | NA | NA | Gm38188 | | 0 | 0 | 0 | 0 | 0 | 0 | 0 | 0 | 0 | 0 | 0 |
| 0 | NA | NA | NA | NA | NA | Gm38161 | | 0 | 0 | 0 | 0 | 0 | 0 | 0 | 0 | 0 | 0 | 0 |

| | | | | | | | | | | | | | | | | | |
|---|---|---|---|---|---|---|---|---|---|---|---|---|---|---|---|---|---|
| 0 | NA | NA | NA | NA | NA | Gm24748 | 0 | 0 | 0 | 0 | 0 | 0 | 0 | 0 | 0 | 0 | 0 |
| 0 | NA | NA | NA | NA | NA | 9430037O13Rik | 0 | 0 | 0 | 0 | 0 | 0 | 0 | 0 | 0 | 0 | 0 |
| 0 | NA | NA | NA | NA | NA | Gm22446 | 0 | 0 | 0 | 0 | 0 | 0 | 0 | 0 | 0 | 0 | 0 |
| 0 | NA | NA | NA | NA | NA | Gm26203 | 0 | 0 | 0 | 0 | 0 | 0 | 0 | 0 | 0 | 0 | 0 |
| 0 | NA | NA | NA | NA | NA | Gm20203 | 0 | 0 | 0 | 0 | 0 | 0 | 0 | 0 | 0 | 0 | 0 |
| 0 | NA | NA | NA | NA | NA | Mir3473c | 0 | 0 | 0 | 0 | 0 | 0 | 0 | 0 | 0 | 0 | 0 |
| 0 | NA | NA | NA | NA | NA | Gm22265 | 0 | 0 | 0 | 0 | 0 | 0 | 0 | 0 | 0 | 0 | 0 |
| 0 | NA | NA | NA | NA | NA | Gm38360 | 0 | 0 | 0 | 0 | 0 | 0 | 0 | 0 | 0 | 0 | 0 |
| 0 | NA | NA | NA | NA | NA | Gm23583 | 0 | 0 | 0 | 0 | 0 | 0 | 0 | 0 | 0 | 0 | 0 |
| 0 | NA | NA | NA | NA | NA | Gm15872 | 0 | 0 | 0 | 0 | 0 | 0 | 0 | 0 | 0 | 0 | 0 |
| 0 | NA | NA | NA | NA | NA | 4930570N18Rik | 0 | 0 | 0 | 0 | 0 | 0 | 0 | 0 | 0 | 0 | 0 |
| 0 | NA | NA | NA | NA | NA | Gm38118 | 0 | 0 | 0 | 0 | 0 | 0 | 0 | 0 | 0 | 0 | 0 |
| 0 | NA | NA | NA | NA | NA | Gm37650 | 0 | 0 | 0 | 0 | 0 | 0 | 0 | 0 | 0 | 0 | 0 |
| 0 | NA | NA | NA | NA | NA | Gm37942 | 0 | 0 | 0 | 0 | 0 | 0 | 0 | 0 | 0 | 0 | 0 |
| 0 | NA | NA | NA | NA | NA | Gm38022 | 0 | 0 | 0 | 0 | 0 | 0 | 0 | 0 | 0 | 0 | 0 |
| 0 | NA | NA | NA | NA | NA | Gm38322 | 0 | 0 | 0 | 0 | 0 | 0 | 0 | 0 | 0 | 0 | 0 |
| 0 | NA | NA | NA | NA | NA | Mir205 | 0 | 0 | 0 | 0 | 0 | 0 | 0 | 0 | 0 | 0 | 0 |
| 0 | NA | NA | NA | NA | NA | Gm21362 | 0 | 0 | 0 | 0 | 0 | 0 | 0 | 0 | 0 | 0 | 0 |
| 0 | NA | NA | NA | NA | NA | 4930503O07Rik | 0 | 0 | 0 | 0 | 0 | 0 | 0 | 0 | 0 | 0 | 0 |
| 0 | NA | NA | NA | NA | NA | 2900035J10Rik | 0 | 0 | 0 | 0 | 0 | 0 | 0 | 0 | 0 | 0 | 0 |
| 0 | NA | NA | NA | NA | NA | Gm19777 | 0 | 0 | 0 | 0 | 0 | 0 | 0 | 0 | 0 | 0 | 0 |
| 0 | NA | NA | NA | NA | NA | Mir3962 | 0 | 0 | 0 | 0 | 0 | 0 | 0 | 0 | 0 | 0 | 0 |
| 0 | NA | NA | NA | NA | NA | Gm23885 | 0 | 0 | 0 | 0 | 0 | 0 | 0 | 0 | 0 | 0 | 0 |
| 0 | NA | NA | NA | NA | NA | Gm32250 | 0 | 0 | 0 | 0 | 0 | 0 | 0 | 0 | 0 | 0 | 0 |
| 0 | NA | NA | NA | NA | NA | Mir29b-2 | 0 | 0 | 0 | 0 | 0 | 0 | 0 | 0 | 0 | 0 | 0 |
| 0 | NA | NA | NA | NA | NA | Mir29c | 0 | 0 | 0 | 0 | 0 | 0 | 0 | 0 | 0 | 0 | 0 |
| 0 | NA | NA | NA | NA | NA | Gm37366 | 0 | 0 | 0 | 0 | 0 | 0 | 0 | 0 | 0 | 0 | 0 |
| 0 | NA | NA | NA | NA | NA | Gm37887 | 0 | 0 | 0 | 0 | 0 | 0 | 0 | 0 | 0 | 0 | 0 |
| 0 | NA | NA | NA | NA | NA | Gm24802 | 0 | 0 | 0 | 0 | 0 | 0 | 0 | 0 | 0 | 0 | 0 |
| 0 | NA | NA | NA | NA | NA | Gm27940 | 0 | 0 | 0 | 0 | 0 | 0 | 0 | 0 | 0 | 0 | 0 |
| 0 | NA | NA | NA | NA | NA | Gm38046 | 0 | 0 | 0 | 0 | 0 | 0 | 0 | 0 | 0 | 0 | 0 |
| 0 | NA | NA | NA | NA | NA | Gm38194 | 0 | 0 | 0 | 0 | 0 | 0 | 0 | 0 | 0 | 0 | 0 |
| 0 | NA | NA | NA | NA | NA | Gm27306 | 0 | 0 | 0 | 0 | 0 | 0 | 0 | 0 | 0 | 0 | 0 |
| 0 | NA | NA | NA | NA | NA | Gm22005 | 0 | 0 | 0 | 0 | 0 | 0 | 0 | 0 | 0 | 0 | 0 |
| 0 | NA | NA | NA | NA | NA | Gm37525 | 0 | 0 | 0 | 0 | 0 | 0 | 0 | 0 | 0 | 0 | 0 |
| 0 | NA | NA | NA | NA | NA | Gm13186 | 0 | 0 | 0 | 0 | 0 | 0 | 0 | 0 | 0 | 0 | 0 |
| 0 | NA | NA | NA | NA | NA | Gm13182 | 0 | 0 | 0 | 0 | 0 | 0 | 0 | 0 | 0 | 0 | 0 |
| 0 | NA | NA | NA | NA | NA | Gm13185 | 0 | 0 | 0 | 0 | 0 | 0 | 0 | 0 | 0 | 0 | 0 |
| 0 | NA | NA | NA | NA | NA | Gm13180 | 0 | 0 | 0 | 0 | 0 | 0 | 0 | 0 | 0 | 0 | 0 |
| 0 | NA | NA | NA | NA | NA | Gm13191 | 0 | 0 | 0 | 0 | 0 | 0 | 0 | 0 | 0 | 0 | 0 |
| 0 | NA | NA | NA | NA | NA | Gm2639 | 0 | 0 | 0 | 0 | 0 | 0 | 0 | 0 | 0 | 0 | 0 |
| 0 | NA | NA | NA | NA | NA | Gm10862 | 0 | 0 | 0 | 0 | 0 | 0 | 0 | 0 | 0 | 0 | 0 |
| 0 | NA | NA | NA | NA | NA | Gm38085 | 0 | 0 | 0 | 0 | 0 | 0 | 0 | 0 | 0 | 0 | 0 |
| 0 | NA | NA | NA | NA | NA | 1700080N15Rik | 0 | 0 | 0 | 0 | 0 | 0 | 0 | 0 | 0 | 0 | 0 |
| 0 | NA | NA | NA | NA | NA | Gm13189 | 0 | 0 | 0 | 0 | 0 | 0 | 0 | 0 | 0 | 0 | 0 |
| 0 | NA | NA | NA | NA | NA | Gm37814 | 0 | 0 | 0 | 0 | 0 | 0 | 0 | 0 | 0 | 0 | 0 |
| 0 | NA | NA | NA | NA | NA | Gm38348 | 0 | 0 | 0 | 0 | 0 | 0 | 0 | 0 | 0 | 0 | 0 |
| 0 | NA | NA | NA | NA | NA | Gm23691 | 0 | 0 | 0 | 0 | 0 | 0 | 0 | 0 | 0 | 0 | 0 |
| 0 | NA | NA | NA | NA | NA | Gm13175 | 0 | 0 | 0 | 0 | 0 | 0 | 0 | 0 | 0 | 0 | 0 |
| 0 | NA | NA | NA | NA | NA | Gm13176 | 0 | 0 | 0 | 0 | 0 | 0 | 0 | 0 | 0 | 0 | 0 |
| 0 | NA | NA | NA | NA | NA | Gm13193 | 0 | 0 | 0 | 0 | 0 | 0 | 0 | 0 | 0 | 0 | 0 |
| 0 | NA | NA | NA | NA | NA | 4930551O13Rik | 0 | 0 | 0 | 0 | 0 | 0 | 0 | 0 | 0 | 0 | 0 |
| 0 | NA | NA | NA | NA | NA | Gm23118 | 0 | 0 | 0 | 0 | 0 | 0 | 0 | 0 | 0 | 0 | 0 |
| 0 | NA | NA | NA | NA | NA | Gm13197 | 0 | 0 | 0 | 0 | 0 | 0 | 0 | 0 | 0 | 0 | 0 |
| 0 | NA | NA | NA | NA | NA | Gm13267 | 0 | 0 | 0 | 0 | 0 | 0 | 0 | 0 | 0 | 0 | 0 |
| 0 | NA | NA | NA | NA | NA | Gm13384 | 0 | 0 | 0 | 0 | 0 | 0 | 0 | 0 | 0 | 0 | 0 |
| 0 | NA | NA | NA | NA | NA | Gm10857 | 0 | 0 | 0 | 0 | 0 | 0 | 0 | 0 | 0 | 0 | 0 |
| 0 | NA | NA | NA | NA | NA | A230108P19Rik | 0 | 0 | 0 | 0 | 0 | 0 | 0 | 0 | 0 | 0 | 0 |
| 0 | NA | NA | NA | NA | NA | Gm13383 | 0 | 0 | 0 | 0 | 0 | 0 | 0 | 0 | 0 | 0 | 0 |
| 0 | NA | NA | NA | NA | NA | Gm13388 | 0 | 0 | 0 | 0 | 0 | 0 | 0 | 0 | 0 | 0 | 0 |

| | | | | | | | | | | | | | | | | | |
|---|---|---|---|---|---|---|---|---|---|---|---|---|---|---|---|---|---|
| 0 | NA | NA | NA | NA | NA | Gm38386 | 0 | 0 | 0 | 0 | 0 | 0 | 0 | 0 | 0 | 0 | 0 |
| 0 | NA | NA | NA | NA | NA | Gm13389 | 0 | 0 | 0 | 0 | 0 | 0 | 0 | 0 | 0 | 0 | 0 |
| 0 | NA | NA | NA | NA | NA | Gm10855 | 0 | 0 | 0 | 0 | 0 | 0 | 0 | 0 | 0 | 0 | 0 |
| 0 | NA | NA | NA | NA | NA | Gm37340 | 0 | 0 | 0 | 0 | 0 | 0 | 0 | 0 | 0 | 0 | 0 |
| 0 | NA | NA | NA | NA | NA | Gm10115 | 0 | 0 | 0 | 0 | 0 | 0 | 0 | 0 | 0 | 0 | 0 |
| 0 | NA | NA | NA | NA | NA | Gm26315 | 0 | 0 | 0 | 0 | 0 | 0 | 0 | 0 | 0 | 0 | 0 |
| 0 | NA | NA | NA | NA | NA | Gm24340 | 0 | 0 | 0 | 0 | 0 | 0 | 0 | 0 | 0 | 0 | 0 |
| 0 | NA | NA | NA | NA | NA | Gm28641 | 0 | 0 | 0 | 0 | 0 | 0 | 0 | 0 | 0 | 0 | 0 |
| 0 | NA | NA | NA | NA | NA | Gm13210 | 0 | 0 | 0 | 0 | 0 | 0 | 0 | 0 | 0 | 0 | 0 |
| 0 | NA | NA | NA | NA | NA | Gm13211 | 0 | 0 | 0 | 0 | 0 | 0 | 0 | 0 | 0 | 0 | 0 |
| 0 | NA | NA | NA | NA | NA | Gm13254 | 0 | 0 | 0 | 0 | 0 | 0 | 0 | 0 | 0 | 0 | 0 |
| 0 | NA | NA | NA | NA | NA | Gm24534 | 0 | 0 | 0 | 0 | 0 | 0 | 0 | 0 | 0 | 0 | 0 |
| 0 | NA | NA | NA | NA | NA | Gm13255 | 0 | 0 | 0 | 0 | 0 | 0 | 0 | 0 | 0 | 0 | 0 |
| 0 | NA | NA | NA | NA | NA | 1700061F12Rik | 0 | 0 | 0 | 0 | 0 | 0 | 0 | 0 | 0 | 0 | 0 |
| 0 | NA | NA | NA | NA | NA | Gm23963 | 0 | 0 | 0 | 0 | 0 | 0 | 0 | 0 | 0 | 0 | 0 |
| 0 | NA | NA | NA | NA | NA | Gm13221 | 0 | 0 | 0 | 0 | 0 | 0 | 0 | 0 | 0 | 0 | 0 |
| 0 | NA | NA | NA | NA | NA | Gm13219 | 0 | 0 | 0 | 0 | 0 | 0 | 0 | 0 | 0 | 0 | 0 |
| 0 | NA | NA | NA | NA | NA | Gm13220 | 0 | 0 | 0 | 0 | 0 | 0 | 0 | 0 | 0 | 0 | 0 |
| 0 | NA | NA | NA | NA | NA | Gm13218 | 0 | 0 | 0 | 0 | 0 | 0 | 0 | 0 | 0 | 0 | 0 |
| 0 | NA | NA | NA | NA | NA | Gm37866 | 0 | 0 | 0 | 0 | 0 | 0 | 0 | 0 | 0 | 0 | 0 |
| 0 | NA | NA | NA | NA | NA | Gm13256 | 0 | 0 | 0 | 0 | 0 | 0 | 0 | 0 | 0 | 0 | 0 |
| 0 | NA | NA | NA | NA | NA | 4930412O13Rik | 0 | 0 | 0 | 0 | 0 | 0 | 0 | 0 | 0 | 0 | 0 |
| 0 | NA | NA | NA | NA | NA | 9230102O04Rik | 0 | 0 | 0 | 0 | 0 | 0 | 0 | 0 | 0 | 0 | 0 |
| 0 | NA | NA | NA | NA | NA | Gm13262 | 0 | 0 | 0 | 0 | 0 | 0 | 0 | 0 | 0 | 0 | 0 |
| 0 | NA | NA | NA | NA | NA | Gm22072 | 0 | 0 | 0 | 0 | 0 | 0 | 0 | 0 | 0 | 0 | 0 |
| 0 | NA | NA | NA | NA | NA | Gm22677 | 0 | 0 | 0 | 0 | 0 | 0 | 0 | 0 | 0 | 0 | 0 |
| 0 | NA | NA | NA | NA | NA | Mir466m | 0 | 0 | 0 | 0 | 0 | 0 | 0 | 0 | 0 | 0 | 0 |
| 0 | NA | NA | NA | NA | NA | Mir466f-1 | 0 | 0 | 0 | 0 | 0 | 0 | 0 | 0 | 0 | 0 | 0 |
| 0 | NA | NA | NA | NA | NA | Mir669f | 0 | 0 | 0 | 0 | 0 | 0 | 0 | 0 | 0 | 0 | 0 |
| 0 | NA | NA | NA | NA | NA | Mir669e | 0 | 0 | 0 | 0 | 0 | 0 | 0 | 0 | 0 | 0 | 0 |
| 0 | NA | NA | NA | NA | NA | Mir669b | 0 | 0 | 0 | 0 | 0 | 0 | 0 | 0 | 0 | 0 | 0 |
| 0 | NA | NA | NA | NA | NA | Gm23578 | 0 | 0 | 0 | 0 | 0 | 0 | 0 | 0 | 0 | 0 | 0 |
| 0 | NA | NA | NA | NA | NA | Mir669d | 0 | 0 | 0 | 0 | 0 | 0 | 0 | 0 | 0 | 0 | 0 |
| 0 | NA | NA | NA | NA | NA | Mir466f-2 | 0 | 0 | 0 | 0 | 0 | 0 | 0 | 0 | 0 | 0 | 0 |
| 0 | NA | NA | NA | NA | NA | Mir669l | 0 | 0 | 0 | 0 | 0 | 0 | 0 | 0 | 0 | 0 | 0 |
| 0 | NA | NA | NA | NA | NA | Mir669d-2 | 0 | 0 | 0 | 0 | 0 | 0 | 0 | 0 | 0 | 0 | 0 |
| 0 | NA | NA | NA | NA | NA | Mir466f-3 | 0 | 0 | 0 | 0 | 0 | 0 | 0 | 0 | 0 | 0 | 0 |
| 0 | NA | NA | NA | NA | NA | Mir297a-2 | 0 | 0 | 0 | 0 | 0 | 0 | 0 | 0 | 0 | 0 | 0 |
| 0 | NA | NA | NA | NA | NA | Mir466o | 0 | 0 | 0 | 0 | 0 | 0 | 0 | 0 | 0 | 0 | 0 |
| 0 | NA | NA | NA | NA | NA | Gm26073 | 0 | 0 | 0 | 0 | 0 | 0 | 0 | 0 | 0 | 0 | 0 |
| 0 | NA | NA | NA | NA | NA | Mir467c | 0 | 0 | 0 | 0 | 0 | 0 | 0 | 0 | 0 | 0 | 0 |
| 0 | NA | NA | NA | NA | NA | Mir466b-1 | 0 | 0 | 0 | 0 | 0 | 0 | 0 | 0 | 0 | 0 | 0 |
| 0 | NA | NA | NA | NA | NA | Mir669a-3 | 0 | 0 | 0 | 0 | 0 | 0 | 0 | 0 | 0 | 0 | 0 |
| 0 | NA | NA | NA | NA | NA | Mir669k | 0 | 0 | 0 | 0 | 0 | 0 | 0 | 0 | 0 | 0 | 0 |
| 0 | NA | NA | NA | NA | NA | Gm26491 | 0 | 0 | 0 | 0 | 0 | 0 | 0 | 0 | 0 | 0 | 0 |
| 0 | NA | NA | NA | NA | NA | Mir467a-1 | 0 | 0 | 0 | 0 | 0 | 0 | 0 | 0 | 0 | 0 | 0 |
| 0 | NA | NA | NA | NA | NA | Mir466b-8 | 0 | 0 | 0 | 0 | 0 | 0 | 0 | 0 | 0 | 0 | 0 |
| 0 | NA | NA | NA | NA | NA | Mir669a-1 | 0 | 0 | 0 | 0 | 0 | 0 | 0 | 0 | 0 | 0 | 0 |
| 0 | NA | NA | NA | NA | NA | Mir669g | 0 | 0 | 0 | 0 | 0 | 0 | 0 | 0 | 0 | 0 | 0 |
| 0 | NA | NA | NA | NA | NA | Gm25701 | 0 | 0 | 0 | 0 | 0 | 0 | 0 | 0 | 0 | 0 | 0 |
| 0 | NA | NA | NA | NA | NA | Mir669j | 0 | 0 | 0 | 0 | 0 | 0 | 0 | 0 | 0 | 0 | 0 |
| 0 | NA | NA | NA | NA | NA | Mir467a-2 | 0 | 0 | 0 | 0 | 0 | 0 | 0 | 0 | 0 | 0 | 0 |
| 0 | NA | NA | NA | NA | NA | Mir466e | 0 | 0 | 0 | 0 | 0 | 0 | 0 | 0 | 0 | 0 | 0 |
| 0 | NA | NA | NA | NA | NA | Mir669a-4 | 0 | 0 | 0 | 0 | 0 | 0 | 0 | 0 | 0 | 0 | 0 |
| 0 | NA | NA | NA | NA | NA | Gm24315 | 0 | 0 | 0 | 0 | 0 | 0 | 0 | 0 | 0 | 0 | 0 |
| 0 | NA | NA | NA | NA | NA | Gm25451 | 0 | 0 | 0 | 0 | 0 | 0 | 0 | 0 | 0 | 0 | 0 |
| 0 | NA | NA | NA | NA | NA | Mir467b | 0 | 0 | 0 | 0 | 0 | 0 | 0 | 0 | 0 | 0 | 0 |
| 0 | NA | NA | NA | NA | NA | Mir466c-1 | 0 | 0 | 0 | 0 | 0 | 0 | 0 | 0 | 0 | 0 | 0 |
| 0 | NA | NA | NA | NA | NA | Mir669a-5 | 0 | 0 | 0 | 0 | 0 | 0 | 0 | 0 | 0 | 0 | 0 |
| 0 | NA | NA | NA | NA | NA | Gm24761 | 0 | 0 | 0 | 0 | 0 | 0 | 0 | 0 | 0 | 0 | 0 |

| | | | | | | | | | | | | | | | | | |
|---|---|---|---|---|---|---|---|---|---|---|---|---|---|---|---|---|---|
| 0 | NA | NA | NA | NA | NA | Gm23651 | 0 | 0 | 0 | 0 | 0 | 0 | 0 | 0 | 0 | 0 | 0 |
| 0 | NA | NA | NA | NA | NA | Mir467a-3 | 0 | 0 | 0 | 0 | 0 | 0 | 0 | 0 | 0 | 0 | 0 |
| 0 | NA | NA | NA | NA | NA | Mir466c-2 | 0 | 0 | 0 | 0 | 0 | 0 | 0 | 0 | 0 | 0 | 0 |
| 0 | NA | NA | NA | NA | NA | Mir669a-6 | 0 | 0 | 0 | 0 | 0 | 0 | 0 | 0 | 0 | 0 | 0 |
| 0 | NA | NA | NA | NA | NA | Gm23108 | 0 | 0 | 0 | 0 | 0 | 0 | 0 | 0 | 0 | 0 | 0 |
| 0 | NA | NA | NA | NA | NA | Gm25680 | 0 | 0 | 0 | 0 | 0 | 0 | 0 | 0 | 0 | 0 | 0 |
| 0 | NA | NA | NA | NA | NA | Gm22775 | 0 | 0 | 0 | 0 | 0 | 0 | 0 | 0 | 0 | 0 | 0 |
| 0 | NA | NA | NA | NA | NA | Mir467a-4 | 0 | 0 | 0 | 0 | 0 | 0 | 0 | 0 | 0 | 0 | 0 |
| 0 | NA | NA | NA | NA | NA | Mir466b-4 | 0 | 0 | 0 | 0 | 0 | 0 | 0 | 0 | 0 | 0 | 0 |
| 0 | NA | NA | NA | NA | NA | Mir669a-7 | 0 | 0 | 0 | 0 | 0 | 0 | 0 | 0 | 0 | 0 | 0 |
| 0 | NA | NA | NA | NA | NA | Gm26060 | 0 | 0 | 0 | 0 | 0 | 0 | 0 | 0 | 0 | 0 | 0 |
| 0 | NA | NA | NA | NA | NA | Gm26431 | 0 | 0 | 0 | 0 | 0 | 0 | 0 | 0 | 0 | 0 | 0 |
| 0 | NA | NA | NA | NA | NA | Gm26229 | 0 | 0 | 0 | 0 | 0 | 0 | 0 | 0 | 0 | 0 | 0 |
| 0 | NA | NA | NA | NA | NA | Mir467a-5 | 0 | 0 | 0 | 0 | 0 | 0 | 0 | 0 | 0 | 0 | 0 |
| 0 | NA | NA | NA | NA | NA | Mir466b-5 | 0 | 0 | 0 | 0 | 0 | 0 | 0 | 0 | 0 | 0 | 0 |
| 0 | NA | NA | NA | NA | NA | Mir669p-1 | 0 | 0 | 0 | 0 | 0 | 0 | 0 | 0 | 0 | 0 | 0 |
| 0 | NA | NA | NA | NA | NA | Gm24167 | 0 | 0 | 0 | 0 | 0 | 0 | 0 | 0 | 0 | 0 | 0 |
| 0 | NA | NA | NA | NA | NA | Gm24640 | 0 | 0 | 0 | 0 | 0 | 0 | 0 | 0 | 0 | 0 | 0 |
| 0 | NA | NA | NA | NA | NA | Gm23288 | 0 | 0 | 0 | 0 | 0 | 0 | 0 | 0 | 0 | 0 | 0 |
| 0 | NA | NA | NA | NA | NA | Mir467a-6 | 0 | 0 | 0 | 0 | 0 | 0 | 0 | 0 | 0 | 0 | 0 |
| 0 | NA | NA | NA | NA | NA | Gm25285 | 0 | 0 | 0 | 0 | 0 | 0 | 0 | 0 | 0 | 0 | 0 |
| 0 | NA | NA | NA | NA | NA | Mir669a-8 | 0 | 0 | 0 | 0 | 0 | 0 | 0 | 0 | 0 | 0 | 0 |
| 0 | NA | NA | NA | NA | NA | Gm24673 | 0 | 0 | 0 | 0 | 0 | 0 | 0 | 0 | 0 | 0 | 0 |
| 0 | NA | NA | NA | NA | NA | Gm22635 | 0 | 0 | 0 | 0 | 0 | 0 | 0 | 0 | 0 | 0 | 0 |
| 0 | NA | NA | NA | NA | NA | Gm23569 | 0 | 0 | 0 | 0 | 0 | 0 | 0 | 0 | 0 | 0 | 0 |
| 0 | NA | NA | NA | NA | NA | Mir467a-7 | 0 | 0 | 0 | 0 | 0 | 0 | 0 | 0 | 0 | 0 | 0 |
| 0 | NA | NA | NA | NA | NA | Mir466b-6 | 0 | 0 | 0 | 0 | 0 | 0 | 0 | 0 | 0 | 0 | 0 |
| 0 | NA | NA | NA | NA | NA | Mir669a-9 | 0 | 0 | 0 | 0 | 0 | 0 | 0 | 0 | 0 | 0 | 0 |
| 0 | NA | NA | NA | NA | NA | Gm26426 | 0 | 0 | 0 | 0 | 0 | 0 | 0 | 0 | 0 | 0 | 0 |
| 0 | NA | NA | NA | NA | NA | Gm24488 | 0 | 0 | 0 | 0 | 0 | 0 | 0 | 0 | 0 | 0 | 0 |
| 0 | NA | NA | NA | NA | NA | Gm25158 | 0 | 0 | 0 | 0 | 0 | 0 | 0 | 0 | 0 | 0 | 0 |
| 0 | NA | NA | NA | NA | NA | Mir467a-8 | 0 | 0 | 0 | 0 | 0 | 0 | 0 | 0 | 0 | 0 | 0 |
| 0 | NA | NA | NA | NA | NA | Mir466b-7 | 0 | 0 | 0 | 0 | 0 | 0 | 0 | 0 | 0 | 0 | 0 |
| 0 | NA | NA | NA | NA | NA | Mir669p-2 | 0 | 0 | 0 | 0 | 0 | 0 | 0 | 0 | 0 | 0 | 0 |
| 0 | NA | NA | NA | NA | NA | Gm24028 | 0 | 0 | 0 | 0 | 0 | 0 | 0 | 0 | 0 | 0 | 0 |
| 0 | NA | NA | NA | NA | NA | Gm22626 | 0 | 0 | 0 | 0 | 0 | 0 | 0 | 0 | 0 | 0 | 0 |
| 0 | NA | NA | NA | NA | NA | Gm22166 | 0 | 0 | 0 | 0 | 0 | 0 | 0 | 0 | 0 | 0 | 0 |
| 0 | NA | NA | NA | NA | NA | Mir467a-9 | 0 | 0 | 0 | 0 | 0 | 0 | 0 | 0 | 0 | 0 | 0 |
| 0 | NA | NA | NA | NA | NA | Mir466b-2 | 0 | 0 | 0 | 0 | 0 | 0 | 0 | 0 | 0 | 0 | 0 |
| 0 | NA | NA | NA | NA | NA | Mir669a-10 | 0 | 0 | 0 | 0 | 0 | 0 | 0 | 0 | 0 | 0 | 0 |
| 0 | NA | NA | NA | NA | NA | Gm26156 | 0 | 0 | 0 | 0 | 0 | 0 | 0 | 0 | 0 | 0 | 0 |
| 0 | NA | NA | NA | NA | NA | Gm23571 | 0 | 0 | 0 | 0 | 0 | 0 | 0 | 0 | 0 | 0 | 0 |
| 0 | NA | NA | NA | NA | NA | Gm25316 | 0 | 0 | 0 | 0 | 0 | 0 | 0 | 0 | 0 | 0 | 0 |
| 0 | NA | NA | NA | NA | NA | Gm25022 | 0 | 0 | 0 | 0 | 0 | 0 | 0 | 0 | 0 | 0 | 0 |
| 0 | NA | NA | NA | NA | NA | Gm25943 | 0 | 0 | 0 | 0 | 0 | 0 | 0 | 0 | 0 | 0 | 0 |
| 0 | NA | NA | NA | NA | NA | Mir669a-11 | 0 | 0 | 0 | 0 | 0 | 0 | 0 | 0 | 0 | 0 | 0 |
| 0 | NA | NA | NA | NA | NA | Gm23988 | 0 | 0 | 0 | 0 | 0 | 0 | 0 | 0 | 0 | 0 | 0 |
| 0 | NA | NA | NA | NA | NA | Gm25673 | 0 | 0 | 0 | 0 | 0 | 0 | 0 | 0 | 0 | 0 | 0 |
| 0 | NA | NA | NA | NA | NA | Gm25753 | 0 | 0 | 0 | 0 | 0 | 0 | 0 | 0 | 0 | 0 | 0 |
| 0 | NA | NA | NA | NA | NA | Mir467a-10 | 0 | 0 | 0 | 0 | 0 | 0 | 0 | 0 | 0 | 0 | 0 |
| 0 | NA | NA | NA | NA | NA | Mir466b-3 | 0 | 0 | 0 | 0 | 0 | 0 | 0 | 0 | 0 | 0 | 0 |
| 0 | NA | NA | NA | NA | NA | Mir669a-12 | 0 | 0 | 0 | 0 | 0 | 0 | 0 | 0 | 0 | 0 | 0 |
| 0 | NA | NA | NA | NA | NA | Gm26398 | 0 | 0 | 0 | 0 | 0 | 0 | 0 | 0 | 0 | 0 | 0 |
| 0 | NA | NA | NA | NA | NA | Mir467e | 0 | 0 | 0 | 0 | 0 | 0 | 0 | 0 | 0 | 0 | 0 |
| 0 | NA | NA | NA | NA | NA | Mir466p | 0 | 0 | 0 | 0 | 0 | 0 | 0 | 0 | 0 | 0 | 0 |
| 0 | NA | NA | NA | NA | NA | Gm22051 | 0 | 0 | 0 | 0 | 0 | 0 | 0 | 0 | 0 | 0 | 0 |
| 0 | NA | NA | NA | NA | NA | Gm22678 | 0 | 0 | 0 | 0 | 0 | 0 | 0 | 0 | 0 | 0 | 0 |
| 0 | NA | NA | NA | NA | NA | Mir467d | 0 | 0 | 0 | 0 | 0 | 0 | 0 | 0 | 0 | 0 | 0 |
| 0 | NA | NA | NA | NA | NA | Mir466 | 0 | 0 | 0 | 0 | 0 | 0 | 0 | 0 | 0 | 0 | 0 |
| 0 | NA | NA | NA | NA | NA | Gm22091 | 0 | 0 | 0 | 0 | 0 | 0 | 0 | 0 | 0 | 0 | 0 |

| | | | | | | | | | | | | | | | | | |
|---|---|---|---|---|---|---|---|---|---|---|---|---|---|---|---|---|---|
| 0 | NA | NA | NA | NA | NA | Mir297c | 0 | 0 | 0 | 0 | 0 | 0 | 0 | 0 | 0 | 0 | 0 |
| 0 | NA | NA | NA | NA | NA | Mir669c | 0 | 0 | 0 | 0 | 0 | 0 | 0 | 0 | 0 | 0 | 0 |
| 0 | NA | NA | NA | NA | NA | Gm24342 | 0 | 0 | 0 | 0 | 0 | 0 | 0 | 0 | 0 | 0 | 0 |
| 0 | NA | NA | NA | NA | NA | Gm25263 | 0 | 0 | 0 | 0 | 0 | 0 | 0 | 0 | 0 | 0 | 0 |
| 0 | NA | NA | NA | NA | NA | Mir669a-2 | 0 | 0 | 0 | 0 | 0 | 0 | 0 | 0 | 0 | 0 | 0 |
| 0 | NA | NA | NA | NA | NA | Gm22239 | 0 | 0 | 0 | 0 | 0 | 0 | 0 | 0 | 0 | 0 | 0 |
| 0 | NA | NA | NA | NA | NA | Gm22925 | 0 | 0 | 0 | 0 | 0 | 0 | 0 | 0 | 0 | 0 | 0 |
| 0 | NA | NA | NA | NA | NA | Gm23542 | 0 | 0 | 0 | 0 | 0 | 0 | 0 | 0 | 0 | 0 | 0 |
| 0 | NA | NA | NA | NA | NA | Mir297b | 0 | 0 | 0 | 0 | 0 | 0 | 0 | 0 | 0 | 0 | 0 |
| 0 | NA | NA | NA | NA | NA | Mir466d | 0 | 0 | 0 | 0 | 0 | 0 | 0 | 0 | 0 | 0 | 0 |
| 0 | NA | NA | NA | NA | NA | Gm26092 | 0 | 0 | 0 | 0 | 0 | 0 | 0 | 0 | 0 | 0 | 0 |
| 0 | NA | NA | NA | NA | NA | Mir669m-1 | 0 | 0 | 0 | 0 | 0 | 0 | 0 | 0 | 0 | 0 | 0 |
| 0 | NA | NA | NA | NA | NA | Mir669m-2 | 0 | 0 | 0 | 0 | 0 | 0 | 0 | 0 | 0 | 0 | 0 |
| 0 | NA | NA | NA | NA | NA | Mir466n | 0 | 0 | 0 | 0 | 0 | 0 | 0 | 0 | 0 | 0 | 0 |
| 0 | NA | NA | NA | NA | NA | Mir669o | 0 | 0 | 0 | 0 | 0 | 0 | 0 | 0 | 0 | 0 | 0 |
| 0 | NA | NA | NA | NA | NA | Mir466g | 0 | 0 | 0 | 0 | 0 | 0 | 0 | 0 | 0 | 0 | 0 |
| 0 | NA | NA | NA | NA | NA | Mir466h | 0 | 0 | 0 | 0 | 0 | 0 | 0 | 0 | 0 | 0 | 0 |
| 0 | NA | NA | NA | NA | NA | Mir297a-3 | 0 | 0 | 0 | 0 | 0 | 0 | 0 | 0 | 0 | 0 | 0 |
| 0 | NA | NA | NA | NA | NA | Mir466l | 0 | 0 | 0 | 0 | 0 | 0 | 0 | 0 | 0 | 0 | 0 |
| 0 | NA | NA | NA | NA | NA | Mir297a-4 | 0 | 0 | 0 | 0 | 0 | 0 | 0 | 0 | 0 | 0 | 0 |
| 0 | NA | NA | NA | NA | NA | Gm24806 | 0 | 0 | 0 | 0 | 0 | 0 | 0 | 0 | 0 | 0 | 0 |
| 0 | NA | NA | NA | NA | NA | Mir669i | 0 | 0 | 0 | 0 | 0 | 0 | 0 | 0 | 0 | 0 | 0 |
| 0 | NA | NA | NA | NA | NA | Mir669h | 0 | 0 | 0 | 0 | 0 | 0 | 0 | 0 | 0 | 0 | 0 |
| 0 | NA | NA | NA | NA | NA | Gm18547 | 0 | 0 | 0 | 0 | 0 | 0 | 0 | 0 | 0 | 0 | 0 |
| 0 | NA | NA | NA | NA | NA | Gm13265 | 0 | 0 | 0 | 0 | 0 | 0 | 0 | 0 | 0 | 0 | 0 |
| 0 | NA | NA | NA | NA | NA | Gm13263 | 0 | 0 | 0 | 0 | 0 | 0 | 0 | 0 | 0 | 0 | 0 |
| 0 | NA | NA | NA | NA | NA | Gm37343 | 0 | 0 | 0 | 0 | 0 | 0 | 0 | 0 | 0 | 0 | 0 |
| 0 | NA | NA | NA | NA | NA | Gm23877 | 0 | 0 | 0 | 0 | 0 | 0 | 0 | 0 | 0 | 0 | 0 |
| 0 | NA | NA | NA | NA | NA | Gm13264 | 0 | 0 | 0 | 0 | 0 | 0 | 0 | 0 | 0 | 0 | 0 |
| 0 | NA | NA | NA | NA | NA | Gm26478 | 0 | 0 | 0 | 0 | 0 | 0 | 0 | 0 | 0 | 0 | 0 |
| 0 | NA | NA | NA | NA | NA | Gm36932 | 0 | 0 | 0 | 0 | 0 | 0 | 0 | 0 | 0 | 0 | 0 |
| 0 | NA | NA | NA | NA | NA | Gm37520 | 0 | 0 | 0 | 0 | 0 | 0 | 0 | 0 | 0 | 0 | 0 |
| 0 | NA | NA | NA | NA | NA | Gm37766 | 0 | 0 | 0 | 0 | 0 | 0 | 0 | 0 | 0 | 0 | 0 |
| 0 | NA | NA | NA | NA | NA | Gm38171 | 0 | 0 | 0 | 0 | 0 | 0 | 0 | 0 | 0 | 0 | 0 |
| 0 | NA | NA | NA | NA | NA | Gm37851 | 0 | 0 | 0 | 0 | 0 | 0 | 0 | 0 | 0 | 0 | 0 |
| 0 | NA | NA | NA | NA | NA | 8030442B05Rik | 0 | 0 | 0 | 0 | 0 | 0 | 0 | 0 | 0 | 0 | 0 |
| 0 | NA | NA | NA | NA | NA | 4933403L11Rik | 0 | 0 | 0 | 0 | 0 | 0 | 0 | 0 | 0 | 0 | 0 |
| 0 | NA | NA | NA | NA | NA | Gm13291 | 0 | 0 | 0 | 0 | 0 | 0 | 0 | 0 | 0 | 0 | 0 |
| 0 | NA | NA | NA | NA | NA | Gm10851 | 0 | 0 | 0 | 0 | 0 | 0 | 0 | 0 | 0 | 0 | 0 |
| 0 | NA | NA | NA | NA | NA | Gm13260 | 0 | 0 | 0 | 0 | 0 | 0 | 0 | 0 | 0 | 0 | 0 |
| 0 | NA | NA | NA | NA | NA | Gm17490 | 0 | 0 | 0 | 0 | 0 | 0 | 0 | 0 | 0 | 0 | 0 |
| 0 | NA | NA | NA | NA | NA | Gm37881 | 0 | 0 | 0 | 0 | 0 | 0 | 0 | 0 | 0 | 0 | 0 |
| 0 | NA | NA | NA | NA | NA | Gm34768 | 0 | 0 | 0 | 0 | 0 | 0 | 0 | 0 | 0 | 0 | 0 |
| 0 | NA | NA | NA | NA | NA | Gm37701 | 0 | 0 | 0 | 0 | 0 | 0 | 0 | 0 | 0 | 0 | 0 |
| 0 | NA | NA | NA | NA | NA | Gm37565 | 0 | 0 | 0 | 0 | 0 | 0 | 0 | 0 | 0 | 0 | 0 |
| 0 | NA | NA | NA | NA | NA | Gm37356 | 0 | 0 | 0 | 0 | 0 | 0 | 0 | 0 | 0 | 0 | 0 |
| 0 | NA | NA | NA | NA | NA | Gm37742 | 0 | 0 | 0 | 0 | 0 | 0 | 0 | 0 | 0 | 0 | 0 |
| 0 | NA | NA | NA | NA | NA | Gm38156 | 0 | 0 | 0 | 0 | 0 | 0 | 0 | 0 | 0 | 0 | 0 |
| 0 | NA | NA | NA | NA | NA | Gm27540 | 0 | 0 | 0 | 0 | 0 | 0 | 0 | 0 | 0 | 0 | 0 |
| 0 | NA | NA | NA | NA | NA | Mir511 | 0 | 0 | 0 | 0 | 0 | 0 | 0 | 0 | 0 | 0 | 0 |
| 0 | NA | NA | NA | NA | NA | Gm37697 | 0 | 0 | 0 | 0 | 0 | 0 | 0 | 0 | 0 | 0 | 0 |
| 0 | NA | NA | NA | NA | NA | Gm13266 | 0 | 0 | 0 | 0 | 0 | 0 | 0 | 0 | 0 | 0 | 0 |
| 0 | NA | NA | NA | NA | NA | Gm13314 | 0 | 0 | 0 | 0 | 0 | 0 | 0 | 0 | 0 | 0 | 0 |
| 0 | NA | NA | NA | NA | NA | Gm10848 | 0 | 0 | 0 | 0 | 0 | 0 | 0 | 0 | 0 | 0 | 0 |
| 0 | NA | NA | NA | NA | NA | Gm13269 | 0 | 0 | 0 | 0 | 0 | 0 | 0 | 0 | 0 | 0 | 0 |
| 0 | NA | NA | NA | NA | NA | Gm13316 | 0 | 0 | 0 | 0 | 0 | 0 | 0 | 0 | 0 | 0 | 0 |
| 0 | NA | NA | NA | NA | NA | Gm36930 | 0 | 0 | 0 | 0 | 0 | 0 | 0 | 0 | 0 | 0 | 0 |
| 0 | NA | NA | NA | NA | NA | Gm13367 | 0 | 0 | 0 | 0 | 0 | 0 | 0 | 0 | 0 | 0 | 0 |
| 0 | NA | NA | NA | NA | NA | Gm13366 | 0 | 0 | 0 | 0 | 0 | 0 | 0 | 0 | 0 | 0 | 0 |
| 0 | NA | NA | NA | NA | NA | Gm13365 | 0 | 0 | 0 | 0 | 0 | 0 | 0 | 0 | 0 | 0 | 0 |

| | | | | | | | | | | | | | | | | | | |
|---|---|---|---|---|---|---|---|---|---|---|---|---|---|---|---|---|---|---|
| 0 | NA | NA | NA | NA | NA | Gm37595 | | 0 | 0 | 0 | 0 | 0 | 0 | 0 | 0 | 0 | 0 | 0 |
| 0 | NA | NA | NA | NA | NA | Gm6587 | | 0 | 0 | 0 | 0 | 0 | 0 | 0 | 0 | 0 | 0 | 0 |
| 0 | NA | NA | NA | NA | NA | Gm37416 | | 0 | 0 | 0 | 0 | 0 | 0 | 0 | 0 | 0 | 0 | 0 |
| 0 | NA | NA | NA | NA | NA | Gm24933 | | 0 | 0 | 0 | 0 | 0 | 0 | 0 | 0 | 0 | 0 | 0 |
| 0 | NA | NA | NA | NA | NA | Gm13325 | | 0 | 0 | 0 | 0 | 0 | 0 | 0 | 0 | 0 | 0 | 0 |
| 0 | NA | NA | NA | NA | NA | Gm13322 | | 0 | 0 | 0 | 0 | 0 | 0 | 0 | 0 | 0 | 0 | 0 |
| 0 | NA | NA | NA | NA | NA | Gm13324 | | 0 | 0 | 0 | 0 | 0 | 0 | 0 | 0 | 0 | 0 | 0 |
| 0 | NA | NA | NA | NA | NA | Mir6419 | | 0 | 0 | 0 | 0 | 0 | 0 | 0 | 0 | 0 | 0 | 0 |
| 0 | NA | NA | NA | NA | NA | Gm13323 | | 0 | 0 | 0 | 0 | 0 | 0 | 0 | 0 | 0 | 0 | 0 |
| 0 | NA | NA | NA | NA | NA | H2al2a | H2A histone family member L2A | 0 | 0 | 0 | 0 | 0 | 0 | 0 | 0 | 0 | 0 | 0 |
| 0 | NA | NA | NA | NA | NA | Gm13329 | | 0 | 0 | 0 | 0 | 0 | 0 | 0 | 0 | 0 | 0 | 0 |
| 0 | NA | NA | NA | NA | NA | Gm23638 | | 0 | 0 | 0 | 0 | 0 | 0 | 0 | 0 | 0 | 0 | 0 |
| 0 | NA | NA | NA | NA | NA | Gm25147 | | 0 | 0 | 0 | 0 | 0 | 0 | 0 | 0 | 0 | 0 | 0 |
| 0 | NA | NA | NA | NA | NA | Gm13353 | | 0 | 0 | 0 | 0 | 0 | 0 | 0 | 0 | 0 | 0 | 0 |
| 0 | NA | NA | NA | NA | NA | Gm13355 | | 0 | 0 | 0 | 0 | 0 | 0 | 0 | 0 | 0 | 0 | 0 |
| 0 | NA | NA | NA | NA | NA | Gm13333 | | 0 | 0 | 0 | 0 | 0 | 0 | 0 | 0 | 0 | 0 | 0 |
| 0 | NA | NA | NA | NA | NA | Carlr | | 0 | 0 | 0 | 0 | 0 | 0 | 0 | 0 | 0 | 0 | 0 |
| 0 | NA | NA | NA | NA | NA | Ptf1aos | | 0 | 0 | 0 | 0 | 0 | 0 | 0 | 0 | 0 | 0 | 0 |
| 0 | NA | NA | NA | NA | NA | 4921504E06Ril | RIKEN cDNA 4921504E06 gene | 0 | 0 | 0 | 0 | 0 | 0 | 0 | 0 | 0 | 0 | 0 |
| 0 | NA | NA | NA | NA | NA | Gm13345 | | 0 | 0 | 0 | 0 | 0 | 0 | 0 | 0 | 0 | 0 | 0 |
| 0 | NA | NA | NA | NA | NA | Gm13348 | | 0 | 0 | 0 | 0 | 0 | 0 | 0 | 0 | 0 | 0 | 0 |
| 0 | NA | NA | NA | NA | NA | Gm13343 | | 0 | 0 | 0 | 0 | 0 | 0 | 0 | 0 | 0 | 0 | 0 |
| 0 | NA | NA | NA | NA | NA | Gm13346 | | 0 | 0 | 0 | 0 | 0 | 0 | 0 | 0 | 0 | 0 | 0 |
| 0 | NA | NA | NA | NA | NA | Gm13347 | | 0 | 0 | 0 | 0 | 0 | 0 | 0 | 0 | 0 | 0 | 0 |
| 0 | NA | NA | NA | NA | NA | Gm24670 | | 0 | 0 | 0 | 0 | 0 | 0 | 0 | 0 | 0 | 0 | 0 |
| 0 | NA | NA | NA | NA | NA | Gm44457 | | 0 | 0 | 0 | 0 | 0 | 0 | 0 | 0 | 0 | 0 | 0 |
| 0 | NA | NA | NA | NA | NA | Gm25859 | | 0 | 0 | 0 | 0 | 0 | 0 | 0 | 0 | 0 | 0 | 0 |
| 0 | NA | NA | NA | NA | NA | Gm13361 | | 0 | 0 | 0 | 0 | 0 | 0 | 0 | 0 | 0 | 0 | 0 |
| 0 | NA | NA | NA | NA | NA | Gm23970 | | 0 | 0 | 0 | 0 | 0 | 0 | 0 | 0 | 0 | 0 | 0 |
| 0 | NA | NA | NA | NA | NA | Gm17171 | | 0 | 0 | 0 | 0 | 0 | 0 | 0 | 0 | 0 | 0 | 0 |
| 0 | NA | NA | NA | NA | NA | Gm13335 | | 0 | 0 | 0 | 0 | 0 | 0 | 0 | 0 | 0 | 0 | 0 |
| 0 | NA | NA | NA | NA | NA | Gm27446 | | 0 | 0 | 0 | 0 | 0 | 0 | 0 | 0 | 0 | 0 | 0 |
| 0 | NA | NA | NA | NA | NA | Gm13377 | | 0 | 0 | 0 | 0 | 0 | 0 | 0 | 0 | 0 | 0 | 0 |
| 0 | NA | NA | NA | NA | NA | Gm13376 | | 0 | 0 | 0 | 0 | 0 | 0 | 0 | 0 | 0 | 0 | 0 |
| 0 | NA | NA | NA | NA | NA | Gm25368 | | 0 | 0 | 0 | 0 | 0 | 0 | 0 | 0 | 0 | 0 | 0 |
| 0 | NA | NA | NA | NA | NA | Gm23007 | | 0 | 0 | 0 | 0 | 0 | 0 | 0 | 0 | 0 | 0 | 0 |
| 0 | NA | NA | NA | NA | NA | Gm24886 | | 0 | 0 | 0 | 0 | 0 | 0 | 0 | 0 | 0 | 0 | 0 |
| 0 | NA | NA | NA | NA | NA | Gm13337 | | 0 | 0 | 0 | 0 | 0 | 0 | 0 | 0 | 0 | 0 | 0 |
| 0 | NA | NA | NA | NA | NA | Mir3967 | | 0 | 0 | 0 | 0 | 0 | 0 | 0 | 0 | 0 | 0 | 0 |
| 0 | NA | NA | NA | NA | NA | Gm23636 | | 0 | 0 | 0 | 0 | 0 | 0 | 0 | 0 | 0 | 0 | 0 |
| 0 | NA | NA | NA | NA | NA | Gm13338 | | 0 | 0 | 0 | 0 | 0 | 0 | 0 | 0 | 0 | 0 | 0 |
| 0 | NA | NA | NA | NA | NA | 1700092C17Rik | | 0 | 0 | 0 | 0 | 0 | 0 | 0 | 0 | 0 | 0 | 0 |
| 0 | NA | NA | NA | NA | NA | Gm13356 | | 0 | 0 | 0 | 0 | 0 | 0 | 0 | 0 | 0 | 0 | 0 |
| 0 | NA | NA | NA | NA | NA | Gm25544 | | 0 | 0 | 0 | 0 | 0 | 0 | 0 | 0 | 0 | 0 | 0 |
| 0 | NA | NA | NA | NA | NA | Gm38309 | | 0 | 0 | 0 | 0 | 0 | 0 | 0 | 0 | 0 | 0 | 0 |
| 0 | NA | NA | NA | NA | NA | Sh3d2c-ps1 | | 0 | 0 | 0 | 0 | 0 | 0 | 0 | 0 | 0 | 0 | 0 |
| 0 | NA | NA | NA | NA | NA | Gm26323 | | 0 | 0 | 0 | 0 | 0 | 0 | 0 | 0 | 0 | 0 | 0 |
| 0 | NA | NA | NA | NA | NA | Tbpl2 | TATA box binding protein like 2 | 0 | 0 | 0 | 0 | 0 | 0 | 0 | 0 | 0 | 0 | 0 |
| 0 | NA | NA | NA | NA | NA | Il1f8 | interleukin 1 family, member 8 | 0 | 0 | 0 | 0 | 0 | 0 | 0 | 0 | 0 | 0 | 0 |
| 0 | NA | NA | NA | NA | NA | Il1f9 | interleukin 1 family, member 9 | 0 | 0 | 0 | 0 | 0 | 0 | 0 | 0 | 0 | 0 | 0 |
| 0 | NA | NA | NA | NA | NA | Il1f6 | interleukin 1 family, member 6 | 0 | 0 | 0 | 0 | 0 | 0 | 0 | 0 | 0 | 0 | 0 |
| 0 | NA | NA | NA | NA | NA | Gm37703 | | 0 | 0 | 0 | 0 | 0 | 0 | 0 | 0 | 0 | 0 | 0 |
| 0 | NA | NA | NA | NA | NA | Gm13409 | | 0 | 0 | 0 | 0 | 0 | 0 | 0 | 0 | 0 | 0 | 0 |
| 0 | NA | NA | NA | NA | NA | Il1f5 | interleukin 1 family, member 5 (delta) | 0 | 0 | 0 | 0 | 0 | 0 | 0 | 0 | 0 | 0 | 0 |
| 0 | NA | NA | NA | NA | NA | Gm13411 | | 0 | 0 | 0 | 0 | 0 | 0 | 0 | 0 | 0 | 0 | 0 |
| 0 | NA | NA | NA | NA | NA | Gm13415 | | 0 | 0 | 0 | 0 | 0 | 0 | 0 | 0 | 0 | 0 | 0 |
| 0 | NA | NA | NA | NA | NA | Gm27193 | | 0 | 0 | 0 | 0 | 0 | 0 | 0 | 0 | 0 | 0 | 0 |
| 0 | NA | NA | NA | NA | NA | Gm13417 | | 0 | 0 | 0 | 0 | 0 | 0 | 0 | 0 | 0 | 0 | 0 |
| 0 | NA | NA | NA | NA | NA | Gm38383 | | 0 | 0 | 0 | 0 | 0 | 0 | 0 | 0 | 0 | 0 | 0 |
| 0 | NA | NA | NA | NA | NA | Gm25455 | | 0 | 0 | 0 | 0 | 0 | 0 | 0 | 0 | 0 | 0 | 0 |

| | | | | | | | | | | | | | | | | | | |
|---|---|---|---|---|---|---|---|---|---|---|---|---|---|---|---|---|---|---|
| 0 | NA | NA | NA | NA | NA | Gm13459 | | 0 | 0 | 0 | 0 | 0 | 0 | 0 | 0 | 0 | 0 | 0 |
| 0 | NA | NA | NA | NA | NA | Gm25518 | | 0 | 0 | 0 | 0 | 0 | 0 | 0 | 0 | 0 | 0 | 0 |
| 0 | NA | NA | NA | NA | NA | Gm24875 | | 0 | 0 | 0 | 0 | 0 | 0 | 0 | 0 | 0 | 0 | 0 |
| 0 | NA | NA | NA | NA | NA | Gm24328 | | 0 | 0 | 0 | 0 | 0 | 0 | 0 | 0 | 0 | 0 | 0 |
| 0 | NA | NA | NA | NA | NA | Mir7664 | | 0 | 0 | 0 | 0 | 0 | 0 | 0 | 0 | 0 | 0 | 0 |
| 0 | NA | NA | NA | NA | NA | Noxa1 | NADPH oxidase activator 1 | 0 | 0 | 0 | 0 | 0 | 0 | 0 | 0 | 0 | 0 | 0 |
| 0 | NA | NA | NA | NA | NA | | | 0 | 0 | 0 | 0 | 0 | 0 | 0 | 0 | 0 | 0 | 0 |
| 0 | NA | NA | NA | NA | NA | Gm22572 | | 0 | 0 | 0 | 0 | 0 | 0 | 0 | 0 | 0 | 0 | 0 |
| 0 | NA | NA | NA | NA | NA | Slc34a3 | solute carrier family 34 (sodium phosphat | 0 | 0 | 0 | 0 | 0 | 0 | 0 | 0 | 0 | 0 | 0 |
| 0 | NA | NA | NA | NA | NA | Rnf224 | ring finger protein 224 | 0 | 0 | 0 | 0 | 0 | 0 | 0 | 0 | 0 | 0 | 0 |
| 0 | NA | NA | NA | NA | NA | Ndor1 | | 0 | 0 | 0 | 0 | 0 | 0 | 0 | 0 | 0 | 0 | 0 |
| 0 | NA | NA | NA | NA | NA | Ndor1 | NADPH dependent diflavin oxidoreducta | 0 | 0 | 0 | 0 | 0 | 0 | 0 | 0 | 0 | 0 | 0 |
| 0 | NA | NA | NA | NA | NA | Tmem210 | transmembrane protein 210 | 0 | 0 | 0 | 0 | 0 | 0 | 0 | 0 | 0 | 0 | 0 |
| 0 | NA | NA | NA | NA | NA | Mir3087 | | 0 | 0 | 0 | 0 | 0 | 0 | 0 | 0 | 0 | 0 | 0 |
| 0 | NA | NA | NA | NA | NA | Lcn12 | lipocalin 12 | 0 | 0 | 0 | 0 | 0 | 0 | 0 | 0 | 0 | 0 | 0 |
| 0 | NA | NA | NA | NA | NA | Gm13432 | | 0 | 0 | 0 | 0 | 0 | 0 | 0 | 0 | 0 | 0 | 0 |
| 0 | NA | NA | NA | NA | NA | Ccdc183 | coiled-coil domain containing 183 | 0 | 0 | 0 | 0 | 0 | 0 | 0 | 0 | 0 | 0 | 0 |
| 0 | NA | NA | NA | NA | NA | Lcn15 | | 0 | 0 | 0 | 0 | 0 | 0 | 0 | 0 | 0 | 0 | 0 |
| 0 | NA | NA | NA | NA | NA | Lcn8 | lipocalin 8 | 0 | 0 | 0 | 0 | 0 | 0 | 0 | 0 | 0 | 0 | 0 |
| 0 | NA | NA | NA | NA | NA | Lcn10 | lipocalin 10 | 0 | 0 | 0 | 0 | 0 | 0 | 0 | 0 | 0 | 0 | 0 |
| 0 | NA | NA | NA | NA | NA | Obp2a | odorant binding protein 2A | 0 | 0 | 0 | 0 | 0 | 0 | 0 | 0 | 0 | 0 | 0 |
| 0 | NA | NA | NA | NA | NA | Lcn3 | lipocalin 3 | 0 | 0 | 0 | 0 | 0 | 0 | 0 | 0 | 0 | 0 | 0 |
| 0 | NA | NA | NA | NA | NA | Gm13539 | | 0 | 0 | 0 | 0 | 0 | 0 | 0 | 0 | 0 | 0 | 0 |
| 0 | NA | NA | NA | NA | NA | Gm13538 | | 0 | 0 | 0 | 0 | 0 | 0 | 0 | 0 | 0 | 0 | 0 |
| 0 | NA | NA | NA | NA | NA | Glt6d1 | glycosyltransferase 6 domain containing | 0 | 0 | 0 | 0 | 0 | 0 | 0 | 0 | 0 | 0 | 0 |
| 0 | NA | NA | NA | NA | NA | Lcn9 | lipocalin 9 | 0 | 0 | 0 | 0 | 0 | 0 | 0 | 0 | 0 | 0 | 0 |
| 0 | NA | NA | NA | NA | NA | Sohlh1 | spermatogenesis and oogenesis specific | 0 | 0 | 0 | 0 | 0 | 0 | 0 | 0 | 0 | 0 | 0 |
| 0 | NA | NA | NA | NA | NA | Gm13542 | | 0 | 0 | 0 | 0 | 0 | 0 | 0 | 0 | 0 | 0 | 0 |
| 0 | NA | NA | NA | NA | NA | Gm13554 | | 0 | 0 | 0 | 0 | 0 | 0 | 0 | 0 | 0 | 0 | 0 |
| 0 | NA | NA | NA | NA | NA | Gm13553 | | 0 | 0 | 0 | 0 | 0 | 0 | 0 | 0 | 0 | 0 | 0 |
| 0 | NA | NA | NA | NA | NA | Gm27196 | | 0 | 0 | 0 | 0 | 0 | 0 | 0 | 0 | 0 | 0 | 0 |
| 0 | NA | NA | NA | NA | NA | Mir6996 | | 0 | 0 | 0 | 0 | 0 | 0 | 0 | 0 | 0 | 0 | 0 |
| 0 | NA | NA | NA | NA | NA | Gm13358 | | 0 | 0 | 0 | 0 | 0 | 0 | 0 | 0 | 0 | 0 | 0 |
| 0 | NA | NA | NA | NA | NA | Gm20532 | | 0 | 0 | 0 | 0 | 0 | 0 | 0 | 0 | 0 | 0 | 0 |
| 0 | NA | NA | NA | NA | NA | Mir126a | | 0 | 0 | 0 | 0 | 0 | 0 | 0 | 0 | 0 | 0 | 0 |
| 0 | NA | NA | NA | NA | NA | Mir126b | | 0 | 0 | 0 | 0 | 0 | 0 | 0 | 0 | 0 | 0 | 0 |
| 0 | NA | NA | NA | NA | NA | Snora43 | | 0 | 0 | 0 | 0 | 0 | 0 | 0 | 0 | 0 | 0 | 0 |
| 0 | NA | NA | NA | NA | NA | Snora43 | | 0 | 0 | 0 | 0 | 0 | 0 | 0 | 0 | 0 | 0 | 0 |
| 0 | NA | NA | NA | NA | NA | Snhg7os | | 0 | 0 | 0 | 0 | 0 | 0 | 0 | 0 | 0 | 0 | 0 |
| 0 | NA | NA | NA | NA | NA | Lcn4 | lipocalin 4 | 0 | 0 | 0 | 0 | 0 | 0 | 0 | 0 | 0 | 0 | 0 |
| 0 | NA | NA | NA | NA | NA | Gm22879 | | 0 | 0 | 0 | 0 | 0 | 0 | 0 | 0 | 0 | 0 | 0 |
| 0 | NA | NA | NA | NA | NA | Gm24134 | | 0 | 0 | 0 | 0 | 0 | 0 | 0 | 0 | 0 | 0 | 0 |
| 0 | NA | NA | NA | NA | NA | Dbhos | | 0 | 0 | 0 | 0 | 0 | 0 | 0 | 0 | 0 | 0 | 0 |
| 0 | NA | NA | NA | NA | NA | Sardhos | | 0 | 0 | 0 | 0 | 0 | 0 | 0 | 0 | 0 | 0 | 0 |
| 0 | NA | NA | NA | NA | NA | AA645442 | | 0 | 0 | 0 | 0 | 0 | 0 | 0 | 0 | 0 | 0 | 0 |
| 0 | NA | NA | NA | NA | NA | Gm24049 | | 0 | 0 | 0 | 0 | 0 | 0 | 0 | 0 | 0 | 0 | 0 |
| 0 | NA | NA | NA | NA | NA | Mir7578 | | 0 | 0 | 0 | 0 | 0 | 0 | 0 | 0 | 0 | 0 | 0 |
| 0 | NA | NA | NA | NA | NA | Gm13372 | | 0 | 0 | 0 | 0 | 0 | 0 | 0 | 0 | 0 | 0 | 0 |
| 0 | NA | NA | NA | NA | NA | Fcnb | ficolin B | 0 | 0 | 0 | 0 | 0 | 0 | 0 | 0 | 0 | 0 | 0 |
| 0 | NA | NA | NA | NA | NA | Gm13371 | | 0 | 0 | 0 | 0 | 0 | 0 | 0 | 0 | 0 | 0 | 0 |
| 0 | NA | NA | NA | NA | NA | Gm22675 | | 0 | 0 | 0 | 0 | 0 | 0 | 0 | 0 | 0 | 0 | 0 |
| 0 | NA | NA | NA | NA | NA | Gm22824 | | 0 | 0 | 0 | 0 | 0 | 0 | 0 | 0 | 0 | 0 | 0 |
| 0 | NA | NA | NA | NA | NA | Gfi1b | growth factor independent 1B | 0 | 0 | 0 | 0 | 0 | 0 | 0 | 0 | 0 | 0 | 0 |
| 0 | NA | NA | NA | NA | NA | Mir3088 | | 0 | 0 | 0 | 0 | 0 | 0 | 0 | 0 | 0 | 0 | 0 |
| 0 | NA | NA | NA | NA | NA | Gm13385 | | 0 | 0 | 0 | 0 | 0 | 0 | 0 | 0 | 0 | 0 | 0 |
| 0 | NA | NA | NA | NA | NA | Gm13386 | | 0 | 0 | 0 | 0 | 0 | 0 | 0 | 0 | 0 | 0 | 0 |
| 0 | NA | NA | NA | NA | NA | Gm13393 | | 0 | 0 | 0 | 0 | 0 | 0 | 0 | 0 | 0 | 0 | 0 |
| 0 | NA | NA | NA | NA | NA | Gm13396 | | 0 | 0 | 0 | 0 | 0 | 0 | 0 | 0 | 0 | 0 | 0 |
| 0 | NA | NA | NA | NA | NA | Gm13402 | | 0 | 0 | 0 | 0 | 0 | 0 | 0 | 0 | 0 | 0 | 0 |
| 0 | NA | NA | NA | NA | NA | Mir133c | | 0 | 0 | 0 | 0 | 0 | 0 | 0 | 0 | 0 | 0 | 0 |

| | | | | | | | | | | | | | | | | | | |
|---|---|---|---|---|---|---|---|---|---|---|---|---|---|---|---|---|---|---|
| 0 | NA | NA | NA | NA | NA | Gm13547 | predicted gene 13547 | 0 | 0 | 0 | 0 | 0 | 0 | 0 | 0 | 0 | 0 | 0 |
| 0 | NA | NA | NA | NA | NA | Mir219a-2 | | 0 | 0 | 0 | 0 | 0 | 0 | 0 | 0 | 0 | 0 | 0 |
| 0 | NA | NA | NA | NA | NA | Mir219b | | 0 | 0 | 0 | 0 | 0 | 0 | 0 | 0 | 0 | 0 | 0 |
| 0 | NA | NA | NA | NA | NA | Gm23865 | | 0 | 0 | 0 | 0 | 0 | 0 | 0 | 0 | 0 | 0 | 0 |
| 0 | NA | NA | NA | NA | NA | Gm28038 | | 0 | 0 | 0 | 0 | 0 | 0 | 0 | 0 | 0 | 0 | 0 |
| 0 | NA | NA | NA | NA | NA | Gm47292 | | 0 | 0 | 0 | 0 | 0 | 0 | 0 | 0 | 0 | 0 | 0 |
| 0 | NA | NA | NA | NA | NA | Gm16323 | | 0 | 0 | 0 | 0 | 0 | 0 | 0 | 0 | 0 | 0 | 0 |
| 0 | NA | NA | NA | NA | NA | 4930527E20Rik | | 0 | 0 | 0 | 0 | 0 | 0 | 0 | 0 | 0 | 0 | 0 |
| 0 | NA | NA | NA | NA | NA | Gm13405 | | 0 | 0 | 0 | 0 | 0 | 0 | 0 | 0 | 0 | 0 | 0 |
| 0 | NA | NA | NA | NA | NA | Gm13426 | | 0 | 0 | 0 | 0 | 0 | 0 | 0 | 0 | 0 | 0 | 0 |
| 0 | NA | NA | NA | NA | NA | Gm13425 | | 0 | 0 | 0 | 0 | 0 | 0 | 0 | 0 | 0 | 0 | 0 |
| 0 | NA | NA | NA | NA | NA | Prdm12 | PR domain containing 12 | 0 | 0 | 0 | 0 | 0 | 0 | 0 | 0 | 0 | 0 | 0 |
| 0 | NA | NA | NA | NA | NA | Gm13424 | | 0 | 0 | 0 | 0 | 0 | 0 | 0 | 0 | 0 | 0 | 0 |
| 0 | NA | NA | NA | NA | NA | | | 0 | 0 | 0 | 0 | 0 | 0 | 0 | 0 | 0 | 0 | 0 |
| 0 | NA | NA | NA | NA | NA | Qrfp | pyroglutamylated RFamide peptide | 0 | 0 | 0 | 0 | 0 | 0 | 0 | 0 | 0 | 0 | 0 |
| 0 | NA | NA | NA | NA | NA | Gm13441 | | 0 | 0 | 0 | 0 | 0 | 0 | 0 | 0 | 0 | 0 | 0 |
| 0 | NA | NA | NA | NA | NA | Gm13442 | | 0 | 0 | 0 | 0 | 0 | 0 | 0 | 0 | 0 | 0 | 0 |
| 0 | NA | NA | NA | NA | NA | Gm16534 | | 0 | 0 | 0 | 0 | 0 | 0 | 0 | 0 | 0 | 0 | 0 |
| 0 | NA | NA | NA | NA | NA | Mir7674 | | 0 | 0 | 0 | 0 | 0 | 0 | 0 | 0 | 0 | 0 | 0 |
| 0 | NA | NA | NA | NA | NA | Gm27805 | | 0 | 0 | 0 | 0 | 0 | 0 | 0 | 0 | 0 | 0 | 0 |
| 0 | NA | NA | NA | NA | NA | Gm23363 | | 0 | 0 | 0 | 0 | 0 | 0 | 0 | 0 | 0 | 0 | 0 |
| 0 | NA | NA | NA | NA | NA | Mir199b | | 0 | 0 | 0 | 0 | 0 | 0 | 0 | 0 | 0 | 0 | 0 |
| 0 | NA | NA | NA | NA | NA | Gm37169 | | 0 | 0 | 0 | 0 | 0 | 0 | 0 | 0 | 0 | 0 | 0 |
| 0 | NA | NA | NA | NA | NA | Gm13412 | | 0 | 0 | 0 | 0 | 0 | 0 | 0 | 0 | 0 | 0 | 0 |
| 0 | NA | NA | NA | NA | NA | Gm23369 | | 0 | 0 | 0 | 0 | 0 | 0 | 0 | 0 | 0 | 0 | 0 |
| 0 | NA | NA | NA | NA | NA | Mir1954 | | 0 | 0 | 0 | 0 | 0 | 0 | 0 | 0 | 0 | 0 | 0 |
| 0 | NA | NA | NA | NA | NA | Gm47291 | | 0 | 0 | 0 | 0 | 0 | 0 | 0 | 0 | 0 | 0 | 0 |
| 0 | NA | NA | NA | NA | NA | Gm26236 | | 0 | 0 | 0 | 0 | 0 | 0 | 0 | 0 | 0 | 0 | 0 |
| 0 | NA | NA | NA | NA | NA | Mir8093 | | 0 | 0 | 0 | 0 | 0 | 0 | 0 | 0 | 0 | 0 | 0 |
| 0 | NA | NA | NA | NA | NA | Mir2861 | | 0 | 0 | 0 | 0 | 0 | 0 | 0 | 0 | 0 | 0 | 0 |
| 0 | NA | NA | NA | NA | NA | Mir3960 | | 0 | 0 | 0 | 0 | 0 | 0 | 0 | 0 | 0 | 0 | 0 |
| 0 | NA | NA | NA | NA | NA | Gm24165 | | 0 | 0 | 0 | 0 | 0 | 0 | 0 | 0 | 0 | 0 | 0 |
| 0 | NA | NA | NA | NA | NA | Snora65 | | 0 | 0 | 0 | 0 | 0 | 0 | 0 | 0 | 0 | 0 | 0 |
| 0 | NA | NA | NA | NA | NA | Gm25998 | | 0 | 0 | 0 | 0 | 0 | 0 | 0 | 0 | 0 | 0 | 0 |
| 0 | NA | NA | NA | NA | NA | Gm13536 | | 0 | 0 | 0 | 0 | 0 | 0 | 0 | 0 | 0 | 0 | 0 |
| 0 | NA | NA | NA | NA | NA | Gm13530 | | 0 | 0 | 0 | 0 | 0 | 0 | 0 | 0 | 0 | 0 | 0 |
| 0 | NA | NA | NA | NA | NA | Gm38011 | | 0 | 0 | 0 | 0 | 0 | 0 | 0 | 0 | 0 | 0 | 0 |
| 0 | NA | NA | NA | NA | NA | Nron | | 0 | 0 | 0 | 0 | 0 | 0 | 0 | 0 | 0 | 0 | 0 |
| 0 | NA | NA | NA | NA | NA | Nron | | 0 | 0 | 0 | 0 | 0 | 0 | 0 | 0 | 0 | 0 | 0 |
| 0 | NA | NA | NA | NA | NA | Gm13403 | | 0 | 0 | 0 | 0 | 0 | 0 | 0 | 0 | 0 | 0 | 0 |
| 0 | NA | NA | NA | NA | NA | Gm13404 | | 0 | 0 | 0 | 0 | 0 | 0 | 0 | 0 | 0 | 0 | 0 |
| 0 | NA | NA | NA | NA | NA | Gm38389 | | 0 | 0 | 0 | 0 | 0 | 0 | 0 | 0 | 0 | 0 | 0 |
| 0 | NA | NA | NA | NA | NA | C79798 | | 0 | 0 | 0 | 0 | 0 | 0 | 0 | 0 | 0 | 0 | 0 |
| 0 | NA | NA | NA | NA | NA | Gm13407 | | 0 | 0 | 0 | 0 | 0 | 0 | 0 | 0 | 0 | 0 | 0 |
| 0 | NA | NA | NA | NA | NA | Gm13406 | | 0 | 0 | 0 | 0 | 0 | 0 | 0 | 0 | 0 | 0 | 0 |
| 0 | NA | NA | NA | NA | NA | Gm13419 | | 0 | 0 | 0 | 0 | 0 | 0 | 0 | 0 | 0 | 0 | 0 |
| 0 | NA | NA | NA | NA | NA | Gm13448 | | 0 | 0 | 0 | 0 | 0 | 0 | 0 | 0 | 0 | 0 | 0 |
| 0 | NA | NA | NA | NA | NA | Gm24905 | | 0 | 0 | 0 | 0 | 0 | 0 | 0 | 0 | 0 | 0 | 0 |
| 0 | NA | NA | NA | NA | NA | Gm13449 | | 0 | 0 | 0 | 0 | 0 | 0 | 0 | 0 | 0 | 0 | 0 |
| 0 | NA | NA | NA | NA | NA | 4930568D16Ri | RIKEN cDNA 4930568D16 gene | 0 | 0 | 0 | 0 | 0 | 0 | 0 | 0 | 0 | 0 | 0 |
| 0 | NA | NA | NA | NA | NA | 4930402F06Ri | RIKEN cDNA 4930402F06 gene | 0 | 0 | 0 | 0 | 0 | 0 | 0 | 0 | 0 | 0 | 0 |
| 0 | NA | NA | NA | NA | NA | Gm43761 | | 0 | 0 | 0 | 0 | 0 | 0 | 0 | 0 | 0 | 0 | 0 |
| 0 | NA | NA | NA | NA | NA | Gm13444 | | 0 | 0 | 0 | 0 | 0 | 0 | 0 | 0 | 0 | 0 | 0 |
| 0 | NA | NA | NA | NA | NA | Gm13445 | | 0 | 0 | 0 | 0 | 0 | 0 | 0 | 0 | 0 | 0 | 0 |
| 0 | NA | NA | NA | NA | NA | Gm35202 | | 0 | 0 | 0 | 0 | 0 | 0 | 0 | 0 | 0 | 0 | 0 |
| 0 | NA | NA | NA | NA | NA | 9030204H09Rik | | 0 | 0 | 0 | 0 | 0 | 0 | 0 | 0 | 0 | 0 | 0 |
| 0 | NA | NA | NA | NA | NA | Gm10829 | | 0 | 0 | 0 | 0 | 0 | 0 | 0 | 0 | 0 | 0 | 0 |
| 0 | NA | NA | NA | NA | NA | Gm43764 | | 0 | 0 | 0 | 0 | 0 | 0 | 0 | 0 | 0 | 0 | 0 |
| 0 | NA | NA | NA | NA | NA | Gm27953 | | 0 | 0 | 0 | 0 | 0 | 0 | 0 | 0 | 0 | 0 | 0 |
| 0 | NA | NA | NA | NA | NA | Gm43621 | | 0 | 0 | 0 | 0 | 0 | 0 | 0 | 0 | 0 | 0 | 0 |

| | | | | | | | | | | | | | | | | | |
|---|---|---|---|---|---|---|---|---|---|---|---|---|---|---|---|---|---|
| 0 | NA | NA | NA | NA | NA | Olfr334-ps1 | | 0 | 0 | 0 | 0 | 0 | 0 | 0 | 0 | 0 | 0 | 0 |
| 0 | NA | NA | NA | NA | NA | Olfr336-ps1 | | 0 | 0 | 0 | 0 | 0 | 0 | 0 | 0 | 0 | 0 | 0 |
| 0 | NA | NA | NA | NA | NA | Olfr337-ps1 | | 0 | 0 | 0 | 0 | 0 | 0 | 0 | 0 | 0 | 0 | 0 |
| 0 | NA | NA | NA | NA | NA | Olfr338 | olfactory receptor 338 | 0 | 0 | 0 | 0 | 0 | 0 | 0 | 0 | 0 | 0 | 0 |
| 0 | NA | NA | NA | NA | NA | Olfr339 | olfactory receptor 339 | 0 | 0 | 0 | 0 | 0 | 0 | 0 | 0 | 0 | 0 | 0 |
| 0 | NA | NA | NA | NA | NA | Olfr340 | olfactory receptor 340 | 0 | 0 | 0 | 0 | 0 | 0 | 0 | 0 | 0 | 0 | 0 |
| 0 | NA | NA | NA | NA | NA | Olfr341 | olfactory receptor 341 | 0 | 0 | 0 | 0 | 0 | 0 | 0 | 0 | 0 | 0 | 0 |
| 0 | NA | NA | NA | NA | NA | Olfr342 | olfactory receptor 342 | 0 | 0 | 0 | 0 | 0 | 0 | 0 | 0 | 0 | 0 | 0 |
| 0 | NA | NA | NA | NA | NA | Gm47783 | | 0 | 0 | 0 | 0 | 0 | 0 | 0 | 0 | 0 | 0 | 0 |
| 0 | NA | NA | NA | NA | NA | Olfr343-ps1 | | 0 | 0 | 0 | 0 | 0 | 0 | 0 | 0 | 0 | 0 | 0 |
| 0 | NA | NA | NA | NA | NA | Olfr344 | olfactory receptor 344 | 0 | 0 | 0 | 0 | 0 | 0 | 0 | 0 | 0 | 0 | 0 |
| 0 | NA | NA | NA | NA | NA | Olfr345 | olfactory receptor 345 | 0 | 0 | 0 | 0 | 0 | 0 | 0 | 0 | 0 | 0 | 0 |
| 0 | NA | NA | NA | NA | NA | Olfr346 | olfactory receptor 346 | 0 | 0 | 0 | 0 | 0 | 0 | 0 | 0 | 0 | 0 | 0 |
| 0 | NA | NA | NA | NA | NA | Olfr347 | olfactory receptor 347 | 0 | 0 | 0 | 0 | 0 | 0 | 0 | 0 | 0 | 0 | 0 |
| 0 | NA | NA | NA | NA | NA | Gm13438 | | 0 | 0 | 0 | 0 | 0 | 0 | 0 | 0 | 0 | 0 | 0 |
| 0 | NA | NA | NA | NA | NA | Olfr348 | olfactory receptor 348 | 0 | 0 | 0 | 0 | 0 | 0 | 0 | 0 | 0 | 0 | 0 |
| 0 | NA | NA | NA | NA | NA | Olfr50 | olfactory receptor 50 | 0 | 0 | 0 | 0 | 0 | 0 | 0 | 0 | 0 | 0 | 0 |
| 0 | NA | NA | NA | NA | NA | Olfr3 | olfactory receptor 3 | 0 | 0 | 0 | 0 | 0 | 0 | 0 | 0 | 0 | 0 | 0 |
| 0 | NA | NA | NA | NA | NA | Olfr349-ps1 | | 0 | 0 | 0 | 0 | 0 | 0 | 0 | 0 | 0 | 0 | 0 |
| 0 | NA | NA | NA | NA | NA | Olfr350 | olfactory receptor 350 | 0 | 0 | 0 | 0 | 0 | 0 | 0 | 0 | 0 | 0 | 0 |
| 0 | NA | NA | NA | NA | NA | Olfr351 | olfactory receptor 351 | 0 | 0 | 0 | 0 | 0 | 0 | 0 | 0 | 0 | 0 | 0 |
| 0 | NA | NA | NA | NA | NA | Olfr352 | olfactory receptor 352 | 0 | 0 | 0 | 0 | 0 | 0 | 0 | 0 | 0 | 0 | 0 |
| 0 | NA | NA | NA | NA | NA | Olfr353 | olfactory receptor 353 | 0 | 0 | 0 | 0 | 0 | 0 | 0 | 0 | 0 | 0 | 0 |
| 0 | NA | NA | NA | NA | NA | Olfr354 | olfactory receptor 354 | 0 | 0 | 0 | 0 | 0 | 0 | 0 | 0 | 0 | 0 | 0 |
| 0 | NA | NA | NA | NA | NA | Gm13440 | | 0 | 0 | 0 | 0 | 0 | 0 | 0 | 0 | 0 | 0 | 0 |
| 0 | NA | NA | NA | NA | NA | Olfr355 | olfactory receptor 355 | 0 | 0 | 0 | 0 | 0 | 0 | 0 | 0 | 0 | 0 | 0 |
| 0 | NA | NA | NA | NA | NA | Olfr356 | olfactory receptor 356 | 0 | 0 | 0 | 0 | 0 | 0 | 0 | 0 | 0 | 0 | 0 |
| 0 | NA | NA | NA | NA | NA | Olfr357 | olfactory receptor 357 | 0 | 0 | 0 | 0 | 0 | 0 | 0 | 0 | 0 | 0 | 0 |
| 0 | NA | NA | NA | NA | NA | Olfr358 | olfactory receptor 358 | 0 | 0 | 0 | 0 | 0 | 0 | 0 | 0 | 0 | 0 | 0 |
| 0 | NA | NA | NA | NA | NA | Olfr359-ps1 | | 0 | 0 | 0 | 0 | 0 | 0 | 0 | 0 | 0 | 0 | 0 |
| 0 | NA | NA | NA | NA | NA | Gm13439 | | 0 | 0 | 0 | 0 | 0 | 0 | 0 | 0 | 0 | 0 | 0 |
| 0 | NA | NA | NA | NA | NA | Olfr360 | olfactory receptor 360 | 0 | 0 | 0 | 0 | 0 | 0 | 0 | 0 | 0 | 0 | 0 |
| 0 | NA | NA | NA | NA | NA | Olfr363-ps | | 0 | 0 | 0 | 0 | 0 | 0 | 0 | 0 | 0 | 0 | 0 |
| 0 | NA | NA | NA | NA | NA | Olfr364-ps1 | | 0 | 0 | 0 | 0 | 0 | 0 | 0 | 0 | 0 | 0 | 0 |
| 0 | NA | NA | NA | NA | NA | Olfr365 | olfactory receptor 365 | 0 | 0 | 0 | 0 | 0 | 0 | 0 | 0 | 0 | 0 | 0 |
| 0 | NA | NA | NA | NA | NA | Olfr366 | olfactory receptor 366 | 0 | 0 | 0 | 0 | 0 | 0 | 0 | 0 | 0 | 0 | 0 |
| 0 | NA | NA | NA | NA | NA | Olfr367-ps | olfactory receptor 367, pseudogene | 0 | 0 | 0 | 0 | 0 | 0 | 0 | 0 | 0 | 0 | 0 |
| 0 | NA | NA | NA | NA | NA | Gm22306 | | 0 | 0 | 0 | 0 | 0 | 0 | 0 | 0 | 0 | 0 | 0 |
| 0 | NA | NA | NA | NA | NA | Olfr368 | olfactory receptor 368 | 0 | 0 | 0 | 0 | 0 | 0 | 0 | 0 | 0 | 0 | 0 |
| 0 | NA | NA | NA | NA | NA | Snord90 | | 0 | 0 | 0 | 0 | 0 | 0 | 0 | 0 | 0 | 0 | 0 |
| 0 | NA | NA | NA | NA | NA | Gm44063 | | 0 | 0 | 0 | 0 | 0 | 0 | 0 | 0 | 0 | 0 | 0 |
| 0 | NA | NA | NA | NA | NA | Gm13543 | | 0 | 0 | 0 | 0 | 0 | 0 | 0 | 0 | 0 | 0 | 0 |
| 0 | NA | NA | NA | NA | NA | Mir5128 | | 0 | 0 | 0 | 0 | 0 | 0 | 0 | 0 | 0 | 0 | 0 |
| 0 | NA | NA | NA | NA | NA | Gm44443 | | 0 | 0 | 0 | 0 | 0 | 0 | 0 | 0 | 0 | 0 | 0 |
| 0 | NA | NA | NA | NA | NA | Gm44455 | | 0 | 0 | 0 | 0 | 0 | 0 | 0 | 0 | 0 | 0 | 0 |
| 0 | NA | NA | NA | NA | NA | Gm13556 | | 0 | 0 | 0 | 0 | 0 | 0 | 0 | 0 | 0 | 0 | 0 |
| 0 | NA | NA | NA | NA | NA | Gm27197 | | 0 | 0 | 0 | 0 | 0 | 0 | 0 | 0 | 0 | 0 | 0 |
| 0 | NA | NA | NA | NA | NA | Gm13584 | | 0 | 0 | 0 | 0 | 0 | 0 | 0 | 0 | 0 | 0 | 0 |
| 0 | NA | NA | NA | NA | NA | Gm13589 | | 0 | 0 | 0 | 0 | 0 | 0 | 0 | 0 | 0 | 0 | 0 |
| 0 | NA | NA | NA | NA | NA | Gm25081 | | 0 | 0 | 0 | 0 | 0 | 0 | 0 | 0 | 0 | 0 | 0 |
| 0 | NA | NA | NA | NA | NA | Gm44062 | | 0 | 0 | 0 | 0 | 0 | 0 | 0 | 0 | 0 | 0 | 0 |
| 0 | NA | NA | NA | NA | NA | Gm35808 | | 0 | 0 | 0 | 0 | 0 | 0 | 0 | 0 | 0 | 0 | 0 |
| 0 | NA | NA | NA | NA | NA | Gm27634 | | 0 | 0 | 0 | 0 | 0 | 0 | 0 | 0 | 0 | 0 | 0 |
| 0 | NA | NA | NA | NA | NA | Adgrd2-ps | | 0 | 0 | 0 | 0 | 0 | 0 | 0 | 0 | 0 | 0 | 0 |
| 0 | NA | NA | NA | NA | NA | Nr5a1os | | 0 | 0 | 0 | 0 | 0 | 0 | 0 | 0 | 0 | 0 | 0 |
| 0 | NA | NA | NA | NA | NA | Rpl15-ps4 | | 0 | 0 | 0 | 0 | 0 | 0 | 0 | 0 | 0 | 0 | 0 |
| 0 | NA | NA | NA | NA | NA | Mir181a-2 | | 0 | 0 | 0 | 0 | 0 | 0 | 0 | 0 | 0 | 0 | 0 |
| 0 | NA | NA | NA | NA | NA | Mir181b-2 | | 0 | 0 | 0 | 0 | 0 | 0 | 0 | 0 | 0 | 0 | 0 |
| 0 | NA | NA | NA | NA | NA | 5730507A11Rik | | 0 | 0 | 0 | 0 | 0 | 0 | 0 | 0 | 0 | 0 | 0 |
| 0 | NA | NA | NA | NA | NA | Gm13473 | | 0 | 0 | 0 | 0 | 0 | 0 | 0 | 0 | 0 | 0 | 0 |

| | | | | | | | | | | | | | | | | | | |
|---|---|---|---|---|---|---|---|---|---|---|---|---|---|---|---|---|---|---|
| 0 | NA | NA | NA | NA | NA | Gm26127 | | 0 | 0 | 0 | 0 | 0 | 0 | 0 | 0 | 0 | 0 | 0 |
| 0 | NA | NA | NA | NA | NA | Gm18791 | | 0 | 0 | 0 | 0 | 0 | 0 | 0 | 0 | 0 | 0 | 0 |
| 0 | NA | NA | NA | NA | NA | Gm13504 | | 0 | 0 | 0 | 0 | 0 | 0 | 0 | 0 | 0 | 0 | 0 |
| 0 | NA | NA | NA | NA | NA | Vmn2r-ps2 | | 0 | 0 | 0 | 0 | 0 | 0 | 0 | 0 | 0 | 0 | 0 |
| 0 | NA | NA | NA | NA | NA | Gm13506 | | 0 | 0 | 0 | 0 | 0 | 0 | 0 | 0 | 0 | 0 | 0 |
| 0 | NA | NA | NA | NA | NA | Gm13454 | | 0 | 0 | 0 | 0 | 0 | 0 | 0 | 0 | 0 | 0 | 0 |
| 0 | NA | NA | NA | NA | NA | Gm13451 | | 0 | 0 | 0 | 0 | 0 | 0 | 0 | 0 | 0 | 0 | 0 |
| 0 | NA | NA | NA | NA | NA | Gm13452 | | 0 | 0 | 0 | 0 | 0 | 0 | 0 | 0 | 0 | 0 | 0 |
| 0 | NA | NA | NA | NA | NA | Gm43944 | | 0 | 0 | 0 | 0 | 0 | 0 | 0 | 0 | 0 | 0 | 0 |
| 0 | NA | NA | NA | NA | NA | Gm43970 | | 0 | 0 | 0 | 0 | 0 | 0 | 0 | 0 | 0 | 0 | 0 |
| 0 | NA | NA | NA | NA | NA | Gm13457 | | 0 | 0 | 0 | 0 | 0 | 0 | 0 | 0 | 0 | 0 | 0 |
| 0 | NA | NA | NA | NA | NA | Gm13458 | | 0 | 0 | 0 | 0 | 0 | 0 | 0 | 0 | 0 | 0 | 0 |
| 0 | NA | NA | NA | NA | NA | Gm13455 | | 0 | 0 | 0 | 0 | 0 | 0 | 0 | 0 | 0 | 0 | 0 |
| 0 | NA | NA | NA | NA | NA | Gm24467 | | 0 | 0 | 0 | 0 | 0 | 0 | 0 | 0 | 0 | 0 | 0 |
| 0 | NA | NA | NA | NA | NA | Gm27348 | | 0 | 0 | 0 | 0 | 0 | 0 | 0 | 0 | 0 | 0 | 0 |
| 0 | NA | NA | NA | NA | NA | Gm13462 | | 0 | 0 | 0 | 0 | 0 | 0 | 0 | 0 | 0 | 0 | 0 |
| 0 | NA | NA | NA | NA | NA | Gm13460 | | 0 | 0 | 0 | 0 | 0 | 0 | 0 | 0 | 0 | 0 | 0 |
| 0 | NA | NA | NA | NA | NA | Gm13463 | | 0 | 0 | 0 | 0 | 0 | 0 | 0 | 0 | 0 | 0 | 0 |
| 0 | NA | NA | NA | NA | NA | Mir6336 | | 0 | 0 | 0 | 0 | 0 | 0 | 0 | 0 | 0 | 0 | 0 |
| 0 | NA | NA | NA | NA | NA | Gm24350 | | 0 | 0 | 0 | 0 | 0 | 0 | 0 | 0 | 0 | 0 | 0 |
| 0 | NA | NA | NA | NA | NA | Gm24350 | | 0 | 0 | 0 | 0 | 0 | 0 | 0 | 0 | 0 | 0 | 0 |
| 0 | NA | NA | NA | NA | NA | Gm22867 | | 0 | 0 | 0 | 0 | 0 | 0 | 0 | 0 | 0 | 0 | 0 |
| 0 | NA | NA | NA | NA | NA | Mir5129 | | 0 | 0 | 0 | 0 | 0 | 0 | 0 | 0 | 0 | 0 | 0 |
| 0 | NA | NA | NA | NA | NA | Gm27761 | | 0 | 0 | 0 | 0 | 0 | 0 | 0 | 0 | 0 | 0 | 0 |
| 0 | NA | NA | NA | NA | NA | Gm27404 | | 0 | 0 | 0 | 0 | 0 | 0 | 0 | 0 | 0 | 0 | 0 |
| 0 | NA | NA | NA | NA | NA | Gm27645 | | 0 | 0 | 0 | 0 | 0 | 0 | 0 | 0 | 0 | 0 | 0 |
| 0 | NA | NA | NA | NA | NA | Gm28643 | | 0 | 0 | 0 | 0 | 0 | 0 | 0 | 0 | 0 | 0 | 0 |
| 0 | NA | NA | NA | NA | NA | Gm13475 | | 0 | 0 | 0 | 0 | 0 | 0 | 0 | 0 | 0 | 0 | 0 |
| 0 | NA | NA | NA | NA | NA | Gm13479 | | 0 | 0 | 0 | 0 | 0 | 0 | 0 | 0 | 0 | 0 | 0 |
| 0 | NA | NA | NA | NA | NA | 1700019E08Rik | | 0 | 0 | 0 | 0 | 0 | 0 | 0 | 0 | 0 | 0 | 0 |
| 0 | NA | NA | NA | NA | NA | Gm13465 | | 0 | 0 | 0 | 0 | 0 | 0 | 0 | 0 | 0 | 0 | 0 |
| 0 | NA | NA | NA | NA | NA | Gm13467 | | 0 | 0 | 0 | 0 | 0 | 0 | 0 | 0 | 0 | 0 | 0 |
| 0 | NA | NA | NA | NA | NA | Gm23072 | | 0 | 0 | 0 | 0 | 0 | 0 | 0 | 0 | 0 | 0 | 0 |
| 0 | NA | NA | NA | NA | NA | Gm13469 | | 0 | 0 | 0 | 0 | 0 | 0 | 0 | 0 | 0 | 0 | 0 |
| 0 | NA | NA | NA | NA | NA | Gm13470 | | 0 | 0 | 0 | 0 | 0 | 0 | 0 | 0 | 0 | 0 | 0 |
| 0 | NA | NA | NA | NA | NA | Gm25264 | | 0 | 0 | 0 | 0 | 0 | 0 | 0 | 0 | 0 | 0 | 0 |
| 0 | NA | NA | NA | NA | NA | Gm25959 | | 0 | 0 | 0 | 0 | 0 | 0 | 0 | 0 | 0 | 0 | 0 |
| 0 | NA | NA | NA | NA | NA | Gm13471 | | 0 | 0 | 0 | 0 | 0 | 0 | 0 | 0 | 0 | 0 | 0 |
| 0 | NA | NA | NA | NA | NA | Gm13482 | | 0 | 0 | 0 | 0 | 0 | 0 | 0 | 0 | 0 | 0 | 0 |
| 0 | NA | NA | NA | NA | NA | Gm25338 | | 0 | 0 | 0 | 0 | 0 | 0 | 0 | 0 | 0 | 0 | 0 |
| 0 | NA | NA | NA | NA | NA | Gm13472 | | 0 | 0 | 0 | 0 | 0 | 0 | 0 | 0 | 0 | 0 | 0 |
| 0 | NA | NA | NA | NA | NA | Gm13510 | | 0 | 0 | 0 | 0 | 0 | 0 | 0 | 0 | 0 | 0 | 0 |
| 0 | NA | NA | NA | NA | NA | Gm25215 | | 0 | 0 | 0 | 0 | 0 | 0 | 0 | 0 | 0 | 0 | 0 |
| 0 | NA | NA | NA | NA | NA | Gm13525 | | 0 | 0 | 0 | 0 | 0 | 0 | 0 | 0 | 0 | 0 | 0 |
| 0 | NA | NA | NA | NA | NA | Gm13480 | | 0 | 0 | 0 | 0 | 0 | 0 | 0 | 0 | 0 | 0 | 0 |
| 0 | NA | NA | NA | NA | NA | Gm25223 | | 0 | 0 | 0 | 0 | 0 | 0 | 0 | 0 | 0 | 0 | 0 |
| 0 | NA | NA | NA | NA | NA | Gm13488 | | 0 | 0 | 0 | 0 | 0 | 0 | 0 | 0 | 0 | 0 | 0 |
| 0 | NA | NA | NA | NA | NA | Gm13485 | | 0 | 0 | 0 | 0 | 0 | 0 | 0 | 0 | 0 | 0 | 0 |
| 0 | NA | NA | NA | NA | NA | Gm13484 | | 0 | 0 | 0 | 0 | 0 | 0 | 0 | 0 | 0 | 0 | 0 |
| 0 | NA | NA | NA | NA | NA | Gm28639 | | 0 | 0 | 0 | 0 | 0 | 0 | 0 | 0 | 0 | 0 | 0 |
| 0 | NA | NA | NA | NA | NA | Gm13500 | | 0 | 0 | 0 | 0 | 0 | 0 | 0 | 0 | 0 | 0 | 0 |
| 0 | NA | NA | NA | NA | NA | Gm13497 | | 0 | 0 | 0 | 0 | 0 | 0 | 0 | 0 | 0 | 0 | 0 |
| 0 | NA | NA | NA | NA | NA | Gm13499 | | 0 | 0 | 0 | 0 | 0 | 0 | 0 | 0 | 0 | 0 | 0 |
| 0 | NA | NA | NA | NA | NA | 1700057H21Rik | | 0 | 0 | 0 | 0 | 0 | 0 | 0 | 0 | 0 | 0 | 0 |
| 0 | NA | NA | NA | NA | NA | Gm23505 | | 0 | 0 | 0 | 0 | 0 | 0 | 0 | 0 | 0 | 0 | 0 |
| 0 | NA | NA | NA | NA | NA | Gm13494 | | 0 | 0 | 0 | 0 | 0 | 0 | 0 | 0 | 0 | 0 | 0 |
| 0 | NA | NA | NA | NA | NA | Gm13491 | | 0 | 0 | 0 | 0 | 0 | 0 | 0 | 0 | 0 | 0 | 0 |
| 0 | NA | NA | NA | NA | NA | Tas2r134 | taste receptor, type 2, member 134 | 0 | 0 | 0 | 0 | 0 | 0 | 0 | 0 | 0 | 0 | 0 |
| 0 | NA | NA | NA | NA | NA | Gm13490 | | 0 | 0 | 0 | 0 | 0 | 0 | 0 | 0 | 0 | 0 | 0 |
| 0 | NA | NA | NA | NA | NA | Gm13492 | | 0 | 0 | 0 | 0 | 0 | 0 | 0 | 0 | 0 | 0 | 0 |

| | | | | | | | | | | | | | | | | | | |
|---|---|---|---|---|---|---|---|---|---|---|---|---|---|---|---|---|---|---|
| 0 | NA | NA | NA | NA | NA | Gm13495 | | 0 | 0 | 0 | 0 | 0 | 0 | 0 | 0 | 0 | 0 | 0 |
| 0 | NA | NA | NA | NA | NA | Gm22095 | | 0 | 0 | 0 | 0 | 0 | 0 | 0 | 0 | 0 | 0 | 0 |
| 0 | NA | NA | NA | NA | NA | Gm13521 | | 0 | 0 | 0 | 0 | 0 | 0 | 0 | 0 | 0 | 0 | 0 |
| 0 | NA | NA | NA | NA | NA | Gm13541 | | 0 | 0 | 0 | 0 | 0 | 0 | 0 | 0 | 0 | 0 | 0 |
| 0 | NA | NA | NA | NA | NA | Gm13548 | | 0 | 0 | 0 | 0 | 0 | 0 | 0 | 0 | 0 | 0 | 0 |
| 0 | NA | NA | NA | NA | NA | Gm25898 | | 0 | 0 | 0 | 0 | 0 | 0 | 0 | 0 | 0 | 0 | 0 |
| 0 | NA | NA | NA | NA | NA | Gm13501 | | 0 | 0 | 0 | 0 | 0 | 0 | 0 | 0 | 0 | 0 | 0 |
| 0 | NA | NA | NA | NA | NA | Gm13503 | | 0 | 0 | 0 | 0 | 0 | 0 | 0 | 0 | 0 | 0 | 0 |
| 0 | NA | NA | NA | NA | NA | Gm23315 | | 0 | 0 | 0 | 0 | 0 | 0 | 0 | 0 | 0 | 0 | 0 |
| 0 | NA | NA | NA | NA | NA | Gm22298 | | 0 | 0 | 0 | 0 | 0 | 0 | 0 | 0 | 0 | 0 | 0 |
| 0 | NA | NA | NA | NA | NA | Gm13505 | | 0 | 0 | 0 | 0 | 0 | 0 | 0 | 0 | 0 | 0 | 0 |
| 0 | NA | NA | NA | NA | NA | Gm14035 | | 0 | 0 | 0 | 0 | 0 | 0 | 0 | 0 | 0 | 0 | 0 |
| 0 | NA | NA | NA | NA | NA | Gm14034 | | 0 | 0 | 0 | 0 | 0 | 0 | 0 | 0 | 0 | 0 | 0 |
| 0 | NA | NA | NA | NA | NA | Gm24549 | | 0 | 0 | 0 | 0 | 0 | 0 | 0 | 0 | 0 | 0 | 0 |
| 0 | NA | NA | NA | NA | NA | Gm25489 | | 0 | 0 | 0 | 0 | 0 | 0 | 0 | 0 | 0 | 0 | 0 |
| 0 | NA | NA | NA | NA | NA | Gm13615 | | 0 | 0 | 0 | 0 | 0 | 0 | 0 | 0 | 0 | 0 | 0 |
| 0 | NA | NA | NA | NA | NA | Gm13518 | | 0 | 0 | 0 | 0 | 0 | 0 | 0 | 0 | 0 | 0 | 0 |
| 0 | NA | NA | NA | NA | NA | Mir195b | | 0 | 0 | 0 | 0 | 0 | 0 | 0 | 0 | 0 | 0 | 0 |
| 0 | NA | NA | NA | NA | NA | Gm13527 | | 0 | 0 | 0 | 0 | 0 | 0 | 0 | 0 | 0 | 0 | 0 |
| 0 | NA | NA | NA | NA | NA | Gm13526 | | 0 | 0 | 0 | 0 | 0 | 0 | 0 | 0 | 0 | 0 | 0 |
| 0 | NA | NA | NA | NA | NA | Gm13535 | | 0 | 0 | 0 | 0 | 0 | 0 | 0 | 0 | 0 | 0 | 0 |
| 0 | NA | NA | NA | NA | NA | Gm13531 | | 0 | 0 | 0 | 0 | 0 | 0 | 0 | 0 | 0 | 0 | 0 |
| 0 | NA | NA | NA | NA | NA | Gm13533 | | 0 | 0 | 0 | 0 | 0 | 0 | 0 | 0 | 0 | 0 | 0 |
| 0 | NA | NA | NA | NA | NA | Gm25388 | | 0 | 0 | 0 | 0 | 0 | 0 | 0 | 0 | 0 | 0 | 0 |
| 0 | NA | NA | NA | NA | NA | Gm33594 | | 0 | 0 | 0 | 0 | 0 | 0 | 0 | 0 | 0 | 0 | 0 |
| 0 | NA | NA | NA | NA | NA | Ermn | ermin, ERM-like protein | 0 | 0 | 0 | 0 | 0 | 0 | 0 | 0 | 0 | 0 | 0 |
| 0 | NA | NA | NA | NA | NA | Gm13546 | | 0 | 0 | 0 | 0 | 0 | 0 | 0 | 0 | 0 | 0 | 0 |
| 0 | NA | NA | NA | NA | NA | Gm17409 | | 0 | 0 | 0 | 0 | 0 | 0 | 0 | 0 | 0 | 0 | 0 |
| 0 | NA | NA | NA | NA | NA | Gm13559 | | 0 | 0 | 0 | 0 | 0 | 0 | 0 | 0 | 0 | 0 | 0 |
| 0 | NA | NA | NA | NA | NA | Gm13560 | | 0 | 0 | 0 | 0 | 0 | 0 | 0 | 0 | 0 | 0 | 0 |
| 0 | NA | NA | NA | NA | NA | Gm13557 | | 0 | 0 | 0 | 0 | 0 | 0 | 0 | 0 | 0 | 0 | 0 |
| 0 | NA | NA | NA | NA | NA | Gm13551 | | 0 | 0 | 0 | 0 | 0 | 0 | 0 | 0 | 0 | 0 | 0 |
| 0 | NA | NA | NA | NA | NA | Gm13549 | | 0 | 0 | 0 | 0 | 0 | 0 | 0 | 0 | 0 | 0 | 0 |
| 0 | NA | NA | NA | NA | NA | Ldha-ps | | 0 | 0 | 0 | 0 | 0 | 0 | 0 | 0 | 0 | 0 | 0 |
| 0 | NA | NA | NA | NA | NA | Gm13571 | | 0 | 0 | 0 | 0 | 0 | 0 | 0 | 0 | 0 | 0 | 0 |
| 0 | NA | NA | NA | NA | NA | Gm13580 | | 0 | 0 | 0 | 0 | 0 | 0 | 0 | 0 | 0 | 0 | 0 |
| 0 | NA | NA | NA | NA | NA | Gm13583 | | 0 | 0 | 0 | 0 | 0 | 0 | 0 | 0 | 0 | 0 | 0 |
| 0 | NA | NA | NA | NA | NA | Gm13567 | | 0 | 0 | 0 | 0 | 0 | 0 | 0 | 0 | 0 | 0 | 0 |
| 0 | NA | NA | NA | NA | NA | Gm13582 | | 0 | 0 | 0 | 0 | 0 | 0 | 0 | 0 | 0 | 0 | 0 |
| 0 | NA | NA | NA | NA | NA | Gm13579 | | 0 | 0 | 0 | 0 | 0 | 0 | 0 | 0 | 0 | 0 | 0 |
| 0 | NA | NA | NA | NA | NA | Gm13581 | | 0 | 0 | 0 | 0 | 0 | 0 | 0 | 0 | 0 | 0 | 0 |
| 0 | NA | NA | NA | NA | NA | Gm26422 | | 0 | 0 | 0 | 0 | 0 | 0 | 0 | 0 | 0 | 0 | 0 |
| 0 | NA | NA | NA | NA | NA | Gm22338 | | 0 | 0 | 0 | 0 | 0 | 0 | 0 | 0 | 0 | 0 | 0 |
| 0 | NA | NA | NA | NA | NA | Gm22403 | | 0 | 0 | 0 | 0 | 0 | 0 | 0 | 0 | 0 | 0 | 0 |
| 0 | NA | NA | NA | NA | NA | Gm13555 | | 0 | 0 | 0 | 0 | 0 | 0 | 0 | 0 | 0 | 0 | 0 |
| 0 | NA | NA | NA | NA | NA | Gm13561 | | 0 | 0 | 0 | 0 | 0 | 0 | 0 | 0 | 0 | 0 | 0 |
| 0 | NA | NA | NA | NA | NA | Gm13590 | | 0 | 0 | 0 | 0 | 0 | 0 | 0 | 0 | 0 | 0 | 0 |
| 0 | NA | NA | NA | NA | NA | Gm22020 | | 0 | 0 | 0 | 0 | 0 | 0 | 0 | 0 | 0 | 0 | 0 |
| 0 | NA | NA | NA | NA | NA | Gm13569 | | 0 | 0 | 0 | 0 | 0 | 0 | 0 | 0 | 0 | 0 | 0 |
| 0 | NA | NA | NA | NA | NA | | | 0 | 0 | 0 | 0 | 0 | 0 | 0 | 0 | 0 | 0 | 0 |
| 0 | NA | NA | NA | NA | NA | Gm23503 | | 0 | 0 | 0 | 0 | 0 | 0 | 0 | 0 | 0 | 0 | 0 |
| 0 | NA | NA | NA | NA | NA | Gm13575 | | 0 | 0 | 0 | 0 | 0 | 0 | 0 | 0 | 0 | 0 | 0 |
| 0 | NA | NA | NA | NA | NA | Gm13576 | | 0 | 0 | 0 | 0 | 0 | 0 | 0 | 0 | 0 | 0 | 0 |
| 0 | NA | NA | NA | NA | NA | Gm13593 | | 0 | 0 | 0 | 0 | 0 | 0 | 0 | 0 | 0 | 0 | 0 |
| 0 | NA | NA | NA | NA | NA | Mir6337 | | 0 | 0 | 0 | 0 | 0 | 0 | 0 | 0 | 0 | 0 | 0 |
| 0 | NA | NA | NA | NA | NA | Gm24138 | | 0 | 0 | 0 | 0 | 0 | 0 | 0 | 0 | 0 | 0 | 0 |
| 0 | NA | NA | NA | NA | NA | Gm13595 | | 0 | 0 | 0 | 0 | 0 | 0 | 0 | 0 | 0 | 0 | 0 |
| 0 | NA | NA | NA | NA | NA | | | 0 | 0 | 0 | 0 | 0 | 0 | 0 | 0 | 0 | 0 | 0 |
| 0 | NA | NA | NA | NA | NA | Gm13619 | | 0 | 0 | 0 | 0 | 0 | 0 | 0 | 0 | 0 | 0 | 0 |
| 0 | NA | NA | NA | NA | NA | Gm13617 | | 0 | 0 | 0 | 0 | 0 | 0 | 0 | 0 | 0 | 0 | 0 |

| | | | | | | | | | | | | | | | | | |
|---|---|---|---|---|---|---|---|---|---|---|---|---|---|---|---|---|---|
| 0 | NA | NA | NA | NA | NA | Gm28638 | | 0 | 0 | 0 | 0 | 0 | 0 | 0 | 0 | 0 | 0 | 0 |
| 0 | NA | NA | NA | NA | NA | Gm13616 | | 0 | 0 | 0 | 0 | 0 | 0 | 0 | 0 | 0 | 0 | 0 |
| 0 | NA | NA | NA | NA | NA | Gm13630 | | 0 | 0 | 0 | 0 | 0 | 0 | 0 | 0 | 0 | 0 | 0 |
| 0 | NA | NA | NA | NA | NA | Gm13629 | | 0 | 0 | 0 | 0 | 0 | 0 | 0 | 0 | 0 | 0 | 0 |
| 0 | NA | NA | NA | NA | NA | Gm23046 | | 0 | 0 | 0 | 0 | 0 | 0 | 0 | 0 | 0 | 0 | 0 |
| 0 | NA | NA | NA | NA | NA | Gm13600 | | 0 | 0 | 0 | 0 | 0 | 0 | 0 | 0 | 0 | 0 | 0 |
| 0 | NA | NA | NA | NA | NA | Gm13598 | | 0 | 0 | 0 | 0 | 0 | 0 | 0 | 0 | 0 | 0 | 0 |
| 0 | NA | NA | NA | NA | NA | Xirp2 | xin actin-binding repeat containing 2 | 0 | 0 | 0 | 0 | 0 | 0 | 0 | 0 | 0 | 0 | 0 |
| 0 | NA | NA | NA | NA | NA | Gm13599 | | 0 | 0 | 0 | 0 | 0 | 0 | 0 | 0 | 0 | 0 | 0 |
| 0 | NA | NA | NA | NA | NA | Gm21830 | | 0 | 0 | 0 | 0 | 0 | 0 | 0 | 0 | 0 | 0 | 0 |
| 0 | NA | NA | NA | NA | NA | Gm23328 | | 0 | 0 | 0 | 0 | 0 | 0 | 0 | 0 | 0 | 0 | 0 |
| 0 | NA | NA | NA | NA | NA | Gm13601 | | 0 | 0 | 0 | 0 | 0 | 0 | 0 | 0 | 0 | 0 | 0 |
| 0 | NA | NA | NA | NA | NA | Mir7224 | | 0 | 0 | 0 | 0 | 0 | 0 | 0 | 0 | 0 | 0 | 0 |
| 0 | NA | NA | NA | NA | NA | Gm26743 | | 0 | 0 | 0 | 0 | 0 | 0 | 0 | 0 | 0 | 0 | 0 |
| 0 | NA | NA | NA | NA | NA | Gm13603 | | 0 | 0 | 0 | 0 | 0 | 0 | 0 | 0 | 0 | 0 | 0 |
| 0 | NA | NA | NA | NA | NA | Mir6335 | | 0 | 0 | 0 | 0 | 0 | 0 | 0 | 0 | 0 | 0 | 0 |
| 0 | NA | NA | NA | NA | NA | Gm37964 | | 0 | 0 | 0 | 0 | 0 | 0 | 0 | 0 | 0 | 0 | 0 |
| 0 | NA | NA | NA | NA | NA | Gm26321 | | 0 | 0 | 0 | 0 | 0 | 0 | 0 | 0 | 0 | 0 | 0 |
| 0 | NA | NA | NA | NA | NA | Gm49461 | | 0 | 0 | 0 | 0 | 0 | 0 | 0 | 0 | 0 | 0 | 0 |
| 0 | NA | NA | NA | NA | NA | Gm49462 | | 0 | 0 | 0 | 0 | 0 | 0 | 0 | 0 | 0 | 0 | 0 |
| 0 | NA | NA | NA | NA | NA | Gm36002 | | 0 | 0 | 0 | 0 | 0 | 0 | 0 | 0 | 0 | 0 | 0 |
| 0 | NA | NA | NA | NA | NA | 4933409G03Ril | RIKEN cDNA 4933409G03 gene | 0 | 0 | 0 | 0 | 0 | 0 | 0 | 0 | 0 | 0 | 0 |
| 0 | NA | NA | NA | NA | NA | 4932414N04Ril | RIKEN cDNA 4932414N04 gene | 0 | 0 | 0 | 0 | 0 | 0 | 0 | 0 | 0 | 0 | 0 |
| 0 | NA | NA | NA | NA | NA | Gm36231 | | 0 | 0 | 0 | 0 | 0 | 0 | 0 | 0 | 0 | 0 | 0 |
| 0 | NA | NA | NA | NA | NA | Gm38236 | | 0 | 0 | 0 | 0 | 0 | 0 | 0 | 0 | 0 | 0 | 0 |
| 0 | NA | NA | NA | NA | NA | Gm37747 | | 0 | 0 | 0 | 0 | 0 | 0 | 0 | 0 | 0 | 0 | 0 |
| 0 | NA | NA | NA | NA | NA | Gm13614 | | 0 | 0 | 0 | 0 | 0 | 0 | 0 | 0 | 0 | 0 | 0 |
| 0 | NA | NA | NA | NA | NA | Gm13631 | | 0 | 0 | 0 | 0 | 0 | 0 | 0 | 0 | 0 | 0 | 0 |
| 0 | NA | NA | NA | NA | NA | Erich2os | | 0 | 0 | 0 | 0 | 0 | 0 | 0 | 0 | 0 | 0 | 0 |
| 0 | NA | NA | NA | NA | NA | Gm37026 | | 0 | 0 | 0 | 0 | 0 | 0 | 0 | 0 | 0 | 0 | 0 |
| 0 | NA | NA | NA | NA | NA | Gm16510 | | 0 | 0 | 0 | 0 | 0 | 0 | 0 | 0 | 0 | 0 | 0 |
| 0 | NA | NA | NA | NA | NA | Gm13633 | | 0 | 0 | 0 | 0 | 0 | 0 | 0 | 0 | 0 | 0 | 0 |
| 0 | NA | NA | NA | NA | NA | Gm13632 | | 0 | 0 | 0 | 0 | 0 | 0 | 0 | 0 | 0 | 0 | 0 |
| 0 | NA | NA | NA | NA | NA | Gm13642 | | 0 | 0 | 0 | 0 | 0 | 0 | 0 | 0 | 0 | 0 | 0 |
| 0 | NA | NA | NA | NA | NA | Gm23166 | | 0 | 0 | 0 | 0 | 0 | 0 | 0 | 0 | 0 | 0 | 0 |
| 0 | NA | NA | NA | NA | NA | Gm23253 | | 0 | 0 | 0 | 0 | 0 | 0 | 0 | 0 | 0 | 0 | 0 |
| 0 | NA | NA | NA | NA | NA | Dlx1as | | 0 | 0 | 0 | 0 | 0 | 0 | 0 | 0 | 0 | 0 | 0 |
| 0 | NA | NA | NA | NA | NA | Dlx2 | distal-less homeobox 2 | 0 | 0 | 0 | 0 | 0 | 0 | 0 | 0 | 0 | 0 | 0 |
| 0 | NA | NA | NA | NA | NA | Gm13647 | | 0 | 0 | 0 | 0 | 0 | 0 | 0 | 0 | 0 | 0 | 0 |
| 0 | NA | NA | NA | NA | NA | Gm13663 | | 0 | 0 | 0 | 0 | 0 | 0 | 0 | 0 | 0 | 0 | 0 |
| 0 | NA | NA | NA | NA | NA | Gm13662 | | 0 | 0 | 0 | 0 | 0 | 0 | 0 | 0 | 0 | 0 | 0 |
| 0 | NA | NA | NA | NA | NA | Gm13739 | | 0 | 0 | 0 | 0 | 0 | 0 | 0 | 0 | 0 | 0 | 0 |
| 0 | NA | NA | NA | NA | NA | Gm13746 | | 0 | 0 | 0 | 0 | 0 | 0 | 0 | 0 | 0 | 0 | 0 |
| 0 | NA | NA | NA | NA | NA | Gm26408 | | 0 | 0 | 0 | 0 | 0 | 0 | 0 | 0 | 0 | 0 | 0 |
| 0 | NA | NA | NA | NA | NA | Rapgef4os1 | | 0 | 0 | 0 | 0 | 0 | 0 | 0 | 0 | 0 | 0 | 0 |
| 0 | NA | NA | NA | NA | NA | Rapgef4os3 | | 0 | 0 | 0 | 0 | 0 | 0 | 0 | 0 | 0 | 0 | 0 |
| 0 | NA | NA | NA | NA | NA | Zakit | | 0 | 0 | 0 | 0 | 0 | 0 | 0 | 0 | 0 | 0 | 0 |
| 0 | NA | NA | NA | NA | NA | Rpl36a-ps4 | | 0 | 0 | 0 | 0 | 0 | 0 | 0 | 0 | 0 | 0 | 0 |
| 0 | NA | NA | NA | NA | NA | Rps13-ps8 | | 0 | 0 | 0 | 0 | 0 | 0 | 0 | 0 | 0 | 0 | 0 |
| 0 | NA | NA | NA | NA | NA | Pex13-ps | | 0 | 0 | 0 | 0 | 0 | 0 | 0 | 0 | 0 | 0 | 0 |
| 0 | NA | NA | NA | NA | NA | 8430437L04Rik | | 0 | 0 | 0 | 0 | 0 | 0 | 0 | 0 | 0 | 0 | 0 |
| 0 | NA | NA | NA | NA | NA | Gm25960 | | 0 | 0 | 0 | 0 | 0 | 0 | 0 | 0 | 0 | 0 | 0 |
| 0 | NA | NA | NA | NA | NA | Hspe1-ps5 | | 0 | 0 | 0 | 0 | 0 | 0 | 0 | 0 | 0 | 0 | 0 |
| 0 | NA | NA | NA | NA | NA | Gm13664 | | 0 | 0 | 0 | 0 | 0 | 0 | 0 | 0 | 0 | 0 | 0 |
| 0 | NA | NA | NA | NA | NA | Sp9 | trans-acting transcription factor 9 | 0 | 0 | 0 | 0 | 0 | 0 | 0 | 0 | 0 | 0 | 0 |
| 0 | NA | NA | NA | NA | NA | Gm13703 | | 0 | 0 | 0 | 0 | 0 | 0 | 0 | 0 | 0 | 0 | 0 |
| 0 | NA | NA | NA | NA | NA | Gm13709 | | 0 | 0 | 0 | 0 | 0 | 0 | 0 | 0 | 0 | 0 | 0 |
| 0 | NA | NA | NA | NA | NA | Gm13708 | | 0 | 0 | 0 | 0 | 0 | 0 | 0 | 0 | 0 | 0 | 0 |
| 0 | NA | NA | NA | NA | NA | Chrna1 | cholinergic receptor, nicotinic, alpha poly | 0 | 0 | 0 | 0 | 0 | 0 | 0 | 0 | 0 | 0 | 0 |
| 0 | NA | NA | NA | NA | NA | Gm13640 | | 0 | 0 | 0 | 0 | 0 | 0 | 0 | 0 | 0 | 0 | 0 |

| | | | | | | | | | | | | | | | | | |
|---|---|---|---|---|---|---|---|---|---|---|---|---|---|---|---|---|---|
| 0 | NA | NA | NA | NA | NA | Gm27421 | | 0 | 0 | 0 | 0 | 0 | 0 | 0 | 0 | 0 | 0 | 0 |
| 0 | NA | NA | NA | NA | NA | A630050E04Rik | | 0 | 0 | 0 | 0 | 0 | 0 | 0 | 0 | 0 | 0 | 0 |
| 0 | NA | NA | NA | NA | NA | Gm28640 | | 0 | 0 | 0 | 0 | 0 | 0 | 0 | 0 | 0 | 0 | 0 |
| 0 | NA | NA | NA | NA | NA | Gm13667 | | 0 | 0 | 0 | 0 | 0 | 0 | 0 | 0 | 0 | 0 | 0 |
| 0 | NA | NA | NA | NA | NA | 4930441J16Rik | | 0 | 0 | 0 | 0 | 0 | 0 | 0 | 0 | 0 | 0 | 0 |
| 0 | NA | NA | NA | NA | NA | 4930445N08Rik | | 0 | 0 | 0 | 0 | 0 | 0 | 0 | 0 | 0 | 0 | 0 |
| 0 | NA | NA | NA | NA | NA | Gm22975 | | 0 | 0 | 0 | 0 | 0 | 0 | 0 | 0 | 0 | 0 | 0 |
| 0 | NA | NA | NA | NA | NA | Gm13672 | | 0 | 0 | 0 | 0 | 0 | 0 | 0 | 0 | 0 | 0 | 0 |
| 0 | NA | NA | NA | NA | NA | Evx2 | even-skipped homeobox 2 | 0 | 0 | 0 | 0 | 0 | 0 | 0 | 0 | 0 | 0 | 0 |
| 0 | NA | NA | NA | NA | NA | Hoxd12 | homeobox D12 | 0 | 0 | 0 | 0 | 0 | 0 | 0 | 0 | 0 | 0 | 0 |
| 0 | NA | NA | NA | NA | NA | Gm28309 | | 0 | 0 | 0 | 0 | 0 | 0 | 0 | 0 | 0 | 0 | 0 |
| 0 | NA | NA | NA | NA | NA | Gm14396 | | 0 | 0 | 0 | 0 | 0 | 0 | 0 | 0 | 0 | 0 | 0 |
| 0 | NA | NA | NA | NA | NA | Gm28793 | | 0 | 0 | 0 | 0 | 0 | 0 | 0 | 0 | 0 | 0 | 0 |
| 0 | NA | NA | NA | NA | NA | Gm44463 | | 0 | 0 | 0 | 0 | 0 | 0 | 0 | 0 | 0 | 0 | 0 |
| 0 | NA | NA | NA | NA | NA | Hoxd3 | homeobox D3 | 0 | 0 | 0 | 0 | 0 | 0 | 0 | 0 | 0 | 0 | 0 |
| 0 | NA | NA | NA | NA | NA | Mir10b | | 0 | 0 | 0 | 0 | 0 | 0 | 0 | 0 | 0 | 0 | 0 |
| 0 | NA | NA | NA | NA | NA | Gm28230 | | 0 | 0 | 0 | 0 | 0 | 0 | 0 | 0 | 0 | 0 | 0 |
| 0 | NA | NA | NA | NA | NA | Haglr | | 0 | 0 | 0 | 0 | 0 | 0 | 0 | 0 | 0 | 0 | 0 |
| 0 | NA | NA | NA | NA | NA | Gm14424 | | 0 | 0 | 0 | 0 | 0 | 0 | 0 | 0 | 0 | 0 | 0 |
| 0 | NA | NA | NA | NA | NA | Gm14426 | | 0 | 0 | 0 | 0 | 0 | 0 | 0 | 0 | 0 | 0 | 0 |
| 0 | NA | NA | NA | NA | NA | Hoxd1 | homeobox D1 | 0 | 0 | 0 | 0 | 0 | 0 | 0 | 0 | 0 | 0 | 0 |
| 0 | NA | NA | NA | NA | NA | Gm14427 | | 0 | 0 | 0 | 0 | 0 | 0 | 0 | 0 | 0 | 0 | 0 |
| 0 | NA | NA | NA | NA | NA | n-R5s198 | | 0 | 0 | 0 | 0 | 0 | 0 | 0 | 0 | 0 | 0 | 0 |
| 0 | NA | NA | NA | NA | NA | Gm18494 | | 0 | 0 | 0 | 0 | 0 | 0 | 0 | 0 | 0 | 0 | 0 |
| 0 | NA | NA | NA | NA | NA | Gm37243 | | 0 | 0 | 0 | 0 | 0 | 0 | 0 | 0 | 0 | 0 | 0 |
| 0 | NA | NA | NA | NA | NA | Gm13658 | | 0 | 0 | 0 | 0 | 0 | 0 | 0 | 0 | 0 | 0 | 0 |
| 0 | NA | NA | NA | NA | NA | Gm13660 | | 0 | 0 | 0 | 0 | 0 | 0 | 0 | 0 | 0 | 0 | 0 |
| 0 | NA | NA | NA | NA | NA | Gm13655 | | 0 | 0 | 0 | 0 | 0 | 0 | 0 | 0 | 0 | 0 | 0 |
| 0 | NA | NA | NA | NA | NA | Gm24574 | | 0 | 0 | 0 | 0 | 0 | 0 | 0 | 0 | 0 | 0 | 0 |
| 0 | NA | NA | NA | NA | NA | Gm24996 | | 0 | 0 | 0 | 0 | 0 | 0 | 0 | 0 | 0 | 0 | 0 |
| 0 | NA | NA | NA | NA | NA | Gm13656 | | 0 | 0 | 0 | 0 | 0 | 0 | 0 | 0 | 0 | 0 | 0 |
| 0 | NA | NA | NA | NA | NA | Gm44337 | | 0 | 0 | 0 | 0 | 0 | 0 | 0 | 0 | 0 | 0 | 0 |
| 0 | NA | NA | NA | NA | NA | Gm23164 | | 0 | 0 | 0 | 0 | 0 | 0 | 0 | 0 | 0 | 0 | 0 |
| 0 | NA | NA | NA | NA | NA | Gm13938 | | 0 | 0 | 0 | 0 | 0 | 0 | 0 | 0 | 0 | 0 | 0 |
| 0 | NA | NA | NA | NA | NA | Gm44360 | | 0 | 0 | 0 | 0 | 0 | 0 | 0 | 0 | 0 | 0 | 0 |
| 0 | NA | NA | NA | NA | NA | Gm13650 | | 0 | 0 | 0 | 0 | 0 | 0 | 0 | 0 | 0 | 0 | 0 |
| 0 | NA | NA | NA | NA | NA | Gm13943 | | 0 | 0 | 0 | 0 | 0 | 0 | 0 | 0 | 0 | 0 | 0 |
| 0 | NA | NA | NA | NA | NA | Gm13649 | | 0 | 0 | 0 | 0 | 0 | 0 | 0 | 0 | 0 | 0 | 0 |
| 0 | NA | NA | NA | NA | NA | Gm13944 | | 0 | 0 | 0 | 0 | 0 | 0 | 0 | 0 | 0 | 0 | 0 |
| 0 | NA | NA | NA | NA | NA | Gm13661 | | 0 | 0 | 0 | 0 | 0 | 0 | 0 | 0 | 0 | 0 | 0 |
| 0 | NA | NA | NA | NA | NA | Gm13726 | | 0 | 0 | 0 | 0 | 0 | 0 | 0 | 0 | 0 | 0 | 0 |
| 0 | NA | NA | NA | NA | NA | 4930440I19Rik | | 0 | 0 | 0 | 0 | 0 | 0 | 0 | 0 | 0 | 0 | 0 |
| 0 | NA | NA | NA | NA | NA | Gm14461 | predicted gene 14461 | 0 | 0 | 0 | 0 | 0 | 0 | 0 | 0 | 0 | 0 | 0 |
| 0 | NA | NA | NA | NA | NA | Gm14460 | | 0 | 0 | 0 | 0 | 0 | 0 | 0 | 0 | 0 | 0 | 0 |
| 0 | NA | NA | NA | NA | NA | Gm23149 | | 0 | 0 | 0 | 0 | 0 | 0 | 0 | 0 | 0 | 0 | 0 |
| 0 | NA | NA | NA | NA | NA | Gm14464 | | 0 | 0 | 0 | 0 | 0 | 0 | 0 | 0 | 0 | 0 | 0 |
| 0 | NA | NA | NA | NA | NA | Gm14463 | | 0 | 0 | 0 | 0 | 0 | 0 | 0 | 0 | 0 | 0 | 0 |
| 0 | NA | NA | NA | NA | NA | Gm17014 | | 0 | 0 | 0 | 0 | 0 | 0 | 0 | 0 | 0 | 0 | 0 |
| 0 | NA | NA | NA | NA | NA | Gm37004 | | 0 | 0 | 0 | 0 | 0 | 0 | 0 | 0 | 0 | 0 | 0 |
| 0 | NA | NA | NA | NA | NA | Gm14466 | | 0 | 0 | 0 | 0 | 0 | 0 | 0 | 0 | 0 | 0 | 0 |
| 0 | NA | NA | NA | NA | NA | Gm14469 | | 0 | 0 | 0 | 0 | 0 | 0 | 0 | 0 | 0 | 0 | 0 |
| 0 | NA | NA | NA | NA | NA | Gm22606 | | 0 | 0 | 0 | 0 | 0 | 0 | 0 | 0 | 0 | 0 | 0 |
| 0 | NA | NA | NA | NA | NA | Gm13673 | | 0 | 0 | 0 | 0 | 0 | 0 | 0 | 0 | 0 | 0 | 0 |
| 0 | NA | NA | NA | NA | NA | Ppp1r1c | protein phosphatase 1, regulatory inhibitc | 0 | 0 | 0 | 0 | 0 | 0 | 0 | 0 | 0 | 0 | 0 |
| 0 | NA | NA | NA | NA | NA | Gm13674 | | 0 | 0 | 0 | 0 | 0 | 0 | 0 | 0 | 0 | 0 | 0 |
| 0 | NA | NA | NA | NA | NA | Gm13752 | | 0 | 0 | 0 | 0 | 0 | 0 | 0 | 0 | 0 | 0 | 0 |
| 0 | NA | NA | NA | NA | NA | Gm13687 | | 0 | 0 | 0 | 0 | 0 | 0 | 0 | 0 | 0 | 0 | 0 |
| 0 | NA | NA | NA | NA | NA | Gm24461 | | 0 | 0 | 0 | 0 | 0 | 0 | 0 | 0 | 0 | 0 | 0 |
| 0 | NA | NA | NA | NA | NA | Gm25420 | | 0 | 0 | 0 | 0 | 0 | 0 | 0 | 0 | 0 | 0 | 0 |
| 0 | NA | NA | NA | NA | NA | Gm22652 | | 0 | 0 | 0 | 0 | 0 | 0 | 0 | 0 | 0 | 0 | 0 |

| | | | | | | | | | | | | | | | | | | |
|---|---|---|---|---|---|---|---|---|---|---|---|---|---|---|---|---|---|---|
| 0 | NA | NA | NA | NA | NA | Gm13700 | | 0 | 0 | 0 | 0 | 0 | 0 | 0 | 0 | 0 | 0 | 0 |
| 0 | NA | NA | NA | NA | NA | Gm13755 | | 0 | 0 | 0 | 0 | 0 | 0 | 0 | 0 | 0 | 0 | 0 |
| 0 | NA | NA | NA | NA | NA | Gm13756 | | 0 | 0 | 0 | 0 | 0 | 0 | 0 | 0 | 0 | 0 | 0 |
| 0 | NA | NA | NA | NA | NA | Gm23452 | | 0 | 0 | 0 | 0 | 0 | 0 | 0 | 0 | 0 | 0 | 0 |
| 0 | NA | NA | NA | NA | NA | Gm23900 | | 0 | 0 | 0 | 0 | 0 | 0 | 0 | 0 | 0 | 0 | 0 |
| 0 | NA | NA | NA | NA | NA | Gm13677 | | 0 | 0 | 0 | 0 | 0 | 0 | 0 | 0 | 0 | 0 | 0 |
| 0 | NA | NA | NA | NA | NA | A130030D18Rik | | 0 | 0 | 0 | 0 | 0 | 0 | 0 | 0 | 0 | 0 | 0 |
| 0 | NA | NA | NA | NA | NA | Gm13676 | | 0 | 0 | 0 | 0 | 0 | 0 | 0 | 0 | 0 | 0 | 0 |
| 0 | NA | NA | NA | NA | NA | Gm13678 | | 0 | 0 | 0 | 0 | 0 | 0 | 0 | 0 | 0 | 0 | 0 |
| 0 | NA | NA | NA | NA | NA | Fsip2 | fibrous sheath-interacting protein 2 | 0 | 0 | 0 | 0 | 0 | 0 | 0 | 0 | 0 | 0 | 0 |
| 0 | NA | NA | NA | NA | NA | Gm28675 | | 0 | 0 | 0 | 0 | 0 | 0 | 0 | 0 | 0 | 0 | 0 |
| 0 | NA | NA | NA | NA | NA | Gm13682 | | 0 | 0 | 0 | 0 | 0 | 0 | 0 | 0 | 0 | 0 | 0 |
| 0 | NA | NA | NA | NA | NA | Gm13681 | | 0 | 0 | 0 | 0 | 0 | 0 | 0 | 0 | 0 | 0 | 0 |
| 0 | NA | NA | NA | NA | NA | Gm13683 | | 0 | 0 | 0 | 0 | 0 | 0 | 0 | 0 | 0 | 0 | 0 |
| 0 | NA | NA | NA | NA | NA | Gm13685 | | 0 | 0 | 0 | 0 | 0 | 0 | 0 | 0 | 0 | 0 | 0 |
| 0 | NA | NA | NA | NA | NA | Gm13686 | | 0 | 0 | 0 | 0 | 0 | 0 | 0 | 0 | 0 | 0 | 0 |
| 0 | NA | NA | NA | NA | NA | Dnmt3l-ps1 | | 0 | 0 | 0 | 0 | 0 | 0 | 0 | 0 | 0 | 0 | 0 |
| 0 | NA | NA | NA | NA | NA | Gm13698 | | 0 | 0 | 0 | 0 | 0 | 0 | 0 | 0 | 0 | 0 | 0 |
| 0 | NA | NA | NA | NA | NA | Gm13693 | | 0 | 0 | 0 | 0 | 0 | 0 | 0 | 0 | 0 | 0 | 0 |
| 0 | NA | NA | NA | NA | NA | Gm13695 | | 0 | 0 | 0 | 0 | 0 | 0 | 0 | 0 | 0 | 0 | 0 |
| 0 | NA | NA | NA | NA | NA | Gm13696 | | 0 | 0 | 0 | 0 | 0 | 0 | 0 | 0 | 0 | 0 | 0 |
| 0 | NA | NA | NA | NA | NA | Gm13694 | | 0 | 0 | 0 | 0 | 0 | 0 | 0 | 0 | 0 | 0 | 0 |
| 0 | NA | NA | NA | NA | NA | Gm13697 | | 0 | 0 | 0 | 0 | 0 | 0 | 0 | 0 | 0 | 0 | 0 |
| 0 | NA | NA | NA | NA | NA | Gm13691 | | 0 | 0 | 0 | 0 | 0 | 0 | 0 | 0 | 0 | 0 | 0 |
| 0 | NA | NA | NA | NA | NA | Gm13692 | | 0 | 0 | 0 | 0 | 0 | 0 | 0 | 0 | 0 | 0 | 0 |
| 0 | NA | NA | NA | NA | NA | Gm13702 | | 0 | 0 | 0 | 0 | 0 | 0 | 0 | 0 | 0 | 0 | 0 |
| 0 | NA | NA | NA | NA | NA | Gm25972 | | 0 | 0 | 0 | 0 | 0 | 0 | 0 | 0 | 0 | 0 | 0 |
| 0 | NA | NA | NA | NA | NA | Gm13711 | | 0 | 0 | 0 | 0 | 0 | 0 | 0 | 0 | 0 | 0 | 0 |
| 0 | NA | NA | NA | NA | NA | | | 0 | 0 | 0 | 0 | 0 | 0 | 0 | 0 | 0 | 0 | 0 |
| 0 | NA | NA | NA | NA | NA | Gm28635 | | 0 | 0 | 0 | 0 | 0 | 0 | 0 | 0 | 0 | 0 | 0 |
| 0 | NA | NA | NA | NA | NA | Gm27942 | | 0 | 0 | 0 | 0 | 0 | 0 | 0 | 0 | 0 | 0 | 0 |
| 0 | NA | NA | NA | NA | NA | Mir130a | | 0 | 0 | 0 | 0 | 0 | 0 | 0 | 0 | 0 | 0 | 0 |
| 0 | NA | NA | NA | NA | NA | Smtnl1 | smoothelin-like 1 | 0 | 0 | 0 | 0 | 0 | 0 | 0 | 0 | 0 | 0 | 0 |
| 0 | NA | NA | NA | NA | NA | Gm24332 | | 0 | 0 | 0 | 0 | 0 | 0 | 0 | 0 | 0 | 0 | 0 |
| 0 | NA | NA | NA | NA | NA | Gm13716 | | 0 | 0 | 0 | 0 | 0 | 0 | 0 | 0 | 0 | 0 | 0 |
| 0 | NA | NA | NA | NA | NA | Gm13715 | | 0 | 0 | 0 | 0 | 0 | 0 | 0 | 0 | 0 | 0 | 0 |
| 0 | NA | NA | NA | NA | NA | Gm13714 | | 0 | 0 | 0 | 0 | 0 | 0 | 0 | 0 | 0 | 0 | 0 |
| 0 | NA | NA | NA | NA | NA | Olfr987 | olfactory receptor 987 | 0 | 0 | 0 | 0 | 0 | 0 | 0 | 0 | 0 | 0 | 0 |
| 0 | NA | NA | NA | NA | NA | Olfr988 | olfactory receptor 988 | 0 | 0 | 0 | 0 | 0 | 0 | 0 | 0 | 0 | 0 | 0 |
| 0 | NA | NA | NA | NA | NA | Olfr989-ps1 | | 0 | 0 | 0 | 0 | 0 | 0 | 0 | 0 | 0 | 0 | 0 |
| 0 | NA | NA | NA | NA | NA | Olfr990-ps1 | | 0 | 0 | 0 | 0 | 0 | 0 | 0 | 0 | 0 | 0 | 0 |
| 0 | NA | NA | NA | NA | NA | Olfr991-ps1 | | 0 | 0 | 0 | 0 | 0 | 0 | 0 | 0 | 0 | 0 | 0 |
| 0 | NA | NA | NA | NA | NA | Olfr992 | olfactory receptor 992 | 0 | 0 | 0 | 0 | 0 | 0 | 0 | 0 | 0 | 0 | 0 |
| 0 | NA | NA | NA | NA | NA | Olfr993 | olfactory receptor 993 | 0 | 0 | 0 | 0 | 0 | 0 | 0 | 0 | 0 | 0 | 0 |
| 0 | NA | NA | NA | NA | NA | Gm23716 | | 0 | 0 | 0 | 0 | 0 | 0 | 0 | 0 | 0 | 0 | 0 |
| 0 | NA | NA | NA | NA | NA | Olfr994 | olfactory receptor 994 | 0 | 0 | 0 | 0 | 0 | 0 | 0 | 0 | 0 | 0 | 0 |
| 0 | NA | NA | NA | NA | NA | Olfr995 | olfactory receptor 995 | 0 | 0 | 0 | 0 | 0 | 0 | 0 | 0 | 0 | 0 | 0 |
| 0 | NA | NA | NA | NA | NA | Fads2b | fatty acid desaturase 2B | 0 | 0 | 0 | 0 | 0 | 0 | 0 | 0 | 0 | 0 | 0 |
| 0 | NA | NA | NA | NA | NA | Olfr996 | olfactory receptor 996 | 0 | 0 | 0 | 0 | 0 | 0 | 0 | 0 | 0 | 0 | 0 |
| 0 | NA | NA | NA | NA | NA | Olfr997-ps1 | | 0 | 0 | 0 | 0 | 0 | 0 | 0 | 0 | 0 | 0 | 0 |
| 0 | NA | NA | NA | NA | NA | Olfr998 | olfactory receptor 998 | 0 | 0 | 0 | 0 | 0 | 0 | 0 | 0 | 0 | 0 | 0 |
| 0 | NA | NA | NA | NA | NA | Olfr999-ps1 | | 0 | 0 | 0 | 0 | 0 | 0 | 0 | 0 | 0 | 0 | 0 |
| 0 | NA | NA | NA | NA | NA | Olfr1000 | olfactory receptor 1000 | 0 | 0 | 0 | 0 | 0 | 0 | 0 | 0 | 0 | 0 | 0 |
| 0 | NA | NA | NA | NA | NA | Olfr1001-ps1 | | 0 | 0 | 0 | 0 | 0 | 0 | 0 | 0 | 0 | 0 | 0 |
| 0 | NA | NA | NA | NA | NA | Olfr1002 | olfactory receptor 1002 | 0 | 0 | 0 | 0 | 0 | 0 | 0 | 0 | 0 | 0 | 0 |
| 0 | NA | NA | NA | NA | NA | Olfr1003-ps1 | | 0 | 0 | 0 | 0 | 0 | 0 | 0 | 0 | 0 | 0 | 0 |
| 0 | NA | NA | NA | NA | NA | Olfr1004-ps1 | | 0 | 0 | 0 | 0 | 0 | 0 | 0 | 0 | 0 | 0 | 0 |
| 0 | NA | NA | NA | NA | NA | Olfr154 | olfactory receptor 154 | 0 | 0 | 0 | 0 | 0 | 0 | 0 | 0 | 0 | 0 | 0 |
| 0 | NA | NA | NA | NA | NA | Olfr1005-ps1 | | 0 | 0 | 0 | 0 | 0 | 0 | 0 | 0 | 0 | 0 | 0 |
| 0 | NA | NA | NA | NA | NA | Olfr1006 | olfactory receptor 1006 | 0 | 0 | 0 | 0 | 0 | 0 | 0 | 0 | 0 | 0 | 0 |

| | | | | | | | | | | | | | | | | | | |
|---|---|---|---|---|---|---|---|---|---|---|---|---|---|---|---|---|---|---|
| 0 | NA | NA | NA | NA | NA | Olfr1007-ps1 | | 0 | 0 | 0 | 0 | 0 | 0 | 0 | 0 | 0 | 0 | 0 |
| 0 | NA | NA | NA | NA | NA | Olfr1008 | olfactory receptor 1008 | 0 | 0 | 0 | 0 | 0 | 0 | 0 | 0 | 0 | 0 | 0 |
| 0 | NA | NA | NA | NA | NA | Olfr1009 | olfactory receptor 1009 | 0 | 0 | 0 | 0 | 0 | 0 | 0 | 0 | 0 | 0 | 0 |
| 0 | NA | NA | NA | NA | NA | Olfr1010 | | 0 | 0 | 0 | 0 | 0 | 0 | 0 | 0 | 0 | 0 | 0 |
| 0 | NA | NA | NA | NA | NA | Olfr1012 | olfactory receptor 1012 | 0 | 0 | 0 | 0 | 0 | 0 | 0 | 0 | 0 | 0 | 0 |
| 0 | NA | NA | NA | NA | NA | Olfr1013 | olfactory receptor 1013 | 0 | 0 | 0 | 0 | 0 | 0 | 0 | 0 | 0 | 0 | 0 |
| 0 | NA | NA | NA | NA | NA | Olfr1014 | olfactory receptor 1014 | 0 | 0 | 0 | 0 | 0 | 0 | 0 | 0 | 0 | 0 | 0 |
| 0 | NA | NA | NA | NA | NA | Gm49332 | | 0 | 0 | 0 | 0 | 0 | 0 | 0 | 0 | 0 | 0 | 0 |
| 0 | NA | NA | NA | NA | NA | Olfr1015 | olfactory receptor 1015 | 0 | 0 | 0 | 0 | 0 | 0 | 0 | 0 | 0 | 0 | 0 |
| 0 | NA | NA | NA | NA | NA | Olfr1016 | olfactory receptor 1016 | 0 | 0 | 0 | 0 | 0 | 0 | 0 | 0 | 0 | 0 | 0 |
| 0 | NA | NA | NA | NA | NA | Olfr1017-ps1 | | 0 | 0 | 0 | 0 | 0 | 0 | 0 | 0 | 0 | 0 | 0 |
| 0 | NA | NA | NA | NA | NA | Gm13720 | | 0 | 0 | 0 | 0 | 0 | 0 | 0 | 0 | 0 | 0 | 0 |
| 0 | NA | NA | NA | NA | NA | Olfr1019 | olfactory receptor 1019 | 0 | 0 | 0 | 0 | 0 | 0 | 0 | 0 | 0 | 0 | 0 |
| 0 | NA | NA | NA | NA | NA | Olfr1020 | olfactory receptor 1020 | 0 | 0 | 0 | 0 | 0 | 0 | 0 | 0 | 0 | 0 | 0 |
| 0 | NA | NA | NA | NA | NA | Olfr1021-ps1 | | 0 | 0 | 0 | 0 | 0 | 0 | 0 | 0 | 0 | 0 | 0 |
| 0 | NA | NA | NA | NA | NA | Olfr1022 | olfactory receptor 1022 | 0 | 0 | 0 | 0 | 0 | 0 | 0 | 0 | 0 | 0 | 0 |
| 0 | NA | NA | NA | NA | NA | Gm24037 | | 0 | 0 | 0 | 0 | 0 | 0 | 0 | 0 | 0 | 0 | 0 |
| 0 | NA | NA | NA | NA | NA | Olfr1023 | olfactory receptor 1023 | 0 | 0 | 0 | 0 | 0 | 0 | 0 | 0 | 0 | 0 | 0 |
| 0 | NA | NA | NA | NA | NA | Gm40020 | | 0 | 0 | 0 | 0 | 0 | 0 | 0 | 0 | 0 | 0 | 0 |
| 0 | NA | NA | NA | NA | NA | Olfr1024 | olfactory receptor 1024 | 0 | 0 | 0 | 0 | 0 | 0 | 0 | 0 | 0 | 0 | 0 |
| 0 | NA | NA | NA | NA | NA | Olfr1025-ps1 | | 0 | 0 | 0 | 0 | 0 | 0 | 0 | 0 | 0 | 0 | 0 |
| 0 | NA | NA | NA | NA | NA | Olfr1026 | olfactory receptor 1026 | 0 | 0 | 0 | 0 | 0 | 0 | 0 | 0 | 0 | 0 | 0 |
| 0 | NA | NA | NA | NA | NA | Olfr1027-ps1 | | 0 | 0 | 0 | 0 | 0 | 0 | 0 | 0 | 0 | 0 | 0 |
| 0 | NA | NA | NA | NA | NA | Olfr1028 | olfactory receptor 1028 | 0 | 0 | 0 | 0 | 0 | 0 | 0 | 0 | 0 | 0 | 0 |
| 0 | NA | NA | NA | NA | NA | Olfr1029 | olfactory receptor 1029 | 0 | 0 | 0 | 0 | 0 | 0 | 0 | 0 | 0 | 0 | 0 |
| 0 | NA | NA | NA | NA | NA | Olfr1030 | olfactory receptor 1030 | 0 | 0 | 0 | 0 | 0 | 0 | 0 | 0 | 0 | 0 | 0 |
| 0 | NA | NA | NA | NA | NA | Olfr1031 | olfactory receptor 1031 | 0 | 0 | 0 | 0 | 0 | 0 | 0 | 0 | 0 | 0 | 0 |
| 0 | NA | NA | NA | NA | NA | Olfr1032 | olfactory receptor 1032 | 0 | 0 | 0 | 0 | 0 | 0 | 0 | 0 | 0 | 0 | 0 |
| 0 | NA | NA | NA | NA | NA | Olfr1035-ps1 | | 0 | 0 | 0 | 0 | 0 | 0 | 0 | 0 | 0 | 0 | 0 |
| 0 | NA | NA | NA | NA | NA | Olfr1036 | olfactory receptor 1036 | 0 | 0 | 0 | 0 | 0 | 0 | 0 | 0 | 0 | 0 | 0 |
| 0 | NA | NA | NA | NA | NA | Olfr1037 | olfactory receptor 1037 | 0 | 0 | 0 | 0 | 0 | 0 | 0 | 0 | 0 | 0 | 0 |
| 0 | NA | NA | NA | NA | NA | Gm13722 | | 0 | 0 | 0 | 0 | 0 | 0 | 0 | 0 | 0 | 0 | 0 |
| 0 | NA | NA | NA | NA | NA | Olfr1038-ps | | 0 | 0 | 0 | 0 | 0 | 0 | 0 | 0 | 0 | 0 | 0 |
| 0 | NA | NA | NA | NA | NA | Olfr1039 | olfactory receptor 1039 | 0 | 0 | 0 | 0 | 0 | 0 | 0 | 0 | 0 | 0 | 0 |
| 0 | NA | NA | NA | NA | NA | Olfr1040 | olfactory receptor 1040 | 0 | 0 | 0 | 0 | 0 | 0 | 0 | 0 | 0 | 0 | 0 |
| 0 | NA | NA | NA | NA | NA | Olfr1041-ps1 | | 0 | 0 | 0 | 0 | 0 | 0 | 0 | 0 | 0 | 0 | 0 |
| 0 | NA | NA | NA | NA | NA | Olfr1042 | olfactory receptor 1042 | 0 | 0 | 0 | 0 | 0 | 0 | 0 | 0 | 0 | 0 | 0 |
| 0 | NA | NA | NA | NA | NA | Olfr1043 | olfactory receptor 1043 | 0 | 0 | 0 | 0 | 0 | 0 | 0 | 0 | 0 | 0 | 0 |
| 0 | NA | NA | NA | NA | NA | Olfr1044 | olfactory receptor 1044 | 0 | 0 | 0 | 0 | 0 | 0 | 0 | 0 | 0 | 0 | 0 |
| 0 | NA | NA | NA | NA | NA | Olfr52 | olfactory receptor 52 | 0 | 0 | 0 | 0 | 0 | 0 | 0 | 0 | 0 | 0 | 0 |
| 0 | NA | NA | NA | NA | NA | Olfr1045 | olfactory receptor 1045 | 0 | 0 | 0 | 0 | 0 | 0 | 0 | 0 | 0 | 0 | 0 |
| 0 | NA | NA | NA | NA | NA | Olfr1046 | olfactory receptor 1046 | 0 | 0 | 0 | 0 | 0 | 0 | 0 | 0 | 0 | 0 | 0 |
| 0 | NA | NA | NA | NA | NA | Olfr1047 | olfactory receptor 1047 | 0 | 0 | 0 | 0 | 0 | 0 | 0 | 0 | 0 | 0 | 0 |
| 0 | NA | NA | NA | NA | NA | Olfr1048 | olfactory receptor 1048 | 0 | 0 | 0 | 0 | 0 | 0 | 0 | 0 | 0 | 0 | 0 |
| 0 | NA | NA | NA | NA | NA | Olfr1049 | olfactory receptor 1049 | 0 | 0 | 0 | 0 | 0 | 0 | 0 | 0 | 0 | 0 | 0 |
| 0 | NA | NA | NA | NA | NA | Olfr1050-ps1 | | 0 | 0 | 0 | 0 | 0 | 0 | 0 | 0 | 0 | 0 | 0 |
| 0 | NA | NA | NA | NA | NA | Olfr1051 | olfactory receptor 1051 | 0 | 0 | 0 | 0 | 0 | 0 | 0 | 0 | 0 | 0 | 0 |
| 0 | NA | NA | NA | NA | NA | Olfr1052 | olfactory receptor 1052 | 0 | 0 | 0 | 0 | 0 | 0 | 0 | 0 | 0 | 0 | 0 |
| 0 | NA | NA | NA | NA | NA | Olfr1053 | olfactory receptor 1053 | 0 | 0 | 0 | 0 | 0 | 0 | 0 | 0 | 0 | 0 | 0 |
| 0 | NA | NA | NA | NA | NA | Olfr1054 | olfactory receptor 1054 | 0 | 0 | 0 | 0 | 0 | 0 | 0 | 0 | 0 | 0 | 0 |
| 0 | NA | NA | NA | NA | NA | Gm48554 | | 0 | 0 | 0 | 0 | 0 | 0 | 0 | 0 | 0 | 0 | 0 |
| 0 | NA | NA | NA | NA | NA | Olfr1055 | olfactory receptor 1055 | 0 | 0 | 0 | 0 | 0 | 0 | 0 | 0 | 0 | 0 | 0 |
| 0 | NA | NA | NA | NA | NA | Olfr1056 | olfactory receptor 1056 | 0 | 0 | 0 | 0 | 0 | 0 | 0 | 0 | 0 | 0 | 0 |
| 0 | NA | NA | NA | NA | NA | Olfr1057 | olfactory receptor 1057 | 0 | 0 | 0 | 0 | 0 | 0 | 0 | 0 | 0 | 0 | 0 |
| 0 | NA | NA | NA | NA | NA | Olfr1058 | olfactory receptor 1058 | 0 | 0 | 0 | 0 | 0 | 0 | 0 | 0 | 0 | 0 | 0 |
| 0 | NA | NA | NA | NA | NA | Olfr1059-ps1 | | 0 | 0 | 0 | 0 | 0 | 0 | 0 | 0 | 0 | 0 | 0 |
| 0 | NA | NA | NA | NA | NA | Gm20734 | | 0 | 0 | 0 | 0 | 0 | 0 | 0 | 0 | 0 | 0 | 0 |
| 0 | NA | NA | NA | NA | NA | Olfr1060-ps1 | | 0 | 0 | 0 | 0 | 0 | 0 | 0 | 0 | 0 | 0 | 0 |
| 0 | NA | NA | NA | NA | NA | Olfr1061 | olfactory receptor 1061 | 0 | 0 | 0 | 0 | 0 | 0 | 0 | 0 | 0 | 0 | 0 |
| 0 | NA | NA | NA | NA | NA | Olfr1062 | olfactory receptor 1062 | 0 | 0 | 0 | 0 | 0 | 0 | 0 | 0 | 0 | 0 | 0 |

| | | | | | | | | | | | | | | | | | | |
|---|---|---|---|---|---|---|---|---|---|---|---|---|---|---|---|---|---|---|
| 0 | NA | NA | NA | NA | NA | Olfr1063-ps1 | | 0 | 0 | 0 | 0 | 0 | 0 | 0 | 0 | 0 | 0 | 0 |
| 0 | NA | NA | NA | NA | NA | Olfr1064-ps1 | | 0 | 0 | 0 | 0 | 0 | 0 | 0 | 0 | 0 | 0 | 0 |
| 0 | NA | NA | NA | NA | NA | Olfr1065 | olfactory receptor 1065 | 0 | 0 | 0 | 0 | 0 | 0 | 0 | 0 | 0 | 0 | 0 |
| 0 | NA | NA | NA | NA | NA | Olfr1066 | olfactory receptor 1066 | 0 | 0 | 0 | 0 | 0 | 0 | 0 | 0 | 0 | 0 | 0 |
| 0 | NA | NA | NA | NA | NA | Olfr1069-ps1 | | 0 | 0 | 0 | 0 | 0 | 0 | 0 | 0 | 0 | 0 | 0 |
| 0 | NA | NA | NA | NA | NA | Gm48628 | | 0 | 0 | 0 | 0 | 0 | 0 | 0 | 0 | 0 | 0 | 0 |
| 0 | NA | NA | NA | NA | NA | Olfr1070-ps1 | | 0 | 0 | 0 | 0 | 0 | 0 | 0 | 0 | 0 | 0 | 0 |
| 0 | NA | NA | NA | NA | NA | Olfr228 | olfactory receptor 228 | 0 | 0 | 0 | 0 | 0 | 0 | 0 | 0 | 0 | 0 | 0 |
| 0 | NA | NA | NA | NA | NA | Olfr1073-ps1 | | 0 | 0 | 0 | 0 | 0 | 0 | 0 | 0 | 0 | 0 | 0 |
| 0 | NA | NA | NA | NA | NA | Olfr1073-ps1 | | 0 | 0 | 0 | 0 | 0 | 0 | 0 | 0 | 0 | 0 | 0 |
| 0 | NA | NA | NA | NA | NA | Olfr1075-ps1 | | 0 | 0 | 0 | 0 | 0 | 0 | 0 | 0 | 0 | 0 | 0 |
| 0 | NA | NA | NA | NA | NA | Gm5023 | | 0 | 0 | 0 | 0 | 0 | 0 | 0 | 0 | 0 | 0 | 0 |
| 0 | NA | NA | NA | NA | NA | Olfr1076 | olfactory receptor 1076 | 0 | 0 | 0 | 0 | 0 | 0 | 0 | 0 | 0 | 0 | 0 |
| 0 | NA | NA | NA | NA | NA | Olfr1077-ps1 | | 0 | 0 | 0 | 0 | 0 | 0 | 0 | 0 | 0 | 0 | 0 |
| 0 | NA | NA | NA | NA | NA | Olfr1078-ps1 | | 0 | 0 | 0 | 0 | 0 | 0 | 0 | 0 | 0 | 0 | 0 |
| 0 | NA | NA | NA | NA | NA | Olfr1079 | olfactory receptor 1079 | 0 | 0 | 0 | 0 | 0 | 0 | 0 | 0 | 0 | 0 | 0 |
| 0 | NA | NA | NA | NA | NA | Olfr1080 | olfactory receptor 1080 | 0 | 0 | 0 | 0 | 0 | 0 | 0 | 0 | 0 | 0 | 0 |
| 0 | NA | NA | NA | NA | NA | Olfr1081-ps1 | | 0 | 0 | 0 | 0 | 0 | 0 | 0 | 0 | 0 | 0 | 0 |
| 0 | NA | NA | NA | NA | NA | Olfr1082 | olfactory receptor 1082 | 0 | 0 | 0 | 0 | 0 | 0 | 0 | 0 | 0 | 0 | 0 |
| 0 | NA | NA | NA | NA | NA | Olfr1083-ps | | 0 | 0 | 0 | 0 | 0 | 0 | 0 | 0 | 0 | 0 | 0 |
| 0 | NA | NA | NA | NA | NA | Olfr1084 | olfactory receptor 1084 | 0 | 0 | 0 | 0 | 0 | 0 | 0 | 0 | 0 | 0 | 0 |
| 0 | NA | NA | NA | NA | NA | Olfr1085 | olfactory receptor 1085 | 0 | 0 | 0 | 0 | 0 | 0 | 0 | 0 | 0 | 0 | 0 |
| 0 | NA | NA | NA | NA | NA | Olfr1086 | olfactory receptor 1086 | 0 | 0 | 0 | 0 | 0 | 0 | 0 | 0 | 0 | 0 | 0 |
| 0 | NA | NA | NA | NA | NA | Olfr1087 | olfactory receptor 1087 | 0 | 0 | 0 | 0 | 0 | 0 | 0 | 0 | 0 | 0 | 0 |
| 0 | NA | NA | NA | NA | NA | Olfr1088-ps1 | | 0 | 0 | 0 | 0 | 0 | 0 | 0 | 0 | 0 | 0 | 0 |
| 0 | NA | NA | NA | NA | NA | Olfr1089 | olfactory receptor 1089 | 0 | 0 | 0 | 0 | 0 | 0 | 0 | 0 | 0 | 0 | 0 |
| 0 | NA | NA | NA | NA | NA | Olfr1090 | olfactory receptor 1090 | 0 | 0 | 0 | 0 | 0 | 0 | 0 | 0 | 0 | 0 | 0 |
| 0 | NA | NA | NA | NA | NA | Olfr1091-ps1 | | 0 | 0 | 0 | 0 | 0 | 0 | 0 | 0 | 0 | 0 | 0 |
| 0 | NA | NA | NA | NA | NA | Olfr1092-ps1 | | 0 | 0 | 0 | 0 | 0 | 0 | 0 | 0 | 0 | 0 | 0 |
| 0 | NA | NA | NA | NA | NA | Olfr1093 | olfactory receptor 1093 | 0 | 0 | 0 | 0 | 0 | 0 | 0 | 0 | 0 | 0 | 0 |
| 0 | NA | NA | NA | NA | NA | Olfr141 | olfactory receptor 141 | 0 | 0 | 0 | 0 | 0 | 0 | 0 | 0 | 0 | 0 | 0 |
| 0 | NA | NA | NA | NA | NA | Olfr1094 | olfactory receptor 1094 | 0 | 0 | 0 | 0 | 0 | 0 | 0 | 0 | 0 | 0 | 0 |
| 0 | NA | NA | NA | NA | NA | Olfr1095 | olfactory receptor 1095 | 0 | 0 | 0 | 0 | 0 | 0 | 0 | 0 | 0 | 0 | 0 |
| 0 | NA | NA | NA | NA | NA | Gm13723 | olfactory receptor 1096, pseudogene 1 | 0 | 0 | 0 | 0 | 0 | 0 | 0 | 0 | 0 | 0 | 0 |
| 0 | NA | NA | NA | NA | NA | Olfr1097 | olfactory receptor 1097 | 0 | 0 | 0 | 0 | 0 | 0 | 0 | 0 | 0 | 0 | 0 |
| 0 | NA | NA | NA | NA | NA | Olfr1098 | olfactory receptor 1098 | 0 | 0 | 0 | 0 | 0 | 0 | 0 | 0 | 0 | 0 | 0 |
| 0 | NA | NA | NA | NA | NA | Gm48002 | | 0 | 0 | 0 | 0 | 0 | 0 | 0 | 0 | 0 | 0 | 0 |
| 0 | NA | NA | NA | NA | NA | Olfr1099 | olfactory receptor 1099 | 0 | 0 | 0 | 0 | 0 | 0 | 0 | 0 | 0 | 0 | 0 |
| 0 | NA | NA | NA | NA | NA | Olfr1100 | olfactory receptor 1100 | 0 | 0 | 0 | 0 | 0 | 0 | 0 | 0 | 0 | 0 | 0 |
| 0 | NA | NA | NA | NA | NA | Olfr1101 | olfactory receptor 1101 | 0 | 0 | 0 | 0 | 0 | 0 | 0 | 0 | 0 | 0 | 0 |
| 0 | NA | NA | NA | NA | NA | Olfr1102 | olfactory receptor 1102 | 0 | 0 | 0 | 0 | 0 | 0 | 0 | 0 | 0 | 0 | 0 |
| 0 | NA | NA | NA | NA | NA | Olfr1103-ps1 | | 0 | 0 | 0 | 0 | 0 | 0 | 0 | 0 | 0 | 0 | 0 |
| 0 | NA | NA | NA | NA | NA | Olfr1104 | olfactory receptor 1104 | 0 | 0 | 0 | 0 | 0 | 0 | 0 | 0 | 0 | 0 | 0 |
| 0 | NA | NA | NA | NA | NA | Olfr1105 | olfactory receptor 1105 | 0 | 0 | 0 | 0 | 0 | 0 | 0 | 0 | 0 | 0 | 0 |
| 0 | NA | NA | NA | NA | NA | Olfr1106 | olfactory receptor 1106 | 0 | 0 | 0 | 0 | 0 | 0 | 0 | 0 | 0 | 0 | 0 |
| 0 | NA | NA | NA | NA | NA | Olfr1107 | olfactory receptor 1107 | 0 | 0 | 0 | 0 | 0 | 0 | 0 | 0 | 0 | 0 | 0 |
| 0 | NA | NA | NA | NA | NA | Olfr1108-ps1 | | 0 | 0 | 0 | 0 | 0 | 0 | 0 | 0 | 0 | 0 | 0 |
| 0 | NA | NA | NA | NA | NA | Olfr1109 | olfactory receptor 1109 | 0 | 0 | 0 | 0 | 0 | 0 | 0 | 0 | 0 | 0 | 0 |
| 0 | NA | NA | NA | NA | NA | Olfr259 | olfactory receptor 259 | 0 | 0 | 0 | 0 | 0 | 0 | 0 | 0 | 0 | 0 | 0 |
| 0 | NA | NA | NA | NA | NA | Olfr1110 | olfactory receptor 1110 | 0 | 0 | 0 | 0 | 0 | 0 | 0 | 0 | 0 | 0 | 0 |
| 0 | NA | NA | NA | NA | NA | Olfr1111 | olfactory receptor 1111 | 0 | 0 | 0 | 0 | 0 | 0 | 0 | 0 | 0 | 0 | 0 |
| 0 | NA | NA | NA | NA | NA | Olfr1112 | olfactory receptor 1112 | 0 | 0 | 0 | 0 | 0 | 0 | 0 | 0 | 0 | 0 | 0 |
| 0 | NA | NA | NA | NA | NA | Olfr1113 | olfactory receptor 1113 | 0 | 0 | 0 | 0 | 0 | 0 | 0 | 0 | 0 | 0 | 0 |
| 0 | NA | NA | NA | NA | NA | Olfr1114-ps1 | | 0 | 0 | 0 | 0 | 0 | 0 | 0 | 0 | 0 | 0 | 0 |
| 0 | NA | NA | NA | NA | NA | Gm13735 | | 0 | 0 | 0 | 0 | 0 | 0 | 0 | 0 | 0 | 0 | 0 |
| 0 | NA | NA | NA | NA | NA | Olfr1115 | olfactory receptor 1115 | 0 | 0 | 0 | 0 | 0 | 0 | 0 | 0 | 0 | 0 | 0 |
| 0 | NA | NA | NA | NA | NA | Gm27438 | | 0 | 0 | 0 | 0 | 0 | 0 | 0 | 0 | 0 | 0 | 0 |
| 0 | NA | NA | NA | NA | NA | Olfr1116 | olfactory receptor 1116 | 0 | 0 | 0 | 0 | 0 | 0 | 0 | 0 | 0 | 0 | 0 |
| 0 | NA | NA | NA | NA | NA | Gm13728 | | 0 | 0 | 0 | 0 | 0 | 0 | 0 | 0 | 0 | 0 | 0 |
| 0 | NA | NA | NA | NA | NA | Olfr1117-ps1 | | 0 | 0 | 0 | 0 | 0 | 0 | 0 | 0 | 0 | 0 | 0 |

| | | | | | | | | | | | | | | | | | | |
|---|---|---|---|---|---|---|---|---|---|---|---|---|---|---|---|---|---|---|
| 0 | NA | NA | NA | NA | NA | Gm13729 | | 0 | 0 | 0 | 0 | 0 | 0 | 0 | 0 | 0 | 0 | 0 |
| 0 | NA | NA | NA | NA | NA | Olfr1118 | olfactory receptor 1118 | 0 | 0 | 0 | 0 | 0 | 0 | 0 | 0 | 0 | 0 | 0 |
| 0 | NA | NA | NA | NA | NA | Gm13737 | | 0 | 0 | 0 | 0 | 0 | 0 | 0 | 0 | 0 | 0 | 0 |
| 0 | NA | NA | NA | NA | NA | Gm13731 | | 0 | 0 | 0 | 0 | 0 | 0 | 0 | 0 | 0 | 0 | 0 |
| 0 | NA | NA | NA | NA | NA | Olfr1119-ps1 | | 0 | 0 | 0 | 0 | 0 | 0 | 0 | 0 | 0 | 0 | 0 |
| 0 | NA | NA | NA | NA | NA | Olfr1120 | olfactory receptor 1120 | 0 | 0 | 0 | 0 | 0 | 0 | 0 | 0 | 0 | 0 | 0 |
| 0 | NA | NA | NA | NA | NA | Olfr1121 | olfactory receptor 1121 | 0 | 0 | 0 | 0 | 0 | 0 | 0 | 0 | 0 | 0 | 0 |
| 0 | NA | NA | NA | NA | NA | Olfr1122 | olfactory receptor 1122 | 0 | 0 | 0 | 0 | 0 | 0 | 0 | 0 | 0 | 0 | 0 |
| 0 | NA | NA | NA | NA | NA | Olfr1123 | olfactory receptor 1123 | 0 | 0 | 0 | 0 | 0 | 0 | 0 | 0 | 0 | 0 | 0 |
| 0 | NA | NA | NA | NA | NA | Olfr1124 | olfactory receptor 1124 | 0 | 0 | 0 | 0 | 0 | 0 | 0 | 0 | 0 | 0 | 0 |
| 0 | NA | NA | NA | NA | NA | Olfr1126 | olfactory receptor 1126 | 0 | 0 | 0 | 0 | 0 | 0 | 0 | 0 | 0 | 0 | 0 |
| 0 | NA | NA | NA | NA | NA | Pramel7 | preferentially expressed antigen in melan | 0 | 0 | 0 | 0 | 0 | 0 | 0 | 0 | 0 | 0 | 0 |
| 0 | NA | NA | NA | NA | NA | Pramel6 | preferentially expressed antigen in melan | 0 | 0 | 0 | 0 | 0 | 0 | 0 | 0 | 0 | 0 | 0 |
| 0 | NA | NA | NA | NA | NA | Gm13738 | | 0 | 0 | 0 | 0 | 0 | 0 | 0 | 0 | 0 | 0 | 0 |
| 0 | NA | NA | NA | NA | NA | Olfr1127-ps1 | | 0 | 0 | 0 | 0 | 0 | 0 | 0 | 0 | 0 | 0 | 0 |
| 0 | NA | NA | NA | NA | NA | Olfr153 | olfactory receptor 153 | 0 | 0 | 0 | 0 | 0 | 0 | 0 | 0 | 0 | 0 | 0 |
| 0 | NA | NA | NA | NA | NA | Olfr1128 | olfactory receptor 1128 | 0 | 0 | 0 | 0 | 0 | 0 | 0 | 0 | 0 | 0 | 0 |
| 0 | NA | NA | NA | NA | NA | Gm13744 | | 0 | 0 | 0 | 0 | 0 | 0 | 0 | 0 | 0 | 0 | 0 |
| 0 | NA | NA | NA | NA | NA | Olfr1129 | olfactory receptor 1129 | 0 | 0 | 0 | 0 | 0 | 0 | 0 | 0 | 0 | 0 | 0 |
| 0 | NA | NA | NA | NA | NA | Olfr1130 | olfactory receptor 1130 | 0 | 0 | 0 | 0 | 0 | 0 | 0 | 0 | 0 | 0 | 0 |
| 0 | NA | NA | NA | NA | NA | Olfr1131 | olfactory receptor 1131 | 0 | 0 | 0 | 0 | 0 | 0 | 0 | 0 | 0 | 0 | 0 |
| 0 | NA | NA | NA | NA | NA | Olfr1132 | olfactory receptor 1132 | 0 | 0 | 0 | 0 | 0 | 0 | 0 | 0 | 0 | 0 | 0 |
| 0 | NA | NA | NA | NA | NA | Gm13742 | | 0 | 0 | 0 | 0 | 0 | 0 | 0 | 0 | 0 | 0 | 0 |
| 0 | NA | NA | NA | NA | NA | Olfr1133 | olfactory receptor 1133 | 0 | 0 | 0 | 0 | 0 | 0 | 0 | 0 | 0 | 0 | 0 |
| 0 | NA | NA | NA | NA | NA | Olfr1134 | olfactory receptor 1134 | 0 | 0 | 0 | 0 | 0 | 0 | 0 | 0 | 0 | 0 | 0 |
| 0 | NA | NA | NA | NA | NA | Olfr1135 | olfactory receptor 1135 | 0 | 0 | 0 | 0 | 0 | 0 | 0 | 0 | 0 | 0 | 0 |
| 0 | NA | NA | NA | NA | NA | Olfr1136 | olfactory receptor 1136 | 0 | 0 | 0 | 0 | 0 | 0 | 0 | 0 | 0 | 0 | 0 |
| 0 | NA | NA | NA | NA | NA | Olfr1137 | olfactory receptor 1137 | 0 | 0 | 0 | 0 | 0 | 0 | 0 | 0 | 0 | 0 | 0 |
| 0 | NA | NA | NA | NA | NA | Olfr1138 | olfactory receptor 1138 | 0 | 0 | 0 | 0 | 0 | 0 | 0 | 0 | 0 | 0 | 0 |
| 0 | NA | NA | NA | NA | NA | Olfr1140 | olfactory receptor 1140 | 0 | 0 | 0 | 0 | 0 | 0 | 0 | 0 | 0 | 0 | 0 |
| 0 | NA | NA | NA | NA | NA | Olfr1139-ps1 | | 0 | 0 | 0 | 0 | 0 | 0 | 0 | 0 | 0 | 0 | 0 |
| 0 | NA | NA | NA | NA | NA | Olfr1141 | olfactory receptor 1141 | 0 | 0 | 0 | 0 | 0 | 0 | 0 | 0 | 0 | 0 | 0 |
| 0 | NA | NA | NA | NA | NA | Olfr152 | olfactory receptor 152 | 0 | 0 | 0 | 0 | 0 | 0 | 0 | 0 | 0 | 0 | 0 |
| 0 | NA | NA | NA | NA | NA | Olfr1142-ps1 | | 0 | 0 | 0 | 0 | 0 | 0 | 0 | 0 | 0 | 0 | 0 |
| 0 | NA | NA | NA | NA | NA | Olfr1143 | olfactory receptor 1143 | 0 | 0 | 0 | 0 | 0 | 0 | 0 | 0 | 0 | 0 | 0 |
| 0 | NA | NA | NA | NA | NA | Olfr1144-ps1 | | 0 | 0 | 0 | 0 | 0 | 0 | 0 | 0 | 0 | 0 | 0 |
| 0 | NA | NA | NA | NA | NA | Olfr1145 | olfactory receptor 1145 | 0 | 0 | 0 | 0 | 0 | 0 | 0 | 0 | 0 | 0 | 0 |
| 0 | NA | NA | NA | NA | NA | Olfr1146-ps1 | | 0 | 0 | 0 | 0 | 0 | 0 | 0 | 0 | 0 | 0 | 0 |
| 0 | NA | NA | NA | NA | NA | Olfr1147-ps1 | | 0 | 0 | 0 | 0 | 0 | 0 | 0 | 0 | 0 | 0 | 0 |
| 0 | NA | NA | NA | NA | NA | Olfr1148 | olfactory receptor 1148 | 0 | 0 | 0 | 0 | 0 | 0 | 0 | 0 | 0 | 0 | 0 |
| 0 | NA | NA | NA | NA | NA | Olfr1149-ps1 | | 0 | 0 | 0 | 0 | 0 | 0 | 0 | 0 | 0 | 0 | 0 |
| 0 | NA | NA | NA | NA | NA | Olfr1150-ps1 | | 0 | 0 | 0 | 0 | 0 | 0 | 0 | 0 | 0 | 0 | 0 |
| 0 | NA | NA | NA | NA | NA | Olfr1151 | olfactory receptor 1151 | 0 | 0 | 0 | 0 | 0 | 0 | 0 | 0 | 0 | 0 | 0 |
| 0 | NA | NA | NA | NA | NA | Olfr1152 | olfactory receptor 1152 | 0 | 0 | 0 | 0 | 0 | 0 | 0 | 0 | 0 | 0 | 0 |
| 0 | NA | NA | NA | NA | NA | Gm13745 | | 0 | 0 | 0 | 0 | 0 | 0 | 0 | 0 | 0 | 0 | 0 |
| 0 | NA | NA | NA | NA | NA | Olfr1153 | olfactory receptor 1153 | 0 | 0 | 0 | 0 | 0 | 0 | 0 | 0 | 0 | 0 | 0 |
| 0 | NA | NA | NA | NA | NA | Olfr1154 | olfactory receptor 1154 | 0 | 0 | 0 | 0 | 0 | 0 | 0 | 0 | 0 | 0 | 0 |
| 0 | NA | NA | NA | NA | NA | Olfr1155 | olfactory receptor 1155 | 0 | 0 | 0 | 0 | 0 | 0 | 0 | 0 | 0 | 0 | 0 |
| 0 | NA | NA | NA | NA | NA | Olfr1156 | olfactory receptor 1156 | 0 | 0 | 0 | 0 | 0 | 0 | 0 | 0 | 0 | 0 | 0 |
| 0 | NA | NA | NA | NA | NA | Olfr1157 | olfactory receptor 1157 | 0 | 0 | 0 | 0 | 0 | 0 | 0 | 0 | 0 | 0 | 0 |
| 0 | NA | NA | NA | NA | NA | Olfr74 | olfactory receptor 74 | 0 | 0 | 0 | 0 | 0 | 0 | 0 | 0 | 0 | 0 | 0 |
| 0 | NA | NA | NA | NA | NA | Olfr1158 | olfactory receptor 1158 | 0 | 0 | 0 | 0 | 0 | 0 | 0 | 0 | 0 | 0 | 0 |
| 0 | NA | NA | NA | NA | NA | Olfr1159-ps1 | | 0 | 0 | 0 | 0 | 0 | 0 | 0 | 0 | 0 | 0 | 0 |
| 0 | NA | NA | NA | NA | NA | Olfr1160 | olfactory receptor 1160 | 0 | 0 | 0 | 0 | 0 | 0 | 0 | 0 | 0 | 0 | 0 |
| 0 | NA | NA | NA | NA | NA | Olfr1161 | olfactory receptor 1161 | 0 | 0 | 0 | 0 | 0 | 0 | 0 | 0 | 0 | 0 | 0 |
| 0 | NA | NA | NA | NA | NA | Olfr73 | olfactory receptor 73 | 0 | 0 | 0 | 0 | 0 | 0 | 0 | 0 | 0 | 0 | 0 |
| 0 | NA | NA | NA | NA | NA | Olfr1162 | olfactory receptor 1162 | 0 | 0 | 0 | 0 | 0 | 0 | 0 | 0 | 0 | 0 | 0 |
| 0 | NA | NA | NA | NA | NA | Olfr1163 | olfactory receptor 1163 | 0 | 0 | 0 | 0 | 0 | 0 | 0 | 0 | 0 | 0 | 0 |
| 0 | NA | NA | NA | NA | NA | Olfr1164 | olfactory receptor 1164 | 0 | 0 | 0 | 0 | 0 | 0 | 0 | 0 | 0 | 0 | 0 |
| 0 | NA | NA | NA | NA | NA | Olfr1165-ps | | 0 | 0 | 0 | 0 | 0 | 0 | 0 | 0 | 0 | 0 | 0 |

| | | | | | | | | | | | | | | | | | | |
|---|---|---|---|---|---|---|---|---|---|---|---|---|---|---|---|---|---|---|
| 0 | NA | NA | NA | NA | NA | Olfr1166 | olfactory receptor 1166 | 0 | 0 | 0 | 0 | 0 | 0 | 0 | 0 | 0 | 0 | 0 |
| 0 | NA | NA | NA | NA | NA | Olfr1167 | olfactory receptor 1167 | 0 | 0 | 0 | 0 | 0 | 0 | 0 | 0 | 0 | 0 | 0 |
| 0 | NA | NA | NA | NA | NA | Olfr1169-ps1 | | 0 | 0 | 0 | 0 | 0 | 0 | 0 | 0 | 0 | 0 | 0 |
| 0 | NA | NA | NA | NA | NA | Olfr1170 | olfactory receptor 1170 | 0 | 0 | 0 | 0 | 0 | 0 | 0 | 0 | 0 | 0 | 0 |
| 0 | NA | NA | NA | NA | NA | Olfr1171-ps1 | | 0 | 0 | 0 | 0 | 0 | 0 | 0 | 0 | 0 | 0 | 0 |
| 0 | NA | NA | NA | NA | NA | Olfr1172-ps1 | | 0 | 0 | 0 | 0 | 0 | 0 | 0 | 0 | 0 | 0 | 0 |
| 0 | NA | NA | NA | NA | NA | Olfr1174 | | 0 | 0 | 0 | 0 | 0 | 0 | 0 | 0 | 0 | 0 | 0 |
| 0 | NA | NA | NA | NA | NA | Olfr1176 | olfactory receptor 1176 | 0 | 0 | 0 | 0 | 0 | 0 | 0 | 0 | 0 | 0 | 0 |
| 0 | NA | NA | NA | NA | NA | Olfr1177-ps | | 0 | 0 | 0 | 0 | 0 | 0 | 0 | 0 | 0 | 0 | 0 |
| 0 | NA | NA | NA | NA | NA | Olfr1178 | olfactory receptor 1178 | 0 | 0 | 0 | 0 | 0 | 0 | 0 | 0 | 0 | 0 | 0 |
| 0 | NA | NA | NA | NA | NA | Olfr1179 | olfactory receptor 1179 | 0 | 0 | 0 | 0 | 0 | 0 | 0 | 0 | 0 | 0 | 0 |
| 0 | NA | NA | NA | NA | NA | Olfr1180 | olfactory receptor 1180 | 0 | 0 | 0 | 0 | 0 | 0 | 0 | 0 | 0 | 0 | 0 |
| 0 | NA | NA | NA | NA | NA | Olfr1181 | olfactory receptor 1181 | 0 | 0 | 0 | 0 | 0 | 0 | 0 | 0 | 0 | 0 | 0 |
| 0 | NA | NA | NA | NA | NA | Olfr1182 | olfactory receptor 1182 | 0 | 0 | 0 | 0 | 0 | 0 | 0 | 0 | 0 | 0 | 0 |
| 0 | NA | NA | NA | NA | NA | Olfr1183 | olfactory receptor 1183 | 0 | 0 | 0 | 0 | 0 | 0 | 0 | 0 | 0 | 0 | 0 |
| 0 | NA | NA | NA | NA | NA | Olfr1184 | olfactory receptor 1184 | 0 | 0 | 0 | 0 | 0 | 0 | 0 | 0 | 0 | 0 | 0 |
| 0 | NA | NA | NA | NA | NA | Olfr1185-ps1 | | 0 | 0 | 0 | 0 | 0 | 0 | 0 | 0 | 0 | 0 | 0 |
| 0 | NA | NA | NA | NA | NA | Olfr1186 | olfactory receptor 1186 | 0 | 0 | 0 | 0 | 0 | 0 | 0 | 0 | 0 | 0 | 0 |
| 0 | NA | NA | NA | NA | NA | Olfr1187-ps1 | | 0 | 0 | 0 | 0 | 0 | 0 | 0 | 0 | 0 | 0 | 0 |
| 0 | NA | NA | NA | NA | NA | Olfr1188 | olfactory receptor 1188 | 0 | 0 | 0 | 0 | 0 | 0 | 0 | 0 | 0 | 0 | 0 |
| 0 | NA | NA | NA | NA | NA | Olfr1189 | olfactory receptor 1189 | 0 | 0 | 0 | 0 | 0 | 0 | 0 | 0 | 0 | 0 | 0 |
| 0 | NA | NA | NA | NA | NA | Gm47581 | | 0 | 0 | 0 | 0 | 0 | 0 | 0 | 0 | 0 | 0 | 0 |
| 0 | NA | NA | NA | NA | NA | Olfr475-ps1 | | 0 | 0 | 0 | 0 | 0 | 0 | 0 | 0 | 0 | 0 | 0 |
| 0 | NA | NA | NA | NA | NA | Olfr1191-ps1 | | 0 | 0 | 0 | 0 | 0 | 0 | 0 | 0 | 0 | 0 | 0 |
| 0 | NA | NA | NA | NA | NA | Olfr1192-ps1 | | 0 | 0 | 0 | 0 | 0 | 0 | 0 | 0 | 0 | 0 | 0 |
| 0 | NA | NA | NA | NA | NA | Olfr1193 | olfactory receptor 1193 | 0 | 0 | 0 | 0 | 0 | 0 | 0 | 0 | 0 | 0 | 0 |
| 0 | NA | NA | NA | NA | NA | Olfr1195 | olfactory receptor 1195 | 0 | 0 | 0 | 0 | 0 | 0 | 0 | 0 | 0 | 0 | 0 |
| 0 | NA | NA | NA | NA | NA | Olfr1196 | olfactory receptor 1196 | 0 | 0 | 0 | 0 | 0 | 0 | 0 | 0 | 0 | 0 | 0 |
| 0 | NA | NA | NA | NA | NA | Olfr1197 | olfactory receptor 1197 | 0 | 0 | 0 | 0 | 0 | 0 | 0 | 0 | 0 | 0 | 0 |
| 0 | NA | NA | NA | NA | NA | Olfr1198 | olfactory receptor 1198 | 0 | 0 | 0 | 0 | 0 | 0 | 0 | 0 | 0 | 0 | 0 |
| 0 | NA | NA | NA | NA | NA | Olfr1199 | olfactory receptor 1199 | 0 | 0 | 0 | 0 | 0 | 0 | 0 | 0 | 0 | 0 | 0 |
| 0 | NA | NA | NA | NA | NA | Olfr1200 | olfactory receptor 1200 | 0 | 0 | 0 | 0 | 0 | 0 | 0 | 0 | 0 | 0 | 0 |
| 0 | NA | NA | NA | NA | NA | Olfr1201 | olfactory receptor 1201 | 0 | 0 | 0 | 0 | 0 | 0 | 0 | 0 | 0 | 0 | 0 |
| 0 | NA | NA | NA | NA | NA | Olfr1202 | olfactory receptor 1202 | 0 | 0 | 0 | 0 | 0 | 0 | 0 | 0 | 0 | 0 | 0 |
| 0 | NA | NA | NA | NA | NA | Olfr1205 | olfactory receptor 1205 | 0 | 0 | 0 | 0 | 0 | 0 | 0 | 0 | 0 | 0 | 0 |
| 0 | NA | NA | NA | NA | NA | Olfr1204 | olfactory receptor 1204 | 0 | 0 | 0 | 0 | 0 | 0 | 0 | 0 | 0 | 0 | 0 |
| 0 | NA | NA | NA | NA | NA | Olfr1206 | olfactory receptor 1206 | 0 | 0 | 0 | 0 | 0 | 0 | 0 | 0 | 0 | 0 | 0 |
| 0 | NA | NA | NA | NA | NA | Olfr1208 | olfactory receptor 1208 | 0 | 0 | 0 | 0 | 0 | 0 | 0 | 0 | 0 | 0 | 0 |
| 0 | NA | NA | NA | NA | NA | Olfr1209 | olfactory receptor 1209 | 0 | 0 | 0 | 0 | 0 | 0 | 0 | 0 | 0 | 0 | 0 |
| 0 | NA | NA | NA | NA | NA | Olfr1211 | olfactory receptor 1211 | 0 | 0 | 0 | 0 | 0 | 0 | 0 | 0 | 0 | 0 | 0 |
| 0 | NA | NA | NA | NA | NA | Olfr1212 | olfactory receptor 1212 | 0 | 0 | 0 | 0 | 0 | 0 | 0 | 0 | 0 | 0 | 0 |
| 0 | NA | NA | NA | NA | NA | Olfr1213 | olfactory receptor 1213 | 0 | 0 | 0 | 0 | 0 | 0 | 0 | 0 | 0 | 0 | 0 |
| 0 | NA | NA | NA | NA | NA | Olfr1214 | olfactory receptor 1214 | 0 | 0 | 0 | 0 | 0 | 0 | 0 | 0 | 0 | 0 | 0 |
| 0 | NA | NA | NA | NA | NA | Olfr1215 | olfactory receptor 1215 | 0 | 0 | 0 | 0 | 0 | 0 | 0 | 0 | 0 | 0 | 0 |
| 0 | NA | NA | NA | NA | NA | Olfr1216 | olfactory receptor 1216 | 0 | 0 | 0 | 0 | 0 | 0 | 0 | 0 | 0 | 0 | 0 |
| 0 | NA | NA | NA | NA | NA | Olfr1217 | olfactory receptor 1217 | 0 | 0 | 0 | 0 | 0 | 0 | 0 | 0 | 0 | 0 | 0 |
| 0 | NA | NA | NA | NA | NA | Olfr1218 | olfactory receptor 1218 | 0 | 0 | 0 | 0 | 0 | 0 | 0 | 0 | 0 | 0 | 0 |
| 0 | NA | NA | NA | NA | NA | Olfr1219 | olfactory receptor 1219 | 0 | 0 | 0 | 0 | 0 | 0 | 0 | 0 | 0 | 0 | 0 |
| 0 | NA | NA | NA | NA | NA | Olfr1220 | olfactory receptor 1220 | 0 | 0 | 0 | 0 | 0 | 0 | 0 | 0 | 0 | 0 | 0 |
| 0 | NA | NA | NA | NA | NA | Olfr1221 | olfactory receptor 1221 | 0 | 0 | 0 | 0 | 0 | 0 | 0 | 0 | 0 | 0 | 0 |
| 0 | NA | NA | NA | NA | NA | Olfr1222 | olfactory receptor 1222 | 0 | 0 | 0 | 0 | 0 | 0 | 0 | 0 | 0 | 0 | 0 |
| 0 | NA | NA | NA | NA | NA | Olfr1223 | olfactory receptor 1223 | 0 | 0 | 0 | 0 | 0 | 0 | 0 | 0 | 0 | 0 | 0 |
| 0 | NA | NA | NA | NA | NA | Olfr1224-ps1 | | 0 | 0 | 0 | 0 | 0 | 0 | 0 | 0 | 0 | 0 | 0 |
| 0 | NA | NA | NA | NA | NA | Olfr1225 | olfactory receptor 1225 | 0 | 0 | 0 | 0 | 0 | 0 | 0 | 0 | 0 | 0 | 0 |
| 0 | NA | NA | NA | NA | NA | Gm13761 | | 0 | 0 | 0 | 0 | 0 | 0 | 0 | 0 | 0 | 0 | 0 |
| 0 | NA | NA | NA | NA | NA | Olfr1226 | olfactory receptor 1226 | 0 | 0 | 0 | 0 | 0 | 0 | 0 | 0 | 0 | 0 | 0 |
| 0 | NA | NA | NA | NA | NA | Gm5035 | | 0 | 0 | 0 | 0 | 0 | 0 | 0 | 0 | 0 | 0 | 0 |
| 0 | NA | NA | NA | NA | NA | Gm18164 | | 0 | 0 | 0 | 0 | 0 | 0 | 0 | 0 | 0 | 0 | 0 |
| 0 | NA | NA | NA | NA | NA | Gm18165 | | 0 | 0 | 0 | 0 | 0 | 0 | 0 | 0 | 0 | 0 | 0 |
| 0 | NA | NA | NA | NA | NA | Gm18166 | | 0 | 0 | 0 | 0 | 0 | 0 | 0 | 0 | 0 | 0 | 0 |

| | | | | | | | | | | | | | | | | | | |
|---|---|---|---|---|---|---|---|---|---|---|---|---|---|---|---|---|---|---|
| 0 | NA | NA | NA | NA | NA | Olfr1227-ps1 | | 0 | 0 | 0 | 0 | 0 | 0 | 0 | 0 | 0 | 0 | 0 |
| 0 | NA | NA | NA | NA | NA | Olfr1228 | olfactory receptor 1228 | 0 | 0 | 0 | 0 | 0 | 0 | 0 | 0 | 0 | 0 | 0 |
| 0 | NA | NA | NA | NA | NA | Olfr1229 | olfactory receptor 1229 | 0 | 0 | 0 | 0 | 0 | 0 | 0 | 0 | 0 | 0 | 0 |
| 0 | NA | NA | NA | NA | NA | Olfr1230 | olfactory receptor 1230 | 0 | 0 | 0 | 0 | 0 | 0 | 0 | 0 | 0 | 0 | 0 |
| 0 | NA | NA | NA | NA | NA | Olfr1231 | olfactory receptor 1231 | 0 | 0 | 0 | 0 | 0 | 0 | 0 | 0 | 0 | 0 | 0 |
| 0 | NA | NA | NA | NA | NA | Olfr1232 | olfactory receptor 1232 | 0 | 0 | 0 | 0 | 0 | 0 | 0 | 0 | 0 | 0 | 0 |
| 0 | NA | NA | NA | NA | NA | Olfr1234 | olfactory receptor 1234 | 0 | 0 | 0 | 0 | 0 | 0 | 0 | 0 | 0 | 0 | 0 |
| 0 | NA | NA | NA | NA | NA | Olfr1235-ps1 | | 0 | 0 | 0 | 0 | 0 | 0 | 0 | 0 | 0 | 0 | 0 |
| 0 | NA | NA | NA | NA | NA | Olfr1239 | olfactory receptor 1239 | 0 | 0 | 0 | 0 | 0 | 0 | 0 | 0 | 0 | 0 | 0 |
| 0 | NA | NA | NA | NA | NA | Olfr1240 | olfactory receptor 1240 | 0 | 0 | 0 | 0 | 0 | 0 | 0 | 0 | 0 | 0 | 0 |
| 0 | NA | NA | NA | NA | NA | Olfr1241 | olfactory receptor 1241 | 0 | 0 | 0 | 0 | 0 | 0 | 0 | 0 | 0 | 0 | 0 |
| 0 | NA | NA | NA | NA | NA | Olfr1242 | olfactory receptor 1242 | 0 | 0 | 0 | 0 | 0 | 0 | 0 | 0 | 0 | 0 | 0 |
| 0 | NA | NA | NA | NA | NA | Olfr1243 | olfactory receptor 1243 | 0 | 0 | 0 | 0 | 0 | 0 | 0 | 0 | 0 | 0 | 0 |
| 0 | NA | NA | NA | NA | NA | Olfr1244-ps1 | | 0 | 0 | 0 | 0 | 0 | 0 | 0 | 0 | 0 | 0 | 0 |
| 0 | NA | NA | NA | NA | NA | Olfr1245 | olfactory receptor 1245 | 0 | 0 | 0 | 0 | 0 | 0 | 0 | 0 | 0 | 0 | 0 |
| 0 | NA | NA | NA | NA | NA | Olfr1246 | olfactory receptor 1246 | 0 | 0 | 0 | 0 | 0 | 0 | 0 | 0 | 0 | 0 | 0 |
| 0 | NA | NA | NA | NA | NA | Olfr1247 | olfactory receptor 1247 | 0 | 0 | 0 | 0 | 0 | 0 | 0 | 0 | 0 | 0 | 0 |
| 0 | NA | NA | NA | NA | NA | Olfr1248 | olfactory receptor 1248 | 0 | 0 | 0 | 0 | 0 | 0 | 0 | 0 | 0 | 0 | 0 |
| 0 | NA | NA | NA | NA | NA | Olfr1249 | olfactory receptor 1249 | 0 | 0 | 0 | 0 | 0 | 0 | 0 | 0 | 0 | 0 | 0 |
| 0 | NA | NA | NA | NA | NA | Olfr1250 | olfactory receptor 1250 | 0 | 0 | 0 | 0 | 0 | 0 | 0 | 0 | 0 | 0 | 0 |
| 0 | NA | NA | NA | NA | NA | Olfr1251 | olfactory receptor 1251 | 0 | 0 | 0 | 0 | 0 | 0 | 0 | 0 | 0 | 0 | 0 |
| 0 | NA | NA | NA | NA | NA | Olfr1253 | olfactory receptor 1253 | 0 | 0 | 0 | 0 | 0 | 0 | 0 | 0 | 0 | 0 | 0 |
| 0 | NA | NA | NA | NA | NA | Olfr1254 | olfactory receptor 1254 | 0 | 0 | 0 | 0 | 0 | 0 | 0 | 0 | 0 | 0 | 0 |
| 0 | NA | NA | NA | NA | NA | Olfr1255 | olfactory receptor 1255 | 0 | 0 | 0 | 0 | 0 | 0 | 0 | 0 | 0 | 0 | 0 |
| 0 | NA | NA | NA | NA | NA | Olfr1256 | olfactory receptor 1256 | 0 | 0 | 0 | 0 | 0 | 0 | 0 | 0 | 0 | 0 | 0 |
| 0 | NA | NA | NA | NA | NA | Olfr48 | olfactory receptor 48 | 0 | 0 | 0 | 0 | 0 | 0 | 0 | 0 | 0 | 0 | 0 |
| 0 | NA | NA | NA | NA | NA | Olfr1257 | olfactory receptor 1257 | 0 | 0 | 0 | 0 | 0 | 0 | 0 | 0 | 0 | 0 | 0 |
| 0 | NA | NA | NA | NA | NA | Olfr1258 | olfactory receptor 1258 | 0 | 0 | 0 | 0 | 0 | 0 | 0 | 0 | 0 | 0 | 0 |
| 0 | NA | NA | NA | NA | NA | Olfr1259 | olfactory receptor 1259 | 0 | 0 | 0 | 0 | 0 | 0 | 0 | 0 | 0 | 0 | 0 |
| 0 | NA | NA | NA | NA | NA | Olfr1260 | olfactory receptor 1260 | 0 | 0 | 0 | 0 | 0 | 0 | 0 | 0 | 0 | 0 | 0 |
| 0 | NA | NA | NA | NA | NA | Olfr1261 | olfactory receptor 1261 | 0 | 0 | 0 | 0 | 0 | 0 | 0 | 0 | 0 | 0 | 0 |
| 0 | NA | NA | NA | NA | NA | Olfr1262 | olfactory receptor 1262 | 0 | 0 | 0 | 0 | 0 | 0 | 0 | 0 | 0 | 0 | 0 |
| 0 | NA | NA | NA | NA | NA | Olfr1263 | olfactory receptor 1263 | 0 | 0 | 0 | 0 | 0 | 0 | 0 | 0 | 0 | 0 | 0 |
| 0 | NA | NA | NA | NA | NA | Olfr1264 | olfactory receptor 1264 | 0 | 0 | 0 | 0 | 0 | 0 | 0 | 0 | 0 | 0 | 0 |
| 0 | NA | NA | NA | NA | NA | Olfr1265 | olfactory receptor 1265 | 0 | 0 | 0 | 0 | 0 | 0 | 0 | 0 | 0 | 0 | 0 |
| 0 | NA | NA | NA | NA | NA | Olfr140 | olfactory receptor 140 | 0 | 0 | 0 | 0 | 0 | 0 | 0 | 0 | 0 | 0 | 0 |
| 0 | NA | NA | NA | NA | NA | Olfr1266-ps1 | | 0 | 0 | 0 | 0 | 0 | 0 | 0 | 0 | 0 | 0 | 0 |
| 0 | NA | NA | NA | NA | NA | Olfr1267-ps1 | | 0 | 0 | 0 | 0 | 0 | 0 | 0 | 0 | 0 | 0 | 0 |
| 0 | NA | NA | NA | NA | NA | Olfr1268-ps1 | | 0 | 0 | 0 | 0 | 0 | 0 | 0 | 0 | 0 | 0 | 0 |
| 0 | NA | NA | NA | NA | NA | Olfr1269 | olfactory receptor 1269 | 0 | 0 | 0 | 0 | 0 | 0 | 0 | 0 | 0 | 0 | 0 |
| 0 | NA | NA | NA | NA | NA | Olfr32 | olfactory receptor 32 | 0 | 0 | 0 | 0 | 0 | 0 | 0 | 0 | 0 | 0 | 0 |
| 0 | NA | NA | NA | NA | NA | Olfr1270 | olfactory receptor 1270 | 0 | 0 | 0 | 0 | 0 | 0 | 0 | 0 | 0 | 0 | 0 |
| 0 | NA | NA | NA | NA | NA | Gm4222 | | 0 | 0 | 0 | 0 | 0 | 0 | 0 | 0 | 0 | 0 | 0 |
| 0 | NA | NA | NA | NA | NA | Gm13768 | | 0 | 0 | 0 | 0 | 0 | 0 | 0 | 0 | 0 | 0 | 0 |
| 0 | NA | NA | NA | NA | NA | Olfr1506 | olfactory receptor 1506 | 0 | 0 | 0 | 0 | 0 | 0 | 0 | 0 | 0 | 0 | 0 |
| 0 | NA | NA | NA | NA | NA | Gm18953 | | 0 | 0 | 0 | 0 | 0 | 0 | 0 | 0 | 0 | 0 | 0 |
| 0 | NA | NA | NA | NA | NA | Olfr142 | olfactory receptor 142 | 0 | 0 | 0 | 0 | 0 | 0 | 0 | 0 | 0 | 0 | 0 |
| 0 | NA | NA | NA | NA | NA | Olfr1271 | olfactory receptor 1271 | 0 | 0 | 0 | 0 | 0 | 0 | 0 | 0 | 0 | 0 | 0 |
| 0 | NA | NA | NA | NA | NA | Olfr1272 | olfactory receptor 1272 | 0 | 0 | 0 | 0 | 0 | 0 | 0 | 0 | 0 | 0 | 0 |
| 0 | NA | NA | NA | NA | NA | Olfr1273-ps | | 0 | 0 | 0 | 0 | 0 | 0 | 0 | 0 | 0 | 0 | 0 |
| 0 | NA | NA | NA | NA | NA | Olfr1565 | olfactory receptor 1506 | 0 | 0 | 0 | 0 | 0 | 0 | 0 | 0 | 0 | 0 | 0 |
| 0 | NA | NA | NA | NA | NA | Gm13767 | | 0 | 0 | 0 | 0 | 0 | 0 | 0 | 0 | 0 | 0 | 0 |
| 0 | NA | NA | NA | NA | NA | Olfr1274 | olfactory receptor 1274 | 0 | 0 | 0 | 0 | 0 | 0 | 0 | 0 | 0 | 0 | 0 |
| 0 | NA | NA | NA | NA | NA | 4933423P22Rik | | 0 | 0 | 0 | 0 | 0 | 0 | 0 | 0 | 0 | 0 | 0 |
| 0 | NA | NA | NA | NA | NA | Gm32514 | | 0 | 0 | 0 | 0 | 0 | 0 | 0 | 0 | 0 | 0 | 0 |
| 0 | NA | NA | NA | NA | NA | Gm23390 | | 0 | 0 | 0 | 0 | 0 | 0 | 0 | 0 | 0 | 0 | 0 |
| 0 | NA | NA | NA | NA | NA | Gm22071 | | 0 | 0 | 0 | 0 | 0 | 0 | 0 | 0 | 0 | 0 | 0 |
| 0 | NA | NA | NA | NA | NA | Snord67 | | 0 | 0 | 0 | 0 | 0 | 0 | 0 | 0 | 0 | 0 | 0 |
| 0 | NA | NA | NA | NA | NA | Gm13776 | | 0 | 0 | 0 | 0 | 0 | 0 | 0 | 0 | 0 | 0 | 0 |
| 0 | NA | NA | NA | NA | NA | Gm9821 | | 0 | 0 | 0 | 0 | 0 | 0 | 0 | 0 | 0 | 0 | 0 |

| | | | | | | | | | | | | | | | | | | |
|---|---|---|---|---|---|---|---|---|---|---|---|---|---|---|---|---|---|---|
| 0 | NA | NA | NA | NA | NA | Gm13780 | | 0 | 0 | 0 | 0 | 0 | 0 | 0 | 0 | 0 | 0 | 0 |
| 0 | NA | NA | NA | NA | NA | Gm13817 | | 0 | 0 | 0 | 0 | 0 | 0 | 0 | 0 | 0 | 0 | 0 |
| 0 | NA | NA | NA | NA | NA | Mir7221 | | 0 | 0 | 0 | 0 | 0 | 0 | 0 | 0 | 0 | 0 | 0 |
| 0 | NA | NA | NA | NA | NA | Mir7222 | | 0 | 0 | 0 | 0 | 0 | 0 | 0 | 0 | 0 | 0 | 0 |
| 0 | NA | NA | NA | NA | NA | 4631405J19Rik | | 0 | 0 | 0 | 0 | 0 | 0 | 0 | 0 | 0 | 0 | 0 |
| 0 | NA | NA | NA | NA | NA | Gm13802 | | 0 | 0 | 0 | 0 | 0 | 0 | 0 | 0 | 0 | 0 | 0 |
| 0 | NA | NA | NA | NA | NA | Gm23380 | | 0 | 0 | 0 | 0 | 0 | 0 | 0 | 0 | 0 | 0 | 0 |
| 0 | NA | NA | NA | NA | NA | Gm13807 | | 0 | 0 | 0 | 0 | 0 | 0 | 0 | 0 | 0 | 0 | 0 |
| 0 | NA | NA | NA | NA | NA | Gm10804 | | 0 | 0 | 0 | 0 | 0 | 0 | 0 | 0 | 0 | 0 | 0 |
| 0 | NA | NA | NA | NA | NA | Gm13818 | | 0 | 0 | 0 | 0 | 0 | 0 | 0 | 0 | 0 | 0 | 0 |
| 0 | NA | NA | NA | NA | NA | Alkbh3os1 | | 0 | 0 | 0 | 0 | 0 | 0 | 0 | 0 | 0 | 0 | 0 |
| 0 | NA | NA | NA | NA | NA | Mir129b | | 0 | 0 | 0 | 0 | 0 | 0 | 0 | 0 | 0 | 0 | 0 |
| 0 | NA | NA | NA | NA | NA | Gm44461 | | 0 | 0 | 0 | 0 | 0 | 0 | 0 | 0 | 0 | 0 | 0 |
| 0 | NA | NA | NA | NA | NA | Gm13788 | | 0 | 0 | 0 | 0 | 0 | 0 | 0 | 0 | 0 | 0 | 0 |
| 0 | NA | NA | NA | NA | NA | Gm24318 | | 0 | 0 | 0 | 0 | 0 | 0 | 0 | 0 | 0 | 0 | 0 |
| 0 | NA | NA | NA | NA | NA | Gm13793 | | 0 | 0 | 0 | 0 | 0 | 0 | 0 | 0 | 0 | 0 | 0 |
| 0 | NA | NA | NA | NA | NA | Gm26396 | | 0 | 0 | 0 | 0 | 0 | 0 | 0 | 0 | 0 | 0 | 0 |
| 0 | NA | NA | NA | NA | NA | Gm13794 | | 0 | 0 | 0 | 0 | 0 | 0 | 0 | 0 | 0 | 0 | 0 |
| 0 | NA | NA | NA | NA | NA | Gm13798 | | 0 | 0 | 0 | 0 | 0 | 0 | 0 | 0 | 0 | 0 | 0 |
| 0 | NA | NA | NA | NA | NA | Gm13797 | | 0 | 0 | 0 | 0 | 0 | 0 | 0 | 0 | 0 | 0 | 0 |
| 0 | NA | NA | NA | NA | NA | Gm13799 | | 0 | 0 | 0 | 0 | 0 | 0 | 0 | 0 | 0 | 0 | 0 |
| 0 | NA | NA | NA | NA | NA | Gm37175 | | 0 | 0 | 0 | 0 | 0 | 0 | 0 | 0 | 0 | 0 | 0 |
| 0 | NA | NA | NA | NA | NA | Gm13801 | | 0 | 0 | 0 | 0 | 0 | 0 | 0 | 0 | 0 | 0 | 0 |
| 0 | NA | NA | NA | NA | NA | 4930445B16Rik | | 0 | 0 | 0 | 0 | 0 | 0 | 0 | 0 | 0 | 0 | 0 |
| 0 | NA | NA | NA | NA | NA | Gm17075 | | 0 | 0 | 0 | 0 | 0 | 0 | 0 | 0 | 0 | 0 | 0 |
| 0 | NA | NA | NA | NA | NA | Gm17076 | | 0 | 0 | 0 | 0 | 0 | 0 | 0 | 0 | 0 | 0 | 0 |
| 0 | NA | NA | NA | NA | NA | Gm22957 | | 0 | 0 | 0 | 0 | 0 | 0 | 0 | 0 | 0 | 0 | 0 |
| 0 | NA | NA | NA | NA | NA | Gm22313 | | 0 | 0 | 0 | 0 | 0 | 0 | 0 | 0 | 0 | 0 | 0 |
| 0 | NA | NA | NA | NA | NA | Gm23317 | | 0 | 0 | 0 | 0 | 0 | 0 | 0 | 0 | 0 | 0 | 0 |
| 0 | NA | NA | NA | NA | NA | Gm44320 | | 0 | 0 | 0 | 0 | 0 | 0 | 0 | 0 | 0 | 0 | 0 |
| 0 | NA | NA | NA | NA | NA | Gm13804 | | 0 | 0 | 0 | 0 | 0 | 0 | 0 | 0 | 0 | 0 | 0 |
| 0 | NA | NA | NA | NA | NA | Gm13806 | | 0 | 0 | 0 | 0 | 0 | 0 | 0 | 0 | 0 | 0 | 0 |
| 0 | NA | NA | NA | NA | NA | Gm29235 | | 0 | 0 | 0 | 0 | 0 | 0 | 0 | 0 | 0 | 0 | 0 |
| 0 | NA | NA | NA | NA | NA | Gm13809 | | 0 | 0 | 0 | 0 | 0 | 0 | 0 | 0 | 0 | 0 | 0 |
| 0 | NA | NA | NA | NA | NA | Gm13808 | | 0 | 0 | 0 | 0 | 0 | 0 | 0 | 0 | 0 | 0 | 0 |
| 0 | NA | NA | NA | NA | NA | Gm13813 | | 0 | 0 | 0 | 0 | 0 | 0 | 0 | 0 | 0 | 0 | 0 |
| 0 | NA | NA | NA | NA | NA | Gm13812 | | 0 | 0 | 0 | 0 | 0 | 0 | 0 | 0 | 0 | 0 | 0 |
| 0 | NA | NA | NA | NA | NA | Gm13810 | | 0 | 0 | 0 | 0 | 0 | 0 | 0 | 0 | 0 | 0 | 0 |
| 0 | NA | NA | NA | NA | NA | Gm20693 | | 0 | 0 | 0 | 0 | 0 | 0 | 0 | 0 | 0 | 0 | 0 |
| 0 | NA | NA | NA | NA | NA | Rag2 | recombination activating gene 2 | 0 | 0 | 0 | 0 | 0 | 0 | 0 | 0 | 0 | 0 | 0 |
| 0 | NA | NA | NA | NA | NA | Rag1 | recombination activating 1 | 0 | 0 | 0 | 0 | 0 | 0 | 0 | 0 | 0 | 0 | 0 |
| 0 | NA | NA | NA | NA | NA | Gm13905 | | 0 | 0 | 0 | 0 | 0 | 0 | 0 | 0 | 0 | 0 | 0 |
| 0 | NA | NA | NA | NA | NA | Gm13920 | | 0 | 0 | 0 | 0 | 0 | 0 | 0 | 0 | 0 | 0 | 0 |
| 0 | NA | NA | NA | NA | NA | Gm13866 | | 0 | 0 | 0 | 0 | 0 | 0 | 0 | 0 | 0 | 0 | 0 |
| 0 | NA | NA | NA | NA | NA | Gm13872 | | 0 | 0 | 0 | 0 | 0 | 0 | 0 | 0 | 0 | 0 | 0 |
| 0 | NA | NA | NA | NA | NA | Gm23731 | | 0 | 0 | 0 | 0 | 0 | 0 | 0 | 0 | 0 | 0 | 0 |
| 0 | NA | NA | NA | NA | NA | Gm13869 | | 0 | 0 | 0 | 0 | 0 | 0 | 0 | 0 | 0 | 0 | 0 |
| 0 | NA | NA | NA | NA | NA | Gm13873 | | 0 | 0 | 0 | 0 | 0 | 0 | 0 | 0 | 0 | 0 | 0 |
| 0 | NA | NA | NA | NA | NA | A930006I01Rik | | 0 | 0 | 0 | 0 | 0 | 0 | 0 | 0 | 0 | 0 | 0 |
| 0 | NA | NA | NA | NA | NA | Gm37220 | | 0 | 0 | 0 | 0 | 0 | 0 | 0 | 0 | 0 | 0 | 0 |
| 0 | NA | NA | NA | NA | NA | Gm13996 | | 0 | 0 | 0 | 0 | 0 | 0 | 0 | 0 | 0 | 0 | 0 |
| 0 | NA | NA | NA | NA | NA | 4930547E08Rik | | 0 | 0 | 0 | 0 | 0 | 0 | 0 | 0 | 0 | 0 | 0 |
| 0 | NA | NA | NA | NA | NA | Gm13878 | | 0 | 0 | 0 | 0 | 0 | 0 | 0 | 0 | 0 | 0 | 0 |
| 0 | NA | NA | NA | NA | NA | Gm13877 | | 0 | 0 | 0 | 0 | 0 | 0 | 0 | 0 | 0 | 0 | 0 |
| 0 | NA | NA | NA | NA | NA | Gm22757 | | 0 | 0 | 0 | 0 | 0 | 0 | 0 | 0 | 0 | 0 | 0 |
| 0 | NA | NA | NA | NA | NA | Gm13879 | | 0 | 0 | 0 | 0 | 0 | 0 | 0 | 0 | 0 | 0 | 0 |
| 0 | NA | NA | NA | NA | NA | Gm22761 | | 0 | 0 | 0 | 0 | 0 | 0 | 0 | 0 | 0 | 0 | 0 |
| 0 | NA | NA | NA | NA | NA | Gm13876 | | 0 | 0 | 0 | 0 | 0 | 0 | 0 | 0 | 0 | 0 | 0 |
| 0 | NA | NA | NA | NA | NA | Gm13881 | | 0 | 0 | 0 | 0 | 0 | 0 | 0 | 0 | 0 | 0 | 0 |
| 0 | NA | NA | NA | NA | NA | Gm10799 | | 0 | 0 | 0 | 0 | 0 | 0 | 0 | 0 | 0 | 0 | 0 |

| | | | | | | | | | | | | | | | | | |
|---|---|---|---|---|---|---|---|---|---|---|---|---|---|---|---|---|---|
| 0 | NA | NA | NA | NA | NA | Gm22501 | | 0 | 0 | 0 | 0 | 0 | 0 | 0 | 0 | 0 | 0 | 0 |
| 0 | NA | NA | NA | NA | NA | Gm13883 | | 0 | 0 | 0 | 0 | 0 | 0 | 0 | 0 | 0 | 0 | 0 |
| 0 | NA | NA | NA | NA | NA | Gm25554 | | 0 | 0 | 0 | 0 | 0 | 0 | 0 | 0 | 0 | 0 | 0 |
| 0 | NA | NA | NA | NA | NA | Mir1902 | | 0 | 0 | 0 | 0 | 0 | 0 | 0 | 0 | 0 | 0 | 0 |
| 0 | NA | NA | NA | NA | NA | Gm13884 | | 0 | 0 | 0 | 0 | 0 | 0 | 0 | 0 | 0 | 0 | 0 |
| 0 | NA | NA | NA | NA | NA | Gm22439 | | 0 | 0 | 0 | 0 | 0 | 0 | 0 | 0 | 0 | 0 | 0 |
| 0 | NA | NA | NA | NA | NA | Gm13885 | | 0 | 0 | 0 | 0 | 0 | 0 | 0 | 0 | 0 | 0 | 0 |
| 0 | NA | NA | NA | NA | NA | Gm13887 | | 0 | 0 | 0 | 0 | 0 | 0 | 0 | 0 | 0 | 0 | 0 |
| 0 | NA | NA | NA | NA | NA | Gm13888 | | 0 | 0 | 0 | 0 | 0 | 0 | 0 | 0 | 0 | 0 | 0 |
| 0 | NA | NA | NA | NA | NA | Gm11060 | | 0 | 0 | 0 | 0 | 0 | 0 | 0 | 0 | 0 | 0 | 0 |
| 0 | NA | NA | NA | NA | NA | Gm27538 | | 0 | 0 | 0 | 0 | 0 | 0 | 0 | 0 | 0 | 0 | 0 |
| 0 | NA | NA | NA | NA | NA | Gm27636 | | 0 | 0 | 0 | 0 | 0 | 0 | 0 | 0 | 0 | 0 | 0 |
| 0 | NA | NA | NA | NA | NA | Gm27308 | | 0 | 0 | 0 | 0 | 0 | 0 | 0 | 0 | 0 | 0 | 0 |
| 0 | NA | NA | NA | NA | NA | Them7 | thioesterase superfamily member 7 | 0 | 0 | 0 | 0 | 0 | 0 | 0 | 0 | 0 | 0 | 0 |
| 0 | NA | NA | NA | NA | NA | Gm22002 | | 0 | 0 | 0 | 0 | 0 | 0 | 0 | 0 | 0 | 0 | 0 |
| 0 | NA | NA | NA | NA | NA | n-R5s201 | | 0 | 0 | 0 | 0 | 0 | 0 | 0 | 0 | 0 | 0 | 0 |
| 0 | NA | NA | NA | NA | NA | Gm13992 | | 0 | 0 | 0 | 0 | 0 | 0 | 0 | 0 | 0 | 0 | 0 |
| 0 | NA | NA | NA | NA | NA | Gm29053 | | 0 | 0 | 0 | 0 | 0 | 0 | 0 | 0 | 0 | 0 | 0 |
| 0 | NA | NA | NA | NA | NA | Gm14015 | | 0 | 0 | 0 | 0 | 0 | 0 | 0 | 0 | 0 | 0 | 0 |
| 0 | NA | NA | NA | NA | NA | Gm22813 | | 0 | 0 | 0 | 0 | 0 | 0 | 0 | 0 | 0 | 0 | 0 |
| 0 | NA | NA | NA | NA | NA | Gm22813 | | 0 | 0 | 0 | 0 | 0 | 0 | 0 | 0 | 0 | 0 | 0 |
| 0 | NA | NA | NA | NA | NA | Gm13901 | | 0 | 0 | 0 | 0 | 0 | 0 | 0 | 0 | 0 | 0 | 0 |
| 0 | NA | NA | NA | NA | NA | Gm13900 | | 0 | 0 | 0 | 0 | 0 | 0 | 0 | 0 | 0 | 0 | 0 |
| 0 | NA | NA | NA | NA | NA | Fshb | follicle stimulating hormone beta | 0 | 0 | 0 | 0 | 0 | 0 | 0 | 0 | 0 | 0 | 0 |
| 0 | NA | NA | NA | NA | NA | Gm24885 | | 0 | 0 | 0 | 0 | 0 | 0 | 0 | 0 | 0 | 0 | 0 |
| 0 | NA | NA | NA | NA | NA | Gm13903 | | 0 | 0 | 0 | 0 | 0 | 0 | 0 | 0 | 0 | 0 | 0 |
| 0 | NA | NA | NA | NA | NA | Gm23439 | | 0 | 0 | 0 | 0 | 0 | 0 | 0 | 0 | 0 | 0 | 0 |
| 0 | NA | NA | NA | NA | NA | Gm23749 | | 0 | 0 | 0 | 0 | 0 | 0 | 0 | 0 | 0 | 0 | 0 |
| 0 | NA | NA | NA | NA | NA | Gm13916 | | 0 | 0 | 0 | 0 | 0 | 0 | 0 | 0 | 0 | 0 | 0 |
| 0 | NA | NA | NA | NA | NA | Gm13915 | | 0 | 0 | 0 | 0 | 0 | 0 | 0 | 0 | 0 | 0 | 0 |
| 0 | NA | NA | NA | NA | NA | Gm13913 | | 0 | 0 | 0 | 0 | 0 | 0 | 0 | 0 | 0 | 0 | 0 |
| 0 | NA | NA | NA | NA | NA | Gm13914 | | 0 | 0 | 0 | 0 | 0 | 0 | 0 | 0 | 0 | 0 | 0 |
| 0 | NA | NA | NA | NA | NA | Gm13923 | | 0 | 0 | 0 | 0 | 0 | 0 | 0 | 0 | 0 | 0 | 0 |
| 0 | NA | NA | NA | NA | NA | Gm13924 | | 0 | 0 | 0 | 0 | 0 | 0 | 0 | 0 | 0 | 0 | 0 |
| 0 | NA | NA | NA | NA | NA | Gm13925 | | 0 | 0 | 0 | 0 | 0 | 0 | 0 | 0 | 0 | 0 | 0 |
| 0 | NA | NA | NA | NA | NA | Gm20638 | | 0 | 0 | 0 | 0 | 0 | 0 | 0 | 0 | 0 | 0 | 0 |
| 0 | NA | NA | NA | NA | NA | Ccdc34os | | 0 | 0 | 0 | 0 | 0 | 0 | 0 | 0 | 0 | 0 | 0 |
| 0 | NA | NA | NA | NA | NA | Gm13936 | | 0 | 0 | 0 | 0 | 0 | 0 | 0 | 0 | 0 | 0 | 0 |
| 0 | NA | NA | NA | NA | NA | Gm13937 | | 0 | 0 | 0 | 0 | 0 | 0 | 0 | 0 | 0 | 0 | 0 |
| 0 | NA | NA | NA | NA | NA | Gm22997 | | 0 | 0 | 0 | 0 | 0 | 0 | 0 | 0 | 0 | 0 | 0 |
| 0 | NA | NA | NA | NA | NA | Gm10794 | | 0 | 0 | 0 | 0 | 0 | 0 | 0 | 0 | 0 | 0 | 0 |
| 0 | NA | NA | NA | NA | NA | Slc5a12 | solute carrier family 5 (sodium/glucose cc | 0 | 0 | 0 | 0 | 0 | 0 | 0 | 0 | 0 | 0 | 0 |
| 0 | NA | NA | NA | NA | NA | Muc15 | mucin 15 | 0 | 0 | 0 | 0 | 0 | 0 | 0 | 0 | 0 | 0 | 0 |
| 0 | NA | NA | NA | NA | NA | Gm13942 | | 0 | 0 | 0 | 0 | 0 | 0 | 0 | 0 | 0 | 0 | 0 |
| 0 | NA | NA | NA | NA | NA | Gm13941 | | 0 | 0 | 0 | 0 | 0 | 0 | 0 | 0 | 0 | 0 | 0 |
| 0 | NA | NA | NA | NA | NA | Gm15130 | | 0 | 0 | 0 | 0 | 0 | 0 | 0 | 0 | 0 | 0 | 0 |
| 0 | NA | NA | NA | NA | NA | A26c3 | ANKRD26-like family C, member 3 | 0 | 0 | 0 | 0 | 0 | 0 | 0 | 0 | 0 | 0 | 0 |
| 0 | NA | NA | NA | NA | NA | Olfr1275 | olfactory receptor 1275 | 0 | 0 | 0 | 0 | 0 | 0 | 0 | 0 | 0 | 0 | 0 |
| 0 | NA | NA | NA | NA | NA | Gm13962 | | 0 | 0 | 0 | 0 | 0 | 0 | 0 | 0 | 0 | 0 | 0 |
| 0 | NA | NA | NA | NA | NA | Olfr1276 | olfactory receptor 1276 | 0 | 0 | 0 | 0 | 0 | 0 | 0 | 0 | 0 | 0 | 0 |
| 0 | NA | NA | NA | NA | NA | Olfr1277 | olfactory receptor 1277 | 0 | 0 | 0 | 0 | 0 | 0 | 0 | 0 | 0 | 0 | 0 |
| 0 | NA | NA | NA | NA | NA | Olfr1278 | olfactory receptor 1278 | 0 | 0 | 0 | 0 | 0 | 0 | 0 | 0 | 0 | 0 | 0 |
| 0 | NA | NA | NA | NA | NA | Olfr1279 | olfactory receptor 1279 | 0 | 0 | 0 | 0 | 0 | 0 | 0 | 0 | 0 | 0 | 0 |
| 0 | NA | NA | NA | NA | NA | Olfr1280 | olfactory receptor 1280 | 0 | 0 | 0 | 0 | 0 | 0 | 0 | 0 | 0 | 0 | 0 |
| 0 | NA | NA | NA | NA | NA | Olfr1281 | olfactory receptor 1281 | 0 | 0 | 0 | 0 | 0 | 0 | 0 | 0 | 0 | 0 | 0 |
| 0 | NA | NA | NA | NA | NA | Olfr1282 | olfactory receptor 1282 | 0 | 0 | 0 | 0 | 0 | 0 | 0 | 0 | 0 | 0 | 0 |
| 0 | NA | NA | NA | NA | NA | Olfr1283 | olfactory receptor 1283 | 0 | 0 | 0 | 0 | 0 | 0 | 0 | 0 | 0 | 0 | 0 |
| 0 | NA | NA | NA | NA | NA | Olfr1284 | olfactory receptor 1284 | 0 | 0 | 0 | 0 | 0 | 0 | 0 | 0 | 0 | 0 | 0 |
| 0 | NA | NA | NA | NA | NA | Olfr1285 | olfactory receptor 1285 | 0 | 0 | 0 | 0 | 0 | 0 | 0 | 0 | 0 | 0 | 0 |
| 0 | NA | NA | NA | NA | NA | Olfr1286 | olfactory receptor 1286 | 0 | 0 | 0 | 0 | 0 | 0 | 0 | 0 | 0 | 0 | 0 |

| | | | | | | | | | | | | | | | | | | |
|---|---|---|---|---|---|---|---|---|---|---|---|---|---|---|---|---|---|---|
| 0 | NA | NA | NA | NA | NA | Olfr1287 | olfactory receptor 1287 | 0 | 0 | 0 | 0 | 0 | 0 | 0 | 0 | 0 | 0 | 0 |
| 0 | NA | NA | NA | NA | NA | Olfr1288 | olfactory receptor 1288 | 0 | 0 | 0 | 0 | 0 | 0 | 0 | 0 | 0 | 0 | 0 |
| 0 | NA | NA | NA | NA | NA | Olfr1289 | olfactory receptor 1289 | 0 | 0 | 0 | 0 | 0 | 0 | 0 | 0 | 0 | 0 | 0 |
| 0 | NA | NA | NA | NA | NA | Olfr1290 | olfactory receptor 1290 | 0 | 0 | 0 | 0 | 0 | 0 | 0 | 0 | 0 | 0 | 0 |
| 0 | NA | NA | NA | NA | NA | Olfr1291-ps1 | | 0 | 0 | 0 | 0 | 0 | 0 | 0 | 0 | 0 | 0 | 0 |
| 0 | NA | NA | NA | NA | NA | Olfr1292-ps1 | | 0 | 0 | 0 | 0 | 0 | 0 | 0 | 0 | 0 | 0 | 0 |
| 0 | NA | NA | NA | NA | NA | Olfr1293-ps | | 0 | 0 | 0 | 0 | 0 | 0 | 0 | 0 | 0 | 0 | 0 |
| 0 | NA | NA | NA | NA | NA | Gm44713 | | 0 | 0 | 0 | 0 | 0 | 0 | 0 | 0 | 0 | 0 | 0 |
| 0 | NA | NA | NA | NA | NA | Olfr1294 | olfactory receptor 1294 | 0 | 0 | 0 | 0 | 0 | 0 | 0 | 0 | 0 | 0 | 0 |
| 0 | NA | NA | NA | NA | NA | Olfr1295 | olfactory receptor 1295 | 0 | 0 | 0 | 0 | 0 | 0 | 0 | 0 | 0 | 0 | 0 |
| 0 | NA | NA | NA | NA | NA | Olfr1296-ps1 | | 0 | 0 | 0 | 0 | 0 | 0 | 0 | 0 | 0 | 0 | 0 |
| 0 | NA | NA | NA | NA | NA | Olfr1297 | olfactory receptor 1297 | 0 | 0 | 0 | 0 | 0 | 0 | 0 | 0 | 0 | 0 | 0 |
| 0 | NA | NA | NA | NA | NA | Olfr1298 | olfactory receptor 1298 | 0 | 0 | 0 | 0 | 0 | 0 | 0 | 0 | 0 | 0 | 0 |
| 0 | NA | NA | NA | NA | NA | Olfr1299 | olfactory receptor 1299 | 0 | 0 | 0 | 0 | 0 | 0 | 0 | 0 | 0 | 0 | 0 |
| 0 | NA | NA | NA | NA | NA | Gm44840 | | 0 | 0 | 0 | 0 | 0 | 0 | 0 | 0 | 0 | 0 | 0 |
| 0 | NA | NA | NA | NA | NA | Olfr1300-ps1 | | 0 | 0 | 0 | 0 | 0 | 0 | 0 | 0 | 0 | 0 | 0 |
| 0 | NA | NA | NA | NA | NA | Gm13922 | | 0 | 0 | 0 | 0 | 0 | 0 | 0 | 0 | 0 | 0 | 0 |
| 0 | NA | NA | NA | NA | NA | Gm13930 | | 0 | 0 | 0 | 0 | 0 | 0 | 0 | 0 | 0 | 0 | 0 |
| 0 | NA | NA | NA | NA | NA | Olfr1301 | olfactory receptor 1301 | 0 | 0 | 0 | 0 | 0 | 0 | 0 | 0 | 0 | 0 | 0 |
| 0 | NA | NA | NA | NA | NA | Olfr1302 | olfactory receptor 1302 | 0 | 0 | 0 | 0 | 0 | 0 | 0 | 0 | 0 | 0 | 0 |
| 0 | NA | NA | NA | NA | NA | Olfr1303 | olfactory receptor 1303 | 0 | 0 | 0 | 0 | 0 | 0 | 0 | 0 | 0 | 0 | 0 |
| 0 | NA | NA | NA | NA | NA | Olfr1304-ps1 | | 0 | 0 | 0 | 0 | 0 | 0 | 0 | 0 | 0 | 0 | 0 |
| 0 | NA | NA | NA | NA | NA | Gm26421 | | 0 | 0 | 0 | 0 | 0 | 0 | 0 | 0 | 0 | 0 | 0 |
| 0 | NA | NA | NA | NA | NA | Olfr268-ps1 | | 0 | 0 | 0 | 0 | 0 | 0 | 0 | 0 | 0 | 0 | 0 |
| 0 | NA | NA | NA | NA | NA | Olfr1305 | olfactory receptor 1305 | 0 | 0 | 0 | 0 | 0 | 0 | 0 | 0 | 0 | 0 | 0 |
| 0 | NA | NA | NA | NA | NA | Olfr1306 | olfactory receptor 1306 | 0 | 0 | 0 | 0 | 0 | 0 | 0 | 0 | 0 | 0 | 0 |
| 0 | NA | NA | NA | NA | NA | Gm13928 | | 0 | 0 | 0 | 0 | 0 | 0 | 0 | 0 | 0 | 0 | 0 |
| 0 | NA | NA | NA | NA | NA | Olfr1307 | olfactory receptor 1307 | 0 | 0 | 0 | 0 | 0 | 0 | 0 | 0 | 0 | 0 | 0 |
| 0 | NA | NA | NA | NA | NA | Olfr1308 | olfactory receptor 1308 | 0 | 0 | 0 | 0 | 0 | 0 | 0 | 0 | 0 | 0 | 0 |
| 0 | NA | NA | NA | NA | NA | Olfr1309 | olfactory receptor 1309 | 0 | 0 | 0 | 0 | 0 | 0 | 0 | 0 | 0 | 0 | 0 |
| 0 | NA | NA | NA | NA | NA | Gm13929 | | 0 | 0 | 0 | 0 | 0 | 0 | 0 | 0 | 0 | 0 | 0 |
| 0 | NA | NA | NA | NA | NA | Olfr1310 | olfactory receptor 1310 | 0 | 0 | 0 | 0 | 0 | 0 | 0 | 0 | 0 | 0 | 0 |
| 0 | NA | NA | NA | NA | NA | Olfr1311 | olfactory receptor 1311 | 0 | 0 | 0 | 0 | 0 | 0 | 0 | 0 | 0 | 0 | 0 |
| 0 | NA | NA | NA | NA | NA | Olfr1312 | olfactory receptor 1312 | 0 | 0 | 0 | 0 | 0 | 0 | 0 | 0 | 0 | 0 | 0 |
| 0 | NA | NA | NA | NA | NA | Olfr1313 | olfactory receptor 1313 | 0 | 0 | 0 | 0 | 0 | 0 | 0 | 0 | 0 | 0 | 0 |
| 0 | NA | NA | NA | NA | NA | Olfr1314 | olfactory receptor 1314 | 0 | 0 | 0 | 0 | 0 | 0 | 0 | 0 | 0 | 0 | 0 |
| 0 | NA | NA | NA | NA | NA | Olfr1315-ps1 | | 0 | 0 | 0 | 0 | 0 | 0 | 0 | 0 | 0 | 0 | 0 |
| 0 | NA | NA | NA | NA | NA | Olfr1316 | olfactory receptor 1316 | 0 | 0 | 0 | 0 | 0 | 0 | 0 | 0 | 0 | 0 | 0 |
| 0 | NA | NA | NA | NA | NA | Olfr1317 | olfactory receptor 1317 | 0 | 0 | 0 | 0 | 0 | 0 | 0 | 0 | 0 | 0 | 0 |
| 0 | NA | NA | NA | NA | NA | Olfr1318 | olfactory receptor 1318 | 0 | 0 | 0 | 0 | 0 | 0 | 0 | 0 | 0 | 0 | 0 |
| 0 | NA | NA | NA | NA | NA | Olfr1319-ps1 | | 0 | 0 | 0 | 0 | 0 | 0 | 0 | 0 | 0 | 0 | 0 |
| 0 | NA | NA | NA | NA | NA | Nutm1 | NUT midline carcinoma, family member 1 | 0 | 0 | 0 | 0 | 0 | 0 | 0 | 0 | 0 | 0 | 0 |
| 0 | NA | NA | NA | NA | NA | Gm21985 | | 0 | 0 | 0 | 0 | 0 | 0 | 0 | 0 | 0 | 0 | 0 |
| 0 | NA | NA | NA | NA | NA | Gm13940 | | 0 | 0 | 0 | 0 | 0 | 0 | 0 | 0 | 0 | 0 | 0 |
| 0 | NA | NA | NA | NA | NA | Gm22676 | | 0 | 0 | 0 | 0 | 0 | 0 | 0 | 0 | 0 | 0 | 0 |
| 0 | NA | NA | NA | NA | NA | Gm44374 | | 0 | 0 | 0 | 0 | 0 | 0 | 0 | 0 | 0 | 0 | 0 |
| 0 | NA | NA | NA | NA | NA | Tmco5b | transmembrane and coiled-coil domains [illegible] | 0 | 0 | 0 | 0 | 0 | 0 | 0 | 0 | 0 | 0 | 0 |
| 0 | NA | NA | NA | NA | NA | Gm26505 | | 0 | 0 | 0 | 0 | 0 | 0 | 0 | 0 | 0 | 0 | 0 |
| 0 | NA | NA | NA | NA | NA | Gm13967 | | 0 | 0 | 0 | 0 | 0 | 0 | 0 | 0 | 0 | 0 | 0 |
| 0 | NA | NA | NA | NA | NA | Gm13963 | | 0 | 0 | 0 | 0 | 0 | 0 | 0 | 0 | 0 | 0 | 0 |
| 0 | NA | NA | NA | NA | NA | Gm13966 | | 0 | 0 | 0 | 0 | 0 | 0 | 0 | 0 | 0 | 0 | 0 |
| 0 | NA | NA | NA | NA | NA | Gm44465 | | 0 | 0 | 0 | 0 | 0 | 0 | 0 | 0 | 0 | 0 | 0 |
| 0 | NA | NA | NA | NA | NA | Gm13964 | | 0 | 0 | 0 | 0 | 0 | 0 | 0 | 0 | 0 | 0 | 0 |
| 0 | NA | NA | NA | NA | NA | 4930533B01Rik | | 0 | 0 | 0 | 0 | 0 | 0 | 0 | 0 | 0 | 0 | 0 |
| 0 | NA | NA | NA | NA | NA | Mir7002 | | 0 | 0 | 0 | 0 | 0 | 0 | 0 | 0 | 0 | 0 | 0 |
| 0 | NA | NA | NA | NA | NA | Gm25103 | | 0 | 0 | 0 | 0 | 0 | 0 | 0 | 0 | 0 | 0 | 0 |
| 0 | NA | NA | NA | NA | NA | Gm29234 | | 0 | 0 | 0 | 0 | 0 | 0 | 0 | 0 | 0 | 0 | 0 |
| 0 | NA | NA | NA | NA | NA | Gm13972 | | 0 | 0 | 0 | 0 | 0 | 0 | 0 | 0 | 0 | 0 | 0 |
| 0 | NA | NA | NA | NA | NA | Gm13975 | | 0 | 0 | 0 | 0 | 0 | 0 | 0 | 0 | 0 | 0 | 0 |
| 0 | NA | NA | NA | NA | NA | Gm13974 | | 0 | 0 | 0 | 0 | 0 | 0 | 0 | 0 | 0 | 0 | 0 |

| | | | | | | | | | | | | | | | | | |
|---|---|---|---|---|---|---|---|---|---|---|---|---|---|---|---|---|---|
| 0 | NA | NA | NA | NA | NA | Gm28494 | | 0 | 0 | 0 | 0 | 0 | 0 | 0 | 0 | 0 | 0 | 0 |
| 0 | NA | NA | NA | NA | NA | Gm28492 | | 0 | 0 | 0 | 0 | 0 | 0 | 0 | 0 | 0 | 0 | 0 |
| 0 | NA | NA | NA | NA | NA | Gm13973 | | 0 | 0 | 0 | 0 | 0 | 0 | 0 | 0 | 0 | 0 | 0 |
| 0 | NA | NA | NA | NA | NA | Gm28493 | | 0 | 0 | 0 | 0 | 0 | 0 | 0 | 0 | 0 | 0 | 0 |
| 0 | NA | NA | NA | NA | NA | 4930528P14Rik | | 0 | 0 | 0 | 0 | 0 | 0 | 0 | 0 | 0 | 0 | 0 |
| 0 | NA | NA | NA | NA | NA | Gm13976 | | 0 | 0 | 0 | 0 | 0 | 0 | 0 | 0 | 0 | 0 | 0 |
| 0 | NA | NA | NA | NA | NA | 3110099E03Rik | | 0 | 0 | 0 | 0 | 0 | 0 | 0 | 0 | 0 | 0 | 0 |
| 0 | NA | NA | NA | NA | NA | Mir1951 | | 0 | 0 | 0 | 0 | 0 | 0 | 0 | 0 | 0 | 0 | 0 |
| 0 | NA | NA | NA | NA | NA | Gm13990 | | 0 | 0 | 0 | 0 | 0 | 0 | 0 | 0 | 0 | 0 | 0 |
| 0 | NA | NA | NA | NA | NA | Tmco5 | transmembrane and coiled-coil domains ! | 0 | 0 | 0 | 0 | 0 | 0 | 0 | 0 | 0 | 0 | 0 |
| 0 | NA | NA | NA | NA | NA | Gm24914 | | 0 | 0 | 0 | 0 | 0 | 0 | 0 | 0 | 0 | 0 | 0 |
| 0 | NA | NA | NA | NA | NA | Gm25514 | | 0 | 0 | 0 | 0 | 0 | 0 | 0 | 0 | 0 | 0 | 0 |
| 0 | NA | NA | NA | NA | NA | Mir674 | | 0 | 0 | 0 | 0 | 0 | 0 | 0 | 0 | 0 | 0 | 0 |
| 0 | NA | NA | NA | NA | NA | Rap1a-ps2 | | 0 | 0 | 0 | 0 | 0 | 0 | 0 | 0 | 0 | 0 | 0 |
| 0 | NA | NA | NA | NA | NA | Gm25189 | | 0 | 0 | 0 | 0 | 0 | 0 | 0 | 0 | 0 | 0 | 0 |
| 0 | NA | NA | NA | NA | NA | Gm13982 | | 0 | 0 | 0 | 0 | 0 | 0 | 0 | 0 | 0 | 0 | 0 |
| 0 | NA | NA | NA | NA | NA | Gm28183 | | 0 | 0 | 0 | 0 | 0 | 0 | 0 | 0 | 0 | 0 | 0 |
| 0 | NA | NA | NA | NA | NA | 4930412B13Rik | | 0 | 0 | 0 | 0 | 0 | 0 | 0 | 0 | 0 | 0 | 0 |
| 0 | NA | NA | NA | NA | NA | Gm13986 | | 0 | 0 | 0 | 0 | 0 | 0 | 0 | 0 | 0 | 0 | 0 |
| 0 | NA | NA | NA | NA | NA | Gm36979 | | 0 | 0 | 0 | 0 | 0 | 0 | 0 | 0 | 0 | 0 | 0 |
| 0 | NA | NA | NA | NA | NA | Gm25308 | | 0 | 0 | 0 | 0 | 0 | 0 | 0 | 0 | 0 | 0 | 0 |
| 0 | NA | NA | NA | NA | NA | Gm29233 | | 0 | 0 | 0 | 0 | 0 | 0 | 0 | 0 | 0 | 0 | 0 |
| 0 | NA | NA | NA | NA | NA | Gm22800 | | 0 | 0 | 0 | 0 | 0 | 0 | 0 | 0 | 0 | 0 | 0 |
| 0 | NA | NA | NA | NA | NA | Gm29232 | | 0 | 0 | 0 | 0 | 0 | 0 | 0 | 0 | 0 | 0 | 0 |
| 0 | NA | NA | NA | NA | NA | A930104D05Rik | | 0 | 0 | 0 | 0 | 0 | 0 | 0 | 0 | 0 | 0 | 0 |
| 0 | NA | NA | NA | NA | NA | 1700054M17Rik | | 0 | 0 | 0 | 0 | 0 | 0 | 0 | 0 | 0 | 0 | 0 |
| 0 | NA | NA | NA | NA | NA | Gm22651 | | 0 | 0 | 0 | 0 | 0 | 0 | 0 | 0 | 0 | 0 | 0 |
| 0 | NA | NA | NA | NA | NA | Gm13983 | | 0 | 0 | 0 | 0 | 0 | 0 | 0 | 0 | 0 | 0 | 0 |
| 0 | NA | NA | NA | NA | NA | Ankrd63 | ankyrin repeat domain 63 | 0 | 0 | 0 | 0 | 0 | 0 | 0 | 0 | 0 | 0 | 0 |
| 0 | NA | NA | NA | NA | NA | Gm14089 | | 0 | 0 | 0 | 0 | 0 | 0 | 0 | 0 | 0 | 0 | 0 |
| 0 | NA | NA | NA | NA | NA | Gm14137 | predicted gene 14137 | 0 | 0 | 0 | 0 | 0 | 0 | 0 | 0 | 0 | 0 | 0 |
| 0 | NA | NA | NA | NA | NA | Gm14175 | | 0 | 0 | 0 | 0 | 0 | 0 | 0 | 0 | 0 | 0 | 0 |
| 0 | NA | NA | NA | NA | NA | Gm14208 | | 0 | 0 | 0 | 0 | 0 | 0 | 0 | 0 | 0 | 0 | 0 |
| 0 | NA | NA | NA | NA | NA | n-R5s204 | | 0 | 0 | 0 | 0 | 0 | 0 | 0 | 0 | 0 | 0 | 0 |
| 0 | NA | NA | NA | NA | NA | Mir7665 | | 0 | 0 | 0 | 0 | 0 | 0 | 0 | 0 | 0 | 0 | 0 |
| 0 | NA | NA | NA | NA | NA | Gm28042 | | 0 | 0 | 0 | 0 | 0 | 0 | 0 | 0 | 0 | 0 | 0 |
| 0 | NA | NA | NA | NA | NA | Pla2g4d | phospholipase A2, group IVD | 0 | 0 | 0 | 0 | 0 | 0 | 0 | 0 | 0 | 0 | 0 |
| 0 | NA | NA | NA | NA | NA | n-R5s205 | | 0 | 0 | 0 | 0 | 0 | 0 | 0 | 0 | 0 | 0 | 0 |
| 0 | NA | NA | NA | NA | NA | Epb42 | erythrocyte membrane protein band 4.2 | 0 | 0 | 0 | 0 | 0 | 0 | 0 | 0 | 0 | 0 | 0 |
| 0 | NA | NA | NA | NA | NA | Gm14213 | | 0 | 0 | 0 | 0 | 0 | 0 | 0 | 0 | 0 | 0 | 0 |
| 0 | NA | NA | NA | NA | NA | Gm23858 | | 0 | 0 | 0 | 0 | 0 | 0 | 0 | 0 | 0 | 0 | 0 |
| 0 | NA | NA | NA | NA | NA | Gm14978 | | 0 | 0 | 0 | 0 | 0 | 0 | 0 | 0 | 0 | 0 | 0 |
| 0 | NA | NA | NA | NA | NA | Gm14018 | | 0 | 0 | 0 | 0 | 0 | 0 | 0 | 0 | 0 | 0 | 0 |
| 0 | NA | NA | NA | NA | NA | Gm14020 | | 0 | 0 | 0 | 0 | 0 | 0 | 0 | 0 | 0 | 0 | 0 |
| 0 | NA | NA | NA | NA | NA | Gm23983 | | 0 | 0 | 0 | 0 | 0 | 0 | 0 | 0 | 0 | 0 | 0 |
| 0 | NA | NA | NA | NA | NA | Gm47314 | | 0 | 0 | 0 | 0 | 0 | 0 | 0 | 0 | 0 | 0 | 0 |
| 0 | NA | NA | NA | NA | NA | Patl2 | protein associated with topoisomerase II I | 0 | 0 | 0 | 0 | 0 | 0 | 0 | 0 | 0 | 0 | 0 |
| 0 | NA | NA | NA | NA | NA | Terb2 | telomere repeat binding bouquet formatic | 0 | 0 | 0 | 0 | 0 | 0 | 0 | 0 | 0 | 0 | 0 |
| 0 | NA | NA | NA | NA | NA | Gm14050 | | 0 | 0 | 0 | 0 | 0 | 0 | 0 | 0 | 0 | 0 | 0 |
| 0 | NA | NA | NA | NA | NA | Gm14058 | | 0 | 0 | 0 | 0 | 0 | 0 | 0 | 0 | 0 | 0 | 0 |
| 0 | NA | NA | NA | NA | NA | Gm24409 | | 0 | 0 | 0 | 0 | 0 | 0 | 0 | 0 | 0 | 0 | 0 |
| 0 | NA | NA | NA | NA | NA | Gm14086 | | 0 | 0 | 0 | 0 | 0 | 0 | 0 | 0 | 0 | 0 | 0 |
| 0 | NA | NA | NA | NA | NA | Mir147 | | 0 | 0 | 0 | 0 | 0 | 0 | 0 | 0 | 0 | 0 | 0 |
| 0 | NA | NA | NA | NA | NA | Gm13988 | | 0 | 0 | 0 | 0 | 0 | 0 | 0 | 0 | 0 | 0 | 0 |
| 0 | NA | NA | NA | NA | NA | 4930517E11Rik | | 0 | 0 | 0 | 0 | 0 | 0 | 0 | 0 | 0 | 0 | 0 |
| 0 | NA | NA | NA | NA | NA | 4930583P06Rik | | 0 | 0 | 0 | 0 | 0 | 0 | 0 | 0 | 0 | 0 | 0 |
| 0 | NA | NA | NA | NA | NA | Gm13994 | | 0 | 0 | 0 | 0 | 0 | 0 | 0 | 0 | 0 | 0 | 0 |
| 0 | NA | NA | NA | NA | NA | Gm22352 | | 0 | 0 | 0 | 0 | 0 | 0 | 0 | 0 | 0 | 0 | 0 |
| 0 | NA | NA | NA | NA | NA | A530010F05Rik | | 0 | 0 | 0 | 0 | 0 | 0 | 0 | 0 | 0 | 0 | 0 |
| 0 | NA | NA | NA | NA | NA | Gm14003 | | 0 | 0 | 0 | 0 | 0 | 0 | 0 | 0 | 0 | 0 | 0 |

| | | | | | | | | | | | | | | | | | | |
|---|---|---|---|---|---|---|---|---|---|---|---|---|---|---|---|---|---|---|
| 0 | NA | NA | NA | NA | NA | Fam227b | family with sequence similarity 227, meml | 0 | 0 | 0 | 0 | 0 | 0 | 0 | 0 | 0 | 0 | 0 |
| 0 | NA | NA | NA | NA | NA | Gm26496 | | 0 | 0 | 0 | 0 | 0 | 0 | 0 | 0 | 0 | 0 | 0 |
| 0 | NA | NA | NA | NA | NA | Gm17555 | | 0 | 0 | 0 | 0 | 0 | 0 | 0 | 0 | 0 | 0 | 0 |
| 0 | NA | NA | NA | NA | NA | Gm24739 | | 0 | 0 | 0 | 0 | 0 | 0 | 0 | 0 | 0 | 0 | 0 |
| 0 | NA | NA | NA | NA | NA | Gm23753 | | 0 | 0 | 0 | 0 | 0 | 0 | 0 | 0 | 0 | 0 | 0 |
| 0 | NA | NA | NA | NA | NA | Gm10774 | | 0 | 0 | 0 | 0 | 0 | 0 | 0 | 0 | 0 | 0 | 0 |
| 0 | NA | NA | NA | NA | NA | Gm14212 | | 0 | 0 | 0 | 0 | 0 | 0 | 0 | 0 | 0 | 0 | 0 |
| 0 | NA | NA | NA | NA | NA | Mir3473g | | 0 | 0 | 0 | 0 | 0 | 0 | 0 | 0 | 0 | 0 | 0 |
| 0 | NA | NA | NA | NA | NA | Astl | astacin-like metalloendopeptidase (M12 f | 0 | 0 | 0 | 0 | 0 | 0 | 0 | 0 | 0 | 0 | 0 |
| 0 | NA | NA | NA | NA | NA | Gm22613 | | 0 | 0 | 0 | 0 | 0 | 0 | 0 | 0 | 0 | 0 | 0 |
| 0 | NA | NA | NA | NA | NA | Gm14229 | | 0 | 0 | 0 | 0 | 0 | 0 | 0 | 0 | 0 | 0 | 0 |
| 0 | NA | NA | NA | NA | NA | Gm22889 | | 0 | 0 | 0 | 0 | 0 | 0 | 0 | 0 | 0 | 0 | 0 |
| 0 | NA | NA | NA | NA | NA | Gm14245 | | 0 | 0 | 0 | 0 | 0 | 0 | 0 | 0 | 0 | 0 | 0 |
| 0 | NA | NA | NA | NA | NA | Gm14244 | | 0 | 0 | 0 | 0 | 0 | 0 | 0 | 0 | 0 | 0 | 0 |
| 0 | NA | NA | NA | NA | NA | Gm22411 | | 0 | 0 | 0 | 0 | 0 | 0 | 0 | 0 | 0 | 0 | 0 |
| 0 | NA | NA | NA | NA | NA | Gm23101 | | 0 | 0 | 0 | 0 | 0 | 0 | 0 | 0 | 0 | 0 | 0 |
| 0 | NA | NA | NA | NA | NA | Gm14010 | | 0 | 0 | 0 | 0 | 0 | 0 | 0 | 0 | 0 | 0 | 0 |
| 0 | NA | NA | NA | NA | NA | Gm29010 | | 0 | 0 | 0 | 0 | 0 | 0 | 0 | 0 | 0 | 0 | 0 |
| 0 | NA | NA | NA | NA | NA | Spdye4c | speedy/RINGO cell cycle regulator family | 0 | 0 | 0 | 0 | 0 | 0 | 0 | 0 | 0 | 0 | 0 |
| 0 | NA | NA | NA | NA | NA | Gm14007 | | 0 | 0 | 0 | 0 | 0 | 0 | 0 | 0 | 0 | 0 | 0 |
| 0 | NA | NA | NA | NA | NA | Gm14011 | | 0 | 0 | 0 | 0 | 0 | 0 | 0 | 0 | 0 | 0 | 0 |
| 0 | NA | NA | NA | NA | NA | Gm23172 | | 0 | 0 | 0 | 0 | 0 | 0 | 0 | 0 | 0 | 0 | 0 |
| 0 | NA | NA | NA | NA | NA | Gm22859 | | 0 | 0 | 0 | 0 | 0 | 0 | 0 | 0 | 0 | 0 | 0 |
| 0 | NA | NA | NA | NA | NA | Gm14028 | | 0 | 0 | 0 | 0 | 0 | 0 | 0 | 0 | 0 | 0 | 0 |
| 0 | NA | NA | NA | NA | NA | 9830144P21Rik | | 0 | 0 | 0 | 0 | 0 | 0 | 0 | 0 | 0 | 0 | 0 |
| 0 | NA | NA | NA | NA | NA | A730036I17Rik | | 0 | 0 | 0 | 0 | 0 | 0 | 0 | 0 | 0 | 0 | 0 |
| 0 | NA | NA | NA | NA | NA | Gm14024 | | 0 | 0 | 0 | 0 | 0 | 0 | 0 | 0 | 0 | 0 | 0 |
| 0 | NA | NA | NA | NA | NA | Gm25703 | | 0 | 0 | 0 | 0 | 0 | 0 | 0 | 0 | 0 | 0 | 0 |
| 0 | NA | NA | NA | NA | NA | Gm14039 | | 0 | 0 | 0 | 0 | 0 | 0 | 0 | 0 | 0 | 0 | 0 |
| 0 | NA | NA | NA | NA | NA | Gm14041 | | 0 | 0 | 0 | 0 | 0 | 0 | 0 | 0 | 0 | 0 | 0 |
| 0 | NA | NA | NA | NA | NA | Gm28196 | | 0 | 0 | 0 | 0 | 0 | 0 | 0 | 0 | 0 | 0 | 0 |
| 0 | NA | NA | NA | NA | NA | Mir6339 | | 0 | 0 | 0 | 0 | 0 | 0 | 0 | 0 | 0 | 0 | 0 |
| 0 | NA | NA | NA | NA | NA | Gm22424 | | 0 | 0 | 0 | 0 | 0 | 0 | 0 | 0 | 0 | 0 | 0 |
| 0 | NA | NA | NA | NA | NA | Gm24665 | | 0 | 0 | 0 | 0 | 0 | 0 | 0 | 0 | 0 | 0 | 0 |
| 0 | NA | NA | NA | NA | NA | AU015228 | | 0 | 0 | 0 | 0 | 0 | 0 | 0 | 0 | 0 | 0 | 0 |
| 0 | NA | NA | NA | NA | NA | Tmc2 | transmembrane channel-like gene family | 0 | 0 | 0 | 0 | 0 | 0 | 0 | 0 | 0 | 0 | 0 |
| 0 | NA | NA | NA | NA | NA | Gm24451 | | 0 | 0 | 0 | 0 | 0 | 0 | 0 | 0 | 0 | 0 | 0 |
| 0 | NA | NA | NA | NA | NA | Gm23187 | | 0 | 0 | 0 | 0 | 0 | 0 | 0 | 0 | 0 | 0 | 0 |
| 0 | NA | NA | NA | NA | NA | Gm23650 | | 0 | 0 | 0 | 0 | 0 | 0 | 0 | 0 | 0 | 0 | 0 |
| 0 | NA | NA | NA | NA | NA | Snord57 | | 0 | 0 | 0 | 0 | 0 | 0 | 0 | 0 | 0 | 0 | 0 |
| 0 | NA | NA | NA | NA | NA | Gm14042 | | 0 | 0 | 0 | 0 | 0 | 0 | 0 | 0 | 0 | 0 | 0 |
| 0 | NA | NA | NA | NA | NA | Gm28372 | | 0 | 0 | 0 | 0 | 0 | 0 | 0 | 0 | 0 | 0 | 0 |
| 0 | NA | NA | NA | NA | NA | 1700020A23Ril | RIKEN cDNA 1700020A23 gene | 0 | 0 | 0 | 0 | 0 | 0 | 0 | 0 | 0 | 0 | 0 |
| 0 | NA | NA | NA | NA | NA | Tmem239 | transmembrane 239 | 0 | 0 | 0 | 0 | 0 | 0 | 0 | 0 | 0 | 0 | 0 |
| 0 | NA | NA | NA | NA | NA | Gm23876 | | 0 | 0 | 0 | 0 | 0 | 0 | 0 | 0 | 0 | 0 | 0 |
| 0 | NA | NA | NA | NA | NA | Oxt | oxytocin | 0 | 0 | 0 | 0 | 0 | 0 | 0 | 0 | 0 | 0 | 0 |
| 0 | NA | NA | NA | NA | NA | Avp | arginine vasopressin | 0 | 0 | 0 | 0 | 0 | 0 | 0 | 0 | 0 | 0 | 0 |
| 0 | NA | NA | NA | NA | NA | Gm24893 | | 0 | 0 | 0 | 0 | 0 | 0 | 0 | 0 | 0 | 0 | 0 |
| 0 | NA | NA | NA | NA | NA | Mir6973b | | 0 | 0 | 0 | 0 | 0 | 0 | 0 | 0 | 0 | 0 | 0 |
| 0 | NA | NA | NA | NA | NA | Gm14233 | | 0 | 0 | 0 | 0 | 0 | 0 | 0 | 0 | 0 | 0 | 0 |
| 0 | NA | NA | NA | NA | NA | Gm22902 | | 0 | 0 | 0 | 0 | 0 | 0 | 0 | 0 | 0 | 0 | 0 |
| 0 | NA | NA | NA | NA | NA | Gm14283 | | 0 | 0 | 0 | 0 | 0 | 0 | 0 | 0 | 0 | 0 | 0 |
| 0 | NA | NA | NA | NA | NA | Gm14304 | | 0 | 0 | 0 | 0 | 0 | 0 | 0 | 0 | 0 | 0 | 0 |
| 0 | NA | NA | NA | NA | NA | Gm14285 | | 0 | 0 | 0 | 0 | 0 | 0 | 0 | 0 | 0 | 0 | 0 |
| 0 | NA | NA | NA | NA | NA | Gm14282 | | 0 | 0 | 0 | 0 | 0 | 0 | 0 | 0 | 0 | 0 | 0 |
| 0 | NA | NA | NA | NA | NA | Gm14281 | | 0 | 0 | 0 | 0 | 0 | 0 | 0 | 0 | 0 | 0 | 0 |
| 0 | NA | NA | NA | NA | NA | Erv3 | endogenous retroviral sequence 3 | 0 | 0 | 0 | 0 | 0 | 0 | 0 | 0 | 0 | 0 | 0 |
| 0 | NA | NA | NA | NA | NA | Lamr1-ps1 | | 0 | 0 | 0 | 0 | 0 | 0 | 0 | 0 | 0 | 0 | 0 |
| 0 | NA | NA | NA | NA | NA | Prn | prion protein readthrough transcript | 0 | 0 | 0 | 0 | 0 | 0 | 0 | 0 | 0 | 0 | 0 |
| 0 | NA | NA | NA | NA | NA | Prnd | prion like protein doppel | 0 | 0 | 0 | 0 | 0 | 0 | 0 | 0 | 0 | 0 | 0 |

| | | | | | | | | | | | | | | | | | | |
|---|---|---|---|---|---|---|---|---|---|---|---|---|---|---|---|---|---|---|
| 0 | NA | NA | NA | NA | NA | Gm14051 | | 0 | 0 | 0 | 0 | 0 | 0 | 0 | 0 | 0 | 0 | 0 |
| 0 | NA | NA | NA | NA | NA | Prokr2 | prokineticin receptor 2 | 0 | 0 | 0 | 0 | 0 | 0 | 0 | 0 | 0 | 0 | 0 |
| 0 | NA | NA | NA | NA | NA | 4921508D12Rik | | 0 | 0 | 0 | 0 | 0 | 0 | 0 | 0 | 0 | 0 | 0 |
| 0 | NA | NA | NA | NA | NA | Gm14095 | | 0 | 0 | 0 | 0 | 0 | 0 | 0 | 0 | 0 | 0 | 0 |
| 0 | NA | NA | NA | NA | NA | 1700026D11Rik | | 0 | 0 | 0 | 0 | 0 | 0 | 0 | 0 | 0 | 0 | 0 |
| 0 | NA | NA | NA | NA | NA | AU019990 | | 0 | 0 | 0 | 0 | 0 | 0 | 0 | 0 | 0 | 0 | 0 |
| 0 | NA | NA | NA | NA | NA | Rpl23a-ps4 | | 0 | 0 | 0 | 0 | 0 | 0 | 0 | 0 | 0 | 0 | 0 |
| 0 | NA | NA | NA | NA | NA | Gm14099 | | 0 | 0 | 0 | 0 | 0 | 0 | 0 | 0 | 0 | 0 | 0 |
| 0 | NA | NA | NA | NA | NA | Gm14098 | | 0 | 0 | 0 | 0 | 0 | 0 | 0 | 0 | 0 | 0 | 0 |
| 0 | NA | NA | NA | NA | NA | Gm14101 | | 0 | 0 | 0 | 0 | 0 | 0 | 0 | 0 | 0 | 0 | 0 |
| 0 | NA | NA | NA | NA | NA | Gm25704 | | 0 | 0 | 0 | 0 | 0 | 0 | 0 | 0 | 0 | 0 | 0 |
| 0 | NA | NA | NA | NA | NA | Gm14103 | | 0 | 0 | 0 | 0 | 0 | 0 | 0 | 0 | 0 | 0 | 0 |
| 0 | NA | NA | NA | NA | NA | Gm27449 | | 0 | 0 | 0 | 0 | 0 | 0 | 0 | 0 | 0 | 0 | 0 |
| 0 | NA | NA | NA | NA | NA | A430048G15Rik | | 0 | 0 | 0 | 0 | 0 | 0 | 0 | 0 | 0 | 0 | 0 |
| 0 | NA | NA | NA | NA | NA | Gm14100 | | 0 | 0 | 0 | 0 | 0 | 0 | 0 | 0 | 0 | 0 | 0 |
| 0 | NA | NA | NA | NA | NA | Mir3090 | | 0 | 0 | 0 | 0 | 0 | 0 | 0 | 0 | 0 | 0 | 0 |
| 0 | NA | NA | NA | NA | NA | Gm25258 | | 0 | 0 | 0 | 0 | 0 | 0 | 0 | 0 | 0 | 0 | 0 |
| 0 | NA | NA | NA | NA | NA | 4930545L23Rik | | 0 | 0 | 0 | 0 | 0 | 0 | 0 | 0 | 0 | 0 | 0 |
| 0 | NA | NA | NA | NA | NA | 9630028H03Rik | | 0 | 0 | 0 | 0 | 0 | 0 | 0 | 0 | 0 | 0 | 0 |
| 0 | NA | NA | NA | NA | NA | Platr3 | | 0 | 0 | 0 | 0 | 0 | 0 | 0 | 0 | 0 | 0 | 0 |
| 0 | NA | NA | NA | NA | NA | Gm14209 | | 0 | 0 | 0 | 0 | 0 | 0 | 0 | 0 | 0 | 0 | 0 |
| 0 | NA | NA | NA | NA | NA | Gm14211 | | 0 | 0 | 0 | 0 | 0 | 0 | 0 | 0 | 0 | 0 | 0 |
| 0 | NA | NA | NA | NA | NA | Gm14210 | | 0 | 0 | 0 | 0 | 0 | 0 | 0 | 0 | 0 | 0 | 0 |
| 0 | NA | NA | NA | NA | NA | Gm26227 | | 0 | 0 | 0 | 0 | 0 | 0 | 0 | 0 | 0 | 0 | 0 |
| 0 | NA | NA | NA | NA | NA | Gm14218 | | 0 | 0 | 0 | 0 | 0 | 0 | 0 | 0 | 0 | 0 | 0 |
| 0 | NA | NA | NA | NA | NA | Gm14054 | | 0 | 0 | 0 | 0 | 0 | 0 | 0 | 0 | 0 | 0 | 0 |
| 0 | NA | NA | NA | NA | NA | Gm36967 | | 0 | 0 | 0 | 0 | 0 | 0 | 0 | 0 | 0 | 0 | 0 |
| 0 | NA | NA | NA | NA | NA | Gm14053 | | 0 | 0 | 0 | 0 | 0 | 0 | 0 | 0 | 0 | 0 | 0 |
| 0 | NA | NA | NA | NA | NA | | | 0 | 0 | 0 | 0 | 0 | 0 | 0 | 0 | 0 | 0 | 0 |
| 0 | NA | NA | NA | NA | NA | | | 0 | 0 | 0 | 0 | 0 | 0 | 0 | 0 | 0 | 0 | 0 |
| 0 | NA | NA | NA | NA | NA | Gm28214 | | 0 | 0 | 0 | 0 | 0 | 0 | 0 | 0 | 0 | 0 | 0 |
| 0 | NA | NA | NA | NA | NA | Gm14055 | | 0 | 0 | 0 | 0 | 0 | 0 | 0 | 0 | 0 | 0 | 0 |
| 0 | NA | NA | NA | NA | NA | Gm14064 | | 0 | 0 | 0 | 0 | 0 | 0 | 0 | 0 | 0 | 0 | 0 |
| 0 | NA | NA | NA | NA | NA | Gm14062 | | 0 | 0 | 0 | 0 | 0 | 0 | 0 | 0 | 0 | 0 | 0 |
| 0 | NA | NA | NA | NA | NA | Gm14061 | | 0 | 0 | 0 | 0 | 0 | 0 | 0 | 0 | 0 | 0 | 0 |
| 0 | NA | NA | NA | NA | NA | Mir1952 | | 0 | 0 | 0 | 0 | 0 | 0 | 0 | 0 | 0 | 0 | 0 |
| 0 | NA | NA | NA | NA | NA | Gm14065 | | 0 | 0 | 0 | 0 | 0 | 0 | 0 | 0 | 0 | 0 | 0 |
| 0 | NA | NA | NA | NA | NA | Gm14067 | | 0 | 0 | 0 | 0 | 0 | 0 | 0 | 0 | 0 | 0 | 0 |
| 0 | NA | NA | NA | NA | NA | Gm14069 | | 0 | 0 | 0 | 0 | 0 | 0 | 0 | 0 | 0 | 0 | 0 |
| 0 | NA | NA | NA | NA | NA | Gm14071 | | 0 | 0 | 0 | 0 | 0 | 0 | 0 | 0 | 0 | 0 | 0 |
| 0 | NA | NA | NA | NA | NA | Sptlc3 | serine palmitoyltransferase, long chain ba | 0 | 0 | 0 | 0 | 0 | 0 | 0 | 0 | 0 | 0 | 0 |
| 0 | NA | NA | NA | NA | NA | Gm27385 | | 0 | 0 | 0 | 0 | 0 | 0 | 0 | 0 | 0 | 0 | 0 |
| 0 | NA | NA | NA | NA | NA | Gm14070 | | 0 | 0 | 0 | 0 | 0 | 0 | 0 | 0 | 0 | 0 | 0 |
| 0 | NA | NA | NA | NA | NA | Gm14073 | | 0 | 0 | 0 | 0 | 0 | 0 | 0 | 0 | 0 | 0 | 0 |
| 0 | NA | NA | NA | NA | NA | Gm17374 | | 0 | 0 | 0 | 0 | 0 | 0 | 0 | 0 | 0 | 0 | 0 |
| 0 | NA | NA | NA | NA | NA | Sel1l2 | sel-1 suppressor of lin-12-like 2 (C. elega | 0 | 0 | 0 | 0 | 0 | 0 | 0 | 0 | 0 | 0 | 0 |
| 0 | NA | NA | NA | NA | NA | Gm23846 | | 0 | 0 | 0 | 0 | 0 | 0 | 0 | 0 | 0 | 0 | 0 |
| 0 | NA | NA | NA | NA | NA | Zcchc9-ps | | 0 | 0 | 0 | 0 | 0 | 0 | 0 | 0 | 0 | 0 | 0 |
| 0 | NA | NA | NA | NA | NA | Macrod2os2 | | 0 | 0 | 0 | 0 | 0 | 0 | 0 | 0 | 0 | 0 | 0 |
| 0 | NA | NA | NA | NA | NA | Macrod2os1 | | 0 | 0 | 0 | 0 | 0 | 0 | 0 | 0 | 0 | 0 | 0 |
| 0 | NA | NA | NA | NA | NA | Gm23179 | | 0 | 0 | 0 | 0 | 0 | 0 | 0 | 0 | 0 | 0 | 0 |
| 0 | NA | NA | NA | NA | NA | Rpl26-ps5 | | 0 | 0 | 0 | 0 | 0 | 0 | 0 | 0 | 0 | 0 | 0 |
| 0 | NA | NA | NA | NA | NA | Gm22310 | | 0 | 0 | 0 | 0 | 0 | 0 | 0 | 0 | 0 | 0 | 0 |
| 0 | NA | NA | NA | NA | NA | Otor | otoraplin | 0 | 0 | 0 | 0 | 0 | 0 | 0 | 0 | 0 | 0 | 0 |
| 0 | NA | NA | NA | NA | NA | Gm14083 | | 0 | 0 | 0 | 0 | 0 | 0 | 0 | 0 | 0 | 0 | 0 |
| 0 | NA | NA | NA | NA | NA | Gm14082 | | 0 | 0 | 0 | 0 | 0 | 0 | 0 | 0 | 0 | 0 | 0 |
| 0 | NA | NA | NA | NA | NA | Gm11644 | | 0 | 0 | 0 | 0 | 0 | 0 | 0 | 0 | 0 | 0 | 0 |
| 0 | NA | NA | NA | NA | NA | Rps13-ps6 | | 0 | 0 | 0 | 0 | 0 | 0 | 0 | 0 | 0 | 0 | 0 |
| 0 | NA | NA | NA | NA | NA | Gm11673 | | 0 | 0 | 0 | 0 | 0 | 0 | 0 | 0 | 0 | 0 | 0 |
| 0 | NA | NA | NA | NA | NA | 4930444E06Rik | | 0 | 0 | 0 | 0 | 0 | 0 | 0 | 0 | 0 | 0 | 0 |

| | | | | | | | | | | | | | | | | | |
|---|---|---|---|---|---|---|---|---|---|---|---|---|---|---|---|---|---|
| 0 | NA | NA | NA | NA | NA | Gm11700 | | 0 | 0 | 0 | 0 | 0 | 0 | 0 | 0 | 0 | 0 | 0 |
| 0 | NA | NA | NA | NA | NA | Pet117 | PET117 homolog | 0 | 0 | 0 | 0 | 0 | 0 | 0 | 0 | 0 | 0 | 0 |
| 0 | NA | NA | NA | NA | NA | Gm24823 | | 0 | 0 | 0 | 0 | 0 | 0 | 0 | 0 | 0 | 0 | 0 |
| 0 | NA | NA | NA | NA | NA | Scp2d1 | SCP2 sterol-binding domain containing 1 | 0 | 0 | 0 | 0 | 0 | 0 | 0 | 0 | 0 | 0 | 0 |
| 0 | NA | NA | NA | NA | NA | Gm11746 | | 0 | 0 | 0 | 0 | 0 | 0 | 0 | 0 | 0 | 0 | 0 |
| 0 | NA | NA | NA | NA | NA | Gm14093 | | 0 | 0 | 0 | 0 | 0 | 0 | 0 | 0 | 0 | 0 | 0 |
| 0 | NA | NA | NA | NA | NA | Gm22731 | | 0 | 0 | 0 | 0 | 0 | 0 | 0 | 0 | 0 | 0 | 0 |
| 0 | NA | NA | NA | NA | NA | Gm22390 | | 0 | 0 | 0 | 0 | 0 | 0 | 0 | 0 | 0 | 0 | 0 |
| 0 | NA | NA | NA | NA | NA | Gm14092 | predicted gene 14092 | 0 | 0 | 0 | 0 | 0 | 0 | 0 | 0 | 0 | 0 | 0 |
| 0 | NA | NA | NA | NA | NA | Gm11763 | | 0 | 0 | 0 | 0 | 0 | 0 | 0 | 0 | 0 | 0 | 0 |
| 0 | NA | NA | NA | NA | NA | Gm25026 | | 0 | 0 | 0 | 0 | 0 | 0 | 0 | 0 | 0 | 0 | 0 |
| 0 | NA | NA | NA | NA | NA | Gm11764 | | 0 | 0 | 0 | 0 | 0 | 0 | 0 | 0 | 0 | 0 | 0 |
| 0 | NA | NA | NA | NA | NA | 4933406D12Rik | | 0 | 0 | 0 | 0 | 0 | 0 | 0 | 0 | 0 | 0 | 0 |
| 0 | NA | NA | NA | NA | NA | Gm14114 | | 0 | 0 | 0 | 0 | 0 | 0 | 0 | 0 | 0 | 0 | 0 |
| 0 | NA | NA | NA | NA | NA | Nkx2-4 | NK2 homeobox 4 | 0 | 0 | 0 | 0 | 0 | 0 | 0 | 0 | 0 | 0 | 0 |
| 0 | NA | NA | NA | NA | NA | Gm14116 | | 0 | 0 | 0 | 0 | 0 | 0 | 0 | 0 | 0 | 0 | 0 |
| 0 | NA | NA | NA | NA | NA | Nkx2-2os | | 0 | 0 | 0 | 0 | 0 | 0 | 0 | 0 | 0 | 0 | 0 |
| 0 | NA | NA | NA | NA | NA | Gm14110 | | 0 | 0 | 0 | 0 | 0 | 0 | 0 | 0 | 0 | 0 | 0 |
| 0 | NA | NA | NA | NA | NA | Gm14109 | | 0 | 0 | 0 | 0 | 0 | 0 | 0 | 0 | 0 | 0 | 0 |
| 0 | NA | NA | NA | NA | NA | Gm14106 | | 0 | 0 | 0 | 0 | 0 | 0 | 0 | 0 | 0 | 0 | 0 |
| 0 | NA | NA | NA | NA | NA | A530006G24Rik | | 0 | 0 | 0 | 0 | 0 | 0 | 0 | 0 | 0 | 0 | 0 |
| 0 | NA | NA | NA | NA | NA | Gm27420 | | 0 | 0 | 0 | 0 | 0 | 0 | 0 | 0 | 0 | 0 | 0 |
| 0 | NA | NA | NA | NA | NA | Gm24221 | | 0 | 0 | 0 | 0 | 0 | 0 | 0 | 0 | 0 | 0 | 0 |
| 0 | NA | NA | NA | NA | NA | Gm14119 | | 0 | 0 | 0 | 0 | 0 | 0 | 0 | 0 | 0 | 0 | 0 |
| 0 | NA | NA | NA | NA | NA | Gm25866 | | 0 | 0 | 0 | 0 | 0 | 0 | 0 | 0 | 0 | 0 | 0 |
| 0 | NA | NA | NA | NA | NA | Cstl1 | cystatin-like 1 | 0 | 0 | 0 | 0 | 0 | 0 | 0 | 0 | 0 | 0 | 0 |
| 0 | NA | NA | NA | NA | NA | Cst11 | cystatin 11 | 0 | 0 | 0 | 0 | 0 | 0 | 0 | 0 | 0 | 0 | 0 |
| 0 | NA | NA | NA | NA | NA | Cstdc1 | cystatin domain containing 1 | 0 | 0 | 0 | 0 | 0 | 0 | 0 | 0 | 0 | 0 | 0 |
| 0 | NA | NA | NA | NA | NA | Cst12 | cystatin 12 | 0 | 0 | 0 | 0 | 0 | 0 | 0 | 0 | 0 | 0 | 0 |
| 0 | NA | NA | NA | NA | NA | Cst8 | cystatin 8 (cystatin-related epididymal spe | 0 | 0 | 0 | 0 | 0 | 0 | 0 | 0 | 0 | 0 | 0 |
| 0 | NA | NA | NA | NA | NA | Cst13 | cystatin 13 | 0 | 0 | 0 | 0 | 0 | 0 | 0 | 0 | 0 | 0 | 0 |
| 0 | NA | NA | NA | NA | NA | Cst9 | cystatin 9 | 0 | 0 | 0 | 0 | 0 | 0 | 0 | 0 | 0 | 0 | 0 |
| 0 | NA | NA | NA | NA | NA | Gm38221 | | 0 | 0 | 0 | 0 | 0 | 0 | 0 | 0 | 0 | 0 | 0 |
| 0 | NA | NA | NA | NA | NA | Gm14129 | | 0 | 0 | 0 | 0 | 0 | 0 | 0 | 0 | 0 | 0 | 0 |
| 0 | NA | NA | NA | NA | NA | Gm1330 | | 0 | 0 | 0 | 0 | 0 | 0 | 0 | 0 | 0 | 0 | 0 |
| 0 | NA | NA | NA | NA | NA | Gm10750 | | 0 | 0 | 0 | 0 | 0 | 0 | 0 | 0 | 0 | 0 | 0 |
| 0 | NA | NA | NA | NA | NA | Gm14135 | | 0 | 0 | 0 | 0 | 0 | 0 | 0 | 0 | 0 | 0 | 0 |
| 0 | NA | NA | NA | NA | NA | Cst10 | cystatin 10 (chondrocytes) | 0 | 0 | 0 | 0 | 0 | 0 | 0 | 0 | 0 | 0 | 0 |
| 0 | NA | NA | NA | NA | NA | Gm24551 | | 0 | 0 | 0 | 0 | 0 | 0 | 0 | 0 | 0 | 0 | 0 |
| 0 | NA | NA | NA | NA | NA | C530025M09Rik | | 0 | 0 | 0 | 0 | 0 | 0 | 0 | 0 | 0 | 0 | 0 |
| 0 | NA | NA | NA | NA | NA | Gm14125 | | 0 | 0 | 0 | 0 | 0 | 0 | 0 | 0 | 0 | 0 | 0 |
| 0 | NA | NA | NA | NA | NA | Gm14126 | | 0 | 0 | 0 | 0 | 0 | 0 | 0 | 0 | 0 | 0 | 0 |
| 0 | NA | NA | NA | NA | NA | Gm14128 | | 0 | 0 | 0 | 0 | 0 | 0 | 0 | 0 | 0 | 0 | 0 |
| 0 | NA | NA | NA | NA | NA | Vsx1 | visual system homeobox 1 | 0 | 0 | 0 | 0 | 0 | 0 | 0 | 0 | 0 | 0 | 0 |
| 0 | NA | NA | NA | NA | NA | Platr15 | | 0 | 0 | 0 | 0 | 0 | 0 | 0 | 0 | 0 | 0 | 0 |
| 0 | NA | NA | NA | NA | NA | Gm28450 | | 0 | 0 | 0 | 0 | 0 | 0 | 0 | 0 | 0 | 0 | 0 |
| 0 | NA | NA | NA | NA | NA | Gm23598 | | 0 | 0 | 0 | 0 | 0 | 0 | 0 | 0 | 0 | 0 | 0 |
| 0 | NA | NA | NA | NA | NA | Gm14143 | | 0 | 0 | 0 | 0 | 0 | 0 | 0 | 0 | 0 | 0 | 0 |
| 0 | NA | NA | NA | NA | NA | Gm14141 | | 0 | 0 | 0 | 0 | 0 | 0 | 0 | 0 | 0 | 0 | 0 |
| 0 | NA | NA | NA | NA | NA | Gm22187 | | 0 | 0 | 0 | 0 | 0 | 0 | 0 | 0 | 0 | 0 | 0 |
| 0 | NA | NA | NA | NA | NA | Gm24199 | | 0 | 0 | 0 | 0 | 0 | 0 | 0 | 0 | 0 | 0 | 0 |
| 0 | NA | NA | NA | NA | NA | Gm14151 | predicted gene 14151 | 0 | 0 | 0 | 0 | 0 | 0 | 0 | 0 | 0 | 0 | 0 |
| 0 | NA | NA | NA | NA | NA | Gm14147 | predicted gene 14147 | 0 | 0 | 0 | 0 | 0 | 0 | 0 | 0 | 0 | 0 | 0 |
| 0 | NA | NA | NA | NA | NA | Gm14145 | | 0 | 0 | 0 | 0 | 0 | 0 | 0 | 0 | 0 | 0 | 0 |
| 0 | NA | NA | NA | NA | NA | Gm22517 | | 0 | 0 | 0 | 0 | 0 | 0 | 0 | 0 | 0 | 0 | 0 |
| 0 | NA | NA | NA | NA | NA | Gm14149 | | 0 | 0 | 0 | 0 | 0 | 0 | 0 | 0 | 0 | 0 | 0 |
| 0 | NA | NA | NA | NA | NA | Gm14144 | | 0 | 0 | 0 | 0 | 0 | 0 | 0 | 0 | 0 | 0 | 0 |
| 0 | NA | NA | NA | NA | NA | Gm23005 | | 0 | 0 | 0 | 0 | 0 | 0 | 0 | 0 | 0 | 0 | 0 |
| 0 | NA | NA | NA | NA | NA | Gm24077 | | 0 | 0 | 0 | 0 | 0 | 0 | 0 | 0 | 0 | 0 | 0 |
| 0 | NA | NA | NA | NA | NA | 4930442J19Rik | | 0 | 0 | 0 | 0 | 0 | 0 | 0 | 0 | 0 | 0 | 0 |

| | | | | | | | | | | | | | | | | | | |
|---|---|---|---|---|---|---|---|---|---|---|---|---|---|---|---|---|---|---|
| 0 | NA | NA | NA | NA | NA | Gm23261 | | 0 | 0 | 0 | 0 | 0 | 0 | 0 | 0 | 0 | 0 | 0 |
| 0 | NA | NA | NA | NA | NA | Gm14152 | | 0 | 0 | 0 | 0 | 0 | 0 | 0 | 0 | 0 | 0 | 0 |
| 0 | NA | NA | NA | NA | NA | Gm24868 | | 0 | 0 | 0 | 0 | 0 | 0 | 0 | 0 | 0 | 0 | 0 |
| 0 | NA | NA | NA | NA | NA | Gm14146 | | 0 | 0 | 0 | 0 | 0 | 0 | 0 | 0 | 0 | 0 | 0 |
| 0 | NA | NA | NA | NA | NA | 4921509C19Ri | RIKEN cDNA 4921509C19 gene | 0 | 0 | 0 | 0 | 0 | 0 | 0 | 0 | 0 | 0 | 0 |
| 0 | NA | NA | NA | NA | NA | Tmem74bos | | 0 | 0 | 0 | 0 | 0 | 0 | 0 | 0 | 0 | 0 | 0 |
| 0 | NA | NA | NA | NA | NA | Gm14159 | | 0 | 0 | 0 | 0 | 0 | 0 | 0 | 0 | 0 | 0 | 0 |
| 0 | NA | NA | NA | NA | NA | 4930556L07Rik | | 0 | 0 | 0 | 0 | 0 | 0 | 0 | 0 | 0 | 0 | 0 |
| 0 | NA | NA | NA | NA | NA | Gm14153 | | 0 | 0 | 0 | 0 | 0 | 0 | 0 | 0 | 0 | 0 | 0 |
| 0 | NA | NA | NA | NA | NA | Gm14155 | | 0 | 0 | 0 | 0 | 0 | 0 | 0 | 0 | 0 | 0 | 0 |
| 0 | NA | NA | NA | NA | NA | Mir1953 | | 0 | 0 | 0 | 0 | 0 | 0 | 0 | 0 | 0 | 0 | 0 |
| 0 | NA | NA | NA | NA | NA | Gm14164 | | 0 | 0 | 0 | 0 | 0 | 0 | 0 | 0 | 0 | 0 | 0 |
| 0 | NA | NA | NA | NA | NA | Nrsn2 | neurensin 2 | 0 | 0 | 0 | 0 | 0 | 0 | 0 | 0 | 0 | 0 | 0 |
| 0 | NA | NA | NA | NA | NA | Gm14158 | | 0 | 0 | 0 | 0 | 0 | 0 | 0 | 0 | 0 | 0 | 0 |
| 0 | NA | NA | NA | NA | NA | Defb23 | defensin beta 23 | 0 | 0 | 0 | 0 | 0 | 0 | 0 | 0 | 0 | 0 | 0 |
| 0 | NA | NA | NA | NA | NA | Defb20 | defensin beta 20 | 0 | 0 | 0 | 0 | 0 | 0 | 0 | 0 | 0 | 0 | 0 |
| 0 | NA | NA | NA | NA | NA | Defb22 | defensin beta 22 | 0 | 0 | 0 | 0 | 0 | 0 | 0 | 0 | 0 | 0 | 0 |
| 0 | NA | NA | NA | NA | NA | Defb26 | defensin beta 26 | 0 | 0 | 0 | 0 | 0 | 0 | 0 | 0 | 0 | 0 | 0 |
| 0 | NA | NA | NA | NA | NA | Defb28 | defensin beta 28 | 0 | 0 | 0 | 0 | 0 | 0 | 0 | 0 | 0 | 0 | 0 |
| 0 | NA | NA | NA | NA | NA | Defb29 | defensin beta 29 | 0 | 0 | 0 | 0 | 0 | 0 | 0 | 0 | 0 | 0 | 0 |
| 0 | NA | NA | NA | NA | NA | Gm14161 | | 0 | 0 | 0 | 0 | 0 | 0 | 0 | 0 | 0 | 0 | 0 |
| 0 | NA | NA | NA | NA | NA | Gm17416 | | 0 | 0 | 0 | 0 | 0 | 0 | 0 | 0 | 0 | 0 | 0 |
| 0 | NA | NA | NA | NA | NA | Defb21 | defensin beta 21 | 0 | 0 | 0 | 0 | 0 | 0 | 0 | 0 | 0 | 0 | 0 |
| 0 | NA | NA | NA | NA | NA | Defb19 | defensin beta 19 | 0 | 0 | 0 | 0 | 0 | 0 | 0 | 0 | 0 | 0 | 0 |
| 0 | NA | NA | NA | NA | NA | | | 0 | 0 | 0 | 0 | 0 | 0 | 0 | 0 | 0 | 0 | 0 |
| 0 | NA | NA | NA | NA | NA | Defb45 | defensin beta 45 | 0 | 0 | 0 | 0 | 0 | 0 | 0 | 0 | 0 | 0 | 0 |
| 0 | NA | NA | NA | NA | NA | Defb36 | defensin beta 36 | 0 | 0 | 0 | 0 | 0 | 0 | 0 | 0 | 0 | 0 | 0 |
| 0 | NA | NA | NA | NA | NA | Gm14163 | | 0 | 0 | 0 | 0 | 0 | 0 | 0 | 0 | 0 | 0 | 0 |
| 0 | NA | NA | NA | NA | NA | Defb25 | defensin beta 25 | 0 | 0 | 0 | 0 | 0 | 0 | 0 | 0 | 0 | 0 | 0 |
| 0 | NA | NA | NA | NA | NA | 1700030C14Rik | | 0 | 0 | 0 | 0 | 0 | 0 | 0 | 0 | 0 | 0 | 0 |
| 0 | NA | NA | NA | NA | NA | Gm14162 | | 0 | 0 | 0 | 0 | 0 | 0 | 0 | 0 | 0 | 0 | 0 |
| 0 | NA | NA | NA | NA | NA | Gm23802 | | 0 | 0 | 0 | 0 | 0 | 0 | 0 | 0 | 0 | 0 | 0 |
| 0 | NA | NA | NA | NA | NA | Mir5622 | | 0 | 0 | 0 | 0 | 0 | 0 | 0 | 0 | 0 | 0 | 0 |
| 0 | NA | NA | NA | NA | NA | Gm14199 | | 0 | 0 | 0 | 0 | 0 | 0 | 0 | 0 | 0 | 0 | 0 |
| 0 | NA | NA | NA | NA | NA | Gm25643 | | 0 | 0 | 0 | 0 | 0 | 0 | 0 | 0 | 0 | 0 | 0 |
| 0 | NA | NA | NA | NA | NA | Sun5 | Sad1 and UNC84 domain containing 5 | 0 | 0 | 0 | 0 | 0 | 0 | 0 | 0 | 0 | 0 | 0 |
| 0 | NA | NA | NA | NA | NA | Bpifb2 | BPI fold containing family B, member 2 | 0 | 0 | 0 | 0 | 0 | 0 | 0 | 0 | 0 | 0 | 0 |
| 0 | NA | NA | NA | NA | NA | Bpifb6 | BPI fold containing family B, member 6 | 0 | 0 | 0 | 0 | 0 | 0 | 0 | 0 | 0 | 0 | 0 |
| 0 | NA | NA | NA | NA | NA | Bpifb4 | BPI fold containing family B, member 4 | 0 | 0 | 0 | 0 | 0 | 0 | 0 | 0 | 0 | 0 | 0 |
| 0 | NA | NA | NA | NA | NA | Bpifa6 | BPI fold containing family A, member 6 | 0 | 0 | 0 | 0 | 0 | 0 | 0 | 0 | 0 | 0 | 0 |
| 0 | NA | NA | NA | NA | NA | Bpifa2 | BPI fold containing family A, member 2 | 0 | 0 | 0 | 0 | 0 | 0 | 0 | 0 | 0 | 0 | 0 |
| 0 | NA | NA | NA | NA | NA | | | 0 | 0 | 0 | 0 | 0 | 0 | 0 | 0 | 0 | 0 | 0 |
| 0 | NA | NA | NA | NA | NA | Gm21850 | | 0 | 0 | 0 | 0 | 0 | 0 | 0 | 0 | 0 | 0 | 0 |
| 0 | NA | NA | NA | NA | NA | Bpifa3 | BPI fold containing family A, member 3 | 0 | 0 | 0 | 0 | 0 | 0 | 0 | 0 | 0 | 0 | 0 |
| 0 | NA | NA | NA | NA | NA | Bpifa1 | BPI fold containing family A, member 1 | 0 | 0 | 0 | 0 | 0 | 0 | 0 | 0 | 0 | 0 | 0 |
| 0 | NA | NA | NA | NA | NA | Bpifa5 | BPI fold containing family A, member 5 | 0 | 0 | 0 | 0 | 0 | 0 | 0 | 0 | 0 | 0 | 0 |
| 0 | NA | NA | NA | NA | NA | Bpifb1 | BPI fold containing family B, member 1 | 0 | 0 | 0 | 0 | 0 | 0 | 0 | 0 | 0 | 0 | 0 |
| 0 | NA | NA | NA | NA | NA | Bpifb5 | BPI fold containing family B, member 5 | 0 | 0 | 0 | 0 | 0 | 0 | 0 | 0 | 0 | 0 | 0 |
| 0 | NA | NA | NA | NA | NA | Bpifb9a | BPI fold containing family B, member 9A | 0 | 0 | 0 | 0 | 0 | 0 | 0 | 0 | 0 | 0 | 0 |
| 0 | NA | NA | NA | NA | NA | Bpifb9b | BPI fold containing family B, member 9B | 0 | 0 | 0 | 0 | 0 | 0 | 0 | 0 | 0 | 0 | 0 |
| 0 | NA | NA | NA | NA | NA | Gm14198 | | 0 | 0 | 0 | 0 | 0 | 0 | 0 | 0 | 0 | 0 | 0 |
| 0 | NA | NA | NA | NA | NA | Gm14215 | | 0 | 0 | 0 | 0 | 0 | 0 | 0 | 0 | 0 | 0 | 0 |
| 0 | NA | NA | NA | NA | NA | Gm14216 | | 0 | 0 | 0 | 0 | 0 | 0 | 0 | 0 | 0 | 0 | 0 |
| 0 | NA | NA | NA | NA | NA | Gm45609 | | 0 | 0 | 0 | 0 | 0 | 0 | 0 | 0 | 0 | 0 | 0 |
| 0 | NA | NA | NA | NA | NA | 2310005A03Rik | | 0 | 0 | 0 | 0 | 0 | 0 | 0 | 0 | 0 | 0 | 0 |
| 0 | NA | NA | NA | NA | NA | Mir695 | | 0 | 0 | 0 | 0 | 0 | 0 | 0 | 0 | 0 | 0 | 0 |
| 0 | NA | NA | NA | NA | NA | Mipep-ps | | 0 | 0 | 0 | 0 | 0 | 0 | 0 | 0 | 0 | 0 | 0 |
| 0 | NA | NA | NA | NA | NA | Acss2os | | 0 | 0 | 0 | 0 | 0 | 0 | 0 | 0 | 0 | 0 | 0 |
| 0 | NA | NA | NA | NA | NA | Gm14257 | | 0 | 0 | 0 | 0 | 0 | 0 | 0 | 0 | 0 | 0 | 0 |
| 0 | NA | NA | NA | NA | NA | Mir499 | | 0 | 0 | 0 | 0 | 0 | 0 | 0 | 0 | 0 | 0 | 0 |

| | | | | | | | | | | | | | | | | | | |
|---|---|---|---|---|---|---|---|---|---|---|---|---|---|---|---|---|---|---|
| 0 | NA | NA | NA | NA | NA | Gm17581 | | 0 | 0 | 0 | 0 | 0 | 0 | 0 | 0 | 0 | 0 | 0 |
| 0 | NA | NA | NA | NA | NA | Fam83c | family with sequence similarity 83, memb | 0 | 0 | 0 | 0 | 0 | 0 | 0 | 0 | 0 | 0 | 0 |
| 0 | NA | NA | NA | NA | NA | Fer1l4 | fer-1-like 4 (C. elegans) | 0 | 0 | 0 | 0 | 0 | 0 | 0 | 0 | 0 | 0 | 0 |
| 0 | NA | NA | NA | NA | NA | Gm28036 | | 0 | 0 | 0 | 0 | 0 | 0 | 0 | 0 | 0 | 0 | 0 |
| 0 | NA | NA | NA | NA | NA | Gm22759 | | 0 | 0 | 0 | 0 | 0 | 0 | 0 | 0 | 0 | 0 | 0 |
| 0 | NA | NA | NA | NA | NA | Gm14252 | | 0 | 0 | 0 | 0 | 0 | 0 | 0 | 0 | 0 | 0 | 0 |
| 0 | NA | NA | NA | NA | NA | Gm14170 | | 0 | 0 | 0 | 0 | 0 | 0 | 0 | 0 | 0 | 0 | 0 |
| 0 | NA | NA | NA | NA | NA | Gm22316 | | 0 | 0 | 0 | 0 | 0 | 0 | 0 | 0 | 0 | 0 | 0 |
| 0 | NA | NA | NA | NA | NA | Gm22758 | | 0 | 0 | 0 | 0 | 0 | 0 | 0 | 0 | 0 | 0 | 0 |
| 0 | NA | NA | NA | NA | NA | Gm14225 | | 0 | 0 | 0 | 0 | 0 | 0 | 0 | 0 | 0 | 0 | 0 |
| 0 | NA | NA | NA | NA | NA | Gm14224 | | 0 | 0 | 0 | 0 | 0 | 0 | 0 | 0 | 0 | 0 | 0 |
| 0 | NA | NA | NA | NA | NA | Gm14169 | | 0 | 0 | 0 | 0 | 0 | 0 | 0 | 0 | 0 | 0 | 0 |
| 0 | NA | NA | NA | NA | NA | Gm14174 | | 0 | 0 | 0 | 0 | 0 | 0 | 0 | 0 | 0 | 0 | 0 |
| 0 | NA | NA | NA | NA | NA | Gm14172 | | 0 | 0 | 0 | 0 | 0 | 0 | 0 | 0 | 0 | 0 | 0 |
| 0 | NA | NA | NA | NA | NA | Gm27651 | | 0 | 0 | 0 | 0 | 0 | 0 | 0 | 0 | 0 | 0 | 0 |
| 0 | NA | NA | NA | NA | NA | Gm14248 | | 0 | 0 | 0 | 0 | 0 | 0 | 0 | 0 | 0 | 0 | 0 |
| 0 | NA | NA | NA | NA | NA | Gm14247 | | 0 | 0 | 0 | 0 | 0 | 0 | 0 | 0 | 0 | 0 | 0 |
| 0 | NA | NA | NA | NA | NA | Gm27653 | | 0 | 0 | 0 | 0 | 0 | 0 | 0 | 0 | 0 | 0 | 0 |
| 0 | NA | NA | NA | NA | NA | Ghrh | growth hormone releasing hormone | 0 | 0 | 0 | 0 | 0 | 0 | 0 | 0 | 0 | 0 | 0 |
| 0 | NA | NA | NA | NA | NA | Gm25407 | | 0 | 0 | 0 | 0 | 0 | 0 | 0 | 0 | 0 | 0 | 0 |
| 0 | NA | NA | NA | NA | NA | 2010009K17Rik | | 0 | 0 | 0 | 0 | 0 | 0 | 0 | 0 | 0 | 0 | 0 |
| 0 | NA | NA | NA | NA | NA | Gm20412 | | 0 | 0 | 0 | 0 | 0 | 0 | 0 | 0 | 0 | 0 | 0 |
| 0 | NA | NA | NA | NA | NA | 1700060C20Rik | | 0 | 0 | 0 | 0 | 0 | 0 | 0 | 0 | 0 | 0 | 0 |
| 0 | NA | NA | NA | NA | NA | Mir7685 | | 0 | 0 | 0 | 0 | 0 | 0 | 0 | 0 | 0 | 0 | 0 |
| 0 | NA | NA | NA | NA | NA | Gm24411 | | 0 | 0 | 0 | 0 | 0 | 0 | 0 | 0 | 0 | 0 | 0 |
| 0 | NA | NA | NA | NA | NA | Gm26003 | | 0 | 0 | 0 | 0 | 0 | 0 | 0 | 0 | 0 | 0 | 0 |
| 0 | NA | NA | NA | NA | NA | Gm23925 | | 0 | 0 | 0 | 0 | 0 | 0 | 0 | 0 | 0 | 0 | 0 |
| 0 | NA | NA | NA | NA | NA | Gm23925 | | 0 | 0 | 0 | 0 | 0 | 0 | 0 | 0 | 0 | 0 | 0 |
| 0 | NA | NA | NA | NA | NA | Gm25187 | | 0 | 0 | 0 | 0 | 0 | 0 | 0 | 0 | 0 | 0 | 0 |
| 0 | NA | NA | NA | NA | NA | Gm25129 | | 0 | 0 | 0 | 0 | 0 | 0 | 0 | 0 | 0 | 0 | 0 |
| 0 | NA | NA | NA | NA | NA | Gm14201 | | 0 | 0 | 0 | 0 | 0 | 0 | 0 | 0 | 0 | 0 | 0 |
| 0 | NA | NA | NA | NA | NA | Arhgap40 | Rho GTPase activating protein 40 | 0 | 0 | 0 | 0 | 0 | 0 | 0 | 0 | 0 | 0 | 0 |
| 0 | NA | NA | NA | NA | NA | Gm14204 | | 0 | 0 | 0 | 0 | 0 | 0 | 0 | 0 | 0 | 0 | 0 |
| 0 | NA | NA | NA | NA | NA | Slc32a1 | solute carrier family 32 (GABA vesicular t | 0 | 0 | 0 | 0 | 0 | 0 | 0 | 0 | 0 | 0 | 0 |
| 0 | NA | NA | NA | NA | NA | Gm14205 | | 0 | 0 | 0 | 0 | 0 | 0 | 0 | 0 | 0 | 0 | 0 |
| 0 | NA | NA | NA | NA | NA | Mir3474 | | 0 | 0 | 0 | 0 | 0 | 0 | 0 | 0 | 0 | 0 | 0 |
| 0 | NA | NA | NA | NA | NA | Gm26141 | | 0 | 0 | 0 | 0 | 0 | 0 | 0 | 0 | 0 | 0 | 0 |
| 0 | NA | NA | NA | NA | NA | Gm44319 | | 0 | 0 | 0 | 0 | 0 | 0 | 0 | 0 | 0 | 0 | 0 |
| 0 | NA | NA | NA | NA | NA | Gm25090 | | 0 | 0 | 0 | 0 | 0 | 0 | 0 | 0 | 0 | 0 | 0 |
| 0 | NA | NA | NA | NA | NA | Gm14219 | | 0 | 0 | 0 | 0 | 0 | 0 | 0 | 0 | 0 | 0 | 0 |
| 0 | NA | NA | NA | NA | NA | Gm11445 | | 0 | 0 | 0 | 0 | 0 | 0 | 0 | 0 | 0 | 0 | 0 |
| 0 | NA | NA | NA | NA | NA | Gm11446 | | 0 | 0 | 0 | 0 | 0 | 0 | 0 | 0 | 0 | 0 | 0 |
| 0 | NA | NA | NA | NA | NA | 9130015L21Rik | | 0 | 0 | 0 | 0 | 0 | 0 | 0 | 0 | 0 | 0 | 0 |
| 0 | NA | NA | NA | NA | NA | Gm14222 | | 0 | 0 | 0 | 0 | 0 | 0 | 0 | 0 | 0 | 0 | 0 |
| 0 | NA | NA | NA | NA | NA | Gm23968 | | 0 | 0 | 0 | 0 | 0 | 0 | 0 | 0 | 0 | 0 | 0 |
| 0 | NA | NA | NA | NA | NA | Gm11447 | | 0 | 0 | 0 | 0 | 0 | 0 | 0 | 0 | 0 | 0 | 0 |
| 0 | NA | NA | NA | NA | NA | Emilin3 | elastin microfibril interfacer 3 | 0 | 0 | 0 | 0 | 0 | 0 | 0 | 0 | 0 | 0 | 0 |
| 0 | NA | NA | NA | NA | NA | Gm14228 | | 0 | 0 | 0 | 0 | 0 | 0 | 0 | 0 | 0 | 0 | 0 |
| 0 | NA | NA | NA | NA | NA | Gm25561 | | 0 | 0 | 0 | 0 | 0 | 0 | 0 | 0 | 0 | 0 | 0 |
| 0 | NA | NA | NA | NA | NA | Gm14243 | | 0 | 0 | 0 | 0 | 0 | 0 | 0 | 0 | 0 | 0 | 0 |
| 0 | NA | NA | NA | NA | NA | Gm14242 | | 0 | 0 | 0 | 0 | 0 | 0 | 0 | 0 | 0 | 0 | 0 |
| 0 | NA | NA | NA | NA | NA | Gm14241 | | 0 | 0 | 0 | 0 | 0 | 0 | 0 | 0 | 0 | 0 | 0 |
| 0 | NA | NA | NA | NA | NA | Ptprtos | | 0 | 0 | 0 | 0 | 0 | 0 | 0 | 0 | 0 | 0 | 0 |
| 0 | NA | NA | NA | NA | NA | Gm22936 | | 0 | 0 | 0 | 0 | 0 | 0 | 0 | 0 | 0 | 0 | 0 |
| 0 | NA | NA | NA | NA | NA | Gm24478 | | 0 | 0 | 0 | 0 | 0 | 0 | 0 | 0 | 0 | 0 | 0 |
| 0 | NA | NA | NA | NA | NA | Gm14255 | | 0 | 0 | 0 | 0 | 0 | 0 | 0 | 0 | 0 | 0 | 0 |
| 0 | NA | NA | NA | NA | NA | Mir7003 | | 0 | 0 | 0 | 0 | 0 | 0 | 0 | 0 | 0 | 0 | 0 |
| 0 | NA | NA | NA | NA | NA | Gtsf1l | gametocyte specific factor 1-like | 0 | 0 | 0 | 0 | 0 | 0 | 0 | 0 | 0 | 0 | 0 |
| 0 | NA | NA | NA | NA | NA | Gm22174 | | 0 | 0 | 0 | 0 | 0 | 0 | 0 | 0 | 0 | 0 | 0 |
| 0 | NA | NA | NA | NA | NA | Gm14254 | | 0 | 0 | 0 | 0 | 0 | 0 | 0 | 0 | 0 | 0 | 0 |

| | | | | | | | | | | | | | | | | | | |
|---|---|---|---|---|---|---|---|---|---|---|---|---|---|---|---|---|---|---|
| 0 | NA | NA | NA | NA | NA | R3hdml | R3H domain containing-like | 0 | 0 | 0 | 0 | 0 | 0 | 0 | 0 | 0 | 0 | 0 |
| 0 | NA | NA | NA | NA | NA | Gm25881 | | 0 | 0 | 0 | 0 | 0 | 0 | 0 | 0 | 0 | 0 | 0 |
| 0 | NA | NA | NA | NA | NA | 0610039K10Rik | | 0 | 0 | 0 | 0 | 0 | 0 | 0 | 0 | 0 | 0 | 0 |
| 0 | NA | NA | NA | NA | NA | Gm16316 | | 0 | 0 | 0 | 0 | 0 | 0 | 0 | 0 | 0 | 0 | 0 |
| 0 | NA | NA | NA | NA | NA | Kcnk15 | potassium channel, subfamily K, member | 0 | 0 | 0 | 0 | 0 | 0 | 0 | 0 | 0 | 0 | 0 |
| 0 | NA | NA | NA | NA | NA | n-R5s207 | | 0 | 0 | 0 | 0 | 0 | 0 | 0 | 0 | 0 | 0 | 0 |
| 0 | NA | NA | NA | NA | NA | Wfdc5 | WAP four-disulfide core domain 5 | 0 | 0 | 0 | 0 | 0 | 0 | 0 | 0 | 0 | 0 | 0 |
| 0 | NA | NA | NA | NA | NA | A730032A03Rik | | 0 | 0 | 0 | 0 | 0 | 0 | 0 | 0 | 0 | 0 | 0 |
| 0 | NA | NA | NA | NA | NA | Wfdc12 | WAP four-disulfide core domain 12 | 0 | 0 | 0 | 0 | 0 | 0 | 0 | 0 | 0 | 0 | 0 |
| 0 | NA | NA | NA | NA | NA | Wfdc15a | WAP four-disulfide core domain 15A | 0 | 0 | 0 | 0 | 0 | 0 | 0 | 0 | 0 | 0 | 0 |
| 0 | NA | NA | NA | NA | NA | Svs2 | seminal vesicle secretory protein 2 | 0 | 0 | 0 | 0 | 0 | 0 | 0 | 0 | 0 | 0 | 0 |
| 0 | NA | NA | NA | NA | NA | Svs3b | seminal vesicle secretory protein 3B | 0 | 0 | 0 | 0 | 0 | 0 | 0 | 0 | 0 | 0 | 0 |
| 0 | NA | NA | NA | NA | NA | Svs4 | seminal vesicle secretory protein 4 | 0 | 0 | 0 | 0 | 0 | 0 | 0 | 0 | 0 | 0 | 0 |
| 0 | NA | NA | NA | NA | NA | Svs3a | seminal vesicle secretory protein 3A | 0 | 0 | 0 | 0 | 0 | 0 | 0 | 0 | 0 | 0 | 0 |
| 0 | NA | NA | NA | NA | NA | Svs6 | seminal vesicle secretory protein 6 | 0 | 0 | 0 | 0 | 0 | 0 | 0 | 0 | 0 | 0 | 0 |
| 0 | NA | NA | NA | NA | NA | Svs5 | seminal vesicle secretory protein 5 | 0 | 0 | 0 | 0 | 0 | 0 | 0 | 0 | 0 | 0 | 0 |
| 0 | NA | NA | NA | NA | NA | Mir7678 | | 0 | 0 | 0 | 0 | 0 | 0 | 0 | 0 | 0 | 0 | 0 |
| 0 | NA | NA | NA | NA | NA | Gm20458 | | 0 | 0 | 0 | 0 | 0 | 0 | 0 | 0 | 0 | 0 | 0 |
| 0 | NA | NA | NA | NA | NA | Trp53tg5 | transformation related protein 53 target 5 | 0 | 0 | 0 | 0 | 0 | 0 | 0 | 0 | 0 | 0 | 0 |
| 0 | NA | NA | NA | NA | NA | Gm14317 | | 0 | 0 | 0 | 0 | 0 | 0 | 0 | 0 | 0 | 0 | 0 |
| 0 | NA | NA | NA | NA | NA | Spint3 | serine peptidase inhibitor, Kunitz type, 3 | 0 | 0 | 0 | 0 | 0 | 0 | 0 | 0 | 0 | 0 | 0 |
| 0 | NA | NA | NA | NA | NA | Eppin | epididymal peptidase inhibitor | 0 | 0 | 0 | 0 | 0 | 0 | 0 | 0 | 0 | 0 | 0 |
| 0 | NA | NA | NA | NA | NA | Wfdc8 | WAP four-disulfide core domain 8 | 0 | 0 | 0 | 0 | 0 | 0 | 0 | 0 | 0 | 0 | 0 |
| 0 | NA | NA | NA | NA | NA | Gm25512 | | 0 | 0 | 0 | 0 | 0 | 0 | 0 | 0 | 0 | 0 | 0 |
| 0 | NA | NA | NA | NA | NA | Gm24473 | | 0 | 0 | 0 | 0 | 0 | 0 | 0 | 0 | 0 | 0 | 0 |
| 0 | NA | NA | NA | NA | NA | Gm11456 | | 0 | 0 | 0 | 0 | 0 | 0 | 0 | 0 | 0 | 0 | 0 |
| 0 | NA | NA | NA | NA | NA | Wfdc9 | WAP four-disulfide core domain 9 | 0 | 0 | 0 | 0 | 0 | 0 | 0 | 0 | 0 | 0 | 0 |
| 0 | NA | NA | NA | NA | NA | Wfdc11 | WAP four-disulfide core domain 11 | 0 | 0 | 0 | 0 | 0 | 0 | 0 | 0 | 0 | 0 | 0 |
| 0 | NA | NA | NA | NA | NA | Wfdc13 | WAP four-disulfide core domain 13 | 0 | 0 | 0 | 0 | 0 | 0 | 0 | 0 | 0 | 0 | 0 |
| 0 | NA | NA | NA | NA | NA | Spint4 | serine protease inhibitor, Kunitz type 4 | 0 | 0 | 0 | 0 | 0 | 0 | 0 | 0 | 0 | 0 | 0 |
| 0 | NA | NA | NA | NA | NA | Spint5 | serine protease inhibitor, Kunitz type 5 | 0 | 0 | 0 | 0 | 0 | 0 | 0 | 0 | 0 | 0 | 0 |
| 0 | NA | NA | NA | NA | NA | Tnnc2 | troponin C2, fast | 0 | 0 | 0 | 0 | 0 | 0 | 0 | 0 | 0 | 0 | 0 |
| 0 | NA | NA | NA | NA | NA | Gm25569 | | 0 | 0 | 0 | 0 | 0 | 0 | 0 | 0 | 0 | 0 | 0 |
| 0 | NA | NA | NA | NA | NA | Gm14437 | | 0 | 0 | 0 | 0 | 0 | 0 | 0 | 0 | 0 | 0 | 0 |
| 0 | NA | NA | NA | NA | NA | Gm28163 | | 0 | 0 | 0 | 0 | 0 | 0 | 0 | 0 | 0 | 0 | 0 |
| 0 | NA | NA | NA | NA | NA | Gm22692 | | 0 | 0 | 0 | 0 | 0 | 0 | 0 | 0 | 0 | 0 | 0 |
| 0 | NA | NA | NA | NA | NA | Gm11461 | | 0 | 0 | 0 | 0 | 0 | 0 | 0 | 0 | 0 | 0 | 0 |
| 0 | NA | NA | NA | NA | NA | Gm22687 | | 0 | 0 | 0 | 0 | 0 | 0 | 0 | 0 | 0 | 0 | 0 |
| 0 | NA | NA | NA | NA | NA | Gm11466 | | 0 | 0 | 0 | 0 | 0 | 0 | 0 | 0 | 0 | 0 | 0 |
| 0 | NA | NA | NA | NA | NA | Gm11467 | | 0 | 0 | 0 | 0 | 0 | 0 | 0 | 0 | 0 | 0 | 0 |
| 0 | NA | NA | NA | NA | NA | Gm11468 | | 0 | 0 | 0 | 0 | 0 | 0 | 0 | 0 | 0 | 0 | 0 |
| 0 | NA | NA | NA | NA | NA | Gm14268 | | 0 | 0 | 0 | 0 | 0 | 0 | 0 | 0 | 0 | 0 | 0 |
| 0 | NA | NA | NA | NA | NA | Gm23152 | | 0 | 0 | 0 | 0 | 0 | 0 | 0 | 0 | 0 | 0 | 0 |
| 0 | NA | NA | NA | NA | NA | Gm25265 | | 0 | 0 | 0 | 0 | 0 | 0 | 0 | 0 | 0 | 0 | 0 |
| 0 | NA | NA | NA | NA | NA | Gm24477 | | 0 | 0 | 0 | 0 | 0 | 0 | 0 | 0 | 0 | 0 | 0 |
| 0 | NA | NA | NA | NA | NA | Gm27494 | | 0 | 0 | 0 | 0 | 0 | 0 | 0 | 0 | 0 | 0 | 0 |
| 0 | NA | NA | NA | NA | NA | Gm25878 | | 0 | 0 | 0 | 0 | 0 | 0 | 0 | 0 | 0 | 0 | 0 |
| 0 | NA | NA | NA | NA | NA | Snord12 | | 0 | 0 | 0 | 0 | 0 | 0 | 0 | 0 | 0 | 0 | 0 |
| 0 | NA | NA | NA | NA | NA | Gm11471 | | 0 | 0 | 0 | 0 | 0 | 0 | 0 | 0 | 0 | 0 | 0 |
| 0 | NA | NA | NA | NA | NA | Gm11473 | | 0 | 0 | 0 | 0 | 0 | 0 | 0 | 0 | 0 | 0 | 0 |
| 0 | NA | NA | NA | NA | NA | Gm11475 | | 0 | 0 | 0 | 0 | 0 | 0 | 0 | 0 | 0 | 0 | 0 |
| 0 | NA | NA | NA | NA | NA | Gm20431 | | 0 | 0 | 0 | 0 | 0 | 0 | 0 | 0 | 0 | 0 | 0 |
| 0 | NA | NA | NA | NA | NA | Gm14319 | | 0 | 0 | 0 | 0 | 0 | 0 | 0 | 0 | 0 | 0 | 0 |
| 0 | NA | NA | NA | NA | NA | Gm14236 | | 0 | 0 | 0 | 0 | 0 | 0 | 0 | 0 | 0 | 0 | 0 |
| 0 | NA | NA | NA | NA | NA | Gm24327 | | 0 | 0 | 0 | 0 | 0 | 0 | 0 | 0 | 0 | 0 | 0 |
| 0 | NA | NA | NA | NA | NA | | | 0 | 0 | 0 | 0 | 0 | 0 | 0 | 0 | 0 | 0 | 0 |
| 0 | NA | NA | NA | NA | NA | Gm24561 | | 0 | 0 | 0 | 0 | 0 | 0 | 0 | 0 | 0 | 0 | 0 |
| 0 | NA | NA | NA | NA | NA | Gm20716 | | 0 | 0 | 0 | 0 | 0 | 0 | 0 | 0 | 0 | 0 | 0 |
| 0 | NA | NA | NA | NA | NA | Gm14237 | | 0 | 0 | 0 | 0 | 0 | 0 | 0 | 0 | 0 | 0 | 0 |
| 0 | NA | NA | NA | NA | NA | Gm14262 | | 0 | 0 | 0 | 0 | 0 | 0 | 0 | 0 | 0 | 0 | 0 |

| | | | | | | | | | | | | | | | | | | |
|---|---|---|---|---|---|---|---|---|---|---|---|---|---|---|---|---|---|---|
| 0 | NA | NA | NA | NA | NA | Gm22704 | | 0 | 0 | 0 | 0 | 0 | 0 | 0 | 0 | 0 | 0 | 0 |
| 0 | NA | NA | NA | NA | NA | Gm14234 | | 0 | 0 | 0 | 0 | 0 | 0 | 0 | 0 | 0 | 0 | 0 |
| 0 | NA | NA | NA | NA | NA | Gm25214 | | 0 | 0 | 0 | 0 | 0 | 0 | 0 | 0 | 0 | 0 | 0 |
| 0 | NA | NA | NA | NA | NA | Gm14260 | | 0 | 0 | 0 | 0 | 0 | 0 | 0 | 0 | 0 | 0 | 0 |
| 0 | NA | NA | NA | NA | NA | Gm26469 | | 0 | 0 | 0 | 0 | 0 | 0 | 0 | 0 | 0 | 0 | 0 |
| 0 | NA | NA | NA | NA | NA | Gm14258 | | 0 | 0 | 0 | 0 | 0 | 0 | 0 | 0 | 0 | 0 | 0 |
| 0 | NA | NA | NA | NA | NA | 9430093N23Rik | | 0 | 0 | 0 | 0 | 0 | 0 | 0 | 0 | 0 | 0 | 0 |
| 0 | NA | NA | NA | NA | NA | Gm14250 | | 0 | 0 | 0 | 0 | 0 | 0 | 0 | 0 | 0 | 0 | 0 |
| 0 | NA | NA | NA | NA | NA | 4930529I22Rik | | 0 | 0 | 0 | 0 | 0 | 0 | 0 | 0 | 0 | 0 | 0 |
| 0 | NA | NA | NA | NA | NA | Gm14249 | | 0 | 0 | 0 | 0 | 0 | 0 | 0 | 0 | 0 | 0 | 0 |
| 0 | NA | NA | NA | NA | NA | Gm11011 | | 0 | 0 | 0 | 0 | 0 | 0 | 0 | 0 | 0 | 0 | 0 |
| 0 | NA | NA | NA | NA | NA | AY702102 | | 0 | 0 | 0 | 0 | 0 | 0 | 0 | 0 | 0 | 0 | 0 |
| 0 | NA | NA | NA | NA | NA | Bcas1os2 | | 0 | 0 | 0 | 0 | 0 | 0 | 0 | 0 | 0 | 0 | 0 |
| 0 | NA | NA | NA | NA | NA | Cyp24a1 | cytochrome P450, family 24, subfamily a, | 0 | 0 | 0 | 0 | 0 | 0 | 0 | 0 | 0 | 0 | 0 |
| 0 | NA | NA | NA | NA | NA | Gm14266 | | 0 | 0 | 0 | 0 | 0 | 0 | 0 | 0 | 0 | 0 | 0 |
| 0 | NA | NA | NA | NA | NA | Gm14640 | | 0 | 0 | 0 | 0 | 0 | 0 | 0 | 0 | 0 | 0 | 0 |
| 0 | NA | NA | NA | NA | NA | 1700007M16Rik | | 0 | 0 | 0 | 0 | 0 | 0 | 0 | 0 | 0 | 0 | 0 |
| 0 | NA | NA | NA | NA | NA | Gm26489 | | 0 | 0 | 0 | 0 | 0 | 0 | 0 | 0 | 0 | 0 | 0 |
| 0 | NA | NA | NA | NA | NA | 1700028P15Rik | | 0 | 0 | 0 | 0 | 0 | 0 | 0 | 0 | 0 | 0 | 0 |
| 0 | NA | NA | NA | NA | NA | Gm14272 | | 0 | 0 | 0 | 0 | 0 | 0 | 0 | 0 | 0 | 0 | 0 |
| 0 | NA | NA | NA | NA | NA | Gm14641 | | 0 | 0 | 0 | 0 | 0 | 0 | 0 | 0 | 0 | 0 | 0 |
| 0 | NA | NA | NA | NA | NA | Gm14271 | | 0 | 0 | 0 | 0 | 0 | 0 | 0 | 0 | 0 | 0 | 0 |
| 0 | NA | NA | NA | NA | NA | Gm44301 | | 0 | 0 | 0 | 0 | 0 | 0 | 0 | 0 | 0 | 0 | 0 |
| 0 | NA | NA | NA | NA | NA | Mc3r | melanocortin 3 receptor | 0 | 0 | 0 | 0 | 0 | 0 | 0 | 0 | 0 | 0 | 0 |
| 0 | NA | NA | NA | NA | NA | Gm14275 | | 0 | 0 | 0 | 0 | 0 | 0 | 0 | 0 | 0 | 0 | 0 |
| 0 | NA | NA | NA | NA | NA | Gm14273 | | 0 | 0 | 0 | 0 | 0 | 0 | 0 | 0 | 0 | 0 | 0 |
| 0 | NA | NA | NA | NA | NA | Gcnt7 | glucosaminyl (N-acetyl) transferase family | 0 | 0 | 0 | 0 | 0 | 0 | 0 | 0 | 0 | 0 | 0 |
| 0 | NA | NA | NA | NA | NA | Tfap2c | transcription factor AP-2, gamma | 0 | 0 | 0 | 0 | 0 | 0 | 0 | 0 | 0 | 0 | 0 |
| 0 | NA | NA | NA | NA | NA | Gm20490 | | 0 | 0 | 0 | 0 | 0 | 0 | 0 | 0 | 0 | 0 | 0 |
| 0 | NA | NA | NA | NA | NA | Gm14453 | | 0 | 0 | 0 | 0 | 0 | 0 | 0 | 0 | 0 | 0 | 0 |
| 0 | NA | NA | NA | NA | NA | Ctcflos | | 0 | 0 | 0 | 0 | 0 | 0 | 0 | 0 | 0 | 0 | 0 |
| 0 | NA | NA | NA | NA | NA | Gm14642 | | 0 | 0 | 0 | 0 | 0 | 0 | 0 | 0 | 0 | 0 | 0 |
| 0 | NA | NA | NA | NA | NA | 1700021F07Rik | RIKEN cDNA 1700021F07 gene | 0 | 0 | 0 | 0 | 0 | 0 | 0 | 0 | 0 | 0 | 0 |
| 0 | NA | NA | NA | NA | NA | Mir6340 | | 0 | 0 | 0 | 0 | 0 | 0 | 0 | 0 | 0 | 0 | 0 |
| 0 | NA | NA | NA | NA | NA | 1700010B08Ril | RIKEN cDNA 1700010B08 gene | 0 | 0 | 0 | 0 | 0 | 0 | 0 | 0 | 0 | 0 | 0 |
| 0 | NA | NA | NA | NA | NA | Gm14371 | | 0 | 0 | 0 | 0 | 0 | 0 | 0 | 0 | 0 | 0 | 0 |
| 0 | NA | NA | NA | NA | NA | Gm10714 | | 0 | 0 | 0 | 0 | 0 | 0 | 0 | 0 | 0 | 0 | 0 |
| 0 | NA | NA | NA | NA | NA | Gm27591 | | 0 | 0 | 0 | 0 | 0 | 0 | 0 | 0 | 0 | 0 | 0 |
| 0 | NA | NA | NA | NA | NA | Mir296 | | 0 | 0 | 0 | 0 | 0 | 0 | 0 | 0 | 0 | 0 | 0 |
| 0 | NA | NA | NA | NA | NA | Mir298 | | 0 | 0 | 0 | 0 | 0 | 0 | 0 | 0 | 0 | 0 | 0 |
| 0 | NA | NA | NA | NA | NA | Gm27869 | | 0 | 0 | 0 | 0 | 0 | 0 | 0 | 0 | 0 | 0 | 0 |
| 0 | NA | NA | NA | NA | NA | Gm27849 | | 0 | 0 | 0 | 0 | 0 | 0 | 0 | 0 | 0 | 0 | 0 |
| 0 | NA | NA | NA | NA | NA | Gm14385 | | 0 | 0 | 0 | 0 | 0 | 0 | 0 | 0 | 0 | 0 | 0 |
| 0 | NA | NA | NA | NA | NA | Vmn1r-ps1 | | 0 | 0 | 0 | 0 | 0 | 0 | 0 | 0 | 0 | 0 | 0 |
| 0 | NA | NA | NA | NA | NA | Gm16354 | | 0 | 0 | 0 | 0 | 0 | 0 | 0 | 0 | 0 | 0 | 0 |
| 0 | NA | NA | NA | NA | NA | Gm14392 | | 0 | 0 | 0 | 0 | 0 | 0 | 0 | 0 | 0 | 0 | 0 |
| 0 | NA | NA | NA | NA | NA | Gm16355 | | 0 | 0 | 0 | 0 | 0 | 0 | 0 | 0 | 0 | 0 | 0 |
| 0 | NA | NA | NA | NA | NA | Gm14442 | | 0 | 0 | 0 | 0 | 0 | 0 | 0 | 0 | 0 | 0 | 0 |
| 0 | NA | NA | NA | NA | NA | Gm14438 | | 0 | 0 | 0 | 0 | 0 | 0 | 0 | 0 | 0 | 0 | 0 |
| 0 | NA | NA | NA | NA | NA | Gm14440 | predicted gene 14440 | 0 | 0 | 0 | 0 | 0 | 0 | 0 | 0 | 0 | 0 | 0 |
| 0 | NA | NA | NA | NA | NA | Gm14298 | | 0 | 0 | 0 | 0 | 0 | 0 | 0 | 0 | 0 | 0 | 0 |
| 0 | NA | NA | NA | NA | NA | Gm4631 | predicted gene 14430 | 0 | 0 | 0 | 0 | 0 | 0 | 0 | 0 | 0 | 0 | 0 |
| 0 | NA | NA | NA | NA | NA | Zfp968 | zinc finger protein 968 | 0 | 0 | 0 | 0 | 0 | 0 | 0 | 0 | 0 | 0 | 0 |
| 0 | NA | NA | NA | NA | NA | Gm14297 | | 0 | 0 | 0 | 0 | 0 | 0 | 0 | 0 | 0 | 0 | 0 |
| 0 | NA | NA | NA | NA | NA | Gm4724 | predicted gene 4724 | 0 | 0 | 0 | 0 | 0 | 0 | 0 | 0 | 0 | 0 | 0 |
| 0 | NA | NA | NA | NA | NA | Zfp965 | zinc finger protein 965 | 0 | 0 | 0 | 0 | 0 | 0 | 0 | 0 | 0 | 0 | 0 |
| 0 | NA | NA | NA | NA | NA | Gm11007 | zinc finger protein 965 | 0 | 0 | 0 | 0 | 0 | 0 | 0 | 0 | 0 | 0 | 0 |
| 0 | NA | NA | NA | NA | NA | Gm16359 | | 0 | 0 | 0 | 0 | 0 | 0 | 0 | 0 | 0 | 0 | 0 |
| 0 | NA | NA | NA | NA | NA | Zfp969 | zinc finger protein 968 | 0 | 0 | 0 | 0 | 0 | 0 | 0 | 0 | 0 | 0 | 0 |
| 0 | NA | NA | NA | NA | NA | Gm2007 | predicted gene 2007 | 0 | 0 | 0 | 0 | 0 | 0 | 0 | 0 | 0 | 0 | 0 |

| | | | | | | | | | | | | | | | | | | |
|---|---|---|---|---|---|---|---|---|---|---|---|---|---|---|---|---|---|---|
| 0 | NA | NA | NA | NA | NA | Gm16358 | | 0 | 0 | 0 | 0 | 0 | 0 | 0 | 0 | 0 | 0 | 0 |
| 0 | NA | NA | NA | NA | NA | Rps8-ps2 | | 0 | 0 | 0 | 0 | 0 | 0 | 0 | 0 | 0 | 0 | 0 |
| 0 | NA | NA | NA | NA | NA | Gm2020 | | 0 | 0 | 0 | 0 | 0 | 0 | 0 | 0 | 0 | 0 | 0 |
| 0 | NA | NA | NA | NA | NA | Gm6710 | predicted gene 6710 | 0 | 0 | 0 | 0 | 0 | 0 | 0 | 0 | 0 | 0 | 0 |
| 0 | NA | NA | NA | NA | NA | Gm16356 | | 0 | 0 | 0 | 0 | 0 | 0 | 0 | 0 | 0 | 0 | 0 |
| 0 | NA | NA | NA | NA | NA | Gm2026 | predicted gene 2026 | 0 | 0 | 0 | 0 | 0 | 0 | 0 | 0 | 0 | 0 | 0 |
| 0 | NA | NA | NA | NA | NA | Gm11008 | | 0 | 0 | 0 | 0 | 0 | 0 | 0 | 0 | 0 | 0 | 0 |
| 0 | NA | NA | NA | NA | NA | Gm16357 | | 0 | 0 | 0 | 0 | 0 | 0 | 0 | 0 | 0 | 0 | 0 |
| 0 | NA | NA | NA | NA | NA | Gm6182 | | 0 | 0 | 0 | 0 | 0 | 0 | 0 | 0 | 0 | 0 | 0 |
| 0 | NA | NA | NA | NA | NA | Gm17258 | | 0 | 0 | 0 | 0 | 0 | 0 | 0 | 0 | 0 | 0 | 0 |
| 0 | NA | NA | NA | NA | NA | Gm1980 | | 0 | 0 | 0 | 0 | 0 | 0 | 0 | 0 | 0 | 0 | 0 |
| 0 | NA | NA | NA | NA | NA | Gm11009 | | 0 | 0 | 0 | 0 | 0 | 0 | 0 | 0 | 0 | 0 | 0 |
| 0 | NA | NA | NA | NA | NA | Zfp968-ps | | 0 | 0 | 0 | 0 | 0 | 0 | 0 | 0 | 0 | 0 | 0 |
| 0 | NA | NA | NA | NA | NA | Gm16362 | | 0 | 0 | 0 | 0 | 0 | 0 | 0 | 0 | 0 | 0 | 0 |
| 0 | NA | NA | NA | NA | NA | Gm14434 | predicted gene 14434 | 0 | 0 | 0 | 0 | 0 | 0 | 0 | 0 | 0 | 0 | 0 |
| 0 | NA | NA | NA | NA | NA | Gm14307 | | 0 | 0 | 0 | 0 | 0 | 0 | 0 | 0 | 0 | 0 | 0 |
| 0 | NA | NA | NA | NA | NA | Gm14308 | predicted gene 14308 | 0 | 0 | 0 | 0 | 0 | 0 | 0 | 0 | 0 | 0 | 0 |
| 0 | NA | NA | NA | NA | NA | Gm14301 | | 0 | 0 | 0 | 0 | 0 | 0 | 0 | 0 | 0 | 0 | 0 |
| 0 | NA | NA | NA | NA | NA | Gm14390 | | 0 | 0 | 0 | 0 | 0 | 0 | 0 | 0 | 0 | 0 | 0 |
| 0 | NA | NA | NA | NA | NA | Gm14408 | | 0 | 0 | 0 | 0 | 0 | 0 | 0 | 0 | 0 | 0 | 0 |
| 0 | NA | NA | NA | NA | NA | Gm14405 | | 0 | 0 | 0 | 0 | 0 | 0 | 0 | 0 | 0 | 0 | 0 |
| 0 | NA | NA | NA | NA | NA | Gm14417 | | 0 | 0 | 0 | 0 | 0 | 0 | 0 | 0 | 0 | 0 | 0 |
| 0 | NA | NA | NA | NA | NA | Gm14402 | | 0 | 0 | 0 | 0 | 0 | 0 | 0 | 0 | 0 | 0 | 0 |
| 0 | NA | NA | NA | NA | NA | Gm14415 | | 0 | 0 | 0 | 0 | 0 | 0 | 0 | 0 | 0 | 0 | 0 |
| 0 | NA | NA | NA | NA | NA | Gm14416 | | 0 | 0 | 0 | 0 | 0 | 0 | 0 | 0 | 0 | 0 | 0 |
| 0 | NA | NA | NA | NA | NA | Gm14404 | | 0 | 0 | 0 | 0 | 0 | 0 | 0 | 0 | 0 | 0 | 0 |
| 0 | NA | NA | NA | NA | NA | Gm16363 | | 0 | 0 | 0 | 0 | 0 | 0 | 0 | 0 | 0 | 0 | 0 |
| 0 | NA | NA | NA | NA | NA | Gm14324 | | 0 | 0 | 0 | 0 | 0 | 0 | 0 | 0 | 0 | 0 | 0 |
| 0 | NA | NA | NA | NA | NA | Gm14428 | | 0 | 0 | 0 | 0 | 0 | 0 | 0 | 0 | 0 | 0 | 0 |
| 0 | NA | NA | NA | NA | NA | Gm14314 | | 0 | 0 | 0 | 0 | 0 | 0 | 0 | 0 | 0 | 0 | 0 |
| 0 | NA | NA | NA | NA | NA | Gm14293 | | 0 | 0 | 0 | 0 | 0 | 0 | 0 | 0 | 0 | 0 | 0 |
| 0 | NA | NA | NA | NA | NA | Gm14294 | | 0 | 0 | 0 | 0 | 0 | 0 | 0 | 0 | 0 | 0 | 0 |
| 0 | NA | NA | NA | NA | NA | 4930591A17Ril | RIKEN cDNA 4930591A17 gene | 0 | 0 | 0 | 0 | 0 | 0 | 0 | 0 | 0 | 0 | 0 |
| 0 | NA | NA | NA | NA | NA | Gm14300 | | 0 | 0 | 0 | 0 | 0 | 0 | 0 | 0 | 0 | 0 | 0 |
| 0 | NA | NA | NA | NA | NA | Gm14299 | | 0 | 0 | 0 | 0 | 0 | 0 | 0 | 0 | 0 | 0 | 0 |
| 0 | NA | NA | NA | NA | NA | Gm17180 | | 0 | 0 | 0 | 0 | 0 | 0 | 0 | 0 | 0 | 0 | 0 |
| 0 | NA | NA | NA | NA | NA | Mir7005 | | 0 | 0 | 0 | 0 | 0 | 0 | 0 | 0 | 0 | 0 | 0 |
| 0 | NA | NA | NA | NA | NA | Gata5os | | 0 | 0 | 0 | 0 | 0 | 0 | 0 | 0 | 0 | 0 | 0 |
| 0 | NA | NA | NA | NA | NA | Mir1a-1 | | 0 | 0 | 0 | 0 | 0 | 0 | 0 | 0 | 0 | 0 | 0 |
| 0 | NA | NA | NA | NA | NA | Mir133a-2 | | 0 | 0 | 0 | 0 | 0 | 0 | 0 | 0 | 0 | 0 | 0 |
| 0 | NA | NA | NA | NA | NA | Gm22502 | | 0 | 0 | 0 | 0 | 0 | 0 | 0 | 0 | 0 | 0 | 0 |
| 0 | NA | NA | NA | NA | NA | Gm14343 | | 0 | 0 | 0 | 0 | 0 | 0 | 0 | 0 | 0 | 0 | 0 |
| 0 | NA | NA | NA | NA | NA | Bhlhe23 | basic helix-loop-helix family, member e23 | 0 | 0 | 0 | 0 | 0 | 0 | 0 | 0 | 0 | 0 | 0 |
| 0 | NA | NA | NA | NA | NA | Gm14340 | | 0 | 0 | 0 | 0 | 0 | 0 | 0 | 0 | 0 | 0 | 0 |
| 0 | NA | NA | NA | NA | NA | Gm14342 | | 0 | 0 | 0 | 0 | 0 | 0 | 0 | 0 | 0 | 0 | 0 |
| 0 | NA | NA | NA | NA | NA | Gm14344 | | 0 | 0 | 0 | 0 | 0 | 0 | 0 | 0 | 0 | 0 | 0 |
| 0 | NA | NA | NA | NA | NA | Gm22842 | | 0 | 0 | 0 | 0 | 0 | 0 | 0 | 0 | 0 | 0 | 0 |
| 0 | NA | NA | NA | NA | NA | Gm14339 | | 0 | 0 | 0 | 0 | 0 | 0 | 0 | 0 | 0 | 0 | 0 |
| 0 | NA | NA | NA | NA | NA | Mir124a-3 | | 0 | 0 | 0 | 0 | 0 | 0 | 0 | 0 | 0 | 0 | 0 |
| 0 | NA | NA | NA | NA | NA | Gm27032 | | 0 | 0 | 0 | 0 | 0 | 0 | 0 | 0 | 0 | 0 | 0 |
| 0 | NA | NA | NA | NA | NA | Birc7 | baculoviral IAP repeat-containing 7 (livin) | 0 | 0 | 0 | 0 | 0 | 0 | 0 | 0 | 0 | 0 | 0 |
| 0 | NA | NA | NA | NA | NA | Gm14341 | | 0 | 0 | 0 | 0 | 0 | 0 | 0 | 0 | 0 | 0 | 0 |
| 0 | NA | NA | NA | NA | NA | Gm24892 | | 0 | 0 | 0 | 0 | 0 | 0 | 0 | 0 | 0 | 0 | 0 |
| 0 | NA | NA | NA | NA | NA | | | 0 | 0 | 0 | 0 | 0 | 0 | 0 | 0 | 0 | 0 | 0 |
| 0 | NA | NA | NA | NA | NA | Gm22708 | | 0 | 0 | 0 | 0 | 0 | 0 | 0 | 0 | 0 | 0 | 0 |
| 0 | NA | NA | NA | NA | NA | Abhd16b | abhydrolase domain containing 16B | 0 | 0 | 0 | 0 | 0 | 0 | 0 | 0 | 0 | 0 | 0 |
| 0 | NA | NA | NA | NA | NA | Gm25879 | | 0 | 0 | 0 | 0 | 0 | 0 | 0 | 0 | 0 | 0 | 0 |
| 0 | NA | NA | NA | NA | NA | Lkaaear1 | LKAAEAR motif containing 1 (IKAAEAR m | 0 | 0 | 0 | 0 | 0 | 0 | 0 | 0 | 0 | 0 | 0 |
| 0 | NA | NA | NA | NA | NA | Gm14489 | | 0 | 0 | 0 | 0 | 0 | 0 | 0 | 0 | 0 | 0 | 0 |
| 0 | NA | NA | NA | NA | NA | Gm14496 | predicted gene 14496 | 0 | 0 | 0 | 0 | 0 | 0 | 0 | 0 | 0 | 0 | 0 |

| | | | | | | | | | | | | | | | | | | |
|---|---|---|---|---|---|---|---|---|---|---|---|---|---|---|---|---|---|---|
| 0 | NA | NA | NA | NA | NA | Gm28002 | | 0 | 0 | 0 | 0 | 0 | 0 | 0 | 0 | 0 | 0 | 0 |
| 0 | NA | NA | NA | NA | NA | Gm27460 | | 0 | 0 | 0 | 0 | 0 | 0 | 0 | 0 | 0 | 0 | 0 |
| 0 | NA | NA | NA | NA | NA | Gm37338 | | 0 | 0 | 0 | 0 | 0 | 0 | 0 | 0 | 0 | 0 | 0 |
| 0 | NA | NA | NA | NA | NA | Gm8747 | | 0 | 0 | 0 | 0 | 0 | 0 | 0 | 0 | 0 | 0 | 0 |
| 0 | NA | NA | NA | NA | NA | Gm37000 | | 0 | 0 | 0 | 0 | 0 | 0 | 0 | 0 | 0 | 0 | 0 |
| 0 | NA | NA | NA | NA | NA | Gm23237 | | 0 | 0 | 0 | 0 | 0 | 0 | 0 | 0 | 0 | 0 | 0 |
| 0 | NA | NA | NA | NA | NA | Gm2071 | | 0 | 0 | 0 | 0 | 0 | 0 | 0 | 0 | 0 | 0 | 0 |
| 0 | NA | NA | NA | NA | NA | Gm2077 | | 0 | 0 | 0 | 0 | 0 | 0 | 0 | 0 | 0 | 0 | 0 |
| 0 | NA | NA | NA | NA | NA | Gm23976 | | 0 | 0 | 0 | 0 | 0 | 0 | 0 | 0 | 0 | 0 | 0 |
| 0 | NA | NA | NA | NA | NA | n-R5s194 | | 0 | 0 | 0 | 0 | 0 | 0 | 0 | 0 | 0 | 0 | 0 |
| 0 | NA | NA | NA | NA | NA | Gm34023 | | 0 | 0 | 0 | 0 | 0 | 0 | 0 | 0 | 0 | 0 | 0 |
| 0 | NA | NA | NA | NA | NA | Gm38131 | | 0 | 0 | 0 | 0 | 0 | 0 | 0 | 0 | 0 | 0 | 0 |
| 0 | NA | NA | NA | NA | NA | Gm37029 | | 0 | 0 | 0 | 0 | 0 | 0 | 0 | 0 | 0 | 0 | 0 |
| 0 | NA | NA | NA | NA | NA | Gm22944 | | 0 | 0 | 0 | 0 | 0 | 0 | 0 | 0 | 0 | 0 | 0 |
| 0 | NA | NA | NA | NA | NA | Gm37763 | | 0 | 0 | 0 | 0 | 0 | 0 | 0 | 0 | 0 | 0 | 0 |
| 0 | NA | NA | NA | NA | NA | Gm37801 | | 0 | 0 | 0 | 0 | 0 | 0 | 0 | 0 | 0 | 0 | 0 |
| 0 | NA | NA | NA | NA | NA | Gm17308 | | 0 | 0 | 0 | 0 | 0 | 0 | 0 | 0 | 0 | 0 | 0 |
| 0 | NA | NA | NA | NA | NA | Gm10748 | | 0 | 0 | 0 | 0 | 0 | 0 | 0 | 0 | 0 | 0 | 0 |
| 0 | NA | NA | NA | NA | NA | 4930555M17Rik | | 0 | 0 | 0 | 0 | 0 | 0 | 0 | 0 | 0 | 0 | 0 |
| 0 | NA | NA | NA | NA | NA | Gm37350 | | 0 | 0 | 0 | 0 | 0 | 0 | 0 | 0 | 0 | 0 | 0 |
| 0 | NA | NA | NA | NA | NA | Gm25352 | | 0 | 0 | 0 | 0 | 0 | 0 | 0 | 0 | 0 | 0 | 0 |
| 0 | NA | NA | NA | NA | NA | Gm25144 | | 0 | 0 | 0 | 0 | 0 | 0 | 0 | 0 | 0 | 0 | 0 |
| 0 | NA | NA | NA | NA | NA | Gm17934 | | 0 | 0 | 0 | 0 | 0 | 0 | 0 | 0 | 0 | 0 | 0 |
| 0 | NA | NA | NA | NA | NA | Gm38298 | | 0 | 0 | 0 | 0 | 0 | 0 | 0 | 0 | 0 | 0 | 0 |
| 0 | NA | NA | NA | NA | NA | Gm23147 | | 0 | 0 | 0 | 0 | 0 | 0 | 0 | 0 | 0 | 0 | 0 |
| 0 | NA | NA | NA | NA | NA | Gm18658 | | 0 | 0 | 0 | 0 | 0 | 0 | 0 | 0 | 0 | 0 | 0 |
| 0 | NA | NA | NA | NA | NA | Gm38291 | | 0 | 0 | 0 | 0 | 0 | 0 | 0 | 0 | 0 | 0 | 0 |
| 0 | NA | NA | NA | NA | NA | 1700008P02Ril | RIKEN cDNA 1700008P02 gene | 0 | 0 | 0 | 0 | 0 | 0 | 0 | 0 | 0 | 0 | 0 |
| 0 | NA | NA | NA | NA | NA | Olfr289-ps1 | | 0 | 0 | 0 | 0 | 0 | 0 | 0 | 0 | 0 | 0 | 0 |
| 0 | NA | NA | NA | NA | NA | Gm22074 | | 0 | 0 | 0 | 0 | 0 | 0 | 0 | 0 | 0 | 0 | 0 |
| 0 | NA | NA | NA | NA | NA | Gm37918 | | 0 | 0 | 0 | 0 | 0 | 0 | 0 | 0 | 0 | 0 | 0 |
| 0 | NA | NA | NA | NA | NA | Gm8826 | | 0 | 0 | 0 | 0 | 0 | 0 | 0 | 0 | 0 | 0 | 0 |
| 0 | NA | NA | NA | NA | NA | Gm23592 | | 0 | 0 | 0 | 0 | 0 | 0 | 0 | 0 | 0 | 0 | 0 |
| 0 | NA | NA | NA | NA | NA | Gm29747 | | 0 | 0 | 0 | 0 | 0 | 0 | 0 | 0 | 0 | 0 | 0 |
| 0 | NA | NA | NA | NA | NA | 1700010I02Rik | | 0 | 0 | 0 | 0 | 0 | 0 | 0 | 0 | 0 | 0 | 0 |
| 0 | NA | NA | NA | NA | NA | Gm7103 | | 0 | 0 | 0 | 0 | 0 | 0 | 0 | 0 | 0 | 0 | 0 |
| 0 | NA | NA | NA | NA | NA | Gm24614 | | 0 | 0 | 0 | 0 | 0 | 0 | 0 | 0 | 0 | 0 | 0 |
| 0 | NA | NA | NA | NA | NA | Gm18822 | | 0 | 0 | 0 | 0 | 0 | 0 | 0 | 0 | 0 | 0 | 0 |
| 0 | NA | NA | NA | NA | NA | Gm32496 | | 0 | 0 | 0 | 0 | 0 | 0 | 0 | 0 | 0 | 0 | 0 |
| 0 | NA | NA | NA | NA | NA | Gm38007 | | 0 | 0 | 0 | 0 | 0 | 0 | 0 | 0 | 0 | 0 | 0 |
| 0 | NA | NA | NA | NA | NA | Gm23670 | | 0 | 0 | 0 | 0 | 0 | 0 | 0 | 0 | 0 | 0 | 0 |
| 0 | NA | NA | NA | NA | NA | C230057A21Rik | | 0 | 0 | 0 | 0 | 0 | 0 | 0 | 0 | 0 | 0 | 0 |
| 0 | NA | NA | NA | NA | NA | Gm37647 | | 0 | 0 | 0 | 0 | 0 | 0 | 0 | 0 | 0 | 0 | 0 |
| 0 | NA | NA | NA | NA | NA | Gm37616 | | 0 | 0 | 0 | 0 | 0 | 0 | 0 | 0 | 0 | 0 | 0 |
| 0 | NA | NA | NA | NA | NA | 4930539M17Rik | | 0 | 0 | 0 | 0 | 0 | 0 | 0 | 0 | 0 | 0 | 0 |
| 0 | NA | NA | NA | NA | NA | Gm22604 | | 0 | 0 | 0 | 0 | 0 | 0 | 0 | 0 | 0 | 0 | 0 |
| 0 | NA | NA | NA | NA | NA | Gm38001 | | 0 | 0 | 0 | 0 | 0 | 0 | 0 | 0 | 0 | 0 | 0 |
| 0 | NA | NA | NA | NA | NA | Gm17806 | | 0 | 0 | 0 | 0 | 0 | 0 | 0 | 0 | 0 | 0 | 0 |
| 0 | NA | NA | NA | NA | NA | Gm16337 | | 0 | 0 | 0 | 0 | 0 | 0 | 0 | 0 | 0 | 0 | 0 |
| 0 | NA | NA | NA | NA | NA | Gm6236 | | 0 | 0 | 0 | 0 | 0 | 0 | 0 | 0 | 0 | 0 | 0 |
| 0 | NA | NA | NA | NA | NA | Gm38335 | | 0 | 0 | 0 | 0 | 0 | 0 | 0 | 0 | 0 | 0 | 0 |
| 0 | NA | NA | NA | NA | NA | Gm37308 | | 0 | 0 | 0 | 0 | 0 | 0 | 0 | 0 | 0 | 0 | 0 |
| 0 | NA | NA | NA | NA | NA | Pmp2 | peripheral myelin protein 2 | 0 | 0 | 0 | 0 | 0 | 0 | 0 | 0 | 0 | 0 | 0 |
| 0 | NA | NA | NA | NA | NA | Gm37389 | | 0 | 0 | 0 | 0 | 0 | 0 | 0 | 0 | 0 | 0 | 0 |
| 0 | NA | NA | NA | NA | NA | Fabp9 | fatty acid binding protein 9, testis | 0 | 0 | 0 | 0 | 0 | 0 | 0 | 0 | 0 | 0 | 0 |
| 0 | NA | NA | NA | NA | NA | 4632432E15Rik | | 0 | 0 | 0 | 0 | 0 | 0 | 0 | 0 | 0 | 0 | 0 |
| 0 | NA | NA | NA | NA | NA | 1700029B24Rik | | 0 | 0 | 0 | 0 | 0 | 0 | 0 | 0 | 0 | 0 | 0 |
| 0 | NA | NA | NA | NA | NA | Fabp12 | fatty acid binding protein 12 | 0 | 0 | 0 | 0 | 0 | 0 | 0 | 0 | 0 | 0 | 0 |
| 0 | NA | NA | NA | NA | NA | A930014E01Rik | | 0 | 0 | 0 | 0 | 0 | 0 | 0 | 0 | 0 | 0 | 0 |
| 0 | NA | NA | NA | NA | NA | Gm38364 | | 0 | 0 | 0 | 0 | 0 | 0 | 0 | 0 | 0 | 0 | 0 |

| | | | | | | | | | | | | | | | | | | |
|---|---|---|---|---|---|---|---|---|---|---|---|---|---|---|---|---|---|---|
| 0 | NA | NA | NA | NA | NA | Gm38303 | | 0 | 0 | 0 | 0 | 0 | 0 | 0 | 0 | 0 | 0 | 0 |
| 0 | NA | NA | NA | NA | NA | Gm37831 | | 0 | 0 | 0 | 0 | 0 | 0 | 0 | 0 | 0 | 0 | 0 |
| 0 | NA | NA | NA | NA | NA | Gm18692 | | 0 | 0 | 0 | 0 | 0 | 0 | 0 | 0 | 0 | 0 | 0 |
| 0 | NA | NA | NA | NA | NA | Gm37244 | | 0 | 0 | 0 | 0 | 0 | 0 | 0 | 0 | 0 | 0 | 0 |
| 0 | NA | NA | NA | NA | NA | Gm22795 | | 0 | 0 | 0 | 0 | 0 | 0 | 0 | 0 | 0 | 0 | 0 |
| 0 | NA | NA | NA | NA | NA | Gm21631 | | 0 | 0 | 0 | 0 | 0 | 0 | 0 | 0 | 0 | 0 | 0 |
| 0 | NA | NA | NA | NA | NA | Gm22547 | | 0 | 0 | 0 | 0 | 0 | 0 | 0 | 0 | 0 | 0 | 0 |
| 0 | NA | NA | NA | NA | NA | Gm8919 | | 0 | 0 | 0 | 0 | 0 | 0 | 0 | 0 | 0 | 0 | 0 |
| 0 | NA | NA | NA | NA | NA | Gm10745 | | 0 | 0 | 0 | 0 | 0 | 0 | 0 | 0 | 0 | 0 | 0 |
| 0 | NA | NA | NA | NA | NA | Gm37996 | | 0 | 0 | 0 | 0 | 0 | 0 | 0 | 0 | 0 | 0 | 0 |
| 0 | NA | NA | NA | NA | NA | Gm2429 | | 0 | 0 | 0 | 0 | 0 | 0 | 0 | 0 | 0 | 0 | 0 |
| 0 | NA | NA | NA | NA | NA | Gm8939 | | 0 | 0 | 0 | 0 | 0 | 0 | 0 | 0 | 0 | 0 | 0 |
| 0 | NA | NA | NA | NA | NA | Gm37813 | | 0 | 0 | 0 | 0 | 0 | 0 | 0 | 0 | 0 | 0 | 0 |
| 0 | NA | NA | NA | NA | NA | Gm37197 | | 0 | 0 | 0 | 0 | 0 | 0 | 0 | 0 | 0 | 0 | 0 |
| 0 | NA | NA | NA | NA | NA | Gm2464 | | 0 | 0 | 0 | 0 | 0 | 0 | 0 | 0 | 0 | 0 | 0 |
| 0 | NA | NA | NA | NA | NA | Gm2474 | | 0 | 0 | 0 | 0 | 0 | 0 | 0 | 0 | 0 | 0 | 0 |
| 0 | NA | NA | NA | NA | NA | Gm19114 | | 0 | 0 | 0 | 0 | 0 | 0 | 0 | 0 | 0 | 0 | 0 |
| 0 | NA | NA | NA | NA | NA | Gm25981 | | 0 | 0 | 0 | 0 | 0 | 0 | 0 | 0 | 0 | 0 | 0 |
| 0 | NA | NA | NA | NA | NA | Gm22026 | | 0 | 0 | 0 | 0 | 0 | 0 | 0 | 0 | 0 | 0 | 0 |
| 0 | NA | NA | NA | NA | NA | Gm23112 | | 0 | 0 | 0 | 0 | 0 | 0 | 0 | 0 | 0 | 0 | 0 |
| 0 | NA | NA | NA | NA | NA | Gm38311 | | 0 | 0 | 0 | 0 | 0 | 0 | 0 | 0 | 0 | 0 | 0 |
| 0 | NA | NA | NA | NA | NA | Gm6300 | | 0 | 0 | 0 | 0 | 0 | 0 | 0 | 0 | 0 | 0 | 0 |
| 0 | NA | NA | NA | NA | NA | Gm8955 | | 0 | 0 | 0 | 0 | 0 | 0 | 0 | 0 | 0 | 0 | 0 |
| 0 | NA | NA | NA | NA | NA | Slc7a12 | solute carrier family 7 (cationic amino aci | 0 | 0 | 0 | 0 | 0 | 0 | 0 | 0 | 0 | 0 | 0 |
| 0 | NA | NA | NA | NA | NA | Gm17119 | | 0 | 0 | 0 | 0 | 0 | 0 | 0 | 0 | 0 | 0 | 0 |
| 0 | NA | NA | NA | NA | NA | Gm38151 | | 0 | 0 | 0 | 0 | 0 | 0 | 0 | 0 | 0 | 0 | 0 |
| 0 | NA | NA | NA | NA | NA | Gm5843 | | 0 | 0 | 0 | 0 | 0 | 0 | 0 | 0 | 0 | 0 | 0 |
| 0 | NA | NA | NA | NA | NA | A930001A20Rik | | 0 | 0 | 0 | 0 | 0 | 0 | 0 | 0 | 0 | 0 | 0 |
| 0 | NA | NA | NA | NA | NA | Gm33819 | | 0 | 0 | 0 | 0 | 0 | 0 | 0 | 0 | 0 | 0 | 0 |
| 0 | NA | NA | NA | NA | NA | Gm37468 | | 0 | 0 | 0 | 0 | 0 | 0 | 0 | 0 | 0 | 0 | 0 |
| 0 | NA | NA | NA | NA | NA | Gm6321 | | 0 | 0 | 0 | 0 | 0 | 0 | 0 | 0 | 0 | 0 | 0 |
| 0 | NA | NA | NA | NA | NA | Gm7343 | | 0 | 0 | 0 | 0 | 0 | 0 | 0 | 0 | 0 | 0 | 0 |
| 0 | NA | NA | NA | NA | NA | Gm24694 | | 0 | 0 | 0 | 0 | 0 | 0 | 0 | 0 | 0 | 0 | 0 |
| 0 | NA | NA | NA | NA | NA | Gm37836 | | 0 | 0 | 0 | 0 | 0 | 0 | 0 | 0 | 0 | 0 | 0 |
| 0 | NA | NA | NA | NA | NA | Gm25863 | | 0 | 0 | 0 | 0 | 0 | 0 | 0 | 0 | 0 | 0 | 0 |
| 0 | NA | NA | NA | NA | NA | Gm26485 | | 0 | 0 | 0 | 0 | 0 | 0 | 0 | 0 | 0 | 0 | 0 |
| 0 | NA | NA | NA | NA | NA | Gm5283 | | 0 | 0 | 0 | 0 | 0 | 0 | 0 | 0 | 0 | 0 | 0 |
| 0 | NA | NA | NA | NA | NA | Gm24963 | | 0 | 0 | 0 | 0 | 0 | 0 | 0 | 0 | 0 | 0 | 0 |
| 0 | NA | NA | NA | NA | NA | Gm6350 | | 0 | 0 | 0 | 0 | 0 | 0 | 0 | 0 | 0 | 0 | 0 |
| 0 | NA | NA | NA | NA | NA | Gm30340 | | 0 | 0 | 0 | 0 | 0 | 0 | 0 | 0 | 0 | 0 | 0 |
| 0 | NA | NA | NA | NA | NA | Gm19694 | | 0 | 0 | 0 | 0 | 0 | 0 | 0 | 0 | 0 | 0 | 0 |
| 0 | NA | NA | NA | NA | NA | Gm38154 | | 0 | 0 | 0 | 0 | 0 | 0 | 0 | 0 | 0 | 0 | 0 |
| 0 | NA | NA | NA | NA | NA | Mir124-2hg | | 0 | 0 | 0 | 0 | 0 | 0 | 0 | 0 | 0 | 0 | 0 |
| 0 | NA | NA | NA | NA | NA | Mir124a-2 | | 0 | 0 | 0 | 0 | 0 | 0 | 0 | 0 | 0 | 0 | 0 |
| 0 | NA | NA | NA | NA | NA | Gm23441 | | 0 | 0 | 0 | 0 | 0 | 0 | 0 | 0 | 0 | 0 | 0 |
| 0 | NA | NA | NA | NA | NA | Gm19175 | | 0 | 0 | 0 | 0 | 0 | 0 | 0 | 0 | 0 | 0 | 0 |
| 0 | NA | NA | NA | NA | NA | Cypt12 | cysteine-rich perinuclear theca 12 | 0 | 0 | 0 | 0 | 0 | 0 | 0 | 0 | 0 | 0 | 0 |
| 0 | NA | NA | NA | NA | NA | Gm23726 | | 0 | 0 | 0 | 0 | 0 | 0 | 0 | 0 | 0 | 0 | 0 |
| 0 | NA | NA | NA | NA | NA | Gm23686 | | 0 | 0 | 0 | 0 | 0 | 0 | 0 | 0 | 0 | 0 | 0 |
| 0 | NA | NA | NA | NA | NA | Gm43834 | | 0 | 0 | 0 | 0 | 0 | 0 | 0 | 0 | 0 | 0 | 0 |
| 0 | NA | NA | NA | NA | NA | Gm30667 | | 0 | 0 | 0 | 0 | 0 | 0 | 0 | 0 | 0 | 0 | 0 |
| 0 | NA | NA | NA | NA | NA | 4930433B08Rik | | 0 | 0 | 0 | 0 | 0 | 0 | 0 | 0 | 0 | 0 | 0 |
| 0 | NA | NA | NA | NA | NA | Gm6369 | | 0 | 0 | 0 | 0 | 0 | 0 | 0 | 0 | 0 | 0 | 0 |
| 0 | NA | NA | NA | NA | NA | Pgk1-ps3 | | 0 | 0 | 0 | 0 | 0 | 0 | 0 | 0 | 0 | 0 | 0 |
| 0 | NA | NA | NA | NA | NA | Gm30341 | | 0 | 0 | 0 | 0 | 0 | 0 | 0 | 0 | 0 | 0 | 0 |
| 0 | NA | NA | NA | NA | NA | Dnajc5b | DnaJ heat shock protein family (Hsp40) m | 0 | 0 | 0 | 0 | 0 | 0 | 0 | 0 | 0 | 0 | 0 |
| 0 | NA | NA | NA | NA | NA | Gm6141 | | 0 | 0 | 0 | 0 | 0 | 0 | 0 | 0 | 0 | 0 | 0 |
| 0 | NA | NA | NA | NA | NA | BC002189 | | 0 | 0 | 0 | 0 | 0 | 0 | 0 | 0 | 0 | 0 | 0 |
| 0 | NA | NA | NA | NA | NA | Crh | corticotropin releasing hormone | 0 | 0 | 0 | 0 | 0 | 0 | 0 | 0 | 0 | 0 | 0 |
| 0 | NA | NA | NA | NA | NA | Gm18864 | | 0 | 0 | 0 | 0 | 0 | 0 | 0 | 0 | 0 | 0 | 0 |

| | | | | | | | | | | | | | | | | | | |
|---|---|---|---|---|---|---|---|---|---|---|---|---|---|---|---|---|---|---|
| 0 | NA | NA | NA | NA | NA | Gm37523 | | 0 | 0 | 0 | 0 | 0 | 0 | 0 | 0 | 0 | 0 | 0 |
| 0 | NA | NA | NA | NA | NA | Mir7007 | | 0 | 0 | 0 | 0 | 0 | 0 | 0 | 0 | 0 | 0 | 0 |
| 0 | NA | NA | NA | NA | NA | Gm18426 | | 0 | 0 | 0 | 0 | 0 | 0 | 0 | 0 | 0 | 0 | 0 |
| 0 | NA | NA | NA | NA | NA | Gm42594 | | 0 | 0 | 0 | 0 | 0 | 0 | 0 | 0 | 0 | 0 | 0 |
| 0 | NA | NA | NA | NA | NA | Gm31466 | | 0 | 0 | 0 | 0 | 0 | 0 | 0 | 0 | 0 | 0 | 0 |
| 0 | NA | NA | NA | NA | NA | Gm18491 | | 0 | 0 | 0 | 0 | 0 | 0 | 0 | 0 | 0 | 0 | 0 |
| 0 | NA | NA | NA | NA | NA | Gm38323 | | 0 | 0 | 0 | 0 | 0 | 0 | 0 | 0 | 0 | 0 | 0 |
| 0 | NA | NA | NA | NA | NA | Gm18493 | | 0 | 0 | 0 | 0 | 0 | 0 | 0 | 0 | 0 | 0 | 0 |
| 0 | NA | NA | NA | NA | NA | Gm37480 | | 0 | 0 | 0 | 0 | 0 | 0 | 0 | 0 | 0 | 0 | 0 |
| 0 | NA | NA | NA | NA | NA | Gm43160 | | 0 | 0 | 0 | 0 | 0 | 0 | 0 | 0 | 0 | 0 | 0 |
| 0 | NA | NA | NA | NA | NA | Gm29137 | | 0 | 0 | 0 | 0 | 0 | 0 | 0 | 0 | 0 | 0 | 0 |
| 0 | NA | NA | NA | NA | NA | Gm43675 | | 0 | 0 | 0 | 0 | 0 | 0 | 0 | 0 | 0 | 0 | 0 |
| 0 | NA | NA | NA | NA | NA | Gm17935 | | 0 | 0 | 0 | 0 | 0 | 0 | 0 | 0 | 0 | 0 | 0 |
| 0 | NA | NA | NA | NA | NA | Gm31693 | | 0 | 0 | 0 | 0 | 0 | 0 | 0 | 0 | 0 | 0 | 0 |
| 0 | NA | NA | NA | NA | NA | Gm43674 | | 0 | 0 | 0 | 0 | 0 | 0 | 0 | 0 | 0 | 0 | 0 |
| 0 | NA | NA | NA | NA | NA | Gm43671 | | 0 | 0 | 0 | 0 | 0 | 0 | 0 | 0 | 0 | 0 | 0 |
| 0 | NA | NA | NA | NA | NA | Rprl2 | | 0 | 0 | 0 | 0 | 0 | 0 | 0 | 0 | 0 | 0 | 0 |
| 0 | NA | NA | NA | NA | NA | Gm38324 | | 0 | 0 | 0 | 0 | 0 | 0 | 0 | 0 | 0 | 0 | 0 |
| 0 | NA | NA | NA | NA | NA | Gm26354 | | 0 | 0 | 0 | 0 | 0 | 0 | 0 | 0 | 0 | 0 | 0 |
| 0 | NA | NA | NA | NA | NA | Gm37889 | | 0 | 0 | 0 | 0 | 0 | 0 | 0 | 0 | 0 | 0 | 0 |
| 0 | NA | NA | NA | NA | NA | Gm37021 | | 0 | 0 | 0 | 0 | 0 | 0 | 0 | 0 | 0 | 0 | 0 |
| 0 | NA | NA | NA | NA | NA | Gm37492 | | 0 | 0 | 0 | 0 | 0 | 0 | 0 | 0 | 0 | 0 | 0 |
| 0 | NA | NA | NA | NA | NA | 4930412E21Rik | | 0 | 0 | 0 | 0 | 0 | 0 | 0 | 0 | 0 | 0 | 0 |
| 0 | NA | NA | NA | NA | NA | Gm24704 | | 0 | 0 | 0 | 0 | 0 | 0 | 0 | 0 | 0 | 0 | 0 |
| 0 | NA | NA | NA | NA | NA | Gm24062 | | 0 | 0 | 0 | 0 | 0 | 0 | 0 | 0 | 0 | 0 | 0 |
| 0 | NA | NA | NA | NA | NA | Gm42774 | | 0 | 0 | 0 | 0 | 0 | 0 | 0 | 0 | 0 | 0 | 0 |
| 0 | NA | NA | NA | NA | NA | Gm37136 | | 0 | 0 | 0 | 0 | 0 | 0 | 0 | 0 | 0 | 0 | 0 |
| 0 | NA | NA | NA | NA | NA | Gm38002 | | 0 | 0 | 0 | 0 | 0 | 0 | 0 | 0 | 0 | 0 | 0 |
| 0 | NA | NA | NA | NA | NA | Gm24250 | | 0 | 0 | 0 | 0 | 0 | 0 | 0 | 0 | 0 | 0 | 0 |
| 0 | NA | NA | NA | NA | NA | A830092H15Rik | | 0 | 0 | 0 | 0 | 0 | 0 | 0 | 0 | 0 | 0 | 0 |
| 0 | NA | NA | NA | NA | NA | Gm4855 | | 0 | 0 | 0 | 0 | 0 | 0 | 0 | 0 | 0 | 0 | 0 |
| 0 | NA | NA | NA | NA | NA | Gm25773 | | 0 | 0 | 0 | 0 | 0 | 0 | 0 | 0 | 0 | 0 | 0 |
| 0 | NA | NA | NA | NA | NA | Spata16 | spermatogenesis associated 16 | 0 | 0 | 0 | 0 | 0 | 0 | 0 | 0 | 0 | 0 | 0 |
| 0 | NA | NA | NA | NA | NA | Gm37659 | | 0 | 0 | 0 | 0 | 0 | 0 | 0 | 0 | 0 | 0 | 0 |
| 0 | NA | NA | NA | NA | NA | Gm44464 | | 0 | 0 | 0 | 0 | 0 | 0 | 0 | 0 | 0 | 0 | 0 |
| 0 | NA | NA | NA | NA | NA | Gm25152 | | 0 | 0 | 0 | 0 | 0 | 0 | 0 | 0 | 0 | 0 | 0 |
| 0 | NA | NA | NA | NA | NA | 1700125G22Rik | | 0 | 0 | 0 | 0 | 0 | 0 | 0 | 0 | 0 | 0 | 0 |
| 0 | NA | NA | NA | NA | NA | Gm43345 | | 0 | 0 | 0 | 0 | 0 | 0 | 0 | 0 | 0 | 0 | 0 |
| 0 | NA | NA | NA | NA | NA | Gm43344 | | 0 | 0 | 0 | 0 | 0 | 0 | 0 | 0 | 0 | 0 | 0 |
| 0 | NA | NA | NA | NA | NA | Mir3092 | | 0 | 0 | 0 | 0 | 0 | 0 | 0 | 0 | 0 | 0 | 0 |
| 0 | NA | NA | NA | NA | NA | Gm26040 | | 0 | 0 | 0 | 0 | 0 | 0 | 0 | 0 | 0 | 0 | 0 |
| 0 | NA | NA | NA | NA | NA | Gm37537 | | 0 | 0 | 0 | 0 | 0 | 0 | 0 | 0 | 0 | 0 | 0 |
| 0 | NA | NA | NA | NA | NA | Mir466q | | 0 | 0 | 0 | 0 | 0 | 0 | 0 | 0 | 0 | 0 | 0 |
| 0 | NA | NA | NA | NA | NA | Gm37867 | | 0 | 0 | 0 | 0 | 0 | 0 | 0 | 0 | 0 | 0 | 0 |
| 0 | NA | NA | NA | NA | NA | Gm37924 | | 0 | 0 | 0 | 0 | 0 | 0 | 0 | 0 | 0 | 0 | 0 |
| 0 | NA | NA | NA | NA | NA | Gm1527 | predicted gene 1527 | 0 | 0 | 0 | 0 | 0 | 0 | 0 | 0 | 0 | 0 | 0 |
| 0 | NA | NA | NA | NA | NA | Gm38029 | | 0 | 0 | 0 | 0 | 0 | 0 | 0 | 0 | 0 | 0 | 0 |
| 0 | NA | NA | NA | NA | NA | Gm37965 | | 0 | 0 | 0 | 0 | 0 | 0 | 0 | 0 | 0 | 0 | 0 |
| 0 | NA | NA | NA | NA | NA | Mir551b | | 0 | 0 | 0 | 0 | 0 | 0 | 0 | 0 | 0 | 0 | 0 |
| 0 | NA | NA | NA | NA | NA | Mir21b | | 0 | 0 | 0 | 0 | 0 | 0 | 0 | 0 | 0 | 0 | 0 |
| 0 | NA | NA | NA | NA | NA | Gm37557 | | 0 | 0 | 0 | 0 | 0 | 0 | 0 | 0 | 0 | 0 | 0 |
| 0 | NA | NA | NA | NA | NA | Gm33206 | | 0 | 0 | 0 | 0 | 0 | 0 | 0 | 0 | 0 | 0 | 0 |
| 0 | NA | NA | NA | NA | NA | Gm37281 | | 0 | 0 | 0 | 0 | 0 | 0 | 0 | 0 | 0 | 0 | 0 |
| 0 | NA | NA | NA | NA | NA | Gm38066 | | 0 | 0 | 0 | 0 | 0 | 0 | 0 | 0 | 0 | 0 | 0 |
| 0 | NA | NA | NA | NA | NA | Gm10258 | | 0 | 0 | 0 | 0 | 0 | 0 | 0 | 0 | 0 | 0 | 0 |
| 0 | NA | NA | NA | NA | NA | Gm38362 | | 0 | 0 | 0 | 0 | 0 | 0 | 0 | 0 | 0 | 0 | 0 |
| 0 | NA | NA | NA | NA | NA | Gm37024 | | 0 | 0 | 0 | 0 | 0 | 0 | 0 | 0 | 0 | 0 | 0 |
| 0 | NA | NA | NA | NA | NA | Gm37652 | | 0 | 0 | 0 | 0 | 0 | 0 | 0 | 0 | 0 | 0 | 0 |
| 0 | NA | NA | NA | NA | NA | Gm38258 | | 0 | 0 | 0 | 0 | 0 | 0 | 0 | 0 | 0 | 0 | 0 |
| 0 | NA | NA | NA | NA | NA | Gm38153 | | 0 | 0 | 0 | 0 | 0 | 0 | 0 | 0 | 0 | 0 | 0 |

| | | | | | | | | | | | | | | | | | | |
|---|---|---|---|---|---|---|---|---|---|---|---|---|---|---|---|---|---|---|
| 0 | NA | NA | NA | NA | NA | 1600017P15Rik | | 0 | 0 | 0 | 0 | 0 | 0 | 0 | 0 | 0 | 0 | 0 |
| 0 | NA | NA | NA | NA | NA | Samd7 | sterile alpha motif domain containing 7 | 0 | 0 | 0 | 0 | 0 | 0 | 0 | 0 | 0 | 0 | 0 |
| 0 | NA | NA | NA | NA | NA | Gm38080 | | 0 | 0 | 0 | 0 | 0 | 0 | 0 | 0 | 0 | 0 | 0 |
| 0 | NA | NA | NA | NA | NA | 9530022L04Rik | | 0 | 0 | 0 | 0 | 0 | 0 | 0 | 0 | 0 | 0 | 0 |
| 0 | NA | NA | NA | NA | NA | Gm2979 | | 0 | 0 | 0 | 0 | 0 | 0 | 0 | 0 | 0 | 0 | 0 |
| 0 | NA | NA | NA | NA | NA | Mir7008 | | 0 | 0 | 0 | 0 | 0 | 0 | 0 | 0 | 0 | 0 | 0 |
| 0 | NA | NA | NA | NA | NA | Gm6558 | | 0 | 0 | 0 | 0 | 0 | 0 | 0 | 0 | 0 | 0 | 0 |
| 0 | NA | NA | NA | NA | NA | Gm37770 | | 0 | 0 | 0 | 0 | 0 | 0 | 0 | 0 | 0 | 0 | 0 |
| 0 | NA | NA | NA | NA | NA | Gm36990 | | 0 | 0 | 0 | 0 | 0 | 0 | 0 | 0 | 0 | 0 | 0 |
| 0 | NA | NA | NA | NA | NA | Kcnmb3 | potassium large conductance calcium-act | 0 | 0 | 0 | 0 | 0 | 0 | 0 | 0 | 0 | 0 | 0 |
| 0 | NA | NA | NA | NA | NA | Gm37592 | | 0 | 0 | 0 | 0 | 0 | 0 | 0 | 0 | 0 | 0 | 0 |
| 0 | NA | NA | NA | NA | NA | Gm37640 | | 0 | 0 | 0 | 0 | 0 | 0 | 0 | 0 | 0 | 0 | 0 |
| 0 | NA | NA | NA | NA | NA | Gm15574 | | 0 | 0 | 0 | 0 | 0 | 0 | 0 | 0 | 0 | 0 | 0 |
| 0 | NA | NA | NA | NA | NA | Gm23214 | | 0 | 0 | 0 | 0 | 0 | 0 | 0 | 0 | 0 | 0 | 0 |
| 0 | NA | NA | NA | NA | NA | 4930419G24Rik | | 0 | 0 | 0 | 0 | 0 | 0 | 0 | 0 | 0 | 0 | 0 |
| 0 | NA | NA | NA | NA | NA | Gm37128 | | 0 | 0 | 0 | 0 | 0 | 0 | 0 | 0 | 0 | 0 | 0 |
| 0 | NA | NA | NA | NA | NA | Gm27974 | | 0 | 0 | 0 | 0 | 0 | 0 | 0 | 0 | 0 | 0 | 0 |
| 0 | NA | NA | NA | NA | NA | Gm24780 | | 0 | 0 | 0 | 0 | 0 | 0 | 0 | 0 | 0 | 0 | 0 |
| 0 | NA | NA | NA | NA | NA | Gm37398 | | 0 | 0 | 0 | 0 | 0 | 0 | 0 | 0 | 0 | 0 | 0 |
| 0 | NA | NA | NA | NA | NA | Gm18611 | | 0 | 0 | 0 | 0 | 0 | 0 | 0 | 0 | 0 | 0 | 0 |
| 0 | NA | NA | NA | NA | NA | Gm43140 | | 0 | 0 | 0 | 0 | 0 | 0 | 0 | 0 | 0 | 0 | 0 |
| 0 | NA | NA | NA | NA | NA | 4930431L21Rik | | 0 | 0 | 0 | 0 | 0 | 0 | 0 | 0 | 0 | 0 | 0 |
| 0 | NA | NA | NA | NA | NA | 4930502C17Rik | | 0 | 0 | 0 | 0 | 0 | 0 | 0 | 0 | 0 | 0 | 0 |
| 0 | NA | NA | NA | NA | NA | Gm23175 | | 0 | 0 | 0 | 0 | 0 | 0 | 0 | 0 | 0 | 0 | 0 |
| 0 | NA | NA | NA | NA | NA | Gm5708 | | 0 | 0 | 0 | 0 | 0 | 0 | 0 | 0 | 0 | 0 | 0 |
| 0 | NA | NA | NA | NA | NA | Gm38505 | | 0 | 0 | 0 | 0 | 0 | 0 | 0 | 0 | 0 | 0 | 0 |
| 0 | NA | NA | NA | NA | NA | Gm34599 | | 0 | 0 | 0 | 0 | 0 | 0 | 0 | 0 | 0 | 0 | 0 |
| 0 | NA | NA | NA | NA | NA | Gm20515 | | 0 | 0 | 0 | 0 | 0 | 0 | 0 | 0 | 0 | 0 | 0 |
| 0 | NA | NA | NA | NA | NA | Gm29135 | | 0 | 0 | 0 | 0 | 0 | 0 | 0 | 0 | 0 | 0 | 0 |
| 0 | NA | NA | NA | NA | NA | Gm27403 | | 0 | 0 | 0 | 0 | 0 | 0 | 0 | 0 | 0 | 0 | 0 |
| 0 | NA | NA | NA | NA | NA | Gm42695 | | 0 | 0 | 0 | 0 | 0 | 0 | 0 | 0 | 0 | 0 | 0 |
| 0 | NA | NA | NA | NA | NA | Gm42205 | | 0 | 0 | 0 | 0 | 0 | 0 | 0 | 0 | 0 | 0 | 0 |
| 0 | NA | NA | NA | NA | NA | | | 0 | 0 | 0 | 0 | 0 | 0 | 0 | 0 | 0 | 0 | 0 |
| 0 | NA | NA | NA | NA | NA | Mir1897 | | 0 | 0 | 0 | 0 | 0 | 0 | 0 | 0 | 0 | 0 | 0 |
| 0 | NA | NA | NA | NA | NA | Gm42692 | | 0 | 0 | 0 | 0 | 0 | 0 | 0 | 0 | 0 | 0 | 0 |
| 0 | NA | NA | NA | NA | NA | Gm27611 | | 0 | 0 | 0 | 0 | 0 | 0 | 0 | 0 | 0 | 0 | 0 |
| 0 | NA | NA | NA | NA | NA | Gm42693 | | 0 | 0 | 0 | 0 | 0 | 0 | 0 | 0 | 0 | 0 | 0 |
| 0 | NA | NA | NA | NA | NA | Gm43206 | | 0 | 0 | 0 | 0 | 0 | 0 | 0 | 0 | 0 | 0 | 0 |
| 0 | NA | NA | NA | NA | NA | Gm3143 | | 0 | 0 | 0 | 0 | 0 | 0 | 0 | 0 | 0 | 0 | 0 |
| 0 | NA | NA | NA | NA | NA | Gm38509 | | 0 | 0 | 0 | 0 | 0 | 0 | 0 | 0 | 0 | 0 | 0 |
| 0 | NA | NA | NA | NA | NA | Gm21388 | | 0 | 0 | 0 | 0 | 0 | 0 | 0 | 0 | 0 | 0 | 0 |
| 0 | NA | NA | NA | NA | NA | Mir6378 | | 0 | 0 | 0 | 0 | 0 | 0 | 0 | 0 | 0 | 0 | 0 |
| 0 | NA | NA | NA | NA | NA | 1700017M07Rik | | 0 | 0 | 0 | 0 | 0 | 0 | 0 | 0 | 0 | 0 | 0 |
| 0 | NA | NA | NA | NA | NA | Gm25442 | | 0 | 0 | 0 | 0 | 0 | 0 | 0 | 0 | 0 | 0 | 0 |
| 0 | NA | NA | NA | NA | NA | Gm43078 | | 0 | 0 | 0 | 0 | 0 | 0 | 0 | 0 | 0 | 0 | 0 |
| 0 | NA | NA | NA | NA | NA | Gm6639 | | 0 | 0 | 0 | 0 | 0 | 0 | 0 | 0 | 0 | 0 | 0 |
| 0 | NA | NA | NA | NA | NA | Gm25696 | | 0 | 0 | 0 | 0 | 0 | 0 | 0 | 0 | 0 | 0 | 0 |
| 0 | NA | NA | NA | NA | NA | B230207O21Rik | | 0 | 0 | 0 | 0 | 0 | 0 | 0 | 0 | 0 | 0 | 0 |
| 0 | NA | NA | NA | NA | NA | Ccdc144b | coiled-coil domain containing 144B | 0 | 0 | 0 | 0 | 0 | 0 | 0 | 0 | 0 | 0 | 0 |
| 0 | NA | NA | NA | NA | NA | Gm43079 | | 0 | 0 | 0 | 0 | 0 | 0 | 0 | 0 | 0 | 0 | 0 |
| 0 | NA | NA | NA | NA | NA | Qrfpr | pyroglutamylated RFamide peptide recep | 0 | 0 | 0 | 0 | 0 | 0 | 0 | 0 | 0 | 0 | 0 |
| 0 | NA | NA | NA | NA | NA | Gm43255 | | 0 | 0 | 0 | 0 | 0 | 0 | 0 | 0 | 0 | 0 | 0 |
| 0 | NA | NA | NA | NA | NA | Gm18893 | | 0 | 0 | 0 | 0 | 0 | 0 | 0 | 0 | 0 | 0 | 0 |
| 0 | NA | NA | NA | NA | NA | Gm42208 | | 0 | 0 | 0 | 0 | 0 | 0 | 0 | 0 | 0 | 0 | 0 |
| 0 | NA | NA | NA | NA | NA | Gm17835 | | 0 | 0 | 0 | 0 | 0 | 0 | 0 | 0 | 0 | 0 | 0 |
| 0 | NA | NA | NA | NA | NA | Mir7009 | | 0 | 0 | 0 | 0 | 0 | 0 | 0 | 0 | 0 | 0 | 0 |
| 0 | NA | NA | NA | NA | NA | Gm11548 | | 0 | 0 | 0 | 0 | 0 | 0 | 0 | 0 | 0 | 0 | 0 |
| 0 | NA | NA | NA | NA | NA | Gm11550 | | 0 | 0 | 0 | 0 | 0 | 0 | 0 | 0 | 0 | 0 | 0 |
| 0 | NA | NA | NA | NA | NA | Gm12377 | | 0 | 0 | 0 | 0 | 0 | 0 | 0 | 0 | 0 | 0 | 0 |
| 0 | NA | NA | NA | NA | NA | Gm12376 | | 0 | 0 | 0 | 0 | 0 | 0 | 0 | 0 | 0 | 0 | 0 |

| | | | | | | | | | | | | | | | | | |
|---|---|---|---|---|---|---|---|---|---|---|---|---|---|---|---|---|---|
| 0 | NA | NA | NA | NA | NA | Gm12584 | 0 | 0 | 0 | 0 | 0 | 0 | 0 | 0 | 0 | 0 | 0 |
| 0 | NA | NA | NA | NA | NA | Gm12532 | 0 | 0 | 0 | 0 | 0 | 0 | 0 | 0 | 0 | 0 | 0 |
| 0 | NA | NA | NA | NA | NA | Gm12534 | 0 | 0 | 0 | 0 | 0 | 0 | 0 | 0 | 0 | 0 | 0 |
| 0 | NA | NA | NA | NA | NA | Gm43822 | 0 | 0 | 0 | 0 | 0 | 0 | 0 | 0 | 0 | 0 | 0 |
| 0 | NA | NA | NA | NA | NA | Gm24169 | 0 | 0 | 0 | 0 | 0 | 0 | 0 | 0 | 0 | 0 | 0 |
| 0 | NA | NA | NA | NA | NA | Gm43439 | 0 | 0 | 0 | 0 | 0 | 0 | 0 | 0 | 0 | 0 | 0 |
| 0 | NA | NA | NA | NA | NA | Fgf2os | 0 | 0 | 0 | 0 | 0 | 0 | 0 | 0 | 0 | 0 | 0 |
| 0 | NA | NA | NA | NA | NA | Gm43488 | 0 | 0 | 0 | 0 | 0 | 0 | 0 | 0 | 0 | 0 | 0 |
| 0 | NA | NA | NA | NA | NA | Gm12564 | 0 | 0 | 0 | 0 | 0 | 0 | 0 | 0 | 0 | 0 | 0 |
| 0 | NA | NA | NA | NA | NA | 4930594O21Rik | 0 | 0 | 0 | 0 | 0 | 0 | 0 | 0 | 0 | 0 | 0 |
| 0 | NA | NA | NA | NA | NA | Gm42921 | 0 | 0 | 0 | 0 | 0 | 0 | 0 | 0 | 0 | 0 | 0 |
| 0 | NA | NA | NA | NA | NA | Gm26404 | 0 | 0 | 0 | 0 | 0 | 0 | 0 | 0 | 0 | 0 | 0 |
| 0 | NA | NA | NA | NA | NA | Gm42753 | 0 | 0 | 0 | 0 | 0 | 0 | 0 | 0 | 0 | 0 | 0 |
| 0 | NA | NA | NA | NA | NA | Gm20755 | 0 | 0 | 0 | 0 | 0 | 0 | 0 | 0 | 0 | 0 | 0 |
| 0 | NA | NA | NA | NA | NA | Gm42751 | 0 | 0 | 0 | 0 | 0 | 0 | 0 | 0 | 0 | 0 | 0 |
| 0 | NA | NA | NA | NA | NA | Gm22899 | 0 | 0 | 0 | 0 | 0 | 0 | 0 | 0 | 0 | 0 | 0 |
| 0 | NA | NA | NA | NA | NA | Gm43821 | 0 | 0 | 0 | 0 | 0 | 0 | 0 | 0 | 0 | 0 | 0 |
| 0 | NA | NA | NA | NA | NA | 5430434I15Rik | 0 | 0 | 0 | 0 | 0 | 0 | 0 | 0 | 0 | 0 | 0 |
| 0 | NA | NA | NA | NA | NA | Gm2965 | 0 | 0 | 0 | 0 | 0 | 0 | 0 | 0 | 0 | 0 | 0 |
| 0 | NA | NA | NA | NA | NA | Gm25574 | 0 | 0 | 0 | 0 | 0 | 0 | 0 | 0 | 0 | 0 | 0 |
| 0 | NA | NA | NA | NA | NA | Gm42920 | 0 | 0 | 0 | 0 | 0 | 0 | 0 | 0 | 0 | 0 | 0 |
| 0 | NA | NA | NA | NA | NA | Gm36823 | 0 | 0 | 0 | 0 | 0 | 0 | 0 | 0 | 0 | 0 | 0 |
| 0 | NA | NA | NA | NA | NA | Gm7815 | 0 | 0 | 0 | 0 | 0 | 0 | 0 | 0 | 0 | 0 | 0 |
| 0 | NA | NA | NA | NA | NA | Gm24462 | 0 | 0 | 0 | 0 | 0 | 0 | 0 | 0 | 0 | 0 | 0 |
| 0 | NA | NA | NA | NA | NA | Gm7824 | 0 | 0 | 0 | 0 | 0 | 0 | 0 | 0 | 0 | 0 | 0 |
| 0 | NA | NA | NA | NA | NA | Gm43538 | 0 | 0 | 0 | 0 | 0 | 0 | 0 | 0 | 0 | 0 | 0 |
| 0 | NA | NA | NA | NA | NA | Gm43539 | 0 | 0 | 0 | 0 | 0 | 0 | 0 | 0 | 0 | 0 | 0 |
| 0 | NA | NA | NA | NA | NA | Gm43308 | 0 | 0 | 0 | 0 | 0 | 0 | 0 | 0 | 0 | 0 | 0 |
| 0 | NA | NA | NA | NA | NA | n-R5s195 | 0 | 0 | 0 | 0 | 0 | 0 | 0 | 0 | 0 | 0 | 0 |
| 0 | NA | NA | NA | NA | NA | Gm42781 | 0 | 0 | 0 | 0 | 0 | 0 | 0 | 0 | 0 | 0 | 0 |
| 0 | NA | NA | NA | NA | NA | Gm23796 | 0 | 0 | 0 | 0 | 0 | 0 | 0 | 0 | 0 | 0 | 0 |
| 0 | NA | NA | NA | NA | NA | Gm23831 | 0 | 0 | 0 | 0 | 0 | 0 | 0 | 0 | 0 | 0 | 0 |
| 0 | NA | NA | NA | NA | NA | Gm42786 | 0 | 0 | 0 | 0 | 0 | 0 | 0 | 0 | 0 | 0 | 0 |
| 0 | NA | NA | NA | NA | NA | Gm42785 | 0 | 0 | 0 | 0 | 0 | 0 | 0 | 0 | 0 | 0 | 0 |
| 0 | NA | NA | NA | NA | NA | 1700017G19Rik | 0 | 0 | 0 | 0 | 0 | 0 | 0 | 0 | 0 | 0 | 0 |
| 0 | NA | NA | NA | NA | NA | Gm23810 | 0 | 0 | 0 | 0 | 0 | 0 | 0 | 0 | 0 | 0 | 0 |
| 0 | NA | NA | NA | NA | NA | Gm16114 | 0 | 0 | 0 | 0 | 0 | 0 | 0 | 0 | 0 | 0 | 0 |
| 0 | NA | NA | NA | NA | NA | Gm10731 | 0 | 0 | 0 | 0 | 0 | 0 | 0 | 0 | 0 | 0 | 0 |
| 0 | NA | NA | NA | NA | NA | Gm40038 | 0 | 0 | 0 | 0 | 0 | 0 | 0 | 0 | 0 | 0 | 0 |
| 0 | NA | NA | NA | NA | NA | Gm16508 | 0 | 0 | 0 | 0 | 0 | 0 | 0 | 0 | 0 | 0 | 0 |
| 0 | NA | NA | NA | NA | NA | 1700052H01Rik | 0 | 0 | 0 | 0 | 0 | 0 | 0 | 0 | 0 | 0 | 0 |
| 0 | NA | NA | NA | NA | NA | Gm25487 | 0 | 0 | 0 | 0 | 0 | 0 | 0 | 0 | 0 | 0 | 0 |
| 0 | NA | NA | NA | NA | NA | Gm37348 | 0 | 0 | 0 | 0 | 0 | 0 | 0 | 0 | 0 | 0 | 0 |
| 0 | NA | NA | NA | NA | NA | Gm37846 | 0 | 0 | 0 | 0 | 0 | 0 | 0 | 0 | 0 | 0 | 0 |
| 0 | NA | NA | NA | NA | NA | Gm38274 | 0 | 0 | 0 | 0 | 0 | 0 | 0 | 0 | 0 | 0 | 0 |
| 0 | NA | NA | NA | NA | NA | Gm37533 | 0 | 0 | 0 | 0 | 0 | 0 | 0 | 0 | 0 | 0 | 0 |
| 0 | NA | NA | NA | NA | NA | Gm38044 | 0 | 0 | 0 | 0 | 0 | 0 | 0 | 0 | 0 | 0 | 0 |
| 0 | NA | NA | NA | NA | NA | Gm18865 | 0 | 0 | 0 | 0 | 0 | 0 | 0 | 0 | 0 | 0 | 0 |
| 0 | NA | NA | NA | NA | NA | Gm25714 | 0 | 0 | 0 | 0 | 0 | 0 | 0 | 0 | 0 | 0 | 0 |
| 0 | NA | NA | NA | NA | NA | Gm32175 | 0 | 0 | 0 | 0 | 0 | 0 | 0 | 0 | 0 | 0 | 0 |
| 0 | NA | NA | NA | NA | NA | Gm37746 | 0 | 0 | 0 | 0 | 0 | 0 | 0 | 0 | 0 | 0 | 0 |
| 0 | NA | NA | NA | NA | NA | Gm37365 | 0 | 0 | 0 | 0 | 0 | 0 | 0 | 0 | 0 | 0 | 0 |
| 0 | NA | NA | NA | NA | NA | Gm38128 | 0 | 0 | 0 | 0 | 0 | 0 | 0 | 0 | 0 | 0 | 0 |
| 0 | NA | NA | NA | NA | NA | Gm37597 | 0 | 0 | 0 | 0 | 0 | 0 | 0 | 0 | 0 | 0 | 0 |
| 0 | NA | NA | NA | NA | NA | Gm24997 | 0 | 0 | 0 | 0 | 0 | 0 | 0 | 0 | 0 | 0 | 0 |
| 0 | NA | NA | NA | NA | NA | Gm27989 | 0 | 0 | 0 | 0 | 0 | 0 | 0 | 0 | 0 | 0 | 0 |
| 0 | NA | NA | NA | NA | NA | Gm36996 | 0 | 0 | 0 | 0 | 0 | 0 | 0 | 0 | 0 | 0 | 0 |
| 0 | NA | NA | NA | NA | NA | 1700027H10Rik | 0 | 0 | 0 | 0 | 0 | 0 | 0 | 0 | 0 | 0 | 0 |
| 0 | NA | NA | NA | NA | NA | Gm37345 | 0 | 0 | 0 | 0 | 0 | 0 | 0 | 0 | 0 | 0 | 0 |
| 0 | NA | NA | NA | NA | NA | Gm37835 | 0 | 0 | 0 | 0 | 0 | 0 | 0 | 0 | 0 | 0 | 0 |

| | | | | | | | | | | | | | | | | | |
|---|---|---|---|---|---|---|---|---|---|---|---|---|---|---|---|---|---|
| 0 | NA | NA | NA | NA | NA | Gm25796 | 0 | 0 | 0 | 0 | 0 | 0 | 0 | 0 | 0 | 0 | 0 |
| 0 | NA | NA | NA | NA | NA | Gm23337 | 0 | 0 | 0 | 0 | 0 | 0 | 0 | 0 | 0 | 0 | 0 |
| 0 | NA | NA | NA | NA | NA | Gm26081 | 0 | 0 | 0 | 0 | 0 | 0 | 0 | 0 | 0 | 0 | 0 |
| 0 | NA | NA | NA | NA | NA | Gm18843 | 0 | 0 | 0 | 0 | 0 | 0 | 0 | 0 | 0 | 0 | 0 |
| 0 | NA | NA | NA | NA | NA | Gm37445 | 0 | 0 | 0 | 0 | 0 | 0 | 0 | 0 | 0 | 0 | 0 |
| 0 | NA | NA | NA | NA | NA | Mir6379 | 0 | 0 | 0 | 0 | 0 | 0 | 0 | 0 | 0 | 0 | 0 |
| 0 | NA | NA | NA | NA | NA | Gm23891 | 0 | 0 | 0 | 0 | 0 | 0 | 0 | 0 | 0 | 0 | 0 |
| 0 | NA | NA | NA | NA | NA | Gm10730 | 0 | 0 | 0 | 0 | 0 | 0 | 0 | 0 | 0 | 0 | 0 |
| 0 | NA | NA | NA | NA | NA | Gm2229 | 0 | 0 | 0 | 0 | 0 | 0 | 0 | 0 | 0 | 0 | 0 |
| 0 | NA | NA | NA | NA | NA | Gm37295 | 0 | 0 | 0 | 0 | 0 | 0 | 0 | 0 | 0 | 0 | 0 |
| 0 | NA | NA | NA | NA | NA | 1700018B24Rik | 0 | 0 | 0 | 0 | 0 | 0 | 0 | 0 | 0 | 0 | 0 |
| 0 | NA | NA | NA | NA | NA | Gm37190 | 0 | 0 | 0 | 0 | 0 | 0 | 0 | 0 | 0 | 0 | 0 |
| 0 | NA | NA | NA | NA | NA | Gm23511 | 0 | 0 | 0 | 0 | 0 | 0 | 0 | 0 | 0 | 0 | 0 |
| 0 | NA | NA | NA | NA | NA | Gm31415 | 0 | 0 | 0 | 0 | 0 | 0 | 0 | 0 | 0 | 0 | 0 |
| 0 | NA | NA | NA | NA | NA | Gm37819 | 0 | 0 | 0 | 0 | 0 | 0 | 0 | 0 | 0 | 0 | 0 |
| 0 | NA | NA | NA | NA | NA | Gm37684 | 0 | 0 | 0 | 0 | 0 | 0 | 0 | 0 | 0 | 0 | 0 |
| 0 | NA | NA | NA | NA | NA | Gm20557 | 0 | 0 | 0 | 0 | 0 | 0 | 0 | 0 | 0 | 0 | 0 |
| 0 | NA | NA | NA | NA | NA | Gm22480 | 0 | 0 | 0 | 0 | 0 | 0 | 0 | 0 | 0 | 0 | 0 |
| 0 | NA | NA | NA | NA | NA | Gm38261 | 0 | 0 | 0 | 0 | 0 | 0 | 0 | 0 | 0 | 0 | 0 |
| 0 | NA | NA | NA | NA | NA | Gm2328 | 0 | 0 | 0 | 0 | 0 | 0 | 0 | 0 | 0 | 0 | 0 |
| 0 | NA | NA | NA | NA | NA | Gm23096 | 0 | 0 | 0 | 0 | 0 | 0 | 0 | 0 | 0 | 0 | 0 |
| 0 | NA | NA | NA | NA | NA | Gm37550 | 0 | 0 | 0 | 0 | 0 | 0 | 0 | 0 | 0 | 0 | 0 |
| 0 | NA | NA | NA | NA | NA | Gm37854 | 0 | 0 | 0 | 0 | 0 | 0 | 0 | 0 | 0 | 0 | 0 |
| 0 | NA | NA | NA | NA | NA | Gm37826 | 0 | 0 | 0 | 0 | 0 | 0 | 0 | 0 | 0 | 0 | 0 |
| 0 | NA | NA | NA | NA | NA | Gm37692 | 0 | 0 | 0 | 0 | 0 | 0 | 0 | 0 | 0 | 0 | 0 |
| 0 | NA | NA | NA | NA | NA | Gm2345 | 0 | 0 | 0 | 0 | 0 | 0 | 0 | 0 | 0 | 0 | 0 |
| 0 | NA | NA | NA | NA | NA | Gm6209 | 0 | 0 | 0 | 0 | 0 | 0 | 0 | 0 | 0 | 0 | 0 |
| 0 | NA | NA | NA | NA | NA | 5830415G21Rik | 0 | 0 | 0 | 0 | 0 | 0 | 0 | 0 | 0 | 0 | 0 |
| 0 | NA | NA | NA | NA | NA | Gm38237 | 0 | 0 | 0 | 0 | 0 | 0 | 0 | 0 | 0 | 0 | 0 |
| 0 | NA | NA | NA | NA | NA | Gm37199 | 0 | 0 | 0 | 0 | 0 | 0 | 0 | 0 | 0 | 0 | 0 |
| 0 | NA | NA | NA | NA | NA | Gm37461 | 0 | 0 | 0 | 0 | 0 | 0 | 0 | 0 | 0 | 0 | 0 |
| 0 | NA | NA | NA | NA | NA | Gm24503 | 0 | 0 | 0 | 0 | 0 | 0 | 0 | 0 | 0 | 0 | 0 |
| 0 | NA | NA | NA | NA | NA | Gm37548 | 0 | 0 | 0 | 0 | 0 | 0 | 0 | 0 | 0 | 0 | 0 |
| 0 | NA | NA | NA | NA | NA | Gm37843 | 0 | 0 | 0 | 0 | 0 | 0 | 0 | 0 | 0 | 0 | 0 |
| 0 | NA | NA | NA | NA | NA | Gm37209 | 0 | 0 | 0 | 0 | 0 | 0 | 0 | 0 | 0 | 0 | 0 |
| 0 | NA | NA | NA | NA | NA | Gm29230 | 0 | 0 | 0 | 0 | 0 | 0 | 0 | 0 | 0 | 0 | 0 |
| 0 | NA | NA | NA | NA | NA | Gm38246 | 0 | 0 | 0 | 0 | 0 | 0 | 0 | 0 | 0 | 0 | 0 |
| 0 | NA | NA | NA | NA | NA | Gm24282 | 0 | 0 | 0 | 0 | 0 | 0 | 0 | 0 | 0 | 0 | 0 |
| 0 | NA | NA | NA | NA | NA | Gm37646 | 0 | 0 | 0 | 0 | 0 | 0 | 0 | 0 | 0 | 0 | 0 |
| 0 | NA | NA | NA | NA | NA | Gm38334 | 0 | 0 | 0 | 0 | 0 | 0 | 0 | 0 | 0 | 0 | 0 |
| 0 | NA | NA | NA | NA | NA | 5430433H01Rik | 0 | 0 | 0 | 0 | 0 | 0 | 0 | 0 | 0 | 0 | 0 |
| 0 | NA | NA | NA | NA | NA | Gm37261 | 0 | 0 | 0 | 0 | 0 | 0 | 0 | 0 | 0 | 0 | 0 |
| 0 | NA | NA | NA | NA | NA | Gm10729 | 0 | 0 | 0 | 0 | 0 | 0 | 0 | 0 | 0 | 0 | 0 |
| 0 | NA | NA | NA | NA | NA | n-R5s196 | 0 | 0 | 0 | 0 | 0 | 0 | 0 | 0 | 0 | 0 | 0 |
| 0 | NA | NA | NA | NA | NA | Gm38000 | 0 | 0 | 0 | 0 | 0 | 0 | 0 | 0 | 0 | 0 | 0 |
| 0 | NA | NA | NA | NA | NA | Gm20089 | 0 | 0 | 0 | 0 | 0 | 0 | 0 | 0 | 0 | 0 | 0 |
| 0 | NA | NA | NA | NA | NA | Gm37526 | 0 | 0 | 0 | 0 | 0 | 0 | 0 | 0 | 0 | 0 | 0 |
| 0 | NA | NA | NA | NA | NA | Gm38132 | 0 | 0 | 0 | 0 | 0 | 0 | 0 | 0 | 0 | 0 | 0 |
| 0 | NA | NA | NA | NA | NA | Olfr1398-ps1 | 0 | 0 | 0 | 0 | 0 | 0 | 0 | 0 | 0 | 0 | 0 |
| 0 | NA | NA | NA | NA | NA | Gm22798 | 0 | 0 | 0 | 0 | 0 | 0 | 0 | 0 | 0 | 0 | 0 |
| 0 | NA | NA | NA | NA | NA | Gm20750 | 0 | 0 | 0 | 0 | 0 | 0 | 0 | 0 | 0 | 0 | 0 |
| 0 | NA | NA | NA | NA | NA | Gm37007 | 0 | 0 | 0 | 0 | 0 | 0 | 0 | 0 | 0 | 0 | 0 |
| 0 | NA | NA | NA | NA | NA | Gm42901 | 0 | 0 | 0 | 0 | 0 | 0 | 0 | 0 | 0 | 0 | 0 |
| 0 | NA | NA | NA | NA | NA | Gm42898 | 0 | 0 | 0 | 0 | 0 | 0 | 0 | 0 | 0 | 0 | 0 |
| 0 | NA | NA | NA | NA | NA | C130075A20Rik | 0 | 0 | 0 | 0 | 0 | 0 | 0 | 0 | 0 | 0 | 0 |
| 0 | NA | NA | NA | NA | NA | Gm43804 | 0 | 0 | 0 | 0 | 0 | 0 | 0 | 0 | 0 | 0 | 0 |
| 0 | NA | NA | NA | NA | NA | Gm43803 | 0 | 0 | 0 | 0 | 0 | 0 | 0 | 0 | 0 | 0 | 0 |
| 0 | NA | NA | NA | NA | NA | Gm43805 | 0 | 0 | 0 | 0 | 0 | 0 | 0 | 0 | 0 | 0 | 0 |
| 0 | NA | NA | NA | NA | NA | Gm43472 | 0 | 0 | 0 | 0 | 0 | 0 | 0 | 0 | 0 | 0 | 0 |
| 0 | NA | NA | NA | NA | NA | Gm16206 | 0 | 0 | 0 | 0 | 0 | 0 | 0 | 0 | 0 | 0 | 0 |

| | | | | | | | | | | | | | | | | | | |
|---|---|---|---|---|---|---|---|---|---|---|---|---|---|---|---|---|---|---|
| 0 | NA | NA | NA | NA | NA | Stoml3 | stomatin (Epb7.2)-like 3 | 0 | 0 | 0 | 0 | 0 | 0 | 0 | 0 | 0 | 0 | 0 |
| 0 | NA | NA | NA | NA | NA | 6430500D05Rik | | 0 | 0 | 0 | 0 | 0 | 0 | 0 | 0 | 0 | 0 | 0 |
| 0 | NA | NA | NA | NA | NA | Gm23675 | | 0 | 0 | 0 | 0 | 0 | 0 | 0 | 0 | 0 | 0 | 0 |
| 0 | NA | NA | NA | NA | NA | Gm6073 | | 0 | 0 | 0 | 0 | 0 | 0 | 0 | 0 | 0 | 0 | 0 |
| 0 | NA | NA | NA | NA | NA | Gm24851 | | 0 | 0 | 0 | 0 | 0 | 0 | 0 | 0 | 0 | 0 | 0 |
| 0 | NA | NA | NA | NA | NA | Gm10985 | | 0 | 0 | 0 | 0 | 0 | 0 | 0 | 0 | 0 | 0 | 0 |
| 0 | NA | NA | NA | NA | NA | Gm31026 | | 0 | 0 | 0 | 0 | 0 | 0 | 0 | 0 | 0 | 0 | 0 |
| 0 | NA | NA | NA | NA | NA | Gm18018 | | 0 | 0 | 0 | 0 | 0 | 0 | 0 | 0 | 0 | 0 | 0 |
| 0 | NA | NA | NA | NA | NA | B020017C02Rik | | 0 | 0 | 0 | 0 | 0 | 0 | 0 | 0 | 0 | 0 | 0 |
| 0 | NA | NA | NA | NA | NA | Gm21958 | | 0 | 0 | 0 | 0 | 0 | 0 | 0 | 0 | 0 | 0 | 0 |
| 0 | NA | NA | NA | NA | NA | Ccna1 | cyclin A1 | 0 | 0 | 0 | 0 | 0 | 0 | 0 | 0 | 0 | 0 | 0 |
| 0 | NA | NA | NA | NA | NA | 4931419H13Rik | | 0 | 0 | 0 | 0 | 0 | 0 | 0 | 0 | 0 | 0 | 0 |
| 0 | NA | NA | NA | NA | NA | Gm22029 | | 0 | 0 | 0 | 0 | 0 | 0 | 0 | 0 | 0 | 0 | 0 |
| 0 | NA | NA | NA | NA | NA | Gm25132 | | 0 | 0 | 0 | 0 | 0 | 0 | 0 | 0 | 0 | 0 | 0 |
| 0 | NA | NA | NA | NA | NA | Gm42609 | | 0 | 0 | 0 | 0 | 0 | 0 | 0 | 0 | 0 | 0 | 0 |
| 0 | NA | NA | NA | NA | NA | Gm21954 | | 0 | 0 | 0 | 0 | 0 | 0 | 0 | 0 | 0 | 0 | 0 |
| 0 | NA | NA | NA | NA | NA | Gm42607 | | 0 | 0 | 0 | 0 | 0 | 0 | 0 | 0 | 0 | 0 | 0 |
| 0 | NA | NA | NA | NA | NA | Gm25404 | | 0 | 0 | 0 | 0 | 0 | 0 | 0 | 0 | 0 | 0 | 0 |
| 0 | NA | NA | NA | NA | NA | Gm43548 | | 0 | 0 | 0 | 0 | 0 | 0 | 0 | 0 | 0 | 0 | 0 |
| 0 | NA | NA | NA | NA | NA | Gm43549 | | 0 | 0 | 0 | 0 | 0 | 0 | 0 | 0 | 0 | 0 | 0 |
| 0 | NA | NA | NA | NA | NA | 4933417G07Rik | | 0 | 0 | 0 | 0 | 0 | 0 | 0 | 0 | 0 | 0 | 0 |
| 0 | NA | NA | NA | NA | NA | Gm43376 | | 0 | 0 | 0 | 0 | 0 | 0 | 0 | 0 | 0 | 0 | 0 |
| 0 | NA | NA | NA | NA | NA | Gm22784 | | 0 | 0 | 0 | 0 | 0 | 0 | 0 | 0 | 0 | 0 | 0 |
| 0 | NA | NA | NA | NA | NA | Gm25727 | | 0 | 0 | 0 | 0 | 0 | 0 | 0 | 0 | 0 | 0 | 0 |
| 0 | NA | NA | NA | NA | NA | Gm22269 | | 0 | 0 | 0 | 0 | 0 | 0 | 0 | 0 | 0 | 0 | 0 |
| 0 | NA | NA | NA | NA | NA | Gm43424 | | 0 | 0 | 0 | 0 | 0 | 0 | 0 | 0 | 0 | 0 | 0 |
| 0 | NA | NA | NA | NA | NA | Gm18660 | | 0 | 0 | 0 | 0 | 0 | 0 | 0 | 0 | 0 | 0 | 0 |
| 0 | NA | NA | NA | NA | NA | Gm16016 | | 0 | 0 | 0 | 0 | 0 | 0 | 0 | 0 | 0 | 0 | 0 |
| 0 | NA | NA | NA | NA | NA | Mir6377 | | 0 | 0 | 0 | 0 | 0 | 0 | 0 | 0 | 0 | 0 | 0 |
| 0 | NA | NA | NA | NA | NA | BB187690 | | 0 | 0 | 0 | 0 | 0 | 0 | 0 | 0 | 0 | 0 | 0 |
| 0 | NA | NA | NA | NA | NA | Gm43098 | | 0 | 0 | 0 | 0 | 0 | 0 | 0 | 0 | 0 | 0 | 0 |
| 0 | NA | NA | NA | NA | NA | Gm24531 | | 0 | 0 | 0 | 0 | 0 | 0 | 0 | 0 | 0 | 0 | 0 |
| 0 | NA | NA | NA | NA | NA | Gm42511 | | 0 | 0 | 0 | 0 | 0 | 0 | 0 | 0 | 0 | 0 | 0 |
| 0 | NA | NA | NA | NA | NA | Gm40055 | | 0 | 0 | 0 | 0 | 0 | 0 | 0 | 0 | 0 | 0 | 0 |
| 0 | NA | NA | NA | NA | NA | Gm42513 | | 0 | 0 | 0 | 0 | 0 | 0 | 0 | 0 | 0 | 0 | 0 |
| 0 | NA | NA | NA | NA | NA | 1700007F19Rik | | 0 | 0 | 0 | 0 | 0 | 0 | 0 | 0 | 0 | 0 | 0 |
| 0 | NA | NA | NA | NA | NA | Gm26166 | | 0 | 0 | 0 | 0 | 0 | 0 | 0 | 0 | 0 | 0 | 0 |
| 0 | NA | NA | NA | NA | NA | Gm43730 | | 0 | 0 | 0 | 0 | 0 | 0 | 0 | 0 | 0 | 0 | 0 |
| 0 | NA | NA | NA | NA | NA | Gm20128 | | 0 | 0 | 0 | 0 | 0 | 0 | 0 | 0 | 0 | 0 | 0 |
| 0 | NA | NA | NA | NA | NA | Gm22491 | | 0 | 0 | 0 | 0 | 0 | 0 | 0 | 0 | 0 | 0 | 0 |
| 0 | NA | NA | NA | NA | NA | Gm25572 | | 0 | 0 | 0 | 0 | 0 | 0 | 0 | 0 | 0 | 0 | 0 |
| 0 | NA | NA | NA | NA | NA | Gm27793 | | 0 | 0 | 0 | 0 | 0 | 0 | 0 | 0 | 0 | 0 | 0 |
| 0 | NA | NA | NA | NA | NA | Gpr87 | G protein-coupled receptor 87 | 0 | 0 | 0 | 0 | 0 | 0 | 0 | 0 | 0 | 0 | 0 |
| 0 | NA | NA | NA | NA | NA | Gm37715 | | 0 | 0 | 0 | 0 | 0 | 0 | 0 | 0 | 0 | 0 | 0 |
| 0 | NA | NA | NA | NA | NA | Gm38288 | | 0 | 0 | 0 | 0 | 0 | 0 | 0 | 0 | 0 | 0 | 0 |
| 0 | NA | NA | NA | NA | NA | Gm37771 | | 0 | 0 | 0 | 0 | 0 | 0 | 0 | 0 | 0 | 0 | 0 |
| 0 | NA | NA | NA | NA | NA | Gm4856 | | 0 | 0 | 0 | 0 | 0 | 0 | 0 | 0 | 0 | 0 | 0 |
| 0 | NA | NA | NA | NA | NA | Gm37926 | | 0 | 0 | 0 | 0 | 0 | 0 | 0 | 0 | 0 | 0 | 0 |
| 0 | NA | NA | NA | NA | NA | Aadacl2fm2 | AADACL2 family member 2 | 0 | 0 | 0 | 0 | 0 | 0 | 0 | 0 | 0 | 0 | 0 |
| 0 | NA | NA | NA | NA | NA | Gm38208 | | 0 | 0 | 0 | 0 | 0 | 0 | 0 | 0 | 0 | 0 | 0 |
| 0 | NA | NA | NA | NA | NA | 4930449A18Rik | | 0 | 0 | 0 | 0 | 0 | 0 | 0 | 0 | 0 | 0 | 0 |
| 0 | NA | NA | NA | NA | NA | Gm2756 | | 0 | 0 | 0 | 0 | 0 | 0 | 0 | 0 | 0 | 0 | 0 |
| 0 | NA | NA | NA | NA | NA | Aadacl2fm3 | AADACL2 family member 3 | 0 | 0 | 0 | 0 | 0 | 0 | 0 | 0 | 0 | 0 | 0 |
| 0 | NA | NA | NA | NA | NA | Aadacl2fm1 | AADACL2 family member 1 | 0 | 0 | 0 | 0 | 0 | 0 | 0 | 0 | 0 | 0 | 0 |
| 0 | NA | NA | NA | NA | NA | Gm9696 | | 0 | 0 | 0 | 0 | 0 | 0 | 0 | 0 | 0 | 0 | 0 |
| 0 | NA | NA | NA | NA | NA | Aadacl2 | arylacetamide deacetylase like 2 | 0 | 0 | 0 | 0 | 0 | 0 | 0 | 0 | 0 | 0 | 0 |
| 0 | NA | NA | NA | NA | NA | Gm18427 | | 0 | 0 | 0 | 0 | 0 | 0 | 0 | 0 | 0 | 0 | 0 |
| 0 | NA | NA | NA | NA | NA | Gm24382 | | 0 | 0 | 0 | 0 | 0 | 0 | 0 | 0 | 0 | 0 | 0 |
| 0 | NA | NA | NA | NA | NA | Tgif1-ps | | 0 | 0 | 0 | 0 | 0 | 0 | 0 | 0 | 0 | 0 | 0 |
| 0 | NA | NA | NA | NA | NA | Gm38280 | | 0 | 0 | 0 | 0 | 0 | 0 | 0 | 0 | 0 | 0 | 0 |

| | | | | | | | | | | | | | | | | | | |
|---|---|---|---|---|---|---|---|---|---|---|---|---|---|---|---|---|---|---|
| 0 | NA | NA | NA | NA | NA | Gm38326 | | 0 | 0 | 0 | 0 | 0 | 0 | 0 | 0 | 0 | 0 | 0 |
| 0 | NA | NA | NA | NA | NA | Gm37719 | | 0 | 0 | 0 | 0 | 0 | 0 | 0 | 0 | 0 | 0 | 0 |
| 0 | NA | NA | NA | NA | NA | Gm19901 | | 0 | 0 | 0 | 0 | 0 | 0 | 0 | 0 | 0 | 0 | 0 |
| 0 | NA | NA | NA | NA | NA | Gm9700 | | 0 | 0 | 0 | 0 | 0 | 0 | 0 | 0 | 0 | 0 | 0 |
| 0 | NA | NA | NA | NA | NA | Gm37404 | | 0 | 0 | 0 | 0 | 0 | 0 | 0 | 0 | 0 | 0 | 0 |
| 0 | NA | NA | NA | NA | NA | 9330121J05Rik | | 0 | 0 | 0 | 0 | 0 | 0 | 0 | 0 | 0 | 0 | 0 |
| 0 | NA | NA | NA | NA | NA | Gpr149 | G protein-coupled receptor 149 | 0 | 0 | 0 | 0 | 0 | 0 | 0 | 0 | 0 | 0 | 0 |
| 0 | NA | NA | NA | NA | NA | Gm18382 | | 0 | 0 | 0 | 0 | 0 | 0 | 0 | 0 | 0 | 0 | 0 |
| 0 | NA | NA | NA | NA | NA | Gm22433 | | 0 | 0 | 0 | 0 | 0 | 0 | 0 | 0 | 0 | 0 | 0 |
| 0 | NA | NA | NA | NA | NA | Gm34240 | | 0 | 0 | 0 | 0 | 0 | 0 | 0 | 0 | 0 | 0 | 0 |
| 0 | NA | NA | NA | NA | NA | Strit1 | small transmembrane regulator of ion trar | 0 | 0 | 0 | 0 | 0 | 0 | 0 | 0 | 0 | 0 | 0 |
| 0 | NA | NA | NA | NA | NA | Gm37279 | | 0 | 0 | 0 | 0 | 0 | 0 | 0 | 0 | 0 | 0 | 0 |
| 0 | NA | NA | NA | NA | NA | Gm24946 | | 0 | 0 | 0 | 0 | 0 | 0 | 0 | 0 | 0 | 0 | 0 |
| 0 | NA | NA | NA | NA | NA | Gm23500 | | 0 | 0 | 0 | 0 | 0 | 0 | 0 | 0 | 0 | 0 | 0 |
| 0 | NA | NA | NA | NA | NA | Gm16162 | | 0 | 0 | 0 | 0 | 0 | 0 | 0 | 0 | 0 | 0 | 0 |
| 0 | NA | NA | NA | NA | NA | Gm16129 | | 0 | 0 | 0 | 0 | 0 | 0 | 0 | 0 | 0 | 0 | 0 |
| 0 | NA | NA | NA | NA | NA | Gm34379 | | 0 | 0 | 0 | 0 | 0 | 0 | 0 | 0 | 0 | 0 | 0 |
| 0 | NA | NA | NA | NA | NA | Vmn2r-ps3 | | 0 | 0 | 0 | 0 | 0 | 0 | 0 | 0 | 0 | 0 | 0 |
| 0 | NA | NA | NA | NA | NA | Gm22165 | | 0 | 0 | 0 | 0 | 0 | 0 | 0 | 0 | 0 | 0 | 0 |
| 0 | NA | NA | NA | NA | NA | Vmn2r2 | vomeronasal 2, receptor 2 | 0 | 0 | 0 | 0 | 0 | 0 | 0 | 0 | 0 | 0 | 0 |
| 0 | NA | NA | NA | NA | NA | Vmn2r3 | vomeronasal 2, receptor 3 | 0 | 0 | 0 | 0 | 0 | 0 | 0 | 0 | 0 | 0 | 0 |
| 0 | NA | NA | NA | NA | NA | Gm23897 | | 0 | 0 | 0 | 0 | 0 | 0 | 0 | 0 | 0 | 0 | 0 |
| 0 | NA | NA | NA | NA | NA | Vmn2r-ps4 | | 0 | 0 | 0 | 0 | 0 | 0 | 0 | 0 | 0 | 0 | 0 |
| 0 | NA | NA | NA | NA | NA | Vmn2r128 | | 0 | 0 | 0 | 0 | 0 | 0 | 0 | 0 | 0 | 0 | 0 |
| 0 | NA | NA | NA | NA | NA | Vmn2r4 | vomeronasal 2, receptor 4 | 0 | 0 | 0 | 0 | 0 | 0 | 0 | 0 | 0 | 0 | 0 |
| 0 | NA | NA | NA | NA | NA | Vmn2r-ps6 | | 0 | 0 | 0 | 0 | 0 | 0 | 0 | 0 | 0 | 0 | 0 |
| 0 | NA | NA | NA | NA | NA | Vmn2r-ps7 | | 0 | 0 | 0 | 0 | 0 | 0 | 0 | 0 | 0 | 0 | 0 |
| 0 | NA | NA | NA | NA | NA | Gm20656 | | 0 | 0 | 0 | 0 | 0 | 0 | 0 | 0 | 0 | 0 | 0 |
| 0 | NA | NA | NA | NA | NA | Vmn2r-ps8 | | 0 | 0 | 0 | 0 | 0 | 0 | 0 | 0 | 0 | 0 | 0 |
| 0 | NA | NA | NA | NA | NA | Vmn2r5 | vomeronasal 2, receptor 5 | 0 | 0 | 0 | 0 | 0 | 0 | 0 | 0 | 0 | 0 | 0 |
| 0 | NA | NA | NA | NA | NA | Vmn2r6 | vomeronasal 2, receptor 6 | 0 | 0 | 0 | 0 | 0 | 0 | 0 | 0 | 0 | 0 | 0 |
| 0 | NA | NA | NA | NA | NA | Vmn2r-ps9 | | 0 | 0 | 0 | 0 | 0 | 0 | 0 | 0 | 0 | 0 | 0 |
| 0 | NA | NA | NA | NA | NA | Vmn2r-ps10 | | 0 | 0 | 0 | 0 | 0 | 0 | 0 | 0 | 0 | 0 | 0 |
| 0 | NA | NA | NA | NA | NA | Vmn2r-ps11 | | 0 | 0 | 0 | 0 | 0 | 0 | 0 | 0 | 0 | 0 | 0 |
| 0 | NA | NA | NA | NA | NA | Gm26939 | | 0 | 0 | 0 | 0 | 0 | 0 | 0 | 0 | 0 | 0 | 0 |
| 0 | NA | NA | NA | NA | NA | Vmn2r7 | vomeronasal 2, receptor 7 | 0 | 0 | 0 | 0 | 0 | 0 | 0 | 0 | 0 | 0 | 0 |
| 0 | NA | NA | NA | NA | NA | Vmn2r-ps12 | | 0 | 0 | 0 | 0 | 0 | 0 | 0 | 0 | 0 | 0 | 0 |
| 0 | NA | NA | NA | NA | NA | Gm25222 | | 0 | 0 | 0 | 0 | 0 | 0 | 0 | 0 | 0 | 0 | 0 |
| 0 | NA | NA | NA | NA | NA | Vmn2r-ps13 | | 0 | 0 | 0 | 0 | 0 | 0 | 0 | 0 | 0 | 0 | 0 |
| 0 | NA | NA | NA | NA | NA | Gm22279 | | 0 | 0 | 0 | 0 | 0 | 0 | 0 | 0 | 0 | 0 | 0 |
| 0 | NA | NA | NA | NA | NA | Gm38233 | | 0 | 0 | 0 | 0 | 0 | 0 | 0 | 0 | 0 | 0 | 0 |
| 0 | NA | NA | NA | NA | NA | Gm25755 | | 0 | 0 | 0 | 0 | 0 | 0 | 0 | 0 | 0 | 0 | 0 |
| 0 | NA | NA | NA | NA | NA | Gm37359 | | 0 | 0 | 0 | 0 | 0 | 0 | 0 | 0 | 0 | 0 | 0 |
| 0 | NA | NA | NA | NA | NA | Gm37037 | | 0 | 0 | 0 | 0 | 0 | 0 | 0 | 0 | 0 | 0 | 0 |
| 0 | NA | NA | NA | NA | NA | Gm19024 | | 0 | 0 | 0 | 0 | 0 | 0 | 0 | 0 | 0 | 0 | 0 |
| 0 | NA | NA | NA | NA | NA | Gm6546 | | 0 | 0 | 0 | 0 | 0 | 0 | 0 | 0 | 0 | 0 | 0 |
| 0 | NA | NA | NA | NA | NA | Gm37822 | | 0 | 0 | 0 | 0 | 0 | 0 | 0 | 0 | 0 | 0 | 0 |
| 0 | NA | NA | NA | NA | NA | Gm26442 | | 0 | 0 | 0 | 0 | 0 | 0 | 0 | 0 | 0 | 0 | 0 |
| 0 | NA | NA | NA | NA | NA | Gm36973 | | 0 | 0 | 0 | 0 | 0 | 0 | 0 | 0 | 0 | 0 | 0 |
| 0 | NA | NA | NA | NA | NA | Tpi-rs2 | | 0 | 0 | 0 | 0 | 0 | 0 | 0 | 0 | 0 | 0 | 0 |
| 0 | NA | NA | NA | NA | NA | Gm37330 | | 0 | 0 | 0 | 0 | 0 | 0 | 0 | 0 | 0 | 0 | 0 |
| 0 | NA | NA | NA | NA | NA | Gm50014 | | 0 | 0 | 0 | 0 | 0 | 0 | 0 | 0 | 0 | 0 | 0 |
| 0 | NA | NA | NA | NA | NA | Gm17952 | | 0 | 0 | 0 | 0 | 0 | 0 | 0 | 0 | 0 | 0 | 0 |
| 0 | NA | NA | NA | NA | NA | Gm6555 | | 0 | 0 | 0 | 0 | 0 | 0 | 0 | 0 | 0 | 0 | 0 |
| 0 | NA | NA | NA | NA | NA | Gm38356 | | 0 | 0 | 0 | 0 | 0 | 0 | 0 | 0 | 0 | 0 | 0 |
| 0 | NA | NA | NA | NA | NA | Gm43516 | | 0 | 0 | 0 | 0 | 0 | 0 | 0 | 0 | 0 | 0 | 0 |
| 0 | NA | NA | NA | NA | NA | Gm22295 | | 0 | 0 | 0 | 0 | 0 | 0 | 0 | 0 | 0 | 0 | 0 |
| 0 | NA | NA | NA | NA | NA | Gm17402 | | 0 | 0 | 0 | 0 | 0 | 0 | 0 | 0 | 0 | 0 | 0 |
| 0 | NA | NA | NA | NA | NA | Gm35299 | | 0 | 0 | 0 | 0 | 0 | 0 | 0 | 0 | 0 | 0 | 0 |
| 0 | NA | NA | NA | NA | NA | 4930402C01Rik | | 0 | 0 | 0 | 0 | 0 | 0 | 0 | 0 | 0 | 0 | 0 |

| | | | | | | | | | | | | | | | | | | |
|---|---|---|---|---|---|---|---|---|---|---|---|---|---|---|---|---|---|---|
| 0 | NA | NA | NA | NA | NA | Gm25919 | | 0 | 0 | 0 | 0 | 0 | 0 | 0 | 0 | 0 | 0 | 0 |
| 0 | NA | NA | NA | NA | NA | Gm412 | | 0 | 0 | 0 | 0 | 0 | 0 | 0 | 0 | 0 | 0 | 0 |
| 0 | NA | NA | NA | NA | NA | Gm37601 | | 0 | 0 | 0 | 0 | 0 | 0 | 0 | 0 | 0 | 0 | 0 |
| 0 | NA | NA | NA | NA | NA | Gm18717 | | 0 | 0 | 0 | 0 | 0 | 0 | 0 | 0 | 0 | 0 | 0 |
| 0 | NA | NA | NA | NA | NA | Iqcj | IQ motif containing J | 0 | 0 | 0 | 0 | 0 | 0 | 0 | 0 | 0 | 0 | 0 |
| 0 | NA | NA | NA | NA | NA | Iqschfp | Iqcj and Schip1 fusion protein | 0 | 0 | 0 | 0 | 0 | 0 | 0 | 0 | 0 | 0 | 0 |
| 0 | NA | NA | NA | NA | NA | Gm10292 | | 0 | 0 | 0 | 0 | 0 | 0 | 0 | 0 | 0 | 0 | 0 |
| 0 | NA | NA | NA | NA | NA | Gm10723 | | 0 | 0 | 0 | 0 | 0 | 0 | 0 | 0 | 0 | 0 | 0 |
| 0 | NA | NA | NA | NA | NA | Gm22073 | | 0 | 0 | 0 | 0 | 0 | 0 | 0 | 0 | 0 | 0 | 0 |
| 0 | NA | NA | NA | NA | NA | Il12a | interleukin 12a | 0 | 0 | 0 | 0 | 0 | 0 | 0 | 0 | 0 | 0 | 0 |
| 0 | NA | NA | NA | NA | NA | Gm10040 | | 0 | 0 | 0 | 0 | 0 | 0 | 0 | 0 | 0 | 0 | 0 |
| 0 | NA | NA | NA | NA | NA | 4930535E02Rik | | 0 | 0 | 0 | 0 | 0 | 0 | 0 | 0 | 0 | 0 | 0 |
| 0 | NA | NA | NA | NA | NA | Gm35584 | | 0 | 0 | 0 | 0 | 0 | 0 | 0 | 0 | 0 | 0 | 0 |
| 0 | NA | NA | NA | NA | NA | Gm7270 | | 0 | 0 | 0 | 0 | 0 | 0 | 0 | 0 | 0 | 0 | 0 |
| 0 | NA | NA | NA | NA | NA | Gm7270 | | 0 | 0 | 0 | 0 | 0 | 0 | 0 | 0 | 0 | 0 | 0 |
| 0 | NA | NA | NA | NA | NA | Gm17641 | | 0 | 0 | 0 | 0 | 0 | 0 | 0 | 0 | 0 | 0 | 0 |
| 0 | NA | NA | NA | NA | NA | Gm20690 | | 0 | 0 | 0 | 0 | 0 | 0 | 0 | 0 | 0 | 0 | 0 |
| 0 | NA | NA | NA | NA | NA | Gm20690 | | 0 | 0 | 0 | 0 | 0 | 0 | 0 | 0 | 0 | 0 | 0 |
| 0 | NA | NA | NA | NA | NA | Mir15b | | 0 | 0 | 0 | 0 | 0 | 0 | 0 | 0 | 0 | 0 | 0 |
| 0 | NA | NA | NA | NA | NA | Mir16-2 | | 0 | 0 | 0 | 0 | 0 | 0 | 0 | 0 | 0 | 0 | 0 |
| 0 | NA | NA | NA | NA | NA | Gm17213 | | 0 | 0 | 0 | 0 | 0 | 0 | 0 | 0 | 0 | 0 | 0 |
| 0 | NA | NA | NA | NA | NA | Gm37380 | | 0 | 0 | 0 | 0 | 0 | 0 | 0 | 0 | 0 | 0 | 0 |
| 0 | NA | NA | NA | NA | NA | Gm25621 | | 0 | 0 | 0 | 0 | 0 | 0 | 0 | 0 | 0 | 0 | 0 |
| 0 | NA | NA | NA | NA | NA | Gm17895 | | 0 | 0 | 0 | 0 | 0 | 0 | 0 | 0 | 0 | 0 | 0 |
| 0 | NA | NA | NA | NA | NA | Gm23484 | | 0 | 0 | 0 | 0 | 0 | 0 | 0 | 0 | 0 | 0 | 0 |
| 0 | NA | NA | NA | NA | NA | Otol1 | otolin 1 | 0 | 0 | 0 | 0 | 0 | 0 | 0 | 0 | 0 | 0 | 0 |
| 0 | NA | NA | NA | NA | NA | Gm37681 | | 0 | 0 | 0 | 0 | 0 | 0 | 0 | 0 | 0 | 0 | 0 |
| 0 | NA | NA | NA | NA | NA | Gm37585 | | 0 | 0 | 0 | 0 | 0 | 0 | 0 | 0 | 0 | 0 | 0 |
| 0 | NA | NA | NA | NA | NA | Gm23477 | | 0 | 0 | 0 | 0 | 0 | 0 | 0 | 0 | 0 | 0 | 0 |
| 0 | NA | NA | NA | NA | NA | Gm6631 | | 0 | 0 | 0 | 0 | 0 | 0 | 0 | 0 | 0 | 0 | 0 |
| 0 | NA | NA | NA | NA | NA | Gm37213 | | 0 | 0 | 0 | 0 | 0 | 0 | 0 | 0 | 0 | 0 | 0 |
| 0 | NA | NA | NA | NA | NA | Gm10780 | | 0 | 0 | 0 | 0 | 0 | 0 | 0 | 0 | 0 | 0 | 0 |
| 0 | NA | NA | NA | NA | NA | Gm19194 | | 0 | 0 | 0 | 0 | 0 | 0 | 0 | 0 | 0 | 0 | 0 |
| 0 | NA | NA | NA | NA | NA | Gm8604 | | 0 | 0 | 0 | 0 | 0 | 0 | 0 | 0 | 0 | 0 | 0 |
| 0 | NA | NA | NA | NA | NA | Gm37617 | | 0 | 0 | 0 | 0 | 0 | 0 | 0 | 0 | 0 | 0 | 0 |
| 0 | NA | NA | NA | NA | NA | Gm3351 | | 0 | 0 | 0 | 0 | 0 | 0 | 0 | 0 | 0 | 0 | 0 |
| 0 | NA | NA | NA | NA | NA | Gm37603 | | 0 | 0 | 0 | 0 | 0 | 0 | 0 | 0 | 0 | 0 | 0 |
| 0 | NA | NA | NA | NA | NA | Gm27465 | | 0 | 0 | 0 | 0 | 0 | 0 | 0 | 0 | 0 | 0 | 0 |
| 0 | NA | NA | NA | NA | NA | Gm37901 | | 0 | 0 | 0 | 0 | 0 | 0 | 0 | 0 | 0 | 0 | 0 |
| 0 | NA | NA | NA | NA | NA | Gm22451 | | 0 | 0 | 0 | 0 | 0 | 0 | 0 | 0 | 0 | 0 | 0 |
| 0 | NA | NA | NA | NA | NA | Gm37056 | | 0 | 0 | 0 | 0 | 0 | 0 | 0 | 0 | 0 | 0 | 0 |
| 0 | NA | NA | NA | NA | NA | Gm24544 | | 0 | 0 | 0 | 0 | 0 | 0 | 0 | 0 | 0 | 0 | 0 |
| 0 | NA | NA | NA | NA | NA | Gm37725 | | 0 | 0 | 0 | 0 | 0 | 0 | 0 | 0 | 0 | 0 | 0 |
| 0 | NA | NA | NA | NA | NA | Gm22538 | | 0 | 0 | 0 | 0 | 0 | 0 | 0 | 0 | 0 | 0 | 0 |
| 0 | NA | NA | NA | NA | NA | 4930509J09Rik | | 0 | 0 | 0 | 0 | 0 | 0 | 0 | 0 | 0 | 0 | 0 |
| 0 | NA | NA | NA | NA | NA | Gm20356 | | 0 | 0 | 0 | 0 | 0 | 0 | 0 | 0 | 0 | 0 | 0 |
| 0 | NA | NA | NA | NA | NA | Gm37050 | | 0 | 0 | 0 | 0 | 0 | 0 | 0 | 0 | 0 | 0 | 0 |
| 0 | NA | NA | NA | NA | NA | Gm37464 | | 0 | 0 | 0 | 0 | 0 | 0 | 0 | 0 | 0 | 0 | 0 |
| 0 | NA | NA | NA | NA | NA | Gm17836 | | 0 | 0 | 0 | 0 | 0 | 0 | 0 | 0 | 0 | 0 | 0 |
| 0 | NA | NA | NA | NA | NA | Platr10 | | 0 | 0 | 0 | 0 | 0 | 0 | 0 | 0 | 0 | 0 | 0 |
| 0 | NA | NA | NA | NA | NA | Gm37193 | | 0 | 0 | 0 | 0 | 0 | 0 | 0 | 0 | 0 | 0 | 0 |
| 0 | NA | NA | NA | NA | NA | Gm25846 | | 0 | 0 | 0 | 0 | 0 | 0 | 0 | 0 | 0 | 0 | 0 |
| 0 | NA | NA | NA | NA | NA | Gm29133 | | 0 | 0 | 0 | 0 | 0 | 0 | 0 | 0 | 0 | 0 | 0 |
| 0 | NA | NA | NA | NA | NA | Gm18428 | | 0 | 0 | 0 | 0 | 0 | 0 | 0 | 0 | 0 | 0 | 0 |
| 0 | NA | NA | NA | NA | NA | Gm37290 | | 0 | 0 | 0 | 0 | 0 | 0 | 0 | 0 | 0 | 0 | 0 |
| 0 | NA | NA | NA | NA | NA | Gm19231 | | 0 | 0 | 0 | 0 | 0 | 0 | 0 | 0 | 0 | 0 | 0 |
| 0 | NA | NA | NA | NA | NA | Gm23644 | | 0 | 0 | 0 | 0 | 0 | 0 | 0 | 0 | 0 | 0 | 0 |
| 0 | NA | NA | NA | NA | NA | Dnmt3a-ps1 | | 0 | 0 | 0 | 0 | 0 | 0 | 0 | 0 | 0 | 0 | 0 |
| 0 | NA | NA | NA | NA | NA | Gm37806 | | 0 | 0 | 0 | 0 | 0 | 0 | 0 | 0 | 0 | 0 | 0 |
| 0 | NA | NA | NA | NA | NA | Gm38195 | | 0 | 0 | 0 | 0 | 0 | 0 | 0 | 0 | 0 | 0 | 0 |

| | | | | | | | | | | | | | | | | | | |
|---|---|---|---|---|---|---|---|---|---|---|---|---|---|---|---|---|---|---|
| 0 | NA | NA | NA | NA | NA | Gm37576 | | 0 | 0 | 0 | 0 | 0 | 0 | 0 | 0 | 0 | 0 | 0 |
| 0 | NA | NA | NA | NA | NA | Gm18719 | | 0 | 0 | 0 | 0 | 0 | 0 | 0 | 0 | 0 | 0 | 0 |
| 0 | NA | NA | NA | NA | NA | Gm6677 | | 0 | 0 | 0 | 0 | 0 | 0 | 0 | 0 | 0 | 0 | 0 |
| 0 | NA | NA | NA | NA | NA | Gm6680 | | 0 | 0 | 0 | 0 | 0 | 0 | 0 | 0 | 0 | 0 | 0 |
| 0 | NA | NA | NA | NA | NA | Gm43326 | | 0 | 0 | 0 | 0 | 0 | 0 | 0 | 0 | 0 | 0 | 0 |
| 0 | NA | NA | NA | NA | NA | Gm15442 | | 0 | 0 | 0 | 0 | 0 | 0 | 0 | 0 | 0 | 0 | 0 |
| 0 | NA | NA | NA | NA | NA | Gm18952 | | 0 | 0 | 0 | 0 | 0 | 0 | 0 | 0 | 0 | 0 | 0 |
| 0 | NA | NA | NA | NA | NA | Gm24749 | | 0 | 0 | 0 | 0 | 0 | 0 | 0 | 0 | 0 | 0 | 0 |
| 0 | NA | NA | NA | NA | NA | Gm17953 | | 0 | 0 | 0 | 0 | 0 | 0 | 0 | 0 | 0 | 0 | 0 |
| 0 | NA | NA | NA | NA | NA | Gm5277 | | 0 | 0 | 0 | 0 | 0 | 0 | 0 | 0 | 0 | 0 | 0 |
| 0 | NA | NA | NA | NA | NA | Gm23440 | | 0 | 0 | 0 | 0 | 0 | 0 | 0 | 0 | 0 | 0 | 0 |
| 0 | NA | NA | NA | NA | NA | Gm24150 | | 0 | 0 | 0 | 0 | 0 | 0 | 0 | 0 | 0 | 0 | 0 |
| 0 | NA | NA | NA | NA | NA | Gm17359 | predicted gene, 17359 | 0 | 0 | 0 | 0 | 0 | 0 | 0 | 0 | 0 | 0 | 0 |
| 0 | NA | NA | NA | NA | NA | 4930589L23Rik | | 0 | 0 | 0 | 0 | 0 | 0 | 0 | 0 | 0 | 0 | 0 |
| 0 | NA | NA | NA | NA | NA | Gm35067 | | 0 | 0 | 0 | 0 | 0 | 0 | 0 | 0 | 0 | 0 | 0 |
| 0 | NA | NA | NA | NA | NA | Gm26984 | | 0 | 0 | 0 | 0 | 0 | 0 | 0 | 0 | 0 | 0 | 0 |
| 0 | NA | NA | NA | NA | NA | Gm26420 | | 0 | 0 | 0 | 0 | 0 | 0 | 0 | 0 | 0 | 0 | 0 |
| 0 | NA | NA | NA | NA | NA | A330069K06Rik | | 0 | 0 | 0 | 0 | 0 | 0 | 0 | 0 | 0 | 0 | 0 |
| 0 | NA | NA | NA | NA | NA | Gm19066 | | 0 | 0 | 0 | 0 | 0 | 0 | 0 | 0 | 0 | 0 | 0 |
| 0 | NA | NA | NA | NA | NA | Gm43839 | | 0 | 0 | 0 | 0 | 0 | 0 | 0 | 0 | 0 | 0 | 0 |
| 0 | NA | NA | NA | NA | NA | Gm38295 | | 0 | 0 | 0 | 0 | 0 | 0 | 0 | 0 | 0 | 0 | 0 |
| 0 | NA | NA | NA | NA | NA | Gm37892 | | 0 | 0 | 0 | 0 | 0 | 0 | 0 | 0 | 0 | 0 | 0 |
| 0 | NA | NA | NA | NA | NA | Gm37971 | | 0 | 0 | 0 | 0 | 0 | 0 | 0 | 0 | 0 | 0 | 0 |
| 0 | NA | NA | NA | NA | NA | Gm25749 | | 0 | 0 | 0 | 0 | 0 | 0 | 0 | 0 | 0 | 0 | 0 |
| 0 | NA | NA | NA | NA | NA | Gm42468 | | 0 | 0 | 0 | 0 | 0 | 0 | 0 | 0 | 0 | 0 | 0 |
| 0 | NA | NA | NA | NA | NA | Gm42476 | | 0 | 0 | 0 | 0 | 0 | 0 | 0 | 0 | 0 | 0 | 0 |
| 0 | NA | NA | NA | NA | NA | Gm16000 | | 0 | 0 | 0 | 0 | 0 | 0 | 0 | 0 | 0 | 0 | 0 |
| 0 | NA | NA | NA | NA | NA | Gm37300 | | 0 | 0 | 0 | 0 | 0 | 0 | 0 | 0 | 0 | 0 | 0 |
| 0 | NA | NA | NA | NA | NA | Gm4857 | | 0 | 0 | 0 | 0 | 0 | 0 | 0 | 0 | 0 | 0 | 0 |
| 0 | NA | NA | NA | NA | NA | Gm43346 | | 0 | 0 | 0 | 0 | 0 | 0 | 0 | 0 | 0 | 0 | 0 |
| 0 | NA | NA | NA | NA | NA | A830029E22Rik | | 0 | 0 | 0 | 0 | 0 | 0 | 0 | 0 | 0 | 0 | 0 |
| 0 | NA | NA | NA | NA | NA | Gm22531 | | 0 | 0 | 0 | 0 | 0 | 0 | 0 | 0 | 0 | 0 | 0 |
| 0 | NA | NA | NA | NA | NA | Gm9989 | | 0 | 0 | 0 | 0 | 0 | 0 | 0 | 0 | 0 | 0 | 0 |
| 0 | NA | NA | NA | NA | NA | Asic5 | acid-sensing (proton-gated) ion channel f | 0 | 0 | 0 | 0 | 0 | 0 | 0 | 0 | 0 | 0 | 0 |
| 0 | NA | NA | NA | NA | NA | Mir7010 | | 0 | 0 | 0 | 0 | 0 | 0 | 0 | 0 | 0 | 0 | 0 |
| 0 | NA | NA | NA | NA | NA | Npy2r | neuropeptide Y receptor Y2 | 0 | 0 | 0 | 0 | 0 | 0 | 0 | 0 | 0 | 0 | 0 |
| 0 | NA | NA | NA | NA | NA | Gm35323 | | 0 | 0 | 0 | 0 | 0 | 0 | 0 | 0 | 0 | 0 | 0 |
| 0 | NA | NA | NA | NA | NA | Gm29999 | | 0 | 0 | 0 | 0 | 0 | 0 | 0 | 0 | 0 | 0 | 0 |
| 0 | NA | NA | NA | NA | NA | Gm43347 | | 0 | 0 | 0 | 0 | 0 | 0 | 0 | 0 | 0 | 0 | 0 |
| 0 | NA | NA | NA | NA | NA | Gm30043 | | 0 | 0 | 0 | 0 | 0 | 0 | 0 | 0 | 0 | 0 | 0 |
| 0 | NA | NA | NA | NA | NA | Gm22038 | | 0 | 0 | 0 | 0 | 0 | 0 | 0 | 0 | 0 | 0 | 0 |
| 0 | NA | NA | NA | NA | NA | Gm43348 | | 0 | 0 | 0 | 0 | 0 | 0 | 0 | 0 | 0 | 0 | 0 |
| 0 | NA | NA | NA | NA | NA | Rbm46os | | 0 | 0 | 0 | 0 | 0 | 0 | 0 | 0 | 0 | 0 | 0 |
| 0 | NA | NA | NA | NA | NA | Gm30097 | | 0 | 0 | 0 | 0 | 0 | 0 | 0 | 0 | 0 | 0 | 0 |
| 0 | NA | NA | NA | NA | NA | Gm38096 | | 0 | 0 | 0 | 0 | 0 | 0 | 0 | 0 | 0 | 0 | 0 |
| 0 | NA | NA | NA | NA | NA | 1700028M03Rik | | 0 | 0 | 0 | 0 | 0 | 0 | 0 | 0 | 0 | 0 | 0 |
| 0 | NA | NA | NA | NA | NA | Gm22342 | | 0 | 0 | 0 | 0 | 0 | 0 | 0 | 0 | 0 | 0 | 0 |
| 0 | NA | NA | NA | NA | NA | Rpl28-ps2 | | 0 | 0 | 0 | 0 | 0 | 0 | 0 | 0 | 0 | 0 | 0 |
| 0 | NA | NA | NA | NA | NA | Gm7115 | | 0 | 0 | 0 | 0 | 0 | 0 | 0 | 0 | 0 | 0 | 0 |
| 0 | NA | NA | NA | NA | NA | Gm36981 | | 0 | 0 | 0 | 0 | 0 | 0 | 0 | 0 | 0 | 0 | 0 |
| 0 | NA | NA | NA | NA | NA | Gm6525 | | 0 | 0 | 0 | 0 | 0 | 0 | 0 | 0 | 0 | 0 | 0 |
| 0 | NA | NA | NA | NA | NA | Gm38041 | | 0 | 0 | 0 | 0 | 0 | 0 | 0 | 0 | 0 | 0 | 0 |
| 0 | NA | NA | NA | NA | NA | Gm38313 | | 0 | 0 | 0 | 0 | 0 | 0 | 0 | 0 | 0 | 0 | 0 |
| 0 | NA | NA | NA | NA | NA | 1700036G14Rik | | 0 | 0 | 0 | 0 | 0 | 0 | 0 | 0 | 0 | 0 | 0 |
| 0 | NA | NA | NA | NA | NA | Gm42813 | | 0 | 0 | 0 | 0 | 0 | 0 | 0 | 0 | 0 | 0 | 0 |
| 0 | NA | NA | NA | NA | NA | Gm45790 | | 0 | 0 | 0 | 0 | 0 | 0 | 0 | 0 | 0 | 0 | 0 |
| 0 | NA | NA | NA | NA | NA | Mir466k | | 0 | 0 | 0 | 0 | 0 | 0 | 0 | 0 | 0 | 0 | 0 |
| 0 | NA | NA | NA | NA | NA | Gm26204 | | 0 | 0 | 0 | 0 | 0 | 0 | 0 | 0 | 0 | 0 | 0 |
| 0 | NA | NA | NA | NA | NA | Gm38030 | | 0 | 0 | 0 | 0 | 0 | 0 | 0 | 0 | 0 | 0 | 0 |
| 0 | NA | NA | NA | NA | NA | Gm25876 | | 0 | 0 | 0 | 0 | 0 | 0 | 0 | 0 | 0 | 0 | 0 |

| | | | | | | | | | | | | | | | | | | |
|---|---|---|---|---|---|---|---|---|---|---|---|---|---|---|---|---|---|---|
| 0 | NA | NA | NA | NA | NA | Gm24501 | | 0 | 0 | 0 | 0 | 0 | 0 | 0 | 0 | 0 | 0 | 0 |
| 0 | NA | NA | NA | NA | NA | Gm27811 | | 0 | 0 | 0 | 0 | 0 | 0 | 0 | 0 | 0 | 0 | 0 |
| 0 | NA | NA | NA | NA | NA | Prss48 | protease, serine 48 | 0 | 0 | 0 | 0 | 0 | 0 | 0 | 0 | 0 | 0 | 0 |
| 0 | NA | NA | NA | NA | NA | Gm25188 | | 0 | 0 | 0 | 0 | 0 | 0 | 0 | 0 | 0 | 0 | 0 |
| 0 | NA | NA | NA | NA | NA | Gm37493 | | 0 | 0 | 0 | 0 | 0 | 0 | 0 | 0 | 0 | 0 | 0 |
| 0 | NA | NA | NA | NA | NA | Gm23025 | | 0 | 0 | 0 | 0 | 0 | 0 | 0 | 0 | 0 | 0 | 0 |
| 0 | NA | NA | NA | NA | NA | Gm37288 | | 0 | 0 | 0 | 0 | 0 | 0 | 0 | 0 | 0 | 0 | 0 |
| 0 | NA | NA | NA | NA | NA | Gm26343 | | 0 | 0 | 0 | 0 | 0 | 0 | 0 | 0 | 0 | 0 | 0 |
| 0 | NA | NA | NA | NA | NA | Gm23850 | | 0 | 0 | 0 | 0 | 0 | 0 | 0 | 0 | 0 | 0 | 0 |
| 0 | NA | NA | NA | NA | NA | Gm25039 | | 0 | 0 | 0 | 0 | 0 | 0 | 0 | 0 | 0 | 0 | 0 |
| 0 | NA | NA | NA | NA | NA | Gm23376 | | 0 | 0 | 0 | 0 | 0 | 0 | 0 | 0 | 0 | 0 | 0 |
| 0 | NA | NA | NA | NA | NA | Gm37207 | | 0 | 0 | 0 | 0 | 0 | 0 | 0 | 0 | 0 | 0 | 0 |
| 0 | NA | NA | NA | NA | NA | Gm18403 | | 0 | 0 | 0 | 0 | 0 | 0 | 0 | 0 | 0 | 0 | 0 |
| 0 | NA | NA | NA | NA | NA | Gm37575 | | 0 | 0 | 0 | 0 | 0 | 0 | 0 | 0 | 0 | 0 | 0 |
| 0 | NA | NA | NA | NA | NA | Gm24472 | | 0 | 0 | 0 | 0 | 0 | 0 | 0 | 0 | 0 | 0 | 0 |
| 0 | NA | NA | NA | NA | NA | Gm37855 | | 0 | 0 | 0 | 0 | 0 | 0 | 0 | 0 | 0 | 0 | 0 |
| 0 | NA | NA | NA | NA | NA | | ets variant 3-like | 0 | 0 | 0 | 0 | 0 | 0 | 0 | 0 | 0 | 0 | 0 |
| 0 | NA | NA | NA | NA | NA | Gm25167 | | 0 | 0 | 0 | 0 | 0 | 0 | 0 | 0 | 0 | 0 | 0 |
| 0 | NA | NA | NA | NA | NA | Gm24109 | | 0 | 0 | 0 | 0 | 0 | 0 | 0 | 0 | 0 | 0 | 0 |
| 0 | NA | NA | NA | NA | NA | AI849053 | | 0 | 0 | 0 | 0 | 0 | 0 | 0 | 0 | 0 | 0 | 0 |
| 0 | NA | NA | NA | NA | NA | Gm3764 | | 0 | 0 | 0 | 0 | 0 | 0 | 0 | 0 | 0 | 0 | 0 |
| 0 | NA | NA | NA | NA | NA | Mir3093 | | 0 | 0 | 0 | 0 | 0 | 0 | 0 | 0 | 0 | 0 | 0 |
| 0 | NA | NA | NA | NA | NA | Gm25641 | | 0 | 0 | 0 | 0 | 0 | 0 | 0 | 0 | 0 | 0 | 0 |
| 0 | NA | NA | NA | NA | NA | Mir9-1 | | 0 | 0 | 0 | 0 | 0 | 0 | 0 | 0 | 0 | 0 | 0 |
| 0 | NA | NA | NA | NA | NA | Rhbg | Rhesus blood group-associated B glycop | 0 | 0 | 0 | 0 | 0 | 0 | 0 | 0 | 0 | 0 | 0 |
| 0 | NA | NA | NA | NA | NA | Gm21956 | | 0 | 0 | 0 | 0 | 0 | 0 | 0 | 0 | 0 | 0 | 0 |
| 0 | NA | NA | NA | NA | NA | Bglap2 | bone gamma-carboxyglutamate protein 2 | 0 | 0 | 0 | 0 | 0 | 0 | 0 | 0 | 0 | 0 | 0 |
| 0 | NA | NA | NA | NA | NA | Gm6821 | | 0 | 0 | 0 | 0 | 0 | 0 | 0 | 0 | 0 | 0 | 0 |
| 0 | NA | NA | NA | NA | NA | Bglap | bone gamma carboxyglutamate protein | 0 | 0 | 0 | 0 | 0 | 0 | 0 | 0 | 0 | 0 | 0 |
| 0 | NA | NA | NA | NA | NA | Mir7011 | | 0 | 0 | 0 | 0 | 0 | 0 | 0 | 0 | 0 | 0 | 0 |
| 0 | NA | NA | NA | NA | NA | Gm5979 | | 0 | 0 | 0 | 0 | 0 | 0 | 0 | 0 | 0 | 0 | 0 |
| 0 | NA | NA | NA | NA | NA | Rxfp4 | relaxin family peptide receptor 4 | 0 | 0 | 0 | 0 | 0 | 0 | 0 | 0 | 0 | 0 | 0 |
| 0 | NA | NA | NA | NA | NA | Gm25820 | | 0 | 0 | 0 | 0 | 0 | 0 | 0 | 0 | 0 | 0 | 0 |
| 0 | NA | NA | NA | NA | NA | Gm43713 | | 0 | 0 | 0 | 0 | 0 | 0 | 0 | 0 | 0 | 0 | 0 |
| 0 | NA | NA | NA | NA | NA | Gm24403 | | 0 | 0 | 0 | 0 | 0 | 0 | 0 | 0 | 0 | 0 | 0 |
| 0 | NA | NA | NA | NA | NA | Gm43738 | | 0 | 0 | 0 | 0 | 0 | 0 | 0 | 0 | 0 | 0 | 0 |
| 0 | NA | NA | NA | NA | NA | Gm29704 | | 0 | 0 | 0 | 0 | 0 | 0 | 0 | 0 | 0 | 0 | 0 |
| 0 | NA | NA | NA | NA | NA | Gm23269 | | 0 | 0 | 0 | 0 | 0 | 0 | 0 | 0 | 0 | 0 | 0 |
| 0 | NA | NA | NA | NA | NA | Mir92b | | 0 | 0 | 0 | 0 | 0 | 0 | 0 | 0 | 0 | 0 | 0 |
| 0 | NA | NA | NA | NA | NA | Gm22642 | | 0 | 0 | 0 | 0 | 0 | 0 | 0 | 0 | 0 | 0 | 0 |
| 0 | NA | NA | NA | NA | NA | 4731419I09Rik | | 0 | 0 | 0 | 0 | 0 | 0 | 0 | 0 | 0 | 0 | 0 |
| 0 | NA | NA | NA | NA | NA | Dcst1 | DC-STAMP domain containing 1 | 0 | 0 | 0 | 0 | 0 | 0 | 0 | 0 | 0 | 0 | 0 |
| 0 | NA | NA | NA | NA | NA | | | 0 | 0 | 0 | 0 | 0 | 0 | 0 | 0 | 0 | 0 | 0 |
| 0 | NA | NA | NA | NA | NA | Gm8969 | | 0 | 0 | 0 | 0 | 0 | 0 | 0 | 0 | 0 | 0 | 0 |
| 0 | NA | NA | NA | NA | NA | Aqp10-ps | | 0 | 0 | 0 | 0 | 0 | 0 | 0 | 0 | 0 | 0 | 0 |
| 0 | NA | NA | NA | NA | NA | Gm24046 | | 0 | 0 | 0 | 0 | 0 | 0 | 0 | 0 | 0 | 0 | 0 |
| 0 | NA | NA | NA | NA | NA | Mir190b | | 0 | 0 | 0 | 0 | 0 | 0 | 0 | 0 | 0 | 0 | 0 |
| 0 | NA | NA | NA | NA | NA | Gm23723 | | 0 | 0 | 0 | 0 | 0 | 0 | 0 | 0 | 0 | 0 | 0 |
| 0 | NA | NA | NA | NA | NA | Gm24185 | | 0 | 0 | 0 | 0 | 0 | 0 | 0 | 0 | 0 | 0 | 0 |
| 0 | NA | NA | NA | NA | NA | Mir7012 | | 0 | 0 | 0 | 0 | 0 | 0 | 0 | 0 | 0 | 0 | 0 |
| 0 | NA | NA | NA | NA | NA | Gm43595 | | 0 | 0 | 0 | 0 | 0 | 0 | 0 | 0 | 0 | 0 | 0 |
| 0 | NA | NA | NA | NA | NA | Gm24593 | | 0 | 0 | 0 | 0 | 0 | 0 | 0 | 0 | 0 | 0 | 0 |
| 0 | NA | NA | NA | NA | NA | Gm23464 | | 0 | 0 | 0 | 0 | 0 | 0 | 0 | 0 | 0 | 0 | 0 |
| 0 | NA | NA | NA | NA | NA | Gm42674 | | 0 | 0 | 0 | 0 | 0 | 0 | 0 | 0 | 0 | 0 | 0 |
| 0 | NA | NA | NA | NA | NA | Gm42579 | | 0 | 0 | 0 | 0 | 0 | 0 | 0 | 0 | 0 | 0 | 0 |
| 0 | NA | NA | NA | NA | NA | Pglyrp4 | peptidoglycan recognition protein 4 | 0 | 0 | 0 | 0 | 0 | 0 | 0 | 0 | 0 | 0 | 0 |
| 0 | NA | NA | NA | NA | NA | Gm5849 | | 0 | 0 | 0 | 0 | 0 | 0 | 0 | 0 | 0 | 0 | 0 |
| 0 | NA | NA | NA | NA | NA | Gm43235 | | 0 | 0 | 0 | 0 | 0 | 0 | 0 | 0 | 0 | 0 | 0 |
| 0 | NA | NA | NA | NA | NA | Gm7166 | | 0 | 0 | 0 | 0 | 0 | 0 | 0 | 0 | 0 | 0 | 0 |
| 0 | NA | NA | NA | NA | NA | S100a7l2 | S100 calcium binding protein A7 like 2 | 0 | 0 | 0 | 0 | 0 | 0 | 0 | 0 | 0 | 0 | 0 |

| | | | | | | | | | | | | | | | | | | |
|---|---|---|---|---|---|---|---|---|---|---|---|---|---|---|---|---|---|---|
| 0 | NA | NA | NA | NA | NA | Gm23697 | | 0 | 0 | 0 | 0 | 0 | 0 | 0 | 0 | 0 | 0 | 0 |
| 0 | NA | NA | NA | NA | NA | Pglyrp3 | peptidoglycan recognition protein 3 | 0 | 0 | 0 | 0 | 0 | 0 | 0 | 0 | 0 | 0 | 0 |
| 0 | NA | NA | NA | NA | NA | Gm25145 | | 0 | 0 | 0 | 0 | 0 | 0 | 0 | 0 | 0 | 0 | 0 |
| 0 | NA | NA | NA | NA | NA | Gm43736 | | 0 | 0 | 0 | 0 | 0 | 0 | 0 | 0 | 0 | 0 | 0 |
| 0 | NA | NA | NA | NA | NA | 4930511M18Rik | | 0 | 0 | 0 | 0 | 0 | 0 | 0 | 0 | 0 | 0 | 0 |
| 0 | NA | NA | NA | NA | NA | Gm43735 | | 0 | 0 | 0 | 0 | 0 | 0 | 0 | 0 | 0 | 0 | 0 |
| 0 | NA | NA | NA | NA | NA | Prr9 | proline rich 9 | 0 | 0 | 0 | 0 | 0 | 0 | 0 | 0 | 0 | 0 | 0 |
| 0 | NA | NA | NA | NA | NA | Lelp1 | late cornified envelope-like proline-rich 1 | 0 | 0 | 0 | 0 | 0 | 0 | 0 | 0 | 0 | 0 | 0 |
| 0 | NA | NA | NA | NA | NA | Gm35439 | | 0 | 0 | 0 | 0 | 0 | 0 | 0 | 0 | 0 | 0 | 0 |
| 0 | NA | NA | NA | NA | NA | | | 0 | 0 | 0 | 0 | 0 | 0 | 0 | 0 | 0 | 0 | 0 |
| 0 | NA | NA | NA | NA | NA | Sprr2a1 | small proline-rich protein 2A1 | 0 | 0 | 0 | 0 | 0 | 0 | 0 | 0 | 0 | 0 | 0 |
| 0 | NA | NA | NA | NA | NA | Sprr2a2 | small proline-rich protein 2A2 | 0 | 0 | 0 | 0 | 0 | 0 | 0 | 0 | 0 | 0 | 0 |
| 0 | NA | NA | NA | NA | NA | Gm5850 | | 0 | 0 | 0 | 0 | 0 | 0 | 0 | 0 | 0 | 0 | 0 |
| 0 | NA | NA | NA | NA | NA | Gm37087 | | 0 | 0 | 0 | 0 | 0 | 0 | 0 | 0 | 0 | 0 | 0 |
| 0 | NA | NA | NA | NA | NA | Gm5851 | | 0 | 0 | 0 | 0 | 0 | 0 | 0 | 0 | 0 | 0 | 0 |
| 0 | NA | NA | NA | NA | NA | Gm37752 | | 0 | 0 | 0 | 0 | 0 | 0 | 0 | 0 | 0 | 0 | 0 |
| 0 | NA | NA | NA | NA | NA | Sprr2a3 | small proline-rich protein 2A3 | 0 | 0 | 0 | 0 | 0 | 0 | 0 | 0 | 0 | 0 | 0 |
| 0 | NA | NA | NA | NA | NA | Sprr2b | small proline-rich protein 2B | 0 | 0 | 0 | 0 | 0 | 0 | 0 | 0 | 0 | 0 | 0 |
| 0 | NA | NA | NA | NA | NA | Gm37428 | | 0 | 0 | 0 | 0 | 0 | 0 | 0 | 0 | 0 | 0 | 0 |
| 0 | NA | NA | NA | NA | NA | Sprr2c-ps | | 0 | 0 | 0 | 0 | 0 | 0 | 0 | 0 | 0 | 0 | 0 |
| 0 | NA | NA | NA | NA | NA | Sprr2d | small proline-rich protein 2D | 0 | 0 | 0 | 0 | 0 | 0 | 0 | 0 | 0 | 0 | 0 |
| 0 | NA | NA | NA | NA | NA | Sprr2e | small proline-rich protein 2E | 0 | 0 | 0 | 0 | 0 | 0 | 0 | 0 | 0 | 0 | 0 |
| 0 | NA | NA | NA | NA | NA | Sprr2f | small proline-rich protein 2F | 0 | 0 | 0 | 0 | 0 | 0 | 0 | 0 | 0 | 0 | 0 |
| 0 | NA | NA | NA | NA | NA | Sprr2g | | 0 | 0 | 0 | 0 | 0 | 0 | 0 | 0 | 0 | 0 | 0 |
| 0 | NA | NA | NA | NA | NA | Sprr2h | small proline-rich protein 2H | 0 | 0 | 0 | 0 | 0 | 0 | 0 | 0 | 0 | 0 | 0 |
| 0 | NA | NA | NA | NA | NA | Sprr2i | small proline-rich protein 2I | 0 | 0 | 0 | 0 | 0 | 0 | 0 | 0 | 0 | 0 | 0 |
| 0 | NA | NA | NA | NA | NA | Sprr2j-ps | | 0 | 0 | 0 | 0 | 0 | 0 | 0 | 0 | 0 | 0 | 0 |
| 0 | NA | NA | NA | NA | NA | Sprr2k | small proline-rich protein 2K | 0 | 0 | 0 | 0 | 0 | 0 | 0 | 0 | 0 | 0 | 0 |
| 0 | NA | NA | NA | NA | NA | Sprr1b | small proline-rich protein 1B | 0 | 0 | 0 | 0 | 0 | 0 | 0 | 0 | 0 | 0 | 0 |
| 0 | NA | NA | NA | NA | NA | Sprr3 | small proline-rich protein 3 | 0 | 0 | 0 | 0 | 0 | 0 | 0 | 0 | 0 | 0 | 0 |
| 0 | NA | NA | NA | NA | NA | Gm37377 | | 0 | 0 | 0 | 0 | 0 | 0 | 0 | 0 | 0 | 0 | 0 |
| 0 | NA | NA | NA | NA | NA | Sprr1a | small proline-rich protein 1A | 0 | 0 | 0 | 0 | 0 | 0 | 0 | 0 | 0 | 0 | 0 |
| 0 | NA | NA | NA | NA | NA | Sprr4 | small proline-rich protein 4 | 0 | 0 | 0 | 0 | 0 | 0 | 0 | 0 | 0 | 0 | 0 |
| 0 | NA | NA | NA | NA | NA | 2310046K23Rik | | 0 | 0 | 0 | 0 | 0 | 0 | 0 | 0 | 0 | 0 | 0 |
| 0 | NA | NA | NA | NA | NA | Ivl | involucrin | 0 | 0 | 0 | 0 | 0 | 0 | 0 | 0 | 0 | 0 | 0 |
| 0 | NA | NA | NA | NA | NA | Smcp | sperm mitochondria-associated cysteine- | 0 | 0 | 0 | 0 | 0 | 0 | 0 | 0 | 0 | 0 | 0 |
| 0 | NA | NA | NA | NA | NA | 2210017I01Rik | | 0 | 0 | 0 | 0 | 0 | 0 | 0 | 0 | 0 | 0 | 0 |
| 0 | NA | NA | NA | NA | NA | Lce6a | late cornified envelope 6A | 0 | 0 | 0 | 0 | 0 | 0 | 0 | 0 | 0 | 0 | 0 |
| 0 | NA | NA | NA | NA | NA | Lce1a1 | late cornified envelope 1A1 | 0 | 0 | 0 | 0 | 0 | 0 | 0 | 0 | 0 | 0 | 0 |
| 0 | NA | NA | NA | NA | NA | Lce1b | late cornified envelope 1B | 0 | 0 | 0 | 0 | 0 | 0 | 0 | 0 | 0 | 0 | 0 |
| 0 | NA | NA | NA | NA | NA | Rpl13-ps4 | | 0 | 0 | 0 | 0 | 0 | 0 | 0 | 0 | 0 | 0 | 0 |
| 0 | NA | NA | NA | NA | NA | Lce1a2 | late cornified envelope 1A2 | 0 | 0 | 0 | 0 | 0 | 0 | 0 | 0 | 0 | 0 | 0 |
| 0 | NA | NA | NA | NA | NA | Lce1c | late cornified envelope 1C | 0 | 0 | 0 | 0 | 0 | 0 | 0 | 0 | 0 | 0 | 0 |
| 0 | NA | NA | NA | NA | NA | Lce1d | late cornified envelope 1D | 0 | 0 | 0 | 0 | 0 | 0 | 0 | 0 | 0 | 0 | 0 |
| 0 | NA | NA | NA | NA | NA | Lce1e | late cornified envelope 1E | 0 | 0 | 0 | 0 | 0 | 0 | 0 | 0 | 0 | 0 | 0 |
| 0 | NA | NA | NA | NA | NA | Lce1f | late cornified envelope 1F | 0 | 0 | 0 | 0 | 0 | 0 | 0 | 0 | 0 | 0 | 0 |
| 0 | NA | NA | NA | NA | NA | Gm38119 | | 0 | 0 | 0 | 0 | 0 | 0 | 0 | 0 | 0 | 0 | 0 |
| 0 | NA | NA | NA | NA | NA | Lce1g | late cornified envelope 1G | 0 | 0 | 0 | 0 | 0 | 0 | 0 | 0 | 0 | 0 | 0 |
| 0 | NA | NA | NA | NA | NA | Lce1h | late cornified envelope 1H | 0 | 0 | 0 | 0 | 0 | 0 | 0 | 0 | 0 | 0 | 0 |
| 0 | NA | NA | NA | NA | NA | Lce1i | late cornified envelope 1I | 0 | 0 | 0 | 0 | 0 | 0 | 0 | 0 | 0 | 0 | 0 |
| 0 | NA | NA | NA | NA | NA | Lce1j | late cornified envelope 1J | 0 | 0 | 0 | 0 | 0 | 0 | 0 | 0 | 0 | 0 | 0 |
| 0 | NA | NA | NA | NA | NA | Lce1k | late cornified envelope 1K | 0 | 0 | 0 | 0 | 0 | 0 | 0 | 0 | 0 | 0 | 0 |
| 0 | NA | NA | NA | NA | NA | Kprp | keratinocyte expressed, proline-rich | 0 | 0 | 0 | 0 | 0 | 0 | 0 | 0 | 0 | 0 | 0 |
| 0 | NA | NA | NA | NA | NA | Gm38201 | | 0 | 0 | 0 | 0 | 0 | 0 | 0 | 0 | 0 | 0 | 0 |
| 0 | NA | NA | NA | NA | NA | Lce1l | late cornified envelope 1L | 0 | 0 | 0 | 0 | 0 | 0 | 0 | 0 | 0 | 0 | 0 |
| 0 | NA | NA | NA | NA | NA | 2310050C09Ril | RIKEN cDNA 2310050C09 gene | 0 | 0 | 0 | 0 | 0 | 0 | 0 | 0 | 0 | 0 | 0 |
| 0 | NA | NA | NA | NA | NA | Lce3a | late cornified envelope 3A | 0 | 0 | 0 | 0 | 0 | 0 | 0 | 0 | 0 | 0 | 0 |
| 0 | NA | NA | NA | NA | NA | Lce3b | late cornified envelope 3B | 0 | 0 | 0 | 0 | 0 | 0 | 0 | 0 | 0 | 0 | 0 |
| 0 | NA | NA | NA | NA | NA | Lce3c | late cornified envelope 3C | 0 | 0 | 0 | 0 | 0 | 0 | 0 | 0 | 0 | 0 | 0 |
| 0 | NA | NA | NA | NA | NA | Gm24481 | | 0 | 0 | 0 | 0 | 0 | 0 | 0 | 0 | 0 | 0 | 0 |

| | | | | | | | | | | | | | | | | | | |
|---|---|---|---|---|---|---|---|---|---|---|---|---|---|---|---|---|---|---|
| 0 | NA | NA | NA | NA | NA | Lce3d | late cornified envelope 3D | 0 | 0 | 0 | 0 | 0 | 0 | 0 | 0 | 0 | 0 | 0 |
| 0 | NA | NA | NA | NA | NA | Gm22577 | | 0 | 0 | 0 | 0 | 0 | 0 | 0 | 0 | 0 | 0 | 0 |
| 0 | NA | NA | NA | NA | NA | Lce3e | late cornified envelope 3E | 0 | 0 | 0 | 0 | 0 | 0 | 0 | 0 | 0 | 0 | 0 |
| 0 | NA | NA | NA | NA | NA | Lce3f | late cornified envelope 3F | 0 | 0 | 0 | 0 | 0 | 0 | 0 | 0 | 0 | 0 | 0 |
| 0 | NA | NA | NA | NA | NA | Crct1 | cysteine-rich C-terminal 1 | 0 | 0 | 0 | 0 | 0 | 0 | 0 | 0 | 0 | 0 | 0 |
| 0 | NA | NA | NA | NA | NA | Lce1m | late cornified envelope 1M | 0 | 0 | 0 | 0 | 0 | 0 | 0 | 0 | 0 | 0 | 0 |
| 0 | NA | NA | NA | NA | NA | Tdpoz8 | TD and POZ domain containing 8 | 0 | 0 | 0 | 0 | 0 | 0 | 0 | 0 | 0 | 0 | 0 |
| 0 | NA | NA | NA | NA | NA | Crnn | cornulin | 0 | 0 | 0 | 0 | 0 | 0 | 0 | 0 | 0 | 0 | 0 |
| 0 | NA | NA | NA | NA | NA | Gm37501 | | 0 | 0 | 0 | 0 | 0 | 0 | 0 | 0 | 0 | 0 | 0 |
| 0 | NA | NA | NA | NA | NA | Flg2 | filaggrin family member 2 | 0 | 0 | 0 | 0 | 0 | 0 | 0 | 0 | 0 | 0 | 0 |
| 0 | NA | NA | NA | NA | NA | Flg | filaggrin | 0 | 0 | 0 | 0 | 0 | 0 | 0 | 0 | 0 | 0 | 0 |
| 0 | NA | NA | NA | NA | NA | Gm37922 | | 0 | 0 | 0 | 0 | 0 | 0 | 0 | 0 | 0 | 0 | 0 |
| 0 | NA | NA | NA | NA | NA | Gm37913 | | 0 | 0 | 0 | 0 | 0 | 0 | 0 | 0 | 0 | 0 | 0 |
| 0 | NA | NA | NA | NA | NA | Gm37245 | | 0 | 0 | 0 | 0 | 0 | 0 | 0 | 0 | 0 | 0 | 0 |
| 0 | NA | NA | NA | NA | NA | Hrnr | hornerin | 0 | 0 | 0 | 0 | 0 | 0 | 0 | 0 | 0 | 0 | 0 |
| 0 | NA | NA | NA | NA | NA | Gm44353 | | 0 | 0 | 0 | 0 | 0 | 0 | 0 | 0 | 0 | 0 | 0 |
| 0 | NA | NA | NA | NA | NA | Gm37857 | | 0 | 0 | 0 | 0 | 0 | 0 | 0 | 0 | 0 | 0 | 0 |
| 0 | NA | NA | NA | NA | NA | Rptn | repetin | 0 | 0 | 0 | 0 | 0 | 0 | 0 | 0 | 0 | 0 | 0 |
| 0 | NA | NA | NA | NA | NA | Gm18296 | | 0 | 0 | 0 | 0 | 0 | 0 | 0 | 0 | 0 | 0 | 0 |
| 0 | NA | NA | NA | NA | NA | Gm22752 | | 0 | 0 | 0 | 0 | 0 | 0 | 0 | 0 | 0 | 0 | 0 |
| 0 | NA | NA | NA | NA | NA | Tchhl1 | trichohyalin-like 1 | 0 | 0 | 0 | 0 | 0 | 0 | 0 | 0 | 0 | 0 | 0 |
| 0 | NA | NA | NA | NA | NA | Gm29950 | | 0 | 0 | 0 | 0 | 0 | 0 | 0 | 0 | 0 | 0 | 0 |
| 0 | NA | NA | NA | NA | NA | Gm5541 | | 0 | 0 | 0 | 0 | 0 | 0 | 0 | 0 | 0 | 0 | 0 |
| 0 | NA | NA | NA | NA | NA | Gm38207 | | 0 | 0 | 0 | 0 | 0 | 0 | 0 | 0 | 0 | 0 | 0 |
| 0 | NA | NA | NA | NA | NA | Tdpoz2 | TD and POZ domain containing 2 | 0 | 0 | 0 | 0 | 0 | 0 | 0 | 0 | 0 | 0 | 0 |
| 0 | NA | NA | NA | NA | NA | Tdpoz1 | TD and POZ domain containing 1 | 0 | 0 | 0 | 0 | 0 | 0 | 0 | 0 | 0 | 0 | 0 |
| 0 | NA | NA | NA | NA | NA | Gm18433 | | 0 | 0 | 0 | 0 | 0 | 0 | 0 | 0 | 0 | 0 | 0 |
| 0 | NA | NA | NA | NA | NA | Tdpoz6 | | 0 | 0 | 0 | 0 | 0 | 0 | 0 | 0 | 0 | 0 | 0 |
| 0 | NA | NA | NA | NA | NA | Gm18433 | | 0 | 0 | 0 | 0 | 0 | 0 | 0 | 0 | 0 | 0 | 0 |
| 0 | NA | NA | NA | NA | NA | Gm38086 | | 0 | 0 | 0 | 0 | 0 | 0 | 0 | 0 | 0 | 0 | 0 |
| 0 | NA | NA | NA | NA | NA | Gm5852 | | 0 | 0 | 0 | 0 | 0 | 0 | 0 | 0 | 0 | 0 | 0 |
| 0 | NA | NA | NA | NA | NA | Gm5773 | | 0 | 0 | 0 | 0 | 0 | 0 | 0 | 0 | 0 | 0 | 0 |
| 0 | NA | NA | NA | NA | NA | Tdpoz4 | TD and POZ domain containing 4 | 0 | 0 | 0 | 0 | 0 | 0 | 0 | 0 | 0 | 0 | 0 |
| 0 | NA | NA | NA | NA | NA | Tdpoz3 | TD and POZ domain containing 3 | 0 | 0 | 0 | 0 | 0 | 0 | 0 | 0 | 0 | 0 | 0 |
| 0 | NA | NA | NA | NA | NA | Gm37430 | | 0 | 0 | 0 | 0 | 0 | 0 | 0 | 0 | 0 | 0 | 0 |
| 0 | NA | NA | NA | NA | NA | Gm9113 | | 0 | 0 | 0 | 0 | 0 | 0 | 0 | 0 | 0 | 0 | 0 |
| 0 | NA | NA | NA | NA | NA | Gm9116 | | 0 | 0 | 0 | 0 | 0 | 0 | 0 | 0 | 0 | 0 | 0 |
| 0 | NA | NA | NA | NA | NA | Gm37185 | | 0 | 0 | 0 | 0 | 0 | 0 | 0 | 0 | 0 | 0 | 0 |
| 0 | NA | NA | NA | NA | NA | Gm9117 | | 0 | 0 | 0 | 0 | 0 | 0 | 0 | 0 | 0 | 0 | 0 |
| 0 | NA | NA | NA | NA | NA | Gm37104 | | 0 | 0 | 0 | 0 | 0 | 0 | 0 | 0 | 0 | 0 | 0 |
| 0 | NA | NA | NA | NA | NA | Tdpoz5 | TD and POZ domain containing 5 | 0 | 0 | 0 | 0 | 0 | 0 | 0 | 0 | 0 | 0 | 0 |
| 0 | NA | NA | NA | NA | NA | Gm37807 | | 0 | 0 | 0 | 0 | 0 | 0 | 0 | 0 | 0 | 0 | 0 |
| 0 | NA | NA | NA | NA | NA | Gm37481 | | 0 | 0 | 0 | 0 | 0 | 0 | 0 | 0 | 0 | 0 | 0 |
| 0 | NA | NA | NA | NA | NA | Gm9121 | | 0 | 0 | 0 | 0 | 0 | 0 | 0 | 0 | 0 | 0 | 0 |
| 0 | NA | NA | NA | NA | NA | Gm9124 | | 0 | 0 | 0 | 0 | 0 | 0 | 0 | 0 | 0 | 0 | 0 |
| 0 | NA | NA | NA | NA | NA | Gm37085 | | 0 | 0 | 0 | 0 | 0 | 0 | 0 | 0 | 0 | 0 | 0 |
| 0 | NA | NA | NA | NA | NA | Tdpoz9 | predicted gene 9125 | 0 | 0 | 0 | 0 | 0 | 0 | 0 | 0 | 0 | 0 | 0 |
| 0 | NA | NA | NA | NA | NA | Gm37081 | | 0 | 0 | 0 | 0 | 0 | 0 | 0 | 0 | 0 | 0 | 0 |
| 0 | NA | NA | NA | NA | NA | Tdpoz7 | Td and POZ domain containing 7 | 0 | 0 | 0 | 0 | 0 | 0 | 0 | 0 | 0 | 0 | 0 |
| 0 | NA | NA | NA | NA | NA | Gm37778 | | 0 | 0 | 0 | 0 | 0 | 0 | 0 | 0 | 0 | 0 | 0 |
| 0 | NA | NA | NA | NA | NA | Gm37155 | | 0 | 0 | 0 | 0 | 0 | 0 | 0 | 0 | 0 | 0 | 0 |
| 0 | NA | NA | NA | NA | NA | Gm5542 | | 0 | 0 | 0 | 0 | 0 | 0 | 0 | 0 | 0 | 0 | 0 |
| 0 | NA | NA | NA | NA | NA | Gm9131 | | 0 | 0 | 0 | 0 | 0 | 0 | 0 | 0 | 0 | 0 | 0 |
| 0 | NA | NA | NA | NA | NA | Gm37882 | | 0 | 0 | 0 | 0 | 0 | 0 | 0 | 0 | 0 | 0 | 0 |
| 0 | NA | NA | NA | NA | NA | Gm5543 | | 0 | 0 | 0 | 0 | 0 | 0 | 0 | 0 | 0 | 0 | 0 |
| 0 | NA | NA | NA | NA | NA | Gm37001 | | 0 | 0 | 0 | 0 | 0 | 0 | 0 | 0 | 0 | 0 | 0 |
| 0 | NA | NA | NA | NA | NA | Gm16560 | | 0 | 0 | 0 | 0 | 0 | 0 | 0 | 0 | 0 | 0 | 0 |
| 0 | NA | NA | NA | NA | NA | Spopfm2 | speckle-type BTB/POZ protein family men | 0 | 0 | 0 | 0 | 0 | 0 | 0 | 0 | 0 | 0 | 0 |
| 0 | NA | NA | NA | NA | NA | Spopfm3 | | 0 | 0 | 0 | 0 | 0 | 0 | 0 | 0 | 0 | 0 | 0 |
| 0 | NA | NA | NA | NA | NA | Gm16584 | | 0 | 0 | 0 | 0 | 0 | 0 | 0 | 0 | 0 | 0 | 0 |

| | | | | | | | | | | | | | | | | | | |
|---|---|---|---|---|---|---|---|---|---|---|---|---|---|---|---|---|---|---|
| 0 | NA | NA | NA | NA | NA | Gm4982 | | 0 | 0 | 0 | 0 | 0 | 0 | 0 | 0 | 0 | 0 | 0 |
| 0 | NA | NA | NA | NA | NA | Gm9141 | | 0 | 0 | 0 | 0 | 0 | 0 | 0 | 0 | 0 | 0 | 0 |
| 0 | NA | NA | NA | NA | NA | Spopfm1 | | 0 | 0 | 0 | 0 | 0 | 0 | 0 | 0 | 0 | 0 | 0 |
| 0 | NA | NA | NA | NA | NA | Gm43743 | | 0 | 0 | 0 | 0 | 0 | 0 | 0 | 0 | 0 | 0 | 0 |
| 0 | NA | NA | NA | NA | NA | Them5 | thioesterase superfamily member 5 | 0 | 0 | 0 | 0 | 0 | 0 | 0 | 0 | 0 | 0 | 0 |
| 0 | NA | NA | NA | NA | NA | Gm38411 | | 0 | 0 | 0 | 0 | 0 | 0 | 0 | 0 | 0 | 0 | 0 |
| 0 | NA | NA | NA | NA | NA | Gm43401 | | 0 | 0 | 0 | 0 | 0 | 0 | 0 | 0 | 0 | 0 | 0 |
| 0 | NA | NA | NA | NA | NA | Gm42464 | | 0 | 0 | 0 | 0 | 0 | 0 | 0 | 0 | 0 | 0 | 0 |
| 0 | NA | NA | NA | NA | NA | Gm30023 | | 0 | 0 | 0 | 0 | 0 | 0 | 0 | 0 | 0 | 0 | 0 |
| 0 | NA | NA | NA | NA | NA | Gm10972 | | 0 | 0 | 0 | 0 | 0 | 0 | 0 | 0 | 0 | 0 | 0 |
| 0 | NA | NA | NA | NA | NA | Gm43711 | | 0 | 0 | 0 | 0 | 0 | 0 | 0 | 0 | 0 | 0 | 0 |
| 0 | NA | NA | NA | NA | NA | Gm15263 | | 0 | 0 | 0 | 0 | 0 | 0 | 0 | 0 | 0 | 0 | 0 |
| 0 | NA | NA | NA | NA | NA | Gm26279 | | 0 | 0 | 0 | 0 | 0 | 0 | 0 | 0 | 0 | 0 | 0 |
| 0 | NA | NA | NA | NA | NA | Gm24920 | | 0 | 0 | 0 | 0 | 0 | 0 | 0 | 0 | 0 | 0 | 0 |
| 0 | NA | NA | NA | NA | NA | Gm22301 | | 0 | 0 | 0 | 0 | 0 | 0 | 0 | 0 | 0 | 0 | 0 |
| 0 | NA | NA | NA | NA | NA | Mir8099-1 | | 0 | 0 | 0 | 0 | 0 | 0 | 0 | 0 | 0 | 0 | 0 |
| 0 | NA | NA | NA | NA | NA | Gm42578 | | 0 | 0 | 0 | 0 | 0 | 0 | 0 | 0 | 0 | 0 | 0 |
| 0 | NA | NA | NA | NA | NA | 4930558C23Rik | | 0 | 0 | 0 | 0 | 0 | 0 | 0 | 0 | 0 | 0 | 0 |
| 0 | NA | NA | NA | NA | NA | Hormad1 | HORMA domain containing 1 | 0 | 0 | 0 | 0 | 0 | 0 | 0 | 0 | 0 | 0 | 0 |
| 0 | NA | NA | NA | NA | NA | Mir7014 | | 0 | 0 | 0 | 0 | 0 | 0 | 0 | 0 | 0 | 0 | 0 |
| 0 | NA | NA | NA | NA | NA | Gm43436 | | 0 | 0 | 0 | 0 | 0 | 0 | 0 | 0 | 0 | 0 | 0 |
| 0 | NA | NA | NA | NA | NA | Gm17815 | | 0 | 0 | 0 | 0 | 0 | 0 | 0 | 0 | 0 | 0 | 0 |
| 0 | NA | NA | NA | NA | NA | Gm26594 | | 0 | 0 | 0 | 0 | 0 | 0 | 0 | 0 | 0 | 0 | 0 |
| 0 | NA | NA | NA | NA | NA | Car14 | carbonic anhydrase 14 | 0 | 0 | 0 | 0 | 0 | 0 | 0 | 0 | 0 | 0 | 0 |
| 0 | NA | NA | NA | NA | NA | Gm20940 | | 0 | 0 | 0 | 0 | 0 | 0 | 0 | 0 | 0 | 0 | 0 |
| 0 | NA | NA | NA | NA | NA | Gm43554 | | 0 | 0 | 0 | 0 | 0 | 0 | 0 | 0 | 0 | 0 | 0 |
| 0 | NA | NA | NA | NA | NA | Gm47311 | | 0 | 0 | 0 | 0 | 0 | 0 | 0 | 0 | 0 | 0 | 0 |
| 0 | NA | NA | NA | NA | NA | H2ac18 | H2A clustered histone 18 | 0 | 0 | 0 | 0 | 0 | 0 | 0 | 0 | 0 | 0 | 0 |
| 0 | NA | NA | NA | NA | NA | Gm47313 | | 0 | 0 | 0 | 0 | 0 | 0 | 0 | 0 | 0 | 0 | 0 |
| 0 | NA | NA | NA | NA | NA | H3c14 | H3 clustered histone 14 | 0 | 0 | 0 | 0 | 0 | 0 | 0 | 0 | 0 | 0 | 0 |
| 0 | NA | NA | NA | NA | NA | Gm24830 | | 0 | 0 | 0 | 0 | 0 | 0 | 0 | 0 | 0 | 0 | 0 |
| 0 | NA | NA | NA | NA | NA | Platr30 | | 0 | 0 | 0 | 0 | 0 | 0 | 0 | 0 | 0 | 0 | 0 |
| 0 | NA | NA | NA | NA | NA | Gm26553 | | 0 | 0 | 0 | 0 | 0 | 0 | 0 | 0 | 0 | 0 | 0 |
| 0 | NA | NA | NA | NA | NA | Rnu1b1 | | 0 | 0 | 0 | 0 | 0 | 0 | 0 | 0 | 0 | 0 | 0 |
| 0 | NA | NA | NA | NA | NA | Rnu1b2 | | 0 | 0 | 0 | 0 | 0 | 0 | 0 | 0 | 0 | 0 | 0 |
| 0 | NA | NA | NA | NA | NA | 4930477E14Rik | | 0 | 0 | 0 | 0 | 0 | 0 | 0 | 0 | 0 | 0 | 0 |
| 0 | NA | NA | NA | NA | NA | Gm26654 | | 0 | 0 | 0 | 0 | 0 | 0 | 0 | 0 | 0 | 0 | 0 |
| 0 | NA | NA | NA | NA | NA | Terc | | 0 | 0 | 0 | 0 | 0 | 0 | 0 | 0 | 0 | 0 | 0 |
| 0 | NA | NA | NA | NA | NA | Terc | | 0 | 0 | 0 | 0 | 0 | 0 | 0 | 0 | 0 | 0 | 0 |
| 0 | NA | NA | NA | NA | NA | Gm24136 | | 0 | 0 | 0 | 0 | 0 | 0 | 0 | 0 | 0 | 0 | 0 |
| 0 | NA | NA | NA | NA | NA | BC107364 | cDNA sequence BC107364 | 0 | 0 | 0 | 0 | 0 | 0 | 0 | 0 | 0 | 0 | 0 |
| 0 | NA | NA | NA | NA | NA | Gm26232 | | 0 | 0 | 0 | 0 | 0 | 0 | 0 | 0 | 0 | 0 | 0 |
| 0 | NA | NA | NA | NA | NA | Gm10685 | | 0 | 0 | 0 | 0 | 0 | 0 | 0 | 0 | 0 | 0 | 0 |
| 0 | NA | NA | NA | NA | NA | Rnu1b6 | | 0 | 0 | 0 | 0 | 0 | 0 | 0 | 0 | 0 | 0 | 0 |
| 0 | NA | NA | NA | NA | NA | Gm43534 | | 0 | 0 | 0 | 0 | 0 | 0 | 0 | 0 | 0 | 0 | 0 |
| 0 | NA | NA | NA | NA | NA | Gm28001 | | 0 | 0 | 0 | 0 | 0 | 0 | 0 | 0 | 0 | 0 | 0 |
| 0 | NA | NA | NA | NA | NA | Gm25890 | | 0 | 0 | 0 | 0 | 0 | 0 | 0 | 0 | 0 | 0 | 0 |
| 0 | NA | NA | NA | NA | NA | Gm22614 | | 0 | 0 | 0 | 0 | 0 | 0 | 0 | 0 | 0 | 0 | 0 |
| 0 | NA | NA | NA | NA | NA | Cd160 | CD160 antigen | 0 | 0 | 0 | 0 | 0 | 0 | 0 | 0 | 0 | 0 | 0 |
| 0 | NA | NA | NA | NA | NA | 4930442L01Rik | | 0 | 0 | 0 | 0 | 0 | 0 | 0 | 0 | 0 | 0 | 0 |
| 0 | NA | NA | NA | NA | NA | Gja8 | gap junction protein, alpha 8 | 0 | 0 | 0 | 0 | 0 | 0 | 0 | 0 | 0 | 0 | 0 |
| 0 | NA | NA | NA | NA | NA | 9230114J08Rik | | 0 | 0 | 0 | 0 | 0 | 0 | 0 | 0 | 0 | 0 | 0 |
| 0 | NA | NA | NA | NA | NA | 4930573H18Rik | | 0 | 0 | 0 | 0 | 0 | 0 | 0 | 0 | 0 | 0 | 0 |
| 0 | NA | NA | NA | NA | NA | Gm43222 | | 0 | 0 | 0 | 0 | 0 | 0 | 0 | 0 | 0 | 0 | 0 |
| 0 | NA | NA | NA | NA | NA | Gm24435 | | 0 | 0 | 0 | 0 | 0 | 0 | 0 | 0 | 0 | 0 | 0 |
| 0 | NA | NA | NA | NA | NA | Gm42508 | | 0 | 0 | 0 | 0 | 0 | 0 | 0 | 0 | 0 | 0 | 0 |
| 0 | NA | NA | NA | NA | NA | Olfr1402 | olfactory receptor 1402 | 0 | 0 | 0 | 0 | 0 | 0 | 0 | 0 | 0 | 0 | 0 |
| 0 | NA | NA | NA | NA | NA | Gm17651 | | 0 | 0 | 0 | 0 | 0 | 0 | 0 | 0 | 0 | 0 | 0 |
| 0 | NA | NA | NA | NA | NA | Mir7225 | | 0 | 0 | 0 | 0 | 0 | 0 | 0 | 0 | 0 | 0 | 0 |
| 0 | NA | NA | NA | NA | NA | Gm31305 | | 0 | 0 | 0 | 0 | 0 | 0 | 0 | 0 | 0 | 0 | 0 |

| | | | | | | | | | | | | | | | | | |
|---|---|---|---|---|---|---|---|---|---|---|---|---|---|---|---|---|---|
| 0 | NA | NA | NA | NA | NA | Gm15999 | | 0 | 0 | 0 | 0 | 0 | 0 | 0 | 0 | 0 | 0 | 0 |
| 0 | NA | NA | NA | NA | NA | Gm42819 | | 0 | 0 | 0 | 0 | 0 | 0 | 0 | 0 | 0 | 0 | 0 |
| 0 | NA | NA | NA | NA | NA | Gm42820 | | 0 | 0 | 0 | 0 | 0 | 0 | 0 | 0 | 0 | 0 | 0 |
| 0 | NA | NA | NA | NA | NA | Adam30 | a disintegrin and metallopeptidase domai | 0 | 0 | 0 | 0 | 0 | 0 | 0 | 0 | 0 | 0 | 0 |
| 0 | NA | NA | NA | NA | NA | Gm9771 | | 0 | 0 | 0 | 0 | 0 | 0 | 0 | 0 | 0 | 0 | 0 |
| 0 | NA | NA | NA | NA | NA | Gm42821 | | 0 | 0 | 0 | 0 | 0 | 0 | 0 | 0 | 0 | 0 | 0 |
| 0 | NA | NA | NA | NA | NA | Gm43189 | | 0 | 0 | 0 | 0 | 0 | 0 | 0 | 0 | 0 | 0 | 0 |
| 0 | NA | NA | NA | NA | NA | Gm4450 | predicted gene 4450 | 0 | 0 | 0 | 0 | 0 | 0 | 0 | 0 | 0 | 0 | 0 |
| 0 | NA | NA | NA | NA | NA | Gm43190 | | 0 | 0 | 0 | 0 | 0 | 0 | 0 | 0 | 0 | 0 | 0 |
| 0 | NA | NA | NA | NA | NA | Gm19210 | | 0 | 0 | 0 | 0 | 0 | 0 | 0 | 0 | 0 | 0 | 0 |
| 0 | NA | NA | NA | NA | NA | Hsd3b4 | hydroxy-delta-5-steroid dehydrogenase, 3 | 0 | 0 | 0 | 0 | 0 | 0 | 0 | 0 | 0 | 0 | 0 |
| 0 | NA | NA | NA | NA | NA | Gm19211 | | 0 | 0 | 0 | 0 | 0 | 0 | 0 | 0 | 0 | 0 | 0 |
| 0 | NA | NA | NA | NA | NA | Gm10681 | predicted gene 10681 | 0 | 0 | 0 | 0 | 0 | 0 | 0 | 0 | 0 | 0 | 0 |
| 0 | NA | NA | NA | NA | NA | Gm9678 | | 0 | 0 | 0 | 0 | 0 | 0 | 0 | 0 | 0 | 0 | 0 |
| 0 | NA | NA | NA | NA | NA | Hsd3b5 | hydroxy-delta-5-steroid dehydrogenase, 3 | 0 | 0 | 0 | 0 | 0 | 0 | 0 | 0 | 0 | 0 | 0 |
| 0 | NA | NA | NA | NA | NA | Gm12399 | | 0 | 0 | 0 | 0 | 0 | 0 | 0 | 0 | 0 | 0 | 0 |
| 0 | NA | NA | NA | NA | NA | Gm12400 | | 0 | 0 | 0 | 0 | 0 | 0 | 0 | 0 | 0 | 0 | 0 |
| 0 | NA | NA | NA | NA | NA | Hsd3b6 | hydroxy-delta-5-steroid dehydrogenase, 3 | 0 | 0 | 0 | 0 | 0 | 0 | 0 | 0 | 0 | 0 | 0 |
| 0 | NA | NA | NA | NA | NA | Gm12427 | | 0 | 0 | 0 | 0 | 0 | 0 | 0 | 0 | 0 | 0 | 0 |
| 0 | NA | NA | NA | NA | NA | Gm12431 | | 0 | 0 | 0 | 0 | 0 | 0 | 0 | 0 | 0 | 0 | 0 |
| 0 | NA | NA | NA | NA | NA | Gm12432 | | 0 | 0 | 0 | 0 | 0 | 0 | 0 | 0 | 0 | 0 | 0 |
| 0 | NA | NA | NA | NA | NA | Gm12433 | | 0 | 0 | 0 | 0 | 0 | 0 | 0 | 0 | 0 | 0 | 0 |
| 0 | NA | NA | NA | NA | NA | Hsd3b1 | hydroxy-delta-5-steroid dehydrogenase, 3 | 0 | 0 | 0 | 0 | 0 | 0 | 0 | 0 | 0 | 0 | 0 |
| 0 | NA | NA | NA | NA | NA | Gm12440 | | 0 | 0 | 0 | 0 | 0 | 0 | 0 | 0 | 0 | 0 | 0 |
| 0 | NA | NA | NA | NA | NA | Gm12441 | | 0 | 0 | 0 | 0 | 0 | 0 | 0 | 0 | 0 | 0 | 0 |
| 0 | NA | NA | NA | NA | NA | Gm22341 | | 0 | 0 | 0 | 0 | 0 | 0 | 0 | 0 | 0 | 0 | 0 |
| 0 | NA | NA | NA | NA | NA | Gm24216 | | 0 | 0 | 0 | 0 | 0 | 0 | 0 | 0 | 0 | 0 | 0 |
| 0 | NA | NA | NA | NA | NA | Gm18982 | | 0 | 0 | 0 | 0 | 0 | 0 | 0 | 0 | 0 | 0 | 0 |
| 0 | NA | NA | NA | NA | NA | Mir6481 | | 0 | 0 | 0 | 0 | 0 | 0 | 0 | 0 | 0 | 0 | 0 |
| 0 | NA | NA | NA | NA | NA | Gm43120 | | 0 | 0 | 0 | 0 | 0 | 0 | 0 | 0 | 0 | 0 | 0 |
| 0 | NA | NA | NA | NA | NA | Gm12449 | | 0 | 0 | 0 | 0 | 0 | 0 | 0 | 0 | 0 | 0 | 0 |
| 0 | NA | NA | NA | NA | NA | Gm12448 | | 0 | 0 | 0 | 0 | 0 | 0 | 0 | 0 | 0 | 0 | 0 |
| 0 | NA | NA | NA | NA | NA | Gm22505 | | 0 | 0 | 0 | 0 | 0 | 0 | 0 | 0 | 0 | 0 | 0 |
| 0 | NA | NA | NA | NA | NA | Rpl36-ps7 | | 0 | 0 | 0 | 0 | 0 | 0 | 0 | 0 | 0 | 0 | 0 |
| 0 | NA | NA | NA | NA | NA | Tra2b-ps | | 0 | 0 | 0 | 0 | 0 | 0 | 0 | 0 | 0 | 0 | 0 |
| 0 | NA | NA | NA | NA | NA | Spag17os | | 0 | 0 | 0 | 0 | 0 | 0 | 0 | 0 | 0 | 0 | 0 |
| 0 | NA | NA | NA | NA | NA | Gm25802 | | 0 | 0 | 0 | 0 | 0 | 0 | 0 | 0 | 0 | 0 | 0 |
| 0 | NA | NA | NA | NA | NA | Gm12485 | | 0 | 0 | 0 | 0 | 0 | 0 | 0 | 0 | 0 | 0 | 0 |
| 0 | NA | NA | NA | NA | NA | Gm23465 | | 0 | 0 | 0 | 0 | 0 | 0 | 0 | 0 | 0 | 0 | 0 |
| 0 | NA | NA | NA | NA | NA | Gm12488 | | 0 | 0 | 0 | 0 | 0 | 0 | 0 | 0 | 0 | 0 | 0 |
| 0 | NA | NA | NA | NA | NA | Gm43465 | | 0 | 0 | 0 | 0 | 0 | 0 | 0 | 0 | 0 | 0 | 0 |
| 0 | NA | NA | NA | NA | NA | Gm12486 | | 0 | 0 | 0 | 0 | 0 | 0 | 0 | 0 | 0 | 0 | 0 |
| 0 | NA | NA | NA | NA | NA | Gm12490 | | 0 | 0 | 0 | 0 | 0 | 0 | 0 | 0 | 0 | 0 | 0 |
| 0 | NA | NA | NA | NA | NA | Gm10355 | | 0 | 0 | 0 | 0 | 0 | 0 | 0 | 0 | 0 | 0 | 0 |
| 0 | NA | NA | NA | NA | NA | Gm43467 | | 0 | 0 | 0 | 0 | 0 | 0 | 0 | 0 | 0 | 0 | 0 |
| 0 | NA | NA | NA | NA | NA | Gm42939 | | 0 | 0 | 0 | 0 | 0 | 0 | 0 | 0 | 0 | 0 | 0 |
| 0 | NA | NA | NA | NA | NA | Gm42938 | | 0 | 0 | 0 | 0 | 0 | 0 | 0 | 0 | 0 | 0 | 0 |
| 0 | NA | NA | NA | NA | NA | Gm42538 | | 0 | 0 | 0 | 0 | 0 | 0 | 0 | 0 | 0 | 0 | 0 |
| 0 | NA | NA | NA | NA | NA | Gm43135 | | 0 | 0 | 0 | 0 | 0 | 0 | 0 | 0 | 0 | 0 | 0 |
| 0 | NA | NA | NA | NA | NA | Gm43134 | | 0 | 0 | 0 | 0 | 0 | 0 | 0 | 0 | 0 | 0 | 0 |
| 0 | NA | NA | NA | NA | NA | Gm43243 | | 0 | 0 | 0 | 0 | 0 | 0 | 0 | 0 | 0 | 0 | 0 |
| 0 | NA | NA | NA | NA | NA | Gm18894 | | 0 | 0 | 0 | 0 | 0 | 0 | 0 | 0 | 0 | 0 | 0 |
| 0 | NA | NA | NA | NA | NA | Gm43244 | | 0 | 0 | 0 | 0 | 0 | 0 | 0 | 0 | 0 | 0 | 0 |
| 0 | NA | NA | NA | NA | NA | Gm16160 | | 0 | 0 | 0 | 0 | 0 | 0 | 0 | 0 | 0 | 0 | 0 |
| 0 | NA | NA | NA | NA | NA | Gm43241 | | 0 | 0 | 0 | 0 | 0 | 0 | 0 | 0 | 0 | 0 | 0 |
| 0 | NA | NA | NA | NA | NA | Gm43242 | | 0 | 0 | 0 | 0 | 0 | 0 | 0 | 0 | 0 | 0 | 0 |
| 0 | NA | NA | NA | NA | NA | Gm42679 | | 0 | 0 | 0 | 0 | 0 | 0 | 0 | 0 | 0 | 0 | 0 |
| 0 | NA | NA | NA | NA | NA | Gm42678 | | 0 | 0 | 0 | 0 | 0 | 0 | 0 | 0 | 0 | 0 | 0 |
| 0 | NA | NA | NA | NA | NA | Gm42677 | | 0 | 0 | 0 | 0 | 0 | 0 | 0 | 0 | 0 | 0 | 0 |
| 0 | NA | NA | NA | NA | NA | Gm42682 | | 0 | 0 | 0 | 0 | 0 | 0 | 0 | 0 | 0 | 0 | 0 |

| | | | | | | | | | | | | | | | | | | |
|---|---|---|---|---|---|---|---|---|---|---|---|---|---|---|---|---|---|---|
| 0 | NA | NA | NA | NA | NA | Tshb | thyroid stimulating hormone, beta subunit | 0 | 0 | 0 | 0 | 0 | 0 | 0 | 0 | 0 | 0 | 0 |
| 0 | NA | NA | NA | NA | NA | Gm42681 | | 0 | 0 | 0 | 0 | 0 | 0 | 0 | 0 | 0 | 0 | 0 |
| 0 | NA | NA | NA | NA | NA | Gm42816 | | 0 | 0 | 0 | 0 | 0 | 0 | 0 | 0 | 0 | 0 | 0 |
| 0 | NA | NA | NA | NA | NA | Gm33952 | | 0 | 0 | 0 | 0 | 0 | 0 | 0 | 0 | 0 | 0 | 0 |
| 0 | NA | NA | NA | NA | NA | Gm33866 | | 0 | 0 | 0 | 0 | 0 | 0 | 0 | 0 | 0 | 0 | 0 |
| 0 | NA | NA | NA | NA | NA | Nr1h5 | nuclear receptor subfamily 1, group H, me | 0 | 0 | 0 | 0 | 0 | 0 | 0 | 0 | 0 | 0 | 0 |
| 0 | NA | NA | NA | NA | NA | Gm22826 | | 0 | 0 | 0 | 0 | 0 | 0 | 0 | 0 | 0 | 0 | 0 |
| 0 | NA | NA | NA | NA | NA | Ampd1 | adenosine monophosphate deaminase 1 | 0 | 0 | 0 | 0 | 0 | 0 | 0 | 0 | 0 | 0 | 0 |
| 0 | NA | NA | NA | NA | NA | Gm23820 | | 0 | 0 | 0 | 0 | 0 | 0 | 0 | 0 | 0 | 0 | 0 |
| 0 | NA | NA | NA | NA | NA | Gm43063 | | 0 | 0 | 0 | 0 | 0 | 0 | 0 | 0 | 0 | 0 | 0 |
| 0 | NA | NA | NA | NA | NA | Gm27024 | | 0 | 0 | 0 | 0 | 0 | 0 | 0 | 0 | 0 | 0 | 0 |
| 0 | NA | NA | NA | NA | NA | 2410024N13Rik | | 0 | 0 | 0 | 0 | 0 | 0 | 0 | 0 | 0 | 0 | 0 |
| 0 | NA | NA | NA | NA | NA | Gm43060 | | 0 | 0 | 0 | 0 | 0 | 0 | 0 | 0 | 0 | 0 | 0 |
| 0 | NA | NA | NA | NA | NA | Gm25009 | | 0 | 0 | 0 | 0 | 0 | 0 | 0 | 0 | 0 | 0 | 0 |
| 0 | NA | NA | NA | NA | NA | Gm25199 | | 0 | 0 | 0 | 0 | 0 | 0 | 0 | 0 | 0 | 0 | 0 |
| 0 | NA | NA | NA | NA | NA | Gm43066 | | 0 | 0 | 0 | 0 | 0 | 0 | 0 | 0 | 0 | 0 | 0 |
| 0 | NA | NA | NA | NA | NA | Gm25325 | | 0 | 0 | 0 | 0 | 0 | 0 | 0 | 0 | 0 | 0 | 0 |
| 0 | NA | NA | NA | NA | NA | Gm15986 | | 0 | 0 | 0 | 0 | 0 | 0 | 0 | 0 | 0 | 0 | 0 |
| 0 | NA | NA | NA | NA | NA | Gm10964 | | 0 | 0 | 0 | 0 | 0 | 0 | 0 | 0 | 0 | 0 | 0 |
| 0 | NA | NA | NA | NA | NA | Gm15472 | | 0 | 0 | 0 | 0 | 0 | 0 | 0 | 0 | 0 | 0 | 0 |
| 0 | NA | NA | NA | NA | NA | Gm43065 | | 0 | 0 | 0 | 0 | 0 | 0 | 0 | 0 | 0 | 0 | 0 |
| 0 | NA | NA | NA | NA | NA | 9530097N15Rik | | 0 | 0 | 0 | 0 | 0 | 0 | 0 | 0 | 0 | 0 | 0 |
| 0 | NA | NA | NA | NA | NA | Gm23830 | | 0 | 0 | 0 | 0 | 0 | 0 | 0 | 0 | 0 | 0 | 0 |
| 0 | NA | NA | NA | NA | NA | Gm42700 | | 0 | 0 | 0 | 0 | 0 | 0 | 0 | 0 | 0 | 0 | 0 |
| 0 | NA | NA | NA | NA | NA | Gm17956 | | 0 | 0 | 0 | 0 | 0 | 0 | 0 | 0 | 0 | 0 | 0 |
| 0 | NA | NA | NA | NA | NA | Gm42702 | | 0 | 0 | 0 | 0 | 0 | 0 | 0 | 0 | 0 | 0 | 0 |
| 0 | NA | NA | NA | NA | NA | Gm42701 | | 0 | 0 | 0 | 0 | 0 | 0 | 0 | 0 | 0 | 0 | 0 |
| 0 | NA | NA | NA | NA | NA | Gm43258 | | 0 | 0 | 0 | 0 | 0 | 0 | 0 | 0 | 0 | 0 | 0 |
| 0 | NA | NA | NA | NA | NA | C030032O16Rik | | 0 | 0 | 0 | 0 | 0 | 0 | 0 | 0 | 0 | 0 | 0 |
| 0 | NA | NA | NA | NA | NA | Gm43259 | | 0 | 0 | 0 | 0 | 0 | 0 | 0 | 0 | 0 | 0 | 0 |
| 0 | NA | NA | NA | NA | NA | Gm43260 | | 0 | 0 | 0 | 0 | 0 | 0 | 0 | 0 | 0 | 0 | 0 |
| 0 | NA | NA | NA | NA | NA | Gm43256 | | 0 | 0 | 0 | 0 | 0 | 0 | 0 | 0 | 0 | 0 | 0 |
| 0 | NA | NA | NA | NA | NA | Gm23077 | | 0 | 0 | 0 | 0 | 0 | 0 | 0 | 0 | 0 | 0 | 0 |
| 0 | NA | NA | NA | NA | NA | Gm5546 | | 0 | 0 | 0 | 0 | 0 | 0 | 0 | 0 | 0 | 0 | 0 |
| 0 | NA | NA | NA | NA | NA | Gm38412 | | 0 | 0 | 0 | 0 | 0 | 0 | 0 | 0 | 0 | 0 | 0 |
| 0 | NA | NA | NA | NA | NA | Gm26091 | | 0 | 0 | 0 | 0 | 0 | 0 | 0 | 0 | 0 | 0 | 0 |
| 0 | NA | NA | NA | NA | NA | Gm43582 | | 0 | 0 | 0 | 0 | 0 | 0 | 0 | 0 | 0 | 0 | 0 |
| 0 | NA | NA | NA | NA | NA | Gm43585 | | 0 | 0 | 0 | 0 | 0 | 0 | 0 | 0 | 0 | 0 | 0 |
| 0 | NA | NA | NA | NA | NA | Gm29561 | | 0 | 0 | 0 | 0 | 0 | 0 | 0 | 0 | 0 | 0 | 0 |
| 0 | NA | NA | NA | NA | NA | Gm29560 | | 0 | 0 | 0 | 0 | 0 | 0 | 0 | 0 | 0 | 0 | 0 |
| 0 | NA | NA | NA | NA | NA | Gm24223 | | 0 | 0 | 0 | 0 | 0 | 0 | 0 | 0 | 0 | 0 | 0 |
| 0 | NA | NA | NA | NA | NA | Gm29572 | | 0 | 0 | 0 | 0 | 0 | 0 | 0 | 0 | 0 | 0 | 0 |
| 0 | NA | NA | NA | NA | NA | Gm43584 | | 0 | 0 | 0 | 0 | 0 | 0 | 0 | 0 | 0 | 0 | 0 |
| 0 | NA | NA | NA | NA | NA | Gm43846 | | 0 | 0 | 0 | 0 | 0 | 0 | 0 | 0 | 0 | 0 | 0 |
| 0 | NA | NA | NA | NA | NA | 4930564D02Ril | RIKEN cDNA 4930564D02 gene | 0 | 0 | 0 | 0 | 0 | 0 | 0 | 0 | 0 | 0 | 0 |
| 0 | NA | NA | NA | NA | NA | Gm43847 | | 0 | 0 | 0 | 0 | 0 | 0 | 0 | 0 | 0 | 0 | 0 |
| 0 | NA | NA | NA | NA | NA | Gm43331 | | 0 | 0 | 0 | 0 | 0 | 0 | 0 | 0 | 0 | 0 | 0 |
| 0 | NA | NA | NA | NA | NA | Tmigd3 | | 0 | 0 | 0 | 0 | 0 | 0 | 0 | 0 | 0 | 0 | 0 |
| 0 | NA | NA | NA | NA | NA | Gm42721 | | 0 | 0 | 0 | 0 | 0 | 0 | 0 | 0 | 0 | 0 | 0 |
| 0 | NA | NA | NA | NA | NA | Gm43709 | | 0 | 0 | 0 | 0 | 0 | 0 | 0 | 0 | 0 | 0 | 0 |
| 0 | NA | NA | NA | NA | NA | Gm6522 | | 0 | 0 | 0 | 0 | 0 | 0 | 0 | 0 | 0 | 0 | 0 |
| 0 | NA | NA | NA | NA | NA | Gm43212 | | 0 | 0 | 0 | 0 | 0 | 0 | 0 | 0 | 0 | 0 | 0 |
| 0 | NA | NA | NA | NA | NA | Gm38244 | | 0 | 0 | 0 | 0 | 0 | 0 | 0 | 0 | 0 | 0 | 0 |
| 0 | NA | NA | NA | NA | NA | Gm43070 | | 0 | 0 | 0 | 0 | 0 | 0 | 0 | 0 | 0 | 0 | 0 |
| 0 | NA | NA | NA | NA | NA | Gm38271 | | 0 | 0 | 0 | 0 | 0 | 0 | 0 | 0 | 0 | 0 | 0 |
| 0 | NA | NA | NA | NA | NA | Olfr266 | olfactory receptor 266 | 0 | 0 | 0 | 0 | 0 | 0 | 0 | 0 | 0 | 0 | 0 |
| 0 | NA | NA | NA | NA | NA | Gm35507 | | 0 | 0 | 0 | 0 | 0 | 0 | 0 | 0 | 0 | 0 | 0 |
| 0 | NA | NA | NA | NA | NA | Gm27008 | | 0 | 0 | 0 | 0 | 0 | 0 | 0 | 0 | 0 | 0 | 0 |
| 0 | NA | NA | NA | NA | NA | A930002I21Rik | | 0 | 0 | 0 | 0 | 0 | 0 | 0 | 0 | 0 | 0 | 0 |
| 0 | NA | NA | NA | NA | NA | Cym | chymosin | 0 | 0 | 0 | 0 | 0 | 0 | 0 | 0 | 0 | 0 | 0 |

| | | | | | | | | | | | | | | | | | | |
|---|---|---|---|---|---|---|---|---|---|---|---|---|---|---|---|---|---|---|
| 0 | NA | NA | NA | NA | NA | Gm5279 | | 0 | 0 | 0 | 0 | 0 | 0 | 0 | 0 | 0 | 0 | 0 |
| 0 | NA | NA | NA | NA | NA | Gm24383 | | 0 | 0 | 0 | 0 | 0 | 0 | 0 | 0 | 0 | 0 | 0 |
| 0 | NA | NA | NA | NA | NA | Ubl4b | ubiquitin-like 4B | 0 | 0 | 0 | 0 | 0 | 0 | 0 | 0 | 0 | 0 | 0 |
| 0 | NA | NA | NA | NA | NA | Alx3 | aristaless-like homeobox 3 | 0 | 0 | 0 | 0 | 0 | 0 | 0 | 0 | 0 | 0 | 0 |
| 0 | NA | NA | NA | NA | NA | Gm43233 | | 0 | 0 | 0 | 0 | 0 | 0 | 0 | 0 | 0 | 0 | 0 |
| 0 | NA | NA | NA | NA | NA | Gm36211 | | 0 | 0 | 0 | 0 | 0 | 0 | 0 | 0 | 0 | 0 | 0 |
| 0 | NA | NA | NA | NA | NA | 4930554G22Rik | | 0 | 0 | 0 | 0 | 0 | 0 | 0 | 0 | 0 | 0 | 0 |
| 0 | NA | NA | NA | NA | NA | Gm43745 | | 0 | 0 | 0 | 0 | 0 | 0 | 0 | 0 | 0 | 0 | 0 |
| 0 | NA | NA | NA | NA | NA | Gm9515 | | 0 | 0 | 0 | 0 | 0 | 0 | 0 | 0 | 0 | 0 | 0 |
| 0 | NA | NA | NA | NA | NA | Gm12489 | | 0 | 0 | 0 | 0 | 0 | 0 | 0 | 0 | 0 | 0 | 0 |
| 0 | NA | NA | NA | NA | NA | Gm12494 | | 0 | 0 | 0 | 0 | 0 | 0 | 0 | 0 | 0 | 0 | 0 |
| 0 | NA | NA | NA | NA | NA | Gm43746 | | 0 | 0 | 0 | 0 | 0 | 0 | 0 | 0 | 0 | 0 | 0 |
| 0 | NA | NA | NA | NA | NA | Gm12498 | | 0 | 0 | 0 | 0 | 0 | 0 | 0 | 0 | 0 | 0 | 0 |
| 0 | NA | NA | NA | NA | NA | Gm12499 | | 0 | 0 | 0 | 0 | 0 | 0 | 0 | 0 | 0 | 0 | 0 |
| 0 | NA | NA | NA | NA | NA | Gm25592 | | 0 | 0 | 0 | 0 | 0 | 0 | 0 | 0 | 0 | 0 | 0 |
| 0 | NA | NA | NA | NA | NA | Gm40123 | | 0 | 0 | 0 | 0 | 0 | 0 | 0 | 0 | 0 | 0 | 0 |
| 0 | NA | NA | NA | NA | NA | Gm22942 | | 0 | 0 | 0 | 0 | 0 | 0 | 0 | 0 | 0 | 0 | 0 |
| 0 | NA | NA | NA | NA | NA | Gm23336 | | 0 | 0 | 0 | 0 | 0 | 0 | 0 | 0 | 0 | 0 | 0 |
| 0 | NA | NA | NA | NA | NA | Gm22860 | | 0 | 0 | 0 | 0 | 0 | 0 | 0 | 0 | 0 | 0 | 0 |
| 0 | NA | NA | NA | NA | NA | Gm27244 | | 0 | 0 | 0 | 0 | 0 | 0 | 0 | 0 | 0 | 0 | 0 |
| 0 | NA | NA | NA | NA | NA | Gm43221 | | 0 | 0 | 0 | 0 | 0 | 0 | 0 | 0 | 0 | 0 | 0 |
| 0 | NA | NA | NA | NA | NA | Gm44384 | | 0 | 0 | 0 | 0 | 0 | 0 | 0 | 0 | 0 | 0 | 0 |
| 0 | NA | NA | NA | NA | NA | 4930408K08Rik | | 0 | 0 | 0 | 0 | 0 | 0 | 0 | 0 | 0 | 0 | 0 |
| 0 | NA | NA | NA | NA | NA | Slc25a54 | solute carrier family 25, member 54 | 0 | 0 | 0 | 0 | 0 | 0 | 0 | 0 | 0 | 0 | 0 |
| 0 | NA | NA | NA | NA | NA | Gm13865 | | 0 | 0 | 0 | 0 | 0 | 0 | 0 | 0 | 0 | 0 | 0 |
| 0 | NA | NA | NA | NA | NA | Gm25681 | | 0 | 0 | 0 | 0 | 0 | 0 | 0 | 0 | 0 | 0 | 0 |
| 0 | NA | NA | NA | NA | NA | Gm12535 | | 0 | 0 | 0 | 0 | 0 | 0 | 0 | 0 | 0 | 0 | 0 |
| 0 | NA | NA | NA | NA | NA | Gm43405 | | 0 | 0 | 0 | 0 | 0 | 0 | 0 | 0 | 0 | 0 | 0 |
| 0 | NA | NA | NA | NA | NA | Gm19185 | | 0 | 0 | 0 | 0 | 0 | 0 | 0 | 0 | 0 | 0 | 0 |
| 0 | NA | NA | NA | NA | NA | Gm43406 | | 0 | 0 | 0 | 0 | 0 | 0 | 0 | 0 | 0 | 0 | 0 |
| 0 | NA | NA | NA | NA | NA | Gm26076 | | 0 | 0 | 0 | 0 | 0 | 0 | 0 | 0 | 0 | 0 | 0 |
| 0 | NA | NA | NA | NA | NA | Gm43404 | | 0 | 0 | 0 | 0 | 0 | 0 | 0 | 0 | 0 | 0 | 0 |
| 0 | NA | NA | NA | NA | NA | Gm43407 | | 0 | 0 | 0 | 0 | 0 | 0 | 0 | 0 | 0 | 0 | 0 |
| 0 | NA | NA | NA | NA | NA | Gm9314 | | 0 | 0 | 0 | 0 | 0 | 0 | 0 | 0 | 0 | 0 | 0 |
| 0 | NA | NA | NA | NA | NA | Gm42905 | | 0 | 0 | 0 | 0 | 0 | 0 | 0 | 0 | 0 | 0 | 0 |
| 0 | NA | NA | NA | NA | NA | Gm43854 | | 0 | 0 | 0 | 0 | 0 | 0 | 0 | 0 | 0 | 0 | 0 |
| 0 | NA | NA | NA | NA | NA | Gm6602 | | 0 | 0 | 0 | 0 | 0 | 0 | 0 | 0 | 0 | 0 | 0 |
| 0 | NA | NA | NA | NA | NA | Gm25519 | | 0 | 0 | 0 | 0 | 0 | 0 | 0 | 0 | 0 | 0 | 0 |
| 0 | NA | NA | NA | NA | NA | Gm43853 | | 0 | 0 | 0 | 0 | 0 | 0 | 0 | 0 | 0 | 0 | 0 |
| 0 | NA | NA | NA | NA | NA | Gm9317 | | 0 | 0 | 0 | 0 | 0 | 0 | 0 | 0 | 0 | 0 | 0 |
| 0 | NA | NA | NA | NA | NA | Gm42526 | | 0 | 0 | 0 | 0 | 0 | 0 | 0 | 0 | 0 | 0 | 0 |
| 0 | NA | NA | NA | NA | NA | Utp14b-ps1 | | 0 | 0 | 0 | 0 | 0 | 0 | 0 | 0 | 0 | 0 | 0 |
| 0 | NA | NA | NA | NA | NA | Gm38395 | | 0 | 0 | 0 | 0 | 0 | 0 | 0 | 0 | 0 | 0 | 0 |
| 0 | NA | NA | NA | NA | NA | Amy2a4 | amylase 2a4 | 0 | 0 | 0 | 0 | 0 | 0 | 0 | 0 | 0 | 0 | 0 |
| 0 | NA | NA | NA | NA | NA | Amy2a3 | amylase 2a3 | 0 | 0 | 0 | 0 | 0 | 0 | 0 | 0 | 0 | 0 | 0 |
| 0 | NA | NA | NA | NA | NA | Amy2a2 | amylase 2a2 | 0 | 0 | 0 | 0 | 0 | 0 | 0 | 0 | 0 | 0 | 0 |
| 0 | NA | NA | NA | NA | NA | Gm18163 | | 0 | 0 | 0 | 0 | 0 | 0 | 0 | 0 | 0 | 0 | 0 |
| 0 | NA | NA | NA | NA | NA | Gm43749 | | 0 | 0 | 0 | 0 | 0 | 0 | 0 | 0 | 0 | 0 | 0 |
| 0 | NA | NA | NA | NA | NA | Gm23750 | | 0 | 0 | 0 | 0 | 0 | 0 | 0 | 0 | 0 | 0 | 0 |
| 0 | NA | NA | NA | NA | NA | Gm19147 | | 0 | 0 | 0 | 0 | 0 | 0 | 0 | 0 | 0 | 0 | 0 |
| 0 | NA | NA | NA | NA | NA | Gm18429 | | 0 | 0 | 0 | 0 | 0 | 0 | 0 | 0 | 0 | 0 | 0 |
| 0 | NA | NA | NA | NA | NA | Gm22337 | | 0 | 0 | 0 | 0 | 0 | 0 | 0 | 0 | 0 | 0 | 0 |
| 0 | NA | NA | NA | NA | NA | Gm25347 | | 0 | 0 | 0 | 0 | 0 | 0 | 0 | 0 | 0 | 0 | 0 |
| 0 | NA | NA | NA | NA | NA | Gm43669 | | 0 | 0 | 0 | 0 | 0 | 0 | 0 | 0 | 0 | 0 | 0 |
| 0 | NA | NA | NA | NA | NA | Gm42687 | | 0 | 0 | 0 | 0 | 0 | 0 | 0 | 0 | 0 | 0 | 0 |
| 0 | NA | NA | NA | NA | NA | Gm43107 | | 0 | 0 | 0 | 0 | 0 | 0 | 0 | 0 | 0 | 0 | 0 |
| 0 | NA | NA | NA | NA | NA | Gm43108 | | 0 | 0 | 0 | 0 | 0 | 0 | 0 | 0 | 0 | 0 | 0 |
| 0 | NA | NA | NA | NA | NA | Mir669n | | 0 | 0 | 0 | 0 | 0 | 0 | 0 | 0 | 0 | 0 | 0 |
| 0 | NA | NA | NA | NA | NA | Gm26344 | | 0 | 0 | 0 | 0 | 0 | 0 | 0 | 0 | 0 | 0 | 0 |
| 0 | NA | NA | NA | NA | NA | Gm43109 | | 0 | 0 | 0 | 0 | 0 | 0 | 0 | 0 | 0 | 0 | 0 |

| | | | | | | | | | | | | | | | | | |
|---|---|---|---|---|---|---|---|---|---|---|---|---|---|---|---|---|---|
| 0 | NA | NA | NA | NA | NA | 5330425B07Rik | 0 | 0 | 0 | 0 | 0 | 0 | 0 | 0 | 0 | 0 | 0 |
| 0 | NA | NA | NA | NA | NA | Gm26544 | 0 | 0 | 0 | 0 | 0 | 0 | 0 | 0 | 0 | 0 | 0 |
| 0 | NA | NA | NA | NA | NA | Gm29151 | 0 | 0 | 0 | 0 | 0 | 0 | 0 | 0 | 0 | 0 | 0 |
| 0 | NA | NA | NA | NA | NA | Gm22361 | 0 | 0 | 0 | 0 | 0 | 0 | 0 | 0 | 0 | 0 | 0 |
| 0 | NA | NA | NA | NA | NA | Gm26358 | 0 | 0 | 0 | 0 | 0 | 0 | 0 | 0 | 0 | 0 | 0 |
| 0 | NA | NA | NA | NA | NA | Gm43191 | 0 | 0 | 0 | 0 | 0 | 0 | 0 | 0 | 0 | 0 | 0 |
| 0 | NA | NA | NA | NA | NA | Gm43520 | 0 | 0 | 0 | 0 | 0 | 0 | 0 | 0 | 0 | 0 | 0 |
| 0 | NA | NA | NA | NA | NA | Gm43521 | 0 | 0 | 0 | 0 | 0 | 0 | 0 | 0 | 0 | 0 | 0 |
| 0 | NA | NA | NA | NA | NA | Gm29736 | 0 | 0 | 0 | 0 | 0 | 0 | 0 | 0 | 0 | 0 | 0 |
| 0 | NA | NA | NA | NA | NA | Gm43542 | 0 | 0 | 0 | 0 | 0 | 0 | 0 | 0 | 0 | 0 | 0 |
| 0 | NA | NA | NA | NA | NA | 4930455H04Rik | 0 | 0 | 0 | 0 | 0 | 0 | 0 | 0 | 0 | 0 | 0 |
| 0 | NA | NA | NA | NA | NA | Gm9332 | 0 | 0 | 0 | 0 | 0 | 0 | 0 | 0 | 0 | 0 | 0 |
| 0 | NA | NA | NA | NA | NA | 1700061I17Rik | 0 | 0 | 0 | 0 | 0 | 0 | 0 | 0 | 0 | 0 | 0 |
| 0 | NA | NA | NA | NA | NA | Gm18430 | 0 | 0 | 0 | 0 | 0 | 0 | 0 | 0 | 0 | 0 | 0 |
| 0 | NA | NA | NA | NA | NA | Gm19243 | 0 | 0 | 0 | 0 | 0 | 0 | 0 | 0 | 0 | 0 | 0 |
| 0 | NA | NA | NA | NA | NA | Gm42891 | 0 | 0 | 0 | 0 | 0 | 0 | 0 | 0 | 0 | 0 | 0 |
| 0 | NA | NA | NA | NA | NA | Gm22408 | 0 | 0 | 0 | 0 | 0 | 0 | 0 | 0 | 0 | 0 | 0 |
| 0 | NA | NA | NA | NA | NA | Gm24811 | 0 | 0 | 0 | 0 | 0 | 0 | 0 | 0 | 0 | 0 | 0 |
| 0 | NA | NA | NA | NA | NA | Gm4321 | 0 | 0 | 0 | 0 | 0 | 0 | 0 | 0 | 0 | 0 | 0 |
| 0 | NA | NA | NA | NA | NA | 4921521D15Rik | 0 | 0 | 0 | 0 | 0 | 0 | 0 | 0 | 0 | 0 | 0 |
| 0 | NA | NA | NA | NA | NA | Mir137 | 0 | 0 | 0 | 0 | 0 | 0 | 0 | 0 | 0 | 0 | 0 |
| 0 | NA | NA | NA | NA | NA | Gm24155 | 0 | 0 | 0 | 0 | 0 | 0 | 0 | 0 | 0 | 0 | 0 |
| 0 | NA | NA | NA | NA | NA | Gm9916 | 0 | 0 | 0 | 0 | 0 | 0 | 0 | 0 | 0 | 0 | 0 |
| 0 | NA | NA | NA | NA | NA | Gm32444 | 0 | 0 | 0 | 0 | 0 | 0 | 0 | 0 | 0 | 0 | 0 |
| 0 | NA | NA | NA | NA | NA | Gm43410 | 0 | 0 | 0 | 0 | 0 | 0 | 0 | 0 | 0 | 0 | 0 |
| 0 | NA | NA | NA | NA | NA | Gm23432 | 0 | 0 | 0 | 0 | 0 | 0 | 0 | 0 | 0 | 0 | 0 |
| 0 | NA | NA | NA | NA | NA | Gm42804 | 0 | 0 | 0 | 0 | 0 | 0 | 0 | 0 | 0 | 0 | 0 |
| 0 | NA | NA | NA | NA | NA | Gm24744 | 0 | 0 | 0 | 0 | 0 | 0 | 0 | 0 | 0 | 0 | 0 |
| 0 | NA | NA | NA | NA | NA | Gm26137 | 0 | 0 | 0 | 0 | 0 | 0 | 0 | 0 | 0 | 0 | 0 |
| 0 | NA | NA | NA | NA | NA | Gm18384 | 0 | 0 | 0 | 0 | 0 | 0 | 0 | 0 | 0 | 0 | 0 |
| 0 | NA | NA | NA | NA | NA | Gm23733 | 0 | 0 | 0 | 0 | 0 | 0 | 0 | 0 | 0 | 0 | 0 |
| 0 | NA | NA | NA | NA | NA | Gm25013 | 0 | 0 | 0 | 0 | 0 | 0 | 0 | 0 | 0 | 0 | 0 |
| 0 | NA | NA | NA | NA | NA | Gm42927 | 0 | 0 | 0 | 0 | 0 | 0 | 0 | 0 | 0 | 0 | 0 |
| 0 | NA | NA | NA | NA | NA | Gm43444 | 0 | 0 | 0 | 0 | 0 | 0 | 0 | 0 | 0 | 0 | 0 |
| 0 | NA | NA | NA | NA | NA | Gm42926 | 0 | 0 | 0 | 0 | 0 | 0 | 0 | 0 | 0 | 0 | 0 |
| 0 | NA | NA | NA | NA | NA | Gm10652 | 0 | 0 | 0 | 0 | 0 | 0 | 0 | 0 | 0 | 0 | 0 |
| 0 | NA | NA | NA | NA | NA | Gm30517 | 0 | 0 | 0 | 0 | 0 | 0 | 0 | 0 | 0 | 0 | 0 |
| 0 | NA | NA | NA | NA | NA | Gm26027 | 0 | 0 | 0 | 0 | 0 | 0 | 0 | 0 | 0 | 0 | 0 |
| 0 | NA | NA | NA | NA | NA | 4930432M17Ri RIKEN cDNA 4930432M17 gene | 0 | 0 | 0 | 0 | 0 | 0 | 0 | 0 | 0 | 0 | 0 |
| 0 | NA | NA | NA | NA | NA | Gm42593 | 0 | 0 | 0 | 0 | 0 | 0 | 0 | 0 | 0 | 0 | 0 |
| 0 | NA | NA | NA | NA | NA | Gm43157 | 0 | 0 | 0 | 0 | 0 | 0 | 0 | 0 | 0 | 0 | 0 |
| 0 | NA | NA | NA | NA | NA | Gm9361 | 0 | 0 | 0 | 0 | 0 | 0 | 0 | 0 | 0 | 0 | 0 |
| 0 | NA | NA | NA | NA | NA | Gm22903 | 0 | 0 | 0 | 0 | 0 | 0 | 0 | 0 | 0 | 0 | 0 |
| 0 | NA | NA | NA | NA | NA | Mir760 | 0 | 0 | 0 | 0 | 0 | 0 | 0 | 0 | 0 | 0 | 0 |
| 0 | NA | NA | NA | NA | NA | Gm42836 | 0 | 0 | 0 | 0 | 0 | 0 | 0 | 0 | 0 | 0 | 0 |
| 0 | NA | NA | NA | NA | NA | Gm25153 | 0 | 0 | 0 | 0 | 0 | 0 | 0 | 0 | 0 | 0 | 0 |
| 0 | NA | NA | NA | NA | NA | Mir7657 | 0 | 0 | 0 | 0 | 0 | 0 | 0 | 0 | 0 | 0 | 0 |
| 0 | NA | NA | NA | NA | NA | Gm43569 | 0 | 0 | 0 | 0 | 0 | 0 | 0 | 0 | 0 | 0 | 0 |
| 0 | NA | NA | NA | NA | NA | Gm42994 | 0 | 0 | 0 | 0 | 0 | 0 | 0 | 0 | 0 | 0 | 0 |
| 0 | NA | NA | NA | NA | NA | Gm34139 | 0 | 0 | 0 | 0 | 0 | 0 | 0 | 0 | 0 | 0 | 0 |
| 0 | NA | NA | NA | NA | NA | Gm10959 | 0 | 0 | 0 | 0 | 0 | 0 | 0 | 0 | 0 | 0 | 0 |
| 0 | NA | NA | NA | NA | NA | Gm43285 | 0 | 0 | 0 | 0 | 0 | 0 | 0 | 0 | 0 | 0 | 0 |
| 0 | NA | NA | NA | NA | NA | Gm22066 | 0 | 0 | 0 | 0 | 0 | 0 | 0 | 0 | 0 | 0 | 0 |
| 0 | NA | NA | NA | NA | NA | Gm43286 | 0 | 0 | 0 | 0 | 0 | 0 | 0 | 0 | 0 | 0 | 0 |
| 0 | NA | NA | NA | NA | NA | Gm43744 | 0 | 0 | 0 | 0 | 0 | 0 | 0 | 0 | 0 | 0 | 0 |
| 0 | NA | NA | NA | NA | NA | Gm43287 | 0 | 0 | 0 | 0 | 0 | 0 | 0 | 0 | 0 | 0 | 0 |
| 0 | NA | NA | NA | NA | NA | Gm43288 | 0 | 0 | 0 | 0 | 0 | 0 | 0 | 0 | 0 | 0 | 0 |
| 0 | NA | NA | NA | NA | NA | Gm43283 | 0 | 0 | 0 | 0 | 0 | 0 | 0 | 0 | 0 | 0 | 0 |
| 0 | NA | NA | NA | NA | NA | Gm22713 | 0 | 0 | 0 | 0 | 0 | 0 | 0 | 0 | 0 | 0 | 0 |
| 0 | NA | NA | NA | NA | NA | Gm35065 | 0 | 0 | 0 | 0 | 0 | 0 | 0 | 0 | 0 | 0 | 0 |

| | | | | | | | | | | | | | | | | | | |
|---|---|---|---|---|---|---|---|---|---|---|---|---|---|---|---|---|---|---|
| 0 | NA | NA | NA | NA | NA | Gm43590 | | 0 | 0 | 0 | 0 | 0 | 0 | 0 | 0 | 0 | 0 | 0 |
| 0 | NA | NA | NA | NA | NA | Snora24 | | 0 | 0 | 0 | 0 | 0 | 0 | 0 | 0 | 0 | 0 | 0 |
| 0 | NA | NA | NA | NA | NA | Gm42510 | | 0 | 0 | 0 | 0 | 0 | 0 | 0 | 0 | 0 | 0 | 0 |
| 0 | NA | NA | NA | NA | NA | Gm24456 | | 0 | 0 | 0 | 0 | 0 | 0 | 0 | 0 | 0 | 0 | 0 |
| 0 | NA | NA | NA | NA | NA | Gm15400 | | 0 | 0 | 0 | 0 | 0 | 0 | 0 | 0 | 0 | 0 | 0 |
| 0 | NA | NA | NA | NA | NA | Gm15399 | | 0 | 0 | 0 | 0 | 0 | 0 | 0 | 0 | 0 | 0 | 0 |
| 0 | NA | NA | NA | NA | NA | Gm43731 | | 0 | 0 | 0 | 0 | 0 | 0 | 0 | 0 | 0 | 0 | 0 |
| 0 | NA | NA | NA | NA | NA | Gm9372 | | 0 | 0 | 0 | 0 | 0 | 0 | 0 | 0 | 0 | 0 | 0 |
| 0 | NA | NA | NA | NA | NA | Gm42824 | | 0 | 0 | 0 | 0 | 0 | 0 | 0 | 0 | 0 | 0 | 0 |
| 0 | NA | NA | NA | NA | NA | 4921527H02Rik | | 0 | 0 | 0 | 0 | 0 | 0 | 0 | 0 | 0 | 0 | 0 |
| 0 | NA | NA | NA | NA | NA | 1700006A11Ril | RIKEN cDNA 1700006A11 gene | 0 | 0 | 0 | 0 | 0 | 0 | 0 | 0 | 0 | 0 | 0 |
| 0 | NA | NA | NA | NA | NA | Gm22575 | | 0 | 0 | 0 | 0 | 0 | 0 | 0 | 0 | 0 | 0 | 0 |
| 0 | NA | NA | NA | NA | NA | Gm42823 | | 0 | 0 | 0 | 0 | 0 | 0 | 0 | 0 | 0 | 0 | 0 |
| 0 | NA | NA | NA | NA | NA | Gm22900 | | 0 | 0 | 0 | 0 | 0 | 0 | 0 | 0 | 0 | 0 | 0 |
| 0 | NA | NA | NA | NA | NA | 1700003H04Ril | RIKEN cDNA 1700003H04 gene | 0 | 0 | 0 | 0 | 0 | 0 | 0 | 0 | 0 | 0 | 0 |
| 0 | NA | NA | NA | NA | NA | Gm42825 | | 0 | 0 | 0 | 0 | 0 | 0 | 0 | 0 | 0 | 0 | 0 |
| 0 | NA | NA | NA | NA | NA | Gm23673 | | 0 | 0 | 0 | 0 | 0 | 0 | 0 | 0 | 0 | 0 | 0 |
| 0 | NA | NA | NA | NA | NA | Gm42826 | | 0 | 0 | 0 | 0 | 0 | 0 | 0 | 0 | 0 | 0 | 0 |
| 0 | NA | NA | NA | NA | NA | Gm26275 | | 0 | 0 | 0 | 0 | 0 | 0 | 0 | 0 | 0 | 0 | 0 |
| 0 | NA | NA | NA | NA | NA | Gm42827 | | 0 | 0 | 0 | 0 | 0 | 0 | 0 | 0 | 0 | 0 | 0 |
| 0 | NA | NA | NA | NA | NA | Gm17225 | | 0 | 0 | 0 | 0 | 0 | 0 | 0 | 0 | 0 | 0 | 0 |
| 0 | NA | NA | NA | NA | NA | Gm43009 | | 0 | 0 | 0 | 0 | 0 | 0 | 0 | 0 | 0 | 0 | 0 |
| 0 | NA | NA | NA | NA | NA | Gm43006 | | 0 | 0 | 0 | 0 | 0 | 0 | 0 | 0 | 0 | 0 | 0 |
| 0 | NA | NA | NA | NA | NA | Gm43005 | | 0 | 0 | 0 | 0 | 0 | 0 | 0 | 0 | 0 | 0 | 0 |
| 0 | NA | NA | NA | NA | NA | Gm15551 | | 0 | 0 | 0 | 0 | 0 | 0 | 0 | 0 | 0 | 0 | 0 |
| 0 | NA | NA | NA | NA | NA | Gm25501 | | 0 | 0 | 0 | 0 | 0 | 0 | 0 | 0 | 0 | 0 | 0 |
| 0 | NA | NA | NA | NA | NA | Gm43355 | | 0 | 0 | 0 | 0 | 0 | 0 | 0 | 0 | 0 | 0 | 0 |
| 0 | NA | NA | NA | NA | NA | Gm42525 | | 0 | 0 | 0 | 0 | 0 | 0 | 0 | 0 | 0 | 0 | 0 |
| 0 | NA | NA | NA | NA | NA | Gm44494 | | 0 | 0 | 0 | 0 | 0 | 0 | 0 | 0 | 0 | 0 | 0 |
| 0 | NA | NA | NA | NA | NA | Gm43354 | | 0 | 0 | 0 | 0 | 0 | 0 | 0 | 0 | 0 | 0 | 0 |
| 0 | NA | NA | NA | NA | NA | Mir302b | | 0 | 0 | 0 | 0 | 0 | 0 | 0 | 0 | 0 | 0 | 0 |
| 0 | NA | NA | NA | NA | NA | Mir302c | | 0 | 0 | 0 | 0 | 0 | 0 | 0 | 0 | 0 | 0 | 0 |
| 0 | NA | NA | NA | NA | NA | Mir302a | | 0 | 0 | 0 | 0 | 0 | 0 | 0 | 0 | 0 | 0 | 0 |
| 0 | NA | NA | NA | NA | NA | Mir302d | | 0 | 0 | 0 | 0 | 0 | 0 | 0 | 0 | 0 | 0 | 0 |
| 0 | NA | NA | NA | NA | NA | Mir367 | | 0 | 0 | 0 | 0 | 0 | 0 | 0 | 0 | 0 | 0 | 0 |
| 0 | NA | NA | NA | NA | NA | Gm43729 | | 0 | 0 | 0 | 0 | 0 | 0 | 0 | 0 | 0 | 0 | 0 |
| 0 | NA | NA | NA | NA | NA | Neurog2 | neurogenin 2 | 0 | 0 | 0 | 0 | 0 | 0 | 0 | 0 | 0 | 0 | 0 |
| 0 | NA | NA | NA | NA | NA | Gm23279 | | 0 | 0 | 0 | 0 | 0 | 0 | 0 | 0 | 0 | 0 | 0 |
| 0 | NA | NA | NA | NA | NA | Gm43652 | | 0 | 0 | 0 | 0 | 0 | 0 | 0 | 0 | 0 | 0 | 0 |
| 0 | NA | NA | NA | NA | NA | 9830132P13Rik | | 0 | 0 | 0 | 0 | 0 | 0 | 0 | 0 | 0 | 0 | 0 |
| 0 | NA | NA | NA | NA | NA | Gm35667 | | 0 | 0 | 0 | 0 | 0 | 0 | 0 | 0 | 0 | 0 | 0 |
| 0 | NA | NA | NA | NA | NA | Gm10650 | | 0 | 0 | 0 | 0 | 0 | 0 | 0 | 0 | 0 | 0 | 0 |
| 0 | NA | NA | NA | NA | NA | D030025E07Rik | | 0 | 0 | 0 | 0 | 0 | 0 | 0 | 0 | 0 | 0 | 0 |
| 0 | NA | NA | NA | NA | NA | Gm43656 | | 0 | 0 | 0 | 0 | 0 | 0 | 0 | 0 | 0 | 0 | 0 |
| 0 | NA | NA | NA | NA | NA | Gm22293 | | 0 | 0 | 0 | 0 | 0 | 0 | 0 | 0 | 0 | 0 | 0 |
| 0 | NA | NA | NA | NA | NA | 4933424H11Rik | | 0 | 0 | 0 | 0 | 0 | 0 | 0 | 0 | 0 | 0 | 0 |
| 0 | NA | NA | NA | NA | NA | Gm4410 | | 0 | 0 | 0 | 0 | 0 | 0 | 0 | 0 | 0 | 0 | 0 |
| 0 | NA | NA | NA | NA | NA | Gm28543 | | 0 | 0 | 0 | 0 | 0 | 0 | 0 | 0 | 0 | 0 | 0 |
| 0 | NA | NA | NA | NA | NA | Gm27195 | | 0 | 0 | 0 | 0 | 0 | 0 | 0 | 0 | 0 | 0 | 0 |
| 0 | NA | NA | NA | NA | NA | Gm43697 | | 0 | 0 | 0 | 0 | 0 | 0 | 0 | 0 | 0 | 0 | 0 |
| 0 | NA | NA | NA | NA | NA | Gm42628 | | 0 | 0 | 0 | 0 | 0 | 0 | 0 | 0 | 0 | 0 | 0 |
| 0 | NA | NA | NA | NA | NA | Gm42630 | | 0 | 0 | 0 | 0 | 0 | 0 | 0 | 0 | 0 | 0 | 0 |
| 0 | NA | NA | NA | NA | NA | Gm5712 | | 0 | 0 | 0 | 0 | 0 | 0 | 0 | 0 | 0 | 0 | 0 |
| 0 | NA | NA | NA | NA | NA | Gm24515 | | 0 | 0 | 0 | 0 | 0 | 0 | 0 | 0 | 0 | 0 | 0 |
| 0 | NA | NA | NA | NA | NA | Gm42650 | | 0 | 0 | 0 | 0 | 0 | 0 | 0 | 0 | 0 | 0 | 0 |
| 0 | NA | NA | NA | NA | NA | Lrit3 | leucine-rich repeat, immunoglobulin-like a | 0 | 0 | 0 | 0 | 0 | 0 | 0 | 0 | 0 | 0 | 0 |
| 0 | NA | NA | NA | NA | NA | Gm42663 | | 0 | 0 | 0 | 0 | 0 | 0 | 0 | 0 | 0 | 0 | 0 |
| 0 | NA | NA | NA | NA | NA | Gm9397 | | 0 | 0 | 0 | 0 | 0 | 0 | 0 | 0 | 0 | 0 | 0 |
| 0 | NA | NA | NA | NA | NA | Gm22682 | | 0 | 0 | 0 | 0 | 0 | 0 | 0 | 0 | 0 | 0 | 0 |
| 0 | NA | NA | NA | NA | NA | Gm43210 | | 0 | 0 | 0 | 0 | 0 | 0 | 0 | 0 | 0 | 0 | 0 |

| | | | | | | | | | | | | | | | | | | |
|---|---|---|---|---|---|---|---|---|---|---|---|---|---|---|---|---|---|---|
| 0 | NA | NA | NA | NA | NA | Gm43211 | | 0 | 0 | 0 | 0 | 0 | 0 | 0 | 0 | 0 | 0 | 0 |
| 0 | NA | NA | NA | NA | NA | Gm42837 | | 0 | 0 | 0 | 0 | 0 | 0 | 0 | 0 | 0 | 0 | 0 |
| 0 | NA | NA | NA | NA | NA | Gm43524 | | 0 | 0 | 0 | 0 | 0 | 0 | 0 | 0 | 0 | 0 | 0 |
| 0 | NA | NA | NA | NA | NA | 2010110G14Rik | | 0 | 0 | 0 | 0 | 0 | 0 | 0 | 0 | 0 | 0 | 0 |
| 0 | NA | NA | NA | NA | NA | Gm5982 | | 0 | 0 | 0 | 0 | 0 | 0 | 0 | 0 | 0 | 0 | 0 |
| 0 | NA | NA | NA | NA | NA | Gm26090 | | 0 | 0 | 0 | 0 | 0 | 0 | 0 | 0 | 0 | 0 | 0 |
| 0 | NA | NA | NA | NA | NA | Gm43350 | | 0 | 0 | 0 | 0 | 0 | 0 | 0 | 0 | 0 | 0 | 0 |
| 0 | NA | NA | NA | NA | NA | 5830437K03Rik | | 0 | 0 | 0 | 0 | 0 | 0 | 0 | 0 | 0 | 0 | 0 |
| 0 | NA | NA | NA | NA | NA | Gm43349 | | 0 | 0 | 0 | 0 | 0 | 0 | 0 | 0 | 0 | 0 | 0 |
| 0 | NA | NA | NA | NA | NA | A430072C10Rik | | 0 | 0 | 0 | 0 | 0 | 0 | 0 | 0 | 0 | 0 | 0 |
| 0 | NA | NA | NA | NA | NA | Lef1os1 | | 0 | 0 | 0 | 0 | 0 | 0 | 0 | 0 | 0 | 0 | 0 |
| 0 | NA | NA | NA | NA | NA | Gm42449 | | 0 | 0 | 0 | 0 | 0 | 0 | 0 | 0 | 0 | 0 | 0 |
| 0 | NA | NA | NA | NA | NA | Gm18492 | | 0 | 0 | 0 | 0 | 0 | 0 | 0 | 0 | 0 | 0 | 0 |
| 0 | NA | NA | NA | NA | NA | Gm42881 | | 0 | 0 | 0 | 0 | 0 | 0 | 0 | 0 | 0 | 0 | 0 |
| 0 | NA | NA | NA | NA | NA | Gm29865 | | 0 | 0 | 0 | 0 | 0 | 0 | 0 | 0 | 0 | 0 | 0 |
| 0 | NA | NA | NA | NA | NA | Olfr375-ps1 | | 0 | 0 | 0 | 0 | 0 | 0 | 0 | 0 | 0 | 0 | 0 |
| 0 | NA | NA | NA | NA | NA | Gm26103 | | 0 | 0 | 0 | 0 | 0 | 0 | 0 | 0 | 0 | 0 | 0 |
| 0 | NA | NA | NA | NA | NA | Gm22421 | | 0 | 0 | 0 | 0 | 0 | 0 | 0 | 0 | 0 | 0 | 0 |
| 0 | NA | NA | NA | NA | NA | Gm42880 | | 0 | 0 | 0 | 0 | 0 | 0 | 0 | 0 | 0 | 0 | 0 |
| 0 | NA | NA | NA | NA | NA | Gm25307 | | 0 | 0 | 0 | 0 | 0 | 0 | 0 | 0 | 0 | 0 | 0 |
| 0 | NA | NA | NA | NA | NA | Gm22322 | | 0 | 0 | 0 | 0 | 0 | 0 | 0 | 0 | 0 | 0 | 0 |
| 0 | NA | NA | NA | NA | NA | Gm23196 | | 0 | 0 | 0 | 0 | 0 | 0 | 0 | 0 | 0 | 0 | 0 |
| 0 | NA | NA | NA | NA | NA | Gm19148 | | 0 | 0 | 0 | 0 | 0 | 0 | 0 | 0 | 0 | 0 | 0 |
| 0 | NA | NA | NA | NA | NA | Gm25042 | | 0 | 0 | 0 | 0 | 0 | 0 | 0 | 0 | 0 | 0 | 0 |
| 0 | NA | NA | NA | NA | NA | Gm49316 | | 0 | 0 | 0 | 0 | 0 | 0 | 0 | 0 | 0 | 0 | 0 |
| 0 | NA | NA | NA | NA | NA | Gm23011 | | 0 | 0 | 0 | 0 | 0 | 0 | 0 | 0 | 0 | 0 | 0 |
| 0 | NA | NA | NA | NA | NA | Gm37228 | | 0 | 0 | 0 | 0 | 0 | 0 | 0 | 0 | 0 | 0 | 0 |
| 0 | NA | NA | NA | NA | NA | Gm40153 | | 0 | 0 | 0 | 0 | 0 | 0 | 0 | 0 | 0 | 0 | 0 |
| 0 | NA | NA | NA | NA | NA | Gm6135 | | 0 | 0 | 0 | 0 | 0 | 0 | 0 | 0 | 0 | 0 | 0 |
| 0 | NA | NA | NA | NA | NA | Gm30484 | | 0 | 0 | 0 | 0 | 0 | 0 | 0 | 0 | 0 | 0 | 0 |
| 0 | NA | NA | NA | NA | NA | Mir1895 | | 0 | 0 | 0 | 0 | 0 | 0 | 0 | 0 | 0 | 0 | 0 |
| 0 | NA | NA | NA | NA | NA | Gm43559 | | 0 | 0 | 0 | 0 | 0 | 0 | 0 | 0 | 0 | 0 | 0 |
| 0 | NA | NA | NA | NA | NA | Gm43558 | | 0 | 0 | 0 | 0 | 0 | 0 | 0 | 0 | 0 | 0 | 0 |
| 0 | NA | NA | NA | NA | NA | Gm6077 | | 0 | 0 | 0 | 0 | 0 | 0 | 0 | 0 | 0 | 0 | 0 |
| 0 | NA | NA | NA | NA | NA | 4930539C22Rik | | 0 | 0 | 0 | 0 | 0 | 0 | 0 | 0 | 0 | 0 | 0 |
| 0 | NA | NA | NA | NA | NA | Gm43036 | | 0 | 0 | 0 | 0 | 0 | 0 | 0 | 0 | 0 | 0 | 0 |
| 0 | NA | NA | NA | NA | NA | Gm43035 | | 0 | 0 | 0 | 0 | 0 | 0 | 0 | 0 | 0 | 0 | 0 |
| 0 | NA | NA | NA | NA | NA | Gm2142 | | 0 | 0 | 0 | 0 | 0 | 0 | 0 | 0 | 0 | 0 | 0 |
| 0 | NA | NA | NA | NA | NA | Gm24048 | | 0 | 0 | 0 | 0 | 0 | 0 | 0 | 0 | 0 | 0 | 0 |
| 0 | NA | NA | NA | NA | NA | Gm18546 | | 0 | 0 | 0 | 0 | 0 | 0 | 0 | 0 | 0 | 0 | 0 |
| 0 | NA | NA | NA | NA | NA | Gm43033 | | 0 | 0 | 0 | 0 | 0 | 0 | 0 | 0 | 0 | 0 | 0 |
| 0 | NA | NA | NA | NA | NA | Slc9b1 | solute carrier family 9, subfamily B (NHA1 | 0 | 0 | 0 | 0 | 0 | 0 | 0 | 0 | 0 | 0 | 0 |
| 0 | NA | NA | NA | NA | NA | Gm42807 | | 0 | 0 | 0 | 0 | 0 | 0 | 0 | 0 | 0 | 0 | 0 |
| 0 | NA | NA | NA | NA | NA | 4933401H06Rik | | 0 | 0 | 0 | 0 | 0 | 0 | 0 | 0 | 0 | 0 | 0 |
| 0 | NA | NA | NA | NA | NA | Gm43429 | | 0 | 0 | 0 | 0 | 0 | 0 | 0 | 0 | 0 | 0 | 0 |
| 0 | NA | NA | NA | NA | NA | Gm18330 | | 0 | 0 | 0 | 0 | 0 | 0 | 0 | 0 | 0 | 0 | 0 |
| 0 | NA | NA | NA | NA | NA | Gm31243 | | 0 | 0 | 0 | 0 | 0 | 0 | 0 | 0 | 0 | 0 | 0 |
| 0 | NA | NA | NA | NA | NA | Gm43602 | | 0 | 0 | 0 | 0 | 0 | 0 | 0 | 0 | 0 | 0 | 0 |
| 0 | NA | NA | NA | NA | NA | Gm5281 | | 0 | 0 | 0 | 0 | 0 | 0 | 0 | 0 | 0 | 0 | 0 |
| 0 | NA | NA | NA | NA | NA | Gm9408 | | 0 | 0 | 0 | 0 | 0 | 0 | 0 | 0 | 0 | 0 | 0 |
| 0 | NA | NA | NA | NA | NA | Gm43402 | | 0 | 0 | 0 | 0 | 0 | 0 | 0 | 0 | 0 | 0 | 0 |
| 0 | NA | NA | NA | NA | NA | 1700030L20Rik | | 0 | 0 | 0 | 0 | 0 | 0 | 0 | 0 | 0 | 0 | 0 |
| 0 | NA | NA | NA | NA | NA | Gm42822 | | 0 | 0 | 0 | 0 | 0 | 0 | 0 | 0 | 0 | 0 | 0 |
| 0 | NA | NA | NA | NA | NA | 4930599N24Rik | | 0 | 0 | 0 | 0 | 0 | 0 | 0 | 0 | 0 | 0 | 0 |
| 0 | NA | NA | NA | NA | NA | Gm44327 | | 0 | 0 | 0 | 0 | 0 | 0 | 0 | 0 | 0 | 0 | 0 |
| 0 | NA | NA | NA | NA | NA | Gm43357 | | 0 | 0 | 0 | 0 | 0 | 0 | 0 | 0 | 0 | 0 | 0 |
| 0 | NA | NA | NA | NA | NA | Gm9409 | | 0 | 0 | 0 | 0 | 0 | 0 | 0 | 0 | 0 | 0 | 0 |
| 0 | NA | NA | NA | NA | NA | Gm43356 | | 0 | 0 | 0 | 0 | 0 | 0 | 0 | 0 | 0 | 0 | 0 |
| 0 | NA | NA | NA | NA | NA | Gm26107 | | 0 | 0 | 0 | 0 | 0 | 0 | 0 | 0 | 0 | 0 | 0 |
| 0 | NA | NA | NA | NA | NA | Gm22468 | | 0 | 0 | 0 | 0 | 0 | 0 | 0 | 0 | 0 | 0 | 0 |

| | | | | | | | | | | | | | | | | | | |
|---|---|---|---|---|---|---|---|---|---|---|---|---|---|---|---|---|---|---|
| 0 | NA | NA | NA | NA | NA | Gm30946 | | 0 | 0 | 0 | 0 | 0 | 0 | 0 | 0 | 0 | 0 | 0 |
| 0 | NA | NA | NA | NA | NA | Gm21962 | | 0 | 0 | 0 | 0 | 0 | 0 | 0 | 0 | 0 | 0 | 0 |
| 0 | NA | NA | NA | NA | NA | Gm42610 | | 0 | 0 | 0 | 0 | 0 | 0 | 0 | 0 | 0 | 0 | 0 |
| 0 | NA | NA | NA | NA | NA | Gm25080 | | 0 | 0 | 0 | 0 | 0 | 0 | 0 | 0 | 0 | 0 | 0 |
| 0 | NA | NA | NA | NA | NA | Mir6380 | | 0 | 0 | 0 | 0 | 0 | 0 | 0 | 0 | 0 | 0 | 0 |
| 0 | NA | NA | NA | NA | NA | Gm31678 | | 0 | 0 | 0 | 0 | 0 | 0 | 0 | 0 | 0 | 0 | 0 |
| 0 | NA | NA | NA | NA | NA | Gm40155 | | 0 | 0 | 0 | 0 | 0 | 0 | 0 | 0 | 0 | 0 | 0 |
| 0 | NA | NA | NA | NA | NA | Gm19708 | | 0 | 0 | 0 | 0 | 0 | 0 | 0 | 0 | 0 | 0 | 0 |
| 0 | NA | NA | NA | NA | NA | Gm43691 | | 0 | 0 | 0 | 0 | 0 | 0 | 0 | 0 | 0 | 0 | 0 |
| 0 | NA | NA | NA | NA | NA | Adh6b | alcohol dehydrogenase 6B (class V) | 0 | 0 | 0 | 0 | 0 | 0 | 0 | 0 | 0 | 0 | 0 |
| 0 | NA | NA | NA | NA | NA | Gm23467 | | 0 | 0 | 0 | 0 | 0 | 0 | 0 | 0 | 0 | 0 | 0 |
| 0 | NA | NA | NA | NA | NA | Adh6-ps1 | | 0 | 0 | 0 | 0 | 0 | 0 | 0 | 0 | 0 | 0 | 0 |
| 0 | NA | NA | NA | NA | NA | Gm43724 | | 0 | 0 | 0 | 0 | 0 | 0 | 0 | 0 | 0 | 0 | 0 |
| 0 | NA | NA | NA | NA | NA | Gm43725 | | 0 | 0 | 0 | 0 | 0 | 0 | 0 | 0 | 0 | 0 | 0 |
| 0 | NA | NA | NA | NA | NA | Mir1956 | | 0 | 0 | 0 | 0 | 0 | 0 | 0 | 0 | 0 | 0 | 0 |
| 0 | NA | NA | NA | NA | NA | Gm9415 | | 0 | 0 | 0 | 0 | 0 | 0 | 0 | 0 | 0 | 0 | 0 |
| 0 | NA | NA | NA | NA | NA | Gm6057 | | 0 | 0 | 0 | 0 | 0 | 0 | 0 | 0 | 0 | 0 | 0 |
| 0 | NA | NA | NA | NA | NA | Gm43723 | | 0 | 0 | 0 | 0 | 0 | 0 | 0 | 0 | 0 | 0 | 0 |
| 0 | NA | NA | NA | NA | NA | 4930425O10Rik | | 0 | 0 | 0 | 0 | 0 | 0 | 0 | 0 | 0 | 0 | 0 |
| 0 | NA | NA | NA | NA | NA | Gm4862 | | 0 | 0 | 0 | 0 | 0 | 0 | 0 | 0 | 0 | 0 | 0 |
| 0 | NA | NA | NA | NA | NA | Gm43163 | | 0 | 0 | 0 | 0 | 0 | 0 | 0 | 0 | 0 | 0 | 0 |
| 0 | NA | NA | NA | NA | NA | Gm32693 | | 0 | 0 | 0 | 0 | 0 | 0 | 0 | 0 | 0 | 0 | 0 |
| 0 | NA | NA | NA | NA | NA | Gm43678 | | 0 | 0 | 0 | 0 | 0 | 0 | 0 | 0 | 0 | 0 | 0 |
| 0 | NA | NA | NA | NA | NA | Mir7j | | 0 | 0 | 0 | 0 | 0 | 0 | 0 | 0 | 0 | 0 | 0 |
| 0 | NA | NA | NA | NA | NA | Gm4863 | | 0 | 0 | 0 | 0 | 0 | 0 | 0 | 0 | 0 | 0 | 0 |
| 0 | NA | NA | NA | NA | NA | Gm19184 | | 0 | 0 | 0 | 0 | 0 | 0 | 0 | 0 | 0 | 0 | 0 |
| 0 | NA | NA | NA | NA | NA | Gm19080 | | 0 | 0 | 0 | 0 | 0 | 0 | 0 | 0 | 0 | 0 | 0 |
| 0 | NA | NA | NA | NA | NA | Gm31211 | | 0 | 0 | 0 | 0 | 0 | 0 | 0 | 0 | 0 | 0 | 0 |
| 0 | NA | NA | NA | NA | NA | Gm43049 | | 0 | 0 | 0 | 0 | 0 | 0 | 0 | 0 | 0 | 0 | 0 |
| 0 | NA | NA | NA | NA | NA | Gm6214 | | 0 | 0 | 0 | 0 | 0 | 0 | 0 | 0 | 0 | 0 | 0 |
| 0 | NA | NA | NA | NA | NA | Gm43276 | | 0 | 0 | 0 | 0 | 0 | 0 | 0 | 0 | 0 | 0 | 0 |
| 0 | NA | NA | NA | NA | NA | Gm32754 | | 0 | 0 | 0 | 0 | 0 | 0 | 0 | 0 | 0 | 0 | 0 |
| 0 | NA | NA | NA | NA | NA | | | 0 | 0 | 0 | 0 | 0 | 0 | 0 | 0 | 0 | 0 | 0 |
| 0 | NA | NA | NA | NA | NA | 4930590L14Rik | | 0 | 0 | 0 | 0 | 0 | 0 | 0 | 0 | 0 | 0 | 0 |
| 0 | NA | NA | NA | NA | NA | Gm43826 | | 0 | 0 | 0 | 0 | 0 | 0 | 0 | 0 | 0 | 0 | 0 |
| 0 | NA | NA | NA | NA | NA | Gm43802 | | 0 | 0 | 0 | 0 | 0 | 0 | 0 | 0 | 0 | 0 | 0 |
| 0 | NA | NA | NA | NA | NA | Mir6381 | | 0 | 0 | 0 | 0 | 0 | 0 | 0 | 0 | 0 | 0 | 0 |
| 0 | NA | NA | NA | NA | NA | Gbp2-ps | | 0 | 0 | 0 | 0 | 0 | 0 | 0 | 0 | 0 | 0 | 0 |
| 0 | NA | NA | NA | NA | NA | Gm15540 | | 0 | 0 | 0 | 0 | 0 | 0 | 0 | 0 | 0 | 0 | 0 |
| 0 | NA | NA | NA | NA | NA | Gm24457 | | 0 | 0 | 0 | 0 | 0 | 0 | 0 | 0 | 0 | 0 | 0 |
| 0 | NA | NA | NA | NA | NA | Gm25965 | | 0 | 0 | 0 | 0 | 0 | 0 | 0 | 0 | 0 | 0 | 0 |
| 0 | NA | NA | NA | NA | NA | Gm43646 | | 0 | 0 | 0 | 0 | 0 | 0 | 0 | 0 | 0 | 0 | 0 |
| 0 | NA | NA | NA | NA | NA | Gm31306 | | 0 | 0 | 0 | 0 | 0 | 0 | 0 | 0 | 0 | 0 | 0 |
| 0 | NA | NA | NA | NA | NA | 4930519L02Rik | RIKEN cDNA 4930519L02 gene | 0 | 0 | 0 | 0 | 0 | 0 | 0 | 0 | 0 | 0 | 0 |
| 0 | NA | NA | NA | NA | NA | Gm43614 | | 0 | 0 | 0 | 0 | 0 | 0 | 0 | 0 | 0 | 0 | 0 |
| 0 | NA | NA | NA | NA | NA | A830019L24Rik | | 0 | 0 | 0 | 0 | 0 | 0 | 0 | 0 | 0 | 0 | 0 |
| 0 | NA | NA | NA | NA | NA | Gm43613 | | 0 | 0 | 0 | 0 | 0 | 0 | 0 | 0 | 0 | 0 | 0 |
| 0 | NA | NA | NA | NA | NA | Gm43612 | | 0 | 0 | 0 | 0 | 0 | 0 | 0 | 0 | 0 | 0 | 0 |
| 0 | NA | NA | NA | NA | NA | 1700001N15Rik | | 0 | 0 | 0 | 0 | 0 | 0 | 0 | 0 | 0 | 0 | 0 |
| 0 | NA | NA | NA | NA | NA | Gm43616 | | 0 | 0 | 0 | 0 | 0 | 0 | 0 | 0 | 0 | 0 | 0 |
| 0 | NA | NA | NA | NA | NA | Gm33651 | | 0 | 0 | 0 | 0 | 0 | 0 | 0 | 0 | 0 | 0 | 0 |
| 0 | NA | NA | NA | NA | NA | Gm33758 | | 0 | 0 | 0 | 0 | 0 | 0 | 0 | 0 | 0 | 0 | 0 |
| 0 | NA | NA | NA | NA | NA | Gm38006 | | 0 | 0 | 0 | 0 | 0 | 0 | 0 | 0 | 0 | 0 | 0 |
| 0 | NA | NA | NA | NA | NA | Gm42704 | | 0 | 0 | 0 | 0 | 0 | 0 | 0 | 0 | 0 | 0 | 0 |
| 0 | NA | NA | NA | NA | NA | Gm6260 | | 0 | 0 | 0 | 0 | 0 | 0 | 0 | 0 | 0 | 0 | 0 |
| 0 | NA | NA | NA | NA | NA | Gm24728 | | 0 | 0 | 0 | 0 | 0 | 0 | 0 | 0 | 0 | 0 | 0 |
| 0 | NA | NA | NA | NA | NA | Gm2505 | | 0 | 0 | 0 | 0 | 0 | 0 | 0 | 0 | 0 | 0 | 0 |
| 0 | NA | NA | NA | NA | NA | Gm42707 | | 0 | 0 | 0 | 0 | 0 | 0 | 0 | 0 | 0 | 0 | 0 |
| 0 | NA | NA | NA | NA | NA | Gm42708 | | 0 | 0 | 0 | 0 | 0 | 0 | 0 | 0 | 0 | 0 | 0 |
| 0 | NA | NA | NA | NA | NA | Gm34078 | | 0 | 0 | 0 | 0 | 0 | 0 | 0 | 0 | 0 | 0 | 0 |

| | | | | | | | | | | | | | | | | | | |
|---|---|---|---|---|---|---|---|---|---|---|---|---|---|---|---|---|---|---|
| 0 | NA | NA | NA | NA | NA | Gm43446 | | 0 | 0 | 0 | 0 | 0 | 0 | 0 | 0 | 0 | 0 | 0 |
| 0 | NA | NA | NA | NA | NA | Gm43447 | | 0 | 0 | 0 | 0 | 0 | 0 | 0 | 0 | 0 | 0 | 0 |
| 0 | NA | NA | NA | NA | NA | Gm43707 | | 0 | 0 | 0 | 0 | 0 | 0 | 0 | 0 | 0 | 0 | 0 |
| 0 | NA | NA | NA | NA | NA | Gm24406 | | 0 | 0 | 0 | 0 | 0 | 0 | 0 | 0 | 0 | 0 | 0 |
| 0 | NA | NA | NA | NA | NA | Gm17743 | | 0 | 0 | 0 | 0 | 0 | 0 | 0 | 0 | 0 | 0 | 0 |
| 0 | NA | NA | NA | NA | NA | Gm43240 | | 0 | 0 | 0 | 0 | 0 | 0 | 0 | 0 | 0 | 0 | 0 |
| 0 | NA | NA | NA | NA | NA | Gm16215 | | 0 | 0 | 0 | 0 | 0 | 0 | 0 | 0 | 0 | 0 | 0 |
| 0 | NA | NA | NA | NA | NA | Clca4c-ps | | 0 | 0 | 0 | 0 | 0 | 0 | 0 | 0 | 0 | 0 | 0 |
| 0 | NA | NA | NA | NA | NA | Gm16233 | | 0 | 0 | 0 | 0 | 0 | 0 | 0 | 0 | 0 | 0 | 0 |
| 0 | NA | NA | NA | NA | NA | Clca4a | chloride channel accessory 4A | 0 | 0 | 0 | 0 | 0 | 0 | 0 | 0 | 0 | 0 | 0 |
| 0 | NA | NA | NA | NA | NA | Gm35066 | | 0 | 0 | 0 | 0 | 0 | 0 | 0 | 0 | 0 | 0 | 0 |
| 0 | NA | NA | NA | NA | NA | 9530034A14Rik | | 0 | 0 | 0 | 0 | 0 | 0 | 0 | 0 | 0 | 0 | 0 |
| 0 | NA | NA | NA | NA | NA | D530037P16Rik | | 0 | 0 | 0 | 0 | 0 | 0 | 0 | 0 | 0 | 0 | 0 |
| 0 | NA | NA | NA | NA | NA | Gm43408 | | 0 | 0 | 0 | 0 | 0 | 0 | 0 | 0 | 0 | 0 | 0 |
| 0 | NA | NA | NA | NA | NA | Gm25627 | | 0 | 0 | 0 | 0 | 0 | 0 | 0 | 0 | 0 | 0 | 0 |
| 0 | NA | NA | NA | NA | NA | Gm43334 | | 0 | 0 | 0 | 0 | 0 | 0 | 0 | 0 | 0 | 0 | 0 |
| 0 | NA | NA | NA | NA | NA | Gm10636 | | 0 | 0 | 0 | 0 | 0 | 0 | 0 | 0 | 0 | 0 | 0 |
| 0 | NA | NA | NA | NA | NA | Gm22078 | | 0 | 0 | 0 | 0 | 0 | 0 | 0 | 0 | 0 | 0 | 0 |
| 0 | NA | NA | NA | NA | NA | Gm22901 | | 0 | 0 | 0 | 0 | 0 | 0 | 0 | 0 | 0 | 0 | 0 |
| 0 | NA | NA | NA | NA | NA | Gm36259 | | 0 | 0 | 0 | 0 | 0 | 0 | 0 | 0 | 0 | 0 | 0 |
| 0 | NA | NA | NA | NA | NA | Gm19104 | | 0 | 0 | 0 | 0 | 0 | 0 | 0 | 0 | 0 | 0 | 0 |
| 0 | NA | NA | NA | NA | NA | Gm29771 | | 0 | 0 | 0 | 0 | 0 | 0 | 0 | 0 | 0 | 0 | 0 |
| 0 | NA | NA | NA | NA | NA | Gm6074 | | 0 | 0 | 0 | 0 | 0 | 0 | 0 | 0 | 0 | 0 | 0 |
| 0 | NA | NA | NA | NA | NA | Gm36831 | | 0 | 0 | 0 | 0 | 0 | 0 | 0 | 0 | 0 | 0 | 0 |
| 0 | NA | NA | NA | NA | NA | Gm37803 | | 0 | 0 | 0 | 0 | 0 | 0 | 0 | 0 | 0 | 0 | 0 |
| 0 | NA | NA | NA | NA | NA | Gm37041 | | 0 | 0 | 0 | 0 | 0 | 0 | 0 | 0 | 0 | 0 | 0 |
| 0 | NA | NA | NA | NA | NA | Gm43574 | | 0 | 0 | 0 | 0 | 0 | 0 | 0 | 0 | 0 | 0 | 0 |
| 0 | NA | NA | NA | NA | NA | Gm23131 | | 0 | 0 | 0 | 0 | 0 | 0 | 0 | 0 | 0 | 0 | 0 |
| 0 | NA | NA | NA | NA | NA | Gm43576 | | 0 | 0 | 0 | 0 | 0 | 0 | 0 | 0 | 0 | 0 | 0 |
| 0 | NA | NA | NA | NA | NA | Gm38068 | | 0 | 0 | 0 | 0 | 0 | 0 | 0 | 0 | 0 | 0 | 0 |
| 0 | NA | NA | NA | NA | NA | Gm43575 | | 0 | 0 | 0 | 0 | 0 | 0 | 0 | 0 | 0 | 0 | 0 |
| 0 | NA | NA | NA | NA | NA | Gm43572 | | 0 | 0 | 0 | 0 | 0 | 0 | 0 | 0 | 0 | 0 | 0 |
| 0 | NA | NA | NA | NA | NA | Gm25127 | | 0 | 0 | 0 | 0 | 0 | 0 | 0 | 0 | 0 | 0 | 0 |
| 0 | NA | NA | NA | NA | NA | Gm10287 | | 0 | 0 | 0 | 0 | 0 | 0 | 0 | 0 | 0 | 0 | 0 |
| 0 | NA | NA | NA | NA | NA | Gm30382 | | 0 | 0 | 0 | 0 | 0 | 0 | 0 | 0 | 0 | 0 | 0 |
| 0 | NA | NA | NA | NA | NA | Gm5149 | | 0 | 0 | 0 | 0 | 0 | 0 | 0 | 0 | 0 | 0 | 0 |
| 0 | NA | NA | NA | NA | NA | Gm26468 | | 0 | 0 | 0 | 0 | 0 | 0 | 0 | 0 | 0 | 0 | 0 |
| 0 | NA | NA | NA | NA | NA | Gm31121 | | 0 | 0 | 0 | 0 | 0 | 0 | 0 | 0 | 0 | 0 | 0 |
| 0 | NA | NA | NA | NA | NA | Gm43470 | | 0 | 0 | 0 | 0 | 0 | 0 | 0 | 0 | 0 | 0 | 0 |
| 0 | NA | NA | NA | NA | NA | Mir3963 | | 0 | 0 | 0 | 0 | 0 | 0 | 0 | 0 | 0 | 0 | 0 |
| 0 | NA | NA | NA | NA | NA | Gm43468 | | 0 | 0 | 0 | 0 | 0 | 0 | 0 | 0 | 0 | 0 | 0 |
| 0 | NA | NA | NA | NA | NA | Gm43469 | | 0 | 0 | 0 | 0 | 0 | 0 | 0 | 0 | 0 | 0 | 0 |
| 0 | NA | NA | NA | NA | NA | Gm43473 | | 0 | 0 | 0 | 0 | 0 | 0 | 0 | 0 | 0 | 0 | 0 |
| 0 | NA | NA | NA | NA | NA | Gm42949 | | 0 | 0 | 0 | 0 | 0 | 0 | 0 | 0 | 0 | 0 | 0 |
| 0 | NA | NA | NA | NA | NA | Gm24458 | | 0 | 0 | 0 | 0 | 0 | 0 | 0 | 0 | 0 | 0 | 0 |
| 0 | NA | NA | NA | NA | NA | Gm43617 | | 0 | 0 | 0 | 0 | 0 | 0 | 0 | 0 | 0 | 0 | 0 |
| 0 | NA | NA | NA | NA | NA | Gm25056 | | 0 | 0 | 0 | 0 | 0 | 0 | 0 | 0 | 0 | 0 | 0 |
| 0 | NA | NA | NA | NA | NA | Gm2939 | | 0 | 0 | 0 | 0 | 0 | 0 | 0 | 0 | 0 | 0 | 0 |
| 0 | NA | NA | NA | NA | NA | Gm2860 | | 0 | 0 | 0 | 0 | 0 | 0 | 0 | 0 | 0 | 0 | 0 |
| 0 | NA | NA | NA | NA | NA | Gm43619 | | 0 | 0 | 0 | 0 | 0 | 0 | 0 | 0 | 0 | 0 | 0 |
| 0 | NA | NA | NA | NA | NA | Gm9682 | | 0 | 0 | 0 | 0 | 0 | 0 | 0 | 0 | 0 | 0 | 0 |
| 0 | NA | NA | NA | NA | NA | D630002J18Rik | | 0 | 0 | 0 | 0 | 0 | 0 | 0 | 0 | 0 | 0 | 0 |
| 0 | NA | NA | NA | NA | NA | Gm43859 | | 0 | 0 | 0 | 0 | 0 | 0 | 0 | 0 | 0 | 0 | 0 |
| 0 | NA | NA | NA | NA | NA | Gm16213 | | 0 | 0 | 0 | 0 | 0 | 0 | 0 | 0 | 0 | 0 | 0 |
| 0 | NA | NA | NA | NA | NA | Gm16231 | | 0 | 0 | 0 | 0 | 0 | 0 | 0 | 0 | 0 | 0 | 0 |
| 0 | NA | NA | NA | NA | NA | Gm42710 | | 0 | 0 | 0 | 0 | 0 | 0 | 0 | 0 | 0 | 0 | 0 |
| 0 | NA | NA | NA | NA | NA | 1700094M23Rik | | 0 | 0 | 0 | 0 | 0 | 0 | 0 | 0 | 0 | 0 | 0 |
| 0 | NA | NA | NA | NA | NA | 1700012D16Rik | | 0 | 0 | 0 | 0 | 0 | 0 | 0 | 0 | 0 | 0 | 0 |
| 0 | NA | NA | NA | NA | NA | A530083M17Rik | | 0 | 0 | 0 | 0 | 0 | 0 | 0 | 0 | 0 | 0 | 0 |
| 0 | NA | NA | NA | NA | NA | Gm27405 | | 0 | 0 | 0 | 0 | 0 | 0 | 0 | 0 | 0 | 0 | 0 |

| | | | | | | | | | | | | | | | | | | |
|---|---|---|---|---|---|---|---|---|---|---|---|---|---|---|---|---|---|---|
| 0 | NA | NA | NA | NA | NA | 1700015C17Rik | | 0 | 0 | 0 | 0 | 0 | 0 | 0 | 0 | 0 | 0 | 0 |
| 0 | NA | NA | NA | NA | NA | 4930597L12Rik | | 0 | 0 | 0 | 0 | 0 | 0 | 0 | 0 | 0 | 0 | 0 |
| 0 | NA | NA | NA | NA | NA | Gm22136 | | 0 | 0 | 0 | 0 | 0 | 0 | 0 | 0 | 0 | 0 | 0 |
| 0 | NA | NA | NA | NA | NA | Gm22206 | | 0 | 0 | 0 | 0 | 0 | 0 | 0 | 0 | 0 | 0 | 0 |
| 0 | NA | NA | NA | NA | NA | Gm43833 | | 0 | 0 | 0 | 0 | 0 | 0 | 0 | 0 | 0 | 0 | 0 |
| 0 | NA | NA | NA | NA | NA | Gm10042 | | 0 | 0 | 0 | 0 | 0 | 0 | 0 | 0 | 0 | 0 | 0 |
| 0 | NA | NA | NA | NA | NA | 5730460C07Rik | | 0 | 0 | 0 | 0 | 0 | 0 | 0 | 0 | 0 | 0 | 0 |
| 0 | NA | NA | NA | NA | NA | Gm25757 | | 0 | 0 | 0 | 0 | 0 | 0 | 0 | 0 | 0 | 0 | 0 |
| 0 | NA | NA | NA | NA | NA | Asb17os | | 0 | 0 | 0 | 0 | 0 | 0 | 0 | 0 | 0 | 0 | 0 |
| 0 | NA | NA | NA | NA | NA | Asb17 | ankyrin repeat and SOCS box-containing | 0 | 0 | 0 | 0 | 0 | 0 | 0 | 0 | 0 | 0 | 0 |
| 0 | NA | NA | NA | NA | NA | Gm24395 | | 0 | 0 | 0 | 0 | 0 | 0 | 0 | 0 | 0 | 0 | 0 |
| 0 | NA | NA | NA | NA | NA | Snord45b | | 0 | 0 | 0 | 0 | 0 | 0 | 0 | 0 | 0 | 0 | 0 |
| 0 | NA | NA | NA | NA | NA | Snord45c | | 0 | 0 | 0 | 0 | 0 | 0 | 0 | 0 | 0 | 0 | 0 |
| 0 | NA | NA | NA | NA | NA | Gm42541 | | 0 | 0 | 0 | 0 | 0 | 0 | 0 | 0 | 0 | 0 | 0 |
| 0 | NA | NA | NA | NA | NA | Gm42540 | | 0 | 0 | 0 | 0 | 0 | 0 | 0 | 0 | 0 | 0 | 0 |
| 0 | NA | NA | NA | NA | NA | Lhx8 | LIM homeobox protein 8 | 0 | 0 | 0 | 0 | 0 | 0 | 0 | 0 | 0 | 0 | 0 |
| 0 | NA | NA | NA | NA | NA | AI606473 | | 0 | 0 | 0 | 0 | 0 | 0 | 0 | 0 | 0 | 0 | 0 |
| 0 | NA | NA | NA | NA | NA | Gm20389 | | 0 | 0 | 0 | 0 | 0 | 0 | 0 | 0 | 0 | 0 | 0 |
| 0 | NA | NA | NA | NA | NA | Gm37890 | | 0 | 0 | 0 | 0 | 0 | 0 | 0 | 0 | 0 | 0 | 0 |
| 0 | NA | NA | NA | NA | NA | Gm26456 | | 0 | 0 | 0 | 0 | 0 | 0 | 0 | 0 | 0 | 0 | 0 |
| 0 | NA | NA | NA | NA | NA | Gm5285 | | 0 | 0 | 0 | 0 | 0 | 0 | 0 | 0 | 0 | 0 | 0 |
| 0 | NA | NA | NA | NA | NA | Gm37449 | | 0 | 0 | 0 | 0 | 0 | 0 | 0 | 0 | 0 | 0 | 0 |
| 0 | NA | NA | NA | NA | NA | Gm37324 | | 0 | 0 | 0 | 0 | 0 | 0 | 0 | 0 | 0 | 0 | 0 |
| 0 | NA | NA | NA | NA | NA | Gm23038 | | 0 | 0 | 0 | 0 | 0 | 0 | 0 | 0 | 0 | 0 | 0 |
| 0 | NA | NA | NA | NA | NA | Gm37102 | | 0 | 0 | 0 | 0 | 0 | 0 | 0 | 0 | 0 | 0 | 0 |
| 0 | NA | NA | NA | NA | NA | Gm37515 | | 0 | 0 | 0 | 0 | 0 | 0 | 0 | 0 | 0 | 0 | 0 |
| 0 | NA | NA | NA | NA | NA | 9330178D15Rik | | 0 | 0 | 0 | 0 | 0 | 0 | 0 | 0 | 0 | 0 | 0 |
| 0 | NA | NA | NA | NA | NA | Gm42947 | | 0 | 0 | 0 | 0 | 0 | 0 | 0 | 0 | 0 | 0 | 0 |
| 0 | NA | NA | NA | NA | NA | Gm6510 | | 0 | 0 | 0 | 0 | 0 | 0 | 0 | 0 | 0 | 0 | 0 |
| 0 | NA | NA | NA | NA | NA | Gm42886 | | 0 | 0 | 0 | 0 | 0 | 0 | 0 | 0 | 0 | 0 | 0 |
| 0 | NA | NA | NA | NA | NA | Gm42946 | | 0 | 0 | 0 | 0 | 0 | 0 | 0 | 0 | 0 | 0 | 0 |
| 0 | NA | NA | NA | NA | NA | Gm42942 | | 0 | 0 | 0 | 0 | 0 | 0 | 0 | 0 | 0 | 0 | 0 |
| 0 | NA | NA | NA | NA | NA | Gm42943 | | 0 | 0 | 0 | 0 | 0 | 0 | 0 | 0 | 0 | 0 | 0 |
| 0 | NA | NA | NA | NA | NA | Gm43528 | | 0 | 0 | 0 | 0 | 0 | 0 | 0 | 0 | 0 | 0 | 0 |
| 0 | NA | NA | NA | NA | NA | Gm15578 | | 0 | 0 | 0 | 0 | 0 | 0 | 0 | 0 | 0 | 0 | 0 |
| 0 | NA | NA | NA | NA | NA | Gm43527 | | 0 | 0 | 0 | 0 | 0 | 0 | 0 | 0 | 0 | 0 | 0 |
| 0 | NA | NA | NA | NA | NA | Gm43526 | | 0 | 0 | 0 | 0 | 0 | 0 | 0 | 0 | 0 | 0 | 0 |
| 0 | NA | NA | NA | NA | NA | 4930566N20Rik | | 0 | 0 | 0 | 0 | 0 | 0 | 0 | 0 | 0 | 0 | 0 |
| 0 | NA | NA | NA | NA | NA | Gm24373 | | 0 | 0 | 0 | 0 | 0 | 0 | 0 | 0 | 0 | 0 | 0 |
| 0 | NA | NA | NA | NA | NA | Gm23725 | | 0 | 0 | 0 | 0 | 0 | 0 | 0 | 0 | 0 | 0 | 0 |
| 0 | NA | NA | NA | NA | NA | Gm22458 | | 0 | 0 | 0 | 0 | 0 | 0 | 0 | 0 | 0 | 0 | 0 |
| 0 | NA | NA | NA | NA | NA | Gm23379 | | 0 | 0 | 0 | 0 | 0 | 0 | 0 | 0 | 0 | 0 | 0 |
| 0 | NA | NA | NA | NA | NA | Gm5983 | | 0 | 0 | 0 | 0 | 0 | 0 | 0 | 0 | 0 | 0 | 0 |
| 0 | NA | NA | NA | NA | NA | Gm43314 | | 0 | 0 | 0 | 0 | 0 | 0 | 0 | 0 | 0 | 0 | 0 |
| 0 | NA | NA | NA | NA | NA | Gm42968 | | 0 | 0 | 0 | 0 | 0 | 0 | 0 | 0 | 0 | 0 | 0 |
| 0 | NA | NA | NA | NA | NA | Gm43485 | | 0 | 0 | 0 | 0 | 0 | 0 | 0 | 0 | 0 | 0 | 0 |
| 0 | NA | NA | NA | NA | NA | Gm43486 | | 0 | 0 | 0 | 0 | 0 | 0 | 0 | 0 | 0 | 0 | 0 |
| 0 | NA | NA | NA | NA | NA | Gm43487 | | 0 | 0 | 0 | 0 | 0 | 0 | 0 | 0 | 0 | 0 | 0 |
| 0 | NA | NA | NA | NA | NA | Gm20752 | | 0 | 0 | 0 | 0 | 0 | 0 | 0 | 0 | 0 | 0 | 0 |
| 0 | NA | NA | NA | NA | NA | Gm43655 | | 0 | 0 | 0 | 0 | 0 | 0 | 0 | 0 | 0 | 0 | 0 |
| 0 | NA | NA | NA | NA | NA | Rpe65 | retinal pigment epithelium 65 | 0 | 0 | 0 | 0 | 0 | 0 | 0 | 0 | 0 | 0 | 0 |
| 0 | NA | NA | NA | NA | NA | Gm23941 | | 0 | 0 | 0 | 0 | 0 | 0 | 0 | 0 | 0 | 0 | 0 |
| 0 | NA | NA | NA | NA | NA | Gm27878 | | 0 | 0 | 0 | 0 | 0 | 0 | 0 | 0 | 0 | 0 | 0 |
| 0 | NA | NA | NA | NA | NA | Vmn1r-ps2 | | 0 | 0 | 0 | 0 | 0 | 0 | 0 | 0 | 0 | 0 | 0 |
| 0 | NA | NA | NA | NA | NA | Vmn1r-ps3 | | 0 | 0 | 0 | 0 | 0 | 0 | 0 | 0 | 0 | 0 | 0 |
| 0 | NA | NA | NA | NA | NA | Vmn1r2 | vomeronasal 1 receptor 2 | 0 | 0 | 0 | 0 | 0 | 0 | 0 | 0 | 0 | 0 | 0 |
| 0 | NA | NA | NA | NA | NA | Vmn1r3 | vomeronasal 1 receptor 3 | 0 | 0 | 0 | 0 | 0 | 0 | 0 | 0 | 0 | 0 | 0 |
| 0 | NA | NA | NA | NA | NA | Gm11783 | | 0 | 0 | 0 | 0 | 0 | 0 | 0 | 0 | 0 | 0 | 0 |
| 0 | NA | NA | NA | NA | NA | Gm11786 | | 0 | 0 | 0 | 0 | 0 | 0 | 0 | 0 | 0 | 0 | 0 |
| 0 | NA | NA | NA | NA | NA | Gm11784 | | 0 | 0 | 0 | 0 | 0 | 0 | 0 | 0 | 0 | 0 | 0 |

| | | | | | | | | | | | | | | | | | | |
|---|---|---|---|---|---|---|---|---|---|---|---|---|---|---|---|---|---|---|
| 0 | NA | NA | NA | NA | NA | Gm11785 | | 0 | 0 | 0 | 0 | 0 | 0 | 0 | 0 | 0 | 0 | 0 |
| 0 | NA | NA | NA | NA | NA | Mir3471-2 | | 0 | 0 | 0 | 0 | 0 | 0 | 0 | 0 | 0 | 0 | 0 |
| 0 | NA | NA | NA | NA | NA | Gm22541 | | 0 | 0 | 0 | 0 | 0 | 0 | 0 | 0 | 0 | 0 | 0 |
| 0 | NA | NA | NA | NA | NA | 2210414B05Rik | | 0 | 0 | 0 | 0 | 0 | 0 | 0 | 0 | 0 | 0 | 0 |
| 0 | NA | NA | NA | NA | NA | Gm11805 | | 0 | 0 | 0 | 0 | 0 | 0 | 0 | 0 | 0 | 0 | 0 |
| 0 | NA | NA | NA | NA | NA | Gm22781 | | 0 | 0 | 0 | 0 | 0 | 0 | 0 | 0 | 0 | 0 | 0 |
| 0 | NA | NA | NA | NA | NA | Gm24016 | | 0 | 0 | 0 | 0 | 0 | 0 | 0 | 0 | 0 | 0 | 0 |
| 0 | NA | NA | NA | NA | NA | Gm24125 | | 0 | 0 | 0 | 0 | 0 | 0 | 0 | 0 | 0 | 0 | 0 |
| 0 | NA | NA | NA | NA | NA | Rps10-ps3 | | 0 | 0 | 0 | 0 | 0 | 0 | 0 | 0 | 0 | 0 | 0 |
| 0 | NA | NA | NA | NA | NA | Sdr16c5 | short chain dehydrogenase/reductase far | 0 | 0 | 0 | 0 | 0 | 0 | 0 | 0 | 0 | 0 | 0 |
| 0 | NA | NA | NA | NA | NA | Sdr16c6 | short chain dehydrogenase/reductase far | 0 | 0 | 0 | 0 | 0 | 0 | 0 | 0 | 0 | 0 | 0 |
| 0 | NA | NA | NA | NA | NA | Gm11780 | | 0 | 0 | 0 | 0 | 0 | 0 | 0 | 0 | 0 | 0 | 0 |
| 0 | NA | NA | NA | NA | NA | Gm26436 | | 0 | 0 | 0 | 0 | 0 | 0 | 0 | 0 | 0 | 0 | 0 |
| 0 | NA | NA | NA | NA | NA | Gm11779 | | 0 | 0 | 0 | 0 | 0 | 0 | 0 | 0 | 0 | 0 | 0 |
| 0 | NA | NA | NA | NA | NA | Gm11782 | | 0 | 0 | 0 | 0 | 0 | 0 | 0 | 0 | 0 | 0 | 0 |
| 0 | NA | NA | NA | NA | NA | Gm11781 | | 0 | 0 | 0 | 0 | 0 | 0 | 0 | 0 | 0 | 0 | 0 |
| 0 | NA | NA | NA | NA | NA | 4930423M02Rik | | 0 | 0 | 0 | 0 | 0 | 0 | 0 | 0 | 0 | 0 | 0 |
| 0 | NA | NA | NA | NA | NA | Gm24337 | | 0 | 0 | 0 | 0 | 0 | 0 | 0 | 0 | 0 | 0 | 0 |
| 0 | NA | NA | NA | NA | NA | Gm11796 | | 0 | 0 | 0 | 0 | 0 | 0 | 0 | 0 | 0 | 0 | 0 |
| 0 | NA | NA | NA | NA | NA | Gm11797 | | 0 | 0 | 0 | 0 | 0 | 0 | 0 | 0 | 0 | 0 | 0 |
| 0 | NA | NA | NA | NA | NA | Gm11798 | | 0 | 0 | 0 | 0 | 0 | 0 | 0 | 0 | 0 | 0 | 0 |
| 0 | NA | NA | NA | NA | NA | Cyp7a1 | cytochrome P450, family 7, subfamily a, p | 0 | 0 | 0 | 0 | 0 | 0 | 0 | 0 | 0 | 0 | 0 |
| 0 | NA | NA | NA | NA | NA | Gm11803 | | 0 | 0 | 0 | 0 | 0 | 0 | 0 | 0 | 0 | 0 | 0 |
| 0 | NA | NA | NA | NA | NA | Gm11801 | | 0 | 0 | 0 | 0 | 0 | 0 | 0 | 0 | 0 | 0 | 0 |
| 0 | NA | NA | NA | NA | NA | Gm22473 | | 0 | 0 | 0 | 0 | 0 | 0 | 0 | 0 | 0 | 0 | 0 |
| 0 | NA | NA | NA | NA | NA | Gm11804 | | 0 | 0 | 0 | 0 | 0 | 0 | 0 | 0 | 0 | 0 | 0 |
| 0 | NA | NA | NA | NA | NA | 8430436N08Rik | | 0 | 0 | 0 | 0 | 0 | 0 | 0 | 0 | 0 | 0 | 0 |
| 0 | NA | NA | NA | NA | NA | Gm11795 | | 0 | 0 | 0 | 0 | 0 | 0 | 0 | 0 | 0 | 0 | 0 |
| 0 | NA | NA | NA | NA | NA | Gm11794 | | 0 | 0 | 0 | 0 | 0 | 0 | 0 | 0 | 0 | 0 | 0 |
| 0 | NA | NA | NA | NA | NA | Gm23860 | | 0 | 0 | 0 | 0 | 0 | 0 | 0 | 0 | 0 | 0 | 0 |
| 0 | NA | NA | NA | NA | NA | Gm11800 | | 0 | 0 | 0 | 0 | 0 | 0 | 0 | 0 | 0 | 0 | 0 |
| 0 | NA | NA | NA | NA | NA | Gm25355 | | 0 | 0 | 0 | 0 | 0 | 0 | 0 | 0 | 0 | 0 | 0 |
| 0 | NA | NA | NA | NA | NA | Gm37386 | | 0 | 0 | 0 | 0 | 0 | 0 | 0 | 0 | 0 | 0 | 0 |
| 0 | NA | NA | NA | NA | NA | Gm11809 | | 0 | 0 | 0 | 0 | 0 | 0 | 0 | 0 | 0 | 0 | 0 |
| 0 | NA | NA | NA | NA | NA | Rps18-ps2 | | 0 | 0 | 0 | 0 | 0 | 0 | 0 | 0 | 0 | 0 | 0 |
| 0 | NA | NA | NA | NA | NA | Gm23423 | | 0 | 0 | 0 | 0 | 0 | 0 | 0 | 0 | 0 | 0 | 0 |
| 0 | NA | NA | NA | NA | NA | Gm11817 | | 0 | 0 | 0 | 0 | 0 | 0 | 0 | 0 | 0 | 0 | 0 |
| 0 | NA | NA | NA | NA | NA | Gm25677 | | 0 | 0 | 0 | 0 | 0 | 0 | 0 | 0 | 0 | 0 | 0 |
| 0 | NA | NA | NA | NA | NA | Gm24152 | | 0 | 0 | 0 | 0 | 0 | 0 | 0 | 0 | 0 | 0 | 0 |
| 0 | NA | NA | NA | NA | NA | 4930448K20Rik | | 0 | 0 | 0 | 0 | 0 | 0 | 0 | 0 | 0 | 0 | 0 |
| 0 | NA | NA | NA | NA | NA | Gm11834 | | 0 | 0 | 0 | 0 | 0 | 0 | 0 | 0 | 0 | 0 | 0 |
| 0 | NA | NA | NA | NA | NA | Gm11835 | | 0 | 0 | 0 | 0 | 0 | 0 | 0 | 0 | 0 | 0 | 0 |
| 0 | NA | NA | NA | NA | NA | 1700123O12Rik | | 0 | 0 | 0 | 0 | 0 | 0 | 0 | 0 | 0 | 0 | 0 |
| 0 | NA | NA | NA | NA | NA | Gm11815 | | 0 | 0 | 0 | 0 | 0 | 0 | 0 | 0 | 0 | 0 | 0 |
| 0 | NA | NA | NA | NA | NA | Gm11813 | | 0 | 0 | 0 | 0 | 0 | 0 | 0 | 0 | 0 | 0 | 0 |
| 0 | NA | NA | NA | NA | NA | Gm12920 | | 0 | 0 | 0 | 0 | 0 | 0 | 0 | 0 | 0 | 0 | 0 |
| 0 | NA | NA | NA | NA | NA | Mir3471-1 | | 0 | 0 | 0 | 0 | 0 | 0 | 0 | 0 | 0 | 0 | 0 |
| 0 | NA | NA | NA | NA | NA | n-TSaga9 | | 0 | 0 | 0 | 0 | 0 | 0 | 0 | 0 | 0 | 0 | 0 |
| 0 | NA | NA | NA | NA | NA | Mir684-2 | | 0 | 0 | 0 | 0 | 0 | 0 | 0 | 0 | 0 | 0 | 0 |
| 0 | NA | NA | NA | NA | NA | Gm11822 | | 0 | 0 | 0 | 0 | 0 | 0 | 0 | 0 | 0 | 0 | 0 |
| 0 | NA | NA | NA | NA | NA | Gm11831 | | 0 | 0 | 0 | 0 | 0 | 0 | 0 | 0 | 0 | 0 | 0 |
| 0 | NA | NA | NA | NA | NA | Gm11832 | | 0 | 0 | 0 | 0 | 0 | 0 | 0 | 0 | 0 | 0 | 0 |
| 0 | NA | NA | NA | NA | NA | Gm11833 | | 0 | 0 | 0 | 0 | 0 | 0 | 0 | 0 | 0 | 0 | 0 |
| 0 | NA | NA | NA | NA | NA | Gm22477 | | 0 | 0 | 0 | 0 | 0 | 0 | 0 | 0 | 0 | 0 | 0 |
| 0 | NA | NA | NA | NA | NA | Gm25002 | | 0 | 0 | 0 | 0 | 0 | 0 | 0 | 0 | 0 | 0 | 0 |
| 0 | NA | NA | NA | NA | NA | Gm11840 | | 0 | 0 | 0 | 0 | 0 | 0 | 0 | 0 | 0 | 0 | 0 |
| 0 | NA | NA | NA | NA | NA | Gm11846 | | 0 | 0 | 0 | 0 | 0 | 0 | 0 | 0 | 0 | 0 | 0 |
| 0 | NA | NA | NA | NA | NA | n-R5s182 | | 0 | 0 | 0 | 0 | 0 | 0 | 0 | 0 | 0 | 0 | 0 |
| 0 | NA | NA | NA | NA | NA | Gm11820 | | 0 | 0 | 0 | 0 | 0 | 0 | 0 | 0 | 0 | 0 | 0 |
| 0 | NA | NA | NA | NA | NA | Gm26250 | | 0 | 0 | 0 | 0 | 0 | 0 | 0 | 0 | 0 | 0 | 0 |

| | | | | | | | | | | | | | | | | | | |
|---|---|---|---|---|---|---|---|---|---|---|---|---|---|---|---|---|---|---|
| 0 | NA | NA | NA | NA | NA | Gm11819 | | 0 | 0 | 0 | 0 | 0 | 0 | 0 | 0 | 0 | 0 | 0 |
| 0 | NA | NA | NA | NA | NA | Gm11825 | | 0 | 0 | 0 | 0 | 0 | 0 | 0 | 0 | 0 | 0 | 0 |
| 0 | NA | NA | NA | NA | NA | Gm23162 | | 0 | 0 | 0 | 0 | 0 | 0 | 0 | 0 | 0 | 0 | 0 |
| 0 | NA | NA | NA | NA | NA | Gm11823 | | 0 | 0 | 0 | 0 | 0 | 0 | 0 | 0 | 0 | 0 | 0 |
| 0 | NA | NA | NA | NA | NA | Gm24068 | | 0 | 0 | 0 | 0 | 0 | 0 | 0 | 0 | 0 | 0 | 0 |
| 0 | NA | NA | NA | NA | NA | 1700120G11Rik | | 0 | 0 | 0 | 0 | 0 | 0 | 0 | 0 | 0 | 0 | 0 |
| 0 | NA | NA | NA | NA | NA | Gm24908 | | 0 | 0 | 0 | 0 | 0 | 0 | 0 | 0 | 0 | 0 | 0 |
| 0 | NA | NA | NA | NA | NA | Lrrc69 | leucine rich repeat containing 69 | 0 | 0 | 0 | 0 | 0 | 0 | 0 | 0 | 0 | 0 | 0 |
| 0 | NA | NA | NA | NA | NA | Gm11844 | | 0 | 0 | 0 | 0 | 0 | 0 | 0 | 0 | 0 | 0 | 0 |
| 0 | NA | NA | NA | NA | NA | Gm11845 | | 0 | 0 | 0 | 0 | 0 | 0 | 0 | 0 | 0 | 0 | 0 |
| 0 | NA | NA | NA | NA | NA | Gm24317 | | 0 | 0 | 0 | 0 | 0 | 0 | 0 | 0 | 0 | 0 | 0 |
| 0 | NA | NA | NA | NA | NA | Gm11856 | | 0 | 0 | 0 | 0 | 0 | 0 | 0 | 0 | 0 | 0 | 0 |
| 0 | NA | NA | NA | NA | NA | Gm11859 | | 0 | 0 | 0 | 0 | 0 | 0 | 0 | 0 | 0 | 0 | 0 |
| 0 | NA | NA | NA | NA | NA | Gm11857 | | 0 | 0 | 0 | 0 | 0 | 0 | 0 | 0 | 0 | 0 | 0 |
| 0 | NA | NA | NA | NA | NA | Gm11858 | | 0 | 0 | 0 | 0 | 0 | 0 | 0 | 0 | 0 | 0 | 0 |
| 0 | NA | NA | NA | NA | NA | Gm24567 | | 0 | 0 | 0 | 0 | 0 | 0 | 0 | 0 | 0 | 0 | 0 |
| 0 | NA | NA | NA | NA | NA | Gm26148 | | 0 | 0 | 0 | 0 | 0 | 0 | 0 | 0 | 0 | 0 | 0 |
| 0 | NA | NA | NA | NA | NA | Gm11869 | | 0 | 0 | 0 | 0 | 0 | 0 | 0 | 0 | 0 | 0 | 0 |
| 0 | NA | NA | NA | NA | NA | Gm37355 | | 0 | 0 | 0 | 0 | 0 | 0 | 0 | 0 | 0 | 0 | 0 |
| 0 | NA | NA | NA | NA | NA | Gm11861 | | 0 | 0 | 0 | 0 | 0 | 0 | 0 | 0 | 0 | 0 | 0 |
| 0 | NA | NA | NA | NA | NA | Gm24978 | | 0 | 0 | 0 | 0 | 0 | 0 | 0 | 0 | 0 | 0 | 0 |
| 0 | NA | NA | NA | NA | NA | Gm11862 | | 0 | 0 | 0 | 0 | 0 | 0 | 0 | 0 | 0 | 0 | 0 |
| 0 | NA | NA | NA | NA | NA | Gm23975 | | 0 | 0 | 0 | 0 | 0 | 0 | 0 | 0 | 0 | 0 | 0 |
| 0 | NA | NA | NA | NA | NA | Gm11848 | | 0 | 0 | 0 | 0 | 0 | 0 | 0 | 0 | 0 | 0 | 0 |
| 0 | NA | NA | NA | NA | NA | Gm11850 | | 0 | 0 | 0 | 0 | 0 | 0 | 0 | 0 | 0 | 0 | 0 |
| 0 | NA | NA | NA | NA | NA | Gm11849 | | 0 | 0 | 0 | 0 | 0 | 0 | 0 | 0 | 0 | 0 | 0 |
| 0 | NA | NA | NA | NA | NA | Gm23384 | | 0 | 0 | 0 | 0 | 0 | 0 | 0 | 0 | 0 | 0 | 0 |
| 0 | NA | NA | NA | NA | NA | Gm11851 | | 0 | 0 | 0 | 0 | 0 | 0 | 0 | 0 | 0 | 0 | 0 |
| 0 | NA | NA | NA | NA | NA | Gm11860 | | 0 | 0 | 0 | 0 | 0 | 0 | 0 | 0 | 0 | 0 | 0 |
| 0 | NA | NA | NA | NA | NA | Gm11852 | | 0 | 0 | 0 | 0 | 0 | 0 | 0 | 0 | 0 | 0 | 0 |
| 0 | NA | NA | NA | NA | NA | Gm11867 | | 0 | 0 | 0 | 0 | 0 | 0 | 0 | 0 | 0 | 0 | 0 |
| 0 | NA | NA | NA | NA | NA | Gm11854 | | 0 | 0 | 0 | 0 | 0 | 0 | 0 | 0 | 0 | 0 | 0 |
| 0 | NA | NA | NA | NA | NA | Gm11853 | | 0 | 0 | 0 | 0 | 0 | 0 | 0 | 0 | 0 | 0 | 0 |
| 0 | NA | NA | NA | NA | NA | Gm25971 | | 0 | 0 | 0 | 0 | 0 | 0 | 0 | 0 | 0 | 0 | 0 |
| 0 | NA | NA | NA | NA | NA | Gm24419 | | 0 | 0 | 0 | 0 | 0 | 0 | 0 | 0 | 0 | 0 | 0 |
| 0 | NA | NA | NA | NA | NA | Cnbd1 | cyclic nucleotide binding domain containi | 0 | 0 | 0 | 0 | 0 | 0 | 0 | 0 | 0 | 0 | 0 |
| 0 | NA | NA | NA | NA | NA | Gm12392 | | 0 | 0 | 0 | 0 | 0 | 0 | 0 | 0 | 0 | 0 | 0 |
| 0 | NA | NA | NA | NA | NA | Gm12393 | | 0 | 0 | 0 | 0 | 0 | 0 | 0 | 0 | 0 | 0 | 0 |
| 0 | NA | NA | NA | NA | NA | Gm25655 | | 0 | 0 | 0 | 0 | 0 | 0 | 0 | 0 | 0 | 0 | 0 |
| 0 | NA | NA | NA | NA | NA | Gm12354 | | 0 | 0 | 0 | 0 | 0 | 0 | 0 | 0 | 0 | 0 | 0 |
| 0 | NA | NA | NA | NA | NA | 4930480G23Rik | | 0 | 0 | 0 | 0 | 0 | 0 | 0 | 0 | 0 | 0 | 0 |
| 0 | NA | NA | NA | NA | NA | Gm11872 | | 0 | 0 | 0 | 0 | 0 | 0 | 0 | 0 | 0 | 0 | 0 |
| 0 | NA | NA | NA | NA | NA | Gm11873 | | 0 | 0 | 0 | 0 | 0 | 0 | 0 | 0 | 0 | 0 | 0 |
| 0 | NA | NA | NA | NA | NA | Gm11871 | | 0 | 0 | 0 | 0 | 0 | 0 | 0 | 0 | 0 | 0 | 0 |
| 0 | NA | NA | NA | NA | NA | Gm22817 | | 0 | 0 | 0 | 0 | 0 | 0 | 0 | 0 | 0 | 0 | 0 |
| 0 | NA | NA | NA | NA | NA | Gm25977 | | 0 | 0 | 0 | 0 | 0 | 0 | 0 | 0 | 0 | 0 | 0 |
| 0 | NA | NA | NA | NA | NA | Gm11875 | | 0 | 0 | 0 | 0 | 0 | 0 | 0 | 0 | 0 | 0 | 0 |
| 0 | NA | NA | NA | NA | NA | Gm11876 | | 0 | 0 | 0 | 0 | 0 | 0 | 0 | 0 | 0 | 0 | 0 |
| 0 | NA | NA | NA | NA | NA | Prdm13 | PR domain containing 13 | 0 | 0 | 0 | 0 | 0 | 0 | 0 | 0 | 0 | 0 | 0 |
| 0 | NA | NA | NA | NA | NA | Gm11885 | | 0 | 0 | 0 | 0 | 0 | 0 | 0 | 0 | 0 | 0 | 0 |
| 0 | NA | NA | NA | NA | NA | Gm11880 | | 0 | 0 | 0 | 0 | 0 | 0 | 0 | 0 | 0 | 0 | 0 |
| 0 | NA | NA | NA | NA | NA | Gm11882 | | 0 | 0 | 0 | 0 | 0 | 0 | 0 | 0 | 0 | 0 | 0 |
| 0 | NA | NA | NA | NA | NA | Pou3f2 | POU domain, class 3, transcription factor | 0 | 0 | 0 | 0 | 0 | 0 | 0 | 0 | 0 | 0 | 0 |
| 0 | NA | NA | NA | NA | NA | Gm30731 | | 0 | 0 | 0 | 0 | 0 | 0 | 0 | 0 | 0 | 0 | 0 |
| 0 | NA | NA | NA | NA | NA | Gm11881 | | 0 | 0 | 0 | 0 | 0 | 0 | 0 | 0 | 0 | 0 | 0 |
| 0 | NA | NA | NA | NA | NA | Gm11879 | | 0 | 0 | 0 | 0 | 0 | 0 | 0 | 0 | 0 | 0 | 0 |
| 0 | NA | NA | NA | NA | NA | Gm24078 | | 0 | 0 | 0 | 0 | 0 | 0 | 0 | 0 | 0 | 0 | 0 |
| 0 | NA | NA | NA | NA | NA | 1700025O08Rik | | 0 | 0 | 0 | 0 | 0 | 0 | 0 | 0 | 0 | 0 | 0 |
| 0 | NA | NA | NA | NA | NA | Gm25978 | | 0 | 0 | 0 | 0 | 0 | 0 | 0 | 0 | 0 | 0 | 0 |
| 0 | NA | NA | NA | NA | NA | Gm11890 | | 0 | 0 | 0 | 0 | 0 | 0 | 0 | 0 | 0 | 0 | 0 |

| | | | | | | | | | | | | | | | | | | |
|---|---|---|---|---|---|---|---|---|---|---|---|---|---|---|---|---|---|---|
| 0 | NA | NA | NA | NA | NA | Gm28448 | | 0 | 0 | 0 | 0 | 0 | 0 | 0 | 0 | 0 | 0 | 0 |
| 0 | NA | NA | NA | NA | NA | Gm11892 | | 0 | 0 | 0 | 0 | 0 | 0 | 0 | 0 | 0 | 0 | 0 |
| 0 | NA | NA | NA | NA | NA | Gm11891 | | 0 | 0 | 0 | 0 | 0 | 0 | 0 | 0 | 0 | 0 | 0 |
| 0 | NA | NA | NA | NA | NA | Gm27243 | | 0 | 0 | 0 | 0 | 0 | 0 | 0 | 0 | 0 | 0 | 0 |
| 0 | NA | NA | NA | NA | NA | Gm11899 | | 0 | 0 | 0 | 0 | 0 | 0 | 0 | 0 | 0 | 0 | 0 |
| 0 | NA | NA | NA | NA | NA | Prdx6-ps2 | | 0 | 0 | 0 | 0 | 0 | 0 | 0 | 0 | 0 | 0 | 0 |
| 0 | NA | NA | NA | NA | NA | Gm11897 | | 0 | 0 | 0 | 0 | 0 | 0 | 0 | 0 | 0 | 0 | 0 |
| 0 | NA | NA | NA | NA | NA | Gm11894 | | 0 | 0 | 0 | 0 | 0 | 0 | 0 | 0 | 0 | 0 | 0 |
| 0 | NA | NA | NA | NA | NA | Gm25975 | | 0 | 0 | 0 | 0 | 0 | 0 | 0 | 0 | 0 | 0 | 0 |
| 0 | NA | NA | NA | NA | NA | Gm11896 | | 0 | 0 | 0 | 0 | 0 | 0 | 0 | 0 | 0 | 0 | 0 |
| 0 | NA | NA | NA | NA | NA | Gm24977 | | 0 | 0 | 0 | 0 | 0 | 0 | 0 | 0 | 0 | 0 | 0 |
| 0 | NA | NA | NA | NA | NA | Fut9 | fucosyltransferase 9 | 0 | 0 | 0 | 0 | 0 | 0 | 0 | 0 | 0 | 0 | 0 |
| 0 | NA | NA | NA | NA | NA | Gm24042 | | 0 | 0 | 0 | 0 | 0 | 0 | 0 | 0 | 0 | 0 | 0 |
| 0 | NA | NA | NA | NA | NA | Gm11898 | | 0 | 0 | 0 | 0 | 0 | 0 | 0 | 0 | 0 | 0 | 0 |
| 0 | NA | NA | NA | NA | NA | Gm26067 | | 0 | 0 | 0 | 0 | 0 | 0 | 0 | 0 | 0 | 0 | 0 |
| 0 | NA | NA | NA | NA | NA | Gm11895 | | 0 | 0 | 0 | 0 | 0 | 0 | 0 | 0 | 0 | 0 | 0 |
| 0 | NA | NA | NA | NA | NA | Gm24965 | | 0 | 0 | 0 | 0 | 0 | 0 | 0 | 0 | 0 | 0 | 0 |
| 0 | NA | NA | NA | NA | NA | Bhmt-ps1 | | 0 | 0 | 0 | 0 | 0 | 0 | 0 | 0 | 0 | 0 | 0 |
| 0 | NA | NA | NA | NA | NA | Gm11903 | | 0 | 0 | 0 | 0 | 0 | 0 | 0 | 0 | 0 | 0 | 0 |
| 0 | NA | NA | NA | NA | NA | Gng2-ps1 | | 0 | 0 | 0 | 0 | 0 | 0 | 0 | 0 | 0 | 0 | 0 |
| 0 | NA | NA | NA | NA | NA | Gm11905 | | 0 | 0 | 0 | 0 | 0 | 0 | 0 | 0 | 0 | 0 | 0 |
| 0 | NA | NA | NA | NA | NA | 4930548K13Rik | | 0 | 0 | 0 | 0 | 0 | 0 | 0 | 0 | 0 | 0 | 0 |
| 0 | NA | NA | NA | NA | NA | Gm11904 | | 0 | 0 | 0 | 0 | 0 | 0 | 0 | 0 | 0 | 0 | 0 |
| 0 | NA | NA | NA | NA | NA | Gm11901 | | 0 | 0 | 0 | 0 | 0 | 0 | 0 | 0 | 0 | 0 | 0 |
| 0 | NA | NA | NA | NA | NA | Gm25173 | | 0 | 0 | 0 | 0 | 0 | 0 | 0 | 0 | 0 | 0 | 0 |
| 0 | NA | NA | NA | NA | NA | Tpm3-rs2 | | 0 | 0 | 0 | 0 | 0 | 0 | 0 | 0 | 0 | 0 | 0 |
| 0 | NA | NA | NA | NA | NA | Gm11911 | | 0 | 0 | 0 | 0 | 0 | 0 | 0 | 0 | 0 | 0 | 0 |
| 0 | NA | NA | NA | NA | NA | Gm11907 | | 0 | 0 | 0 | 0 | 0 | 0 | 0 | 0 | 0 | 0 | 0 |
| 0 | NA | NA | NA | NA | NA | Gm25378 | | 0 | 0 | 0 | 0 | 0 | 0 | 0 | 0 | 0 | 0 | 0 |
| 0 | NA | NA | NA | NA | NA | Gm11908 | | 0 | 0 | 0 | 0 | 0 | 0 | 0 | 0 | 0 | 0 | 0 |
| 0 | NA | NA | NA | NA | NA | Gm11909 | | 0 | 0 | 0 | 0 | 0 | 0 | 0 | 0 | 0 | 0 | 0 |
| 0 | NA | NA | NA | NA | NA | Gm11910 | | 0 | 0 | 0 | 0 | 0 | 0 | 0 | 0 | 0 | 0 | 0 |
| 0 | NA | NA | NA | NA | NA | Gm11917 | | 0 | 0 | 0 | 0 | 0 | 0 | 0 | 0 | 0 | 0 | 0 |
| 0 | NA | NA | NA | NA | NA | Gm11916 | | 0 | 0 | 0 | 0 | 0 | 0 | 0 | 0 | 0 | 0 | 0 |
| 0 | NA | NA | NA | NA | NA | Gm11918 | | 0 | 0 | 0 | 0 | 0 | 0 | 0 | 0 | 0 | 0 | 0 |
| 0 | NA | NA | NA | NA | NA | Gm11919 | | 0 | 0 | 0 | 0 | 0 | 0 | 0 | 0 | 0 | 0 | 0 |
| 0 | NA | NA | NA | NA | NA | Gm11923 | | 0 | 0 | 0 | 0 | 0 | 0 | 0 | 0 | 0 | 0 | 0 |
| 0 | NA | NA | NA | NA | NA | Gm11925 | | 0 | 0 | 0 | 0 | 0 | 0 | 0 | 0 | 0 | 0 | 0 |
| 0 | NA | NA | NA | NA | NA | Gm11924 | | 0 | 0 | 0 | 0 | 0 | 0 | 0 | 0 | 0 | 0 | 0 |
| 0 | NA | NA | NA | NA | NA | 4930556G01Rik | | 0 | 0 | 0 | 0 | 0 | 0 | 0 | 0 | 0 | 0 | 0 |
| 0 | NA | NA | NA | NA | NA | Gm22831 | | 0 | 0 | 0 | 0 | 0 | 0 | 0 | 0 | 0 | 0 | 0 |
| 0 | NA | NA | NA | NA | NA | Gm11920 | | 0 | 0 | 0 | 0 | 0 | 0 | 0 | 0 | 0 | 0 | 0 |
| 0 | NA | NA | NA | NA | NA | Gm25705 | | 0 | 0 | 0 | 0 | 0 | 0 | 0 | 0 | 0 | 0 | 0 |
| 0 | NA | NA | NA | NA | NA | Gm11922 | | 0 | 0 | 0 | 0 | 0 | 0 | 0 | 0 | 0 | 0 | 0 |
| 0 | NA | NA | NA | NA | NA | Gm11926 | | 0 | 0 | 0 | 0 | 0 | 0 | 0 | 0 | 0 | 0 | 0 |
| 0 | NA | NA | NA | NA | NA | Gm26254 | | 0 | 0 | 0 | 0 | 0 | 0 | 0 | 0 | 0 | 0 | 0 |
| 0 | NA | NA | NA | NA | NA | Gm11927 | | 0 | 0 | 0 | 0 | 0 | 0 | 0 | 0 | 0 | 0 | 0 |
| 0 | NA | NA | NA | NA | NA | Gm11928 | | 0 | 0 | 0 | 0 | 0 | 0 | 0 | 0 | 0 | 0 | 0 |
| 0 | NA | NA | NA | NA | NA | Bach2it1 | | 0 | 0 | 0 | 0 | 0 | 0 | 0 | 0 | 0 | 0 | 0 |
| 0 | NA | NA | NA | NA | NA | Gm24371 | | 0 | 0 | 0 | 0 | 0 | 0 | 0 | 0 | 0 | 0 | 0 |
| 0 | NA | NA | NA | NA | NA | Gm11933 | | 0 | 0 | 0 | 0 | 0 | 0 | 0 | 0 | 0 | 0 | 0 |
| 0 | NA | NA | NA | NA | NA | Gm11936 | | 0 | 0 | 0 | 0 | 0 | 0 | 0 | 0 | 0 | 0 | 0 |
| 0 | NA | NA | NA | NA | NA | Gm11941 | | 0 | 0 | 0 | 0 | 0 | 0 | 0 | 0 | 0 | 0 | 0 |
| 0 | NA | NA | NA | NA | NA | Gm11943 | | 0 | 0 | 0 | 0 | 0 | 0 | 0 | 0 | 0 | 0 | 0 |
| 0 | NA | NA | NA | NA | NA | Gm11931 | | 0 | 0 | 0 | 0 | 0 | 0 | 0 | 0 | 0 | 0 | 0 |
| 0 | NA | NA | NA | NA | NA | Mir8118 | | 0 | 0 | 0 | 0 | 0 | 0 | 0 | 0 | 0 | 0 | 0 |
| 0 | NA | NA | NA | NA | NA | Gm11935 | | 0 | 0 | 0 | 0 | 0 | 0 | 0 | 0 | 0 | 0 | 0 |
| 0 | NA | NA | NA | NA | NA | Gm23304 | | 0 | 0 | 0 | 0 | 0 | 0 | 0 | 0 | 0 | 0 | 0 |
| 0 | NA | NA | NA | NA | NA | Gm24341 | | 0 | 0 | 0 | 0 | 0 | 0 | 0 | 0 | 0 | 0 | 0 |
| 0 | NA | NA | NA | NA | NA | Gm22010 | | 0 | 0 | 0 | 0 | 0 | 0 | 0 | 0 | 0 | 0 | 0 |

| | | | | | | | | | | | | | | | | | |
|---|---|---|---|---|---|---|---|---|---|---|---|---|---|---|---|---|---|
| 0 | NA | NA | NA | NA | NA | Gm25623 | | 0 | 0 | 0 | 0 | 0 | 0 | 0 | 0 | 0 | 0 | 0 |
| 0 | NA | NA | NA | NA | NA | Gm12753 | | 0 | 0 | 0 | 0 | 0 | 0 | 0 | 0 | 0 | 0 | 0 |
| 0 | NA | NA | NA | NA | NA | Gm12752 | | 0 | 0 | 0 | 0 | 0 | 0 | 0 | 0 | 0 | 0 | 0 |
| 0 | NA | NA | NA | NA | NA | Gm12754 | | 0 | 0 | 0 | 0 | 0 | 0 | 0 | 0 | 0 | 0 | 0 |
| 0 | NA | NA | NA | NA | NA | Gm23466 | | 0 | 0 | 0 | 0 | 0 | 0 | 0 | 0 | 0 | 0 | 0 |
| 0 | NA | NA | NA | NA | NA | Gm12351 | | 0 | 0 | 0 | 0 | 0 | 0 | 0 | 0 | 0 | 0 | 0 |
| 0 | NA | NA | NA | NA | NA | Gm22849 | | 0 | 0 | 0 | 0 | 0 | 0 | 0 | 0 | 0 | 0 | 0 |
| 0 | NA | NA | NA | NA | NA | Gm136 | predicted gene 136 | 0 | 0 | 0 | 0 | 0 | 0 | 0 | 0 | 0 | 0 | 0 |
| 0 | NA | NA | NA | NA | NA | Gm12363 | | 0 | 0 | 0 | 0 | 0 | 0 | 0 | 0 | 0 | 0 | 0 |
| 0 | NA | NA | NA | NA | NA | Cga | glycoprotein hormones, alpha subunit | 0 | 0 | 0 | 0 | 0 | 0 | 0 | 0 | 0 | 0 | 0 |
| 0 | NA | NA | NA | NA | NA | Platr9 | | 0 | 0 | 0 | 0 | 0 | 0 | 0 | 0 | 0 | 0 | 0 |
| 0 | NA | NA | NA | NA | NA | Gm12361 | | 0 | 0 | 0 | 0 | 0 | 0 | 0 | 0 | 0 | 0 | 0 |
| 0 | NA | NA | NA | NA | NA | Gm12364 | | 0 | 0 | 0 | 0 | 0 | 0 | 0 | 0 | 0 | 0 | 0 |
| 0 | NA | NA | NA | NA | NA | Ifnk | interferon kappa | 0 | 0 | 0 | 0 | 0 | 0 | 0 | 0 | 0 | 0 | 0 |
| 0 | NA | NA | NA | NA | NA | Gm12368 | | 0 | 0 | 0 | 0 | 0 | 0 | 0 | 0 | 0 | 0 | 0 |
| 0 | NA | NA | NA | NA | NA | Gm12365 | | 0 | 0 | 0 | 0 | 0 | 0 | 0 | 0 | 0 | 0 | 0 |
| 0 | NA | NA | NA | NA | NA | Lingo2 | leucine rich repeat and Ig domain contain | 0 | 0 | 0 | 0 | 0 | 0 | 0 | 0 | 0 | 0 | 0 |
| 0 | NA | NA | NA | NA | NA | Gm12369 | | 0 | 0 | 0 | 0 | 0 | 0 | 0 | 0 | 0 | 0 | 0 |
| 0 | NA | NA | NA | NA | NA | Gm12370 | | 0 | 0 | 0 | 0 | 0 | 0 | 0 | 0 | 0 | 0 | 0 |
| 0 | NA | NA | NA | NA | NA | Mir876 | | 0 | 0 | 0 | 0 | 0 | 0 | 0 | 0 | 0 | 0 | 0 |
| 0 | NA | NA | NA | NA | NA | Mir873a | | 0 | 0 | 0 | 0 | 0 | 0 | 0 | 0 | 0 | 0 | 0 |
| 0 | NA | NA | NA | NA | NA | Mir873b | | 0 | 0 | 0 | 0 | 0 | 0 | 0 | 0 | 0 | 0 | 0 |
| 0 | NA | NA | NA | NA | NA | Gm23314 | | 0 | 0 | 0 | 0 | 0 | 0 | 0 | 0 | 0 | 0 | 0 |
| 0 | NA | NA | NA | NA | NA | Gm12371 | | 0 | 0 | 0 | 0 | 0 | 0 | 0 | 0 | 0 | 0 | 0 |
| 0 | NA | NA | NA | NA | NA | Rpl23a-ps6 | | 0 | 0 | 0 | 0 | 0 | 0 | 0 | 0 | 0 | 0 | 0 |
| 0 | NA | NA | NA | NA | NA | Gm12374 | | 0 | 0 | 0 | 0 | 0 | 0 | 0 | 0 | 0 | 0 | 0 |
| 0 | NA | NA | NA | NA | NA | Gm22732 | | 0 | 0 | 0 | 0 | 0 | 0 | 0 | 0 | 0 | 0 | 0 |
| 0 | NA | NA | NA | NA | NA | Gm12375 | | 0 | 0 | 0 | 0 | 0 | 0 | 0 | 0 | 0 | 0 | 0 |
| 0 | NA | NA | NA | NA | NA | Gm12378 | | 0 | 0 | 0 | 0 | 0 | 0 | 0 | 0 | 0 | 0 | 0 |
| 0 | NA | NA | NA | NA | NA | Gm12379 | | 0 | 0 | 0 | 0 | 0 | 0 | 0 | 0 | 0 | 0 | 0 |
| 0 | NA | NA | NA | NA | NA | Gm12381 | | 0 | 0 | 0 | 0 | 0 | 0 | 0 | 0 | 0 | 0 | 0 |
| 0 | NA | NA | NA | NA | NA | Gm12382 | | 0 | 0 | 0 | 0 | 0 | 0 | 0 | 0 | 0 | 0 | 0 |
| 0 | NA | NA | NA | NA | NA | Gm25581 | | 0 | 0 | 0 | 0 | 0 | 0 | 0 | 0 | 0 | 0 | 0 |
| 0 | NA | NA | NA | NA | NA | Gm12383 | | 0 | 0 | 0 | 0 | 0 | 0 | 0 | 0 | 0 | 0 | 0 |
| 0 | NA | NA | NA | NA | NA | Gm23607 | | 0 | 0 | 0 | 0 | 0 | 0 | 0 | 0 | 0 | 0 | 0 |
| 0 | NA | NA | NA | NA | NA | Gm17094 | | 0 | 0 | 0 | 0 | 0 | 0 | 0 | 0 | 0 | 0 | 0 |
| 0 | NA | NA | NA | NA | NA | 1700009N14Ril | RIKEN cDNA 1700009N14 gene | 0 | 0 | 0 | 0 | 0 | 0 | 0 | 0 | 0 | 0 | 0 |
| 0 | NA | NA | NA | NA | NA | Gm12385 | | 0 | 0 | 0 | 0 | 0 | 0 | 0 | 0 | 0 | 0 | 0 |
| 0 | NA | NA | NA | NA | NA | Gm12384 | | 0 | 0 | 0 | 0 | 0 | 0 | 0 | 0 | 0 | 0 | 0 |
| 0 | NA | NA | NA | NA | NA | Gm12386 | | 0 | 0 | 0 | 0 | 0 | 0 | 0 | 0 | 0 | 0 | 0 |
| 0 | NA | NA | NA | NA | NA | Gm22850 | | 0 | 0 | 0 | 0 | 0 | 0 | 0 | 0 | 0 | 0 | 0 |
| 0 | NA | NA | NA | NA | NA | Gm12387 | | 0 | 0 | 0 | 0 | 0 | 0 | 0 | 0 | 0 | 0 | 0 |
| 0 | NA | NA | NA | NA | NA | Gm26087 | | 0 | 0 | 0 | 0 | 0 | 0 | 0 | 0 | 0 | 0 | 0 |
| 0 | NA | NA | NA | NA | NA | Gm12388 | | 0 | 0 | 0 | 0 | 0 | 0 | 0 | 0 | 0 | 0 | 0 |
| 0 | NA | NA | NA | NA | NA | 4930509K18Rik | | 0 | 0 | 0 | 0 | 0 | 0 | 0 | 0 | 0 | 0 | 0 |
| 0 | NA | NA | NA | NA | NA | Gm12391 | | 0 | 0 | 0 | 0 | 0 | 0 | 0 | 0 | 0 | 0 | 0 |
| 0 | NA | NA | NA | NA | NA | Mir5123 | | 0 | 0 | 0 | 0 | 0 | 0 | 0 | 0 | 0 | 0 | 0 |
| 0 | NA | NA | NA | NA | NA | Gm24112 | | 0 | 0 | 0 | 0 | 0 | 0 | 0 | 0 | 0 | 0 | 0 |
| 0 | NA | NA | NA | NA | NA | Gm25931 | | 0 | 0 | 0 | 0 | 0 | 0 | 0 | 0 | 0 | 0 | 0 |
| 0 | NA | NA | NA | NA | NA | Gm12402 | | 0 | 0 | 0 | 0 | 0 | 0 | 0 | 0 | 0 | 0 | 0 |
| 0 | NA | NA | NA | NA | NA | Gm24650 | | 0 | 0 | 0 | 0 | 0 | 0 | 0 | 0 | 0 | 0 | 0 |
| 0 | NA | NA | NA | NA | NA | Gm23090 | | 0 | 0 | 0 | 0 | 0 | 0 | 0 | 0 | 0 | 0 | 0 |
| 0 | NA | NA | NA | NA | NA | Gm12401 | | 0 | 0 | 0 | 0 | 0 | 0 | 0 | 0 | 0 | 0 | 0 |
| 0 | NA | NA | NA | NA | NA | n-R5s183 | | 0 | 0 | 0 | 0 | 0 | 0 | 0 | 0 | 0 | 0 | 0 |
| 0 | NA | NA | NA | NA | NA | Gm26084 | | 0 | 0 | 0 | 0 | 0 | 0 | 0 | 0 | 0 | 0 | 0 |
| 0 | NA | NA | NA | NA | NA | Fam219aos | | 0 | 0 | 0 | 0 | 0 | 0 | 0 | 0 | 0 | 0 | 0 |
| 0 | NA | NA | NA | NA | NA | Gm10601 | | 0 | 0 | 0 | 0 | 0 | 0 | 0 | 0 | 0 | 0 | 0 |
| 0 | NA | NA | NA | NA | NA | Gm13302 | predicted gene 13302 | 0 | 0 | 0 | 0 | 0 | 0 | 0 | 0 | 0 | 0 | 0 |
| 0 | NA | NA | NA | NA | NA | Ccl21d | C-C motif chemokine 21c | 0 | 0 | 0 | 0 | 0 | 0 | 0 | 0 | 0 | 0 | 0 |
| 0 | NA | NA | NA | NA | NA | Ccl19-ps1 | | 0 | 0 | 0 | 0 | 0 | 0 | 0 | 0 | 0 | 0 | 0 |

| | | | | | | | | | | | | | | | | | |
|---|---|---|---|---|---|---|---|---|---|---|---|---|---|---|---|---|---|
| 0 | NA | NA | NA | NA | NA | Gm20878 | predicted gene, 20878 | 0 | 0 | 0 | 0 | 0 | 0 | 0 | 0 | 0 | 0 | 0 |
| 0 | NA | NA | NA | NA | NA | Fam205a4 | family with sequence similarity 205, meml | 0 | 0 | 0 | 0 | 0 | 0 | 0 | 0 | 0 | 0 | 0 |
| 0 | NA | NA | NA | NA | NA | Gm13303 | | 0 | 0 | 0 | 0 | 0 | 0 | 0 | 0 | 0 | 0 | 0 |
| 0 | NA | NA | NA | NA | NA | Gm21093 | | 0 | 0 | 0 | 0 | 0 | 0 | 0 | 0 | 0 | 0 | 0 |
| 0 | NA | NA | NA | NA | NA | Gm37530 | | 0 | 0 | 0 | 0 | 0 | 0 | 0 | 0 | 0 | 0 | 0 |
| 0 | NA | NA | NA | NA | NA | Gm3892 | | 0 | 0 | 0 | 0 | 0 | 0 | 0 | 0 | 0 | 0 | 0 |
| 0 | NA | NA | NA | NA | NA | Gm3893 | | 0 | 0 | 0 | 0 | 0 | 0 | 0 | 0 | 0 | 0 | 0 |
| 0 | NA | NA | NA | NA | NA | Gm13304 | C-C motif chemokine 21c | 0 | 0 | 0 | 0 | 0 | 0 | 0 | 0 | 0 | 0 | 0 |
| 0 | NA | NA | NA | NA | NA | Gm13299 | | 0 | 0 | 0 | 0 | 0 | 0 | 0 | 0 | 0 | 0 | 0 |
| 0 | NA | NA | NA | NA | NA | Ccl19-ps3 | | 0 | 0 | 0 | 0 | 0 | 0 | 0 | 0 | 0 | 0 | 0 |
| 0 | NA | NA | NA | NA | NA | Gm13305 | predicted gene 13305 | 0 | 0 | 0 | 0 | 0 | 0 | 0 | 0 | 0 | 0 | 0 |
| 0 | NA | NA | NA | NA | NA | 1700045I11Rik | | 0 | 0 | 0 | 0 | 0 | 0 | 0 | 0 | 0 | 0 | 0 |
| 0 | NA | NA | NA | NA | NA | Gm13301 | | 0 | 0 | 0 | 0 | 0 | 0 | 0 | 0 | 0 | 0 | 0 |
| 0 | NA | NA | NA | NA | NA | Gm10600 | | 0 | 0 | 0 | 0 | 0 | 0 | 0 | 0 | 0 | 0 | 0 |
| 0 | NA | NA | NA | NA | NA | Gm13301 | | 0 | 0 | 0 | 0 | 0 | 0 | 0 | 0 | 0 | 0 | 0 |
| 0 | NA | NA | NA | NA | NA | Gm5859 | | 0 | 0 | 0 | 0 | 0 | 0 | 0 | 0 | 0 | 0 | 0 |
| 0 | NA | NA | NA | NA | NA | Gm17081 | | 0 | 0 | 0 | 0 | 0 | 0 | 0 | 0 | 0 | 0 | 0 |
| 0 | NA | NA | NA | NA | NA | Gm10599 | | 0 | 0 | 0 | 0 | 0 | 0 | 0 | 0 | 0 | 0 | 0 |
| 0 | NA | NA | NA | NA | NA | Gm50470 | | 0 | 0 | 0 | 0 | 0 | 0 | 0 | 0 | 0 | 0 | 0 |
| 0 | NA | NA | NA | NA | NA | Ccl21b | C-C motif chemokine 21c | 0 | 0 | 0 | 0 | 0 | 0 | 0 | 0 | 0 | 0 | 0 |
| 0 | NA | NA | NA | NA | NA | Gm50471 | | 0 | 0 | 0 | 0 | 0 | 0 | 0 | 0 | 0 | 0 | 0 |
| 0 | NA | NA | NA | NA | NA | Gm21980 | | 0 | 0 | 0 | 0 | 0 | 0 | 0 | 0 | 0 | 0 | 0 |
| 0 | NA | NA | NA | NA | NA | Gm21953 | predicted gene, 21953 | 0 | 0 | 0 | 0 | 0 | 0 | 0 | 0 | 0 | 0 | 0 |
| 0 | NA | NA | NA | NA | NA | Fam205a3 | family with sequence similarity 205, meml | 0 | 0 | 0 | 0 | 0 | 0 | 0 | 0 | 0 | 0 | 0 |
| 0 | NA | NA | NA | NA | NA | Gm21586 | predicted gene, 21586 | 0 | 0 | 0 | 0 | 0 | 0 | 0 | 0 | 0 | 0 | 0 |
| 0 | NA | NA | NA | NA | NA | Gm3883 | | 0 | 0 | 0 | 0 | 0 | 0 | 0 | 0 | 0 | 0 | 0 |
| 0 | NA | NA | NA | NA | NA | Gm2163 | | 0 | 0 | 0 | 0 | 0 | 0 | 0 | 0 | 0 | 0 | 0 |
| 0 | NA | NA | NA | NA | NA | Fam205a2 | family with sequence similarity 205, meml | 0 | 0 | 0 | 0 | 0 | 0 | 0 | 0 | 0 | 0 | 0 |
| 0 | NA | NA | NA | NA | NA | Gm10593 | | 0 | 0 | 0 | 0 | 0 | 0 | 0 | 0 | 0 | 0 | 0 |
| 0 | NA | NA | NA | NA | NA | | protein FAM205A-2-like | 0 | 0 | 0 | 0 | 0 | 0 | 0 | 0 | 0 | 0 | 0 |
| 0 | NA | NA | NA | NA | NA | Gm10591 | C-C motif chemokine 21c | 0 | 0 | 0 | 0 | 0 | 0 | 0 | 0 | 0 | 0 | 0 |
| 0 | NA | NA | NA | NA | NA | Gm2564 | | 0 | 0 | 0 | 0 | 0 | 0 | 0 | 0 | 0 | 0 | 0 |
| 0 | NA | NA | NA | NA | NA | Ccl27b | chemokine (C-C motif) ligand 27b | 0 | 0 | 0 | 0 | 0 | 0 | 0 | 0 | 0 | 0 | 0 |
| 0 | NA | NA | NA | NA | NA | Il11ra2 | interleukin 11 receptor, alpha chain 2 | 0 | 0 | 0 | 0 | 0 | 0 | 0 | 0 | 0 | 0 | 0 |
| 0 | NA | NA | NA | NA | NA | 4930578G10Rik | | 0 | 0 | 0 | 0 | 0 | 0 | 0 | 0 | 0 | 0 | 0 |
| 0 | NA | NA | NA | NA | NA | Gm12395 | | 0 | 0 | 0 | 0 | 0 | 0 | 0 | 0 | 0 | 0 | 0 |
| 0 | NA | NA | NA | NA | NA | Gm12394 | | 0 | 0 | 0 | 0 | 0 | 0 | 0 | 0 | 0 | 0 | 0 |
| 0 | NA | NA | NA | NA | NA | Fam205a1 | family with sequence similarity 205, meml | 0 | 0 | 0 | 0 | 0 | 0 | 0 | 0 | 0 | 0 | 0 |
| 0 | NA | NA | NA | NA | NA | Gm23709 | | 0 | 0 | 0 | 0 | 0 | 0 | 0 | 0 | 0 | 0 | 0 |
| 0 | NA | NA | NA | NA | NA | Gm25010 | | 0 | 0 | 0 | 0 | 0 | 0 | 0 | 0 | 0 | 0 | 0 |
| 0 | NA | NA | NA | NA | NA | Rmrp | | 0 | 0 | 0 | 0 | 0 | 0 | 0 | 0 | 0 | 0 | 0 |
| 0 | NA | NA | NA | NA | NA | Rmrp | | 0 | 0 | 0 | 0 | 0 | 0 | 0 | 0 | 0 | 0 | 0 |
| 0 | NA | NA | NA | NA | NA | Olfr70 | olfactory receptor 70 | 0 | 0 | 0 | 0 | 0 | 0 | 0 | 0 | 0 | 0 | 0 |
| 0 | NA | NA | NA | NA | NA | Olfr71 | olfactory receptor 71 | 0 | 0 | 0 | 0 | 0 | 0 | 0 | 0 | 0 | 0 | 0 |
| 0 | NA | NA | NA | NA | NA | Olfr269-ps1 | | 0 | 0 | 0 | 0 | 0 | 0 | 0 | 0 | 0 | 0 | 0 |
| 0 | NA | NA | NA | NA | NA | Olfr159 | olfactory receptor 159 | 0 | 0 | 0 | 0 | 0 | 0 | 0 | 0 | 0 | 0 | 0 |
| 0 | NA | NA | NA | NA | NA | Olfr29-ps1 | | 0 | 0 | 0 | 0 | 0 | 0 | 0 | 0 | 0 | 0 | 0 |
| 0 | NA | NA | NA | NA | NA | Olfr156 | olfactory receptor 156 | 0 | 0 | 0 | 0 | 0 | 0 | 0 | 0 | 0 | 0 | 0 |
| 0 | NA | NA | NA | NA | NA | Olfr157 | olfactory receptor 157 | 0 | 0 | 0 | 0 | 0 | 0 | 0 | 0 | 0 | 0 | 0 |
| 0 | NA | NA | NA | NA | NA | Olfr155 | olfactory receptor 155 | 0 | 0 | 0 | 0 | 0 | 0 | 0 | 0 | 0 | 0 | 0 |
| 0 | NA | NA | NA | NA | NA | Gm27265 | | 0 | 0 | 0 | 0 | 0 | 0 | 0 | 0 | 0 | 0 | 0 |
| 0 | NA | NA | NA | NA | NA | Ccin | calicin | 0 | 0 | 0 | 0 | 0 | 0 | 0 | 0 | 0 | 0 | 0 |
| 0 | NA | NA | NA | NA | NA | Gm22706 | | 0 | 0 | 0 | 0 | 0 | 0 | 0 | 0 | 0 | 0 | 0 |
| 0 | NA | NA | NA | NA | NA | Gm12503 | | 0 | 0 | 0 | 0 | 0 | 0 | 0 | 0 | 0 | 0 | 0 |
| 0 | NA | NA | NA | NA | NA | Gm12677 | | 0 | 0 | 0 | 0 | 0 | 0 | 0 | 0 | 0 | 0 | 0 |
| 0 | NA | NA | NA | NA | NA | Gm12455 | | 0 | 0 | 0 | 0 | 0 | 0 | 0 | 0 | 0 | 0 | 0 |
| 0 | NA | NA | NA | NA | NA | Gm24376 | | 0 | 0 | 0 | 0 | 0 | 0 | 0 | 0 | 0 | 0 | 0 |
| 0 | NA | NA | NA | NA | NA | Gm12462 | | 0 | 0 | 0 | 0 | 0 | 0 | 0 | 0 | 0 | 0 | 0 |
| 0 | NA | NA | NA | NA | NA | 4930517J16Rik | | 0 | 0 | 0 | 0 | 0 | 0 | 0 | 0 | 0 | 0 | 0 |
| 0 | NA | NA | NA | NA | NA | 5730488B01Rik | | 0 | 0 | 0 | 0 | 0 | 0 | 0 | 0 | 0 | 0 | 0 |

| | | | | | | | | | | | | | | | | | |
|---|---|---|---|---|---|---|---|---|---|---|---|---|---|---|---|---|---|
| 0 | NA | NA | NA | NA | NA | Gm22639 | | 0 | 0 | 0 | 0 | 0 | 0 | 0 | 0 | 0 | 0 | 0 |
| 0 | NA | NA | NA | NA | NA | Gm12493 | | 0 | 0 | 0 | 0 | 0 | 0 | 0 | 0 | 0 | 0 | 0 |
| 0 | NA | NA | NA | NA | NA | Gm12566 | | 0 | 0 | 0 | 0 | 0 | 0 | 0 | 0 | 0 | 0 | 0 |
| 0 | NA | NA | NA | NA | NA | Frmpd1os | | 0 | 0 | 0 | 0 | 0 | 0 | 0 | 0 | 0 | 0 | 0 |
| 0 | NA | NA | NA | NA | NA | Gm12585 | | 0 | 0 | 0 | 0 | 0 | 0 | 0 | 0 | 0 | 0 | 0 |
| 0 | NA | NA | NA | NA | NA | Gm25692 | | 0 | 0 | 0 | 0 | 0 | 0 | 0 | 0 | 0 | 0 | 0 |
| 0 | NA | NA | NA | NA | NA | Gm22055 | | 0 | 0 | 0 | 0 | 0 | 0 | 0 | 0 | 0 | 0 | 0 |
| 0 | NA | NA | NA | NA | NA | Gm12410 | | 0 | 0 | 0 | 0 | 0 | 0 | 0 | 0 | 0 | 0 | 0 |
| 0 | NA | NA | NA | NA | NA | Gm23250 | | 0 | 0 | 0 | 0 | 0 | 0 | 0 | 0 | 0 | 0 | 0 |
| 0 | NA | NA | NA | NA | NA | Gm12445 | | 0 | 0 | 0 | 0 | 0 | 0 | 0 | 0 | 0 | 0 | 0 |
| 0 | NA | NA | NA | NA | NA | Gm12446 | | 0 | 0 | 0 | 0 | 0 | 0 | 0 | 0 | 0 | 0 | 0 |
| 0 | NA | NA | NA | NA | NA | Foxe1 | forkhead box E1 | 0 | 0 | 0 | 0 | 0 | 0 | 0 | 0 | 0 | 0 | 0 |
| 0 | NA | NA | NA | NA | NA | Gm12443 | | 0 | 0 | 0 | 0 | 0 | 0 | 0 | 0 | 0 | 0 | 0 |
| 0 | NA | NA | NA | NA | NA | Gm22247 | | 0 | 0 | 0 | 0 | 0 | 0 | 0 | 0 | 0 | 0 | 0 |
| 0 | NA | NA | NA | NA | NA | Gm12425 | | 0 | 0 | 0 | 0 | 0 | 0 | 0 | 0 | 0 | 0 | 0 |
| 0 | NA | NA | NA | NA | NA | Gm12423 | | 0 | 0 | 0 | 0 | 0 | 0 | 0 | 0 | 0 | 0 | 0 |
| 0 | NA | NA | NA | NA | NA | Gm27601 | | 0 | 0 | 0 | 0 | 0 | 0 | 0 | 0 | 0 | 0 | 0 |
| 0 | NA | NA | NA | NA | NA | Gm27668 | | 0 | 0 | 0 | 0 | 0 | 0 | 0 | 0 | 0 | 0 | 0 |
| 0 | NA | NA | NA | NA | NA | Gm38343 | | 0 | 0 | 0 | 0 | 0 | 0 | 0 | 0 | 0 | 0 | 0 |
| 0 | NA | NA | NA | NA | NA | Gm22670 | | 0 | 0 | 0 | 0 | 0 | 0 | 0 | 0 | 0 | 0 | 0 |
| 0 | NA | NA | NA | NA | NA | Gm12434 | | 0 | 0 | 0 | 0 | 0 | 0 | 0 | 0 | 0 | 0 | 0 |
| 0 | NA | NA | NA | NA | NA | Mir1958 | | 0 | 0 | 0 | 0 | 0 | 0 | 0 | 0 | 0 | 0 | 0 |
| 0 | NA | NA | NA | NA | NA | Gm12438 | | 0 | 0 | 0 | 0 | 0 | 0 | 0 | 0 | 0 | 0 | 0 |
| 0 | NA | NA | NA | NA | NA | Gm24573 | | 0 | 0 | 0 | 0 | 0 | 0 | 0 | 0 | 0 | 0 | 0 |
| 0 | NA | NA | NA | NA | NA | Gm23746 | | 0 | 0 | 0 | 0 | 0 | 0 | 0 | 0 | 0 | 0 | 0 |
| 0 | NA | NA | NA | NA | NA | Gm12436 | | 0 | 0 | 0 | 0 | 0 | 0 | 0 | 0 | 0 | 0 | 0 |
| 0 | NA | NA | NA | NA | NA | Gm12442 | | 0 | 0 | 0 | 0 | 0 | 0 | 0 | 0 | 0 | 0 | 0 |
| 0 | NA | NA | NA | NA | NA | Gm22685 | | 0 | 0 | 0 | 0 | 0 | 0 | 0 | 0 | 0 | 0 | 0 |
| 0 | NA | NA | NA | NA | NA | Rbpj-ps1 | | 0 | 0 | 0 | 0 | 0 | 0 | 0 | 0 | 0 | 0 | 0 |
| 0 | NA | NA | NA | NA | NA | Acnat2 | acyl-coenzyme A amino acid N-acyltransf | 0 | 0 | 0 | 0 | 0 | 0 | 0 | 0 | 0 | 0 | 0 |
| 0 | NA | NA | NA | NA | NA | Gm12451 | | 0 | 0 | 0 | 0 | 0 | 0 | 0 | 0 | 0 | 0 | 0 |
| 0 | NA | NA | NA | NA | NA | Gm12452 | | 0 | 0 | 0 | 0 | 0 | 0 | 0 | 0 | 0 | 0 | 0 |
| 0 | NA | NA | NA | NA | NA | Gm26424 | | 0 | 0 | 0 | 0 | 0 | 0 | 0 | 0 | 0 | 0 | 0 |
| 0 | NA | NA | NA | NA | NA | Ppp3r2 | protein phosphatase 3, regulatory subuni | 0 | 0 | 0 | 0 | 0 | 0 | 0 | 0 | 0 | 0 | 0 |
| 0 | NA | NA | NA | NA | NA | Gm12459 | | 0 | 0 | 0 | 0 | 0 | 0 | 0 | 0 | 0 | 0 | 0 |
| 0 | NA | NA | NA | NA | NA | Gm25485 | | 0 | 0 | 0 | 0 | 0 | 0 | 0 | 0 | 0 | 0 | 0 |
| 0 | NA | NA | NA | NA | NA | Cylc2 | cylicin, basic protein of sperm head cytosl | 0 | 0 | 0 | 0 | 0 | 0 | 0 | 0 | 0 | 0 | 0 |
| 0 | NA | NA | NA | NA | NA | Gm22563 | | 0 | 0 | 0 | 0 | 0 | 0 | 0 | 0 | 0 | 0 | 0 |
| 0 | NA | NA | NA | NA | NA | Gm23043 | | 0 | 0 | 0 | 0 | 0 | 0 | 0 | 0 | 0 | 0 | 0 |
| 0 | NA | NA | NA | NA | NA | Gm24943 | | 0 | 0 | 0 | 0 | 0 | 0 | 0 | 0 | 0 | 0 | 0 |
| 0 | NA | NA | NA | NA | NA | Gm22042 | | 0 | 0 | 0 | 0 | 0 | 0 | 0 | 0 | 0 | 0 | 0 |
| 0 | NA | NA | NA | NA | NA | Gm12465 | | 0 | 0 | 0 | 0 | 0 | 0 | 0 | 0 | 0 | 0 | 0 |
| 0 | NA | NA | NA | NA | NA | Gm12467 | | 0 | 0 | 0 | 0 | 0 | 0 | 0 | 0 | 0 | 0 | 0 |
| 0 | NA | NA | NA | NA | NA | Tmc4-ps | | 0 | 0 | 0 | 0 | 0 | 0 | 0 | 0 | 0 | 0 | 0 |
| 0 | NA | NA | NA | NA | NA | Toporsl | topoisomerase I binding, arginine/serine- | 0 | 0 | 0 | 0 | 0 | 0 | 0 | 0 | 0 | 0 | 0 |
| 0 | NA | NA | NA | NA | NA | Olfr275 | olfactory receptor 275 | 0 | 0 | 0 | 0 | 0 | 0 | 0 | 0 | 0 | 0 | 0 |
| 0 | NA | NA | NA | NA | NA | Olfr274-ps1 | | 0 | 0 | 0 | 0 | 0 | 0 | 0 | 0 | 0 | 0 | 0 |
| 0 | NA | NA | NA | NA | NA | Gm47786 | | 0 | 0 | 0 | 0 | 0 | 0 | 0 | 0 | 0 | 0 | 0 |
| 0 | NA | NA | NA | NA | NA | Olfr272 | olfactory receptor 272 | 0 | 0 | 0 | 0 | 0 | 0 | 0 | 0 | 0 | 0 | 0 |
| 0 | NA | NA | NA | NA | NA | Olfr271-ps1 | | 0 | 0 | 0 | 0 | 0 | 0 | 0 | 0 | 0 | 0 | 0 |
| 0 | NA | NA | NA | NA | NA | Olfr270 | olfactory receptor 270 | 0 | 0 | 0 | 0 | 0 | 0 | 0 | 0 | 0 | 0 | 0 |
| 0 | NA | NA | NA | NA | NA | Nipsnap3a | nipsnap homolog 3A | 0 | 0 | 0 | 0 | 0 | 0 | 0 | 0 | 0 | 0 | 0 |
| 0 | NA | NA | NA | NA | NA | 4930522O17Rik | | 0 | 0 | 0 | 0 | 0 | 0 | 0 | 0 | 0 | 0 | 0 |
| 0 | NA | NA | NA | NA | NA | Gm12495 | | 0 | 0 | 0 | 0 | 0 | 0 | 0 | 0 | 0 | 0 | 0 |
| 0 | NA | NA | NA | NA | NA | Gm12496 | | 0 | 0 | 0 | 0 | 0 | 0 | 0 | 0 | 0 | 0 | 0 |
| 0 | NA | NA | NA | NA | NA | Gm25445 | | 0 | 0 | 0 | 0 | 0 | 0 | 0 | 0 | 0 | 0 | 0 |
| 0 | NA | NA | NA | NA | NA | n-R5s185 | | 0 | 0 | 0 | 0 | 0 | 0 | 0 | 0 | 0 | 0 | 0 |
| 0 | NA | NA | NA | NA | NA | Gm23745 | | 0 | 0 | 0 | 0 | 0 | 0 | 0 | 0 | 0 | 0 | 0 |
| 0 | NA | NA | NA | NA | NA | Gm24177 | | 0 | 0 | 0 | 0 | 0 | 0 | 0 | 0 | 0 | 0 | 0 |
| 0 | NA | NA | NA | NA | NA | Gm12484 | | 0 | 0 | 0 | 0 | 0 | 0 | 0 | 0 | 0 | 0 | 0 |

| | | | | | | | | | | | | | | | | | | |
|---|---|---|---|---|---|---|---|---|---|---|---|---|---|---|---|---|---|---|
| 0 | NA | NA | NA | NA | NA | Gm25687 | | 0 | 0 | 0 | 0 | 0 | 0 | 0 | 0 | 0 | 0 | 0 |
| 0 | NA | NA | NA | NA | NA | Gm29067 | | 0 | 0 | 0 | 0 | 0 | 0 | 0 | 0 | 0 | 0 | 0 |
| 0 | NA | NA | NA | NA | NA | Gm12480 | | 0 | 0 | 0 | 0 | 0 | 0 | 0 | 0 | 0 | 0 | 0 |
| 0 | NA | NA | NA | NA | NA | Gm12477 | | 0 | 0 | 0 | 0 | 0 | 0 | 0 | 0 | 0 | 0 | 0 |
| 0 | NA | NA | NA | NA | NA | Gm24628 | | 0 | 0 | 0 | 0 | 0 | 0 | 0 | 0 | 0 | 0 | 0 |
| 0 | NA | NA | NA | NA | NA | Gm12479 | | 0 | 0 | 0 | 0 | 0 | 0 | 0 | 0 | 0 | 0 | 0 |
| 0 | NA | NA | NA | NA | NA | Gm12515 | | 0 | 0 | 0 | 0 | 0 | 0 | 0 | 0 | 0 | 0 | 0 |
| 0 | NA | NA | NA | NA | NA | Gm12514 | | 0 | 0 | 0 | 0 | 0 | 0 | 0 | 0 | 0 | 0 | 0 |
| 0 | NA | NA | NA | NA | NA | Gm12509 | | 0 | 0 | 0 | 0 | 0 | 0 | 0 | 0 | 0 | 0 | 0 |
| 0 | NA | NA | NA | NA | NA | Gm12512 | | 0 | 0 | 0 | 0 | 0 | 0 | 0 | 0 | 0 | 0 | 0 |
| 0 | NA | NA | NA | NA | NA | Gm25419 | | 0 | 0 | 0 | 0 | 0 | 0 | 0 | 0 | 0 | 0 | 0 |
| 0 | NA | NA | NA | NA | NA | Gm12505 | | 0 | 0 | 0 | 0 | 0 | 0 | 0 | 0 | 0 | 0 | 0 |
| 0 | NA | NA | NA | NA | NA | Gm12510 | | 0 | 0 | 0 | 0 | 0 | 0 | 0 | 0 | 0 | 0 | 0 |
| 0 | NA | NA | NA | NA | NA | Gm22150 | | 0 | 0 | 0 | 0 | 0 | 0 | 0 | 0 | 0 | 0 | 0 |
| 0 | NA | NA | NA | NA | NA | Gm12507 | | 0 | 0 | 0 | 0 | 0 | 0 | 0 | 0 | 0 | 0 | 0 |
| 0 | NA | NA | NA | NA | NA | Gm12506 | | 0 | 0 | 0 | 0 | 0 | 0 | 0 | 0 | 0 | 0 | 0 |
| 0 | NA | NA | NA | NA | NA | Gm12520 | | 0 | 0 | 0 | 0 | 0 | 0 | 0 | 0 | 0 | 0 | 0 |
| 0 | NA | NA | NA | NA | NA | Gm12518 | | 0 | 0 | 0 | 0 | 0 | 0 | 0 | 0 | 0 | 0 | 0 |
| 0 | NA | NA | NA | NA | NA | Gm26144 | | 0 | 0 | 0 | 0 | 0 | 0 | 0 | 0 | 0 | 0 | 0 |
| 0 | NA | NA | NA | NA | NA | Actl7b | actin-like 7b | 0 | 0 | 0 | 0 | 0 | 0 | 0 | 0 | 0 | 0 | 0 |
| 0 | NA | NA | NA | NA | NA | Gm26657 | | 0 | 0 | 0 | 0 | 0 | 0 | 0 | 0 | 0 | 0 | 0 |
| 0 | NA | NA | NA | NA | NA | Actl7a | actin-like 7a | 0 | 0 | 0 | 0 | 0 | 0 | 0 | 0 | 0 | 0 | 0 |
| 0 | NA | NA | NA | NA | NA | Gm25053 | | 0 | 0 | 0 | 0 | 0 | 0 | 0 | 0 | 0 | 0 | 0 |
| 0 | NA | NA | NA | NA | NA | Gm25053 | | 0 | 0 | 0 | 0 | 0 | 0 | 0 | 0 | 0 | 0 | 0 |
| 0 | NA | NA | NA | NA | NA | Gm22300 | | 0 | 0 | 0 | 0 | 0 | 0 | 0 | 0 | 0 | 0 | 0 |
| 0 | NA | NA | NA | NA | NA | 1700042G15Rik | | 0 | 0 | 0 | 0 | 0 | 0 | 0 | 0 | 0 | 0 | 0 |
| 0 | NA | NA | NA | NA | NA | Gm12539 | | 0 | 0 | 0 | 0 | 0 | 0 | 0 | 0 | 0 | 0 | 0 |
| 0 | NA | NA | NA | NA | NA | Pakap | paralemmin A kinase anchor protein | 0 | 0 | 0 | 0 | 0 | 0 | 0 | 0 | 0 | 0 | 0 |
| 0 | NA | NA | NA | NA | NA | Gm12526 | | 0 | 0 | 0 | 0 | 0 | 0 | 0 | 0 | 0 | 0 | 0 |
| 0 | NA | NA | NA | NA | NA | Gm24277 | | 0 | 0 | 0 | 0 | 0 | 0 | 0 | 0 | 0 | 0 | 0 |
| 0 | NA | NA | NA | NA | NA | Gm26138 | | 0 | 0 | 0 | 0 | 0 | 0 | 0 | 0 | 0 | 0 | 0 |
| 0 | NA | NA | NA | NA | NA | Txndc8 | thioredoxin domain containing 8 | 0 | 0 | 0 | 0 | 0 | 0 | 0 | 0 | 0 | 0 | 0 |
| 0 | NA | NA | NA | NA | NA | Mir3095 | | 0 | 0 | 0 | 0 | 0 | 0 | 0 | 0 | 0 | 0 | 0 |
| 0 | NA | NA | NA | NA | NA | Gm12578 | | 0 | 0 | 0 | 0 | 0 | 0 | 0 | 0 | 0 | 0 | 0 |
| 0 | NA | NA | NA | NA | NA | Olfr267 | olfactory receptor 267 | 0 | 0 | 0 | 0 | 0 | 0 | 0 | 0 | 0 | 0 | 0 |
| 0 | NA | NA | NA | NA | NA | Gm26400 | | 0 | 0 | 0 | 0 | 0 | 0 | 0 | 0 | 0 | 0 | 0 |
| 0 | NA | NA | NA | NA | NA | Gm20503 | | 0 | 0 | 0 | 0 | 0 | 0 | 0 | 0 | 0 | 0 | 0 |
| 0 | NA | NA | NA | NA | NA | Shoc1 | shortage in chiasmata 1 | 0 | 0 | 0 | 0 | 0 | 0 | 0 | 0 | 0 | 0 | 0 |
| 0 | NA | NA | NA | NA | NA | Gm12593 | | 0 | 0 | 0 | 0 | 0 | 0 | 0 | 0 | 0 | 0 | 0 |
| 0 | NA | NA | NA | NA | NA | Gm12594 | | 0 | 0 | 0 | 0 | 0 | 0 | 0 | 0 | 0 | 0 | 0 |
| 0 | NA | NA | NA | NA | NA | Gm12596 | | 0 | 0 | 0 | 0 | 0 | 0 | 0 | 0 | 0 | 0 | 0 |
| 0 | NA | NA | NA | NA | NA | Gm12529 | | 0 | 0 | 0 | 0 | 0 | 0 | 0 | 0 | 0 | 0 | 0 |
| 0 | NA | NA | NA | NA | NA | Gm12542 | | 0 | 0 | 0 | 0 | 0 | 0 | 0 | 0 | 0 | 0 | 0 |
| 0 | NA | NA | NA | NA | NA | Mup4 | major urinary protein 4 | 0 | 0 | 0 | 0 | 0 | 0 | 0 | 0 | 0 | 0 | 0 |
| 0 | NA | NA | NA | NA | NA | Mup-ps1 | | 0 | 0 | 0 | 0 | 0 | 0 | 0 | 0 | 0 | 0 | 0 |
| 0 | NA | NA | NA | NA | NA | Mup2 | major urinary protein 2 | 0 | 0 | 0 | 0 | 0 | 0 | 0 | 0 | 0 | 0 | 0 |
| 0 | NA | NA | NA | NA | NA | Mup-ps2 | | 0 | 0 | 0 | 0 | 0 | 0 | 0 | 0 | 0 | 0 | 0 |
| 0 | NA | NA | NA | NA | NA | Mup8 | major urinary protein 8 | 0 | 0 | 0 | 0 | 0 | 0 | 0 | 0 | 0 | 0 | 0 |
| 0 | NA | NA | NA | NA | NA | Mup-ps3 | | 0 | 0 | 0 | 0 | 0 | 0 | 0 | 0 | 0 | 0 | 0 |
| 0 | NA | NA | NA | NA | NA | Mup-ps29 | | 0 | 0 | 0 | 0 | 0 | 0 | 0 | 0 | 0 | 0 | 0 |
| 0 | NA | NA | NA | NA | NA | Mup-ps28 | | 0 | 0 | 0 | 0 | 0 | 0 | 0 | 0 | 0 | 0 | 0 |
| 0 | NA | NA | NA | NA | NA | Mup9 | major urinary protein 9 | 0 | 0 | 0 | 0 | 0 | 0 | 0 | 0 | 0 | 0 | 0 |
| 0 | NA | NA | NA | NA | NA | Mup-ps4 | | 0 | 0 | 0 | 0 | 0 | 0 | 0 | 0 | 0 | 0 | 0 |
| 0 | NA | NA | NA | NA | NA | Mup-ps26 | | 0 | 0 | 0 | 0 | 0 | 0 | 0 | 0 | 0 | 0 | 0 |
| 0 | NA | NA | NA | NA | NA | Mup1 | major urinary protein 1 | 0 | 0 | 0 | 0 | 0 | 0 | 0 | 0 | 0 | 0 | 0 |
| 0 | NA | NA | NA | NA | NA | Gm13773 | | 0 | 0 | 0 | 0 | 0 | 0 | 0 | 0 | 0 | 0 | 0 |
| 0 | NA | NA | NA | NA | NA | Mup-ps6 | | 0 | 0 | 0 | 0 | 0 | 0 | 0 | 0 | 0 | 0 | 0 |
| 0 | NA | NA | NA | NA | NA | Mup-ps24 | | 0 | 0 | 0 | 0 | 0 | 0 | 0 | 0 | 0 | 0 | 0 |
| 0 | NA | NA | NA | NA | NA | Mup10 | major urinary protein 10 | 0 | 0 | 0 | 0 | 0 | 0 | 0 | 0 | 0 | 0 | 0 |
| 0 | NA | NA | NA | NA | NA | Mup-ps5 | | 0 | 0 | 0 | 0 | 0 | 0 | 0 | 0 | 0 | 0 | 0 |

| | | | | | | | | | | | | | | | | | | |
|---|---|---|---|---|---|---|---|---|---|---|---|---|---|---|---|---|---|---|
| 0 | NA | NA | NA | NA | NA | Mup11 | major urinary protein 11 | 0 | 0 | 0 | 0 | 0 | 0 | 0 | 0 | 0 | 0 | 0 |
| 0 | NA | NA | NA | NA | NA | Mup-ps7 | | 0 | 0 | 0 | 0 | 0 | 0 | 0 | 0 | 0 | 0 | 0 |
| 0 | NA | NA | NA | NA | NA | Mup12 | major urinary protein 12 | 0 | 0 | 0 | 0 | 0 | 0 | 0 | 0 | 0 | 0 | 0 |
| 0 | NA | NA | NA | NA | NA | Mup-ps8 | | 0 | 0 | 0 | 0 | 0 | 0 | 0 | 0 | 0 | 0 | 0 |
| 0 | NA | NA | NA | NA | NA | Gm21286 | | 0 | 0 | 0 | 0 | 0 | 0 | 0 | 0 | 0 | 0 | 0 |
| 0 | NA | NA | NA | NA | NA | Mup-ps9 | | 0 | 0 | 0 | 0 | 0 | 0 | 0 | 0 | 0 | 0 | 0 |
| 0 | NA | NA | NA | NA | NA | Mup13 | major urinary protein 13 | 0 | 0 | 0 | 0 | 0 | 0 | 0 | 0 | 0 | 0 | 0 |
| 0 | NA | NA | NA | NA | NA | Mup-ps10 | | 0 | 0 | 0 | 0 | 0 | 0 | 0 | 0 | 0 | 0 | 0 |
| 0 | NA | NA | NA | NA | NA | Mup-ps11 | | 0 | 0 | 0 | 0 | 0 | 0 | 0 | 0 | 0 | 0 | 0 |
| 0 | NA | NA | NA | NA | NA | Mup15 | major urinary protein 15 | 0 | 0 | 0 | 0 | 0 | 0 | 0 | 0 | 0 | 0 | 0 |
| 0 | NA | NA | NA | NA | NA | Mup-ps12 | | 0 | 0 | 0 | 0 | 0 | 0 | 0 | 0 | 0 | 0 | 0 |
| 0 | NA | NA | NA | NA | NA | Mup16 | major urinary protein 13 | 0 | 0 | 0 | 0 | 0 | 0 | 0 | 0 | 0 | 0 | 0 |
| 0 | NA | NA | NA | NA | NA | Mup-ps13 | | 0 | 0 | 0 | 0 | 0 | 0 | 0 | 0 | 0 | 0 | 0 |
| 0 | NA | NA | NA | NA | NA | Mup17 | major urinary protein 13 | 0 | 0 | 0 | 0 | 0 | 0 | 0 | 0 | 0 | 0 | 0 |
| 0 | NA | NA | NA | NA | NA | Mup-ps14 | | 0 | 0 | 0 | 0 | 0 | 0 | 0 | 0 | 0 | 0 | 0 |
| 0 | NA | NA | NA | NA | NA | Mup18 | major urinary protein 18 | 0 | 0 | 0 | 0 | 0 | 0 | 0 | 0 | 0 | 0 | 0 |
| 0 | NA | NA | NA | NA | NA | Mup-ps15 | | 0 | 0 | 0 | 0 | 0 | 0 | 0 | 0 | 0 | 0 | 0 |
| 0 | NA | NA | NA | NA | NA | Gm12573 | | 0 | 0 | 0 | 0 | 0 | 0 | 0 | 0 | 0 | 0 | 0 |
| 0 | NA | NA | NA | NA | NA | Mup19 | major urinary protein 13 | 0 | 0 | 0 | 0 | 0 | 0 | 0 | 0 | 0 | 0 | 0 |
| 0 | NA | NA | NA | NA | NA | Gm13775 | major urinary protein, pseudogene 16 | 0 | 0 | 0 | 0 | 0 | 0 | 0 | 0 | 0 | 0 | 0 |
| 0 | NA | NA | NA | NA | NA | Mup-ps16 | | 0 | 0 | 0 | 0 | 0 | 0 | 0 | 0 | 0 | 0 | 0 |
| 0 | NA | NA | NA | NA | NA | Mup5 | major urinary protein 5 | 0 | 0 | 0 | 0 | 0 | 0 | 0 | 0 | 0 | 0 | 0 |
| 0 | NA | NA | NA | NA | NA | Mup-ps23 | | 0 | 0 | 0 | 0 | 0 | 0 | 0 | 0 | 0 | 0 | 0 |
| 0 | NA | NA | NA | NA | NA | Mup-ps17 | | 0 | 0 | 0 | 0 | 0 | 0 | 0 | 0 | 0 | 0 | 0 |
| 0 | NA | NA | NA | NA | NA | Mup-ps18 | | 0 | 0 | 0 | 0 | 0 | 0 | 0 | 0 | 0 | 0 | 0 |
| 0 | NA | NA | NA | NA | NA | Mup-ps19 | | 0 | 0 | 0 | 0 | 0 | 0 | 0 | 0 | 0 | 0 | 0 |
| 0 | NA | NA | NA | NA | NA | Mup-ps20 | | 0 | 0 | 0 | 0 | 0 | 0 | 0 | 0 | 0 | 0 | 0 |
| 0 | NA | NA | NA | NA | NA | Mup-ps21 | | 0 | 0 | 0 | 0 | 0 | 0 | 0 | 0 | 0 | 0 | 0 |
| 0 | NA | NA | NA | NA | NA | Gm12909 | | 0 | 0 | 0 | 0 | 0 | 0 | 0 | 0 | 0 | 0 | 0 |
| 0 | NA | NA | NA | NA | NA | Gm12910 | | 0 | 0 | 0 | 0 | 0 | 0 | 0 | 0 | 0 | 0 | 0 |
| 0 | NA | NA | NA | NA | NA | Mup21 | major urinary protein 21 | 0 | 0 | 0 | 0 | 0 | 0 | 0 | 0 | 0 | 0 | 0 |
| 0 | NA | NA | NA | NA | NA | Gm11209 | | 0 | 0 | 0 | 0 | 0 | 0 | 0 | 0 | 0 | 0 | 0 |
| 0 | NA | NA | NA | NA | NA | Gm11211 | | 0 | 0 | 0 | 0 | 0 | 0 | 0 | 0 | 0 | 0 | 0 |
| 0 | NA | NA | NA | NA | NA | Gm24117 | | 0 | 0 | 0 | 0 | 0 | 0 | 0 | 0 | 0 | 0 | 0 |
| 0 | NA | NA | NA | NA | NA | Gm25391 | | 0 | 0 | 0 | 0 | 0 | 0 | 0 | 0 | 0 | 0 | 0 |
| 0 | NA | NA | NA | NA | NA | Gm11481 | | 0 | 0 | 0 | 0 | 0 | 0 | 0 | 0 | 0 | 0 | 0 |
| 0 | NA | NA | NA | NA | NA | Gm11480 | | 0 | 0 | 0 | 0 | 0 | 0 | 0 | 0 | 0 | 0 | 0 |
| 0 | NA | NA | NA | NA | NA | Mir455 | | 0 | 0 | 0 | 0 | 0 | 0 | 0 | 0 | 0 | 0 | 0 |
| 0 | NA | NA | NA | NA | NA | Gm11212 | | 0 | 0 | 0 | 0 | 0 | 0 | 0 | 0 | 0 | 0 | 0 |
| 0 | NA | NA | NA | NA | NA | Orm3 | orosomucoid 3 | 0 | 0 | 0 | 0 | 0 | 0 | 0 | 0 | 0 | 0 | 0 |
| 0 | NA | NA | NA | NA | NA | Orm2 | orosomucoid 2 | 0 | 0 | 0 | 0 | 0 | 0 | 0 | 0 | 0 | 0 | 0 |
| 0 | NA | NA | NA | NA | NA | Aknaos | | 0 | 0 | 0 | 0 | 0 | 0 | 0 | 0 | 0 | 0 | 0 |
| 0 | NA | NA | NA | NA | NA | Gm11483 | | 0 | 0 | 0 | 0 | 0 | 0 | 0 | 0 | 0 | 0 | 0 |
| 0 | NA | NA | NA | NA | NA | Gm24448 | | 0 | 0 | 0 | 0 | 0 | 0 | 0 | 0 | 0 | 0 | 0 |
| 0 | NA | NA | NA | NA | NA | Gm11213 | predicted gene 11213 | 0 | 0 | 0 | 0 | 0 | 0 | 0 | 0 | 0 | 0 | 0 |
| 0 | NA | NA | NA | NA | NA | Tex48 | testis expressed 48 | 0 | 0 | 0 | 0 | 0 | 0 | 0 | 0 | 0 | 0 | 0 |
| 0 | NA | NA | NA | NA | NA | Gm24234 | | 0 | 0 | 0 | 0 | 0 | 0 | 0 | 0 | 0 | 0 | 0 |
| 0 | NA | NA | NA | NA | NA | Gm11482 | | 0 | 0 | 0 | 0 | 0 | 0 | 0 | 0 | 0 | 0 | 0 |
| 0 | NA | NA | NA | NA | NA | Gm11214 | | 0 | 0 | 0 | 0 | 0 | 0 | 0 | 0 | 0 | 0 | 0 |
| 0 | NA | NA | NA | NA | NA | Gm11216 | | 0 | 0 | 0 | 0 | 0 | 0 | 0 | 0 | 0 | 0 | 0 |
| 0 | NA | NA | NA | NA | NA | Gm11217 | | 0 | 0 | 0 | 0 | 0 | 0 | 0 | 0 | 0 | 0 | 0 |
| 0 | NA | NA | NA | NA | NA | Gm23950 | | 0 | 0 | 0 | 0 | 0 | 0 | 0 | 0 | 0 | 0 | 0 |
| 0 | NA | NA | NA | NA | NA | Gm11218 | | 0 | 0 | 0 | 0 | 0 | 0 | 0 | 0 | 0 | 0 | 0 |
| 0 | NA | NA | NA | NA | NA | Gm11219 | | 0 | 0 | 0 | 0 | 0 | 0 | 0 | 0 | 0 | 0 | 0 |
| 0 | NA | NA | NA | NA | NA | Gm11484 | | 0 | 0 | 0 | 0 | 0 | 0 | 0 | 0 | 0 | 0 | 0 |
| 0 | NA | NA | NA | NA | NA | Gm25480 | | 0 | 0 | 0 | 0 | 0 | 0 | 0 | 0 | 0 | 0 | 0 |
| 0 | NA | NA | NA | NA | NA | Gm11220 | | 0 | 0 | 0 | 0 | 0 | 0 | 0 | 0 | 0 | 0 | 0 |
| 0 | NA | NA | NA | NA | NA | Gm11403 | | 0 | 0 | 0 | 0 | 0 | 0 | 0 | 0 | 0 | 0 | 0 |
| 0 | NA | NA | NA | NA | NA | Gm44497 | | 0 | 0 | 0 | 0 | 0 | 0 | 0 | 0 | 0 | 0 | 0 |
| 0 | NA | NA | NA | NA | NA | Gm11751 | | 0 | 0 | 0 | 0 | 0 | 0 | 0 | 0 | 0 | 0 | 0 |

| | | | | | | | | | | | | | | | | | | |
|---|---|---|---|---|---|---|---|---|---|---|---|---|---|---|---|---|---|---|
| 0 | NA | NA | NA | NA | NA | Gm23738 | | 0 | 0 | 0 | 0 | 0 | 0 | 0 | 0 | 0 | 0 | 0 |
| 0 | NA | NA | NA | NA | NA | Tcp1-ps1 | | 0 | 0 | 0 | 0 | 0 | 0 | 0 | 0 | 0 | 0 | 0 |
| 0 | NA | NA | NA | NA | NA | Gm12911 | | 0 | 0 | 0 | 0 | 0 | 0 | 0 | 0 | 0 | 0 | 0 |
| 0 | NA | NA | NA | NA | NA | Gm24697 | | 0 | 0 | 0 | 0 | 0 | 0 | 0 | 0 | 0 | 0 | 0 |
| 0 | NA | NA | NA | NA | NA | Gm11404 | | 0 | 0 | 0 | 0 | 0 | 0 | 0 | 0 | 0 | 0 | 0 |
| 0 | NA | NA | NA | NA | NA | Gm26320 | | 0 | 0 | 0 | 0 | 0 | 0 | 0 | 0 | 0 | 0 | 0 |
| 0 | NA | NA | NA | NA | NA | Gm11227 | | 0 | 0 | 0 | 0 | 0 | 0 | 0 | 0 | 0 | 0 | 0 |
| 0 | NA | NA | NA | NA | NA | Gm11228 | | 0 | 0 | 0 | 0 | 0 | 0 | 0 | 0 | 0 | 0 | 0 |
| 0 | NA | NA | NA | NA | NA | Gm11229 | | 0 | 0 | 0 | 0 | 0 | 0 | 0 | 0 | 0 | 0 | 0 |
| 0 | NA | NA | NA | NA | NA | Gm11231 | | 0 | 0 | 0 | 0 | 0 | 0 | 0 | 0 | 0 | 0 | 0 |
| 0 | NA | NA | NA | NA | NA | Gm11232 | predicted gene 11232 | 0 | 0 | 0 | 0 | 0 | 0 | 0 | 0 | 0 | 0 | 0 |
| 0 | NA | NA | NA | NA | NA | Gm11233 | | 0 | 0 | 0 | 0 | 0 | 0 | 0 | 0 | 0 | 0 | 0 |
| 0 | NA | NA | NA | NA | NA | Gm11234 | | 0 | 0 | 0 | 0 | 0 | 0 | 0 | 0 | 0 | 0 | 0 |
| 0 | NA | NA | NA | NA | NA | Gm11250 | | 0 | 0 | 0 | 0 | 0 | 0 | 0 | 0 | 0 | 0 | 0 |
| 0 | NA | NA | NA | NA | NA | Gm5287 | | 0 | 0 | 0 | 0 | 0 | 0 | 0 | 0 | 0 | 0 | 0 |
| 0 | NA | NA | NA | NA | NA | Gm25653 | | 0 | 0 | 0 | 0 | 0 | 0 | 0 | 0 | 0 | 0 | 0 |
| 0 | NA | NA | NA | NA | NA | Gm11251 | | 0 | 0 | 0 | 0 | 0 | 0 | 0 | 0 | 0 | 0 | 0 |
| 0 | NA | NA | NA | NA | NA | Gm25769 | | 0 | 0 | 0 | 0 | 0 | 0 | 0 | 0 | 0 | 0 | 0 |
| 0 | NA | NA | NA | NA | NA | Gm11487 | predicted gene 11487 | 0 | 0 | 0 | 0 | 0 | 0 | 0 | 0 | 0 | 0 | 0 |
| 0 | NA | NA | NA | NA | NA | Gm11486 | | 0 | 0 | 0 | 0 | 0 | 0 | 0 | 0 | 0 | 0 | 0 |
| 0 | NA | NA | NA | NA | NA | Gm11485 | | 0 | 0 | 0 | 0 | 0 | 0 | 0 | 0 | 0 | 0 | 0 |
| 0 | NA | NA | NA | NA | NA | Gm24170 | | 0 | 0 | 0 | 0 | 0 | 0 | 0 | 0 | 0 | 0 | 0 |
| 0 | NA | NA | NA | NA | NA | Gm11238 | predicted gene 11238 | 0 | 0 | 0 | 0 | 0 | 0 | 0 | 0 | 0 | 0 | 0 |
| 0 | NA | NA | NA | NA | NA | Gm11237 | predicted gene 11237 | 0 | 0 | 0 | 0 | 0 | 0 | 0 | 0 | 0 | 0 | 0 |
| 0 | NA | NA | NA | NA | NA | Gm11236 | predicted gene 11236 | 0 | 0 | 0 | 0 | 0 | 0 | 0 | 0 | 0 | 0 | 0 |
| 0 | NA | NA | NA | NA | NA | Gm11239 | predicted gene 11239 | 0 | 0 | 0 | 0 | 0 | 0 | 0 | 0 | 0 | 0 | 0 |
| 0 | NA | NA | NA | NA | NA | Gm428 | predicted gene 428 | 0 | 0 | 0 | 0 | 0 | 0 | 0 | 0 | 0 | 0 | 0 |
| 0 | NA | NA | NA | NA | NA | Gm11240 | | 0 | 0 | 0 | 0 | 0 | 0 | 0 | 0 | 0 | 0 | 0 |
| 0 | NA | NA | NA | NA | NA | Gm13867 | | 0 | 0 | 0 | 0 | 0 | 0 | 0 | 0 | 0 | 0 | 0 |
| 0 | NA | NA | NA | NA | NA | Gm11758 | predicted gene 11758 | 0 | 0 | 0 | 0 | 0 | 0 | 0 | 0 | 0 | 0 | 0 |
| 0 | NA | NA | NA | NA | NA | Gm11757 | predicted gene 11757 | 0 | 0 | 0 | 0 | 0 | 0 | 0 | 0 | 0 | 0 | 0 |
| 0 | NA | NA | NA | NA | NA | Gm13871 | predicted gene 13871 | 0 | 0 | 0 | 0 | 0 | 0 | 0 | 0 | 0 | 0 | 0 |
| 0 | NA | NA | NA | NA | NA | Gm11756 | predicted gene 11756 | 0 | 0 | 0 | 0 | 0 | 0 | 0 | 0 | 0 | 0 | 0 |
| 0 | NA | NA | NA | NA | NA | 2310002L09Ri | RIKEN cDNA 2310002L09 gene | 0 | 0 | 0 | 0 | 0 | 0 | 0 | 0 | 0 | 0 | 0 |
| 0 | NA | NA | NA | NA | NA | Gm26379 | | 0 | 0 | 0 | 0 | 0 | 0 | 0 | 0 | 0 | 0 | 0 |
| 0 | NA | NA | NA | NA | NA | Gm26379 | | 0 | 0 | 0 | 0 | 0 | 0 | 0 | 0 | 0 | 0 | 0 |
| 0 | NA | NA | NA | NA | NA | Gm11254 | | 0 | 0 | 0 | 0 | 0 | 0 | 0 | 0 | 0 | 0 | 0 |
| 0 | NA | NA | NA | NA | NA | Gm11258 | | 0 | 0 | 0 | 0 | 0 | 0 | 0 | 0 | 0 | 0 | 0 |
| 0 | NA | NA | NA | NA | NA | Gm11253 | | 0 | 0 | 0 | 0 | 0 | 0 | 0 | 0 | 0 | 0 | 0 |
| 0 | NA | NA | NA | NA | NA | Gm11257 | | 0 | 0 | 0 | 0 | 0 | 0 | 0 | 0 | 0 | 0 | 0 |
| 0 | NA | NA | NA | NA | NA | Gm22864 | | 0 | 0 | 0 | 0 | 0 | 0 | 0 | 0 | 0 | 0 | 0 |
| 0 | NA | NA | NA | NA | NA | Gm11259 | | 0 | 0 | 0 | 0 | 0 | 0 | 0 | 0 | 0 | 0 | 0 |
| 0 | NA | NA | NA | NA | NA | Gm11256 | | 0 | 0 | 0 | 0 | 0 | 0 | 0 | 0 | 0 | 0 | 0 |
| 0 | NA | NA | NA | NA | NA | Gm11241 | | 0 | 0 | 0 | 0 | 0 | 0 | 0 | 0 | 0 | 0 | 0 |
| 0 | NA | NA | NA | NA | NA | Gm11252 | | 0 | 0 | 0 | 0 | 0 | 0 | 0 | 0 | 0 | 0 | 0 |
| 0 | NA | NA | NA | NA | NA | Gm11242 | | 0 | 0 | 0 | 0 | 0 | 0 | 0 | 0 | 0 | 0 | 0 |
| 0 | NA | NA | NA | NA | NA | Gm11243 | | 0 | 0 | 0 | 0 | 0 | 0 | 0 | 0 | 0 | 0 | 0 |
| 0 | NA | NA | NA | NA | NA | Gm11244 | | 0 | 0 | 0 | 0 | 0 | 0 | 0 | 0 | 0 | 0 | 0 |
| 0 | NA | NA | NA | NA | NA | Gm11245 | | 0 | 0 | 0 | 0 | 0 | 0 | 0 | 0 | 0 | 0 | 0 |
| 0 | NA | NA | NA | NA | NA | Gm11246 | | 0 | 0 | 0 | 0 | 0 | 0 | 0 | 0 | 0 | 0 | 0 |
| 0 | NA | NA | NA | NA | NA | Gm23159 | | 0 | 0 | 0 | 0 | 0 | 0 | 0 | 0 | 0 | 0 | 0 |
| 0 | NA | NA | NA | NA | NA | Gm11260 | | 0 | 0 | 0 | 0 | 0 | 0 | 0 | 0 | 0 | 0 | 0 |
| 0 | NA | NA | NA | NA | NA | Gm11261 | | 0 | 0 | 0 | 0 | 0 | 0 | 0 | 0 | 0 | 0 | 0 |
| 0 | NA | NA | NA | NA | NA | Gm22262 | | 0 | 0 | 0 | 0 | 0 | 0 | 0 | 0 | 0 | 0 | 0 |
| 0 | NA | NA | NA | NA | NA | Gm11262 | | 0 | 0 | 0 | 0 | 0 | 0 | 0 | 0 | 0 | 0 | 0 |
| 0 | NA | NA | NA | NA | NA | Gm25836 | | 0 | 0 | 0 | 0 | 0 | 0 | 0 | 0 | 0 | 0 | 0 |
| 0 | NA | NA | NA | NA | NA | Gm11409 | | 0 | 0 | 0 | 0 | 0 | 0 | 0 | 0 | 0 | 0 | 0 |
| 0 | NA | NA | NA | NA | NA | Gm11408 | | 0 | 0 | 0 | 0 | 0 | 0 | 0 | 0 | 0 | 0 | 0 |
| 0 | NA | NA | NA | NA | NA | Gm11406 | | 0 | 0 | 0 | 0 | 0 | 0 | 0 | 0 | 0 | 0 | 0 |
| 0 | NA | NA | NA | NA | NA | Tyrp1 | tyrosinase-related protein 1 | 0 | 0 | 0 | 0 | 0 | 0 | 0 | 0 | 0 | 0 | 0 |

| | | | | | | | | | | | | | | | | | | |
|---|---|---|---|---|---|---|---|---|---|---|---|---|---|---|---|---|---|---|
| 0 | NA | NA | NA | NA | NA | Gm27452 | | 0 | 0 | 0 | 0 | 0 | 0 | 0 | 0 | 0 | 0 | 0 |
| 0 | NA | NA | NA | NA | NA | Gm25803 | | 0 | 0 | 0 | 0 | 0 | 0 | 0 | 0 | 0 | 0 | 0 |
| 0 | NA | NA | NA | NA | NA | Gm11411 | | 0 | 0 | 0 | 0 | 0 | 0 | 0 | 0 | 0 | 0 | 0 |
| 0 | NA | NA | NA | NA | NA | n-R5s187 | | 0 | 0 | 0 | 0 | 0 | 0 | 0 | 0 | 0 | 0 | 0 |
| 0 | NA | NA | NA | NA | NA | Gm12914 | | 0 | 0 | 0 | 0 | 0 | 0 | 0 | 0 | 0 | 0 | 0 |
| 0 | NA | NA | NA | NA | NA | Gm22402 | | 0 | 0 | 0 | 0 | 0 | 0 | 0 | 0 | 0 | 0 | 0 |
| 0 | NA | NA | NA | NA | NA | n-R5s188 | | 0 | 0 | 0 | 0 | 0 | 0 | 0 | 0 | 0 | 0 | 0 |
| 0 | NA | NA | NA | NA | NA | Gm11267 | | 0 | 0 | 0 | 0 | 0 | 0 | 0 | 0 | 0 | 0 | 0 |
| 0 | NA | NA | NA | NA | NA | Gm11268 | | 0 | 0 | 0 | 0 | 0 | 0 | 0 | 0 | 0 | 0 | 0 |
| 0 | NA | NA | NA | NA | NA | Gm11269 | | 0 | 0 | 0 | 0 | 0 | 0 | 0 | 0 | 0 | 0 | 0 |
| 0 | NA | NA | NA | NA | NA | Gm24884 | | 0 | 0 | 0 | 0 | 0 | 0 | 0 | 0 | 0 | 0 | 0 |
| 0 | NA | NA | NA | NA | NA | Gm11248 | | 0 | 0 | 0 | 0 | 0 | 0 | 0 | 0 | 0 | 0 | 0 |
| 0 | NA | NA | NA | NA | NA | Gm23412 | | 0 | 0 | 0 | 0 | 0 | 0 | 0 | 0 | 0 | 0 | 0 |
| 0 | NA | NA | NA | NA | NA | Gm11184 | | 0 | 0 | 0 | 0 | 0 | 0 | 0 | 0 | 0 | 0 | 0 |
| 0 | NA | NA | NA | NA | NA | Gm11414 | | 0 | 0 | 0 | 0 | 0 | 0 | 0 | 0 | 0 | 0 | 0 |
| 0 | NA | NA | NA | NA | NA | Gm10154 | | 0 | 0 | 0 | 0 | 0 | 0 | 0 | 0 | 0 | 0 | 0 |
| 0 | NA | NA | NA | NA | NA | Gm25899 | | 0 | 0 | 0 | 0 | 0 | 0 | 0 | 0 | 0 | 0 | 0 |
| 0 | NA | NA | NA | NA | NA | Gm26968 | | 0 | 0 | 0 | 0 | 0 | 0 | 0 | 0 | 0 | 0 | 0 |
| 0 | NA | NA | NA | NA | NA | Gm11415 | | 0 | 0 | 0 | 0 | 0 | 0 | 0 | 0 | 0 | 0 | 0 |
| 0 | NA | NA | NA | NA | NA | 6030471H07Rik | | 0 | 0 | 0 | 0 | 0 | 0 | 0 | 0 | 0 | 0 | 0 |
| 0 | NA | NA | NA | NA | NA | Gm12414 | | 0 | 0 | 0 | 0 | 0 | 0 | 0 | 0 | 0 | 0 | 0 |
| 0 | NA | NA | NA | NA | NA | Gm12415 | | 0 | 0 | 0 | 0 | 0 | 0 | 0 | 0 | 0 | 0 | 0 |
| 0 | NA | NA | NA | NA | NA | Gm12417 | | 0 | 0 | 0 | 0 | 0 | 0 | 0 | 0 | 0 | 0 | 0 |
| 0 | NA | NA | NA | NA | NA | Gm12413 | | 0 | 0 | 0 | 0 | 0 | 0 | 0 | 0 | 0 | 0 | 0 |
| 0 | NA | NA | NA | NA | NA | Gm26069 | | 0 | 0 | 0 | 0 | 0 | 0 | 0 | 0 | 0 | 0 | 0 |
| 0 | NA | NA | NA | NA | NA | Gm25811 | | 0 | 0 | 0 | 0 | 0 | 0 | 0 | 0 | 0 | 0 | 0 |
| 0 | NA | NA | NA | NA | NA | Gm12550 | | 0 | 0 | 0 | 0 | 0 | 0 | 0 | 0 | 0 | 0 | 0 |
| 0 | NA | NA | NA | NA | NA | Saxo1 | stabilizer of axonemal microtubules 1 | 0 | 0 | 0 | 0 | 0 | 0 | 0 | 0 | 0 | 0 | 0 |
| 0 | NA | NA | NA | NA | NA | Gm26017 | | 0 | 0 | 0 | 0 | 0 | 0 | 0 | 0 | 0 | 0 | 0 |
| 0 | NA | NA | NA | NA | NA | Saxo1os | | 0 | 0 | 0 | 0 | 0 | 0 | 0 | 0 | 0 | 0 | 0 |
| 0 | NA | NA | NA | NA | NA | Scarna8 | | 0 | 0 | 0 | 0 | 0 | 0 | 0 | 0 | 0 | 0 | 0 |
| 0 | NA | NA | NA | NA | NA | Gm12600 | | 0 | 0 | 0 | 0 | 0 | 0 | 0 | 0 | 0 | 0 | 0 |
| 0 | NA | NA | NA | NA | NA | Gm23154 | | 0 | 0 | 0 | 0 | 0 | 0 | 0 | 0 | 0 | 0 | 0 |
| 0 | NA | NA | NA | NA | NA | Gm12604 | | 0 | 0 | 0 | 0 | 0 | 0 | 0 | 0 | 0 | 0 | 0 |
| 0 | NA | NA | NA | NA | NA | Gm12646 | | 0 | 0 | 0 | 0 | 0 | 0 | 0 | 0 | 0 | 0 | 0 |
| 0 | NA | NA | NA | NA | NA | Mir491 | | 0 | 0 | 0 | 0 | 0 | 0 | 0 | 0 | 0 | 0 | 0 |
| 0 | NA | NA | NA | NA | NA | Gm12645 | | 0 | 0 | 0 | 0 | 0 | 0 | 0 | 0 | 0 | 0 | 0 |
| 0 | NA | NA | NA | NA | NA | Ifna15 | interferon alpha 15 | 0 | 0 | 0 | 0 | 0 | 0 | 0 | 0 | 0 | 0 | 0 |
| 0 | NA | NA | NA | NA | NA | Ifna14 | interferon alpha 14 | 0 | 0 | 0 | 0 | 0 | 0 | 0 | 0 | 0 | 0 | 0 |
| 0 | NA | NA | NA | NA | NA | Ifna-ps1 | | 0 | 0 | 0 | 0 | 0 | 0 | 0 | 0 | 0 | 0 | 0 |
| 0 | NA | NA | NA | NA | NA | Ifna9 | interferon alpha 9 | 0 | 0 | 0 | 0 | 0 | 0 | 0 | 0 | 0 | 0 | 0 |
| 0 | NA | NA | NA | NA | NA | Ifna12 | interferon alpha 12 | 0 | 0 | 0 | 0 | 0 | 0 | 0 | 0 | 0 | 0 | 0 |
| 0 | NA | NA | NA | NA | NA | Gm12601 | | 0 | 0 | 0 | 0 | 0 | 0 | 0 | 0 | 0 | 0 | 0 |
| 0 | NA | NA | NA | NA | NA | C87499 | expressed sequence C87499 | 0 | 0 | 0 | 0 | 0 | 0 | 0 | 0 | 0 | 0 | 0 |
| 0 | NA | NA | NA | NA | NA | Ifna13 | interferon alpha 13 | 0 | 0 | 0 | 0 | 0 | 0 | 0 | 0 | 0 | 0 | 0 |
| 0 | NA | NA | NA | NA | NA | Gm13281 | | 0 | 0 | 0 | 0 | 0 | 0 | 0 | 0 | 0 | 0 | 0 |
| 0 | NA | NA | NA | NA | NA | Ifna16 | interferon alpha 16 | 0 | 0 | 0 | 0 | 0 | 0 | 0 | 0 | 0 | 0 | 0 |
| 0 | NA | NA | NA | NA | NA | Ifna2 | interferon alpha 2 | 0 | 0 | 0 | 0 | 0 | 0 | 0 | 0 | 0 | 0 | 0 |
| 0 | NA | NA | NA | NA | NA | Ifnab | interferon alpha B | 0 | 0 | 0 | 0 | 0 | 0 | 0 | 0 | 0 | 0 | 0 |
| 0 | NA | NA | NA | NA | NA | Gm13273 | | 0 | 0 | 0 | 0 | 0 | 0 | 0 | 0 | 0 | 0 | 0 |
| 0 | NA | NA | NA | NA | NA | Gm13282 | | 0 | 0 | 0 | 0 | 0 | 0 | 0 | 0 | 0 | 0 | 0 |
| 0 | NA | NA | NA | NA | NA | Gm49884 | | 0 | 0 | 0 | 0 | 0 | 0 | 0 | 0 | 0 | 0 | 0 |
| 0 | NA | NA | NA | NA | NA | Gm13271 | predicted gene 13271 | 0 | 0 | 0 | 0 | 0 | 0 | 0 | 0 | 0 | 0 | 0 |
| 0 | NA | NA | NA | NA | NA | Gm13286 | predicted gene 13286 | 0 | 0 | 0 | 0 | 0 | 0 | 0 | 0 | 0 | 0 | 0 |
| 0 | NA | NA | NA | NA | NA | Gm10583 | | 0 | 0 | 0 | 0 | 0 | 0 | 0 | 0 | 0 | 0 | 0 |
| 0 | NA | NA | NA | NA | NA | Gm13283 | predicted gene 13283 | 0 | 0 | 0 | 0 | 0 | 0 | 0 | 0 | 0 | 0 | 0 |
| 0 | NA | NA | NA | NA | NA | Gm13284 | | 0 | 0 | 0 | 0 | 0 | 0 | 0 | 0 | 0 | 0 | 0 |
| 0 | NA | NA | NA | NA | NA | Gm10581 | | 0 | 0 | 0 | 0 | 0 | 0 | 0 | 0 | 0 | 0 | 0 |
| 0 | NA | NA | NA | NA | NA | Gm13290 | predicted gene 13290 | 0 | 0 | 0 | 0 | 0 | 0 | 0 | 0 | 0 | 0 | 0 |
| 0 | NA | NA | NA | NA | NA | Gm10580 | | 0 | 0 | 0 | 0 | 0 | 0 | 0 | 0 | 0 | 0 | 0 |

| | | | | | | | | | | | | | | | | | | |
|---|---|---|---|---|---|---|---|---|---|---|---|---|---|---|---|---|---|---|
| 0 | NA | NA | NA | NA | NA | Gm13289 | predicted gene 13289 | 0 | 0 | 0 | 0 | 0 | 0 | 0 | 0 | 0 | 0 | 0 |
| 0 | NA | NA | NA | NA | NA | Gm13272 | predicted gene 13272 | 0 | 0 | 0 | 0 | 0 | 0 | 0 | 0 | 0 | 0 | 0 |
| 0 | NA | NA | NA | NA | NA | Ifnz | interferon zeta | 0 | 0 | 0 | 0 | 0 | 0 | 0 | 0 | 0 | 0 | 0 |
| 0 | NA | NA | NA | NA | NA | Gm13276 | predicted gene 13276 | 0 | 0 | 0 | 0 | 0 | 0 | 0 | 0 | 0 | 0 | 0 |
| 0 | NA | NA | NA | NA | NA | Gm13277 | predicted gene 13277 | 0 | 0 | 0 | 0 | 0 | 0 | 0 | 0 | 0 | 0 | 0 |
| 0 | NA | NA | NA | NA | NA | Gm13278 | predicted gene 13278 | 0 | 0 | 0 | 0 | 0 | 0 | 0 | 0 | 0 | 0 | 0 |
| 0 | NA | NA | NA | NA | NA | Gm13275 | predicted gene 13275 | 0 | 0 | 0 | 0 | 0 | 0 | 0 | 0 | 0 | 0 | 0 |
| 0 | NA | NA | NA | NA | NA | Gm13279 | predicted gene 13279 | 0 | 0 | 0 | 0 | 0 | 0 | 0 | 0 | 0 | 0 | 0 |
| 0 | NA | NA | NA | NA | NA | Gm13285 | predicted gene 13285 | 0 | 0 | 0 | 0 | 0 | 0 | 0 | 0 | 0 | 0 | 0 |
| 0 | NA | NA | NA | NA | NA | Gm13287 | predicted gene 13287 | 0 | 0 | 0 | 0 | 0 | 0 | 0 | 0 | 0 | 0 | 0 |
| 0 | NA | NA | NA | NA | NA | Gm13288 | predicted gene 13288 | 0 | 0 | 0 | 0 | 0 | 0 | 0 | 0 | 0 | 0 | 0 |
| 0 | NA | NA | NA | NA | NA | Ifna7 | interferon alpha 7 | 0 | 0 | 0 | 0 | 0 | 0 | 0 | 0 | 0 | 0 | 0 |
| 0 | NA | NA | NA | NA | NA | Ifna11 | interferon alpha 11 | 0 | 0 | 0 | 0 | 0 | 0 | 0 | 0 | 0 | 0 | 0 |
| 0 | NA | NA | NA | NA | NA | Ifna6 | interferon alpha 6 | 0 | 0 | 0 | 0 | 0 | 0 | 0 | 0 | 0 | 0 | 0 |
| 0 | NA | NA | NA | NA | NA | Ifna5 | interferon alpha 5 | 0 | 0 | 0 | 0 | 0 | 0 | 0 | 0 | 0 | 0 | 0 |
| 0 | NA | NA | NA | NA | NA | Ifna4 | interferon alpha 4 | 0 | 0 | 0 | 0 | 0 | 0 | 0 | 0 | 0 | 0 | 0 |
| 0 | NA | NA | NA | NA | NA | Ifna1 | interferon alpha 1 | 0 | 0 | 0 | 0 | 0 | 0 | 0 | 0 | 0 | 0 | 0 |
| 0 | NA | NA | NA | NA | NA | 4930553M12Rik | | 0 | 0 | 0 | 0 | 0 | 0 | 0 | 0 | 0 | 0 | 0 |
| 0 | NA | NA | NA | NA | NA | Ifne | interferon epsilon | 0 | 0 | 0 | 0 | 0 | 0 | 0 | 0 | 0 | 0 | 0 |
| 0 | NA | NA | NA | NA | NA | Mir31 | | 0 | 0 | 0 | 0 | 0 | 0 | 0 | 0 | 0 | 0 | 0 |
| 0 | NA | NA | NA | NA | NA | Gm49891 | | 0 | 0 | 0 | 0 | 0 | 0 | 0 | 0 | 0 | 0 | 0 |
| 0 | NA | NA | NA | NA | NA | Wincr1 | | 0 | 0 | 0 | 0 | 0 | 0 | 0 | 0 | 0 | 0 | 0 |
| 0 | NA | NA | NA | NA | NA | Gm12602 | | 0 | 0 | 0 | 0 | 0 | 0 | 0 | 0 | 0 | 0 | 0 |
| 0 | NA | NA | NA | NA | NA | Gm12607 | | 0 | 0 | 0 | 0 | 0 | 0 | 0 | 0 | 0 | 0 | 0 |
| 0 | NA | NA | NA | NA | NA | Gm26490 | | 0 | 0 | 0 | 0 | 0 | 0 | 0 | 0 | 0 | 0 | 0 |
| 0 | NA | NA | NA | NA | NA | Tgif2-ps1 | | 0 | 0 | 0 | 0 | 0 | 0 | 0 | 0 | 0 | 0 | 0 |
| 0 | NA | NA | NA | NA | NA | Gm12606 | | 0 | 0 | 0 | 0 | 0 | 0 | 0 | 0 | 0 | 0 | 0 |
| 0 | NA | NA | NA | NA | NA | Gm27580 | | 0 | 0 | 0 | 0 | 0 | 0 | 0 | 0 | 0 | 0 | 0 |
| 0 | NA | NA | NA | NA | NA | Gm12629 | | 0 | 0 | 0 | 0 | 0 | 0 | 0 | 0 | 0 | 0 | 0 |
| 0 | NA | NA | NA | NA | NA | Zfp352 | zinc finger protein 352 | 0 | 0 | 0 | 0 | 0 | 0 | 0 | 0 | 0 | 0 | 0 |
| 0 | NA | NA | NA | NA | NA | Gm12633 | | 0 | 0 | 0 | 0 | 0 | 0 | 0 | 0 | 0 | 0 | 0 |
| 0 | NA | NA | NA | NA | NA | Gm12634 | | 0 | 0 | 0 | 0 | 0 | 0 | 0 | 0 | 0 | 0 | 0 |
| 0 | NA | NA | NA | NA | NA | Gm12635 | | 0 | 0 | 0 | 0 | 0 | 0 | 0 | 0 | 0 | 0 | 0 |
| 0 | NA | NA | NA | NA | NA | Gm12636 | | 0 | 0 | 0 | 0 | 0 | 0 | 0 | 0 | 0 | 0 | 0 |
| 0 | NA | NA | NA | NA | NA | Gm12632 | | 0 | 0 | 0 | 0 | 0 | 0 | 0 | 0 | 0 | 0 | 0 |
| 0 | NA | NA | NA | NA | NA | Gm25244 | | 0 | 0 | 0 | 0 | 0 | 0 | 0 | 0 | 0 | 0 | 0 |
| 0 | NA | NA | NA | NA | NA | Gm12644 | | 0 | 0 | 0 | 0 | 0 | 0 | 0 | 0 | 0 | 0 | 0 |
| 0 | NA | NA | NA | NA | NA | Gm12653 | | 0 | 0 | 0 | 0 | 0 | 0 | 0 | 0 | 0 | 0 | 0 |
| 0 | NA | NA | NA | NA | NA | Mir6402 | | 0 | 0 | 0 | 0 | 0 | 0 | 0 | 0 | 0 | 0 | 0 |
| 0 | NA | NA | NA | NA | NA | Gm12668 | | 0 | 0 | 0 | 0 | 0 | 0 | 0 | 0 | 0 | 0 | 0 |
| 0 | NA | NA | NA | NA | NA | Gm12670 | | 0 | 0 | 0 | 0 | 0 | 0 | 0 | 0 | 0 | 0 | 0 |
| 0 | NA | NA | NA | NA | NA | Gm12667 | | 0 | 0 | 0 | 0 | 0 | 0 | 0 | 0 | 0 | 0 | 0 |
| 0 | NA | NA | NA | NA | NA | 4930577H14Rik | | 0 | 0 | 0 | 0 | 0 | 0 | 0 | 0 | 0 | 0 | 0 |
| 0 | NA | NA | NA | NA | NA | Izumo3 | IZUMO family member 3 | 0 | 0 | 0 | 0 | 0 | 0 | 0 | 0 | 0 | 0 | 0 |
| 0 | NA | NA | NA | NA | NA | Gm12666 | | 0 | 0 | 0 | 0 | 0 | 0 | 0 | 0 | 0 | 0 | 0 |
| 0 | NA | NA | NA | NA | NA | Gm12638 | | 0 | 0 | 0 | 0 | 0 | 0 | 0 | 0 | 0 | 0 | 0 |
| 0 | NA | NA | NA | NA | NA | Gm12637 | | 0 | 0 | 0 | 0 | 0 | 0 | 0 | 0 | 0 | 0 | 0 |
| 0 | NA | NA | NA | NA | NA | Gm22605 | | 0 | 0 | 0 | 0 | 0 | 0 | 0 | 0 | 0 | 0 | 0 |
| 0 | NA | NA | NA | NA | NA | Gm12639 | | 0 | 0 | 0 | 0 | 0 | 0 | 0 | 0 | 0 | 0 | 0 |
| 0 | NA | NA | NA | NA | NA | Gm12641 | | 0 | 0 | 0 | 0 | 0 | 0 | 0 | 0 | 0 | 0 | 0 |
| 0 | NA | NA | NA | NA | NA | Gm12649 | | 0 | 0 | 0 | 0 | 0 | 0 | 0 | 0 | 0 | 0 | 0 |
| 0 | NA | NA | NA | NA | NA | Gm23443 | | 0 | 0 | 0 | 0 | 0 | 0 | 0 | 0 | 0 | 0 | 0 |
| 0 | NA | NA | NA | NA | NA | Gm12640 | | 0 | 0 | 0 | 0 | 0 | 0 | 0 | 0 | 0 | 0 | 0 |
| 0 | NA | NA | NA | NA | NA | Gm24796 | | 0 | 0 | 0 | 0 | 0 | 0 | 0 | 0 | 0 | 0 | 0 |
| 0 | NA | NA | NA | NA | NA | Gm22619 | | 0 | 0 | 0 | 0 | 0 | 0 | 0 | 0 | 0 | 0 | 0 |
| 0 | NA | NA | NA | NA | NA | Gm12647 | | 0 | 0 | 0 | 0 | 0 | 0 | 0 | 0 | 0 | 0 | 0 |
| 0 | NA | NA | NA | NA | NA | Gm12648 | | 0 | 0 | 0 | 0 | 0 | 0 | 0 | 0 | 0 | 0 | 0 |
| 0 | NA | NA | NA | NA | NA | Gm23836 | | 0 | 0 | 0 | 0 | 0 | 0 | 0 | 0 | 0 | 0 | 0 |
| 0 | NA | NA | NA | NA | NA | Gm12654 | | 0 | 0 | 0 | 0 | 0 | 0 | 0 | 0 | 0 | 0 | 0 |
| 0 | NA | NA | NA | NA | NA | Gm26410 | | 0 | 0 | 0 | 0 | 0 | 0 | 0 | 0 | 0 | 0 | 0 |

| | | | | | | | | | | | | | | | | | | |
|---|---|---|---|---|---|---|---|---|---|---|---|---|---|---|---|---|---|---|
| 0 | NA | NA | NA | NA | NA | Gm22804 | | 0 | 0 | 0 | 0 | 0 | 0 | 0 | 0 | 0 | 0 | 0 |
| 0 | NA | NA | NA | NA | NA | Gm12656 | | 0 | 0 | 0 | 0 | 0 | 0 | 0 | 0 | 0 | 0 | 0 |
| 0 | NA | NA | NA | NA | NA | Mir872 | | 0 | 0 | 0 | 0 | 0 | 0 | 0 | 0 | 0 | 0 | 0 |
| 0 | NA | NA | NA | NA | NA | Gm12694 | | 0 | 0 | 0 | 0 | 0 | 0 | 0 | 0 | 0 | 0 | 0 |
| 0 | NA | NA | NA | NA | NA | Gm12703 | | 0 | 0 | 0 | 0 | 0 | 0 | 0 | 0 | 0 | 0 | 0 |
| 0 | NA | NA | NA | NA | NA | Gm28096 | | 0 | 0 | 0 | 0 | 0 | 0 | 0 | 0 | 0 | 0 | 0 |
| 0 | NA | NA | NA | NA | NA | Gm12708 | | 0 | 0 | 0 | 0 | 0 | 0 | 0 | 0 | 0 | 0 | 0 |
| 0 | NA | NA | NA | NA | NA | Gm29064 | | 0 | 0 | 0 | 0 | 0 | 0 | 0 | 0 | 0 | 0 | 0 |
| 0 | NA | NA | NA | NA | NA | Gm12707 | | 0 | 0 | 0 | 0 | 0 | 0 | 0 | 0 | 0 | 0 | 0 |
| 0 | NA | NA | NA | NA | NA | Gm25004 | | 0 | 0 | 0 | 0 | 0 | 0 | 0 | 0 | 0 | 0 | 0 |
| 0 | NA | NA | NA | NA | NA | Gm20651 | | 0 | 0 | 0 | 0 | 0 | 0 | 0 | 0 | 0 | 0 | 0 |
| 0 | NA | NA | NA | NA | NA | Cyp2j13 | cytochrome P450, family 2, subfamily j, pc | 0 | 0 | 0 | 0 | 0 | 0 | 0 | 0 | 0 | 0 | 0 |
| 0 | NA | NA | NA | NA | NA | | | 0 | 0 | 0 | 0 | 0 | 0 | 0 | 0 | 0 | 0 | 0 |
| 0 | NA | NA | NA | NA | NA | Hspe1-ps6 | | 0 | 0 | 0 | 0 | 0 | 0 | 0 | 0 | 0 | 0 | 0 |
| 0 | NA | NA | NA | NA | NA | Cyp2j12 | cytochrome P450, family 2, subfamily j, pc | 0 | 0 | 0 | 0 | 0 | 0 | 0 | 0 | 0 | 0 | 0 |
| 0 | NA | NA | NA | NA | NA | Cyp2j7-ps3 | | 0 | 0 | 0 | 0 | 0 | 0 | 0 | 0 | 0 | 0 | 0 |
| 0 | NA | NA | NA | NA | NA | Cyp2j7-ps2 | | 0 | 0 | 0 | 0 | 0 | 0 | 0 | 0 | 0 | 0 | 0 |
| 0 | NA | NA | NA | NA | NA | Cyp2j7-ps1 | | 0 | 0 | 0 | 0 | 0 | 0 | 0 | 0 | 0 | 0 | 0 |
| 0 | NA | NA | NA | NA | NA | Cyp2j7 | | 0 | 0 | 0 | 0 | 0 | 0 | 0 | 0 | 0 | 0 | 0 |
| 0 | NA | NA | NA | NA | NA | Cyp2j15-ps | | 0 | 0 | 0 | 0 | 0 | 0 | 0 | 0 | 0 | 0 | 0 |
| 0 | NA | NA | NA | NA | NA | Cyp2j11 | cytochrome P450, family 2, subfamily j, pc | 0 | 0 | 0 | 0 | 0 | 0 | 0 | 0 | 0 | 0 | 0 |
| 0 | NA | NA | NA | NA | NA | Cyp2j14-ps | | 0 | 0 | 0 | 0 | 0 | 0 | 0 | 0 | 0 | 0 | 0 |
| 0 | NA | NA | NA | NA | NA | Gm12695 | predicted gene 12695 | 0 | 0 | 0 | 0 | 0 | 0 | 0 | 0 | 0 | 0 | 0 |
| 0 | NA | NA | NA | NA | NA | Gm27579 | | 0 | 0 | 0 | 0 | 0 | 0 | 0 | 0 | 0 | 0 | 0 |
| 0 | NA | NA | NA | NA | NA | Gm27521 | | 0 | 0 | 0 | 0 | 0 | 0 | 0 | 0 | 0 | 0 | 0 |
| 0 | NA | NA | NA | NA | NA | Gm12697 | | 0 | 0 | 0 | 0 | 0 | 0 | 0 | 0 | 0 | 0 | 0 |
| 0 | NA | NA | NA | NA | NA | Gm12692 | | 0 | 0 | 0 | 0 | 0 | 0 | 0 | 0 | 0 | 0 | 0 |
| 0 | NA | NA | NA | NA | NA | Gm12787 | | 0 | 0 | 0 | 0 | 0 | 0 | 0 | 0 | 0 | 0 | 0 |
| 0 | NA | NA | NA | NA | NA | Gm24203 | | 0 | 0 | 0 | 0 | 0 | 0 | 0 | 0 | 0 | 0 | 0 |
| 0 | NA | NA | NA | NA | NA | Gm12788 | | 0 | 0 | 0 | 0 | 0 | 0 | 0 | 0 | 0 | 0 | 0 |
| 0 | NA | NA | NA | NA | NA | Gm24960 | | 0 | 0 | 0 | 0 | 0 | 0 | 0 | 0 | 0 | 0 | 0 |
| 0 | NA | NA | NA | NA | NA | Gm22625 | | 0 | 0 | 0 | 0 | 0 | 0 | 0 | 0 | 0 | 0 | 0 |
| 0 | NA | NA | NA | NA | NA | Gm26154 | | 0 | 0 | 0 | 0 | 0 | 0 | 0 | 0 | 0 | 0 | 0 |
| 0 | NA | NA | NA | NA | NA | Gm17376 | | 0 | 0 | 0 | 0 | 0 | 0 | 0 | 0 | 0 | 0 | 0 |
| 0 | NA | NA | NA | NA | NA | Kank4os | | 0 | 0 | 0 | 0 | 0 | 0 | 0 | 0 | 0 | 0 | 0 |
| 0 | NA | NA | NA | NA | NA | I0C0044D17Rik | | 0 | 0 | 0 | 0 | 0 | 0 | 0 | 0 | 0 | 0 | 0 |
| 0 | NA | NA | NA | NA | NA | Gm12851 | | 0 | 0 | 0 | 0 | 0 | 0 | 0 | 0 | 0 | 0 | 0 |
| 0 | NA | NA | NA | NA | NA | Gm12705 | | 0 | 0 | 0 | 0 | 0 | 0 | 0 | 0 | 0 | 0 | 0 |
| 0 | NA | NA | NA | NA | NA | Gm12681 | | 0 | 0 | 0 | 0 | 0 | 0 | 0 | 0 | 0 | 0 | 0 |
| 0 | NA | NA | NA | NA | NA | Gm37556 | | 0 | 0 | 0 | 0 | 0 | 0 | 0 | 0 | 0 | 0 | 0 |
| 0 | NA | NA | NA | NA | NA | Gm12685 | | 0 | 0 | 0 | 0 | 0 | 0 | 0 | 0 | 0 | 0 | 0 |
| 0 | NA | NA | NA | NA | NA | Gm12686 | | 0 | 0 | 0 | 0 | 0 | 0 | 0 | 0 | 0 | 0 | 0 |
| 0 | NA | NA | NA | NA | NA | Gm12684 | | 0 | 0 | 0 | 0 | 0 | 0 | 0 | 0 | 0 | 0 | 0 |
| 0 | NA | NA | NA | NA | NA | Gm12683 | | 0 | 0 | 0 | 0 | 0 | 0 | 0 | 0 | 0 | 0 | 0 |
| 0 | NA | NA | NA | NA | NA | Gm37117 | | 0 | 0 | 0 | 0 | 0 | 0 | 0 | 0 | 0 | 0 | 0 |
| 0 | NA | NA | NA | NA | NA | Gm12690 | | 0 | 0 | 0 | 0 | 0 | 0 | 0 | 0 | 0 | 0 | 0 |
| 0 | NA | NA | NA | NA | NA | Gm12687 | | 0 | 0 | 0 | 0 | 0 | 0 | 0 | 0 | 0 | 0 | 0 |
| 0 | NA | NA | NA | NA | NA | Gm23366 | | 0 | 0 | 0 | 0 | 0 | 0 | 0 | 0 | 0 | 0 | 0 |
| 0 | NA | NA | NA | NA | NA | Gm12682 | | 0 | 0 | 0 | 0 | 0 | 0 | 0 | 0 | 0 | 0 | 0 |
| 0 | NA | NA | NA | NA | NA | Gm12702 | | 0 | 0 | 0 | 0 | 0 | 0 | 0 | 0 | 0 | 0 | 0 |
| 0 | NA | NA | NA | NA | NA | Gm12701 | | 0 | 0 | 0 | 0 | 0 | 0 | 0 | 0 | 0 | 0 | 0 |
| 0 | NA | NA | NA | NA | NA | Ube2uos | | 0 | 0 | 0 | 0 | 0 | 0 | 0 | 0 | 0 | 0 | 0 |
| 0 | NA | NA | NA | NA | NA | Gm10577 | | 0 | 0 | 0 | 0 | 0 | 0 | 0 | 0 | 0 | 0 | 0 |
| 0 | NA | NA | NA | NA | NA | Gm10576 | | 0 | 0 | 0 | 0 | 0 | 0 | 0 | 0 | 0 | 0 | 0 |
| 0 | NA | NA | NA | NA | NA | Gm22100 | | 0 | 0 | 0 | 0 | 0 | 0 | 0 | 0 | 0 | 0 | 0 |
| 0 | NA | NA | NA | NA | NA | Gm24468 | | 0 | 0 | 0 | 0 | 0 | 0 | 0 | 0 | 0 | 0 | 0 |
| 0 | NA | NA | NA | NA | NA | Gm12785 | | 0 | 0 | 0 | 0 | 0 | 0 | 0 | 0 | 0 | 0 | 0 |
| 0 | NA | NA | NA | NA | NA | Gm25124 | | 0 | 0 | 0 | 0 | 0 | 0 | 0 | 0 | 0 | 0 | 0 |
| 0 | NA | NA | NA | NA | NA | Gm12801 | | 0 | 0 | 0 | 0 | 0 | 0 | 0 | 0 | 0 | 0 | 0 |
| 0 | NA | NA | NA | NA | NA | Gm12793 | | 0 | 0 | 0 | 0 | 0 | 0 | 0 | 0 | 0 | 0 | 0 |

| | | | | | | | | | | | | | | | | | | |
|---|---|---|---|---|---|---|---|---|---|---|---|---|---|---|---|---|---|---|
| 0 | NA | NA | NA | NA | NA | Mir101a | | 0 | 0 | 0 | 0 | 0 | 0 | 0 | 0 | 0 | 0 | 0 |
| 0 | NA | NA | NA | NA | NA | Gm12795 | | 0 | 0 | 0 | 0 | 0 | 0 | 0 | 0 | 0 | 0 | 0 |
| 0 | NA | NA | NA | NA | NA | Gm22533 | | 0 | 0 | 0 | 0 | 0 | 0 | 0 | 0 | 0 | 0 | 0 |
| 0 | NA | NA | NA | NA | NA | B020004J07Ril | RIKEN cDNA B020004J07 gene | 0 | 0 | 0 | 0 | 0 | 0 | 0 | 0 | 0 | 0 | 0 |
| 0 | NA | NA | NA | NA | NA | Gm12797 | | 0 | 0 | 0 | 0 | 0 | 0 | 0 | 0 | 0 | 0 | 0 |
| 0 | NA | NA | NA | NA | NA | C130073F10Ril | RIKEN cDNA C130073F10 gene | 0 | 0 | 0 | 0 | 0 | 0 | 0 | 0 | 0 | 0 | 0 |
| 0 | NA | NA | NA | NA | NA | Gm12800 | predicted gene 12800 | 0 | 0 | 0 | 0 | 0 | 0 | 0 | 0 | 0 | 0 | 0 |
| 0 | NA | NA | NA | NA | NA | Gm12794 | predicted gene 12794 | 0 | 0 | 0 | 0 | 0 | 0 | 0 | 0 | 0 | 0 | 0 |
| 0 | NA | NA | NA | NA | NA | Gm12790 | oogenesin-2-like | 0 | 0 | 0 | 0 | 0 | 0 | 0 | 0 | 0 | 0 | 0 |
| 0 | NA | NA | NA | NA | NA | Gm12789 | predicted gene 12789 | 0 | 0 | 0 | 0 | 0 | 0 | 0 | 0 | 0 | 0 | 0 |
| 0 | NA | NA | NA | NA | NA | Gm24869 | | 0 | 0 | 0 | 0 | 0 | 0 | 0 | 0 | 0 | 0 | 0 |
| 0 | NA | NA | NA | NA | NA | Tctex1d1 | Tctex1 domain containing 1 | 0 | 0 | 0 | 0 | 0 | 0 | 0 | 0 | 0 | 0 | 0 |
| 0 | NA | NA | NA | NA | NA | BB031773 | | 0 | 0 | 0 | 0 | 0 | 0 | 0 | 0 | 0 | 0 | 0 |
| 0 | NA | NA | NA | NA | NA | Insl5 | insulin-like 5 | 0 | 0 | 0 | 0 | 0 | 0 | 0 | 0 | 0 | 0 | 0 |
| 0 | NA | NA | NA | NA | NA | Gm12710 | | 0 | 0 | 0 | 0 | 0 | 0 | 0 | 0 | 0 | 0 | 0 |
| 0 | NA | NA | NA | NA | NA | 4921539E11Ril | RIKEN cDNA 4921539E11 gene | 0 | 0 | 0 | 0 | 0 | 0 | 0 | 0 | 0 | 0 | 0 |
| 0 | NA | NA | NA | NA | NA | 4930456L15Rik | | 0 | 0 | 0 | 0 | 0 | 0 | 0 | 0 | 0 | 0 | 0 |
| 0 | NA | NA | NA | NA | NA | Gm12718 | | 0 | 0 | 0 | 0 | 0 | 0 | 0 | 0 | 0 | 0 | 0 |
| 0 | NA | NA | NA | NA | NA | Gm23064 | | 0 | 0 | 0 | 0 | 0 | 0 | 0 | 0 | 0 | 0 | 0 |
| 0 | NA | NA | NA | NA | NA | Gm12720 | | 0 | 0 | 0 | 0 | 0 | 0 | 0 | 0 | 0 | 0 | 0 |
| 0 | NA | NA | NA | NA | NA | Gm22801 | | 0 | 0 | 0 | 0 | 0 | 0 | 0 | 0 | 0 | 0 | 0 |
| 0 | NA | NA | NA | NA | NA | Gm25877 | | 0 | 0 | 0 | 0 | 0 | 0 | 0 | 0 | 0 | 0 | 0 |
| 0 | NA | NA | NA | NA | NA | Gm12719 | | 0 | 0 | 0 | 0 | 0 | 0 | 0 | 0 | 0 | 0 | 0 |
| 0 | NA | NA | NA | NA | NA | Gm12717 | | 0 | 0 | 0 | 0 | 0 | 0 | 0 | 0 | 0 | 0 | 0 |
| 0 | NA | NA | NA | NA | NA | C8a | complement component 8, alpha polypep | 0 | 0 | 0 | 0 | 0 | 0 | 0 | 0 | 0 | 0 | 0 |
| 0 | NA | NA | NA | NA | NA | Gm17662 | | 0 | 0 | 0 | 0 | 0 | 0 | 0 | 0 | 0 | 0 | 0 |
| 0 | NA | NA | NA | NA | NA | Gm12721 | | 0 | 0 | 0 | 0 | 0 | 0 | 0 | 0 | 0 | 0 | 0 |
| 0 | NA | NA | NA | NA | NA | Gm12722 | | 0 | 0 | 0 | 0 | 0 | 0 | 0 | 0 | 0 | 0 | 0 |
| 0 | NA | NA | NA | NA | NA | Gm12723 | | 0 | 0 | 0 | 0 | 0 | 0 | 0 | 0 | 0 | 0 | 0 |
| 0 | NA | NA | NA | NA | NA | Gm24363 | | 0 | 0 | 0 | 0 | 0 | 0 | 0 | 0 | 0 | 0 | 0 |
| 0 | NA | NA | NA | NA | NA | Gm12729 | | 0 | 0 | 0 | 0 | 0 | 0 | 0 | 0 | 0 | 0 | 0 |
| 0 | NA | NA | NA | NA | NA | Gm26047 | | 0 | 0 | 0 | 0 | 0 | 0 | 0 | 0 | 0 | 0 | 0 |
| 0 | NA | NA | NA | NA | NA | Gm26047 | | 0 | 0 | 0 | 0 | 0 | 0 | 0 | 0 | 0 | 0 | 0 |
| 0 | NA | NA | NA | NA | NA | Gm12724 | | 0 | 0 | 0 | 0 | 0 | 0 | 0 | 0 | 0 | 0 | 0 |
| 0 | NA | NA | NA | NA | NA | Gm12731 | | 0 | 0 | 0 | 0 | 0 | 0 | 0 | 0 | 0 | 0 | 0 |
| 0 | NA | NA | NA | NA | NA | Gm37253 | | 0 | 0 | 0 | 0 | 0 | 0 | 0 | 0 | 0 | 0 | 0 |
| 0 | NA | NA | NA | NA | NA | Bsnd | barttin CLCNK type accessory beta subur | 0 | 0 | 0 | 0 | 0 | 0 | 0 | 0 | 0 | 0 | 0 |
| 0 | NA | NA | NA | NA | NA | | | 0 | 0 | 0 | 0 | 0 | 0 | 0 | 0 | 0 | 0 | 0 |
| 0 | NA | NA | NA | NA | NA | Tmem61 | transmembrane protein 61 | 0 | 0 | 0 | 0 | 0 | 0 | 0 | 0 | 0 | 0 | 0 |
| 0 | NA | NA | NA | NA | NA | Gm12744 | | 0 | 0 | 0 | 0 | 0 | 0 | 0 | 0 | 0 | 0 | 0 |
| 0 | NA | NA | NA | NA | NA | Gm12746 | | 0 | 0 | 0 | 0 | 0 | 0 | 0 | 0 | 0 | 0 | 0 |
| 0 | NA | NA | NA | NA | NA | Gm24804 | | 0 | 0 | 0 | 0 | 0 | 0 | 0 | 0 | 0 | 0 | 0 |
| 0 | NA | NA | NA | NA | NA | Dmrtb1 | DMRT-like family B with proline-rich C-ter | 0 | 0 | 0 | 0 | 0 | 0 | 0 | 0 | 0 | 0 | 0 |
| 0 | NA | NA | NA | NA | NA | Gm12899 | | 0 | 0 | 0 | 0 | 0 | 0 | 0 | 0 | 0 | 0 | 0 |
| 0 | NA | NA | NA | NA | NA | 1700047F07Rik | | 0 | 0 | 0 | 0 | 0 | 0 | 0 | 0 | 0 | 0 | 0 |
| 0 | NA | NA | NA | NA | NA | Gm25123 | | 0 | 0 | 0 | 0 | 0 | 0 | 0 | 0 | 0 | 0 | 0 |
| 0 | NA | NA | NA | NA | NA | Gm12907 | | 0 | 0 | 0 | 0 | 0 | 0 | 0 | 0 | 0 | 0 | 0 |
| 0 | NA | NA | NA | NA | NA | Gm23354 | | 0 | 0 | 0 | 0 | 0 | 0 | 0 | 0 | 0 | 0 | 0 |
| 0 | NA | NA | NA | NA | NA | Mir6397 | | 0 | 0 | 0 | 0 | 0 | 0 | 0 | 0 | 0 | 0 | 0 |
| 0 | NA | NA | NA | NA | NA | Gm12742 | | 0 | 0 | 0 | 0 | 0 | 0 | 0 | 0 | 0 | 0 | 0 |
| 0 | NA | NA | NA | NA | NA | Gm12740 | | 0 | 0 | 0 | 0 | 0 | 0 | 0 | 0 | 0 | 0 | 0 |
| 0 | NA | NA | NA | NA | NA | Gm12738 | | 0 | 0 | 0 | 0 | 0 | 0 | 0 | 0 | 0 | 0 | 0 |
| 0 | NA | NA | NA | NA | NA | Gm23138 | | 0 | 0 | 0 | 0 | 0 | 0 | 0 | 0 | 0 | 0 | 0 |
| 0 | NA | NA | NA | NA | NA | Gm25827 | | 0 | 0 | 0 | 0 | 0 | 0 | 0 | 0 | 0 | 0 | 0 |
| 0 | NA | NA | NA | NA | NA | Gm22345 | | 0 | 0 | 0 | 0 | 0 | 0 | 0 | 0 | 0 | 0 | 0 |
| 0 | NA | NA | NA | NA | NA | Gm12741 | | 0 | 0 | 0 | 0 | 0 | 0 | 0 | 0 | 0 | 0 | 0 |
| 0 | NA | NA | NA | NA | NA | Gm23589 | | 0 | 0 | 0 | 0 | 0 | 0 | 0 | 0 | 0 | 0 | 0 |
| 0 | NA | NA | NA | NA | NA | Mir761 | | 0 | 0 | 0 | 0 | 0 | 0 | 0 | 0 | 0 | 0 | 0 |
| 0 | NA | NA | NA | NA | NA | Gm23743 | | 0 | 0 | 0 | 0 | 0 | 0 | 0 | 0 | 0 | 0 | 0 |
| 0 | NA | NA | NA | NA | NA | 4930522H14Ril | RIKEN cDNA 4930522H14 gene | 0 | 0 | 0 | 0 | 0 | 0 | 0 | 0 | 0 | 0 | 0 |

| | | | | | | | | | | | | | | | | | |
|---|---|---|---|---|---|---|---|---|---|---|---|---|---|---|---|---|---|
| 0 | NA | NA | NA | NA | NA | Gm22497 | | 0 | 0 | 0 | 0 | 0 | 0 | 0 | 0 | 0 | 0 |
| 0 | NA | NA | NA | NA | NA | Gm12803 | | 0 | 0 | 0 | 0 | 0 | 0 | 0 | 0 | 0 | 0 |
| 0 | NA | NA | NA | NA | NA | Gm12808 | | 0 | 0 | 0 | 0 | 0 | 0 | 0 | 0 | 0 | 0 |
| 0 | NA | NA | NA | NA | NA | Dmrta2os | | 0 | 0 | 0 | 0 | 0 | 0 | 0 | 0 | 0 | 0 |
| 0 | NA | NA | NA | NA | NA | Gm12806 | | 0 | 0 | 0 | 0 | 0 | 0 | 0 | 0 | 0 | 0 |
| 0 | NA | NA | NA | NA | NA | Gm12805 | | 0 | 0 | 0 | 0 | 0 | 0 | 0 | 0 | 0 | 0 |
| 0 | NA | NA | NA | NA | NA | n-R5s191 | | 0 | 0 | 0 | 0 | 0 | 0 | 0 | 0 | 0 | 0 |
| 0 | NA | NA | NA | NA | NA | Skint8 | selection and upkeep of intraepithelial T c | 0 | 0 | 0 | 0 | 0 | 0 | 0 | 0 | 0 | 0 |
| 0 | NA | NA | NA | NA | NA | Skint7 | selection and upkeep of intraepithelial T c | 0 | 0 | 0 | 0 | 0 | 0 | 0 | 0 | 0 | 0 |
| 0 | NA | NA | NA | NA | NA | Gm25844 | | 0 | 0 | 0 | 0 | 0 | 0 | 0 | 0 | 0 | 0 |
| 0 | NA | NA | NA | NA | NA | Skint1 | selection and upkeep of intraepithelial T c | 0 | 0 | 0 | 0 | 0 | 0 | 0 | 0 | 0 | 0 |
| 0 | NA | NA | NA | NA | NA | Skint4 | selection and upkeep of intraepithelial T c | 0 | 0 | 0 | 0 | 0 | 0 | 0 | 0 | 0 | 0 |
| 0 | NA | NA | NA | NA | NA | Gm12820 | | 0 | 0 | 0 | 0 | 0 | 0 | 0 | 0 | 0 | 0 |
| 0 | NA | NA | NA | NA | NA | Gm12815 | | 0 | 0 | 0 | 0 | 0 | 0 | 0 | 0 | 0 | 0 |
| 0 | NA | NA | NA | NA | NA | Skint9 | selection and upkeep of intraepithelial T c | 0 | 0 | 0 | 0 | 0 | 0 | 0 | 0 | 0 | 0 |
| 0 | NA | NA | NA | NA | NA | Gm12819 | | 0 | 0 | 0 | 0 | 0 | 0 | 0 | 0 | 0 | 0 |
| 0 | NA | NA | NA | NA | NA | Gm12814 | | 0 | 0 | 0 | 0 | 0 | 0 | 0 | 0 | 0 | 0 |
| 0 | NA | NA | NA | NA | NA | Gm24099 | | 0 | 0 | 0 | 0 | 0 | 0 | 0 | 0 | 0 | 0 |
| 0 | NA | NA | NA | NA | NA | Gm12817 | | 0 | 0 | 0 | 0 | 0 | 0 | 0 | 0 | 0 | 0 |
| 0 | NA | NA | NA | NA | NA | Skint10 | selection and upkeep of intraepithelial T c | 0 | 0 | 0 | 0 | 0 | 0 | 0 | 0 | 0 | 0 |
| 0 | NA | NA | NA | NA | NA | Skint6 | selection and upkeep of intraepithelial T c | 0 | 0 | 0 | 0 | 0 | 0 | 0 | 0 | 0 | 0 |
| 0 | NA | NA | NA | NA | NA | Gm12823 | | 0 | 0 | 0 | 0 | 0 | 0 | 0 | 0 | 0 | 0 |
| 0 | NA | NA | NA | NA | NA | Gm12813 | | 0 | 0 | 0 | 0 | 0 | 0 | 0 | 0 | 0 | 0 |
| 0 | NA | NA | NA | NA | NA | Gm12825 | | 0 | 0 | 0 | 0 | 0 | 0 | 0 | 0 | 0 | 0 |
| 0 | NA | NA | NA | NA | NA | | | 0 | 0 | 0 | 0 | 0 | 0 | 0 | 0 | 0 | 0 |
| 0 | NA | NA | NA | NA | NA | Skint11 | selection and upkeep of intraepithelial T c | 0 | 0 | 0 | 0 | 0 | 0 | 0 | 0 | 0 | 0 |
| 0 | NA | NA | NA | NA | NA | Gm15741 | | 0 | 0 | 0 | 0 | 0 | 0 | 0 | 0 | 0 | 0 |
| 0 | NA | NA | NA | NA | NA | Gm28864 | | 0 | 0 | 0 | 0 | 0 | 0 | 0 | 0 | 0 | 0 |
| 0 | NA | NA | NA | NA | NA | Gm12829 | | 0 | 0 | 0 | 0 | 0 | 0 | 0 | 0 | 0 | 0 |
| 0 | NA | NA | NA | NA | NA | Gm23230 | | 0 | 0 | 0 | 0 | 0 | 0 | 0 | 0 | 0 | 0 |
| 0 | NA | NA | NA | NA | NA | Foxe3 | forkhead box E3 | 0 | 0 | 0 | 0 | 0 | 0 | 0 | 0 | 0 | 0 |
| 0 | NA | NA | NA | NA | NA | Cyp4a28-ps | | 0 | 0 | 0 | 0 | 0 | 0 | 0 | 0 | 0 | 0 |
| 0 | NA | NA | NA | NA | NA | Cyp4a29-ps1 | | 0 | 0 | 0 | 0 | 0 | 0 | 0 | 0 | 0 | 0 |
| 0 | NA | NA | NA | NA | NA | | | 0 | 0 | 0 | 0 | 0 | 0 | 0 | 0 | 0 | 0 |
| 0 | NA | NA | NA | NA | NA | Cyp4a29 | cytochrome P450, family 4, subfamily a, p | 0 | 0 | 0 | 0 | 0 | 0 | 0 | 0 | 0 | 0 |
| 0 | NA | NA | NA | NA | NA | Cyp4a12a | cytochrome P450, family 4, subfamily a, p | 0 | 0 | 0 | 0 | 0 | 0 | 0 | 0 | 0 | 0 |
| 0 | NA | NA | NA | NA | NA | Cyp4a30-ps | | 0 | 0 | 0 | 0 | 0 | 0 | 0 | 0 | 0 | 0 |
| 0 | NA | NA | NA | NA | NA | Cyp4a12b | cytochrome P450, family 4, subfamily a, p | 0 | 0 | 0 | 0 | 0 | 0 | 0 | 0 | 0 | 0 |
| 0 | NA | NA | NA | NA | NA | Gm12837 | | 0 | 0 | 0 | 0 | 0 | 0 | 0 | 0 | 0 | 0 |
| 0 | NA | NA | NA | NA | NA | Cyp4a30b | cytochrome P450, family 4, subfamily a, p | 0 | 0 | 0 | 0 | 0 | 0 | 0 | 0 | 0 | 0 |
| 0 | NA | NA | NA | NA | NA | Gm12833 | | 0 | 0 | 0 | 0 | 0 | 0 | 0 | 0 | 0 | 0 |
| 0 | NA | NA | NA | NA | NA | Cyp4b1-ps1 | | 0 | 0 | 0 | 0 | 0 | 0 | 0 | 0 | 0 | 0 |
| 0 | NA | NA | NA | NA | NA | Cyp4a31 | cytochrome P450, family 4, subfamily a, p | 0 | 0 | 0 | 0 | 0 | 0 | 0 | 0 | 0 | 0 |
| 0 | NA | NA | NA | NA | NA | Cyp4b1-ps2 | | 0 | 0 | 0 | 0 | 0 | 0 | 0 | 0 | 0 | 0 |
| 0 | NA | NA | NA | NA | NA | Gm12849 | | 0 | 0 | 0 | 0 | 0 | 0 | 0 | 0 | 0 | 0 |
| 0 | NA | NA | NA | NA | NA | 6430628N08Ril | RIKEN cDNA 6430628N08 gene | 0 | 0 | 0 | 0 | 0 | 0 | 0 | 0 | 0 | 0 |
| 0 | NA | NA | NA | NA | NA | Dmbx1 | diencephalon/mesencephalon homeobox | 0 | 0 | 0 | 0 | 0 | 0 | 0 | 0 | 0 | 0 |
| 0 | NA | NA | NA | NA | NA | 1700042G07Ril | RIKEN cDNA 1700042G07 gene | 0 | 0 | 0 | 0 | 0 | 0 | 0 | 0 | 0 | 0 |
| 0 | NA | NA | NA | NA | NA | Gm49337 | | 0 | 0 | 0 | 0 | 0 | 0 | 0 | 0 | 0 | 0 |
| 0 | NA | NA | NA | NA | NA | Gm12951 | | 0 | 0 | 0 | 0 | 0 | 0 | 0 | 0 | 0 | 0 |
| 0 | NA | NA | NA | NA | NA | Gm25137 | | 0 | 0 | 0 | 0 | 0 | 0 | 0 | 0 | 0 | 0 |
| 0 | NA | NA | NA | NA | NA | Gm27688 | | 0 | 0 | 0 | 0 | 0 | 0 | 0 | 0 | 0 | 0 |
| 0 | NA | NA | NA | NA | NA | Rpl36-ps8 | | 0 | 0 | 0 | 0 | 0 | 0 | 0 | 0 | 0 | 0 |
| 0 | NA | NA | NA | NA | NA | Gm12994 | | 0 | 0 | 0 | 0 | 0 | 0 | 0 | 0 | 0 | 0 |
| 0 | NA | NA | NA | NA | NA | Gm26355 | | 0 | 0 | 0 | 0 | 0 | 0 | 0 | 0 | 0 | 0 |
| 0 | NA | NA | NA | NA | NA | Gm25435 | | 0 | 0 | 0 | 0 | 0 | 0 | 0 | 0 | 0 | 0 |
| 0 | NA | NA | NA | NA | NA | Gm22092 | | 0 | 0 | 0 | 0 | 0 | 0 | 0 | 0 | 0 | 0 |
| 0 | NA | NA | NA | NA | NA | Gm12997 | | 0 | 0 | 0 | 0 | 0 | 0 | 0 | 0 | 0 | 0 |
| 0 | NA | NA | NA | NA | NA | Best4-ps | | 0 | 0 | 0 | 0 | 0 | 0 | 0 | 0 | 0 | 0 |
| 0 | NA | NA | NA | NA | NA | Snord38a | | 0 | 0 | 0 | 0 | 0 | 0 | 0 | 0 | 0 | 0 |

| | | | | | | | | | | | | | | | | | |
|---|---|---|---|---|---|---|---|---|---|---|---|---|---|---|---|---|---|
| 0 | NA | NA | NA | NA | NA | Gm26330 | | 0 | 0 | 0 | 0 | 0 | 0 | 0 | 0 | 0 | 0 | 0 |
| 0 | NA | NA | NA | NA | NA | Gm22398 | | 0 | 0 | 0 | 0 | 0 | 0 | 0 | 0 | 0 | 0 | 0 |
| 0 | NA | NA | NA | NA | NA | Gm25099 | | 0 | 0 | 0 | 0 | 0 | 0 | 0 | 0 | 0 | 0 | 0 |
| 0 | NA | NA | NA | NA | NA | Gm23143 | | 0 | 0 | 0 | 0 | 0 | 0 | 0 | 0 | 0 | 0 | 0 |
| 0 | NA | NA | NA | NA | NA | Gm23287 | | 0 | 0 | 0 | 0 | 0 | 0 | 0 | 0 | 0 | 0 | 0 |
| 0 | NA | NA | NA | NA | NA | 1700012C08Rik | | 0 | 0 | 0 | 0 | 0 | 0 | 0 | 0 | 0 | 0 | 0 |
| 0 | NA | NA | NA | NA | NA | Gm12827 | | 0 | 0 | 0 | 0 | 0 | 0 | 0 | 0 | 0 | 0 | 0 |
| 0 | NA | NA | NA | NA | NA | Gm12826 | | 0 | 0 | 0 | 0 | 0 | 0 | 0 | 0 | 0 | 0 | 0 |
| 0 | NA | NA | NA | NA | NA | Gm25559 | | 0 | 0 | 0 | 0 | 0 | 0 | 0 | 0 | 0 | 0 | 0 |
| 0 | NA | NA | NA | NA | NA | Gm12845 | | 0 | 0 | 0 | 0 | 0 | 0 | 0 | 0 | 0 | 0 | 0 |
| 0 | NA | NA | NA | NA | NA | Klf17 | Kruppel-like factor 17 | 0 | 0 | 0 | 0 | 0 | 0 | 0 | 0 | 0 | 0 | 0 |
| 0 | NA | NA | NA | NA | NA | Gm23382 | | 0 | 0 | 0 | 0 | 0 | 0 | 0 | 0 | 0 | 0 | 0 |
| 0 | NA | NA | NA | NA | NA | Mir7226 | | 0 | 0 | 0 | 0 | 0 | 0 | 0 | 0 | 0 | 0 | 0 |
| 0 | NA | NA | NA | NA | NA | Gm12858 | | 0 | 0 | 0 | 0 | 0 | 0 | 0 | 0 | 0 | 0 | 0 |
| 0 | NA | NA | NA | NA | NA | Gm12859 | | 0 | 0 | 0 | 0 | 0 | 0 | 0 | 0 | 0 | 0 | 0 |
| 0 | NA | NA | NA | NA | NA | Gm12857 | | 0 | 0 | 0 | 0 | 0 | 0 | 0 | 0 | 0 | 0 | 0 |
| 0 | NA | NA | NA | NA | NA | Gm12853 | | 0 | 0 | 0 | 0 | 0 | 0 | 0 | 0 | 0 | 0 | 0 |
| 0 | NA | NA | NA | NA | NA | Olfr62 | olfactory receptor 62 | 0 | 0 | 0 | 0 | 0 | 0 | 0 | 0 | 0 | 0 | 0 |
| 0 | NA | NA | NA | NA | NA | Olfr1343-ps1 | | 0 | 0 | 0 | 0 | 0 | 0 | 0 | 0 | 0 | 0 | 0 |
| 0 | NA | NA | NA | NA | NA | Olfr1341 | olfactory receptor 1341 | 0 | 0 | 0 | 0 | 0 | 0 | 0 | 0 | 0 | 0 | 0 |
| 0 | NA | NA | NA | NA | NA | Olfr1340 | olfactory receptor 1340 | 0 | 0 | 0 | 0 | 0 | 0 | 0 | 0 | 0 | 0 | 0 |
| 0 | NA | NA | NA | NA | NA | Olfr1339 | olfactory receptor 1339 | 0 | 0 | 0 | 0 | 0 | 0 | 0 | 0 | 0 | 0 | 0 |
| 0 | NA | NA | NA | NA | NA | Olfr1338 | olfactory receptor 1338 | 0 | 0 | 0 | 0 | 0 | 0 | 0 | 0 | 0 | 0 | 0 |
| 0 | NA | NA | NA | NA | NA | Olfr1337 | olfactory receptor 1337 | 0 | 0 | 0 | 0 | 0 | 0 | 0 | 0 | 0 | 0 | 0 |
| 0 | NA | NA | NA | NA | NA | Gm12855 | | 0 | 0 | 0 | 0 | 0 | 0 | 0 | 0 | 0 | 0 | 0 |
| 0 | NA | NA | NA | NA | NA | Olfr1335 | olfactory receptor 1335 | 0 | 0 | 0 | 0 | 0 | 0 | 0 | 0 | 0 | 0 | 0 |
| 0 | NA | NA | NA | NA | NA | Olfr1333 | olfactory receptor 1333 | 0 | 0 | 0 | 0 | 0 | 0 | 0 | 0 | 0 | 0 | 0 |
| 0 | NA | NA | NA | NA | NA | Olfr1332-ps1 | | 0 | 0 | 0 | 0 | 0 | 0 | 0 | 0 | 0 | 0 | 0 |
| 0 | NA | NA | NA | NA | NA | Olfr1331 | olfactory receptor 1331 | 0 | 0 | 0 | 0 | 0 | 0 | 0 | 0 | 0 | 0 | 0 |
| 0 | NA | NA | NA | NA | NA | Olfr1330 | olfactory receptor 1330 | 0 | 0 | 0 | 0 | 0 | 0 | 0 | 0 | 0 | 0 | 0 |
| 0 | NA | NA | NA | NA | NA | Olfr1329 | olfactory receptor 1329 | 0 | 0 | 0 | 0 | 0 | 0 | 0 | 0 | 0 | 0 | 0 |
| 0 | NA | NA | NA | NA | NA | Olfr1328 | olfactory receptor 1328 | 0 | 0 | 0 | 0 | 0 | 0 | 0 | 0 | 0 | 0 | 0 |
| 0 | NA | NA | NA | NA | NA | Gm12856 | | 0 | 0 | 0 | 0 | 0 | 0 | 0 | 0 | 0 | 0 | 0 |
| 0 | NA | NA | NA | NA | NA | Gm12861 | | 0 | 0 | 0 | 0 | 0 | 0 | 0 | 0 | 0 | 0 | 0 |
| 0 | NA | NA | NA | NA | NA | Gm24352 | | 0 | 0 | 0 | 0 | 0 | 0 | 0 | 0 | 0 | 0 | 0 |
| 0 | NA | NA | NA | NA | NA | Gm12864 | | 0 | 0 | 0 | 0 | 0 | 0 | 0 | 0 | 0 | 0 | 0 |
| 0 | NA | NA | NA | NA | NA | Gm12862 | | 0 | 0 | 0 | 0 | 0 | 0 | 0 | 0 | 0 | 0 | 0 |
| 0 | NA | NA | NA | NA | NA | Gm12866 | | 0 | 0 | 0 | 0 | 0 | 0 | 0 | 0 | 0 | 0 | 0 |
| 0 | NA | NA | NA | NA | NA | Mir1957a | | 0 | 0 | 0 | 0 | 0 | 0 | 0 | 0 | 0 | 0 | 0 |
| 0 | NA | NA | NA | NA | NA | Gm12898 | | 0 | 0 | 0 | 0 | 0 | 0 | 0 | 0 | 0 | 0 | 0 |
| 0 | NA | NA | NA | NA | NA | Gm12927 | | 0 | 0 | 0 | 0 | 0 | 0 | 0 | 0 | 0 | 0 | 0 |
| 0 | NA | NA | NA | NA | NA | Cldn19 | claudin 19 | 0 | 0 | 0 | 0 | 0 | 0 | 0 | 0 | 0 | 0 | 0 |
| 0 | NA | NA | NA | NA | NA | Gm37817 | | 0 | 0 | 0 | 0 | 0 | 0 | 0 | 0 | 0 | 0 | 0 |
| 0 | NA | NA | NA | NA | NA | Gm12954 | | 0 | 0 | 0 | 0 | 0 | 0 | 0 | 0 | 0 | 0 | 0 |
| 0 | NA | NA | NA | NA | NA | Gm12956 | | 0 | 0 | 0 | 0 | 0 | 0 | 0 | 0 | 0 | 0 | 0 |
| 0 | NA | NA | NA | NA | NA | Gm12957 | | 0 | 0 | 0 | 0 | 0 | 0 | 0 | 0 | 0 | 0 | 0 |
| 0 | NA | NA | NA | NA | NA | Rnu6-ps2 | | 0 | 0 | 0 | 0 | 0 | 0 | 0 | 0 | 0 | 0 | 0 |
| 0 | NA | NA | NA | NA | NA | Gm12959 | | 0 | 0 | 0 | 0 | 0 | 0 | 0 | 0 | 0 | 0 | 0 |
| 0 | NA | NA | NA | NA | NA | Guca2b | guanylate cyclase activator 2b (retina) | 0 | 0 | 0 | 0 | 0 | 0 | 0 | 0 | 0 | 0 | 0 |
| 0 | NA | NA | NA | NA | NA | | | 0 | 0 | 0 | 0 | 0 | 0 | 0 | 0 | 0 | 0 | 0 |
| 0 | NA | NA | NA | NA | NA | Gm8439 | predicted gene 8439 | 0 | 0 | 0 | 0 | 0 | 0 | 0 | 0 | 0 | 0 | 0 |
| 0 | NA | NA | NA | NA | NA | Mir30e | | 0 | 0 | 0 | 0 | 0 | 0 | 0 | 0 | 0 | 0 | 0 |
| 0 | NA | NA | NA | NA | NA | Mir30f | | 0 | 0 | 0 | 0 | 0 | 0 | 0 | 0 | 0 | 0 | 0 |
| 0 | NA | NA | NA | NA | NA | Gm23936 | | 0 | 0 | 0 | 0 | 0 | 0 | 0 | 0 | 0 | 0 | 0 |
| 0 | NA | NA | NA | NA | NA | Gm12871 | | 0 | 0 | 0 | 0 | 0 | 0 | 0 | 0 | 0 | 0 | 0 |
| 0 | NA | NA | NA | NA | NA | Gm24678 | | 0 | 0 | 0 | 0 | 0 | 0 | 0 | 0 | 0 | 0 | 0 |
| 0 | NA | NA | NA | NA | NA | Gm26186 | | 0 | 0 | 0 | 0 | 0 | 0 | 0 | 0 | 0 | 0 | 0 |
| 0 | NA | NA | NA | NA | NA | Gm23540 | | 0 | 0 | 0 | 0 | 0 | 0 | 0 | 0 | 0 | 0 | 0 |
| 0 | NA | NA | NA | NA | NA | Tmco2 | transmembrane and coiled-coil domains 2 | 0 | 0 | 0 | 0 | 0 | 0 | 0 | 0 | 0 | 0 | 0 |
| 0 | NA | NA | NA | NA | NA | Gm12882 | | 0 | 0 | 0 | 0 | 0 | 0 | 0 | 0 | 0 | 0 | 0 |

| | | | | | | | | | | | | | | | | | | |
|---|---|---|---|---|---|---|---|---|---|---|---|---|---|---|---|---|---|---|
| 0 | NA | NA | NA | NA | NA | Gm12881 | | 0 | 0 | 0 | 0 | 0 | 0 | 0 | 0 | 0 | 0 | 0 |
| 0 | NA | NA | NA | NA | NA | Gm12879 | | 0 | 0 | 0 | 0 | 0 | 0 | 0 | 0 | 0 | 0 | 0 |
| 0 | NA | NA | NA | NA | NA | Gm22982 | | 0 | 0 | 0 | 0 | 0 | 0 | 0 | 0 | 0 | 0 | 0 |
| 0 | NA | NA | NA | NA | NA | Gm12890 | | 0 | 0 | 0 | 0 | 0 | 0 | 0 | 0 | 0 | 0 | 0 |
| 0 | NA | NA | NA | NA | NA | Gm12888 | predicted gene 12888 | 0 | 0 | 0 | 0 | 0 | 0 | 0 | 0 | 0 | 0 | 0 |
| 0 | NA | NA | NA | NA | NA | Gm12889 | | 0 | 0 | 0 | 0 | 0 | 0 | 0 | 0 | 0 | 0 | 0 |
| 0 | NA | NA | NA | NA | NA | Gm12885 | | 0 | 0 | 0 | 0 | 0 | 0 | 0 | 0 | 0 | 0 | 0 |
| 0 | NA | NA | NA | NA | NA | Gm12884 | | 0 | 0 | 0 | 0 | 0 | 0 | 0 | 0 | 0 | 0 | 0 |
| 0 | NA | NA | NA | NA | NA | Gm12883 | | 0 | 0 | 0 | 0 | 0 | 0 | 0 | 0 | 0 | 0 | 0 |
| 0 | NA | NA | NA | NA | NA | Gm12886 | predicted gene 12886 | 0 | 0 | 0 | 0 | 0 | 0 | 0 | 0 | 0 | 0 | 0 |
| 0 | NA | NA | NA | NA | NA | Gm12880 | | 0 | 0 | 0 | 0 | 0 | 0 | 0 | 0 | 0 | 0 | 0 |
| 0 | NA | NA | NA | NA | NA | Gm26162 | | 0 | 0 | 0 | 0 | 0 | 0 | 0 | 0 | 0 | 0 | 0 |
| 0 | NA | NA | NA | NA | NA | Gm12887 | predicted gene 12887 | 0 | 0 | 0 | 0 | 0 | 0 | 0 | 0 | 0 | 0 | 0 |
| 0 | NA | NA | NA | NA | NA | Gm12878 | | 0 | 0 | 0 | 0 | 0 | 0 | 0 | 0 | 0 | 0 | 0 |
| 0 | NA | NA | NA | NA | NA | Gm12896 | | 0 | 0 | 0 | 0 | 0 | 0 | 0 | 0 | 0 | 0 | 0 |
| 0 | NA | NA | NA | NA | NA | Gm12897 | | 0 | 0 | 0 | 0 | 0 | 0 | 0 | 0 | 0 | 0 | 0 |
| 0 | NA | NA | NA | NA | NA | Gm12893 | | 0 | 0 | 0 | 0 | 0 | 0 | 0 | 0 | 0 | 0 | 0 |
| 0 | NA | NA | NA | NA | NA | Gm12895 | | 0 | 0 | 0 | 0 | 0 | 0 | 0 | 0 | 0 | 0 | 0 |
| 0 | NA | NA | NA | NA | NA | Gm12877 | | 0 | 0 | 0 | 0 | 0 | 0 | 0 | 0 | 0 | 0 | 0 |
| 0 | NA | NA | NA | NA | NA | Gm12874 | | 0 | 0 | 0 | 0 | 0 | 0 | 0 | 0 | 0 | 0 | 0 |
| 0 | NA | NA | NA | NA | NA | Gm12875 | | 0 | 0 | 0 | 0 | 0 | 0 | 0 | 0 | 0 | 0 | 0 |
| 0 | NA | NA | NA | NA | NA | Gm12876 | | 0 | 0 | 0 | 0 | 0 | 0 | 0 | 0 | 0 | 0 | 0 |
| 0 | NA | NA | NA | NA | NA | 9530002B09Ril | RIKEN cDNA 9530002B09 gene | 0 | 0 | 0 | 0 | 0 | 0 | 0 | 0 | 0 | 0 | 0 |
| 0 | NA | NA | NA | NA | NA | Gm12872 | | 0 | 0 | 0 | 0 | 0 | 0 | 0 | 0 | 0 | 0 | 0 |
| 0 | NA | NA | NA | NA | NA | Gm12873 | predicted gene 12873 | 0 | 0 | 0 | 0 | 0 | 0 | 0 | 0 | 0 | 0 | 0 |
| 0 | NA | NA | NA | NA | NA | Gm12891 | | 0 | 0 | 0 | 0 | 0 | 0 | 0 | 0 | 0 | 0 | 0 |
| 0 | NA | NA | NA | NA | NA | Gm12923 | | 0 | 0 | 0 | 0 | 0 | 0 | 0 | 0 | 0 | 0 | 0 |
| 0 | NA | NA | NA | NA | NA | Gm17244 | | 0 | 0 | 0 | 0 | 0 | 0 | 0 | 0 | 0 | 0 | 0 |
| 0 | NA | NA | NA | NA | NA | Oxct2b | 3-oxoacid CoA transferase 2B | 0 | 0 | 0 | 0 | 0 | 0 | 0 | 0 | 0 | 0 | 0 |
| 0 | NA | NA | NA | NA | NA | Nt5c1a | 5'-nucleotidase, cytosolic IA | 0 | 0 | 0 | 0 | 0 | 0 | 0 | 0 | 0 | 0 | 0 |
| 0 | NA | NA | NA | NA | NA | | | 0 | 0 | 0 | 0 | 0 | 0 | 0 | 0 | 0 | 0 | 0 |
| 0 | NA | NA | NA | NA | NA | Oxct2a | 3-oxoacid CoA transferase 2A | 0 | 0 | 0 | 0 | 0 | 0 | 0 | 0 | 0 | 0 | 0 |
| 0 | NA | NA | NA | NA | NA | Gm12924 | | 0 | 0 | 0 | 0 | 0 | 0 | 0 | 0 | 0 | 0 | 0 |
| 0 | NA | NA | NA | NA | NA | Gm23548 | | 0 | 0 | 0 | 0 | 0 | 0 | 0 | 0 | 0 | 0 | 0 |
| 0 | NA | NA | NA | NA | NA | Mir6398 | | 0 | 0 | 0 | 0 | 0 | 0 | 0 | 0 | 0 | 0 | 0 |
| 0 | NA | NA | NA | NA | NA | Gm12926 | | 0 | 0 | 0 | 0 | 0 | 0 | 0 | 0 | 0 | 0 | 0 |
| 0 | NA | NA | NA | NA | NA | Gm12903 | | 0 | 0 | 0 | 0 | 0 | 0 | 0 | 0 | 0 | 0 | 0 |
| 0 | NA | NA | NA | NA | NA | Gm37667 | | 0 | 0 | 0 | 0 | 0 | 0 | 0 | 0 | 0 | 0 | 0 |
| 0 | NA | NA | NA | NA | NA | Gm12916 | | 0 | 0 | 0 | 0 | 0 | 0 | 0 | 0 | 0 | 0 | 0 |
| 0 | NA | NA | NA | NA | NA | 1700021L23Rik | | 0 | 0 | 0 | 0 | 0 | 0 | 0 | 0 | 0 | 0 | 0 |
| 0 | NA | NA | NA | NA | NA | 4933435F18Rik | | 0 | 0 | 0 | 0 | 0 | 0 | 0 | 0 | 0 | 0 | 0 |
| 0 | NA | NA | NA | NA | NA | Gm22983 | | 0 | 0 | 0 | 0 | 0 | 0 | 0 | 0 | 0 | 0 | 0 |
| 0 | NA | NA | NA | NA | NA | 1700057H15Rik | | 0 | 0 | 0 | 0 | 0 | 0 | 0 | 0 | 0 | 0 | 0 |
| 0 | NA | NA | NA | NA | NA | Gm24480 | | 0 | 0 | 0 | 0 | 0 | 0 | 0 | 0 | 0 | 0 | 0 |
| 0 | NA | NA | NA | NA | NA | Gm24721 | | 0 | 0 | 0 | 0 | 0 | 0 | 0 | 0 | 0 | 0 | 0 |
| 0 | NA | NA | NA | NA | NA | Mir698 | | 0 | 0 | 0 | 0 | 0 | 0 | 0 | 0 | 0 | 0 | 0 |
| 0 | NA | NA | NA | NA | NA | 1700125G02Rik | | 0 | 0 | 0 | 0 | 0 | 0 | 0 | 0 | 0 | 0 | 0 |
| 0 | NA | NA | NA | NA | NA | Gm12930 | | 0 | 0 | 0 | 0 | 0 | 0 | 0 | 0 | 0 | 0 | 0 |
| 0 | NA | NA | NA | NA | NA | Gm12931 | | 0 | 0 | 0 | 0 | 0 | 0 | 0 | 0 | 0 | 0 | 0 |
| 0 | NA | NA | NA | NA | NA | 1700041M05Rik | | 0 | 0 | 0 | 0 | 0 | 0 | 0 | 0 | 0 | 0 | 0 |
| 0 | NA | NA | NA | NA | NA | Mir692-2 | | 0 | 0 | 0 | 0 | 0 | 0 | 0 | 0 | 0 | 0 | 0 |
| 0 | NA | NA | NA | NA | NA | Gm12946 | | 0 | 0 | 0 | 0 | 0 | 0 | 0 | 0 | 0 | 0 | 0 |
| 0 | NA | NA | NA | NA | NA | Gm12935 | | 0 | 0 | 0 | 0 | 0 | 0 | 0 | 0 | 0 | 0 | 0 |
| 0 | NA | NA | NA | NA | NA | Mir7119 | | 0 | 0 | 0 | 0 | 0 | 0 | 0 | 0 | 0 | 0 | 0 |
| 0 | NA | NA | NA | NA | NA | Gm12937 | | 0 | 0 | 0 | 0 | 0 | 0 | 0 | 0 | 0 | 0 | 0 |
| 0 | NA | NA | NA | NA | NA | Gm24161 | | 0 | 0 | 0 | 0 | 0 | 0 | 0 | 0 | 0 | 0 | 0 |
| 0 | NA | NA | NA | NA | NA | Gm12939 | | 0 | 0 | 0 | 0 | 0 | 0 | 0 | 0 | 0 | 0 | 0 |
| 0 | NA | NA | NA | NA | NA | Gm25600 | | 0 | 0 | 0 | 0 | 0 | 0 | 0 | 0 | 0 | 0 | 0 |
| 0 | NA | NA | NA | NA | NA | Gm22221 | | 0 | 0 | 0 | 0 | 0 | 0 | 0 | 0 | 0 | 0 | 0 |
| 0 | NA | NA | NA | NA | NA | Gm12948 | | 0 | 0 | 0 | 0 | 0 | 0 | 0 | 0 | 0 | 0 | 0 |

| | | | | | | | | | | | | | | | | | | |
|---|---|---|---|---|---|---|---|---|---|---|---|---|---|---|---|---|---|---|
| 0 | NA | NA | NA | NA | NA | 1700112K13Rik | | 0 | 0 | 0 | 0 | 0 | 0 | 0 | 0 | 0 | 0 | 0 |
| 0 | NA | NA | NA | NA | NA | Hmgb4 | high-mobility group box 4 | 0 | 0 | 0 | 0 | 0 | 0 | 0 | 0 | 0 | 0 | 0 |
| 0 | NA | NA | NA | NA | NA | Hmgb4os | | 0 | 0 | 0 | 0 | 0 | 0 | 0 | 0 | 0 | 0 | 0 |
| 0 | NA | NA | NA | NA | NA | Gm12968 | | 0 | 0 | 0 | 0 | 0 | 0 | 0 | 0 | 0 | 0 | 0 |
| 0 | NA | NA | NA | NA | NA | Gm15904 | | 0 | 0 | 0 | 0 | 0 | 0 | 0 | 0 | 0 | 0 | 0 |
| 0 | NA | NA | NA | NA | NA | Gm22994 | | 0 | 0 | 0 | 0 | 0 | 0 | 0 | 0 | 0 | 0 | 0 |
| 0 | NA | NA | NA | NA | NA | Gm12980 | | 0 | 0 | 0 | 0 | 0 | 0 | 0 | 0 | 0 | 0 | 0 |
| 0 | NA | NA | NA | NA | NA | Gm24621 | | 0 | 0 | 0 | 0 | 0 | 0 | 0 | 0 | 0 | 0 | 0 |
| 0 | NA | NA | NA | NA | NA | Gm12979 | | 0 | 0 | 0 | 0 | 0 | 0 | 0 | 0 | 0 | 0 | 0 |
| 0 | NA | NA | NA | NA | NA | Mir8119 | | 0 | 0 | 0 | 0 | 0 | 0 | 0 | 0 | 0 | 0 | 0 |
| 0 | NA | NA | NA | NA | NA | Gm12967 | | 0 | 0 | 0 | 0 | 0 | 0 | 0 | 0 | 0 | 0 | 0 |
| 0 | NA | NA | NA | NA | NA | Gm28017 | | 0 | 0 | 0 | 0 | 0 | 0 | 0 | 0 | 0 | 0 | 0 |
| 0 | NA | NA | NA | NA | NA | Gm23845 | | 0 | 0 | 0 | 0 | 0 | 0 | 0 | 0 | 0 | 0 | 0 |
| 0 | NA | NA | NA | NA | NA | E330017L17Rik | | 0 | 0 | 0 | 0 | 0 | 0 | 0 | 0 | 0 | 0 | 0 |
| 0 | NA | NA | NA | NA | NA | Ldc1 | leucine decarboxylase 1 | 0 | 0 | 0 | 0 | 0 | 0 | 0 | 0 | 0 | 0 | 0 |
| 0 | NA | NA | NA | NA | NA | Snord85 | | 0 | 0 | 0 | 0 | 0 | 0 | 0 | 0 | 0 | 0 | 0 |
| 0 | NA | NA | NA | NA | NA | Gm12972 | | 0 | 0 | 0 | 0 | 0 | 0 | 0 | 0 | 0 | 0 | 0 |
| 0 | NA | NA | NA | NA | NA | Gm12970 | | 0 | 0 | 0 | 0 | 0 | 0 | 0 | 0 | 0 | 0 | 0 |
| 0 | NA | NA | NA | NA | NA | Gm22660 | | 0 | 0 | 0 | 0 | 0 | 0 | 0 | 0 | 0 | 0 | 0 |
| 0 | NA | NA | NA | NA | NA | A930031H19Rik | | 0 | 0 | 0 | 0 | 0 | 0 | 0 | 0 | 0 | 0 | 0 |
| 0 | NA | NA | NA | NA | NA | Gm25261 | | 0 | 0 | 0 | 0 | 0 | 0 | 0 | 0 | 0 | 0 | 0 |
| 0 | NA | NA | NA | NA | NA | Gm16080 | | 0 | 0 | 0 | 0 | 0 | 0 | 0 | 0 | 0 | 0 | 0 |
| 0 | NA | NA | NA | NA | NA | Gm12978 | | 0 | 0 | 0 | 0 | 0 | 0 | 0 | 0 | 0 | 0 | 0 |
| 0 | NA | NA | NA | NA | NA | Gm27445 | | 0 | 0 | 0 | 0 | 0 | 0 | 0 | 0 | 0 | 0 | 0 |
| 0 | NA | NA | NA | NA | NA | Gm13214 | | 0 | 0 | 0 | 0 | 0 | 0 | 0 | 0 | 0 | 0 | 0 |
| 0 | NA | NA | NA | NA | NA | Gm22033 | | 0 | 0 | 0 | 0 | 0 | 0 | 0 | 0 | 0 | 0 | 0 |
| 0 | NA | NA | NA | NA | NA | Gm28872 | | 0 | 0 | 0 | 0 | 0 | 0 | 0 | 0 | 0 | 0 | 0 |
| 0 | NA | NA | NA | NA | NA | Snora16a | | 0 | 0 | 0 | 0 | 0 | 0 | 0 | 0 | 0 | 0 | 0 |
| 0 | NA | NA | NA | NA | NA | Snora16a | | 0 | 0 | 0 | 0 | 0 | 0 | 0 | 0 | 0 | 0 | 0 |
| 0 | NA | NA | NA | NA | NA | Snora44 | | 0 | 0 | 0 | 0 | 0 | 0 | 0 | 0 | 0 | 0 | 0 |
| 0 | NA | NA | NA | NA | NA | Snora61 | | 0 | 0 | 0 | 0 | 0 | 0 | 0 | 0 | 0 | 0 | 0 |
| 0 | NA | NA | NA | NA | NA | Snord99 | | 0 | 0 | 0 | 0 | 0 | 0 | 0 | 0 | 0 | 0 | 0 |
| 0 | NA | NA | NA | NA | NA | Gm17300 | | 0 | 0 | 0 | 0 | 0 | 0 | 0 | 0 | 0 | 0 | 0 |
| 0 | NA | NA | NA | NA | NA | Gm25707 | | 0 | 0 | 0 | 0 | 0 | 0 | 0 | 0 | 0 | 0 | 0 |
| 0 | NA | NA | NA | NA | NA | Gm23055 | | 0 | 0 | 0 | 0 | 0 | 0 | 0 | 0 | 0 | 0 | 0 |
| 0 | NA | NA | NA | NA | NA | Gm24762 | | 0 | 0 | 0 | 0 | 0 | 0 | 0 | 0 | 0 | 0 | 0 |
| 0 | NA | NA | NA | NA | NA | Gm24913 | | 0 | 0 | 0 | 0 | 0 | 0 | 0 | 0 | 0 | 0 | 0 |
| 0 | NA | NA | NA | NA | NA | Gm24636 | | 0 | 0 | 0 | 0 | 0 | 0 | 0 | 0 | 0 | 0 | 0 |
| 0 | NA | NA | NA | NA | NA | Mir7017 | | 0 | 0 | 0 | 0 | 0 | 0 | 0 | 0 | 0 | 0 | 0 |
| 0 | NA | NA | NA | NA | NA | Gm13259 | | 0 | 0 | 0 | 0 | 0 | 0 | 0 | 0 | 0 | 0 | 0 |
| 0 | NA | NA | NA | NA | NA | Mir5122 | | 0 | 0 | 0 | 0 | 0 | 0 | 0 | 0 | 0 | 0 | 0 |
| 0 | NA | NA | NA | NA | NA | Gm23158 | | 0 | 0 | 0 | 0 | 0 | 0 | 0 | 0 | 0 | 0 | 0 |
| 0 | NA | NA | NA | NA | NA | Gm22662 | | 0 | 0 | 0 | 0 | 0 | 0 | 0 | 0 | 0 | 0 | 0 |
| 0 | NA | NA | NA | NA | NA | Mir7227 | | 0 | 0 | 0 | 0 | 0 | 0 | 0 | 0 | 0 | 0 | 0 |
| 0 | NA | NA | NA | NA | NA | Gm25270 | | 0 | 0 | 0 | 0 | 0 | 0 | 0 | 0 | 0 | 0 | 0 |
| 0 | NA | NA | NA | NA | NA | Gm12977 | | 0 | 0 | 0 | 0 | 0 | 0 | 0 | 0 | 0 | 0 | 0 |
| 0 | NA | NA | NA | NA | NA | Gm12985 | | 0 | 0 | 0 | 0 | 0 | 0 | 0 | 0 | 0 | 0 | 0 |
| 0 | NA | NA | NA | NA | NA | Gm22523 | | 0 | 0 | 0 | 0 | 0 | 0 | 0 | 0 | 0 | 0 | 0 |
| 0 | NA | NA | NA | NA | NA | Gm27619 | | 0 | 0 | 0 | 0 | 0 | 0 | 0 | 0 | 0 | 0 | 0 |
| 0 | NA | NA | NA | NA | NA | Zpld2 | zona pellucida like domain containing 2 | 0 | 0 | 0 | 0 | 0 | 0 | 0 | 0 | 0 | 0 | 0 |
| 0 | NA | NA | NA | NA | NA | E130218I03Rik | | 0 | 0 | 0 | 0 | 0 | 0 | 0 | 0 | 0 | 0 | 0 |
| 0 | NA | NA | NA | NA | NA | Gm30191 | predicted gene, 30191 | 0 | 0 | 0 | 0 | 0 | 0 | 0 | 0 | 0 | 0 | 0 |
| 0 | NA | NA | NA | NA | NA | Gm13195 | | 0 | 0 | 0 | 0 | 0 | 0 | 0 | 0 | 0 | 0 | 0 |
| 0 | NA | NA | NA | NA | NA | 1700021N21Rik | | 0 | 0 | 0 | 0 | 0 | 0 | 0 | 0 | 0 | 0 | 0 |
| 0 | NA | NA | NA | NA | NA | Gm13250 | | 0 | 0 | 0 | 0 | 0 | 0 | 0 | 0 | 0 | 0 | 0 |
| 0 | NA | NA | NA | NA | NA | Mir6403 | | 0 | 0 | 0 | 0 | 0 | 0 | 0 | 0 | 0 | 0 | 0 |
| 0 | NA | NA | NA | NA | NA | Gm16225 | | 0 | 0 | 0 | 0 | 0 | 0 | 0 | 0 | 0 | 0 | 0 |
| 0 | NA | NA | NA | NA | NA | Gm16224 | | 0 | 0 | 0 | 0 | 0 | 0 | 0 | 0 | 0 | 0 | 0 |
| 0 | NA | NA | NA | NA | NA | Gm12983 | | 0 | 0 | 0 | 0 | 0 | 0 | 0 | 0 | 0 | 0 | 0 |
| 0 | NA | NA | NA | NA | NA | Gm12984 | | 0 | 0 | 0 | 0 | 0 | 0 | 0 | 0 | 0 | 0 | 0 |

| | | | | | | | | | | | | | | | | | |
|---|---|---|---|---|---|---|---|---|---|---|---|---|---|---|---|---|---|
| 0 | NA | NA | NA | NA | NA | Gm12982 | | 0 | 0 | 0 | 0 | 0 | 0 | 0 | 0 | 0 | 0 | 0 |
| 0 | NA | NA | NA | NA | NA | Gm12991 | | 0 | 0 | 0 | 0 | 0 | 0 | 0 | 0 | 0 | 0 | 0 |
| 0 | NA | NA | NA | NA | NA | Gm25317 | | 0 | 0 | 0 | 0 | 0 | 0 | 0 | 0 | 0 | 0 | 0 |
| 0 | NA | NA | NA | NA | NA | Gm23106 | | 0 | 0 | 0 | 0 | 0 | 0 | 0 | 0 | 0 | 0 | 0 |
| 0 | NA | NA | NA | NA | NA | 1700029M20Rik | | 0 | 0 | 0 | 0 | 0 | 0 | 0 | 0 | 0 | 0 | 0 |
| 0 | NA | NA | NA | NA | NA | Gm13000 | | 0 | 0 | 0 | 0 | 0 | 0 | 0 | 0 | 0 | 0 | 0 |
| 0 | NA | NA | NA | NA | NA | Gm13006 | | 0 | 0 | 0 | 0 | 0 | 0 | 0 | 0 | 0 | 0 | 0 |
| 0 | NA | NA | NA | NA | NA | Gm26001 | | 0 | 0 | 0 | 0 | 0 | 0 | 0 | 0 | 0 | 0 | 0 |
| 0 | NA | NA | NA | NA | NA | Gm27022 | | 0 | 0 | 0 | 0 | 0 | 0 | 0 | 0 | 0 | 0 | 0 |
| 0 | NA | NA | NA | NA | NA | Gm22180 | | 0 | 0 | 0 | 0 | 0 | 0 | 0 | 0 | 0 | 0 | 0 |
| 0 | NA | NA | NA | NA | NA | Gm37583 | | 0 | 0 | 0 | 0 | 0 | 0 | 0 | 0 | 0 | 0 | 0 |
| 0 | NA | NA | NA | NA | NA | Gm23834 | | 0 | 0 | 0 | 0 | 0 | 0 | 0 | 0 | 0 | 0 | 0 |
| 0 | NA | NA | NA | NA | NA | Gm25399 | | 0 | 0 | 0 | 0 | 0 | 0 | 0 | 0 | 0 | 0 | 0 |
| 0 | NA | NA | NA | NA | NA | Gm23263 | | 0 | 0 | 0 | 0 | 0 | 0 | 0 | 0 | 0 | 0 | 0 |
| 0 | NA | NA | NA | NA | NA | Gm13005 | | 0 | 0 | 0 | 0 | 0 | 0 | 0 | 0 | 0 | 0 | 0 |
| 0 | NA | NA | NA | NA | NA | Gm13002 | | 0 | 0 | 0 | 0 | 0 | 0 | 0 | 0 | 0 | 0 | 0 |
| 0 | NA | NA | NA | NA | NA | Gm25772 | | 0 | 0 | 0 | 0 | 0 | 0 | 0 | 0 | 0 | 0 | 0 |
| 0 | NA | NA | NA | NA | NA | 1700013G24Ril | RIKEN cDNA 1700013G24 gene | 0 | 0 | 0 | 0 | 0 | 0 | 0 | 0 | 0 | 0 | 0 |
| 0 | NA | NA | NA | NA | NA | Mir7018 | | 0 | 0 | 0 | 0 | 0 | 0 | 0 | 0 | 0 | 0 | 0 |
| 0 | NA | NA | NA | NA | NA | 1700095J12Rik | | 0 | 0 | 0 | 0 | 0 | 0 | 0 | 0 | 0 | 0 | 0 |
| 0 | NA | NA | NA | NA | NA | Mir7019 | | 0 | 0 | 0 | 0 | 0 | 0 | 0 | 0 | 0 | 0 | 0 |
| 0 | NA | NA | NA | NA | NA | Gm27646 | | 0 | 0 | 0 | 0 | 0 | 0 | 0 | 0 | 0 | 0 | 0 |
| 0 | NA | NA | NA | NA | NA | AB041806 | | 0 | 0 | 0 | 0 | 0 | 0 | 0 | 0 | 0 | 0 | 0 |
| 0 | NA | NA | NA | NA | NA | Gm25280 | | 0 | 0 | 0 | 0 | 0 | 0 | 0 | 0 | 0 | 0 | 0 |
| 0 | NA | NA | NA | NA | NA | Pla2g2a | phospholipase A2, group IIA (platelets, sy | 0 | 0 | 0 | 0 | 0 | 0 | 0 | 0 | 0 | 0 | 0 |
| 0 | NA | NA | NA | NA | NA | Gm13030 | predicted gene 13030 | 0 | 0 | 0 | 0 | 0 | 0 | 0 | 0 | 0 | 0 | 0 |
| 0 | NA | NA | NA | NA | NA | Pla2g2e | phospholipase A2, group IIE | 0 | 0 | 0 | 0 | 0 | 0 | 0 | 0 | 0 | 0 | 0 |
| 0 | NA | NA | NA | NA | NA | Gm45533 | | 0 | 0 | 0 | 0 | 0 | 0 | 0 | 0 | 0 | 0 | 0 |
| 0 | NA | NA | NA | NA | NA | Hspe1-ps4 | | 0 | 0 | 0 | 0 | 0 | 0 | 0 | 0 | 0 | 0 | 0 |
| 0 | NA | NA | NA | NA | NA | | | 0 | 0 | 0 | 0 | 0 | 0 | 0 | 0 | 0 | 0 | 0 |
| 0 | NA | NA | NA | NA | NA | Gm13019 | | 0 | 0 | 0 | 0 | 0 | 0 | 0 | 0 | 0 | 0 | 0 |
| 0 | NA | NA | NA | NA | NA | Gm25571 | | 0 | 0 | 0 | 0 | 0 | 0 | 0 | 0 | 0 | 0 | 0 |
| 0 | NA | NA | NA | NA | NA | Gm21969 | | 0 | 0 | 0 | 0 | 0 | 0 | 0 | 0 | 0 | 0 | 0 |
| 0 | NA | NA | NA | NA | NA | Mir7020 | | 0 | 0 | 0 | 0 | 0 | 0 | 0 | 0 | 0 | 0 | 0 |
| 0 | NA | NA | NA | NA | NA | Pax7 | paired box 7 | 0 | 0 | 0 | 0 | 0 | 0 | 0 | 0 | 0 | 0 | 0 |
| 0 | NA | NA | NA | NA | NA | Gm1667 | | 0 | 0 | 0 | 0 | 0 | 0 | 0 | 0 | 0 | 0 | 0 |
| 0 | NA | NA | NA | NA | NA | Gm13028 | | 0 | 0 | 0 | 0 | 0 | 0 | 0 | 0 | 0 | 0 | 0 |
| 0 | NA | NA | NA | NA | NA | Mir2139 | | 0 | 0 | 0 | 0 | 0 | 0 | 0 | 0 | 0 | 0 | 0 |
| 0 | NA | NA | NA | NA | NA | Gm9867 | | 0 | 0 | 0 | 0 | 0 | 0 | 0 | 0 | 0 | 0 | 0 |
| 0 | NA | NA | NA | NA | NA | Gm13016 | | 0 | 0 | 0 | 0 | 0 | 0 | 0 | 0 | 0 | 0 | 0 |
| 0 | NA | NA | NA | NA | NA | Gm13017 | | 0 | 0 | 0 | 0 | 0 | 0 | 0 | 0 | 0 | 0 | 0 |
| 0 | NA | NA | NA | NA | NA | Gm13021 | | 0 | 0 | 0 | 0 | 0 | 0 | 0 | 0 | 0 | 0 | 0 |
| 0 | NA | NA | NA | NA | NA | Gm13026 | | 0 | 0 | 0 | 0 | 0 | 0 | 0 | 0 | 0 | 0 | 0 |
| 0 | NA | NA | NA | NA | NA | Gm50453 | | 0 | 0 | 0 | 0 | 0 | 0 | 0 | 0 | 0 | 0 | 0 |
| 0 | NA | NA | NA | NA | NA | Gm13025 | | 0 | 0 | 0 | 0 | 0 | 0 | 0 | 0 | 0 | 0 | 0 |
| 0 | NA | NA | NA | NA | NA | Gm23045 | | 0 | 0 | 0 | 0 | 0 | 0 | 0 | 0 | 0 | 0 | 0 |
| 0 | NA | NA | NA | NA | NA | Gm25951 | | 0 | 0 | 0 | 0 | 0 | 0 | 0 | 0 | 0 | 0 | 0 |
| 0 | NA | NA | NA | NA | NA | Padi6 | peptidyl arginine deiminase, type VI | 0 | 0 | 0 | 0 | 0 | 0 | 0 | 0 | 0 | 0 | 0 |
| 0 | NA | NA | NA | NA | NA | 4930515B02Rik | | 0 | 0 | 0 | 0 | 0 | 0 | 0 | 0 | 0 | 0 | 0 |
| 0 | NA | NA | NA | NA | NA | Spata21 | spermatogenesis associated 21 | 0 | 0 | 0 | 0 | 0 | 0 | 0 | 0 | 0 | 0 | 0 |
| 0 | NA | NA | NA | NA | NA | Gm13055 | | 0 | 0 | 0 | 0 | 0 | 0 | 0 | 0 | 0 | 0 | 0 |
| 0 | NA | NA | NA | NA | NA | Gm25690 | | 0 | 0 | 0 | 0 | 0 | 0 | 0 | 0 | 0 | 0 | 0 |
| 0 | NA | NA | NA | NA | NA | Gm13074 | | 0 | 0 | 0 | 0 | 0 | 0 | 0 | 0 | 0 | 0 | 0 |
| 0 | NA | NA | NA | NA | NA | Gm13076 | | 0 | 0 | 0 | 0 | 0 | 0 | 0 | 0 | 0 | 0 | 0 |
| 0 | NA | NA | NA | NA | NA | Clcnka | chloride channel, voltage-sensitive Ka | 0 | 0 | 0 | 0 | 0 | 0 | 0 | 0 | 0 | 0 | 0 |
| 0 | NA | NA | NA | NA | NA | Clcnkb | chloride channel, voltage-sensitive Kb | 0 | 0 | 0 | 0 | 0 | 0 | 0 | 0 | 0 | 0 | 0 |
| 0 | NA | NA | NA | NA | NA | Rsc1a1 | regulatory solute carrier protein, family 1, | 0 | 0 | 0 | 0 | 0 | 0 | 0 | 0 | 0 | 0 | 0 |
| 0 | NA | NA | NA | NA | NA | Agmat | agmatine ureohydrolase (agmatinase) | 0 | 0 | 0 | 0 | 0 | 0 | 0 | 0 | 0 | 0 | 0 |
| 0 | NA | NA | NA | NA | NA | Fhad1os2 | | 0 | 0 | 0 | 0 | 0 | 0 | 0 | 0 | 0 | 0 | 0 |
| 0 | NA | NA | NA | NA | NA | Gm13059 | | 0 | 0 | 0 | 0 | 0 | 0 | 0 | 0 | 0 | 0 | 0 |

| | | | | | | | | | | | | | | | | | | |
|---|---|---|---|---|---|---|---|---|---|---|---|---|---|---|---|---|---|---|
| 0 | NA | NA | NA | NA | NA | Fhad1os1 | | 0 | 0 | 0 | 0 | 0 | 0 | 0 | 0 | 0 | 0 | 0 |
| 0 | NA | NA | NA | NA | NA | Gm13062 | | 0 | 0 | 0 | 0 | 0 | 0 | 0 | 0 | 0 | 0 | 0 |
| 0 | NA | NA | NA | NA | NA | Gm37624 | | 0 | 0 | 0 | 0 | 0 | 0 | 0 | 0 | 0 | 0 | 0 |
| 0 | NA | NA | NA | NA | NA | Mir7021 | | 0 | 0 | 0 | 0 | 0 | 0 | 0 | 0 | 0 | 0 | 0 |
| 0 | NA | NA | NA | NA | NA | Gm13038 | | 0 | 0 | 0 | 0 | 0 | 0 | 0 | 0 | 0 | 0 | 0 |
| 0 | NA | NA | NA | NA | NA | Gm22039 | | 0 | 0 | 0 | 0 | 0 | 0 | 0 | 0 | 0 | 0 | 0 |
| 0 | NA | NA | NA | NA | NA | Gm26313 | | 0 | 0 | 0 | 0 | 0 | 0 | 0 | 0 | 0 | 0 | 0 |
| 0 | NA | NA | NA | NA | NA | Pramel1 | preferentially expressed antigen in melan | 0 | 0 | 0 | 0 | 0 | 0 | 0 | 0 | 0 | 0 | 0 |
| 0 | NA | NA | NA | NA | NA | Gm13045 | | 0 | 0 | 0 | 0 | 0 | 0 | 0 | 0 | 0 | 0 | 0 |
| 0 | NA | NA | NA | NA | NA | Oog4 | oogenesin 4 | 0 | 0 | 0 | 0 | 0 | 0 | 0 | 0 | 0 | 0 | 0 |
| 0 | NA | NA | NA | NA | NA | Gm13046 | | 0 | 0 | 0 | 0 | 0 | 0 | 0 | 0 | 0 | 0 | 0 |
| 0 | NA | NA | NA | NA | NA | Gm13041 | | 0 | 0 | 0 | 0 | 0 | 0 | 0 | 0 | 0 | 0 | 0 |
| 0 | NA | NA | NA | NA | NA | Gm13043 | predicted gene 13043 | 0 | 0 | 0 | 0 | 0 | 0 | 0 | 0 | 0 | 0 | 0 |
| 0 | NA | NA | NA | NA | NA | Gm13040 | predicted gene 13040 | 0 | 0 | 0 | 0 | 0 | 0 | 0 | 0 | 0 | 0 | 0 |
| 0 | NA | NA | NA | NA | NA | BC080695 | cDNA sequence BC080695 | 0 | 0 | 0 | 0 | 0 | 0 | 0 | 0 | 0 | 0 | 0 |
| 0 | NA | NA | NA | NA | NA | Gm13057 | predicted gene 13057 | 0 | 0 | 0 | 0 | 0 | 0 | 0 | 0 | 0 | 0 | 0 |
| 0 | NA | NA | NA | NA | NA | Gm13058 | | 0 | 0 | 0 | 0 | 0 | 0 | 0 | 0 | 0 | 0 | 0 |
| 0 | NA | NA | NA | NA | NA | Gm13083 | predicted gene 13083 | 0 | 0 | 0 | 0 | 0 | 0 | 0 | 0 | 0 | 0 | 0 |
| 0 | NA | NA | NA | NA | NA | Gm13088 | predicted gene 13088 | 0 | 0 | 0 | 0 | 0 | 0 | 0 | 0 | 0 | 0 | 0 |
| 0 | NA | NA | NA | NA | NA | Gm13085 | | 0 | 0 | 0 | 0 | 0 | 0 | 0 | 0 | 0 | 0 | 0 |
| 0 | NA | NA | NA | NA | NA | Gm13089 | predicted gene 13089 | 0 | 0 | 0 | 0 | 0 | 0 | 0 | 0 | 0 | 0 | 0 |
| 0 | NA | NA | NA | NA | NA | Gm13078 | predicted gene 13078 | 0 | 0 | 0 | 0 | 0 | 0 | 0 | 0 | 0 | 0 | 0 |
| 0 | NA | NA | NA | NA | NA | Gm13087 | | 0 | 0 | 0 | 0 | 0 | 0 | 0 | 0 | 0 | 0 | 0 |
| 0 | NA | NA | NA | NA | NA | Gm13081 | | 0 | 0 | 0 | 0 | 0 | 0 | 0 | 0 | 0 | 0 | 0 |
| 0 | NA | NA | NA | NA | NA | Gm13023 | predicted gene 13023 | 0 | 0 | 0 | 0 | 0 | 0 | 0 | 0 | 0 | 0 | 0 |
| 0 | NA | NA | NA | NA | NA | Gm13084 | predicted gene 13084 | 0 | 0 | 0 | 0 | 0 | 0 | 0 | 0 | 0 | 0 | 0 |
| 0 | NA | NA | NA | NA | NA | Gm13103 | predicted gene 13103 | 0 | 0 | 0 | 0 | 0 | 0 | 0 | 0 | 0 | 0 | 0 |
| 0 | NA | NA | NA | NA | NA | Pramef6 | PRAME family member 6 | 0 | 0 | 0 | 0 | 0 | 0 | 0 | 0 | 0 | 0 | 0 |
| 0 | NA | NA | NA | NA | NA | Gm13100 | | 0 | 0 | 0 | 0 | 0 | 0 | 0 | 0 | 0 | 0 | 0 |
| 0 | NA | NA | NA | NA | NA | Gm13106 | | 0 | 0 | 0 | 0 | 0 | 0 | 0 | 0 | 0 | 0 | 0 |
| 0 | NA | NA | NA | NA | NA | Pramef25 | PRAME family member 25 | 0 | 0 | 0 | 0 | 0 | 0 | 0 | 0 | 0 | 0 | 0 |
| 0 | NA | NA | NA | NA | NA | Gm13101 | | 0 | 0 | 0 | 0 | 0 | 0 | 0 | 0 | 0 | 0 | 0 |
| 0 | NA | NA | NA | NA | NA | Pramef17 | PRAME family member 17 | 0 | 0 | 0 | 0 | 0 | 0 | 0 | 0 | 0 | 0 | 0 |
| 0 | NA | NA | NA | NA | NA | Gm13108 | | 0 | 0 | 0 | 0 | 0 | 0 | 0 | 0 | 0 | 0 | 0 |
| 0 | NA | NA | NA | NA | NA | Gm13104 | | 0 | 0 | 0 | 0 | 0 | 0 | 0 | 0 | 0 | 0 | 0 |
| 0 | NA | NA | NA | NA | NA | Pramel4 | preferentially expressed antigen in melan | 0 | 0 | 0 | 0 | 0 | 0 | 0 | 0 | 0 | 0 | 0 |
| 0 | NA | NA | NA | NA | NA | Gm13102 | predicted gene 13102 | 0 | 0 | 0 | 0 | 0 | 0 | 0 | 0 | 0 | 0 | 0 |
| 0 | NA | NA | NA | NA | NA | Oog3 | oogenesin 3 | 0 | 0 | 0 | 0 | 0 | 0 | 0 | 0 | 0 | 0 | 0 |
| 0 | NA | NA | NA | NA | NA | Oog2 | oogenesin 2 | 0 | 0 | 0 | 0 | 0 | 0 | 0 | 0 | 0 | 0 | 0 |
| 0 | NA | NA | NA | NA | NA | C87977 | expressed sequence C87977 | 0 | 0 | 0 | 0 | 0 | 0 | 0 | 0 | 0 | 0 | 0 |
| 0 | NA | NA | NA | NA | NA | Pramel5 | preferentially expressed antigen in melan | 0 | 0 | 0 | 0 | 0 | 0 | 0 | 0 | 0 | 0 | 0 |
| 0 | NA | NA | NA | NA | NA | Gm13120 | | 0 | 0 | 0 | 0 | 0 | 0 | 0 | 0 | 0 | 0 | 0 |
| 0 | NA | NA | NA | NA | NA | Gm13130 | | 0 | 0 | 0 | 0 | 0 | 0 | 0 | 0 | 0 | 0 | 0 |
| 0 | NA | NA | NA | NA | NA | Gm13128 | predicted gene 13128 | 0 | 0 | 0 | 0 | 0 | 0 | 0 | 0 | 0 | 0 | 0 |
| 0 | NA | NA | NA | NA | NA | Gm13122 | | 0 | 0 | 0 | 0 | 0 | 0 | 0 | 0 | 0 | 0 | 0 |
| 0 | NA | NA | NA | NA | NA | Gm13119 | predicted gene 13119 | 0 | 0 | 0 | 0 | 0 | 0 | 0 | 0 | 0 | 0 | 0 |
| 0 | NA | NA | NA | NA | NA | Pramef20 | PRAME family member 20 | 0 | 0 | 0 | 0 | 0 | 0 | 0 | 0 | 0 | 0 | 0 |
| 0 | NA | NA | NA | NA | NA | Pramef12os | | 0 | 0 | 0 | 0 | 0 | 0 | 0 | 0 | 0 | 0 | 0 |
| 0 | NA | NA | NA | NA | NA | 1700012P22Ril | RIKEN cDNA 1700012P22 gene | 0 | 0 | 0 | 0 | 0 | 0 | 0 | 0 | 0 | 0 | 0 |
| 0 | NA | NA | NA | NA | NA | Gm9944 | | 0 | 0 | 0 | 0 | 0 | 0 | 0 | 0 | 0 | 0 | 0 |
| 0 | NA | NA | NA | NA | NA | Aadacl3 | arylacetamide deacetylase like 3 | 0 | 0 | 0 | 0 | 0 | 0 | 0 | 0 | 0 | 0 | 0 |
| 0 | NA | NA | NA | NA | NA | | | 0 | 0 | 0 | 0 | 0 | 0 | 0 | 0 | 0 | 0 | 0 |
| 0 | NA | NA | NA | NA | NA | Aadacl4fm1 | AADACL4 family member 1 | 0 | 0 | 0 | 0 | 0 | 0 | 0 | 0 | 0 | 0 | 0 |
| 0 | NA | NA | NA | NA | NA | Gm13126 | | 0 | 0 | 0 | 0 | 0 | 0 | 0 | 0 | 0 | 0 | 0 |
| 0 | NA | NA | NA | NA | NA | Aadacl4fm2 | AADACL4 family member 2 | 0 | 0 | 0 | 0 | 0 | 0 | 0 | 0 | 0 | 0 | 0 |
| 0 | NA | NA | NA | NA | NA | Aadacl4 | arylacetamide deacetylase like 4 | 0 | 0 | 0 | 0 | 0 | 0 | 0 | 0 | 0 | 0 | 0 |
| 0 | NA | NA | NA | NA | NA | Aadacl4fm4 | AADACL4 family member 4 | 0 | 0 | 0 | 0 | 0 | 0 | 0 | 0 | 0 | 0 | 0 |
| 0 | NA | NA | NA | NA | NA | Gm25938 | | 0 | 0 | 0 | 0 | 0 | 0 | 0 | 0 | 0 | 0 | 0 |
| 0 | NA | NA | NA | NA | NA | Aadacl4fm5 | AADACL4 family member 5 | 0 | 0 | 0 | 0 | 0 | 0 | 0 | 0 | 0 | 0 | 0 |
| 0 | NA | NA | NA | NA | NA | Gm38074 | | 0 | 0 | 0 | 0 | 0 | 0 | 0 | 0 | 0 | 0 | 0 |

| | | | | | | | | | | | | | | | | | | |
|---|---|---|---|---|---|---|---|---|---|---|---|---|---|---|---|---|---|---|
| 0 | NA | NA | NA | NA | NA | Gm13024 | | 0 | 0 | 0 | 0 | 0 | 0 | 0 | 0 | 0 | 0 | 0 |
| 0 | NA | NA | NA | NA | NA | Gm13224 | | 0 | 0 | 0 | 0 | 0 | 0 | 0 | 0 | 0 | 0 | 0 |
| 0 | NA | NA | NA | NA | NA | Gm13230 | | 0 | 0 | 0 | 0 | 0 | 0 | 0 | 0 | 0 | 0 | 0 |
| 0 | NA | NA | NA | NA | NA | Gm13229 | | 0 | 0 | 0 | 0 | 0 | 0 | 0 | 0 | 0 | 0 | 0 |
| 0 | NA | NA | NA | NA | NA | Gm21978 | | 0 | 0 | 0 | 0 | 0 | 0 | 0 | 0 | 0 | 0 | 0 |
| 0 | NA | NA | NA | NA | NA | Gm13228 | | 0 | 0 | 0 | 0 | 0 | 0 | 0 | 0 | 0 | 0 | 0 |
| 0 | NA | NA | NA | NA | NA | Gm13222 | | 0 | 0 | 0 | 0 | 0 | 0 | 0 | 0 | 0 | 0 | 0 |
| 0 | NA | NA | NA | NA | NA | Gm13234 | | 0 | 0 | 0 | 0 | 0 | 0 | 0 | 0 | 0 | 0 | 0 |
| 0 | NA | NA | NA | NA | NA | Gm13249 | | 0 | 0 | 0 | 0 | 0 | 0 | 0 | 0 | 0 | 0 | 0 |
| 0 | NA | NA | NA | NA | NA | Gm13239 | | 0 | 0 | 0 | 0 | 0 | 0 | 0 | 0 | 0 | 0 | 0 |
| 0 | NA | NA | NA | NA | NA | Gm13244 | | 0 | 0 | 0 | 0 | 0 | 0 | 0 | 0 | 0 | 0 | 0 |
| 0 | NA | NA | NA | NA | NA | Gm13236 | | 0 | 0 | 0 | 0 | 0 | 0 | 0 | 0 | 0 | 0 | 0 |
| 0 | NA | NA | NA | NA | NA | Vmn2r-ps15 | | 0 | 0 | 0 | 0 | 0 | 0 | 0 | 0 | 0 | 0 | 0 |
| 0 | NA | NA | NA | NA | NA | Gm13246 | | 0 | 0 | 0 | 0 | 0 | 0 | 0 | 0 | 0 | 0 | 0 |
| 0 | NA | NA | NA | NA | NA | Gm13035 | predicted gene 13035 | 0 | 0 | 0 | 0 | 0 | 0 | 0 | 0 | 0 | 0 | 0 |
| 0 | NA | NA | NA | NA | NA | Zfp987 | zinc finger protein 987 | 0 | 0 | 0 | 0 | 0 | 0 | 0 | 0 | 0 | 0 | 0 |
| 0 | NA | NA | NA | NA | NA | Gm13165 | | 0 | 0 | 0 | 0 | 0 | 0 | 0 | 0 | 0 | 0 | 0 |
| 0 | NA | NA | NA | NA | NA | Gm13170 | | 0 | 0 | 0 | 0 | 0 | 0 | 0 | 0 | 0 | 0 | 0 |
| 0 | NA | NA | NA | NA | NA | Gm13167 | | 0 | 0 | 0 | 0 | 0 | 0 | 0 | 0 | 0 | 0 | 0 |
| 0 | NA | NA | NA | NA | NA | Vmn2r-ps17 | | 0 | 0 | 0 | 0 | 0 | 0 | 0 | 0 | 0 | 0 | 0 |
| 0 | NA | NA | NA | NA | NA | 1700095A21Rik | | 0 | 0 | 0 | 0 | 0 | 0 | 0 | 0 | 0 | 0 | 0 |
| 0 | NA | NA | NA | NA | NA | Gm13237 | | 0 | 0 | 0 | 0 | 0 | 0 | 0 | 0 | 0 | 0 | 0 |
| 0 | NA | NA | NA | NA | NA | Gm13233 | | 0 | 0 | 0 | 0 | 0 | 0 | 0 | 0 | 0 | 0 | 0 |
| 0 | NA | NA | NA | NA | NA | Gm13240 | | 0 | 0 | 0 | 0 | 0 | 0 | 0 | 0 | 0 | 0 | 0 |
| 0 | NA | NA | NA | NA | NA | Vmn2r-ps16 | | 0 | 0 | 0 | 0 | 0 | 0 | 0 | 0 | 0 | 0 | 0 |
| 0 | NA | NA | NA | NA | NA | Gm20760 | | 0 | 0 | 0 | 0 | 0 | 0 | 0 | 0 | 0 | 0 | 0 |
| 0 | NA | NA | NA | NA | NA | Gm13149 | | 0 | 0 | 0 | 0 | 0 | 0 | 0 | 0 | 0 | 0 | 0 |
| 0 | NA | NA | NA | NA | NA | Gm13153 | | 0 | 0 | 0 | 0 | 0 | 0 | 0 | 0 | 0 | 0 | 0 |
| 0 | NA | NA | NA | NA | NA | Gm13163 | | 0 | 0 | 0 | 0 | 0 | 0 | 0 | 0 | 0 | 0 | 0 |
| 0 | NA | NA | NA | NA | NA | Gm13147 | | 0 | 0 | 0 | 0 | 0 | 0 | 0 | 0 | 0 | 0 | 0 |
| 0 | NA | NA | NA | NA | NA | Gm13158 | | 0 | 0 | 0 | 0 | 0 | 0 | 0 | 0 | 0 | 0 | 0 |
| 0 | NA | NA | NA | NA | NA | Gm13144 | | 0 | 0 | 0 | 0 | 0 | 0 | 0 | 0 | 0 | 0 | 0 |
| 0 | NA | NA | NA | NA | NA | Gm8178 | | 0 | 0 | 0 | 0 | 0 | 0 | 0 | 0 | 0 | 0 | 0 |
| 0 | NA | NA | NA | NA | NA | Gm15951 | | 0 | 0 | 0 | 0 | 0 | 0 | 0 | 0 | 0 | 0 | 0 |
| 0 | NA | NA | NA | NA | NA | Gm13141 | | 0 | 0 | 0 | 0 | 0 | 0 | 0 | 0 | 0 | 0 | 0 |
| 0 | NA | NA | NA | NA | NA | Gm13155 | | 0 | 0 | 0 | 0 | 0 | 0 | 0 | 0 | 0 | 0 | 0 |
| 0 | NA | NA | NA | NA | NA | Gm16211 | | 0 | 0 | 0 | 0 | 0 | 0 | 0 | 0 | 0 | 0 | 0 |
| 0 | NA | NA | NA | NA | NA | Zfp534 | zinc finger protein 534 | 0 | 0 | 0 | 0 | 0 | 0 | 0 | 0 | 0 | 0 | 0 |
| 0 | NA | NA | NA | NA | NA | Gm13137 | | 0 | 0 | 0 | 0 | 0 | 0 | 0 | 0 | 0 | 0 | 0 |
| 0 | NA | NA | NA | NA | NA | Gm13148 | | 0 | 0 | 0 | 0 | 0 | 0 | 0 | 0 | 0 | 0 | 0 |
| 0 | NA | NA | NA | NA | NA | Gm13156 | | 0 | 0 | 0 | 0 | 0 | 0 | 0 | 0 | 0 | 0 | 0 |
| 0 | NA | NA | NA | NA | NA | Nppb | natriuretic peptide type B | 0 | 0 | 0 | 0 | 0 | 0 | 0 | 0 | 0 | 0 | 0 |
| 0 | NA | NA | NA | NA | NA | Gm13207 | | 0 | 0 | 0 | 0 | 0 | 0 | 0 | 0 | 0 | 0 | 0 |
| 0 | NA | NA | NA | NA | NA | Mir7022 | | 0 | 0 | 0 | 0 | 0 | 0 | 0 | 0 | 0 | 0 | 0 |
| 0 | NA | NA | NA | NA | NA | Gm13206 | | 0 | 0 | 0 | 0 | 0 | 0 | 0 | 0 | 0 | 0 | 0 |
| 0 | NA | NA | NA | NA | NA | Gm13200 | | 0 | 0 | 0 | 0 | 0 | 0 | 0 | 0 | 0 | 0 | 0 |
| 0 | NA | NA | NA | NA | NA | Gm13204 | | 0 | 0 | 0 | 0 | 0 | 0 | 0 | 0 | 0 | 0 | 0 |
| 0 | NA | NA | NA | NA | NA | Gm13203 | | 0 | 0 | 0 | 0 | 0 | 0 | 0 | 0 | 0 | 0 | 0 |
| 0 | NA | NA | NA | NA | NA | Gm15969 | | 0 | 0 | 0 | 0 | 0 | 0 | 0 | 0 | 0 | 0 | 0 |
| 0 | NA | NA | NA | NA | NA | Ube4bos2 | | 0 | 0 | 0 | 0 | 0 | 0 | 0 | 0 | 0 | 0 | 0 |
| 0 | NA | NA | NA | NA | NA | Tmem274 | transmembrame protein 274 | 0 | 0 | 0 | 0 | 0 | 0 | 0 | 0 | 0 | 0 | 0 |
| 0 | NA | NA | NA | NA | NA | Gm24381 | | 0 | 0 | 0 | 0 | 0 | 0 | 0 | 0 | 0 | 0 | 0 |
| 0 | NA | NA | NA | NA | NA | Mir5616 | | 0 | 0 | 0 | 0 | 0 | 0 | 0 | 0 | 0 | 0 | 0 |
| 0 | NA | NA | NA | NA | NA | Mir7023 | | 0 | 0 | 0 | 0 | 0 | 0 | 0 | 0 | 0 | 0 | 0 |
| 0 | NA | NA | NA | NA | NA | Gm13068 | | 0 | 0 | 0 | 0 | 0 | 0 | 0 | 0 | 0 | 0 | 0 |
| 0 | NA | NA | NA | NA | NA | Gm13073 | | 0 | 0 | 0 | 0 | 0 | 0 | 0 | 0 | 0 | 0 | 0 |
| 0 | NA | NA | NA | NA | NA | Gm13065 | | 0 | 0 | 0 | 0 | 0 | 0 | 0 | 0 | 0 | 0 | 0 |
| 0 | NA | NA | NA | NA | NA | Gm13064 | | 0 | 0 | 0 | 0 | 0 | 0 | 0 | 0 | 0 | 0 | 0 |
| 0 | NA | NA | NA | NA | NA | n-R5s193 | | 0 | 0 | 0 | 0 | 0 | 0 | 0 | 0 | 0 | 0 | 0 |
| 0 | NA | NA | NA | NA | NA | Gm47301 | | 0 | 0 | 0 | 0 | 0 | 0 | 0 | 0 | 0 | 0 | 0 |

| | | | | | | | | | | | | | | | | | |
|---|---|---|---|---|---|---|---|---|---|---|---|---|---|---|---|---|---|
| 0 | NA | NA | NA | NA | NA | Gm13070 | | 0 | 0 | 0 | 0 | 0 | 0 | 0 | 0 | 0 | 0 | 0 |
| 0 | NA | NA | NA | NA | NA | Mir34a | | 0 | 0 | 0 | 0 | 0 | 0 | 0 | 0 | 0 | 0 | 0 |
| 0 | NA | NA | NA | NA | NA | | | 0 | 0 | 0 | 0 | 0 | 0 | 0 | 0 | 0 | 0 | 0 |
| 0 | NA | NA | NA | NA | NA | Gm23209 | | 0 | 0 | 0 | 0 | 0 | 0 | 0 | 0 | 0 | 0 | 0 |
| 0 | NA | NA | NA | NA | NA | Gm16079 | | 0 | 0 | 0 | 0 | 0 | 0 | 0 | 0 | 0 | 0 | 0 |
| 0 | NA | NA | NA | NA | NA | Gm13049 | | 0 | 0 | 0 | 0 | 0 | 0 | 0 | 0 | 0 | 0 | 0 |
| 0 | NA | NA | NA | NA | NA | Gm23405 | | 0 | 0 | 0 | 0 | 0 | 0 | 0 | 0 | 0 | 0 | 0 |
| 0 | NA | NA | NA | NA | NA | Uts2 | urotensin 2 | 0 | 0 | 0 | 0 | 0 | 0 | 0 | 0 | 0 | 0 | 0 |
| 0 | NA | NA | NA | NA | NA | Gm13090 | predicted gene 13090 | 0 | 0 | 0 | 0 | 0 | 0 | 0 | 0 | 0 | 0 | 0 |
| 0 | NA | NA | NA | NA | NA | 4930589P08Rik | | 0 | 0 | 0 | 0 | 0 | 0 | 0 | 0 | 0 | 0 | 0 |
| 0 | NA | NA | NA | NA | NA | Gm37141 | | 0 | 0 | 0 | 0 | 0 | 0 | 0 | 0 | 0 | 0 | 0 |
| 0 | NA | NA | NA | NA | NA | Gm27500 | | 0 | 0 | 0 | 0 | 0 | 0 | 0 | 0 | 0 | 0 | 0 |
| 0 | NA | NA | NA | NA | NA | Gm13096 | | 0 | 0 | 0 | 0 | 0 | 0 | 0 | 0 | 0 | 0 | 0 |
| 0 | NA | NA | NA | NA | NA | Hes3 | hes family bHLH transcription factor 3 | 0 | 0 | 0 | 0 | 0 | 0 | 0 | 0 | 0 | 0 | 0 |
| 0 | NA | NA | NA | NA | NA | Gm13645 | | 0 | 0 | 0 | 0 | 0 | 0 | 0 | 0 | 0 | 0 | 0 |
| 0 | NA | NA | NA | NA | NA | Gm16333 | | 0 | 0 | 0 | 0 | 0 | 0 | 0 | 0 | 0 | 0 | 0 |
| 0 | NA | NA | NA | NA | NA | Gm25779 | | 0 | 0 | 0 | 0 | 0 | 0 | 0 | 0 | 0 | 0 | 0 |
| 0 | NA | NA | NA | NA | NA | Gm13173 | | 0 | 0 | 0 | 0 | 0 | 0 | 0 | 0 | 0 | 0 | 0 |
| 0 | NA | NA | NA | NA | NA | Gm13174 | | 0 | 0 | 0 | 0 | 0 | 0 | 0 | 0 | 0 | 0 | 0 |
| 0 | NA | NA | NA | NA | NA | Gm25880 | | 0 | 0 | 0 | 0 | 0 | 0 | 0 | 0 | 0 | 0 | 0 |
| 0 | NA | NA | NA | NA | NA | Gm13117 | | 0 | 0 | 0 | 0 | 0 | 0 | 0 | 0 | 0 | 0 | 0 |
| 0 | NA | NA | NA | NA | NA | Gm13114 | | 0 | 0 | 0 | 0 | 0 | 0 | 0 | 0 | 0 | 0 | 0 |
| 0 | NA | NA | NA | NA | NA | Trp73os | | 0 | 0 | 0 | 0 | 0 | 0 | 0 | 0 | 0 | 0 | 0 |
| 0 | NA | NA | NA | NA | NA | Gm26036 | | 0 | 0 | 0 | 0 | 0 | 0 | 0 | 0 | 0 | 0 | 0 |
| 0 | NA | NA | NA | NA | NA | B230104I21Rik | | 0 | 0 | 0 | 0 | 0 | 0 | 0 | 0 | 0 | 0 | 0 |
| 0 | NA | NA | NA | NA | NA | Gm13133 | | 0 | 0 | 0 | 0 | 0 | 0 | 0 | 0 | 0 | 0 | 0 |
| 0 | NA | NA | NA | NA | NA | Gm13134 | | 0 | 0 | 0 | 0 | 0 | 0 | 0 | 0 | 0 | 0 | 0 |
| 0 | NA | NA | NA | NA | NA | Gm27202 | | 0 | 0 | 0 | 0 | 0 | 0 | 0 | 0 | 0 | 0 | 0 |
| 0 | NA | NA | NA | NA | NA | Actrt2 | actin-related protein T2 | 0 | 0 | 0 | 0 | 0 | 0 | 0 | 0 | 0 | 0 | 0 |
| 0 | NA | NA | NA | NA | NA | Gm13112 | | 0 | 0 | 0 | 0 | 0 | 0 | 0 | 0 | 0 | 0 | 0 |
| 0 | NA | NA | NA | NA | NA | Hes5 | hes family bHLH transcription factor 5 | 0 | 0 | 0 | 0 | 0 | 0 | 0 | 0 | 0 | 0 | 0 |
| 0 | NA | NA | NA | NA | NA | Gm10564 | | 0 | 0 | 0 | 0 | 0 | 0 | 0 | 0 | 0 | 0 | 0 |
| 0 | NA | NA | NA | NA | NA | Gm27200 | | 0 | 0 | 0 | 0 | 0 | 0 | 0 | 0 | 0 | 0 | 0 |
| 0 | NA | NA | NA | NA | NA | | | 0 | 0 | 0 | 0 | 0 | 0 | 0 | 0 | 0 | 0 | 0 |
| 0 | NA | NA | NA | NA | NA | Gm13171 | | 0 | 0 | 0 | 0 | 0 | 0 | 0 | 0 | 0 | 0 | 0 |
| 0 | NA | NA | NA | NA | NA | Gm22573 | | 0 | 0 | 0 | 0 | 0 | 0 | 0 | 0 | 0 | 0 | 0 |
| 0 | NA | NA | NA | NA | NA | Gm27743 | | 0 | 0 | 0 | 0 | 0 | 0 | 0 | 0 | 0 | 0 | 0 |
| 0 | NA | NA | NA | NA | NA | Gm22991 | | 0 | 0 | 0 | 0 | 0 | 0 | 0 | 0 | 0 | 0 | 0 |
| 0 | NA | NA | NA | NA | NA | Mir429 | | 0 | 0 | 0 | 0 | 0 | 0 | 0 | 0 | 0 | 0 | 0 |
| 0 | NA | NA | NA | NA | NA | Mir200a | | 0 | 0 | 0 | 0 | 0 | 0 | 0 | 0 | 0 | 0 | 0 |
| 0 | NA | NA | NA | NA | NA | Mir200b | | 0 | 0 | 0 | 0 | 0 | 0 | 0 | 0 | 0 | 0 | 0 |
| 0 | NA | NA | NA | NA | NA | Mir7658 | | 0 | 0 | 0 | 0 | 0 | 0 | 0 | 0 | 0 | 0 | 0 |
| 0 | NA | NA | NA | NA | NA | Gm47252 | | 0 | 0 | 0 | 0 | 0 | 0 | 0 | 0 | 0 | 0 | 0 |
| 0 | NA | NA | NA | NA | NA | Vmn2r-ps159 | vomeronasal 2, receptor 123 | 0 | 0 | 0 | 0 | 0 | 0 | 0 | 0 | 0 | 0 | 0 |
| 0 | NA | NA | NA | NA | NA | Vmn2r125 | | 0 | 0 | 0 | 0 | 0 | 0 | 0 | 0 | 0 | 0 | 0 |
| 0 | NA | NA | NA | NA | NA | V1rg10 | | 0 | 0 | 0 | 0 | 0 | 0 | 0 | 0 | 0 | 0 | 0 |
| 0 | NA | NA | NA | NA | NA | Gm3027 | | 0 | 0 | 0 | 0 | 0 | 0 | 0 | 0 | 0 | 0 | 0 |
| 0 | NA | NA | NA | NA | NA | Gm18187 | | 0 | 0 | 0 | 0 | 0 | 0 | 0 | 0 | 0 | 0 | 0 |
| 0 | NA | NA | NA | NA | NA | Gm8706 | | 0 | 0 | 0 | 0 | 0 | 0 | 0 | 0 | 0 | 0 | 0 |
| 0 | NA | NA | NA | NA | NA | Gm43529 | | 0 | 0 | 0 | 0 | 0 | 0 | 0 | 0 | 0 | 0 | 0 |
| 0 | NA | NA | NA | NA | NA | Gm6486 | | 0 | 0 | 0 | 0 | 0 | 0 | 0 | 0 | 0 | 0 | 0 |
| 0 | NA | NA | NA | NA | NA | Gm3053 | | 0 | 0 | 0 | 0 | 0 | 0 | 0 | 0 | 0 | 0 | 0 |
| 0 | NA | NA | NA | NA | NA | Gm43530 | | 0 | 0 | 0 | 0 | 0 | 0 | 0 | 0 | 0 | 0 | 0 |
| 0 | NA | NA | NA | NA | NA | Gm8715 | | 0 | 0 | 0 | 0 | 0 | 0 | 0 | 0 | 0 | 0 | 0 |
| 0 | NA | NA | NA | NA | NA | Gm18985 | | 0 | 0 | 0 | 0 | 0 | 0 | 0 | 0 | 0 | 0 | 0 |
| 0 | NA | NA | NA | NA | NA | Gm17590 | | 0 | 0 | 0 | 0 | 0 | 0 | 0 | 0 | 0 | 0 | 0 |
| 0 | NA | NA | NA | NA | NA | Gm43017 | | 0 | 0 | 0 | 0 | 0 | 0 | 0 | 0 | 0 | 0 | 0 |
| 0 | NA | NA | NA | NA | NA | n-R5s169 | | 0 | 0 | 0 | 0 | 0 | 0 | 0 | 0 | 0 | 0 | 0 |
| 0 | NA | NA | NA | NA | NA | Tmbim7 | transmembrane BAX inhibitor motif contai | 0 | 0 | 0 | 0 | 0 | 0 | 0 | 0 | 0 | 0 | 0 |
| 0 | NA | NA | NA | NA | NA | Gm26238 | | 0 | 0 | 0 | 0 | 0 | 0 | 0 | 0 | 0 | 0 | 0 |

| | | | | | | | | | | | | | | | | | | |
|---|---|---|---|---|---|---|---|---|---|---|---|---|---|---|---|---|---|---|
| 0 | NA | NA | NA | NA | NA | Gm22926 | | 0 | 0 | 0 | 0 | 0 | 0 | 0 | 0 | 0 | 0 | 0 |
| 0 | NA | NA | NA | NA | NA | Gm44483 | | 0 | 0 | 0 | 0 | 0 | 0 | 0 | 0 | 0 | 0 | 0 |
| 0 | NA | NA | NA | NA | NA | Gm25037 | | 0 | 0 | 0 | 0 | 0 | 0 | 0 | 0 | 0 | 0 | 0 |
| 0 | NA | NA | NA | NA | NA | Gm22897 | | 0 | 0 | 0 | 0 | 0 | 0 | 0 | 0 | 0 | 0 | 0 |
| 0 | NA | NA | NA | NA | NA | Gm22897 | | 0 | 0 | 0 | 0 | 0 | 0 | 0 | 0 | 0 | 0 | 0 |
| 0 | NA | NA | NA | NA | NA | Gm43623 | | 0 | 0 | 0 | 0 | 0 | 0 | 0 | 0 | 0 | 0 | 0 |
| 0 | NA | NA | NA | NA | NA | Gm10484 | | 0 | 0 | 0 | 0 | 0 | 0 | 0 | 0 | 0 | 0 | 0 |
| 0 | NA | NA | NA | NA | NA | Gm43728 | | 0 | 0 | 0 | 0 | 0 | 0 | 0 | 0 | 0 | 0 | 0 |
| 0 | NA | NA | NA | NA | NA | Gm43219 | | 0 | 0 | 0 | 0 | 0 | 0 | 0 | 0 | 0 | 0 | 0 |
| 0 | NA | NA | NA | NA | NA | Gm38697 | | 0 | 0 | 0 | 0 | 0 | 0 | 0 | 0 | 0 | 0 | 0 |
| 0 | NA | NA | NA | NA | NA | Gm23838 | | 0 | 0 | 0 | 0 | 0 | 0 | 0 | 0 | 0 | 0 | 0 |
| 0 | NA | NA | NA | NA | NA | Gm42718 | | 0 | 0 | 0 | 0 | 0 | 0 | 0 | 0 | 0 | 0 | 0 |
| 0 | NA | NA | NA | NA | NA | Gm42719 | | 0 | 0 | 0 | 0 | 0 | 0 | 0 | 0 | 0 | 0 | 0 |
| 0 | NA | NA | NA | NA | NA | Gm24686 | | 0 | 0 | 0 | 0 | 0 | 0 | 0 | 0 | 0 | 0 | 0 |
| 0 | NA | NA | NA | NA | NA | Gm40263 | | 0 | 0 | 0 | 0 | 0 | 0 | 0 | 0 | 0 | 0 | 0 |
| 0 | NA | NA | NA | NA | NA | Gm3203 | | 0 | 0 | 0 | 0 | 0 | 0 | 0 | 0 | 0 | 0 | 0 |
| 0 | NA | NA | NA | NA | NA | Gm24525 | | 0 | 0 | 0 | 0 | 0 | 0 | 0 | 0 | 0 | 0 | 0 |
| 0 | NA | NA | NA | NA | NA | Gm42598 | | 0 | 0 | 0 | 0 | 0 | 0 | 0 | 0 | 0 | 0 | 0 |
| 0 | NA | NA | NA | NA | NA | Gm43841 | | 0 | 0 | 0 | 0 | 0 | 0 | 0 | 0 | 0 | 0 | 0 |
| 0 | NA | NA | NA | NA | NA | Tex47 | testis expressed 47 | 0 | 0 | 0 | 0 | 0 | 0 | 0 | 0 | 0 | 0 | 0 |
| 0 | NA | NA | NA | NA | NA | Gm30835 | | 0 | 0 | 0 | 0 | 0 | 0 | 0 | 0 | 0 | 0 | 0 |
| 0 | NA | NA | NA | NA | NA | Gm22196 | | 0 | 0 | 0 | 0 | 0 | 0 | 0 | 0 | 0 | 0 | 0 |
| 0 | NA | NA | NA | NA | NA | Gm43478 | | 0 | 0 | 0 | 0 | 0 | 0 | 0 | 0 | 0 | 0 | 0 |
| 0 | NA | NA | NA | NA | NA | Gm21759 | | 0 | 0 | 0 | 0 | 0 | 0 | 0 | 0 | 0 | 0 | 0 |
| 0 | NA | NA | NA | NA | NA | Gm23993 | | 0 | 0 | 0 | 0 | 0 | 0 | 0 | 0 | 0 | 0 | 0 |
| 0 | NA | NA | NA | NA | NA | Gm42684 | | 0 | 0 | 0 | 0 | 0 | 0 | 0 | 0 | 0 | 0 | 0 |
| 0 | NA | NA | NA | NA | NA | Gm42686 | | 0 | 0 | 0 | 0 | 0 | 0 | 0 | 0 | 0 | 0 | 0 |
| 0 | NA | NA | NA | NA | NA | 7330423F06Rik | | 0 | 0 | 0 | 0 | 0 | 0 | 0 | 0 | 0 | 0 | 0 |
| 0 | NA | NA | NA | NA | NA | Gm37153 | | 0 | 0 | 0 | 0 | 0 | 0 | 0 | 0 | 0 | 0 | 0 |
| 0 | NA | NA | NA | NA | NA | Gm27941 | | 0 | 0 | 0 | 0 | 0 | 0 | 0 | 0 | 0 | 0 | 0 |
| 0 | NA | NA | NA | NA | NA | Gm15611 | | 0 | 0 | 0 | 0 | 0 | 0 | 0 | 0 | 0 | 0 | 0 |
| 0 | NA | NA | NA | NA | NA | Gm24666 | | 0 | 0 | 0 | 0 | 0 | 0 | 0 | 0 | 0 | 0 | 0 |
| 0 | NA | NA | NA | NA | NA | Gm15733 | | 0 | 0 | 0 | 0 | 0 | 0 | 0 | 0 | 0 | 0 | 0 |
| 0 | NA | NA | NA | NA | NA | Mir879 | | 0 | 0 | 0 | 0 | 0 | 0 | 0 | 0 | 0 | 0 | 0 |
| 0 | NA | NA | NA | NA | NA | Gm42455 | | 0 | 0 | 0 | 0 | 0 | 0 | 0 | 0 | 0 | 0 | 0 |
| 0 | NA | NA | NA | NA | NA | Gm23312 | | 0 | 0 | 0 | 0 | 0 | 0 | 0 | 0 | 0 | 0 | 0 |
| 0 | NA | NA | NA | NA | NA | Gm35212 | | 0 | 0 | 0 | 0 | 0 | 0 | 0 | 0 | 0 | 0 | 0 |
| 0 | NA | NA | NA | NA | NA | Gm42832 | | 0 | 0 | 0 | 0 | 0 | 0 | 0 | 0 | 0 | 0 | 0 |
| 0 | NA | NA | NA | NA | NA | Gm6586 | | 0 | 0 | 0 | 0 | 0 | 0 | 0 | 0 | 0 | 0 | 0 |
| 0 | NA | NA | NA | NA | NA | Gm5715 | | 0 | 0 | 0 | 0 | 0 | 0 | 0 | 0 | 0 | 0 | 0 |
| 0 | NA | NA | NA | NA | NA | Gm10482 | | 0 | 0 | 0 | 0 | 0 | 0 | 0 | 0 | 0 | 0 | 0 |
| 0 | NA | NA | NA | NA | NA | Gm5152 | | 0 | 0 | 0 | 0 | 0 | 0 | 0 | 0 | 0 | 0 | 0 |
| 0 | NA | NA | NA | NA | NA | Gm31831 | | 0 | 0 | 0 | 0 | 0 | 0 | 0 | 0 | 0 | 0 | 0 |
| 0 | NA | NA | NA | NA | NA | Gm17091 | | 0 | 0 | 0 | 0 | 0 | 0 | 0 | 0 | 0 | 0 | 0 |
| 0 | NA | NA | NA | NA | NA | Gm42828 | | 0 | 0 | 0 | 0 | 0 | 0 | 0 | 0 | 0 | 0 | 0 |
| 0 | NA | NA | NA | NA | NA | Gm42833 | | 0 | 0 | 0 | 0 | 0 | 0 | 0 | 0 | 0 | 0 | 0 |
| 0 | NA | NA | NA | NA | NA | Gm6455 | | 0 | 0 | 0 | 0 | 0 | 0 | 0 | 0 | 0 | 0 | 0 |
| 0 | NA | NA | NA | NA | NA | Gm3321 | | 0 | 0 | 0 | 0 | 0 | 0 | 0 | 0 | 0 | 0 | 0 |
| 0 | NA | NA | NA | NA | NA | Gm43358 | | 0 | 0 | 0 | 0 | 0 | 0 | 0 | 0 | 0 | 0 | 0 |
| 0 | NA | NA | NA | NA | NA | Gm8871 | | 0 | 0 | 0 | 0 | 0 | 0 | 0 | 0 | 0 | 0 | 0 |
| 0 | NA | NA | NA | NA | NA | Gm8857 | | 0 | 0 | 0 | 0 | 0 | 0 | 0 | 0 | 0 | 0 | 0 |
| 0 | NA | NA | NA | NA | NA | Gm8861 | | 0 | 0 | 0 | 0 | 0 | 0 | 0 | 0 | 0 | 0 | 0 |
| 0 | NA | NA | NA | NA | NA | Gm43492 | | 0 | 0 | 0 | 0 | 0 | 0 | 0 | 0 | 0 | 0 | 0 |
| 0 | NA | NA | NA | NA | NA | Gm8874 | | 0 | 0 | 0 | 0 | 0 | 0 | 0 | 0 | 0 | 0 | 0 |
| 0 | NA | NA | NA | NA | NA | 4930558F17Rik | | 0 | 0 | 0 | 0 | 0 | 0 | 0 | 0 | 0 | 0 | 0 |
| 0 | NA | NA | NA | NA | NA | | | 0 | 0 | 0 | 0 | 0 | 0 | 0 | 0 | 0 | 0 | 0 |
| 0 | NA | NA | NA | NA | NA | Gm32554 | | 0 | 0 | 0 | 0 | 0 | 0 | 0 | 0 | 0 | 0 | 0 |
| 0 | NA | NA | NA | NA | NA | Gm33093 | | 0 | 0 | 0 | 0 | 0 | 0 | 0 | 0 | 0 | 0 | 0 |
| 0 | NA | NA | NA | NA | NA | Gm8890 | | 0 | 0 | 0 | 0 | 0 | 0 | 0 | 0 | 0 | 0 | 0 |
| 0 | NA | NA | NA | NA | NA | Gm8891 | | 0 | 0 | 0 | 0 | 0 | 0 | 0 | 0 | 0 | 0 | 0 |

| | | | | | | | | | | | | | | | | | | |
|---|---|---|---|---|---|---|---|---|---|---|---|---|---|---|---|---|---|---|
| 0 | NA | NA | NA | NA | NA | Gm32877 | | 0 | 0 | 0 | 0 | 0 | 0 | 0 | 0 | 0 | 0 | 0 |
| 0 | NA | NA | NA | NA | NA | Speer1 | spermatogenesis associated glutamate (E | 0 | 0 | 0 | 0 | 0 | 0 | 0 | 0 | 0 | 0 | 0 |
| 0 | NA | NA | NA | NA | NA | Gm8896 | | 0 | 0 | 0 | 0 | 0 | 0 | 0 | 0 | 0 | 0 | 0 |
| 0 | NA | NA | NA | NA | NA | 1700108N06Rik | | 0 | 0 | 0 | 0 | 0 | 0 | 0 | 0 | 0 | 0 | 0 |
| 0 | NA | NA | NA | NA | NA | Gm8897 | | 0 | 0 | 0 | 0 | 0 | 0 | 0 | 0 | 0 | 0 | 0 |
| 0 | NA | NA | NA | NA | NA | Gm29745 | | 0 | 0 | 0 | 0 | 0 | 0 | 0 | 0 | 0 | 0 | 0 |
| 0 | NA | NA | NA | NA | NA | Gm43540 | | 0 | 0 | 0 | 0 | 0 | 0 | 0 | 0 | 0 | 0 | 0 |
| 0 | NA | NA | NA | NA | NA | 4930437M23Rik | | 0 | 0 | 0 | 0 | 0 | 0 | 0 | 0 | 0 | 0 | 0 |
| 0 | NA | NA | NA | NA | NA | Gm21040 | | 0 | 0 | 0 | 0 | 0 | 0 | 0 | 0 | 0 | 0 | 0 |
| 0 | NA | NA | NA | NA | NA | Gm43640 | | 0 | 0 | 0 | 0 | 0 | 0 | 0 | 0 | 0 | 0 | 0 |
| 0 | NA | NA | NA | NA | NA | Gm6460 | predicted gene 6460 | 0 | 0 | 0 | 0 | 0 | 0 | 0 | 0 | 0 | 0 | 0 |
| 0 | NA | NA | NA | NA | NA | Gm8920 | | 0 | 0 | 0 | 0 | 0 | 0 | 0 | 0 | 0 | 0 | 0 |
| 0 | NA | NA | NA | NA | NA | Gm33847 | | 0 | 0 | 0 | 0 | 0 | 0 | 0 | 0 | 0 | 0 | 0 |
| 0 | NA | NA | NA | NA | NA | Gm8922 | | 0 | 0 | 0 | 0 | 0 | 0 | 0 | 0 | 0 | 0 | 0 |
| 0 | NA | NA | NA | NA | NA | 1700003C15Rik | | 0 | 0 | 0 | 0 | 0 | 0 | 0 | 0 | 0 | 0 | 0 |
| 0 | NA | NA | NA | NA | NA | Gm8924 | | 0 | 0 | 0 | 0 | 0 | 0 | 0 | 0 | 0 | 0 | 0 |
| 0 | NA | NA | NA | NA | NA | Gm8926 | | 0 | 0 | 0 | 0 | 0 | 0 | 0 | 0 | 0 | 0 | 0 |
| 0 | NA | NA | NA | NA | NA | Gm43132 | | 0 | 0 | 0 | 0 | 0 | 0 | 0 | 0 | 0 | 0 | 0 |
| 0 | NA | NA | NA | NA | NA | Gm6465 | | 0 | 0 | 0 | 0 | 0 | 0 | 0 | 0 | 0 | 0 | 0 |
| 0 | NA | NA | NA | NA | NA | Gm29755 | | 0 | 0 | 0 | 0 | 0 | 0 | 0 | 0 | 0 | 0 | 0 |
| 0 | NA | NA | NA | NA | NA | Gm43131 | | 0 | 0 | 0 | 0 | 0 | 0 | 0 | 0 | 0 | 0 | 0 |
| 0 | NA | NA | NA | NA | NA | 4933402N22Ril | RIKEN cDNA 4933402N22 gene | 0 | 0 | 0 | 0 | 0 | 0 | 0 | 0 | 0 | 0 | 0 |
| 0 | NA | NA | NA | NA | NA | Gm5288 | | 0 | 0 | 0 | 0 | 0 | 0 | 0 | 0 | 0 | 0 | 0 |
| 0 | NA | NA | NA | NA | NA | Gm29066 | | 0 | 0 | 0 | 0 | 0 | 0 | 0 | 0 | 0 | 0 | 0 |
| 0 | NA | NA | NA | NA | NA | Gm38242 | | 0 | 0 | 0 | 0 | 0 | 0 | 0 | 0 | 0 | 0 | 0 |
| 0 | NA | NA | NA | NA | NA | Gm42910 | | 0 | 0 | 0 | 0 | 0 | 0 | 0 | 0 | 0 | 0 | 0 |
| 0 | NA | NA | NA | NA | NA | Gm8953 | | 0 | 0 | 0 | 0 | 0 | 0 | 0 | 0 | 0 | 0 | 0 |
| 0 | NA | NA | NA | NA | NA | Gm42909 | | 0 | 0 | 0 | 0 | 0 | 0 | 0 | 0 | 0 | 0 | 0 |
| 0 | NA | NA | NA | NA | NA | Gm42580 | | 0 | 0 | 0 | 0 | 0 | 0 | 0 | 0 | 0 | 0 | 0 |
| 0 | NA | NA | NA | NA | NA | Gm23243 | | 0 | 0 | 0 | 0 | 0 | 0 | 0 | 0 | 0 | 0 | 0 |
| 0 | NA | NA | NA | NA | NA | Gm10108 | | 0 | 0 | 0 | 0 | 0 | 0 | 0 | 0 | 0 | 0 | 0 |
| 0 | NA | NA | NA | NA | NA | Gm42680 | | 0 | 0 | 0 | 0 | 0 | 0 | 0 | 0 | 0 | 0 | 0 |
| 0 | NA | NA | NA | NA | NA | Gm43130 | | 0 | 0 | 0 | 0 | 0 | 0 | 0 | 0 | 0 | 0 | 0 |
| 0 | NA | NA | NA | NA | NA | Gm42676 | | 0 | 0 | 0 | 0 | 0 | 0 | 0 | 0 | 0 | 0 | 0 |
| 0 | NA | NA | NA | NA | NA | Gm42675 | | 0 | 0 | 0 | 0 | 0 | 0 | 0 | 0 | 0 | 0 | 0 |
| 0 | NA | NA | NA | NA | NA | Gm43718 | | 0 | 0 | 0 | 0 | 0 | 0 | 0 | 0 | 0 | 0 | 0 |
| 0 | NA | NA | NA | NA | NA | 4930568A13Rik | | 0 | 0 | 0 | 0 | 0 | 0 | 0 | 0 | 0 | 0 | 0 |
| 0 | NA | NA | NA | NA | NA | Gm6647 | | 0 | 0 | 0 | 0 | 0 | 0 | 0 | 0 | 0 | 0 | 0 |
| 0 | NA | NA | NA | NA | NA | Speer3 | spermatogenesis associated glutamate (E | 0 | 0 | 0 | 0 | 0 | 0 | 0 | 0 | 0 | 0 | 0 |
| 0 | NA | NA | NA | NA | NA | Gm6650 | | 0 | 0 | 0 | 0 | 0 | 0 | 0 | 0 | 0 | 0 | 0 |
| 0 | NA | NA | NA | NA | NA | Gm8968 | | 0 | 0 | 0 | 0 | 0 | 0 | 0 | 0 | 0 | 0 | 0 |
| 0 | NA | NA | NA | NA | NA | Gm35702 | | 0 | 0 | 0 | 0 | 0 | 0 | 0 | 0 | 0 | 0 | 0 |
| 0 | NA | NA | NA | NA | NA | Dnd1-ps | | 0 | 0 | 0 | 0 | 0 | 0 | 0 | 0 | 0 | 0 | 0 |
| 0 | NA | NA | NA | NA | NA | Gm24054 | | 0 | 0 | 0 | 0 | 0 | 0 | 0 | 0 | 0 | 0 | 0 |
| 0 | NA | NA | NA | NA | NA | Gm4128 | | 0 | 0 | 0 | 0 | 0 | 0 | 0 | 0 | 0 | 0 | 0 |
| 0 | NA | NA | NA | NA | NA | Gm43523 | | 0 | 0 | 0 | 0 | 0 | 0 | 0 | 0 | 0 | 0 | 0 |
| 0 | NA | NA | NA | NA | NA | Gm26918 | | 0 | 0 | 0 | 0 | 0 | 0 | 0 | 0 | 0 | 0 | 0 |
| 0 | NA | NA | NA | NA | NA | Gm43663 | | 0 | 0 | 0 | 0 | 0 | 0 | 0 | 0 | 0 | 0 | 0 |
| 0 | NA | NA | NA | NA | NA | Gm9758 | predicted gene 9758 | 0 | 0 | 0 | 0 | 0 | 0 | 0 | 0 | 0 | 0 | 0 |
| 0 | NA | NA | NA | NA | NA | Speer4e | spermatogenesis associated glutamate (E | 0 | 0 | 0 | 0 | 0 | 0 | 0 | 0 | 0 | 0 | 0 |
| 0 | NA | NA | NA | NA | NA | Gm10354 | predicted gene 10354 | 0 | 0 | 0 | 0 | 0 | 0 | 0 | 0 | 0 | 0 | 0 |
| 0 | NA | NA | NA | NA | NA | Gm17019 | predicted gene 17019 | 0 | 0 | 0 | 0 | 0 | 0 | 0 | 0 | 0 | 0 | 0 |
| 0 | NA | NA | NA | NA | NA | Gm43391 | | 0 | 0 | 0 | 0 | 0 | 0 | 0 | 0 | 0 | 0 | 0 |
| 0 | NA | NA | NA | NA | NA | Gm21149 | predicted gene, 21149 | 0 | 0 | 0 | 0 | 0 | 0 | 0 | 0 | 0 | 0 | 0 |
| 0 | NA | NA | NA | NA | NA | Gm21847 | | 0 | 0 | 0 | 0 | 0 | 0 | 0 | 0 | 0 | 0 | 0 |
| 0 | NA | NA | NA | NA | NA | Gm21190 | predicted gene, 21190 | 0 | 0 | 0 | 0 | 0 | 0 | 0 | 0 | 0 | 0 | 0 |
| 0 | NA | NA | NA | NA | NA | Gm30613 | | 0 | 0 | 0 | 0 | 0 | 0 | 0 | 0 | 0 | 0 | 0 |
| 0 | NA | NA | NA | NA | NA | Gm21083 | predicted gene, 21083 | 0 | 0 | 0 | 0 | 0 | 0 | 0 | 0 | 0 | 0 | 0 |
| 0 | NA | NA | NA | NA | NA | Speer4d | spermatogenesis associated glutamate (E | 0 | 0 | 0 | 0 | 0 | 0 | 0 | 0 | 0 | 0 | 0 |
| 0 | NA | NA | NA | NA | NA | 4930572O03Rik | | 0 | 0 | 0 | 0 | 0 | 0 | 0 | 0 | 0 | 0 | 0 |

| | | | | | | | | | | | | | | | | | | |
|---|---|---|---|---|---|---|---|---|---|---|---|---|---|---|---|---|---|---|
| 0 | NA | NA | NA | NA | NA | Speer4cos | | 0 | 0 | 0 | 0 | 0 | 0 | 0 | 0 | 0 | 0 | 0 |
| 0 | NA | NA | NA | NA | NA | Speer4c | spermatogenesis associated glutamate (E | 0 | 0 | 0 | 0 | 0 | 0 | 0 | 0 | 0 | 0 | 0 |
| 0 | NA | NA | NA | NA | NA | Gm42452 | | 0 | 0 | 0 | 0 | 0 | 0 | 0 | 0 | 0 | 0 | 0 |
| 0 | NA | NA | NA | NA | NA | Gm42454 | | 0 | 0 | 0 | 0 | 0 | 0 | 0 | 0 | 0 | 0 | 0 |
| 0 | NA | NA | NA | NA | NA | 4930519H02Rik | | 0 | 0 | 0 | 0 | 0 | 0 | 0 | 0 | 0 | 0 | 0 |
| 0 | NA | NA | NA | NA | NA | Gm42453 | | 0 | 0 | 0 | 0 | 0 | 0 | 0 | 0 | 0 | 0 | 0 |
| 0 | NA | NA | NA | NA | NA | Gm42451 | | 0 | 0 | 0 | 0 | 0 | 0 | 0 | 0 | 0 | 0 | 0 |
| 0 | NA | NA | NA | NA | NA | Gm43491 | | 0 | 0 | 0 | 0 | 0 | 0 | 0 | 0 | 0 | 0 | 0 |
| 0 | NA | NA | NA | NA | NA | Gm28710 | | 0 | 0 | 0 | 0 | 0 | 0 | 0 | 0 | 0 | 0 | 0 |
| 0 | NA | NA | NA | NA | NA | Gm28710 | | 0 | 0 | 0 | 0 | 0 | 0 | 0 | 0 | 0 | 0 | 0 |
| 0 | NA | NA | NA | NA | NA | Gm42855 | | 0 | 0 | 0 | 0 | 0 | 0 | 0 | 0 | 0 | 0 | 0 |
| 0 | NA | NA | NA | NA | NA | Gm6673 | | 0 | 0 | 0 | 0 | 0 | 0 | 0 | 0 | 0 | 0 | 0 |
| 0 | NA | NA | NA | NA | NA | Speer4f2 | spermatogenesis associated glutamate (E | 0 | 0 | 0 | 0 | 0 | 0 | 0 | 0 | 0 | 0 | 0 |
| 0 | NA | NA | NA | NA | NA | Gm21999 | | 0 | 0 | 0 | 0 | 0 | 0 | 0 | 0 | 0 | 0 | 0 |
| 0 | NA | NA | NA | NA | NA | Speer4f1 | spermatogenesis associated glutamate (E | 0 | 0 | 0 | 0 | 0 | 0 | 0 | 0 | 0 | 0 | 0 |
| 0 | NA | NA | NA | NA | NA | Gm3527 | | 0 | 0 | 0 | 0 | 0 | 0 | 0 | 0 | 0 | 0 | 0 |
| 0 | NA | NA | NA | NA | NA | Gm31752 | | 0 | 0 | 0 | 0 | 0 | 0 | 0 | 0 | 0 | 0 | 0 |
| 0 | NA | NA | NA | NA | NA | Gm17889 | | 0 | 0 | 0 | 0 | 0 | 0 | 0 | 0 | 0 | 0 | 0 |
| 0 | NA | NA | NA | NA | NA | Gm23822 | | 0 | 0 | 0 | 0 | 0 | 0 | 0 | 0 | 0 | 0 | 0 |
| 0 | NA | NA | NA | NA | NA | Gm32102 | | 0 | 0 | 0 | 0 | 0 | 0 | 0 | 0 | 0 | 0 | 0 |
| 0 | NA | NA | NA | NA | NA | Gm21009 | | 0 | 0 | 0 | 0 | 0 | 0 | 0 | 0 | 0 | 0 | 0 |
| 0 | NA | NA | NA | NA | NA | Gm25335 | | 0 | 0 | 0 | 0 | 0 | 0 | 0 | 0 | 0 | 0 | 0 |
| 0 | NA | NA | NA | NA | NA | Gm23570 | | 0 | 0 | 0 | 0 | 0 | 0 | 0 | 0 | 0 | 0 | 0 |
| 0 | NA | NA | NA | NA | NA | Gm25761 | | 0 | 0 | 0 | 0 | 0 | 0 | 0 | 0 | 0 | 0 | 0 |
| 0 | NA | NA | NA | NA | NA | Gm43680 | | 0 | 0 | 0 | 0 | 0 | 0 | 0 | 0 | 0 | 0 | 0 |
| 0 | NA | NA | NA | NA | NA | Gm22755 | | 0 | 0 | 0 | 0 | 0 | 0 | 0 | 0 | 0 | 0 | 0 |
| 0 | NA | NA | NA | NA | NA | Gm3544 | | 0 | 0 | 0 | 0 | 0 | 0 | 0 | 0 | 0 | 0 | 0 |
| 0 | NA | NA | NA | NA | NA | 1110060G06Rik | | 0 | 0 | 0 | 0 | 0 | 0 | 0 | 0 | 0 | 0 | 0 |
| 0 | NA | NA | NA | NA | NA | Rpl31-ps5 | | 0 | 0 | 0 | 0 | 0 | 0 | 0 | 0 | 0 | 0 | 0 |
| 0 | NA | NA | NA | NA | NA | Gm43216 | | 0 | 0 | 0 | 0 | 0 | 0 | 0 | 0 | 0 | 0 | 0 |
| 0 | NA | NA | NA | NA | NA | Gm22490 | | 0 | 0 | 0 | 0 | 0 | 0 | 0 | 0 | 0 | 0 | 0 |
| 0 | NA | NA | NA | NA | NA | Gm18830 | | 0 | 0 | 0 | 0 | 0 | 0 | 0 | 0 | 0 | 0 | 0 |
| 0 | NA | NA | NA | NA | NA | Gm43214 | | 0 | 0 | 0 | 0 | 0 | 0 | 0 | 0 | 0 | 0 | 0 |
| 0 | NA | NA | NA | NA | NA | Gm15719 | | 0 | 0 | 0 | 0 | 0 | 0 | 0 | 0 | 0 | 0 | 0 |
| 0 | NA | NA | NA | NA | NA | Gm15715 | | 0 | 0 | 0 | 0 | 0 | 0 | 0 | 0 | 0 | 0 | 0 |
| 0 | NA | NA | NA | NA | NA | Gm16113 | | 0 | 0 | 0 | 0 | 0 | 0 | 0 | 0 | 0 | 0 | 0 |
| 0 | NA | NA | NA | NA | NA | Gm16112 | | 0 | 0 | 0 | 0 | 0 | 0 | 0 | 0 | 0 | 0 | 0 |
| 0 | NA | NA | NA | NA | NA | Gm16110 | | 0 | 0 | 0 | 0 | 0 | 0 | 0 | 0 | 0 | 0 | 0 |
| 0 | NA | NA | NA | NA | NA | Gm25459 | | 0 | 0 | 0 | 0 | 0 | 0 | 0 | 0 | 0 | 0 | 0 |
| 0 | NA | NA | NA | NA | NA | Gm43657 | | 0 | 0 | 0 | 0 | 0 | 0 | 0 | 0 | 0 | 0 | 0 |
| 0 | NA | NA | NA | NA | NA | n-R5s170 | | 0 | 0 | 0 | 0 | 0 | 0 | 0 | 0 | 0 | 0 | 0 |
| 0 | NA | NA | NA | NA | NA | Gm24512 | | 0 | 0 | 0 | 0 | 0 | 0 | 0 | 0 | 0 | 0 | 0 |
| 0 | NA | NA | NA | NA | NA | Gm21846 | | 0 | 0 | 0 | 0 | 0 | 0 | 0 | 0 | 0 | 0 | 0 |
| 0 | NA | NA | NA | NA | NA | Gm43169 | | 0 | 0 | 0 | 0 | 0 | 0 | 0 | 0 | 0 | 0 | 0 |
| 0 | NA | NA | NA | NA | NA | Gm42948 | | 0 | 0 | 0 | 0 | 0 | 0 | 0 | 0 | 0 | 0 | 0 |
| 0 | NA | NA | NA | NA | NA | Gm28022 | | 0 | 0 | 0 | 0 | 0 | 0 | 0 | 0 | 0 | 0 | 0 |
| 0 | NA | NA | NA | NA | NA | Gm42951 | | 0 | 0 | 0 | 0 | 0 | 0 | 0 | 0 | 0 | 0 | 0 |
| 0 | NA | NA | NA | NA | NA | Gm43722 | | 0 | 0 | 0 | 0 | 0 | 0 | 0 | 0 | 0 | 0 | 0 |
| 0 | NA | NA | NA | NA | NA | 4930580E04Rik | | 0 | 0 | 0 | 0 | 0 | 0 | 0 | 0 | 0 | 0 | 0 |
| 0 | NA | NA | NA | NA | NA | Gm43058 | | 0 | 0 | 0 | 0 | 0 | 0 | 0 | 0 | 0 | 0 | 0 |
| 0 | NA | NA | NA | NA | NA | Gm42509 | | 0 | 0 | 0 | 0 | 0 | 0 | 0 | 0 | 0 | 0 | 0 |
| 0 | NA | NA | NA | NA | NA | Gm24009 | | 0 | 0 | 0 | 0 | 0 | 0 | 0 | 0 | 0 | 0 | 0 |
| 0 | NA | NA | NA | NA | NA | Gm50451 | | 0 | 0 | 0 | 0 | 0 | 0 | 0 | 0 | 0 | 0 | 0 |
| 0 | NA | NA | NA | NA | NA | | | 0 | 0 | 0 | 0 | 0 | 0 | 0 | 0 | 0 | 0 | 0 |
| 0 | NA | NA | NA | NA | NA | | | 0 | 0 | 0 | 0 | 0 | 0 | 0 | 0 | 0 | 0 | 0 |
| 0 | NA | NA | NA | NA | NA | Gm22054 | | 0 | 0 | 0 | 0 | 0 | 0 | 0 | 0 | 0 | 0 | 0 |
| 0 | NA | NA | NA | NA | NA | Snord93 | | 0 | 0 | 0 | 0 | 0 | 0 | 0 | 0 | 0 | 0 | 0 |
| 0 | NA | NA | NA | NA | NA | | | 0 | 0 | 0 | 0 | 0 | 0 | 0 | 0 | 0 | 0 | 0 |
| 0 | NA | NA | NA | NA | NA | Gm18038 | | 0 | 0 | 0 | 0 | 0 | 0 | 0 | 0 | 0 | 0 | 0 |
| 0 | NA | NA | NA | NA | NA | Gm43828 | | 0 | 0 | 0 | 0 | 0 | 0 | 0 | 0 | 0 | 0 | 0 |

| | | | | | | | | | | | | | | | | | | |
|---|---|---|---|---|---|---|---|---|---|---|---|---|---|---|---|---|---|---|
| 0 | NA | NA | NA | NA | NA | Gm43827 | | 0 | 0 | 0 | 0 | 0 | 0 | 0 | 0 | 0 | 0 | 0 |
| 0 | NA | NA | NA | NA | NA | Gm15589 | | 0 | 0 | 0 | 0 | 0 | 0 | 0 | 0 | 0 | 0 | 0 |
| 0 | NA | NA | NA | NA | NA | Asic3 | acid-sensing (proton-gated) ion channel 3 | 0 | 0 | 0 | 0 | 0 | 0 | 0 | 0 | 0 | 0 | 0 |
| 0 | NA | NA | NA | NA | NA | Gm22333 | | 0 | 0 | 0 | 0 | 0 | 0 | 0 | 0 | 0 | 0 | 0 |
| 0 | NA | NA | NA | NA | NA | Mir671 | | 0 | 0 | 0 | 0 | 0 | 0 | 0 | 0 | 0 | 0 | 0 |
| 0 | NA | NA | NA | NA | NA | 1700022A21Rik | | 0 | 0 | 0 | 0 | 0 | 0 | 0 | 0 | 0 | 0 | 0 |
| 0 | NA | NA | NA | NA | NA | Prkag2os1 | | 0 | 0 | 0 | 0 | 0 | 0 | 0 | 0 | 0 | 0 | 0 |
| 0 | NA | NA | NA | NA | NA | Gm19088 | | 0 | 0 | 0 | 0 | 0 | 0 | 0 | 0 | 0 | 0 | 0 |
| 0 | NA | NA | NA | NA | NA | Gm26051 | | 0 | 0 | 0 | 0 | 0 | 0 | 0 | 0 | 0 | 0 | 0 |
| 0 | NA | NA | NA | NA | NA | Galntl5 | UDP-N-acetyl-alpha-D-galactosamine:po | 0 | 0 | 0 | 0 | 0 | 0 | 0 | 0 | 0 | 0 | 0 |
| 0 | NA | NA | NA | NA | NA | Gm26356 | | 0 | 0 | 0 | 0 | 0 | 0 | 0 | 0 | 0 | 0 | 0 |
| 0 | NA | NA | NA | NA | NA | Gm20432 | | 0 | 0 | 0 | 0 | 0 | 0 | 0 | 0 | 0 | 0 | 0 |
| 0 | NA | NA | NA | NA | NA | Gm18343 | | 0 | 0 | 0 | 0 | 0 | 0 | 0 | 0 | 0 | 0 | 0 |
| 0 | NA | NA | NA | NA | NA | Cct8l1 | chaperonin containing TCP1, subunit 8 (t | 0 | 0 | 0 | 0 | 0 | 0 | 0 | 0 | 0 | 0 | 0 |
| 0 | NA | NA | NA | NA | NA | Gm43141 | | 0 | 0 | 0 | 0 | 0 | 0 | 0 | 0 | 0 | 0 | 0 |
| 0 | NA | NA | NA | NA | NA | Gm10062 | | 0 | 0 | 0 | 0 | 0 | 0 | 0 | 0 | 0 | 0 | 0 |
| 0 | NA | NA | NA | NA | NA | Gm43972 | | 0 | 0 | 0 | 0 | 0 | 0 | 0 | 0 | 0 | 0 | 0 |
| 0 | NA | NA | NA | NA | NA | Gm21655 | | 0 | 0 | 0 | 0 | 0 | 0 | 0 | 0 | 0 | 0 | 0 |
| 0 | NA | NA | NA | NA | NA | Gm21663 | predicted gene, 21663 | 0 | 0 | 0 | 0 | 0 | 0 | 0 | 0 | 0 | 0 | 0 |
| 0 | NA | NA | NA | NA | NA | Gm21671 | predicted gene, 21671 | 0 | 0 | 0 | 0 | 0 | 0 | 0 | 0 | 0 | 0 | 0 |
| 0 | NA | NA | NA | NA | NA | Gm21680 | predicted gene, 21680 | 0 | 0 | 0 | 0 | 0 | 0 | 0 | 0 | 0 | 0 | 0 |
| 0 | NA | NA | NA | NA | NA | Gm21698 | predicted gene, 21698 | 0 | 0 | 0 | 0 | 0 | 0 | 0 | 0 | 0 | 0 | 0 |
| 0 | NA | NA | NA | NA | NA | Gm1979 | predicted gene 1979 | 0 | 0 | 0 | 0 | 0 | 0 | 0 | 0 | 0 | 0 | 0 |
| 0 | NA | NA | NA | NA | NA | Gm5067 | | 0 | 0 | 0 | 0 | 0 | 0 | 0 | 0 | 0 | 0 | 0 |
| 0 | NA | NA | NA | NA | NA | Gm5862 | predicted gene 5862 | 0 | 0 | 0 | 0 | 0 | 0 | 0 | 0 | 0 | 0 | 0 |
| 0 | NA | NA | NA | NA | NA | Speer4a | spermatogenesis associated glutamate (E | 0 | 0 | 0 | 0 | 0 | 0 | 0 | 0 | 0 | 0 | 0 |
| 0 | NA | NA | NA | NA | NA | Gm7347 | predicted gene 7347 | 0 | 0 | 0 | 0 | 0 | 0 | 0 | 0 | 0 | 0 | 0 |
| 0 | NA | NA | NA | NA | NA | Gm6089 | | 0 | 0 | 0 | 0 | 0 | 0 | 0 | 0 | 0 | 0 | 0 |
| 0 | NA | NA | NA | NA | NA | 5031410I06Rik | RIKEN cDNA 5031410I06 gene | 0 | 0 | 0 | 0 | 0 | 0 | 0 | 0 | 0 | 0 | 0 |
| 0 | NA | NA | NA | NA | NA | Gm10220 | predicted gene 10220 | 0 | 0 | 0 | 0 | 0 | 0 | 0 | 0 | 0 | 0 | 0 |
| 0 | NA | NA | NA | NA | NA | Gm18957 | | 0 | 0 | 0 | 0 | 0 | 0 | 0 | 0 | 0 | 0 | 0 |
| 0 | NA | NA | NA | NA | NA | 4930584F24Rik | | 0 | 0 | 0 | 0 | 0 | 0 | 0 | 0 | 0 | 0 | 0 |
| 0 | NA | NA | NA | NA | NA | Gm42844 | | 0 | 0 | 0 | 0 | 0 | 0 | 0 | 0 | 0 | 0 | 0 |
| 0 | NA | NA | NA | NA | NA | Gm29784 | | 0 | 0 | 0 | 0 | 0 | 0 | 0 | 0 | 0 | 0 | 0 |
| 0 | NA | NA | NA | NA | NA | Gm16056 | | 0 | 0 | 0 | 0 | 0 | 0 | 0 | 0 | 0 | 0 | 0 |
| 0 | NA | NA | NA | NA | NA | Gm16057 | | 0 | 0 | 0 | 0 | 0 | 0 | 0 | 0 | 0 | 0 | 0 |
| 0 | NA | NA | NA | NA | NA | Gm43776 | | 0 | 0 | 0 | 0 | 0 | 0 | 0 | 0 | 0 | 0 | 0 |
| 0 | NA | NA | NA | NA | NA | Gm43502 | | 0 | 0 | 0 | 0 | 0 | 0 | 0 | 0 | 0 | 0 | 0 |
| 0 | NA | NA | NA | NA | NA | Speer4b | spermatogenesis associated glutamate (E | 0 | 0 | 0 | 0 | 0 | 0 | 0 | 0 | 0 | 0 | 0 |
| 0 | NA | NA | NA | NA | NA | Gm16058 | | 0 | 0 | 0 | 0 | 0 | 0 | 0 | 0 | 0 | 0 | 0 |
| 0 | NA | NA | NA | NA | NA | Gm43433 | | 0 | 0 | 0 | 0 | 0 | 0 | 0 | 0 | 0 | 0 | 0 |
| 0 | NA | NA | NA | NA | NA | Gm20239 | | 0 | 0 | 0 | 0 | 0 | 0 | 0 | 0 | 0 | 0 | 0 |
| 0 | NA | NA | NA | NA | NA | | | 0 | 0 | 0 | 0 | 0 | 0 | 0 | 0 | 0 | 0 | 0 |
| 0 | NA | NA | NA | NA | NA | Gm43611 | | 0 | 0 | 0 | 0 | 0 | 0 | 0 | 0 | 0 | 0 | 0 |
| 0 | NA | NA | NA | NA | NA | Gm43610 | | 0 | 0 | 0 | 0 | 0 | 0 | 0 | 0 | 0 | 0 | 0 |
| 0 | NA | NA | NA | NA | NA | Cnpy1 | canopy FGF signaling regulator 1 | 0 | 0 | 0 | 0 | 0 | 0 | 0 | 0 | 0 | 0 | 0 |
| 0 | NA | NA | NA | NA | NA | Gm43161 | | 0 | 0 | 0 | 0 | 0 | 0 | 0 | 0 | 0 | 0 | 0 |
| 0 | NA | NA | NA | NA | NA | Gm23801 | | 0 | 0 | 0 | 0 | 0 | 0 | 0 | 0 | 0 | 0 | 0 |
| 0 | NA | NA | NA | NA | NA | Gm24538 | | 0 | 0 | 0 | 0 | 0 | 0 | 0 | 0 | 0 | 0 | 0 |
| 0 | NA | NA | NA | NA | NA | Gm42570 | | 0 | 0 | 0 | 0 | 0 | 0 | 0 | 0 | 0 | 0 | 0 |
| 0 | NA | NA | NA | NA | NA | Gm26346 | | 0 | 0 | 0 | 0 | 0 | 0 | 0 | 0 | 0 | 0 | 0 |
| 0 | NA | NA | NA | NA | NA | Gm42450 | | 0 | 0 | 0 | 0 | 0 | 0 | 0 | 0 | 0 | 0 | 0 |
| 0 | NA | NA | NA | NA | NA | Gm43386 | | 0 | 0 | 0 | 0 | 0 | 0 | 0 | 0 | 0 | 0 | 0 |
| 0 | NA | NA | NA | NA | NA | A230098N10Rik | | 0 | 0 | 0 | 0 | 0 | 0 | 0 | 0 | 0 | 0 | 0 |
| 0 | NA | NA | NA | NA | NA | 3110082J24Rik | RIKEN cDNA 3110082J24 gene | 0 | 0 | 0 | 0 | 0 | 0 | 0 | 0 | 0 | 0 | 0 |
| 0 | NA | NA | NA | NA | NA | Gm43755 | | 0 | 0 | 0 | 0 | 0 | 0 | 0 | 0 | 0 | 0 | 0 |
| 0 | NA | NA | NA | NA | NA | Gm25666 | | 0 | 0 | 0 | 0 | 0 | 0 | 0 | 0 | 0 | 0 | 0 |
| 0 | NA | NA | NA | NA | NA | Mir5625 | | 0 | 0 | 0 | 0 | 0 | 0 | 0 | 0 | 0 | 0 | 0 |
| 0 | NA | NA | NA | NA | NA | Gm42764 | | 0 | 0 | 0 | 0 | 0 | 0 | 0 | 0 | 0 | 0 | 0 |
| 0 | NA | NA | NA | NA | NA | Gm15461 | | 0 | 0 | 0 | 0 | 0 | 0 | 0 | 0 | 0 | 0 | 0 |

| | | | | | | | | | | | | | | | | | | |
|---|---|---|---|---|---|---|---|---|---|---|---|---|---|---|---|---|---|---|
| 0 | NA | NA | NA | NA | NA | Gm9924 | | 0 | 0 | 0 | 0 | 0 | 0 | 0 | 0 | 0 | 0 | 0 |
| 0 | NA | NA | NA | NA | NA | Dnajc5g | DnaJ heat shock protein family (Hsp40) m | 0 | 0 | 0 | 0 | 0 | 0 | 0 | 0 | 0 | 0 | 0 |
| 0 | NA | NA | NA | NA | NA | Trim54 | tripartite motif-containing 54 | 0 | 0 | 0 | 0 | 0 | 0 | 0 | 0 | 0 | 0 | 0 |
| 0 | NA | NA | NA | NA | NA | Ucn | urocortin | 0 | 0 | 0 | 0 | 0 | 0 | 0 | 0 | 0 | 0 | 0 |
| 0 | NA | NA | NA | NA | NA | Gm43659 | | 0 | 0 | 0 | 0 | 0 | 0 | 0 | 0 | 0 | 0 | 0 |
| 0 | NA | NA | NA | NA | NA | 4930478M09Rik | | 0 | 0 | 0 | 0 | 0 | 0 | 0 | 0 | 0 | 0 | 0 |
| 0 | NA | NA | NA | NA | NA | 4930566F21Rik | | 0 | 0 | 0 | 0 | 0 | 0 | 0 | 0 | 0 | 0 | 0 |
| 0 | NA | NA | NA | NA | NA | Gm43809 | | 0 | 0 | 0 | 0 | 0 | 0 | 0 | 0 | 0 | 0 | 0 |
| 0 | NA | NA | NA | NA | NA | Gm24436 | | 0 | 0 | 0 | 0 | 0 | 0 | 0 | 0 | 0 | 0 | 0 |
| 0 | NA | NA | NA | NA | NA | Gm43811 | | 0 | 0 | 0 | 0 | 0 | 0 | 0 | 0 | 0 | 0 | 0 |
| 0 | NA | NA | NA | NA | NA | Gm26335 | | 0 | 0 | 0 | 0 | 0 | 0 | 0 | 0 | 0 | 0 | 0 |
| 0 | NA | NA | NA | NA | NA | Gm9555 | | 0 | 0 | 0 | 0 | 0 | 0 | 0 | 0 | 0 | 0 | 0 |
| 0 | NA | NA | NA | NA | NA | Gm15615 | | 0 | 0 | 0 | 0 | 0 | 0 | 0 | 0 | 0 | 0 | 0 |
| 0 | NA | NA | NA | NA | NA | Gm36840 | | 0 | 0 | 0 | 0 | 0 | 0 | 0 | 0 | 0 | 0 | 0 |
| 0 | NA | NA | NA | NA | NA | Gm15613 | | 0 | 0 | 0 | 0 | 0 | 0 | 0 | 0 | 0 | 0 | 0 |
| 0 | NA | NA | NA | NA | NA | Gm15614 | | 0 | 0 | 0 | 0 | 0 | 0 | 0 | 0 | 0 | 0 | 0 |
| 0 | NA | NA | NA | NA | NA | Gm42964 | | 0 | 0 | 0 | 0 | 0 | 0 | 0 | 0 | 0 | 0 | 0 |
| 0 | NA | NA | NA | NA | NA | Gm43812 | | 0 | 0 | 0 | 0 | 0 | 0 | 0 | 0 | 0 | 0 | 0 |
| 0 | NA | NA | NA | NA | NA | Gm20671 | | 0 | 0 | 0 | 0 | 0 | 0 | 0 | 0 | 0 | 0 | 0 |
| 0 | NA | NA | NA | NA | NA | Gm43695 | | 0 | 0 | 0 | 0 | 0 | 0 | 0 | 0 | 0 | 0 | 0 |
| 0 | NA | NA | NA | NA | NA | Gm24003 | | 0 | 0 | 0 | 0 | 0 | 0 | 0 | 0 | 0 | 0 | 0 |
| 0 | NA | NA | NA | NA | NA | Gm43849 | | 0 | 0 | 0 | 0 | 0 | 0 | 0 | 0 | 0 | 0 | 0 |
| 0 | NA | NA | NA | NA | NA | Gm15936 | | 0 | 0 | 0 | 0 | 0 | 0 | 0 | 0 | 0 | 0 | 0 |
| 0 | NA | NA | NA | NA | NA | 5033423K11Rik | | 0 | 0 | 0 | 0 | 0 | 0 | 0 | 0 | 0 | 0 | 0 |
| 0 | NA | NA | NA | NA | NA | Gm20465 | | 0 | 0 | 0 | 0 | 0 | 0 | 0 | 0 | 0 | 0 | 0 |
| 0 | NA | NA | NA | NA | NA | Gm43851 | | 0 | 0 | 0 | 0 | 0 | 0 | 0 | 0 | 0 | 0 | 0 |
| 0 | NA | NA | NA | NA | NA | Gm42965 | | 0 | 0 | 0 | 0 | 0 | 0 | 0 | 0 | 0 | 0 | 0 |
| 0 | NA | NA | NA | NA | NA | Gm30708 | | 0 | 0 | 0 | 0 | 0 | 0 | 0 | 0 | 0 | 0 | 0 |
| 0 | NA | NA | NA | NA | NA | Mir7024 | | 0 | 0 | 0 | 0 | 0 | 0 | 0 | 0 | 0 | 0 | 0 |
| 0 | NA | NA | NA | NA | NA | Gm43459 | | 0 | 0 | 0 | 0 | 0 | 0 | 0 | 0 | 0 | 0 | 0 |
| 0 | NA | NA | NA | NA | NA | 4930557J02Rik | | 0 | 0 | 0 | 0 | 0 | 0 | 0 | 0 | 0 | 0 | 0 |
| 0 | NA | NA | NA | NA | NA | Gm25918 | | 0 | 0 | 0 | 0 | 0 | 0 | 0 | 0 | 0 | 0 | 0 |
| 0 | NA | NA | NA | NA | NA | Gm42848 | | 0 | 0 | 0 | 0 | 0 | 0 | 0 | 0 | 0 | 0 | 0 |
| 0 | NA | NA | NA | NA | NA | Cfap99 | cilia and flagella associated protein 99 | 0 | 0 | 0 | 0 | 0 | 0 | 0 | 0 | 0 | 0 | 0 |
| 0 | NA | NA | NA | NA | NA | Gm43458 | | 0 | 0 | 0 | 0 | 0 | 0 | 0 | 0 | 0 | 0 | 0 |
| 0 | NA | NA | NA | NA | NA | Gm22847 | | 0 | 0 | 0 | 0 | 0 | 0 | 0 | 0 | 0 | 0 | 0 |
| 0 | NA | NA | NA | NA | NA | Mir7036b | | 0 | 0 | 0 | 0 | 0 | 0 | 0 | 0 | 0 | 0 | 0 |
| 0 | NA | NA | NA | NA | NA | Gm42560 | | 0 | 0 | 0 | 0 | 0 | 0 | 0 | 0 | 0 | 0 | 0 |
| 0 | NA | NA | NA | NA | NA | Mir467g | | 0 | 0 | 0 | 0 | 0 | 0 | 0 | 0 | 0 | 0 | 0 |
| 0 | NA | NA | NA | NA | NA | Mir3097 | | 0 | 0 | 0 | 0 | 0 | 0 | 0 | 0 | 0 | 0 | 0 |
| 0 | NA | NA | NA | NA | NA | Gm43791 | | 0 | 0 | 0 | 0 | 0 | 0 | 0 | 0 | 0 | 0 | 0 |
| 0 | NA | NA | NA | NA | NA | 4930478P22Rik | | 0 | 0 | 0 | 0 | 0 | 0 | 0 | 0 | 0 | 0 | 0 |
| 0 | NA | NA | NA | NA | NA | E130018O15Rik | | 0 | 0 | 0 | 0 | 0 | 0 | 0 | 0 | 0 | 0 | 0 |
| 0 | NA | NA | NA | NA | NA | Hmx1 | H6 homeobox 1 | 0 | 0 | 0 | 0 | 0 | 0 | 0 | 0 | 0 | 0 | 0 |
| 0 | NA | NA | NA | NA | NA | 4930442P19Rik | | 0 | 0 | 0 | 0 | 0 | 0 | 0 | 0 | 0 | 0 | 0 |
| 0 | NA | NA | NA | NA | NA | Gm43377 | | 0 | 0 | 0 | 0 | 0 | 0 | 0 | 0 | 0 | 0 | 0 |
| 0 | NA | NA | NA | NA | NA | Plk-ps1 | | 0 | 0 | 0 | 0 | 0 | 0 | 0 | 0 | 0 | 0 | 0 |
| 0 | NA | NA | NA | NA | NA | Gm15650 | | 0 | 0 | 0 | 0 | 0 | 0 | 0 | 0 | 0 | 0 | 0 |
| 0 | NA | NA | NA | NA | NA | H2af-ps | | 0 | 0 | 0 | 0 | 0 | 0 | 0 | 0 | 0 | 0 | 0 |
| 0 | NA | NA | NA | NA | NA | Psapl1 | prosaposin-like 1 | 0 | 0 | 0 | 0 | 0 | 0 | 0 | 0 | 0 | 0 | 0 |
| 0 | NA | NA | NA | NA | NA | Mir7689 | | 0 | 0 | 0 | 0 | 0 | 0 | 0 | 0 | 0 | 0 | 0 |
| 0 | NA | NA | NA | NA | NA | Gm42936 | | 0 | 0 | 0 | 0 | 0 | 0 | 0 | 0 | 0 | 0 | 0 |
| 0 | NA | NA | NA | NA | NA | Gm42507 | | 0 | 0 | 0 | 0 | 0 | 0 | 0 | 0 | 0 | 0 | 0 |
| 0 | NA | NA | NA | NA | NA | Gm25767 | | 0 | 0 | 0 | 0 | 0 | 0 | 0 | 0 | 0 | 0 | 0 |
| 0 | NA | NA | NA | NA | NA | Gm43701 | | 0 | 0 | 0 | 0 | 0 | 0 | 0 | 0 | 0 | 0 | 0 |
| 0 | NA | NA | NA | NA | NA | Gm40318 | | 0 | 0 | 0 | 0 | 0 | 0 | 0 | 0 | 0 | 0 | 0 |
| 0 | NA | NA | NA | NA | NA | Ppp2r2cos | | 0 | 0 | 0 | 0 | 0 | 0 | 0 | 0 | 0 | 0 | 0 |
| 0 | NA | NA | NA | NA | NA | Gm42506 | | 0 | 0 | 0 | 0 | 0 | 0 | 0 | 0 | 0 | 0 | 0 |
| 0 | NA | NA | NA | NA | NA | Gm25551 | | 0 | 0 | 0 | 0 | 0 | 0 | 0 | 0 | 0 | 0 | 0 |
| 0 | NA | NA | NA | NA | NA | Jakmip1 | | 0 | 0 | 0 | 0 | 0 | 0 | 0 | 0 | 0 | 0 | 0 |

| | | | | | | | | | | | | | | | | | | |
|---|---|---|---|---|---|---|---|---|---|---|---|---|---|---|---|---|---|---|
| 0 | NA | NA | NA | NA | NA | Gm42935 | | 0 | 0 | 0 | 0 | 0 | 0 | 0 | 0 | 0 | 0 | 0 |
| 0 | NA | NA | NA | NA | NA | Gm43763 | | 0 | 0 | 0 | 0 | 0 | 0 | 0 | 0 | 0 | 0 | 0 |
| 0 | NA | NA | NA | NA | NA | Gm20052 | | 0 | 0 | 0 | 0 | 0 | 0 | 0 | 0 | 0 | 0 | 0 |
| 0 | NA | NA | NA | NA | NA | Gm40289 | | 0 | 0 | 0 | 0 | 0 | 0 | 0 | 0 | 0 | 0 | 0 |
| 0 | NA | NA | NA | NA | NA | Gm43762 | | 0 | 0 | 0 | 0 | 0 | 0 | 0 | 0 | 0 | 0 | 0 |
| 0 | NA | NA | NA | NA | NA | Otop1 | otopetrin 1 | 0 | 0 | 0 | 0 | 0 | 0 | 0 | 0 | 0 | 0 | 0 |
| 0 | NA | NA | NA | NA | NA | 4930487D11Rik | | 0 | 0 | 0 | 0 | 0 | 0 | 0 | 0 | 0 | 0 | 0 |
| 0 | NA | NA | NA | NA | NA | Gm22249 | | 0 | 0 | 0 | 0 | 0 | 0 | 0 | 0 | 0 | 0 | 0 |
| 0 | NA | NA | NA | NA | NA | 4930421P07Rik | | 0 | 0 | 0 | 0 | 0 | 0 | 0 | 0 | 0 | 0 | 0 |
| 0 | NA | NA | NA | NA | NA | Gm22779 | | 0 | 0 | 0 | 0 | 0 | 0 | 0 | 0 | 0 | 0 | 0 |
| 0 | NA | NA | NA | NA | NA | Clnkos | | 0 | 0 | 0 | 0 | 0 | 0 | 0 | 0 | 0 | 0 | 0 |
| 0 | NA | NA | NA | NA | NA | Gm42857 | | 0 | 0 | 0 | 0 | 0 | 0 | 0 | 0 | 0 | 0 | 0 |
| 0 | NA | NA | NA | NA | NA | Gm40293 | | 0 | 0 | 0 | 0 | 0 | 0 | 0 | 0 | 0 | 0 | 0 |
| 0 | NA | NA | NA | NA | NA | Gm40292 | | 0 | 0 | 0 | 0 | 0 | 0 | 0 | 0 | 0 | 0 | 0 |
| 0 | NA | NA | NA | NA | NA | Gm43456 | | 0 | 0 | 0 | 0 | 0 | 0 | 0 | 0 | 0 | 0 | 0 |
| 0 | NA | NA | NA | NA | NA | Gm25505 | | 0 | 0 | 0 | 0 | 0 | 0 | 0 | 0 | 0 | 0 | 0 |
| 0 | NA | NA | NA | NA | NA | 4930421C12Rik | | 0 | 0 | 0 | 0 | 0 | 0 | 0 | 0 | 0 | 0 | 0 |
| 0 | NA | NA | NA | NA | NA | Gm42524 | | 0 | 0 | 0 | 0 | 0 | 0 | 0 | 0 | 0 | 0 | 0 |
| 0 | NA | NA | NA | NA | NA | 4930513D17Rik | | 0 | 0 | 0 | 0 | 0 | 0 | 0 | 0 | 0 | 0 | 0 |
| 0 | NA | NA | NA | NA | NA | Gm25377 | | 0 | 0 | 0 | 0 | 0 | 0 | 0 | 0 | 0 | 0 | 0 |
| 0 | NA | NA | NA | NA | NA | Gm35191 | | 0 | 0 | 0 | 0 | 0 | 0 | 0 | 0 | 0 | 0 | 0 |
| 0 | NA | NA | NA | NA | NA | Gm23022 | | 0 | 0 | 0 | 0 | 0 | 0 | 0 | 0 | 0 | 0 | 0 |
| 0 | NA | NA | NA | NA | NA | 4930519E07Rik | | 0 | 0 | 0 | 0 | 0 | 0 | 0 | 0 | 0 | 0 | 0 |
| 0 | NA | NA | NA | NA | NA | Gm43806 | | 0 | 0 | 0 | 0 | 0 | 0 | 0 | 0 | 0 | 0 | 0 |
| 0 | NA | NA | NA | NA | NA | Mir6414 | | 0 | 0 | 0 | 0 | 0 | 0 | 0 | 0 | 0 | 0 | 0 |
| 0 | NA | NA | NA | NA | NA | Gm16223 | | 0 | 0 | 0 | 0 | 0 | 0 | 0 | 0 | 0 | 0 | 0 |
| 0 | NA | NA | NA | NA | NA | Gm5554 | | 0 | 0 | 0 | 0 | 0 | 0 | 0 | 0 | 0 | 0 | 0 |
| 0 | NA | NA | NA | NA | NA | Gm43700 | | 0 | 0 | 0 | 0 | 0 | 0 | 0 | 0 | 0 | 0 | 0 |
| 0 | NA | NA | NA | NA | NA | Gm42711 | | 0 | 0 | 0 | 0 | 0 | 0 | 0 | 0 | 0 | 0 | 0 |
| 0 | NA | NA | NA | NA | NA | Gm42553 | | 0 | 0 | 0 | 0 | 0 | 0 | 0 | 0 | 0 | 0 | 0 |
| 0 | NA | NA | NA | NA | NA | Gm42552 | | 0 | 0 | 0 | 0 | 0 | 0 | 0 | 0 | 0 | 0 | 0 |
| 0 | NA | NA | NA | NA | NA | Gm43699 | | 0 | 0 | 0 | 0 | 0 | 0 | 0 | 0 | 0 | 0 | 0 |
| 0 | NA | NA | NA | NA | NA | Gm43021 | | 0 | 0 | 0 | 0 | 0 | 0 | 0 | 0 | 0 | 0 | 0 |
| 0 | NA | NA | NA | NA | NA | Gm43022 | | 0 | 0 | 0 | 0 | 0 | 0 | 0 | 0 | 0 | 0 | 0 |
| 0 | NA | NA | NA | NA | NA | Gm43023 | | 0 | 0 | 0 | 0 | 0 | 0 | 0 | 0 | 0 | 0 | 0 |
| 0 | NA | NA | NA | NA | NA | Gm43181 | | 0 | 0 | 0 | 0 | 0 | 0 | 0 | 0 | 0 | 0 | 0 |
| 0 | NA | NA | NA | NA | NA | Gm42555 | | 0 | 0 | 0 | 0 | 0 | 0 | 0 | 0 | 0 | 0 | 0 |
| 0 | NA | NA | NA | NA | NA | Gm16014 | | 0 | 0 | 0 | 0 | 0 | 0 | 0 | 0 | 0 | 0 | 0 |
| 0 | NA | NA | NA | NA | NA | Gm16015 | | 0 | 0 | 0 | 0 | 0 | 0 | 0 | 0 | 0 | 0 | 0 |
| 0 | NA | NA | NA | NA | NA | Gm7882 | | 0 | 0 | 0 | 0 | 0 | 0 | 0 | 0 | 0 | 0 | 0 |
| 0 | NA | NA | NA | NA | NA | Gm43184 | | 0 | 0 | 0 | 0 | 0 | 0 | 0 | 0 | 0 | 0 | 0 |
| 0 | NA | NA | NA | NA | NA | Gm42429 | | 0 | 0 | 0 | 0 | 0 | 0 | 0 | 0 | 0 | 0 | 0 |
| 0 | NA | NA | NA | NA | NA | Gm42982 | | 0 | 0 | 0 | 0 | 0 | 0 | 0 | 0 | 0 | 0 | 0 |
| 0 | NA | NA | NA | NA | NA | Gm42983 | | 0 | 0 | 0 | 0 | 0 | 0 | 0 | 0 | 0 | 0 | 0 |
| 0 | NA | NA | NA | NA | NA | Gm42698 | | 0 | 0 | 0 | 0 | 0 | 0 | 0 | 0 | 0 | 0 | 0 |
| 0 | NA | NA | NA | NA | NA | Gm43114 | | 0 | 0 | 0 | 0 | 0 | 0 | 0 | 0 | 0 | 0 | 0 |
| 0 | NA | NA | NA | NA | NA | Gm36017 | | 0 | 0 | 0 | 0 | 0 | 0 | 0 | 0 | 0 | 0 | 0 |
| 0 | NA | NA | NA | NA | NA | 8030487O14Rik | | 0 | 0 | 0 | 0 | 0 | 0 | 0 | 0 | 0 | 0 | 0 |
| 0 | NA | NA | NA | NA | NA | Gm43506 | | 0 | 0 | 0 | 0 | 0 | 0 | 0 | 0 | 0 | 0 | 0 |
| 0 | NA | NA | NA | NA | NA | Gm43505 | | 0 | 0 | 0 | 0 | 0 | 0 | 0 | 0 | 0 | 0 | 0 |
| 0 | NA | NA | NA | NA | NA | Gm43508 | | 0 | 0 | 0 | 0 | 0 | 0 | 0 | 0 | 0 | 0 | 0 |
| 0 | NA | NA | NA | NA | NA | Gm43507 | | 0 | 0 | 0 | 0 | 0 | 0 | 0 | 0 | 0 | 0 | 0 |
| 0 | NA | NA | NA | NA | NA | Gm43504 | | 0 | 0 | 0 | 0 | 0 | 0 | 0 | 0 | 0 | 0 | 0 |
| 0 | NA | NA | NA | NA | NA | Gm43503 | | 0 | 0 | 0 | 0 | 0 | 0 | 0 | 0 | 0 | 0 | 0 |
| 0 | NA | NA | NA | NA | NA | E030026E10Rik | | 0 | 0 | 0 | 0 | 0 | 0 | 0 | 0 | 0 | 0 | 0 |
| 0 | NA | NA | NA | NA | NA | Gm43509 | | 0 | 0 | 0 | 0 | 0 | 0 | 0 | 0 | 0 | 0 | 0 |
| 0 | NA | NA | NA | NA | NA | Gm42750 | | 0 | 0 | 0 | 0 | 0 | 0 | 0 | 0 | 0 | 0 | 0 |
| 0 | NA | NA | NA | NA | NA | 4930431F12Rik | | 0 | 0 | 0 | 0 | 0 | 0 | 0 | 0 | 0 | 0 | 0 |
| 0 | NA | NA | NA | NA | NA | Gm10205 | | 0 | 0 | 0 | 0 | 0 | 0 | 0 | 0 | 0 | 0 | 0 |
| 0 | NA | NA | NA | NA | NA | Gm43303 | | 0 | 0 | 0 | 0 | 0 | 0 | 0 | 0 | 0 | 0 | 0 |

| | | | | | | | | | | | | | | | | | | |
|---|---|---|---|---|---|---|---|---|---|---|---|---|---|---|---|---|---|---|
| 0 | NA | NA | NA | NA | NA | Gm42519 | | 0 | 0 | 0 | 0 | 0 | 0 | 0 | 0 | 0 | 0 | 0 |
| 0 | NA | NA | NA | NA | NA | Clrn2 | clarin 2 | 0 | 0 | 0 | 0 | 0 | 0 | 0 | 0 | 0 | 0 | 0 |
| 0 | NA | NA | NA | NA | NA | Gm42520 | | 0 | 0 | 0 | 0 | 0 | 0 | 0 | 0 | 0 | 0 | 0 |
| 0 | NA | NA | NA | NA | NA | Gm43367 | predicted gene 43367 | 0 | 0 | 0 | 0 | 0 | 0 | 0 | 0 | 0 | 0 | 0 |
| 0 | NA | NA | NA | NA | NA | Gm43366 | | 0 | 0 | 0 | 0 | 0 | 0 | 0 | 0 | 0 | 0 | 0 |
| 0 | NA | NA | NA | NA | NA | 4930405L22Rik | | 0 | 0 | 0 | 0 | 0 | 0 | 0 | 0 | 0 | 0 | 0 |
| 0 | NA | NA | NA | NA | NA | Gm43093 | | 0 | 0 | 0 | 0 | 0 | 0 | 0 | 0 | 0 | 0 | 0 |
| 0 | NA | NA | NA | NA | NA | Gm43092 | | 0 | 0 | 0 | 0 | 0 | 0 | 0 | 0 | 0 | 0 | 0 |
| 0 | NA | NA | NA | NA | NA | Gm43601 | | 0 | 0 | 0 | 0 | 0 | 0 | 0 | 0 | 0 | 0 | 0 |
| 0 | NA | NA | NA | NA | NA | Gm17824 | | 0 | 0 | 0 | 0 | 0 | 0 | 0 | 0 | 0 | 0 | 0 |
| 0 | NA | NA | NA | NA | NA | Gm3010 | | 0 | 0 | 0 | 0 | 0 | 0 | 0 | 0 | 0 | 0 | 0 |
| 0 | NA | NA | NA | NA | NA | Gm43427 | | 0 | 0 | 0 | 0 | 0 | 0 | 0 | 0 | 0 | 0 | 0 |
| 0 | NA | NA | NA | NA | NA | 4930518C09Rik | | 0 | 0 | 0 | 0 | 0 | 0 | 0 | 0 | 0 | 0 | 0 |
| 0 | NA | NA | NA | NA | NA | Gm20475 | | 0 | 0 | 0 | 0 | 0 | 0 | 0 | 0 | 0 | 0 | 0 |
| 0 | NA | NA | NA | NA | NA | Gm43426 | | 0 | 0 | 0 | 0 | 0 | 0 | 0 | 0 | 0 | 0 | 0 |
| 0 | NA | NA | NA | NA | NA | Gm42536 | | 0 | 0 | 0 | 0 | 0 | 0 | 0 | 0 | 0 | 0 | 0 |
| 0 | NA | NA | NA | NA | NA | Gm42537 | | 0 | 0 | 0 | 0 | 0 | 0 | 0 | 0 | 0 | 0 | 0 |
| 0 | NA | NA | NA | NA | NA | Mir218-1 | | 0 | 0 | 0 | 0 | 0 | 0 | 0 | 0 | 0 | 0 | 0 |
| 0 | NA | NA | NA | NA | NA | Gm42534 | | 0 | 0 | 0 | 0 | 0 | 0 | 0 | 0 | 0 | 0 | 0 |
| 0 | NA | NA | NA | NA | NA | Gm42535 | | 0 | 0 | 0 | 0 | 0 | 0 | 0 | 0 | 0 | 0 | 0 |
| 0 | NA | NA | NA | NA | NA | Gm43831 | | 0 | 0 | 0 | 0 | 0 | 0 | 0 | 0 | 0 | 0 | 0 |
| 0 | NA | NA | NA | NA | NA | Gm20700 | | 0 | 0 | 0 | 0 | 0 | 0 | 0 | 0 | 0 | 0 | 0 |
| 0 | NA | NA | NA | NA | NA | Gm43648 | | 0 | 0 | 0 | 0 | 0 | 0 | 0 | 0 | 0 | 0 | 0 |
| 0 | NA | NA | NA | NA | NA | Gm19031 | | 0 | 0 | 0 | 0 | 0 | 0 | 0 | 0 | 0 | 0 | 0 |
| 0 | NA | NA | NA | NA | NA | Gm43829 | | 0 | 0 | 0 | 0 | 0 | 0 | 0 | 0 | 0 | 0 | 0 |
| 0 | NA | NA | NA | NA | NA | Gm43830 | | 0 | 0 | 0 | 0 | 0 | 0 | 0 | 0 | 0 | 0 | 0 |
| 0 | NA | NA | NA | NA | NA | Gm42919 | | 0 | 0 | 0 | 0 | 0 | 0 | 0 | 0 | 0 | 0 | 0 |
| 0 | NA | NA | NA | NA | NA | Gm10025 | | 0 | 0 | 0 | 0 | 0 | 0 | 0 | 0 | 0 | 0 | 0 |
| 0 | NA | NA | NA | NA | NA | Gm43341 | | 0 | 0 | 0 | 0 | 0 | 0 | 0 | 0 | 0 | 0 | 0 |
| 0 | NA | NA | NA | NA | NA | C030015E24Rik | | 0 | 0 | 0 | 0 | 0 | 0 | 0 | 0 | 0 | 0 | 0 |
| 0 | NA | NA | NA | NA | NA | Gm43047 | | 0 | 0 | 0 | 0 | 0 | 0 | 0 | 0 | 0 | 0 | 0 |
| 0 | NA | NA | NA | NA | NA | Gm42767 | | 0 | 0 | 0 | 0 | 0 | 0 | 0 | 0 | 0 | 0 | 0 |
| 0 | NA | NA | NA | NA | NA | Gm19932 | | 0 | 0 | 0 | 0 | 0 | 0 | 0 | 0 | 0 | 0 | 0 |
| 0 | NA | NA | NA | NA | NA | Gm43670 | | 0 | 0 | 0 | 0 | 0 | 0 | 0 | 0 | 0 | 0 | 0 |
| 0 | NA | NA | NA | NA | NA | 2900064K03Rik | | 0 | 0 | 0 | 0 | 0 | 0 | 0 | 0 | 0 | 0 | 0 |
| 0 | NA | NA | NA | NA | NA | Gm7988 | | 0 | 0 | 0 | 0 | 0 | 0 | 0 | 0 | 0 | 0 | 0 |
| 0 | NA | NA | NA | NA | NA | 4930448I18Rik | | 0 | 0 | 0 | 0 | 0 | 0 | 0 | 0 | 0 | 0 | 0 |
| 0 | NA | NA | NA | NA | NA | Mir8117 | | 0 | 0 | 0 | 0 | 0 | 0 | 0 | 0 | 0 | 0 | 0 |
| 0 | NA | NA | NA | NA | NA | Gm22618 | | 0 | 0 | 0 | 0 | 0 | 0 | 0 | 0 | 0 | 0 | 0 |
| 0 | NA | NA | NA | NA | NA | Gm36814 | | 0 | 0 | 0 | 0 | 0 | 0 | 0 | 0 | 0 | 0 | 0 |
| 0 | NA | NA | NA | NA | NA | Gm42459 | | 0 | 0 | 0 | 0 | 0 | 0 | 0 | 0 | 0 | 0 | 0 |
| 0 | NA | NA | NA | NA | NA | Gm4962 | | 0 | 0 | 0 | 0 | 0 | 0 | 0 | 0 | 0 | 0 | 0 |
| 0 | NA | NA | NA | NA | NA | Gm43014 | | 0 | 0 | 0 | 0 | 0 | 0 | 0 | 0 | 0 | 0 | 0 |
| 0 | NA | NA | NA | NA | NA | Gm43015 | | 0 | 0 | 0 | 0 | 0 | 0 | 0 | 0 | 0 | 0 | 0 |
| 0 | NA | NA | NA | NA | NA | Gm44377 | | 0 | 0 | 0 | 0 | 0 | 0 | 0 | 0 | 0 | 0 | 0 |
| 0 | NA | NA | NA | NA | NA | Gm42613 | | 0 | 0 | 0 | 0 | 0 | 0 | 0 | 0 | 0 | 0 | 0 |
| 0 | NA | NA | NA | NA | NA | Gm43605 | | 0 | 0 | 0 | 0 | 0 | 0 | 0 | 0 | 0 | 0 | 0 |
| 0 | NA | NA | NA | NA | NA | Gm43606 | | 0 | 0 | 0 | 0 | 0 | 0 | 0 | 0 | 0 | 0 | 0 |
| 0 | NA | NA | NA | NA | NA | Gm42616 | | 0 | 0 | 0 | 0 | 0 | 0 | 0 | 0 | 0 | 0 | 0 |
| 0 | NA | NA | NA | NA | NA | Gm42617 | | 0 | 0 | 0 | 0 | 0 | 0 | 0 | 0 | 0 | 0 | 0 |
| 0 | NA | NA | NA | NA | NA | Gm43175 | | 0 | 0 | 0 | 0 | 0 | 0 | 0 | 0 | 0 | 0 | 0 |
| 0 | NA | NA | NA | NA | NA | Gm43174 | | 0 | 0 | 0 | 0 | 0 | 0 | 0 | 0 | 0 | 0 | 0 |
| 0 | NA | NA | NA | NA | NA | Gm43177 | | 0 | 0 | 0 | 0 | 0 | 0 | 0 | 0 | 0 | 0 | 0 |
| 0 | NA | NA | NA | NA | NA | Mir6417 | | 0 | 0 | 0 | 0 | 0 | 0 | 0 | 0 | 0 | 0 | 0 |
| 0 | NA | NA | NA | NA | NA | Gm43176 | | 0 | 0 | 0 | 0 | 0 | 0 | 0 | 0 | 0 | 0 | 0 |
| 0 | NA | NA | NA | NA | NA | Gm43178 | | 0 | 0 | 0 | 0 | 0 | 0 | 0 | 0 | 0 | 0 | 0 |
| 0 | NA | NA | NA | NA | NA | Gm43179 | | 0 | 0 | 0 | 0 | 0 | 0 | 0 | 0 | 0 | 0 | 0 |
| 0 | NA | NA | NA | NA | NA | Gm25601 | | 0 | 0 | 0 | 0 | 0 | 0 | 0 | 0 | 0 | 0 | 0 |
| 0 | NA | NA | NA | NA | NA | Gm3519 | | 0 | 0 | 0 | 0 | 0 | 0 | 0 | 0 | 0 | 0 | 0 |
| 0 | NA | NA | NA | NA | NA | Gm43664 | | 0 | 0 | 0 | 0 | 0 | 0 | 0 | 0 | 0 | 0 | 0 |

| | | | | | | | | | | | | | | | | | |
|---|---|---|---|---|---|---|---|---|---|---|---|---|---|---|---|---|---|
| 0 | NA | NA | NA | NA | NA | Gm23532 | 0 | 0 | 0 | 0 | 0 | 0 | 0 | 0 | 0 | 0 | 0 |
| 0 | NA | NA | NA | NA | NA | Gm30301 | 0 | 0 | 0 | 0 | 0 | 0 | 0 | 0 | 0 | 0 | 0 |
| 0 | NA | NA | NA | NA | NA | Gm17182 | 0 | 0 | 0 | 0 | 0 | 0 | 0 | 0 | 0 | 0 | 0 |
| 0 | NA | NA | NA | NA | NA | n-R5s171 | 0 | 0 | 0 | 0 | 0 | 0 | 0 | 0 | 0 | 0 | 0 |
| 0 | NA | NA | NA | NA | NA | n-R5s172 | 0 | 0 | 0 | 0 | 0 | 0 | 0 | 0 | 0 | 0 | 0 |
| 0 | NA | NA | NA | NA | NA | Gm42745 | 0 | 0 | 0 | 0 | 0 | 0 | 0 | 0 | 0 | 0 | 0 |
| 0 | NA | NA | NA | NA | NA | Gm26486 | 0 | 0 | 0 | 0 | 0 | 0 | 0 | 0 | 0 | 0 | 0 |
| 0 | NA | NA | NA | NA | NA | Gm10441 | 0 | 0 | 0 | 0 | 0 | 0 | 0 | 0 | 0 | 0 | 0 |
| 0 | NA | NA | NA | NA | NA | Gm40304 | 0 | 0 | 0 | 0 | 0 | 0 | 0 | 0 | 0 | 0 | 0 |
| 0 | NA | NA | NA | NA | NA | Gm15820 | 0 | 0 | 0 | 0 | 0 | 0 | 0 | 0 | 0 | 0 | 0 |
| 0 | NA | NA | NA | NA | NA | Gm10440 | 0 | 0 | 0 | 0 | 0 | 0 | 0 | 0 | 0 | 0 | 0 |
| 0 | NA | NA | NA | NA | NA | Gm18451 | 0 | 0 | 0 | 0 | 0 | 0 | 0 | 0 | 0 | 0 | 0 |
| 0 | NA | NA | NA | NA | NA | Gm43781 | 0 | 0 | 0 | 0 | 0 | 0 | 0 | 0 | 0 | 0 | 0 |
| 0 | NA | NA | NA | NA | NA | Gm24235 | 0 | 0 | 0 | 0 | 0 | 0 | 0 | 0 | 0 | 0 | 0 |
| 0 | NA | NA | NA | NA | NA | Gm43779 | 0 | 0 | 0 | 0 | 0 | 0 | 0 | 0 | 0 | 0 | 0 |
| 0 | NA | NA | NA | NA | NA | Gm25513 | 0 | 0 | 0 | 0 | 0 | 0 | 0 | 0 | 0 | 0 | 0 |
| 0 | NA | NA | NA | NA | NA | Gm43780 | 0 | 0 | 0 | 0 | 0 | 0 | 0 | 0 | 0 | 0 | 0 |
| 0 | NA | NA | NA | NA | NA | Gm43396 | 0 | 0 | 0 | 0 | 0 | 0 | 0 | 0 | 0 | 0 | 0 |
| 0 | NA | NA | NA | NA | NA | Gm43199 | 0 | 0 | 0 | 0 | 0 | 0 | 0 | 0 | 0 | 0 | 0 |
| 0 | NA | NA | NA | NA | NA | Gm43778 | 0 | 0 | 0 | 0 | 0 | 0 | 0 | 0 | 0 | 0 | 0 |
| 0 | NA | NA | NA | NA | NA | Gm20223 | 0 | 0 | 0 | 0 | 0 | 0 | 0 | 0 | 0 | 0 | 0 |
| 0 | NA | NA | NA | NA | NA | Gm43705 | 0 | 0 | 0 | 0 | 0 | 0 | 0 | 0 | 0 | 0 | 0 |
| 0 | NA | NA | NA | NA | NA | Gm43315 | 0 | 0 | 0 | 0 | 0 | 0 | 0 | 0 | 0 | 0 | 0 |
| 0 | NA | NA | NA | NA | NA | Gm42977 | 0 | 0 | 0 | 0 | 0 | 0 | 0 | 0 | 0 | 0 | 0 |
| 0 | NA | NA | NA | NA | NA | Gm23896 | 0 | 0 | 0 | 0 | 0 | 0 | 0 | 0 | 0 | 0 | 0 |
| 0 | NA | NA | NA | NA | NA | Gm24902 | 0 | 0 | 0 | 0 | 0 | 0 | 0 | 0 | 0 | 0 | 0 |
| 0 | NA | NA | NA | NA | NA | Gm23015 | 0 | 0 | 0 | 0 | 0 | 0 | 0 | 0 | 0 | 0 | 0 |
| 0 | NA | NA | NA | NA | NA | Gm3283 | 0 | 0 | 0 | 0 | 0 | 0 | 0 | 0 | 0 | 0 | 0 |
| 0 | NA | NA | NA | NA | NA | Gm18901 | 0 | 0 | 0 | 0 | 0 | 0 | 0 | 0 | 0 | 0 | 0 |
| 0 | NA | NA | NA | NA | NA | Gm43706 | 0 | 0 | 0 | 0 | 0 | 0 | 0 | 0 | 0 | 0 | 0 |
| 0 | NA | NA | NA | NA | NA | Gm6615 | 0 | 0 | 0 | 0 | 0 | 0 | 0 | 0 | 0 | 0 | 0 |
| 0 | NA | NA | NA | NA | NA | Gm43002 | 0 | 0 | 0 | 0 | 0 | 0 | 0 | 0 | 0 | 0 | 0 |
| 0 | NA | NA | NA | NA | NA | Gm42634 | 0 | 0 | 0 | 0 | 0 | 0 | 0 | 0 | 0 | 0 | 0 |
| 0 | NA | NA | NA | NA | NA | Gm42635 | 0 | 0 | 0 | 0 | 0 | 0 | 0 | 0 | 0 | 0 | 0 |
| 0 | NA | NA | NA | NA | NA | Gm42481 | 0 | 0 | 0 | 0 | 0 | 0 | 0 | 0 | 0 | 0 | 0 |
| 0 | NA | NA | NA | NA | NA | Gm42480 | 0 | 0 | 0 | 0 | 0 | 0 | 0 | 0 | 0 | 0 | 0 |
| 0 | NA | NA | NA | NA | NA | Gm42482 | 0 | 0 | 0 | 0 | 0 | 0 | 0 | 0 | 0 | 0 | 0 |
| 0 | NA | NA | NA | NA | NA | Gm17977 | 0 | 0 | 0 | 0 | 0 | 0 | 0 | 0 | 0 | 0 | 0 |
| 0 | NA | NA | NA | NA | NA | Gm43715 | 0 | 0 | 0 | 0 | 0 | 0 | 0 | 0 | 0 | 0 | 0 |
| 0 | NA | NA | NA | NA | NA | Gm42479 | 0 | 0 | 0 | 0 | 0 | 0 | 0 | 0 | 0 | 0 | 0 |
| 0 | NA | NA | NA | NA | NA | Gm42478 | 0 | 0 | 0 | 0 | 0 | 0 | 0 | 0 | 0 | 0 | 0 |
| 0 | NA | NA | NA | NA | NA | Gm42640 | 0 | 0 | 0 | 0 | 0 | 0 | 0 | 0 | 0 | 0 | 0 |
| 0 | NA | NA | NA | NA | NA | Gm42484 | 0 | 0 | 0 | 0 | 0 | 0 | 0 | 0 | 0 | 0 | 0 |
| 0 | NA | NA | NA | NA | NA | Gm37720 | 0 | 0 | 0 | 0 | 0 | 0 | 0 | 0 | 0 | 0 | 0 |
| 0 | NA | NA | NA | NA | NA | Gm42483 | 0 | 0 | 0 | 0 | 0 | 0 | 0 | 0 | 0 | 0 | 0 |
| 0 | NA | NA | NA | NA | NA | Gm42486 | 0 | 0 | 0 | 0 | 0 | 0 | 0 | 0 | 0 | 0 | 0 |
| 0 | NA | NA | NA | NA | NA | Gm42976 | 0 | 0 | 0 | 0 | 0 | 0 | 0 | 0 | 0 | 0 | 0 |
| 0 | NA | NA | NA | NA | NA | Gm43045 | 0 | 0 | 0 | 0 | 0 | 0 | 0 | 0 | 0 | 0 | 0 |
| 0 | NA | NA | NA | NA | NA | Gm42999 | 0 | 0 | 0 | 0 | 0 | 0 | 0 | 0 | 0 | 0 | 0 |
| 0 | NA | NA | NA | NA | NA | Gm22536 | 0 | 0 | 0 | 0 | 0 | 0 | 0 | 0 | 0 | 0 | 0 |
| 0 | NA | NA | NA | NA | NA | 4930459L07Rik | 0 | 0 | 0 | 0 | 0 | 0 | 0 | 0 | 0 | 0 | 0 |
| 0 | NA | NA | NA | NA | NA | 1700029E06Rik | 0 | 0 | 0 | 0 | 0 | 0 | 0 | 0 | 0 | 0 | 0 |
| 0 | NA | NA | NA | NA | NA | Gm43394 | 0 | 0 | 0 | 0 | 0 | 0 | 0 | 0 | 0 | 0 | 0 |
| 0 | NA | NA | NA | NA | NA | Gm6632 | 0 | 0 | 0 | 0 | 0 | 0 | 0 | 0 | 0 | 0 | 0 |
| 0 | NA | NA | NA | NA | NA | Gm43395 | 0 | 0 | 0 | 0 | 0 | 0 | 0 | 0 | 0 | 0 | 0 |
| 0 | NA | NA | NA | NA | NA | Gm22308 | 0 | 0 | 0 | 0 | 0 | 0 | 0 | 0 | 0 | 0 | 0 |
| 0 | NA | NA | NA | NA | NA | Gm43044 | 0 | 0 | 0 | 0 | 0 | 0 | 0 | 0 | 0 | 0 | 0 |
| 0 | NA | NA | NA | NA | NA | Gm43043 | 0 | 0 | 0 | 0 | 0 | 0 | 0 | 0 | 0 | 0 | 0 |
| 0 | NA | NA | NA | NA | NA | Gm43041 | 0 | 0 | 0 | 0 | 0 | 0 | 0 | 0 | 0 | 0 | 0 |
| 0 | NA | NA | NA | NA | NA | 4933402J10Rik | 0 | 0 | 0 | 0 | 0 | 0 | 0 | 0 | 0 | 0 | 0 |

| | | | | | | | | | | | | | | | | | | |
|---|---|---|---|---|---|---|---|---|---|---|---|---|---|---|---|---|---|---|
| 0 | NA | NA | NA | NA | NA | Gm42862 | | 0 | 0 | 0 | 0 | 0 | 0 | 0 | 0 | 0 | 0 | 0 |
| 0 | NA | NA | NA | NA | NA | Gm43392 | | 0 | 0 | 0 | 0 | 0 | 0 | 0 | 0 | 0 | 0 | 0 |
| 0 | NA | NA | NA | NA | NA | Gm43393 | | 0 | 0 | 0 | 0 | 0 | 0 | 0 | 0 | 0 | 0 | 0 |
| 0 | NA | NA | NA | NA | NA | Gm42861 | | 0 | 0 | 0 | 0 | 0 | 0 | 0 | 0 | 0 | 0 | 0 |
| 0 | NA | NA | NA | NA | NA | Gm23331 | | 0 | 0 | 0 | 0 | 0 | 0 | 0 | 0 | 0 | 0 | 0 |
| 0 | NA | NA | NA | NA | NA | Gm43390 | | 0 | 0 | 0 | 0 | 0 | 0 | 0 | 0 | 0 | 0 | 0 |
| 0 | NA | NA | NA | NA | NA | Gm3372 | | 0 | 0 | 0 | 0 | 0 | 0 | 0 | 0 | 0 | 0 | 0 |
| 0 | NA | NA | NA | NA | NA | Gm23708 | | 0 | 0 | 0 | 0 | 0 | 0 | 0 | 0 | 0 | 0 | 0 |
| 0 | NA | NA | NA | NA | NA | 1700014F14Rik | | 0 | 0 | 0 | 0 | 0 | 0 | 0 | 0 | 0 | 0 | 0 |
| 0 | NA | NA | NA | NA | NA | Gm43382 | | 0 | 0 | 0 | 0 | 0 | 0 | 0 | 0 | 0 | 0 | 0 |
| 0 | NA | NA | NA | NA | NA | Gm43383 | | 0 | 0 | 0 | 0 | 0 | 0 | 0 | 0 | 0 | 0 | 0 |
| 0 | NA | NA | NA | NA | NA | Gm43384 | | 0 | 0 | 0 | 0 | 0 | 0 | 0 | 0 | 0 | 0 | 0 |
| 0 | NA | NA | NA | NA | NA | Gm42430 | | 0 | 0 | 0 | 0 | 0 | 0 | 0 | 0 | 0 | 0 | 0 |
| 0 | NA | NA | NA | NA | NA | Gm22273 | | 0 | 0 | 0 | 0 | 0 | 0 | 0 | 0 | 0 | 0 | 0 |
| 0 | NA | NA | NA | NA | NA | Gm43238 | | 0 | 0 | 0 | 0 | 0 | 0 | 0 | 0 | 0 | 0 | 0 |
| 0 | NA | NA | NA | NA | NA | Gm24079 | | 0 | 0 | 0 | 0 | 0 | 0 | 0 | 0 | 0 | 0 | 0 |
| 0 | NA | NA | NA | NA | NA | Gm43237 | | 0 | 0 | 0 | 0 | 0 | 0 | 0 | 0 | 0 | 0 | 0 |
| 0 | NA | NA | NA | NA | NA | Gm43753 | | 0 | 0 | 0 | 0 | 0 | 0 | 0 | 0 | 0 | 0 | 0 |
| 0 | NA | NA | NA | NA | NA | Dthd1 | death domain containing 1 | 0 | 0 | 0 | 0 | 0 | 0 | 0 | 0 | 0 | 0 | 0 |
| 0 | NA | NA | NA | NA | NA | Gm3687 | | 0 | 0 | 0 | 0 | 0 | 0 | 0 | 0 | 0 | 0 | 0 |
| 0 | NA | NA | NA | NA | NA | Gm15819 | | 0 | 0 | 0 | 0 | 0 | 0 | 0 | 0 | 0 | 0 | 0 |
| 0 | NA | NA | NA | NA | NA | Gm43840 | | 0 | 0 | 0 | 0 | 0 | 0 | 0 | 0 | 0 | 0 | 0 |
| 0 | NA | NA | NA | NA | NA | Gm25608 | | 0 | 0 | 0 | 0 | 0 | 0 | 0 | 0 | 0 | 0 | 0 |
| 0 | NA | NA | NA | NA | NA | Gm23967 | | 0 | 0 | 0 | 0 | 0 | 0 | 0 | 0 | 0 | 0 | 0 |
| 0 | NA | NA | NA | NA | NA | Gm22395 | | 0 | 0 | 0 | 0 | 0 | 0 | 0 | 0 | 0 | 0 | 0 |
| 0 | NA | NA | NA | NA | NA | Gm22559 | | 0 | 0 | 0 | 0 | 0 | 0 | 0 | 0 | 0 | 0 | 0 |
| 0 | NA | NA | NA | NA | NA | Gm15824 | | 0 | 0 | 0 | 0 | 0 | 0 | 0 | 0 | 0 | 0 | 0 |
| 0 | NA | NA | NA | NA | NA | Gm43838 | | 0 | 0 | 0 | 0 | 0 | 0 | 0 | 0 | 0 | 0 | 0 |
| 0 | NA | NA | NA | NA | NA | Gm31363 | | 0 | 0 | 0 | 0 | 0 | 0 | 0 | 0 | 0 | 0 | 0 |
| 0 | NA | NA | NA | NA | NA | 1700027F09Rik | | 0 | 0 | 0 | 0 | 0 | 0 | 0 | 0 | 0 | 0 | 0 |
| 0 | NA | NA | NA | NA | NA | 4930526M16Rik | | 0 | 0 | 0 | 0 | 0 | 0 | 0 | 0 | 0 | 0 | 0 |
| 0 | NA | NA | NA | NA | NA | C030018K13Rik | | 0 | 0 | 0 | 0 | 0 | 0 | 0 | 0 | 0 | 0 | 0 |
| 0 | NA | NA | NA | NA | NA | Gm43836 | | 0 | 0 | 0 | 0 | 0 | 0 | 0 | 0 | 0 | 0 | 0 |
| 0 | NA | NA | NA | NA | NA | Gm43835 | | 0 | 0 | 0 | 0 | 0 | 0 | 0 | 0 | 0 | 0 | 0 |
| 0 | NA | NA | NA | NA | NA | Gm42564 | | 0 | 0 | 0 | 0 | 0 | 0 | 0 | 0 | 0 | 0 | 0 |
| 0 | NA | NA | NA | NA | NA | Gm42565 | | 0 | 0 | 0 | 0 | 0 | 0 | 0 | 0 | 0 | 0 | 0 |
| 0 | NA | NA | NA | NA | NA | Gm25306 | | 0 | 0 | 0 | 0 | 0 | 0 | 0 | 0 | 0 | 0 | 0 |
| 0 | NA | NA | NA | NA | NA | Mir574 | | 0 | 0 | 0 | 0 | 0 | 0 | 0 | 0 | 0 | 0 | 0 |
| 0 | NA | NA | NA | NA | NA | Gm25154 | | 0 | 0 | 0 | 0 | 0 | 0 | 0 | 0 | 0 | 0 | 0 |
| 0 | NA | NA | NA | NA | NA | Gm43552 | | 0 | 0 | 0 | 0 | 0 | 0 | 0 | 0 | 0 | 0 | 0 |
| 0 | NA | NA | NA | NA | NA | Gm43289 | | 0 | 0 | 0 | 0 | 0 | 0 | 0 | 0 | 0 | 0 | 0 |
| 0 | NA | NA | NA | NA | NA | Gm42726 | | 0 | 0 | 0 | 0 | 0 | 0 | 0 | 0 | 0 | 0 | 0 |
| 0 | NA | NA | NA | NA | NA | Gm24399 | | 0 | 0 | 0 | 0 | 0 | 0 | 0 | 0 | 0 | 0 | 0 |
| 0 | NA | NA | NA | NA | NA | Gm42645 | | 0 | 0 | 0 | 0 | 0 | 0 | 0 | 0 | 0 | 0 | 0 |
| 0 | NA | NA | NA | NA | NA | Gm42646 | | 0 | 0 | 0 | 0 | 0 | 0 | 0 | 0 | 0 | 0 | 0 |
| 0 | NA | NA | NA | NA | NA | Gm43004 | | 0 | 0 | 0 | 0 | 0 | 0 | 0 | 0 | 0 | 0 | 0 |
| 0 | NA | NA | NA | NA | NA | Gm43769 | | 0 | 0 | 0 | 0 | 0 | 0 | 0 | 0 | 0 | 0 | 0 |
| 0 | NA | NA | NA | NA | NA | Chrna9 | cholinergic receptor, nicotinic, alpha poly | 0 | 0 | 0 | 0 | 0 | 0 | 0 | 0 | 0 | 0 | 0 |
| 0 | NA | NA | NA | NA | NA | 4930480C01Rik | | 0 | 0 | 0 | 0 | 0 | 0 | 0 | 0 | 0 | 0 | 0 |
| 0 | NA | NA | NA | NA | NA | Gm43771 | | 0 | 0 | 0 | 0 | 0 | 0 | 0 | 0 | 0 | 0 | 0 |
| 0 | NA | NA | NA | NA | NA | Gm25952 | | 0 | 0 | 0 | 0 | 0 | 0 | 0 | 0 | 0 | 0 | 0 |
| 0 | NA | NA | NA | NA | NA | Gm43323 | | 0 | 0 | 0 | 0 | 0 | 0 | 0 | 0 | 0 | 0 | 0 |
| 0 | NA | NA | NA | NA | NA | Gm22929 | | 0 | 0 | 0 | 0 | 0 | 0 | 0 | 0 | 0 | 0 | 0 |
| 0 | NA | NA | NA | NA | NA | 1700126H18Rik | | 0 | 0 | 0 | 0 | 0 | 0 | 0 | 0 | 0 | 0 | 0 |
| 0 | NA | NA | NA | NA | NA | Gm15794 | | 0 | 0 | 0 | 0 | 0 | 0 | 0 | 0 | 0 | 0 | 0 |
| 0 | NA | NA | NA | NA | NA | Gm3822 | | 0 | 0 | 0 | 0 | 0 | 0 | 0 | 0 | 0 | 0 | 0 |
| 0 | NA | NA | NA | NA | NA | Gm16273 | | 0 | 0 | 0 | 0 | 0 | 0 | 0 | 0 | 0 | 0 | 0 |
| 0 | NA | NA | NA | NA | NA | Gm43280 | | 0 | 0 | 0 | 0 | 0 | 0 | 0 | 0 | 0 | 0 | 0 |
| 0 | NA | NA | NA | NA | NA | Gm43281 | | 0 | 0 | 0 | 0 | 0 | 0 | 0 | 0 | 0 | 0 | 0 |
| 0 | NA | NA | NA | NA | NA | Gm43282 | | 0 | 0 | 0 | 0 | 0 | 0 | 0 | 0 | 0 | 0 | 0 |

| | | | | | | | | | | | | | | | | | | |
|---|---|---|---|---|---|---|---|---|---|---|---|---|---|---|---|---|---|---|
| 0 | NA | NA | NA | NA | NA | Gm15949 | | 0 | 0 | 0 | 0 | 0 | 0 | 0 | 0 | 0 | 0 | 0 |
| 0 | NA | NA | NA | NA | NA | Gm15948 | | 0 | 0 | 0 | 0 | 0 | 0 | 0 | 0 | 0 | 0 | 0 |
| 0 | NA | NA | NA | NA | NA | Gm42713 | | 0 | 0 | 0 | 0 | 0 | 0 | 0 | 0 | 0 | 0 | 0 |
| 0 | NA | NA | NA | NA | NA | Gm23841 | | 0 | 0 | 0 | 0 | 0 | 0 | 0 | 0 | 0 | 0 | 0 |
| 0 | NA | NA | NA | NA | NA | Gm33167 | | 0 | 0 | 0 | 0 | 0 | 0 | 0 | 0 | 0 | 0 | 0 |
| 0 | NA | NA | NA | NA | NA | 1700025A08Rik | | 0 | 0 | 0 | 0 | 0 | 0 | 0 | 0 | 0 | 0 | 0 |
| 0 | NA | NA | NA | NA | NA | Gm23617 | | 0 | 0 | 0 | 0 | 0 | 0 | 0 | 0 | 0 | 0 | 0 |
| 0 | NA | NA | NA | NA | NA | Gm15478 | | 0 | 0 | 0 | 0 | 0 | 0 | 0 | 0 | 0 | 0 | 0 |
| 0 | NA | NA | NA | NA | NA | Gm42737 | | 0 | 0 | 0 | 0 | 0 | 0 | 0 | 0 | 0 | 0 | 0 |
| 0 | NA | NA | NA | NA | NA | Gm42736 | | 0 | 0 | 0 | 0 | 0 | 0 | 0 | 0 | 0 | 0 | 0 |
| 0 | NA | NA | NA | NA | NA | Gm21905 | | 0 | 0 | 0 | 0 | 0 | 0 | 0 | 0 | 0 | 0 | 0 |
| 0 | NA | NA | NA | NA | NA | 4930425K10Rik | | 0 | 0 | 0 | 0 | 0 | 0 | 0 | 0 | 0 | 0 | 0 |
| 0 | NA | NA | NA | NA | NA | Gm43027 | | 0 | 0 | 0 | 0 | 0 | 0 | 0 | 0 | 0 | 0 | 0 |
| 0 | NA | NA | NA | NA | NA | Grxcr1 | glutaredoxin, cysteine rich 1 | 0 | 0 | 0 | 0 | 0 | 0 | 0 | 0 | 0 | 0 | 0 |
| 0 | NA | NA | NA | NA | NA | Gm43025 | | 0 | 0 | 0 | 0 | 0 | 0 | 0 | 0 | 0 | 0 | 0 |
| 0 | NA | NA | NA | NA | NA | Gm43026 | | 0 | 0 | 0 | 0 | 0 | 0 | 0 | 0 | 0 | 0 | 0 |
| 0 | NA | NA | NA | NA | NA | Gm23605 | | 0 | 0 | 0 | 0 | 0 | 0 | 0 | 0 | 0 | 0 | 0 |
| 0 | NA | NA | NA | NA | NA | Gm42850 | | 0 | 0 | 0 | 0 | 0 | 0 | 0 | 0 | 0 | 0 | 0 |
| 0 | NA | NA | NA | NA | NA | Gm43842 | | 0 | 0 | 0 | 0 | 0 | 0 | 0 | 0 | 0 | 0 | 0 |
| 0 | NA | NA | NA | NA | NA | Gm43373 | | 0 | 0 | 0 | 0 | 0 | 0 | 0 | 0 | 0 | 0 | 0 |
| 0 | NA | NA | NA | NA | NA | Gm43374 | | 0 | 0 | 0 | 0 | 0 | 0 | 0 | 0 | 0 | 0 | 0 |
| 0 | NA | NA | NA | NA | NA | Gm23932 | | 0 | 0 | 0 | 0 | 0 | 0 | 0 | 0 | 0 | 0 | 0 |
| 0 | NA | NA | NA | NA | NA | Gm42887 | | 0 | 0 | 0 | 0 | 0 | 0 | 0 | 0 | 0 | 0 | 0 |
| 0 | NA | NA | NA | NA | NA | Gm26044 | | 0 | 0 | 0 | 0 | 0 | 0 | 0 | 0 | 0 | 0 | 0 |
| 0 | NA | NA | NA | NA | NA | Gm23661 | | 0 | 0 | 0 | 0 | 0 | 0 | 0 | 0 | 0 | 0 | 0 |
| 0 | NA | NA | NA | NA | NA | Gm43139 | | 0 | 0 | 0 | 0 | 0 | 0 | 0 | 0 | 0 | 0 | 0 |
| 0 | NA | NA | NA | NA | NA | Gm23067 | | 0 | 0 | 0 | 0 | 0 | 0 | 0 | 0 | 0 | 0 | 0 |
| 0 | NA | NA | NA | NA | NA | Gm26072 | | 0 | 0 | 0 | 0 | 0 | 0 | 0 | 0 | 0 | 0 | 0 |
| 0 | NA | NA | NA | NA | NA | Gm42652 | | 0 | 0 | 0 | 0 | 0 | 0 | 0 | 0 | 0 | 0 | 0 |
| 0 | NA | NA | NA | NA | NA | Gm42653 | | 0 | 0 | 0 | 0 | 0 | 0 | 0 | 0 | 0 | 0 | 0 |
| 0 | NA | NA | NA | NA | NA | Gm38562 | | 0 | 0 | 0 | 0 | 0 | 0 | 0 | 0 | 0 | 0 | 0 |
| 0 | NA | NA | NA | NA | NA | Gm15617 | | 0 | 0 | 0 | 0 | 0 | 0 | 0 | 0 | 0 | 0 | 0 |
| 0 | NA | NA | NA | NA | NA | Gm42569 | | 0 | 0 | 0 | 0 | 0 | 0 | 0 | 0 | 0 | 0 | 0 |
| 0 | NA | NA | NA | NA | NA | Gm25768 | | 0 | 0 | 0 | 0 | 0 | 0 | 0 | 0 | 0 | 0 | 0 |
| 0 | NA | NA | NA | NA | NA | Gm20647 | | 0 | 0 | 0 | 0 | 0 | 0 | 0 | 0 | 0 | 0 | 0 |
| 0 | NA | NA | NA | NA | NA | Gm19560 | | 0 | 0 | 0 | 0 | 0 | 0 | 0 | 0 | 0 | 0 | 0 |
| 0 | NA | NA | NA | NA | NA | Gm23712 | | 0 | 0 | 0 | 0 | 0 | 0 | 0 | 0 | 0 | 0 | 0 |
| 0 | NA | NA | NA | NA | NA | 4933408A14Rik | | 0 | 0 | 0 | 0 | 0 | 0 | 0 | 0 | 0 | 0 | 0 |
| 0 | NA | NA | NA | NA | NA | Gm43717 | | 0 | 0 | 0 | 0 | 0 | 0 | 0 | 0 | 0 | 0 | 0 |
| 0 | NA | NA | NA | NA | NA | Gm43499 | | 0 | 0 | 0 | 0 | 0 | 0 | 0 | 0 | 0 | 0 | 0 |
| 0 | NA | NA | NA | NA | NA | Gm24776 | | 0 | 0 | 0 | 0 | 0 | 0 | 0 | 0 | 0 | 0 | 0 |
| 0 | NA | NA | NA | NA | NA | Gm15847 | | 0 | 0 | 0 | 0 | 0 | 0 | 0 | 0 | 0 | 0 | 0 |
| 0 | NA | NA | NA | NA | NA | Gm42668 | | 0 | 0 | 0 | 0 | 0 | 0 | 0 | 0 | 0 | 0 | 0 |
| 0 | NA | NA | NA | NA | NA | 1700071G01Rik | | 0 | 0 | 0 | 0 | 0 | 0 | 0 | 0 | 0 | 0 | 0 |
| 0 | NA | NA | NA | NA | NA | Gm42571 | | 0 | 0 | 0 | 0 | 0 | 0 | 0 | 0 | 0 | 0 | 0 |
| 0 | NA | NA | NA | NA | NA | Gm34411 | | 0 | 0 | 0 | 0 | 0 | 0 | 0 | 0 | 0 | 0 | 0 |
| 0 | NA | NA | NA | NA | NA | 1700025M24Rik | | 0 | 0 | 0 | 0 | 0 | 0 | 0 | 0 | 0 | 0 | 0 |
| 0 | NA | NA | NA | NA | NA | Gm42734 | | 0 | 0 | 0 | 0 | 0 | 0 | 0 | 0 | 0 | 0 | 0 |
| 0 | NA | NA | NA | NA | NA | Gm42733 | | 0 | 0 | 0 | 0 | 0 | 0 | 0 | 0 | 0 | 0 | 0 |
| 0 | NA | NA | NA | NA | NA | Gm34583 | | 0 | 0 | 0 | 0 | 0 | 0 | 0 | 0 | 0 | 0 | 0 |
| 0 | NA | NA | NA | NA | NA | Gm35960 | | 0 | 0 | 0 | 0 | 0 | 0 | 0 | 0 | 0 | 0 | 0 |
| 0 | NA | NA | NA | NA | NA | Gm43798 | | 0 | 0 | 0 | 0 | 0 | 0 | 0 | 0 | 0 | 0 | 0 |
| 0 | NA | NA | NA | NA | NA | Spata18 | spermatogenesis associated 18 | 0 | 0 | 0 | 0 | 0 | 0 | 0 | 0 | 0 | 0 | 0 |
| 0 | NA | NA | NA | NA | NA | 1700019F05Rik | | 0 | 0 | 0 | 0 | 0 | 0 | 0 | 0 | 0 | 0 | 0 |
| 0 | NA | NA | NA | NA | NA | C78283 | | 0 | 0 | 0 | 0 | 0 | 0 | 0 | 0 | 0 | 0 | 0 |
| 0 | NA | NA | NA | NA | NA | Usp46os1 | | 0 | 0 | 0 | 0 | 0 | 0 | 0 | 0 | 0 | 0 | 0 |
| 0 | NA | NA | NA | NA | NA | Gm43417 | | 0 | 0 | 0 | 0 | 0 | 0 | 0 | 0 | 0 | 0 | 0 |
| 0 | NA | NA | NA | NA | NA | Gm43415 | | 0 | 0 | 0 | 0 | 0 | 0 | 0 | 0 | 0 | 0 | 0 |
| 0 | NA | NA | NA | NA | NA | Gm43416 | | 0 | 0 | 0 | 0 | 0 | 0 | 0 | 0 | 0 | 0 | 0 |
| 0 | NA | NA | NA | NA | NA | Snora26 | | 0 | 0 | 0 | 0 | 0 | 0 | 0 | 0 | 0 | 0 | 0 |

| | | | | | | | | | | | | | | | | | | |
|---|---|---|---|---|---|---|---|---|---|---|---|---|---|---|---|---|---|---|
| 0 | NA | NA | NA | NA | NA | A330058E17Rik | | 0 | 0 | 0 | 0 | 0 | 0 | 0 | 0 | 0 | 0 | 0 |
| 0 | NA | NA | NA | NA | NA | Gm42599 | | 0 | 0 | 0 | 0 | 0 | 0 | 0 | 0 | 0 | 0 | 0 |
| 0 | NA | NA | NA | NA | NA | Gm15985 | | 0 | 0 | 0 | 0 | 0 | 0 | 0 | 0 | 0 | 0 | 0 |
| 0 | NA | NA | NA | NA | NA | Gm42575 | | 0 | 0 | 0 | 0 | 0 | 0 | 0 | 0 | 0 | 0 | 0 |
| 0 | NA | NA | NA | NA | NA | Gm17906 | | 0 | 0 | 0 | 0 | 0 | 0 | 0 | 0 | 0 | 0 | 0 |
| 0 | NA | NA | NA | NA | NA | Gm6116 | | 0 | 0 | 0 | 0 | 0 | 0 | 0 | 0 | 0 | 0 | 0 |
| 0 | NA | NA | NA | NA | NA | Gm18593 | | 0 | 0 | 0 | 0 | 0 | 0 | 0 | 0 | 0 | 0 | 0 |
| 0 | NA | NA | NA | NA | NA | Gsx2 | GS homeobox 2 | 0 | 0 | 0 | 0 | 0 | 0 | 0 | 0 | 0 | 0 | 0 |
| 0 | NA | NA | NA | NA | NA | Gm2040 | | 0 | 0 | 0 | 0 | 0 | 0 | 0 | 0 | 0 | 0 | 0 |
| 0 | NA | NA | NA | NA | NA | Gm19583 | | 0 | 0 | 0 | 0 | 0 | 0 | 0 | 0 | 0 | 0 | 0 |
| 0 | NA | NA | NA | NA | NA | Olfr1401-ps1 | | 0 | 0 | 0 | 0 | 0 | 0 | 0 | 0 | 0 | 0 | 0 |
| 0 | NA | NA | NA | NA | NA | Gm42803 | | 0 | 0 | 0 | 0 | 0 | 0 | 0 | 0 | 0 | 0 | 0 |
| 0 | NA | NA | NA | NA | NA | Mir7025 | | 0 | 0 | 0 | 0 | 0 | 0 | 0 | 0 | 0 | 0 | 0 |
| 0 | NA | NA | NA | NA | NA | Gm42799 | | 0 | 0 | 0 | 0 | 0 | 0 | 0 | 0 | 0 | 0 | 0 |
| 0 | NA | NA | NA | NA | NA | Gm22084 | | 0 | 0 | 0 | 0 | 0 | 0 | 0 | 0 | 0 | 0 | 0 |
| 0 | NA | NA | NA | NA | NA | Gm24502 | | 0 | 0 | 0 | 0 | 0 | 0 | 0 | 0 | 0 | 0 | 0 |
| 0 | NA | NA | NA | NA | NA | Gm43100 | | 0 | 0 | 0 | 0 | 0 | 0 | 0 | 0 | 0 | 0 | 0 |
| 0 | NA | NA | NA | NA | NA | Gm38180 | | 0 | 0 | 0 | 0 | 0 | 0 | 0 | 0 | 0 | 0 | 0 |
| 0 | NA | NA | NA | NA | NA | Gm28865 | | 0 | 0 | 0 | 0 | 0 | 0 | 0 | 0 | 0 | 0 | 0 |
| 0 | NA | NA | NA | NA | NA | Gm25374 | | 0 | 0 | 0 | 0 | 0 | 0 | 0 | 0 | 0 | 0 | 0 |
| 0 | NA | NA | NA | NA | NA | Gm24738 | | 0 | 0 | 0 | 0 | 0 | 0 | 0 | 0 | 0 | 0 | 0 |
| 0 | NA | NA | NA | NA | NA | Gm32780 | | 0 | 0 | 0 | 0 | 0 | 0 | 0 | 0 | 0 | 0 | 0 |
| 0 | NA | NA | NA | NA | NA | Pdcl2 | phosducin-like 2 | 0 | 0 | 0 | 0 | 0 | 0 | 0 | 0 | 0 | 0 | 0 |
| 0 | NA | NA | NA | NA | NA | Nmu | neuromedin U | 0 | 0 | 0 | 0 | 0 | 0 | 0 | 0 | 0 | 0 | 0 |
| 0 | NA | NA | NA | NA | NA | Gm42665 | | 0 | 0 | 0 | 0 | 0 | 0 | 0 | 0 | 0 | 0 | 0 |
| 0 | NA | NA | NA | NA | NA | Gm7271 | | 0 | 0 | 0 | 0 | 0 | 0 | 0 | 0 | 0 | 0 | 0 |
| 0 | NA | NA | NA | NA | NA | Gm7494 | | 0 | 0 | 0 | 0 | 0 | 0 | 0 | 0 | 0 | 0 | 0 |
| 0 | NA | NA | NA | NA | NA | 1700112J05Rik | | 0 | 0 | 0 | 0 | 0 | 0 | 0 | 0 | 0 | 0 | 0 |
| 0 | NA | NA | NA | NA | NA | Gm15831 | | 0 | 0 | 0 | 0 | 0 | 0 | 0 | 0 | 0 | 0 | 0 |
| 0 | NA | NA | NA | NA | NA | Chaer1 | | 0 | 0 | 0 | 0 | 0 | 0 | 0 | 0 | 0 | 0 | 0 |
| 0 | NA | NA | NA | NA | NA | Gm19060 | | 0 | 0 | 0 | 0 | 0 | 0 | 0 | 0 | 0 | 0 | 0 |
| 0 | NA | NA | NA | NA | NA | Mir5098 | | 0 | 0 | 0 | 0 | 0 | 0 | 0 | 0 | 0 | 0 | 0 |
| 0 | NA | NA | NA | NA | NA | Gm42758 | | 0 | 0 | 0 | 0 | 0 | 0 | 0 | 0 | 0 | 0 | 0 |
| 0 | NA | NA | NA | NA | NA | Gm34648 | | 0 | 0 | 0 | 0 | 0 | 0 | 0 | 0 | 0 | 0 | 0 |
| 0 | NA | NA | NA | NA | NA | Gm34728 | | 0 | 0 | 0 | 0 | 0 | 0 | 0 | 0 | 0 | 0 | 0 |
| 0 | NA | NA | NA | NA | NA | 1700017L05Rik | | 0 | 0 | 0 | 0 | 0 | 0 | 0 | 0 | 0 | 0 | 0 |
| 0 | NA | NA | NA | NA | NA | Gm42761 | | 0 | 0 | 0 | 0 | 0 | 0 | 0 | 0 | 0 | 0 | 0 |
| 0 | NA | NA | NA | NA | NA | Pea15b | | 0 | 0 | 0 | 0 | 0 | 0 | 0 | 0 | 0 | 0 | 0 |
| 0 | NA | NA | NA | NA | NA | Gm22011 | | 0 | 0 | 0 | 0 | 0 | 0 | 0 | 0 | 0 | 0 | 0 |
| 0 | NA | NA | NA | NA | NA | Gm31049 | | 0 | 0 | 0 | 0 | 0 | 0 | 0 | 0 | 0 | 0 | 0 |
| 0 | NA | NA | NA | NA | NA | Gm42673 | | 0 | 0 | 0 | 0 | 0 | 0 | 0 | 0 | 0 | 0 | 0 |
| 0 | NA | NA | NA | NA | NA | Mthfr-ps1 | | 0 | 0 | 0 | 0 | 0 | 0 | 0 | 0 | 0 | 0 | 0 |
| 0 | NA | NA | NA | NA | NA | Gm43232 | | 0 | 0 | 0 | 0 | 0 | 0 | 0 | 0 | 0 | 0 | 0 |
| 0 | NA | NA | NA | NA | NA | Gm25756 | | 0 | 0 | 0 | 0 | 0 | 0 | 0 | 0 | 0 | 0 | 0 |
| 0 | NA | NA | NA | NA | NA | Gm43234 | | 0 | 0 | 0 | 0 | 0 | 0 | 0 | 0 | 0 | 0 | 0 |
| 0 | NA | NA | NA | NA | NA | Gm18452 | | 0 | 0 | 0 | 0 | 0 | 0 | 0 | 0 | 0 | 0 | 0 |
| 0 | NA | NA | NA | NA | NA | Gm22553 | | 0 | 0 | 0 | 0 | 0 | 0 | 0 | 0 | 0 | 0 | 0 |
| 0 | NA | NA | NA | NA | NA | Gm23429 | | 0 | 0 | 0 | 0 | 0 | 0 | 0 | 0 | 0 | 0 | 0 |
| 0 | NA | NA | NA | NA | NA | Gm24304 | | 0 | 0 | 0 | 0 | 0 | 0 | 0 | 0 | 0 | 0 | 0 |
| 0 | NA | NA | NA | NA | NA | Gm22974 | | 0 | 0 | 0 | 0 | 0 | 0 | 0 | 0 | 0 | 0 | 0 |
| 0 | NA | NA | NA | NA | NA | Gm43594 | | 0 | 0 | 0 | 0 | 0 | 0 | 0 | 0 | 0 | 0 | 0 |
| 0 | NA | NA | NA | NA | NA | Gm22048 | | 0 | 0 | 0 | 0 | 0 | 0 | 0 | 0 | 0 | 0 | 0 |
| 0 | NA | NA | NA | NA | NA | 1700008H02Rik | | 0 | 0 | 0 | 0 | 0 | 0 | 0 | 0 | 0 | 0 | 0 |
| 0 | NA | NA | NA | NA | NA | 1700031L13Rik | | 0 | 0 | 0 | 0 | 0 | 0 | 0 | 0 | 0 | 0 | 0 |
| 0 | NA | NA | NA | NA | NA | Mir1187 | | 0 | 0 | 0 | 0 | 0 | 0 | 0 | 0 | 0 | 0 | 0 |
| 0 | NA | NA | NA | NA | NA | Gm24129 | | 0 | 0 | 0 | 0 | 0 | 0 | 0 | 0 | 0 | 0 | 0 |
| 0 | NA | NA | NA | NA | NA | Gm43217 | | 0 | 0 | 0 | 0 | 0 | 0 | 0 | 0 | 0 | 0 | 0 |
| 0 | NA | NA | NA | NA | NA | Gm15626 | | 0 | 0 | 0 | 0 | 0 | 0 | 0 | 0 | 0 | 0 | 0 |
| 0 | NA | NA | NA | NA | NA | Gm44303 | | 0 | 0 | 0 | 0 | 0 | 0 | 0 | 0 | 0 | 0 | 0 |
| 0 | NA | NA | NA | NA | NA | Gm25765 | | 0 | 0 | 0 | 0 | 0 | 0 | 0 | 0 | 0 | 0 | 0 |

| | | | | | | | | | | | | | | | | | |
|---|---|---|---|---|---|---|---|---|---|---|---|---|---|---|---|---|---|
| 0 | NA | NA | NA | NA | NA | Gm25806 | | 0 | 0 | 0 | 0 | 0 | 0 | 0 | 0 | 0 | 0 | 0 |
| 0 | NA | NA | NA | NA | NA | Gm24792 | | 0 | 0 | 0 | 0 | 0 | 0 | 0 | 0 | 0 | 0 | 0 |
| 0 | NA | NA | NA | NA | NA | Gm43532 | | 0 | 0 | 0 | 0 | 0 | 0 | 0 | 0 | 0 | 0 | 0 |
| 0 | NA | NA | NA | NA | NA | Gm43531 | | 0 | 0 | 0 | 0 | 0 | 0 | 0 | 0 | 0 | 0 | 0 |
| 0 | NA | NA | NA | NA | NA | Gm21006 | | 0 | 0 | 0 | 0 | 0 | 0 | 0 | 0 | 0 | 0 | 0 |
| 0 | NA | NA | NA | NA | NA | Gm43567 | | 0 | 0 | 0 | 0 | 0 | 0 | 0 | 0 | 0 | 0 | 0 |
| 0 | NA | NA | NA | NA | NA | Gm24626 | | 0 | 0 | 0 | 0 | 0 | 0 | 0 | 0 | 0 | 0 | 0 |
| 0 | NA | NA | NA | NA | NA | Gm43564 | | 0 | 0 | 0 | 0 | 0 | 0 | 0 | 0 | 0 | 0 | 0 |
| 0 | NA | NA | NA | NA | NA | Gm24524 | | 0 | 0 | 0 | 0 | 0 | 0 | 0 | 0 | 0 | 0 | 0 |
| 0 | NA | NA | NA | NA | NA | Tmprss11c | transmembrane protease, serine 11c | 0 | 0 | 0 | 0 | 0 | 0 | 0 | 0 | 0 | 0 | 0 |
| 0 | NA | NA | NA | NA | NA | Gm25473 | | 0 | 0 | 0 | 0 | 0 | 0 | 0 | 0 | 0 | 0 | 0 |
| 0 | NA | NA | NA | NA | NA | Tmprss11d | transmembrane protease, serine 11d | 0 | 0 | 0 | 0 | 0 | 0 | 0 | 0 | 0 | 0 | 0 |
| 0 | NA | NA | NA | NA | NA | Tmprss11a | transmembrane protease, serine 11a | 0 | 0 | 0 | 0 | 0 | 0 | 0 | 0 | 0 | 0 | 0 |
| 0 | NA | NA | NA | NA | NA | Tmprss11g | transmembrane protease, serine 11g | 0 | 0 | 0 | 0 | 0 | 0 | 0 | 0 | 0 | 0 | 0 |
| 0 | NA | NA | NA | NA | NA | Tmprss11f | transmembrane protease, serine 11f | 0 | 0 | 0 | 0 | 0 | 0 | 0 | 0 | 0 | 0 | 0 |
| 0 | NA | NA | NA | NA | NA | Tmprss11b | transmembrane protease, serine 11B | 0 | 0 | 0 | 0 | 0 | 0 | 0 | 0 | 0 | 0 | 0 |
| 0 | NA | NA | NA | NA | NA | Gm43056 | | 0 | 0 | 0 | 0 | 0 | 0 | 0 | 0 | 0 | 0 | 0 |
| 0 | NA | NA | NA | NA | NA | Tmprss11e | transmembrane protease, serine 11e | 0 | 0 | 0 | 0 | 0 | 0 | 0 | 0 | 0 | 0 | 0 |
| 0 | NA | NA | NA | NA | NA | Gm16579 | | 0 | 0 | 0 | 0 | 0 | 0 | 0 | 0 | 0 | 0 | 0 |
| 0 | NA | NA | NA | NA | NA | Gm43055 | | 0 | 0 | 0 | 0 | 0 | 0 | 0 | 0 | 0 | 0 | 0 |
| 0 | NA | NA | NA | NA | NA | Gm43057 | | 0 | 0 | 0 | 0 | 0 | 0 | 0 | 0 | 0 | 0 | 0 |
| 0 | NA | NA | NA | NA | NA | Gm25211 | | 0 | 0 | 0 | 0 | 0 | 0 | 0 | 0 | 0 | 0 | 0 |
| 0 | NA | NA | NA | NA | NA | Ugt2b1 | UDP glucuronosyltransferase 2 family, pc | 0 | 0 | 0 | 0 | 0 | 0 | 0 | 0 | 0 | 0 | 0 |
| 0 | NA | NA | NA | NA | NA | Gm24121 | | 0 | 0 | 0 | 0 | 0 | 0 | 0 | 0 | 0 | 0 | 0 |
| 0 | NA | NA | NA | NA | NA | Gm20121 | | 0 | 0 | 0 | 0 | 0 | 0 | 0 | 0 | 0 | 0 | 0 |
| 0 | NA | NA | NA | NA | NA | | | 0 | 0 | 0 | 0 | 0 | 0 | 0 | 0 | 0 | 0 | 0 |
| 0 | NA | NA | NA | NA | NA | Gm7631 | | 0 | 0 | 0 | 0 | 0 | 0 | 0 | 0 | 0 | 0 | 0 |
| 0 | NA | NA | NA | NA | NA | Gm18635 | | 0 | 0 | 0 | 0 | 0 | 0 | 0 | 0 | 0 | 0 | 0 |
| 0 | NA | NA | NA | NA | NA | Gm42465 | | 0 | 0 | 0 | 0 | 0 | 0 | 0 | 0 | 0 | 0 | 0 |
| 0 | NA | NA | NA | NA | NA | Gm31230 | | 0 | 0 | 0 | 0 | 0 | 0 | 0 | 0 | 0 | 0 | 0 |
| 0 | NA | NA | NA | NA | NA | Gm42795 | | 0 | 0 | 0 | 0 | 0 | 0 | 0 | 0 | 0 | 0 | 0 |
| 0 | NA | NA | NA | NA | NA | Ugt2b5 | UDP glucuronosyltransferase 2 family, pc | 0 | 0 | 0 | 0 | 0 | 0 | 0 | 0 | 0 | 0 | 0 |
| 0 | NA | NA | NA | NA | NA | Gm5717 | | 0 | 0 | 0 | 0 | 0 | 0 | 0 | 0 | 0 | 0 | 0 |
| 0 | NA | NA | NA | NA | NA | Gm43369 | | 0 | 0 | 0 | 0 | 0 | 0 | 0 | 0 | 0 | 0 | 0 |
| 0 | NA | NA | NA | NA | NA | Gm19418 | | 0 | 0 | 0 | 0 | 0 | 0 | 0 | 0 | 0 | 0 | 0 |
| 0 | NA | NA | NA | NA | NA | Ugt2b37 | UDP glucuronosyltransferase 2 family, pc | 0 | 0 | 0 | 0 | 0 | 0 | 0 | 0 | 0 | 0 | 0 |
| 0 | NA | NA | NA | NA | NA | Gm7646 | | 0 | 0 | 0 | 0 | 0 | 0 | 0 | 0 | 0 | 0 | 0 |
| 0 | NA | NA | NA | NA | NA | Gm7652 | | 0 | 0 | 0 | 0 | 0 | 0 | 0 | 0 | 0 | 0 | 0 |
| 0 | NA | NA | NA | NA | NA | Gm42796 | | 0 | 0 | 0 | 0 | 0 | 0 | 0 | 0 | 0 | 0 | 0 |
| 0 | NA | NA | NA | NA | NA | Gm35570 | | 0 | 0 | 0 | 0 | 0 | 0 | 0 | 0 | 0 | 0 | 0 |
| 0 | NA | NA | NA | NA | NA | Gm21049 | | 0 | 0 | 0 | 0 | 0 | 0 | 0 | 0 | 0 | 0 | 0 |
| 0 | NA | NA | NA | NA | NA | Gm43370 | | 0 | 0 | 0 | 0 | 0 | 0 | 0 | 0 | 0 | 0 | 0 |
| 0 | NA | NA | NA | NA | NA | Ugt2b38 | UDP glucuronosyltransferase 2 family, pc | 0 | 0 | 0 | 0 | 0 | 0 | 0 | 0 | 0 | 0 | 0 |
| 0 | NA | NA | NA | NA | NA | Ugt2a2 | UDP glucuronosyltransferase 2 family, pc | 0 | 0 | 0 | 0 | 0 | 0 | 0 | 0 | 0 | 0 | 0 |
| 0 | NA | NA | NA | NA | NA | Ugt2a1 | UDP glucuronosyltransferase 2 family, pc | 0 | 0 | 0 | 0 | 0 | 0 | 0 | 0 | 0 | 0 | 0 |
| 0 | NA | NA | NA | NA | NA | Gm43638 | | 0 | 0 | 0 | 0 | 0 | 0 | 0 | 0 | 0 | 0 | 0 |
| 0 | NA | NA | NA | NA | NA | Mir3969 | | 0 | 0 | 0 | 0 | 0 | 0 | 0 | 0 | 0 | 0 | 0 |
| 0 | NA | NA | NA | NA | NA | Gm21043 | | 0 | 0 | 0 | 0 | 0 | 0 | 0 | 0 | 0 | 0 | 0 |
| 0 | NA | NA | NA | NA | NA | Csn1s1 | casein alpha s1 | 0 | 0 | 0 | 0 | 0 | 0 | 0 | 0 | 0 | 0 | 0 |
| 0 | NA | NA | NA | NA | NA | Csn2 | casein beta | 0 | 0 | 0 | 0 | 0 | 0 | 0 | 0 | 0 | 0 | 0 |
| 0 | NA | NA | NA | NA | NA | Gm25066 | | 0 | 0 | 0 | 0 | 0 | 0 | 0 | 0 | 0 | 0 | 0 |
| 0 | NA | NA | NA | NA | NA | Gm42845 | | 0 | 0 | 0 | 0 | 0 | 0 | 0 | 0 | 0 | 0 | 0 |
| 0 | NA | NA | NA | NA | NA | Gm24314 | | 0 | 0 | 0 | 0 | 0 | 0 | 0 | 0 | 0 | 0 | 0 |
| 0 | NA | NA | NA | NA | NA | Csn1s2a | casein alpha s2-like A | 0 | 0 | 0 | 0 | 0 | 0 | 0 | 0 | 0 | 0 | 0 |
| 0 | NA | NA | NA | NA | NA | Prr27 | proline rich 27 | 0 | 0 | 0 | 0 | 0 | 0 | 0 | 0 | 0 | 0 | 0 |
| 0 | NA | NA | NA | NA | NA | Gm42794 | | 0 | 0 | 0 | 0 | 0 | 0 | 0 | 0 | 0 | 0 | 0 |
| 0 | NA | NA | NA | NA | NA | Gm7337 | | 0 | 0 | 0 | 0 | 0 | 0 | 0 | 0 | 0 | 0 | 0 |
| 0 | NA | NA | NA | NA | NA | Odam | odontogenic, ameloblast asssociated | 0 | 0 | 0 | 0 | 0 | 0 | 0 | 0 | 0 | 0 | 0 |
| 0 | NA | NA | NA | NA | NA | 1700066N21Rik | | 0 | 0 | 0 | 0 | 0 | 0 | 0 | 0 | 0 | 0 | 0 |
| 0 | NA | NA | NA | NA | NA | BC037156 | cDNA sequence BC037156 | 0 | 0 | 0 | 0 | 0 | 0 | 0 | 0 | 0 | 0 | 0 |

| | | | | | | | | | | | | | | | | | |
|---|---|---|---|---|---|---|---|---|---|---|---|---|---|---|---|---|---|
| 0 | NA | NA | NA | NA | NA | BC051076 | | 0 | 0 | 0 | 0 | 0 | 0 | 0 | 0 | 0 | 0 | 0 |
| 0 | NA | NA | NA | NA | NA | 2310003L06Rik | | 0 | 0 | 0 | 0 | 0 | 0 | 0 | 0 | 0 | 0 | 0 |
| 0 | NA | NA | NA | NA | NA | Gm28434 | | 0 | 0 | 0 | 0 | 0 | 0 | 0 | 0 | 0 | 0 | 0 |
| 0 | NA | NA | NA | NA | NA | Cabs1 | calcium binding protein, spermatid specifi | 0 | 0 | 0 | 0 | 0 | 0 | 0 | 0 | 0 | 0 | 0 |
| 0 | NA | NA | NA | NA | NA | Smr3a | submaxillary gland androgen regulated p | 0 | 0 | 0 | 0 | 0 | 0 | 0 | 0 | 0 | 0 | 0 |
| 0 | NA | NA | NA | NA | NA | Gm7709 | | 0 | 0 | 0 | 0 | 0 | 0 | 0 | 0 | 0 | 0 | 0 |
| 0 | NA | NA | NA | NA | NA | Gm42623 | | 0 | 0 | 0 | 0 | 0 | 0 | 0 | 0 | 0 | 0 | 0 |
| 0 | NA | NA | NA | NA | NA | Smr2 | submaxillary gland androgen regulated p | 0 | 0 | 0 | 0 | 0 | 0 | 0 | 0 | 0 | 0 | 0 |
| 0 | NA | NA | NA | NA | NA | Gm11116 | | 0 | 0 | 0 | 0 | 0 | 0 | 0 | 0 | 0 | 0 | 0 |
| 0 | NA | NA | NA | NA | NA | Gm42622 | | 0 | 0 | 0 | 0 | 0 | 0 | 0 | 0 | 0 | 0 | 0 |
| 0 | NA | NA | NA | NA | NA | Gm11115 | | 0 | 0 | 0 | 0 | 0 | 0 | 0 | 0 | 0 | 0 | 0 |
| 0 | NA | NA | NA | NA | NA | Gm42621 | | 0 | 0 | 0 | 0 | 0 | 0 | 0 | 0 | 0 | 0 | 0 |
| 0 | NA | NA | NA | NA | NA | Gm42164 | | 0 | 0 | 0 | 0 | 0 | 0 | 0 | 0 | 0 | 0 | 0 |
| 0 | NA | NA | NA | NA | NA | Gm7714 | predicted gene 7714 | 0 | 0 | 0 | 0 | 0 | 0 | 0 | 0 | 0 | 0 | 0 |
| 0 | NA | NA | NA | NA | NA | Prol1 | proline rich, lacrimal 1 | 0 | 0 | 0 | 0 | 0 | 0 | 0 | 0 | 0 | 0 | 0 |
| 0 | NA | NA | NA | NA | NA | Amtn | amelotin | 0 | 0 | 0 | 0 | 0 | 0 | 0 | 0 | 0 | 0 | 0 |
| 0 | NA | NA | NA | NA | NA | Gm43431 | | 0 | 0 | 0 | 0 | 0 | 0 | 0 | 0 | 0 | 0 | 0 |
| 0 | NA | NA | NA | NA | NA | Ambn | ameloblastin | 0 | 0 | 0 | 0 | 0 | 0 | 0 | 0 | 0 | 0 | 0 |
| 0 | NA | NA | NA | NA | NA | Enam | enamelin | 0 | 0 | 0 | 0 | 0 | 0 | 0 | 0 | 0 | 0 | 0 |
| 0 | NA | NA | NA | NA | NA | Gm42912 | | 0 | 0 | 0 | 0 | 0 | 0 | 0 | 0 | 0 | 0 | 0 |
| 0 | NA | NA | NA | NA | NA | Gm42911 | | 0 | 0 | 0 | 0 | 0 | 0 | 0 | 0 | 0 | 0 | 0 |
| 0 | NA | NA | NA | NA | NA | Gm43515 | | 0 | 0 | 0 | 0 | 0 | 0 | 0 | 0 | 0 | 0 | 0 |
| 0 | NA | NA | NA | NA | NA | Gm25320 | | 0 | 0 | 0 | 0 | 0 | 0 | 0 | 0 | 0 | 0 | 0 |
| 0 | NA | NA | NA | NA | NA | Gm25758 | | 0 | 0 | 0 | 0 | 0 | 0 | 0 | 0 | 0 | 0 | 0 |
| 0 | NA | NA | NA | NA | NA | Gm43363 | | 0 | 0 | 0 | 0 | 0 | 0 | 0 | 0 | 0 | 0 | 0 |
| 0 | NA | NA | NA | NA | NA | Gm43449 | | 0 | 0 | 0 | 0 | 0 | 0 | 0 | 0 | 0 | 0 | 0 |
| 0 | NA | NA | NA | NA | NA | Gm42109 | | 0 | 0 | 0 | 0 | 0 | 0 | 0 | 0 | 0 | 0 | 0 |
| 0 | NA | NA | NA | NA | NA | Gm2602 | | 0 | 0 | 0 | 0 | 0 | 0 | 0 | 0 | 0 | 0 | 0 |
| 0 | NA | NA | NA | NA | NA | Gm43090 | | 0 | 0 | 0 | 0 | 0 | 0 | 0 | 0 | 0 | 0 | 0 |
| 0 | NA | NA | NA | NA | NA | Cxcl5 | chemokine (C-X-C motif) ligand 5 | 0 | 0 | 0 | 0 | 0 | 0 | 0 | 0 | 0 | 0 | 0 |
| 0 | NA | NA | NA | NA | NA | Cxcl15 | chemokine (C-X-C motif) ligand 15 | 0 | 0 | 0 | 0 | 0 | 0 | 0 | 0 | 0 | 0 | 0 |
| 0 | NA | NA | NA | NA | NA | Gm24322 | | 0 | 0 | 0 | 0 | 0 | 0 | 0 | 0 | 0 | 0 | 0 |
| 0 | NA | NA | NA | NA | NA | Gm43085 | | 0 | 0 | 0 | 0 | 0 | 0 | 0 | 0 | 0 | 0 | 0 |
| 0 | NA | NA | NA | NA | NA | Gm43086 | | 0 | 0 | 0 | 0 | 0 | 0 | 0 | 0 | 0 | 0 | 0 |
| 0 | NA | NA | NA | NA | NA | Gm22816 | | 0 | 0 | 0 | 0 | 0 | 0 | 0 | 0 | 0 | 0 | 0 |
| 0 | NA | NA | NA | NA | NA | Gm42530 | | 0 | 0 | 0 | 0 | 0 | 0 | 0 | 0 | 0 | 0 | 0 |
| 0 | NA | NA | NA | NA | NA | Epgn | epithelial mitogen | 0 | 0 | 0 | 0 | 0 | 0 | 0 | 0 | 0 | 0 | 0 |
| 0 | NA | NA | NA | NA | NA | Gm19610 | | 0 | 0 | 0 | 0 | 0 | 0 | 0 | 0 | 0 | 0 | 0 |
| 0 | NA | NA | NA | NA | NA | Gm23092 | | 0 | 0 | 0 | 0 | 0 | 0 | 0 | 0 | 0 | 0 | 0 |
| 0 | NA | NA | NA | NA | NA | Gm19619 | | 0 | 0 | 0 | 0 | 0 | 0 | 0 | 0 | 0 | 0 | 0 |
| 0 | NA | NA | NA | NA | NA | Gm28271 | | 0 | 0 | 0 | 0 | 0 | 0 | 0 | 0 | 0 | 0 | 0 |
| 0 | NA | NA | NA | NA | NA | 1700063O14Rik | | 0 | 0 | 0 | 0 | 0 | 0 | 0 | 0 | 0 | 0 | 0 |
| 0 | NA | NA | NA | NA | NA | Gm19649 | | 0 | 0 | 0 | 0 | 0 | 0 | 0 | 0 | 0 | 0 | 0 |
| 0 | NA | NA | NA | NA | NA | Gm43596 | | 0 | 0 | 0 | 0 | 0 | 0 | 0 | 0 | 0 | 0 | 0 |
| 0 | NA | NA | NA | NA | NA | Gm43688 | | 0 | 0 | 0 | 0 | 0 | 0 | 0 | 0 | 0 | 0 | 0 |
| 0 | NA | NA | NA | NA | NA | Gm24113 | | 0 | 0 | 0 | 0 | 0 | 0 | 0 | 0 | 0 | 0 | 0 |
| 0 | NA | NA | NA | NA | NA | Gm43040 | | 0 | 0 | 0 | 0 | 0 | 0 | 0 | 0 | 0 | 0 | 0 |
| 0 | NA | NA | NA | NA | NA | U90926 | | 0 | 0 | 0 | 0 | 0 | 0 | 0 | 0 | 0 | 0 | 0 |
| 0 | NA | NA | NA | NA | NA | Gm23031 | | 0 | 0 | 0 | 0 | 0 | 0 | 0 | 0 | 0 | 0 | 0 |
| 0 | NA | NA | NA | NA | NA | Gm2673 | | 0 | 0 | 0 | 0 | 0 | 0 | 0 | 0 | 0 | 0 | 0 |
| 0 | NA | NA | NA | NA | NA | Gm26039 | | 0 | 0 | 0 | 0 | 0 | 0 | 0 | 0 | 0 | 0 | 0 |
| 0 | NA | NA | NA | NA | NA | Mir6415 | | 0 | 0 | 0 | 0 | 0 | 0 | 0 | 0 | 0 | 0 | 0 |
| 0 | NA | NA | NA | NA | NA | Gm20500 | | 0 | 0 | 0 | 0 | 0 | 0 | 0 | 0 | 0 | 0 | 0 |
| 0 | NA | NA | NA | NA | NA | Mir1961 | | 0 | 0 | 0 | 0 | 0 | 0 | 0 | 0 | 0 | 0 | 0 |
| 0 | NA | NA | NA | NA | NA | Gm25521 | | 0 | 0 | 0 | 0 | 0 | 0 | 0 | 0 | 0 | 0 | 0 |
| 0 | NA | NA | NA | NA | NA | Gm22915 | | 0 | 0 | 0 | 0 | 0 | 0 | 0 | 0 | 0 | 0 | 0 |
| 0 | NA | NA | NA | NA | NA | Gm18118 | | 0 | 0 | 0 | 0 | 0 | 0 | 0 | 0 | 0 | 0 | 0 |
| 0 | NA | NA | NA | NA | NA | Gm43224 | | 0 | 0 | 0 | 0 | 0 | 0 | 0 | 0 | 0 | 0 | 0 |
| 0 | NA | NA | NA | NA | NA | Gm16427 | predicted gene 16427 | 0 | 0 | 0 | 0 | 0 | 0 | 0 | 0 | 0 | 0 | 0 |
| 0 | NA | NA | NA | NA | NA | C87414 | expressed sequence C87414 | 0 | 0 | 0 | 0 | 0 | 0 | 0 | 0 | 0 | 0 | 0 |

| | | | | | | | | | | | | | | | | | | |
|---|---|---|---|---|---|---|---|---|---|---|---|---|---|---|---|---|---|---|
| 0 | NA | NA | NA | NA | NA | Gm43225 | | 0 | 0 | 0 | 0 | 0 | 0 | 0 | 0 | 0 | 0 | 0 |
| 0 | NA | NA | NA | NA | NA | Gm43226 | | 0 | 0 | 0 | 0 | 0 | 0 | 0 | 0 | 0 | 0 | 0 |
| 0 | NA | NA | NA | NA | NA | Gm7919 | | 0 | 0 | 0 | 0 | 0 | 0 | 0 | 0 | 0 | 0 | 0 |
| 0 | NA | NA | NA | NA | NA | Gm43227 | | 0 | 0 | 0 | 0 | 0 | 0 | 0 | 0 | 0 | 0 | 0 |
| 0 | NA | NA | NA | NA | NA | Gm7792 | predicted gene 7792 | 0 | 0 | 0 | 0 | 0 | 0 | 0 | 0 | 0 | 0 | 0 |
| 0 | NA | NA | NA | NA | NA | Gm16429 | predicted gene 16429 | 0 | 0 | 0 | 0 | 0 | 0 | 0 | 0 | 0 | 0 | 0 |
| 0 | NA | NA | NA | NA | NA | Gm6346 | predicted gene 6346 | 0 | 0 | 0 | 0 | 0 | 0 | 0 | 0 | 0 | 0 | 0 |
| 0 | NA | NA | NA | NA | NA | Gm3089 | | 0 | 0 | 0 | 0 | 0 | 0 | 0 | 0 | 0 | 0 | 0 |
| 0 | NA | NA | NA | NA | NA | BC061212 | cDNA sequence BC061212 | 0 | 0 | 0 | 0 | 0 | 0 | 0 | 0 | 0 | 0 | 0 |
| 0 | NA | NA | NA | NA | NA | Gm6351 | predicted gene 6351 | 0 | 0 | 0 | 0 | 0 | 0 | 0 | 0 | 0 | 0 | 0 |
| 0 | NA | NA | NA | NA | NA | Gm3106 | predicted gene 3106 | 0 | 0 | 0 | 0 | 0 | 0 | 0 | 0 | 0 | 0 | 0 |
| 0 | NA | NA | NA | NA | NA | A430089I19Rik | RIKEN cDNA A430089I19 gene | 0 | 0 | 0 | 0 | 0 | 0 | 0 | 0 | 0 | 0 | 0 |
| 0 | NA | NA | NA | NA | NA | Gm6502 | predicted gene 6502 | 0 | 0 | 0 | 0 | 0 | 0 | 0 | 0 | 0 | 0 | 0 |
| 0 | NA | NA | NA | NA | NA | AA792892 | expressed sequence AA792892 | 0 | 0 | 0 | 0 | 0 | 0 | 0 | 0 | 0 | 0 | 0 |
| 0 | NA | NA | NA | NA | NA | BC080696 | | 0 | 0 | 0 | 0 | 0 | 0 | 0 | 0 | 0 | 0 | 0 |
| 0 | NA | NA | NA | NA | NA | Gm7682 | PRAME family member 8-like | 0 | 0 | 0 | 0 | 0 | 0 | 0 | 0 | 0 | 0 | 0 |
| 0 | NA | NA | NA | NA | NA | Gm3139 | predicted gene 3139 | 0 | 0 | 0 | 0 | 0 | 0 | 0 | 0 | 0 | 0 | 0 |
| 0 | NA | NA | NA | NA | NA | Gm3147 | predicted gene 3147 | 0 | 0 | 0 | 0 | 0 | 0 | 0 | 0 | 0 | 0 | 0 |
| 0 | NA | NA | NA | NA | NA | Gm6509 | predicted gene 6509 | 0 | 0 | 0 | 0 | 0 | 0 | 0 | 0 | 0 | 0 | 0 |
| 0 | NA | NA | NA | NA | NA | Gm6205 | | 0 | 0 | 0 | 0 | 0 | 0 | 0 | 0 | 0 | 0 | 0 |
| 0 | NA | NA | NA | NA | NA | Gm42561 | | 0 | 0 | 0 | 0 | 0 | 0 | 0 | 0 | 0 | 0 | 0 |
| 0 | NA | NA | NA | NA | NA | Gm7663 | | 0 | 0 | 0 | 0 | 0 | 0 | 0 | 0 | 0 | 0 | 0 |
| 0 | NA | NA | NA | NA | NA | Gm3183 | predicted gene 3183 | 0 | 0 | 0 | 0 | 0 | 0 | 0 | 0 | 0 | 0 | 0 |
| 0 | NA | NA | NA | NA | NA | Gm3176 | | 0 | 0 | 0 | 0 | 0 | 0 | 0 | 0 | 0 | 0 | 0 |
| 0 | NA | NA | NA | NA | NA | Gm7647 | | 0 | 0 | 0 | 0 | 0 | 0 | 0 | 0 | 0 | 0 | 0 |
| 0 | NA | NA | NA | NA | NA | Gm6367 | | 0 | 0 | 0 | 0 | 0 | 0 | 0 | 0 | 0 | 0 | 0 |
| 0 | NA | NA | NA | NA | NA | Gm42661 | | 0 | 0 | 0 | 0 | 0 | 0 | 0 | 0 | 0 | 0 | 0 |
| 0 | NA | NA | NA | NA | NA | Gm7939 | | 0 | 0 | 0 | 0 | 0 | 0 | 0 | 0 | 0 | 0 | 0 |
| 0 | NA | NA | NA | NA | NA | Gm7942 | PRAME family member 8-like | 0 | 0 | 0 | 0 | 0 | 0 | 0 | 0 | 0 | 0 | 0 |
| 0 | NA | NA | NA | NA | NA | Gm16513 | predicted gene, Gm16513 | 0 | 0 | 0 | 0 | 0 | 0 | 0 | 0 | 0 | 0 | 0 |
| 0 | NA | NA | NA | NA | NA | Gm10424 | predicted gene 10424 | 0 | 0 | 0 | 0 | 0 | 0 | 0 | 0 | 0 | 0 | 0 |
| 0 | NA | NA | NA | NA | NA | Gm6468 | predicted gene 6468 | 0 | 0 | 0 | 0 | 0 | 0 | 0 | 0 | 0 | 0 | 0 |
| 0 | NA | NA | NA | NA | NA | Gm3259 | predicted gene 3259 | 0 | 0 | 0 | 0 | 0 | 0 | 0 | 0 | 0 | 0 | 0 |
| 0 | NA | NA | NA | NA | NA | Gm32072 | | 0 | 0 | 0 | 0 | 0 | 0 | 0 | 0 | 0 | 0 | 0 |
| 0 | NA | NA | NA | NA | NA | Gm7961 | | 0 | 0 | 0 | 0 | 0 | 0 | 0 | 0 | 0 | 0 | 0 |
| 0 | NA | NA | NA | NA | NA | Gm42662 | | 0 | 0 | 0 | 0 | 0 | 0 | 0 | 0 | 0 | 0 | 0 |
| 0 | NA | NA | NA | NA | NA | D5Ertd577e | DNA segment, Chr 5, ERATO Doi 577, ex | 0 | 0 | 0 | 0 | 0 | 0 | 0 | 0 | 0 | 0 | 0 |
| 0 | NA | NA | NA | NA | NA | Gm3286 | predicted gene 3286 | 0 | 0 | 0 | 0 | 0 | 0 | 0 | 0 | 0 | 0 | 0 |
| 0 | NA | NA | NA | NA | NA | Gm7971 | predicted gene 7971 | 0 | 0 | 0 | 0 | 0 | 0 | 0 | 0 | 0 | 0 | 0 |
| 0 | NA | NA | NA | NA | NA | Gm6348 | predicted gene 6348 | 0 | 0 | 0 | 0 | 0 | 0 | 0 | 0 | 0 | 0 | 0 |
| 0 | NA | NA | NA | NA | NA | Gm3302 | | 0 | 0 | 0 | 0 | 0 | 0 | 0 | 0 | 0 | 0 | 0 |
| 0 | NA | NA | NA | NA | NA | Gm7978 | predicted gene 7978 | 0 | 0 | 0 | 0 | 0 | 0 | 0 | 0 | 0 | 0 | 0 |
| 0 | NA | NA | NA | NA | NA | Gm7982 | predicted gene 7982 | 0 | 0 | 0 | 0 | 0 | 0 | 0 | 0 | 0 | 0 | 0 |
| 0 | NA | NA | NA | NA | NA | E330014E10Ril | RIKEN cDNA E330014E10 gene | 0 | 0 | 0 | 0 | 0 | 0 | 0 | 0 | 0 | 0 | 0 |
| 0 | NA | NA | NA | NA | NA | Gm42714 | | 0 | 0 | 0 | 0 | 0 | 0 | 0 | 0 | 0 | 0 | 0 |
| 0 | NA | NA | NA | NA | NA | Gm25937 | | 0 | 0 | 0 | 0 | 0 | 0 | 0 | 0 | 0 | 0 | 0 |
| 0 | NA | NA | NA | NA | NA | Gm2840 | | 0 | 0 | 0 | 0 | 0 | 0 | 0 | 0 | 0 | 0 | 0 |
| 0 | NA | NA | NA | NA | NA | Gm42605 | | 0 | 0 | 0 | 0 | 0 | 0 | 0 | 0 | 0 | 0 | 0 |
| 0 | NA | NA | NA | NA | NA | Gm42603 | | 0 | 0 | 0 | 0 | 0 | 0 | 0 | 0 | 0 | 0 | 0 |
| 0 | NA | NA | NA | NA | NA | 1700016F12Rik | | 0 | 0 | 0 | 0 | 0 | 0 | 0 | 0 | 0 | 0 | 0 |
| 0 | NA | NA | NA | NA | NA | Gm42602 | | 0 | 0 | 0 | 0 | 0 | 0 | 0 | 0 | 0 | 0 | 0 |
| 0 | NA | NA | NA | NA | NA | Gm24623 | | 0 | 0 | 0 | 0 | 0 | 0 | 0 | 0 | 0 | 0 | 0 |
| 0 | NA | NA | NA | NA | NA | Gm43147 | | 0 | 0 | 0 | 0 | 0 | 0 | 0 | 0 | 0 | 0 | 0 |
| 0 | NA | NA | NA | NA | NA | Gm43148 | | 0 | 0 | 0 | 0 | 0 | 0 | 0 | 0 | 0 | 0 | 0 |
| 0 | NA | NA | NA | NA | NA | Gm43144 | | 0 | 0 | 0 | 0 | 0 | 0 | 0 | 0 | 0 | 0 | 0 |
| 0 | NA | NA | NA | NA | NA | Gm43145 | | 0 | 0 | 0 | 0 | 0 | 0 | 0 | 0 | 0 | 0 | 0 |
| 0 | NA | NA | NA | NA | NA | Gm33969 | | 0 | 0 | 0 | 0 | 0 | 0 | 0 | 0 | 0 | 0 | 0 |
| 0 | NA | NA | NA | NA | NA | Naa11 | N(alpha)-acetyltransferase 11, NatA catal | 0 | 0 | 0 | 0 | 0 | 0 | 0 | 0 | 0 | 0 | 0 |
| 0 | NA | NA | NA | NA | NA | 4930467D21Rik | | 0 | 0 | 0 | 0 | 0 | 0 | 0 | 0 | 0 | 0 | 0 |
| 0 | NA | NA | NA | NA | NA | Gk2 | glycerol kinase 2 | 0 | 0 | 0 | 0 | 0 | 0 | 0 | 0 | 0 | 0 | 0 |

| | | | | | | | | | | | | | | | | | | |
|---|---|---|---|---|---|---|---|---|---|---|---|---|---|---|---|---|---|---|
| 0 | NA | NA | NA | NA | NA | C430019N01Rik | | 0 | 0 | 0 | 0 | 0 | 0 | 0 | 0 | 0 | 0 | 0 |
| 0 | NA | NA | NA | NA | NA | Gm42759 | | 0 | 0 | 0 | 0 | 0 | 0 | 0 | 0 | 0 | 0 | 0 |
| 0 | NA | NA | NA | NA | NA | Gm42760 | | 0 | 0 | 0 | 0 | 0 | 0 | 0 | 0 | 0 | 0 | 0 |
| 0 | NA | NA | NA | NA | NA | E030032P16Rik | | 0 | 0 | 0 | 0 | 0 | 0 | 0 | 0 | 0 | 0 | 0 |
| 0 | NA | NA | NA | NA | NA | Gm24279 | | 0 | 0 | 0 | 0 | 0 | 0 | 0 | 0 | 0 | 0 | 0 |
| 0 | NA | NA | NA | NA | NA | Gm11111 | | 0 | 0 | 0 | 0 | 0 | 0 | 0 | 0 | 0 | 0 | 0 |
| 0 | NA | NA | NA | NA | NA | 1700010H22Rik | | 0 | 0 | 0 | 0 | 0 | 0 | 0 | 0 | 0 | 0 | 0 |
| 0 | NA | NA | NA | NA | NA | Gm3470 | | 0 | 0 | 0 | 0 | 0 | 0 | 0 | 0 | 0 | 0 | 0 |
| 0 | NA | NA | NA | NA | NA | Gm43251 | | 0 | 0 | 0 | 0 | 0 | 0 | 0 | 0 | 0 | 0 | 0 |
| 0 | NA | NA | NA | NA | NA | Gm35394 | | 0 | 0 | 0 | 0 | 0 | 0 | 0 | 0 | 0 | 0 | 0 |
| 0 | NA | NA | NA | NA | NA | Gm16227 | | 0 | 0 | 0 | 0 | 0 | 0 | 0 | 0 | 0 | 0 | 0 |
| 0 | NA | NA | NA | NA | NA | Gm16228 | | 0 | 0 | 0 | 0 | 0 | 0 | 0 | 0 | 0 | 0 | 0 |
| 0 | NA | NA | NA | NA | NA | 4930405H06Rik | | 0 | 0 | 0 | 0 | 0 | 0 | 0 | 0 | 0 | 0 | 0 |
| 0 | NA | NA | NA | NA | NA | Gm25969 | | 0 | 0 | 0 | 0 | 0 | 0 | 0 | 0 | 0 | 0 | 0 |
| 0 | NA | NA | NA | NA | NA | Gm16226 | | 0 | 0 | 0 | 0 | 0 | 0 | 0 | 0 | 0 | 0 | 0 |
| 0 | NA | NA | NA | NA | NA | 4930522N08Rik | | 0 | 0 | 0 | 0 | 0 | 0 | 0 | 0 | 0 | 0 | 0 |
| 0 | NA | NA | NA | NA | NA | Gm43249 | | 0 | 0 | 0 | 0 | 0 | 0 | 0 | 0 | 0 | 0 | 0 |
| 0 | NA | NA | NA | NA | NA | Gm26252 | | 0 | 0 | 0 | 0 | 0 | 0 | 0 | 0 | 0 | 0 | 0 |
| 0 | NA | NA | NA | NA | NA | Gm22838 | | 0 | 0 | 0 | 0 | 0 | 0 | 0 | 0 | 0 | 0 | 0 |
| 0 | NA | NA | NA | NA | NA | Gm26363 | | 0 | 0 | 0 | 0 | 0 | 0 | 0 | 0 | 0 | 0 | 0 |
| 0 | NA | NA | NA | NA | NA | Gm9932 | | 0 | 0 | 0 | 0 | 0 | 0 | 0 | 0 | 0 | 0 | 0 |
| 0 | NA | NA | NA | NA | NA | 2310034O05Rik | | 0 | 0 | 0 | 0 | 0 | 0 | 0 | 0 | 0 | 0 | 0 |
| 0 | NA | NA | NA | NA | NA | Gm8091 | | 0 | 0 | 0 | 0 | 0 | 0 | 0 | 0 | 0 | 0 | 0 |
| 0 | NA | NA | NA | NA | NA | Gm27997 | | 0 | 0 | 0 | 0 | 0 | 0 | 0 | 0 | 0 | 0 | 0 |
| 0 | NA | NA | NA | NA | NA | Gm28030 | | 0 | 0 | 0 | 0 | 0 | 0 | 0 | 0 | 0 | 0 | 0 |
| 0 | NA | NA | NA | NA | NA | Gm27286 | | 0 | 0 | 0 | 0 | 0 | 0 | 0 | 0 | 0 | 0 | 0 |
| 0 | NA | NA | NA | NA | NA | Gm23222 | | 0 | 0 | 0 | 0 | 0 | 0 | 0 | 0 | 0 | 0 | 0 |
| 0 | NA | NA | NA | NA | NA | Gm23173 | | 0 | 0 | 0 | 0 | 0 | 0 | 0 | 0 | 0 | 0 | 0 |
| 0 | NA | NA | NA | NA | NA | Gm15777 | | 0 | 0 | 0 | 0 | 0 | 0 | 0 | 0 | 0 | 0 | 0 |
| 0 | NA | NA | NA | NA | NA | Gm26286 | | 0 | 0 | 0 | 0 | 0 | 0 | 0 | 0 | 0 | 0 | 0 |
| 0 | NA | NA | NA | NA | NA | Gm42987 | | 0 | 0 | 0 | 0 | 0 | 0 | 0 | 0 | 0 | 0 | 0 |
| 0 | NA | NA | NA | NA | NA | Gm36793 | | 0 | 0 | 0 | 0 | 0 | 0 | 0 | 0 | 0 | 0 | 0 |
| 0 | NA | NA | NA | NA | NA | Gm42934 | | 0 | 0 | 0 | 0 | 0 | 0 | 0 | 0 | 0 | 0 | 0 |
| 0 | NA | NA | NA | NA | NA | Gm17978 | | 0 | 0 | 0 | 0 | 0 | 0 | 0 | 0 | 0 | 0 | 0 |
| 0 | NA | NA | NA | NA | NA | Gm42932 | | 0 | 0 | 0 | 0 | 0 | 0 | 0 | 0 | 0 | 0 | 0 |
| 0 | NA | NA | NA | NA | NA | 1700013M08Rik | | 0 | 0 | 0 | 0 | 0 | 0 | 0 | 0 | 0 | 0 | 0 |
| 0 | NA | NA | NA | NA | NA | 4930429D17Rik | | 0 | 0 | 0 | 0 | 0 | 0 | 0 | 0 | 0 | 0 | 0 |
| 0 | NA | NA | NA | NA | NA | Gm18702 | | 0 | 0 | 0 | 0 | 0 | 0 | 0 | 0 | 0 | 0 | 0 |
| 0 | NA | NA | NA | NA | NA | Gm42619 | | 0 | 0 | 0 | 0 | 0 | 0 | 0 | 0 | 0 | 0 | 0 |
| 0 | NA | NA | NA | NA | NA | 1700021F02Rik | | 0 | 0 | 0 | 0 | 0 | 0 | 0 | 0 | 0 | 0 | 0 |
| 0 | NA | NA | NA | NA | NA | Gm23085 | | 0 | 0 | 0 | 0 | 0 | 0 | 0 | 0 | 0 | 0 | 0 |
| 0 | NA | NA | NA | NA | NA | Gm15844 | | 0 | 0 | 0 | 0 | 0 | 0 | 0 | 0 | 0 | 0 | 0 |
| 0 | NA | NA | NA | NA | NA | Gm43546 | | 0 | 0 | 0 | 0 | 0 | 0 | 0 | 0 | 0 | 0 | 0 |
| 0 | NA | NA | NA | NA | NA | Gm25721 | | 0 | 0 | 0 | 0 | 0 | 0 | 0 | 0 | 0 | 0 | 0 |
| 0 | NA | NA | NA | NA | NA | Gm24897 | | 0 | 0 | 0 | 0 | 0 | 0 | 0 | 0 | 0 | 0 | 0 |
| 0 | NA | NA | NA | NA | NA | Gm8200 | | 0 | 0 | 0 | 0 | 0 | 0 | 0 | 0 | 0 | 0 | 0 |
| 0 | NA | NA | NA | NA | NA | Gm43333 | | 0 | 0 | 0 | 0 | 0 | 0 | 0 | 0 | 0 | 0 | 0 |
| 0 | NA | NA | NA | NA | NA | Dspp | dentin sialophosphoprotein | 0 | 0 | 0 | 0 | 0 | 0 | 0 | 0 | 0 | 0 | 0 |
| 0 | NA | NA | NA | NA | NA | Gm31048 | | 0 | 0 | 0 | 0 | 0 | 0 | 0 | 0 | 0 | 0 | 0 |
| 0 | NA | NA | NA | NA | NA | Gm23616 | | 0 | 0 | 0 | 0 | 0 | 0 | 0 | 0 | 0 | 0 | 0 |
| 0 | NA | NA | NA | NA | NA | Ibsp | integrin binding sialoprotein | 0 | 0 | 0 | 0 | 0 | 0 | 0 | 0 | 0 | 0 | 0 |
| 0 | NA | NA | NA | NA | NA | Mepe | matrix extracellular phosphoglycoprotein | 0 | 0 | 0 | 0 | 0 | 0 | 0 | 0 | 0 | 0 | 0 |
| 0 | NA | NA | NA | NA | NA | Gm24097 | | 0 | 0 | 0 | 0 | 0 | 0 | 0 | 0 | 0 | 0 | 0 |
| 0 | NA | NA | NA | NA | NA | Gm43856 | | 0 | 0 | 0 | 0 | 0 | 0 | 0 | 0 | 0 | 0 | 0 |
| 0 | NA | NA | NA | NA | NA | Gm43855 | | 0 | 0 | 0 | 0 | 0 | 0 | 0 | 0 | 0 | 0 | 0 |
| 0 | NA | NA | NA | NA | NA | Gm43302 | | 0 | 0 | 0 | 0 | 0 | 0 | 0 | 0 | 0 | 0 | 0 |
| 0 | NA | NA | NA | NA | NA | Gm26519 | | 0 | 0 | 0 | 0 | 0 | 0 | 0 | 0 | 0 | 0 | 0 |
| 0 | NA | NA | NA | NA | NA | Gm32051 | | 0 | 0 | 0 | 0 | 0 | 0 | 0 | 0 | 0 | 0 | 0 |
| 0 | NA | NA | NA | NA | NA | C230066G23Rik | | 0 | 0 | 0 | 0 | 0 | 0 | 0 | 0 | 0 | 0 | 0 |
| 0 | NA | NA | NA | NA | NA | Lrrc8dos | | 0 | 0 | 0 | 0 | 0 | 0 | 0 | 0 | 0 | 0 | 0 |

| | | | | | | | | | | | | | | | | | | |
|---|---|---|---|---|---|---|---|---|---|---|---|---|---|---|---|---|---|---|
| 0 | NA | NA | NA | NA | NA | Gm43818 | | 0 | 0 | 0 | 0 | 0 | 0 | 0 | 0 | 0 | 0 | 0 |
| 0 | NA | NA | NA | NA | NA | Gm43817 | | 0 | 0 | 0 | 0 | 0 | 0 | 0 | 0 | 0 | 0 | 0 |
| 0 | NA | NA | NA | NA | NA | Gm19566 | | 0 | 0 | 0 | 0 | 0 | 0 | 0 | 0 | 0 | 0 | 0 |
| 0 | NA | NA | NA | NA | NA | Gm25672 | | 0 | 0 | 0 | 0 | 0 | 0 | 0 | 0 | 0 | 0 | 0 |
| 0 | NA | NA | NA | NA | NA | Gm32736 | | 0 | 0 | 0 | 0 | 0 | 0 | 0 | 0 | 0 | 0 | 0 |
| 0 | NA | NA | NA | NA | NA | Gm5987 | | 0 | 0 | 0 | 0 | 0 | 0 | 0 | 0 | 0 | 0 | 0 |
| 0 | NA | NA | NA | NA | NA | Gm38433 | | 0 | 0 | 0 | 0 | 0 | 0 | 0 | 0 | 0 | 0 | 0 |
| 0 | NA | NA | NA | NA | NA | Gm32921 | | 0 | 0 | 0 | 0 | 0 | 0 | 0 | 0 | 0 | 0 | 0 |
| 0 | NA | NA | NA | NA | NA | 4930458A03Rik | | 0 | 0 | 0 | 0 | 0 | 0 | 0 | 0 | 0 | 0 | 0 |
| 0 | NA | NA | NA | NA | NA | Gm42915 | | 0 | 0 | 0 | 0 | 0 | 0 | 0 | 0 | 0 | 0 | 0 |
| 0 | NA | NA | NA | NA | NA | 4930432H08Rik | | 0 | 0 | 0 | 0 | 0 | 0 | 0 | 0 | 0 | 0 | 0 |
| 0 | NA | NA | NA | NA | NA | Gm42916 | | 0 | 0 | 0 | 0 | 0 | 0 | 0 | 0 | 0 | 0 | 0 |
| 0 | NA | NA | NA | NA | NA | Gm26872 | | 0 | 0 | 0 | 0 | 0 | 0 | 0 | 0 | 0 | 0 | 0 |
| 0 | NA | NA | NA | NA | NA | Gm28050 | | 0 | 0 | 0 | 0 | 0 | 0 | 0 | 0 | 0 | 0 | 0 |
| 0 | NA | NA | NA | NA | NA | Gm28039 | | 0 | 0 | 0 | 0 | 0 | 0 | 0 | 0 | 0 | 0 | 0 |
| 0 | NA | NA | NA | NA | NA | Gm29464 | | 0 | 0 | 0 | 0 | 0 | 0 | 0 | 0 | 0 | 0 | 0 |
| 0 | NA | NA | NA | NA | NA | Gm17304 | | 0 | 0 | 0 | 0 | 0 | 0 | 0 | 0 | 0 | 0 | 0 |
| 0 | NA | NA | NA | NA | NA | Gm17937 | | 0 | 0 | 0 | 0 | 0 | 0 | 0 | 0 | 0 | 0 | 0 |
| 0 | NA | NA | NA | NA | NA | Gm24191 | | 0 | 0 | 0 | 0 | 0 | 0 | 0 | 0 | 0 | 0 | 0 |
| 0 | NA | NA | NA | NA | NA | Gm27527 | | 0 | 0 | 0 | 0 | 0 | 0 | 0 | 0 | 0 | 0 | 0 |
| 0 | NA | NA | NA | NA | NA | Olfr719-ps | | 0 | 0 | 0 | 0 | 0 | 0 | 0 | 0 | 0 | 0 | 0 |
| 0 | NA | NA | NA | NA | NA | Gm8145 | | 0 | 0 | 0 | 0 | 0 | 0 | 0 | 0 | 0 | 0 | 0 |
| 0 | NA | NA | NA | NA | NA | Gm42900 | | 0 | 0 | 0 | 0 | 0 | 0 | 0 | 0 | 0 | 0 | 0 |
| 0 | NA | NA | NA | NA | NA | Gm22557 | | 0 | 0 | 0 | 0 | 0 | 0 | 0 | 0 | 0 | 0 | 0 |
| 0 | NA | NA | NA | NA | NA | n-R5s173 | | 0 | 0 | 0 | 0 | 0 | 0 | 0 | 0 | 0 | 0 | 0 |
| 0 | NA | NA | NA | NA | NA | Lpcat2b | lysophosphatidylcholine acyltransferase 2 | 0 | 0 | 0 | 0 | 0 | 0 | 0 | 0 | 0 | 0 | 0 |
| 0 | NA | NA | NA | NA | NA | 5830411K02Rik | | 0 | 0 | 0 | 0 | 0 | 0 | 0 | 0 | 0 | 0 | 0 |
| 0 | NA | NA | NA | NA | NA | Ube2d2b | ubiquitin-conjugating enzyme E2D 2B | 0 | 0 | 0 | 0 | 0 | 0 | 0 | 0 | 0 | 0 | 0 |
| 0 | NA | NA | NA | NA | NA | Gm16007 | | 0 | 0 | 0 | 0 | 0 | 0 | 0 | 0 | 0 | 0 | 0 |
| 0 | NA | NA | NA | NA | NA | Snord21 | | 0 | 0 | 0 | 0 | 0 | 0 | 0 | 0 | 0 | 0 | 0 |
| 0 | NA | NA | NA | NA | NA | Rps19-ps8 | | 0 | 0 | 0 | 0 | 0 | 0 | 0 | 0 | 0 | 0 | 0 |
| 0 | NA | NA | NA | NA | NA | Gm8190 | | 0 | 0 | 0 | 0 | 0 | 0 | 0 | 0 | 0 | 0 | 0 |
| 0 | NA | NA | NA | NA | NA | Gm43083 | | 0 | 0 | 0 | 0 | 0 | 0 | 0 | 0 | 0 | 0 | 0 |
| 0 | NA | NA | NA | NA | NA | Gm42518 | | 0 | 0 | 0 | 0 | 0 | 0 | 0 | 0 | 0 | 0 | 0 |
| 0 | NA | NA | NA | NA | NA | | | 0 | 0 | 0 | 0 | 0 | 0 | 0 | 0 | 0 | 0 | 0 |
| 0 | NA | NA | NA | NA | NA | 1700047L14Rik | | 0 | 0 | 0 | 0 | 0 | 0 | 0 | 0 | 0 | 0 | 0 |
| 0 | NA | NA | NA | NA | NA | Vmn2r8 | vomeronasal 2, receptor 8 | 0 | 0 | 0 | 0 | 0 | 0 | 0 | 0 | 0 | 0 | 0 |
| 0 | NA | NA | NA | NA | NA | Gm18594 | | 0 | 0 | 0 | 0 | 0 | 0 | 0 | 0 | 0 | 0 | 0 |
| 0 | NA | NA | NA | NA | NA | Vmn2r9 | vomeronasal 2, receptor 9 | 0 | 0 | 0 | 0 | 0 | 0 | 0 | 0 | 0 | 0 | 0 |
| 0 | NA | NA | NA | NA | NA | Gm20629 | | 0 | 0 | 0 | 0 | 0 | 0 | 0 | 0 | 0 | 0 | 0 |
| 0 | NA | NA | NA | NA | NA | Gm10417 | | 0 | 0 | 0 | 0 | 0 | 0 | 0 | 0 | 0 | 0 | 0 |
| 0 | NA | NA | NA | NA | NA | Vmn2r-ps21 | | 0 | 0 | 0 | 0 | 0 | 0 | 0 | 0 | 0 | 0 | 0 |
| 0 | NA | NA | NA | NA | NA | Gm18971 | | 0 | 0 | 0 | 0 | 0 | 0 | 0 | 0 | 0 | 0 | 0 |
| 0 | NA | NA | NA | NA | NA | Gm18595 | | 0 | 0 | 0 | 0 | 0 | 0 | 0 | 0 | 0 | 0 | 0 |
| 0 | NA | NA | NA | NA | NA | Vmn2r10 | vomeronasal 2, receptor 10 | 0 | 0 | 0 | 0 | 0 | 0 | 0 | 0 | 0 | 0 | 0 |
| 0 | NA | NA | NA | NA | NA | Vmn2r11 | vomeronasal 2, receptor 11 | 0 | 0 | 0 | 0 | 0 | 0 | 0 | 0 | 0 | 0 | 0 |
| 0 | NA | NA | NA | NA | NA | Vmn2r12 | vomeronasal 2, receptor 12 | 0 | 0 | 0 | 0 | 0 | 0 | 0 | 0 | 0 | 0 | 0 |
| 0 | NA | NA | NA | NA | NA | Vmn2r13 | vomeronasal 2, receptor 13 | 0 | 0 | 0 | 0 | 0 | 0 | 0 | 0 | 0 | 0 | 0 |
| 0 | NA | NA | NA | NA | NA | Vmn2r14 | vomeronasal 2, receptor 14 | 0 | 0 | 0 | 0 | 0 | 0 | 0 | 0 | 0 | 0 | 0 |
| 0 | NA | NA | NA | NA | NA | Gm8473 | | 0 | 0 | 0 | 0 | 0 | 0 | 0 | 0 | 0 | 0 | 0 |
| 0 | NA | NA | NA | NA | NA | Vmn2r15 | vomeronasal 2, receptor 15 | 0 | 0 | 0 | 0 | 0 | 0 | 0 | 0 | 0 | 0 | 0 |
| 0 | NA | NA | NA | NA | NA | Vmn2r16 | vomeronasal 2, receptor 16 | 0 | 0 | 0 | 0 | 0 | 0 | 0 | 0 | 0 | 0 | 0 |
| 0 | NA | NA | NA | NA | NA | Gm7171 | | 0 | 0 | 0 | 0 | 0 | 0 | 0 | 0 | 0 | 0 | 0 |
| 0 | NA | NA | NA | NA | NA | Vmn2r17 | vomeronasal 2, receptor 17 | 0 | 0 | 0 | 0 | 0 | 0 | 0 | 0 | 0 | 0 | 0 |
| 0 | NA | NA | NA | NA | NA | Gm8488 | | 0 | 0 | 0 | 0 | 0 | 0 | 0 | 0 | 0 | 0 | 0 |
| 0 | NA | NA | NA | NA | NA | Vmn2r-ps23 | | 0 | 0 | 0 | 0 | 0 | 0 | 0 | 0 | 0 | 0 | 0 |
| 0 | NA | NA | NA | NA | NA | Gm28000 | | 0 | 0 | 0 | 0 | 0 | 0 | 0 | 0 | 0 | 0 | 0 |
| 0 | NA | NA | NA | NA | NA | Gm26277 | | 0 | 0 | 0 | 0 | 0 | 0 | 0 | 0 | 0 | 0 | 0 |
| 0 | NA | NA | NA | NA | NA | Gm15791 | | 0 | 0 | 0 | 0 | 0 | 0 | 0 | 0 | 0 | 0 | 0 |
| 0 | NA | NA | NA | NA | NA | Gm15792 | | 0 | 0 | 0 | 0 | 0 | 0 | 0 | 0 | 0 | 0 | 0 |

| | | | | | | | | | | | | | | | | | |
|---|---|---|---|---|---|---|---|---|---|---|---|---|---|---|---|---|---|
| 0 | NA | NA | NA | NA | NA | Lrcol1 | leucine rich colipase-like 1 | 0 | 0 | 0 | 0 | 0 | 0 | 0 | 0 | 0 | 0 | 0 |
| 0 | NA | NA | NA | NA | NA | Gm32996 | | 0 | 0 | 0 | 0 | 0 | 0 | 0 | 0 | 0 | 0 | 0 |
| 0 | NA | NA | NA | NA | NA | Gm26079 | | 0 | 0 | 0 | 0 | 0 | 0 | 0 | 0 | 0 | 0 | 0 |
| 0 | NA | NA | NA | NA | NA | Gm22583 | | 0 | 0 | 0 | 0 | 0 | 0 | 0 | 0 | 0 | 0 | 0 |
| 0 | NA | NA | NA | NA | NA | Gm42779 | | 0 | 0 | 0 | 0 | 0 | 0 | 0 | 0 | 0 | 0 | 0 |
| 0 | NA | NA | NA | NA | NA | Gm15988 | | 0 | 0 | 0 | 0 | 0 | 0 | 0 | 0 | 0 | 0 | 0 |
| 0 | NA | NA | NA | NA | NA | Gm43677 | | 0 | 0 | 0 | 0 | 0 | 0 | 0 | 0 | 0 | 0 | 0 |
| 0 | NA | NA | NA | NA | NA | Gm27602 | | 0 | 0 | 0 | 0 | 0 | 0 | 0 | 0 | 0 | 0 | 0 |
| 0 | NA | NA | NA | NA | NA | Gm27680 | | 0 | 0 | 0 | 0 | 0 | 0 | 0 | 0 | 0 | 0 | 0 |
| 0 | NA | NA | NA | NA | NA | Gm27680 | | 0 | 0 | 0 | 0 | 0 | 0 | 0 | 0 | 0 | 0 | 0 |
| 0 | NA | NA | NA | NA | NA | Gm43119 | | 0 | 0 | 0 | 0 | 0 | 0 | 0 | 0 | 0 | 0 | 0 |
| 0 | NA | NA | NA | NA | NA | Gm42489 | | 0 | 0 | 0 | 0 | 0 | 0 | 0 | 0 | 0 | 0 | 0 |
| 0 | NA | NA | NA | NA | NA | Gm26897 | | 0 | 0 | 0 | 0 | 0 | 0 | 0 | 0 | 0 | 0 | 0 |
| 0 | NA | NA | NA | NA | NA | E130006D01Rik | | 0 | 0 | 0 | 0 | 0 | 0 | 0 | 0 | 0 | 0 | 0 |
| 0 | NA | NA | NA | NA | NA | Gm16019 | | 0 | 0 | 0 | 0 | 0 | 0 | 0 | 0 | 0 | 0 | 0 |
| 0 | NA | NA | NA | NA | NA | Gm42488 | | 0 | 0 | 0 | 0 | 0 | 0 | 0 | 0 | 0 | 0 | 0 |
| 0 | NA | NA | NA | NA | NA | 1700016B01Rik | | 0 | 0 | 0 | 0 | 0 | 0 | 0 | 0 | 0 | 0 | 0 |
| 0 | NA | NA | NA | NA | NA | 1700028D13Rik | | 0 | 0 | 0 | 0 | 0 | 0 | 0 | 0 | 0 | 0 | 0 |
| 0 | NA | NA | NA | NA | NA | Gm27998 | | 0 | 0 | 0 | 0 | 0 | 0 | 0 | 0 | 0 | 0 | 0 |
| 0 | NA | NA | NA | NA | NA | Gm27882 | | 0 | 0 | 0 | 0 | 0 | 0 | 0 | 0 | 0 | 0 | 0 |
| 0 | NA | NA | NA | NA | NA | Gm27627 | | 0 | 0 | 0 | 0 | 0 | 0 | 0 | 0 | 0 | 0 | 0 |
| 0 | NA | NA | NA | NA | NA | Gm27357 | | 0 | 0 | 0 | 0 | 0 | 0 | 0 | 0 | 0 | 0 | 0 |
| 0 | NA | NA | NA | NA | NA | Gm43758 | | 0 | 0 | 0 | 0 | 0 | 0 | 0 | 0 | 0 | 0 | 0 |
| 0 | NA | NA | NA | NA | NA | Gm20636 | | 0 | 0 | 0 | 0 | 0 | 0 | 0 | 0 | 0 | 0 | 0 |
| 0 | NA | NA | NA | NA | NA | Gm6588 | predicted gene 6588 | 0 | 0 | 0 | 0 | 0 | 0 | 0 | 0 | 0 | 0 | 0 |
| 0 | NA | NA | NA | NA | NA | 4933415J04Rik | | 0 | 0 | 0 | 0 | 0 | 0 | 0 | 0 | 0 | 0 | 0 |
| 0 | NA | NA | NA | NA | NA | Gm42864 | | 0 | 0 | 0 | 0 | 0 | 0 | 0 | 0 | 0 | 0 | 0 |
| 0 | NA | NA | NA | NA | NA | Gm42865 | | 0 | 0 | 0 | 0 | 0 | 0 | 0 | 0 | 0 | 0 | 0 |
| 0 | NA | NA | NA | NA | NA | 1700095B10Rik | | 0 | 0 | 0 | 0 | 0 | 0 | 0 | 0 | 0 | 0 | 0 |
| 0 | NA | NA | NA | NA | NA | Crybb2 | crystallin, beta B2 | 0 | 0 | 0 | 0 | 0 | 0 | 0 | 0 | 0 | 0 | 0 |
| 0 | NA | NA | NA | NA | NA | Gm43246 | | 0 | 0 | 0 | 0 | 0 | 0 | 0 | 0 | 0 | 0 | 0 |
| 0 | NA | NA | NA | NA | NA | 4930557B06Rik | | 0 | 0 | 0 | 0 | 0 | 0 | 0 | 0 | 0 | 0 | 0 |
| 0 | NA | NA | NA | NA | NA | Gm42161 | | 0 | 0 | 0 | 0 | 0 | 0 | 0 | 0 | 0 | 0 | 0 |
| 0 | NA | NA | NA | NA | NA | Tmem211 | transmembrane protein 211 | 0 | 0 | 0 | 0 | 0 | 0 | 0 | 0 | 0 | 0 | 0 |
| 0 | NA | NA | NA | NA | NA | Mir7229 | | 0 | 0 | 0 | 0 | 0 | 0 | 0 | 0 | 0 | 0 | 0 |
| 0 | NA | NA | NA | NA | NA | Mir7230 | | 0 | 0 | 0 | 0 | 0 | 0 | 0 | 0 | 0 | 0 | 0 |
| 0 | NA | NA | NA | NA | NA | Aym1 | activator of yeast meiotic promoters 1 | 0 | 0 | 0 | 0 | 0 | 0 | 0 | 0 | 0 | 0 | 0 |
| 0 | NA | NA | NA | NA | NA | Gm15736 | | 0 | 0 | 0 | 0 | 0 | 0 | 0 | 0 | 0 | 0 | 0 |
| 0 | NA | NA | NA | NA | NA | 1700069L16Rik | | 0 | 0 | 0 | 0 | 0 | 0 | 0 | 0 | 0 | 0 | 0 |
| 0 | NA | NA | NA | NA | NA | Gm19931 | | 0 | 0 | 0 | 0 | 0 | 0 | 0 | 0 | 0 | 0 | 0 |
| 0 | NA | NA | NA | NA | NA | Gm43335 | | 0 | 0 | 0 | 0 | 0 | 0 | 0 | 0 | 0 | 0 | 0 |
| 0 | NA | NA | NA | NA | NA | Gm22056 | | 0 | 0 | 0 | 0 | 0 | 0 | 0 | 0 | 0 | 0 | 0 |
| 0 | NA | NA | NA | NA | NA | Dao | D-amino acid oxidase | 0 | 0 | 0 | 0 | 0 | 0 | 0 | 0 | 0 | 0 | 0 |
| 0 | NA | NA | NA | NA | NA | Gm13784 | | 0 | 0 | 0 | 0 | 0 | 0 | 0 | 0 | 0 | 0 | 0 |
| 0 | NA | NA | NA | NA | NA | Foxn4 | forkhead box N4 | 0 | 0 | 0 | 0 | 0 | 0 | 0 | 0 | 0 | 0 | 0 |
| 0 | NA | NA | NA | NA | NA | Gm13821 | | 0 | 0 | 0 | 0 | 0 | 0 | 0 | 0 | 0 | 0 | 0 |
| 0 | NA | NA | NA | NA | NA | Mir6240 | | 0 | 0 | 0 | 0 | 0 | 0 | 0 | 0 | 0 | 0 | 0 |
| 0 | NA | NA | NA | NA | NA | 4930519G04Ril | RIKEN cDNA 4930519G04 gene | 0 | 0 | 0 | 0 | 0 | 0 | 0 | 0 | 0 | 0 | 0 |
| 0 | NA | NA | NA | NA | NA | Gm13823 | | 0 | 0 | 0 | 0 | 0 | 0 | 0 | 0 | 0 | 0 | 0 |
| 0 | NA | NA | NA | NA | NA | Gm13829 | | 0 | 0 | 0 | 0 | 0 | 0 | 0 | 0 | 0 | 0 | 0 |
| 0 | NA | NA | NA | NA | NA | 4930401G09Rik | | 0 | 0 | 0 | 0 | 0 | 0 | 0 | 0 | 0 | 0 | 0 |
| 0 | NA | NA | NA | NA | NA | Gm13832 | | 0 | 0 | 0 | 0 | 0 | 0 | 0 | 0 | 0 | 0 | 0 |
| 0 | NA | NA | NA | NA | NA | Gm24407 | | 0 | 0 | 0 | 0 | 0 | 0 | 0 | 0 | 0 | 0 | 0 |
| 0 | NA | NA | NA | NA | NA | Mir7029 | | 0 | 0 | 0 | 0 | 0 | 0 | 0 | 0 | 0 | 0 | 0 |
| 0 | NA | NA | NA | NA | NA | Gm42473 | | 0 | 0 | 0 | 0 | 0 | 0 | 0 | 0 | 0 | 0 | 0 |
| 0 | NA | NA | NA | NA | NA | Gm42471 | | 0 | 0 | 0 | 0 | 0 | 0 | 0 | 0 | 0 | 0 | 0 |
| 0 | NA | NA | NA | NA | NA | Gm13842 | | 0 | 0 | 0 | 0 | 0 | 0 | 0 | 0 | 0 | 0 | 0 |
| 0 | NA | NA | NA | NA | NA | Gm17420 | | 0 | 0 | 0 | 0 | 0 | 0 | 0 | 0 | 0 | 0 | 0 |
| 0 | NA | NA | NA | NA | NA | Gm14507 | | 0 | 0 | 0 | 0 | 0 | 0 | 0 | 0 | 0 | 0 | 0 |
| 0 | NA | NA | NA | NA | NA | Gm13837 | | 0 | 0 | 0 | 0 | 0 | 0 | 0 | 0 | 0 | 0 | 0 |

| | | | | | | | | | | | | | | | | | | |
|---|---|---|---|---|---|---|---|---|---|---|---|---|---|---|---|---|---|---|
| 0 | NA | NA | NA | NA | NA | Gm13839 | | 0 | 0 | 0 | 0 | 0 | 0 | 0 | 0 | 0 | 0 | 0 |
| 0 | NA | NA | NA | NA | NA | Gm13838 | | 0 | 0 | 0 | 0 | 0 | 0 | 0 | 0 | 0 | 0 | 0 |
| 0 | NA | NA | NA | NA | NA | B230112J18Rik | | 0 | 0 | 0 | 0 | 0 | 0 | 0 | 0 | 0 | 0 | 0 |
| 0 | NA | NA | NA | NA | NA | 4933424N20Rik | | 0 | 0 | 0 | 0 | 0 | 0 | 0 | 0 | 0 | 0 | 0 |
| 0 | NA | NA | NA | NA | NA | 4930562A09Rik | | 0 | 0 | 0 | 0 | 0 | 0 | 0 | 0 | 0 | 0 | 0 |
| 0 | NA | NA | NA | NA | NA | 4933430H06Rik | | 0 | 0 | 0 | 0 | 0 | 0 | 0 | 0 | 0 | 0 | 0 |
| 0 | NA | NA | NA | NA | NA | Gm42854 | | 0 | 0 | 0 | 0 | 0 | 0 | 0 | 0 | 0 | 0 | 0 |
| 0 | NA | NA | NA | NA | NA | Gm43122 | | 0 | 0 | 0 | 0 | 0 | 0 | 0 | 0 | 0 | 0 | 0 |
| 0 | NA | NA | NA | NA | NA | Gm29926 | | 0 | 0 | 0 | 0 | 0 | 0 | 0 | 0 | 0 | 0 | 0 |
| 0 | NA | NA | NA | NA | NA | 4930569F06Rik | | 0 | 0 | 0 | 0 | 0 | 0 | 0 | 0 | 0 | 0 | 0 |
| 0 | NA | NA | NA | NA | NA | n-R5s174 | | 0 | 0 | 0 | 0 | 0 | 0 | 0 | 0 | 0 | 0 | 0 |
| 0 | NA | NA | NA | NA | NA | Gm3786 | | 0 | 0 | 0 | 0 | 0 | 0 | 0 | 0 | 0 | 0 | 0 |
| 0 | NA | NA | NA | NA | NA | Gm42546 | | 0 | 0 | 0 | 0 | 0 | 0 | 0 | 0 | 0 | 0 | 0 |
| 0 | NA | NA | NA | NA | NA | Gm42550 | | 0 | 0 | 0 | 0 | 0 | 0 | 0 | 0 | 0 | 0 | 0 |
| 0 | NA | NA | NA | NA | NA | Gm37028 | | 0 | 0 | 0 | 0 | 0 | 0 | 0 | 0 | 0 | 0 | 0 |
| 0 | NA | NA | NA | NA | NA | Gm25343 | | 0 | 0 | 0 | 0 | 0 | 0 | 0 | 0 | 0 | 0 | 0 |
| 0 | NA | NA | NA | NA | NA | Gm42545 | | 0 | 0 | 0 | 0 | 0 | 0 | 0 | 0 | 0 | 0 | 0 |
| 0 | NA | NA | NA | NA | NA | Gm26411 | | 0 | 0 | 0 | 0 | 0 | 0 | 0 | 0 | 0 | 0 | 0 |
| 0 | NA | NA | NA | NA | NA | Gm28563 | | 0 | 0 | 0 | 0 | 0 | 0 | 0 | 0 | 0 | 0 | 0 |
| 0 | NA | NA | NA | NA | NA | Gm26455 | | 0 | 0 | 0 | 0 | 0 | 0 | 0 | 0 | 0 | 0 | 0 |
| 0 | NA | NA | NA | NA | NA | Gm15754 | | 0 | 0 | 0 | 0 | 0 | 0 | 0 | 0 | 0 | 0 | 0 |
| 0 | NA | NA | NA | NA | NA | Gm43785 | | 0 | 0 | 0 | 0 | 0 | 0 | 0 | 0 | 0 | 0 | 0 |
| 0 | NA | NA | NA | NA | NA | Gm43052 | | 0 | 0 | 0 | 0 | 0 | 0 | 0 | 0 | 0 | 0 | 0 |
| 0 | NA | NA | NA | NA | NA | Gm43782 | | 0 | 0 | 0 | 0 | 0 | 0 | 0 | 0 | 0 | 0 | 0 |
| 0 | NA | NA | NA | NA | NA | Gm43342 | | 0 | 0 | 0 | 0 | 0 | 0 | 0 | 0 | 0 | 0 | 0 |
| 0 | NA | NA | NA | NA | NA | 4930413E15Rik | | 0 | 0 | 0 | 0 | 0 | 0 | 0 | 0 | 0 | 0 | 0 |
| 0 | NA | NA | NA | NA | NA | Gm43784 | | 0 | 0 | 0 | 0 | 0 | 0 | 0 | 0 | 0 | 0 | 0 |
| 0 | NA | NA | NA | NA | NA | 1700081H04Rik | | 0 | 0 | 0 | 0 | 0 | 0 | 0 | 0 | 0 | 0 | 0 |
| 0 | NA | NA | NA | NA | NA | n-R5s175 | | 0 | 0 | 0 | 0 | 0 | 0 | 0 | 0 | 0 | 0 | 0 |
| 0 | NA | NA | NA | NA | NA | Gm31314 | | 0 | 0 | 0 | 0 | 0 | 0 | 0 | 0 | 0 | 0 | 0 |
| 0 | NA | NA | NA | NA | NA | Gm16063 | | 0 | 0 | 0 | 0 | 0 | 0 | 0 | 0 | 0 | 0 | 0 |
| 0 | NA | NA | NA | NA | NA | Gm16064 | | 0 | 0 | 0 | 0 | 0 | 0 | 0 | 0 | 0 | 0 | 0 |
| 0 | NA | NA | NA | NA | NA | 1700021F13Rik | | 0 | 0 | 0 | 0 | 0 | 0 | 0 | 0 | 0 | 0 | 0 |
| 0 | NA | NA | NA | NA | NA | Gm43050 | | 0 | 0 | 0 | 0 | 0 | 0 | 0 | 0 | 0 | 0 | 0 |
| 0 | NA | NA | NA | NA | NA | Tbx5 | T-box 5 | 0 | 0 | 0 | 0 | 0 | 0 | 0 | 0 | 0 | 0 | 0 |
| 0 | NA | NA | NA | NA | NA | Gm43269 | | 0 | 0 | 0 | 0 | 0 | 0 | 0 | 0 | 0 | 0 | 0 |
| 0 | NA | NA | NA | NA | NA | Gm38554 | | 0 | 0 | 0 | 0 | 0 | 0 | 0 | 0 | 0 | 0 | 0 |
| 0 | NA | NA | NA | NA | NA | n-R5s176 | | 0 | 0 | 0 | 0 | 0 | 0 | 0 | 0 | 0 | 0 | 0 |
| 0 | NA | NA | NA | NA | NA | Gm10390 | | 0 | 0 | 0 | 0 | 0 | 0 | 0 | 0 | 0 | 0 | 0 |
| 0 | NA | NA | NA | NA | NA | Gm26474 | | 0 | 0 | 0 | 0 | 0 | 0 | 0 | 0 | 0 | 0 | 0 |
| 0 | NA | NA | NA | NA | NA | Gm42654 | | 0 | 0 | 0 | 0 | 0 | 0 | 0 | 0 | 0 | 0 | 0 |
| 0 | NA | NA | NA | NA | NA | Gm27199 | | 0 | 0 | 0 | 0 | 0 | 0 | 0 | 0 | 0 | 0 | 0 |
| 0 | NA | NA | NA | NA | NA | Lhx5 | LIM homeobox protein 5 | 0 | 0 | 0 | 0 | 0 | 0 | 0 | 0 | 0 | 0 | 0 |
| 0 | NA | NA | NA | NA | NA | Sds | serine dehydratase | 0 | 0 | 0 | 0 | 0 | 0 | 0 | 0 | 0 | 0 | 0 |
| 0 | NA | NA | NA | NA | NA | Gm42657 | | 0 | 0 | 0 | 0 | 0 | 0 | 0 | 0 | 0 | 0 | 0 |
| 0 | NA | NA | NA | NA | NA | Gm43068 | | 0 | 0 | 0 | 0 | 0 | 0 | 0 | 0 | 0 | 0 | 0 |
| 0 | NA | NA | NA | NA | NA | Gm43069 | | 0 | 0 | 0 | 0 | 0 | 0 | 0 | 0 | 0 | 0 | 0 |
| 0 | NA | NA | NA | NA | NA | Oas1f | 2'-5' oligoadenylate synthetase 1F | 0 | 0 | 0 | 0 | 0 | 0 | 0 | 0 | 0 | 0 | 0 |
| 0 | NA | NA | NA | NA | NA | Gm22907 | | 0 | 0 | 0 | 0 | 0 | 0 | 0 | 0 | 0 | 0 | 0 |
| 0 | NA | NA | NA | NA | NA | Oas1h | 2'-5' oligoadenylate synthetase 1H | 0 | 0 | 0 | 0 | 0 | 0 | 0 | 0 | 0 | 0 | 0 |
| 0 | NA | NA | NA | NA | NA | Oas1d | 2'-5' oligoadenylate synthetase 1D | 0 | 0 | 0 | 0 | 0 | 0 | 0 | 0 | 0 | 0 | 0 |
| 0 | NA | NA | NA | NA | NA | | | 0 | 0 | 0 | 0 | 0 | 0 | 0 | 0 | 0 | 0 | 0 |
| 0 | NA | NA | NA | NA | NA | Gm42656 | | 0 | 0 | 0 | 0 | 0 | 0 | 0 | 0 | 0 | 0 | 0 |
| 0 | NA | NA | NA | NA | NA | Gm26205 | | 0 | 0 | 0 | 0 | 0 | 0 | 0 | 0 | 0 | 0 | 0 |
| 0 | NA | NA | NA | NA | NA | Gm24671 | | 0 | 0 | 0 | 0 | 0 | 0 | 0 | 0 | 0 | 0 | 0 |
| 0 | NA | NA | NA | NA | NA | Gm43420 | | 0 | 0 | 0 | 0 | 0 | 0 | 0 | 0 | 0 | 0 | 0 |
| 0 | NA | NA | NA | NA | NA | Adam1b | a disintegrin and metallopeptidase domai | 0 | 0 | 0 | 0 | 0 | 0 | 0 | 0 | 0 | 0 | 0 |
| 0 | NA | NA | NA | NA | NA | Gm16552 | | 0 | 0 | 0 | 0 | 0 | 0 | 0 | 0 | 0 | 0 | 0 |
| 0 | NA | NA | NA | NA | NA | Gm3970 | | 0 | 0 | 0 | 0 | 0 | 0 | 0 | 0 | 0 | 0 | 0 |
| 0 | NA | NA | NA | NA | NA | Gm15637 | | 0 | 0 | 0 | 0 | 0 | 0 | 0 | 0 | 0 | 0 | 0 |

| | | | | | | | | | | | | | | | | | |
|---|---|---|---|---|---|---|---|---|---|---|---|---|---|---|---|---|---|
| 0 | NA | NA | NA | NA | NA | Myl2 | myosin, light polypeptide 2, regulatory, ca | 0 | 0 | 0 | 0 | 0 | 0 | 0 | 0 | 0 | 0 | 0 |
| 0 | NA | NA | NA | NA | NA | Gm15481 | | 0 | 0 | 0 | 0 | 0 | 0 | 0 | 0 | 0 | 0 | 0 |
| 0 | NA | NA | NA | NA | NA | Gm15846 | | 0 | 0 | 0 | 0 | 0 | 0 | 0 | 0 | 0 | 0 | 0 |
| 0 | NA | NA | NA | NA | NA | Mir8115 | | 0 | 0 | 0 | 0 | 0 | 0 | 0 | 0 | 0 | 0 | 0 |
| 0 | NA | NA | NA | NA | NA | Gm10064 | | 0 | 0 | 0 | 0 | 0 | 0 | 0 | 0 | 0 | 0 | 0 |
| 0 | NA | NA | NA | NA | NA | Gm43411 | | 0 | 0 | 0 | 0 | 0 | 0 | 0 | 0 | 0 | 0 | 0 |
| 0 | NA | NA | NA | NA | NA | Gm44574 | | 0 | 0 | 0 | 0 | 0 | 0 | 0 | 0 | 0 | 0 | 0 |
| 0 | NA | NA | NA | NA | NA | Gm2479 | | 0 | 0 | 0 | 0 | 0 | 0 | 0 | 0 | 0 | 0 | 0 |
| 0 | NA | NA | NA | NA | NA | Gm43409 | | 0 | 0 | 0 | 0 | 0 | 0 | 0 | 0 | 0 | 0 | 0 |
| 0 | NA | NA | NA | NA | NA | Il31 | interleukin 31 | 0 | 0 | 0 | 0 | 0 | 0 | 0 | 0 | 0 | 0 | 0 |
| 0 | NA | NA | NA | NA | NA | Lrrc43 | leucine rich repeat containing 43 | 0 | 0 | 0 | 0 | 0 | 0 | 0 | 0 | 0 | 0 | 0 |
| 0 | NA | NA | NA | NA | NA | Gm24682 | | 0 | 0 | 0 | 0 | 0 | 0 | 0 | 0 | 0 | 0 | 0 |
| 0 | NA | NA | NA | NA | NA | Gm24840 | | 0 | 0 | 0 | 0 | 0 | 0 | 0 | 0 | 0 | 0 | 0 |
| 0 | NA | NA | NA | NA | NA | Mir7032 | | 0 | 0 | 0 | 0 | 0 | 0 | 0 | 0 | 0 | 0 | 0 |
| 0 | NA | NA | NA | NA | NA | Gm37939 | | 0 | 0 | 0 | 0 | 0 | 0 | 0 | 0 | 0 | 0 | 0 |
| 0 | NA | NA | NA | NA | NA | Pitpnm2os2 | | 0 | 0 | 0 | 0 | 0 | 0 | 0 | 0 | 0 | 0 | 0 |
| 0 | NA | NA | NA | NA | NA | Gm23171 | | 0 | 0 | 0 | 0 | 0 | 0 | 0 | 0 | 0 | 0 | 0 |
| 0 | NA | NA | NA | NA | NA | Gm27486 | | 0 | 0 | 0 | 0 | 0 | 0 | 0 | 0 | 0 | 0 | 0 |
| 0 | NA | NA | NA | NA | NA | Gm24854 | | 0 | 0 | 0 | 0 | 0 | 0 | 0 | 0 | 0 | 0 | 0 |
| 0 | NA | NA | NA | NA | NA | 4930404A12Rik | | 0 | 0 | 0 | 0 | 0 | 0 | 0 | 0 | 0 | 0 | 0 |
| 0 | NA | NA | NA | NA | NA | Gm43777 | | 0 | 0 | 0 | 0 | 0 | 0 | 0 | 0 | 0 | 0 | 0 |
| 0 | NA | NA | NA | NA | NA | Gm42838 | | 0 | 0 | 0 | 0 | 0 | 0 | 0 | 0 | 0 | 0 | 0 |
| 0 | NA | NA | NA | NA | NA | Gm32585 | | 0 | 0 | 0 | 0 | 0 | 0 | 0 | 0 | 0 | 0 | 0 |
| 0 | NA | NA | NA | NA | NA | Gm42633 | | 0 | 0 | 0 | 0 | 0 | 0 | 0 | 0 | 0 | 0 | 0 |
| 0 | NA | NA | NA | NA | NA | Gm43756 | | 0 | 0 | 0 | 0 | 0 | 0 | 0 | 0 | 0 | 0 | 0 |
| 0 | NA | NA | NA | NA | NA | Gm27551 | | 0 | 0 | 0 | 0 | 0 | 0 | 0 | 0 | 0 | 0 | 0 |
| 0 | NA | NA | NA | NA | NA | Gm4868 | | 0 | 0 | 0 | 0 | 0 | 0 | 0 | 0 | 0 | 0 | 0 |
| 0 | NA | NA | NA | NA | NA | 1700048F04Rik | | 0 | 0 | 0 | 0 | 0 | 0 | 0 | 0 | 0 | 0 | 0 |
| 0 | NA | NA | NA | NA | NA | Gm42945 | | 0 | 0 | 0 | 0 | 0 | 0 | 0 | 0 | 0 | 0 | 0 |
| 0 | NA | NA | NA | NA | NA | Gm42952 | | 0 | 0 | 0 | 0 | 0 | 0 | 0 | 0 | 0 | 0 | 0 |
| 0 | NA | NA | NA | NA | NA | Gm23151 | | 0 | 0 | 0 | 0 | 0 | 0 | 0 | 0 | 0 | 0 | 0 |
| 0 | NA | NA | NA | NA | NA | Gm37700 | | 0 | 0 | 0 | 0 | 0 | 0 | 0 | 0 | 0 | 0 | 0 |
| 0 | NA | NA | NA | NA | NA | Gm40330 | | 0 | 0 | 0 | 0 | 0 | 0 | 0 | 0 | 0 | 0 | 0 |
| 0 | NA | NA | NA | NA | NA | Gm24839 | | 0 | 0 | 0 | 0 | 0 | 0 | 0 | 0 | 0 | 0 | 0 |
| 0 | NA | NA | NA | NA | NA | 1700081B01Rik | | 0 | 0 | 0 | 0 | 0 | 0 | 0 | 0 | 0 | 0 | 0 |
| 0 | NA | NA | NA | NA | NA | Gm33347 | | 0 | 0 | 0 | 0 | 0 | 0 | 0 | 0 | 0 | 0 | 0 |
| 0 | NA | NA | NA | NA | NA | Gm42953 | | 0 | 0 | 0 | 0 | 0 | 0 | 0 | 0 | 0 | 0 | 0 |
| 0 | NA | NA | NA | NA | NA | Gm40331 | | 0 | 0 | 0 | 0 | 0 | 0 | 0 | 0 | 0 | 0 | 0 |
| 0 | NA | NA | NA | NA | NA | Gm42955 | | 0 | 0 | 0 | 0 | 0 | 0 | 0 | 0 | 0 | 0 | 0 |
| 0 | NA | NA | NA | NA | NA | 4933438B17Rik | | 0 | 0 | 0 | 0 | 0 | 0 | 0 | 0 | 0 | 0 | 0 |
| 0 | NA | NA | NA | NA | NA | Gm43475 | | 0 | 0 | 0 | 0 | 0 | 0 | 0 | 0 | 0 | 0 | 0 |
| 0 | NA | NA | NA | NA | NA | 4930572K03Rik | | 0 | 0 | 0 | 0 | 0 | 0 | 0 | 0 | 0 | 0 | 0 |
| 0 | NA | NA | NA | NA | NA | Gm43476 | | 0 | 0 | 0 | 0 | 0 | 0 | 0 | 0 | 0 | 0 | 0 |
| 0 | NA | NA | NA | NA | NA | Tmem132cos | | 0 | 0 | 0 | 0 | 0 | 0 | 0 | 0 | 0 | 0 | 0 |
| 0 | NA | NA | NA | NA | NA | 9430087B13Rik | | 0 | 0 | 0 | 0 | 0 | 0 | 0 | 0 | 0 | 0 | 0 |
| 0 | NA | NA | NA | NA | NA | Gm42980 | | 0 | 0 | 0 | 0 | 0 | 0 | 0 | 0 | 0 | 0 | 0 |
| 0 | NA | NA | NA | NA | NA | Gm17132 | | 0 | 0 | 0 | 0 | 0 | 0 | 0 | 0 | 0 | 0 | 0 |
| 0 | NA | NA | NA | NA | NA | Gm17131 | | 0 | 0 | 0 | 0 | 0 | 0 | 0 | 0 | 0 | 0 | 0 |
| 0 | NA | NA | NA | NA | NA | Gm42497 | | 0 | 0 | 0 | 0 | 0 | 0 | 0 | 0 | 0 | 0 | 0 |
| 0 | NA | NA | NA | NA | NA | Gm42496 | | 0 | 0 | 0 | 0 | 0 | 0 | 0 | 0 | 0 | 0 | 0 |
| 0 | NA | NA | NA | NA | NA | Gm42817 | | 0 | 0 | 0 | 0 | 0 | 0 | 0 | 0 | 0 | 0 | 0 |
| 0 | NA | NA | NA | NA | NA | Gm42495 | | 0 | 0 | 0 | 0 | 0 | 0 | 0 | 0 | 0 | 0 | 0 |
| 0 | NA | NA | NA | NA | NA | Gm42500 | | 0 | 0 | 0 | 0 | 0 | 0 | 0 | 0 | 0 | 0 | 0 |
| 0 | NA | NA | NA | NA | NA | 4930573I07Rik | | 0 | 0 | 0 | 0 | 0 | 0 | 0 | 0 | 0 | 0 | 0 |
| 0 | NA | NA | NA | NA | NA | Gm42499 | | 0 | 0 | 0 | 0 | 0 | 0 | 0 | 0 | 0 | 0 | 0 |
| 0 | NA | NA | NA | NA | NA | Gm42498 | | 0 | 0 | 0 | 0 | 0 | 0 | 0 | 0 | 0 | 0 | 0 |
| 0 | NA | NA | NA | NA | NA | Piwil1 | piwi-like RNA-mediated gene silencing 1 | 0 | 0 | 0 | 0 | 0 | 0 | 0 | 0 | 0 | 0 | 0 |
| 0 | NA | NA | NA | NA | NA | Gm43001 | | 0 | 0 | 0 | 0 | 0 | 0 | 0 | 0 | 0 | 0 | 0 |
| 0 | NA | NA | NA | NA | NA | Septin14 | septin 14 | 0 | 0 | 0 | 0 | 0 | 0 | 0 | 0 | 0 | 0 | 0 |
| 0 | NA | NA | NA | NA | NA | Snora15 | | 0 | 0 | 0 | 0 | 0 | 0 | 0 | 0 | 0 | 0 | 0 |

| | | | | | | | | | | | | | | | | | | |
|---|---|---|---|---|---|---|---|---|---|---|---|---|---|---|---|---|---|---|
| 0 | NA | NA | NA | NA | NA | A330070K13Ri | RIKEN cDNA A330070K13 gene | 0 | 0 | 0 | 0 | 0 | 0 | 0 | 0 | 0 | 0 | 0 |
| 0 | NA | NA | NA | NA | NA | Gm43418 | | 0 | 0 | 0 | 0 | 0 | 0 | 0 | 0 | 0 | 0 | 0 |
| 0 | NA | NA | NA | NA | NA | Gm23761 | | 0 | 0 | 0 | 0 | 0 | 0 | 0 | 0 | 0 | 0 | 0 |
| 0 | NA | NA | NA | NA | NA | Gm42894 | | 0 | 0 | 0 | 0 | 0 | 0 | 0 | 0 | 0 | 0 | 0 |
| 0 | NA | NA | NA | NA | NA | Gm42897 | | 0 | 0 | 0 | 0 | 0 | 0 | 0 | 0 | 0 | 0 | 0 |
| 0 | NA | NA | NA | NA | NA | Gm42896 | | 0 | 0 | 0 | 0 | 0 | 0 | 0 | 0 | 0 | 0 | 0 |
| 0 | NA | NA | NA | NA | NA | Gm43165 | | 0 | 0 | 0 | 0 | 0 | 0 | 0 | 0 | 0 | 0 | 0 |
| 0 | NA | NA | NA | NA | NA | Gm43166 | | 0 | 0 | 0 | 0 | 0 | 0 | 0 | 0 | 0 | 0 | 0 |
| 0 | NA | NA | NA | NA | NA | Gm27266 | | 0 | 0 | 0 | 0 | 0 | 0 | 0 | 0 | 0 | 0 | 0 |
| 0 | NA | NA | NA | NA | NA | Gm42591 | | 0 | 0 | 0 | 0 | 0 | 0 | 0 | 0 | 0 | 0 | 0 |
| 0 | NA | NA | NA | NA | NA | Gm43483 | | 0 | 0 | 0 | 0 | 0 | 0 | 0 | 0 | 0 | 0 | 0 |
| 0 | NA | NA | NA | NA | NA | 4930563F08Rik | | 0 | 0 | 0 | 0 | 0 | 0 | 0 | 0 | 0 | 0 | 0 |
| 0 | NA | NA | NA | NA | NA | Gm42441 | | 0 | 0 | 0 | 0 | 0 | 0 | 0 | 0 | 0 | 0 | 0 |
| 0 | NA | NA | NA | NA | NA | Gm42440 | | 0 | 0 | 0 | 0 | 0 | 0 | 0 | 0 | 0 | 0 | 0 |
| 0 | NA | NA | NA | NA | NA | Gm42439 | | 0 | 0 | 0 | 0 | 0 | 0 | 0 | 0 | 0 | 0 | 0 |
| 0 | NA | NA | NA | NA | NA | Gm42995 | | 0 | 0 | 0 | 0 | 0 | 0 | 0 | 0 | 0 | 0 | 0 |
| 0 | NA | NA | NA | NA | NA | Gm42996 | | 0 | 0 | 0 | 0 | 0 | 0 | 0 | 0 | 0 | 0 | 0 |
| 0 | NA | NA | NA | NA | NA | Gm43019 | | 0 | 0 | 0 | 0 | 0 | 0 | 0 | 0 | 0 | 0 | 0 |
| 0 | NA | NA | NA | NA | NA | Gm43187 | | 0 | 0 | 0 | 0 | 0 | 0 | 0 | 0 | 0 | 0 | 0 |
| 0 | NA | NA | NA | NA | NA | Gm43186 | | 0 | 0 | 0 | 0 | 0 | 0 | 0 | 0 | 0 | 0 | 0 |
| 0 | NA | NA | NA | NA | NA | Gm28616 | | 0 | 0 | 0 | 0 | 0 | 0 | 0 | 0 | 0 | 0 | 0 |
| 0 | NA | NA | NA | NA | NA | Gm42626 | | 0 | 0 | 0 | 0 | 0 | 0 | 0 | 0 | 0 | 0 | 0 |
| 0 | NA | NA | NA | NA | NA | Gm38373 | | 0 | 0 | 0 | 0 | 0 | 0 | 0 | 0 | 0 | 0 | 0 |
| 0 | NA | NA | NA | NA | NA | Gm25722 | | 0 | 0 | 0 | 0 | 0 | 0 | 0 | 0 | 0 | 0 | 0 |
| 0 | NA | NA | NA | NA | NA | Gm42625 | | 0 | 0 | 0 | 0 | 0 | 0 | 0 | 0 | 0 | 0 | 0 |
| 0 | NA | NA | NA | NA | NA | Gm17979 | | 0 | 0 | 0 | 0 | 0 | 0 | 0 | 0 | 0 | 0 | 0 |
| 0 | NA | NA | NA | NA | NA | Gm36667 | | 0 | 0 | 0 | 0 | 0 | 0 | 0 | 0 | 0 | 0 | 0 |
| 0 | NA | NA | NA | NA | NA | Gm2404 | | 0 | 0 | 0 | 0 | 0 | 0 | 0 | 0 | 0 | 0 | 0 |
| 0 | NA | NA | NA | NA | NA | 1700030N18Rik | | 0 | 0 | 0 | 0 | 0 | 0 | 0 | 0 | 0 | 0 | 0 |
| 0 | NA | NA | NA | NA | NA | Mir3965 | | 0 | 0 | 0 | 0 | 0 | 0 | 0 | 0 | 0 | 0 | 0 |
| 0 | NA | NA | NA | NA | NA | n-R5s178 | | 0 | 0 | 0 | 0 | 0 | 0 | 0 | 0 | 0 | 0 | 0 |
| 0 | NA | NA | NA | NA | NA | Gm26340 | | 0 | 0 | 0 | 0 | 0 | 0 | 0 | 0 | 0 | 0 | 0 |
| 0 | NA | NA | NA | NA | NA | Gm42885 | | 0 | 0 | 0 | 0 | 0 | 0 | 0 | 0 | 0 | 0 | 0 |
| 0 | NA | NA | NA | NA | NA | Mir7228 | | 0 | 0 | 0 | 0 | 0 | 0 | 0 | 0 | 0 | 0 | 0 |
| 0 | NA | NA | NA | NA | NA | Tmem270 | transmembrane protein 270 | 0 | 0 | 0 | 0 | 0 | 0 | 0 | 0 | 0 | 0 | 0 |
| 0 | NA | NA | NA | NA | NA | Cldn13 | claudin 13 | 0 | 0 | 0 | 0 | 0 | 0 | 0 | 0 | 0 | 0 | 0 |
| 0 | NA | NA | NA | NA | NA | Gm50473 | | 0 | 0 | 0 | 0 | 0 | 0 | 0 | 0 | 0 | 0 | 0 |
| 0 | NA | NA | NA | NA | NA | 4933439J24Rik | | 0 | 0 | 0 | 0 | 0 | 0 | 0 | 0 | 0 | 0 | 0 |
| 0 | NA | NA | NA | NA | NA | Gm50474 | | 0 | 0 | 0 | 0 | 0 | 0 | 0 | 0 | 0 | 0 | 0 |
| 0 | NA | NA | NA | NA | NA | 9030607J07Rik | | 0 | 0 | 0 | 0 | 0 | 0 | 0 | 0 | 0 | 0 | 0 |
| 0 | NA | NA | NA | NA | NA | Rps29-ps | | 0 | 0 | 0 | 0 | 0 | 0 | 0 | 0 | 0 | 0 | 0 |
| 0 | NA | NA | NA | NA | NA | Gm25315 | | 0 | 0 | 0 | 0 | 0 | 0 | 0 | 0 | 0 | 0 | 0 |
| 0 | NA | NA | NA | NA | NA | 9430007M09Rik | | 0 | 0 | 0 | 0 | 0 | 0 | 0 | 0 | 0 | 0 | 0 |
| 0 | NA | NA | NA | NA | NA | Mir7033 | | 0 | 0 | 0 | 0 | 0 | 0 | 0 | 0 | 0 | 0 | 0 |
| 0 | NA | NA | NA | NA | NA | Ccl26 | chemokine (C-C motif) ligand 26 | 0 | 0 | 0 | 0 | 0 | 0 | 0 | 0 | 0 | 0 | 0 |
| 0 | NA | NA | NA | NA | NA | Gm43123 | | 0 | 0 | 0 | 0 | 0 | 0 | 0 | 0 | 0 | 0 | 0 |
| 0 | NA | NA | NA | NA | NA | Gm43091 | | 0 | 0 | 0 | 0 | 0 | 0 | 0 | 0 | 0 | 0 | 0 |
| 0 | NA | NA | NA | NA | NA | Mir7034 | | 0 | 0 | 0 | 0 | 0 | 0 | 0 | 0 | 0 | 0 | 0 |
| 0 | NA | NA | NA | NA | NA | Gm23268 | | 0 | 0 | 0 | 0 | 0 | 0 | 0 | 0 | 0 | 0 | 0 |
| 0 | NA | NA | NA | NA | NA | Srrm3os | | 0 | 0 | 0 | 0 | 0 | 0 | 0 | 0 | 0 | 0 | 0 |
| 0 | NA | NA | NA | NA | NA | Zp3 | zona pellucida glycoprotein 3 | 0 | 0 | 0 | 0 | 0 | 0 | 0 | 0 | 0 | 0 | 0 |
| 0 | NA | NA | NA | NA | NA | Gm22214 | | 0 | 0 | 0 | 0 | 0 | 0 | 0 | 0 | 0 | 0 | 0 |
| 0 | NA | NA | NA | NA | NA | Gm15701 | | 0 | 0 | 0 | 0 | 0 | 0 | 0 | 0 | 0 | 0 | 0 |
| 0 | NA | NA | NA | NA | NA | Mir7035 | | 0 | 0 | 0 | 0 | 0 | 0 | 0 | 0 | 0 | 0 | 0 |
| 0 | NA | NA | NA | NA | NA | Mir721 | | 0 | 0 | 0 | 0 | 0 | 0 | 0 | 0 | 0 | 0 | 0 |
| 0 | NA | NA | NA | NA | NA | Myl10 | myosin, light chain 10, regulatory | 0 | 0 | 0 | 0 | 0 | 0 | 0 | 0 | 0 | 0 | 0 |
| 0 | NA | NA | NA | NA | NA | Mir702 | | 0 | 0 | 0 | 0 | 0 | 0 | 0 | 0 | 0 | 0 | 0 |
| 0 | NA | NA | NA | NA | NA | Gm7284 | | 0 | 0 | 0 | 0 | 0 | 0 | 0 | 0 | 0 | 0 | 0 |
| 0 | NA | NA | NA | NA | NA | Gm26989 | | 0 | 0 | 0 | 0 | 0 | 0 | 0 | 0 | 0 | 0 | 0 |
| 0 | NA | NA | NA | NA | NA | Gm40348 | | 0 | 0 | 0 | 0 | 0 | 0 | 0 | 0 | 0 | 0 | 0 |

| | | | | | | | | | | | | | | | | | |
|---|---|---|---|---|---|---|---|---|---|---|---|---|---|---|---|---|---|
| 0 | NA | NA | NA | NA | NA | Mir7036 | | 0 | 0 | 0 | 0 | 0 | 0 | 0 | 0 | 0 | 0 | 0 |
| 0 | NA | NA | NA | NA | NA | Rpl36-ps9 | | 0 | 0 | 0 | 0 | 0 | 0 | 0 | 0 | 0 | 0 | 0 |
| 0 | NA | NA | NA | NA | NA | Epo | erythropoietin | 0 | 0 | 0 | 0 | 0 | 0 | 0 | 0 | 0 | 0 | 0 |
| 0 | NA | NA | NA | NA | NA | Mir6418 | | 0 | 0 | 0 | 0 | 0 | 0 | 0 | 0 | 0 | 0 | 0 |
| 0 | NA | NA | NA | NA | NA | Sap25 | sin3 associated polypeptide | 0 | 0 | 0 | 0 | 0 | 0 | 0 | 0 | 0 | 0 | 0 |
| 0 | NA | NA | NA | NA | NA | Gm15498 | | 0 | 0 | 0 | 0 | 0 | 0 | 0 | 0 | 0 | 0 | 0 |
| 0 | NA | NA | NA | NA | NA | Gm43495 | | 0 | 0 | 0 | 0 | 0 | 0 | 0 | 0 | 0 | 0 | 0 |
| 0 | NA | NA | NA | NA | NA | Gm43494 | | 0 | 0 | 0 | 0 | 0 | 0 | 0 | 0 | 0 | 0 | 0 |
| 0 | NA | NA | NA | NA | NA | Gm3103 | | 0 | 0 | 0 | 0 | 0 | 0 | 0 | 0 | 0 | 0 | 0 |
| 0 | NA | NA | NA | NA | NA | Gm29127 | | 0 | 0 | 0 | 0 | 0 | 0 | 0 | 0 | 0 | 0 | 0 |
| 0 | NA | NA | NA | NA | NA | Smok3a | sperm motility kinase 3A | 0 | 0 | 0 | 0 | 0 | 0 | 0 | 0 | 0 | 0 | 0 |
| 0 | NA | NA | NA | NA | NA | Smok3b | sperm motility kinase 3B | 0 | 0 | 0 | 0 | 0 | 0 | 0 | 0 | 0 | 0 | 0 |
| 0 | NA | NA | NA | NA | NA | Smok3c | sperm motility kinase 3C | 0 | 0 | 0 | 0 | 0 | 0 | 0 | 0 | 0 | 0 | 0 |
| 0 | NA | NA | NA | NA | NA | Gm43493 | | 0 | 0 | 0 | 0 | 0 | 0 | 0 | 0 | 0 | 0 | 0 |
| 0 | NA | NA | NA | NA | NA | Mir25 | | 0 | 0 | 0 | 0 | 0 | 0 | 0 | 0 | 0 | 0 | 0 |
| 0 | NA | NA | NA | NA | NA | Mir93 | | 0 | 0 | 0 | 0 | 0 | 0 | 0 | 0 | 0 | 0 | 0 |
| 0 | NA | NA | NA | NA | NA | Mir106b | | 0 | 0 | 0 | 0 | 0 | 0 | 0 | 0 | 0 | 0 | 0 |
| 0 | NA | NA | NA | NA | NA | Gm454 | | 0 | 0 | 0 | 0 | 0 | 0 | 0 | 0 | 0 | 0 | 0 |
| 0 | NA | NA | NA | NA | NA | Gm43645 | | 0 | 0 | 0 | 0 | 0 | 0 | 0 | 0 | 0 | 0 | 0 |
| 0 | NA | NA | NA | NA | NA | Gm22956 | | 0 | 0 | 0 | 0 | 0 | 0 | 0 | 0 | 0 | 0 | 0 |
| 0 | NA | NA | NA | NA | NA | Gal3st4 | galactose-3-O-sulfotransferase 4 | 0 | 0 | 0 | 0 | 0 | 0 | 0 | 0 | 0 | 0 | 0 |
| 0 | NA | NA | NA | NA | NA | Gm43720 | | 0 | 0 | 0 | 0 | 0 | 0 | 0 | 0 | 0 | 0 | 0 |
| 0 | NA | NA | NA | NA | NA | Pvrig-ps | | 0 | 0 | 0 | 0 | 0 | 0 | 0 | 0 | 0 | 0 | 0 |
| 0 | NA | NA | NA | NA | NA | Gm10874 | | 0 | 0 | 0 | 0 | 0 | 0 | 0 | 0 | 0 | 0 | 0 |
| 0 | NA | NA | NA | NA | NA | Gm25525 | | 0 | 0 | 0 | 0 | 0 | 0 | 0 | 0 | 0 | 0 | 0 |
| 0 | NA | NA | NA | NA | NA | Cyp3a61-ps1 | | 0 | 0 | 0 | 0 | 0 | 0 | 0 | 0 | 0 | 0 | 0 |
| 0 | NA | NA | NA | NA | NA | Gm48029 | | 0 | 0 | 0 | 0 | 0 | 0 | 0 | 0 | 0 | 0 | 0 |
| 0 | NA | NA | NA | NA | NA | Vmn2r-ps24 | | 0 | 0 | 0 | 0 | 0 | 0 | 0 | 0 | 0 | 0 | 0 |
| 0 | NA | NA | NA | NA | NA | 1700123K08Ril | RIKEN cDNA 1700123K08 gene | 0 | 0 | 0 | 0 | 0 | 0 | 0 | 0 | 0 | 0 | 0 |
| 0 | NA | NA | NA | NA | NA | Gm38232 | | 0 | 0 | 0 | 0 | 0 | 0 | 0 | 0 | 0 | 0 | 0 |
| 0 | NA | NA | NA | NA | NA | 4930520M14Rik | | 0 | 0 | 0 | 0 | 0 | 0 | 0 | 0 | 0 | 0 | 0 |
| 0 | NA | NA | NA | NA | NA | Gm27316 | | 0 | 0 | 0 | 0 | 0 | 0 | 0 | 0 | 0 | 0 | 0 |
| 0 | NA | NA | NA | NA | NA | Cyp2w1 | cytochrome P450, family 2, subfamily w, p | 0 | 0 | 0 | 0 | 0 | 0 | 0 | 0 | 0 | 0 | 0 |
| 0 | NA | NA | NA | NA | NA | Gm26285 | | 0 | 0 | 0 | 0 | 0 | 0 | 0 | 0 | 0 | 0 | 0 |
| 0 | NA | NA | NA | NA | NA | 4930500L23Rik | | 0 | 0 | 0 | 0 | 0 | 0 | 0 | 0 | 0 | 0 | 0 |
| 0 | NA | NA | NA | NA | NA | Uncx | UNC homeobox | 0 | 0 | 0 | 0 | 0 | 0 | 0 | 0 | 0 | 0 | 0 |
| 0 | NA | NA | NA | NA | NA | Gm42424 | | 0 | 0 | 0 | 0 | 0 | 0 | 0 | 0 | 0 | 0 | 0 |
| 0 | NA | NA | NA | NA | NA | Mir7037 | | 0 | 0 | 0 | 0 | 0 | 0 | 0 | 0 | 0 | 0 | 0 |
| 0 | NA | NA | NA | NA | NA | Gm26938 | | 0 | 0 | 0 | 0 | 0 | 0 | 0 | 0 | 0 | 0 | 0 |
| 0 | NA | NA | NA | NA | NA | Gm24146 | | 0 | 0 | 0 | 0 | 0 | 0 | 0 | 0 | 0 | 0 | 0 |
| 0 | NA | NA | NA | NA | NA | Gm42423 | | 0 | 0 | 0 | 0 | 0 | 0 | 0 | 0 | 0 | 0 | 0 |
| 0 | NA | NA | NA | NA | NA | Gm16122 | | 0 | 0 | 0 | 0 | 0 | 0 | 0 | 0 | 0 | 0 | 0 |
| 0 | NA | NA | NA | NA | NA | Gm16121 | | 0 | 0 | 0 | 0 | 0 | 0 | 0 | 0 | 0 | 0 | 0 |
| 0 | NA | NA | NA | NA | NA | Gm16120 | | 0 | 0 | 0 | 0 | 0 | 0 | 0 | 0 | 0 | 0 | 0 |
| 0 | NA | NA | NA | NA | NA | Gm43338 | | 0 | 0 | 0 | 0 | 0 | 0 | 0 | 0 | 0 | 0 | 0 |
| 0 | NA | NA | NA | NA | NA | Mir7038 | | 0 | 0 | 0 | 0 | 0 | 0 | 0 | 0 | 0 | 0 | 0 |
| 0 | NA | NA | NA | NA | NA | Gm42805 | | 0 | 0 | 0 | 0 | 0 | 0 | 0 | 0 | 0 | 0 | 0 |
| 0 | NA | NA | NA | NA | NA | Grifin | galectin-related inter-fiber protein | 0 | 0 | 0 | 0 | 0 | 0 | 0 | 0 | 0 | 0 | 0 |
| 0 | NA | NA | NA | NA | NA | Gm44385 | | 0 | 0 | 0 | 0 | 0 | 0 | 0 | 0 | 0 | 0 | 0 |
| 0 | NA | NA | NA | NA | NA | Gm26113 | | 0 | 0 | 0 | 0 | 0 | 0 | 0 | 0 | 0 | 0 | 0 |
| 0 | NA | NA | NA | NA | NA | Gm43708 | | 0 | 0 | 0 | 0 | 0 | 0 | 0 | 0 | 0 | 0 | 0 |
| 0 | NA | NA | NA | NA | NA | Gm43556 | | 0 | 0 | 0 | 0 | 0 | 0 | 0 | 0 | 0 | 0 | 0 |
| 0 | NA | NA | NA | NA | NA | Gm43557 | | 0 | 0 | 0 | 0 | 0 | 0 | 0 | 0 | 0 | 0 | 0 |
| 0 | NA | NA | NA | NA | NA | Gm24131 | | 0 | 0 | 0 | 0 | 0 | 0 | 0 | 0 | 0 | 0 | 0 |
| 0 | NA | NA | NA | NA | NA | Gm16036 | | 0 | 0 | 0 | 0 | 0 | 0 | 0 | 0 | 0 | 0 | 0 |
| 0 | NA | NA | NA | NA | NA | Gm43037 | | 0 | 0 | 0 | 0 | 0 | 0 | 0 | 0 | 0 | 0 | 0 |
| 0 | NA | NA | NA | NA | NA | Gm15770 | | 0 | 0 | 0 | 0 | 0 | 0 | 0 | 0 | 0 | 0 | 0 |
| 0 | NA | NA | NA | NA | NA | Olfr718-ps1 | | 0 | 0 | 0 | 0 | 0 | 0 | 0 | 0 | 0 | 0 | 0 |
| 0 | NA | NA | NA | NA | NA | Gm7291 | | 0 | 0 | 0 | 0 | 0 | 0 | 0 | 0 | 0 | 0 | 0 |
| 0 | NA | NA | NA | NA | NA | Gm44405 | | 0 | 0 | 0 | 0 | 0 | 0 | 0 | 0 | 0 | 0 | 0 |

| | | | | | | | | | | | | | | | | | | |
|---|---|---|---|---|---|---|---|---|---|---|---|---|---|---|---|---|---|---|
| 0 | NA | NA | NA | NA | NA | Rbakdn | | 0 | 0 | 0 | 0 | 0 | 0 | 0 | 0 | 0 | 0 | 0 |
| 0 | NA | NA | NA | NA | NA | Spdye4a | speedy/RINGO cell cycle regulator family | 0 | 0 | 0 | 0 | 0 | 0 | 0 | 0 | 0 | 0 | 0 |
| 0 | NA | NA | NA | NA | NA | 4930448H16Rik | | 0 | 0 | 0 | 0 | 0 | 0 | 0 | 0 | 0 | 0 | 0 |
| 0 | NA | NA | NA | NA | NA | Gm42504 | | 0 | 0 | 0 | 0 | 0 | 0 | 0 | 0 | 0 | 0 | 0 |
| 0 | NA | NA | NA | NA | NA | Gm4870 | | 0 | 0 | 0 | 0 | 0 | 0 | 0 | 0 | 0 | 0 | 0 |
| 0 | NA | NA | NA | NA | NA | Gm42502 | | 0 | 0 | 0 | 0 | 0 | 0 | 0 | 0 | 0 | 0 | 0 |
| 0 | NA | NA | NA | NA | NA | Fam220a | family with sequence similarity 220, mem | 0 | 0 | 0 | 0 | 0 | 0 | 0 | 0 | 0 | 0 | 0 |
| 0 | NA | NA | NA | NA | NA | Gm17135 | | 0 | 0 | 0 | 0 | 0 | 0 | 0 | 0 | 0 | 0 | 0 |
| 0 | NA | NA | NA | NA | NA | Gm42421 | | 0 | 0 | 0 | 0 | 0 | 0 | 0 | 0 | 0 | 0 | 0 |
| 0 | NA | NA | NA | NA | NA | Gm24311 | | 0 | 0 | 0 | 0 | 0 | 0 | 0 | 0 | 0 | 0 | 0 |
| 0 | NA | NA | NA | NA | NA | Ocm | oncomodulin | 0 | 0 | 0 | 0 | 0 | 0 | 0 | 0 | 0 | 0 | 0 |
| 0 | NA | NA | NA | NA | NA | Gm43615 | | 0 | 0 | 0 | 0 | 0 | 0 | 0 | 0 | 0 | 0 | 0 |
| 0 | NA | NA | NA | NA | NA | Gm26925 | | 0 | 0 | 0 | 0 | 0 | 0 | 0 | 0 | 0 | 0 | 0 |
| 0 | NA | NA | NA | NA | NA | Dmrt1i | | 0 | 0 | 0 | 0 | 0 | 0 | 0 | 0 | 0 | 0 | 0 |
| 0 | NA | NA | NA | NA | NA | Gm15709 | | 0 | 0 | 0 | 0 | 0 | 0 | 0 | 0 | 0 | 0 | 0 |
| 0 | NA | NA | NA | NA | NA | Gm6272 | | 0 | 0 | 0 | 0 | 0 | 0 | 0 | 0 | 0 | 0 | 0 |
| 0 | NA | NA | NA | NA | NA | Mir7039 | | 0 | 0 | 0 | 0 | 0 | 0 | 0 | 0 | 0 | 0 | 0 |
| 0 | NA | NA | NA | NA | NA | Gm43563 | | 0 | 0 | 0 | 0 | 0 | 0 | 0 | 0 | 0 | 0 | 0 |
| 0 | NA | NA | NA | NA | NA | Gm4871 | predicted gene 4871 | 0 | 0 | 0 | 0 | 0 | 0 | 0 | 0 | 0 | 0 | 0 |
| 0 | NA | NA | NA | NA | NA | Gm43562 | | 0 | 0 | 0 | 0 | 0 | 0 | 0 | 0 | 0 | 0 | 0 |
| 0 | NA | NA | NA | NA | NA | 1700018F24Ri | RIKEN cDNA 1700018F24 gene | 0 | 0 | 0 | 0 | 0 | 0 | 0 | 0 | 0 | 0 | 0 |
| 0 | NA | NA | NA | NA | NA | Gm50458 | | 0 | 0 | 0 | 0 | 0 | 0 | 0 | 0 | 0 | 0 | 0 |
| 0 | NA | NA | NA | NA | NA | Gm9710 | | 0 | 0 | 0 | 0 | 0 | 0 | 0 | 0 | 0 | 0 | 0 |
| 0 | NA | NA | NA | NA | NA | Cyp3a57 | cytochrome P450, family 3, subfamily a, p | 0 | 0 | 0 | 0 | 0 | 0 | 0 | 0 | 0 | 0 | 0 |
| 0 | NA | NA | NA | NA | NA | Gm2314 | | 0 | 0 | 0 | 0 | 0 | 0 | 0 | 0 | 0 | 0 | 0 |
| 0 | NA | NA | NA | NA | NA | Gm42859 | | 0 | 0 | 0 | 0 | 0 | 0 | 0 | 0 | 0 | 0 | 0 |
| 0 | NA | NA | NA | NA | NA | Cyp3a16 | cytochrome P450, family 3, subfamily a, p | 0 | 0 | 0 | 0 | 0 | 0 | 0 | 0 | 0 | 0 | 0 |
| 0 | NA | NA | NA | NA | NA | Gm42863 | | 0 | 0 | 0 | 0 | 0 | 0 | 0 | 0 | 0 | 0 | 0 |
| 0 | NA | NA | NA | NA | NA | Cyp3a41b | cytochrome P450, family 3, subfamily a, p | 0 | 0 | 0 | 0 | 0 | 0 | 0 | 0 | 0 | 0 | 0 |
| 0 | NA | NA | NA | NA | NA | Gm43381 | | 0 | 0 | 0 | 0 | 0 | 0 | 0 | 0 | 0 | 0 | 0 |
| 0 | NA | NA | NA | NA | NA | Cyp3a63-ps | | 0 | 0 | 0 | 0 | 0 | 0 | 0 | 0 | 0 | 0 | 0 |
| 0 | NA | NA | NA | NA | NA | Cyp3a41a | cytochrome P450, family 3, subfamily a, p | 0 | 0 | 0 | 0 | 0 | 0 | 0 | 0 | 0 | 0 | 0 |
| 0 | NA | NA | NA | NA | NA | Gm43626 | | 0 | 0 | 0 | 0 | 0 | 0 | 0 | 0 | 0 | 0 | 0 |
| 0 | NA | NA | NA | NA | NA | Cyp3a44 | cytochrome P450, family 3, subfamily a, p | 0 | 0 | 0 | 0 | 0 | 0 | 0 | 0 | 0 | 0 | 0 |
| 0 | NA | NA | NA | NA | NA | Gm19089 | | 0 | 0 | 0 | 0 | 0 | 0 | 0 | 0 | 0 | 0 | 0 |
| 0 | NA | NA | NA | NA | NA | Gm42431 | | 0 | 0 | 0 | 0 | 0 | 0 | 0 | 0 | 0 | 0 | 0 |
| 0 | NA | NA | NA | NA | NA | Gm43624 | | 0 | 0 | 0 | 0 | 0 | 0 | 0 | 0 | 0 | 0 | 0 |
| 0 | NA | NA | NA | NA | NA | Gm43115 | | 0 | 0 | 0 | 0 | 0 | 0 | 0 | 0 | 0 | 0 | 0 |
| 0 | NA | NA | NA | NA | NA | Cyp3a25 | cytochrome P450, family 3, subfamily a, p | 0 | 0 | 0 | 0 | 0 | 0 | 0 | 0 | 0 | 0 | 0 |
| 0 | NA | NA | NA | NA | NA | Gm8454 | | 0 | 0 | 0 | 0 | 0 | 0 | 0 | 0 | 0 | 0 | 0 |
| 0 | NA | NA | NA | NA | NA | Gm43239 | | 0 | 0 | 0 | 0 | 0 | 0 | 0 | 0 | 0 | 0 | 0 |
| 0 | NA | NA | NA | NA | NA | Gm8458 | | 0 | 0 | 0 | 0 | 0 | 0 | 0 | 0 | 0 | 0 | 0 |
| 0 | NA | NA | NA | NA | NA | Cyp3a59 | cytochrome P450, family 3, subfamily a, p | 0 | 0 | 0 | 0 | 0 | 0 | 0 | 0 | 0 | 0 | 0 |
| 0 | NA | NA | NA | NA | NA | Gm5565 | predicted gene 5565 | 0 | 0 | 0 | 0 | 0 | 0 | 0 | 0 | 0 | 0 | 0 |
| 0 | NA | NA | NA | NA | NA | Gm6309 | predicted gene 6309 | 0 | 0 | 0 | 0 | 0 | 0 | 0 | 0 | 0 | 0 | 0 |
| 0 | NA | NA | NA | NA | NA | Gm6408 | predicted gene 6408 | 0 | 0 | 0 | 0 | 0 | 0 | 0 | 0 | 0 | 0 | 0 |
| 0 | NA | NA | NA | NA | NA | Gm6370 | predicted gene 6370 | 0 | 0 | 0 | 0 | 0 | 0 | 0 | 0 | 0 | 0 | 0 |
| 0 | NA | NA | NA | NA | NA | 4930449I24Rik | RIKEN cDNA 4930449I24 gene | 0 | 0 | 0 | 0 | 0 | 0 | 0 | 0 | 0 | 0 | 0 |
| 0 | NA | NA | NA | NA | NA | Gm3402 | predicted gene 3402 | 0 | 0 | 0 | 0 | 0 | 0 | 0 | 0 | 0 | 0 | 0 |
| 0 | NA | NA | NA | NA | NA | Gm3404 | predicted gene 3404 | 0 | 0 | 0 | 0 | 0 | 0 | 0 | 0 | 0 | 0 | 0 |
| 0 | NA | NA | NA | NA | NA | Gm3409 | predicted gene 3409 | 0 | 0 | 0 | 0 | 0 | 0 | 0 | 0 | 0 | 0 | 0 |
| 0 | NA | NA | NA | NA | NA | Gm3415 | predicted gene 3415 | 0 | 0 | 0 | 0 | 0 | 0 | 0 | 0 | 0 | 0 | 0 |
| 0 | NA | NA | NA | NA | NA | Gpr12 | G-protein coupled receptor 12 | 0 | 0 | 0 | 0 | 0 | 0 | 0 | 0 | 0 | 0 | 0 |
| 0 | NA | NA | NA | NA | NA | Gm34333 | | 0 | 0 | 0 | 0 | 0 | 0 | 0 | 0 | 0 | 0 | 0 |
| 0 | NA | NA | NA | NA | NA | 4930573C15Rik | | 0 | 0 | 0 | 0 | 0 | 0 | 0 | 0 | 0 | 0 | 0 |
| 0 | NA | NA | NA | NA | NA | 1700041I07Rik | | 0 | 0 | 0 | 0 | 0 | 0 | 0 | 0 | 0 | 0 | 0 |
| 0 | NA | NA | NA | NA | NA | Gm25091 | | 0 | 0 | 0 | 0 | 0 | 0 | 0 | 0 | 0 | 0 | 0 |
| 0 | NA | NA | NA | NA | NA | Gm23202 | | 0 | 0 | 0 | 0 | 0 | 0 | 0 | 0 | 0 | 0 | 0 |
| 0 | NA | NA | NA | NA | NA | Gm24556 | | 0 | 0 | 0 | 0 | 0 | 0 | 0 | 0 | 0 | 0 | 0 |
| 0 | NA | NA | NA | NA | NA | Mir7k | | 0 | 0 | 0 | 0 | 0 | 0 | 0 | 0 | 0 | 0 | 0 |

| | | | | | | | | | | | | | | | | | | |
|---|---|---|---|---|---|---|---|---|---|---|---|---|---|---|---|---|---|---|
| 0 | NA | NA | NA | NA | NA | Urad | ureidoimidazoline (2-oxo-4-hydroxy-4-ca | 0 | 0 | 0 | 0 | 0 | 0 | 0 | 0 | 0 | 0 | 0 |
| 0 | NA | NA | NA | NA | NA | Gm29778 | | 0 | 0 | 0 | 0 | 0 | 0 | 0 | 0 | 0 | 0 | 0 |
| 0 | NA | NA | NA | NA | NA | Gm26597 | | 0 | 0 | 0 | 0 | 0 | 0 | 0 | 0 | 0 | 0 | 0 |
| 0 | NA | NA | NA | NA | NA | Gm43460 | | 0 | 0 | 0 | 0 | 0 | 0 | 0 | 0 | 0 | 0 | 0 |
| 0 | NA | NA | NA | NA | NA | Gm42883 | | 0 | 0 | 0 | 0 | 0 | 0 | 0 | 0 | 0 | 0 | 0 |
| 0 | NA | NA | NA | NA | NA | Gm43156 | | 0 | 0 | 0 | 0 | 0 | 0 | 0 | 0 | 0 | 0 | 0 |
| 0 | NA | NA | NA | NA | NA | Gm15721 | | 0 | 0 | 0 | 0 | 0 | 0 | 0 | 0 | 0 | 0 | 0 |
| 0 | NA | NA | NA | NA | NA | Gm43553 | | 0 | 0 | 0 | 0 | 0 | 0 | 0 | 0 | 0 | 0 | 0 |
| 0 | NA | NA | NA | NA | NA | Gm19617 | | 0 | 0 | 0 | 0 | 0 | 0 | 0 | 0 | 0 | 0 | 0 |
| 0 | NA | NA | NA | NA | NA | 1700013A02Rik | | 0 | 0 | 0 | 0 | 0 | 0 | 0 | 0 | 0 | 0 | 0 |
| 0 | NA | NA | NA | NA | NA | Gm29815 | | 0 | 0 | 0 | 0 | 0 | 0 | 0 | 0 | 0 | 0 | 0 |
| 0 | NA | NA | NA | NA | NA | Gm36186 | | 0 | 0 | 0 | 0 | 0 | 0 | 0 | 0 | 0 | 0 | 0 |
| 0 | NA | NA | NA | NA | NA | n-R5s179 | | 0 | 0 | 0 | 0 | 0 | 0 | 0 | 0 | 0 | 0 | 0 |
| 0 | NA | NA | NA | NA | NA | 4930505K14Rik | | 0 | 0 | 0 | 0 | 0 | 0 | 0 | 0 | 0 | 0 | 0 |
| 0 | NA | NA | NA | NA | NA | Gm43600 | | 0 | 0 | 0 | 0 | 0 | 0 | 0 | 0 | 0 | 0 | 0 |
| 0 | NA | NA | NA | NA | NA | Gm15406 | | 0 | 0 | 0 | 0 | 0 | 0 | 0 | 0 | 0 | 0 | 0 |
| 0 | NA | NA | NA | NA | NA | 8430423G03Rik | | 0 | 0 | 0 | 0 | 0 | 0 | 0 | 0 | 0 | 0 | 0 |
| 0 | NA | NA | NA | NA | NA | Gm42787 | | 0 | 0 | 0 | 0 | 0 | 0 | 0 | 0 | 0 | 0 | 0 |
| 0 | NA | NA | NA | NA | NA | Gm42788 | | 0 | 0 | 0 | 0 | 0 | 0 | 0 | 0 | 0 | 0 | 0 |
| 0 | NA | NA | NA | NA | NA | Gm23362 | | 0 | 0 | 0 | 0 | 0 | 0 | 0 | 0 | 0 | 0 | 0 |
| 0 | NA | NA | NA | NA | NA | Gm24105 | | 0 | 0 | 0 | 0 | 0 | 0 | 0 | 0 | 0 | 0 | 0 |
| 0 | NA | NA | NA | NA | NA | n-R5s180 | | 0 | 0 | 0 | 0 | 0 | 0 | 0 | 0 | 0 | 0 | 0 |
| 0 | NA | NA | NA | NA | NA | Gm20488 | | 0 | 0 | 0 | 0 | 0 | 0 | 0 | 0 | 0 | 0 | 0 |
| 0 | NA | NA | NA | NA | NA | Gm43151 | | 0 | 0 | 0 | 0 | 0 | 0 | 0 | 0 | 0 | 0 | 0 |
| 0 | NA | NA | NA | NA | NA | Gm15997 | | 0 | 0 | 0 | 0 | 0 | 0 | 0 | 0 | 0 | 0 | 0 |
| 0 | NA | NA | NA | NA | NA | Gm2566 | | 0 | 0 | 0 | 0 | 0 | 0 | 0 | 0 | 0 | 0 | 0 |
| 0 | NA | NA | NA | NA | NA | Wdr95 | WD40 repeat domain 95 | 0 | 0 | 0 | 0 | 0 | 0 | 0 | 0 | 0 | 0 | 0 |
| 0 | NA | NA | NA | NA | NA | Gm21221 | | 0 | 0 | 0 | 0 | 0 | 0 | 0 | 0 | 0 | 0 | 0 |
| 0 | NA | NA | NA | NA | NA | 4930500F04Rik | | 0 | 0 | 0 | 0 | 0 | 0 | 0 | 0 | 0 | 0 | 0 |
| 0 | NA | NA | NA | NA | NA | Gm43450 | | 0 | 0 | 0 | 0 | 0 | 0 | 0 | 0 | 0 | 0 | 0 |
| 0 | NA | NA | NA | NA | NA | Gm43452 | | 0 | 0 | 0 | 0 | 0 | 0 | 0 | 0 | 0 | 0 | 0 |
| 0 | NA | NA | NA | NA | NA | Gm21048 | | 0 | 0 | 0 | 0 | 0 | 0 | 0 | 0 | 0 | 0 | 0 |
| 0 | NA | NA | NA | NA | NA | Gm42558 | | 0 | 0 | 0 | 0 | 0 | 0 | 0 | 0 | 0 | 0 | 0 |
| 0 | NA | NA | NA | NA | NA | Gm42557 | | 0 | 0 | 0 | 0 | 0 | 0 | 0 | 0 | 0 | 0 | 0 |
| 0 | NA | NA | NA | NA | NA | Gm42556 | | 0 | 0 | 0 | 0 | 0 | 0 | 0 | 0 | 0 | 0 | 0 |
| 0 | NA | NA | NA | NA | NA | | | 0 | 0 | 0 | 0 | 0 | 0 | 0 | 0 | 0 | 0 | 0 |
| 0 | NA | NA | NA | NA | NA | Gm5 | | 0 | 0 | 0 | 0 | 0 | 0 | 0 | 0 | 0 | 0 | 0 |
| 0 | NA | NA | NA | NA | NA | Gm15933 | | 0 | 0 | 0 | 0 | 0 | 0 | 0 | 0 | 0 | 0 | 0 |
| 0 | NA | NA | NA | NA | NA | Gm25284 | | 0 | 0 | 0 | 0 | 0 | 0 | 0 | 0 | 0 | 0 | 0 |
| 0 | NA | NA | NA | NA | NA | Gm23641 | | 0 | 0 | 0 | 0 | 0 | 0 | 0 | 0 | 0 | 0 | 0 |
| 0 | NA | NA | NA | NA | NA | Gm42529 | | 0 | 0 | 0 | 0 | 0 | 0 | 0 | 0 | 0 | 0 | 0 |
| 0 | NA | NA | NA | NA | NA | Gm24630 | | 0 | 0 | 0 | 0 | 0 | 0 | 0 | 0 | 0 | 0 | 0 |
| 0 | NA | NA | NA | NA | NA | Gm26013 | | 0 | 0 | 0 | 0 | 0 | 0 | 0 | 0 | 0 | 0 | 0 |
| 0 | NA | NA | NA | NA | NA | Gm43807 | | 0 | 0 | 0 | 0 | 0 | 0 | 0 | 0 | 0 | 0 | 0 |
| 0 | NA | NA | NA | NA | NA | Gm43298 | | 0 | 0 | 0 | 0 | 0 | 0 | 0 | 0 | 0 | 0 | 0 |
| 0 | NA | NA | NA | NA | NA | Gm42906 | | 0 | 0 | 0 | 0 | 0 | 0 | 0 | 0 | 0 | 0 | 0 |
| 0 | NA | NA | NA | NA | NA | 5430435K18Rik | | 0 | 0 | 0 | 0 | 0 | 0 | 0 | 0 | 0 | 0 | 0 |
| 0 | NA | NA | NA | NA | NA | Gm7041 | | 0 | 0 | 0 | 0 | 0 | 0 | 0 | 0 | 0 | 0 | 0 |
| 0 | NA | NA | NA | NA | NA | Vmn2r-ps25 | | 0 | 0 | 0 | 0 | 0 | 0 | 0 | 0 | 0 | 0 | 0 |
| 0 | NA | NA | NA | NA | NA | Vmn2r-ps26 | | 0 | 0 | 0 | 0 | 0 | 0 | 0 | 0 | 0 | 0 | 0 |
| 0 | NA | NA | NA | NA | NA | Vmn2r18 | vomeronasal 2, receptor 18 | 0 | 0 | 0 | 0 | 0 | 0 | 0 | 0 | 0 | 0 | 0 |
| 0 | NA | NA | NA | NA | NA | | | 0 | 0 | 0 | 0 | 0 | 0 | 0 | 0 | 0 | 0 | 0 |
| 0 | NA | NA | NA | NA | NA | Gm43038 | | 0 | 0 | 0 | 0 | 0 | 0 | 0 | 0 | 0 | 0 | 0 |
| 0 | NA | NA | NA | NA | NA | Gm43129 | | 0 | 0 | 0 | 0 | 0 | 0 | 0 | 0 | 0 | 0 | 0 |
| 0 | NA | NA | NA | NA | NA | Gm24242 | | 0 | 0 | 0 | 0 | 0 | 0 | 0 | 0 | 0 | 0 | 0 |
| 0 | NA | NA | NA | NA | NA | Gm26463 | | 0 | 0 | 0 | 0 | 0 | 0 | 0 | 0 | 0 | 0 | 0 |
| 0 | NA | NA | NA | NA | NA | Gm23551 | | 0 | 0 | 0 | 0 | 0 | 0 | 0 | 0 | 0 | 0 | 0 |
| 0 | NA | NA | NA | NA | NA | Vmn1r-ps5 | | 0 | 0 | 0 | 0 | 0 | 0 | 0 | 0 | 0 | 0 | 0 |
| 0 | NA | NA | NA | NA | NA | Vmn1r-ps6 | | 0 | 0 | 0 | 0 | 0 | 0 | 0 | 0 | 0 | 0 | 0 |
| 0 | NA | NA | NA | NA | NA | Vmn1r261 | | 0 | 0 | 0 | 0 | 0 | 0 | 0 | 0 | 0 | 0 | 0 |

| | | | | | | | | | | | | | | | | | | |
|---|---|---|---|---|---|---|---|---|---|---|---|---|---|---|---|---|---|---|
| 0 | NA | NA | NA | NA | NA | Gm42642 | | 0 | 0 | 0 | 0 | 0 | 0 | 0 | 0 | 0 | 0 | 0 |
| 0 | NA | NA | NA | NA | NA | Gm2651 | | 0 | 0 | 0 | 0 | 0 | 0 | 0 | 0 | 0 | 0 | 0 |
| 0 | NA | NA | NA | NA | NA | Gm42960 | | 0 | 0 | 0 | 0 | 0 | 0 | 0 | 0 | 0 | 0 | 0 |
| 0 | NA | NA | NA | NA | NA | Gm42961 | | 0 | 0 | 0 | 0 | 0 | 0 | 0 | 0 | 0 | 0 | 0 |
| 0 | NA | NA | NA | NA | NA | Gm18222 | | 0 | 0 | 0 | 0 | 0 | 0 | 0 | 0 | 0 | 0 | 0 |
| 0 | NA | NA | NA | NA | NA | Gm43198 | | 0 | 0 | 0 | 0 | 0 | 0 | 0 | 0 | 0 | 0 | 0 |
| 0 | NA | NA | NA | NA | NA | Mir653 | | 0 | 0 | 0 | 0 | 0 | 0 | 0 | 0 | 0 | 0 | 0 |
| 0 | NA | NA | NA | NA | NA | Mir489 | | 0 | 0 | 0 | 0 | 0 | 0 | 0 | 0 | 0 | 0 | 0 |
| 0 | NA | NA | NA | NA | NA | Gm44221 | | 0 | 0 | 0 | 0 | 0 | 0 | 0 | 0 | 0 | 0 | 0 |
| 0 | NA | NA | NA | NA | NA | Gngt1 | guanine nucleotide binding protein (G prc | 0 | 0 | 0 | 0 | 0 | 0 | 0 | 0 | 0 | 0 | 0 |
| 0 | NA | NA | NA | NA | NA | 4930528H21Rik | | 0 | 0 | 0 | 0 | 0 | 0 | 0 | 0 | 0 | 0 | 0 |
| 0 | NA | NA | NA | NA | NA | Gm44055 | | 0 | 0 | 0 | 0 | 0 | 0 | 0 | 0 | 0 | 0 | 0 |
| 0 | NA | NA | NA | NA | NA | Gm44057 | | 0 | 0 | 0 | 0 | 0 | 0 | 0 | 0 | 0 | 0 | 0 |
| 0 | NA | NA | NA | NA | NA | Gm44379 | | 0 | 0 | 0 | 0 | 0 | 0 | 0 | 0 | 0 | 0 | 0 |
| 0 | NA | NA | NA | NA | NA | Gm44380 | | 0 | 0 | 0 | 0 | 0 | 0 | 0 | 0 | 0 | 0 | 0 |
| 0 | NA | NA | NA | NA | NA | Gm24442 | | 0 | 0 | 0 | 0 | 0 | 0 | 0 | 0 | 0 | 0 | 0 |
| 0 | NA | NA | NA | NA | NA | Gm2522 | | 0 | 0 | 0 | 0 | 0 | 0 | 0 | 0 | 0 | 0 | 0 |
| 0 | NA | NA | NA | NA | NA | Gm37883 | | 0 | 0 | 0 | 0 | 0 | 0 | 0 | 0 | 0 | 0 | 0 |
| 0 | NA | NA | NA | NA | NA | Gm18289 | | 0 | 0 | 0 | 0 | 0 | 0 | 0 | 0 | 0 | 0 | 0 |
| 0 | NA | NA | NA | NA | NA | Gm44121 | | 0 | 0 | 0 | 0 | 0 | 0 | 0 | 0 | 0 | 0 | 0 |
| 0 | NA | NA | NA | NA | NA | Gm20714 | | 0 | 0 | 0 | 0 | 0 | 0 | 0 | 0 | 0 | 0 | 0 |
| 0 | NA | NA | NA | NA | NA | Gm26828 | | 0 | 0 | 0 | 0 | 0 | 0 | 0 | 0 | 0 | 0 | 0 |
| 0 | NA | NA | NA | NA | NA | Gm26556 | | 0 | 0 | 0 | 0 | 0 | 0 | 0 | 0 | 0 | 0 | 0 |
| 0 | NA | NA | NA | NA | NA | Pon1 | paraoxonase 1 | 0 | 0 | 0 | 0 | 0 | 0 | 0 | 0 | 0 | 0 | 0 |
| 0 | NA | NA | NA | NA | NA | Gm44424 | | 0 | 0 | 0 | 0 | 0 | 0 | 0 | 0 | 0 | 0 | 0 |
| 0 | NA | NA | NA | NA | NA | 1700019G24Rik | | 0 | 0 | 0 | 0 | 0 | 0 | 0 | 0 | 0 | 0 | 0 |
| 0 | NA | NA | NA | NA | NA | Gm8649 | | 0 | 0 | 0 | 0 | 0 | 0 | 0 | 0 | 0 | 0 | 0 |
| 0 | NA | NA | NA | NA | NA | Gm20617 | | 0 | 0 | 0 | 0 | 0 | 0 | 0 | 0 | 0 | 0 | 0 |
| 0 | NA | NA | NA | NA | NA | Gm20685 | | 0 | 0 | 0 | 0 | 0 | 0 | 0 | 0 | 0 | 0 | 0 |
| 0 | NA | NA | NA | NA | NA | Gm20618 | | 0 | 0 | 0 | 0 | 0 | 0 | 0 | 0 | 0 | 0 | 0 |
| 0 | NA | NA | NA | NA | NA | Gm18006 | | 0 | 0 | 0 | 0 | 0 | 0 | 0 | 0 | 0 | 0 | 0 |
| 0 | NA | NA | NA | NA | NA | Dlx6os1 | | 0 | 0 | 0 | 0 | 0 | 0 | 0 | 0 | 0 | 0 | 0 |
| 0 | NA | NA | NA | NA | NA | Gm27699 | | 0 | 0 | 0 | 0 | 0 | 0 | 0 | 0 | 0 | 0 | 0 |
| 0 | NA | NA | NA | NA | NA | Gm27905 | | 0 | 0 | 0 | 0 | 0 | 0 | 0 | 0 | 0 | 0 | 0 |
| 0 | NA | NA | NA | NA | NA | Gm44094 | | 0 | 0 | 0 | 0 | 0 | 0 | 0 | 0 | 0 | 0 | 0 |
| 0 | NA | NA | NA | NA | NA | Dlx6 | distal-less homeobox 6 | 0 | 0 | 0 | 0 | 0 | 0 | 0 | 0 | 0 | 0 | 0 |
| 0 | NA | NA | NA | NA | NA | Dlx6os2 | | 0 | 0 | 0 | 0 | 0 | 0 | 0 | 0 | 0 | 0 | 0 |
| 0 | NA | NA | NA | NA | NA | Dlx6os1 | | 0 | 0 | 0 | 0 | 0 | 0 | 0 | 0 | 0 | 0 | 0 |
| 0 | NA | NA | NA | NA | NA | Gm22155 | | 0 | 0 | 0 | 0 | 0 | 0 | 0 | 0 | 0 | 0 | 0 |
| 0 | NA | NA | NA | NA | NA | Gm26774 | | 0 | 0 | 0 | 0 | 0 | 0 | 0 | 0 | 0 | 0 | 0 |
| 0 | NA | NA | NA | NA | NA | Gm27190 | | 0 | 0 | 0 | 0 | 0 | 0 | 0 | 0 | 0 | 0 | 0 |
| 0 | NA | NA | NA | NA | NA | Gm8669 | | 0 | 0 | 0 | 0 | 0 | 0 | 0 | 0 | 0 | 0 | 0 |
| 0 | NA | NA | NA | NA | NA | Gm19272 | | 0 | 0 | 0 | 0 | 0 | 0 | 0 | 0 | 0 | 0 | 0 |
| 0 | NA | NA | NA | NA | NA | Gm44046 | | 0 | 0 | 0 | 0 | 0 | 0 | 0 | 0 | 0 | 0 | 0 |
| 0 | NA | NA | NA | NA | NA | Gm44048 | | 0 | 0 | 0 | 0 | 0 | 0 | 0 | 0 | 0 | 0 | 0 |
| 0 | NA | NA | NA | NA | NA | Gm8676 | | 0 | 0 | 0 | 0 | 0 | 0 | 0 | 0 | 0 | 0 | 0 |
| 0 | NA | NA | NA | NA | NA | 1700012J15Rik | | 0 | 0 | 0 | 0 | 0 | 0 | 0 | 0 | 0 | 0 | 0 |
| 0 | NA | NA | NA | NA | NA | Gm43962 | | 0 | 0 | 0 | 0 | 0 | 0 | 0 | 0 | 0 | 0 | 0 |
| 0 | NA | NA | NA | NA | NA | Gm22276 | | 0 | 0 | 0 | 0 | 0 | 0 | 0 | 0 | 0 | 0 | 0 |
| 0 | NA | NA | NA | NA | NA | Gm45062 | | 0 | 0 | 0 | 0 | 0 | 0 | 0 | 0 | 0 | 0 | 0 |
| 0 | NA | NA | NA | NA | NA | Gm8428 | | 0 | 0 | 0 | 0 | 0 | 0 | 0 | 0 | 0 | 0 | 0 |
| 0 | NA | NA | NA | NA | NA | Gm24059 | | 0 | 0 | 0 | 0 | 0 | 0 | 0 | 0 | 0 | 0 | 0 |
| 0 | NA | NA | NA | NA | NA | Gm26075 | | 0 | 0 | 0 | 0 | 0 | 0 | 0 | 0 | 0 | 0 | 0 |
| 0 | NA | NA | NA | NA | NA | Gm16042 | | 0 | 0 | 0 | 0 | 0 | 0 | 0 | 0 | 0 | 0 | 0 |
| 0 | NA | NA | NA | NA | NA | Gm35736 | | 0 | 0 | 0 | 0 | 0 | 0 | 0 | 0 | 0 | 0 | 0 |
| 0 | NA | NA | NA | NA | NA | Gm18421 | | 0 | 0 | 0 | 0 | 0 | 0 | 0 | 0 | 0 | 0 | 0 |
| 0 | NA | NA | NA | NA | NA | Gm2691 | | 0 | 0 | 0 | 0 | 0 | 0 | 0 | 0 | 0 | 0 | 0 |
| 0 | NA | NA | NA | NA | NA | Gm43960 | | 0 | 0 | 0 | 0 | 0 | 0 | 0 | 0 | 0 | 0 | 0 |
| 0 | NA | NA | NA | NA | NA | Gm27894 | | 0 | 0 | 0 | 0 | 0 | 0 | 0 | 0 | 0 | 0 | 0 |
| 0 | NA | NA | NA | NA | NA | Gm8744 | | 0 | 0 | 0 | 0 | 0 | 0 | 0 | 0 | 0 | 0 | 0 |

| | | | | | | | | | | | | | | | | | |
|---|---|---|---|---|---|---|---|---|---|---|---|---|---|---|---|---|---|
| 0 | NA | NA | NA | NA | NA | Gm35822 | 0 | 0 | 0 | 0 | 0 | 0 | 0 | 0 | 0 | 0 | 0 |
| 0 | NA | NA | NA | NA | NA | AA545190 | 0 | 0 | 0 | 0 | 0 | 0 | 0 | 0 | 0 | 0 | 0 |
| 0 | NA | NA | NA | NA | NA | Gm6578 | 0 | 0 | 0 | 0 | 0 | 0 | 0 | 0 | 0 | 0 | 0 |
| 0 | NA | NA | NA | NA | NA | Gm32222 | 0 | 0 | 0 | 0 | 0 | 0 | 0 | 0 | 0 | 0 | 0 |
| 0 | NA | NA | NA | NA | NA | Gm16123 | 0 | 0 | 0 | 0 | 0 | 0 | 0 | 0 | 0 | 0 | 0 |
| 0 | NA | NA | NA | NA | NA | Gm44129 | 0 | 0 | 0 | 0 | 0 | 0 | 0 | 0 | 0 | 0 | 0 |
| 0 | NA | NA | NA | NA | NA | n-R5s160 | 0 | 0 | 0 | 0 | 0 | 0 | 0 | 0 | 0 | 0 | 0 |
| 0 | NA | NA | NA | NA | NA | Gm15529 | 0 | 0 | 0 | 0 | 0 | 0 | 0 | 0 | 0 | 0 | 0 |
| 0 | NA | NA | NA | NA | NA | Gm44472 | 0 | 0 | 0 | 0 | 0 | 0 | 0 | 0 | 0 | 0 | 0 |
| 0 | NA | NA | NA | NA | NA | Gm24985 | 0 | 0 | 0 | 0 | 0 | 0 | 0 | 0 | 0 | 0 | 0 |
| 0 | NA | NA | NA | NA | NA | predicted gene 725 | 0 | 0 | 0 | 0 | 0 | 0 | 0 | 0 | 0 | 0 | 0 |
| 0 | NA | NA | NA | NA | NA | 1700016P04Rik | 0 | 0 | 0 | 0 | 0 | 0 | 0 | 0 | 0 | 0 | 0 |
| 0 | NA | NA | NA | NA | NA | Gm25737 | 0 | 0 | 0 | 0 | 0 | 0 | 0 | 0 | 0 | 0 | 0 |
| 0 | NA | NA | NA | NA | NA | Gm44412 | 0 | 0 | 0 | 0 | 0 | 0 | 0 | 0 | 0 | 0 | 0 |
| 0 | NA | NA | NA | NA | NA | Gm19252 | 0 | 0 | 0 | 0 | 0 | 0 | 0 | 0 | 0 | 0 | 0 |
| 0 | NA | NA | NA | NA | NA | Gm23760 | 0 | 0 | 0 | 0 | 0 | 0 | 0 | 0 | 0 | 0 | 0 |
| 0 | NA | NA | NA | NA | NA | Gm8820 | 0 | 0 | 0 | 0 | 0 | 0 | 0 | 0 | 0 | 0 | 0 |
| 0 | NA | NA | NA | NA | NA | Gm23013 | 0 | 0 | 0 | 0 | 0 | 0 | 0 | 0 | 0 | 0 | 0 |
| 0 | NA | NA | NA | NA | NA | Gm44446 | 0 | 0 | 0 | 0 | 0 | 0 | 0 | 0 | 0 | 0 | 0 |
| 0 | NA | NA | NA | NA | NA | Gm25470 | 0 | 0 | 0 | 0 | 0 | 0 | 0 | 0 | 0 | 0 | 0 |
| 0 | NA | NA | NA | NA | NA | Gm44039 | 0 | 0 | 0 | 0 | 0 | 0 | 0 | 0 | 0 | 0 | 0 |
| 0 | NA | NA | NA | NA | NA | Gm43924 | 0 | 0 | 0 | 0 | 0 | 0 | 0 | 0 | 0 | 0 | 0 |
| 0 | NA | NA | NA | NA | NA | Gm43990 | 0 | 0 | 0 | 0 | 0 | 0 | 0 | 0 | 0 | 0 | 0 |
| 0 | NA | NA | NA | NA | NA | Gm43991 | 0 | 0 | 0 | 0 | 0 | 0 | 0 | 0 | 0 | 0 | 0 |
| 0 | NA | NA | NA | NA | NA | Gm43993 | 0 | 0 | 0 | 0 | 0 | 0 | 0 | 0 | 0 | 0 | 0 |
| 0 | NA | NA | NA | NA | NA | Gm3148 | 0 | 0 | 0 | 0 | 0 | 0 | 0 | 0 | 0 | 0 | 0 |
| 0 | NA | NA | NA | NA | NA | Gm36503 | 0 | 0 | 0 | 0 | 0 | 0 | 0 | 0 | 0 | 0 | 0 |
| 0 | NA | NA | NA | NA | NA | Gm36669 | 0 | 0 | 0 | 0 | 0 | 0 | 0 | 0 | 0 | 0 | 0 |
| 0 | NA | NA | NA | NA | NA | Gm5721 | 0 | 0 | 0 | 0 | 0 | 0 | 0 | 0 | 0 | 0 | 0 |
| 0 | NA | NA | NA | NA | NA | Gm15581 | 0 | 0 | 0 | 0 | 0 | 0 | 0 | 0 | 0 | 0 | 0 |
| 0 | NA | NA | NA | NA | NA | Gm29591 | 0 | 0 | 0 | 0 | 0 | 0 | 0 | 0 | 0 | 0 | 0 |
| 0 | NA | NA | NA | NA | NA | Gm25021 | 0 | 0 | 0 | 0 | 0 | 0 | 0 | 0 | 0 | 0 | 0 |
| 0 | NA | NA | NA | NA | NA | Gm27755 | 0 | 0 | 0 | 0 | 0 | 0 | 0 | 0 | 0 | 0 | 0 |
| 0 | NA | NA | NA | NA | NA | Gm27762 | 0 | 0 | 0 | 0 | 0 | 0 | 0 | 0 | 0 | 0 | 0 |
| 0 | NA | NA | NA | NA | NA | Gm27960 | 0 | 0 | 0 | 0 | 0 | 0 | 0 | 0 | 0 | 0 | 0 |
| 0 | NA | NA | NA | NA | NA | Gm27365 | 0 | 0 | 0 | 0 | 0 | 0 | 0 | 0 | 0 | 0 | 0 |
| 0 | NA | NA | NA | NA | NA | Gm27542 | 0 | 0 | 0 | 0 | 0 | 0 | 0 | 0 | 0 | 0 | 0 |
| 0 | NA | NA | NA | NA | NA | Gm13898 | 0 | 0 | 0 | 0 | 0 | 0 | 0 | 0 | 0 | 0 | 0 |
| 0 | NA | NA | NA | NA | NA | Gm13899 | 0 | 0 | 0 | 0 | 0 | 0 | 0 | 0 | 0 | 0 | 0 |
| 0 | NA | NA | NA | NA | NA | Gm43533 | 0 | 0 | 0 | 0 | 0 | 0 | 0 | 0 | 0 | 0 | 0 |
| 0 | NA | NA | NA | NA | NA | Gm42852 | 0 | 0 | 0 | 0 | 0 | 0 | 0 | 0 | 0 | 0 | 0 |
| 0 | NA | NA | NA | NA | NA | Gm27444 | 0 | 0 | 0 | 0 | 0 | 0 | 0 | 0 | 0 | 0 | 0 |
| 0 | NA | NA | NA | NA | NA | Gm27402 | 0 | 0 | 0 | 0 | 0 | 0 | 0 | 0 | 0 | 0 | 0 |
| 0 | NA | NA | NA | NA | NA | Gm43845 | 0 | 0 | 0 | 0 | 0 | 0 | 0 | 0 | 0 | 0 | 0 |
| 0 | NA | NA | NA | NA | NA | Gm15592 | 0 | 0 | 0 | 0 | 0 | 0 | 0 | 0 | 0 | 0 | 0 |
| 0 | NA | NA | NA | NA | NA | Gm15595 | 0 | 0 | 0 | 0 | 0 | 0 | 0 | 0 | 0 | 0 | 0 |
| 0 | NA | NA | NA | NA | NA | Gm15594 | 0 | 0 | 0 | 0 | 0 | 0 | 0 | 0 | 0 | 0 | 0 |
| 0 | NA | NA | NA | NA | NA | Gm26233 | 0 | 0 | 0 | 0 | 0 | 0 | 0 | 0 | 0 | 0 | 0 |
| 0 | NA | NA | NA | NA | NA | Gm23648 | 0 | 0 | 0 | 0 | 0 | 0 | 0 | 0 | 0 | 0 | 0 |
| 0 | NA | NA | NA | NA | NA | Gm43228 | 0 | 0 | 0 | 0 | 0 | 0 | 0 | 0 | 0 | 0 | 0 |
| 0 | NA | NA | NA | NA | NA | Gm43498 | 0 | 0 | 0 | 0 | 0 | 0 | 0 | 0 | 0 | 0 | 0 |
| 0 | NA | NA | NA | NA | NA | Gm22718 | 0 | 0 | 0 | 0 | 0 | 0 | 0 | 0 | 0 | 0 | 0 |
| 0 | NA | NA | NA | NA | NA | 4930524B17Rik | 0 | 0 | 0 | 0 | 0 | 0 | 0 | 0 | 0 | 0 | 0 |
| 0 | NA | NA | NA | NA | NA | Gm24213 | 0 | 0 | 0 | 0 | 0 | 0 | 0 | 0 | 0 | 0 | 0 |
| 0 | NA | NA | NA | NA | NA | Mir6370 | 0 | 0 | 0 | 0 | 0 | 0 | 0 | 0 | 0 | 0 | 0 |
| 0 | NA | NA | NA | NA | NA | Gm22828 | 0 | 0 | 0 | 0 | 0 | 0 | 0 | 0 | 0 | 0 | 0 |
| 0 | NA | NA | NA | NA | NA | Gm18482 | 0 | 0 | 0 | 0 | 0 | 0 | 0 | 0 | 0 | 0 | 0 |
| 0 | NA | NA | NA | NA | NA | Gm43501 | 0 | 0 | 0 | 0 | 0 | 0 | 0 | 0 | 0 | 0 | 0 |
| 0 | NA | NA | NA | NA | NA | Gm42581 | 0 | 0 | 0 | 0 | 0 | 0 | 0 | 0 | 0 | 0 | 0 |
| 0 | NA | NA | NA | NA | NA | Gm19102 | 0 | 0 | 0 | 0 | 0 | 0 | 0 | 0 | 0 | 0 | 0 |

| | | | | | | | | | | | | | | | | | | |
|---|---|---|---|---|---|---|---|---|---|---|---|---|---|---|---|---|---|---|
| 0 | NA | NA | NA | NA | NA | Gm26373 | | 0 | 0 | 0 | 0 | 0 | 0 | 0 | 0 | 0 | 0 | 0 |
| 0 | NA | NA | NA | NA | NA | Gm42583 | | 0 | 0 | 0 | 0 | 0 | 0 | 0 | 0 | 0 | 0 | 0 |
| 0 | NA | NA | NA | NA | NA | Gm42584 | | 0 | 0 | 0 | 0 | 0 | 0 | 0 | 0 | 0 | 0 | 0 |
| 0 | NA | NA | NA | NA | NA | Gm43630 | | 0 | 0 | 0 | 0 | 0 | 0 | 0 | 0 | 0 | 0 | 0 |
| 0 | NA | NA | NA | NA | NA | Gm18861 | | 0 | 0 | 0 | 0 | 0 | 0 | 0 | 0 | 0 | 0 | 0 |
| 0 | NA | NA | NA | NA | NA | Gm23961 | | 0 | 0 | 0 | 0 | 0 | 0 | 0 | 0 | 0 | 0 | 0 |
| 0 | NA | NA | NA | NA | NA | Gm25942 | | 0 | 0 | 0 | 0 | 0 | 0 | 0 | 0 | 0 | 0 | 0 |
| 0 | NA | NA | NA | NA | NA | Gm8945 | | 0 | 0 | 0 | 0 | 0 | 0 | 0 | 0 | 0 | 0 | 0 |
| 0 | NA | NA | NA | NA | NA | Gm5989 | | 0 | 0 | 0 | 0 | 0 | 0 | 0 | 0 | 0 | 0 | 0 |
| 0 | NA | NA | NA | NA | NA | Gm8950 | | 0 | 0 | 0 | 0 | 0 | 0 | 0 | 0 | 0 | 0 | 0 |
| 0 | NA | NA | NA | NA | NA | Gm5301 | | 0 | 0 | 0 | 0 | 0 | 0 | 0 | 0 | 0 | 0 | 0 |
| 0 | NA | NA | NA | NA | NA | C130093G08Rik | | 0 | 0 | 0 | 0 | 0 | 0 | 0 | 0 | 0 | 0 | 0 |
| 0 | NA | NA | NA | NA | NA | Gm27959 | | 0 | 0 | 0 | 0 | 0 | 0 | 0 | 0 | 0 | 0 | 0 |
| 0 | NA | NA | NA | NA | NA | Gm24217 | | 0 | 0 | 0 | 0 | 0 | 0 | 0 | 0 | 0 | 0 | 0 |
| 0 | NA | NA | NA | NA | NA | Rnf133 | ring finger protein 133 | 0 | 0 | 0 | 0 | 0 | 0 | 0 | 0 | 0 | 0 | 0 |
| 0 | NA | NA | NA | NA | NA | Rnf148 | ring finger protein 148 | 0 | 0 | 0 | 0 | 0 | 0 | 0 | 0 | 0 | 0 | 0 |
| 0 | NA | NA | NA | NA | NA | Tas2r118 | taste receptor, type 2, member 118 | 0 | 0 | 0 | 0 | 0 | 0 | 0 | 0 | 0 | 0 | 0 |
| 0 | NA | NA | NA | NA | NA | Slc13a1 | solute carrier family 13 (sodium/sulfate sy | 0 | 0 | 0 | 0 | 0 | 0 | 0 | 0 | 0 | 0 | 0 |
| 0 | NA | NA | NA | NA | NA | Gm20706 | | 0 | 0 | 0 | 0 | 0 | 0 | 0 | 0 | 0 | 0 | 0 |
| 0 | NA | NA | NA | NA | NA | Gm26386 | | 0 | 0 | 0 | 0 | 0 | 0 | 0 | 0 | 0 | 0 | 0 |
| 0 | NA | NA | NA | NA | NA | Gm42689 | | 0 | 0 | 0 | 0 | 0 | 0 | 0 | 0 | 0 | 0 | 0 |
| 0 | NA | NA | NA | NA | NA | Lmod2 | leiomodin 2 (cardiac) | 0 | 0 | 0 | 0 | 0 | 0 | 0 | 0 | 0 | 0 | 0 |
| 0 | NA | NA | NA | NA | NA | Hyal6 | hyaluronoglucosaminidase 6 | 0 | 0 | 0 | 0 | 0 | 0 | 0 | 0 | 0 | 0 | 0 |
| 0 | NA | NA | NA | NA | NA | Hyal4 | hyaluronoglucosaminidase 4 | 0 | 0 | 0 | 0 | 0 | 0 | 0 | 0 | 0 | 0 | 0 |
| 0 | NA | NA | NA | NA | NA | Spam1 | sperm adhesion molecule 1 | 0 | 0 | 0 | 0 | 0 | 0 | 0 | 0 | 0 | 0 | 0 |
| 0 | NA | NA | NA | NA | NA | Gm18327 | | 0 | 0 | 0 | 0 | 0 | 0 | 0 | 0 | 0 | 0 | 0 |
| 0 | NA | NA | NA | NA | NA | Hyal5 | hyaluronoglucosaminidase 5 | 0 | 0 | 0 | 0 | 0 | 0 | 0 | 0 | 0 | 0 | 0 |
| 0 | NA | NA | NA | NA | NA | Gm22267 | | 0 | 0 | 0 | 0 | 0 | 0 | 0 | 0 | 0 | 0 | 0 |
| 0 | NA | NA | NA | NA | NA | Gm24443 | | 0 | 0 | 0 | 0 | 0 | 0 | 0 | 0 | 0 | 0 | 0 |
| 0 | NA | NA | NA | NA | NA | Gm22529 | | 0 | 0 | 0 | 0 | 0 | 0 | 0 | 0 | 0 | 0 | 0 |
| 0 | NA | NA | NA | NA | NA | Gm20756 | | 0 | 0 | 0 | 0 | 0 | 0 | 0 | 0 | 0 | 0 | 0 |
| 0 | NA | NA | NA | NA | NA | Gm6719 | | 0 | 0 | 0 | 0 | 0 | 0 | 0 | 0 | 0 | 0 | 0 |
| 0 | NA | NA | NA | NA | NA | Gm24691 | | 0 | 0 | 0 | 0 | 0 | 0 | 0 | 0 | 0 | 0 | 0 |
| 0 | NA | NA | NA | NA | NA | Gm42474 | | 0 | 0 | 0 | 0 | 0 | 0 | 0 | 0 | 0 | 0 | 0 |
| 0 | NA | NA | NA | NA | NA | Gm42801 | | 0 | 0 | 0 | 0 | 0 | 0 | 0 | 0 | 0 | 0 | 0 |
| 0 | NA | NA | NA | NA | NA | Gm26310 | | 0 | 0 | 0 | 0 | 0 | 0 | 0 | 0 | 0 | 0 | 0 |
| 0 | NA | NA | NA | NA | NA | Mir592 | | 0 | 0 | 0 | 0 | 0 | 0 | 0 | 0 | 0 | 0 | 0 |
| 0 | NA | NA | NA | NA | NA | Pax4 | paired box 4 | 0 | 0 | 0 | 0 | 0 | 0 | 0 | 0 | 0 | 0 | 0 |
| 0 | NA | NA | NA | NA | NA | Gm43264 | | 0 | 0 | 0 | 0 | 0 | 0 | 0 | 0 | 0 | 0 | 0 |
| 0 | NA | NA | NA | NA | NA | Mir129-1 | | 0 | 0 | 0 | 0 | 0 | 0 | 0 | 0 | 0 | 0 | 0 |
| 0 | NA | NA | NA | NA | NA | Gm25589 | | 0 | 0 | 0 | 0 | 0 | 0 | 0 | 0 | 0 | 0 | 0 |
| 0 | NA | NA | NA | NA | NA | Gm24303 | | 0 | 0 | 0 | 0 | 0 | 0 | 0 | 0 | 0 | 0 | 0 |
| 0 | NA | NA | NA | NA | NA | Gm26021 | | 0 | 0 | 0 | 0 | 0 | 0 | 0 | 0 | 0 | 0 | 0 |
| 0 | NA | NA | NA | NA | NA | Fam71f1 | family with sequence similarity 71, membe | 0 | 0 | 0 | 0 | 0 | 0 | 0 | 0 | 0 | 0 | 0 |
| 0 | NA | NA | NA | NA | NA | Gm17172 | | 0 | 0 | 0 | 0 | 0 | 0 | 0 | 0 | 0 | 0 | 0 |
| 0 | NA | NA | NA | NA | NA | Gm26220 | | 0 | 0 | 0 | 0 | 0 | 0 | 0 | 0 | 0 | 0 | 0 |
| 0 | NA | NA | NA | NA | NA | Gm44354 | | 0 | 0 | 0 | 0 | 0 | 0 | 0 | 0 | 0 | 0 | 0 |
| 0 | NA | NA | NA | NA | NA | Gm26484 | | 0 | 0 | 0 | 0 | 0 | 0 | 0 | 0 | 0 | 0 | 0 |
| 0 | NA | NA | NA | NA | NA | Gm26627 | | 0 | 0 | 0 | 0 | 0 | 0 | 0 | 0 | 0 | 0 | 0 |
| 0 | NA | NA | NA | NA | NA | 4930528J11Rik | | 0 | 0 | 0 | 0 | 0 | 0 | 0 | 0 | 0 | 0 | 0 |
| 0 | NA | NA | NA | NA | NA | Gm13724 | | 0 | 0 | 0 | 0 | 0 | 0 | 0 | 0 | 0 | 0 | 0 |
| 0 | NA | NA | NA | NA | NA | Gm24536 | | 0 | 0 | 0 | 0 | 0 | 0 | 0 | 0 | 0 | 0 | 0 |
| 0 | NA | NA | NA | NA | NA | Gm13725 | | 0 | 0 | 0 | 0 | 0 | 0 | 0 | 0 | 0 | 0 | 0 |
| 0 | NA | NA | NA | NA | NA | Gm23049 | | 0 | 0 | 0 | 0 | 0 | 0 | 0 | 0 | 0 | 0 | 0 |
| 0 | NA | NA | NA | NA | NA | Gm25580 | | 0 | 0 | 0 | 0 | 0 | 0 | 0 | 0 | 0 | 0 | 0 |
| 0 | NA | NA | NA | NA | NA | Mir182 | | 0 | 0 | 0 | 0 | 0 | 0 | 0 | 0 | 0 | 0 | 0 |
| 0 | NA | NA | NA | NA | NA | Mir96 | | 0 | 0 | 0 | 0 | 0 | 0 | 0 | 0 | 0 | 0 | 0 |
| 0 | NA | NA | NA | NA | NA | Mir183 | | 0 | 0 | 0 | 0 | 0 | 0 | 0 | 0 | 0 | 0 | 0 |
| 0 | NA | NA | NA | NA | NA | 1700095J07Rik | | 0 | 0 | 0 | 0 | 0 | 0 | 0 | 0 | 0 | 0 | 0 |
| 0 | NA | NA | NA | NA | NA | Gm25625 | | 0 | 0 | 0 | 0 | 0 | 0 | 0 | 0 | 0 | 0 | 0 |

| | | | | | | | | | | | | | | | | | | |
|---|---|---|---|---|---|---|---|---|---|---|---|---|---|---|---|---|---|---|
| 0 | NA | NA | NA | NA | NA | n-R5s161 | | 0 | 0 | 0 | 0 | 0 | 0 | 0 | 0 | 0 | 0 | 0 |
| 0 | NA | NA | NA | NA | NA | Gm13781 | | 0 | 0 | 0 | 0 | 0 | 0 | 0 | 0 | 0 | 0 | 0 |
| 0 | NA | NA | NA | NA | NA | Gm13782 | | 0 | 0 | 0 | 0 | 0 | 0 | 0 | 0 | 0 | 0 | 0 |
| 0 | NA | NA | NA | NA | NA | Gm14528 | | 0 | 0 | 0 | 0 | 0 | 0 | 0 | 0 | 0 | 0 | 0 |
| 0 | NA | NA | NA | NA | NA | Gm27522 | | 0 | 0 | 0 | 0 | 0 | 0 | 0 | 0 | 0 | 0 | 0 |
| 0 | NA | NA | NA | NA | NA | Gm44296 | | 0 | 0 | 0 | 0 | 0 | 0 | 0 | 0 | 0 | 0 | 0 |
| 0 | NA | NA | NA | NA | NA | Gm27568 | | 0 | 0 | 0 | 0 | 0 | 0 | 0 | 0 | 0 | 0 | 0 |
| 0 | NA | NA | NA | NA | NA | Mir335 | | 0 | 0 | 0 | 0 | 0 | 0 | 0 | 0 | 0 | 0 | 0 |
| 0 | NA | NA | NA | NA | NA | Tsga13 | testis specific gene A13 | 0 | 0 | 0 | 0 | 0 | 0 | 0 | 0 | 0 | 0 | 0 |
| 0 | NA | NA | NA | NA | NA | Mir29b-1 | | 0 | 0 | 0 | 0 | 0 | 0 | 0 | 0 | 0 | 0 | 0 |
| 0 | NA | NA | NA | NA | NA | Gm13833 | | 0 | 0 | 0 | 0 | 0 | 0 | 0 | 0 | 0 | 0 | 0 |
| 0 | NA | NA | NA | NA | NA | Gm14532 | | 0 | 0 | 0 | 0 | 0 | 0 | 0 | 0 | 0 | 0 | 0 |
| 0 | NA | NA | NA | NA | NA | Gm6117 | | 0 | 0 | 0 | 0 | 0 | 0 | 0 | 0 | 0 | 0 | 0 |
| 0 | NA | NA | NA | NA | NA | Gm13848 | | 0 | 0 | 0 | 0 | 0 | 0 | 0 | 0 | 0 | 0 | 0 |
| 0 | NA | NA | NA | NA | NA | Gm43293 | | 0 | 0 | 0 | 0 | 0 | 0 | 0 | 0 | 0 | 0 | 0 |
| 0 | NA | NA | NA | NA | NA | Gm13849 | | 0 | 0 | 0 | 0 | 0 | 0 | 0 | 0 | 0 | 0 | 0 |
| 0 | NA | NA | NA | NA | NA | 1700012A03Ril | RIKEN cDNA 1700012A03 gene | 0 | 0 | 0 | 0 | 0 | 0 | 0 | 0 | 0 | 0 | 0 |
| 0 | NA | NA | NA | NA | NA | Gm13850 | | 0 | 0 | 0 | 0 | 0 | 0 | 0 | 0 | 0 | 0 | 0 |
| 0 | NA | NA | NA | NA | NA | Plxna4os3 | | 0 | 0 | 0 | 0 | 0 | 0 | 0 | 0 | 0 | 0 | 0 |
| 0 | NA | NA | NA | NA | NA | Plxna4os2 | | 0 | 0 | 0 | 0 | 0 | 0 | 0 | 0 | 0 | 0 | 0 |
| 0 | NA | NA | NA | NA | NA | Gm13852 | | 0 | 0 | 0 | 0 | 0 | 0 | 0 | 0 | 0 | 0 | 0 |
| 0 | NA | NA | NA | NA | NA | BC049739 | | 0 | 0 | 0 | 0 | 0 | 0 | 0 | 0 | 0 | 0 | 0 |
| 0 | NA | NA | NA | NA | NA | Gm14540 | | 0 | 0 | 0 | 0 | 0 | 0 | 0 | 0 | 0 | 0 | 0 |
| 0 | NA | NA | NA | NA | NA | Gm22321 | | 0 | 0 | 0 | 0 | 0 | 0 | 0 | 0 | 0 | 0 | 0 |
| 0 | NA | NA | NA | NA | NA | Gm1401 | | 0 | 0 | 0 | 0 | 0 | 0 | 0 | 0 | 0 | 0 | 0 |
| 0 | NA | NA | NA | NA | NA | Gm13854 | | 0 | 0 | 0 | 0 | 0 | 0 | 0 | 0 | 0 | 0 | 0 |
| 0 | NA | NA | NA | NA | NA | Gm43168 | | 0 | 0 | 0 | 0 | 0 | 0 | 0 | 0 | 0 | 0 | 0 |
| 0 | NA | NA | NA | NA | NA | Gm43164 | | 0 | 0 | 0 | 0 | 0 | 0 | 0 | 0 | 0 | 0 | 0 |
| 0 | NA | NA | NA | NA | NA | Gm13853 | | 0 | 0 | 0 | 0 | 0 | 0 | 0 | 0 | 0 | 0 | 0 |
| 0 | NA | NA | NA | NA | NA | Gm13857 | | 0 | 0 | 0 | 0 | 0 | 0 | 0 | 0 | 0 | 0 | 0 |
| 0 | NA | NA | NA | NA | NA | Gm13860 | | 0 | 0 | 0 | 0 | 0 | 0 | 0 | 0 | 0 | 0 | 0 |
| 0 | NA | NA | NA | NA | NA | Npn2 | | 0 | 0 | 0 | 0 | 0 | 0 | 0 | 0 | 0 | 0 | 0 |
| 0 | NA | NA | NA | NA | NA | Gm14547 | | 0 | 0 | 0 | 0 | 0 | 0 | 0 | 0 | 0 | 0 | 0 |
| 0 | NA | NA | NA | NA | NA | Gm13862 | | 0 | 0 | 0 | 0 | 0 | 0 | 0 | 0 | 0 | 0 | 0 |
| 0 | NA | NA | NA | NA | NA | Gm13863 | | 0 | 0 | 0 | 0 | 0 | 0 | 0 | 0 | 0 | 0 | 0 |
| 0 | NA | NA | NA | NA | NA | Stra8 | stimulated by retinoic acid gene 8 | 0 | 0 | 0 | 0 | 0 | 0 | 0 | 0 | 0 | 0 | 0 |
| 0 | NA | NA | NA | NA | NA | Gm23435 | | 0 | 0 | 0 | 0 | 0 | 0 | 0 | 0 | 0 | 0 | 0 |
| 0 | NA | NA | NA | NA | NA | Slc13a4 | solute carrier family 13 (sodium/sulfate sy | 0 | 0 | 0 | 0 | 0 | 0 | 0 | 0 | 0 | 0 | 0 |
| 0 | NA | NA | NA | NA | NA | 1700065J11Rik | | 0 | 0 | 0 | 0 | 0 | 0 | 0 | 0 | 0 | 0 | 0 |
| 0 | NA | NA | NA | NA | NA | Gm23053 | | 0 | 0 | 0 | 0 | 0 | 0 | 0 | 0 | 0 | 0 | 0 |
| 0 | NA | NA | NA | NA | NA | Gm42931 | | 0 | 0 | 0 | 0 | 0 | 0 | 0 | 0 | 0 | 0 | 0 |
| 0 | NA | NA | NA | NA | NA | Gm23273 | | 0 | 0 | 0 | 0 | 0 | 0 | 0 | 0 | 0 | 0 | 0 |
| 0 | NA | NA | NA | NA | NA | Gm43443 | | 0 | 0 | 0 | 0 | 0 | 0 | 0 | 0 | 0 | 0 | 0 |
| 0 | NA | NA | NA | NA | NA | Mir490 | | 0 | 0 | 0 | 0 | 0 | 0 | 0 | 0 | 0 | 0 | 0 |
| 0 | NA | NA | NA | NA | NA | Gm43441 | | 0 | 0 | 0 | 0 | 0 | 0 | 0 | 0 | 0 | 0 | 0 |
| 0 | NA | NA | NA | NA | NA | Gm2189 | | 0 | 0 | 0 | 0 | 0 | 0 | 0 | 0 | 0 | 0 | 0 |
| 0 | NA | NA | NA | NA | NA | Gm25111 | | 0 | 0 | 0 | 0 | 0 | 0 | 0 | 0 | 0 | 0 | 0 |
| 0 | NA | NA | NA | NA | NA | 1700111E14Rik | | 0 | 0 | 0 | 0 | 0 | 0 | 0 | 0 | 0 | 0 | 0 |
| 0 | NA | NA | NA | NA | NA | Gm7492 | | 0 | 0 | 0 | 0 | 0 | 0 | 0 | 0 | 0 | 0 | 0 |
| 0 | NA | NA | NA | NA | NA | Tmem213 | transmembrane protein 213 | 0 | 0 | 0 | 0 | 0 | 0 | 0 | 0 | 0 | 0 | 0 |
| 0 | NA | NA | NA | NA | NA | Gm23231 | | 0 | 0 | 0 | 0 | 0 | 0 | 0 | 0 | 0 | 0 | 0 |
| 0 | NA | NA | NA | NA | NA | Mir7670 | | 0 | 0 | 0 | 0 | 0 | 0 | 0 | 0 | 0 | 0 | 0 |
| 0 | NA | NA | NA | NA | NA | Gm23010 | | 0 | 0 | 0 | 0 | 0 | 0 | 0 | 0 | 0 | 0 | 0 |
| 0 | NA | NA | NA | NA | NA | 1700025N23Rik | | 0 | 0 | 0 | 0 | 0 | 0 | 0 | 0 | 0 | 0 | 0 |
| 0 | NA | NA | NA | NA | NA | Gm42963 | | 0 | 0 | 0 | 0 | 0 | 0 | 0 | 0 | 0 | 0 | 0 |
| 0 | NA | NA | NA | NA | NA | Gm43479 | | 0 | 0 | 0 | 0 | 0 | 0 | 0 | 0 | 0 | 0 | 0 |
| 0 | NA | NA | NA | NA | NA | Gm26008 | | 0 | 0 | 0 | 0 | 0 | 0 | 0 | 0 | 0 | 0 | 0 |
| 0 | NA | NA | NA | NA | NA | Gm25402 | | 0 | 0 | 0 | 0 | 0 | 0 | 0 | 0 | 0 | 0 | 0 |
| 0 | NA | NA | NA | NA | NA | Gm42420 | | 0 | 0 | 0 | 0 | 0 | 0 | 0 | 0 | 0 | 0 | 0 |
| 0 | NA | NA | NA | NA | NA | Gm7554 | | 0 | 0 | 0 | 0 | 0 | 0 | 0 | 0 | 0 | 0 | 0 |

| | | | | | | | | | | | | | | | | | | |
|---|---|---|---|---|---|---|---|---|---|---|---|---|---|---|---|---|---|---|
| 0 | NA | NA | NA | NA | NA | Gm37995 | | 0 | 0 | 0 | 0 | 0 | 0 | 0 | 0 | 0 | 0 | 0 |
| 0 | NA | NA | NA | NA | NA | Gm26638 | | 0 | 0 | 0 | 0 | 0 | 0 | 0 | 0 | 0 | 0 | 0 |
| 0 | NA | NA | NA | NA | NA | Gm26807 | | 0 | 0 | 0 | 0 | 0 | 0 | 0 | 0 | 0 | 0 | 0 |
| 0 | NA | NA | NA | NA | NA | Tas2r137 | taste receptor, type 2, member 137 | 0 | 0 | 0 | 0 | 0 | 0 | 0 | 0 | 0 | 0 | 0 |
| 0 | NA | NA | NA | NA | NA | Prss37 | protease, serine 37 | 0 | 0 | 0 | 0 | 0 | 0 | 0 | 0 | 0 | 0 | 0 |
| 0 | NA | NA | NA | NA | NA | Olfr461 | olfactory receptor 461 | 0 | 0 | 0 | 0 | 0 | 0 | 0 | 0 | 0 | 0 | 0 |
| 0 | NA | NA | NA | NA | NA | Olfr460 | olfactory receptor 460 | 0 | 0 | 0 | 0 | 0 | 0 | 0 | 0 | 0 | 0 | 0 |
| 0 | NA | NA | NA | NA | NA | Tas2r138 | taste receptor, type 2, member 138 | 0 | 0 | 0 | 0 | 0 | 0 | 0 | 0 | 0 | 0 | 0 |
| 0 | NA | NA | NA | NA | NA | Moxd2 | monooxygenase, DBH-like 2 | 0 | 0 | 0 | 0 | 0 | 0 | 0 | 0 | 0 | 0 | 0 |
| 0 | NA | NA | NA | NA | NA | Gm38276 | | 0 | 0 | 0 | 0 | 0 | 0 | 0 | 0 | 0 | 0 | 0 |
| 0 | NA | NA | NA | NA | NA | Prss58 | protease, serine 58 | 0 | 0 | 0 | 0 | 0 | 0 | 0 | 0 | 0 | 0 | 0 |
| 0 | NA | NA | NA | NA | NA | 1700074P13Ril | RIKEN cDNA 1700074P13 gene | 0 | 0 | 0 | 0 | 0 | 0 | 0 | 0 | 0 | 0 | 0 |
| 0 | NA | NA | NA | NA | NA | Gm29588 | | 0 | 0 | 0 | 0 | 0 | 0 | 0 | 0 | 0 | 0 | 0 |
| 0 | NA | NA | NA | NA | NA | Gm2663 | predicted gene 2663 | 0 | 0 | 0 | 0 | 0 | 0 | 0 | 0 | 0 | 0 | 0 |
| 0 | NA | NA | NA | NA | NA | Gm38015 | | 0 | 0 | 0 | 0 | 0 | 0 | 0 | 0 | 0 | 0 | 0 |
| 0 | NA | NA | NA | NA | NA | Gm37878 | | 0 | 0 | 0 | 0 | 0 | 0 | 0 | 0 | 0 | 0 | 0 |
| 0 | NA | NA | NA | NA | NA | Gm38294 | | 0 | 0 | 0 | 0 | 0 | 0 | 0 | 0 | 0 | 0 | 0 |
| 0 | NA | NA | NA | NA | NA | 5830405F06Rik | | 0 | 0 | 0 | 0 | 0 | 0 | 0 | 0 | 0 | 0 | 0 |
| 0 | NA | NA | NA | NA | NA | Trbv4 | | 0 | 0 | 0 | 0 | 0 | 0 | 0 | 0 | 0 | 0 | 0 |
| 0 | NA | NA | NA | NA | NA | Trbv5 | | 0 | 0 | 0 | 0 | 0 | 0 | 0 | 0 | 0 | 0 | 0 |
| 0 | NA | NA | NA | NA | NA | Trbv6 | | 0 | 0 | 0 | 0 | 0 | 0 | 0 | 0 | 0 | 0 | 0 |
| 0 | NA | NA | NA | NA | NA | Trbv7 | | 0 | 0 | 0 | 0 | 0 | 0 | 0 | 0 | 0 | 0 | 0 |
| 0 | NA | NA | NA | NA | NA | Gm49885 | | 0 | 0 | 0 | 0 | 0 | 0 | 0 | 0 | 0 | 0 | 0 |
| 0 | NA | NA | NA | NA | NA | Trbv8 | | 0 | 0 | 0 | 0 | 0 | 0 | 0 | 0 | 0 | 0 | 0 |
| 0 | NA | NA | NA | NA | NA | Trbv9 | | 0 | 0 | 0 | 0 | 0 | 0 | 0 | 0 | 0 | 0 | 0 |
| 0 | NA | NA | NA | NA | NA | Trbv10 | | 0 | 0 | 0 | 0 | 0 | 0 | 0 | 0 | 0 | 0 | 0 |
| 0 | NA | NA | NA | NA | NA | Trbv11 | | 0 | 0 | 0 | 0 | 0 | 0 | 0 | 0 | 0 | 0 | 0 |
| 0 | NA | NA | NA | NA | NA | Trbv12-1 | | 0 | 0 | 0 | 0 | 0 | 0 | 0 | 0 | 0 | 0 | 0 |
| 0 | NA | NA | NA | NA | NA | Trbv12-3 | | 0 | 0 | 0 | 0 | 0 | 0 | 0 | 0 | 0 | 0 | 0 |
| 0 | NA | NA | NA | NA | NA | Trbv14 | | 0 | 0 | 0 | 0 | 0 | 0 | 0 | 0 | 0 | 0 | 0 |
| 0 | NA | NA | NA | NA | NA | Trbv15 | | 0 | 0 | 0 | 0 | 0 | 0 | 0 | 0 | 0 | 0 | 0 |
| 0 | NA | NA | NA | NA | NA | Trbv17 | | 0 | 0 | 0 | 0 | 0 | 0 | 0 | 0 | 0 | 0 | 0 |
| 0 | NA | NA | NA | NA | NA | Trbv18 | | 0 | 0 | 0 | 0 | 0 | 0 | 0 | 0 | 0 | 0 | 0 |
| 0 | NA | NA | NA | NA | NA | Trbv20 | | 0 | 0 | 0 | 0 | 0 | 0 | 0 | 0 | 0 | 0 | 0 |
| 0 | NA | NA | NA | NA | NA | Trbv21 | | 0 | 0 | 0 | 0 | 0 | 0 | 0 | 0 | 0 | 0 | 0 |
| 0 | NA | NA | NA | NA | NA | Trbv22 | | 0 | 0 | 0 | 0 | 0 | 0 | 0 | 0 | 0 | 0 | 0 |
| 0 | NA | NA | NA | NA | NA | Trbv23 | | 0 | 0 | 0 | 0 | 0 | 0 | 0 | 0 | 0 | 0 | 0 |
| 0 | NA | NA | NA | NA | NA | Trbv24 | | 0 | 0 | 0 | 0 | 0 | 0 | 0 | 0 | 0 | 0 | 0 |
| 0 | NA | NA | NA | NA | NA | Trbv25 | | 0 | 0 | 0 | 0 | 0 | 0 | 0 | 0 | 0 | 0 | 0 |
| 0 | NA | NA | NA | NA | NA | Trbv26 | | 0 | 0 | 0 | 0 | 0 | 0 | 0 | 0 | 0 | 0 | 0 |
| 0 | NA | NA | NA | NA | NA | Trbv27 | | 0 | 0 | 0 | 0 | 0 | 0 | 0 | 0 | 0 | 0 | 0 |
| 0 | NA | NA | NA | NA | NA | Trbv28 | | 0 | 0 | 0 | 0 | 0 | 0 | 0 | 0 | 0 | 0 | 0 |
| 0 | NA | NA | NA | NA | NA | Trbv30 | | 0 | 0 | 0 | 0 | 0 | 0 | 0 | 0 | 0 | 0 | 0 |
| 0 | NA | NA | NA | NA | NA | Gm18964 | | 0 | 0 | 0 | 0 | 0 | 0 | 0 | 0 | 0 | 0 | 0 |
| 0 | NA | NA | NA | NA | NA | Gm38306 | | 0 | 0 | 0 | 0 | 0 | 0 | 0 | 0 | 0 | 0 | 0 |
| 0 | NA | NA | NA | NA | NA | Gm29813 | | 0 | 0 | 0 | 0 | 0 | 0 | 0 | 0 | 0 | 0 | 0 |
| 0 | NA | NA | NA | NA | NA | Gm37841 | | 0 | 0 | 0 | 0 | 0 | 0 | 0 | 0 | 0 | 0 | 0 |
| 0 | NA | NA | NA | NA | NA | Gm38018 | | 0 | 0 | 0 | 0 | 0 | 0 | 0 | 0 | 0 | 0 | 0 |
| 0 | NA | NA | NA | NA | NA | Trbd1 | | 0 | 0 | 0 | 0 | 0 | 0 | 0 | 0 | 0 | 0 | 0 |
| 0 | NA | NA | NA | NA | NA | Trbj1-1 | | 0 | 0 | 0 | 0 | 0 | 0 | 0 | 0 | 0 | 0 | 0 |
| 0 | NA | NA | NA | NA | NA | Trbj1-2 | | 0 | 0 | 0 | 0 | 0 | 0 | 0 | 0 | 0 | 0 | 0 |
| 0 | NA | NA | NA | NA | NA | Trbj1-3 | | 0 | 0 | 0 | 0 | 0 | 0 | 0 | 0 | 0 | 0 | 0 |
| 0 | NA | NA | NA | NA | NA | Trbj1-4 | | 0 | 0 | 0 | 0 | 0 | 0 | 0 | 0 | 0 | 0 | 0 |
| 0 | NA | NA | NA | NA | NA | Trbj1-5 | | 0 | 0 | 0 | 0 | 0 | 0 | 0 | 0 | 0 | 0 | 0 |
| 0 | NA | NA | NA | NA | NA | Trbj1-6 | | 0 | 0 | 0 | 0 | 0 | 0 | 0 | 0 | 0 | 0 | 0 |
| 0 | NA | NA | NA | NA | NA | Trbj1-7 | | 0 | 0 | 0 | 0 | 0 | 0 | 0 | 0 | 0 | 0 | 0 |
| 0 | NA | NA | NA | NA | NA | Trbd2 | | 0 | 0 | 0 | 0 | 0 | 0 | 0 | 0 | 0 | 0 | 0 |
| 0 | NA | NA | NA | NA | NA | Trbj2-1 | | 0 | 0 | 0 | 0 | 0 | 0 | 0 | 0 | 0 | 0 | 0 |
| 0 | NA | NA | NA | NA | NA | Trbj2-2 | | 0 | 0 | 0 | 0 | 0 | 0 | 0 | 0 | 0 | 0 | 0 |
| 0 | NA | NA | NA | NA | NA | Trbj2-3 | | 0 | 0 | 0 | 0 | 0 | 0 | 0 | 0 | 0 | 0 | 0 |

| | | | | | | | | | | | | | | | | | |
|---|---|---|---|---|---|---|---|---|---|---|---|---|---|---|---|---|---|
| 0 | NA | NA | NA | NA | NA | Trbj2-4 | | 0 | 0 | 0 | 0 | 0 | 0 | 0 | 0 | 0 | 0 | 0 |
| 0 | NA | NA | NA | NA | NA | Trbj2-5 | | 0 | 0 | 0 | 0 | 0 | 0 | 0 | 0 | 0 | 0 | 0 |
| 0 | NA | NA | NA | NA | NA | Trbj2-6 | | 0 | 0 | 0 | 0 | 0 | 0 | 0 | 0 | 0 | 0 | 0 |
| 0 | NA | NA | NA | NA | NA | Trbj2-7 | | 0 | 0 | 0 | 0 | 0 | 0 | 0 | 0 | 0 | 0 | 0 |
| 0 | NA | NA | NA | NA | NA | Gm27549 | | 0 | 0 | 0 | 0 | 0 | 0 | 0 | 0 | 0 | 0 | 0 |
| 0 | NA | NA | NA | NA | NA | Trpv5 | transient receptor potential cation channe | 0 | 0 | 0 | 0 | 0 | 0 | 0 | 0 | 0 | 0 | 0 |
| 0 | NA | NA | NA | NA | NA | Llcfc1 | LLLL and CFNLAS motif containing 1 | 0 | 0 | 0 | 0 | 0 | 0 | 0 | 0 | 0 | 0 | 0 |
| 0 | NA | NA | NA | NA | NA | Kel | Kell blood group | 0 | 0 | 0 | 0 | 0 | 0 | 0 | 0 | 0 | 0 | 0 |
| 0 | NA | NA | NA | NA | NA | Gm37527 | | 0 | 0 | 0 | 0 | 0 | 0 | 0 | 0 | 0 | 0 | 0 |
| 0 | NA | NA | NA | NA | NA | Olfr459 | olfactory receptor 459 | 0 | 0 | 0 | 0 | 0 | 0 | 0 | 0 | 0 | 0 | 0 |
| 0 | NA | NA | NA | NA | NA | Pip | prolactin induced protein | 0 | 0 | 0 | 0 | 0 | 0 | 0 | 0 | 0 | 0 | 0 |
| 0 | NA | NA | NA | NA | NA | Sval3 | seminal vesicle antigen-like 3 | 0 | 0 | 0 | 0 | 0 | 0 | 0 | 0 | 0 | 0 | 0 |
| 0 | NA | NA | NA | NA | NA | Sva | seminal vesicle antigen | 0 | 0 | 0 | 0 | 0 | 0 | 0 | 0 | 0 | 0 | 0 |
| 0 | NA | NA | NA | NA | NA | Gm34701 | | 0 | 0 | 0 | 0 | 0 | 0 | 0 | 0 | 0 | 0 | 0 |
| 0 | NA | NA | NA | NA | NA | Tas2r139 | taste receptor, type 2, member 139 | 0 | 0 | 0 | 0 | 0 | 0 | 0 | 0 | 0 | 0 | 0 |
| 0 | NA | NA | NA | NA | NA | Tas2r144 | taste receptor, type 2, member 144 | 0 | 0 | 0 | 0 | 0 | 0 | 0 | 0 | 0 | 0 | 0 |
| 0 | NA | NA | NA | NA | NA | Gm44284 | | 0 | 0 | 0 | 0 | 0 | 0 | 0 | 0 | 0 | 0 | 0 |
| 0 | NA | NA | NA | NA | NA | Gm24611 | | 0 | 0 | 0 | 0 | 0 | 0 | 0 | 0 | 0 | 0 | 0 |
| 0 | NA | NA | NA | NA | NA | Tas2r143 | taste receptor, type 2, member 143 | 0 | 0 | 0 | 0 | 0 | 0 | 0 | 0 | 0 | 0 | 0 |
| 0 | NA | NA | NA | NA | NA | Tas2r135 | taste receptor, type 2, member 135 | 0 | 0 | 0 | 0 | 0 | 0 | 0 | 0 | 0 | 0 | 0 |
| 0 | NA | NA | NA | NA | NA | Tas2r126 | taste receptor, type 2, member 126 | 0 | 0 | 0 | 0 | 0 | 0 | 0 | 0 | 0 | 0 | 0 |
| 0 | NA | NA | NA | NA | NA | Gm44470 | | 0 | 0 | 0 | 0 | 0 | 0 | 0 | 0 | 0 | 0 | 0 |
| 0 | NA | NA | NA | NA | NA | Olfr458 | olfactory receptor 458 | 0 | 0 | 0 | 0 | 0 | 0 | 0 | 0 | 0 | 0 | 0 |
| 0 | NA | NA | NA | NA | NA | Olfr457 | olfactory receptor 457 | 0 | 0 | 0 | 0 | 0 | 0 | 0 | 0 | 0 | 0 | 0 |
| 0 | NA | NA | NA | NA | NA | Olfr456 | olfactory receptor 456 | 0 | 0 | 0 | 0 | 0 | 0 | 0 | 0 | 0 | 0 | 0 |
| 0 | NA | NA | NA | NA | NA | Tcaf3 | TRPM8 channel-associated factor 3 | 0 | 0 | 0 | 0 | 0 | 0 | 0 | 0 | 0 | 0 | 0 |
| 0 | NA | NA | NA | NA | NA | Olfr454-ps1 | | 0 | 0 | 0 | 0 | 0 | 0 | 0 | 0 | 0 | 0 | 0 |
| 0 | NA | NA | NA | NA | NA | Gm44030 | | 0 | 0 | 0 | 0 | 0 | 0 | 0 | 0 | 0 | 0 | 0 |
| 0 | NA | NA | NA | NA | NA | Gm44032 | | 0 | 0 | 0 | 0 | 0 | 0 | 0 | 0 | 0 | 0 | 0 |
| 0 | NA | NA | NA | NA | NA | Olfr453 | olfactory receptor 453 | 0 | 0 | 0 | 0 | 0 | 0 | 0 | 0 | 0 | 0 | 0 |
| 0 | NA | NA | NA | NA | NA | Gm44034 | | 0 | 0 | 0 | 0 | 0 | 0 | 0 | 0 | 0 | 0 | 0 |
| 0 | NA | NA | NA | NA | NA | Olfr38 | olfactory receptor 38 | 0 | 0 | 0 | 0 | 0 | 0 | 0 | 0 | 0 | 0 | 0 |
| 0 | NA | NA | NA | NA | NA | Gm22363 | | 0 | 0 | 0 | 0 | 0 | 0 | 0 | 0 | 0 | 0 | 0 |
| 0 | NA | NA | NA | NA | NA | Olfr452 | olfactory receptor 452 | 0 | 0 | 0 | 0 | 0 | 0 | 0 | 0 | 0 | 0 | 0 |
| 0 | NA | NA | NA | NA | NA | Olfr451-ps1 | | 0 | 0 | 0 | 0 | 0 | 0 | 0 | 0 | 0 | 0 | 0 |
| 0 | NA | NA | NA | NA | NA | Olfr450 | olfactory receptor 450 | 0 | 0 | 0 | 0 | 0 | 0 | 0 | 0 | 0 | 0 | 0 |
| 0 | NA | NA | NA | NA | NA | Gm23982 | | 0 | 0 | 0 | 0 | 0 | 0 | 0 | 0 | 0 | 0 | 0 |
| 0 | NA | NA | NA | NA | NA | Olfr449 | olfactory receptor 449 | 0 | 0 | 0 | 0 | 0 | 0 | 0 | 0 | 0 | 0 | 0 |
| 0 | NA | NA | NA | NA | NA | Olfr448 | olfactory receptor 448 | 0 | 0 | 0 | 0 | 0 | 0 | 0 | 0 | 0 | 0 | 0 |
| 0 | NA | NA | NA | NA | NA | Olfr447 | olfactory receptor 447 | 0 | 0 | 0 | 0 | 0 | 0 | 0 | 0 | 0 | 0 | 0 |
| 0 | NA | NA | NA | NA | NA | Olfr446 | olfactory receptor 446 | 0 | 0 | 0 | 0 | 0 | 0 | 0 | 0 | 0 | 0 | 0 |
| 0 | NA | NA | NA | NA | NA | Olrf445-ps1 | | 0 | 0 | 0 | 0 | 0 | 0 | 0 | 0 | 0 | 0 | 0 |
| 0 | NA | NA | NA | NA | NA | Olfr444 | olfactory receptor 444 | 0 | 0 | 0 | 0 | 0 | 0 | 0 | 0 | 0 | 0 | 0 |
| 0 | NA | NA | NA | NA | NA | Gm17472 | | 0 | 0 | 0 | 0 | 0 | 0 | 0 | 0 | 0 | 0 | 0 |
| 0 | NA | NA | NA | NA | NA | Olfr443-ps1 | | 0 | 0 | 0 | 0 | 0 | 0 | 0 | 0 | 0 | 0 | 0 |
| 0 | NA | NA | NA | NA | NA | Olfr442-ps1 | | 0 | 0 | 0 | 0 | 0 | 0 | 0 | 0 | 0 | 0 | 0 |
| 0 | NA | NA | NA | NA | NA | Olfr441 | olfactory receptor 441 | 0 | 0 | 0 | 0 | 0 | 0 | 0 | 0 | 0 | 0 | 0 |
| 0 | NA | NA | NA | NA | NA | Olfr440-ps1 | | 0 | 0 | 0 | 0 | 0 | 0 | 0 | 0 | 0 | 0 | 0 |
| 0 | NA | NA | NA | NA | NA | Olfr439-ps1 | | 0 | 0 | 0 | 0 | 0 | 0 | 0 | 0 | 0 | 0 | 0 |
| 0 | NA | NA | NA | NA | NA | Olfr237 | olfactory receptor 237 | 0 | 0 | 0 | 0 | 0 | 0 | 0 | 0 | 0 | 0 | 0 |
| 0 | NA | NA | NA | NA | NA | Olfr437 | olfactory receptor 437 | 0 | 0 | 0 | 0 | 0 | 0 | 0 | 0 | 0 | 0 | 0 |
| 0 | NA | NA | NA | NA | NA | Olfr13 | olfactory receptor 13 | 0 | 0 | 0 | 0 | 0 | 0 | 0 | 0 | 0 | 0 | 0 |
| 0 | NA | NA | NA | NA | NA | Olfr436-ps1 | | 0 | 0 | 0 | 0 | 0 | 0 | 0 | 0 | 0 | 0 | 0 |
| 0 | NA | NA | NA | NA | NA | Olfr435 | olfactory receptor 435 | 0 | 0 | 0 | 0 | 0 | 0 | 0 | 0 | 0 | 0 | 0 |
| 0 | NA | NA | NA | NA | NA | Gm21827 | | 0 | 0 | 0 | 0 | 0 | 0 | 0 | 0 | 0 | 0 | 0 |
| 0 | NA | NA | NA | NA | NA | Olfr434 | olfactory receptor 434 | 0 | 0 | 0 | 0 | 0 | 0 | 0 | 0 | 0 | 0 | 0 |
| 0 | NA | NA | NA | NA | NA | Olfr47 | olfactory receptor 47 | 0 | 0 | 0 | 0 | 0 | 0 | 0 | 0 | 0 | 0 | 0 |
| 0 | NA | NA | NA | NA | NA | Nobox | NOBOX oogenesis homeobox | 0 | 0 | 0 | 0 | 0 | 0 | 0 | 0 | 0 | 0 | 0 |
| 0 | NA | NA | NA | NA | NA | 9430018G01Rik | | 0 | 0 | 0 | 0 | 0 | 0 | 0 | 0 | 0 | 0 | 0 |
| 0 | NA | NA | NA | NA | NA | Gm22939 | | 0 | 0 | 0 | 0 | 0 | 0 | 0 | 0 | 0 | 0 | 0 |

| | | | | | | | | | | | | | | | | | | |
|---|---|---|---|---|---|---|---|---|---|---|---|---|---|---|---|---|---|---|
| 0 | NA | NA | NA | NA | NA | Gm35216 | | 0 | 0 | 0 | 0 | 0 | 0 | 0 | 0 | 0 | 0 | 0 |
| 0 | NA | NA | NA | NA | NA | Gm44211 | | 0 | 0 | 0 | 0 | 0 | 0 | 0 | 0 | 0 | 0 | 0 |
| 0 | NA | NA | NA | NA | NA | Gm44212 | | 0 | 0 | 0 | 0 | 0 | 0 | 0 | 0 | 0 | 0 | 0 |
| 0 | NA | NA | NA | NA | NA | Gm44213 | | 0 | 0 | 0 | 0 | 0 | 0 | 0 | 0 | 0 | 0 | 0 |
| 0 | NA | NA | NA | NA | NA | Gm43884 | | 0 | 0 | 0 | 0 | 0 | 0 | 0 | 0 | 0 | 0 | 0 |
| 0 | NA | NA | NA | NA | NA | Gm22839 | | 0 | 0 | 0 | 0 | 0 | 0 | 0 | 0 | 0 | 0 | 0 |
| 0 | NA | NA | NA | NA | NA | Gm43888 | | 0 | 0 | 0 | 0 | 0 | 0 | 0 | 0 | 0 | 0 | 0 |
| 0 | NA | NA | NA | NA | NA | Gm25161 | | 0 | 0 | 0 | 0 | 0 | 0 | 0 | 0 | 0 | 0 | 0 |
| 0 | NA | NA | NA | NA | NA | Gm18484 | | 0 | 0 | 0 | 0 | 0 | 0 | 0 | 0 | 0 | 0 | 0 |
| 0 | NA | NA | NA | NA | NA | Gm35931 | | 0 | 0 | 0 | 0 | 0 | 0 | 0 | 0 | 0 | 0 | 0 |
| 0 | NA | NA | NA | NA | NA | Gm44282 | | 0 | 0 | 0 | 0 | 0 | 0 | 0 | 0 | 0 | 0 | 0 |
| 0 | NA | NA | NA | NA | NA | Gm44141 | | 0 | 0 | 0 | 0 | 0 | 0 | 0 | 0 | 0 | 0 | 0 |
| 0 | NA | NA | NA | NA | NA | Gm25283 | | 0 | 0 | 0 | 0 | 0 | 0 | 0 | 0 | 0 | 0 | 0 |
| 0 | NA | NA | NA | NA | NA | Rny3 | | 0 | 0 | 0 | 0 | 0 | 0 | 0 | 0 | 0 | 0 | 0 |
| 0 | NA | NA | NA | NA | NA | Gm16630 | | 0 | 0 | 0 | 0 | 0 | 0 | 0 | 0 | 0 | 0 | 0 |
| 0 | NA | NA | NA | NA | NA | Gm22122 | | 0 | 0 | 0 | 0 | 0 | 0 | 0 | 0 | 0 | 0 | 0 |
| 0 | NA | NA | NA | NA | NA | Gm7880 | | 0 | 0 | 0 | 0 | 0 | 0 | 0 | 0 | 0 | 0 | 0 |
| 0 | NA | NA | NA | NA | NA | Gm5111 | predicted gene 5111 | 0 | 0 | 0 | 0 | 0 | 0 | 0 | 0 | 0 | 0 | 0 |
| 0 | NA | NA | NA | NA | NA | Gm45021 | | 0 | 0 | 0 | 0 | 0 | 0 | 0 | 0 | 0 | 0 | 0 |
| 0 | NA | NA | NA | NA | NA | Gm44965 | | 0 | 0 | 0 | 0 | 0 | 0 | 0 | 0 | 0 | 0 | 0 |
| 0 | NA | NA | NA | NA | NA | Gm44262 | | 0 | 0 | 0 | 0 | 0 | 0 | 0 | 0 | 0 | 0 | 0 |
| 0 | NA | NA | NA | NA | NA | Gm44226 | | 0 | 0 | 0 | 0 | 0 | 0 | 0 | 0 | 0 | 0 | 0 |
| 0 | NA | NA | NA | NA | NA | Gm30781 | | 0 | 0 | 0 | 0 | 0 | 0 | 0 | 0 | 0 | 0 | 0 |
| 0 | NA | NA | NA | NA | NA | Gm17834 | | 0 | 0 | 0 | 0 | 0 | 0 | 0 | 0 | 0 | 0 | 0 |
| 0 | NA | NA | NA | NA | NA | Gm24348 | | 0 | 0 | 0 | 0 | 0 | 0 | 0 | 0 | 0 | 0 | 0 |
| 0 | NA | NA | NA | NA | NA | Gm10242 | | 0 | 0 | 0 | 0 | 0 | 0 | 0 | 0 | 0 | 0 | 0 |
| 0 | NA | NA | NA | NA | NA | Gm18538 | | 0 | 0 | 0 | 0 | 0 | 0 | 0 | 0 | 0 | 0 | 0 |
| 0 | NA | NA | NA | NA | NA | Gm19172 | | 0 | 0 | 0 | 0 | 0 | 0 | 0 | 0 | 0 | 0 | 0 |
| 0 | NA | NA | NA | NA | NA | Doxl2 | diamine oxidase-like protein 2 | 0 | 0 | 0 | 0 | 0 | 0 | 0 | 0 | 0 | 0 | 0 |
| 0 | NA | NA | NA | NA | NA | Gm18539 | | 0 | 0 | 0 | 0 | 0 | 0 | 0 | 0 | 0 | 0 | 0 |
| 0 | NA | NA | NA | NA | NA | 4930563H07Rik | | 0 | 0 | 0 | 0 | 0 | 0 | 0 | 0 | 0 | 0 | 0 |
| 0 | NA | NA | NA | NA | NA | Svs1 | seminal vesicle secretory protein 1 | 0 | 0 | 0 | 0 | 0 | 0 | 0 | 0 | 0 | 0 | 0 |
| 0 | NA | NA | NA | NA | NA | Gm19173 | | 0 | 0 | 0 | 0 | 0 | 0 | 0 | 0 | 0 | 0 | 0 |
| 0 | NA | NA | NA | NA | NA | Gm19253 | | 0 | 0 | 0 | 0 | 0 | 0 | 0 | 0 | 0 | 0 | 0 |
| 0 | NA | NA | NA | NA | NA | Gm18010 | | 0 | 0 | 0 | 0 | 0 | 0 | 0 | 0 | 0 | 0 | 0 |
| 0 | NA | NA | NA | NA | NA | Gm18011 | | 0 | 0 | 0 | 0 | 0 | 0 | 0 | 0 | 0 | 0 | 0 |
| 0 | NA | NA | NA | NA | NA | Gm44219 | | 0 | 0 | 0 | 0 | 0 | 0 | 0 | 0 | 0 | 0 | 0 |
| 0 | NA | NA | NA | NA | NA | Gm18007 | | 0 | 0 | 0 | 0 | 0 | 0 | 0 | 0 | 0 | 0 | 0 |
| 0 | NA | NA | NA | NA | NA | Gm3455 | | 0 | 0 | 0 | 0 | 0 | 0 | 0 | 0 | 0 | 0 | 0 |
| 0 | NA | NA | NA | NA | NA | Gm26289 | | 0 | 0 | 0 | 0 | 0 | 0 | 0 | 0 | 0 | 0 | 0 |
| 0 | NA | NA | NA | NA | NA | Mir6371 | | 0 | 0 | 0 | 0 | 0 | 0 | 0 | 0 | 0 | 0 | 0 |
| 0 | NA | NA | NA | NA | NA | Gm44054 | | 0 | 0 | 0 | 0 | 0 | 0 | 0 | 0 | 0 | 0 | 0 |
| 0 | NA | NA | NA | NA | NA | Gm4782 | | 0 | 0 | 0 | 0 | 0 | 0 | 0 | 0 | 0 | 0 | 0 |
| 0 | NA | NA | NA | NA | NA | Gm8035 | | 0 | 0 | 0 | 0 | 0 | 0 | 0 | 0 | 0 | 0 | 0 |
| 0 | NA | NA | NA | NA | NA | Gm8038 | | 0 | 0 | 0 | 0 | 0 | 0 | 0 | 0 | 0 | 0 | 0 |
| 0 | NA | NA | NA | NA | NA | Gm8039 | | 0 | 0 | 0 | 0 | 0 | 0 | 0 | 0 | 0 | 0 | 0 |
| 0 | NA | NA | NA | NA | NA | Gm18765 | | 0 | 0 | 0 | 0 | 0 | 0 | 0 | 0 | 0 | 0 | 0 |
| 0 | NA | NA | NA | NA | NA | Npvf | neuropeptide VF precursor | 0 | 0 | 0 | 0 | 0 | 0 | 0 | 0 | 0 | 0 | 0 |
| 0 | NA | NA | NA | NA | NA | Gm3017 | | 0 | 0 | 0 | 0 | 0 | 0 | 0 | 0 | 0 | 0 | 0 |
| 0 | NA | NA | NA | NA | NA | | | 0 | 0 | 0 | 0 | 0 | 0 | 0 | 0 | 0 | 0 | 0 |
| 0 | NA | NA | NA | NA | NA | Gm31579 | | 0 | 0 | 0 | 0 | 0 | 0 | 0 | 0 | 0 | 0 | 0 |
| 0 | NA | NA | NA | NA | NA | Gm44402 | | 0 | 0 | 0 | 0 | 0 | 0 | 0 | 0 | 0 | 0 | 0 |
| 0 | NA | NA | NA | NA | NA | Gm22914 | | 0 | 0 | 0 | 0 | 0 | 0 | 0 | 0 | 0 | 0 | 0 |
| 0 | NA | NA | NA | NA | NA | Gm38811 | | 0 | 0 | 0 | 0 | 0 | 0 | 0 | 0 | 0 | 0 | 0 |
| 0 | NA | NA | NA | NA | NA | Halr1 | | 0 | 0 | 0 | 0 | 0 | 0 | 0 | 0 | 0 | 0 | 0 |
| 0 | NA | NA | NA | NA | NA | Rps19-ps14 | | 0 | 0 | 0 | 0 | 0 | 0 | 0 | 0 | 0 | 0 | 0 |
| 0 | NA | NA | NA | NA | NA | Gm27867 | | 0 | 0 | 0 | 0 | 0 | 0 | 0 | 0 | 0 | 0 | 0 |
| 0 | NA | NA | NA | NA | NA | Gm27861 | | 0 | 0 | 0 | 0 | 0 | 0 | 0 | 0 | 0 | 0 | 0 |
| 0 | NA | NA | NA | NA | NA | Gm27868 | | 0 | 0 | 0 | 0 | 0 | 0 | 0 | 0 | 0 | 0 | 0 |
| 0 | NA | NA | NA | NA | NA | Gm28308 | | 0 | 0 | 0 | 0 | 0 | 0 | 0 | 0 | 0 | 0 | 0 |

| | | | | | | | | | | | | | | | | | | |
|---|---|---|---|---|---|---|---|---|---|---|---|---|---|---|---|---|---|---|
| 0 | NA | NA | NA | NA | NA | Mir196b | | 0 | 0 | 0 | 0 | 0 | 0 | 0 | 0 | 0 | 0 | 0 |
| 0 | NA | NA | NA | NA | NA | Gm28032 | | 0 | 0 | 0 | 0 | 0 | 0 | 0 | 0 | 0 | 0 | 0 |
| 0 | NA | NA | NA | NA | NA | Gm27492 | | 0 | 0 | 0 | 0 | 0 | 0 | 0 | 0 | 0 | 0 | 0 |
| 0 | NA | NA | NA | NA | NA | Gm27695 | | 0 | 0 | 0 | 0 | 0 | 0 | 0 | 0 | 0 | 0 | 0 |
| 0 | NA | NA | NA | NA | NA | Gm27817 | | 0 | 0 | 0 | 0 | 0 | 0 | 0 | 0 | 0 | 0 | 0 |
| 0 | NA | NA | NA | NA | NA | Gm27454 | | 0 | 0 | 0 | 0 | 0 | 0 | 0 | 0 | 0 | 0 | 0 |
| 0 | NA | NA | NA | NA | NA | Gm27543 | | 0 | 0 | 0 | 0 | 0 | 0 | 0 | 0 | 0 | 0 | 0 |
| 0 | NA | NA | NA | NA | NA | Gm27280 | | 0 | 0 | 0 | 0 | 0 | 0 | 0 | 0 | 0 | 0 | 0 |
| 0 | NA | NA | NA | NA | NA | Gm27985 | | 0 | 0 | 0 | 0 | 0 | 0 | 0 | 0 | 0 | 0 | 0 |
| 0 | NA | NA | NA | NA | NA | Hottip | | 0 | 0 | 0 | 0 | 0 | 0 | 0 | 0 | 0 | 0 | 0 |
| 0 | NA | NA | NA | NA | NA | Gm27383 | | 0 | 0 | 0 | 0 | 0 | 0 | 0 | 0 | 0 | 0 | 0 |
| 0 | NA | NA | NA | NA | NA | Gm27866 | | 0 | 0 | 0 | 0 | 0 | 0 | 0 | 0 | 0 | 0 | 0 |
| 0 | NA | NA | NA | NA | NA | Gm44294 | | 0 | 0 | 0 | 0 | 0 | 0 | 0 | 0 | 0 | 0 | 0 |
| 0 | NA | NA | NA | NA | NA | Evx1os | | 0 | 0 | 0 | 0 | 0 | 0 | 0 | 0 | 0 | 0 | 0 |
| 0 | NA | NA | NA | NA | NA | Evx1 | even-skipped homeobox 1 | 0 | 0 | 0 | 0 | 0 | 0 | 0 | 0 | 0 | 0 | 0 |
| 0 | NA | NA | NA | NA | NA | Gm15054 | | 0 | 0 | 0 | 0 | 0 | 0 | 0 | 0 | 0 | 0 | 0 |
| 0 | NA | NA | NA | NA | NA | 1700094M24Rik | | 0 | 0 | 0 | 0 | 0 | 0 | 0 | 0 | 0 | 0 | 0 |
| 0 | NA | NA | NA | NA | NA | Gm44028 | | 0 | 0 | 0 | 0 | 0 | 0 | 0 | 0 | 0 | 0 | 0 |
| 0 | NA | NA | NA | NA | NA | Gm44445 | | 0 | 0 | 0 | 0 | 0 | 0 | 0 | 0 | 0 | 0 | 0 |
| 0 | NA | NA | NA | NA | NA | Gm24355 | | 0 | 0 | 0 | 0 | 0 | 0 | 0 | 0 | 0 | 0 | 0 |
| 0 | NA | NA | NA | NA | NA | Gm43967 | | 0 | 0 | 0 | 0 | 0 | 0 | 0 | 0 | 0 | 0 | 0 |
| 0 | NA | NA | NA | NA | NA | Gm26215 | | 0 | 0 | 0 | 0 | 0 | 0 | 0 | 0 | 0 | 0 | 0 |
| 0 | NA | NA | NA | NA | NA | Gm8143 | | 0 | 0 | 0 | 0 | 0 | 0 | 0 | 0 | 0 | 0 | 0 |
| 0 | NA | NA | NA | NA | NA | Gm4872 | | 0 | 0 | 0 | 0 | 0 | 0 | 0 | 0 | 0 | 0 | 0 |
| 0 | NA | NA | NA | NA | NA | Gm25818 | | 0 | 0 | 0 | 0 | 0 | 0 | 0 | 0 | 0 | 0 | 0 |
| 0 | NA | NA | NA | NA | NA | Gm44080 | | 0 | 0 | 0 | 0 | 0 | 0 | 0 | 0 | 0 | 0 | 0 |
| 0 | NA | NA | NA | NA | NA | Gm5306 | | 0 | 0 | 0 | 0 | 0 | 0 | 0 | 0 | 0 | 0 | 0 |
| 0 | NA | NA | NA | NA | NA | Gm16499 | | 0 | 0 | 0 | 0 | 0 | 0 | 0 | 0 | 0 | 0 | 0 |
| 0 | NA | NA | NA | NA | NA | Gm22910 | | 0 | 0 | 0 | 0 | 0 | 0 | 0 | 0 | 0 | 0 | 0 |
| 0 | NA | NA | NA | NA | NA | Cpvl | carboxypeptidase, vitellogenic-like | 0 | 0 | 0 | 0 | 0 | 0 | 0 | 0 | 0 | 0 | 0 |
| 0 | NA | NA | NA | NA | NA | 4921529L05Rik | | 0 | 0 | 0 | 0 | 0 | 0 | 0 | 0 | 0 | 0 | 0 |
| 0 | NA | NA | NA | NA | NA | Gm15527 | | 0 | 0 | 0 | 0 | 0 | 0 | 0 | 0 | 0 | 0 | 0 |
| 0 | NA | NA | NA | NA | NA | Gm24230 | | 0 | 0 | 0 | 0 | 0 | 0 | 0 | 0 | 0 | 0 | 0 |
| 0 | NA | NA | NA | NA | NA | Gm34358 | | 0 | 0 | 0 | 0 | 0 | 0 | 0 | 0 | 0 | 0 | 0 |
| 0 | NA | NA | NA | NA | NA | Gm44073 | | 0 | 0 | 0 | 0 | 0 | 0 | 0 | 0 | 0 | 0 | 0 |
| 0 | NA | NA | NA | NA | NA | Gm25458 | | 0 | 0 | 0 | 0 | 0 | 0 | 0 | 0 | 0 | 0 | 0 |
| 0 | NA | NA | NA | NA | NA | Ghrhr | growth hormone releasing hormone rece[illegible] | 0 | 0 | 0 | 0 | 0 | 0 | 0 | 0 | 0 | 0 | 0 |
| 0 | NA | NA | NA | NA | NA | Gm44352 | | 0 | 0 | 0 | 0 | 0 | 0 | 0 | 0 | 0 | 0 | 0 |
| 0 | NA | NA | NA | NA | NA | Gm3279 | | 0 | 0 | 0 | 0 | 0 | 0 | 0 | 0 | 0 | 0 | 0 |
| 0 | NA | NA | NA | NA | NA | Itprid1 | ITPR interacting domain containing 1 | 0 | 0 | 0 | 0 | 0 | 0 | 0 | 0 | 0 | 0 | 0 |
| 0 | NA | NA | NA | NA | NA | Gm44413 | | 0 | 0 | 0 | 0 | 0 | 0 | 0 | 0 | 0 | 0 | 0 |
| 0 | NA | NA | NA | NA | NA | Gm24709 | | 0 | 0 | 0 | 0 | 0 | 0 | 0 | 0 | 0 | 0 | 0 |
| 0 | NA | NA | NA | NA | NA | Gm22493 | | 0 | 0 | 0 | 0 | 0 | 0 | 0 | 0 | 0 | 0 | 0 |
| 0 | NA | NA | NA | NA | NA | Gm44049 | | 0 | 0 | 0 | 0 | 0 | 0 | 0 | 0 | 0 | 0 | 0 |
| 0 | NA | NA | NA | NA | NA | Vmn1r5 | vomeronasal 1 receptor 5 | 0 | 0 | 0 | 0 | 0 | 0 | 0 | 0 | 0 | 0 | 0 |
| 0 | NA | NA | NA | NA | NA | Vmn1r6 | vomeronasal 1 receptor 6 | 0 | 0 | 0 | 0 | 0 | 0 | 0 | 0 | 0 | 0 | 0 |
| 0 | NA | NA | NA | NA | NA | Vmn1r7 | vomeronasal 1 receptor 7 | 0 | 0 | 0 | 0 | 0 | 0 | 0 | 0 | 0 | 0 | 0 |
| 0 | NA | NA | NA | NA | NA | Vmn1r8 | vomeronasal 1 receptor 8 | 0 | 0 | 0 | 0 | 0 | 0 | 0 | 0 | 0 | 0 | 0 |
| 0 | NA | NA | NA | NA | NA | Vmn1r9 | vomeronasal 1 receptor 9 | 0 | 0 | 0 | 0 | 0 | 0 | 0 | 0 | 0 | 0 | 0 |
| 0 | NA | NA | NA | NA | NA | Vmn1r10 | vomeronasal 1 receptor 10 | 0 | 0 | 0 | 0 | 0 | 0 | 0 | 0 | 0 | 0 | 0 |
| 0 | NA | NA | NA | NA | NA | Vmn1r11 | vomeronasal 1 receptor 11 | 0 | 0 | 0 | 0 | 0 | 0 | 0 | 0 | 0 | 0 | 0 |
| 0 | NA | NA | NA | NA | NA | Vmn1r12 | vomeronasal 1 receptor 12 | 0 | 0 | 0 | 0 | 0 | 0 | 0 | 0 | 0 | 0 | 0 |
| 0 | NA | NA | NA | NA | NA | Vmn1r-ps8 | | 0 | 0 | 0 | 0 | 0 | 0 | 0 | 0 | 0 | 0 | 0 |
| 0 | NA | NA | NA | NA | NA | Vmn1r13 | vomeronasal 1 receptor 13 | 0 | 0 | 0 | 0 | 0 | 0 | 0 | 0 | 0 | 0 | 0 |
| 0 | NA | NA | NA | NA | NA | Vmn1r14 | vomeronasal 1 receptor 14 | 0 | 0 | 0 | 0 | 0 | 0 | 0 | 0 | 0 | 0 | 0 |
| 0 | NA | NA | NA | NA | NA | Vmn1r15 | vomeronasal 1 receptor 15 | 0 | 0 | 0 | 0 | 0 | 0 | 0 | 0 | 0 | 0 | 0 |
| 0 | NA | NA | NA | NA | NA | Vmn1r-ps9 | | 0 | 0 | 0 | 0 | 0 | 0 | 0 | 0 | 0 | 0 | 0 |
| 0 | NA | NA | NA | NA | NA | Vmn1r-ps10 | | 0 | 0 | 0 | 0 | 0 | 0 | 0 | 0 | 0 | 0 | 0 |
| 0 | NA | NA | NA | NA | NA | Vmn1r17 | vomeronasal 1 receptor 17 | 0 | 0 | 0 | 0 | 0 | 0 | 0 | 0 | 0 | 0 | 0 |
| 0 | NA | NA | NA | NA | NA | Vmn1r18 | vomeronasal 1 receptor 18 | 0 | 0 | 0 | 0 | 0 | 0 | 0 | 0 | 0 | 0 | 0 |

| | | | | | | | | | | | | | | | | | | |
|---|---|---|---|---|---|---|---|---|---|---|---|---|---|---|---|---|---|---|
| 0 | NA | NA | NA | NA | NA | Vmn1r19 | vomeronasal 1 receptor 19 | 0 | 0 | 0 | 0 | 0 | 0 | 0 | 0 | 0 | 0 | 0 |
| 0 | NA | NA | NA | NA | NA | Vmn1r20 | vomeronasal 1 receptor 20 | 0 | 0 | 0 | 0 | 0 | 0 | 0 | 0 | 0 | 0 | 0 |
| 0 | NA | NA | NA | NA | NA | Vmn1r-ps11 | | 0 | 0 | 0 | 0 | 0 | 0 | 0 | 0 | 0 | 0 | 0 |
| 0 | NA | NA | NA | NA | NA | 4930533I22Rik | | 0 | 0 | 0 | 0 | 0 | 0 | 0 | 0 | 0 | 0 | 0 |
| 0 | NA | NA | NA | NA | NA | Gm45193 | | 0 | 0 | 0 | 0 | 0 | 0 | 0 | 0 | 0 | 0 | 0 |
| 0 | NA | NA | NA | NA | NA | Gm19244 | | 0 | 0 | 0 | 0 | 0 | 0 | 0 | 0 | 0 | 0 | 0 |
| 0 | NA | NA | NA | NA | NA | Gm26712 | | 0 | 0 | 0 | 0 | 0 | 0 | 0 | 0 | 0 | 0 | 0 |
| 0 | NA | NA | NA | NA | NA | Vmn1r21 | vomeronasal 1 receptor 21 | 0 | 0 | 0 | 0 | 0 | 0 | 0 | 0 | 0 | 0 | 0 |
| 0 | NA | NA | NA | NA | NA | Gm44114 | | 0 | 0 | 0 | 0 | 0 | 0 | 0 | 0 | 0 | 0 | 0 |
| 0 | NA | NA | NA | NA | NA | Gm25726 | | 0 | 0 | 0 | 0 | 0 | 0 | 0 | 0 | 0 | 0 | 0 |
| 0 | NA | NA | NA | NA | NA | Vmn1r22 | vomeronasal 1 receptor 22 | 0 | 0 | 0 | 0 | 0 | 0 | 0 | 0 | 0 | 0 | 0 |
| 0 | NA | NA | NA | NA | NA | Vmn1r23 | vomeronasal 1 receptor 23 | 0 | 0 | 0 | 0 | 0 | 0 | 0 | 0 | 0 | 0 | 0 |
| 0 | NA | NA | NA | NA | NA | Gm20660 | | 0 | 0 | 0 | 0 | 0 | 0 | 0 | 0 | 0 | 0 | 0 |
| 0 | NA | NA | NA | NA | NA | Vmn1r24 | vomeronasal 1 receptor 24 | 0 | 0 | 0 | 0 | 0 | 0 | 0 | 0 | 0 | 0 | 0 |
| 0 | NA | NA | NA | NA | NA | Vmn1r25 | vomeronasal 1 receptor 25 | 0 | 0 | 0 | 0 | 0 | 0 | 0 | 0 | 0 | 0 | 0 |
| 0 | NA | NA | NA | NA | NA | Vmn1r-ps13 | | 0 | 0 | 0 | 0 | 0 | 0 | 0 | 0 | 0 | 0 | 0 |
| 0 | NA | NA | NA | NA | NA | Vmn1r26 | vomeronasal 1 receptor 26 | 0 | 0 | 0 | 0 | 0 | 0 | 0 | 0 | 0 | 0 | 0 |
| 0 | NA | NA | NA | NA | NA | Vmn1r-ps14 | | 0 | 0 | 0 | 0 | 0 | 0 | 0 | 0 | 0 | 0 | 0 |
| 0 | NA | NA | NA | NA | NA | Vmn1r-ps15 | | 0 | 0 | 0 | 0 | 0 | 0 | 0 | 0 | 0 | 0 | 0 |
| 0 | NA | NA | NA | NA | NA | Vmn1r-ps16 | | 0 | 0 | 0 | 0 | 0 | 0 | 0 | 0 | 0 | 0 | 0 |
| 0 | NA | NA | NA | NA | NA | Vmn1r-ps17 | | 0 | 0 | 0 | 0 | 0 | 0 | 0 | 0 | 0 | 0 | 0 |
| 0 | NA | NA | NA | NA | NA | Gm20718 | | 0 | 0 | 0 | 0 | 0 | 0 | 0 | 0 | 0 | 0 | 0 |
| 0 | NA | NA | NA | NA | NA | Gm43994 | | 0 | 0 | 0 | 0 | 0 | 0 | 0 | 0 | 0 | 0 | 0 |
| 0 | NA | NA | NA | NA | NA | Vmn1r-ps18 | | 0 | 0 | 0 | 0 | 0 | 0 | 0 | 0 | 0 | 0 | 0 |
| 0 | NA | NA | NA | NA | NA | Vmn1r27 | vomeronasal 1 receptor 27 | 0 | 0 | 0 | 0 | 0 | 0 | 0 | 0 | 0 | 0 | 0 |
| 0 | NA | NA | NA | NA | NA | Vmn1r-ps19 | | 0 | 0 | 0 | 0 | 0 | 0 | 0 | 0 | 0 | 0 | 0 |
| 0 | NA | NA | NA | NA | NA | Vmn1r28 | vomeronasal 1 receptor 28 | 0 | 0 | 0 | 0 | 0 | 0 | 0 | 0 | 0 | 0 | 0 |
| 0 | NA | NA | NA | NA | NA | Vmn1r29 | vomeronasal 1 receptor 29 | 0 | 0 | 0 | 0 | 0 | 0 | 0 | 0 | 0 | 0 | 0 |
| 0 | NA | NA | NA | NA | NA | Gm20674 | | 0 | 0 | 0 | 0 | 0 | 0 | 0 | 0 | 0 | 0 | 0 |
| 0 | NA | NA | NA | NA | NA | Vmn1r-ps20 | | 0 | 0 | 0 | 0 | 0 | 0 | 0 | 0 | 0 | 0 | 0 |
| 0 | NA | NA | NA | NA | NA | Gm44016 | | 0 | 0 | 0 | 0 | 0 | 0 | 0 | 0 | 0 | 0 | 0 |
| 0 | NA | NA | NA | NA | NA | Vmn1r-ps21 | | 0 | 0 | 0 | 0 | 0 | 0 | 0 | 0 | 0 | 0 | 0 |
| 0 | NA | NA | NA | NA | NA | Gm35077 | | 0 | 0 | 0 | 0 | 0 | 0 | 0 | 0 | 0 | 0 | 0 |
| 0 | NA | NA | NA | NA | NA | Vmn1r31 | vomeronasal 1 receptor 31 | 0 | 0 | 0 | 0 | 0 | 0 | 0 | 0 | 0 | 0 | 0 |
| 0 | NA | NA | NA | NA | NA | Vmn1r-ps22 | | 0 | 0 | 0 | 0 | 0 | 0 | 0 | 0 | 0 | 0 | 0 |
| 0 | NA | NA | NA | NA | NA | Vmn1r-ps23 | | 0 | 0 | 0 | 0 | 0 | 0 | 0 | 0 | 0 | 0 | 0 |
| 0 | NA | NA | NA | NA | NA | Gm20672 | | 0 | 0 | 0 | 0 | 0 | 0 | 0 | 0 | 0 | 0 | 0 |
| 0 | NA | NA | NA | NA | NA | Gm43899 | | 0 | 0 | 0 | 0 | 0 | 0 | 0 | 0 | 0 | 0 | 0 |
| 0 | NA | NA | NA | NA | NA | BB365896 | | 0 | 0 | 0 | 0 | 0 | 0 | 0 | 0 | 0 | 0 | 0 |
| 0 | NA | NA | NA | NA | NA | Gm45061 | | 0 | 0 | 0 | 0 | 0 | 0 | 0 | 0 | 0 | 0 | 0 |
| 0 | NA | NA | NA | NA | NA | Gm19165 | | 0 | 0 | 0 | 0 | 0 | 0 | 0 | 0 | 0 | 0 | 0 |
| 0 | NA | NA | NA | NA | NA | Gm22367 | | 0 | 0 | 0 | 0 | 0 | 0 | 0 | 0 | 0 | 0 | 0 |
| 0 | NA | NA | NA | NA | NA | Gm43905 | | 0 | 0 | 0 | 0 | 0 | 0 | 0 | 0 | 0 | 0 | 0 |
| 0 | NA | NA | NA | NA | NA | Gm22622 | | 0 | 0 | 0 | 0 | 0 | 0 | 0 | 0 | 0 | 0 | 0 |
| 0 | NA | NA | NA | NA | NA | Gm18012 | | 0 | 0 | 0 | 0 | 0 | 0 | 0 | 0 | 0 | 0 | 0 |
| 0 | NA | NA | NA | NA | NA | Gm5570 | | 0 | 0 | 0 | 0 | 0 | 0 | 0 | 0 | 0 | 0 | 0 |
| 0 | NA | NA | NA | NA | NA | Gm22723 | | 0 | 0 | 0 | 0 | 0 | 0 | 0 | 0 | 0 | 0 | 0 |
| 0 | NA | NA | NA | NA | NA | n-R5s162 | | 0 | 0 | 0 | 0 | 0 | 0 | 0 | 0 | 0 | 0 | 0 |
| 0 | NA | NA | NA | NA | NA | A530053G22Rik | | 0 | 0 | 0 | 0 | 0 | 0 | 0 | 0 | 0 | 0 | 0 |
| 0 | NA | NA | NA | NA | NA | Gm44410 | | 0 | 0 | 0 | 0 | 0 | 0 | 0 | 0 | 0 | 0 | 0 |
| 0 | NA | NA | NA | NA | NA | Gm35386 | | 0 | 0 | 0 | 0 | 0 | 0 | 0 | 0 | 0 | 0 | 0 |
| 0 | NA | NA | NA | NA | NA | Mageb16-ps2 | | 0 | 0 | 0 | 0 | 0 | 0 | 0 | 0 | 0 | 0 | 0 |
| 0 | NA | NA | NA | NA | NA | Gm18838 | | 0 | 0 | 0 | 0 | 0 | 0 | 0 | 0 | 0 | 0 | 0 |
| 0 | NA | NA | NA | NA | NA | Gm43892 | | 0 | 0 | 0 | 0 | 0 | 0 | 0 | 0 | 0 | 0 | 0 |
| 0 | NA | NA | NA | NA | NA | Gm43893 | | 0 | 0 | 0 | 0 | 0 | 0 | 0 | 0 | 0 | 0 | 0 |
| 0 | NA | NA | NA | NA | NA | Gm43894 | | 0 | 0 | 0 | 0 | 0 | 0 | 0 | 0 | 0 | 0 | 0 |
| 0 | NA | NA | NA | NA | NA | Gm43898 | | 0 | 0 | 0 | 0 | 0 | 0 | 0 | 0 | 0 | 0 | 0 |
| 0 | NA | NA | NA | NA | NA | Gm43897 | | 0 | 0 | 0 | 0 | 0 | 0 | 0 | 0 | 0 | 0 | 0 |
| 0 | NA | NA | NA | NA | NA | Gm43895 | | 0 | 0 | 0 | 0 | 0 | 0 | 0 | 0 | 0 | 0 | 0 |
| 0 | NA | NA | NA | NA | NA | Gm19237 | | 0 | 0 | 0 | 0 | 0 | 0 | 0 | 0 | 0 | 0 | 0 |

| | | | | | | | | | | | | | | | | | | |
|---|---|---|---|---|---|---|---|---|---|---|---|---|---|---|---|---|---|---|
| 0 | NA | NA | NA | NA | NA | Gm44088 | | 0 | 0 | 0 | 0 | 0 | 0 | 0 | 0 | 0 | 0 | 0 |
| 0 | NA | NA | NA | NA | NA | Gm44092 | | 0 | 0 | 0 | 0 | 0 | 0 | 0 | 0 | 0 | 0 | 0 |
| 0 | NA | NA | NA | NA | NA | Gm18540 | | 0 | 0 | 0 | 0 | 0 | 0 | 0 | 0 | 0 | 0 | 0 |
| 0 | NA | NA | NA | NA | NA | Gm5001 | | 0 | 0 | 0 | 0 | 0 | 0 | 0 | 0 | 0 | 0 | 0 |
| 0 | NA | NA | NA | NA | NA | Gm19131 | | 0 | 0 | 0 | 0 | 0 | 0 | 0 | 0 | 0 | 0 | 0 |
| 0 | NA | NA | NA | NA | NA | Nasp-ps1 | | 0 | 0 | 0 | 0 | 0 | 0 | 0 | 0 | 0 | 0 | 0 |
| 0 | NA | NA | NA | NA | NA | Gm22212 | | 0 | 0 | 0 | 0 | 0 | 0 | 0 | 0 | 0 | 0 | 0 |
| 0 | NA | NA | NA | NA | NA | 9330118I20Rik | | 0 | 0 | 0 | 0 | 0 | 0 | 0 | 0 | 0 | 0 | 0 |
| 0 | NA | NA | NA | NA | NA | Gm44071 | | 0 | 0 | 0 | 0 | 0 | 0 | 0 | 0 | 0 | 0 | 0 |
| 0 | NA | NA | NA | NA | NA | Gm44072 | | 0 | 0 | 0 | 0 | 0 | 0 | 0 | 0 | 0 | 0 | 0 |
| 0 | NA | NA | NA | NA | NA | Gm20144 | | 0 | 0 | 0 | 0 | 0 | 0 | 0 | 0 | 0 | 0 | 0 |
| 0 | NA | NA | NA | NA | NA | Gm25205 | | 0 | 0 | 0 | 0 | 0 | 0 | 0 | 0 | 0 | 0 | 0 |
| 0 | NA | NA | NA | NA | NA | Atoh1 | atonal bHLH transcription factor 1 | 0 | 0 | 0 | 0 | 0 | 0 | 0 | 0 | 0 | 0 | 0 |
| 0 | NA | NA | NA | NA | NA | Gm18839 | | 0 | 0 | 0 | 0 | 0 | 0 | 0 | 0 | 0 | 0 | 0 |
| 0 | NA | NA | NA | NA | NA | Gm44038 | | 0 | 0 | 0 | 0 | 0 | 0 | 0 | 0 | 0 | 0 | 0 |
| 0 | NA | NA | NA | NA | NA | Gm17147 | | 0 | 0 | 0 | 0 | 0 | 0 | 0 | 0 | 0 | 0 | 0 |
| 0 | NA | NA | NA | NA | NA | Gm3549 | | 0 | 0 | 0 | 0 | 0 | 0 | 0 | 0 | 0 | 0 | 0 |
| 0 | NA | NA | NA | NA | NA | Gm44058 | | 0 | 0 | 0 | 0 | 0 | 0 | 0 | 0 | 0 | 0 | 0 |
| 0 | NA | NA | NA | NA | NA | Gm44059 | | 0 | 0 | 0 | 0 | 0 | 0 | 0 | 0 | 0 | 0 | 0 |
| 0 | NA | NA | NA | NA | NA | Gm20177 | | 0 | 0 | 0 | 0 | 0 | 0 | 0 | 0 | 0 | 0 | 0 |
| 0 | NA | NA | NA | NA | NA | Gm44060 | | 0 | 0 | 0 | 0 | 0 | 0 | 0 | 0 | 0 | 0 | 0 |
| 0 | NA | NA | NA | NA | NA | Gm44061 | | 0 | 0 | 0 | 0 | 0 | 0 | 0 | 0 | 0 | 0 | 0 |
| 0 | NA | NA | NA | NA | NA | Gm8479 | | 0 | 0 | 0 | 0 | 0 | 0 | 0 | 0 | 0 | 0 | 0 |
| 0 | NA | NA | NA | NA | NA | Qrfprl | pyroglutamylated RFamide peptide recep | 0 | 0 | 0 | 0 | 0 | 0 | 0 | 0 | 0 | 0 | 0 |
| 0 | NA | NA | NA | NA | NA | Gm18716 | | 0 | 0 | 0 | 0 | 0 | 0 | 0 | 0 | 0 | 0 | 0 |
| 0 | NA | NA | NA | NA | NA | Gm15490 | | 0 | 0 | 0 | 0 | 0 | 0 | 0 | 0 | 0 | 0 | 0 |
| 0 | NA | NA | NA | NA | NA | Gm16838 | | 0 | 0 | 0 | 0 | 0 | 0 | 0 | 0 | 0 | 0 | 0 |
| 0 | NA | NA | NA | NA | NA | Gm23543 | | 0 | 0 | 0 | 0 | 0 | 0 | 0 | 0 | 0 | 0 | 0 |
| 0 | NA | NA | NA | NA | NA | Gm20445 | | 0 | 0 | 0 | 0 | 0 | 0 | 0 | 0 | 0 | 0 | 0 |
| 0 | NA | NA | NA | NA | NA | 4930544G11Ril | RIKEN cDNA 4930544G11 gene | 0 | 0 | 0 | 0 | 0 | 0 | 0 | 0 | 0 | 0 | 0 |
| 0 | NA | NA | NA | NA | NA | Gm44404 | | 0 | 0 | 0 | 0 | 0 | 0 | 0 | 0 | 0 | 0 | 0 |
| 0 | NA | NA | NA | NA | NA | Gm23625 | | 0 | 0 | 0 | 0 | 0 | 0 | 0 | 0 | 0 | 0 | 0 |
| 0 | NA | NA | NA | NA | NA | Gm44289 | | 0 | 0 | 0 | 0 | 0 | 0 | 0 | 0 | 0 | 0 | 0 |
| 0 | NA | NA | NA | NA | NA | Gm31520 | | 0 | 0 | 0 | 0 | 0 | 0 | 0 | 0 | 0 | 0 | 0 |
| 0 | NA | NA | NA | NA | NA | Gm44281 | | 0 | 0 | 0 | 0 | 0 | 0 | 0 | 0 | 0 | 0 | 0 |
| 0 | NA | NA | NA | NA | NA | Gm44233 | | 0 | 0 | 0 | 0 | 0 | 0 | 0 | 0 | 0 | 0 | 0 |
| 0 | NA | NA | NA | NA | NA | 2610300M13Rik | | 0 | 0 | 0 | 0 | 0 | 0 | 0 | 0 | 0 | 0 | 0 |
| 0 | NA | NA | NA | NA | NA | Gm19764 | | 0 | 0 | 0 | 0 | 0 | 0 | 0 | 0 | 0 | 0 | 0 |
| 0 | NA | NA | NA | NA | NA | Vmn1r32 | vomeronasal 1 receptor 32 | 0 | 0 | 0 | 0 | 0 | 0 | 0 | 0 | 0 | 0 | 0 |
| 0 | NA | NA | NA | NA | NA | Gm5307 | | 0 | 0 | 0 | 0 | 0 | 0 | 0 | 0 | 0 | 0 | 0 |
| 0 | NA | NA | NA | NA | NA | Vmn1r33 | vomeronasal 1 receptor 33 | 0 | 0 | 0 | 0 | 0 | 0 | 0 | 0 | 0 | 0 | 0 |
| 0 | NA | NA | NA | NA | NA | Vmn1r34 | vomeronasal 1 receptor 34 | 0 | 0 | 0 | 0 | 0 | 0 | 0 | 0 | 0 | 0 | 0 |
| 0 | NA | NA | NA | NA | NA | Vmn1r35 | vomeronasal 1 receptor 35 | 0 | 0 | 0 | 0 | 0 | 0 | 0 | 0 | 0 | 0 | 0 |
| 0 | NA | NA | NA | NA | NA | Vmn1r36 | vomeronasal 1 receptor 36 | 0 | 0 | 0 | 0 | 0 | 0 | 0 | 0 | 0 | 0 | 0 |
| 0 | NA | NA | NA | NA | NA | Vmn1r37 | vomeronasal 1 receptor 37 | 0 | 0 | 0 | 0 | 0 | 0 | 0 | 0 | 0 | 0 | 0 |
| 0 | NA | NA | NA | NA | NA | Gm44261 | | 0 | 0 | 0 | 0 | 0 | 0 | 0 | 0 | 0 | 0 | 0 |
| 0 | NA | NA | NA | NA | NA | Vmn1r-ps24 | | 0 | 0 | 0 | 0 | 0 | 0 | 0 | 0 | 0 | 0 | 0 |
| 0 | NA | NA | NA | NA | NA | Vmn1r38 | vomeronasal 1 receptor 38 | 0 | 0 | 0 | 0 | 0 | 0 | 0 | 0 | 0 | 0 | 0 |
| 0 | NA | NA | NA | NA | NA | Vmn1r39 | vomeronasal 1 receptor 39 | 0 | 0 | 0 | 0 | 0 | 0 | 0 | 0 | 0 | 0 | 0 |
| 0 | NA | NA | NA | NA | NA | Gm36816 | | 0 | 0 | 0 | 0 | 0 | 0 | 0 | 0 | 0 | 0 | 0 |
| 0 | NA | NA | NA | NA | NA | Gm38825 | | 0 | 0 | 0 | 0 | 0 | 0 | 0 | 0 | 0 | 0 | 0 |
| 0 | NA | NA | NA | NA | NA | Gm44052 | | 0 | 0 | 0 | 0 | 0 | 0 | 0 | 0 | 0 | 0 | 0 |
| 0 | NA | NA | NA | NA | NA | Gm8574 | | 0 | 0 | 0 | 0 | 0 | 0 | 0 | 0 | 0 | 0 | 0 |
| 0 | NA | NA | NA | NA | NA | Gm44208 | | 0 | 0 | 0 | 0 | 0 | 0 | 0 | 0 | 0 | 0 | 0 |
| 0 | NA | NA | NA | NA | NA | Gm44083 | | 0 | 0 | 0 | 0 | 0 | 0 | 0 | 0 | 0 | 0 | 0 |
| 0 | NA | NA | NA | NA | NA | Il23r | interleukin 23 receptor | 0 | 0 | 0 | 0 | 0 | 0 | 0 | 0 | 0 | 0 | 0 |
| 0 | NA | NA | NA | NA | NA | Gm42889 | | 0 | 0 | 0 | 0 | 0 | 0 | 0 | 0 | 0 | 0 | 0 |
| 0 | NA | NA | NA | NA | NA | Igkv2-137 | | 0 | 0 | 0 | 0 | 0 | 0 | 0 | 0 | 0 | 0 | 0 |
| 0 | NA | NA | NA | NA | NA | 4930515G16Rik | | 0 | 0 | 0 | 0 | 0 | 0 | 0 | 0 | 0 | 0 | 0 |
| 0 | NA | NA | NA | NA | NA | Igkv1-136 | | 0 | 0 | 0 | 0 | 0 | 0 | 0 | 0 | 0 | 0 | 0 |

| | | | | | | | | | | | | | | | | | |
|---|---|---|---|---|---|---|---|---|---|---|---|---|---|---|---|---|---|
| 0 | NA | NA | NA | NA | NA | Gm9728 | 0 | 0 | 0 | 0 | 0 | 0 | 0 | 0 | 0 | 0 | 0 |
| 0 | NA | NA | NA | NA | NA | Rpl18a-ps2 | 0 | 0 | 0 | 0 | 0 | 0 | 0 | 0 | 0 | 0 | 0 |
| 0 | NA | NA | NA | NA | NA | Igkv14-134-1 | 0 | 0 | 0 | 0 | 0 | 0 | 0 | 0 | 0 | 0 | 0 |
| 0 | NA | NA | NA | NA | NA | Igkv17-134 | 0 | 0 | 0 | 0 | 0 | 0 | 0 | 0 | 0 | 0 | 0 |
| 0 | NA | NA | NA | NA | NA | Rpl18a-ps3 | 0 | 0 | 0 | 0 | 0 | 0 | 0 | 0 | 0 | 0 | 0 |
| 0 | NA | NA | NA | NA | NA | Gm43284 | 0 | 0 | 0 | 0 | 0 | 0 | 0 | 0 | 0 | 0 | 0 |
| 0 | NA | NA | NA | NA | NA | Gm18980 | 0 | 0 | 0 | 0 | 0 | 0 | 0 | 0 | 0 | 0 | 0 |
| 0 | NA | NA | NA | NA | NA | Gm22425 | 0 | 0 | 0 | 0 | 0 | 0 | 0 | 0 | 0 | 0 | 0 |
| 0 | NA | NA | NA | NA | NA | Gm24911 | 0 | 0 | 0 | 0 | 0 | 0 | 0 | 0 | 0 | 0 | 0 |
| 0 | NA | NA | NA | NA | NA | Igkv9-128 | 0 | 0 | 0 | 0 | 0 | 0 | 0 | 0 | 0 | 0 | 0 |
| 0 | NA | NA | NA | NA | NA | Igkv14-126-1 | 0 | 0 | 0 | 0 | 0 | 0 | 0 | 0 | 0 | 0 | 0 |
| 0 | NA | NA | NA | NA | NA | Gm25910 | 0 | 0 | 0 | 0 | 0 | 0 | 0 | 0 | 0 | 0 | 0 |
| 0 | NA | NA | NA | NA | NA | Igkv1-122 | 0 | 0 | 0 | 0 | 0 | 0 | 0 | 0 | 0 | 0 | 0 |
| 0 | NA | NA | NA | NA | NA | Igkv14-118-2 | 0 | 0 | 0 | 0 | 0 | 0 | 0 | 0 | 0 | 0 | 0 |
| 0 | NA | NA | NA | NA | NA | Igkv14-118-1 | 0 | 0 | 0 | 0 | 0 | 0 | 0 | 0 | 0 | 0 | 0 |
| 0 | NA | NA | NA | NA | NA | Gm43739 | 0 | 0 | 0 | 0 | 0 | 0 | 0 | 0 | 0 | 0 | 0 |
| 0 | NA | NA | NA | NA | NA | Igkv11-118 | 0 | 0 | 0 | 0 | 0 | 0 | 0 | 0 | 0 | 0 | 0 |
| 0 | NA | NA | NA | NA | NA | Gm43171 | 0 | 0 | 0 | 0 | 0 | 0 | 0 | 0 | 0 | 0 | 0 |
| 0 | NA | NA | NA | NA | NA | Gm43170 | 0 | 0 | 0 | 0 | 0 | 0 | 0 | 0 | 0 | 0 | 0 |
| 0 | NA | NA | NA | NA | NA | Igkv1-115 | 0 | 0 | 0 | 0 | 0 | 0 | 0 | 0 | 0 | 0 | 0 |
| 0 | NA | NA | NA | NA | NA | Igkv2-113 | 0 | 0 | 0 | 0 | 0 | 0 | 0 | 0 | 0 | 0 | 0 |
| 0 | NA | NA | NA | NA | NA | Gm25436 | 0 | 0 | 0 | 0 | 0 | 0 | 0 | 0 | 0 | 0 | 0 |
| 0 | NA | NA | NA | NA | NA | Gm43007 | 0 | 0 | 0 | 0 | 0 | 0 | 0 | 0 | 0 | 0 | 0 |
| 0 | NA | NA | NA | NA | NA | Gm42612 | 0 | 0 | 0 | 0 | 0 | 0 | 0 | 0 | 0 | 0 | 0 |
| 0 | NA | NA | NA | NA | NA | Igkv1-108 | 0 | 0 | 0 | 0 | 0 | 0 | 0 | 0 | 0 | 0 | 0 |
| 0 | NA | NA | NA | NA | NA | Igkv2-107 | 0 | 0 | 0 | 0 | 0 | 0 | 0 | 0 | 0 | 0 | 0 |
| 0 | NA | NA | NA | NA | NA | Igkv11-106 | 0 | 0 | 0 | 0 | 0 | 0 | 0 | 0 | 0 | 0 | 0 |
| 0 | NA | NA | NA | NA | NA | Igkv2-105 | 0 | 0 | 0 | 0 | 0 | 0 | 0 | 0 | 0 | 0 | 0 |
| 0 | NA | NA | NA | NA | NA | Gm5310 | 0 | 0 | 0 | 0 | 0 | 0 | 0 | 0 | 0 | 0 | 0 |
| 0 | NA | NA | NA | NA | NA | Igkv15-102 | 0 | 0 | 0 | 0 | 0 | 0 | 0 | 0 | 0 | 0 | 0 |
| 0 | NA | NA | NA | NA | NA | Igkv20-101-2 | 0 | 0 | 0 | 0 | 0 | 0 | 0 | 0 | 0 | 0 | 0 |
| 0 | NA | NA | NA | NA | NA | Igkv15-101-1 | 0 | 0 | 0 | 0 | 0 | 0 | 0 | 0 | 0 | 0 | 0 |
| 0 | NA | NA | NA | NA | NA | Igkv15-101 | 0 | 0 | 0 | 0 | 0 | 0 | 0 | 0 | 0 | 0 | 0 |
| 0 | NA | NA | NA | NA | NA | Gm25833 | 0 | 0 | 0 | 0 | 0 | 0 | 0 | 0 | 0 | 0 | 0 |
| 0 | NA | NA | NA | NA | NA | Igkv12-98 | 0 | 0 | 0 | 0 | 0 | 0 | 0 | 0 | 0 | 0 | 0 |
| 0 | NA | NA | NA | NA | NA | Igkv15-97 | 0 | 0 | 0 | 0 | 0 | 0 | 0 | 0 | 0 | 0 | 0 |
| 0 | NA | NA | NA | NA | NA | Igkv2-95-2 | 0 | 0 | 0 | 0 | 0 | 0 | 0 | 0 | 0 | 0 | 0 |
| 0 | NA | NA | NA | NA | NA | Igkv2-95-1 | 0 | 0 | 0 | 0 | 0 | 0 | 0 | 0 | 0 | 0 | 0 |
| 0 | NA | NA | NA | NA | NA | Igkv10-95 | 0 | 0 | 0 | 0 | 0 | 0 | 0 | 0 | 0 | 0 | 0 |
| 0 | NA | NA | NA | NA | NA | Igkv2-93-1 | 0 | 0 | 0 | 0 | 0 | 0 | 0 | 0 | 0 | 0 | 0 |
| 0 | NA | NA | NA | NA | NA | Gm42925 | 0 | 0 | 0 | 0 | 0 | 0 | 0 | 0 | 0 | 0 | 0 |
| 0 | NA | NA | NA | NA | NA | Igkv4-90 | 0 | 0 | 0 | 0 | 0 | 0 | 0 | 0 | 0 | 0 | 0 |
| 0 | NA | NA | NA | NA | NA | Gm42924 | 0 | 0 | 0 | 0 | 0 | 0 | 0 | 0 | 0 | 0 | 0 |
| 0 | NA | NA | NA | NA | NA | Igkv12-89 | 0 | 0 | 0 | 0 | 0 | 0 | 0 | 0 | 0 | 0 | 0 |
| 0 | NA | NA | NA | NA | NA | Gm42539 | 0 | 0 | 0 | 0 | 0 | 0 | 0 | 0 | 0 | 0 | 0 |
| 0 | NA | NA | NA | NA | NA | Gm42542 | 0 | 0 | 0 | 0 | 0 | 0 | 0 | 0 | 0 | 0 | 0 |
| 0 | NA | NA | NA | NA | NA | Igkv13-87 | 0 | 0 | 0 | 0 | 0 | 0 | 0 | 0 | 0 | 0 | 0 |
| 0 | NA | NA | NA | NA | NA | Gm43710 | 0 | 0 | 0 | 0 | 0 | 0 | 0 | 0 | 0 | 0 | 0 |
| 0 | NA | NA | NA | NA | NA | Igkv13-82 | 0 | 0 | 0 | 0 | 0 | 0 | 0 | 0 | 0 | 0 | 0 |
| 0 | NA | NA | NA | NA | NA | Igkv13-80-1 | 0 | 0 | 0 | 0 | 0 | 0 | 0 | 0 | 0 | 0 | 0 |
| 0 | NA | NA | NA | NA | NA | Igkv13-78-1 | 0 | 0 | 0 | 0 | 0 | 0 | 0 | 0 | 0 | 0 | 0 |
| 0 | NA | NA | NA | NA | NA | Gm43583 | 0 | 0 | 0 | 0 | 0 | 0 | 0 | 0 | 0 | 0 | 0 |
| 0 | NA | NA | NA | NA | NA | Igkv13-76 | 0 | 0 | 0 | 0 | 0 | 0 | 0 | 0 | 0 | 0 | 0 |
| 0 | NA | NA | NA | NA | NA | Igkv13-74-1 | 0 | 0 | 0 | 0 | 0 | 0 | 0 | 0 | 0 | 0 | 0 |
| 0 | NA | NA | NA | NA | NA | Igkv13-73-1 | 0 | 0 | 0 | 0 | 0 | 0 | 0 | 0 | 0 | 0 | 0 |
| 0 | NA | NA | NA | NA | NA | Igkv4-72 | 0 | 0 | 0 | 0 | 0 | 0 | 0 | 0 | 0 | 0 | 0 |
| 0 | NA | NA | NA | NA | NA | Igkv13-71-1 | 0 | 0 | 0 | 0 | 0 | 0 | 0 | 0 | 0 | 0 | 0 |
| 0 | NA | NA | NA | NA | NA | Igkv12-67 | 0 | 0 | 0 | 0 | 0 | 0 | 0 | 0 | 0 | 0 | 0 |
| 0 | NA | NA | NA | NA | NA | Rprl1 | 0 | 0 | 0 | 0 | 0 | 0 | 0 | 0 | 0 | 0 | 0 |
| 0 | NA | NA | NA | NA | NA | Rprl1 | 0 | 0 | 0 | 0 | 0 | 0 | 0 | 0 | 0 | 0 | 0 |

| | | | | | | | | | | | | | | | | | | |
|---|---|---|---|---|---|---|---|---|---|---|---|---|---|---|---|---|---|---|
| 0 | NA | NA | NA | NA | NA | Igkv12-66 | | 0 | 0 | 0 | 0 | 0 | 0 | 0 | 0 | 0 | 0 | 0 |
| 0 | NA | NA | NA | NA | NA | Igkv4-65 | | 0 | 0 | 0 | 0 | 0 | 0 | 0 | 0 | 0 | 0 | 0 |
| 0 | NA | NA | NA | NA | NA | Igkv13-64 | | 0 | 0 | 0 | 0 | 0 | 0 | 0 | 0 | 0 | 0 | 0 |
| 0 | NA | NA | NA | NA | NA | Igkv13-62-1 | | 0 | 0 | 0 | 0 | 0 | 0 | 0 | 0 | 0 | 0 | 0 |
| 0 | NA | NA | NA | NA | NA | Igkv13-61-1 | | 0 | 0 | 0 | 0 | 0 | 0 | 0 | 0 | 0 | 0 | 0 |
| 0 | NA | NA | NA | NA | NA | Igkv4-60 | | 0 | 0 | 0 | 0 | 0 | 0 | 0 | 0 | 0 | 0 | 0 |
| 0 | NA | NA | NA | NA | NA | Igkv4-58 | | 0 | 0 | 0 | 0 | 0 | 0 | 0 | 0 | 0 | 0 | 0 |
| 0 | NA | NA | NA | NA | NA | Rn7s6 | | 0 | 0 | 0 | 0 | 0 | 0 | 0 | 0 | 0 | 0 | 0 |
| 0 | NA | NA | NA | NA | NA | Gm47304 | | 0 | 0 | 0 | 0 | 0 | 0 | 0 | 0 | 0 | 0 | 0 |
| 0 | NA | NA | NA | NA | NA | Igkv13-57-2 | | 0 | 0 | 0 | 0 | 0 | 0 | 0 | 0 | 0 | 0 | 0 |
| 0 | NA | NA | NA | NA | NA | Igkv13-57-1 | | 0 | 0 | 0 | 0 | 0 | 0 | 0 | 0 | 0 | 0 | 0 |
| 0 | NA | NA | NA | NA | NA | Igkv13-56-1 | | 0 | 0 | 0 | 0 | 0 | 0 | 0 | 0 | 0 | 0 | 0 |
| 0 | NA | NA | NA | NA | NA | Igkv13-55-1 | | 0 | 0 | 0 | 0 | 0 | 0 | 0 | 0 | 0 | 0 | 0 |
| 0 | NA | NA | NA | NA | NA | Igkv13-54-1 | | 0 | 0 | 0 | 0 | 0 | 0 | 0 | 0 | 0 | 0 | 0 |
| 0 | NA | NA | NA | NA | NA | Igkv4-54 | | 0 | 0 | 0 | 0 | 0 | 0 | 0 | 0 | 0 | 0 | 0 |
| 0 | NA | NA | NA | NA | NA | Igkv4-53 | | 0 | 0 | 0 | 0 | 0 | 0 | 0 | 0 | 0 | 0 | 0 |
| 0 | NA | NA | NA | NA | NA | Gm42606 | | 0 | 0 | 0 | 0 | 0 | 0 | 0 | 0 | 0 | 0 | 0 |
| 0 | NA | NA | NA | NA | NA | Igkv4-51 | | 0 | 0 | 0 | 0 | 0 | 0 | 0 | 0 | 0 | 0 | 0 |
| 0 | NA | NA | NA | NA | NA | Gm42958 | | 0 | 0 | 0 | 0 | 0 | 0 | 0 | 0 | 0 | 0 | 0 |
| 0 | NA | NA | NA | NA | NA | Igkv12-49 | | 0 | 0 | 0 | 0 | 0 | 0 | 0 | 0 | 0 | 0 | 0 |
| 0 | NA | NA | NA | NA | NA | Igkv12-47 | | 0 | 0 | 0 | 0 | 0 | 0 | 0 | 0 | 0 | 0 | 0 |
| 0 | NA | NA | NA | NA | NA | Igkv12-42 | | 0 | 0 | 0 | 0 | 0 | 0 | 0 | 0 | 0 | 0 | 0 |
| 0 | NA | NA | NA | NA | NA | Igkv5-40-1 | | 0 | 0 | 0 | 0 | 0 | 0 | 0 | 0 | 0 | 0 | 0 |
| 0 | NA | NA | NA | NA | NA | Igkv12-40 | | 0 | 0 | 0 | 0 | 0 | 0 | 0 | 0 | 0 | 0 | 0 |
| 0 | NA | NA | NA | NA | NA | Gm43220 | | 0 | 0 | 0 | 0 | 0 | 0 | 0 | 0 | 0 | 0 | 0 |
| 0 | NA | NA | NA | NA | NA | Igkv5-37 | | 0 | 0 | 0 | 0 | 0 | 0 | 0 | 0 | 0 | 0 | 0 |
| 0 | NA | NA | NA | NA | NA | Igkv18-36 | | 0 | 0 | 0 | 0 | 0 | 0 | 0 | 0 | 0 | 0 | 0 |
| 0 | NA | NA | NA | NA | NA | Igkv1-35 | | 0 | 0 | 0 | 0 | 0 | 0 | 0 | 0 | 0 | 0 | 0 |
| 0 | NA | NA | NA | NA | NA | Gm23694 | | 0 | 0 | 0 | 0 | 0 | 0 | 0 | 0 | 0 | 0 | 0 |
| 0 | NA | NA | NA | NA | NA | Gm43586 | | 0 | 0 | 0 | 0 | 0 | 0 | 0 | 0 | 0 | 0 | 0 |
| 0 | NA | NA | NA | NA | NA | Igkv8-34 | | 0 | 0 | 0 | 0 | 0 | 0 | 0 | 0 | 0 | 0 | 0 |
| 0 | NA | NA | NA | NA | NA | Igkv7-33 | | 0 | 0 | 0 | 0 | 0 | 0 | 0 | 0 | 0 | 0 | 0 |
| 0 | NA | NA | NA | NA | NA | Igkv8-31 | | 0 | 0 | 0 | 0 | 0 | 0 | 0 | 0 | 0 | 0 | 0 |
| 0 | NA | NA | NA | NA | NA | Igkv6-29 | | 0 | 0 | 0 | 0 | 0 | 0 | 0 | 0 | 0 | 0 | 0 |
| 0 | NA | NA | NA | NA | NA | Gm24096 | | 0 | 0 | 0 | 0 | 0 | 0 | 0 | 0 | 0 | 0 | 0 |
| 0 | NA | NA | NA | NA | NA | Igkv8-26 | | 0 | 0 | 0 | 0 | 0 | 0 | 0 | 0 | 0 | 0 | 0 |
| 0 | NA | NA | NA | NA | NA | Gm8828 | | 0 | 0 | 0 | 0 | 0 | 0 | 0 | 0 | 0 | 0 | 0 |
| 0 | NA | NA | NA | NA | NA | Gm11065 | | 0 | 0 | 0 | 0 | 0 | 0 | 0 | 0 | 0 | 0 | 0 |
| 0 | NA | NA | NA | NA | NA | Igkv8-22 | | 0 | 0 | 0 | 0 | 0 | 0 | 0 | 0 | 0 | 0 | 0 |
| 0 | NA | NA | NA | NA | NA | Igkv8-21 | | 0 | 0 | 0 | 0 | 0 | 0 | 0 | 0 | 0 | 0 | 0 |
| 0 | NA | NA | NA | NA | NA | Amd-ps5 | | 0 | 0 | 0 | 0 | 0 | 0 | 0 | 0 | 0 | 0 | 0 |
| 0 | NA | NA | NA | NA | NA | Igkv8-18 | | 0 | 0 | 0 | 0 | 0 | 0 | 0 | 0 | 0 | 0 | 0 |
| 0 | NA | NA | NA | NA | NA | Gm42667 | | 0 | 0 | 0 | 0 | 0 | 0 | 0 | 0 | 0 | 0 | 0 |
| 0 | NA | NA | NA | NA | NA | Gm8853 | | 0 | 0 | 0 | 0 | 0 | 0 | 0 | 0 | 0 | 0 | 0 |
| 0 | NA | NA | NA | NA | NA | Gm10360 | | 0 | 0 | 0 | 0 | 0 | 0 | 0 | 0 | 0 | 0 | 0 |
| 0 | NA | NA | NA | NA | NA | Igkv6-14 | | 0 | 0 | 0 | 0 | 0 | 0 | 0 | 0 | 0 | 0 | 0 |
| 0 | NA | NA | NA | NA | NA | Gm8872 | | 0 | 0 | 0 | 0 | 0 | 0 | 0 | 0 | 0 | 0 | 0 |
| 0 | NA | NA | NA | NA | NA | Gm42720 | | 0 | 0 | 0 | 0 | 0 | 0 | 0 | 0 | 0 | 0 | 0 |
| 0 | NA | NA | NA | NA | NA | Gm42773 | | 0 | 0 | 0 | 0 | 0 | 0 | 0 | 0 | 0 | 0 | 0 |
| 0 | NA | NA | NA | NA | NA | Igkv3-12-1 | | 0 | 0 | 0 | 0 | 0 | 0 | 0 | 0 | 0 | 0 | 0 |
| 0 | NA | NA | NA | NA | NA | Igkv3-11 | | 0 | 0 | 0 | 0 | 0 | 0 | 0 | 0 | 0 | 0 | 0 |
| 0 | NA | NA | NA | NA | NA | Igkv3-8 | | 0 | 0 | 0 | 0 | 0 | 0 | 0 | 0 | 0 | 0 | 0 |
| 0 | NA | NA | NA | NA | NA | Igkv3-6 | | 0 | 0 | 0 | 0 | 0 | 0 | 0 | 0 | 0 | 0 | 0 |
| 0 | NA | NA | NA | NA | NA | Tex37 | testis expressed 37 | 0 | 0 | 0 | 0 | 0 | 0 | 0 | 0 | 0 | 0 | 0 |
| 0 | NA | NA | NA | NA | NA | Foxi3 | forkhead box I3 | 0 | 0 | 0 | 0 | 0 | 0 | 0 | 0 | 0 | 0 | 0 |
| 0 | NA | NA | NA | NA | NA | Gm23485 | | 0 | 0 | 0 | 0 | 0 | 0 | 0 | 0 | 0 | 0 | 0 |
| 0 | NA | NA | NA | NA | NA | Gm43906 | | 0 | 0 | 0 | 0 | 0 | 0 | 0 | 0 | 0 | 0 | 0 |
| 0 | NA | NA | NA | NA | NA | Mir8112 | | 0 | 0 | 0 | 0 | 0 | 0 | 0 | 0 | 0 | 0 | 0 |
| 0 | NA | NA | NA | NA | NA | Gm44110 | | 0 | 0 | 0 | 0 | 0 | 0 | 0 | 0 | 0 | 0 | 0 |
| 0 | NA | NA | NA | NA | NA | Gm44111 | | 0 | 0 | 0 | 0 | 0 | 0 | 0 | 0 | 0 | 0 | 0 |

| | | | | | | | | | | | | | | | | | | |
|---|---|---|---|---|---|---|---|---|---|---|---|---|---|---|---|---|---|---|
| 0 | NA | NA | NA | NA | NA | Gm44174 | | 0 | 0 | 0 | 0 | 0 | 0 | 0 | 0 | 0 | 0 | 0 |
| 0 | NA | NA | NA | NA | NA | Gm44172 | | 0 | 0 | 0 | 0 | 0 | 0 | 0 | 0 | 0 | 0 | 0 |
| 0 | NA | NA | NA | NA | NA | Gm44186 | | 0 | 0 | 0 | 0 | 0 | 0 | 0 | 0 | 0 | 0 | 0 |
| 0 | NA | NA | NA | NA | NA | Gm44194 | | 0 | 0 | 0 | 0 | 0 | 0 | 0 | 0 | 0 | 0 | 0 |
| 0 | NA | NA | NA | NA | NA | Gm44769 | | 0 | 0 | 0 | 0 | 0 | 0 | 0 | 0 | 0 | 0 | 0 |
| 0 | NA | NA | NA | NA | NA | Gm19078 | | 0 | 0 | 0 | 0 | 0 | 0 | 0 | 0 | 0 | 0 | 0 |
| 0 | NA | NA | NA | NA | NA | Gm26628 | | 0 | 0 | 0 | 0 | 0 | 0 | 0 | 0 | 0 | 0 | 0 |
| 0 | NA | NA | NA | NA | NA | 9130221F21Rik | | 0 | 0 | 0 | 0 | 0 | 0 | 0 | 0 | 0 | 0 | 0 |
| 0 | NA | NA | NA | NA | NA | Gm8952 | | 0 | 0 | 0 | 0 | 0 | 0 | 0 | 0 | 0 | 0 | 0 |
| 0 | NA | NA | NA | NA | NA | Gm44604 | | 0 | 0 | 0 | 0 | 0 | 0 | 0 | 0 | 0 | 0 | 0 |
| 0 | NA | NA | NA | NA | NA | 4933431G14Rik | | 0 | 0 | 0 | 0 | 0 | 0 | 0 | 0 | 0 | 0 | 0 |
| 0 | NA | NA | NA | NA | NA | Sftpb | surfactant associated protein B | 0 | 0 | 0 | 0 | 0 | 0 | 0 | 0 | 0 | 0 | 0 |
| 0 | NA | NA | NA | NA | NA | Gm38320 | | 0 | 0 | 0 | 0 | 0 | 0 | 0 | 0 | 0 | 0 | 0 |
| 0 | NA | NA | NA | NA | NA | Gm15402 | | 0 | 0 | 0 | 0 | 0 | 0 | 0 | 0 | 0 | 0 | 0 |
| 0 | NA | NA | NA | NA | NA | Gm37736 | | 0 | 0 | 0 | 0 | 0 | 0 | 0 | 0 | 0 | 0 | 0 |
| 0 | NA | NA | NA | NA | NA | Gm18402 | | 0 | 0 | 0 | 0 | 0 | 0 | 0 | 0 | 0 | 0 | 0 |
| 0 | NA | NA | NA | NA | NA | Gm20560 | | 0 | 0 | 0 | 0 | 0 | 0 | 0 | 0 | 0 | 0 | 0 |
| 0 | NA | NA | NA | NA | NA | Gm24533 | | 0 | 0 | 0 | 0 | 0 | 0 | 0 | 0 | 0 | 0 | 0 |
| 0 | NA | NA | NA | NA | NA | Gm44475 | | 0 | 0 | 0 | 0 | 0 | 0 | 0 | 0 | 0 | 0 | 0 |
| 0 | NA | NA | NA | NA | NA | Gm24370 | | 0 | 0 | 0 | 0 | 0 | 0 | 0 | 0 | 0 | 0 | 0 |
| 0 | NA | NA | NA | NA | NA | Olfr210-ps1 | | 0 | 0 | 0 | 0 | 0 | 0 | 0 | 0 | 0 | 0 | 0 |
| 0 | NA | NA | NA | NA | NA | 1700065L07Rik | | 0 | 0 | 0 | 0 | 0 | 0 | 0 | 0 | 0 | 0 | 0 |
| 0 | NA | NA | NA | NA | NA | 4931417E11Ril | RIKEN cDNA 4931417E11 gene | 0 | 0 | 0 | 0 | 0 | 0 | 0 | 0 | 0 | 0 | 0 |
| 0 | NA | NA | NA | NA | NA | Gm44225 | | 0 | 0 | 0 | 0 | 0 | 0 | 0 | 0 | 0 | 0 | 0 |
| 0 | NA | NA | NA | NA | NA | Gm26179 | | 0 | 0 | 0 | 0 | 0 | 0 | 0 | 0 | 0 | 0 | 0 |
| 0 | NA | NA | NA | NA | NA | Gm23924 | | 0 | 0 | 0 | 0 | 0 | 0 | 0 | 0 | 0 | 0 | 0 |
| 0 | NA | NA | NA | NA | NA | Gm22254 | | 0 | 0 | 0 | 0 | 0 | 0 | 0 | 0 | 0 | 0 | 0 |
| 0 | NA | NA | NA | NA | NA | Gm8992 | | 0 | 0 | 0 | 0 | 0 | 0 | 0 | 0 | 0 | 0 | 0 |
| 0 | NA | NA | NA | NA | NA | Gm8993 | | 0 | 0 | 0 | 0 | 0 | 0 | 0 | 0 | 0 | 0 | 0 |
| 0 | NA | NA | NA | NA | NA | Gm8999 | | 0 | 0 | 0 | 0 | 0 | 0 | 0 | 0 | 0 | 0 | 0 |
| 0 | NA | NA | NA | NA | NA | 4933439N06Rik | | 0 | 0 | 0 | 0 | 0 | 0 | 0 | 0 | 0 | 0 | 0 |
| 0 | NA | NA | NA | NA | NA | Gm44113 | | 0 | 0 | 0 | 0 | 0 | 0 | 0 | 0 | 0 | 0 | 0 |
| 0 | NA | NA | NA | NA | NA | Gm5995 | | 0 | 0 | 0 | 0 | 0 | 0 | 0 | 0 | 0 | 0 | 0 |
| 0 | NA | NA | NA | NA | NA | Gm44334 | | 0 | 0 | 0 | 0 | 0 | 0 | 0 | 0 | 0 | 0 | 0 |
| 0 | NA | NA | NA | NA | NA | Gm22283 | | 0 | 0 | 0 | 0 | 0 | 0 | 0 | 0 | 0 | 0 | 0 |
| 0 | NA | NA | NA | NA | NA | Gm44166 | | 0 | 0 | 0 | 0 | 0 | 0 | 0 | 0 | 0 | 0 | 0 |
| 0 | NA | NA | NA | NA | NA | Gm44435 | | 0 | 0 | 0 | 0 | 0 | 0 | 0 | 0 | 0 | 0 | 0 |
| 0 | NA | NA | NA | NA | NA | Gm44235 | | 0 | 0 | 0 | 0 | 0 | 0 | 0 | 0 | 0 | 0 | 0 |
| 0 | NA | NA | NA | NA | NA | Gm44286 | | 0 | 0 | 0 | 0 | 0 | 0 | 0 | 0 | 0 | 0 | 0 |
| 0 | NA | NA | NA | NA | NA | Gm44255 | | 0 | 0 | 0 | 0 | 0 | 0 | 0 | 0 | 0 | 0 | 0 |
| 0 | NA | NA | NA | NA | NA | Gm21045 | | 0 | 0 | 0 | 0 | 0 | 0 | 0 | 0 | 0 | 0 | 0 |
| 0 | NA | NA | NA | NA | NA | Gm21392 | | 0 | 0 | 0 | 0 | 0 | 0 | 0 | 0 | 0 | 0 | 0 |
| 0 | NA | NA | NA | NA | NA | Gm6072 | | 0 | 0 | 0 | 0 | 0 | 0 | 0 | 0 | 0 | 0 | 0 |
| 0 | NA | NA | NA | NA | NA | Gm23727 | | 0 | 0 | 0 | 0 | 0 | 0 | 0 | 0 | 0 | 0 | 0 |
| 0 | NA | NA | NA | NA | NA | Gm29005 | | 0 | 0 | 0 | 0 | 0 | 0 | 0 | 0 | 0 | 0 | 0 |
| 0 | NA | NA | NA | NA | NA | Gm20362 | | 0 | 0 | 0 | 0 | 0 | 0 | 0 | 0 | 0 | 0 | 0 |
| 0 | NA | NA | NA | NA | NA | Gm6261 | | 0 | 0 | 0 | 0 | 0 | 0 | 0 | 0 | 0 | 0 | 0 |
| 0 | NA | NA | NA | NA | NA | Gm44271 | | 0 | 0 | 0 | 0 | 0 | 0 | 0 | 0 | 0 | 0 | 0 |
| 0 | NA | NA | NA | NA | NA | Gm20594 | predicted gene, 20594 | 0 | 0 | 0 | 0 | 0 | 0 | 0 | 0 | 0 | 0 | 0 |
| 0 | NA | NA | NA | NA | NA | Gm4409 | | 0 | 0 | 0 | 0 | 0 | 0 | 0 | 0 | 0 | 0 | 0 |
| 0 | NA | NA | NA | NA | NA | Gm44240 | | 0 | 0 | 0 | 0 | 0 | 0 | 0 | 0 | 0 | 0 | 0 |
| 0 | NA | NA | NA | NA | NA | Gm44228 | | 0 | 0 | 0 | 0 | 0 | 0 | 0 | 0 | 0 | 0 | 0 |
| 0 | NA | NA | NA | NA | NA | Gm44230 | | 0 | 0 | 0 | 0 | 0 | 0 | 0 | 0 | 0 | 0 | 0 |
| 0 | NA | NA | NA | NA | NA | Gm20371 | | 0 | 0 | 0 | 0 | 0 | 0 | 0 | 0 | 0 | 0 | 0 |
| 0 | NA | NA | NA | NA | NA | Gm44241 | | 0 | 0 | 0 | 0 | 0 | 0 | 0 | 0 | 0 | 0 | 0 |
| 0 | NA | NA | NA | NA | NA | Gm44229 | | 0 | 0 | 0 | 0 | 0 | 0 | 0 | 0 | 0 | 0 | 0 |
| 0 | NA | NA | NA | NA | NA | Gm44242 | | 0 | 0 | 0 | 0 | 0 | 0 | 0 | 0 | 0 | 0 | 0 |
| 0 | NA | NA | NA | NA | NA | Gm44231 | | 0 | 0 | 0 | 0 | 0 | 0 | 0 | 0 | 0 | 0 | 0 |
| 0 | NA | NA | NA | NA | NA | Gm43900 | | 0 | 0 | 0 | 0 | 0 | 0 | 0 | 0 | 0 | 0 | 0 |
| 0 | NA | NA | NA | NA | NA | Gm43901 | | 0 | 0 | 0 | 0 | 0 | 0 | 0 | 0 | 0 | 0 | 0 |

| | | | | | | | | | | | | | | | | | | |
|---|---|---|---|---|---|---|---|---|---|---|---|---|---|---|---|---|---|---|
| 0 | NA | NA | NA | NA | NA | Gm26264 | | 0 | 0 | 0 | 0 | 0 | 0 | 0 | 0 | 0 | 0 | 0 |
| 0 | NA | NA | NA | NA | NA | Gm38839 | | 0 | 0 | 0 | 0 | 0 | 0 | 0 | 0 | 0 | 0 | 0 |
| 0 | NA | NA | NA | NA | NA | Gm22530 | | 0 | 0 | 0 | 0 | 0 | 0 | 0 | 0 | 0 | 0 | 0 |
| 0 | NA | NA | NA | NA | NA | Gm4874 | | 0 | 0 | 0 | 0 | 0 | 0 | 0 | 0 | 0 | 0 | 0 |
| 0 | NA | NA | NA | NA | NA | Gm38840 | | 0 | 0 | 0 | 0 | 0 | 0 | 0 | 0 | 0 | 0 | 0 |
| 0 | NA | NA | NA | NA | NA | 1700097M23Rik | | 0 | 0 | 0 | 0 | 0 | 0 | 0 | 0 | 0 | 0 | 0 |
| 0 | NA | NA | NA | NA | NA | Mir7232 | | 0 | 0 | 0 | 0 | 0 | 0 | 0 | 0 | 0 | 0 | 0 |
| 0 | NA | NA | NA | NA | NA | Mir468 | | 0 | 0 | 0 | 0 | 0 | 0 | 0 | 0 | 0 | 0 | 0 |
| 0 | NA | NA | NA | NA | NA | Gm44202 | | 0 | 0 | 0 | 0 | 0 | 0 | 0 | 0 | 0 | 0 | 0 |
| 0 | NA | NA | NA | NA | NA | Gm15864 | | 0 | 0 | 0 | 0 | 0 | 0 | 0 | 0 | 0 | 0 | 0 |
| 0 | NA | NA | NA | NA | NA | Gm31861 | | 0 | 0 | 0 | 0 | 0 | 0 | 0 | 0 | 0 | 0 | 0 |
| 0 | NA | NA | NA | NA | NA | Gm44209 | | 0 | 0 | 0 | 0 | 0 | 0 | 0 | 0 | 0 | 0 | 0 |
| 0 | NA | NA | NA | NA | NA | Gm44145 | | 0 | 0 | 0 | 0 | 0 | 0 | 0 | 0 | 0 | 0 | 0 |
| 0 | NA | NA | NA | NA | NA | Gm17034 | | 0 | 0 | 0 | 0 | 0 | 0 | 0 | 0 | 0 | 0 | 0 |
| 0 | NA | NA | NA | NA | NA | Gm32591 | | 0 | 0 | 0 | 0 | 0 | 0 | 0 | 0 | 0 | 0 | 0 |
| 0 | NA | NA | NA | NA | NA | 2310069B03Rik | | 0 | 0 | 0 | 0 | 0 | 0 | 0 | 0 | 0 | 0 | 0 |
| 0 | NA | NA | NA | NA | NA | Gm6312 | | 0 | 0 | 0 | 0 | 0 | 0 | 0 | 0 | 0 | 0 | 0 |
| 0 | NA | NA | NA | NA | NA | Gm45145 | | 0 | 0 | 0 | 0 | 0 | 0 | 0 | 0 | 0 | 0 | 0 |
| 0 | NA | NA | NA | NA | NA | Gm43985 | | 0 | 0 | 0 | 0 | 0 | 0 | 0 | 0 | 0 | 0 | 0 |
| 0 | NA | NA | NA | NA | NA | Gm43981 | | 0 | 0 | 0 | 0 | 0 | 0 | 0 | 0 | 0 | 0 | 0 |
| 0 | NA | NA | NA | NA | NA | Mir7040 | | 0 | 0 | 0 | 0 | 0 | 0 | 0 | 0 | 0 | 0 | 0 |
| 0 | NA | NA | NA | NA | NA | Mir3470a | | 0 | 0 | 0 | 0 | 0 | 0 | 0 | 0 | 0 | 0 | 0 |
| 0 | NA | NA | NA | NA | NA | Mir6374 | | 0 | 0 | 0 | 0 | 0 | 0 | 0 | 0 | 0 | 0 | 0 |
| 0 | NA | NA | NA | NA | NA | Gm24990 | | 0 | 0 | 0 | 0 | 0 | 0 | 0 | 0 | 0 | 0 | 0 |
| 0 | NA | NA | NA | NA | NA | Clec4f | C-type lectin domain family 4, member f | 0 | 0 | 0 | 0 | 0 | 0 | 0 | 0 | 0 | 0 | 0 |
| 0 | NA | NA | NA | NA | NA | Gm27772 | | 0 | 0 | 0 | 0 | 0 | 0 | 0 | 0 | 0 | 0 | 0 |
| 0 | NA | NA | NA | NA | NA | Gm27631 | | 0 | 0 | 0 | 0 | 0 | 0 | 0 | 0 | 0 | 0 | 0 |
| 0 | NA | NA | NA | NA | NA | Gm27893 | | 0 | 0 | 0 | 0 | 0 | 0 | 0 | 0 | 0 | 0 | 0 |
| 0 | NA | NA | NA | NA | NA | Vax2 | ventral anterior homeobox 2 | 0 | 0 | 0 | 0 | 0 | 0 | 0 | 0 | 0 | 0 | 0 |
| 0 | NA | NA | NA | NA | NA | Gm25054 | | 0 | 0 | 0 | 0 | 0 | 0 | 0 | 0 | 0 | 0 | 0 |
| 0 | NA | NA | NA | NA | NA | Atp6v1b1 | ATPase, H+ transporting, lysosomal V1 su | 0 | 0 | 0 | 0 | 0 | 0 | 0 | 0 | 0 | 0 | 0 |
| 0 | NA | NA | NA | NA | NA | Ankrd53 | ankyrin repeat domain 53 | 0 | 0 | 0 | 0 | 0 | 0 | 0 | 0 | 0 | 0 | 0 |
| 0 | NA | NA | NA | NA | NA | Gm23014 | | 0 | 0 | 0 | 0 | 0 | 0 | 0 | 0 | 0 | 0 | 0 |
| 0 | NA | NA | NA | NA | NA | Gm6342 | | 0 | 0 | 0 | 0 | 0 | 0 | 0 | 0 | 0 | 0 | 0 |
| 0 | NA | NA | NA | NA | NA | Gm44750 | | 0 | 0 | 0 | 0 | 0 | 0 | 0 | 0 | 0 | 0 | 0 |
| 0 | NA | NA | NA | NA | NA | Gm43919 | | 0 | 0 | 0 | 0 | 0 | 0 | 0 | 0 | 0 | 0 | 0 |
| 0 | NA | NA | NA | NA | NA | Gm33024 | | 0 | 0 | 0 | 0 | 0 | 0 | 0 | 0 | 0 | 0 | 0 |
| 0 | NA | NA | NA | NA | NA | Gm15475 | | 0 | 0 | 0 | 0 | 0 | 0 | 0 | 0 | 0 | 0 | 0 |
| 0 | NA | NA | NA | NA | NA | Gm44127 | | 0 | 0 | 0 | 0 | 0 | 0 | 0 | 0 | 0 | 0 | 0 |
| 0 | NA | NA | NA | NA | NA | Gm10445 | | 0 | 0 | 0 | 0 | 0 | 0 | 0 | 0 | 0 | 0 | 0 |
| 0 | NA | NA | NA | NA | NA | 4930504D19Rik | | 0 | 0 | 0 | 0 | 0 | 0 | 0 | 0 | 0 | 0 | 0 |
| 0 | NA | NA | NA | NA | NA | Gm44012 | | 0 | 0 | 0 | 0 | 0 | 0 | 0 | 0 | 0 | 0 | 0 |
| 0 | NA | NA | NA | NA | NA | Gm44011 | | 0 | 0 | 0 | 0 | 0 | 0 | 0 | 0 | 0 | 0 | 0 |
| 0 | NA | NA | NA | NA | NA | Gm26353 | | 0 | 0 | 0 | 0 | 0 | 0 | 0 | 0 | 0 | 0 | 0 |
| 0 | NA | NA | NA | NA | NA | Gm43954 | | 0 | 0 | 0 | 0 | 0 | 0 | 0 | 0 | 0 | 0 | 0 |
| 0 | NA | NA | NA | NA | NA | Gm43955 | | 0 | 0 | 0 | 0 | 0 | 0 | 0 | 0 | 0 | 0 | 0 |
| 0 | NA | NA | NA | NA | NA | Gm43957 | | 0 | 0 | 0 | 0 | 0 | 0 | 0 | 0 | 0 | 0 | 0 |
| 0 | NA | NA | NA | NA | NA | Gm5878 | | 0 | 0 | 0 | 0 | 0 | 0 | 0 | 0 | 0 | 0 | 0 |
| 0 | NA | NA | NA | NA | NA | Gm43959 | | 0 | 0 | 0 | 0 | 0 | 0 | 0 | 0 | 0 | 0 | 0 |
| 0 | NA | NA | NA | NA | NA | Gm44285 | | 0 | 0 | 0 | 0 | 0 | 0 | 0 | 0 | 0 | 0 | 0 |
| 0 | NA | NA | NA | NA | NA | Gm22893 | | 0 | 0 | 0 | 0 | 0 | 0 | 0 | 0 | 0 | 0 | 0 |
| 0 | NA | NA | NA | NA | NA | Mir705 | | 0 | 0 | 0 | 0 | 0 | 0 | 0 | 0 | 0 | 0 | 0 |
| 0 | NA | NA | NA | NA | NA | Noto | notochord homeobox | 0 | 0 | 0 | 0 | 0 | 0 | 0 | 0 | 0 | 0 | 0 |
| 0 | NA | NA | NA | NA | NA | Gm44023 | | 0 | 0 | 0 | 0 | 0 | 0 | 0 | 0 | 0 | 0 | 0 |
| 0 | NA | NA | NA | NA | NA | Gm27317 | | 0 | 0 | 0 | 0 | 0 | 0 | 0 | 0 | 0 | 0 | 0 |
| 0 | NA | NA | NA | NA | NA | Alms1-ps2 | | 0 | 0 | 0 | 0 | 0 | 0 | 0 | 0 | 0 | 0 | 0 |
| 0 | NA | NA | NA | NA | NA | Mir6375 | | 0 | 0 | 0 | 0 | 0 | 0 | 0 | 0 | 0 | 0 | 0 |
| 0 | NA | NA | NA | NA | NA | Gm43309 | | 0 | 0 | 0 | 0 | 0 | 0 | 0 | 0 | 0 | 0 | 0 |
| 0 | NA | NA | NA | NA | NA | Gm18290 | | 0 | 0 | 0 | 0 | 0 | 0 | 0 | 0 | 0 | 0 | 0 |
| 0 | NA | NA | NA | NA | NA | Nat8b-ps | | 0 | 0 | 0 | 0 | 0 | 0 | 0 | 0 | 0 | 0 | 0 |

| | | | | | | | | | | | | | | | | | |
|---|---|---|---|---|---|---|---|---|---|---|---|---|---|---|---|---|---|
| 0 | NA | NA | NA | NA | NA | Figla | folliculogenesis specific basic helix-loop-l | 0 | 0 | 0 | 0 | 0 | 0 | 0 | 0 | 0 | 0 | 0 |
| 0 | NA | NA | NA | NA | NA | Gm19596 | | 0 | 0 | 0 | 0 | 0 | 0 | 0 | 0 | 0 | 0 | 0 |
| 0 | NA | NA | NA | NA | NA | Gm43917 | | 0 | 0 | 0 | 0 | 0 | 0 | 0 | 0 | 0 | 0 | 0 |
| 0 | NA | NA | NA | NA | NA | Gm18862 | | 0 | 0 | 0 | 0 | 0 | 0 | 0 | 0 | 0 | 0 | 0 |
| 0 | NA | NA | NA | NA | NA | Gm44941 | | 0 | 0 | 0 | 0 | 0 | 0 | 0 | 0 | 0 | 0 | 0 |
| 0 | NA | NA | NA | NA | NA | Gm44790 | | 0 | 0 | 0 | 0 | 0 | 0 | 0 | 0 | 0 | 0 | 0 |
| 0 | NA | NA | NA | NA | NA | Gm28720 | | 0 | 0 | 0 | 0 | 0 | 0 | 0 | 0 | 0 | 0 | 0 |
| 0 | NA | NA | NA | NA | NA | Gm22184 | | 0 | 0 | 0 | 0 | 0 | 0 | 0 | 0 | 0 | 0 | 0 |
| 0 | NA | NA | NA | NA | NA | Gm44152 | | 0 | 0 | 0 | 0 | 0 | 0 | 0 | 0 | 0 | 0 | 0 |
| 0 | NA | NA | NA | NA | NA | Gm44415 | | 0 | 0 | 0 | 0 | 0 | 0 | 0 | 0 | 0 | 0 | 0 |
| 0 | NA | NA | NA | NA | NA | Gm44416 | | 0 | 0 | 0 | 0 | 0 | 0 | 0 | 0 | 0 | 0 | 0 |
| 0 | NA | NA | NA | NA | NA | Gm44417 | | 0 | 0 | 0 | 0 | 0 | 0 | 0 | 0 | 0 | 0 | 0 |
| 0 | NA | NA | NA | NA | NA | 9530013L04Rik | | 0 | 0 | 0 | 0 | 0 | 0 | 0 | 0 | 0 | 0 | 0 |
| 0 | NA | NA | NA | NA | NA | Gm44414 | | 0 | 0 | 0 | 0 | 0 | 0 | 0 | 0 | 0 | 0 | 0 |
| 0 | NA | NA | NA | NA | NA | Gkn1 | gastrokine 1 | 0 | 0 | 0 | 0 | 0 | 0 | 0 | 0 | 0 | 0 | 0 |
| 0 | NA | NA | NA | NA | NA | Gkn2 | gastrokine 2 | 0 | 0 | 0 | 0 | 0 | 0 | 0 | 0 | 0 | 0 | 0 |
| 0 | NA | NA | NA | NA | NA | Bmp10 | bone morphogenetic protein 10 | 0 | 0 | 0 | 0 | 0 | 0 | 0 | 0 | 0 | 0 | 0 |
| 0 | NA | NA | NA | NA | NA | Gm44198 | | 0 | 0 | 0 | 0 | 0 | 0 | 0 | 0 | 0 | 0 | 0 |
| 0 | NA | NA | NA | NA | NA | Gm34312 | | 0 | 0 | 0 | 0 | 0 | 0 | 0 | 0 | 0 | 0 | 0 |
| 0 | NA | NA | NA | NA | NA | Gm44097 | | 0 | 0 | 0 | 0 | 0 | 0 | 0 | 0 | 0 | 0 | 0 |
| 0 | NA | NA | NA | NA | NA | Gm43904 | | 0 | 0 | 0 | 0 | 0 | 0 | 0 | 0 | 0 | 0 | 0 |
| 0 | NA | NA | NA | NA | NA | Gm47286 | | 0 | 0 | 0 | 0 | 0 | 0 | 0 | 0 | 0 | 0 | 0 |
| 0 | NA | NA | NA | NA | NA | 4933412L11Rik | | 0 | 0 | 0 | 0 | 0 | 0 | 0 | 0 | 0 | 0 | 0 |
| 0 | NA | NA | NA | NA | NA | Mir6376 | | 0 | 0 | 0 | 0 | 0 | 0 | 0 | 0 | 0 | 0 | 0 |
| 0 | NA | NA | NA | NA | NA | Gm44388 | | 0 | 0 | 0 | 0 | 0 | 0 | 0 | 0 | 0 | 0 | 0 |
| 0 | NA | NA | NA | NA | NA | Dnajb8 | DnaJ heat shock protein family (Hsp40) m | 0 | 0 | 0 | 0 | 0 | 0 | 0 | 0 | 0 | 0 | 0 |
| 0 | NA | NA | NA | NA | NA | 1700031F10Rik | | 0 | 0 | 0 | 0 | 0 | 0 | 0 | 0 | 0 | 0 | 0 |
| 0 | NA | NA | NA | NA | NA | Gm44265 | | 0 | 0 | 0 | 0 | 0 | 0 | 0 | 0 | 0 | 0 | 0 |
| 0 | NA | NA | NA | NA | NA | Gm25178 | | 0 | 0 | 0 | 0 | 0 | 0 | 0 | 0 | 0 | 0 | 0 |
| 0 | NA | NA | NA | NA | NA | Gm44264 | | 0 | 0 | 0 | 0 | 0 | 0 | 0 | 0 | 0 | 0 | 0 |
| 0 | NA | NA | NA | NA | NA | Gm43999 | | 0 | 0 | 0 | 0 | 0 | 0 | 0 | 0 | 0 | 0 | 0 |
| 0 | NA | NA | NA | NA | NA | Gm43937 | | 0 | 0 | 0 | 0 | 0 | 0 | 0 | 0 | 0 | 0 | 0 |
| 0 | NA | NA | NA | NA | NA | Gm44001 | | 0 | 0 | 0 | 0 | 0 | 0 | 0 | 0 | 0 | 0 | 0 |
| 0 | NA | NA | NA | NA | NA | Gm44178 | | 0 | 0 | 0 | 0 | 0 | 0 | 0 | 0 | 0 | 0 | 0 |
| 0 | NA | NA | NA | NA | NA | 4933427D06Rik | | 0 | 0 | 0 | 0 | 0 | 0 | 0 | 0 | 0 | 0 | 0 |
| 0 | NA | NA | NA | NA | NA | Gm6507 | | 0 | 0 | 0 | 0 | 0 | 0 | 0 | 0 | 0 | 0 | 0 |
| 0 | NA | NA | NA | NA | NA | Gm44203 | | 0 | 0 | 0 | 0 | 0 | 0 | 0 | 0 | 0 | 0 | 0 |
| 0 | NA | NA | NA | NA | NA | Gm839 | | 0 | 0 | 0 | 0 | 0 | 0 | 0 | 0 | 0 | 0 | 0 |
| 0 | NA | NA | NA | NA | NA | Gm25961 | | 0 | 0 | 0 | 0 | 0 | 0 | 0 | 0 | 0 | 0 | 0 |
| 0 | NA | NA | NA | NA | NA | Gm44204 | | 0 | 0 | 0 | 0 | 0 | 0 | 0 | 0 | 0 | 0 | 0 |
| 0 | NA | NA | NA | NA | NA | 4930512J16Rik | | 0 | 0 | 0 | 0 | 0 | 0 | 0 | 0 | 0 | 0 | 0 |
| 0 | NA | NA | NA | NA | NA | Gm26811 | | 0 | 0 | 0 | 0 | 0 | 0 | 0 | 0 | 0 | 0 | 0 |
| 0 | NA | NA | NA | NA | NA | Gm44207 | | 0 | 0 | 0 | 0 | 0 | 0 | 0 | 0 | 0 | 0 | 0 |
| 0 | NA | NA | NA | NA | NA | Gm43903 | | 0 | 0 | 0 | 0 | 0 | 0 | 0 | 0 | 0 | 0 | 0 |
| 0 | NA | NA | NA | NA | NA | Vmn1r-ps25 | | 0 | 0 | 0 | 0 | 0 | 0 | 0 | 0 | 0 | 0 | 0 |
| 0 | NA | NA | NA | NA | NA | Vmn1r40 | vomeronasal 1 receptor 40 | 0 | 0 | 0 | 0 | 0 | 0 | 0 | 0 | 0 | 0 | 0 |
| 0 | NA | NA | NA | NA | NA | Vmn1r41 | vomeronasal 1 receptor 41 | 0 | 0 | 0 | 0 | 0 | 0 | 0 | 0 | 0 | 0 | 0 |
| 0 | NA | NA | NA | NA | NA | Vmn1r-ps26 | | 0 | 0 | 0 | 0 | 0 | 0 | 0 | 0 | 0 | 0 | 0 |
| 0 | NA | NA | NA | NA | NA | Vmn1r-ps27 | | 0 | 0 | 0 | 0 | 0 | 0 | 0 | 0 | 0 | 0 | 0 |
| 0 | NA | NA | NA | NA | NA | Vmn1r42 | vomeronasal 1 receptor 42 | 0 | 0 | 0 | 0 | 0 | 0 | 0 | 0 | 0 | 0 | 0 |
| 0 | NA | NA | NA | NA | NA | Vmn1r43 | vomeronasal 1 receptor 43 | 0 | 0 | 0 | 0 | 0 | 0 | 0 | 0 | 0 | 0 | 0 |
| 0 | NA | NA | NA | NA | NA | Vmn1r44 | vomeronasal 1 receptor 44 | 0 | 0 | 0 | 0 | 0 | 0 | 0 | 0 | 0 | 0 | 0 |
| 0 | NA | NA | NA | NA | NA | Vmn1r-ps28 | | 0 | 0 | 0 | 0 | 0 | 0 | 0 | 0 | 0 | 0 | 0 |
| 0 | NA | NA | NA | NA | NA | Vmn1r45 | vomeronasal 1 receptor 45 | 0 | 0 | 0 | 0 | 0 | 0 | 0 | 0 | 0 | 0 | 0 |
| 0 | NA | NA | NA | NA | NA | Vmn1r46 | vomeronasal 1 receptor 46 | 0 | 0 | 0 | 0 | 0 | 0 | 0 | 0 | 0 | 0 | 0 |
| 0 | NA | NA | NA | NA | NA | Vmn1r-ps29 | | 0 | 0 | 0 | 0 | 0 | 0 | 0 | 0 | 0 | 0 | 0 |
| 0 | NA | NA | NA | NA | NA | Vmn1r47 | vomeronasal 1 receptor 47 | 0 | 0 | 0 | 0 | 0 | 0 | 0 | 0 | 0 | 0 | 0 |
| 0 | NA | NA | NA | NA | NA | Vmn1r48 | vomeronasal 1 receptor 48 | 0 | 0 | 0 | 0 | 0 | 0 | 0 | 0 | 0 | 0 | 0 |
| 0 | NA | NA | NA | NA | NA | Vmn1r-ps31 | | 0 | 0 | 0 | 0 | 0 | 0 | 0 | 0 | 0 | 0 | 0 |
| 0 | NA | NA | NA | NA | NA | Gm20534 | | 0 | 0 | 0 | 0 | 0 | 0 | 0 | 0 | 0 | 0 | 0 |

| | | | | | | | | | | | | | | | | | | |
|---|---|---|---|---|---|---|---|---|---|---|---|---|---|---|---|---|---|---|
| 0 | NA | NA | NA | NA | NA | Vmn1r49 | vomeronasal 1, receptor 49 | 0 | 0 | 0 | 0 | 0 | 0 | 0 | 0 | 0 | 0 | 0 |
| 0 | NA | NA | NA | NA | NA | Vmn1r50 | vomeronasal 1 receptor 50 | 0 | 0 | 0 | 0 | 0 | 0 | 0 | 0 | 0 | 0 | 0 |
| 0 | NA | NA | NA | NA | NA | Vmn1r51 | vomeronasal 1 receptor 51 | 0 | 0 | 0 | 0 | 0 | 0 | 0 | 0 | 0 | 0 | 0 |
| 0 | NA | NA | NA | NA | NA | Vmn1r-ps32 | | 0 | 0 | 0 | 0 | 0 | 0 | 0 | 0 | 0 | 0 | 0 |
| 0 | NA | NA | NA | NA | NA | Vmn1r52 | vomeronasal 1 receptor 52 | 0 | 0 | 0 | 0 | 0 | 0 | 0 | 0 | 0 | 0 | 0 |
| 0 | NA | NA | NA | NA | NA | V1ra8 | vomeronasal 1 receptor, A8 | 0 | 0 | 0 | 0 | 0 | 0 | 0 | 0 | 0 | 0 | 0 |
| 0 | NA | NA | NA | NA | NA | Vmn1r-ps34 | | 0 | 0 | 0 | 0 | 0 | 0 | 0 | 0 | 0 | 0 | 0 |
| 0 | NA | NA | NA | NA | NA | Vmn1r54 | vomeronasal 1 receptor 54 | 0 | 0 | 0 | 0 | 0 | 0 | 0 | 0 | 0 | 0 | 0 |
| 0 | NA | NA | NA | NA | NA | Gm43891 | | 0 | 0 | 0 | 0 | 0 | 0 | 0 | 0 | 0 | 0 | 0 |
| 0 | NA | NA | NA | NA | NA | BC048671 | cDNA sequence BC048671 | 0 | 0 | 0 | 0 | 0 | 0 | 0 | 0 | 0 | 0 | 0 |
| 0 | NA | NA | NA | NA | NA | Gm45901 | | 0 | 0 | 0 | 0 | 0 | 0 | 0 | 0 | 0 | 0 | 0 |
| 0 | NA | NA | NA | NA | NA | Gm44236 | | 0 | 0 | 0 | 0 | 0 | 0 | 0 | 0 | 0 | 0 | 0 |
| 0 | NA | NA | NA | NA | NA | Gm15756 | | 0 | 0 | 0 | 0 | 0 | 0 | 0 | 0 | 0 | 0 | 0 |
| 0 | NA | NA | NA | NA | NA | Gm24816 | | 0 | 0 | 0 | 0 | 0 | 0 | 0 | 0 | 0 | 0 | 0 |
| 0 | NA | NA | NA | NA | NA | Gm44123 | | 0 | 0 | 0 | 0 | 0 | 0 | 0 | 0 | 0 | 0 | 0 |
| 0 | NA | NA | NA | NA | NA | Gm25185 | | 0 | 0 | 0 | 0 | 0 | 0 | 0 | 0 | 0 | 0 | 0 |
| 0 | NA | NA | NA | NA | NA | Gm44278 | | 0 | 0 | 0 | 0 | 0 | 0 | 0 | 0 | 0 | 0 | 0 |
| 0 | NA | NA | NA | NA | NA | 4930402H05Rik | | 0 | 0 | 0 | 0 | 0 | 0 | 0 | 0 | 0 | 0 | 0 |
| 0 | NA | NA | NA | NA | NA | Gm30437 | | 0 | 0 | 0 | 0 | 0 | 0 | 0 | 0 | 0 | 0 | 0 |
| 0 | NA | NA | NA | NA | NA | Gm44629 | | 0 | 0 | 0 | 0 | 0 | 0 | 0 | 0 | 0 | 0 | 0 |
| 0 | NA | NA | NA | NA | NA | Gm45215 | | 0 | 0 | 0 | 0 | 0 | 0 | 0 | 0 | 0 | 0 | 0 |
| 0 | NA | NA | NA | NA | NA | Gm45217 | | 0 | 0 | 0 | 0 | 0 | 0 | 0 | 0 | 0 | 0 | 0 |
| 0 | NA | NA | NA | NA | NA | 4930517G19Rik | | 0 | 0 | 0 | 0 | 0 | 0 | 0 | 0 | 0 | 0 | 0 |
| 0 | NA | NA | NA | NA | NA | Gm44981 | | 0 | 0 | 0 | 0 | 0 | 0 | 0 | 0 | 0 | 0 | 0 |
| 0 | NA | NA | NA | NA | NA | Gm44711 | | 0 | 0 | 0 | 0 | 0 | 0 | 0 | 0 | 0 | 0 | 0 |
| 0 | NA | NA | NA | NA | NA | Gm44095 | | 0 | 0 | 0 | 0 | 0 | 0 | 0 | 0 | 0 | 0 | 0 |
| 0 | NA | NA | NA | NA | NA | 4930466I24Rik | | 0 | 0 | 0 | 0 | 0 | 0 | 0 | 0 | 0 | 0 | 0 |
| 0 | NA | NA | NA | NA | NA | Gm44803 | | 0 | 0 | 0 | 0 | 0 | 0 | 0 | 0 | 0 | 0 | 0 |
| 0 | NA | NA | NA | NA | NA | Gm44024 | | 0 | 0 | 0 | 0 | 0 | 0 | 0 | 0 | 0 | 0 | 0 |
| 0 | NA | NA | NA | NA | NA | Gm17797 | | 0 | 0 | 0 | 0 | 0 | 0 | 0 | 0 | 0 | 0 | 0 |
| 0 | NA | NA | NA | NA | NA | Gm44224 | | 0 | 0 | 0 | 0 | 0 | 0 | 0 | 0 | 0 | 0 | 0 |
| 0 | NA | NA | NA | NA | NA | Gm5313 | | 0 | 0 | 0 | 0 | 0 | 0 | 0 | 0 | 0 | 0 | 0 |
| 0 | NA | NA | NA | NA | NA | Gm25094 | | 0 | 0 | 0 | 0 | 0 | 0 | 0 | 0 | 0 | 0 | 0 |
| 0 | NA | NA | NA | NA | NA | Gm44181 | | 0 | 0 | 0 | 0 | 0 | 0 | 0 | 0 | 0 | 0 | 0 |
| 0 | NA | NA | NA | NA | NA | Gm22312 | | 0 | 0 | 0 | 0 | 0 | 0 | 0 | 0 | 0 | 0 | 0 |
| 0 | NA | NA | NA | NA | NA | Gm7812 | | 0 | 0 | 0 | 0 | 0 | 0 | 0 | 0 | 0 | 0 | 0 |
| 0 | NA | NA | NA | NA | NA | Gm23035 | | 0 | 0 | 0 | 0 | 0 | 0 | 0 | 0 | 0 | 0 | 0 |
| 0 | NA | NA | NA | NA | NA | Gm22840 | | 0 | 0 | 0 | 0 | 0 | 0 | 0 | 0 | 0 | 0 | 0 |
| 0 | NA | NA | NA | NA | NA | Gm44307 | | 0 | 0 | 0 | 0 | 0 | 0 | 0 | 0 | 0 | 0 | 0 |
| 0 | NA | NA | NA | NA | NA | Gm19908 | | 0 | 0 | 0 | 0 | 0 | 0 | 0 | 0 | 0 | 0 | 0 |
| 0 | NA | NA | NA | NA | NA | Gm7825 | | 0 | 0 | 0 | 0 | 0 | 0 | 0 | 0 | 0 | 0 | 0 |
| 0 | NA | NA | NA | NA | NA | Gm44315 | | 0 | 0 | 0 | 0 | 0 | 0 | 0 | 0 | 0 | 0 | 0 |
| 0 | NA | NA | NA | NA | NA | Mir7041 | | 0 | 0 | 0 | 0 | 0 | 0 | 0 | 0 | 0 | 0 | 0 |
| 0 | NA | NA | NA | NA | NA | Gm43997 | | 0 | 0 | 0 | 0 | 0 | 0 | 0 | 0 | 0 | 0 | 0 |
| 0 | NA | NA | NA | NA | NA | Gm23364 | | 0 | 0 | 0 | 0 | 0 | 0 | 0 | 0 | 0 | 0 | 0 |
| 0 | NA | NA | NA | NA | NA | Gm44100 | | 0 | 0 | 0 | 0 | 0 | 0 | 0 | 0 | 0 | 0 | 0 |
| 0 | NA | NA | NA | NA | NA | 4930511E03Rik | | 0 | 0 | 0 | 0 | 0 | 0 | 0 | 0 | 0 | 0 | 0 |
| 0 | NA | NA | NA | NA | NA | Gm26501 | | 0 | 0 | 0 | 0 | 0 | 0 | 0 | 0 | 0 | 0 | 0 |
| 0 | NA | NA | NA | NA | NA | Gm17822 | | 0 | 0 | 0 | 0 | 0 | 0 | 0 | 0 | 0 | 0 | 0 |
| 0 | NA | NA | NA | NA | NA | Gm5723 | | 0 | 0 | 0 | 0 | 0 | 0 | 0 | 0 | 0 | 0 | 0 |
| 0 | NA | NA | NA | NA | NA | AY512915 | | 0 | 0 | 0 | 0 | 0 | 0 | 0 | 0 | 0 | 0 | 0 |
| 0 | NA | NA | NA | NA | NA | Gm10234 | | 0 | 0 | 0 | 0 | 0 | 0 | 0 | 0 | 0 | 0 | 0 |
| 0 | NA | NA | NA | NA | NA | 1700123L14Rik | | 0 | 0 | 0 | 0 | 0 | 0 | 0 | 0 | 0 | 0 | 0 |
| 0 | NA | NA | NA | NA | NA | Gm22971 | | 0 | 0 | 0 | 0 | 0 | 0 | 0 | 0 | 0 | 0 | 0 |
| 0 | NA | NA | NA | NA | NA | Gm26011 | | 0 | 0 | 0 | 0 | 0 | 0 | 0 | 0 | 0 | 0 | 0 |
| 0 | NA | NA | NA | NA | NA | Gm19653 | | 0 | 0 | 0 | 0 | 0 | 0 | 0 | 0 | 0 | 0 | 0 |
| 0 | NA | NA | NA | NA | NA | Gm23510 | | 0 | 0 | 0 | 0 | 0 | 0 | 0 | 0 | 0 | 0 | 0 |
| 0 | NA | NA | NA | NA | NA | Gm24565 | | 0 | 0 | 0 | 0 | 0 | 0 | 0 | 0 | 0 | 0 | 0 |
| 0 | NA | NA | NA | NA | NA | Gm23703 | | 0 | 0 | 0 | 0 | 0 | 0 | 0 | 0 | 0 | 0 | 0 |
| 0 | NA | NA | NA | NA | NA | Gm32592 | | 0 | 0 | 0 | 0 | 0 | 0 | 0 | 0 | 0 | 0 | 0 |

| | | | | | | | | | | | | | | | | | |
|---|---|---|---|---|---|---|---|---|---|---|---|---|---|---|---|---|---|
| 0 | NA | NA | NA | NA | NA | Gm44196 | 0 | 0 | 0 | 0 | 0 | 0 | 0 | 0 | 0 | 0 | 0 |
| 0 | NA | NA | NA | NA | NA | 4930595L18Rik | 0 | 0 | 0 | 0 | 0 | 0 | 0 | 0 | 0 | 0 | 0 |
| 0 | NA | NA | NA | NA | NA | Gm32308 | 0 | 0 | 0 | 0 | 0 | 0 | 0 | 0 | 0 | 0 | 0 |
| 0 | NA | NA | NA | NA | NA | Gm32381 | 0 | 0 | 0 | 0 | 0 | 0 | 0 | 0 | 0 | 0 | 0 |
| 0 | NA | NA | NA | NA | NA | Gm49348 | 0 | 0 | 0 | 0 | 0 | 0 | 0 | 0 | 0 | 0 | 0 |
| 0 | NA | NA | NA | NA | NA | Gm22328 | 0 | 0 | 0 | 0 | 0 | 0 | 0 | 0 | 0 | 0 | 0 |
| 0 | NA | NA | NA | NA | NA | Gm24936 | 0 | 0 | 0 | 0 | 0 | 0 | 0 | 0 | 0 | 0 | 0 |
| 0 | NA | NA | NA | NA | NA | Gm26748 | 0 | 0 | 0 | 0 | 0 | 0 | 0 | 0 | 0 | 0 | 0 |
| 0 | NA | NA | NA | NA | NA | Gm5314 | 0 | 0 | 0 | 0 | 0 | 0 | 0 | 0 | 0 | 0 | 0 |
| 0 | NA | NA | NA | NA | NA | Gm24248 | 0 | 0 | 0 | 0 | 0 | 0 | 0 | 0 | 0 | 0 | 0 |
| 0 | NA | NA | NA | NA | NA | Gm18422 | 0 | 0 | 0 | 0 | 0 | 0 | 0 | 0 | 0 | 0 | 0 |
| 0 | NA | NA | NA | NA | NA | Gm44044 | 0 | 0 | 0 | 0 | 0 | 0 | 0 | 0 | 0 | 0 | 0 |
| 0 | NA | NA | NA | NA | NA | Gm23234 | 0 | 0 | 0 | 0 | 0 | 0 | 0 | 0 | 0 | 0 | 0 |
| 0 | NA | NA | NA | NA | NA | Gm44107 | 0 | 0 | 0 | 0 | 0 | 0 | 0 | 0 | 0 | 0 | 0 |
| 0 | NA | NA | NA | NA | NA | 1700049E22Rik | 0 | 0 | 0 | 0 | 0 | 0 | 0 | 0 | 0 | 0 | 0 |
| 0 | NA | NA | NA | NA | NA | Gm15576 | 0 | 0 | 0 | 0 | 0 | 0 | 0 | 0 | 0 | 0 | 0 |
| 0 | NA | NA | NA | NA | NA | Gm19692 | 0 | 0 | 0 | 0 | 0 | 0 | 0 | 0 | 0 | 0 | 0 |
| 0 | NA | NA | NA | NA | NA | Gm43941 | 0 | 0 | 0 | 0 | 0 | 0 | 0 | 0 | 0 | 0 | 0 |
| 0 | NA | NA | NA | NA | NA | Gm43950 | 0 | 0 | 0 | 0 | 0 | 0 | 0 | 0 | 0 | 0 | 0 |
| 0 | NA | NA | NA | NA | NA | Gm43942 | 0 | 0 | 0 | 0 | 0 | 0 | 0 | 0 | 0 | 0 | 0 |
| 0 | NA | NA | NA | NA | NA | Gm26911 | 0 | 0 | 0 | 0 | 0 | 0 | 0 | 0 | 0 | 0 | 0 |
| 0 | NA | NA | NA | NA | NA | 9530086O07Rik | 0 | 0 | 0 | 0 | 0 | 0 | 0 | 0 | 0 | 0 | 0 |
| 0 | NA | NA | NA | NA | NA | Gm43953 | 0 | 0 | 0 | 0 | 0 | 0 | 0 | 0 | 0 | 0 | 0 |
| 0 | NA | NA | NA | NA | NA | Gm43947 | 0 | 0 | 0 | 0 | 0 | 0 | 0 | 0 | 0 | 0 | 0 |
| 0 | NA | NA | NA | NA | NA | Gm44169 | 0 | 0 | 0 | 0 | 0 | 0 | 0 | 0 | 0 | 0 | 0 |
| 0 | NA | NA | NA | NA | NA | Gm44171 | 0 | 0 | 0 | 0 | 0 | 0 | 0 | 0 | 0 | 0 | 0 |
| 0 | NA | NA | NA | NA | NA | Gm32453 | 0 | 0 | 0 | 0 | 0 | 0 | 0 | 0 | 0 | 0 | 0 |
| 0 | NA | NA | NA | NA | NA | Gm9871 | 0 | 0 | 0 | 0 | 0 | 0 | 0 | 0 | 0 | 0 | 0 |
| 0 | NA | NA | NA | NA | NA | Gm44176 | 0 | 0 | 0 | 0 | 0 | 0 | 0 | 0 | 0 | 0 | 0 |
| 0 | NA | NA | NA | NA | NA | Gm44177 | 0 | 0 | 0 | 0 | 0 | 0 | 0 | 0 | 0 | 0 | 0 |
| 0 | NA | NA | NA | NA | NA | 4930587E11Rik | 0 | 0 | 0 | 0 | 0 | 0 | 0 | 0 | 0 | 0 | 0 |
| 0 | NA | NA | NA | NA | NA | Gm44422 | 0 | 0 | 0 | 0 | 0 | 0 | 0 | 0 | 0 | 0 | 0 |
| 0 | NA | NA | NA | NA | NA | Mir6373 | 0 | 0 | 0 | 0 | 0 | 0 | 0 | 0 | 0 | 0 | 0 |
| 0 | NA | NA | NA | NA | NA | Gm26022 | 0 | 0 | 0 | 0 | 0 | 0 | 0 | 0 | 0 | 0 | 0 |
| 0 | NA | NA | NA | NA | NA | Gm44429 | 0 | 0 | 0 | 0 | 0 | 0 | 0 | 0 | 0 | 0 | 0 |
| 0 | NA | NA | NA | NA | NA | Gm44449 | 0 | 0 | 0 | 0 | 0 | 0 | 0 | 0 | 0 | 0 | 0 |
| 0 | NA | NA | NA | NA | NA | Gm44295 | 0 | 0 | 0 | 0 | 0 | 0 | 0 | 0 | 0 | 0 | 0 |
| 0 | NA | NA | NA | NA | NA | Gm44439 | 0 | 0 | 0 | 0 | 0 | 0 | 0 | 0 | 0 | 0 | 0 |
| 0 | NA | NA | NA | NA | NA | Gm7946 | 0 | 0 | 0 | 0 | 0 | 0 | 0 | 0 | 0 | 0 | 0 |
| 0 | NA | NA | NA | NA | NA | Gm44441 | 0 | 0 | 0 | 0 | 0 | 0 | 0 | 0 | 0 | 0 | 0 |
| 0 | NA | NA | NA | NA | NA | Gm24784 | 0 | 0 | 0 | 0 | 0 | 0 | 0 | 0 | 0 | 0 | 0 |
| 0 | NA | NA | NA | NA | NA | Gm44440 | 0 | 0 | 0 | 0 | 0 | 0 | 0 | 0 | 0 | 0 | 0 |
| 0 | NA | NA | NA | NA | NA | Gm44436 | 0 | 0 | 0 | 0 | 0 | 0 | 0 | 0 | 0 | 0 | 0 |
| 0 | NA | NA | NA | NA | NA | Gm25656 | 0 | 0 | 0 | 0 | 0 | 0 | 0 | 0 | 0 | 0 | 0 |
| 0 | NA | NA | NA | NA | NA | Gm21054 | 0 | 0 | 0 | 0 | 0 | 0 | 0 | 0 | 0 | 0 | 0 |
| 0 | NA | NA | NA | NA | NA | Gm18100 | 0 | 0 | 0 | 0 | 0 | 0 | 0 | 0 | 0 | 0 | 0 |
| 0 | NA | NA | NA | NA | NA | Gm23339 | 0 | 0 | 0 | 0 | 0 | 0 | 0 | 0 | 0 | 0 | 0 |
| 0 | NA | NA | NA | NA | NA | Gm25570 | 0 | 0 | 0 | 0 | 0 | 0 | 0 | 0 | 0 | 0 | 0 |
| 0 | NA | NA | NA | NA | NA | Gm19757 | 0 | 0 | 0 | 0 | 0 | 0 | 0 | 0 | 0 | 0 | 0 |
| 0 | NA | NA | NA | NA | NA | Gm15631 | 0 | 0 | 0 | 0 | 0 | 0 | 0 | 0 | 0 | 0 | 0 |
| 0 | NA | NA | NA | NA | NA | Gm22418 | 0 | 0 | 0 | 0 | 0 | 0 | 0 | 0 | 0 | 0 | 0 |
| 0 | NA | NA | NA | NA | NA | Gm6064 | 0 | 0 | 0 | 0 | 0 | 0 | 0 | 0 | 0 | 0 | 0 |
| 0 | NA | NA | NA | NA | NA | Gm34933 | 0 | 0 | 0 | 0 | 0 | 0 | 0 | 0 | 0 | 0 | 0 |
| 0 | NA | NA | NA | NA | NA | Gm44399 | 0 | 0 | 0 | 0 | 0 | 0 | 0 | 0 | 0 | 0 | 0 |
| 0 | NA | NA | NA | NA | NA | Gm27690 | 0 | 0 | 0 | 0 | 0 | 0 | 0 | 0 | 0 | 0 | 0 |
| 0 | NA | NA | NA | NA | NA | Gm44277 | 0 | 0 | 0 | 0 | 0 | 0 | 0 | 0 | 0 | 0 | 0 |
| 0 | NA | NA | NA | NA | NA | Gm19177 | 0 | 0 | 0 | 0 | 0 | 0 | 0 | 0 | 0 | 0 | 0 |
| 0 | NA | NA | NA | NA | NA | Gm44279 | 0 | 0 | 0 | 0 | 0 | 0 | 0 | 0 | 0 | 0 | 0 |
| 0 | NA | NA | NA | NA | NA | Gm27813 | 0 | 0 | 0 | 0 | 0 | 0 | 0 | 0 | 0 | 0 | 0 |
| 0 | NA | NA | NA | NA | NA | Mir7661 | 0 | 0 | 0 | 0 | 0 | 0 | 0 | 0 | 0 | 0 | 0 |

| | | | | | | | | | | | | | | | | | | |
|---|---|---|---|---|---|---|---|---|---|---|---|---|---|---|---|---|---|---|
| 0 | NA | NA | NA | NA | NA | Gm44040 | | 0 | 0 | 0 | 0 | 0 | 0 | 0 | 0 | 0 | 0 | 0 |
| 0 | NA | NA | NA | NA | NA | Gm38167 | | 0 | 0 | 0 | 0 | 0 | 0 | 0 | 0 | 0 | 0 | 0 |
| 0 | NA | NA | NA | NA | NA | Gm32858 | | 0 | 0 | 0 | 0 | 0 | 0 | 0 | 0 | 0 | 0 | 0 |
| 0 | NA | NA | NA | NA | NA | Gm44125 | | 0 | 0 | 0 | 0 | 0 | 0 | 0 | 0 | 0 | 0 | 0 |
| 0 | NA | NA | NA | NA | NA | Gm4605 | | 0 | 0 | 0 | 0 | 0 | 0 | 0 | 0 | 0 | 0 | 0 |
| 0 | NA | NA | NA | NA | NA | Gm44161 | | 0 | 0 | 0 | 0 | 0 | 0 | 0 | 0 | 0 | 0 | 0 |
| 0 | NA | NA | NA | NA | NA | Gm44162 | | 0 | 0 | 0 | 0 | 0 | 0 | 0 | 0 | 0 | 0 | 0 |
| 0 | NA | NA | NA | NA | NA | Gm19039 | | 0 | 0 | 0 | 0 | 0 | 0 | 0 | 0 | 0 | 0 | 0 |
| 0 | NA | NA | NA | NA | NA | Gm20387 | | 0 | 0 | 0 | 0 | 0 | 0 | 0 | 0 | 0 | 0 | 0 |
| 0 | NA | NA | NA | NA | NA | Grm7 | glutamate receptor, metabotropic 7 | 0 | 0 | 0 | 0 | 0 | 0 | 0 | 0 | 0 | 0 | 0 |
| 0 | NA | NA | NA | NA | NA | Gm20386 | | 0 | 0 | 0 | 0 | 0 | 0 | 0 | 0 | 0 | 0 | 0 |
| 0 | NA | NA | NA | NA | NA | Gm22093 | | 0 | 0 | 0 | 0 | 0 | 0 | 0 | 0 | 0 | 0 | 0 |
| 0 | NA | NA | NA | NA | NA | Gm24005 | | 0 | 0 | 0 | 0 | 0 | 0 | 0 | 0 | 0 | 0 | 0 |
| 0 | NA | NA | NA | NA | NA | Gm26362 | | 0 | 0 | 0 | 0 | 0 | 0 | 0 | 0 | 0 | 0 | 0 |
| 0 | NA | NA | NA | NA | NA | Gm25454 | | 0 | 0 | 0 | 0 | 0 | 0 | 0 | 0 | 0 | 0 | 0 |
| 0 | NA | NA | NA | NA | NA | Gm44192 | | 0 | 0 | 0 | 0 | 0 | 0 | 0 | 0 | 0 | 0 | 0 |
| 0 | NA | NA | NA | NA | NA | n-R5s165 | | 0 | 0 | 0 | 0 | 0 | 0 | 0 | 0 | 0 | 0 | 0 |
| 0 | NA | NA | NA | NA | NA | 5031434C07Rik | | 0 | 0 | 0 | 0 | 0 | 0 | 0 | 0 | 0 | 0 | 0 |
| 0 | NA | NA | NA | NA | NA | Gm44336 | | 0 | 0 | 0 | 0 | 0 | 0 | 0 | 0 | 0 | 0 | 0 |
| 0 | NA | NA | NA | NA | NA | Gm27944 | | 0 | 0 | 0 | 0 | 0 | 0 | 0 | 0 | 0 | 0 | 0 |
| 0 | NA | NA | NA | NA | NA | Gm23002 | | 0 | 0 | 0 | 0 | 0 | 0 | 0 | 0 | 0 | 0 | 0 |
| 0 | NA | NA | NA | NA | NA | Gm43865 | | 0 | 0 | 0 | 0 | 0 | 0 | 0 | 0 | 0 | 0 | 0 |
| 0 | NA | NA | NA | NA | NA | Gm8083 | | 0 | 0 | 0 | 0 | 0 | 0 | 0 | 0 | 0 | 0 | 0 |
| 0 | NA | NA | NA | NA | NA | Gm23244 | | 0 | 0 | 0 | 0 | 0 | 0 | 0 | 0 | 0 | 0 | 0 |
| 0 | NA | NA | NA | NA | NA | Gm22591 | | 0 | 0 | 0 | 0 | 0 | 0 | 0 | 0 | 0 | 0 | 0 |
| 0 | NA | NA | NA | NA | NA | Gm16161 | | 0 | 0 | 0 | 0 | 0 | 0 | 0 | 0 | 0 | 0 | 0 |
| 0 | NA | NA | NA | NA | NA | Gm20589 | | 0 | 0 | 0 | 0 | 0 | 0 | 0 | 0 | 0 | 0 | 0 |
| 0 | NA | NA | NA | NA | NA | Gm24387 | | 0 | 0 | 0 | 0 | 0 | 0 | 0 | 0 | 0 | 0 | 0 |
| 0 | NA | NA | NA | NA | NA | Gm43928 | | 0 | 0 | 0 | 0 | 0 | 0 | 0 | 0 | 0 | 0 | 0 |
| 0 | NA | NA | NA | NA | NA | Gm23365 | | 0 | 0 | 0 | 0 | 0 | 0 | 0 | 0 | 0 | 0 | 0 |
| 0 | NA | NA | NA | NA | NA | Gm44232 | | 0 | 0 | 0 | 0 | 0 | 0 | 0 | 0 | 0 | 0 | 0 |
| 0 | NA | NA | NA | NA | NA | Fancd2os | Fancd2 opposite strand | 0 | 0 | 0 | 0 | 0 | 0 | 0 | 0 | 0 | 0 | 0 |
| 0 | NA | NA | NA | NA | NA | 1700015O11Rik | | 0 | 0 | 0 | 0 | 0 | 0 | 0 | 0 | 0 | 0 | 0 |
| 0 | NA | NA | NA | NA | NA | Gm43964 | | 0 | 0 | 0 | 0 | 0 | 0 | 0 | 0 | 0 | 0 | 0 |
| 0 | NA | NA | NA | NA | NA | Gm43963 | | 0 | 0 | 0 | 0 | 0 | 0 | 0 | 0 | 0 | 0 | 0 |
| 0 | NA | NA | NA | NA | NA | Gm15082 | | 0 | 0 | 0 | 0 | 0 | 0 | 0 | 0 | 0 | 0 | 0 |
| 0 | NA | NA | NA | NA | NA | Gm43911 | | 0 | 0 | 0 | 0 | 0 | 0 | 0 | 0 | 0 | 0 | 0 |
| 0 | NA | NA | NA | NA | NA | Gm43932 | | 0 | 0 | 0 | 0 | 0 | 0 | 0 | 0 | 0 | 0 | 0 |
| 0 | NA | NA | NA | NA | NA | | | 0 | 0 | 0 | 0 | 0 | 0 | 0 | 0 | 0 | 0 | 0 |
| 0 | NA | NA | NA | NA | NA | Gm44332 | | 0 | 0 | 0 | 0 | 0 | 0 | 0 | 0 | 0 | 0 | 0 |
| 0 | NA | NA | NA | NA | NA | Gm44331 | | 0 | 0 | 0 | 0 | 0 | 0 | 0 | 0 | 0 | 0 | 0 |
| 0 | NA | NA | NA | NA | NA | Gm24767 | | 0 | 0 | 0 | 0 | 0 | 0 | 0 | 0 | 0 | 0 | 0 |
| 0 | NA | NA | NA | NA | NA | Gm44010 | | 0 | 0 | 0 | 0 | 0 | 0 | 0 | 0 | 0 | 0 | 0 |
| 0 | NA | NA | NA | NA | NA | Gm18325 | | 0 | 0 | 0 | 0 | 0 | 0 | 0 | 0 | 0 | 0 | 0 |
| 0 | NA | NA | NA | NA | NA | Gm44079 | | 0 | 0 | 0 | 0 | 0 | 0 | 0 | 0 | 0 | 0 | 0 |
| 0 | NA | NA | NA | NA | NA | Gm24008 | | 0 | 0 | 0 | 0 | 0 | 0 | 0 | 0 | 0 | 0 | 0 |
| 0 | NA | NA | NA | NA | NA | Gm27853 | | 0 | 0 | 0 | 0 | 0 | 0 | 0 | 0 | 0 | 0 | 0 |
| 0 | NA | NA | NA | NA | NA | Mir7660 | | 0 | 0 | 0 | 0 | 0 | 0 | 0 | 0 | 0 | 0 | 0 |
| 0 | NA | NA | NA | NA | NA | Snora7a | | 0 | 0 | 0 | 0 | 0 | 0 | 0 | 0 | 0 | 0 | 0 |
| 0 | NA | NA | NA | NA | NA | Gm44206 | | 0 | 0 | 0 | 0 | 0 | 0 | 0 | 0 | 0 | 0 | 0 |
| 0 | NA | NA | NA | NA | NA | Gm43866 | | 0 | 0 | 0 | 0 | 0 | 0 | 0 | 0 | 0 | 0 | 0 |
| 0 | NA | NA | NA | NA | NA | H1f8 | H1.8 linker histone | 0 | 0 | 0 | 0 | 0 | 0 | 0 | 0 | 0 | 0 | 0 |
| 0 | NA | NA | NA | NA | NA | Gm20404 | | 0 | 0 | 0 | 0 | 0 | 0 | 0 | 0 | 0 | 0 | 0 |
| 0 | NA | NA | NA | NA | NA | Gm23527 | | 0 | 0 | 0 | 0 | 0 | 0 | 0 | 0 | 0 | 0 | 0 |
| 0 | NA | NA | NA | NA | NA | Gm44076 | | 0 | 0 | 0 | 0 | 0 | 0 | 0 | 0 | 0 | 0 | 0 |
| 0 | NA | NA | NA | NA | NA | Olfr211 | olfactory receptor 211 | 0 | 0 | 0 | 0 | 0 | 0 | 0 | 0 | 0 | 0 | 0 |
| 0 | NA | NA | NA | NA | NA | Olfr212 | olfactory receptor 212 | 0 | 0 | 0 | 0 | 0 | 0 | 0 | 0 | 0 | 0 | 0 |
| 0 | NA | NA | NA | NA | NA | Olfr213 | olfactory receptor 213 | 0 | 0 | 0 | 0 | 0 | 0 | 0 | 0 | 0 | 0 | 0 |
| 0 | NA | NA | NA | NA | NA | Olfr214 | olfactory receptor 214 | 0 | 0 | 0 | 0 | 0 | 0 | 0 | 0 | 0 | 0 | 0 |
| 0 | NA | NA | NA | NA | NA | Olfr215 | olfactory receptor 215 | 0 | 0 | 0 | 0 | 0 | 0 | 0 | 0 | 0 | 0 | 0 |

| | | | | | | | | | | | | | | | | | |
|---|---|---|---|---|---|---|---|---|---|---|---|---|---|---|---|---|---|
| 0 | NA | NA | NA | NA | NA | Gm18485 | | 0 | 0 | 0 | 0 | 0 | 0 | 0 | 0 | 0 | 0 | 0 |
| 0 | NA | NA | NA | NA | NA | Gm22882 | | 0 | 0 | 0 | 0 | 0 | 0 | 0 | 0 | 0 | 0 | 0 |
| 0 | NA | NA | NA | NA | NA | Gm43926 | | 0 | 0 | 0 | 0 | 0 | 0 | 0 | 0 | 0 | 0 | 0 |
| 0 | NA | NA | NA | NA | NA | Gm43929 | | 0 | 0 | 0 | 0 | 0 | 0 | 0 | 0 | 0 | 0 | 0 |
| 0 | NA | NA | NA | NA | NA | Gm43930 | | 0 | 0 | 0 | 0 | 0 | 0 | 0 | 0 | 0 | 0 | 0 |
| 0 | NA | NA | NA | NA | NA | D030044L04Rik | | 0 | 0 | 0 | 0 | 0 | 0 | 0 | 0 | 0 | 0 | 0 |
| 0 | NA | NA | NA | NA | NA | Rpl28-ps4 | | 0 | 0 | 0 | 0 | 0 | 0 | 0 | 0 | 0 | 0 | 0 |
| 0 | NA | NA | NA | NA | NA | Gm4640 | | 0 | 0 | 0 | 0 | 0 | 0 | 0 | 0 | 0 | 0 | 0 |
| 0 | NA | NA | NA | NA | NA | Gm6227 | | 0 | 0 | 0 | 0 | 0 | 0 | 0 | 0 | 0 | 0 | 0 |
| 0 | NA | NA | NA | NA | NA | 6820426E19Rik | | 0 | 0 | 0 | 0 | 0 | 0 | 0 | 0 | 0 | 0 | 0 |
| 0 | NA | NA | NA | NA | NA | Gm9946 | | 0 | 0 | 0 | 0 | 0 | 0 | 0 | 0 | 0 | 0 | 0 |
| 0 | NA | NA | NA | NA | NA | 1700030F04Rik | | 0 | 0 | 0 | 0 | 0 | 0 | 0 | 0 | 0 | 0 | 0 |
| 0 | NA | NA | NA | NA | NA | Gemin6-ps | | 0 | 0 | 0 | 0 | 0 | 0 | 0 | 0 | 0 | 0 | 0 |
| 0 | NA | NA | NA | NA | NA | Gm29509 | | 0 | 0 | 0 | 0 | 0 | 0 | 0 | 0 | 0 | 0 | 0 |
| 0 | NA | NA | NA | NA | NA | Fxyd4 | FXYD domain-containing ion transport re | 0 | 0 | 0 | 0 | 0 | 0 | 0 | 0 | 0 | 0 | 0 |
| 0 | NA | NA | NA | NA | NA | Gm25535 | | 0 | 0 | 0 | 0 | 0 | 0 | 0 | 0 | 0 | 0 | 0 |
| 0 | NA | NA | NA | NA | NA | Mir7044 | | 0 | 0 | 0 | 0 | 0 | 0 | 0 | 0 | 0 | 0 | 0 |
| 0 | NA | NA | NA | NA | NA | 1700069P05Rik | | 0 | 0 | 0 | 0 | 0 | 0 | 0 | 0 | 0 | 0 | 0 |
| 0 | NA | NA | NA | NA | NA | Gm30557 | | 0 | 0 | 0 | 0 | 0 | 0 | 0 | 0 | 0 | 0 | 0 |
| 0 | NA | NA | NA | NA | NA | Gm30498 | | 0 | 0 | 0 | 0 | 0 | 0 | 0 | 0 | 0 | 0 | 0 |
| 0 | NA | NA | NA | NA | NA | Gm25953 | | 0 | 0 | 0 | 0 | 0 | 0 | 0 | 0 | 0 | 0 | 0 |
| 0 | NA | NA | NA | NA | NA | | | 0 | 0 | 0 | 0 | 0 | 0 | 0 | 0 | 0 | 0 | 0 |
| 0 | NA | NA | NA | NA | NA | Gm25905 | | 0 | 0 | 0 | 0 | 0 | 0 | 0 | 0 | 0 | 0 | 0 |
| 0 | NA | NA | NA | NA | NA | Gm15532 | | 0 | 0 | 0 | 0 | 0 | 0 | 0 | 0 | 0 | 0 | 0 |
| 0 | NA | NA | NA | NA | NA | Mir706 | | 0 | 0 | 0 | 0 | 0 | 0 | 0 | 0 | 0 | 0 | 0 |
| 0 | NA | NA | NA | NA | NA | Gm25327 | | 0 | 0 | 0 | 0 | 0 | 0 | 0 | 0 | 0 | 0 | 0 |
| 0 | NA | NA | NA | NA | NA | Gm24592 | | 0 | 0 | 0 | 0 | 0 | 0 | 0 | 0 | 0 | 0 | 0 |
| 0 | NA | NA | NA | NA | NA | Gm24391 | | 0 | 0 | 0 | 0 | 0 | 0 | 0 | 0 | 0 | 0 | 0 |
| 0 | NA | NA | NA | NA | NA | 1700072O05Rik | | 0 | 0 | 0 | 0 | 0 | 0 | 0 | 0 | 0 | 0 | 0 |
| 0 | NA | NA | NA | NA | NA | Gm44074 | | 0 | 0 | 0 | 0 | 0 | 0 | 0 | 0 | 0 | 0 | 0 |
| 0 | NA | NA | NA | NA | NA | Gm44124 | | 0 | 0 | 0 | 0 | 0 | 0 | 0 | 0 | 0 | 0 | 0 |
| 0 | NA | NA | NA | NA | NA | Gm17812 | | 0 | 0 | 0 | 0 | 0 | 0 | 0 | 0 | 0 | 0 | 0 |
| 0 | NA | NA | NA | NA | NA | Gm43916 | | 0 | 0 | 0 | 0 | 0 | 0 | 0 | 0 | 0 | 0 | 0 |
| 0 | NA | NA | NA | NA | NA | Gm43915 | | 0 | 0 | 0 | 0 | 0 | 0 | 0 | 0 | 0 | 0 | 0 |
| 0 | NA | NA | NA | NA | NA | Gm44014 | | 0 | 0 | 0 | 0 | 0 | 0 | 0 | 0 | 0 | 0 | 0 |
| 0 | NA | NA | NA | NA | NA | Gm24855 | | 0 | 0 | 0 | 0 | 0 | 0 | 0 | 0 | 0 | 0 | 0 |
| 0 | NA | NA | NA | NA | NA | Gm43922 | | 0 | 0 | 0 | 0 | 0 | 0 | 0 | 0 | 0 | 0 | 0 |
| 0 | NA | NA | NA | NA | NA | Gm43933 | | 0 | 0 | 0 | 0 | 0 | 0 | 0 | 0 | 0 | 0 | 0 |
| 0 | NA | NA | NA | NA | NA | Gm43934 | | 0 | 0 | 0 | 0 | 0 | 0 | 0 | 0 | 0 | 0 | 0 |
| 0 | NA | NA | NA | NA | NA | Gm43945 | | 0 | 0 | 0 | 0 | 0 | 0 | 0 | 0 | 0 | 0 | 0 |
| 0 | NA | NA | NA | NA | NA | Gm44018 | | 0 | 0 | 0 | 0 | 0 | 0 | 0 | 0 | 0 | 0 | 0 |
| 0 | NA | NA | NA | NA | NA | Gm44173 | | 0 | 0 | 0 | 0 | 0 | 0 | 0 | 0 | 0 | 0 | 0 |
| 0 | NA | NA | NA | NA | NA | Mug4-ps | | 0 | 0 | 0 | 0 | 0 | 0 | 0 | 0 | 0 | 0 | 0 |
| 0 | NA | NA | NA | NA | NA | Mug2 | murinoglobulin 2 | 0 | 0 | 0 | 0 | 0 | 0 | 0 | 0 | 0 | 0 | 0 |
| 0 | NA | NA | NA | NA | NA | Gm44184 | | 0 | 0 | 0 | 0 | 0 | 0 | 0 | 0 | 0 | 0 | 0 |
| 0 | NA | NA | NA | NA | NA | Gm10319 | | 0 | 0 | 0 | 0 | 0 | 0 | 0 | 0 | 0 | 0 | 0 |
| 0 | NA | NA | NA | NA | NA | Gm27703 | | 0 | 0 | 0 | 0 | 0 | 0 | 0 | 0 | 0 | 0 | 0 |
| 0 | NA | NA | NA | NA | NA | Gm27826 | | 0 | 0 | 0 | 0 | 0 | 0 | 0 | 0 | 0 | 0 | 0 |
| 0 | NA | NA | NA | NA | NA | Gm44227 | | 0 | 0 | 0 | 0 | 0 | 0 | 0 | 0 | 0 | 0 | 0 |
| 0 | NA | NA | NA | NA | NA | 1700063H04Rik | | 0 | 0 | 0 | 0 | 0 | 0 | 0 | 0 | 0 | 0 | 0 |
| 0 | NA | NA | NA | NA | NA | Gm19254 | | 0 | 0 | 0 | 0 | 0 | 0 | 0 | 0 | 0 | 0 | 0 |
| 0 | NA | NA | NA | NA | NA | Gm16557 | | 0 | 0 | 0 | 0 | 0 | 0 | 0 | 0 | 0 | 0 | 0 |
| 0 | NA | NA | NA | NA | NA | Gm26168 | | 0 | 0 | 0 | 0 | 0 | 0 | 0 | 0 | 0 | 0 | 0 |
| 0 | NA | NA | NA | NA | NA | Mir7231 | | 0 | 0 | 0 | 0 | 0 | 0 | 0 | 0 | 0 | 0 | 0 |
| 0 | NA | NA | NA | NA | NA | Gm5316 | | 0 | 0 | 0 | 0 | 0 | 0 | 0 | 0 | 0 | 0 | 0 |
| 0 | NA | NA | NA | NA | NA | Gm44132 | | 0 | 0 | 0 | 0 | 0 | 0 | 0 | 0 | 0 | 0 | 0 |
| 0 | NA | NA | NA | NA | NA | Gm44147 | | 0 | 0 | 0 | 0 | 0 | 0 | 0 | 0 | 0 | 0 | 0 |
| 0 | NA | NA | NA | NA | NA | Clec4b2 | C-type lectin domain family 4, member b2 | 0 | 0 | 0 | 0 | 0 | 0 | 0 | 0 | 0 | 0 | 0 |
| 0 | NA | NA | NA | NA | NA | Gm22946 | | 0 | 0 | 0 | 0 | 0 | 0 | 0 | 0 | 0 | 0 | 0 |
| 0 | NA | NA | NA | NA | NA | Vmn2r19 | vomeronasal 2, receptor 19 | 0 | 0 | 0 | 0 | 0 | 0 | 0 | 0 | 0 | 0 | 0 |

| | | | | | | | | | | | | | | | | | | |
|---|---|---|---|---|---|---|---|---|---|---|---|---|---|---|---|---|---|---|
| 0 | NA | NA | NA | NA | NA | Vmn2r20 | vomeronasal 2, receptor 20 | 0 | 0 | 0 | 0 | 0 | 0 | 0 | 0 | 0 | 0 | 0 |
| 0 | NA | NA | NA | NA | NA | Gm22833 | | 0 | 0 | 0 | 0 | 0 | 0 | 0 | 0 | 0 | 0 | 0 |
| 0 | NA | NA | NA | NA | NA | Vmn2r-ps31 | | 0 | 0 | 0 | 0 | 0 | 0 | 0 | 0 | 0 | 0 | 0 |
| 0 | NA | NA | NA | NA | NA | Vmn2r21 | vomeronasal 2, receptor 21 | 0 | 0 | 0 | 0 | 0 | 0 | 0 | 0 | 0 | 0 | 0 |
| 0 | NA | NA | NA | NA | NA | Gm25891 | | 0 | 0 | 0 | 0 | 0 | 0 | 0 | 0 | 0 | 0 | 0 |
| 0 | NA | NA | NA | NA | NA | Gm22722 | | 0 | 0 | 0 | 0 | 0 | 0 | 0 | 0 | 0 | 0 | 0 |
| 0 | NA | NA | NA | NA | NA | Vmn2r-ps32 | | 0 | 0 | 0 | 0 | 0 | 0 | 0 | 0 | 0 | 0 | 0 |
| 0 | NA | NA | NA | NA | NA | Vmn2r22 | vomeronasal 2, receptor 22 | 0 | 0 | 0 | 0 | 0 | 0 | 0 | 0 | 0 | 0 | 0 |
| 0 | NA | NA | NA | NA | NA | Gm25065 | | 0 | 0 | 0 | 0 | 0 | 0 | 0 | 0 | 0 | 0 | 0 |
| 0 | NA | NA | NA | NA | NA | Vmn2r23 | vomeronasal 2, receptor 23 | 0 | 0 | 0 | 0 | 0 | 0 | 0 | 0 | 0 | 0 | 0 |
| 0 | NA | NA | NA | NA | NA | Gm22724 | | 0 | 0 | 0 | 0 | 0 | 0 | 0 | 0 | 0 | 0 | 0 |
| 0 | NA | NA | NA | NA | NA | Vmn2r24 | vomeronasal 2, receptor 24 | 0 | 0 | 0 | 0 | 0 | 0 | 0 | 0 | 0 | 0 | 0 |
| 0 | NA | NA | NA | NA | NA | Gm23079 | | 0 | 0 | 0 | 0 | 0 | 0 | 0 | 0 | 0 | 0 | 0 |
| 0 | NA | NA | NA | NA | NA | Gm5317 | | 0 | 0 | 0 | 0 | 0 | 0 | 0 | 0 | 0 | 0 | 0 |
| 0 | NA | NA | NA | NA | NA | Gm44823 | | 0 | 0 | 0 | 0 | 0 | 0 | 0 | 0 | 0 | 0 | 0 |
| 0 | NA | NA | NA | NA | NA | A330044P14Rik | | 0 | 0 | 0 | 0 | 0 | 0 | 0 | 0 | 0 | 0 | 0 |
| 0 | NA | NA | NA | NA | NA | Gm22763 | | 0 | 0 | 0 | 0 | 0 | 0 | 0 | 0 | 0 | 0 | 0 |
| 0 | NA | NA | NA | NA | NA | Gm24843 | | 0 | 0 | 0 | 0 | 0 | 0 | 0 | 0 | 0 | 0 | 0 |
| 0 | NA | NA | NA | NA | NA | Vmn2r-ps33 | | 0 | 0 | 0 | 0 | 0 | 0 | 0 | 0 | 0 | 0 | 0 |
| 0 | NA | NA | NA | NA | NA | Gm22916 | | 0 | 0 | 0 | 0 | 0 | 0 | 0 | 0 | 0 | 0 | 0 |
| 0 | NA | NA | NA | NA | NA | Vmn2r-ps34 | | 0 | 0 | 0 | 0 | 0 | 0 | 0 | 0 | 0 | 0 | 0 |
| 0 | NA | NA | NA | NA | NA | Gm23554 | | 0 | 0 | 0 | 0 | 0 | 0 | 0 | 0 | 0 | 0 | 0 |
| 0 | NA | NA | NA | NA | NA | Vmn2r27 | vomeronasal 2, receptor27 | 0 | 0 | 0 | 0 | 0 | 0 | 0 | 0 | 0 | 0 | 0 |
| 0 | NA | NA | NA | NA | NA | Gm43971 | | 0 | 0 | 0 | 0 | 0 | 0 | 0 | 0 | 0 | 0 | 0 |
| 0 | NA | NA | NA | NA | NA | Gm25311 | | 0 | 0 | 0 | 0 | 0 | 0 | 0 | 0 | 0 | 0 | 0 |
| 0 | NA | NA | NA | NA | NA | 1700013D24Rik | | 0 | 0 | 0 | 0 | 0 | 0 | 0 | 0 | 0 | 0 | 0 |
| 0 | NA | NA | NA | NA | NA | 1700027F06Rik | | 0 | 0 | 0 | 0 | 0 | 0 | 0 | 0 | 0 | 0 | 0 |
| 0 | NA | NA | NA | NA | NA | Gm44006 | | 0 | 0 | 0 | 0 | 0 | 0 | 0 | 0 | 0 | 0 | 0 |
| 0 | NA | NA | NA | NA | NA | Gm44007 | | 0 | 0 | 0 | 0 | 0 | 0 | 0 | 0 | 0 | 0 | 0 |
| 0 | NA | NA | NA | NA | NA | Gm50463 | | 0 | 0 | 0 | 0 | 0 | 0 | 0 | 0 | 0 | 0 | 0 |
| 0 | NA | NA | NA | NA | NA | Gm44096 | | 0 | 0 | 0 | 0 | 0 | 0 | 0 | 0 | 0 | 0 | 0 |
| 0 | NA | NA | NA | NA | NA | Gm18840 | | 0 | 0 | 0 | 0 | 0 | 0 | 0 | 0 | 0 | 0 | 0 |
| 0 | NA | NA | NA | NA | NA | Gm23547 | | 0 | 0 | 0 | 0 | 0 | 0 | 0 | 0 | 0 | 0 | 0 |
| 0 | NA | NA | NA | NA | NA | Mir141 | | 0 | 0 | 0 | 0 | 0 | 0 | 0 | 0 | 0 | 0 | 0 |
| 0 | NA | NA | NA | NA | NA | Mir200c | | 0 | 0 | 0 | 0 | 0 | 0 | 0 | 0 | 0 | 0 | 0 |
| 0 | NA | NA | NA | NA | NA | Rnu7 | | 0 | 0 | 0 | 0 | 0 | 0 | 0 | 0 | 0 | 0 | 0 |
| 0 | NA | NA | NA | NA | NA | Gm23851 | | 0 | 0 | 0 | 0 | 0 | 0 | 0 | 0 | 0 | 0 | 0 |
| 0 | NA | NA | NA | NA | NA | 4930557K07Rik | | 0 | 0 | 0 | 0 | 0 | 0 | 0 | 0 | 0 | 0 | 0 |
| 0 | NA | NA | NA | NA | NA | Mir7045 | | 0 | 0 | 0 | 0 | 0 | 0 | 0 | 0 | 0 | 0 | 0 |
| 0 | NA | NA | NA | NA | NA | Gm23751 | | 0 | 0 | 0 | 0 | 0 | 0 | 0 | 0 | 0 | 0 | 0 |
| 0 | NA | NA | NA | NA | NA | Scarna10 | | 0 | 0 | 0 | 0 | 0 | 0 | 0 | 0 | 0 | 0 | 0 |
| 0 | NA | NA | NA | NA | NA | Gm28967 | | 0 | 0 | 0 | 0 | 0 | 0 | 0 | 0 | 0 | 0 | 0 |
| 0 | NA | NA | NA | NA | NA | Mir3098 | | 0 | 0 | 0 | 0 | 0 | 0 | 0 | 0 | 0 | 0 | 0 |
| 0 | NA | NA | NA | NA | NA | Gm44487 | | 0 | 0 | 0 | 0 | 0 | 0 | 0 | 0 | 0 | 0 | 0 |
| 0 | NA | NA | NA | NA | NA | Mir8113 | | 0 | 0 | 0 | 0 | 0 | 0 | 0 | 0 | 0 | 0 | 0 |
| 0 | NA | NA | NA | NA | NA | Gm44259 | | 0 | 0 | 0 | 0 | 0 | 0 | 0 | 0 | 0 | 0 | 0 |
| 0 | NA | NA | NA | NA | NA | Gm24175 | | 0 | 0 | 0 | 0 | 0 | 0 | 0 | 0 | 0 | 0 | 0 |
| 0 | NA | NA | NA | NA | NA | Gm16303 | | 0 | 0 | 0 | 0 | 0 | 0 | 0 | 0 | 0 | 0 | 0 |
| 0 | NA | NA | NA | NA | NA | Gm24947 | | 0 | 0 | 0 | 0 | 0 | 0 | 0 | 0 | 0 | 0 | 0 |
| 0 | NA | NA | NA | NA | NA | Gm10415 | | 0 | 0 | 0 | 0 | 0 | 0 | 0 | 0 | 0 | 0 | 0 |
| 0 | NA | NA | NA | NA | NA | Gm29009 | | 0 | 0 | 0 | 0 | 0 | 0 | 0 | 0 | 0 | 0 | 0 |
| 0 | NA | NA | NA | NA | NA | Gm44431 | | 0 | 0 | 0 | 0 | 0 | 0 | 0 | 0 | 0 | 0 | 0 |
| 0 | NA | NA | NA | NA | NA | Gm33320 | | 0 | 0 | 0 | 0 | 0 | 0 | 0 | 0 | 0 | 0 | 0 |
| 0 | NA | NA | NA | NA | NA | Gm26673 | | 0 | 0 | 0 | 0 | 0 | 0 | 0 | 0 | 0 | 0 | 0 |
| 0 | NA | NA | NA | NA | NA | Gm29788 | | 0 | 0 | 0 | 0 | 0 | 0 | 0 | 0 | 0 | 0 | 0 |
| 0 | NA | NA | NA | NA | NA | Gm17808 | | 0 | 0 | 0 | 0 | 0 | 0 | 0 | 0 | 0 | 0 | 0 |
| 0 | NA | NA | NA | NA | NA | Gm43125 | | 0 | 0 | 0 | 0 | 0 | 0 | 0 | 0 | 0 | 0 | 0 |
| 0 | NA | NA | NA | NA | NA | Fgf6 | fibroblast growth factor 6 | 0 | 0 | 0 | 0 | 0 | 0 | 0 | 0 | 0 | 0 | 0 |
| 0 | NA | NA | NA | NA | NA | Gm18995 | | 0 | 0 | 0 | 0 | 0 | 0 | 0 | 0 | 0 | 0 | 0 |
| 0 | NA | NA | NA | NA | NA | Gm38404 | | 0 | 0 | 0 | 0 | 0 | 0 | 0 | 0 | 0 | 0 | 0 |

| | | | | | | | | | | | | | | | | | |
|---|---|---|---|---|---|---|---|---|---|---|---|---|---|---|---|---|---|
| 0 | NA | NA | NA | NA | NA | Gm42458 | | 0 | 0 | 0 | 0 | 0 | 0 | 0 | 0 | 0 | 0 | 0 |
| 0 | NA | NA | NA | NA | NA | Gm43631 | | 0 | 0 | 0 | 0 | 0 | 0 | 0 | 0 | 0 | 0 | 0 |
| 0 | NA | NA | NA | NA | NA | Gm43634 | | 0 | 0 | 0 | 0 | 0 | 0 | 0 | 0 | 0 | 0 | 0 |
| 0 | NA | NA | NA | NA | NA | Gm42739 | | 0 | 0 | 0 | 0 | 0 | 0 | 0 | 0 | 0 | 0 | 0 |
| 0 | NA | NA | NA | NA | NA | Gm42738 | | 0 | 0 | 0 | 0 | 0 | 0 | 0 | 0 | 0 | 0 | 0 |
| 0 | NA | NA | NA | NA | NA | Gm42741 | | 0 | 0 | 0 | 0 | 0 | 0 | 0 | 0 | 0 | 0 | 0 |
| 0 | NA | NA | NA | NA | NA | 4921513H07Rik | | 0 | 0 | 0 | 0 | 0 | 0 | 0 | 0 | 0 | 0 | 0 |
| 0 | NA | NA | NA | NA | NA | Mir7233 | | 0 | 0 | 0 | 0 | 0 | 0 | 0 | 0 | 0 | 0 | 0 |
| 0 | NA | NA | NA | NA | NA | Gm42811 | | 0 | 0 | 0 | 0 | 0 | 0 | 0 | 0 | 0 | 0 | 0 |
| 0 | NA | NA | NA | NA | NA | Gm23098 | | 0 | 0 | 0 | 0 | 0 | 0 | 0 | 0 | 0 | 0 | 0 |
| 0 | NA | NA | NA | NA | NA | Gm15870 | | 0 | 0 | 0 | 0 | 0 | 0 | 0 | 0 | 0 | 0 | 0 |
| 0 | NA | NA | NA | NA | NA | Gm26338 | | 0 | 0 | 0 | 0 | 0 | 0 | 0 | 0 | 0 | 0 | 0 |
| 0 | NA | NA | NA | NA | NA | Gm44056 | | 0 | 0 | 0 | 0 | 0 | 0 | 0 | 0 | 0 | 0 | 0 |
| 0 | NA | NA | NA | NA | NA | Gm18609 | | 0 | 0 | 0 | 0 | 0 | 0 | 0 | 0 | 0 | 0 | 0 |
| 0 | NA | NA | NA | NA | NA | Gm19065 | | 0 | 0 | 0 | 0 | 0 | 0 | 0 | 0 | 0 | 0 | 0 |
| 0 | NA | NA | NA | NA | NA | Gm8703 | | 0 | 0 | 0 | 0 | 0 | 0 | 0 | 0 | 0 | 0 | 0 |
| 0 | NA | NA | NA | NA | NA | Gm21893 | | 0 | 0 | 0 | 0 | 0 | 0 | 0 | 0 | 0 | 0 | 0 |
| 0 | NA | NA | NA | NA | NA | Gm26770 | | 0 | 0 | 0 | 0 | 0 | 0 | 0 | 0 | 0 | 0 | 0 |
| 0 | NA | NA | NA | NA | NA | Gm44371 | | 0 | 0 | 0 | 0 | 0 | 0 | 0 | 0 | 0 | 0 | 0 |
| 0 | NA | NA | NA | NA | NA | Tex52 | testis expressed 52 | 0 | 0 | 0 | 0 | 0 | 0 | 0 | 0 | 0 | 0 | 0 |
| 0 | NA | NA | NA | NA | NA | Gm44596 | | 0 | 0 | 0 | 0 | 0 | 0 | 0 | 0 | 0 | 0 | 0 |
| 0 | NA | NA | NA | NA | NA | Gm5318 | | 0 | 0 | 0 | 0 | 0 | 0 | 0 | 0 | 0 | 0 | 0 |
| 0 | NA | NA | NA | NA | NA | Gm44009 | | 0 | 0 | 0 | 0 | 0 | 0 | 0 | 0 | 0 | 0 | 0 |
| 0 | NA | NA | NA | NA | NA | Gm44478 | | 0 | 0 | 0 | 0 | 0 | 0 | 0 | 0 | 0 | 0 | 0 |
| 0 | NA | NA | NA | NA | NA | Gm44420 | | 0 | 0 | 0 | 0 | 0 | 0 | 0 | 0 | 0 | 0 | 0 |
| 0 | NA | NA | NA | NA | NA | Clec2j | | 0 | 0 | 0 | 0 | 0 | 0 | 0 | 0 | 0 | 0 | 0 |
| 0 | NA | NA | NA | NA | NA | Gm44216 | | 0 | 0 | 0 | 0 | 0 | 0 | 0 | 0 | 0 | 0 | 0 |
| 0 | NA | NA | NA | NA | NA | Klrb1 | killer cell lectin-like receptor subfamily B r | 0 | 0 | 0 | 0 | 0 | 0 | 0 | 0 | 0 | 0 | 0 |
| 0 | NA | NA | NA | NA | NA | Gm43907 | | 0 | 0 | 0 | 0 | 0 | 0 | 0 | 0 | 0 | 0 | 0 |
| 0 | NA | NA | NA | NA | NA | Gm44511 | | 0 | 0 | 0 | 0 | 0 | 0 | 0 | 0 | 0 | 0 | 0 |
| 0 | NA | NA | NA | NA | NA | Gm44078 | | 0 | 0 | 0 | 0 | 0 | 0 | 0 | 0 | 0 | 0 | 0 |
| 0 | NA | NA | NA | NA | NA | Gm8774 | | 0 | 0 | 0 | 0 | 0 | 0 | 0 | 0 | 0 | 0 | 0 |
| 0 | NA | NA | NA | NA | NA | Gm44223 | | 0 | 0 | 0 | 0 | 0 | 0 | 0 | 0 | 0 | 0 | 0 |
| 0 | NA | NA | NA | NA | NA | Gm27336 | | 0 | 0 | 0 | 0 | 0 | 0 | 0 | 0 | 0 | 0 | 0 |
| 0 | NA | NA | NA | NA | NA | Mir7649 | | 0 | 0 | 0 | 0 | 0 | 0 | 0 | 0 | 0 | 0 | 0 |
| 0 | NA | NA | NA | NA | NA | Clec2f | C-type lectin domain family 2, member f | 0 | 0 | 0 | 0 | 0 | 0 | 0 | 0 | 0 | 0 | 0 |
| 0 | NA | NA | NA | NA | NA | Gm16186 | | 0 | 0 | 0 | 0 | 0 | 0 | 0 | 0 | 0 | 0 | 0 |
| 0 | NA | NA | NA | NA | NA | Gm43992 | | 0 | 0 | 0 | 0 | 0 | 0 | 0 | 0 | 0 | 0 | 0 |
| 0 | NA | NA | NA | NA | NA | Clec2e | C-type lectin domain family 2, member e | 0 | 0 | 0 | 0 | 0 | 0 | 0 | 0 | 0 | 0 | 0 |
| 0 | NA | NA | NA | NA | NA | Klrb1-ps1 | | 0 | 0 | 0 | 0 | 0 | 0 | 0 | 0 | 0 | 0 | 0 |
| 0 | NA | NA | NA | NA | NA | Gm26160 | | 0 | 0 | 0 | 0 | 0 | 0 | 0 | 0 | 0 | 0 | 0 |
| 0 | NA | NA | NA | NA | NA | Gm27514 | | 0 | 0 | 0 | 0 | 0 | 0 | 0 | 0 | 0 | 0 | 0 |
| 0 | NA | NA | NA | NA | NA | Gm47861 | | 0 | 0 | 0 | 0 | 0 | 0 | 0 | 0 | 0 | 0 | 0 |
| 0 | NA | NA | NA | NA | NA | Clec2m | C-type lectin domain family 2, member m | 0 | 0 | 0 | 0 | 0 | 0 | 0 | 0 | 0 | 0 | 0 |
| 0 | NA | NA | NA | NA | NA | Gm44120 | | 0 | 0 | 0 | 0 | 0 | 0 | 0 | 0 | 0 | 0 | 0 |
| 0 | NA | NA | NA | NA | NA | Gm44251 | | 0 | 0 | 0 | 0 | 0 | 0 | 0 | 0 | 0 | 0 | 0 |
| 0 | NA | NA | NA | NA | NA | Klrc3 | killer cell lectin-like receptor subfamily C, | 0 | 0 | 0 | 0 | 0 | 0 | 0 | 0 | 0 | 0 | 0 |
| 0 | NA | NA | NA | NA | NA | Klrc2 | killer cell lectin-like receptor subfamily C, | 0 | 0 | 0 | 0 | 0 | 0 | 0 | 0 | 0 | 0 | 0 |
| 0 | NA | NA | NA | NA | NA | Gm7309 | | 0 | 0 | 0 | 0 | 0 | 0 | 0 | 0 | 0 | 0 | 0 |
| 0 | NA | NA | NA | NA | NA | Mir680-1 | | 0 | 0 | 0 | 0 | 0 | 0 | 0 | 0 | 0 | 0 | 0 |
| 0 | NA | NA | NA | NA | NA | Klrh1 | killer cell lectin-like receptor subfamily H, | 0 | 0 | 0 | 0 | 0 | 0 | 0 | 0 | 0 | 0 | 0 |
| 0 | NA | NA | NA | NA | NA | Gm16242 | | 0 | 0 | 0 | 0 | 0 | 0 | 0 | 0 | 0 | 0 | 0 |
| 0 | NA | NA | NA | NA | NA | Gm44248 | | 0 | 0 | 0 | 0 | 0 | 0 | 0 | 0 | 0 | 0 | 0 |
| 0 | NA | NA | NA | NA | NA | Gm17631 | | 0 | 0 | 0 | 0 | 0 | 0 | 0 | 0 | 0 | 0 | 0 |
| 0 | NA | NA | NA | NA | NA | Gm23977 | | 0 | 0 | 0 | 0 | 0 | 0 | 0 | 0 | 0 | 0 | 0 |
| 0 | NA | NA | NA | NA | NA | Klra5 | killer cell lectin-like receptor, subfamily A, | 0 | 0 | 0 | 0 | 0 | 0 | 0 | 0 | 0 | 0 | 0 |
| 0 | NA | NA | NA | NA | NA | Gm15854 | killer cell lectin-like receptor subfamily A, | 0 | 0 | 0 | 0 | 0 | 0 | 0 | 0 | 0 | 0 | 0 |
| 0 | NA | NA | NA | NA | NA | Gm22555 | | 0 | 0 | 0 | 0 | 0 | 0 | 0 | 0 | 0 | 0 | 0 |
| 0 | NA | NA | NA | NA | NA | Klra6 | killer cell lectin-like receptor, subfamily A, | 0 | 0 | 0 | 0 | 0 | 0 | 0 | 0 | 0 | 0 | 0 |
| 0 | NA | NA | NA | NA | NA | Klra4 | killer cell lectin-like receptor, subfamily A, | 0 | 0 | 0 | 0 | 0 | 0 | 0 | 0 | 0 | 0 | 0 |

| | | | | | | | | | | | | | | | | | | |
|---|---|---|---|---|---|---|---|---|---|---|---|---|---|---|---|---|---|---|
| 0 | NA | NA | NA | NA | NA | Gm24676 | | 0 | 0 | 0 | 0 | 0 | 0 | 0 | 0 | 0 | 0 | 0 |
| 0 | NA | NA | NA | NA | NA | Klra11-ps | | 0 | 0 | 0 | 0 | 0 | 0 | 0 | 0 | 0 | 0 | 0 |
| 0 | NA | NA | NA | NA | NA | Gm43988 | | 0 | 0 | 0 | 0 | 0 | 0 | 0 | 0 | 0 | 0 | 0 |
| 0 | NA | NA | NA | NA | NA | Gm22621 | | 0 | 0 | 0 | 0 | 0 | 0 | 0 | 0 | 0 | 0 | 0 |
| 0 | NA | NA | NA | NA | NA | Klra8 | killer cell lectin-like receptor, subfamily A, | 0 | 0 | 0 | 0 | 0 | 0 | 0 | 0 | 0 | 0 | 0 |
| 0 | NA | NA | NA | NA | NA | Gm24712 | | 0 | 0 | 0 | 0 | 0 | 0 | 0 | 0 | 0 | 0 | 0 |
| 0 | NA | NA | NA | NA | NA | Klra14-ps | | 0 | 0 | 0 | 0 | 0 | 0 | 0 | 0 | 0 | 0 | 0 |
| 0 | NA | NA | NA | NA | NA | Gm23552 | | 0 | 0 | 0 | 0 | 0 | 0 | 0 | 0 | 0 | 0 | 0 |
| 0 | NA | NA | NA | NA | NA | Klra9 | killer cell lectin-like receptor subfamily A, | 0 | 0 | 0 | 0 | 0 | 0 | 0 | 0 | 0 | 0 | 0 |
| 0 | NA | NA | NA | NA | NA | Klra7 | killer cell lectin-like receptor, subfamily A, | 0 | 0 | 0 | 0 | 0 | 0 | 0 | 0 | 0 | 0 | 0 |
| 0 | NA | NA | NA | NA | NA | Klra10 | killer cell lectin-like receptor subfamily A, | 0 | 0 | 0 | 0 | 0 | 0 | 0 | 0 | 0 | 0 | 0 |
| 0 | NA | NA | NA | NA | NA | Klra13-ps | | 0 | 0 | 0 | 0 | 0 | 0 | 0 | 0 | 0 | 0 | 0 |
| 0 | NA | NA | NA | NA | NA | Gm24072 | | 0 | 0 | 0 | 0 | 0 | 0 | 0 | 0 | 0 | 0 | 0 |
| 0 | NA | NA | NA | NA | NA | Klra3 | killer cell lectin-like receptor, subfamily A, | 0 | 0 | 0 | 0 | 0 | 0 | 0 | 0 | 0 | 0 | 0 |
| 0 | NA | NA | NA | NA | NA | Klra1 | killer cell lectin-like receptor, subfamily A, | 0 | 0 | 0 | 0 | 0 | 0 | 0 | 0 | 0 | 0 | 0 |
| 0 | NA | NA | NA | NA | NA | Gm6584 | | 0 | 0 | 0 | 0 | 0 | 0 | 0 | 0 | 0 | 0 | 0 |
| 0 | NA | NA | NA | NA | NA | Gm15861 | | 0 | 0 | 0 | 0 | 0 | 0 | 0 | 0 | 0 | 0 | 0 |
| 0 | NA | NA | NA | NA | NA | Gm6590 | predicted gene 6590 | 0 | 0 | 0 | 0 | 0 | 0 | 0 | 0 | 0 | 0 | 0 |
| 0 | NA | NA | NA | NA | NA | Gm38706 | | 0 | 0 | 0 | 0 | 0 | 0 | 0 | 0 | 0 | 0 | 0 |
| 0 | NA | NA | NA | NA | NA | Gm27013 | | 0 | 0 | 0 | 0 | 0 | 0 | 0 | 0 | 0 | 0 | 0 |
| 0 | NA | NA | NA | NA | NA | Gm44526 | | 0 | 0 | 0 | 0 | 0 | 0 | 0 | 0 | 0 | 0 | 0 |
| 0 | NA | NA | NA | NA | NA | Gm8816 | | 0 | 0 | 0 | 0 | 0 | 0 | 0 | 0 | 0 | 0 | 0 |
| 0 | NA | NA | NA | NA | NA | Gm26996 | | 0 | 0 | 0 | 0 | 0 | 0 | 0 | 0 | 0 | 0 | 0 |
| 0 | NA | NA | NA | NA | NA | Gm26914 | | 0 | 0 | 0 | 0 | 0 | 0 | 0 | 0 | 0 | 0 | 0 |
| 0 | NA | NA | NA | NA | NA | Gm27047 | | 0 | 0 | 0 | 0 | 0 | 0 | 0 | 0 | 0 | 0 | 0 |
| 0 | NA | NA | NA | NA | NA | Gm27013 | | 0 | 0 | 0 | 0 | 0 | 0 | 0 | 0 | 0 | 0 | 0 |
| 0 | NA | NA | NA | NA | NA | Gm20997 | | 0 | 0 | 0 | 0 | 0 | 0 | 0 | 0 | 0 | 0 | 0 |
| 0 | NA | NA | NA | NA | NA | Gm44527 | | 0 | 0 | 0 | 0 | 0 | 0 | 0 | 0 | 0 | 0 | 0 |
| 0 | NA | NA | NA | NA | NA | Gm18423 | | 0 | 0 | 0 | 0 | 0 | 0 | 0 | 0 | 0 | 0 | 0 |
| 0 | NA | NA | NA | NA | NA | Gm44525 | | 0 | 0 | 0 | 0 | 0 | 0 | 0 | 0 | 0 | 0 | 0 |
| 0 | NA | NA | NA | NA | NA | Gm44118 | | 0 | 0 | 0 | 0 | 0 | 0 | 0 | 0 | 0 | 0 | 0 |
| 0 | NA | NA | NA | NA | NA | Gm21940 | | 0 | 0 | 0 | 0 | 0 | 0 | 0 | 0 | 0 | 0 | 0 |
| 0 | NA | NA | NA | NA | NA | Gm33962 | | 0 | 0 | 0 | 0 | 0 | 0 | 0 | 0 | 0 | 0 | 0 |
| 0 | NA | NA | NA | NA | NA | Gm44128 | | 0 | 0 | 0 | 0 | 0 | 0 | 0 | 0 | 0 | 0 | 0 |
| 0 | NA | NA | NA | NA | NA | Gm44988 | | 0 | 0 | 0 | 0 | 0 | 0 | 0 | 0 | 0 | 0 | 0 |
| 0 | NA | NA | NA | NA | NA | Gm44828 | | 0 | 0 | 0 | 0 | 0 | 0 | 0 | 0 | 0 | 0 | 0 |
| 0 | NA | NA | NA | NA | NA | Gm44989 | | 0 | 0 | 0 | 0 | 0 | 0 | 0 | 0 | 0 | 0 | 0 |
| 0 | NA | NA | NA | NA | NA | Gm44133 | | 0 | 0 | 0 | 0 | 0 | 0 | 0 | 0 | 0 | 0 | 0 |
| 0 | NA | NA | NA | NA | NA | Gm44134 | | 0 | 0 | 0 | 0 | 0 | 0 | 0 | 0 | 0 | 0 | 0 |
| 0 | NA | NA | NA | NA | NA | Gm44149 | | 0 | 0 | 0 | 0 | 0 | 0 | 0 | 0 | 0 | 0 | 0 |
| 0 | NA | NA | NA | NA | NA | Gm44150 | | 0 | 0 | 0 | 0 | 0 | 0 | 0 | 0 | 0 | 0 | 0 |
| 0 | NA | NA | NA | NA | NA | Gm8837 | | 0 | 0 | 0 | 0 | 0 | 0 | 0 | 0 | 0 | 0 | 0 |
| 0 | NA | NA | NA | NA | NA | Gm5581 | | 0 | 0 | 0 | 0 | 0 | 0 | 0 | 0 | 0 | 0 | 0 |
| 0 | NA | NA | NA | NA | NA | Gm5582 | | 0 | 0 | 0 | 0 | 0 | 0 | 0 | 0 | 0 | 0 | 0 |
| 0 | NA | NA | NA | NA | NA | Gm44182 | | 0 | 0 | 0 | 0 | 0 | 0 | 0 | 0 | 0 | 0 | 0 |
| 0 | NA | NA | NA | NA | NA | Gm18688 | | 0 | 0 | 0 | 0 | 0 | 0 | 0 | 0 | 0 | 0 | 0 |
| 0 | NA | NA | NA | NA | NA | Gm6619 | predicted gene 6619 | 0 | 0 | 0 | 0 | 0 | 0 | 0 | 0 | 0 | 0 | 0 |
| 0 | NA | NA | NA | NA | NA | 5430401F13Rik | RIKEN cDNA 5430401F13 gene | 0 | 0 | 0 | 0 | 0 | 0 | 0 | 0 | 0 | 0 | 0 |
| 0 | NA | NA | NA | NA | NA | Gm22190 | | 0 | 0 | 0 | 0 | 0 | 0 | 0 | 0 | 0 | 0 | 0 |
| 0 | NA | NA | NA | NA | NA | Gm16571 | | 0 | 0 | 0 | 0 | 0 | 0 | 0 | 0 | 0 | 0 | 0 |
| 0 | NA | NA | NA | NA | NA | Hmgb1l | | 0 | 0 | 0 | 0 | 0 | 0 | 0 | 0 | 0 | 0 | 0 |
| 0 | NA | NA | NA | NA | NA | Gm43943 | | 0 | 0 | 0 | 0 | 0 | 0 | 0 | 0 | 0 | 0 | 0 |
| 0 | NA | NA | NA | NA | NA | Tas2r130 | taste receptor, type 2, member 130 | 0 | 0 | 0 | 0 | 0 | 0 | 0 | 0 | 0 | 0 | 0 |
| 0 | NA | NA | NA | NA | NA | Tas2r107 | taste receptor, type 2, member 107 | 0 | 0 | 0 | 0 | 0 | 0 | 0 | 0 | 0 | 0 | 0 |
| 0 | NA | NA | NA | NA | NA | Tas2r106 | taste receptor, type 2, member 106 | 0 | 0 | 0 | 0 | 0 | 0 | 0 | 0 | 0 | 0 | 0 |
| 0 | NA | NA | NA | NA | NA | Tas2r104 | taste receptor, type 2, member 104 | 0 | 0 | 0 | 0 | 0 | 0 | 0 | 0 | 0 | 0 | 0 |
| 0 | NA | NA | NA | NA | NA | Tas2r105 | taste receptor, type 2, member 105 | 0 | 0 | 0 | 0 | 0 | 0 | 0 | 0 | 0 | 0 | 0 |
| 0 | NA | NA | NA | NA | NA | Tas2r114 | taste receptor, type 2, member 114 | 0 | 0 | 0 | 0 | 0 | 0 | 0 | 0 | 0 | 0 | 0 |
| 0 | NA | NA | NA | NA | NA | Gm43973 | | 0 | 0 | 0 | 0 | 0 | 0 | 0 | 0 | 0 | 0 | 0 |
| 0 | NA | NA | NA | NA | NA | Gm43974 | | 0 | 0 | 0 | 0 | 0 | 0 | 0 | 0 | 0 | 0 | 0 |

| | | | | | | | | | | | | | | | | | |
|---|---|---|---|---|---|---|---|---|---|---|---|---|---|---|---|---|---|
| 0 | NA | NA | NA | NA | NA | Prb3-ps | | 0 | 0 | 0 | 0 | 0 | 0 | 0 | 0 | 0 | 0 | 0 |
| 0 | NA | NA | NA | NA | NA | Gm43975 | | 0 | 0 | 0 | 0 | 0 | 0 | 0 | 0 | 0 | 0 | 0 |
| 0 | NA | NA | NA | NA | NA | Gm4736 | | 0 | 0 | 0 | 0 | 0 | 0 | 0 | 0 | 0 | 0 | 0 |
| 0 | NA | NA | NA | NA | NA | Gm43976 | | 0 | 0 | 0 | 0 | 0 | 0 | 0 | 0 | 0 | 0 | 0 |
| 0 | NA | NA | NA | NA | NA | Gm43978 | | 0 | 0 | 0 | 0 | 0 | 0 | 0 | 0 | 0 | 0 | 0 |
| 0 | NA | NA | NA | NA | NA | Prb1 | proline-rich protein BstNI subfamily 1 | 0 | 0 | 0 | 0 | 0 | 0 | 0 | 0 | 0 | 0 | 0 |
| 0 | NA | NA | NA | NA | NA | Gm44687 | | 0 | 0 | 0 | 0 | 0 | 0 | 0 | 0 | 0 | 0 | 0 |
| 0 | NA | NA | NA | NA | NA | Gm43979 | | 0 | 0 | 0 | 0 | 0 | 0 | 0 | 0 | 0 | 0 | 0 |
| 0 | NA | NA | NA | NA | NA | Prpmp5 | proline-rich protein MP5 | 0 | 0 | 0 | 0 | 0 | 0 | 0 | 0 | 0 | 0 | 0 |
| 0 | NA | NA | NA | NA | NA | Gm44990 | | 0 | 0 | 0 | 0 | 0 | 0 | 0 | 0 | 0 | 0 | 0 |
| 0 | NA | NA | NA | NA | NA | Gm44727 | | 0 | 0 | 0 | 0 | 0 | 0 | 0 | 0 | 0 | 0 | 0 |
| 0 | NA | NA | NA | NA | NA | Gm8882 | predicted gene 8882 | 0 | 0 | 0 | 0 | 0 | 0 | 0 | 0 | 0 | 0 | 0 |
| 0 | NA | NA | NA | NA | NA | Gm44726 | | 0 | 0 | 0 | 0 | 0 | 0 | 0 | 0 | 0 | 0 | 0 |
| 0 | NA | NA | NA | NA | NA | Gm44003 | | 0 | 0 | 0 | 0 | 0 | 0 | 0 | 0 | 0 | 0 | 0 |
| 0 | NA | NA | NA | NA | NA | Gm5154 | | 0 | 0 | 0 | 0 | 0 | 0 | 0 | 0 | 0 | 0 | 0 |
| 0 | NA | NA | NA | NA | NA | Gm45063 | | 0 | 0 | 0 | 0 | 0 | 0 | 0 | 0 | 0 | 0 | 0 |
| 0 | NA | NA | NA | NA | NA | Gm44952 | | 0 | 0 | 0 | 0 | 0 | 0 | 0 | 0 | 0 | 0 | 0 |
| 0 | NA | NA | NA | NA | NA | Gm44070 | | 0 | 0 | 0 | 0 | 0 | 0 | 0 | 0 | 0 | 0 | 0 |
| 0 | NA | NA | NA | NA | NA | n-R5s166 | | 0 | 0 | 0 | 0 | 0 | 0 | 0 | 0 | 0 | 0 | 0 |
| 0 | NA | NA | NA | NA | NA | Prh1 | proline rich protein HaeIII subfamily 1 | 0 | 0 | 0 | 0 | 0 | 0 | 0 | 0 | 0 | 0 | 0 |
| 0 | NA | NA | NA | NA | NA | n-R5s167 | | 0 | 0 | 0 | 0 | 0 | 0 | 0 | 0 | 0 | 0 | 0 |
| 0 | NA | NA | NA | NA | NA | Prp2 | proline rich protein 2 | 0 | 0 | 0 | 0 | 0 | 0 | 0 | 0 | 0 | 0 | 0 |
| 0 | NA | NA | NA | NA | NA | A630073D07Ri | RIKEN cDNA A630073D07 gene | 0 | 0 | 0 | 0 | 0 | 0 | 0 | 0 | 0 | 0 | 0 |
| 0 | NA | NA | NA | NA | NA | Tas2r120 | taste receptor, type 2, member 120 | 0 | 0 | 0 | 0 | 0 | 0 | 0 | 0 | 0 | 0 | 0 |
| 0 | NA | NA | NA | NA | NA | Tas2r121 | taste receptor, type 2, member 121 | 0 | 0 | 0 | 0 | 0 | 0 | 0 | 0 | 0 | 0 | 0 |
| 0 | NA | NA | NA | NA | NA | Tas2r122 | taste receptor, type 2, member 122 | 0 | 0 | 0 | 0 | 0 | 0 | 0 | 0 | 0 | 0 | 0 |
| 0 | NA | NA | NA | NA | NA | Tas2r115 | taste receptor, type 2, member 115 | 0 | 0 | 0 | 0 | 0 | 0 | 0 | 0 | 0 | 0 | 0 |
| 0 | NA | NA | NA | NA | NA | Gm44137 | | 0 | 0 | 0 | 0 | 0 | 0 | 0 | 0 | 0 | 0 | 0 |
| 0 | NA | NA | NA | NA | NA | Tas2r124 | taste receptor, type 2, member 124 | 0 | 0 | 0 | 0 | 0 | 0 | 0 | 0 | 0 | 0 | 0 |
| 0 | NA | NA | NA | NA | NA | Gm44138 | | 0 | 0 | 0 | 0 | 0 | 0 | 0 | 0 | 0 | 0 | 0 |
| 0 | NA | NA | NA | NA | NA | Tas2r102 | taste receptor, type 2, member 102 | 0 | 0 | 0 | 0 | 0 | 0 | 0 | 0 | 0 | 0 | 0 |
| 0 | NA | NA | NA | NA | NA | Tas2r136 | taste receptor, type 2, member 136 | 0 | 0 | 0 | 0 | 0 | 0 | 0 | 0 | 0 | 0 | 0 |
| 0 | NA | NA | NA | NA | NA | Tas2r145-ps3 | | 0 | 0 | 0 | 0 | 0 | 0 | 0 | 0 | 0 | 0 | 0 |
| 0 | NA | NA | NA | NA | NA | Tas2r117 | taste receptor, type 2, member 117 | 0 | 0 | 0 | 0 | 0 | 0 | 0 | 0 | 0 | 0 | 0 |
| 0 | NA | NA | NA | NA | NA | Tas2r111-ps2 | | 0 | 0 | 0 | 0 | 0 | 0 | 0 | 0 | 0 | 0 | 0 |
| 0 | NA | NA | NA | NA | NA | Gm44139 | | 0 | 0 | 0 | 0 | 0 | 0 | 0 | 0 | 0 | 0 | 0 |
| 0 | NA | NA | NA | NA | NA | Tas2r123 | taste receptor, type 2, member 123 | 0 | 0 | 0 | 0 | 0 | 0 | 0 | 0 | 0 | 0 | 0 |
| 0 | NA | NA | NA | NA | NA | Tas2r116 | taste receptor, type 2, member 116 | 0 | 0 | 0 | 0 | 0 | 0 | 0 | 0 | 0 | 0 | 0 |
| 0 | NA | NA | NA | NA | NA | Gm44142 | | 0 | 0 | 0 | 0 | 0 | 0 | 0 | 0 | 0 | 0 | 0 |
| 0 | NA | NA | NA | NA | NA | Tas2r110 | taste receptor, type 2, member 110 | 0 | 0 | 0 | 0 | 0 | 0 | 0 | 0 | 0 | 0 | 0 |
| 0 | NA | NA | NA | NA | NA | Tas2r113 | taste receptor, type 2, member 113 | 0 | 0 | 0 | 0 | 0 | 0 | 0 | 0 | 0 | 0 | 0 |
| 0 | NA | NA | NA | NA | NA | Tas2r142-ps5 | | 0 | 0 | 0 | 0 | 0 | 0 | 0 | 0 | 0 | 0 | 0 |
| 0 | NA | NA | NA | NA | NA | Tas2r125 | taste receptor, type 2, member 125 | 0 | 0 | 0 | 0 | 0 | 0 | 0 | 0 | 0 | 0 | 0 |
| 0 | NA | NA | NA | NA | NA | Tas2r146-ps1 | | 0 | 0 | 0 | 0 | 0 | 0 | 0 | 0 | 0 | 0 | 0 |
| 0 | NA | NA | NA | NA | NA | Tas2r129 | taste receptor, type 2, member 129 | 0 | 0 | 0 | 0 | 0 | 0 | 0 | 0 | 0 | 0 | 0 |
| 0 | NA | NA | NA | NA | NA | Tas2r131 | taste receptor, type 2, member 131 | 0 | 0 | 0 | 0 | 0 | 0 | 0 | 0 | 0 | 0 | 0 |
| 0 | NA | NA | NA | NA | NA | Gm44425 | | 0 | 0 | 0 | 0 | 0 | 0 | 0 | 0 | 0 | 0 | 0 |
| 0 | NA | NA | NA | NA | NA | Tas2r109 | taste receptor, type 2, member 109 | 0 | 0 | 0 | 0 | 0 | 0 | 0 | 0 | 0 | 0 | 0 |
| 0 | NA | NA | NA | NA | NA | Gm44426 | | 0 | 0 | 0 | 0 | 0 | 0 | 0 | 0 | 0 | 0 | 0 |
| 0 | NA | NA | NA | NA | NA | Gm18689 | | 0 | 0 | 0 | 0 | 0 | 0 | 0 | 0 | 0 | 0 | 0 |
| 0 | NA | NA | NA | NA | NA | Gm18690 | | 0 | 0 | 0 | 0 | 0 | 0 | 0 | 0 | 0 | 0 | 0 |
| 0 | NA | NA | NA | NA | NA | Tas2r103 | taste receptor, type 2, member 103 | 0 | 0 | 0 | 0 | 0 | 0 | 0 | 0 | 0 | 0 | 0 |
| 0 | NA | NA | NA | NA | NA | Tas2r140 | taste receptor, type 2, member 140 | 0 | 0 | 0 | 0 | 0 | 0 | 0 | 0 | 0 | 0 | 0 |
| 0 | NA | NA | NA | NA | NA | Gm18691 | | 0 | 0 | 0 | 0 | 0 | 0 | 0 | 0 | 0 | 0 | 0 |
| 0 | NA | NA | NA | NA | NA | Gm8921 | | 0 | 0 | 0 | 0 | 0 | 0 | 0 | 0 | 0 | 0 | 0 |
| 0 | NA | NA | NA | NA | NA | Tas2r141-ps4 | | 0 | 0 | 0 | 0 | 0 | 0 | 0 | 0 | 0 | 0 | 0 |
| 0 | NA | NA | NA | NA | NA | Gm7039 | | 0 | 0 | 0 | 0 | 0 | 0 | 0 | 0 | 0 | 0 | 0 |
| 0 | NA | NA | NA | NA | NA | Gm23403 | | 0 | 0 | 0 | 0 | 0 | 0 | 0 | 0 | 0 | 0 | 0 |
| 0 | NA | NA | NA | NA | NA | Gm25019 | | 0 | 0 | 0 | 0 | 0 | 0 | 0 | 0 | 0 | 0 | 0 |
| 0 | NA | NA | NA | NA | NA | 5530400C23Ril | RIKEN cDNA 5530400C23 gene | 0 | 0 | 0 | 0 | 0 | 0 | 0 | 0 | 0 | 0 | 0 |

| | | | | | | | | | | | | | | | | | | |
|---|---|---|---|---|---|---|---|---|---|---|---|---|---|---|---|---|---|---|
| 0 | NA | NA | NA | NA | NA | Gm34149 | | 0 | 0 | 0 | 0 | 0 | 0 | 0 | 0 | 0 | 0 | 0 |
| 0 | NA | NA | NA | NA | NA | Gm44239 | | 0 | 0 | 0 | 0 | 0 | 0 | 0 | 0 | 0 | 0 | 0 |
| 0 | NA | NA | NA | NA | NA | Gm5885 | predicted gene 5885 | 0 | 0 | 0 | 0 | 0 | 0 | 0 | 0 | 0 | 0 | 0 |
| 0 | NA | NA | NA | NA | NA | Gm44245 | | 0 | 0 | 0 | 0 | 0 | 0 | 0 | 0 | 0 | 0 | 0 |
| 0 | NA | NA | NA | NA | NA | Gm34224 | | 0 | 0 | 0 | 0 | 0 | 0 | 0 | 0 | 0 | 0 | 0 |
| 0 | NA | NA | NA | NA | NA | Gm34314 | | 0 | 0 | 0 | 0 | 0 | 0 | 0 | 0 | 0 | 0 | 0 |
| 0 | NA | NA | NA | NA | NA | Gm44247 | | 0 | 0 | 0 | 0 | 0 | 0 | 0 | 0 | 0 | 0 | 0 |
| 0 | NA | NA | NA | NA | NA | Gm44252 | | 0 | 0 | 0 | 0 | 0 | 0 | 0 | 0 | 0 | 0 | 0 |
| 0 | NA | NA | NA | NA | NA | Gm23375 | | 0 | 0 | 0 | 0 | 0 | 0 | 0 | 0 | 0 | 0 | 0 |
| 0 | NA | NA | NA | NA | NA | Gm5886 | | 0 | 0 | 0 | 0 | 0 | 0 | 0 | 0 | 0 | 0 | 0 |
| 0 | NA | NA | NA | NA | NA | Kap | kidney androgen regulated protein | 0 | 0 | 0 | 0 | 0 | 0 | 0 | 0 | 0 | 0 | 0 |
| 0 | NA | NA | NA | NA | NA | Gm43968 | | 0 | 0 | 0 | 0 | 0 | 0 | 0 | 0 | 0 | 0 | 0 |
| 0 | NA | NA | NA | NA | NA | Gm8956 | | 0 | 0 | 0 | 0 | 0 | 0 | 0 | 0 | 0 | 0 | 0 |
| 0 | NA | NA | NA | NA | NA | Gm43984 | | 0 | 0 | 0 | 0 | 0 | 0 | 0 | 0 | 0 | 0 | 0 |
| 0 | NA | NA | NA | NA | NA | Gm17088 | | 0 | 0 | 0 | 0 | 0 | 0 | 0 | 0 | 0 | 0 | 0 |
| 0 | NA | NA | NA | NA | NA | Gm17089 | | 0 | 0 | 0 | 0 | 0 | 0 | 0 | 0 | 0 | 0 | 0 |
| 0 | NA | NA | NA | NA | NA | 1700051K13Rik | | 0 | 0 | 0 | 0 | 0 | 0 | 0 | 0 | 0 | 0 | 0 |
| 0 | NA | NA | NA | NA | NA | 4933406J09Rik | | 0 | 0 | 0 | 0 | 0 | 0 | 0 | 0 | 0 | 0 | 0 |
| 0 | NA | NA | NA | NA | NA | Gm44448 | | 0 | 0 | 0 | 0 | 0 | 0 | 0 | 0 | 0 | 0 | 0 |
| 0 | NA | NA | NA | NA | NA | Gm43952 | | 0 | 0 | 0 | 0 | 0 | 0 | 0 | 0 | 0 | 0 | 0 |
| 0 | NA | NA | NA | NA | NA | Gm36328 | | 0 | 0 | 0 | 0 | 0 | 0 | 0 | 0 | 0 | 0 | 0 |
| 0 | NA | NA | NA | NA | NA | Gm44238 | | 0 | 0 | 0 | 0 | 0 | 0 | 0 | 0 | 0 | 0 | 0 |
| 0 | NA | NA | NA | NA | NA | Gm29803 | | 0 | 0 | 0 | 0 | 0 | 0 | 0 | 0 | 0 | 0 | 0 |
| 0 | NA | NA | NA | NA | NA | Gm22881 | | 0 | 0 | 0 | 0 | 0 | 0 | 0 | 0 | 0 | 0 | 0 |
| 0 | NA | NA | NA | NA | NA | Gm34391 | | 0 | 0 | 0 | 0 | 0 | 0 | 0 | 0 | 0 | 0 | 0 |
| 0 | NA | NA | NA | NA | NA | Pbp2 | phosphatidylethanolamine binding protei | 0 | 0 | 0 | 0 | 0 | 0 | 0 | 0 | 0 | 0 | 0 |
| 0 | NA | NA | NA | NA | NA | Gm44274 | | 0 | 0 | 0 | 0 | 0 | 0 | 0 | 0 | 0 | 0 | 0 |
| 0 | NA | NA | NA | NA | NA | Gm44275 | | 0 | 0 | 0 | 0 | 0 | 0 | 0 | 0 | 0 | 0 | 0 |
| 0 | NA | NA | NA | NA | NA | Gm25136 | | 0 | 0 | 0 | 0 | 0 | 0 | 0 | 0 | 0 | 0 | 0 |
| 0 | NA | NA | NA | NA | NA | 4930425L21Rik | | 0 | 0 | 0 | 0 | 0 | 0 | 0 | 0 | 0 | 0 | 0 |
| 0 | NA | NA | NA | NA | NA | Gm14329 | | 0 | 0 | 0 | 0 | 0 | 0 | 0 | 0 | 0 | 0 | 0 |
| 0 | NA | NA | NA | NA | NA | Gm22892 | | 0 | 0 | 0 | 0 | 0 | 0 | 0 | 0 | 0 | 0 | 0 |
| 0 | NA | NA | NA | NA | NA | Gm14330 | | 0 | 0 | 0 | 0 | 0 | 0 | 0 | 0 | 0 | 0 | 0 |
| 0 | NA | NA | NA | NA | NA | Gm26653 | | 0 | 0 | 0 | 0 | 0 | 0 | 0 | 0 | 0 | 0 | 0 |
| 0 | NA | NA | NA | NA | NA | Gm26105 | | 0 | 0 | 0 | 0 | 0 | 0 | 0 | 0 | 0 | 0 | 0 |
| 0 | NA | NA | NA | NA | NA | Gm15476 | | 0 | 0 | 0 | 0 | 0 | 0 | 0 | 0 | 0 | 0 | 0 |
| 0 | NA | NA | NA | NA | NA | E330021D16Ri | RIKEN cDNA E330021D16 gene | 0 | 0 | 0 | 0 | 0 | 0 | 0 | 0 | 0 | 0 | 0 |
| 0 | NA | NA | NA | NA | NA | Gm25882 | | 0 | 0 | 0 | 0 | 0 | 0 | 0 | 0 | 0 | 0 | 0 |
| 0 | NA | NA | NA | NA | NA | Gm44131 | | 0 | 0 | 0 | 0 | 0 | 0 | 0 | 0 | 0 | 0 | 0 |
| 0 | NA | NA | NA | NA | NA | Gm44401 | | 0 | 0 | 0 | 0 | 0 | 0 | 0 | 0 | 0 | 0 | 0 |
| 0 | NA | NA | NA | NA | NA | Gm43969 | | 0 | 0 | 0 | 0 | 0 | 0 | 0 | 0 | 0 | 0 | 0 |
| 0 | NA | NA | NA | NA | NA | Gm44069 | | 0 | 0 | 0 | 0 | 0 | 0 | 0 | 0 | 0 | 0 | 0 |
| 0 | NA | NA | NA | NA | NA | 4930480K02Rik | | 0 | 0 | 0 | 0 | 0 | 0 | 0 | 0 | 0 | 0 | 0 |
| 0 | NA | NA | NA | NA | NA | Gm44258 | | 0 | 0 | 0 | 0 | 0 | 0 | 0 | 0 | 0 | 0 | 0 |
| 0 | NA | NA | NA | NA | NA | Gm44106 | | 0 | 0 | 0 | 0 | 0 | 0 | 0 | 0 | 0 | 0 | 0 |
| 0 | NA | NA | NA | NA | NA | Gm24308 | | 0 | 0 | 0 | 0 | 0 | 0 | 0 | 0 | 0 | 0 | 0 |
| 0 | NA | NA | NA | NA | NA | Slc15a5 | solute carrier family 15, member 5 | 0 | 0 | 0 | 0 | 0 | 0 | 0 | 0 | 0 | 0 | 0 |
| 0 | NA | NA | NA | NA | NA | Gm30332 | | 0 | 0 | 0 | 0 | 0 | 0 | 0 | 0 | 0 | 0 | 0 |
| 0 | NA | NA | NA | NA | NA | B230110G15Rik | | 0 | 0 | 0 | 0 | 0 | 0 | 0 | 0 | 0 | 0 | 0 |
| 0 | NA | NA | NA | NA | NA | Gm44266 | | 0 | 0 | 0 | 0 | 0 | 0 | 0 | 0 | 0 | 0 | 0 |
| 0 | NA | NA | NA | NA | NA | Igbp1b | immunoglobulin (CD79A) binding protein | 0 | 0 | 0 | 0 | 0 | 0 | 0 | 0 | 0 | 0 | 0 |
| 0 | NA | NA | NA | NA | NA | Gm9038 | | 0 | 0 | 0 | 0 | 0 | 0 | 0 | 0 | 0 | 0 | 0 |
| 0 | NA | NA | NA | NA | NA | Gm44267 | | 0 | 0 | 0 | 0 | 0 | 0 | 0 | 0 | 0 | 0 | 0 |
| 0 | NA | NA | NA | NA | NA | Gm44268 | | 0 | 0 | 0 | 0 | 0 | 0 | 0 | 0 | 0 | 0 | 0 |
| 0 | NA | NA | NA | NA | NA | Gm25434 | | 0 | 0 | 0 | 0 | 0 | 0 | 0 | 0 | 0 | 0 | 0 |
| 0 | NA | NA | NA | NA | NA | 4922502N22Rik | | 0 | 0 | 0 | 0 | 0 | 0 | 0 | 0 | 0 | 0 | 0 |
| 0 | NA | NA | NA | NA | NA | 4930404I20Rik | | 0 | 0 | 0 | 0 | 0 | 0 | 0 | 0 | 0 | 0 | 0 |
| 0 | NA | NA | NA | NA | NA | 4930443G03Rik | | 0 | 0 | 0 | 0 | 0 | 0 | 0 | 0 | 0 | 0 | 0 |
| 0 | NA | NA | NA | NA | NA | Gm43940 | | 0 | 0 | 0 | 0 | 0 | 0 | 0 | 0 | 0 | 0 | 0 |
| 0 | NA | NA | NA | NA | NA | Gm44112 | | 0 | 0 | 0 | 0 | 0 | 0 | 0 | 0 | 0 | 0 | 0 |

| | | | | | | | | | | | | | | | | | | |
|---|---|---|---|---|---|---|---|---|---|---|---|---|---|---|---|---|---|---|
| 0 | NA | NA | NA | NA | NA | Capza3 | capping protein (actin filament) muscle Z- | 0 | 0 | 0 | 0 | 0 | 0 | 0 | 0 | 0 | 0 | 0 |
| 0 | NA | NA | NA | NA | NA | Gm44156 | | 0 | 0 | 0 | 0 | 0 | 0 | 0 | 0 | 0 | 0 | 0 |
| 0 | NA | NA | NA | NA | NA | Gm44155 | | 0 | 0 | 0 | 0 | 0 | 0 | 0 | 0 | 0 | 0 | 0 |
| 0 | NA | NA | NA | NA | NA | Gm44158 | | 0 | 0 | 0 | 0 | 0 | 0 | 0 | 0 | 0 | 0 | 0 |
| 0 | NA | NA | NA | NA | NA | Gm44159 | | 0 | 0 | 0 | 0 | 0 | 0 | 0 | 0 | 0 | 0 | 0 |
| 0 | NA | NA | NA | NA | NA | Gm24174 | | 0 | 0 | 0 | 0 | 0 | 0 | 0 | 0 | 0 | 0 | 0 |
| 0 | NA | NA | NA | NA | NA | Gm3961 | | 0 | 0 | 0 | 0 | 0 | 0 | 0 | 0 | 0 | 0 | 0 |
| 0 | NA | NA | NA | NA | NA | 1700102F20Rik | | 0 | 0 | 0 | 0 | 0 | 0 | 0 | 0 | 0 | 0 | 0 |
| 0 | NA | NA | NA | NA | NA | n-R5s168 | | 0 | 0 | 0 | 0 | 0 | 0 | 0 | 0 | 0 | 0 | 0 |
| 0 | NA | NA | NA | NA | NA | Gm11077 | | 0 | 0 | 0 | 0 | 0 | 0 | 0 | 0 | 0 | 0 | 0 |
| 0 | NA | NA | NA | NA | NA | Gm43925 | | 0 | 0 | 0 | 0 | 0 | 0 | 0 | 0 | 0 | 0 | 0 |
| 0 | NA | NA | NA | NA | NA | Gm43927 | | 0 | 0 | 0 | 0 | 0 | 0 | 0 | 0 | 0 | 0 | 0 |
| 0 | NA | NA | NA | NA | NA | Gm10400 | | 0 | 0 | 0 | 0 | 0 | 0 | 0 | 0 | 0 | 0 | 0 |
| 0 | NA | NA | NA | NA | NA | Gm43958 | | 0 | 0 | 0 | 0 | 0 | 0 | 0 | 0 | 0 | 0 | 0 |
| 0 | NA | NA | NA | NA | NA | Gm23069 | | 0 | 0 | 0 | 0 | 0 | 0 | 0 | 0 | 0 | 0 | 0 |
| 0 | NA | NA | NA | NA | NA | Gm30784 | | 0 | 0 | 0 | 0 | 0 | 0 | 0 | 0 | 0 | 0 | 0 |
| 0 | NA | NA | NA | NA | NA | Gm22036 | | 0 | 0 | 0 | 0 | 0 | 0 | 0 | 0 | 0 | 0 | 0 |
| 0 | NA | NA | NA | NA | NA | Gm23775 | | 0 | 0 | 0 | 0 | 0 | 0 | 0 | 0 | 0 | 0 | 0 |
| 0 | NA | NA | NA | NA | NA | Gm18159 | | 0 | 0 | 0 | 0 | 0 | 0 | 0 | 0 | 0 | 0 | 0 |
| 0 | NA | NA | NA | NA | NA | 4930434O05Rik | | 0 | 0 | 0 | 0 | 0 | 0 | 0 | 0 | 0 | 0 | 0 |
| 0 | NA | NA | NA | NA | NA | Gm31108 | | 0 | 0 | 0 | 0 | 0 | 0 | 0 | 0 | 0 | 0 | 0 |
| 0 | NA | NA | NA | NA | NA | Gm44312 | | 0 | 0 | 0 | 0 | 0 | 0 | 0 | 0 | 0 | 0 | 0 |
| 0 | NA | NA | NA | NA | NA | Gm44306 | | 0 | 0 | 0 | 0 | 0 | 0 | 0 | 0 | 0 | 0 | 0 |
| 0 | NA | NA | NA | NA | NA | Gm23272 | | 0 | 0 | 0 | 0 | 0 | 0 | 0 | 0 | 0 | 0 | 0 |
| 0 | NA | NA | NA | NA | NA | 4930579D09Rik | | 0 | 0 | 0 | 0 | 0 | 0 | 0 | 0 | 0 | 0 | 0 |
| 0 | NA | NA | NA | NA | NA | Gm44068 | | 0 | 0 | 0 | 0 | 0 | 0 | 0 | 0 | 0 | 0 | 0 |
| 0 | NA | NA | NA | NA | NA | Sox5os1 | | 0 | 0 | 0 | 0 | 0 | 0 | 0 | 0 | 0 | 0 | 0 |
| 0 | NA | NA | NA | NA | NA | Sox5os2 | | 0 | 0 | 0 | 0 | 0 | 0 | 0 | 0 | 0 | 0 | 0 |
| 0 | NA | NA | NA | NA | NA | Sox5it | | 0 | 0 | 0 | 0 | 0 | 0 | 0 | 0 | 0 | 0 | 0 |
| 0 | NA | NA | NA | NA | NA | Sox5os3 | | 0 | 0 | 0 | 0 | 0 | 0 | 0 | 0 | 0 | 0 | 0 |
| 0 | NA | NA | NA | NA | NA | Sox5os4 | | 0 | 0 | 0 | 0 | 0 | 0 | 0 | 0 | 0 | 0 | 0 |
| 0 | NA | NA | NA | NA | NA | Sox5os5 | | 0 | 0 | 0 | 0 | 0 | 0 | 0 | 0 | 0 | 0 | 0 |
| 0 | NA | NA | NA | NA | NA | Gm22792 | | 0 | 0 | 0 | 0 | 0 | 0 | 0 | 0 | 0 | 0 | 0 |
| 0 | NA | NA | NA | NA | NA | Gm23071 | | 0 | 0 | 0 | 0 | 0 | 0 | 0 | 0 | 0 | 0 | 0 |
| 0 | NA | NA | NA | NA | NA | Gm26666 | | 0 | 0 | 0 | 0 | 0 | 0 | 0 | 0 | 0 | 0 | 0 |
| 0 | NA | NA | NA | NA | NA | Gm15687 | | 0 | 0 | 0 | 0 | 0 | 0 | 0 | 0 | 0 | 0 | 0 |
| 0 | NA | NA | NA | NA | NA | Gm43910 | | 0 | 0 | 0 | 0 | 0 | 0 | 0 | 0 | 0 | 0 | 0 |
| 0 | NA | NA | NA | NA | NA | Gm15543 | | 0 | 0 | 0 | 0 | 0 | 0 | 0 | 0 | 0 | 0 | 0 |
| 0 | NA | NA | NA | NA | NA | Gm22793 | | 0 | 0 | 0 | 0 | 0 | 0 | 0 | 0 | 0 | 0 | 0 |
| 0 | NA | NA | NA | NA | NA | 2010013B24Rik | | 0 | 0 | 0 | 0 | 0 | 0 | 0 | 0 | 0 | 0 | 0 |
| 0 | NA | NA | NA | NA | NA | Gm15707 | | 0 | 0 | 0 | 0 | 0 | 0 | 0 | 0 | 0 | 0 | 0 |
| 0 | NA | NA | NA | NA | NA | 1700073E17Rik | | 0 | 0 | 0 | 0 | 0 | 0 | 0 | 0 | 0 | 0 | 0 |
| 0 | NA | NA | NA | NA | NA | Gm15499 | | 0 | 0 | 0 | 0 | 0 | 0 | 0 | 0 | 0 | 0 | 0 |
| 0 | NA | NA | NA | NA | NA | Gm25373 | | 0 | 0 | 0 | 0 | 0 | 0 | 0 | 0 | 0 | 0 | 0 |
| 0 | NA | NA | NA | NA | NA | Tuba3b | tubulin, alpha 3B | 0 | 0 | 0 | 0 | 0 | 0 | 0 | 0 | 0 | 0 | 0 |
| 0 | NA | NA | NA | NA | NA | Gm44087 | | 0 | 0 | 0 | 0 | 0 | 0 | 0 | 0 | 0 | 0 | 0 |
| 0 | NA | NA | NA | NA | NA | 4930447C11Rik | | 0 | 0 | 0 | 0 | 0 | 0 | 0 | 0 | 0 | 0 | 0 |
| 0 | NA | NA | NA | NA | NA | Smco2 | single-pass membrane protein with coilec | 0 | 0 | 0 | 0 | 0 | 0 | 0 | 0 | 0 | 0 | 0 |
| 0 | NA | NA | NA | NA | NA | 1700034J05Rik | RIKEN cDNA 1700034J05 gene | 0 | 0 | 0 | 0 | 0 | 0 | 0 | 0 | 0 | 0 | 0 |
| 0 | NA | NA | NA | NA | NA | Gm7571 | | 0 | 0 | 0 | 0 | 0 | 0 | 0 | 0 | 0 | 0 | 0 |
| 0 | NA | NA | NA | NA | NA | E330012B07Rik | | 0 | 0 | 0 | 0 | 0 | 0 | 0 | 0 | 0 | 0 | 0 |
| 0 | NA | NA | NA | NA | NA | 4933406L23Rik | | 0 | 0 | 0 | 0 | 0 | 0 | 0 | 0 | 0 | 0 | 0 |
| 0 | NA | NA | NA | NA | NA | Far2os2 | | 0 | 0 | 0 | 0 | 0 | 0 | 0 | 0 | 0 | 0 | 0 |
| 0 | NA | NA | NA | NA | NA | Far2os1 | | 0 | 0 | 0 | 0 | 0 | 0 | 0 | 0 | 0 | 0 | 0 |
| 0 | NA | NA | NA | NA | NA | Gm6313 | | 0 | 0 | 0 | 0 | 0 | 0 | 0 | 0 | 0 | 0 | 0 |
| 0 | NA | NA | NA | NA | NA | Gm23456 | | 0 | 0 | 0 | 0 | 0 | 0 | 0 | 0 | 0 | 0 | 0 |
| 0 | NA | NA | NA | NA | NA | 4930528G23Rik | | 0 | 0 | 0 | 0 | 0 | 0 | 0 | 0 | 0 | 0 | 0 |
| 0 | NA | NA | NA | NA | NA | Gm15778 | | 0 | 0 | 0 | 0 | 0 | 0 | 0 | 0 | 0 | 0 | 0 |
| 0 | NA | NA | NA | NA | NA | Gm44115 | | 0 | 0 | 0 | 0 | 0 | 0 | 0 | 0 | 0 | 0 | 0 |
| 0 | NA | NA | NA | NA | NA | Gm43908 | | 0 | 0 | 0 | 0 | 0 | 0 | 0 | 0 | 0 | 0 | 0 |

| | | | | | | | | | | | | | | | | | | |
|---|---|---|---|---|---|---|---|---|---|---|---|---|---|---|---|---|---|---|
| 0 | NA | NA | NA | NA | NA | Gm43913 | | 0 | 0 | 0 | 0 | 0 | 0 | 0 | 0 | 0 | 0 | 0 |
| 0 | NA | NA | NA | NA | NA | Gm25539 | | 0 | 0 | 0 | 0 | 0 | 0 | 0 | 0 | 0 | 0 | 0 |
| 0 | NA | NA | NA | NA | NA | Gm15781 | | 0 | 0 | 0 | 0 | 0 | 0 | 0 | 0 | 0 | 0 | 0 |
| 0 | NA | NA | NA | NA | NA | Gm22578 | | 0 | 0 | 0 | 0 | 0 | 0 | 0 | 0 | 0 | 0 | 0 |
| 0 | NA | NA | NA | NA | NA | Gm15780 | | 0 | 0 | 0 | 0 | 0 | 0 | 0 | 0 | 0 | 0 | 0 |
| 0 | NA | NA | NA | NA | NA | Gm23462 | | 0 | 0 | 0 | 0 | 0 | 0 | 0 | 0 | 0 | 0 | 0 |
| 0 | NA | NA | NA | NA | NA | Gm10203 | | 0 | 0 | 0 | 0 | 0 | 0 | 0 | 0 | 0 | 0 | 0 |
| 0 | NA | NA | NA | NA | NA | 1700003I16Rik | | 0 | 0 | 0 | 0 | 0 | 0 | 0 | 0 | 0 | 0 | 0 |
| 0 | NA | NA | NA | NA | NA | Gm10388 | | 0 | 0 | 0 | 0 | 0 | 0 | 0 | 0 | 0 | 0 | 0 |
| 0 | NA | NA | NA | NA | NA | Gm15786 | | 0 | 0 | 0 | 0 | 0 | 0 | 0 | 0 | 0 | 0 | 0 |
| 0 | NA | NA | NA | NA | NA | Gm15783 | | 0 | 0 | 0 | 0 | 0 | 0 | 0 | 0 | 0 | 0 | 0 |
| 0 | NA | NA | NA | NA | NA | Gm44473 | | 0 | 0 | 0 | 0 | 0 | 0 | 0 | 0 | 0 | 0 | 0 |
| 0 | NA | NA | NA | NA | NA | Gm3104 | | 0 | 0 | 0 | 0 | 0 | 0 | 0 | 0 | 0 | 0 | 0 |
| 0 | NA | NA | NA | NA | NA | Gm18198 | | 0 | 0 | 0 | 0 | 0 | 0 | 0 | 0 | 0 | 0 | 0 |
| 0 | NA | NA | NA | NA | NA | Gm6727 | | 0 | 0 | 0 | 0 | 0 | 0 | 0 | 0 | 0 | 0 | 0 |
| 0 | NA | NA | NA | NA | NA | Gm44451 | | 0 | 0 | 0 | 0 | 0 | 0 | 0 | 0 | 0 | 0 | 0 |
| 0 | NA | NA | NA | NA | NA | Gm44459 | | 0 | 0 | 0 | 0 | 0 | 0 | 0 | 0 | 0 | 0 | 0 |
| 0 | NA | NA | NA | NA | NA | Speer9-ps1 | | 0 | 0 | 0 | 0 | 0 | 0 | 0 | 0 | 0 | 0 | 0 |
| 0 | NA | NA | NA | NA | NA | Gm29708 | | 0 | 0 | 0 | 0 | 0 | 0 | 0 | 0 | 0 | 0 | 0 |
| 0 | NA | NA | NA | NA | NA | AU018091 | expressed sequence AU018091 | 0 | 0 | 0 | 0 | 0 | 0 | 0 | 0 | 0 | 0 | 0 |
| 0 | NA | NA | NA | NA | NA | Mir290a | | 0 | 0 | 0 | 0 | 0 | 0 | 0 | 0 | 0 | 0 | 0 |
| 0 | NA | NA | NA | NA | NA | Mir290b | | 0 | 0 | 0 | 0 | 0 | 0 | 0 | 0 | 0 | 0 | 0 |
| 0 | NA | NA | NA | NA | NA | Mir291a | | 0 | 0 | 0 | 0 | 0 | 0 | 0 | 0 | 0 | 0 | 0 |
| 0 | NA | NA | NA | NA | NA | Mir292 | | 0 | 0 | 0 | 0 | 0 | 0 | 0 | 0 | 0 | 0 | 0 |
| 0 | NA | NA | NA | NA | NA | Mir292b | | 0 | 0 | 0 | 0 | 0 | 0 | 0 | 0 | 0 | 0 | 0 |
| 0 | NA | NA | NA | NA | NA | Mir291b | | 0 | 0 | 0 | 0 | 0 | 0 | 0 | 0 | 0 | 0 | 0 |
| 0 | NA | NA | NA | NA | NA | Mir293 | | 0 | 0 | 0 | 0 | 0 | 0 | 0 | 0 | 0 | 0 | 0 |
| 0 | NA | NA | NA | NA | NA | Mir294 | | 0 | 0 | 0 | 0 | 0 | 0 | 0 | 0 | 0 | 0 | 0 |
| 0 | NA | NA | NA | NA | NA | Mir295 | | 0 | 0 | 0 | 0 | 0 | 0 | 0 | 0 | 0 | 0 | 0 |
| 0 | NA | NA | NA | NA | NA | Gm20553 | | 0 | 0 | 0 | 0 | 0 | 0 | 0 | 0 | 0 | 0 | 0 |
| 0 | NA | NA | NA | NA | NA | Gm44256 | | 0 | 0 | 0 | 0 | 0 | 0 | 0 | 0 | 0 | 0 | 0 |
| 0 | NA | NA | NA | NA | NA | Gm44257 | | 0 | 0 | 0 | 0 | 0 | 0 | 0 | 0 | 0 | 0 | 0 |
| 0 | NA | NA | NA | NA | NA | Gm44151 | | 0 | 0 | 0 | 0 | 0 | 0 | 0 | 0 | 0 | 0 | 0 |
| 0 | NA | NA | NA | NA | NA | Gm23450 | | 0 | 0 | 0 | 0 | 0 | 0 | 0 | 0 | 0 | 0 | 0 |
| 0 | NA | NA | NA | NA | NA | 3300002P13Rik | | 0 | 0 | 0 | 0 | 0 | 0 | 0 | 0 | 0 | 0 | 0 |
| 0 | NA | NA | NA | NA | NA | Gm44165 | | 0 | 0 | 0 | 0 | 0 | 0 | 0 | 0 | 0 | 0 | 0 |
| 0 | NA | NA | NA | NA | NA | | | 0 | 0 | 0 | 0 | 0 | 0 | 0 | 0 | 0 | 0 | 0 |
| 0 | NA | NA | NA | NA | NA | Gm44921 | | 0 | 0 | 0 | 0 | 0 | 0 | 0 | 0 | 0 | 0 | 0 |
| 0 | NA | NA | NA | NA | NA | Gm25100 | | 0 | 0 | 0 | 0 | 0 | 0 | 0 | 0 | 0 | 0 | 0 |
| 0 | NA | NA | NA | NA | NA | Gm44920 | | 0 | 0 | 0 | 0 | 0 | 0 | 0 | 0 | 0 | 0 | 0 |
| 0 | NA | NA | NA | NA | NA | Mir3572 | | 0 | 0 | 0 | 0 | 0 | 0 | 0 | 0 | 0 | 0 | 0 |
| 0 | NA | NA | NA | NA | NA | Gm15930 | | 0 | 0 | 0 | 0 | 0 | 0 | 0 | 0 | 0 | 0 | 0 |
| 0 | NA | NA | NA | NA | NA | Gm15925 | | 0 | 0 | 0 | 0 | 0 | 0 | 0 | 0 | 0 | 0 | 0 |
| 0 | NA | NA | NA | NA | NA | Gm44581 | | 0 | 0 | 0 | 0 | 0 | 0 | 0 | 0 | 0 | 0 | 0 |
| 0 | NA | NA | NA | NA | NA | Gm44580 | | 0 | 0 | 0 | 0 | 0 | 0 | 0 | 0 | 0 | 0 | 0 |
| 0 | NA | NA | NA | NA | NA | Gm3189 | | 0 | 0 | 0 | 0 | 0 | 0 | 0 | 0 | 0 | 0 | 0 |
| 0 | NA | NA | NA | NA | NA | 1700095K22Rik | | 0 | 0 | 0 | 0 | 0 | 0 | 0 | 0 | 0 | 0 | 0 |
| 0 | NA | NA | NA | NA | NA | Gm15928 | | 0 | 0 | 0 | 0 | 0 | 0 | 0 | 0 | 0 | 0 | 0 |
| 0 | NA | NA | NA | NA | NA | Gm23741 | | 0 | 0 | 0 | 0 | 0 | 0 | 0 | 0 | 0 | 0 | 0 |
| 0 | NA | NA | NA | NA | NA | Gm44807 | | 0 | 0 | 0 | 0 | 0 | 0 | 0 | 0 | 0 | 0 | 0 |
| 0 | NA | NA | NA | NA | NA | Gm15926 | | 0 | 0 | 0 | 0 | 0 | 0 | 0 | 0 | 0 | 0 | 0 |
| 0 | NA | NA | NA | NA | NA | Gm15923 | | 0 | 0 | 0 | 0 | 0 | 0 | 0 | 0 | 0 | 0 | 0 |
| 0 | NA | NA | NA | NA | NA | Gm44881 | | 0 | 0 | 0 | 0 | 0 | 0 | 0 | 0 | 0 | 0 | 0 |
| 0 | NA | NA | NA | NA | NA | Gm45013 | | 0 | 0 | 0 | 0 | 0 | 0 | 0 | 0 | 0 | 0 | 0 |
| 0 | NA | NA | NA | NA | NA | Gp6 | glycoprotein 6 (platelet) | 0 | 0 | 0 | 0 | 0 | 0 | 0 | 0 | 0 | 0 | 0 |
| 0 | NA | NA | NA | NA | NA | Gm25349 | | 0 | 0 | 0 | 0 | 0 | 0 | 0 | 0 | 0 | 0 | 0 |
| 0 | NA | NA | NA | NA | NA | Gm24752 | | 0 | 0 | 0 | 0 | 0 | 0 | 0 | 0 | 0 | 0 | 0 |
| 0 | NA | NA | NA | NA | NA | Gm24605 | | 0 | 0 | 0 | 0 | 0 | 0 | 0 | 0 | 0 | 0 | 0 |
| 0 | NA | NA | NA | NA | NA | Gm23577 | | 0 | 0 | 0 | 0 | 0 | 0 | 0 | 0 | 0 | 0 | 0 |
| 0 | NA | NA | NA | NA | NA | Gm9463 | | 0 | 0 | 0 | 0 | 0 | 0 | 0 | 0 | 0 | 0 | 0 |

| | | | | | | | | | | | | | | | | | | |
|---|---|---|---|---|---|---|---|---|---|---|---|---|---|---|---|---|---|---|
| 0 | NA | NA | NA | NA | NA | Gm18973 | | 0 | 0 | 0 | 0 | 0 | 0 | 0 | 0 | 0 | 0 | 0 |
| 0 | NA | NA | NA | NA | NA | Gm44877 | | 0 | 0 | 0 | 0 | 0 | 0 | 0 | 0 | 0 | 0 | 0 |
| 0 | NA | NA | NA | NA | NA | Gm24537 | | 0 | 0 | 0 | 0 | 0 | 0 | 0 | 0 | 0 | 0 | 0 |
| 0 | NA | NA | NA | NA | NA | Fam71e2 | family with sequence similarity 71, membe | 0 | 0 | 0 | 0 | 0 | 0 | 0 | 0 | 0 | 0 | 0 |
| 0 | NA | NA | NA | NA | NA | Tmem190 | transmembrane protein 190 | 0 | 0 | 0 | 0 | 0 | 0 | 0 | 0 | 0 | 0 | 0 |
| 0 | NA | NA | NA | NA | NA | Gm44973 | | 0 | 0 | 0 | 0 | 0 | 0 | 0 | 0 | 0 | 0 | 0 |
| 0 | NA | NA | NA | NA | NA | 4632433K11Rik | | 0 | 0 | 0 | 0 | 0 | 0 | 0 | 0 | 0 | 0 | 0 |
| 0 | NA | NA | NA | NA | NA | Gm38941 | | 0 | 0 | 0 | 0 | 0 | 0 | 0 | 0 | 0 | 0 | 0 |
| 0 | NA | NA | NA | NA | NA | Rfpl4 | ret finger protein-like 4 | 0 | 0 | 0 | 0 | 0 | 0 | 0 | 0 | 0 | 0 | 0 |
| 0 | NA | NA | NA | NA | NA | Gm26217 | | 0 | 0 | 0 | 0 | 0 | 0 | 0 | 0 | 0 | 0 | 0 |
| 0 | NA | NA | NA | NA | NA | Vmn1r56 | vomeronasal 1 receptor 56 | 0 | 0 | 0 | 0 | 0 | 0 | 0 | 0 | 0 | 0 | 0 |
| 0 | NA | NA | NA | NA | NA | Vmn1r57 | vomeronasal 1 receptor 57 | 0 | 0 | 0 | 0 | 0 | 0 | 0 | 0 | 0 | 0 | 0 |
| 0 | NA | NA | NA | NA | NA | Gm45146 | | 0 | 0 | 0 | 0 | 0 | 0 | 0 | 0 | 0 | 0 | 0 |
| 0 | NA | NA | NA | NA | NA | Gm23942 | | 0 | 0 | 0 | 0 | 0 | 0 | 0 | 0 | 0 | 0 | 0 |
| 0 | NA | NA | NA | NA | NA | Nlrp2 | NLR family, pyrin domain containing 2 | 0 | 0 | 0 | 0 | 0 | 0 | 0 | 0 | 0 | 0 | 0 |
| 0 | NA | NA | NA | NA | NA | Gm18757 | | 0 | 0 | 0 | 0 | 0 | 0 | 0 | 0 | 0 | 0 | 0 |
| 0 | NA | NA | NA | NA | NA | Gm5065 | | 0 | 0 | 0 | 0 | 0 | 0 | 0 | 0 | 0 | 0 | 0 |
| 0 | NA | NA | NA | NA | NA | Gm45147 | | 0 | 0 | 0 | 0 | 0 | 0 | 0 | 0 | 0 | 0 | 0 |
| 0 | NA | NA | NA | NA | NA | Vmn1r58 | vomeronasal 1 receptor 58 | 0 | 0 | 0 | 0 | 0 | 0 | 0 | 0 | 0 | 0 | 0 |
| 0 | NA | NA | NA | NA | NA | Gm44543 | | 0 | 0 | 0 | 0 | 0 | 0 | 0 | 0 | 0 | 0 | 0 |
| 0 | NA | NA | NA | NA | NA | Vmn1r59 | vomeronasal 1 receptor 59 | 0 | 0 | 0 | 0 | 0 | 0 | 0 | 0 | 0 | 0 | 0 |
| 0 | NA | NA | NA | NA | NA | Vmn2r28 | vomeronasal 2, receptor 28 | 0 | 0 | 0 | 0 | 0 | 0 | 0 | 0 | 0 | 0 | 0 |
| 0 | NA | NA | NA | NA | NA | Vmn1r60 | vomeronasal 1 receptor 60 | 0 | 0 | 0 | 0 | 0 | 0 | 0 | 0 | 0 | 0 | 0 |
| 0 | NA | NA | NA | NA | NA | Vmn1r-ps40 | | 0 | 0 | 0 | 0 | 0 | 0 | 0 | 0 | 0 | 0 | 0 |
| 0 | NA | NA | NA | NA | NA | Vmn1r61 | vomeronasal 1 receptor 61 | 0 | 0 | 0 | 0 | 0 | 0 | 0 | 0 | 0 | 0 | 0 |
| 0 | NA | NA | NA | NA | NA | Vmn1r62 | vomeronasal 1 receptor 62 | 0 | 0 | 0 | 0 | 0 | 0 | 0 | 0 | 0 | 0 | 0 |
| 0 | NA | NA | NA | NA | NA | Gm44821 | | 0 | 0 | 0 | 0 | 0 | 0 | 0 | 0 | 0 | 0 | 0 |
| 0 | NA | NA | NA | NA | NA | Vmn1r63 | vomeronasal 1 receptor 63 | 0 | 0 | 0 | 0 | 0 | 0 | 0 | 0 | 0 | 0 | 0 |
| 0 | NA | NA | NA | NA | NA | Gm44820 | | 0 | 0 | 0 | 0 | 0 | 0 | 0 | 0 | 0 | 0 | 0 |
| 0 | NA | NA | NA | NA | NA | Vmn1r64 | vomeronasal 1 receptor 64 | 0 | 0 | 0 | 0 | 0 | 0 | 0 | 0 | 0 | 0 | 0 |
| 0 | NA | NA | NA | NA | NA | Gm20399 | | 0 | 0 | 0 | 0 | 0 | 0 | 0 | 0 | 0 | 0 | 0 |
| 0 | NA | NA | NA | NA | NA | Vmn1r-ps45 | | 0 | 0 | 0 | 0 | 0 | 0 | 0 | 0 | 0 | 0 | 0 |
| 0 | NA | NA | NA | NA | NA | Vmn1r-ps46 | | 0 | 0 | 0 | 0 | 0 | 0 | 0 | 0 | 0 | 0 | 0 |
| 0 | NA | NA | NA | NA | NA | Vmn1r65 | vomeronasal 1 receptor 65 | 0 | 0 | 0 | 0 | 0 | 0 | 0 | 0 | 0 | 0 | 0 |
| 0 | NA | NA | NA | NA | NA | 4933402C05Rik | | 0 | 0 | 0 | 0 | 0 | 0 | 0 | 0 | 0 | 0 | 0 |
| 0 | NA | NA | NA | NA | NA | Gm44733 | | 0 | 0 | 0 | 0 | 0 | 0 | 0 | 0 | 0 | 0 | 0 |
| 0 | NA | NA | NA | NA | NA | Nlrp4c | NLR family, pyrin domain containing 4C | 0 | 0 | 0 | 0 | 0 | 0 | 0 | 0 | 0 | 0 | 0 |
| 0 | NA | NA | NA | NA | NA | 4933431C10Rik | | 0 | 0 | 0 | 0 | 0 | 0 | 0 | 0 | 0 | 0 | 0 |
| 0 | NA | NA | NA | NA | NA | Gm45841 | | 0 | 0 | 0 | 0 | 0 | 0 | 0 | 0 | 0 | 0 | 0 |
| 0 | NA | NA | NA | NA | NA | Galp | galanin-like peptide | 0 | 0 | 0 | 0 | 0 | 0 | 0 | 0 | 0 | 0 | 0 |
| 0 | NA | NA | NA | NA | NA | Zscan5b | zinc finger and SCAN domain containing | 0 | 0 | 0 | 0 | 0 | 0 | 0 | 0 | 0 | 0 | 0 |
| 0 | NA | NA | NA | NA | NA | Epp13 | epididymal protein 13 | 0 | 0 | 0 | 0 | 0 | 0 | 0 | 0 | 0 | 0 | 0 |
| 0 | NA | NA | NA | NA | NA | Gm16307 | | 0 | 0 | 0 | 0 | 0 | 0 | 0 | 0 | 0 | 0 | 0 |
| 0 | NA | NA | NA | NA | NA | Olfr1344 | olfactory receptor 1344 | 0 | 0 | 0 | 0 | 0 | 0 | 0 | 0 | 0 | 0 | 0 |
| 0 | NA | NA | NA | NA | NA | Olfr1345-ps1 | | 0 | 0 | 0 | 0 | 0 | 0 | 0 | 0 | 0 | 0 | 0 |
| 0 | NA | NA | NA | NA | NA | Olfr1336 | olfactory receptor 1336 | 0 | 0 | 0 | 0 | 0 | 0 | 0 | 0 | 0 | 0 | 0 |
| 0 | NA | NA | NA | NA | NA | Olfr1346 | olfactory receptor 1346 | 0 | 0 | 0 | 0 | 0 | 0 | 0 | 0 | 0 | 0 | 0 |
| 0 | NA | NA | NA | NA | NA | Olfr5 | olfactory receptor 5 | 0 | 0 | 0 | 0 | 0 | 0 | 0 | 0 | 0 | 0 | 0 |
| 0 | NA | NA | NA | NA | NA | n-R5s150 | | 0 | 0 | 0 | 0 | 0 | 0 | 0 | 0 | 0 | 0 | 0 |
| 0 | NA | NA | NA | NA | NA | Gm7587 | | 0 | 0 | 0 | 0 | 0 | 0 | 0 | 0 | 0 | 0 | 0 |
| 0 | NA | NA | NA | NA | NA | Olfr1350 | olfactory receptor 1350 | 0 | 0 | 0 | 0 | 0 | 0 | 0 | 0 | 0 | 0 | 0 |
| 0 | NA | NA | NA | NA | NA | Zim2 | | 0 | 0 | 0 | 0 | 0 | 0 | 0 | 0 | 0 | 0 | 0 |
| 0 | NA | NA | NA | NA | NA | Gm27675 | | 0 | 0 | 0 | 0 | 0 | 0 | 0 | 0 | 0 | 0 | 0 |
| 0 | NA | NA | NA | NA | NA | Mir3099 | | 0 | 0 | 0 | 0 | 0 | 0 | 0 | 0 | 0 | 0 | 0 |
| 0 | NA | NA | NA | NA | NA | Gm44585 | | 0 | 0 | 0 | 0 | 0 | 0 | 0 | 0 | 0 | 0 | 0 |
| 0 | NA | NA | NA | NA | NA | Gm22649 | | 0 | 0 | 0 | 0 | 0 | 0 | 0 | 0 | 0 | 0 | 0 |
| 0 | NA | NA | NA | NA | NA | Zim3 | | 0 | 0 | 0 | 0 | 0 | 0 | 0 | 0 | 0 | 0 | 0 |
| 0 | NA | NA | NA | NA | NA | | | 0 | 0 | 0 | 0 | 0 | 0 | 0 | 0 | 0 | 0 | 0 |
| 0 | NA | NA | NA | NA | NA | Gm44583 | | 0 | 0 | 0 | 0 | 0 | 0 | 0 | 0 | 0 | 0 | 0 |
| 0 | NA | NA | NA | NA | NA | Vmn2r-ps35 | | 0 | 0 | 0 | 0 | 0 | 0 | 0 | 0 | 0 | 0 | 0 |

| | | | | | | | | | | | | | | | | | | |
|---|---|---|---|---|---|---|---|---|---|---|---|---|---|---|---|---|---|---|
| 0 | NA | NA | NA | NA | NA | Vmn2r30 | vomeronasal 2, receptor 30 | 0 | 0 | 0 | 0 | 0 | 0 | 0 | 0 | 0 | 0 | 0 |
| 0 | NA | NA | NA | NA | NA | Gm45783 | | 0 | 0 | 0 | 0 | 0 | 0 | 0 | 0 | 0 | 0 | 0 |
| 0 | NA | NA | NA | NA | NA | Vmn2r31 | vomeronasal 2, receptor 31 | 0 | 0 | 0 | 0 | 0 | 0 | 0 | 0 | 0 | 0 | 0 |
| 0 | NA | NA | NA | NA | NA | Vmn2r-ps36 | | 0 | 0 | 0 | 0 | 0 | 0 | 0 | 0 | 0 | 0 | 0 |
| 0 | NA | NA | NA | NA | NA | Vmn2r32 | vomeronasal 2, receptor 32 | 0 | 0 | 0 | 0 | 0 | 0 | 0 | 0 | 0 | 0 | 0 |
| 0 | NA | NA | NA | NA | NA | Gm3563 | | 0 | 0 | 0 | 0 | 0 | 0 | 0 | 0 | 0 | 0 | 0 |
| 0 | NA | NA | NA | NA | NA | Gm6842 | | 0 | 0 | 0 | 0 | 0 | 0 | 0 | 0 | 0 | 0 | 0 |
| 0 | NA | NA | NA | NA | NA | Vmn2r33 | vomeronasal 2, receptor 33 | 0 | 0 | 0 | 0 | 0 | 0 | 0 | 0 | 0 | 0 | 0 |
| 0 | NA | NA | NA | NA | NA | Gm7648 | | 0 | 0 | 0 | 0 | 0 | 0 | 0 | 0 | 0 | 0 | 0 |
| 0 | NA | NA | NA | NA | NA | Gm7651 | | 0 | 0 | 0 | 0 | 0 | 0 | 0 | 0 | 0 | 0 | 0 |
| 0 | NA | NA | NA | NA | NA | Vmn2r-ps37 | | 0 | 0 | 0 | 0 | 0 | 0 | 0 | 0 | 0 | 0 | 0 |
| 0 | NA | NA | NA | NA | NA | Vmn2r34 | vomeronasal 2, receptor 34 | 0 | 0 | 0 | 0 | 0 | 0 | 0 | 0 | 0 | 0 | 0 |
| 0 | NA | NA | NA | NA | NA | Vmn2r-ps38 | | 0 | 0 | 0 | 0 | 0 | 0 | 0 | 0 | 0 | 0 | 0 |
| 0 | NA | NA | NA | NA | NA | Vmn2r35 | vomeronasal 2, receptor 35 | 0 | 0 | 0 | 0 | 0 | 0 | 0 | 0 | 0 | 0 | 0 |
| 0 | NA | NA | NA | NA | NA | Vmn2r-ps40 | | 0 | 0 | 0 | 0 | 0 | 0 | 0 | 0 | 0 | 0 | 0 |
| 0 | NA | NA | NA | NA | NA | Vmn2r36 | vomeronasal 2, receptor 36 | 0 | 0 | 0 | 0 | 0 | 0 | 0 | 0 | 0 | 0 | 0 |
| 0 | NA | NA | NA | NA | NA | Gm7230 | | 0 | 0 | 0 | 0 | 0 | 0 | 0 | 0 | 0 | 0 | 0 |
| 0 | NA | NA | NA | NA | NA | Gm3978 | | 0 | 0 | 0 | 0 | 0 | 0 | 0 | 0 | 0 | 0 | 0 |
| 0 | NA | NA | NA | NA | NA | Gm20438 | | 0 | 0 | 0 | 0 | 0 | 0 | 0 | 0 | 0 | 0 | 0 |
| 0 | NA | NA | NA | NA | NA | Gm18200 | | 0 | 0 | 0 | 0 | 0 | 0 | 0 | 0 | 0 | 0 | 0 |
| 0 | NA | NA | NA | NA | NA | Gm4073 | | 0 | 0 | 0 | 0 | 0 | 0 | 0 | 0 | 0 | 0 | 0 |
| 0 | NA | NA | NA | NA | NA | Gm3675 | | 0 | 0 | 0 | 0 | 0 | 0 | 0 | 0 | 0 | 0 | 0 |
| 0 | NA | NA | NA | NA | NA | 2810047C21Rik1 | | 0 | 0 | 0 | 0 | 0 | 0 | 0 | 0 | 0 | 0 | 0 |
| 0 | NA | NA | NA | NA | NA | Vmn2r41 | vomeronasal 2, receptor 41 | 0 | 0 | 0 | 0 | 0 | 0 | 0 | 0 | 0 | 0 | 0 |
| 0 | NA | NA | NA | NA | NA | Vmn2r42 | vomeronasal 2, receptor 42 | 0 | 0 | 0 | 0 | 0 | 0 | 0 | 0 | 0 | 0 | 0 |
| 0 | NA | NA | NA | NA | NA | Vmn2r43 | vomeronasal 2, receptor 43 | 0 | 0 | 0 | 0 | 0 | 0 | 0 | 0 | 0 | 0 | 0 |
| 0 | NA | NA | NA | NA | NA | Vmn2r-ps45 | | 0 | 0 | 0 | 0 | 0 | 0 | 0 | 0 | 0 | 0 | 0 |
| 0 | NA | NA | NA | NA | NA | Gm7696 | | 0 | 0 | 0 | 0 | 0 | 0 | 0 | 0 | 0 | 0 | 0 |
| 0 | NA | NA | NA | NA | NA | Vmn2r44 | vomeronasal 2, receptor 44 | 0 | 0 | 0 | 0 | 0 | 0 | 0 | 0 | 0 | 0 | 0 |
| 0 | NA | NA | NA | NA | NA | Vmn2r-ps46 | | 0 | 0 | 0 | 0 | 0 | 0 | 0 | 0 | 0 | 0 | 0 |
| 0 | NA | NA | NA | NA | NA | Vmn2r45 | vomeronasal 2, receptor 45 | 0 | 0 | 0 | 0 | 0 | 0 | 0 | 0 | 0 | 0 | 0 |
| 0 | NA | NA | NA | NA | NA | Vmn2r-ps47 | | 0 | 0 | 0 | 0 | 0 | 0 | 0 | 0 | 0 | 0 | 0 |
| 0 | NA | NA | NA | NA | NA | Vmn2r-ps49 | | 0 | 0 | 0 | 0 | 0 | 0 | 0 | 0 | 0 | 0 | 0 |
| 0 | NA | NA | NA | NA | NA | Gm3584 | | 0 | 0 | 0 | 0 | 0 | 0 | 0 | 0 | 0 | 0 | 0 |
| 0 | NA | NA | NA | NA | NA | Gm3956 | | 0 | 0 | 0 | 0 | 0 | 0 | 0 | 0 | 0 | 0 | 0 |
| 0 | NA | NA | NA | NA | NA | Gm3585 | | 0 | 0 | 0 | 0 | 0 | 0 | 0 | 0 | 0 | 0 | 0 |
| 0 | NA | NA | NA | NA | NA | Gm3962 | | 0 | 0 | 0 | 0 | 0 | 0 | 0 | 0 | 0 | 0 | 0 |
| 0 | NA | NA | NA | NA | NA | Gm18191 | | 0 | 0 | 0 | 0 | 0 | 0 | 0 | 0 | 0 | 0 | 0 |
| 0 | NA | NA | NA | NA | NA | Vmn2r-ps44 | | 0 | 0 | 0 | 0 | 0 | 0 | 0 | 0 | 0 | 0 | 0 |
| 0 | NA | NA | NA | NA | NA | Gm48598 | | 0 | 0 | 0 | 0 | 0 | 0 | 0 | 0 | 0 | 0 | 0 |
| 0 | NA | NA | NA | NA | NA | Vmn2r-ps43 | | 0 | 0 | 0 | 0 | 0 | 0 | 0 | 0 | 0 | 0 | 0 |
| 0 | NA | NA | NA | NA | NA | Vmn2r40 | vomeronasal 2, receptor 40 | 0 | 0 | 0 | 0 | 0 | 0 | 0 | 0 | 0 | 0 | 0 |
| 0 | NA | NA | NA | NA | NA | Gm45530 | | 0 | 0 | 0 | 0 | 0 | 0 | 0 | 0 | 0 | 0 | 0 |
| 0 | NA | NA | NA | NA | NA | Vmn2r-ps42 | | 0 | 0 | 0 | 0 | 0 | 0 | 0 | 0 | 0 | 0 | 0 |
| 0 | NA | NA | NA | NA | NA | Vmn2r39 | vomeronasal 2, receptor 39 | 0 | 0 | 0 | 0 | 0 | 0 | 0 | 0 | 0 | 0 | 0 |
| 0 | NA | NA | NA | NA | NA | Zfp419 | | 0 | 0 | 0 | 0 | 0 | 0 | 0 | 0 | 0 | 0 | 0 |
| 0 | NA | NA | NA | NA | NA | Vmn2r38 | vomeronasal 2, receptor 38 | 0 | 0 | 0 | 0 | 0 | 0 | 0 | 0 | 0 | 0 | 0 |
| 0 | NA | NA | NA | NA | NA | Gm5583 | | 0 | 0 | 0 | 0 | 0 | 0 | 0 | 0 | 0 | 0 | 0 |
| 0 | NA | NA | NA | NA | NA | Gm18201 | | 0 | 0 | 0 | 0 | 0 | 0 | 0 | 0 | 0 | 0 | 0 |
| 0 | NA | NA | NA | NA | NA | Vmn2r-ps41 | | 0 | 0 | 0 | 0 | 0 | 0 | 0 | 0 | 0 | 0 | 0 |
| 0 | NA | NA | NA | NA | NA | Vmn2r37 | vomeronasal 2, receptor 37 | 0 | 0 | 0 | 0 | 0 | 0 | 0 | 0 | 0 | 0 | 0 |
| 0 | NA | NA | NA | NA | NA | Gm4011 | | 0 | 0 | 0 | 0 | 0 | 0 | 0 | 0 | 0 | 0 | 0 |
| 0 | NA | NA | NA | NA | NA | Gm3623 | | 0 | 0 | 0 | 0 | 0 | 0 | 0 | 0 | 0 | 0 | 0 |
| 0 | NA | NA | NA | NA | NA | Gm3616 | | 0 | 0 | 0 | 0 | 0 | 0 | 0 | 0 | 0 | 0 | 0 |
| 0 | NA | NA | NA | NA | NA | Gm20287 | | 0 | 0 | 0 | 0 | 0 | 0 | 0 | 0 | 0 | 0 | 0 |
| 0 | NA | NA | NA | NA | NA | Gm3611 | | 0 | 0 | 0 | 0 | 0 | 0 | 0 | 0 | 0 | 0 | 0 |
| 0 | NA | NA | NA | NA | NA | Gm3972 | | 0 | 0 | 0 | 0 | 0 | 0 | 0 | 0 | 0 | 0 | 0 |
| 0 | NA | NA | NA | NA | NA | Gm18199 | | 0 | 0 | 0 | 0 | 0 | 0 | 0 | 0 | 0 | 0 | 0 |
| 0 | NA | NA | NA | NA | NA | Gm10302 | predicted gene 10302 | 0 | 0 | 0 | 0 | 0 | 0 | 0 | 0 | 0 | 0 | 0 |
| 0 | NA | NA | NA | NA | NA | Gm3683 | | 0 | 0 | 0 | 0 | 0 | 0 | 0 | 0 | 0 | 0 | 0 |

| | | | | | | | | | | | | | | | | | |
|---|---|---|---|---|---|---|---|---|---|---|---|---|---|---|---|---|---|
| 0 | NA | NA | NA | NA | NA | Gm20345 | | 0 | 0 | 0 | 0 | 0 | 0 | 0 | 0 | 0 | 0 | 0 |
| 0 | NA | NA | NA | NA | NA | Gm3690 | | 0 | 0 | 0 | 0 | 0 | 0 | 0 | 0 | 0 | 0 | 0 |
| 0 | NA | NA | NA | NA | NA | Vmn2r46 | vomeronasal 2, receptor 46 | 0 | 0 | 0 | 0 | 0 | 0 | 0 | 0 | 0 | 0 | 0 |
| 0 | NA | NA | NA | NA | NA | Gm18192 | | 0 | 0 | 0 | 0 | 0 | 0 | 0 | 0 | 0 | 0 | 0 |
| 0 | NA | NA | NA | NA | NA | Vmn2r47 | vomeronasal 2, receptor 47 | 0 | 0 | 0 | 0 | 0 | 0 | 0 | 0 | 0 | 0 | 0 |
| 0 | NA | NA | NA | NA | NA | Vmn2r-ps51 | | 0 | 0 | 0 | 0 | 0 | 0 | 0 | 0 | 0 | 0 | 0 |
| 0 | NA | NA | NA | NA | NA | Vmn2r-ps52 | | 0 | 0 | 0 | 0 | 0 | 0 | 0 | 0 | 0 | 0 | 0 |
| 0 | NA | NA | NA | NA | NA | Gm45403 | | 0 | 0 | 0 | 0 | 0 | 0 | 0 | 0 | 0 | 0 | 0 |
| 0 | NA | NA | NA | NA | NA | Vmn2r48 | vomeronasal 2, receptor 48 | 0 | 0 | 0 | 0 | 0 | 0 | 0 | 0 | 0 | 0 | 0 |
| 0 | NA | NA | NA | NA | NA | Gm3702 | | 0 | 0 | 0 | 0 | 0 | 0 | 0 | 0 | 0 | 0 | 0 |
| 0 | NA | NA | NA | NA | NA | Vmn2r49 | vomeronasal 2, receptor 49 | 0 | 0 | 0 | 0 | 0 | 0 | 0 | 0 | 0 | 0 | 0 |
| 0 | NA | NA | NA | NA | NA | Vmn2r50 | vomeronasal 2, receptor 50 | 0 | 0 | 0 | 0 | 0 | 0 | 0 | 0 | 0 | 0 | 0 |
| 0 | NA | NA | NA | NA | NA | Gm6850 | | 0 | 0 | 0 | 0 | 0 | 0 | 0 | 0 | 0 | 0 | 0 |
| 0 | NA | NA | NA | NA | NA | Vmn2r51 | vomeronasal 2, receptor 51 | 0 | 0 | 0 | 0 | 0 | 0 | 0 | 0 | 0 | 0 | 0 |
| 0 | NA | NA | NA | NA | NA | Vmn2r52 | vomeronasal 2, receptor 52 | 0 | 0 | 0 | 0 | 0 | 0 | 0 | 0 | 0 | 0 | 0 |
| 0 | NA | NA | NA | NA | NA | Gm20679 | | 0 | 0 | 0 | 0 | 0 | 0 | 0 | 0 | 0 | 0 | 0 |
| 0 | NA | NA | NA | NA | NA | Vmn1r66 | vomeronasal 1 receptor 66 | 0 | 0 | 0 | 0 | 0 | 0 | 0 | 0 | 0 | 0 | 0 |
| 0 | NA | NA | NA | NA | NA | Vmn1r-ps47 | | 0 | 0 | 0 | 0 | 0 | 0 | 0 | 0 | 0 | 0 | 0 |
| 0 | NA | NA | NA | NA | NA | Vmn1r-ps48 | | 0 | 0 | 0 | 0 | 0 | 0 | 0 | 0 | 0 | 0 | 0 |
| 0 | NA | NA | NA | NA | NA | Gm45402 | | 0 | 0 | 0 | 0 | 0 | 0 | 0 | 0 | 0 | 0 | 0 |
| 0 | NA | NA | NA | NA | NA | Gm45400 | | 0 | 0 | 0 | 0 | 0 | 0 | 0 | 0 | 0 | 0 | 0 |
| 0 | NA | NA | NA | NA | NA | Gm45401 | | 0 | 0 | 0 | 0 | 0 | 0 | 0 | 0 | 0 | 0 | 0 |
| 0 | NA | NA | NA | NA | NA | Nlrp4d | | 0 | 0 | 0 | 0 | 0 | 0 | 0 | 0 | 0 | 0 | 0 |
| 0 | NA | NA | NA | NA | NA | Gm3726 | | 0 | 0 | 0 | 0 | 0 | 0 | 0 | 0 | 0 | 0 | 0 |
| 0 | NA | NA | NA | NA | NA | Vmn1r67 | vomeronasal 1 receptor 67 | 0 | 0 | 0 | 0 | 0 | 0 | 0 | 0 | 0 | 0 | 0 |
| 0 | NA | NA | NA | NA | NA | Gm30645 | | 0 | 0 | 0 | 0 | 0 | 0 | 0 | 0 | 0 | 0 | 0 |
| 0 | NA | NA | NA | NA | NA | Gm18193 | | 0 | 0 | 0 | 0 | 0 | 0 | 0 | 0 | 0 | 0 | 0 |
| 0 | NA | NA | NA | NA | NA | Vmn1r68 | vomeronasal 1 receptor 68 | 0 | 0 | 0 | 0 | 0 | 0 | 0 | 0 | 0 | 0 | 0 |
| 0 | NA | NA | NA | NA | NA | Vmn1r-ps49 | | 0 | 0 | 0 | 0 | 0 | 0 | 0 | 0 | 0 | 0 | 0 |
| 0 | NA | NA | NA | NA | NA | Vmn1r69 | vomeronasal 1 receptor 69 | 0 | 0 | 0 | 0 | 0 | 0 | 0 | 0 | 0 | 0 | 0 |
| 0 | NA | NA | NA | NA | NA | Vmn1r-ps50 | | 0 | 0 | 0 | 0 | 0 | 0 | 0 | 0 | 0 | 0 | 0 |
| 0 | NA | NA | NA | NA | NA | Vmn1r70 | vomeronasal 1 receptor 70 | 0 | 0 | 0 | 0 | 0 | 0 | 0 | 0 | 0 | 0 | 0 |
| 0 | NA | NA | NA | NA | NA | Vmn1r-ps51 | | 0 | 0 | 0 | 0 | 0 | 0 | 0 | 0 | 0 | 0 | 0 |
| 0 | NA | NA | NA | NA | NA | Vmn1r-ps52 | | 0 | 0 | 0 | 0 | 0 | 0 | 0 | 0 | 0 | 0 | 0 |
| 0 | NA | NA | NA | NA | NA | Nlrp4b | NLR family, pyrin domain containing 4B | 0 | 0 | 0 | 0 | 0 | 0 | 0 | 0 | 0 | 0 | 0 |
| 0 | NA | NA | NA | NA | NA | Gm45379 | | 0 | 0 | 0 | 0 | 0 | 0 | 0 | 0 | 0 | 0 | 0 |
| 0 | NA | NA | NA | NA | NA | Vmn1r71 | vomeronasal 1 receptor 71 | 0 | 0 | 0 | 0 | 0 | 0 | 0 | 0 | 0 | 0 | 0 |
| 0 | NA | NA | NA | NA | NA | Gm45438 | | 0 | 0 | 0 | 0 | 0 | 0 | 0 | 0 | 0 | 0 | 0 |
| 0 | NA | NA | NA | NA | NA | Zscan4a | | 0 | 0 | 0 | 0 | 0 | 0 | 0 | 0 | 0 | 0 | 0 |
| 0 | NA | NA | NA | NA | NA | Gm18781 | | 0 | 0 | 0 | 0 | 0 | 0 | 0 | 0 | 0 | 0 | 0 |
| 0 | NA | NA | NA | NA | NA | Gm7203 | | 0 | 0 | 0 | 0 | 0 | 0 | 0 | 0 | 0 | 0 | 0 |
| 0 | NA | NA | NA | NA | NA | Gm4903 | | 0 | 0 | 0 | 0 | 0 | 0 | 0 | 0 | 0 | 0 | 0 |
| 0 | NA | NA | NA | NA | NA | Gm7836 | | 0 | 0 | 0 | 0 | 0 | 0 | 0 | 0 | 0 | 0 | 0 |
| 0 | NA | NA | NA | NA | NA | Gm18638 | | 0 | 0 | 0 | 0 | 0 | 0 | 0 | 0 | 0 | 0 | 0 |
| 0 | NA | NA | NA | NA | NA | Zscan4b | zinc finger and SCAN domain containing | 0 | 0 | 0 | 0 | 0 | 0 | 0 | 0 | 0 | 0 | 0 |
| 0 | NA | NA | NA | NA | NA | Gm45279 | | 0 | 0 | 0 | 0 | 0 | 0 | 0 | 0 | 0 | 0 | 0 |
| 0 | NA | NA | NA | NA | NA | Mir297-1 | | 0 | 0 | 0 | 0 | 0 | 0 | 0 | 0 | 0 | 0 | 0 |
| 0 | NA | NA | NA | NA | NA | Gm7786 | | 0 | 0 | 0 | 0 | 0 | 0 | 0 | 0 | 0 | 0 | 0 |
| 0 | NA | NA | NA | NA | NA | Gm7789 | | 0 | 0 | 0 | 0 | 0 | 0 | 0 | 0 | 0 | 0 | 0 |
| 0 | NA | NA | NA | NA | NA | Gm18122 | | 0 | 0 | 0 | 0 | 0 | 0 | 0 | 0 | 0 | 0 | 0 |
| 0 | NA | NA | NA | NA | NA | Zscan4c | zinc finger and SCAN domain containing | 0 | 0 | 0 | 0 | 0 | 0 | 0 | 0 | 0 | 0 | 0 |
| 0 | NA | NA | NA | NA | NA | Gm45327 | | 0 | 0 | 0 | 0 | 0 | 0 | 0 | 0 | 0 | 0 | 0 |
| 0 | NA | NA | NA | NA | NA | Gm45395 | | 0 | 0 | 0 | 0 | 0 | 0 | 0 | 0 | 0 | 0 | 0 |
| 0 | NA | NA | NA | NA | NA | Zscan4-ps1 | | 0 | 0 | 0 | 0 | 0 | 0 | 0 | 0 | 0 | 0 | 0 |
| 0 | NA | NA | NA | NA | NA | Gm33601 | | 0 | 0 | 0 | 0 | 0 | 0 | 0 | 0 | 0 | 0 | 0 |
| 0 | NA | NA | NA | NA | NA | Gm7151 | | 0 | 0 | 0 | 0 | 0 | 0 | 0 | 0 | 0 | 0 | 0 |
| 0 | NA | NA | NA | NA | NA | Gm4157 | | 0 | 0 | 0 | 0 | 0 | 0 | 0 | 0 | 0 | 0 | 0 |
| 0 | NA | NA | NA | NA | NA | Gm3774 | | 0 | 0 | 0 | 0 | 0 | 0 | 0 | 0 | 0 | 0 | 0 |
| 0 | NA | NA | NA | NA | NA | Gm38430 | | 0 | 0 | 0 | 0 | 0 | 0 | 0 | 0 | 0 | 0 | 0 |
| 0 | NA | NA | NA | NA | NA | Zscan4d | zinc finger and SCAN domain containing | 0 | 0 | 0 | 0 | 0 | 0 | 0 | 0 | 0 | 0 | 0 |

| | | | | | | | | | | | | | | | | | | |
|---|---|---|---|---|---|---|---|---|---|---|---|---|---|---|---|---|---|---|
| 0 | NA | NA | NA | NA | NA | Gm18204 | | 0 | 0 | 0 | 0 | 0 | 0 | 0 | 0 | 0 | 0 | 0 |
| 0 | NA | NA | NA | NA | NA | Gm6909 | | 0 | 0 | 0 | 0 | 0 | 0 | 0 | 0 | 0 | 0 | 0 |
| 0 | NA | NA | NA | NA | NA | Gm9689 | | 0 | 0 | 0 | 0 | 0 | 0 | 0 | 0 | 0 | 0 | 0 |
| 0 | NA | NA | NA | NA | NA | Gm39481 | | 0 | 0 | 0 | 0 | 0 | 0 | 0 | 0 | 0 | 0 | 0 |
| 0 | NA | NA | NA | NA | NA | Gm5996 | | 0 | 0 | 0 | 0 | 0 | 0 | 0 | 0 | 0 | 0 | 0 |
| 0 | NA | NA | NA | NA | NA | Gm9652 | | 0 | 0 | 0 | 0 | 0 | 0 | 0 | 0 | 0 | 0 | 0 |
| 0 | NA | NA | NA | NA | NA | Gm19683 | | 0 | 0 | 0 | 0 | 0 | 0 | 0 | 0 | 0 | 0 | 0 |
| 0 | NA | NA | NA | NA | NA | Gm26248 | | 0 | 0 | 0 | 0 | 0 | 0 | 0 | 0 | 0 | 0 | 0 |
| 0 | NA | NA | NA | NA | NA | Zscan4e | | 0 | 0 | 0 | 0 | 0 | 0 | 0 | 0 | 0 | 0 | 0 |
| 0 | NA | NA | NA | NA | NA | Gm45870 | | 0 | 0 | 0 | 0 | 0 | 0 | 0 | 0 | 0 | 0 | 0 |
| 0 | NA | NA | NA | NA | NA | Gm23087 | | 0 | 0 | 0 | 0 | 0 | 0 | 0 | 0 | 0 | 0 | 0 |
| 0 | NA | NA | NA | NA | NA | Gm45596 | | 0 | 0 | 0 | 0 | 0 | 0 | 0 | 0 | 0 | 0 | 0 |
| 0 | NA | NA | NA | NA | NA | Gm45598 | | 0 | 0 | 0 | 0 | 0 | 0 | 0 | 0 | 0 | 0 | 0 |
| 0 | NA | NA | NA | NA | NA | Gm18123 | | 0 | 0 | 0 | 0 | 0 | 0 | 0 | 0 | 0 | 0 | 0 |
| 0 | NA | NA | NA | NA | NA | Zscan4f | zinc finger and SCAN domain containing | 0 | 0 | 0 | 0 | 0 | 0 | 0 | 0 | 0 | 0 | 0 |
| 0 | NA | NA | NA | NA | NA | Gm18639 | | 0 | 0 | 0 | 0 | 0 | 0 | 0 | 0 | 0 | 0 | 0 |
| 0 | NA | NA | NA | NA | NA | Gm9604 | | 0 | 0 | 0 | 0 | 0 | 0 | 0 | 0 | 0 | 0 | 0 |
| 0 | NA | NA | NA | NA | NA | Gm7839 | | 0 | 0 | 0 | 0 | 0 | 0 | 0 | 0 | 0 | 0 | 0 |
| 0 | NA | NA | NA | NA | NA | Gm45382 | | 0 | 0 | 0 | 0 | 0 | 0 | 0 | 0 | 0 | 0 | 0 |
| 0 | NA | NA | NA | NA | NA | Gm7207 | | 0 | 0 | 0 | 0 | 0 | 0 | 0 | 0 | 0 | 0 | 0 |
| 0 | NA | NA | NA | NA | NA | Gm45550 | | 0 | 0 | 0 | 0 | 0 | 0 | 0 | 0 | 0 | 0 | 0 |
| 0 | NA | NA | NA | NA | NA | Gm45551 | | 0 | 0 | 0 | 0 | 0 | 0 | 0 | 0 | 0 | 0 | 0 |
| 0 | NA | NA | NA | NA | NA | Gm45549 | | 0 | 0 | 0 | 0 | 0 | 0 | 0 | 0 | 0 | 0 | 0 |
| 0 | NA | NA | NA | NA | NA | Gm18124 | | 0 | 0 | 0 | 0 | 0 | 0 | 0 | 0 | 0 | 0 | 0 |
| 0 | NA | NA | NA | NA | NA | Zscan4-ps2 | | 0 | 0 | 0 | 0 | 0 | 0 | 0 | 0 | 0 | 0 | 0 |
| 0 | NA | NA | NA | NA | NA | Gm18640 | | 0 | 0 | 0 | 0 | 0 | 0 | 0 | 0 | 0 | 0 | 0 |
| 0 | NA | NA | NA | NA | NA | Gm3816 | | 0 | 0 | 0 | 0 | 0 | 0 | 0 | 0 | 0 | 0 | 0 |
| 0 | NA | NA | NA | NA | NA | Gm7848 | | 0 | 0 | 0 | 0 | 0 | 0 | 0 | 0 | 0 | 0 | 0 |
| 0 | NA | NA | NA | NA | NA | Gm7210 | | 0 | 0 | 0 | 0 | 0 | 0 | 0 | 0 | 0 | 0 | 0 |
| 0 | NA | NA | NA | NA | NA | Gm45500 | | 0 | 0 | 0 | 0 | 0 | 0 | 0 | 0 | 0 | 0 | 0 |
| 0 | NA | NA | NA | NA | NA | Zscan4-ps3 | | 0 | 0 | 0 | 0 | 0 | 0 | 0 | 0 | 0 | 0 | 0 |
| 0 | NA | NA | NA | NA | NA | Gm18641 | | 0 | 0 | 0 | 0 | 0 | 0 | 0 | 0 | 0 | 0 | 0 |
| 0 | NA | NA | NA | NA | NA | Vmn1r-ps53 | | 0 | 0 | 0 | 0 | 0 | 0 | 0 | 0 | 0 | 0 | 0 |
| 0 | NA | NA | NA | NA | NA | Gm45283 | | 0 | 0 | 0 | 0 | 0 | 0 | 0 | 0 | 0 | 0 | 0 |
| 0 | NA | NA | NA | NA | NA | Gm18850 | | 0 | 0 | 0 | 0 | 0 | 0 | 0 | 0 | 0 | 0 | 0 |
| 0 | NA | NA | NA | NA | NA | Gm19093 | | 0 | 0 | 0 | 0 | 0 | 0 | 0 | 0 | 0 | 0 | 0 |
| 0 | NA | NA | NA | NA | NA | Vmn1r73 | vomeronasal 1 receptor 73 | 0 | 0 | 0 | 0 | 0 | 0 | 0 | 0 | 0 | 0 | 0 |
| 0 | NA | NA | NA | NA | NA | Gm45616 | | 0 | 0 | 0 | 0 | 0 | 0 | 0 | 0 | 0 | 0 | 0 |
| 0 | NA | NA | NA | NA | NA | Gm45617 | | 0 | 0 | 0 | 0 | 0 | 0 | 0 | 0 | 0 | 0 | 0 |
| 0 | NA | NA | NA | NA | NA | Vmn1r74 | vomeronasal 1 receptor 74 | 0 | 0 | 0 | 0 | 0 | 0 | 0 | 0 | 0 | 0 | 0 |
| 0 | NA | NA | NA | NA | NA | Vmn1r-ps55 | | 0 | 0 | 0 | 0 | 0 | 0 | 0 | 0 | 0 | 0 | 0 |
| 0 | NA | NA | NA | NA | NA | Gm34779 | | 0 | 0 | 0 | 0 | 0 | 0 | 0 | 0 | 0 | 0 | 0 |
| 0 | NA | NA | NA | NA | NA | Vmn1r75 | vomeronasal 1 receptor 75 | 0 | 0 | 0 | 0 | 0 | 0 | 0 | 0 | 0 | 0 | 0 |
| 0 | NA | NA | NA | NA | NA | Gm18851 | | 0 | 0 | 0 | 0 | 0 | 0 | 0 | 0 | 0 | 0 | 0 |
| 0 | NA | NA | NA | NA | NA | Vmn1r-ps56 | | 0 | 0 | 0 | 0 | 0 | 0 | 0 | 0 | 0 | 0 | 0 |
| 0 | NA | NA | NA | NA | NA | Vmn1r76 | vomeronasal 1 receptor 76 | 0 | 0 | 0 | 0 | 0 | 0 | 0 | 0 | 0 | 0 | 0 |
| 0 | NA | NA | NA | NA | NA | Gm45387 | | 0 | 0 | 0 | 0 | 0 | 0 | 0 | 0 | 0 | 0 | 0 |
| 0 | NA | NA | NA | NA | NA | Vmn1r-ps57 | | 0 | 0 | 0 | 0 | 0 | 0 | 0 | 0 | 0 | 0 | 0 |
| 0 | NA | NA | NA | NA | NA | Gm3836 | | 0 | 0 | 0 | 0 | 0 | 0 | 0 | 0 | 0 | 0 | 0 |
| 0 | NA | NA | NA | NA | NA | Vmn1r-ps58 | | 0 | 0 | 0 | 0 | 0 | 0 | 0 | 0 | 0 | 0 | 0 |
| 0 | NA | NA | NA | NA | NA | Gm17737 | | 0 | 0 | 0 | 0 | 0 | 0 | 0 | 0 | 0 | 0 | 0 |
| 0 | NA | NA | NA | NA | NA | Gm45429 | | 0 | 0 | 0 | 0 | 0 | 0 | 0 | 0 | 0 | 0 | 0 |
| 0 | NA | NA | NA | NA | NA | Vmn1r-ps59 | | 0 | 0 | 0 | 0 | 0 | 0 | 0 | 0 | 0 | 0 | 0 |
| 0 | NA | NA | NA | NA | NA | Gm18597 | | 0 | 0 | 0 | 0 | 0 | 0 | 0 | 0 | 0 | 0 | 0 |
| 0 | NA | NA | NA | NA | NA | Vmn1r-ps60 | | 0 | 0 | 0 | 0 | 0 | 0 | 0 | 0 | 0 | 0 | 0 |
| 0 | NA | NA | NA | NA | NA | Gm18854 | | 0 | 0 | 0 | 0 | 0 | 0 | 0 | 0 | 0 | 0 | 0 |
| 0 | NA | NA | NA | NA | NA | Vmn1r77 | vomeronasal 1 receptor 77 | 0 | 0 | 0 | 0 | 0 | 0 | 0 | 0 | 0 | 0 | 0 |
| 0 | NA | NA | NA | NA | NA | Vmn1r-ps61 | | 0 | 0 | 0 | 0 | 0 | 0 | 0 | 0 | 0 | 0 | 0 |
| 0 | NA | NA | NA | NA | NA | Gm45562 | | 0 | 0 | 0 | 0 | 0 | 0 | 0 | 0 | 0 | 0 | 0 |
| 0 | NA | NA | NA | NA | NA | Vmn1r-ps62 | | 0 | 0 | 0 | 0 | 0 | 0 | 0 | 0 | 0 | 0 | 0 |

| | | | | | | | | | | | | | | | | | | |
|---|---|---|---|---|---|---|---|---|---|---|---|---|---|---|---|---|---|---|
| 0 | NA | NA | NA | NA | NA | Gm45498 | | 0 | 0 | 0 | 0 | 0 | 0 | 0 | 0 | 0 | 0 | 0 |
| 0 | NA | NA | NA | NA | NA | Vmn1r-ps63 | | 0 | 0 | 0 | 0 | 0 | 0 | 0 | 0 | 0 | 0 | 0 |
| 0 | NA | NA | NA | NA | NA | Vmn1r78 | vomeronasal 1 receptor 78 | 0 | 0 | 0 | 0 | 0 | 0 | 0 | 0 | 0 | 0 | 0 |
| 0 | NA | NA | NA | NA | NA | Gm45365 | | 0 | 0 | 0 | 0 | 0 | 0 | 0 | 0 | 0 | 0 | 0 |
| 0 | NA | NA | NA | NA | NA | Vmn1r79 | vomeronasal 1 receptor 79 | 0 | 0 | 0 | 0 | 0 | 0 | 0 | 0 | 0 | 0 | 0 |
| 0 | NA | NA | NA | NA | NA | Gm19201 | | 0 | 0 | 0 | 0 | 0 | 0 | 0 | 0 | 0 | 0 | 0 |
| 0 | NA | NA | NA | NA | NA | Vmn1r80 | vomeronasal 1 receptor 80 | 0 | 0 | 0 | 0 | 0 | 0 | 0 | 0 | 0 | 0 | 0 |
| 0 | NA | NA | NA | NA | NA | Vmn1r-ps64 | | 0 | 0 | 0 | 0 | 0 | 0 | 0 | 0 | 0 | 0 | 0 |
| 0 | NA | NA | NA | NA | NA | Gm45364 | | 0 | 0 | 0 | 0 | 0 | 0 | 0 | 0 | 0 | 0 | 0 |
| 0 | NA | NA | NA | NA | NA | Vmn1r-ps65 | | 0 | 0 | 0 | 0 | 0 | 0 | 0 | 0 | 0 | 0 | 0 |
| 0 | NA | NA | NA | NA | NA | Gm45367 | | 0 | 0 | 0 | 0 | 0 | 0 | 0 | 0 | 0 | 0 | 0 |
| 0 | NA | NA | NA | NA | NA | Gm4224 | | 0 | 0 | 0 | 0 | 0 | 0 | 0 | 0 | 0 | 0 | 0 |
| 0 | NA | NA | NA | NA | NA | Vmn1r81 | vomeronasal 1 receptor 81 | 0 | 0 | 0 | 0 | 0 | 0 | 0 | 0 | 0 | 0 | 0 |
| 0 | NA | NA | NA | NA | NA | Vmn1r82 | vomeronasal 1 receptor 82 | 0 | 0 | 0 | 0 | 0 | 0 | 0 | 0 | 0 | 0 | 0 |
| 0 | NA | NA | NA | NA | NA | Vmn1r83 | vomeronasal 1 receptor 83 | 0 | 0 | 0 | 0 | 0 | 0 | 0 | 0 | 0 | 0 | 0 |
| 0 | NA | NA | NA | NA | NA | Gm23521 | | 0 | 0 | 0 | 0 | 0 | 0 | 0 | 0 | 0 | 0 | 0 |
| 0 | NA | NA | NA | NA | NA | Vmn1r84 | vomeronasal 1 receptor 84 | 0 | 0 | 0 | 0 | 0 | 0 | 0 | 0 | 0 | 0 | 0 |
| 0 | NA | NA | NA | NA | NA | Vmn1r-ps66 | | 0 | 0 | 0 | 0 | 0 | 0 | 0 | 0 | 0 | 0 | 0 |
| 0 | NA | NA | NA | NA | NA | Vmn1r-ps67 | | 0 | 0 | 0 | 0 | 0 | 0 | 0 | 0 | 0 | 0 | 0 |
| 0 | NA | NA | NA | NA | NA | Vmn1r-ps68 | | 0 | 0 | 0 | 0 | 0 | 0 | 0 | 0 | 0 | 0 | 0 |
| 0 | NA | NA | NA | NA | NA | Vmn2r-ps53 | | 0 | 0 | 0 | 0 | 0 | 0 | 0 | 0 | 0 | 0 | 0 |
| 0 | NA | NA | NA | NA | NA | 2900092C05Ril | RIKEN cDNA 2900092C05 gene | 0 | 0 | 0 | 0 | 0 | 0 | 0 | 0 | 0 | 0 | 0 |
| 0 | NA | NA | NA | NA | NA | Gm45475 | | 0 | 0 | 0 | 0 | 0 | 0 | 0 | 0 | 0 | 0 | 0 |
| 0 | NA | NA | NA | NA | NA | Vmn2r53 | vomeronasal 2, receptor 53 | 0 | 0 | 0 | 0 | 0 | 0 | 0 | 0 | 0 | 0 | 0 |
| 0 | NA | NA | NA | NA | NA | Vmn2r54 | vomeronasal 2, receptor 54 | 0 | 0 | 0 | 0 | 0 | 0 | 0 | 0 | 0 | 0 | 0 |
| 0 | NA | NA | NA | NA | NA | Vmn2r55 | vomeronasal 2, receptor 55 | 0 | 0 | 0 | 0 | 0 | 0 | 0 | 0 | 0 | 0 | 0 |
| 0 | NA | NA | NA | NA | NA | Vmn2r56 | vomeronasal 2, receptor 56 | 0 | 0 | 0 | 0 | 0 | 0 | 0 | 0 | 0 | 0 | 0 |
| 0 | NA | NA | NA | NA | NA | Gm18205 | | 0 | 0 | 0 | 0 | 0 | 0 | 0 | 0 | 0 | 0 | 0 |
| 0 | NA | NA | NA | NA | NA | 1700047O18Rik | | 0 | 0 | 0 | 0 | 0 | 0 | 0 | 0 | 0 | 0 | 0 |
| 0 | NA | NA | NA | NA | NA | Gm45593 | | 0 | 0 | 0 | 0 | 0 | 0 | 0 | 0 | 0 | 0 | 0 |
| 0 | NA | NA | NA | NA | NA | Gm26325 | | 0 | 0 | 0 | 0 | 0 | 0 | 0 | 0 | 0 | 0 | 0 |
| 0 | NA | NA | NA | NA | NA | Vmn1r85 | vomeronasal 1 receptor 85 | 0 | 0 | 0 | 0 | 0 | 0 | 0 | 0 | 0 | 0 | 0 |
| 0 | NA | NA | NA | NA | NA | Gm45436 | | 0 | 0 | 0 | 0 | 0 | 0 | 0 | 0 | 0 | 0 | 0 |
| 0 | NA | NA | NA | NA | NA | Gm4270 | | 0 | 0 | 0 | 0 | 0 | 0 | 0 | 0 | 0 | 0 | 0 |
| 0 | NA | NA | NA | NA | NA | Vmn1r86 | vomeronasal 1 receptor 86 | 0 | 0 | 0 | 0 | 0 | 0 | 0 | 0 | 0 | 0 | 0 |
| 0 | NA | NA | NA | NA | NA | Vmn1r87 | vomeronasal 1 receptor 87 | 0 | 0 | 0 | 0 | 0 | 0 | 0 | 0 | 0 | 0 | 0 |
| 0 | NA | NA | NA | NA | NA | Vmn1r88 | vomeronasal 1 receptor, 88 | 0 | 0 | 0 | 0 | 0 | 0 | 0 | 0 | 0 | 0 | 0 |
| 0 | NA | NA | NA | NA | NA | Gm4879 | | 0 | 0 | 0 | 0 | 0 | 0 | 0 | 0 | 0 | 0 | 0 |
| 0 | NA | NA | NA | NA | NA | Gm45525 | | 0 | 0 | 0 | 0 | 0 | 0 | 0 | 0 | 0 | 0 | 0 |
| 0 | NA | NA | NA | NA | NA | Vmn1r89 | vomeronasal 1 receptor 89 | 0 | 0 | 0 | 0 | 0 | 0 | 0 | 0 | 0 | 0 | 0 |
| 0 | NA | NA | NA | NA | NA | Vmn1r-ps69 | | 0 | 0 | 0 | 0 | 0 | 0 | 0 | 0 | 0 | 0 | 0 |
| 0 | NA | NA | NA | NA | NA | Gm24909 | | 0 | 0 | 0 | 0 | 0 | 0 | 0 | 0 | 0 | 0 | 0 |
| 0 | NA | NA | NA | NA | NA | Pla2g4c-ps | | 0 | 0 | 0 | 0 | 0 | 0 | 0 | 0 | 0 | 0 | 0 |
| 0 | NA | NA | NA | NA | NA | Gm45268 | | 0 | 0 | 0 | 0 | 0 | 0 | 0 | 0 | 0 | 0 | 0 |
| 0 | NA | NA | NA | NA | NA | Gm23499 | | 0 | 0 | 0 | 0 | 0 | 0 | 0 | 0 | 0 | 0 | 0 |
| 0 | NA | NA | NA | NA | NA | Rpl7a-ps14 | | 0 | 0 | 0 | 0 | 0 | 0 | 0 | 0 | 0 | 0 | 0 |
| 0 | NA | NA | NA | NA | NA | Bsph1 | binder of sperm protein homolog 1 | 0 | 0 | 0 | 0 | 0 | 0 | 0 | 0 | 0 | 0 | 0 |
| 0 | NA | NA | NA | NA | NA | Gm44254 | | 0 | 0 | 0 | 0 | 0 | 0 | 0 | 0 | 0 | 0 | 0 |
| 0 | NA | NA | NA | NA | NA | Bsph2-ps | | 0 | 0 | 0 | 0 | 0 | 0 | 0 | 0 | 0 | 0 | 0 |
| 0 | NA | NA | NA | NA | NA | Bsph2 | binder of sperm protein homolog 2 | 0 | 0 | 0 | 0 | 0 | 0 | 0 | 0 | 0 | 0 | 0 |
| 0 | NA | NA | NA | NA | NA | Sult2a5 | sulfotransferase family 2A, dehydroepian | 0 | 0 | 0 | 0 | 0 | 0 | 0 | 0 | 0 | 0 | 0 |
| 0 | NA | NA | NA | NA | NA | Kat6b-ps1 | | 0 | 0 | 0 | 0 | 0 | 0 | 0 | 0 | 0 | 0 | 0 |
| 0 | NA | NA | NA | NA | NA | Tmem167-ps1 | | 0 | 0 | 0 | 0 | 0 | 0 | 0 | 0 | 0 | 0 | 0 |
| 0 | NA | NA | NA | NA | NA | Sult2a-ps1 | | 0 | 0 | 0 | 0 | 0 | 0 | 0 | 0 | 0 | 0 | 0 |
| 0 | NA | NA | NA | NA | NA | Sult2a2 | sulfotransferase family 2A, dehydroepian | 0 | 0 | 0 | 0 | 0 | 0 | 0 | 0 | 0 | 0 | 0 |
| 0 | NA | NA | NA | NA | NA | Zbed4-ps2 | | 0 | 0 | 0 | 0 | 0 | 0 | 0 | 0 | 0 | 0 | 0 |
| 0 | NA | NA | NA | NA | NA | Sult2a1 | sulfotransferase family 2A, dehydroepian | 0 | 0 | 0 | 0 | 0 | 0 | 0 | 0 | 0 | 0 | 0 |
| 0 | NA | NA | NA | NA | NA | Kat6b-ps2 | | 0 | 0 | 0 | 0 | 0 | 0 | 0 | 0 | 0 | 0 | 0 |
| 0 | NA | NA | NA | NA | NA | Sult2a-ps2 | | 0 | 0 | 0 | 0 | 0 | 0 | 0 | 0 | 0 | 0 | 0 |
| 0 | NA | NA | NA | NA | NA | Tmem167-ps2 | | 0 | 0 | 0 | 0 | 0 | 0 | 0 | 0 | 0 | 0 | 0 |

| | | | | | | | | | | | | | | | | | | |
|---|---|---|---|---|---|---|---|---|---|---|---|---|---|---|---|---|---|---|
| 0 | NA | NA | NA | NA | NA | Sult2a4 | sulfotransferase family 2A, dehydroepian | 0 | 0 | 0 | 0 | 0 | 0 | 0 | 0 | 0 | 0 | 0 |
| 0 | NA | NA | NA | NA | NA | Tmem167-ps3 | | 0 | 0 | 0 | 0 | 0 | 0 | 0 | 0 | 0 | 0 | 0 |
| 0 | NA | NA | NA | NA | NA | Sult2a3 | sulfotransferase family 2A, dehydroepian | 0 | 0 | 0 | 0 | 0 | 0 | 0 | 0 | 0 | 0 | 0 |
| 0 | NA | NA | NA | NA | NA | Sult2a-ps3 | | 0 | 0 | 0 | 0 | 0 | 0 | 0 | 0 | 0 | 0 | 0 |
| 0 | NA | NA | NA | NA | NA | Sult2a-ps4 | | 0 | 0 | 0 | 0 | 0 | 0 | 0 | 0 | 0 | 0 | 0 |
| 0 | NA | NA | NA | NA | NA | Rpap3-ps1 | | 0 | 0 | 0 | 0 | 0 | 0 | 0 | 0 | 0 | 0 | 0 |
| 0 | NA | NA | NA | NA | NA | Sult2a6 | sulfotransferase family 2A, dehydroepian | 0 | 0 | 0 | 0 | 0 | 0 | 0 | 0 | 0 | 0 | 0 |
| 0 | NA | NA | NA | NA | NA | Rpap3-ps2 | | 0 | 0 | 0 | 0 | 0 | 0 | 0 | 0 | 0 | 0 | 0 |
| 0 | NA | NA | NA | NA | NA | Obox8 | | 0 | 0 | 0 | 0 | 0 | 0 | 0 | 0 | 0 | 0 | 0 |
| 0 | NA | NA | NA | NA | NA | Mrps36-ps3 | | 0 | 0 | 0 | 0 | 0 | 0 | 0 | 0 | 0 | 0 | 0 |
| 0 | NA | NA | NA | NA | NA | Crs-ps | | 0 | 0 | 0 | 0 | 0 | 0 | 0 | 0 | 0 | 0 | 0 |
| 0 | NA | NA | NA | NA | NA | Sult2a7 | sulfotransferase family 2A, dehydroepian | 0 | 0 | 0 | 0 | 0 | 0 | 0 | 0 | 0 | 0 | 0 |
| 0 | NA | NA | NA | NA | NA | Obox4-ps1 | | 0 | 0 | 0 | 0 | 0 | 0 | 0 | 0 | 0 | 0 | 0 |
| 0 | NA | NA | NA | NA | NA | Obox4-ps2 | | 0 | 0 | 0 | 0 | 0 | 0 | 0 | 0 | 0 | 0 | 0 |
| 0 | NA | NA | NA | NA | NA | Obox4-ps3 | | 0 | 0 | 0 | 0 | 0 | 0 | 0 | 0 | 0 | 0 | 0 |
| 0 | NA | NA | NA | NA | NA | Zfp-ps | | 0 | 0 | 0 | 0 | 0 | 0 | 0 | 0 | 0 | 0 | 0 |
| 0 | NA | NA | NA | NA | NA | Ranbp2-ps1 | | 0 | 0 | 0 | 0 | 0 | 0 | 0 | 0 | 0 | 0 | 0 |
| 0 | NA | NA | NA | NA | NA | Obox3-ps7 | | 0 | 0 | 0 | 0 | 0 | 0 | 0 | 0 | 0 | 0 | 0 |
| 0 | NA | NA | NA | NA | NA | Obox7 | oocyte specific homeobox 7 | 0 | 0 | 0 | 0 | 0 | 0 | 0 | 0 | 0 | 0 | 0 |
| 0 | NA | NA | NA | NA | NA | Obox3-ps1 | | 0 | 0 | 0 | 0 | 0 | 0 | 0 | 0 | 0 | 0 | 0 |
| 0 | NA | NA | NA | NA | NA | Obox4-ps4 | | 0 | 0 | 0 | 0 | 0 | 0 | 0 | 0 | 0 | 0 | 0 |
| 0 | NA | NA | NA | NA | NA | Gtpbp4-ps1 | | 0 | 0 | 0 | 0 | 0 | 0 | 0 | 0 | 0 | 0 | 0 |
| 0 | NA | NA | NA | NA | NA | Ranbp2-ps2 | | 0 | 0 | 0 | 0 | 0 | 0 | 0 | 0 | 0 | 0 | 0 |
| 0 | NA | NA | NA | NA | NA | Obox4-ps36 | | 0 | 0 | 0 | 0 | 0 | 0 | 0 | 0 | 0 | 0 | 0 |
| 0 | NA | NA | NA | NA | NA | Obox4-ps37 | | 0 | 0 | 0 | 0 | 0 | 0 | 0 | 0 | 0 | 0 | 0 |
| 0 | NA | NA | NA | NA | NA | Gm26056 | | 0 | 0 | 0 | 0 | 0 | 0 | 0 | 0 | 0 | 0 | 0 |
| 0 | NA | NA | NA | NA | NA | Obox3-ps2 | | 0 | 0 | 0 | 0 | 0 | 0 | 0 | 0 | 0 | 0 | 0 |
| 0 | NA | NA | NA | NA | NA | Obox4-ps5 | | 0 | 0 | 0 | 0 | 0 | 0 | 0 | 0 | 0 | 0 | 0 |
| 0 | NA | NA | NA | NA | NA | Gtpbp4-ps2 | | 0 | 0 | 0 | 0 | 0 | 0 | 0 | 0 | 0 | 0 | 0 |
| 0 | NA | NA | NA | NA | NA | Ranbp2-ps3 | | 0 | 0 | 0 | 0 | 0 | 0 | 0 | 0 | 0 | 0 | 0 |
| 0 | NA | NA | NA | NA | NA | Obox4-ps38 | | 0 | 0 | 0 | 0 | 0 | 0 | 0 | 0 | 0 | 0 | 0 |
| 0 | NA | NA | NA | NA | NA | Obox4-ps39 | | 0 | 0 | 0 | 0 | 0 | 0 | 0 | 0 | 0 | 0 | 0 |
| 0 | NA | NA | NA | NA | NA | Ranbp2-ps4 | | 0 | 0 | 0 | 0 | 0 | 0 | 0 | 0 | 0 | 0 | 0 |
| 0 | NA | NA | NA | NA | NA | Obox4-ps6 | | 0 | 0 | 0 | 0 | 0 | 0 | 0 | 0 | 0 | 0 | 0 |
| 0 | NA | NA | NA | NA | NA | Gm24701 | | 0 | 0 | 0 | 0 | 0 | 0 | 0 | 0 | 0 | 0 | 0 |
| 0 | NA | NA | NA | NA | NA | Obox3-ps3 | | 0 | 0 | 0 | 0 | 0 | 0 | 0 | 0 | 0 | 0 | 0 |
| 0 | NA | NA | NA | NA | NA | Obox4-ps7 | | 0 | 0 | 0 | 0 | 0 | 0 | 0 | 0 | 0 | 0 | 0 |
| 0 | NA | NA | NA | NA | NA | Gtpbp4-ps3 | | 0 | 0 | 0 | 0 | 0 | 0 | 0 | 0 | 0 | 0 | 0 |
| 0 | NA | NA | NA | NA | NA | Ranbp2-ps5 | | 0 | 0 | 0 | 0 | 0 | 0 | 0 | 0 | 0 | 0 | 0 |
| 0 | NA | NA | NA | NA | NA | Obox4-ps8 | | 0 | 0 | 0 | 0 | 0 | 0 | 0 | 0 | 0 | 0 | 0 |
| 0 | NA | NA | NA | NA | NA | Obox4-ps9 | | 0 | 0 | 0 | 0 | 0 | 0 | 0 | 0 | 0 | 0 | 0 |
| 0 | NA | NA | NA | NA | NA | Obox3-ps4 | | 0 | 0 | 0 | 0 | 0 | 0 | 0 | 0 | 0 | 0 | 0 |
| 0 | NA | NA | NA | NA | NA | Obox4-ps10 | | 0 | 0 | 0 | 0 | 0 | 0 | 0 | 0 | 0 | 0 | 0 |
| 0 | NA | NA | NA | NA | NA | Gtpbp4-ps4 | | 0 | 0 | 0 | 0 | 0 | 0 | 0 | 0 | 0 | 0 | 0 |
| 0 | NA | NA | NA | NA | NA | Ranbp2-ps6 | | 0 | 0 | 0 | 0 | 0 | 0 | 0 | 0 | 0 | 0 | 0 |
| 0 | NA | NA | NA | NA | NA | Obox4-ps11 | | 0 | 0 | 0 | 0 | 0 | 0 | 0 | 0 | 0 | 0 | 0 |
| 0 | NA | NA | NA | NA | NA | Obox4-ps12 | | 0 | 0 | 0 | 0 | 0 | 0 | 0 | 0 | 0 | 0 | 0 |
| 0 | NA | NA | NA | NA | NA | Obox4-ps13 | | 0 | 0 | 0 | 0 | 0 | 0 | 0 | 0 | 0 | 0 | 0 |
| 0 | NA | NA | NA | NA | NA | Obox4-ps14 | | 0 | 0 | 0 | 0 | 0 | 0 | 0 | 0 | 0 | 0 | 0 |
| 0 | NA | NA | NA | NA | NA | Obox4-ps15 | | 0 | 0 | 0 | 0 | 0 | 0 | 0 | 0 | 0 | 0 | 0 |
| 0 | NA | NA | NA | NA | NA | Obox4-ps16 | | 0 | 0 | 0 | 0 | 0 | 0 | 0 | 0 | 0 | 0 | 0 |
| 0 | NA | NA | NA | NA | NA | Obox4-ps17 | | 0 | 0 | 0 | 0 | 0 | 0 | 0 | 0 | 0 | 0 | 0 |
| 0 | NA | NA | NA | NA | NA | Obox4-ps18 | | 0 | 0 | 0 | 0 | 0 | 0 | 0 | 0 | 0 | 0 | 0 |
| 0 | NA | NA | NA | NA | NA | Obox4-ps19 | | 0 | 0 | 0 | 0 | 0 | 0 | 0 | 0 | 0 | 0 | 0 |
| 0 | NA | NA | NA | NA | NA | Obox4-ps20 | | 0 | 0 | 0 | 0 | 0 | 0 | 0 | 0 | 0 | 0 | 0 |
| 0 | NA | NA | NA | NA | NA | Obox4-ps21 | | 0 | 0 | 0 | 0 | 0 | 0 | 0 | 0 | 0 | 0 | 0 |
| 0 | NA | NA | NA | NA | NA | Obox4-ps22 | | 0 | 0 | 0 | 0 | 0 | 0 | 0 | 0 | 0 | 0 | 0 |
| 0 | NA | NA | NA | NA | NA | Obox4-ps23 | | 0 | 0 | 0 | 0 | 0 | 0 | 0 | 0 | 0 | 0 | 0 |
| 0 | NA | NA | NA | NA | NA | Obox3-ps5 | | 0 | 0 | 0 | 0 | 0 | 0 | 0 | 0 | 0 | 0 | 0 |
| 0 | NA | NA | NA | NA | NA | Obox4-ps24 | | 0 | 0 | 0 | 0 | 0 | 0 | 0 | 0 | 0 | 0 | 0 |

| | | | | | | | | | | | | | | | | | |
|---|---|---|---|---|---|---|---|---|---|---|---|---|---|---|---|---|---|
| 0 | NA | NA | NA | NA | NA | Gtpbp4-ps5 | | 0 | 0 | 0 | 0 | 0 | 0 | 0 | 0 | 0 | 0 | 0 |
| 0 | NA | NA | NA | NA | NA | Obox4-ps25 | | 0 | 0 | 0 | 0 | 0 | 0 | 0 | 0 | 0 | 0 | 0 |
| 0 | NA | NA | NA | NA | NA | Obox4-ps26 | | 0 | 0 | 0 | 0 | 0 | 0 | 0 | 0 | 0 | 0 | 0 |
| 0 | NA | NA | NA | NA | NA | Ranbp2-ps7 | | 0 | 0 | 0 | 0 | 0 | 0 | 0 | 0 | 0 | 0 | 0 |
| 0 | NA | NA | NA | NA | NA | Obox2 | oocyte specific homeobox 2 | 0 | 0 | 0 | 0 | 0 | 0 | 0 | 0 | 0 | 0 | 0 |
| 0 | NA | NA | NA | NA | NA | Obox1 | oocyte specific homeobox 1 | 0 | 0 | 0 | 0 | 0 | 0 | 0 | 0 | 0 | 0 | 0 |
| 0 | NA | NA | NA | NA | NA | Ranbp2-ps8 | | 0 | 0 | 0 | 0 | 0 | 0 | 0 | 0 | 0 | 0 | 0 |
| 0 | NA | NA | NA | NA | NA | Obox4-ps27 | | 0 | 0 | 0 | 0 | 0 | 0 | 0 | 0 | 0 | 0 | 0 |
| 0 | NA | NA | NA | NA | NA | Obox4-ps28 | | 0 | 0 | 0 | 0 | 0 | 0 | 0 | 0 | 0 | 0 | 0 |
| 0 | NA | NA | NA | NA | NA | Obox4-ps29 | | 0 | 0 | 0 | 0 | 0 | 0 | 0 | 0 | 0 | 0 | 0 |
| 0 | NA | NA | NA | NA | NA | Obox3 | oocyte specific homeobox 3 | 0 | 0 | 0 | 0 | 0 | 0 | 0 | 0 | 0 | 0 | 0 |
| 0 | NA | NA | NA | NA | NA | Obox3-ps6 | | 0 | 0 | 0 | 0 | 0 | 0 | 0 | 0 | 0 | 0 | 0 |
| 0 | NA | NA | NA | NA | NA | Obox4-ps30 | | 0 | 0 | 0 | 0 | 0 | 0 | 0 | 0 | 0 | 0 | 0 |
| 0 | NA | NA | NA | NA | NA | Ranbp2-ps9 | | 0 | 0 | 0 | 0 | 0 | 0 | 0 | 0 | 0 | 0 | 0 |
| 0 | NA | NA | NA | NA | NA | Ranbp2-ps10 | | 0 | 0 | 0 | 0 | 0 | 0 | 0 | 0 | 0 | 0 | 0 |
| 0 | NA | NA | NA | NA | NA | Obox4-ps31 | | 0 | 0 | 0 | 0 | 0 | 0 | 0 | 0 | 0 | 0 | 0 |
| 0 | NA | NA | NA | NA | NA | Gm25354 | | 0 | 0 | 0 | 0 | 0 | 0 | 0 | 0 | 0 | 0 | 0 |
| 0 | NA | NA | NA | NA | NA | Obox4-ps32 | | 0 | 0 | 0 | 0 | 0 | 0 | 0 | 0 | 0 | 0 | 0 |
| 0 | NA | NA | NA | NA | NA | Gtpbp4-ps6 | | 0 | 0 | 0 | 0 | 0 | 0 | 0 | 0 | 0 | 0 | 0 |
| 0 | NA | NA | NA | NA | NA | Obox4-ps33 | | 0 | 0 | 0 | 0 | 0 | 0 | 0 | 0 | 0 | 0 | 0 |
| 0 | NA | NA | NA | NA | NA | Gm23714 | | 0 | 0 | 0 | 0 | 0 | 0 | 0 | 0 | 0 | 0 | 0 |
| 0 | NA | NA | NA | NA | NA | Obox4-ps34 | | 0 | 0 | 0 | 0 | 0 | 0 | 0 | 0 | 0 | 0 | 0 |
| 0 | NA | NA | NA | NA | NA | Ranbp2-ps11 | | 0 | 0 | 0 | 0 | 0 | 0 | 0 | 0 | 0 | 0 | 0 |
| 0 | NA | NA | NA | NA | NA | Obox5 | oocyte specific homeobox 5 | 0 | 0 | 0 | 0 | 0 | 0 | 0 | 0 | 0 | 0 | 0 |
| 0 | NA | NA | NA | NA | NA | Ranbp2-ps12 | | 0 | 0 | 0 | 0 | 0 | 0 | 0 | 0 | 0 | 0 | 0 |
| 0 | NA | NA | NA | NA | NA | Gtpbp4-ps7 | | 0 | 0 | 0 | 0 | 0 | 0 | 0 | 0 | 0 | 0 | 0 |
| 0 | NA | NA | NA | NA | NA | Obox4-ps35 | | 0 | 0 | 0 | 0 | 0 | 0 | 0 | 0 | 0 | 0 | 0 |
| 0 | NA | NA | NA | NA | NA | Obox6 | oocyte specific homeobox 6 | 0 | 0 | 0 | 0 | 0 | 0 | 0 | 0 | 0 | 0 | 0 |
| 0 | NA | NA | NA | NA | NA | Crx | cone-rod homeobox | 0 | 0 | 0 | 0 | 0 | 0 | 0 | 0 | 0 | 0 | 0 |
| 0 | NA | NA | NA | NA | NA | Crxos | cone-rod homeobox, opposite strand | 0 | 0 | 0 | 0 | 0 | 0 | 0 | 0 | 0 | 0 | 0 |
| 0 | NA | NA | NA | NA | NA | Ass-ps1 | | 0 | 0 | 0 | 0 | 0 | 0 | 0 | 0 | 0 | 0 | 0 |
| 0 | NA | NA | NA | NA | NA | Gm24576 | | 0 | 0 | 0 | 0 | 0 | 0 | 0 | 0 | 0 | 0 | 0 |
| 0 | NA | NA | NA | NA | NA | Gm38948 | | 0 | 0 | 0 | 0 | 0 | 0 | 0 | 0 | 0 | 0 | 0 |
| 0 | NA | NA | NA | NA | NA | Gm45409 | | 0 | 0 | 0 | 0 | 0 | 0 | 0 | 0 | 0 | 0 | 0 |
| 0 | NA | NA | NA | NA | NA | Gm26473 | | 0 | 0 | 0 | 0 | 0 | 0 | 0 | 0 | 0 | 0 | 0 |
| 0 | NA | NA | NA | NA | NA | Gm45508 | | 0 | 0 | 0 | 0 | 0 | 0 | 0 | 0 | 0 | 0 | 0 |
| 0 | NA | NA | NA | NA | NA | Gm45509 | | 0 | 0 | 0 | 0 | 0 | 0 | 0 | 0 | 0 | 0 | 0 |
| 0 | NA | NA | NA | NA | NA | Gm45510 | | 0 | 0 | 0 | 0 | 0 | 0 | 0 | 0 | 0 | 0 | 0 |
| 0 | NA | NA | NA | NA | NA | Npas1 | neuronal PAS domain protein 1 | 0 | 0 | 0 | 0 | 0 | 0 | 0 | 0 | 0 | 0 | 0 |
| 0 | NA | NA | NA | NA | NA | Gm29442 | | 0 | 0 | 0 | 0 | 0 | 0 | 0 | 0 | 0 | 0 | 0 |
| 0 | NA | NA | NA | NA | NA | Gm24708 | | 0 | 0 | 0 | 0 | 0 | 0 | 0 | 0 | 0 | 0 | 0 |
| 0 | NA | NA | NA | NA | NA | Ceacam-ps1 | | 0 | 0 | 0 | 0 | 0 | 0 | 0 | 0 | 0 | 0 | 0 |
| 0 | NA | NA | NA | NA | NA | 2700080J24Rik | | 0 | 0 | 0 | 0 | 0 | 0 | 0 | 0 | 0 | 0 | 0 |
| 0 | NA | NA | NA | NA | NA | Gm25815 | | 0 | 0 | 0 | 0 | 0 | 0 | 0 | 0 | 0 | 0 | 0 |
| 0 | NA | NA | NA | NA | NA | 4833404L02Rik | | 0 | 0 | 0 | 0 | 0 | 0 | 0 | 0 | 0 | 0 | 0 |
| 0 | NA | NA | NA | NA | NA | Gm45219 | | 0 | 0 | 0 | 0 | 0 | 0 | 0 | 0 | 0 | 0 | 0 |
| 0 | NA | NA | NA | NA | NA | Ceacam3 | carcinoembryonic antigen-related cell adl | 0 | 0 | 0 | 0 | 0 | 0 | 0 | 0 | 0 | 0 | 0 |
| 0 | NA | NA | NA | NA | NA | Gm45032 | | 0 | 0 | 0 | 0 | 0 | 0 | 0 | 0 | 0 | 0 | 0 |
| 0 | NA | NA | NA | NA | NA | Gm32772 | | 0 | 0 | 0 | 0 | 0 | 0 | 0 | 0 | 0 | 0 | 0 |
| 0 | NA | NA | NA | NA | NA | Gm32846 | | 0 | 0 | 0 | 0 | 0 | 0 | 0 | 0 | 0 | 0 | 0 |
| 0 | NA | NA | NA | NA | NA | Gm24520 | | 0 | 0 | 0 | 0 | 0 | 0 | 0 | 0 | 0 | 0 | 0 |
| 0 | NA | NA | NA | NA | NA | Gm25068 | | 0 | 0 | 0 | 0 | 0 | 0 | 0 | 0 | 0 | 0 | 0 |
| 0 | NA | NA | NA | NA | NA | Gm5418 | | 0 | 0 | 0 | 0 | 0 | 0 | 0 | 0 | 0 | 0 | 0 |
| 0 | NA | NA | NA | NA | NA | Gm9519 | | 0 | 0 | 0 | 0 | 0 | 0 | 0 | 0 | 0 | 0 | 0 |
| 0 | NA | NA | NA | NA | NA | Psg-ps1 | | 0 | 0 | 0 | 0 | 0 | 0 | 0 | 0 | 0 | 0 | 0 |
| 0 | NA | NA | NA | NA | NA | Gm45150 | | 0 | 0 | 0 | 0 | 0 | 0 | 0 | 0 | 0 | 0 | 0 |
| 0 | NA | NA | NA | NA | NA | Ceacam5 | carcinoembryonic antigen-related cell adl | 0 | 0 | 0 | 0 | 0 | 0 | 0 | 0 | 0 | 0 | 0 |
| 0 | NA | NA | NA | NA | NA | Gm45149 | | 0 | 0 | 0 | 0 | 0 | 0 | 0 | 0 | 0 | 0 | 0 |
| 0 | NA | NA | NA | NA | NA | Ceacam14 | carcinoembryonic antigen-related cell adl | 0 | 0 | 0 | 0 | 0 | 0 | 0 | 0 | 0 | 0 | 0 |
| 0 | NA | NA | NA | NA | NA | Gm5155 | predicted gene 5155 | 0 | 0 | 0 | 0 | 0 | 0 | 0 | 0 | 0 | 0 | 0 |

| | | | | | | | | | | | | | | | | | |
|---|---|---|---|---|---|---|---|---|---|---|---|---|---|---|---|---|---|
| 0 | NA | NA | NA | NA | NA | Ceacam11 | carcinoembryonic antigen-related cell adl | 0 | 0 | 0 | 0 | 0 | 0 | 0 | 0 | 0 | 0 | 0 |
| 0 | NA | NA | NA | NA | NA | Ceacam13 | carcinoembryonic antigen-related cell adl | 0 | 0 | 0 | 0 | 0 | 0 | 0 | 0 | 0 | 0 | 0 |
| 0 | NA | NA | NA | NA | NA | Gm22414 | | 0 | 0 | 0 | 0 | 0 | 0 | 0 | 0 | 0 | 0 | 0 |
| 0 | NA | NA | NA | NA | NA | Igfl3 | IGF-like family member 3 | 0 | 0 | 0 | 0 | 0 | 0 | 0 | 0 | 0 | 0 | 0 |
| 0 | NA | NA | NA | NA | NA | Mill1 | MHC I like leukocyte 1 | 0 | 0 | 0 | 0 | 0 | 0 | 0 | 0 | 0 | 0 | 0 |
| 0 | NA | NA | NA | NA | NA | Gm33626 | | 0 | 0 | 0 | 0 | 0 | 0 | 0 | 0 | 0 | 0 | 0 |
| 0 | NA | NA | NA | NA | NA | Psg18 | pregnancy specific glycoprotein 18 | 0 | 0 | 0 | 0 | 0 | 0 | 0 | 0 | 0 | 0 | 0 |
| 0 | NA | NA | NA | NA | NA | Gm24715 | | 0 | 0 | 0 | 0 | 0 | 0 | 0 | 0 | 0 | 0 | 0 |
| 0 | NA | NA | NA | NA | NA | Psg28 | pregnancy-specific glycoprotein 28 | 0 | 0 | 0 | 0 | 0 | 0 | 0 | 0 | 0 | 0 | 0 |
| 0 | NA | NA | NA | NA | NA | Gm44871 | | 0 | 0 | 0 | 0 | 0 | 0 | 0 | 0 | 0 | 0 | 0 |
| 0 | NA | NA | NA | NA | NA | Psg26 | pregnancy-specific glycoprotein 26 | 0 | 0 | 0 | 0 | 0 | 0 | 0 | 0 | 0 | 0 | 0 |
| 0 | NA | NA | NA | NA | NA | Gm45156 | | 0 | 0 | 0 | 0 | 0 | 0 | 0 | 0 | 0 | 0 | 0 |
| 0 | NA | NA | NA | NA | NA | Psg25 | pregnancy-specific glycoprotein 25 | 0 | 0 | 0 | 0 | 0 | 0 | 0 | 0 | 0 | 0 | 0 |
| 0 | NA | NA | NA | NA | NA | Gm26062 | | 0 | 0 | 0 | 0 | 0 | 0 | 0 | 0 | 0 | 0 | 0 |
| 0 | NA | NA | NA | NA | NA | Psg27 | pregnancy-specific glycoprotein 27 | 0 | 0 | 0 | 0 | 0 | 0 | 0 | 0 | 0 | 0 | 0 |
| 0 | NA | NA | NA | NA | NA | Gm19091 | | 0 | 0 | 0 | 0 | 0 | 0 | 0 | 0 | 0 | 0 | 0 |
| 0 | NA | NA | NA | NA | NA | Gm44657 | | 0 | 0 | 0 | 0 | 0 | 0 | 0 | 0 | 0 | 0 | 0 |
| 0 | NA | NA | NA | NA | NA | Gm19092 | | 0 | 0 | 0 | 0 | 0 | 0 | 0 | 0 | 0 | 0 | 0 |
| 0 | NA | NA | NA | NA | NA | Gm44656 | | 0 | 0 | 0 | 0 | 0 | 0 | 0 | 0 | 0 | 0 | 0 |
| 0 | NA | NA | NA | NA | NA | Psg21 | pregnancy-specific glycoprotein 21 | 0 | 0 | 0 | 0 | 0 | 0 | 0 | 0 | 0 | 0 | 0 |
| 0 | NA | NA | NA | NA | NA | Gm44654 | | 0 | 0 | 0 | 0 | 0 | 0 | 0 | 0 | 0 | 0 | 0 |
| 0 | NA | NA | NA | NA | NA | Psg20 | pregnancy-specific glycoprotein 20 | 0 | 0 | 0 | 0 | 0 | 0 | 0 | 0 | 0 | 0 | 0 |
| 0 | NA | NA | NA | NA | NA | Gm44655 | | 0 | 0 | 0 | 0 | 0 | 0 | 0 | 0 | 0 | 0 | 0 |
| 0 | NA | NA | NA | NA | NA | Gm44563 | | 0 | 0 | 0 | 0 | 0 | 0 | 0 | 0 | 0 | 0 | 0 |
| 0 | NA | NA | NA | NA | NA | Gm44564 | | 0 | 0 | 0 | 0 | 0 | 0 | 0 | 0 | 0 | 0 | 0 |
| 0 | NA | NA | NA | NA | NA | Gm44566 | | 0 | 0 | 0 | 0 | 0 | 0 | 0 | 0 | 0 | 0 | 0 |
| 0 | NA | NA | NA | NA | NA | Gm44565 | | 0 | 0 | 0 | 0 | 0 | 0 | 0 | 0 | 0 | 0 | 0 |
| 0 | NA | NA | NA | NA | NA | Gm24464 | | 0 | 0 | 0 | 0 | 0 | 0 | 0 | 0 | 0 | 0 | 0 |
| 0 | NA | NA | NA | NA | NA | Gm44561 | | 0 | 0 | 0 | 0 | 0 | 0 | 0 | 0 | 0 | 0 | 0 |
| 0 | NA | NA | NA | NA | NA | Sycp1-ps1 | | 0 | 0 | 0 | 0 | 0 | 0 | 0 | 0 | 0 | 0 | 0 |
| 0 | NA | NA | NA | NA | NA | Psg19 | pregnancy specific glycoprotein 19 | 0 | 0 | 0 | 0 | 0 | 0 | 0 | 0 | 0 | 0 | 0 |
| 0 | NA | NA | NA | NA | NA | Gm19126 | | 0 | 0 | 0 | 0 | 0 | 0 | 0 | 0 | 0 | 0 | 0 |
| 0 | NA | NA | NA | NA | NA | Psg17 | pregnancy specific glycoprotein 17 | 0 | 0 | 0 | 0 | 0 | 0 | 0 | 0 | 0 | 0 | 0 |
| 0 | NA | NA | NA | NA | NA | Gm18855 | | 0 | 0 | 0 | 0 | 0 | 0 | 0 | 0 | 0 | 0 | 0 |
| 0 | NA | NA | NA | NA | NA | Gm45058 | | 0 | 0 | 0 | 0 | 0 | 0 | 0 | 0 | 0 | 0 | 0 |
| 0 | NA | NA | NA | NA | NA | Rsph6a | radial spoke head 6 homolog A (Chlamyc | 0 | 0 | 0 | 0 | 0 | 0 | 0 | 0 | 0 | 0 | 0 |
| 0 | NA | NA | NA | NA | NA | Mir3100 | | 0 | 0 | 0 | 0 | 0 | 0 | 0 | 0 | 0 | 0 | 0 |
| 0 | NA | NA | NA | NA | NA | Mir330 | | 0 | 0 | 0 | 0 | 0 | 0 | 0 | 0 | 0 | 0 | 0 |
| 0 | NA | NA | NA | NA | NA | 1700058P15Rik | | 0 | 0 | 0 | 0 | 0 | 0 | 0 | 0 | 0 | 0 | 0 |
| 0 | NA | NA | NA | NA | NA | Ppm1n | protein phosphatase, Mg2+/Mn2+ depenc | 0 | 0 | 0 | 0 | 0 | 0 | 0 | 0 | 0 | 0 | 0 |
| 0 | NA | NA | NA | NA | NA | Gm23099 | | 0 | 0 | 0 | 0 | 0 | 0 | 0 | 0 | 0 | 0 | 0 |
| 0 | NA | NA | NA | NA | NA | Gm45167 | | 0 | 0 | 0 | 0 | 0 | 0 | 0 | 0 | 0 | 0 | 0 |
| 0 | NA | NA | NA | NA | NA | Apoc2 | apolipoprotein C-II | 0 | 0 | 0 | 0 | 0 | 0 | 0 | 0 | 0 | 0 | 0 |
| 0 | NA | NA | NA | NA | NA | Gm44805 | predicted gene 44805 | 0 | 0 | 0 | 0 | 0 | 0 | 0 | 0 | 0 | 0 | 0 |
| 0 | NA | NA | NA | NA | NA | Apoc4 | apolipoprotein C-IV | 0 | 0 | 0 | 0 | 0 | 0 | 0 | 0 | 0 | 0 | 0 |
| 0 | NA | NA | NA | NA | NA | Gm26600 | | 0 | 0 | 0 | 0 | 0 | 0 | 0 | 0 | 0 | 0 | 0 |
| 0 | NA | NA | NA | NA | NA | Ceacam16 | carcinoembryonic antigen-related cell adl | 0 | 0 | 0 | 0 | 0 | 0 | 0 | 0 | 0 | 0 | 0 |
| 0 | NA | NA | NA | NA | NA | Gm44659 | | 0 | 0 | 0 | 0 | 0 | 0 | 0 | 0 | 0 | 0 | 0 |
| 0 | NA | NA | NA | NA | NA | Gm44660 | | 0 | 0 | 0 | 0 | 0 | 0 | 0 | 0 | 0 | 0 | 0 |
| 0 | NA | NA | NA | NA | NA | Gm16184 | | 0 | 0 | 0 | 0 | 0 | 0 | 0 | 0 | 0 | 0 | 0 |
| 0 | NA | NA | NA | NA | NA | Gm44651 | | 0 | 0 | 0 | 0 | 0 | 0 | 0 | 0 | 0 | 0 | 0 |
| 0 | NA | NA | NA | NA | NA | Nlrp9b | NLR family, pyrin domain containing 9B | 0 | 0 | 0 | 0 | 0 | 0 | 0 | 0 | 0 | 0 | 0 |
| 0 | NA | NA | NA | NA | NA | n-R5s152 | | 0 | 0 | 0 | 0 | 0 | 0 | 0 | 0 | 0 | 0 | 0 |
| 0 | NA | NA | NA | NA | NA | Gm18877 | | 0 | 0 | 0 | 0 | 0 | 0 | 0 | 0 | 0 | 0 | 0 |
| 0 | NA | NA | NA | NA | NA | Vmn1r91 | vomeronasal 1 receptor 91 | 0 | 0 | 0 | 0 | 0 | 0 | 0 | 0 | 0 | 0 | 0 |
| 0 | NA | NA | NA | NA | NA | Vmn1r240 | vomeronasal 1 receptor 240 | 0 | 0 | 0 | 0 | 0 | 0 | 0 | 0 | 0 | 0 | 0 |
| 0 | NA | NA | NA | NA | NA | Vmn1r93 | vomeronasal 1 receptor 93 | 0 | 0 | 0 | 0 | 0 | 0 | 0 | 0 | 0 | 0 | 0 |
| 0 | NA | NA | NA | NA | NA | Vmn1r94 | vomeronasal 1 receptor 94 | 0 | 0 | 0 | 0 | 0 | 0 | 0 | 0 | 0 | 0 | 0 |
| 0 | NA | NA | NA | NA | NA | Gm44598 | | 0 | 0 | 0 | 0 | 0 | 0 | 0 | 0 | 0 | 0 | 0 |
| 0 | NA | NA | NA | NA | NA | Vmn1r95 | vomeronasal 1 receptor, 95 | 0 | 0 | 0 | 0 | 0 | 0 | 0 | 0 | 0 | 0 | 0 |

| | | | | | | | | | | | | | | | | | | |
|---|---|---|---|---|---|---|---|---|---|---|---|---|---|---|---|---|---|---|
| 0 | NA | NA | NA | NA | NA | Vmn1r241 | vomeronasal 1 receptor 241 | 0 | 0 | 0 | 0 | 0 | 0 | 0 | 0 | 0 | 0 | 0 |
| 0 | NA | NA | NA | NA | NA | Vmn1r242 | vomeronasal 1 receptor 242 | 0 | 0 | 0 | 0 | 0 | 0 | 0 | 0 | 0 | 0 | 0 |
| 0 | NA | NA | NA | NA | NA | Gm18049 | | 0 | 0 | 0 | 0 | 0 | 0 | 0 | 0 | 0 | 0 | 0 |
| 0 | NA | NA | NA | NA | NA | Gm5890 | | 0 | 0 | 0 | 0 | 0 | 0 | 0 | 0 | 0 | 0 | 0 |
| 0 | NA | NA | NA | NA | NA | Gm18455 | | 0 | 0 | 0 | 0 | 0 | 0 | 0 | 0 | 0 | 0 | 0 |
| 0 | NA | NA | NA | NA | NA | Gm4501 | | 0 | 0 | 0 | 0 | 0 | 0 | 0 | 0 | 0 | 0 | 0 |
| 0 | NA | NA | NA | NA | NA | Vmn1r243 | vomeronasal 1 receptor 243 | 0 | 0 | 0 | 0 | 0 | 0 | 0 | 0 | 0 | 0 | 0 |
| 0 | NA | NA | NA | NA | NA | Vmn1r-ps79 | | 0 | 0 | 0 | 0 | 0 | 0 | 0 | 0 | 0 | 0 | 0 |
| 0 | NA | NA | NA | NA | NA | Vmn1r244 | vomeronasal 1 receptor 244 | 0 | 0 | 0 | 0 | 0 | 0 | 0 | 0 | 0 | 0 | 0 |
| 0 | NA | NA | NA | NA | NA | Vmn1r100 | vomeronasal 1 receptor 100 | 0 | 0 | 0 | 0 | 0 | 0 | 0 | 0 | 0 | 0 | 0 |
| 0 | NA | NA | NA | NA | NA | Vmn1r101 | vomeronasal 1 receptor 101 | 0 | 0 | 0 | 0 | 0 | 0 | 0 | 0 | 0 | 0 | 0 |
| 0 | NA | NA | NA | NA | NA | Gm10670 | predicted gene 10670 | 0 | 0 | 0 | 0 | 0 | 0 | 0 | 0 | 0 | 0 | 0 |
| 0 | NA | NA | NA | NA | NA | Vmn1r103 | vomeronasal 1 receptor 103 | 0 | 0 | 0 | 0 | 0 | 0 | 0 | 0 | 0 | 0 | 0 |
| 0 | NA | NA | NA | NA | NA | Vmn1r104 | vomeronasal 1 receptor 104 | 0 | 0 | 0 | 0 | 0 | 0 | 0 | 0 | 0 | 0 | 0 |
| 0 | NA | NA | NA | NA | NA | Vmn1r245 | vomeronasal 1 receptor 245 | 0 | 0 | 0 | 0 | 0 | 0 | 0 | 0 | 0 | 0 | 0 |
| 0 | NA | NA | NA | NA | NA | Vmn1r246 | vomeronasal 1 receptor 246 | 0 | 0 | 0 | 0 | 0 | 0 | 0 | 0 | 0 | 0 | 0 |
| 0 | NA | NA | NA | NA | NA | Gm4513 | serine/threonine-protein kinase MARK2-li | 0 | 0 | 0 | 0 | 0 | 0 | 0 | 0 | 0 | 0 | 0 |
| 0 | NA | NA | NA | NA | NA | Gm8486 | | 0 | 0 | 0 | 0 | 0 | 0 | 0 | 0 | 0 | 0 | 0 |
| 0 | NA | NA | NA | NA | NA | Vmn1r107 | vomeronasal 1 receptor 107 | 0 | 0 | 0 | 0 | 0 | 0 | 0 | 0 | 0 | 0 | 0 |
| 0 | NA | NA | NA | NA | NA | Vmn1r-ps71 | | 0 | 0 | 0 | 0 | 0 | 0 | 0 | 0 | 0 | 0 | 0 |
| 0 | NA | NA | NA | NA | NA | Vmn1r111 | vomeronasal 1 receptor 111 | 0 | 0 | 0 | 0 | 0 | 0 | 0 | 0 | 0 | 0 | 0 |
| 0 | NA | NA | NA | NA | NA | Gm18052 | | 0 | 0 | 0 | 0 | 0 | 0 | 0 | 0 | 0 | 0 | 0 |
| 0 | NA | NA | NA | NA | NA | Gm10668 | predicted gene 10668 | 0 | 0 | 0 | 0 | 0 | 0 | 0 | 0 | 0 | 0 | 0 |
| 0 | NA | NA | NA | NA | NA | Gm18458 | | 0 | 0 | 0 | 0 | 0 | 0 | 0 | 0 | 0 | 0 | 0 |
| 0 | NA | NA | NA | NA | NA | Vmn1r112 | vomeronasal 1 receptor 112 | 0 | 0 | 0 | 0 | 0 | 0 | 0 | 0 | 0 | 0 | 0 |
| 0 | NA | NA | NA | NA | NA | Vmn1r113 | vomeronasal 1 receptor 113 | 0 | 0 | 0 | 0 | 0 | 0 | 0 | 0 | 0 | 0 | 0 |
| 0 | NA | NA | NA | NA | NA | Vmn1r114 | vomeronasal 1 receptor 114 | 0 | 0 | 0 | 0 | 0 | 0 | 0 | 0 | 0 | 0 | 0 |
| 0 | NA | NA | NA | NA | NA | Vmn1r115 | vomeronasal 1 receptor 115 | 0 | 0 | 0 | 0 | 0 | 0 | 0 | 0 | 0 | 0 | 0 |
| 0 | NA | NA | NA | NA | NA | Vmn1r116 | vomeronasal 1 receptor 116 | 0 | 0 | 0 | 0 | 0 | 0 | 0 | 0 | 0 | 0 | 0 |
| 0 | NA | NA | NA | NA | NA | Vmn1r117 | vomeronasal 1 receptor 117 | 0 | 0 | 0 | 0 | 0 | 0 | 0 | 0 | 0 | 0 | 0 |
| 0 | NA | NA | NA | NA | NA | Vmn1r118 | vomeronasal 1 receptor 118 | 0 | 0 | 0 | 0 | 0 | 0 | 0 | 0 | 0 | 0 | 0 |
| 0 | NA | NA | NA | NA | NA | Gm4526 | | 0 | 0 | 0 | 0 | 0 | 0 | 0 | 0 | 0 | 0 | 0 |
| 0 | NA | NA | NA | NA | NA | Gm4528 | | 0 | 0 | 0 | 0 | 0 | 0 | 0 | 0 | 0 | 0 | 0 |
| 0 | NA | NA | NA | NA | NA | Vmn1r119 | vomeronasal 1 receptor 119 | 0 | 0 | 0 | 0 | 0 | 0 | 0 | 0 | 0 | 0 | 0 |
| 0 | NA | NA | NA | NA | NA | Vmn1r-ps73 | | 0 | 0 | 0 | 0 | 0 | 0 | 0 | 0 | 0 | 0 | 0 |
| 0 | NA | NA | NA | NA | NA | Vmn1r120 | vomeronasal 1 receptor 120 | 0 | 0 | 0 | 0 | 0 | 0 | 0 | 0 | 0 | 0 | 0 |
| 0 | NA | NA | NA | NA | NA | Vmn1r-ps74 | | 0 | 0 | 0 | 0 | 0 | 0 | 0 | 0 | 0 | 0 | 0 |
| 0 | NA | NA | NA | NA | NA | Vmn1r121 | vomeronasal 1 receptor 121 | 0 | 0 | 0 | 0 | 0 | 0 | 0 | 0 | 0 | 0 | 0 |
| 0 | NA | NA | NA | NA | NA | Vmn1r122 | vomeronasal 1 receptor 122 | 0 | 0 | 0 | 0 | 0 | 0 | 0 | 0 | 0 | 0 | 0 |
| 0 | NA | NA | NA | NA | NA | Vmn1r123 | vomeronasal 1 receptor 123 | 0 | 0 | 0 | 0 | 0 | 0 | 0 | 0 | 0 | 0 | 0 |
| 0 | NA | NA | NA | NA | NA | Gm18054 | | 0 | 0 | 0 | 0 | 0 | 0 | 0 | 0 | 0 | 0 | 0 |
| 0 | NA | NA | NA | NA | NA | Gm5157 | predicted gene 5157 | 0 | 0 | 0 | 0 | 0 | 0 | 0 | 0 | 0 | 0 | 0 |
| 0 | NA | NA | NA | NA | NA | Gm18460 | | 0 | 0 | 0 | 0 | 0 | 0 | 0 | 0 | 0 | 0 | 0 |
| 0 | NA | NA | NA | NA | NA | Gm4536 | | 0 | 0 | 0 | 0 | 0 | 0 | 0 | 0 | 0 | 0 | 0 |
| 0 | NA | NA | NA | NA | NA | Vmn1r124 | vomeronasal 1 receptor 124 | 0 | 0 | 0 | 0 | 0 | 0 | 0 | 0 | 0 | 0 | 0 |
| 0 | NA | NA | NA | NA | NA | Vmn1r125 | vomeronasal 1 receptor 125 | 0 | 0 | 0 | 0 | 0 | 0 | 0 | 0 | 0 | 0 | 0 |
| 0 | NA | NA | NA | NA | NA | Vmn1r126 | vomeronasal 1 receptor 126 | 0 | 0 | 0 | 0 | 0 | 0 | 0 | 0 | 0 | 0 | 0 |
| 0 | NA | NA | NA | NA | NA | Vmn1r127 | vomeronasal 1 receptor 127 | 0 | 0 | 0 | 0 | 0 | 0 | 0 | 0 | 0 | 0 | 0 |
| 0 | NA | NA | NA | NA | NA | Vmn1r128 | vomeronasal 1 receptor 128 | 0 | 0 | 0 | 0 | 0 | 0 | 0 | 0 | 0 | 0 | 0 |
| 0 | NA | NA | NA | NA | NA | Vmn1r129 | vomeronasal 1 receptor 129 | 0 | 0 | 0 | 0 | 0 | 0 | 0 | 0 | 0 | 0 | 0 |
| 0 | NA | NA | NA | NA | NA | Vmn1r-ps75 | | 0 | 0 | 0 | 0 | 0 | 0 | 0 | 0 | 0 | 0 | 0 |
| 0 | NA | NA | NA | NA | NA | Gm18055 | | 0 | 0 | 0 | 0 | 0 | 0 | 0 | 0 | 0 | 0 | 0 |
| 0 | NA | NA | NA | NA | NA | Gm6882 | predicted gene 6882 | 0 | 0 | 0 | 0 | 0 | 0 | 0 | 0 | 0 | 0 | 0 |
| 0 | NA | NA | NA | NA | NA | Gm18461 | | 0 | 0 | 0 | 0 | 0 | 0 | 0 | 0 | 0 | 0 | 0 |
| 0 | NA | NA | NA | NA | NA | Gm4541 | | 0 | 0 | 0 | 0 | 0 | 0 | 0 | 0 | 0 | 0 | 0 |
| 0 | NA | NA | NA | NA | NA | Vmn1r130 | vomeronasal 1 receptor 130 | 0 | 0 | 0 | 0 | 0 | 0 | 0 | 0 | 0 | 0 | 0 |
| 0 | NA | NA | NA | NA | NA | Vmn1r-ps76 | | 0 | 0 | 0 | 0 | 0 | 0 | 0 | 0 | 0 | 0 | 0 |
| 0 | NA | NA | NA | NA | NA | Vmn1r131 | vomeronasal 1 receptor 131 | 0 | 0 | 0 | 0 | 0 | 0 | 0 | 0 | 0 | 0 | 0 |
| 0 | NA | NA | NA | NA | NA | Gm18056 | | 0 | 0 | 0 | 0 | 0 | 0 | 0 | 0 | 0 | 0 | 0 |
| 0 | NA | NA | NA | NA | NA | Gm4545 | | 0 | 0 | 0 | 0 | 0 | 0 | 0 | 0 | 0 | 0 | 0 |

| | | | | | | | | | | | | | | | | | | |
|---|---|---|---|---|---|---|---|---|---|---|---|---|---|---|---|---|---|---|
| 0 | NA | NA | NA | NA | NA | Gm18462 | | 0 | 0 | 0 | 0 | 0 | 0 | 0 | 0 | 0 | 0 | 0 |
| 0 | NA | NA | NA | NA | NA | Vmn1r132 | vomeronasal 1 receptor 132 | 0 | 0 | 0 | 0 | 0 | 0 | 0 | 0 | 0 | 0 | 0 |
| 0 | NA | NA | NA | NA | NA | Gm8600 | predicted gene 8600 | 0 | 0 | 0 | 0 | 0 | 0 | 0 | 0 | 0 | 0 | 0 |
| 0 | NA | NA | NA | NA | NA | Gm18463 | | 0 | 0 | 0 | 0 | 0 | 0 | 0 | 0 | 0 | 0 | 0 |
| 0 | NA | NA | NA | NA | NA | Gm4548 | | 0 | 0 | 0 | 0 | 0 | 0 | 0 | 0 | 0 | 0 | 0 |
| 0 | NA | NA | NA | NA | NA | Vmn1r247 | vomeronasal 1 receptor 247 | 0 | 0 | 0 | 0 | 0 | 0 | 0 | 0 | 0 | 0 | 0 |
| 0 | NA | NA | NA | NA | NA | Vmn1r248 | vomeronasal 1 receptor 248 | 0 | 0 | 0 | 0 | 0 | 0 | 0 | 0 | 0 | 0 | 0 |
| 0 | NA | NA | NA | NA | NA | Gm44554 | | 0 | 0 | 0 | 0 | 0 | 0 | 0 | 0 | 0 | 0 | 0 |
| 0 | NA | NA | NA | NA | NA | Vmn1r135 | vomeronasal 1 receptor 135 | 0 | 0 | 0 | 0 | 0 | 0 | 0 | 0 | 0 | 0 | 0 |
| 0 | NA | NA | NA | NA | NA | Vmn1r249 | vomeronasal 1 receptor 249 | 0 | 0 | 0 | 0 | 0 | 0 | 0 | 0 | 0 | 0 | 0 |
| 0 | NA | NA | NA | NA | NA | Vmn1r-ps77 | | 0 | 0 | 0 | 0 | 0 | 0 | 0 | 0 | 0 | 0 | 0 |
| 0 | NA | NA | NA | NA | NA | Gm18057 | | 0 | 0 | 0 | 0 | 0 | 0 | 0 | 0 | 0 | 0 | 0 |
| 0 | NA | NA | NA | NA | NA | Gm5891 | predicted gene 5891 | 0 | 0 | 0 | 0 | 0 | 0 | 0 | 0 | 0 | 0 | 0 |
| 0 | NA | NA | NA | NA | NA | Gm18464 | | 0 | 0 | 0 | 0 | 0 | 0 | 0 | 0 | 0 | 0 | 0 |
| 0 | NA | NA | NA | NA | NA | Gm16443 | | 0 | 0 | 0 | 0 | 0 | 0 | 0 | 0 | 0 | 0 | 0 |
| 0 | NA | NA | NA | NA | NA | Vmn1r137 | vomeronasal 1 receptor 137 | 0 | 0 | 0 | 0 | 0 | 0 | 0 | 0 | 0 | 0 | 0 |
| 0 | NA | NA | NA | NA | NA | Vmn1r-ps78 | | 0 | 0 | 0 | 0 | 0 | 0 | 0 | 0 | 0 | 0 | 0 |
| 0 | NA | NA | NA | NA | NA | Vmn1r138 | vomeronasal 1 receptor 138 | 0 | 0 | 0 | 0 | 0 | 0 | 0 | 0 | 0 | 0 | 0 |
| 0 | NA | NA | NA | NA | NA | Gm18058 | | 0 | 0 | 0 | 0 | 0 | 0 | 0 | 0 | 0 | 0 | 0 |
| 0 | NA | NA | NA | NA | NA | Gm6176 | predicted gene 6176 | 0 | 0 | 0 | 0 | 0 | 0 | 0 | 0 | 0 | 0 | 0 |
| 0 | NA | NA | NA | NA | NA | Gm18465 | | 0 | 0 | 0 | 0 | 0 | 0 | 0 | 0 | 0 | 0 | 0 |
| 0 | NA | NA | NA | NA | NA | Vmn1r139 | vomeronasal 1 receptor 139 | 0 | 0 | 0 | 0 | 0 | 0 | 0 | 0 | 0 | 0 | 0 |
| 0 | NA | NA | NA | NA | NA | Gm16451 | predicted pseudogene 16451 | 0 | 0 | 0 | 0 | 0 | 0 | 0 | 0 | 0 | 0 | 0 |
| 0 | NA | NA | NA | NA | NA | Vmn1r250 | vomeronasal 1 receptor 250 | 0 | 0 | 0 | 0 | 0 | 0 | 0 | 0 | 0 | 0 | 0 |
| 0 | NA | NA | NA | NA | NA | Vmn1r142 | vomeronasal 1 receptor 142 | 0 | 0 | 0 | 0 | 0 | 0 | 0 | 0 | 0 | 0 | 0 |
| 0 | NA | NA | NA | NA | NA | Vmn1r143 | vomeronasal 1 receptor 143 | 0 | 0 | 0 | 0 | 0 | 0 | 0 | 0 | 0 | 0 | 0 |
| 0 | NA | NA | NA | NA | NA | Vmn1r251 | vomeronasal 1 receptor 251 | 0 | 0 | 0 | 0 | 0 | 0 | 0 | 0 | 0 | 0 | 0 |
| 0 | NA | NA | NA | NA | NA | Vmn1r252 | vomeronasal 1 receptor 252 | 0 | 0 | 0 | 0 | 0 | 0 | 0 | 0 | 0 | 0 | 0 |
| 0 | NA | NA | NA | NA | NA | Gm18059 | | 0 | 0 | 0 | 0 | 0 | 0 | 0 | 0 | 0 | 0 | 0 |
| 0 | NA | NA | NA | NA | NA | Gm4557 | | 0 | 0 | 0 | 0 | 0 | 0 | 0 | 0 | 0 | 0 | 0 |
| 0 | NA | NA | NA | NA | NA | Gm18466 | | 0 | 0 | 0 | 0 | 0 | 0 | 0 | 0 | 0 | 0 | 0 |
| 0 | NA | NA | NA | NA | NA | Gm4558 | | 0 | 0 | 0 | 0 | 0 | 0 | 0 | 0 | 0 | 0 | 0 |
| 0 | NA | NA | NA | NA | NA | Vmn1r253 | vomeronasal 1 receptor 253 | 0 | 0 | 0 | 0 | 0 | 0 | 0 | 0 | 0 | 0 | 0 |
| 0 | NA | NA | NA | NA | NA | Vmn1r-ps70 | | 0 | 0 | 0 | 0 | 0 | 0 | 0 | 0 | 0 | 0 | 0 |
| 0 | NA | NA | NA | NA | NA | Vmn1r254 | vomeronasal 1 receptor 254 | 0 | 0 | 0 | 0 | 0 | 0 | 0 | 0 | 0 | 0 | 0 |
| 0 | NA | NA | NA | NA | NA | Vmn1r148 | vomeronasal 1 receptor 148 | 0 | 0 | 0 | 0 | 0 | 0 | 0 | 0 | 0 | 0 | 0 |
| 0 | NA | NA | NA | NA | NA | Vmn1r149 | vomeronasal 1 receptor 149 | 0 | 0 | 0 | 0 | 0 | 0 | 0 | 0 | 0 | 0 | 0 |
| 0 | NA | NA | NA | NA | NA | Gm10665 | predicted gene 10665 | 0 | 0 | 0 | 0 | 0 | 0 | 0 | 0 | 0 | 0 | 0 |
| 0 | NA | NA | NA | NA | NA | Vmn1r151 | vomeronasal 1 receptor 151 | 0 | 0 | 0 | 0 | 0 | 0 | 0 | 0 | 0 | 0 | 0 |
| 0 | NA | NA | NA | NA | NA | Vmn1r152 | vomeronasal 1 receptor 152 | 0 | 0 | 0 | 0 | 0 | 0 | 0 | 0 | 0 | 0 | 0 |
| 0 | NA | NA | NA | NA | NA | Vmn1r255 | vomeronasal 1 receptor 255 | 0 | 0 | 0 | 0 | 0 | 0 | 0 | 0 | 0 | 0 | 0 |
| 0 | NA | NA | NA | NA | NA | Vmn1r256 | vomeronasal 1 receptor 256 | 0 | 0 | 0 | 0 | 0 | 0 | 0 | 0 | 0 | 0 | 0 |
| 0 | NA | NA | NA | NA | NA | Gm18060 | | 0 | 0 | 0 | 0 | 0 | 0 | 0 | 0 | 0 | 0 | 0 |
| 0 | NA | NA | NA | NA | NA | Gm4565 | predicted gene 4565 | 0 | 0 | 0 | 0 | 0 | 0 | 0 | 0 | 0 | 0 | 0 |
| 0 | NA | NA | NA | NA | NA | Gm18467 | | 0 | 0 | 0 | 0 | 0 | 0 | 0 | 0 | 0 | 0 | 0 |
| 0 | NA | NA | NA | NA | NA | Gm8685 | | 0 | 0 | 0 | 0 | 0 | 0 | 0 | 0 | 0 | 0 | 0 |
| 0 | NA | NA | NA | NA | NA | Vmn1r155 | vomeronasal 1 receptor 155 | 0 | 0 | 0 | 0 | 0 | 0 | 0 | 0 | 0 | 0 | 0 |
| 0 | NA | NA | NA | NA | NA | Vmn1r-ps80 | | 0 | 0 | 0 | 0 | 0 | 0 | 0 | 0 | 0 | 0 | 0 |
| 0 | NA | NA | NA | NA | NA | Vmn1r257 | vomeronasal 1 receptor 257 | 0 | 0 | 0 | 0 | 0 | 0 | 0 | 0 | 0 | 0 | 0 |
| 0 | NA | NA | NA | NA | NA | Gm18061 | | 0 | 0 | 0 | 0 | 0 | 0 | 0 | 0 | 0 | 0 | 0 |
| 0 | NA | NA | NA | NA | NA | Gm4567 | predicted gene 4567 | 0 | 0 | 0 | 0 | 0 | 0 | 0 | 0 | 0 | 0 | 0 |
| 0 | NA | NA | NA | NA | NA | Gm18468 | | 0 | 0 | 0 | 0 | 0 | 0 | 0 | 0 | 0 | 0 | 0 |
| 0 | NA | NA | NA | NA | NA | Vmn1r157 | vomeronasal 1 receptor 157 | 0 | 0 | 0 | 0 | 0 | 0 | 0 | 0 | 0 | 0 | 0 |
| 0 | NA | NA | NA | NA | NA | Vmn1r158 | vomeronasal 1 receptor 158 | 0 | 0 | 0 | 0 | 0 | 0 | 0 | 0 | 0 | 0 | 0 |
| 0 | NA | NA | NA | NA | NA | Gm4207 | | 0 | 0 | 0 | 0 | 0 | 0 | 0 | 0 | 0 | 0 | 0 |
| 0 | NA | NA | NA | NA | NA | Vmn1r159 | vomeronasal 1 receptor 159 | 0 | 0 | 0 | 0 | 0 | 0 | 0 | 0 | 0 | 0 | 0 |
| 0 | NA | NA | NA | NA | NA | Gm44825 | | 0 | 0 | 0 | 0 | 0 | 0 | 0 | 0 | 0 | 0 | 0 |
| 0 | NA | NA | NA | NA | NA | Vmn1r160 | vomeronasal 1 receptor 160 | 0 | 0 | 0 | 0 | 0 | 0 | 0 | 0 | 0 | 0 | 0 |
| 0 | NA | NA | NA | NA | NA | Gm19900 | | 0 | 0 | 0 | 0 | 0 | 0 | 0 | 0 | 0 | 0 | 0 |
| 0 | NA | NA | NA | NA | NA | Gm8708 | predicted gene 8708 | 0 | 0 | 0 | 0 | 0 | 0 | 0 | 0 | 0 | 0 | 0 |

| | | | | | | | | | | | | | | | | | | |
|---|---|---|---|---|---|---|---|---|---|---|---|---|---|---|---|---|---|---|
| 0 | NA | NA | NA | NA | NA | Gm18469 | | 0 | 0 | 0 | 0 | 0 | 0 | 0 | 0 | 0 | 0 | 0 |
| 0 | NA | NA | NA | NA | NA | Gm4574 | | 0 | 0 | 0 | 0 | 0 | 0 | 0 | 0 | 0 | 0 | 0 |
| 0 | NA | NA | NA | NA | NA | Vmn1r258 | vomeronasal 1 receptor 258 | 0 | 0 | 0 | 0 | 0 | 0 | 0 | 0 | 0 | 0 | 0 |
| 0 | NA | NA | NA | NA | NA | Vmn1r259 | vomeronasal 1 receptor 259 | 0 | 0 | 0 | 0 | 0 | 0 | 0 | 0 | 0 | 0 | 0 |
| 0 | NA | NA | NA | NA | NA | Vmn1r163 | vomeronasal 1 receptor 163 | 0 | 0 | 0 | 0 | 0 | 0 | 0 | 0 | 0 | 0 | 0 |
| 0 | NA | NA | NA | NA | NA | Vmn1r260 | vomeronasal 1 receptor 260 | 0 | 0 | 0 | 0 | 0 | 0 | 0 | 0 | 0 | 0 | 0 |
| 0 | NA | NA | NA | NA | NA | Gm16456 | | 0 | 0 | 0 | 0 | 0 | 0 | 0 | 0 | 0 | 0 | 0 |
| 0 | NA | NA | NA | NA | NA | Gm18062 | | 0 | 0 | 0 | 0 | 0 | 0 | 0 | 0 | 0 | 0 | 0 |
| 0 | NA | NA | NA | NA | NA | Gm10662 | predicted gene 10662 | 0 | 0 | 0 | 0 | 0 | 0 | 0 | 0 | 0 | 0 | 0 |
| 0 | NA | NA | NA | NA | NA | Gm18470 | | 0 | 0 | 0 | 0 | 0 | 0 | 0 | 0 | 0 | 0 | 0 |
| 0 | NA | NA | NA | NA | NA | Gm8728 | | 0 | 0 | 0 | 0 | 0 | 0 | 0 | 0 | 0 | 0 | 0 |
| 0 | NA | NA | NA | NA | NA | Vmn1r165 | vomeronasal 1 receptor 165 | 0 | 0 | 0 | 0 | 0 | 0 | 0 | 0 | 0 | 0 | 0 |
| 0 | NA | NA | NA | NA | NA | Vmn1r-ps83 | | 0 | 0 | 0 | 0 | 0 | 0 | 0 | 0 | 0 | 0 | 0 |
| 0 | NA | NA | NA | NA | NA | Vmn1r166 | vomeronasal 1 receptor 166 | 0 | 0 | 0 | 0 | 0 | 0 | 0 | 0 | 0 | 0 | 0 |
| 0 | NA | NA | NA | NA | NA | Gm18063 | | 0 | 0 | 0 | 0 | 0 | 0 | 0 | 0 | 0 | 0 | 0 |
| 0 | NA | NA | NA | NA | NA | Gm6902 | predicted gene 6902 | 0 | 0 | 0 | 0 | 0 | 0 | 0 | 0 | 0 | 0 | 0 |
| 0 | NA | NA | NA | NA | NA | Gm18471 | | 0 | 0 | 0 | 0 | 0 | 0 | 0 | 0 | 0 | 0 | 0 |
| 0 | NA | NA | NA | NA | NA | Nlrp4e | NLR family, pyrin domain containing 4E | 0 | 0 | 0 | 0 | 0 | 0 | 0 | 0 | 0 | 0 | 0 |
| 0 | NA | NA | NA | NA | NA | Nlrp5 | NLR family, pyrin domain containing 5 | 0 | 0 | 0 | 0 | 0 | 0 | 0 | 0 | 0 | 0 | 0 |
| 0 | NA | NA | NA | NA | NA | Gm25196 | | 0 | 0 | 0 | 0 | 0 | 0 | 0 | 0 | 0 | 0 | 0 |
| 0 | NA | NA | NA | NA | NA | Gm18598 | | 0 | 0 | 0 | 0 | 0 | 0 | 0 | 0 | 0 | 0 | 0 |
| 0 | NA | NA | NA | NA | NA | Gm18306 | | 0 | 0 | 0 | 0 | 0 | 0 | 0 | 0 | 0 | 0 | 0 |
| 0 | NA | NA | NA | NA | NA | Vmn1r-ps84 | | 0 | 0 | 0 | 0 | 0 | 0 | 0 | 0 | 0 | 0 | 0 |
| 0 | NA | NA | NA | NA | NA | Vmn1r167 | vomeronasal 1 receptor 167 | 0 | 0 | 0 | 0 | 0 | 0 | 0 | 0 | 0 | 0 | 0 |
| 0 | NA | NA | NA | NA | NA | Gm8745 | | 0 | 0 | 0 | 0 | 0 | 0 | 0 | 0 | 0 | 0 | 0 |
| 0 | NA | NA | NA | NA | NA | Gm18195 | | 0 | 0 | 0 | 0 | 0 | 0 | 0 | 0 | 0 | 0 | 0 |
| 0 | NA | NA | NA | NA | NA | Vmn1r168 | vomeronasal 1 receptor 168 | 0 | 0 | 0 | 0 | 0 | 0 | 0 | 0 | 0 | 0 | 0 |
| 0 | NA | NA | NA | NA | NA | Vmn1r169 | vomeronasal 1 receptor 169 | 0 | 0 | 0 | 0 | 0 | 0 | 0 | 0 | 0 | 0 | 0 |
| 0 | NA | NA | NA | NA | NA | Vmn1r170 | vomeronasal 1 receptor 170 | 0 | 0 | 0 | 0 | 0 | 0 | 0 | 0 | 0 | 0 | 0 |
| 0 | NA | NA | NA | NA | NA | Vmn1r171 | vomeronasal 1 receptor 171 | 0 | 0 | 0 | 0 | 0 | 0 | 0 | 0 | 0 | 0 | 0 |
| 0 | NA | NA | NA | NA | NA | Vmn1r172 | vomeronasal 1 receptor 172 | 0 | 0 | 0 | 0 | 0 | 0 | 0 | 0 | 0 | 0 | 0 |
| 0 | NA | NA | NA | NA | NA | Vmn1r173 | vomeronasal 1 receptor 173 | 0 | 0 | 0 | 0 | 0 | 0 | 0 | 0 | 0 | 0 | 0 |
| 0 | NA | NA | NA | NA | NA | Vmn1r174 | vomeronasal 1 receptor 174 | 0 | 0 | 0 | 0 | 0 | 0 | 0 | 0 | 0 | 0 | 0 |
| 0 | NA | NA | NA | NA | NA | Vmn1r-ps85 | | 0 | 0 | 0 | 0 | 0 | 0 | 0 | 0 | 0 | 0 | 0 |
| 0 | NA | NA | NA | NA | NA | Gm17780 | | 0 | 0 | 0 | 0 | 0 | 0 | 0 | 0 | 0 | 0 | 0 |
| 0 | NA | NA | NA | NA | NA | Gm44576 | | 0 | 0 | 0 | 0 | 0 | 0 | 0 | 0 | 0 | 0 | 0 |
| 0 | NA | NA | NA | NA | NA | Vmn1r175 | vomeronasal 1 receptor 175 | 0 | 0 | 0 | 0 | 0 | 0 | 0 | 0 | 0 | 0 | 0 |
| 0 | NA | NA | NA | NA | NA | Vmn1r176 | vomeronasal 1 receptor 176 | 0 | 0 | 0 | 0 | 0 | 0 | 0 | 0 | 0 | 0 | 0 |
| 0 | NA | NA | NA | NA | NA | Vmn1r177 | vomeronasal 1 receptor 177 | 0 | 0 | 0 | 0 | 0 | 0 | 0 | 0 | 0 | 0 | 0 |
| 0 | NA | NA | NA | NA | NA | Gm18206 | | 0 | 0 | 0 | 0 | 0 | 0 | 0 | 0 | 0 | 0 | 0 |
| 0 | NA | NA | NA | NA | NA | Vmn1r178 | vomeronasal 1 receptor 178 | 0 | 0 | 0 | 0 | 0 | 0 | 0 | 0 | 0 | 0 | 0 |
| 0 | NA | NA | NA | NA | NA | Vmn1r179 | vomeronasal 1 receptor 179 | 0 | 0 | 0 | 0 | 0 | 0 | 0 | 0 | 0 | 0 | 0 |
| 0 | NA | NA | NA | NA | NA | Vmn1r180 | vomeronasal 1 receptor 180 | 0 | 0 | 0 | 0 | 0 | 0 | 0 | 0 | 0 | 0 | 0 |
| 0 | NA | NA | NA | NA | NA | Gm5324 | | 0 | 0 | 0 | 0 | 0 | 0 | 0 | 0 | 0 | 0 | 0 |
| 0 | NA | NA | NA | NA | NA | Vmn1r181 | vomeronasal 1 receptor 181 | 0 | 0 | 0 | 0 | 0 | 0 | 0 | 0 | 0 | 0 | 0 |
| 0 | NA | NA | NA | NA | NA | V1rd19 | vomeronasal 1 receptor, D19 | 0 | 0 | 0 | 0 | 0 | 0 | 0 | 0 | 0 | 0 | 0 |
| 0 | NA | NA | NA | NA | NA | Vmn1r183 | vomeronasal 1 receptor 183 | 0 | 0 | 0 | 0 | 0 | 0 | 0 | 0 | 0 | 0 | 0 |
| 0 | NA | NA | NA | NA | NA | Gm36694 | | 0 | 0 | 0 | 0 | 0 | 0 | 0 | 0 | 0 | 0 | 0 |
| 0 | NA | NA | NA | NA | NA | Ptma-ps1 | | 0 | 0 | 0 | 0 | 0 | 0 | 0 | 0 | 0 | 0 | 0 |
| 0 | NA | NA | NA | NA | NA | Gm44943 | | 0 | 0 | 0 | 0 | 0 | 0 | 0 | 0 | 0 | 0 | 0 |
| 0 | NA | NA | NA | NA | NA | Gm44796 | | 0 | 0 | 0 | 0 | 0 | 0 | 0 | 0 | 0 | 0 | 0 |
| 0 | NA | NA | NA | NA | NA | Gm44513 | | 0 | 0 | 0 | 0 | 0 | 0 | 0 | 0 | 0 | 0 | 0 |
| 0 | NA | NA | NA | NA | NA | Gm15465 | | 0 | 0 | 0 | 0 | 0 | 0 | 0 | 0 | 0 | 0 | 0 |
| 0 | NA | NA | NA | NA | NA | | | 0 | 0 | 0 | 0 | 0 | 0 | 0 | 0 | 0 | 0 | 0 |
| 0 | NA | NA | NA | NA | NA | Gm27592 | | 0 | 0 | 0 | 0 | 0 | 0 | 0 | 0 | 0 | 0 | 0 |
| 0 | NA | NA | NA | NA | NA | Lypd5 | Ly6/Plaur domain containing 5 | 0 | 0 | 0 | 0 | 0 | 0 | 0 | 0 | 0 | 0 | 0 |
| 0 | NA | NA | NA | NA | NA | Gm36159 | | 0 | 0 | 0 | 0 | 0 | 0 | 0 | 0 | 0 | 0 | 0 |
| 0 | NA | NA | NA | NA | NA | Gm50092 | | 0 | 0 | 0 | 0 | 0 | 0 | 0 | 0 | 0 | 0 | 0 |
| 0 | NA | NA | NA | NA | NA | Gm26550 | | 0 | 0 | 0 | 0 | 0 | 0 | 0 | 0 | 0 | 0 | 0 |
| 0 | NA | NA | NA | NA | NA | 2200007N16Rik | | 0 | 0 | 0 | 0 | 0 | 0 | 0 | 0 | 0 | 0 | 0 |

| | | | | | | | | | | | | | | | | | | |
|---|---|---|---|---|---|---|---|---|---|---|---|---|---|---|---|---|---|---|
| 0 | NA | NA | NA | NA | NA | Tex101 | testis expressed gene 101 | 0 | 0 | 0 | 0 | 0 | 0 | 0 | 0 | 0 | 0 | 0 |
| 0 | NA | NA | NA | NA | NA | Gm36776 | | 0 | 0 | 0 | 0 | 0 | 0 | 0 | 0 | 0 | 0 | 0 |
| 0 | NA | NA | NA | NA | NA | Lypd11 | Ly6/PLAUR domain containing 11 | 0 | 0 | 0 | 0 | 0 | 0 | 0 | 0 | 0 | 0 | 0 |
| 0 | NA | NA | NA | NA | NA | Gm44972 | | 0 | 0 | 0 | 0 | 0 | 0 | 0 | 0 | 0 | 0 | 0 |
| 0 | NA | NA | NA | NA | NA | Gm5893 | | 0 | 0 | 0 | 0 | 0 | 0 | 0 | 0 | 0 | 0 | 0 |
| 0 | NA | NA | NA | NA | NA | Gm18207 | | 0 | 0 | 0 | 0 | 0 | 0 | 0 | 0 | 0 | 0 | 0 |
| 0 | NA | NA | NA | NA | NA | Lypd4 | Ly6/Plaur domain containing 4 | 0 | 0 | 0 | 0 | 0 | 0 | 0 | 0 | 0 | 0 | 0 |
| 0 | NA | NA | NA | NA | NA | Dmrtc2 | doublesex and mab-3 related transcriptio | 0 | 0 | 0 | 0 | 0 | 0 | 0 | 0 | 0 | 0 | 0 |
| 0 | NA | NA | NA | NA | NA | Erfl | ETS repressor factor like | 0 | 0 | 0 | 0 | 0 | 0 | 0 | 0 | 0 | 0 | 0 |
| 0 | NA | NA | NA | NA | NA | Gm44970 | | 0 | 0 | 0 | 0 | 0 | 0 | 0 | 0 | 0 | 0 | 0 |
| 0 | NA | NA | NA | NA | NA | Mir7046 | | 0 | 0 | 0 | 0 | 0 | 0 | 0 | 0 | 0 | 0 | 0 |
| 0 | NA | NA | NA | NA | NA | Mir7047 | | 0 | 0 | 0 | 0 | 0 | 0 | 0 | 0 | 0 | 0 | 0 |
| 0 | NA | NA | NA | NA | NA | Mir6537 | | 0 | 0 | 0 | 0 | 0 | 0 | 0 | 0 | 0 | 0 | 0 |
| 0 | NA | NA | NA | NA | NA | D930028M14Rik | | 0 | 0 | 0 | 0 | 0 | 0 | 0 | 0 | 0 | 0 | 0 |
| 0 | NA | NA | NA | NA | NA | Cnfn | cornifelin | 0 | 0 | 0 | 0 | 0 | 0 | 0 | 0 | 0 | 0 | 0 |
| 0 | NA | NA | NA | NA | NA | Gm20949 | | 0 | 0 | 0 | 0 | 0 | 0 | 0 | 0 | 0 | 0 | 0 |
| 0 | NA | NA | NA | NA | NA | Gm29918 | | 0 | 0 | 0 | 0 | 0 | 0 | 0 | 0 | 0 | 0 | 0 |
| 0 | NA | NA | NA | NA | NA | Gm38569 | | 0 | 0 | 0 | 0 | 0 | 0 | 0 | 0 | 0 | 0 | 0 |
| 0 | NA | NA | NA | NA | NA | Gm45226 | | 0 | 0 | 0 | 0 | 0 | 0 | 0 | 0 | 0 | 0 | 0 |
| 0 | NA | NA | NA | NA | NA | Gm24865 | | 0 | 0 | 0 | 0 | 0 | 0 | 0 | 0 | 0 | 0 | 0 |
| 0 | NA | NA | NA | NA | NA | Gm30146 | | 0 | 0 | 0 | 0 | 0 | 0 | 0 | 0 | 0 | 0 | 0 |
| 0 | NA | NA | NA | NA | NA | Gm23921 | | 0 | 0 | 0 | 0 | 0 | 0 | 0 | 0 | 0 | 0 | 0 |
| 0 | NA | NA | NA | NA | NA | Gm45228 | | 0 | 0 | 0 | 0 | 0 | 0 | 0 | 0 | 0 | 0 | 0 |
| 0 | NA | NA | NA | NA | NA | Gm45227 | | 0 | 0 | 0 | 0 | 0 | 0 | 0 | 0 | 0 | 0 | 0 |
| 0 | NA | NA | NA | NA | NA | Gm45225 | | 0 | 0 | 0 | 0 | 0 | 0 | 0 | 0 | 0 | 0 | 0 |
| 0 | NA | NA | NA | NA | NA | Cyp2b27-ps | | 0 | 0 | 0 | 0 | 0 | 0 | 0 | 0 | 0 | 0 | 0 |
| 0 | NA | NA | NA | NA | NA | BC026762 | | 0 | 0 | 0 | 0 | 0 | 0 | 0 | 0 | 0 | 0 | 0 |
| 0 | NA | NA | NA | NA | NA | Cyp2b13 | cytochrome P450, family 2, subfamily b, p | 0 | 0 | 0 | 0 | 0 | 0 | 0 | 0 | 0 | 0 | 0 |
| 0 | NA | NA | NA | NA | NA | Gm29920 | | 0 | 0 | 0 | 0 | 0 | 0 | 0 | 0 | 0 | 0 | 0 |
| 0 | NA | NA | NA | NA | NA | Cyp2b26-ps | | 0 | 0 | 0 | 0 | 0 | 0 | 0 | 0 | 0 | 0 | 0 |
| 0 | NA | NA | NA | NA | NA | Cyp2b9 | cytochrome P450, family 2, subfamily b, p | 0 | 0 | 0 | 0 | 0 | 0 | 0 | 0 | 0 | 0 | 0 |
| 0 | NA | NA | NA | NA | NA | Vmn1r-ps86 | | 0 | 0 | 0 | 0 | 0 | 0 | 0 | 0 | 0 | 0 | 0 |
| 0 | NA | NA | NA | NA | NA | Vmn1r184 | vomeronasal 1 receptor, 184 | 0 | 0 | 0 | 0 | 0 | 0 | 0 | 0 | 0 | 0 | 0 |
| 0 | NA | NA | NA | NA | NA | Gm44964 | | 0 | 0 | 0 | 0 | 0 | 0 | 0 | 0 | 0 | 0 | 0 |
| 0 | NA | NA | NA | NA | NA | Gm42375 | | 0 | 0 | 0 | 0 | 0 | 0 | 0 | 0 | 0 | 0 | 0 |
| 0 | NA | NA | NA | NA | NA | Cyp2a4 | cytochrome P450, family 2, subfamily a, p | 0 | 0 | 0 | 0 | 0 | 0 | 0 | 0 | 0 | 0 | 0 |
| 0 | NA | NA | NA | NA | NA | Nlrp9c | NLR family, pyrin domain containing 9C | 0 | 0 | 0 | 0 | 0 | 0 | 0 | 0 | 0 | 0 | 0 |
| 0 | NA | NA | NA | NA | NA | Nlrp4a | NLR family, pyrin domain containing 4A | 0 | 0 | 0 | 0 | 0 | 0 | 0 | 0 | 0 | 0 | 0 |
| 0 | NA | NA | NA | NA | NA | Gm25947 | | 0 | 0 | 0 | 0 | 0 | 0 | 0 | 0 | 0 | 0 | 0 |
| 0 | NA | NA | NA | NA | NA | Nlrp9a | NLR family, pyrin domain containing 9A | 0 | 0 | 0 | 0 | 0 | 0 | 0 | 0 | 0 | 0 | 0 |
| 0 | NA | NA | NA | NA | NA | Vmn1r185 | vomeronasal 1 receptor 185 | 0 | 0 | 0 | 0 | 0 | 0 | 0 | 0 | 0 | 0 | 0 |
| 0 | NA | NA | NA | NA | NA | | | 0 | 0 | 0 | 0 | 0 | 0 | 0 | 0 | 0 | 0 | 0 |
| 0 | NA | NA | NA | NA | NA | Cyp2b23 | cytochrome P450, family 2, subfamily b, p | 0 | 0 | 0 | 0 | 0 | 0 | 0 | 0 | 0 | 0 | 0 |
| 0 | NA | NA | NA | NA | NA | Cyp2b19 | cytochrome P450, family 2, subfamily b, p | 0 | 0 | 0 | 0 | 0 | 0 | 0 | 0 | 0 | 0 | 0 |
| 0 | NA | NA | NA | NA | NA | Cyp2g1 | cytochrome P450, family 2, subfamily g, p | 0 | 0 | 0 | 0 | 0 | 0 | 0 | 0 | 0 | 0 | 0 |
| 0 | NA | NA | NA | NA | NA | Cyp2a5 | cytochrome P450, family 2, subfamily a, p | 0 | 0 | 0 | 0 | 0 | 0 | 0 | 0 | 0 | 0 | 0 |
| 0 | NA | NA | NA | NA | NA | Cyp2a21-ps | | 0 | 0 | 0 | 0 | 0 | 0 | 0 | 0 | 0 | 0 | 0 |
| 0 | NA | NA | NA | NA | NA | Api5-ps | | 0 | 0 | 0 | 0 | 0 | 0 | 0 | 0 | 0 | 0 | 0 |
| 0 | NA | NA | NA | NA | NA | Cyp2a22 | cytochrome P450, family 2, subfamily a, p | 0 | 0 | 0 | 0 | 0 | 0 | 0 | 0 | 0 | 0 | 0 |
| 0 | NA | NA | NA | NA | NA | Cyp2a12 | cytochrome P450, family 2, subfamily a, p | 0 | 0 | 0 | 0 | 0 | 0 | 0 | 0 | 0 | 0 | 0 |
| 0 | NA | NA | NA | NA | NA | Gm4290 | | 0 | 0 | 0 | 0 | 0 | 0 | 0 | 0 | 0 | 0 | 0 |
| 0 | NA | NA | NA | NA | NA | Gm50480 | | 0 | 0 | 0 | 0 | 0 | 0 | 0 | 0 | 0 | 0 | 0 |
| 0 | NA | NA | NA | NA | NA | Gm29754 | | 0 | 0 | 0 | 0 | 0 | 0 | 0 | 0 | 0 | 0 | 0 |
| 0 | NA | NA | NA | NA | NA | Gm15883 | | 0 | 0 | 0 | 0 | 0 | 0 | 0 | 0 | 0 | 0 | 0 |
| 0 | NA | NA | NA | NA | NA | Cyp2t4 | cytochrome P450, family 2, subfamily t, po | 0 | 0 | 0 | 0 | 0 | 0 | 0 | 0 | 0 | 0 | 0 |
| 0 | NA | NA | NA | NA | NA | Gm21983 | | 0 | 0 | 0 | 0 | 0 | 0 | 0 | 0 | 0 | 0 | 0 |
| 0 | NA | NA | NA | NA | NA | Mir3101 | | 0 | 0 | 0 | 0 | 0 | 0 | 0 | 0 | 0 | 0 | 0 |
| 0 | NA | NA | NA | NA | NA | Mir1191 | | 0 | 0 | 0 | 0 | 0 | 0 | 0 | 0 | 0 | 0 | 0 |
| 0 | NA | NA | NA | NA | NA | Gm44641 | | 0 | 0 | 0 | 0 | 0 | 0 | 0 | 0 | 0 | 0 | 0 |
| 0 | NA | NA | NA | NA | NA | Gm15570 | | 0 | 0 | 0 | 0 | 0 | 0 | 0 | 0 | 0 | 0 | 0 |

| | | | | | | | | | | | | | | | | | |
|---|---|---|---|---|---|---|---|---|---|---|---|---|---|---|---|---|---|
| 0 | NA | NA | NA | NA | NA | Gm20479 | | 0 | 0 | 0 | 0 | 0 | 0 | 0 | 0 | 0 | 0 | 0 |
| 0 | NA | NA | NA | NA | NA | Gm44924 | | 0 | 0 | 0 | 0 | 0 | 0 | 0 | 0 | 0 | 0 | 0 |
| 0 | NA | NA | NA | NA | NA | | | 0 | 0 | 0 | 0 | 0 | 0 | 0 | 0 | 0 | 0 | 0 |
| 0 | NA | NA | NA | NA | NA | | | 0 | 0 | 0 | 0 | 0 | 0 | 0 | 0 | 0 | 0 | 0 |
| 0 | NA | NA | NA | NA | NA | Gm24356 | | 0 | 0 | 0 | 0 | 0 | 0 | 0 | 0 | 0 | 0 | 0 |
| 0 | NA | NA | NA | NA | NA | Gm45011 | | 0 | 0 | 0 | 0 | 0 | 0 | 0 | 0 | 0 | 0 | 0 |
| 0 | NA | NA | NA | NA | NA | Gm18672 | | 0 | 0 | 0 | 0 | 0 | 0 | 0 | 0 | 0 | 0 | 0 |
| 0 | NA | NA | NA | NA | NA | Selenov | selenoprotein V | 0 | 0 | 0 | 0 | 0 | 0 | 0 | 0 | 0 | 0 | 0 |
| 0 | NA | NA | NA | NA | NA | 1700028B04Rik | | 0 | 0 | 0 | 0 | 0 | 0 | 0 | 0 | 0 | 0 | 0 |
| 0 | NA | NA | NA | NA | NA | Ifnl2 | interferon lambda 2 | 0 | 0 | 0 | 0 | 0 | 0 | 0 | 0 | 0 | 0 | 0 |
| 0 | NA | NA | NA | NA | NA | | | 0 | 0 | 0 | 0 | 0 | 0 | 0 | 0 | 0 | 0 | 0 |
| 0 | NA | NA | NA | NA | NA | Ifnl3 | interferon lambda 3 | 0 | 0 | 0 | 0 | 0 | 0 | 0 | 0 | 0 | 0 | 0 |
| 0 | NA | NA | NA | NA | NA | Acp7 | acid phosphatase 7, tartrate resistant | 0 | 0 | 0 | 0 | 0 | 0 | 0 | 0 | 0 | 0 | 0 |
| 0 | NA | NA | NA | NA | NA | Gm44618 | | 0 | 0 | 0 | 0 | 0 | 0 | 0 | 0 | 0 | 0 | 0 |
| 0 | NA | NA | NA | NA | NA | Gm44617 | | 0 | 0 | 0 | 0 | 0 | 0 | 0 | 0 | 0 | 0 | 0 |
| 0 | NA | NA | NA | NA | NA | Gm44700 | | 0 | 0 | 0 | 0 | 0 | 0 | 0 | 0 | 0 | 0 | 0 |
| 0 | NA | NA | NA | NA | NA | Gm38979 | | 0 | 0 | 0 | 0 | 0 | 0 | 0 | 0 | 0 | 0 | 0 |
| 0 | NA | NA | NA | NA | NA | Gm44408 | | 0 | 0 | 0 | 0 | 0 | 0 | 0 | 0 | 0 | 0 | 0 |
| 0 | NA | NA | NA | NA | NA | Mir1963 | | 0 | 0 | 0 | 0 | 0 | 0 | 0 | 0 | 0 | 0 | 0 |
| 0 | NA | NA | NA | NA | NA | Gm16282 | | 0 | 0 | 0 | 0 | 0 | 0 | 0 | 0 | 0 | 0 | 0 |
| 0 | NA | NA | NA | NA | NA | Gm17113 | | 0 | 0 | 0 | 0 | 0 | 0 | 0 | 0 | 0 | 0 | 0 |
| 0 | NA | NA | NA | NA | NA | Gm44632 | | 0 | 0 | 0 | 0 | 0 | 0 | 0 | 0 | 0 | 0 | 0 |
| 0 | NA | NA | NA | NA | NA | 4930432E11Rik | | 0 | 0 | 0 | 0 | 0 | 0 | 0 | 0 | 0 | 0 | 0 |
| 0 | NA | NA | NA | NA | NA | Gm29326 | | 0 | 0 | 0 | 0 | 0 | 0 | 0 | 0 | 0 | 0 | 0 |
| 0 | NA | NA | NA | NA | NA | A330087D11Rik | | 0 | 0 | 0 | 0 | 0 | 0 | 0 | 0 | 0 | 0 | 0 |
| 0 | NA | NA | NA | NA | NA | n-R5s153 | | 0 | 0 | 0 | 0 | 0 | 0 | 0 | 0 | 0 | 0 | 0 |
| 0 | NA | NA | NA | NA | NA | Gm26920 | | 0 | 0 | 0 | 0 | 0 | 0 | 0 | 0 | 0 | 0 | 0 |
| 0 | NA | NA | NA | NA | NA | Gm44872 | | 0 | 0 | 0 | 0 | 0 | 0 | 0 | 0 | 0 | 0 | 0 |
| 0 | NA | NA | NA | NA | NA | Gm44600 | | 0 | 0 | 0 | 0 | 0 | 0 | 0 | 0 | 0 | 0 | 0 |
| 0 | NA | NA | NA | NA | NA | Gm24757 | | 0 | 0 | 0 | 0 | 0 | 0 | 0 | 0 | 0 | 0 | 0 |
| 0 | NA | NA | NA | NA | NA | Gm30646 | | 0 | 0 | 0 | 0 | 0 | 0 | 0 | 0 | 0 | 0 | 0 |
| 0 | NA | NA | NA | NA | NA | Nphs1os | | 0 | 0 | 0 | 0 | 0 | 0 | 0 | 0 | 0 | 0 | 0 |
| 0 | NA | NA | NA | NA | NA | Arhgap33os | | 0 | 0 | 0 | 0 | 0 | 0 | 0 | 0 | 0 | 0 | 0 |
| 0 | NA | NA | NA | NA | NA | 1700019A23Rik | | 0 | 0 | 0 | 0 | 0 | 0 | 0 | 0 | 0 | 0 | 0 |
| 0 | NA | NA | NA | NA | NA | Gm38670 | | 0 | 0 | 0 | 0 | 0 | 0 | 0 | 0 | 0 | 0 | 0 |
| 0 | NA | NA | NA | NA | NA | Igflr1 | IGF-like family receptor 1 | 0 | 0 | 0 | 0 | 0 | 0 | 0 | 0 | 0 | 0 | 0 |
| 0 | NA | NA | NA | NA | NA | Etv2 | ets variant 2 | 0 | 0 | 0 | 0 | 0 | 0 | 0 | 0 | 0 | 0 | 0 |
| 0 | NA | NA | NA | NA | NA | Gm21982 | | 0 | 0 | 0 | 0 | 0 | 0 | 0 | 0 | 0 | 0 | 0 |
| 0 | NA | NA | NA | NA | NA | Pmis2 | | 0 | 0 | 0 | 0 | 0 | 0 | 0 | 0 | 0 | 0 | 0 |
| 0 | NA | NA | NA | NA | NA | Tmem147os | | 0 | 0 | 0 | 0 | 0 | 0 | 0 | 0 | 0 | 0 | 0 |
| 0 | NA | NA | NA | NA | NA | Mir7050 | | 0 | 0 | 0 | 0 | 0 | 0 | 0 | 0 | 0 | 0 | 0 |
| 0 | NA | NA | NA | NA | NA | Gm9569 | | 0 | 0 | 0 | 0 | 0 | 0 | 0 | 0 | 0 | 0 | 0 |
| 0 | NA | NA | NA | NA | NA | Scgb1b1-ps | | 0 | 0 | 0 | 0 | 0 | 0 | 0 | 0 | 0 | 0 | 0 |
| 0 | NA | NA | NA | NA | NA | Scgb2b1 | secretoglobin, family 2B, member 1 | 0 | 0 | 0 | 0 | 0 | 0 | 0 | 0 | 0 | 0 | 0 |
| 0 | NA | NA | NA | NA | NA | Scgb2b2 | secretoglobin, family 2B, member 2 | 0 | 0 | 0 | 0 | 0 | 0 | 0 | 0 | 0 | 0 | 0 |
| 0 | NA | NA | NA | NA | NA | Scgb1b2 | secretoglobin, family 1B, member 2 | 0 | 0 | 0 | 0 | 0 | 0 | 0 | 0 | 0 | 0 | 0 |
| 0 | NA | NA | NA | NA | NA | Scgb2b3 | secretoglobin, family 2B, member 3 | 0 | 0 | 0 | 0 | 0 | 0 | 0 | 0 | 0 | 0 | 0 |
| 0 | NA | NA | NA | NA | NA | Scgb1b3 | secretoglobin, family 1B, member 3 | 0 | 0 | 0 | 0 | 0 | 0 | 0 | 0 | 0 | 0 | 0 |
| 0 | NA | NA | NA | NA | NA | Scgb2b4-ps | secretoglobin, family 2B, member 4, pseu | 0 | 0 | 0 | 0 | 0 | 0 | 0 | 0 | 0 | 0 | 0 |
| 0 | NA | NA | NA | NA | NA | Scgb1b4-ps | | 0 | 0 | 0 | 0 | 0 | 0 | 0 | 0 | 0 | 0 | 0 |
| 0 | NA | NA | NA | NA | NA | Scgb1b28-ps | | 0 | 0 | 0 | 0 | 0 | 0 | 0 | 0 | 0 | 0 | 0 |
| 0 | NA | NA | NA | NA | NA | Gm29492 | | 0 | 0 | 0 | 0 | 0 | 0 | 0 | 0 | 0 | 0 | 0 |
| 0 | NA | NA | NA | NA | NA | Scgb2b5-ps | | 0 | 0 | 0 | 0 | 0 | 0 | 0 | 0 | 0 | 0 | 0 |
| 0 | NA | NA | NA | NA | NA | Scgb1b5-ps | | 0 | 0 | 0 | 0 | 0 | 0 | 0 | 0 | 0 | 0 | 0 |
| 0 | NA | NA | NA | NA | NA | Scgb2b28-ps | | 0 | 0 | 0 | 0 | 0 | 0 | 0 | 0 | 0 | 0 | 0 |
| 0 | NA | NA | NA | NA | NA | Rpl23a-ps8 | | 0 | 0 | 0 | 0 | 0 | 0 | 0 | 0 | 0 | 0 | 0 |
| 0 | NA | NA | NA | NA | NA | Scgb2b6 | secretoglobin, family 2B, member 6 | 0 | 0 | 0 | 0 | 0 | 0 | 0 | 0 | 0 | 0 | 0 |
| 0 | NA | NA | NA | NA | NA | Scgb1b6-ps | | 0 | 0 | 0 | 0 | 0 | 0 | 0 | 0 | 0 | 0 | 0 |
| 0 | NA | NA | NA | NA | NA | Gm4373 | | 0 | 0 | 0 | 0 | 0 | 0 | 0 | 0 | 0 | 0 | 0 |
| 0 | NA | NA | NA | NA | NA | Rpl23a-ps9 | | 0 | 0 | 0 | 0 | 0 | 0 | 0 | 0 | 0 | 0 | 0 |

| | | | | | | | | | | | | | | | | | | |
|---|---|---|---|---|---|---|---|---|---|---|---|---|---|---|---|---|---|---|
| 0 | NA | NA | NA | NA | NA | Scgb2b7 | secretoglobin, family 2B, member 7 | 0 | 0 | 0 | 0 | 0 | 0 | 0 | 0 | 0 | 0 | 0 |
| 0 | NA | NA | NA | NA | NA | Scgb1b7 | secretoglobin, family 1B, member 7 | 0 | 0 | 0 | 0 | 0 | 0 | 0 | 0 | 0 | 0 | 0 |
| 0 | NA | NA | NA | NA | NA | Gm9114 | | 0 | 0 | 0 | 0 | 0 | 0 | 0 | 0 | 0 | 0 | 0 |
| 0 | NA | NA | NA | NA | NA | Gm29627 | | 0 | 0 | 0 | 0 | 0 | 0 | 0 | 0 | 0 | 0 | 0 |
| 0 | NA | NA | NA | NA | NA | Scgb2b8-ps | | 0 | 0 | 0 | 0 | 0 | 0 | 0 | 0 | 0 | 0 | 0 |
| 0 | NA | NA | NA | NA | NA | Scgb1b8-ps | | 0 | 0 | 0 | 0 | 0 | 0 | 0 | 0 | 0 | 0 | 0 |
| 0 | NA | NA | NA | NA | NA | Gm6669 | | 0 | 0 | 0 | 0 | 0 | 0 | 0 | 0 | 0 | 0 | 0 |
| 0 | NA | NA | NA | NA | NA | Gm9120 | | 0 | 0 | 0 | 0 | 0 | 0 | 0 | 0 | 0 | 0 | 0 |
| 0 | NA | NA | NA | NA | NA | Gm21673 | | 0 | 0 | 0 | 0 | 0 | 0 | 0 | 0 | 0 | 0 | 0 |
| 0 | NA | NA | NA | NA | NA | Scgb2b9-ps | | 0 | 0 | 0 | 0 | 0 | 0 | 0 | 0 | 0 | 0 | 0 |
| 0 | NA | NA | NA | NA | NA | Scgb1b9-ps | | 0 | 0 | 0 | 0 | 0 | 0 | 0 | 0 | 0 | 0 | 0 |
| 0 | NA | NA | NA | NA | NA | Gm5329 | | 0 | 0 | 0 | 0 | 0 | 0 | 0 | 0 | 0 | 0 | 0 |
| 0 | NA | NA | NA | NA | NA | Scgb2b29-ps | | 0 | 0 | 0 | 0 | 0 | 0 | 0 | 0 | 0 | 0 | 0 |
| 0 | NA | NA | NA | NA | NA | Gm29262 | | 0 | 0 | 0 | 0 | 0 | 0 | 0 | 0 | 0 | 0 | 0 |
| 0 | NA | NA | NA | NA | NA | Gm29263 | | 0 | 0 | 0 | 0 | 0 | 0 | 0 | 0 | 0 | 0 | 0 |
| 0 | NA | NA | NA | NA | NA | Scgb2b10 | secretoglobin, family 2B, member 10 | 0 | 0 | 0 | 0 | 0 | 0 | 0 | 0 | 0 | 0 | 0 |
| 0 | NA | NA | NA | NA | NA | Scgb1b10 | secretoglobin, family 1B, member 10 | 0 | 0 | 0 | 0 | 0 | 0 | 0 | 0 | 0 | 0 | 0 |
| 0 | NA | NA | NA | NA | NA | Rpl23a-ps10 | | 0 | 0 | 0 | 0 | 0 | 0 | 0 | 0 | 0 | 0 | 0 |
| 0 | NA | NA | NA | NA | NA | Scgb2b30 | secretoglobin, family 2B, member 30 | 0 | 0 | 0 | 0 | 0 | 0 | 0 | 0 | 0 | 0 | 0 |
| 0 | NA | NA | NA | NA | NA | Gm9135 | | 0 | 0 | 0 | 0 | 0 | 0 | 0 | 0 | 0 | 0 | 0 |
| 0 | NA | NA | NA | NA | NA | Gm28256 | | 0 | 0 | 0 | 0 | 0 | 0 | 0 | 0 | 0 | 0 | 0 |
| 0 | NA | NA | NA | NA | NA | Gm28257 | | 0 | 0 | 0 | 0 | 0 | 0 | 0 | 0 | 0 | 0 | 0 |
| 0 | NA | NA | NA | NA | NA | Scgb2b11 | secretoglobin, family 2B, member 11 | 0 | 0 | 0 | 0 | 0 | 0 | 0 | 0 | 0 | 0 | 0 |
| 0 | NA | NA | NA | NA | NA | Scgb1b11 | | 0 | 0 | 0 | 0 | 0 | 0 | 0 | 0 | 0 | 0 | 0 |
| 0 | NA | NA | NA | NA | NA | Scgb2b12 | secretoglobin, family 2B, member 12 | 0 | 0 | 0 | 0 | 0 | 0 | 0 | 0 | 0 | 0 | 0 |
| 0 | NA | NA | NA | NA | NA | Scgb1b12 | secretoglobin, family 1B, member 12 | 0 | 0 | 0 | 0 | 0 | 0 | 0 | 0 | 0 | 0 | 0 |
| 0 | NA | NA | NA | NA | NA | Gm28862 | | 0 | 0 | 0 | 0 | 0 | 0 | 0 | 0 | 0 | 0 | 0 |
| 0 | NA | NA | NA | NA | NA | Rpl23a-ps11 | | 0 | 0 | 0 | 0 | 0 | 0 | 0 | 0 | 0 | 0 | 0 |
| 0 | NA | NA | NA | NA | NA | Scgb1b29 | secretoglobin, family 1B, member 29 | 0 | 0 | 0 | 0 | 0 | 0 | 0 | 0 | 0 | 0 | 0 |
| 0 | NA | NA | NA | NA | NA | Gm5116 | | 0 | 0 | 0 | 0 | 0 | 0 | 0 | 0 | 0 | 0 | 0 |
| 0 | NA | NA | NA | NA | NA | Scgb2b13-ps | secretoglobin, family 2B, member 13, pse | 0 | 0 | 0 | 0 | 0 | 0 | 0 | 0 | 0 | 0 | 0 |
| 0 | NA | NA | NA | NA | NA | Scgb1b13-ps | | 0 | 0 | 0 | 0 | 0 | 0 | 0 | 0 | 0 | 0 | 0 |
| 0 | NA | NA | NA | NA | NA | Gm5327 | | 0 | 0 | 0 | 0 | 0 | 0 | 0 | 0 | 0 | 0 | 0 |
| 0 | NA | NA | NA | NA | NA | Gm9149 | | 0 | 0 | 0 | 0 | 0 | 0 | 0 | 0 | 0 | 0 | 0 |
| 0 | NA | NA | NA | NA | NA | Gm21135 | | 0 | 0 | 0 | 0 | 0 | 0 | 0 | 0 | 0 | 0 | 0 |
| 0 | NA | NA | NA | NA | NA | Scgb2b14-ps | | 0 | 0 | 0 | 0 | 0 | 0 | 0 | 0 | 0 | 0 | 0 |
| 0 | NA | NA | NA | NA | NA | Scgb1b14-ps | | 0 | 0 | 0 | 0 | 0 | 0 | 0 | 0 | 0 | 0 | 0 |
| 0 | NA | NA | NA | NA | NA | Gm24766 | | 0 | 0 | 0 | 0 | 0 | 0 | 0 | 0 | 0 | 0 | 0 |
| 0 | NA | NA | NA | NA | NA | Gm5058 | | 0 | 0 | 0 | 0 | 0 | 0 | 0 | 0 | 0 | 0 | 0 |
| 0 | NA | NA | NA | NA | NA | Gm29429 | | 0 | 0 | 0 | 0 | 0 | 0 | 0 | 0 | 0 | 0 | 0 |
| 0 | NA | NA | NA | NA | NA | Scgb2b31-ps | | 0 | 0 | 0 | 0 | 0 | 0 | 0 | 0 | 0 | 0 | 0 |
| 0 | NA | NA | NA | NA | NA | Gm4398 | | 0 | 0 | 0 | 0 | 0 | 0 | 0 | 0 | 0 | 0 | 0 |
| 0 | NA | NA | NA | NA | NA | Gm29432 | | 0 | 0 | 0 | 0 | 0 | 0 | 0 | 0 | 0 | 0 | 0 |
| 0 | NA | NA | NA | NA | NA | Gm28408 | | 0 | 0 | 0 | 0 | 0 | 0 | 0 | 0 | 0 | 0 | 0 |
| 0 | NA | NA | NA | NA | NA | Scgb2b15 | secretoglobin, family 2B, member 15 | 0 | 0 | 0 | 0 | 0 | 0 | 0 | 0 | 0 | 0 | 0 |
| 0 | NA | NA | NA | NA | NA | Scgb1b15 | secretoglobin, family 1B, member 15 | 0 | 0 | 0 | 0 | 0 | 0 | 0 | 0 | 0 | 0 | 0 |
| 0 | NA | NA | NA | NA | NA | Scgb2b16-ps | | 0 | 0 | 0 | 0 | 0 | 0 | 0 | 0 | 0 | 0 | 0 |
| 0 | NA | NA | NA | NA | NA | Scgb1b16-ps | | 0 | 0 | 0 | 0 | 0 | 0 | 0 | 0 | 0 | 0 | 0 |
| 0 | NA | NA | NA | NA | NA | Gm25817 | | 0 | 0 | 0 | 0 | 0 | 0 | 0 | 0 | 0 | 0 | 0 |
| 0 | NA | NA | NA | NA | NA | Gm9156 | | 0 | 0 | 0 | 0 | 0 | 0 | 0 | 0 | 0 | 0 | 0 |
| 0 | NA | NA | NA | NA | NA | Gm28409 | | 0 | 0 | 0 | 0 | 0 | 0 | 0 | 0 | 0 | 0 | 0 |
| 0 | NA | NA | NA | NA | NA | Scgb2b32-ps | | 0 | 0 | 0 | 0 | 0 | 0 | 0 | 0 | 0 | 0 | 0 |
| 0 | NA | NA | NA | NA | NA | Gm4405 | | 0 | 0 | 0 | 0 | 0 | 0 | 0 | 0 | 0 | 0 | 0 |
| 0 | NA | NA | NA | NA | NA | Gm28383 | | 0 | 0 | 0 | 0 | 0 | 0 | 0 | 0 | 0 | 0 | 0 |
| 0 | NA | NA | NA | NA | NA | Gm28384 | | 0 | 0 | 0 | 0 | 0 | 0 | 0 | 0 | 0 | 0 | 0 |
| 0 | NA | NA | NA | NA | NA | Scgb2b17 | secretoglobin, family 2B, member 10 | 0 | 0 | 0 | 0 | 0 | 0 | 0 | 0 | 0 | 0 | 0 |
| 0 | NA | NA | NA | NA | NA | Scgb1b17 | secretoglobin, family 1B, member 17 | 0 | 0 | 0 | 0 | 0 | 0 | 0 | 0 | 0 | 0 | 0 |
| 0 | NA | NA | NA | NA | NA | Gm28992 | | 0 | 0 | 0 | 0 | 0 | 0 | 0 | 0 | 0 | 0 | 0 |
| 0 | NA | NA | NA | NA | NA | Rpl23a-ps7 | | 0 | 0 | 0 | 0 | 0 | 0 | 0 | 0 | 0 | 0 | 0 |
| 0 | NA | NA | NA | NA | NA | Scgb2b33 | secretoglobin, family 2B, member 33 | 0 | 0 | 0 | 0 | 0 | 0 | 0 | 0 | 0 | 0 | 0 |

| | | | | | | | | | | | | | | | | | |
|---|---|---|---|---|---|---|---|---|---|---|---|---|---|---|---|---|---|
| 0 | NA | NA | NA | NA | NA | Gm9162 | | 0 | 0 | 0 | 0 | 0 | 0 | 0 | 0 | 0 | 0 | 0 |
| 0 | NA | NA | NA | NA | NA | Gm28559 | | 0 | 0 | 0 | 0 | 0 | 0 | 0 | 0 | 0 | 0 | 0 |
| 0 | NA | NA | NA | NA | NA | Gm6003 | | 0 | 0 | 0 | 0 | 0 | 0 | 0 | 0 | 0 | 0 | 0 |
| 0 | NA | NA | NA | NA | NA | Scgb2b18 | secretoglobin, family 2B, member 18 | 0 | 0 | 0 | 0 | 0 | 0 | 0 | 0 | 0 | 0 | 0 |
| 0 | NA | NA | NA | NA | NA | Scgb1b18 | | 0 | 0 | 0 | 0 | 0 | 0 | 0 | 0 | 0 | 0 | 0 |
| 0 | NA | NA | NA | NA | NA | Scgb2b19 | secretoglobin, family 2B, member 19 | 0 | 0 | 0 | 0 | 0 | 0 | 0 | 0 | 0 | 0 | 0 |
| 0 | NA | NA | NA | NA | NA | Scgb1b19 | secretoglobin, family 1B, member 19 | 0 | 0 | 0 | 0 | 0 | 0 | 0 | 0 | 0 | 0 | 0 |
| 0 | NA | NA | NA | NA | NA | Scgb2b20 | secretoglobin, family 2B, member 20 | 0 | 0 | 0 | 0 | 0 | 0 | 0 | 0 | 0 | 0 | 0 |
| 0 | NA | NA | NA | NA | NA | Scgb1b20 | secretoglobin, family 1B, member 20 | 0 | 0 | 0 | 0 | 0 | 0 | 0 | 0 | 0 | 0 | 0 |
| 0 | NA | NA | NA | NA | NA | Gm28794 | | 0 | 0 | 0 | 0 | 0 | 0 | 0 | 0 | 0 | 0 | 0 |
| 0 | NA | NA | NA | NA | NA | Gm5731 | | 0 | 0 | 0 | 0 | 0 | 0 | 0 | 0 | 0 | 0 | 0 |
| 0 | NA | NA | NA | NA | NA | Scgb1b34-ps | | 0 | 0 | 0 | 0 | 0 | 0 | 0 | 0 | 0 | 0 | 0 |
| 0 | NA | NA | NA | NA | NA | Scgb2b21 | secretoglobin, family 2B, member 21 | 0 | 0 | 0 | 0 | 0 | 0 | 0 | 0 | 0 | 0 | 0 |
| 0 | NA | NA | NA | NA | NA | Scgb1b21 | | 0 | 0 | 0 | 0 | 0 | 0 | 0 | 0 | 0 | 0 | 0 |
| 0 | NA | NA | NA | NA | NA | Gm12770 | | 0 | 0 | 0 | 0 | 0 | 0 | 0 | 0 | 0 | 0 | 0 |
| 0 | NA | NA | NA | NA | NA | Gm12771 | | 0 | 0 | 0 | 0 | 0 | 0 | 0 | 0 | 0 | 0 | 0 |
| 0 | NA | NA | NA | NA | NA | Scgb2b22-ps | | 0 | 0 | 0 | 0 | 0 | 0 | 0 | 0 | 0 | 0 | 0 |
| 0 | NA | NA | NA | NA | NA | Scgb1b22-ps | | 0 | 0 | 0 | 0 | 0 | 0 | 0 | 0 | 0 | 0 | 0 |
| 0 | NA | NA | NA | NA | NA | Scgb2b23-ps | | 0 | 0 | 0 | 0 | 0 | 0 | 0 | 0 | 0 | 0 | 0 |
| 0 | NA | NA | NA | NA | NA | Scgb1b23-ps | | 0 | 0 | 0 | 0 | 0 | 0 | 0 | 0 | 0 | 0 | 0 |
| 0 | NA | NA | NA | NA | NA | Gm12763 | | 0 | 0 | 0 | 0 | 0 | 0 | 0 | 0 | 0 | 0 | 0 |
| 0 | NA | NA | NA | NA | NA | Rpl23a-ps12 | | 0 | 0 | 0 | 0 | 0 | 0 | 0 | 0 | 0 | 0 | 0 |
| 0 | NA | NA | NA | NA | NA | Scgb2b24 | secretoglobin, family 2B, member 24 | 0 | 0 | 0 | 0 | 0 | 0 | 0 | 0 | 0 | 0 | 0 |
| 0 | NA | NA | NA | NA | NA | Scgb1b24 | secretoglobin, family 1B, member 24 | 0 | 0 | 0 | 0 | 0 | 0 | 0 | 0 | 0 | 0 | 0 |
| 0 | NA | NA | NA | NA | NA | Gm12782 | | 0 | 0 | 0 | 0 | 0 | 0 | 0 | 0 | 0 | 0 | 0 |
| 0 | NA | NA | NA | NA | NA | Gm28614 | | 0 | 0 | 0 | 0 | 0 | 0 | 0 | 0 | 0 | 0 | 0 |
| 0 | NA | NA | NA | NA | NA | Gm26443 | | 0 | 0 | 0 | 0 | 0 | 0 | 0 | 0 | 0 | 0 | 0 |
| 0 | NA | NA | NA | NA | NA | Gm12768 | | 0 | 0 | 0 | 0 | 0 | 0 | 0 | 0 | 0 | 0 | 0 |
| 0 | NA | NA | NA | NA | NA | Scgb2b25-ps | | 0 | 0 | 0 | 0 | 0 | 0 | 0 | 0 | 0 | 0 | 0 |
| 0 | NA | NA | NA | NA | NA | Scgb1b25-ps | | 0 | 0 | 0 | 0 | 0 | 0 | 0 | 0 | 0 | 0 | 0 |
| 0 | NA | NA | NA | NA | NA | Gm28615 | | 0 | 0 | 0 | 0 | 0 | 0 | 0 | 0 | 0 | 0 | 0 |
| 0 | NA | NA | NA | NA | NA | Gm12765 | | 0 | 0 | 0 | 0 | 0 | 0 | 0 | 0 | 0 | 0 | 0 |
| 0 | NA | NA | NA | NA | NA | Scgb2b26 | secretoglobin, family 2B, member 26 | 0 | 0 | 0 | 0 | 0 | 0 | 0 | 0 | 0 | 0 | 0 |
| 0 | NA | NA | NA | NA | NA | Scgb1b26-ps | secretoglobin, family 1B, member 26, pse | 0 | 0 | 0 | 0 | 0 | 0 | 0 | 0 | 0 | 0 | 0 |
| 0 | NA | NA | NA | NA | NA | Gm28184 | | 0 | 0 | 0 | 0 | 0 | 0 | 0 | 0 | 0 | 0 | 0 |
| 0 | NA | NA | NA | NA | NA | Gm12773 | | 0 | 0 | 0 | 0 | 0 | 0 | 0 | 0 | 0 | 0 | 0 |
| 0 | NA | NA | NA | NA | NA | Scgb2b27 | secretoglobin, family 2B, member 27 | 0 | 0 | 0 | 0 | 0 | 0 | 0 | 0 | 0 | 0 | 0 |
| 0 | NA | NA | NA | NA | NA | Scgb1b27 | secretoglobin, family 1B, member 27 | 0 | 0 | 0 | 0 | 0 | 0 | 0 | 0 | 0 | 0 | 0 |
| 0 | NA | NA | NA | NA | NA | Gm12779 | | 0 | 0 | 0 | 0 | 0 | 0 | 0 | 0 | 0 | 0 | 0 |
| 0 | NA | NA | NA | NA | NA | Scgb1b30 | secretoglobin, family 1B, member 30 | 0 | 0 | 0 | 0 | 0 | 0 | 0 | 0 | 0 | 0 | 0 |
| 0 | NA | NA | NA | NA | NA | Gm12783 | | 0 | 0 | 0 | 0 | 0 | 0 | 0 | 0 | 0 | 0 | 0 |
| 0 | NA | NA | NA | NA | NA | Gm6096 | | 0 | 0 | 0 | 0 | 0 | 0 | 0 | 0 | 0 | 0 | 0 |
| 0 | NA | NA | NA | NA | NA | Gm12774 | | 0 | 0 | 0 | 0 | 0 | 0 | 0 | 0 | 0 | 0 | 0 |
| 0 | NA | NA | NA | NA | NA | Gm12780 | | 0 | 0 | 0 | 0 | 0 | 0 | 0 | 0 | 0 | 0 | 0 |
| 0 | NA | NA | NA | NA | NA | Gm12756 | | 0 | 0 | 0 | 0 | 0 | 0 | 0 | 0 | 0 | 0 | 0 |
| 0 | NA | NA | NA | NA | NA | Gm25922 | | 0 | 0 | 0 | 0 | 0 | 0 | 0 | 0 | 0 | 0 | 0 |
| 0 | NA | NA | NA | NA | NA | Gm45096 | | 0 | 0 | 0 | 0 | 0 | 0 | 0 | 0 | 0 | 0 | 0 |
| 0 | NA | NA | NA | NA | NA | Gm25033 | | 0 | 0 | 0 | 0 | 0 | 0 | 0 | 0 | 0 | 0 | 0 |
| 0 | NA | NA | NA | NA | NA | Gm22535 | | 0 | 0 | 0 | 0 | 0 | 0 | 0 | 0 | 0 | 0 | 0 |
| 0 | NA | NA | NA | NA | NA | Gm35665 | | 0 | 0 | 0 | 0 | 0 | 0 | 0 | 0 | 0 | 0 | 0 |
| 0 | NA | NA | NA | NA | NA | Gm24505 | | 0 | 0 | 0 | 0 | 0 | 0 | 0 | 0 | 0 | 0 | 0 |
| 0 | NA | NA | NA | NA | NA | Gm44636 | | 0 | 0 | 0 | 0 | 0 | 0 | 0 | 0 | 0 | 0 | 0 |
| 0 | NA | NA | NA | NA | NA | Gm18254 | | 0 | 0 | 0 | 0 | 0 | 0 | 0 | 0 | 0 | 0 | 0 |
| 0 | NA | NA | NA | NA | NA | Slc7a9 | solute carrier family 7 (cationic amino acid | 0 | 0 | 0 | 0 | 0 | 0 | 0 | 0 | 0 | 0 | 0 |
| 0 | NA | NA | NA | NA | NA | Gm18558 | | 0 | 0 | 0 | 0 | 0 | 0 | 0 | 0 | 0 | 0 | 0 |
| 0 | NA | NA | NA | NA | NA | Gm28514 | | 0 | 0 | 0 | 0 | 0 | 0 | 0 | 0 | 0 | 0 | 0 |
| 0 | NA | NA | NA | NA | NA | Gm38991 | | 0 | 0 | 0 | 0 | 0 | 0 | 0 | 0 | 0 | 0 | 0 |
| 0 | NA | NA | NA | NA | NA | Gm23292 | | 0 | 0 | 0 | 0 | 0 | 0 | 0 | 0 | 0 | 0 | 0 |
| 0 | NA | NA | NA | NA | NA | Gm25247 | | 0 | 0 | 0 | 0 | 0 | 0 | 0 | 0 | 0 | 0 | 0 |
| 0 | NA | NA | NA | NA | NA | Gm29129 | | 0 | 0 | 0 | 0 | 0 | 0 | 0 | 0 | 0 | 0 | 0 |

| | | | | | | | | | | | | | | | | | | |
|---|---|---|---|---|---|---|---|---|---|---|---|---|---|---|---|---|---|---|
| 0 | NA | NA | NA | NA | NA | Gm24550 | | 0 | 0 | 0 | 0 | 0 | 0 | 0 | 0 | 0 | 0 | 0 |
| 0 | NA | NA | NA | NA | NA | Gm28078 | | 0 | 0 | 0 | 0 | 0 | 0 | 0 | 0 | 0 | 0 | 0 |
| 0 | NA | NA | NA | NA | NA | Gm28076 | | 0 | 0 | 0 | 0 | 0 | 0 | 0 | 0 | 0 | 0 | 0 |
| 0 | NA | NA | NA | NA | NA | Gm28075 | | 0 | 0 | 0 | 0 | 0 | 0 | 0 | 0 | 0 | 0 | 0 |
| 0 | NA | NA | NA | NA | NA | Gm28077 | | 0 | 0 | 0 | 0 | 0 | 0 | 0 | 0 | 0 | 0 | 0 |
| 0 | NA | NA | NA | NA | NA | 6720469O03Rik | | 0 | 0 | 0 | 0 | 0 | 0 | 0 | 0 | 0 | 0 | 0 |
| 0 | NA | NA | NA | NA | NA | Gm44806 | | 0 | 0 | 0 | 0 | 0 | 0 | 0 | 0 | 0 | 0 | 0 |
| 0 | NA | NA | NA | NA | NA | Gm44882 | | 0 | 0 | 0 | 0 | 0 | 0 | 0 | 0 | 0 | 0 | 0 |
| 0 | NA | NA | NA | NA | NA | Gm44883 | | 0 | 0 | 0 | 0 | 0 | 0 | 0 | 0 | 0 | 0 | 0 |
| 0 | NA | NA | NA | NA | NA | 4930505M18Rik | | 0 | 0 | 0 | 0 | 0 | 0 | 0 | 0 | 0 | 0 | 0 |
| 0 | NA | NA | NA | NA | NA | Gm22302 | | 0 | 0 | 0 | 0 | 0 | 0 | 0 | 0 | 0 | 0 | 0 |
| 0 | NA | NA | NA | NA | NA | Gm30684 | | 0 | 0 | 0 | 0 | 0 | 0 | 0 | 0 | 0 | 0 | 0 |
| 0 | NA | NA | NA | NA | NA | Gm30771 | | 0 | 0 | 0 | 0 | 0 | 0 | 0 | 0 | 0 | 0 | 0 |
| 0 | NA | NA | NA | NA | NA | Gm22203 | | 0 | 0 | 0 | 0 | 0 | 0 | 0 | 0 | 0 | 0 | 0 |
| 0 | NA | NA | NA | NA | NA | Gm44767 | | 0 | 0 | 0 | 0 | 0 | 0 | 0 | 0 | 0 | 0 | 0 |
| 0 | NA | NA | NA | NA | NA | Gm28581 | | 0 | 0 | 0 | 0 | 0 | 0 | 0 | 0 | 0 | 0 | 0 |
| 0 | NA | NA | NA | NA | NA | Gm9220 | | 0 | 0 | 0 | 0 | 0 | 0 | 0 | 0 | 0 | 0 | 0 |
| 0 | NA | NA | NA | NA | NA | Gm4451 | | 0 | 0 | 0 | 0 | 0 | 0 | 0 | 0 | 0 | 0 | 0 |
| 0 | NA | NA | NA | NA | NA | Gm28453 | | 0 | 0 | 0 | 0 | 0 | 0 | 0 | 0 | 0 | 0 | 0 |
| 0 | NA | NA | NA | NA | NA | Gm22638 | | 0 | 0 | 0 | 0 | 0 | 0 | 0 | 0 | 0 | 0 | 0 |
| 0 | NA | NA | NA | NA | NA | Gm6818 | | 0 | 0 | 0 | 0 | 0 | 0 | 0 | 0 | 0 | 0 | 0 |
| 0 | NA | NA | NA | NA | NA | Gm28583 | | 0 | 0 | 0 | 0 | 0 | 0 | 0 | 0 | 0 | 0 | 0 |
| 0 | NA | NA | NA | NA | NA | Gm6605 | | 0 | 0 | 0 | 0 | 0 | 0 | 0 | 0 | 0 | 0 | 0 |
| 0 | NA | NA | NA | NA | NA | Gm6819 | | 0 | 0 | 0 | 0 | 0 | 0 | 0 | 0 | 0 | 0 | 0 |
| 0 | NA | NA | NA | NA | NA | Gm29087 | | 0 | 0 | 0 | 0 | 0 | 0 | 0 | 0 | 0 | 0 | 0 |
| 0 | NA | NA | NA | NA | NA | Gm5591 | predicted gene 5591 | 0 | 0 | 0 | 0 | 0 | 0 | 0 | 0 | 0 | 0 | 0 |
| 0 | NA | NA | NA | NA | NA | Gm4454 | | 0 | 0 | 0 | 0 | 0 | 0 | 0 | 0 | 0 | 0 | 0 |
| 0 | NA | NA | NA | NA | NA | Gm20749 | | 0 | 0 | 0 | 0 | 0 | 0 | 0 | 0 | 0 | 0 | 0 |
| 0 | NA | NA | NA | NA | NA | Gm29086 | | 0 | 0 | 0 | 0 | 0 | 0 | 0 | 0 | 0 | 0 | 0 |
| 0 | NA | NA | NA | NA | NA | Gm25286 | | 0 | 0 | 0 | 0 | 0 | 0 | 0 | 0 | 0 | 0 | 0 |
| 0 | NA | NA | NA | NA | NA | Gm21115 | | 0 | 0 | 0 | 0 | 0 | 0 | 0 | 0 | 0 | 0 | 0 |
| 0 | NA | NA | NA | NA | NA | Gm21276 | | 0 | 0 | 0 | 0 | 0 | 0 | 0 | 0 | 0 | 0 | 0 |
| 0 | NA | NA | NA | NA | NA | Gm21122 | | 0 | 0 | 0 | 0 | 0 | 0 | 0 | 0 | 0 | 0 | 0 |
| 0 | NA | NA | NA | NA | NA | Gm29085 | | 0 | 0 | 0 | 0 | 0 | 0 | 0 | 0 | 0 | 0 | 0 |
| 0 | NA | NA | NA | NA | NA | 1700010N08Rik | | 0 | 0 | 0 | 0 | 0 | 0 | 0 | 0 | 0 | 0 | 0 |
| 0 | NA | NA | NA | NA | NA | Gm24797 | | 0 | 0 | 0 | 0 | 0 | 0 | 0 | 0 | 0 | 0 | 0 |
| 0 | NA | NA | NA | NA | NA | Gm21129 | | 0 | 0 | 0 | 0 | 0 | 0 | 0 | 0 | 0 | 0 | 0 |
| 0 | NA | NA | NA | NA | NA | Gm21136 | | 0 | 0 | 0 | 0 | 0 | 0 | 0 | 0 | 0 | 0 | 0 |
| 0 | NA | NA | NA | NA | NA | Gm21142 | | 0 | 0 | 0 | 0 | 0 | 0 | 0 | 0 | 0 | 0 | 0 |
| 0 | NA | NA | NA | NA | NA | Gm21152 | | 0 | 0 | 0 | 0 | 0 | 0 | 0 | 0 | 0 | 0 | 0 |
| 0 | NA | NA | NA | NA | NA | Gm25671 | | 0 | 0 | 0 | 0 | 0 | 0 | 0 | 0 | 0 | 0 | 0 |
| 0 | NA | NA | NA | NA | NA | Gm28451 | | 0 | 0 | 0 | 0 | 0 | 0 | 0 | 0 | 0 | 0 | 0 |
| 0 | NA | NA | NA | NA | NA | Gm1982 | | 0 | 0 | 0 | 0 | 0 | 0 | 0 | 0 | 0 | 0 | 0 |
| 0 | NA | NA | NA | NA | NA | Gm1988 | | 0 | 0 | 0 | 0 | 0 | 0 | 0 | 0 | 0 | 0 | 0 |
| 0 | NA | NA | NA | NA | NA | Gm6124 | | 0 | 0 | 0 | 0 | 0 | 0 | 0 | 0 | 0 | 0 | 0 |
| 0 | NA | NA | NA | NA | NA | Gm16387 | | 0 | 0 | 0 | 0 | 0 | 0 | 0 | 0 | 0 | 0 | 0 |
| 0 | NA | NA | NA | NA | NA | Gm9226 | | 0 | 0 | 0 | 0 | 0 | 0 | 0 | 0 | 0 | 0 | 0 |
| 0 | NA | NA | NA | NA | NA | Gm23307 | | 0 | 0 | 0 | 0 | 0 | 0 | 0 | 0 | 0 | 0 | 0 |
| 0 | NA | NA | NA | NA | NA | Gm29258 | | 0 | 0 | 0 | 0 | 0 | 0 | 0 | 0 | 0 | 0 | 0 |
| 0 | NA | NA | NA | NA | NA | Gm9271 | | 0 | 0 | 0 | 0 | 0 | 0 | 0 | 0 | 0 | 0 | 0 |
| 0 | NA | NA | NA | NA | NA | Gm5114 | predicted gene 5114 | 0 | 0 | 0 | 0 | 0 | 0 | 0 | 0 | 0 | 0 | 0 |
| 0 | NA | NA | NA | NA | NA | Gm27437 | | 0 | 0 | 0 | 0 | 0 | 0 | 0 | 0 | 0 | 0 | 0 |
| 0 | NA | NA | NA | NA | NA | Gm17983 | | 0 | 0 | 0 | 0 | 0 | 0 | 0 | 0 | 0 | 0 | 0 |
| 0 | NA | NA | NA | NA | NA | Gm26327 | | 0 | 0 | 0 | 0 | 0 | 0 | 0 | 0 | 0 | 0 | 0 |
| 0 | NA | NA | NA | NA | NA | Gm2021 | | 0 | 0 | 0 | 0 | 0 | 0 | 0 | 0 | 0 | 0 | 0 |
| 0 | NA | NA | NA | NA | NA | Gm9230 | | 0 | 0 | 0 | 0 | 0 | 0 | 0 | 0 | 0 | 0 | 0 |
| 0 | NA | NA | NA | NA | NA | Gm45839 | | 0 | 0 | 0 | 0 | 0 | 0 | 0 | 0 | 0 | 0 | 0 |
| 0 | NA | NA | NA | NA | NA | 4930435C17Rik | | 0 | 0 | 0 | 0 | 0 | 0 | 0 | 0 | 0 | 0 | 0 |
| 0 | NA | NA | NA | NA | NA | 4930558N11Rik | | 0 | 0 | 0 | 0 | 0 | 0 | 0 | 0 | 0 | 0 | 0 |
| 0 | NA | NA | NA | NA | NA | Gm37302 | | 0 | 0 | 0 | 0 | 0 | 0 | 0 | 0 | 0 | 0 | 0 |

| | | | | | | | | | | | | | | | | | | |
|---|---|---|---|---|---|---|---|---|---|---|---|---|---|---|---|---|---|---|
| 0 | NA | NA | NA | NA | NA | Gm44992 | | 0 | 0 | 0 | 0 | 0 | 0 | 0 | 0 | 0 | 0 | 0 |
| 0 | NA | NA | NA | NA | NA | Gm38199 | | 0 | 0 | 0 | 0 | 0 | 0 | 0 | 0 | 0 | 0 | 0 |
| 0 | NA | NA | NA | NA | NA | 4933404I11Rik | | 0 | 0 | 0 | 0 | 0 | 0 | 0 | 0 | 0 | 0 | 0 |
| 0 | NA | NA | NA | NA | NA | Gm24910 | | 0 | 0 | 0 | 0 | 0 | 0 | 0 | 0 | 0 | 0 | 0 |
| 0 | NA | NA | NA | NA | NA | Gm28807 | | 0 | 0 | 0 | 0 | 0 | 0 | 0 | 0 | 0 | 0 | 0 |
| 0 | NA | NA | NA | NA | NA | Gm50466 | | 0 | 0 | 0 | 0 | 0 | 0 | 0 | 0 | 0 | 0 | 0 |
| 0 | NA | NA | NA | NA | NA | 4933402C06Rik | | 0 | 0 | 0 | 0 | 0 | 0 | 0 | 0 | 0 | 0 | 0 |
| 0 | NA | NA | NA | NA | NA | A230077H06Rik | | 0 | 0 | 0 | 0 | 0 | 0 | 0 | 0 | 0 | 0 | 0 |
| 0 | NA | NA | NA | NA | NA | Gm45008 | | 0 | 0 | 0 | 0 | 0 | 0 | 0 | 0 | 0 | 0 | 0 |
| 0 | NA | NA | NA | NA | NA | Gm28307 | | 0 | 0 | 0 | 0 | 0 | 0 | 0 | 0 | 0 | 0 | 0 |
| 0 | NA | NA | NA | NA | NA | Gm25784 | | 0 | 0 | 0 | 0 | 0 | 0 | 0 | 0 | 0 | 0 | 0 |
| 0 | NA | NA | NA | NA | NA | Gm29293 | | 0 | 0 | 0 | 0 | 0 | 0 | 0 | 0 | 0 | 0 | 0 |
| 0 | NA | NA | NA | NA | NA | 4930433I11Rik | RIKEN cDNA 4930433I11 gene | 0 | 0 | 0 | 0 | 0 | 0 | 0 | 0 | 0 | 0 | 0 |
| 0 | NA | NA | NA | NA | NA | Gm45171 | | 0 | 0 | 0 | 0 | 0 | 0 | 0 | 0 | 0 | 0 | 0 |
| 0 | NA | NA | NA | NA | NA | Gm38997 | | 0 | 0 | 0 | 0 | 0 | 0 | 0 | 0 | 0 | 0 | 0 |
| 0 | NA | NA | NA | NA | NA | Gm4884 | predicted gene 4884 | 0 | 0 | 0 | 0 | 0 | 0 | 0 | 0 | 0 | 0 | 0 |
| 0 | NA | NA | NA | NA | NA | Topbp1-ps2 | | 0 | 0 | 0 | 0 | 0 | 0 | 0 | 0 | 0 | 0 | 0 |
| 0 | NA | NA | NA | NA | NA | Gm45172 | | 0 | 0 | 0 | 0 | 0 | 0 | 0 | 0 | 0 | 0 | 0 |
| 0 | NA | NA | NA | NA | NA | 1700110I07Rik | | 0 | 0 | 0 | 0 | 0 | 0 | 0 | 0 | 0 | 0 | 0 |
| 0 | NA | NA | NA | NA | NA | Gm28061 | | 0 | 0 | 0 | 0 | 0 | 0 | 0 | 0 | 0 | 0 | 0 |
| 0 | NA | NA | NA | NA | NA | Gm44839 | | 0 | 0 | 0 | 0 | 0 | 0 | 0 | 0 | 0 | 0 | 0 |
| 0 | NA | NA | NA | NA | NA | Gm9239 | | 0 | 0 | 0 | 0 | 0 | 0 | 0 | 0 | 0 | 0 | 0 |
| 0 | NA | NA | NA | NA | NA | Gm6820 | | 0 | 0 | 0 | 0 | 0 | 0 | 0 | 0 | 0 | 0 | 0 |
| 0 | NA | NA | NA | NA | NA | Gm44748 | | 0 | 0 | 0 | 0 | 0 | 0 | 0 | 0 | 0 | 0 | 0 |
| 0 | NA | NA | NA | NA | NA | Gm5592 | predicted gene 5592 | 0 | 0 | 0 | 0 | 0 | 0 | 0 | 0 | 0 | 0 | 0 |
| 0 | NA | NA | NA | NA | NA | Gm2128 | | 0 | 0 | 0 | 0 | 0 | 0 | 0 | 0 | 0 | 0 | 0 |
| 0 | NA | NA | NA | NA | NA | Gm28060 | | 0 | 0 | 0 | 0 | 0 | 0 | 0 | 0 | 0 | 0 | 0 |
| 0 | NA | NA | NA | NA | NA | Gm28667 | | 0 | 0 | 0 | 0 | 0 | 0 | 0 | 0 | 0 | 0 | 0 |
| 0 | NA | NA | NA | NA | NA | Gm34811 | | 0 | 0 | 0 | 0 | 0 | 0 | 0 | 0 | 0 | 0 | 0 |
| 0 | NA | NA | NA | NA | NA | Gm6866 | predicted gene 6866 | 0 | 0 | 0 | 0 | 0 | 0 | 0 | 0 | 0 | 0 | 0 |
| 0 | NA | NA | NA | NA | NA | Gm45691 | | 0 | 0 | 0 | 0 | 0 | 0 | 0 | 0 | 0 | 0 | 0 |
| 0 | NA | NA | NA | NA | NA | Gm9244 | | 0 | 0 | 0 | 0 | 0 | 0 | 0 | 0 | 0 | 0 | 0 |
| 0 | NA | NA | NA | NA | NA | Gm44520 | | 0 | 0 | 0 | 0 | 0 | 0 | 0 | 0 | 0 | 0 | 0 |
| 0 | NA | NA | NA | NA | NA | Topbp1-ps1 | | 0 | 0 | 0 | 0 | 0 | 0 | 0 | 0 | 0 | 0 | 0 |
| 0 | NA | NA | NA | NA | NA | Gm2108 | | 0 | 0 | 0 | 0 | 0 | 0 | 0 | 0 | 0 | 0 | 0 |
| 0 | NA | NA | NA | NA | NA | Gm17791 | | 0 | 0 | 0 | 0 | 0 | 0 | 0 | 0 | 0 | 0 | 0 |
| 0 | NA | NA | NA | NA | NA | Gm17217 | | 0 | 0 | 0 | 0 | 0 | 0 | 0 | 0 | 0 | 0 | 0 |
| 0 | NA | NA | NA | NA | NA | Gm45078 | | 0 | 0 | 0 | 0 | 0 | 0 | 0 | 0 | 0 | 0 | 0 |
| 0 | NA | NA | NA | NA | NA | Gm9252 | | 0 | 0 | 0 | 0 | 0 | 0 | 0 | 0 | 0 | 0 | 0 |
| 0 | NA | NA | NA | NA | NA | Vmn2r-ps54 | | 0 | 0 | 0 | 0 | 0 | 0 | 0 | 0 | 0 | 0 | 0 |
| 0 | NA | NA | NA | NA | NA | Vmn2r-ps55 | | 0 | 0 | 0 | 0 | 0 | 0 | 0 | 0 | 0 | 0 | 0 |
| 0 | NA | NA | NA | NA | NA | Vmn2r-ps56 | | 0 | 0 | 0 | 0 | 0 | 0 | 0 | 0 | 0 | 0 | 0 |
| 0 | NA | NA | NA | NA | NA | Vmn2r58 | vomeronasal 2, receptor 58 | 0 | 0 | 0 | 0 | 0 | 0 | 0 | 0 | 0 | 0 | 0 |
| 0 | NA | NA | NA | NA | NA | Vmn2r-ps57 | | 0 | 0 | 0 | 0 | 0 | 0 | 0 | 0 | 0 | 0 | 0 |
| 0 | NA | NA | NA | NA | NA | Vmn2r60 | vomeronasal 2, receptor 60 | 0 | 0 | 0 | 0 | 0 | 0 | 0 | 0 | 0 | 0 | 0 |
| 0 | NA | NA | NA | NA | NA | Gm44519 | | 0 | 0 | 0 | 0 | 0 | 0 | 0 | 0 | 0 | 0 | 0 |
| 0 | NA | NA | NA | NA | NA | Vmn2r61 | vomeronasal 2, receptor 61 | 0 | 0 | 0 | 0 | 0 | 0 | 0 | 0 | 0 | 0 | 0 |
| 0 | NA | NA | NA | NA | NA | Vmn2r-ps58 | | 0 | 0 | 0 | 0 | 0 | 0 | 0 | 0 | 0 | 0 | 0 |
| 0 | NA | NA | NA | NA | NA | Gm5594 | | 0 | 0 | 0 | 0 | 0 | 0 | 0 | 0 | 0 | 0 | 0 |
| 0 | NA | NA | NA | NA | NA | Vmn2r-ps59 | | 0 | 0 | 0 | 0 | 0 | 0 | 0 | 0 | 0 | 0 | 0 |
| 0 | NA | NA | NA | NA | NA | Vmn2r-ps60 | | 0 | 0 | 0 | 0 | 0 | 0 | 0 | 0 | 0 | 0 | 0 |
| 0 | NA | NA | NA | NA | NA | 4933421I07Rik | RIKEN cDNA 4933421I07 gene | 0 | 0 | 0 | 0 | 0 | 0 | 0 | 0 | 0 | 0 | 0 |
| 0 | NA | NA | NA | NA | NA | Gm44523 | | 0 | 0 | 0 | 0 | 0 | 0 | 0 | 0 | 0 | 0 | 0 |
| 0 | NA | NA | NA | NA | NA | Gm17067 | | 0 | 0 | 0 | 0 | 0 | 0 | 0 | 0 | 0 | 0 | 0 |
| 0 | NA | NA | NA | NA | NA | Gm18128 | | 0 | 0 | 0 | 0 | 0 | 0 | 0 | 0 | 0 | 0 | 0 |
| 0 | NA | NA | NA | NA | NA | Gm6004 | | 0 | 0 | 0 | 0 | 0 | 0 | 0 | 0 | 0 | 0 | 0 |
| 0 | NA | NA | NA | NA | NA | Vmn2r62 | vomeronasal 2, receptor 62 | 0 | 0 | 0 | 0 | 0 | 0 | 0 | 0 | 0 | 0 | 0 |
| 0 | NA | NA | NA | NA | NA | Gm2381 | | 0 | 0 | 0 | 0 | 0 | 0 | 0 | 0 | 0 | 0 | 0 |
| 0 | NA | NA | NA | NA | NA | Gm7211 | | 0 | 0 | 0 | 0 | 0 | 0 | 0 | 0 | 0 | 0 | 0 |
| 0 | NA | NA | NA | NA | NA | Vmn2r63 | vomeronasal 2, receptor 63 | 0 | 0 | 0 | 0 | 0 | 0 | 0 | 0 | 0 | 0 | 0 |

| | | | | | | | | | | | | | | | | | | |
|---|---|---|---|---|---|---|---|---|---|---|---|---|---|---|---|---|---|---|
| 0 | NA | NA | NA | NA | NA | Gm9266 | | 0 | 0 | 0 | 0 | 0 | 0 | 0 | 0 | 0 | 0 | 0 |
| 0 | NA | NA | NA | NA | NA | Gm23256 | | 0 | 0 | 0 | 0 | 0 | 0 | 0 | 0 | 0 | 0 | 0 |
| 0 | NA | NA | NA | NA | NA | Vmn2r126 | vomeronasal 2, receptor 126 | 0 | 0 | 0 | 0 | 0 | 0 | 0 | 0 | 0 | 0 | 0 |
| 0 | NA | NA | NA | NA | NA | Gm44538 | | 0 | 0 | 0 | 0 | 0 | 0 | 0 | 0 | 0 | 0 | 0 |
| 0 | NA | NA | NA | NA | NA | Gm17768 | | 0 | 0 | 0 | 0 | 0 | 0 | 0 | 0 | 0 | 0 | 0 |
| 0 | NA | NA | NA | NA | NA | Rpl18a-ps4 | | 0 | 0 | 0 | 0 | 0 | 0 | 0 | 0 | 0 | 0 | 0 |
| 0 | NA | NA | NA | NA | NA | Gm10351 | | 0 | 0 | 0 | 0 | 0 | 0 | 0 | 0 | 0 | 0 | 0 |
| 0 | NA | NA | NA | NA | NA | Gm22707 | | 0 | 0 | 0 | 0 | 0 | 0 | 0 | 0 | 0 | 0 | 0 |
| 0 | NA | NA | NA | NA | NA | Gm44539 | | 0 | 0 | 0 | 0 | 0 | 0 | 0 | 0 | 0 | 0 | 0 |
| 0 | NA | NA | NA | NA | NA | Gm10298 | | 0 | 0 | 0 | 0 | 0 | 0 | 0 | 0 | 0 | 0 | 0 |
| 0 | NA | NA | NA | NA | NA | Gm44540 | | 0 | 0 | 0 | 0 | 0 | 0 | 0 | 0 | 0 | 0 | 0 |
| 0 | NA | NA | NA | NA | NA | Lim2 | | 0 | 0 | 0 | 0 | 0 | 0 | 0 | 0 | 0 | 0 | 0 |
| 0 | NA | NA | NA | NA | NA | | predicted gene, 38999 | 0 | 0 | 0 | 0 | 0 | 0 | 0 | 0 | 0 | 0 | 0 |
| 0 | NA | NA | NA | NA | NA | Lim2 | lens intrinsic membrane protein 2 | 0 | 0 | 0 | 0 | 0 | 0 | 0 | 0 | 0 | 0 | 0 |
| 0 | NA | NA | NA | NA | NA | Cldnd2 | claudin domain containing 2 | 0 | 0 | 0 | 0 | 0 | 0 | 0 | 0 | 0 | 0 | 0 |
| 0 | NA | NA | NA | NA | NA | Etfbl | | 0 | 0 | 0 | 0 | 0 | 0 | 0 | 0 | 0 | 0 | 0 |
| 0 | NA | NA | NA | NA | NA | 4931406B18Ril | RIKEN cDNA 4931406B18 gene | 0 | 0 | 0 | 0 | 0 | 0 | 0 | 0 | 0 | 0 | 0 |
| 0 | NA | NA | NA | NA | NA | Gm2511 | predicted gene 2511 | 0 | 0 | 0 | 0 | 0 | 0 | 0 | 0 | 0 | 0 | 0 |
| 0 | NA | NA | NA | NA | NA | Gm44818 | | 0 | 0 | 0 | 0 | 0 | 0 | 0 | 0 | 0 | 0 | 0 |
| 0 | NA | NA | NA | NA | NA | Zfp819 | zinc finger protein 819 | 0 | 0 | 0 | 0 | 0 | 0 | 0 | 0 | 0 | 0 | 0 |
| 0 | NA | NA | NA | NA | NA | Ceacam18 | carcinoembryonic antigen-related cell adl | 0 | 0 | 0 | 0 | 0 | 0 | 0 | 0 | 0 | 0 | 0 |
| 0 | NA | NA | NA | NA | NA | Klk14 | kallikrein related-peptidase 14 | 0 | 0 | 0 | 0 | 0 | 0 | 0 | 0 | 0 | 0 | 0 |
| 0 | NA | NA | NA | NA | NA | Gm44817 | | 0 | 0 | 0 | 0 | 0 | 0 | 0 | 0 | 0 | 0 | 0 |
| 0 | NA | NA | NA | NA | NA | 2310002F09Rik | | 0 | 0 | 0 | 0 | 0 | 0 | 0 | 0 | 0 | 0 | 0 |
| 0 | NA | NA | NA | NA | NA | Gm44814 | | 0 | 0 | 0 | 0 | 0 | 0 | 0 | 0 | 0 | 0 | 0 |
| 0 | NA | NA | NA | NA | NA | Klk12 | kallikrein related-peptidase 12 | 0 | 0 | 0 | 0 | 0 | 0 | 0 | 0 | 0 | 0 | 0 |
| 0 | NA | NA | NA | NA | NA | Klk11 | kallikrein related-peptidase 11 | 0 | 0 | 0 | 0 | 0 | 0 | 0 | 0 | 0 | 0 | 0 |
| 0 | NA | NA | NA | NA | NA | Klk9 | kallikrein related-peptidase 9 | 0 | 0 | 0 | 0 | 0 | 0 | 0 | 0 | 0 | 0 | 0 |
| 0 | NA | NA | NA | NA | NA | Klk7 | kallikrein related-peptidase 7 (chymotrypt | 0 | 0 | 0 | 0 | 0 | 0 | 0 | 0 | 0 | 0 | 0 |
| 0 | NA | NA | NA | NA | NA | Klk5 | kallikrein related-peptidase 5 | 0 | 0 | 0 | 0 | 0 | 0 | 0 | 0 | 0 | 0 | 0 |
| 0 | NA | NA | NA | NA | NA | Gm44707 | | 0 | 0 | 0 | 0 | 0 | 0 | 0 | 0 | 0 | 0 | 0 |
| 0 | NA | NA | NA | NA | NA | Klk2-ps | | 0 | 0 | 0 | 0 | 0 | 0 | 0 | 0 | 0 | 0 | 0 |
| 0 | NA | NA | NA | NA | NA | Klk15 | kallikrein related-peptidase 15 | 0 | 0 | 0 | 0 | 0 | 0 | 0 | 0 | 0 | 0 | 0 |
| 0 | NA | NA | NA | NA | NA | Gm44756 | | 0 | 0 | 0 | 0 | 0 | 0 | 0 | 0 | 0 | 0 | 0 |
| 0 | NA | NA | NA | NA | NA | Klk1b7-ps | | 0 | 0 | 0 | 0 | 0 | 0 | 0 | 0 | 0 | 0 | 0 |
| 0 | NA | NA | NA | NA | NA | Gm44945 | | 0 | 0 | 0 | 0 | 0 | 0 | 0 | 0 | 0 | 0 | 0 |
| 0 | NA | NA | NA | NA | NA | Klk1b2-ps | | 0 | 0 | 0 | 0 | 0 | 0 | 0 | 0 | 0 | 0 | 0 |
| 0 | NA | NA | NA | NA | NA | Klk1b1 | kallikrein 1-related peptidase b1 | 0 | 0 | 0 | 0 | 0 | 0 | 0 | 0 | 0 | 0 | 0 |
| 0 | NA | NA | NA | NA | NA | Gm45198 | | 0 | 0 | 0 | 0 | 0 | 0 | 0 | 0 | 0 | 0 | 0 |
| 0 | NA | NA | NA | NA | NA | Klk1b9 | kallikrein 1-related peptidase b9 | 0 | 0 | 0 | 0 | 0 | 0 | 0 | 0 | 0 | 0 | 0 |
| 0 | NA | NA | NA | NA | NA | Gm45135 | | 0 | 0 | 0 | 0 | 0 | 0 | 0 | 0 | 0 | 0 | 0 |
| 0 | NA | NA | NA | NA | NA | Klk1b10-ps | | 0 | 0 | 0 | 0 | 0 | 0 | 0 | 0 | 0 | 0 | 0 |
| 0 | NA | NA | NA | NA | NA | Gm45195 | | 0 | 0 | 0 | 0 | 0 | 0 | 0 | 0 | 0 | 0 | 0 |
| 0 | NA | NA | NA | NA | NA | Klk1b12-ps | | 0 | 0 | 0 | 0 | 0 | 0 | 0 | 0 | 0 | 0 | 0 |
| 0 | NA | NA | NA | NA | NA | Klk1b26 | kallikrein 1-related petidase b26 | 0 | 0 | 0 | 0 | 0 | 0 | 0 | 0 | 0 | 0 | 0 |
| 0 | NA | NA | NA | NA | NA | Klk1b18-ps | | 0 | 0 | 0 | 0 | 0 | 0 | 0 | 0 | 0 | 0 | 0 |
| 0 | NA | NA | NA | NA | NA | Klk1b28-ps | | 0 | 0 | 0 | 0 | 0 | 0 | 0 | 0 | 0 | 0 | 0 |
| 0 | NA | NA | NA | NA | NA | Gm45196 | | 0 | 0 | 0 | 0 | 0 | 0 | 0 | 0 | 0 | 0 | 0 |
| 0 | NA | NA | NA | NA | NA | Klk1b19-ps | | 0 | 0 | 0 | 0 | 0 | 0 | 0 | 0 | 0 | 0 | 0 |
| 0 | NA | NA | NA | NA | NA | Klk1b27 | kallikrein 1-related peptidase b27 | 0 | 0 | 0 | 0 | 0 | 0 | 0 | 0 | 0 | 0 | 0 |
| 0 | NA | NA | NA | NA | NA | Gm45197 | | 0 | 0 | 0 | 0 | 0 | 0 | 0 | 0 | 0 | 0 | 0 |
| 0 | NA | NA | NA | NA | NA | Klk1b14-ps | | 0 | 0 | 0 | 0 | 0 | 0 | 0 | 0 | 0 | 0 | 0 |
| 0 | NA | NA | NA | NA | NA | Gm45026 | | 0 | 0 | 0 | 0 | 0 | 0 | 0 | 0 | 0 | 0 | 0 |
| 0 | NA | NA | NA | NA | NA | Klk1b15-ps | | 0 | 0 | 0 | 0 | 0 | 0 | 0 | 0 | 0 | 0 | 0 |
| 0 | NA | NA | NA | NA | NA | Gm45125 | | 0 | 0 | 0 | 0 | 0 | 0 | 0 | 0 | 0 | 0 | 0 |
| 0 | NA | NA | NA | NA | NA | Klk1b21 | kallikrein 1-related peptidase b21 | 0 | 0 | 0 | 0 | 0 | 0 | 0 | 0 | 0 | 0 | 0 |
| 0 | NA | NA | NA | NA | NA | Gm45127 | | 0 | 0 | 0 | 0 | 0 | 0 | 0 | 0 | 0 | 0 | 0 |
| 0 | NA | NA | NA | NA | NA | Klk1b22 | kallikrein 1-related peptidase b22 | 0 | 0 | 0 | 0 | 0 | 0 | 0 | 0 | 0 | 0 | 0 |
| 0 | NA | NA | NA | NA | NA | Gm45126 | | 0 | 0 | 0 | 0 | 0 | 0 | 0 | 0 | 0 | 0 | 0 |
| 0 | NA | NA | NA | NA | NA | Gm45122 | | 0 | 0 | 0 | 0 | 0 | 0 | 0 | 0 | 0 | 0 | 0 |

| | | | | | | | | | | | | | | | | | | |
|---|---|---|---|---|---|---|---|---|---|---|---|---|---|---|---|---|---|---|
| 0 | NA | NA | NA | NA | NA | Klk1b23-ps | | 0 | 0 | 0 | 0 | 0 | 0 | 0 | 0 | 0 | 0 | 0 |
| 0 | NA | NA | NA | NA | NA | Klk1b16 | kallikrein 1-related peptidase b16 | 0 | 0 | 0 | 0 | 0 | 0 | 0 | 0 | 0 | 0 | 0 |
| 0 | NA | NA | NA | NA | NA | Gm45136 | | 0 | 0 | 0 | 0 | 0 | 0 | 0 | 0 | 0 | 0 | 0 |
| 0 | NA | NA | NA | NA | NA | Gm44779 | | 0 | 0 | 0 | 0 | 0 | 0 | 0 | 0 | 0 | 0 | 0 |
| 0 | NA | NA | NA | NA | NA | Gm45134 | | 0 | 0 | 0 | 0 | 0 | 0 | 0 | 0 | 0 | 0 | 0 |
| 0 | NA | NA | NA | NA | NA | Gm25386 | | 0 | 0 | 0 | 0 | 0 | 0 | 0 | 0 | 0 | 0 | 0 |
| 0 | NA | NA | NA | NA | NA | Gm26247 | | 0 | 0 | 0 | 0 | 0 | 0 | 0 | 0 | 0 | 0 | 0 |
| 0 | NA | NA | NA | NA | NA | Snord88a | | 0 | 0 | 0 | 0 | 0 | 0 | 0 | 0 | 0 | 0 | 0 |
| 0 | NA | NA | NA | NA | NA | Gm23991 | | 0 | 0 | 0 | 0 | 0 | 0 | 0 | 0 | 0 | 0 | 0 |
| 0 | NA | NA | NA | NA | NA | Acp4 | acid phosphatase 4 | 0 | 0 | 0 | 0 | 0 | 0 | 0 | 0 | 0 | 0 | 0 |
| 0 | NA | NA | NA | NA | NA | Gm15517 | | 0 | 0 | 0 | 0 | 0 | 0 | 0 | 0 | 0 | 0 | 0 |
| 0 | NA | NA | NA | NA | NA | Gm18905 | | 0 | 0 | 0 | 0 | 0 | 0 | 0 | 0 | 0 | 0 | 0 |
| 0 | NA | NA | NA | NA | NA | Gm18307 | | 0 | 0 | 0 | 0 | 0 | 0 | 0 | 0 | 0 | 0 | 0 |
| 0 | NA | NA | NA | NA | NA | Gm7238 | | 0 | 0 | 0 | 0 | 0 | 0 | 0 | 0 | 0 | 0 | 0 |
| 0 | NA | NA | NA | NA | NA | Gm44757 | | 0 | 0 | 0 | 0 | 0 | 0 | 0 | 0 | 0 | 0 | 0 |
| 0 | NA | NA | NA | NA | NA | Aspdh | aspartate dehydrogenase domain contair | 0 | 0 | 0 | 0 | 0 | 0 | 0 | 0 | 0 | 0 | 0 |
| 0 | NA | NA | NA | NA | NA | 5430431A17Rik | | 0 | 0 | 0 | 0 | 0 | 0 | 0 | 0 | 0 | 0 | 0 |
| 0 | NA | NA | NA | NA | NA | Mir7053 | | 0 | 0 | 0 | 0 | 0 | 0 | 0 | 0 | 0 | 0 | 0 |
| 0 | NA | NA | NA | NA | NA | Gm44833 | | 0 | 0 | 0 | 0 | 0 | 0 | 0 | 0 | 0 | 0 | 0 |
| 0 | NA | NA | NA | NA | NA | Gm18256 | | 0 | 0 | 0 | 0 | 0 | 0 | 0 | 0 | 0 | 0 | 0 |
| 0 | NA | NA | NA | NA | NA | Izumo2 | IZUMO family member 2 | 0 | 0 | 0 | 0 | 0 | 0 | 0 | 0 | 0 | 0 | 0 |
| 0 | NA | NA | NA | NA | NA | Mir707 | | 0 | 0 | 0 | 0 | 0 | 0 | 0 | 0 | 0 | 0 | 0 |
| 0 | NA | NA | NA | NA | NA | Mir150 | | 0 | 0 | 0 | 0 | 0 | 0 | 0 | 0 | 0 | 0 | 0 |
| 0 | NA | NA | NA | NA | NA | Snord35b | | 0 | 0 | 0 | 0 | 0 | 0 | 0 | 0 | 0 | 0 | 0 |
| 0 | NA | NA | NA | NA | NA | Gm45713 | | 0 | 0 | 0 | 0 | 0 | 0 | 0 | 0 | 0 | 0 | 0 |
| 0 | NA | NA | NA | NA | NA | Snord33 | | 0 | 0 | 0 | 0 | 0 | 0 | 0 | 0 | 0 | 0 | 0 |
| 0 | NA | NA | NA | NA | NA | Mir5121 | | 0 | 0 | 0 | 0 | 0 | 0 | 0 | 0 | 0 | 0 | 0 |
| 0 | NA | NA | NA | NA | NA | Mir7055 | | 0 | 0 | 0 | 0 | 0 | 0 | 0 | 0 | 0 | 0 | 0 |
| 0 | NA | NA | NA | NA | NA | Pth2 | parathyroid hormone 2 | 0 | 0 | 0 | 0 | 0 | 0 | 0 | 0 | 0 | 0 | 0 |
| 0 | NA | NA | NA | NA | NA | Slc6a16 | solute carrier family 6, member 16 | 0 | 0 | 0 | 0 | 0 | 0 | 0 | 0 | 0 | 0 | 0 |
| 0 | NA | NA | NA | NA | NA | Gm23585 | | 0 | 0 | 0 | 0 | 0 | 0 | 0 | 0 | 0 | 0 | 0 |
| 0 | NA | NA | NA | NA | NA | Slc6a21 | solute carrier family 6 member 21 | 0 | 0 | 0 | 0 | 0 | 0 | 0 | 0 | 0 | 0 | 0 |
| 0 | NA | NA | NA | NA | NA | Gm22478 | | 0 | 0 | 0 | 0 | 0 | 0 | 0 | 0 | 0 | 0 | 0 |
| 0 | NA | NA | NA | NA | NA | Kcna7 | potassium voltage-gated channel, shaker | 0 | 0 | 0 | 0 | 0 | 0 | 0 | 0 | 0 | 0 | 0 |
| 0 | NA | NA | NA | NA | NA | | luteinizing hormone beta | 0 | 0 | 0 | 0 | 0 | 0 | 0 | 0 | 0 | 0 | 0 |
| 0 | NA | NA | NA | NA | NA | Lhb | luteinizing hormone beta | 0 | 0 | 0 | 0 | 0 | 0 | 0 | 0 | 0 | 0 | 0 |
| 0 | NA | NA | NA | NA | NA | Gm22133 | | 0 | 0 | 0 | 0 | 0 | 0 | 0 | 0 | 0 | 0 | 0 |
| 0 | NA | NA | NA | NA | NA | Gm45808 | | 0 | 0 | 0 | 0 | 0 | 0 | 0 | 0 | 0 | 0 | 0 |
| 0 | NA | NA | NA | NA | NA | Gm45478 | | 0 | 0 | 0 | 0 | 0 | 0 | 0 | 0 | 0 | 0 | 0 |
| 0 | NA | NA | NA | NA | NA | Gm45564 | | 0 | 0 | 0 | 0 | 0 | 0 | 0 | 0 | 0 | 0 | 0 |
| 0 | NA | NA | NA | NA | NA | Hsd17b14 | hydroxysteroid (17-beta) dehydrogenase | 0 | 0 | 0 | 0 | 0 | 0 | 0 | 0 | 0 | 0 | 0 |
| 0 | NA | NA | NA | NA | NA | Ntn5 | netrin 5 | 0 | 0 | 0 | 0 | 0 | 0 | 0 | 0 | 0 | 0 | 0 |
| 0 | NA | NA | NA | NA | NA | Gm22098 | | 0 | 0 | 0 | 0 | 0 | 0 | 0 | 0 | 0 | 0 | 0 |
| 0 | NA | NA | NA | NA | NA | Gm45674 | | 0 | 0 | 0 | 0 | 0 | 0 | 0 | 0 | 0 | 0 | 0 |
| 0 | NA | NA | NA | NA | NA | 1700025L06Rik | | 0 | 0 | 0 | 0 | 0 | 0 | 0 | 0 | 0 | 0 | 0 |
| 0 | NA | NA | NA | NA | NA | Gm15700 | | 0 | 0 | 0 | 0 | 0 | 0 | 0 | 0 | 0 | 0 | 0 |
| 0 | NA | NA | NA | NA | NA | Gm45310 | | 0 | 0 | 0 | 0 | 0 | 0 | 0 | 0 | 0 | 0 | 0 |
| 0 | NA | NA | NA | NA | NA | Saa4 | serum amyloid A 4 | 0 | 0 | 0 | 0 | 0 | 0 | 0 | 0 | 0 | 0 | 0 |
| 0 | NA | NA | NA | NA | NA | Saa-ps | | 0 | 0 | 0 | 0 | 0 | 0 | 0 | 0 | 0 | 0 | 0 |
| 0 | NA | NA | NA | NA | NA | Ldhc | lactate dehydrogenase C | 0 | 0 | 0 | 0 | 0 | 0 | 0 | 0 | 0 | 0 | 0 |
| 0 | NA | NA | NA | NA | NA | Gm45628 | | 0 | 0 | 0 | 0 | 0 | 0 | 0 | 0 | 0 | 0 | 0 |
| 0 | NA | NA | NA | NA | NA | C86187 | | 0 | 0 | 0 | 0 | 0 | 0 | 0 | 0 | 0 | 0 | 0 |
| 0 | NA | NA | NA | NA | NA | Gm31941 | | 0 | 0 | 0 | 0 | 0 | 0 | 0 | 0 | 0 | 0 | 0 |
| 0 | NA | NA | NA | NA | NA | Gm9999 | | 0 | 0 | 0 | 0 | 0 | 0 | 0 | 0 | 0 | 0 | 0 |
| 0 | NA | NA | NA | NA | NA | LOC102637012 | | 0 | 0 | 0 | 0 | 0 | 0 | 0 | 0 | 0 | 0 | 0 |
| 0 | NA | NA | NA | NA | NA | Ptpn5 | protein tyrosine phosphatase, non-recept | 0 | 0 | 0 | 0 | 0 | 0 | 0 | 0 | 0 | 0 | 0 |
| 0 | NA | NA | NA | NA | NA | Mir7056 | | 0 | 0 | 0 | 0 | 0 | 0 | 0 | 0 | 0 | 0 | 0 |
| 0 | NA | NA | NA | NA | NA | Gm14377 | | 0 | 0 | 0 | 0 | 0 | 0 | 0 | 0 | 0 | 0 | 0 |
| 0 | NA | NA | NA | NA | NA | Gm44969 | | 0 | 0 | 0 | 0 | 0 | 0 | 0 | 0 | 0 | 0 | 0 |
| 0 | NA | NA | NA | NA | NA | Mrgpra6 | MAS-related GPR, member A6 | 0 | 0 | 0 | 0 | 0 | 0 | 0 | 0 | 0 | 0 | 0 |

| | | | | | | | | | | | | | | | | | | |
|---|---|---|---|---|---|---|---|---|---|---|---|---|---|---|---|---|---|---|
| 0 | NA | NA | NA | NA | NA | Mrgpra9 | MAS-related GPR, member A9 | 0 | 0 | 0 | 0 | 0 | 0 | 0 | 0 | 0 | 0 | 0 |
| 0 | NA | NA | NA | NA | NA | Mrgprx3-ps | | 0 | 0 | 0 | 0 | 0 | 0 | 0 | 0 | 0 | 0 | 0 |
| 0 | NA | NA | NA | NA | NA | Mrgpra1 | MAS-related GPR, member A1 | 0 | 0 | 0 | 0 | 0 | 0 | 0 | 0 | 0 | 0 | 0 |
| 0 | NA | NA | NA | NA | NA | Gm45085 | | 0 | 0 | 0 | 0 | 0 | 0 | 0 | 0 | 0 | 0 | 0 |
| 0 | NA | NA | NA | NA | NA | Gm29479 | | 0 | 0 | 0 | 0 | 0 | 0 | 0 | 0 | 0 | 0 | 0 |
| 0 | NA | NA | NA | NA | NA | Gm25326 | | 0 | 0 | 0 | 0 | 0 | 0 | 0 | 0 | 0 | 0 | 0 |
| 0 | NA | NA | NA | NA | NA | Mrgpra2a | MAS-related GPR, member A2A | 0 | 0 | 0 | 0 | 0 | 0 | 0 | 0 | 0 | 0 | 0 |
| 0 | NA | NA | NA | NA | NA | Mrgpra2b | MAS-related GPR, member A2B | 0 | 0 | 0 | 0 | 0 | 0 | 0 | 0 | 0 | 0 | 0 |
| 0 | NA | NA | NA | NA | NA | Gm7255 | | 0 | 0 | 0 | 0 | 0 | 0 | 0 | 0 | 0 | 0 | 0 |
| 0 | NA | NA | NA | NA | NA | Mrgprc4-ps | | 0 | 0 | 0 | 0 | 0 | 0 | 0 | 0 | 0 | 0 | 0 |
| 0 | NA | NA | NA | NA | NA | Mrgpra14 | | 0 | 0 | 0 | 0 | 0 | 0 | 0 | 0 | 0 | 0 | 0 |
| 0 | NA | NA | NA | NA | NA | Gm9322 | | 0 | 0 | 0 | 0 | 0 | 0 | 0 | 0 | 0 | 0 | 0 |
| 0 | NA | NA | NA | NA | NA | Mrgpra3 | MAS-related GPR, member A3 | 0 | 0 | 0 | 0 | 0 | 0 | 0 | 0 | 0 | 0 | 0 |
| 0 | NA | NA | NA | NA | NA | Mrgprc1-ps | | 0 | 0 | 0 | 0 | 0 | 0 | 0 | 0 | 0 | 0 | 0 |
| 0 | NA | NA | NA | NA | NA | Gm2707 | | 0 | 0 | 0 | 0 | 0 | 0 | 0 | 0 | 0 | 0 | 0 |
| 0 | NA | NA | NA | NA | NA | Gm28288 | | 0 | 0 | 0 | 0 | 0 | 0 | 0 | 0 | 0 | 0 | 0 |
| 0 | NA | NA | NA | NA | NA | Mrgprc3-ps | | 0 | 0 | 0 | 0 | 0 | 0 | 0 | 0 | 0 | 0 | 0 |
| 0 | NA | NA | NA | NA | NA | Gm5733 | | 0 | 0 | 0 | 0 | 0 | 0 | 0 | 0 | 0 | 0 | 0 |
| 0 | NA | NA | NA | NA | NA | Gm22427 | | 0 | 0 | 0 | 0 | 0 | 0 | 0 | 0 | 0 | 0 | 0 |
| 0 | NA | NA | NA | NA | NA | Gm18348 | | 0 | 0 | 0 | 0 | 0 | 0 | 0 | 0 | 0 | 0 | 0 |
| 0 | NA | NA | NA | NA | NA | Gm29563 | | 0 | 0 | 0 | 0 | 0 | 0 | 0 | 0 | 0 | 0 | 0 |
| 0 | NA | NA | NA | NA | NA | Mrgprc7-ps | | 0 | 0 | 0 | 0 | 0 | 0 | 0 | 0 | 0 | 0 | 0 |
| 0 | NA | NA | NA | NA | NA | Gm18349 | | 0 | 0 | 0 | 0 | 0 | 0 | 0 | 0 | 0 | 0 | 0 |
| 0 | NA | NA | NA | NA | NA | Mrgpra18-ps | | 0 | 0 | 0 | 0 | 0 | 0 | 0 | 0 | 0 | 0 | 0 |
| 0 | NA | NA | NA | NA | NA | Gm18350 | | 0 | 0 | 0 | 0 | 0 | 0 | 0 | 0 | 0 | 0 | 0 |
| 0 | NA | NA | NA | NA | NA | Gm5734 | | 0 | 0 | 0 | 0 | 0 | 0 | 0 | 0 | 0 | 0 | 0 |
| 0 | NA | NA | NA | NA | NA | Mrgprc2-ps | | 0 | 0 | 0 | 0 | 0 | 0 | 0 | 0 | 0 | 0 | 0 |
| 0 | NA | NA | NA | NA | NA | Mrgpra4 | MAS-related GPR, member A4 | 0 | 0 | 0 | 0 | 0 | 0 | 0 | 0 | 0 | 0 | 0 |
| 0 | NA | NA | NA | NA | NA | Mrgprx1 | MAS-related GPR, member X1 | 0 | 0 | 0 | 0 | 0 | 0 | 0 | 0 | 0 | 0 | 0 |
| 0 | NA | NA | NA | NA | NA | Mrgprb7-ps | | 0 | 0 | 0 | 0 | 0 | 0 | 0 | 0 | 0 | 0 | 0 |
| 0 | NA | NA | NA | NA | NA | Gm28666 | | 0 | 0 | 0 | 0 | 0 | 0 | 0 | 0 | 0 | 0 | 0 |
| 0 | NA | NA | NA | NA | NA | Mrgprb12-ps | | 0 | 0 | 0 | 0 | 0 | 0 | 0 | 0 | 0 | 0 | 0 |
| 0 | NA | NA | NA | NA | NA | Gm18351 | | 0 | 0 | 0 | 0 | 0 | 0 | 0 | 0 | 0 | 0 | 0 |
| 0 | NA | NA | NA | NA | NA | Gm28665 | | 0 | 0 | 0 | 0 | 0 | 0 | 0 | 0 | 0 | 0 | 0 |
| 0 | NA | NA | NA | NA | NA | Mrgprb5 | MAS-related GPR, member B5 | 0 | 0 | 0 | 0 | 0 | 0 | 0 | 0 | 0 | 0 | 0 |
| 0 | NA | NA | NA | NA | NA | Gm18352 | | 0 | 0 | 0 | 0 | 0 | 0 | 0 | 0 | 0 | 0 | 0 |
| 0 | NA | NA | NA | NA | NA | Mrgprb4 | MAS-related GPR, member B4 | 0 | 0 | 0 | 0 | 0 | 0 | 0 | 0 | 0 | 0 | 0 |
| 0 | NA | NA | NA | NA | NA | Gm26950 | | 0 | 0 | 0 | 0 | 0 | 0 | 0 | 0 | 0 | 0 | 0 |
| 0 | NA | NA | NA | NA | NA | Gm18356 | | 0 | 0 | 0 | 0 | 0 | 0 | 0 | 0 | 0 | 0 | 0 |
| 0 | NA | NA | NA | NA | NA | Mrgprb13 | MAS-related GPR, member B13 | 0 | 0 | 0 | 0 | 0 | 0 | 0 | 0 | 0 | 0 | 0 |
| 0 | NA | NA | NA | NA | NA | Gm18357 | | 0 | 0 | 0 | 0 | 0 | 0 | 0 | 0 | 0 | 0 | 0 |
| 0 | NA | NA | NA | NA | NA | Mrgprb8 | MAS-related GPR, member B8 | 0 | 0 | 0 | 0 | 0 | 0 | 0 | 0 | 0 | 0 | 0 |
| 0 | NA | NA | NA | NA | NA | Mrgprb9-ps | | 0 | 0 | 0 | 0 | 0 | 0 | 0 | 0 | 0 | 0 | 0 |
| 0 | NA | NA | NA | NA | NA | Gm18353 | | 0 | 0 | 0 | 0 | 0 | 0 | 0 | 0 | 0 | 0 | 0 |
| 0 | NA | NA | NA | NA | NA | Mrgprb3 | MAS-related GPR, member B3 | 0 | 0 | 0 | 0 | 0 | 0 | 0 | 0 | 0 | 0 | 0 |
| 0 | NA | NA | NA | NA | NA | Mrgprb11-ps | | 0 | 0 | 0 | 0 | 0 | 0 | 0 | 0 | 0 | 0 | 0 |
| 0 | NA | NA | NA | NA | NA | Gm45086 | | 0 | 0 | 0 | 0 | 0 | 0 | 0 | 0 | 0 | 0 | 0 |
| 0 | NA | NA | NA | NA | NA | Csrp3 | cysteine and glycine-rich protein 3 | 0 | 0 | 0 | 0 | 0 | 0 | 0 | 0 | 0 | 0 | 0 |
| 0 | NA | NA | NA | NA | NA | Gm18559 | | 0 | 0 | 0 | 0 | 0 | 0 | 0 | 0 | 0 | 0 | 0 |
| 0 | NA | NA | NA | NA | NA | Gm45207 | | 0 | 0 | 0 | 0 | 0 | 0 | 0 | 0 | 0 | 0 | 0 |
| 0 | NA | NA | NA | NA | NA | Gm38059 | | 0 | 0 | 0 | 0 | 0 | 0 | 0 | 0 | 0 | 0 | 0 |
| 0 | NA | NA | NA | NA | NA | Dbx1 | developing brain homeobox 1 | 0 | 0 | 0 | 0 | 0 | 0 | 0 | 0 | 0 | 0 | 0 |
| 0 | NA | NA | NA | NA | NA | Nell1os | | 0 | 0 | 0 | 0 | 0 | 0 | 0 | 0 | 0 | 0 | 0 |
| 0 | NA | NA | NA | NA | NA | Gm9343 | | 0 | 0 | 0 | 0 | 0 | 0 | 0 | 0 | 0 | 0 | 0 |
| 0 | NA | NA | NA | NA | NA | Gm21032 | | 0 | 0 | 0 | 0 | 0 | 0 | 0 | 0 | 0 | 0 | 0 |
| 0 | NA | NA | NA | NA | NA | Gm21033 | | 0 | 0 | 0 | 0 | 0 | 0 | 0 | 0 | 0 | 0 | 0 |
| 0 | NA | NA | NA | NA | NA | Gm21034 | | 0 | 0 | 0 | 0 | 0 | 0 | 0 | 0 | 0 | 0 | 0 |
| 0 | NA | NA | NA | NA | NA | Gm21035 | | 0 | 0 | 0 | 0 | 0 | 0 | 0 | 0 | 0 | 0 | 0 |
| 0 | NA | NA | NA | NA | NA | Gm21036 | | 0 | 0 | 0 | 0 | 0 | 0 | 0 | 0 | 0 | 0 | 0 |
| 0 | NA | NA | NA | NA | NA | Gm21037 | | 0 | 0 | 0 | 0 | 0 | 0 | 0 | 0 | 0 | 0 | 0 |

| | | | | | | | | | | | | | | | | | |
|---|---|---|---|---|---|---|---|---|---|---|---|---|---|---|---|---|---|
| 0 | NA | NA | NA | NA | NA | Gm16478 | | 0 | 0 | 0 | 0 | 0 | 0 | 0 | 0 | 0 | 0 | 0 |
| 0 | NA | NA | NA | NA | NA | Gm45002 | | 0 | 0 | 0 | 0 | 0 | 0 | 0 | 0 | 0 | 0 | 0 |
| 0 | NA | NA | NA | NA | NA | Ano5 | anoctamin 5 | 0 | 0 | 0 | 0 | 0 | 0 | 0 | 0 | 0 | 0 | 0 |
| 0 | NA | NA | NA | NA | NA | Gm45072 | | 0 | 0 | 0 | 0 | 0 | 0 | 0 | 0 | 0 | 0 | 0 |
| 0 | NA | NA | NA | NA | NA | Gm29296 | | 0 | 0 | 0 | 0 | 0 | 0 | 0 | 0 | 0 | 0 | 0 |
| 0 | NA | NA | NA | NA | NA | Gm45001 | | 0 | 0 | 0 | 0 | 0 | 0 | 0 | 0 | 0 | 0 | 0 |
| 0 | NA | NA | NA | NA | NA | Gm45235 | | 0 | 0 | 0 | 0 | 0 | 0 | 0 | 0 | 0 | 0 | 0 |
| 0 | NA | NA | NA | NA | NA | Gm9357 | | 0 | 0 | 0 | 0 | 0 | 0 | 0 | 0 | 0 | 0 | 0 |
| 0 | NA | NA | NA | NA | NA | Gm22211 | | 0 | 0 | 0 | 0 | 0 | 0 | 0 | 0 | 0 | 0 | 0 |
| 0 | NA | NA | NA | NA | NA | Gm44683 | | 0 | 0 | 0 | 0 | 0 | 0 | 0 | 0 | 0 | 0 | 0 |
| 0 | NA | NA | NA | NA | NA | Gm20074 | | 0 | 0 | 0 | 0 | 0 | 0 | 0 | 0 | 0 | 0 | 0 |
| 0 | NA | NA | NA | NA | NA | Gm22444 | | 0 | 0 | 0 | 0 | 0 | 0 | 0 | 0 | 0 | 0 | 0 |
| 0 | NA | NA | NA | NA | NA | Mir6238 | | 0 | 0 | 0 | 0 | 0 | 0 | 0 | 0 | 0 | 0 | 0 |
| 0 | NA | NA | NA | NA | NA | Gm44682 | | 0 | 0 | 0 | 0 | 0 | 0 | 0 | 0 | 0 | 0 | 0 |
| 0 | NA | NA | NA | NA | NA | Gm27921 | | 0 | 0 | 0 | 0 | 0 | 0 | 0 | 0 | 0 | 0 | 0 |
| 0 | NA | NA | NA | NA | NA | Gm25956 | | 0 | 0 | 0 | 0 | 0 | 0 | 0 | 0 | 0 | 0 | 0 |
| 0 | NA | NA | NA | NA | NA | Gm44895 | | 0 | 0 | 0 | 0 | 0 | 0 | 0 | 0 | 0 | 0 | 0 |
| 0 | NA | NA | NA | NA | NA | Gm44897 | | 0 | 0 | 0 | 0 | 0 | 0 | 0 | 0 | 0 | 0 | 0 |
| 0 | NA | NA | NA | NA | NA | Gm44896 | | 0 | 0 | 0 | 0 | 0 | 0 | 0 | 0 | 0 | 0 | 0 |
| 0 | NA | NA | NA | NA | NA | 4833421K07Rik | | 0 | 0 | 0 | 0 | 0 | 0 | 0 | 0 | 0 | 0 | 0 |
| 0 | NA | NA | NA | NA | NA | Gm17838 | | 0 | 0 | 0 | 0 | 0 | 0 | 0 | 0 | 0 | 0 | 0 |
| 0 | NA | NA | NA | NA | NA | Gm24116 | | 0 | 0 | 0 | 0 | 0 | 0 | 0 | 0 | 0 | 0 | 0 |
| 0 | NA | NA | NA | NA | NA | Gm44898 | | 0 | 0 | 0 | 0 | 0 | 0 | 0 | 0 | 0 | 0 | 0 |
| 0 | NA | NA | NA | NA | NA | Gm9367 | | 0 | 0 | 0 | 0 | 0 | 0 | 0 | 0 | 0 | 0 | 0 |
| 0 | NA | NA | NA | NA | NA | Gm44615 | | 0 | 0 | 0 | 0 | 0 | 0 | 0 | 0 | 0 | 0 | 0 |
| 0 | NA | NA | NA | NA | NA | Gm26482 | | 0 | 0 | 0 | 0 | 0 | 0 | 0 | 0 | 0 | 0 | 0 |
| 0 | NA | NA | NA | NA | NA | Gm17907 | | 0 | 0 | 0 | 0 | 0 | 0 | 0 | 0 | 0 | 0 | 0 |
| 0 | NA | NA | NA | NA | NA | Gm15888 | | 0 | 0 | 0 | 0 | 0 | 0 | 0 | 0 | 0 | 0 | 0 |
| 0 | NA | NA | NA | NA | NA | Gm19187 | | 0 | 0 | 0 | 0 | 0 | 0 | 0 | 0 | 0 | 0 | 0 |
| 0 | NA | NA | NA | NA | NA | Oca2 | oculocutaneous albinism II | 0 | 0 | 0 | 0 | 0 | 0 | 0 | 0 | 0 | 0 | 0 |
| 0 | NA | NA | NA | NA | NA | Gm44506 | | 0 | 0 | 0 | 0 | 0 | 0 | 0 | 0 | 0 | 0 | 0 |
| 0 | NA | NA | NA | NA | NA | Gm44979 | | 0 | 0 | 0 | 0 | 0 | 0 | 0 | 0 | 0 | 0 | 0 |
| 0 | NA | NA | NA | NA | NA | Gm17984 | | 0 | 0 | 0 | 0 | 0 | 0 | 0 | 0 | 0 | 0 | 0 |
| 0 | NA | NA | NA | NA | NA | Gm44976 | | 0 | 0 | 0 | 0 | 0 | 0 | 0 | 0 | 0 | 0 | 0 |
| 0 | NA | NA | NA | NA | NA | Gm44977 | | 0 | 0 | 0 | 0 | 0 | 0 | 0 | 0 | 0 | 0 | 0 |
| 0 | NA | NA | NA | NA | NA | Gm17987 | | 0 | 0 | 0 | 0 | 0 | 0 | 0 | 0 | 0 | 0 | 0 |
| 0 | NA | NA | NA | NA | NA | Gm18642 | | 0 | 0 | 0 | 0 | 0 | 0 | 0 | 0 | 0 | 0 | 0 |
| 0 | NA | NA | NA | NA | NA | Mir6389 | | 0 | 0 | 0 | 0 | 0 | 0 | 0 | 0 | 0 | 0 | 0 |
| 0 | NA | NA | NA | NA | NA | Gm44853 | | 0 | 0 | 0 | 0 | 0 | 0 | 0 | 0 | 0 | 0 | 0 |
| 0 | NA | NA | NA | NA | NA | Gm34272 | | 0 | 0 | 0 | 0 | 0 | 0 | 0 | 0 | 0 | 0 | 0 |
| 0 | NA | NA | NA | NA | NA | Gm18127 | | 0 | 0 | 0 | 0 | 0 | 0 | 0 | 0 | 0 | 0 | 0 |
| 0 | NA | NA | NA | NA | NA | Gm23911 | | 0 | 0 | 0 | 0 | 0 | 0 | 0 | 0 | 0 | 0 | 0 |
| 0 | NA | NA | NA | NA | NA | Gm26288 | | 0 | 0 | 0 | 0 | 0 | 0 | 0 | 0 | 0 | 0 | 0 |
| 0 | NA | NA | NA | NA | NA | Gm44852 | | 0 | 0 | 0 | 0 | 0 | 0 | 0 | 0 | 0 | 0 | 0 |
| 0 | NA | NA | NA | NA | NA | Gm45157 | | 0 | 0 | 0 | 0 | 0 | 0 | 0 | 0 | 0 | 0 | 0 |
| 0 | NA | NA | NA | NA | NA | Gm44936 | | 0 | 0 | 0 | 0 | 0 | 0 | 0 | 0 | 0 | 0 | 0 |
| 0 | NA | NA | NA | NA | NA | Gm44937 | | 0 | 0 | 0 | 0 | 0 | 0 | 0 | 0 | 0 | 0 | 0 |
| 0 | NA | NA | NA | NA | NA | Gm44938 | | 0 | 0 | 0 | 0 | 0 | 0 | 0 | 0 | 0 | 0 | 0 |
| 0 | NA | NA | NA | NA | NA | Gm18453 | | 0 | 0 | 0 | 0 | 0 | 0 | 0 | 0 | 0 | 0 | 0 |
| 0 | NA | NA | NA | NA | NA | Gm9373 | | 0 | 0 | 0 | 0 | 0 | 0 | 0 | 0 | 0 | 0 | 0 |
| 0 | NA | NA | NA | NA | NA | Gm45189 | | 0 | 0 | 0 | 0 | 0 | 0 | 0 | 0 | 0 | 0 | 0 |
| 0 | NA | NA | NA | NA | NA | Gm9377 | | 0 | 0 | 0 | 0 | 0 | 0 | 0 | 0 | 0 | 0 | 0 |
| 0 | NA | NA | NA | NA | NA | Gm44743 | | 0 | 0 | 0 | 0 | 0 | 0 | 0 | 0 | 0 | 0 | 0 |
| 0 | NA | NA | NA | NA | NA | Gm44562 | | 0 | 0 | 0 | 0 | 0 | 0 | 0 | 0 | 0 | 0 | 0 |
| 0 | NA | NA | NA | NA | NA | Gm44559 | | 0 | 0 | 0 | 0 | 0 | 0 | 0 | 0 | 0 | 0 | 0 |
| 0 | NA | NA | NA | NA | NA | A230103L15Rik | | 0 | 0 | 0 | 0 | 0 | 0 | 0 | 0 | 0 | 0 | 0 |
| 0 | NA | NA | NA | NA | NA | Gm25499 | | 0 | 0 | 0 | 0 | 0 | 0 | 0 | 0 | 0 | 0 | 0 |
| 0 | NA | NA | NA | NA | NA | Gm24966 | | 0 | 0 | 0 | 0 | 0 | 0 | 0 | 0 | 0 | 0 | 0 |
| 0 | NA | NA | NA | NA | NA | Gm22496 | | 0 | 0 | 0 | 0 | 0 | 0 | 0 | 0 | 0 | 0 | 0 |
| 0 | NA | NA | NA | NA | NA | Gm25648 | | 0 | 0 | 0 | 0 | 0 | 0 | 0 | 0 | 0 | 0 | 0 |

| | | | | | | | | | | | | | | | | | |
|---|---|---|---|---|---|---|---|---|---|---|---|---|---|---|---|---|---|
| 0 | NA | NA | NA | NA | NA | Gm24306 | 0 | 0 | 0 | 0 | 0 | 0 | 0 | 0 | 0 | 0 | 0 |
| 0 | NA | NA | NA | NA | NA | Gm25934 | 0 | 0 | 0 | 0 | 0 | 0 | 0 | 0 | 0 | 0 | 0 |
| 0 | NA | NA | NA | NA | NA | Gm25156 | 0 | 0 | 0 | 0 | 0 | 0 | 0 | 0 | 0 | 0 | 0 |
| 0 | NA | NA | NA | NA | NA | Gm26488 | 0 | 0 | 0 | 0 | 0 | 0 | 0 | 0 | 0 | 0 | 0 |
| 0 | NA | NA | NA | NA | NA | Gm24220 | 0 | 0 | 0 | 0 | 0 | 0 | 0 | 0 | 0 | 0 | 0 |
| 0 | NA | NA | NA | NA | NA | Gm24653 | 0 | 0 | 0 | 0 | 0 | 0 | 0 | 0 | 0 | 0 | 0 |
| 0 | NA | NA | NA | NA | NA | Gm24969 | 0 | 0 | 0 | 0 | 0 | 0 | 0 | 0 | 0 | 0 | 0 |
| 0 | NA | NA | NA | NA | NA | Gm24417 | 0 | 0 | 0 | 0 | 0 | 0 | 0 | 0 | 0 | 0 | 0 |
| 0 | NA | NA | NA | NA | NA | Gm25121 | 0 | 0 | 0 | 0 | 0 | 0 | 0 | 0 | 0 | 0 | 0 |
| 0 | NA | NA | NA | NA | NA | Gm24657 | 0 | 0 | 0 | 0 | 0 | 0 | 0 | 0 | 0 | 0 | 0 |
| 0 | NA | NA | NA | NA | NA | Gm26230 | 0 | 0 | 0 | 0 | 0 | 0 | 0 | 0 | 0 | 0 | 0 |
| 0 | NA | NA | NA | NA | NA | Gm23922 | 0 | 0 | 0 | 0 | 0 | 0 | 0 | 0 | 0 | 0 | 0 |
| 0 | NA | NA | NA | NA | NA | Gm25314 | 0 | 0 | 0 | 0 | 0 | 0 | 0 | 0 | 0 | 0 | 0 |
| 0 | NA | NA | NA | NA | NA | Gm26096 | 0 | 0 | 0 | 0 | 0 | 0 | 0 | 0 | 0 | 0 | 0 |
| 0 | NA | NA | NA | NA | NA | Gm22996 | 0 | 0 | 0 | 0 | 0 | 0 | 0 | 0 | 0 | 0 | 0 |
| 0 | NA | NA | NA | NA | NA | Gm26499 | 0 | 0 | 0 | 0 | 0 | 0 | 0 | 0 | 0 | 0 | 0 |
| 0 | NA | NA | NA | NA | NA | Gm25646 | 0 | 0 | 0 | 0 | 0 | 0 | 0 | 0 | 0 | 0 | 0 |
| 0 | NA | NA | NA | NA | NA | Gm26498 | 0 | 0 | 0 | 0 | 0 | 0 | 0 | 0 | 0 | 0 | 0 |
| 0 | NA | NA | NA | NA | NA | Gm23660 | 0 | 0 | 0 | 0 | 0 | 0 | 0 | 0 | 0 | 0 | 0 |
| 0 | NA | NA | NA | NA | NA | Gm23933 | 0 | 0 | 0 | 0 | 0 | 0 | 0 | 0 | 0 | 0 | 0 |
| 0 | NA | NA | NA | NA | NA | Gm24862 | 0 | 0 | 0 | 0 | 0 | 0 | 0 | 0 | 0 | 0 | 0 |
| 0 | NA | NA | NA | NA | NA | Gm24866 | 0 | 0 | 0 | 0 | 0 | 0 | 0 | 0 | 0 | 0 | 0 |
| 0 | NA | NA | NA | NA | NA | Gm22348 | 0 | 0 | 0 | 0 | 0 | 0 | 0 | 0 | 0 | 0 | 0 |
| 0 | NA | NA | NA | NA | NA | Gm26466 | 0 | 0 | 0 | 0 | 0 | 0 | 0 | 0 | 0 | 0 | 0 |
| 0 | NA | NA | NA | NA | NA | Gm24799 | 0 | 0 | 0 | 0 | 0 | 0 | 0 | 0 | 0 | 0 | 0 |
| 0 | NA | NA | NA | NA | NA | Gm24088 | 0 | 0 | 0 | 0 | 0 | 0 | 0 | 0 | 0 | 0 | 0 |
| 0 | NA | NA | NA | NA | NA | Gm25463 | 0 | 0 | 0 | 0 | 0 | 0 | 0 | 0 | 0 | 0 | 0 |
| 0 | NA | NA | NA | NA | NA | Gm25229 | 0 | 0 | 0 | 0 | 0 | 0 | 0 | 0 | 0 | 0 | 0 |
| 0 | NA | NA | NA | NA | NA | Gm24528 | 0 | 0 | 0 | 0 | 0 | 0 | 0 | 0 | 0 | 0 | 0 |
| 0 | NA | NA | NA | NA | NA | Gm23356 | 0 | 0 | 0 | 0 | 0 | 0 | 0 | 0 | 0 | 0 | 0 |
| 0 | NA | NA | NA | NA | NA | Gm25449 | 0 | 0 | 0 | 0 | 0 | 0 | 0 | 0 | 0 | 0 | 0 |
| 0 | NA | NA | NA | NA | NA | Gm24872 | 0 | 0 | 0 | 0 | 0 | 0 | 0 | 0 | 0 | 0 | 0 |
| 0 | NA | NA | NA | NA | NA | Gm26337 | 0 | 0 | 0 | 0 | 0 | 0 | 0 | 0 | 0 | 0 | 0 |
| 0 | NA | NA | NA | NA | NA | Gm23145 | 0 | 0 | 0 | 0 | 0 | 0 | 0 | 0 | 0 | 0 | 0 |
| 0 | NA | NA | NA | NA | NA | Gm22130 | 0 | 0 | 0 | 0 | 0 | 0 | 0 | 0 | 0 | 0 | 0 |
| 0 | NA | NA | NA | NA | NA | Gm25988 | 0 | 0 | 0 | 0 | 0 | 0 | 0 | 0 | 0 | 0 | 0 |
| 0 | NA | NA | NA | NA | NA | Gm25221 | 0 | 0 | 0 | 0 | 0 | 0 | 0 | 0 | 0 | 0 | 0 |
| 0 | NA | NA | NA | NA | NA | Gm26389 | 0 | 0 | 0 | 0 | 0 | 0 | 0 | 0 | 0 | 0 | 0 |
| 0 | NA | NA | NA | NA | NA | Gm23471 | 0 | 0 | 0 | 0 | 0 | 0 | 0 | 0 | 0 | 0 | 0 |
| 0 | NA | NA | NA | NA | NA | Gm25120 | 0 | 0 | 0 | 0 | 0 | 0 | 0 | 0 | 0 | 0 | 0 |
| 0 | NA | NA | NA | NA | NA | Gm24418 | 0 | 0 | 0 | 0 | 0 | 0 | 0 | 0 | 0 | 0 | 0 |
| 0 | NA | NA | NA | NA | NA | Gm23687 | 0 | 0 | 0 | 0 | 0 | 0 | 0 | 0 | 0 | 0 | 0 |
| 0 | NA | NA | NA | NA | NA | Gm22393 | 0 | 0 | 0 | 0 | 0 | 0 | 0 | 0 | 0 | 0 | 0 |
| 0 | NA | NA | NA | NA | NA | Gm24639 | 0 | 0 | 0 | 0 | 0 | 0 | 0 | 0 | 0 | 0 | 0 |
| 0 | NA | NA | NA | NA | NA | Gm24566 | 0 | 0 | 0 | 0 | 0 | 0 | 0 | 0 | 0 | 0 | 0 |
| 0 | NA | NA | NA | NA | NA | Gm22111 | 0 | 0 | 0 | 0 | 0 | 0 | 0 | 0 | 0 | 0 | 0 |
| 0 | NA | NA | NA | NA | NA | Gm23076 | 0 | 0 | 0 | 0 | 0 | 0 | 0 | 0 | 0 | 0 | 0 |
| 0 | NA | NA | NA | NA | NA | Gm25874 | 0 | 0 | 0 | 0 | 0 | 0 | 0 | 0 | 0 | 0 | 0 |
| 0 | NA | NA | NA | NA | NA | Gm25088 | 0 | 0 | 0 | 0 | 0 | 0 | 0 | 0 | 0 | 0 | 0 |
| 0 | NA | NA | NA | NA | NA | Gm22510 | 0 | 0 | 0 | 0 | 0 | 0 | 0 | 0 | 0 | 0 | 0 |
| 0 | NA | NA | NA | NA | NA | Gm24983 | 0 | 0 | 0 | 0 | 0 | 0 | 0 | 0 | 0 | 0 | 0 |
| 0 | NA | NA | NA | NA | NA | Gm24027 | 0 | 0 | 0 | 0 | 0 | 0 | 0 | 0 | 0 | 0 | 0 |
| 0 | NA | NA | NA | NA | NA | Gm22524 | 0 | 0 | 0 | 0 | 0 | 0 | 0 | 0 | 0 | 0 | 0 |
| 0 | NA | NA | NA | NA | NA | Gm22630 | 0 | 0 | 0 | 0 | 0 | 0 | 0 | 0 | 0 | 0 | 0 |
| 0 | NA | NA | NA | NA | NA | Gm25741 | 0 | 0 | 0 | 0 | 0 | 0 | 0 | 0 | 0 | 0 | 0 |
| 0 | NA | NA | NA | NA | NA | Gm26059 | 0 | 0 | 0 | 0 | 0 | 0 | 0 | 0 | 0 | 0 | 0 |
| 0 | NA | NA | NA | NA | NA | Gm25175 | 0 | 0 | 0 | 0 | 0 | 0 | 0 | 0 | 0 | 0 | 0 |
| 0 | NA | NA | NA | NA | NA | Gm22511 | 0 | 0 | 0 | 0 | 0 | 0 | 0 | 0 | 0 | 0 | 0 |
| 0 | NA | NA | NA | NA | NA | Gm25077 | 0 | 0 | 0 | 0 | 0 | 0 | 0 | 0 | 0 | 0 | 0 |
| 0 | NA | NA | NA | NA | NA | Gm22834 | 0 | 0 | 0 | 0 | 0 | 0 | 0 | 0 | 0 | 0 | 0 |

| | | | | | | | | | | | | | | | | | |
|---|---|---|---|---|---|---|---|---|---|---|---|---|---|---|---|---|---|
| 0 | NA | NA | NA | NA | NA | Gm24926 | 0 | 0 | 0 | 0 | 0 | 0 | 0 | 0 | 0 | 0 | 0 |
| 0 | NA | NA | NA | NA | NA | Gm26200 | 0 | 0 | 0 | 0 | 0 | 0 | 0 | 0 | 0 | 0 | 0 |
| 0 | NA | NA | NA | NA | NA | Gm22632 | 0 | 0 | 0 | 0 | 0 | 0 | 0 | 0 | 0 | 0 | 0 |
| 0 | NA | NA | NA | NA | NA | Gm23254 | 0 | 0 | 0 | 0 | 0 | 0 | 0 | 0 | 0 | 0 | 0 |
| 0 | NA | NA | NA | NA | NA | Gm26390 | 0 | 0 | 0 | 0 | 0 | 0 | 0 | 0 | 0 | 0 | 0 |
| 0 | NA | NA | NA | NA | NA | Gm26366 | 0 | 0 | 0 | 0 | 0 | 0 | 0 | 0 | 0 | 0 | 0 |
| 0 | NA | NA | NA | NA | NA | Gm26336 | 0 | 0 | 0 | 0 | 0 | 0 | 0 | 0 | 0 | 0 | 0 |
| 0 | NA | NA | NA | NA | NA | Gm26283 | 0 | 0 | 0 | 0 | 0 | 0 | 0 | 0 | 0 | 0 | 0 |
| 0 | NA | NA | NA | NA | NA | Gm25098 | 0 | 0 | 0 | 0 | 0 | 0 | 0 | 0 | 0 | 0 | 0 |
| 0 | NA | NA | NA | NA | NA | Gm24702 | 0 | 0 | 0 | 0 | 0 | 0 | 0 | 0 | 0 | 0 | 0 |
| 0 | NA | NA | NA | NA | NA | Gm26432 | 0 | 0 | 0 | 0 | 0 | 0 | 0 | 0 | 0 | 0 | 0 |
| 0 | NA | NA | NA | NA | NA | Gm25462 | 0 | 0 | 0 | 0 | 0 | 0 | 0 | 0 | 0 | 0 | 0 |
| 0 | NA | NA | NA | NA | NA | Gm24495 | 0 | 0 | 0 | 0 | 0 | 0 | 0 | 0 | 0 | 0 | 0 |
| 0 | NA | NA | NA | NA | NA | Gm25523 | 0 | 0 | 0 | 0 | 0 | 0 | 0 | 0 | 0 | 0 | 0 |
| 0 | NA | NA | NA | NA | NA | Gm22255 | 0 | 0 | 0 | 0 | 0 | 0 | 0 | 0 | 0 | 0 | 0 |
| 0 | NA | NA | NA | NA | NA | Gm23449 | 0 | 0 | 0 | 0 | 0 | 0 | 0 | 0 | 0 | 0 | 0 |
| 0 | NA | NA | NA | NA | NA | Gm23310 | 0 | 0 | 0 | 0 | 0 | 0 | 0 | 0 | 0 | 0 | 0 |
| 0 | NA | NA | NA | NA | NA | Gm25461 | 0 | 0 | 0 | 0 | 0 | 0 | 0 | 0 | 0 | 0 | 0 |
| 0 | NA | NA | NA | NA | NA | Gm24310 | 0 | 0 | 0 | 0 | 0 | 0 | 0 | 0 | 0 | 0 | 0 |
| 0 | NA | NA | NA | NA | NA | Gm22909 | 0 | 0 | 0 | 0 | 0 | 0 | 0 | 0 | 0 | 0 | 0 |
| 0 | NA | NA | NA | NA | NA | Gm25823 | 0 | 0 | 0 | 0 | 0 | 0 | 0 | 0 | 0 | 0 | 0 |
| 0 | NA | NA | NA | NA | NA | Gm25647 | 0 | 0 | 0 | 0 | 0 | 0 | 0 | 0 | 0 | 0 | 0 |
| 0 | NA | NA | NA | NA | NA | Gm22252 | 0 | 0 | 0 | 0 | 0 | 0 | 0 | 0 | 0 | 0 | 0 |
| 0 | NA | NA | NA | NA | NA | Gm26392 | 0 | 0 | 0 | 0 | 0 | 0 | 0 | 0 | 0 | 0 | 0 |
| 0 | NA | NA | NA | NA | NA | Gm24219 | 0 | 0 | 0 | 0 | 0 | 0 | 0 | 0 | 0 | 0 | 0 |
| 0 | NA | NA | NA | NA | NA | Gm25230 | 0 | 0 | 0 | 0 | 0 | 0 | 0 | 0 | 0 | 0 | 0 |
| 0 | NA | NA | NA | NA | NA | Gm22627 | 0 | 0 | 0 | 0 | 0 | 0 | 0 | 0 | 0 | 0 | 0 |
| 0 | NA | NA | NA | NA | NA | Gm24585 | 0 | 0 | 0 | 0 | 0 | 0 | 0 | 0 | 0 | 0 | 0 |
| 0 | NA | NA | NA | NA | NA | Gm23305 | 0 | 0 | 0 | 0 | 0 | 0 | 0 | 0 | 0 | 0 | 0 |
| 0 | NA | NA | NA | NA | NA | Gm24206 | 0 | 0 | 0 | 0 | 0 | 0 | 0 | 0 | 0 | 0 | 0 |
| 0 | NA | NA | NA | NA | NA | Gm25085 | 0 | 0 | 0 | 0 | 0 | 0 | 0 | 0 | 0 | 0 | 0 |
| 0 | NA | NA | NA | NA | NA | Gm25209 | 0 | 0 | 0 | 0 | 0 | 0 | 0 | 0 | 0 | 0 | 0 |
| 0 | NA | NA | NA | NA | NA | Gm23682 | 0 | 0 | 0 | 0 | 0 | 0 | 0 | 0 | 0 | 0 | 0 |
| 0 | NA | NA | NA | NA | NA | Gm24654 | 0 | 0 | 0 | 0 | 0 | 0 | 0 | 0 | 0 | 0 | 0 |
| 0 | NA | NA | NA | NA | NA | Gm26334 | 0 | 0 | 0 | 0 | 0 | 0 | 0 | 0 | 0 | 0 | 0 |
| 0 | NA | NA | NA | NA | NA | Gm25742 | 0 | 0 | 0 | 0 | 0 | 0 | 0 | 0 | 0 | 0 | 0 |
| 0 | NA | NA | NA | NA | NA | Gm25585 | 0 | 0 | 0 | 0 | 0 | 0 | 0 | 0 | 0 | 0 | 0 |
| 0 | NA | NA | NA | NA | NA | Gm25840 | 0 | 0 | 0 | 0 | 0 | 0 | 0 | 0 | 0 | 0 | 0 |
| 0 | NA | NA | NA | NA | NA | Gm25089 | 0 | 0 | 0 | 0 | 0 | 0 | 0 | 0 | 0 | 0 | 0 |
| 0 | NA | NA | NA | NA | NA | Gm24040 | 0 | 0 | 0 | 0 | 0 | 0 | 0 | 0 | 0 | 0 | 0 |
| 0 | NA | NA | NA | NA | NA | Gm22050 | 0 | 0 | 0 | 0 | 0 | 0 | 0 | 0 | 0 | 0 | 0 |
| 0 | NA | NA | NA | NA | NA | Gm23286 | 0 | 0 | 0 | 0 | 0 | 0 | 0 | 0 | 0 | 0 | 0 |
| 0 | NA | NA | NA | NA | NA | Gm24021 | 0 | 0 | 0 | 0 | 0 | 0 | 0 | 0 | 0 | 0 | 0 |
| 0 | NA | NA | NA | NA | NA | Gm22274 | 0 | 0 | 0 | 0 | 0 | 0 | 0 | 0 | 0 | 0 | 0 |
| 0 | NA | NA | NA | NA | NA | Gm24652 | 0 | 0 | 0 | 0 | 0 | 0 | 0 | 0 | 0 | 0 | 0 |
| 0 | NA | NA | NA | NA | NA | Gm23618 | 0 | 0 | 0 | 0 | 0 | 0 | 0 | 0 | 0 | 0 | 0 |
| 0 | NA | NA | NA | NA | NA | Gm23688 | 0 | 0 | 0 | 0 | 0 | 0 | 0 | 0 | 0 | 0 | 0 |
| 0 | NA | NA | NA | NA | NA | Gm24570 | 0 | 0 | 0 | 0 | 0 | 0 | 0 | 0 | 0 | 0 | 0 |
| 0 | NA | NA | NA | NA | NA | Gm26467 | 0 | 0 | 0 | 0 | 0 | 0 | 0 | 0 | 0 | 0 | 0 |
| 0 | NA | NA | NA | NA | NA | Gm25194 | 0 | 0 | 0 | 0 | 0 | 0 | 0 | 0 | 0 | 0 | 0 |
| 0 | NA | NA | NA | NA | NA | Gm22640 | 0 | 0 | 0 | 0 | 0 | 0 | 0 | 0 | 0 | 0 | 0 |
| 0 | NA | NA | NA | NA | NA | Gm24952 | 0 | 0 | 0 | 0 | 0 | 0 | 0 | 0 | 0 | 0 | 0 |
| 0 | NA | NA | NA | NA | NA | Gm25087 | 0 | 0 | 0 | 0 | 0 | 0 | 0 | 0 | 0 | 0 | 0 |
| 0 | NA | NA | NA | NA | NA | Gm23944 | 0 | 0 | 0 | 0 | 0 | 0 | 0 | 0 | 0 | 0 | 0 |
| 0 | NA | NA | NA | NA | NA | Gm22629 | 0 | 0 | 0 | 0 | 0 | 0 | 0 | 0 | 0 | 0 | 0 |
| 0 | NA | NA | NA | NA | NA | Gm26374 | 0 | 0 | 0 | 0 | 0 | 0 | 0 | 0 | 0 | 0 | 0 |
| 0 | NA | NA | NA | NA | NA | Gm26284 | 0 | 0 | 0 | 0 | 0 | 0 | 0 | 0 | 0 | 0 | 0 |
| 0 | NA | NA | NA | NA | NA | Gm25452 | 0 | 0 | 0 | 0 | 0 | 0 | 0 | 0 | 0 | 0 | 0 |
| 0 | NA | NA | NA | NA | NA | Gm25984 | 0 | 0 | 0 | 0 | 0 | 0 | 0 | 0 | 0 | 0 | 0 |
| 0 | NA | NA | NA | NA | NA | Gm25146 | 0 | 0 | 0 | 0 | 0 | 0 | 0 | 0 | 0 | 0 | 0 |

| | | | | | | | | | | | | | | | | | |
|---|---|---|---|---|---|---|---|---|---|---|---|---|---|---|---|---|---|
| 0 | NA | NA | NA | NA | NA | Gm24759 | 0 | 0 | 0 | 0 | 0 | 0 | 0 | 0 | 0 | 0 | 0 |
| 0 | NA | NA | NA | NA | NA | Gm24100 | 0 | 0 | 0 | 0 | 0 | 0 | 0 | 0 | 0 | 0 | 0 |
| 0 | NA | NA | NA | NA | NA | Gm22253 | 0 | 0 | 0 | 0 | 0 | 0 | 0 | 0 | 0 | 0 | 0 |
| 0 | NA | NA | NA | NA | NA | Gm25017 | 0 | 0 | 0 | 0 | 0 | 0 | 0 | 0 | 0 | 0 | 0 |
| 0 | NA | NA | NA | NA | NA | Gm23575 | 0 | 0 | 0 | 0 | 0 | 0 | 0 | 0 | 0 | 0 | 0 |
| 0 | NA | NA | NA | NA | NA | Gm22391 | 0 | 0 | 0 | 0 | 0 | 0 | 0 | 0 | 0 | 0 | 0 |
| 0 | NA | NA | NA | NA | NA | Gm22449 | 0 | 0 | 0 | 0 | 0 | 0 | 0 | 0 | 0 | 0 | 0 |
| 0 | NA | NA | NA | NA | NA | Gm24137 | 0 | 0 | 0 | 0 | 0 | 0 | 0 | 0 | 0 | 0 | 0 |
| 0 | NA | NA | NA | NA | NA | Gm22912 | 0 | 0 | 0 | 0 | 0 | 0 | 0 | 0 | 0 | 0 | 0 |
| 0 | NA | NA | NA | NA | NA | Gm25710 | 0 | 0 | 0 | 0 | 0 | 0 | 0 | 0 | 0 | 0 | 0 |
| 0 | NA | NA | NA | NA | NA | Gm26434 | 0 | 0 | 0 | 0 | 0 | 0 | 0 | 0 | 0 | 0 | 0 |
| 0 | NA | NA | NA | NA | NA | Gm23560 | 0 | 0 | 0 | 0 | 0 | 0 | 0 | 0 | 0 | 0 | 0 |
| 0 | NA | NA | NA | NA | NA | Gm25870 | 0 | 0 | 0 | 0 | 0 | 0 | 0 | 0 | 0 | 0 | 0 |
| 0 | NA | NA | NA | NA | NA | Gm44831 | 0 | 0 | 0 | 0 | 0 | 0 | 0 | 0 | 0 | 0 | 0 |
| 0 | NA | NA | NA | NA | NA | Gm23862 | 0 | 0 | 0 | 0 | 0 | 0 | 0 | 0 | 0 | 0 | 0 |
| 0 | NA | NA | NA | NA | NA | Gm26504 | 0 | 0 | 0 | 0 | 0 | 0 | 0 | 0 | 0 | 0 | 0 |
| 0 | NA | NA | NA | NA | NA | Gm22285 | 0 | 0 | 0 | 0 | 0 | 0 | 0 | 0 | 0 | 0 | 0 |
| 0 | NA | NA | NA | NA | NA | Gm22131 | 0 | 0 | 0 | 0 | 0 | 0 | 0 | 0 | 0 | 0 | 0 |
| 0 | NA | NA | NA | NA | NA | Gm24153 | 0 | 0 | 0 | 0 | 0 | 0 | 0 | 0 | 0 | 0 | 0 |
| 0 | NA | NA | NA | NA | NA | Gm26365 | 0 | 0 | 0 | 0 | 0 | 0 | 0 | 0 | 0 | 0 | 0 |
| 0 | NA | NA | NA | NA | NA | Gm24609 | 0 | 0 | 0 | 0 | 0 | 0 | 0 | 0 | 0 | 0 | 0 |
| 0 | NA | NA | NA | NA | NA | Gm24264 | 0 | 0 | 0 | 0 | 0 | 0 | 0 | 0 | 0 | 0 | 0 |
| 0 | NA | NA | NA | NA | NA | Gm25155 | 0 | 0 | 0 | 0 | 0 | 0 | 0 | 0 | 0 | 0 | 0 |
| 0 | NA | NA | NA | NA | NA | Gm25816 | 0 | 0 | 0 | 0 | 0 | 0 | 0 | 0 | 0 | 0 | 0 |
| 0 | NA | NA | NA | NA | NA | Gm23141 | 0 | 0 | 0 | 0 | 0 | 0 | 0 | 0 | 0 | 0 | 0 |
| 0 | NA | NA | NA | NA | NA | Gm23767 | 0 | 0 | 0 | 0 | 0 | 0 | 0 | 0 | 0 | 0 | 0 |
| 0 | NA | NA | NA | NA | NA | Gm22417 | 0 | 0 | 0 | 0 | 0 | 0 | 0 | 0 | 0 | 0 | 0 |
| 0 | NA | NA | NA | NA | NA | Gm22584 | 0 | 0 | 0 | 0 | 0 | 0 | 0 | 0 | 0 | 0 | 0 |
| 0 | NA | NA | NA | NA | NA | Gm26270 | 0 | 0 | 0 | 0 | 0 | 0 | 0 | 0 | 0 | 0 | 0 |
| 0 | NA | NA | NA | NA | NA | Gm23089 | 0 | 0 | 0 | 0 | 0 | 0 | 0 | 0 | 0 | 0 | 0 |
| 0 | NA | NA | NA | NA | NA | Gm24711 | 0 | 0 | 0 | 0 | 0 | 0 | 0 | 0 | 0 | 0 | 0 |
| 0 | NA | NA | NA | NA | NA | Gm26188 | 0 | 0 | 0 | 0 | 0 | 0 | 0 | 0 | 0 | 0 | 0 |
| 0 | NA | NA | NA | NA | NA | Gm23265 | 0 | 0 | 0 | 0 | 0 | 0 | 0 | 0 | 0 | 0 | 0 |
| 0 | NA | NA | NA | NA | NA | Gm24518 | 0 | 0 | 0 | 0 | 0 | 0 | 0 | 0 | 0 | 0 | 0 |
| 0 | NA | NA | NA | NA | NA | Gm25944 | 0 | 0 | 0 | 0 | 0 | 0 | 0 | 0 | 0 | 0 | 0 |
| 0 | NA | NA | NA | NA | NA | Gm23047 | 0 | 0 | 0 | 0 | 0 | 0 | 0 | 0 | 0 | 0 | 0 |
| 0 | NA | NA | NA | NA | NA | Gm23953 | 0 | 0 | 0 | 0 | 0 | 0 | 0 | 0 | 0 | 0 | 0 |
| 0 | NA | NA | NA | NA | NA | Gm26332 | 0 | 0 | 0 | 0 | 0 | 0 | 0 | 0 | 0 | 0 | 0 |
| 0 | NA | NA | NA | NA | NA | Gm26032 | 0 | 0 | 0 | 0 | 0 | 0 | 0 | 0 | 0 | 0 | 0 |
| 0 | NA | NA | NA | NA | NA | Gm22863 | 0 | 0 | 0 | 0 | 0 | 0 | 0 | 0 | 0 | 0 | 0 |
| 0 | NA | NA | NA | NA | NA | Gm24618 | 0 | 0 | 0 | 0 | 0 | 0 | 0 | 0 | 0 | 0 | 0 |
| 0 | NA | NA | NA | NA | NA | Gm22046 | 0 | 0 | 0 | 0 | 0 | 0 | 0 | 0 | 0 | 0 | 0 |
| 0 | NA | NA | NA | NA | NA | Gm24760 | 0 | 0 | 0 | 0 | 0 | 0 | 0 | 0 | 0 | 0 | 0 |
| 0 | NA | NA | NA | NA | NA | Gm25350 | 0 | 0 | 0 | 0 | 0 | 0 | 0 | 0 | 0 | 0 | 0 |
| 0 | NA | NA | NA | NA | NA | Gm26502 | 0 | 0 | 0 | 0 | 0 | 0 | 0 | 0 | 0 | 0 | 0 |
| 0 | NA | NA | NA | NA | NA | Gm23313 | 0 | 0 | 0 | 0 | 0 | 0 | 0 | 0 | 0 | 0 | 0 |
| 0 | NA | NA | NA | NA | NA | Gm25471 | 0 | 0 | 0 | 0 | 0 | 0 | 0 | 0 | 0 | 0 | 0 |
| 0 | NA | NA | NA | NA | NA | Gm22776 | 0 | 0 | 0 | 0 | 0 | 0 | 0 | 0 | 0 | 0 | 0 |
| 0 | NA | NA | NA | NA | NA | Gm23446 | 0 | 0 | 0 | 0 | 0 | 0 | 0 | 0 | 0 | 0 | 0 |
| 0 | NA | NA | NA | NA | NA | Gm25210 | 0 | 0 | 0 | 0 | 0 | 0 | 0 | 0 | 0 | 0 | 0 |
| 0 | NA | NA | NA | NA | NA | Gm22128 | 0 | 0 | 0 | 0 | 0 | 0 | 0 | 0 | 0 | 0 | 0 |
| 0 | NA | NA | NA | NA | NA | Gm22851 | 0 | 0 | 0 | 0 | 0 | 0 | 0 | 0 | 0 | 0 | 0 |
| 0 | NA | NA | NA | NA | NA | Gm25074 | 0 | 0 | 0 | 0 | 0 | 0 | 0 | 0 | 0 | 0 | 0 |
| 0 | NA | NA | NA | NA | NA | Gm25157 | 0 | 0 | 0 | 0 | 0 | 0 | 0 | 0 | 0 | 0 | 0 |
| 0 | NA | NA | NA | NA | NA | Gm22631 | 0 | 0 | 0 | 0 | 0 | 0 | 0 | 0 | 0 | 0 | 0 |
| 0 | NA | NA | NA | NA | NA | Gm25474 | 0 | 0 | 0 | 0 | 0 | 0 | 0 | 0 | 0 | 0 | 0 |
| 0 | NA | NA | NA | NA | NA | Gm22941 | 0 | 0 | 0 | 0 | 0 | 0 | 0 | 0 | 0 | 0 | 0 |
| 0 | NA | NA | NA | NA | NA | Gm26246 | 0 | 0 | 0 | 0 | 0 | 0 | 0 | 0 | 0 | 0 | 0 |
| 0 | NA | NA | NA | NA | NA | Gm22047 | 0 | 0 | 0 | 0 | 0 | 0 | 0 | 0 | 0 | 0 | 0 |
| 0 | NA | NA | NA | NA | NA | Gm25597 | 0 | 0 | 0 | 0 | 0 | 0 | 0 | 0 | 0 | 0 | 0 |

| | | | | | | | | | | | | | | | | | | |
|---|---|---|---|---|---|---|---|---|---|---|---|---|---|---|---|---|---|---|
| 0 | NA | NA | NA | NA | NA | Gm26097 | | 0 | 0 | 0 | 0 | 0 | 0 | 0 | 0 | 0 | 0 | 0 |
| 0 | NA | NA | NA | NA | NA | Gm22258 | | 0 | 0 | 0 | 0 | 0 | 0 | 0 | 0 | 0 | 0 | 0 |
| 0 | NA | NA | NA | NA | NA | Gm26201 | | 0 | 0 | 0 | 0 | 0 | 0 | 0 | 0 | 0 | 0 | 0 |
| 0 | NA | NA | NA | NA | NA | Gm23619 | | 0 | 0 | 0 | 0 | 0 | 0 | 0 | 0 | 0 | 0 | 0 |
| 0 | NA | NA | NA | NA | NA | Gm22812 | | 0 | 0 | 0 | 0 | 0 | 0 | 0 | 0 | 0 | 0 | 0 |
| 0 | NA | NA | NA | NA | NA | Gm26433 | | 0 | 0 | 0 | 0 | 0 | 0 | 0 | 0 | 0 | 0 | 0 |
| 0 | NA | NA | NA | NA | NA | Snord116l17 | | 0 | 0 | 0 | 0 | 0 | 0 | 0 | 0 | 0 | 0 | 0 |
| 0 | NA | NA | NA | NA | NA | Gm25615 | | 0 | 0 | 0 | 0 | 0 | 0 | 0 | 0 | 0 | 0 | 0 |
| 0 | NA | NA | NA | NA | NA | Snord116l16 | | 0 | 0 | 0 | 0 | 0 | 0 | 0 | 0 | 0 | 0 | 0 |
| 0 | NA | NA | NA | NA | NA | Snord116l15 | | 0 | 0 | 0 | 0 | 0 | 0 | 0 | 0 | 0 | 0 | 0 |
| 0 | NA | NA | NA | NA | NA | Snord116l14 | | 0 | 0 | 0 | 0 | 0 | 0 | 0 | 0 | 0 | 0 | 0 |
| 0 | NA | NA | NA | NA | NA | Snord116l13 | | 0 | 0 | 0 | 0 | 0 | 0 | 0 | 0 | 0 | 0 | 0 |
| 0 | NA | NA | NA | NA | NA | Snord116l2 | | 0 | 0 | 0 | 0 | 0 | 0 | 0 | 0 | 0 | 0 | 0 |
| 0 | NA | NA | NA | NA | NA | Snord116l8 | | 0 | 0 | 0 | 0 | 0 | 0 | 0 | 0 | 0 | 0 | 0 |
| 0 | NA | NA | NA | NA | NA | Snord116l7 | | 0 | 0 | 0 | 0 | 0 | 0 | 0 | 0 | 0 | 0 | 0 |
| 0 | NA | NA | NA | NA | NA | Snord116l6 | | 0 | 0 | 0 | 0 | 0 | 0 | 0 | 0 | 0 | 0 | 0 |
| 0 | NA | NA | NA | NA | NA | Snord116l5 | | 0 | 0 | 0 | 0 | 0 | 0 | 0 | 0 | 0 | 0 | 0 |
| 0 | NA | NA | NA | NA | NA | Snord116l4 | | 0 | 0 | 0 | 0 | 0 | 0 | 0 | 0 | 0 | 0 | 0 |
| 0 | NA | NA | NA | NA | NA | Snord116l3 | | 0 | 0 | 0 | 0 | 0 | 0 | 0 | 0 | 0 | 0 | 0 |
| 0 | NA | NA | NA | NA | NA | Snord116l1 | | 0 | 0 | 0 | 0 | 0 | 0 | 0 | 0 | 0 | 0 | 0 |
| 0 | NA | NA | NA | NA | NA | Snord116l9 | | 0 | 0 | 0 | 0 | 0 | 0 | 0 | 0 | 0 | 0 | 0 |
| 0 | NA | NA | NA | NA | NA | Snord116l10 | | 0 | 0 | 0 | 0 | 0 | 0 | 0 | 0 | 0 | 0 | 0 |
| 0 | NA | NA | NA | NA | NA | Snord116l11 | | 0 | 0 | 0 | 0 | 0 | 0 | 0 | 0 | 0 | 0 | 0 |
| 0 | NA | NA | NA | NA | NA | Snord116l12 | | 0 | 0 | 0 | 0 | 0 | 0 | 0 | 0 | 0 | 0 | 0 |
| 0 | NA | NA | NA | NA | NA | Gm22373 | | 0 | 0 | 0 | 0 | 0 | 0 | 0 | 0 | 0 | 0 | 0 |
| 0 | NA | NA | NA | NA | NA | Snord64 | | 0 | 0 | 0 | 0 | 0 | 0 | 0 | 0 | 0 | 0 | 0 |
| 0 | NA | NA | NA | NA | NA | Snord107 | | 0 | 0 | 0 | 0 | 0 | 0 | 0 | 0 | 0 | 0 | 0 |
| 0 | NA | NA | NA | NA | NA | Snurf | SNRPN upstream reading frame | 0 | 0 | 0 | 0 | 0 | 0 | 0 | 0 | 0 | 0 | 0 |
| 0 | NA | NA | NA | NA | NA | Gm18908 | | 0 | 0 | 0 | 0 | 0 | 0 | 0 | 0 | 0 | 0 | 0 |
| 0 | NA | NA | NA | NA | NA | Gm18129 | | 0 | 0 | 0 | 0 | 0 | 0 | 0 | 0 | 0 | 0 | 0 |
| 0 | NA | NA | NA | NA | NA | Gm30196 | | 0 | 0 | 0 | 0 | 0 | 0 | 0 | 0 | 0 | 0 | 0 |
| 0 | NA | NA | NA | NA | NA | Gm3198 | | 0 | 0 | 0 | 0 | 0 | 0 | 0 | 0 | 0 | 0 | 0 |
| 0 | NA | NA | NA | NA | NA | Gm7394 | | 0 | 0 | 0 | 0 | 0 | 0 | 0 | 0 | 0 | 0 | 0 |
| 0 | NA | NA | NA | NA | NA | Gm44644 | | 0 | 0 | 0 | 0 | 0 | 0 | 0 | 0 | 0 | 0 | 0 |
| 0 | NA | NA | NA | NA | NA | Gm44643 | | 0 | 0 | 0 | 0 | 0 | 0 | 0 | 0 | 0 | 0 | 0 |
| 0 | NA | NA | NA | NA | NA | A230006K03Rik | | 0 | 0 | 0 | 0 | 0 | 0 | 0 | 0 | 0 | 0 | 0 |
| 0 | NA | NA | NA | NA | NA | Gm38451 | | 0 | 0 | 0 | 0 | 0 | 0 | 0 | 0 | 0 | 0 | 0 |
| 0 | NA | NA | NA | NA | NA | B230209E15Rik | | 0 | 0 | 0 | 0 | 0 | 0 | 0 | 0 | 0 | 0 | 0 |
| 0 | NA | NA | NA | NA | NA | Gm25533 | | 0 | 0 | 0 | 0 | 0 | 0 | 0 | 0 | 0 | 0 | 0 |
| 0 | NA | NA | NA | NA | NA | Gm23490 | | 0 | 0 | 0 | 0 | 0 | 0 | 0 | 0 | 0 | 0 | 0 |
| 0 | NA | NA | NA | NA | NA | Gm22466 | | 0 | 0 | 0 | 0 | 0 | 0 | 0 | 0 | 0 | 0 | 0 |
| 0 | NA | NA | NA | NA | NA | Mir344d-3 | | 0 | 0 | 0 | 0 | 0 | 0 | 0 | 0 | 0 | 0 | 0 |
| 0 | NA | NA | NA | NA | NA | Mir344e | | 0 | 0 | 0 | 0 | 0 | 0 | 0 | 0 | 0 | 0 | 0 |
| 0 | NA | NA | NA | NA | NA | Mir344h-1 | | 0 | 0 | 0 | 0 | 0 | 0 | 0 | 0 | 0 | 0 | 0 |
| 0 | NA | NA | NA | NA | NA | Mir344h-2 | | 0 | 0 | 0 | 0 | 0 | 0 | 0 | 0 | 0 | 0 | 0 |
| 0 | NA | NA | NA | NA | NA | Mir344b | | 0 | 0 | 0 | 0 | 0 | 0 | 0 | 0 | 0 | 0 | 0 |
| 0 | NA | NA | NA | NA | NA | Gm19032 | | 0 | 0 | 0 | 0 | 0 | 0 | 0 | 0 | 0 | 0 | 0 |
| 0 | NA | NA | NA | NA | NA | Gm3257 | | 0 | 0 | 0 | 0 | 0 | 0 | 0 | 0 | 0 | 0 | 0 |
| 0 | NA | NA | NA | NA | NA | Gm32061 | | 0 | 0 | 0 | 0 | 0 | 0 | 0 | 0 | 0 | 0 | 0 |
| 0 | NA | NA | NA | NA | NA | Vmn2r-ps61 | | 0 | 0 | 0 | 0 | 0 | 0 | 0 | 0 | 0 | 0 | 0 |
| 0 | NA | NA | NA | NA | NA | Gm22494 | | 0 | 0 | 0 | 0 | 0 | 0 | 0 | 0 | 0 | 0 | 0 |
| 0 | NA | NA | NA | NA | NA | Gm9445 | | 0 | 0 | 0 | 0 | 0 | 0 | 0 | 0 | 0 | 0 | 0 |
| 0 | NA | NA | NA | NA | NA | A26c2 | ANKRD26-like family C, member 2 | 0 | 0 | 0 | 0 | 0 | 0 | 0 | 0 | 0 | 0 | 0 |
| 0 | NA | NA | NA | NA | NA | Gm32111 | | 0 | 0 | 0 | 0 | 0 | 0 | 0 | 0 | 0 | 0 | 0 |
| 0 | NA | NA | NA | NA | NA | 4930554H23Rik | | 0 | 0 | 0 | 0 | 0 | 0 | 0 | 0 | 0 | 0 | 0 |
| 0 | NA | NA | NA | NA | NA | Gm22228 | | 0 | 0 | 0 | 0 | 0 | 0 | 0 | 0 | 0 | 0 | 0 |
| 0 | NA | NA | NA | NA | NA | Gm44720 | | 0 | 0 | 0 | 0 | 0 | 0 | 0 | 0 | 0 | 0 | 0 |
| 0 | NA | NA | NA | NA | NA | Gm45052 | | 0 | 0 | 0 | 0 | 0 | 0 | 0 | 0 | 0 | 0 | 0 |
| 0 | NA | NA | NA | NA | NA | Gm45054 | | 0 | 0 | 0 | 0 | 0 | 0 | 0 | 0 | 0 | 0 | 0 |
| 0 | NA | NA | NA | NA | NA | Gm20670 | | 0 | 0 | 0 | 0 | 0 | 0 | 0 | 0 | 0 | 0 | 0 |

| | | | | | | | | | | | | | | | | | |
|---|---|---|---|---|---|---|---|---|---|---|---|---|---|---|---|---|---|
| 0 | NA | NA | NA | NA | NA | Gm44722 | 0 | 0 | 0 | 0 | 0 | 0 | 0 | 0 | 0 | 0 | 0 |
| 0 | NA | NA | NA | NA | NA | Mir211 | 0 | 0 | 0 | 0 | 0 | 0 | 0 | 0 | 0 | 0 | 0 |
| 0 | NA | NA | NA | NA | NA | Gm44719 | 0 | 0 | 0 | 0 | 0 | 0 | 0 | 0 | 0 | 0 | 0 |
| 0 | NA | NA | NA | NA | NA | Mir6416 | 0 | 0 | 0 | 0 | 0 | 0 | 0 | 0 | 0 | 0 | 0 |
| 0 | NA | NA | NA | NA | NA | Gm44765 | 0 | 0 | 0 | 0 | 0 | 0 | 0 | 0 | 0 | 0 | 0 |
| 0 | NA | NA | NA | NA | NA | Gm45082 | 0 | 0 | 0 | 0 | 0 | 0 | 0 | 0 | 0 | 0 | 0 |
| 0 | NA | NA | NA | NA | NA | Gm44721 | 0 | 0 | 0 | 0 | 0 | 0 | 0 | 0 | 0 | 0 | 0 |
| 0 | NA | NA | NA | NA | NA | Gm25249 | 0 | 0 | 0 | 0 | 0 | 0 | 0 | 0 | 0 | 0 | 0 |
| 0 | NA | NA | NA | NA | NA | Gm17908 | 0 | 0 | 0 | 0 | 0 | 0 | 0 | 0 | 0 | 0 | 0 |
| 0 | NA | NA | NA | NA | NA | Gm17918 | 0 | 0 | 0 | 0 | 0 | 0 | 0 | 0 | 0 | 0 | 0 |
| 0 | NA | NA | NA | NA | NA | 1810049I09Rik | 0 | 0 | 0 | 0 | 0 | 0 | 0 | 0 | 0 | 0 | 0 |
| 0 | NA | NA | NA | NA | NA | Gm44794 | 0 | 0 | 0 | 0 | 0 | 0 | 0 | 0 | 0 | 0 | 0 |
| 0 | NA | NA | NA | NA | NA | Gm44795 | 0 | 0 | 0 | 0 | 0 | 0 | 0 | 0 | 0 | 0 | 0 |
| 0 | NA | NA | NA | NA | NA | Gm44792 | 0 | 0 | 0 | 0 | 0 | 0 | 0 | 0 | 0 | 0 | 0 |
| 0 | NA | NA | NA | NA | NA | Gm45213 | 0 | 0 | 0 | 0 | 0 | 0 | 0 | 0 | 0 | 0 | 0 |
| 0 | NA | NA | NA | NA | NA | Gm26827 | 0 | 0 | 0 | 0 | 0 | 0 | 0 | 0 | 0 | 0 | 0 |
| 0 | NA | NA | NA | NA | NA | Gm23042 | 0 | 0 | 0 | 0 | 0 | 0 | 0 | 0 | 0 | 0 | 0 |
| 0 | NA | NA | NA | NA | NA | Gm44752 | 0 | 0 | 0 | 0 | 0 | 0 | 0 | 0 | 0 | 0 | 0 |
| 0 | NA | NA | NA | NA | NA | Gm10974 | 0 | 0 | 0 | 0 | 0 | 0 | 0 | 0 | 0 | 0 | 0 |
| 0 | NA | NA | NA | NA | NA | Gm23957 | 0 | 0 | 0 | 0 | 0 | 0 | 0 | 0 | 0 | 0 | 0 |
| 0 | NA | NA | NA | NA | NA | Mir7057 | 0 | 0 | 0 | 0 | 0 | 0 | 0 | 0 | 0 | 0 | 0 |
| 0 | NA | NA | NA | NA | NA | Gm44753 | 0 | 0 | 0 | 0 | 0 | 0 | 0 | 0 | 0 | 0 | 0 |
| 0 | NA | NA | NA | NA | NA | Gm44754 | 0 | 0 | 0 | 0 | 0 | 0 | 0 | 0 | 0 | 0 | 0 |
| 0 | NA | NA | NA | NA | NA | Gm33570 | 0 | 0 | 0 | 0 | 0 | 0 | 0 | 0 | 0 | 0 | 0 |
| 0 | NA | NA | NA | NA | NA | Gm44610 | 0 | 0 | 0 | 0 | 0 | 0 | 0 | 0 | 0 | 0 | 0 |
| 0 | NA | NA | NA | NA | NA | Gm45098 | 0 | 0 | 0 | 0 | 0 | 0 | 0 | 0 | 0 | 0 | 0 |
| 0 | NA | NA | NA | NA | NA | Gm10623 | 0 | 0 | 0 | 0 | 0 | 0 | 0 | 0 | 0 | 0 | 0 |
| 0 | NA | NA | NA | NA | NA | Gm45101 | 0 | 0 | 0 | 0 | 0 | 0 | 0 | 0 | 0 | 0 | 0 |
| 0 | NA | NA | NA | NA | NA | Gm44668 | 0 | 0 | 0 | 0 | 0 | 0 | 0 | 0 | 0 | 0 | 0 |
| 0 | NA | NA | NA | NA | NA | Gm33926 | 0 | 0 | 0 | 0 | 0 | 0 | 0 | 0 | 0 | 0 | 0 |
| 0 | NA | NA | NA | NA | NA | Gm23233 | 0 | 0 | 0 | 0 | 0 | 0 | 0 | 0 | 0 | 0 | 0 |
| 0 | NA | NA | NA | NA | NA | Gm45100 | 0 | 0 | 0 | 0 | 0 | 0 | 0 | 0 | 0 | 0 | 0 |
| 0 | NA | NA | NA | NA | NA | Gm44669 | 0 | 0 | 0 | 0 | 0 | 0 | 0 | 0 | 0 | 0 | 0 |
| 0 | NA | NA | NA | NA | NA | Gm45102 | 0 | 0 | 0 | 0 | 0 | 0 | 0 | 0 | 0 | 0 | 0 |
| 0 | NA | NA | NA | NA | NA | Gm23844 | 0 | 0 | 0 | 0 | 0 | 0 | 0 | 0 | 0 | 0 | 0 |
| 0 | NA | NA | NA | NA | NA | Gm44666 | 0 | 0 | 0 | 0 | 0 | 0 | 0 | 0 | 0 | 0 | 0 |
| 0 | NA | NA | NA | NA | NA | Gm16157 | 0 | 0 | 0 | 0 | 0 | 0 | 0 | 0 | 0 | 0 | 0 |
| 0 | NA | NA | NA | NA | NA | 4930405G09Rik | 0 | 0 | 0 | 0 | 0 | 0 | 0 | 0 | 0 | 0 | 0 |
| 0 | NA | NA | NA | NA | NA | Gm44888 | 0 | 0 | 0 | 0 | 0 | 0 | 0 | 0 | 0 | 0 | 0 |
| 0 | NA | NA | NA | NA | NA | Gm16158 | 0 | 0 | 0 | 0 | 0 | 0 | 0 | 0 | 0 | 0 | 0 |
| 0 | NA | NA | NA | NA | NA | Gm44889 | 0 | 0 | 0 | 0 | 0 | 0 | 0 | 0 | 0 | 0 | 0 |
| 0 | NA | NA | NA | NA | NA | 4933436H12Rik | 0 | 0 | 0 | 0 | 0 | 0 | 0 | 0 | 0 | 0 | 0 |
| 0 | NA | NA | NA | NA | NA | Gm34549 | 0 | 0 | 0 | 0 | 0 | 0 | 0 | 0 | 0 | 0 | 0 |
| 0 | NA | NA | NA | NA | NA | Gm44886 | 0 | 0 | 0 | 0 | 0 | 0 | 0 | 0 | 0 | 0 | 0 |
| 0 | NA | NA | NA | NA | NA | Gm44887 | 0 | 0 | 0 | 0 | 0 | 0 | 0 | 0 | 0 | 0 | 0 |
| 0 | NA | NA | NA | NA | NA | Gm5334 | 0 | 0 | 0 | 0 | 0 | 0 | 0 | 0 | 0 | 0 | 0 |
| 0 | NA | NA | NA | NA | NA | Gm44692 | 0 | 0 | 0 | 0 | 0 | 0 | 0 | 0 | 0 | 0 | 0 |
| 0 | NA | NA | NA | NA | NA | Gm34664 | 0 | 0 | 0 | 0 | 0 | 0 | 0 | 0 | 0 | 0 | 0 |
| 0 | NA | NA | NA | NA | NA | Gm23795 | 0 | 0 | 0 | 0 | 0 | 0 | 0 | 0 | 0 | 0 | 0 |
| 0 | NA | NA | NA | NA | NA | Gm44691 | 0 | 0 | 0 | 0 | 0 | 0 | 0 | 0 | 0 | 0 | 0 |
| 0 | NA | NA | NA | NA | NA | Gm5342 | 0 | 0 | 0 | 0 | 0 | 0 | 0 | 0 | 0 | 0 | 0 |
| 0 | NA | NA | NA | NA | NA | Gm44533 | 0 | 0 | 0 | 0 | 0 | 0 | 0 | 0 | 0 | 0 | 0 |
| 0 | NA | NA | NA | NA | NA | Gm34838 | 0 | 0 | 0 | 0 | 0 | 0 | 0 | 0 | 0 | 0 | 0 |
| 0 | NA | NA | NA | NA | NA | Gm38584 | 0 | 0 | 0 | 0 | 0 | 0 | 0 | 0 | 0 | 0 | 0 |
| 0 | NA | NA | NA | NA | NA | 4930402F11Rik | 0 | 0 | 0 | 0 | 0 | 0 | 0 | 0 | 0 | 0 | 0 |
| 0 | NA | NA | NA | NA | NA | Gm23815 | 0 | 0 | 0 | 0 | 0 | 0 | 0 | 0 | 0 | 0 | 0 |
| 0 | NA | NA | NA | NA | NA | Gm44535 | 0 | 0 | 0 | 0 | 0 | 0 | 0 | 0 | 0 | 0 | 0 |
| 0 | NA | NA | NA | NA | NA | Gm7627 | 0 | 0 | 0 | 0 | 0 | 0 | 0 | 0 | 0 | 0 | 0 |
| 0 | NA | NA | NA | NA | NA | Gm44690 | 0 | 0 | 0 | 0 | 0 | 0 | 0 | 0 | 0 | 0 | 0 |
| 0 | NA | NA | NA | NA | NA | 4933423L19Rik | 0 | 0 | 0 | 0 | 0 | 0 | 0 | 0 | 0 | 0 | 0 |

| | | | | | | | | | | | | | | | | | |
|---|---|---|---|---|---|---|---|---|---|---|---|---|---|---|---|---|---|
| 0 | NA | NA | NA | NA | NA | Gm24120 | 0 | 0 | 0 | 0 | 0 | 0 | 0 | 0 | 0 | 0 | 0 |
| 0 | NA | NA | NA | NA | NA | Gm35325 | 0 | 0 | 0 | 0 | 0 | 0 | 0 | 0 | 0 | 0 | 0 |
| 0 | NA | NA | NA | NA | NA | Gm29327 | 0 | 0 | 0 | 0 | 0 | 0 | 0 | 0 | 0 | 0 | 0 |
| 0 | NA | NA | NA | NA | NA | Gm44946 | 0 | 0 | 0 | 0 | 0 | 0 | 0 | 0 | 0 | 0 | 0 |
| 0 | NA | NA | NA | NA | NA | Gm44948 | 0 | 0 | 0 | 0 | 0 | 0 | 0 | 0 | 0 | 0 | 0 |
| 0 | NA | NA | NA | NA | NA | Gm28258 | 0 | 0 | 0 | 0 | 0 | 0 | 0 | 0 | 0 | 0 | 0 |
| 0 | NA | NA | NA | NA | NA | Gm44532 | 0 | 0 | 0 | 0 | 0 | 0 | 0 | 0 | 0 | 0 | 0 |
| 0 | NA | NA | NA | NA | NA | Gm7656 | 0 | 0 | 0 | 0 | 0 | 0 | 0 | 0 | 0 | 0 | 0 |
| 0 | NA | NA | NA | NA | NA | Gm44811 | 0 | 0 | 0 | 0 | 0 | 0 | 0 | 0 | 0 | 0 | 0 |
| 0 | NA | NA | NA | NA | NA | Gm35842 | 0 | 0 | 0 | 0 | 0 | 0 | 0 | 0 | 0 | 0 | 0 |
| 0 | NA | NA | NA | NA | NA | Gm44813 | 0 | 0 | 0 | 0 | 0 | 0 | 0 | 0 | 0 | 0 | 0 |
| 0 | NA | NA | NA | NA | NA | Gm44809 | 0 | 0 | 0 | 0 | 0 | 0 | 0 | 0 | 0 | 0 | 0 |
| 0 | NA | NA | NA | NA | NA | Gm44808 | 0 | 0 | 0 | 0 | 0 | 0 | 0 | 0 | 0 | 0 | 0 |
| 0 | NA | NA | NA | NA | NA | Gm24880 | 0 | 0 | 0 | 0 | 0 | 0 | 0 | 0 | 0 | 0 | 0 |
| 0 | NA | NA | NA | NA | NA | Gm42397 | 0 | 0 | 0 | 0 | 0 | 0 | 0 | 0 | 0 | 0 | 0 |
| 0 | NA | NA | NA | NA | NA | Gm34783 | 0 | 0 | 0 | 0 | 0 | 0 | 0 | 0 | 0 | 0 | 0 |
| 0 | NA | NA | NA | NA | NA | Gm36011 | 0 | 0 | 0 | 0 | 0 | 0 | 0 | 0 | 0 | 0 | 0 |
| 0 | NA | NA | NA | NA | NA | Gm39033 | 0 | 0 | 0 | 0 | 0 | 0 | 0 | 0 | 0 | 0 | 0 |
| 0 | NA | NA | NA | NA | NA | 6030442E23Rik | 0 | 0 | 0 | 0 | 0 | 0 | 0 | 0 | 0 | 0 | 0 |
| 0 | NA | NA | NA | NA | NA | Gm29328 | 0 | 0 | 0 | 0 | 0 | 0 | 0 | 0 | 0 | 0 | 0 |
| 0 | NA | NA | NA | NA | NA | Gm45066 | 0 | 0 | 0 | 0 | 0 | 0 | 0 | 0 | 0 | 0 | 0 |
| 0 | NA | NA | NA | NA | NA | Gm10172 | 0 | 0 | 0 | 0 | 0 | 0 | 0 | 0 | 0 | 0 | 0 |
| 0 | NA | NA | NA | NA | NA | Gm29295 | 0 | 0 | 0 | 0 | 0 | 0 | 0 | 0 | 0 | 0 | 0 |
| 0 | NA | NA | NA | NA | NA | Gm44688 | 0 | 0 | 0 | 0 | 0 | 0 | 0 | 0 | 0 | 0 | 0 |
| 0 | NA | NA | NA | NA | NA | Gm17988 | 0 | 0 | 0 | 0 | 0 | 0 | 0 | 0 | 0 | 0 | 0 |
| 0 | NA | NA | NA | NA | NA | 4930441H08Rik | 0 | 0 | 0 | 0 | 0 | 0 | 0 | 0 | 0 | 0 | 0 |
| 0 | NA | NA | NA | NA | NA | Gm36633 | 0 | 0 | 0 | 0 | 0 | 0 | 0 | 0 | 0 | 0 | 0 |
| 0 | NA | NA | NA | NA | NA | 1700011C11Rik | 0 | 0 | 0 | 0 | 0 | 0 | 0 | 0 | 0 | 0 | 0 |
| 0 | NA | NA | NA | NA | NA | Gm36696 | 0 | 0 | 0 | 0 | 0 | 0 | 0 | 0 | 0 | 0 | 0 |
| 0 | NA | NA | NA | NA | NA | Gm45715 | 0 | 0 | 0 | 0 | 0 | 0 | 0 | 0 | 0 | 0 | 0 |
| 0 | NA | NA | NA | NA | NA | Gm37620 | 0 | 0 | 0 | 0 | 0 | 0 | 0 | 0 | 0 | 0 | 0 |
| 0 | NA | NA | NA | NA | NA | Gm28745 | 0 | 0 | 0 | 0 | 0 | 0 | 0 | 0 | 0 | 0 | 0 |
| 0 | NA | NA | NA | NA | NA | Gm28744 | 0 | 0 | 0 | 0 | 0 | 0 | 0 | 0 | 0 | 0 | 0 |
| 0 | NA | NA | NA | NA | NA | Gm7693 | 0 | 0 | 0 | 0 | 0 | 0 | 0 | 0 | 0 | 0 | 0 |
| 0 | NA | NA | NA | NA | NA | Gm44725 | 0 | 0 | 0 | 0 | 0 | 0 | 0 | 0 | 0 | 0 | 0 |
| 0 | NA | NA | NA | NA | NA | Gm6567 | 0 | 0 | 0 | 0 | 0 | 0 | 0 | 0 | 0 | 0 | 0 |
| 0 | NA | NA | NA | NA | NA | 4933435G04Rik | 0 | 0 | 0 | 0 | 0 | 0 | 0 | 0 | 0 | 0 | 0 |
| 0 | NA | NA | NA | NA | NA | Gm28746 | 0 | 0 | 0 | 0 | 0 | 0 | 0 | 0 | 0 | 0 | 0 |
| 0 | NA | NA | NA | NA | NA | Gm30075 | 0 | 0 | 0 | 0 | 0 | 0 | 0 | 0 | 0 | 0 | 0 |
| 0 | NA | NA | NA | NA | NA | 4930429H19Rik | 0 | 0 | 0 | 0 | 0 | 0 | 0 | 0 | 0 | 0 | 0 |
| 0 | NA | NA | NA | NA | NA | Gm44552 | 0 | 0 | 0 | 0 | 0 | 0 | 0 | 0 | 0 | 0 | 0 |
| 0 | NA | NA | NA | NA | NA | Gm44734 | 0 | 0 | 0 | 0 | 0 | 0 | 0 | 0 | 0 | 0 | 0 |
| 0 | NA | NA | NA | NA | NA | Gm44738 | 0 | 0 | 0 | 0 | 0 | 0 | 0 | 0 | 0 | 0 | 0 |
| 0 | NA | NA | NA | NA | NA | Gm26176 | 0 | 0 | 0 | 0 | 0 | 0 | 0 | 0 | 0 | 0 | 0 |
| 0 | NA | NA | NA | NA | NA | Gm23020 | 0 | 0 | 0 | 0 | 0 | 0 | 0 | 0 | 0 | 0 | 0 |
| 0 | NA | NA | NA | NA | NA | Gm44736 | 0 | 0 | 0 | 0 | 0 | 0 | 0 | 0 | 0 | 0 | 0 |
| 0 | NA | NA | NA | NA | NA | Gm44739 | 0 | 0 | 0 | 0 | 0 | 0 | 0 | 0 | 0 | 0 | 0 |
| 0 | NA | NA | NA | NA | NA | Gm45005 | 0 | 0 | 0 | 0 | 0 | 0 | 0 | 0 | 0 | 0 | 0 |
| 0 | NA | NA | NA | NA | NA | Gm45006 | 0 | 0 | 0 | 0 | 0 | 0 | 0 | 0 | 0 | 0 | 0 |
| 0 | NA | NA | NA | NA | NA | Mir7234 | 0 | 0 | 0 | 0 | 0 | 0 | 0 | 0 | 0 | 0 | 0 |
| 0 | NA | NA | NA | NA | NA | Gm45003 | 0 | 0 | 0 | 0 | 0 | 0 | 0 | 0 | 0 | 0 | 0 |
| 0 | NA | NA | NA | NA | NA | Gm38143 | 0 | 0 | 0 | 0 | 0 | 0 | 0 | 0 | 0 | 0 | 0 |
| 0 | NA | NA | NA | NA | NA | Gm45004 | 0 | 0 | 0 | 0 | 0 | 0 | 0 | 0 | 0 | 0 | 0 |
| 0 | NA | NA | NA | NA | NA | Gm27459 | 0 | 0 | 0 | 0 | 0 | 0 | 0 | 0 | 0 | 0 | 0 |
| 0 | NA | NA | NA | NA | NA | Gm7726 | 0 | 0 | 0 | 0 | 0 | 0 | 0 | 0 | 0 | 0 | 0 |
| 0 | NA | NA | NA | NA | NA | Gm30459 | 0 | 0 | 0 | 0 | 0 | 0 | 0 | 0 | 0 | 0 | 0 |
| 0 | NA | NA | NA | NA | NA | 0610006L08Rik | 0 | 0 | 0 | 0 | 0 | 0 | 0 | 0 | 0 | 0 | 0 |
| 0 | NA | NA | NA | NA | NA | Gm44903 | 0 | 0 | 0 | 0 | 0 | 0 | 0 | 0 | 0 | 0 | 0 |
| 0 | NA | NA | NA | NA | NA | Gm44902 | 0 | 0 | 0 | 0 | 0 | 0 | 0 | 0 | 0 | 0 | 0 |
| 0 | NA | NA | NA | NA | NA | Gm22326 | 0 | 0 | 0 | 0 | 0 | 0 | 0 | 0 | 0 | 0 | 0 |

| | | | | | | | | | | | | | | | | | | |
|---|---|---|---|---|---|---|---|---|---|---|---|---|---|---|---|---|---|---|
| 0 | NA | NA | NA | NA | NA | 4930533N22Rik | | 0 | 0 | 0 | 0 | 0 | 0 | 0 | 0 | 0 | 0 | 0 |
| 0 | NA | NA | NA | NA | NA | Gm24858 | | 0 | 0 | 0 | 0 | 0 | 0 | 0 | 0 | 0 | 0 | 0 |
| 0 | NA | NA | NA | NA | NA | Gm44962 | | 0 | 0 | 0 | 0 | 0 | 0 | 0 | 0 | 0 | 0 | 0 |
| 0 | NA | NA | NA | NA | NA | Gm44834 | | 0 | 0 | 0 | 0 | 0 | 0 | 0 | 0 | 0 | 0 | 0 |
| 0 | NA | NA | NA | NA | NA | Gm44835 | | 0 | 0 | 0 | 0 | 0 | 0 | 0 | 0 | 0 | 0 | 0 |
| 0 | NA | NA | NA | NA | NA | Gm25282 | | 0 | 0 | 0 | 0 | 0 | 0 | 0 | 0 | 0 | 0 | 0 |
| 0 | NA | NA | NA | NA | NA | Gm23251 | | 0 | 0 | 0 | 0 | 0 | 0 | 0 | 0 | 0 | 0 | 0 |
| 0 | NA | NA | NA | NA | NA | Gm26729 | | 0 | 0 | 0 | 0 | 0 | 0 | 0 | 0 | 0 | 0 | 0 |
| 0 | NA | NA | NA | NA | NA | Gm45139 | | 0 | 0 | 0 | 0 | 0 | 0 | 0 | 0 | 0 | 0 | 0 |
| 0 | NA | NA | NA | NA | NA | Gm45211 | | 0 | 0 | 0 | 0 | 0 | 0 | 0 | 0 | 0 | 0 | 0 |
| 0 | NA | NA | NA | NA | NA | Agbl1 | ATP/GTP binding protein-like 1 | 0 | 0 | 0 | 0 | 0 | 0 | 0 | 0 | 0 | 0 | 0 |
| 0 | NA | NA | NA | NA | NA | 2310001K20Rik | | 0 | 0 | 0 | 0 | 0 | 0 | 0 | 0 | 0 | 0 | 0 |
| 0 | NA | NA | NA | NA | NA | Gm45210 | | 0 | 0 | 0 | 0 | 0 | 0 | 0 | 0 | 0 | 0 | 0 |
| 0 | NA | NA | NA | NA | NA | Gm44879 | | 0 | 0 | 0 | 0 | 0 | 0 | 0 | 0 | 0 | 0 | 0 |
| 0 | NA | NA | NA | NA | NA | Gm44608 | | 0 | 0 | 0 | 0 | 0 | 0 | 0 | 0 | 0 | 0 | 0 |
| 0 | NA | NA | NA | NA | NA | Gm23239 | | 0 | 0 | 0 | 0 | 0 | 0 | 0 | 0 | 0 | 0 | 0 |
| 0 | NA | NA | NA | NA | NA | Gm44358 | | 0 | 0 | 0 | 0 | 0 | 0 | 0 | 0 | 0 | 0 | 0 |
| 0 | NA | NA | NA | NA | NA | Gm44607 | | 0 | 0 | 0 | 0 | 0 | 0 | 0 | 0 | 0 | 0 | 0 |
| 0 | NA | NA | NA | NA | NA | 4921513I08Rik | | 0 | 0 | 0 | 0 | 0 | 0 | 0 | 0 | 0 | 0 | 0 |
| 0 | NA | NA | NA | NA | NA | Gm39038 | | 0 | 0 | 0 | 0 | 0 | 0 | 0 | 0 | 0 | 0 | 0 |
| 0 | NA | NA | NA | NA | NA | Gm44730 | | 0 | 0 | 0 | 0 | 0 | 0 | 0 | 0 | 0 | 0 | 0 |
| 0 | NA | NA | NA | NA | NA | Gm44815 | | 0 | 0 | 0 | 0 | 0 | 0 | 0 | 0 | 0 | 0 | 0 |
| 0 | NA | NA | NA | NA | NA | Acan | aggrecan | 0 | 0 | 0 | 0 | 0 | 0 | 0 | 0 | 0 | 0 | 0 |
| 0 | NA | NA | NA | NA | NA | Gm24224 | | 0 | 0 | 0 | 0 | 0 | 0 | 0 | 0 | 0 | 0 | 0 |
| 0 | NA | NA | NA | NA | NA | Gm16017 | | 0 | 0 | 0 | 0 | 0 | 0 | 0 | 0 | 0 | 0 | 0 |
| 0 | NA | NA | NA | NA | NA | Mir9-3hg | | 0 | 0 | 0 | 0 | 0 | 0 | 0 | 0 | 0 | 0 | 0 |
| 0 | NA | NA | NA | NA | NA | Mir9-3 | | 0 | 0 | 0 | 0 | 0 | 0 | 0 | 0 | 0 | 0 | 0 |
| 0 | NA | NA | NA | NA | NA | 2900037B21Rik | | 0 | 0 | 0 | 0 | 0 | 0 | 0 | 0 | 0 | 0 | 0 |
| 0 | NA | NA | NA | NA | NA | Gm37608 | | 0 | 0 | 0 | 0 | 0 | 0 | 0 | 0 | 0 | 0 | 0 |
| 0 | NA | NA | NA | NA | NA | Gm35040 | | 0 | 0 | 0 | 0 | 0 | 0 | 0 | 0 | 0 | 0 | 0 |
| 0 | NA | NA | NA | NA | NA | Gm45168 | | 0 | 0 | 0 | 0 | 0 | 0 | 0 | 0 | 0 | 0 | 0 |
| 0 | NA | NA | NA | NA | NA | Rhcg | Rhesus blood group-associated C glycop | 0 | 0 | 0 | 0 | 0 | 0 | 0 | 0 | 0 | 0 | 0 |
| 0 | NA | NA | NA | NA | NA | Gm45170 | | 0 | 0 | 0 | 0 | 0 | 0 | 0 | 0 | 0 | 0 | 0 |
| 0 | NA | NA | NA | NA | NA | 9330171B17Rik | | 0 | 0 | 0 | 0 | 0 | 0 | 0 | 0 | 0 | 0 | 0 |
| 0 | NA | NA | NA | NA | NA | Gm44706 | | 0 | 0 | 0 | 0 | 0 | 0 | 0 | 0 | 0 | 0 | 0 |
| 0 | NA | NA | NA | NA | NA | Gm45202 | | 0 | 0 | 0 | 0 | 0 | 0 | 0 | 0 | 0 | 0 | 0 |
| 0 | NA | NA | NA | NA | NA | Gm15504 | | 0 | 0 | 0 | 0 | 0 | 0 | 0 | 0 | 0 | 0 | 0 |
| 0 | NA | NA | NA | NA | NA | Gm44649 | | 0 | 0 | 0 | 0 | 0 | 0 | 0 | 0 | 0 | 0 | 0 |
| 0 | NA | NA | NA | NA | NA | Gm16171 | | 0 | 0 | 0 | 0 | 0 | 0 | 0 | 0 | 0 | 0 | 0 |
| 0 | NA | NA | NA | NA | NA | Gm18782 | | 0 | 0 | 0 | 0 | 0 | 0 | 0 | 0 | 0 | 0 | 0 |
| 0 | NA | NA | NA | NA | NA | Platr32 | | 0 | 0 | 0 | 0 | 0 | 0 | 0 | 0 | 0 | 0 | 0 |
| 0 | NA | NA | NA | NA | NA | Gm44968 | | 0 | 0 | 0 | 0 | 0 | 0 | 0 | 0 | 0 | 0 | 0 |
| 0 | NA | NA | NA | NA | NA | BC048679 | cDNA sequence BC048679 | 0 | 0 | 0 | 0 | 0 | 0 | 0 | 0 | 0 | 0 | 0 |
| 0 | NA | NA | NA | NA | NA | Mir1839 | | 0 | 0 | 0 | 0 | 0 | 0 | 0 | 0 | 0 | 0 | 0 |
| 0 | NA | NA | NA | NA | NA | Mir1839 | | 0 | 0 | 0 | 0 | 0 | 0 | 0 | 0 | 0 | 0 | 0 |
| 0 | NA | NA | NA | NA | NA | Gm44570 | | 0 | 0 | 0 | 0 | 0 | 0 | 0 | 0 | 0 | 0 | 0 |
| 0 | NA | NA | NA | NA | NA | Gm26149 | | 0 | 0 | 0 | 0 | 0 | 0 | 0 | 0 | 0 | 0 | 0 |
| 0 | NA | NA | NA | NA | NA | Gm18806 | | 0 | 0 | 0 | 0 | 0 | 0 | 0 | 0 | 0 | 0 | 0 |
| 0 | NA | NA | NA | NA | NA | Gm25907 | | 0 | 0 | 0 | 0 | 0 | 0 | 0 | 0 | 0 | 0 | 0 |
| 0 | NA | NA | NA | NA | NA | 4833418N17Rik | | 0 | 0 | 0 | 0 | 0 | 0 | 0 | 0 | 0 | 0 | 0 |
| 0 | NA | NA | NA | NA | NA | Gm20744 | | 0 | 0 | 0 | 0 | 0 | 0 | 0 | 0 | 0 | 0 | 0 |
| 0 | NA | NA | NA | NA | NA | Gm45056 | | 0 | 0 | 0 | 0 | 0 | 0 | 0 | 0 | 0 | 0 | 0 |
| 0 | NA | NA | NA | NA | NA | Gm32850 | | 0 | 0 | 0 | 0 | 0 | 0 | 0 | 0 | 0 | 0 | 0 |
| 0 | NA | NA | NA | NA | NA | Gm45090 | | 0 | 0 | 0 | 0 | 0 | 0 | 0 | 0 | 0 | 0 | 0 |
| 0 | NA | NA | NA | NA | NA | Gm45089 | | 0 | 0 | 0 | 0 | 0 | 0 | 0 | 0 | 0 | 0 | 0 |
| 0 | NA | NA | NA | NA | NA | Gm44919 | | 0 | 0 | 0 | 0 | 0 | 0 | 0 | 0 | 0 | 0 | 0 |
| 0 | NA | NA | NA | NA | NA | 4933406J10Rik | | 0 | 0 | 0 | 0 | 0 | 0 | 0 | 0 | 0 | 0 | 0 |
| 0 | NA | NA | NA | NA | NA | n-R5s154 | | 0 | 0 | 0 | 0 | 0 | 0 | 0 | 0 | 0 | 0 | 0 |
| 0 | NA | NA | NA | NA | NA | 1700010L04Rik | | 0 | 0 | 0 | 0 | 0 | 0 | 0 | 0 | 0 | 0 | 0 |
| 0 | NA | NA | NA | NA | NA | 4933430H16Rik | | 0 | 0 | 0 | 0 | 0 | 0 | 0 | 0 | 0 | 0 | 0 |

| | | | | | | | | | | | | | | | | | |
|---|---|---|---|---|---|---|---|---|---|---|---|---|---|---|---|---|---|
| 0 | NA | NA | NA | NA | NA | Gm22934 | | 0 | 0 | 0 | 0 | 0 | 0 | 0 | 0 | 0 | 0 | 0 |
| 0 | NA | NA | NA | NA | NA | 5930435M05Rik | | 0 | 0 | 0 | 0 | 0 | 0 | 0 | 0 | 0 | 0 | 0 |
| 0 | NA | NA | NA | NA | NA | Gm44845 | | 0 | 0 | 0 | 0 | 0 | 0 | 0 | 0 | 0 | 0 | 0 |
| 0 | NA | NA | NA | NA | NA | Gm44724 | | 0 | 0 | 0 | 0 | 0 | 0 | 0 | 0 | 0 | 0 | 0 |
| 0 | NA | NA | NA | NA | NA | Gm44723 | | 0 | 0 | 0 | 0 | 0 | 0 | 0 | 0 | 0 | 0 | 0 |
| 0 | NA | NA | NA | NA | NA | Gm7957 | | 0 | 0 | 0 | 0 | 0 | 0 | 0 | 0 | 0 | 0 | 0 |
| 0 | NA | NA | NA | NA | NA | Gm44991 | | 0 | 0 | 0 | 0 | 0 | 0 | 0 | 0 | 0 | 0 | 0 |
| 0 | NA | NA | NA | NA | NA | Gm10610 | | 0 | 0 | 0 | 0 | 0 | 0 | 0 | 0 | 0 | 0 | 0 |
| 0 | NA | NA | NA | NA | NA | 2310044K18Rik | | 0 | 0 | 0 | 0 | 0 | 0 | 0 | 0 | 0 | 0 | 0 |
| 0 | NA | NA | NA | NA | NA | Phgdh-ps1 | | 0 | 0 | 0 | 0 | 0 | 0 | 0 | 0 | 0 | 0 | 0 |
| 0 | NA | NA | NA | NA | NA | Cfap161 | cilia and flagella associated protein 161 | 0 | 0 | 0 | 0 | 0 | 0 | 0 | 0 | 0 | 0 | 0 |
| 0 | NA | NA | NA | NA | NA | Gm44993 | | 0 | 0 | 0 | 0 | 0 | 0 | 0 | 0 | 0 | 0 | 0 |
| 0 | NA | NA | NA | NA | NA | Gm39043 | | 0 | 0 | 0 | 0 | 0 | 0 | 0 | 0 | 0 | 0 | 0 |
| 0 | NA | NA | NA | NA | NA | 3110009M11Rik | | 0 | 0 | 0 | 0 | 0 | 0 | 0 | 0 | 0 | 0 | 0 |
| 0 | NA | NA | NA | NA | NA | Gm44531 | | 0 | 0 | 0 | 0 | 0 | 0 | 0 | 0 | 0 | 0 | 0 |
| 0 | NA | NA | NA | NA | NA | Gm44530 | | 0 | 0 | 0 | 0 | 0 | 0 | 0 | 0 | 0 | 0 | 0 |
| 0 | NA | NA | NA | NA | NA | Gm22177 | | 0 | 0 | 0 | 0 | 0 | 0 | 0 | 0 | 0 | 0 | 0 |
| 0 | NA | NA | NA | NA | NA | Gm44826 | | 0 | 0 | 0 | 0 | 0 | 0 | 0 | 0 | 0 | 0 | 0 |
| 0 | NA | NA | NA | NA | NA | Gm44827 | | 0 | 0 | 0 | 0 | 0 | 0 | 0 | 0 | 0 | 0 | 0 |
| 0 | NA | NA | NA | NA | NA | Gm25790 | | 0 | 0 | 0 | 0 | 0 | 0 | 0 | 0 | 0 | 0 | 0 |
| 0 | NA | NA | NA | NA | NA | Gm44933 | | 0 | 0 | 0 | 0 | 0 | 0 | 0 | 0 | 0 | 0 | 0 |
| 0 | NA | NA | NA | NA | NA | Gm45175 | | 0 | 0 | 0 | 0 | 0 | 0 | 0 | 0 | 0 | 0 | 0 |
| 0 | NA | NA | NA | NA | NA | Gm44928 | | 0 | 0 | 0 | 0 | 0 | 0 | 0 | 0 | 0 | 0 | 0 |
| 0 | NA | NA | NA | NA | NA | Vmn2r-ps62 | | 0 | 0 | 0 | 0 | 0 | 0 | 0 | 0 | 0 | 0 | 0 |
| 0 | NA | NA | NA | NA | NA | Olfr291 | olfactory receptor 291 | 0 | 0 | 0 | 0 | 0 | 0 | 0 | 0 | 0 | 0 | 0 |
| 0 | NA | NA | NA | NA | NA | Gm8018 | | 0 | 0 | 0 | 0 | 0 | 0 | 0 | 0 | 0 | 0 | 0 |
| 0 | NA | NA | NA | NA | NA | Olfr290 | | 0 | 0 | 0 | 0 | 0 | 0 | 0 | 0 | 0 | 0 | 0 |
| 0 | NA | NA | NA | NA | NA | Olfr290 | olfactory receptor 290 | 0 | 0 | 0 | 0 | 0 | 0 | 0 | 0 | 0 | 0 | 0 |
| 0 | NA | NA | NA | NA | NA | Gm18560 | | 0 | 0 | 0 | 0 | 0 | 0 | 0 | 0 | 0 | 0 | 0 |
| 0 | NA | NA | NA | NA | NA | Vmn2r65 | vomeronasal 2, receptor 65 | 0 | 0 | 0 | 0 | 0 | 0 | 0 | 0 | 0 | 0 | 0 |
| 0 | NA | NA | NA | NA | NA | Vmn2r-ps64 | | 0 | 0 | 0 | 0 | 0 | 0 | 0 | 0 | 0 | 0 | 0 |
| 0 | NA | NA | NA | NA | NA | Vmn2r-ps65 | | 0 | 0 | 0 | 0 | 0 | 0 | 0 | 0 | 0 | 0 | 0 |
| 0 | NA | NA | NA | NA | NA | Vmn2r-ps66 | | 0 | 0 | 0 | 0 | 0 | 0 | 0 | 0 | 0 | 0 | 0 |
| 0 | NA | NA | NA | NA | NA | Vmn2r-ps67 | | 0 | 0 | 0 | 0 | 0 | 0 | 0 | 0 | 0 | 0 | 0 |
| 0 | NA | NA | NA | NA | NA | Gm18358 | | 0 | 0 | 0 | 0 | 0 | 0 | 0 | 0 | 0 | 0 | 0 |
| 0 | NA | NA | NA | NA | NA | Vmn2r67 | vomeronasal 2, receptor 67 | 0 | 0 | 0 | 0 | 0 | 0 | 0 | 0 | 0 | 0 | 0 |
| 0 | NA | NA | NA | NA | NA | Vmn2r-ps68 | | 0 | 0 | 0 | 0 | 0 | 0 | 0 | 0 | 0 | 0 | 0 |
| 0 | NA | NA | NA | NA | NA | Vmn2r68 | vomeronasal 2, receptor 68 | 0 | 0 | 0 | 0 | 0 | 0 | 0 | 0 | 0 | 0 | 0 |
| 0 | NA | NA | NA | NA | NA | Gm20414 | | 0 | 0 | 0 | 0 | 0 | 0 | 0 | 0 | 0 | 0 | 0 |
| 0 | NA | NA | NA | NA | NA | Gm20415 | | 0 | 0 | 0 | 0 | 0 | 0 | 0 | 0 | 0 | 0 | 0 |
| 0 | NA | NA | NA | NA | NA | Vmn2r-ps69 | | 0 | 0 | 0 | 0 | 0 | 0 | 0 | 0 | 0 | 0 | 0 |
| 0 | NA | NA | NA | NA | NA | Vmn2r69 | vomeronasal 2, receptor 69 | 0 | 0 | 0 | 0 | 0 | 0 | 0 | 0 | 0 | 0 | 0 |
| 0 | NA | NA | NA | NA | NA | Gm18359 | | 0 | 0 | 0 | 0 | 0 | 0 | 0 | 0 | 0 | 0 | 0 |
| 0 | NA | NA | NA | NA | NA | Vmn2r-ps70 | | 0 | 0 | 0 | 0 | 0 | 0 | 0 | 0 | 0 | 0 | 0 |
| 0 | NA | NA | NA | NA | NA | Gm18360 | | 0 | 0 | 0 | 0 | 0 | 0 | 0 | 0 | 0 | 0 | 0 |
| 0 | NA | NA | NA | NA | NA | Vmn2r70 | vomeronasal 2, receptor 70 | 0 | 0 | 0 | 0 | 0 | 0 | 0 | 0 | 0 | 0 | 0 |
| 0 | NA | NA | NA | NA | NA | Vmn2r71 | vomeronasal 2, receptor 71 | 0 | 0 | 0 | 0 | 0 | 0 | 0 | 0 | 0 | 0 | 0 |
| 0 | NA | NA | NA | NA | NA | Vmn2r-ps72 | | 0 | 0 | 0 | 0 | 0 | 0 | 0 | 0 | 0 | 0 | 0 |
| 0 | NA | NA | NA | NA | NA | Vmn2r-ps73 | | 0 | 0 | 0 | 0 | 0 | 0 | 0 | 0 | 0 | 0 | 0 |
| 0 | NA | NA | NA | NA | NA | Vmn2r72 | vomeronasal 2, receptor 72 | 0 | 0 | 0 | 0 | 0 | 0 | 0 | 0 | 0 | 0 | 0 |
| 0 | NA | NA | NA | NA | NA | Gm18779 | | 0 | 0 | 0 | 0 | 0 | 0 | 0 | 0 | 0 | 0 | 0 |
| 0 | NA | NA | NA | NA | NA | Gm18354 | | 0 | 0 | 0 | 0 | 0 | 0 | 0 | 0 | 0 | 0 | 0 |
| 0 | NA | NA | NA | NA | NA | Vmn2r-ps75 | | 0 | 0 | 0 | 0 | 0 | 0 | 0 | 0 | 0 | 0 | 0 |
| 0 | NA | NA | NA | NA | NA | Vmn2r-ps76 | | 0 | 0 | 0 | 0 | 0 | 0 | 0 | 0 | 0 | 0 | 0 |
| 0 | NA | NA | NA | NA | NA | Vmn2r-ps77 | | 0 | 0 | 0 | 0 | 0 | 0 | 0 | 0 | 0 | 0 | 0 |
| 0 | NA | NA | NA | NA | NA | Vmn2r73 | vomeronasal 2, receptor 73 | 0 | 0 | 0 | 0 | 0 | 0 | 0 | 0 | 0 | 0 | 0 |
| 0 | NA | NA | NA | NA | NA | Vmn2r-ps78 | | 0 | 0 | 0 | 0 | 0 | 0 | 0 | 0 | 0 | 0 | 0 |
| 0 | NA | NA | NA | NA | NA | Vmn2r74 | vomeronasal 2, receptor 74 | 0 | 0 | 0 | 0 | 0 | 0 | 0 | 0 | 0 | 0 | 0 |
| 0 | NA | NA | NA | NA | NA | Vmn2r-ps79 | | 0 | 0 | 0 | 0 | 0 | 0 | 0 | 0 | 0 | 0 | 0 |
| 0 | NA | NA | NA | NA | NA | Vmn2r-ps80 | | 0 | 0 | 0 | 0 | 0 | 0 | 0 | 0 | 0 | 0 | 0 |

| | | | | | | | | | | | | | | | | | |
|---|---|---|---|---|---|---|---|---|---|---|---|---|---|---|---|---|---|
| 0 | NA | NA | NA | NA | NA | Vmn2r-ps81 | | 0 | 0 | 0 | 0 | 0 | 0 | 0 | 0 | 0 | 0 | 0 |
| 0 | NA | NA | NA | NA | NA | Vmn2r-ps82 | | 0 | 0 | 0 | 0 | 0 | 0 | 0 | 0 | 0 | 0 | 0 |
| 0 | NA | NA | NA | NA | NA | Vmn2r-ps83 | | 0 | 0 | 0 | 0 | 0 | 0 | 0 | 0 | 0 | 0 | 0 |
| 0 | NA | NA | NA | NA | NA | Vmn2r-ps84 | | 0 | 0 | 0 | 0 | 0 | 0 | 0 | 0 | 0 | 0 | 0 |
| 0 | NA | NA | NA | NA | NA | Vmn2r75 | vomeronasal 2, receptor 75 | 0 | 0 | 0 | 0 | 0 | 0 | 0 | 0 | 0 | 0 | 0 |
| 0 | NA | NA | NA | NA | NA | Vmn2r-ps85 | | 0 | 0 | 0 | 0 | 0 | 0 | 0 | 0 | 0 | 0 | 0 |
| 0 | NA | NA | NA | NA | NA | Vmn2r-ps86 | | 0 | 0 | 0 | 0 | 0 | 0 | 0 | 0 | 0 | 0 | 0 |
| 0 | NA | NA | NA | NA | NA | Vmn2r76 | vomeronasal 2, receptor 76 | 0 | 0 | 0 | 0 | 0 | 0 | 0 | 0 | 0 | 0 | 0 |
| 0 | NA | NA | NA | NA | NA | Olfr310 | olfactory receptor 310 | 0 | 0 | 0 | 0 | 0 | 0 | 0 | 0 | 0 | 0 | 0 |
| 0 | NA | NA | NA | NA | NA | Gm20480 | | 0 | 0 | 0 | 0 | 0 | 0 | 0 | 0 | 0 | 0 | 0 |
| 0 | NA | NA | NA | NA | NA | Gm23192 | | 0 | 0 | 0 | 0 | 0 | 0 | 0 | 0 | 0 | 0 | 0 |
| 0 | NA | NA | NA | NA | NA | Olfr309 | olfactory receptor 309 | 0 | 0 | 0 | 0 | 0 | 0 | 0 | 0 | 0 | 0 | 0 |
| 0 | NA | NA | NA | NA | NA | Olfr308 | olfactory receptor 308 | 0 | 0 | 0 | 0 | 0 | 0 | 0 | 0 | 0 | 0 | 0 |
| 0 | NA | NA | NA | NA | NA | Olfr307 | olfactory receptor 307 | 0 | 0 | 0 | 0 | 0 | 0 | 0 | 0 | 0 | 0 | 0 |
| 0 | NA | NA | NA | NA | NA | Olfr306-ps1 | | 0 | 0 | 0 | 0 | 0 | 0 | 0 | 0 | 0 | 0 | 0 |
| 0 | NA | NA | NA | NA | NA | Olfr305 | olfactory receptor 305 | 0 | 0 | 0 | 0 | 0 | 0 | 0 | 0 | 0 | 0 | 0 |
| 0 | NA | NA | NA | NA | NA | Olfr304 | olfactory receptor 304 | 0 | 0 | 0 | 0 | 0 | 0 | 0 | 0 | 0 | 0 | 0 |
| 0 | NA | NA | NA | NA | NA | Olfr303 | olfactory receptor 303 | 0 | 0 | 0 | 0 | 0 | 0 | 0 | 0 | 0 | 0 | 0 |
| 0 | NA | NA | NA | NA | NA | Gm45541 | | 0 | 0 | 0 | 0 | 0 | 0 | 0 | 0 | 0 | 0 | 0 |
| 0 | NA | NA | NA | NA | NA | Olfr301 | olfactory receptor 301 | 0 | 0 | 0 | 0 | 0 | 0 | 0 | 0 | 0 | 0 | 0 |
| 0 | NA | NA | NA | NA | NA | Olfr300-ps1 | | 0 | 0 | 0 | 0 | 0 | 0 | 0 | 0 | 0 | 0 | 0 |
| 0 | NA | NA | NA | NA | NA | Olfr299 | olfactory receptor 299 | 0 | 0 | 0 | 0 | 0 | 0 | 0 | 0 | 0 | 0 | 0 |
| 0 | NA | NA | NA | NA | NA | Olfr298 | olfactory receptor 298 | 0 | 0 | 0 | 0 | 0 | 0 | 0 | 0 | 0 | 0 | 0 |
| 0 | NA | NA | NA | NA | NA | Olfr297 | olfactory receptor 297 | 0 | 0 | 0 | 0 | 0 | 0 | 0 | 0 | 0 | 0 | 0 |
| 0 | NA | NA | NA | NA | NA | Gm17919 | | 0 | 0 | 0 | 0 | 0 | 0 | 0 | 0 | 0 | 0 | 0 |
| 0 | NA | NA | NA | NA | NA | Olfr296-ps1 | | 0 | 0 | 0 | 0 | 0 | 0 | 0 | 0 | 0 | 0 | 0 |
| 0 | NA | NA | NA | NA | NA | Olfr295 | olfactory receptor 295 | 0 | 0 | 0 | 0 | 0 | 0 | 0 | 0 | 0 | 0 | 0 |
| 0 | NA | NA | NA | NA | NA | Olfr294 | olfactory receptor 294 | 0 | 0 | 0 | 0 | 0 | 0 | 0 | 0 | 0 | 0 | 0 |
| 0 | NA | NA | NA | NA | NA | Olfr293 | olfactory receptor 293 | 0 | 0 | 0 | 0 | 0 | 0 | 0 | 0 | 0 | 0 | 0 |
| 0 | NA | NA | NA | NA | NA | Olfr292 | olfactory receptor 292 | 0 | 0 | 0 | 0 | 0 | 0 | 0 | 0 | 0 | 0 | 0 |
| 0 | NA | NA | NA | NA | NA | Vmn2r77 | vomeronasal 2, receptor 77 | 0 | 0 | 0 | 0 | 0 | 0 | 0 | 0 | 0 | 0 | 0 |
| 0 | NA | NA | NA | NA | NA | Vmn2r-ps87 | | 0 | 0 | 0 | 0 | 0 | 0 | 0 | 0 | 0 | 0 | 0 |
| 0 | NA | NA | NA | NA | NA | Gm19203 | | 0 | 0 | 0 | 0 | 0 | 0 | 0 | 0 | 0 | 0 | 0 |
| 0 | NA | NA | NA | NA | NA | Vmn2r78 | vomeronasal 2, receptor 78 | 0 | 0 | 0 | 0 | 0 | 0 | 0 | 0 | 0 | 0 | 0 |
| 0 | NA | NA | NA | NA | NA | Gm44925 | | 0 | 0 | 0 | 0 | 0 | 0 | 0 | 0 | 0 | 0 | 0 |
| 0 | NA | NA | NA | NA | NA | Gm6230 | | 0 | 0 | 0 | 0 | 0 | 0 | 0 | 0 | 0 | 0 | 0 |
| 0 | NA | NA | NA | NA | NA | Tyr | tyrosinase | 0 | 0 | 0 | 0 | 0 | 0 | 0 | 0 | 0 | 0 | 0 |
| 0 | NA | NA | NA | NA | NA | Grm5 | glutamate receptor, metabotropic 5 | 0 | 0 | 0 | 0 | 0 | 0 | 0 | 0 | 0 | 0 | 0 |
| 0 | NA | NA | NA | NA | NA | Gm44926 | | 0 | 0 | 0 | 0 | 0 | 0 | 0 | 0 | 0 | 0 | 0 |
| 0 | NA | NA | NA | NA | NA | Gm44927 | | 0 | 0 | 0 | 0 | 0 | 0 | 0 | 0 | 0 | 0 | 0 |
| 0 | NA | NA | NA | NA | NA | Gm44848 | | 0 | 0 | 0 | 0 | 0 | 0 | 0 | 0 | 0 | 0 | 0 |
| 0 | NA | NA | NA | NA | NA | Gm44847 | | 0 | 0 | 0 | 0 | 0 | 0 | 0 | 0 | 0 | 0 | 0 |
| 0 | NA | NA | NA | NA | NA | Gm24249 | | 0 | 0 | 0 | 0 | 0 | 0 | 0 | 0 | 0 | 0 | 0 |
| 0 | NA | NA | NA | NA | NA | Gm44846 | | 0 | 0 | 0 | 0 | 0 | 0 | 0 | 0 | 0 | 0 | 0 |
| 0 | NA | NA | NA | NA | NA | Gm25345 | | 0 | 0 | 0 | 0 | 0 | 0 | 0 | 0 | 0 | 0 | 0 |
| 0 | NA | NA | NA | NA | NA | Gm8183 | | 0 | 0 | 0 | 0 | 0 | 0 | 0 | 0 | 0 | 0 | 0 |
| 0 | NA | NA | NA | NA | NA | Gm6240 | | 0 | 0 | 0 | 0 | 0 | 0 | 0 | 0 | 0 | 0 | 0 |
| 0 | NA | NA | NA | NA | NA | Gm45144 | | 0 | 0 | 0 | 0 | 0 | 0 | 0 | 0 | 0 | 0 | 0 |
| 0 | NA | NA | NA | NA | NA | Gm24659 | | 0 | 0 | 0 | 0 | 0 | 0 | 0 | 0 | 0 | 0 | 0 |
| 0 | NA | NA | NA | NA | NA | Gm22171 | | 0 | 0 | 0 | 0 | 0 | 0 | 0 | 0 | 0 | 0 | 0 |
| 0 | NA | NA | NA | NA | NA | Gm45064 | | 0 | 0 | 0 | 0 | 0 | 0 | 0 | 0 | 0 | 0 | 0 |
| 0 | NA | NA | NA | NA | NA | Ccdc81 | coiled-coil domain containing 81 | 0 | 0 | 0 | 0 | 0 | 0 | 0 | 0 | 0 | 0 | 0 |
| 0 | NA | NA | NA | NA | NA | Gm5341 | | 0 | 0 | 0 | 0 | 0 | 0 | 0 | 0 | 0 | 0 | 0 |
| 0 | NA | NA | NA | NA | NA | Ccdc83 | coiled-coil domain containing 83 | 0 | 0 | 0 | 0 | 0 | 0 | 0 | 0 | 0 | 0 | 0 |
| 0 | NA | NA | NA | NA | NA | n-R5s155 | | 0 | 0 | 0 | 0 | 0 | 0 | 0 | 0 | 0 | 0 | 0 |
| 0 | NA | NA | NA | NA | NA | A930002H02Rik | | 0 | 0 | 0 | 0 | 0 | 0 | 0 | 0 | 0 | 0 | 0 |
| 0 | NA | NA | NA | NA | NA | Gm45161 | | 0 | 0 | 0 | 0 | 0 | 0 | 0 | 0 | 0 | 0 | 0 |
| 0 | NA | NA | NA | NA | NA | Gm45178 | | 0 | 0 | 0 | 0 | 0 | 0 | 0 | 0 | 0 | 0 | 0 |
| 0 | NA | NA | NA | NA | NA | Gm27470 | | 0 | 0 | 0 | 0 | 0 | 0 | 0 | 0 | 0 | 0 | 0 |
| 0 | NA | NA | NA | NA | NA | Gm27660 | | 0 | 0 | 0 | 0 | 0 | 0 | 0 | 0 | 0 | 0 | 0 |

| | | | | | | | | | | | | | | | | | | |
|---|---|---|---|---|---|---|---|---|---|---|---|---|---|---|---|---|---|---|
| 0 | NA | NA | NA | NA | NA | Gm45176 | | 0 | 0 | 0 | 0 | 0 | 0 | 0 | 0 | 0 | 0 | 0 |
| 0 | NA | NA | NA | NA | NA | Gm45177 | | 0 | 0 | 0 | 0 | 0 | 0 | 0 | 0 | 0 | 0 | 0 |
| 0 | NA | NA | NA | NA | NA | Gm45182 | | 0 | 0 | 0 | 0 | 0 | 0 | 0 | 0 | 0 | 0 | 0 |
| 0 | NA | NA | NA | NA | NA | Gm24552 | | 0 | 0 | 0 | 0 | 0 | 0 | 0 | 0 | 0 | 0 | 0 |
| 0 | NA | NA | NA | NA | NA | Gm45183 | | 0 | 0 | 0 | 0 | 0 | 0 | 0 | 0 | 0 | 0 | 0 |
| 0 | NA | NA | NA | NA | NA | Gm44678 | | 0 | 0 | 0 | 0 | 0 | 0 | 0 | 0 | 0 | 0 | 0 |
| 0 | NA | NA | NA | NA | NA | Gm22080 | | 0 | 0 | 0 | 0 | 0 | 0 | 0 | 0 | 0 | 0 | 0 |
| 0 | NA | NA | NA | NA | NA | Gm44679 | | 0 | 0 | 0 | 0 | 0 | 0 | 0 | 0 | 0 | 0 | 0 |
| 0 | NA | NA | NA | NA | NA | Gm44675 | | 0 | 0 | 0 | 0 | 0 | 0 | 0 | 0 | 0 | 0 | 0 |
| 0 | NA | NA | NA | NA | NA | Gm44676 | | 0 | 0 | 0 | 0 | 0 | 0 | 0 | 0 | 0 | 0 | 0 |
| 0 | NA | NA | NA | NA | NA | Gm44677 | | 0 | 0 | 0 | 0 | 0 | 0 | 0 | 0 | 0 | 0 | 0 |
| 0 | NA | NA | NA | NA | NA | Gm44680 | | 0 | 0 | 0 | 0 | 0 | 0 | 0 | 0 | 0 | 0 | 0 |
| 0 | NA | NA | NA | NA | NA | Gm44681 | | 0 | 0 | 0 | 0 | 0 | 0 | 0 | 0 | 0 | 0 | 0 |
| 0 | NA | NA | NA | NA | NA | Gm23928 | | 0 | 0 | 0 | 0 | 0 | 0 | 0 | 0 | 0 | 0 | 0 |
| 0 | NA | NA | NA | NA | NA | C030038I04Rik | | 0 | 0 | 0 | 0 | 0 | 0 | 0 | 0 | 0 | 0 | 0 |
| 0 | NA | NA | NA | NA | NA | Gm45130 | | 0 | 0 | 0 | 0 | 0 | 0 | 0 | 0 | 0 | 0 | 0 |
| 0 | NA | NA | NA | NA | NA | Gm45131 | | 0 | 0 | 0 | 0 | 0 | 0 | 0 | 0 | 0 | 0 | 0 |
| 0 | NA | NA | NA | NA | NA | 4931412I15Rik | | 0 | 0 | 0 | 0 | 0 | 0 | 0 | 0 | 0 | 0 | 0 |
| 0 | NA | NA | NA | NA | NA | 4930567K12Rik | | 0 | 0 | 0 | 0 | 0 | 0 | 0 | 0 | 0 | 0 | 0 |
| 0 | NA | NA | NA | NA | NA | B230206I08Rik | | 0 | 0 | 0 | 0 | 0 | 0 | 0 | 0 | 0 | 0 | 0 |
| 0 | NA | NA | NA | NA | NA | Gm45199 | | 0 | 0 | 0 | 0 | 0 | 0 | 0 | 0 | 0 | 0 | 0 |
| 0 | NA | NA | NA | NA | NA | Gm26981 | | 0 | 0 | 0 | 0 | 0 | 0 | 0 | 0 | 0 | 0 | 0 |
| 0 | NA | NA | NA | NA | NA | Gm45200 | | 0 | 0 | 0 | 0 | 0 | 0 | 0 | 0 | 0 | 0 | 0 |
| 0 | NA | NA | NA | NA | NA | 9530078K11Rik | | 0 | 0 | 0 | 0 | 0 | 0 | 0 | 0 | 0 | 0 | 0 |
| 0 | NA | NA | NA | NA | NA | Gm31663 | | 0 | 0 | 0 | 0 | 0 | 0 | 0 | 0 | 0 | 0 | 0 |
| 0 | NA | NA | NA | NA | NA | Gm9934 | | 0 | 0 | 0 | 0 | 0 | 0 | 0 | 0 | 0 | 0 | 0 |
| 0 | NA | NA | NA | NA | NA | Gm25860 | | 0 | 0 | 0 | 0 | 0 | 0 | 0 | 0 | 0 | 0 | 0 |
| 0 | NA | NA | NA | NA | NA | Gm44715 | | 0 | 0 | 0 | 0 | 0 | 0 | 0 | 0 | 0 | 0 | 0 |
| 0 | NA | NA | NA | NA | NA | Gm44947 | | 0 | 0 | 0 | 0 | 0 | 0 | 0 | 0 | 0 | 0 | 0 |
| 0 | NA | NA | NA | NA | NA | Gm45037 | | 0 | 0 | 0 | 0 | 0 | 0 | 0 | 0 | 0 | 0 | 0 |
| 0 | NA | NA | NA | NA | NA | Gm45038 | | 0 | 0 | 0 | 0 | 0 | 0 | 0 | 0 | 0 | 0 | 0 |
| 0 | NA | NA | NA | NA | NA | Gm8285 | | 0 | 0 | 0 | 0 | 0 | 0 | 0 | 0 | 0 | 0 | 0 |
| 0 | NA | NA | NA | NA | NA | Gm45039 | | 0 | 0 | 0 | 0 | 0 | 0 | 0 | 0 | 0 | 0 | 0 |
| 0 | NA | NA | NA | NA | NA | Bc1-ps1 | | 0 | 0 | 0 | 0 | 0 | 0 | 0 | 0 | 0 | 0 | 0 |
| 0 | NA | NA | NA | NA | NA | | | 0 | 0 | 0 | 0 | 0 | 0 | 0 | 0 | 0 | 0 | 0 |
| 0 | NA | NA | NA | NA | NA | Gm5899 | | 0 | 0 | 0 | 0 | 0 | 0 | 0 | 0 | 0 | 0 | 0 |
| 0 | NA | NA | NA | NA | NA | Gm44601 | | 0 | 0 | 0 | 0 | 0 | 0 | 0 | 0 | 0 | 0 | 0 |
| 0 | NA | NA | NA | NA | NA | Gm44602 | | 0 | 0 | 0 | 0 | 0 | 0 | 0 | 0 | 0 | 0 | 0 |
| 0 | NA | NA | NA | NA | NA | Gm19761 | | 0 | 0 | 0 | 0 | 0 | 0 | 0 | 0 | 0 | 0 | 0 |
| 0 | NA | NA | NA | NA | NA | Gm23884 | | 0 | 0 | 0 | 0 | 0 | 0 | 0 | 0 | 0 | 0 | 0 |
| 0 | NA | NA | NA | NA | NA | Gm25684 | | 0 | 0 | 0 | 0 | 0 | 0 | 0 | 0 | 0 | 0 | 0 |
| 0 | NA | NA | NA | NA | NA | Gm8309 | | 0 | 0 | 0 | 0 | 0 | 0 | 0 | 0 | 0 | 0 | 0 |
| 0 | NA | NA | NA | NA | NA | Gm44603 | | 0 | 0 | 0 | 0 | 0 | 0 | 0 | 0 | 0 | 0 | 0 |
| 0 | NA | NA | NA | NA | NA | Gm5037 | | 0 | 0 | 0 | 0 | 0 | 0 | 0 | 0 | 0 | 0 | 0 |
| 0 | NA | NA | NA | NA | NA | Mir708 | | 0 | 0 | 0 | 0 | 0 | 0 | 0 | 0 | 0 | 0 | 0 |
| 0 | NA | NA | NA | NA | NA | Mir6394 | | 0 | 0 | 0 | 0 | 0 | 0 | 0 | 0 | 0 | 0 | 0 |
| 0 | NA | NA | NA | NA | NA | Gm15412 | | 0 | 0 | 0 | 0 | 0 | 0 | 0 | 0 | 0 | 0 | 0 |
| 0 | NA | NA | NA | NA | NA | Gm15414 | | 0 | 0 | 0 | 0 | 0 | 0 | 0 | 0 | 0 | 0 | 0 |
| 0 | NA | NA | NA | NA | NA | Gm15416 | | 0 | 0 | 0 | 0 | 0 | 0 | 0 | 0 | 0 | 0 | 0 |
| 0 | NA | NA | NA | NA | NA | Gm44633 | | 0 | 0 | 0 | 0 | 0 | 0 | 0 | 0 | 0 | 0 | 0 |
| 0 | NA | NA | NA | NA | NA | Gm25712 | | 0 | 0 | 0 | 0 | 0 | 0 | 0 | 0 | 0 | 0 | 0 |
| 0 | NA | NA | NA | NA | NA | Gm22226 | | 0 | 0 | 0 | 0 | 0 | 0 | 0 | 0 | 0 | 0 | 0 |
| 0 | NA | NA | NA | NA | NA | Ywhaq-ps1 | | 0 | 0 | 0 | 0 | 0 | 0 | 0 | 0 | 0 | 0 | 0 |
| 0 | NA | NA | NA | NA | NA | Rsf1os1 | | 0 | 0 | 0 | 0 | 0 | 0 | 0 | 0 | 0 | 0 | 0 |
| 0 | NA | NA | NA | NA | NA | Gm45186 | | 0 | 0 | 0 | 0 | 0 | 0 | 0 | 0 | 0 | 0 | 0 |
| 0 | NA | NA | NA | NA | NA | Gm44590 | | 0 | 0 | 0 | 0 | 0 | 0 | 0 | 0 | 0 | 0 | 0 |
| 0 | NA | NA | NA | NA | NA | Gdpd4 | glycerophosphodiester phosphodiesteras | 0 | 0 | 0 | 0 | 0 | 0 | 0 | 0 | 0 | 0 | 0 |
| 0 | NA | NA | NA | NA | NA | Gm44507 | | 0 | 0 | 0 | 0 | 0 | 0 | 0 | 0 | 0 | 0 | 0 |
| 0 | NA | NA | NA | NA | NA | Gm44980 | | 0 | 0 | 0 | 0 | 0 | 0 | 0 | 0 | 0 | 0 | 0 |
| 0 | NA | NA | NA | NA | NA | Gucy2d | guanylate cyclase 2d | 0 | 0 | 0 | 0 | 0 | 0 | 0 | 0 | 0 | 0 | 0 |

| | | | | | | | | | | | | | | | | | | |
|---|---|---|---|---|---|---|---|---|---|---|---|---|---|---|---|---|---|---|
| 0 | NA | NA | NA | NA | NA | Gm23040 | | 0 | 0 | 0 | 0 | 0 | 0 | 0 | 0 | 0 | 0 | 0 |
| 0 | NA | NA | NA | NA | NA | A630091E08Rik | | 0 | 0 | 0 | 0 | 0 | 0 | 0 | 0 | 0 | 0 | 0 |
| 0 | NA | NA | NA | NA | NA | Gm45132 | | 0 | 0 | 0 | 0 | 0 | 0 | 0 | 0 | 0 | 0 | 0 |
| 0 | NA | NA | NA | NA | NA | Gm44934 | | 0 | 0 | 0 | 0 | 0 | 0 | 0 | 0 | 0 | 0 | 0 |
| 0 | NA | NA | NA | NA | NA | Gm8398 | | 0 | 0 | 0 | 0 | 0 | 0 | 0 | 0 | 0 | 0 | 0 |
| 0 | NA | NA | NA | NA | NA | Gm45188 | | 0 | 0 | 0 | 0 | 0 | 0 | 0 | 0 | 0 | 0 | 0 |
| 0 | NA | NA | NA | NA | NA | Gm23479 | | 0 | 0 | 0 | 0 | 0 | 0 | 0 | 0 | 0 | 0 | 0 |
| 0 | NA | NA | NA | NA | NA | Gm44975 | | 0 | 0 | 0 | 0 | 0 | 0 | 0 | 0 | 0 | 0 | 0 |
| 0 | NA | NA | NA | NA | NA | Gm45012 | | 0 | 0 | 0 | 0 | 0 | 0 | 0 | 0 | 0 | 0 | 0 |
| 0 | NA | NA | NA | NA | NA | Snord15b | | 0 | 0 | 0 | 0 | 0 | 0 | 0 | 0 | 0 | 0 | 0 |
| 0 | NA | NA | NA | NA | NA | n-R5s156 | | 0 | 0 | 0 | 0 | 0 | 0 | 0 | 0 | 0 | 0 | 0 |
| 0 | NA | NA | NA | NA | NA | Mir326 | | 0 | 0 | 0 | 0 | 0 | 0 | 0 | 0 | 0 | 0 | 0 |
| 0 | NA | NA | NA | NA | NA | Gm25409 | | 0 | 0 | 0 | 0 | 0 | 0 | 0 | 0 | 0 | 0 | 0 |
| 0 | NA | NA | NA | NA | NA | Gm15635 | | 0 | 0 | 0 | 0 | 0 | 0 | 0 | 0 | 0 | 0 | 0 |
| 0 | NA | NA | NA | NA | NA | Gm44551 | | 0 | 0 | 0 | 0 | 0 | 0 | 0 | 0 | 0 | 0 | 0 |
| 0 | NA | NA | NA | NA | NA | Olfr520 | olfactory receptor 520 | 0 | 0 | 0 | 0 | 0 | 0 | 0 | 0 | 0 | 0 | 0 |
| 0 | NA | NA | NA | NA | NA | Gm19182 | | 0 | 0 | 0 | 0 | 0 | 0 | 0 | 0 | 0 | 0 | 0 |
| 0 | NA | NA | NA | NA | NA | Olfr521 | olfactory receptor 521 | 0 | 0 | 0 | 0 | 0 | 0 | 0 | 0 | 0 | 0 | 0 |
| 0 | NA | NA | NA | NA | NA | F730035P03Rik | | 0 | 0 | 0 | 0 | 0 | 0 | 0 | 0 | 0 | 0 | 0 |
| 0 | NA | NA | NA | NA | NA | Gm17931 | | 0 | 0 | 0 | 0 | 0 | 0 | 0 | 0 | 0 | 0 | 0 |
| 0 | NA | NA | NA | NA | NA | Gm7067 | | 0 | 0 | 0 | 0 | 0 | 0 | 0 | 0 | 0 | 0 | 0 |
| 0 | NA | NA | NA | NA | NA | Gm44648 | | 0 | 0 | 0 | 0 | 0 | 0 | 0 | 0 | 0 | 0 | 0 |
| 0 | NA | NA | NA | NA | NA | Gm45209 | | 0 | 0 | 0 | 0 | 0 | 0 | 0 | 0 | 0 | 0 | 0 |
| 0 | NA | NA | NA | NA | NA | Mir3102 | | 0 | 0 | 0 | 0 | 0 | 0 | 0 | 0 | 0 | 0 | 0 |
| 0 | NA | NA | NA | NA | NA | Gm45070 | | 0 | 0 | 0 | 0 | 0 | 0 | 0 | 0 | 0 | 0 | 0 |
| 0 | NA | NA | NA | NA | NA | Gm47324 | | 0 | 0 | 0 | 0 | 0 | 0 | 0 | 0 | 0 | 0 | 0 |
| 0 | NA | NA | NA | NA | NA | Gm45548 | | 0 | 0 | 0 | 0 | 0 | 0 | 0 | 0 | 0 | 0 | 0 |
| 0 | NA | NA | NA | NA | NA | Pde2a | phosphodiesterase 2A, cGMP-stimulated | 0 | 0 | 0 | 0 | 0 | 0 | 0 | 0 | 0 | 0 | 0 |
| 0 | NA | NA | NA | NA | NA | Mir139 | | 0 | 0 | 0 | 0 | 0 | 0 | 0 | 0 | 0 | 0 | 0 |
| 0 | NA | NA | NA | NA | NA | Gm35363 | | 0 | 0 | 0 | 0 | 0 | 0 | 0 | 0 | 0 | 0 | 0 |
| 0 | NA | NA | NA | NA | NA | Gm22655 | | 0 | 0 | 0 | 0 | 0 | 0 | 0 | 0 | 0 | 0 | 0 |
| 0 | NA | NA | NA | NA | NA | Gm8523 | | 0 | 0 | 0 | 0 | 0 | 0 | 0 | 0 | 0 | 0 | 0 |
| 0 | NA | NA | NA | NA | NA | Gm45312 | | 0 | 0 | 0 | 0 | 0 | 0 | 0 | 0 | 0 | 0 | 0 |
| 0 | NA | NA | NA | NA | NA | Gm45313 | | 0 | 0 | 0 | 0 | 0 | 0 | 0 | 0 | 0 | 0 | 0 |
| 0 | NA | NA | NA | NA | NA | Trpc2 | transient receptor potential cation channe | 0 | 0 | 0 | 0 | 0 | 0 | 0 | 0 | 0 | 0 | 0 |
| 0 | NA | NA | NA | NA | NA | | | 0 | 0 | 0 | 0 | 0 | 0 | 0 | 0 | 0 | 0 | 0 |
| 0 | NA | NA | NA | NA | NA | Gm18255 | | 0 | 0 | 0 | 0 | 0 | 0 | 0 | 0 | 0 | 0 | 0 |
| 0 | NA | NA | NA | NA | NA | Olfr543 | olfactory receptor 543 | 0 | 0 | 0 | 0 | 0 | 0 | 0 | 0 | 0 | 0 | 0 |
| 0 | NA | NA | NA | NA | NA | Olfr544 | olfactory receptor 544 | 0 | 0 | 0 | 0 | 0 | 0 | 0 | 0 | 0 | 0 | 0 |
| 0 | NA | NA | NA | NA | NA | Olfr545 | olfactory receptor 545 | 0 | 0 | 0 | 0 | 0 | 0 | 0 | 0 | 0 | 0 | 0 |
| 0 | NA | NA | NA | NA | NA | Olfr546-ps1 | | 0 | 0 | 0 | 0 | 0 | 0 | 0 | 0 | 0 | 0 | 0 |
| 0 | NA | NA | NA | NA | NA | Olfr547 | olfactory receptor 547 | 0 | 0 | 0 | 0 | 0 | 0 | 0 | 0 | 0 | 0 | 0 |
| 0 | NA | NA | NA | NA | NA | Olfr548-ps1 | | 0 | 0 | 0 | 0 | 0 | 0 | 0 | 0 | 0 | 0 | 0 |
| 0 | NA | NA | NA | NA | NA | Olfr549 | olfactory receptor 549 | 0 | 0 | 0 | 0 | 0 | 0 | 0 | 0 | 0 | 0 | 0 |
| 0 | NA | NA | NA | NA | NA | Olfr550 | olfactory receptor 550 | 0 | 0 | 0 | 0 | 0 | 0 | 0 | 0 | 0 | 0 | 0 |
| 0 | NA | NA | NA | NA | NA | Olfr551 | olfactory receptor 551 | 0 | 0 | 0 | 0 | 0 | 0 | 0 | 0 | 0 | 0 | 0 |
| 0 | NA | NA | NA | NA | NA | Olfr552 | olfactory receptor 552 | 0 | 0 | 0 | 0 | 0 | 0 | 0 | 0 | 0 | 0 | 0 |
| 0 | NA | NA | NA | NA | NA | Olfr553 | olfactory receptor 553 | 0 | 0 | 0 | 0 | 0 | 0 | 0 | 0 | 0 | 0 | 0 |
| 0 | NA | NA | NA | NA | NA | Olfr554 | olfactory receptor 554 | 0 | 0 | 0 | 0 | 0 | 0 | 0 | 0 | 0 | 0 | 0 |
| 0 | NA | NA | NA | NA | NA | Olfr555 | olfactory receptor 555 | 0 | 0 | 0 | 0 | 0 | 0 | 0 | 0 | 0 | 0 | 0 |
| 0 | NA | NA | NA | NA | NA | Olfr556 | olfactory receptor 556 | 0 | 0 | 0 | 0 | 0 | 0 | 0 | 0 | 0 | 0 | 0 |
| 0 | NA | NA | NA | NA | NA | Gm45687 | | 0 | 0 | 0 | 0 | 0 | 0 | 0 | 0 | 0 | 0 | 0 |
| 0 | NA | NA | NA | NA | NA | Gm45688 | | 0 | 0 | 0 | 0 | 0 | 0 | 0 | 0 | 0 | 0 | 0 |
| 0 | NA | NA | NA | NA | NA | Olfr33 | olfactory receptor 33 | 0 | 0 | 0 | 0 | 0 | 0 | 0 | 0 | 0 | 0 | 0 |
| 0 | NA | NA | NA | NA | NA | Olfr559 | olfactory receptor 559 | 0 | 0 | 0 | 0 | 0 | 0 | 0 | 0 | 0 | 0 | 0 |
| 0 | NA | NA | NA | NA | NA | Gm24210 | | 0 | 0 | 0 | 0 | 0 | 0 | 0 | 0 | 0 | 0 | 0 |
| 0 | NA | NA | NA | NA | NA | Olfr560 | olfactory receptor 560 | 0 | 0 | 0 | 0 | 0 | 0 | 0 | 0 | 0 | 0 | 0 |
| 0 | NA | NA | NA | NA | NA | Olfr561 | olfactory receptor 561 | 0 | 0 | 0 | 0 | 0 | 0 | 0 | 0 | 0 | 0 | 0 |
| 0 | NA | NA | NA | NA | NA | Olfr562-ps1 | | 0 | 0 | 0 | 0 | 0 | 0 | 0 | 0 | 0 | 0 | 0 |
| 0 | NA | NA | NA | NA | NA | Olfr563-ps1 | | 0 | 0 | 0 | 0 | 0 | 0 | 0 | 0 | 0 | 0 | 0 |

| | | | | | | | | | | | | | | | | | |
|---|---|---|---|---|---|---|---|---|---|---|---|---|---|---|---|---|---|
| 0 | NA | NA | NA | NA | NA | Olfr564 | olfactory receptor 564 | 0 | 0 | 0 | 0 | 0 | 0 | 0 | 0 | 0 | 0 | 0 |
| 0 | NA | NA | NA | NA | NA | Olfr565-ps1 | | 0 | 0 | 0 | 0 | 0 | 0 | 0 | 0 | 0 | 0 | 0 |
| 0 | NA | NA | NA | NA | NA | Olfr566 | olfactory receptor 566 | 0 | 0 | 0 | 0 | 0 | 0 | 0 | 0 | 0 | 0 | 0 |
| 0 | NA | NA | NA | NA | NA | Gm45689 | | 0 | 0 | 0 | 0 | 0 | 0 | 0 | 0 | 0 | 0 | 0 |
| 0 | NA | NA | NA | NA | NA | Olfr567-ps1 | | 0 | 0 | 0 | 0 | 0 | 0 | 0 | 0 | 0 | 0 | 0 |
| 0 | NA | NA | NA | NA | NA | Olfr568 | olfactory receptor 568 | 0 | 0 | 0 | 0 | 0 | 0 | 0 | 0 | 0 | 0 | 0 |
| 0 | NA | NA | NA | NA | NA | Olfr569 | olfactory receptor 569 | 0 | 0 | 0 | 0 | 0 | 0 | 0 | 0 | 0 | 0 | 0 |
| 0 | NA | NA | NA | NA | NA | Olfr570 | olfactory receptor 570 | 0 | 0 | 0 | 0 | 0 | 0 | 0 | 0 | 0 | 0 | 0 |
| 0 | NA | NA | NA | NA | NA | Olfr571 | olfactory receptor 571 | 0 | 0 | 0 | 0 | 0 | 0 | 0 | 0 | 0 | 0 | 0 |
| 0 | NA | NA | NA | NA | NA | Olfr572 | olfactory receptor 572 | 0 | 0 | 0 | 0 | 0 | 0 | 0 | 0 | 0 | 0 | 0 |
| 0 | NA | NA | NA | NA | NA | Olfr573-ps1 | | 0 | 0 | 0 | 0 | 0 | 0 | 0 | 0 | 0 | 0 | 0 |
| 0 | NA | NA | NA | NA | NA | Olfr574 | olfactory receptor 574 | 0 | 0 | 0 | 0 | 0 | 0 | 0 | 0 | 0 | 0 | 0 |
| 0 | NA | NA | NA | NA | NA | Olfr575 | olfactory receptor 575 | 0 | 0 | 0 | 0 | 0 | 0 | 0 | 0 | 0 | 0 | 0 |
| 0 | NA | NA | NA | NA | NA | Olfr576 | olfactory receptor 576 | 0 | 0 | 0 | 0 | 0 | 0 | 0 | 0 | 0 | 0 | 0 |
| 0 | NA | NA | NA | NA | NA | Olfr577 | olfactory receptor 577 | 0 | 0 | 0 | 0 | 0 | 0 | 0 | 0 | 0 | 0 | 0 |
| 0 | NA | NA | NA | NA | NA | Olfr578 | olfactory receptor 578 | 0 | 0 | 0 | 0 | 0 | 0 | 0 | 0 | 0 | 0 | 0 |
| 0 | NA | NA | NA | NA | NA | | | 0 | 0 | 0 | 0 | 0 | 0 | 0 | 0 | 0 | 0 | 0 |
| 0 | NA | NA | NA | NA | NA | Olfr580-ps1 | | 0 | 0 | 0 | 0 | 0 | 0 | 0 | 0 | 0 | 0 | 0 |
| 0 | NA | NA | NA | NA | NA | Olfr581-ps1 | | 0 | 0 | 0 | 0 | 0 | 0 | 0 | 0 | 0 | 0 | 0 |
| 0 | NA | NA | NA | NA | NA | Olfr582 | olfactory receptor 582 | 0 | 0 | 0 | 0 | 0 | 0 | 0 | 0 | 0 | 0 | 0 |
| 0 | NA | NA | NA | NA | NA | Olfr583 | olfactory receptor 583 | 0 | 0 | 0 | 0 | 0 | 0 | 0 | 0 | 0 | 0 | 0 |
| 0 | NA | NA | NA | NA | NA | Olfr584 | olfactory receptor 584 | 0 | 0 | 0 | 0 | 0 | 0 | 0 | 0 | 0 | 0 | 0 |
| 0 | NA | NA | NA | NA | NA | Olfr585 | olfactory receptor 585 | 0 | 0 | 0 | 0 | 0 | 0 | 0 | 0 | 0 | 0 | 0 |
| 0 | NA | NA | NA | NA | NA | Olfr586 | olfactory receptor 586 | 0 | 0 | 0 | 0 | 0 | 0 | 0 | 0 | 0 | 0 | 0 |
| 0 | NA | NA | NA | NA | NA | Olfr587-ps1 | | 0 | 0 | 0 | 0 | 0 | 0 | 0 | 0 | 0 | 0 | 0 |
| 0 | NA | NA | NA | NA | NA | Gm15120 | | 0 | 0 | 0 | 0 | 0 | 0 | 0 | 0 | 0 | 0 | 0 |
| 0 | NA | NA | NA | NA | NA | Olfr588-ps1 | | 0 | 0 | 0 | 0 | 0 | 0 | 0 | 0 | 0 | 0 | 0 |
| 0 | NA | NA | NA | NA | NA | Gm15116 | | 0 | 0 | 0 | 0 | 0 | 0 | 0 | 0 | 0 | 0 | 0 |
| 0 | NA | NA | NA | NA | NA | Olfr589 | olfactory receptor 589 | 0 | 0 | 0 | 0 | 0 | 0 | 0 | 0 | 0 | 0 | 0 |
| 0 | NA | NA | NA | NA | NA | Olfr590-ps1 | | 0 | 0 | 0 | 0 | 0 | 0 | 0 | 0 | 0 | 0 | 0 |
| 0 | NA | NA | NA | NA | NA | Olfr591 | olfactory receptor 591 | 0 | 0 | 0 | 0 | 0 | 0 | 0 | 0 | 0 | 0 | 0 |
| 0 | NA | NA | NA | NA | NA | Olfr592 | olfactory receptor 592 | 0 | 0 | 0 | 0 | 0 | 0 | 0 | 0 | 0 | 0 | 0 |
| 0 | NA | NA | NA | NA | NA | Olfr593 | olfactory receptor 593 | 0 | 0 | 0 | 0 | 0 | 0 | 0 | 0 | 0 | 0 | 0 |
| 0 | NA | NA | NA | NA | NA | Olfr594 | olfactory receptor 594 | 0 | 0 | 0 | 0 | 0 | 0 | 0 | 0 | 0 | 0 | 0 |
| 0 | NA | NA | NA | NA | NA | Olfr595-ps1 | | 0 | 0 | 0 | 0 | 0 | 0 | 0 | 0 | 0 | 0 | 0 |
| 0 | NA | NA | NA | NA | NA | Usp17ld | ubiquitin specific peptidase 17-like D | 0 | 0 | 0 | 0 | 0 | 0 | 0 | 0 | 0 | 0 | 0 |
| 0 | NA | NA | NA | NA | NA | Gm25171 | | 0 | 0 | 0 | 0 | 0 | 0 | 0 | 0 | 0 | 0 | 0 |
| 0 | NA | NA | NA | NA | NA | Gm23550 | | 0 | 0 | 0 | 0 | 0 | 0 | 0 | 0 | 0 | 0 | 0 |
| 0 | NA | NA | NA | NA | NA | Gm23973 | | 0 | 0 | 0 | 0 | 0 | 0 | 0 | 0 | 0 | 0 | 0 |
| 0 | NA | NA | NA | NA | NA | Olfr596 | olfactory receptor 596 | 0 | 0 | 0 | 0 | 0 | 0 | 0 | 0 | 0 | 0 | 0 |
| 0 | NA | NA | NA | NA | NA | Olfr597 | olfactory receptor 597 | 0 | 0 | 0 | 0 | 0 | 0 | 0 | 0 | 0 | 0 | 0 |
| 0 | NA | NA | NA | NA | NA | Olfr598 | olfactory receptor 598 | 0 | 0 | 0 | 0 | 0 | 0 | 0 | 0 | 0 | 0 | 0 |
| 0 | NA | NA | NA | NA | NA | Olfr599 | olfactory receptor 599 | 0 | 0 | 0 | 0 | 0 | 0 | 0 | 0 | 0 | 0 | 0 |
| 0 | NA | NA | NA | NA | NA | Olfr600 | olfactory receptor 600 | 0 | 0 | 0 | 0 | 0 | 0 | 0 | 0 | 0 | 0 | 0 |
| 0 | NA | NA | NA | NA | NA | Olfr601 | olfactory receptor 601 | 0 | 0 | 0 | 0 | 0 | 0 | 0 | 0 | 0 | 0 | 0 |
| 0 | NA | NA | NA | NA | NA | Olfr602-ps1 | | 0 | 0 | 0 | 0 | 0 | 0 | 0 | 0 | 0 | 0 | 0 |
| 0 | NA | NA | NA | NA | NA | Olfr603 | olfactory receptor 603 | 0 | 0 | 0 | 0 | 0 | 0 | 0 | 0 | 0 | 0 | 0 |
| 0 | NA | NA | NA | NA | NA | Usp17lc | ubiquitin specific peptidase 17-like C | 0 | 0 | 0 | 0 | 0 | 0 | 0 | 0 | 0 | 0 | 0 |
| 0 | NA | NA | NA | NA | NA | Gm25774 | | 0 | 0 | 0 | 0 | 0 | 0 | 0 | 0 | 0 | 0 | 0 |
| 0 | NA | NA | NA | NA | NA | Olfr605 | olfactory receptor 605 | 0 | 0 | 0 | 0 | 0 | 0 | 0 | 0 | 0 | 0 | 0 |
| 0 | NA | NA | NA | NA | NA | Olfr606 | olfactory receptor 606 | 0 | 0 | 0 | 0 | 0 | 0 | 0 | 0 | 0 | 0 | 0 |
| 0 | NA | NA | NA | NA | NA | Olfr607 | olfactory receptor 607 | 0 | 0 | 0 | 0 | 0 | 0 | 0 | 0 | 0 | 0 | 0 |
| 0 | NA | NA | NA | NA | NA | Olfr608 | olfactory receptor 608 | 0 | 0 | 0 | 0 | 0 | 0 | 0 | 0 | 0 | 0 | 0 |
| 0 | NA | NA | NA | NA | NA | Olfr609 | olfactory receptor 609 | 0 | 0 | 0 | 0 | 0 | 0 | 0 | 0 | 0 | 0 | 0 |
| 0 | NA | NA | NA | NA | NA | Olfr610 | olfactory receptor 610 | 0 | 0 | 0 | 0 | 0 | 0 | 0 | 0 | 0 | 0 | 0 |
| 0 | NA | NA | NA | NA | NA | Olfr611 | olfactory receptor 611 | 0 | 0 | 0 | 0 | 0 | 0 | 0 | 0 | 0 | 0 | 0 |
| 0 | NA | NA | NA | NA | NA | Olfr612 | olfactory receptor 612 | 0 | 0 | 0 | 0 | 0 | 0 | 0 | 0 | 0 | 0 | 0 |
| 0 | NA | NA | NA | NA | NA | Olfr613 | olfactory receptor 613 | 0 | 0 | 0 | 0 | 0 | 0 | 0 | 0 | 0 | 0 | 0 |
| 0 | NA | NA | NA | NA | NA | Olfr615 | olfactory receptor 615 | 0 | 0 | 0 | 0 | 0 | 0 | 0 | 0 | 0 | 0 | 0 |
| 0 | NA | NA | NA | NA | NA | Olfr616 | olfactory receptor 616 | 0 | 0 | 0 | 0 | 0 | 0 | 0 | 0 | 0 | 0 | 0 |

| | | | | | | | | | | | | | | | | | | |
|---|---|---|---|---|---|---|---|---|---|---|---|---|---|---|---|---|---|---|
| 0 | NA | NA | NA | NA | NA | Olfr617 | olfactory receptor 617 | 0 | 0 | 0 | 0 | 0 | 0 | 0 | 0 | 0 | 0 | 0 |
| 0 | NA | NA | NA | NA | NA | Gm26506 | | 0 | 0 | 0 | 0 | 0 | 0 | 0 | 0 | 0 | 0 | 0 |
| 0 | NA | NA | NA | NA | NA | Olfr618 | olfactory receptor 618 | 0 | 0 | 0 | 0 | 0 | 0 | 0 | 0 | 0 | 0 | 0 |
| 0 | NA | NA | NA | NA | NA | Olfr619 | olfactory receptor 619 | 0 | 0 | 0 | 0 | 0 | 0 | 0 | 0 | 0 | 0 | 0 |
| 0 | NA | NA | NA | NA | NA | Olfr620 | olfactory receptor 620 | 0 | 0 | 0 | 0 | 0 | 0 | 0 | 0 | 0 | 0 | 0 |
| 0 | NA | NA | NA | NA | NA | Olfr621-ps1 | | 0 | 0 | 0 | 0 | 0 | 0 | 0 | 0 | 0 | 0 | 0 |
| 0 | NA | NA | NA | NA | NA | Olfr622 | olfactory receptor 622 | 0 | 0 | 0 | 0 | 0 | 0 | 0 | 0 | 0 | 0 | 0 |
| 0 | NA | NA | NA | NA | NA | Olfr623 | olfactory receptor 623 | 0 | 0 | 0 | 0 | 0 | 0 | 0 | 0 | 0 | 0 | 0 |
| 0 | NA | NA | NA | NA | NA | Olfr624 | olfactory receptor 624 | 0 | 0 | 0 | 0 | 0 | 0 | 0 | 0 | 0 | 0 | 0 |
| 0 | NA | NA | NA | NA | NA | Olfr625-ps1 | | 0 | 0 | 0 | 0 | 0 | 0 | 0 | 0 | 0 | 0 | 0 |
| 0 | NA | NA | NA | NA | NA | Olfr626-ps1 | | 0 | 0 | 0 | 0 | 0 | 0 | 0 | 0 | 0 | 0 | 0 |
| 0 | NA | NA | NA | NA | NA | Olfr260-ps1 | | 0 | 0 | 0 | 0 | 0 | 0 | 0 | 0 | 0 | 0 | 0 |
| 0 | NA | NA | NA | NA | NA | Olfr243 | olfactory receptor 243 | 0 | 0 | 0 | 0 | 0 | 0 | 0 | 0 | 0 | 0 | 0 |
| 0 | NA | NA | NA | NA | NA | Olfr629 | olfactory receptor 629 | 0 | 0 | 0 | 0 | 0 | 0 | 0 | 0 | 0 | 0 | 0 |
| 0 | NA | NA | NA | NA | NA | Olfr630 | olfactory receptor 630 | 0 | 0 | 0 | 0 | 0 | 0 | 0 | 0 | 0 | 0 | 0 |
| 0 | NA | NA | NA | NA | NA | Olfr69 | olfactory receptor 69 | 0 | 0 | 0 | 0 | 0 | 0 | 0 | 0 | 0 | 0 | 0 |
| 0 | NA | NA | NA | NA | NA | Olfr68 | olfactory receptor 68 | 0 | 0 | 0 | 0 | 0 | 0 | 0 | 0 | 0 | 0 | 0 |
| 0 | NA | NA | NA | NA | NA | Olfr67 | olfactory receptor 67 | 0 | 0 | 0 | 0 | 0 | 0 | 0 | 0 | 0 | 0 | 0 |
| 0 | NA | NA | NA | NA | NA | Gm15115 | | 0 | 0 | 0 | 0 | 0 | 0 | 0 | 0 | 0 | 0 | 0 |
| 0 | NA | NA | NA | NA | NA | Hbb-bh3 | | 0 | 0 | 0 | 0 | 0 | 0 | 0 | 0 | 0 | 0 | 0 |
| 0 | NA | NA | NA | NA | NA | Hbb-bh2 | hemoglobin beta, bh2 | 0 | 0 | 0 | 0 | 0 | 0 | 0 | 0 | 0 | 0 | 0 |
| 0 | NA | NA | NA | NA | NA | Hbb-bh1 | hemoglobin Z, beta-like embryonic chain | 0 | 0 | 0 | 0 | 0 | 0 | 0 | 0 | 0 | 0 | 0 |
| 0 | NA | NA | NA | NA | NA | Hbb-bh0 | | 0 | 0 | 0 | 0 | 0 | 0 | 0 | 0 | 0 | 0 | 0 |
| 0 | NA | NA | NA | NA | NA | Hbb-y | hemoglobin Y, beta-like embryonic chain | 0 | 0 | 0 | 0 | 0 | 0 | 0 | 0 | 0 | 0 | 0 |
| 0 | NA | NA | NA | NA | NA | Olfr66 | olfactory receptor 66 | 0 | 0 | 0 | 0 | 0 | 0 | 0 | 0 | 0 | 0 | 0 |
| 0 | NA | NA | NA | NA | NA | Olfr64 | olfactory receptor 64 | 0 | 0 | 0 | 0 | 0 | 0 | 0 | 0 | 0 | 0 | 0 |
| 0 | NA | NA | NA | NA | NA | Olfr65 | olfactory receptor 65 | 0 | 0 | 0 | 0 | 0 | 0 | 0 | 0 | 0 | 0 | 0 |
| 0 | NA | NA | NA | NA | NA | Olfr631 | olfactory receptor 631 | 0 | 0 | 0 | 0 | 0 | 0 | 0 | 0 | 0 | 0 | 0 |
| 0 | NA | NA | NA | NA | NA | Olfr632 | olfactory receptor 632 | 0 | 0 | 0 | 0 | 0 | 0 | 0 | 0 | 0 | 0 | 0 |
| 0 | NA | NA | NA | NA | NA | Gm45655 | | 0 | 0 | 0 | 0 | 0 | 0 | 0 | 0 | 0 | 0 | 0 |
| 0 | NA | NA | NA | NA | NA | Olfr633 | olfactory receptor 633 | 0 | 0 | 0 | 0 | 0 | 0 | 0 | 0 | 0 | 0 | 0 |
| 0 | NA | NA | NA | NA | NA | Olfr634-ps1 | | 0 | 0 | 0 | 0 | 0 | 0 | 0 | 0 | 0 | 0 | 0 |
| 0 | NA | NA | NA | NA | NA | Olfr635 | olfactory receptor 635 | 0 | 0 | 0 | 0 | 0 | 0 | 0 | 0 | 0 | 0 | 0 |
| 0 | NA | NA | NA | NA | NA | Gm4886 | | 0 | 0 | 0 | 0 | 0 | 0 | 0 | 0 | 0 | 0 | 0 |
| 0 | NA | NA | NA | NA | NA | Olfr636-ps1 | | 0 | 0 | 0 | 0 | 0 | 0 | 0 | 0 | 0 | 0 | 0 |
| 0 | NA | NA | NA | NA | NA | Olfr637-ps1 | | 0 | 0 | 0 | 0 | 0 | 0 | 0 | 0 | 0 | 0 | 0 |
| 0 | NA | NA | NA | NA | NA | Olfr638 | olfactory receptor 638 | 0 | 0 | 0 | 0 | 0 | 0 | 0 | 0 | 0 | 0 | 0 |
| 0 | NA | NA | NA | NA | NA | Olfr643 | | 0 | 0 | 0 | 0 | 0 | 0 | 0 | 0 | 0 | 0 | 0 |
| 0 | NA | NA | NA | NA | NA | Olfr640 | olfactory receptor 640 | 0 | 0 | 0 | 0 | 0 | 0 | 0 | 0 | 0 | 0 | 0 |
| 0 | NA | NA | NA | NA | NA | Olfr641 | olfactory receptor 641 | 0 | 0 | 0 | 0 | 0 | 0 | 0 | 0 | 0 | 0 | 0 |
| 0 | NA | NA | NA | NA | NA | Olfr642 | olfactory receptor 642 | 0 | 0 | 0 | 0 | 0 | 0 | 0 | 0 | 0 | 0 | 0 |
| 0 | NA | NA | NA | NA | NA | 4930516K23Ril | olfactory receptor 643 | 0 | 0 | 0 | 0 | 0 | 0 | 0 | 0 | 0 | 0 | 0 |
| 0 | NA | NA | NA | NA | NA | Olfr644 | olfactory receptor 644 | 0 | 0 | 0 | 0 | 0 | 0 | 0 | 0 | 0 | 0 | 0 |
| 0 | NA | NA | NA | NA | NA | Olfr645 | olfactory receptor 645 | 0 | 0 | 0 | 0 | 0 | 0 | 0 | 0 | 0 | 0 | 0 |
| 0 | NA | NA | NA | NA | NA | Olfr646 | olfactory receptor 646 | 0 | 0 | 0 | 0 | 0 | 0 | 0 | 0 | 0 | 0 | 0 |
| 0 | NA | NA | NA | NA | NA | Ubqln5 | ubiquilin 5 | 0 | 0 | 0 | 0 | 0 | 0 | 0 | 0 | 0 | 0 | 0 |
| 0 | NA | NA | NA | NA | NA | Ubqln3 | ubiquilin 3 | 0 | 0 | 0 | 0 | 0 | 0 | 0 | 0 | 0 | 0 | 0 |
| 0 | NA | NA | NA | NA | NA | Ubqlnl | ubiquilin-like | 0 | 0 | 0 | 0 | 0 | 0 | 0 | 0 | 0 | 0 | 0 |
| 0 | NA | NA | NA | NA | NA | Olfm5 | olfactomedin 5 | 0 | 0 | 0 | 0 | 0 | 0 | 0 | 0 | 0 | 0 | 0 |
| 0 | NA | NA | NA | NA | NA | Olfr647-ps1 | | 0 | 0 | 0 | 0 | 0 | 0 | 0 | 0 | 0 | 0 | 0 |
| 0 | NA | NA | NA | NA | NA | Olfr648 | olfactory receptor 648 | 0 | 0 | 0 | 0 | 0 | 0 | 0 | 0 | 0 | 0 | 0 |
| 0 | NA | NA | NA | NA | NA | Olfr649 | olfactory receptor 649 | 0 | 0 | 0 | 0 | 0 | 0 | 0 | 0 | 0 | 0 | 0 |
| 0 | NA | NA | NA | NA | NA | Olfr650-ps1 | | 0 | 0 | 0 | 0 | 0 | 0 | 0 | 0 | 0 | 0 | 0 |
| 0 | NA | NA | NA | NA | NA | Gm47248 | | 0 | 0 | 0 | 0 | 0 | 0 | 0 | 0 | 0 | 0 | 0 |
| 0 | NA | NA | NA | NA | NA | Gm24846 | | 0 | 0 | 0 | 0 | 0 | 0 | 0 | 0 | 0 | 0 | 0 |
| 0 | NA | NA | NA | NA | NA | Gm27995 | | 0 | 0 | 0 | 0 | 0 | 0 | 0 | 0 | 0 | 0 | 0 |
| 0 | NA | NA | NA | NA | NA | Gm45471 | | 0 | 0 | 0 | 0 | 0 | 0 | 0 | 0 | 0 | 0 | 0 |
| 0 | NA | NA | NA | NA | NA | Gm25405 | | 0 | 0 | 0 | 0 | 0 | 0 | 0 | 0 | 0 | 0 | 0 |
| 0 | NA | NA | NA | NA | NA | Gm6574 | | 0 | 0 | 0 | 0 | 0 | 0 | 0 | 0 | 0 | 0 | 0 |
| 0 | NA | NA | NA | NA | NA | Olfr651 | olfactory receptor 651 | 0 | 0 | 0 | 0 | 0 | 0 | 0 | 0 | 0 | 0 | 0 |

| | | | | | | | | | | | | | | | | | | |
|---|---|---|---|---|---|---|---|---|---|---|---|---|---|---|---|---|---|---|
| 0 | NA | NA | NA | NA | NA | Olfr652 | olfactory receptor 652 | 0 | 0 | 0 | 0 | 0 | 0 | 0 | 0 | 0 | 0 | 0 |
| 0 | NA | NA | NA | NA | NA | Olfr653 | olfactory receptor 653 | 0 | 0 | 0 | 0 | 0 | 0 | 0 | 0 | 0 | 0 | 0 |
| 0 | NA | NA | NA | NA | NA | Olfr654 | olfactory receptor 654 | 0 | 0 | 0 | 0 | 0 | 0 | 0 | 0 | 0 | 0 | 0 |
| 0 | NA | NA | NA | NA | NA | Olfr655 | olfactory receptor 655 | 0 | 0 | 0 | 0 | 0 | 0 | 0 | 0 | 0 | 0 | 0 |
| 0 | NA | NA | NA | NA | NA | Olfr656 | olfactory receptor 656 | 0 | 0 | 0 | 0 | 0 | 0 | 0 | 0 | 0 | 0 | 0 |
| 0 | NA | NA | NA | NA | NA | Olfr657 | olfactory receptor 657 | 0 | 0 | 0 | 0 | 0 | 0 | 0 | 0 | 0 | 0 | 0 |
| 0 | NA | NA | NA | NA | NA | Olfr658 | olfactory receptor 658 | 0 | 0 | 0 | 0 | 0 | 0 | 0 | 0 | 0 | 0 | 0 |
| 0 | NA | NA | NA | NA | NA | Usp17le-ps | | 0 | 0 | 0 | 0 | 0 | 0 | 0 | 0 | 0 | 0 | 0 |
| 0 | NA | NA | NA | NA | NA | Olfr659 | olfactory receptor 659 | 0 | 0 | 0 | 0 | 0 | 0 | 0 | 0 | 0 | 0 | 0 |
| 0 | NA | NA | NA | NA | NA | Olfr660-ps1 | | 0 | 0 | 0 | 0 | 0 | 0 | 0 | 0 | 0 | 0 | 0 |
| 0 | NA | NA | NA | NA | NA | Olfr661 | olfactory receptor 661 | 0 | 0 | 0 | 0 | 0 | 0 | 0 | 0 | 0 | 0 | 0 |
| 0 | NA | NA | NA | NA | NA | Gm6593 | | 0 | 0 | 0 | 0 | 0 | 0 | 0 | 0 | 0 | 0 | 0 |
| 0 | NA | NA | NA | NA | NA | Olfr662-ps1 | | 0 | 0 | 0 | 0 | 0 | 0 | 0 | 0 | 0 | 0 | 0 |
| 0 | NA | NA | NA | NA | NA | Olfr663 | olfactory receptor 663 | 0 | 0 | 0 | 0 | 0 | 0 | 0 | 0 | 0 | 0 | 0 |
| 0 | NA | NA | NA | NA | NA | Olfr664 | | 0 | 0 | 0 | 0 | 0 | 0 | 0 | 0 | 0 | 0 | 0 |
| 0 | NA | NA | NA | NA | NA | Usp17le | ubiquitin specific peptidase 17-like E | 0 | 0 | 0 | 0 | 0 | 0 | 0 | 0 | 0 | 0 | 0 |
| 0 | NA | NA | NA | NA | NA | Usp17lb | ubiquitin specific peptidase 17-like B | 0 | 0 | 0 | 0 | 0 | 0 | 0 | 0 | 0 | 0 | 0 |
| 0 | NA | NA | NA | NA | NA | Usp17la | ubiquitin specific peptidase 17-like A | 0 | 0 | 0 | 0 | 0 | 0 | 0 | 0 | 0 | 0 | 0 |
| 0 | NA | NA | NA | NA | NA | Gm45556 | | 0 | 0 | 0 | 0 | 0 | 0 | 0 | 0 | 0 | 0 | 0 |
| 0 | NA | NA | NA | NA | NA | Olfr665 | olfactory receptor 665 | 0 | 0 | 0 | 0 | 0 | 0 | 0 | 0 | 0 | 0 | 0 |
| 0 | NA | NA | NA | NA | NA | Olfr666 | olfactory receptor 666 | 0 | 0 | 0 | 0 | 0 | 0 | 0 | 0 | 0 | 0 | 0 |
| 0 | NA | NA | NA | NA | NA | Gm45272 | | 0 | 0 | 0 | 0 | 0 | 0 | 0 | 0 | 0 | 0 | 0 |
| 0 | NA | NA | NA | NA | NA | Gm45275 | | 0 | 0 | 0 | 0 | 0 | 0 | 0 | 0 | 0 | 0 | 0 |
| 0 | NA | NA | NA | NA | NA | Olfr667 | olfactory receptor 667 | 0 | 0 | 0 | 0 | 0 | 0 | 0 | 0 | 0 | 0 | 0 |
| 0 | NA | NA | NA | NA | NA | Olfr668 | olfactory receptor 668 | 0 | 0 | 0 | 0 | 0 | 0 | 0 | 0 | 0 | 0 | 0 |
| 0 | NA | NA | NA | NA | NA | Olfr669 | olfactory receptor 669 | 0 | 0 | 0 | 0 | 0 | 0 | 0 | 0 | 0 | 0 | 0 |
| 0 | NA | NA | NA | NA | NA | Gm45273 | | 0 | 0 | 0 | 0 | 0 | 0 | 0 | 0 | 0 | 0 | 0 |
| 0 | NA | NA | NA | NA | NA | Olfr670 | olfactory receptor 670 | 0 | 0 | 0 | 0 | 0 | 0 | 0 | 0 | 0 | 0 | 0 |
| 0 | NA | NA | NA | NA | NA | Olfr671 | olfactory receptor 671 | 0 | 0 | 0 | 0 | 0 | 0 | 0 | 0 | 0 | 0 | 0 |
| 0 | NA | NA | NA | NA | NA | Olfr672 | olfactory receptor 672 | 0 | 0 | 0 | 0 | 0 | 0 | 0 | 0 | 0 | 0 | 0 |
| 0 | NA | NA | NA | NA | NA | Olfr673-ps1 | | 0 | 0 | 0 | 0 | 0 | 0 | 0 | 0 | 0 | 0 | 0 |
| 0 | NA | NA | NA | NA | NA | Olfr674-ps1 | | 0 | 0 | 0 | 0 | 0 | 0 | 0 | 0 | 0 | 0 | 0 |
| 0 | NA | NA | NA | NA | NA | Olfr675 | olfactory receptor 675 | 0 | 0 | 0 | 0 | 0 | 0 | 0 | 0 | 0 | 0 | 0 |
| 0 | NA | NA | NA | NA | NA | Olfr676 | olfactory receptor 676 | 0 | 0 | 0 | 0 | 0 | 0 | 0 | 0 | 0 | 0 | 0 |
| 0 | NA | NA | NA | NA | NA | Olfr677 | olfactory receptor 677 | 0 | 0 | 0 | 0 | 0 | 0 | 0 | 0 | 0 | 0 | 0 |
| 0 | NA | NA | NA | NA | NA | Gm8913 | | 0 | 0 | 0 | 0 | 0 | 0 | 0 | 0 | 0 | 0 | 0 |
| 0 | NA | NA | NA | NA | NA | Olfr678 | olfactory receptor 678 | 0 | 0 | 0 | 0 | 0 | 0 | 0 | 0 | 0 | 0 | 0 |
| 0 | NA | NA | NA | NA | NA | Gm17989 | | 0 | 0 | 0 | 0 | 0 | 0 | 0 | 0 | 0 | 0 | 0 |
| 0 | NA | NA | NA | NA | NA | Gm45557 | | 0 | 0 | 0 | 0 | 0 | 0 | 0 | 0 | 0 | 0 | 0 |
| 0 | NA | NA | NA | NA | NA | Olfr679 | olfactory receptor 679 | 0 | 0 | 0 | 0 | 0 | 0 | 0 | 0 | 0 | 0 | 0 |
| 0 | NA | NA | NA | NA | NA | Olfr680-ps1 | | 0 | 0 | 0 | 0 | 0 | 0 | 0 | 0 | 0 | 0 | 0 |
| 0 | NA | NA | NA | NA | NA | Gm17990 | | 0 | 0 | 0 | 0 | 0 | 0 | 0 | 0 | 0 | 0 | 0 |
| 0 | NA | NA | NA | NA | NA | Gm45558 | | 0 | 0 | 0 | 0 | 0 | 0 | 0 | 0 | 0 | 0 | 0 |
| 0 | NA | NA | NA | NA | NA | Olfr681 | olfactory receptor 681 | 0 | 0 | 0 | 0 | 0 | 0 | 0 | 0 | 0 | 0 | 0 |
| 0 | NA | NA | NA | NA | NA | Olfr682-ps1 | | 0 | 0 | 0 | 0 | 0 | 0 | 0 | 0 | 0 | 0 | 0 |
| 0 | NA | NA | NA | NA | NA | Olfr683 | olfactory receptor 683 | 0 | 0 | 0 | 0 | 0 | 0 | 0 | 0 | 0 | 0 | 0 |
| 0 | NA | NA | NA | NA | NA | Gm45374 | | 0 | 0 | 0 | 0 | 0 | 0 | 0 | 0 | 0 | 0 | 0 |
| 0 | NA | NA | NA | NA | NA | Olfr684 | olfactory receptor 684 | 0 | 0 | 0 | 0 | 0 | 0 | 0 | 0 | 0 | 0 | 0 |
| 0 | NA | NA | NA | NA | NA | Olfr685 | olfactory receptor 685 | 0 | 0 | 0 | 0 | 0 | 0 | 0 | 0 | 0 | 0 | 0 |
| 0 | NA | NA | NA | NA | NA | Olfr686 | olfactory receptor 686 | 0 | 0 | 0 | 0 | 0 | 0 | 0 | 0 | 0 | 0 | 0 |
| 0 | NA | NA | NA | NA | NA | Olfr687 | | 0 | 0 | 0 | 0 | 0 | 0 | 0 | 0 | 0 | 0 | 0 |
| 0 | NA | NA | NA | NA | NA | Olfr688 | olfactory receptor 688 | 0 | 0 | 0 | 0 | 0 | 0 | 0 | 0 | 0 | 0 | 0 |
| 0 | NA | NA | NA | NA | NA | Olfr689 | olfactory receptor 689 | 0 | 0 | 0 | 0 | 0 | 0 | 0 | 0 | 0 | 0 | 0 |
| 0 | NA | NA | NA | NA | NA | Gm45331 | | 0 | 0 | 0 | 0 | 0 | 0 | 0 | 0 | 0 | 0 | 0 |
| 0 | NA | NA | NA | NA | NA | Olfr690 | olfactory receptor 690 | 0 | 0 | 0 | 0 | 0 | 0 | 0 | 0 | 0 | 0 | 0 |
| 0 | NA | NA | NA | NA | NA | Olfr691 | olfactory receptor 691 | 0 | 0 | 0 | 0 | 0 | 0 | 0 | 0 | 0 | 0 | 0 |
| 0 | NA | NA | NA | NA | NA | Olfr692 | olfactory receptor 692 | 0 | 0 | 0 | 0 | 0 | 0 | 0 | 0 | 0 | 0 | 0 |
| 0 | NA | NA | NA | NA | NA | Gm5901 | predicted gene 5901 | 0 | 0 | 0 | 0 | 0 | 0 | 0 | 0 | 0 | 0 | 0 |
| 0 | NA | NA | NA | NA | NA | Gm22504 | | 0 | 0 | 0 | 0 | 0 | 0 | 0 | 0 | 0 | 0 | 0 |
| 0 | NA | NA | NA | NA | NA | Gm45684 | | 0 | 0 | 0 | 0 | 0 | 0 | 0 | 0 | 0 | 0 | 0 |

| | | | | | | | | | | | | | | | | | | |
|---|---|---|---|---|---|---|---|---|---|---|---|---|---|---|---|---|---|---|
| 0 | NA | NA | NA | NA | NA | Gm20663 | | 0 | 0 | 0 | 0 | 0 | 0 | 0 | 0 | 0 | 0 | 0 |
| 0 | NA | NA | NA | NA | NA | Gm27228 | | 0 | 0 | 0 | 0 | 0 | 0 | 0 | 0 | 0 | 0 | 0 |
| 0 | NA | NA | NA | NA | NA | Gm27149 | | 0 | 0 | 0 | 0 | 0 | 0 | 0 | 0 | 0 | 0 | 0 |
| 0 | NA | NA | NA | NA | NA | Gm8982 | | 0 | 0 | 0 | 0 | 0 | 0 | 0 | 0 | 0 | 0 | 0 |
| 0 | NA | NA | NA | NA | NA | Gm27150 | | 0 | 0 | 0 | 0 | 0 | 0 | 0 | 0 | 0 | 0 | 0 |
| 0 | NA | NA | NA | NA | NA | Gm22372 | | 0 | 0 | 0 | 0 | 0 | 0 | 0 | 0 | 0 | 0 | 0 |
| 0 | NA | NA | NA | NA | NA | Olfr693 | olfactory receptor 693 | 0 | 0 | 0 | 0 | 0 | 0 | 0 | 0 | 0 | 0 | 0 |
| 0 | NA | NA | NA | NA | NA | Olfr694 | olfactory receptor 694 | 0 | 0 | 0 | 0 | 0 | 0 | 0 | 0 | 0 | 0 | 0 |
| 0 | NA | NA | NA | NA | NA | Gm44857 | | 0 | 0 | 0 | 0 | 0 | 0 | 0 | 0 | 0 | 0 | 0 |
| 0 | NA | NA | NA | NA | NA | Olfr695 | olfactory receptor 695 | 0 | 0 | 0 | 0 | 0 | 0 | 0 | 0 | 0 | 0 | 0 |
| 0 | NA | NA | NA | NA | NA | Olfr696-ps1 | | 0 | 0 | 0 | 0 | 0 | 0 | 0 | 0 | 0 | 0 | 0 |
| 0 | NA | NA | NA | NA | NA | Olfr697 | olfactory receptor 697 | 0 | 0 | 0 | 0 | 0 | 0 | 0 | 0 | 0 | 0 | 0 |
| 0 | NA | NA | NA | NA | NA | Olfr698 | olfactory receptor 698 | 0 | 0 | 0 | 0 | 0 | 0 | 0 | 0 | 0 | 0 | 0 |
| 0 | NA | NA | NA | NA | NA | Olfr699 | olfactory receptor 699 | 0 | 0 | 0 | 0 | 0 | 0 | 0 | 0 | 0 | 0 | 0 |
| 0 | NA | NA | NA | NA | NA | Olfr700 | olfactory receptor 700 | 0 | 0 | 0 | 0 | 0 | 0 | 0 | 0 | 0 | 0 | 0 |
| 0 | NA | NA | NA | NA | NA | 4930458B22Rik | | 0 | 0 | 0 | 0 | 0 | 0 | 0 | 0 | 0 | 0 | 0 |
| 0 | NA | NA | NA | NA | NA | Olfr702 | olfactory receptor 702 | 0 | 0 | 0 | 0 | 0 | 0 | 0 | 0 | 0 | 0 | 0 |
| 0 | NA | NA | NA | NA | NA | Olfr703 | olfactory receptor 703 | 0 | 0 | 0 | 0 | 0 | 0 | 0 | 0 | 0 | 0 | 0 |
| 0 | NA | NA | NA | NA | NA | Olfr704 | olfactory receptor 704 | 0 | 0 | 0 | 0 | 0 | 0 | 0 | 0 | 0 | 0 | 0 |
| 0 | NA | NA | NA | NA | NA | Olfr705 | olfactory receptor 705 | 0 | 0 | 0 | 0 | 0 | 0 | 0 | 0 | 0 | 0 | 0 |
| 0 | NA | NA | NA | NA | NA | Olfr706 | olfactory receptor 706 | 0 | 0 | 0 | 0 | 0 | 0 | 0 | 0 | 0 | 0 | 0 |
| 0 | NA | NA | NA | NA | NA | Olfr707 | olfactory receptor 707 | 0 | 0 | 0 | 0 | 0 | 0 | 0 | 0 | 0 | 0 | 0 |
| 0 | NA | NA | NA | NA | NA | Olfr1532-ps1 | olfactory receptor 1532, pseudogene 1 | 0 | 0 | 0 | 0 | 0 | 0 | 0 | 0 | 0 | 0 | 0 |
| 0 | NA | NA | NA | NA | NA | Olfr709-ps1 | | 0 | 0 | 0 | 0 | 0 | 0 | 0 | 0 | 0 | 0 | 0 |
| 0 | NA | NA | NA | NA | NA | Olfr710 | olfactory receptor 710 | 0 | 0 | 0 | 0 | 0 | 0 | 0 | 0 | 0 | 0 | 0 |
| 0 | NA | NA | NA | NA | NA | Gm44582 | | 0 | 0 | 0 | 0 | 0 | 0 | 0 | 0 | 0 | 0 | 0 |
| 0 | NA | NA | NA | NA | NA | Olfr6 | olfactory receptor 6 | 0 | 0 | 0 | 0 | 0 | 0 | 0 | 0 | 0 | 0 | 0 |
| 0 | NA | NA | NA | NA | NA | Olfr711 | olfactory receptor 711 | 0 | 0 | 0 | 0 | 0 | 0 | 0 | 0 | 0 | 0 | 0 |
| 0 | NA | NA | NA | NA | NA | Olfr712-ps1 | | 0 | 0 | 0 | 0 | 0 | 0 | 0 | 0 | 0 | 0 | 0 |
| 0 | NA | NA | NA | NA | NA | Olfr713 | olfactory receptor 713 | 0 | 0 | 0 | 0 | 0 | 0 | 0 | 0 | 0 | 0 | 0 |
| 0 | NA | NA | NA | NA | NA | Gm17991 | | 0 | 0 | 0 | 0 | 0 | 0 | 0 | 0 | 0 | 0 | 0 |
| 0 | NA | NA | NA | NA | NA | Olfr714 | olfactory receptor 714 | 0 | 0 | 0 | 0 | 0 | 0 | 0 | 0 | 0 | 0 | 0 |
| 0 | NA | NA | NA | NA | NA | Gm45023 | | 0 | 0 | 0 | 0 | 0 | 0 | 0 | 0 | 0 | 0 | 0 |
| 0 | NA | NA | NA | NA | NA | Olfr17 | olfactory receptor 17 | 0 | 0 | 0 | 0 | 0 | 0 | 0 | 0 | 0 | 0 | 0 |
| 0 | NA | NA | NA | NA | NA | Olfr715b | olfactory receptor 715B | 0 | 0 | 0 | 0 | 0 | 0 | 0 | 0 | 0 | 0 | 0 |
| 0 | NA | NA | NA | NA | NA | Gm45868 | | 0 | 0 | 0 | 0 | 0 | 0 | 0 | 0 | 0 | 0 | 0 |
| 0 | NA | NA | NA | NA | NA | Olfr715 | olfactory receptor 715 | 0 | 0 | 0 | 0 | 0 | 0 | 0 | 0 | 0 | 0 | 0 |
| 0 | NA | NA | NA | NA | NA | Olfr716 | olfactory receptor 716 | 0 | 0 | 0 | 0 | 0 | 0 | 0 | 0 | 0 | 0 | 0 |
| 0 | NA | NA | NA | NA | NA | Nlrp14 | NLR family, pyrin domain containing 14 | 0 | 0 | 0 | 0 | 0 | 0 | 0 | 0 | 0 | 0 | 0 |
| 0 | NA | NA | NA | NA | NA | Gm3765 | | 0 | 0 | 0 | 0 | 0 | 0 | 0 | 0 | 0 | 0 | 0 |
| 0 | NA | NA | NA | NA | NA | Rbmxl2 | RNA binding motif protein, X-linked-like 2 | 0 | 0 | 0 | 0 | 0 | 0 | 0 | 0 | 0 | 0 | 0 |
| 0 | NA | NA | NA | NA | NA | Platr28 | | 0 | 0 | 0 | 0 | 0 | 0 | 0 | 0 | 0 | 0 | 0 |
| 0 | NA | NA | NA | NA | NA | Gm24932 | | 0 | 0 | 0 | 0 | 0 | 0 | 0 | 0 | 0 | 0 | 0 |
| 0 | NA | NA | NA | NA | NA | Gm19033 | | 0 | 0 | 0 | 0 | 0 | 0 | 0 | 0 | 0 | 0 | 0 |
| 0 | NA | NA | NA | NA | NA | Gm44773 | | 0 | 0 | 0 | 0 | 0 | 0 | 0 | 0 | 0 | 0 | 0 |
| 0 | NA | NA | NA | NA | NA | 5330417H12Rik | | 0 | 0 | 0 | 0 | 0 | 0 | 0 | 0 | 0 | 0 | 0 |
| 0 | NA | NA | NA | NA | NA | Gm45040 | | 0 | 0 | 0 | 0 | 0 | 0 | 0 | 0 | 0 | 0 | 0 |
| 0 | NA | NA | NA | NA | NA | Ovch2 | ovochymase 2 | 0 | 0 | 0 | 0 | 0 | 0 | 0 | 0 | 0 | 0 | 0 |
| 0 | NA | NA | NA | NA | NA | Olfr467 | olfactory receptor 467 | 0 | 0 | 0 | 0 | 0 | 0 | 0 | 0 | 0 | 0 | 0 |
| 0 | NA | NA | NA | NA | NA | Olfr468-ps1 | | 0 | 0 | 0 | 0 | 0 | 0 | 0 | 0 | 0 | 0 | 0 |
| 0 | NA | NA | NA | NA | NA | Olfr469 | olfactory receptor 469 | 0 | 0 | 0 | 0 | 0 | 0 | 0 | 0 | 0 | 0 | 0 |
| 0 | NA | NA | NA | NA | NA | Gm5002 | | 0 | 0 | 0 | 0 | 0 | 0 | 0 | 0 | 0 | 0 | 0 |
| 0 | NA | NA | NA | NA | NA | Olfr470 | olfactory receptor 470 | 0 | 0 | 0 | 0 | 0 | 0 | 0 | 0 | 0 | 0 | 0 |
| 0 | NA | NA | NA | NA | NA | Olfr471 | | 0 | 0 | 0 | 0 | 0 | 0 | 0 | 0 | 0 | 0 | 0 |
| 0 | NA | NA | NA | NA | NA | Olfr473 | olfactory receptor 473 | 0 | 0 | 0 | 0 | 0 | 0 | 0 | 0 | 0 | 0 | 0 |
| 0 | NA | NA | NA | NA | NA | Olfr474 | olfactory receptor 474 | 0 | 0 | 0 | 0 | 0 | 0 | 0 | 0 | 0 | 0 | 0 |
| 0 | NA | NA | NA | NA | NA | Olfr476 | olfactory receptor 476 | 0 | 0 | 0 | 0 | 0 | 0 | 0 | 0 | 0 | 0 | 0 |
| 0 | NA | NA | NA | NA | NA | Olfr477 | olfactory receptor 477 | 0 | 0 | 0 | 0 | 0 | 0 | 0 | 0 | 0 | 0 | 0 |
| 0 | NA | NA | NA | NA | NA | Olfr478 | olfactory receptor 478 | 0 | 0 | 0 | 0 | 0 | 0 | 0 | 0 | 0 | 0 | 0 |
| 0 | NA | NA | NA | NA | NA | Olfr479 | olfactory receptor 479 | 0 | 0 | 0 | 0 | 0 | 0 | 0 | 0 | 0 | 0 | 0 |

| | | | | | | | | | | | | | | | | | | |
|---|---|---|---|---|---|---|---|---|---|---|---|---|---|---|---|---|---|---|
| 0 | NA | NA | NA | NA | NA | Olfr480 | olfactory receptor 480 | 0 | 0 | 0 | 0 | 0 | 0 | 0 | 0 | 0 | 0 | 0 |
| 0 | NA | NA | NA | NA | NA | Olfr481 | olfactory receptor 481 | 0 | 0 | 0 | 0 | 0 | 0 | 0 | 0 | 0 | 0 | 0 |
| 0 | NA | NA | NA | NA | NA | Gm45375 | | 0 | 0 | 0 | 0 | 0 | 0 | 0 | 0 | 0 | 0 | 0 |
| 0 | NA | NA | NA | NA | NA | Olfr482 | olfactory receptor 482 | 0 | 0 | 0 | 0 | 0 | 0 | 0 | 0 | 0 | 0 | 0 |
| 0 | NA | NA | NA | NA | NA | Gm44870 | | 0 | 0 | 0 | 0 | 0 | 0 | 0 | 0 | 0 | 0 | 0 |
| 0 | NA | NA | NA | NA | NA | Olfr483 | olfactory receptor 483 | 0 | 0 | 0 | 0 | 0 | 0 | 0 | 0 | 0 | 0 | 0 |
| 0 | NA | NA | NA | NA | NA | Gm25424 | | 0 | 0 | 0 | 0 | 0 | 0 | 0 | 0 | 0 | 0 | 0 |
| 0 | NA | NA | NA | NA | NA | Olfr484 | olfactory receptor 484 | 0 | 0 | 0 | 0 | 0 | 0 | 0 | 0 | 0 | 0 | 0 |
| 0 | NA | NA | NA | NA | NA | Olfr485 | olfactory receptor 485 | 0 | 0 | 0 | 0 | 0 | 0 | 0 | 0 | 0 | 0 | 0 |
| 0 | NA | NA | NA | NA | NA | Olfr486 | olfactory receptor 486 | 0 | 0 | 0 | 0 | 0 | 0 | 0 | 0 | 0 | 0 | 0 |
| 0 | NA | NA | NA | NA | NA | Gm18944 | | 0 | 0 | 0 | 0 | 0 | 0 | 0 | 0 | 0 | 0 | 0 |
| 0 | NA | NA | NA | NA | NA | Olfr487 | olfactory receptor 487 | 0 | 0 | 0 | 0 | 0 | 0 | 0 | 0 | 0 | 0 | 0 |
| 0 | NA | NA | NA | NA | NA | Olfr489-ps1 | | 0 | 0 | 0 | 0 | 0 | 0 | 0 | 0 | 0 | 0 | 0 |
| 0 | NA | NA | NA | NA | NA | Olfr490 | olfactory receptor 490 | 0 | 0 | 0 | 0 | 0 | 0 | 0 | 0 | 0 | 0 | 0 |
| 0 | NA | NA | NA | NA | NA | Gm44594 | | 0 | 0 | 0 | 0 | 0 | 0 | 0 | 0 | 0 | 0 | 0 |
| 0 | NA | NA | NA | NA | NA | Olfr491 | olfactory receptor 491 | 0 | 0 | 0 | 0 | 0 | 0 | 0 | 0 | 0 | 0 | 0 |
| 0 | NA | NA | NA | NA | NA | Olfr492 | olfactory receptor 492 | 0 | 0 | 0 | 0 | 0 | 0 | 0 | 0 | 0 | 0 | 0 |
| 0 | NA | NA | NA | NA | NA | Olfr493 | olfactory receptor 493 | 0 | 0 | 0 | 0 | 0 | 0 | 0 | 0 | 0 | 0 | 0 |
| 0 | NA | NA | NA | NA | NA | Olfr494 | olfactory receptor 494 | 0 | 0 | 0 | 0 | 0 | 0 | 0 | 0 | 0 | 0 | 0 |
| 0 | NA | NA | NA | NA | NA | Gm45392 | | 0 | 0 | 0 | 0 | 0 | 0 | 0 | 0 | 0 | 0 | 0 |
| 0 | NA | NA | NA | NA | NA | Olfr495 | olfactory receptor 495 | 0 | 0 | 0 | 0 | 0 | 0 | 0 | 0 | 0 | 0 | 0 |
| 0 | NA | NA | NA | NA | NA | Olfr496-ps1 | | 0 | 0 | 0 | 0 | 0 | 0 | 0 | 0 | 0 | 0 | 0 |
| 0 | NA | NA | NA | NA | NA | Olfr497 | olfactory receptor 497 | 0 | 0 | 0 | 0 | 0 | 0 | 0 | 0 | 0 | 0 | 0 |
| 0 | NA | NA | NA | NA | NA | Gm44611 | | 0 | 0 | 0 | 0 | 0 | 0 | 0 | 0 | 0 | 0 | 0 |
| 0 | NA | NA | NA | NA | NA | Olfr498 | olfactory receptor 498 | 0 | 0 | 0 | 0 | 0 | 0 | 0 | 0 | 0 | 0 | 0 |
| 0 | NA | NA | NA | NA | NA | Olfr499-ps1 | | 0 | 0 | 0 | 0 | 0 | 0 | 0 | 0 | 0 | 0 | 0 |
| 0 | NA | NA | NA | NA | NA | Olfr500-ps1 | | 0 | 0 | 0 | 0 | 0 | 0 | 0 | 0 | 0 | 0 | 0 |
| 0 | NA | NA | NA | NA | NA | Olfr501-ps1 | | 0 | 0 | 0 | 0 | 0 | 0 | 0 | 0 | 0 | 0 | 0 |
| 0 | NA | NA | NA | NA | NA | Olfr502 | olfactory receptor 502 | 0 | 0 | 0 | 0 | 0 | 0 | 0 | 0 | 0 | 0 | 0 |
| 0 | NA | NA | NA | NA | NA | Gm5053 | | 0 | 0 | 0 | 0 | 0 | 0 | 0 | 0 | 0 | 0 | 0 |
| 0 | NA | NA | NA | NA | NA | Olfr503 | olfactory receptor 503 | 0 | 0 | 0 | 0 | 0 | 0 | 0 | 0 | 0 | 0 | 0 |
| 0 | NA | NA | NA | NA | NA | Olfr504 | olfactory receptor 504 | 0 | 0 | 0 | 0 | 0 | 0 | 0 | 0 | 0 | 0 | 0 |
| 0 | NA | NA | NA | NA | NA | Gm18410 | | 0 | 0 | 0 | 0 | 0 | 0 | 0 | 0 | 0 | 0 | 0 |
| 0 | NA | NA | NA | NA | NA | Gm44865 | | 0 | 0 | 0 | 0 | 0 | 0 | 0 | 0 | 0 | 0 | 0 |
| 0 | NA | NA | NA | NA | NA | Olfr505-ps1 | | 0 | 0 | 0 | 0 | 0 | 0 | 0 | 0 | 0 | 0 | 0 |
| 0 | NA | NA | NA | NA | NA | Gm19127 | | 0 | 0 | 0 | 0 | 0 | 0 | 0 | 0 | 0 | 0 | 0 |
| 0 | NA | NA | NA | NA | NA | Olfr506 | olfactory receptor 506 | 0 | 0 | 0 | 0 | 0 | 0 | 0 | 0 | 0 | 0 | 0 |
| 0 | NA | NA | NA | NA | NA | Olfr507 | olfactory receptor 507 | 0 | 0 | 0 | 0 | 0 | 0 | 0 | 0 | 0 | 0 | 0 |
| 0 | NA | NA | NA | NA | NA | Olfr508 | olfactory receptor 508 | 0 | 0 | 0 | 0 | 0 | 0 | 0 | 0 | 0 | 0 | 0 |
| 0 | NA | NA | NA | NA | NA | Olfr509 | olfactory receptor 509 | 0 | 0 | 0 | 0 | 0 | 0 | 0 | 0 | 0 | 0 | 0 |
| 0 | NA | NA | NA | NA | NA | Olfr510 | olfactory receptor 510 | 0 | 0 | 0 | 0 | 0 | 0 | 0 | 0 | 0 | 0 | 0 |
| 0 | NA | NA | NA | NA | NA | Olfr511-ps1 | | 0 | 0 | 0 | 0 | 0 | 0 | 0 | 0 | 0 | 0 | 0 |
| 0 | NA | NA | NA | NA | NA | Olfr512 | olfactory receptor 512 | 0 | 0 | 0 | 0 | 0 | 0 | 0 | 0 | 0 | 0 | 0 |
| 0 | NA | NA | NA | NA | NA | Olfr513 | olfactory receptor 513 | 0 | 0 | 0 | 0 | 0 | 0 | 0 | 0 | 0 | 0 | 0 |
| 0 | NA | NA | NA | NA | NA | Gm45232 | | 0 | 0 | 0 | 0 | 0 | 0 | 0 | 0 | 0 | 0 | 0 |
| 0 | NA | NA | NA | NA | NA | Olfr514 | olfactory receptor 514 | 0 | 0 | 0 | 0 | 0 | 0 | 0 | 0 | 0 | 0 | 0 |
| 0 | NA | NA | NA | NA | NA | Olfr515-ps1 | | 0 | 0 | 0 | 0 | 0 | 0 | 0 | 0 | 0 | 0 | 0 |
| 0 | NA | NA | NA | NA | NA | Olfr516 | olfactory receptor 516 | 0 | 0 | 0 | 0 | 0 | 0 | 0 | 0 | 0 | 0 | 0 |
| 0 | NA | NA | NA | NA | NA | Gm17992 | | 0 | 0 | 0 | 0 | 0 | 0 | 0 | 0 | 0 | 0 | 0 |
| 0 | NA | NA | NA | NA | NA | Olfr517 | olfactory receptor 517 | 0 | 0 | 0 | 0 | 0 | 0 | 0 | 0 | 0 | 0 | 0 |
| 0 | NA | NA | NA | NA | NA | Olfr518 | olfactory receptor 518 | 0 | 0 | 0 | 0 | 0 | 0 | 0 | 0 | 0 | 0 | 0 |
| 0 | NA | NA | NA | NA | NA | Gm44781 | | 0 | 0 | 0 | 0 | 0 | 0 | 0 | 0 | 0 | 0 | 0 |
| 0 | NA | NA | NA | NA | NA | Snora3 | | 0 | 0 | 0 | 0 | 0 | 0 | 0 | 0 | 0 | 0 | 0 |
| 0 | NA | NA | NA | NA | NA | Gm44864 | | 0 | 0 | 0 | 0 | 0 | 0 | 0 | 0 | 0 | 0 | 0 |
| 0 | NA | NA | NA | NA | NA | Gm22822 | | 0 | 0 | 0 | 0 | 0 | 0 | 0 | 0 | 0 | 0 | 0 |
| 0 | NA | NA | NA | NA | NA | Gm24842 | | 0 | 0 | 0 | 0 | 0 | 0 | 0 | 0 | 0 | 0 | 0 |
| 0 | NA | NA | NA | NA | NA | Gm23100 | | 0 | 0 | 0 | 0 | 0 | 0 | 0 | 0 | 0 | 0 | 0 |
| 0 | NA | NA | NA | NA | NA | Gm25636 | | 0 | 0 | 0 | 0 | 0 | 0 | 0 | 0 | 0 | 0 | 0 |
| 0 | NA | NA | NA | NA | NA | Gm45819 | | 0 | 0 | 0 | 0 | 0 | 0 | 0 | 0 | 0 | 0 | 0 |
| 0 | NA | NA | NA | NA | NA | Gm45515 | | 0 | 0 | 0 | 0 | 0 | 0 | 0 | 0 | 0 | 0 | 0 |

| | | | | | | | | | | | | | | | | | | |
|---|---|---|---|---|---|---|---|---|---|---|---|---|---|---|---|---|---|---|
| 0 | NA | NA | NA | NA | NA | Gm9064 | | 0 | 0 | 0 | 0 | 0 | 0 | 0 | 0 | 0 | 0 | 0 |
| 0 | NA | NA | NA | NA | NA | Gm17219 | | 0 | 0 | 0 | 0 | 0 | 0 | 0 | 0 | 0 | 0 | 0 |
| 0 | NA | NA | NA | NA | NA | Gm10087 | | 0 | 0 | 0 | 0 | 0 | 0 | 0 | 0 | 0 | 0 | 0 |
| 0 | NA | NA | NA | NA | NA | Gm28863 | | 0 | 0 | 0 | 0 | 0 | 0 | 0 | 0 | 0 | 0 | 0 |
| 0 | NA | NA | NA | NA | NA | Gm21123 | | 0 | 0 | 0 | 0 | 0 | 0 | 0 | 0 | 0 | 0 | 0 |
| 0 | NA | NA | NA | NA | NA | Gm18907 | | 0 | 0 | 0 | 0 | 0 | 0 | 0 | 0 | 0 | 0 | 0 |
| 0 | NA | NA | NA | NA | NA | B430319F04Rik | | 0 | 0 | 0 | 0 | 0 | 0 | 0 | 0 | 0 | 0 | 0 |
| 0 | NA | NA | NA | NA | NA | Gm47282 | | 0 | 0 | 0 | 0 | 0 | 0 | 0 | 0 | 0 | 0 | 0 |
| 0 | NA | NA | NA | NA | NA | Gm16336 | | 0 | 0 | 0 | 0 | 0 | 0 | 0 | 0 | 0 | 0 | 0 |
| 0 | NA | NA | NA | NA | NA | 5430402P08Rik | | 0 | 0 | 0 | 0 | 0 | 0 | 0 | 0 | 0 | 0 | 0 |
| 0 | NA | NA | NA | NA | NA | Gm25787 | | 0 | 0 | 0 | 0 | 0 | 0 | 0 | 0 | 0 | 0 | 0 |
| 0 | NA | NA | NA | NA | NA | Gm45588 | | 0 | 0 | 0 | 0 | 0 | 0 | 0 | 0 | 0 | 0 | 0 |
| 0 | NA | NA | NA | NA | NA | Gm23345 | | 0 | 0 | 0 | 0 | 0 | 0 | 0 | 0 | 0 | 0 | 0 |
| 0 | NA | NA | NA | NA | NA | Gm24154 | | 0 | 0 | 0 | 0 | 0 | 0 | 0 | 0 | 0 | 0 | 0 |
| 0 | NA | NA | NA | NA | NA | Gm44462 | | 0 | 0 | 0 | 0 | 0 | 0 | 0 | 0 | 0 | 0 | 0 |
| 0 | NA | NA | NA | NA | NA | 4930543E12Rik | | 0 | 0 | 0 | 0 | 0 | 0 | 0 | 0 | 0 | 0 | 0 |
| 0 | NA | NA | NA | NA | NA | Gm23662 | | 0 | 0 | 0 | 0 | 0 | 0 | 0 | 0 | 0 | 0 | 0 |
| 0 | NA | NA | NA | NA | NA | Gm45355 | | 0 | 0 | 0 | 0 | 0 | 0 | 0 | 0 | 0 | 0 | 0 |
| 0 | NA | NA | NA | NA | NA | Gm15994 | | 0 | 0 | 0 | 0 | 0 | 0 | 0 | 0 | 0 | 0 | 0 |
| 0 | NA | NA | NA | NA | NA | Pth | parathyroid hormone | 0 | 0 | 0 | 0 | 0 | 0 | 0 | 0 | 0 | 0 | 0 |
| 0 | NA | NA | NA | NA | NA | Gm45614 | | 0 | 0 | 0 | 0 | 0 | 0 | 0 | 0 | 0 | 0 | 0 |
| 0 | NA | NA | NA | NA | NA | Gm18780 | | 0 | 0 | 0 | 0 | 0 | 0 | 0 | 0 | 0 | 0 | 0 |
| 0 | NA | NA | NA | NA | NA | Gm45612 | | 0 | 0 | 0 | 0 | 0 | 0 | 0 | 0 | 0 | 0 | 0 |
| 0 | NA | NA | NA | NA | NA | Gm23700 | | 0 | 0 | 0 | 0 | 0 | 0 | 0 | 0 | 0 | 0 | 0 |
| 0 | NA | NA | NA | NA | NA | Gm5600 | | 0 | 0 | 0 | 0 | 0 | 0 | 0 | 0 | 0 | 0 | 0 |
| 0 | NA | NA | NA | NA | NA | Gm29507 | | 0 | 0 | 0 | 0 | 0 | 0 | 0 | 0 | 0 | 0 | 0 |
| 0 | NA | NA | NA | NA | NA | Gm45615 | | 0 | 0 | 0 | 0 | 0 | 0 | 0 | 0 | 0 | 0 | 0 |
| 0 | NA | NA | NA | NA | NA | Gm18599 | | 0 | 0 | 0 | 0 | 0 | 0 | 0 | 0 | 0 | 0 | 0 |
| 0 | NA | NA | NA | NA | NA | Gm24982 | | 0 | 0 | 0 | 0 | 0 | 0 | 0 | 0 | 0 | 0 | 0 |
| 0 | NA | NA | NA | NA | NA | Gm45017 | | 0 | 0 | 0 | 0 | 0 | 0 | 0 | 0 | 0 | 0 | 0 |
| 0 | NA | NA | NA | NA | NA | Gm22669 | | 0 | 0 | 0 | 0 | 0 | 0 | 0 | 0 | 0 | 0 | 0 |
| 0 | NA | NA | NA | NA | NA | Gm45020 | | 0 | 0 | 0 | 0 | 0 | 0 | 0 | 0 | 0 | 0 | 0 |
| 0 | NA | NA | NA | NA | NA | Gm27684 | | 0 | 0 | 0 | 0 | 0 | 0 | 0 | 0 | 0 | 0 | 0 |
| 0 | NA | NA | NA | NA | NA | Gm45025 | | 0 | 0 | 0 | 0 | 0 | 0 | 0 | 0 | 0 | 0 | 0 |
| 0 | NA | NA | NA | NA | NA | Sox6os | | 0 | 0 | 0 | 0 | 0 | 0 | 0 | 0 | 0 | 0 | 0 |
| 0 | NA | NA | NA | NA | NA | Gm44867 | | 0 | 0 | 0 | 0 | 0 | 0 | 0 | 0 | 0 | 0 | 0 |
| 0 | NA | NA | NA | NA | NA | Snord14a | | 0 | 0 | 0 | 0 | 0 | 0 | 0 | 0 | 0 | 0 | 0 |
| 0 | NA | NA | NA | NA | NA | Gm45031 | | 0 | 0 | 0 | 0 | 0 | 0 | 0 | 0 | 0 | 0 | 0 |
| 0 | NA | NA | NA | NA | NA | n-R5s157 | | 0 | 0 | 0 | 0 | 0 | 0 | 0 | 0 | 0 | 0 | 0 |
| 0 | NA | NA | NA | NA | NA | Gm25423 | | 0 | 0 | 0 | 0 | 0 | 0 | 0 | 0 | 0 | 0 | 0 |
| 0 | NA | NA | NA | NA | NA | Gm6008 | | 0 | 0 | 0 | 0 | 0 | 0 | 0 | 0 | 0 | 0 | 0 |
| 0 | NA | NA | NA | NA | NA | Gm26054 | | 0 | 0 | 0 | 0 | 0 | 0 | 0 | 0 | 0 | 0 | 0 |
| 0 | NA | NA | NA | NA | NA | Gm39075 | | 0 | 0 | 0 | 0 | 0 | 0 | 0 | 0 | 0 | 0 | 0 |
| 0 | NA | NA | NA | NA | NA | Gm45153 | | 0 | 0 | 0 | 0 | 0 | 0 | 0 | 0 | 0 | 0 | 0 |
| 0 | NA | NA | NA | NA | NA | Gm24063 | | 0 | 0 | 0 | 0 | 0 | 0 | 0 | 0 | 0 | 0 | 0 |
| 0 | NA | NA | NA | NA | NA | Gm25683 | | 0 | 0 | 0 | 0 | 0 | 0 | 0 | 0 | 0 | 0 | 0 |
| 0 | NA | NA | NA | NA | NA | Gm23229 | | 0 | 0 | 0 | 0 | 0 | 0 | 0 | 0 | 0 | 0 | 0 |
| 0 | NA | NA | NA | NA | NA | Gm35309 | | 0 | 0 | 0 | 0 | 0 | 0 | 0 | 0 | 0 | 0 | 0 |
| 0 | NA | NA | NA | NA | NA | Gm32766 | | 0 | 0 | 0 | 0 | 0 | 0 | 0 | 0 | 0 | 0 | 0 |
| 0 | NA | NA | NA | NA | NA | Gm44652 | | 0 | 0 | 0 | 0 | 0 | 0 | 0 | 0 | 0 | 0 | 0 |
| 0 | NA | NA | NA | NA | NA | B230311B06Ri | RIKEN cDNA B230311B06 gene | 0 | 0 | 0 | 0 | 0 | 0 | 0 | 0 | 0 | 0 | 0 |
| 0 | NA | NA | NA | NA | NA | Gm45236 | | 0 | 0 | 0 | 0 | 0 | 0 | 0 | 0 | 0 | 0 | 0 |
| 0 | NA | NA | NA | NA | NA | 1700016B15Rik | | 0 | 0 | 0 | 0 | 0 | 0 | 0 | 0 | 0 | 0 | 0 |
| 0 | NA | NA | NA | NA | NA | Gpr139 | G protein-coupled receptor 139 | 0 | 0 | 0 | 0 | 0 | 0 | 0 | 0 | 0 | 0 | 0 |
| 0 | NA | NA | NA | NA | NA | Gm4083 | | 0 | 0 | 0 | 0 | 0 | 0 | 0 | 0 | 0 | 0 | 0 |
| 0 | NA | NA | NA | NA | NA | Gm23321 | | 0 | 0 | 0 | 0 | 0 | 0 | 0 | 0 | 0 | 0 | 0 |
| 0 | NA | NA | NA | NA | NA | Gm44588 | | 0 | 0 | 0 | 0 | 0 | 0 | 0 | 0 | 0 | 0 | 0 |
| 0 | NA | NA | NA | NA | NA | Pdilt | protein disulfide isomerase-like, testis exp | 0 | 0 | 0 | 0 | 0 | 0 | 0 | 0 | 0 | 0 | 0 |
| 0 | NA | NA | NA | NA | NA | Gm37518 | | 0 | 0 | 0 | 0 | 0 | 0 | 0 | 0 | 0 | 0 | 0 |
| 0 | NA | NA | NA | NA | NA | Gm26967 | | 0 | 0 | 0 | 0 | 0 | 0 | 0 | 0 | 0 | 0 | 0 |

| | | | | | | | | | | | | | | | | | | |
|---|---|---|---|---|---|---|---|---|---|---|---|---|---|---|---|---|---|---|
| 0 | NA | NA | NA | NA | NA | Acsm2 | acyl-CoA synthetase medium-chain famil | 0 | 0 | 0 | 0 | 0 | 0 | 0 | 0 | 0 | 0 | 0 |
| 0 | NA | NA | NA | NA | NA | Gm39078 | | 0 | 0 | 0 | 0 | 0 | 0 | 0 | 0 | 0 | 0 | 0 |
| 0 | NA | NA | NA | NA | NA | Gm19950 | | 0 | 0 | 0 | 0 | 0 | 0 | 0 | 0 | 0 | 0 | 0 |
| 0 | NA | NA | NA | NA | NA | Gm45792 | | 0 | 0 | 0 | 0 | 0 | 0 | 0 | 0 | 0 | 0 | 0 |
| 0 | NA | NA | NA | NA | NA | Acsm4 | acyl-CoA synthetase medium-chain famil | 0 | 0 | 0 | 0 | 0 | 0 | 0 | 0 | 0 | 0 | 0 |
| 0 | NA | NA | NA | NA | NA | Gm44866 | | 0 | 0 | 0 | 0 | 0 | 0 | 0 | 0 | 0 | 0 | 0 |
| 0 | NA | NA | NA | NA | NA | Gm19963 | | 0 | 0 | 0 | 0 | 0 | 0 | 0 | 0 | 0 | 0 | 0 |
| 0 | NA | NA | NA | NA | NA | Gm44966 | | 0 | 0 | 0 | 0 | 0 | 0 | 0 | 0 | 0 | 0 | 0 |
| 0 | NA | NA | NA | NA | NA | Gm25217 | | 0 | 0 | 0 | 0 | 0 | 0 | 0 | 0 | 0 | 0 | 0 |
| 0 | NA | NA | NA | NA | NA | 4930505K13Rik | | 0 | 0 | 0 | 0 | 0 | 0 | 0 | 0 | 0 | 0 | 0 |
| 0 | NA | NA | NA | NA | NA | Abca15 | ATP-binding cassette, sub-family A (ABC | 0 | 0 | 0 | 0 | 0 | 0 | 0 | 0 | 0 | 0 | 0 |
| 0 | NA | NA | NA | NA | NA | Abca16 | ATP-binding cassette, sub-family A (ABC | 0 | 0 | 0 | 0 | 0 | 0 | 0 | 0 | 0 | 0 | 0 |
| 0 | NA | NA | NA | NA | NA | Gm16475 | | 0 | 0 | 0 | 0 | 0 | 0 | 0 | 0 | 0 | 0 | 0 |
| 0 | NA | NA | NA | NA | NA | Gm32916 | ankyrin repeat domain-containing protein | 0 | 0 | 0 | 0 | 0 | 0 | 0 | 0 | 0 | 0 | 0 |
| 0 | NA | NA | NA | NA | NA | 4930588G17Rik | | 0 | 0 | 0 | 0 | 0 | 0 | 0 | 0 | 0 | 0 | 0 |
| 0 | NA | NA | NA | NA | NA | Gm15774 | | 0 | 0 | 0 | 0 | 0 | 0 | 0 | 0 | 0 | 0 | 0 |
| 0 | NA | NA | NA | NA | NA | Gm25924 | | 0 | 0 | 0 | 0 | 0 | 0 | 0 | 0 | 0 | 0 | 0 |
| 0 | NA | NA | NA | NA | NA | Gm27991 | | 0 | 0 | 0 | 0 | 0 | 0 | 0 | 0 | 0 | 0 | 0 |
| 0 | NA | NA | NA | NA | NA | Gm45079 | | 0 | 0 | 0 | 0 | 0 | 0 | 0 | 0 | 0 | 0 | 0 |
| 0 | NA | NA | NA | NA | NA | 4930486N12Rik | | 0 | 0 | 0 | 0 | 0 | 0 | 0 | 0 | 0 | 0 | 0 |
| 0 | NA | NA | NA | NA | NA | Gm36736 | | 0 | 0 | 0 | 0 | 0 | 0 | 0 | 0 | 0 | 0 | 0 |
| 0 | NA | NA | NA | NA | NA | Gm44740 | | 0 | 0 | 0 | 0 | 0 | 0 | 0 | 0 | 0 | 0 | 0 |
| 0 | NA | NA | NA | NA | NA | Gm16326 | | 0 | 0 | 0 | 0 | 0 | 0 | 0 | 0 | 0 | 0 | 0 |
| 0 | NA | NA | NA | NA | NA | Gm29977 | | 0 | 0 | 0 | 0 | 0 | 0 | 0 | 0 | 0 | 0 | 0 |
| 0 | NA | NA | NA | NA | NA | Gm23614 | | 0 | 0 | 0 | 0 | 0 | 0 | 0 | 0 | 0 | 0 | 0 |
| 0 | NA | NA | NA | NA | NA | Gm44745 | | 0 | 0 | 0 | 0 | 0 | 0 | 0 | 0 | 0 | 0 | 0 |
| 0 | NA | NA | NA | NA | NA | Gm14389 | | 0 | 0 | 0 | 0 | 0 | 0 | 0 | 0 | 0 | 0 | 0 |
| 0 | NA | NA | NA | NA | NA | Gm44749 | | 0 | 0 | 0 | 0 | 0 | 0 | 0 | 0 | 0 | 0 | 0 |
| 0 | NA | NA | NA | NA | NA | Gm29759 | | 0 | 0 | 0 | 0 | 0 | 0 | 0 | 0 | 0 | 0 | 0 |
| 0 | NA | NA | NA | NA | NA | Slc5a11 | solute carrier family 5 (sodium/glucose cc | 0 | 0 | 0 | 0 | 0 | 0 | 0 | 0 | 0 | 0 | 0 |
| 0 | NA | NA | NA | NA | NA | Gm23559 | | 0 | 0 | 0 | 0 | 0 | 0 | 0 | 0 | 0 | 0 | 0 |
| 0 | NA | NA | NA | NA | NA | Gm23788 | | 0 | 0 | 0 | 0 | 0 | 0 | 0 | 0 | 0 | 0 | 0 |
| 0 | NA | NA | NA | NA | NA | Gm26974 | | 0 | 0 | 0 | 0 | 0 | 0 | 0 | 0 | 0 | 0 | 0 |
| 0 | NA | NA | NA | NA | NA | Gm27040 | | 0 | 0 | 0 | 0 | 0 | 0 | 0 | 0 | 0 | 0 | 0 |
| 0 | NA | NA | NA | NA | NA | Gm25759 | | 0 | 0 | 0 | 0 | 0 | 0 | 0 | 0 | 0 | 0 | 0 |
| 0 | NA | NA | NA | NA | NA | Gm15338 | | 0 | 0 | 0 | 0 | 0 | 0 | 0 | 0 | 0 | 0 | 0 |
| 0 | NA | NA | NA | NA | NA | Gm15339 | | 0 | 0 | 0 | 0 | 0 | 0 | 0 | 0 | 0 | 0 | 0 |
| 0 | NA | NA | NA | NA | NA | Gm23168 | | 0 | 0 | 0 | 0 | 0 | 0 | 0 | 0 | 0 | 0 | 0 |
| 0 | NA | NA | NA | NA | NA | 3100003L05Rik | | 0 | 0 | 0 | 0 | 0 | 0 | 0 | 0 | 0 | 0 | 0 |
| 0 | NA | NA | NA | NA | NA | Gm45092 | | 0 | 0 | 0 | 0 | 0 | 0 | 0 | 0 | 0 | 0 | 0 |
| 0 | NA | NA | NA | NA | NA | Gm45093 | | 0 | 0 | 0 | 0 | 0 | 0 | 0 | 0 | 0 | 0 | 0 |
| 0 | NA | NA | NA | NA | NA | Gm21957 | | 0 | 0 | 0 | 0 | 0 | 0 | 0 | 0 | 0 | 0 | 0 |
| 0 | NA | NA | NA | NA | NA | 4930533L02Rik | | 0 | 0 | 0 | 0 | 0 | 0 | 0 | 0 | 0 | 0 | 0 |
| 0 | NA | NA | NA | NA | NA | 4930571K23Ril | RIKEN cDNA 4930571K23 gene | 0 | 0 | 0 | 0 | 0 | 0 | 0 | 0 | 0 | 0 | 0 |
| 0 | NA | NA | NA | NA | NA | Gm44876 | | 0 | 0 | 0 | 0 | 0 | 0 | 0 | 0 | 0 | 0 | 0 |
| 0 | NA | NA | NA | NA | NA | Gm44874 | | 0 | 0 | 0 | 0 | 0 | 0 | 0 | 0 | 0 | 0 | 0 |
| 0 | NA | NA | NA | NA | NA | Gm17137 | | 0 | 0 | 0 | 0 | 0 | 0 | 0 | 0 | 0 | 0 | 0 |
| 0 | NA | NA | NA | NA | NA | 1700123J17Rik | | 0 | 0 | 0 | 0 | 0 | 0 | 0 | 0 | 0 | 0 | 0 |
| 0 | NA | NA | NA | NA | NA | Gm44703 | | 0 | 0 | 0 | 0 | 0 | 0 | 0 | 0 | 0 | 0 | 0 |
| 0 | NA | NA | NA | NA | NA | Gm30928 | | 0 | 0 | 0 | 0 | 0 | 0 | 0 | 0 | 0 | 0 | 0 |
| 0 | NA | NA | NA | NA | NA | Mir7058 | | 0 | 0 | 0 | 0 | 0 | 0 | 0 | 0 | 0 | 0 | 0 |
| 0 | NA | NA | NA | NA | NA | Gm45141 | | 0 | 0 | 0 | 0 | 0 | 0 | 0 | 0 | 0 | 0 | 0 |
| 0 | NA | NA | NA | NA | NA | Gm25579 | | 0 | 0 | 0 | 0 | 0 | 0 | 0 | 0 | 0 | 0 | 0 |
| 0 | NA | NA | NA | NA | NA | 4930448A20Rik | | 0 | 0 | 0 | 0 | 0 | 0 | 0 | 0 | 0 | 0 | 0 |
| 0 | NA | NA | NA | NA | NA | Gm44824 | | 0 | 0 | 0 | 0 | 0 | 0 | 0 | 0 | 0 | 0 | 0 |
| 0 | NA | NA | NA | NA | NA | Gm44856 | | 0 | 0 | 0 | 0 | 0 | 0 | 0 | 0 | 0 | 0 | 0 |
| 0 | NA | NA | NA | NA | NA | 4930451I11Rik | RIKEN cDNA 4930451I11 gene | 0 | 0 | 0 | 0 | 0 | 0 | 0 | 0 | 0 | 0 | 0 |
| 0 | NA | NA | NA | NA | NA | Gm44854 | | 0 | 0 | 0 | 0 | 0 | 0 | 0 | 0 | 0 | 0 | 0 |
| 0 | NA | NA | NA | NA | NA | Pagr1b | | 0 | 0 | 0 | 0 | 0 | 0 | 0 | 0 | 0 | 0 | 0 |
| 0 | NA | NA | NA | NA | NA | Pagr1a | | 0 | 0 | 0 | 0 | 0 | 0 | 0 | 0 | 0 | 0 | 0 |

| | | | | | | | | | | | | | | | | | | |
|---|---|---|---|---|---|---|---|---|---|---|---|---|---|---|---|---|---|---|
| 0 | NA | NA | NA | NA | NA | Gm25333 | | 0 | 0 | 0 | 0 | 0 | 0 | 0 | 0 | 0 | 0 | 0 |
| 0 | NA | NA | NA | NA | NA | Gm44343 | | 0 | 0 | 0 | 0 | 0 | 0 | 0 | 0 | 0 | 0 | 0 |
| 0 | NA | NA | NA | NA | NA | Gm44939 | | 0 | 0 | 0 | 0 | 0 | 0 | 0 | 0 | 0 | 0 | 0 |
| 0 | NA | NA | NA | NA | NA | Gm22747 | | 0 | 0 | 0 | 0 | 0 | 0 | 0 | 0 | 0 | 0 | 0 |
| 0 | NA | NA | NA | NA | NA | Gm44940 | | 0 | 0 | 0 | 0 | 0 | 0 | 0 | 0 | 0 | 0 | 0 |
| 0 | NA | NA | NA | NA | NA | Gm25735 | | 0 | 0 | 0 | 0 | 0 | 0 | 0 | 0 | 0 | 0 | 0 |
| 0 | NA | NA | NA | NA | NA | Gm33080 | | 0 | 0 | 0 | 0 | 0 | 0 | 0 | 0 | 0 | 0 | 0 |
| 0 | NA | NA | NA | NA | NA | Mir7060 | | 0 | 0 | 0 | 0 | 0 | 0 | 0 | 0 | 0 | 0 | 0 |
| 0 | NA | NA | NA | NA | NA | Gm47278 | | 0 | 0 | 0 | 0 | 0 | 0 | 0 | 0 | 0 | 0 | 0 |
| 0 | NA | NA | NA | NA | NA | Tmem265 | transmembrane protein 265 | 0 | 0 | 0 | 0 | 0 | 0 | 0 | 0 | 0 | 0 | 0 |
| 0 | NA | NA | NA | NA | NA | Ctf2 | cardiotrophin 2 | 0 | 0 | 0 | 0 | 0 | 0 | 0 | 0 | 0 | 0 | 0 |
| 0 | NA | NA | NA | NA | NA | Gm49388 | | 0 | 0 | 0 | 0 | 0 | 0 | 0 | 0 | 0 | 0 | 0 |
| 0 | NA | NA | NA | NA | NA | Gm21974 | | 0 | 0 | 0 | 0 | 0 | 0 | 0 | 0 | 0 | 0 | 0 |
| 0 | NA | NA | NA | NA | NA | 2200002A13Rik | | 0 | 0 | 0 | 0 | 0 | 0 | 0 | 0 | 0 | 0 | 0 |
| 0 | NA | NA | NA | NA | NA | Gm15533 | | 0 | 0 | 0 | 0 | 0 | 0 | 0 | 0 | 0 | 0 | 0 |
| 0 | NA | NA | NA | NA | NA | Gm22809 | | 0 | 0 | 0 | 0 | 0 | 0 | 0 | 0 | 0 | 0 | 0 |
| 0 | NA | NA | NA | NA | NA | Gm49368 | | 0 | 0 | 0 | 0 | 0 | 0 | 0 | 0 | 0 | 0 | 0 |
| 0 | NA | NA | NA | NA | NA | Gm45700 | | 0 | 0 | 0 | 0 | 0 | 0 | 0 | 0 | 0 | 0 | 0 |
| 0 | NA | NA | NA | NA | NA | Gm17831 | | 0 | 0 | 0 | 0 | 0 | 0 | 0 | 0 | 0 | 0 | 0 |
| 0 | NA | NA | NA | NA | NA | Gm39091 | | 0 | 0 | 0 | 0 | 0 | 0 | 0 | 0 | 0 | 0 | 0 |
| 0 | NA | NA | NA | NA | NA | Mir3103 | | 0 | 0 | 0 | 0 | 0 | 0 | 0 | 0 | 0 | 0 | 0 |
| 0 | NA | NA | NA | NA | NA | Gm44650 | | 0 | 0 | 0 | 0 | 0 | 0 | 0 | 0 | 0 | 0 | 0 |
| 0 | NA | NA | NA | NA | NA | Gm24365 | | 0 | 0 | 0 | 0 | 0 | 0 | 0 | 0 | 0 | 0 | 0 |
| 0 | NA | NA | NA | NA | NA | Gm40457 | | 0 | 0 | 0 | 0 | 0 | 0 | 0 | 0 | 0 | 0 | 0 |
| 0 | NA | NA | NA | NA | NA | Gm7258 | | 0 | 0 | 0 | 0 | 0 | 0 | 0 | 0 | 0 | 0 | 0 |
| 0 | NA | NA | NA | NA | NA | Gm43580 | | 0 | 0 | 0 | 0 | 0 | 0 | 0 | 0 | 0 | 0 | 0 |
| 0 | NA | NA | NA | NA | NA | Gm44674 | | 0 | 0 | 0 | 0 | 0 | 0 | 0 | 0 | 0 | 0 | 0 |
| 0 | NA | NA | NA | NA | NA | Gm23783 | | 0 | 0 | 0 | 0 | 0 | 0 | 0 | 0 | 0 | 0 | 0 |
| 0 | NA | NA | NA | NA | NA | n-R5s158 | | 0 | 0 | 0 | 0 | 0 | 0 | 0 | 0 | 0 | 0 | 0 |
| 0 | NA | NA | NA | NA | NA | Gm32884 | | 0 | 0 | 0 | 0 | 0 | 0 | 0 | 0 | 0 | 0 | 0 |
| 0 | NA | NA | NA | NA | NA | Gm25778 | | 0 | 0 | 0 | 0 | 0 | 0 | 0 | 0 | 0 | 0 | 0 |
| 0 | NA | NA | NA | NA | NA | Gm4259 | | 0 | 0 | 0 | 0 | 0 | 0 | 0 | 0 | 0 | 0 | 0 |
| 0 | NA | NA | NA | NA | NA | Gm33027 | | 0 | 0 | 0 | 0 | 0 | 0 | 0 | 0 | 0 | 0 | 0 |
| 0 | NA | NA | NA | NA | NA | Gm39094 | | 0 | 0 | 0 | 0 | 0 | 0 | 0 | 0 | 0 | 0 | 0 |
| 0 | NA | NA | NA | NA | NA | Gm33248 | | 0 | 0 | 0 | 0 | 0 | 0 | 0 | 0 | 0 | 0 | 0 |
| 0 | NA | NA | NA | NA | NA | Gm44778 | | 0 | 0 | 0 | 0 | 0 | 0 | 0 | 0 | 0 | 0 | 0 |
| 0 | NA | NA | NA | NA | NA | Gm4265 | | 0 | 0 | 0 | 0 | 0 | 0 | 0 | 0 | 0 | 0 | 0 |
| 0 | NA | NA | NA | NA | NA | Gm23847 | | 0 | 0 | 0 | 0 | 0 | 0 | 0 | 0 | 0 | 0 | 0 |
| 0 | NA | NA | NA | NA | NA | Gm27884 | | 0 | 0 | 0 | 0 | 0 | 0 | 0 | 0 | 0 | 0 | 0 |
| 0 | NA | NA | NA | NA | NA | 4930513N20Rik | | 0 | 0 | 0 | 0 | 0 | 0 | 0 | 0 | 0 | 0 | 0 |
| 0 | NA | NA | NA | NA | NA | Etos1 | | 0 | 0 | 0 | 0 | 0 | 0 | 0 | 0 | 0 | 0 | 0 |
| 0 | NA | NA | NA | NA | NA | 4933402N03Ril | RIKEN cDNA 4933402N03 gene | 0 | 0 | 0 | 0 | 0 | 0 | 0 | 0 | 0 | 0 | 0 |
| 0 | NA | NA | NA | NA | NA | 1700029B22Rik | | 0 | 0 | 0 | 0 | 0 | 0 | 0 | 0 | 0 | 0 | 0 |
| 0 | NA | NA | NA | NA | NA | | | 0 | 0 | 0 | 0 | 0 | 0 | 0 | 0 | 0 | 0 | 0 |
| 0 | NA | NA | NA | NA | NA | Fam24b | family with sequence similarity 24 membe | 0 | 0 | 0 | 0 | 0 | 0 | 0 | 0 | 0 | 0 | 0 |
| 0 | NA | NA | NA | NA | NA | Fam24a | family with sequence similarity 24, memb | 0 | 0 | 0 | 0 | 0 | 0 | 0 | 0 | 0 | 0 | 0 |
| 0 | NA | NA | NA | NA | NA | Gm44984 | | 0 | 0 | 0 | 0 | 0 | 0 | 0 | 0 | 0 | 0 | 0 |
| 0 | NA | NA | NA | NA | NA | Gm45105 | | 0 | 0 | 0 | 0 | 0 | 0 | 0 | 0 | 0 | 0 | 0 |
| 0 | NA | NA | NA | NA | NA | Hmx3 | H6 homeobox 3 | 0 | 0 | 0 | 0 | 0 | 0 | 0 | 0 | 0 | 0 | 0 |
| 0 | NA | NA | NA | NA | NA | Gm4585 | | 0 | 0 | 0 | 0 | 0 | 0 | 0 | 0 | 0 | 0 | 0 |
| 0 | NA | NA | NA | NA | NA | Gm44547 | | 0 | 0 | 0 | 0 | 0 | 0 | 0 | 0 | 0 | 0 | 0 |
| 0 | NA | NA | NA | NA | NA | Gm44546 | | 0 | 0 | 0 | 0 | 0 | 0 | 0 | 0 | 0 | 0 | 0 |
| 0 | NA | NA | NA | NA | NA | Gm44548 | | 0 | 0 | 0 | 0 | 0 | 0 | 0 | 0 | 0 | 0 | 0 |
| 0 | NA | NA | NA | NA | NA | Gpr26 | G protein-coupled receptor 26 | 0 | 0 | 0 | 0 | 0 | 0 | 0 | 0 | 0 | 0 | 0 |
| 0 | NA | NA | NA | NA | NA | n-R5s159 | | 0 | 0 | 0 | 0 | 0 | 0 | 0 | 0 | 0 | 0 | 0 |
| 0 | NA | NA | NA | NA | NA | Gm15677 | | 0 | 0 | 0 | 0 | 0 | 0 | 0 | 0 | 0 | 0 | 0 |
| 0 | NA | NA | NA | NA | NA | Gm44892 | | 0 | 0 | 0 | 0 | 0 | 0 | 0 | 0 | 0 | 0 | 0 |
| 0 | NA | NA | NA | NA | NA | Gm34908 | | 0 | 0 | 0 | 0 | 0 | 0 | 0 | 0 | 0 | 0 | 0 |
| 0 | NA | NA | NA | NA | NA | Gm44893 | | 0 | 0 | 0 | 0 | 0 | 0 | 0 | 0 | 0 | 0 | 0 |
| 0 | NA | NA | NA | NA | NA | Gm35147 | | 0 | 0 | 0 | 0 | 0 | 0 | 0 | 0 | 0 | 0 | 0 |

| | | | | | | | | | | | | | | | | | | |
|---|---|---|---|---|---|---|---|---|---|---|---|---|---|---|---|---|---|---|
| 0 | NA | NA | NA | NA | NA | Gm4587 | | 0 | 0 | 0 | 0 | 0 | 0 | 0 | 0 | 0 | 0 | 0 |
| 0 | NA | NA | NA | NA | NA | Nkx1-2 | NK1 homeobox 2 | 0 | 0 | 0 | 0 | 0 | 0 | 0 | 0 | 0 | 0 | 0 |
| 0 | NA | NA | NA | NA | NA | Gm16764 | | 0 | 0 | 0 | 0 | 0 | 0 | 0 | 0 | 0 | 0 | 0 |
| 0 | NA | NA | NA | NA | NA | Gm15582 | | 0 | 0 | 0 | 0 | 0 | 0 | 0 | 0 | 0 | 0 | 0 |
| 0 | NA | NA | NA | NA | NA | Gm45502 | | 0 | 0 | 0 | 0 | 0 | 0 | 0 | 0 | 0 | 0 | 0 |
| 0 | NA | NA | NA | NA | NA | Gm45672 | | 0 | 0 | 0 | 0 | 0 | 0 | 0 | 0 | 0 | 0 | 0 |
| 0 | NA | NA | NA | NA | NA | Gm35625 | | 0 | 0 | 0 | 0 | 0 | 0 | 0 | 0 | 0 | 0 | 0 |
| 0 | NA | NA | NA | NA | NA | Gm45671 | | 0 | 0 | 0 | 0 | 0 | 0 | 0 | 0 | 0 | 0 | 0 |
| 0 | NA | NA | NA | NA | NA | Gm28747 | | 0 | 0 | 0 | 0 | 0 | 0 | 0 | 0 | 0 | 0 | 0 |
| 0 | NA | NA | NA | NA | NA | Mmp21 | matrix metallopeptidase 21 | 0 | 0 | 0 | 0 | 0 | 0 | 0 | 0 | 0 | 0 | 0 |
| 0 | NA | NA | NA | NA | NA | Gm23631 | | 0 | 0 | 0 | 0 | 0 | 0 | 0 | 0 | 0 | 0 | 0 |
| 0 | NA | NA | NA | NA | NA | A130023I24Rik | | 0 | 0 | 0 | 0 | 0 | 0 | 0 | 0 | 0 | 0 | 0 |
| 0 | NA | NA | NA | NA | NA | Gm24581 | | 0 | 0 | 0 | 0 | 0 | 0 | 0 | 0 | 0 | 0 | 0 |
| 0 | NA | NA | NA | NA | NA | 1700120G07Rik | | 0 | 0 | 0 | 0 | 0 | 0 | 0 | 0 | 0 | 0 | 0 |
| 0 | NA | NA | NA | NA | NA | Nps | neuropeptide S | 0 | 0 | 0 | 0 | 0 | 0 | 0 | 0 | 0 | 0 | 0 |
| 0 | NA | NA | NA | NA | NA | Gm45339 | | 0 | 0 | 0 | 0 | 0 | 0 | 0 | 0 | 0 | 0 | 0 |
| 0 | NA | NA | NA | NA | NA | Foxi2 | forkhead box I2 | 0 | 0 | 0 | 0 | 0 | 0 | 0 | 0 | 0 | 0 | 0 |
| 0 | NA | NA | NA | NA | NA | Gm36356 | | 0 | 0 | 0 | 0 | 0 | 0 | 0 | 0 | 0 | 0 | 0 |
| 0 | NA | NA | NA | NA | NA | Mir1962 | | 0 | 0 | 0 | 0 | 0 | 0 | 0 | 0 | 0 | 0 | 0 |
| 0 | NA | NA | NA | NA | NA | Gm45240 | | 0 | 0 | 0 | 0 | 0 | 0 | 0 | 0 | 0 | 0 | 0 |
| 0 | NA | NA | NA | NA | NA | Gm36431 | | 0 | 0 | 0 | 0 | 0 | 0 | 0 | 0 | 0 | 0 | 0 |
| 0 | NA | NA | NA | NA | NA | Gm45241 | | 0 | 0 | 0 | 0 | 0 | 0 | 0 | 0 | 0 | 0 | 0 |
| 0 | NA | NA | NA | NA | NA | 6330420H09Rik | | 0 | 0 | 0 | 0 | 0 | 0 | 0 | 0 | 0 | 0 | 0 |
| 0 | NA | NA | NA | NA | NA | Gm45440 | | 0 | 0 | 0 | 0 | 0 | 0 | 0 | 0 | 0 | 0 | 0 |
| 0 | NA | NA | NA | NA | NA | Gm45439 | | 0 | 0 | 0 | 0 | 0 | 0 | 0 | 0 | 0 | 0 | 0 |
| 0 | NA | NA | NA | NA | NA | Gm36737 | | 0 | 0 | 0 | 0 | 0 | 0 | 0 | 0 | 0 | 0 | 0 |
| 0 | NA | NA | NA | NA | NA | C230079O03Rik | | 0 | 0 | 0 | 0 | 0 | 0 | 0 | 0 | 0 | 0 | 0 |
| 0 | NA | NA | NA | NA | NA | Gm36849 | | 0 | 0 | 0 | 0 | 0 | 0 | 0 | 0 | 0 | 0 | 0 |
| 0 | NA | NA | NA | NA | NA | Gm24444 | | 0 | 0 | 0 | 0 | 0 | 0 | 0 | 0 | 0 | 0 | 0 |
| 0 | NA | NA | NA | NA | NA | Gm6249 | | 0 | 0 | 0 | 0 | 0 | 0 | 0 | 0 | 0 | 0 | 0 |
| 0 | NA | NA | NA | NA | NA | Gm10578 | | 0 | 0 | 0 | 0 | 0 | 0 | 0 | 0 | 0 | 0 | 0 |
| 0 | NA | NA | NA | NA | NA | Gm45514 | | 0 | 0 | 0 | 0 | 0 | 0 | 0 | 0 | 0 | 0 | 0 |
| 0 | NA | NA | NA | NA | NA | Gm17841 | | 0 | 0 | 0 | 0 | 0 | 0 | 0 | 0 | 0 | 0 | 0 |
| 0 | NA | NA | NA | NA | NA | Gm45683 | | 0 | 0 | 0 | 0 | 0 | 0 | 0 | 0 | 0 | 0 | 0 |
| 0 | NA | NA | NA | NA | NA | Gm45682 | | 0 | 0 | 0 | 0 | 0 | 0 | 0 | 0 | 0 | 0 | 0 |
| 0 | NA | NA | NA | NA | NA | Gm5650 | | 0 | 0 | 0 | 0 | 0 | 0 | 0 | 0 | 0 | 0 | 0 |
| 0 | NA | NA | NA | NA | NA | Gm25798 | | 0 | 0 | 0 | 0 | 0 | 0 | 0 | 0 | 0 | 0 | 0 |
| 0 | NA | NA | NA | NA | NA | Gm45686 | | 0 | 0 | 0 | 0 | 0 | 0 | 0 | 0 | 0 | 0 | 0 |
| 0 | NA | NA | NA | NA | NA | Gm30808 | | 0 | 0 | 0 | 0 | 0 | 0 | 0 | 0 | 0 | 0 | 0 |
| 0 | NA | NA | NA | NA | NA | Gm45506 | | 0 | 0 | 0 | 0 | 0 | 0 | 0 | 0 | 0 | 0 | 0 |
| 0 | NA | NA | NA | NA | NA | Gm45668 | | 0 | 0 | 0 | 0 | 0 | 0 | 0 | 0 | 0 | 0 | 0 |
| 0 | NA | NA | NA | NA | NA | Gm45613 | | 0 | 0 | 0 | 0 | 0 | 0 | 0 | 0 | 0 | 0 | 0 |
| 0 | NA | NA | NA | NA | NA | Gm27975 | | 0 | 0 | 0 | 0 | 0 | 0 | 0 | 0 | 0 | 0 | 0 |
| 0 | NA | NA | NA | NA | NA | Gm32486 | | 0 | 0 | 0 | 0 | 0 | 0 | 0 | 0 | 0 | 0 | 0 |
| 0 | NA | NA | NA | NA | NA | Mir6401 | | 0 | 0 | 0 | 0 | 0 | 0 | 0 | 0 | 0 | 0 | 0 |
| 0 | NA | NA | NA | NA | NA | Gm2044 | | 0 | 0 | 0 | 0 | 0 | 0 | 0 | 0 | 0 | 0 | 0 |
| 0 | NA | NA | NA | NA | NA | Mir7686 | | 0 | 0 | 0 | 0 | 0 | 0 | 0 | 0 | 0 | 0 | 0 |
| 0 | NA | NA | NA | NA | NA | Mir202 | | 0 | 0 | 0 | 0 | 0 | 0 | 0 | 0 | 0 | 0 | 0 |
| 0 | NA | NA | NA | NA | NA | Mir7062 | | 0 | 0 | 0 | 0 | 0 | 0 | 0 | 0 | 0 | 0 | 0 |
| 0 | NA | NA | NA | NA | NA | Msx3 | msh homeobox 3 | 0 | 0 | 0 | 0 | 0 | 0 | 0 | 0 | 0 | 0 | 0 |
| 0 | NA | NA | NA | NA | NA | Prap1 | proline-rich acidic protein 1 | 0 | 0 | 0 | 0 | 0 | 0 | 0 | 0 | 0 | 0 | 0 |
| 0 | NA | NA | NA | NA | NA | Olfr522 | olfactory receptor 522 | 0 | 0 | 0 | 0 | 0 | 0 | 0 | 0 | 0 | 0 | 0 |
| 0 | NA | NA | NA | NA | NA | Olfr523 | olfactory receptor 523 | 0 | 0 | 0 | 0 | 0 | 0 | 0 | 0 | 0 | 0 | 0 |
| 0 | NA | NA | NA | NA | NA | Olfr524 | olfactory receptor 524 | 0 | 0 | 0 | 0 | 0 | 0 | 0 | 0 | 0 | 0 | 0 |
| 0 | NA | NA | NA | NA | NA | 5830411N06Ri | RIKEN cDNA 5830411N06 gene | 0 | 0 | 0 | 0 | 0 | 0 | 0 | 0 | 0 | 0 | 0 |
| 0 | NA | NA | NA | NA | NA | Olfr527 | olfactory receptor 527 | 0 | 0 | 0 | 0 | 0 | 0 | 0 | 0 | 0 | 0 | 0 |
| 0 | NA | NA | NA | NA | NA | Olfr528-ps1 | | 0 | 0 | 0 | 0 | 0 | 0 | 0 | 0 | 0 | 0 | 0 |
| 0 | NA | NA | NA | NA | NA | Olfr60 | olfactory receptor 60 | 0 | 0 | 0 | 0 | 0 | 0 | 0 | 0 | 0 | 0 | 0 |
| 0 | NA | NA | NA | NA | NA | Olfr529-ps1 | | 0 | 0 | 0 | 0 | 0 | 0 | 0 | 0 | 0 | 0 | 0 |
| 0 | NA | NA | NA | NA | NA | Olfr530 | olfactory receptor 530 | 0 | 0 | 0 | 0 | 0 | 0 | 0 | 0 | 0 | 0 | 0 |

| | | | | | | | | | | | | | | | | | |
|---|---|---|---|---|---|---|---|---|---|---|---|---|---|---|---|---|---|
| 0 | NA | NA | NA | NA | NA | Gm6376 | | 0 | 0 | 0 | 0 | 0 | 0 | 0 | 0 | 0 | 0 | 0 |
| 0 | NA | NA | NA | NA | NA | Olfr531 | olfactory receptor 531 | 0 | 0 | 0 | 0 | 0 | 0 | 0 | 0 | 0 | 0 | 0 |
| 0 | NA | NA | NA | NA | NA | Olfr532 | olfactory receptor 532 | 0 | 0 | 0 | 0 | 0 | 0 | 0 | 0 | 0 | 0 | 0 |
| 0 | NA | NA | NA | NA | NA | Olfr533 | olfactory receptor 533 | 0 | 0 | 0 | 0 | 0 | 0 | 0 | 0 | 0 | 0 | 0 |
| 0 | NA | NA | NA | NA | NA | Olfr534-ps1 | | 0 | 0 | 0 | 0 | 0 | 0 | 0 | 0 | 0 | 0 | 0 |
| 0 | NA | NA | NA | NA | NA | Olfr535 | olfactory receptor 535 | 0 | 0 | 0 | 0 | 0 | 0 | 0 | 0 | 0 | 0 | 0 |
| 0 | NA | NA | NA | NA | NA | Olfr536 | olfactory receptor 536 | 0 | 0 | 0 | 0 | 0 | 0 | 0 | 0 | 0 | 0 | 0 |
| 0 | NA | NA | NA | NA | NA | Olfr537-ps1 | | 0 | 0 | 0 | 0 | 0 | 0 | 0 | 0 | 0 | 0 | 0 |
| 0 | NA | NA | NA | NA | NA | Olfr538 | olfactory receptor 538 | 0 | 0 | 0 | 0 | 0 | 0 | 0 | 0 | 0 | 0 | 0 |
| 0 | NA | NA | NA | NA | NA | Olfr46 | olfactory receptor 46 | 0 | 0 | 0 | 0 | 0 | 0 | 0 | 0 | 0 | 0 | 0 |
| 0 | NA | NA | NA | NA | NA | Olfr61 | olfactory receptor 61 | 0 | 0 | 0 | 0 | 0 | 0 | 0 | 0 | 0 | 0 | 0 |
| 0 | NA | NA | NA | NA | NA | Olfr53 | olfactory receptor 53 | 0 | 0 | 0 | 0 | 0 | 0 | 0 | 0 | 0 | 0 | 0 |
| 0 | NA | NA | NA | NA | NA | Olfr540-ps1 | | 0 | 0 | 0 | 0 | 0 | 0 | 0 | 0 | 0 | 0 | 0 |
| 0 | NA | NA | NA | NA | NA | Olfr45 | olfactory receptor 45 | 0 | 0 | 0 | 0 | 0 | 0 | 0 | 0 | 0 | 0 | 0 |
| 0 | NA | NA | NA | NA | NA | Olfr541 | olfactory receptor 541 | 0 | 0 | 0 | 0 | 0 | 0 | 0 | 0 | 0 | 0 | 0 |
| 0 | NA | NA | NA | NA | NA | Olfr542-ps1 | | 0 | 0 | 0 | 0 | 0 | 0 | 0 | 0 | 0 | 0 | 0 |
| 0 | NA | NA | NA | NA | NA | Gm29799 | | 0 | 0 | 0 | 0 | 0 | 0 | 0 | 0 | 0 | 0 | 0 |
| 0 | NA | NA | NA | NA | NA | Scgb1c1 | secretoglobin, family 1C, member 1 | 0 | 0 | 0 | 0 | 0 | 0 | 0 | 0 | 0 | 0 | 0 |
| 0 | NA | NA | NA | NA | NA | Odf3 | outer dense fiber of sperm tails 3 | 0 | 0 | 0 | 0 | 0 | 0 | 0 | 0 | 0 | 0 | 0 |
| 0 | NA | NA | NA | NA | NA | Cox8b | cytochrome c oxidase subunit 8B | 0 | 0 | 0 | 0 | 0 | 0 | 0 | 0 | 0 | 0 | 0 |
| 0 | NA | NA | NA | NA | NA | BC024386 | | 0 | 0 | 0 | 0 | 0 | 0 | 0 | 0 | 0 | 0 | 0 |
| 0 | NA | NA | NA | NA | NA | Ifitm5 | interferon induced transmembrane protei | 0 | 0 | 0 | 0 | 0 | 0 | 0 | 0 | 0 | 0 | 0 |
| 0 | NA | NA | NA | NA | NA | Gm45717 | | 0 | 0 | 0 | 0 | 0 | 0 | 0 | 0 | 0 | 0 | 0 |
| 0 | NA | NA | NA | NA | NA | Gm17387 | | 0 | 0 | 0 | 0 | 0 | 0 | 0 | 0 | 0 | 0 | 0 |
| 0 | NA | NA | NA | NA | NA | Gm45299 | | 0 | 0 | 0 | 0 | 0 | 0 | 0 | 0 | 0 | 0 | 0 |
| 0 | NA | NA | NA | NA | NA | Gm24962 | | 0 | 0 | 0 | 0 | 0 | 0 | 0 | 0 | 0 | 0 | 0 |
| 0 | NA | NA | NA | NA | NA | Gm22019 | | 0 | 0 | 0 | 0 | 0 | 0 | 0 | 0 | 0 | 0 | 0 |
| 0 | NA | NA | NA | NA | NA | Mir210 | | 0 | 0 | 0 | 0 | 0 | 0 | 0 | 0 | 0 | 0 | 0 |
| 0 | NA | NA | NA | NA | NA | Drd4 | dopamine receptor D4 | 0 | 0 | 0 | 0 | 0 | 0 | 0 | 0 | 0 | 0 | 0 |
| 0 | NA | NA | NA | NA | NA | Gm25725 | | 0 | 0 | 0 | 0 | 0 | 0 | 0 | 0 | 0 | 0 | 0 |
| 0 | NA | NA | NA | NA | NA | Snora52 | | 0 | 0 | 0 | 0 | 0 | 0 | 0 | 0 | 0 | 0 | 0 |
| 0 | NA | NA | NA | NA | NA | Muc6 | mucin 6, gastric | 0 | 0 | 0 | 0 | 0 | 0 | 0 | 0 | 0 | 0 | 0 |
| 0 | NA | NA | NA | NA | NA | Muc5ac | mucin 5, subtypes A and C, tracheobronc | 0 | 0 | 0 | 0 | 0 | 0 | 0 | 0 | 0 | 0 | 0 |
| 0 | NA | NA | NA | NA | NA | Muc5b | mucin 5, subtype B, tracheobronchial | 0 | 0 | 0 | 0 | 0 | 0 | 0 | 0 | 0 | 0 | 0 |
| 0 | NA | NA | NA | NA | NA | Mir3104 | | 0 | 0 | 0 | 0 | 0 | 0 | 0 | 0 | 0 | 0 | 0 |
| 0 | NA | NA | NA | NA | NA | Gm25416 | | 0 | 0 | 0 | 0 | 0 | 0 | 0 | 0 | 0 | 0 | 0 |
| 0 | NA | NA | NA | NA | NA | Gm45417 | | 0 | 0 | 0 | 0 | 0 | 0 | 0 | 0 | 0 | 0 | 0 |
| 0 | NA | NA | NA | NA | NA | Gm45618 | predicted gene 45618 | 0 | 0 | 0 | 0 | 0 | 0 | 0 | 0 | 0 | 0 | 0 |
| 0 | NA | NA | NA | NA | NA | Gm39115 | predicted gene, 39115 | 0 | 0 | 0 | 0 | 0 | 0 | 0 | 0 | 0 | 0 | 0 |
| 0 | NA | NA | NA | NA | NA | Gm45337 | predicted gene 45337 | 0 | 0 | 0 | 0 | 0 | 0 | 0 | 0 | 0 | 0 | 0 |
| 0 | NA | NA | NA | NA | NA | Gm25370 | | 0 | 0 | 0 | 0 | 0 | 0 | 0 | 0 | 0 | 0 | 0 |
| 0 | NA | NA | NA | NA | NA | Gm29735 | predicted gene, 29735 | 0 | 0 | 0 | 0 | 0 | 0 | 0 | 0 | 0 | 0 | 0 |
| 0 | NA | NA | NA | NA | NA | Gm4553 | predicted gene 4553 | 0 | 0 | 0 | 0 | 0 | 0 | 0 | 0 | 0 | 0 | 0 |
| 0 | NA | NA | NA | NA | NA | Krtap5-2 | keratin associated protein 5-2 | 0 | 0 | 0 | 0 | 0 | 0 | 0 | 0 | 0 | 0 | 0 |
| 0 | NA | NA | NA | NA | NA | Gm10153 | | 0 | 0 | 0 | 0 | 0 | 0 | 0 | 0 | 0 | 0 | 0 |
| 0 | NA | NA | NA | NA | NA | Krtap5-3 | keratin associated protein 5-3 | 0 | 0 | 0 | 0 | 0 | 0 | 0 | 0 | 0 | 0 | 0 |
| 0 | NA | NA | NA | NA | NA | Gm7579 | predicted gene 7579 | 0 | 0 | 0 | 0 | 0 | 0 | 0 | 0 | 0 | 0 | 0 |
| 0 | NA | NA | NA | NA | NA | Gm19094 | | 0 | 0 | 0 | 0 | 0 | 0 | 0 | 0 | 0 | 0 | 0 |
| 0 | NA | NA | NA | NA | NA | Krtap5-5 | keratin associated protein 5-5 | 0 | 0 | 0 | 0 | 0 | 0 | 0 | 0 | 0 | 0 | 0 |
| 0 | NA | NA | NA | NA | NA | Gm40460 | predicted gene, 40460 | 0 | 0 | 0 | 0 | 0 | 0 | 0 | 0 | 0 | 0 | 0 |
| 0 | NA | NA | NA | NA | NA | Gm47762 | | 0 | 0 | 0 | 0 | 0 | 0 | 0 | 0 | 0 | 0 | 0 |
| 0 | NA | NA | NA | NA | NA | Gm2431 | | 0 | 0 | 0 | 0 | 0 | 0 | 0 | 0 | 0 | 0 | 0 |
| 0 | NA | NA | NA | NA | NA | Gm4559 | predicted gene 4559 | 0 | 0 | 0 | 0 | 0 | 0 | 0 | 0 | 0 | 0 | 0 |
| 0 | NA | NA | NA | NA | NA | Gm10013 | | 0 | 0 | 0 | 0 | 0 | 0 | 0 | 0 | 0 | 0 | 0 |
| 0 | NA | NA | NA | NA | NA | Krtap5-1 | keratin associated protein 5-1 | 0 | 0 | 0 | 0 | 0 | 0 | 0 | 0 | 0 | 0 | 0 |
| 0 | NA | NA | NA | NA | NA | Krtap5-4 | keratin associated protein 5-4 | 0 | 0 | 0 | 0 | 0 | 0 | 0 | 0 | 0 | 0 | 0 |
| 0 | NA | NA | NA | NA | NA | Gm49369 | | 0 | 0 | 0 | 0 | 0 | 0 | 0 | 0 | 0 | 0 | 0 |
| 0 | NA | NA | NA | NA | NA | Gm26143 | | 0 | 0 | 0 | 0 | 0 | 0 | 0 | 0 | 0 | 0 | 0 |
| 0 | NA | NA | NA | NA | NA | Syt8 | synaptotagmin VIII | 0 | 0 | 0 | 0 | 0 | 0 | 0 | 0 | 0 | 0 | 0 |
| 0 | NA | NA | NA | NA | NA | Prr33 | | 0 | 0 | 0 | 0 | 0 | 0 | 0 | 0 | 0 | 0 | 0 |

| | | | | | | | | | | | | | | | | | | |
|---|---|---|---|---|---|---|---|---|---|---|---|---|---|---|---|---|---|---|
| 0 | NA | NA | NA | NA | NA | Tnnt3 | troponin T3, skeletal, fast | 0 | 0 | 0 | 0 | 0 | 0 | 0 | 0 | 0 | 0 | 0 |
| 0 | NA | NA | NA | NA | NA | Nctc1 | | 0 | 0 | 0 | 0 | 0 | 0 | 0 | 0 | 0 | 0 | 0 |
| 0 | NA | NA | NA | NA | NA | Gm27483 | | 0 | 0 | 0 | 0 | 0 | 0 | 0 | 0 | 0 | 0 | 0 |
| 0 | NA | NA | NA | NA | NA | Mir675 | | 0 | 0 | 0 | 0 | 0 | 0 | 0 | 0 | 0 | 0 | 0 |
| 0 | NA | NA | NA | NA | NA | Gm27786 | | 0 | 0 | 0 | 0 | 0 | 0 | 0 | 0 | 0 | 0 | 0 |
| 0 | NA | NA | NA | NA | NA | Gm33148 | | 0 | 0 | 0 | 0 | 0 | 0 | 0 | 0 | 0 | 0 | 0 |
| 0 | NA | NA | NA | NA | NA | Mir483 | | 0 | 0 | 0 | 0 | 0 | 0 | 0 | 0 | 0 | 0 | 0 |
| 0 | NA | NA | NA | NA | NA | Gm6471 | | 0 | 0 | 0 | 0 | 0 | 0 | 0 | 0 | 0 | 0 | 0 |
| 0 | NA | NA | NA | NA | NA | Gm39117 | | 0 | 0 | 0 | 0 | 0 | 0 | 0 | 0 | 0 | 0 | 0 |
| 0 | NA | NA | NA | NA | NA | Gm38321 | | 0 | 0 | 0 | 0 | 0 | 0 | 0 | 0 | 0 | 0 | 0 |
| 0 | NA | NA | NA | NA | NA | Gm37235 | | 0 | 0 | 0 | 0 | 0 | 0 | 0 | 0 | 0 | 0 | 0 |
| 0 | NA | NA | NA | NA | NA | Gm28821 | | 0 | 0 | 0 | 0 | 0 | 0 | 0 | 0 | 0 | 0 | 0 |
| 0 | NA | NA | NA | NA | NA | Gm27395 | | 0 | 0 | 0 | 0 | 0 | 0 | 0 | 0 | 0 | 0 | 0 |
| 0 | NA | NA | NA | NA | NA | Gm27389 | | 0 | 0 | 0 | 0 | 0 | 0 | 0 | 0 | 0 | 0 | 0 |
| 0 | NA | NA | NA | NA | NA | Gm27803 | | 0 | 0 | 0 | 0 | 0 | 0 | 0 | 0 | 0 | 0 | 0 |
| 0 | NA | NA | NA | NA | NA | Gm27539 | | 0 | 0 | 0 | 0 | 0 | 0 | 0 | 0 | 0 | 0 | 0 |
| 0 | NA | NA | NA | NA | NA | 4933417O13Rik | | 0 | 0 | 0 | 0 | 0 | 0 | 0 | 0 | 0 | 0 | 0 |
| 0 | NA | NA | NA | NA | NA | Gm38095 | | 0 | 0 | 0 | 0 | 0 | 0 | 0 | 0 | 0 | 0 | 0 |
| 0 | NA | NA | NA | NA | NA | Gm38065 | | 0 | 0 | 0 | 0 | 0 | 0 | 0 | 0 | 0 | 0 | 0 |
| 0 | NA | NA | NA | NA | NA | Gm27901 | | 0 | 0 | 0 | 0 | 0 | 0 | 0 | 0 | 0 | 0 | 0 |
| 0 | NA | NA | NA | NA | NA | Gm23297 | | 0 | 0 | 0 | 0 | 0 | 0 | 0 | 0 | 0 | 0 | 0 |
| 0 | NA | NA | NA | NA | NA | Gm15579 | | 0 | 0 | 0 | 0 | 0 | 0 | 0 | 0 | 0 | 0 | 0 |
| 0 | NA | NA | NA | NA | NA | E230032D23Rik | | 0 | 0 | 0 | 0 | 0 | 0 | 0 | 0 | 0 | 0 | 0 |
| 0 | NA | NA | NA | NA | NA | Mrgprg | MAS-related GPR, member G | 0 | 0 | 0 | 0 | 0 | 0 | 0 | 0 | 0 | 0 | 0 |
| 0 | NA | NA | NA | NA | NA | Gm44998 | | 0 | 0 | 0 | 0 | 0 | 0 | 0 | 0 | 0 | 0 | 0 |
| 0 | NA | NA | NA | NA | NA | Gm22064 | | 0 | 0 | 0 | 0 | 0 | 0 | 0 | 0 | 0 | 0 | 0 |
| 0 | NA | NA | NA | NA | NA | Acte1 | actin, epsilon 1 | 0 | 0 | 0 | 0 | 0 | 0 | 0 | 0 | 0 | 0 | 0 |
| 0 | NA | NA | NA | NA | NA | Mir6404 | | 0 | 0 | 0 | 0 | 0 | 0 | 0 | 0 | 0 | 0 | 0 |
| 0 | NA | NA | NA | NA | NA | Mir3105 | | 0 | 0 | 0 | 0 | 0 | 0 | 0 | 0 | 0 | 0 | 0 |
| 0 | NA | NA | NA | NA | NA | Gm44999 | | 0 | 0 | 0 | 0 | 0 | 0 | 0 | 0 | 0 | 0 | 0 |
| 0 | NA | NA | NA | NA | NA | Gm14372 | | 0 | 0 | 0 | 0 | 0 | 0 | 0 | 0 | 0 | 0 | 0 |
| 0 | NA | NA | NA | NA | NA | Faddos | | 0 | 0 | 0 | 0 | 0 | 0 | 0 | 0 | 0 | 0 | 0 |
| 0 | NA | NA | NA | NA | NA | Gm10152 | | 0 | 0 | 0 | 0 | 0 | 0 | 0 | 0 | 0 | 0 | 0 |
| 0 | NA | NA | NA | NA | NA | Gm34964 | | 0 | 0 | 0 | 0 | 0 | 0 | 0 | 0 | 0 | 0 | 0 |
| 0 | NA | NA | NA | NA | NA | Fgf3 | fibroblast growth factor 3 | 0 | 0 | 0 | 0 | 0 | 0 | 0 | 0 | 0 | 0 | 0 |
| 0 | NA | NA | NA | NA | NA | Fgf4 | fibroblast growth factor 4 | 0 | 0 | 0 | 0 | 0 | 0 | 0 | 0 | 0 | 0 | 0 |
| 0 | NA | NA | NA | NA | NA | Gm26793 | | 0 | 0 | 0 | 0 | 0 | 0 | 0 | 0 | 0 | 0 | 0 |
| 0 | NA | NA | NA | NA | NA | Fgf15 | fibroblast growth factor 15 | 0 | 0 | 0 | 0 | 0 | 0 | 0 | 0 | 0 | 0 | 0 |
| 0 | NA | NA | NA | NA | NA | Gm45181 | | 0 | 0 | 0 | 0 | 0 | 0 | 0 | 0 | 0 | 0 | 0 |
| 0 | NA | NA | NA | NA | NA | Mrgprd | MAS-related GPR, member D | 0 | 0 | 0 | 0 | 0 | 0 | 0 | 0 | 0 | 0 | 0 |
| 0 | NA | NA | NA | NA | NA | Gm44634 | | 0 | 0 | 0 | 0 | 0 | 0 | 0 | 0 | 0 | 0 | 0 |
| 0 | NA | NA | NA | NA | NA | Gm7346 | | 0 | 0 | 0 | 0 | 0 | 0 | 0 | 0 | 0 | 0 | 0 |
| 0 | NA | NA | NA | NA | NA | Gm25775 | | 0 | 0 | 0 | 0 | 0 | 0 | 0 | 0 | 0 | 0 | 0 |
| 0 | NA | NA | NA | NA | NA | Gm18210 | | 0 | 0 | 0 | 0 | 0 | 0 | 0 | 0 | 0 | 0 | 0 |
| 0 | NA | NA | NA | NA | NA | Vmn2r-ps88 | | 0 | 0 | 0 | 0 | 0 | 0 | 0 | 0 | 0 | 0 | 0 |
| 0 | NA | NA | NA | NA | NA | Gm16180 | | 0 | 0 | 0 | 0 | 0 | 0 | 0 | 0 | 0 | 0 | 0 |
| 0 | NA | NA | NA | NA | NA | Gm22204 | | 0 | 0 | 0 | 0 | 0 | 0 | 0 | 0 | 0 | 0 | 0 |
| 0 | NA | NA | NA | NA | NA | Pcp2 | Purkinje cell protein 2 (L7) | 0 | 0 | 0 | 0 | 0 | 0 | 0 | 0 | 0 | 0 | 0 |
| 0 | NA | NA | NA | NA | NA | Retn | resistin | 0 | 0 | 0 | 0 | 0 | 0 | 0 | 0 | 0 | 0 | 0 |
| 0 | NA | NA | NA | NA | NA | Gm44776 | | 0 | 0 | 0 | 0 | 0 | 0 | 0 | 0 | 0 | 0 | 0 |
| 0 | NA | NA | NA | NA | NA | Gm23886 | | 0 | 0 | 0 | 0 | 0 | 0 | 0 | 0 | 0 | 0 | 0 |
| 0 | NA | NA | NA | NA | NA | Gm24934 | | 0 | 0 | 0 | 0 | 0 | 0 | 0 | 0 | 0 | 0 | 0 |
| 0 | NA | NA | NA | NA | NA | Rprl3 | | 0 | 0 | 0 | 0 | 0 | 0 | 0 | 0 | 0 | 0 | 0 |
| 0 | NA | NA | NA | NA | NA | Gm45048 | | 0 | 0 | 0 | 0 | 0 | 0 | 0 | 0 | 0 | 0 | 0 |
| 0 | NA | NA | NA | NA | NA | Gm7397 | | 0 | 0 | 0 | 0 | 0 | 0 | 0 | 0 | 0 | 0 | 0 |
| 0 | NA | NA | NA | NA | NA | Gm23888 | | 0 | 0 | 0 | 0 | 0 | 0 | 0 | 0 | 0 | 0 | 0 |
| 0 | NA | NA | NA | NA | NA | Gm49320 | | 0 | 0 | 0 | 0 | 0 | 0 | 0 | 0 | 0 | 0 | 0 |
| 0 | NA | NA | NA | NA | NA | Gm26102 | | 0 | 0 | 0 | 0 | 0 | 0 | 0 | 0 | 0 | 0 | 0 |
| 0 | NA | NA | NA | NA | NA | Gm44869 | | 0 | 0 | 0 | 0 | 0 | 0 | 0 | 0 | 0 | 0 | 0 |
| 0 | NA | NA | NA | NA | NA | Gm45118 | | 0 | 0 | 0 | 0 | 0 | 0 | 0 | 0 | 0 | 0 | 0 |

| | | | | | | | | | | | | | | | | | |
|---|---|---|---|---|---|---|---|---|---|---|---|---|---|---|---|---|---|
| 0 | NA | NA | NA | NA | NA | Gm45117 | 0 | 0 | 0 | 0 | 0 | 0 | 0 | 0 | 0 | 0 | 0 |
| 0 | NA | NA | NA | NA | NA | Gm36956 | 0 | 0 | 0 | 0 | 0 | 0 | 0 | 0 | 0 | 0 | 0 |
| 0 | NA | NA | NA | NA | NA | Vmn2r-ps89 | 0 | 0 | 0 | 0 | 0 | 0 | 0 | 0 | 0 | 0 | 0 |
| 0 | NA | NA | NA | NA | NA | Gm44959 | 0 | 0 | 0 | 0 | 0 | 0 | 0 | 0 | 0 | 0 | 0 |
| 0 | NA | NA | NA | NA | NA | Gm44960 | 0 | 0 | 0 | 0 | 0 | 0 | 0 | 0 | 0 | 0 | 0 |
| 0 | NA | NA | NA | NA | NA | Gm44961 | 0 | 0 | 0 | 0 | 0 | 0 | 0 | 0 | 0 | 0 | 0 |
| 0 | NA | NA | NA | NA | NA | n-R5s93 | 0 | 0 | 0 | 0 | 0 | 0 | 0 | 0 | 0 | 0 | 0 |
| 0 | NA | NA | NA | NA | NA | Gm5739 | 0 | 0 | 0 | 0 | 0 | 0 | 0 | 0 | 0 | 0 | 0 |
| 0 | NA | NA | NA | NA | NA | n-R5s94 | 0 | 0 | 0 | 0 | 0 | 0 | 0 | 0 | 0 | 0 | 0 |
| 0 | NA | NA | NA | NA | NA | Gm44693 | 0 | 0 | 0 | 0 | 0 | 0 | 0 | 0 | 0 | 0 | 0 |
| 0 | NA | NA | NA | NA | NA | Gm22565 | 0 | 0 | 0 | 0 | 0 | 0 | 0 | 0 | 0 | 0 | 0 |
| 0 | NA | NA | NA | NA | NA | Gm2734 | 0 | 0 | 0 | 0 | 0 | 0 | 0 | 0 | 0 | 0 | 0 |
| 0 | NA | NA | NA | NA | NA | Gm44843 | 0 | 0 | 0 | 0 | 0 | 0 | 0 | 0 | 0 | 0 | 0 |
| 0 | NA | NA | NA | NA | NA | Gm44842 | 0 | 0 | 0 | 0 | 0 | 0 | 0 | 0 | 0 | 0 | 0 |
| 0 | NA | NA | NA | NA | NA | Gm24288 | 0 | 0 | 0 | 0 | 0 | 0 | 0 | 0 | 0 | 0 | 0 |
| 0 | NA | NA | NA | NA | NA | Gm44844 | 0 | 0 | 0 | 0 | 0 | 0 | 0 | 0 | 0 | 0 | 0 |
| 0 | NA | NA | NA | NA | NA | Gm44909 | 0 | 0 | 0 | 0 | 0 | 0 | 0 | 0 | 0 | 0 | 0 |
| 0 | NA | NA | NA | NA | NA | Gm7506 | 0 | 0 | 0 | 0 | 0 | 0 | 0 | 0 | 0 | 0 | 0 |
| 0 | NA | NA | NA | NA | NA | Gm44910 | 0 | 0 | 0 | 0 | 0 | 0 | 0 | 0 | 0 | 0 | 0 |
| 0 | NA | NA | NA | NA | NA | Gm44908 | 0 | 0 | 0 | 0 | 0 | 0 | 0 | 0 | 0 | 0 | 0 |
| 0 | NA | NA | NA | NA | NA | Gm17215 | 0 | 0 | 0 | 0 | 0 | 0 | 0 | 0 | 0 | 0 | 0 |
| 0 | NA | NA | NA | NA | NA | Gm26427 | 0 | 0 | 0 | 0 | 0 | 0 | 0 | 0 | 0 | 0 | 0 |
| 0 | NA | NA | NA | NA | NA | Gm30954 | 0 | 0 | 0 | 0 | 0 | 0 | 0 | 0 | 0 | 0 | 0 |
| 0 | NA | NA | NA | NA | NA | Gm27585 | 0 | 0 | 0 | 0 | 0 | 0 | 0 | 0 | 0 | 0 | 0 |
| 0 | NA | NA | NA | NA | NA | Gm27545 | 0 | 0 | 0 | 0 | 0 | 0 | 0 | 0 | 0 | 0 | 0 |
| 0 | NA | NA | NA | NA | NA | Gm25169 | 0 | 0 | 0 | 0 | 0 | 0 | 0 | 0 | 0 | 0 | 0 |
| 0 | NA | NA | NA | NA | NA | Gm28262 | 0 | 0 | 0 | 0 | 0 | 0 | 0 | 0 | 0 | 0 | 0 |
| 0 | NA | NA | NA | NA | NA | Gm31135 | 0 | 0 | 0 | 0 | 0 | 0 | 0 | 0 | 0 | 0 | 0 |
| 0 | NA | NA | NA | NA | NA | Gm31401 | 0 | 0 | 0 | 0 | 0 | 0 | 0 | 0 | 0 | 0 | 0 |
| 0 | NA | NA | NA | NA | NA | Gm45076 | 0 | 0 | 0 | 0 | 0 | 0 | 0 | 0 | 0 | 0 | 0 |
| 0 | NA | NA | NA | NA | NA | Gm31343 | 0 | 0 | 0 | 0 | 0 | 0 | 0 | 0 | 0 | 0 | 0 |
| 0 | NA | NA | NA | NA | NA | Gm2448 | 0 | 0 | 0 | 0 | 0 | 0 | 0 | 0 | 0 | 0 | 0 |
| 0 | NA | NA | NA | NA | NA | D530014G21Rik | 0 | 0 | 0 | 0 | 0 | 0 | 0 | 0 | 0 | 0 | 0 |
| 0 | NA | NA | NA | NA | NA | Gm31463 | 0 | 0 | 0 | 0 | 0 | 0 | 0 | 0 | 0 | 0 | 0 |
| 0 | NA | NA | NA | NA | NA | Gm45075 | 0 | 0 | 0 | 0 | 0 | 0 | 0 | 0 | 0 | 0 | 0 |
| 0 | NA | NA | NA | NA | NA | 4921522P10Rik | 0 | 0 | 0 | 0 | 0 | 0 | 0 | 0 | 0 | 0 | 0 |
| 0 | NA | NA | NA | NA | NA | Mir7654 | 0 | 0 | 0 | 0 | 0 | 0 | 0 | 0 | 0 | 0 | 0 |
| 0 | NA | NA | NA | NA | NA | Gm18393 | 0 | 0 | 0 | 0 | 0 | 0 | 0 | 0 | 0 | 0 | 0 |
| 0 | NA | NA | NA | NA | NA | Gm44621 | 0 | 0 | 0 | 0 | 0 | 0 | 0 | 0 | 0 | 0 | 0 |
| 0 | NA | NA | NA | NA | NA | Gm44515 | 0 | 0 | 0 | 0 | 0 | 0 | 0 | 0 | 0 | 0 | 0 |
| 0 | NA | NA | NA | NA | NA | 4933430N04Rik | 0 | 0 | 0 | 0 | 0 | 0 | 0 | 0 | 0 | 0 | 0 |
| 0 | NA | NA | NA | NA | NA | 4930453L07Rik | 0 | 0 | 0 | 0 | 0 | 0 | 0 | 0 | 0 | 0 | 0 |
| 0 | NA | NA | NA | NA | NA | Gm44516 | 0 | 0 | 0 | 0 | 0 | 0 | 0 | 0 | 0 | 0 | 0 |
| 0 | NA | NA | NA | NA | NA | 4930435N07Rik | 0 | 0 | 0 | 0 | 0 | 0 | 0 | 0 | 0 | 0 | 0 |
| 0 | NA | NA | NA | NA | NA | Gm47312 | 0 | 0 | 0 | 0 | 0 | 0 | 0 | 0 | 0 | 0 | 0 |
| 0 | NA | NA | NA | NA | NA | Gm44761 | 0 | 0 | 0 | 0 | 0 | 0 | 0 | 0 | 0 | 0 | 0 |
| 0 | NA | NA | NA | NA | NA | Gm10217 | 0 | 0 | 0 | 0 | 0 | 0 | 0 | 0 | 0 | 0 | 0 |
| 0 | NA | NA | NA | NA | NA | Gm10067 | 0 | 0 | 0 | 0 | 0 | 0 | 0 | 0 | 0 | 0 | 0 |
| 0 | NA | NA | NA | NA | NA | Gm44789 | 0 | 0 | 0 | 0 | 0 | 0 | 0 | 0 | 0 | 0 | 0 |
| 0 | NA | NA | NA | NA | NA | Gm44788 | 0 | 0 | 0 | 0 | 0 | 0 | 0 | 0 | 0 | 0 | 0 |
| 0 | NA | NA | NA | NA | NA | Gm44786 | 0 | 0 | 0 | 0 | 0 | 0 | 0 | 0 | 0 | 0 | 0 |
| 0 | NA | NA | NA | NA | NA | Gm44785 | 0 | 0 | 0 | 0 | 0 | 0 | 0 | 0 | 0 | 0 | 0 |
| 0 | NA | NA | NA | NA | NA | Gm44784 | 0 | 0 | 0 | 0 | 0 | 0 | 0 | 0 | 0 | 0 | 0 |
| 0 | NA | NA | NA | NA | NA | Gm45045 | 0 | 0 | 0 | 0 | 0 | 0 | 0 | 0 | 0 | 0 | 0 |
| 0 | NA | NA | NA | NA | NA | Gm39129 | 0 | 0 | 0 | 0 | 0 | 0 | 0 | 0 | 0 | 0 | 0 |
| 0 | NA | NA | NA | NA | NA | Rps16-ps3 | 0 | 0 | 0 | 0 | 0 | 0 | 0 | 0 | 0 | 0 | 0 |
| 0 | NA | NA | NA | NA | NA | Gm32540 | 0 | 0 | 0 | 0 | 0 | 0 | 0 | 0 | 0 | 0 | 0 |
| 0 | NA | NA | NA | NA | NA | Gm45044 | 0 | 0 | 0 | 0 | 0 | 0 | 0 | 0 | 0 | 0 | 0 |
| 0 | NA | NA | NA | NA | NA | Gm45042 | 0 | 0 | 0 | 0 | 0 | 0 | 0 | 0 | 0 | 0 | 0 |
| 0 | NA | NA | NA | NA | NA | Gm44955 | 0 | 0 | 0 | 0 | 0 | 0 | 0 | 0 | 0 | 0 | 0 |

| | | | | | | | | | | | | | | | | | |
|---|---|---|---|---|---|---|---|---|---|---|---|---|---|---|---|---|---|
| 0 | NA | NA | NA | NA | NA | Gm44716 | | 0 | 0 | 0 | 0 | 0 | 0 | 0 | 0 | 0 | 0 | 0 |
| 0 | NA | NA | NA | NA | NA | Mir7238 | | 0 | 0 | 0 | 0 | 0 | 0 | 0 | 0 | 0 | 0 | 0 |
| 0 | NA | NA | NA | NA | NA | 1700016D06Ril | RIKEN cDNA 1700016D06 gene | 0 | 0 | 0 | 0 | 0 | 0 | 0 | 0 | 0 | 0 | 0 |
| 0 | NA | NA | NA | NA | NA | Gm7562 | | 0 | 0 | 0 | 0 | 0 | 0 | 0 | 0 | 0 | 0 | 0 |
| 0 | NA | NA | NA | NA | NA | 4930465I24Rik | | 0 | 0 | 0 | 0 | 0 | 0 | 0 | 0 | 0 | 0 | 0 |
| 0 | NA | NA | NA | NA | NA | Gm27854 | | 0 | 0 | 0 | 0 | 0 | 0 | 0 | 0 | 0 | 0 | 0 |
| 0 | NA | NA | NA | NA | NA | Gm45420 | | 0 | 0 | 0 | 0 | 0 | 0 | 0 | 0 | 0 | 0 | 0 |
| 0 | NA | NA | NA | NA | NA | Gm18991 | | 0 | 0 | 0 | 0 | 0 | 0 | 0 | 0 | 0 | 0 | 0 |
| 0 | NA | NA | NA | NA | NA | Gm15875 | | 0 | 0 | 0 | 0 | 0 | 0 | 0 | 0 | 0 | 0 | 0 |
| 0 | NA | NA | NA | NA | NA | Gm39132 | | 0 | 0 | 0 | 0 | 0 | 0 | 0 | 0 | 0 | 0 | 0 |
| 0 | NA | NA | NA | NA | NA | Gm45680 | | 0 | 0 | 0 | 0 | 0 | 0 | 0 | 0 | 0 | 0 | 0 |
| 0 | NA | NA | NA | NA | NA | B020031H02Rik | | 0 | 0 | 0 | 0 | 0 | 0 | 0 | 0 | 0 | 0 | 0 |
| 0 | NA | NA | NA | NA | NA | A230072I06Rik | RIKEN cDNA A230072I06 gene | 0 | 0 | 0 | 0 | 0 | 0 | 0 | 0 | 0 | 0 | 0 |
| 0 | NA | NA | NA | NA | NA | Gm29175 | | 0 | 0 | 0 | 0 | 0 | 0 | 0 | 0 | 0 | 0 | 0 |
| 0 | NA | NA | NA | NA | NA | Gm33175 | | 0 | 0 | 0 | 0 | 0 | 0 | 0 | 0 | 0 | 0 | 0 |
| 0 | NA | NA | NA | NA | NA | Gm22207 | | 0 | 0 | 0 | 0 | 0 | 0 | 0 | 0 | 0 | 0 | 0 |
| 0 | NA | NA | NA | NA | NA | Sox1ot | | 0 | 0 | 0 | 0 | 0 | 0 | 0 | 0 | 0 | 0 | 0 |
| 0 | NA | NA | NA | NA | NA | Gm45560 | | 0 | 0 | 0 | 0 | 0 | 0 | 0 | 0 | 0 | 0 | 0 |
| 0 | NA | NA | NA | NA | NA | Sox1 | SRY (sex determining region Y)-box 1 | 0 | 0 | 0 | 0 | 0 | 0 | 0 | 0 | 0 | 0 | 0 |
| 0 | NA | NA | NA | NA | NA | Gm25239 | | 0 | 0 | 0 | 0 | 0 | 0 | 0 | 0 | 0 | 0 | 0 |
| 0 | NA | NA | NA | NA | NA | Gm33326 | | 0 | 0 | 0 | 0 | 0 | 0 | 0 | 0 | 0 | 0 | 0 |
| 0 | NA | NA | NA | NA | NA | 4933439N14Rik | | 0 | 0 | 0 | 0 | 0 | 0 | 0 | 0 | 0 | 0 | 0 |
| 0 | NA | NA | NA | NA | NA | Gm26483 | | 0 | 0 | 0 | 0 | 0 | 0 | 0 | 0 | 0 | 0 | 0 |
| 0 | NA | NA | NA | NA | NA | Gm15347 | | 0 | 0 | 0 | 0 | 0 | 0 | 0 | 0 | 0 | 0 | 0 |
| 0 | NA | NA | NA | NA | NA | Gm38671 | | 0 | 0 | 0 | 0 | 0 | 0 | 0 | 0 | 0 | 0 | 0 |
| 0 | NA | NA | NA | NA | NA | Gm15349 | | 0 | 0 | 0 | 0 | 0 | 0 | 0 | 0 | 0 | 0 | 0 |
| 0 | NA | NA | NA | NA | NA | Gm17023 | | 0 | 0 | 0 | 0 | 0 | 0 | 0 | 0 | 0 | 0 | 0 |
| 0 | NA | NA | NA | NA | NA | Gm17022 | | 0 | 0 | 0 | 0 | 0 | 0 | 0 | 0 | 0 | 0 | 0 |
| 0 | NA | NA | NA | NA | NA | Mir7065 | | 0 | 0 | 0 | 0 | 0 | 0 | 0 | 0 | 0 | 0 | 0 |
| 0 | NA | NA | NA | NA | NA | Atp4b | ATPase, H+/K+ exchanging, beta polypep | 0 | 0 | 0 | 0 | 0 | 0 | 0 | 0 | 0 | 0 | 0 |
| 0 | NA | NA | NA | NA | NA | Gm45410 | | 0 | 0 | 0 | 0 | 0 | 0 | 0 | 0 | 0 | 0 | 0 |
| 0 | NA | NA | NA | NA | NA | Gm5608 | | 0 | 0 | 0 | 0 | 0 | 0 | 0 | 0 | 0 | 0 | 0 |
| 0 | NA | NA | NA | NA | NA | 1700029H14Ril | RIKEN cDNA 1700029H14 gene | 0 | 0 | 0 | 0 | 0 | 0 | 0 | 0 | 0 | 0 | 0 |
| 0 | NA | NA | NA | NA | NA | Gm7669 | | 0 | 0 | 0 | 0 | 0 | 0 | 0 | 0 | 0 | 0 | 0 |
| 0 | NA | NA | NA | NA | NA | AF366264 | cDNA sequence AF366264 | 0 | 0 | 0 | 0 | 0 | 0 | 0 | 0 | 0 | 0 | 0 |
| 0 | NA | NA | NA | NA | NA | Gm47277 | | 0 | 0 | 0 | 0 | 0 | 0 | 0 | 0 | 0 | 0 | 0 |
| 0 | NA | NA | NA | NA | NA | Gm24698 | | 0 | 0 | 0 | 0 | 0 | 0 | 0 | 0 | 0 | 0 | 0 |
| 0 | NA | NA | NA | NA | NA | Gm7676 | | 0 | 0 | 0 | 0 | 0 | 0 | 0 | 0 | 0 | 0 | 0 |
| 0 | NA | NA | NA | NA | NA | Gm5907 | | 0 | 0 | 0 | 0 | 0 | 0 | 0 | 0 | 0 | 0 | 0 |
| 0 | NA | NA | NA | NA | NA | Gm45520 | | 0 | 0 | 0 | 0 | 0 | 0 | 0 | 0 | 0 | 0 | 0 |
| 0 | NA | NA | NA | NA | NA | Gm26184 | | 0 | 0 | 0 | 0 | 0 | 0 | 0 | 0 | 0 | 0 | 0 |
| 0 | NA | NA | NA | NA | NA | Gm44347 | | 0 | 0 | 0 | 0 | 0 | 0 | 0 | 0 | 0 | 0 | 0 |
| 0 | NA | NA | NA | NA | NA | C030037F17Rik | | 0 | 0 | 0 | 0 | 0 | 0 | 0 | 0 | 0 | 0 | 0 |
| 0 | NA | NA | NA | NA | NA | BB014433 | | 0 | 0 | 0 | 0 | 0 | 0 | 0 | 0 | 0 | 0 | 0 |
| 0 | NA | NA | NA | NA | NA | Gm45276 | | 0 | 0 | 0 | 0 | 0 | 0 | 0 | 0 | 0 | 0 | 0 |
| 0 | NA | NA | NA | NA | NA | Gm45274 | | 0 | 0 | 0 | 0 | 0 | 0 | 0 | 0 | 0 | 0 | 0 |
| 0 | NA | NA | NA | NA | NA | Myom2 | myomesin 2 | 0 | 0 | 0 | 0 | 0 | 0 | 0 | 0 | 0 | 0 | 0 |
| 0 | NA | NA | NA | NA | NA | Gm15678 | | 0 | 0 | 0 | 0 | 0 | 0 | 0 | 0 | 0 | 0 | 0 |
| 0 | NA | NA | NA | NA | NA | Gm7706 | | 0 | 0 | 0 | 0 | 0 | 0 | 0 | 0 | 0 | 0 | 0 |
| 0 | NA | NA | NA | NA | NA | Gm45271 | | 0 | 0 | 0 | 0 | 0 | 0 | 0 | 0 | 0 | 0 | 0 |
| 0 | NA | NA | NA | NA | NA | Gm24231 | | 0 | 0 | 0 | 0 | 0 | 0 | 0 | 0 | 0 | 0 | 0 |
| 0 | NA | NA | NA | NA | NA | Gm35392 | | 0 | 0 | 0 | 0 | 0 | 0 | 0 | 0 | 0 | 0 | 0 |
| 0 | NA | NA | NA | NA | NA | Gm24222 | | 0 | 0 | 0 | 0 | 0 | 0 | 0 | 0 | 0 | 0 | 0 |
| 0 | NA | NA | NA | NA | NA | Gm25871 | | 0 | 0 | 0 | 0 | 0 | 0 | 0 | 0 | 0 | 0 | 0 |
| 0 | NA | NA | NA | NA | NA | Gm25255 | | 0 | 0 | 0 | 0 | 0 | 0 | 0 | 0 | 0 | 0 | 0 |
| 0 | NA | NA | NA | NA | NA | Gm45677 | | 0 | 0 | 0 | 0 | 0 | 0 | 0 | 0 | 0 | 0 | 0 |
| 0 | NA | NA | NA | NA | NA | Gm35453 | | 0 | 0 | 0 | 0 | 0 | 0 | 0 | 0 | 0 | 0 | 0 |
| 0 | NA | NA | NA | NA | NA | Gm45676 | | 0 | 0 | 0 | 0 | 0 | 0 | 0 | 0 | 0 | 0 | 0 |
| 0 | NA | NA | NA | NA | NA | Gm45408 | | 0 | 0 | 0 | 0 | 0 | 0 | 0 | 0 | 0 | 0 | 0 |
| 0 | NA | NA | NA | NA | NA | Gm45675 | | 0 | 0 | 0 | 0 | 0 | 0 | 0 | 0 | 0 | 0 | 0 |

| | | | | | | | | | | | | | | | | | | |
|---|---|---|---|---|---|---|---|---|---|---|---|---|---|---|---|---|---|---|
| 0 | NA | NA | NA | NA | NA | Gm7128 | | 0 | 0 | 0 | 0 | 0 | 0 | 0 | 0 | 0 | 0 | 0 |
| 0 | NA | NA | NA | NA | NA | Mir3106 | | 0 | 0 | 0 | 0 | 0 | 0 | 0 | 0 | 0 | 0 | 0 |
| 0 | NA | NA | NA | NA | NA | Gm16346 | | 0 | 0 | 0 | 0 | 0 | 0 | 0 | 0 | 0 | 0 | 0 |
| 0 | NA | NA | NA | NA | NA | Gm16347 | | 0 | 0 | 0 | 0 | 0 | 0 | 0 | 0 | 0 | 0 | 0 |
| 0 | NA | NA | NA | NA | NA | Gm45393 | | 0 | 0 | 0 | 0 | 0 | 0 | 0 | 0 | 0 | 0 | 0 |
| 0 | NA | NA | NA | NA | NA | Gm45394 | | 0 | 0 | 0 | 0 | 0 | 0 | 0 | 0 | 0 | 0 | 0 |
| 0 | NA | NA | NA | NA | NA | Gm25278 | | 0 | 0 | 0 | 0 | 0 | 0 | 0 | 0 | 0 | 0 | 0 |
| 0 | NA | NA | NA | NA | NA | 3110080E11Rik | | 0 | 0 | 0 | 0 | 0 | 0 | 0 | 0 | 0 | 0 | 0 |
| 0 | NA | NA | NA | NA | NA | Gm25665 | | 0 | 0 | 0 | 0 | 0 | 0 | 0 | 0 | 0 | 0 | 0 |
| 0 | NA | NA | NA | NA | NA | 1700014L14Rik | | 0 | 0 | 0 | 0 | 0 | 0 | 0 | 0 | 0 | 0 | 0 |
| 0 | NA | NA | NA | NA | NA | Gm24469 | | 0 | 0 | 0 | 0 | 0 | 0 | 0 | 0 | 0 | 0 | 0 |
| 0 | NA | NA | NA | NA | NA | Gm25763 | | 0 | 0 | 0 | 0 | 0 | 0 | 0 | 0 | 0 | 0 | 0 |
| 0 | NA | NA | NA | NA | NA | Gm16725 | | 0 | 0 | 0 | 0 | 0 | 0 | 0 | 0 | 0 | 0 | 0 |
| 0 | NA | NA | NA | NA | NA | Gm35934 | | 0 | 0 | 0 | 0 | 0 | 0 | 0 | 0 | 0 | 0 | 0 |
| 0 | NA | NA | NA | NA | NA | Gm45770 | | 0 | 0 | 0 | 0 | 0 | 0 | 0 | 0 | 0 | 0 | 0 |
| 0 | NA | NA | NA | NA | NA | Gm22551 | | 0 | 0 | 0 | 0 | 0 | 0 | 0 | 0 | 0 | 0 | 0 |
| 0 | NA | NA | NA | NA | NA | Defb40 | defensin beta 40 | 0 | 0 | 0 | 0 | 0 | 0 | 0 | 0 | 0 | 0 | 0 |
| 0 | NA | NA | NA | NA | NA | Defb37 | defensin beta 37 | 0 | 0 | 0 | 0 | 0 | 0 | 0 | 0 | 0 | 0 | 0 |
| 0 | NA | NA | NA | NA | NA | Defb38 | defensin beta 38 | 0 | 0 | 0 | 0 | 0 | 0 | 0 | 0 | 0 | 0 | 0 |
| 0 | NA | NA | NA | NA | NA | Gm15366 | | 0 | 0 | 0 | 0 | 0 | 0 | 0 | 0 | 0 | 0 | 0 |
| 0 | NA | NA | NA | NA | NA | Defb39 | defensin beta 39 | 0 | 0 | 0 | 0 | 0 | 0 | 0 | 0 | 0 | 0 | 0 |
| 0 | NA | NA | NA | NA | NA | Defb12 | defensin beta 12 | 0 | 0 | 0 | 0 | 0 | 0 | 0 | 0 | 0 | 0 | 0 |
| 0 | NA | NA | NA | NA | NA | Defb34 | defensin beta 34 | 0 | 0 | 0 | 0 | 0 | 0 | 0 | 0 | 0 | 0 | 0 |
| 0 | NA | NA | NA | NA | NA | Spag11b | sperm associated antigen 11B | 0 | 0 | 0 | 0 | 0 | 0 | 0 | 0 | 0 | 0 | 0 |
| 0 | NA | NA | NA | NA | NA | Gm45826 | | 0 | 0 | 0 | 0 | 0 | 0 | 0 | 0 | 0 | 0 | 0 |
| 0 | NA | NA | NA | NA | NA | Spag11a | sperm associated antigen 11A | 0 | 0 | 0 | 0 | 0 | 0 | 0 | 0 | 0 | 0 | 0 |
| 0 | NA | NA | NA | NA | NA | Gm39139 | | 0 | 0 | 0 | 0 | 0 | 0 | 0 | 0 | 0 | 0 | 0 |
| 0 | NA | NA | NA | NA | NA | Defb4 | defensin beta 4 | 0 | 0 | 0 | 0 | 0 | 0 | 0 | 0 | 0 | 0 | 0 |
| 0 | NA | NA | NA | NA | NA | Defa-ps12 | | 0 | 0 | 0 | 0 | 0 | 0 | 0 | 0 | 0 | 0 | 0 |
| 0 | NA | NA | NA | NA | NA | Defb6 | defensin beta 6 | 0 | 0 | 0 | 0 | 0 | 0 | 0 | 0 | 0 | 0 | 0 |
| 0 | NA | NA | NA | NA | NA | Defb46 | defensin beta 46 | 0 | 0 | 0 | 0 | 0 | 0 | 0 | 0 | 0 | 0 | 0 |
| 0 | NA | NA | NA | NA | NA | Defb5 | defensin beta 5 | 0 | 0 | 0 | 0 | 0 | 0 | 0 | 0 | 0 | 0 | 0 |
| 0 | NA | NA | NA | NA | NA | Gm15357 | | 0 | 0 | 0 | 0 | 0 | 0 | 0 | 0 | 0 | 0 | 0 |
| 0 | NA | NA | NA | NA | NA | Defb3 | defensin beta 3 | 0 | 0 | 0 | 0 | 0 | 0 | 0 | 0 | 0 | 0 | 0 |
| 0 | NA | NA | NA | NA | NA | Defa-ps13 | | 0 | 0 | 0 | 0 | 0 | 0 | 0 | 0 | 0 | 0 | 0 |
| 0 | NA | NA | NA | NA | NA | Defa-ps14 | | 0 | 0 | 0 | 0 | 0 | 0 | 0 | 0 | 0 | 0 | 0 |
| 0 | NA | NA | NA | NA | NA | Defb8 | defensin beta 8 | 0 | 0 | 0 | 0 | 0 | 0 | 0 | 0 | 0 | 0 | 0 |
| 0 | NA | NA | NA | NA | NA | Defa-ps15 | | 0 | 0 | 0 | 0 | 0 | 0 | 0 | 0 | 0 | 0 | 0 |
| 0 | NA | NA | NA | NA | NA | Gm27288 | | 0 | 0 | 0 | 0 | 0 | 0 | 0 | 0 | 0 | 0 | 0 |
| 0 | NA | NA | NA | NA | NA | Defb7 | defensin beta 7 | 0 | 0 | 0 | 0 | 0 | 0 | 0 | 0 | 0 | 0 | 0 |
| 0 | NA | NA | NA | NA | NA | Gm15358 | | 0 | 0 | 0 | 0 | 0 | 0 | 0 | 0 | 0 | 0 | 0 |
| 0 | NA | NA | NA | NA | NA | Gm27852 | | 0 | 0 | 0 | 0 | 0 | 0 | 0 | 0 | 0 | 0 | 0 |
| 0 | NA | NA | NA | NA | NA | Gm35998 | | 0 | 0 | 0 | 0 | 0 | 0 | 0 | 0 | 0 | 0 | 0 |
| 0 | NA | NA | NA | NA | NA | 4930467E23Ril | RIKEN cDNA 4930467E23 gene | 0 | 0 | 0 | 0 | 0 | 0 | 0 | 0 | 0 | 0 | 0 |
| 0 | NA | NA | NA | NA | NA | Gm31371 | RIKEN cDNA 9530057J20 gene | 0 | 0 | 0 | 0 | 0 | 0 | 0 | 0 | 0 | 0 | 0 |
| 0 | NA | NA | NA | NA | NA | Gm45754 | | 0 | 0 | 0 | 0 | 0 | 0 | 0 | 0 | 0 | 0 | 0 |
| 0 | NA | NA | NA | NA | NA | Gm15319 | predicted gene 15319 | 0 | 0 | 0 | 0 | 0 | 0 | 0 | 0 | 0 | 0 | 0 |
| 0 | NA | NA | NA | NA | NA | Gm7807 | | 0 | 0 | 0 | 0 | 0 | 0 | 0 | 0 | 0 | 0 | 0 |
| 0 | NA | NA | NA | NA | NA | Gm21182 | | 0 | 0 | 0 | 0 | 0 | 0 | 0 | 0 | 0 | 0 | 0 |
| 0 | NA | NA | NA | NA | NA | Gm21112 | | 0 | 0 | 0 | 0 | 0 | 0 | 0 | 0 | 0 | 0 | 0 |
| 0 | NA | NA | NA | NA | NA | Gm21119 | predicted gene, 21119 | 0 | 0 | 0 | 0 | 0 | 0 | 0 | 0 | 0 | 0 | 0 |
| 0 | NA | NA | NA | NA | NA | Gm20797 | | 0 | 0 | 0 | 0 | 0 | 0 | 0 | 0 | 0 | 0 | 0 |
| 0 | NA | NA | NA | NA | NA | Gm20945 | | 0 | 0 | 0 | 0 | 0 | 0 | 0 | 0 | 0 | 0 | 0 |
| 0 | NA | NA | NA | NA | NA | Defb33 | defensin beta 33 | 0 | 0 | 0 | 0 | 0 | 0 | 0 | 0 | 0 | 0 | 0 |
| 0 | NA | NA | NA | NA | NA | Gm6040 | predicted gene 6040 | 0 | 0 | 0 | 0 | 0 | 0 | 0 | 0 | 0 | 0 | 0 |
| 0 | NA | NA | NA | NA | NA | AY761185 | cDNA sequence AY761185 | 0 | 0 | 0 | 0 | 0 | 0 | 0 | 0 | 0 | 0 | 0 |
| 0 | NA | NA | NA | NA | NA | Defa21 | defensin, alpha, 21 | 0 | 0 | 0 | 0 | 0 | 0 | 0 | 0 | 0 | 0 | 0 |
| 0 | NA | NA | NA | NA | NA | Gm23142 | | 0 | 0 | 0 | 0 | 0 | 0 | 0 | 0 | 0 | 0 | 0 |
| 0 | NA | NA | NA | NA | NA | Defa23 | defensin, alpha, 23 | 0 | 0 | 0 | 0 | 0 | 0 | 0 | 0 | 0 | 0 | 0 |
| 0 | NA | NA | NA | NA | NA | Defa35 | defensin, alpha, 35 | 0 | 0 | 0 | 0 | 0 | 0 | 0 | 0 | 0 | 0 | 0 |

| | | | | | | | | | | | | | | | | | |
|---|---|---|---|---|---|---|---|---|---|---|---|---|---|---|---|---|---|
| 0 | NA | NA | NA | NA | NA | Defa-ps3 | | 0 | 0 | 0 | 0 | 0 | 0 | 0 | 0 | 0 | 0 | 0 |
| 0 | NA | NA | NA | NA | NA | Defa25 | defensin, alpha, 25 | 0 | 0 | 0 | 0 | 0 | 0 | 0 | 0 | 0 | 0 | 0 |
| 0 | NA | NA | NA | NA | NA | Defa38 | defensin, alpha, 38 | 0 | 0 | 0 | 0 | 0 | 0 | 0 | 0 | 0 | 0 | 0 |
| 0 | NA | NA | NA | NA | NA | Gm24568 | | 0 | 0 | 0 | 0 | 0 | 0 | 0 | 0 | 0 | 0 | 0 |
| 0 | NA | NA | NA | NA | NA | Gm2869 | | 0 | 0 | 0 | 0 | 0 | 0 | 0 | 0 | 0 | 0 | 0 |
| 0 | NA | NA | NA | NA | NA | Defa-ps4 | | 0 | 0 | 0 | 0 | 0 | 0 | 0 | 0 | 0 | 0 | 0 |
| 0 | NA | NA | NA | NA | NA | Defa30 | defensin, alpha, 30 | 0 | 0 | 0 | 0 | 0 | 0 | 0 | 0 | 0 | 0 | 0 |
| 0 | NA | NA | NA | NA | NA | Defa22 | defensin, alpha, 22 | 0 | 0 | 0 | 0 | 0 | 0 | 0 | 0 | 0 | 0 | 0 |
| 0 | NA | NA | NA | NA | NA | Gm26240 | | 0 | 0 | 0 | 0 | 0 | 0 | 0 | 0 | 0 | 0 | 0 |
| 0 | NA | NA | NA | NA | NA | Defa31 | defensin, alpha, 31 | 0 | 0 | 0 | 0 | 0 | 0 | 0 | 0 | 0 | 0 | 0 |
| 0 | NA | NA | NA | NA | NA | Defa41 | defensin, alpha, 41 | 0 | 0 | 0 | 0 | 0 | 0 | 0 | 0 | 0 | 0 | 0 |
| 0 | NA | NA | NA | NA | NA | Defa-ps5 | | 0 | 0 | 0 | 0 | 0 | 0 | 0 | 0 | 0 | 0 | 0 |
| 0 | NA | NA | NA | NA | NA | Defa-ps6 | | 0 | 0 | 0 | 0 | 0 | 0 | 0 | 0 | 0 | 0 | 0 |
| 0 | NA | NA | NA | NA | NA | Defa-ps7 | | 0 | 0 | 0 | 0 | 0 | 0 | 0 | 0 | 0 | 0 | 0 |
| 0 | NA | NA | NA | NA | NA | Gm15303 | | 0 | 0 | 0 | 0 | 0 | 0 | 0 | 0 | 0 | 0 | 0 |
| 0 | NA | NA | NA | NA | NA | Defa40 | defensin, alpha, 40 | 0 | 0 | 0 | 0 | 0 | 0 | 0 | 0 | 0 | 0 | 0 |
| 0 | NA | NA | NA | NA | NA | Defa-ps8 | defensin, alpha, pseudogene 8 | 0 | 0 | 0 | 0 | 0 | 0 | 0 | 0 | 0 | 0 | 0 |
| 0 | NA | NA | NA | NA | NA | Defa3 | defensin, alpha, 3 | 0 | 0 | 0 | 0 | 0 | 0 | 0 | 0 | 0 | 0 | 0 |
| 0 | NA | NA | NA | NA | NA | Defa5 | defensin, alpha, 5 | 0 | 0 | 0 | 0 | 0 | 0 | 0 | 0 | 0 | 0 | 0 |
| 0 | NA | NA | NA | NA | NA | Gm15300 | | 0 | 0 | 0 | 0 | 0 | 0 | 0 | 0 | 0 | 0 | 0 |
| 0 | NA | NA | NA | NA | NA | Defa27 | defensin, alpha, 27 | 0 | 0 | 0 | 0 | 0 | 0 | 0 | 0 | 0 | 0 | 0 |
| 0 | NA | NA | NA | NA | NA | Defa29 | defensin, alpha, 29 | 0 | 0 | 0 | 0 | 0 | 0 | 0 | 0 | 0 | 0 | 0 |
| 0 | NA | NA | NA | NA | NA | Gm23561 | | 0 | 0 | 0 | 0 | 0 | 0 | 0 | 0 | 0 | 0 | 0 |
| 0 | NA | NA | NA | NA | NA | Defa-ps16 | | 0 | 0 | 0 | 0 | 0 | 0 | 0 | 0 | 0 | 0 | 0 |
| 0 | NA | NA | NA | NA | NA | Defa2 | defensin, alpha, 2 | 0 | 0 | 0 | 0 | 0 | 0 | 0 | 0 | 0 | 0 | 0 |
| 0 | NA | NA | NA | NA | NA | Defa33 | defensin, alpha, 33 | 0 | 0 | 0 | 0 | 0 | 0 | 0 | 0 | 0 | 0 | 0 |
| 0 | NA | NA | NA | NA | NA | Defa36 | defensin, alpha, 36 | 0 | 0 | 0 | 0 | 0 | 0 | 0 | 0 | 0 | 0 | 0 |
| 0 | NA | NA | NA | NA | NA | Gm15305 | | 0 | 0 | 0 | 0 | 0 | 0 | 0 | 0 | 0 | 0 | 0 |
| 0 | NA | NA | NA | NA | NA | Defa-ps18 | | 0 | 0 | 0 | 0 | 0 | 0 | 0 | 0 | 0 | 0 | 0 |
| 0 | NA | NA | NA | NA | NA | Gm15309 | | 0 | 0 | 0 | 0 | 0 | 0 | 0 | 0 | 0 | 0 | 0 |
| 0 | NA | NA | NA | NA | NA | Defa42 | defensin, alpha, 42 | 0 | 0 | 0 | 0 | 0 | 0 | 0 | 0 | 0 | 0 | 0 |
| 0 | NA | NA | NA | NA | NA | Gm23568 | | 0 | 0 | 0 | 0 | 0 | 0 | 0 | 0 | 0 | 0 | 0 |
| 0 | NA | NA | NA | NA | NA | Gm15304 | | 0 | 0 | 0 | 0 | 0 | 0 | 0 | 0 | 0 | 0 | 0 |
| 0 | NA | NA | NA | NA | NA | Defa-ps17 | | 0 | 0 | 0 | 0 | 0 | 0 | 0 | 0 | 0 | 0 | 0 |
| 0 | NA | NA | NA | NA | NA | Defa20 | defensin, alpha, 20 | 0 | 0 | 0 | 0 | 0 | 0 | 0 | 0 | 0 | 0 | 0 |
| 0 | NA | NA | NA | NA | NA | Defa32 | defensin, alpha, 32 | 0 | 0 | 0 | 0 | 0 | 0 | 0 | 0 | 0 | 0 | 0 |
| 0 | NA | NA | NA | NA | NA | Gm24423 | | 0 | 0 | 0 | 0 | 0 | 0 | 0 | 0 | 0 | 0 | 0 |
| 0 | NA | NA | NA | NA | NA | Defa37 | defensin, alpha, 37 | 0 | 0 | 0 | 0 | 0 | 0 | 0 | 0 | 0 | 0 | 0 |
| 0 | NA | NA | NA | NA | NA | Gm15311 | | 0 | 0 | 0 | 0 | 0 | 0 | 0 | 0 | 0 | 0 | 0 |
| 0 | NA | NA | NA | NA | NA | Defa28 | defensin, alpha, 28 | 0 | 0 | 0 | 0 | 0 | 0 | 0 | 0 | 0 | 0 | 0 |
| 0 | NA | NA | NA | NA | NA | Gm15312 | | 0 | 0 | 0 | 0 | 0 | 0 | 0 | 0 | 0 | 0 | 0 |
| 0 | NA | NA | NA | NA | NA | Defa43 | defensin, alpha, 43 | 0 | 0 | 0 | 0 | 0 | 0 | 0 | 0 | 0 | 0 | 0 |
| 0 | NA | NA | NA | NA | NA | Gm24829 | | 0 | 0 | 0 | 0 | 0 | 0 | 0 | 0 | 0 | 0 | 0 |
| 0 | NA | NA | NA | NA | NA | Gm15314 | | 0 | 0 | 0 | 0 | 0 | 0 | 0 | 0 | 0 | 0 | 0 |
| 0 | NA | NA | NA | NA | NA | Defa26 | defensin, alpha, 26 | 0 | 0 | 0 | 0 | 0 | 0 | 0 | 0 | 0 | 0 | 0 |
| 0 | NA | NA | NA | NA | NA | Gm15313 | predicted gene 15313 | 0 | 0 | 0 | 0 | 0 | 0 | 0 | 0 | 0 | 0 | 0 |
| 0 | NA | NA | NA | NA | NA | Defa17 | defensin, alpha, 17 | 0 | 0 | 0 | 0 | 0 | 0 | 0 | 0 | 0 | 0 | 0 |
| 0 | NA | NA | NA | NA | NA | Defa34 | defensin, alpha, 34 | 0 | 0 | 0 | 0 | 0 | 0 | 0 | 0 | 0 | 0 | 0 |
| 0 | NA | NA | NA | NA | NA | Defa-ps9 | | 0 | 0 | 0 | 0 | 0 | 0 | 0 | 0 | 0 | 0 | 0 |
| 0 | NA | NA | NA | NA | NA | Defa-ps1 | | 0 | 0 | 0 | 0 | 0 | 0 | 0 | 0 | 0 | 0 | 0 |
| 0 | NA | NA | NA | NA | NA | Gm22447 | | 0 | 0 | 0 | 0 | 0 | 0 | 0 | 0 | 0 | 0 | 0 |
| 0 | NA | NA | NA | NA | NA | Gm7869 | | 0 | 0 | 0 | 0 | 0 | 0 | 0 | 0 | 0 | 0 | 0 |
| 0 | NA | NA | NA | NA | NA | Defa-ps10 | | 0 | 0 | 0 | 0 | 0 | 0 | 0 | 0 | 0 | 0 | 0 |
| 0 | NA | NA | NA | NA | NA | Defa24 | defensin, alpha, 24 | 0 | 0 | 0 | 0 | 0 | 0 | 0 | 0 | 0 | 0 | 0 |
| 0 | NA | NA | NA | NA | NA | Defa-ps11 | | 0 | 0 | 0 | 0 | 0 | 0 | 0 | 0 | 0 | 0 | 0 |
| 0 | NA | NA | NA | NA | NA | Gm15316 | | 0 | 0 | 0 | 0 | 0 | 0 | 0 | 0 | 0 | 0 | 0 |
| 0 | NA | NA | NA | NA | NA | Defb50 | defensin beta 50 | 0 | 0 | 0 | 0 | 0 | 0 | 0 | 0 | 0 | 0 | 0 |
| 0 | NA | NA | NA | NA | NA | Defb2 | defensin beta 2 | 0 | 0 | 0 | 0 | 0 | 0 | 0 | 0 | 0 | 0 | 0 |
| 0 | NA | NA | NA | NA | NA | Defb10 | defensin beta 10 | 0 | 0 | 0 | 0 | 0 | 0 | 0 | 0 | 0 | 0 | 0 |
| 0 | NA | NA | NA | NA | NA | Defb9 | defensin beta 9 | 0 | 0 | 0 | 0 | 0 | 0 | 0 | 0 | 0 | 0 | 0 |

| | | | | | | | | | | | | | | | | | | |
|---|---|---|---|---|---|---|---|---|---|---|---|---|---|---|---|---|---|---|
| 0 | NA | NA | NA | NA | NA | Defb11 | defensin beta 11 | 0 | 0 | 0 | 0 | 0 | 0 | 0 | 0 | 0 | 0 | 0 |
| 0 | NA | NA | NA | NA | NA | Gm36879 | | 0 | 0 | 0 | 0 | 0 | 0 | 0 | 0 | 0 | 0 | 0 |
| 0 | NA | NA | NA | NA | NA | Defb15 | defensin beta 15 | 0 | 0 | 0 | 0 | 0 | 0 | 0 | 0 | 0 | 0 | 0 |
| 0 | NA | NA | NA | NA | NA | Gm45796 | | 0 | 0 | 0 | 0 | 0 | 0 | 0 | 0 | 0 | 0 | 0 |
| 0 | NA | NA | NA | NA | NA | Defb35 | defensin beta 35 | 0 | 0 | 0 | 0 | 0 | 0 | 0 | 0 | 0 | 0 | 0 |
| 0 | NA | NA | NA | NA | NA | Defb13 | defensin beta 13 | 0 | 0 | 0 | 0 | 0 | 0 | 0 | 0 | 0 | 0 | 0 |
| 0 | NA | NA | NA | NA | NA | Fam90a1a | family with sequence similarity 90, membe | 0 | 0 | 0 | 0 | 0 | 0 | 0 | 0 | 0 | 0 | 0 |
| 0 | NA | NA | NA | NA | NA | Ccdc70 | coiled-coil domain containing 70 | 0 | 0 | 0 | 0 | 0 | 0 | 0 | 0 | 0 | 0 | 0 |
| 0 | NA | NA | NA | NA | NA | Gm45452 | | 0 | 0 | 0 | 0 | 0 | 0 | 0 | 0 | 0 | 0 | 0 |
| 0 | NA | NA | NA | NA | NA | Gm45453 | | 0 | 0 | 0 | 0 | 0 | 0 | 0 | 0 | 0 | 0 | 0 |
| 0 | NA | NA | NA | NA | NA | Dkk4 | dickkopf WNT signaling pathway inhibitor | 0 | 0 | 0 | 0 | 0 | 0 | 0 | 0 | 0 | 0 | 0 |
| 0 | NA | NA | NA | NA | NA | Gm15346 | | 0 | 0 | 0 | 0 | 0 | 0 | 0 | 0 | 0 | 0 | 0 |
| 0 | NA | NA | NA | NA | NA | Gm24735 | | 0 | 0 | 0 | 0 | 0 | 0 | 0 | 0 | 0 | 0 | 0 |
| 0 | NA | NA | NA | NA | NA | Gm44365 | | 0 | 0 | 0 | 0 | 0 | 0 | 0 | 0 | 0 | 0 | 0 |
| 0 | NA | NA | NA | NA | NA | Gm15815 | | 0 | 0 | 0 | 0 | 0 | 0 | 0 | 0 | 0 | 0 | 0 |
| 0 | NA | NA | NA | NA | NA | Mir486 | | 0 | 0 | 0 | 0 | 0 | 0 | 0 | 0 | 0 | 0 | 0 |
| 0 | NA | NA | NA | NA | NA | Mir3107 | | 0 | 0 | 0 | 0 | 0 | 0 | 0 | 0 | 0 | 0 | 0 |
| 0 | NA | NA | NA | NA | NA | Gm24335 | | 0 | 0 | 0 | 0 | 0 | 0 | 0 | 0 | 0 | 0 | 0 |
| 0 | NA | NA | NA | NA | NA | Gm45165 | | 0 | 0 | 0 | 0 | 0 | 0 | 0 | 0 | 0 | 0 | 0 |
| 0 | NA | NA | NA | NA | NA | Gm45166 | | 0 | 0 | 0 | 0 | 0 | 0 | 0 | 0 | 0 | 0 | 0 |
| 0 | NA | NA | NA | NA | NA | A730045E13Rik | | 0 | 0 | 0 | 0 | 0 | 0 | 0 | 0 | 0 | 0 | 0 |
| 0 | NA | NA | NA | NA | NA | Gm45163 | | 0 | 0 | 0 | 0 | 0 | 0 | 0 | 0 | 0 | 0 | 0 |
| 0 | NA | NA | NA | NA | NA | Gm45164 | | 0 | 0 | 0 | 0 | 0 | 0 | 0 | 0 | 0 | 0 | 0 |
| 0 | NA | NA | NA | NA | NA | Gm30692 | | 0 | 0 | 0 | 0 | 0 | 0 | 0 | 0 | 0 | 0 | 0 |
| 0 | NA | NA | NA | NA | NA | Gm44620 | | 0 | 0 | 0 | 0 | 0 | 0 | 0 | 0 | 0 | 0 | 0 |
| 0 | NA | NA | NA | NA | NA | Ido2 | indoleamine 2,3-dioxygenase 2 | 0 | 0 | 0 | 0 | 0 | 0 | 0 | 0 | 0 | 0 | 0 |
| 0 | NA | NA | NA | NA | NA | Adam18 | a disintegrin and metallopeptidase domai | 0 | 0 | 0 | 0 | 0 | 0 | 0 | 0 | 0 | 0 | 0 |
| 0 | NA | NA | NA | NA | NA | Adam3 | a disintegrin and metallopeptidase domai | 0 | 0 | 0 | 0 | 0 | 0 | 0 | 0 | 0 | 0 | 0 |
| 0 | NA | NA | NA | NA | NA | Gm26208 | | 0 | 0 | 0 | 0 | 0 | 0 | 0 | 0 | 0 | 0 | 0 |
| 0 | NA | NA | NA | NA | NA | Gm45528 | | 0 | 0 | 0 | 0 | 0 | 0 | 0 | 0 | 0 | 0 | 0 |
| 0 | NA | NA | NA | NA | NA | Gm23898 | | 0 | 0 | 0 | 0 | 0 | 0 | 0 | 0 | 0 | 0 | 0 |
| 0 | NA | NA | NA | NA | NA | Gm45745 | | 0 | 0 | 0 | 0 | 0 | 0 | 0 | 0 | 0 | 0 | 0 |
| 0 | NA | NA | NA | NA | NA | Mir8108 | | 0 | 0 | 0 | 0 | 0 | 0 | 0 | 0 | 0 | 0 | 0 |
| 0 | NA | NA | NA | NA | NA | Gm23184 | | 0 | 0 | 0 | 0 | 0 | 0 | 0 | 0 | 0 | 0 | 0 |
| 0 | NA | NA | NA | NA | NA | Gm24992 | | 0 | 0 | 0 | 0 | 0 | 0 | 0 | 0 | 0 | 0 | 0 |
| 0 | NA | NA | NA | NA | NA | Gm45411 | | 0 | 0 | 0 | 0 | 0 | 0 | 0 | 0 | 0 | 0 | 0 |
| 0 | NA | NA | NA | NA | NA | Gm45692 | | 0 | 0 | 0 | 0 | 0 | 0 | 0 | 0 | 0 | 0 | 0 |
| 0 | NA | NA | NA | NA | NA | Gm44299 | | 0 | 0 | 0 | 0 | 0 | 0 | 0 | 0 | 0 | 0 | 0 |
| 0 | NA | NA | NA | NA | NA | Gm50479 | | 0 | 0 | 0 | 0 | 0 | 0 | 0 | 0 | 0 | 0 | 0 |
| 0 | NA | NA | NA | NA | NA | Gm22227 | | 0 | 0 | 0 | 0 | 0 | 0 | 0 | 0 | 0 | 0 | 0 |
| 0 | NA | NA | NA | NA | NA | Gm45305 | | 0 | 0 | 0 | 0 | 0 | 0 | 0 | 0 | 0 | 0 | 0 |
| 0 | NA | NA | NA | NA | NA | Gm31898 | | 0 | 0 | 0 | 0 | 0 | 0 | 0 | 0 | 0 | 0 | 0 |
| 0 | NA | NA | NA | NA | NA | Gm45580 | | 0 | 0 | 0 | 0 | 0 | 0 | 0 | 0 | 0 | 0 | 0 |
| 0 | NA | NA | NA | NA | NA | Gm31983 | | 0 | 0 | 0 | 0 | 0 | 0 | 0 | 0 | 0 | 0 | 0 |
| 0 | NA | NA | NA | NA | NA | Gm39149 | | 0 | 0 | 0 | 0 | 0 | 0 | 0 | 0 | 0 | 0 | 0 |
| 0 | NA | NA | NA | NA | NA | Gm32050 | | 0 | 0 | 0 | 0 | 0 | 0 | 0 | 0 | 0 | 0 | 0 |
| 0 | NA | NA | NA | NA | NA | Gm45584 | | 0 | 0 | 0 | 0 | 0 | 0 | 0 | 0 | 0 | 0 | 0 |
| 0 | NA | NA | NA | NA | NA | Gm45646 | | 0 | 0 | 0 | 0 | 0 | 0 | 0 | 0 | 0 | 0 | 0 |
| 0 | NA | NA | NA | NA | NA | Gm45647 | | 0 | 0 | 0 | 0 | 0 | 0 | 0 | 0 | 0 | 0 | 0 |
| 0 | NA | NA | NA | NA | NA | 2310008N11Rik | | 0 | 0 | 0 | 0 | 0 | 0 | 0 | 0 | 0 | 0 | 0 |
| 0 | NA | NA | NA | NA | NA | Gm45572 | | 0 | 0 | 0 | 0 | 0 | 0 | 0 | 0 | 0 | 0 | 0 |
| 0 | NA | NA | NA | NA | NA | Gm45370 | | 0 | 0 | 0 | 0 | 0 | 0 | 0 | 0 | 0 | 0 | 0 |
| 0 | NA | NA | NA | NA | NA | Gm19164 | | 0 | 0 | 0 | 0 | 0 | 0 | 0 | 0 | 0 | 0 | 0 |
| 0 | NA | NA | NA | NA | NA | 4933416M07Rik | | 0 | 0 | 0 | 0 | 0 | 0 | 0 | 0 | 0 | 0 | 0 |
| 0 | NA | NA | NA | NA | NA | Gm45267 | | 0 | 0 | 0 | 0 | 0 | 0 | 0 | 0 | 0 | 0 | 0 |
| 0 | NA | NA | NA | NA | NA | Adrb3 | adrenergic receptor, beta 3 | 0 | 0 | 0 | 0 | 0 | 0 | 0 | 0 | 0 | 0 | 0 |
| 0 | NA | NA | NA | NA | NA | Gm9731 | | 0 | 0 | 0 | 0 | 0 | 0 | 0 | 0 | 0 | 0 | 0 |
| 0 | NA | NA | NA | NA | NA | Tex24 | testis expressed gene 24 | 0 | 0 | 0 | 0 | 0 | 0 | 0 | 0 | 0 | 0 | 0 |
| 0 | NA | NA | NA | NA | NA | Chrnb3 | cholinergic receptor, nicotinic, beta polype | 0 | 0 | 0 | 0 | 0 | 0 | 0 | 0 | 0 | 0 | 0 |
| 0 | NA | NA | NA | NA | NA | Chrna6 | cholinergic receptor, nicotinic, alpha poly | 0 | 0 | 0 | 0 | 0 | 0 | 0 | 0 | 0 | 0 | 0 |

| | | | | | | | | | | | | | | | | | | |
|---|---|---|---|---|---|---|---|---|---|---|---|---|---|---|---|---|---|---|
| 0 | NA | NA | NA | NA | NA | Gm24410 | | 0 | 0 | 0 | 0 | 0 | 0 | 0 | 0 | 0 | 0 | 0 |
| 0 | NA | NA | NA | NA | NA | Gm45861 | POTE ankyrin domain family, member G | 0 | 0 | 0 | 0 | 0 | 0 | 0 | 0 | 0 | 0 | 0 |
| 0 | NA | NA | NA | NA | NA | Poteg | POTE ankyrin domain family, member G | 0 | 0 | 0 | 0 | 0 | 0 | 0 | 0 | 0 | 0 | 0 |
| 0 | NA | NA | NA | NA | NA | Gm1698 | | 0 | 0 | 0 | 0 | 0 | 0 | 0 | 0 | 0 | 0 | 0 |
| 0 | NA | NA | NA | NA | NA | Gm44493 | | 0 | 0 | 0 | 0 | 0 | 0 | 0 | 0 | 0 | 0 | 0 |
| 0 | NA | NA | NA | NA | NA | Gm24202 | | 0 | 0 | 0 | 0 | 0 | 0 | 0 | 0 | 0 | 0 | 0 |
| 0 | NA | NA | NA | NA | NA | Gm45751 | | 0 | 0 | 0 | 0 | 0 | 0 | 0 | 0 | 0 | 0 | 0 |
| 0 | NA | NA | NA | NA | NA | Gm8096 | | 0 | 0 | 0 | 0 | 0 | 0 | 0 | 0 | 0 | 0 | 0 |
| 0 | NA | NA | NA | NA | NA | 1700008N11Rik | | 0 | 0 | 0 | 0 | 0 | 0 | 0 | 0 | 0 | 0 | 0 |
| 0 | NA | NA | NA | NA | NA | Gm8110 | | 0 | 0 | 0 | 0 | 0 | 0 | 0 | 0 | 0 | 0 | 0 |
| 0 | NA | NA | NA | NA | NA | Gm45571 | | 0 | 0 | 0 | 0 | 0 | 0 | 0 | 0 | 0 | 0 | 0 |
| 0 | NA | NA | NA | NA | NA | Gm26795 | | 0 | 0 | 0 | 0 | 0 | 0 | 0 | 0 | 0 | 0 | 0 |
| 0 | NA | NA | NA | NA | NA | Gm9911 | | 0 | 0 | 0 | 0 | 0 | 0 | 0 | 0 | 0 | 0 | 0 |
| 0 | NA | NA | NA | NA | NA | Gm45652 | | 0 | 0 | 0 | 0 | 0 | 0 | 0 | 0 | 0 | 0 | 0 |
| 0 | NA | NA | NA | NA | NA | Svet1 | | 0 | 0 | 0 | 0 | 0 | 0 | 0 | 0 | 0 | 0 | 0 |
| 0 | NA | NA | NA | NA | NA | Gm32567 | | 0 | 0 | 0 | 0 | 0 | 0 | 0 | 0 | 0 | 0 | 0 |
| 0 | NA | NA | NA | NA | NA | Gm45492 | | 0 | 0 | 0 | 0 | 0 | 0 | 0 | 0 | 0 | 0 | 0 |
| 0 | NA | NA | NA | NA | NA | Gm39154 | | 0 | 0 | 0 | 0 | 0 | 0 | 0 | 0 | 0 | 0 | 0 |
| 0 | NA | NA | NA | NA | NA | Gm6853 | | 0 | 0 | 0 | 0 | 0 | 0 | 0 | 0 | 0 | 0 | 0 |
| 0 | NA | NA | NA | NA | NA | Gm45489 | | 0 | 0 | 0 | 0 | 0 | 0 | 0 | 0 | 0 | 0 | 0 |
| 0 | NA | NA | NA | NA | NA | 4933433F19Rik | | 0 | 0 | 0 | 0 | 0 | 0 | 0 | 0 | 0 | 0 | 0 |
| 0 | NA | NA | NA | NA | NA | Gm45252 | | 0 | 0 | 0 | 0 | 0 | 0 | 0 | 0 | 0 | 0 | 0 |
| 0 | NA | NA | NA | NA | NA | Gm8051 | | 0 | 0 | 0 | 0 | 0 | 0 | 0 | 0 | 0 | 0 | 0 |
| 0 | NA | NA | NA | NA | NA | 7420700N18Rik | | 0 | 0 | 0 | 0 | 0 | 0 | 0 | 0 | 0 | 0 | 0 |
| 0 | NA | NA | NA | NA | NA | Gm19096 | | 0 | 0 | 0 | 0 | 0 | 0 | 0 | 0 | 0 | 0 | 0 |
| 0 | NA | NA | NA | NA | NA | | | 0 | 0 | 0 | 0 | 0 | 0 | 0 | 0 | 0 | 0 | 0 |
| 0 | NA | NA | NA | NA | NA | Gm45303 | | 0 | 0 | 0 | 0 | 0 | 0 | 0 | 0 | 0 | 0 | 0 |
| 0 | NA | NA | NA | NA | NA | Gm23539 | | 0 | 0 | 0 | 0 | 0 | 0 | 0 | 0 | 0 | 0 | 0 |
| 0 | NA | NA | NA | NA | NA | Gm45304 | | 0 | 0 | 0 | 0 | 0 | 0 | 0 | 0 | 0 | 0 | 0 |
| 0 | NA | NA | NA | NA | NA | Gm24184 | | 0 | 0 | 0 | 0 | 0 | 0 | 0 | 0 | 0 | 0 | 0 |
| 0 | NA | NA | NA | NA | NA | | | 0 | 0 | 0 | 0 | 0 | 0 | 0 | 0 | 0 | 0 | 0 |
| 0 | NA | NA | NA | NA | NA | Gm22140 | | 0 | 0 | 0 | 0 | 0 | 0 | 0 | 0 | 0 | 0 | 0 |
| 0 | NA | NA | NA | NA | NA | Gm3985 | predicted gene 3985 | 0 | 0 | 0 | 0 | 0 | 0 | 0 | 0 | 0 | 0 | 0 |
| 0 | NA | NA | NA | NA | NA | Gm45587 | | 0 | 0 | 0 | 0 | 0 | 0 | 0 | 0 | 0 | 0 | 0 |
| 0 | NA | NA | NA | NA | NA | Gm26632 | | 0 | 0 | 0 | 0 | 0 | 0 | 0 | 0 | 0 | 0 | 0 |
| 0 | NA | NA | NA | NA | NA | Gm26978 | | 0 | 0 | 0 | 0 | 0 | 0 | 0 | 0 | 0 | 0 | 0 |
| 0 | NA | NA | NA | NA | NA | Gm39158 | | 0 | 0 | 0 | 0 | 0 | 0 | 0 | 0 | 0 | 0 | 0 |
| 0 | NA | NA | NA | NA | NA | Mir6395 | | 0 | 0 | 0 | 0 | 0 | 0 | 0 | 0 | 0 | 0 | 0 |
| 0 | NA | NA | NA | NA | NA | Gm29243 | | 0 | 0 | 0 | 0 | 0 | 0 | 0 | 0 | 0 | 0 | 0 |
| 0 | NA | NA | NA | NA | NA | Gm23177 | | 0 | 0 | 0 | 0 | 0 | 0 | 0 | 0 | 0 | 0 | 0 |
| 0 | NA | NA | NA | NA | NA | Gm6100 | | 0 | 0 | 0 | 0 | 0 | 0 | 0 | 0 | 0 | 0 | 0 |
| 0 | NA | NA | NA | NA | NA | Gm8254 | | 0 | 0 | 0 | 0 | 0 | 0 | 0 | 0 | 0 | 0 | 0 |
| 0 | NA | NA | NA | NA | NA | Gm45350 | | 0 | 0 | 0 | 0 | 0 | 0 | 0 | 0 | 0 | 0 | 0 |
| 0 | NA | NA | NA | NA | NA | Gm33831 | | 0 | 0 | 0 | 0 | 0 | 0 | 0 | 0 | 0 | 0 | 0 |
| 0 | NA | NA | NA | NA | NA | Gm33968 | | 0 | 0 | 0 | 0 | 0 | 0 | 0 | 0 | 0 | 0 | 0 |
| 0 | NA | NA | NA | NA | NA | Gm34096 | | 0 | 0 | 0 | 0 | 0 | 0 | 0 | 0 | 0 | 0 | 0 |
| 0 | NA | NA | NA | NA | NA | Gm45627 | | 0 | 0 | 0 | 0 | 0 | 0 | 0 | 0 | 0 | 0 | 0 |
| 0 | NA | NA | NA | NA | NA | Gm34368 | | 0 | 0 | 0 | 0 | 0 | 0 | 0 | 0 | 0 | 0 | 0 |
| 0 | NA | NA | NA | NA | NA | Gm34419 | | 0 | 0 | 0 | 0 | 0 | 0 | 0 | 0 | 0 | 0 | 0 |
| 0 | NA | NA | NA | NA | NA | Rpl31-ps23 | | 0 | 0 | 0 | 0 | 0 | 0 | 0 | 0 | 0 | 0 | 0 |
| 0 | NA | NA | NA | NA | NA | Gm45626 | | 0 | 0 | 0 | 0 | 0 | 0 | 0 | 0 | 0 | 0 | 0 |
| 0 | NA | NA | NA | NA | NA | Gm45625 | | 0 | 0 | 0 | 0 | 0 | 0 | 0 | 0 | 0 | 0 | 0 |
| 0 | NA | NA | NA | NA | NA | Gm34474 | | 0 | 0 | 0 | 0 | 0 | 0 | 0 | 0 | 0 | 0 | 0 |
| 0 | NA | NA | NA | NA | NA | Gm34597 | | 0 | 0 | 0 | 0 | 0 | 0 | 0 | 0 | 0 | 0 | 0 |
| 0 | NA | NA | NA | NA | NA | Gm45301 | | 0 | 0 | 0 | 0 | 0 | 0 | 0 | 0 | 0 | 0 | 0 |
| 0 | NA | NA | NA | NA | NA | Gm34853 | | 0 | 0 | 0 | 0 | 0 | 0 | 0 | 0 | 0 | 0 | 0 |
| 0 | NA | NA | NA | NA | NA | 1700015I17Rik | | 0 | 0 | 0 | 0 | 0 | 0 | 0 | 0 | 0 | 0 | 0 |
| 0 | NA | NA | NA | NA | NA | Gm22216 | | 0 | 0 | 0 | 0 | 0 | 0 | 0 | 0 | 0 | 0 | 0 |
| 0 | NA | NA | NA | NA | NA | Gm44474 | | 0 | 0 | 0 | 0 | 0 | 0 | 0 | 0 | 0 | 0 | 0 |
| 0 | NA | NA | NA | NA | NA | Gm45566 | | 0 | 0 | 0 | 0 | 0 | 0 | 0 | 0 | 0 | 0 | 0 |

| | | | | | | | | | | | | | | | | | |
|---|---|---|---|---|---|---|---|---|---|---|---|---|---|---|---|---|---|
| 0 | NA | NA | NA | NA | NA | Gm22030 | | 0 | 0 | 0 | 0 | 0 | 0 | 0 | 0 | 0 | 0 | 0 |
| 0 | NA | NA | NA | NA | NA | Gm39164 | | 0 | 0 | 0 | 0 | 0 | 0 | 0 | 0 | 0 | 0 | 0 |
| 0 | NA | NA | NA | NA | NA | 4933430A20Rik | | 0 | 0 | 0 | 0 | 0 | 0 | 0 | 0 | 0 | 0 | 0 |
| 0 | NA | NA | NA | NA | NA | Gm35520 | | 0 | 0 | 0 | 0 | 0 | 0 | 0 | 0 | 0 | 0 | 0 |
| 0 | NA | NA | NA | NA | NA | Gm26150 | | 0 | 0 | 0 | 0 | 0 | 0 | 0 | 0 | 0 | 0 | 0 |
| 0 | NA | NA | NA | NA | NA | Gm19140 | | 0 | 0 | 0 | 0 | 0 | 0 | 0 | 0 | 0 | 0 | 0 |
| 0 | NA | NA | NA | NA | NA | G630064G18Rik | | 0 | 0 | 0 | 0 | 0 | 0 | 0 | 0 | 0 | 0 | 0 |
| 0 | NA | NA | NA | NA | NA | Gm45690 | | 0 | 0 | 0 | 0 | 0 | 0 | 0 | 0 | 0 | 0 | 0 |
| 0 | NA | NA | NA | NA | NA | Gm10683 | | 0 | 0 | 0 | 0 | 0 | 0 | 0 | 0 | 0 | 0 | 0 |
| 0 | NA | NA | NA | NA | NA | Gm8342 | | 0 | 0 | 0 | 0 | 0 | 0 | 0 | 0 | 0 | 0 | 0 |
| 0 | NA | NA | NA | NA | NA | Gm45524 | | 0 | 0 | 0 | 0 | 0 | 0 | 0 | 0 | 0 | 0 | 0 |
| 0 | NA | NA | NA | NA | NA | Gm3433 | | 0 | 0 | 0 | 0 | 0 | 0 | 0 | 0 | 0 | 0 | 0 |
| 0 | NA | NA | NA | NA | NA | Gm7319 | | 0 | 0 | 0 | 0 | 0 | 0 | 0 | 0 | 0 | 0 | 0 |
| 0 | NA | NA | NA | NA | NA | Gm45523 | | 0 | 0 | 0 | 0 | 0 | 0 | 0 | 0 | 0 | 0 | 0 |
| 0 | NA | NA | NA | NA | NA | Gm25126 | | 0 | 0 | 0 | 0 | 0 | 0 | 0 | 0 | 0 | 0 | 0 |
| 0 | NA | NA | NA | NA | NA | Gm45368 | | 0 | 0 | 0 | 0 | 0 | 0 | 0 | 0 | 0 | 0 | 0 |
| 0 | NA | NA | NA | NA | NA | Gm6203 | | 0 | 0 | 0 | 0 | 0 | 0 | 0 | 0 | 0 | 0 | 0 |
| 0 | NA | NA | NA | NA | NA | Gm16205 | | 0 | 0 | 0 | 0 | 0 | 0 | 0 | 0 | 0 | 0 | 0 |
| 0 | NA | NA | NA | NA | NA | Mir383 | | 0 | 0 | 0 | 0 | 0 | 0 | 0 | 0 | 0 | 0 | 0 |
| 0 | NA | NA | NA | NA | NA | | | 0 | 0 | 0 | 0 | 0 | 0 | 0 | 0 | 0 | 0 | 0 |
| 0 | NA | NA | NA | NA | NA | Gm23128 | | 0 | 0 | 0 | 0 | 0 | 0 | 0 | 0 | 0 | 0 | 0 |
| 0 | NA | NA | NA | NA | NA | Gm23128 | | 0 | 0 | 0 | 0 | 0 | 0 | 0 | 0 | 0 | 0 | 0 |
| 0 | NA | NA | NA | NA | NA | Gm20948 | | 0 | 0 | 0 | 0 | 0 | 0 | 0 | 0 | 0 | 0 | 0 |
| 0 | NA | NA | NA | NA | NA | Gm17766 | | 0 | 0 | 0 | 0 | 0 | 0 | 0 | 0 | 0 | 0 | 0 |
| 0 | NA | NA | NA | NA | NA | Gm45398 | | 0 | 0 | 0 | 0 | 0 | 0 | 0 | 0 | 0 | 0 | 0 |
| 0 | NA | NA | NA | NA | NA | Zfp148-ps1 | | 0 | 0 | 0 | 0 | 0 | 0 | 0 | 0 | 0 | 0 | 0 |
| 0 | NA | NA | NA | NA | NA | Gm18644 | | 0 | 0 | 0 | 0 | 0 | 0 | 0 | 0 | 0 | 0 | 0 |
| 0 | NA | NA | NA | NA | NA | Gm5609 | | 0 | 0 | 0 | 0 | 0 | 0 | 0 | 0 | 0 | 0 | 0 |
| 0 | NA | NA | NA | NA | NA | Adam24 | a disintegrin and metallopeptidase domai | 0 | 0 | 0 | 0 | 0 | 0 | 0 | 0 | 0 | 0 | 0 |
| 0 | NA | NA | NA | NA | NA | Gm2080 | | 0 | 0 | 0 | 0 | 0 | 0 | 0 | 0 | 0 | 0 | 0 |
| 0 | NA | NA | NA | NA | NA | Adam25 | a disintegrin and metallopeptidase domai | 0 | 0 | 0 | 0 | 0 | 0 | 0 | 0 | 0 | 0 | 0 |
| 0 | NA | NA | NA | NA | NA | Adam20 | a disintegrin and metallopeptidase domai | 0 | 0 | 0 | 0 | 0 | 0 | 0 | 0 | 0 | 0 | 0 |
| 0 | NA | NA | NA | NA | NA | Adam39 | a disintegrin and metallopeptidase domai | 0 | 0 | 0 | 0 | 0 | 0 | 0 | 0 | 0 | 0 | 0 |
| 0 | NA | NA | NA | NA | NA | Gm16193 | | 0 | 0 | 0 | 0 | 0 | 0 | 0 | 0 | 0 | 0 | 0 |
| 0 | NA | NA | NA | NA | NA | 2810404M03Rik | | 0 | 0 | 0 | 0 | 0 | 0 | 0 | 0 | 0 | 0 | 0 |
| 0 | NA | NA | NA | NA | NA | Hspd1-ps1 | | 0 | 0 | 0 | 0 | 0 | 0 | 0 | 0 | 0 | 0 | 0 |
| 0 | NA | NA | NA | NA | NA | Zfp353-ps | | 0 | 0 | 0 | 0 | 0 | 0 | 0 | 0 | 0 | 0 | 0 |
| 0 | NA | NA | NA | NA | NA | Gm23531 | | 0 | 0 | 0 | 0 | 0 | 0 | 0 | 0 | 0 | 0 | 0 |
| 0 | NA | NA | NA | NA | NA | Gm26181 | | 0 | 0 | 0 | 0 | 0 | 0 | 0 | 0 | 0 | 0 | 0 |
| 0 | NA | NA | NA | NA | NA | Gm18414 | | 0 | 0 | 0 | 0 | 0 | 0 | 0 | 0 | 0 | 0 | 0 |
| 0 | NA | NA | NA | NA | NA | Gm45590 | | 0 | 0 | 0 | 0 | 0 | 0 | 0 | 0 | 0 | 0 | 0 |
| 0 | NA | NA | NA | NA | NA | Triml1 | tripartite motif family-like 1 | 0 | 0 | 0 | 0 | 0 | 0 | 0 | 0 | 0 | 0 | 0 |
| 0 | NA | NA | NA | NA | NA | Gm6284 | | 0 | 0 | 0 | 0 | 0 | 0 | 0 | 0 | 0 | 0 | 0 |
| 0 | NA | NA | NA | NA | NA | Triml2 | tripartite motif family-like 2 | 0 | 0 | 0 | 0 | 0 | 0 | 0 | 0 | 0 | 0 | 0 |
| 0 | NA | NA | NA | NA | NA | Gm2221 | | 0 | 0 | 0 | 0 | 0 | 0 | 0 | 0 | 0 | 0 | 0 |
| 0 | NA | NA | NA | NA | NA | Zfp42 | zinc finger protein 42 | 0 | 0 | 0 | 0 | 0 | 0 | 0 | 0 | 0 | 0 | 0 |
| 0 | NA | NA | NA | NA | NA | Gm22986 | | 0 | 0 | 0 | 0 | 0 | 0 | 0 | 0 | 0 | 0 | 0 |
| 0 | NA | NA | NA | NA | NA | Adam26b | a disintegrin and metallopeptidase domai | 0 | 0 | 0 | 0 | 0 | 0 | 0 | 0 | 0 | 0 | 0 |
| 0 | NA | NA | NA | NA | NA | Adam26a | a disintegrin and metallopeptidase domai | 0 | 0 | 0 | 0 | 0 | 0 | 0 | 0 | 0 | 0 | 0 |
| 0 | NA | NA | NA | NA | NA | Adam34l | a disintegrin and metallopeptidase domai | 0 | 0 | 0 | 0 | 0 | 0 | 0 | 0 | 0 | 0 | 0 |
| 0 | NA | NA | NA | NA | NA | Adam34 | a disintegrin and metallopeptidase domai | 0 | 0 | 0 | 0 | 0 | 0 | 0 | 0 | 0 | 0 | 0 |
| 0 | NA | NA | NA | NA | NA | Gm5347 | | 0 | 0 | 0 | 0 | 0 | 0 | 0 | 0 | 0 | 0 | 0 |
| 0 | NA | NA | NA | NA | NA | Gm35629 | | 0 | 0 | 0 | 0 | 0 | 0 | 0 | 0 | 0 | 0 | 0 |
| 0 | NA | NA | NA | NA | NA | Gm45488 | | 0 | 0 | 0 | 0 | 0 | 0 | 0 | 0 | 0 | 0 | 0 |
| 0 | NA | NA | NA | NA | NA | Gm6011 | | 0 | 0 | 0 | 0 | 0 | 0 | 0 | 0 | 0 | 0 | 0 |
| 0 | NA | NA | NA | NA | NA | Gm6303 | | 0 | 0 | 0 | 0 | 0 | 0 | 0 | 0 | 0 | 0 | 0 |
| 0 | NA | NA | NA | NA | NA | Gm45657 | | 0 | 0 | 0 | 0 | 0 | 0 | 0 | 0 | 0 | 0 | 0 |
| 0 | NA | NA | NA | NA | NA | Gm45658 | | 0 | 0 | 0 | 0 | 0 | 0 | 0 | 0 | 0 | 0 | 0 |
| 0 | NA | NA | NA | NA | NA | Gm45659 | | 0 | 0 | 0 | 0 | 0 | 0 | 0 | 0 | 0 | 0 | 0 |
| 0 | NA | NA | NA | NA | NA | Gm24591 | | 0 | 0 | 0 | 0 | 0 | 0 | 0 | 0 | 0 | 0 | 0 |

| | | | | | | | | | | | | | | | | | | |
|---|---|---|---|---|---|---|---|---|---|---|---|---|---|---|---|---|---|---|
| 0 | NA | NA | NA | NA | NA | Gm19855 | | 0 | 0 | 0 | 0 | 0 | 0 | 0 | 0 | 0 | 0 | 0 |
| 0 | NA | NA | NA | NA | NA | Gm37972 | | 0 | 0 | 0 | 0 | 0 | 0 | 0 | 0 | 0 | 0 | 0 |
| 0 | NA | NA | NA | NA | NA | Gm26241 | | 0 | 0 | 0 | 0 | 0 | 0 | 0 | 0 | 0 | 0 | 0 |
| 0 | NA | NA | NA | NA | NA | Gm26089 | | 0 | 0 | 0 | 0 | 0 | 0 | 0 | 0 | 0 | 0 | 0 |
| 0 | NA | NA | NA | NA | NA | Gm2366 | | 0 | 0 | 0 | 0 | 0 | 0 | 0 | 0 | 0 | 0 | 0 |
| 0 | NA | NA | NA | NA | NA | Mtnr1a | melatonin receptor 1A | 0 | 0 | 0 | 0 | 0 | 0 | 0 | 0 | 0 | 0 | 0 |
| 0 | NA | NA | NA | NA | NA | Gm30504 | | 0 | 0 | 0 | 0 | 0 | 0 | 0 | 0 | 0 | 0 | 0 |
| 0 | NA | NA | NA | NA | NA | Gm6329 | | 0 | 0 | 0 | 0 | 0 | 0 | 0 | 0 | 0 | 0 | 0 |
| 0 | NA | NA | NA | NA | NA | Gm45753 | | 0 | 0 | 0 | 0 | 0 | 0 | 0 | 0 | 0 | 0 | 0 |
| 0 | NA | NA | NA | NA | NA | Gm23604 | | 0 | 0 | 0 | 0 | 0 | 0 | 0 | 0 | 0 | 0 | 0 |
| 0 | NA | NA | NA | NA | NA | Gm23604 | | 0 | 0 | 0 | 0 | 0 | 0 | 0 | 0 | 0 | 0 | 0 |
| 0 | NA | NA | NA | NA | NA | Sorbs2os | | 0 | 0 | 0 | 0 | 0 | 0 | 0 | 0 | 0 | 0 | 0 |
| 0 | NA | NA | NA | NA | NA | Gm25309 | | 0 | 0 | 0 | 0 | 0 | 0 | 0 | 0 | 0 | 0 | 0 |
| 0 | NA | NA | NA | NA | NA | Gm15878 | | 0 | 0 | 0 | 0 | 0 | 0 | 0 | 0 | 0 | 0 | 0 |
| 0 | NA | NA | NA | NA | NA | Gm23030 | | 0 | 0 | 0 | 0 | 0 | 0 | 0 | 0 | 0 | 0 | 0 |
| 0 | NA | NA | NA | NA | NA | Gm23812 | | 0 | 0 | 0 | 0 | 0 | 0 | 0 | 0 | 0 | 0 | 0 |
| 0 | NA | NA | NA | NA | NA | Gm24669 | | 0 | 0 | 0 | 0 | 0 | 0 | 0 | 0 | 0 | 0 | 0 |
| 0 | NA | NA | NA | NA | NA | Gm8575 | | 0 | 0 | 0 | 0 | 0 | 0 | 0 | 0 | 0 | 0 | 0 |
| 0 | NA | NA | NA | NA | NA | Helt | helt bHLH transcription factor | 0 | 0 | 0 | 0 | 0 | 0 | 0 | 0 | 0 | 0 | 0 |
| 0 | NA | NA | NA | NA | NA | Gm39168 | | 0 | 0 | 0 | 0 | 0 | 0 | 0 | 0 | 0 | 0 | 0 |
| 0 | NA | NA | NA | NA | NA | Gm45255 | | 0 | 0 | 0 | 0 | 0 | 0 | 0 | 0 | 0 | 0 | 0 |
| 0 | NA | NA | NA | NA | NA | Gm30931 | | 0 | 0 | 0 | 0 | 0 | 0 | 0 | 0 | 0 | 0 | 0 |
| 0 | NA | NA | NA | NA | NA | Gm45253 | | 0 | 0 | 0 | 0 | 0 | 0 | 0 | 0 | 0 | 0 | 0 |
| 0 | NA | NA | NA | NA | NA | Gm45245 | | 0 | 0 | 0 | 0 | 0 | 0 | 0 | 0 | 0 | 0 | 0 |
| 0 | NA | NA | NA | NA | NA | Gm45244 | | 0 | 0 | 0 | 0 | 0 | 0 | 0 | 0 | 0 | 0 | 0 |
| 0 | NA | NA | NA | NA | NA | Gm45607 | | 0 | 0 | 0 | 0 | 0 | 0 | 0 | 0 | 0 | 0 | 0 |
| 0 | NA | NA | NA | NA | NA | Gm45479 | | 0 | 0 | 0 | 0 | 0 | 0 | 0 | 0 | 0 | 0 | 0 |
| 0 | NA | NA | NA | NA | NA | Gm31172 | | 0 | 0 | 0 | 0 | 0 | 0 | 0 | 0 | 0 | 0 | 0 |
| 0 | NA | NA | NA | NA | NA | Gm45481 | | 0 | 0 | 0 | 0 | 0 | 0 | 0 | 0 | 0 | 0 | 0 |
| 0 | NA | NA | NA | NA | NA | Gm45600 | | 0 | 0 | 0 | 0 | 0 | 0 | 0 | 0 | 0 | 0 | 0 |
| 0 | NA | NA | NA | NA | NA | 4930579M01Rik | | 0 | 0 | 0 | 0 | 0 | 0 | 0 | 0 | 0 | 0 | 0 |
| 0 | NA | NA | NA | NA | NA | E330018M18Rik | | 0 | 0 | 0 | 0 | 0 | 0 | 0 | 0 | 0 | 0 | 0 |
| 0 | NA | NA | NA | NA | NA | Gm22857 | | 0 | 0 | 0 | 0 | 0 | 0 | 0 | 0 | 0 | 0 | 0 |
| 0 | NA | NA | NA | NA | NA | Gm45576 | | 0 | 0 | 0 | 0 | 0 | 0 | 0 | 0 | 0 | 0 | 0 |
| 0 | NA | NA | NA | NA | NA | Gm45824 | | 0 | 0 | 0 | 0 | 0 | 0 | 0 | 0 | 0 | 0 | 0 |
| 0 | NA | NA | NA | NA | NA | Gm45864 | | 0 | 0 | 0 | 0 | 0 | 0 | 0 | 0 | 0 | 0 | 0 |
| 0 | NA | NA | NA | NA | NA | 1700061N14Rik | | 0 | 0 | 0 | 0 | 0 | 0 | 0 | 0 | 0 | 0 | 0 |
| 0 | NA | NA | NA | NA | NA | Cldn24 | claudin 24 | 0 | 0 | 0 | 0 | 0 | 0 | 0 | 0 | 0 | 0 | 0 |
| 0 | NA | NA | NA | NA | NA | n-R5s97 | | 0 | 0 | 0 | 0 | 0 | 0 | 0 | 0 | 0 | 0 | 0 |
| 0 | NA | NA | NA | NA | NA | Gm32052 | | 0 | 0 | 0 | 0 | 0 | 0 | 0 | 0 | 0 | 0 | 0 |
| 0 | NA | NA | NA | NA | NA | Gm45772 | | 0 | 0 | 0 | 0 | 0 | 0 | 0 | 0 | 0 | 0 | 0 |
| 0 | NA | NA | NA | NA | NA | Gm25830 | | 0 | 0 | 0 | 0 | 0 | 0 | 0 | 0 | 0 | 0 | 0 |
| 0 | NA | NA | NA | NA | NA | Gm45830 | | 0 | 0 | 0 | 0 | 0 | 0 | 0 | 0 | 0 | 0 | 0 |
| 0 | NA | NA | NA | NA | NA | Gm45829 | | 0 | 0 | 0 | 0 | 0 | 0 | 0 | 0 | 0 | 0 | 0 |
| 0 | NA | NA | NA | NA | NA | Gm45832 | | 0 | 0 | 0 | 0 | 0 | 0 | 0 | 0 | 0 | 0 | 0 |
| 0 | NA | NA | NA | NA | NA | Gm45577 | | 0 | 0 | 0 | 0 | 0 | 0 | 0 | 0 | 0 | 0 | 0 |
| 0 | NA | NA | NA | NA | NA | Gm45697 | | 0 | 0 | 0 | 0 | 0 | 0 | 0 | 0 | 0 | 0 | 0 |
| 0 | NA | NA | NA | NA | NA | Gm45322 | | 0 | 0 | 0 | 0 | 0 | 0 | 0 | 0 | 0 | 0 | 0 |
| 0 | NA | NA | NA | NA | NA | Gm45323 | | 0 | 0 | 0 | 0 | 0 | 0 | 0 | 0 | 0 | 0 | 0 |
| 0 | NA | NA | NA | NA | NA | Gm45324 | | 0 | 0 | 0 | 0 | 0 | 0 | 0 | 0 | 0 | 0 | 0 |
| 0 | NA | NA | NA | NA | NA | Gm45341 | | 0 | 0 | 0 | 0 | 0 | 0 | 0 | 0 | 0 | 0 | 0 |
| 0 | NA | NA | NA | NA | NA | Gm45455 | | 0 | 0 | 0 | 0 | 0 | 0 | 0 | 0 | 0 | 0 | 0 |
| 0 | NA | NA | NA | NA | NA | Gm2516 | | 0 | 0 | 0 | 0 | 0 | 0 | 0 | 0 | 0 | 0 | 0 |
| 0 | NA | NA | NA | NA | NA | Gm45259 | | 0 | 0 | 0 | 0 | 0 | 0 | 0 | 0 | 0 | 0 | 0 |
| 0 | NA | NA | NA | NA | NA | Gm23986 | | 0 | 0 | 0 | 0 | 0 | 0 | 0 | 0 | 0 | 0 | 0 |
| 0 | NA | NA | NA | NA | NA | Gm45386 | | 0 | 0 | 0 | 0 | 0 | 0 | 0 | 0 | 0 | 0 | 0 |
| 0 | NA | NA | NA | NA | NA | Gm32975 | | 0 | 0 | 0 | 0 | 0 | 0 | 0 | 0 | 0 | 0 | 0 |
| 0 | NA | NA | NA | NA | NA | Gm8679 | | 0 | 0 | 0 | 0 | 0 | 0 | 0 | 0 | 0 | 0 | 0 |
| 0 | NA | NA | NA | NA | NA | Gm17911 | | 0 | 0 | 0 | 0 | 0 | 0 | 0 | 0 | 0 | 0 | 0 |
| 0 | NA | NA | NA | NA | NA | Gm45389 | | 0 | 0 | 0 | 0 | 0 | 0 | 0 | 0 | 0 | 0 | 0 |

| | | | | | | | | | | | | | | | | | | |
|---|---|---|---|---|---|---|---|---|---|---|---|---|---|---|---|---|---|---|
| 0 | NA | NA | NA | NA | NA | Gm44387 | | 0 | 0 | 0 | 0 | 0 | 0 | 0 | 0 | 0 | 0 | 0 |
| 0 | NA | NA | NA | NA | NA | Gm24929 | | 0 | 0 | 0 | 0 | 0 | 0 | 0 | 0 | 0 | 0 | 0 |
| 0 | NA | NA | NA | NA | NA | Gm45388 | | 0 | 0 | 0 | 0 | 0 | 0 | 0 | 0 | 0 | 0 | 0 |
| 0 | NA | NA | NA | NA | NA | Gm25813 | | 0 | 0 | 0 | 0 | 0 | 0 | 0 | 0 | 0 | 0 | 0 |
| 0 | NA | NA | NA | NA | NA | 1190028D05Rik | | 0 | 0 | 0 | 0 | 0 | 0 | 0 | 0 | 0 | 0 | 0 |
| 0 | NA | NA | NA | NA | NA | Gm45673 | | 0 | 0 | 0 | 0 | 0 | 0 | 0 | 0 | 0 | 0 | 0 |
| 0 | NA | NA | NA | NA | NA | Gm9685 | | 0 | 0 | 0 | 0 | 0 | 0 | 0 | 0 | 0 | 0 | 0 |
| 0 | NA | NA | NA | NA | NA | Gm23666 | | 0 | 0 | 0 | 0 | 0 | 0 | 0 | 0 | 0 | 0 | 0 |
| 0 | NA | NA | NA | NA | NA | Gm45554 | | 0 | 0 | 0 | 0 | 0 | 0 | 0 | 0 | 0 | 0 | 0 |
| 0 | NA | NA | NA | NA | NA | Gm27964 | | 0 | 0 | 0 | 0 | 0 | 0 | 0 | 0 | 0 | 0 | 0 |
| 0 | NA | NA | NA | NA | NA | Gm22841 | | 0 | 0 | 0 | 0 | 0 | 0 | 0 | 0 | 0 | 0 | 0 |
| 0 | NA | NA | NA | NA | NA | Gm19921 | | 0 | 0 | 0 | 0 | 0 | 0 | 0 | 0 | 0 | 0 | 0 |
| 0 | NA | NA | NA | NA | NA | Gm45553 | | 0 | 0 | 0 | 0 | 0 | 0 | 0 | 0 | 0 | 0 | 0 |
| 0 | NA | NA | NA | NA | NA | Gm15758 | | 0 | 0 | 0 | 0 | 0 | 0 | 0 | 0 | 0 | 0 | 0 |
| 0 | NA | NA | NA | NA | NA | n-R5s98 | | 0 | 0 | 0 | 0 | 0 | 0 | 0 | 0 | 0 | 0 | 0 |
| 0 | NA | NA | NA | NA | NA | Gm45263 | | 0 | 0 | 0 | 0 | 0 | 0 | 0 | 0 | 0 | 0 | 0 |
| 0 | NA | NA | NA | NA | NA | Gm45265 | | 0 | 0 | 0 | 0 | 0 | 0 | 0 | 0 | 0 | 0 | 0 |
| 0 | NA | NA | NA | NA | NA | Gm45262 | | 0 | 0 | 0 | 0 | 0 | 0 | 0 | 0 | 0 | 0 | 0 |
| 0 | NA | NA | NA | NA | NA | A230085B16Rik | | 0 | 0 | 0 | 0 | 0 | 0 | 0 | 0 | 0 | 0 | 0 |
| 0 | NA | NA | NA | NA | NA | 9330121K16Rik | | 0 | 0 | 0 | 0 | 0 | 0 | 0 | 0 | 0 | 0 | 0 |
| 0 | NA | NA | NA | NA | NA | Gm8734 | | 0 | 0 | 0 | 0 | 0 | 0 | 0 | 0 | 0 | 0 | 0 |
| 0 | NA | NA | NA | NA | NA | Gm20586 | | 0 | 0 | 0 | 0 | 0 | 0 | 0 | 0 | 0 | 0 | 0 |
| 0 | NA | NA | NA | NA | NA | Gm18320 | | 0 | 0 | 0 | 0 | 0 | 0 | 0 | 0 | 0 | 0 | 0 |
| 0 | NA | NA | NA | NA | NA | Gm45610 | | 0 | 0 | 0 | 0 | 0 | 0 | 0 | 0 | 0 | 0 | 0 |
| 0 | NA | NA | NA | NA | NA | Adam29 | a disintegrin and metallopeptidase domai | 0 | 0 | 0 | 0 | 0 | 0 | 0 | 0 | 0 | 0 | 0 |
| 0 | NA | NA | NA | NA | NA | Gm45366 | | 0 | 0 | 0 | 0 | 0 | 0 | 0 | 0 | 0 | 0 | 0 |
| 0 | NA | NA | NA | NA | NA | Gm7419 | | 0 | 0 | 0 | 0 | 0 | 0 | 0 | 0 | 0 | 0 | 0 |
| 0 | NA | NA | NA | NA | NA | 4930518J21Rik | | 0 | 0 | 0 | 0 | 0 | 0 | 0 | 0 | 0 | 0 | 0 |
| 0 | NA | NA | NA | NA | NA | Gm24375 | | 0 | 0 | 0 | 0 | 0 | 0 | 0 | 0 | 0 | 0 | 0 |
| 0 | NA | NA | NA | NA | NA | Gm2767 | | 0 | 0 | 0 | 0 | 0 | 0 | 0 | 0 | 0 | 0 | 0 |
| 0 | NA | NA | NA | NA | NA | Gm25992 | | 0 | 0 | 0 | 0 | 0 | 0 | 0 | 0 | 0 | 0 | 0 |
| 0 | NA | NA | NA | NA | NA | Gm6503 | | 0 | 0 | 0 | 0 | 0 | 0 | 0 | 0 | 0 | 0 | 0 |
| 0 | NA | NA | NA | NA | NA | Gm33501 | | 0 | 0 | 0 | 0 | 0 | 0 | 0 | 0 | 0 | 0 | 0 |
| 0 | NA | NA | NA | NA | NA | Gm17994 | | 0 | 0 | 0 | 0 | 0 | 0 | 0 | 0 | 0 | 0 | 0 |
| 0 | NA | NA | NA | NA | NA | Gm34099 | | 0 | 0 | 0 | 0 | 0 | 0 | 0 | 0 | 0 | 0 | 0 |
| 0 | NA | NA | NA | NA | NA | Scrg1 | scrapie responsive gene 1 | 0 | 0 | 0 | 0 | 0 | 0 | 0 | 0 | 0 | 0 | 0 |
| 0 | NA | NA | NA | NA | NA | Gm25379 | | 0 | 0 | 0 | 0 | 0 | 0 | 0 | 0 | 0 | 0 | 0 |
| 0 | NA | NA | NA | NA | NA | Gm45359 | | 0 | 0 | 0 | 0 | 0 | 0 | 0 | 0 | 0 | 0 | 0 |
| 0 | NA | NA | NA | NA | NA | Gm45644 | | 0 | 0 | 0 | 0 | 0 | 0 | 0 | 0 | 0 | 0 | 0 |
| 0 | NA | NA | NA | NA | NA | Gm45356 | | 0 | 0 | 0 | 0 | 0 | 0 | 0 | 0 | 0 | 0 | 0 |
| 0 | NA | NA | NA | NA | NA | Galntl6 | UDP-N-acetyl-alpha-D-galactosamine:po | 0 | 0 | 0 | 0 | 0 | 0 | 0 | 0 | 0 | 0 | 0 |
| 0 | NA | NA | NA | NA | NA | Gm45635 | | 0 | 0 | 0 | 0 | 0 | 0 | 0 | 0 | 0 | 0 | 0 |
| 0 | NA | NA | NA | NA | NA | Gm15882 | | 0 | 0 | 0 | 0 | 0 | 0 | 0 | 0 | 0 | 0 | 0 |
| 0 | NA | NA | NA | NA | NA | Gm45634 | | 0 | 0 | 0 | 0 | 0 | 0 | 0 | 0 | 0 | 0 | 0 |
| 0 | NA | NA | NA | NA | NA | Gm45636 | | 0 | 0 | 0 | 0 | 0 | 0 | 0 | 0 | 0 | 0 | 0 |
| 0 | NA | NA | NA | NA | NA | Gm15881 | | 0 | 0 | 0 | 0 | 0 | 0 | 0 | 0 | 0 | 0 | 0 |
| 0 | NA | NA | NA | NA | NA | Gm7499 | | 0 | 0 | 0 | 0 | 0 | 0 | 0 | 0 | 0 | 0 | 0 |
| 0 | NA | NA | NA | NA | NA | Gm5740 | | 0 | 0 | 0 | 0 | 0 | 0 | 0 | 0 | 0 | 0 | 0 |
| 0 | NA | NA | NA | NA | NA | Gm34370 | | 0 | 0 | 0 | 0 | 0 | 0 | 0 | 0 | 0 | 0 | 0 |
| 0 | NA | NA | NA | NA | NA | Gm24847 | | 0 | 0 | 0 | 0 | 0 | 0 | 0 | 0 | 0 | 0 | 0 |
| 0 | NA | NA | NA | NA | NA | 4930470O06Rik | | 0 | 0 | 0 | 0 | 0 | 0 | 0 | 0 | 0 | 0 | 0 |
| 0 | NA | NA | NA | NA | NA | Gm26164 | | 0 | 0 | 0 | 0 | 0 | 0 | 0 | 0 | 0 | 0 | 0 |
| 0 | NA | NA | NA | NA | NA | Gm34623 | | 0 | 0 | 0 | 0 | 0 | 0 | 0 | 0 | 0 | 0 | 0 |
| 0 | NA | NA | NA | NA | NA | Gm10283 | | 0 | 0 | 0 | 0 | 0 | 0 | 0 | 0 | 0 | 0 | 0 |
| 0 | NA | NA | NA | NA | NA | Gm34730 | | 0 | 0 | 0 | 0 | 0 | 0 | 0 | 0 | 0 | 0 | 0 |
| 0 | NA | NA | NA | NA | NA | Gm45650 | | 0 | 0 | 0 | 0 | 0 | 0 | 0 | 0 | 0 | 0 | 0 |
| 0 | NA | NA | NA | NA | NA | Gm45651 | | 0 | 0 | 0 | 0 | 0 | 0 | 0 | 0 | 0 | 0 | 0 |
| 0 | NA | NA | NA | NA | NA | Stambp-ps2 | | 0 | 0 | 0 | 0 | 0 | 0 | 0 | 0 | 0 | 0 | 0 |
| 0 | NA | NA | NA | NA | NA | Gm23329 | | 0 | 0 | 0 | 0 | 0 | 0 | 0 | 0 | 0 | 0 | 0 |
| 0 | NA | NA | NA | NA | NA | Gm45485 | | 0 | 0 | 0 | 0 | 0 | 0 | 0 | 0 | 0 | 0 | 0 |

| | | | | | | | | | | | | | | | | | | |
|---|---|---|---|---|---|---|---|---|---|---|---|---|---|---|---|---|---|---|
| 0 | NA | NA | NA | NA | NA | Gm16179 | | 0 | 0 | 0 | 0 | 0 | 0 | 0 | 0 | 0 | 0 | 0 |
| 0 | NA | NA | NA | NA | NA | Gm16178 | | 0 | 0 | 0 | 0 | 0 | 0 | 0 | 0 | 0 | 0 | 0 |
| 0 | NA | NA | NA | NA | NA | Gm18066 | | 0 | 0 | 0 | 0 | 0 | 0 | 0 | 0 | 0 | 0 | 0 |
| 0 | NA | NA | NA | NA | NA | Gm2961 | | 0 | 0 | 0 | 0 | 0 | 0 | 0 | 0 | 0 | 0 | 0 |
| 0 | NA | NA | NA | NA | NA | Gm2975 | | 0 | 0 | 0 | 0 | 0 | 0 | 0 | 0 | 0 | 0 | 0 |
| 0 | NA | NA | NA | NA | NA | Gm45460 | | 0 | 0 | 0 | 0 | 0 | 0 | 0 | 0 | 0 | 0 | 0 |
| 0 | NA | NA | NA | NA | NA | Gm45459 | | 0 | 0 | 0 | 0 | 0 | 0 | 0 | 0 | 0 | 0 | 0 |
| 0 | NA | NA | NA | NA | NA | Gm45462 | | 0 | 0 | 0 | 0 | 0 | 0 | 0 | 0 | 0 | 0 | 0 |
| 0 | NA | NA | NA | NA | NA | Gm22324 | | 0 | 0 | 0 | 0 | 0 | 0 | 0 | 0 | 0 | 0 | 0 |
| 0 | NA | NA | NA | NA | NA | Gm45461 | | 0 | 0 | 0 | 0 | 0 | 0 | 0 | 0 | 0 | 0 | 0 |
| 0 | NA | NA | NA | NA | NA | Gm45516 | | 0 | 0 | 0 | 0 | 0 | 0 | 0 | 0 | 0 | 0 | 0 |
| 0 | NA | NA | NA | NA | NA | Gm24236 | | 0 | 0 | 0 | 0 | 0 | 0 | 0 | 0 | 0 | 0 | 0 |
| 0 | NA | NA | NA | NA | NA | Gm45517 | | 0 | 0 | 0 | 0 | 0 | 0 | 0 | 0 | 0 | 0 | 0 |
| 0 | NA | NA | NA | NA | NA | Gm18645 | | 0 | 0 | 0 | 0 | 0 | 0 | 0 | 0 | 0 | 0 | 0 |
| 0 | NA | NA | NA | NA | NA | Gm18645 | | 0 | 0 | 0 | 0 | 0 | 0 | 0 | 0 | 0 | 0 | 0 |
| 0 | NA | NA | NA | NA | NA | Gm35368 | | 0 | 0 | 0 | 0 | 0 | 0 | 0 | 0 | 0 | 0 | 0 |
| 0 | NA | NA | NA | NA | NA | Gm45589 | | 0 | 0 | 0 | 0 | 0 | 0 | 0 | 0 | 0 | 0 | 0 |
| 0 | NA | NA | NA | NA | NA | Gm24580 | | 0 | 0 | 0 | 0 | 0 | 0 | 0 | 0 | 0 | 0 | 0 |
| 0 | NA | NA | NA | NA | NA | Sgo2b | shugoshin 2B | 0 | 0 | 0 | 0 | 0 | 0 | 0 | 0 | 0 | 0 | 0 |
| 0 | NA | NA | NA | NA | NA | Gm35713 | | 0 | 0 | 0 | 0 | 0 | 0 | 0 | 0 | 0 | 0 | 0 |
| 0 | NA | NA | NA | NA | NA | Gm45513 | | 0 | 0 | 0 | 0 | 0 | 0 | 0 | 0 | 0 | 0 | 0 |
| 0 | NA | NA | NA | NA | NA | Gm22586 | | 0 | 0 | 0 | 0 | 0 | 0 | 0 | 0 | 0 | 0 | 0 |
| 0 | NA | NA | NA | NA | NA | Gm35521 | | 0 | 0 | 0 | 0 | 0 | 0 | 0 | 0 | 0 | 0 | 0 |
| 0 | NA | NA | NA | NA | NA | Mir710 | | 0 | 0 | 0 | 0 | 0 | 0 | 0 | 0 | 0 | 0 | 0 |
| 0 | NA | NA | NA | NA | NA | Gm45345 | | 0 | 0 | 0 | 0 | 0 | 0 | 0 | 0 | 0 | 0 | 0 |
| 0 | NA | NA | NA | NA | NA | Gm26210 | | 0 | 0 | 0 | 0 | 0 | 0 | 0 | 0 | 0 | 0 | 0 |
| 0 | NA | NA | NA | NA | NA | Gm32507 | | 0 | 0 | 0 | 0 | 0 | 0 | 0 | 0 | 0 | 0 | 0 |
| 0 | NA | NA | NA | NA | NA | Trim75 | tripartite motif-containing 75 | 0 | 0 | 0 | 0 | 0 | 0 | 0 | 0 | 0 | 0 | 0 |
| 0 | NA | NA | NA | NA | NA | Trim60 | tripartite motif-containing 60 | 0 | 0 | 0 | 0 | 0 | 0 | 0 | 0 | 0 | 0 | 0 |
| 0 | NA | NA | NA | NA | NA | Trim61 | tripartite motif-containing 61 | 0 | 0 | 0 | 0 | 0 | 0 | 0 | 0 | 0 | 0 | 0 |
| 0 | NA | NA | NA | NA | NA | Gm45886 | | 0 | 0 | 0 | 0 | 0 | 0 | 0 | 0 | 0 | 0 | 0 |
| 0 | NA | NA | NA | NA | NA | Gm45448 | | 0 | 0 | 0 | 0 | 0 | 0 | 0 | 0 | 0 | 0 | 0 |
| 0 | NA | NA | NA | NA | NA | Apela | apelin receptor early endogenous ligand | 0 | 0 | 0 | 0 | 0 | 0 | 0 | 0 | 0 | 0 | 0 |
| 0 | NA | NA | NA | NA | NA | Gm10663 | | 0 | 0 | 0 | 0 | 0 | 0 | 0 | 0 | 0 | 0 | 0 |
| 0 | NA | NA | NA | NA | NA | Gm45445 | | 0 | 0 | 0 | 0 | 0 | 0 | 0 | 0 | 0 | 0 | 0 |
| 0 | NA | NA | NA | NA | NA | Gm48810 | | 0 | 0 | 0 | 0 | 0 | 0 | 0 | 0 | 0 | 0 | 0 |
| 0 | NA | NA | NA | NA | NA | Gm24333 | | 0 | 0 | 0 | 0 | 0 | 0 | 0 | 0 | 0 | 0 | 0 |
| 0 | NA | NA | NA | NA | NA | Gm45249 | | 0 | 0 | 0 | 0 | 0 | 0 | 0 | 0 | 0 | 0 | 0 |
| 0 | NA | NA | NA | NA | NA | Gm7624 | | 0 | 0 | 0 | 0 | 0 | 0 | 0 | 0 | 0 | 0 | 0 |
| 0 | NA | NA | NA | NA | NA | Gm25403 | | 0 | 0 | 0 | 0 | 0 | 0 | 0 | 0 | 0 | 0 | 0 |
| 0 | NA | NA | NA | NA | NA | Gm23515 | | 0 | 0 | 0 | 0 | 0 | 0 | 0 | 0 | 0 | 0 | 0 |
| 0 | NA | NA | NA | NA | NA | Gm15355 | | 0 | 0 | 0 | 0 | 0 | 0 | 0 | 0 | 0 | 0 | 0 |
| 0 | NA | NA | NA | NA | NA | Gm26123 | | 0 | 0 | 0 | 0 | 0 | 0 | 0 | 0 | 0 | 0 | 0 |
| 0 | NA | NA | NA | NA | NA | Gm24205 | | 0 | 0 | 0 | 0 | 0 | 0 | 0 | 0 | 0 | 0 | 0 |
| 0 | NA | NA | NA | NA | NA | Gm15354 | | 0 | 0 | 0 | 0 | 0 | 0 | 0 | 0 | 0 | 0 | 0 |
| 0 | NA | NA | NA | NA | NA | Gm15356 | | 0 | 0 | 0 | 0 | 0 | 0 | 0 | 0 | 0 | 0 | 0 |
| 0 | NA | NA | NA | NA | NA | Gm32568 | | 0 | 0 | 0 | 0 | 0 | 0 | 0 | 0 | 0 | 0 | 0 |
| 0 | NA | NA | NA | NA | NA | Tktl2 | transketolase-like 2 | 0 | 0 | 0 | 0 | 0 | 0 | 0 | 0 | 0 | 0 | 0 |
| 0 | NA | NA | NA | NA | NA | Gm16330 | | 0 | 0 | 0 | 0 | 0 | 0 | 0 | 0 | 0 | 0 | 0 |
| 0 | NA | NA | NA | NA | NA | Npy5r | neuropeptide Y receptor Y5 | 0 | 0 | 0 | 0 | 0 | 0 | 0 | 0 | 0 | 0 | 0 |
| 0 | NA | NA | NA | NA | NA | Gm45911 | | 0 | 0 | 0 | 0 | 0 | 0 | 0 | 0 | 0 | 0 | 0 |
| 0 | NA | NA | NA | NA | NA | Gm39185 | | 0 | 0 | 0 | 0 | 0 | 0 | 0 | 0 | 0 | 0 | 0 |
| 0 | NA | NA | NA | NA | NA | Gm25275 | | 0 | 0 | 0 | 0 | 0 | 0 | 0 | 0 | 0 | 0 | 0 |
| 0 | NA | NA | NA | NA | NA | Gm7639 | | 0 | 0 | 0 | 0 | 0 | 0 | 0 | 0 | 0 | 0 | 0 |
| 0 | NA | NA | NA | NA | NA | Gm45827 | | 0 | 0 | 0 | 0 | 0 | 0 | 0 | 0 | 0 | 0 | 0 |
| 0 | NA | NA | NA | NA | NA | Gm24274 | | 0 | 0 | 0 | 0 | 0 | 0 | 0 | 0 | 0 | 0 | 0 |
| 0 | NA | NA | NA | NA | NA | Gm24677 | | 0 | 0 | 0 | 0 | 0 | 0 | 0 | 0 | 0 | 0 | 0 |
| 0 | NA | NA | NA | NA | NA | Gm45880 | | 0 | 0 | 0 | 0 | 0 | 0 | 0 | 0 | 0 | 0 | 0 |
| 0 | NA | NA | NA | NA | NA | Gm10660 | | 0 | 0 | 0 | 0 | 0 | 0 | 0 | 0 | 0 | 0 | 0 |
| 0 | NA | NA | NA | NA | NA | Nat3 | N-acetyltransferase 3 | 0 | 0 | 0 | 0 | 0 | 0 | 0 | 0 | 0 | 0 | 0 |

| | | | | | | | | | | | | | | | | | | |
|---|---|---|---|---|---|---|---|---|---|---|---|---|---|---|---|---|---|---|
| 0 | NA | NA | NA | NA | NA | Gm36247 | | 0 | 0 | 0 | 0 | 0 | 0 | 0 | 0 | 0 | 0 | 0 |
| 0 | NA | NA | NA | NA | NA | Gm44391 | | 0 | 0 | 0 | 0 | 0 | 0 | 0 | 0 | 0 | 0 | 0 |
| 0 | NA | NA | NA | NA | NA | Gm22018 | | 0 | 0 | 0 | 0 | 0 | 0 | 0 | 0 | 0 | 0 | 0 |
| 0 | NA | NA | NA | NA | NA | Gm45739 | | 0 | 0 | 0 | 0 | 0 | 0 | 0 | 0 | 0 | 0 | 0 |
| 0 | NA | NA | NA | NA | NA | Gm19170 | | 0 | 0 | 0 | 0 | 0 | 0 | 0 | 0 | 0 | 0 | 0 |
| 0 | NA | NA | NA | NA | NA | Gm15656 | | 0 | 0 | 0 | 0 | 0 | 0 | 0 | 0 | 0 | 0 | 0 |
| 0 | NA | NA | NA | NA | NA | Gm15654 | | 0 | 0 | 0 | 0 | 0 | 0 | 0 | 0 | 0 | 0 | 0 |
| 0 | NA | NA | NA | NA | NA | Gm45780 | | 0 | 0 | 0 | 0 | 0 | 0 | 0 | 0 | 0 | 0 | 0 |
| 0 | NA | NA | NA | NA | NA | Gm33103 | | 0 | 0 | 0 | 0 | 0 | 0 | 0 | 0 | 0 | 0 | 0 |
| 0 | NA | NA | NA | NA | NA | Gm15716 | | 0 | 0 | 0 | 0 | 0 | 0 | 0 | 0 | 0 | 0 | 0 |
| 0 | NA | NA | NA | NA | NA | Gm33178 | | 0 | 0 | 0 | 0 | 0 | 0 | 0 | 0 | 0 | 0 | 0 |
| 0 | NA | NA | NA | NA | NA | Gm45848 | | 0 | 0 | 0 | 0 | 0 | 0 | 0 | 0 | 0 | 0 | 0 |
| 0 | NA | NA | NA | NA | NA | Gm9495 | predicted gene 9495 | 0 | 0 | 0 | 0 | 0 | 0 | 0 | 0 | 0 | 0 | 0 |
| 0 | NA | NA | NA | NA | NA | Gm45850 | | 0 | 0 | 0 | 0 | 0 | 0 | 0 | 0 | 0 | 0 | 0 |
| 0 | NA | NA | NA | NA | NA | Gm7697 | predicted gene 7697 | 0 | 0 | 0 | 0 | 0 | 0 | 0 | 0 | 0 | 0 | 0 |
| 0 | NA | NA | NA | NA | NA | Rps7-ps1 | | 0 | 0 | 0 | 0 | 0 | 0 | 0 | 0 | 0 | 0 | 0 |
| 0 | NA | NA | NA | NA | NA | Gm20422 | | 0 | 0 | 0 | 0 | 0 | 0 | 0 | 0 | 0 | 0 | 0 |
| 0 | NA | NA | NA | NA | NA | Mir7066 | | 0 | 0 | 0 | 0 | 0 | 0 | 0 | 0 | 0 | 0 | 0 |
| 0 | NA | NA | NA | NA | NA | Gm25906 | | 0 | 0 | 0 | 0 | 0 | 0 | 0 | 0 | 0 | 0 | 0 |
| 0 | NA | NA | NA | NA | NA | Gdf1 | growth differentiation factor 1 | 0 | 0 | 0 | 0 | 0 | 0 | 0 | 0 | 0 | 0 | 0 |
| 0 | NA | NA | NA | NA | NA | Mir7240 | | 0 | 0 | 0 | 0 | 0 | 0 | 0 | 0 | 0 | 0 | 0 |
| 0 | NA | NA | NA | NA | NA | Mir7067 | | 0 | 0 | 0 | 0 | 0 | 0 | 0 | 0 | 0 | 0 | 0 |
| 0 | NA | NA | NA | NA | NA | Snora68 | | 0 | 0 | 0 | 0 | 0 | 0 | 0 | 0 | 0 | 0 | 0 |
| 0 | NA | NA | NA | NA | NA | Mir1969 | | 0 | 0 | 0 | 0 | 0 | 0 | 0 | 0 | 0 | 0 | 0 |
| 0 | NA | NA | NA | NA | NA | Gm27628 | | 0 | 0 | 0 | 0 | 0 | 0 | 0 | 0 | 0 | 0 | 0 |
| 0 | NA | NA | NA | NA | NA | Mir6769b | | 0 | 0 | 0 | 0 | 0 | 0 | 0 | 0 | 0 | 0 | 0 |
| 0 | NA | NA | NA | NA | NA | Gm29705 | | 0 | 0 | 0 | 0 | 0 | 0 | 0 | 0 | 0 | 0 | 0 |
| 0 | NA | NA | NA | NA | NA | Vmn2r-ps91 | | 0 | 0 | 0 | 0 | 0 | 0 | 0 | 0 | 0 | 0 | 0 |
| 0 | NA | NA | NA | NA | NA | Gm18924 | | 0 | 0 | 0 | 0 | 0 | 0 | 0 | 0 | 0 | 0 | 0 |
| 0 | NA | NA | NA | NA | NA | Gm25052 | | 0 | 0 | 0 | 0 | 0 | 0 | 0 | 0 | 0 | 0 | 0 |
| 0 | NA | NA | NA | NA | NA | Olfr372 | olfactory receptor 372 | 0 | 0 | 0 | 0 | 0 | 0 | 0 | 0 | 0 | 0 | 0 |
| 0 | NA | NA | NA | NA | NA | Gm45544 | | 0 | 0 | 0 | 0 | 0 | 0 | 0 | 0 | 0 | 0 | 0 |
| 0 | NA | NA | NA | NA | NA | Gm45735 | | 0 | 0 | 0 | 0 | 0 | 0 | 0 | 0 | 0 | 0 | 0 |
| 0 | NA | NA | NA | NA | NA | Olfr373 | olfactory receptor 373 | 0 | 0 | 0 | 0 | 0 | 0 | 0 | 0 | 0 | 0 | 0 |
| 0 | NA | NA | NA | NA | NA | Gm45543 | | 0 | 0 | 0 | 0 | 0 | 0 | 0 | 0 | 0 | 0 | 0 |
| 0 | NA | NA | NA | NA | NA | Gm24747 | | 0 | 0 | 0 | 0 | 0 | 0 | 0 | 0 | 0 | 0 | 0 |
| 0 | NA | NA | NA | NA | NA | Gm11034 | | 0 | 0 | 0 | 0 | 0 | 0 | 0 | 0 | 0 | 0 | 0 |
| 0 | NA | NA | NA | NA | NA | Mir7068 | | 0 | 0 | 0 | 0 | 0 | 0 | 0 | 0 | 0 | 0 | 0 |
| 0 | NA | NA | NA | NA | NA | Gm28019 | | 0 | 0 | 0 | 0 | 0 | 0 | 0 | 0 | 0 | 0 | 0 |
| 0 | NA | NA | NA | NA | NA | Gm26237 | | 0 | 0 | 0 | 0 | 0 | 0 | 0 | 0 | 0 | 0 | 0 |
| 0 | NA | NA | NA | NA | NA | Gm45704 | | 0 | 0 | 0 | 0 | 0 | 0 | 0 | 0 | 0 | 0 | 0 |
| 0 | NA | NA | NA | NA | NA | Gm45791 | | 0 | 0 | 0 | 0 | 0 | 0 | 0 | 0 | 0 | 0 | 0 |
| 0 | NA | NA | NA | NA | NA | Mir28b | | 0 | 0 | 0 | 0 | 0 | 0 | 0 | 0 | 0 | 0 | 0 |
| 0 | NA | NA | NA | NA | NA | Gm22532 | | 0 | 0 | 0 | 0 | 0 | 0 | 0 | 0 | 0 | 0 | 0 |
| 0 | NA | NA | NA | NA | NA | Gm7943 | | 0 | 0 | 0 | 0 | 0 | 0 | 0 | 0 | 0 | 0 | 0 |
| 0 | NA | NA | NA | NA | NA | Gm23255 | | 0 | 0 | 0 | 0 | 0 | 0 | 0 | 0 | 0 | 0 | 0 |
| 0 | NA | NA | NA | NA | NA | Gm7948 | | 0 | 0 | 0 | 0 | 0 | 0 | 0 | 0 | 0 | 0 | 0 |
| 0 | NA | NA | NA | NA | NA | Gm45821 | | 0 | 0 | 0 | 0 | 0 | 0 | 0 | 0 | 0 | 0 | 0 |
| 0 | NA | NA | NA | NA | NA | Gm27864 | | 0 | 0 | 0 | 0 | 0 | 0 | 0 | 0 | 0 | 0 | 0 |
| 0 | NA | NA | NA | NA | NA | Gm11033 | | 0 | 0 | 0 | 0 | 0 | 0 | 0 | 0 | 0 | 0 | 0 |
| 0 | NA | NA | NA | NA | NA | Gm45822 | | 0 | 0 | 0 | 0 | 0 | 0 | 0 | 0 | 0 | 0 | 0 |
| 0 | NA | NA | NA | NA | NA | Gm45820 | | 0 | 0 | 0 | 0 | 0 | 0 | 0 | 0 | 0 | 0 | 0 |
| 0 | NA | NA | NA | NA | NA | Gm2059 | | 0 | 0 | 0 | 0 | 0 | 0 | 0 | 0 | 0 | 0 | 0 |
| 0 | NA | NA | NA | NA | NA | Gm45899 | | 0 | 0 | 0 | 0 | 0 | 0 | 0 | 0 | 0 | 0 | 0 |
| 0 | NA | NA | NA | NA | NA | Gm45903 | | 0 | 0 | 0 | 0 | 0 | 0 | 0 | 0 | 0 | 0 | 0 |
| 0 | NA | NA | NA | NA | NA | Gm45907 | | 0 | 0 | 0 | 0 | 0 | 0 | 0 | 0 | 0 | 0 | 0 |
| 0 | NA | NA | NA | NA | NA | Umpk-ps | | 0 | 0 | 0 | 0 | 0 | 0 | 0 | 0 | 0 | 0 | 0 |
| 0 | NA | NA | NA | NA | NA | Gm19885 | | 0 | 0 | 0 | 0 | 0 | 0 | 0 | 0 | 0 | 0 | 0 |
| 0 | NA | NA | NA | NA | NA | Gm38270 | | 0 | 0 | 0 | 0 | 0 | 0 | 0 | 0 | 0 | 0 | 0 |
| 0 | NA | NA | NA | NA | NA | Gm45740 | | 0 | 0 | 0 | 0 | 0 | 0 | 0 | 0 | 0 | 0 | 0 |

| | | | | | | | | | | | | | | | | | | |
|---|---|---|---|---|---|---|---|---|---|---|---|---|---|---|---|---|---|---|
| 0 | NA | NA | NA | NA | NA | Gm45741 | | 0 | 0 | 0 | 0 | 0 | 0 | 0 | 0 | 0 | 0 | 0 |
| 0 | NA | NA | NA | NA | NA | Gm22544 | | 0 | 0 | 0 | 0 | 0 | 0 | 0 | 0 | 0 | 0 | 0 |
| 0 | NA | NA | NA | NA | NA | Gm45742 | | 0 | 0 | 0 | 0 | 0 | 0 | 0 | 0 | 0 | 0 | 0 |
| 0 | NA | NA | NA | NA | NA | Gm22509 | | 0 | 0 | 0 | 0 | 0 | 0 | 0 | 0 | 0 | 0 | 0 |
| 0 | NA | NA | NA | NA | NA | Gm45798 | | 0 | 0 | 0 | 0 | 0 | 0 | 0 | 0 | 0 | 0 | 0 |
| 0 | NA | NA | NA | NA | NA | Gm7990 | | 0 | 0 | 0 | 0 | 0 | 0 | 0 | 0 | 0 | 0 | 0 |
| 0 | NA | NA | NA | NA | NA | Gm27355 | | 0 | 0 | 0 | 0 | 0 | 0 | 0 | 0 | 0 | 0 | 0 |
| 0 | NA | NA | NA | NA | NA | Gm24838 | | 0 | 0 | 0 | 0 | 0 | 0 | 0 | 0 | 0 | 0 | 0 |
| 0 | NA | NA | NA | NA | NA | Gm23260 | | 0 | 0 | 0 | 0 | 0 | 0 | 0 | 0 | 0 | 0 | 0 |
| 0 | NA | NA | NA | NA | NA | Gm45406 | | 0 | 0 | 0 | 0 | 0 | 0 | 0 | 0 | 0 | 0 | 0 |
| 0 | NA | NA | NA | NA | NA | 4933421D24Rik | | 0 | 0 | 0 | 0 | 0 | 0 | 0 | 0 | 0 | 0 | 0 |
| 0 | NA | NA | NA | NA | NA | Gm45285 | | 0 | 0 | 0 | 0 | 0 | 0 | 0 | 0 | 0 | 0 | 0 |
| 0 | NA | NA | NA | NA | NA | Gm45407 | | 0 | 0 | 0 | 0 | 0 | 0 | 0 | 0 | 0 | 0 | 0 |
| 0 | NA | NA | NA | NA | NA | Gm23468 | | 0 | 0 | 0 | 0 | 0 | 0 | 0 | 0 | 0 | 0 | 0 |
| 0 | NA | NA | NA | NA | NA | 1700092C02Rik | | 0 | 0 | 0 | 0 | 0 | 0 | 0 | 0 | 0 | 0 | 0 |
| 0 | NA | NA | NA | NA | NA | Gm45714 | | 0 | 0 | 0 | 0 | 0 | 0 | 0 | 0 | 0 | 0 | 0 |
| 0 | NA | NA | NA | NA | NA | Gm23981 | | 0 | 0 | 0 | 0 | 0 | 0 | 0 | 0 | 0 | 0 | 0 |
| 0 | NA | NA | NA | NA | NA | Gm29895 | | 0 | 0 | 0 | 0 | 0 | 0 | 0 | 0 | 0 | 0 | 0 |
| 0 | NA | NA | NA | NA | NA | Gm45288 | | 0 | 0 | 0 | 0 | 0 | 0 | 0 | 0 | 0 | 0 | 0 |
| 0 | NA | NA | NA | NA | NA | Gm45574 | | 0 | 0 | 0 | 0 | 0 | 0 | 0 | 0 | 0 | 0 | 0 |
| 0 | NA | NA | NA | NA | NA | Ttc29 | tetratricopeptide repeat domain 29 | 0 | 0 | 0 | 0 | 0 | 0 | 0 | 0 | 0 | 0 | 0 |
| 0 | NA | NA | NA | NA | NA | Pou4f2 | POU domain, class 4, transcription factor | 0 | 0 | 0 | 0 | 0 | 0 | 0 | 0 | 0 | 0 | 0 |
| 0 | NA | NA | NA | NA | NA | Gm47209 | | 0 | 0 | 0 | 0 | 0 | 0 | 0 | 0 | 0 | 0 | 0 |
| 0 | NA | NA | NA | NA | NA | Gm45446 | | 0 | 0 | 0 | 0 | 0 | 0 | 0 | 0 | 0 | 0 | 0 |
| 0 | NA | NA | NA | NA | NA | Gm45326 | | 0 | 0 | 0 | 0 | 0 | 0 | 0 | 0 | 0 | 0 | 0 |
| 0 | NA | NA | NA | NA | NA | Gm39204 | | 0 | 0 | 0 | 0 | 0 | 0 | 0 | 0 | 0 | 0 | 0 |
| 0 | NA | NA | NA | NA | NA | Gm22044 | | 0 | 0 | 0 | 0 | 0 | 0 | 0 | 0 | 0 | 0 | 0 |
| 0 | NA | NA | NA | NA | NA | Gm39288 | | 0 | 0 | 0 | 0 | 0 | 0 | 0 | 0 | 0 | 0 | 0 |
| 0 | NA | NA | NA | NA | NA | Gm45602 | | 0 | 0 | 0 | 0 | 0 | 0 | 0 | 0 | 0 | 0 | 0 |
| 0 | NA | NA | NA | NA | NA | Gm8077 | | 0 | 0 | 0 | 0 | 0 | 0 | 0 | 0 | 0 | 0 | 0 |
| 0 | NA | NA | NA | NA | NA | Gm27006 | | 0 | 0 | 0 | 0 | 0 | 0 | 0 | 0 | 0 | 0 | 0 |
| 0 | NA | NA | NA | NA | NA | Gm45435 | | 0 | 0 | 0 | 0 | 0 | 0 | 0 | 0 | 0 | 0 | 0 |
| 0 | NA | NA | NA | NA | NA | Gm45434 | | 0 | 0 | 0 | 0 | 0 | 0 | 0 | 0 | 0 | 0 | 0 |
| 0 | NA | NA | NA | NA | NA | Gm45433 | | 0 | 0 | 0 | 0 | 0 | 0 | 0 | 0 | 0 | 0 | 0 |
| 0 | NA | NA | NA | NA | NA | Gm45432 | | 0 | 0 | 0 | 0 | 0 | 0 | 0 | 0 | 0 | 0 | 0 |
| 0 | NA | NA | NA | NA | NA | Gm27820 | | 0 | 0 | 0 | 0 | 0 | 0 | 0 | 0 | 0 | 0 | 0 |
| 0 | NA | NA | NA | NA | NA | Gm33096 | | 0 | 0 | 0 | 0 | 0 | 0 | 0 | 0 | 0 | 0 | 0 |
| 0 | NA | NA | NA | NA | NA | Gm45431 | | 0 | 0 | 0 | 0 | 0 | 0 | 0 | 0 | 0 | 0 | 0 |
| 0 | NA | NA | NA | NA | NA | Gm45430 | | 0 | 0 | 0 | 0 | 0 | 0 | 0 | 0 | 0 | 0 | 0 |
| 0 | NA | NA | NA | NA | NA | 4930505O20Rik | | 0 | 0 | 0 | 0 | 0 | 0 | 0 | 0 | 0 | 0 | 0 |
| 0 | NA | NA | NA | NA | NA | Frem3 | Fras1 related extracellular matrix protein 3 | 0 | 0 | 0 | 0 | 0 | 0 | 0 | 0 | 0 | 0 | 0 |
| 0 | NA | NA | NA | NA | NA | Gm45872 | | 0 | 0 | 0 | 0 | 0 | 0 | 0 | 0 | 0 | 0 | 0 |
| 0 | NA | NA | NA | NA | NA | Gm7997 | | 0 | 0 | 0 | 0 | 0 | 0 | 0 | 0 | 0 | 0 | 0 |
| 0 | NA | NA | NA | NA | NA | Gm27048 | | 0 | 0 | 0 | 0 | 0 | 0 | 0 | 0 | 0 | 0 | 0 |
| 0 | NA | NA | NA | NA | NA | Gm45538 | | 0 | 0 | 0 | 0 | 0 | 0 | 0 | 0 | 0 | 0 | 0 |
| 0 | NA | NA | NA | NA | NA | Gm4899 | | 0 | 0 | 0 | 0 | 0 | 0 | 0 | 0 | 0 | 0 | 0 |
| 0 | NA | NA | NA | NA | NA | Gm45539 | | 0 | 0 | 0 | 0 | 0 | 0 | 0 | 0 | 0 | 0 | 0 |
| 0 | NA | NA | NA | NA | NA | Gm5354 | | 0 | 0 | 0 | 0 | 0 | 0 | 0 | 0 | 0 | 0 | 0 |
| 0 | NA | NA | NA | NA | NA | 4930579O11Rik | | 0 | 0 | 0 | 0 | 0 | 0 | 0 | 0 | 0 | 0 | 0 |
| 0 | NA | NA | NA | NA | NA | Gm17072 | | 0 | 0 | 0 | 0 | 0 | 0 | 0 | 0 | 0 | 0 | 0 |
| 0 | NA | NA | NA | NA | NA | Gm25267 | | 0 | 0 | 0 | 0 | 0 | 0 | 0 | 0 | 0 | 0 | 0 |
| 0 | NA | NA | NA | NA | NA | Gm9520 | | 0 | 0 | 0 | 0 | 0 | 0 | 0 | 0 | 0 | 0 | 0 |
| 0 | NA | NA | NA | NA | NA | Gm38590 | | 0 | 0 | 0 | 0 | 0 | 0 | 0 | 0 | 0 | 0 | 0 |
| 0 | NA | NA | NA | NA | NA | Gm25138 | | 0 | 0 | 0 | 0 | 0 | 0 | 0 | 0 | 0 | 0 | 0 |
| 0 | NA | NA | NA | NA | NA | Gm8167 | | 0 | 0 | 0 | 0 | 0 | 0 | 0 | 0 | 0 | 0 | 0 |
| 0 | NA | NA | NA | NA | NA | Mgat4d | MGAT4 family, member C | 0 | 0 | 0 | 0 | 0 | 0 | 0 | 0 | 0 | 0 | 0 |
| 0 | NA | NA | NA | NA | NA | Gm33242 | | 0 | 0 | 0 | 0 | 0 | 0 | 0 | 0 | 0 | 0 | 0 |
| 0 | NA | NA | NA | NA | NA | Olfr370 | olfactory receptor 370 | 0 | 0 | 0 | 0 | 0 | 0 | 0 | 0 | 0 | 0 | 0 |
| 0 | NA | NA | NA | NA | NA | Gm18001 | | 0 | 0 | 0 | 0 | 0 | 0 | 0 | 0 | 0 | 0 | 0 |
| 0 | NA | NA | NA | NA | NA | Mir1668 | | 0 | 0 | 0 | 0 | 0 | 0 | 0 | 0 | 0 | 0 | 0 |

| | | | | | | | | | | | | | | | | | | |
|---|---|---|---|---|---|---|---|---|---|---|---|---|---|---|---|---|---|---|
| 0 | NA | NA | NA | NA | NA | Gm22024 | | 0 | 0 | 0 | 0 | 0 | 0 | 0 | 0 | 0 | 0 | 0 |
| 0 | NA | NA | NA | NA | NA | Gm44330 | | 0 | 0 | 0 | 0 | 0 | 0 | 0 | 0 | 0 | 0 | 0 |
| 0 | NA | NA | NA | NA | NA | Mir8111 | | 0 | 0 | 0 | 0 | 0 | 0 | 0 | 0 | 0 | 0 | 0 |
| 0 | NA | NA | NA | NA | NA | Mir1199 | | 0 | 0 | 0 | 0 | 0 | 0 | 0 | 0 | 0 | 0 | 0 |
| 0 | NA | NA | NA | NA | NA | Gm25110 | | 0 | 0 | 0 | 0 | 0 | 0 | 0 | 0 | 0 | 0 | 0 |
| 0 | NA | NA | NA | NA | NA | Mir181c | | 0 | 0 | 0 | 0 | 0 | 0 | 0 | 0 | 0 | 0 | 0 |
| 0 | NA | NA | NA | NA | NA | Mir27a | | 0 | 0 | 0 | 0 | 0 | 0 | 0 | 0 | 0 | 0 | 0 |
| 0 | NA | NA | NA | NA | NA | Mir3074-2 | | 0 | 0 | 0 | 0 | 0 | 0 | 0 | 0 | 0 | 0 | 0 |
| 0 | NA | NA | NA | NA | NA | Gm23626 | | 0 | 0 | 0 | 0 | 0 | 0 | 0 | 0 | 0 | 0 | 0 |
| 0 | NA | NA | NA | NA | NA | Mir7069 | | 0 | 0 | 0 | 0 | 0 | 0 | 0 | 0 | 0 | 0 | 0 |
| 0 | NA | NA | NA | NA | NA | Gm24197 | | 0 | 0 | 0 | 0 | 0 | 0 | 0 | 0 | 0 | 0 | 0 |
| 0 | NA | NA | NA | NA | NA | Gm49661 | | 0 | 0 | 0 | 0 | 0 | 0 | 0 | 0 | 0 | 0 | 0 |
| 0 | NA | NA | NA | NA | NA | Gm25506 | | 0 | 0 | 0 | 0 | 0 | 0 | 0 | 0 | 0 | 0 | 0 |
| 0 | NA | NA | NA | NA | NA | Mir7070 | | 0 | 0 | 0 | 0 | 0 | 0 | 0 | 0 | 0 | 0 | 0 |
| 0 | NA | NA | NA | NA | NA | Gm45317 | | 0 | 0 | 0 | 0 | 0 | 0 | 0 | 0 | 0 | 0 | 0 |
| 0 | NA | NA | NA | NA | NA | Gm45316 | | 0 | 0 | 0 | 0 | 0 | 0 | 0 | 0 | 0 | 0 | 0 |
| 0 | NA | NA | NA | NA | NA | Cks1brt | CDC28 protein kinase 1b, retrogene | 0 | 0 | 0 | 0 | 0 | 0 | 0 | 0 | 0 | 0 | 0 |
| 0 | NA | NA | NA | NA | NA | Olfr371 | olfactory receptor 371 | 0 | 0 | 0 | 0 | 0 | 0 | 0 | 0 | 0 | 0 | 0 |
| 0 | NA | NA | NA | NA | NA | Gm22336 | | 0 | 0 | 0 | 0 | 0 | 0 | 0 | 0 | 0 | 0 | 0 |
| 0 | NA | NA | NA | NA | NA | AA672651 | | 0 | 0 | 0 | 0 | 0 | 0 | 0 | 0 | 0 | 0 | 0 |
| 0 | NA | NA | NA | NA | NA | 1700051O22Rik | | 0 | 0 | 0 | 0 | 0 | 0 | 0 | 0 | 0 | 0 | 0 |
| 0 | NA | NA | NA | NA | NA | Gm45425 | | 0 | 0 | 0 | 0 | 0 | 0 | 0 | 0 | 0 | 0 | 0 |
| 0 | NA | NA | NA | NA | NA | Mir8109 | | 0 | 0 | 0 | 0 | 0 | 0 | 0 | 0 | 0 | 0 | 0 |
| 0 | NA | NA | NA | NA | NA | Gm16167 | | 0 | 0 | 0 | 0 | 0 | 0 | 0 | 0 | 0 | 0 | 0 |
| 0 | NA | NA | NA | NA | NA | Gm23904 | | 0 | 0 | 0 | 0 | 0 | 0 | 0 | 0 | 0 | 0 | 0 |
| 0 | NA | NA | NA | NA | NA | Gm24490 | | 0 | 0 | 0 | 0 | 0 | 0 | 0 | 0 | 0 | 0 | 0 |
| 0 | NA | NA | NA | NA | NA | Gm44348 | | 0 | 0 | 0 | 0 | 0 | 0 | 0 | 0 | 0 | 0 | 0 |
| 0 | NA | NA | NA | NA | NA | Gm22305 | | 0 | 0 | 0 | 0 | 0 | 0 | 0 | 0 | 0 | 0 | 0 |
| 0 | NA | NA | NA | NA | NA | Gm24781 | | 0 | 0 | 0 | 0 | 0 | 0 | 0 | 0 | 0 | 0 | 0 |
| 0 | NA | NA | NA | NA | NA | Rps13-ps1 | | 0 | 0 | 0 | 0 | 0 | 0 | 0 | 0 | 0 | 0 | 0 |
| 0 | NA | NA | NA | NA | NA | Gm18602 | | 0 | 0 | 0 | 0 | 0 | 0 | 0 | 0 | 0 | 0 | 0 |
| 0 | NA | NA | NA | NA | NA | Gm5118 | | 0 | 0 | 0 | 0 | 0 | 0 | 0 | 0 | 0 | 0 | 0 |
| 0 | NA | NA | NA | NA | NA | Gm27246 | | 0 | 0 | 0 | 0 | 0 | 0 | 0 | 0 | 0 | 0 | 0 |
| 0 | NA | NA | NA | NA | NA | Gm27207 | | 0 | 0 | 0 | 0 | 0 | 0 | 0 | 0 | 0 | 0 | 0 |
| 0 | NA | NA | NA | NA | NA | Gm27208 | | 0 | 0 | 0 | 0 | 0 | 0 | 0 | 0 | 0 | 0 | 0 |
| 0 | NA | NA | NA | NA | NA | 4930535O05Rik | | 0 | 0 | 0 | 0 | 0 | 0 | 0 | 0 | 0 | 0 | 0 |
| 0 | NA | NA | NA | NA | NA | Gm19872 | | 0 | 0 | 0 | 0 | 0 | 0 | 0 | 0 | 0 | 0 | 0 |
| 0 | NA | NA | NA | NA | NA | Gm27016 | | 0 | 0 | 0 | 0 | 0 | 0 | 0 | 0 | 0 | 0 | 0 |
| 0 | NA | NA | NA | NA | NA | 1700120C18Rik | | 0 | 0 | 0 | 0 | 0 | 0 | 0 | 0 | 0 | 0 | 0 |
| 0 | NA | NA | NA | NA | NA | 4933402J07Rik | RIKEN cDNA 4933402J07 gene | 0 | 0 | 0 | 0 | 0 | 0 | 0 | 0 | 0 | 0 | 0 |
| 0 | NA | NA | NA | NA | NA | Gm24841 | | 0 | 0 | 0 | 0 | 0 | 0 | 0 | 0 | 0 | 0 | 0 |
| 0 | NA | NA | NA | NA | NA | Gm2716 | | 0 | 0 | 0 | 0 | 0 | 0 | 0 | 0 | 0 | 0 | 0 |
| 0 | NA | NA | NA | NA | NA | | | 0 | 0 | 0 | 0 | 0 | 0 | 0 | 0 | 0 | 0 | 0 |
| 0 | NA | NA | NA | NA | NA | Mir7071 | | 0 | 0 | 0 | 0 | 0 | 0 | 0 | 0 | 0 | 0 | 0 |
| 0 | NA | NA | NA | NA | NA | Gm45497 | | 0 | 0 | 0 | 0 | 0 | 0 | 0 | 0 | 0 | 0 | 0 |
| 0 | NA | NA | NA | NA | NA | Gm45496 | | 0 | 0 | 0 | 0 | 0 | 0 | 0 | 0 | 0 | 0 | 0 |
| 0 | NA | NA | NA | NA | NA | Gm17995 | | 0 | 0 | 0 | 0 | 0 | 0 | 0 | 0 | 0 | 0 | 0 |
| 0 | NA | NA | NA | NA | NA | Gm17910 | | 0 | 0 | 0 | 0 | 0 | 0 | 0 | 0 | 0 | 0 | 0 |
| 0 | NA | NA | NA | NA | NA | Gm45504 | | 0 | 0 | 0 | 0 | 0 | 0 | 0 | 0 | 0 | 0 | 0 |
| 0 | NA | NA | NA | NA | NA | Mir8110 | | 0 | 0 | 0 | 0 | 0 | 0 | 0 | 0 | 0 | 0 | 0 |
| 0 | NA | NA | NA | NA | NA | Gm26331 | | 0 | 0 | 0 | 0 | 0 | 0 | 0 | 0 | 0 | 0 | 0 |
| 0 | NA | NA | NA | NA | NA | Gm24212 | | 0 | 0 | 0 | 0 | 0 | 0 | 0 | 0 | 0 | 0 | 0 |
| 0 | NA | NA | NA | NA | NA | Gm45639 | | 0 | 0 | 0 | 0 | 0 | 0 | 0 | 0 | 0 | 0 | 0 |
| 0 | NA | NA | NA | NA | NA | Gm45638 | | 0 | 0 | 0 | 0 | 0 | 0 | 0 | 0 | 0 | 0 | 0 |
| 0 | NA | NA | NA | NA | NA | Gm35963 | | 0 | 0 | 0 | 0 | 0 | 0 | 0 | 0 | 0 | 0 | 0 |
| 0 | NA | NA | NA | NA | NA | Gm44382 | | 0 | 0 | 0 | 0 | 0 | 0 | 0 | 0 | 0 | 0 | 0 |
| 0 | NA | NA | NA | NA | NA | Gm45641 | | 0 | 0 | 0 | 0 | 0 | 0 | 0 | 0 | 0 | 0 | 0 |
| 0 | NA | NA | NA | NA | NA | Gm22428 | | 0 | 0 | 0 | 0 | 0 | 0 | 0 | 0 | 0 | 0 | 0 |
| 0 | NA | NA | NA | NA | NA | Gm45665 | | 0 | 0 | 0 | 0 | 0 | 0 | 0 | 0 | 0 | 0 | 0 |
| 0 | NA | NA | NA | NA | NA | Gm8489 | | 0 | 0 | 0 | 0 | 0 | 0 | 0 | 0 | 0 | 0 | 0 |

| | | | | | | | | | | | | | | | | | | |
|---|---|---|---|---|---|---|---|---|---|---|---|---|---|---|---|---|---|---|
| 0 | NA | NA | NA | NA | NA | Gm45666 | | 0 | 0 | 0 | 0 | 0 | 0 | 0 | 0 | 0 | 0 | 0 |
| 0 | NA | NA | NA | NA | NA | Gm23408 | | 0 | 0 | 0 | 0 | 0 | 0 | 0 | 0 | 0 | 0 | 0 |
| 0 | NA | NA | NA | NA | NA | Gm36243 | | 0 | 0 | 0 | 0 | 0 | 0 | 0 | 0 | 0 | 0 | 0 |
| 0 | NA | NA | NA | NA | NA | Gm45662 | | 0 | 0 | 0 | 0 | 0 | 0 | 0 | 0 | 0 | 0 | 0 |
| 0 | NA | NA | NA | NA | NA | Gm45427 | | 0 | 0 | 0 | 0 | 0 | 0 | 0 | 0 | 0 | 0 | 0 |
| 0 | NA | NA | NA | NA | NA | 4831440D22Rik | | 0 | 0 | 0 | 0 | 0 | 0 | 0 | 0 | 0 | 0 | 0 |
| 0 | NA | NA | NA | NA | NA | Gm45290 | | 0 | 0 | 0 | 0 | 0 | 0 | 0 | 0 | 0 | 0 | 0 |
| 0 | NA | NA | NA | NA | NA | Gm45294 | | 0 | 0 | 0 | 0 | 0 | 0 | 0 | 0 | 0 | 0 | 0 |
| 0 | NA | NA | NA | NA | NA | Gm3235 | | 0 | 0 | 0 | 0 | 0 | 0 | 0 | 0 | 0 | 0 | 0 |
| 0 | NA | NA | NA | NA | NA | Gm45295 | | 0 | 0 | 0 | 0 | 0 | 0 | 0 | 0 | 0 | 0 | 0 |
| 0 | NA | NA | NA | NA | NA | Gm2875 | | 0 | 0 | 0 | 0 | 0 | 0 | 0 | 0 | 0 | 0 | 0 |
| 0 | NA | NA | NA | NA | NA | Gm36325 | | 0 | 0 | 0 | 0 | 0 | 0 | 0 | 0 | 0 | 0 | 0 |
| 0 | NA | NA | NA | NA | NA | Gm36380 | | 0 | 0 | 0 | 0 | 0 | 0 | 0 | 0 | 0 | 0 | 0 |
| 0 | NA | NA | NA | NA | NA | Irx3os | | 0 | 0 | 0 | 0 | 0 | 0 | 0 | 0 | 0 | 0 | 0 |
| 0 | NA | NA | NA | NA | NA | Gm28511 | | 0 | 0 | 0 | 0 | 0 | 0 | 0 | 0 | 0 | 0 | 0 |
| 0 | NA | NA | NA | NA | NA | Gm28515 | | 0 | 0 | 0 | 0 | 0 | 0 | 0 | 0 | 0 | 0 | 0 |
| 0 | NA | NA | NA | NA | NA | Gm36670 | | 0 | 0 | 0 | 0 | 0 | 0 | 0 | 0 | 0 | 0 | 0 |
| 0 | NA | NA | NA | NA | NA | Gm36843 | | 0 | 0 | 0 | 0 | 0 | 0 | 0 | 0 | 0 | 0 | 0 |
| 0 | NA | NA | NA | NA | NA | Gm28516 | | 0 | 0 | 0 | 0 | 0 | 0 | 0 | 0 | 0 | 0 | 0 |
| 0 | NA | NA | NA | NA | NA | Gm24254 | | 0 | 0 | 0 | 0 | 0 | 0 | 0 | 0 | 0 | 0 | 0 |
| 0 | NA | NA | NA | NA | NA | Gm45332 | | 0 | 0 | 0 | 0 | 0 | 0 | 0 | 0 | 0 | 0 | 0 |
| 0 | NA | NA | NA | NA | NA | Gm45335 | | 0 | 0 | 0 | 0 | 0 | 0 | 0 | 0 | 0 | 0 | 0 |
| 0 | NA | NA | NA | NA | NA | Gm45334 | | 0 | 0 | 0 | 0 | 0 | 0 | 0 | 0 | 0 | 0 | 0 |
| 0 | NA | NA | NA | NA | NA | Gm21817 | | 0 | 0 | 0 | 0 | 0 | 0 | 0 | 0 | 0 | 0 | 0 |
| 0 | NA | NA | NA | NA | NA | Irx5 | Iroquois homeobox 5 | 0 | 0 | 0 | 0 | 0 | 0 | 0 | 0 | 0 | 0 | 0 |
| 0 | NA | NA | NA | NA | NA | Gm45336 | | 0 | 0 | 0 | 0 | 0 | 0 | 0 | 0 | 0 | 0 | 0 |
| 0 | NA | NA | NA | NA | NA | Irx6 | Iroquois homeobox 6 | 0 | 0 | 0 | 0 | 0 | 0 | 0 | 0 | 0 | 0 | 0 |
| 0 | NA | NA | NA | NA | NA | Gm45505 | | 0 | 0 | 0 | 0 | 0 | 0 | 0 | 0 | 0 | 0 | 0 |
| 0 | NA | NA | NA | NA | NA | Gm45472 | | 0 | 0 | 0 | 0 | 0 | 0 | 0 | 0 | 0 | 0 | 0 |
| 0 | NA | NA | NA | NA | NA | Capns2 | calpain, small subunit 2 | 0 | 0 | 0 | 0 | 0 | 0 | 0 | 0 | 0 | 0 | 0 |
| 0 | NA | NA | NA | NA | NA | Gm22151 | | 0 | 0 | 0 | 0 | 0 | 0 | 0 | 0 | 0 | 0 | 0 |
| 0 | NA | NA | NA | NA | NA | Gm45463 | | 0 | 0 | 0 | 0 | 0 | 0 | 0 | 0 | 0 | 0 | 0 |
| 0 | NA | NA | NA | NA | NA | Ces1a | carboxylesterase 1A | 0 | 0 | 0 | 0 | 0 | 0 | 0 | 0 | 0 | 0 | 0 |
| 0 | NA | NA | NA | NA | NA | Gm45910 | | 0 | 0 | 0 | 0 | 0 | 0 | 0 | 0 | 0 | 0 | 0 |
| 0 | NA | NA | NA | NA | NA | Gm45909 | | 0 | 0 | 0 | 0 | 0 | 0 | 0 | 0 | 0 | 0 | 0 |
| 0 | NA | NA | NA | NA | NA | Gm45727 | | 0 | 0 | 0 | 0 | 0 | 0 | 0 | 0 | 0 | 0 | 0 |
| 0 | NA | NA | NA | NA | NA | Ces1f | carboxylesterase 1F | 0 | 0 | 0 | 0 | 0 | 0 | 0 | 0 | 0 | 0 | 0 |
| 0 | NA | NA | NA | NA | NA | Ces1g | carboxylesterase 1G | 0 | 0 | 0 | 0 | 0 | 0 | 0 | 0 | 0 | 0 | 0 |
| 0 | NA | NA | NA | NA | NA | Ces1h | carboxylesterase 1H | 0 | 0 | 0 | 0 | 0 | 0 | 0 | 0 | 0 | 0 | 0 |
| 0 | NA | NA | NA | NA | NA | Gm45765 | | 0 | 0 | 0 | 0 | 0 | 0 | 0 | 0 | 0 | 0 | 0 |
| 0 | NA | NA | NA | NA | NA | Gm45764 | | 0 | 0 | 0 | 0 | 0 | 0 | 0 | 0 | 0 | 0 | 0 |
| 0 | NA | NA | NA | NA | NA | Gm30132 | | 0 | 0 | 0 | 0 | 0 | 0 | 0 | 0 | 0 | 0 | 0 |
| 0 | NA | NA | NA | NA | NA | Gm45766 | | 0 | 0 | 0 | 0 | 0 | 0 | 0 | 0 | 0 | 0 | 0 |
| 0 | NA | NA | NA | NA | NA | | | 0 | 0 | 0 | 0 | 0 | 0 | 0 | 0 | 0 | 0 | 0 |
| 0 | NA | NA | NA | NA | NA | Gm24159 | | 0 | 0 | 0 | 0 | 0 | 0 | 0 | 0 | 0 | 0 | 0 |
| 0 | NA | NA | NA | NA | NA | Mt4 | metallothionein 4 | 0 | 0 | 0 | 0 | 0 | 0 | 0 | 0 | 0 | 0 | 0 |
| 0 | NA | NA | NA | NA | NA | Mir138-2 | | 0 | 0 | 0 | 0 | 0 | 0 | 0 | 0 | 0 | 0 | 0 |
| 0 | NA | NA | NA | NA | NA | Mir7072 | | 0 | 0 | 0 | 0 | 0 | 0 | 0 | 0 | 0 | 0 | 0 |
| 0 | NA | NA | NA | NA | NA | Gm22846 | | 0 | 0 | 0 | 0 | 0 | 0 | 0 | 0 | 0 | 0 | 0 |
| 0 | NA | NA | NA | NA | NA | Gnb1-ps3 | | 0 | 0 | 0 | 0 | 0 | 0 | 0 | 0 | 0 | 0 | 0 |
| 0 | NA | NA | NA | NA | NA | Gm31224 | | 0 | 0 | 0 | 0 | 0 | 0 | 0 | 0 | 0 | 0 | 0 |
| 0 | NA | NA | NA | NA | NA | Gm45812 | | 0 | 0 | 0 | 0 | 0 | 0 | 0 | 0 | 0 | 0 | 0 |
| 0 | NA | NA | NA | NA | NA | Gm45854 | | 0 | 0 | 0 | 0 | 0 | 0 | 0 | 0 | 0 | 0 | 0 |
| 0 | NA | NA | NA | NA | NA | Prss54 | protease, serine 54 | 0 | 0 | 0 | 0 | 0 | 0 | 0 | 0 | 0 | 0 | 0 |
| 0 | NA | NA | NA | NA | NA | Gm26265 | | 0 | 0 | 0 | 0 | 0 | 0 | 0 | 0 | 0 | 0 | 0 |
| 0 | NA | NA | NA | NA | NA | Gm26265 | | 0 | 0 | 0 | 0 | 0 | 0 | 0 | 0 | 0 | 0 | 0 |
| 0 | NA | NA | NA | NA | NA | Gm45762 | | 0 | 0 | 0 | 0 | 0 | 0 | 0 | 0 | 0 | 0 | 0 |
| 0 | NA | NA | NA | NA | NA | Gm31659 | | 0 | 0 | 0 | 0 | 0 | 0 | 0 | 0 | 0 | 0 | 0 |
| 0 | NA | NA | NA | NA | NA | Gm8637 | | 0 | 0 | 0 | 0 | 0 | 0 | 0 | 0 | 0 | 0 | 0 |
| 0 | NA | NA | NA | NA | NA | Gm32005 | | 0 | 0 | 0 | 0 | 0 | 0 | 0 | 0 | 0 | 0 | 0 |

| | | | | | | | | | | | | | | | | | | |
|---|---|---|---|---|---|---|---|---|---|---|---|---|---|---|---|---|---|---|
| 0 | NA | NA | NA | NA | NA | Gm22780 | | 0 | 0 | 0 | 0 | 0 | 0 | 0 | 0 | 0 | 0 | 0 |
| 0 | NA | NA | NA | NA | NA | Gm32122 | | 0 | 0 | 0 | 0 | 0 | 0 | 0 | 0 | 0 | 0 | 0 |
| 0 | NA | NA | NA | NA | NA | 1700047G07Rik | | 0 | 0 | 0 | 0 | 0 | 0 | 0 | 0 | 0 | 0 | 0 |
| 0 | NA | NA | NA | NA | NA | Gm24132 | | 0 | 0 | 0 | 0 | 0 | 0 | 0 | 0 | 0 | 0 | 0 |
| 0 | NA | NA | NA | NA | NA | Gm45703 | | 0 | 0 | 0 | 0 | 0 | 0 | 0 | 0 | 0 | 0 | 0 |
| 0 | NA | NA | NA | NA | NA | Gm45788 | | 0 | 0 | 0 | 0 | 0 | 0 | 0 | 0 | 0 | 0 | 0 |
| 0 | NA | NA | NA | NA | NA | Gm25160 | | 0 | 0 | 0 | 0 | 0 | 0 | 0 | 0 | 0 | 0 | 0 |
| 0 | NA | NA | NA | NA | NA | Gm26399 | | 0 | 0 | 0 | 0 | 0 | 0 | 0 | 0 | 0 | 0 | 0 |
| 0 | NA | NA | NA | NA | NA | Gm45247 | | 0 | 0 | 0 | 0 | 0 | 0 | 0 | 0 | 0 | 0 | 0 |
| 0 | NA | NA | NA | NA | NA | Gm39231 | | 0 | 0 | 0 | 0 | 0 | 0 | 0 | 0 | 0 | 0 | 0 |
| 0 | NA | NA | NA | NA | NA | Gm17997 | | 0 | 0 | 0 | 0 | 0 | 0 | 0 | 0 | 0 | 0 | 0 |
| 0 | NA | NA | NA | NA | NA | Gm23494 | | 0 | 0 | 0 | 0 | 0 | 0 | 0 | 0 | 0 | 0 | 0 |
| 0 | NA | NA | NA | NA | NA | Gm29817 | | 0 | 0 | 0 | 0 | 0 | 0 | 0 | 0 | 0 | 0 | 0 |
| 0 | NA | NA | NA | NA | NA | Gm15679 | | 0 | 0 | 0 | 0 | 0 | 0 | 0 | 0 | 0 | 0 | 0 |
| 0 | NA | NA | NA | NA | NA | Gm19232 | | 0 | 0 | 0 | 0 | 0 | 0 | 0 | 0 | 0 | 0 | 0 |
| 0 | NA | NA | NA | NA | NA | Gm5742 | | 0 | 0 | 0 | 0 | 0 | 0 | 0 | 0 | 0 | 0 | 0 |
| 0 | NA | NA | NA | NA | NA | Gm45296 | | 0 | 0 | 0 | 0 | 0 | 0 | 0 | 0 | 0 | 0 | 0 |
| 0 | NA | NA | NA | NA | NA | Gm5361 | | 0 | 0 | 0 | 0 | 0 | 0 | 0 | 0 | 0 | 0 | 0 |
| 0 | NA | NA | NA | NA | NA | Gm39232 | | 0 | 0 | 0 | 0 | 0 | 0 | 0 | 0 | 0 | 0 | 0 |
| 0 | NA | NA | NA | NA | NA | Gm22223 | | 0 | 0 | 0 | 0 | 0 | 0 | 0 | 0 | 0 | 0 | 0 |
| 0 | NA | NA | NA | NA | NA | Gm24602 | | 0 | 0 | 0 | 0 | 0 | 0 | 0 | 0 | 0 | 0 | 0 |
| 0 | NA | NA | NA | NA | NA | Gm45257 | | 0 | 0 | 0 | 0 | 0 | 0 | 0 | 0 | 0 | 0 | 0 |
| 0 | NA | NA | NA | NA | NA | 9430099M06Rik | | 0 | 0 | 0 | 0 | 0 | 0 | 0 | 0 | 0 | 0 | 0 |
| 0 | NA | NA | NA | NA | NA | Gm26301 | | 0 | 0 | 0 | 0 | 0 | 0 | 0 | 0 | 0 | 0 | 0 |
| 0 | NA | NA | NA | NA | NA | Gm45258 | | 0 | 0 | 0 | 0 | 0 | 0 | 0 | 0 | 0 | 0 | 0 |
| 0 | NA | NA | NA | NA | NA | Gm45391 | | 0 | 0 | 0 | 0 | 0 | 0 | 0 | 0 | 0 | 0 | 0 |
| 0 | NA | NA | NA | NA | NA | Gm32531 | | 0 | 0 | 0 | 0 | 0 | 0 | 0 | 0 | 0 | 0 | 0 |
| 0 | NA | NA | NA | NA | NA | 1600027J07Rik | | 0 | 0 | 0 | 0 | 0 | 0 | 0 | 0 | 0 | 0 | 0 |
| 0 | NA | NA | NA | NA | NA | Gm45277 | | 0 | 0 | 0 | 0 | 0 | 0 | 0 | 0 | 0 | 0 | 0 |
| 0 | NA | NA | NA | NA | NA | 4933400L20Rik | | 0 | 0 | 0 | 0 | 0 | 0 | 0 | 0 | 0 | 0 | 0 |
| 0 | NA | NA | NA | NA | NA | Gm29682 | | 0 | 0 | 0 | 0 | 0 | 0 | 0 | 0 | 0 | 0 | 0 |
| 0 | NA | NA | NA | NA | NA | Gm10631 | | 0 | 0 | 0 | 0 | 0 | 0 | 0 | 0 | 0 | 0 | 0 |
| 0 | NA | NA | NA | NA | NA | Gm45892 | | 0 | 0 | 0 | 0 | 0 | 0 | 0 | 0 | 0 | 0 | 0 |
| 0 | NA | NA | NA | NA | NA | Gm45887 | | 0 | 0 | 0 | 0 | 0 | 0 | 0 | 0 | 0 | 0 | 0 |
| 0 | NA | NA | NA | NA | NA | Gm45750 | | 0 | 0 | 0 | 0 | 0 | 0 | 0 | 0 | 0 | 0 | 0 |
| 0 | NA | NA | NA | NA | NA | Cmtm2a | CKLF-like MARVEL transmembrane dom | 0 | 0 | 0 | 0 | 0 | 0 | 0 | 0 | 0 | 0 | 0 |
| 0 | NA | NA | NA | NA | NA | Gm45711 | | 0 | 0 | 0 | 0 | 0 | 0 | 0 | 0 | 0 | 0 | 0 |
| 0 | NA | NA | NA | NA | NA | Cmtm1 | CKLF-like MARVEL transmembrane dom | 0 | 0 | 0 | 0 | 0 | 0 | 0 | 0 | 0 | 0 | 0 |
| 0 | NA | NA | NA | NA | NA | Gm45710 | | 0 | 0 | 0 | 0 | 0 | 0 | 0 | 0 | 0 | 0 | 0 |
| 0 | NA | NA | NA | NA | NA | Gm11020 | | 0 | 0 | 0 | 0 | 0 | 0 | 0 | 0 | 0 | 0 | 0 |
| 0 | NA | NA | NA | NA | NA | Cmtm2b | CKLF-like MARVEL transmembrane dom | 0 | 0 | 0 | 0 | 0 | 0 | 0 | 0 | 0 | 0 | 0 |
| 0 | NA | NA | NA | NA | NA | Gm45878 | | 0 | 0 | 0 | 0 | 0 | 0 | 0 | 0 | 0 | 0 | 0 |
| 0 | NA | NA | NA | NA | NA | Gm45731 | | 0 | 0 | 0 | 0 | 0 | 0 | 0 | 0 | 0 | 0 | 0 |
| 0 | NA | NA | NA | NA | NA | Gm22148 | | 0 | 0 | 0 | 0 | 0 | 0 | 0 | 0 | 0 | 0 | 0 |
| 0 | NA | NA | NA | NA | NA | Gm45843 | | 0 | 0 | 0 | 0 | 0 | 0 | 0 | 0 | 0 | 0 | 0 |
| 0 | NA | NA | NA | NA | NA | Gm7208 | | 0 | 0 | 0 | 0 | 0 | 0 | 0 | 0 | 0 | 0 | 0 |
| 0 | NA | NA | NA | NA | NA | Gm5159 | | 0 | 0 | 0 | 0 | 0 | 0 | 0 | 0 | 0 | 0 | 0 |
| 0 | NA | NA | NA | NA | NA | Gm44328 | | 0 | 0 | 0 | 0 | 0 | 0 | 0 | 0 | 0 | 0 | 0 |
| 0 | NA | NA | NA | NA | NA | | | 0 | 0 | 0 | 0 | 0 | 0 | 0 | 0 | 0 | 0 | 0 |
| 0 | NA | NA | NA | NA | NA | 9230110F11Rik | | 0 | 0 | 0 | 0 | 0 | 0 | 0 | 0 | 0 | 0 | 0 |
| 0 | NA | NA | NA | NA | NA | Gm6831 | | 0 | 0 | 0 | 0 | 0 | 0 | 0 | 0 | 0 | 0 | 0 |
| 0 | NA | NA | NA | NA | NA | Gm45782 | | 0 | 0 | 0 | 0 | 0 | 0 | 0 | 0 | 0 | 0 | 0 |
| 0 | NA | NA | NA | NA | NA | 4932416K20Rik | | 0 | 0 | 0 | 0 | 0 | 0 | 0 | 0 | 0 | 0 | 0 |
| 0 | NA | NA | NA | NA | NA | Matr3-ps2 | | 0 | 0 | 0 | 0 | 0 | 0 | 0 | 0 | 0 | 0 | 0 |
| 0 | NA | NA | NA | NA | NA | Ces2d-ps | | 0 | 0 | 0 | 0 | 0 | 0 | 0 | 0 | 0 | 0 | 0 |
| 0 | NA | NA | NA | NA | NA | Gm8798 | | 0 | 0 | 0 | 0 | 0 | 0 | 0 | 0 | 0 | 0 | 0 |
| 0 | NA | NA | NA | NA | NA | | | 0 | 0 | 0 | 0 | 0 | 0 | 0 | 0 | 0 | 0 | 0 |
| 0 | NA | NA | NA | NA | NA | Ces2f | carboxylesterase 2F | 0 | 0 | 0 | 0 | 0 | 0 | 0 | 0 | 0 | 0 | 0 |
| 0 | NA | NA | NA | NA | NA | Gm5743 | | 0 | 0 | 0 | 0 | 0 | 0 | 0 | 0 | 0 | 0 | 0 |
| 0 | NA | NA | NA | NA | NA | Ces2h | carboxylesterase 2H | 0 | 0 | 0 | 0 | 0 | 0 | 0 | 0 | 0 | 0 | 0 |

| | | | | | | | | | | | | | | | | | | |
|---|---|---|---|---|---|---|---|---|---|---|---|---|---|---|---|---|---|---|
| 0 | NA | NA | NA | NA | NA | Gm45725 | | 0 | 0 | 0 | 0 | 0 | 0 | 0 | 0 | 0 | 0 | 0 |
| 0 | NA | NA | NA | NA | NA | Gm45724 | | 0 | 0 | 0 | 0 | 0 | 0 | 0 | 0 | 0 | 0 | 0 |
| 0 | NA | NA | NA | NA | NA | Ces3b | carboxylesterase 3B | 0 | 0 | 0 | 0 | 0 | 0 | 0 | 0 | 0 | 0 | 0 |
| 0 | NA | NA | NA | NA | NA | Gm8804 | | 0 | 0 | 0 | 0 | 0 | 0 | 0 | 0 | 0 | 0 | 0 |
| 0 | NA | NA | NA | NA | NA | Gm3830 | | 0 | 0 | 0 | 0 | 0 | 0 | 0 | 0 | 0 | 0 | 0 |
| 0 | NA | NA | NA | NA | NA | Gm24629 | | 0 | 0 | 0 | 0 | 0 | 0 | 0 | 0 | 0 | 0 | 0 |
| 0 | NA | NA | NA | NA | NA | Gm8838 | | 0 | 0 | 0 | 0 | 0 | 0 | 0 | 0 | 0 | 0 | 0 |
| 0 | NA | NA | NA | NA | NA | Mir1966 | | 0 | 0 | 0 | 0 | 0 | 0 | 0 | 0 | 0 | 0 | 0 |
| 0 | NA | NA | NA | NA | NA | Nrn1l | neuritin 1-like | 0 | 0 | 0 | 0 | 0 | 0 | 0 | 0 | 0 | 0 | 0 |
| 0 | NA | NA | NA | NA | NA | Mir7074 | | 0 | 0 | 0 | 0 | 0 | 0 | 0 | 0 | 0 | 0 | 0 |
| 0 | NA | NA | NA | NA | NA | Gm45752 | | 0 | 0 | 0 | 0 | 0 | 0 | 0 | 0 | 0 | 0 | 0 |
| 0 | NA | NA | NA | NA | NA | Dpep3 | dipeptidase 3 | 0 | 0 | 0 | 0 | 0 | 0 | 0 | 0 | 0 | 0 | 0 |
| 0 | NA | NA | NA | NA | NA | Dpep2 | dipeptidase 2 | 0 | 0 | 0 | 0 | 0 | 0 | 0 | 0 | 0 | 0 | 0 |
| 0 | NA | NA | NA | NA | NA | Dpep2 | dipeptidase 2 | 0 | 0 | 0 | 0 | 0 | 0 | 0 | 0 | 0 | 0 | 0 |
| 0 | NA | NA | NA | NA | NA | Gm22972 | | 0 | 0 | 0 | 0 | 0 | 0 | 0 | 0 | 0 | 0 | 0 |
| 0 | NA | NA | NA | NA | NA | Dpep2nb | | 0 | 0 | 0 | 0 | 0 | 0 | 0 | 0 | 0 | 0 | 0 |
| 0 | NA | NA | NA | NA | NA | Gm27956 | | 0 | 0 | 0 | 0 | 0 | 0 | 0 | 0 | 0 | 0 | 0 |
| 0 | NA | NA | NA | NA | NA | Gm10629 | | 0 | 0 | 0 | 0 | 0 | 0 | 0 | 0 | 0 | 0 | 0 |
| 0 | NA | NA | NA | NA | NA | Gm50241 | | 0 | 0 | 0 | 0 | 0 | 0 | 0 | 0 | 0 | 0 | 0 |
| 0 | NA | NA | NA | NA | NA | Gm22085 | | 0 | 0 | 0 | 0 | 0 | 0 | 0 | 0 | 0 | 0 | 0 |
| 0 | NA | NA | NA | NA | NA | Mir7075 | | 0 | 0 | 0 | 0 | 0 | 0 | 0 | 0 | 0 | 0 | 0 |
| 0 | NA | NA | NA | NA | NA | C630050I24Rik | | 0 | 0 | 0 | 0 | 0 | 0 | 0 | 0 | 0 | 0 | 0 |
| 0 | NA | NA | NA | NA | NA | Gm45900 | | 0 | 0 | 0 | 0 | 0 | 0 | 0 | 0 | 0 | 0 | 0 |
| 0 | NA | NA | NA | NA | NA | Mir140 | | 0 | 0 | 0 | 0 | 0 | 0 | 0 | 0 | 0 | 0 | 0 |
| 0 | NA | NA | NA | NA | NA | Gm23646 | | 0 | 0 | 0 | 0 | 0 | 0 | 0 | 0 | 0 | 0 | 0 |
| 0 | NA | NA | NA | NA | NA | Gm22825 | | 0 | 0 | 0 | 0 | 0 | 0 | 0 | 0 | 0 | 0 | 0 |
| 0 | NA | NA | NA | NA | NA | 5033426E14Rik | | 0 | 0 | 0 | 0 | 0 | 0 | 0 | 0 | 0 | 0 | 0 |
| 0 | NA | NA | NA | NA | NA | Gm39244 | | 0 | 0 | 0 | 0 | 0 | 0 | 0 | 0 | 0 | 0 | 0 |
| 0 | NA | NA | NA | NA | NA | Lncbate1 | | 0 | 0 | 0 | 0 | 0 | 0 | 0 | 0 | 0 | 0 | 0 |
| 0 | NA | NA | NA | NA | NA | Gm20686 | | 0 | 0 | 0 | 0 | 0 | 0 | 0 | 0 | 0 | 0 | 0 |
| 0 | NA | NA | NA | NA | NA | Mir3108 | | 0 | 0 | 0 | 0 | 0 | 0 | 0 | 0 | 0 | 0 | 0 |
| 0 | NA | NA | NA | NA | NA | D030068K23Rik | | 0 | 0 | 0 | 0 | 0 | 0 | 0 | 0 | 0 | 0 | 0 |
| 0 | NA | NA | NA | NA | NA | Gm23163 | | 0 | 0 | 0 | 0 | 0 | 0 | 0 | 0 | 0 | 0 | 0 |
| 0 | NA | NA | NA | NA | NA | Gm25321 | | 0 | 0 | 0 | 0 | 0 | 0 | 0 | 0 | 0 | 0 | 0 |
| 0 | NA | NA | NA | NA | NA | 8030455M16Rik | | 0 | 0 | 0 | 0 | 0 | 0 | 0 | 0 | 0 | 0 | 0 |
| 0 | NA | NA | NA | NA | NA | Gm17720 | | 0 | 0 | 0 | 0 | 0 | 0 | 0 | 0 | 0 | 0 | 0 |
| 0 | NA | NA | NA | NA | NA | Tle7 | TLE family member 7 | 0 | 0 | 0 | 0 | 0 | 0 | 0 | 0 | 0 | 0 | 0 |
| 0 | NA | NA | NA | NA | NA | Gm26832 | | 0 | 0 | 0 | 0 | 0 | 0 | 0 | 0 | 0 | 0 | 0 |
| 0 | NA | NA | NA | NA | NA | Gm23627 | | 0 | 0 | 0 | 0 | 0 | 0 | 0 | 0 | 0 | 0 | 0 |
| 0 | NA | NA | NA | NA | NA | Gm22701 | | 0 | 0 | 0 | 0 | 0 | 0 | 0 | 0 | 0 | 0 | 0 |
| 0 | NA | NA | NA | NA | NA | Gm47306 | | 0 | 0 | 0 | 0 | 0 | 0 | 0 | 0 | 0 | 0 | 0 |
| 0 | NA | NA | NA | NA | NA | Gm44489 | | 0 | 0 | 0 | 0 | 0 | 0 | 0 | 0 | 0 | 0 | 0 |
| 0 | NA | NA | NA | NA | NA | Gm15894 | | 0 | 0 | 0 | 0 | 0 | 0 | 0 | 0 | 0 | 0 | 0 |
| 0 | NA | NA | NA | NA | NA | Snord111 | | 0 | 0 | 0 | 0 | 0 | 0 | 0 | 0 | 0 | 0 | 0 |
| 0 | NA | NA | NA | NA | NA | Gm22193 | | 0 | 0 | 0 | 0 | 0 | 0 | 0 | 0 | 0 | 0 | 0 |
| 0 | NA | NA | NA | NA | NA | Gm26132 | | 0 | 0 | 0 | 0 | 0 | 0 | 0 | 0 | 0 | 0 | 0 |
| 0 | NA | NA | NA | NA | NA | | | 0 | 0 | 0 | 0 | 0 | 0 | 0 | 0 | 0 | 0 | 0 |
| 0 | NA | NA | NA | NA | NA | Gm24103 | | 0 | 0 | 0 | 0 | 0 | 0 | 0 | 0 | 0 | 0 | 0 |
| 0 | NA | NA | NA | NA | NA | Exosc6 | exosome component 6 | 0 | 0 | 0 | 0 | 0 | 0 | 0 | 0 | 0 | 0 | 0 |
| 0 | NA | NA | NA | NA | NA | Clec18a | C-type lectin domain family 18, member A | 0 | 0 | 0 | 0 | 0 | 0 | 0 | 0 | 0 | 0 | 0 |
| 0 | NA | NA | NA | NA | NA | | | 0 | 0 | 0 | 0 | 0 | 0 | 0 | 0 | 0 | 0 | 0 |
| 0 | NA | NA | NA | NA | NA | Gm23193 | | 0 | 0 | 0 | 0 | 0 | 0 | 0 | 0 | 0 | 0 | 0 |
| 0 | NA | NA | NA | NA | NA | Gm45712 | | 0 | 0 | 0 | 0 | 0 | 0 | 0 | 0 | 0 | 0 | 0 |
| 0 | NA | NA | NA | NA | NA | Gm24793 | | 0 | 0 | 0 | 0 | 0 | 0 | 0 | 0 | 0 | 0 | 0 |
| 0 | NA | NA | NA | NA | NA | Mir1957b | | 0 | 0 | 0 | 0 | 0 | 0 | 0 | 0 | 0 | 0 | 0 |
| 0 | NA | NA | NA | NA | NA | Mir7076 | | 0 | 0 | 0 | 0 | 0 | 0 | 0 | 0 | 0 | 0 | 0 |
| 0 | NA | NA | NA | NA | NA | Gm23853 | | 0 | 0 | 0 | 0 | 0 | 0 | 0 | 0 | 0 | 0 | 0 |
| 0 | NA | NA | NA | NA | NA | Chst5 | carbohydrate (N-acetylglucosamine 6-O) | 0 | 0 | 0 | 0 | 0 | 0 | 0 | 0 | 0 | 0 | 0 |
| 0 | NA | NA | NA | NA | NA | Gm22827 | | 0 | 0 | 0 | 0 | 0 | 0 | 0 | 0 | 0 | 0 | 0 |
| 0 | NA | NA | NA | NA | NA | Gm26994 | | 0 | 0 | 0 | 0 | 0 | 0 | 0 | 0 | 0 | 0 | 0 |

| | | | | | | | | | | | | | | | | | | |
|---|---|---|---|---|---|---|---|---|---|---|---|---|---|---|---|---|---|---|
| 0 | NA | NA | NA | NA | NA | Gm22664 | | 0 | 0 | 0 | 0 | 0 | 0 | 0 | 0 | 0 | 0 | 0 |
| 0 | NA | NA | NA | NA | NA | Gm45904 | | 0 | 0 | 0 | 0 | 0 | 0 | 0 | 0 | 0 | 0 | 0 |
| 0 | NA | NA | NA | NA | NA | Gm10280 | | 0 | 0 | 0 | 0 | 0 | 0 | 0 | 0 | 0 | 0 | 0 |
| 0 | NA | NA | NA | NA | NA | Gm25766 | | 0 | 0 | 0 | 0 | 0 | 0 | 0 | 0 | 0 | 0 | 0 |
| 0 | NA | NA | NA | NA | NA | Gm24291 | | 0 | 0 | 0 | 0 | 0 | 0 | 0 | 0 | 0 | 0 | 0 |
| 0 | NA | NA | NA | NA | NA | Gm45789 | | 0 | 0 | 0 | 0 | 0 | 0 | 0 | 0 | 0 | 0 | 0 |
| 0 | NA | NA | NA | NA | NA | Gm30052 | | 0 | 0 | 0 | 0 | 0 | 0 | 0 | 0 | 0 | 0 | 0 |
| 0 | NA | NA | NA | NA | NA | Clec3a | C-type lectin domain family 3, member a | 0 | 0 | 0 | 0 | 0 | 0 | 0 | 0 | 0 | 0 | 0 |
| 0 | NA | NA | NA | NA | NA | Gm16117 | | 0 | 0 | 0 | 0 | 0 | 0 | 0 | 0 | 0 | 0 | 0 |
| 0 | NA | NA | NA | NA | NA | Gm22556 | | 0 | 0 | 0 | 0 | 0 | 0 | 0 | 0 | 0 | 0 | 0 |
| 0 | NA | NA | NA | NA | NA | 4930488N15Rik | | 0 | 0 | 0 | 0 | 0 | 0 | 0 | 0 | 0 | 0 | 0 |
| 0 | NA | NA | NA | NA | NA | Gm23677 | | 0 | 0 | 0 | 0 | 0 | 0 | 0 | 0 | 0 | 0 | 0 |
| 0 | NA | NA | NA | NA | NA | 4930422C21Rik | | 0 | 0 | 0 | 0 | 0 | 0 | 0 | 0 | 0 | 0 | 0 |
| 0 | NA | NA | NA | NA | NA | 1700018P08Rik | | 0 | 0 | 0 | 0 | 0 | 0 | 0 | 0 | 0 | 0 | 0 |
| 0 | NA | NA | NA | NA | NA | Gm5357 | | 0 | 0 | 0 | 0 | 0 | 0 | 0 | 0 | 0 | 0 | 0 |
| 0 | NA | NA | NA | NA | NA | Gm10620 | | 0 | 0 | 0 | 0 | 0 | 0 | 0 | 0 | 0 | 0 | 0 |
| 0 | NA | NA | NA | NA | NA | Gm15395 | | 0 | 0 | 0 | 0 | 0 | 0 | 0 | 0 | 0 | 0 | 0 |
| 0 | NA | NA | NA | NA | NA | Mir7077 | | 0 | 0 | 0 | 0 | 0 | 0 | 0 | 0 | 0 | 0 | 0 |
| 0 | NA | NA | NA | NA | NA | Mir7078 | | 0 | 0 | 0 | 0 | 0 | 0 | 0 | 0 | 0 | 0 | 0 |
| 0 | NA | NA | NA | NA | NA | Gm27361 | | 0 | 0 | 0 | 0 | 0 | 0 | 0 | 0 | 0 | 0 | 0 |
| 0 | NA | NA | NA | NA | NA | Gm31717 | | 0 | 0 | 0 | 0 | 0 | 0 | 0 | 0 | 0 | 0 | 0 |
| 0 | NA | NA | NA | NA | NA | Gm38416 | | 0 | 0 | 0 | 0 | 0 | 0 | 0 | 0 | 0 | 0 | 0 |
| 0 | NA | NA | NA | NA | NA | Gm45776 | | 0 | 0 | 0 | 0 | 0 | 0 | 0 | 0 | 0 | 0 | 0 |
| 0 | NA | NA | NA | NA | NA | Gm31774 | | 0 | 0 | 0 | 0 | 0 | 0 | 0 | 0 | 0 | 0 | 0 |
| 0 | NA | NA | NA | NA | NA | Gm25200 | | 0 | 0 | 0 | 0 | 0 | 0 | 0 | 0 | 0 | 0 | 0 |
| 0 | NA | NA | NA | NA | NA | Gm45722 | | 0 | 0 | 0 | 0 | 0 | 0 | 0 | 0 | 0 | 0 | 0 |
| 0 | NA | NA | NA | NA | NA | Gm24035 | | 0 | 0 | 0 | 0 | 0 | 0 | 0 | 0 | 0 | 0 | 0 |
| 0 | NA | NA | NA | NA | NA | Gm45721 | | 0 | 0 | 0 | 0 | 0 | 0 | 0 | 0 | 0 | 0 | 0 |
| 0 | NA | NA | NA | NA | NA | Gm45720 | | 0 | 0 | 0 | 0 | 0 | 0 | 0 | 0 | 0 | 0 | 0 |
| 0 | NA | NA | NA | NA | NA | Gm22999 | | 0 | 0 | 0 | 0 | 0 | 0 | 0 | 0 | 0 | 0 | 0 |
| 0 | NA | NA | NA | NA | NA | Gm45723 | | 0 | 0 | 0 | 0 | 0 | 0 | 0 | 0 | 0 | 0 | 0 |
| 0 | NA | NA | NA | NA | NA | Adad2 | adenosine deaminase domain containing | 0 | 0 | 0 | 0 | 0 | 0 | 0 | 0 | 0 | 0 | 0 |
| 0 | NA | NA | NA | NA | NA | Gm45747 | | 0 | 0 | 0 | 0 | 0 | 0 | 0 | 0 | 0 | 0 | 0 |
| 0 | NA | NA | NA | NA | NA | Galnt2l | | 0 | 0 | 0 | 0 | 0 | 0 | 0 | 0 | 0 | 0 | 0 |
| 0 | NA | NA | NA | NA | NA | Gm15684 | | 0 | 0 | 0 | 0 | 0 | 0 | 0 | 0 | 0 | 0 | 0 |
| 0 | NA | NA | NA | NA | NA | Gm22715 | | 0 | 0 | 0 | 0 | 0 | 0 | 0 | 0 | 0 | 0 | 0 |
| 0 | NA | NA | NA | NA | NA | Mir7687 | | 0 | 0 | 0 | 0 | 0 | 0 | 0 | 0 | 0 | 0 | 0 |
| 0 | NA | NA | NA | NA | NA | Gm45834 | | 0 | 0 | 0 | 0 | 0 | 0 | 0 | 0 | 0 | 0 | 0 |
| 0 | NA | NA | NA | NA | NA | Gm27021 | predicted gene, 27021 | 0 | 0 | 0 | 0 | 0 | 0 | 0 | 0 | 0 | 0 | 0 |
| 0 | NA | NA | NA | NA | NA | Gm26537 | | 0 | 0 | 0 | 0 | 0 | 0 | 0 | 0 | 0 | 0 | 0 |
| 0 | NA | NA | NA | NA | NA | Gm33142 | | 0 | 0 | 0 | 0 | 0 | 0 | 0 | 0 | 0 | 0 | 0 |
| 0 | NA | NA | NA | NA | NA | Gm24239 | | 0 | 0 | 0 | 0 | 0 | 0 | 0 | 0 | 0 | 0 | 0 |
| 0 | NA | NA | NA | NA | NA | Gm26878 | | 0 | 0 | 0 | 0 | 0 | 0 | 0 | 0 | 0 | 0 | 0 |
| 0 | NA | NA | NA | NA | NA | Gm28324 | | 0 | 0 | 0 | 0 | 0 | 0 | 0 | 0 | 0 | 0 | 0 |
| 0 | NA | NA | NA | NA | NA | Gm37609 | | 0 | 0 | 0 | 0 | 0 | 0 | 0 | 0 | 0 | 0 | 0 |
| 0 | NA | NA | NA | NA | NA | 5033426O07Rik | | 0 | 0 | 0 | 0 | 0 | 0 | 0 | 0 | 0 | 0 | 0 |
| 0 | NA | NA | NA | NA | NA | Gm26784 | | 0 | 0 | 0 | 0 | 0 | 0 | 0 | 0 | 0 | 0 | 0 |
| 0 | NA | NA | NA | NA | NA | 1700018B08Ril | RIKEN cDNA 1700018B08 gene | 0 | 0 | 0 | 0 | 0 | 0 | 0 | 0 | 0 | 0 | 0 |
| 0 | NA | NA | NA | NA | NA | Gm23299 | | 0 | 0 | 0 | 0 | 0 | 0 | 0 | 0 | 0 | 0 | 0 |
| 0 | NA | NA | NA | NA | NA | Gm45833 | | 0 | 0 | 0 | 0 | 0 | 0 | 0 | 0 | 0 | 0 | 0 |
| 0 | NA | NA | NA | NA | NA | Gm20406 | | 0 | 0 | 0 | 0 | 0 | 0 | 0 | 0 | 0 | 0 | 0 |
| 0 | NA | NA | NA | NA | NA | Gm24283 | | 0 | 0 | 0 | 0 | 0 | 0 | 0 | 0 | 0 | 0 | 0 |
| 0 | NA | NA | NA | NA | NA | 9530085P06Rik | | 0 | 0 | 0 | 0 | 0 | 0 | 0 | 0 | 0 | 0 | 0 |
| 0 | NA | NA | NA | NA | NA | Car5a | carbonic anhydrase 5a, mitochondrial | 0 | 0 | 0 | 0 | 0 | 0 | 0 | 0 | 0 | 0 | 0 |
| 0 | NA | NA | NA | NA | NA | Trhr2 | thyrotropin releasing hormone receptor 2 | 0 | 0 | 0 | 0 | 0 | 0 | 0 | 0 | 0 | 0 | 0 |
| 0 | NA | NA | NA | NA | NA | Il17c | interleukin 17C | 0 | 0 | 0 | 0 | 0 | 0 | 0 | 0 | 0 | 0 | 0 |
| 0 | NA | NA | NA | NA | NA | Gm26497 | | 0 | 0 | 0 | 0 | 0 | 0 | 0 | 0 | 0 | 0 | 0 |
| 0 | NA | NA | NA | NA | NA | Mir6396 | | 0 | 0 | 0 | 0 | 0 | 0 | 0 | 0 | 0 | 0 | 0 |
| 0 | NA | NA | NA | NA | NA | Gm10612 | | 0 | 0 | 0 | 0 | 0 | 0 | 0 | 0 | 0 | 0 | 0 |
| 0 | NA | NA | NA | NA | NA | Gm24445 | | 0 | 0 | 0 | 0 | 0 | 0 | 0 | 0 | 0 | 0 | 0 |

| | | | | | | | | | | | | | | | | | | |
|---|---|---|---|---|---|---|---|---|---|---|---|---|---|---|---|---|---|---|
| 0 | NA | NA | NA | NA | NA | Snord68 | | 0 | 0 | 0 | 0 | 0 | 0 | 0 | 0 | 0 | 0 | 0 |
| 0 | NA | NA | NA | NA | NA | Mir7079 | | 0 | 0 | 0 | 0 | 0 | 0 | 0 | 0 | 0 | 0 | 0 |
| 0 | NA | NA | NA | NA | NA | Mir7080 | | 0 | 0 | 0 | 0 | 0 | 0 | 0 | 0 | 0 | 0 | 0 |
| 0 | NA | NA | NA | NA | NA | Gm45842 | | 0 | 0 | 0 | 0 | 0 | 0 | 0 | 0 | 0 | 0 | 0 |
| 0 | NA | NA | NA | NA | NA | Mc1r | melanocortin 1 receptor | 0 | 0 | 0 | 0 | 0 | 0 | 0 | 0 | 0 | 0 | 0 |
| 0 | NA | NA | NA | NA | NA | Gm38092 | | 0 | 0 | 0 | 0 | 0 | 0 | 0 | 0 | 0 | 0 | 0 |
| 0 | NA | NA | NA | NA | NA | Gm25428 | | 0 | 0 | 0 | 0 | 0 | 0 | 0 | 0 | 0 | 0 | 0 |
| 0 | NA | NA | NA | NA | NA | n-R5s100 | | 0 | 0 | 0 | 0 | 0 | 0 | 0 | 0 | 0 | 0 | 0 |
| 0 | NA | NA | NA | NA | NA | n-R5s101 | | 0 | 0 | 0 | 0 | 0 | 0 | 0 | 0 | 0 | 0 | 0 |
| 0 | NA | NA | NA | NA | NA | n-R5s102 | | 0 | 0 | 0 | 0 | 0 | 0 | 0 | 0 | 0 | 0 | 0 |
| 0 | NA | NA | NA | NA | NA | n-R5s103 | | 0 | 0 | 0 | 0 | 0 | 0 | 0 | 0 | 0 | 0 | 0 |
| 0 | NA | NA | NA | NA | NA | n-R5s104 | | 0 | 0 | 0 | 0 | 0 | 0 | 0 | 0 | 0 | 0 | 0 |
| 0 | NA | NA | NA | NA | NA | n-R5s105 | | 0 | 0 | 0 | 0 | 0 | 0 | 0 | 0 | 0 | 0 | 0 |
| 0 | NA | NA | NA | NA | NA | n-R5s106 | | 0 | 0 | 0 | 0 | 0 | 0 | 0 | 0 | 0 | 0 | 0 |
| 0 | NA | NA | NA | NA | NA | n-R5s107 | | 0 | 0 | 0 | 0 | 0 | 0 | 0 | 0 | 0 | 0 | 0 |
| 0 | NA | NA | NA | NA | NA | n-R5s108 | | 0 | 0 | 0 | 0 | 0 | 0 | 0 | 0 | 0 | 0 | 0 |
| 0 | NA | NA | NA | NA | NA | n-R5s109 | | 0 | 0 | 0 | 0 | 0 | 0 | 0 | 0 | 0 | 0 | 0 |
| 0 | NA | NA | NA | NA | NA | n-R5s110 | | 0 | 0 | 0 | 0 | 0 | 0 | 0 | 0 | 0 | 0 | 0 |
| 0 | NA | NA | NA | NA | NA | n-R5s111 | | 0 | 0 | 0 | 0 | 0 | 0 | 0 | 0 | 0 | 0 | 0 |
| 0 | NA | NA | NA | NA | NA | n-R5s112 | | 0 | 0 | 0 | 0 | 0 | 0 | 0 | 0 | 0 | 0 | 0 |
| 0 | NA | NA | NA | NA | NA | n-R5s113 | | 0 | 0 | 0 | 0 | 0 | 0 | 0 | 0 | 0 | 0 | 0 |
| 0 | NA | NA | NA | NA | NA | n-R5s114 | | 0 | 0 | 0 | 0 | 0 | 0 | 0 | 0 | 0 | 0 | 0 |
| 0 | NA | NA | NA | NA | NA | n-R5s115 | | 0 | 0 | 0 | 0 | 0 | 0 | 0 | 0 | 0 | 0 | 0 |
| 0 | NA | NA | NA | NA | NA | Gm25018 | | 0 | 0 | 0 | 0 | 0 | 0 | 0 | 0 | 0 | 0 | 0 |
| 0 | NA | NA | NA | NA | NA | n-R5s117 | | 0 | 0 | 0 | 0 | 0 | 0 | 0 | 0 | 0 | 0 | 0 |
| 0 | NA | NA | NA | NA | NA | n-R5s118 | | 0 | 0 | 0 | 0 | 0 | 0 | 0 | 0 | 0 | 0 | 0 |
| 0 | NA | NA | NA | NA | NA | n-R5s119 | | 0 | 0 | 0 | 0 | 0 | 0 | 0 | 0 | 0 | 0 | 0 |
| 0 | NA | NA | NA | NA | NA | n-R5s120 | | 0 | 0 | 0 | 0 | 0 | 0 | 0 | 0 | 0 | 0 | 0 |
| 0 | NA | NA | NA | NA | NA | n-R5s121 | | 0 | 0 | 0 | 0 | 0 | 0 | 0 | 0 | 0 | 0 | 0 |
| 0 | NA | NA | NA | NA | NA | n-R5s122 | | 0 | 0 | 0 | 0 | 0 | 0 | 0 | 0 | 0 | 0 | 0 |
| 0 | NA | NA | NA | NA | NA | n-R5s123 | | 0 | 0 | 0 | 0 | 0 | 0 | 0 | 0 | 0 | 0 | 0 |
| 0 | NA | NA | NA | NA | NA | n-R5s124 | | 0 | 0 | 0 | 0 | 0 | 0 | 0 | 0 | 0 | 0 | 0 |
| 0 | NA | NA | NA | NA | NA | n-R5s125 | | 0 | 0 | 0 | 0 | 0 | 0 | 0 | 0 | 0 | 0 | 0 |
| 0 | NA | NA | NA | NA | NA | Gm24312 | | 0 | 0 | 0 | 0 | 0 | 0 | 0 | 0 | 0 | 0 | 0 |
| 0 | NA | NA | NA | NA | NA | n-R5s127 | | 0 | 0 | 0 | 0 | 0 | 0 | 0 | 0 | 0 | 0 | 0 |
| 0 | NA | NA | NA | NA | NA | n-R5s128 | | 0 | 0 | 0 | 0 | 0 | 0 | 0 | 0 | 0 | 0 | 0 |
| 0 | NA | NA | NA | NA | NA | n-R5s129 | | 0 | 0 | 0 | 0 | 0 | 0 | 0 | 0 | 0 | 0 | 0 |
| 0 | NA | NA | NA | NA | NA | n-R5s130 | | 0 | 0 | 0 | 0 | 0 | 0 | 0 | 0 | 0 | 0 | 0 |
| 0 | NA | NA | NA | NA | NA | n-R5s131 | | 0 | 0 | 0 | 0 | 0 | 0 | 0 | 0 | 0 | 0 | 0 |
| 0 | NA | NA | NA | NA | NA | Gm26391 | | 0 | 0 | 0 | 0 | 0 | 0 | 0 | 0 | 0 | 0 | 0 |
| 0 | NA | NA | NA | NA | NA | n-R5s133 | | 0 | 0 | 0 | 0 | 0 | 0 | 0 | 0 | 0 | 0 | 0 |
| 0 | NA | NA | NA | NA | NA | n-R5s134 | | 0 | 0 | 0 | 0 | 0 | 0 | 0 | 0 | 0 | 0 | 0 |
| 0 | NA | NA | NA | NA | NA | Rn5s | | 0 | 0 | 0 | 0 | 0 | 0 | 0 | 0 | 0 | 0 | 0 |
| 0 | NA | NA | NA | NA | NA | n-R5s136 | | 0 | 0 | 0 | 0 | 0 | 0 | 0 | 0 | 0 | 0 | 0 |
| 0 | NA | NA | NA | NA | NA | Gm24089 | | 0 | 0 | 0 | 0 | 0 | 0 | 0 | 0 | 0 | 0 | 0 |
| 0 | NA | NA | NA | NA | NA | n-R5s138 | | 0 | 0 | 0 | 0 | 0 | 0 | 0 | 0 | 0 | 0 | 0 |
| 0 | NA | NA | NA | NA | NA | n-R5s139 | | 0 | 0 | 0 | 0 | 0 | 0 | 0 | 0 | 0 | 0 | 0 |
| 0 | NA | NA | NA | NA | NA | Gm22109 | | 0 | 0 | 0 | 0 | 0 | 0 | 0 | 0 | 0 | 0 | 0 |
| 0 | NA | NA | NA | NA | NA | n-R5s141 | | 0 | 0 | 0 | 0 | 0 | 0 | 0 | 0 | 0 | 0 | 0 |
| 0 | NA | NA | NA | NA | NA | n-R5s142 | | 0 | 0 | 0 | 0 | 0 | 0 | 0 | 0 | 0 | 0 | 0 |
| 0 | NA | NA | NA | NA | NA | n-R5s143 | | 0 | 0 | 0 | 0 | 0 | 0 | 0 | 0 | 0 | 0 | 0 |
| 0 | NA | NA | NA | NA | NA | n-R5s144 | | 0 | 0 | 0 | 0 | 0 | 0 | 0 | 0 | 0 | 0 | 0 |
| 0 | NA | NA | NA | NA | NA | Gm25212 | | 0 | 0 | 0 | 0 | 0 | 0 | 0 | 0 | 0 | 0 | 0 |
| 0 | NA | NA | NA | NA | NA | n-R5s146 | | 0 | 0 | 0 | 0 | 0 | 0 | 0 | 0 | 0 | 0 | 0 |
| 0 | NA | NA | NA | NA | NA | Gm22291 | | 0 | 0 | 0 | 0 | 0 | 0 | 0 | 0 | 0 | 0 | 0 |
| 0 | NA | NA | NA | NA | NA | Gm23284 | | 0 | 0 | 0 | 0 | 0 | 0 | 0 | 0 | 0 | 0 | 0 |
| 0 | NA | NA | NA | NA | NA | n-R5s149 | | 0 | 0 | 0 | 0 | 0 | 0 | 0 | 0 | 0 | 0 | 0 |
| 0 | NA | NA | NA | NA | NA | Gm23647 | | 0 | 0 | 0 | 0 | 0 | 0 | 0 | 0 | 0 | 0 | 0 |
| 0 | NA | NA | NA | NA | NA | Gm45781 | | 0 | 0 | 0 | 0 | 0 | 0 | 0 | 0 | 0 | 0 | 0 |
| 0 | NA | NA | NA | NA | NA | Mir1967 | | 0 | 0 | 0 | 0 | 0 | 0 | 0 | 0 | 0 | 0 | 0 |

| | | | | | | | | | | | | | | | | | |
|---|---|---|---|---|---|---|---|---|---|---|---|---|---|---|---|---|---|
| 0 | NA | NA | NA | NA | NA | Gm45732 | | 0 | 0 | 0 | 0 | 0 | 0 | 0 | 0 | 0 | 0 | 0 |
| 0 | NA | NA | NA | NA | NA | Gm3889 | | 0 | 0 | 0 | 0 | 0 | 0 | 0 | 0 | 0 | 0 | 0 |
| 0 | NA | NA | NA | NA | NA | Gm15775 | | 0 | 0 | 0 | 0 | 0 | 0 | 0 | 0 | 0 | 0 | 0 |
| 0 | NA | NA | NA | NA | NA | Gm24459 | | 0 | 0 | 0 | 0 | 0 | 0 | 0 | 0 | 0 | 0 | 0 |
| 0 | NA | NA | NA | NA | NA | Gm9172 | | 0 | 0 | 0 | 0 | 0 | 0 | 0 | 0 | 0 | 0 | 0 |
| 0 | NA | NA | NA | NA | NA | Gm45856 | | 0 | 0 | 0 | 0 | 0 | 0 | 0 | 0 | 0 | 0 | 0 |
| 0 | NA | NA | NA | NA | NA | Gm45874 | | 0 | 0 | 0 | 0 | 0 | 0 | 0 | 0 | 0 | 0 | 0 |
| 0 | NA | NA | NA | NA | NA | Gm22197 | | 0 | 0 | 0 | 0 | 0 | 0 | 0 | 0 | 0 | 0 | 0 |
| 0 | NA | NA | NA | NA | NA | Gm16237 | | 0 | 0 | 0 | 0 | 0 | 0 | 0 | 0 | 0 | 0 | 0 |
| 0 | NA | NA | NA | NA | NA | A730098A19Rik | | 0 | 0 | 0 | 0 | 0 | 0 | 0 | 0 | 0 | 0 | 0 |
| 0 | NA | NA | NA | NA | NA | Gm45875 | | 0 | 0 | 0 | 0 | 0 | 0 | 0 | 0 | 0 | 0 | 0 |
| 0 | NA | NA | NA | NA | NA | Gm45876 | | 0 | 0 | 0 | 0 | 0 | 0 | 0 | 0 | 0 | 0 | 0 |
| 0 | NA | NA | NA | NA | NA | Gm6091 | | 0 | 0 | 0 | 0 | 0 | 0 | 0 | 0 | 0 | 0 | 0 |
| 0 | NA | NA | NA | NA | NA | Gm5358 | | 0 | 0 | 0 | 0 | 0 | 0 | 0 | 0 | 0 | 0 | 0 |
| 0 | NA | NA | NA | NA | NA | Gm22730 | | 0 | 0 | 0 | 0 | 0 | 0 | 0 | 0 | 0 | 0 | 0 |
| 0 | NA | NA | NA | NA | NA | Gm22968 | | 0 | 0 | 0 | 0 | 0 | 0 | 0 | 0 | 0 | 0 | 0 |
| 0 | NA | NA | NA | NA | NA | Mir21c | | 0 | 0 | 0 | 0 | 0 | 0 | 0 | 0 | 0 | 0 | 0 |
| 0 | NA | NA | NA | NA | NA | Mir1903 | | 0 | 0 | 0 | 0 | 0 | 0 | 0 | 0 | 0 | 0 | 0 |
| 0 | NA | NA | NA | NA | NA | Gm26002 | | 0 | 0 | 0 | 0 | 0 | 0 | 0 | 0 | 0 | 0 | 0 |
| 0 | NA | NA | NA | NA | NA | Gm45746 | | 0 | 0 | 0 | 0 | 0 | 0 | 0 | 0 | 0 | 0 | 0 |
| 0 | NA | NA | NA | NA | NA | Ccdc7a | coiled-coil domain containing 7A | 0 | 0 | 0 | 0 | 0 | 0 | 0 | 0 | 0 | 0 | 0 |
| 0 | NA | NA | NA | NA | NA | Gm3952 | | 0 | 0 | 0 | 0 | 0 | 0 | 0 | 0 | 0 | 0 | 0 |
| 0 | NA | NA | NA | NA | NA | Gm26100 | | 0 | 0 | 0 | 0 | 0 | 0 | 0 | 0 | 0 | 0 | 0 |
| 0 | NA | NA | NA | NA | NA | Gm22545 | | 0 | 0 | 0 | 0 | 0 | 0 | 0 | 0 | 0 | 0 | 0 |
| 0 | NA | NA | NA | NA | NA | Ccdc7b | coiled-coil domain containing 7B | 0 | 0 | 0 | 0 | 0 | 0 | 0 | 0 | 0 | 0 | 0 |
| 0 | NA | NA | NA | NA | NA | Gm22374 | | 0 | 0 | 0 | 0 | 0 | 0 | 0 | 0 | 0 | 0 | 0 |
| 0 | NA | NA | NA | NA | NA | Gm10999 | | 0 | 0 | 0 | 0 | 0 | 0 | 0 | 0 | 0 | 0 | 0 |
| 0 | NA | NA | NA | NA | NA | Gm10722 | | 0 | 0 | 0 | 0 | 0 | 0 | 0 | 0 | 0 | 0 | 0 |
| 0 | NA | NA | NA | NA | NA | Gm11168 | | 0 | 0 | 0 | 0 | 0 | 0 | 0 | 0 | 0 | 0 | 0 |
| 0 | NA | NA | NA | NA | NA | Gm10721 | | 0 | 0 | 0 | 0 | 0 | 0 | 0 | 0 | 0 | 0 | 0 |
| 0 | NA | NA | NA | NA | NA | Gm10720 | | 0 | 0 | 0 | 0 | 0 | 0 | 0 | 0 | 0 | 0 | 0 |
| 0 | NA | NA | NA | NA | NA | Gm10719 | | 0 | 0 | 0 | 0 | 0 | 0 | 0 | 0 | 0 | 0 | 0 |
| 0 | NA | NA | NA | NA | NA | Gm10718 | | 0 | 0 | 0 | 0 | 0 | 0 | 0 | 0 | 0 | 0 | 0 |
| 0 | NA | NA | NA | NA | NA | Gm17535 | | 0 | 0 | 0 | 0 | 0 | 0 | 0 | 0 | 0 | 0 | 0 |
| 0 | NA | NA | NA | NA | NA | Gm10715 | | 0 | 0 | 0 | 0 | 0 | 0 | 0 | 0 | 0 | 0 | 0 |
| 0 | NA | NA | NA | NA | NA | Mir101c | | 0 | 0 | 0 | 0 | 0 | 0 | 0 | 0 | 0 | 0 | 0 |
| 0 | NA | NA | NA | NA | NA | Gm32390 | | 0 | 0 | 0 | 0 | 0 | 0 | 0 | 0 | 0 | 0 | 0 |
| 0 | NA | NA | NA | NA | NA | Gm7344 | | 0 | 0 | 0 | 0 | 0 | 0 | 0 | 0 | 0 | 0 | 0 |
| 0 | NA | NA | NA | NA | NA | Olfr369-ps1 | | 0 | 0 | 0 | 0 | 0 | 0 | 0 | 0 | 0 | 0 | 0 |
| 0 | NA | NA | NA | NA | NA | Gm27679 | | 0 | 0 | 0 | 0 | 0 | 0 | 0 | 0 | 0 | 0 | 0 |
| 0 | NA | NA | NA | NA | NA | Gm24918 | | 0 | 0 | 0 | 0 | 0 | 0 | 0 | 0 | 0 | 0 | 0 |
| 0 | NA | NA | NA | NA | NA | Gm23275 | | 0 | 0 | 0 | 0 | 0 | 0 | 0 | 0 | 0 | 0 | 0 |
| 0 | NA | NA | NA | NA | NA | Gm5363 | | 0 | 0 | 0 | 0 | 0 | 0 | 0 | 0 | 0 | 0 | 0 |
| 0 | NA | NA | NA | NA | NA | Gm22195 | | 0 | 0 | 0 | 0 | 0 | 0 | 0 | 0 | 0 | 0 | 0 |
| 0 | NA | NA | NA | NA | NA | Gm23992 | | 0 | 0 | 0 | 0 | 0 | 0 | 0 | 0 | 0 | 0 | 0 |
| 0 | NA | NA | NA | NA | NA | Gm24896 | | 0 | 0 | 0 | 0 | 0 | 0 | 0 | 0 | 0 | 0 | 0 |
| 0 | NA | NA | NA | NA | NA | Gm48506 | | 0 | 0 | 0 | 0 | 0 | 0 | 0 | 0 | 0 | 0 | 0 |
| 0 | NA | NA | NA | NA | NA | Gm9662 | | 0 | 0 | 0 | 0 | 0 | 0 | 0 | 0 | 0 | 0 | 0 |
| 0 | NA | NA | NA | NA | NA | Ddi1 | DNA-damage inducible 1 | 0 | 0 | 0 | 0 | 0 | 0 | 0 | 0 | 0 | 0 | 0 |
| 0 | NA | NA | NA | NA | NA | Gm48674 | | 0 | 0 | 0 | 0 | 0 | 0 | 0 | 0 | 0 | 0 | 0 |
| 0 | NA | NA | NA | NA | NA | Gm23097 | | 0 | 0 | 0 | 0 | 0 | 0 | 0 | 0 | 0 | 0 | 0 |
| 0 | NA | NA | NA | NA | NA | Gm48675 | | 0 | 0 | 0 | 0 | 0 | 0 | 0 | 0 | 0 | 0 | 0 |
| 0 | NA | NA | NA | NA | NA | Gm47741 | | 0 | 0 | 0 | 0 | 0 | 0 | 0 | 0 | 0 | 0 | 0 |
| 0 | NA | NA | NA | NA | NA | Mmp1b | matrix metallopeptidase 1b (interstitial col | 0 | 0 | 0 | 0 | 0 | 0 | 0 | 0 | 0 | 0 | 0 |
| 0 | NA | NA | NA | NA | NA | Mmp1a | matrix metallopeptidase 1a (interstitial col | 0 | 0 | 0 | 0 | 0 | 0 | 0 | 0 | 0 | 0 | 0 |
| 0 | NA | NA | NA | NA | NA | Mmp10 | matrix metallopeptidase 10 | 0 | 0 | 0 | 0 | 0 | 0 | 0 | 0 | 0 | 0 | 0 |
| 0 | NA | NA | NA | NA | NA | Gm47616 | | 0 | 0 | 0 | 0 | 0 | 0 | 0 | 0 | 0 | 0 | 0 |
| 0 | NA | NA | NA | NA | NA | Mmp20 | matrix metallopeptidase 20 (enamelysin) | 0 | 0 | 0 | 0 | 0 | 0 | 0 | 0 | 0 | 0 | 0 |
| 0 | NA | NA | NA | NA | NA | AV064505 | | 0 | 0 | 0 | 0 | 0 | 0 | 0 | 0 | 0 | 0 | 0 |
| 0 | NA | NA | NA | NA | NA | Gm19212 | | 0 | 0 | 0 | 0 | 0 | 0 | 0 | 0 | 0 | 0 | 0 |

| | | | | | | | | | | | | | | | | |
|---|---|---|---|---|---|---|---|---|---|---|---|---|---|---|---|---|
| 0 | NA | NA | NA | NA | NA | Cbx3-ps8 | 0 | 0 | 0 | 0 | 0 | 0 | 0 | 0 | 0 | 0 | 0 |
| 0 | NA | NA | NA | NA | NA | Mir5618 | 0 | 0 | 0 | 0 | 0 | 0 | 0 | 0 | 0 | 0 | 0 |
| 0 | NA | NA | NA | NA | NA | Gm20416 | 0 | 0 | 0 | 0 | 0 | 0 | 0 | 0 | 0 | 0 | 0 |
| 0 | NA | NA | NA | NA | NA | 1700128F08Rik | 0 | 0 | 0 | 0 | 0 | 0 | 0 | 0 | 0 | 0 | 0 |
| 0 | NA | NA | NA | NA | NA | Mir1899 | 0 | 0 | 0 | 0 | 0 | 0 | 0 | 0 | 0 | 0 | 0 |
| 0 | NA | NA | NA | NA | NA | Gm47517 | 0 | 0 | 0 | 0 | 0 | 0 | 0 | 0 | 0 | 0 | 0 |
| 0 | NA | NA | NA | NA | NA | Gm32014 | 0 | 0 | 0 | 0 | 0 | 0 | 0 | 0 | 0 | 0 | 0 |
| 0 | NA | NA | NA | NA | NA | Gm16833 | 0 | 0 | 0 | 0 | 0 | 0 | 0 | 0 | 0 | 0 | 0 |
| 0 | NA | NA | NA | NA | NA | Gm22815 | 0 | 0 | 0 | 0 | 0 | 0 | 0 | 0 | 0 | 0 | 0 |
| 0 | NA | NA | NA | NA | NA | Gm46102 | 0 | 0 | 0 | 0 | 0 | 0 | 0 | 0 | 0 | 0 | 0 |
| 0 | NA | NA | NA | NA | NA | Gm47333 | 0 | 0 | 0 | 0 | 0 | 0 | 0 | 0 | 0 | 0 | 0 |
| 0 | NA | NA | NA | NA | NA | Gm47334 | 0 | 0 | 0 | 0 | 0 | 0 | 0 | 0 | 0 | 0 | 0 |
| 0 | NA | NA | NA | NA | NA | Gm26195 | 0 | 0 | 0 | 0 | 0 | 0 | 0 | 0 | 0 | 0 | 0 |
| 0 | NA | NA | NA | NA | NA | Mir6237 | 0 | 0 | 0 | 0 | 0 | 0 | 0 | 0 | 0 | 0 | 0 |
| 0 | NA | NA | NA | NA | NA | Gm24496 | 0 | 0 | 0 | 0 | 0 | 0 | 0 | 0 | 0 | 0 | 0 |
| 0 | NA | NA | NA | NA | NA | Gm25086 | 0 | 0 | 0 | 0 | 0 | 0 | 0 | 0 | 0 | 0 | 0 |
| 0 | NA | NA | NA | NA | NA | Gm46103 | 0 | 0 | 0 | 0 | 0 | 0 | 0 | 0 | 0 | 0 | 0 |
| 0 | NA | NA | NA | NA | NA | Gm7529 | 0 | 0 | 0 | 0 | 0 | 0 | 0 | 0 | 0 | 0 | 0 |
| 0 | NA | NA | NA | NA | NA | Gm32710 | 0 | 0 | 0 | 0 | 0 | 0 | 0 | 0 | 0 | 0 | 0 |
| 0 | NA | NA | NA | NA | NA | Gm18488 | 0 | 0 | 0 | 0 | 0 | 0 | 0 | 0 | 0 | 0 | 0 |
| 0 | NA | NA | NA | NA | NA | Gm47001 | 0 | 0 | 0 | 0 | 0 | 0 | 0 | 0 | 0 | 0 | 0 |
| 0 | NA | NA | NA | NA | NA | Gm47002 | 0 | 0 | 0 | 0 | 0 | 0 | 0 | 0 | 0 | 0 | 0 |
| 0 | NA | NA | NA | NA | NA | Gm47029 | 0 | 0 | 0 | 0 | 0 | 0 | 0 | 0 | 0 | 0 | 0 |
| 0 | NA | NA | NA | NA | NA | Gm18934 | 0 | 0 | 0 | 0 | 0 | 0 | 0 | 0 | 0 | 0 | 0 |
| 0 | NA | NA | NA | NA | NA | Gm47034 | 0 | 0 | 0 | 0 | 0 | 0 | 0 | 0 | 0 | 0 | 0 |
| 0 | NA | NA | NA | NA | NA | Gm18160 | 0 | 0 | 0 | 0 | 0 | 0 | 0 | 0 | 0 | 0 | 0 |
| 0 | NA | NA | NA | NA | NA | Gm47035 | 0 | 0 | 0 | 0 | 0 | 0 | 0 | 0 | 0 | 0 | 0 |
| 0 | NA | NA | NA | NA | NA | Gm47037 | 0 | 0 | 0 | 0 | 0 | 0 | 0 | 0 | 0 | 0 | 0 |
| 0 | NA | NA | NA | NA | NA | Gm29785 | 0 | 0 | 0 | 0 | 0 | 0 | 0 | 0 | 0 | 0 | 0 |
| 0 | NA | NA | NA | NA | NA | Gm25365 | 0 | 0 | 0 | 0 | 0 | 0 | 0 | 0 | 0 | 0 | 0 |
| 0 | NA | NA | NA | NA | NA | Gm7570 | 0 | 0 | 0 | 0 | 0 | 0 | 0 | 0 | 0 | 0 | 0 |
| 0 | NA | NA | NA | NA | NA | Gm21070 | 0 | 0 | 0 | 0 | 0 | 0 | 0 | 0 | 0 | 0 | 0 |
| 0 | NA | NA | NA | NA | NA | 4930568E12Rik | 0 | 0 | 0 | 0 | 0 | 0 | 0 | 0 | 0 | 0 | 0 |
| 0 | NA | NA | NA | NA | NA | Gm47051 | 0 | 0 | 0 | 0 | 0 | 0 | 0 | 0 | 0 | 0 | 0 |
| 0 | NA | NA | NA | NA | NA | Phxr4 | 0 | 0 | 0 | 0 | 0 | 0 | 0 | 0 | 0 | 0 | 0 |
| 0 | NA | NA | NA | NA | NA | Gm17571 | 0 | 0 | 0 | 0 | 0 | 0 | 0 | 0 | 0 | 0 | 0 |
| 0 | NA | NA | NA | NA | NA | Gm47089 | 0 | 0 | 0 | 0 | 0 | 0 | 0 | 0 | 0 | 0 | 0 |
| 0 | NA | NA | NA | NA | NA | Gm22643 | 0 | 0 | 0 | 0 | 0 | 0 | 0 | 0 | 0 | 0 | 0 |
| 0 | NA | NA | NA | NA | NA | Gm47105 | 0 | 0 | 0 | 0 | 0 | 0 | 0 | 0 | 0 | 0 | 0 |
| 0 | NA | NA | NA | NA | NA | Gm47535 | 0 | 0 | 0 | 0 | 0 | 0 | 0 | 0 | 0 | 0 | 0 |
| 0 | NA | NA | NA | NA | NA | Gm47557 | 0 | 0 | 0 | 0 | 0 | 0 | 0 | 0 | 0 | 0 | 0 |
| 0 | NA | NA | NA | NA | NA | 1700019J19Rik | 0 | 0 | 0 | 0 | 0 | 0 | 0 | 0 | 0 | 0 | 0 |
| 0 | NA | NA | NA | NA | NA | Gm47559 | 0 | 0 | 0 | 0 | 0 | 0 | 0 | 0 | 0 | 0 | 0 |
| 0 | NA | NA | NA | NA | NA | Gm47564 | 0 | 0 | 0 | 0 | 0 | 0 | 0 | 0 | 0 | 0 | 0 |
| 0 | NA | NA | NA | NA | NA | Gm47565 | 0 | 0 | 0 | 0 | 0 | 0 | 0 | 0 | 0 | 0 | 0 |
| 0 | NA | NA | NA | NA | NA | Gm47326 | 0 | 0 | 0 | 0 | 0 | 0 | 0 | 0 | 0 | 0 | 0 |
| 0 | NA | NA | NA | NA | NA | Gm47321 | 0 | 0 | 0 | 0 | 0 | 0 | 0 | 0 | 0 | 0 | 0 |
| 0 | NA | NA | NA | NA | NA | Gm47319 | 0 | 0 | 0 | 0 | 0 | 0 | 0 | 0 | 0 | 0 | 0 |
| 0 | NA | NA | NA | NA | NA | Gm47318 | 0 | 0 | 0 | 0 | 0 | 0 | 0 | 0 | 0 | 0 | 0 |
| 0 | NA | NA | NA | NA | NA | Gm47317 | 0 | 0 | 0 | 0 | 0 | 0 | 0 | 0 | 0 | 0 | 0 |
| 0 | NA | NA | NA | NA | NA | Gm47090 | 0 | 0 | 0 | 0 | 0 | 0 | 0 | 0 | 0 | 0 | 0 |
| 0 | NA | NA | NA | NA | NA | 4930584E12Rik | 0 | 0 | 0 | 0 | 0 | 0 | 0 | 0 | 0 | 0 | 0 |
| 0 | NA | NA | NA | NA | NA | Gm34263 | 0 | 0 | 0 | 0 | 0 | 0 | 0 | 0 | 0 | 0 | 0 |
| 0 | NA | NA | NA | NA | NA | Gm19790 | 0 | 0 | 0 | 0 | 0 | 0 | 0 | 0 | 0 | 0 | 0 |
| 0 | NA | NA | NA | NA | NA | Gm18997 | 0 | 0 | 0 | 0 | 0 | 0 | 0 | 0 | 0 | 0 | 0 |
| 0 | NA | NA | NA | NA | NA | Gm10706 | 0 | 0 | 0 | 0 | 0 | 0 | 0 | 0 | 0 | 0 | 0 |
| 0 | NA | NA | NA | NA | NA | Gm15586 | 0 | 0 | 0 | 0 | 0 | 0 | 0 | 0 | 0 | 0 | 0 |
| 0 | NA | NA | NA | NA | NA | 1700012B09Ril RIKEN cDNA 1700012B09 gene | 0 | 0 | 0 | 0 | 0 | 0 | 0 | 0 | 0 | 0 | 0 |
| 0 | NA | NA | NA | NA | NA | Gm47154 | 0 | 0 | 0 | 0 | 0 | 0 | 0 | 0 | 0 | 0 | 0 |
| 0 | NA | NA | NA | NA | NA | Gm18789 | 0 | 0 | 0 | 0 | 0 | 0 | 0 | 0 | 0 | 0 | 0 |

| | | | | | | | | | | | | | | | | | | |
|---|---|---|---|---|---|---|---|---|---|---|---|---|---|---|---|---|---|---|
| 0 | NA | NA | NA | NA | NA | Gm47214 | | 0 | 0 | 0 | 0 | 0 | 0 | 0 | 0 | 0 | 0 | 0 |
| 0 | NA | NA | NA | NA | NA | Hephl1 | hephaestin-like 1 | 0 | 0 | 0 | 0 | 0 | 0 | 0 | 0 | 0 | 0 | 0 |
| 0 | NA | NA | NA | NA | NA | Gm22658 | | 0 | 0 | 0 | 0 | 0 | 0 | 0 | 0 | 0 | 0 | 0 |
| 0 | NA | NA | NA | NA | NA | Gm25500 | | 0 | 0 | 0 | 0 | 0 | 0 | 0 | 0 | 0 | 0 | 0 |
| 0 | NA | NA | NA | NA | NA | Mir7650 | | 0 | 0 | 0 | 0 | 0 | 0 | 0 | 0 | 0 | 0 | 0 |
| 0 | NA | NA | NA | NA | NA | Gm24455 | | 0 | 0 | 0 | 0 | 0 | 0 | 0 | 0 | 0 | 0 | 0 |
| 0 | NA | NA | NA | NA | NA | Gm24299 | | 0 | 0 | 0 | 0 | 0 | 0 | 0 | 0 | 0 | 0 | 0 |
| 0 | NA | NA | NA | NA | NA | Gm49349 | | 0 | 0 | 0 | 0 | 0 | 0 | 0 | 0 | 0 | 0 | 0 |
| 0 | NA | NA | NA | NA | NA | Gm25791 | | 0 | 0 | 0 | 0 | 0 | 0 | 0 | 0 | 0 | 0 | 0 |
| 0 | NA | NA | NA | NA | NA | Gm22620 | | 0 | 0 | 0 | 0 | 0 | 0 | 0 | 0 | 0 | 0 | 0 |
| 0 | NA | NA | NA | NA | NA | Gm24357 | | 0 | 0 | 0 | 0 | 0 | 0 | 0 | 0 | 0 | 0 | 0 |
| 0 | NA | NA | NA | NA | NA | Gm23455 | | 0 | 0 | 0 | 0 | 0 | 0 | 0 | 0 | 0 | 0 | 0 |
| 0 | NA | NA | NA | NA | NA | Gm22579 | | 0 | 0 | 0 | 0 | 0 | 0 | 0 | 0 | 0 | 0 | 0 |
| 0 | NA | NA | NA | NA | NA | Gm48142 | | 0 | 0 | 0 | 0 | 0 | 0 | 0 | 0 | 0 | 0 | 0 |
| 0 | NA | NA | NA | NA | NA | Gm48141 | | 0 | 0 | 0 | 0 | 0 | 0 | 0 | 0 | 0 | 0 | 0 |
| 0 | NA | NA | NA | NA | NA | Scarna9 | | 0 | 0 | 0 | 0 | 0 | 0 | 0 | 0 | 0 | 0 | 0 |
| 0 | NA | NA | NA | NA | NA | Gm16568 | | 0 | 0 | 0 | 0 | 0 | 0 | 0 | 0 | 0 | 0 | 0 |
| 0 | NA | NA | NA | NA | NA | Gm34973 | | 0 | 0 | 0 | 0 | 0 | 0 | 0 | 0 | 0 | 0 | 0 |
| 0 | NA | NA | NA | NA | NA | Gm2517 | | 0 | 0 | 0 | 0 | 0 | 0 | 0 | 0 | 0 | 0 | 0 |
| 0 | NA | NA | NA | NA | NA | Gm48634 | | 0 | 0 | 0 | 0 | 0 | 0 | 0 | 0 | 0 | 0 | 0 |
| 0 | NA | NA | NA | NA | NA | 4930540M03Rik | | 0 | 0 | 0 | 0 | 0 | 0 | 0 | 0 | 0 | 0 | 0 |
| 0 | NA | NA | NA | NA | NA | Gm5119 | | 0 | 0 | 0 | 0 | 0 | 0 | 0 | 0 | 0 | 0 | 0 |
| 0 | NA | NA | NA | NA | NA | Gm48711 | | 0 | 0 | 0 | 0 | 0 | 0 | 0 | 0 | 0 | 0 | 0 |
| 0 | NA | NA | NA | NA | NA | Mtnr1b | melatonin receptor 1B | 0 | 0 | 0 | 0 | 0 | 0 | 0 | 0 | 0 | 0 | 0 |
| 0 | NA | NA | NA | NA | NA | Gm48726 | | 0 | 0 | 0 | 0 | 0 | 0 | 0 | 0 | 0 | 0 | 0 |
| 0 | NA | NA | NA | NA | NA | Gm2594 | | 0 | 0 | 0 | 0 | 0 | 0 | 0 | 0 | 0 | 0 | 0 |
| 0 | NA | NA | NA | NA | NA | Gm48846 | | 0 | 0 | 0 | 0 | 0 | 0 | 0 | 0 | 0 | 0 | 0 |
| 0 | NA | NA | NA | NA | NA | Gm48848 | | 0 | 0 | 0 | 0 | 0 | 0 | 0 | 0 | 0 | 0 | 0 |
| 0 | NA | NA | NA | NA | NA | Gm48850 | | 0 | 0 | 0 | 0 | 0 | 0 | 0 | 0 | 0 | 0 | 0 |
| 0 | NA | NA | NA | NA | NA | Gm44498 | | 0 | 0 | 0 | 0 | 0 | 0 | 0 | 0 | 0 | 0 | 0 |
| 0 | NA | NA | NA | NA | NA | Gm7692 | | 0 | 0 | 0 | 0 | 0 | 0 | 0 | 0 | 0 | 0 | 0 |
| 0 | NA | NA | NA | NA | NA | Gm19174 | | 0 | 0 | 0 | 0 | 0 | 0 | 0 | 0 | 0 | 0 | 0 |
| 0 | NA | NA | NA | NA | NA | Ubtfl1 | upstream binding transcription factor, RN/ | 0 | 0 | 0 | 0 | 0 | 0 | 0 | 0 | 0 | 0 | 0 |
| 0 | NA | NA | NA | NA | NA | Mbd3l2 | methyl-CpG binding domain protein 3-like | 0 | 0 | 0 | 0 | 0 | 0 | 0 | 0 | 0 | 0 | 0 |
| 0 | NA | NA | NA | NA | NA | Gm22854 | | 0 | 0 | 0 | 0 | 0 | 0 | 0 | 0 | 0 | 0 | 0 |
| 0 | NA | NA | NA | NA | NA | Mbd3l1 | methyl-CpG binding domain protein 3-like | 0 | 0 | 0 | 0 | 0 | 0 | 0 | 0 | 0 | 0 | 0 |
| 0 | NA | NA | NA | NA | NA | Gm25295 | | 0 | 0 | 0 | 0 | 0 | 0 | 0 | 0 | 0 | 0 | 0 |
| 0 | NA | NA | NA | NA | NA | Gm24546 | | 0 | 0 | 0 | 0 | 0 | 0 | 0 | 0 | 0 | 0 | 0 |
| 0 | NA | NA | NA | NA | NA | Olfr24 | olfactory receptor 24 | 0 | 0 | 0 | 0 | 0 | 0 | 0 | 0 | 0 | 0 | 0 |
| 0 | NA | NA | NA | NA | NA | Olfr828 | olfactory receptor 828 | 0 | 0 | 0 | 0 | 0 | 0 | 0 | 0 | 0 | 0 | 0 |
| 0 | NA | NA | NA | NA | NA | Olfr829 | olfactory receptor 829 | 0 | 0 | 0 | 0 | 0 | 0 | 0 | 0 | 0 | 0 | 0 |
| 0 | NA | NA | NA | NA | NA | Olfr830 | olfactory receptor 830 | 0 | 0 | 0 | 0 | 0 | 0 | 0 | 0 | 0 | 0 | 0 |
| 0 | NA | NA | NA | NA | NA | Olfr831-ps1 | | 0 | 0 | 0 | 0 | 0 | 0 | 0 | 0 | 0 | 0 | 0 |
| 0 | NA | NA | NA | NA | NA | Olfr832 | olfactory receptor 832 | 0 | 0 | 0 | 0 | 0 | 0 | 0 | 0 | 0 | 0 | 0 |
| 0 | NA | NA | NA | NA | NA | Olfr833-ps1 | | 0 | 0 | 0 | 0 | 0 | 0 | 0 | 0 | 0 | 0 | 0 |
| 0 | NA | NA | NA | NA | NA | Olfr834 | olfactory receptor 834 | 0 | 0 | 0 | 0 | 0 | 0 | 0 | 0 | 0 | 0 | 0 |
| 0 | NA | NA | NA | NA | NA | Gm45814 | | 0 | 0 | 0 | 0 | 0 | 0 | 0 | 0 | 0 | 0 | 0 |
| 0 | NA | NA | NA | NA | NA | Olfr835 | olfactory receptor 835 | 0 | 0 | 0 | 0 | 0 | 0 | 0 | 0 | 0 | 0 | 0 |
| 0 | NA | NA | NA | NA | NA | Olfr836 | olfactory receptor 836 | 0 | 0 | 0 | 0 | 0 | 0 | 0 | 0 | 0 | 0 | 0 |
| 0 | NA | NA | NA | NA | NA | Olfr837 | olfactory receptor 837 | 0 | 0 | 0 | 0 | 0 | 0 | 0 | 0 | 0 | 0 | 0 |
| 0 | NA | NA | NA | NA | NA | Olfr838-ps1 | | 0 | 0 | 0 | 0 | 0 | 0 | 0 | 0 | 0 | 0 | 0 |
| 0 | NA | NA | NA | NA | NA | Olfr839-ps1 | | 0 | 0 | 0 | 0 | 0 | 0 | 0 | 0 | 0 | 0 | 0 |
| 0 | NA | NA | NA | NA | NA | Olfr840-ps1 | | 0 | 0 | 0 | 0 | 0 | 0 | 0 | 0 | 0 | 0 | 0 |
| 0 | NA | NA | NA | NA | NA | | | 0 | 0 | 0 | 0 | 0 | 0 | 0 | 0 | 0 | 0 | 0 |
| 0 | NA | NA | NA | NA | NA | Olfr843 | olfactory receptor 843 | 0 | 0 | 0 | 0 | 0 | 0 | 0 | 0 | 0 | 0 | 0 |
| 0 | NA | NA | NA | NA | NA | Olfr844 | | 0 | 0 | 0 | 0 | 0 | 0 | 0 | 0 | 0 | 0 | 0 |
| 0 | NA | NA | NA | NA | NA | Olfr845 | olfactory receptor 845 | 0 | 0 | 0 | 0 | 0 | 0 | 0 | 0 | 0 | 0 | 0 |
| 0 | NA | NA | NA | NA | NA | Olfr846 | olfactory receptor 846 | 0 | 0 | 0 | 0 | 0 | 0 | 0 | 0 | 0 | 0 | 0 |
| 0 | NA | NA | NA | NA | NA | Gm7751 | | 0 | 0 | 0 | 0 | 0 | 0 | 0 | 0 | 0 | 0 | 0 |
| 0 | NA | NA | NA | NA | NA | Olfr847 | olfactory receptor 847 | 0 | 0 | 0 | 0 | 0 | 0 | 0 | 0 | 0 | 0 | 0 |

| | | | | | | | | | | | | | | | | | | |
|---|---|---|---|---|---|---|---|---|---|---|---|---|---|---|---|---|---|---|
| 0 | NA | NA | NA | NA | NA | Olfr848-ps1 | | 0 | 0 | 0 | 0 | 0 | 0 | 0 | 0 | 0 | 0 | 0 |
| 0 | NA | NA | NA | NA | NA | Gm23048 | | 0 | 0 | 0 | 0 | 0 | 0 | 0 | 0 | 0 | 0 | 0 |
| 0 | NA | NA | NA | NA | NA | Olfr849 | olfactory receptor 849 | 0 | 0 | 0 | 0 | 0 | 0 | 0 | 0 | 0 | 0 | 0 |
| 0 | NA | NA | NA | NA | NA | Olfr850 | olfactory receptor 850 | 0 | 0 | 0 | 0 | 0 | 0 | 0 | 0 | 0 | 0 | 0 |
| 0 | NA | NA | NA | NA | NA | Olfr851 | olfactory receptor 851 | 0 | 0 | 0 | 0 | 0 | 0 | 0 | 0 | 0 | 0 | 0 |
| 0 | NA | NA | NA | NA | NA | Olfr852-ps1 | | 0 | 0 | 0 | 0 | 0 | 0 | 0 | 0 | 0 | 0 | 0 |
| 0 | NA | NA | NA | NA | NA | Olfr853 | olfactory receptor 853 | 0 | 0 | 0 | 0 | 0 | 0 | 0 | 0 | 0 | 0 | 0 |
| 0 | NA | NA | NA | NA | NA | Olfr854 | olfactory receptor 854 | 0 | 0 | 0 | 0 | 0 | 0 | 0 | 0 | 0 | 0 | 0 |
| 0 | NA | NA | NA | NA | NA | Olfr855 | olfactory receptor 855 | 0 | 0 | 0 | 0 | 0 | 0 | 0 | 0 | 0 | 0 | 0 |
| 0 | NA | NA | NA | NA | NA | Gm45730 | | 0 | 0 | 0 | 0 | 0 | 0 | 0 | 0 | 0 | 0 | 0 |
| 0 | NA | NA | NA | NA | NA | Olfr856-ps1 | | 0 | 0 | 0 | 0 | 0 | 0 | 0 | 0 | 0 | 0 | 0 |
| 0 | NA | NA | NA | NA | NA | Gm7769 | | 0 | 0 | 0 | 0 | 0 | 0 | 0 | 0 | 0 | 0 | 0 |
| 0 | NA | NA | NA | NA | NA | Olfr857 | olfactory receptor 857 | 0 | 0 | 0 | 0 | 0 | 0 | 0 | 0 | 0 | 0 | 0 |
| 0 | NA | NA | NA | NA | NA | Olfr858-ps1 | | 0 | 0 | 0 | 0 | 0 | 0 | 0 | 0 | 0 | 0 | 0 |
| 0 | NA | NA | NA | NA | NA | Olfr58 | olfactory receptor 58 | 0 | 0 | 0 | 0 | 0 | 0 | 0 | 0 | 0 | 0 | 0 |
| 0 | NA | NA | NA | NA | NA | Olfr859 | olfactory receptor 859 | 0 | 0 | 0 | 0 | 0 | 0 | 0 | 0 | 0 | 0 | 0 |
| 0 | NA | NA | NA | NA | NA | Gm45699 | | 0 | 0 | 0 | 0 | 0 | 0 | 0 | 0 | 0 | 0 | 0 |
| 0 | NA | NA | NA | NA | NA | Olfr860 | olfactory receptor 860 | 0 | 0 | 0 | 0 | 0 | 0 | 0 | 0 | 0 | 0 | 0 |
| 0 | NA | NA | NA | NA | NA | Olfr861-ps1 | | 0 | 0 | 0 | 0 | 0 | 0 | 0 | 0 | 0 | 0 | 0 |
| 0 | NA | NA | NA | NA | NA | Olfr862 | olfactory receptor 862 | 0 | 0 | 0 | 0 | 0 | 0 | 0 | 0 | 0 | 0 | 0 |
| 0 | NA | NA | NA | NA | NA | Olfr77 | olfactory receptor 77 | 0 | 0 | 0 | 0 | 0 | 0 | 0 | 0 | 0 | 0 | 0 |
| 0 | NA | NA | NA | NA | NA | Olfr863-ps1 | | 0 | 0 | 0 | 0 | 0 | 0 | 0 | 0 | 0 | 0 | 0 |
| 0 | NA | NA | NA | NA | NA | Olfr864-ps1 | | 0 | 0 | 0 | 0 | 0 | 0 | 0 | 0 | 0 | 0 | 0 |
| 0 | NA | NA | NA | NA | NA | Olfr865-ps1 | | 0 | 0 | 0 | 0 | 0 | 0 | 0 | 0 | 0 | 0 | 0 |
| 0 | NA | NA | NA | NA | NA | Olfr866 | olfactory receptor 866 | 0 | 0 | 0 | 0 | 0 | 0 | 0 | 0 | 0 | 0 | 0 |
| 0 | NA | NA | NA | NA | NA | Gm44395 | | 0 | 0 | 0 | 0 | 0 | 0 | 0 | 0 | 0 | 0 | 0 |
| 0 | NA | NA | NA | NA | NA | Gm26231 | | 0 | 0 | 0 | 0 | 0 | 0 | 0 | 0 | 0 | 0 | 0 |
| 0 | NA | NA | NA | NA | NA | Olfr867 | olfactory receptor 867 | 0 | 0 | 0 | 0 | 0 | 0 | 0 | 0 | 0 | 0 | 0 |
| 0 | NA | NA | NA | NA | NA | Olfr868 | olfactory receptor 868 | 0 | 0 | 0 | 0 | 0 | 0 | 0 | 0 | 0 | 0 | 0 |
| 0 | NA | NA | NA | NA | NA | Olfr869 | olfactory receptor 869 | 0 | 0 | 0 | 0 | 0 | 0 | 0 | 0 | 0 | 0 | 0 |
| 0 | NA | NA | NA | NA | NA | Olfr870 | olfactory receptor 870 | 0 | 0 | 0 | 0 | 0 | 0 | 0 | 0 | 0 | 0 | 0 |
| 0 | NA | NA | NA | NA | NA | Gm25914 | | 0 | 0 | 0 | 0 | 0 | 0 | 0 | 0 | 0 | 0 | 0 |
| 0 | NA | NA | NA | NA | NA | Gm26290 | | 0 | 0 | 0 | 0 | 0 | 0 | 0 | 0 | 0 | 0 | 0 |
| 0 | NA | NA | NA | NA | NA | Olfr871 | olfactory receptor 871 | 0 | 0 | 0 | 0 | 0 | 0 | 0 | 0 | 0 | 0 | 0 |
| 0 | NA | NA | NA | NA | NA | Olfr872 | olfactory receptor 872 | 0 | 0 | 0 | 0 | 0 | 0 | 0 | 0 | 0 | 0 | 0 |
| 0 | NA | NA | NA | NA | NA | Gm25425 | | 0 | 0 | 0 | 0 | 0 | 0 | 0 | 0 | 0 | 0 | 0 |
| 0 | NA | NA | NA | NA | NA | Olfr39 | olfactory receptor 39 | 0 | 0 | 0 | 0 | 0 | 0 | 0 | 0 | 0 | 0 | 0 |
| 0 | NA | NA | NA | NA | NA | Olfr873 | olfactory receptor 873 | 0 | 0 | 0 | 0 | 0 | 0 | 0 | 0 | 0 | 0 | 0 |
| 0 | NA | NA | NA | NA | NA | Olfr18 | olfactory receptor 18 | 0 | 0 | 0 | 0 | 0 | 0 | 0 | 0 | 0 | 0 | 0 |
| 0 | NA | NA | NA | NA | NA | Gm45815 | | 0 | 0 | 0 | 0 | 0 | 0 | 0 | 0 | 0 | 0 | 0 |
| 0 | NA | NA | NA | NA | NA | Gm29761 | | 0 | 0 | 0 | 0 | 0 | 0 | 0 | 0 | 0 | 0 | 0 |
| 0 | NA | NA | NA | NA | NA | Gm48729 | | 0 | 0 | 0 | 0 | 0 | 0 | 0 | 0 | 0 | 0 | 0 |
| 0 | NA | NA | NA | NA | NA | Gm48738 | | 0 | 0 | 0 | 0 | 0 | 0 | 0 | 0 | 0 | 0 | 0 |
| 0 | NA | NA | NA | NA | NA | Gm23316 | | 0 | 0 | 0 | 0 | 0 | 0 | 0 | 0 | 0 | 0 | 0 |
| 0 | NA | NA | NA | NA | NA | Gm26274 | | 0 | 0 | 0 | 0 | 0 | 0 | 0 | 0 | 0 | 0 | 0 |
| 0 | NA | NA | NA | NA | NA | Rdh8 | retinol dehydrogenase 8 | 0 | 0 | 0 | 0 | 0 | 0 | 0 | 0 | 0 | 0 | 0 |
| 0 | NA | NA | NA | NA | NA | | | 0 | 0 | 0 | 0 | 0 | 0 | 0 | 0 | 0 | 0 | 0 |
| 0 | NA | NA | NA | NA | NA | Gm24067 | | 0 | 0 | 0 | 0 | 0 | 0 | 0 | 0 | 0 | 0 | 0 |
| 0 | NA | NA | NA | NA | NA | Gm23008 | | 0 | 0 | 0 | 0 | 0 | 0 | 0 | 0 | 0 | 0 | 0 |
| 0 | NA | NA | NA | NA | NA | Gm50491 | | 0 | 0 | 0 | 0 | 0 | 0 | 0 | 0 | 0 | 0 | 0 |
| 0 | NA | NA | NA | NA | NA | Mir1900 | | 0 | 0 | 0 | 0 | 0 | 0 | 0 | 0 | 0 | 0 | 0 |
| 0 | NA | NA | NA | NA | NA | Zglp1 | zinc finger, GATA-like protein 1 | 0 | 0 | 0 | 0 | 0 | 0 | 0 | 0 | 0 | 0 | 0 |
| 0 | NA | NA | NA | NA | NA | Raver1 | ribonucleoprotein, PTB-binding 1 | 0 | 0 | 0 | 0 | 0 | 0 | 0 | 0 | 0 | 0 | 0 |
| 0 | NA | NA | NA | NA | NA | Mir7082 | | 0 | 0 | 0 | 0 | 0 | 0 | 0 | 0 | 0 | 0 | 0 |
| 0 | NA | NA | NA | NA | NA | Gm48303 | | 0 | 0 | 0 | 0 | 0 | 0 | 0 | 0 | 0 | 0 | 0 |
| 0 | NA | NA | NA | NA | NA | Mir199a-1 | | 0 | 0 | 0 | 0 | 0 | 0 | 0 | 0 | 0 | 0 | 0 |
| 0 | NA | NA | NA | NA | NA | Gm7904 | | 0 | 0 | 0 | 0 | 0 | 0 | 0 | 0 | 0 | 0 | 0 |
| 0 | NA | NA | NA | NA | NA | Gm27373 | | 0 | 0 | 0 | 0 | 0 | 0 | 0 | 0 | 0 | 0 | 0 |
| 0 | NA | NA | NA | NA | NA | Mir7083 | | 0 | 0 | 0 | 0 | 0 | 0 | 0 | 0 | 0 | 0 | 0 |
| 0 | NA | NA | NA | NA | NA | Gm47873 | | 0 | 0 | 0 | 0 | 0 | 0 | 0 | 0 | 0 | 0 | 0 |

| | | | | | | | | | | | | | | | | | |
|---|---|---|---|---|---|---|---|---|---|---|---|---|---|---|---|---|---|
| 0 | NA | NA | NA | NA | NA | Gm49318 | | 0 | 0 | 0 | 0 | 0 | 0 | 0 | 0 | 0 | 0 | 0 |
| 0 | NA | NA | NA | NA | NA | Mir7084 | | 0 | 0 | 0 | 0 | 0 | 0 | 0 | 0 | 0 | 0 | 0 |
| 0 | NA | NA | NA | NA | NA | Rpl15-ps3 | | 0 | 0 | 0 | 0 | 0 | 0 | 0 | 0 | 0 | 0 | 0 |
| 0 | NA | NA | NA | NA | NA | Gm18228 | | 0 | 0 | 0 | 0 | 0 | 0 | 0 | 0 | 0 | 0 | 0 |
| 0 | NA | NA | NA | NA | NA | Gm48652 | | 0 | 0 | 0 | 0 | 0 | 0 | 0 | 0 | 0 | 0 | 0 |
| 0 | NA | NA | NA | NA | NA | Gm16165 | | 0 | 0 | 0 | 0 | 0 | 0 | 0 | 0 | 0 | 0 | 0 |
| 0 | NA | NA | NA | NA | NA | Gm16164 | | 0 | 0 | 0 | 0 | 0 | 0 | 0 | 0 | 0 | 0 | 0 |
| 0 | NA | NA | NA | NA | NA | Gm27639 | | 0 | 0 | 0 | 0 | 0 | 0 | 0 | 0 | 0 | 0 | 0 |
| 0 | NA | NA | NA | NA | NA | Gm3011 | | 0 | 0 | 0 | 0 | 0 | 0 | 0 | 0 | 0 | 0 | 0 |
| 0 | NA | NA | NA | NA | NA | Gm47638 | | 0 | 0 | 0 | 0 | 0 | 0 | 0 | 0 | 0 | 0 | 0 |
| 0 | NA | NA | NA | NA | NA | Gm3028 | | 0 | 0 | 0 | 0 | 0 | 0 | 0 | 0 | 0 | 0 | 0 |
| 0 | NA | NA | NA | NA | NA | Gm47663 | | 0 | 0 | 0 | 0 | 0 | 0 | 0 | 0 | 0 | 0 | 0 |
| 0 | NA | NA | NA | NA | NA | Gm47668 | | 0 | 0 | 0 | 0 | 0 | 0 | 0 | 0 | 0 | 0 | 0 |
| 0 | NA | NA | NA | NA | NA | Gm47670 | | 0 | 0 | 0 | 0 | 0 | 0 | 0 | 0 | 0 | 0 | 0 |
| 0 | NA | NA | NA | NA | NA | Npsr1 | neuropeptide S receptor 1 | 0 | 0 | 0 | 0 | 0 | 0 | 0 | 0 | 0 | 0 | 0 |
| 0 | NA | NA | NA | NA | NA | Gm47674 | | 0 | 0 | 0 | 0 | 0 | 0 | 0 | 0 | 0 | 0 | 0 |
| 0 | NA | NA | NA | NA | NA | Gm15600 | | 0 | 0 | 0 | 0 | 0 | 0 | 0 | 0 | 0 | 0 | 0 |
| 0 | NA | NA | NA | NA | NA | Gm10701 | | 0 | 0 | 0 | 0 | 0 | 0 | 0 | 0 | 0 | 0 | 0 |
| 0 | NA | NA | NA | NA | NA | Gm22380 | | 0 | 0 | 0 | 0 | 0 | 0 | 0 | 0 | 0 | 0 | 0 |
| 0 | NA | NA | NA | NA | NA | Dpy19l2 | dpy-19-like 2 (C. elegans) | 0 | 0 | 0 | 0 | 0 | 0 | 0 | 0 | 0 | 0 | 0 |
| 0 | NA | NA | NA | NA | NA | Cypt4 | cysteine-rich perinuclear theca 4 | 0 | 0 | 0 | 0 | 0 | 0 | 0 | 0 | 0 | 0 | 0 |
| 0 | NA | NA | NA | NA | NA | Gm29824 | | 0 | 0 | 0 | 0 | 0 | 0 | 0 | 0 | 0 | 0 | 0 |
| 0 | NA | NA | NA | NA | NA | Amd-ps6 | | 0 | 0 | 0 | 0 | 0 | 0 | 0 | 0 | 0 | 0 | 0 |
| 0 | NA | NA | NA | NA | NA | Gm48255 | | 0 | 0 | 0 | 0 | 0 | 0 | 0 | 0 | 0 | 0 | 0 |
| 0 | NA | NA | NA | NA | NA | Gm18377 | | 0 | 0 | 0 | 0 | 0 | 0 | 0 | 0 | 0 | 0 | 0 |
| 0 | NA | NA | NA | NA | NA | Gm23893 | | 0 | 0 | 0 | 0 | 0 | 0 | 0 | 0 | 0 | 0 | 0 |
| 0 | NA | NA | NA | NA | NA | E130101E03Rik | | 0 | 0 | 0 | 0 | 0 | 0 | 0 | 0 | 0 | 0 | 0 |
| 0 | NA | NA | NA | NA | NA | Gm48256 | | 0 | 0 | 0 | 0 | 0 | 0 | 0 | 0 | 0 | 0 | 0 |
| 0 | NA | NA | NA | NA | NA | Gm8031 | | 0 | 0 | 0 | 0 | 0 | 0 | 0 | 0 | 0 | 0 | 0 |
| 0 | NA | NA | NA | NA | NA | Gm48272 | | 0 | 0 | 0 | 0 | 0 | 0 | 0 | 0 | 0 | 0 | 0 |
| 0 | NA | NA | NA | NA | NA | Gm18891 | | 0 | 0 | 0 | 0 | 0 | 0 | 0 | 0 | 0 | 0 | 0 |
| 0 | NA | NA | NA | NA | NA | Gm25861 | | 0 | 0 | 0 | 0 | 0 | 0 | 0 | 0 | 0 | 0 | 0 |
| 0 | NA | NA | NA | NA | NA | Gm25346 | | 0 | 0 | 0 | 0 | 0 | 0 | 0 | 0 | 0 | 0 | 0 |
| 0 | NA | NA | NA | NA | NA | Gm48330 | | 0 | 0 | 0 | 0 | 0 | 0 | 0 | 0 | 0 | 0 | 0 |
| 0 | NA | NA | NA | NA | NA | Gm48331 | | 0 | 0 | 0 | 0 | 0 | 0 | 0 | 0 | 0 | 0 | 0 |
| 0 | NA | NA | NA | NA | NA | Gm48337 | | 0 | 0 | 0 | 0 | 0 | 0 | 0 | 0 | 0 | 0 | 0 |
| 0 | NA | NA | NA | NA | NA | Gm26190 | | 0 | 0 | 0 | 0 | 0 | 0 | 0 | 0 | 0 | 0 | 0 |
| 0 | NA | NA | NA | NA | NA | Gm39312 | | 0 | 0 | 0 | 0 | 0 | 0 | 0 | 0 | 0 | 0 | 0 |
| 0 | NA | NA | NA | NA | NA | Gm19113 | | 0 | 0 | 0 | 0 | 0 | 0 | 0 | 0 | 0 | 0 | 0 |
| 0 | NA | NA | NA | NA | NA | Gm48373 | | 0 | 0 | 0 | 0 | 0 | 0 | 0 | 0 | 0 | 0 | 0 |
| 0 | NA | NA | NA | NA | NA | Gm30313 | | 0 | 0 | 0 | 0 | 0 | 0 | 0 | 0 | 0 | 0 | 0 |
| 0 | NA | NA | NA | NA | NA | Gm48393 | | 0 | 0 | 0 | 0 | 0 | 0 | 0 | 0 | 0 | 0 | 0 |
| 0 | NA | NA | NA | NA | NA | Gm1110 | predicted gene 1110 | 0 | 0 | 0 | 0 | 0 | 0 | 0 | 0 | 0 | 0 | 0 |
| 0 | NA | NA | NA | NA | NA | Gm48803 | | 0 | 0 | 0 | 0 | 0 | 0 | 0 | 0 | 0 | 0 | 0 |
| 0 | NA | NA | NA | NA | NA | Gm48858 | | 0 | 0 | 0 | 0 | 0 | 0 | 0 | 0 | 0 | 0 | 0 |
| 0 | NA | NA | NA | NA | NA | Gm48901 | | 0 | 0 | 0 | 0 | 0 | 0 | 0 | 0 | 0 | 0 | 0 |
| 0 | NA | NA | NA | NA | NA | Gm18585 | | 0 | 0 | 0 | 0 | 0 | 0 | 0 | 0 | 0 | 0 | 0 |
| 0 | NA | NA | NA | NA | NA | Gm8106 | | 0 | 0 | 0 | 0 | 0 | 0 | 0 | 0 | 0 | 0 | 0 |
| 0 | NA | NA | NA | NA | NA | Gm48902 | | 0 | 0 | 0 | 0 | 0 | 0 | 0 | 0 | 0 | 0 | 0 |
| 0 | NA | NA | NA | NA | NA | Gm15606 | | 0 | 0 | 0 | 0 | 0 | 0 | 0 | 0 | 0 | 0 | 0 |
| 0 | NA | NA | NA | NA | NA | Gm44316 | | 0 | 0 | 0 | 0 | 0 | 0 | 0 | 0 | 0 | 0 | 0 |
| 0 | NA | NA | NA | NA | NA | Gm48091 | | 0 | 0 | 0 | 0 | 0 | 0 | 0 | 0 | 0 | 0 | 0 |
| 0 | NA | NA | NA | NA | NA | Gm15520 | | 0 | 0 | 0 | 0 | 0 | 0 | 0 | 0 | 0 | 0 | 0 |
| 0 | NA | NA | NA | NA | NA | Gm19640 | | 0 | 0 | 0 | 0 | 0 | 0 | 0 | 0 | 0 | 0 | 0 |
| 0 | NA | NA | NA | NA | NA | Gm47677 | | 0 | 0 | 0 | 0 | 0 | 0 | 0 | 0 | 0 | 0 | 0 |
| 0 | NA | NA | NA | NA | NA | Gm26435 | | 0 | 0 | 0 | 0 | 0 | 0 | 0 | 0 | 0 | 0 | 0 |
| 0 | NA | NA | NA | NA | NA | Gm30933 | | 0 | 0 | 0 | 0 | 0 | 0 | 0 | 0 | 0 | 0 | 0 |
| 0 | NA | NA | NA | NA | NA | Gm47714 | | 0 | 0 | 0 | 0 | 0 | 0 | 0 | 0 | 0 | 0 | 0 |
| 0 | NA | NA | NA | NA | NA | Rps2-ps12 | | 0 | 0 | 0 | 0 | 0 | 0 | 0 | 0 | 0 | 0 | 0 |
| 0 | NA | NA | NA | NA | NA | Gm47716 | | 0 | 0 | 0 | 0 | 0 | 0 | 0 | 0 | 0 | 0 | 0 |

| | | | | | | | | | | | | | | | | | | |
|---|---|---|---|---|---|---|---|---|---|---|---|---|---|---|---|---|---|---|
| 0 | NA | NA | NA | NA | NA | Gm31013 | | 0 | 0 | 0 | 0 | 0 | 0 | 0 | 0 | 0 | 0 | 0 |
| 0 | NA | NA | NA | NA | NA | Gm47717 | | 0 | 0 | 0 | 0 | 0 | 0 | 0 | 0 | 0 | 0 | 0 |
| 0 | NA | NA | NA | NA | NA | Gm17508 | | 0 | 0 | 0 | 0 | 0 | 0 | 0 | 0 | 0 | 0 | 0 |
| 0 | NA | NA | NA | NA | NA | Gm7244 | | 0 | 0 | 0 | 0 | 0 | 0 | 0 | 0 | 0 | 0 | 0 |
| 0 | NA | NA | NA | NA | NA | 5033425B01Rik | | 0 | 0 | 0 | 0 | 0 | 0 | 0 | 0 | 0 | 0 | 0 |
| 0 | NA | NA | NA | NA | NA | 4930434F21Rik | | 0 | 0 | 0 | 0 | 0 | 0 | 0 | 0 | 0 | 0 | 0 |
| 0 | NA | NA | NA | NA | NA | Gm31408 | | 0 | 0 | 0 | 0 | 0 | 0 | 0 | 0 | 0 | 0 | 0 |
| 0 | NA | NA | NA | NA | NA | Gm18226 | | 0 | 0 | 0 | 0 | 0 | 0 | 0 | 0 | 0 | 0 | 0 |
| 0 | NA | NA | NA | NA | NA | Gm18820 | | 0 | 0 | 0 | 0 | 0 | 0 | 0 | 0 | 0 | 0 | 0 |
| 0 | NA | NA | NA | NA | NA | Gm47436 | | 0 | 0 | 0 | 0 | 0 | 0 | 0 | 0 | 0 | 0 | 0 |
| 0 | NA | NA | NA | NA | NA | Gm31497 | | 0 | 0 | 0 | 0 | 0 | 0 | 0 | 0 | 0 | 0 | 0 |
| 0 | NA | NA | NA | NA | NA | Gm47465 | | 0 | 0 | 0 | 0 | 0 | 0 | 0 | 0 | 0 | 0 | 0 |
| 0 | NA | NA | NA | NA | NA | Tpi-rs4 | | 0 | 0 | 0 | 0 | 0 | 0 | 0 | 0 | 0 | 0 | 0 |
| 0 | NA | NA | NA | NA | NA | Gm44453 | | 0 | 0 | 0 | 0 | 0 | 0 | 0 | 0 | 0 | 0 | 0 |
| 0 | NA | NA | NA | NA | NA | Gm47496 | | 0 | 0 | 0 | 0 | 0 | 0 | 0 | 0 | 0 | 0 | 0 |
| 0 | NA | NA | NA | NA | NA | Gm47536 | | 0 | 0 | 0 | 0 | 0 | 0 | 0 | 0 | 0 | 0 | 0 |
| 0 | NA | NA | NA | NA | NA | Kcnj1 | potassium inwardly-rectifying channel, su | 0 | 0 | 0 | 0 | 0 | 0 | 0 | 0 | 0 | 0 | 0 |
| 0 | NA | NA | NA | NA | NA | Gm22060 | | 0 | 0 | 0 | 0 | 0 | 0 | 0 | 0 | 0 | 0 | 0 |
| 0 | NA | NA | NA | NA | NA | Gm38292 | | 0 | 0 | 0 | 0 | 0 | 0 | 0 | 0 | 0 | 0 | 0 |
| 0 | NA | NA | NA | NA | NA | Gm38346 | | 0 | 0 | 0 | 0 | 0 | 0 | 0 | 0 | 0 | 0 | 0 |
| 0 | NA | NA | NA | NA | NA | Gm27201 | | 0 | 0 | 0 | 0 | 0 | 0 | 0 | 0 | 0 | 0 | 0 |
| 0 | NA | NA | NA | NA | NA | Gm37474 | | 0 | 0 | 0 | 0 | 0 | 0 | 0 | 0 | 0 | 0 | 0 |
| 0 | NA | NA | NA | NA | NA | Gm47680 | | 0 | 0 | 0 | 0 | 0 | 0 | 0 | 0 | 0 | 0 | 0 |
| 0 | NA | NA | NA | NA | NA | Gm27163 | | 0 | 0 | 0 | 0 | 0 | 0 | 0 | 0 | 0 | 0 | 0 |
| 0 | NA | NA | NA | NA | NA | Gm27240 | | 0 | 0 | 0 | 0 | 0 | 0 | 0 | 0 | 0 | 0 | 0 |
| 0 | NA | NA | NA | NA | NA | Gm27162 | | 0 | 0 | 0 | 0 | 0 | 0 | 0 | 0 | 0 | 0 | 0 |
| 0 | NA | NA | NA | NA | NA | Gm37192 | | 0 | 0 | 0 | 0 | 0 | 0 | 0 | 0 | 0 | 0 | 0 |
| 0 | NA | NA | NA | NA | NA | Gm27166 | | 0 | 0 | 0 | 0 | 0 | 0 | 0 | 0 | 0 | 0 | 0 |
| 0 | NA | NA | NA | NA | NA | Gm32171 | | 0 | 0 | 0 | 0 | 0 | 0 | 0 | 0 | 0 | 0 | 0 |
| 0 | NA | NA | NA | NA | NA | Gm37758 | | 0 | 0 | 0 | 0 | 0 | 0 | 0 | 0 | 0 | 0 | 0 |
| 0 | NA | NA | NA | NA | NA | Gm47775 | | 0 | 0 | 0 | 0 | 0 | 0 | 0 | 0 | 0 | 0 | 0 |
| 0 | NA | NA | NA | NA | NA | Gm47777 | | 0 | 0 | 0 | 0 | 0 | 0 | 0 | 0 | 0 | 0 | 0 |
| 0 | NA | NA | NA | NA | NA | Gm47778 | | 0 | 0 | 0 | 0 | 0 | 0 | 0 | 0 | 0 | 0 | 0 |
| 0 | NA | NA | NA | NA | NA | Gm47784 | | 0 | 0 | 0 | 0 | 0 | 0 | 0 | 0 | 0 | 0 | 0 |
| 0 | NA | NA | NA | NA | NA | Gm47785 | | 0 | 0 | 0 | 0 | 0 | 0 | 0 | 0 | 0 | 0 | 0 |
| 0 | NA | NA | NA | NA | NA | Gm25439 | | 0 | 0 | 0 | 0 | 0 | 0 | 0 | 0 | 0 | 0 | 0 |
| 0 | NA | NA | NA | NA | NA | Gm27165 | | 0 | 0 | 0 | 0 | 0 | 0 | 0 | 0 | 0 | 0 | 0 |
| 0 | NA | NA | NA | NA | NA | Gm27164 | | 0 | 0 | 0 | 0 | 0 | 0 | 0 | 0 | 0 | 0 | 0 |
| 0 | NA | NA | NA | NA | NA | Gm47858 | | 0 | 0 | 0 | 0 | 0 | 0 | 0 | 0 | 0 | 0 | 0 |
| 0 | NA | NA | NA | NA | NA | Gm38217 | | 0 | 0 | 0 | 0 | 0 | 0 | 0 | 0 | 0 | 0 | 0 |
| 0 | NA | NA | NA | NA | NA | Gm23702 | | 0 | 0 | 0 | 0 | 0 | 0 | 0 | 0 | 0 | 0 | 0 |
| 0 | NA | NA | NA | NA | NA | Gm24262 | | 0 | 0 | 0 | 0 | 0 | 0 | 0 | 0 | 0 | 0 | 0 |
| 0 | NA | NA | NA | NA | NA | Gm33838 | | 0 | 0 | 0 | 0 | 0 | 0 | 0 | 0 | 0 | 0 | 0 |
| 0 | NA | NA | NA | NA | NA | 2610105M22Rik | | 0 | 0 | 0 | 0 | 0 | 0 | 0 | 0 | 0 | 0 | 0 |
| 0 | NA | NA | NA | NA | NA | Gm24188 | | 0 | 0 | 0 | 0 | 0 | 0 | 0 | 0 | 0 | 0 | 0 |
| 0 | NA | NA | NA | NA | NA | Gm1113 | predicted gene 1113 | 0 | 0 | 0 | 0 | 0 | 0 | 0 | 0 | 0 | 0 | 0 |
| 0 | NA | NA | NA | NA | NA | Gm6762 | | 0 | 0 | 0 | 0 | 0 | 0 | 0 | 0 | 0 | 0 | 0 |
| 0 | NA | NA | NA | NA | NA | Gm10105 | | 0 | 0 | 0 | 0 | 0 | 0 | 0 | 0 | 0 | 0 | 0 |
| 0 | NA | NA | NA | NA | NA | Pate4 | prostate and testis expressed 4 | 0 | 0 | 0 | 0 | 0 | 0 | 0 | 0 | 0 | 0 | 0 |
| 0 | NA | NA | NA | NA | NA | Pate3 | prostate and testis expressed 3 | 0 | 0 | 0 | 0 | 0 | 0 | 0 | 0 | 0 | 0 | 0 |
| 0 | NA | NA | NA | NA | NA | Pate1 | prostate and testis expressed 1 | 0 | 0 | 0 | 0 | 0 | 0 | 0 | 0 | 0 | 0 | 0 |
| 0 | NA | NA | NA | NA | NA | Pate10 | prostate and testis expressed 10 | 0 | 0 | 0 | 0 | 0 | 0 | 0 | 0 | 0 | 0 | 0 |
| 0 | NA | NA | NA | NA | NA | Pate7 | prostate and testis expressed 7 | 0 | 0 | 0 | 0 | 0 | 0 | 0 | 0 | 0 | 0 | 0 |
| 0 | NA | NA | NA | NA | NA | Gm27235 | | 0 | 0 | 0 | 0 | 0 | 0 | 0 | 0 | 0 | 0 | 0 |
| 0 | NA | NA | NA | NA | NA | Pate6 | prostate and testis expressed 6 | 0 | 0 | 0 | 0 | 0 | 0 | 0 | 0 | 0 | 0 | 0 |
| 0 | NA | NA | NA | NA | NA | Gm36974 | | 0 | 0 | 0 | 0 | 0 | 0 | 0 | 0 | 0 | 0 | 0 |
| 0 | NA | NA | NA | NA | NA | Pate5 | prostate and testis expressed 5 | 0 | 0 | 0 | 0 | 0 | 0 | 0 | 0 | 0 | 0 | 0 |
| 0 | NA | NA | NA | NA | NA | Gm3428 | | 0 | 0 | 0 | 0 | 0 | 0 | 0 | 0 | 0 | 0 | 0 |
| 0 | NA | NA | NA | NA | NA | Pate13 | prostate and testis expressed 13 | 0 | 0 | 0 | 0 | 0 | 0 | 0 | 0 | 0 | 0 | 0 |
| 0 | NA | NA | NA | NA | NA | Gm27187 | | 0 | 0 | 0 | 0 | 0 | 0 | 0 | 0 | 0 | 0 | 0 |

| | | | | | | | | | | | | | | | | | | |
|---|---|---|---|---|---|---|---|---|---|---|---|---|---|---|---|---|---|---|
| 0 | NA | NA | NA | NA | NA | Gm48348 | | 0 | 0 | 0 | 0 | 0 | 0 | 0 | 0 | 0 | 0 | 0 |
| 0 | NA | NA | NA | NA | NA | Gm5916 | predicted gene 5916 | 0 | 0 | 0 | 0 | 0 | 0 | 0 | 0 | 0 | 0 | 0 |
| 0 | NA | NA | NA | NA | NA | Gm3434 | | 0 | 0 | 0 | 0 | 0 | 0 | 0 | 0 | 0 | 0 | 0 |
| 0 | NA | NA | NA | NA | NA | Gm48391 | | 0 | 0 | 0 | 0 | 0 | 0 | 0 | 0 | 0 | 0 | 0 |
| 0 | NA | NA | NA | NA | NA | Gm3867 | | 0 | 0 | 0 | 0 | 0 | 0 | 0 | 0 | 0 | 0 | 0 |
| 0 | NA | NA | NA | NA | NA | Gm48401 | | 0 | 0 | 0 | 0 | 0 | 0 | 0 | 0 | 0 | 0 | 0 |
| 0 | NA | NA | NA | NA | NA | Gm48407 | | 0 | 0 | 0 | 0 | 0 | 0 | 0 | 0 | 0 | 0 | 0 |
| 0 | NA | NA | NA | NA | NA | Pate12 | prostate and testis expressed 12 | 0 | 0 | 0 | 0 | 0 | 0 | 0 | 0 | 0 | 0 | 0 |
| 0 | NA | NA | NA | NA | NA | Pate11 | prostate and testis expressed 11 | 0 | 0 | 0 | 0 | 0 | 0 | 0 | 0 | 0 | 0 | 0 |
| 0 | NA | NA | NA | NA | NA | Gm27160 | | 0 | 0 | 0 | 0 | 0 | 0 | 0 | 0 | 0 | 0 | 0 |
| 0 | NA | NA | NA | NA | NA | Pate9 | prostate and testis expressed 9 | 0 | 0 | 0 | 0 | 0 | 0 | 0 | 0 | 0 | 0 | 0 |
| 0 | NA | NA | NA | NA | NA | Gm27218 | | 0 | 0 | 0 | 0 | 0 | 0 | 0 | 0 | 0 | 0 | 0 |
| 0 | NA | NA | NA | NA | NA | Gm48431 | | 0 | 0 | 0 | 0 | 0 | 0 | 0 | 0 | 0 | 0 | 0 |
| 0 | NA | NA | NA | NA | NA | Pate8 | prostate and testis expressed 8 | 0 | 0 | 0 | 0 | 0 | 0 | 0 | 0 | 0 | 0 | 0 |
| 0 | NA | NA | NA | NA | NA | Gm49367 | | 0 | 0 | 0 | 0 | 0 | 0 | 0 | 0 | 0 | 0 | 0 |
| 0 | NA | NA | NA | NA | NA | Pate14 | prostate and testis expressed 14 | 0 | 0 | 0 | 0 | 0 | 0 | 0 | 0 | 0 | 0 | 0 |
| 0 | NA | NA | NA | NA | NA | 1700027I24Rik | | 0 | 0 | 0 | 0 | 0 | 0 | 0 | 0 | 0 | 0 | 0 |
| 0 | NA | NA | NA | NA | NA | Gm48443 | | 0 | 0 | 0 | 0 | 0 | 0 | 0 | 0 | 0 | 0 | 0 |
| 0 | NA | NA | NA | NA | NA | Acrv1 | acrosomal vesicle protein 1 | 0 | 0 | 0 | 0 | 0 | 0 | 0 | 0 | 0 | 0 | 0 |
| 0 | NA | NA | NA | NA | NA | Gm3896 | | 0 | 0 | 0 | 0 | 0 | 0 | 0 | 0 | 0 | 0 | 0 |
| 0 | NA | NA | NA | NA | NA | n-R5s82 | | 0 | 0 | 0 | 0 | 0 | 0 | 0 | 0 | 0 | 0 | 0 |
| 0 | NA | NA | NA | NA | NA | Gm48716 | | 0 | 0 | 0 | 0 | 0 | 0 | 0 | 0 | 0 | 0 | 0 |
| 0 | NA | NA | NA | NA | NA | Gm48717 | | 0 | 0 | 0 | 0 | 0 | 0 | 0 | 0 | 0 | 0 | 0 |
| 0 | NA | NA | NA | NA | NA | Gm25273 | | 0 | 0 | 0 | 0 | 0 | 0 | 0 | 0 | 0 | 0 | 0 |
| 0 | NA | NA | NA | NA | NA | Gm47633 | | 0 | 0 | 0 | 0 | 0 | 0 | 0 | 0 | 0 | 0 | 0 |
| 0 | NA | NA | NA | NA | NA | Gm47640 | | 0 | 0 | 0 | 0 | 0 | 0 | 0 | 0 | 0 | 0 | 0 |
| 0 | NA | NA | NA | NA | NA | Gm18329 | | 0 | 0 | 0 | 0 | 0 | 0 | 0 | 0 | 0 | 0 | 0 |
| 0 | NA | NA | NA | NA | NA | Gm34885 | | 0 | 0 | 0 | 0 | 0 | 0 | 0 | 0 | 0 | 0 | 0 |
| 0 | NA | NA | NA | NA | NA | Gm47945 | | 0 | 0 | 0 | 0 | 0 | 0 | 0 | 0 | 0 | 0 | 0 |
| 0 | NA | NA | NA | NA | NA | Panx3 | pannexin 3 | 0 | 0 | 0 | 0 | 0 | 0 | 0 | 0 | 0 | 0 | 0 |
| 0 | NA | NA | NA | NA | NA | Olfr160 | olfactory receptor 160 | 0 | 0 | 0 | 0 | 0 | 0 | 0 | 0 | 0 | 0 | 0 |
| 0 | NA | NA | NA | NA | NA | Olfr151 | olfactory receptor 151 | 0 | 0 | 0 | 0 | 0 | 0 | 0 | 0 | 0 | 0 | 0 |
| 0 | NA | NA | NA | NA | NA | Olfr874 | olfactory receptor 874 | 0 | 0 | 0 | 0 | 0 | 0 | 0 | 0 | 0 | 0 | 0 |
| 0 | NA | NA | NA | NA | NA | Olfr875 | olfactory receptor 875 | 0 | 0 | 0 | 0 | 0 | 0 | 0 | 0 | 0 | 0 | 0 |
| 0 | NA | NA | NA | NA | NA | Olfr876 | olfactory receptor 876 | 0 | 0 | 0 | 0 | 0 | 0 | 0 | 0 | 0 | 0 | 0 |
| 0 | NA | NA | NA | NA | NA | Gm24764 | | 0 | 0 | 0 | 0 | 0 | 0 | 0 | 0 | 0 | 0 | 0 |
| 0 | NA | NA | NA | NA | NA | Gm47005 | | 0 | 0 | 0 | 0 | 0 | 0 | 0 | 0 | 0 | 0 | 0 |
| 0 | NA | NA | NA | NA | NA | Olfr877 | olfactory receptor 877 | 0 | 0 | 0 | 0 | 0 | 0 | 0 | 0 | 0 | 0 | 0 |
| 0 | NA | NA | NA | NA | NA | Olfr145 | olfactory receptor 145 | 0 | 0 | 0 | 0 | 0 | 0 | 0 | 0 | 0 | 0 | 0 |
| 0 | NA | NA | NA | NA | NA | Gm6796 | | 0 | 0 | 0 | 0 | 0 | 0 | 0 | 0 | 0 | 0 | 0 |
| 0 | NA | NA | NA | NA | NA | Olfr878 | olfactory receptor 878 | 0 | 0 | 0 | 0 | 0 | 0 | 0 | 0 | 0 | 0 | 0 |
| 0 | NA | NA | NA | NA | NA | Olfr879-ps1 | | 0 | 0 | 0 | 0 | 0 | 0 | 0 | 0 | 0 | 0 | 0 |
| 0 | NA | NA | NA | NA | NA | Olfr880-ps1 | | 0 | 0 | 0 | 0 | 0 | 0 | 0 | 0 | 0 | 0 | 0 |
| 0 | NA | NA | NA | NA | NA | Olfr881 | olfactory receptor 881 | 0 | 0 | 0 | 0 | 0 | 0 | 0 | 0 | 0 | 0 | 0 |
| 0 | NA | NA | NA | NA | NA | Gm45701 | | 0 | 0 | 0 | 0 | 0 | 0 | 0 | 0 | 0 | 0 | 0 |
| 0 | NA | NA | NA | NA | NA | Olfr882-ps1 | | 0 | 0 | 0 | 0 | 0 | 0 | 0 | 0 | 0 | 0 | 0 |
| 0 | NA | NA | NA | NA | NA | Olfr883 | olfactory receptor 883 | 0 | 0 | 0 | 0 | 0 | 0 | 0 | 0 | 0 | 0 | 0 |
| 0 | NA | NA | NA | NA | NA | Olfr884 | olfactory receptor 884 | 0 | 0 | 0 | 0 | 0 | 0 | 0 | 0 | 0 | 0 | 0 |
| 0 | NA | NA | NA | NA | NA | Olfr885 | olfactory receptor 885 | 0 | 0 | 0 | 0 | 0 | 0 | 0 | 0 | 0 | 0 | 0 |
| 0 | NA | NA | NA | NA | NA | Olfr886-ps1 | | 0 | 0 | 0 | 0 | 0 | 0 | 0 | 0 | 0 | 0 | 0 |
| 0 | NA | NA | NA | NA | NA | Olfr887 | olfactory receptor 887 | 0 | 0 | 0 | 0 | 0 | 0 | 0 | 0 | 0 | 0 | 0 |
| 0 | NA | NA | NA | NA | NA | Olfr888 | olfactory receptor 888 | 0 | 0 | 0 | 0 | 0 | 0 | 0 | 0 | 0 | 0 | 0 |
| 0 | NA | NA | NA | NA | NA | Olfr889 | olfactory receptor 889 | 0 | 0 | 0 | 0 | 0 | 0 | 0 | 0 | 0 | 0 | 0 |
| 0 | NA | NA | NA | NA | NA | Olfr890 | olfactory receptor 890 | 0 | 0 | 0 | 0 | 0 | 0 | 0 | 0 | 0 | 0 | 0 |
| 0 | NA | NA | NA | NA | NA | Olfr891 | olfactory receptor 891 | 0 | 0 | 0 | 0 | 0 | 0 | 0 | 0 | 0 | 0 | 0 |
| 0 | NA | NA | NA | NA | NA | Gm17798 | | 0 | 0 | 0 | 0 | 0 | 0 | 0 | 0 | 0 | 0 | 0 |
| 0 | NA | NA | NA | NA | NA | Olfr892-ps1 | | 0 | 0 | 0 | 0 | 0 | 0 | 0 | 0 | 0 | 0 | 0 |
| 0 | NA | NA | NA | NA | NA | Olfr893 | olfactory receptor 893 | 0 | 0 | 0 | 0 | 0 | 0 | 0 | 0 | 0 | 0 | 0 |
| 0 | NA | NA | NA | NA | NA | Olfr894 | olfactory receptor 894 | 0 | 0 | 0 | 0 | 0 | 0 | 0 | 0 | 0 | 0 | 0 |
| 0 | NA | NA | NA | NA | NA | Olfr143 | olfactory receptor 143 | 0 | 0 | 0 | 0 | 0 | 0 | 0 | 0 | 0 | 0 | 0 |

| | | | | | | | | | | | | | | | | | | |
|---|---|---|---|---|---|---|---|---|---|---|---|---|---|---|---|---|---|---|
| 0 | NA | NA | NA | NA | NA | Olfr895 | olfactory receptor 895 | 0 | 0 | 0 | 0 | 0 | 0 | 0 | 0 | 0 | 0 | 0 |
| 0 | NA | NA | NA | NA | NA | Olfr896-ps1 | | 0 | 0 | 0 | 0 | 0 | 0 | 0 | 0 | 0 | 0 | 0 |
| 0 | NA | NA | NA | NA | NA | Olfr897-ps1 | | 0 | 0 | 0 | 0 | 0 | 0 | 0 | 0 | 0 | 0 | 0 |
| 0 | NA | NA | NA | NA | NA | Olfr25 | olfactory receptor 25 | 0 | 0 | 0 | 0 | 0 | 0 | 0 | 0 | 0 | 0 | 0 |
| 0 | NA | NA | NA | NA | NA | Olfr898 | olfactory receptor 898 | 0 | 0 | 0 | 0 | 0 | 0 | 0 | 0 | 0 | 0 | 0 |
| 0 | NA | NA | NA | NA | NA | Olfr250 | olfactory receptor 899 | 0 | 0 | 0 | 0 | 0 | 0 | 0 | 0 | 0 | 0 | 0 |
| 0 | NA | NA | NA | NA | NA | Olfr251 | olfactory receptor 900 | 0 | 0 | 0 | 0 | 0 | 0 | 0 | 0 | 0 | 0 | 0 |
| 0 | NA | NA | NA | NA | NA | Olfr147 | olfactory receptor 147 | 0 | 0 | 0 | 0 | 0 | 0 | 0 | 0 | 0 | 0 | 0 |
| 0 | NA | NA | NA | NA | NA | Olfr901 | olfactory receptor 901 | 0 | 0 | 0 | 0 | 0 | 0 | 0 | 0 | 0 | 0 | 0 |
| 0 | NA | NA | NA | NA | NA | Olfr902 | olfactory receptor 902 | 0 | 0 | 0 | 0 | 0 | 0 | 0 | 0 | 0 | 0 | 0 |
| 0 | NA | NA | NA | NA | NA | Olfr1543-ps1 | | 0 | 0 | 0 | 0 | 0 | 0 | 0 | 0 | 0 | 0 | 0 |
| 0 | NA | NA | NA | NA | NA | Olfr904 | olfactory receptor 904 | 0 | 0 | 0 | 0 | 0 | 0 | 0 | 0 | 0 | 0 | 0 |
| 0 | NA | NA | NA | NA | NA | Olfr905 | olfactory receptor 905 | 0 | 0 | 0 | 0 | 0 | 0 | 0 | 0 | 0 | 0 | 0 |
| 0 | NA | NA | NA | NA | NA | Olfr906 | olfactory receptor 906 | 0 | 0 | 0 | 0 | 0 | 0 | 0 | 0 | 0 | 0 | 0 |
| 0 | NA | NA | NA | NA | NA | Olfr907 | olfactory receptor 907 | 0 | 0 | 0 | 0 | 0 | 0 | 0 | 0 | 0 | 0 | 0 |
| 0 | NA | NA | NA | NA | NA | Gm23271 | | 0 | 0 | 0 | 0 | 0 | 0 | 0 | 0 | 0 | 0 | 0 |
| 0 | NA | NA | NA | NA | NA | Olfr908 | olfactory receptor 908 | 0 | 0 | 0 | 0 | 0 | 0 | 0 | 0 | 0 | 0 | 0 |
| 0 | NA | NA | NA | NA | NA | Olfr911-ps1 | olfactory receptor 911, pseudogene 1 | 0 | 0 | 0 | 0 | 0 | 0 | 0 | 0 | 0 | 0 | 0 |
| 0 | NA | NA | NA | NA | NA | Olfr910 | olfactory receptor 910 | 0 | 0 | 0 | 0 | 0 | 0 | 0 | 0 | 0 | 0 | 0 |
| 0 | NA | NA | NA | NA | NA | Olfr913 | olfactory receptor 913 | 0 | 0 | 0 | 0 | 0 | 0 | 0 | 0 | 0 | 0 | 0 |
| 0 | NA | NA | NA | NA | NA | Olfr914 | olfactory receptor 914 | 0 | 0 | 0 | 0 | 0 | 0 | 0 | 0 | 0 | 0 | 0 |
| 0 | NA | NA | NA | NA | NA | Olfr915 | olfactory receptor 915 | 0 | 0 | 0 | 0 | 0 | 0 | 0 | 0 | 0 | 0 | 0 |
| 0 | NA | NA | NA | NA | NA | Gm49385 | | 0 | 0 | 0 | 0 | 0 | 0 | 0 | 0 | 0 | 0 | 0 |
| 0 | NA | NA | NA | NA | NA | Olfr916 | olfactory receptor 916 | 0 | 0 | 0 | 0 | 0 | 0 | 0 | 0 | 0 | 0 | 0 |
| 0 | NA | NA | NA | NA | NA | Gm47449 | | 0 | 0 | 0 | 0 | 0 | 0 | 0 | 0 | 0 | 0 | 0 |
| 0 | NA | NA | NA | NA | NA | Olfr917 | olfactory receptor 917 | 0 | 0 | 0 | 0 | 0 | 0 | 0 | 0 | 0 | 0 | 0 |
| 0 | NA | NA | NA | NA | NA | Gm47458 | | 0 | 0 | 0 | 0 | 0 | 0 | 0 | 0 | 0 | 0 | 0 |
| 0 | NA | NA | NA | NA | NA | Olfr918 | olfactory receptor 918 | 0 | 0 | 0 | 0 | 0 | 0 | 0 | 0 | 0 | 0 | 0 |
| 0 | NA | NA | NA | NA | NA | Gm5010 | | 0 | 0 | 0 | 0 | 0 | 0 | 0 | 0 | 0 | 0 | 0 |
| 0 | NA | NA | NA | NA | NA | Olfr919 | olfactory receptor 919 | 0 | 0 | 0 | 0 | 0 | 0 | 0 | 0 | 0 | 0 | 0 |
| 0 | NA | NA | NA | NA | NA | Gm19709 | | 0 | 0 | 0 | 0 | 0 | 0 | 0 | 0 | 0 | 0 | 0 |
| 0 | NA | NA | NA | NA | NA | Olfr922 | olfactory receptor 922 | 0 | 0 | 0 | 0 | 0 | 0 | 0 | 0 | 0 | 0 | 0 |
| 0 | NA | NA | NA | NA | NA | Olfr923 | olfactory receptor 923 | 0 | 0 | 0 | 0 | 0 | 0 | 0 | 0 | 0 | 0 | 0 |
| 0 | NA | NA | NA | NA | NA | Olfr924 | olfactory receptor 924 | 0 | 0 | 0 | 0 | 0 | 0 | 0 | 0 | 0 | 0 | 0 |
| 0 | NA | NA | NA | NA | NA | Olfr26 | olfactory receptor 26 | 0 | 0 | 0 | 0 | 0 | 0 | 0 | 0 | 0 | 0 | 0 |
| 0 | NA | NA | NA | NA | NA | Olfr925-ps1 | | 0 | 0 | 0 | 0 | 0 | 0 | 0 | 0 | 0 | 0 | 0 |
| 0 | NA | NA | NA | NA | NA | Olfr926 | olfactory receptor 926 | 0 | 0 | 0 | 0 | 0 | 0 | 0 | 0 | 0 | 0 | 0 |
| 0 | NA | NA | NA | NA | NA | Olfr927-ps1 | | 0 | 0 | 0 | 0 | 0 | 0 | 0 | 0 | 0 | 0 | 0 |
| 0 | NA | NA | NA | NA | NA | Olfr928-ps1 | | 0 | 0 | 0 | 0 | 0 | 0 | 0 | 0 | 0 | 0 | 0 |
| 0 | NA | NA | NA | NA | NA | Olfr929-ps1 | | 0 | 0 | 0 | 0 | 0 | 0 | 0 | 0 | 0 | 0 | 0 |
| 0 | NA | NA | NA | NA | NA | Olfr930 | olfactory receptor 930 | 0 | 0 | 0 | 0 | 0 | 0 | 0 | 0 | 0 | 0 | 0 |
| 0 | NA | NA | NA | NA | NA | Gm20953 | | 0 | 0 | 0 | 0 | 0 | 0 | 0 | 0 | 0 | 0 | 0 |
| 0 | NA | NA | NA | NA | NA | Olfr931-ps1 | | 0 | 0 | 0 | 0 | 0 | 0 | 0 | 0 | 0 | 0 | 0 |
| 0 | NA | NA | NA | NA | NA | | | 0 | 0 | 0 | 0 | 0 | 0 | 0 | 0 | 0 | 0 | 0 |
| 0 | NA | NA | NA | NA | NA | Olfr933 | olfactory receptor 933 | 0 | 0 | 0 | 0 | 0 | 0 | 0 | 0 | 0 | 0 | 0 |
| 0 | NA | NA | NA | NA | NA | Olfr934 | olfactory receptor 934 | 0 | 0 | 0 | 0 | 0 | 0 | 0 | 0 | 0 | 0 | 0 |
| 0 | NA | NA | NA | NA | NA | Olfr935 | olfactory receptor 935 | 0 | 0 | 0 | 0 | 0 | 0 | 0 | 0 | 0 | 0 | 0 |
| 0 | NA | NA | NA | NA | NA | Olfr146 | olfactory receptor 146 | 0 | 0 | 0 | 0 | 0 | 0 | 0 | 0 | 0 | 0 | 0 |
| 0 | NA | NA | NA | NA | NA | Olfr936 | olfactory receptor 936 | 0 | 0 | 0 | 0 | 0 | 0 | 0 | 0 | 0 | 0 | 0 |
| 0 | NA | NA | NA | NA | NA | Gm25529 | | 0 | 0 | 0 | 0 | 0 | 0 | 0 | 0 | 0 | 0 | 0 |
| 0 | NA | NA | NA | NA | NA | Olfr937 | olfactory receptor 937 | 0 | 0 | 0 | 0 | 0 | 0 | 0 | 0 | 0 | 0 | 0 |
| 0 | NA | NA | NA | NA | NA | Gm25032 | | 0 | 0 | 0 | 0 | 0 | 0 | 0 | 0 | 0 | 0 | 0 |
| 0 | NA | NA | NA | NA | NA | Olfr938 | olfactory receptor 938 | 0 | 0 | 0 | 0 | 0 | 0 | 0 | 0 | 0 | 0 | 0 |
| 0 | NA | NA | NA | NA | NA | | | 0 | 0 | 0 | 0 | 0 | 0 | 0 | 0 | 0 | 0 | 0 |
| 0 | NA | NA | NA | NA | NA | Gm47085 | | 0 | 0 | 0 | 0 | 0 | 0 | 0 | 0 | 0 | 0 | 0 |
| 0 | NA | NA | NA | NA | NA | Olfr941-ps1 | | 0 | 0 | 0 | 0 | 0 | 0 | 0 | 0 | 0 | 0 | 0 |
| 0 | NA | NA | NA | NA | NA | Olfr27 | olfactory receptor 27 | 0 | 0 | 0 | 0 | 0 | 0 | 0 | 0 | 0 | 0 | 0 |
| 0 | NA | NA | NA | NA | NA | Olfr942-ps1 | | 0 | 0 | 0 | 0 | 0 | 0 | 0 | 0 | 0 | 0 | 0 |
| 0 | NA | NA | NA | NA | NA | Olfr943 | olfactory receptor 943 | 0 | 0 | 0 | 0 | 0 | 0 | 0 | 0 | 0 | 0 | 0 |
| 0 | NA | NA | NA | NA | NA | Gm47100 | | 0 | 0 | 0 | 0 | 0 | 0 | 0 | 0 | 0 | 0 | 0 |

| | | | | | | | | | | | | | | | | | | |
|---|---|---|---|---|---|---|---|---|---|---|---|---|---|---|---|---|---|---|
| 0 | NA | NA | NA | NA | NA | Olfr944 | olfactory receptor 944 | 0 | 0 | 0 | 0 | 0 | 0 | 0 | 0 | 0 | 0 | 0 |
| 0 | NA | NA | NA | NA | NA | Olfr1537 | olfactory receptor 1537 | 0 | 0 | 0 | 0 | 0 | 0 | 0 | 0 | 0 | 0 | 0 |
| 0 | NA | NA | NA | NA | NA | Olfr945 | olfactory receptor 945 | 0 | 0 | 0 | 0 | 0 | 0 | 0 | 0 | 0 | 0 | 0 |
| 0 | NA | NA | NA | NA | NA | Olfr946-ps1 | | 0 | 0 | 0 | 0 | 0 | 0 | 0 | 0 | 0 | 0 | 0 |
| 0 | NA | NA | NA | NA | NA | Olfr947-ps1 | | 0 | 0 | 0 | 0 | 0 | 0 | 0 | 0 | 0 | 0 | 0 |
| 0 | NA | NA | NA | NA | NA | Olfr948 | olfactory receptor 948 | 0 | 0 | 0 | 0 | 0 | 0 | 0 | 0 | 0 | 0 | 0 |
| 0 | NA | NA | NA | NA | NA | Gm21041 | | 0 | 0 | 0 | 0 | 0 | 0 | 0 | 0 | 0 | 0 | 0 |
| 0 | NA | NA | NA | NA | NA | Olfr949-ps1 | | 0 | 0 | 0 | 0 | 0 | 0 | 0 | 0 | 0 | 0 | 0 |
| 0 | NA | NA | NA | NA | NA | Olfr950-ps1 | | 0 | 0 | 0 | 0 | 0 | 0 | 0 | 0 | 0 | 0 | 0 |
| 0 | NA | NA | NA | NA | NA | Olfr951 | olfactory receptor 951 | 0 | 0 | 0 | 0 | 0 | 0 | 0 | 0 | 0 | 0 | 0 |
| 0 | NA | NA | NA | NA | NA | Olfr952 | olfactory receptor 952 | 0 | 0 | 0 | 0 | 0 | 0 | 0 | 0 | 0 | 0 | 0 |
| 0 | NA | NA | NA | NA | NA | Olfr953-ps1 | | 0 | 0 | 0 | 0 | 0 | 0 | 0 | 0 | 0 | 0 | 0 |
| 0 | NA | NA | NA | NA | NA | Olfr954 | olfactory receptor 954 | 0 | 0 | 0 | 0 | 0 | 0 | 0 | 0 | 0 | 0 | 0 |
| 0 | NA | NA | NA | NA | NA | Olfr955 | olfactory receptor 955 | 0 | 0 | 0 | 0 | 0 | 0 | 0 | 0 | 0 | 0 | 0 |
| 0 | NA | NA | NA | NA | NA | Olfr956-ps1 | | 0 | 0 | 0 | 0 | 0 | 0 | 0 | 0 | 0 | 0 | 0 |
| 0 | NA | NA | NA | NA | NA | Olfr44 | olfactory receptor 44 | 0 | 0 | 0 | 0 | 0 | 0 | 0 | 0 | 0 | 0 | 0 |
| 0 | NA | NA | NA | NA | NA | Olfr957 | olfactory receptor 957 | 0 | 0 | 0 | 0 | 0 | 0 | 0 | 0 | 0 | 0 | 0 |
| 0 | NA | NA | NA | NA | NA | Gm24278 | | 0 | 0 | 0 | 0 | 0 | 0 | 0 | 0 | 0 | 0 | 0 |
| 0 | NA | NA | NA | NA | NA | Olfr958 | olfactory receptor 958 | 0 | 0 | 0 | 0 | 0 | 0 | 0 | 0 | 0 | 0 | 0 |
| 0 | NA | NA | NA | NA | NA | Olfr959 | olfactory receptor 959 | 0 | 0 | 0 | 0 | 0 | 0 | 0 | 0 | 0 | 0 | 0 |
| 0 | NA | NA | NA | NA | NA | Olfr148 | olfactory receptor 148 | 0 | 0 | 0 | 0 | 0 | 0 | 0 | 0 | 0 | 0 | 0 |
| 0 | NA | NA | NA | NA | NA | Olfr960 | olfactory receptor 960 | 0 | 0 | 0 | 0 | 0 | 0 | 0 | 0 | 0 | 0 | 0 |
| 0 | NA | NA | NA | NA | NA | Olfr961 | olfactory receptor 961 | 0 | 0 | 0 | 0 | 0 | 0 | 0 | 0 | 0 | 0 | 0 |
| 0 | NA | NA | NA | NA | NA | Olfr962-ps1 | | 0 | 0 | 0 | 0 | 0 | 0 | 0 | 0 | 0 | 0 | 0 |
| 0 | NA | NA | NA | NA | NA | Olfr963 | olfactory receptor 963 | 0 | 0 | 0 | 0 | 0 | 0 | 0 | 0 | 0 | 0 | 0 |
| 0 | NA | NA | NA | NA | NA | Olfr964-ps1 | | 0 | 0 | 0 | 0 | 0 | 0 | 0 | 0 | 0 | 0 | 0 |
| 0 | NA | NA | NA | NA | NA | Olfr149 | olfactory receptor 149 | 0 | 0 | 0 | 0 | 0 | 0 | 0 | 0 | 0 | 0 | 0 |
| 0 | NA | NA | NA | NA | NA | Olfr965 | olfactory receptor 965 | 0 | 0 | 0 | 0 | 0 | 0 | 0 | 0 | 0 | 0 | 0 |
| 0 | NA | NA | NA | NA | NA | Olfr966-ps1 | | 0 | 0 | 0 | 0 | 0 | 0 | 0 | 0 | 0 | 0 | 0 |
| 0 | NA | NA | NA | NA | NA | Olfr150 | olfactory receptor 150 | 0 | 0 | 0 | 0 | 0 | 0 | 0 | 0 | 0 | 0 | 0 |
| 0 | NA | NA | NA | NA | NA | Olfr967 | olfactory receptor 967 | 0 | 0 | 0 | 0 | 0 | 0 | 0 | 0 | 0 | 0 | 0 |
| 0 | NA | NA | NA | NA | NA | Olfr968 | olfactory receptor 968 | 0 | 0 | 0 | 0 | 0 | 0 | 0 | 0 | 0 | 0 | 0 |
| 0 | NA | NA | NA | NA | NA | Olfr969 | olfactory receptor 969 | 0 | 0 | 0 | 0 | 0 | 0 | 0 | 0 | 0 | 0 | 0 |
| 0 | NA | NA | NA | NA | NA | Olfr970 | olfactory receptor 970 | 0 | 0 | 0 | 0 | 0 | 0 | 0 | 0 | 0 | 0 | 0 |
| 0 | NA | NA | NA | NA | NA | Gm47219 | | 0 | 0 | 0 | 0 | 0 | 0 | 0 | 0 | 0 | 0 | 0 |
| 0 | NA | NA | NA | NA | NA | Olfr971 | olfactory receptor 971 | 0 | 0 | 0 | 0 | 0 | 0 | 0 | 0 | 0 | 0 | 0 |
| 0 | NA | NA | NA | NA | NA | Olfr972 | olfactory receptor 972 | 0 | 0 | 0 | 0 | 0 | 0 | 0 | 0 | 0 | 0 | 0 |
| 0 | NA | NA | NA | NA | NA | Olfr229 | olfactory receptor 229 | 0 | 0 | 0 | 0 | 0 | 0 | 0 | 0 | 0 | 0 | 0 |
| 0 | NA | NA | NA | NA | NA | Gm47229 | | 0 | 0 | 0 | 0 | 0 | 0 | 0 | 0 | 0 | 0 | 0 |
| 0 | NA | NA | NA | NA | NA | Olfr974 | olfactory receptor 974 | 0 | 0 | 0 | 0 | 0 | 0 | 0 | 0 | 0 | 0 | 0 |
| 0 | NA | NA | NA | NA | NA | Olfr975 | olfactory receptor 975 | 0 | 0 | 0 | 0 | 0 | 0 | 0 | 0 | 0 | 0 | 0 |
| 0 | NA | NA | NA | NA | NA | Olfr976 | olfactory receptor 976 | 0 | 0 | 0 | 0 | 0 | 0 | 0 | 0 | 0 | 0 | 0 |
| 0 | NA | NA | NA | NA | NA | Olfr977-ps1 | | 0 | 0 | 0 | 0 | 0 | 0 | 0 | 0 | 0 | 0 | 0 |
| 0 | NA | NA | NA | NA | NA | Olfr978 | olfactory receptor 978 | 0 | 0 | 0 | 0 | 0 | 0 | 0 | 0 | 0 | 0 | 0 |
| 0 | NA | NA | NA | NA | NA | Olfr979 | olfactory receptor 979 | 0 | 0 | 0 | 0 | 0 | 0 | 0 | 0 | 0 | 0 | 0 |
| 0 | NA | NA | NA | NA | NA | Olfr980 | olfactory receptor 980 | 0 | 0 | 0 | 0 | 0 | 0 | 0 | 0 | 0 | 0 | 0 |
| 0 | NA | NA | NA | NA | NA | Olfr981 | olfactory receptor 981 | 0 | 0 | 0 | 0 | 0 | 0 | 0 | 0 | 0 | 0 | 0 |
| 0 | NA | NA | NA | NA | NA | Gm18892 | | 0 | 0 | 0 | 0 | 0 | 0 | 0 | 0 | 0 | 0 | 0 |
| 0 | NA | NA | NA | NA | NA | 1700001J11Rik | | 0 | 0 | 0 | 0 | 0 | 0 | 0 | 0 | 0 | 0 | 0 |
| 0 | NA | NA | NA | NA | NA | Olfr982 | olfactory receptor 982 | 0 | 0 | 0 | 0 | 0 | 0 | 0 | 0 | 0 | 0 | 0 |
| 0 | NA | NA | NA | NA | NA | Olfr983 | olfactory receptor 983 | 0 | 0 | 0 | 0 | 0 | 0 | 0 | 0 | 0 | 0 | 0 |
| 0 | NA | NA | NA | NA | NA | Olfr984 | olfactory receptor 984 | 0 | 0 | 0 | 0 | 0 | 0 | 0 | 0 | 0 | 0 | 0 |
| 0 | NA | NA | NA | NA | NA | Olfr985 | olfactory receptor 985 | 0 | 0 | 0 | 0 | 0 | 0 | 0 | 0 | 0 | 0 | 0 |
| 0 | NA | NA | NA | NA | NA | Tmem225 | transmembrane protein 225 | 0 | 0 | 0 | 0 | 0 | 0 | 0 | 0 | 0 | 0 | 0 |
| 0 | NA | NA | NA | NA | NA | Olfr986 | olfactory receptor 986 | 0 | 0 | 0 | 0 | 0 | 0 | 0 | 0 | 0 | 0 | 0 |
| 0 | NA | NA | NA | NA | NA | Gm47266 | | 0 | 0 | 0 | 0 | 0 | 0 | 0 | 0 | 0 | 0 | 0 |
| 0 | NA | NA | NA | NA | NA | Gm48127 | | 0 | 0 | 0 | 0 | 0 | 0 | 0 | 0 | 0 | 0 | 0 |
| 0 | NA | NA | NA | NA | NA | Gm17540 | | 0 | 0 | 0 | 0 | 0 | 0 | 0 | 0 | 0 | 0 | 0 |
| 0 | NA | NA | NA | NA | NA | Gm35177 | | 0 | 0 | 0 | 0 | 0 | 0 | 0 | 0 | 0 | 0 | 0 |
| 0 | NA | NA | NA | NA | NA | Gm48277 | | 0 | 0 | 0 | 0 | 0 | 0 | 0 | 0 | 0 | 0 | 0 |

| | | | | | | | | | | | | | | | | | | |
|---|---|---|---|---|---|---|---|---|---|---|---|---|---|---|---|---|---|---|
| 0 | NA | NA | NA | NA | NA | Gm48284 | | 0 | 0 | 0 | 0 | 0 | 0 | 0 | 0 | 0 | 0 | 0 |
| 0 | NA | NA | NA | NA | NA | Gm35371 | | 0 | 0 | 0 | 0 | 0 | 0 | 0 | 0 | 0 | 0 | 0 |
| 0 | NA | NA | NA | NA | NA | Gm21915 | | 0 | 0 | 0 | 0 | 0 | 0 | 0 | 0 | 0 | 0 | 0 |
| 0 | NA | NA | NA | NA | NA | Gm48288 | | 0 | 0 | 0 | 0 | 0 | 0 | 0 | 0 | 0 | 0 | 0 |
| 0 | NA | NA | NA | NA | NA | Gm48294 | | 0 | 0 | 0 | 0 | 0 | 0 | 0 | 0 | 0 | 0 | 0 |
| 0 | NA | NA | NA | NA | NA | Gm16095 | | 0 | 0 | 0 | 0 | 0 | 0 | 0 | 0 | 0 | 0 | 0 |
| 0 | NA | NA | NA | NA | NA | Snord14c | | 0 | 0 | 0 | 0 | 0 | 0 | 0 | 0 | 0 | 0 | 0 |
| 0 | NA | NA | NA | NA | NA | Snord14d | | 0 | 0 | 0 | 0 | 0 | 0 | 0 | 0 | 0 | 0 | 0 |
| 0 | NA | NA | NA | NA | NA | Snord14e | | 0 | 0 | 0 | 0 | 0 | 0 | 0 | 0 | 0 | 0 | 0 |
| 0 | NA | NA | NA | NA | NA | Bsx | brain specific homeobox | 0 | 0 | 0 | 0 | 0 | 0 | 0 | 0 | 0 | 0 | 0 |
| 0 | NA | NA | NA | NA | NA | Gm18586 | | 0 | 0 | 0 | 0 | 0 | 0 | 0 | 0 | 0 | 0 | 0 |
| 0 | NA | NA | NA | NA | NA | Gm48611 | | 0 | 0 | 0 | 0 | 0 | 0 | 0 | 0 | 0 | 0 | 0 |
| 0 | NA | NA | NA | NA | NA | Gm6885 | | 0 | 0 | 0 | 0 | 0 | 0 | 0 | 0 | 0 | 0 | 0 |
| 0 | NA | NA | NA | NA | NA | Gm48710 | | 0 | 0 | 0 | 0 | 0 | 0 | 0 | 0 | 0 | 0 | 0 |
| 0 | NA | NA | NA | NA | NA | Gm25401 | | 0 | 0 | 0 | 0 | 0 | 0 | 0 | 0 | 0 | 0 | 0 |
| 0 | NA | NA | NA | NA | NA | Gm48742 | | 0 | 0 | 0 | 0 | 0 | 0 | 0 | 0 | 0 | 0 | 0 |
| 0 | NA | NA | NA | NA | NA | Gm48739 | | 0 | 0 | 0 | 0 | 0 | 0 | 0 | 0 | 0 | 0 | 0 |
| 0 | NA | NA | NA | NA | NA | Gm49317 | | 0 | 0 | 0 | 0 | 0 | 0 | 0 | 0 | 0 | 0 | 0 |
| 0 | NA | NA | NA | NA | NA | Gm48736 | | 0 | 0 | 0 | 0 | 0 | 0 | 0 | 0 | 0 | 0 | 0 |
| 0 | NA | NA | NA | NA | NA | Mir100 | | 0 | 0 | 0 | 0 | 0 | 0 | 0 | 0 | 0 | 0 | 0 |
| 0 | NA | NA | NA | NA | NA | Mirlet7a-2 | | 0 | 0 | 0 | 0 | 0 | 0 | 0 | 0 | 0 | 0 | 0 |
| 0 | NA | NA | NA | NA | NA | Gm48740 | | 0 | 0 | 0 | 0 | 0 | 0 | 0 | 0 | 0 | 0 | 0 |
| 0 | NA | NA | NA | NA | NA | Mir125b-1 | | 0 | 0 | 0 | 0 | 0 | 0 | 0 | 0 | 0 | 0 | 0 |
| 0 | NA | NA | NA | NA | NA | Gm48737 | | 0 | 0 | 0 | 0 | 0 | 0 | 0 | 0 | 0 | 0 | 0 |
| 0 | NA | NA | NA | NA | NA | Gm8707 | | 0 | 0 | 0 | 0 | 0 | 0 | 0 | 0 | 0 | 0 | 0 |
| 0 | NA | NA | NA | NA | NA | Gm48784 | | 0 | 0 | 0 | 0 | 0 | 0 | 0 | 0 | 0 | 0 | 0 |
| 0 | NA | NA | NA | NA | NA | Gm35835 | | 0 | 0 | 0 | 0 | 0 | 0 | 0 | 0 | 0 | 0 | 0 |
| 0 | NA | NA | NA | NA | NA | | | 0 | 0 | 0 | 0 | 0 | 0 | 0 | 0 | 0 | 0 | 0 |
| 0 | NA | NA | NA | NA | NA | Gm35940 | | 0 | 0 | 0 | 0 | 0 | 0 | 0 | 0 | 0 | 0 | 0 |
| 0 | NA | NA | NA | NA | NA | Gm40513 | | 0 | 0 | 0 | 0 | 0 | 0 | 0 | 0 | 0 | 0 | 0 |
| 0 | NA | NA | NA | NA | NA | Gm16214 | | 0 | 0 | 0 | 0 | 0 | 0 | 0 | 0 | 0 | 0 | 0 |
| 0 | NA | NA | NA | NA | NA | Gm39321 | | 0 | 0 | 0 | 0 | 0 | 0 | 0 | 0 | 0 | 0 | 0 |
| 0 | NA | NA | NA | NA | NA | 4930546K05Rik | | 0 | 0 | 0 | 0 | 0 | 0 | 0 | 0 | 0 | 0 | 0 |
| 0 | NA | NA | NA | NA | NA | Gm36435 | | 0 | 0 | 0 | 0 | 0 | 0 | 0 | 0 | 0 | 0 | 0 |
| 0 | NA | NA | NA | NA | NA | Gm26272 | | 0 | 0 | 0 | 0 | 0 | 0 | 0 | 0 | 0 | 0 | 0 |
| 0 | NA | NA | NA | NA | NA | Gm47629 | | 0 | 0 | 0 | 0 | 0 | 0 | 0 | 0 | 0 | 0 | 0 |
| 0 | NA | NA | NA | NA | NA | Gm44311 | | 0 | 0 | 0 | 0 | 0 | 0 | 0 | 0 | 0 | 0 | 0 |
| 0 | NA | NA | NA | NA | NA | Gm24420 | | 0 | 0 | 0 | 0 | 0 | 0 | 0 | 0 | 0 | 0 | 0 |
| 0 | NA | NA | NA | NA | NA | Gm27509 | | 0 | 0 | 0 | 0 | 0 | 0 | 0 | 0 | 0 | 0 | 0 |
| 0 | NA | NA | NA | NA | NA | Gm47931 | | 0 | 0 | 0 | 0 | 0 | 0 | 0 | 0 | 0 | 0 | 0 |
| 0 | NA | NA | NA | NA | NA | Rpl26-ps6 | | 0 | 0 | 0 | 0 | 0 | 0 | 0 | 0 | 0 | 0 | 0 |
| 0 | NA | NA | NA | NA | NA | Gm36799 | | 0 | 0 | 0 | 0 | 0 | 0 | 0 | 0 | 0 | 0 | 0 |
| 0 | NA | NA | NA | NA | NA | Gm47475 | | 0 | 0 | 0 | 0 | 0 | 0 | 0 | 0 | 0 | 0 | 0 |
| 0 | NA | NA | NA | NA | NA | Gm28215 | | 0 | 0 | 0 | 0 | 0 | 0 | 0 | 0 | 0 | 0 | 0 |
| 0 | NA | NA | NA | NA | NA | Gm36855 | | 0 | 0 | 0 | 0 | 0 | 0 | 0 | 0 | 0 | 0 | 0 |
| 0 | NA | NA | NA | NA | NA | Gm29909 | | 0 | 0 | 0 | 0 | 0 | 0 | 0 | 0 | 0 | 0 | 0 |
| 0 | NA | NA | NA | NA | NA | Gm29961 | | 0 | 0 | 0 | 0 | 0 | 0 | 0 | 0 | 0 | 0 | 0 |
| 0 | NA | NA | NA | NA | NA | Gm23326 | | 0 | 0 | 0 | 0 | 0 | 0 | 0 | 0 | 0 | 0 | 0 |
| 0 | NA | NA | NA | NA | NA | Gm10688 | | 0 | 0 | 0 | 0 | 0 | 0 | 0 | 0 | 0 | 0 | 0 |
| 0 | NA | NA | NA | NA | NA | E230034D01Rik | | 0 | 0 | 0 | 0 | 0 | 0 | 0 | 0 | 0 | 0 | 0 |
| 0 | NA | NA | NA | NA | NA | Gm47345 | | 0 | 0 | 0 | 0 | 0 | 0 | 0 | 0 | 0 | 0 | 0 |
| 0 | NA | NA | NA | NA | NA | Gm39325 | | 0 | 0 | 0 | 0 | 0 | 0 | 0 | 0 | 0 | 0 | 0 |
| 0 | NA | NA | NA | NA | NA | C1qtnf5 | C1q and tumor necrosis factor related prc | 0 | 0 | 0 | 0 | 0 | 0 | 0 | 0 | 0 | 0 | 0 |
| 0 | NA | NA | NA | NA | NA | Gm47327 | | 0 | 0 | 0 | 0 | 0 | 0 | 0 | 0 | 0 | 0 | 0 |
| 0 | NA | NA | NA | NA | NA | Gm47255 | | 0 | 0 | 0 | 0 | 0 | 0 | 0 | 0 | 0 | 0 | 0 |
| 0 | NA | NA | NA | NA | NA | Ccdc153 | coiled-coil domain containing 153 | 0 | 0 | 0 | 0 | 0 | 0 | 0 | 0 | 0 | 0 | 0 |
| 0 | NA | NA | NA | NA | NA | Gm10686 | | 0 | 0 | 0 | 0 | 0 | 0 | 0 | 0 | 0 | 0 | 0 |
| 0 | NA | NA | NA | NA | NA | Gm26306 | | 0 | 0 | 0 | 0 | 0 | 0 | 0 | 0 | 0 | 0 | 0 |
| 0 | NA | NA | NA | NA | NA | Gm47186 | | 0 | 0 | 0 | 0 | 0 | 0 | 0 | 0 | 0 | 0 | 0 |
| 0 | NA | NA | NA | NA | NA | Upk2 | uroplakin 2 | 0 | 0 | 0 | 0 | 0 | 0 | 0 | 0 | 0 | 0 | 0 |

| | | | | | | | | | | | | | | | | | | |
|---|---|---|---|---|---|---|---|---|---|---|---|---|---|---|---|---|---|---|
| 0 | NA | NA | NA | NA | NA | Gm30671 | | 0 | 0 | 0 | 0 | 0 | 0 | 0 | 0 | 0 | 0 | 0 |
| 0 | NA | NA | NA | NA | NA | Gm47238 | | 0 | 0 | 0 | 0 | 0 | 0 | 0 | 0 | 0 | 0 | 0 |
| 0 | NA | NA | NA | NA | NA | Gm3953 | | 0 | 0 | 0 | 0 | 0 | 0 | 0 | 0 | 0 | 0 | 0 |
| 0 | NA | NA | NA | NA | NA | BC049987 | | 0 | 0 | 0 | 0 | 0 | 0 | 0 | 0 | 0 | 0 | 0 |
| 0 | NA | NA | NA | NA | NA | Treh | trehalase (brush-border membrane glyco | 0 | 0 | 0 | 0 | 0 | 0 | 0 | 0 | 0 | 0 | 0 |
| 0 | NA | NA | NA | NA | NA | Gm47239 | | 0 | 0 | 0 | 0 | 0 | 0 | 0 | 0 | 0 | 0 | 0 |
| 0 | NA | NA | NA | NA | NA | Gm24166 | | 0 | 0 | 0 | 0 | 0 | 0 | 0 | 0 | 0 | 0 | 0 |
| 0 | NA | NA | NA | NA | NA | Gm26249 | | 0 | 0 | 0 | 0 | 0 | 0 | 0 | 0 | 0 | 0 | 0 |
| 0 | NA | NA | NA | NA | NA | Gm19121 | | 0 | 0 | 0 | 0 | 0 | 0 | 0 | 0 | 0 | 0 | 0 |
| 0 | NA | NA | NA | NA | NA | Gm48719 | | 0 | 0 | 0 | 0 | 0 | 0 | 0 | 0 | 0 | 0 | 0 |
| 0 | NA | NA | NA | NA | NA | Gm48720 | | 0 | 0 | 0 | 0 | 0 | 0 | 0 | 0 | 0 | 0 | 0 |
| 0 | NA | NA | NA | NA | NA | Gm47596 | | 0 | 0 | 0 | 0 | 0 | 0 | 0 | 0 | 0 | 0 | 0 |
| 0 | NA | NA | NA | NA | NA | Gm20612 | | 0 | 0 | 0 | 0 | 0 | 0 | 0 | 0 | 0 | 0 | 0 |
| 0 | NA | NA | NA | NA | NA | Mir7086 | | 0 | 0 | 0 | 0 | 0 | 0 | 0 | 0 | 0 | 0 | 0 |
| 0 | NA | NA | NA | NA | NA | Gm47964 | | 0 | 0 | 0 | 0 | 0 | 0 | 0 | 0 | 0 | 0 | 0 |
| 0 | NA | NA | NA | NA | NA | Lncbate3 | | 0 | 0 | 0 | 0 | 0 | 0 | 0 | 0 | 0 | 0 | 0 |
| 0 | NA | NA | NA | NA | NA | Gm48135 | | 0 | 0 | 0 | 0 | 0 | 0 | 0 | 0 | 0 | 0 | 0 |
| 0 | NA | NA | NA | NA | NA | 4833428L15Rik | | 0 | 0 | 0 | 0 | 0 | 0 | 0 | 0 | 0 | 0 | 0 |
| 0 | NA | NA | NA | NA | NA | Gm22069 | | 0 | 0 | 0 | 0 | 0 | 0 | 0 | 0 | 0 | 0 | 0 |
| 0 | NA | NA | NA | NA | NA | Gm47528 | | 0 | 0 | 0 | 0 | 0 | 0 | 0 | 0 | 0 | 0 | 0 |
| 0 | NA | NA | NA | NA | NA | Apoa4 | apolipoprotein A-IV | 0 | 0 | 0 | 0 | 0 | 0 | 0 | 0 | 0 | 0 | 0 |
| 0 | NA | NA | NA | NA | NA | Gm10680 | | 0 | 0 | 0 | 0 | 0 | 0 | 0 | 0 | 0 | 0 | 0 |
| 0 | NA | NA | NA | NA | NA | Apoa5 | apolipoprotein A-V | 0 | 0 | 0 | 0 | 0 | 0 | 0 | 0 | 0 | 0 | 0 |
| 0 | NA | NA | NA | NA | NA | Gm47544 | | 0 | 0 | 0 | 0 | 0 | 0 | 0 | 0 | 0 | 0 | 0 |
| 0 | NA | NA | NA | NA | NA | 4931429L15Ri | RIKEN cDNA 4931429L15 gene | 0 | 0 | 0 | 0 | 0 | 0 | 0 | 0 | 0 | 0 | 0 |
| 0 | NA | NA | NA | NA | NA | Gm39329 | | 0 | 0 | 0 | 0 | 0 | 0 | 0 | 0 | 0 | 0 | 0 |
| 0 | NA | NA | NA | NA | NA | Gm47144 | | 0 | 0 | 0 | 0 | 0 | 0 | 0 | 0 | 0 | 0 | 0 |
| 0 | NA | NA | NA | NA | NA | Gm22805 | | 0 | 0 | 0 | 0 | 0 | 0 | 0 | 0 | 0 | 0 | 0 |
| 0 | NA | NA | NA | NA | NA | Gm8863 | | 0 | 0 | 0 | 0 | 0 | 0 | 0 | 0 | 0 | 0 | 0 |
| 0 | NA | NA | NA | NA | NA | Gm47153 | | 0 | 0 | 0 | 0 | 0 | 0 | 0 | 0 | 0 | 0 | 0 |
| 0 | NA | NA | NA | NA | NA | Gm47161 | | 0 | 0 | 0 | 0 | 0 | 0 | 0 | 0 | 0 | 0 | 0 |
| 0 | NA | NA | NA | NA | NA | Gm47198 | | 0 | 0 | 0 | 0 | 0 | 0 | 0 | 0 | 0 | 0 | 0 |
| 0 | NA | NA | NA | NA | NA | Gm47197 | | 0 | 0 | 0 | 0 | 0 | 0 | 0 | 0 | 0 | 0 | 0 |
| 0 | NA | NA | NA | NA | NA | Gm47213 | | 0 | 0 | 0 | 0 | 0 | 0 | 0 | 0 | 0 | 0 | 0 |
| 0 | NA | NA | NA | NA | NA | Gm47215 | | 0 | 0 | 0 | 0 | 0 | 0 | 0 | 0 | 0 | 0 | 0 |
| 0 | NA | NA | NA | NA | NA | Gm47236 | | 0 | 0 | 0 | 0 | 0 | 0 | 0 | 0 | 0 | 0 | 0 |
| 0 | NA | NA | NA | NA | NA | Gm40518 | | 0 | 0 | 0 | 0 | 0 | 0 | 0 | 0 | 0 | 0 | 0 |
| 0 | NA | NA | NA | NA | NA | Gm23653 | | 0 | 0 | 0 | 0 | 0 | 0 | 0 | 0 | 0 | 0 | 0 |
| 0 | NA | NA | NA | NA | NA | Gm32335 | | 0 | 0 | 0 | 0 | 0 | 0 | 0 | 0 | 0 | 0 | 0 |
| 0 | NA | NA | NA | NA | NA | Cldn25-ps | | 0 | 0 | 0 | 0 | 0 | 0 | 0 | 0 | 0 | 0 | 0 |
| 0 | NA | NA | NA | NA | NA | Gm4894 | predicted gene 4894 | 0 | 0 | 0 | 0 | 0 | 0 | 0 | 0 | 0 | 0 | 0 |
| 0 | NA | NA | NA | NA | NA | Ankk1 | ankyrin repeat and kinase domain contai | 0 | 0 | 0 | 0 | 0 | 0 | 0 | 0 | 0 | 0 | 0 |
| 0 | NA | NA | NA | NA | NA | Gm25633 | | 0 | 0 | 0 | 0 | 0 | 0 | 0 | 0 | 0 | 0 | 0 |
| 0 | NA | NA | NA | NA | NA | Gm47543 | | 0 | 0 | 0 | 0 | 0 | 0 | 0 | 0 | 0 | 0 | 0 |
| 0 | NA | NA | NA | NA | NA | Gm8907 | | 0 | 0 | 0 | 0 | 0 | 0 | 0 | 0 | 0 | 0 | 0 |
| 0 | NA | NA | NA | NA | NA | Gm4042 | | 0 | 0 | 0 | 0 | 0 | 0 | 0 | 0 | 0 | 0 | 0 |
| 0 | NA | NA | NA | NA | NA | 2310003N18Rik | | 0 | 0 | 0 | 0 | 0 | 0 | 0 | 0 | 0 | 0 | 0 |
| 0 | NA | NA | NA | NA | NA | Gm39336 | | 0 | 0 | 0 | 0 | 0 | 0 | 0 | 0 | 0 | 0 | 0 |
| 0 | NA | NA | NA | NA | NA | Tex12 | testis expressed 12 | 0 | 0 | 0 | 0 | 0 | 0 | 0 | 0 | 0 | 0 | 0 |
| 0 | NA | NA | NA | NA | NA | Gm25558 | | 0 | 0 | 0 | 0 | 0 | 0 | 0 | 0 | 0 | 0 | 0 |
| 0 | NA | NA | NA | NA | NA | Btg4 | BTG anti-proliferation factor 4 | 0 | 0 | 0 | 0 | 0 | 0 | 0 | 0 | 0 | 0 | 0 |
| 0 | NA | NA | NA | NA | NA | Mir34c | | 0 | 0 | 0 | 0 | 0 | 0 | 0 | 0 | 0 | 0 | 0 |
| 0 | NA | NA | NA | NA | NA | Mir34b | | 0 | 0 | 0 | 0 | 0 | 0 | 0 | 0 | 0 | 0 | 0 |
| 0 | NA | NA | NA | NA | NA | Gm32742 | predicted gene, 32742 | 0 | 0 | 0 | 0 | 0 | 0 | 0 | 0 | 0 | 0 | 0 |
| 0 | NA | NA | NA | NA | NA | Gm47828 | | 0 | 0 | 0 | 0 | 0 | 0 | 0 | 0 | 0 | 0 | 0 |
| 0 | NA | NA | NA | NA | NA | Gm47854 | | 0 | 0 | 0 | 0 | 0 | 0 | 0 | 0 | 0 | 0 | 0 |
| 0 | NA | NA | NA | NA | NA | Arhgap20os | | 0 | 0 | 0 | 0 | 0 | 0 | 0 | 0 | 0 | 0 | 0 |
| 0 | NA | NA | NA | NA | NA | Gm6980 | | 0 | 0 | 0 | 0 | 0 | 0 | 0 | 0 | 0 | 0 | 0 |
| 0 | NA | NA | NA | NA | NA | 4933407I05Rik | | 0 | 0 | 0 | 0 | 0 | 0 | 0 | 0 | 0 | 0 | 0 |
| 0 | NA | NA | NA | NA | NA | Gm27686 | | 0 | 0 | 0 | 0 | 0 | 0 | 0 | 0 | 0 | 0 | 0 |

| | | | | | | | | | | | | | | | | | | |
|---|---|---|---|---|---|---|---|---|---|---|---|---|---|---|---|---|---|---|
| 0 | NA | NA | NA | NA | NA | Mir6244 | | 0 | 0 | 0 | 0 | 0 | 0 | 0 | 0 | 0 | 0 | 0 |
| 0 | NA | NA | NA | NA | NA | Gm25562 | | 0 | 0 | 0 | 0 | 0 | 0 | 0 | 0 | 0 | 0 | 0 |
| 0 | NA | NA | NA | NA | NA | Gm47957 | | 0 | 0 | 0 | 0 | 0 | 0 | 0 | 0 | 0 | 0 | 0 |
| 0 | NA | NA | NA | NA | NA | Gm47960 | | 0 | 0 | 0 | 0 | 0 | 0 | 0 | 0 | 0 | 0 | 0 |
| 0 | NA | NA | NA | NA | NA | Gm33125 | | 0 | 0 | 0 | 0 | 0 | 0 | 0 | 0 | 0 | 0 | 0 |
| 0 | NA | NA | NA | NA | NA | Gm1715 | | 0 | 0 | 0 | 0 | 0 | 0 | 0 | 0 | 0 | 0 | 0 |
| 0 | NA | NA | NA | NA | NA | Gm22242 | | 0 | 0 | 0 | 0 | 0 | 0 | 0 | 0 | 0 | 0 | 0 |
| 0 | NA | NA | NA | NA | NA | Gm47981 | | 0 | 0 | 0 | 0 | 0 | 0 | 0 | 0 | 0 | 0 | 0 |
| 0 | NA | NA | NA | NA | NA | 4930510E17Rik | | 0 | 0 | 0 | 0 | 0 | 0 | 0 | 0 | 0 | 0 | 0 |
| 0 | NA | NA | NA | NA | NA | Mir130c | | 0 | 0 | 0 | 0 | 0 | 0 | 0 | 0 | 0 | 0 | 0 |
| 0 | NA | NA | NA | NA | NA | Gm22304 | | 0 | 0 | 0 | 0 | 0 | 0 | 0 | 0 | 0 | 0 | 0 |
| 0 | NA | NA | NA | NA | NA | Gm48292 | | 0 | 0 | 0 | 0 | 0 | 0 | 0 | 0 | 0 | 0 | 0 |
| 0 | NA | NA | NA | NA | NA | Gm48346 | | 0 | 0 | 0 | 0 | 0 | 0 | 0 | 0 | 0 | 0 | 0 |
| 0 | NA | NA | NA | NA | NA | Gm48345 | | 0 | 0 | 0 | 0 | 0 | 0 | 0 | 0 | 0 | 0 | 0 |
| 0 | NA | NA | NA | NA | NA | Cyp19a1 | cytochrome P450, family 19, subfamily a, | 0 | 0 | 0 | 0 | 0 | 0 | 0 | 0 | 0 | 0 | 0 |
| 0 | NA | NA | NA | NA | NA | Gldnos | | 0 | 0 | 0 | 0 | 0 | 0 | 0 | 0 | 0 | 0 | 0 |
| 0 | NA | NA | NA | NA | NA | Sh2d7 | SH2 domain containing 7 | 0 | 0 | 0 | 0 | 0 | 0 | 0 | 0 | 0 | 0 | 0 |
| 0 | NA | NA | NA | NA | NA | Gm25980 | | 0 | 0 | 0 | 0 | 0 | 0 | 0 | 0 | 0 | 0 | 0 |
| 0 | NA | NA | NA | NA | NA | AY074887 | cDNA sequence AY074887 | 0 | 0 | 0 | 0 | 0 | 0 | 0 | 0 | 0 | 0 | 0 |
| 0 | NA | NA | NA | NA | NA | Chrna5 | cholinergic receptor, nicotinic, alpha poly | 0 | 0 | 0 | 0 | 0 | 0 | 0 | 0 | 0 | 0 | 0 |
| 0 | NA | NA | NA | NA | NA | Gm47877 | | 0 | 0 | 0 | 0 | 0 | 0 | 0 | 0 | 0 | 0 | 0 |
| 0 | NA | NA | NA | NA | NA | Gm47886 | | 0 | 0 | 0 | 0 | 0 | 0 | 0 | 0 | 0 | 0 | 0 |
| 0 | NA | NA | NA | NA | NA | Gm7395 | | 0 | 0 | 0 | 0 | 0 | 0 | 0 | 0 | 0 | 0 | 0 |
| 0 | NA | NA | NA | NA | NA | Gm15625 | | 0 | 0 | 0 | 0 | 0 | 0 | 0 | 0 | 0 | 0 | 0 |
| 0 | NA | NA | NA | NA | NA | Gm17226 | | 0 | 0 | 0 | 0 | 0 | 0 | 0 | 0 | 0 | 0 | 0 |
| 0 | NA | NA | NA | NA | NA | Gm23407 | | 0 | 0 | 0 | 0 | 0 | 0 | 0 | 0 | 0 | 0 | 0 |
| 0 | NA | NA | NA | NA | NA | 4930563M21Ri | RIKEN cDNA 4930563M21 gene | 0 | 0 | 0 | 0 | 0 | 0 | 0 | 0 | 0 | 0 | 0 |
| 0 | NA | NA | NA | NA | NA | Gm29322 | | 0 | 0 | 0 | 0 | 0 | 0 | 0 | 0 | 0 | 0 | 0 |
| 0 | NA | NA | NA | NA | NA | Gm37842 | | 0 | 0 | 0 | 0 | 0 | 0 | 0 | 0 | 0 | 0 | 0 |
| 0 | NA | NA | NA | NA | NA | Gm47176 | | 0 | 0 | 0 | 0 | 0 | 0 | 0 | 0 | 0 | 0 | 0 |
| 0 | NA | NA | NA | NA | NA | Gm47178 | | 0 | 0 | 0 | 0 | 0 | 0 | 0 | 0 | 0 | 0 | 0 |
| 0 | NA | NA | NA | NA | NA | Odf3l1 | outer dense fiber of sperm tails 3-like 1 | 0 | 0 | 0 | 0 | 0 | 0 | 0 | 0 | 0 | 0 | 0 |
| 0 | NA | NA | NA | NA | NA | Gm47237 | | 0 | 0 | 0 | 0 | 0 | 0 | 0 | 0 | 0 | 0 | 0 |
| 0 | NA | NA | NA | NA | NA | 4930442G15Rik | | 0 | 0 | 0 | 0 | 0 | 0 | 0 | 0 | 0 | 0 | 0 |
| 0 | NA | NA | NA | NA | NA | 1700041C23Rik | | 0 | 0 | 0 | 0 | 0 | 0 | 0 | 0 | 0 | 0 | 0 |
| 0 | NA | NA | NA | NA | NA | Trcg1 | taste receptor cell gene 1 | 0 | 0 | 0 | 0 | 0 | 0 | 0 | 0 | 0 | 0 | 0 |
| 0 | NA | NA | NA | NA | NA | Gm18103 | | 0 | 0 | 0 | 0 | 0 | 0 | 0 | 0 | 0 | 0 | 0 |
| 0 | NA | NA | NA | NA | NA | Cyp1a2 | cytochrome P450, family 1, subfamily a, p | 0 | 0 | 0 | 0 | 0 | 0 | 0 | 0 | 0 | 0 | 0 |
| 0 | NA | NA | NA | NA | NA | Gm16131 | | 0 | 0 | 0 | 0 | 0 | 0 | 0 | 0 | 0 | 0 | 0 |
| 0 | NA | NA | NA | NA | NA | Gm47174 | | 0 | 0 | 0 | 0 | 0 | 0 | 0 | 0 | 0 | 0 | 0 |
| 0 | NA | NA | NA | NA | NA | 1600029O15Rik | | 0 | 0 | 0 | 0 | 0 | 0 | 0 | 0 | 0 | 0 | 0 |
| 0 | NA | NA | NA | NA | NA | Mir6385 | | 0 | 0 | 0 | 0 | 0 | 0 | 0 | 0 | 0 | 0 | 0 |
| 0 | NA | NA | NA | NA | NA | Tbc1d21 | TBC1 domain family, member 21 | 0 | 0 | 0 | 0 | 0 | 0 | 0 | 0 | 0 | 0 | 0 |
| 0 | NA | NA | NA | NA | NA | 1700072B07Rik | | 0 | 0 | 0 | 0 | 0 | 0 | 0 | 0 | 0 | 0 | 0 |
| 0 | NA | NA | NA | NA | NA | 4930461G14Rik | | 0 | 0 | 0 | 0 | 0 | 0 | 0 | 0 | 0 | 0 | 0 |
| 0 | NA | NA | NA | NA | NA | Gm47271 | | 0 | 0 | 0 | 0 | 0 | 0 | 0 | 0 | 0 | 0 | 0 |
| 0 | NA | NA | NA | NA | NA | Gm33180 | | 0 | 0 | 0 | 0 | 0 | 0 | 0 | 0 | 0 | 0 | 0 |
| 0 | NA | NA | NA | NA | NA | Nr2e3 | nuclear receptor subfamily 2, group E, me | 0 | 0 | 0 | 0 | 0 | 0 | 0 | 0 | 0 | 0 | 0 |
| 0 | NA | NA | NA | NA | NA | Gm15507 | | 0 | 0 | 0 | 0 | 0 | 0 | 0 | 0 | 0 | 0 | 0 |
| 0 | NA | NA | NA | NA | NA | 2010001M07Rik | | 0 | 0 | 0 | 0 | 0 | 0 | 0 | 0 | 0 | 0 | 0 |
| 0 | NA | NA | NA | NA | NA | Gm22099 | | 0 | 0 | 0 | 0 | 0 | 0 | 0 | 0 | 0 | 0 | 0 |
| 0 | NA | NA | NA | NA | NA | Gm47740 | | 0 | 0 | 0 | 0 | 0 | 0 | 0 | 0 | 0 | 0 | 0 |
| 0 | NA | NA | NA | NA | NA | 1700036A12Rik | | 0 | 0 | 0 | 0 | 0 | 0 | 0 | 0 | 0 | 0 | 0 |
| 0 | NA | NA | NA | NA | NA | Gm47923 | | 0 | 0 | 0 | 0 | 0 | 0 | 0 | 0 | 0 | 0 | 0 |
| 0 | NA | NA | NA | NA | NA | Gm5122 | | 0 | 0 | 0 | 0 | 0 | 0 | 0 | 0 | 0 | 0 | 0 |
| 0 | NA | NA | NA | NA | NA | Gm39348 | | 0 | 0 | 0 | 0 | 0 | 0 | 0 | 0 | 0 | 0 | 0 |
| 0 | NA | NA | NA | NA | NA | Gm34004 | | 0 | 0 | 0 | 0 | 0 | 0 | 0 | 0 | 0 | 0 | 0 |
| 0 | NA | NA | NA | NA | NA | Gm33914 | | 0 | 0 | 0 | 0 | 0 | 0 | 0 | 0 | 0 | 0 | 0 |
| 0 | NA | NA | NA | NA | NA | 4933433G08Rik | | 0 | 0 | 0 | 0 | 0 | 0 | 0 | 0 | 0 | 0 | 0 |
| 0 | NA | NA | NA | NA | NA | I730028E13Rik | | 0 | 0 | 0 | 0 | 0 | 0 | 0 | 0 | 0 | 0 | 0 |

| | | | | | | | | | | | | | | | | | |
|---|---|---|---|---|---|---|---|---|---|---|---|---|---|---|---|---|---|
| 0 | NA | NA | NA | NA | NA | Gm34105 | | 0 | 0 | 0 | 0 | 0 | 0 | 0 | 0 | 0 | 0 | 0 |
| 0 | NA | NA | NA | NA | NA | B930092H01Rik | | 0 | 0 | 0 | 0 | 0 | 0 | 0 | 0 | 0 | 0 | 0 |
| 0 | NA | NA | NA | NA | NA | Gm34322 | | 0 | 0 | 0 | 0 | 0 | 0 | 0 | 0 | 0 | 0 | 0 |
| 0 | NA | NA | NA | NA | NA | Gm34424 | | 0 | 0 | 0 | 0 | 0 | 0 | 0 | 0 | 0 | 0 | 0 |
| 0 | NA | NA | NA | NA | NA | Gm47241 | | 0 | 0 | 0 | 0 | 0 | 0 | 0 | 0 | 0 | 0 | 0 |
| 0 | NA | NA | NA | NA | NA | Gm19208 | | 0 | 0 | 0 | 0 | 0 | 0 | 0 | 0 | 0 | 0 | 0 |
| 0 | NA | NA | NA | NA | NA | Gm23221 | | 0 | 0 | 0 | 0 | 0 | 0 | 0 | 0 | 0 | 0 | 0 |
| 0 | NA | NA | NA | NA | NA | Mir5133 | | 0 | 0 | 0 | 0 | 0 | 0 | 0 | 0 | 0 | 0 | 0 |
| 0 | NA | NA | NA | NA | NA | Gm47262 | | 0 | 0 | 0 | 0 | 0 | 0 | 0 | 0 | 0 | 0 | 0 |
| 0 | NA | NA | NA | NA | NA | Gm25603 | | 0 | 0 | 0 | 0 | 0 | 0 | 0 | 0 | 0 | 0 | 0 |
| 0 | NA | NA | NA | NA | NA | Spesp1 | sperm equatorial segment protein 1 | 0 | 0 | 0 | 0 | 0 | 0 | 0 | 0 | 0 | 0 | 0 |
| 0 | NA | NA | NA | NA | NA | Gm35288 | | 0 | 0 | 0 | 0 | 0 | 0 | 0 | 0 | 0 | 0 | 0 |
| 0 | NA | NA | NA | NA | NA | n-R5s84 | | 0 | 0 | 0 | 0 | 0 | 0 | 0 | 0 | 0 | 0 | 0 |
| 0 | NA | NA | NA | NA | NA | Gm7653 | | 0 | 0 | 0 | 0 | 0 | 0 | 0 | 0 | 0 | 0 | 0 |
| 0 | NA | NA | NA | NA | NA | Gm47274 | | 0 | 0 | 0 | 0 | 0 | 0 | 0 | 0 | 0 | 0 | 0 |
| 0 | NA | NA | NA | NA | NA | Gm24526 | | 0 | 0 | 0 | 0 | 0 | 0 | 0 | 0 | 0 | 0 | 0 |
| 0 | NA | NA | NA | NA | NA | Gm48193 | | 0 | 0 | 0 | 0 | 0 | 0 | 0 | 0 | 0 | 0 | 0 |
| 0 | NA | NA | NA | NA | NA | Gm6082 | | 0 | 0 | 0 | 0 | 0 | 0 | 0 | 0 | 0 | 0 | 0 |
| 0 | NA | NA | NA | NA | NA | Gm25995 | | 0 | 0 | 0 | 0 | 0 | 0 | 0 | 0 | 0 | 0 | 0 |
| 0 | NA | NA | NA | NA | NA | Gm2553 | | 0 | 0 | 0 | 0 | 0 | 0 | 0 | 0 | 0 | 0 | 0 |
| 0 | NA | NA | NA | NA | NA | Gm36033 | | 0 | 0 | 0 | 0 | 0 | 0 | 0 | 0 | 0 | 0 | 0 |
| 0 | NA | NA | NA | NA | NA | 1700055C04Rik | | 0 | 0 | 0 | 0 | 0 | 0 | 0 | 0 | 0 | 0 | 0 |
| 0 | NA | NA | NA | NA | NA | Gm25606 | | 0 | 0 | 0 | 0 | 0 | 0 | 0 | 0 | 0 | 0 | 0 |
| 0 | NA | NA | NA | NA | NA | Scarletltr | | 0 | 0 | 0 | 0 | 0 | 0 | 0 | 0 | 0 | 0 | 0 |
| 0 | NA | NA | NA | NA | NA | Lctl | lactase-like | 0 | 0 | 0 | 0 | 0 | 0 | 0 | 0 | 0 | 0 | 0 |
| 0 | NA | NA | NA | NA | NA | Gm23344 | | 0 | 0 | 0 | 0 | 0 | 0 | 0 | 0 | 0 | 0 | 0 |
| 0 | NA | NA | NA | NA | NA | Snord16a | | 0 | 0 | 0 | 0 | 0 | 0 | 0 | 0 | 0 | 0 | 0 |
| 0 | NA | NA | NA | NA | NA | Gm22455 | | 0 | 0 | 0 | 0 | 0 | 0 | 0 | 0 | 0 | 0 | 0 |
| 0 | NA | NA | NA | NA | NA | Gm24106 | | 0 | 0 | 0 | 0 | 0 | 0 | 0 | 0 | 0 | 0 | 0 |
| 0 | NA | NA | NA | NA | NA | Gm47308 | | 0 | 0 | 0 | 0 | 0 | 0 | 0 | 0 | 0 | 0 | 0 |
| 0 | NA | NA | NA | NA | NA | Gm24347 | | 0 | 0 | 0 | 0 | 0 | 0 | 0 | 0 | 0 | 0 | 0 |
| 0 | NA | NA | NA | NA | NA | Gm25313 | | 0 | 0 | 0 | 0 | 0 | 0 | 0 | 0 | 0 | 0 | 0 |
| 0 | NA | NA | NA | NA | NA | Gm16218 | | 0 | 0 | 0 | 0 | 0 | 0 | 0 | 0 | 0 | 0 | 0 |
| 0 | NA | NA | NA | NA | NA | Gm39363 | | 0 | 0 | 0 | 0 | 0 | 0 | 0 | 0 | 0 | 0 | 0 |
| 0 | NA | NA | NA | NA | NA | Gm19219 | | 0 | 0 | 0 | 0 | 0 | 0 | 0 | 0 | 0 | 0 | 0 |
| 0 | NA | NA | NA | NA | NA | Gm47192 | | 0 | 0 | 0 | 0 | 0 | 0 | 0 | 0 | 0 | 0 | 0 |
| 0 | NA | NA | NA | NA | NA | Ankdd1a | ankyrin repeat and death domain contain | 0 | 0 | 0 | 0 | 0 | 0 | 0 | 0 | 0 | 0 | 0 |
| 0 | NA | NA | NA | NA | NA | n-R5s85 | | 0 | 0 | 0 | 0 | 0 | 0 | 0 | 0 | 0 | 0 | 0 |
| 0 | NA | NA | NA | NA | NA | Gm23248 | | 0 | 0 | 0 | 0 | 0 | 0 | 0 | 0 | 0 | 0 | 0 |
| 0 | NA | NA | NA | NA | NA | Gm22657 | | 0 | 0 | 0 | 0 | 0 | 0 | 0 | 0 | 0 | 0 | 0 |
| 0 | NA | NA | NA | NA | NA | Gm23428 | | 0 | 0 | 0 | 0 | 0 | 0 | 0 | 0 | 0 | 0 | 0 |
| 0 | NA | NA | NA | NA | NA | Gm15563 | | 0 | 0 | 0 | 0 | 0 | 0 | 0 | 0 | 0 | 0 | 0 |
| 0 | NA | NA | NA | NA | NA | Gm24225 | | 0 | 0 | 0 | 0 | 0 | 0 | 0 | 0 | 0 | 0 | 0 |
| 0 | NA | NA | NA | NA | NA | Gm10646 | | 0 | 0 | 0 | 0 | 0 | 0 | 0 | 0 | 0 | 0 | 0 |
| 0 | NA | NA | NA | NA | NA | Gm22145 | | 0 | 0 | 0 | 0 | 0 | 0 | 0 | 0 | 0 | 0 | 0 |
| 0 | NA | NA | NA | NA | NA | Gm19299 | | 0 | 0 | 0 | 0 | 0 | 0 | 0 | 0 | 0 | 0 | 0 |
| 0 | NA | NA | NA | NA | NA | Mir190a | | 0 | 0 | 0 | 0 | 0 | 0 | 0 | 0 | 0 | 0 | 0 |
| 0 | NA | NA | NA | NA | NA | Gm47047 | | 0 | 0 | 0 | 0 | 0 | 0 | 0 | 0 | 0 | 0 | 0 |
| 0 | NA | NA | NA | NA | NA | Gm47110 | | 0 | 0 | 0 | 0 | 0 | 0 | 0 | 0 | 0 | 0 | 0 |
| 0 | NA | NA | NA | NA | NA | Mir7241 | | 0 | 0 | 0 | 0 | 0 | 0 | 0 | 0 | 0 | 0 | 0 |
| 0 | NA | NA | NA | NA | NA | Mir7242 | | 0 | 0 | 0 | 0 | 0 | 0 | 0 | 0 | 0 | 0 | 0 |
| 0 | NA | NA | NA | NA | NA | Gm20745 | | 0 | 0 | 0 | 0 | 0 | 0 | 0 | 0 | 0 | 0 | 0 |
| 0 | NA | NA | NA | NA | NA | Gm28195 | | 0 | 0 | 0 | 0 | 0 | 0 | 0 | 0 | 0 | 0 | 0 |
| 0 | NA | NA | NA | NA | NA | Gm22382 | | 0 | 0 | 0 | 0 | 0 | 0 | 0 | 0 | 0 | 0 | 0 |
| 0 | NA | NA | NA | NA | NA | Gm19987 | | 0 | 0 | 0 | 0 | 0 | 0 | 0 | 0 | 0 | 0 | 0 |
| 0 | NA | NA | NA | NA | NA | Gm22962 | | 0 | 0 | 0 | 0 | 0 | 0 | 0 | 0 | 0 | 0 | 0 |
| 0 | NA | NA | NA | NA | NA | Gm7802 | | 0 | 0 | 0 | 0 | 0 | 0 | 0 | 0 | 0 | 0 | 0 |
| 0 | NA | NA | NA | NA | NA | Gm18587 | | 0 | 0 | 0 | 0 | 0 | 0 | 0 | 0 | 0 | 0 | 0 |
| 0 | NA | NA | NA | NA | NA | Gm16144 | | 0 | 0 | 0 | 0 | 0 | 0 | 0 | 0 | 0 | 0 | 0 |
| 0 | NA | NA | NA | NA | NA | Gm47203 | | 0 | 0 | 0 | 0 | 0 | 0 | 0 | 0 | 0 | 0 | 0 |

| | | | | | | | | | | | | | | | | | | |
|---|---|---|---|---|---|---|---|---|---|---|---|---|---|---|---|---|---|---|
| 0 | NA | NA | NA | NA | NA | Foxb1 | forkhead box B1 | 0 | 0 | 0 | 0 | 0 | 0 | 0 | 0 | 0 | 0 | 0 |
| 0 | NA | NA | NA | NA | NA | B230323A14Rik | | 0 | 0 | 0 | 0 | 0 | 0 | 0 | 0 | 0 | 0 | 0 |
| 0 | NA | NA | NA | NA | NA | Gm18745 | | 0 | 0 | 0 | 0 | 0 | 0 | 0 | 0 | 0 | 0 | 0 |
| 0 | NA | NA | NA | NA | NA | Gcnt3 | glucosaminyl (N-acetyl) transferase 3, mu | 0 | 0 | 0 | 0 | 0 | 0 | 0 | 0 | 0 | 0 | 0 |
| 0 | NA | NA | NA | NA | NA | Gm47233 | | 0 | 0 | 0 | 0 | 0 | 0 | 0 | 0 | 0 | 0 | 0 |
| 0 | NA | NA | NA | NA | NA | 2210414F02Rik | | 0 | 0 | 0 | 0 | 0 | 0 | 0 | 0 | 0 | 0 | 0 |
| 0 | NA | NA | NA | NA | NA | Gm24257 | | 0 | 0 | 0 | 0 | 0 | 0 | 0 | 0 | 0 | 0 | 0 |
| 0 | NA | NA | NA | NA | NA | Mir5626 | | 0 | 0 | 0 | 0 | 0 | 0 | 0 | 0 | 0 | 0 | 0 |
| 0 | NA | NA | NA | NA | NA | Gm24615 | | 0 | 0 | 0 | 0 | 0 | 0 | 0 | 0 | 0 | 0 | 0 |
| 0 | NA | NA | NA | NA | NA | Gm23588 | | 0 | 0 | 0 | 0 | 0 | 0 | 0 | 0 | 0 | 0 | 0 |
| 0 | NA | NA | NA | NA | NA | Gm23730 | | 0 | 0 | 0 | 0 | 0 | 0 | 0 | 0 | 0 | 0 | 0 |
| 0 | NA | NA | NA | NA | NA | Gm22229 | | 0 | 0 | 0 | 0 | 0 | 0 | 0 | 0 | 0 | 0 | 0 |
| 0 | NA | NA | NA | NA | NA | Gm25149 | | 0 | 0 | 0 | 0 | 0 | 0 | 0 | 0 | 0 | 0 | 0 |
| 0 | NA | NA | NA | NA | NA | Gm32017 | | 0 | 0 | 0 | 0 | 0 | 0 | 0 | 0 | 0 | 0 | 0 |
| 0 | NA | NA | NA | NA | NA | Gm47268 | | 0 | 0 | 0 | 0 | 0 | 0 | 0 | 0 | 0 | 0 | 0 |
| 0 | NA | NA | NA | NA | NA | Gm32511 | | 0 | 0 | 0 | 0 | 0 | 0 | 0 | 0 | 0 | 0 | 0 |
| 0 | NA | NA | NA | NA | NA | Gm48124 | | 0 | 0 | 0 | 0 | 0 | 0 | 0 | 0 | 0 | 0 | 0 |
| 0 | NA | NA | NA | NA | NA | Gm6018 | | 0 | 0 | 0 | 0 | 0 | 0 | 0 | 0 | 0 | 0 | 0 |
| 0 | NA | NA | NA | NA | NA | Gm27220 | | 0 | 0 | 0 | 0 | 0 | 0 | 0 | 0 | 0 | 0 | 0 |
| 0 | NA | NA | NA | NA | NA | Gm38111 | | 0 | 0 | 0 | 0 | 0 | 0 | 0 | 0 | 0 | 0 | 0 |
| 0 | NA | NA | NA | NA | NA | Gm27152 | | 0 | 0 | 0 | 0 | 0 | 0 | 0 | 0 | 0 | 0 | 0 |
| 0 | NA | NA | NA | NA | NA | Gm37023 | | 0 | 0 | 0 | 0 | 0 | 0 | 0 | 0 | 0 | 0 | 0 |
| 0 | NA | NA | NA | NA | NA | Gm27450 | | 0 | 0 | 0 | 0 | 0 | 0 | 0 | 0 | 0 | 0 | 0 |
| 0 | NA | NA | NA | NA | NA | Gm25045 | | 0 | 0 | 0 | 0 | 0 | 0 | 0 | 0 | 0 | 0 | 0 |
| 0 | NA | NA | NA | NA | NA | Gm7863 | | 0 | 0 | 0 | 0 | 0 | 0 | 0 | 0 | 0 | 0 | 0 |
| 0 | NA | NA | NA | NA | NA | Gm27159 | | 0 | 0 | 0 | 0 | 0 | 0 | 0 | 0 | 0 | 0 | 0 |
| 0 | NA | NA | NA | NA | NA | Gm27188 | | 0 | 0 | 0 | 0 | 0 | 0 | 0 | 0 | 0 | 0 | 0 |
| 0 | NA | NA | NA | NA | NA | Gm25163 | | 0 | 0 | 0 | 0 | 0 | 0 | 0 | 0 | 0 | 0 | 0 |
| 0 | NA | NA | NA | NA | NA | Gm37150 | | 0 | 0 | 0 | 0 | 0 | 0 | 0 | 0 | 0 | 0 | 0 |
| 0 | NA | NA | NA | NA | NA | Gm27230 | | 0 | 0 | 0 | 0 | 0 | 0 | 0 | 0 | 0 | 0 | 0 |
| 0 | NA | NA | NA | NA | NA | Gm27255 | | 0 | 0 | 0 | 0 | 0 | 0 | 0 | 0 | 0 | 0 | 0 |
| 0 | NA | NA | NA | NA | NA | Gm27254 | | 0 | 0 | 0 | 0 | 0 | 0 | 0 | 0 | 0 | 0 | 0 |
| 0 | NA | NA | NA | NA | NA | Gm28010 | | 0 | 0 | 0 | 0 | 0 | 0 | 0 | 0 | 0 | 0 | 0 |
| 0 | NA | NA | NA | NA | NA | Gm18014 | | 0 | 0 | 0 | 0 | 0 | 0 | 0 | 0 | 0 | 0 | 0 |
| 0 | NA | NA | NA | NA | NA | Gm27241 | | 0 | 0 | 0 | 0 | 0 | 0 | 0 | 0 | 0 | 0 | 0 |
| 0 | NA | NA | NA | NA | NA | Gm27232 | | 0 | 0 | 0 | 0 | 0 | 0 | 0 | 0 | 0 | 0 | 0 |
| 0 | NA | NA | NA | NA | NA | Gm23057 | | 0 | 0 | 0 | 0 | 0 | 0 | 0 | 0 | 0 | 0 | 0 |
| 0 | NA | NA | NA | NA | NA | Gm27204 | | 0 | 0 | 0 | 0 | 0 | 0 | 0 | 0 | 0 | 0 | 0 |
| 0 | NA | NA | NA | NA | NA | Gm27203 | | 0 | 0 | 0 | 0 | 0 | 0 | 0 | 0 | 0 | 0 | 0 |
| 0 | NA | NA | NA | NA | NA | A730062M13Rik | | 0 | 0 | 0 | 0 | 0 | 0 | 0 | 0 | 0 | 0 | 0 |
| 0 | NA | NA | NA | NA | NA | Gm37234 | | 0 | 0 | 0 | 0 | 0 | 0 | 0 | 0 | 0 | 0 | 0 |
| 0 | NA | NA | NA | NA | NA | Gm44503 | | 0 | 0 | 0 | 0 | 0 | 0 | 0 | 0 | 0 | 0 | 0 |
| 0 | NA | NA | NA | NA | NA | Gm20509 | | 0 | 0 | 0 | 0 | 0 | 0 | 0 | 0 | 0 | 0 | 0 |
| 0 | NA | NA | NA | NA | NA | Gm27211 | | 0 | 0 | 0 | 0 | 0 | 0 | 0 | 0 | 0 | 0 | 0 |
| 0 | NA | NA | NA | NA | NA | Gm37611 | | 0 | 0 | 0 | 0 | 0 | 0 | 0 | 0 | 0 | 0 | 0 |
| 0 | NA | NA | NA | NA | NA | Gm27148 | | 0 | 0 | 0 | 0 | 0 | 0 | 0 | 0 | 0 | 0 | 0 |
| 0 | NA | NA | NA | NA | NA | Gm25105 | | 0 | 0 | 0 | 0 | 0 | 0 | 0 | 0 | 0 | 0 | 0 |
| 0 | NA | NA | NA | NA | NA | Gm37326 | | 0 | 0 | 0 | 0 | 0 | 0 | 0 | 0 | 0 | 0 | 0 |
| 0 | NA | NA | NA | NA | NA | n-R5s86 | | 0 | 0 | 0 | 0 | 0 | 0 | 0 | 0 | 0 | 0 | 0 |
| 0 | NA | NA | NA | NA | NA | Gm24141 | | 0 | 0 | 0 | 0 | 0 | 0 | 0 | 0 | 0 | 0 | 0 |
| 0 | NA | NA | NA | NA | NA | Nr1h2-ps | | 0 | 0 | 0 | 0 | 0 | 0 | 0 | 0 | 0 | 0 | 0 |
| 0 | NA | NA | NA | NA | NA | Gm28622 | | 0 | 0 | 0 | 0 | 0 | 0 | 0 | 0 | 0 | 0 | 0 |
| 0 | NA | NA | NA | NA | NA | Gm27189 | | 0 | 0 | 0 | 0 | 0 | 0 | 0 | 0 | 0 | 0 | 0 |
| 0 | NA | NA | NA | NA | NA | Gm27233 | | 0 | 0 | 0 | 0 | 0 | 0 | 0 | 0 | 0 | 0 | 0 |
| 0 | NA | NA | NA | NA | NA | Gm22315 | | 0 | 0 | 0 | 0 | 0 | 0 | 0 | 0 | 0 | 0 | 0 |
| 0 | NA | NA | NA | NA | NA | Gm20649 | | 0 | 0 | 0 | 0 | 0 | 0 | 0 | 0 | 0 | 0 | 0 |
| 0 | NA | NA | NA | NA | NA | Gm28182 | | 0 | 0 | 0 | 0 | 0 | 0 | 0 | 0 | 0 | 0 | 0 |
| 0 | NA | NA | NA | NA | NA | Gm16353 | | 0 | 0 | 0 | 0 | 0 | 0 | 0 | 0 | 0 | 0 | 0 |
| 0 | NA | NA | NA | NA | NA | Gm48677 | | 0 | 0 | 0 | 0 | 0 | 0 | 0 | 0 | 0 | 0 | 0 |
| 0 | NA | NA | NA | NA | NA | Gm18015 | | 0 | 0 | 0 | 0 | 0 | 0 | 0 | 0 | 0 | 0 | 0 |

| | | | | | | | | | | | | | | | | | | |
|---|---|---|---|---|---|---|---|---|---|---|---|---|---|---|---|---|---|---|
| 0 | NA | NA | NA | NA | NA | Gm5366 | | 0 | 0 | 0 | 0 | 0 | 0 | 0 | 0 | 0 | 0 | 0 |
| 0 | NA | NA | NA | NA | NA | Gm48789 | | 0 | 0 | 0 | 0 | 0 | 0 | 0 | 0 | 0 | 0 | 0 |
| 0 | NA | NA | NA | NA | NA | Gm47723 | | 0 | 0 | 0 | 0 | 0 | 0 | 0 | 0 | 0 | 0 | 0 |
| 0 | NA | NA | NA | NA | NA | Gm44355 | | 0 | 0 | 0 | 0 | 0 | 0 | 0 | 0 | 0 | 0 | 0 |
| 0 | NA | NA | NA | NA | NA | Gm34005 | | 0 | 0 | 0 | 0 | 0 | 0 | 0 | 0 | 0 | 0 | 0 |
| 0 | NA | NA | NA | NA | NA | Gm25300 | | 0 | 0 | 0 | 0 | 0 | 0 | 0 | 0 | 0 | 0 | 0 |
| 0 | NA | NA | NA | NA | NA | Gfral | GDNF family receptor alpha like | 0 | 0 | 0 | 0 | 0 | 0 | 0 | 0 | 0 | 0 | 0 |
| 0 | NA | NA | NA | NA | NA | Hcrtr2 | hypocretin (orexin) receptor 2 | 0 | 0 | 0 | 0 | 0 | 0 | 0 | 0 | 0 | 0 | 0 |
| 0 | NA | NA | NA | NA | NA | Gm27234 | | 0 | 0 | 0 | 0 | 0 | 0 | 0 | 0 | 0 | 0 | 0 |
| 0 | NA | NA | NA | NA | NA | Gm27156 | | 0 | 0 | 0 | 0 | 0 | 0 | 0 | 0 | 0 | 0 | 0 |
| 0 | NA | NA | NA | NA | NA | Gm27245 | | 0 | 0 | 0 | 0 | 0 | 0 | 0 | 0 | 0 | 0 | 0 |
| 0 | NA | NA | NA | NA | NA | Gm22938 | | 0 | 0 | 0 | 0 | 0 | 0 | 0 | 0 | 0 | 0 | 0 |
| 0 | NA | NA | NA | NA | NA | Gm34158 | | 0 | 0 | 0 | 0 | 0 | 0 | 0 | 0 | 0 | 0 | 0 |
| 0 | NA | NA | NA | NA | NA | Gm27155 | | 0 | 0 | 0 | 0 | 0 | 0 | 0 | 0 | 0 | 0 | 0 |
| 0 | NA | NA | NA | NA | NA | Gm23791 | | 0 | 0 | 0 | 0 | 0 | 0 | 0 | 0 | 0 | 0 | 0 |
| 0 | NA | NA | NA | NA | NA | Gm47800 | | 0 | 0 | 0 | 0 | 0 | 0 | 0 | 0 | 0 | 0 | 0 |
| 0 | NA | NA | NA | NA | NA | Gm18486 | | 0 | 0 | 0 | 0 | 0 | 0 | 0 | 0 | 0 | 0 | 0 |
| 0 | NA | NA | NA | NA | NA | Gm27158 | | 0 | 0 | 0 | 0 | 0 | 0 | 0 | 0 | 0 | 0 | 0 |
| 0 | NA | NA | NA | NA | NA | Gm19541 | | 0 | 0 | 0 | 0 | 0 | 0 | 0 | 0 | 0 | 0 | 0 |
| 0 | NA | NA | NA | NA | NA | Gm5746 | | 0 | 0 | 0 | 0 | 0 | 0 | 0 | 0 | 0 | 0 | 0 |
| 0 | NA | NA | NA | NA | NA | Gm27210 | | 0 | 0 | 0 | 0 | 0 | 0 | 0 | 0 | 0 | 0 | 0 |
| 0 | NA | NA | NA | NA | NA | Gm38254 | | 0 | 0 | 0 | 0 | 0 | 0 | 0 | 0 | 0 | 0 | 0 |
| 0 | NA | NA | NA | NA | NA | 4930590H14Rik | | 0 | 0 | 0 | 0 | 0 | 0 | 0 | 0 | 0 | 0 | 0 |
| 0 | NA | NA | NA | NA | NA | Gm25250 | | 0 | 0 | 0 | 0 | 0 | 0 | 0 | 0 | 0 | 0 | 0 |
| 0 | NA | NA | NA | NA | NA | Gm47790 | | 0 | 0 | 0 | 0 | 0 | 0 | 0 | 0 | 0 | 0 | 0 |
| 0 | NA | NA | NA | NA | NA | Gm47789 | | 0 | 0 | 0 | 0 | 0 | 0 | 0 | 0 | 0 | 0 | 0 |
| 0 | NA | NA | NA | NA | NA | Gm47795 | | 0 | 0 | 0 | 0 | 0 | 0 | 0 | 0 | 0 | 0 | 0 |
| 0 | NA | NA | NA | NA | NA | Gm47818 | | 0 | 0 | 0 | 0 | 0 | 0 | 0 | 0 | 0 | 0 | 0 |
| 0 | NA | NA | NA | NA | NA | Gm19572 | | 0 | 0 | 0 | 0 | 0 | 0 | 0 | 0 | 0 | 0 | 0 |
| 0 | NA | NA | NA | NA | NA | Gm23609 | | 0 | 0 | 0 | 0 | 0 | 0 | 0 | 0 | 0 | 0 | 0 |
| 0 | NA | NA | NA | NA | NA | Gm39375 | | 0 | 0 | 0 | 0 | 0 | 0 | 0 | 0 | 0 | 0 | 0 |
| 0 | NA | NA | NA | NA | NA | Gm47829 | | 0 | 0 | 0 | 0 | 0 | 0 | 0 | 0 | 0 | 0 | 0 |
| 0 | NA | NA | NA | NA | NA | Gcm1 | glial cells missing homolog 1 | 0 | 0 | 0 | 0 | 0 | 0 | 0 | 0 | 0 | 0 | 0 |
| 0 | NA | NA | NA | NA | NA | Gm3126 | | 0 | 0 | 0 | 0 | 0 | 0 | 0 | 0 | 0 | 0 | 0 |
| 0 | NA | NA | NA | NA | NA | Gsta1 | glutathione S-transferase, alpha 1 (Ya) | 0 | 0 | 0 | 0 | 0 | 0 | 0 | 0 | 0 | 0 | 0 |
| 0 | NA | NA | NA | NA | NA | Dppa5b | | 0 | 0 | 0 | 0 | 0 | 0 | 0 | 0 | 0 | 0 | 0 |
| 0 | NA | NA | NA | NA | NA | Gm3776 | predicted gene 3776 | 0 | 0 | 0 | 0 | 0 | 0 | 0 | 0 | 0 | 0 | 0 |
| 0 | NA | NA | NA | NA | NA | Dppa5c | | 0 | 0 | 0 | 0 | 0 | 0 | 0 | 0 | 0 | 0 | 0 |
| 0 | NA | NA | NA | NA | NA | Gm10639 | predicted gene 10639 | 0 | 0 | 0 | 0 | 0 | 0 | 0 | 0 | 0 | 0 | 0 |
| 0 | NA | NA | NA | NA | NA | Omt2a | oocyte maturation, alpha | 0 | 0 | 0 | 0 | 0 | 0 | 0 | 0 | 0 | 0 | 0 |
| 0 | NA | NA | NA | NA | NA | Gm8074 | | 0 | 0 | 0 | 0 | 0 | 0 | 0 | 0 | 0 | 0 | 0 |
| 0 | NA | NA | NA | NA | NA | Omt2b | oocyte maturation, beta | 0 | 0 | 0 | 0 | 0 | 0 | 0 | 0 | 0 | 0 | 0 |
| 0 | NA | NA | NA | NA | NA | Gsta2 | glutathione S-transferase, alpha 2 (Yc2) | 0 | 0 | 0 | 0 | 0 | 0 | 0 | 0 | 0 | 0 | 0 |
| 0 | NA | NA | NA | NA | NA | 4930542C12Rik | | 0 | 0 | 0 | 0 | 0 | 0 | 0 | 0 | 0 | 0 | 0 |
| 0 | NA | NA | NA | NA | NA | Gm26377 | | 0 | 0 | 0 | 0 | 0 | 0 | 0 | 0 | 0 | 0 | 0 |
| 0 | NA | NA | NA | NA | NA | 4921509A06Rik | | 0 | 0 | 0 | 0 | 0 | 0 | 0 | 0 | 0 | 0 | 0 |
| 0 | NA | NA | NA | NA | NA | Gm47497 | | 0 | 0 | 0 | 0 | 0 | 0 | 0 | 0 | 0 | 0 | 0 |
| 0 | NA | NA | NA | NA | NA | Gm46138 | | 0 | 0 | 0 | 0 | 0 | 0 | 0 | 0 | 0 | 0 | 0 |
| 0 | NA | NA | NA | NA | NA | Gm47498 | | 0 | 0 | 0 | 0 | 0 | 0 | 0 | 0 | 0 | 0 | 0 |
| 0 | NA | NA | NA | NA | NA | Gm18863 | | 0 | 0 | 0 | 0 | 0 | 0 | 0 | 0 | 0 | 0 | 0 |
| 0 | NA | NA | NA | NA | NA | Gm35024 | | 0 | 0 | 0 | 0 | 0 | 0 | 0 | 0 | 0 | 0 | 0 |
| 0 | NA | NA | NA | NA | NA | Gm47499 | | 0 | 0 | 0 | 0 | 0 | 0 | 0 | 0 | 0 | 0 | 0 |
| 0 | NA | NA | NA | NA | NA | Gm47501 | | 0 | 0 | 0 | 0 | 0 | 0 | 0 | 0 | 0 | 0 | 0 |
| 0 | NA | NA | NA | NA | NA | Gm28620 | | 0 | 0 | 0 | 0 | 0 | 0 | 0 | 0 | 0 | 0 | 0 |
| 0 | NA | NA | NA | NA | NA | Gm47500 | | 0 | 0 | 0 | 0 | 0 | 0 | 0 | 0 | 0 | 0 | 0 |
| 0 | NA | NA | NA | NA | NA | Gm49264 | | 0 | 0 | 0 | 0 | 0 | 0 | 0 | 0 | 0 | 0 | 0 |
| 0 | NA | NA | NA | NA | NA | Aldoa-ps3 | | 0 | 0 | 0 | 0 | 0 | 0 | 0 | 0 | 0 | 0 | 0 |
| 0 | NA | NA | NA | NA | NA | Gm39377 | | 0 | 0 | 0 | 0 | 0 | 0 | 0 | 0 | 0 | 0 | 0 |
| 0 | NA | NA | NA | NA | NA | Gm35262 | | 0 | 0 | 0 | 0 | 0 | 0 | 0 | 0 | 0 | 0 | 0 |
| 0 | NA | NA | NA | NA | NA | Gm10635 | | 0 | 0 | 0 | 0 | 0 | 0 | 0 | 0 | 0 | 0 | 0 |

| | | | | | | | | | | | | | | | | | | |
|---|---|---|---|---|---|---|---|---|---|---|---|---|---|---|---|---|---|---|
| 0 | NA | NA | NA | NA | NA | Gm8125 | | 0 | 0 | 0 | 0 | 0 | 0 | 0 | 0 | 0 | 0 | 0 |
| 0 | NA | NA | NA | NA | NA | 4930562D21Rik | | 0 | 0 | 0 | 0 | 0 | 0 | 0 | 0 | 0 | 0 | 0 |
| 0 | NA | NA | NA | NA | NA | Gm20477 | | 0 | 0 | 0 | 0 | 0 | 0 | 0 | 0 | 0 | 0 | 0 |
| 0 | NA | NA | NA | NA | NA | Gm25433 | | 0 | 0 | 0 | 0 | 0 | 0 | 0 | 0 | 0 | 0 | 0 |
| 0 | NA | NA | NA | NA | NA | Gm46146 | | 0 | 0 | 0 | 0 | 0 | 0 | 0 | 0 | 0 | 0 | 0 |
| 0 | NA | NA | NA | NA | NA | Gm20648 | | 0 | 0 | 0 | 0 | 0 | 0 | 0 | 0 | 0 | 0 | 0 |
| 0 | NA | NA | NA | NA | NA | Gm27227 | | 0 | 0 | 0 | 0 | 0 | 0 | 0 | 0 | 0 | 0 | 0 |
| 0 | NA | NA | NA | NA | NA | Gm18292 | | 0 | 0 | 0 | 0 | 0 | 0 | 0 | 0 | 0 | 0 | 0 |
| 0 | NA | NA | NA | NA | NA | Gm48387 | | 0 | 0 | 0 | 0 | 0 | 0 | 0 | 0 | 0 | 0 | 0 |
| 0 | NA | NA | NA | NA | NA | Gm39380 | | 0 | 0 | 0 | 0 | 0 | 0 | 0 | 0 | 0 | 0 | 0 |
| 0 | NA | NA | NA | NA | NA | Gm27224 | | 0 | 0 | 0 | 0 | 0 | 0 | 0 | 0 | 0 | 0 | 0 |
| 0 | NA | NA | NA | NA | NA | Gm27225 | | 0 | 0 | 0 | 0 | 0 | 0 | 0 | 0 | 0 | 0 | 0 |
| 0 | NA | NA | NA | NA | NA | Gm26713 | | 0 | 0 | 0 | 0 | 0 | 0 | 0 | 0 | 0 | 0 | 0 |
| 0 | NA | NA | NA | NA | NA | Gm8151 | | 0 | 0 | 0 | 0 | 0 | 0 | 0 | 0 | 0 | 0 | 0 |
| 0 | NA | NA | NA | NA | NA | Gm27226 | | 0 | 0 | 0 | 0 | 0 | 0 | 0 | 0 | 0 | 0 | 0 |
| 0 | NA | NA | NA | NA | NA | Gm26907 | | 0 | 0 | 0 | 0 | 0 | 0 | 0 | 0 | 0 | 0 | 0 |
| 0 | NA | NA | NA | NA | NA | Gm37837 | | 0 | 0 | 0 | 0 | 0 | 0 | 0 | 0 | 0 | 0 | 0 |
| 0 | NA | NA | NA | NA | NA | Gm19228 | | 0 | 0 | 0 | 0 | 0 | 0 | 0 | 0 | 0 | 0 | 0 |
| 0 | NA | NA | NA | NA | NA | Gm23748 | | 0 | 0 | 0 | 0 | 0 | 0 | 0 | 0 | 0 | 0 | 0 |
| 0 | NA | NA | NA | NA | NA | Gm35548 | | 0 | 0 | 0 | 0 | 0 | 0 | 0 | 0 | 0 | 0 | 0 |
| 0 | NA | NA | NA | NA | NA | 1700063H06Rik | | 0 | 0 | 0 | 0 | 0 | 0 | 0 | 0 | 0 | 0 | 0 |
| 0 | NA | NA | NA | NA | NA | Gm22836 | | 0 | 0 | 0 | 0 | 0 | 0 | 0 | 0 | 0 | 0 | 0 |
| 0 | NA | NA | NA | NA | NA | Gm24393 | | 0 | 0 | 0 | 0 | 0 | 0 | 0 | 0 | 0 | 0 | 0 |
| 0 | NA | NA | NA | NA | NA | Gm27249 | | 0 | 0 | 0 | 0 | 0 | 0 | 0 | 0 | 0 | 0 | 0 |
| 0 | NA | NA | NA | NA | NA | Gm27248 | | 0 | 0 | 0 | 0 | 0 | 0 | 0 | 0 | 0 | 0 | 0 |
| 0 | NA | NA | NA | NA | NA | Gm38398 | | 0 | 0 | 0 | 0 | 0 | 0 | 0 | 0 | 0 | 0 | 0 |
| 0 | NA | NA | NA | NA | NA | Gm27216 | | 0 | 0 | 0 | 0 | 0 | 0 | 0 | 0 | 0 | 0 | 0 |
| 0 | NA | NA | NA | NA | NA | Gm28552 | | 0 | 0 | 0 | 0 | 0 | 0 | 0 | 0 | 0 | 0 | 0 |
| 0 | NA | NA | NA | NA | NA | Gm28477 | | 0 | 0 | 0 | 0 | 0 | 0 | 0 | 0 | 0 | 0 | 0 |
| 0 | NA | NA | NA | NA | NA | Gm29163 | | 0 | 0 | 0 | 0 | 0 | 0 | 0 | 0 | 0 | 0 | 0 |
| 0 | NA | NA | NA | NA | NA | Gm46124 | | 0 | 0 | 0 | 0 | 0 | 0 | 0 | 0 | 0 | 0 | 0 |
| 0 | NA | NA | NA | NA | NA | Gm26146 | | 0 | 0 | 0 | 0 | 0 | 0 | 0 | 0 | 0 | 0 | 0 |
| 0 | NA | NA | NA | NA | NA | Gm18487 | | 0 | 0 | 0 | 0 | 0 | 0 | 0 | 0 | 0 | 0 | 0 |
| 0 | NA | NA | NA | NA | NA | Gm26126 | | 0 | 0 | 0 | 0 | 0 | 0 | 0 | 0 | 0 | 0 | 0 |
| 0 | NA | NA | NA | NA | NA | Gm48811 | | 0 | 0 | 0 | 0 | 0 | 0 | 0 | 0 | 0 | 0 | 0 |
| 0 | NA | NA | NA | NA | NA | 4930554C24Rik | | 0 | 0 | 0 | 0 | 0 | 0 | 0 | 0 | 0 | 0 | 0 |
| 0 | NA | NA | NA | NA | NA | Gm48820 | | 0 | 0 | 0 | 0 | 0 | 0 | 0 | 0 | 0 | 0 | 0 |
| 0 | NA | NA | NA | NA | NA | Gm28625 | | 0 | 0 | 0 | 0 | 0 | 0 | 0 | 0 | 0 | 0 | 0 |
| 0 | NA | NA | NA | NA | NA | Gm48821 | | 0 | 0 | 0 | 0 | 0 | 0 | 0 | 0 | 0 | 0 | 0 |
| 0 | NA | NA | NA | NA | NA | Gm28070 | | 0 | 0 | 0 | 0 | 0 | 0 | 0 | 0 | 0 | 0 | 0 |
| 0 | NA | NA | NA | NA | NA | Gm48830 | | 0 | 0 | 0 | 0 | 0 | 0 | 0 | 0 | 0 | 0 | 0 |
| 0 | NA | NA | NA | NA | NA | Gm48831 | | 0 | 0 | 0 | 0 | 0 | 0 | 0 | 0 | 0 | 0 | 0 |
| 0 | NA | NA | NA | NA | NA | Gm11114 | | 0 | 0 | 0 | 0 | 0 | 0 | 0 | 0 | 0 | 0 | 0 |
| 0 | NA | NA | NA | NA | NA | Gm48832 | | 0 | 0 | 0 | 0 | 0 | 0 | 0 | 0 | 0 | 0 | 0 |
| 0 | NA | NA | NA | NA | NA | Gm36927 | | 0 | 0 | 0 | 0 | 0 | 0 | 0 | 0 | 0 | 0 | 0 |
| 0 | NA | NA | NA | NA | NA | Gm48833 | | 0 | 0 | 0 | 0 | 0 | 0 | 0 | 0 | 0 | 0 | 0 |
| 0 | NA | NA | NA | NA | NA | Gm25125 | | 0 | 0 | 0 | 0 | 0 | 0 | 0 | 0 | 0 | 0 | 0 |
| 0 | NA | NA | NA | NA | NA | Gm48834 | | 0 | 0 | 0 | 0 | 0 | 0 | 0 | 0 | 0 | 0 | 0 |
| 0 | NA | NA | NA | NA | NA | Gm48842 | | 0 | 0 | 0 | 0 | 0 | 0 | 0 | 0 | 0 | 0 | 0 |
| 0 | NA | NA | NA | NA | NA | Gm22830 | | 0 | 0 | 0 | 0 | 0 | 0 | 0 | 0 | 0 | 0 | 0 |
| 0 | NA | NA | NA | NA | NA | Gm18766 | | 0 | 0 | 0 | 0 | 0 | 0 | 0 | 0 | 0 | 0 | 0 |
| 0 | NA | NA | NA | NA | NA | Gm47604 | | 0 | 0 | 0 | 0 | 0 | 0 | 0 | 0 | 0 | 0 | 0 |
| 0 | NA | NA | NA | NA | NA | Gm18016 | | 0 | 0 | 0 | 0 | 0 | 0 | 0 | 0 | 0 | 0 | 0 |
| 0 | NA | NA | NA | NA | NA | Gm38230 | | 0 | 0 | 0 | 0 | 0 | 0 | 0 | 0 | 0 | 0 | 0 |
| 0 | NA | NA | NA | NA | NA | Gm25949 | | 0 | 0 | 0 | 0 | 0 | 0 | 0 | 0 | 0 | 0 | 0 |
| 0 | NA | NA | NA | NA | NA | Gm22138 | | 0 | 0 | 0 | 0 | 0 | 0 | 0 | 0 | 0 | 0 | 0 |
| 0 | NA | NA | NA | NA | NA | Gm36539 | | 0 | 0 | 0 | 0 | 0 | 0 | 0 | 0 | 0 | 0 | 0 |
| 0 | NA | NA | NA | NA | NA | Ripply2 | ripply transcriptional repressor 2 | 0 | 0 | 0 | 0 | 0 | 0 | 0 | 0 | 0 | 0 | 0 |
| 0 | NA | NA | NA | NA | NA | Gm8282 | | 0 | 0 | 0 | 0 | 0 | 0 | 0 | 0 | 0 | 0 | 0 |
| 0 | NA | NA | NA | NA | NA | Gm9540 | | 0 | 0 | 0 | 0 | 0 | 0 | 0 | 0 | 0 | 0 | 0 |

| | | | | | | | | | | | | | | | | | | |
|---|---|---|---|---|---|---|---|---|---|---|---|---|---|---|---|---|---|---|
| 0 | NA | NA | NA | NA | NA | Gm37375 | | 0 | 0 | 0 | 0 | 0 | 0 | 0 | 0 | 0 | 0 | 0 |
| 0 | NA | NA | NA | NA | NA | D030062O11Rik | | 0 | 0 | 0 | 0 | 0 | 0 | 0 | 0 | 0 | 0 | 0 |
| 0 | NA | NA | NA | NA | NA | Gm37374 | | 0 | 0 | 0 | 0 | 0 | 0 | 0 | 0 | 0 | 0 | 0 |
| 0 | NA | NA | NA | NA | NA | Gm9541 | | 0 | 0 | 0 | 0 | 0 | 0 | 0 | 0 | 0 | 0 | 0 |
| 0 | NA | NA | NA | NA | NA | Gm25528 | | 0 | 0 | 0 | 0 | 0 | 0 | 0 | 0 | 0 | 0 | 0 |
| 0 | NA | NA | NA | NA | NA | Gm23176 | | 0 | 0 | 0 | 0 | 0 | 0 | 0 | 0 | 0 | 0 | 0 |
| 0 | NA | NA | NA | NA | NA | Gm28877 | | 0 | 0 | 0 | 0 | 0 | 0 | 0 | 0 | 0 | 0 | 0 |
| 0 | NA | NA | NA | NA | NA | Gm2324 | | 0 | 0 | 0 | 0 | 0 | 0 | 0 | 0 | 0 | 0 | 0 |
| 0 | NA | NA | NA | NA | NA | Gm5368 | | 0 | 0 | 0 | 0 | 0 | 0 | 0 | 0 | 0 | 0 | 0 |
| 0 | NA | NA | NA | NA | NA | Gm28231 | | 0 | 0 | 0 | 0 | 0 | 0 | 0 | 0 | 0 | 0 | 0 |
| 0 | NA | NA | NA | NA | NA | Gm10163 | | 0 | 0 | 0 | 0 | 0 | 0 | 0 | 0 | 0 | 0 | 0 |
| 0 | NA | NA | NA | NA | NA | Snord50a | | 0 | 0 | 0 | 0 | 0 | 0 | 0 | 0 | 0 | 0 | 0 |
| 0 | NA | NA | NA | NA | NA | Snord50b | | 0 | 0 | 0 | 0 | 0 | 0 | 0 | 0 | 0 | 0 | 0 |
| 0 | NA | NA | NA | NA | NA | Gm22985 | | 0 | 0 | 0 | 0 | 0 | 0 | 0 | 0 | 0 | 0 | 0 |
| 0 | NA | NA | NA | NA | NA | Gm29603 | | 0 | 0 | 0 | 0 | 0 | 0 | 0 | 0 | 0 | 0 | 0 |
| 0 | NA | NA | NA | NA | NA | Gm23017 | | 0 | 0 | 0 | 0 | 0 | 0 | 0 | 0 | 0 | 0 | 0 |
| 0 | NA | NA | NA | NA | NA | Gm37560 | | 0 | 0 | 0 | 0 | 0 | 0 | 0 | 0 | 0 | 0 | 0 |
| 0 | NA | NA | NA | NA | NA | Gm26082 | | 0 | 0 | 0 | 0 | 0 | 0 | 0 | 0 | 0 | 0 | 0 |
| 0 | NA | NA | NA | NA | NA | Gm37614 | | 0 | 0 | 0 | 0 | 0 | 0 | 0 | 0 | 0 | 0 | 0 |
| 0 | NA | NA | NA | NA | NA | Gm9775 | | 0 | 0 | 0 | 0 | 0 | 0 | 0 | 0 | 0 | 0 | 0 |
| 0 | NA | NA | NA | NA | NA | Gm26135 | | 0 | 0 | 0 | 0 | 0 | 0 | 0 | 0 | 0 | 0 | 0 |
| 0 | NA | NA | NA | NA | NA | Gm2396 | predicted gene 2396 | 0 | 0 | 0 | 0 | 0 | 0 | 0 | 0 | 0 | 0 | 0 |
| 0 | NA | NA | NA | NA | NA | Gm23707 | | 0 | 0 | 0 | 0 | 0 | 0 | 0 | 0 | 0 | 0 | 0 |
| 0 | NA | NA | NA | NA | NA | Gm37833 | | 0 | 0 | 0 | 0 | 0 | 0 | 0 | 0 | 0 | 0 | 0 |
| 0 | NA | NA | NA | NA | NA | Gm37993 | | 0 | 0 | 0 | 0 | 0 | 0 | 0 | 0 | 0 | 0 | 0 |
| 0 | NA | NA | NA | NA | NA | Gm24463 | | 0 | 0 | 0 | 0 | 0 | 0 | 0 | 0 | 0 | 0 | 0 |
| 0 | NA | NA | NA | NA | NA | Gm19498 | | 0 | 0 | 0 | 0 | 0 | 0 | 0 | 0 | 0 | 0 | 0 |
| 0 | NA | NA | NA | NA | NA | Trim43b | tripartite motif-containing 43B | 0 | 0 | 0 | 0 | 0 | 0 | 0 | 0 | 0 | 0 | 0 |
| 0 | NA | NA | NA | NA | NA | 4930579C12Rik | | 0 | 0 | 0 | 0 | 0 | 0 | 0 | 0 | 0 | 0 | 0 |
| 0 | NA | NA | NA | NA | NA | Gm26423 | | 0 | 0 | 0 | 0 | 0 | 0 | 0 | 0 | 0 | 0 | 0 |
| 0 | NA | NA | NA | NA | NA | Gm29094 | | 0 | 0 | 0 | 0 | 0 | 0 | 0 | 0 | 0 | 0 | 0 |
| 0 | NA | NA | NA | NA | NA | Gm38091 | | 0 | 0 | 0 | 0 | 0 | 0 | 0 | 0 | 0 | 0 | 0 |
| 0 | NA | NA | NA | NA | NA | Gm47403 | | 0 | 0 | 0 | 0 | 0 | 0 | 0 | 0 | 0 | 0 | 0 |
| 0 | NA | NA | NA | NA | NA | Gm47410 | | 0 | 0 | 0 | 0 | 0 | 0 | 0 | 0 | 0 | 0 | 0 |
| 0 | NA | NA | NA | NA | NA | Gm29608 | | 0 | 0 | 0 | 0 | 0 | 0 | 0 | 0 | 0 | 0 | 0 |
| 0 | NA | NA | NA | NA | NA | Mir184 | | 0 | 0 | 0 | 0 | 0 | 0 | 0 | 0 | 0 | 0 | 0 |
| 0 | NA | NA | NA | NA | NA | 1700017I07Rik | | 0 | 0 | 0 | 0 | 0 | 0 | 0 | 0 | 0 | 0 | 0 |
| 0 | NA | NA | NA | NA | NA | Gm23759 | | 0 | 0 | 0 | 0 | 0 | 0 | 0 | 0 | 0 | 0 | 0 |
| 0 | NA | NA | NA | NA | NA | Gm17944 | | 0 | 0 | 0 | 0 | 0 | 0 | 0 | 0 | 0 | 0 | 0 |
| 0 | NA | NA | NA | NA | NA | Gm30849 | | 0 | 0 | 0 | 0 | 0 | 0 | 0 | 0 | 0 | 0 | 0 |
| 0 | NA | NA | NA | NA | NA | Gm22717 | | 0 | 0 | 0 | 0 | 0 | 0 | 0 | 0 | 0 | 0 | 0 |
| 0 | NA | NA | NA | NA | NA | Gm22866 | | 0 | 0 | 0 | 0 | 0 | 0 | 0 | 0 | 0 | 0 | 0 |
| 0 | NA | NA | NA | NA | NA | Gm2497 | | 0 | 0 | 0 | 0 | 0 | 0 | 0 | 0 | 0 | 0 | 0 |
| 0 | NA | NA | NA | NA | NA | Gm29605 | | 0 | 0 | 0 | 0 | 0 | 0 | 0 | 0 | 0 | 0 | 0 |
| 0 | NA | NA | NA | NA | NA | Gm29604 | | 0 | 0 | 0 | 0 | 0 | 0 | 0 | 0 | 0 | 0 | 0 |
| 0 | NA | NA | NA | NA | NA | Gm28703 | | 0 | 0 | 0 | 0 | 0 | 0 | 0 | 0 | 0 | 0 | 0 |
| 0 | NA | NA | NA | NA | NA | Gm37665 | | 0 | 0 | 0 | 0 | 0 | 0 | 0 | 0 | 0 | 0 | 0 |
| 0 | NA | NA | NA | NA | NA | Gm20561 | | 0 | 0 | 0 | 0 | 0 | 0 | 0 | 0 | 0 | 0 | 0 |
| 0 | NA | NA | NA | NA | NA | Gm29408 | | 0 | 0 | 0 | 0 | 0 | 0 | 0 | 0 | 0 | 0 | 0 |
| 0 | NA | NA | NA | NA | NA | Gm36960 | | 0 | 0 | 0 | 0 | 0 | 0 | 0 | 0 | 0 | 0 | 0 |
| 0 | NA | NA | NA | NA | NA | Gm29602 | | 0 | 0 | 0 | 0 | 0 | 0 | 0 | 0 | 0 | 0 | 0 |
| 0 | NA | NA | NA | NA | NA | Gm37812 | | 0 | 0 | 0 | 0 | 0 | 0 | 0 | 0 | 0 | 0 | 0 |
| 0 | NA | NA | NA | NA | NA | Gm18378 | | 0 | 0 | 0 | 0 | 0 | 0 | 0 | 0 | 0 | 0 | 0 |
| 0 | NA | NA | NA | NA | NA | Gm37457 | | 0 | 0 | 0 | 0 | 0 | 0 | 0 | 0 | 0 | 0 | 0 |
| 0 | NA | NA | NA | NA | NA | Gm37040 | | 0 | 0 | 0 | 0 | 0 | 0 | 0 | 0 | 0 | 0 | 0 |
| 0 | NA | NA | NA | NA | NA | Gm28652 | | 0 | 0 | 0 | 0 | 0 | 0 | 0 | 0 | 0 | 0 | 0 |
| 0 | NA | NA | NA | NA | NA | A730094K22Rik | | 0 | 0 | 0 | 0 | 0 | 0 | 0 | 0 | 0 | 0 | 0 |
| 0 | NA | NA | NA | NA | NA | Zic1 | zinc finger protein of the cerebellum 1 | 0 | 0 | 0 | 0 | 0 | 0 | 0 | 0 | 0 | 0 | 0 |
| 0 | NA | NA | NA | NA | NA | Gm37477 | | 0 | 0 | 0 | 0 | 0 | 0 | 0 | 0 | 0 | 0 | 0 |
| 0 | NA | NA | NA | NA | NA | Gm29478 | | 0 | 0 | 0 | 0 | 0 | 0 | 0 | 0 | 0 | 0 | 0 |

| | | | | | | | | | | | | | | | | | | |
|---|---|---|---|---|---|---|---|---|---|---|---|---|---|---|---|---|---|---|
| 0 | NA | NA | NA | NA | NA | Gm37987 | | 0 | 0 | 0 | 0 | 0 | 0 | 0 | 0 | 0 | 0 | 0 |
| 0 | NA | NA | NA | NA | NA | Gm26101 | | 0 | 0 | 0 | 0 | 0 | 0 | 0 | 0 | 0 | 0 | 0 |
| 0 | NA | NA | NA | NA | NA | Gm31409 | | 0 | 0 | 0 | 0 | 0 | 0 | 0 | 0 | 0 | 0 | 0 |
| 0 | NA | NA | NA | NA | NA | Plscr5 | phospholipid scramblase family, member | 0 | 0 | 0 | 0 | 0 | 0 | 0 | 0 | 0 | 0 | 0 |
| 0 | NA | NA | NA | NA | NA | Mir6386 | | 0 | 0 | 0 | 0 | 0 | 0 | 0 | 0 | 0 | 0 | 0 |
| 0 | NA | NA | NA | NA | NA | Gm29139 | | 0 | 0 | 0 | 0 | 0 | 0 | 0 | 0 | 0 | 0 | 0 |
| 0 | NA | NA | NA | NA | NA | 1700057G04Ri | RIKEN cDNA 1700057G04 gene | 0 | 0 | 0 | 0 | 0 | 0 | 0 | 0 | 0 | 0 | 0 |
| 0 | NA | NA | NA | NA | NA | Gm15493 | | 0 | 0 | 0 | 0 | 0 | 0 | 0 | 0 | 0 | 0 | 0 |
| 0 | NA | NA | NA | NA | NA | Gm25417 | | 0 | 0 | 0 | 0 | 0 | 0 | 0 | 0 | 0 | 0 | 0 |
| 0 | NA | NA | NA | NA | NA | Gm37545 | | 0 | 0 | 0 | 0 | 0 | 0 | 0 | 0 | 0 | 0 | 0 |
| 0 | NA | NA | NA | NA | NA | Gm2495 | | 0 | 0 | 0 | 0 | 0 | 0 | 0 | 0 | 0 | 0 | 0 |
| 0 | NA | NA | NA | NA | NA | Gm29396 | | 0 | 0 | 0 | 0 | 0 | 0 | 0 | 0 | 0 | 0 | 0 |
| 0 | NA | NA | NA | NA | NA | 4933400C23Rik | | 0 | 0 | 0 | 0 | 0 | 0 | 0 | 0 | 0 | 0 | 0 |
| 0 | NA | NA | NA | NA | NA | Gm28054 | | 0 | 0 | 0 | 0 | 0 | 0 | 0 | 0 | 0 | 0 | 0 |
| 0 | NA | NA | NA | NA | NA | 1700034K08Rik | | 0 | 0 | 0 | 0 | 0 | 0 | 0 | 0 | 0 | 0 | 0 |
| 0 | NA | NA | NA | NA | NA | Gm9621 | | 0 | 0 | 0 | 0 | 0 | 0 | 0 | 0 | 0 | 0 | 0 |
| 0 | NA | NA | NA | NA | NA | Gm47165 | | 0 | 0 | 0 | 0 | 0 | 0 | 0 | 0 | 0 | 0 | 0 |
| 0 | NA | NA | NA | NA | NA | Gm5369 | | 0 | 0 | 0 | 0 | 0 | 0 | 0 | 0 | 0 | 0 | 0 |
| 0 | NA | NA | NA | NA | NA | Gm24200 | | 0 | 0 | 0 | 0 | 0 | 0 | 0 | 0 | 0 | 0 | 0 |
| 0 | NA | NA | NA | NA | NA | Gm28934 | | 0 | 0 | 0 | 0 | 0 | 0 | 0 | 0 | 0 | 0 | 0 |
| 0 | NA | NA | NA | NA | NA | Gm39404 | | 0 | 0 | 0 | 0 | 0 | 0 | 0 | 0 | 0 | 0 | 0 |
| 0 | NA | NA | NA | NA | NA | Gm24178 | | 0 | 0 | 0 | 0 | 0 | 0 | 0 | 0 | 0 | 0 | 0 |
| 0 | NA | NA | NA | NA | NA | Gm16262 | | 0 | 0 | 0 | 0 | 0 | 0 | 0 | 0 | 0 | 0 | 0 |
| 0 | NA | NA | NA | NA | NA | Mir7656 | | 0 | 0 | 0 | 0 | 0 | 0 | 0 | 0 | 0 | 0 | 0 |
| 0 | NA | NA | NA | NA | NA | Gm47168 | | 0 | 0 | 0 | 0 | 0 | 0 | 0 | 0 | 0 | 0 | 0 |
| 0 | NA | NA | NA | NA | NA | Gm8449 | | 0 | 0 | 0 | 0 | 0 | 0 | 0 | 0 | 0 | 0 | 0 |
| 0 | NA | NA | NA | NA | NA | Gm19193 | | 0 | 0 | 0 | 0 | 0 | 0 | 0 | 0 | 0 | 0 | 0 |
| 0 | NA | NA | NA | NA | NA | Gm32281 | | 0 | 0 | 0 | 0 | 0 | 0 | 0 | 0 | 0 | 0 | 0 |
| 0 | NA | NA | NA | NA | NA | Gm16126 | | 0 | 0 | 0 | 0 | 0 | 0 | 0 | 0 | 0 | 0 | 0 |
| 0 | NA | NA | NA | NA | NA | Gm28164 | | 0 | 0 | 0 | 0 | 0 | 0 | 0 | 0 | 0 | 0 | 0 |
| 0 | NA | NA | NA | NA | NA | Gm25582 | | 0 | 0 | 0 | 0 | 0 | 0 | 0 | 0 | 0 | 0 | 0 |
| 0 | NA | NA | NA | NA | NA | Gm16127 | | 0 | 0 | 0 | 0 | 0 | 0 | 0 | 0 | 0 | 0 | 0 |
| 0 | NA | NA | NA | NA | NA | Itpa-ps2 | | 0 | 0 | 0 | 0 | 0 | 0 | 0 | 0 | 0 | 0 | 0 |
| 0 | NA | NA | NA | NA | NA | Gm24004 | | 0 | 0 | 0 | 0 | 0 | 0 | 0 | 0 | 0 | 0 | 0 |
| 0 | NA | NA | NA | NA | NA | C78334 | | 0 | 0 | 0 | 0 | 0 | 0 | 0 | 0 | 0 | 0 | 0 |
| 0 | NA | NA | NA | NA | NA | Gm28085 | | 0 | 0 | 0 | 0 | 0 | 0 | 0 | 0 | 0 | 0 | 0 |
| 0 | NA | NA | NA | NA | NA | Gm28924 | | 0 | 0 | 0 | 0 | 0 | 0 | 0 | 0 | 0 | 0 | 0 |
| 0 | NA | NA | NA | NA | NA | Gm50488 | | 0 | 0 | 0 | 0 | 0 | 0 | 0 | 0 | 0 | 0 | 0 |
| 0 | NA | NA | NA | NA | NA | Gm27289 | | 0 | 0 | 0 | 0 | 0 | 0 | 0 | 0 | 0 | 0 | 0 |
| 0 | NA | NA | NA | NA | NA | n-R5s87 | | 0 | 0 | 0 | 0 | 0 | 0 | 0 | 0 | 0 | 0 | 0 |
| 0 | NA | NA | NA | NA | NA | Gm19280 | | 0 | 0 | 0 | 0 | 0 | 0 | 0 | 0 | 0 | 0 | 0 |
| 0 | NA | NA | NA | NA | NA | C430002N11Rik | | 0 | 0 | 0 | 0 | 0 | 0 | 0 | 0 | 0 | 0 | 0 |
| 0 | NA | NA | NA | NA | NA | Gm28688 | | 0 | 0 | 0 | 0 | 0 | 0 | 0 | 0 | 0 | 0 | 0 |
| 0 | NA | NA | NA | NA | NA | Gm5371 | | 0 | 0 | 0 | 0 | 0 | 0 | 0 | 0 | 0 | 0 | 0 |
| 0 | NA | NA | NA | NA | NA | Gm29241 | | 0 | 0 | 0 | 0 | 0 | 0 | 0 | 0 | 0 | 0 | 0 |
| 0 | NA | NA | NA | NA | NA | Gm29240 | | 0 | 0 | 0 | 0 | 0 | 0 | 0 | 0 | 0 | 0 | 0 |
| 0 | NA | NA | NA | NA | NA | Gm15891 | | 0 | 0 | 0 | 0 | 0 | 0 | 0 | 0 | 0 | 0 | 0 |
| 0 | NA | NA | NA | NA | NA | Gm25607 | | 0 | 0 | 0 | 0 | 0 | 0 | 0 | 0 | 0 | 0 | 0 |
| 0 | NA | NA | NA | NA | NA | Gm37661 | | 0 | 0 | 0 | 0 | 0 | 0 | 0 | 0 | 0 | 0 | 0 |
| 0 | NA | NA | NA | NA | NA | 4921534H16Rik | | 0 | 0 | 0 | 0 | 0 | 0 | 0 | 0 | 0 | 0 | 0 |
| 0 | NA | NA | NA | NA | NA | 2610303G11Rik | | 0 | 0 | 0 | 0 | 0 | 0 | 0 | 0 | 0 | 0 | 0 |
| 0 | NA | NA | NA | NA | NA | Gm37113 | | 0 | 0 | 0 | 0 | 0 | 0 | 0 | 0 | 0 | 0 | 0 |
| 0 | NA | NA | NA | NA | NA | Gm47147 | | 0 | 0 | 0 | 0 | 0 | 0 | 0 | 0 | 0 | 0 | 0 |
| 0 | NA | NA | NA | NA | NA | Gm27919 | | 0 | 0 | 0 | 0 | 0 | 0 | 0 | 0 | 0 | 0 | 0 |
| 0 | NA | NA | NA | NA | NA | Pisrt1 | | 0 | 0 | 0 | 0 | 0 | 0 | 0 | 0 | 0 | 0 | 0 |
| 0 | NA | NA | NA | NA | NA | E330023G01Rik | | 0 | 0 | 0 | 0 | 0 | 0 | 0 | 0 | 0 | 0 | 0 |
| 0 | NA | NA | NA | NA | NA | Gm37282 | | 0 | 0 | 0 | 0 | 0 | 0 | 0 | 0 | 0 | 0 | 0 |
| 0 | NA | NA | NA | NA | NA | Gm37917 | | 0 | 0 | 0 | 0 | 0 | 0 | 0 | 0 | 0 | 0 | 0 |
| 0 | NA | NA | NA | NA | NA | Gm38226 | | 0 | 0 | 0 | 0 | 0 | 0 | 0 | 0 | 0 | 0 | 0 |
| 0 | NA | NA | NA | NA | NA | Prr23a1 | proline rich 23A, member 1 | 0 | 0 | 0 | 0 | 0 | 0 | 0 | 0 | 0 | 0 | 0 |

| | | | | | | | | | | | | | | | | | | |
|---|---|---|---|---|---|---|---|---|---|---|---|---|---|---|---|---|---|---|
| 0 | NA | NA | NA | NA | NA | Prr23a2 | proline rich 23A, member 2 | 0 | 0 | 0 | 0 | 0 | 0 | 0 | 0 | 0 | 0 | 0 |
| 0 | NA | NA | NA | NA | NA | Gm33700 | predicted gene, 33700 | 0 | 0 | 0 | 0 | 0 | 0 | 0 | 0 | 0 | 0 | 0 |
| 0 | NA | NA | NA | NA | NA | 7420426K07Ril | RIKEN cDNA 7420426K07 gene | 0 | 0 | 0 | 0 | 0 | 0 | 0 | 0 | 0 | 0 | 0 |
| 0 | NA | NA | NA | NA | NA | Gm33778 | | 0 | 0 | 0 | 0 | 0 | 0 | 0 | 0 | 0 | 0 | 0 |
| 0 | NA | NA | NA | NA | NA | Gm28266 | | 0 | 0 | 0 | 0 | 0 | 0 | 0 | 0 | 0 | 0 | 0 |
| 0 | NA | NA | NA | NA | NA | Foxl2os | | 0 | 0 | 0 | 0 | 0 | 0 | 0 | 0 | 0 | 0 | 0 |
| 0 | NA | NA | NA | NA | NA | Foxl2 | forkhead box L2 | 0 | 0 | 0 | 0 | 0 | 0 | 0 | 0 | 0 | 0 | 0 |
| 0 | NA | NA | NA | NA | NA | Gm2950 | | 0 | 0 | 0 | 0 | 0 | 0 | 0 | 0 | 0 | 0 | 0 |
| 0 | NA | NA | NA | NA | NA | Faiml | Fas apoptotic inhibitory molecule like | 0 | 0 | 0 | 0 | 0 | 0 | 0 | 0 | 0 | 0 | 0 |
| 0 | NA | NA | NA | NA | NA | Gm47199 | | 0 | 0 | 0 | 0 | 0 | 0 | 0 | 0 | 0 | 0 | 0 |
| 0 | NA | NA | NA | NA | NA | Gm18914 | | 0 | 0 | 0 | 0 | 0 | 0 | 0 | 0 | 0 | 0 | 0 |
| 0 | NA | NA | NA | NA | NA | Gm28087 | | 0 | 0 | 0 | 0 | 0 | 0 | 0 | 0 | 0 | 0 | 0 |
| 0 | NA | NA | NA | NA | NA | A4gnt | alpha-1,4-N-acetylglucosaminyltransferas | 0 | 0 | 0 | 0 | 0 | 0 | 0 | 0 | 0 | 0 | 0 |
| 0 | NA | NA | NA | NA | NA | Cldn18 | claudin 18 | 0 | 0 | 0 | 0 | 0 | 0 | 0 | 0 | 0 | 0 | 0 |
| 0 | NA | NA | NA | NA | NA | Gm16004 | | 0 | 0 | 0 | 0 | 0 | 0 | 0 | 0 | 0 | 0 | 0 |
| 0 | NA | NA | NA | NA | NA | Gm23949 | | 0 | 0 | 0 | 0 | 0 | 0 | 0 | 0 | 0 | 0 | 0 |
| 0 | NA | NA | NA | NA | NA | Gm47757 | | 0 | 0 | 0 | 0 | 0 | 0 | 0 | 0 | 0 | 0 | 0 |
| 0 | NA | NA | NA | NA | NA | Sox14 | SRY (sex determining region Y)-box 14 | 0 | 0 | 0 | 0 | 0 | 0 | 0 | 0 | 0 | 0 | 0 |
| 0 | NA | NA | NA | NA | NA | Gm23594 | | 0 | 0 | 0 | 0 | 0 | 0 | 0 | 0 | 0 | 0 | 0 |
| 0 | NA | NA | NA | NA | NA | 4930519F24Rik | | 0 | 0 | 0 | 0 | 0 | 0 | 0 | 0 | 0 | 0 | 0 |
| 0 | NA | NA | NA | NA | NA | 4930540E01Rik | | 0 | 0 | 0 | 0 | 0 | 0 | 0 | 0 | 0 | 0 | 0 |
| 0 | NA | NA | NA | NA | NA | Gm28167 | | 0 | 0 | 0 | 0 | 0 | 0 | 0 | 0 | 0 | 0 | 0 |
| 0 | NA | NA | NA | NA | NA | Gm34397 | | 0 | 0 | 0 | 0 | 0 | 0 | 0 | 0 | 0 | 0 | 0 |
| 0 | NA | NA | NA | NA | NA | Gm28267 | | 0 | 0 | 0 | 0 | 0 | 0 | 0 | 0 | 0 | 0 | 0 |
| 0 | NA | NA | NA | NA | NA | Gm28166 | | 0 | 0 | 0 | 0 | 0 | 0 | 0 | 0 | 0 | 0 | 0 |
| 0 | NA | NA | NA | NA | NA | Gm47803 | | 0 | 0 | 0 | 0 | 0 | 0 | 0 | 0 | 0 | 0 | 0 |
| 0 | NA | NA | NA | NA | NA | Gm24338 | | 0 | 0 | 0 | 0 | 0 | 0 | 0 | 0 | 0 | 0 | 0 |
| 0 | NA | NA | NA | NA | NA | Gm3121 | | 0 | 0 | 0 | 0 | 0 | 0 | 0 | 0 | 0 | 0 | 0 |
| 0 | NA | NA | NA | NA | NA | Gm28660 | | 0 | 0 | 0 | 0 | 0 | 0 | 0 | 0 | 0 | 0 | 0 |
| 0 | NA | NA | NA | NA | NA | Gm24638 | | 0 | 0 | 0 | 0 | 0 | 0 | 0 | 0 | 0 | 0 | 0 |
| 0 | NA | NA | NA | NA | NA | Gm29385 | | 0 | 0 | 0 | 0 | 0 | 0 | 0 | 0 | 0 | 0 | 0 |
| 0 | NA | NA | NA | NA | NA | Gm47932 | | 0 | 0 | 0 | 0 | 0 | 0 | 0 | 0 | 0 | 0 | 0 |
| 0 | NA | NA | NA | NA | NA | Gm47933 | | 0 | 0 | 0 | 0 | 0 | 0 | 0 | 0 | 0 | 0 | 0 |
| 0 | NA | NA | NA | NA | NA | Gm47934 | | 0 | 0 | 0 | 0 | 0 | 0 | 0 | 0 | 0 | 0 | 0 |
| 0 | NA | NA | NA | NA | NA | Gm37460 | | 0 | 0 | 0 | 0 | 0 | 0 | 0 | 0 | 0 | 0 | 0 |
| 0 | NA | NA | NA | NA | NA | Gm31326 | | 0 | 0 | 0 | 0 | 0 | 0 | 0 | 0 | 0 | 0 | 0 |
| 0 | NA | NA | NA | NA | NA | Gm37562 | | 0 | 0 | 0 | 0 | 0 | 0 | 0 | 0 | 0 | 0 | 0 |
| 0 | NA | NA | NA | NA | NA | n-R5s88 | | 0 | 0 | 0 | 0 | 0 | 0 | 0 | 0 | 0 | 0 | 0 |
| 0 | NA | NA | NA | NA | NA | Gm29388 | | 0 | 0 | 0 | 0 | 0 | 0 | 0 | 0 | 0 | 0 | 0 |
| 0 | NA | NA | NA | NA | NA | Gm37945 | | 0 | 0 | 0 | 0 | 0 | 0 | 0 | 0 | 0 | 0 | 0 |
| 0 | NA | NA | NA | NA | NA | Gm37260 | | 0 | 0 | 0 | 0 | 0 | 0 | 0 | 0 | 0 | 0 | 0 |
| 0 | NA | NA | NA | NA | NA | Mir7243 | | 0 | 0 | 0 | 0 | 0 | 0 | 0 | 0 | 0 | 0 | 0 |
| 0 | NA | NA | NA | NA | NA | Gm22894 | | 0 | 0 | 0 | 0 | 0 | 0 | 0 | 0 | 0 | 0 | 0 |
| 0 | NA | NA | NA | NA | NA | Gm47983 | | 0 | 0 | 0 | 0 | 0 | 0 | 0 | 0 | 0 | 0 | 0 |
| 0 | NA | NA | NA | NA | NA | Gm25669 | | 0 | 0 | 0 | 0 | 0 | 0 | 0 | 0 | 0 | 0 | 0 |
| 0 | NA | NA | NA | NA | NA | Gm29143 | | 0 | 0 | 0 | 0 | 0 | 0 | 0 | 0 | 0 | 0 | 0 |
| 0 | NA | NA | NA | NA | NA | Gm17950 | | 0 | 0 | 0 | 0 | 0 | 0 | 0 | 0 | 0 | 0 | 0 |
| 0 | NA | NA | NA | NA | NA | Gm36957 | | 0 | 0 | 0 | 0 | 0 | 0 | 0 | 0 | 0 | 0 | 0 |
| 0 | NA | NA | NA | NA | NA | Gm5627 | | 0 | 0 | 0 | 0 | 0 | 0 | 0 | 0 | 0 | 0 | 0 |
| 0 | NA | NA | NA | NA | NA | Gm36958 | | 0 | 0 | 0 | 0 | 0 | 0 | 0 | 0 | 0 | 0 | 0 |
| 0 | NA | NA | NA | NA | NA | Gm47416 | | 0 | 0 | 0 | 0 | 0 | 0 | 0 | 0 | 0 | 0 | 0 |
| 0 | NA | NA | NA | NA | NA | Gm25842 | | 0 | 0 | 0 | 0 | 0 | 0 | 0 | 0 | 0 | 0 | 0 |
| 0 | NA | NA | NA | NA | NA | n-R5s89 | | 0 | 0 | 0 | 0 | 0 | 0 | 0 | 0 | 0 | 0 | 0 |
| 0 | NA | NA | NA | NA | NA | Gm22595 | | 0 | 0 | 0 | 0 | 0 | 0 | 0 | 0 | 0 | 0 | 0 |
| 0 | NA | NA | NA | NA | NA | Gm47468 | | 0 | 0 | 0 | 0 | 0 | 0 | 0 | 0 | 0 | 0 | 0 |
| 0 | NA | NA | NA | NA | NA | Gm29600 | | 0 | 0 | 0 | 0 | 0 | 0 | 0 | 0 | 0 | 0 | 0 |
| 0 | NA | NA | NA | NA | NA | Gm47328 | | 0 | 0 | 0 | 0 | 0 | 0 | 0 | 0 | 0 | 0 | 0 |
| 0 | NA | NA | NA | NA | NA | Gm26360 | | 0 | 0 | 0 | 0 | 0 | 0 | 0 | 0 | 0 | 0 | 0 |
| 0 | NA | NA | NA | NA | NA | Gm38032 | | 0 | 0 | 0 | 0 | 0 | 0 | 0 | 0 | 0 | 0 | 0 |
| 0 | NA | NA | NA | NA | NA | Gm5372 | | 0 | 0 | 0 | 0 | 0 | 0 | 0 | 0 | 0 | 0 | 0 |

| | | | | | | | | | | | | | | | | | | |
|---|---|---|---|---|---|---|---|---|---|---|---|---|---|---|---|---|---|---|
| 0 | NA | NA | NA | NA | NA | Gm28305 | | 0 | 0 | 0 | 0 | 0 | 0 | 0 | 0 | 0 | 0 | 0 |
| 0 | NA | NA | NA | NA | NA | Gm37750 | | 0 | 0 | 0 | 0 | 0 | 0 | 0 | 0 | 0 | 0 | 0 |
| 0 | NA | NA | NA | NA | NA | Gm46999 | | 0 | 0 | 0 | 0 | 0 | 0 | 0 | 0 | 0 | 0 | 0 |
| 0 | NA | NA | NA | NA | NA | Gm28996 | | 0 | 0 | 0 | 0 | 0 | 0 | 0 | 0 | 0 | 0 | 0 |
| 0 | NA | NA | NA | NA | NA | Gm28548 | | 0 | 0 | 0 | 0 | 0 | 0 | 0 | 0 | 0 | 0 | 0 |
| 0 | NA | NA | NA | NA | NA | Mir2136 | | 0 | 0 | 0 | 0 | 0 | 0 | 0 | 0 | 0 | 0 | 0 |
| 0 | NA | NA | NA | NA | NA | Gm25904 | | 0 | 0 | 0 | 0 | 0 | 0 | 0 | 0 | 0 | 0 | 0 |
| 0 | NA | NA | NA | NA | NA | Gm23391 | | 0 | 0 | 0 | 0 | 0 | 0 | 0 | 0 | 0 | 0 | 0 |
| 0 | NA | NA | NA | NA | NA | Gm47106 | | 0 | 0 | 0 | 0 | 0 | 0 | 0 | 0 | 0 | 0 | 0 |
| 0 | NA | NA | NA | NA | NA | Gm47103 | | 0 | 0 | 0 | 0 | 0 | 0 | 0 | 0 | 0 | 0 | 0 |
| 0 | NA | NA | NA | NA | NA | Gm47102 | | 0 | 0 | 0 | 0 | 0 | 0 | 0 | 0 | 0 | 0 | 0 |
| 0 | NA | NA | NA | NA | NA | Nudt16l2 | nudix (nucleoside diphosphate linked mo | 0 | 0 | 0 | 0 | 0 | 0 | 0 | 0 | 0 | 0 | 0 |
| 0 | NA | NA | NA | NA | NA | Gm37582 | | 0 | 0 | 0 | 0 | 0 | 0 | 0 | 0 | 0 | 0 | 0 |
| 0 | NA | NA | NA | NA | NA | Gm20643 | | 0 | 0 | 0 | 0 | 0 | 0 | 0 | 0 | 0 | 0 | 0 |
| 0 | NA | NA | NA | NA | NA | Gm15619 | | 0 | 0 | 0 | 0 | 0 | 0 | 0 | 0 | 0 | 0 | 0 |
| 0 | NA | NA | NA | NA | NA | Gm19667 | | 0 | 0 | 0 | 0 | 0 | 0 | 0 | 0 | 0 | 0 | 0 |
| 0 | NA | NA | NA | NA | NA | Gm22720 | | 0 | 0 | 0 | 0 | 0 | 0 | 0 | 0 | 0 | 0 | 0 |
| 0 | NA | NA | NA | NA | NA | Gm24631 | | 0 | 0 | 0 | 0 | 0 | 0 | 0 | 0 | 0 | 0 | 0 |
| 0 | NA | NA | NA | NA | NA | Gm47575 | | 0 | 0 | 0 | 0 | 0 | 0 | 0 | 0 | 0 | 0 | 0 |
| 0 | NA | NA | NA | NA | NA | Gm9451 | | 0 | 0 | 0 | 0 | 0 | 0 | 0 | 0 | 0 | 0 | 0 |
| 0 | NA | NA | NA | NA | NA | Gm29208 | | 0 | 0 | 0 | 0 | 0 | 0 | 0 | 0 | 0 | 0 | 0 |
| 0 | NA | NA | NA | NA | NA | Mir135a-1 | | 0 | 0 | 0 | 0 | 0 | 0 | 0 | 0 | 0 | 0 | 0 |
| 0 | NA | NA | NA | NA | NA | Mirlet7g | | 0 | 0 | 0 | 0 | 0 | 0 | 0 | 0 | 0 | 0 | 0 |
| 0 | NA | NA | NA | NA | NA | Gm29123 | | 0 | 0 | 0 | 0 | 0 | 0 | 0 | 0 | 0 | 0 | 0 |
| 0 | NA | NA | NA | NA | NA | Iqcf1 | IQ motif containing F1 | 0 | 0 | 0 | 0 | 0 | 0 | 0 | 0 | 0 | 0 | 0 |
| 0 | NA | NA | NA | NA | NA | Iqcf2 | IQ motif containing F3 | 0 | 0 | 0 | 0 | 0 | 0 | 0 | 0 | 0 | 0 | 0 |
| 0 | NA | NA | NA | NA | NA | Iqcf4 | IQ motif containing F4 | 0 | 0 | 0 | 0 | 0 | 0 | 0 | 0 | 0 | 0 | 0 |
| 0 | NA | NA | NA | NA | NA | Iqcf6 | IQ motif containing F6 | 0 | 0 | 0 | 0 | 0 | 0 | 0 | 0 | 0 | 0 | 0 |
| 0 | NA | NA | NA | NA | NA | Gm26185 | | 0 | 0 | 0 | 0 | 0 | 0 | 0 | 0 | 0 | 0 | 0 |
| 0 | NA | NA | NA | NA | NA | Gm28111 | | 0 | 0 | 0 | 0 | 0 | 0 | 0 | 0 | 0 | 0 | 0 |
| 0 | NA | NA | NA | NA | NA | Gm22384 | | 0 | 0 | 0 | 0 | 0 | 0 | 0 | 0 | 0 | 0 | 0 |
| 0 | NA | NA | NA | NA | NA | Gm17205 | | 0 | 0 | 0 | 0 | 0 | 0 | 0 | 0 | 0 | 0 | 0 |
| 0 | NA | NA | NA | NA | NA | Gm22454 | | 0 | 0 | 0 | 0 | 0 | 0 | 0 | 0 | 0 | 0 | 0 |
| 0 | NA | NA | NA | NA | NA | Gm17117 | | 0 | 0 | 0 | 0 | 0 | 0 | 0 | 0 | 0 | 0 | 0 |
| 0 | NA | NA | NA | NA | NA | Gm23595 | | 0 | 0 | 0 | 0 | 0 | 0 | 0 | 0 | 0 | 0 | 0 |
| 0 | NA | NA | NA | NA | NA | AA543401 | | 0 | 0 | 0 | 0 | 0 | 0 | 0 | 0 | 0 | 0 | 0 |
| 0 | NA | NA | NA | NA | NA | Gm22238 | | 0 | 0 | 0 | 0 | 0 | 0 | 0 | 0 | 0 | 0 | 0 |
| 0 | NA | NA | NA | NA | NA | 1700039M15Rik | | 0 | 0 | 0 | 0 | 0 | 0 | 0 | 0 | 0 | 0 | 0 |
| 0 | NA | NA | NA | NA | NA | 4930429P21Rik | | 0 | 0 | 0 | 0 | 0 | 0 | 0 | 0 | 0 | 0 | 0 |
| 0 | NA | NA | NA | NA | NA | Gm17141 | | 0 | 0 | 0 | 0 | 0 | 0 | 0 | 0 | 0 | 0 | 0 |
| 0 | NA | NA | NA | NA | NA | Gm17118 | | 0 | 0 | 0 | 0 | 0 | 0 | 0 | 0 | 0 | 0 | 0 |
| 0 | NA | NA | NA | NA | NA | A930036K24Rik | | 0 | 0 | 0 | 0 | 0 | 0 | 0 | 0 | 0 | 0 | 0 |
| 0 | NA | NA | NA | NA | NA | Gm23856 | | 0 | 0 | 0 | 0 | 0 | 0 | 0 | 0 | 0 | 0 | 0 |
| 0 | NA | NA | NA | NA | NA | Actl11 | actin-like 11 | 0 | 0 | 0 | 0 | 0 | 0 | 0 | 0 | 0 | 0 | 0 |
| 0 | NA | NA | NA | NA | NA | Gm37974 | | 0 | 0 | 0 | 0 | 0 | 0 | 0 | 0 | 0 | 0 | 0 |
| 0 | NA | NA | NA | NA | NA | Camkv | CaM kinase-like vesicle-associated | 0 | 0 | 0 | 0 | 0 | 0 | 0 | 0 | 0 | 0 | 0 |
| 0 | NA | NA | NA | NA | NA | Gm20661 | | 0 | 0 | 0 | 0 | 0 | 0 | 0 | 0 | 0 | 0 | 0 |
| 0 | NA | NA | NA | NA | NA | Gm24944 | | 0 | 0 | 0 | 0 | 0 | 0 | 0 | 0 | 0 | 0 | 0 |
| 0 | NA | NA | NA | NA | NA | 4921523L03Rik | | 0 | 0 | 0 | 0 | 0 | 0 | 0 | 0 | 0 | 0 | 0 |
| 0 | NA | NA | NA | NA | NA | Gm20662 | | 0 | 0 | 0 | 0 | 0 | 0 | 0 | 0 | 0 | 0 | 0 |
| 0 | NA | NA | NA | NA | NA | Gm38134 | | 0 | 0 | 0 | 0 | 0 | 0 | 0 | 0 | 0 | 0 | 0 |
| 0 | NA | NA | NA | NA | NA | Gm20529 | | 0 | 0 | 0 | 0 | 0 | 0 | 0 | 0 | 0 | 0 | 0 |
| 0 | NA | NA | NA | NA | NA | Mir7088 | | 0 | 0 | 0 | 0 | 0 | 0 | 0 | 0 | 0 | 0 | 0 |
| 0 | NA | NA | NA | NA | NA | Gm37230 | | 0 | 0 | 0 | 0 | 0 | 0 | 0 | 0 | 0 | 0 | 0 |
| 0 | NA | NA | NA | NA | NA | Gm25456 | | 0 | 0 | 0 | 0 | 0 | 0 | 0 | 0 | 0 | 0 | 0 |
| 0 | NA | NA | NA | NA | NA | BC048562 | cDNA sequence BC048562 | 0 | 0 | 0 | 0 | 0 | 0 | 0 | 0 | 0 | 0 | 0 |
| 0 | NA | NA | NA | NA | NA | Gm24083 | | 0 | 0 | 0 | 0 | 0 | 0 | 0 | 0 | 0 | 0 | 0 |
| 0 | NA | NA | NA | NA | NA | Mir191 | | 0 | 0 | 0 | 0 | 0 | 0 | 0 | 0 | 0 | 0 | 0 |
| 0 | NA | NA | NA | NA | NA | Mir425 | | 0 | 0 | 0 | 0 | 0 | 0 | 0 | 0 | 0 | 0 | 0 |
| 0 | NA | NA | NA | NA | NA | Gm49209 | | 0 | 0 | 0 | 0 | 0 | 0 | 0 | 0 | 0 | 0 | 0 |

| | | | | | | | | | | | | | | | | | | |
|---|---|---|---|---|---|---|---|---|---|---|---|---|---|---|---|---|---|---|
| 0 | NA | NA | NA | NA | NA | Gm26298 | | 0 | 0 | 0 | 0 | 0 | 0 | 0 | 0 | 0 | 0 | 0 |
| 0 | NA | NA | NA | NA | NA | Gm24259 | | 0 | 0 | 0 | 0 | 0 | 0 | 0 | 0 | 0 | 0 | 0 |
| 0 | NA | NA | NA | NA | NA | Gm23034 | | 0 | 0 | 0 | 0 | 0 | 0 | 0 | 0 | 0 | 0 | 0 |
| 0 | NA | NA | NA | NA | NA | Gm37284 | | 0 | 0 | 0 | 0 | 0 | 0 | 0 | 0 | 0 | 0 | 0 |
| 0 | NA | NA | NA | NA | NA | Mir711 | | 0 | 0 | 0 | 0 | 0 | 0 | 0 | 0 | 0 | 0 | 0 |
| 0 | NA | NA | NA | NA | NA | Trex1 | three prime repair exonuclease 1 | 0 | 0 | 0 | 0 | 0 | 0 | 0 | 0 | 0 | 0 | 0 |
| 0 | NA | NA | NA | NA | NA | Fbxw21 | F-box and WD-40 domain protein 21 | 0 | 0 | 0 | 0 | 0 | 0 | 0 | 0 | 0 | 0 | 0 |
| 0 | NA | NA | NA | NA | NA | Fbxw13 | F-box and WD-40 domain protein 13 | 0 | 0 | 0 | 0 | 0 | 0 | 0 | 0 | 0 | 0 | 0 |
| 0 | NA | NA | NA | NA | NA | Fbxw20 | F-box and WD-40 domain protein 20 | 0 | 0 | 0 | 0 | 0 | 0 | 0 | 0 | 0 | 0 | 0 |
| 0 | NA | NA | NA | NA | NA | Fbxw14 | F-box and WD-40 domain protein 14 | 0 | 0 | 0 | 0 | 0 | 0 | 0 | 0 | 0 | 0 | 0 |
| 0 | NA | NA | NA | NA | NA | Gm22821 | | 0 | 0 | 0 | 0 | 0 | 0 | 0 | 0 | 0 | 0 | 0 |
| 0 | NA | NA | NA | NA | NA | Fbxw28 | F-box and WD-40 domain protein 28 | 0 | 0 | 0 | 0 | 0 | 0 | 0 | 0 | 0 | 0 | 0 |
| 0 | NA | NA | NA | NA | NA | Gm25484 | | 0 | 0 | 0 | 0 | 0 | 0 | 0 | 0 | 0 | 0 | 0 |
| 0 | NA | NA | NA | NA | NA | Gm24641 | | 0 | 0 | 0 | 0 | 0 | 0 | 0 | 0 | 0 | 0 | 0 |
| 0 | NA | NA | NA | NA | NA | Fbxw22 | F-box and WD-40 domain protein 22 | 0 | 0 | 0 | 0 | 0 | 0 | 0 | 0 | 0 | 0 | 0 |
| 0 | NA | NA | NA | NA | NA | Fbxw16 | F-box and WD-40 domain protein 16 | 0 | 0 | 0 | 0 | 0 | 0 | 0 | 0 | 0 | 0 | 0 |
| 0 | NA | NA | NA | NA | NA | Gm42494 | | 0 | 0 | 0 | 0 | 0 | 0 | 0 | 0 | 0 | 0 | 0 |
| 0 | NA | NA | NA | NA | NA | Gm42493 | | 0 | 0 | 0 | 0 | 0 | 0 | 0 | 0 | 0 | 0 | 0 |
| 0 | NA | NA | NA | NA | NA | Fbxw19 | F-box and WD-40 domain protein 19 | 0 | 0 | 0 | 0 | 0 | 0 | 0 | 0 | 0 | 0 | 0 |
| 0 | NA | NA | NA | NA | NA | Fbxw23 | | 0 | 0 | 0 | 0 | 0 | 0 | 0 | 0 | 0 | 0 | 0 |
| 0 | NA | NA | NA | NA | NA | Gm42756 | | 0 | 0 | 0 | 0 | 0 | 0 | 0 | 0 | 0 | 0 | 0 |
| 0 | NA | NA | NA | NA | NA | Gm43089 | | 0 | 0 | 0 | 0 | 0 | 0 | 0 | 0 | 0 | 0 | 0 |
| 0 | NA | NA | NA | NA | NA | Fbxw25 | F-box and WD-40 domain protein 25 | 0 | 0 | 0 | 0 | 0 | 0 | 0 | 0 | 0 | 0 | 0 |
| 0 | NA | NA | NA | NA | NA | Fbxw18 | F-box and WD-40 domain protein 18 | 0 | 0 | 0 | 0 | 0 | 0 | 0 | 0 | 0 | 0 | 0 |
| 0 | NA | NA | NA | NA | NA | Gm43209 | | 0 | 0 | 0 | 0 | 0 | 0 | 0 | 0 | 0 | 0 | 0 |
| 0 | NA | NA | NA | NA | NA | Fbxw26 | F-box and WD-40 domain protein 26 | 0 | 0 | 0 | 0 | 0 | 0 | 0 | 0 | 0 | 0 | 0 |
| 0 | NA | NA | NA | NA | NA | Fbxw27 | F-box and WD-40 domain protein 27 | 0 | 0 | 0 | 0 | 0 | 0 | 0 | 0 | 0 | 0 | 0 |
| 0 | NA | NA | NA | NA | NA | Spink8 | serine peptidase inhibitor, Kazal type 8 | 0 | 0 | 0 | 0 | 0 | 0 | 0 | 0 | 0 | 0 | 0 |
| 0 | NA | NA | NA | NA | NA | Camp | cathelicidin antimicrobial peptide | 0 | 0 | 0 | 0 | 0 | 0 | 0 | 0 | 0 | 0 | 0 |
| 0 | NA | NA | NA | NA | NA | Gm4644 | | 0 | 0 | 0 | 0 | 0 | 0 | 0 | 0 | 0 | 0 | 0 |
| 0 | NA | NA | NA | NA | NA | Gm43732 | | 0 | 0 | 0 | 0 | 0 | 0 | 0 | 0 | 0 | 0 | 0 |
| 0 | NA | NA | NA | NA | NA | Gm42433 | | 0 | 0 | 0 | 0 | 0 | 0 | 0 | 0 | 0 | 0 | 0 |
| 0 | NA | NA | NA | NA | NA | Gm27855 | | 0 | 0 | 0 | 0 | 0 | 0 | 0 | 0 | 0 | 0 | 0 |
| 0 | NA | NA | NA | NA | NA | Gm23377 | | 0 | 0 | 0 | 0 | 0 | 0 | 0 | 0 | 0 | 0 | 0 |
| 0 | NA | NA | NA | NA | NA | Gm23377 | | 0 | 0 | 0 | 0 | 0 | 0 | 0 | 0 | 0 | 0 | 0 |
| 0 | NA | NA | NA | NA | NA | Gm43622 | | 0 | 0 | 0 | 0 | 0 | 0 | 0 | 0 | 0 | 0 | 0 |
| 0 | NA | NA | NA | NA | NA | Gm42775 | | 0 | 0 | 0 | 0 | 0 | 0 | 0 | 0 | 0 | 0 | 0 |
| 0 | NA | NA | NA | NA | NA | Gm42638 | | 0 | 0 | 0 | 0 | 0 | 0 | 0 | 0 | 0 | 0 | 0 |
| 0 | NA | NA | NA | NA | NA | Ngp | neutrophilic granule protein | 0 | 0 | 0 | 0 | 0 | 0 | 0 | 0 | 0 | 0 | 0 |
| 0 | NA | NA | NA | NA | NA | Mir8107 | | 0 | 0 | 0 | 0 | 0 | 0 | 0 | 0 | 0 | 0 | 0 |
| 0 | NA | NA | NA | NA | NA | Gm35715 | | 0 | 0 | 0 | 0 | 0 | 0 | 0 | 0 | 0 | 0 | 0 |
| 0 | NA | NA | NA | NA | NA | Myl3 | myosin, light polypeptide 3 | 0 | 0 | 0 | 0 | 0 | 0 | 0 | 0 | 0 | 0 | 0 |
| 0 | NA | NA | NA | NA | NA | Prss42 | protease, serine 42 | 0 | 0 | 0 | 0 | 0 | 0 | 0 | 0 | 0 | 0 | 0 |
| 0 | NA | NA | NA | NA | NA | Prss43 | protease, serine 43 | 0 | 0 | 0 | 0 | 0 | 0 | 0 | 0 | 0 | 0 | 0 |
| 0 | NA | NA | NA | NA | NA | Prss46 | protease, serine 46 | 0 | 0 | 0 | 0 | 0 | 0 | 0 | 0 | 0 | 0 | 0 |
| 0 | NA | NA | NA | NA | NA | Gm43127 | | 0 | 0 | 0 | 0 | 0 | 0 | 0 | 0 | 0 | 0 | 0 |
| 0 | NA | NA | NA | NA | NA | Fam240a | family with sequence similarity 240 memb | 0 | 0 | 0 | 0 | 0 | 0 | 0 | 0 | 0 | 0 | 0 |
| 0 | NA | NA | NA | NA | NA | Gm24269 | | 0 | 0 | 0 | 0 | 0 | 0 | 0 | 0 | 0 | 0 | 0 |
| 0 | NA | NA | NA | NA | NA | Gm43535 | | 0 | 0 | 0 | 0 | 0 | 0 | 0 | 0 | 0 | 0 | 0 |
| 0 | NA | NA | NA | NA | NA | 1700061E17Rik | | 0 | 0 | 0 | 0 | 0 | 0 | 0 | 0 | 0 | 0 | 0 |
| 0 | NA | NA | NA | NA | NA | Gm26409 | | 0 | 0 | 0 | 0 | 0 | 0 | 0 | 0 | 0 | 0 | 0 |
| 0 | NA | NA | NA | NA | NA | Gm42527 | | 0 | 0 | 0 | 0 | 0 | 0 | 0 | 0 | 0 | 0 | 0 |
| 0 | NA | NA | NA | NA | NA | Gm42523 | | 0 | 0 | 0 | 0 | 0 | 0 | 0 | 0 | 0 | 0 | 0 |
| 0 | NA | NA | NA | NA | NA | Gm43236 | | 0 | 0 | 0 | 0 | 0 | 0 | 0 | 0 | 0 | 0 | 0 |
| 0 | NA | NA | NA | NA | NA | n-R5s90 | | 0 | 0 | 0 | 0 | 0 | 0 | 0 | 0 | 0 | 0 | 0 |
| 0 | NA | NA | NA | NA | NA | 4930545L08Rik | | 0 | 0 | 0 | 0 | 0 | 0 | 0 | 0 | 0 | 0 | 0 |
| 0 | NA | NA | NA | NA | NA | Mir128-2 | | 0 | 0 | 0 | 0 | 0 | 0 | 0 | 0 | 0 | 0 | 0 |
| 0 | NA | NA | NA | NA | NA | 2310075C17Rik | | 0 | 0 | 0 | 0 | 0 | 0 | 0 | 0 | 0 | 0 | 0 |
| 0 | NA | NA | NA | NA | NA | 2900079G21Rik | | 0 | 0 | 0 | 0 | 0 | 0 | 0 | 0 | 0 | 0 | 0 |
| 0 | NA | NA | NA | NA | NA | 4933411E02Rik | | 0 | 0 | 0 | 0 | 0 | 0 | 0 | 0 | 0 | 0 | 0 |

| | | | | | | | | | | | | | | | | | | |
|---|---|---|---|---|---|---|---|---|---|---|---|---|---|---|---|---|---|---|
| 0 | NA | NA | NA | NA | NA | Gm27847 | | 0 | 0 | 0 | 0 | 0 | 0 | 0 | 0 | 0 | 0 | 0 |
| 0 | NA | NA | NA | NA | NA | Gm24957 | | 0 | 0 | 0 | 0 | 0 | 0 | 0 | 0 | 0 | 0 | 0 |
| 0 | NA | NA | NA | NA | NA | Gm36251 | | 0 | 0 | 0 | 0 | 0 | 0 | 0 | 0 | 0 | 0 | 0 |
| 0 | NA | NA | NA | NA | NA | Gm4657 | | 0 | 0 | 0 | 0 | 0 | 0 | 0 | 0 | 0 | 0 | 0 |
| 0 | NA | NA | NA | NA | NA | Gm36367 | | 0 | 0 | 0 | 0 | 0 | 0 | 0 | 0 | 0 | 0 | 0 |
| 0 | NA | NA | NA | NA | NA | Gm39436 | | 0 | 0 | 0 | 0 | 0 | 0 | 0 | 0 | 0 | 0 | 0 |
| 0 | NA | NA | NA | NA | NA | Gm22492 | | 0 | 0 | 0 | 0 | 0 | 0 | 0 | 0 | 0 | 0 | 0 |
| 0 | NA | NA | NA | NA | NA | Gm36485 | | 0 | 0 | 0 | 0 | 0 | 0 | 0 | 0 | 0 | 0 | 0 |
| 0 | NA | NA | NA | NA | NA | Gm47875 | | 0 | 0 | 0 | 0 | 0 | 0 | 0 | 0 | 0 | 0 | 0 |
| 0 | NA | NA | NA | NA | NA | AU023762 | | 0 | 0 | 0 | 0 | 0 | 0 | 0 | 0 | 0 | 0 | 0 |
| 0 | NA | NA | NA | NA | NA | Gm25059 | | 0 | 0 | 0 | 0 | 0 | 0 | 0 | 0 | 0 | 0 | 0 |
| 0 | NA | NA | NA | NA | NA | | | 0 | 0 | 0 | 0 | 0 | 0 | 0 | 0 | 0 | 0 | 0 |
| 0 | NA | NA | NA | NA | NA | 4921528I07Rik | | 0 | 0 | 0 | 0 | 0 | 0 | 0 | 0 | 0 | 0 | 0 |
| 0 | NA | NA | NA | NA | NA | Bcl2a1c | B cell leukemia/lymphoma 2 related prote | 0 | 0 | 0 | 0 | 0 | 0 | 0 | 0 | 0 | 0 | 0 |
| 0 | NA | NA | NA | NA | NA | C130032M10Rik | | 0 | 0 | 0 | 0 | 0 | 0 | 0 | 0 | 0 | 0 | 0 |
| 0 | NA | NA | NA | NA | NA | Gm23232 | | 0 | 0 | 0 | 0 | 0 | 0 | 0 | 0 | 0 | 0 | 0 |
| 0 | NA | NA | NA | NA | NA | Gm23889 | | 0 | 0 | 0 | 0 | 0 | 0 | 0 | 0 | 0 | 0 | 0 |
| 0 | NA | NA | NA | NA | NA | Gm4665 | | 0 | 0 | 0 | 0 | 0 | 0 | 0 | 0 | 0 | 0 | 0 |
| 0 | NA | NA | NA | NA | NA | Gm26962 | | 0 | 0 | 0 | 0 | 0 | 0 | 0 | 0 | 0 | 0 | 0 |
| 0 | NA | NA | NA | NA | NA | Stmn1-rs1 | | 0 | 0 | 0 | 0 | 0 | 0 | 0 | 0 | 0 | 0 | 0 |
| 0 | NA | NA | NA | NA | NA | Mir467h | | 0 | 0 | 0 | 0 | 0 | 0 | 0 | 0 | 0 | 0 | 0 |
| 0 | NA | NA | NA | NA | NA | Gm26303 | | 0 | 0 | 0 | 0 | 0 | 0 | 0 | 0 | 0 | 0 | 0 |
| 0 | NA | NA | NA | NA | NA | Gm25510 | | 0 | 0 | 0 | 0 | 0 | 0 | 0 | 0 | 0 | 0 | 0 |
| 0 | NA | NA | NA | NA | NA | Gm22243 | | 0 | 0 | 0 | 0 | 0 | 0 | 0 | 0 | 0 | 0 | 0 |
| 0 | NA | NA | NA | NA | NA | n-R5s92 | | 0 | 0 | 0 | 0 | 0 | 0 | 0 | 0 | 0 | 0 | 0 |
| 0 | NA | NA | NA | NA | NA | D330037F02Rik | | 0 | 0 | 0 | 0 | 0 | 0 | 0 | 0 | 0 | 0 | 0 |
| 0 | NA | NA | NA | NA | NA | Gm48007 | | 0 | 0 | 0 | 0 | 0 | 0 | 0 | 0 | 0 | 0 | 0 |
| 0 | NA | NA | NA | NA | NA | Gm16142 | | 0 | 0 | 0 | 0 | 0 | 0 | 0 | 0 | 0 | 0 | 0 |
| 0 | NA | NA | NA | NA | NA | Gm4668 | | 0 | 0 | 0 | 0 | 0 | 0 | 0 | 0 | 0 | 0 | 0 |
| 0 | NA | NA | NA | NA | NA | Gm48038 | | 0 | 0 | 0 | 0 | 0 | 0 | 0 | 0 | 0 | 0 | 0 |
| 0 | NA | NA | NA | NA | NA | Gm31410 | | 0 | 0 | 0 | 0 | 0 | 0 | 0 | 0 | 0 | 0 | 0 |
| 0 | NA | NA | NA | NA | NA | D730003K21Rik | | 0 | 0 | 0 | 0 | 0 | 0 | 0 | 0 | 0 | 0 | 0 |
| 0 | NA | NA | NA | NA | NA | Gm18489 | | 0 | 0 | 0 | 0 | 0 | 0 | 0 | 0 | 0 | 0 | 0 |
| 0 | NA | NA | NA | NA | NA | Gm20397 | | 0 | 0 | 0 | 0 | 0 | 0 | 0 | 0 | 0 | 0 | 0 |
| 0 | NA | NA | NA | NA | NA | Gm20396 | | 0 | 0 | 0 | 0 | 0 | 0 | 0 | 0 | 0 | 0 | 0 |
| 0 | NA | NA | NA | NA | NA | Gm22008 | | 0 | 0 | 0 | 0 | 0 | 0 | 0 | 0 | 0 | 0 | 0 |
| 0 | NA | NA | NA | NA | NA | Gm22198 | | 0 | 0 | 0 | 0 | 0 | 0 | 0 | 0 | 0 | 0 | 0 |
| 0 | NA | NA | NA | NA | NA | 2610509F24Rik | | 0 | 0 | 0 | 0 | 0 | 0 | 0 | 0 | 0 | 0 | 0 |
| 0 | NA | NA | NA | NA | NA | Platr11 | | 0 | 0 | 0 | 0 | 0 | 0 | 0 | 0 | 0 | 0 | 0 |
| 0 | NA | NA | NA | NA | NA | 4933432G23Rik | | 0 | 0 | 0 | 0 | 0 | 0 | 0 | 0 | 0 | 0 | 0 |
| 0 | NA | NA | NA | NA | NA | Gm22479 | | 0 | 0 | 0 | 0 | 0 | 0 | 0 | 0 | 0 | 0 | 0 |
| 0 | NA | NA | NA | NA | NA | Gm33460 | | 0 | 0 | 0 | 0 | 0 | 0 | 0 | 0 | 0 | 0 | 0 |
| 0 | NA | NA | NA | NA | NA | Gm18935 | | 0 | 0 | 0 | 0 | 0 | 0 | 0 | 0 | 0 | 0 | 0 |
| 0 | NA | NA | NA | NA | NA | Gm45897 | | 0 | 0 | 0 | 0 | 0 | 0 | 0 | 0 | 0 | 0 | 0 |
| 0 | NA | NA | NA | NA | NA | Gm47289 | | 0 | 0 | 0 | 0 | 0 | 0 | 0 | 0 | 0 | 0 | 0 |
| 0 | NA | NA | NA | NA | NA | Gm22729 | | 0 | 0 | 0 | 0 | 0 | 0 | 0 | 0 | 0 | 0 | 0 |
| 0 | NA | NA | NA | NA | NA | Gm47320 | | 0 | 0 | 0 | 0 | 0 | 0 | 0 | 0 | 0 | 0 | 0 |
| 0 | NA | NA | NA | NA | NA | Scn10a | sodium channel, voltage-gated, type X, al | 0 | 0 | 0 | 0 | 0 | 0 | 0 | 0 | 0 | 0 | 0 |
| 0 | NA | NA | NA | NA | NA | Gm47043 | | 0 | 0 | 0 | 0 | 0 | 0 | 0 | 0 | 0 | 0 | 0 |
| 0 | NA | NA | NA | NA | NA | Gm49427 | | 0 | 0 | 0 | 0 | 0 | 0 | 0 | 0 | 0 | 0 | 0 |
| 0 | NA | NA | NA | NA | NA | Gm33858 | | 0 | 0 | 0 | 0 | 0 | 0 | 0 | 0 | 0 | 0 | 0 |
| 0 | NA | NA | NA | NA | NA | Ccr8 | chemokine (C-C motif) receptor 8 | 0 | 0 | 0 | 0 | 0 | 0 | 0 | 0 | 0 | 0 | 0 |
| 0 | NA | NA | NA | NA | NA | Gm24044 | | 0 | 0 | 0 | 0 | 0 | 0 | 0 | 0 | 0 | 0 | 0 |
| 0 | NA | NA | NA | NA | NA | Gm5922 | | 0 | 0 | 0 | 0 | 0 | 0 | 0 | 0 | 0 | 0 | 0 |
| 0 | NA | NA | NA | NA | NA | Gm38642 | | 0 | 0 | 0 | 0 | 0 | 0 | 0 | 0 | 0 | 0 | 0 |
| 0 | NA | NA | NA | NA | NA | Gm38642 | | 0 | 0 | 0 | 0 | 0 | 0 | 0 | 0 | 0 | 0 | 0 |
| 0 | NA | NA | NA | NA | NA | Gm47059 | | 0 | 0 | 0 | 0 | 0 | 0 | 0 | 0 | 0 | 0 | 0 |
| 0 | NA | NA | NA | NA | NA | Gm15566 | | 0 | 0 | 0 | 0 | 0 | 0 | 0 | 0 | 0 | 0 | 0 |
| 0 | NA | NA | NA | NA | NA | Gm15565 | | 0 | 0 | 0 | 0 | 0 | 0 | 0 | 0 | 0 | 0 | 0 |
| 0 | NA | NA | NA | NA | NA | Gm47062 | | 0 | 0 | 0 | 0 | 0 | 0 | 0 | 0 | 0 | 0 | 0 |

| | | | | | | | | | | | | | | | | | | |
|---|---|---|---|---|---|---|---|---|---|---|---|---|---|---|---|---|---|---|
| 0 | NA | NA | NA | NA | NA | Gm39456 | | 0 | 0 | 0 | 0 | 0 | 0 | 0 | 0 | 0 | 0 | 0 |
| 0 | NA | NA | NA | NA | NA | Gm39458 | | 0 | 0 | 0 | 0 | 0 | 0 | 0 | 0 | 0 | 0 | 0 |
| 0 | NA | NA | NA | NA | NA | Gm47063 | | 0 | 0 | 0 | 0 | 0 | 0 | 0 | 0 | 0 | 0 | 0 |
| 0 | NA | NA | NA | NA | NA | Gm47064 | | 0 | 0 | 0 | 0 | 0 | 0 | 0 | 0 | 0 | 0 | 0 |
| 0 | NA | NA | NA | NA | NA | Gm23624 | | 0 | 0 | 0 | 0 | 0 | 0 | 0 | 0 | 0 | 0 | 0 |
| 0 | NA | NA | NA | NA | NA | Gm39459 | | 0 | 0 | 0 | 0 | 0 | 0 | 0 | 0 | 0 | 0 | 0 |
| 0 | NA | NA | NA | NA | NA | Gm47068 | | 0 | 0 | 0 | 0 | 0 | 0 | 0 | 0 | 0 | 0 | 0 |
| 0 | NA | NA | NA | NA | NA | Gm47069 | | 0 | 0 | 0 | 0 | 0 | 0 | 0 | 0 | 0 | 0 | 0 |
| 0 | NA | NA | NA | NA | NA | Gm47070 | | 0 | 0 | 0 | 0 | 0 | 0 | 0 | 0 | 0 | 0 | 0 |
| 0 | NA | NA | NA | NA | NA | 1700020M21Rik | | 0 | 0 | 0 | 0 | 0 | 0 | 0 | 0 | 0 | 0 | 0 |
| 0 | NA | NA | NA | NA | NA | Gm22987 | | 0 | 0 | 0 | 0 | 0 | 0 | 0 | 0 | 0 | 0 | 0 |
| 0 | NA | NA | NA | NA | NA | Gm17157 | | 0 | 0 | 0 | 0 | 0 | 0 | 0 | 0 | 0 | 0 | 0 |
| 0 | NA | NA | NA | NA | NA | Gm47092 | | 0 | 0 | 0 | 0 | 0 | 0 | 0 | 0 | 0 | 0 | 0 |
| 0 | NA | NA | NA | NA | NA | Gm47095 | | 0 | 0 | 0 | 0 | 0 | 0 | 0 | 0 | 0 | 0 | 0 |
| 0 | NA | NA | NA | NA | NA | Gm38661 | | 0 | 0 | 0 | 0 | 0 | 0 | 0 | 0 | 0 | 0 | 0 |
| 0 | NA | NA | NA | NA | NA | Lyzl4os | | 0 | 0 | 0 | 0 | 0 | 0 | 0 | 0 | 0 | 0 | 0 |
| 0 | NA | NA | NA | NA | NA | Gm17163 | | 0 | 0 | 0 | 0 | 0 | 0 | 0 | 0 | 0 | 0 | 0 |
| 0 | NA | NA | NA | NA | NA | Gm47111 | | 0 | 0 | 0 | 0 | 0 | 0 | 0 | 0 | 0 | 0 | 0 |
| 0 | NA | NA | NA | NA | NA | Gm47115 | | 0 | 0 | 0 | 0 | 0 | 0 | 0 | 0 | 0 | 0 | 0 |
| 0 | NA | NA | NA | NA | NA | Gm49343 | | 0 | 0 | 0 | 0 | 0 | 0 | 0 | 0 | 0 | 0 | 0 |
| 0 | NA | NA | NA | NA | NA | Gm47117 | | 0 | 0 | 0 | 0 | 0 | 0 | 0 | 0 | 0 | 0 | 0 |
| 0 | NA | NA | NA | NA | NA | Mir8106 | | 0 | 0 | 0 | 0 | 0 | 0 | 0 | 0 | 0 | 0 | 0 |
| 0 | NA | NA | NA | NA | NA | Gm47126 | | 0 | 0 | 0 | 0 | 0 | 0 | 0 | 0 | 0 | 0 | 0 |
| 0 | NA | NA | NA | NA | NA | Gm47128 | | 0 | 0 | 0 | 0 | 0 | 0 | 0 | 0 | 0 | 0 | 0 |
| 0 | NA | NA | NA | NA | NA | Gm35454 | | 0 | 0 | 0 | 0 | 0 | 0 | 0 | 0 | 0 | 0 | 0 |
| 0 | NA | NA | NA | NA | NA | Gm47129 | | 0 | 0 | 0 | 0 | 0 | 0 | 0 | 0 | 0 | 0 | 0 |
| 0 | NA | NA | NA | NA | NA | Gm47130 | | 0 | 0 | 0 | 0 | 0 | 0 | 0 | 0 | 0 | 0 | 0 |
| 0 | NA | NA | NA | NA | NA | Gm47131 | | 0 | 0 | 0 | 0 | 0 | 0 | 0 | 0 | 0 | 0 | 0 |
| 0 | NA | NA | NA | NA | NA | Gm47133 | | 0 | 0 | 0 | 0 | 0 | 0 | 0 | 0 | 0 | 0 | 0 |
| 0 | NA | NA | NA | NA | NA | 9530059O14Rik | | 0 | 0 | 0 | 0 | 0 | 0 | 0 | 0 | 0 | 0 | 0 |
| 0 | NA | NA | NA | NA | NA | Gm47134 | | 0 | 0 | 0 | 0 | 0 | 0 | 0 | 0 | 0 | 0 | 0 |
| 0 | NA | NA | NA | NA | NA | Mir138-1 | | 0 | 0 | 0 | 0 | 0 | 0 | 0 | 0 | 0 | 0 | 0 |
| 0 | NA | NA | NA | NA | NA | Gm35501 | | 0 | 0 | 0 | 0 | 0 | 0 | 0 | 0 | 0 | 0 | 0 |
| 0 | NA | NA | NA | NA | NA | Gm47137 | | 0 | 0 | 0 | 0 | 0 | 0 | 0 | 0 | 0 | 0 | 0 |
| 0 | NA | NA | NA | NA | NA | Topaz1 | testis and ovary specific PAZ domain cont | 0 | 0 | 0 | 0 | 0 | 0 | 0 | 0 | 0 | 0 | 0 |
| 0 | NA | NA | NA | NA | NA | Gm35549 | predicted gene, 35549 | 0 | 0 | 0 | 0 | 0 | 0 | 0 | 0 | 0 | 0 | 0 |
| 0 | NA | NA | NA | NA | NA | Gm47145 | | 0 | 0 | 0 | 0 | 0 | 0 | 0 | 0 | 0 | 0 | 0 |
| 0 | NA | NA | NA | NA | NA | Gm47146 | | 0 | 0 | 0 | 0 | 0 | 0 | 0 | 0 | 0 | 0 | 0 |
| 0 | NA | NA | NA | NA | NA | Gm49323 | | 0 | 0 | 0 | 0 | 0 | 0 | 0 | 0 | 0 | 0 | 0 |
| 0 | NA | NA | NA | NA | NA | Gm47159 | | 0 | 0 | 0 | 0 | 0 | 0 | 0 | 0 | 0 | 0 | 0 |
| 0 | NA | NA | NA | NA | NA | Gm17120 | | 0 | 0 | 0 | 0 | 0 | 0 | 0 | 0 | 0 | 0 | 0 |
| 0 | NA | NA | NA | NA | NA | Gm47173 | | 0 | 0 | 0 | 0 | 0 | 0 | 0 | 0 | 0 | 0 | 0 |
| 0 | NA | NA | NA | NA | NA | Gm17020 | | 0 | 0 | 0 | 0 | 0 | 0 | 0 | 0 | 0 | 0 | 0 |
| 0 | NA | NA | NA | NA | NA | Gm2774 | | 0 | 0 | 0 | 0 | 0 | 0 | 0 | 0 | 0 | 0 | 0 |
| 0 | NA | NA | NA | NA | NA | Ccr1l1 | chemokine (C-C motif) receptor 1-like 1 | 0 | 0 | 0 | 0 | 0 | 0 | 0 | 0 | 0 | 0 | 0 |
| 0 | NA | NA | NA | NA | NA | Gm47180 | | 0 | 0 | 0 | 0 | 0 | 0 | 0 | 0 | 0 | 0 | 0 |
| 0 | NA | NA | NA | NA | NA | Gm47179 | | 0 | 0 | 0 | 0 | 0 | 0 | 0 | 0 | 0 | 0 | 0 |
| 0 | NA | NA | NA | NA | NA | Gm47181 | | 0 | 0 | 0 | 0 | 0 | 0 | 0 | 0 | 0 | 0 | 0 |
| 0 | NA | NA | NA | NA | NA | Nlrp4g | NLR family, pyrin domain containing 4G | 0 | 0 | 0 | 0 | 0 | 0 | 0 | 0 | 0 | 0 | 0 |
| 0 | NA | NA | NA | NA | NA | Gm47182 | | 0 | 0 | 0 | 0 | 0 | 0 | 0 | 0 | 0 | 0 | 0 |
| 0 | NA | NA | NA | NA | NA | Gm47185 | | 0 | 0 | 0 | 0 | 0 | 0 | 0 | 0 | 0 | 0 | 0 |
| 0 | NA | NA | NA | NA | NA | Gm47183 | | 0 | 0 | 0 | 0 | 0 | 0 | 0 | 0 | 0 | 0 | 0 |
| 0 | NA | NA | NA | NA | NA | Gm21836 | | 0 | 0 | 0 | 0 | 0 | 0 | 0 | 0 | 0 | 0 | 0 |
| 0 | NA | NA | NA | NA | NA | Gm30674 | | 0 | 0 | 0 | 0 | 0 | 0 | 0 | 0 | 0 | 0 | 0 |
| 0 | NA | NA | NA | NA | NA | Gm3318 | | 0 | 0 | 0 | 0 | 0 | 0 | 0 | 0 | 0 | 0 | 0 |
| 0 | NA | NA | NA | NA | NA | 9230019H11Ril | RIKEN cDNA 9230019H11 gene | 0 | 0 | 0 | 0 | 0 | 0 | 0 | 0 | 0 | 0 | 0 |
| 0 | NA | NA | NA | NA | NA | B020014A21Rik | | 0 | 0 | 0 | 0 | 0 | 0 | 0 | 0 | 0 | 0 | 0 |
| 0 | NA | NA | NA | NA | NA | Gm8155 | | 0 | 0 | 0 | 0 | 0 | 0 | 0 | 0 | 0 | 0 | 0 |
| 0 | NA | NA | NA | NA | NA | Gm48097 | | 0 | 0 | 0 | 0 | 0 | 0 | 0 | 0 | 0 | 0 | 0 |
| 0 | NA | NA | NA | NA | NA | Gm6667 | | 0 | 0 | 0 | 0 | 0 | 0 | 0 | 0 | 0 | 0 | 0 |

| | | | | | | | | | | | | | | | | | | |
|---|---|---|---|---|---|---|---|---|---|---|---|---|---|---|---|---|---|---|
| 0 | NA | NA | NA | NA | NA | H60c | histocompatibility 60c | 0 | 0 | 0 | 0 | 0 | 0 | 0 | 0 | 0 | 0 | 0 |
| 0 | NA | NA | NA | NA | NA | Gm18703 | | 0 | 0 | 0 | 0 | 0 | 0 | 0 | 0 | 0 | 0 | 0 |
| 0 | NA | NA | NA | NA | NA | Gm24482 | | 0 | 0 | 0 | 0 | 0 | 0 | 0 | 0 | 0 | 0 | 0 |
| 0 | NA | NA | NA | NA | NA | Gm16149 | | 0 | 0 | 0 | 0 | 0 | 0 | 0 | 0 | 0 | 0 | 0 |
| 0 | NA | NA | NA | NA | NA | Gm23023 | | 0 | 0 | 0 | 0 | 0 | 0 | 0 | 0 | 0 | 0 | 0 |
| 0 | NA | NA | NA | NA | NA | Gm25515 | | 0 | 0 | 0 | 0 | 0 | 0 | 0 | 0 | 0 | 0 | 0 |
| 0 | NA | NA | NA | NA | NA | Gm25369 | | 0 | 0 | 0 | 0 | 0 | 0 | 0 | 0 | 0 | 0 | 0 |
| 0 | NA | NA | NA | NA | NA | Gm30570 | | 0 | 0 | 0 | 0 | 0 | 0 | 0 | 0 | 0 | 0 | 0 |
| 0 | NA | NA | NA | NA | NA | Gm16153 | | 0 | 0 | 0 | 0 | 0 | 0 | 0 | 0 | 0 | 0 | 0 |
| 0 | NA | NA | NA | NA | NA | Gm22739 | | 0 | 0 | 0 | 0 | 0 | 0 | 0 | 0 | 0 | 0 | 0 |
| 0 | NA | NA | NA | NA | NA | Gm3266 | | 0 | 0 | 0 | 0 | 0 | 0 | 0 | 0 | 0 | 0 | 0 |
| 0 | NA | NA | NA | NA | NA | Gm10097 | | 0 | 0 | 0 | 0 | 0 | 0 | 0 | 0 | 0 | 0 | 0 |
| 0 | NA | NA | NA | NA | NA | Gm48836 | | 0 | 0 | 0 | 0 | 0 | 0 | 0 | 0 | 0 | 0 | 0 |
| 0 | NA | NA | NA | NA | NA | Gm23573 | | 0 | 0 | 0 | 0 | 0 | 0 | 0 | 0 | 0 | 0 | 0 |
| 0 | NA | NA | NA | NA | NA | Gm47679 | | 0 | 0 | 0 | 0 | 0 | 0 | 0 | 0 | 0 | 0 | 0 |
| 0 | NA | NA | NA | NA | NA | Gm3213 | | 0 | 0 | 0 | 0 | 0 | 0 | 0 | 0 | 0 | 0 | 0 |
| 0 | NA | NA | NA | NA | NA | Gm47755 | | 0 | 0 | 0 | 0 | 0 | 0 | 0 | 0 | 0 | 0 | 0 |
| 0 | NA | NA | NA | NA | NA | Gm4921 | | 0 | 0 | 0 | 0 | 0 | 0 | 0 | 0 | 0 | 0 | 0 |
| 0 | NA | NA | NA | NA | NA | Gm10945 | | 0 | 0 | 0 | 0 | 0 | 0 | 0 | 0 | 0 | 0 | 0 |
| 0 | NA | NA | NA | NA | NA | Gm21321 | | 0 | 0 | 0 | 0 | 0 | 0 | 0 | 0 | 0 | 0 | 0 |
| 0 | NA | NA | NA | NA | NA | Gm48121 | | 0 | 0 | 0 | 0 | 0 | 0 | 0 | 0 | 0 | 0 | 0 |
| 0 | NA | NA | NA | NA | NA | Gm16254 | | 0 | 0 | 0 | 0 | 0 | 0 | 0 | 0 | 0 | 0 | 0 |
| 0 | NA | NA | NA | NA | NA | Gm15560 | | 0 | 0 | 0 | 0 | 0 | 0 | 0 | 0 | 0 | 0 | 0 |
| 0 | NA | NA | NA | NA | NA | Mir5104 | | 0 | 0 | 0 | 0 | 0 | 0 | 0 | 0 | 0 | 0 | 0 |
| 0 | NA | NA | NA | NA | NA | Gm24726 | | 0 | 0 | 0 | 0 | 0 | 0 | 0 | 0 | 0 | 0 | 0 |
| 0 | NA | NA | NA | NA | NA | Gm48614 | | 0 | 0 | 0 | 0 | 0 | 0 | 0 | 0 | 0 | 0 | 0 |
| 0 | NA | NA | NA | NA | NA | Gm30906 | | 0 | 0 | 0 | 0 | 0 | 0 | 0 | 0 | 0 | 0 | 0 |
| 0 | NA | NA | NA | NA | NA | Gm24374 | | 0 | 0 | 0 | 0 | 0 | 0 | 0 | 0 | 0 | 0 | 0 |
| 0 | NA | NA | NA | NA | NA | Gm48654 | | 0 | 0 | 0 | 0 | 0 | 0 | 0 | 0 | 0 | 0 | 0 |
| 0 | NA | NA | NA | NA | NA | Gm25410 | | 0 | 0 | 0 | 0 | 0 | 0 | 0 | 0 | 0 | 0 | 0 |
| 0 | NA | NA | NA | NA | NA | 4930553I21Rik | | 0 | 0 | 0 | 0 | 0 | 0 | 0 | 0 | 0 | 0 | 0 |
| 0 | NA | NA | NA | NA | NA | Gm48728 | | 0 | 0 | 0 | 0 | 0 | 0 | 0 | 0 | 0 | 0 | 0 |
| 0 | NA | NA | NA | NA | NA | Gm48744 | | 0 | 0 | 0 | 0 | 0 | 0 | 0 | 0 | 0 | 0 | 0 |
| 0 | NA | NA | NA | NA | NA | Gm48748 | | 0 | 0 | 0 | 0 | 0 | 0 | 0 | 0 | 0 | 0 | 0 |
| 0 | NA | NA | NA | NA | NA | Gm48749 | | 0 | 0 | 0 | 0 | 0 | 0 | 0 | 0 | 0 | 0 | 0 |
| 0 | NA | NA | NA | NA | NA | Gm48751 | | 0 | 0 | 0 | 0 | 0 | 0 | 0 | 0 | 0 | 0 | 0 |
| 0 | NA | NA | NA | NA | NA | Gm48753 | | 0 | 0 | 0 | 0 | 0 | 0 | 0 | 0 | 0 | 0 | 0 |
| 0 | NA | NA | NA | NA | NA | Gm9930 | | 0 | 0 | 0 | 0 | 0 | 0 | 0 | 0 | 0 | 0 | 0 |
| 0 | NA | NA | NA | NA | NA | Gm48755 | | 0 | 0 | 0 | 0 | 0 | 0 | 0 | 0 | 0 | 0 | 0 |
| 0 | NA | NA | NA | NA | NA | Gm48757 | | 0 | 0 | 0 | 0 | 0 | 0 | 0 | 0 | 0 | 0 | 0 |
| 0 | NA | NA | NA | NA | NA | Gm48758 | | 0 | 0 | 0 | 0 | 0 | 0 | 0 | 0 | 0 | 0 | 0 |
| 0 | NA | NA | NA | NA | NA | Gm48759 | | 0 | 0 | 0 | 0 | 0 | 0 | 0 | 0 | 0 | 0 | 0 |
| 0 | NA | NA | NA | NA | NA | Gm48760 | | 0 | 0 | 0 | 0 | 0 | 0 | 0 | 0 | 0 | 0 | 0 |
| 0 | NA | NA | NA | NA | NA | Gm6150 | | 0 | 0 | 0 | 0 | 0 | 0 | 0 | 0 | 0 | 0 | 0 |
| 0 | NA | NA | NA | NA | NA | Gm19161 | | 0 | 0 | 0 | 0 | 0 | 0 | 0 | 0 | 0 | 0 | 0 |
| 0 | NA | NA | NA | NA | NA | Gm25635 | | 0 | 0 | 0 | 0 | 0 | 0 | 0 | 0 | 0 | 0 | 0 |
| 0 | NA | NA | NA | NA | NA | Gm48282 | | 0 | 0 | 0 | 0 | 0 | 0 | 0 | 0 | 0 | 0 | 0 |
| 0 | NA | NA | NA | NA | NA | Gm48281 | | 0 | 0 | 0 | 0 | 0 | 0 | 0 | 0 | 0 | 0 | 0 |
| 0 | NA | NA | NA | NA | NA | Gm25682 | | 0 | 0 | 0 | 0 | 0 | 0 | 0 | 0 | 0 | 0 | 0 |
| 0 | NA | NA | NA | NA | NA | Gm48279 | | 0 | 0 | 0 | 0 | 0 | 0 | 0 | 0 | 0 | 0 | 0 |
| 0 | NA | NA | NA | NA | NA | Gm48280 | | 0 | 0 | 0 | 0 | 0 | 0 | 0 | 0 | 0 | 0 | 0 |
| 0 | NA | NA | NA | NA | NA | Gm31904 | | 0 | 0 | 0 | 0 | 0 | 0 | 0 | 0 | 0 | 0 | 0 |
| 0 | NA | NA | NA | NA | NA | Gm48324 | | 0 | 0 | 0 | 0 | 0 | 0 | 0 | 0 | 0 | 0 | 0 |
| 0 | NA | NA | NA | NA | NA | Gm48326 | | 0 | 0 | 0 | 0 | 0 | 0 | 0 | 0 | 0 | 0 | 0 |
| 0 | NA | NA | NA | NA | NA | Gm23601 | | 0 | 0 | 0 | 0 | 0 | 0 | 0 | 0 | 0 | 0 | 0 |
| 0 | NA | NA | NA | NA | NA | Gm10944 | | 0 | 0 | 0 | 0 | 0 | 0 | 0 | 0 | 0 | 0 | 0 |
| 0 | NA | NA | NA | NA | NA | Gm48400 | | 0 | 0 | 0 | 0 | 0 | 0 | 0 | 0 | 0 | 0 | 0 |
| 0 | NA | NA | NA | NA | NA | 4930567K20Rik | | 0 | 0 | 0 | 0 | 0 | 0 | 0 | 0 | 0 | 0 | 0 |
| 0 | NA | NA | NA | NA | NA | Gm48406 | | 0 | 0 | 0 | 0 | 0 | 0 | 0 | 0 | 0 | 0 | 0 |
| 0 | NA | NA | NA | NA | NA | Gm48666 | | 0 | 0 | 0 | 0 | 0 | 0 | 0 | 0 | 0 | 0 | 0 |

| | | | | | | | | | | | | | | | | | | |
|---|---|---|---|---|---|---|---|---|---|---|---|---|---|---|---|---|---|---|
| 0 | NA | NA | NA | NA | NA | Gm25979 | | 0 | 0 | 0 | 0 | 0 | 0 | 0 | 0 | 0 | 0 | 0 |
| 0 | NA | NA | NA | NA | NA | Rpl17-ps11 | | 0 | 0 | 0 | 0 | 0 | 0 | 0 | 0 | 0 | 0 | 0 |
| 0 | NA | NA | NA | NA | NA | Gm48679 | | 0 | 0 | 0 | 0 | 0 | 0 | 0 | 0 | 0 | 0 | 0 |
| 0 | NA | NA | NA | NA | NA | Gm48680 | | 0 | 0 | 0 | 0 | 0 | 0 | 0 | 0 | 0 | 0 | 0 |
| 0 | NA | NA | NA | NA | NA | Gm22546 | | 0 | 0 | 0 | 0 | 0 | 0 | 0 | 0 | 0 | 0 | 0 |
| 0 | NA | NA | NA | NA | NA | Gm48697 | | 0 | 0 | 0 | 0 | 0 | 0 | 0 | 0 | 0 | 0 | 0 |
| 0 | NA | NA | NA | NA | NA | Gm48698 | | 0 | 0 | 0 | 0 | 0 | 0 | 0 | 0 | 0 | 0 | 0 |
| 0 | NA | NA | NA | NA | NA | Gm22194 | | 0 | 0 | 0 | 0 | 0 | 0 | 0 | 0 | 0 | 0 | 0 |
| 0 | NA | NA | NA | NA | NA | Gm25011 | | 0 | 0 | 0 | 0 | 0 | 0 | 0 | 0 | 0 | 0 | 0 |
| 0 | NA | NA | NA | NA | NA | Gm48721 | | 0 | 0 | 0 | 0 | 0 | 0 | 0 | 0 | 0 | 0 | 0 |
| 0 | NA | NA | NA | NA | NA | Gm48722 | | 0 | 0 | 0 | 0 | 0 | 0 | 0 | 0 | 0 | 0 | 0 |
| 0 | NA | NA | NA | NA | NA | Gm48723 | | 0 | 0 | 0 | 0 | 0 | 0 | 0 | 0 | 0 | 0 | 0 |
| 0 | NA | NA | NA | NA | NA | Gm48724 | | 0 | 0 | 0 | 0 | 0 | 0 | 0 | 0 | 0 | 0 | 0 |
| 0 | NA | NA | NA | NA | NA | Gm23680 | | 0 | 0 | 0 | 0 | 0 | 0 | 0 | 0 | 0 | 0 | 0 |
| 0 | NA | NA | NA | NA | NA | Gm25139 | | 0 | 0 | 0 | 0 | 0 | 0 | 0 | 0 | 0 | 0 | 0 |
| 0 | NA | NA | NA | NA | NA | Gm7525 | | 0 | 0 | 0 | 0 | 0 | 0 | 0 | 0 | 0 | 0 | 0 |
| 0 | NA | NA | NA | NA | NA | Gm47691 | | 0 | 0 | 0 | 0 | 0 | 0 | 0 | 0 | 0 | 0 | 0 |
| 0 | NA | NA | NA | NA | NA | Gm27741 | | 0 | 0 | 0 | 0 | 0 | 0 | 0 | 0 | 0 | 0 | 0 |
| 0 | NA | NA | NA | NA | NA | Zc2hc1b | zinc finger, C2HC-type containing 1B | 0 | 0 | 0 | 0 | 0 | 0 | 0 | 0 | 0 | 0 | 0 |
| 0 | NA | NA | NA | NA | NA | Gm47402 | | 0 | 0 | 0 | 0 | 0 | 0 | 0 | 0 | 0 | 0 | 0 |
| 0 | NA | NA | NA | NA | NA | Gm48773 | | 0 | 0 | 0 | 0 | 0 | 0 | 0 | 0 | 0 | 0 | 0 |
| 0 | NA | NA | NA | NA | NA | Gm32172 | | 0 | 0 | 0 | 0 | 0 | 0 | 0 | 0 | 0 | 0 | 0 |
| 0 | NA | NA | NA | NA | NA | Gm32105 | | 0 | 0 | 0 | 0 | 0 | 0 | 0 | 0 | 0 | 0 | 0 |
| 0 | NA | NA | NA | NA | NA | Gm22995 | | 0 | 0 | 0 | 0 | 0 | 0 | 0 | 0 | 0 | 0 | 0 |
| 0 | NA | NA | NA | NA | NA | 4930444K16Rik | | 0 | 0 | 0 | 0 | 0 | 0 | 0 | 0 | 0 | 0 | 0 |
| 0 | NA | NA | NA | NA | NA | Gm28289 | | 0 | 0 | 0 | 0 | 0 | 0 | 0 | 0 | 0 | 0 | 0 |
| 0 | NA | NA | NA | NA | NA | Gm48843 | | 0 | 0 | 0 | 0 | 0 | 0 | 0 | 0 | 0 | 0 | 0 |
| 0 | NA | NA | NA | NA | NA | Gm48845 | | 0 | 0 | 0 | 0 | 0 | 0 | 0 | 0 | 0 | 0 | 0 |
| 0 | NA | NA | NA | NA | NA | Gm48844 | | 0 | 0 | 0 | 0 | 0 | 0 | 0 | 0 | 0 | 0 | 0 |
| 0 | NA | NA | NA | NA | NA | Gm24343 | | 0 | 0 | 0 | 0 | 0 | 0 | 0 | 0 | 0 | 0 | 0 |
| 0 | NA | NA | NA | NA | NA | Gm40603 | | 0 | 0 | 0 | 0 | 0 | 0 | 0 | 0 | 0 | 0 | 0 |
| 0 | NA | NA | NA | NA | NA | Gm37299 | | 0 | 0 | 0 | 0 | 0 | 0 | 0 | 0 | 0 | 0 | 0 |
| 0 | NA | NA | NA | NA | NA | Gm48854 | | 0 | 0 | 0 | 0 | 0 | 0 | 0 | 0 | 0 | 0 | 0 |
| 0 | NA | NA | NA | NA | NA | B230364G03Rik | | 0 | 0 | 0 | 0 | 0 | 0 | 0 | 0 | 0 | 0 | 0 |
| 0 | NA | NA | NA | NA | NA | Gm47698 | | 0 | 0 | 0 | 0 | 0 | 0 | 0 | 0 | 0 | 0 | 0 |
| 0 | NA | NA | NA | NA | NA | Gm47699 | | 0 | 0 | 0 | 0 | 0 | 0 | 0 | 0 | 0 | 0 | 0 |
| 0 | NA | NA | NA | NA | NA | Gm23892 | | 0 | 0 | 0 | 0 | 0 | 0 | 0 | 0 | 0 | 0 | 0 |
| 0 | NA | NA | NA | NA | NA | Gm47703 | | 0 | 0 | 0 | 0 | 0 | 0 | 0 | 0 | 0 | 0 | 0 |
| 0 | NA | NA | NA | NA | NA | Gm18188 | | 0 | 0 | 0 | 0 | 0 | 0 | 0 | 0 | 0 | 0 | 0 |
| 0 | NA | NA | NA | NA | NA | Gm47710 | | 0 | 0 | 0 | 0 | 0 | 0 | 0 | 0 | 0 | 0 | 0 |
| 0 | NA | NA | NA | NA | NA | Gm47712 | | 0 | 0 | 0 | 0 | 0 | 0 | 0 | 0 | 0 | 0 | 0 |
| 0 | NA | NA | NA | NA | NA | Gm46188 | | 0 | 0 | 0 | 0 | 0 | 0 | 0 | 0 | 0 | 0 | 0 |
| 0 | NA | NA | NA | NA | NA | Gm32283 | | 0 | 0 | 0 | 0 | 0 | 0 | 0 | 0 | 0 | 0 | 0 |
| 0 | NA | NA | NA | NA | NA | C330004P14Rik | | 0 | 0 | 0 | 0 | 0 | 0 | 0 | 0 | 0 | 0 | 0 |
| 0 | NA | NA | NA | NA | NA | Gm18557 | | 0 | 0 | 0 | 0 | 0 | 0 | 0 | 0 | 0 | 0 | 0 |
| 0 | NA | NA | NA | NA | NA | Gm47729 | | 0 | 0 | 0 | 0 | 0 | 0 | 0 | 0 | 0 | 0 | 0 |
| 0 | NA | NA | NA | NA | NA | Gm5948 | | 0 | 0 | 0 | 0 | 0 | 0 | 0 | 0 | 0 | 0 | 0 |
| 0 | NA | NA | NA | NA | NA | Gm18039 | | 0 | 0 | 0 | 0 | 0 | 0 | 0 | 0 | 0 | 0 | 0 |
| 0 | NA | NA | NA | NA | NA | Gm47736 | | 0 | 0 | 0 | 0 | 0 | 0 | 0 | 0 | 0 | 0 | 0 |
| 0 | NA | NA | NA | NA | NA | Gm25382 | | 0 | 0 | 0 | 0 | 0 | 0 | 0 | 0 | 0 | 0 | 0 |
| 0 | NA | NA | NA | NA | NA | Gm47760 | | 0 | 0 | 0 | 0 | 0 | 0 | 0 | 0 | 0 | 0 | 0 |
| 0 | NA | NA | NA | NA | NA | Gm47768 | | 0 | 0 | 0 | 0 | 0 | 0 | 0 | 0 | 0 | 0 | 0 |
| 0 | NA | NA | NA | NA | NA | Gm47771 | | 0 | 0 | 0 | 0 | 0 | 0 | 0 | 0 | 0 | 0 | 0 |
| 0 | NA | NA | NA | NA | NA | Gm7584 | | 0 | 0 | 0 | 0 | 0 | 0 | 0 | 0 | 0 | 0 | 0 |
| 0 | NA | NA | NA | NA | NA | Gm17711 | | 0 | 0 | 0 | 0 | 0 | 0 | 0 | 0 | 0 | 0 | 0 |
| 0 | NA | NA | NA | NA | NA | Gm48229 | | 0 | 0 | 0 | 0 | 0 | 0 | 0 | 0 | 0 | 0 | 0 |
| 0 | NA | NA | NA | NA | NA | Gm18942 | | 0 | 0 | 0 | 0 | 0 | 0 | 0 | 0 | 0 | 0 | 0 |
| 0 | NA | NA | NA | NA | NA | Gm4922 | predicted gene 4922 | 0 | 0 | 0 | 0 | 0 | 0 | 0 | 0 | 0 | 0 | 0 |
| 0 | NA | NA | NA | NA | NA | 4930444F02Rik | | 0 | 0 | 0 | 0 | 0 | 0 | 0 | 0 | 0 | 0 | 0 |
| 0 | NA | NA | NA | NA | NA | Gm48483 | | 0 | 0 | 0 | 0 | 0 | 0 | 0 | 0 | 0 | 0 | 0 |

| | | | | | | | | | | | | | | | | | |
|---|---|---|---|---|---|---|---|---|---|---|---|---|---|---|---|---|---|
| 0 | NA | NA | NA | NA | NA | 1700124M09Rik | | 0 | 0 | 0 | 0 | 0 | 0 | 0 | 0 | 0 | 0 | 0 |
| 0 | NA | NA | NA | NA | NA | Gm48545 | | 0 | 0 | 0 | 0 | 0 | 0 | 0 | 0 | 0 | 0 | 0 |
| 0 | NA | NA | NA | NA | NA | Gm20139 | | 0 | 0 | 0 | 0 | 0 | 0 | 0 | 0 | 0 | 0 | 0 |
| 0 | NA | NA | NA | NA | NA | Gm33056 | | 0 | 0 | 0 | 0 | 0 | 0 | 0 | 0 | 0 | 0 | 0 |
| 0 | NA | NA | NA | NA | NA | Olig3 | oligodendrocyte transcription factor 3 | 0 | 0 | 0 | 0 | 0 | 0 | 0 | 0 | 0 | 0 | 0 |
| 0 | NA | NA | NA | NA | NA | Gm48563 | | 0 | 0 | 0 | 0 | 0 | 0 | 0 | 0 | 0 | 0 | 0 |
| 0 | NA | NA | NA | NA | NA | 9230106D20Rik | | 0 | 0 | 0 | 0 | 0 | 0 | 0 | 0 | 0 | 0 | 0 |
| 0 | NA | NA | NA | NA | NA | Gm33332 | | 0 | 0 | 0 | 0 | 0 | 0 | 0 | 0 | 0 | 0 | 0 |
| 0 | NA | NA | NA | NA | NA | Gm25256 | | 0 | 0 | 0 | 0 | 0 | 0 | 0 | 0 | 0 | 0 | 0 |
| 0 | NA | NA | NA | NA | NA | 4930405J17Rik | | 0 | 0 | 0 | 0 | 0 | 0 | 0 | 0 | 0 | 0 | 0 |
| 0 | NA | NA | NA | NA | NA | Gm30285 | | 0 | 0 | 0 | 0 | 0 | 0 | 0 | 0 | 0 | 0 | 0 |
| 0 | NA | NA | NA | NA | NA | Mir6413 | | 0 | 0 | 0 | 0 | 0 | 0 | 0 | 0 | 0 | 0 | 0 |
| 0 | NA | NA | NA | NA | NA | Gm48537 | | 0 | 0 | 0 | 0 | 0 | 0 | 0 | 0 | 0 | 0 | 0 |
| 0 | NA | NA | NA | NA | NA | Gm48651 | | 0 | 0 | 0 | 0 | 0 | 0 | 0 | 0 | 0 | 0 | 0 |
| 0 | NA | NA | NA | NA | NA | Rps2-ps3 | | 0 | 0 | 0 | 0 | 0 | 0 | 0 | 0 | 0 | 0 | 0 |
| 0 | NA | NA | NA | NA | NA | Gm10826 | | 0 | 0 | 0 | 0 | 0 | 0 | 0 | 0 | 0 | 0 | 0 |
| 0 | NA | NA | NA | NA | NA | Gm20149 | | 0 | 0 | 0 | 0 | 0 | 0 | 0 | 0 | 0 | 0 | 0 |
| 0 | NA | NA | NA | NA | NA | Gm48189 | | 0 | 0 | 0 | 0 | 0 | 0 | 0 | 0 | 0 | 0 | 0 |
| 0 | NA | NA | NA | NA | NA | Gm40608 | | 0 | 0 | 0 | 0 | 0 | 0 | 0 | 0 | 0 | 0 | 0 |
| 0 | NA | NA | NA | NA | NA | Gm33619 | | 0 | 0 | 0 | 0 | 0 | 0 | 0 | 0 | 0 | 0 | 0 |
| 0 | NA | NA | NA | NA | NA | Gm33728 | | 0 | 0 | 0 | 0 | 0 | 0 | 0 | 0 | 0 | 0 | 0 |
| 0 | NA | NA | NA | NA | NA | Aldh8a1 | aldehyde dehydrogenase 8 family, memb | 0 | 0 | 0 | 0 | 0 | 0 | 0 | 0 | 0 | 0 | 0 |
| 0 | NA | NA | NA | NA | NA | Fkbp1a-ps2 | | 0 | 0 | 0 | 0 | 0 | 0 | 0 | 0 | 0 | 0 | 0 |
| 0 | NA | NA | NA | NA | NA | 1700021A07Rik | | 0 | 0 | 0 | 0 | 0 | 0 | 0 | 0 | 0 | 0 | 0 |
| 0 | NA | NA | NA | NA | NA | Gm48386 | | 0 | 0 | 0 | 0 | 0 | 0 | 0 | 0 | 0 | 0 | 0 |
| 0 | NA | NA | NA | NA | NA | 1700020N01Rik | | 0 | 0 | 0 | 0 | 0 | 0 | 0 | 0 | 0 | 0 | 0 |
| 0 | NA | NA | NA | NA | NA | Gm29323 | | 0 | 0 | 0 | 0 | 0 | 0 | 0 | 0 | 0 | 0 | 0 |
| 0 | NA | NA | NA | NA | NA | Platr6 | | 0 | 0 | 0 | 0 | 0 | 0 | 0 | 0 | 0 | 0 | 0 |
| 0 | NA | NA | NA | NA | NA | Gm23839 | | 0 | 0 | 0 | 0 | 0 | 0 | 0 | 0 | 0 | 0 | 0 |
| 0 | NA | NA | NA | NA | NA | Gm18637 | | 0 | 0 | 0 | 0 | 0 | 0 | 0 | 0 | 0 | 0 | 0 |
| 0 | NA | NA | NA | NA | NA | 4930444G20Ril | RIKEN cDNA 4930444G20 gene | 0 | 0 | 0 | 0 | 0 | 0 | 0 | 0 | 0 | 0 | 0 |
| 0 | NA | NA | NA | NA | NA | Gm2539 | | 0 | 0 | 0 | 0 | 0 | 0 | 0 | 0 | 0 | 0 | 0 |
| 0 | NA | NA | NA | NA | NA | Gm19791 | | 0 | 0 | 0 | 0 | 0 | 0 | 0 | 0 | 0 | 0 | 0 |
| 0 | NA | NA | NA | NA | NA | Gm7678 | | 0 | 0 | 0 | 0 | 0 | 0 | 0 | 0 | 0 | 0 | 0 |
| 0 | NA | NA | NA | NA | NA | Gm26581 | | 0 | 0 | 0 | 0 | 0 | 0 | 0 | 0 | 0 | 0 | 0 |
| 0 | NA | NA | NA | NA | NA | Gm1972 | | 0 | 0 | 0 | 0 | 0 | 0 | 0 | 0 | 0 | 0 | 0 |
| 0 | NA | NA | NA | NA | NA | Gm16531 | | 0 | 0 | 0 | 0 | 0 | 0 | 0 | 0 | 0 | 0 | 0 |
| 0 | NA | NA | NA | NA | NA | | | 0 | 0 | 0 | 0 | 0 | 0 | 0 | 0 | 0 | 0 | 0 |
| 0 | NA | NA | NA | NA | NA | Gm47634 | | 0 | 0 | 0 | 0 | 0 | 0 | 0 | 0 | 0 | 0 | 0 |
| 0 | NA | NA | NA | NA | NA | Gm47787 | | 0 | 0 | 0 | 0 | 0 | 0 | 0 | 0 | 0 | 0 | 0 |
| 0 | NA | NA | NA | NA | NA | Gm24655 | | 0 | 0 | 0 | 0 | 0 | 0 | 0 | 0 | 0 | 0 | 0 |
| 0 | NA | NA | NA | NA | NA | Gm47843 | | 0 | 0 | 0 | 0 | 0 | 0 | 0 | 0 | 0 | 0 | 0 |
| 0 | NA | NA | NA | NA | NA | 4930520K02Rik | | 0 | 0 | 0 | 0 | 0 | 0 | 0 | 0 | 0 | 0 | 0 |
| 0 | NA | NA | NA | NA | NA | Snora33 | | 0 | 0 | 0 | 0 | 0 | 0 | 0 | 0 | 0 | 0 | 0 |
| 0 | NA | NA | NA | NA | NA | Snord100 | | 0 | 0 | 0 | 0 | 0 | 0 | 0 | 0 | 0 | 0 | 0 |
| 0 | NA | NA | NA | NA | NA | Gm23130 | | 0 | 0 | 0 | 0 | 0 | 0 | 0 | 0 | 0 | 0 | 0 |
| 0 | NA | NA | NA | NA | NA | Taar5 | trace amine-associated receptor 5 | 0 | 0 | 0 | 0 | 0 | 0 | 0 | 0 | 0 | 0 | 0 |
| 0 | NA | NA | NA | NA | NA | Taar6 | trace amine-associated receptor 6 | 0 | 0 | 0 | 0 | 0 | 0 | 0 | 0 | 0 | 0 | 0 |
| 0 | NA | NA | NA | NA | NA | Taar7a | trace amine-associated receptor 7A | 0 | 0 | 0 | 0 | 0 | 0 | 0 | 0 | 0 | 0 | 0 |
| 0 | NA | NA | NA | NA | NA | Taar7b | trace amine-associated receptor 7B | 0 | 0 | 0 | 0 | 0 | 0 | 0 | 0 | 0 | 0 | 0 |
| 0 | NA | NA | NA | NA | NA | Taar7c-ps | | 0 | 0 | 0 | 0 | 0 | 0 | 0 | 0 | 0 | 0 | 0 |
| 0 | NA | NA | NA | NA | NA | Gm15136 | | 0 | 0 | 0 | 0 | 0 | 0 | 0 | 0 | 0 | 0 | 0 |
| 0 | NA | NA | NA | NA | NA | Taar7d | trace amine-associated receptor 7D | 0 | 0 | 0 | 0 | 0 | 0 | 0 | 0 | 0 | 0 | 0 |
| 0 | NA | NA | NA | NA | NA | Taar7e | trace amine-associated receptor 7E | 0 | 0 | 0 | 0 | 0 | 0 | 0 | 0 | 0 | 0 | 0 |
| 0 | NA | NA | NA | NA | NA | Gm6884 | | 0 | 0 | 0 | 0 | 0 | 0 | 0 | 0 | 0 | 0 | 0 |
| 0 | NA | NA | NA | NA | NA | Taar7f | trace amine-associated receptor 7F | 0 | 0 | 0 | 0 | 0 | 0 | 0 | 0 | 0 | 0 | 0 |
| 0 | NA | NA | NA | NA | NA | | | 0 | 0 | 0 | 0 | 0 | 0 | 0 | 0 | 0 | 0 | 0 |
| 0 | NA | NA | NA | NA | NA | Taar8a | trace amine-associated receptor 8A | 0 | 0 | 0 | 0 | 0 | 0 | 0 | 0 | 0 | 0 | 0 |
| 0 | NA | NA | NA | NA | NA | Taar8b | trace amine-associated receptor 8B | 0 | 0 | 0 | 0 | 0 | 0 | 0 | 0 | 0 | 0 | 0 |
| 0 | NA | NA | NA | NA | NA | Taar8c | trace amine-associated receptor 8C | 0 | 0 | 0 | 0 | 0 | 0 | 0 | 0 | 0 | 0 | 0 |

| | | | | | | | | | | | | | | | | | |
|---|---|---|---|---|---|---|---|---|---|---|---|---|---|---|---|---|---|
| 0 | NA | NA | NA | NA | NA | Taar9 | trace amine-associated receptor 9 | 0 | 0 | 0 | 0 | 0 | 0 | 0 | 0 | 0 | 0 | 0 |
| 0 | NA | NA | NA | NA | NA | Gm15271 | | 0 | 0 | 0 | 0 | 0 | 0 | 0 | 0 | 0 | 0 | 0 |
| 0 | NA | NA | NA | NA | NA | Gm6893 | | 0 | 0 | 0 | 0 | 0 | 0 | 0 | 0 | 0 | 0 | 0 |
| 0 | NA | NA | NA | NA | NA | Gm23410 | | 0 | 0 | 0 | 0 | 0 | 0 | 0 | 0 | 0 | 0 | 0 |
| 0 | NA | NA | NA | NA | NA | Gm47715 | | 0 | 0 | 0 | 0 | 0 | 0 | 0 | 0 | 0 | 0 | 0 |
| 0 | NA | NA | NA | NA | NA | Gm22566 | | 0 | 0 | 0 | 0 | 0 | 0 | 0 | 0 | 0 | 0 | 0 |
| 0 | NA | NA | NA | NA | NA | Gm7791 | | 0 | 0 | 0 | 0 | 0 | 0 | 0 | 0 | 0 | 0 | 0 |
| 0 | NA | NA | NA | NA | NA | Gm25016 | | 0 | 0 | 0 | 0 | 0 | 0 | 0 | 0 | 0 | 0 | 0 |
| 0 | NA | NA | NA | NA | NA | Gm47830 | | 0 | 0 | 0 | 0 | 0 | 0 | 0 | 0 | 0 | 0 | 0 |
| 0 | NA | NA | NA | NA | NA | Smlr1 | small leucine-rich protein 1 | 0 | 0 | 0 | 0 | 0 | 0 | 0 | 0 | 0 | 0 | 0 |
| 0 | NA | NA | NA | NA | NA | Gm29571 | | 0 | 0 | 0 | 0 | 0 | 0 | 0 | 0 | 0 | 0 | 0 |
| 0 | NA | NA | NA | NA | NA | Gm36908 | | 0 | 0 | 0 | 0 | 0 | 0 | 0 | 0 | 0 | 0 | 0 |
| 0 | NA | NA | NA | NA | NA | Gm47924 | | 0 | 0 | 0 | 0 | 0 | 0 | 0 | 0 | 0 | 0 | 0 |
| 0 | NA | NA | NA | NA | NA | 4930401C15Rik | | 0 | 0 | 0 | 0 | 0 | 0 | 0 | 0 | 0 | 0 | 0 |
| 0 | NA | NA | NA | NA | NA | Gm8681 | | 0 | 0 | 0 | 0 | 0 | 0 | 0 | 0 | 0 | 0 | 0 |
| 0 | NA | NA | NA | NA | NA | Gm47930 | | 0 | 0 | 0 | 0 | 0 | 0 | 0 | 0 | 0 | 0 | 0 |
| 0 | NA | NA | NA | NA | NA | Gm47939 | | 0 | 0 | 0 | 0 | 0 | 0 | 0 | 0 | 0 | 0 | 0 |
| 0 | NA | NA | NA | NA | NA | Gm27754 | | 0 | 0 | 0 | 0 | 0 | 0 | 0 | 0 | 0 | 0 | 0 |
| 0 | NA | NA | NA | NA | NA | Gm23757 | | 0 | 0 | 0 | 0 | 0 | 0 | 0 | 0 | 0 | 0 | 0 |
| 0 | NA | NA | NA | NA | NA | Gm5649 | | 0 | 0 | 0 | 0 | 0 | 0 | 0 | 0 | 0 | 0 | 0 |
| 0 | NA | NA | NA | NA | NA | Gm30034 | | 0 | 0 | 0 | 0 | 0 | 0 | 0 | 0 | 0 | 0 | 0 |
| 0 | NA | NA | NA | NA | NA | Gm48864 | | 0 | 0 | 0 | 0 | 0 | 0 | 0 | 0 | 0 | 0 | 0 |
| 0 | NA | NA | NA | NA | NA | Gm8709 | | 0 | 0 | 0 | 0 | 0 | 0 | 0 | 0 | 0 | 0 | 0 |
| 0 | NA | NA | NA | NA | NA | Gm48893 | | 0 | 0 | 0 | 0 | 0 | 0 | 0 | 0 | 0 | 0 | 0 |
| 0 | NA | NA | NA | NA | NA | Gm48891 | | 0 | 0 | 0 | 0 | 0 | 0 | 0 | 0 | 0 | 0 | 0 |
| 0 | NA | NA | NA | NA | NA | Gm48892 | | 0 | 0 | 0 | 0 | 0 | 0 | 0 | 0 | 0 | 0 | 0 |
| 0 | NA | NA | NA | NA | NA | Gm48084 | | 0 | 0 | 0 | 0 | 0 | 0 | 0 | 0 | 0 | 0 | 0 |
| 0 | NA | NA | NA | NA | NA | Gm30534 | | 0 | 0 | 0 | 0 | 0 | 0 | 0 | 0 | 0 | 0 | 0 |
| 0 | NA | NA | NA | NA | NA | Gm17917 | | 0 | 0 | 0 | 0 | 0 | 0 | 0 | 0 | 0 | 0 | 0 |
| 0 | NA | NA | NA | NA | NA | Gm48083 | | 0 | 0 | 0 | 0 | 0 | 0 | 0 | 0 | 0 | 0 | 0 |
| 0 | NA | NA | NA | NA | NA | 4930579H20Rik | | 0 | 0 | 0 | 0 | 0 | 0 | 0 | 0 | 0 | 0 | 0 |
| 0 | NA | NA | NA | NA | NA | Gm23353 | | 0 | 0 | 0 | 0 | 0 | 0 | 0 | 0 | 0 | 0 | 0 |
| 0 | NA | NA | NA | NA | NA | Gm36962 | | 0 | 0 | 0 | 0 | 0 | 0 | 0 | 0 | 0 | 0 | 0 |
| 0 | NA | NA | NA | NA | NA | Gm30165 | | 0 | 0 | 0 | 0 | 0 | 0 | 0 | 0 | 0 | 0 | 0 |
| 0 | NA | NA | NA | NA | NA | Gm18041 | | 0 | 0 | 0 | 0 | 0 | 0 | 0 | 0 | 0 | 0 | 0 |
| 0 | NA | NA | NA | NA | NA | Gm47696 | | 0 | 0 | 0 | 0 | 0 | 0 | 0 | 0 | 0 | 0 | 0 |
| 0 | NA | NA | NA | NA | NA | Gm18972 | | 0 | 0 | 0 | 0 | 0 | 0 | 0 | 0 | 0 | 0 | 0 |
| 0 | NA | NA | NA | NA | NA | Gm18507 | | 0 | 0 | 0 | 0 | 0 | 0 | 0 | 0 | 0 | 0 | 0 |
| 0 | NA | NA | NA | NA | NA | Gm22370 | | 0 | 0 | 0 | 0 | 0 | 0 | 0 | 0 | 0 | 0 | 0 |
| 0 | NA | NA | NA | NA | NA | Gm47834 | | 0 | 0 | 0 | 0 | 0 | 0 | 0 | 0 | 0 | 0 | 0 |
| 0 | NA | NA | NA | NA | NA | 4930519F09Rik | | 0 | 0 | 0 | 0 | 0 | 0 | 0 | 0 | 0 | 0 | 0 |
| 0 | NA | NA | NA | NA | NA | Gm22811 | | 0 | 0 | 0 | 0 | 0 | 0 | 0 | 0 | 0 | 0 | 0 |
| 0 | NA | NA | NA | NA | NA | Gm18508 | | 0 | 0 | 0 | 0 | 0 | 0 | 0 | 0 | 0 | 0 | 0 |
| 0 | NA | NA | NA | NA | NA | Gm30619 | | 0 | 0 | 0 | 0 | 0 | 0 | 0 | 0 | 0 | 0 | 0 |
| 0 | NA | NA | NA | NA | NA | Gm49353 | | 0 | 0 | 0 | 0 | 0 | 0 | 0 | 0 | 0 | 0 | 0 |
| 0 | NA | NA | NA | NA | NA | Gm40621 | | 0 | 0 | 0 | 0 | 0 | 0 | 0 | 0 | 0 | 0 | 0 |
| 0 | NA | NA | NA | NA | NA | Gm24286 | | 0 | 0 | 0 | 0 | 0 | 0 | 0 | 0 | 0 | 0 | 0 |
| 0 | NA | NA | NA | NA | NA | Gm18903 | | 0 | 0 | 0 | 0 | 0 | 0 | 0 | 0 | 0 | 0 | 0 |
| 0 | NA | NA | NA | NA | NA | Gm20276 | | 0 | 0 | 0 | 0 | 0 | 0 | 0 | 0 | 0 | 0 | 0 |
| 0 | NA | NA | NA | NA | NA | Gm48159 | | 0 | 0 | 0 | 0 | 0 | 0 | 0 | 0 | 0 | 0 | 0 |
| 0 | NA | NA | NA | NA | NA | Gm46197 | | 0 | 0 | 0 | 0 | 0 | 0 | 0 | 0 | 0 | 0 | 0 |
| 0 | NA | NA | NA | NA | NA | Gm6390 | | 0 | 0 | 0 | 0 | 0 | 0 | 0 | 0 | 0 | 0 | 0 |
| 0 | NA | NA | NA | NA | NA | Gm2829 | | 0 | 0 | 0 | 0 | 0 | 0 | 0 | 0 | 0 | 0 | 0 |
| 0 | NA | NA | NA | NA | NA | Gm4780 | | 0 | 0 | 0 | 0 | 0 | 0 | 0 | 0 | 0 | 0 | 0 |
| 0 | NA | NA | NA | NA | NA | Gm22623 | | 0 | 0 | 0 | 0 | 0 | 0 | 0 | 0 | 0 | 0 | 0 |
| 0 | NA | NA | NA | NA | NA | Gm48232 | | 0 | 0 | 0 | 0 | 0 | 0 | 0 | 0 | 0 | 0 | 0 |
| 0 | NA | NA | NA | NA | NA | Gm48233 | | 0 | 0 | 0 | 0 | 0 | 0 | 0 | 0 | 0 | 0 | 0 |
| 0 | NA | NA | NA | NA | NA | Gm48862 | | 0 | 0 | 0 | 0 | 0 | 0 | 0 | 0 | 0 | 0 | 0 |
| 0 | NA | NA | NA | NA | NA | Gm24168 | | 0 | 0 | 0 | 0 | 0 | 0 | 0 | 0 | 0 | 0 | 0 |
| 0 | NA | NA | NA | NA | NA | Gm30676 | | 0 | 0 | 0 | 0 | 0 | 0 | 0 | 0 | 0 | 0 | 0 |

| | | | | | | | | | | | | | | | | | | |
|---|---|---|---|---|---|---|---|---|---|---|---|---|---|---|---|---|---|---|
| 0 | NA | NA | NA | NA | NA | Gm23790 | | 0 | 0 | 0 | 0 | 0 | 0 | 0 | 0 | 0 | 0 | 0 |
| 0 | NA | NA | NA | NA | NA | Gm7333 | | 0 | 0 | 0 | 0 | 0 | 0 | 0 | 0 | 0 | 0 | 0 |
| 0 | NA | NA | NA | NA | NA | n-R5s76 | | 0 | 0 | 0 | 0 | 0 | 0 | 0 | 0 | 0 | 0 | 0 |
| 0 | NA | NA | NA | NA | NA | Gm48094 | | 0 | 0 | 0 | 0 | 0 | 0 | 0 | 0 | 0 | 0 | 0 |
| 0 | NA | NA | NA | NA | NA | Gm48093 | | 0 | 0 | 0 | 0 | 0 | 0 | 0 | 0 | 0 | 0 | 0 |
| 0 | NA | NA | NA | NA | NA | Gm18189 | | 0 | 0 | 0 | 0 | 0 | 0 | 0 | 0 | 0 | 0 | 0 |
| 0 | NA | NA | NA | NA | NA | A230081H15Rik | | 0 | 0 | 0 | 0 | 0 | 0 | 0 | 0 | 0 | 0 | 0 |
| 0 | NA | NA | NA | NA | NA | C030006N10Rik | | 0 | 0 | 0 | 0 | 0 | 0 | 0 | 0 | 0 | 0 | 0 |
| 0 | NA | NA | NA | NA | NA | Gm18704 | | 0 | 0 | 0 | 0 | 0 | 0 | 0 | 0 | 0 | 0 | 0 |
| 0 | NA | NA | NA | NA | NA | Gm47700 | | 0 | 0 | 0 | 0 | 0 | 0 | 0 | 0 | 0 | 0 | 0 |
| 0 | NA | NA | NA | NA | NA | Gm47702 | | 0 | 0 | 0 | 0 | 0 | 0 | 0 | 0 | 0 | 0 | 0 |
| 0 | NA | NA | NA | NA | NA | Gm47823 | | 0 | 0 | 0 | 0 | 0 | 0 | 0 | 0 | 0 | 0 | 0 |
| 0 | NA | NA | NA | NA | NA | Trdn | triadin | 0 | 0 | 0 | 0 | 0 | 0 | 0 | 0 | 0 | 0 | 0 |
| 0 | NA | NA | NA | NA | NA | Gm47831 | | 0 | 0 | 0 | 0 | 0 | 0 | 0 | 0 | 0 | 0 | 0 |
| 0 | NA | NA | NA | NA | NA | Gm47845 | | 0 | 0 | 0 | 0 | 0 | 0 | 0 | 0 | 0 | 0 | 0 |
| 0 | NA | NA | NA | NA | NA | D830005E20Rik | | 0 | 0 | 0 | 0 | 0 | 0 | 0 | 0 | 0 | 0 | 0 |
| 0 | NA | NA | NA | NA | NA | Gm19073 | | 0 | 0 | 0 | 0 | 0 | 0 | 0 | 0 | 0 | 0 | 0 |
| 0 | NA | NA | NA | NA | NA | Gm47842 | | 0 | 0 | 0 | 0 | 0 | 0 | 0 | 0 | 0 | 0 | 0 |
| 0 | NA | NA | NA | NA | NA | Gm29884 | | 0 | 0 | 0 | 0 | 0 | 0 | 0 | 0 | 0 | 0 | 0 |
| 0 | NA | NA | NA | NA | NA | Gm47844 | | 0 | 0 | 0 | 0 | 0 | 0 | 0 | 0 | 0 | 0 | 0 |
| 0 | NA | NA | NA | NA | NA | Gm20304 | | 0 | 0 | 0 | 0 | 0 | 0 | 0 | 0 | 0 | 0 | 0 |
| 0 | NA | NA | NA | NA | NA | Gm47908 | | 0 | 0 | 0 | 0 | 0 | 0 | 0 | 0 | 0 | 0 | 0 |
| 0 | NA | NA | NA | NA | NA | Clvs2 | clavesin 2 | 0 | 0 | 0 | 0 | 0 | 0 | 0 | 0 | 0 | 0 | 0 |
| 0 | NA | NA | NA | NA | NA | Gm47910 | | 0 | 0 | 0 | 0 | 0 | 0 | 0 | 0 | 0 | 0 | 0 |
| 0 | NA | NA | NA | NA | NA | Gm47911 | | 0 | 0 | 0 | 0 | 0 | 0 | 0 | 0 | 0 | 0 | 0 |
| 0 | NA | NA | NA | NA | NA | Gm15939 | | 0 | 0 | 0 | 0 | 0 | 0 | 0 | 0 | 0 | 0 | 0 |
| 0 | NA | NA | NA | NA | NA | Gm47929 | | 0 | 0 | 0 | 0 | 0 | 0 | 0 | 0 | 0 | 0 | 0 |
| 0 | NA | NA | NA | NA | NA | Sult3a2 | sulfotransferase family 3A, member 2 | 0 | 0 | 0 | 0 | 0 | 0 | 0 | 0 | 0 | 0 | 0 |
| 0 | NA | NA | NA | NA | NA | Sult3a1 | sulfotransferase family 3A, member 1 | 0 | 0 | 0 | 0 | 0 | 0 | 0 | 0 | 0 | 0 | 0 |
| 0 | NA | NA | NA | NA | NA | Gm47935 | | 0 | 0 | 0 | 0 | 0 | 0 | 0 | 0 | 0 | 0 | 0 |
| 0 | NA | NA | NA | NA | NA | Gm48018 | | 0 | 0 | 0 | 0 | 0 | 0 | 0 | 0 | 0 | 0 | 0 |
| 0 | NA | NA | NA | NA | NA | Gm48034 | | 0 | 0 | 0 | 0 | 0 | 0 | 0 | 0 | 0 | 0 | 0 |
| 0 | NA | NA | NA | NA | NA | Trappc3l | trafficking protein particle complex 3 like | 0 | 0 | 0 | 0 | 0 | 0 | 0 | 0 | 0 | 0 | 0 |
| 0 | NA | NA | NA | NA | NA | Calhm4 | calcium homeostasis modulator family me | 0 | 0 | 0 | 0 | 0 | 0 | 0 | 0 | 0 | 0 | 0 |
| 0 | NA | NA | NA | NA | NA | Gm18346 | | 0 | 0 | 0 | 0 | 0 | 0 | 0 | 0 | 0 | 0 | 0 |
| 0 | NA | NA | NA | NA | NA | Gm26564 | | 0 | 0 | 0 | 0 | 0 | 0 | 0 | 0 | 0 | 0 | 0 |
| 0 | NA | NA | NA | NA | NA | Gm47485 | | 0 | 0 | 0 | 0 | 0 | 0 | 0 | 0 | 0 | 0 | 0 |
| 0 | NA | NA | NA | NA | NA | Gm18252 | | 0 | 0 | 0 | 0 | 0 | 0 | 0 | 0 | 0 | 0 | 0 |
| 0 | NA | NA | NA | NA | NA | Gm26341 | | 0 | 0 | 0 | 0 | 0 | 0 | 0 | 0 | 0 | 0 | 0 |
| 0 | NA | NA | NA | NA | NA | Gm47044 | | 0 | 0 | 0 | 0 | 0 | 0 | 0 | 0 | 0 | 0 | 0 |
| 0 | NA | NA | NA | NA | NA | Gm18729 | | 0 | 0 | 0 | 0 | 0 | 0 | 0 | 0 | 0 | 0 | 0 |
| 0 | NA | NA | NA | NA | NA | 1700054O05Rik | | 0 | 0 | 0 | 0 | 0 | 0 | 0 | 0 | 0 | 0 | 0 |
| 0 | NA | NA | NA | NA | NA | Gm49446 | | 0 | 0 | 0 | 0 | 0 | 0 | 0 | 0 | 0 | 0 | 0 |
| 0 | NA | NA | NA | NA | NA | Gm47046 | | 0 | 0 | 0 | 0 | 0 | 0 | 0 | 0 | 0 | 0 | 0 |
| 0 | NA | NA | NA | NA | NA | Gm47048 | | 0 | 0 | 0 | 0 | 0 | 0 | 0 | 0 | 0 | 0 | 0 |
| 0 | NA | NA | NA | NA | NA | Gm47049 | | 0 | 0 | 0 | 0 | 0 | 0 | 0 | 0 | 0 | 0 | 0 |
| 0 | NA | NA | NA | NA | NA | Gm47054 | | 0 | 0 | 0 | 0 | 0 | 0 | 0 | 0 | 0 | 0 | 0 |
| 0 | NA | NA | NA | NA | NA | Gm22554 | | 0 | 0 | 0 | 0 | 0 | 0 | 0 | 0 | 0 | 0 | 0 |
| 0 | NA | NA | NA | NA | NA | Gm38231 | | 0 | 0 | 0 | 0 | 0 | 0 | 0 | 0 | 0 | 0 | 0 |
| 0 | NA | NA | NA | NA | NA | Gm26535 | | 0 | 0 | 0 | 0 | 0 | 0 | 0 | 0 | 0 | 0 | 0 |
| 0 | NA | NA | NA | NA | NA | Gm48172 | | 0 | 0 | 0 | 0 | 0 | 0 | 0 | 0 | 0 | 0 | 0 |
| 0 | NA | NA | NA | NA | NA | Gm24710 | | 0 | 0 | 0 | 0 | 0 | 0 | 0 | 0 | 0 | 0 | 0 |
| 0 | NA | NA | NA | NA | NA | 4930591E09Rik | | 0 | 0 | 0 | 0 | 0 | 0 | 0 | 0 | 0 | 0 | 0 |
| 0 | NA | NA | NA | NA | NA | Gm24889 | | 0 | 0 | 0 | 0 | 0 | 0 | 0 | 0 | 0 | 0 | 0 |
| 0 | NA | NA | NA | NA | NA | Gm24297 | | 0 | 0 | 0 | 0 | 0 | 0 | 0 | 0 | 0 | 0 | 0 |
| 0 | NA | NA | NA | NA | NA | Gm48173 | | 0 | 0 | 0 | 0 | 0 | 0 | 0 | 0 | 0 | 0 | 0 |
| 0 | NA | NA | NA | NA | NA | Gm48197 | | 0 | 0 | 0 | 0 | 0 | 0 | 0 | 0 | 0 | 0 | 0 |
| 0 | NA | NA | NA | NA | NA | Gm48198 | | 0 | 0 | 0 | 0 | 0 | 0 | 0 | 0 | 0 | 0 | 0 |
| 0 | NA | NA | NA | NA | NA | Gm22911 | | 0 | 0 | 0 | 0 | 0 | 0 | 0 | 0 | 0 | 0 | 0 |
| 0 | NA | NA | NA | NA | NA | Rfpl4b | ret finger protein-like 4B | 0 | 0 | 0 | 0 | 0 | 0 | 0 | 0 | 0 | 0 | 0 |

| | | | | | | | | | | | | | | | | | |
|---|---|---|---|---|---|---|---|---|---|---|---|---|---|---|---|---|---|
| 0 | NA | NA | NA | NA | NA | Gm31562 | | 0 | 0 | 0 | 0 | 0 | 0 | 0 | 0 | 0 | 0 | 0 |
| 0 | NA | NA | NA | NA | NA | Gm48486 | | 0 | 0 | 0 | 0 | 0 | 0 | 0 | 0 | 0 | 0 | 0 |
| 0 | NA | NA | NA | NA | NA | Gm6963 | | 0 | 0 | 0 | 0 | 0 | 0 | 0 | 0 | 0 | 0 | 0 |
| 0 | NA | NA | NA | NA | NA | Gm48839 | | 0 | 0 | 0 | 0 | 0 | 0 | 0 | 0 | 0 | 0 | 0 |
| 0 | NA | NA | NA | NA | NA | Gm25613 | | 0 | 0 | 0 | 0 | 0 | 0 | 0 | 0 | 0 | 0 | 0 |
| 0 | NA | NA | NA | NA | NA | Rnu3a | | 0 | 0 | 0 | 0 | 0 | 0 | 0 | 0 | 0 | 0 | 0 |
| 0 | NA | NA | NA | NA | NA | Rnu3a | | 0 | 0 | 0 | 0 | 0 | 0 | 0 | 0 | 0 | 0 | 0 |
| 0 | NA | NA | NA | NA | NA | Gm47867 | | 0 | 0 | 0 | 0 | 0 | 0 | 0 | 0 | 0 | 0 | 0 |
| 0 | NA | NA | NA | NA | NA | Gm8948 | | 0 | 0 | 0 | 0 | 0 | 0 | 0 | 0 | 0 | 0 | 0 |
| 0 | NA | NA | NA | NA | NA | Gm18671 | | 0 | 0 | 0 | 0 | 0 | 0 | 0 | 0 | 0 | 0 | 0 |
| 0 | NA | NA | NA | NA | NA | Slc22a16 | solute carrier family 22 (organic cation tra | 0 | 0 | 0 | 0 | 0 | 0 | 0 | 0 | 0 | 0 | 0 |
| 0 | NA | NA | NA | NA | NA | Gm22948 | | 0 | 0 | 0 | 0 | 0 | 0 | 0 | 0 | 0 | 0 | 0 |
| 0 | NA | NA | NA | NA | NA | Gm48057 | | 0 | 0 | 0 | 0 | 0 | 0 | 0 | 0 | 0 | 0 | 0 |
| 0 | NA | NA | NA | NA | NA | Gm19611 | | 0 | 0 | 0 | 0 | 0 | 0 | 0 | 0 | 0 | 0 | 0 |
| 0 | NA | NA | NA | NA | NA | Gm32284 | | 0 | 0 | 0 | 0 | 0 | 0 | 0 | 0 | 0 | 0 | 0 |
| 0 | NA | NA | NA | NA | NA | Gm32338 | | 0 | 0 | 0 | 0 | 0 | 0 | 0 | 0 | 0 | 0 | 0 |
| 0 | NA | NA | NA | NA | NA | Gm25526 | | 0 | 0 | 0 | 0 | 0 | 0 | 0 | 0 | 0 | 0 | 0 |
| 0 | NA | NA | NA | NA | NA | Gm18305 | | 0 | 0 | 0 | 0 | 0 | 0 | 0 | 0 | 0 | 0 | 0 |
| 0 | NA | NA | NA | NA | NA | Gm18986 | | 0 | 0 | 0 | 0 | 0 | 0 | 0 | 0 | 0 | 0 | 0 |
| 0 | NA | NA | NA | NA | NA | Gm25626 | | 0 | 0 | 0 | 0 | 0 | 0 | 0 | 0 | 0 | 0 | 0 |
| 0 | NA | NA | NA | NA | NA | Gm47473 | | 0 | 0 | 0 | 0 | 0 | 0 | 0 | 0 | 0 | 0 | 0 |
| 0 | NA | NA | NA | NA | NA | Gm27783 | | 0 | 0 | 0 | 0 | 0 | 0 | 0 | 0 | 0 | 0 | 0 |
| 0 | NA | NA | NA | NA | NA | Mir3473b | | 0 | 0 | 0 | 0 | 0 | 0 | 0 | 0 | 0 | 0 | 0 |
| 0 | NA | NA | NA | NA | NA | Gm23533 | | 0 | 0 | 0 | 0 | 0 | 0 | 0 | 0 | 0 | 0 | 0 |
| 0 | NA | NA | NA | NA | NA | Gm15198 | | 0 | 0 | 0 | 0 | 0 | 0 | 0 | 0 | 0 | 0 | 0 |
| 0 | NA | NA | NA | NA | NA | Gm25277 | | 0 | 0 | 0 | 0 | 0 | 0 | 0 | 0 | 0 | 0 | 0 |
| 0 | NA | NA | NA | NA | NA | Nr2e1 | nuclear receptor subfamily 2, group E, me | 0 | 0 | 0 | 0 | 0 | 0 | 0 | 0 | 0 | 0 | 0 |
| 0 | NA | NA | NA | NA | NA | Gm15200 | | 0 | 0 | 0 | 0 | 0 | 0 | 0 | 0 | 0 | 0 | 0 |
| 0 | NA | NA | NA | NA | NA | Gm15199 | | 0 | 0 | 0 | 0 | 0 | 0 | 0 | 0 | 0 | 0 | 0 |
| 0 | NA | NA | NA | NA | NA | Gm26464 | | 0 | 0 | 0 | 0 | 0 | 0 | 0 | 0 | 0 | 0 | 0 |
| 0 | NA | NA | NA | NA | NA | Gm48671 | | 0 | 0 | 0 | 0 | 0 | 0 | 0 | 0 | 0 | 0 | 0 |
| 0 | NA | NA | NA | NA | NA | Gm47815 | | 0 | 0 | 0 | 0 | 0 | 0 | 0 | 0 | 0 | 0 | 0 |
| 0 | NA | NA | NA | NA | NA | Gm29245 | | 0 | 0 | 0 | 0 | 0 | 0 | 0 | 0 | 0 | 0 | 0 |
| 0 | NA | NA | NA | NA | NA | Rps19-ps11 | | 0 | 0 | 0 | 0 | 0 | 0 | 0 | 0 | 0 | 0 | 0 |
| 0 | NA | NA | NA | NA | NA | Rpl26-ps1 | | 0 | 0 | 0 | 0 | 0 | 0 | 0 | 0 | 0 | 0 | 0 |
| 0 | NA | NA | NA | NA | NA | Gm34481 | | 0 | 0 | 0 | 0 | 0 | 0 | 0 | 0 | 0 | 0 | 0 |
| 0 | NA | NA | NA | NA | NA | Gm47927 | | 0 | 0 | 0 | 0 | 0 | 0 | 0 | 0 | 0 | 0 | 0 |
| 0 | NA | NA | NA | NA | NA | 1700027J07Rik | | 0 | 0 | 0 | 0 | 0 | 0 | 0 | 0 | 0 | 0 | 0 |
| 0 | NA | NA | NA | NA | NA | Speer5-ps1 | | 0 | 0 | 0 | 0 | 0 | 0 | 0 | 0 | 0 | 0 | 0 |
| 0 | NA | NA | NA | NA | NA | Gm35028 | | 0 | 0 | 0 | 0 | 0 | 0 | 0 | 0 | 0 | 0 | 0 |
| 0 | NA | NA | NA | NA | NA | Gm48061 | | 0 | 0 | 0 | 0 | 0 | 0 | 0 | 0 | 0 | 0 | 0 |
| 0 | NA | NA | NA | NA | NA | Gm22087 | | 0 | 0 | 0 | 0 | 0 | 0 | 0 | 0 | 0 | 0 | 0 |
| 0 | NA | NA | NA | NA | NA | Gm48066 | | 0 | 0 | 0 | 0 | 0 | 0 | 0 | 0 | 0 | 0 | 0 |
| 0 | NA | NA | NA | NA | NA | Gm27977 | | 0 | 0 | 0 | 0 | 0 | 0 | 0 | 0 | 0 | 0 | 0 |
| 0 | NA | NA | NA | NA | NA | 4930431F10Rik | | 0 | 0 | 0 | 0 | 0 | 0 | 0 | 0 | 0 | 0 | 0 |
| 0 | NA | NA | NA | NA | NA | Gm40638 | | 0 | 0 | 0 | 0 | 0 | 0 | 0 | 0 | 0 | 0 | 0 |
| 0 | NA | NA | NA | NA | NA | Gm24271 | | 0 | 0 | 0 | 0 | 0 | 0 | 0 | 0 | 0 | 0 | 0 |
| 0 | NA | NA | NA | NA | NA | Gm40639 | | 0 | 0 | 0 | 0 | 0 | 0 | 0 | 0 | 0 | 0 | 0 |
| 0 | NA | NA | NA | NA | NA | Gm23766 | | 0 | 0 | 0 | 0 | 0 | 0 | 0 | 0 | 0 | 0 | 0 |
| 0 | NA | NA | NA | NA | NA | Gm27723 | | 0 | 0 | 0 | 0 | 0 | 0 | 0 | 0 | 0 | 0 | 0 |
| 0 | NA | NA | NA | NA | NA | Gm26016 | | 0 | 0 | 0 | 0 | 0 | 0 | 0 | 0 | 0 | 0 | 0 |
| 0 | NA | NA | NA | NA | NA | Gm15934 | | 0 | 0 | 0 | 0 | 0 | 0 | 0 | 0 | 0 | 0 | 0 |
| 0 | NA | NA | NA | NA | NA | Gm47433 | | 0 | 0 | 0 | 0 | 0 | 0 | 0 | 0 | 0 | 0 | 0 |
| 0 | NA | NA | NA | NA | NA | Bves | blood vessel epicardial substance | 0 | 0 | 0 | 0 | 0 | 0 | 0 | 0 | 0 | 0 | 0 |
| 0 | NA | NA | NA | NA | NA | Gm27731 | | 0 | 0 | 0 | 0 | 0 | 0 | 0 | 0 | 0 | 0 | 0 |
| 0 | NA | NA | NA | NA | NA | Gm47502 | | 0 | 0 | 0 | 0 | 0 | 0 | 0 | 0 | 0 | 0 | 0 |
| 0 | NA | NA | NA | NA | NA | Gm27410 | | 0 | 0 | 0 | 0 | 0 | 0 | 0 | 0 | 0 | 0 | 0 |
| 0 | NA | NA | NA | NA | NA | Gm35430 | | 0 | 0 | 0 | 0 | 0 | 0 | 0 | 0 | 0 | 0 | 0 |
| 0 | NA | NA | NA | NA | NA | Gm25740 | | 0 | 0 | 0 | 0 | 0 | 0 | 0 | 0 | 0 | 0 | 0 |
| 0 | NA | NA | NA | NA | NA | Gm23481 | | 0 | 0 | 0 | 0 | 0 | 0 | 0 | 0 | 0 | 0 | 0 |

| | | | | | | | | | | | | | | | | | |
|---|---|---|---|---|---|---|---|---|---|---|---|---|---|---|---|---|---|
| 0 | NA | NA | NA | NA | NA | Gm19994 | 0 | 0 | 0 | 0 | 0 | 0 | 0 | 0 | 0 | 0 | 0 |
| 0 | NA | NA | NA | NA | NA | Gm25650 | 0 | 0 | 0 | 0 | 0 | 0 | 0 | 0 | 0 | 0 | 0 |
| 0 | NA | NA | NA | NA | NA | Gm47562 | 0 | 0 | 0 | 0 | 0 | 0 | 0 | 0 | 0 | 0 | 0 |
| 0 | NA | NA | NA | NA | NA | Gm35552 | 0 | 0 | 0 | 0 | 0 | 0 | 0 | 0 | 0 | 0 | 0 |
| 0 | NA | NA | NA | NA | NA | Gm47568 | 0 | 0 | 0 | 0 | 0 | 0 | 0 | 0 | 0 | 0 | 0 |
| 0 | NA | NA | NA | NA | NA | Gm3442 | 0 | 0 | 0 | 0 | 0 | 0 | 0 | 0 | 0 | 0 | 0 |
| 0 | NA | NA | NA | NA | NA | Gm19137 | 0 | 0 | 0 | 0 | 0 | 0 | 0 | 0 | 0 | 0 | 0 |
| 0 | NA | NA | NA | NA | NA | Gm9068 | 0 | 0 | 0 | 0 | 0 | 0 | 0 | 0 | 0 | 0 | 0 |
| 0 | NA | NA | NA | NA | NA | Gm47571 | 0 | 0 | 0 | 0 | 0 | 0 | 0 | 0 | 0 | 0 | 0 |
| 0 | NA | NA | NA | NA | NA | Gm47331 | 0 | 0 | 0 | 0 | 0 | 0 | 0 | 0 | 0 | 0 | 0 |
| 0 | NA | NA | NA | NA | NA | Gm47332 | 0 | 0 | 0 | 0 | 0 | 0 | 0 | 0 | 0 | 0 | 0 |
| 0 | NA | NA | NA | NA | NA | Gm47353 | 0 | 0 | 0 | 0 | 0 | 0 | 0 | 0 | 0 | 0 | 0 |
| 0 | NA | NA | NA | NA | NA | Gm23584 | 0 | 0 | 0 | 0 | 0 | 0 | 0 | 0 | 0 | 0 | 0 |
| 0 | NA | NA | NA | NA | NA | Gm47354 | 0 | 0 | 0 | 0 | 0 | 0 | 0 | 0 | 0 | 0 | 0 |
| 0 | NA | NA | NA | NA | NA | Gm47355 | 0 | 0 | 0 | 0 | 0 | 0 | 0 | 0 | 0 | 0 | 0 |
| 0 | NA | NA | NA | NA | NA | Gm24256 | 0 | 0 | 0 | 0 | 0 | 0 | 0 | 0 | 0 | 0 | 0 |
| 0 | NA | NA | NA | NA | NA | Gm47356 | 0 | 0 | 0 | 0 | 0 | 0 | 0 | 0 | 0 | 0 | 0 |
| 0 | NA | NA | NA | NA | NA | Gm47364 | 0 | 0 | 0 | 0 | 0 | 0 | 0 | 0 | 0 | 0 | 0 |
| 0 | NA | NA | NA | NA | NA | Gm47377 | 0 | 0 | 0 | 0 | 0 | 0 | 0 | 0 | 0 | 0 | 0 |
| 0 | NA | NA | NA | NA | NA | Gm47379 | 0 | 0 | 0 | 0 | 0 | 0 | 0 | 0 | 0 | 0 | 0 |
| 0 | NA | NA | NA | NA | NA | Gm3473 | 0 | 0 | 0 | 0 | 0 | 0 | 0 | 0 | 0 | 0 | 0 |
| 0 | NA | NA | NA | NA | NA | Gm47378 | 0 | 0 | 0 | 0 | 0 | 0 | 0 | 0 | 0 | 0 | 0 |
| 0 | NA | NA | NA | NA | NA | Gm47380 | 0 | 0 | 0 | 0 | 0 | 0 | 0 | 0 | 0 | 0 | 0 |
| 0 | NA | NA | NA | NA | NA | Gm47017 | 0 | 0 | 0 | 0 | 0 | 0 | 0 | 0 | 0 | 0 | 0 |
| 0 | NA | NA | NA | NA | NA | Gm26543 | 0 | 0 | 0 | 0 | 0 | 0 | 0 | 0 | 0 | 0 | 0 |
| 0 | NA | NA | NA | NA | NA | Gm47018 | 0 | 0 | 0 | 0 | 0 | 0 | 0 | 0 | 0 | 0 | 0 |
| 0 | NA | NA | NA | NA | NA | Gm47023 | 0 | 0 | 0 | 0 | 0 | 0 | 0 | 0 | 0 | 0 | 0 |
| 0 | NA | NA | NA | NA | NA | Gm47016 | 0 | 0 | 0 | 0 | 0 | 0 | 0 | 0 | 0 | 0 | 0 |
| 0 | NA | NA | NA | NA | NA | Gm47022 | 0 | 0 | 0 | 0 | 0 | 0 | 0 | 0 | 0 | 0 | 0 |
| 0 | NA | NA | NA | NA | NA | Gm48123 | 0 | 0 | 0 | 0 | 0 | 0 | 0 | 0 | 0 | 0 | 0 |
| 0 | NA | NA | NA | NA | NA | Gm48125 | 0 | 0 | 0 | 0 | 0 | 0 | 0 | 0 | 0 | 0 | 0 |
| 0 | NA | NA | NA | NA | NA | Gm48128 | 0 | 0 | 0 | 0 | 0 | 0 | 0 | 0 | 0 | 0 | 0 |
| 0 | NA | NA | NA | NA | NA | Gm48130 | 0 | 0 | 0 | 0 | 0 | 0 | 0 | 0 | 0 | 0 | 0 |
| 0 | NA | NA | NA | NA | NA | Gm48131 | 0 | 0 | 0 | 0 | 0 | 0 | 0 | 0 | 0 | 0 | 0 |
| 0 | NA | NA | NA | NA | NA | Gm48132 | 0 | 0 | 0 | 0 | 0 | 0 | 0 | 0 | 0 | 0 | 0 |
| 0 | NA | NA | NA | NA | NA | Gm48269 | 0 | 0 | 0 | 0 | 0 | 0 | 0 | 0 | 0 | 0 | 0 |
| 0 | NA | NA | NA | NA | NA | Gm48270 | 0 | 0 | 0 | 0 | 0 | 0 | 0 | 0 | 0 | 0 | 0 |
| 0 | NA | NA | NA | NA | NA | Gm48273 | 0 | 0 | 0 | 0 | 0 | 0 | 0 | 0 | 0 | 0 | 0 |
| 0 | NA | NA | NA | NA | NA | Gm46223 | 0 | 0 | 0 | 0 | 0 | 0 | 0 | 0 | 0 | 0 | 0 |
| 0 | NA | NA | NA | NA | NA | Gm48274 | 0 | 0 | 0 | 0 | 0 | 0 | 0 | 0 | 0 | 0 | 0 |
| 0 | NA | NA | NA | NA | NA | Gm48275 | 0 | 0 | 0 | 0 | 0 | 0 | 0 | 0 | 0 | 0 | 0 |
| 0 | NA | NA | NA | NA | NA | Gm6991 | 0 | 0 | 0 | 0 | 0 | 0 | 0 | 0 | 0 | 0 | 0 |
| 0 | NA | NA | NA | NA | NA | Gm46199 | 0 | 0 | 0 | 0 | 0 | 0 | 0 | 0 | 0 | 0 | 0 |
| 0 | NA | NA | NA | NA | NA | Gm48332 | 0 | 0 | 0 | 0 | 0 | 0 | 0 | 0 | 0 | 0 | 0 |
| 0 | NA | NA | NA | NA | NA | Gm18120 | 0 | 0 | 0 | 0 | 0 | 0 | 0 | 0 | 0 | 0 | 0 |
| 0 | NA | NA | NA | NA | NA | Gm48333 | 0 | 0 | 0 | 0 | 0 | 0 | 0 | 0 | 0 | 0 | 0 |
| 0 | NA | NA | NA | NA | NA | Gm23883 | 0 | 0 | 0 | 0 | 0 | 0 | 0 | 0 | 0 | 0 | 0 |
| 0 | NA | NA | NA | NA | NA | Gm26257 | 0 | 0 | 0 | 0 | 0 | 0 | 0 | 0 | 0 | 0 | 0 |
| 0 | NA | NA | NA | NA | NA | Gm46189 | 0 | 0 | 0 | 0 | 0 | 0 | 0 | 0 | 0 | 0 | 0 |
| 0 | NA | NA | NA | NA | NA | Gm49339 | 0 | 0 | 0 | 0 | 0 | 0 | 0 | 0 | 0 | 0 | 0 |
| 0 | NA | NA | NA | NA | NA | Gm40645 | 0 | 0 | 0 | 0 | 0 | 0 | 0 | 0 | 0 | 0 | 0 |
| 0 | NA | NA | NA | NA | NA | Gm48484 | 0 | 0 | 0 | 0 | 0 | 0 | 0 | 0 | 0 | 0 | 0 |
| 0 | NA | NA | NA | NA | NA | Gm48786 | 0 | 0 | 0 | 0 | 0 | 0 | 0 | 0 | 0 | 0 | 0 |
| 0 | NA | NA | NA | NA | NA | Gm48787 | 0 | 0 | 0 | 0 | 0 | 0 | 0 | 0 | 0 | 0 | 0 |
| 0 | NA | NA | NA | NA | NA | Gm48788 | 0 | 0 | 0 | 0 | 0 | 0 | 0 | 0 | 0 | 0 | 0 |
| 0 | NA | NA | NA | NA | NA | 1700042O05Rik | 0 | 0 | 0 | 0 | 0 | 0 | 0 | 0 | 0 | 0 | 0 |
| 0 | NA | NA | NA | NA | NA | Gm47613 | 0 | 0 | 0 | 0 | 0 | 0 | 0 | 0 | 0 | 0 | 0 |
| 0 | NA | NA | NA | NA | NA | Gm47614 | 0 | 0 | 0 | 0 | 0 | 0 | 0 | 0 | 0 | 0 | 0 |
| 0 | NA | NA | NA | NA | NA | Gm40646 | 0 | 0 | 0 | 0 | 0 | 0 | 0 | 0 | 0 | 0 | 0 |
| 0 | NA | NA | NA | NA | NA | Gm23906 | 0 | 0 | 0 | 0 | 0 | 0 | 0 | 0 | 0 | 0 | 0 |

| | | | | | | | | | | | | | | | | | | |
|---|---|---|---|---|---|---|---|---|---|---|---|---|---|---|---|---|---|---|
| 0 | NA | NA | NA | NA | NA | Gm47626 | | 0 | 0 | 0 | 0 | 0 | 0 | 0 | 0 | 0 | 0 | 0 |
| 0 | NA | NA | NA | NA | NA | Vgll2 | vestigial like family member 2 | 0 | 0 | 0 | 0 | 0 | 0 | 0 | 0 | 0 | 0 | 0 |
| 0 | NA | NA | NA | NA | NA | Gm24084 | | 0 | 0 | 0 | 0 | 0 | 0 | 0 | 0 | 0 | 0 | 0 |
| 0 | NA | NA | NA | NA | NA | Gm20630 | | 0 | 0 | 0 | 0 | 0 | 0 | 0 | 0 | 0 | 0 | 0 |
| 0 | NA | NA | NA | NA | NA | Gm18513 | | 0 | 0 | 0 | 0 | 0 | 0 | 0 | 0 | 0 | 0 | 0 |
| 0 | NA | NA | NA | NA | NA | Nepn | nephrocan | 0 | 0 | 0 | 0 | 0 | 0 | 0 | 0 | 0 | 0 | 0 |
| 0 | NA | NA | NA | NA | NA | Gm47608 | | 0 | 0 | 0 | 0 | 0 | 0 | 0 | 0 | 0 | 0 | 0 |
| 0 | NA | NA | NA | NA | NA | Gm47607 | | 0 | 0 | 0 | 0 | 0 | 0 | 0 | 0 | 0 | 0 | 0 |
| 0 | NA | NA | NA | NA | NA | Gm47609 | | 0 | 0 | 0 | 0 | 0 | 0 | 0 | 0 | 0 | 0 | 0 |
| 0 | NA | NA | NA | NA | NA | Gm47610 | | 0 | 0 | 0 | 0 | 0 | 0 | 0 | 0 | 0 | 0 | 0 |
| 0 | NA | NA | NA | NA | NA | Gm47612 | | 0 | 0 | 0 | 0 | 0 | 0 | 0 | 0 | 0 | 0 | 0 |
| 0 | NA | NA | NA | NA | NA | Gm36065 | | 0 | 0 | 0 | 0 | 0 | 0 | 0 | 0 | 0 | 0 | 0 |
| 0 | NA | NA | NA | NA | NA | Gm25664 | | 0 | 0 | 0 | 0 | 0 | 0 | 0 | 0 | 0 | 0 | 0 |
| 0 | NA | NA | NA | NA | NA | Gm40649 | | 0 | 0 | 0 | 0 | 0 | 0 | 0 | 0 | 0 | 0 | 0 |
| 0 | NA | NA | NA | NA | NA | Gm47637 | | 0 | 0 | 0 | 0 | 0 | 0 | 0 | 0 | 0 | 0 | 0 |
| 0 | NA | NA | NA | NA | NA | Gm36173 | | 0 | 0 | 0 | 0 | 0 | 0 | 0 | 0 | 0 | 0 | 0 |
| 0 | NA | NA | NA | NA | NA | Gm19395 | | 0 | 0 | 0 | 0 | 0 | 0 | 0 | 0 | 0 | 0 | 0 |
| 0 | NA | NA | NA | NA | NA | Gm47641 | | 0 | 0 | 0 | 0 | 0 | 0 | 0 | 0 | 0 | 0 | 0 |
| 0 | NA | NA | NA | NA | NA | Gm36229 | | 0 | 0 | 0 | 0 | 0 | 0 | 0 | 0 | 0 | 0 | 0 |
| 0 | NA | NA | NA | NA | NA | Gm40650 | | 0 | 0 | 0 | 0 | 0 | 0 | 0 | 0 | 0 | 0 | 0 |
| 0 | NA | NA | NA | NA | NA | Gm47722 | | 0 | 0 | 0 | 0 | 0 | 0 | 0 | 0 | 0 | 0 | 0 |
| 0 | NA | NA | NA | NA | NA | Gm47738 | | 0 | 0 | 0 | 0 | 0 | 0 | 0 | 0 | 0 | 0 | 0 |
| 0 | NA | NA | NA | NA | NA | Gm47739 | | 0 | 0 | 0 | 0 | 0 | 0 | 0 | 0 | 0 | 0 | 0 |
| 0 | NA | NA | NA | NA | NA | Gm47737 | | 0 | 0 | 0 | 0 | 0 | 0 | 0 | 0 | 0 | 0 | 0 |
| 0 | NA | NA | NA | NA | NA | Gm47915 | | 0 | 0 | 0 | 0 | 0 | 0 | 0 | 0 | 0 | 0 | 0 |
| 0 | NA | NA | NA | NA | NA | Gm47917 | | 0 | 0 | 0 | 0 | 0 | 0 | 0 | 0 | 0 | 0 | 0 |
| 0 | NA | NA | NA | NA | NA | Gm47967 | | 0 | 0 | 0 | 0 | 0 | 0 | 0 | 0 | 0 | 0 | 0 |
| 0 | NA | NA | NA | NA | NA | Gm47968 | | 0 | 0 | 0 | 0 | 0 | 0 | 0 | 0 | 0 | 0 | 0 |
| 0 | NA | NA | NA | NA | NA | Gm26177 | | 0 | 0 | 0 | 0 | 0 | 0 | 0 | 0 | 0 | 0 | 0 |
| 0 | NA | NA | NA | NA | NA | Gm47970 | | 0 | 0 | 0 | 0 | 0 | 0 | 0 | 0 | 0 | 0 | 0 |
| 0 | NA | NA | NA | NA | NA | Gm25602 | | 0 | 0 | 0 | 0 | 0 | 0 | 0 | 0 | 0 | 0 | 0 |
| 0 | NA | NA | NA | NA | NA | Gm31849 | | 0 | 0 | 0 | 0 | 0 | 0 | 0 | 0 | 0 | 0 | 0 |
| 0 | NA | NA | NA | NA | NA | Gm46190 | | 0 | 0 | 0 | 0 | 0 | 0 | 0 | 0 | 0 | 0 | 0 |
| 0 | NA | NA | NA | NA | NA | Gm36595 | | 0 | 0 | 0 | 0 | 0 | 0 | 0 | 0 | 0 | 0 | 0 |
| 0 | NA | NA | NA | NA | NA | 4930452L12Rik | | 0 | 0 | 0 | 0 | 0 | 0 | 0 | 0 | 0 | 0 | 0 |
| 0 | NA | NA | NA | NA | NA | Gm31931 | | 0 | 0 | 0 | 0 | 0 | 0 | 0 | 0 | 0 | 0 | 0 |
| 0 | NA | NA | NA | NA | NA | Gm48053 | | 0 | 0 | 0 | 0 | 0 | 0 | 0 | 0 | 0 | 0 | 0 |
| 0 | NA | NA | NA | NA | NA | Gm36827 | | 0 | 0 | 0 | 0 | 0 | 0 | 0 | 0 | 0 | 0 | 0 |
| 0 | NA | NA | NA | NA | NA | Gm40652 | | 0 | 0 | 0 | 0 | 0 | 0 | 0 | 0 | 0 | 0 | 0 |
| 0 | NA | NA | NA | NA | NA | Gm24285 | | 0 | 0 | 0 | 0 | 0 | 0 | 0 | 0 | 0 | 0 | 0 |
| 0 | NA | NA | NA | NA | NA | Gm48055 | | 0 | 0 | 0 | 0 | 0 | 0 | 0 | 0 | 0 | 0 | 0 |
| 0 | NA | NA | NA | NA | NA | Gm9129 | | 0 | 0 | 0 | 0 | 0 | 0 | 0 | 0 | 0 | 0 | 0 |
| 0 | NA | NA | NA | NA | NA | Gm19256 | | 0 | 0 | 0 | 0 | 0 | 0 | 0 | 0 | 0 | 0 | 0 |
| 0 | NA | NA | NA | NA | NA | Rpl48-ps1 | | 0 | 0 | 0 | 0 | 0 | 0 | 0 | 0 | 0 | 0 | 0 |
| 0 | NA | NA | NA | NA | NA | AW822073 | double homeobox protein 4-like | 0 | 0 | 0 | 0 | 0 | 0 | 0 | 0 | 0 | 0 | 0 |
| 0 | NA | NA | NA | NA | NA | Gm10807 | | 0 | 0 | 0 | 0 | 0 | 0 | 0 | 0 | 0 | 0 | 0 |
| 0 | NA | NA | NA | NA | NA | Duxf3 | double homeobox family member 3 | 0 | 0 | 0 | 0 | 0 | 0 | 0 | 0 | 0 | 0 | 0 |
| 0 | NA | NA | NA | NA | NA | Gm4981 | predicted gene 4981 | 0 | 0 | 0 | 0 | 0 | 0 | 0 | 0 | 0 | 0 | 0 |
| 0 | NA | NA | NA | NA | NA | BB019430 | | 0 | 0 | 0 | 0 | 0 | 0 | 0 | 0 | 0 | 0 | 0 |
| 0 | NA | NA | NA | NA | NA | Gm25055 | | 0 | 0 | 0 | 0 | 0 | 0 | 0 | 0 | 0 | 0 | 0 |
| 0 | NA | NA | NA | NA | NA | Gm17542 | | 0 | 0 | 0 | 0 | 0 | 0 | 0 | 0 | 0 | 0 | 0 |
| 0 | NA | NA | NA | NA | NA | Gm9987 | | 0 | 0 | 0 | 0 | 0 | 0 | 0 | 0 | 0 | 0 | 0 |
| 0 | NA | NA | NA | NA | NA | Gm27672 | | 0 | 0 | 0 | 0 | 0 | 0 | 0 | 0 | 0 | 0 | 0 |
| 0 | NA | NA | NA | NA | NA | Gm23223 | | 0 | 0 | 0 | 0 | 0 | 0 | 0 | 0 | 0 | 0 | 0 |
| 0 | NA | NA | NA | NA | NA | Gm10273 | | 0 | 0 | 0 | 0 | 0 | 0 | 0 | 0 | 0 | 0 | 0 |
| 0 | NA | NA | NA | NA | NA | Gm17148 | | 0 | 0 | 0 | 0 | 0 | 0 | 0 | 0 | 0 | 0 | 0 |
| 0 | NA | NA | NA | NA | NA | Gm17059 | | 0 | 0 | 0 | 0 | 0 | 0 | 0 | 0 | 0 | 0 | 0 |
| 0 | NA | NA | NA | NA | NA | Gm23803 | | 0 | 0 | 0 | 0 | 0 | 0 | 0 | 0 | 0 | 0 | 0 |
| 0 | NA | NA | NA | NA | NA | Gm22117 | | 0 | 0 | 0 | 0 | 0 | 0 | 0 | 0 | 0 | 0 | 0 |
| 0 | NA | NA | NA | NA | NA | Gm17455 | predicted gene, 17455 | 0 | 0 | 0 | 0 | 0 | 0 | 0 | 0 | 0 | 0 | 0 |

| | | | | | | | | | | | | | | | | | | |
|---|---|---|---|---|---|---|---|---|---|---|---|---|---|---|---|---|---|---|
| 0 | NA | NA | NA | NA | NA | Gm19972 | | 0 | 0 | 0 | 0 | 0 | 0 | 0 | 0 | 0 | 0 | 0 |
| 0 | NA | NA | NA | NA | NA | Mir6408 | | 0 | 0 | 0 | 0 | 0 | 0 | 0 | 0 | 0 | 0 | 0 |
| 0 | NA | NA | NA | NA | NA | Pldi | | 0 | 0 | 0 | 0 | 0 | 0 | 0 | 0 | 0 | 0 | 0 |
| 0 | NA | NA | NA | NA | NA | Mir466j | | 0 | 0 | 0 | 0 | 0 | 0 | 0 | 0 | 0 | 0 | 0 |
| 0 | NA | NA | NA | NA | NA | Gm20611 | | 0 | 0 | 0 | 0 | 0 | 0 | 0 | 0 | 0 | 0 | 0 |
| 0 | NA | NA | NA | NA | NA | Gm20625 | | 0 | 0 | 0 | 0 | 0 | 0 | 0 | 0 | 0 | 0 | 0 |
| 0 | NA | NA | NA | NA | NA | Gm20610 | | 0 | 0 | 0 | 0 | 0 | 0 | 0 | 0 | 0 | 0 | 0 |
| 0 | NA | NA | NA | NA | NA | Gm44308 | | 0 | 0 | 0 | 0 | 0 | 0 | 0 | 0 | 0 | 0 | 0 |
| 0 | NA | NA | NA | NA | NA | Gm48077 | | 0 | 0 | 0 | 0 | 0 | 0 | 0 | 0 | 0 | 0 | 0 |
| 0 | NA | NA | NA | NA | NA | Gm48074 | | 0 | 0 | 0 | 0 | 0 | 0 | 0 | 0 | 0 | 0 | 0 |
| 0 | NA | NA | NA | NA | NA | Gm47595 | | 0 | 0 | 0 | 0 | 0 | 0 | 0 | 0 | 0 | 0 | 0 |
| 0 | NA | NA | NA | NA | NA | Gm47593 | | 0 | 0 | 0 | 0 | 0 | 0 | 0 | 0 | 0 | 0 | 0 |
| 0 | NA | NA | NA | NA | NA | Gm28447 | | 0 | 0 | 0 | 0 | 0 | 0 | 0 | 0 | 0 | 0 | 0 |
| 0 | NA | NA | NA | NA | NA | Mir5108 | | 0 | 0 | 0 | 0 | 0 | 0 | 0 | 0 | 0 | 0 | 0 |
| 0 | NA | NA | NA | NA | NA | Gm47919 | | 0 | 0 | 0 | 0 | 0 | 0 | 0 | 0 | 0 | 0 | 0 |
| 0 | NA | NA | NA | NA | NA | Mir7662 | | 0 | 0 | 0 | 0 | 0 | 0 | 0 | 0 | 0 | 0 | 0 |
| 0 | NA | NA | NA | NA | NA | Tacr2 | tachykinin receptor 2 | 0 | 0 | 0 | 0 | 0 | 0 | 0 | 0 | 0 | 0 | 0 |
| 0 | NA | NA | NA | NA | NA | Mir7215 | | 0 | 0 | 0 | 0 | 0 | 0 | 0 | 0 | 0 | 0 | 0 |
| 0 | NA | NA | NA | NA | NA | Gm47261 | | 0 | 0 | 0 | 0 | 0 | 0 | 0 | 0 | 0 | 0 | 0 |
| 0 | NA | NA | NA | NA | NA | Gm26479 | | 0 | 0 | 0 | 0 | 0 | 0 | 0 | 0 | 0 | 0 | 0 |
| 0 | NA | NA | NA | NA | NA | Gm22803 | | 0 | 0 | 0 | 0 | 0 | 0 | 0 | 0 | 0 | 0 | 0 |
| 0 | NA | NA | NA | NA | NA | Gm25862 | | 0 | 0 | 0 | 0 | 0 | 0 | 0 | 0 | 0 | 0 | 0 |
| 0 | NA | NA | NA | NA | NA | Gm16143 | | 0 | 0 | 0 | 0 | 0 | 0 | 0 | 0 | 0 | 0 | 0 |
| 0 | NA | NA | NA | NA | NA | Gm48072 | | 0 | 0 | 0 | 0 | 0 | 0 | 0 | 0 | 0 | 0 | 0 |
| 0 | NA | NA | NA | NA | NA | Mypn | myopalladin | 0 | 0 | 0 | 0 | 0 | 0 | 0 | 0 | 0 | 0 | 0 |
| 0 | NA | NA | NA | NA | NA | Gm26789 | | 0 | 0 | 0 | 0 | 0 | 0 | 0 | 0 | 0 | 0 | 0 |
| 0 | NA | NA | NA | NA | NA | Gm16145 | | 0 | 0 | 0 | 0 | 0 | 0 | 0 | 0 | 0 | 0 | 0 |
| 0 | NA | NA | NA | NA | NA | Gm23854 | | 0 | 0 | 0 | 0 | 0 | 0 | 0 | 0 | 0 | 0 | 0 |
| 0 | NA | NA | NA | NA | NA | Gm28881 | | 0 | 0 | 0 | 0 | 0 | 0 | 0 | 0 | 0 | 0 | 0 |
| 0 | NA | NA | NA | NA | NA | Gm7540 | | 0 | 0 | 0 | 0 | 0 | 0 | 0 | 0 | 0 | 0 | 0 |
| 0 | NA | NA | NA | NA | NA | Gm19116 | | 0 | 0 | 0 | 0 | 0 | 0 | 0 | 0 | 0 | 0 | 0 |
| 0 | NA | NA | NA | NA | NA | Gm30853 | | 0 | 0 | 0 | 0 | 0 | 0 | 0 | 0 | 0 | 0 | 0 |
| 0 | NA | NA | NA | NA | NA | 1700023F02Rik | | 0 | 0 | 0 | 0 | 0 | 0 | 0 | 0 | 0 | 0 | 0 |
| 0 | NA | NA | NA | NA | NA | 4930407I19Rik | | 0 | 0 | 0 | 0 | 0 | 0 | 0 | 0 | 0 | 0 | 0 |
| 0 | NA | NA | NA | NA | NA | Gm22829 | | 0 | 0 | 0 | 0 | 0 | 0 | 0 | 0 | 0 | 0 | 0 |
| 0 | NA | NA | NA | NA | NA | Gm22594 | | 0 | 0 | 0 | 0 | 0 | 0 | 0 | 0 | 0 | 0 | 0 |
| 0 | NA | NA | NA | NA | NA | Gm28139 | | 0 | 0 | 0 | 0 | 0 | 0 | 0 | 0 | 0 | 0 | 0 |
| 0 | NA | NA | NA | NA | NA | Gm31181 | | 0 | 0 | 0 | 0 | 0 | 0 | 0 | 0 | 0 | 0 | 0 |
| 0 | NA | NA | NA | NA | NA | Gm18777 | | 0 | 0 | 0 | 0 | 0 | 0 | 0 | 0 | 0 | 0 | 0 |
| 0 | NA | NA | NA | NA | NA | Gm47895 | | 0 | 0 | 0 | 0 | 0 | 0 | 0 | 0 | 0 | 0 | 0 |
| 0 | NA | NA | NA | NA | NA | Gm47896 | | 0 | 0 | 0 | 0 | 0 | 0 | 0 | 0 | 0 | 0 | 0 |
| 0 | NA | NA | NA | NA | NA | Gm47897 | | 0 | 0 | 0 | 0 | 0 | 0 | 0 | 0 | 0 | 0 | 0 |
| 0 | NA | NA | NA | NA | NA | Gm47900 | | 0 | 0 | 0 | 0 | 0 | 0 | 0 | 0 | 0 | 0 | 0 |
| 0 | NA | NA | NA | NA | NA | Gm47903 | | 0 | 0 | 0 | 0 | 0 | 0 | 0 | 0 | 0 | 0 | 0 |
| 0 | NA | NA | NA | NA | NA | Gm26359 | | 0 | 0 | 0 | 0 | 0 | 0 | 0 | 0 | 0 | 0 | 0 |
| 0 | NA | NA | NA | NA | NA | Gm49454 | | 0 | 0 | 0 | 0 | 0 | 0 | 0 | 0 | 0 | 0 | 0 |
| 0 | NA | NA | NA | NA | NA | Gm47516 | | 0 | 0 | 0 | 0 | 0 | 0 | 0 | 0 | 0 | 0 | 0 |
| 0 | NA | NA | NA | NA | NA | Gm32023 | | 0 | 0 | 0 | 0 | 0 | 0 | 0 | 0 | 0 | 0 | 0 |
| 0 | NA | NA | NA | NA | NA | Gm47531 | | 0 | 0 | 0 | 0 | 0 | 0 | 0 | 0 | 0 | 0 | 0 |
| 0 | NA | NA | NA | NA | NA | 4930563J15Rik | | 0 | 0 | 0 | 0 | 0 | 0 | 0 | 0 | 0 | 0 | 0 |
| 0 | NA | NA | NA | NA | NA | Gm10797 | | 0 | 0 | 0 | 0 | 0 | 0 | 0 | 0 | 0 | 0 | 0 |
| 0 | NA | NA | NA | NA | NA | Gm47574 | | 0 | 0 | 0 | 0 | 0 | 0 | 0 | 0 | 0 | 0 | 0 |
| 0 | NA | NA | NA | NA | NA | 1700030E10Rik | | 0 | 0 | 0 | 0 | 0 | 0 | 0 | 0 | 0 | 0 | 0 |
| 0 | NA | NA | NA | NA | NA | 4930543I11Rik | | 0 | 0 | 0 | 0 | 0 | 0 | 0 | 0 | 0 | 0 | 0 |
| 0 | NA | NA | NA | NA | NA | Gm49766 | | 0 | 0 | 0 | 0 | 0 | 0 | 0 | 0 | 0 | 0 | 0 |
| 0 | NA | NA | NA | NA | NA | Gm29774 | | 0 | 0 | 0 | 0 | 0 | 0 | 0 | 0 | 0 | 0 | 0 |
| 0 | NA | NA | NA | NA | NA | 4930521O17Rik | | 0 | 0 | 0 | 0 | 0 | 0 | 0 | 0 | 0 | 0 | 0 |
| 0 | NA | NA | NA | NA | NA | Gm16212 | | 0 | 0 | 0 | 0 | 0 | 0 | 0 | 0 | 0 | 0 | 0 |
| 0 | NA | NA | NA | NA | NA | Gm47078 | | 0 | 0 | 0 | 0 | 0 | 0 | 0 | 0 | 0 | 0 | 0 |
| 0 | NA | NA | NA | NA | NA | 4930545H06Rik | | 0 | 0 | 0 | 0 | 0 | 0 | 0 | 0 | 0 | 0 | 0 |

| | | | | | | | | | | | | | | | | | | |
|---|---|---|---|---|---|---|---|---|---|---|---|---|---|---|---|---|---|---|
| 0 | NA | NA | NA | NA | NA | Gm32872 | | 0 | 0 | 0 | 0 | 0 | 0 | 0 | 0 | 0 | 0 | 0 |
| 0 | NA | NA | NA | NA | NA | Gm7594 | | 0 | 0 | 0 | 0 | 0 | 0 | 0 | 0 | 0 | 0 | 0 |
| 0 | NA | NA | NA | NA | NA | 1700048P04Rik | | 0 | 0 | 0 | 0 | 0 | 0 | 0 | 0 | 0 | 0 | 0 |
| 0 | NA | NA | NA | NA | NA | Gm24172 | | 0 | 0 | 0 | 0 | 0 | 0 | 0 | 0 | 0 | 0 | 0 |
| 0 | NA | NA | NA | NA | NA | Gm40685 | | 0 | 0 | 0 | 0 | 0 | 0 | 0 | 0 | 0 | 0 | 0 |
| 0 | NA | NA | NA | NA | NA | Gm47211 | | 0 | 0 | 0 | 0 | 0 | 0 | 0 | 0 | 0 | 0 | 0 |
| 0 | NA | NA | NA | NA | NA | Gm18636 | | 0 | 0 | 0 | 0 | 0 | 0 | 0 | 0 | 0 | 0 | 0 |
| 0 | NA | NA | NA | NA | NA | Gm46231 | | 0 | 0 | 0 | 0 | 0 | 0 | 0 | 0 | 0 | 0 | 0 |
| 0 | NA | NA | NA | NA | NA | Gm20108 | | 0 | 0 | 0 | 0 | 0 | 0 | 0 | 0 | 0 | 0 | 0 |
| 0 | NA | NA | NA | NA | NA | Gm47946 | | 0 | 0 | 0 | 0 | 0 | 0 | 0 | 0 | 0 | 0 | 0 |
| 0 | NA | NA | NA | NA | NA | Gm22320 | | 0 | 0 | 0 | 0 | 0 | 0 | 0 | 0 | 0 | 0 | 0 |
| 0 | NA | NA | NA | NA | NA | Gm33979 | | 0 | 0 | 0 | 0 | 0 | 0 | 0 | 0 | 0 | 0 | 0 |
| 0 | NA | NA | NA | NA | NA | Gm48021 | | 0 | 0 | 0 | 0 | 0 | 0 | 0 | 0 | 0 | 0 | 0 |
| 0 | NA | NA | NA | NA | NA | Gm48022 | | 0 | 0 | 0 | 0 | 0 | 0 | 0 | 0 | 0 | 0 | 0 |
| 0 | NA | NA | NA | NA | NA | Gm40689 | | 0 | 0 | 0 | 0 | 0 | 0 | 0 | 0 | 0 | 0 | 0 |
| 0 | NA | NA | NA | NA | NA | Gm48023 | | 0 | 0 | 0 | 0 | 0 | 0 | 0 | 0 | 0 | 0 | 0 |
| 0 | NA | NA | NA | NA | NA | Gm48024 | | 0 | 0 | 0 | 0 | 0 | 0 | 0 | 0 | 0 | 0 | 0 |
| 0 | NA | NA | NA | NA | NA | Gm48036 | | 0 | 0 | 0 | 0 | 0 | 0 | 0 | 0 | 0 | 0 | 0 |
| 0 | NA | NA | NA | NA | NA | Gm24486 | | 0 | 0 | 0 | 0 | 0 | 0 | 0 | 0 | 0 | 0 | 0 |
| 0 | NA | NA | NA | NA | NA | Gm48025 | | 0 | 0 | 0 | 0 | 0 | 0 | 0 | 0 | 0 | 0 | 0 |
| 0 | NA | NA | NA | NA | NA | Gm15647 | | 0 | 0 | 0 | 0 | 0 | 0 | 0 | 0 | 0 | 0 | 0 |
| 0 | NA | NA | NA | NA | NA | Gm32025 | | 0 | 0 | 0 | 0 | 0 | 0 | 0 | 0 | 0 | 0 | 0 |
| 0 | NA | NA | NA | NA | NA | 1700113B09Rik | | 0 | 0 | 0 | 0 | 0 | 0 | 0 | 0 | 0 | 0 | 0 |
| 0 | NA | NA | NA | NA | NA | Gm20609 | | 0 | 0 | 0 | 0 | 0 | 0 | 0 | 0 | 0 | 0 | 0 |
| 0 | NA | NA | NA | NA | NA | Gm47028 | | 0 | 0 | 0 | 0 | 0 | 0 | 0 | 0 | 0 | 0 | 0 |
| 0 | NA | NA | NA | NA | NA | Gm19074 | | 0 | 0 | 0 | 0 | 0 | 0 | 0 | 0 | 0 | 0 | 0 |
| 0 | NA | NA | NA | NA | NA | Gm47045 | | 0 | 0 | 0 | 0 | 0 | 0 | 0 | 0 | 0 | 0 | 0 |
| 0 | NA | NA | NA | NA | NA | Gm6362 | | 0 | 0 | 0 | 0 | 0 | 0 | 0 | 0 | 0 | 0 | 0 |
| 0 | NA | NA | NA | NA | NA | Gm5080 | | 0 | 0 | 0 | 0 | 0 | 0 | 0 | 0 | 0 | 0 | 0 |
| 0 | NA | NA | NA | NA | NA | Gm22937 | | 0 | 0 | 0 | 0 | 0 | 0 | 0 | 0 | 0 | 0 | 0 |
| 0 | NA | NA | NA | NA | NA | 1700049L16Rik | | 0 | 0 | 0 | 0 | 0 | 0 | 0 | 0 | 0 | 0 | 0 |
| 0 | NA | NA | NA | NA | NA | Gm47060 | | 0 | 0 | 0 | 0 | 0 | 0 | 0 | 0 | 0 | 0 | 0 |
| 0 | NA | NA | NA | NA | NA | Gm24984 | | 0 | 0 | 0 | 0 | 0 | 0 | 0 | 0 | 0 | 0 | 0 |
| 0 | NA | NA | NA | NA | NA | Gm34776 | | 0 | 0 | 0 | 0 | 0 | 0 | 0 | 0 | 0 | 0 | 0 |
| 0 | NA | NA | NA | NA | NA | A330049N07Rik | | 0 | 0 | 0 | 0 | 0 | 0 | 0 | 0 | 0 | 0 | 0 |
| 0 | NA | NA | NA | NA | NA | Gm37438 | | 0 | 0 | 0 | 0 | 0 | 0 | 0 | 0 | 0 | 0 | 0 |
| 0 | NA | NA | NA | NA | NA | Gm15398 | | 0 | 0 | 0 | 0 | 0 | 0 | 0 | 0 | 0 | 0 | 0 |
| 0 | NA | NA | NA | NA | NA | Gm47098 | | 0 | 0 | 0 | 0 | 0 | 0 | 0 | 0 | 0 | 0 | 0 |
| 0 | NA | NA | NA | NA | NA | Gm19168 | | 0 | 0 | 0 | 0 | 0 | 0 | 0 | 0 | 0 | 0 | 0 |
| 0 | NA | NA | NA | NA | NA | Gm15397 | | 0 | 0 | 0 | 0 | 0 | 0 | 0 | 0 | 0 | 0 | 0 |
| 0 | NA | NA | NA | NA | NA | Gm7034 | | 0 | 0 | 0 | 0 | 0 | 0 | 0 | 0 | 0 | 0 | 0 |
| 0 | NA | NA | NA | NA | NA | Gm46234 | | 0 | 0 | 0 | 0 | 0 | 0 | 0 | 0 | 0 | 0 | 0 |
| 0 | NA | NA | NA | NA | NA | Gm48158 | | 0 | 0 | 0 | 0 | 0 | 0 | 0 | 0 | 0 | 0 | 0 |
| 0 | NA | NA | NA | NA | NA | Gm48148 | | 0 | 0 | 0 | 0 | 0 | 0 | 0 | 0 | 0 | 0 | 0 |
| 0 | NA | NA | NA | NA | NA | Gm9985 | | 0 | 0 | 0 | 0 | 0 | 0 | 0 | 0 | 0 | 0 | 0 |
| 0 | NA | NA | NA | NA | NA | Gm26402 | | 0 | 0 | 0 | 0 | 0 | 0 | 0 | 0 | 0 | 0 | 0 |
| 0 | NA | NA | NA | NA | NA | Gm19326 | | 0 | 0 | 0 | 0 | 0 | 0 | 0 | 0 | 0 | 0 | 0 |
| 0 | NA | NA | NA | NA | NA | Gm20441 | | 0 | 0 | 0 | 0 | 0 | 0 | 0 | 0 | 0 | 0 | 0 |
| 0 | NA | NA | NA | NA | NA | Gm47744 | | 0 | 0 | 0 | 0 | 0 | 0 | 0 | 0 | 0 | 0 | 0 |
| 0 | NA | NA | NA | NA | NA | Gm16221 | | 0 | 0 | 0 | 0 | 0 | 0 | 0 | 0 | 0 | 0 | 0 |
| 0 | NA | NA | NA | NA | NA | Gm16219 | | 0 | 0 | 0 | 0 | 0 | 0 | 0 | 0 | 0 | 0 | 0 |
| 0 | NA | NA | NA | NA | NA | Gm16220 | | 0 | 0 | 0 | 0 | 0 | 0 | 0 | 0 | 0 | 0 | 0 |
| 0 | NA | NA | NA | NA | NA | Mir6409 | | 0 | 0 | 0 | 0 | 0 | 0 | 0 | 0 | 0 | 0 | 0 |
| 0 | NA | NA | NA | NA | NA | Slc5a4b | solute carrier family 5 (neutral amino acid | 0 | 0 | 0 | 0 | 0 | 0 | 0 | 0 | 0 | 0 | 0 |
| 0 | NA | NA | NA | NA | NA | Gm19217 | | 0 | 0 | 0 | 0 | 0 | 0 | 0 | 0 | 0 | 0 | 0 |
| 0 | NA | NA | NA | NA | NA | Slc5a4a | solute carrier family 5, member 4a | 0 | 0 | 0 | 0 | 0 | 0 | 0 | 0 | 0 | 0 | 0 |
| 0 | NA | NA | NA | NA | NA | Mir678 | | 0 | 0 | 0 | 0 | 0 | 0 | 0 | 0 | 0 | 0 | 0 |
| 0 | NA | NA | NA | NA | NA | Gm16240 | | 0 | 0 | 0 | 0 | 0 | 0 | 0 | 0 | 0 | 0 | 0 |
| 0 | NA | NA | NA | NA | NA | | | 0 | 0 | 0 | 0 | 0 | 0 | 0 | 0 | 0 | 0 | 0 |
| 0 | NA | NA | NA | NA | NA | Gm35608 | | 0 | 0 | 0 | 0 | 0 | 0 | 0 | 0 | 0 | 0 | 0 |

| | | | | | | | | | | | | | | | | | |
|---|---|---|---|---|---|---|---|---|---|---|---|---|---|---|---|---|---|
| 0 | NA | NA | NA | NA | NA | Gm37782 | | 0 | 0 | 0 | 0 | 0 | 0 | 0 | 0 | 0 | 0 | 0 |
| 0 | NA | NA | NA | NA | NA | Gm3137 | | 0 | 0 | 0 | 0 | 0 | 0 | 0 | 0 | 0 | 0 | 0 |
| 0 | NA | NA | NA | NA | NA | Gm35920 | | 0 | 0 | 0 | 0 | 0 | 0 | 0 | 0 | 0 | 0 | 0 |
| 0 | NA | NA | NA | NA | NA | Gm49325 | | 0 | 0 | 0 | 0 | 0 | 0 | 0 | 0 | 0 | 0 | 0 |
| 0 | NA | NA | NA | NA | NA | Mir1930 | | 0 | 0 | 0 | 0 | 0 | 0 | 0 | 0 | 0 | 0 | 0 |
| 0 | NA | NA | NA | NA | NA | Gm48160 | | 0 | 0 | 0 | 0 | 0 | 0 | 0 | 0 | 0 | 0 | 0 |
| 0 | NA | NA | NA | NA | NA | Gm19402 | predicted gene, 19402 | 0 | 0 | 0 | 0 | 0 | 0 | 0 | 0 | 0 | 0 | 0 |
| 0 | NA | NA | NA | NA | NA | Gm9508 | predicted gene 9508 | 0 | 0 | 0 | 0 | 0 | 0 | 0 | 0 | 0 | 0 | 0 |
| 0 | NA | NA | NA | NA | NA | Gm10272 | | 0 | 0 | 0 | 0 | 0 | 0 | 0 | 0 | 0 | 0 | 0 |
| 0 | NA | NA | NA | NA | NA | Gm10024 | predicted gene 10024 | 0 | 0 | 0 | 0 | 0 | 0 | 0 | 0 | 0 | 0 | 0 |
| 0 | NA | NA | NA | NA | NA | Gm10142 | predicted gene 10142 | 0 | 0 | 0 | 0 | 0 | 0 | 0 | 0 | 0 | 0 | 0 |
| 0 | NA | NA | NA | NA | NA | Gm48179 | | 0 | 0 | 0 | 0 | 0 | 0 | 0 | 0 | 0 | 0 | 0 |
| 0 | NA | NA | NA | NA | NA | Krtap12-1 | keratin associated protein 12-1 | 0 | 0 | 0 | 0 | 0 | 0 | 0 | 0 | 0 | 0 | 0 |
| 0 | NA | NA | NA | NA | NA | Gm10100 | predicted gene 10100 | 0 | 0 | 0 | 0 | 0 | 0 | 0 | 0 | 0 | 0 | 0 |
| 0 | NA | NA | NA | NA | NA | Gm48180 | | 0 | 0 | 0 | 0 | 0 | 0 | 0 | 0 | 0 | 0 | 0 |
| 0 | NA | NA | NA | NA | NA | Gm18596 | | 0 | 0 | 0 | 0 | 0 | 0 | 0 | 0 | 0 | 0 | 0 |
| 0 | NA | NA | NA | NA | NA | Gm48178 | | 0 | 0 | 0 | 0 | 0 | 0 | 0 | 0 | 0 | 0 | 0 |
| 0 | NA | NA | NA | NA | NA | Gm9736 | predicted gene 9736 | 0 | 0 | 0 | 0 | 0 | 0 | 0 | 0 | 0 | 0 | 0 |
| 0 | NA | NA | NA | NA | NA | Gm3233 | predicted gene 3233 | 0 | 0 | 0 | 0 | 0 | 0 | 0 | 0 | 0 | 0 | 0 |
| 0 | NA | NA | NA | NA | NA | Gm3238 | predicted gene 3238 | 0 | 0 | 0 | 0 | 0 | 0 | 0 | 0 | 0 | 0 | 0 |
| 0 | NA | NA | NA | NA | NA | Gm7138 | predicted gene 7138 | 0 | 0 | 0 | 0 | 0 | 0 | 0 | 0 | 0 | 0 | 0 |
| 0 | NA | NA | NA | NA | NA | Gm3250 | predicted gene 3250 | 0 | 0 | 0 | 0 | 0 | 0 | 0 | 0 | 0 | 0 | 0 |
| 0 | NA | NA | NA | NA | NA | Gm7137 | predicted gene 7137 | 0 | 0 | 0 | 0 | 0 | 0 | 0 | 0 | 0 | 0 | 0 |
| 0 | NA | NA | NA | NA | NA | Gm9639 | predicted gene 9639 | 0 | 0 | 0 | 0 | 0 | 0 | 0 | 0 | 0 | 0 | 0 |
| 0 | NA | NA | NA | NA | NA | Gm19668 | predicted gene, 19668 | 0 | 0 | 0 | 0 | 0 | 0 | 0 | 0 | 0 | 0 | 0 |
| 0 | NA | NA | NA | NA | NA | Gm9507 | predicted gene 9507 | 0 | 0 | 0 | 0 | 0 | 0 | 0 | 0 | 0 | 0 | 0 |
| 0 | NA | NA | NA | NA | NA | Gm49918 | predicted gene 2696 | 0 | 0 | 0 | 0 | 0 | 0 | 0 | 0 | 0 | 0 | 0 |
| 0 | NA | NA | NA | NA | NA | Krtap10-4 | keratin associated protein 10-4 | 0 | 0 | 0 | 0 | 0 | 0 | 0 | 0 | 0 | 0 | 0 |
| 0 | NA | NA | NA | NA | NA | Krtap10-10 | keratin associated protein 10-10 | 0 | 0 | 0 | 0 | 0 | 0 | 0 | 0 | 0 | 0 | 0 |
| 0 | NA | NA | NA | NA | NA | Gm36176 | predicted gene, 36176 | 0 | 0 | 0 | 0 | 0 | 0 | 0 | 0 | 0 | 0 | 0 |
| 0 | NA | NA | NA | NA | NA | Gm10318 | predicted gene 10318 | 0 | 0 | 0 | 0 | 0 | 0 | 0 | 0 | 0 | 0 | 0 |
| 0 | NA | NA | NA | NA | NA | Gm3285 | predicted gene 3285 | 0 | 0 | 0 | 0 | 0 | 0 | 0 | 0 | 0 | 0 | 0 |
| 0 | NA | NA | NA | NA | NA | Gm22394 | | 0 | 0 | 0 | 0 | 0 | 0 | 0 | 0 | 0 | 0 | 0 |
| 0 | NA | NA | NA | NA | NA | Gm22871 | | 0 | 0 | 0 | 0 | 0 | 0 | 0 | 0 | 0 | 0 | 0 |
| 0 | NA | NA | NA | NA | NA | | | 0 | 0 | 0 | 0 | 0 | 0 | 0 | 0 | 0 | 0 | 0 |
| 0 | NA | NA | NA | NA | NA | n-R5s77 | | 0 | 0 | 0 | 0 | 0 | 0 | 0 | 0 | 0 | 0 | 0 |
| 0 | NA | NA | NA | NA | NA | Mir7659 | | 0 | 0 | 0 | 0 | 0 | 0 | 0 | 0 | 0 | 0 | 0 |
| 0 | NA | NA | NA | NA | NA | Gm22514 | | 0 | 0 | 0 | 0 | 0 | 0 | 0 | 0 | 0 | 0 | 0 |
| 0 | NA | NA | NA | NA | NA | Mir6410 | | 0 | 0 | 0 | 0 | 0 | 0 | 0 | 0 | 0 | 0 | 0 |
| 0 | NA | NA | NA | NA | NA | Mir6908 | | 0 | 0 | 0 | 0 | 0 | 0 | 0 | 0 | 0 | 0 | 0 |
| 0 | NA | NA | NA | NA | NA | 4933436P19Rik | | 0 | 0 | 0 | 0 | 0 | 0 | 0 | 0 | 0 | 0 | 0 |
| 0 | NA | NA | NA | NA | NA | Gm30232 | | 0 | 0 | 0 | 0 | 0 | 0 | 0 | 0 | 0 | 0 | 0 |
| 0 | NA | NA | NA | NA | NA | Gm9978 | | 0 | 0 | 0 | 0 | 0 | 0 | 0 | 0 | 0 | 0 | 0 |
| 0 | NA | NA | NA | NA | NA | Gm47944 | | 0 | 0 | 0 | 0 | 0 | 0 | 0 | 0 | 0 | 0 | 0 |
| 0 | NA | NA | NA | NA | NA | Olfr1358 | olfactory receptor 1358 | 0 | 0 | 0 | 0 | 0 | 0 | 0 | 0 | 0 | 0 | 0 |
| 0 | NA | NA | NA | NA | NA | Gm25164 | | 0 | 0 | 0 | 0 | 0 | 0 | 0 | 0 | 0 | 0 | 0 |
| 0 | NA | NA | NA | NA | NA | Gm27541 | | 0 | 0 | 0 | 0 | 0 | 0 | 0 | 0 | 0 | 0 | 0 |
| 0 | NA | NA | NA | NA | NA | Gm5951 | | 0 | 0 | 0 | 0 | 0 | 0 | 0 | 0 | 0 | 0 | 0 |
| 0 | NA | NA | NA | NA | NA | Vmn2r-ps92 | | 0 | 0 | 0 | 0 | 0 | 0 | 0 | 0 | 0 | 0 | 0 |
| 0 | NA | NA | NA | NA | NA | Gm30346 | | 0 | 0 | 0 | 0 | 0 | 0 | 0 | 0 | 0 | 0 | 0 |
| 0 | NA | NA | NA | NA | NA | Gm37514 | | 0 | 0 | 0 | 0 | 0 | 0 | 0 | 0 | 0 | 0 | 0 |
| 0 | NA | NA | NA | NA | NA | Olfr1357 | olfactory receptor 1357 | 0 | 0 | 0 | 0 | 0 | 0 | 0 | 0 | 0 | 0 | 0 |
| 0 | NA | NA | NA | NA | NA | Gm47476 | | 0 | 0 | 0 | 0 | 0 | 0 | 0 | 0 | 0 | 0 | 0 |
| 0 | NA | NA | NA | NA | NA | Gm47495 | | 0 | 0 | 0 | 0 | 0 | 0 | 0 | 0 | 0 | 0 | 0 |
| 0 | NA | NA | NA | NA | NA | Gm30400 | | 0 | 0 | 0 | 0 | 0 | 0 | 0 | 0 | 0 | 0 | 0 |
| 0 | NA | NA | NA | NA | NA | Olfr1355 | olfactory receptor 1355 | 0 | 0 | 0 | 0 | 0 | 0 | 0 | 0 | 0 | 0 | 0 |
| 0 | NA | NA | NA | NA | NA | Olfr1354 | olfactory receptor 1354 | 0 | 0 | 0 | 0 | 0 | 0 | 0 | 0 | 0 | 0 | 0 |
| 0 | NA | NA | NA | NA | NA | Olfr8 | olfactory receptor 8 | 0 | 0 | 0 | 0 | 0 | 0 | 0 | 0 | 0 | 0 | 0 |
| 0 | NA | NA | NA | NA | NA | Olfr1353 | olfactory receptor 1353 | 0 | 0 | 0 | 0 | 0 | 0 | 0 | 0 | 0 | 0 | 0 |
| 0 | NA | NA | NA | NA | NA | Gm47343 | | 0 | 0 | 0 | 0 | 0 | 0 | 0 | 0 | 0 | 0 | 0 |

| | | | | | | | | | | | | | | | | | | |
|---|---|---|---|---|---|---|---|---|---|---|---|---|---|---|---|---|---|---|
| 0 | NA | NA | NA | NA | NA | Olfr1352 | olfactory receptor 1352 | 0 | 0 | 0 | 0 | 0 | 0 | 0 | 0 | 0 | 0 | 0 |
| 0 | NA | NA | NA | NA | NA | Olfr1351 | olfactory receptor 1351 | 0 | 0 | 0 | 0 | 0 | 0 | 0 | 0 | 0 | 0 | 0 |
| 0 | NA | NA | NA | NA | NA | Olfr57 | olfactory receptor 57 | 0 | 0 | 0 | 0 | 0 | 0 | 0 | 0 | 0 | 0 | 0 |
| 0 | NA | NA | NA | NA | NA | Vmn2r80 | vomeronasal 2, receptor 80 | 0 | 0 | 0 | 0 | 0 | 0 | 0 | 0 | 0 | 0 | 0 |
| 0 | NA | NA | NA | NA | NA | Vmn2r81 | vomeronasal 2, receptor 81 | 0 | 0 | 0 | 0 | 0 | 0 | 0 | 0 | 0 | 0 | 0 |
| 0 | NA | NA | NA | NA | NA | Vmn2r82 | vomeronasal 2, receptor 82 | 0 | 0 | 0 | 0 | 0 | 0 | 0 | 0 | 0 | 0 | 0 |
| 0 | NA | NA | NA | NA | NA | Vmn2r83 | vomeronasal 2, receptor 83 | 0 | 0 | 0 | 0 | 0 | 0 | 0 | 0 | 0 | 0 | 0 |
| 0 | NA | NA | NA | NA | NA | Mir6909 | | 0 | 0 | 0 | 0 | 0 | 0 | 0 | 0 | 0 | 0 | 0 |
| 0 | NA | NA | NA | NA | NA | Theg | testicular haploid expressed gene | 0 | 0 | 0 | 0 | 0 | 0 | 0 | 0 | 0 | 0 | 0 |
| 0 | NA | NA | NA | NA | NA | Mir6910 | | 0 | 0 | 0 | 0 | 0 | 0 | 0 | 0 | 0 | 0 | 0 |
| 0 | NA | NA | NA | NA | NA | Elane | elastase, neutrophil expressed | 0 | 0 | 0 | 0 | 0 | 0 | 0 | 0 | 0 | 0 | 0 |
| 0 | NA | NA | NA | NA | NA | Gm24019 | | 0 | 0 | 0 | 0 | 0 | 0 | 0 | 0 | 0 | 0 | 0 |
| 0 | NA | NA | NA | NA | NA | Gm15122 | | 0 | 0 | 0 | 0 | 0 | 0 | 0 | 0 | 0 | 0 | 0 |
| 0 | NA | NA | NA | NA | NA | Gm15123 | | 0 | 0 | 0 | 0 | 0 | 0 | 0 | 0 | 0 | 0 | 0 |
| 0 | NA | NA | NA | NA | NA | Gm47793 | | 0 | 0 | 0 | 0 | 0 | 0 | 0 | 0 | 0 | 0 | 0 |
| 0 | NA | NA | NA | NA | NA | Gm22721 | | 0 | 0 | 0 | 0 | 0 | 0 | 0 | 0 | 0 | 0 | 0 |
| 0 | NA | NA | NA | NA | NA | Gm25044 | | 0 | 0 | 0 | 0 | 0 | 0 | 0 | 0 | 0 | 0 | 0 |
| 0 | NA | NA | NA | NA | NA | 2610034O05Rik | | 0 | 0 | 0 | 0 | 0 | 0 | 0 | 0 | 0 | 0 | 0 |
| 0 | NA | NA | NA | NA | NA | Adat3 | adenosine deaminase, tRNA-specific 3 | 0 | 0 | 0 | 0 | 0 | 0 | 0 | 0 | 0 | 0 | 0 |
| 0 | NA | NA | NA | NA | NA | Mir6912 | | 0 | 0 | 0 | 0 | 0 | 0 | 0 | 0 | 0 | 0 | 0 |
| 0 | NA | NA | NA | NA | NA | Gm17151 | | 0 | 0 | 0 | 0 | 0 | 0 | 0 | 0 | 0 | 0 | 0 |
| 0 | NA | NA | NA | NA | NA | Mir1982 | | 0 | 0 | 0 | 0 | 0 | 0 | 0 | 0 | 0 | 0 | 0 |
| 0 | NA | NA | NA | NA | NA | Mir5615-2 | | 0 | 0 | 0 | 0 | 0 | 0 | 0 | 0 | 0 | 0 | 0 |
| 0 | NA | NA | NA | NA | NA | Nmrk2 | nicotinamide riboside kinase 2 | 0 | 0 | 0 | 0 | 0 | 0 | 0 | 0 | 0 | 0 | 0 |
| 0 | NA | NA | NA | NA | NA | Mir3057 | | 0 | 0 | 0 | 0 | 0 | 0 | 0 | 0 | 0 | 0 | 0 |
| 0 | NA | NA | NA | NA | NA | Gm16105 | | 0 | 0 | 0 | 0 | 0 | 0 | 0 | 0 | 0 | 0 | 0 |
| 0 | NA | NA | NA | NA | NA | Gm25595 | | 0 | 0 | 0 | 0 | 0 | 0 | 0 | 0 | 0 | 0 | 0 |
| 0 | NA | NA | NA | NA | NA | Gm47527 | | 0 | 0 | 0 | 0 | 0 | 0 | 0 | 0 | 0 | 0 | 0 |
| 0 | NA | NA | NA | NA | NA | Gm8112 | | 0 | 0 | 0 | 0 | 0 | 0 | 0 | 0 | 0 | 0 | 0 |
| 0 | NA | NA | NA | NA | NA | Gm47529 | | 0 | 0 | 0 | 0 | 0 | 0 | 0 | 0 | 0 | 0 | 0 |
| 0 | NA | NA | NA | NA | NA | Gm47537 | | 0 | 0 | 0 | 0 | 0 | 0 | 0 | 0 | 0 | 0 | 0 |
| 0 | NA | NA | NA | NA | NA | Gm8264 | | 0 | 0 | 0 | 0 | 0 | 0 | 0 | 0 | 0 | 0 | 0 |
| 0 | NA | NA | NA | NA | NA | Gm8137 | | 0 | 0 | 0 | 0 | 0 | 0 | 0 | 0 | 0 | 0 | 0 |
| 0 | NA | NA | NA | NA | NA | Gm8274 | | 0 | 0 | 0 | 0 | 0 | 0 | 0 | 0 | 0 | 0 | 0 |
| 0 | NA | NA | NA | NA | NA | Gm47539 | | 0 | 0 | 0 | 0 | 0 | 0 | 0 | 0 | 0 | 0 | 0 |
| 0 | NA | NA | NA | NA | NA | Gm47546 | | 0 | 0 | 0 | 0 | 0 | 0 | 0 | 0 | 0 | 0 | 0 |
| 0 | NA | NA | NA | NA | NA | Gm8158 | | 0 | 0 | 0 | 0 | 0 | 0 | 0 | 0 | 0 | 0 | 0 |
| 0 | NA | NA | NA | NA | NA | Gm46194 | | 0 | 0 | 0 | 0 | 0 | 0 | 0 | 0 | 0 | 0 | 0 |
| 0 | NA | NA | NA | NA | NA | Gm18556 | | 0 | 0 | 0 | 0 | 0 | 0 | 0 | 0 | 0 | 0 | 0 |
| 0 | NA | NA | NA | NA | NA | Gm47382 | | 0 | 0 | 0 | 0 | 0 | 0 | 0 | 0 | 0 | 0 | 0 |
| 0 | NA | NA | NA | NA | NA | Gm1553 | predicted gene 1553 | 0 | 0 | 0 | 0 | 0 | 0 | 0 | 0 | 0 | 0 | 0 |
| 0 | NA | NA | NA | NA | NA | Gm19186 | | 0 | 0 | 0 | 0 | 0 | 0 | 0 | 0 | 0 | 0 | 0 |
| 0 | NA | NA | NA | NA | NA | Gm47560 | | 0 | 0 | 0 | 0 | 0 | 0 | 0 | 0 | 0 | 0 | 0 |
| 0 | NA | NA | NA | NA | NA | Gm47561 | | 0 | 0 | 0 | 0 | 0 | 0 | 0 | 0 | 0 | 0 | 0 |
| 0 | NA | NA | NA | NA | NA | Gm10773 | | 0 | 0 | 0 | 0 | 0 | 0 | 0 | 0 | 0 | 0 | 0 |
| 0 | NA | NA | NA | NA | NA | 1700025N21Rik | | 0 | 0 | 0 | 0 | 0 | 0 | 0 | 0 | 0 | 0 | 0 |
| 0 | NA | NA | NA | NA | NA | Gm23122 | | 0 | 0 | 0 | 0 | 0 | 0 | 0 | 0 | 0 | 0 | 0 |
| 0 | NA | NA | NA | NA | NA | Gm47170 | | 0 | 0 | 0 | 0 | 0 | 0 | 0 | 0 | 0 | 0 | 0 |
| 0 | NA | NA | NA | NA | NA | Gm47244 | | 0 | 0 | 0 | 0 | 0 | 0 | 0 | 0 | 0 | 0 | 0 |
| 0 | NA | NA | NA | NA | NA | Gm47245 | | 0 | 0 | 0 | 0 | 0 | 0 | 0 | 0 | 0 | 0 | 0 |
| 0 | NA | NA | NA | NA | NA | C230099D08Rik | | 0 | 0 | 0 | 0 | 0 | 0 | 0 | 0 | 0 | 0 | 0 |
| 0 | NA | NA | NA | NA | NA | Gm37908 | | 0 | 0 | 0 | 0 | 0 | 0 | 0 | 0 | 0 | 0 | 0 |
| 0 | NA | NA | NA | NA | NA | Gm34113 | | 0 | 0 | 0 | 0 | 0 | 0 | 0 | 0 | 0 | 0 | 0 |
| 0 | NA | NA | NA | NA | NA | Gm47253 | | 0 | 0 | 0 | 0 | 0 | 0 | 0 | 0 | 0 | 0 | 0 |
| 0 | NA | NA | NA | NA | NA | 4930463O16Rik | | 0 | 0 | 0 | 0 | 0 | 0 | 0 | 0 | 0 | 0 | 0 |
| 0 | NA | NA | NA | NA | NA | 1700092E16Rik | | 0 | 0 | 0 | 0 | 0 | 0 | 0 | 0 | 0 | 0 | 0 |
| 0 | NA | NA | NA | NA | NA | Gm18251 | | 0 | 0 | 0 | 0 | 0 | 0 | 0 | 0 | 0 | 0 | 0 |
| 0 | NA | NA | NA | NA | NA | Gm25657 | | 0 | 0 | 0 | 0 | 0 | 0 | 0 | 0 | 0 | 0 | 0 |
| 0 | NA | NA | NA | NA | NA | Gm24226 | | 0 | 0 | 0 | 0 | 0 | 0 | 0 | 0 | 0 | 0 | 0 |
| 0 | NA | NA | NA | NA | NA | Gm38575 | | 0 | 0 | 0 | 0 | 0 | 0 | 0 | 0 | 0 | 0 | 0 |

| | | | | | | | | | | | | | | | | | | |
|---|---|---|---|---|---|---|---|---|---|---|---|---|---|---|---|---|---|---|
| 0 | NA | NA | NA | NA | NA | Gm27392 | | 0 | 0 | 0 | 0 | 0 | 0 | 0 | 0 | 0 | 0 | 0 |
| 0 | NA | NA | NA | NA | NA | Gm25281 | | 0 | 0 | 0 | 0 | 0 | 0 | 0 | 0 | 0 | 0 | 0 |
| 0 | NA | NA | NA | NA | NA | Gm47358 | | 0 | 0 | 0 | 0 | 0 | 0 | 0 | 0 | 0 | 0 | 0 |
| 0 | NA | NA | NA | NA | NA | Gm6713 | | 0 | 0 | 0 | 0 | 0 | 0 | 0 | 0 | 0 | 0 | 0 |
| 0 | NA | NA | NA | NA | NA | Gm6714 | | 0 | 0 | 0 | 0 | 0 | 0 | 0 | 0 | 0 | 0 | 0 |
| 0 | NA | NA | NA | NA | NA | Gm21026 | | 0 | 0 | 0 | 0 | 0 | 0 | 0 | 0 | 0 | 0 | 0 |
| 0 | NA | NA | NA | NA | NA | Gm47362 | | 0 | 0 | 0 | 0 | 0 | 0 | 0 | 0 | 0 | 0 | 0 |
| 0 | NA | NA | NA | NA | NA | Gm47363 | | 0 | 0 | 0 | 0 | 0 | 0 | 0 | 0 | 0 | 0 | 0 |
| 0 | NA | NA | NA | NA | NA | Gm47366 | | 0 | 0 | 0 | 0 | 0 | 0 | 0 | 0 | 0 | 0 | 0 |
| 0 | NA | NA | NA | NA | NA | Gm47365 | | 0 | 0 | 0 | 0 | 0 | 0 | 0 | 0 | 0 | 0 | 0 |
| 0 | NA | NA | NA | NA | NA | Gm47367 | | 0 | 0 | 0 | 0 | 0 | 0 | 0 | 0 | 0 | 0 | 0 |
| 0 | NA | NA | NA | NA | NA | Gm9466 | | 0 | 0 | 0 | 0 | 0 | 0 | 0 | 0 | 0 | 0 | 0 |
| 0 | NA | NA | NA | NA | NA | Gm19612 | | 0 | 0 | 0 | 0 | 0 | 0 | 0 | 0 | 0 | 0 | 0 |
| 0 | NA | NA | NA | NA | NA | Gm1559 | | 0 | 0 | 0 | 0 | 0 | 0 | 0 | 0 | 0 | 0 | 0 |
| 0 | NA | NA | NA | NA | NA | Gm6721 | | 0 | 0 | 0 | 0 | 0 | 0 | 0 | 0 | 0 | 0 | 0 |
| 0 | NA | NA | NA | NA | NA | Gm48175 | | 0 | 0 | 0 | 0 | 0 | 0 | 0 | 0 | 0 | 0 | 0 |
| 0 | NA | NA | NA | NA | NA | 4930486F22Rik | | 0 | 0 | 0 | 0 | 0 | 0 | 0 | 0 | 0 | 0 | 0 |
| 0 | NA | NA | NA | NA | NA | Gm15990 | | 0 | 0 | 0 | 0 | 0 | 0 | 0 | 0 | 0 | 0 | 0 |
| 0 | NA | NA | NA | NA | NA | Gm6729 | predicted gene 6729 | 0 | 0 | 0 | 0 | 0 | 0 | 0 | 0 | 0 | 0 | 0 |
| 0 | NA | NA | NA | NA | NA | Gm6729 | | 0 | 0 | 0 | 0 | 0 | 0 | 0 | 0 | 0 | 0 | 0 |
| 0 | NA | NA | NA | NA | NA | Gm9583 | | 0 | 0 | 0 | 0 | 0 | 0 | 0 | 0 | 0 | 0 | 0 |
| 0 | NA | NA | NA | NA | NA | Gm21027 | | 0 | 0 | 0 | 0 | 0 | 0 | 0 | 0 | 0 | 0 | 0 |
| 0 | NA | NA | NA | NA | NA | Gm48448 | | 0 | 0 | 0 | 0 | 0 | 0 | 0 | 0 | 0 | 0 | 0 |
| 0 | NA | NA | NA | NA | NA | Gm48450 | | 0 | 0 | 0 | 0 | 0 | 0 | 0 | 0 | 0 | 0 | 0 |
| 0 | NA | NA | NA | NA | NA | Gm48451 | | 0 | 0 | 0 | 0 | 0 | 0 | 0 | 0 | 0 | 0 | 0 |
| 0 | NA | NA | NA | NA | NA | Gm48454 | | 0 | 0 | 0 | 0 | 0 | 0 | 0 | 0 | 0 | 0 | 0 |
| 0 | NA | NA | NA | NA | NA | Gm48460 | | 0 | 0 | 0 | 0 | 0 | 0 | 0 | 0 | 0 | 0 | 0 |
| 0 | NA | NA | NA | NA | NA | Gm46191 | | 0 | 0 | 0 | 0 | 0 | 0 | 0 | 0 | 0 | 0 | 0 |
| 0 | NA | NA | NA | NA | NA | Gm48461 | | 0 | 0 | 0 | 0 | 0 | 0 | 0 | 0 | 0 | 0 | 0 |
| 0 | NA | NA | NA | NA | NA | Gm48463 | | 0 | 0 | 0 | 0 | 0 | 0 | 0 | 0 | 0 | 0 | 0 |
| 0 | NA | NA | NA | NA | NA | Gm48464 | | 0 | 0 | 0 | 0 | 0 | 0 | 0 | 0 | 0 | 0 | 0 |
| 0 | NA | NA | NA | NA | NA | Gm48465 | | 0 | 0 | 0 | 0 | 0 | 0 | 0 | 0 | 0 | 0 | 0 |
| 0 | NA | NA | NA | NA | NA | Gm48466 | | 0 | 0 | 0 | 0 | 0 | 0 | 0 | 0 | 0 | 0 | 0 |
| 0 | NA | NA | NA | NA | NA | Gm48473 | | 0 | 0 | 0 | 0 | 0 | 0 | 0 | 0 | 0 | 0 | 0 |
| 0 | NA | NA | NA | NA | NA | Gm48474 | | 0 | 0 | 0 | 0 | 0 | 0 | 0 | 0 | 0 | 0 | 0 |
| 0 | NA | NA | NA | NA | NA | Gm48475 | | 0 | 0 | 0 | 0 | 0 | 0 | 0 | 0 | 0 | 0 | 0 |
| 0 | NA | NA | NA | NA | NA | Gm19989 | | 0 | 0 | 0 | 0 | 0 | 0 | 0 | 0 | 0 | 0 | 0 |
| 0 | NA | NA | NA | NA | NA | Gm18876 | | 0 | 0 | 0 | 0 | 0 | 0 | 0 | 0 | 0 | 0 | 0 |
| 0 | NA | NA | NA | NA | NA | Gm5174 | | 0 | 0 | 0 | 0 | 0 | 0 | 0 | 0 | 0 | 0 | 0 |
| 0 | NA | NA | NA | NA | NA | Gm48485 | predicted gene 5174 | 0 | 0 | 0 | 0 | 0 | 0 | 0 | 0 | 0 | 0 | 0 |
| 0 | NA | NA | NA | NA | NA | Gm25745 | | 0 | 0 | 0 | 0 | 0 | 0 | 0 | 0 | 0 | 0 | 0 |
| 0 | NA | NA | NA | NA | NA | Gm49358 | | 0 | 0 | 0 | 0 | 0 | 0 | 0 | 0 | 0 | 0 | 0 |
| 0 | NA | NA | NA | NA | NA | Gm16280 | | 0 | 0 | 0 | 0 | 0 | 0 | 0 | 0 | 0 | 0 | 0 |
| 0 | NA | NA | NA | NA | NA | Gm16270 | | 0 | 0 | 0 | 0 | 0 | 0 | 0 | 0 | 0 | 0 | 0 |
| 0 | NA | NA | NA | NA | NA | Gm16269 | | 0 | 0 | 0 | 0 | 0 | 0 | 0 | 0 | 0 | 0 | 0 |
| 0 | NA | NA | NA | NA | NA | 4930555G07Rik | | 0 | 0 | 0 | 0 | 0 | 0 | 0 | 0 | 0 | 0 | 0 |
| 0 | NA | NA | NA | NA | NA | Gm10764 | | 0 | 0 | 0 | 0 | 0 | 0 | 0 | 0 | 0 | 0 | 0 |
| 0 | NA | NA | NA | NA | NA | Gm23191 | | 0 | 0 | 0 | 0 | 0 | 0 | 0 | 0 | 0 | 0 | 0 |
| 0 | NA | NA | NA | NA | NA | Gm48120 | | 0 | 0 | 0 | 0 | 0 | 0 | 0 | 0 | 0 | 0 | 0 |
| 0 | NA | NA | NA | NA | NA | Gm48195 | | 0 | 0 | 0 | 0 | 0 | 0 | 0 | 0 | 0 | 0 | 0 |
| 0 | NA | NA | NA | NA | NA | Gm48306 | | 0 | 0 | 0 | 0 | 0 | 0 | 0 | 0 | 0 | 0 | 0 |
| 0 | NA | NA | NA | NA | NA | Gm48307 | | 0 | 0 | 0 | 0 | 0 | 0 | 0 | 0 | 0 | 0 | 0 |
| 0 | NA | NA | NA | NA | NA | Gm48308 | | 0 | 0 | 0 | 0 | 0 | 0 | 0 | 0 | 0 | 0 | 0 |
| 0 | NA | NA | NA | NA | NA | Pmch | pro-melanin-concentrating hormone | 0 | 0 | 0 | 0 | 0 | 0 | 0 | 0 | 0 | 0 | 0 |
| 0 | NA | NA | NA | NA | NA | Gm6653 | | 0 | 0 | 0 | 0 | 0 | 0 | 0 | 0 | 0 | 0 | 0 |
| 0 | NA | NA | NA | NA | NA | 4921515L22Rik | | 0 | 0 | 0 | 0 | 0 | 0 | 0 | 0 | 0 | 0 | 0 |
| 0 | NA | NA | NA | NA | NA | Gm26357 | | 0 | 0 | 0 | 0 | 0 | 0 | 0 | 0 | 0 | 0 | 0 |
| 0 | NA | NA | NA | NA | NA | Gm17980 | | 0 | 0 | 0 | 0 | 0 | 0 | 0 | 0 | 0 | 0 | 0 |
| 0 | NA | NA | NA | NA | NA | Gm48476 | | 0 | 0 | 0 | 0 | 0 | 0 | 0 | 0 | 0 | 0 | 0 |
| 0 | NA | NA | NA | NA | NA | Gm48752 | | 0 | 0 | 0 | 0 | 0 | 0 | 0 | 0 | 0 | 0 | 0 |

| | | | | | | | | | | | | | | | | | | |
|---|---|---|---|---|---|---|---|---|---|---|---|---|---|---|---|---|---|---|
| 0 | NA | NA | NA | NA | NA | Gm8435 | | 0 | 0 | 0 | 0 | 0 | 0 | 0 | 0 | 0 | 0 | 0 |
| 0 | NA | NA | NA | NA | NA | Gm48108 | | 0 | 0 | 0 | 0 | 0 | 0 | 0 | 0 | 0 | 0 | 0 |
| 0 | NA | NA | NA | NA | NA | Gm29807 | | 0 | 0 | 0 | 0 | 0 | 0 | 0 | 0 | 0 | 0 | 0 |
| 0 | NA | NA | NA | NA | NA | n-R5s79 | | 0 | 0 | 0 | 0 | 0 | 0 | 0 | 0 | 0 | 0 | 0 |
| 0 | NA | NA | NA | NA | NA | Gm48144 | | 0 | 0 | 0 | 0 | 0 | 0 | 0 | 0 | 0 | 0 | 0 |
| 0 | NA | NA | NA | NA | NA | Ano4 | anoctamin 4 | 0 | 0 | 0 | 0 | 0 | 0 | 0 | 0 | 0 | 0 | 0 |
| 0 | NA | NA | NA | NA | NA | Gm48087 | | 0 | 0 | 0 | 0 | 0 | 0 | 0 | 0 | 0 | 0 | 0 |
| 0 | NA | NA | NA | NA | NA | Gm26180 | | 0 | 0 | 0 | 0 | 0 | 0 | 0 | 0 | 0 | 0 | 0 |
| 0 | NA | NA | NA | NA | NA | Gm22602 | | 0 | 0 | 0 | 0 | 0 | 0 | 0 | 0 | 0 | 0 | 0 |
| 0 | NA | NA | NA | NA | NA | Gm47591 | | 0 | 0 | 0 | 0 | 0 | 0 | 0 | 0 | 0 | 0 | 0 |
| 0 | NA | NA | NA | NA | NA | Gm18670 | | 0 | 0 | 0 | 0 | 0 | 0 | 0 | 0 | 0 | 0 | 0 |
| 0 | NA | NA | NA | NA | NA | Gm47589 | | 0 | 0 | 0 | 0 | 0 | 0 | 0 | 0 | 0 | 0 | 0 |
| 0 | NA | NA | NA | NA | NA | Gm47592 | | 0 | 0 | 0 | 0 | 0 | 0 | 0 | 0 | 0 | 0 | 0 |
| 0 | NA | NA | NA | NA | NA | Gm25390 | | 0 | 0 | 0 | 0 | 0 | 0 | 0 | 0 | 0 | 0 | 0 |
| 0 | NA | NA | NA | NA | NA | Gm47081 | | 0 | 0 | 0 | 0 | 0 | 0 | 0 | 0 | 0 | 0 | 0 |
| 0 | NA | NA | NA | NA | NA | Gm47083 | | 0 | 0 | 0 | 0 | 0 | 0 | 0 | 0 | 0 | 0 | 0 |
| 0 | NA | NA | NA | NA | NA | Gm47084 | | 0 | 0 | 0 | 0 | 0 | 0 | 0 | 0 | 0 | 0 | 0 |
| 0 | NA | NA | NA | NA | NA | Gm47091 | | 0 | 0 | 0 | 0 | 0 | 0 | 0 | 0 | 0 | 0 | 0 |
| 0 | NA | NA | NA | NA | NA | Gm31592 | | 0 | 0 | 0 | 0 | 0 | 0 | 0 | 0 | 0 | 0 | 0 |
| 0 | NA | NA | NA | NA | NA | Mir135a-2 | | 0 | 0 | 0 | 0 | 0 | 0 | 0 | 0 | 0 | 0 | 0 |
| 0 | NA | NA | NA | NA | NA | Gm27999 | | 0 | 0 | 0 | 0 | 0 | 0 | 0 | 0 | 0 | 0 | 0 |
| 0 | NA | NA | NA | NA | NA | Gm27608 | | 0 | 0 | 0 | 0 | 0 | 0 | 0 | 0 | 0 | 0 | 0 |
| 0 | NA | NA | NA | NA | NA | Gm27588 | | 0 | 0 | 0 | 0 | 0 | 0 | 0 | 0 | 0 | 0 | 0 |
| 0 | NA | NA | NA | NA | NA | Gm27753 | | 0 | 0 | 0 | 0 | 0 | 0 | 0 | 0 | 0 | 0 | 0 |
| 0 | NA | NA | NA | NA | NA | Gm27582 | | 0 | 0 | 0 | 0 | 0 | 0 | 0 | 0 | 0 | 0 | 0 |
| 0 | NA | NA | NA | NA | NA | Gm27366 | | 0 | 0 | 0 | 0 | 0 | 0 | 0 | 0 | 0 | 0 | 0 |
| 0 | NA | NA | NA | NA | NA | Gm27419 | | 0 | 0 | 0 | 0 | 0 | 0 | 0 | 0 | 0 | 0 | 0 |
| 0 | NA | NA | NA | NA | NA | Gm20757 | | 0 | 0 | 0 | 0 | 0 | 0 | 0 | 0 | 0 | 0 | 0 |
| 0 | NA | NA | NA | NA | NA | Gm47124 | | 0 | 0 | 0 | 0 | 0 | 0 | 0 | 0 | 0 | 0 | 0 |
| 0 | NA | NA | NA | NA | NA | 4930401A07Rik | | 0 | 0 | 0 | 0 | 0 | 0 | 0 | 0 | 0 | 0 | 0 |
| 0 | NA | NA | NA | NA | NA | Gm18043 | | 0 | 0 | 0 | 0 | 0 | 0 | 0 | 0 | 0 | 0 | 0 |
| 0 | NA | NA | NA | NA | NA | Gm25884 | | 0 | 0 | 0 | 0 | 0 | 0 | 0 | 0 | 0 | 0 | 0 |
| 0 | NA | NA | NA | NA | NA | Gm3523 | | 0 | 0 | 0 | 0 | 0 | 0 | 0 | 0 | 0 | 0 | 0 |
| 0 | NA | NA | NA | NA | NA | Gm22582 | | 0 | 0 | 0 | 0 | 0 | 0 | 0 | 0 | 0 | 0 | 0 |
| 0 | NA | NA | NA | NA | NA | Gm31822 | | 0 | 0 | 0 | 0 | 0 | 0 | 0 | 0 | 0 | 0 | 0 |
| 0 | NA | NA | NA | NA | NA | Gm47993 | | 0 | 0 | 0 | 0 | 0 | 0 | 0 | 0 | 0 | 0 | 0 |
| 0 | NA | NA | NA | NA | NA | Gm32468 | | 0 | 0 | 0 | 0 | 0 | 0 | 0 | 0 | 0 | 0 | 0 |
| 0 | NA | NA | NA | NA | NA | Mir1931 | | 0 | 0 | 0 | 0 | 0 | 0 | 0 | 0 | 0 | 0 | 0 |
| 0 | NA | NA | NA | NA | NA | Gm47344 | | 0 | 0 | 0 | 0 | 0 | 0 | 0 | 0 | 0 | 0 | 0 |
| 0 | NA | NA | NA | NA | NA | Ccdc38 | coiled-coil domain containing 38 | 0 | 0 | 0 | 0 | 0 | 0 | 0 | 0 | 0 | 0 | 0 |
| 0 | NA | NA | NA | NA | NA | Gm47156 | | 0 | 0 | 0 | 0 | 0 | 0 | 0 | 0 | 0 | 0 | 0 |
| 0 | NA | NA | NA | NA | NA | Gm15963 | | 0 | 0 | 0 | 0 | 0 | 0 | 0 | 0 | 0 | 0 | 0 |
| 0 | NA | NA | NA | NA | NA | Gm26239 | | 0 | 0 | 0 | 0 | 0 | 0 | 0 | 0 | 0 | 0 | 0 |
| 0 | NA | NA | NA | NA | NA | Gm25962 | | 0 | 0 | 0 | 0 | 0 | 0 | 0 | 0 | 0 | 0 | 0 |
| 0 | NA | NA | NA | NA | NA | Gm48126 | | 0 | 0 | 0 | 0 | 0 | 0 | 0 | 0 | 0 | 0 | 0 |
| 0 | NA | NA | NA | NA | NA | Mir331 | | 0 | 0 | 0 | 0 | 0 | 0 | 0 | 0 | 0 | 0 | 0 |
| 0 | NA | NA | NA | NA | NA | Gm48191 | | 0 | 0 | 0 | 0 | 0 | 0 | 0 | 0 | 0 | 0 | 0 |
| 0 | NA | NA | NA | NA | NA | Gm18391 | | 0 | 0 | 0 | 0 | 0 | 0 | 0 | 0 | 0 | 0 | 0 |
| 0 | NA | NA | NA | NA | NA | Gm26122 | | 0 | 0 | 0 | 0 | 0 | 0 | 0 | 0 | 0 | 0 | 0 |
| 0 | NA | NA | NA | NA | NA | Tmcc3os | | 0 | 0 | 0 | 0 | 0 | 0 | 0 | 0 | 0 | 0 | 0 |
| 0 | NA | NA | NA | NA | NA | Mir7211 | | 0 | 0 | 0 | 0 | 0 | 0 | 0 | 0 | 0 | 0 | 0 |
| 0 | NA | NA | NA | NA | NA | Gm48657 | | 0 | 0 | 0 | 0 | 0 | 0 | 0 | 0 | 0 | 0 | 0 |
| 0 | NA | NA | NA | NA | NA | Gm48658 | | 0 | 0 | 0 | 0 | 0 | 0 | 0 | 0 | 0 | 0 | 0 |
| 0 | NA | NA | NA | NA | NA | Gm49167 | | 0 | 0 | 0 | 0 | 0 | 0 | 0 | 0 | 0 | 0 | 0 |
| 0 | NA | NA | NA | NA | NA | Gm48689 | | 0 | 0 | 0 | 0 | 0 | 0 | 0 | 0 | 0 | 0 | 0 |
| 0 | NA | NA | NA | NA | NA | Gm24186 | | 0 | 0 | 0 | 0 | 0 | 0 | 0 | 0 | 0 | 0 | 0 |
| 0 | NA | NA | NA | NA | NA | Gm48718 | | 0 | 0 | 0 | 0 | 0 | 0 | 0 | 0 | 0 | 0 | 0 |
| 0 | NA | NA | NA | NA | NA | Gm25266 | | 0 | 0 | 0 | 0 | 0 | 0 | 0 | 0 | 0 | 0 | 0 |
| 0 | NA | NA | NA | NA | NA | Gm48867 | | 0 | 0 | 0 | 0 | 0 | 0 | 0 | 0 | 0 | 0 | 0 |
| 0 | NA | NA | NA | NA | NA | Gm48868 | | 0 | 0 | 0 | 0 | 0 | 0 | 0 | 0 | 0 | 0 | 0 |

| | | | | | | | | | | | | | | | | | | |
|---|---|---|---|---|---|---|---|---|---|---|---|---|---|---|---|---|---|---|
| 0 | NA | NA | NA | NA | NA | Gm29684 | | 0 | 0 | 0 | 0 | 0 | 0 | 0 | 0 | 0 | 0 | 0 |
| 0 | NA | NA | NA | NA | NA | Gm33091 | | 0 | 0 | 0 | 0 | 0 | 0 | 0 | 0 | 0 | 0 | 0 |
| 0 | NA | NA | NA | NA | NA | Gm48874 | | 0 | 0 | 0 | 0 | 0 | 0 | 0 | 0 | 0 | 0 | 0 |
| 0 | NA | NA | NA | NA | NA | 2310039L15Rik | | 0 | 0 | 0 | 0 | 0 | 0 | 0 | 0 | 0 | 0 | 0 |
| 0 | NA | NA | NA | NA | NA | Gm47600 | | 0 | 0 | 0 | 0 | 0 | 0 | 0 | 0 | 0 | 0 | 0 |
| 0 | NA | NA | NA | NA | NA | Gm23776 | | 0 | 0 | 0 | 0 | 0 | 0 | 0 | 0 | 0 | 0 | 0 |
| 0 | NA | NA | NA | NA | NA | Gm33336 | | 0 | 0 | 0 | 0 | 0 | 0 | 0 | 0 | 0 | 0 | 0 |
| 0 | NA | NA | NA | NA | NA | Gm47667 | | 0 | 0 | 0 | 0 | 0 | 0 | 0 | 0 | 0 | 0 | 0 |
| 0 | NA | NA | NA | NA | NA | Gm47669 | | 0 | 0 | 0 | 0 | 0 | 0 | 0 | 0 | 0 | 0 | 0 |
| 0 | NA | NA | NA | NA | NA | Gm47671 | | 0 | 0 | 0 | 0 | 0 | 0 | 0 | 0 | 0 | 0 | 0 |
| 0 | NA | NA | NA | NA | NA | 1110019B22Rik | | 0 | 0 | 0 | 0 | 0 | 0 | 0 | 0 | 0 | 0 | 0 |
| 0 | NA | NA | NA | NA | NA | Gm47708 | | 0 | 0 | 0 | 0 | 0 | 0 | 0 | 0 | 0 | 0 | 0 |
| 0 | NA | NA | NA | NA | NA | Gm47718 | | 0 | 0 | 0 | 0 | 0 | 0 | 0 | 0 | 0 | 0 | 0 |
| 0 | NA | NA | NA | NA | NA | Gm47719 | | 0 | 0 | 0 | 0 | 0 | 0 | 0 | 0 | 0 | 0 | 0 |
| 0 | NA | NA | NA | NA | NA | Gm24433 | | 0 | 0 | 0 | 0 | 0 | 0 | 0 | 0 | 0 | 0 | 0 |
| 0 | NA | NA | NA | NA | NA | Gm49817 | | 0 | 0 | 0 | 0 | 0 | 0 | 0 | 0 | 0 | 0 | 0 |
| 0 | NA | NA | NA | NA | NA | Gm48425 | | 0 | 0 | 0 | 0 | 0 | 0 | 0 | 0 | 0 | 0 | 0 |
| 0 | NA | NA | NA | NA | NA | Gm22913 | | 0 | 0 | 0 | 0 | 0 | 0 | 0 | 0 | 0 | 0 | 0 |
| 0 | NA | NA | NA | NA | NA | 4930459C07Rik | | 0 | 0 | 0 | 0 | 0 | 0 | 0 | 0 | 0 | 0 | 0 |
| 0 | NA | NA | NA | NA | NA | Gm48428 | | 0 | 0 | 0 | 0 | 0 | 0 | 0 | 0 | 0 | 0 | 0 |
| 0 | NA | NA | NA | NA | NA | Gm8601 | | 0 | 0 | 0 | 0 | 0 | 0 | 0 | 0 | 0 | 0 | 0 |
| 0 | NA | NA | NA | NA | NA | Gm48507 | | 0 | 0 | 0 | 0 | 0 | 0 | 0 | 0 | 0 | 0 | 0 |
| 0 | NA | NA | NA | NA | NA | Gm6859 | | 0 | 0 | 0 | 0 | 0 | 0 | 0 | 0 | 0 | 0 | 0 |
| 0 | NA | NA | NA | NA | NA | Gm48509 | | 0 | 0 | 0 | 0 | 0 | 0 | 0 | 0 | 0 | 0 | 0 |
| 0 | NA | NA | NA | NA | NA | Gm48521 | | 0 | 0 | 0 | 0 | 0 | 0 | 0 | 0 | 0 | 0 | 0 |
| 0 | NA | NA | NA | NA | NA | Gm24587 | | 0 | 0 | 0 | 0 | 0 | 0 | 0 | 0 | 0 | 0 | 0 |
| 0 | NA | NA | NA | NA | NA | Gm33981 | | 0 | 0 | 0 | 0 | 0 | 0 | 0 | 0 | 0 | 0 | 0 |
| 0 | NA | NA | NA | NA | NA | 4930556N09Rik | | 0 | 0 | 0 | 0 | 0 | 0 | 0 | 0 | 0 | 0 | 0 |
| 0 | NA | NA | NA | NA | NA | Gm48555 | | 0 | 0 | 0 | 0 | 0 | 0 | 0 | 0 | 0 | 0 | 0 |
| 0 | NA | NA | NA | NA | NA | Gm8613 | | 0 | 0 | 0 | 0 | 0 | 0 | 0 | 0 | 0 | 0 | 0 |
| 0 | NA | NA | NA | NA | NA | Gm18515 | | 0 | 0 | 0 | 0 | 0 | 0 | 0 | 0 | 0 | 0 | 0 |
| 0 | NA | NA | NA | NA | NA | Mir3966 | | 0 | 0 | 0 | 0 | 0 | 0 | 0 | 0 | 0 | 0 | 0 |
| 0 | NA | NA | NA | NA | NA | 4930473O22Rik | | 0 | 0 | 0 | 0 | 0 | 0 | 0 | 0 | 0 | 0 | 0 |
| 0 | NA | NA | NA | NA | NA | Kera | keratocan | 0 | 0 | 0 | 0 | 0 | 0 | 0 | 0 | 0 | 0 | 0 |
| 0 | NA | NA | NA | NA | NA | Gm10754 | | 0 | 0 | 0 | 0 | 0 | 0 | 0 | 0 | 0 | 0 | 0 |
| 0 | NA | NA | NA | NA | NA | Ccer1 | coiled-coil glutamate-rich protein 1 | 0 | 0 | 0 | 0 | 0 | 0 | 0 | 0 | 0 | 0 | 0 |
| 0 | NA | NA | NA | NA | NA | 4930525C09Rik | | 0 | 0 | 0 | 0 | 0 | 0 | 0 | 0 | 0 | 0 | 0 |
| 0 | NA | NA | NA | NA | NA | C030005K15Rik | | 0 | 0 | 0 | 0 | 0 | 0 | 0 | 0 | 0 | 0 | 0 |
| 0 | NA | NA | NA | NA | NA | Gm25658 | | 0 | 0 | 0 | 0 | 0 | 0 | 0 | 0 | 0 | 0 | 0 |
| 0 | NA | NA | NA | NA | NA | Gm8633 | | 0 | 0 | 0 | 0 | 0 | 0 | 0 | 0 | 0 | 0 | 0 |
| 0 | NA | NA | NA | NA | NA | Gm48687 | | 0 | 0 | 0 | 0 | 0 | 0 | 0 | 0 | 0 | 0 | 0 |
| 0 | NA | NA | NA | NA | NA | Gm37631 | | 0 | 0 | 0 | 0 | 0 | 0 | 0 | 0 | 0 | 0 | 0 |
| 0 | NA | NA | NA | NA | NA | Gm34297 | | 0 | 0 | 0 | 0 | 0 | 0 | 0 | 0 | 0 | 0 | 0 |
| 0 | NA | NA | NA | NA | NA | Gm5427 | | 0 | 0 | 0 | 0 | 0 | 0 | 0 | 0 | 0 | 0 | 0 |
| 0 | NA | NA | NA | NA | NA | Gm48761 | | 0 | 0 | 0 | 0 | 0 | 0 | 0 | 0 | 0 | 0 | 0 |
| 0 | NA | NA | NA | NA | NA | Gm16239 | | 0 | 0 | 0 | 0 | 0 | 0 | 0 | 0 | 0 | 0 | 0 |
| 0 | NA | NA | NA | NA | NA | Gm48884 | | 0 | 0 | 0 | 0 | 0 | 0 | 0 | 0 | 0 | 0 | 0 |
| 0 | NA | NA | NA | NA | NA | Gm34777 | | 0 | 0 | 0 | 0 | 0 | 0 | 0 | 0 | 0 | 0 | 0 |
| 0 | NA | NA | NA | NA | NA | Gm34921 | | 0 | 0 | 0 | 0 | 0 | 0 | 0 | 0 | 0 | 0 | 0 |
| 0 | NA | NA | NA | NA | NA | Gm48089 | | 0 | 0 | 0 | 0 | 0 | 0 | 0 | 0 | 0 | 0 | 0 |
| 0 | NA | NA | NA | NA | NA | Gm35035 | | 0 | 0 | 0 | 0 | 0 | 0 | 0 | 0 | 0 | 0 | 0 |
| 0 | NA | NA | NA | NA | NA | Gm18409 | | 0 | 0 | 0 | 0 | 0 | 0 | 0 | 0 | 0 | 0 | 0 |
| 0 | NA | NA | NA | NA | NA | Gm35101 | | 0 | 0 | 0 | 0 | 0 | 0 | 0 | 0 | 0 | 0 | 0 |
| 0 | NA | NA | NA | NA | NA | Gm47578 | | 0 | 0 | 0 | 0 | 0 | 0 | 0 | 0 | 0 | 0 | 0 |
| 0 | NA | NA | NA | NA | NA | Gm35206 | | 0 | 0 | 0 | 0 | 0 | 0 | 0 | 0 | 0 | 0 | 0 |
| 0 | NA | NA | NA | NA | NA | Gm47579 | | 0 | 0 | 0 | 0 | 0 | 0 | 0 | 0 | 0 | 0 | 0 |
| 0 | NA | NA | NA | NA | NA | Gm17870 | | 0 | 0 | 0 | 0 | 0 | 0 | 0 | 0 | 0 | 0 | 0 |
| 0 | NA | NA | NA | NA | NA | Gm35533 | | 0 | 0 | 0 | 0 | 0 | 0 | 0 | 0 | 0 | 0 | 0 |
| 0 | NA | NA | NA | NA | NA | Gm22918 | | 0 | 0 | 0 | 0 | 0 | 0 | 0 | 0 | 0 | 0 | 0 |
| 0 | NA | NA | NA | NA | NA | Gm25287 | | 0 | 0 | 0 | 0 | 0 | 0 | 0 | 0 | 0 | 0 | 0 |

| | | | | | | | | | | | | | | | | | | |
|---|---|---|---|---|---|---|---|---|---|---|---|---|---|---|---|---|---|---|
| 0 | NA | NA | NA | NA | NA | Gm47627 | | 0 | 0 | 0 | 0 | 0 | 0 | 0 | 0 | 0 | 0 | 0 |
| 0 | NA | NA | NA | NA | NA | Gm8689 | | 0 | 0 | 0 | 0 | 0 | 0 | 0 | 0 | 0 | 0 | 0 |
| 0 | NA | NA | NA | NA | NA | Gm9476 | | 0 | 0 | 0 | 0 | 0 | 0 | 0 | 0 | 0 | 0 | 0 |
| 0 | NA | NA | NA | NA | NA | Gm47632 | | 0 | 0 | 0 | 0 | 0 | 0 | 0 | 0 | 0 | 0 | 0 |
| 0 | NA | NA | NA | NA | NA | Gm47628 | | 0 | 0 | 0 | 0 | 0 | 0 | 0 | 0 | 0 | 0 | 0 |
| 0 | NA | NA | NA | NA | NA | Gm47630 | | 0 | 0 | 0 | 0 | 0 | 0 | 0 | 0 | 0 | 0 | 0 |
| 0 | NA | NA | NA | NA | NA | Gm8701 | | 0 | 0 | 0 | 0 | 0 | 0 | 0 | 0 | 0 | 0 | 0 |
| 0 | NA | NA | NA | NA | NA | Gm4301 | predicted gene 4301 | 0 | 0 | 0 | 0 | 0 | 0 | 0 | 0 | 0 | 0 | 0 |
| 0 | NA | NA | NA | NA | NA | Gm4302 | predicted gene 4302 | 0 | 0 | 0 | 0 | 0 | 0 | 0 | 0 | 0 | 0 | 0 |
| 0 | NA | NA | NA | NA | NA | Gm4303 | predicted gene 4303 | 0 | 0 | 0 | 0 | 0 | 0 | 0 | 0 | 0 | 0 | 0 |
| 0 | NA | NA | NA | NA | NA | Gm4305 | predicted gene 4305 | 0 | 0 | 0 | 0 | 0 | 0 | 0 | 0 | 0 | 0 | 0 |
| 0 | NA | NA | NA | NA | NA | Gm8711 | | 0 | 0 | 0 | 0 | 0 | 0 | 0 | 0 | 0 | 0 | 0 |
| 0 | NA | NA | NA | NA | NA | Gm4307 | predicted gene 4307 | 0 | 0 | 0 | 0 | 0 | 0 | 0 | 0 | 0 | 0 | 0 |
| 0 | NA | NA | NA | NA | NA | Gm4308 | | 0 | 0 | 0 | 0 | 0 | 0 | 0 | 0 | 0 | 0 | 0 |
| 0 | NA | NA | NA | NA | NA | Gm4310 | | 0 | 0 | 0 | 0 | 0 | 0 | 0 | 0 | 0 | 0 | 0 |
| 0 | NA | NA | NA | NA | NA | Gm47631 | | 0 | 0 | 0 | 0 | 0 | 0 | 0 | 0 | 0 | 0 | 0 |
| 0 | NA | NA | NA | NA | NA | Gm8723 | | 0 | 0 | 0 | 0 | 0 | 0 | 0 | 0 | 0 | 0 | 0 |
| 0 | NA | NA | NA | NA | NA | Gm4312 | predicted gene 4312 | 0 | 0 | 0 | 0 | 0 | 0 | 0 | 0 | 0 | 0 | 0 |
| 0 | NA | NA | NA | NA | NA | Gm4781 | | 0 | 0 | 0 | 0 | 0 | 0 | 0 | 0 | 0 | 0 | 0 |
| 0 | NA | NA | NA | NA | NA | 1700017N19Ril | RIKEN cDNA 1700017N19 gene | 0 | 0 | 0 | 0 | 0 | 0 | 0 | 0 | 0 | 0 | 0 |
| 0 | NA | NA | NA | NA | NA | Gm47325 | | 0 | 0 | 0 | 0 | 0 | 0 | 0 | 0 | 0 | 0 | 0 |
| 0 | NA | NA | NA | NA | NA | Gm35722 | | 0 | 0 | 0 | 0 | 0 | 0 | 0 | 0 | 0 | 0 | 0 |
| 0 | NA | NA | NA | NA | NA | Gm19233 | | 0 | 0 | 0 | 0 | 0 | 0 | 0 | 0 | 0 | 0 | 0 |
| 0 | NA | NA | NA | NA | NA | Gm26278 | | 0 | 0 | 0 | 0 | 0 | 0 | 0 | 0 | 0 | 0 | 0 |
| 0 | NA | NA | NA | NA | NA | Gm47032 | | 0 | 0 | 0 | 0 | 0 | 0 | 0 | 0 | 0 | 0 | 0 |
| 0 | NA | NA | NA | NA | NA | Mir3059 | | 0 | 0 | 0 | 0 | 0 | 0 | 0 | 0 | 0 | 0 | 0 |
| 0 | NA | NA | NA | NA | NA | Gm47031 | | 0 | 0 | 0 | 0 | 0 | 0 | 0 | 0 | 0 | 0 | 0 |
| 0 | NA | NA | NA | NA | NA | Gm47033 | | 0 | 0 | 0 | 0 | 0 | 0 | 0 | 0 | 0 | 0 | 0 |
| 0 | NA | NA | NA | NA | NA | Gm47025 | | 0 | 0 | 0 | 0 | 0 | 0 | 0 | 0 | 0 | 0 | 0 |
| 0 | NA | NA | NA | NA | NA | Gm47101 | | 0 | 0 | 0 | 0 | 0 | 0 | 0 | 0 | 0 | 0 | 0 |
| 0 | NA | NA | NA | NA | NA | Gm26988 | | 0 | 0 | 0 | 0 | 0 | 0 | 0 | 0 | 0 | 0 | 0 |
| 0 | NA | NA | NA | NA | NA | Gm47142 | | 0 | 0 | 0 | 0 | 0 | 0 | 0 | 0 | 0 | 0 | 0 |
| 0 | NA | NA | NA | NA | NA | Gm47143 | | 0 | 0 | 0 | 0 | 0 | 0 | 0 | 0 | 0 | 0 | 0 |
| 0 | NA | NA | NA | NA | NA | Alx1 | ALX homeobox 1 | 0 | 0 | 0 | 0 | 0 | 0 | 0 | 0 | 0 | 0 | 0 |
| 0 | NA | NA | NA | NA | NA | Gm17028 | | 0 | 0 | 0 | 0 | 0 | 0 | 0 | 0 | 0 | 0 | 0 |
| 0 | NA | NA | NA | NA | NA | Gm25376 | | 0 | 0 | 0 | 0 | 0 | 0 | 0 | 0 | 0 | 0 | 0 |
| 0 | NA | NA | NA | NA | NA | Gm47164 | | 0 | 0 | 0 | 0 | 0 | 0 | 0 | 0 | 0 | 0 | 0 |
| 0 | NA | NA | NA | NA | NA | Gm8768 | | 0 | 0 | 0 | 0 | 0 | 0 | 0 | 0 | 0 | 0 | 0 |
| 0 | NA | NA | NA | NA | NA | Gm47223 | | 0 | 0 | 0 | 0 | 0 | 0 | 0 | 0 | 0 | 0 | 0 |
| 0 | NA | NA | NA | NA | NA | Gm47221 | | 0 | 0 | 0 | 0 | 0 | 0 | 0 | 0 | 0 | 0 | 0 |
| 0 | NA | NA | NA | NA | NA | Gm47220 | | 0 | 0 | 0 | 0 | 0 | 0 | 0 | 0 | 0 | 0 | 0 |
| 0 | NA | NA | NA | NA | NA | Gm47222 | | 0 | 0 | 0 | 0 | 0 | 0 | 0 | 0 | 0 | 0 | 0 |
| 0 | NA | NA | NA | NA | NA | Gm26923 | | 0 | 0 | 0 | 0 | 0 | 0 | 0 | 0 | 0 | 0 | 0 |
| 0 | NA | NA | NA | NA | NA | Gm47224 | | 0 | 0 | 0 | 0 | 0 | 0 | 0 | 0 | 0 | 0 | 0 |
| 0 | NA | NA | NA | NA | NA | Gm47225 | | 0 | 0 | 0 | 0 | 0 | 0 | 0 | 0 | 0 | 0 | 0 |
| 0 | NA | NA | NA | NA | NA | Gm25143 | | 0 | 0 | 0 | 0 | 0 | 0 | 0 | 0 | 0 | 0 | 0 |
| 0 | NA | NA | NA | NA | NA | Gm47226 | | 0 | 0 | 0 | 0 | 0 | 0 | 0 | 0 | 0 | 0 | 0 |
| 0 | NA | NA | NA | NA | NA | Gm47227 | | 0 | 0 | 0 | 0 | 0 | 0 | 0 | 0 | 0 | 0 | 0 |
| 0 | NA | NA | NA | NA | NA | Gm47228 | | 0 | 0 | 0 | 0 | 0 | 0 | 0 | 0 | 0 | 0 | 0 |
| 0 | NA | NA | NA | NA | NA | Gm21293 | predicted gene, 21293 | 0 | 0 | 0 | 0 | 0 | 0 | 0 | 0 | 0 | 0 | 0 |
| 0 | NA | NA | NA | NA | NA | Gm6763 | predicted gene 6763 | 0 | 0 | 0 | 0 | 0 | 0 | 0 | 0 | 0 | 0 | 0 |
| 0 | NA | NA | NA | NA | NA | Gm8764 | predicted gene 8764 | 0 | 0 | 0 | 0 | 0 | 0 | 0 | 0 | 0 | 0 | 0 |
| 0 | NA | NA | NA | NA | NA | Gm21304 | predicted gene, 21304 | 0 | 0 | 0 | 0 | 0 | 0 | 0 | 0 | 0 | 0 | 0 |
| 0 | NA | NA | NA | NA | NA | Gm21312 | predicted gene, 21312 | 0 | 0 | 0 | 0 | 0 | 0 | 0 | 0 | 0 | 0 | 0 |
| 0 | NA | NA | NA | NA | NA | Gm20765 | predicted gene, 20765 | 0 | 0 | 0 | 0 | 0 | 0 | 0 | 0 | 0 | 0 | 0 |
| 0 | NA | NA | NA | NA | NA | Gm4340 | predicted gene 4340 | 0 | 0 | 0 | 0 | 0 | 0 | 0 | 0 | 0 | 0 | 0 |
| 0 | NA | NA | NA | NA | NA | Gm22339 | | 0 | 0 | 0 | 0 | 0 | 0 | 0 | 0 | 0 | 0 | 0 |
| 0 | NA | NA | NA | NA | NA | Mir6411 | | 0 | 0 | 0 | 0 | 0 | 0 | 0 | 0 | 0 | 0 | 0 |
| 0 | NA | NA | NA | NA | NA | Gm22945 | | 0 | 0 | 0 | 0 | 0 | 0 | 0 | 0 | 0 | 0 | 0 |
| 0 | NA | NA | NA | NA | NA | Gm25522 | | 0 | 0 | 0 | 0 | 0 | 0 | 0 | 0 | 0 | 0 | 0 |

| | | | | | | | | | | | | | | | | | |
|---|---|---|---|---|---|---|---|---|---|---|---|---|---|---|---|---|---|
| 0 | NA | NA | NA | NA | NA | Gm48184 | | 0 | 0 | 0 | 0 | 0 | 0 | 0 | 0 | 0 | 0 | 0 |
| 0 | NA | NA | NA | NA | NA | Gm25587 | | 0 | 0 | 0 | 0 | 0 | 0 | 0 | 0 | 0 | 0 | 0 |
| 0 | NA | NA | NA | NA | NA | Gm25422 | | 0 | 0 | 0 | 0 | 0 | 0 | 0 | 0 | 0 | 0 | 0 |
| 0 | NA | NA | NA | NA | NA | Gm36997 | | 0 | 0 | 0 | 0 | 0 | 0 | 0 | 0 | 0 | 0 | 0 |
| 0 | NA | NA | NA | NA | NA | Gm48206 | | 0 | 0 | 0 | 0 | 0 | 0 | 0 | 0 | 0 | 0 | 0 |
| 0 | NA | NA | NA | NA | NA | Gm48205 | | 0 | 0 | 0 | 0 | 0 | 0 | 0 | 0 | 0 | 0 | 0 |
| 0 | NA | NA | NA | NA | NA | Gm15664 | | 0 | 0 | 0 | 0 | 0 | 0 | 0 | 0 | 0 | 0 | 0 |
| 0 | NA | NA | NA | NA | NA | n-R5s80 | | 0 | 0 | 0 | 0 | 0 | 0 | 0 | 0 | 0 | 0 | 0 |
| 0 | NA | NA | NA | NA | NA | Gm24237 | | 0 | 0 | 0 | 0 | 0 | 0 | 0 | 0 | 0 | 0 | 0 |
| 0 | NA | NA | NA | NA | NA | Gm25342 | | 0 | 0 | 0 | 0 | 0 | 0 | 0 | 0 | 0 | 0 | 0 |
| 0 | NA | NA | NA | NA | NA | Gm48363 | | 0 | 0 | 0 | 0 | 0 | 0 | 0 | 0 | 0 | 0 | 0 |
| 0 | NA | NA | NA | NA | NA | Gm48376 | | 0 | 0 | 0 | 0 | 0 | 0 | 0 | 0 | 0 | 0 | 0 |
| 0 | NA | NA | NA | NA | NA | Gm48377 | | 0 | 0 | 0 | 0 | 0 | 0 | 0 | 0 | 0 | 0 | 0 |
| 0 | NA | NA | NA | NA | NA | Gm48379 | | 0 | 0 | 0 | 0 | 0 | 0 | 0 | 0 | 0 | 0 | 0 |
| 0 | NA | NA | NA | NA | NA | Gm48380 | | 0 | 0 | 0 | 0 | 0 | 0 | 0 | 0 | 0 | 0 | 0 |
| 0 | NA | NA | NA | NA | NA | Olfr1556-ps1 | | 0 | 0 | 0 | 0 | 0 | 0 | 0 | 0 | 0 | 0 | 0 |
| 0 | NA | NA | NA | NA | NA | 4930532I03Rik | | 0 | 0 | 0 | 0 | 0 | 0 | 0 | 0 | 0 | 0 | 0 |
| 0 | NA | NA | NA | NA | NA | Gm19007 | | 0 | 0 | 0 | 0 | 0 | 0 | 0 | 0 | 0 | 0 | 0 |
| 0 | NA | NA | NA | NA | NA | Gm18040 | | 0 | 0 | 0 | 0 | 0 | 0 | 0 | 0 | 0 | 0 | 0 |
| 0 | NA | NA | NA | NA | NA | Gm19181 | | 0 | 0 | 0 | 0 | 0 | 0 | 0 | 0 | 0 | 0 | 0 |
| 0 | NA | NA | NA | NA | NA | Gm24745 | | 0 | 0 | 0 | 0 | 0 | 0 | 0 | 0 | 0 | 0 | 0 |
| 0 | NA | NA | NA | NA | NA | Gm29674 | | 0 | 0 | 0 | 0 | 0 | 0 | 0 | 0 | 0 | 0 | 0 |
| 0 | NA | NA | NA | NA | NA | Gm38117 | | 0 | 0 | 0 | 0 | 0 | 0 | 0 | 0 | 0 | 0 | 0 |
| 0 | NA | NA | NA | NA | NA | Gm44403 | | 0 | 0 | 0 | 0 | 0 | 0 | 0 | 0 | 0 | 0 | 0 |
| 0 | NA | NA | NA | NA | NA | Myf5 | myogenic factor 5 | 0 | 0 | 0 | 0 | 0 | 0 | 0 | 0 | 0 | 0 | 0 |
| 0 | NA | NA | NA | NA | NA | Myf6 | myogenic factor 6 | 0 | 0 | 0 | 0 | 0 | 0 | 0 | 0 | 0 | 0 | 0 |
| 0 | NA | NA | NA | NA | NA | Ptprq | protein tyrosine phosphatase, receptor typ | 0 | 0 | 0 | 0 | 0 | 0 | 0 | 0 | 0 | 0 | 0 |
| 0 | NA | NA | NA | NA | NA | Gm29685 | | 0 | 0 | 0 | 0 | 0 | 0 | 0 | 0 | 0 | 0 | 0 |
| 0 | NA | NA | NA | NA | NA | Gm47999 | | 0 | 0 | 0 | 0 | 0 | 0 | 0 | 0 | 0 | 0 | 0 |
| 0 | NA | NA | NA | NA | NA | Gm25591 | | 0 | 0 | 0 | 0 | 0 | 0 | 0 | 0 | 0 | 0 | 0 |
| 0 | NA | NA | NA | NA | NA | Gm47466 | | 0 | 0 | 0 | 0 | 0 | 0 | 0 | 0 | 0 | 0 | 0 |
| 0 | NA | NA | NA | NA | NA | Gm5136 | predicted gene 5136 | 0 | 0 | 0 | 0 | 0 | 0 | 0 | 0 | 0 | 0 | 0 |
| 0 | NA | NA | NA | NA | NA | Gm47472 | | 0 | 0 | 0 | 0 | 0 | 0 | 0 | 0 | 0 | 0 | 0 |
| 0 | NA | NA | NA | NA | NA | Gm47477 | | 0 | 0 | 0 | 0 | 0 | 0 | 0 | 0 | 0 | 0 | 0 |
| 0 | NA | NA | NA | NA | NA | Gm22097 | | 0 | 0 | 0 | 0 | 0 | 0 | 0 | 0 | 0 | 0 | 0 |
| 0 | NA | NA | NA | NA | NA | Gm44495 | | 0 | 0 | 0 | 0 | 0 | 0 | 0 | 0 | 0 | 0 | 0 |
| 0 | NA | NA | NA | NA | NA | Gm23041 | | 0 | 0 | 0 | 0 | 0 | 0 | 0 | 0 | 0 | 0 | 0 |
| 0 | NA | NA | NA | NA | NA | Gm47329 | | 0 | 0 | 0 | 0 | 0 | 0 | 0 | 0 | 0 | 0 | 0 |
| 0 | NA | NA | NA | NA | NA | 3110043J17Rik | | 0 | 0 | 0 | 0 | 0 | 0 | 0 | 0 | 0 | 0 | 0 |
| 0 | NA | NA | NA | NA | NA | Gm36935 | | 0 | 0 | 0 | 0 | 0 | 0 | 0 | 0 | 0 | 0 | 0 |
| 0 | NA | NA | NA | NA | NA | Gm47337 | | 0 | 0 | 0 | 0 | 0 | 0 | 0 | 0 | 0 | 0 | 0 |
| 0 | NA | NA | NA | NA | NA | Gm47339 | | 0 | 0 | 0 | 0 | 0 | 0 | 0 | 0 | 0 | 0 | 0 |
| 0 | NA | NA | NA | NA | NA | Gm47338 | | 0 | 0 | 0 | 0 | 0 | 0 | 0 | 0 | 0 | 0 | 0 |
| 0 | NA | NA | NA | NA | NA | Gm47055 | | 0 | 0 | 0 | 0 | 0 | 0 | 0 | 0 | 0 | 0 | 0 |
| 0 | NA | NA | NA | NA | NA | 9230102K24Rik | | 0 | 0 | 0 | 0 | 0 | 0 | 0 | 0 | 0 | 0 | 0 |
| 0 | NA | NA | NA | NA | NA | Gm47058 | | 0 | 0 | 0 | 0 | 0 | 0 | 0 | 0 | 0 | 0 | 0 |
| 0 | NA | NA | NA | NA | NA | Gm23698 | | 0 | 0 | 0 | 0 | 0 | 0 | 0 | 0 | 0 | 0 | 0 |
| 0 | NA | NA | NA | NA | NA | Gm22186 | | 0 | 0 | 0 | 0 | 0 | 0 | 0 | 0 | 0 | 0 | 0 |
| 0 | NA | NA | NA | NA | NA | Gm22236 | | 0 | 0 | 0 | 0 | 0 | 0 | 0 | 0 | 0 | 0 | 0 |
| 0 | NA | NA | NA | NA | NA | Gm22101 | | 0 | 0 | 0 | 0 | 0 | 0 | 0 | 0 | 0 | 0 | 0 |
| 0 | NA | NA | NA | NA | NA | Gm17849 | | 0 | 0 | 0 | 0 | 0 | 0 | 0 | 0 | 0 | 0 | 0 |
| 0 | NA | NA | NA | NA | NA | Gm40761 | | 0 | 0 | 0 | 0 | 0 | 0 | 0 | 0 | 0 | 0 | 0 |
| 0 | NA | NA | NA | NA | NA | 4933440J02Rik | | 0 | 0 | 0 | 0 | 0 | 0 | 0 | 0 | 0 | 0 | 0 |
| 0 | NA | NA | NA | NA | NA | Gm47864 | | 0 | 0 | 0 | 0 | 0 | 0 | 0 | 0 | 0 | 0 | 0 |
| 0 | NA | NA | NA | NA | NA | Gm30624 | | 0 | 0 | 0 | 0 | 0 | 0 | 0 | 0 | 0 | 0 | 0 |
| 0 | NA | NA | NA | NA | NA | Gm47865 | | 0 | 0 | 0 | 0 | 0 | 0 | 0 | 0 | 0 | 0 | 0 |
| 0 | NA | NA | NA | NA | NA | Gm46193 | | 0 | 0 | 0 | 0 | 0 | 0 | 0 | 0 | 0 | 0 | 0 |
| 0 | NA | NA | NA | NA | NA | Gm19005 | | 0 | 0 | 0 | 0 | 0 | 0 | 0 | 0 | 0 | 0 | 0 |
| 0 | NA | NA | NA | NA | NA | Gm37189 | | 0 | 0 | 0 | 0 | 0 | 0 | 0 | 0 | 0 | 0 | 0 |
| 0 | NA | NA | NA | NA | NA | Gm47866 | | 0 | 0 | 0 | 0 | 0 | 0 | 0 | 0 | 0 | 0 | 0 |

| | | | | | | | | | | | | | | | | | |
|---|---|---|---|---|---|---|---|---|---|---|---|---|---|---|---|---|---|
| 0 | NA | NA | NA | NA | NA | Gm20758 | | 0 | 0 | 0 | 0 | 0 | 0 | 0 | 0 | 0 | 0 | 0 |
| 0 | NA | NA | NA | NA | NA | Gm47880 | | 0 | 0 | 0 | 0 | 0 | 0 | 0 | 0 | 0 | 0 | 0 |
| 0 | NA | NA | NA | NA | NA | Gm7269 | | 0 | 0 | 0 | 0 | 0 | 0 | 0 | 0 | 0 | 0 | 0 |
| 0 | NA | NA | NA | NA | NA | Glipr1l1 | GLI pathogenesis-related 1 like 1 | 0 | 0 | 0 | 0 | 0 | 0 | 0 | 0 | 0 | 0 | 0 |
| 0 | NA | NA | NA | NA | NA | Glipr1l2 | GLI pathogenesis-related 1 like 2 | 0 | 0 | 0 | 0 | 0 | 0 | 0 | 0 | 0 | 0 | 0 |
| 0 | NA | NA | NA | NA | NA | Gm6808 | | 0 | 0 | 0 | 0 | 0 | 0 | 0 | 0 | 0 | 0 | 0 |
| 0 | NA | NA | NA | NA | NA | Gm15914 | | 0 | 0 | 0 | 0 | 0 | 0 | 0 | 0 | 0 | 0 | 0 |
| 0 | NA | NA | NA | NA | NA | Gm48047 | | 0 | 0 | 0 | 0 | 0 | 0 | 0 | 0 | 0 | 0 | 0 |
| 0 | NA | NA | NA | NA | NA | Gm47419 | | 0 | 0 | 0 | 0 | 0 | 0 | 0 | 0 | 0 | 0 | 0 |
| 0 | NA | NA | NA | NA | NA | Gm47492 | | 0 | 0 | 0 | 0 | 0 | 0 | 0 | 0 | 0 | 0 | 0 |
| 0 | NA | NA | NA | NA | NA | 1700010J16Rik | | 0 | 0 | 0 | 0 | 0 | 0 | 0 | 0 | 0 | 0 | 0 |
| 0 | NA | NA | NA | NA | NA | Gm3942 | | 0 | 0 | 0 | 0 | 0 | 0 | 0 | 0 | 0 | 0 | 0 |
| 0 | NA | NA | NA | NA | NA | Gm47519 | | 0 | 0 | 0 | 0 | 0 | 0 | 0 | 0 | 0 | 0 | 0 |
| 0 | NA | NA | NA | NA | NA | Gm47522 | | 0 | 0 | 0 | 0 | 0 | 0 | 0 | 0 | 0 | 0 | 0 |
| 0 | NA | NA | NA | NA | NA | Gm47524 | | 0 | 0 | 0 | 0 | 0 | 0 | 0 | 0 | 0 | 0 | 0 |
| 0 | NA | NA | NA | NA | NA | Gm47532 | | 0 | 0 | 0 | 0 | 0 | 0 | 0 | 0 | 0 | 0 | 0 |
| 0 | NA | NA | NA | NA | NA | Gm40765 | | 0 | 0 | 0 | 0 | 0 | 0 | 0 | 0 | 0 | 0 | 0 |
| 0 | NA | NA | NA | NA | NA | Gm40810 | | 0 | 0 | 0 | 0 | 0 | 0 | 0 | 0 | 0 | 0 | 0 |
| 0 | NA | NA | NA | NA | NA | Gm47538 | | 0 | 0 | 0 | 0 | 0 | 0 | 0 | 0 | 0 | 0 | 0 |
| 0 | NA | NA | NA | NA | NA | Gm38333 | | 0 | 0 | 0 | 0 | 0 | 0 | 0 | 0 | 0 | 0 | 0 |
| 0 | NA | NA | NA | NA | NA | Gm48104 | | 0 | 0 | 0 | 0 | 0 | 0 | 0 | 0 | 0 | 0 | 0 |
| 0 | NA | NA | NA | NA | NA | Gm38281 | | 0 | 0 | 0 | 0 | 0 | 0 | 0 | 0 | 0 | 0 | 0 |
| 0 | NA | NA | NA | NA | NA | 4930473D10Rik | | 0 | 0 | 0 | 0 | 0 | 0 | 0 | 0 | 0 | 0 | 0 |
| 0 | NA | NA | NA | NA | NA | Gm31182 | | 0 | 0 | 0 | 0 | 0 | 0 | 0 | 0 | 0 | 0 | 0 |
| 0 | NA | NA | NA | NA | NA | Gm25319 | | 0 | 0 | 0 | 0 | 0 | 0 | 0 | 0 | 0 | 0 | 0 |
| 0 | NA | NA | NA | NA | NA | Gm31267 | | 0 | 0 | 0 | 0 | 0 | 0 | 0 | 0 | 0 | 0 | 0 |
| 0 | NA | NA | NA | NA | NA | Gm22070 | | 0 | 0 | 0 | 0 | 0 | 0 | 0 | 0 | 0 | 0 | 0 |
| 0 | NA | NA | NA | NA | NA | Gm22794 | | 0 | 0 | 0 | 0 | 0 | 0 | 0 | 0 | 0 | 0 | 0 |
| 0 | NA | NA | NA | NA | NA | A930009A15Ril | RIKEN cDNA A930009A15 gene | 0 | 0 | 0 | 0 | 0 | 0 | 0 | 0 | 0 | 0 | 0 |
| 0 | NA | NA | NA | NA | NA | Taf7l2 | Taf7l2 | 0 | 0 | 0 | 0 | 0 | 0 | 0 | 0 | 0 | 0 | 0 |
| 0 | NA | NA | NA | NA | NA | Gm26495 | | 0 | 0 | 0 | 0 | 0 | 0 | 0 | 0 | 0 | 0 | 0 |
| 0 | NA | NA | NA | NA | NA | Kcnmb4os1 | | 0 | 0 | 0 | 0 | 0 | 0 | 0 | 0 | 0 | 0 | 0 |
| 0 | NA | NA | NA | NA | NA | Gm49344 | | 0 | 0 | 0 | 0 | 0 | 0 | 0 | 0 | 0 | 0 | 0 |
| 0 | NA | NA | NA | NA | NA | Gm48403 | | 0 | 0 | 0 | 0 | 0 | 0 | 0 | 0 | 0 | 0 | 0 |
| 0 | NA | NA | NA | NA | NA | Gm26579 | | 0 | 0 | 0 | 0 | 0 | 0 | 0 | 0 | 0 | 0 | 0 |
| 0 | NA | NA | NA | NA | NA | 1700030O20Rik | | 0 | 0 | 0 | 0 | 0 | 0 | 0 | 0 | 0 | 0 | 0 |
| 0 | NA | NA | NA | NA | NA | 4930579P08Rik | | 0 | 0 | 0 | 0 | 0 | 0 | 0 | 0 | 0 | 0 | 0 |
| 0 | NA | NA | NA | NA | NA | Gm10271 | | 0 | 0 | 0 | 0 | 0 | 0 | 0 | 0 | 0 | 0 | 0 |
| 0 | NA | NA | NA | NA | NA | 4930423D24Rik | | 0 | 0 | 0 | 0 | 0 | 0 | 0 | 0 | 0 | 0 | 0 |
| 0 | NA | NA | NA | NA | NA | Gm26311 | | 0 | 0 | 0 | 0 | 0 | 0 | 0 | 0 | 0 | 0 | 0 |
| 0 | NA | NA | NA | NA | NA | 9530003J23Ril | RIKEN cDNA 9530003J23 gene | 0 | 0 | 0 | 0 | 0 | 0 | 0 | 0 | 0 | 0 | 0 |
| 0 | NA | NA | NA | NA | NA | Gm10936 | | 0 | 0 | 0 | 0 | 0 | 0 | 0 | 0 | 0 | 0 | 0 |
| 0 | NA | NA | NA | NA | NA | Gm48903 | | 0 | 0 | 0 | 0 | 0 | 0 | 0 | 0 | 0 | 0 | 0 |
| 0 | NA | NA | NA | NA | NA | Gm9002 | | 0 | 0 | 0 | 0 | 0 | 0 | 0 | 0 | 0 | 0 | 0 |
| 0 | NA | NA | NA | NA | NA | Gm32141 | | 0 | 0 | 0 | 0 | 0 | 0 | 0 | 0 | 0 | 0 | 0 |
| 0 | NA | NA | NA | NA | NA | Gm32235 | | 0 | 0 | 0 | 0 | 0 | 0 | 0 | 0 | 0 | 0 | 0 |
| 0 | NA | NA | NA | NA | NA | Gm9004 | | 0 | 0 | 0 | 0 | 0 | 0 | 0 | 0 | 0 | 0 | 0 |
| 0 | NA | NA | NA | NA | NA | Kifc5c-ps | | 0 | 0 | 0 | 0 | 0 | 0 | 0 | 0 | 0 | 0 | 0 |
| 0 | NA | NA | NA | NA | NA | Gm25709 | | 0 | 0 | 0 | 0 | 0 | 0 | 0 | 0 | 0 | 0 | 0 |
| 0 | NA | NA | NA | NA | NA | Gm47899 | | 0 | 0 | 0 | 0 | 0 | 0 | 0 | 0 | 0 | 0 | 0 |
| 0 | NA | NA | NA | NA | NA | Gm37852 | | 0 | 0 | 0 | 0 | 0 | 0 | 0 | 0 | 0 | 0 | 0 |
| 0 | NA | NA | NA | NA | NA | Gm9585 | | 0 | 0 | 0 | 0 | 0 | 0 | 0 | 0 | 0 | 0 | 0 |
| 0 | NA | NA | NA | NA | NA | Gm32717 | | 0 | 0 | 0 | 0 | 0 | 0 | 0 | 0 | 0 | 0 | 0 |
| 0 | NA | NA | NA | NA | NA | Gm9029 | | 0 | 0 | 0 | 0 | 0 | 0 | 0 | 0 | 0 | 0 | 0 |
| 0 | NA | NA | NA | NA | NA | Gm9030 | | 0 | 0 | 0 | 0 | 0 | 0 | 0 | 0 | 0 | 0 | 0 |
| 0 | NA | NA | NA | NA | NA | Gm32802 | | 0 | 0 | 0 | 0 | 0 | 0 | 0 | 0 | 0 | 0 | 0 |
| 0 | NA | NA | NA | NA | NA | Gm9035 | | 0 | 0 | 0 | 0 | 0 | 0 | 0 | 0 | 0 | 0 | 0 |
| 0 | NA | NA | NA | NA | NA | Gm9040 | | 0 | 0 | 0 | 0 | 0 | 0 | 0 | 0 | 0 | 0 | 0 |
| 0 | NA | NA | NA | NA | NA | Gm9044 | | 0 | 0 | 0 | 0 | 0 | 0 | 0 | 0 | 0 | 0 | 0 |
| 0 | NA | NA | NA | NA | NA | Gm9045 | | 0 | 0 | 0 | 0 | 0 | 0 | 0 | 0 | 0 | 0 | 0 |

| | | | | | | | | | | | | | | | | | | |
|---|---|---|---|---|---|---|---|---|---|---|---|---|---|---|---|---|---|---|
| 0 | NA | NA | NA | NA | NA | Gm9046 | | 0 | 0 | 0 | 0 | 0 | 0 | 0 | 0 | 0 | 0 | 0 |
| 0 | NA | NA | NA | NA | NA | Gm9048 | | 0 | 0 | 0 | 0 | 0 | 0 | 0 | 0 | 0 | 0 | 0 |
| 0 | NA | NA | NA | NA | NA | Gm9049 | | 0 | 0 | 0 | 0 | 0 | 0 | 0 | 0 | 0 | 0 | 0 |
| 0 | NA | NA | NA | NA | NA | Gm49751 | | 0 | 0 | 0 | 0 | 0 | 0 | 0 | 0 | 0 | 0 | 0 |
| 0 | NA | NA | NA | NA | NA | Ifngas1 | | 0 | 0 | 0 | 0 | 0 | 0 | 0 | 0 | 0 | 0 | 0 |
| 0 | NA | NA | NA | NA | NA | Gm33337 | | 0 | 0 | 0 | 0 | 0 | 0 | 0 | 0 | 0 | 0 | 0 |
| 0 | NA | NA | NA | NA | NA | Gm47425 | | 0 | 0 | 0 | 0 | 0 | 0 | 0 | 0 | 0 | 0 | 0 |
| 0 | NA | NA | NA | NA | NA | Gm47434 | | 0 | 0 | 0 | 0 | 0 | 0 | 0 | 0 | 0 | 0 | 0 |
| 0 | NA | NA | NA | NA | NA | 4932442E05Rik | | 0 | 0 | 0 | 0 | 0 | 0 | 0 | 0 | 0 | 0 | 0 |
| 0 | NA | NA | NA | NA | NA | Gm38403 | | 0 | 0 | 0 | 0 | 0 | 0 | 0 | 0 | 0 | 0 | 0 |
| 0 | NA | NA | NA | NA | NA | Gm47461 | | 0 | 0 | 0 | 0 | 0 | 0 | 0 | 0 | 0 | 0 | 0 |
| 0 | NA | NA | NA | NA | NA | 1700064J06Rik | | 0 | 0 | 0 | 0 | 0 | 0 | 0 | 0 | 0 | 0 | 0 |
| 0 | NA | NA | NA | NA | NA | 4930477N07Rik | | 0 | 0 | 0 | 0 | 0 | 0 | 0 | 0 | 0 | 0 | 0 |
| 0 | NA | NA | NA | NA | NA | Gm19090 | | 0 | 0 | 0 | 0 | 0 | 0 | 0 | 0 | 0 | 0 | 0 |
| 0 | NA | NA | NA | NA | NA | Gm40778 | | 0 | 0 | 0 | 0 | 0 | 0 | 0 | 0 | 0 | 0 | 0 |
| 0 | NA | NA | NA | NA | NA | Grip1os3 | | 0 | 0 | 0 | 0 | 0 | 0 | 0 | 0 | 0 | 0 | 0 |
| 0 | NA | NA | NA | NA | NA | Gm16321 | | 0 | 0 | 0 | 0 | 0 | 0 | 0 | 0 | 0 | 0 | 0 |
| 0 | NA | NA | NA | NA | NA | Gm47341 | | 0 | 0 | 0 | 0 | 0 | 0 | 0 | 0 | 0 | 0 | 0 |
| 0 | NA | NA | NA | NA | NA | Gm47024 | | 0 | 0 | 0 | 0 | 0 | 0 | 0 | 0 | 0 | 0 | 0 |
| 0 | NA | NA | NA | NA | NA | Gm47342 | | 0 | 0 | 0 | 0 | 0 | 0 | 0 | 0 | 0 | 0 | 0 |
| 0 | NA | NA | NA | NA | NA | Gm47026 | | 0 | 0 | 0 | 0 | 0 | 0 | 0 | 0 | 0 | 0 | 0 |
| 0 | NA | NA | NA | NA | NA | Grip1os2 | | 0 | 0 | 0 | 0 | 0 | 0 | 0 | 0 | 0 | 0 | 0 |
| 0 | NA | NA | NA | NA | NA | Grip1os1 | | 0 | 0 | 0 | 0 | 0 | 0 | 0 | 0 | 0 | 0 | 0 |
| 0 | NA | NA | NA | NA | NA | Gm48115 | | 0 | 0 | 0 | 0 | 0 | 0 | 0 | 0 | 0 | 0 | 0 |
| 0 | NA | NA | NA | NA | NA | Gm48118 | | 0 | 0 | 0 | 0 | 0 | 0 | 0 | 0 | 0 | 0 | 0 |
| 0 | NA | NA | NA | NA | NA | Gm38141 | | 0 | 0 | 0 | 0 | 0 | 0 | 0 | 0 | 0 | 0 | 0 |
| 0 | NA | NA | NA | NA | NA | Hmga2 | high mobility group AT-hook 2 | 0 | 0 | 0 | 0 | 0 | 0 | 0 | 0 | 0 | 0 | 0 |
| 0 | NA | NA | NA | NA | NA | 9230105E05Rik | | 0 | 0 | 0 | 0 | 0 | 0 | 0 | 0 | 0 | 0 | 0 |
| 0 | NA | NA | NA | NA | NA | Mir763 | | 0 | 0 | 0 | 0 | 0 | 0 | 0 | 0 | 0 | 0 | 0 |
| 0 | NA | NA | NA | NA | NA | Gm24298 | | 0 | 0 | 0 | 0 | 0 | 0 | 0 | 0 | 0 | 0 | 0 |
| 0 | NA | NA | NA | NA | NA | 4930471E19Rik | | 0 | 0 | 0 | 0 | 0 | 0 | 0 | 0 | 0 | 0 | 0 |
| 0 | NA | NA | NA | NA | NA | Gm38289 | | 0 | 0 | 0 | 0 | 0 | 0 | 0 | 0 | 0 | 0 | 0 |
| 0 | NA | NA | NA | NA | NA | Gm37505 | | 0 | 0 | 0 | 0 | 0 | 0 | 0 | 0 | 0 | 0 | 0 |
| 0 | NA | NA | NA | NA | NA | Gm10743 | | 0 | 0 | 0 | 0 | 0 | 0 | 0 | 0 | 0 | 0 | 0 |
| 0 | NA | NA | NA | NA | NA | Gm48202 | | 0 | 0 | 0 | 0 | 0 | 0 | 0 | 0 | 0 | 0 | 0 |
| 0 | NA | NA | NA | NA | NA | 4921513I03Rik | | 0 | 0 | 0 | 0 | 0 | 0 | 0 | 0 | 0 | 0 | 0 |
| 0 | NA | NA | NA | NA | NA | Gm15961 | | 0 | 0 | 0 | 0 | 0 | 0 | 0 | 0 | 0 | 0 | 0 |
| 0 | NA | NA | NA | NA | NA | Gm4473 | | 0 | 0 | 0 | 0 | 0 | 0 | 0 | 0 | 0 | 0 | 0 |
| 0 | NA | NA | NA | NA | NA | Gm23632 | | 0 | 0 | 0 | 0 | 0 | 0 | 0 | 0 | 0 | 0 | 0 |
| 0 | NA | NA | NA | NA | NA | Gm48341 | | 0 | 0 | 0 | 0 | 0 | 0 | 0 | 0 | 0 | 0 | 0 |
| 0 | NA | NA | NA | NA | NA | Gm48410 | | 0 | 0 | 0 | 0 | 0 | 0 | 0 | 0 | 0 | 0 | 0 |
| 0 | NA | NA | NA | NA | NA | Gm48426 | | 0 | 0 | 0 | 0 | 0 | 0 | 0 | 0 | 0 | 0 | 0 |
| 0 | NA | NA | NA | NA | NA | Gm18730 | | 0 | 0 | 0 | 0 | 0 | 0 | 0 | 0 | 0 | 0 | 0 |
| 0 | NA | NA | NA | NA | NA | Gm40787 | | 0 | 0 | 0 | 0 | 0 | 0 | 0 | 0 | 0 | 0 | 0 |
| 0 | NA | NA | NA | NA | NA | Gm50454 | | 0 | 0 | 0 | 0 | 0 | 0 | 0 | 0 | 0 | 0 | 0 |
| 0 | NA | NA | NA | NA | NA | Gm24330 | | 0 | 0 | 0 | 0 | 0 | 0 | 0 | 0 | 0 | 0 | 0 |
| 0 | NA | NA | NA | NA | NA | Gm35865 | | 0 | 0 | 0 | 0 | 0 | 0 | 0 | 0 | 0 | 0 | 0 |
| 0 | NA | NA | NA | NA | NA | Gm48826 | | 0 | 0 | 0 | 0 | 0 | 0 | 0 | 0 | 0 | 0 | 0 |
| 0 | NA | NA | NA | NA | NA | Gm48828 | | 0 | 0 | 0 | 0 | 0 | 0 | 0 | 0 | 0 | 0 | 0 |
| 0 | NA | NA | NA | NA | NA | Gm48875 | | 0 | 0 | 0 | 0 | 0 | 0 | 0 | 0 | 0 | 0 | 0 |
| 0 | NA | NA | NA | NA | NA | Gm36041 | | 0 | 0 | 0 | 0 | 0 | 0 | 0 | 0 | 0 | 0 | 0 |
| 0 | NA | NA | NA | NA | NA | Gm48877 | | 0 | 0 | 0 | 0 | 0 | 0 | 0 | 0 | 0 | 0 | 0 |
| 0 | NA | NA | NA | NA | NA | Mir6412 | | 0 | 0 | 0 | 0 | 0 | 0 | 0 | 0 | 0 | 0 | 0 |
| 0 | NA | NA | NA | NA | NA | Mir8104 | | 0 | 0 | 0 | 0 | 0 | 0 | 0 | 0 | 0 | 0 | 0 |
| 0 | NA | NA | NA | NA | NA | Gm47582 | | 0 | 0 | 0 | 0 | 0 | 0 | 0 | 0 | 0 | 0 | 0 |
| 0 | NA | NA | NA | NA | NA | Gm36208 | | 0 | 0 | 0 | 0 | 0 | 0 | 0 | 0 | 0 | 0 | 0 |
| 0 | NA | NA | NA | NA | NA | Mirlet7i | | 0 | 0 | 0 | 0 | 0 | 0 | 0 | 0 | 0 | 0 | 0 |
| 0 | NA | NA | NA | NA | NA | Gm27920 | | 0 | 0 | 0 | 0 | 0 | 0 | 0 | 0 | 0 | 0 | 0 |
| 0 | NA | NA | NA | NA | NA | Gm22958 | | 0 | 0 | 0 | 0 | 0 | 0 | 0 | 0 | 0 | 0 | 0 |
| 0 | NA | NA | NA | NA | NA | Gm47801 | | 0 | 0 | 0 | 0 | 0 | 0 | 0 | 0 | 0 | 0 | 0 |

| | | | | | | | | | | | | | | | | | |
|---|---|---|---|---|---|---|---|---|---|---|---|---|---|---|---|---|---|
| 0 | NA | NA | NA | NA | NA | Gm19169 | | 0 | 0 | 0 | 0 | 0 | 0 | 0 | 0 | 0 | 0 | 0 |
| 0 | NA | NA | NA | NA | NA | Gm18510 | | 0 | 0 | 0 | 0 | 0 | 0 | 0 | 0 | 0 | 0 | 0 |
| 0 | NA | NA | NA | NA | NA | Gm47837 | | 0 | 0 | 0 | 0 | 0 | 0 | 0 | 0 | 0 | 0 | 0 |
| 0 | NA | NA | NA | NA | NA | Gm23777 | | 0 | 0 | 0 | 0 | 0 | 0 | 0 | 0 | 0 | 0 | 0 |
| 0 | NA | NA | NA | NA | NA | 4930503E24Rik | | 0 | 0 | 0 | 0 | 0 | 0 | 0 | 0 | 0 | 0 | 0 |
| 0 | NA | NA | NA | NA | NA | Gm47839 | | 0 | 0 | 0 | 0 | 0 | 0 | 0 | 0 | 0 | 0 | 0 |
| 0 | NA | NA | NA | NA | NA | Gm47840 | | 0 | 0 | 0 | 0 | 0 | 0 | 0 | 0 | 0 | 0 | 0 |
| 0 | NA | NA | NA | NA | NA | Gm36719 | | 0 | 0 | 0 | 0 | 0 | 0 | 0 | 0 | 0 | 0 | 0 |
| 0 | NA | NA | NA | NA | NA | Gm45604 | | 0 | 0 | 0 | 0 | 0 | 0 | 0 | 0 | 0 | 0 | 0 |
| 0 | NA | NA | NA | NA | NA | Gm9102 | | 0 | 0 | 0 | 0 | 0 | 0 | 0 | 0 | 0 | 0 | 0 |
| 0 | NA | NA | NA | NA | NA | Gm22183 | | 0 | 0 | 0 | 0 | 0 | 0 | 0 | 0 | 0 | 0 | 0 |
| 0 | NA | NA | NA | NA | NA | Gm4510 | | 0 | 0 | 0 | 0 | 0 | 0 | 0 | 0 | 0 | 0 | 0 |
| 0 | NA | NA | NA | NA | NA | 1700021G15Rik | | 0 | 0 | 0 | 0 | 0 | 0 | 0 | 0 | 0 | 0 | 0 |
| 0 | NA | NA | NA | NA | NA | Gm47965 | | 0 | 0 | 0 | 0 | 0 | 0 | 0 | 0 | 0 | 0 | 0 |
| 0 | NA | NA | NA | NA | NA | Gm40797 | | 0 | 0 | 0 | 0 | 0 | 0 | 0 | 0 | 0 | 0 | 0 |
| 0 | NA | NA | NA | NA | NA | Mir378d | | 0 | 0 | 0 | 0 | 0 | 0 | 0 | 0 | 0 | 0 | 0 |
| 0 | NA | NA | NA | NA | NA | Gm47966 | | 0 | 0 | 0 | 0 | 0 | 0 | 0 | 0 | 0 | 0 | 0 |
| 0 | NA | NA | NA | NA | NA | 4930484H19Rik | | 0 | 0 | 0 | 0 | 0 | 0 | 0 | 0 | 0 | 0 | 0 |
| 0 | NA | NA | NA | NA | NA | Gm9111 | | 0 | 0 | 0 | 0 | 0 | 0 | 0 | 0 | 0 | 0 | 0 |
| 0 | NA | NA | NA | NA | NA | Gm24179 | | 0 | 0 | 0 | 0 | 0 | 0 | 0 | 0 | 0 | 0 | 0 |
| 0 | NA | NA | NA | NA | NA | Mir546 | | 0 | 0 | 0 | 0 | 0 | 0 | 0 | 0 | 0 | 0 | 0 |
| 0 | NA | NA | NA | NA | NA | Gm4178 | | 0 | 0 | 0 | 0 | 0 | 0 | 0 | 0 | 0 | 0 | 0 |
| 0 | NA | NA | NA | NA | NA | Gm23664 | | 0 | 0 | 0 | 0 | 0 | 0 | 0 | 0 | 0 | 0 | 0 |
| 0 | NA | NA | NA | NA | NA | Ddit3 | | 0 | 0 | 0 | 0 | 0 | 0 | 0 | 0 | 0 | 0 | 0 |
| 0 | NA | NA | NA | NA | NA | Inhbe | inhibin beta-E | 0 | 0 | 0 | 0 | 0 | 0 | 0 | 0 | 0 | 0 | 0 |
| 0 | NA | NA | NA | NA | NA | Inhbc | inhibin beta-C | 0 | 0 | 0 | 0 | 0 | 0 | 0 | 0 | 0 | 0 | 0 |
| 0 | NA | NA | NA | NA | NA | Gm23819 | | 0 | 0 | 0 | 0 | 0 | 0 | 0 | 0 | 0 | 0 | 0 |
| 0 | NA | NA | NA | NA | NA | Gm16217 | | 0 | 0 | 0 | 0 | 0 | 0 | 0 | 0 | 0 | 0 | 0 |
| 0 | NA | NA | NA | NA | NA | Gm48815 | | 0 | 0 | 0 | 0 | 0 | 0 | 0 | 0 | 0 | 0 | 0 |
| 0 | NA | NA | NA | NA | NA | Rdh16f1 | | 0 | 0 | 0 | 0 | 0 | 0 | 0 | 0 | 0 | 0 | 0 |
| 0 | NA | NA | NA | NA | NA | Rdh18-ps | | 0 | 0 | 0 | 0 | 0 | 0 | 0 | 0 | 0 | 0 | 0 |
| 0 | NA | NA | NA | NA | NA | Rdh19 | retinol dehydrogenase 19 | 0 | 0 | 0 | 0 | 0 | 0 | 0 | 0 | 0 | 0 | 0 |
| 0 | NA | NA | NA | NA | NA | Gm15905 | | 0 | 0 | 0 | 0 | 0 | 0 | 0 | 0 | 0 | 0 | 0 |
| 0 | NA | NA | NA | NA | NA | Gm23996 | | 0 | 0 | 0 | 0 | 0 | 0 | 0 | 0 | 0 | 0 | 0 |
| 0 | NA | NA | NA | NA | NA | Gm47949 | | 0 | 0 | 0 | 0 | 0 | 0 | 0 | 0 | 0 | 0 | 0 |
| 0 | NA | NA | NA | NA | NA | Hsd17b6 | hydroxysteroid (17-beta) dehydrogenase | 0 | 0 | 0 | 0 | 0 | 0 | 0 | 0 | 0 | 0 | 0 |
| 0 | NA | NA | NA | NA | NA | Gm50449 | | 0 | 0 | 0 | 0 | 0 | 0 | 0 | 0 | 0 | 0 | 0 |
| 0 | NA | NA | NA | NA | NA | Mip | major intrinsic protein of lens fiber | 0 | 0 | 0 | 0 | 0 | 0 | 0 | 0 | 0 | 0 | 0 |
| 0 | NA | NA | NA | NA | NA | Gm23241 | | 0 | 0 | 0 | 0 | 0 | 0 | 0 | 0 | 0 | 0 | 0 |
| 0 | NA | NA | NA | NA | NA | Gm26308 | | 0 | 0 | 0 | 0 | 0 | 0 | 0 | 0 | 0 | 0 | 0 |
| 0 | NA | NA | NA | NA | NA | Mir6914 | | 0 | 0 | 0 | 0 | 0 | 0 | 0 | 0 | 0 | 0 | 0 |
| 0 | NA | NA | NA | NA | NA | Mir6915 | | 0 | 0 | 0 | 0 | 0 | 0 | 0 | 0 | 0 | 0 | 0 |
| 0 | NA | NA | NA | NA | NA | Gm25361 | | 0 | 0 | 0 | 0 | 0 | 0 | 0 | 0 | 0 | 0 | 0 |
| 0 | NA | NA | NA | NA | NA | Mir8105 | | 0 | 0 | 0 | 0 | 0 | 0 | 0 | 0 | 0 | 0 | 0 |
| 0 | NA | NA | NA | NA | NA | Gm47275 | | 0 | 0 | 0 | 0 | 0 | 0 | 0 | 0 | 0 | 0 | 0 |
| 0 | NA | NA | NA | NA | NA | Mir6917 | | 0 | 0 | 0 | 0 | 0 | 0 | 0 | 0 | 0 | 0 | 0 |
| 0 | NA | NA | NA | NA | NA | Gm25494 | | 0 | 0 | 0 | 0 | 0 | 0 | 0 | 0 | 0 | 0 | 0 |
| 0 | NA | NA | NA | NA | NA | 9030616G12Rik | | 0 | 0 | 0 | 0 | 0 | 0 | 0 | 0 | 0 | 0 | 0 |
| 0 | NA | NA | NA | NA | NA | Olfr9 | olfactory receptor 9 | 0 | 0 | 0 | 0 | 0 | 0 | 0 | 0 | 0 | 0 | 0 |
| 0 | NA | NA | NA | NA | NA | Olfr763 | olfactory receptor 763 | 0 | 0 | 0 | 0 | 0 | 0 | 0 | 0 | 0 | 0 | 0 |
| 0 | NA | NA | NA | NA | NA | Olfr764-ps1 | | 0 | 0 | 0 | 0 | 0 | 0 | 0 | 0 | 0 | 0 | 0 |
| 0 | NA | NA | NA | NA | NA | Olfr765 | olfactory receptor 765 | 0 | 0 | 0 | 0 | 0 | 0 | 0 | 0 | 0 | 0 | 0 |
| 0 | NA | NA | NA | NA | NA | Olfr766-ps1 | | 0 | 0 | 0 | 0 | 0 | 0 | 0 | 0 | 0 | 0 | 0 |
| 0 | NA | NA | NA | NA | NA | Olfr767 | olfactory receptor 767 | 0 | 0 | 0 | 0 | 0 | 0 | 0 | 0 | 0 | 0 | 0 |
| 0 | NA | NA | NA | NA | NA | Olfr768 | olfactory receptor 768 | 0 | 0 | 0 | 0 | 0 | 0 | 0 | 0 | 0 | 0 | 0 |
| 0 | NA | NA | NA | NA | NA | Olfr769 | olfactory receptor 769 | 0 | 0 | 0 | 0 | 0 | 0 | 0 | 0 | 0 | 0 | 0 |
| 0 | NA | NA | NA | NA | NA | Olfr770 | olfactory receptor 770 | 0 | 0 | 0 | 0 | 0 | 0 | 0 | 0 | 0 | 0 | 0 |
| 0 | NA | NA | NA | NA | NA | Olfr771 | olfactory receptor 771 | 0 | 0 | 0 | 0 | 0 | 0 | 0 | 0 | 0 | 0 | 0 |
| 0 | NA | NA | NA | NA | NA | Gm25807 | | 0 | 0 | 0 | 0 | 0 | 0 | 0 | 0 | 0 | 0 | 0 |
| 0 | NA | NA | NA | NA | NA | Olfr772 | olfactory receptor 772 | 0 | 0 | 0 | 0 | 0 | 0 | 0 | 0 | 0 | 0 | 0 |

| | | | | | | | | | | | | | | | | | | |
|---|---|---|---|---|---|---|---|---|---|---|---|---|---|---|---|---|---|---|
| 0 | NA | NA | NA | NA | NA | Olfr773 | olfactory receptor 773 | 0 | 0 | 0 | 0 | 0 | 0 | 0 | 0 | 0 | 0 | 0 |
| 0 | NA | NA | NA | NA | NA | Olfr774 | olfactory receptor 774 | 0 | 0 | 0 | 0 | 0 | 0 | 0 | 0 | 0 | 0 | 0 |
| 0 | NA | NA | NA | NA | NA | Olfr775 | olfactory receptor 775 | 0 | 0 | 0 | 0 | 0 | 0 | 0 | 0 | 0 | 0 | 0 |
| 0 | NA | NA | NA | NA | NA | Olfr776 | olfactory receptor 776 | 0 | 0 | 0 | 0 | 0 | 0 | 0 | 0 | 0 | 0 | 0 |
| 0 | NA | NA | NA | NA | NA | Olfr777 | olfactory receptor 777 | 0 | 0 | 0 | 0 | 0 | 0 | 0 | 0 | 0 | 0 | 0 |
| 0 | NA | NA | NA | NA | NA | Olfr778-ps1 | | 0 | 0 | 0 | 0 | 0 | 0 | 0 | 0 | 0 | 0 | 0 |
| 0 | NA | NA | NA | NA | NA | Gm25764 | | 0 | 0 | 0 | 0 | 0 | 0 | 0 | 0 | 0 | 0 | 0 |
| 0 | NA | NA | NA | NA | NA | Olfr779 | | 0 | 0 | 0 | 0 | 0 | 0 | 0 | 0 | 0 | 0 | 0 |
| 0 | NA | NA | NA | NA | NA | Olfr780 | olfactory receptor 780 | 0 | 0 | 0 | 0 | 0 | 0 | 0 | 0 | 0 | 0 | 0 |
| 0 | NA | NA | NA | NA | NA | Olfr781 | olfactory receptor 781 | 0 | 0 | 0 | 0 | 0 | 0 | 0 | 0 | 0 | 0 | 0 |
| 0 | NA | NA | NA | NA | NA | Olfr782 | olfactory receptor 782 | 0 | 0 | 0 | 0 | 0 | 0 | 0 | 0 | 0 | 0 | 0 |
| 0 | NA | NA | NA | NA | NA | Olfr783-ps1 | | 0 | 0 | 0 | 0 | 0 | 0 | 0 | 0 | 0 | 0 | 0 |
| 0 | NA | NA | NA | NA | NA | Olfr784 | olfactory receptor 784 | 0 | 0 | 0 | 0 | 0 | 0 | 0 | 0 | 0 | 0 | 0 |
| 0 | NA | NA | NA | NA | NA | Olfr785 | olfactory receptor 785 | 0 | 0 | 0 | 0 | 0 | 0 | 0 | 0 | 0 | 0 | 0 |
| 0 | NA | NA | NA | NA | NA | Olfr786 | olfactory receptor 786 | 0 | 0 | 0 | 0 | 0 | 0 | 0 | 0 | 0 | 0 | 0 |
| 0 | NA | NA | NA | NA | NA | Olfr787 | olfactory receptor 787 | 0 | 0 | 0 | 0 | 0 | 0 | 0 | 0 | 0 | 0 | 0 |
| 0 | NA | NA | NA | NA | NA | Olfr788 | olfactory receptor 788 | 0 | 0 | 0 | 0 | 0 | 0 | 0 | 0 | 0 | 0 | 0 |
| 0 | NA | NA | NA | NA | NA | Olfr789 | olfactory receptor 789 | 0 | 0 | 0 | 0 | 0 | 0 | 0 | 0 | 0 | 0 | 0 |
| 0 | NA | NA | NA | NA | NA | Olfr790 | olfactory receptor 790 | 0 | 0 | 0 | 0 | 0 | 0 | 0 | 0 | 0 | 0 | 0 |
| 0 | NA | NA | NA | NA | NA | Olfr791 | olfactory receptor 791 | 0 | 0 | 0 | 0 | 0 | 0 | 0 | 0 | 0 | 0 | 0 |
| 0 | NA | NA | NA | NA | NA | Olfr792 | olfactory receptor 792 | 0 | 0 | 0 | 0 | 0 | 0 | 0 | 0 | 0 | 0 | 0 |
| 0 | NA | NA | NA | NA | NA | Olfr793-ps1 | | 0 | 0 | 0 | 0 | 0 | 0 | 0 | 0 | 0 | 0 | 0 |
| 0 | NA | NA | NA | NA | NA | Olfr794 | olfactory receptor 794 | 0 | 0 | 0 | 0 | 0 | 0 | 0 | 0 | 0 | 0 | 0 |
| 0 | NA | NA | NA | NA | NA | Olfr795-ps1 | | 0 | 0 | 0 | 0 | 0 | 0 | 0 | 0 | 0 | 0 | 0 |
| 0 | NA | NA | NA | NA | NA | Gm23252 | | 0 | 0 | 0 | 0 | 0 | 0 | 0 | 0 | 0 | 0 | 0 |
| 0 | NA | NA | NA | NA | NA | Olfr796 | olfactory receptor 796 | 0 | 0 | 0 | 0 | 0 | 0 | 0 | 0 | 0 | 0 | 0 |
| 0 | NA | NA | NA | NA | NA | Olfr797-ps1 | | 0 | 0 | 0 | 0 | 0 | 0 | 0 | 0 | 0 | 0 | 0 |
| 0 | NA | NA | NA | NA | NA | Olfr798 | olfactory receptor 798 | 0 | 0 | 0 | 0 | 0 | 0 | 0 | 0 | 0 | 0 | 0 |
| 0 | NA | NA | NA | NA | NA | Olfr799 | olfactory receptor 799 | 0 | 0 | 0 | 0 | 0 | 0 | 0 | 0 | 0 | 0 | 0 |
| 0 | NA | NA | NA | NA | NA | Olfr800 | olfactory receptor 800 | 0 | 0 | 0 | 0 | 0 | 0 | 0 | 0 | 0 | 0 | 0 |
| 0 | NA | NA | NA | NA | NA | Olfr801 | olfactory receptor 801 | 0 | 0 | 0 | 0 | 0 | 0 | 0 | 0 | 0 | 0 | 0 |
| 0 | NA | NA | NA | NA | NA | Olfr802 | olfactory receptor 802 | 0 | 0 | 0 | 0 | 0 | 0 | 0 | 0 | 0 | 0 | 0 |
| 0 | NA | NA | NA | NA | NA | Olfr803 | olfactory receptor 803 | 0 | 0 | 0 | 0 | 0 | 0 | 0 | 0 | 0 | 0 | 0 |
| 0 | NA | NA | NA | NA | NA | Gm24717 | | 0 | 0 | 0 | 0 | 0 | 0 | 0 | 0 | 0 | 0 | 0 |
| 0 | NA | NA | NA | NA | NA | Olfr804 | olfactory receptor 804 | 0 | 0 | 0 | 0 | 0 | 0 | 0 | 0 | 0 | 0 | 0 |
| 0 | NA | NA | NA | NA | NA | Olfr805 | olfactory receptor 805 | 0 | 0 | 0 | 0 | 0 | 0 | 0 | 0 | 0 | 0 | 0 |
| 0 | NA | NA | NA | NA | NA | Gm24485 | | 0 | 0 | 0 | 0 | 0 | 0 | 0 | 0 | 0 | 0 | 0 |
| 0 | NA | NA | NA | NA | NA | Olfr806 | olfactory receptor 806 | 0 | 0 | 0 | 0 | 0 | 0 | 0 | 0 | 0 | 0 | 0 |
| 0 | NA | NA | NA | NA | NA | Olfr807 | olfactory receptor 807 | 0 | 0 | 0 | 0 | 0 | 0 | 0 | 0 | 0 | 0 | 0 |
| 0 | NA | NA | NA | NA | NA | Olfr808 | olfactory receptor 808 | 0 | 0 | 0 | 0 | 0 | 0 | 0 | 0 | 0 | 0 | 0 |
| 0 | NA | NA | NA | NA | NA | Olfr809 | olfactory receptor 809 | 0 | 0 | 0 | 0 | 0 | 0 | 0 | 0 | 0 | 0 | 0 |
| 0 | NA | NA | NA | NA | NA | Olfr810 | olfactory receptor 810 | 0 | 0 | 0 | 0 | 0 | 0 | 0 | 0 | 0 | 0 | 0 |
| 0 | NA | NA | NA | NA | NA | Olfr811 | olfactory receptor 811 | 0 | 0 | 0 | 0 | 0 | 0 | 0 | 0 | 0 | 0 | 0 |
| 0 | NA | NA | NA | NA | NA | Olfr812 | olfactory receptor 812 | 0 | 0 | 0 | 0 | 0 | 0 | 0 | 0 | 0 | 0 | 0 |
| 0 | NA | NA | NA | NA | NA | Olfr813 | olfactory receptor 813 | 0 | 0 | 0 | 0 | 0 | 0 | 0 | 0 | 0 | 0 | 0 |
| 0 | NA | NA | NA | NA | NA | Olfr815 | olfactory receptor 815 | 0 | 0 | 0 | 0 | 0 | 0 | 0 | 0 | 0 | 0 | 0 |
| 0 | NA | NA | NA | NA | NA | Olfr816 | olfactory receptor 816 | 0 | 0 | 0 | 0 | 0 | 0 | 0 | 0 | 0 | 0 | 0 |
| 0 | NA | NA | NA | NA | NA | | | 0 | 0 | 0 | 0 | 0 | 0 | 0 | 0 | 0 | 0 | 0 |
| 0 | NA | NA | NA | NA | NA | Olfr818 | olfactory receptor 818 | 0 | 0 | 0 | 0 | 0 | 0 | 0 | 0 | 0 | 0 | 0 |
| 0 | NA | NA | NA | NA | NA | Olfr819 | olfactory receptor 819 | 0 | 0 | 0 | 0 | 0 | 0 | 0 | 0 | 0 | 0 | 0 |
| 0 | NA | NA | NA | NA | NA | Olfr247 | olfactory receptor 247 | 0 | 0 | 0 | 0 | 0 | 0 | 0 | 0 | 0 | 0 | 0 |
| 0 | NA | NA | NA | NA | NA | Gm10310 | predicted pseudogene 10310 | 0 | 0 | 0 | 0 | 0 | 0 | 0 | 0 | 0 | 0 | 0 |
| 0 | NA | NA | NA | NA | NA | Gm43977 | | 0 | 0 | 0 | 0 | 0 | 0 | 0 | 0 | 0 | 0 | 0 |
| 0 | NA | NA | NA | NA | NA | Olfr820 | olfactory receptor 820 | 0 | 0 | 0 | 0 | 0 | 0 | 0 | 0 | 0 | 0 | 0 |
| 0 | NA | NA | NA | NA | NA | Olfr821 | olfactory receptor 821 | 0 | 0 | 0 | 0 | 0 | 0 | 0 | 0 | 0 | 0 | 0 |
| 0 | NA | NA | NA | NA | NA | Olfr822 | olfactory receptor 822 | 0 | 0 | 0 | 0 | 0 | 0 | 0 | 0 | 0 | 0 | 0 |
| 0 | NA | NA | NA | NA | NA | Olfr1542-ps1 | | 0 | 0 | 0 | 0 | 0 | 0 | 0 | 0 | 0 | 0 | 0 |
| 0 | NA | NA | NA | NA | NA | Olfr823 | olfactory receptor 823 | 0 | 0 | 0 | 0 | 0 | 0 | 0 | 0 | 0 | 0 | 0 |
| 0 | NA | NA | NA | NA | NA | Gm5181 | | 0 | 0 | 0 | 0 | 0 | 0 | 0 | 0 | 0 | 0 | 0 |
| 0 | NA | NA | NA | NA | NA | Olfr824 | olfactory receptor 824 | 0 | 0 | 0 | 0 | 0 | 0 | 0 | 0 | 0 | 0 | 0 |

| | | | | | | | | | | | | | | | | | |
|---|---|---|---|---|---|---|---|---|---|---|---|---|---|---|---|---|---|
| 0 | NA | NA | NA | NA | NA | Gm9560 | | 0 | 0 | 0 | 0 | 0 | 0 | 0 | 0 | 0 | 0 | 0 |
| 0 | NA | NA | NA | NA | NA | Olfr825 | olfactory receptor 825 | 0 | 0 | 0 | 0 | 0 | 0 | 0 | 0 | 0 | 0 | 0 |
| 0 | NA | NA | NA | NA | NA | Gm48514 | | 0 | 0 | 0 | 0 | 0 | 0 | 0 | 0 | 0 | 0 | 0 |
| 0 | NA | NA | NA | NA | NA | Olfr826 | olfactory receptor 826 | 0 | 0 | 0 | 0 | 0 | 0 | 0 | 0 | 0 | 0 | 0 |
| 0 | NA | NA | NA | NA | NA | Olfr827 | olfactory receptor 827 | 0 | 0 | 0 | 0 | 0 | 0 | 0 | 0 | 0 | 0 | 0 |
| 0 | NA | NA | NA | NA | NA | Vmn2r84 | vomeronasal 2, receptor 84 | 0 | 0 | 0 | 0 | 0 | 0 | 0 | 0 | 0 | 0 | 0 |
| 0 | NA | NA | NA | NA | NA | Vmn2r85 | vomeronasal 2, receptor 85 | 0 | 0 | 0 | 0 | 0 | 0 | 0 | 0 | 0 | 0 | 0 |
| 0 | NA | NA | NA | NA | NA | Vmn2r86 | vomeronasal 2, receptor 86 | 0 | 0 | 0 | 0 | 0 | 0 | 0 | 0 | 0 | 0 | 0 |
| 0 | NA | NA | NA | NA | NA | Vmn2r87 | vomeronasal 2, receptor 87 | 0 | 0 | 0 | 0 | 0 | 0 | 0 | 0 | 0 | 0 | 0 |
| 0 | NA | NA | NA | NA | NA | Gm23793 | | 0 | 0 | 0 | 0 | 0 | 0 | 0 | 0 | 0 | 0 | 0 |
| 0 | NA | NA | NA | NA | NA | Gm11399 | | 0 | 0 | 0 | 0 | 0 | 0 | 0 | 0 | 0 | 0 | 0 |
| 0 | NA | NA | NA | NA | NA | Fau-ps2 | | 0 | 0 | 0 | 0 | 0 | 0 | 0 | 0 | 0 | 0 | 0 |
| 0 | NA | NA | NA | NA | NA | Gm24439 | | 0 | 0 | 0 | 0 | 0 | 0 | 0 | 0 | 0 | 0 | 0 |
| 0 | NA | NA | NA | NA | NA | Gm11946 | | 0 | 0 | 0 | 0 | 0 | 0 | 0 | 0 | 0 | 0 | 0 |
| 0 | NA | NA | NA | NA | NA | Gm11947 | | 0 | 0 | 0 | 0 | 0 | 0 | 0 | 0 | 0 | 0 | 0 |
| 0 | NA | NA | NA | NA | NA | Gm27624 | | 0 | 0 | 0 | 0 | 0 | 0 | 0 | 0 | 0 | 0 | 0 |
| 0 | NA | NA | NA | NA | NA | Gm27517 | | 0 | 0 | 0 | 0 | 0 | 0 | 0 | 0 | 0 | 0 | 0 |
| 0 | NA | NA | NA | NA | NA | Gm27937 | | 0 | 0 | 0 | 0 | 0 | 0 | 0 | 0 | 0 | 0 | 0 |
| 0 | NA | NA | NA | NA | NA | Gm27640 | | 0 | 0 | 0 | 0 | 0 | 0 | 0 | 0 | 0 | 0 | 0 |
| 0 | NA | NA | NA | NA | NA | Gm26393 | | 0 | 0 | 0 | 0 | 0 | 0 | 0 | 0 | 0 | 0 | 0 |
| 0 | NA | NA | NA | NA | NA | Gm24013 | | 0 | 0 | 0 | 0 | 0 | 0 | 0 | 0 | 0 | 0 | 0 |
| 0 | NA | NA | NA | NA | NA | Gm11952 | | 0 | 0 | 0 | 0 | 0 | 0 | 0 | 0 | 0 | 0 | 0 |
| 0 | NA | NA | NA | NA | NA | Gm11954 | | 0 | 0 | 0 | 0 | 0 | 0 | 0 | 0 | 0 | 0 | 0 |
| 0 | NA | NA | NA | NA | NA | Gm11955 | | 0 | 0 | 0 | 0 | 0 | 0 | 0 | 0 | 0 | 0 | 0 |
| 0 | NA | NA | NA | NA | NA | Gm11957 | | 0 | 0 | 0 | 0 | 0 | 0 | 0 | 0 | 0 | 0 | 0 |
| 0 | NA | NA | NA | NA | NA | Gm11956 | | 0 | 0 | 0 | 0 | 0 | 0 | 0 | 0 | 0 | 0 | 0 |
| 0 | NA | NA | NA | NA | NA | Gm11959 | | 0 | 0 | 0 | 0 | 0 | 0 | 0 | 0 | 0 | 0 | 0 |
| 0 | NA | NA | NA | NA | NA | Gm11958 | | 0 | 0 | 0 | 0 | 0 | 0 | 0 | 0 | 0 | 0 | 0 |
| 0 | NA | NA | NA | NA | NA | Gm24803 | | 0 | 0 | 0 | 0 | 0 | 0 | 0 | 0 | 0 | 0 | 0 |
| 0 | NA | NA | NA | NA | NA | Gm11032 | | 0 | 0 | 0 | 0 | 0 | 0 | 0 | 0 | 0 | 0 | 0 |
| 0 | NA | NA | NA | NA | NA | Gm24958 | | 0 | 0 | 0 | 0 | 0 | 0 | 0 | 0 | 0 | 0 | 0 |
| 0 | NA | NA | NA | NA | NA | Gm25142 | | 0 | 0 | 0 | 0 | 0 | 0 | 0 | 0 | 0 | 0 | 0 |
| 0 | NA | NA | NA | NA | NA | Gm30172 | | 0 | 0 | 0 | 0 | 0 | 0 | 0 | 0 | 0 | 0 | 0 |
| 0 | NA | NA | NA | NA | NA | Gm11962 | | 0 | 0 | 0 | 0 | 0 | 0 | 0 | 0 | 0 | 0 | 0 |
| 0 | NA | NA | NA | NA | NA | Gm22471 | | 0 | 0 | 0 | 0 | 0 | 0 | 0 | 0 | 0 | 0 | 0 |
| 0 | NA | NA | NA | NA | NA | Ankrd36 | ankyrin repeat domain 36 | 0 | 0 | 0 | 0 | 0 | 0 | 0 | 0 | 0 | 0 | 0 |
| 0 | NA | NA | NA | NA | NA | n-R5s67 | | 0 | 0 | 0 | 0 | 0 | 0 | 0 | 0 | 0 | 0 | 0 |
| 0 | NA | NA | NA | NA | NA | Gm22123 | | 0 | 0 | 0 | 0 | 0 | 0 | 0 | 0 | 0 | 0 | 0 |
| 0 | NA | NA | NA | NA | NA | Snora9 | | 0 | 0 | 0 | 0 | 0 | 0 | 0 | 0 | 0 | 0 | 0 |
| 0 | NA | NA | NA | NA | NA | Gm24000 | | 0 | 0 | 0 | 0 | 0 | 0 | 0 | 0 | 0 | 0 | 0 |
| 0 | NA | NA | NA | NA | NA | Gm24313 | | 0 | 0 | 0 | 0 | 0 | 0 | 0 | 0 | 0 | 0 | 0 |
| 0 | NA | NA | NA | NA | NA | Wap | whey acidic protein | 0 | 0 | 0 | 0 | 0 | 0 | 0 | 0 | 0 | 0 | 0 |
| 0 | NA | NA | NA | NA | NA | Gm11978 | | 0 | 0 | 0 | 0 | 0 | 0 | 0 | 0 | 0 | 0 | 0 |
| 0 | NA | NA | NA | NA | NA | Gm11980 | | 0 | 0 | 0 | 0 | 0 | 0 | 0 | 0 | 0 | 0 | 0 |
| 0 | NA | NA | NA | NA | NA | Gm11981 | | 0 | 0 | 0 | 0 | 0 | 0 | 0 | 0 | 0 | 0 | 0 |
| 0 | NA | NA | NA | NA | NA | Gm11976 | | 0 | 0 | 0 | 0 | 0 | 0 | 0 | 0 | 0 | 0 | 0 |
| 0 | NA | NA | NA | NA | NA | Gm11982 | | 0 | 0 | 0 | 0 | 0 | 0 | 0 | 0 | 0 | 0 | 0 |
| 0 | NA | NA | NA | NA | NA | Gm11983 | | 0 | 0 | 0 | 0 | 0 | 0 | 0 | 0 | 0 | 0 | 0 |
| 0 | NA | NA | NA | NA | NA | Ccdc201 | | 0 | 0 | 0 | 0 | 0 | 0 | 0 | 0 | 0 | 0 | 0 |
| 0 | NA | NA | NA | NA | NA | Gm11986 | | 0 | 0 | 0 | 0 | 0 | 0 | 0 | 0 | 0 | 0 | 0 |
| 0 | NA | NA | NA | NA | NA | Gm27393 | | 0 | 0 | 0 | 0 | 0 | 0 | 0 | 0 | 0 | 0 | 0 |
| 0 | NA | NA | NA | NA | NA | Gm11987 | | 0 | 0 | 0 | 0 | 0 | 0 | 0 | 0 | 0 | 0 | 0 |
| 0 | NA | NA | NA | NA | NA | Gm24051 | | 0 | 0 | 0 | 0 | 0 | 0 | 0 | 0 | 0 | 0 | 0 |
| 0 | NA | NA | NA | NA | NA | Gm11988 | | 0 | 0 | 0 | 0 | 0 | 0 | 0 | 0 | 0 | 0 | 0 |
| 0 | NA | NA | NA | NA | NA | Gm11990 | | 0 | 0 | 0 | 0 | 0 | 0 | 0 | 0 | 0 | 0 | 0 |
| 0 | NA | NA | NA | NA | NA | Gm11991 | | 0 | 0 | 0 | 0 | 0 | 0 | 0 | 0 | 0 | 0 | 0 |
| 0 | NA | NA | NA | NA | NA | Gm36954 | | 0 | 0 | 0 | 0 | 0 | 0 | 0 | 0 | 0 | 0 | 0 |
| 0 | NA | NA | NA | NA | NA | Gm11996 | | 0 | 0 | 0 | 0 | 0 | 0 | 0 | 0 | 0 | 0 | 0 |
| 0 | NA | NA | NA | NA | NA | Gm11997 | | 0 | 0 | 0 | 0 | 0 | 0 | 0 | 0 | 0 | 0 | 0 |
| 0 | NA | NA | NA | NA | NA | Gm23065 | | 0 | 0 | 0 | 0 | 0 | 0 | 0 | 0 | 0 | 0 | 0 |

| | | | | | | | | | | | | | | | | | | |
|---|---|---|---|---|---|---|---|---|---|---|---|---|---|---|---|---|---|---|
| 0 | NA | NA | NA | NA | NA | Gm12598 | | 0 | 0 | 0 | 0 | 0 | 0 | 0 | 0 | 0 | 0 | 0 |
| 0 | NA | NA | NA | NA | NA | Gm26458 | | 0 | 0 | 0 | 0 | 0 | 0 | 0 | 0 | 0 | 0 | 0 |
| 0 | NA | NA | NA | NA | NA | 4930512M02Ri | RIKEN cDNA 4930512M02 gene | 0 | 0 | 0 | 0 | 0 | 0 | 0 | 0 | 0 | 0 | 0 |
| 0 | NA | NA | NA | NA | NA | Gm11999 | | 0 | 0 | 0 | 0 | 0 | 0 | 0 | 0 | 0 | 0 | 0 |
| 0 | NA | NA | NA | NA | NA | Gm12000 | | 0 | 0 | 0 | 0 | 0 | 0 | 0 | 0 | 0 | 0 | 0 |
| 0 | NA | NA | NA | NA | NA | Gm25058 | | 0 | 0 | 0 | 0 | 0 | 0 | 0 | 0 | 0 | 0 | 0 |
| 0 | NA | NA | NA | NA | NA | Gm24036 | | 0 | 0 | 0 | 0 | 0 | 0 | 0 | 0 | 0 | 0 | 0 |
| 0 | NA | NA | NA | NA | NA | Gm12001 | | 0 | 0 | 0 | 0 | 0 | 0 | 0 | 0 | 0 | 0 | 0 |
| 0 | NA | NA | NA | NA | NA | Gm22156 | | 0 | 0 | 0 | 0 | 0 | 0 | 0 | 0 | 0 | 0 | 0 |
| 0 | NA | NA | NA | NA | NA | Gm22789 | | 0 | 0 | 0 | 0 | 0 | 0 | 0 | 0 | 0 | 0 | 0 |
| 0 | NA | NA | NA | NA | NA | Gm12004 | | 0 | 0 | 0 | 0 | 0 | 0 | 0 | 0 | 0 | 0 | 0 |
| 0 | NA | NA | NA | NA | NA | 4930554G24Rik | | 0 | 0 | 0 | 0 | 0 | 0 | 0 | 0 | 0 | 0 | 0 |
| 0 | NA | NA | NA | NA | NA | Gm12006 | | 0 | 0 | 0 | 0 | 0 | 0 | 0 | 0 | 0 | 0 | 0 |
| 0 | NA | NA | NA | NA | NA | 1700046C09Rik | | 0 | 0 | 0 | 0 | 0 | 0 | 0 | 0 | 0 | 0 | 0 |
| 0 | NA | NA | NA | NA | NA | Pom121l12 | POM121 membrane glycoprotein-like 12 | 0 | 0 | 0 | 0 | 0 | 0 | 0 | 0 | 0 | 0 | 0 |
| 0 | NA | NA | NA | NA | NA | Gm12007 | | 0 | 0 | 0 | 0 | 0 | 0 | 0 | 0 | 0 | 0 | 0 |
| 0 | NA | NA | NA | NA | NA | Gm12008 | | 0 | 0 | 0 | 0 | 0 | 0 | 0 | 0 | 0 | 0 | 0 |
| 0 | NA | NA | NA | NA | NA | Gm24707 | | 0 | 0 | 0 | 0 | 0 | 0 | 0 | 0 | 0 | 0 | 0 |
| 0 | NA | NA | NA | NA | NA | Gm25490 | | 0 | 0 | 0 | 0 | 0 | 0 | 0 | 0 | 0 | 0 | 0 |
| 0 | NA | NA | NA | NA | NA | Gm12010 | | 0 | 0 | 0 | 0 | 0 | 0 | 0 | 0 | 0 | 0 | 0 |
| 0 | NA | NA | NA | NA | NA | Gm37319 | | 0 | 0 | 0 | 0 | 0 | 0 | 0 | 0 | 0 | 0 | 0 |
| 0 | NA | NA | NA | NA | NA | Gm12011 | | 0 | 0 | 0 | 0 | 0 | 0 | 0 | 0 | 0 | 0 | 0 |
| 0 | NA | NA | NA | NA | NA | Gm12012 | | 0 | 0 | 0 | 0 | 0 | 0 | 0 | 0 | 0 | 0 | 0 |
| 0 | NA | NA | NA | NA | NA | Gm24714 | | 0 | 0 | 0 | 0 | 0 | 0 | 0 | 0 | 0 | 0 | 0 |
| 0 | NA | NA | NA | NA | NA | Gm25698 | | 0 | 0 | 0 | 0 | 0 | 0 | 0 | 0 | 0 | 0 | 0 |
| 0 | NA | NA | NA | NA | NA | Egfros | | 0 | 0 | 0 | 0 | 0 | 0 | 0 | 0 | 0 | 0 | 0 |
| 0 | NA | NA | NA | NA | NA | Gm12014 | | 0 | 0 | 0 | 0 | 0 | 0 | 0 | 0 | 0 | 0 | 0 |
| 0 | NA | NA | NA | NA | NA | Gm12015 | | 0 | 0 | 0 | 0 | 0 | 0 | 0 | 0 | 0 | 0 | 0 |
| 0 | NA | NA | NA | NA | NA | Gm23959 | | 0 | 0 | 0 | 0 | 0 | 0 | 0 | 0 | 0 | 0 | 0 |
| 0 | NA | NA | NA | NA | NA | Gm12016 | | 0 | 0 | 0 | 0 | 0 | 0 | 0 | 0 | 0 | 0 | 0 |
| 0 | NA | NA | NA | NA | NA | Gm12018 | | 0 | 0 | 0 | 0 | 0 | 0 | 0 | 0 | 0 | 0 | 0 |
| 0 | NA | NA | NA | NA | NA | Gm12019 | | 0 | 0 | 0 | 0 | 0 | 0 | 0 | 0 | 0 | 0 | 0 |
| 0 | NA | NA | NA | NA | NA | Gm37869 | | 0 | 0 | 0 | 0 | 0 | 0 | 0 | 0 | 0 | 0 | 0 |
| 0 | NA | NA | NA | NA | NA | Gm12021 | | 0 | 0 | 0 | 0 | 0 | 0 | 0 | 0 | 0 | 0 | 0 |
| 0 | NA | NA | NA | NA | NA | Gm12024 | | 0 | 0 | 0 | 0 | 0 | 0 | 0 | 0 | 0 | 0 | 0 |
| 0 | NA | NA | NA | NA | NA | Gm12023 | | 0 | 0 | 0 | 0 | 0 | 0 | 0 | 0 | 0 | 0 | 0 |
| 0 | NA | NA | NA | NA | NA | Gm23093 | | 0 | 0 | 0 | 0 | 0 | 0 | 0 | 0 | 0 | 0 | 0 |
| 0 | NA | NA | NA | NA | NA | Gm23021 | | 0 | 0 | 0 | 0 | 0 | 0 | 0 | 0 | 0 | 0 | 0 |
| 0 | NA | NA | NA | NA | NA | 4933406G16Rik | | 0 | 0 | 0 | 0 | 0 | 0 | 0 | 0 | 0 | 0 | 0 |
| 0 | NA | NA | NA | NA | NA | Gm12025 | | 0 | 0 | 0 | 0 | 0 | 0 | 0 | 0 | 0 | 0 | 0 |
| 0 | NA | NA | NA | NA | NA | Gm12027 | | 0 | 0 | 0 | 0 | 0 | 0 | 0 | 0 | 0 | 0 | 0 |
| 0 | NA | NA | NA | NA | NA | Gm23386 | | 0 | 0 | 0 | 0 | 0 | 0 | 0 | 0 | 0 | 0 | 0 |
| 0 | NA | NA | NA | NA | NA | Gm12028 | | 0 | 0 | 0 | 0 | 0 | 0 | 0 | 0 | 0 | 0 | 0 |
| 0 | NA | NA | NA | NA | NA | Gm12029 | | 0 | 0 | 0 | 0 | 0 | 0 | 0 | 0 | 0 | 0 | 0 |
| 0 | NA | NA | NA | NA | NA | Gm12030 | | 0 | 0 | 0 | 0 | 0 | 0 | 0 | 0 | 0 | 0 | 0 |
| 0 | NA | NA | NA | NA | NA | Gm26253 | | 0 | 0 | 0 | 0 | 0 | 0 | 0 | 0 | 0 | 0 | 0 |
| 0 | NA | NA | NA | NA | NA | Rpl12-ps2 | | 0 | 0 | 0 | 0 | 0 | 0 | 0 | 0 | 0 | 0 | 0 |
| 0 | NA | NA | NA | NA | NA | Gm12036 | | 0 | 0 | 0 | 0 | 0 | 0 | 0 | 0 | 0 | 0 | 0 |
| 0 | NA | NA | NA | NA | NA | Gm12037 | | 0 | 0 | 0 | 0 | 0 | 0 | 0 | 0 | 0 | 0 | 0 |
| 0 | NA | NA | NA | NA | NA | Gm12038 | | 0 | 0 | 0 | 0 | 0 | 0 | 0 | 0 | 0 | 0 | 0 |
| 0 | NA | NA | NA | NA | NA | Gm22807 | | 0 | 0 | 0 | 0 | 0 | 0 | 0 | 0 | 0 | 0 | 0 |
| 0 | NA | NA | NA | NA | NA | Gm23681 | | 0 | 0 | 0 | 0 | 0 | 0 | 0 | 0 | 0 | 0 | 0 |
| 0 | NA | NA | NA | NA | NA | 2610024D14Rik | | 0 | 0 | 0 | 0 | 0 | 0 | 0 | 0 | 0 | 0 | 0 |
| 0 | NA | NA | NA | NA | NA | Gm22600 | | 0 | 0 | 0 | 0 | 0 | 0 | 0 | 0 | 0 | 0 | 0 |
| 0 | NA | NA | NA | NA | NA | Gm12041 | | 0 | 0 | 0 | 0 | 0 | 0 | 0 | 0 | 0 | 0 | 0 |
| 0 | NA | NA | NA | NA | NA | Gm23582 | | 0 | 0 | 0 | 0 | 0 | 0 | 0 | 0 | 0 | 0 | 0 |
| 0 | NA | NA | NA | NA | NA | Gm12039 | | 0 | 0 | 0 | 0 | 0 | 0 | 0 | 0 | 0 | 0 | 0 |
| 0 | NA | NA | NA | NA | NA | Mir1933 | | 0 | 0 | 0 | 0 | 0 | 0 | 0 | 0 | 0 | 0 | 0 |
| 0 | NA | NA | NA | NA | NA | Gm12044 | | 0 | 0 | 0 | 0 | 0 | 0 | 0 | 0 | 0 | 0 | 0 |
| 0 | NA | NA | NA | NA | NA | Gm12046 | | 0 | 0 | 0 | 0 | 0 | 0 | 0 | 0 | 0 | 0 | 0 |

| | | | | | | | | | | | | | | | | |
|---|---|---|---|---|---|---|---|---|---|---|---|---|---|---|---|---|
| 0 | NA | NA | NA | NA | NA | 4932414J04Rik | 0 | 0 | 0 | 0 | 0 | 0 | 0 | 0 | 0 | 0 | 0 |
| 0 | NA | NA | NA | NA | NA | Gm12047 | 0 | 0 | 0 | 0 | 0 | 0 | 0 | 0 | 0 | 0 | 0 |
| 0 | NA | NA | NA | NA | NA | Gm12050 | 0 | 0 | 0 | 0 | 0 | 0 | 0 | 0 | 0 | 0 | 0 |
| 0 | NA | NA | NA | NA | NA | Gm12051 | 0 | 0 | 0 | 0 | 0 | 0 | 0 | 0 | 0 | 0 | 0 |
| 0 | NA | NA | NA | NA | NA | Gm12053 | 0 | 0 | 0 | 0 | 0 | 0 | 0 | 0 | 0 | 0 | 0 |
| 0 | NA | NA | NA | NA | NA | Gm12055 | 0 | 0 | 0 | 0 | 0 | 0 | 0 | 0 | 0 | 0 | 0 |
| 0 | NA | NA | NA | NA | NA | Gm12058 | 0 | 0 | 0 | 0 | 0 | 0 | 0 | 0 | 0 | 0 | 0 |
| 0 | NA | NA | NA | NA | NA | Commd1b | 0 | 0 | 0 | 0 | 0 | 0 | 0 | 0 | 0 | 0 | 0 |
| 0 | NA | NA | NA | NA | NA | Gm20456 | 0 | 0 | 0 | 0 | 0 | 0 | 0 | 0 | 0 | 0 | 0 |
| 0 | NA | NA | NA | NA | NA | Gm23772 | 0 | 0 | 0 | 0 | 0 | 0 | 0 | 0 | 0 | 0 | 0 |
| 0 | NA | NA | NA | NA | NA | Gm24917 | 0 | 0 | 0 | 0 | 0 | 0 | 0 | 0 | 0 | 0 | 0 |
| 0 | NA | NA | NA | NA | NA | Gm23740 | 0 | 0 | 0 | 0 | 0 | 0 | 0 | 0 | 0 | 0 | 0 |
| 0 | NA | NA | NA | NA | NA | 1700061J23Rik | 0 | 0 | 0 | 0 | 0 | 0 | 0 | 0 | 0 | 0 | 0 |
| 0 | NA | NA | NA | NA | NA | Gm24398 | 0 | 0 | 0 | 0 | 0 | 0 | 0 | 0 | 0 | 0 | 0 |
| 0 | NA | NA | NA | NA | NA | Gm23114 | 0 | 0 | 0 | 0 | 0 | 0 | 0 | 0 | 0 | 0 | 0 |
| 0 | NA | NA | NA | NA | NA | Gm23827 | 0 | 0 | 0 | 0 | 0 | 0 | 0 | 0 | 0 | 0 | 0 |
| 0 | NA | NA | NA | NA | NA | Gm12061 | 0 | 0 | 0 | 0 | 0 | 0 | 0 | 0 | 0 | 0 | 0 |
| 0 | NA | NA | NA | NA | NA | Gm12062 | 0 | 0 | 0 | 0 | 0 | 0 | 0 | 0 | 0 | 0 | 0 |
| 0 | NA | NA | NA | NA | NA | Gm44381 | 0 | 0 | 0 | 0 | 0 | 0 | 0 | 0 | 0 | 0 | 0 |
| 0 | NA | NA | NA | NA | NA | A830031A19Rik | 0 | 0 | 0 | 0 | 0 | 0 | 0 | 0 | 0 | 0 | 0 |
| 0 | NA | NA | NA | NA | NA | Gm12063 | 0 | 0 | 0 | 0 | 0 | 0 | 0 | 0 | 0 | 0 | 0 |
| 0 | NA | NA | NA | NA | NA | Gm12064 | 0 | 0 | 0 | 0 | 0 | 0 | 0 | 0 | 0 | 0 | 0 |
| 0 | NA | NA | NA | NA | NA | 4930538E20Rik | 0 | 0 | 0 | 0 | 0 | 0 | 0 | 0 | 0 | 0 | 0 |
| 0 | NA | NA | NA | NA | NA | Gm12068 | 0 | 0 | 0 | 0 | 0 | 0 | 0 | 0 | 0 | 0 | 0 |
| 0 | NA | NA | NA | NA | NA | 4933430M04Rik | 0 | 0 | 0 | 0 | 0 | 0 | 0 | 0 | 0 | 0 | 0 |
| 0 | NA | NA | NA | NA | NA | Gm12067 | 0 | 0 | 0 | 0 | 0 | 0 | 0 | 0 | 0 | 0 | 0 |
| 0 | NA | NA | NA | NA | NA | Gm22787 | 0 | 0 | 0 | 0 | 0 | 0 | 0 | 0 | 0 | 0 | 0 |
| 0 | NA | NA | NA | NA | NA | Gm25507 | 0 | 0 | 0 | 0 | 0 | 0 | 0 | 0 | 0 | 0 | 0 |
| 0 | NA | NA | NA | NA | NA | Gm10466 | 0 | 0 | 0 | 0 | 0 | 0 | 0 | 0 | 0 | 0 | 0 |
| 0 | NA | NA | NA | NA | NA | Gm12665 | 0 | 0 | 0 | 0 | 0 | 0 | 0 | 0 | 0 | 0 | 0 |
| 0 | NA | NA | NA | NA | NA | 4933427E13Rik | 0 | 0 | 0 | 0 | 0 | 0 | 0 | 0 | 0 | 0 | 0 |
| 0 | NA | NA | NA | NA | NA | Gm24255 | 0 | 0 | 0 | 0 | 0 | 0 | 0 | 0 | 0 | 0 | 0 |
| 0 | NA | NA | NA | NA | NA | Gm12069 | 0 | 0 | 0 | 0 | 0 | 0 | 0 | 0 | 0 | 0 | 0 |
| 0 | NA | NA | NA | NA | NA | Gm23514 | 0 | 0 | 0 | 0 | 0 | 0 | 0 | 0 | 0 | 0 | 0 |
| 0 | NA | NA | NA | NA | NA | Mir6372 | 0 | 0 | 0 | 0 | 0 | 0 | 0 | 0 | 0 | 0 | 0 |
| 0 | NA | NA | NA | NA | NA | Gm22868 | 0 | 0 | 0 | 0 | 0 | 0 | 0 | 0 | 0 | 0 | 0 |
| 0 | NA | NA | NA | NA | NA | Gm12713 | 0 | 0 | 0 | 0 | 0 | 0 | 0 | 0 | 0 | 0 | 0 |
| 0 | NA | NA | NA | NA | NA | Gm12071 | 0 | 0 | 0 | 0 | 0 | 0 | 0 | 0 | 0 | 0 | 0 |
| 0 | NA | NA | NA | NA | NA | Gm12072 | 0 | 0 | 0 | 0 | 0 | 0 | 0 | 0 | 0 | 0 | 0 |
| 0 | NA | NA | NA | NA | NA | Gm12073 | 0 | 0 | 0 | 0 | 0 | 0 | 0 | 0 | 0 | 0 | 0 |
| 0 | NA | NA | NA | NA | NA | Gm12074 | 0 | 0 | 0 | 0 | 0 | 0 | 0 | 0 | 0 | 0 | 0 |
| 0 | NA | NA | NA | NA | NA | n-R5s68 | 0 | 0 | 0 | 0 | 0 | 0 | 0 | 0 | 0 | 0 | 0 |
| 0 | NA | NA | NA | NA | NA | Gm24850 | 0 | 0 | 0 | 0 | 0 | 0 | 0 | 0 | 0 | 0 | 0 |
| 0 | NA | NA | NA | NA | NA | Gm25387 | 0 | 0 | 0 | 0 | 0 | 0 | 0 | 0 | 0 | 0 | 0 |
| 0 | NA | NA | NA | NA | NA | Gm12075 | 0 | 0 | 0 | 0 | 0 | 0 | 0 | 0 | 0 | 0 | 0 |
| 0 | NA | NA | NA | NA | NA | Gm12077 | 0 | 0 | 0 | 0 | 0 | 0 | 0 | 0 | 0 | 0 | 0 |
| 0 | NA | NA | NA | NA | NA | Gm3810 | 0 | 0 | 0 | 0 | 0 | 0 | 0 | 0 | 0 | 0 | 0 |
| 0 | NA | NA | NA | NA | NA | Gm12080 | 0 | 0 | 0 | 0 | 0 | 0 | 0 | 0 | 0 | 0 | 0 |
| 0 | NA | NA | NA | NA | NA | Gm12081 | 0 | 0 | 0 | 0 | 0 | 0 | 0 | 0 | 0 | 0 | 0 |
| 0 | NA | NA | NA | NA | NA | 2810471M01Rik | 0 | 0 | 0 | 0 | 0 | 0 | 0 | 0 | 0 | 0 | 0 |
| 0 | NA | NA | NA | NA | NA | Mir216b | 0 | 0 | 0 | 0 | 0 | 0 | 0 | 0 | 0 | 0 | 0 |
| 0 | NA | NA | NA | NA | NA | Mir216c | 0 | 0 | 0 | 0 | 0 | 0 | 0 | 0 | 0 | 0 | 0 |
| 0 | NA | NA | NA | NA | NA | Mir217hg | 0 | 0 | 0 | 0 | 0 | 0 | 0 | 0 | 0 | 0 | 0 |
| 0 | NA | NA | NA | NA | NA | Mir216a | 0 | 0 | 0 | 0 | 0 | 0 | 0 | 0 | 0 | 0 | 0 |
| 0 | NA | NA | NA | NA | NA | Mir217 | 0 | 0 | 0 | 0 | 0 | 0 | 0 | 0 | 0 | 0 | 0 |
| 0 | NA | NA | NA | NA | NA | Gm12085 | 0 | 0 | 0 | 0 | 0 | 0 | 0 | 0 | 0 | 0 | 0 |
| 0 | NA | NA | NA | NA | NA | Gm23109 | 0 | 0 | 0 | 0 | 0 | 0 | 0 | 0 | 0 | 0 | 0 |
| 0 | NA | NA | NA | NA | NA | Gm12086 | 0 | 0 | 0 | 0 | 0 | 0 | 0 | 0 | 0 | 0 | 0 |
| 0 | NA | NA | NA | NA | NA | Gm12087 | 0 | 0 | 0 | 0 | 0 | 0 | 0 | 0 | 0 | 0 | 0 |
| 0 | NA | NA | NA | NA | NA | Gm12092 | 0 | 0 | 0 | 0 | 0 | 0 | 0 | 0 | 0 | 0 | 0 |

| | | | | | | | | | | | | | | | | | | |
|---|---|---|---|---|---|---|---|---|---|---|---|---|---|---|---|---|---|---|
| 0 | NA | NA | NA | NA | NA | Gm12093 | | 0 | 0 | 0 | 0 | 0 | 0 | 0 | 0 | 0 | 0 | 0 |
| 0 | NA | NA | NA | NA | NA | Gm22588 | | 0 | 0 | 0 | 0 | 0 | 0 | 0 | 0 | 0 | 0 | 0 |
| 0 | NA | NA | NA | NA | NA | Gm12095 | | 0 | 0 | 0 | 0 | 0 | 0 | 0 | 0 | 0 | 0 | 0 |
| 0 | NA | NA | NA | NA | NA | Gm29237 | | 0 | 0 | 0 | 0 | 0 | 0 | 0 | 0 | 0 | 0 | 0 |
| 0 | NA | NA | NA | NA | NA | Gm12103 | | 0 | 0 | 0 | 0 | 0 | 0 | 0 | 0 | 0 | 0 | 0 |
| 0 | NA | NA | NA | NA | NA | Gm12102 | | 0 | 0 | 0 | 0 | 0 | 0 | 0 | 0 | 0 | 0 | 0 |
| 0 | NA | NA | NA | NA | NA | Gm12106 | | 0 | 0 | 0 | 0 | 0 | 0 | 0 | 0 | 0 | 0 | 0 |
| 0 | NA | NA | NA | NA | NA | Gm17332 | | 0 | 0 | 0 | 0 | 0 | 0 | 0 | 0 | 0 | 0 | 0 |
| 0 | NA | NA | NA | NA | NA | Gm38061 | | 0 | 0 | 0 | 0 | 0 | 0 | 0 | 0 | 0 | 0 | 0 |
| 0 | NA | NA | NA | NA | NA | Gm12107 | | 0 | 0 | 0 | 0 | 0 | 0 | 0 | 0 | 0 | 0 | 0 |
| 0 | NA | NA | NA | NA | NA | Gm8192 | | 0 | 0 | 0 | 0 | 0 | 0 | 0 | 0 | 0 | 0 | 0 |
| 0 | NA | NA | NA | NA | NA | 4930555O08Rik | | 0 | 0 | 0 | 0 | 0 | 0 | 0 | 0 | 0 | 0 | 0 |
| 0 | NA | NA | NA | NA | NA | Smim23 | small integral membrane protein 23 | 0 | 0 | 0 | 0 | 0 | 0 | 0 | 0 | 0 | 0 | 0 |
| 0 | NA | NA | NA | NA | NA | Gm12110 | | 0 | 0 | 0 | 0 | 0 | 0 | 0 | 0 | 0 | 0 | 0 |
| 0 | NA | NA | NA | NA | NA | Gm26052 | | 0 | 0 | 0 | 0 | 0 | 0 | 0 | 0 | 0 | 0 | 0 |
| 0 | NA | NA | NA | NA | NA | Gm12114 | | 0 | 0 | 0 | 0 | 0 | 0 | 0 | 0 | 0 | 0 | 0 |
| 0 | NA | NA | NA | NA | NA | Tlx3 | T cell leukemia, homeobox 3 | 0 | 0 | 0 | 0 | 0 | 0 | 0 | 0 | 0 | 0 | 0 |
| 0 | NA | NA | NA | NA | NA | Gm12116 | | 0 | 0 | 0 | 0 | 0 | 0 | 0 | 0 | 0 | 0 | 0 |
| 0 | NA | NA | NA | NA | NA | Gm12117 | | 0 | 0 | 0 | 0 | 0 | 0 | 0 | 0 | 0 | 0 | 0 |
| 0 | NA | NA | NA | NA | NA | Gm12118 | | 0 | 0 | 0 | 0 | 0 | 0 | 0 | 0 | 0 | 0 | 0 |
| 0 | NA | NA | NA | NA | NA | D130052B06Rik | | 0 | 0 | 0 | 0 | 0 | 0 | 0 | 0 | 0 | 0 | 0 |
| 0 | NA | NA | NA | NA | NA | Gm12119 | | 0 | 0 | 0 | 0 | 0 | 0 | 0 | 0 | 0 | 0 | 0 |
| 0 | NA | NA | NA | NA | NA | Gm12121 | | 0 | 0 | 0 | 0 | 0 | 0 | 0 | 0 | 0 | 0 | 0 |
| 0 | NA | NA | NA | NA | NA | Foxi1 | forkhead box I1 | 0 | 0 | 0 | 0 | 0 | 0 | 0 | 0 | 0 | 0 | 0 |
| 0 | NA | NA | NA | NA | NA | 4930403D09Rik | | 0 | 0 | 0 | 0 | 0 | 0 | 0 | 0 | 0 | 0 | 0 |
| 0 | NA | NA | NA | NA | NA | Gm22022 | | 0 | 0 | 0 | 0 | 0 | 0 | 0 | 0 | 0 | 0 | 0 |
| 0 | NA | NA | NA | NA | NA | Gm12711 | | 0 | 0 | 0 | 0 | 0 | 0 | 0 | 0 | 0 | 0 | 0 |
| 0 | NA | NA | NA | NA | NA | Gm25799 | | 0 | 0 | 0 | 0 | 0 | 0 | 0 | 0 | 0 | 0 | 0 |
| 0 | NA | NA | NA | NA | NA | Gm26070 | | 0 | 0 | 0 | 0 | 0 | 0 | 0 | 0 | 0 | 0 | 0 |
| 0 | NA | NA | NA | NA | NA | Gm12122 | | 0 | 0 | 0 | 0 | 0 | 0 | 0 | 0 | 0 | 0 | 0 |
| 0 | NA | NA | NA | NA | NA | Mir218-2 | | 0 | 0 | 0 | 0 | 0 | 0 | 0 | 0 | 0 | 0 | 0 |
| 0 | NA | NA | NA | NA | NA | Mir103-1 | | 0 | 0 | 0 | 0 | 0 | 0 | 0 | 0 | 0 | 0 | 0 |
| 0 | NA | NA | NA | NA | NA | Gm12124 | | 0 | 0 | 0 | 0 | 0 | 0 | 0 | 0 | 0 | 0 | 0 |
| 0 | NA | NA | NA | NA | NA | Gm12125 | | 0 | 0 | 0 | 0 | 0 | 0 | 0 | 0 | 0 | 0 | 0 |
| 0 | NA | NA | NA | NA | NA | Gm12126 | | 0 | 0 | 0 | 0 | 0 | 0 | 0 | 0 | 0 | 0 | 0 |
| 0 | NA | NA | NA | NA | NA | Gm12127 | | 0 | 0 | 0 | 0 | 0 | 0 | 0 | 0 | 0 | 0 | 0 |
| 0 | NA | NA | NA | NA | NA | Gm22127 | | 0 | 0 | 0 | 0 | 0 | 0 | 0 | 0 | 0 | 0 | 0 |
| 0 | NA | NA | NA | NA | NA | Gm12128 | | 0 | 0 | 0 | 0 | 0 | 0 | 0 | 0 | 0 | 0 | 0 |
| 0 | NA | NA | NA | NA | NA | Gm25693 | | 0 | 0 | 0 | 0 | 0 | 0 | 0 | 0 | 0 | 0 | 0 |
| 0 | NA | NA | NA | NA | NA | Gm12130 | | 0 | 0 | 0 | 0 | 0 | 0 | 0 | 0 | 0 | 0 | 0 |
| 0 | NA | NA | NA | NA | NA | Gm23520 | | 0 | 0 | 0 | 0 | 0 | 0 | 0 | 0 | 0 | 0 | 0 |
| 0 | NA | NA | NA | NA | NA | Gm12131 | | 0 | 0 | 0 | 0 | 0 | 0 | 0 | 0 | 0 | 0 | 0 |
| 0 | NA | NA | NA | NA | NA | Gm24192 | | 0 | 0 | 0 | 0 | 0 | 0 | 0 | 0 | 0 | 0 | 0 |
| 0 | NA | NA | NA | NA | NA | Gm12132 | | 0 | 0 | 0 | 0 | 0 | 0 | 0 | 0 | 0 | 0 | 0 |
| 0 | NA | NA | NA | NA | NA | Gm12133 | | 0 | 0 | 0 | 0 | 0 | 0 | 0 | 0 | 0 | 0 | 0 |
| 0 | NA | NA | NA | NA | NA | Gm12134 | | 0 | 0 | 0 | 0 | 0 | 0 | 0 | 0 | 0 | 0 | 0 |
| 0 | NA | NA | NA | NA | NA | 3110004A20Rik | | 0 | 0 | 0 | 0 | 0 | 0 | 0 | 0 | 0 | 0 | 0 |
| 0 | NA | NA | NA | NA | NA | Gm12135 | | 0 | 0 | 0 | 0 | 0 | 0 | 0 | 0 | 0 | 0 | 0 |
| 0 | NA | NA | NA | NA | NA | Gm12137 | | 0 | 0 | 0 | 0 | 0 | 0 | 0 | 0 | 0 | 0 | 0 |
| 0 | NA | NA | NA | NA | NA | Gm24446 | | 0 | 0 | 0 | 0 | 0 | 0 | 0 | 0 | 0 | 0 | 0 |
| 0 | NA | NA | NA | NA | NA | Platr13 | | 0 | 0 | 0 | 0 | 0 | 0 | 0 | 0 | 0 | 0 | 0 |
| 0 | NA | NA | NA | NA | NA | Gm12142 | | 0 | 0 | 0 | 0 | 0 | 0 | 0 | 0 | 0 | 0 | 0 |
| 0 | NA | NA | NA | NA | NA | A230004M16Rik | | 0 | 0 | 0 | 0 | 0 | 0 | 0 | 0 | 0 | 0 | 0 |
| 0 | NA | NA | NA | NA | NA | Gm12143 | | 0 | 0 | 0 | 0 | 0 | 0 | 0 | 0 | 0 | 0 | 0 |
| 0 | NA | NA | NA | NA | NA | Gabra6 | gamma-aminobutyric acid (GABA) A rece | 0 | 0 | 0 | 0 | 0 | 0 | 0 | 0 | 0 | 0 | 0 |
| 0 | NA | NA | NA | NA | NA | Gm25543 | | 0 | 0 | 0 | 0 | 0 | 0 | 0 | 0 | 0 | 0 | 0 |
| 0 | NA | NA | NA | NA | NA | Gm9972 | | 0 | 0 | 0 | 0 | 0 | 0 | 0 | 0 | 0 | 0 | 0 |
| 0 | NA | NA | NA | NA | NA | Gm12145 | | 0 | 0 | 0 | 0 | 0 | 0 | 0 | 0 | 0 | 0 | 0 |
| 0 | NA | NA | NA | NA | NA | Gm12146 | | 0 | 0 | 0 | 0 | 0 | 0 | 0 | 0 | 0 | 0 | 0 |
| 0 | NA | NA | NA | NA | NA | Gm12144 | | 0 | 0 | 0 | 0 | 0 | 0 | 0 | 0 | 0 | 0 | 0 |

| | | | | | | | | | | | | | | | | | | |
|---|---|---|---|---|---|---|---|---|---|---|---|---|---|---|---|---|---|---|
| 0 | NA | NA | NA | NA | NA | Gm12147 | | 0 | 0 | 0 | 0 | 0 | 0 | 0 | 0 | 0 | 0 | 0 |
| 0 | NA | NA | NA | NA | NA | Mir146 | | 0 | 0 | 0 | 0 | 0 | 0 | 0 | 0 | 0 | 0 | 0 |
| 0 | NA | NA | NA | NA | NA | Gm12148 | | 0 | 0 | 0 | 0 | 0 | 0 | 0 | 0 | 0 | 0 | 0 |
| 0 | NA | NA | NA | NA | NA | Gm12151 | | 0 | 0 | 0 | 0 | 0 | 0 | 0 | 0 | 0 | 0 | 0 |
| 0 | NA | NA | NA | NA | NA | Gm12150 | | 0 | 0 | 0 | 0 | 0 | 0 | 0 | 0 | 0 | 0 | 0 |
| 0 | NA | NA | NA | NA | NA | Fabp6 | fatty acid binding protein 6 | 0 | 0 | 0 | 0 | 0 | 0 | 0 | 0 | 0 | 0 | 0 |
| 0 | NA | NA | NA | NA | NA | Gm12149 | | 0 | 0 | 0 | 0 | 0 | 0 | 0 | 0 | 0 | 0 | 0 |
| 0 | NA | NA | NA | NA | NA | Gm12152 | | 0 | 0 | 0 | 0 | 0 | 0 | 0 | 0 | 0 | 0 | 0 |
| 0 | NA | NA | NA | NA | NA | Gm12153 | | 0 | 0 | 0 | 0 | 0 | 0 | 0 | 0 | 0 | 0 | 0 |
| 0 | NA | NA | NA | NA | NA | Gm12154 | | 0 | 0 | 0 | 0 | 0 | 0 | 0 | 0 | 0 | 0 | 0 |
| 0 | NA | NA | NA | NA | NA | Gm12155 | | 0 | 0 | 0 | 0 | 0 | 0 | 0 | 0 | 0 | 0 | 0 |
| 0 | NA | NA | NA | NA | NA | Gm22121 | | 0 | 0 | 0 | 0 | 0 | 0 | 0 | 0 | 0 | 0 | 0 |
| 0 | NA | NA | NA | NA | NA | Gm12156 | | 0 | 0 | 0 | 0 | 0 | 0 | 0 | 0 | 0 | 0 | 0 |
| 0 | NA | NA | NA | NA | NA | 4930597A21Rik | | 0 | 0 | 0 | 0 | 0 | 0 | 0 | 0 | 0 | 0 | 0 |
| 0 | NA | NA | NA | NA | NA | Gm37695 | | 0 | 0 | 0 | 0 | 0 | 0 | 0 | 0 | 0 | 0 | 0 |
| 0 | NA | NA | NA | NA | NA | Gm12158 | | 0 | 0 | 0 | 0 | 0 | 0 | 0 | 0 | 0 | 0 | 0 |
| 0 | NA | NA | NA | NA | NA | Gm12160 | | 0 | 0 | 0 | 0 | 0 | 0 | 0 | 0 | 0 | 0 | 0 |
| 0 | NA | NA | NA | NA | NA | Gm38087 | | 0 | 0 | 0 | 0 | 0 | 0 | 0 | 0 | 0 | 0 | 0 |
| 0 | NA | NA | NA | NA | NA | Gm37312 | | 0 | 0 | 0 | 0 | 0 | 0 | 0 | 0 | 0 | 0 | 0 |
| 0 | NA | NA | NA | NA | NA | Gm22284 | | 0 | 0 | 0 | 0 | 0 | 0 | 0 | 0 | 0 | 0 | 0 |
| 0 | NA | NA | NA | NA | NA | F630206G17Rik | | 0 | 0 | 0 | 0 | 0 | 0 | 0 | 0 | 0 | 0 | 0 |
| 0 | NA | NA | NA | NA | NA | Gm12163 | | 0 | 0 | 0 | 0 | 0 | 0 | 0 | 0 | 0 | 0 | 0 |
| 0 | NA | NA | NA | NA | NA | Gm22751 | | 0 | 0 | 0 | 0 | 0 | 0 | 0 | 0 | 0 | 0 | 0 |
| 0 | NA | NA | NA | NA | NA | Gm12165 | | 0 | 0 | 0 | 0 | 0 | 0 | 0 | 0 | 0 | 0 | 0 |
| 0 | NA | NA | NA | NA | NA | Mir8100 | | 0 | 0 | 0 | 0 | 0 | 0 | 0 | 0 | 0 | 0 | 0 |
| 0 | NA | NA | NA | NA | NA | Gm16033 | | 0 | 0 | 0 | 0 | 0 | 0 | 0 | 0 | 0 | 0 | 0 |
| 0 | NA | NA | NA | NA | NA | Gm16034 | | 0 | 0 | 0 | 0 | 0 | 0 | 0 | 0 | 0 | 0 | 0 |
| 0 | NA | NA | NA | NA | NA | Mir7239 | | 0 | 0 | 0 | 0 | 0 | 0 | 0 | 0 | 0 | 0 | 0 |
| 0 | NA | NA | NA | NA | NA | Fndc9 | fibronectin type III domain containing 9 | 0 | 0 | 0 | 0 | 0 | 0 | 0 | 0 | 0 | 0 | 0 |
| 0 | NA | NA | NA | NA | NA | Gm12167 | | 0 | 0 | 0 | 0 | 0 | 0 | 0 | 0 | 0 | 0 | 0 |
| 0 | NA | NA | NA | NA | NA | Gm12169 | predicted gene 12169 | 0 | 0 | 0 | 0 | 0 | 0 | 0 | 0 | 0 | 0 | 0 |
| 0 | NA | NA | NA | NA | NA | Gm12170 | | 0 | 0 | 0 | 0 | 0 | 0 | 0 | 0 | 0 | 0 | 0 |
| 0 | NA | NA | NA | NA | NA | Gm12171 | predicted gene 12171 | 0 | 0 | 0 | 0 | 0 | 0 | 0 | 0 | 0 | 0 | 0 |
| 0 | NA | NA | NA | NA | NA | Rpl9-ps2 | | 0 | 0 | 0 | 0 | 0 | 0 | 0 | 0 | 0 | 0 | 0 |
| 0 | NA | NA | NA | NA | NA | BC053393 | cDNA sequence BC053393 | 0 | 0 | 0 | 0 | 0 | 0 | 0 | 0 | 0 | 0 | 0 |
| 0 | NA | NA | NA | NA | NA | Dppa1 | developmental pluripotency associated 1 | 0 | 0 | 0 | 0 | 0 | 0 | 0 | 0 | 0 | 0 | 0 |
| 0 | NA | NA | NA | NA | NA | Gm4926 | | 0 | 0 | 0 | 0 | 0 | 0 | 0 | 0 | 0 | 0 | 0 |
| 0 | NA | NA | NA | NA | NA | Gm12173 | | 0 | 0 | 0 | 0 | 0 | 0 | 0 | 0 | 0 | 0 | 0 |
| 0 | NA | NA | NA | NA | NA | Gm12175 | | 0 | 0 | 0 | 0 | 0 | 0 | 0 | 0 | 0 | 0 | 0 |
| 0 | NA | NA | NA | NA | NA | Gm23039 | | 0 | 0 | 0 | 0 | 0 | 0 | 0 | 0 | 0 | 0 | 0 |
| 0 | NA | NA | NA | NA | NA | Gm12179 | | 0 | 0 | 0 | 0 | 0 | 0 | 0 | 0 | 0 | 0 | 0 |
| 0 | NA | NA | NA | NA | NA | Gm12180 | | 0 | 0 | 0 | 0 | 0 | 0 | 0 | 0 | 0 | 0 | 0 |
| 0 | NA | NA | NA | NA | NA | Gm24422 | | 0 | 0 | 0 | 0 | 0 | 0 | 0 | 0 | 0 | 0 | 0 |
| 0 | NA | NA | NA | NA | NA | Gm25296 | | 0 | 0 | 0 | 0 | 0 | 0 | 0 | 0 | 0 | 0 | 0 |
| 0 | NA | NA | NA | NA | NA | Snord96a | | 0 | 0 | 0 | 0 | 0 | 0 | 0 | 0 | 0 | 0 | 0 |
| 0 | NA | NA | NA | NA | NA | Snord95 | | 0 | 0 | 0 | 0 | 0 | 0 | 0 | 0 | 0 | 0 | 0 |
| 0 | NA | NA | NA | NA | NA | Gm12187 | | 0 | 0 | 0 | 0 | 0 | 0 | 0 | 0 | 0 | 0 | 0 |
| 0 | NA | NA | NA | NA | NA | Gm12188 | | 0 | 0 | 0 | 0 | 0 | 0 | 0 | 0 | 0 | 0 | 0 |
| 0 | NA | NA | NA | NA | NA | Olfr1395 | olfactory receptor 1395 | 0 | 0 | 0 | 0 | 0 | 0 | 0 | 0 | 0 | 0 | 0 |
| 0 | NA | NA | NA | NA | NA | Olfr1394 | olfactory receptor 1394 | 0 | 0 | 0 | 0 | 0 | 0 | 0 | 0 | 0 | 0 | 0 |
| 0 | NA | NA | NA | NA | NA | Olfr1393 | olfactory receptor 1393 | 0 | 0 | 0 | 0 | 0 | 0 | 0 | 0 | 0 | 0 | 0 |
| 0 | NA | NA | NA | NA | NA | Olfr1392 | olfactory receptor 1392 | 0 | 0 | 0 | 0 | 0 | 0 | 0 | 0 | 0 | 0 | 0 |
| 0 | NA | NA | NA | NA | NA | Olfr10 | olfactory receptor 10 | 0 | 0 | 0 | 0 | 0 | 0 | 0 | 0 | 0 | 0 | 0 |
| 0 | NA | NA | NA | NA | NA | Olfr1391 | olfactory receptor 1391 | 0 | 0 | 0 | 0 | 0 | 0 | 0 | 0 | 0 | 0 | 0 |
| 0 | NA | NA | NA | NA | NA | Olfr1390 | olfactory receptor 1390 | 0 | 0 | 0 | 0 | 0 | 0 | 0 | 0 | 0 | 0 | 0 |
| 0 | NA | NA | NA | NA | NA | Gm12189 | | 0 | 0 | 0 | 0 | 0 | 0 | 0 | 0 | 0 | 0 | 0 |
| 0 | NA | NA | NA | NA | NA | Olfr1389 | olfactory receptor 1389 | 0 | 0 | 0 | 0 | 0 | 0 | 0 | 0 | 0 | 0 | 0 |
| 0 | NA | NA | NA | NA | NA | Olfr1388 | olfactory receptor 1388 | 0 | 0 | 0 | 0 | 0 | 0 | 0 | 0 | 0 | 0 | 0 |
| 0 | NA | NA | NA | NA | NA | Olfr1387 | olfactory receptor 1387 | 0 | 0 | 0 | 0 | 0 | 0 | 0 | 0 | 0 | 0 | 0 |
| 0 | NA | NA | NA | NA | NA | Olfr1386 | olfactory receptor 1386 | 0 | 0 | 0 | 0 | 0 | 0 | 0 | 0 | 0 | 0 | 0 |

| | | | | | | | | | | | | | | | | | | |
|---|---|---|---|---|---|---|---|---|---|---|---|---|---|---|---|---|---|---|
| 0 | NA | NA | NA | NA | NA | Olfr1385 | olfactory receptor 1385 | 0 | 0 | 0 | 0 | 0 | 0 | 0 | 0 | 0 | 0 | 0 |
| 0 | NA | NA | NA | NA | NA | Olfr1384 | olfactory receptor 1384 | 0 | 0 | 0 | 0 | 0 | 0 | 0 | 0 | 0 | 0 | 0 |
| 0 | NA | NA | NA | NA | NA | Olfr1383 | olfactory receptor 1383 | 0 | 0 | 0 | 0 | 0 | 0 | 0 | 0 | 0 | 0 | 0 |
| 0 | NA | NA | NA | NA | NA | Olfr1382 | olfactory receptor 1382 | 0 | 0 | 0 | 0 | 0 | 0 | 0 | 0 | 0 | 0 | 0 |
| 0 | NA | NA | NA | NA | NA | Olfr1381 | olfactory receptor 1381 | 0 | 0 | 0 | 0 | 0 | 0 | 0 | 0 | 0 | 0 | 0 |
| 0 | NA | NA | NA | NA | NA | Olfr1380 | olfactory receptor 1380 | 0 | 0 | 0 | 0 | 0 | 0 | 0 | 0 | 0 | 0 | 0 |
| 0 | NA | NA | NA | NA | NA | 1700024J04Rik | | 0 | 0 | 0 | 0 | 0 | 0 | 0 | 0 | 0 | 0 | 0 |
| 0 | NA | NA | NA | NA | NA | Gm12190 | | 0 | 0 | 0 | 0 | 0 | 0 | 0 | 0 | 0 | 0 | 0 |
| 0 | NA | NA | NA | NA | NA | Gm12192 | | 0 | 0 | 0 | 0 | 0 | 0 | 0 | 0 | 0 | 0 | 0 |
| 0 | NA | NA | NA | NA | NA | Gm23813 | | 0 | 0 | 0 | 0 | 0 | 0 | 0 | 0 | 0 | 0 | 0 |
| 0 | NA | NA | NA | NA | NA | Gm12193 | | 0 | 0 | 0 | 0 | 0 | 0 | 0 | 0 | 0 | 0 | 0 |
| 0 | NA | NA | NA | NA | NA | Gm23492 | | 0 | 0 | 0 | 0 | 0 | 0 | 0 | 0 | 0 | 0 | 0 |
| 0 | NA | NA | NA | NA | NA | Gm12195 | | 0 | 0 | 0 | 0 | 0 | 0 | 0 | 0 | 0 | 0 | 0 |
| 0 | NA | NA | NA | NA | NA | Mir6919 | | 0 | 0 | 0 | 0 | 0 | 0 | 0 | 0 | 0 | 0 | 0 |
| 0 | NA | NA | NA | NA | NA | Gm23824 | | 0 | 0 | 0 | 0 | 0 | 0 | 0 | 0 | 0 | 0 | 0 |
| 0 | NA | NA | NA | NA | NA | Gm12199 | | 0 | 0 | 0 | 0 | 0 | 0 | 0 | 0 | 0 | 0 | 0 |
| 0 | NA | NA | NA | NA | NA | Gm12200 | | 0 | 0 | 0 | 0 | 0 | 0 | 0 | 0 | 0 | 0 | 0 |
| 0 | NA | NA | NA | NA | NA | Gm23392 | | 0 | 0 | 0 | 0 | 0 | 0 | 0 | 0 | 0 | 0 | 0 |
| 0 | NA | NA | NA | NA | NA | Grm6 | glutamate receptor, metabotropic 6 | 0 | 0 | 0 | 0 | 0 | 0 | 0 | 0 | 0 | 0 | 0 |
| 0 | NA | NA | NA | NA | NA | Gm12660 | | 0 | 0 | 0 | 0 | 0 | 0 | 0 | 0 | 0 | 0 | 0 |
| 0 | NA | NA | NA | NA | NA | 4933414I15Rik | | 0 | 0 | 0 | 0 | 0 | 0 | 0 | 0 | 0 | 0 | 0 |
| 0 | NA | NA | NA | NA | NA | Prop1 | paired like homeodomain factor 1 | 0 | 0 | 0 | 0 | 0 | 0 | 0 | 0 | 0 | 0 | 0 |
| 0 | NA | NA | NA | NA | NA | Olfr1379-ps1 | | 0 | 0 | 0 | 0 | 0 | 0 | 0 | 0 | 0 | 0 | 0 |
| 0 | NA | NA | NA | NA | NA | Olfr1378 | olfactory receptor 1378 | 0 | 0 | 0 | 0 | 0 | 0 | 0 | 0 | 0 | 0 | 0 |
| 0 | NA | NA | NA | NA | NA | Gm12661 | | 0 | 0 | 0 | 0 | 0 | 0 | 0 | 0 | 0 | 0 | 0 |
| 0 | NA | NA | NA | NA | NA | Olfr1377 | olfactory receptor 1377 | 0 | 0 | 0 | 0 | 0 | 0 | 0 | 0 | 0 | 0 | 0 |
| 0 | NA | NA | NA | NA | NA | Olfr1376-ps1 | | 0 | 0 | 0 | 0 | 0 | 0 | 0 | 0 | 0 | 0 | 0 |
| 0 | NA | NA | NA | NA | NA | Olfr51 | olfactory receptor 51 | 0 | 0 | 0 | 0 | 0 | 0 | 0 | 0 | 0 | 0 | 0 |
| 0 | NA | NA | NA | NA | NA | Olfr54 | olfactory receptor 54 | 0 | 0 | 0 | 0 | 0 | 0 | 0 | 0 | 0 | 0 | 0 |
| 0 | NA | NA | NA | NA | NA | Olfr1375 | olfactory receptor 1375 | 0 | 0 | 0 | 0 | 0 | 0 | 0 | 0 | 0 | 0 | 0 |
| 0 | NA | NA | NA | NA | NA | 9230009I02Rik | | 0 | 0 | 0 | 0 | 0 | 0 | 0 | 0 | 0 | 0 | 0 |
| 0 | NA | NA | NA | NA | NA | Gm12202 | | 0 | 0 | 0 | 0 | 0 | 0 | 0 | 0 | 0 | 0 | 0 |
| 0 | NA | NA | NA | NA | NA | Gm12201 | | 0 | 0 | 0 | 0 | 0 | 0 | 0 | 0 | 0 | 0 | 0 |
| 0 | NA | NA | NA | NA | NA | Olfr1374-ps1 | | 0 | 0 | 0 | 0 | 0 | 0 | 0 | 0 | 0 | 0 | 0 |
| 0 | NA | NA | NA | NA | NA | Gm12203 | | 0 | 0 | 0 | 0 | 0 | 0 | 0 | 0 | 0 | 0 | 0 |
| 0 | NA | NA | NA | NA | NA | Platr20 | | 0 | 0 | 0 | 0 | 0 | 0 | 0 | 0 | 0 | 0 | 0 |
| 0 | NA | NA | NA | NA | NA | Gm12569 | predicted gene 12569 | 0 | 0 | 0 | 0 | 0 | 0 | 0 | 0 | 0 | 0 | 0 |
| 0 | NA | NA | NA | NA | NA | Gm26061 | | 0 | 0 | 0 | 0 | 0 | 0 | 0 | 0 | 0 | 0 | 0 |
| 0 | NA | NA | NA | NA | NA | Gm12205 | | 0 | 0 | 0 | 0 | 0 | 0 | 0 | 0 | 0 | 0 | 0 |
| 0 | NA | NA | NA | NA | NA | Mir3061 | | 0 | 0 | 0 | 0 | 0 | 0 | 0 | 0 | 0 | 0 | 0 |
| 0 | NA | NA | NA | NA | NA | Olfr1373 | olfactory receptor 1373 | 0 | 0 | 0 | 0 | 0 | 0 | 0 | 0 | 0 | 0 | 0 |
| 0 | NA | NA | NA | NA | NA | Gm12206 | | 0 | 0 | 0 | 0 | 0 | 0 | 0 | 0 | 0 | 0 | 0 |
| 0 | NA | NA | NA | NA | NA | Olfr1371 | olfactory receptor 1371 | 0 | 0 | 0 | 0 | 0 | 0 | 0 | 0 | 0 | 0 | 0 |
| 0 | NA | NA | NA | NA | NA | Gm12209 | | 0 | 0 | 0 | 0 | 0 | 0 | 0 | 0 | 0 | 0 | 0 |
| 0 | NA | NA | NA | NA | NA | Gm12210 | | 0 | 0 | 0 | 0 | 0 | 0 | 0 | 0 | 0 | 0 | 0 |
| 0 | NA | NA | NA | NA | NA | Gm11186 | | 0 | 0 | 0 | 0 | 0 | 0 | 0 | 0 | 0 | 0 | 0 |
| 0 | NA | NA | NA | NA | NA | Gm11188 | | 0 | 0 | 0 | 0 | 0 | 0 | 0 | 0 | 0 | 0 | 0 |
| 0 | NA | NA | NA | NA | NA | Gm25930 | | 0 | 0 | 0 | 0 | 0 | 0 | 0 | 0 | 0 | 0 | 0 |
| 0 | NA | NA | NA | NA | NA | Gm10447 | | 0 | 0 | 0 | 0 | 0 | 0 | 0 | 0 | 0 | 0 | 0 |
| 0 | NA | NA | NA | NA | NA | Gm9837 | | 0 | 0 | 0 | 0 | 0 | 0 | 0 | 0 | 0 | 0 | 0 |
| 0 | NA | NA | NA | NA | NA | Il4 | interleukin 4 | 0 | 0 | 0 | 0 | 0 | 0 | 0 | 0 | 0 | 0 | 0 |
| 0 | NA | NA | NA | NA | NA | Il13 | interleukin 13 | 0 | 0 | 0 | 0 | 0 | 0 | 0 | 0 | 0 | 0 | 0 |
| 0 | NA | NA | NA | NA | NA | Gm22275 | | 0 | 0 | 0 | 0 | 0 | 0 | 0 | 0 | 0 | 0 | 0 |
| 0 | NA | NA | NA | NA | NA | Gm12214 | | 0 | 0 | 0 | 0 | 0 | 0 | 0 | 0 | 0 | 0 | 0 |
| 0 | NA | NA | NA | NA | NA | Gm12213 | | 0 | 0 | 0 | 0 | 0 | 0 | 0 | 0 | 0 | 0 | 0 |
| 0 | NA | NA | NA | NA | NA | Gm50459 | | 0 | 0 | 0 | 0 | 0 | 0 | 0 | 0 | 0 | 0 | 0 |
| 0 | NA | NA | NA | NA | NA | Gm12220 | | 0 | 0 | 0 | 0 | 0 | 0 | 0 | 0 | 0 | 0 | 0 |
| 0 | NA | NA | NA | NA | NA | 4933405E24Rik | | 0 | 0 | 0 | 0 | 0 | 0 | 0 | 0 | 0 | 0 | 0 |
| 0 | NA | NA | NA | NA | NA | Gm12222 | | 0 | 0 | 0 | 0 | 0 | 0 | 0 | 0 | 0 | 0 | 0 |
| 0 | NA | NA | NA | NA | NA | Csf2 | colony stimulating factor 2 (granulocyte-m | 0 | 0 | 0 | 0 | 0 | 0 | 0 | 0 | 0 | 0 | 0 |

| | | | | | | | | | | | | | | | | | |
|---|---|---|---|---|---|---|---|---|---|---|---|---|---|---|---|---|---|
| 0 | NA | NA | NA | NA | NA | Gm12225 | | 0 | 0 | 0 | 0 | 0 | 0 | 0 | 0 | 0 | 0 | 0 |
| 0 | NA | NA | NA | NA | NA | Gm24198 | | 0 | 0 | 0 | 0 | 0 | 0 | 0 | 0 | 0 | 0 | 0 |
| 0 | NA | NA | NA | NA | NA | Gm26116 | | 0 | 0 | 0 | 0 | 0 | 0 | 0 | 0 | 0 | 0 | 0 |
| 0 | NA | NA | NA | NA | NA | Gm12227 | | 0 | 0 | 0 | 0 | 0 | 0 | 0 | 0 | 0 | 0 | 0 |
| 0 | NA | NA | NA | NA | NA | Lyrm7os | | 0 | 0 | 0 | 0 | 0 | 0 | 0 | 0 | 0 | 0 | 0 |
| 0 | NA | NA | NA | NA | NA | Gm25426 | | 0 | 0 | 0 | 0 | 0 | 0 | 0 | 0 | 0 | 0 | 0 |
| 0 | NA | NA | NA | NA | NA | Mup-ps22 | | 0 | 0 | 0 | 0 | 0 | 0 | 0 | 0 | 0 | 0 | 0 |
| 0 | NA | NA | NA | NA | NA | Slc36a3 | solute carrier family 36 (proton/amino acid | 0 | 0 | 0 | 0 | 0 | 0 | 0 | 0 | 0 | 0 | 0 |
| 0 | NA | NA | NA | NA | NA | Slc36a1os | | 0 | 0 | 0 | 0 | 0 | 0 | 0 | 0 | 0 | 0 | 0 |
| 0 | NA | NA | NA | NA | NA | Gm12235 | | 0 | 0 | 0 | 0 | 0 | 0 | 0 | 0 | 0 | 0 | 0 |
| 0 | NA | NA | NA | NA | NA | Gm22884 | | 0 | 0 | 0 | 0 | 0 | 0 | 0 | 0 | 0 | 0 | 0 |
| 0 | NA | NA | NA | NA | NA | Gm12238 | | 0 | 0 | 0 | 0 | 0 | 0 | 0 | 0 | 0 | 0 | 0 |
| 0 | NA | NA | NA | NA | NA | Mir7652 | | 0 | 0 | 0 | 0 | 0 | 0 | 0 | 0 | 0 | 0 | 0 |
| 0 | NA | NA | NA | NA | NA | Gm12236 | | 0 | 0 | 0 | 0 | 0 | 0 | 0 | 0 | 0 | 0 | 0 |
| 0 | NA | NA | NA | NA | NA | Gm12237 | | 0 | 0 | 0 | 0 | 0 | 0 | 0 | 0 | 0 | 0 | 0 |
| 0 | NA | NA | NA | NA | NA | Gm12240 | | 0 | 0 | 0 | 0 | 0 | 0 | 0 | 0 | 0 | 0 | 0 |
| 0 | NA | NA | NA | NA | NA | Gm24253 | | 0 | 0 | 0 | 0 | 0 | 0 | 0 | 0 | 0 | 0 | 0 |
| 0 | NA | NA | NA | NA | NA | Gm12241 | | 0 | 0 | 0 | 0 | 0 | 0 | 0 | 0 | 0 | 0 | 0 |
| 0 | NA | NA | NA | NA | NA | Gm22603 | | 0 | 0 | 0 | 0 | 0 | 0 | 0 | 0 | 0 | 0 | 0 |
| 0 | NA | NA | NA | NA | NA | 4933424L21Rik | | 0 | 0 | 0 | 0 | 0 | 0 | 0 | 0 | 0 | 0 | 0 |
| 0 | NA | NA | NA | NA | NA | 4933426K07Rik | | 0 | 0 | 0 | 0 | 0 | 0 | 0 | 0 | 0 | 0 | 0 |
| 0 | NA | NA | NA | NA | NA | n-R5s69 | | 0 | 0 | 0 | 0 | 0 | 0 | 0 | 0 | 0 | 0 | 0 |
| 0 | NA | NA | NA | NA | NA | Hand1 | heart and neural crest derivatives express | 0 | 0 | 0 | 0 | 0 | 0 | 0 | 0 | 0 | 0 | 0 |
| 0 | NA | NA | NA | NA | NA | Gm12244 | | 0 | 0 | 0 | 0 | 0 | 0 | 0 | 0 | 0 | 0 | 0 |
| 0 | NA | NA | NA | NA | NA | Gm26025 | | 0 | 0 | 0 | 0 | 0 | 0 | 0 | 0 | 0 | 0 | 0 |
| 0 | NA | NA | NA | NA | NA | Gm5038 | | 0 | 0 | 0 | 0 | 0 | 0 | 0 | 0 | 0 | 0 | 0 |
| 0 | NA | NA | NA | NA | NA | 4930438A08Ril | RIKEN cDNA 4930438A08 gene | 0 | 0 | 0 | 0 | 0 | 0 | 0 | 0 | 0 | 0 | 0 |
| 0 | NA | NA | NA | NA | NA | Gm12252 | | 0 | 0 | 0 | 0 | 0 | 0 | 0 | 0 | 0 | 0 | 0 |
| 0 | NA | NA | NA | NA | NA | Lypd9 | LY6/PLAUR domain containing 9 | 0 | 0 | 0 | 0 | 0 | 0 | 0 | 0 | 0 | 0 | 0 |
| 0 | NA | NA | NA | NA | NA | Olfr30 | olfactory receptor 30 | 0 | 0 | 0 | 0 | 0 | 0 | 0 | 0 | 0 | 0 | 0 |
| 0 | NA | NA | NA | NA | NA | Olfr333-ps1 | | 0 | 0 | 0 | 0 | 0 | 0 | 0 | 0 | 0 | 0 | 0 |
| 0 | NA | NA | NA | NA | NA | Olfr332 | olfactory receptor 332 | 0 | 0 | 0 | 0 | 0 | 0 | 0 | 0 | 0 | 0 | 0 |
| 0 | NA | NA | NA | NA | NA | Olfr331 | olfactory receptor 331 | 0 | 0 | 0 | 0 | 0 | 0 | 0 | 0 | 0 | 0 | 0 |
| 0 | NA | NA | NA | NA | NA | Gm12255 | | 0 | 0 | 0 | 0 | 0 | 0 | 0 | 0 | 0 | 0 | 0 |
| 0 | NA | NA | NA | NA | NA | Olfr329 | olfactory receptor 329 | 0 | 0 | 0 | 0 | 0 | 0 | 0 | 0 | 0 | 0 | 0 |
| 0 | NA | NA | NA | NA | NA | Olfr328 | olfactory receptor 328 | 0 | 0 | 0 | 0 | 0 | 0 | 0 | 0 | 0 | 0 | 0 |
| 0 | NA | NA | NA | NA | NA | Olfr224 | olfactory receptor 224 | 0 | 0 | 0 | 0 | 0 | 0 | 0 | 0 | 0 | 0 | 0 |
| 0 | NA | NA | NA | NA | NA | Olfr326-ps1 | | 0 | 0 | 0 | 0 | 0 | 0 | 0 | 0 | 0 | 0 | 0 |
| 0 | NA | NA | NA | NA | NA | Olfr325 | olfactory receptor 325 | 0 | 0 | 0 | 0 | 0 | 0 | 0 | 0 | 0 | 0 | 0 |
| 0 | NA | NA | NA | NA | NA | Olfr324 | olfactory receptor 324 | 0 | 0 | 0 | 0 | 0 | 0 | 0 | 0 | 0 | 0 | 0 |
| 0 | NA | NA | NA | NA | NA | Olfr323 | olfactory receptor 323 | 0 | 0 | 0 | 0 | 0 | 0 | 0 | 0 | 0 | 0 | 0 |
| 0 | NA | NA | NA | NA | NA | Olfr322 | olfactory receptor 322 | 0 | 0 | 0 | 0 | 0 | 0 | 0 | 0 | 0 | 0 | 0 |
| 0 | NA | NA | NA | NA | NA | Gm22599 | | 0 | 0 | 0 | 0 | 0 | 0 | 0 | 0 | 0 | 0 | 0 |
| 0 | NA | NA | NA | NA | NA | Olfr320 | olfactory receptor 320 | 0 | 0 | 0 | 0 | 0 | 0 | 0 | 0 | 0 | 0 | 0 |
| 0 | NA | NA | NA | NA | NA | Gm12570 | | 0 | 0 | 0 | 0 | 0 | 0 | 0 | 0 | 0 | 0 | 0 |
| 0 | NA | NA | NA | NA | NA | Olfr319 | olfactory receptor 319 | 0 | 0 | 0 | 0 | 0 | 0 | 0 | 0 | 0 | 0 | 0 |
| 0 | NA | NA | NA | NA | NA | Olfr318 | olfactory receptor 318 | 0 | 0 | 0 | 0 | 0 | 0 | 0 | 0 | 0 | 0 | 0 |
| 0 | NA | NA | NA | NA | NA | Olfr315 | olfactory receptor 315 | 0 | 0 | 0 | 0 | 0 | 0 | 0 | 0 | 0 | 0 | 0 |
| 0 | NA | NA | NA | NA | NA | Olfr314 | olfactory receptor 314 | 0 | 0 | 0 | 0 | 0 | 0 | 0 | 0 | 0 | 0 | 0 |
| 0 | NA | NA | NA | NA | NA | Olfr313 | olfactory receptor 313 | 0 | 0 | 0 | 0 | 0 | 0 | 0 | 0 | 0 | 0 | 0 |
| 0 | NA | NA | NA | NA | NA | Olfr311 | olfactory receptor 311 | 0 | 0 | 0 | 0 | 0 | 0 | 0 | 0 | 0 | 0 | 0 |
| 0 | NA | NA | NA | NA | NA | Gm12257 | | 0 | 0 | 0 | 0 | 0 | 0 | 0 | 0 | 0 | 0 | 0 |
| 0 | NA | NA | NA | NA | NA | Btnl10 | butyrophilin-like 10 | 0 | 0 | 0 | 0 | 0 | 0 | 0 | 0 | 0 | 0 | 0 |
| 0 | NA | NA | NA | NA | NA | Gm10435 | | 0 | 0 | 0 | 0 | 0 | 0 | 0 | 0 | 0 | 0 | 0 |
| 0 | NA | NA | NA | NA | NA | Gm24756 | | 0 | 0 | 0 | 0 | 0 | 0 | 0 | 0 | 0 | 0 | 0 |
| 0 | NA | NA | NA | NA | NA | Gm15755 | | 0 | 0 | 0 | 0 | 0 | 0 | 0 | 0 | 0 | 0 | 0 |
| 0 | NA | NA | NA | NA | NA | Gm12262 | | 0 | 0 | 0 | 0 | 0 | 0 | 0 | 0 | 0 | 0 | 0 |
| 0 | NA | NA | NA | NA | NA | Olfr222 | olfactory receptor 222 | 0 | 0 | 0 | 0 | 0 | 0 | 0 | 0 | 0 | 0 | 0 |
| 0 | NA | NA | NA | NA | NA | Olfr223 | olfactory receptor 223 | 0 | 0 | 0 | 0 | 0 | 0 | 0 | 0 | 0 | 0 | 0 |
| 0 | NA | NA | NA | NA | NA | Gm5027 | | 0 | 0 | 0 | 0 | 0 | 0 | 0 | 0 | 0 | 0 | 0 |

| | | | | | | | | | | | | | | | | | | |
|---|---|---|---|---|---|---|---|---|---|---|---|---|---|---|---|---|---|---|
| 0 | NA | NA | NA | NA | NA | Olfr225 | olfactory receptor 225 | 0 | 0 | 0 | 0 | 0 | 0 | 0 | 0 | 0 | 0 | 0 |
| 0 | NA | NA | NA | NA | NA | 1700007J10Rik | | 0 | 0 | 0 | 0 | 0 | 0 | 0 | 0 | 0 | 0 | 0 |
| 0 | NA | NA | NA | NA | NA | Gm24939 | | 0 | 0 | 0 | 0 | 0 | 0 | 0 | 0 | 0 | 0 | 0 |
| 0 | NA | NA | NA | NA | NA | Gm12714 | | 0 | 0 | 0 | 0 | 0 | 0 | 0 | 0 | 0 | 0 | 0 |
| 0 | NA | NA | NA | NA | NA | Gm27511 | | 0 | 0 | 0 | 0 | 0 | 0 | 0 | 0 | 0 | 0 | 0 |
| 0 | NA | NA | NA | NA | NA | Mir6921 | | 0 | 0 | 0 | 0 | 0 | 0 | 0 | 0 | 0 | 0 | 0 |
| 0 | NA | NA | NA | NA | NA | Mir6922 | | 0 | 0 | 0 | 0 | 0 | 0 | 0 | 0 | 0 | 0 | 0 |
| 0 | NA | NA | NA | NA | NA | Gm12268 | | 0 | 0 | 0 | 0 | 0 | 0 | 0 | 0 | 0 | 0 | 0 |
| 0 | NA | NA | NA | NA | NA | Gm12622 | | 0 | 0 | 0 | 0 | 0 | 0 | 0 | 0 | 0 | 0 | 0 |
| 0 | NA | NA | NA | NA | NA | Gm12623 | | 0 | 0 | 0 | 0 | 0 | 0 | 0 | 0 | 0 | 0 | 0 |
| 0 | NA | NA | NA | NA | NA | Gm12621 | | 0 | 0 | 0 | 0 | 0 | 0 | 0 | 0 | 0 | 0 | 0 |
| 0 | NA | NA | NA | NA | NA | Gm12627 | | 0 | 0 | 0 | 0 | 0 | 0 | 0 | 0 | 0 | 0 | 0 |
| 0 | NA | NA | NA | NA | NA | Gm12613 | | 0 | 0 | 0 | 0 | 0 | 0 | 0 | 0 | 0 | 0 | 0 |
| 0 | NA | NA | NA | NA | NA | Gm12614 | | 0 | 0 | 0 | 0 | 0 | 0 | 0 | 0 | 0 | 0 | 0 |
| 0 | NA | NA | NA | NA | NA | Gm12615 | | 0 | 0 | 0 | 0 | 0 | 0 | 0 | 0 | 0 | 0 | 0 |
| 0 | NA | NA | NA | NA | NA | Gm12625 | | 0 | 0 | 0 | 0 | 0 | 0 | 0 | 0 | 0 | 0 | 0 |
| 0 | NA | NA | NA | NA | NA | Gm12619 | | 0 | 0 | 0 | 0 | 0 | 0 | 0 | 0 | 0 | 0 | 0 |
| 0 | NA | NA | NA | NA | NA | Gm12624 | | 0 | 0 | 0 | 0 | 0 | 0 | 0 | 0 | 0 | 0 | 0 |
| 0 | NA | NA | NA | NA | NA | Gm12616 | | 0 | 0 | 0 | 0 | 0 | 0 | 0 | 0 | 0 | 0 | 0 |
| 0 | NA | NA | NA | NA | NA | Gm12620 | | 0 | 0 | 0 | 0 | 0 | 0 | 0 | 0 | 0 | 0 | 0 |
| 0 | NA | NA | NA | NA | NA | Gm12617 | | 0 | 0 | 0 | 0 | 0 | 0 | 0 | 0 | 0 | 0 | 0 |
| 0 | NA | NA | NA | NA | NA | Gm23341 | | 0 | 0 | 0 | 0 | 0 | 0 | 0 | 0 | 0 | 0 | 0 |
| 0 | NA | NA | NA | NA | NA | Gm24945 | | 0 | 0 | 0 | 0 | 0 | 0 | 0 | 0 | 0 | 0 | 0 |
| 0 | NA | NA | NA | NA | NA | Gm12269 | | 0 | 0 | 0 | 0 | 0 | 0 | 0 | 0 | 0 | 0 | 0 |
| 0 | NA | NA | NA | NA | NA | Gm25929 | | 0 | 0 | 0 | 0 | 0 | 0 | 0 | 0 | 0 | 0 | 0 |
| 0 | NA | NA | NA | NA | NA | Gm12272 | | 0 | 0 | 0 | 0 | 0 | 0 | 0 | 0 | 0 | 0 | 0 |
| 0 | NA | NA | NA | NA | NA | Gm26964 | | 0 | 0 | 0 | 0 | 0 | 0 | 0 | 0 | 0 | 0 | 0 |
| 0 | NA | NA | NA | NA | NA | Gm23721 | | 0 | 0 | 0 | 0 | 0 | 0 | 0 | 0 | 0 | 0 | 0 |
| 0 | NA | NA | NA | NA | NA | Gm12273 | | 0 | 0 | 0 | 0 | 0 | 0 | 0 | 0 | 0 | 0 | 0 |
| 0 | NA | NA | NA | NA | NA | Gm12274 | | 0 | 0 | 0 | 0 | 0 | 0 | 0 | 0 | 0 | 0 | 0 |
| 0 | NA | NA | NA | NA | NA | | | 0 | 0 | 0 | 0 | 0 | 0 | 0 | 0 | 0 | 0 | 0 |
| 0 | NA | NA | NA | NA | NA | Gm12280 | | 0 | 0 | 0 | 0 | 0 | 0 | 0 | 0 | 0 | 0 | 0 |
| 0 | NA | NA | NA | NA | NA | Lrrc75aos1 | | 0 | 0 | 0 | 0 | 0 | 0 | 0 | 0 | 0 | 0 | 0 |
| 0 | NA | NA | NA | NA | NA | Gm12281 | | 0 | 0 | 0 | 0 | 0 | 0 | 0 | 0 | 0 | 0 | 0 |
| 0 | NA | NA | NA | NA | NA | Gm27194 | | 0 | 0 | 0 | 0 | 0 | 0 | 0 | 0 | 0 | 0 | 0 |
| 0 | NA | NA | NA | NA | NA | Gm10428 | | 0 | 0 | 0 | 0 | 0 | 0 | 0 | 0 | 0 | 0 | 0 |
| 0 | NA | NA | NA | NA | NA | Zfp286os | | 0 | 0 | 0 | 0 | 0 | 0 | 0 | 0 | 0 | 0 | 0 |
| 0 | NA | NA | NA | NA | NA | Gm24151 | | 0 | 0 | 0 | 0 | 0 | 0 | 0 | 0 | 0 | 0 | 0 |
| 0 | NA | NA | NA | NA | NA | Cdrt4os2 | | 0 | 0 | 0 | 0 | 0 | 0 | 0 | 0 | 0 | 0 | 0 |
| 0 | NA | NA | NA | NA | NA | Cdrt4 | CMT1A duplicated region transcript 4 | 0 | 0 | 0 | 0 | 0 | 0 | 0 | 0 | 0 | 0 | 0 |
| 0 | NA | NA | NA | NA | NA | Cdrt4os1 | | 0 | 0 | 0 | 0 | 0 | 0 | 0 | 0 | 0 | 0 | 0 |
| 0 | NA | NA | NA | NA | NA | Tekt3 | tektin 3 | 0 | 0 | 0 | 0 | 0 | 0 | 0 | 0 | 0 | 0 | 0 |
| 0 | NA | NA | NA | NA | NA | B430319H21Rik | | 0 | 0 | 0 | 0 | 0 | 0 | 0 | 0 | 0 | 0 | 0 |
| 0 | NA | NA | NA | NA | NA | B130011K05Rik | | 0 | 0 | 0 | 0 | 0 | 0 | 0 | 0 | 0 | 0 | 0 |
| 0 | NA | NA | NA | NA | NA | Gm12287 | | 0 | 0 | 0 | 0 | 0 | 0 | 0 | 0 | 0 | 0 | 0 |
| 0 | NA | NA | NA | NA | NA | Gm12288 | | 0 | 0 | 0 | 0 | 0 | 0 | 0 | 0 | 0 | 0 | 0 |
| 0 | NA | NA | NA | NA | NA | 9630013K17Rik | | 0 | 0 | 0 | 0 | 0 | 0 | 0 | 0 | 0 | 0 | 0 |
| 0 | NA | NA | NA | NA | NA | Gm12289 | | 0 | 0 | 0 | 0 | 0 | 0 | 0 | 0 | 0 | 0 | 0 |
| 0 | NA | NA | NA | NA | NA | Gm12290 | | 0 | 0 | 0 | 0 | 0 | 0 | 0 | 0 | 0 | 0 | 0 |
| 0 | NA | NA | NA | NA | NA | Gm24275 | | 0 | 0 | 0 | 0 | 0 | 0 | 0 | 0 | 0 | 0 | 0 |
| 0 | NA | NA | NA | NA | NA | Gm12291 | | 0 | 0 | 0 | 0 | 0 | 0 | 0 | 0 | 0 | 0 | 0 |
| 0 | NA | NA | NA | NA | NA | Gm25991 | | 0 | 0 | 0 | 0 | 0 | 0 | 0 | 0 | 0 | 0 | 0 |
| 0 | NA | NA | NA | NA | NA | | | 0 | 0 | 0 | 0 | 0 | 0 | 0 | 0 | 0 | 0 | 0 |
| 0 | NA | NA | NA | NA | NA | Gm12293 | | 0 | 0 | 0 | 0 | 0 | 0 | 0 | 0 | 0 | 0 | 0 |
| 0 | NA | NA | NA | NA | NA | Gm12295 | | 0 | 0 | 0 | 0 | 0 | 0 | 0 | 0 | 0 | 0 | 0 |
| 0 | NA | NA | NA | NA | NA | Gm26128 | | 0 | 0 | 0 | 0 | 0 | 0 | 0 | 0 | 0 | 0 | 0 |
| 0 | NA | NA | NA | NA | NA | Mir744 | | 0 | 0 | 0 | 0 | 0 | 0 | 0 | 0 | 0 | 0 | 0 |
| 0 | NA | NA | NA | NA | NA | Gm12297 | | 0 | 0 | 0 | 0 | 0 | 0 | 0 | 0 | 0 | 0 | 0 |
| 0 | NA | NA | NA | NA | NA | Mir6923 | | 0 | 0 | 0 | 0 | 0 | 0 | 0 | 0 | 0 | 0 | 0 |
| 0 | NA | NA | NA | NA | NA | 2310065F04Rik | | 0 | 0 | 0 | 0 | 0 | 0 | 0 | 0 | 0 | 0 | 0 |

| | | | | | | | | | | | | | | | | | | |
|---|---|---|---|---|---|---|---|---|---|---|---|---|---|---|---|---|---|---|
| 0 | NA | NA | NA | NA | NA | Myh2 | myosin, heavy polypeptide 2, skeletal mu | 0 | 0 | 0 | 0 | 0 | 0 | 0 | 0 | 0 | 0 | 0 |
| 0 | NA | NA | NA | NA | NA | 9430073C21Rik | | 0 | 0 | 0 | 0 | 0 | 0 | 0 | 0 | 0 | 0 | 0 |
| 0 | NA | NA | NA | NA | NA | Myh4 | myosin, heavy polypeptide 4, skeletal mu | 0 | 0 | 0 | 0 | 0 | 0 | 0 | 0 | 0 | 0 | 0 |
| 0 | NA | NA | NA | NA | NA | Myh8 | myosin, heavy polypeptide 8, skeletal mu | 0 | 0 | 0 | 0 | 0 | 0 | 0 | 0 | 0 | 0 | 0 |
| 0 | NA | NA | NA | NA | NA | Gm26476 | | 0 | 0 | 0 | 0 | 0 | 0 | 0 | 0 | 0 | 0 | 0 |
| 0 | NA | NA | NA | NA | NA | Gm12301 | | 0 | 0 | 0 | 0 | 0 | 0 | 0 | 0 | 0 | 0 | 0 |
| 0 | NA | NA | NA | NA | NA | Rcvrn | recoverin | 0 | 0 | 0 | 0 | 0 | 0 | 0 | 0 | 0 | 0 | 0 |
| 0 | NA | NA | NA | NA | NA | Gsg1l2 | GSG1-like 2 | 0 | 0 | 0 | 0 | 0 | 0 | 0 | 0 | 0 | 0 | 0 |
| 0 | NA | NA | NA | NA | NA | Gm12303 | | 0 | 0 | 0 | 0 | 0 | 0 | 0 | 0 | 0 | 0 | 0 |
| 0 | NA | NA | NA | NA | NA | Gm26157 | | 0 | 0 | 0 | 0 | 0 | 0 | 0 | 0 | 0 | 0 | 0 |
| 0 | NA | NA | NA | NA | NA | Gm12304 | | 0 | 0 | 0 | 0 | 0 | 0 | 0 | 0 | 0 | 0 | 0 |
| 0 | NA | NA | NA | NA | NA | Mir6406 | | 0 | 0 | 0 | 0 | 0 | 0 | 0 | 0 | 0 | 0 | 0 |
| 0 | NA | NA | NA | NA | NA | Gm12305 | | 0 | 0 | 0 | 0 | 0 | 0 | 0 | 0 | 0 | 0 | 0 |
| 0 | NA | NA | NA | NA | NA | Ccdc42os | | 0 | 0 | 0 | 0 | 0 | 0 | 0 | 0 | 0 | 0 | 0 |
| 0 | NA | NA | NA | NA | NA | Ccdc42 | coiled-coil domain containing 42 | 0 | 0 | 0 | 0 | 0 | 0 | 0 | 0 | 0 | 0 | 0 |
| 0 | NA | NA | NA | NA | NA | Gm22608 | | 0 | 0 | 0 | 0 | 0 | 0 | 0 | 0 | 0 | 0 | 0 |
| 0 | NA | NA | NA | NA | NA | Odf4 | outer dense fiber of sperm tails 4 | 0 | 0 | 0 | 0 | 0 | 0 | 0 | 0 | 0 | 0 | 0 |
| 0 | NA | NA | NA | NA | NA | Mir3062 | | 0 | 0 | 0 | 0 | 0 | 0 | 0 | 0 | 0 | 0 | 0 |
| 0 | NA | NA | NA | NA | NA | Gm25819 | | 0 | 0 | 0 | 0 | 0 | 0 | 0 | 0 | 0 | 0 | 0 |
| 0 | NA | NA | NA | NA | NA | Snord118 | | 0 | 0 | 0 | 0 | 0 | 0 | 0 | 0 | 0 | 0 | 0 |
| 0 | NA | NA | NA | NA | NA | Gm12306 | | 0 | 0 | 0 | 0 | 0 | 0 | 0 | 0 | 0 | 0 | 0 |
| 0 | NA | NA | NA | NA | NA | Gm25371 | | 0 | 0 | 0 | 0 | 0 | 0 | 0 | 0 | 0 | 0 | 0 |
| 0 | NA | NA | NA | NA | NA | Gm23194 | | 0 | 0 | 0 | 0 | 0 | 0 | 0 | 0 | 0 | 0 | 0 |
| 0 | NA | NA | NA | NA | NA | Cntrobos | | 0 | 0 | 0 | 0 | 0 | 0 | 0 | 0 | 0 | 0 | 0 |
| 0 | NA | NA | NA | NA | NA | Shbg | sex hormone binding globulin | 0 | 0 | 0 | 0 | 0 | 0 | 0 | 0 | 0 | 0 | 0 |
| 0 | NA | NA | NA | NA | NA | Gm12308 | | 0 | 0 | 0 | 0 | 0 | 0 | 0 | 0 | 0 | 0 | 0 |
| 0 | NA | NA | NA | NA | NA | Gm12309 | | 0 | 0 | 0 | 0 | 0 | 0 | 0 | 0 | 0 | 0 | 0 |
| 0 | NA | NA | NA | NA | NA | Mir467f | | 0 | 0 | 0 | 0 | 0 | 0 | 0 | 0 | 0 | 0 | 0 |
| 0 | NA | NA | NA | NA | NA | Mir1934 | | 0 | 0 | 0 | 0 | 0 | 0 | 0 | 0 | 0 | 0 | 0 |
| 0 | NA | NA | NA | NA | NA | Gm24029 | | 0 | 0 | 0 | 0 | 0 | 0 | 0 | 0 | 0 | 0 | 0 |
| 0 | NA | NA | NA | NA | NA | Gm22988 | | 0 | 0 | 0 | 0 | 0 | 0 | 0 | 0 | 0 | 0 | 0 |
| 0 | NA | NA | NA | NA | NA | Gm12307 | | 0 | 0 | 0 | 0 | 0 | 0 | 0 | 0 | 0 | 0 | 0 |
| 0 | NA | NA | NA | NA | NA | Gm39566 | predicted gene, 39566 | 0 | 0 | 0 | 0 | 0 | 0 | 0 | 0 | 0 | 0 | 0 |
| 0 | NA | NA | NA | NA | NA | Spem2 | SPEM family member 2 | 0 | 0 | 0 | 0 | 0 | 0 | 0 | 0 | 0 | 0 | 0 |
| 0 | NA | NA | NA | NA | NA | Mir6924 | | 0 | 0 | 0 | 0 | 0 | 0 | 0 | 0 | 0 | 0 | 0 |
| 0 | NA | NA | NA | NA | NA | Gm12310 | | 0 | 0 | 0 | 0 | 0 | 0 | 0 | 0 | 0 | 0 | 0 |
| 0 | NA | NA | NA | NA | NA | Mir324 | | 0 | 0 | 0 | 0 | 0 | 0 | 0 | 0 | 0 | 0 | 0 |
| 0 | NA | NA | NA | NA | NA | Gm12311 | | 0 | 0 | 0 | 0 | 0 | 0 | 0 | 0 | 0 | 0 | 0 |
| 0 | NA | NA | NA | NA | NA | Mir497b | | 0 | 0 | 0 | 0 | 0 | 0 | 0 | 0 | 0 | 0 | 0 |
| 0 | NA | NA | NA | NA | NA | Mir497 | | 0 | 0 | 0 | 0 | 0 | 0 | 0 | 0 | 0 | 0 | 0 |
| 0 | NA | NA | NA | NA | NA | Mir195a | | 0 | 0 | 0 | 0 | 0 | 0 | 0 | 0 | 0 | 0 | 0 |
| 0 | NA | NA | NA | NA | NA | Gm12313 | | 0 | 0 | 0 | 0 | 0 | 0 | 0 | 0 | 0 | 0 | 0 |
| 0 | NA | NA | NA | NA | NA | Gm12314 | | 0 | 0 | 0 | 0 | 0 | 0 | 0 | 0 | 0 | 0 | 0 |
| 0 | NA | NA | NA | NA | NA | Mir7115 | | 0 | 0 | 0 | 0 | 0 | 0 | 0 | 0 | 0 | 0 | 0 |
| 0 | NA | NA | NA | NA | NA | Gm12317 | | 0 | 0 | 0 | 0 | 0 | 0 | 0 | 0 | 0 | 0 | 0 |
| 0 | NA | NA | NA | NA | NA | Gm40193 | | 0 | 0 | 0 | 0 | 0 | 0 | 0 | 0 | 0 | 0 | 0 |
| 0 | NA | NA | NA | NA | NA | Gm10418 | | 0 | 0 | 0 | 0 | 0 | 0 | 0 | 0 | 0 | 0 | 0 |
| 0 | NA | NA | NA | NA | NA | Chrne | cholinergic receptor, nicotinic, epsilon pol | 0 | 0 | 0 | 0 | 0 | 0 | 0 | 0 | 0 | 0 | 0 |
| 0 | NA | NA | NA | NA | NA | 4930544D05Ril | RIKEN cDNA 4930544D05 gene | 0 | 0 | 0 | 0 | 0 | 0 | 0 | 0 | 0 | 0 | 0 |
| 0 | NA | NA | NA | NA | NA | Gm12315 | | 0 | 0 | 0 | 0 | 0 | 0 | 0 | 0 | 0 | 0 | 0 |
| 0 | NA | NA | NA | NA | NA | Mir6925 | | 0 | 0 | 0 | 0 | 0 | 0 | 0 | 0 | 0 | 0 | 0 |
| 0 | NA | NA | NA | NA | NA | Gm12320 | | 0 | 0 | 0 | 0 | 0 | 0 | 0 | 0 | 0 | 0 | 0 |
| 0 | NA | NA | NA | NA | NA | Gm22760 | | 0 | 0 | 0 | 0 | 0 | 0 | 0 | 0 | 0 | 0 | 0 |
| 0 | NA | NA | NA | NA | NA | Rpl23a-ps2 | | 0 | 0 | 0 | 0 | 0 | 0 | 0 | 0 | 0 | 0 | 0 |
| 0 | NA | NA | NA | NA | NA | Gm12321 | | 0 | 0 | 0 | 0 | 0 | 0 | 0 | 0 | 0 | 0 | 0 |
| 0 | NA | NA | NA | NA | NA | Gm15377 | | 0 | 0 | 0 | 0 | 0 | 0 | 0 | 0 | 0 | 0 | 0 |
| 0 | NA | NA | NA | NA | NA | Gm23266 | | 0 | 0 | 0 | 0 | 0 | 0 | 0 | 0 | 0 | 0 | 0 |
| 0 | NA | NA | NA | NA | NA | Gm23311 | | 0 | 0 | 0 | 0 | 0 | 0 | 0 | 0 | 0 | 0 | 0 |
| 0 | NA | NA | NA | NA | NA | Gm16013 | | 0 | 0 | 0 | 0 | 0 | 0 | 0 | 0 | 0 | 0 | 0 |
| 0 | NA | NA | NA | NA | NA | Gm33351 | | 0 | 0 | 0 | 0 | 0 | 0 | 0 | 0 | 0 | 0 | 0 |

| | | | | | | | | | | | | | | | | | | |
|---|---|---|---|---|---|---|---|---|---|---|---|---|---|---|---|---|---|---|
| 0 | NA | NA | NA | NA | NA | Gm12324 | | 0 | 0 | 0 | 0 | 0 | 0 | 0 | 0 | 0 | 0 | 0 |
| 0 | NA | NA | NA | NA | NA | Gm48818 | | 0 | 0 | 0 | 0 | 0 | 0 | 0 | 0 | 0 | 0 | 0 |
| 0 | NA | NA | NA | NA | NA | Gm12325 | | 0 | 0 | 0 | 0 | 0 | 0 | 0 | 0 | 0 | 0 | 0 |
| 0 | NA | NA | NA | NA | NA | Gm12326 | | 0 | 0 | 0 | 0 | 0 | 0 | 0 | 0 | 0 | 0 | 0 |
| 0 | NA | NA | NA | NA | NA | Gm22927 | | 0 | 0 | 0 | 0 | 0 | 0 | 0 | 0 | 0 | 0 | 0 |
| 0 | NA | NA | NA | NA | NA | 4930401O10Rik | | 0 | 0 | 0 | 0 | 0 | 0 | 0 | 0 | 0 | 0 | 0 |
| 0 | NA | NA | NA | NA | NA | Gm23226 | | 0 | 0 | 0 | 0 | 0 | 0 | 0 | 0 | 0 | 0 | 0 |
| 0 | NA | NA | NA | NA | NA | 1700051A21Rik | | 0 | 0 | 0 | 0 | 0 | 0 | 0 | 0 | 0 | 0 | 0 |
| 0 | NA | NA | NA | NA | NA | n-R5s70 | | 0 | 0 | 0 | 0 | 0 | 0 | 0 | 0 | 0 | 0 | 0 |
| 0 | NA | NA | NA | NA | NA | Fbxo39 | F-box protein 39 | 0 | 0 | 0 | 0 | 0 | 0 | 0 | 0 | 0 | 0 | 0 |
| 0 | NA | NA | NA | NA | NA | Gm23224 | | 0 | 0 | 0 | 0 | 0 | 0 | 0 | 0 | 0 | 0 | 0 |
| 0 | NA | NA | NA | NA | NA | Mir6338 | | 0 | 0 | 0 | 0 | 0 | 0 | 0 | 0 | 0 | 0 | 0 |
| 0 | NA | NA | NA | NA | NA | Gm24143 | | 0 | 0 | 0 | 0 | 0 | 0 | 0 | 0 | 0 | 0 | 0 |
| 0 | NA | NA | NA | NA | NA | Gm25678 | | 0 | 0 | 0 | 0 | 0 | 0 | 0 | 0 | 0 | 0 | 0 |
| 0 | NA | NA | NA | NA | NA | Gm22075 | | 0 | 0 | 0 | 0 | 0 | 0 | 0 | 0 | 0 | 0 | 0 |
| 0 | NA | NA | NA | NA | NA | Gm44467 | | 0 | 0 | 0 | 0 | 0 | 0 | 0 | 0 | 0 | 0 | 0 |
| 0 | NA | NA | NA | NA | NA | Gm16021 | | 0 | 0 | 0 | 0 | 0 | 0 | 0 | 0 | 0 | 0 | 0 |
| 0 | NA | NA | NA | NA | NA | Spata22 | spermatogenesis associated 22 | 0 | 0 | 0 | 0 | 0 | 0 | 0 | 0 | 0 | 0 | 0 |
| 0 | NA | NA | NA | NA | NA | Gm49340 | | 0 | 0 | 0 | 0 | 0 | 0 | 0 | 0 | 0 | 0 | 0 |
| 0 | NA | NA | NA | NA | NA | Olfr20 | olfactory receptor 20 | 0 | 0 | 0 | 0 | 0 | 0 | 0 | 0 | 0 | 0 | 0 |
| 0 | NA | NA | NA | NA | NA | Olfr376 | olfactory receptor 376 | 0 | 0 | 0 | 0 | 0 | 0 | 0 | 0 | 0 | 0 | 0 |
| 0 | NA | NA | NA | NA | NA | Olfr377-ps1 | | 0 | 0 | 0 | 0 | 0 | 0 | 0 | 0 | 0 | 0 | 0 |
| 0 | NA | NA | NA | NA | NA | Gm17811 | | 0 | 0 | 0 | 0 | 0 | 0 | 0 | 0 | 0 | 0 | 0 |
| 0 | NA | NA | NA | NA | NA | Olfr1 | olfactory receptor 1 | 0 | 0 | 0 | 0 | 0 | 0 | 0 | 0 | 0 | 0 | 0 |
| 0 | NA | NA | NA | NA | NA | Olfr75-ps1 | | 0 | 0 | 0 | 0 | 0 | 0 | 0 | 0 | 0 | 0 | 0 |
| 0 | NA | NA | NA | NA | NA | Gm44922 | | 0 | 0 | 0 | 0 | 0 | 0 | 0 | 0 | 0 | 0 | 0 |
| 0 | NA | NA | NA | NA | NA | Olfr378 | olfactory receptor 378 | 0 | 0 | 0 | 0 | 0 | 0 | 0 | 0 | 0 | 0 | 0 |
| 0 | NA | NA | NA | NA | NA | Olfr379-ps1 | | 0 | 0 | 0 | 0 | 0 | 0 | 0 | 0 | 0 | 0 | 0 |
| 0 | NA | NA | NA | NA | NA | Olfr380 | olfactory receptor 380 | 0 | 0 | 0 | 0 | 0 | 0 | 0 | 0 | 0 | 0 | 0 |
| 0 | NA | NA | NA | NA | NA | Gm44923 | | 0 | 0 | 0 | 0 | 0 | 0 | 0 | 0 | 0 | 0 | 0 |
| 0 | NA | NA | NA | NA | NA | Olfr381 | olfactory receptor 381 | 0 | 0 | 0 | 0 | 0 | 0 | 0 | 0 | 0 | 0 | 0 |
| 0 | NA | NA | NA | NA | NA | Olfr382 | olfactory receptor 382 | 0 | 0 | 0 | 0 | 0 | 0 | 0 | 0 | 0 | 0 | 0 |
| 0 | NA | NA | NA | NA | NA | Olfr383-ps1 | | 0 | 0 | 0 | 0 | 0 | 0 | 0 | 0 | 0 | 0 | 0 |
| 0 | NA | NA | NA | NA | NA | Olfr385 | olfactory receptor 385 | 0 | 0 | 0 | 0 | 0 | 0 | 0 | 0 | 0 | 0 | 0 |
| 0 | NA | NA | NA | NA | NA | Olfr384 | olfactory receptor 384 | 0 | 0 | 0 | 0 | 0 | 0 | 0 | 0 | 0 | 0 | 0 |
| 0 | NA | NA | NA | NA | NA | Gm12327 | | 0 | 0 | 0 | 0 | 0 | 0 | 0 | 0 | 0 | 0 | 0 |
| 0 | NA | NA | NA | NA | NA | Olfr387-ps1 | | 0 | 0 | 0 | 0 | 0 | 0 | 0 | 0 | 0 | 0 | 0 |
| 0 | NA | NA | NA | NA | NA | Zfp735 | zinc finger protein 735 | 0 | 0 | 0 | 0 | 0 | 0 | 0 | 0 | 0 | 0 | 0 |
| 0 | NA | NA | NA | NA | NA | Gm44665 | | 0 | 0 | 0 | 0 | 0 | 0 | 0 | 0 | 0 | 0 | 0 |
| 0 | NA | NA | NA | NA | NA | Olfr388-ps1 | | 0 | 0 | 0 | 0 | 0 | 0 | 0 | 0 | 0 | 0 | 0 |
| 0 | NA | NA | NA | NA | NA | Gm22381 | | 0 | 0 | 0 | 0 | 0 | 0 | 0 | 0 | 0 | 0 | 0 |
| 0 | NA | NA | NA | NA | NA | Olfr389 | olfactory receptor 389 | 0 | 0 | 0 | 0 | 0 | 0 | 0 | 0 | 0 | 0 | 0 |
| 0 | NA | NA | NA | NA | NA | Olfr390 | olfactory receptor 390 | 0 | 0 | 0 | 0 | 0 | 0 | 0 | 0 | 0 | 0 | 0 |
| 0 | NA | NA | NA | NA | NA | Olfr391 | olfactory receptor 391 | 0 | 0 | 0 | 0 | 0 | 0 | 0 | 0 | 0 | 0 | 0 |
| 0 | NA | NA | NA | NA | NA | Olfr392 | olfactory receptor 392 | 0 | 0 | 0 | 0 | 0 | 0 | 0 | 0 | 0 | 0 | 0 |
| 0 | NA | NA | NA | NA | NA | Gm12329 | | 0 | 0 | 0 | 0 | 0 | 0 | 0 | 0 | 0 | 0 | 0 |
| 0 | NA | NA | NA | NA | NA | Olfr393 | olfactory receptor 393 | 0 | 0 | 0 | 0 | 0 | 0 | 0 | 0 | 0 | 0 | 0 |
| 0 | NA | NA | NA | NA | NA | Gm44638 | | 0 | 0 | 0 | 0 | 0 | 0 | 0 | 0 | 0 | 0 | 0 |
| 0 | NA | NA | NA | NA | NA | Olfr394 | olfactory receptor 394 | 0 | 0 | 0 | 0 | 0 | 0 | 0 | 0 | 0 | 0 | 0 |
| 0 | NA | NA | NA | NA | NA | Gm44575 | | 0 | 0 | 0 | 0 | 0 | 0 | 0 | 0 | 0 | 0 | 0 |
| 0 | NA | NA | NA | NA | NA | Olfr395 | olfactory receptor 395 | 0 | 0 | 0 | 0 | 0 | 0 | 0 | 0 | 0 | 0 | 0 |
| 0 | NA | NA | NA | NA | NA | Olfr396-ps1 | | 0 | 0 | 0 | 0 | 0 | 0 | 0 | 0 | 0 | 0 | 0 |
| 0 | NA | NA | NA | NA | NA | Olfr23 | olfactory receptor 23 | 0 | 0 | 0 | 0 | 0 | 0 | 0 | 0 | 0 | 0 | 0 |
| 0 | NA | NA | NA | NA | NA | Olfr22-ps1 | | 0 | 0 | 0 | 0 | 0 | 0 | 0 | 0 | 0 | 0 | 0 |
| 0 | NA | NA | NA | NA | NA | Olfr397 | olfactory receptor 397 | 0 | 0 | 0 | 0 | 0 | 0 | 0 | 0 | 0 | 0 | 0 |
| 0 | NA | NA | NA | NA | NA | Olfr398 | olfactory receptor 398 | 0 | 0 | 0 | 0 | 0 | 0 | 0 | 0 | 0 | 0 | 0 |
| 0 | NA | NA | NA | NA | NA | Olfr139 | olfactory receptor 139 | 0 | 0 | 0 | 0 | 0 | 0 | 0 | 0 | 0 | 0 | 0 |
| 0 | NA | NA | NA | NA | NA | Olfr399 | olfactory receptor 399 | 0 | 0 | 0 | 0 | 0 | 0 | 0 | 0 | 0 | 0 | 0 |
| 0 | NA | NA | NA | NA | NA | Zfp616 | zinc finger protein 616 | 0 | 0 | 0 | 0 | 0 | 0 | 0 | 0 | 0 | 0 | 0 |
| 0 | NA | NA | NA | NA | NA | Olfr400-ps1 | | 0 | 0 | 0 | 0 | 0 | 0 | 0 | 0 | 0 | 0 | 0 |

| | | | | | | | | | | | | | | | | | | |
|---|---|---|---|---|---|---|---|---|---|---|---|---|---|---|---|---|---|---|
| 0 | NA | NA | NA | NA | NA | Olfr401 | olfactory receptor 401 | 0 | 0 | 0 | 0 | 0 | 0 | 0 | 0 | 0 | 0 | 0 |
| 0 | NA | NA | NA | NA | NA | Olfr402 | olfactory receptor 402 | 0 | 0 | 0 | 0 | 0 | 0 | 0 | 0 | 0 | 0 | 0 |
| 0 | NA | NA | NA | NA | NA | Olfr403 | olfactory receptor 403 | 0 | 0 | 0 | 0 | 0 | 0 | 0 | 0 | 0 | 0 | 0 |
| 0 | NA | NA | NA | NA | NA | Olfr43 | olfactory receptor 43 | 0 | 0 | 0 | 0 | 0 | 0 | 0 | 0 | 0 | 0 | 0 |
| 0 | NA | NA | NA | NA | NA | Olfr404-ps1 | | 0 | 0 | 0 | 0 | 0 | 0 | 0 | 0 | 0 | 0 | 0 |
| 0 | NA | NA | NA | NA | NA | Olfr405-ps1 | | 0 | 0 | 0 | 0 | 0 | 0 | 0 | 0 | 0 | 0 | 0 |
| 0 | NA | NA | NA | NA | NA | Olfr406 | olfactory receptor 406 | 0 | 0 | 0 | 0 | 0 | 0 | 0 | 0 | 0 | 0 | 0 |
| 0 | NA | NA | NA | NA | NA | Olfr407-ps1 | | 0 | 0 | 0 | 0 | 0 | 0 | 0 | 0 | 0 | 0 | 0 |
| 0 | NA | NA | NA | NA | NA | Olfr59 | olfactory receptor 59 | 0 | 0 | 0 | 0 | 0 | 0 | 0 | 0 | 0 | 0 | 0 |
| 0 | NA | NA | NA | NA | NA | Olfr408-ps1 | | 0 | 0 | 0 | 0 | 0 | 0 | 0 | 0 | 0 | 0 | 0 |
| 0 | NA | NA | NA | NA | NA | Olfr409-ps1 | | 0 | 0 | 0 | 0 | 0 | 0 | 0 | 0 | 0 | 0 | 0 |
| 0 | NA | NA | NA | NA | NA | n-R5s71 | | 0 | 0 | 0 | 0 | 0 | 0 | 0 | 0 | 0 | 0 | 0 |
| 0 | NA | NA | NA | NA | NA | Gm12332 | | 0 | 0 | 0 | 0 | 0 | 0 | 0 | 0 | 0 | 0 | 0 |
| 0 | NA | NA | NA | NA | NA | Olfr410 | olfactory receptor 410 | 0 | 0 | 0 | 0 | 0 | 0 | 0 | 0 | 0 | 0 | 0 |
| 0 | NA | NA | NA | NA | NA | Olfr411 | olfactory receptor 411 | 0 | 0 | 0 | 0 | 0 | 0 | 0 | 0 | 0 | 0 | 0 |
| 0 | NA | NA | NA | NA | NA | Olfr412 | olfactory receptor 412 | 0 | 0 | 0 | 0 | 0 | 0 | 0 | 0 | 0 | 0 | 0 |
| 0 | NA | NA | NA | NA | NA | Gm22993 | | 0 | 0 | 0 | 0 | 0 | 0 | 0 | 0 | 0 | 0 | 0 |
| 0 | NA | NA | NA | NA | NA | Gm24211 | | 0 | 0 | 0 | 0 | 0 | 0 | 0 | 0 | 0 | 0 | 0 |
| 0 | NA | NA | NA | NA | NA | Gm16032 | | 0 | 0 | 0 | 0 | 0 | 0 | 0 | 0 | 0 | 0 | 0 |
| 0 | NA | NA | NA | NA | NA | Mir6926 | | 0 | 0 | 0 | 0 | 0 | 0 | 0 | 0 | 0 | 0 | 0 |
| 0 | NA | NA | NA | NA | NA | Gm22733 | | 0 | 0 | 0 | 0 | 0 | 0 | 0 | 0 | 0 | 0 | 0 |
| 0 | NA | NA | NA | NA | NA | Gm28180 | | 0 | 0 | 0 | 0 | 0 | 0 | 0 | 0 | 0 | 0 | 0 |
| 0 | NA | NA | NA | NA | NA | Gm12333 | | 0 | 0 | 0 | 0 | 0 | 0 | 0 | 0 | 0 | 0 | 0 |
| 0 | NA | NA | NA | NA | NA | Gm47300 | | 0 | 0 | 0 | 0 | 0 | 0 | 0 | 0 | 0 | 0 | 0 |
| 0 | NA | NA | NA | NA | NA | Gm12336 | | 0 | 0 | 0 | 0 | 0 | 0 | 0 | 0 | 0 | 0 | 0 |
| 0 | NA | NA | NA | NA | NA | Mir22 | | 0 | 0 | 0 | 0 | 0 | 0 | 0 | 0 | 0 | 0 | 0 |
| 0 | NA | NA | NA | NA | NA | 4931413K12Rik | | 0 | 0 | 0 | 0 | 0 | 0 | 0 | 0 | 0 | 0 | 0 |
| 0 | NA | NA | NA | NA | NA | Gm16075 | | 0 | 0 | 0 | 0 | 0 | 0 | 0 | 0 | 0 | 0 | 0 |
| 0 | NA | NA | NA | NA | NA | Gm12340 | | 0 | 0 | 0 | 0 | 0 | 0 | 0 | 0 | 0 | 0 | 0 |
| 0 | NA | NA | NA | NA | NA | Gm12343 | | 0 | 0 | 0 | 0 | 0 | 0 | 0 | 0 | 0 | 0 | 0 |
| 0 | NA | NA | NA | NA | NA | Gm22772 | | 0 | 0 | 0 | 0 | 0 | 0 | 0 | 0 | 0 | 0 | 0 |
| 0 | NA | NA | NA | NA | NA | Gm24989 | | 0 | 0 | 0 | 0 | 0 | 0 | 0 | 0 | 0 | 0 | 0 |
| 0 | NA | NA | NA | NA | NA | Gm23132 | | 0 | 0 | 0 | 0 | 0 | 0 | 0 | 0 | 0 | 0 | 0 |
| 0 | NA | NA | NA | NA | NA | Cryba1 | crystallin, beta A1 | 0 | 0 | 0 | 0 | 0 | 0 | 0 | 0 | 0 | 0 | 0 |
| 0 | NA | NA | NA | NA | NA | Gm10277 | predicted gene 10277 | 0 | 0 | 0 | 0 | 0 | 0 | 0 | 0 | 0 | 0 | 0 |
| 0 | NA | NA | NA | NA | NA | Gm11190 | | 0 | 0 | 0 | 0 | 0 | 0 | 0 | 0 | 0 | 0 | 0 |
| 0 | NA | NA | NA | NA | NA | Mir144 | | 0 | 0 | 0 | 0 | 0 | 0 | 0 | 0 | 0 | 0 | 0 |
| 0 | NA | NA | NA | NA | NA | Mir451b | | 0 | 0 | 0 | 0 | 0 | 0 | 0 | 0 | 0 | 0 | 0 |
| 0 | NA | NA | NA | NA | NA | Mir451a | | 0 | 0 | 0 | 0 | 0 | 0 | 0 | 0 | 0 | 0 | 0 |
| 0 | NA | NA | NA | NA | NA | Mir7653 | | 0 | 0 | 0 | 0 | 0 | 0 | 0 | 0 | 0 | 0 | 0 |
| 0 | NA | NA | NA | NA | NA | Snord42a | | 0 | 0 | 0 | 0 | 0 | 0 | 0 | 0 | 0 | 0 | 0 |
| 0 | NA | NA | NA | NA | NA | Snord4a | | 0 | 0 | 0 | 0 | 0 | 0 | 0 | 0 | 0 | 0 | 0 |
| 0 | NA | NA | NA | NA | NA | Gm11192 | | 0 | 0 | 0 | 0 | 0 | 0 | 0 | 0 | 0 | 0 | 0 |
| 0 | NA | NA | NA | NA | NA | Slc13a2os | | 0 | 0 | 0 | 0 | 0 | 0 | 0 | 0 | 0 | 0 | 0 |
| 0 | NA | NA | NA | NA | NA | Gm11195 | | 0 | 0 | 0 | 0 | 0 | 0 | 0 | 0 | 0 | 0 | 0 |
| 0 | NA | NA | NA | NA | NA | D11Bhm181e | | 0 | 0 | 0 | 0 | 0 | 0 | 0 | 0 | 0 | 0 | 0 |
| 0 | NA | NA | NA | NA | NA | Gm22947 | | 0 | 0 | 0 | 0 | 0 | 0 | 0 | 0 | 0 | 0 | 0 |
| 0 | NA | NA | NA | NA | NA | Gm17235 | | 0 | 0 | 0 | 0 | 0 | 0 | 0 | 0 | 0 | 0 | 0 |
| 0 | NA | NA | NA | NA | NA | Gm11197 | | 0 | 0 | 0 | 0 | 0 | 0 | 0 | 0 | 0 | 0 | 0 |
| 0 | NA | NA | NA | NA | NA | Gm23840 | | 0 | 0 | 0 | 0 | 0 | 0 | 0 | 0 | 0 | 0 | 0 |
| 0 | NA | NA | NA | NA | NA | Gm23219 | | 0 | 0 | 0 | 0 | 0 | 0 | 0 | 0 | 0 | 0 | 0 |
| 0 | NA | NA | NA | NA | NA | Gm11201 | | 0 | 0 | 0 | 0 | 0 | 0 | 0 | 0 | 0 | 0 | 0 |
| 0 | NA | NA | NA | NA | NA | Gm11200 | | 0 | 0 | 0 | 0 | 0 | 0 | 0 | 0 | 0 | 0 | 0 |
| 0 | NA | NA | NA | NA | NA | Gm9964 | | 0 | 0 | 0 | 0 | 0 | 0 | 0 | 0 | 0 | 0 | 0 |
| 0 | NA | NA | NA | NA | NA | Gm11199 | | 0 | 0 | 0 | 0 | 0 | 0 | 0 | 0 | 0 | 0 | 0 |
| 0 | NA | NA | NA | NA | NA | AU040972 | | 0 | 0 | 0 | 0 | 0 | 0 | 0 | 0 | 0 | 0 | 0 |
| 0 | NA | NA | NA | NA | NA | Evi2b | ecotropic viral integration site 2b | 0 | 0 | 0 | 0 | 0 | 0 | 0 | 0 | 0 | 0 | 0 |
| 0 | NA | NA | NA | NA | NA | Gm23293 | | 0 | 0 | 0 | 0 | 0 | 0 | 0 | 0 | 0 | 0 | 0 |
| 0 | NA | NA | NA | NA | NA | Gm25867 | | 0 | 0 | 0 | 0 | 0 | 0 | 0 | 0 | 0 | 0 | 0 |
| 0 | NA | NA | NA | NA | NA | Gm24887 | | 0 | 0 | 0 | 0 | 0 | 0 | 0 | 0 | 0 | 0 | 0 |

| | | | | | | | | | | | | | | | | | | |
|---|---|---|---|---|---|---|---|---|---|---|---|---|---|---|---|---|---|---|
| 0 | NA | NA | NA | NA | NA | Rab11fip4os2 | | 0 | 0 | 0 | 0 | 0 | 0 | 0 | 0 | 0 | 0 | 0 |
| 0 | NA | NA | NA | NA | NA | Mir193a | | 0 | 0 | 0 | 0 | 0 | 0 | 0 | 0 | 0 | 0 | 0 |
| 0 | NA | NA | NA | NA | NA | Mir365-2 | | 0 | 0 | 0 | 0 | 0 | 0 | 0 | 0 | 0 | 0 | 0 |
| 0 | NA | NA | NA | NA | NA | 9130204K15Rik | | 0 | 0 | 0 | 0 | 0 | 0 | 0 | 0 | 0 | 0 | 0 |
| 0 | NA | NA | NA | NA | NA | Gm24454 | | 0 | 0 | 0 | 0 | 0 | 0 | 0 | 0 | 0 | 0 | 0 |
| 0 | NA | NA | NA | NA | NA | Gm11203 | | 0 | 0 | 0 | 0 | 0 | 0 | 0 | 0 | 0 | 0 | 0 |
| 0 | NA | NA | NA | NA | NA | Adap2os | | 0 | 0 | 0 | 0 | 0 | 0 | 0 | 0 | 0 | 0 | 0 |
| 0 | NA | NA | NA | NA | NA | Gm11207 | | 0 | 0 | 0 | 0 | 0 | 0 | 0 | 0 | 0 | 0 | 0 |
| 0 | NA | NA | NA | NA | NA | 4930507D10Rik | | 0 | 0 | 0 | 0 | 0 | 0 | 0 | 0 | 0 | 0 | 0 |
| 0 | NA | NA | NA | NA | NA | Spaca3 | sperm acrosome associated 3 | 0 | 0 | 0 | 0 | 0 | 0 | 0 | 0 | 0 | 0 | 0 |
| 0 | NA | NA | NA | NA | NA | Gm11416 | | 0 | 0 | 0 | 0 | 0 | 0 | 0 | 0 | 0 | 0 | 0 |
| 0 | NA | NA | NA | NA | NA | 4930527B05Rik | | 0 | 0 | 0 | 0 | 0 | 0 | 0 | 0 | 0 | 0 | 0 |
| 0 | NA | NA | NA | NA | NA | 1700071K01Ril | RIKEN cDNA 1700071K01 gene | 0 | 0 | 0 | 0 | 0 | 0 | 0 | 0 | 0 | 0 | 0 |
| 0 | NA | NA | NA | NA | NA | Gm11418 | | 0 | 0 | 0 | 0 | 0 | 0 | 0 | 0 | 0 | 0 | 0 |
| 0 | NA | NA | NA | NA | NA | Gm31522 | | 0 | 0 | 0 | 0 | 0 | 0 | 0 | 0 | 0 | 0 | 0 |
| 0 | NA | NA | NA | NA | NA | Gm17268 | | 0 | 0 | 0 | 0 | 0 | 0 | 0 | 0 | 0 | 0 | 0 |
| 0 | NA | NA | NA | NA | NA | Gm38577 | | 0 | 0 | 0 | 0 | 0 | 0 | 0 | 0 | 0 | 0 | 0 |
| 0 | NA | NA | NA | NA | NA | Gm24612 | | 0 | 0 | 0 | 0 | 0 | 0 | 0 | 0 | 0 | 0 | 0 |
| 0 | NA | NA | NA | NA | NA | Gm11426 | | 0 | 0 | 0 | 0 | 0 | 0 | 0 | 0 | 0 | 0 | 0 |
| 0 | NA | NA | NA | NA | NA | Gm11421 | | 0 | 0 | 0 | 0 | 0 | 0 | 0 | 0 | 0 | 0 | 0 |
| 0 | NA | NA | NA | NA | NA | Gm11422 | | 0 | 0 | 0 | 0 | 0 | 0 | 0 | 0 | 0 | 0 | 0 |
| 0 | NA | NA | NA | NA | NA | Mir7679 | | 0 | 0 | 0 | 0 | 0 | 0 | 0 | 0 | 0 | 0 | 0 |
| 0 | NA | NA | NA | NA | NA | Gm27598 | | 0 | 0 | 0 | 0 | 0 | 0 | 0 | 0 | 0 | 0 | 0 |
| 0 | NA | NA | NA | NA | NA | Gm48434 | | 0 | 0 | 0 | 0 | 0 | 0 | 0 | 0 | 0 | 0 | 0 |
| 0 | NA | NA | NA | NA | NA | Gm20234 | | 0 | 0 | 0 | 0 | 0 | 0 | 0 | 0 | 0 | 0 | 0 |
| 0 | NA | NA | NA | NA | NA | Gm48462 | | 0 | 0 | 0 | 0 | 0 | 0 | 0 | 0 | 0 | 0 | 0 |
| 0 | NA | NA | NA | NA | NA | Gm48467 | | 0 | 0 | 0 | 0 | 0 | 0 | 0 | 0 | 0 | 0 | 0 |
| 0 | NA | NA | NA | NA | NA | Gm48468 | | 0 | 0 | 0 | 0 | 0 | 0 | 0 | 0 | 0 | 0 | 0 |
| 0 | NA | NA | NA | NA | NA | Gm48469 | | 0 | 0 | 0 | 0 | 0 | 0 | 0 | 0 | 0 | 0 | 0 |
| 0 | NA | NA | NA | NA | NA | Gm48477 | | 0 | 0 | 0 | 0 | 0 | 0 | 0 | 0 | 0 | 0 | 0 |
| 0 | NA | NA | NA | NA | NA | Gm48490 | | 0 | 0 | 0 | 0 | 0 | 0 | 0 | 0 | 0 | 0 | 0 |
| 0 | NA | NA | NA | NA | NA | Gm23951 | | 0 | 0 | 0 | 0 | 0 | 0 | 0 | 0 | 0 | 0 | 0 |
| 0 | NA | NA | NA | NA | NA | Gas2l2 | growth arrest-specific 2 like 2 | 0 | 0 | 0 | 0 | 0 | 0 | 0 | 0 | 0 | 0 | 0 |
| 0 | NA | NA | NA | NA | NA | Heatr9 | HEAT repeat containing 9 | 0 | 0 | 0 | 0 | 0 | 0 | 0 | 0 | 0 | 0 | 0 |
| 0 | NA | NA | NA | NA | NA | Gm23247 | | 0 | 0 | 0 | 0 | 0 | 0 | 0 | 0 | 0 | 0 | 0 |
| 0 | NA | NA | NA | NA | NA | Gm11433 | | 0 | 0 | 0 | 0 | 0 | 0 | 0 | 0 | 0 | 0 | 0 |
| 0 | NA | NA | NA | NA | NA | Gm11438 | | 0 | 0 | 0 | 0 | 0 | 0 | 0 | 0 | 0 | 0 | 0 |
| 0 | NA | NA | NA | NA | NA | Gm11432 | | 0 | 0 | 0 | 0 | 0 | 0 | 0 | 0 | 0 | 0 | 0 |
| 0 | NA | NA | NA | NA | NA | Gm11430 | | 0 | 0 | 0 | 0 | 0 | 0 | 0 | 0 | 0 | 0 | 0 |
| 0 | NA | NA | NA | NA | NA | Gm11429 | | 0 | 0 | 0 | 0 | 0 | 0 | 0 | 0 | 0 | 0 | 0 |
| 0 | NA | NA | NA | NA | NA | Wfdc21 | WAP four-disulfide core domain 21 | 0 | 0 | 0 | 0 | 0 | 0 | 0 | 0 | 0 | 0 | 0 |
| 0 | NA | NA | NA | NA | NA | Gm22311 | | 0 | 0 | 0 | 0 | 0 | 0 | 0 | 0 | 0 | 0 | 0 |
| 0 | NA | NA | NA | NA | NA | Gm23211 | | 0 | 0 | 0 | 0 | 0 | 0 | 0 | 0 | 0 | 0 | 0 |
| 0 | NA | NA | NA | NA | NA | Gm37391 | | 0 | 0 | 0 | 0 | 0 | 0 | 0 | 0 | 0 | 0 | 0 |
| 0 | NA | NA | NA | NA | NA | 1700109G15Rik | | 0 | 0 | 0 | 0 | 0 | 0 | 0 | 0 | 0 | 0 | 0 |
| 0 | NA | NA | NA | NA | NA | Gm23506 | | 0 | 0 | 0 | 0 | 0 | 0 | 0 | 0 | 0 | 0 | 0 |
| 0 | NA | NA | NA | NA | NA | 1700125H20Ril | RIKEN cDNA 1700125H20 gene | 0 | 0 | 0 | 0 | 0 | 0 | 0 | 0 | 0 | 0 | 0 |
| 0 | NA | NA | NA | NA | NA | Gm22587 | | 0 | 0 | 0 | 0 | 0 | 0 | 0 | 0 | 0 | 0 | 0 |
| 0 | NA | NA | NA | NA | NA | Gm23765 | | 0 | 0 | 0 | 0 | 0 | 0 | 0 | 0 | 0 | 0 | 0 |
| 0 | NA | NA | NA | NA | NA | Bcas3os1 | | 0 | 0 | 0 | 0 | 0 | 0 | 0 | 0 | 0 | 0 | 0 |
| 0 | NA | NA | NA | NA | NA | Mir5110 | | 0 | 0 | 0 | 0 | 0 | 0 | 0 | 0 | 0 | 0 | 0 |
| 0 | NA | NA | NA | NA | NA | Gm23473 | | 0 | 0 | 0 | 0 | 0 | 0 | 0 | 0 | 0 | 0 | 0 |
| 0 | NA | NA | NA | NA | NA | Gm11444 | | 0 | 0 | 0 | 0 | 0 | 0 | 0 | 0 | 0 | 0 | 0 |
| 0 | NA | NA | NA | NA | NA | Gm25427 | | 0 | 0 | 0 | 0 | 0 | 0 | 0 | 0 | 0 | 0 | 0 |
| 0 | NA | NA | NA | NA | NA | Gm23667 | | 0 | 0 | 0 | 0 | 0 | 0 | 0 | 0 | 0 | 0 | 0 |
| 0 | NA | NA | NA | NA | NA | Gm26439 | | 0 | 0 | 0 | 0 | 0 | 0 | 0 | 0 | 0 | 0 | 0 |
| 0 | NA | NA | NA | NA | NA | Mir21a | | 0 | 0 | 0 | 0 | 0 | 0 | 0 | 0 | 0 | 0 | 0 |
| 0 | NA | NA | NA | NA | NA | Gm22387 | | 0 | 0 | 0 | 0 | 0 | 0 | 0 | 0 | 0 | 0 | 0 |
| 0 | NA | NA | NA | NA | NA | Gm11479 | | 0 | 0 | 0 | 0 | 0 | 0 | 0 | 0 | 0 | 0 | 0 |
| 0 | NA | NA | NA | NA | NA | Gm11490 | | 0 | 0 | 0 | 0 | 0 | 0 | 0 | 0 | 0 | 0 | 0 |

| | | | | | | | | | | | | | | | | | | |
|---|---|---|---|---|---|---|---|---|---|---|---|---|---|---|---|---|---|---|
| 0 | NA | NA | NA | NA | NA | Gm22883 | | 0 | 0 | 0 | 0 | 0 | 0 | 0 | 0 | 0 | 0 | 0 |
| 0 | NA | NA | NA | NA | NA | Gm22068 | | 0 | 0 | 0 | 0 | 0 | 0 | 0 | 0 | 0 | 0 | 0 |
| 0 | NA | NA | NA | NA | NA | Rnu3b1 | | 0 | 0 | 0 | 0 | 0 | 0 | 0 | 0 | 0 | 0 | 0 |
| 0 | NA | NA | NA | NA | NA | Rnu3b1 | | 0 | 0 | 0 | 0 | 0 | 0 | 0 | 0 | 0 | 0 | 0 |
| 0 | NA | NA | NA | NA | NA | Gm23137 | | 0 | 0 | 0 | 0 | 0 | 0 | 0 | 0 | 0 | 0 | 0 |
| 0 | NA | NA | NA | NA | NA | Rnu3b3 | | 0 | 0 | 0 | 0 | 0 | 0 | 0 | 0 | 0 | 0 | 0 |
| 0 | NA | NA | NA | NA | NA | Rnu3b3 | | 0 | 0 | 0 | 0 | 0 | 0 | 0 | 0 | 0 | 0 | 0 |
| 0 | NA | NA | NA | NA | NA | Rnu3b4 | | 0 | 0 | 0 | 0 | 0 | 0 | 0 | 0 | 0 | 0 | 0 |
| 0 | NA | NA | NA | NA | NA | Rnu3b4 | | 0 | 0 | 0 | 0 | 0 | 0 | 0 | 0 | 0 | 0 | 0 |
| 0 | NA | NA | NA | NA | NA | Gm11489 | | 0 | 0 | 0 | 0 | 0 | 0 | 0 | 0 | 0 | 0 | 0 |
| 0 | NA | NA | NA | NA | NA | Rnu3b2 | | 0 | 0 | 0 | 0 | 0 | 0 | 0 | 0 | 0 | 0 | 0 |
| 0 | NA | NA | NA | NA | NA | Rnu3b2 | | 0 | 0 | 0 | 0 | 0 | 0 | 0 | 0 | 0 | 0 | 0 |
| 0 | NA | NA | NA | NA | NA | Gm25968 | | 0 | 0 | 0 | 0 | 0 | 0 | 0 | 0 | 0 | 0 | 0 |
| 0 | NA | NA | NA | NA | NA | 1110028F11Rik | | 0 | 0 | 0 | 0 | 0 | 0 | 0 | 0 | 0 | 0 | 0 |
| 0 | NA | NA | NA | NA | NA | Mir142b | | 0 | 0 | 0 | 0 | 0 | 0 | 0 | 0 | 0 | 0 | 0 |
| 0 | NA | NA | NA | NA | NA | Mpo | myeloperoxidase | 0 | 0 | 0 | 0 | 0 | 0 | 0 | 0 | 0 | 0 | 0 |
| 0 | NA | NA | NA | NA | NA | Gm11505 | | 0 | 0 | 0 | 0 | 0 | 0 | 0 | 0 | 0 | 0 | 0 |
| 0 | NA | NA | NA | NA | NA | Epx | eosinophil peroxidase | 0 | 0 | 0 | 0 | 0 | 0 | 0 | 0 | 0 | 0 | 0 |
| 0 | NA | NA | NA | NA | NA | Olfr462 | olfactory receptor 462 | 0 | 0 | 0 | 0 | 0 | 0 | 0 | 0 | 0 | 0 | 0 |
| 0 | NA | NA | NA | NA | NA | Olfr463 | olfactory receptor 463 | 0 | 0 | 0 | 0 | 0 | 0 | 0 | 0 | 0 | 0 | 0 |
| 0 | NA | NA | NA | NA | NA | Olfr464 | olfactory receptor 464 | 0 | 0 | 0 | 0 | 0 | 0 | 0 | 0 | 0 | 0 | 0 |
| 0 | NA | NA | NA | NA | NA | Gm11504 | | 0 | 0 | 0 | 0 | 0 | 0 | 0 | 0 | 0 | 0 | 0 |
| 0 | NA | NA | NA | NA | NA | Gm11507 | | 0 | 0 | 0 | 0 | 0 | 0 | 0 | 0 | 0 | 0 | 0 |
| 0 | NA | NA | NA | NA | NA | Gm38534 | | 0 | 0 | 0 | 0 | 0 | 0 | 0 | 0 | 0 | 0 | 0 |
| 0 | NA | NA | NA | NA | NA | Ccdc182 | coiled-coil domain containing 182 | 0 | 0 | 0 | 0 | 0 | 0 | 0 | 0 | 0 | 0 | 0 |
| 0 | NA | NA | NA | NA | NA | Mir378b | | 0 | 0 | 0 | 0 | 0 | 0 | 0 | 0 | 0 | 0 | 0 |
| 0 | NA | NA | NA | NA | NA | Gm11493 | | 0 | 0 | 0 | 0 | 0 | 0 | 0 | 0 | 0 | 0 | 0 |
| 0 | NA | NA | NA | NA | NA | 4930556N13Rik | | 0 | 0 | 0 | 0 | 0 | 0 | 0 | 0 | 0 | 0 | 0 |
| 0 | NA | NA | NA | NA | NA | Gm23507 | | 0 | 0 | 0 | 0 | 0 | 0 | 0 | 0 | 0 | 0 | 0 |
| 0 | NA | NA | NA | NA | NA | 2210409E12Rik | | 0 | 0 | 0 | 0 | 0 | 0 | 0 | 0 | 0 | 0 | 0 |
| 0 | NA | NA | NA | NA | NA | Gm22534 | | 0 | 0 | 0 | 0 | 0 | 0 | 0 | 0 | 0 | 0 | 0 |
| 0 | NA | NA | NA | NA | NA | Gm11497 | | 0 | 0 | 0 | 0 | 0 | 0 | 0 | 0 | 0 | 0 | 0 |
| 0 | NA | NA | NA | NA | NA | Gm24734 | | 0 | 0 | 0 | 0 | 0 | 0 | 0 | 0 | 0 | 0 | 0 |
| 0 | NA | NA | NA | NA | NA | Gm2018 | | 0 | 0 | 0 | 0 | 0 | 0 | 0 | 0 | 0 | 0 | 0 |
| 0 | NA | NA | NA | NA | NA | n-R5s72 | | 0 | 0 | 0 | 0 | 0 | 0 | 0 | 0 | 0 | 0 | 0 |
| 0 | NA | NA | NA | NA | NA | Gm45883 | | 0 | 0 | 0 | 0 | 0 | 0 | 0 | 0 | 0 | 0 | 0 |
| 0 | NA | NA | NA | NA | NA | Gm22762 | | 0 | 0 | 0 | 0 | 0 | 0 | 0 | 0 | 0 | 0 | 0 |
| 0 | NA | NA | NA | NA | NA | 4930405D11Rik | | 0 | 0 | 0 | 0 | 0 | 0 | 0 | 0 | 0 | 0 | 0 |
| 0 | NA | NA | NA | NA | NA | Kif2b | kinesin family member 2B | 0 | 0 | 0 | 0 | 0 | 0 | 0 | 0 | 0 | 0 | 0 |
| 0 | NA | NA | NA | NA | NA | Gm11498 | | 0 | 0 | 0 | 0 | 0 | 0 | 0 | 0 | 0 | 0 | 0 |
| 0 | NA | NA | NA | NA | NA | Gm22702 | | 0 | 0 | 0 | 0 | 0 | 0 | 0 | 0 | 0 | 0 | 0 |
| 0 | NA | NA | NA | NA | NA | Gm11500 | | 0 | 0 | 0 | 0 | 0 | 0 | 0 | 0 | 0 | 0 | 0 |
| 0 | NA | NA | NA | NA | NA | Gm11501 | | 0 | 0 | 0 | 0 | 0 | 0 | 0 | 0 | 0 | 0 | 0 |
| 0 | NA | NA | NA | NA | NA | Gm25113 | | 0 | 0 | 0 | 0 | 0 | 0 | 0 | 0 | 0 | 0 | 0 |
| 0 | NA | NA | NA | NA | NA | Gm24856 | | 0 | 0 | 0 | 0 | 0 | 0 | 0 | 0 | 0 | 0 | 0 |
| 0 | NA | NA | NA | NA | NA | Gm11506 | | 0 | 0 | 0 | 0 | 0 | 0 | 0 | 0 | 0 | 0 | 0 |
| 0 | NA | NA | NA | NA | NA | Gm20390 | | 0 | 0 | 0 | 0 | 0 | 0 | 0 | 0 | 0 | 0 | 0 |
| 0 | NA | NA | NA | NA | NA | Gm21885 | predicted gene, 21885 | 0 | 0 | 0 | 0 | 0 | 0 | 0 | 0 | 0 | 0 | 0 |
| 0 | NA | NA | NA | NA | NA | Gm11540 | | 0 | 0 | 0 | 0 | 0 | 0 | 0 | 0 | 0 | 0 | 0 |
| 0 | NA | NA | NA | NA | NA | Gm11541 | predicted gene 11541 | 0 | 0 | 0 | 0 | 0 | 0 | 0 | 0 | 0 | 0 | 0 |
| 0 | NA | NA | NA | NA | NA | Tmem92 | transmembrane protein 92 | 0 | 0 | 0 | 0 | 0 | 0 | 0 | 0 | 0 | 0 | 0 |
| 0 | NA | NA | NA | NA | NA | Gm11546 | | 0 | 0 | 0 | 0 | 0 | 0 | 0 | 0 | 0 | 0 | 0 |
| 0 | NA | NA | NA | NA | NA | AA623943 | | 0 | 0 | 0 | 0 | 0 | 0 | 0 | 0 | 0 | 0 | 0 |
| 0 | NA | NA | NA | NA | NA | Gm11543 | | 0 | 0 | 0 | 0 | 0 | 0 | 0 | 0 | 0 | 0 | 0 |
| 0 | NA | NA | NA | NA | NA | Gm11544 | | 0 | 0 | 0 | 0 | 0 | 0 | 0 | 0 | 0 | 0 | 0 |
| 0 | NA | NA | NA | NA | NA | A430060F13Rik | | 0 | 0 | 0 | 0 | 0 | 0 | 0 | 0 | 0 | 0 | 0 |
| 0 | NA | NA | NA | NA | NA | Gm22456 | | 0 | 0 | 0 | 0 | 0 | 0 | 0 | 0 | 0 | 0 | 0 |
| 0 | NA | NA | NA | NA | NA | Gm24877 | | 0 | 0 | 0 | 0 | 0 | 0 | 0 | 0 | 0 | 0 | 0 |
| 0 | NA | NA | NA | NA | NA | Gm11513 | | 0 | 0 | 0 | 0 | 0 | 0 | 0 | 0 | 0 | 0 | 0 |
| 0 | NA | NA | NA | NA | NA | Dlx4 | distal-less homeobox 4 | 0 | 0 | 0 | 0 | 0 | 0 | 0 | 0 | 0 | 0 | 0 |

| | | | | | | | | | | | | | | | | | | |
|---|---|---|---|---|---|---|---|---|---|---|---|---|---|---|---|---|---|---|
| 0 | NA | NA | NA | NA | NA | Dlx4os | | 0 | 0 | 0 | 0 | 0 | 0 | 0 | 0 | 0 | 0 | 0 |
| 0 | NA | NA | NA | NA | NA | Gm11514 | | 0 | 0 | 0 | 0 | 0 | 0 | 0 | 0 | 0 | 0 | 0 |
| 0 | NA | NA | NA | NA | NA | Tac4 | tachykinin 4 | 0 | 0 | 0 | 0 | 0 | 0 | 0 | 0 | 0 | 0 | 0 |
| 0 | NA | NA | NA | NA | NA | Gm29477 | | 0 | 0 | 0 | 0 | 0 | 0 | 0 | 0 | 0 | 0 | 0 |
| 0 | NA | NA | NA | NA | NA | Gm11522 | | 0 | 0 | 0 | 0 | 0 | 0 | 0 | 0 | 0 | 0 | 0 |
| 0 | NA | NA | NA | NA | NA | Gm11528 | | 0 | 0 | 0 | 0 | 0 | 0 | 0 | 0 | 0 | 0 | 0 |
| 0 | NA | NA | NA | NA | NA | Gm24725 | | 0 | 0 | 0 | 0 | 0 | 0 | 0 | 0 | 0 | 0 | 0 |
| 0 | NA | NA | NA | NA | NA | Mir3063 | | 0 | 0 | 0 | 0 | 0 | 0 | 0 | 0 | 0 | 0 | 0 |
| 0 | NA | NA | NA | NA | NA | Gm29202 | | 0 | 0 | 0 | 0 | 0 | 0 | 0 | 0 | 0 | 0 | 0 |
| 0 | NA | NA | NA | NA | NA | Gm37179 | | 0 | 0 | 0 | 0 | 0 | 0 | 0 | 0 | 0 | 0 | 0 |
| 0 | NA | NA | NA | NA | NA | Gip | gastric inhibitory polypeptide | 0 | 0 | 0 | 0 | 0 | 0 | 0 | 0 | 0 | 0 | 0 |
| 0 | NA | NA | NA | NA | NA | Gm24949 | | 0 | 0 | 0 | 0 | 0 | 0 | 0 | 0 | 0 | 0 | 0 |
| 0 | NA | NA | NA | NA | NA | Gm22423 | | 0 | 0 | 0 | 0 | 0 | 0 | 0 | 0 | 0 | 0 | 0 |
| 0 | NA | NA | NA | NA | NA | Calcoco2 | calcium binding and coiled-coil domain 2 | 0 | 0 | 0 | 0 | 0 | 0 | 0 | 0 | 0 | 0 | 0 |
| 0 | NA | NA | NA | NA | NA | Gm11535 | | 0 | 0 | 0 | 0 | 0 | 0 | 0 | 0 | 0 | 0 | 0 |
| 0 | NA | NA | NA | NA | NA | Gm27898 | | 0 | 0 | 0 | 0 | 0 | 0 | 0 | 0 | 0 | 0 | 0 |
| 0 | NA | NA | NA | NA | NA | Gm27605 | | 0 | 0 | 0 | 0 | 0 | 0 | 0 | 0 | 0 | 0 | 0 |
| 0 | NA | NA | NA | NA | NA | Mir196a-1 | | 0 | 0 | 0 | 0 | 0 | 0 | 0 | 0 | 0 | 0 | 0 |
| 0 | NA | NA | NA | NA | NA | Mir10a | | 0 | 0 | 0 | 0 | 0 | 0 | 0 | 0 | 0 | 0 | 0 |
| 0 | NA | NA | NA | NA | NA | Hoxb1 | homeobox B1 | 0 | 0 | 0 | 0 | 0 | 0 | 0 | 0 | 0 | 0 | 0 |
| 0 | NA | NA | NA | NA | NA | Gm11524 | | 0 | 0 | 0 | 0 | 0 | 0 | 0 | 0 | 0 | 0 | 0 |
| 0 | NA | NA | NA | NA | NA | Gm11525 | | 0 | 0 | 0 | 0 | 0 | 0 | 0 | 0 | 0 | 0 | 0 |
| 0 | NA | NA | NA | NA | NA | Gm11573 | | 0 | 0 | 0 | 0 | 0 | 0 | 0 | 0 | 0 | 0 | 0 |
| 0 | NA | NA | NA | NA | NA | Gm11572 | | 0 | 0 | 0 | 0 | 0 | 0 | 0 | 0 | 0 | 0 | 0 |
| 0 | NA | NA | NA | NA | NA | Mir7235 | | 0 | 0 | 0 | 0 | 0 | 0 | 0 | 0 | 0 | 0 | 0 |
| 0 | NA | NA | NA | NA | NA | Gm11594 | | 0 | 0 | 0 | 0 | 0 | 0 | 0 | 0 | 0 | 0 | 0 |
| 0 | NA | NA | NA | NA | NA | 4933428G20Ril | RIKEN cDNA 4933428G20 gene | 0 | 0 | 0 | 0 | 0 | 0 | 0 | 0 | 0 | 0 | 0 |
| 0 | NA | NA | NA | NA | NA | 2410003L11Rik | | 0 | 0 | 0 | 0 | 0 | 0 | 0 | 0 | 0 | 0 | 0 |
| 0 | NA | NA | NA | NA | NA | Gm24158 | | 0 | 0 | 0 | 0 | 0 | 0 | 0 | 0 | 0 | 0 | 0 |
| 0 | NA | NA | NA | NA | NA | Mir6927 | | 0 | 0 | 0 | 0 | 0 | 0 | 0 | 0 | 0 | 0 | 0 |
| 0 | NA | NA | NA | NA | NA | Gm11614 | | 0 | 0 | 0 | 0 | 0 | 0 | 0 | 0 | 0 | 0 | 0 |
| 0 | NA | NA | NA | NA | NA | Gm22461 | | 0 | 0 | 0 | 0 | 0 | 0 | 0 | 0 | 0 | 0 | 0 |
| 0 | NA | NA | NA | NA | NA | Gm27597 | | 0 | 0 | 0 | 0 | 0 | 0 | 0 | 0 | 0 | 0 | 0 |
| 0 | NA | NA | NA | NA | NA | Gm11632 | | 0 | 0 | 0 | 0 | 0 | 0 | 0 | 0 | 0 | 0 | 0 |
| 0 | NA | NA | NA | NA | NA | Gm27399 | | 0 | 0 | 0 | 0 | 0 | 0 | 0 | 0 | 0 | 0 | 0 |
| 0 | NA | NA | NA | NA | NA | n-R5s73 | | 0 | 0 | 0 | 0 | 0 | 0 | 0 | 0 | 0 | 0 | 0 |
| 0 | NA | NA | NA | NA | NA | 1700003D09Rik | | 0 | 0 | 0 | 0 | 0 | 0 | 0 | 0 | 0 | 0 | 0 |
| 0 | NA | NA | NA | NA | NA | Pnmt | phenylethanolamine-N-methyltransferase | 0 | 0 | 0 | 0 | 0 | 0 | 0 | 0 | 0 | 0 | 0 |
| 0 | NA | NA | NA | NA | NA | Gm25048 | | 0 | 0 | 0 | 0 | 0 | 0 | 0 | 0 | 0 | 0 | 0 |
| 0 | NA | NA | NA | NA | NA | Gm25106 | | 0 | 0 | 0 | 0 | 0 | 0 | 0 | 0 | 0 | 0 | 0 |
| 0 | NA | NA | NA | NA | NA | Lrrc3c | leucine rich repeat containing 3C | 0 | 0 | 0 | 0 | 0 | 0 | 0 | 0 | 0 | 0 | 0 |
| 0 | NA | NA | NA | NA | NA | Gm23640 | | 0 | 0 | 0 | 0 | 0 | 0 | 0 | 0 | 0 | 0 | 0 |
| 0 | NA | NA | NA | NA | NA | Gm22061 | | 0 | 0 | 0 | 0 | 0 | 0 | 0 | 0 | 0 | 0 | 0 |
| 0 | NA | NA | NA | NA | NA | Gm26394 | | 0 | 0 | 0 | 0 | 0 | 0 | 0 | 0 | 0 | 0 | 0 |
| 0 | NA | NA | NA | NA | NA | Gm23451 | | 0 | 0 | 0 | 0 | 0 | 0 | 0 | 0 | 0 | 0 | 0 |
| 0 | NA | NA | NA | NA | NA | Krt24 | keratin 24 | 0 | 0 | 0 | 0 | 0 | 0 | 0 | 0 | 0 | 0 | 0 |
| 0 | NA | NA | NA | NA | NA | Gm12358 | | 0 | 0 | 0 | 0 | 0 | 0 | 0 | 0 | 0 | 0 | 0 |
| 0 | NA | NA | NA | NA | NA | Krt25 | keratin 25 | 0 | 0 | 0 | 0 | 0 | 0 | 0 | 0 | 0 | 0 | 0 |
| 0 | NA | NA | NA | NA | NA | Krt26 | keratin 26 | 0 | 0 | 0 | 0 | 0 | 0 | 0 | 0 | 0 | 0 | 0 |
| 0 | NA | NA | NA | NA | NA | Krt12 | keratin 12 | 0 | 0 | 0 | 0 | 0 | 0 | 0 | 0 | 0 | 0 | 0 |
| 0 | NA | NA | NA | NA | NA | A830036E02Rik | | 0 | 0 | 0 | 0 | 0 | 0 | 0 | 0 | 0 | 0 | 0 |
| 0 | NA | NA | NA | NA | NA | Krt39 | keratin 39 | 0 | 0 | 0 | 0 | 0 | 0 | 0 | 0 | 0 | 0 | 0 |
| 0 | NA | NA | NA | NA | NA | Krt40 | keratin 40 | 0 | 0 | 0 | 0 | 0 | 0 | 0 | 0 | 0 | 0 | 0 |
| 0 | NA | NA | NA | NA | NA | Krtap3-3 | keratin associated protein 3-3 | 0 | 0 | 0 | 0 | 0 | 0 | 0 | 0 | 0 | 0 | 0 |
| 0 | NA | NA | NA | NA | NA | Krtap3-2 | keratin associated protein 3-2 | 0 | 0 | 0 | 0 | 0 | 0 | 0 | 0 | 0 | 0 | 0 |
| 0 | NA | NA | NA | NA | NA | Gm11939 | | 0 | 0 | 0 | 0 | 0 | 0 | 0 | 0 | 0 | 0 | 0 |
| 0 | NA | NA | NA | NA | NA | Krtap3-1 | keratin associated protein 3-1 | 0 | 0 | 0 | 0 | 0 | 0 | 0 | 0 | 0 | 0 | 0 |
| 0 | NA | NA | NA | NA | NA | Krtap1-5 | keratin associated protein 1-5 | 0 | 0 | 0 | 0 | 0 | 0 | 0 | 0 | 0 | 0 | 0 |
| 0 | NA | NA | NA | NA | NA | Krtap1-4 | keratin associated protein 1-4 | 0 | 0 | 0 | 0 | 0 | 0 | 0 | 0 | 0 | 0 | 0 |
| 0 | NA | NA | NA | NA | NA | Krtap1-3 | keratin associated protein 1-3 | 0 | 0 | 0 | 0 | 0 | 0 | 0 | 0 | 0 | 0 | 0 |

| | | | | | | | | | | | | | | | | | | |
|---|---|---|---|---|---|---|---|---|---|---|---|---|---|---|---|---|---|---|
| 0 | NA | NA | NA | NA | NA | Krtap9-3 | keratin associated protein 9-3 | 0 | 0 | 0 | 0 | 0 | 0 | 0 | 0 | 0 | 0 | 0 |
| 0 | NA | NA | NA | NA | NA | Gm11938 | predicted gene 11938 | 0 | 0 | 0 | 0 | 0 | 0 | 0 | 0 | 0 | 0 | 0 |
| 0 | NA | NA | NA | NA | NA | Gm11937 | predicted gene 11937 | 0 | 0 | 0 | 0 | 0 | 0 | 0 | 0 | 0 | 0 | 0 |
| 0 | NA | NA | NA | NA | NA | Krtap2-4 | keratin associated protein 2-4 | 0 | 0 | 0 | 0 | 0 | 0 | 0 | 0 | 0 | 0 | 0 |
| 0 | NA | NA | NA | NA | NA | Gm11562 | predicted gene 11562 | 0 | 0 | 0 | 0 | 0 | 0 | 0 | 0 | 0 | 0 | 0 |
| 0 | NA | NA | NA | NA | NA | Krtap4-1 | keratin associated protein 4-1 | 0 | 0 | 0 | 0 | 0 | 0 | 0 | 0 | 0 | 0 | 0 |
| 0 | NA | NA | NA | NA | NA | Krtap4-2 | keratin associated protein 4-2 | 0 | 0 | 0 | 0 | 0 | 0 | 0 | 0 | 0 | 0 | 0 |
| 0 | NA | NA | NA | NA | NA | Krtap4-7 | keratin associated protein 4-7 | 0 | 0 | 0 | 0 | 0 | 0 | 0 | 0 | 0 | 0 | 0 |
| 0 | NA | NA | NA | NA | NA | Gm11555 | predicted gene 11555 | 0 | 0 | 0 | 0 | 0 | 0 | 0 | 0 | 0 | 0 | 0 |
| 0 | NA | NA | NA | NA | NA | Gm11563 | predicted gene 11563 | 0 | 0 | 0 | 0 | 0 | 0 | 0 | 0 | 0 | 0 | 0 |
| 0 | NA | NA | NA | NA | NA | Gm14182 | | 0 | 0 | 0 | 0 | 0 | 0 | 0 | 0 | 0 | 0 | 0 |
| 0 | NA | NA | NA | NA | NA | Krtap4-6 | keratin associated protein 4-6 | 0 | 0 | 0 | 0 | 0 | 0 | 0 | 0 | 0 | 0 | 0 |
| 0 | NA | NA | NA | NA | NA | 4930415H17Rik | | 0 | 0 | 0 | 0 | 0 | 0 | 0 | 0 | 0 | 0 | 0 |
| 0 | NA | NA | NA | NA | NA | Gm14190 | | 0 | 0 | 0 | 0 | 0 | 0 | 0 | 0 | 0 | 0 | 0 |
| 0 | NA | NA | NA | NA | NA | | | 0 | 0 | 0 | 0 | 0 | 0 | 0 | 0 | 0 | 0 | 0 |
| 0 | NA | NA | NA | NA | NA | | | 0 | 0 | 0 | 0 | 0 | 0 | 0 | 0 | 0 | 0 | 0 |
| 0 | NA | NA | NA | NA | NA | | | 0 | 0 | 0 | 0 | 0 | 0 | 0 | 0 | 0 | 0 | 0 |
| 0 | NA | NA | NA | NA | NA | Gm14186 | | 0 | 0 | 0 | 0 | 0 | 0 | 0 | 0 | 0 | 0 | 0 |
| 0 | NA | NA | NA | NA | NA | Gm14193 | | 0 | 0 | 0 | 0 | 0 | 0 | 0 | 0 | 0 | 0 | 0 |
| 0 | NA | NA | NA | NA | NA | Gm14180 | | 0 | 0 | 0 | 0 | 0 | 0 | 0 | 0 | 0 | 0 | 0 |
| 0 | NA | NA | NA | NA | NA | | | 0 | 0 | 0 | 0 | 0 | 0 | 0 | 0 | 0 | 0 | 0 |
| 0 | NA | NA | NA | NA | NA | Gm14188 | | 0 | 0 | 0 | 0 | 0 | 0 | 0 | 0 | 0 | 0 | 0 |
| 0 | NA | NA | NA | NA | NA | Gm11595 | predicted gene 11595 | 0 | 0 | 0 | 0 | 0 | 0 | 0 | 0 | 0 | 0 | 0 |
| 0 | NA | NA | NA | NA | NA | Krtap4-8 | keratin associated protein 4-8 | 0 | 0 | 0 | 0 | 0 | 0 | 0 | 0 | 0 | 0 | 0 |
| 0 | NA | NA | NA | NA | NA | Krtap4-9 | keratin associated protein 4-9 | 0 | 0 | 0 | 0 | 0 | 0 | 0 | 0 | 0 | 0 | 0 |
| 0 | NA | NA | NA | NA | NA | Gm11596 | predicted gene 11596 | 0 | 0 | 0 | 0 | 0 | 0 | 0 | 0 | 0 | 0 | 0 |
| 0 | NA | NA | NA | NA | NA | Gm11569 | predicted gene 11569 | 0 | 0 | 0 | 0 | 0 | 0 | 0 | 0 | 0 | 0 | 0 |
| 0 | NA | NA | NA | NA | NA | Gm11554 | predicted gene 11554 | 0 | 0 | 0 | 0 | 0 | 0 | 0 | 0 | 0 | 0 | 0 |
| 0 | NA | NA | NA | NA | NA | Krtap4-13 | keratin associated protein 4-13 | 0 | 0 | 0 | 0 | 0 | 0 | 0 | 0 | 0 | 0 | 0 |
| 0 | NA | NA | NA | NA | NA | Gm11564 | predicted gene 11564 | 0 | 0 | 0 | 0 | 0 | 0 | 0 | 0 | 0 | 0 | 0 |
| 0 | NA | NA | NA | NA | NA | 2300003K06Ril | RIKEN cDNA 2300003K06 gene | 0 | 0 | 0 | 0 | 0 | 0 | 0 | 0 | 0 | 0 | 0 |
| 0 | NA | NA | NA | NA | NA | Gm11556 | | 0 | 0 | 0 | 0 | 0 | 0 | 0 | 0 | 0 | 0 | 0 |
| 0 | NA | NA | NA | NA | NA | Krtap4-16 | keratin associated protein 4-16 | 0 | 0 | 0 | 0 | 0 | 0 | 0 | 0 | 0 | 0 | 0 |
| 0 | NA | NA | NA | NA | NA | Gm26324 | | 0 | 0 | 0 | 0 | 0 | 0 | 0 | 0 | 0 | 0 | 0 |
| 0 | NA | NA | NA | NA | NA | Gm11568 | predicted gene 11568 | 0 | 0 | 0 | 0 | 0 | 0 | 0 | 0 | 0 | 0 | 0 |
| 0 | NA | NA | NA | NA | NA | Gm11559 | predicted gene 11559 | 0 | 0 | 0 | 0 | 0 | 0 | 0 | 0 | 0 | 0 | 0 |
| 0 | NA | NA | NA | NA | NA | Krtap9-1 | keratin associated protein 9-1 | 0 | 0 | 0 | 0 | 0 | 0 | 0 | 0 | 0 | 0 | 0 |
| 0 | NA | NA | NA | NA | NA | Gm11567 | predicted gene 11567 | 0 | 0 | 0 | 0 | 0 | 0 | 0 | 0 | 0 | 0 | 0 |
| 0 | NA | NA | NA | NA | NA | Gm11566 | | 0 | 0 | 0 | 0 | 0 | 0 | 0 | 0 | 0 | 0 | 0 |
| 0 | NA | NA | NA | NA | NA | Krtap31-1 | keratin associated protein 31-1 | 0 | 0 | 0 | 0 | 0 | 0 | 0 | 0 | 0 | 0 | 0 |
| 0 | NA | NA | NA | NA | NA | Gm11565 | predicted gene 11565 | 0 | 0 | 0 | 0 | 0 | 0 | 0 | 0 | 0 | 0 | 0 |
| 0 | NA | NA | NA | NA | NA | Krtap31-2 | keratin associated protein 31-2 | 0 | 0 | 0 | 0 | 0 | 0 | 0 | 0 | 0 | 0 | 0 |
| 0 | NA | NA | NA | NA | NA | Krtap9-5 | keratin associated protein 9-5 | 0 | 0 | 0 | 0 | 0 | 0 | 0 | 0 | 0 | 0 | 0 |
| 0 | NA | NA | NA | NA | NA | Gm11561 | | 0 | 0 | 0 | 0 | 0 | 0 | 0 | 0 | 0 | 0 | 0 |
| 0 | NA | NA | NA | NA | NA | Gm11553 | | 0 | 0 | 0 | 0 | 0 | 0 | 0 | 0 | 0 | 0 | 0 |
| 0 | NA | NA | NA | NA | NA | Gm11558 | | 0 | 0 | 0 | 0 | 0 | 0 | 0 | 0 | 0 | 0 | 0 |
| 0 | NA | NA | NA | NA | NA | Krt33a | keratin 33A | 0 | 0 | 0 | 0 | 0 | 0 | 0 | 0 | 0 | 0 | 0 |
| 0 | NA | NA | NA | NA | NA | Krt33b | keratin 33B | 0 | 0 | 0 | 0 | 0 | 0 | 0 | 0 | 0 | 0 | 0 |
| 0 | NA | NA | NA | NA | NA | Krt34 | keratin 34 | 0 | 0 | 0 | 0 | 0 | 0 | 0 | 0 | 0 | 0 | 0 |
| 0 | NA | NA | NA | NA | NA | Krt31 | keratin 31 | 0 | 0 | 0 | 0 | 0 | 0 | 0 | 0 | 0 | 0 | 0 |
| 0 | NA | NA | NA | NA | NA | Gm11571 | | 0 | 0 | 0 | 0 | 0 | 0 | 0 | 0 | 0 | 0 | 0 |
| 0 | NA | NA | NA | NA | NA | | | 0 | 0 | 0 | 0 | 0 | 0 | 0 | 0 | 0 | 0 | 0 |
| 0 | NA | NA | NA | NA | NA | Krt32 | keratin 32 | 0 | 0 | 0 | 0 | 0 | 0 | 0 | 0 | 0 | 0 | 0 |
| 0 | NA | NA | NA | NA | NA | Gm12347 | | 0 | 0 | 0 | 0 | 0 | 0 | 0 | 0 | 0 | 0 | 0 |
| 0 | NA | NA | NA | NA | NA | Krt13 | keratin 13 | 0 | 0 | 0 | 0 | 0 | 0 | 0 | 0 | 0 | 0 | 0 |
| 0 | NA | NA | NA | NA | NA | Krt9 | keratin 9 | 0 | 0 | 0 | 0 | 0 | 0 | 0 | 0 | 0 | 0 | 0 |
| 0 | NA | NA | NA | NA | NA | Gm11597 | | 0 | 0 | 0 | 0 | 0 | 0 | 0 | 0 | 0 | 0 | 0 |
| 0 | NA | NA | NA | NA | NA | Krt16 | keratin 16 | 0 | 0 | 0 | 0 | 0 | 0 | 0 | 0 | 0 | 0 | 0 |
| 0 | NA | NA | NA | NA | NA | Krt42 | keratin 42 | 0 | 0 | 0 | 0 | 0 | 0 | 0 | 0 | 0 | 0 | 0 |
| 0 | NA | NA | NA | NA | NA | Gm14206 | | 0 | 0 | 0 | 0 | 0 | 0 | 0 | 0 | 0 | 0 | 0 |

| | | | | | | | | | | | | | | | | | |
|---|---|---|---|---|---|---|---|---|---|---|---|---|---|---|---|---|---|
| 0 | NA | NA | NA | NA | NA | Gm27374 | | 0 | 0 | 0 | 0 | 0 | 0 | 0 | 0 | 0 | 0 | 0 |
| 0 | NA | NA | NA | NA | NA | Gm44544 | | 0 | 0 | 0 | 0 | 0 | 0 | 0 | 0 | 0 | 0 | 0 |
| 0 | NA | NA | NA | NA | NA | Hcrt | hypocretin | 0 | 0 | 0 | 0 | 0 | 0 | 0 | 0 | 0 | 0 | 0 |
| 0 | NA | NA | NA | NA | NA | Gm24358 | | 0 | 0 | 0 | 0 | 0 | 0 | 0 | 0 | 0 | 0 | 0 |
| 0 | NA | NA | NA | NA | NA | Mir6928 | | 0 | 0 | 0 | 0 | 0 | 0 | 0 | 0 | 0 | 0 | 0 |
| 0 | NA | NA | NA | NA | NA | Gm11617 | | 0 | 0 | 0 | 0 | 0 | 0 | 0 | 0 | 0 | 0 | 0 |
| 0 | NA | NA | NA | NA | NA | Gm28156 | | 0 | 0 | 0 | 0 | 0 | 0 | 0 | 0 | 0 | 0 | 0 |
| 0 | NA | NA | NA | NA | NA | Gm11618 | | 0 | 0 | 0 | 0 | 0 | 0 | 0 | 0 | 0 | 0 | 0 |
| 0 | NA | NA | NA | NA | NA | Gm11619 | | 0 | 0 | 0 | 0 | 0 | 0 | 0 | 0 | 0 | 0 | 0 |
| 0 | NA | NA | NA | NA | NA | Gm25593 | | 0 | 0 | 0 | 0 | 0 | 0 | 0 | 0 | 0 | 0 | 0 |
| 0 | NA | NA | NA | NA | NA | | | 0 | 0 | 0 | 0 | 0 | 0 | 0 | 0 | 0 | 0 | 0 |
| 0 | NA | NA | NA | NA | NA | Gm25475 | | 0 | 0 | 0 | 0 | 0 | 0 | 0 | 0 | 0 | 0 | 0 |
| 0 | NA | NA | NA | NA | NA | Mir6929 | | 0 | 0 | 0 | 0 | 0 | 0 | 0 | 0 | 0 | 0 | 0 |
| 0 | NA | NA | NA | NA | NA | Gm11626 | | 0 | 0 | 0 | 0 | 0 | 0 | 0 | 0 | 0 | 0 | 0 |
| 0 | NA | NA | NA | NA | NA | Gm11634 | predicted gene 11634 | 0 | 0 | 0 | 0 | 0 | 0 | 0 | 0 | 0 | 0 | 0 |
| 0 | NA | NA | NA | NA | NA | Gm26316 | | 0 | 0 | 0 | 0 | 0 | 0 | 0 | 0 | 0 | 0 | 0 |
| 0 | NA | NA | NA | NA | NA | Gm11635 | uncharacterized LOC666331 | 0 | 0 | 0 | 0 | 0 | 0 | 0 | 0 | 0 | 0 | 0 |
| 0 | NA | NA | NA | NA | NA | Gm24950 | | 0 | 0 | 0 | 0 | 0 | 0 | 0 | 0 | 0 | 0 | 0 |
| 0 | NA | NA | NA | NA | NA | Gm23971 | | 0 | 0 | 0 | 0 | 0 | 0 | 0 | 0 | 0 | 0 | 0 |
| 0 | NA | NA | NA | NA | NA | 4930417O22Rik | | 0 | 0 | 0 | 0 | 0 | 0 | 0 | 0 | 0 | 0 | 0 |
| 0 | NA | NA | NA | NA | NA | Cfap97d1 | CFAP97 domain containing 1 | 0 | 0 | 0 | 0 | 0 | 0 | 0 | 0 | 0 | 0 | 0 |
| 0 | NA | NA | NA | NA | NA | E130111B04Rik | | 0 | 0 | 0 | 0 | 0 | 0 | 0 | 0 | 0 | 0 | 0 |
| 0 | NA | NA | NA | NA | NA | Gm11584 | | 0 | 0 | 0 | 0 | 0 | 0 | 0 | 0 | 0 | 0 | 0 |
| 0 | NA | NA | NA | NA | NA | Gm11586 | | 0 | 0 | 0 | 0 | 0 | 0 | 0 | 0 | 0 | 0 | 0 |
| 0 | NA | NA | NA | NA | NA | Mir8101 | | 0 | 0 | 0 | 0 | 0 | 0 | 0 | 0 | 0 | 0 | 0 |
| 0 | NA | NA | NA | NA | NA | Mir6930 | | 0 | 0 | 0 | 0 | 0 | 0 | 0 | 0 | 0 | 0 | 0 |
| 0 | NA | NA | NA | NA | NA | Gm23563 | | 0 | 0 | 0 | 0 | 0 | 0 | 0 | 0 | 0 | 0 | 0 |
| 0 | NA | NA | NA | NA | NA | Mdk-ps1 | | 0 | 0 | 0 | 0 | 0 | 0 | 0 | 0 | 0 | 0 | 0 |
| 0 | NA | NA | NA | NA | NA | Gm11636 | | 0 | 0 | 0 | 0 | 0 | 0 | 0 | 0 | 0 | 0 | 0 |
| 0 | NA | NA | NA | NA | NA | Gm22920 | | 0 | 0 | 0 | 0 | 0 | 0 | 0 | 0 | 0 | 0 | 0 |
| 0 | NA | NA | NA | NA | NA | Meioc | meiosis specific with coiled-coil domain | 0 | 0 | 0 | 0 | 0 | 0 | 0 | 0 | 0 | 0 | 0 |
| 0 | NA | NA | NA | NA | NA | Gm11637 | | 0 | 0 | 0 | 0 | 0 | 0 | 0 | 0 | 0 | 0 | 0 |
| 0 | NA | NA | NA | NA | NA | Gm22683 | | 0 | 0 | 0 | 0 | 0 | 0 | 0 | 0 | 0 | 0 | 0 |
| 0 | NA | NA | NA | NA | NA | Fam187a | family with sequence similarity 187, meml | 0 | 0 | 0 | 0 | 0 | 0 | 0 | 0 | 0 | 0 | 0 |
| 0 | NA | NA | NA | NA | NA | Mir6931 | | 0 | 0 | 0 | 0 | 0 | 0 | 0 | 0 | 0 | 0 | 0 |
| 0 | NA | NA | NA | NA | NA | Lrrc37a | leucine rich repeat containing 37A | 0 | 0 | 0 | 0 | 0 | 0 | 0 | 0 | 0 | 0 | 0 |
| 0 | NA | NA | NA | NA | NA | Gm884 | predicted gene 884 | 0 | 0 | 0 | 0 | 0 | 0 | 0 | 0 | 0 | 0 | 0 |
| 0 | NA | NA | NA | NA | NA | Lyzl6 | lysozyme-like 6 | 0 | 0 | 0 | 0 | 0 | 0 | 0 | 0 | 0 | 0 | 0 |
| 0 | NA | NA | NA | NA | NA | Sppl2c | signal peptide peptidase 2C | 0 | 0 | 0 | 0 | 0 | 0 | 0 | 0 | 0 | 0 | 0 |
| 0 | NA | NA | NA | NA | NA | Gm11659 | | 0 | 0 | 0 | 0 | 0 | 0 | 0 | 0 | 0 | 0 | 0 |
| 0 | NA | NA | NA | NA | NA | Gm22743 | | 0 | 0 | 0 | 0 | 0 | 0 | 0 | 0 | 0 | 0 | 0 |
| 0 | NA | NA | NA | NA | NA | Gm47310 | | 0 | 0 | 0 | 0 | 0 | 0 | 0 | 0 | 0 | 0 | 0 |
| 0 | NA | NA | NA | NA | NA | Gm47315 | | 0 | 0 | 0 | 0 | 0 | 0 | 0 | 0 | 0 | 0 | 0 |
| 0 | NA | NA | NA | NA | NA | Gm11660 | | 0 | 0 | 0 | 0 | 0 | 0 | 0 | 0 | 0 | 0 | 0 |
| 0 | NA | NA | NA | NA | NA | Gm11665 | | 0 | 0 | 0 | 0 | 0 | 0 | 0 | 0 | 0 | 0 | 0 |
| 0 | NA | NA | NA | NA | NA | Gm11661 | | 0 | 0 | 0 | 0 | 0 | 0 | 0 | 0 | 0 | 0 | 0 |
| 0 | NA | NA | NA | NA | NA | Gm11639 | | 0 | 0 | 0 | 0 | 0 | 0 | 0 | 0 | 0 | 0 | 0 |
| 0 | NA | NA | NA | NA | NA | Gm11623 | | 0 | 0 | 0 | 0 | 0 | 0 | 0 | 0 | 0 | 0 | 0 |
| 0 | NA | NA | NA | NA | NA | Gm11621 | | 0 | 0 | 0 | 0 | 0 | 0 | 0 | 0 | 0 | 0 | 0 |
| 0 | NA | NA | NA | NA | NA | Gm11622 | | 0 | 0 | 0 | 0 | 0 | 0 | 0 | 0 | 0 | 0 | 0 |
| 0 | NA | NA | NA | NA | NA | Gm11624 | | 0 | 0 | 0 | 0 | 0 | 0 | 0 | 0 | 0 | 0 | 0 |
| 0 | NA | NA | NA | NA | NA | Efcab3 | EF-hand calcium binding domain 3 | 0 | 0 | 0 | 0 | 0 | 0 | 0 | 0 | 0 | 0 | 0 |
| 0 | NA | NA | NA | NA | NA | Gm25362 | | 0 | 0 | 0 | 0 | 0 | 0 | 0 | 0 | 0 | 0 | 0 |
| 0 | NA | NA | NA | NA | NA | Gm24290 | | 0 | 0 | 0 | 0 | 0 | 0 | 0 | 0 | 0 | 0 | 0 |
| 0 | NA | NA | NA | NA | NA | Gm11640 | | 0 | 0 | 0 | 0 | 0 | 0 | 0 | 0 | 0 | 0 | 0 |
| 0 | NA | NA | NA | NA | NA | Gm23990 | | 0 | 0 | 0 | 0 | 0 | 0 | 0 | 0 | 0 | 0 | 0 |
| 0 | NA | NA | NA | NA | NA | Gm11651 | | 0 | 0 | 0 | 0 | 0 | 0 | 0 | 0 | 0 | 0 | 0 |
| 0 | NA | NA | NA | NA | NA | Gm11672 | | 0 | 0 | 0 | 0 | 0 | 0 | 0 | 0 | 0 | 0 | 0 |
| 0 | NA | NA | NA | NA | NA | Gm25556 | | 0 | 0 | 0 | 0 | 0 | 0 | 0 | 0 | 0 | 0 | 0 |
| 0 | NA | NA | NA | NA | NA | Gh | growth hormone | 0 | 0 | 0 | 0 | 0 | 0 | 0 | 0 | 0 | 0 | 0 |

| | | | | | | | | | | | | | | | | | | |
|---|---|---|---|---|---|---|---|---|---|---|---|---|---|---|---|---|---|---|
| 0 | NA | NA | NA | NA | NA | Gm22711 | | 0 | 0 | 0 | 0 | 0 | 0 | 0 | 0 | 0 | 0 | 0 |
| 0 | NA | NA | NA | NA | NA | Gm22711 | | 0 | 0 | 0 | 0 | 0 | 0 | 0 | 0 | 0 | 0 | 0 |
| 0 | NA | NA | NA | NA | NA | Gm25889 | | 0 | 0 | 0 | 0 | 0 | 0 | 0 | 0 | 0 | 0 | 0 |
| 0 | NA | NA | NA | NA | NA | Mir3064 | | 0 | 0 | 0 | 0 | 0 | 0 | 0 | 0 | 0 | 0 | 0 |
| 0 | NA | NA | NA | NA | NA | Gm25994 | | 0 | 0 | 0 | 0 | 0 | 0 | 0 | 0 | 0 | 0 | 0 |
| 0 | NA | NA | NA | NA | NA | Gm27322 | | 0 | 0 | 0 | 0 | 0 | 0 | 0 | 0 | 0 | 0 | 0 |
| 0 | NA | NA | NA | NA | NA | Gm11708 | | 0 | 0 | 0 | 0 | 0 | 0 | 0 | 0 | 0 | 0 | 0 |
| 0 | NA | NA | NA | NA | NA | Gm11715 | predicted gene 11715 | 0 | 0 | 0 | 0 | 0 | 0 | 0 | 0 | 0 | 0 | 0 |
| 0 | NA | NA | NA | NA | NA | Gm11718 | | 0 | 0 | 0 | 0 | 0 | 0 | 0 | 0 | 0 | 0 | 0 |
| 0 | NA | NA | NA | NA | NA | Gm11719 | | 0 | 0 | 0 | 0 | 0 | 0 | 0 | 0 | 0 | 0 | 0 |
| 0 | NA | NA | NA | NA | NA | Gm11712 | | 0 | 0 | 0 | 0 | 0 | 0 | 0 | 0 | 0 | 0 | 0 |
| 0 | NA | NA | NA | NA | NA | C330019F10Rik | | 0 | 0 | 0 | 0 | 0 | 0 | 0 | 0 | 0 | 0 | 0 |
| 0 | NA | NA | NA | NA | NA | A830035A12Rik | | 0 | 0 | 0 | 0 | 0 | 0 | 0 | 0 | 0 | 0 | 0 |
| 0 | NA | NA | NA | NA | NA | Gm11649 | | 0 | 0 | 0 | 0 | 0 | 0 | 0 | 0 | 0 | 0 | 0 |
| 0 | NA | NA | NA | NA | NA | Mir6932 | | 0 | 0 | 0 | 0 | 0 | 0 | 0 | 0 | 0 | 0 | 0 |
| 0 | NA | NA | NA | NA | NA | Gm23120 | | 0 | 0 | 0 | 0 | 0 | 0 | 0 | 0 | 0 | 0 | 0 |
| 0 | NA | NA | NA | NA | NA | Gm27595 | | 0 | 0 | 0 | 0 | 0 | 0 | 0 | 0 | 0 | 0 | 0 |
| 0 | NA | NA | NA | NA | NA | Gm11657 | | 0 | 0 | 0 | 0 | 0 | 0 | 0 | 0 | 0 | 0 | 0 |
| 0 | NA | NA | NA | NA | NA | Mir7223 | | 0 | 0 | 0 | 0 | 0 | 0 | 0 | 0 | 0 | 0 | 0 |
| 0 | NA | NA | NA | NA | NA | Gm11655 | | 0 | 0 | 0 | 0 | 0 | 0 | 0 | 0 | 0 | 0 | 0 |
| 0 | NA | NA | NA | NA | NA | Gm11656 | | 0 | 0 | 0 | 0 | 0 | 0 | 0 | 0 | 0 | 0 | 0 |
| 0 | NA | NA | NA | NA | NA | Gm11667 | | 0 | 0 | 0 | 0 | 0 | 0 | 0 | 0 | 0 | 0 | 0 |
| 0 | NA | NA | NA | NA | NA | Cep112os1 | | 0 | 0 | 0 | 0 | 0 | 0 | 0 | 0 | 0 | 0 | 0 |
| 0 | NA | NA | NA | NA | NA | Olfr1397-ps1 | | 0 | 0 | 0 | 0 | 0 | 0 | 0 | 0 | 0 | 0 | 0 |
| 0 | NA | NA | NA | NA | NA | Gm11670 | | 0 | 0 | 0 | 0 | 0 | 0 | 0 | 0 | 0 | 0 | 0 |
| 0 | NA | NA | NA | NA | NA | Gm11668 | | 0 | 0 | 0 | 0 | 0 | 0 | 0 | 0 | 0 | 0 | 0 |
| 0 | NA | NA | NA | NA | NA | Gm24149 | | 0 | 0 | 0 | 0 | 0 | 0 | 0 | 0 | 0 | 0 | 0 |
| 0 | NA | NA | NA | NA | NA | E030025P04Rik | | 0 | 0 | 0 | 0 | 0 | 0 | 0 | 0 | 0 | 0 | 0 |
| 0 | NA | NA | NA | NA | NA | 1700096J18Rik | | 0 | 0 | 0 | 0 | 0 | 0 | 0 | 0 | 0 | 0 | 0 |
| 0 | NA | NA | NA | NA | NA | Gm25540 | | 0 | 0 | 0 | 0 | 0 | 0 | 0 | 0 | 0 | 0 | 0 |
| 0 | NA | NA | NA | NA | NA | Gm22378 | | 0 | 0 | 0 | 0 | 0 | 0 | 0 | 0 | 0 | 0 | 0 |
| 0 | NA | NA | NA | NA | NA | Gm11684 | | 0 | 0 | 0 | 0 | 0 | 0 | 0 | 0 | 0 | 0 | 0 |
| 0 | NA | NA | NA | NA | NA | Gm11697 | | 0 | 0 | 0 | 0 | 0 | 0 | 0 | 0 | 0 | 0 | 0 |
| 0 | NA | NA | NA | NA | NA | Gm11676 | | 0 | 0 | 0 | 0 | 0 | 0 | 0 | 0 | 0 | 0 | 0 |
| 0 | NA | NA | NA | NA | NA | Gm11677 | | 0 | 0 | 0 | 0 | 0 | 0 | 0 | 0 | 0 | 0 | 0 |
| 0 | NA | NA | NA | NA | NA | Gm11674 | | 0 | 0 | 0 | 0 | 0 | 0 | 0 | 0 | 0 | 0 | 0 |
| 0 | NA | NA | NA | NA | NA | Gm11679 | | 0 | 0 | 0 | 0 | 0 | 0 | 0 | 0 | 0 | 0 | 0 |
| 0 | NA | NA | NA | NA | NA | Gm11680 | | 0 | 0 | 0 | 0 | 0 | 0 | 0 | 0 | 0 | 0 | 0 |
| 0 | NA | NA | NA | NA | NA | Gm11681 | | 0 | 0 | 0 | 0 | 0 | 0 | 0 | 0 | 0 | 0 | 0 |
| 0 | NA | NA | NA | NA | NA | 4933434M16Rik | | 0 | 0 | 0 | 0 | 0 | 0 | 0 | 0 | 0 | 0 | 0 |
| 0 | NA | NA | NA | NA | NA | Gm11693 | | 0 | 0 | 0 | 0 | 0 | 0 | 0 | 0 | 0 | 0 | 0 |
| 0 | NA | NA | NA | NA | NA | Cpsf4l | cleavage and polyadenylation specific fac | 0 | 0 | 0 | 0 | 0 | 0 | 0 | 0 | 0 | 0 | 0 |
| 0 | NA | NA | NA | NA | NA | Gm11736 | | 0 | 0 | 0 | 0 | 0 | 0 | 0 | 0 | 0 | 0 | 0 |
| 0 | NA | NA | NA | NA | NA | 1700025D23Rik | | 0 | 0 | 0 | 0 | 0 | 0 | 0 | 0 | 0 | 0 | 0 |
| 0 | NA | NA | NA | NA | NA | 1700001J04Rik | | 0 | 0 | 0 | 0 | 0 | 0 | 0 | 0 | 0 | 0 | 0 |
| 0 | NA | NA | NA | NA | NA | 1700092K14Rik | | 0 | 0 | 0 | 0 | 0 | 0 | 0 | 0 | 0 | 0 | 0 |
| 0 | NA | NA | NA | NA | NA | Gm11690 | | 0 | 0 | 0 | 0 | 0 | 0 | 0 | 0 | 0 | 0 | 0 |
| 0 | NA | NA | NA | NA | NA | Gm11691 | | 0 | 0 | 0 | 0 | 0 | 0 | 0 | 0 | 0 | 0 | 0 |
| 0 | NA | NA | NA | NA | NA | Gm11692 | | 0 | 0 | 0 | 0 | 0 | 0 | 0 | 0 | 0 | 0 | 0 |
| 0 | NA | NA | NA | NA | NA | Gm11698 | | 0 | 0 | 0 | 0 | 0 | 0 | 0 | 0 | 0 | 0 | 0 |
| 0 | NA | NA | NA | NA | NA | Gm11699 | | 0 | 0 | 0 | 0 | 0 | 0 | 0 | 0 | 0 | 0 | 0 |
| 0 | NA | NA | NA | NA | NA | Cd300ld4 | CD300 molecule like family member D4 | 0 | 0 | 0 | 0 | 0 | 0 | 0 | 0 | 0 | 0 | 0 |
| 0 | NA | NA | NA | NA | NA | Cd300ld5 | CD300 molecule like family member D5 | 0 | 0 | 0 | 0 | 0 | 0 | 0 | 0 | 0 | 0 | 0 |
| 0 | NA | NA | NA | NA | NA | Gm25837 | | 0 | 0 | 0 | 0 | 0 | 0 | 0 | 0 | 0 | 0 | 0 |
| 0 | NA | NA | NA | NA | NA | Otop2 | otopetrin 2 | 0 | 0 | 0 | 0 | 0 | 0 | 0 | 0 | 0 | 0 | 0 |
| 0 | NA | NA | NA | NA | NA | Mir3968 | | 0 | 0 | 0 | 0 | 0 | 0 | 0 | 0 | 0 | 0 | 0 |
| 0 | NA | NA | NA | NA | NA | Gm25364 | | 0 | 0 | 0 | 0 | 0 | 0 | 0 | 0 | 0 | 0 | 0 |
| 0 | NA | NA | NA | NA | NA | Gm23699 | | 0 | 0 | 0 | 0 | 0 | 0 | 0 | 0 | 0 | 0 | 0 |
| 0 | NA | NA | NA | NA | NA | Gm11704 | | 0 | 0 | 0 | 0 | 0 | 0 | 0 | 0 | 0 | 0 | 0 |
| 0 | NA | NA | NA | NA | NA | Recql5os1 | | 0 | 0 | 0 | 0 | 0 | 0 | 0 | 0 | 0 | 0 | 0 |

| | | | | | | | | | | | | | | | | | |
|---|---|---|---|---|---|---|---|---|---|---|---|---|---|---|---|---|---|
| 0 | NA | NA | NA | NA | NA | Gm26413 | | 0 | 0 | 0 | 0 | 0 | 0 | 0 | 0 | 0 | 0 | 0 |
| 0 | NA | NA | NA | NA | NA | Gm26413 | | 0 | 0 | 0 | 0 | 0 | 0 | 0 | 0 | 0 | 0 | 0 |
| 0 | NA | NA | NA | NA | NA | | | 0 | 0 | 0 | 0 | 0 | 0 | 0 | 0 | 0 | 0 | 0 |
| 0 | NA | NA | NA | NA | NA | n-R5s74 | | 0 | 0 | 0 | 0 | 0 | 0 | 0 | 0 | 0 | 0 | 0 |
| 0 | NA | NA | NA | NA | NA | Rps11-ps2 | | 0 | 0 | 0 | 0 | 0 | 0 | 0 | 0 | 0 | 0 | 0 |
| 0 | NA | NA | NA | NA | NA | Gm11739 | | 0 | 0 | 0 | 0 | 0 | 0 | 0 | 0 | 0 | 0 | 0 |
| 0 | NA | NA | NA | NA | NA | Snord1c | | 0 | 0 | 0 | 0 | 0 | 0 | 0 | 0 | 0 | 0 | 0 |
| 0 | NA | NA | NA | NA | NA | Snord1b | | 0 | 0 | 0 | 0 | 0 | 0 | 0 | 0 | 0 | 0 | 0 |
| 0 | NA | NA | NA | NA | NA | Snord1a | | 0 | 0 | 0 | 0 | 0 | 0 | 0 | 0 | 0 | 0 | 0 |
| 0 | NA | NA | NA | NA | NA | Gm11735 | | 0 | 0 | 0 | 0 | 0 | 0 | 0 | 0 | 0 | 0 | 0 |
| 0 | NA | NA | NA | NA | NA | BC018473 | | 0 | 0 | 0 | 0 | 0 | 0 | 0 | 0 | 0 | 0 | 0 |
| 0 | NA | NA | NA | NA | NA | St6galnac1 | ST6 (alpha-N-acetyl-neuraminyl-2,3-beta | 0 | 0 | 0 | 0 | 0 | 0 | 0 | 0 | 0 | 0 | 0 |
| 0 | NA | NA | NA | NA | NA | Gm27401 | | 0 | 0 | 0 | 0 | 0 | 0 | 0 | 0 | 0 | 0 | 0 |
| 0 | NA | NA | NA | NA | NA | Gm11728 | | 0 | 0 | 0 | 0 | 0 | 0 | 0 | 0 | 0 | 0 | 0 |
| 0 | NA | NA | NA | NA | NA | Gm11731 | | 0 | 0 | 0 | 0 | 0 | 0 | 0 | 0 | 0 | 0 | 0 |
| 0 | NA | NA | NA | NA | NA | Gm26083 | | 0 | 0 | 0 | 0 | 0 | 0 | 0 | 0 | 0 | 0 | 0 |
| 0 | NA | NA | NA | NA | NA | Mir6516 | | 0 | 0 | 0 | 0 | 0 | 0 | 0 | 0 | 0 | 0 | 0 |
| 0 | NA | NA | NA | NA | NA | Gm24323 | | 0 | 0 | 0 | 0 | 0 | 0 | 0 | 0 | 0 | 0 | 0 |
| 0 | NA | NA | NA | NA | NA | Gm11729 | | 0 | 0 | 0 | 0 | 0 | 0 | 0 | 0 | 0 | 0 | 0 |
| 0 | NA | NA | NA | NA | NA | 6030468B19Ril | RIKEN cDNA 6030468B19 gene | 0 | 0 | 0 | 0 | 0 | 0 | 0 | 0 | 0 | 0 | 0 |
| 0 | NA | NA | NA | NA | NA | Gm20708 | | 0 | 0 | 0 | 0 | 0 | 0 | 0 | 0 | 0 | 0 | 0 |
| 0 | NA | NA | NA | NA | NA | Tmem235 | transmembrane protein 235 | 0 | 0 | 0 | 0 | 0 | 0 | 0 | 0 | 0 | 0 | 0 |
| 0 | NA | NA | NA | NA | NA | Gm11726 | | 0 | 0 | 0 | 0 | 0 | 0 | 0 | 0 | 0 | 0 | 0 |
| 0 | NA | NA | NA | NA | NA | Mir6933 | | 0 | 0 | 0 | 0 | 0 | 0 | 0 | 0 | 0 | 0 | 0 |
| 0 | NA | NA | NA | NA | NA | Gm11738 | | 0 | 0 | 0 | 0 | 0 | 0 | 0 | 0 | 0 | 0 | 0 |
| 0 | NA | NA | NA | NA | NA | Gm11747 | | 0 | 0 | 0 | 0 | 0 | 0 | 0 | 0 | 0 | 0 | 0 |
| 0 | NA | NA | NA | NA | NA | Gm25040 | | 0 | 0 | 0 | 0 | 0 | 0 | 0 | 0 | 0 | 0 | 0 |
| 0 | NA | NA | NA | NA | NA | Gm11749 | | 0 | 0 | 0 | 0 | 0 | 0 | 0 | 0 | 0 | 0 | 0 |
| 0 | NA | NA | NA | NA | NA | Enpp7 | ectonucleotide pyrophosphatase/phosph | 0 | 0 | 0 | 0 | 0 | 0 | 0 | 0 | 0 | 0 | 0 |
| 0 | NA | NA | NA | NA | NA | Gm11754 | | 0 | 0 | 0 | 0 | 0 | 0 | 0 | 0 | 0 | 0 | 0 |
| 0 | NA | NA | NA | NA | NA | Mir6934 | | 0 | 0 | 0 | 0 | 0 | 0 | 0 | 0 | 0 | 0 | 0 |
| 0 | NA | NA | NA | NA | NA | Mir1932 | | 0 | 0 | 0 | 0 | 0 | 0 | 0 | 0 | 0 | 0 | 0 |
| 0 | NA | NA | NA | NA | NA | Gm11761 | | 0 | 0 | 0 | 0 | 0 | 0 | 0 | 0 | 0 | 0 | 0 |
| 0 | NA | NA | NA | NA | NA | Gm23663 | | 0 | 0 | 0 | 0 | 0 | 0 | 0 | 0 | 0 | 0 | 0 |
| 0 | NA | NA | NA | NA | NA | Mir338 | | 0 | 0 | 0 | 0 | 0 | 0 | 0 | 0 | 0 | 0 | 0 |
| 0 | NA | NA | NA | NA | NA | Mir3065 | | 0 | 0 | 0 | 0 | 0 | 0 | 0 | 0 | 0 | 0 | 0 |
| 0 | NA | NA | NA | NA | NA | Gm47846 | | 0 | 0 | 0 | 0 | 0 | 0 | 0 | 0 | 0 | 0 | 0 |
| 0 | NA | NA | NA | NA | NA | Gm47297 | | 0 | 0 | 0 | 0 | 0 | 0 | 0 | 0 | 0 | 0 | 0 |
| 0 | NA | NA | NA | NA | NA | Gm23419 | | 0 | 0 | 0 | 0 | 0 | 0 | 0 | 0 | 0 | 0 | 0 |
| 0 | NA | NA | NA | NA | NA | Mir6935 | | 0 | 0 | 0 | 0 | 0 | 0 | 0 | 0 | 0 | 0 | 0 |
| 0 | NA | NA | NA | NA | NA | Mir6936 | | 0 | 0 | 0 | 0 | 0 | 0 | 0 | 0 | 0 | 0 | 0 |
| 0 | NA | NA | NA | NA | NA | Myadml2os | | 0 | 0 | 0 | 0 | 0 | 0 | 0 | 0 | 0 | 0 | 0 |
| 0 | NA | NA | NA | NA | NA | Notumos | | 0 | 0 | 0 | 0 | 0 | 0 | 0 | 0 | 0 | 0 | 0 |
| 0 | NA | NA | NA | NA | NA | Gm17178 | | 0 | 0 | 0 | 0 | 0 | 0 | 0 | 0 | 0 | 0 | 0 |
| 0 | NA | NA | NA | NA | NA | Gm11771 | | 0 | 0 | 0 | 0 | 0 | 0 | 0 | 0 | 0 | 0 | 0 |
| 0 | NA | NA | NA | NA | NA | Gm11773 | | 0 | 0 | 0 | 0 | 0 | 0 | 0 | 0 | 0 | 0 | 0 |
| 0 | NA | NA | NA | NA | NA | Gm28192 | | 0 | 0 | 0 | 0 | 0 | 0 | 0 | 0 | 0 | 0 | 0 |
| 0 | NA | NA | NA | NA | NA | Gm11774 | | 0 | 0 | 0 | 0 | 0 | 0 | 0 | 0 | 0 | 0 | 0 |
| 0 | NA | NA | NA | NA | NA | Gm11775 | | 0 | 0 | 0 | 0 | 0 | 0 | 0 | 0 | 0 | 0 | 0 |
| 0 | NA | NA | NA | NA | NA | Gm22152 | | 0 | 0 | 0 | 0 | 0 | 0 | 0 | 0 | 0 | 0 | 0 |
| 0 | NA | NA | NA | NA | NA | Gm11792 | | 0 | 0 | 0 | 0 | 0 | 0 | 0 | 0 | 0 | 0 | 0 |
| 0 | NA | NA | NA | NA | NA | Gm11791 | | 0 | 0 | 0 | 0 | 0 | 0 | 0 | 0 | 0 | 0 | 0 |
| 0 | NA | NA | NA | NA | NA | Tex19.2 | testis expressed gene 19.2 | 0 | 0 | 0 | 0 | 0 | 0 | 0 | 0 | 0 | 0 | 0 |
| 0 | NA | NA | NA | NA | NA | Tex19.1 | testis expressed gene 19.1 | 0 | 0 | 0 | 0 | 0 | 0 | 0 | 0 | 0 | 0 | 0 |
| 0 | NA | NA | NA | NA | NA | Uts2r | urotensin 2 receptor | 0 | 0 | 0 | 0 | 0 | 0 | 0 | 0 | 0 | 0 | 0 |
| 0 | NA | NA | NA | NA | NA | Gm11790 | | 0 | 0 | 0 | 0 | 0 | 0 | 0 | 0 | 0 | 0 | 0 |
| 0 | NA | NA | NA | NA | NA | Mir7236 | | 0 | 0 | 0 | 0 | 0 | 0 | 0 | 0 | 0 | 0 | 0 |
| 0 | NA | NA | NA | NA | NA | Gm11866 | | 0 | 0 | 0 | 0 | 0 | 0 | 0 | 0 | 0 | 0 | 0 |
| 0 | NA | NA | NA | NA | NA | Gm12589 | | 0 | 0 | 0 | 0 | 0 | 0 | 0 | 0 | 0 | 0 | 0 |
| 0 | NA | NA | NA | NA | NA | Gm29513 | | 0 | 0 | 0 | 0 | 0 | 0 | 0 | 0 | 0 | 0 | 0 |

| | | | | | | | | | | | | | | | | | | |
|---|---|---|---|---|---|---|---|---|---|---|---|---|---|---|---|---|---|---|
| 0 | NA | NA | NA | NA | NA | Gm15643 | | 0 | 0 | 0 | 0 | 0 | 0 | 0 | 0 | 0 | 0 | 0 |
| 0 | NA | NA | NA | NA | NA | Ptchd3 | patched domain containing 3 | 0 | 0 | 0 | 0 | 0 | 0 | 0 | 0 | 0 | 0 | 0 |
| 0 | NA | NA | NA | NA | NA | Gm12590 | | 0 | 0 | 0 | 0 | 0 | 0 | 0 | 0 | 0 | 0 | 0 |
| 0 | NA | NA | NA | NA | NA | Gm12591 | | 0 | 0 | 0 | 0 | 0 | 0 | 0 | 0 | 0 | 0 | 0 |
| 0 | NA | NA | NA | NA | NA | Gm9071 | | 0 | 0 | 0 | 0 | 0 | 0 | 0 | 0 | 0 | 0 | 0 |
| 0 | NA | NA | NA | NA | NA | Gm18244 | | 0 | 0 | 0 | 0 | 0 | 0 | 0 | 0 | 0 | 0 | 0 |
| 0 | NA | NA | NA | NA | NA | Gm9072 | | 0 | 0 | 0 | 0 | 0 | 0 | 0 | 0 | 0 | 0 | 0 |
| 0 | NA | NA | NA | NA | NA | Gm48310 | | 0 | 0 | 0 | 0 | 0 | 0 | 0 | 0 | 0 | 0 | 0 |
| 0 | NA | NA | NA | NA | NA | Gm9075 | | 0 | 0 | 0 | 0 | 0 | 0 | 0 | 0 | 0 | 0 | 0 |
| 0 | NA | NA | NA | NA | NA | Gm48512 | | 0 | 0 | 0 | 0 | 0 | 0 | 0 | 0 | 0 | 0 | 0 |
| 0 | NA | NA | NA | NA | NA | Gm48511 | | 0 | 0 | 0 | 0 | 0 | 0 | 0 | 0 | 0 | 0 | 0 |
| 0 | NA | NA | NA | NA | NA | Gm26050 | | 0 | 0 | 0 | 0 | 0 | 0 | 0 | 0 | 0 | 0 | 0 |
| 0 | NA | NA | NA | NA | NA | Gm9088 | | 0 | 0 | 0 | 0 | 0 | 0 | 0 | 0 | 0 | 0 | 0 |
| 0 | NA | NA | NA | NA | NA | Gm48001 | | 0 | 0 | 0 | 0 | 0 | 0 | 0 | 0 | 0 | 0 | 0 |
| 0 | NA | NA | NA | NA | NA | Gm48122 | | 0 | 0 | 0 | 0 | 0 | 0 | 0 | 0 | 0 | 0 | 0 |
| 0 | NA | NA | NA | NA | NA | Gm48210 | | 0 | 0 | 0 | 0 | 0 | 0 | 0 | 0 | 0 | 0 | 0 |
| 0 | NA | NA | NA | NA | NA | Gm29696 | | 0 | 0 | 0 | 0 | 0 | 0 | 0 | 0 | 0 | 0 | 0 |
| 0 | NA | NA | NA | NA | NA | Gm23756 | | 0 | 0 | 0 | 0 | 0 | 0 | 0 | 0 | 0 | 0 | 0 |
| 0 | NA | NA | NA | NA | NA | Gm22343 | | 0 | 0 | 0 | 0 | 0 | 0 | 0 | 0 | 0 | 0 | 0 |
| 0 | NA | NA | NA | NA | NA | Gm48678 | | 0 | 0 | 0 | 0 | 0 | 0 | 0 | 0 | 0 | 0 | 0 |
| 0 | NA | NA | NA | NA | NA | Gm23676 | | 0 | 0 | 0 | 0 | 0 | 0 | 0 | 0 | 0 | 0 | 0 |
| 0 | NA | NA | NA | NA | NA | Gm22003 | | 0 | 0 | 0 | 0 | 0 | 0 | 0 | 0 | 0 | 0 | 0 |
| 0 | NA | NA | NA | NA | NA | | | 0 | 0 | 0 | 0 | 0 | 0 | 0 | 0 | 0 | 0 | 0 |
| 0 | NA | NA | NA | NA | NA | Gm23167 | | 0 | 0 | 0 | 0 | 0 | 0 | 0 | 0 | 0 | 0 | 0 |
| 0 | NA | NA | NA | NA | NA | Gm9110 | | 0 | 0 | 0 | 0 | 0 | 0 | 0 | 0 | 0 | 0 | 0 |
| 0 | NA | NA | NA | NA | NA | Gm48532 | | 0 | 0 | 0 | 0 | 0 | 0 | 0 | 0 | 0 | 0 | 0 |
| 0 | NA | NA | NA | NA | NA | Gm48599 | | 0 | 0 | 0 | 0 | 0 | 0 | 0 | 0 | 0 | 0 | 0 |
| 0 | NA | NA | NA | NA | NA | Gm32442 | | 0 | 0 | 0 | 0 | 0 | 0 | 0 | 0 | 0 | 0 | 0 |
| 0 | NA | NA | NA | NA | NA | Gm23752 | | 0 | 0 | 0 | 0 | 0 | 0 | 0 | 0 | 0 | 0 | 0 |
| 0 | NA | NA | NA | NA | NA | Gm48608 | | 0 | 0 | 0 | 0 | 0 | 0 | 0 | 0 | 0 | 0 | 0 |
| 0 | NA | NA | NA | NA | NA | Gm4927 | | 0 | 0 | 0 | 0 | 0 | 0 | 0 | 0 | 0 | 0 | 0 |
| 0 | NA | NA | NA | NA | NA | Gm3678 | | 0 | 0 | 0 | 0 | 0 | 0 | 0 | 0 | 0 | 0 | 0 |
| 0 | NA | NA | NA | NA | NA | Gm48619 | | 0 | 0 | 0 | 0 | 0 | 0 | 0 | 0 | 0 | 0 | 0 |
| 0 | NA | NA | NA | NA | NA | 1700101O22Rik | | 0 | 0 | 0 | 0 | 0 | 0 | 0 | 0 | 0 | 0 | 0 |
| 0 | NA | NA | NA | NA | NA | Gm9139 | | 0 | 0 | 0 | 0 | 0 | 0 | 0 | 0 | 0 | 0 | 0 |
| 0 | NA | NA | NA | NA | NA | Gm18387 | | 0 | 0 | 0 | 0 | 0 | 0 | 0 | 0 | 0 | 0 | 0 |
| 0 | NA | NA | NA | NA | NA | Gdf7 | growth differentiation factor 7 | 0 | 0 | 0 | 0 | 0 | 0 | 0 | 0 | 0 | 0 | 0 |
| 0 | NA | NA | NA | NA | NA | Gm48076 | | 0 | 0 | 0 | 0 | 0 | 0 | 0 | 0 | 0 | 0 | 0 |
| 0 | NA | NA | NA | NA | NA | Gm47391 | | 0 | 0 | 0 | 0 | 0 | 0 | 0 | 0 | 0 | 0 | 0 |
| 0 | NA | NA | NA | NA | NA | Matn3 | matrilin 3 | 0 | 0 | 0 | 0 | 0 | 0 | 0 | 0 | 0 | 0 | 0 |
| 0 | NA | NA | NA | NA | NA | Gm18696 | | 0 | 0 | 0 | 0 | 0 | 0 | 0 | 0 | 0 | 0 | 0 |
| 0 | NA | NA | NA | NA | NA | Gm48762 | | 0 | 0 | 0 | 0 | 0 | 0 | 0 | 0 | 0 | 0 | 0 |
| 0 | NA | NA | NA | NA | NA | Gm48764 | | 0 | 0 | 0 | 0 | 0 | 0 | 0 | 0 | 0 | 0 | 0 |
| 0 | NA | NA | NA | NA | NA | 1700022H16Rik | | 0 | 0 | 0 | 0 | 0 | 0 | 0 | 0 | 0 | 0 | 0 |
| 0 | NA | NA | NA | NA | NA | Gm23979 | | 0 | 0 | 0 | 0 | 0 | 0 | 0 | 0 | 0 | 0 | 0 |
| 0 | NA | NA | NA | NA | NA | Gm48774 | | 0 | 0 | 0 | 0 | 0 | 0 | 0 | 0 | 0 | 0 | 0 |
| 0 | NA | NA | NA | NA | NA | Gm48790 | | 0 | 0 | 0 | 0 | 0 | 0 | 0 | 0 | 0 | 0 | 0 |
| 0 | NA | NA | NA | NA | NA | Gm22845 | | 0 | 0 | 0 | 0 | 0 | 0 | 0 | 0 | 0 | 0 | 0 |
| 0 | NA | NA | NA | NA | NA | Gm48791 | | 0 | 0 | 0 | 0 | 0 | 0 | 0 | 0 | 0 | 0 | 0 |
| 0 | NA | NA | NA | NA | NA | Gm48792 | | 0 | 0 | 0 | 0 | 0 | 0 | 0 | 0 | 0 | 0 | 0 |
| 0 | NA | NA | NA | NA | NA | Gm48793 | | 0 | 0 | 0 | 0 | 0 | 0 | 0 | 0 | 0 | 0 | 0 |
| 0 | NA | NA | NA | NA | NA | Gm48794 | | 0 | 0 | 0 | 0 | 0 | 0 | 0 | 0 | 0 | 0 | 0 |
| 0 | NA | NA | NA | NA | NA | Gm48886 | | 0 | 0 | 0 | 0 | 0 | 0 | 0 | 0 | 0 | 0 | 0 |
| 0 | NA | NA | NA | NA | NA | Gm48082 | | 0 | 0 | 0 | 0 | 0 | 0 | 0 | 0 | 0 | 0 | 0 |
| 0 | NA | NA | NA | NA | NA | 7420701I03Rik | | 0 | 0 | 0 | 0 | 0 | 0 | 0 | 0 | 0 | 0 | 0 |
| 0 | NA | NA | NA | NA | NA | 9530020I12Rik | | 0 | 0 | 0 | 0 | 0 | 0 | 0 | 0 | 0 | 0 | 0 |
| 0 | NA | NA | NA | NA | NA | Msgn1 | mesogenin 1 | 0 | 0 | 0 | 0 | 0 | 0 | 0 | 0 | 0 | 0 | 0 |
| 0 | NA | NA | NA | NA | NA | Gm4803 | | 0 | 0 | 0 | 0 | 0 | 0 | 0 | 0 | 0 | 0 | 0 |
| 0 | NA | NA | NA | NA | NA | Gm23781 | | 0 | 0 | 0 | 0 | 0 | 0 | 0 | 0 | 0 | 0 | 0 |
| 0 | NA | NA | NA | NA | NA | Gm47821 | | 0 | 0 | 0 | 0 | 0 | 0 | 0 | 0 | 0 | 0 | 0 |

| | | | | | | | | | | | | | | | | | | |
|---|---|---|---|---|---|---|---|---|---|---|---|---|---|---|---|---|---|---|
| 0 | NA | NA | NA | NA | NA | Gm47822 | | 0 | 0 | 0 | 0 | 0 | 0 | 0 | 0 | 0 | 0 | 0 |
| 0 | NA | NA | NA | NA | NA | Gm25610 | | 0 | 0 | 0 | 0 | 0 | 0 | 0 | 0 | 0 | 0 | 0 |
| 0 | NA | NA | NA | NA | NA | Gm26171 | | 0 | 0 | 0 | 0 | 0 | 0 | 0 | 0 | 0 | 0 | 0 |
| 0 | NA | NA | NA | NA | NA | 4921511I17Rik | | 0 | 0 | 0 | 0 | 0 | 0 | 0 | 0 | 0 | 0 | 0 |
| 0 | NA | NA | NA | NA | NA | | | 0 | 0 | 0 | 0 | 0 | 0 | 0 | 0 | 0 | 0 | 0 |
| 0 | NA | NA | NA | NA | NA | Gm27952 | | 0 | 0 | 0 | 0 | 0 | 0 | 0 | 0 | 0 | 0 | 0 |
| 0 | NA | NA | NA | NA | NA | Gm25538 | | 0 | 0 | 0 | 0 | 0 | 0 | 0 | 0 | 0 | 0 | 0 |
| 0 | NA | NA | NA | NA | NA | 3732407C23Rik | | 0 | 0 | 0 | 0 | 0 | 0 | 0 | 0 | 0 | 0 | 0 |
| 0 | NA | NA | NA | NA | NA | Platr19 | | 0 | 0 | 0 | 0 | 0 | 0 | 0 | 0 | 0 | 0 | 0 |
| 0 | NA | NA | NA | NA | NA | Gm48187 | | 0 | 0 | 0 | 0 | 0 | 0 | 0 | 0 | 0 | 0 | 0 |
| 0 | NA | NA | NA | NA | NA | Gm40271 | | 0 | 0 | 0 | 0 | 0 | 0 | 0 | 0 | 0 | 0 | 0 |
| 0 | NA | NA | NA | NA | NA | Gm48201 | | 0 | 0 | 0 | 0 | 0 | 0 | 0 | 0 | 0 | 0 | 0 |
| 0 | NA | NA | NA | NA | NA | Gm35208 | | 0 | 0 | 0 | 0 | 0 | 0 | 0 | 0 | 0 | 0 | 0 |
| 0 | NA | NA | NA | NA | NA | Gm48209 | | 0 | 0 | 0 | 0 | 0 | 0 | 0 | 0 | 0 | 0 | 0 |
| 0 | NA | NA | NA | NA | NA | Gm35298 | | 0 | 0 | 0 | 0 | 0 | 0 | 0 | 0 | 0 | 0 | 0 |
| 0 | NA | NA | NA | NA | NA | Gm48213 | | 0 | 0 | 0 | 0 | 0 | 0 | 0 | 0 | 0 | 0 | 0 |
| 0 | NA | NA | NA | NA | NA | Rpl21-ps2 | | 0 | 0 | 0 | 0 | 0 | 0 | 0 | 0 | 0 | 0 | 0 |
| 0 | NA | NA | NA | NA | NA | Gm24208 | | 0 | 0 | 0 | 0 | 0 | 0 | 0 | 0 | 0 | 0 | 0 |
| 0 | NA | NA | NA | NA | NA | Gm48309 | | 0 | 0 | 0 | 0 | 0 | 0 | 0 | 0 | 0 | 0 | 0 |
| 0 | NA | NA | NA | NA | NA | Gm48311 | | 0 | 0 | 0 | 0 | 0 | 0 | 0 | 0 | 0 | 0 | 0 |
| 0 | NA | NA | NA | NA | NA | Gm24142 | | 0 | 0 | 0 | 0 | 0 | 0 | 0 | 0 | 0 | 0 | 0 |
| 0 | NA | NA | NA | NA | NA | Gm22170 | | 0 | 0 | 0 | 0 | 0 | 0 | 0 | 0 | 0 | 0 | 0 |
| 0 | NA | NA | NA | NA | NA | Gm35725 | | 0 | 0 | 0 | 0 | 0 | 0 | 0 | 0 | 0 | 0 | 0 |
| 0 | NA | NA | NA | NA | NA | Gm35890 | | 0 | 0 | 0 | 0 | 0 | 0 | 0 | 0 | 0 | 0 | 0 |
| 0 | NA | NA | NA | NA | NA | Gm49374 | | 0 | 0 | 0 | 0 | 0 | 0 | 0 | 0 | 0 | 0 | 0 |
| 0 | NA | NA | NA | NA | NA | Gm7126 | | 0 | 0 | 0 | 0 | 0 | 0 | 0 | 0 | 0 | 0 | 0 |
| 0 | NA | NA | NA | NA | NA | Gm9194 | | 0 | 0 | 0 | 0 | 0 | 0 | 0 | 0 | 0 | 0 | 0 |
| 0 | NA | NA | NA | NA | NA | Gm48479 | | 0 | 0 | 0 | 0 | 0 | 0 | 0 | 0 | 0 | 0 | 0 |
| 0 | NA | NA | NA | NA | NA | Gm48480 | | 0 | 0 | 0 | 0 | 0 | 0 | 0 | 0 | 0 | 0 | 0 |
| 0 | NA | NA | NA | NA | NA | Gm16497 | | 0 | 0 | 0 | 0 | 0 | 0 | 0 | 0 | 0 | 0 | 0 |
| 0 | NA | NA | NA | NA | NA | Gm48498 | | 0 | 0 | 0 | 0 | 0 | 0 | 0 | 0 | 0 | 0 | 0 |
| 0 | NA | NA | NA | NA | NA | Gm25414 | | 0 | 0 | 0 | 0 | 0 | 0 | 0 | 0 | 0 | 0 | 0 |
| 0 | NA | NA | NA | NA | NA | Gm48504 | | 0 | 0 | 0 | 0 | 0 | 0 | 0 | 0 | 0 | 0 | 0 |
| 0 | NA | NA | NA | NA | NA | Gm26449 | | 0 | 0 | 0 | 0 | 0 | 0 | 0 | 0 | 0 | 0 | 0 |
| 0 | NA | NA | NA | NA | NA | 4930448C13Rik | | 0 | 0 | 0 | 0 | 0 | 0 | 0 | 0 | 0 | 0 | 0 |
| 0 | NA | NA | NA | NA | NA | Gm27682 | | 0 | 0 | 0 | 0 | 0 | 0 | 0 | 0 | 0 | 0 | 0 |
| 0 | NA | NA | NA | NA | NA | Gm36128 | | 0 | 0 | 0 | 0 | 0 | 0 | 0 | 0 | 0 | 0 | 0 |
| 0 | NA | NA | NA | NA | NA | Gm48539 | | 0 | 0 | 0 | 0 | 0 | 0 | 0 | 0 | 0 | 0 | 0 |
| 0 | NA | NA | NA | NA | NA | Gm48541 | | 0 | 0 | 0 | 0 | 0 | 0 | 0 | 0 | 0 | 0 | 0 |
| 0 | NA | NA | NA | NA | NA | Gm17915 | | 0 | 0 | 0 | 0 | 0 | 0 | 0 | 0 | 0 | 0 | 0 |
| 0 | NA | NA | NA | NA | NA | Gm48584 | | 0 | 0 | 0 | 0 | 0 | 0 | 0 | 0 | 0 | 0 | 0 |
| 0 | NA | NA | NA | NA | NA | Gm18114 | | 0 | 0 | 0 | 0 | 0 | 0 | 0 | 0 | 0 | 0 | 0 |
| 0 | NA | NA | NA | NA | NA | Gm48607 | | 0 | 0 | 0 | 0 | 0 | 0 | 0 | 0 | 0 | 0 | 0 |
| 0 | NA | NA | NA | NA | NA | Gm36372 | | 0 | 0 | 0 | 0 | 0 | 0 | 0 | 0 | 0 | 0 | 0 |
| 0 | NA | NA | NA | NA | NA | Ntsr2 | neurotensin receptor 2 | 0 | 0 | 0 | 0 | 0 | 0 | 0 | 0 | 0 | 0 | 0 |
| 0 | NA | NA | NA | NA | NA | Gm48140 | | 0 | 0 | 0 | 0 | 0 | 0 | 0 | 0 | 0 | 0 | 0 |
| 0 | NA | NA | NA | NA | NA | Gm48227 | | 0 | 0 | 0 | 0 | 0 | 0 | 0 | 0 | 0 | 0 | 0 |
| 0 | NA | NA | NA | NA | NA | Gm48291 | | 0 | 0 | 0 | 0 | 0 | 0 | 0 | 0 | 0 | 0 | 0 |
| 0 | NA | NA | NA | NA | NA | Gm22303 | | 0 | 0 | 0 | 0 | 0 | 0 | 0 | 0 | 0 | 0 | 0 |
| 0 | NA | NA | NA | NA | NA | Gm48538 | | 0 | 0 | 0 | 0 | 0 | 0 | 0 | 0 | 0 | 0 | 0 |
| 0 | NA | NA | NA | NA | NA | Mir3066 | | 0 | 0 | 0 | 0 | 0 | 0 | 0 | 0 | 0 | 0 | 0 |
| 0 | NA | NA | NA | NA | NA | Gm48313 | | 0 | 0 | 0 | 0 | 0 | 0 | 0 | 0 | 0 | 0 | 0 |
| 0 | NA | NA | NA | NA | NA | Gm25925 | | 0 | 0 | 0 | 0 | 0 | 0 | 0 | 0 | 0 | 0 | 0 |
| 0 | NA | NA | NA | NA | NA | Gm48315 | | 0 | 0 | 0 | 0 | 0 | 0 | 0 | 0 | 0 | 0 | 0 |
| 0 | NA | NA | NA | NA | NA | Gm48317 | | 0 | 0 | 0 | 0 | 0 | 0 | 0 | 0 | 0 | 0 | 0 |
| 0 | NA | NA | NA | NA | NA | Gm48319 | | 0 | 0 | 0 | 0 | 0 | 0 | 0 | 0 | 0 | 0 | 0 |
| 0 | NA | NA | NA | NA | NA | Gm48318 | | 0 | 0 | 0 | 0 | 0 | 0 | 0 | 0 | 0 | 0 | 0 |
| 0 | NA | NA | NA | NA | NA | Gm48347 | | 0 | 0 | 0 | 0 | 0 | 0 | 0 | 0 | 0 | 0 | 0 |
| 0 | NA | NA | NA | NA | NA | Gm48354 | | 0 | 0 | 0 | 0 | 0 | 0 | 0 | 0 | 0 | 0 | 0 |
| 0 | NA | NA | NA | NA | NA | Gm48355 | | 0 | 0 | 0 | 0 | 0 | 0 | 0 | 0 | 0 | 0 | 0 |

| | | | | | | | | | | | | | | | | | |
|---|---|---|---|---|---|---|---|---|---|---|---|---|---|---|---|---|---|
| 0 | NA | NA | NA | NA | NA | Gm40879 | 0 | 0 | 0 | 0 | 0 | 0 | 0 | 0 | 0 | 0 | 0 |
| 0 | NA | NA | NA | NA | NA | Gm19368 | 0 | 0 | 0 | 0 | 0 | 0 | 0 | 0 | 0 | 0 | 0 |
| 0 | NA | NA | NA | NA | NA | Gm7093 | 0 | 0 | 0 | 0 | 0 | 0 | 0 | 0 | 0 | 0 | 0 |
| 0 | NA | NA | NA | NA | NA | Gm40848 | 0 | 0 | 0 | 0 | 0 | 0 | 0 | 0 | 0 | 0 | 0 |
| 0 | NA | NA | NA | NA | NA | Gm48392 | 0 | 0 | 0 | 0 | 0 | 0 | 0 | 0 | 0 | 0 | 0 |
| 0 | NA | NA | NA | NA | NA | Gm19786 | 0 | 0 | 0 | 0 | 0 | 0 | 0 | 0 | 0 | 0 | 0 |
| 0 | NA | NA | NA | NA | NA | Gm48409 | 0 | 0 | 0 | 0 | 0 | 0 | 0 | 0 | 0 | 0 | 0 |
| 0 | NA | NA | NA | NA | NA | Gm9237 | 0 | 0 | 0 | 0 | 0 | 0 | 0 | 0 | 0 | 0 | 0 |
| 0 | NA | NA | NA | NA | NA | Gm48411 | 0 | 0 | 0 | 0 | 0 | 0 | 0 | 0 | 0 | 0 | 0 |
| 0 | NA | NA | NA | NA | NA | Gm48446 | 0 | 0 | 0 | 0 | 0 | 0 | 0 | 0 | 0 | 0 | 0 |
| 0 | NA | NA | NA | NA | NA | Gm49370 | 0 | 0 | 0 | 0 | 0 | 0 | 0 | 0 | 0 | 0 | 0 |
| 0 | NA | NA | NA | NA | NA | Gm46338 | 0 | 0 | 0 | 0 | 0 | 0 | 0 | 0 | 0 | 0 | 0 |
| 0 | NA | NA | NA | NA | NA | Gm49764 | 0 | 0 | 0 | 0 | 0 | 0 | 0 | 0 | 0 | 0 | 0 |
| 0 | NA | NA | NA | NA | NA | Gm48472 | 0 | 0 | 0 | 0 | 0 | 0 | 0 | 0 | 0 | 0 | 0 |
| 0 | NA | NA | NA | NA | NA | Gm9249 | 0 | 0 | 0 | 0 | 0 | 0 | 0 | 0 | 0 | 0 | 0 |
| 0 | NA | NA | NA | NA | NA | Gm48516 | 0 | 0 | 0 | 0 | 0 | 0 | 0 | 0 | 0 | 0 | 0 |
| 0 | NA | NA | NA | NA | NA | Gm40849 | 0 | 0 | 0 | 0 | 0 | 0 | 0 | 0 | 0 | 0 | 0 |
| 0 | NA | NA | NA | NA | NA | Gm9158 | 0 | 0 | 0 | 0 | 0 | 0 | 0 | 0 | 0 | 0 | 0 |
| 0 | NA | NA | NA | NA | NA | Gm48523 | 0 | 0 | 0 | 0 | 0 | 0 | 0 | 0 | 0 | 0 | 0 |
| 0 | NA | NA | NA | NA | NA | Gm48525 | 0 | 0 | 0 | 0 | 0 | 0 | 0 | 0 | 0 | 0 | 0 |
| 0 | NA | NA | NA | NA | NA | Gm48528 | 0 | 0 | 0 | 0 | 0 | 0 | 0 | 0 | 0 | 0 | 0 |
| 0 | NA | NA | NA | NA | NA | Gm49957 | 0 | 0 | 0 | 0 | 0 | 0 | 0 | 0 | 0 | 0 | 0 |
| 0 | NA | NA | NA | NA | NA | Gm48540 | 0 | 0 | 0 | 0 | 0 | 0 | 0 | 0 | 0 | 0 | 0 |
| 0 | NA | NA | NA | NA | NA | Gm9255 | 0 | 0 | 0 | 0 | 0 | 0 | 0 | 0 | 0 | 0 | 0 |
| 0 | NA | NA | NA | NA | NA | Gm48576 | 0 | 0 | 0 | 0 | 0 | 0 | 0 | 0 | 0 | 0 | 0 |
| 0 | NA | NA | NA | NA | NA | Gm6802 | 0 | 0 | 0 | 0 | 0 | 0 | 0 | 0 | 0 | 0 | 0 |
| 0 | NA | NA | NA | NA | NA | Gm6943 | 0 | 0 | 0 | 0 | 0 | 0 | 0 | 0 | 0 | 0 | 0 |
| 0 | NA | NA | NA | NA | NA | Gm49372 | 0 | 0 | 0 | 0 | 0 | 0 | 0 | 0 | 0 | 0 | 0 |
| 0 | NA | NA | NA | NA | NA | Gm49371 | 0 | 0 | 0 | 0 | 0 | 0 | 0 | 0 | 0 | 0 | 0 |
| 0 | NA | NA | NA | NA | NA | Gm48625 | 0 | 0 | 0 | 0 | 0 | 0 | 0 | 0 | 0 | 0 | 0 |
| 0 | NA | NA | NA | NA | NA | Gm9257 | 0 | 0 | 0 | 0 | 0 | 0 | 0 | 0 | 0 | 0 | 0 |
| 0 | NA | NA | NA | NA | NA | Gm48635 | 0 | 0 | 0 | 0 | 0 | 0 | 0 | 0 | 0 | 0 | 0 |
| 0 | NA | NA | NA | NA | NA | Gm48637 | 0 | 0 | 0 | 0 | 0 | 0 | 0 | 0 | 0 | 0 | 0 |
| 0 | NA | NA | NA | NA | NA | Gm48638 | 0 | 0 | 0 | 0 | 0 | 0 | 0 | 0 | 0 | 0 | 0 |
| 0 | NA | NA | NA | NA | NA | Gm48641 | 0 | 0 | 0 | 0 | 0 | 0 | 0 | 0 | 0 | 0 | 0 |
| 0 | NA | NA | NA | NA | NA | Gm9259 | 0 | 0 | 0 | 0 | 0 | 0 | 0 | 0 | 0 | 0 | 0 |
| 0 | NA | NA | NA | NA | NA | Gm49379 | 0 | 0 | 0 | 0 | 0 | 0 | 0 | 0 | 0 | 0 | 0 |
| 0 | NA | NA | NA | NA | NA | Gm22189 | 0 | 0 | 0 | 0 | 0 | 0 | 0 | 0 | 0 | 0 | 0 |
| 0 | NA | NA | NA | NA | NA | Gm48644 | 0 | 0 | 0 | 0 | 0 | 0 | 0 | 0 | 0 | 0 | 0 |
| 0 | NA | NA | NA | NA | NA | Gm21863 | 0 | 0 | 0 | 0 | 0 | 0 | 0 | 0 | 0 | 0 | 0 |
| 0 | NA | NA | NA | NA | NA | Gm48645 | 0 | 0 | 0 | 0 | 0 | 0 | 0 | 0 | 0 | 0 | 0 |
| 0 | NA | NA | NA | NA | NA | Gm40853 | 0 | 0 | 0 | 0 | 0 | 0 | 0 | 0 | 0 | 0 | 0 |
| 0 | NA | NA | NA | NA | NA | Gm48647 | 0 | 0 | 0 | 0 | 0 | 0 | 0 | 0 | 0 | 0 | 0 |
| 0 | NA | NA | NA | NA | NA | Gm49381 | 0 | 0 | 0 | 0 | 0 | 0 | 0 | 0 | 0 | 0 | 0 |
| 0 | NA | NA | NA | NA | NA | Gm9265 | 0 | 0 | 0 | 0 | 0 | 0 | 0 | 0 | 0 | 0 | 0 |
| 0 | NA | NA | NA | NA | NA | Gm49382 | 0 | 0 | 0 | 0 | 0 | 0 | 0 | 0 | 0 | 0 | 0 |
| 0 | NA | NA | NA | NA | NA | Gm49692 | 0 | 0 | 0 | 0 | 0 | 0 | 0 | 0 | 0 | 0 | 0 |
| 0 | NA | NA | NA | NA | NA | Gm48668 | 0 | 0 | 0 | 0 | 0 | 0 | 0 | 0 | 0 | 0 | 0 |
| 0 | NA | NA | NA | NA | NA | Gm49690 | 0 | 0 | 0 | 0 | 0 | 0 | 0 | 0 | 0 | 0 | 0 |
| 0 | NA | NA | NA | NA | NA | Gm49697 | 0 | 0 | 0 | 0 | 0 | 0 | 0 | 0 | 0 | 0 | 0 |
| 0 | NA | NA | NA | NA | NA | Gm40881 | 0 | 0 | 0 | 0 | 0 | 0 | 0 | 0 | 0 | 0 | 0 |
| 0 | NA | NA | NA | NA | NA | Gm49929 | 0 | 0 | 0 | 0 | 0 | 0 | 0 | 0 | 0 | 0 | 0 |
| 0 | NA | NA | NA | NA | NA | Mrto4-ps1 | 0 | 0 | 0 | 0 | 0 | 0 | 0 | 0 | 0 | 0 | 0 |
| 0 | NA | NA | NA | NA | NA | Gm10479 | 0 | 0 | 0 | 0 | 0 | 0 | 0 | 0 | 0 | 0 | 0 |
| 0 | NA | NA | NA | NA | NA | Gm49698 | 0 | 0 | 0 | 0 | 0 | 0 | 0 | 0 | 0 | 0 | 0 |
| 0 | NA | NA | NA | NA | NA | Gm48669 | 0 | 0 | 0 | 0 | 0 | 0 | 0 | 0 | 0 | 0 | 0 |
| 0 | NA | NA | NA | NA | NA | Gm4012 | 0 | 0 | 0 | 0 | 0 | 0 | 0 | 0 | 0 | 0 | 0 |
| 0 | NA | NA | NA | NA | NA | Gm48688 | 0 | 0 | 0 | 0 | 0 | 0 | 0 | 0 | 0 | 0 | 0 |
| 0 | NA | NA | NA | NA | NA | Gm48693 | 0 | 0 | 0 | 0 | 0 | 0 | 0 | 0 | 0 | 0 | 0 |
| 0 | NA | NA | NA | NA | NA | Gm16476 | 0 | 0 | 0 | 0 | 0 | 0 | 0 | 0 | 0 | 0 | 0 |

| | | | | | | | | | | | | | | | | | |
|---|---|---|---|---|---|---|---|---|---|---|---|---|---|---|---|---|---|
| 0 | NA | NA | NA | NA | NA | Gm28503 | 0 | 0 | 0 | 0 | 0 | 0 | 0 | 0 | 0 | 0 | 0 |
| 0 | NA | NA | NA | NA | NA | Gm21989 | 0 | 0 | 0 | 0 | 0 | 0 | 0 | 0 | 0 | 0 | 0 |
| 0 | NA | NA | NA | NA | NA | | 0 | 0 | 0 | 0 | 0 | 0 | 0 | 0 | 0 | 0 | 0 |
| 0 | NA | NA | NA | NA | NA | Gm22766 | 0 | 0 | 0 | 0 | 0 | 0 | 0 | 0 | 0 | 0 | 0 |
| 0 | NA | NA | NA | NA | NA | Gm4036 | 0 | 0 | 0 | 0 | 0 | 0 | 0 | 0 | 0 | 0 | 0 |
| 0 | NA | NA | NA | NA | NA | Mrto4-ps2 | 0 | 0 | 0 | 0 | 0 | 0 | 0 | 0 | 0 | 0 | 0 |
| 0 | NA | NA | NA | NA | NA | Gm22908 | 0 | 0 | 0 | 0 | 0 | 0 | 0 | 0 | 0 | 0 | 0 |
| 0 | NA | NA | NA | NA | NA | Gm9289 | 0 | 0 | 0 | 0 | 0 | 0 | 0 | 0 | 0 | 0 | 0 |
| 0 | NA | NA | NA | NA | NA | Gm20486 | 0 | 0 | 0 | 0 | 0 | 0 | 0 | 0 | 0 | 0 | 0 |
| 0 | NA | NA | NA | NA | NA | Gm6838 | 0 | 0 | 0 | 0 | 0 | 0 | 0 | 0 | 0 | 0 | 0 |
| 0 | NA | NA | NA | NA | NA | Gm49248 | 0 | 0 | 0 | 0 | 0 | 0 | 0 | 0 | 0 | 0 | 0 |
| 0 | NA | NA | NA | NA | NA | Gm49251 | 0 | 0 | 0 | 0 | 0 | 0 | 0 | 0 | 0 | 0 | 0 |
| 0 | NA | NA | NA | NA | NA | Gm47997 | 0 | 0 | 0 | 0 | 0 | 0 | 0 | 0 | 0 | 0 | 0 |
| 0 | NA | NA | NA | NA | NA | Gm48012 | 0 | 0 | 0 | 0 | 0 | 0 | 0 | 0 | 0 | 0 | 0 |
| 0 | NA | NA | NA | NA | NA | Gm48016 | 0 | 0 | 0 | 0 | 0 | 0 | 0 | 0 | 0 | 0 | 0 |
| 0 | NA | NA | NA | NA | NA | Gm40864 | 0 | 0 | 0 | 0 | 0 | 0 | 0 | 0 | 0 | 0 | 0 |
| 0 | NA | NA | NA | NA | NA | Gm48019 | 0 | 0 | 0 | 0 | 0 | 0 | 0 | 0 | 0 | 0 | 0 |
| 0 | NA | NA | NA | NA | NA | Gm48020 | 0 | 0 | 0 | 0 | 0 | 0 | 0 | 0 | 0 | 0 | 0 |
| 0 | NA | NA | NA | NA | NA | Gm17391 | 0 | 0 | 0 | 0 | 0 | 0 | 0 | 0 | 0 | 0 | 0 |
| 0 | NA | NA | NA | NA | NA | Gm48030 | 0 | 0 | 0 | 0 | 0 | 0 | 0 | 0 | 0 | 0 | 0 |
| 0 | NA | NA | NA | NA | NA | Gm48031 | 0 | 0 | 0 | 0 | 0 | 0 | 0 | 0 | 0 | 0 | 0 |
| 0 | NA | NA | NA | NA | NA | Gm48032 | 0 | 0 | 0 | 0 | 0 | 0 | 0 | 0 | 0 | 0 | 0 |
| 0 | NA | NA | NA | NA | NA | Gm48033 | 0 | 0 | 0 | 0 | 0 | 0 | 0 | 0 | 0 | 0 | 0 |
| 0 | NA | NA | NA | NA | NA | Gm48037 | 0 | 0 | 0 | 0 | 0 | 0 | 0 | 0 | 0 | 0 | 0 |
| 0 | NA | NA | NA | NA | NA | Gm20336 | 0 | 0 | 0 | 0 | 0 | 0 | 0 | 0 | 0 | 0 | 0 |
| 0 | NA | NA | NA | NA | NA | Gm48039 | 0 | 0 | 0 | 0 | 0 | 0 | 0 | 0 | 0 | 0 | 0 |
| 0 | NA | NA | NA | NA | NA | Gm48041 | 0 | 0 | 0 | 0 | 0 | 0 | 0 | 0 | 0 | 0 | 0 |
| 0 | NA | NA | NA | NA | NA | Gm48042 | 0 | 0 | 0 | 0 | 0 | 0 | 0 | 0 | 0 | 0 | 0 |
| 0 | NA | NA | NA | NA | NA | Gm48043 | 0 | 0 | 0 | 0 | 0 | 0 | 0 | 0 | 0 | 0 | 0 |
| 0 | NA | NA | NA | NA | NA | Gm49686 | 0 | 0 | 0 | 0 | 0 | 0 | 0 | 0 | 0 | 0 | 0 |
| 0 | NA | NA | NA | NA | NA | Gm48044 | 0 | 0 | 0 | 0 | 0 | 0 | 0 | 0 | 0 | 0 | 0 |
| 0 | NA | NA | NA | NA | NA | Gm48046 | 0 | 0 | 0 | 0 | 0 | 0 | 0 | 0 | 0 | 0 | 0 |
| 0 | NA | NA | NA | NA | NA | Gm48064 | 0 | 0 | 0 | 0 | 0 | 0 | 0 | 0 | 0 | 0 | 0 |
| 0 | NA | NA | NA | NA | NA | Gm9293 | 0 | 0 | 0 | 0 | 0 | 0 | 0 | 0 | 0 | 0 | 0 |
| 0 | NA | NA | NA | NA | NA | Gm47383 | 0 | 0 | 0 | 0 | 0 | 0 | 0 | 0 | 0 | 0 | 0 |
| 0 | NA | NA | NA | NA | NA | Gm47384 | 0 | 0 | 0 | 0 | 0 | 0 | 0 | 0 | 0 | 0 | 0 |
| 0 | NA | NA | NA | NA | NA | Gm47385 | 0 | 0 | 0 | 0 | 0 | 0 | 0 | 0 | 0 | 0 | 0 |
| 0 | NA | NA | NA | NA | NA | Gm47386 | 0 | 0 | 0 | 0 | 0 | 0 | 0 | 0 | 0 | 0 | 0 |
| 0 | NA | NA | NA | NA | NA | Gm26140 | 0 | 0 | 0 | 0 | 0 | 0 | 0 | 0 | 0 | 0 | 0 |
| 0 | NA | NA | NA | NA | NA | Gm20472 | 0 | 0 | 0 | 0 | 0 | 0 | 0 | 0 | 0 | 0 | 0 |
| 0 | NA | NA | NA | NA | NA | Gm20473 | 0 | 0 | 0 | 0 | 0 | 0 | 0 | 0 | 0 | 0 | 0 |
| 0 | NA | NA | NA | NA | NA | Gm9285 | 0 | 0 | 0 | 0 | 0 | 0 | 0 | 0 | 0 | 0 | 0 |
| 0 | NA | NA | NA | NA | NA | Gm47393 | 0 | 0 | 0 | 0 | 0 | 0 | 0 | 0 | 0 | 0 | 0 |
| 0 | NA | NA | NA | NA | NA | Gm47420 | 0 | 0 | 0 | 0 | 0 | 0 | 0 | 0 | 0 | 0 | 0 |
| 0 | NA | NA | NA | NA | NA | Gm47421 | 0 | 0 | 0 | 0 | 0 | 0 | 0 | 0 | 0 | 0 | 0 |
| 0 | NA | NA | NA | NA | NA | Gm9286 | 0 | 0 | 0 | 0 | 0 | 0 | 0 | 0 | 0 | 0 | 0 |
| 0 | NA | NA | NA | NA | NA | Gm47422 | 0 | 0 | 0 | 0 | 0 | 0 | 0 | 0 | 0 | 0 | 0 |
| 0 | NA | NA | NA | NA | NA | Gm20474 | 0 | 0 | 0 | 0 | 0 | 0 | 0 | 0 | 0 | 0 | 0 |
| 0 | NA | NA | NA | NA | NA | Gm49386 | 0 | 0 | 0 | 0 | 0 | 0 | 0 | 0 | 0 | 0 | 0 |
| 0 | NA | NA | NA | NA | NA | Gm9305 | 0 | 0 | 0 | 0 | 0 | 0 | 0 | 0 | 0 | 0 | 0 |
| 0 | NA | NA | NA | NA | NA | Gm9309 | 0 | 0 | 0 | 0 | 0 | 0 | 0 | 0 | 0 | 0 | 0 |
| 0 | NA | NA | NA | NA | NA | Gm10330 | 0 | 0 | 0 | 0 | 0 | 0 | 0 | 0 | 0 | 0 | 0 |
| 0 | NA | NA | NA | NA | NA | Gm47437 | 0 | 0 | 0 | 0 | 0 | 0 | 0 | 0 | 0 | 0 | 0 |
| 0 | NA | NA | NA | NA | NA | Gm47440 | 0 | 0 | 0 | 0 | 0 | 0 | 0 | 0 | 0 | 0 | 0 |
| 0 | NA | NA | NA | NA | NA | Gm46342 | 0 | 0 | 0 | 0 | 0 | 0 | 0 | 0 | 0 | 0 | 0 |
| 0 | NA | NA | NA | NA | NA | Gm47442 | 0 | 0 | 0 | 0 | 0 | 0 | 0 | 0 | 0 | 0 | 0 |
| 0 | NA | NA | NA | NA | NA | Gm47443 | 0 | 0 | 0 | 0 | 0 | 0 | 0 | 0 | 0 | 0 | 0 |
| 0 | NA | NA | NA | NA | NA | Gm17330 | 0 | 0 | 0 | 0 | 0 | 0 | 0 | 0 | 0 | 0 | 0 |
| 0 | NA | NA | NA | NA | NA | Gm47444 | 0 | 0 | 0 | 0 | 0 | 0 | 0 | 0 | 0 | 0 | 0 |
| 0 | NA | NA | NA | NA | NA | Gm5954 | 0 | 0 | 0 | 0 | 0 | 0 | 0 | 0 | 0 | 0 | 0 |

| | | | | | | | | | | | | | | | | | | |
|---|---|---|---|---|---|---|---|---|---|---|---|---|---|---|---|---|---|---|
| 0 | NA | NA | NA | NA | NA | Gm4120 | | 0 | 0 | 0 | 0 | 0 | 0 | 0 | 0 | 0 | 0 | 0 |
| 0 | NA | NA | NA | NA | NA | Gm47478 | | 0 | 0 | 0 | 0 | 0 | 0 | 0 | 0 | 0 | 0 | 0 |
| 0 | NA | NA | NA | NA | NA | Gm47482 | | 0 | 0 | 0 | 0 | 0 | 0 | 0 | 0 | 0 | 0 | 0 |
| 0 | NA | NA | NA | NA | NA | Gm48895 | | 0 | 0 | 0 | 0 | 0 | 0 | 0 | 0 | 0 | 0 | 0 |
| 0 | NA | NA | NA | NA | NA | Gm48896 | | 0 | 0 | 0 | 0 | 0 | 0 | 0 | 0 | 0 | 0 | 0 |
| 0 | NA | NA | NA | NA | NA | Gm9318 | | 0 | 0 | 0 | 0 | 0 | 0 | 0 | 0 | 0 | 0 | 0 |
| 0 | NA | NA | NA | NA | NA | Gm26211 | | 0 | 0 | 0 | 0 | 0 | 0 | 0 | 0 | 0 | 0 | 0 |
| 0 | NA | NA | NA | NA | NA | Gm48897 | | 0 | 0 | 0 | 0 | 0 | 0 | 0 | 0 | 0 | 0 | 0 |
| 0 | NA | NA | NA | NA | NA | Gm17746 | | 0 | 0 | 0 | 0 | 0 | 0 | 0 | 0 | 0 | 0 | 0 |
| 0 | NA | NA | NA | NA | NA | Gm19340 | | 0 | 0 | 0 | 0 | 0 | 0 | 0 | 0 | 0 | 0 | 0 |
| 0 | NA | NA | NA | NA | NA | Gm22527 | | 0 | 0 | 0 | 0 | 0 | 0 | 0 | 0 | 0 | 0 | 0 |
| 0 | NA | NA | NA | NA | NA | Gm9321 | | 0 | 0 | 0 | 0 | 0 | 0 | 0 | 0 | 0 | 0 | 0 |
| 0 | NA | NA | NA | NA | NA | Gm47733 | | 0 | 0 | 0 | 0 | 0 | 0 | 0 | 0 | 0 | 0 | 0 |
| 0 | NA | NA | NA | NA | NA | Gm29687 | | 0 | 0 | 0 | 0 | 0 | 0 | 0 | 0 | 0 | 0 | 0 |
| 0 | NA | NA | NA | NA | NA | 4930480M12Rik | | 0 | 0 | 0 | 0 | 0 | 0 | 0 | 0 | 0 | 0 | 0 |
| 0 | NA | NA | NA | NA | NA | Gm29968 | | 0 | 0 | 0 | 0 | 0 | 0 | 0 | 0 | 0 | 0 | 0 |
| 0 | NA | NA | NA | NA | NA | 4930549C15Rik | | 0 | 0 | 0 | 0 | 0 | 0 | 0 | 0 | 0 | 0 | 0 |
| 0 | NA | NA | NA | NA | NA | Mir6538 | | 0 | 0 | 0 | 0 | 0 | 0 | 0 | 0 | 0 | 0 | 0 |
| 0 | NA | NA | NA | NA | NA | Gm47799 | | 0 | 0 | 0 | 0 | 0 | 0 | 0 | 0 | 0 | 0 | 0 |
| 0 | NA | NA | NA | NA | NA | 1700020D12Rik | | 0 | 0 | 0 | 0 | 0 | 0 | 0 | 0 | 0 | 0 | 0 |
| 0 | NA | NA | NA | NA | NA | Gm47847 | | 0 | 0 | 0 | 0 | 0 | 0 | 0 | 0 | 0 | 0 | 0 |
| 0 | NA | NA | NA | NA | NA | 4933409F18Rik | | 0 | 0 | 0 | 0 | 0 | 0 | 0 | 0 | 0 | 0 | 0 |
| 0 | NA | NA | NA | NA | NA | Gm47860 | | 0 | 0 | 0 | 0 | 0 | 0 | 0 | 0 | 0 | 0 | 0 |
| 0 | NA | NA | NA | NA | NA | Gm4166 | | 0 | 0 | 0 | 0 | 0 | 0 | 0 | 0 | 0 | 0 | 0 |
| 0 | NA | NA | NA | NA | NA | Gm24326 | | 0 | 0 | 0 | 0 | 0 | 0 | 0 | 0 | 0 | 0 | 0 |
| 0 | NA | NA | NA | NA | NA | Gm45941 | | 0 | 0 | 0 | 0 | 0 | 0 | 0 | 0 | 0 | 0 | 0 |
| 0 | NA | NA | NA | NA | NA | Gm31025 | | 0 | 0 | 0 | 0 | 0 | 0 | 0 | 0 | 0 | 0 | 0 |
| 0 | NA | NA | NA | NA | NA | Gm25923 | | 0 | 0 | 0 | 0 | 0 | 0 | 0 | 0 | 0 | 0 | 0 |
| 0 | NA | NA | NA | NA | NA | Gm47872 | | 0 | 0 | 0 | 0 | 0 | 0 | 0 | 0 | 0 | 0 | 0 |
| 0 | NA | NA | NA | NA | NA | Gm47871 | | 0 | 0 | 0 | 0 | 0 | 0 | 0 | 0 | 0 | 0 | 0 |
| 0 | NA | NA | NA | NA | NA | Gm25564 | | 0 | 0 | 0 | 0 | 0 | 0 | 0 | 0 | 0 | 0 | 0 |
| 0 | NA | NA | NA | NA | NA | Dcdc2c | doublecortin domain containing 2C | 0 | 0 | 0 | 0 | 0 | 0 | 0 | 0 | 0 | 0 | 0 |
| 0 | NA | NA | NA | NA | NA | Gm47878 | | 0 | 0 | 0 | 0 | 0 | 0 | 0 | 0 | 0 | 0 | 0 |
| 0 | NA | NA | NA | NA | NA | Allc | allantoicase | 0 | 0 | 0 | 0 | 0 | 0 | 0 | 0 | 0 | 0 | 0 |
| 0 | NA | NA | NA | NA | NA | Gm48904 | | 0 | 0 | 0 | 0 | 0 | 0 | 0 | 0 | 0 | 0 | 0 |
| 0 | NA | NA | NA | NA | NA | Gm48080 | | 0 | 0 | 0 | 0 | 0 | 0 | 0 | 0 | 0 | 0 | 0 |
| 0 | NA | NA | NA | NA | NA | Gm23342 | | 0 | 0 | 0 | 0 | 0 | 0 | 0 | 0 | 0 | 0 | 0 |
| 0 | NA | NA | NA | NA | NA | Gm6989 | | 0 | 0 | 0 | 0 | 0 | 0 | 0 | 0 | 0 | 0 | 0 |
| 0 | NA | NA | NA | NA | NA | C630031E19Rik | | 0 | 0 | 0 | 0 | 0 | 0 | 0 | 0 | 0 | 0 | 0 |
| 0 | NA | NA | NA | NA | NA | Gm15691 | | 0 | 0 | 0 | 0 | 0 | 0 | 0 | 0 | 0 | 0 | 0 |
| 0 | NA | NA | NA | NA | NA | 2310016D03Rik | | 0 | 0 | 0 | 0 | 0 | 0 | 0 | 0 | 0 | 0 | 0 |
| 0 | NA | NA | NA | NA | NA | Gm25865 | | 0 | 0 | 0 | 0 | 0 | 0 | 0 | 0 | 0 | 0 | 0 |
| 0 | NA | NA | NA | NA | NA | Gm28806 | | 0 | 0 | 0 | 0 | 0 | 0 | 0 | 0 | 0 | 0 | 0 |
| 0 | NA | NA | NA | NA | NA | Ryk-ps1 | | 0 | 0 | 0 | 0 | 0 | 0 | 0 | 0 | 0 | 0 | 0 |
| 0 | NA | NA | NA | NA | NA | Gm25243 | | 0 | 0 | 0 | 0 | 0 | 0 | 0 | 0 | 0 | 0 | 0 |
| 0 | NA | NA | NA | NA | NA | Gm18502 | | 0 | 0 | 0 | 0 | 0 | 0 | 0 | 0 | 0 | 0 | 0 |
| 0 | NA | NA | NA | NA | NA | Gm25408 | | 0 | 0 | 0 | 0 | 0 | 0 | 0 | 0 | 0 | 0 | 0 |
| 0 | NA | NA | NA | NA | NA | Gm32443 | | 0 | 0 | 0 | 0 | 0 | 0 | 0 | 0 | 0 | 0 | 0 |
| 0 | NA | NA | NA | NA | NA | Gm18024 | | 0 | 0 | 0 | 0 | 0 | 0 | 0 | 0 | 0 | 0 | 0 |
| 0 | NA | NA | NA | NA | NA | Gm48117 | | 0 | 0 | 0 | 0 | 0 | 0 | 0 | 0 | 0 | 0 | 0 |
| 0 | NA | NA | NA | NA | NA | Gm6993 | | 0 | 0 | 0 | 0 | 0 | 0 | 0 | 0 | 0 | 0 | 0 |
| 0 | NA | NA | NA | NA | NA | Gm24613 | | 0 | 0 | 0 | 0 | 0 | 0 | 0 | 0 | 0 | 0 | 0 |
| 0 | NA | NA | NA | NA | NA | Gm9359 | | 0 | 0 | 0 | 0 | 0 | 0 | 0 | 0 | 0 | 0 | 0 |
| 0 | NA | NA | NA | NA | NA | Gm29541 | | 0 | 0 | 0 | 0 | 0 | 0 | 0 | 0 | 0 | 0 | 0 |
| 0 | NA | NA | NA | NA | NA | Gm48524 | | 0 | 0 | 0 | 0 | 0 | 0 | 0 | 0 | 0 | 0 | 0 |
| 0 | NA | NA | NA | NA | NA | Gm48808 | | 0 | 0 | 0 | 0 | 0 | 0 | 0 | 0 | 0 | 0 | 0 |
| 0 | NA | NA | NA | NA | NA | Gm18029 | | 0 | 0 | 0 | 0 | 0 | 0 | 0 | 0 | 0 | 0 | 0 |
| 0 | NA | NA | NA | NA | NA | Gm48809 | | 0 | 0 | 0 | 0 | 0 | 0 | 0 | 0 | 0 | 0 | 0 |
| 0 | NA | NA | NA | NA | NA | Gm33111 | | 0 | 0 | 0 | 0 | 0 | 0 | 0 | 0 | 0 | 0 | 0 |
| 0 | NA | NA | NA | NA | NA | 5430401H09Rik | | 0 | 0 | 0 | 0 | 0 | 0 | 0 | 0 | 0 | 0 | 0 |

| | | | | | | | | | | | | | | | | | | |
|---|---|---|---|---|---|---|---|---|---|---|---|---|---|---|---|---|---|---|
| 0 | NA | NA | NA | NA | NA | Gm47928 | | 0 | 0 | 0 | 0 | 0 | 0 | 0 | 0 | 0 | 0 | 0 |
| 0 | NA | NA | NA | NA | NA | Gm33308 | | 0 | 0 | 0 | 0 | 0 | 0 | 0 | 0 | 0 | 0 | 0 |
| 0 | NA | NA | NA | NA | NA | Gm17820 | | 0 | 0 | 0 | 0 | 0 | 0 | 0 | 0 | 0 | 0 | 0 |
| 0 | NA | NA | NA | NA | NA | Gm47937 | | 0 | 0 | 0 | 0 | 0 | 0 | 0 | 0 | 0 | 0 | 0 |
| 0 | NA | NA | NA | NA | NA | Gm47948 | | 0 | 0 | 0 | 0 | 0 | 0 | 0 | 0 | 0 | 0 | 0 |
| 0 | NA | NA | NA | NA | NA | Gm18622 | | 0 | 0 | 0 | 0 | 0 | 0 | 0 | 0 | 0 | 0 | 0 |
| 0 | NA | NA | NA | NA | NA | Gm47961 | | 0 | 0 | 0 | 0 | 0 | 0 | 0 | 0 | 0 | 0 | 0 |
| 0 | NA | NA | NA | NA | NA | Gm24922 | | 0 | 0 | 0 | 0 | 0 | 0 | 0 | 0 | 0 | 0 | 0 |
| 0 | NA | NA | NA | NA | NA | Gm22904 | | 0 | 0 | 0 | 0 | 0 | 0 | 0 | 0 | 0 | 0 | 0 |
| 0 | NA | NA | NA | NA | NA | Gm16267 | | 0 | 0 | 0 | 0 | 0 | 0 | 0 | 0 | 0 | 0 | 0 |
| 0 | NA | NA | NA | NA | NA | Atxn7l1os1 | | 0 | 0 | 0 | 0 | 0 | 0 | 0 | 0 | 0 | 0 | 0 |
| 0 | NA | NA | NA | NA | NA | Atxn7l1os2 | | 0 | 0 | 0 | 0 | 0 | 0 | 0 | 0 | 0 | 0 | 0 |
| 0 | NA | NA | NA | NA | NA | Gm23529 | | 0 | 0 | 0 | 0 | 0 | 0 | 0 | 0 | 0 | 0 | 0 |
| 0 | NA | NA | NA | NA | NA | Gm9368 | | 0 | 0 | 0 | 0 | 0 | 0 | 0 | 0 | 0 | 0 | 0 |
| 0 | NA | NA | NA | NA | NA | Gm40383 | | 0 | 0 | 0 | 0 | 0 | 0 | 0 | 0 | 0 | 0 | 0 |
| 0 | NA | NA | NA | NA | NA | Ferd3l | Fer3 like bHLH transcription factor | 0 | 0 | 0 | 0 | 0 | 0 | 0 | 0 | 0 | 0 | 0 |
| 0 | NA | NA | NA | NA | NA | Gm47357 | | 0 | 0 | 0 | 0 | 0 | 0 | 0 | 0 | 0 | 0 | 0 |
| 0 | NA | NA | NA | NA | NA | 4921508M14Rik | | 0 | 0 | 0 | 0 | 0 | 0 | 0 | 0 | 0 | 0 | 0 |
| 0 | NA | NA | NA | NA | NA | Gm24429 | | 0 | 0 | 0 | 0 | 0 | 0 | 0 | 0 | 0 | 0 | 0 |
| 0 | NA | NA | NA | NA | NA | Prps1l1 | phosphoribosyl pyrophosphate synthetas | 0 | 0 | 0 | 0 | 0 | 0 | 0 | 0 | 0 | 0 | 0 |
| 0 | NA | NA | NA | NA | NA | Gm24359 | | 0 | 0 | 0 | 0 | 0 | 0 | 0 | 0 | 0 | 0 | 0 |
| 0 | NA | NA | NA | NA | NA | Mir680-3 | | 0 | 0 | 0 | 0 | 0 | 0 | 0 | 0 | 0 | 0 | 0 |
| 0 | NA | NA | NA | NA | NA | Gm44396 | | 0 | 0 | 0 | 0 | 0 | 0 | 0 | 0 | 0 | 0 | 0 |
| 0 | NA | NA | NA | NA | NA | Gm48237 | | 0 | 0 | 0 | 0 | 0 | 0 | 0 | 0 | 0 | 0 | 0 |
| 0 | NA | NA | NA | NA | NA | Gm34047 | | 0 | 0 | 0 | 0 | 0 | 0 | 0 | 0 | 0 | 0 | 0 |
| 0 | NA | NA | NA | NA | NA | Gm48264 | | 0 | 0 | 0 | 0 | 0 | 0 | 0 | 0 | 0 | 0 | 0 |
| 0 | NA | NA | NA | NA | NA | Gm9472 | | 0 | 0 | 0 | 0 | 0 | 0 | 0 | 0 | 0 | 0 | 0 |
| 0 | NA | NA | NA | NA | NA | Gm18115 | | 0 | 0 | 0 | 0 | 0 | 0 | 0 | 0 | 0 | 0 | 0 |
| 0 | NA | NA | NA | NA | NA | Gm40392 | | 0 | 0 | 0 | 0 | 0 | 0 | 0 | 0 | 0 | 0 | 0 |
| 0 | NA | NA | NA | NA | NA | Gm25438 | | 0 | 0 | 0 | 0 | 0 | 0 | 0 | 0 | 0 | 0 | 0 |
| 0 | NA | NA | NA | NA | NA | Mir5099 | | 0 | 0 | 0 | 0 | 0 | 0 | 0 | 0 | 0 | 0 | 0 |
| 0 | NA | NA | NA | NA | NA | Gm48613 | | 0 | 0 | 0 | 0 | 0 | 0 | 0 | 0 | 0 | 0 | 0 |
| 0 | NA | NA | NA | NA | NA | Gm18026 | | 0 | 0 | 0 | 0 | 0 | 0 | 0 | 0 | 0 | 0 | 0 |
| 0 | NA | NA | NA | NA | NA | Gm44326 | | 0 | 0 | 0 | 0 | 0 | 0 | 0 | 0 | 0 | 0 | 0 |
| 0 | NA | NA | NA | NA | NA | Gm48616 | | 0 | 0 | 0 | 0 | 0 | 0 | 0 | 0 | 0 | 0 | 0 |
| 0 | NA | NA | NA | NA | NA | Gm48617 | | 0 | 0 | 0 | 0 | 0 | 0 | 0 | 0 | 0 | 0 | 0 |
| 0 | NA | NA | NA | NA | NA | Gm29007 | | 0 | 0 | 0 | 0 | 0 | 0 | 0 | 0 | 0 | 0 | 0 |
| 0 | NA | NA | NA | NA | NA | Gm34408 | | 0 | 0 | 0 | 0 | 0 | 0 | 0 | 0 | 0 | 0 | 0 |
| 0 | NA | NA | NA | NA | NA | Gm48660 | | 0 | 0 | 0 | 0 | 0 | 0 | 0 | 0 | 0 | 0 | 0 |
| 0 | NA | NA | NA | NA | NA | Gm48715 | | 0 | 0 | 0 | 0 | 0 | 0 | 0 | 0 | 0 | 0 | 0 |
| 0 | NA | NA | NA | NA | NA | Gm18338 | | 0 | 0 | 0 | 0 | 0 | 0 | 0 | 0 | 0 | 0 | 0 |
| 0 | NA | NA | NA | NA | NA | 4930428E07Rik | | 0 | 0 | 0 | 0 | 0 | 0 | 0 | 0 | 0 | 0 | 0 |
| 0 | NA | NA | NA | NA | NA | Gm48813 | | 0 | 0 | 0 | 0 | 0 | 0 | 0 | 0 | 0 | 0 | 0 |
| 0 | NA | NA | NA | NA | NA | Gm4257 | | 0 | 0 | 0 | 0 | 0 | 0 | 0 | 0 | 0 | 0 | 0 |
| 0 | NA | NA | NA | NA | NA | Gm18442 | | 0 | 0 | 0 | 0 | 0 | 0 | 0 | 0 | 0 | 0 | 0 |
| 0 | NA | NA | NA | NA | NA | Gm47832 | | 0 | 0 | 0 | 0 | 0 | 0 | 0 | 0 | 0 | 0 | 0 |
| 0 | NA | NA | NA | NA | NA | Gm34662 | | 0 | 0 | 0 | 0 | 0 | 0 | 0 | 0 | 0 | 0 | 0 |
| 0 | NA | NA | NA | NA | NA | Gm19563 | | 0 | 0 | 0 | 0 | 0 | 0 | 0 | 0 | 0 | 0 | 0 |
| 0 | NA | NA | NA | NA | NA | Gm34809 | | 0 | 0 | 0 | 0 | 0 | 0 | 0 | 0 | 0 | 0 | 0 |
| 0 | NA | NA | NA | NA | NA | Gm18591 | | 0 | 0 | 0 | 0 | 0 | 0 | 0 | 0 | 0 | 0 | 0 |
| 0 | NA | NA | NA | NA | NA | Gm47855 | | 0 | 0 | 0 | 0 | 0 | 0 | 0 | 0 | 0 | 0 | 0 |
| 0 | NA | NA | NA | NA | NA | Gm24069 | | 0 | 0 | 0 | 0 | 0 | 0 | 0 | 0 | 0 | 0 | 0 |
| 0 | NA | NA | NA | NA | NA | Gm34923 | | 0 | 0 | 0 | 0 | 0 | 0 | 0 | 0 | 0 | 0 | 0 |
| 0 | NA | NA | NA | NA | NA | Gm47859 | | 0 | 0 | 0 | 0 | 0 | 0 | 0 | 0 | 0 | 0 | 0 |
| 0 | NA | NA | NA | NA | NA | Gm24434 | | 0 | 0 | 0 | 0 | 0 | 0 | 0 | 0 | 0 | 0 | 0 |
| 0 | NA | NA | NA | NA | NA | Gm4263 | | 0 | 0 | 0 | 0 | 0 | 0 | 0 | 0 | 0 | 0 | 0 |
| 0 | NA | NA | NA | NA | NA | Gm47868 | | 0 | 0 | 0 | 0 | 0 | 0 | 0 | 0 | 0 | 0 | 0 |
| 0 | NA | NA | NA | NA | NA | Gm47954 | | 0 | 0 | 0 | 0 | 0 | 0 | 0 | 0 | 0 | 0 | 0 |
| 0 | NA | NA | NA | NA | NA | Gm19441 | | 0 | 0 | 0 | 0 | 0 | 0 | 0 | 0 | 0 | 0 | 0 |
| 0 | NA | NA | NA | NA | NA | Gm19220 | | 0 | 0 | 0 | 0 | 0 | 0 | 0 | 0 | 0 | 0 | 0 |

| | | | | | | | | | | | | | | | | | | |
|---|---|---|---|---|---|---|---|---|---|---|---|---|---|---|---|---|---|---|
| 0 | NA | NA | NA | NA | NA | Gm18246 | | 0 | 0 | 0 | 0 | 0 | 0 | 0 | 0 | 0 | 0 | 0 |
| 0 | NA | NA | NA | NA | NA | Gm18954 | | 0 | 0 | 0 | 0 | 0 | 0 | 0 | 0 | 0 | 0 | 0 |
| 0 | NA | NA | NA | NA | NA | Gm47375 | | 0 | 0 | 0 | 0 | 0 | 0 | 0 | 0 | 0 | 0 | 0 |
| 0 | NA | NA | NA | NA | NA | Gm47376 | | 0 | 0 | 0 | 0 | 0 | 0 | 0 | 0 | 0 | 0 | 0 |
| 0 | NA | NA | NA | NA | NA | Gm25497 | | 0 | 0 | 0 | 0 | 0 | 0 | 0 | 0 | 0 | 0 | 0 |
| 0 | NA | NA | NA | NA | NA | Gm47373 | | 0 | 0 | 0 | 0 | 0 | 0 | 0 | 0 | 0 | 0 | 0 |
| 0 | NA | NA | NA | NA | NA | Gm47368 | | 0 | 0 | 0 | 0 | 0 | 0 | 0 | 0 | 0 | 0 | 0 |
| 0 | NA | NA | NA | NA | NA | Gm47374 | | 0 | 0 | 0 | 0 | 0 | 0 | 0 | 0 | 0 | 0 | 0 |
| 0 | NA | NA | NA | NA | NA | Gm47369 | | 0 | 0 | 0 | 0 | 0 | 0 | 0 | 0 | 0 | 0 | 0 |
| 0 | NA | NA | NA | NA | NA | Gm47371 | | 0 | 0 | 0 | 0 | 0 | 0 | 0 | 0 | 0 | 0 | 0 |
| 0 | NA | NA | NA | NA | NA | Gm46348 | | 0 | 0 | 0 | 0 | 0 | 0 | 0 | 0 | 0 | 0 | 0 |
| 0 | NA | NA | NA | NA | NA | Gm47012 | | 0 | 0 | 0 | 0 | 0 | 0 | 0 | 0 | 0 | 0 | 0 |
| 0 | NA | NA | NA | NA | NA | Gm7242 | | 0 | 0 | 0 | 0 | 0 | 0 | 0 | 0 | 0 | 0 | 0 |
| 0 | NA | NA | NA | NA | NA | Gm47013 | | 0 | 0 | 0 | 0 | 0 | 0 | 0 | 0 | 0 | 0 | 0 |
| 0 | NA | NA | NA | NA | NA | Gm47020 | | 0 | 0 | 0 | 0 | 0 | 0 | 0 | 0 | 0 | 0 | 0 |
| 0 | NA | NA | NA | NA | NA | Gm25248 | | 0 | 0 | 0 | 0 | 0 | 0 | 0 | 0 | 0 | 0 | 0 |
| 0 | NA | NA | NA | NA | NA | Gm33340 | | 0 | 0 | 0 | 0 | 0 | 0 | 0 | 0 | 0 | 0 | 0 |
| 0 | NA | NA | NA | NA | NA | Gm47030 | | 0 | 0 | 0 | 0 | 0 | 0 | 0 | 0 | 0 | 0 | 0 |
| 0 | NA | NA | NA | NA | NA | Gm26326 | | 0 | 0 | 0 | 0 | 0 | 0 | 0 | 0 | 0 | 0 | 0 |
| 0 | NA | NA | NA | NA | NA | Gm47036 | | 0 | 0 | 0 | 0 | 0 | 0 | 0 | 0 | 0 | 0 | 0 |
| 0 | NA | NA | NA | NA | NA | Gm10165 | | 0 | 0 | 0 | 0 | 0 | 0 | 0 | 0 | 0 | 0 | 0 |
| 0 | NA | NA | NA | NA | NA | Gm46349 | | 0 | 0 | 0 | 0 | 0 | 0 | 0 | 0 | 0 | 0 | 0 |
| 0 | NA | NA | NA | NA | NA | Gm47056 | | 0 | 0 | 0 | 0 | 0 | 0 | 0 | 0 | 0 | 0 | 0 |
| 0 | NA | NA | NA | NA | NA | Mir5627 | | 0 | 0 | 0 | 0 | 0 | 0 | 0 | 0 | 0 | 0 | 0 |
| 0 | NA | NA | NA | NA | NA | Gm48181 | | 0 | 0 | 0 | 0 | 0 | 0 | 0 | 0 | 0 | 0 | 0 |
| 0 | NA | NA | NA | NA | NA | Gm48182 | | 0 | 0 | 0 | 0 | 0 | 0 | 0 | 0 | 0 | 0 | 0 |
| 0 | NA | NA | NA | NA | NA | Gm15901 | | 0 | 0 | 0 | 0 | 0 | 0 | 0 | 0 | 0 | 0 | 0 |
| 0 | NA | NA | NA | NA | NA | Gm15902 | | 0 | 0 | 0 | 0 | 0 | 0 | 0 | 0 | 0 | 0 | 0 |
| 0 | NA | NA | NA | NA | NA | Gm48508 | | 0 | 0 | 0 | 0 | 0 | 0 | 0 | 0 | 0 | 0 | 0 |
| 0 | NA | NA | NA | NA | NA | Gm24731 | | 0 | 0 | 0 | 0 | 0 | 0 | 0 | 0 | 0 | 0 | 0 |
| 0 | NA | NA | NA | NA | NA | Gm22076 | | 0 | 0 | 0 | 0 | 0 | 0 | 0 | 0 | 0 | 0 | 0 |
| 0 | NA | NA | NA | NA | NA | Gm48517 | | 0 | 0 | 0 | 0 | 0 | 0 | 0 | 0 | 0 | 0 | 0 |
| 0 | NA | NA | NA | NA | NA | Gm48518 | | 0 | 0 | 0 | 0 | 0 | 0 | 0 | 0 | 0 | 0 | 0 |
| 0 | NA | NA | NA | NA | NA | Gm48522 | | 0 | 0 | 0 | 0 | 0 | 0 | 0 | 0 | 0 | 0 | 0 |
| 0 | NA | NA | NA | NA | NA | Gm24948 | | 0 | 0 | 0 | 0 | 0 | 0 | 0 | 0 | 0 | 0 | 0 |
| 0 | NA | NA | NA | NA | NA | Gm18112 | | 0 | 0 | 0 | 0 | 0 | 0 | 0 | 0 | 0 | 0 | 0 |
| 0 | NA | NA | NA | NA | NA | Gm48531 | | 0 | 0 | 0 | 0 | 0 | 0 | 0 | 0 | 0 | 0 | 0 |
| 0 | NA | NA | NA | NA | NA | B230217J21Rik | | 0 | 0 | 0 | 0 | 0 | 0 | 0 | 0 | 0 | 0 | 0 |
| 0 | NA | NA | NA | NA | NA | Gm48543 | | 0 | 0 | 0 | 0 | 0 | 0 | 0 | 0 | 0 | 0 | 0 |
| 0 | NA | NA | NA | NA | NA | Gm25051 | | 0 | 0 | 0 | 0 | 0 | 0 | 0 | 0 | 0 | 0 | 0 |
| 0 | NA | NA | NA | NA | NA | Gm36971 | | 0 | 0 | 0 | 0 | 0 | 0 | 0 | 0 | 0 | 0 | 0 |
| 0 | NA | NA | NA | NA | NA | Gm2089 | | 0 | 0 | 0 | 0 | 0 | 0 | 0 | 0 | 0 | 0 | 0 |
| 0 | NA | NA | NA | NA | NA | Gm48578 | | 0 | 0 | 0 | 0 | 0 | 0 | 0 | 0 | 0 | 0 | 0 |
| 0 | NA | NA | NA | NA | NA | Gm46327 | | 0 | 0 | 0 | 0 | 0 | 0 | 0 | 0 | 0 | 0 | 0 |
| 0 | NA | NA | NA | NA | NA | Gm48587 | | 0 | 0 | 0 | 0 | 0 | 0 | 0 | 0 | 0 | 0 | 0 |
| 0 | NA | NA | NA | NA | NA | Gm29818 | | 0 | 0 | 0 | 0 | 0 | 0 | 0 | 0 | 0 | 0 | 0 |
| 0 | NA | NA | NA | NA | NA | | | 0 | 0 | 0 | 0 | 0 | 0 | 0 | 0 | 0 | 0 | 0 |
| 0 | NA | NA | NA | NA | NA | Gm26454 | | 0 | 0 | 0 | 0 | 0 | 0 | 0 | 0 | 0 | 0 | 0 |
| 0 | NA | NA | NA | NA | NA | Gm18623 | | 0 | 0 | 0 | 0 | 0 | 0 | 0 | 0 | 0 | 0 | 0 |
| 0 | NA | NA | NA | NA | NA | Gm34304 | | 0 | 0 | 0 | 0 | 0 | 0 | 0 | 0 | 0 | 0 | 0 |
| 0 | NA | NA | NA | NA | NA | Foxg1 | forkhead box G1 | 0 | 0 | 0 | 0 | 0 | 0 | 0 | 0 | 0 | 0 | 0 |
| 0 | NA | NA | NA | NA | NA | Gm43517 | | 0 | 0 | 0 | 0 | 0 | 0 | 0 | 0 | 0 | 0 | 0 |
| 0 | NA | NA | NA | NA | NA | Gm48659 | | 0 | 0 | 0 | 0 | 0 | 0 | 0 | 0 | 0 | 0 | 0 |
| 0 | NA | NA | NA | NA | NA | Gm19599 | | 0 | 0 | 0 | 0 | 0 | 0 | 0 | 0 | 0 | 0 | 0 |
| 0 | NA | NA | NA | NA | NA | Gm25015 | | 0 | 0 | 0 | 0 | 0 | 0 | 0 | 0 | 0 | 0 | 0 |
| 0 | NA | NA | NA | NA | NA | Gm40418 | | 0 | 0 | 0 | 0 | 0 | 0 | 0 | 0 | 0 | 0 | 0 |
| 0 | NA | NA | NA | NA | NA | Gm48777 | | 0 | 0 | 0 | 0 | 0 | 0 | 0 | 0 | 0 | 0 | 0 |
| 0 | NA | NA | NA | NA | NA | Gm40421 | | 0 | 0 | 0 | 0 | 0 | 0 | 0 | 0 | 0 | 0 | 0 |
| 0 | NA | NA | NA | NA | NA | 1700008C04Rik | | 0 | 0 | 0 | 0 | 0 | 0 | 0 | 0 | 0 | 0 | 0 |
| 0 | NA | NA | NA | NA | NA | Gm22088 | | 0 | 0 | 0 | 0 | 0 | 0 | 0 | 0 | 0 | 0 | 0 |

| | | | | | | | | | | | | | | | | | |
|---|---|---|---|---|---|---|---|---|---|---|---|---|---|---|---|---|---|
| 0 | NA | NA | NA | NA | NA | Gm18503 | | 0 | 0 | 0 | 0 | 0 | 0 | 0 | 0 | 0 | 0 | 0 |
| 0 | NA | NA | NA | NA | NA | Gm22663 | | 0 | 0 | 0 | 0 | 0 | 0 | 0 | 0 | 0 | 0 | 0 |
| 0 | NA | NA | NA | NA | NA | Gm19309 | | 0 | 0 | 0 | 0 | 0 | 0 | 0 | 0 | 0 | 0 | 0 |
| 0 | NA | NA | NA | NA | NA | 1700031P21Rik | | 0 | 0 | 0 | 0 | 0 | 0 | 0 | 0 | 0 | 0 | 0 |
| 0 | NA | NA | NA | NA | NA | Gm47488 | | 0 | 0 | 0 | 0 | 0 | 0 | 0 | 0 | 0 | 0 | 0 |
| 0 | NA | NA | NA | NA | NA | n-R5s58 | | 0 | 0 | 0 | 0 | 0 | 0 | 0 | 0 | 0 | 0 | 0 |
| 0 | NA | NA | NA | NA | NA | 1700030L22Rik | | 0 | 0 | 0 | 0 | 0 | 0 | 0 | 0 | 0 | 0 | 0 |
| 0 | NA | NA | NA | NA | NA | 1700060O08Rik | | 0 | 0 | 0 | 0 | 0 | 0 | 0 | 0 | 0 | 0 | 0 |
| 0 | NA | NA | NA | NA | NA | Gm35818 | | 0 | 0 | 0 | 0 | 0 | 0 | 0 | 0 | 0 | 0 | 0 |
| 0 | NA | NA | NA | NA | NA | 1700104L18Rik | | 0 | 0 | 0 | 0 | 0 | 0 | 0 | 0 | 0 | 0 | 0 |
| 0 | NA | NA | NA | NA | NA | Gm22220 | | 0 | 0 | 0 | 0 | 0 | 0 | 0 | 0 | 0 | 0 | 0 |
| 0 | NA | NA | NA | NA | NA | Gm24692 | | 0 | 0 | 0 | 0 | 0 | 0 | 0 | 0 | 0 | 0 | 0 |
| 0 | NA | NA | NA | NA | NA | Gm47552 | | 0 | 0 | 0 | 0 | 0 | 0 | 0 | 0 | 0 | 0 | 0 |
| 0 | NA | NA | NA | NA | NA | Gm7550 | | 0 | 0 | 0 | 0 | 0 | 0 | 0 | 0 | 0 | 0 | 0 |
| 0 | NA | NA | NA | NA | NA | Gm46328 | | 0 | 0 | 0 | 0 | 0 | 0 | 0 | 0 | 0 | 0 | 0 |
| 0 | NA | NA | NA | NA | NA | Gm7557 | | 0 | 0 | 0 | 0 | 0 | 0 | 0 | 0 | 0 | 0 | 0 |
| 0 | NA | NA | NA | NA | NA | Gm10465 | | 0 | 0 | 0 | 0 | 0 | 0 | 0 | 0 | 0 | 0 | 0 |
| 0 | NA | NA | NA | NA | NA | Mir1892 | | 0 | 0 | 0 | 0 | 0 | 0 | 0 | 0 | 0 | 0 | 0 |
| 0 | NA | NA | NA | NA | NA | Gm24971 | | 0 | 0 | 0 | 0 | 0 | 0 | 0 | 0 | 0 | 0 | 0 |
| 0 | NA | NA | NA | NA | NA | Gm22634 | | 0 | 0 | 0 | 0 | 0 | 0 | 0 | 0 | 0 | 0 | 0 |
| 0 | NA | NA | NA | NA | NA | Gm23804 | | 0 | 0 | 0 | 0 | 0 | 0 | 0 | 0 | 0 | 0 | 0 |
| 0 | NA | NA | NA | NA | NA | Gm2395 | | 0 | 0 | 0 | 0 | 0 | 0 | 0 | 0 | 0 | 0 | 0 |
| 0 | NA | NA | NA | NA | NA | Gm25679 | | 0 | 0 | 0 | 0 | 0 | 0 | 0 | 0 | 0 | 0 | 0 |
| 0 | NA | NA | NA | NA | NA | Gm26444 | | 0 | 0 | 0 | 0 | 0 | 0 | 0 | 0 | 0 | 0 | 0 |
| 0 | NA | NA | NA | NA | NA | Gm2412 | | 0 | 0 | 0 | 0 | 0 | 0 | 0 | 0 | 0 | 0 | 0 |
| 0 | NA | NA | NA | NA | NA | Gm24305 | | 0 | 0 | 0 | 0 | 0 | 0 | 0 | 0 | 0 | 0 | 0 |
| 0 | NA | NA | NA | NA | NA | Gm22317 | | 0 | 0 | 0 | 0 | 0 | 0 | 0 | 0 | 0 | 0 | 0 |
| 0 | NA | NA | NA | NA | NA | Gm27014 | | 0 | 0 | 0 | 0 | 0 | 0 | 0 | 0 | 0 | 0 | 0 |
| 0 | NA | NA | NA | NA | NA | Gm20403 | | 0 | 0 | 0 | 0 | 0 | 0 | 0 | 0 | 0 | 0 | 0 |
| 0 | NA | NA | NA | NA | NA | Gm24296 | | 0 | 0 | 0 | 0 | 0 | 0 | 0 | 0 | 0 | 0 | 0 |
| 0 | NA | NA | NA | NA | NA | Fam177a | family with sequence similarity 177, meml | 0 | 0 | 0 | 0 | 0 | 0 | 0 | 0 | 0 | 0 | 0 |
| 0 | NA | NA | NA | NA | NA | Gm2436 | predicted gene 2436 | 0 | 0 | 0 | 0 | 0 | 0 | 0 | 0 | 0 | 0 | 0 |
| 0 | NA | NA | NA | NA | NA | Gm48340 | | 0 | 0 | 0 | 0 | 0 | 0 | 0 | 0 | 0 | 0 | 0 |
| 0 | NA | NA | NA | NA | NA | Srp54b | signal recognition particle 54B | 0 | 0 | 0 | 0 | 0 | 0 | 0 | 0 | 0 | 0 | 0 |
| 0 | NA | NA | NA | NA | NA | Fam177a2 | family with sequence similarity 177 memb | 0 | 0 | 0 | 0 | 0 | 0 | 0 | 0 | 0 | 0 | 0 |
| 0 | NA | NA | NA | NA | NA | Gm2446 | predicted gene 2446 | 0 | 0 | 0 | 0 | 0 | 0 | 0 | 0 | 0 | 0 | 0 |
| 0 | NA | NA | NA | NA | NA | Gm48352 | | 0 | 0 | 0 | 0 | 0 | 0 | 0 | 0 | 0 | 0 | 0 |
| 0 | NA | NA | NA | NA | NA | Gm23394 | | 0 | 0 | 0 | 0 | 0 | 0 | 0 | 0 | 0 | 0 | 0 |
| 0 | NA | NA | NA | NA | NA | Gm22296 | | 0 | 0 | 0 | 0 | 0 | 0 | 0 | 0 | 0 | 0 | 0 |
| 0 | NA | NA | NA | NA | NA | Gm5183 | | 0 | 0 | 0 | 0 | 0 | 0 | 0 | 0 | 0 | 0 | 0 |
| 0 | NA | NA | NA | NA | NA | Gm23799 | | 0 | 0 | 0 | 0 | 0 | 0 | 0 | 0 | 0 | 0 | 0 |
| 0 | NA | NA | NA | NA | NA | Gm47682 | | 0 | 0 | 0 | 0 | 0 | 0 | 0 | 0 | 0 | 0 | 0 |
| 0 | NA | NA | NA | NA | NA | Gm25917 | | 0 | 0 | 0 | 0 | 0 | 0 | 0 | 0 | 0 | 0 | 0 |
| 0 | NA | NA | NA | NA | NA | Gm19990 | | 0 | 0 | 0 | 0 | 0 | 0 | 0 | 0 | 0 | 0 | 0 |
| 0 | NA | NA | NA | NA | NA | Gm24685 | | 0 | 0 | 0 | 0 | 0 | 0 | 0 | 0 | 0 | 0 | 0 |
| 0 | NA | NA | NA | NA | NA | Gm18027 | | 0 | 0 | 0 | 0 | 0 | 0 | 0 | 0 | 0 | 0 | 0 |
| 0 | NA | NA | NA | NA | NA | Sfta3-ps | | 0 | 0 | 0 | 0 | 0 | 0 | 0 | 0 | 0 | 0 | 0 |
| 0 | NA | NA | NA | NA | NA | Gm47720 | | 0 | 0 | 0 | 0 | 0 | 0 | 0 | 0 | 0 | 0 | 0 |
| 0 | NA | NA | NA | NA | NA | Gm26973 | | 0 | 0 | 0 | 0 | 0 | 0 | 0 | 0 | 0 | 0 | 0 |
| 0 | NA | NA | NA | NA | NA | Gm25760 | | 0 | 0 | 0 | 0 | 0 | 0 | 0 | 0 | 0 | 0 | 0 |
| 0 | NA | NA | NA | NA | NA | C87198 | | 0 | 0 | 0 | 0 | 0 | 0 | 0 | 0 | 0 | 0 | 0 |
| 0 | NA | NA | NA | NA | NA | Gm38103 | | 0 | 0 | 0 | 0 | 0 | 0 | 0 | 0 | 0 | 0 | 0 |
| 0 | NA | NA | NA | NA | NA | Gm48835 | | 0 | 0 | 0 | 0 | 0 | 0 | 0 | 0 | 0 | 0 | 0 |
| 0 | NA | NA | NA | NA | NA | Gm18339 | | 0 | 0 | 0 | 0 | 0 | 0 | 0 | 0 | 0 | 0 | 0 |
| 0 | NA | NA | NA | NA | NA | Gm15524 | | 0 | 0 | 0 | 0 | 0 | 0 | 0 | 0 | 0 | 0 | 0 |
| 0 | NA | NA | NA | NA | NA | Pax9 | paired box 9 | 0 | 0 | 0 | 0 | 0 | 0 | 0 | 0 | 0 | 0 | 0 |
| 0 | NA | NA | NA | NA | NA | Gm11025 | | 0 | 0 | 0 | 0 | 0 | 0 | 0 | 0 | 0 | 0 | 0 |
| 0 | NA | NA | NA | NA | NA | Gm16246 | | 0 | 0 | 0 | 0 | 0 | 0 | 0 | 0 | 0 | 0 | 0 |
| 0 | NA | NA | NA | NA | NA | Gm2568 | | 0 | 0 | 0 | 0 | 0 | 0 | 0 | 0 | 0 | 0 | 0 |
| 0 | NA | NA | NA | NA | NA | Gm37291 | | 0 | 0 | 0 | 0 | 0 | 0 | 0 | 0 | 0 | 0 | 0 |

| | | | | | | | | | | | | | | | | | |
|---|---|---|---|---|---|---|---|---|---|---|---|---|---|---|---|---|---|
| 0 | NA | NA | NA | NA | NA | 4921518K17Rik | | 0 | 0 | 0 | 0 | 0 | 0 | 0 | 0 | 0 | 0 | 0 |
| 0 | NA | NA | NA | NA | NA | BC042761 | | 0 | 0 | 0 | 0 | 0 | 0 | 0 | 0 | 0 | 0 | 0 |
| 0 | NA | NA | NA | NA | NA | Gm47617 | | 0 | 0 | 0 | 0 | 0 | 0 | 0 | 0 | 0 | 0 | 0 |
| 0 | NA | NA | NA | NA | NA | Gm18873 | | 0 | 0 | 0 | 0 | 0 | 0 | 0 | 0 | 0 | 0 | 0 |
| 0 | NA | NA | NA | NA | NA | Gm47645 | | 0 | 0 | 0 | 0 | 0 | 0 | 0 | 0 | 0 | 0 | 0 |
| 0 | NA | NA | NA | NA | NA | Gm2649 | | 0 | 0 | 0 | 0 | 0 | 0 | 0 | 0 | 0 | 0 | 0 |
| 0 | NA | NA | NA | NA | NA | Gm47647 | | 0 | 0 | 0 | 0 | 0 | 0 | 0 | 0 | 0 | 0 | 0 |
| 0 | NA | NA | NA | NA | NA | Gm17529 | | 0 | 0 | 0 | 0 | 0 | 0 | 0 | 0 | 0 | 0 | 0 |
| 0 | NA | NA | NA | NA | NA | Gm22973 | | 0 | 0 | 0 | 0 | 0 | 0 | 0 | 0 | 0 | 0 | 0 |
| 0 | NA | NA | NA | NA | NA | Gm27284 | | 0 | 0 | 0 | 0 | 0 | 0 | 0 | 0 | 0 | 0 | 0 |
| 0 | NA | NA | NA | NA | NA | Gm48258 | | 0 | 0 | 0 | 0 | 0 | 0 | 0 | 0 | 0 | 0 | 0 |
| 0 | NA | NA | NA | NA | NA | Gm7719 | | 0 | 0 | 0 | 0 | 0 | 0 | 0 | 0 | 0 | 0 | 0 |
| 0 | NA | NA | NA | NA | NA | Gm4930 | | 0 | 0 | 0 | 0 | 0 | 0 | 0 | 0 | 0 | 0 | 0 |
| 0 | NA | NA | NA | NA | NA | Gm48263 | | 0 | 0 | 0 | 0 | 0 | 0 | 0 | 0 | 0 | 0 | 0 |
| 0 | NA | NA | NA | NA | NA | Gm18242 | | 0 | 0 | 0 | 0 | 0 | 0 | 0 | 0 | 0 | 0 | 0 |
| 0 | NA | NA | NA | NA | NA | Gm48268 | | 0 | 0 | 0 | 0 | 0 | 0 | 0 | 0 | 0 | 0 | 0 |
| 0 | NA | NA | NA | NA | NA | Gm24377 | | 0 | 0 | 0 | 0 | 0 | 0 | 0 | 0 | 0 | 0 | 0 |
| 0 | NA | NA | NA | NA | NA | Gm22673 | | 0 | 0 | 0 | 0 | 0 | 0 | 0 | 0 | 0 | 0 | 0 |
| 0 | NA | NA | NA | NA | NA | Gm48285 | | 0 | 0 | 0 | 0 | 0 | 0 | 0 | 0 | 0 | 0 | 0 |
| 0 | NA | NA | NA | NA | NA | Gm29809 | | 0 | 0 | 0 | 0 | 0 | 0 | 0 | 0 | 0 | 0 | 0 |
| 0 | NA | NA | NA | NA | NA | Gm48289 | | 0 | 0 | 0 | 0 | 0 | 0 | 0 | 0 | 0 | 0 | 0 |
| 0 | NA | NA | NA | NA | NA | Gm32936 | predicted gene, 32936 | 0 | 0 | 0 | 0 | 0 | 0 | 0 | 0 | 0 | 0 | 0 |
| 0 | NA | NA | NA | NA | NA | Gm18871 | | 0 | 0 | 0 | 0 | 0 | 0 | 0 | 0 | 0 | 0 | 0 |
| 0 | NA | NA | NA | NA | NA | Gm20049 | | 0 | 0 | 0 | 0 | 0 | 0 | 0 | 0 | 0 | 0 | 0 |
| 0 | NA | NA | NA | NA | NA | Gm7739 | | 0 | 0 | 0 | 0 | 0 | 0 | 0 | 0 | 0 | 0 | 0 |
| 0 | NA | NA | NA | NA | NA | Gm23059 | | 0 | 0 | 0 | 0 | 0 | 0 | 0 | 0 | 0 | 0 | 0 |
| 0 | NA | NA | NA | NA | NA | Gm38172 | | 0 | 0 | 0 | 0 | 0 | 0 | 0 | 0 | 0 | 0 | 0 |
| 0 | NA | NA | NA | NA | NA | Gm48301 | | 0 | 0 | 0 | 0 | 0 | 0 | 0 | 0 | 0 | 0 | 0 |
| 0 | NA | NA | NA | NA | NA | Gm48304 | | 0 | 0 | 0 | 0 | 0 | 0 | 0 | 0 | 0 | 0 | 0 |
| 0 | NA | NA | NA | NA | NA | Gm48305 | | 0 | 0 | 0 | 0 | 0 | 0 | 0 | 0 | 0 | 0 | 0 |
| 0 | NA | NA | NA | NA | NA | Gm7745 | | 0 | 0 | 0 | 0 | 0 | 0 | 0 | 0 | 0 | 0 | 0 |
| 0 | NA | NA | NA | NA | NA | Nanog-ps2 | | 0 | 0 | 0 | 0 | 0 | 0 | 0 | 0 | 0 | 0 | 0 |
| 0 | NA | NA | NA | NA | NA | Gm20063 | | 0 | 0 | 0 | 0 | 0 | 0 | 0 | 0 | 0 | 0 | 0 |
| 0 | NA | NA | NA | NA | NA | Gm48328 | | 0 | 0 | 0 | 0 | 0 | 0 | 0 | 0 | 0 | 0 | 0 |
| 0 | NA | NA | NA | NA | NA | Gm23213 | | 0 | 0 | 0 | 0 | 0 | 0 | 0 | 0 | 0 | 0 | 0 |
| 0 | NA | NA | NA | NA | NA | Gm7753 | | 0 | 0 | 0 | 0 | 0 | 0 | 0 | 0 | 0 | 0 | 0 |
| 0 | NA | NA | NA | NA | NA | Gm24750 | | 0 | 0 | 0 | 0 | 0 | 0 | 0 | 0 | 0 | 0 | 0 |
| 0 | NA | NA | NA | NA | NA | Gm48358 | | 0 | 0 | 0 | 0 | 0 | 0 | 0 | 0 | 0 | 0 | 0 |
| 0 | NA | NA | NA | NA | NA | Gm2754 | | 0 | 0 | 0 | 0 | 0 | 0 | 0 | 0 | 0 | 0 | 0 |
| 0 | NA | NA | NA | NA | NA | Gm48359 | | 0 | 0 | 0 | 0 | 0 | 0 | 0 | 0 | 0 | 0 | 0 |
| 0 | NA | NA | NA | NA | NA | Gm7754 | | 0 | 0 | 0 | 0 | 0 | 0 | 0 | 0 | 0 | 0 | 0 |
| 0 | NA | NA | NA | NA | NA | Gm6353 | | 0 | 0 | 0 | 0 | 0 | 0 | 0 | 0 | 0 | 0 | 0 |
| 0 | NA | NA | NA | NA | NA | Gm48367 | | 0 | 0 | 0 | 0 | 0 | 0 | 0 | 0 | 0 | 0 | 0 |
| 0 | NA | NA | NA | NA | NA | Gm48368 | | 0 | 0 | 0 | 0 | 0 | 0 | 0 | 0 | 0 | 0 | 0 |
| 0 | NA | NA | NA | NA | NA | Gm24590 | | 0 | 0 | 0 | 0 | 0 | 0 | 0 | 0 | 0 | 0 | 0 |
| 0 | NA | NA | NA | NA | NA | Spanxn4 | | 0 | 0 | 0 | 0 | 0 | 0 | 0 | 0 | 0 | 0 | 0 |
| 0 | NA | NA | NA | NA | NA | Gm19134 | | 0 | 0 | 0 | 0 | 0 | 0 | 0 | 0 | 0 | 0 | 0 |
| 0 | NA | NA | NA | NA | NA | Tspyl-ps | | 0 | 0 | 0 | 0 | 0 | 0 | 0 | 0 | 0 | 0 | 0 |
| 0 | NA | NA | NA | NA | NA | Gm48378 | | 0 | 0 | 0 | 0 | 0 | 0 | 0 | 0 | 0 | 0 | 0 |
| 0 | NA | NA | NA | NA | NA | Gm7763 | | 0 | 0 | 0 | 0 | 0 | 0 | 0 | 0 | 0 | 0 | 0 |
| 0 | NA | NA | NA | NA | NA | Gm5185 | | 0 | 0 | 0 | 0 | 0 | 0 | 0 | 0 | 0 | 0 | 0 |
| 0 | NA | NA | NA | NA | NA | Gm48415 | | 0 | 0 | 0 | 0 | 0 | 0 | 0 | 0 | 0 | 0 | 0 |
| 0 | NA | NA | NA | NA | NA | Gm48416 | | 0 | 0 | 0 | 0 | 0 | 0 | 0 | 0 | 0 | 0 | 0 |
| 0 | NA | NA | NA | NA | NA | Gm48417 | | 0 | 0 | 0 | 0 | 0 | 0 | 0 | 0 | 0 | 0 | 0 |
| 0 | NA | NA | NA | NA | NA | Gm24921 | | 0 | 0 | 0 | 0 | 0 | 0 | 0 | 0 | 0 | 0 | 0 |
| 0 | NA | NA | NA | NA | NA | Gm48418 | | 0 | 0 | 0 | 0 | 0 | 0 | 0 | 0 | 0 | 0 | 0 |
| 0 | NA | NA | NA | NA | NA | Gm5653 | | 0 | 0 | 0 | 0 | 0 | 0 | 0 | 0 | 0 | 0 | 0 |
| 0 | NA | NA | NA | NA | NA | Gm48421 | | 0 | 0 | 0 | 0 | 0 | 0 | 0 | 0 | 0 | 0 | 0 |
| 0 | NA | NA | NA | NA | NA | Gm48422 | | 0 | 0 | 0 | 0 | 0 | 0 | 0 | 0 | 0 | 0 | 0 |
| 0 | NA | NA | NA | NA | NA | Fscb | fibrous sheath CABYR binding protein | 0 | 0 | 0 | 0 | 0 | 0 | 0 | 0 | 0 | 0 | 0 |

| | | | | | | | | | | | | | | | | | | |
|---|---|---|---|---|---|---|---|---|---|---|---|---|---|---|---|---|---|---|
| 0 | NA | NA | NA | NA | NA | Gm24066 | | 0 | 0 | 0 | 0 | 0 | 0 | 0 | 0 | 0 | 0 | 0 |
| 0 | NA | NA | NA | NA | NA | Gm4806 | | 0 | 0 | 0 | 0 | 0 | 0 | 0 | 0 | 0 | 0 | 0 |
| 0 | NA | NA | NA | NA | NA | Gm47869 | | 0 | 0 | 0 | 0 | 0 | 0 | 0 | 0 | 0 | 0 | 0 |
| 0 | NA | NA | NA | NA | NA | 4930471E15Rik | | 0 | 0 | 0 | 0 | 0 | 0 | 0 | 0 | 0 | 0 | 0 |
| 0 | NA | NA | NA | NA | NA | Gm31063 | | 0 | 0 | 0 | 0 | 0 | 0 | 0 | 0 | 0 | 0 | 0 |
| 0 | NA | NA | NA | NA | NA | Gm47973 | | 0 | 0 | 0 | 0 | 0 | 0 | 0 | 0 | 0 | 0 | 0 |
| 0 | NA | NA | NA | NA | NA | Gm26015 | | 0 | 0 | 0 | 0 | 0 | 0 | 0 | 0 | 0 | 0 | 0 |
| 0 | NA | NA | NA | NA | NA | Gm25599 | | 0 | 0 | 0 | 0 | 0 | 0 | 0 | 0 | 0 | 0 | 0 |
| 0 | NA | NA | NA | NA | NA | Gm23948 | | 0 | 0 | 0 | 0 | 0 | 0 | 0 | 0 | 0 | 0 | 0 |
| 0 | NA | NA | NA | NA | NA | Gm22782 | | 0 | 0 | 0 | 0 | 0 | 0 | 0 | 0 | 0 | 0 | 0 |
| 0 | NA | NA | NA | NA | NA | Gm47975 | | 0 | 0 | 0 | 0 | 0 | 0 | 0 | 0 | 0 | 0 | 0 |
| 0 | NA | NA | NA | NA | NA | Gm2915 | | 0 | 0 | 0 | 0 | 0 | 0 | 0 | 0 | 0 | 0 | 0 |
| 0 | NA | NA | NA | NA | NA | Rpl10l | ribosomal protein L10-like | 0 | 0 | 0 | 0 | 0 | 0 | 0 | 0 | 0 | 0 | 0 |
| 0 | NA | NA | NA | NA | NA | Gm47979 | | 0 | 0 | 0 | 0 | 0 | 0 | 0 | 0 | 0 | 0 | 0 |
| 0 | NA | NA | NA | NA | NA | Gm47982 | | 0 | 0 | 0 | 0 | 0 | 0 | 0 | 0 | 0 | 0 | 0 |
| 0 | NA | NA | NA | NA | NA | Gm19001 | | 0 | 0 | 0 | 0 | 0 | 0 | 0 | 0 | 0 | 0 | 0 |
| 0 | NA | NA | NA | NA | NA | Gm47988 | | 0 | 0 | 0 | 0 | 0 | 0 | 0 | 0 | 0 | 0 | 0 |
| 0 | NA | NA | NA | NA | NA | Gm23826 | | 0 | 0 | 0 | 0 | 0 | 0 | 0 | 0 | 0 | 0 | 0 |
| 0 | NA | NA | NA | NA | NA | Gm18940 | | 0 | 0 | 0 | 0 | 0 | 0 | 0 | 0 | 0 | 0 | 0 |
| 0 | NA | NA | NA | NA | NA | Gm47989 | | 0 | 0 | 0 | 0 | 0 | 0 | 0 | 0 | 0 | 0 | 0 |
| 0 | NA | NA | NA | NA | NA | Gm47987 | | 0 | 0 | 0 | 0 | 0 | 0 | 0 | 0 | 0 | 0 | 0 |
| 0 | NA | NA | NA | NA | NA | Gm48059 | | 0 | 0 | 0 | 0 | 0 | 0 | 0 | 0 | 0 | 0 | 0 |
| 0 | NA | NA | NA | NA | NA | Gm48060 | | 0 | 0 | 0 | 0 | 0 | 0 | 0 | 0 | 0 | 0 | 0 |
| 0 | NA | NA | NA | NA | NA | n-R5s60 | | 0 | 0 | 0 | 0 | 0 | 0 | 0 | 0 | 0 | 0 | 0 |
| 0 | NA | NA | NA | NA | NA | Gm26256 | | 0 | 0 | 0 | 0 | 0 | 0 | 0 | 0 | 0 | 0 | 0 |
| 0 | NA | NA | NA | NA | NA | Gm47432 | | 0 | 0 | 0 | 0 | 0 | 0 | 0 | 0 | 0 | 0 | 0 |
| 0 | NA | NA | NA | NA | NA | Gm17821 | | 0 | 0 | 0 | 0 | 0 | 0 | 0 | 0 | 0 | 0 | 0 |
| 0 | NA | NA | NA | NA | NA | Gm47452 | | 0 | 0 | 0 | 0 | 0 | 0 | 0 | 0 | 0 | 0 | 0 |
| 0 | NA | NA | NA | NA | NA | Gm20102 | | 0 | 0 | 0 | 0 | 0 | 0 | 0 | 0 | 0 | 0 | 0 |
| 0 | NA | NA | NA | NA | NA | Gm47453 | | 0 | 0 | 0 | 0 | 0 | 0 | 0 | 0 | 0 | 0 | 0 |
| 0 | NA | NA | NA | NA | NA | Gm47454 | | 0 | 0 | 0 | 0 | 0 | 0 | 0 | 0 | 0 | 0 | 0 |
| 0 | NA | NA | NA | NA | NA | Gm47457 | | 0 | 0 | 0 | 0 | 0 | 0 | 0 | 0 | 0 | 0 | 0 |
| 0 | NA | NA | NA | NA | NA | Gm19176 | | 0 | 0 | 0 | 0 | 0 | 0 | 0 | 0 | 0 | 0 | 0 |
| 0 | NA | NA | NA | NA | NA | Gm47489 | | 0 | 0 | 0 | 0 | 0 | 0 | 0 | 0 | 0 | 0 | 0 |
| 0 | NA | NA | NA | NA | NA | Gm19047 | | 0 | 0 | 0 | 0 | 0 | 0 | 0 | 0 | 0 | 0 | 0 |
| 0 | NA | NA | NA | NA | NA | Gm47490 | | 0 | 0 | 0 | 0 | 0 | 0 | 0 | 0 | 0 | 0 | 0 |
| 0 | NA | NA | NA | NA | NA | n-R5s61 | | 0 | 0 | 0 | 0 | 0 | 0 | 0 | 0 | 0 | 0 | 0 |
| 0 | NA | NA | NA | NA | NA | Gm47491 | | 0 | 0 | 0 | 0 | 0 | 0 | 0 | 0 | 0 | 0 | 0 |
| 0 | NA | NA | NA | NA | NA | Gm47515 | | 0 | 0 | 0 | 0 | 0 | 0 | 0 | 0 | 0 | 0 | 0 |
| 0 | NA | NA | NA | NA | NA | Gm2912 | | 0 | 0 | 0 | 0 | 0 | 0 | 0 | 0 | 0 | 0 | 0 |
| 0 | NA | NA | NA | NA | NA | Gm47518 | | 0 | 0 | 0 | 0 | 0 | 0 | 0 | 0 | 0 | 0 | 0 |
| 0 | NA | NA | NA | NA | NA | Gm31447 | | 0 | 0 | 0 | 0 | 0 | 0 | 0 | 0 | 0 | 0 | 0 |
| 0 | NA | NA | NA | NA | NA | Gm15571 | | 0 | 0 | 0 | 0 | 0 | 0 | 0 | 0 | 0 | 0 | 0 |
| 0 | NA | NA | NA | NA | NA | Gm24499 | | 0 | 0 | 0 | 0 | 0 | 0 | 0 | 0 | 0 | 0 | 0 |
| 0 | NA | NA | NA | NA | NA | Gm47287 | | 0 | 0 | 0 | 0 | 0 | 0 | 0 | 0 | 0 | 0 | 0 |
| 0 | NA | NA | NA | NA | NA | Gm47295 | | 0 | 0 | 0 | 0 | 0 | 0 | 0 | 0 | 0 | 0 | 0 |
| 0 | NA | NA | NA | NA | NA | Gm49383 | | 0 | 0 | 0 | 0 | 0 | 0 | 0 | 0 | 0 | 0 | 0 |
| 0 | NA | NA | NA | NA | NA | Gm49384 | | 0 | 0 | 0 | 0 | 0 | 0 | 0 | 0 | 0 | 0 | 0 |
| 0 | NA | NA | NA | NA | NA | Gm47545 | | 0 | 0 | 0 | 0 | 0 | 0 | 0 | 0 | 0 | 0 | 0 |
| 0 | NA | NA | NA | NA | NA | Gm15561 | | 0 | 0 | 0 | 0 | 0 | 0 | 0 | 0 | 0 | 0 | 0 |
| 0 | NA | NA | NA | NA | NA | Gm47299 | | 0 | 0 | 0 | 0 | 0 | 0 | 0 | 0 | 0 | 0 | 0 |
| 0 | NA | NA | NA | NA | NA | Rn7s2 | | 0 | 0 | 0 | 0 | 0 | 0 | 0 | 0 | 0 | 0 | 0 |
| 0 | NA | NA | NA | NA | NA | Gm47305 | | 0 | 0 | 0 | 0 | 0 | 0 | 0 | 0 | 0 | 0 | 0 |
| 0 | NA | NA | NA | NA | NA | Gm23018 | | 0 | 0 | 0 | 0 | 0 | 0 | 0 | 0 | 0 | 0 | 0 |
| 0 | NA | NA | NA | NA | NA | Gm24449 | | 0 | 0 | 0 | 0 | 0 | 0 | 0 | 0 | 0 | 0 | 0 |
| 0 | NA | NA | NA | NA | NA | Gm18030 | | 0 | 0 | 0 | 0 | 0 | 0 | 0 | 0 | 0 | 0 | 0 |
| 0 | NA | NA | NA | NA | NA | Gm22058 | | 0 | 0 | 0 | 0 | 0 | 0 | 0 | 0 | 0 | 0 | 0 |
| 0 | NA | NA | NA | NA | NA | Gm19144 | | 0 | 0 | 0 | 0 | 0 | 0 | 0 | 0 | 0 | 0 | 0 |
| 0 | NA | NA | NA | NA | NA | Gm23579 | | 0 | 0 | 0 | 0 | 0 | 0 | 0 | 0 | 0 | 0 | 0 |
| 0 | NA | NA | NA | NA | NA | Gm48695 | | 0 | 0 | 0 | 0 | 0 | 0 | 0 | 0 | 0 | 0 | 0 |

| | | | | | | | | | | | | | | | | | | |
|---|---|---|---|---|---|---|---|---|---|---|---|---|---|---|---|---|---|---|
| 0 | NA | NA | NA | NA | NA | Abhd12b | abhydrolase domain containing 12B | 0 | 0 | 0 | 0 | 0 | 0 | 0 | 0 | 0 | 0 | 0 |
| 0 | NA | NA | NA | NA | NA | Gm48847 | | 0 | 0 | 0 | 0 | 0 | 0 | 0 | 0 | 0 | 0 | 0 |
| 0 | NA | NA | NA | NA | NA | Gm24474 | | 0 | 0 | 0 | 0 | 0 | 0 | 0 | 0 | 0 | 0 | 0 |
| 0 | NA | NA | NA | NA | NA | Gm47290 | | 0 | 0 | 0 | 0 | 0 | 0 | 0 | 0 | 0 | 0 | 0 |
| 0 | NA | NA | NA | NA | NA | Gm48866 | | 0 | 0 | 0 | 0 | 0 | 0 | 0 | 0 | 0 | 0 | 0 |
| 0 | NA | NA | NA | NA | NA | Gm32369 | | 0 | 0 | 0 | 0 | 0 | 0 | 0 | 0 | 0 | 0 | 0 |
| 0 | NA | NA | NA | NA | NA | Gm40437 | | 0 | 0 | 0 | 0 | 0 | 0 | 0 | 0 | 0 | 0 | 0 |
| 0 | NA | NA | NA | NA | NA | Gm46355 | | 0 | 0 | 0 | 0 | 0 | 0 | 0 | 0 | 0 | 0 | 0 |
| 0 | NA | NA | NA | NA | NA | Gm10457 | | 0 | 0 | 0 | 0 | 0 | 0 | 0 | 0 | 0 | 0 | 0 |
| 0 | NA | NA | NA | NA | NA | Gm24857 | | 0 | 0 | 0 | 0 | 0 | 0 | 0 | 0 | 0 | 0 | 0 |
| 0 | NA | NA | NA | NA | NA | Tomm20l | translocase of outer mitochondrial membr | 0 | 0 | 0 | 0 | 0 | 0 | 0 | 0 | 0 | 0 | 0 |
| 0 | NA | NA | NA | NA | NA | Gm19045 | | 0 | 0 | 0 | 0 | 0 | 0 | 0 | 0 | 0 | 0 | 0 |
| 0 | NA | NA | NA | NA | NA | Gm7959 | | 0 | 0 | 0 | 0 | 0 | 0 | 0 | 0 | 0 | 0 | 0 |
| 0 | NA | NA | NA | NA | NA | 1700083H02Rik | | 0 | 0 | 0 | 0 | 0 | 0 | 0 | 0 | 0 | 0 | 0 |
| 0 | NA | NA | NA | NA | NA | Gm22959 | | 0 | 0 | 0 | 0 | 0 | 0 | 0 | 0 | 0 | 0 | 0 |
| 0 | NA | NA | NA | NA | NA | Gm47555 | | 0 | 0 | 0 | 0 | 0 | 0 | 0 | 0 | 0 | 0 | 0 |
| 0 | NA | NA | NA | NA | NA | Gm18243 | | 0 | 0 | 0 | 0 | 0 | 0 | 0 | 0 | 0 | 0 | 0 |
| 0 | NA | NA | NA | NA | NA | Gm24466 | | 0 | 0 | 0 | 0 | 0 | 0 | 0 | 0 | 0 | 0 | 0 |
| 0 | NA | NA | NA | NA | NA | Gm44392 | | 0 | 0 | 0 | 0 | 0 | 0 | 0 | 0 | 0 | 0 | 0 |
| 0 | NA | NA | NA | NA | NA | Gm4756 | | 0 | 0 | 0 | 0 | 0 | 0 | 0 | 0 | 0 | 0 | 0 |
| 0 | NA | NA | NA | NA | NA | Gm48143 | | 0 | 0 | 0 | 0 | 0 | 0 | 0 | 0 | 0 | 0 | 0 |
| 0 | NA | NA | NA | NA | NA | Six6 | sine oculis-related homeobox 6 | 0 | 0 | 0 | 0 | 0 | 0 | 0 | 0 | 0 | 0 | 0 |
| 0 | NA | NA | NA | NA | NA | Gm26709 | | 0 | 0 | 0 | 0 | 0 | 0 | 0 | 0 | 0 | 0 | 0 |
| 0 | NA | NA | NA | NA | NA | Gm38170 | | 0 | 0 | 0 | 0 | 0 | 0 | 0 | 0 | 0 | 0 | 0 |
| 0 | NA | NA | NA | NA | NA | Gm23910 | | 0 | 0 | 0 | 0 | 0 | 0 | 0 | 0 | 0 | 0 | 0 |
| 0 | NA | NA | NA | NA | NA | Gm33929 | | 0 | 0 | 0 | 0 | 0 | 0 | 0 | 0 | 0 | 0 | 0 |
| 0 | NA | NA | NA | NA | NA | Gm48656 | | 0 | 0 | 0 | 0 | 0 | 0 | 0 | 0 | 0 | 0 | 0 |
| 0 | NA | NA | NA | NA | NA | Gm34016 | | 0 | 0 | 0 | 0 | 0 | 0 | 0 | 0 | 0 | 0 | 0 |
| 0 | NA | NA | NA | NA | NA | Gm8075 | | 0 | 0 | 0 | 0 | 0 | 0 | 0 | 0 | 0 | 0 | 0 |
| 0 | NA | NA | NA | NA | NA | Gm48823 | | 0 | 0 | 0 | 0 | 0 | 0 | 0 | 0 | 0 | 0 | 0 |
| 0 | NA | NA | NA | NA | NA | Snapc1l | | 0 | 0 | 0 | 0 | 0 | 0 | 0 | 0 | 0 | 0 | 0 |
| 0 | NA | NA | NA | NA | NA | S100a11-ps | | 0 | 0 | 0 | 0 | 0 | 0 | 0 | 0 | 0 | 0 | 0 |
| 0 | NA | NA | NA | NA | NA | Gm9544 | | 0 | 0 | 0 | 0 | 0 | 0 | 0 | 0 | 0 | 0 | 0 |
| 0 | NA | NA | NA | NA | NA | Gm47618 | | 0 | 0 | 0 | 0 | 0 | 0 | 0 | 0 | 0 | 0 | 0 |
| 0 | NA | NA | NA | NA | NA | Gm47619 | | 0 | 0 | 0 | 0 | 0 | 0 | 0 | 0 | 0 | 0 | 0 |
| 0 | NA | NA | NA | NA | NA | Gm39473 | | 0 | 0 | 0 | 0 | 0 | 0 | 0 | 0 | 0 | 0 | 0 |
| 0 | NA | NA | NA | NA | NA | 4930442G10Rik | | 0 | 0 | 0 | 0 | 0 | 0 | 0 | 0 | 0 | 0 | 0 |
| 0 | NA | NA | NA | NA | NA | Gphb5 | glycoprotein hormone beta 5 | 0 | 0 | 0 | 0 | 0 | 0 | 0 | 0 | 0 | 0 | 0 |
| 0 | NA | NA | NA | NA | NA | Gm47689 | | 0 | 0 | 0 | 0 | 0 | 0 | 0 | 0 | 0 | 0 | 0 |
| 0 | NA | NA | NA | NA | NA | | | 0 | 0 | 0 | 0 | 0 | 0 | 0 | 0 | 0 | 0 | 0 |
| 0 | NA | NA | NA | NA | NA | Mir5101 | | 0 | 0 | 0 | 0 | 0 | 0 | 0 | 0 | 0 | 0 | 0 |
| 0 | NA | NA | NA | NA | NA | Tex21 | testis expressed gene 21 | 0 | 0 | 0 | 0 | 0 | 0 | 0 | 0 | 0 | 0 | 0 |
| 0 | NA | NA | NA | NA | NA | Gm47526 | | 0 | 0 | 0 | 0 | 0 | 0 | 0 | 0 | 0 | 0 | 0 |
| 0 | NA | NA | NA | NA | NA | Gm34868 | | 0 | 0 | 0 | 0 | 0 | 0 | 0 | 0 | 0 | 0 | 0 |
| 0 | NA | NA | NA | NA | NA | Gm25563 | | 0 | 0 | 0 | 0 | 0 | 0 | 0 | 0 | 0 | 0 | 0 |
| 0 | NA | NA | NA | NA | NA | Gm47053 | | 0 | 0 | 0 | 0 | 0 | 0 | 0 | 0 | 0 | 0 | 0 |
| 0 | NA | NA | NA | NA | NA | Gm24010 | | 0 | 0 | 0 | 0 | 0 | 0 | 0 | 0 | 0 | 0 | 0 |
| 0 | NA | NA | NA | NA | NA | Gm35041 | | 0 | 0 | 0 | 0 | 0 | 0 | 0 | 0 | 0 | 0 | 0 |
| 0 | NA | NA | NA | NA | NA | Gm35189 | | 0 | 0 | 0 | 0 | 0 | 0 | 0 | 0 | 0 | 0 | 0 |
| 0 | NA | NA | NA | NA | NA | Gm22696 | | 0 | 0 | 0 | 0 | 0 | 0 | 0 | 0 | 0 | 0 | 0 |
| 0 | NA | NA | NA | NA | NA | Gm24775 | | 0 | 0 | 0 | 0 | 0 | 0 | 0 | 0 | 0 | 0 | 0 |
| 0 | NA | NA | NA | NA | NA | Gm48177 | | 0 | 0 | 0 | 0 | 0 | 0 | 0 | 0 | 0 | 0 | 0 |
| 0 | NA | NA | NA | NA | NA | Gm48196 | | 0 | 0 | 0 | 0 | 0 | 0 | 0 | 0 | 0 | 0 | 0 |
| 0 | NA | NA | NA | NA | NA | Gm24070 | | 0 | 0 | 0 | 0 | 0 | 0 | 0 | 0 | 0 | 0 | 0 |
| 0 | NA | NA | NA | NA | NA | Gm24994 | | 0 | 0 | 0 | 0 | 0 | 0 | 0 | 0 | 0 | 0 | 0 |
| 0 | NA | NA | NA | NA | NA | Gm6657 | predicted gene 6657 | 0 | 0 | 0 | 0 | 0 | 0 | 0 | 0 | 0 | 0 | 0 |
| 0 | NA | NA | NA | NA | NA | Gm48224 | | 0 | 0 | 0 | 0 | 0 | 0 | 0 | 0 | 0 | 0 | 0 |
| 0 | NA | NA | NA | NA | NA | Gm25225 | | 0 | 0 | 0 | 0 | 0 | 0 | 0 | 0 | 0 | 0 | 0 |
| 0 | NA | NA | NA | NA | NA | Gm27297 | | 0 | 0 | 0 | 0 | 0 | 0 | 0 | 0 | 0 | 0 | 0 |
| 0 | NA | NA | NA | NA | NA | Gm8255 | | 0 | 0 | 0 | 0 | 0 | 0 | 0 | 0 | 0 | 0 | 0 |

| | | | | | | | | | | | | | | | | | | |
|---|---|---|---|---|---|---|---|---|---|---|---|---|---|---|---|---|---|---|
| 0 | NA | NA | NA | NA | NA | Gm17194 | | 0 | 0 | 0 | 0 | 0 | 0 | 0 | 0 | 0 | 0 | 0 |
| 0 | NA | NA | NA | NA | NA | Gm5654 | | 0 | 0 | 0 | 0 | 0 | 0 | 0 | 0 | 0 | 0 | 0 |
| 0 | NA | NA | NA | NA | NA | Gm8275 | | 0 | 0 | 0 | 0 | 0 | 0 | 0 | 0 | 0 | 0 | 0 |
| 0 | NA | NA | NA | NA | NA | Gm47752 | | 0 | 0 | 0 | 0 | 0 | 0 | 0 | 0 | 0 | 0 | 0 |
| 0 | NA | NA | NA | NA | NA | Mir1843a | | 0 | 0 | 0 | 0 | 0 | 0 | 0 | 0 | 0 | 0 | 0 |
| 0 | NA | NA | NA | NA | NA | Scarna3b | | 0 | 0 | 0 | 0 | 0 | 0 | 0 | 0 | 0 | 0 | 0 |
| 0 | NA | NA | NA | NA | NA | Gm50278 | | 0 | 0 | 0 | 0 | 0 | 0 | 0 | 0 | 0 | 0 | 0 |
| 0 | NA | NA | NA | NA | NA | 2310002D06Rik | | 0 | 0 | 0 | 0 | 0 | 0 | 0 | 0 | 0 | 0 | 0 |
| 0 | NA | NA | NA | NA | NA | 4933406B15Rik | | 0 | 0 | 0 | 0 | 0 | 0 | 0 | 0 | 0 | 0 | 0 |
| 0 | NA | NA | NA | NA | NA | Gm46364 | | 0 | 0 | 0 | 0 | 0 | 0 | 0 | 0 | 0 | 0 | 0 |
| 0 | NA | NA | NA | NA | NA | Gm23817 | | 0 | 0 | 0 | 0 | 0 | 0 | 0 | 0 | 0 | 0 | 0 |
| 0 | NA | NA | NA | NA | NA | Gm47852 | | 0 | 0 | 0 | 0 | 0 | 0 | 0 | 0 | 0 | 0 | 0 |
| 0 | NA | NA | NA | NA | NA | Gm47879 | | 0 | 0 | 0 | 0 | 0 | 0 | 0 | 0 | 0 | 0 | 0 |
| 0 | NA | NA | NA | NA | NA | Gm47874 | | 0 | 0 | 0 | 0 | 0 | 0 | 0 | 0 | 0 | 0 | 0 |
| 0 | NA | NA | NA | NA | NA | Gm23298 | | 0 | 0 | 0 | 0 | 0 | 0 | 0 | 0 | 0 | 0 | 0 |
| 0 | NA | NA | NA | NA | NA | 1700052I22Rik | | 0 | 0 | 0 | 0 | 0 | 0 | 0 | 0 | 0 | 0 | 0 |
| 0 | NA | NA | NA | NA | NA | Slc10a1 | solute carrier family 10 (sodium/bile acid [illegible] | 0 | 0 | 0 | 0 | 0 | 0 | 0 | 0 | 0 | 0 | 0 |
| 0 | NA | NA | NA | NA | NA | Mir3067 | | 0 | 0 | 0 | 0 | 0 | 0 | 0 | 0 | 0 | 0 | 0 |
| 0 | NA | NA | NA | NA | NA | 6530401F13Rik | | 0 | 0 | 0 | 0 | 0 | 0 | 0 | 0 | 0 | 0 | 0 |
| 0 | NA | NA | NA | NA | NA | Gm16572 | | 0 | 0 | 0 | 0 | 0 | 0 | 0 | 0 | 0 | 0 | 0 |
| 0 | NA | NA | NA | NA | NA | Gm49749 | | 0 | 0 | 0 | 0 | 0 | 0 | 0 | 0 | 0 | 0 | 0 |
| 0 | NA | NA | NA | NA | NA | n-R5s62 | | 0 | 0 | 0 | 0 | 0 | 0 | 0 | 0 | 0 | 0 | 0 |
| 0 | NA | NA | NA | NA | NA | Gm8358 | | 0 | 0 | 0 | 0 | 0 | 0 | 0 | 0 | 0 | 0 | 0 |
| 0 | NA | NA | NA | NA | NA | n-R5s63 | | 0 | 0 | 0 | 0 | 0 | 0 | 0 | 0 | 0 | 0 | 0 |
| 0 | NA | NA | NA | NA | NA | Gm22149 | | 0 | 0 | 0 | 0 | 0 | 0 | 0 | 0 | 0 | 0 | 0 |
| 0 | NA | NA | NA | NA | NA | Gm22149 | | 0 | 0 | 0 | 0 | 0 | 0 | 0 | 0 | 0 | 0 | 0 |
| 0 | NA | NA | NA | NA | NA | 1700085C21Rik | | 0 | 0 | 0 | 0 | 0 | 0 | 0 | 0 | 0 | 0 | 0 |
| 0 | NA | NA | NA | NA | NA | 4930423C22Rik | | 0 | 0 | 0 | 0 | 0 | 0 | 0 | 0 | 0 | 0 | 0 |
| 0 | NA | NA | NA | NA | NA | Gm29530 | | 0 | 0 | 0 | 0 | 0 | 0 | 0 | 0 | 0 | 0 | 0 |
| 0 | NA | NA | NA | NA | NA | 4933436F18Rik | | 0 | 0 | 0 | 0 | 0 | 0 | 0 | 0 | 0 | 0 | 0 |
| 0 | NA | NA | NA | NA | NA | 1700030I03Rik | | 0 | 0 | 0 | 0 | 0 | 0 | 0 | 0 | 0 | 0 | 0 |
| 0 | NA | NA | NA | NA | NA | Gm47184 | | 0 | 0 | 0 | 0 | 0 | 0 | 0 | 0 | 0 | 0 | 0 |
| 0 | NA | NA | NA | NA | NA | Gm47666 | | 0 | 0 | 0 | 0 | 0 | 0 | 0 | 0 | 0 | 0 | 0 |
| 0 | NA | NA | NA | NA | NA | Gm18592 | | 0 | 0 | 0 | 0 | 0 | 0 | 0 | 0 | 0 | 0 | 0 |
| 0 | NA | NA | NA | NA | NA | Gm46367 | | 0 | 0 | 0 | 0 | 0 | 0 | 0 | 0 | 0 | 0 | 0 |
| 0 | NA | NA | NA | NA | NA | Gm47748 | | 0 | 0 | 0 | 0 | 0 | 0 | 0 | 0 | 0 | 0 | 0 |
| 0 | NA | NA | NA | NA | NA | Gm47943 | | 0 | 0 | 0 | 0 | 0 | 0 | 0 | 0 | 0 | 0 | 0 |
| 0 | NA | NA | NA | NA | NA | Gm49366 | | 0 | 0 | 0 | 0 | 0 | 0 | 0 | 0 | 0 | 0 | 0 |
| 0 | NA | NA | NA | NA | NA | Gm8385 | | 0 | 0 | 0 | 0 | 0 | 0 | 0 | 0 | 0 | 0 | 0 |
| 0 | NA | NA | NA | NA | NA | Gm47447 | | 0 | 0 | 0 | 0 | 0 | 0 | 0 | 0 | 0 | 0 | 0 |
| 0 | NA | NA | NA | NA | NA | Vrtn | vertebrae development associated | 0 | 0 | 0 | 0 | 0 | 0 | 0 | 0 | 0 | 0 | 0 |
| 0 | NA | NA | NA | NA | NA | Gm48573 | | 0 | 0 | 0 | 0 | 0 | 0 | 0 | 0 | 0 | 0 | 0 |
| 0 | NA | NA | NA | NA | NA | Gm48709 | | 0 | 0 | 0 | 0 | 0 | 0 | 0 | 0 | 0 | 0 | 0 |
| 0 | NA | NA | NA | NA | NA | Gm32296 | | 0 | 0 | 0 | 0 | 0 | 0 | 0 | 0 | 0 | 0 | 0 |
| 0 | NA | NA | NA | NA | NA | n-R5s64 | | 0 | 0 | 0 | 0 | 0 | 0 | 0 | 0 | 0 | 0 | 0 |
| 0 | NA | NA | NA | NA | NA | Gm47285 | | 0 | 0 | 0 | 0 | 0 | 0 | 0 | 0 | 0 | 0 | 0 |
| 0 | NA | NA | NA | NA | NA | Gm47901 | | 0 | 0 | 0 | 0 | 0 | 0 | 0 | 0 | 0 | 0 | 0 |
| 0 | NA | NA | NA | NA | NA | Gm26531 | | 0 | 0 | 0 | 0 | 0 | 0 | 0 | 0 | 0 | 0 | 0 |
| 0 | NA | NA | NA | NA | NA | Gm37804 | | 0 | 0 | 0 | 0 | 0 | 0 | 0 | 0 | 0 | 0 | 0 |
| 0 | NA | NA | NA | NA | NA | Gm805 | | 0 | 0 | 0 | 0 | 0 | 0 | 0 | 0 | 0 | 0 | 0 |
| 0 | NA | NA | NA | NA | NA | Gm48101 | | 0 | 0 | 0 | 0 | 0 | 0 | 0 | 0 | 0 | 0 | 0 |
| 0 | NA | NA | NA | NA | NA | Gm8504 | | 0 | 0 | 0 | 0 | 0 | 0 | 0 | 0 | 0 | 0 | 0 |
| 0 | NA | NA | NA | NA | NA | Gm27993 | | 0 | 0 | 0 | 0 | 0 | 0 | 0 | 0 | 0 | 0 | 0 |
| 0 | NA | NA | NA | NA | NA | Gm22004 | | 0 | 0 | 0 | 0 | 0 | 0 | 0 | 0 | 0 | 0 | 0 |
| 0 | NA | NA | NA | NA | NA | Gm17138 | | 0 | 0 | 0 | 0 | 0 | 0 | 0 | 0 | 0 | 0 | 0 |
| 0 | NA | NA | NA | NA | NA | Lrrc74a | leucine rich repeat containing 74A | 0 | 0 | 0 | 0 | 0 | 0 | 0 | 0 | 0 | 0 | 0 |
| 0 | NA | NA | NA | NA | NA | Gm25888 | | 0 | 0 | 0 | 0 | 0 | 0 | 0 | 0 | 0 | 0 | 0 |
| 0 | NA | NA | NA | NA | NA | Gm25847 | | 0 | 0 | 0 | 0 | 0 | 0 | 0 | 0 | 0 | 0 | 0 |
| 0 | NA | NA | NA | NA | NA | Gm18499 | | 0 | 0 | 0 | 0 | 0 | 0 | 0 | 0 | 0 | 0 | 0 |
| 0 | NA | NA | NA | NA | NA | Gm47293 | | 0 | 0 | 0 | 0 | 0 | 0 | 0 | 0 | 0 | 0 | 0 |

| | | | | | | | | | | | | | | | | | | |
|---|---|---|---|---|---|---|---|---|---|---|---|---|---|---|---|---|---|---|
| 0 | NA | NA | NA | NA | NA | Gm48806 | | 0 | 0 | 0 | 0 | 0 | 0 | 0 | 0 | 0 | 0 | 0 |
| 0 | NA | NA | NA | NA | NA | Gm48873 | | 0 | 0 | 0 | 0 | 0 | 0 | 0 | 0 | 0 | 0 | 0 |
| 0 | NA | NA | NA | NA | NA | Mir3068 | | 0 | 0 | 0 | 0 | 0 | 0 | 0 | 0 | 0 | 0 | 0 |
| 0 | NA | NA | NA | NA | NA | Gm50450 | | 0 | 0 | 0 | 0 | 0 | 0 | 0 | 0 | 0 | 0 | 0 |
| 0 | NA | NA | NA | NA | NA | Gm29362 | | 0 | 0 | 0 | 0 | 0 | 0 | 0 | 0 | 0 | 0 | 0 |
| 0 | NA | NA | NA | NA | NA | Gm18983 | | 0 | 0 | 0 | 0 | 0 | 0 | 0 | 0 | 0 | 0 | 0 |
| 0 | NA | NA | NA | NA | NA | Gm18624 | | 0 | 0 | 0 | 0 | 0 | 0 | 0 | 0 | 0 | 0 | 0 |
| 0 | NA | NA | NA | NA | NA | Gm5788 | | 0 | 0 | 0 | 0 | 0 | 0 | 0 | 0 | 0 | 0 | 0 |
| 0 | NA | NA | NA | NA | NA | Gm18625 | | 0 | 0 | 0 | 0 | 0 | 0 | 0 | 0 | 0 | 0 | 0 |
| 0 | NA | NA | NA | NA | NA | Gm49362 | | 0 | 0 | 0 | 0 | 0 | 0 | 0 | 0 | 0 | 0 | 0 |
| 0 | NA | NA | NA | NA | NA | Gm8300 | predicted gene 8300 | 0 | 0 | 0 | 0 | 0 | 0 | 0 | 0 | 0 | 0 | 0 |
| 0 | NA | NA | NA | NA | NA | Gm18626 | | 0 | 0 | 0 | 0 | 0 | 0 | 0 | 0 | 0 | 0 | 0 |
| 0 | NA | NA | NA | NA | NA | Gm18615 | | 0 | 0 | 0 | 0 | 0 | 0 | 0 | 0 | 0 | 0 | 0 |
| 0 | NA | NA | NA | NA | NA | Oog1 | oogenesin 1 | 0 | 0 | 0 | 0 | 0 | 0 | 0 | 0 | 0 | 0 | 0 |
| 0 | NA | NA | NA | NA | NA | Gm4027 | predicted gene 4027 | 0 | 0 | 0 | 0 | 0 | 0 | 0 | 0 | 0 | 0 | 0 |
| 0 | NA | NA | NA | NA | NA | Gm3742 | | 0 | 0 | 0 | 0 | 0 | 0 | 0 | 0 | 0 | 0 | 0 |
| 0 | NA | NA | NA | NA | NA | Gm16381 | predicted gene 16381 | 0 | 0 | 0 | 0 | 0 | 0 | 0 | 0 | 0 | 0 | 0 |
| 0 | NA | NA | NA | NA | NA | Gm21937 | | 0 | 0 | 0 | 0 | 0 | 0 | 0 | 0 | 0 | 0 | 0 |
| 0 | NA | NA | NA | NA | NA | | | 0 | 0 | 0 | 0 | 0 | 0 | 0 | 0 | 0 | 0 | 0 |
| 0 | NA | NA | NA | NA | NA | Gm48150 | | 0 | 0 | 0 | 0 | 0 | 0 | 0 | 0 | 0 | 0 | 0 |
| 0 | NA | NA | NA | NA | NA | Gm21559 | | 0 | 0 | 0 | 0 | 0 | 0 | 0 | 0 | 0 | 0 | 0 |
| 0 | NA | NA | NA | NA | NA | Gm21569 | | 0 | 0 | 0 | 0 | 0 | 0 | 0 | 0 | 0 | 0 | 0 |
| 0 | NA | NA | NA | NA | NA | Gm21319 | predicted gene, 21319 | 0 | 0 | 0 | 0 | 0 | 0 | 0 | 0 | 0 | 0 | 0 |
| 0 | NA | NA | NA | NA | NA | Gm48152 | | 0 | 0 | 0 | 0 | 0 | 0 | 0 | 0 | 0 | 0 | 0 |
| 0 | NA | NA | NA | NA | NA | Gm21936 | predicted gene, 21936 | 0 | 0 | 0 | 0 | 0 | 0 | 0 | 0 | 0 | 0 | 0 |
| 0 | NA | NA | NA | NA | NA | Gm48149 | | 0 | 0 | 0 | 0 | 0 | 0 | 0 | 0 | 0 | 0 | 0 |
| 0 | NA | NA | NA | NA | NA | Gm48154 | | 0 | 0 | 0 | 0 | 0 | 0 | 0 | 0 | 0 | 0 | 0 |
| 0 | NA | NA | NA | NA | NA | BB287469 | expressed sequence BB287469 | 0 | 0 | 0 | 0 | 0 | 0 | 0 | 0 | 0 | 0 | 0 |
| 0 | NA | NA | NA | NA | NA | Gm21038 | | 0 | 0 | 0 | 0 | 0 | 0 | 0 | 0 | 0 | 0 | 0 |
| 0 | NA | NA | NA | NA | NA | Gm2001 | predicted gene 2001 | 0 | 0 | 0 | 0 | 0 | 0 | 0 | 0 | 0 | 0 | 0 |
| 0 | NA | NA | NA | NA | NA | Gm1995 | | 0 | 0 | 0 | 0 | 0 | 0 | 0 | 0 | 0 | 0 | 0 |
| 0 | NA | NA | NA | NA | NA | Gm18627 | | 0 | 0 | 0 | 0 | 0 | 0 | 0 | 0 | 0 | 0 | 0 |
| 0 | NA | NA | NA | NA | NA | Gm2016 | predicted gene 2016 | 0 | 0 | 0 | 0 | 0 | 0 | 0 | 0 | 0 | 0 | 0 |
| 0 | NA | NA | NA | NA | NA | Gm49363 | | 0 | 0 | 0 | 0 | 0 | 0 | 0 | 0 | 0 | 0 | 0 |
| 0 | NA | NA | NA | NA | NA | Gm2022 | predicted pseudogene 2022 | 0 | 0 | 0 | 0 | 0 | 0 | 0 | 0 | 0 | 0 | 0 |
| 0 | NA | NA | NA | NA | NA | Gm18616 | | 0 | 0 | 0 | 0 | 0 | 0 | 0 | 0 | 0 | 0 | 0 |
| 0 | NA | NA | NA | NA | NA | Gm2035 | | 0 | 0 | 0 | 0 | 0 | 0 | 0 | 0 | 0 | 0 | 0 |
| 0 | NA | NA | NA | NA | NA | Gm18628 | | 0 | 0 | 0 | 0 | 0 | 0 | 0 | 0 | 0 | 0 | 0 |
| 0 | NA | NA | NA | NA | NA | Gm2042 | predicted gene 2042 | 0 | 0 | 0 | 0 | 0 | 0 | 0 | 0 | 0 | 0 | 0 |
| 0 | NA | NA | NA | NA | NA | Gm48389 | | 0 | 0 | 0 | 0 | 0 | 0 | 0 | 0 | 0 | 0 | 0 |
| 0 | NA | NA | NA | NA | NA | Gm48388 | | 0 | 0 | 0 | 0 | 0 | 0 | 0 | 0 | 0 | 0 | 0 |
| 0 | NA | NA | NA | NA | NA | Gm46992 | | 0 | 0 | 0 | 0 | 0 | 0 | 0 | 0 | 0 | 0 | 0 |
| 0 | NA | NA | NA | NA | NA | Gm2075 | | 0 | 0 | 0 | 0 | 0 | 0 | 0 | 0 | 0 | 0 | 0 |
| 0 | NA | NA | NA | NA | NA | Gm6803 | predicted gene 6803 | 0 | 0 | 0 | 0 | 0 | 0 | 0 | 0 | 0 | 0 | 0 |
| 0 | NA | NA | NA | NA | NA | Gm2056 | | 0 | 0 | 0 | 0 | 0 | 0 | 0 | 0 | 0 | 0 | 0 |
| 0 | NA | NA | NA | NA | NA | Gm18629 | | 0 | 0 | 0 | 0 | 0 | 0 | 0 | 0 | 0 | 0 | 0 |
| 0 | NA | NA | NA | NA | NA | Gm46993 | | 0 | 0 | 0 | 0 | 0 | 0 | 0 | 0 | 0 | 0 | 0 |
| 0 | NA | NA | NA | NA | NA | Gm18247 | | 0 | 0 | 0 | 0 | 0 | 0 | 0 | 0 | 0 | 0 | 0 |
| 0 | NA | NA | NA | NA | NA | Gm46998 | | 0 | 0 | 0 | 0 | 0 | 0 | 0 | 0 | 0 | 0 | 0 |
| 0 | NA | NA | NA | NA | NA | Gm46997 | | 0 | 0 | 0 | 0 | 0 | 0 | 0 | 0 | 0 | 0 | 0 |
| 0 | NA | NA | NA | NA | NA | Gm46994 | | 0 | 0 | 0 | 0 | 0 | 0 | 0 | 0 | 0 | 0 | 0 |
| 0 | NA | NA | NA | NA | NA | Gm46995 | | 0 | 0 | 0 | 0 | 0 | 0 | 0 | 0 | 0 | 0 | 0 |
| 0 | NA | NA | NA | NA | NA | Gm46996 | | 0 | 0 | 0 | 0 | 0 | 0 | 0 | 0 | 0 | 0 | 0 |
| 0 | NA | NA | NA | NA | NA | Gm18617 | | 0 | 0 | 0 | 0 | 0 | 0 | 0 | 0 | 0 | 0 | 0 |
| 0 | NA | NA | NA | NA | NA | Gm16368 | | 0 | 0 | 0 | 0 | 0 | 0 | 0 | 0 | 0 | 0 | 0 |
| 0 | NA | NA | NA | NA | NA | Gm5442 | | 0 | 0 | 0 | 0 | 0 | 0 | 0 | 0 | 0 | 0 | 0 |
| 0 | NA | NA | NA | NA | NA | Gm8587 | | 0 | 0 | 0 | 0 | 0 | 0 | 0 | 0 | 0 | 0 | 0 |
| 0 | NA | NA | NA | NA | NA | Gm48459 | | 0 | 0 | 0 | 0 | 0 | 0 | 0 | 0 | 0 | 0 | 0 |
| 0 | NA | NA | NA | NA | NA | Gm48487 | | 0 | 0 | 0 | 0 | 0 | 0 | 0 | 0 | 0 | 0 | 0 |
| 0 | NA | NA | NA | NA | NA | Gm33654 | | 0 | 0 | 0 | 0 | 0 | 0 | 0 | 0 | 0 | 0 | 0 |

| | | | | | | | | | | | | | | | | | | |
|---|---|---|---|---|---|---|---|---|---|---|---|---|---|---|---|---|---|---|
| 0 | NA | NA | NA | NA | NA | Gm5955 | predicted gene 5955 | 0 | 0 | 0 | 0 | 0 | 0 | 0 | 0 | 0 | 0 | 0 |
| 0 | NA | NA | NA | NA | NA | Gm8605 | | 0 | 0 | 0 | 0 | 0 | 0 | 0 | 0 | 0 | 0 | 0 |
| 0 | NA | NA | NA | NA | NA | Gm8607 | | 0 | 0 | 0 | 0 | 0 | 0 | 0 | 0 | 0 | 0 | 0 |
| 0 | NA | NA | NA | NA | NA | Gm18630 | | 0 | 0 | 0 | 0 | 0 | 0 | 0 | 0 | 0 | 0 | 0 |
| 0 | NA | NA | NA | NA | NA | Gm10436 | predicted gene 10436 | 0 | 0 | 0 | 0 | 0 | 0 | 0 | 0 | 0 | 0 | 0 |
| 0 | NA | NA | NA | NA | NA | Gm18631 | | 0 | 0 | 0 | 0 | 0 | 0 | 0 | 0 | 0 | 0 | 0 |
| 0 | NA | NA | NA | NA | NA | Gm48560 | | 0 | 0 | 0 | 0 | 0 | 0 | 0 | 0 | 0 | 0 | 0 |
| 0 | NA | NA | NA | NA | NA | Gm8332 | | 0 | 0 | 0 | 0 | 0 | 0 | 0 | 0 | 0 | 0 | 0 |
| 0 | NA | NA | NA | NA | NA | Gm18618 | | 0 | 0 | 0 | 0 | 0 | 0 | 0 | 0 | 0 | 0 | 0 |
| 0 | NA | NA | NA | NA | NA | Gm48561 | | 0 | 0 | 0 | 0 | 0 | 0 | 0 | 0 | 0 | 0 | 0 |
| 0 | NA | NA | NA | NA | NA | Gm5662 | predicted gene 5662 | 0 | 0 | 0 | 0 | 0 | 0 | 0 | 0 | 0 | 0 | 0 |
| 0 | NA | NA | NA | NA | NA | Gm21039 | | 0 | 0 | 0 | 0 | 0 | 0 | 0 | 0 | 0 | 0 | 0 |
| 0 | NA | NA | NA | NA | NA | Gm7104 | | 0 | 0 | 0 | 0 | 0 | 0 | 0 | 0 | 0 | 0 | 0 |
| 0 | NA | NA | NA | NA | NA | Gm8634 | | 0 | 0 | 0 | 0 | 0 | 0 | 0 | 0 | 0 | 0 | 0 |
| 0 | NA | NA | NA | NA | NA | Gm48564 | | 0 | 0 | 0 | 0 | 0 | 0 | 0 | 0 | 0 | 0 | 0 |
| 0 | NA | NA | NA | NA | NA | Gm18619 | | 0 | 0 | 0 | 0 | 0 | 0 | 0 | 0 | 0 | 0 | 0 |
| 0 | NA | NA | NA | NA | NA | Gm5039 | | 0 | 0 | 0 | 0 | 0 | 0 | 0 | 0 | 0 | 0 | 0 |
| 0 | NA | NA | NA | NA | NA | Gm10264 | | 0 | 0 | 0 | 0 | 0 | 0 | 0 | 0 | 0 | 0 | 0 |
| 0 | NA | NA | NA | NA | NA | Gm18632 | | 0 | 0 | 0 | 0 | 0 | 0 | 0 | 0 | 0 | 0 | 0 |
| 0 | NA | NA | NA | NA | NA | Gm48572 | | 0 | 0 | 0 | 0 | 0 | 0 | 0 | 0 | 0 | 0 | 0 |
| 0 | NA | NA | NA | NA | NA | Gm18620 | | 0 | 0 | 0 | 0 | 0 | 0 | 0 | 0 | 0 | 0 | 0 |
| 0 | NA | NA | NA | NA | NA | | | 0 | 0 | 0 | 0 | 0 | 0 | 0 | 0 | 0 | 0 | 0 |
| 0 | NA | NA | NA | NA | NA | Gm48577 | | 0 | 0 | 0 | 0 | 0 | 0 | 0 | 0 | 0 | 0 | 0 |
| 0 | NA | NA | NA | NA | NA | 3200001D21Rik | | 0 | 0 | 0 | 0 | 0 | 0 | 0 | 0 | 0 | 0 | 0 |
| 0 | NA | NA | NA | NA | NA | 4930473H19Rik | | 0 | 0 | 0 | 0 | 0 | 0 | 0 | 0 | 0 | 0 | 0 |
| 0 | NA | NA | NA | NA | NA | Gm48642 | | 0 | 0 | 0 | 0 | 0 | 0 | 0 | 0 | 0 | 0 | 0 |
| 0 | NA | NA | NA | NA | NA | Gm19540 | | 0 | 0 | 0 | 0 | 0 | 0 | 0 | 0 | 0 | 0 | 0 |
| 0 | NA | NA | NA | NA | NA | Gm48665 | | 0 | 0 | 0 | 0 | 0 | 0 | 0 | 0 | 0 | 0 | 0 |
| 0 | NA | NA | NA | NA | NA | Gm48664 | | 0 | 0 | 0 | 0 | 0 | 0 | 0 | 0 | 0 | 0 | 0 |
| 0 | NA | NA | NA | NA | NA | 1700040E09Rik | | 0 | 0 | 0 | 0 | 0 | 0 | 0 | 0 | 0 | 0 | 0 |
| 0 | NA | NA | NA | NA | NA | Gm37980 | | 0 | 0 | 0 | 0 | 0 | 0 | 0 | 0 | 0 | 0 | 0 |
| 0 | NA | NA | NA | NA | NA | Gm27313 | | 0 | 0 | 0 | 0 | 0 | 0 | 0 | 0 | 0 | 0 | 0 |
| 0 | NA | NA | NA | NA | NA | Gm23989 | | 0 | 0 | 0 | 0 | 0 | 0 | 0 | 0 | 0 | 0 | 0 |
| 0 | NA | NA | NA | NA | NA | Gm48692 | | 0 | 0 | 0 | 0 | 0 | 0 | 0 | 0 | 0 | 0 | 0 |
| 0 | NA | NA | NA | NA | NA | Gm48700 | | 0 | 0 | 0 | 0 | 0 | 0 | 0 | 0 | 0 | 0 | 0 |
| 0 | NA | NA | NA | NA | NA | Gm2270 | | 0 | 0 | 0 | 0 | 0 | 0 | 0 | 0 | 0 | 0 | 0 |
| 0 | NA | NA | NA | NA | NA | Gm47684 | | 0 | 0 | 0 | 0 | 0 | 0 | 0 | 0 | 0 | 0 | 0 |
| 0 | NA | NA | NA | NA | NA | Gm40538 | | 0 | 0 | 0 | 0 | 0 | 0 | 0 | 0 | 0 | 0 | 0 |
| 0 | NA | NA | NA | NA | NA | Gm47686 | | 0 | 0 | 0 | 0 | 0 | 0 | 0 | 0 | 0 | 0 | 0 |
| 0 | NA | NA | NA | NA | NA | Gm2139 | | 0 | 0 | 0 | 0 | 0 | 0 | 0 | 0 | 0 | 0 | 0 |
| 0 | NA | NA | NA | NA | NA | 4930544I03Rik | | 0 | 0 | 0 | 0 | 0 | 0 | 0 | 0 | 0 | 0 | 0 |
| 0 | NA | NA | NA | NA | NA | n-R5s65 | | 0 | 0 | 0 | 0 | 0 | 0 | 0 | 0 | 0 | 0 | 0 |
| 0 | NA | NA | NA | NA | NA | Gm23195 | | 0 | 0 | 0 | 0 | 0 | 0 | 0 | 0 | 0 | 0 | 0 |
| 0 | NA | NA | NA | NA | NA | Gm47688 | | 0 | 0 | 0 | 0 | 0 | 0 | 0 | 0 | 0 | 0 | 0 |
| 0 | NA | NA | NA | NA | NA | 1700105G05Rik | | 0 | 0 | 0 | 0 | 0 | 0 | 0 | 0 | 0 | 0 | 0 |
| 0 | NA | NA | NA | NA | NA | Gm26477 | | 0 | 0 | 0 | 0 | 0 | 0 | 0 | 0 | 0 | 0 | 0 |
| 0 | NA | NA | NA | NA | NA | 5430427M07Rik | | 0 | 0 | 0 | 0 | 0 | 0 | 0 | 0 | 0 | 0 | 0 |
| 0 | NA | NA | NA | NA | NA | Gm44370 | | 0 | 0 | 0 | 0 | 0 | 0 | 0 | 0 | 0 | 0 | 0 |
| 0 | NA | NA | NA | NA | NA | Gm18968 | | 0 | 0 | 0 | 0 | 0 | 0 | 0 | 0 | 0 | 0 | 0 |
| 0 | NA | NA | NA | NA | NA | Gm47742 | | 0 | 0 | 0 | 0 | 0 | 0 | 0 | 0 | 0 | 0 | 0 |
| 0 | NA | NA | NA | NA | NA | Gm18388 | | 0 | 0 | 0 | 0 | 0 | 0 | 0 | 0 | 0 | 0 | 0 |
| 0 | NA | NA | NA | NA | NA | Gm18827 | | 0 | 0 | 0 | 0 | 0 | 0 | 0 | 0 | 0 | 0 | 0 |
| 0 | NA | NA | NA | NA | NA | Gm47804 | | 0 | 0 | 0 | 0 | 0 | 0 | 0 | 0 | 0 | 0 | 0 |
| 0 | NA | NA | NA | NA | NA | Gm8378 | | 0 | 0 | 0 | 0 | 0 | 0 | 0 | 0 | 0 | 0 | 0 |
| 0 | NA | NA | NA | NA | NA | Gm16876 | | 0 | 0 | 0 | 0 | 0 | 0 | 0 | 0 | 0 | 0 | 0 |
| 0 | NA | NA | NA | NA | NA | Gm22015 | | 0 | 0 | 0 | 0 | 0 | 0 | 0 | 0 | 0 | 0 | 0 |
| 0 | NA | NA | NA | NA | NA | Gm21051 | | 0 | 0 | 0 | 0 | 0 | 0 | 0 | 0 | 0 | 0 | 0 |
| 0 | NA | NA | NA | NA | NA | Gm18621 | | 0 | 0 | 0 | 0 | 0 | 0 | 0 | 0 | 0 | 0 | 0 |
| 0 | NA | NA | NA | NA | NA | Gm21614 | | 0 | 0 | 0 | 0 | 0 | 0 | 0 | 0 | 0 | 0 | 0 |
| 0 | NA | NA | NA | NA | NA | Gm47984 | | 0 | 0 | 0 | 0 | 0 | 0 | 0 | 0 | 0 | 0 | 0 |

| | | | | | | | | | | | | | | | | | |
|---|---|---|---|---|---|---|---|---|---|---|---|---|---|---|---|---|---|
| 0 | NA | NA | NA | NA | NA | Gm6841 | 0 | 0 | 0 | 0 | 0 | 0 | 0 | 0 | 0 | 0 | 0 |
| 0 | NA | NA | NA | NA | NA | Gm18500 | 0 | 0 | 0 | 0 | 0 | 0 | 0 | 0 | 0 | 0 | 0 |
| 0 | NA | NA | NA | NA | NA | Gm47998 | 0 | 0 | 0 | 0 | 0 | 0 | 0 | 0 | 0 | 0 | 0 |
| 0 | NA | NA | NA | NA | NA | 4930559C10Rik | 0 | 0 | 0 | 0 | 0 | 0 | 0 | 0 | 0 | 0 | 0 |
| 0 | NA | NA | NA | NA | NA | Gm21007 | 0 | 0 | 0 | 0 | 0 | 0 | 0 | 0 | 0 | 0 | 0 |
| 0 | NA | NA | NA | NA | NA | Gm23249 | 0 | 0 | 0 | 0 | 0 | 0 | 0 | 0 | 0 | 0 | 0 |
| 0 | NA | NA | NA | NA | NA | Mir8099-2 | 0 | 0 | 0 | 0 | 0 | 0 | 0 | 0 | 0 | 0 | 0 |
| 0 | NA | NA | NA | NA | NA | Gm7119 | 0 | 0 | 0 | 0 | 0 | 0 | 0 | 0 | 0 | 0 | 0 |
| 0 | NA | NA | NA | NA | NA | Gm18749 | 0 | 0 | 0 | 0 | 0 | 0 | 0 | 0 | 0 | 0 | 0 |
| 0 | NA | NA | NA | NA | NA | Gm18504 | 0 | 0 | 0 | 0 | 0 | 0 | 0 | 0 | 0 | 0 | 0 |
| 0 | NA | NA | NA | NA | NA | Gm48011 | 0 | 0 | 0 | 0 | 0 | 0 | 0 | 0 | 0 | 0 | 0 |
| 0 | NA | NA | NA | NA | NA | Gm4808 | 0 | 0 | 0 | 0 | 0 | 0 | 0 | 0 | 0 | 0 | 0 |
| 0 | NA | NA | NA | NA | NA | Gm48013 | 0 | 0 | 0 | 0 | 0 | 0 | 0 | 0 | 0 | 0 | 0 |
| 0 | NA | NA | NA | NA | NA | Gm26055 | 0 | 0 | 0 | 0 | 0 | 0 | 0 | 0 | 0 | 0 | 0 |
| 0 | NA | NA | NA | NA | NA | Gm22246 | 0 | 0 | 0 | 0 | 0 | 0 | 0 | 0 | 0 | 0 | 0 |
| 0 | NA | NA | NA | NA | NA | Rpl17-ps2 | 0 | 0 | 0 | 0 | 0 | 0 | 0 | 0 | 0 | 0 | 0 |
| 0 | NA | NA | NA | NA | NA | Gm48014 | 0 | 0 | 0 | 0 | 0 | 0 | 0 | 0 | 0 | 0 | 0 |
| 0 | NA | NA | NA | NA | NA | Gm48015 | 0 | 0 | 0 | 0 | 0 | 0 | 0 | 0 | 0 | 0 | 0 |
| 0 | NA | NA | NA | NA | NA | 1700019M22Rik | 0 | 0 | 0 | 0 | 0 | 0 | 0 | 0 | 0 | 0 | 0 |
| 0 | NA | NA | NA | NA | NA | Gm47394 | 0 | 0 | 0 | 0 | 0 | 0 | 0 | 0 | 0 | 0 | 0 |
| 0 | NA | NA | NA | NA | NA | Gm26480 | 0 | 0 | 0 | 0 | 0 | 0 | 0 | 0 | 0 | 0 | 0 |
| 0 | NA | NA | NA | NA | NA | Gm47395 | 0 | 0 | 0 | 0 | 0 | 0 | 0 | 0 | 0 | 0 | 0 |
| 0 | NA | NA | NA | NA | NA | Gm47396 | 0 | 0 | 0 | 0 | 0 | 0 | 0 | 0 | 0 | 0 | 0 |
| 0 | NA | NA | NA | NA | NA | Gm47397 | 0 | 0 | 0 | 0 | 0 | 0 | 0 | 0 | 0 | 0 | 0 |
| 0 | NA | NA | NA | NA | NA | Gm40552 | 0 | 0 | 0 | 0 | 0 | 0 | 0 | 0 | 0 | 0 | 0 |
| 0 | NA | NA | NA | NA | NA | Gm23558 | 0 | 0 | 0 | 0 | 0 | 0 | 0 | 0 | 0 | 0 | 0 |
| 0 | NA | NA | NA | NA | NA | Gm5186 | 0 | 0 | 0 | 0 | 0 | 0 | 0 | 0 | 0 | 0 | 0 |
| 0 | NA | NA | NA | NA | NA | Gm2398 | 0 | 0 | 0 | 0 | 0 | 0 | 0 | 0 | 0 | 0 | 0 |
| 0 | NA | NA | NA | NA | NA | Gm8785 | 0 | 0 | 0 | 0 | 0 | 0 | 0 | 0 | 0 | 0 | 0 |
| 0 | NA | NA | NA | NA | NA | Gm35274 | 0 | 0 | 0 | 0 | 0 | 0 | 0 | 0 | 0 | 0 | 0 |
| 0 | NA | NA | NA | NA | NA | Gm35326 | 0 | 0 | 0 | 0 | 0 | 0 | 0 | 0 | 0 | 0 | 0 |
| 0 | NA | NA | NA | NA | NA | Gm47415 | 0 | 0 | 0 | 0 | 0 | 0 | 0 | 0 | 0 | 0 | 0 |
| 0 | NA | NA | NA | NA | NA | Gm47566 | 0 | 0 | 0 | 0 | 0 | 0 | 0 | 0 | 0 | 0 | 0 |
| 0 | NA | NA | NA | NA | NA | 5330409N07Rik | 0 | 0 | 0 | 0 | 0 | 0 | 0 | 0 | 0 | 0 | 0 |
| 0 | NA | NA | NA | NA | NA | Gm47346 | 0 | 0 | 0 | 0 | 0 | 0 | 0 | 0 | 0 | 0 | 0 |
| 0 | NA | NA | NA | NA | NA | Gm25486 | 0 | 0 | 0 | 0 | 0 | 0 | 0 | 0 | 0 | 0 | 0 |
| 0 | NA | NA | NA | NA | NA | Gm47372 | 0 | 0 | 0 | 0 | 0 | 0 | 0 | 0 | 0 | 0 | 0 |
| 0 | NA | NA | NA | NA | NA | A930040O22Rik | 0 | 0 | 0 | 0 | 0 | 0 | 0 | 0 | 0 | 0 | 0 |
| 0 | NA | NA | NA | NA | NA | Gm47116 | 0 | 0 | 0 | 0 | 0 | 0 | 0 | 0 | 0 | 0 | 0 |
| 0 | NA | NA | NA | NA | NA | Gm19898 | 0 | 0 | 0 | 0 | 0 | 0 | 0 | 0 | 0 | 0 | 0 |
| 0 | NA | NA | NA | NA | NA | Gm47138 | 0 | 0 | 0 | 0 | 0 | 0 | 0 | 0 | 0 | 0 | 0 |
| 0 | NA | NA | NA | NA | NA | Gm47177 | 0 | 0 | 0 | 0 | 0 | 0 | 0 | 0 | 0 | 0 | 0 |
| 0 | NA | NA | NA | NA | NA | Gm47196 | 0 | 0 | 0 | 0 | 0 | 0 | 0 | 0 | 0 | 0 | 0 |
| 0 | NA | NA | NA | NA | NA | Gm19951 | 0 | 0 | 0 | 0 | 0 | 0 | 0 | 0 | 0 | 0 | 0 |
| 0 | NA | NA | NA | NA | NA | Gm47208 | 0 | 0 | 0 | 0 | 0 | 0 | 0 | 0 | 0 | 0 | 0 |
| 0 | NA | NA | NA | NA | NA | Gm19238 | 0 | 0 | 0 | 0 | 0 | 0 | 0 | 0 | 0 | 0 | 0 |
| 0 | NA | NA | NA | NA | NA | Gm47207 | 0 | 0 | 0 | 0 | 0 | 0 | 0 | 0 | 0 | 0 | 0 |
| 0 | NA | NA | NA | NA | NA | Gm47206 | 0 | 0 | 0 | 0 | 0 | 0 | 0 | 0 | 0 | 0 | 0 |
| 0 | NA | NA | NA | NA | NA | Gm23291 | 0 | 0 | 0 | 0 | 0 | 0 | 0 | 0 | 0 | 0 | 0 |
| 0 | NA | NA | NA | NA | NA | 4930477G07Rik | 0 | 0 | 0 | 0 | 0 | 0 | 0 | 0 | 0 | 0 | 0 |
| 0 | NA | NA | NA | NA | NA | Gm48381 | 0 | 0 | 0 | 0 | 0 | 0 | 0 | 0 | 0 | 0 | 0 |
| 0 | NA | NA | NA | NA | NA | Gm48383 | 0 | 0 | 0 | 0 | 0 | 0 | 0 | 0 | 0 | 0 | 0 |
| 0 | NA | NA | NA | NA | NA | Gm25801 | 0 | 0 | 0 | 0 | 0 | 0 | 0 | 0 | 0 | 0 | 0 |
| 0 | NA | NA | NA | NA | NA | Gm8482 | 0 | 0 | 0 | 0 | 0 | 0 | 0 | 0 | 0 | 0 | 0 |
| 0 | NA | NA | NA | NA | NA | | 0 | 0 | 0 | 0 | 0 | 0 | 0 | 0 | 0 | 0 | 0 |
| 0 | NA | NA | NA | NA | NA | A630072L19Rik | 0 | 0 | 0 | 0 | 0 | 0 | 0 | 0 | 0 | 0 | 0 |
| 0 | NA | NA | NA | NA | NA | Mir1190 | 0 | 0 | 0 | 0 | 0 | 0 | 0 | 0 | 0 | 0 | 0 |
| 0 | NA | NA | NA | NA | NA | Kif4-ps | 0 | 0 | 0 | 0 | 0 | 0 | 0 | 0 | 0 | 0 | 0 |
| 0 | NA | NA | NA | NA | NA | Gm18795 | 0 | 0 | 0 | 0 | 0 | 0 | 0 | 0 | 0 | 0 | 0 |
| 0 | NA | NA | NA | NA | NA | Mylf-ps | 0 | 0 | 0 | 0 | 0 | 0 | 0 | 0 | 0 | 0 | 0 |

| | | | | | | | | | | | | | | | | | |
|---|---|---|---|---|---|---|---|---|---|---|---|---|---|---|---|---|---|
| 0 | NA | NA | NA | NA | NA | Gm22749 | | 0 | 0 | 0 | 0 | 0 | 0 | 0 | 0 | 0 | 0 | 0 |
| 0 | NA | NA | NA | NA | NA | Catsperb | cation channel sperm associated auxiliar | 0 | 0 | 0 | 0 | 0 | 0 | 0 | 0 | 0 | 0 | 0 |
| 0 | NA | NA | NA | NA | NA | Gm24542 | | 0 | 0 | 0 | 0 | 0 | 0 | 0 | 0 | 0 | 0 | 0 |
| 0 | NA | NA | NA | NA | NA | Gm16285 | | 0 | 0 | 0 | 0 | 0 | 0 | 0 | 0 | 0 | 0 | 0 |
| 0 | NA | NA | NA | NA | NA | Gm20701 | | 0 | 0 | 0 | 0 | 0 | 0 | 0 | 0 | 0 | 0 | 0 |
| 0 | NA | NA | NA | NA | NA | Gm16339 | | 0 | 0 | 0 | 0 | 0 | 0 | 0 | 0 | 0 | 0 | 0 |
| 0 | NA | NA | NA | NA | NA | Gm25203 | | 0 | 0 | 0 | 0 | 0 | 0 | 0 | 0 | 0 | 0 | 0 |
| 0 | NA | NA | NA | NA | NA | Gm25203 | | 0 | 0 | 0 | 0 | 0 | 0 | 0 | 0 | 0 | 0 | 0 |
| 0 | NA | NA | NA | NA | NA | Gm30198 | | 0 | 0 | 0 | 0 | 0 | 0 | 0 | 0 | 0 | 0 | 0 |
| 0 | NA | NA | NA | NA | NA | Gm20069 | | 0 | 0 | 0 | 0 | 0 | 0 | 0 | 0 | 0 | 0 | 0 |
| 0 | NA | NA | NA | NA | NA | Mir1936 | | 0 | 0 | 0 | 0 | 0 | 0 | 0 | 0 | 0 | 0 | 0 |
| 0 | NA | NA | NA | NA | NA | Gm28051 | | 0 | 0 | 0 | 0 | 0 | 0 | 0 | 0 | 0 | 0 | 0 |
| 0 | NA | NA | NA | NA | NA | Gm21971 | | 0 | 0 | 0 | 0 | 0 | 0 | 0 | 0 | 0 | 0 | 0 |
| 0 | NA | NA | NA | NA | NA | Gm47042 | | 0 | 0 | 0 | 0 | 0 | 0 | 0 | 0 | 0 | 0 | 0 |
| 0 | NA | NA | NA | NA | NA | Gm47167 | | 0 | 0 | 0 | 0 | 0 | 0 | 0 | 0 | 0 | 0 | 0 |
| 0 | NA | NA | NA | NA | NA | Cox8c | cytochrome c oxidase subunit 8C | 0 | 0 | 0 | 0 | 0 | 0 | 0 | 0 | 0 | 0 | 0 |
| 0 | NA | NA | NA | NA | NA | Gm27976 | | 0 | 0 | 0 | 0 | 0 | 0 | 0 | 0 | 0 | 0 | 0 |
| 0 | NA | NA | NA | NA | NA | Fam181a | family with sequence similarity 181, meml | 0 | 0 | 0 | 0 | 0 | 0 | 0 | 0 | 0 | 0 | 0 |
| 0 | NA | NA | NA | NA | NA | Gm29508 | | 0 | 0 | 0 | 0 | 0 | 0 | 0 | 0 | 0 | 0 | 0 |
| 0 | NA | NA | NA | NA | NA | Gm45777 | | 0 | 0 | 0 | 0 | 0 | 0 | 0 | 0 | 0 | 0 | 0 |
| 0 | NA | NA | NA | NA | NA | Gm47267 | | 0 | 0 | 0 | 0 | 0 | 0 | 0 | 0 | 0 | 0 | 0 |
| 0 | NA | NA | NA | NA | NA | Gm18501 | | 0 | 0 | 0 | 0 | 0 | 0 | 0 | 0 | 0 | 0 | 0 |
| 0 | NA | NA | NA | NA | NA | Serpina16 | serine (or cysteine) peptidase inhibitor, cl | 0 | 0 | 0 | 0 | 0 | 0 | 0 | 0 | 0 | 0 | 0 |
| 0 | NA | NA | NA | NA | NA | Serpina1f | serine (or cysteine) peptidase inhibitor, cl | 0 | 0 | 0 | 0 | 0 | 0 | 0 | 0 | 0 | 0 | 0 |
| 0 | NA | NA | NA | NA | NA | Gm5656 | | 0 | 0 | 0 | 0 | 0 | 0 | 0 | 0 | 0 | 0 | 0 |
| 0 | NA | NA | NA | NA | NA | Gm17142 | | 0 | 0 | 0 | 0 | 0 | 0 | 0 | 0 | 0 | 0 | 0 |
| 0 | NA | NA | NA | NA | NA | Gm17042 | | 0 | 0 | 0 | 0 | 0 | 0 | 0 | 0 | 0 | 0 | 0 |
| 0 | NA | NA | NA | NA | NA | Gm17043 | | 0 | 0 | 0 | 0 | 0 | 0 | 0 | 0 | 0 | 0 | 0 |
| 0 | NA | NA | NA | NA | NA | Gm6918 | | 0 | 0 | 0 | 0 | 0 | 0 | 0 | 0 | 0 | 0 | 0 |
| 0 | NA | NA | NA | NA | NA | Gm8895 | | 0 | 0 | 0 | 0 | 0 | 0 | 0 | 0 | 0 | 0 | 0 |
| 0 | NA | NA | NA | NA | NA | Gm17198 | | 0 | 0 | 0 | 0 | 0 | 0 | 0 | 0 | 0 | 0 | 0 |
| 0 | NA | NA | NA | NA | NA | Serpina11 | serine (or cysteine) peptidase inhibitor, cl | 0 | 0 | 0 | 0 | 0 | 0 | 0 | 0 | 0 | 0 | 0 |
| 0 | NA | NA | NA | NA | NA | Gm28577 | | 0 | 0 | 0 | 0 | 0 | 0 | 0 | 0 | 0 | 0 | 0 |
| 0 | NA | NA | NA | NA | NA | Gm37496 | | 0 | 0 | 0 | 0 | 0 | 0 | 0 | 0 | 0 | 0 | 0 |
| 0 | NA | NA | NA | NA | NA | Serpina12 | serine (or cysteine) peptidase inhibitor, cl | 0 | 0 | 0 | 0 | 0 | 0 | 0 | 0 | 0 | 0 | 0 |
| 0 | NA | NA | NA | NA | NA | Serpina4-ps1 | | 0 | 0 | 0 | 0 | 0 | 0 | 0 | 0 | 0 | 0 | 0 |
| 0 | NA | NA | NA | NA | NA | Serpina5 | serine (or cysteine) peptidase inhibitor, cl | 0 | 0 | 0 | 0 | 0 | 0 | 0 | 0 | 0 | 0 | 0 |
| 0 | NA | NA | NA | NA | NA | Gm23289 | | 0 | 0 | 0 | 0 | 0 | 0 | 0 | 0 | 0 | 0 | 0 |
| 0 | NA | NA | NA | NA | NA | Serpina3a | serine (or cysteine) peptidase inhibitor, cl | 0 | 0 | 0 | 0 | 0 | 0 | 0 | 0 | 0 | 0 | 0 |
| 0 | NA | NA | NA | NA | NA | Serpina3b | serine (or cysteine) peptidase inhibitor, cl | 0 | 0 | 0 | 0 | 0 | 0 | 0 | 0 | 0 | 0 | 0 |
| 0 | NA | NA | NA | NA | NA | Serpina3d-ps | | 0 | 0 | 0 | 0 | 0 | 0 | 0 | 0 | 0 | 0 | 0 |
| 0 | NA | NA | NA | NA | NA | | | 0 | 0 | 0 | 0 | 0 | 0 | 0 | 0 | 0 | 0 | 0 |
| 0 | NA | NA | NA | NA | NA | Serpina3j | serine (or cysteine) peptidase inhibitor, cl | 0 | 0 | 0 | 0 | 0 | 0 | 0 | 0 | 0 | 0 | 0 |
| 0 | NA | NA | NA | NA | NA | Serpina3l-ps | | 0 | 0 | 0 | 0 | 0 | 0 | 0 | 0 | 0 | 0 | 0 |
| 0 | NA | NA | NA | NA | NA | Gm24334 | | 0 | 0 | 0 | 0 | 0 | 0 | 0 | 0 | 0 | 0 | 0 |
| 0 | NA | NA | NA | NA | NA | Gm10000 | | 0 | 0 | 0 | 0 | 0 | 0 | 0 | 0 | 0 | 0 | 0 |
| 0 | NA | NA | NA | NA | NA | Gm49732 | | 0 | 0 | 0 | 0 | 0 | 0 | 0 | 0 | 0 | 0 | 0 |
| 0 | NA | NA | NA | NA | NA | 4930408O17Rik | | 0 | 0 | 0 | 0 | 0 | 0 | 0 | 0 | 0 | 0 | 0 |
| 0 | NA | NA | NA | NA | NA | Gm47650 | | 0 | 0 | 0 | 0 | 0 | 0 | 0 | 0 | 0 | 0 | 0 |
| 0 | NA | NA | NA | NA | NA | Mir3069 | | 0 | 0 | 0 | 0 | 0 | 0 | 0 | 0 | 0 | 0 | 0 |
| 0 | NA | NA | NA | NA | NA | Gm47672 | | 0 | 0 | 0 | 0 | 0 | 0 | 0 | 0 | 0 | 0 | 0 |
| 0 | NA | NA | NA | NA | NA | Tcl1b2 | T cell leukemia/lymphoma 1B, 2 | 0 | 0 | 0 | 0 | 0 | 0 | 0 | 0 | 0 | 0 | 0 |
| 0 | NA | NA | NA | NA | NA | Tcl1b1 | T cell leukemia/lymphoma 1B, 1 | 0 | 0 | 0 | 0 | 0 | 0 | 0 | 0 | 0 | 0 | 0 |
| 0 | NA | NA | NA | NA | NA | Tcl1b5 | T cell leukemia/lymphoma 1B, 5 | 0 | 0 | 0 | 0 | 0 | 0 | 0 | 0 | 0 | 0 | 0 |
| 0 | NA | NA | NA | NA | NA | Tcl1b3 | T cell leukemia/lymphoma 1B, 3 | 0 | 0 | 0 | 0 | 0 | 0 | 0 | 0 | 0 | 0 | 0 |
| 0 | NA | NA | NA | NA | NA | Tcl1b4 | T cell leukemia/lymphoma 1B, 4 | 0 | 0 | 0 | 0 | 0 | 0 | 0 | 0 | 0 | 0 | 0 |
| 0 | NA | NA | NA | NA | NA | Tcl1 | T cell lymphoma breakpoint 1 | 0 | 0 | 0 | 0 | 0 | 0 | 0 | 0 | 0 | 0 | 0 |
| 0 | NA | NA | NA | NA | NA | Gm2721 | | 0 | 0 | 0 | 0 | 0 | 0 | 0 | 0 | 0 | 0 | 0 |
| 0 | NA | NA | NA | NA | NA | Gm46376 | | 0 | 0 | 0 | 0 | 0 | 0 | 0 | 0 | 0 | 0 | 0 |
| 0 | NA | NA | NA | NA | NA | AU015791 | | 0 | 0 | 0 | 0 | 0 | 0 | 0 | 0 | 0 | 0 | 0 |

| | | | | | | | | | | | | | | | | | |
|---|---|---|---|---|---|---|---|---|---|---|---|---|---|---|---|---|---|
| 0 | NA | NA | NA | NA | NA | Gm22790 | 0 | 0 | 0 | 0 | 0 | 0 | 0 | 0 | 0 | 0 | 0 |
| 0 | NA | NA | NA | NA | NA | Gm48807 | 0 | 0 | 0 | 0 | 0 | 0 | 0 | 0 | 0 | 0 | 0 |
| 0 | NA | NA | NA | NA | NA | Gm22120 | 0 | 0 | 0 | 0 | 0 | 0 | 0 | 0 | 0 | 0 | 0 |
| 0 | NA | NA | NA | NA | NA | Gm32635 | 0 | 0 | 0 | 0 | 0 | 0 | 0 | 0 | 0 | 0 | 0 |
| 0 | NA | NA | NA | NA | NA | Gm47646 | 0 | 0 | 0 | 0 | 0 | 0 | 0 | 0 | 0 | 0 | 0 |
| 0 | NA | NA | NA | NA | NA | Gm24986 | 0 | 0 | 0 | 0 | 0 | 0 | 0 | 0 | 0 | 0 | 0 |
| 0 | NA | NA | NA | NA | NA | Gm17032 | 0 | 0 | 0 | 0 | 0 | 0 | 0 | 0 | 0 | 0 | 0 |
| 0 | NA | NA | NA | NA | NA | 1700121N20Rik | 0 | 0 | 0 | 0 | 0 | 0 | 0 | 0 | 0 | 0 | 0 |
| 0 | NA | NA | NA | NA | NA | Gm17031 | 0 | 0 | 0 | 0 | 0 | 0 | 0 | 0 | 0 | 0 | 0 |
| 0 | NA | NA | NA | NA | NA | Gm17053 | 0 | 0 | 0 | 0 | 0 | 0 | 0 | 0 | 0 | 0 | 0 |
| 0 | NA | NA | NA | NA | NA | Gm17033 | 0 | 0 | 0 | 0 | 0 | 0 | 0 | 0 | 0 | 0 | 0 |
| 0 | NA | NA | NA | NA | NA | 4933406K04Rik | 0 | 0 | 0 | 0 | 0 | 0 | 0 | 0 | 0 | 0 | 0 |
| 0 | NA | NA | NA | NA | NA | Gm9599 | 0 | 0 | 0 | 0 | 0 | 0 | 0 | 0 | 0 | 0 | 0 |
| 0 | NA | NA | NA | NA | NA | Gm16087 | 0 | 0 | 0 | 0 | 0 | 0 | 0 | 0 | 0 | 0 | 0 |
| 0 | NA | NA | NA | NA | NA | Gm16085 | 0 | 0 | 0 | 0 | 0 | 0 | 0 | 0 | 0 | 0 | 0 |
| 0 | NA | NA | NA | NA | NA | Gm16084 | 0 | 0 | 0 | 0 | 0 | 0 | 0 | 0 | 0 | 0 | 0 |
| 0 | NA | NA | NA | NA | NA | Gm2800 | 0 | 0 | 0 | 0 | 0 | 0 | 0 | 0 | 0 | 0 | 0 |
| 0 | NA | NA | NA | NA | NA | Gm17052 | 0 | 0 | 0 | 0 | 0 | 0 | 0 | 0 | 0 | 0 | 0 |
| 0 | NA | NA | NA | NA | NA | Gm3234 | 0 | 0 | 0 | 0 | 0 | 0 | 0 | 0 | 0 | 0 | 0 |
| 0 | NA | NA | NA | NA | NA | 3110018I06Rik | 0 | 0 | 0 | 0 | 0 | 0 | 0 | 0 | 0 | 0 | 0 |
| 0 | NA | NA | NA | NA | NA | Gm9006 | 0 | 0 | 0 | 0 | 0 | 0 | 0 | 0 | 0 | 0 | 0 |
| 0 | NA | NA | NA | NA | NA | 4930465M20Rik | 0 | 0 | 0 | 0 | 0 | 0 | 0 | 0 | 0 | 0 | 0 |
| 0 | NA | NA | NA | NA | NA | Gm15208 | 0 | 0 | 0 | 0 | 0 | 0 | 0 | 0 | 0 | 0 | 0 |
| 0 | NA | NA | NA | NA | NA | 4930478K11Rik | 0 | 0 | 0 | 0 | 0 | 0 | 0 | 0 | 0 | 0 | 0 |
| 0 | NA | NA | NA | NA | NA | Gm15636 | 0 | 0 | 0 | 0 | 0 | 0 | 0 | 0 | 0 | 0 | 0 |
| 0 | NA | NA | NA | NA | NA | Mir342 | 0 | 0 | 0 | 0 | 0 | 0 | 0 | 0 | 0 | 0 | 0 |
| 0 | NA | NA | NA | NA | NA | Gm23872 | 0 | 0 | 0 | 0 | 0 | 0 | 0 | 0 | 0 | 0 | 0 |
| 0 | NA | NA | NA | NA | NA | Gm33682 | 0 | 0 | 0 | 0 | 0 | 0 | 0 | 0 | 0 | 0 | 0 |
| 0 | NA | NA | NA | NA | NA | Mir345 | 0 | 0 | 0 | 0 | 0 | 0 | 0 | 0 | 0 | 0 | 0 |
| 0 | NA | NA | NA | NA | NA | Gm18405 | 0 | 0 | 0 | 0 | 0 | 0 | 0 | 0 | 0 | 0 | 0 |
| 0 | NA | NA | NA | NA | NA | Mif-ps7 | 0 | 0 | 0 | 0 | 0 | 0 | 0 | 0 | 0 | 0 | 0 |
| 0 | NA | NA | NA | NA | NA | Gm22234 | 0 | 0 | 0 | 0 | 0 | 0 | 0 | 0 | 0 | 0 | 0 |
| 0 | NA | NA | NA | NA | NA | Gm25297 | 0 | 0 | 0 | 0 | 0 | 0 | 0 | 0 | 0 | 0 | 0 |
| 0 | NA | NA | NA | NA | NA | Gm27528 | 0 | 0 | 0 | 0 | 0 | 0 | 0 | 0 | 0 | 0 | 0 |
| 0 | NA | NA | NA | NA | NA | Gm27528 | 0 | 0 | 0 | 0 | 0 | 0 | 0 | 0 | 0 | 0 | 0 |
| 0 | NA | NA | NA | NA | NA | Gm27300 | 0 | 0 | 0 | 0 | 0 | 0 | 0 | 0 | 0 | 0 | 0 |
| 0 | NA | NA | NA | NA | NA | Mir1906-1 | 0 | 0 | 0 | 0 | 0 | 0 | 0 | 0 | 0 | 0 | 0 |
| 0 | NA | NA | NA | NA | NA | Mir770 | 0 | 0 | 0 | 0 | 0 | 0 | 0 | 0 | 0 | 0 | 0 |
| 0 | NA | NA | NA | NA | NA | Mir673 | 0 | 0 | 0 | 0 | 0 | 0 | 0 | 0 | 0 | 0 | 0 |
| 0 | NA | NA | NA | NA | NA | Mir493 | 0 | 0 | 0 | 0 | 0 | 0 | 0 | 0 | 0 | 0 | 0 |
| 0 | NA | NA | NA | NA | NA | Mir3544 | 0 | 0 | 0 | 0 | 0 | 0 | 0 | 0 | 0 | 0 | 0 |
| 0 | NA | NA | NA | NA | NA | Mir540 | 0 | 0 | 0 | 0 | 0 | 0 | 0 | 0 | 0 | 0 | 0 |
| 0 | NA | NA | NA | NA | NA | Mir665 | 0 | 0 | 0 | 0 | 0 | 0 | 0 | 0 | 0 | 0 | 0 |
| 0 | NA | NA | NA | NA | NA | Mir3070a | 0 | 0 | 0 | 0 | 0 | 0 | 0 | 0 | 0 | 0 | 0 |
| 0 | NA | NA | NA | NA | NA | Mir127 | 0 | 0 | 0 | 0 | 0 | 0 | 0 | 0 | 0 | 0 | 0 |
| 0 | NA | NA | NA | NA | NA | Mir432 | 0 | 0 | 0 | 0 | 0 | 0 | 0 | 0 | 0 | 0 | 0 |
| 0 | NA | NA | NA | NA | NA | Mir3071 | 0 | 0 | 0 | 0 | 0 | 0 | 0 | 0 | 0 | 0 | 0 |
| 0 | NA | NA | NA | NA | NA | Mir136 | 0 | 0 | 0 | 0 | 0 | 0 | 0 | 0 | 0 | 0 | 0 |
| 0 | NA | NA | NA | NA | NA | Gm27350 | 0 | 0 | 0 | 0 | 0 | 0 | 0 | 0 | 0 | 0 | 0 |
| 0 | NA | NA | NA | NA | NA | Gm23912 | 0 | 0 | 0 | 0 | 0 | 0 | 0 | 0 | 0 | 0 | 0 |
| 0 | NA | NA | NA | NA | NA | Mir370 | 0 | 0 | 0 | 0 | 0 | 0 | 0 | 0 | 0 | 0 | 0 |
| 0 | NA | NA | NA | NA | NA | Gm23474 | 0 | 0 | 0 | 0 | 0 | 0 | 0 | 0 | 0 | 0 | 0 |
| 0 | NA | NA | NA | NA | NA | AF357425 | 0 | 0 | 0 | 0 | 0 | 0 | 0 | 0 | 0 | 0 | 0 |
| 0 | NA | NA | NA | NA | NA | Gm24899 | 0 | 0 | 0 | 0 | 0 | 0 | 0 | 0 | 0 | 0 | 0 |
| 0 | NA | NA | NA | NA | NA | Gm23600 | 0 | 0 | 0 | 0 | 0 | 0 | 0 | 0 | 0 | 0 | 0 |
| 0 | NA | NA | NA | NA | NA | Gm24895 | 0 | 0 | 0 | 0 | 0 | 0 | 0 | 0 | 0 | 0 | 0 |
| 0 | NA | NA | NA | NA | NA | DQ267100 | 0 | 0 | 0 | 0 | 0 | 0 | 0 | 0 | 0 | 0 | 0 |
| 0 | NA | NA | NA | NA | NA | DQ267101 | 0 | 0 | 0 | 0 | 0 | 0 | 0 | 0 | 0 | 0 | 0 |
| 0 | NA | NA | NA | NA | NA | Gm23508 | 0 | 0 | 0 | 0 | 0 | 0 | 0 | 0 | 0 | 0 | 0 |
| 0 | NA | NA | NA | NA | NA | Gm26922 | 0 | 0 | 0 | 0 | 0 | 0 | 0 | 0 | 0 | 0 | 0 |

| | | | | | | | | | | | | | | | | | | |
|---|---|---|---|---|---|---|---|---|---|---|---|---|---|---|---|---|---|---|
| 0 | NA | NA | NA | NA | NA | DQ267102 | | 0 | 0 | 0 | 0 | 0 | 0 | 0 | 0 | 0 | 0 | 0 |
| 0 | NA | NA | NA | NA | NA | Gm25357 | | 0 | 0 | 0 | 0 | 0 | 0 | 0 | 0 | 0 | 0 | 0 |
| 0 | NA | NA | NA | NA | NA | Gm27604 | | 0 | 0 | 0 | 0 | 0 | 0 | 0 | 0 | 0 | 0 | 0 |
| 0 | NA | NA | NA | NA | NA | Gm23347 | | 0 | 0 | 0 | 0 | 0 | 0 | 0 | 0 | 0 | 0 | 0 |
| 0 | NA | NA | NA | NA | NA | AF357428 | | 0 | 0 | 0 | 0 | 0 | 0 | 0 | 0 | 0 | 0 | 0 |
| 0 | NA | NA | NA | NA | NA | Gm23736 | | 0 | 0 | 0 | 0 | 0 | 0 | 0 | 0 | 0 | 0 | 0 |
| 0 | NA | NA | NA | NA | NA | AF357341 | | 0 | 0 | 0 | 0 | 0 | 0 | 0 | 0 | 0 | 0 | 0 |
| 0 | NA | NA | NA | NA | NA | Mir882 | | 0 | 0 | 0 | 0 | 0 | 0 | 0 | 0 | 0 | 0 | 0 |
| 0 | NA | NA | NA | NA | NA | Gm22981 | | 0 | 0 | 0 | 0 | 0 | 0 | 0 | 0 | 0 | 0 | 0 |
| 0 | NA | NA | NA | NA | NA | Gm23787 | | 0 | 0 | 0 | 0 | 0 | 0 | 0 | 0 | 0 | 0 | 0 |
| 0 | NA | NA | NA | NA | NA | Gm25854 | | 0 | 0 | 0 | 0 | 0 | 0 | 0 | 0 | 0 | 0 | 0 |
| 0 | NA | NA | NA | NA | NA | Gm22205 | | 0 | 0 | 0 | 0 | 0 | 0 | 0 | 0 | 0 | 0 | 0 |
| 0 | NA | NA | NA | NA | NA | Gm25224 | | 0 | 0 | 0 | 0 | 0 | 0 | 0 | 0 | 0 | 0 | 0 |
| 0 | NA | NA | NA | NA | NA | Mir379 | | 0 | 0 | 0 | 0 | 0 | 0 | 0 | 0 | 0 | 0 | 0 |
| 0 | NA | NA | NA | NA | NA | Mir411 | | 0 | 0 | 0 | 0 | 0 | 0 | 0 | 0 | 0 | 0 | 0 |
| 0 | NA | NA | NA | NA | NA | Mir299a | | 0 | 0 | 0 | 0 | 0 | 0 | 0 | 0 | 0 | 0 | 0 |
| 0 | NA | NA | NA | NA | NA | Mir299b | | 0 | 0 | 0 | 0 | 0 | 0 | 0 | 0 | 0 | 0 | 0 |
| 0 | NA | NA | NA | NA | NA | Gm25750 | | 0 | 0 | 0 | 0 | 0 | 0 | 0 | 0 | 0 | 0 | 0 |
| 0 | NA | NA | NA | NA | NA | Mir380 | | 0 | 0 | 0 | 0 | 0 | 0 | 0 | 0 | 0 | 0 | 0 |
| 0 | NA | NA | NA | NA | NA | Mir323 | | 0 | 0 | 0 | 0 | 0 | 0 | 0 | 0 | 0 | 0 | 0 |
| 0 | NA | NA | NA | NA | NA | Mir758 | | 0 | 0 | 0 | 0 | 0 | 0 | 0 | 0 | 0 | 0 | 0 |
| 0 | NA | NA | NA | NA | NA | Mir329 | | 0 | 0 | 0 | 0 | 0 | 0 | 0 | 0 | 0 | 0 | 0 |
| 0 | NA | NA | NA | NA | NA | Mir666 | | 0 | 0 | 0 | 0 | 0 | 0 | 0 | 0 | 0 | 0 | 0 |
| 0 | NA | NA | NA | NA | NA | Mir543 | | 0 | 0 | 0 | 0 | 0 | 0 | 0 | 0 | 0 | 0 | 0 |
| 0 | NA | NA | NA | NA | NA | Mir495 | | 0 | 0 | 0 | 0 | 0 | 0 | 0 | 0 | 0 | 0 | 0 |
| 0 | NA | NA | NA | NA | NA | Mir667 | | 0 | 0 | 0 | 0 | 0 | 0 | 0 | 0 | 0 | 0 | 0 |
| 0 | NA | NA | NA | NA | NA | Mir376c | | 0 | 0 | 0 | 0 | 0 | 0 | 0 | 0 | 0 | 0 | 0 |
| 0 | NA | NA | NA | NA | NA | Mir654 | | 0 | 0 | 0 | 0 | 0 | 0 | 0 | 0 | 0 | 0 | 0 |
| 0 | NA | NA | NA | NA | NA | Mir376b | | 0 | 0 | 0 | 0 | 0 | 0 | 0 | 0 | 0 | 0 | 0 |
| 0 | NA | NA | NA | NA | NA | Gm25252 | | 0 | 0 | 0 | 0 | 0 | 0 | 0 | 0 | 0 | 0 | 0 |
| 0 | NA | NA | NA | NA | NA | Mir376a | | 0 | 0 | 0 | 0 | 0 | 0 | 0 | 0 | 0 | 0 | 0 |
| 0 | NA | NA | NA | NA | NA | Mir300 | | 0 | 0 | 0 | 0 | 0 | 0 | 0 | 0 | 0 | 0 | 0 |
| 0 | NA | NA | NA | NA | NA | Mir487b | | 0 | 0 | 0 | 0 | 0 | 0 | 0 | 0 | 0 | 0 | 0 |
| 0 | NA | NA | NA | NA | NA | Gm44322 | | 0 | 0 | 0 | 0 | 0 | 0 | 0 | 0 | 0 | 0 | 0 |
| 0 | NA | NA | NA | NA | NA | Mir539 | | 0 | 0 | 0 | 0 | 0 | 0 | 0 | 0 | 0 | 0 | 0 |
| 0 | NA | NA | NA | NA | NA | Gm27379 | | 0 | 0 | 0 | 0 | 0 | 0 | 0 | 0 | 0 | 0 | 0 |
| 0 | NA | NA | NA | NA | NA | Mir544 | | 0 | 0 | 0 | 0 | 0 | 0 | 0 | 0 | 0 | 0 | 0 |
| 0 | NA | NA | NA | NA | NA | Mir134 | | 0 | 0 | 0 | 0 | 0 | 0 | 0 | 0 | 0 | 0 | 0 |
| 0 | NA | NA | NA | NA | NA | Mir668 | | 0 | 0 | 0 | 0 | 0 | 0 | 0 | 0 | 0 | 0 | 0 |
| 0 | NA | NA | NA | NA | NA | Mir485 | | 0 | 0 | 0 | 0 | 0 | 0 | 0 | 0 | 0 | 0 | 0 |
| 0 | NA | NA | NA | NA | NA | Mir377 | | 0 | 0 | 0 | 0 | 0 | 0 | 0 | 0 | 0 | 0 | 0 |
| 0 | NA | NA | NA | NA | NA | Gm24321 | | 0 | 0 | 0 | 0 | 0 | 0 | 0 | 0 | 0 | 0 | 0 |
| 0 | NA | NA | NA | NA | NA | Mir410 | | 0 | 0 | 0 | 0 | 0 | 0 | 0 | 0 | 0 | 0 | 0 |
| 0 | NA | NA | NA | NA | NA | Mir3072 | | 0 | 0 | 0 | 0 | 0 | 0 | 0 | 0 | 0 | 0 | 0 |
| 0 | NA | NA | NA | NA | NA | Gm9661 | | 0 | 0 | 0 | 0 | 0 | 0 | 0 | 0 | 0 | 0 | 0 |
| 0 | NA | NA | NA | NA | NA | 4930511J24Rik | | 0 | 0 | 0 | 0 | 0 | 0 | 0 | 0 | 0 | 0 | 0 |
| 0 | NA | NA | NA | NA | NA | Gm34667 | | 0 | 0 | 0 | 0 | 0 | 0 | 0 | 0 | 0 | 0 | 0 |
| 0 | NA | NA | NA | NA | NA | Gm34719 | | 0 | 0 | 0 | 0 | 0 | 0 | 0 | 0 | 0 | 0 | 0 |
| 0 | NA | NA | NA | NA | NA | Gm47187 | | 0 | 0 | 0 | 0 | 0 | 0 | 0 | 0 | 0 | 0 | 0 |
| 0 | NA | NA | NA | NA | NA | Gm47188 | | 0 | 0 | 0 | 0 | 0 | 0 | 0 | 0 | 0 | 0 | 0 |
| 0 | NA | NA | NA | NA | NA | Gm34785 | | 0 | 0 | 0 | 0 | 0 | 0 | 0 | 0 | 0 | 0 | 0 |
| 0 | NA | NA | NA | NA | NA | Gm40576 | | 0 | 0 | 0 | 0 | 0 | 0 | 0 | 0 | 0 | 0 | 0 |
| 0 | NA | NA | NA | NA | NA | 4930425K24Rik | | 0 | 0 | 0 | 0 | 0 | 0 | 0 | 0 | 0 | 0 | 0 |
| 0 | NA | NA | NA | NA | NA | Mir1247 | | 0 | 0 | 0 | 0 | 0 | 0 | 0 | 0 | 0 | 0 | 0 |
| 0 | NA | NA | NA | NA | NA | 1700001K19Ril | RIKEN cDNA 1700001K19 gene | 0 | 0 | 0 | 0 | 0 | 0 | 0 | 0 | 0 | 0 | 0 |
| 0 | NA | NA | NA | NA | NA | Gm48630 | | 0 | 0 | 0 | 0 | 0 | 0 | 0 | 0 | 0 | 0 | 0 |
| 0 | NA | NA | NA | NA | NA | 6030440G07Rik | | 0 | 0 | 0 | 0 | 0 | 0 | 0 | 0 | 0 | 0 | 0 |
| 0 | NA | NA | NA | NA | NA | Gm48631 | | 0 | 0 | 0 | 0 | 0 | 0 | 0 | 0 | 0 | 0 | 0 |
| 0 | NA | NA | NA | NA | NA | Lbhd2 | LBH domain containing 2 | 0 | 0 | 0 | 0 | 0 | 0 | 0 | 0 | 0 | 0 | 0 |
| 0 | NA | NA | NA | NA | NA | Gm40578 | | 0 | 0 | 0 | 0 | 0 | 0 | 0 | 0 | 0 | 0 | 0 |

| | | | | | | | | | | | | | | | | | |
|---|---|---|---|---|---|---|---|---|---|---|---|---|---|---|---|---|---|
| 0 | NA | NA | NA | NA | NA | Snora28 | | 0 | 0 | 0 | 0 | 0 | 0 | 0 | 0 | 0 | 0 | 0 |
| 0 | NA | NA | NA | NA | NA | Gm15995 | | 0 | 0 | 0 | 0 | 0 | 0 | 0 | 0 | 0 | 0 | 0 |
| 0 | NA | NA | NA | NA | NA | Gm15996 | | 0 | 0 | 0 | 0 | 0 | 0 | 0 | 0 | 0 | 0 | 0 |
| 0 | NA | NA | NA | NA | NA | Rd3l | retinal degeneration 3-like | 0 | 0 | 0 | 0 | 0 | 0 | 0 | 0 | 0 | 0 | 0 |
| 0 | NA | NA | NA | NA | NA | Mir3073a | | 0 | 0 | 0 | 0 | 0 | 0 | 0 | 0 | 0 | 0 | 0 |
| 0 | NA | NA | NA | NA | NA | Mir3073b | | 0 | 0 | 0 | 0 | 0 | 0 | 0 | 0 | 0 | 0 | 0 |
| 0 | NA | NA | NA | NA | NA | Mir203 | | 0 | 0 | 0 | 0 | 0 | 0 | 0 | 0 | 0 | 0 | 0 |
| 0 | NA | NA | NA | NA | NA | Gm20368 | | 0 | 0 | 0 | 0 | 0 | 0 | 0 | 0 | 0 | 0 | 0 |
| 0 | NA | NA | NA | NA | NA | Gm38123 | | 0 | 0 | 0 | 0 | 0 | 0 | 0 | 0 | 0 | 0 | 0 |
| 0 | NA | NA | NA | NA | NA | E330035G20Rik | | 0 | 0 | 0 | 0 | 0 | 0 | 0 | 0 | 0 | 0 | 0 |
| 0 | NA | NA | NA | NA | NA | A730018C14Rik | | 0 | 0 | 0 | 0 | 0 | 0 | 0 | 0 | 0 | 0 | 0 |
| 0 | NA | NA | NA | NA | NA | B020018J22Rik | | 0 | 0 | 0 | 0 | 0 | 0 | 0 | 0 | 0 | 0 | 0 |
| 0 | NA | NA | NA | NA | NA | Gm5189 | | 0 | 0 | 0 | 0 | 0 | 0 | 0 | 0 | 0 | 0 | 0 |
| 0 | NA | NA | NA | NA | NA | Gm47474 | | 0 | 0 | 0 | 0 | 0 | 0 | 0 | 0 | 0 | 0 | 0 |
| 0 | NA | NA | NA | NA | NA | 1700127F24Rik | | 0 | 0 | 0 | 0 | 0 | 0 | 0 | 0 | 0 | 0 | 0 |
| 0 | NA | NA | NA | NA | NA | Gm44471 | | 0 | 0 | 0 | 0 | 0 | 0 | 0 | 0 | 0 | 0 | 0 |
| 0 | NA | NA | NA | NA | NA | Gm43310 | | 0 | 0 | 0 | 0 | 0 | 0 | 0 | 0 | 0 | 0 | 0 |
| 0 | NA | NA | NA | NA | NA | Gm25622 | | 0 | 0 | 0 | 0 | 0 | 0 | 0 | 0 | 0 | 0 | 0 |
| 0 | NA | NA | NA | NA | NA | Gm37944 | | 0 | 0 | 0 | 0 | 0 | 0 | 0 | 0 | 0 | 0 | 0 |
| 0 | NA | NA | NA | NA | NA | Gm17421 | | 0 | 0 | 0 | 0 | 0 | 0 | 0 | 0 | 0 | 0 | 0 |
| 0 | NA | NA | NA | NA | NA | Gm9136 | | 0 | 0 | 0 | 0 | 0 | 0 | 0 | 0 | 0 | 0 | 0 |
| 0 | NA | NA | NA | NA | NA | Gm5190 | | 0 | 0 | 0 | 0 | 0 | 0 | 0 | 0 | 0 | 0 | 0 |
| 0 | NA | NA | NA | NA | NA | Gm25747 | | 0 | 0 | 0 | 0 | 0 | 0 | 0 | 0 | 0 | 0 | 0 |
| 0 | NA | NA | NA | NA | NA | Gm24022 | | 0 | 0 | 0 | 0 | 0 | 0 | 0 | 0 | 0 | 0 | 0 |
| 0 | NA | NA | NA | NA | NA | Gm24022 | | 0 | 0 | 0 | 0 | 0 | 0 | 0 | 0 | 0 | 0 | 0 |
| 0 | NA | NA | NA | NA | NA | Gm22175 | | 0 | 0 | 0 | 0 | 0 | 0 | 0 | 0 | 0 | 0 | 0 |
| 0 | NA | NA | NA | NA | NA | Ighj2 | | 0 | 0 | 0 | 0 | 0 | 0 | 0 | 0 | 0 | 0 | 0 |
| 0 | NA | NA | NA | NA | NA | Ighd3-2 | | 0 | 0 | 0 | 0 | 0 | 0 | 0 | 0 | 0 | 0 | 0 |
| 0 | NA | NA | NA | NA | NA | Ighd5-6 | | 0 | 0 | 0 | 0 | 0 | 0 | 0 | 0 | 0 | 0 | 0 |
| 0 | NA | NA | NA | NA | NA | Ighd2-8 | | 0 | 0 | 0 | 0 | 0 | 0 | 0 | 0 | 0 | 0 | 0 |
| 0 | NA | NA | NA | NA | NA | Ighd5-5 | | 0 | 0 | 0 | 0 | 0 | 0 | 0 | 0 | 0 | 0 | 0 |
| 0 | NA | NA | NA | NA | NA | Ighd2-7 | | 0 | 0 | 0 | 0 | 0 | 0 | 0 | 0 | 0 | 0 | 0 |
| 0 | NA | NA | NA | NA | NA | Ighd5-8 | | 0 | 0 | 0 | 0 | 0 | 0 | 0 | 0 | 0 | 0 | 0 |
| 0 | NA | NA | NA | NA | NA | Ighd5-4 | | 0 | 0 | 0 | 0 | 0 | 0 | 0 | 0 | 0 | 0 | 0 |
| 0 | NA | NA | NA | NA | NA | Ighd2-6 | | 0 | 0 | 0 | 0 | 0 | 0 | 0 | 0 | 0 | 0 | 0 |
| 0 | NA | NA | NA | NA | NA | Ighd5-7 | | 0 | 0 | 0 | 0 | 0 | 0 | 0 | 0 | 0 | 0 | 0 |
| 0 | NA | NA | NA | NA | NA | Ighd5-3 | | 0 | 0 | 0 | 0 | 0 | 0 | 0 | 0 | 0 | 0 | 0 |
| 0 | NA | NA | NA | NA | NA | Ighd2-5 | | 0 | 0 | 0 | 0 | 0 | 0 | 0 | 0 | 0 | 0 | 0 |
| 0 | NA | NA | NA | NA | NA | Gm37327 | | 0 | 0 | 0 | 0 | 0 | 0 | 0 | 0 | 0 | 0 | 0 |
| 0 | NA | NA | NA | NA | NA | Ighd5-2 | | 0 | 0 | 0 | 0 | 0 | 0 | 0 | 0 | 0 | 0 | 0 |
| 0 | NA | NA | NA | NA | NA | Ighd2-4 | | 0 | 0 | 0 | 0 | 0 | 0 | 0 | 0 | 0 | 0 | 0 |
| 0 | NA | NA | NA | NA | NA | Ighd6-2 | | 0 | 0 | 0 | 0 | 0 | 0 | 0 | 0 | 0 | 0 | 0 |
| 0 | NA | NA | NA | NA | NA | Gm37227 | | 0 | 0 | 0 | 0 | 0 | 0 | 0 | 0 | 0 | 0 | 0 |
| 0 | NA | NA | NA | NA | NA | Ighd2-3 | | 0 | 0 | 0 | 0 | 0 | 0 | 0 | 0 | 0 | 0 | 0 |
| 0 | NA | NA | NA | NA | NA | Ighd6-1 | | 0 | 0 | 0 | 0 | 0 | 0 | 0 | 0 | 0 | 0 | 0 |
| 0 | NA | NA | NA | NA | NA | Ighd1-1 | | 0 | 0 | 0 | 0 | 0 | 0 | 0 | 0 | 0 | 0 | 0 |
| 0 | NA | NA | NA | NA | NA | Ighd3-1 | | 0 | 0 | 0 | 0 | 0 | 0 | 0 | 0 | 0 | 0 | 0 |
| 0 | NA | NA | NA | NA | NA | Ighd5-1 | | 0 | 0 | 0 | 0 | 0 | 0 | 0 | 0 | 0 | 0 | 0 |
| 0 | NA | NA | NA | NA | NA | Gm16968 | | 0 | 0 | 0 | 0 | 0 | 0 | 0 | 0 | 0 | 0 | 0 |
| 0 | NA | NA | NA | NA | NA | Adam6a | a disintegrin and metallopeptidase domai | 0 | 0 | 0 | 0 | 0 | 0 | 0 | 0 | 0 | 0 | 0 |
| 0 | NA | NA | NA | NA | NA | Ighv2-1 | | 0 | 0 | 0 | 0 | 0 | 0 | 0 | 0 | 0 | 0 | 0 |
| 0 | NA | NA | NA | NA | NA | Ighv5-2 | | 0 | 0 | 0 | 0 | 0 | 0 | 0 | 0 | 0 | 0 | 0 |
| 0 | NA | NA | NA | NA | NA | Ighv5-3 | | 0 | 0 | 0 | 0 | 0 | 0 | 0 | 0 | 0 | 0 | 0 |
| 0 | NA | NA | NA | NA | NA | Ighv6-1 | | 0 | 0 | 0 | 0 | 0 | 0 | 0 | 0 | 0 | 0 | 0 |
| 0 | NA | NA | NA | NA | NA | Ighv5-5 | | 0 | 0 | 0 | 0 | 0 | 0 | 0 | 0 | 0 | 0 | 0 |
| 0 | NA | NA | NA | NA | NA | Ighv5-7 | | 0 | 0 | 0 | 0 | 0 | 0 | 0 | 0 | 0 | 0 | 0 |
| 0 | NA | NA | NA | NA | NA | Ighv5-8 | | 0 | 0 | 0 | 0 | 0 | 0 | 0 | 0 | 0 | 0 | 0 |
| 0 | NA | NA | NA | NA | NA | Ighv5-8 | | 0 | 0 | 0 | 0 | 0 | 0 | 0 | 0 | 0 | 0 | 0 |
| 0 | NA | NA | NA | NA | NA | Ighv5-10 | | 0 | 0 | 0 | 0 | 0 | 0 | 0 | 0 | 0 | 0 | 0 |
| 0 | NA | NA | NA | NA | NA | Gm37187 | | 0 | 0 | 0 | 0 | 0 | 0 | 0 | 0 | 0 | 0 | 0 |

| | | | | | | | | | | | | | | | | | |
|---|---|---|---|---|---|---|---|---|---|---|---|---|---|---|---|---|---|
| 0 | NA | NA | NA | NA | NA | Ighv5-11 | 0 | 0 | 0 | 0 | 0 | 0 | 0 | 0 | 0 | 0 | 0 |
| 0 | NA | NA | NA | NA | NA | Gm37976 | 0 | 0 | 0 | 0 | 0 | 0 | 0 | 0 | 0 | 0 | 0 |
| 0 | NA | NA | NA | NA | NA | Gm37418 | 0 | 0 | 0 | 0 | 0 | 0 | 0 | 0 | 0 | 0 | 0 |
| 0 | NA | NA | NA | NA | NA | Gm38203 | 0 | 0 | 0 | 0 | 0 | 0 | 0 | 0 | 0 | 0 | 0 |
| 0 | NA | NA | NA | NA | NA | Ighv5-12-4 | 0 | 0 | 0 | 0 | 0 | 0 | 0 | 0 | 0 | 0 | 0 |
| 0 | NA | NA | NA | NA | NA | Gm38205 | 0 | 0 | 0 | 0 | 0 | 0 | 0 | 0 | 0 | 0 | 0 |
| 0 | NA | NA | NA | NA | NA | Gm37722 | 0 | 0 | 0 | 0 | 0 | 0 | 0 | 0 | 0 | 0 | 0 |
| 0 | NA | NA | NA | NA | NA | Ighv5-13 | 0 | 0 | 0 | 0 | 0 | 0 | 0 | 0 | 0 | 0 | 0 |
| 0 | NA | NA | NA | NA | NA | Gm7003 | 0 | 0 | 0 | 0 | 0 | 0 | 0 | 0 | 0 | 0 | 0 |
| 0 | NA | NA | NA | NA | NA | Rpl26-ps3 | 0 | 0 | 0 | 0 | 0 | 0 | 0 | 0 | 0 | 0 | 0 |
| 0 | NA | NA | NA | NA | NA | Ighv5-16 | 0 | 0 | 0 | 0 | 0 | 0 | 0 | 0 | 0 | 0 | 0 |
| 0 | NA | NA | NA | NA | NA | Ighv2-8 | 0 | 0 | 0 | 0 | 0 | 0 | 0 | 0 | 0 | 0 | 0 |
| 0 | NA | NA | NA | NA | NA | Ighv5-19 | 0 | 0 | 0 | 0 | 0 | 0 | 0 | 0 | 0 | 0 | 0 |
| 0 | NA | NA | NA | NA | NA | Ighv7-2 | 0 | 0 | 0 | 0 | 0 | 0 | 0 | 0 | 0 | 0 | 0 |
| 0 | NA | NA | NA | NA | NA | Gm37961 | 0 | 0 | 0 | 0 | 0 | 0 | 0 | 0 | 0 | 0 | 0 |
| 0 | NA | NA | NA | NA | NA | Ighv16-1 | 0 | 0 | 0 | 0 | 0 | 0 | 0 | 0 | 0 | 0 | 0 |
| 0 | NA | NA | NA | NA | NA | Ighv6-2 | 0 | 0 | 0 | 0 | 0 | 0 | 0 | 0 | 0 | 0 | 0 |
| 0 | NA | NA | NA | NA | NA | Ighv12-1 | 0 | 0 | 0 | 0 | 0 | 0 | 0 | 0 | 0 | 0 | 0 |
| 0 | NA | NA | NA | NA | NA | Ighv12-2 | 0 | 0 | 0 | 0 | 0 | 0 | 0 | 0 | 0 | 0 | 0 |
| 0 | NA | NA | NA | NA | NA | Ighv7-3 | 0 | 0 | 0 | 0 | 0 | 0 | 0 | 0 | 0 | 0 | 0 |
| 0 | NA | NA | NA | NA | NA | Ighv15-1 | 0 | 0 | 0 | 0 | 0 | 0 | 0 | 0 | 0 | 0 | 0 |
| 0 | NA | NA | NA | NA | NA | Gm37838 | 0 | 0 | 0 | 0 | 0 | 0 | 0 | 0 | 0 | 0 | 0 |
| 0 | NA | NA | NA | NA | NA | Ighv3-3 | 0 | 0 | 0 | 0 | 0 | 0 | 0 | 0 | 0 | 0 | 0 |
| 0 | NA | NA | NA | NA | NA | Ighv7-4 | 0 | 0 | 0 | 0 | 0 | 0 | 0 | 0 | 0 | 0 | 0 |
| 0 | NA | NA | NA | NA | NA | Gm18551 | 0 | 0 | 0 | 0 | 0 | 0 | 0 | 0 | 0 | 0 | 0 |
| 0 | NA | NA | NA | NA | NA | Ighv13-1 | 0 | 0 | 0 | 0 | 0 | 0 | 0 | 0 | 0 | 0 | 0 |
| 0 | NA | NA | NA | NA | NA | Ighv5-21 | 0 | 0 | 0 | 0 | 0 | 0 | 0 | 0 | 0 | 0 | 0 |
| 0 | NA | NA | NA | NA | NA | Ighv8-1 | 0 | 0 | 0 | 0 | 0 | 0 | 0 | 0 | 0 | 0 | 0 |
| 0 | NA | NA | NA | NA | NA | Ighv1-1 | 0 | 0 | 0 | 0 | 0 | 0 | 0 | 0 | 0 | 0 | 0 |
| 0 | NA | NA | NA | NA | NA | Ighv13-2 | 0 | 0 | 0 | 0 | 0 | 0 | 0 | 0 | 0 | 0 | 0 |
| 0 | NA | NA | NA | NA | NA | Ighv6-5 | 0 | 0 | 0 | 0 | 0 | 0 | 0 | 0 | 0 | 0 | 0 |
| 0 | NA | NA | NA | NA | NA | Ighv8-2 | 0 | 0 | 0 | 0 | 0 | 0 | 0 | 0 | 0 | 0 | 0 |
| 0 | NA | NA | NA | NA | NA | Mir7094-1 | 0 | 0 | 0 | 0 | 0 | 0 | 0 | 0 | 0 | 0 | 0 |
| 0 | NA | NA | NA | NA | NA | Ighv1-3 | 0 | 0 | 0 | 0 | 0 | 0 | 0 | 0 | 0 | 0 | 0 |
| 0 | NA | NA | NA | NA | NA | Ighv10-2 | 0 | 0 | 0 | 0 | 0 | 0 | 0 | 0 | 0 | 0 | 0 |
| 0 | NA | NA | NA | NA | NA | Mir7094-2 | 0 | 0 | 0 | 0 | 0 | 0 | 0 | 0 | 0 | 0 | 0 |
| 0 | NA | NA | NA | NA | NA | Ighv1-6 | 0 | 0 | 0 | 0 | 0 | 0 | 0 | 0 | 0 | 0 | 0 |
| 0 | NA | NA | NA | NA | NA | Ighv10-4 | 0 | 0 | 0 | 0 | 0 | 0 | 0 | 0 | 0 | 0 | 0 |
| 0 | NA | NA | NA | NA | NA | Ighv1-8 | 0 | 0 | 0 | 0 | 0 | 0 | 0 | 0 | 0 | 0 | 0 |
| 0 | NA | NA | NA | NA | NA | Ighv1-10 | 0 | 0 | 0 | 0 | 0 | 0 | 0 | 0 | 0 | 0 | 0 |
| 0 | NA | NA | NA | NA | NA | Ighv1-11 | 0 | 0 | 0 | 0 | 0 | 0 | 0 | 0 | 0 | 0 | 0 |
| 0 | NA | NA | NA | NA | NA | Ighv1-12 | 0 | 0 | 0 | 0 | 0 | 0 | 0 | 0 | 0 | 0 | 0 |
| 0 | NA | NA | NA | NA | NA | Ighv1-13 | 0 | 0 | 0 | 0 | 0 | 0 | 0 | 0 | 0 | 0 | 0 |
| 0 | NA | NA | NA | NA | NA | Ighv1-13 | 0 | 0 | 0 | 0 | 0 | 0 | 0 | 0 | 0 | 0 | 0 |
| 0 | NA | NA | NA | NA | NA | Ighv1-16 | 0 | 0 | 0 | 0 | 0 | 0 | 0 | 0 | 0 | 0 | 0 |
| 0 | NA | NA | NA | NA | NA | Ighv1-17-1 | 0 | 0 | 0 | 0 | 0 | 0 | 0 | 0 | 0 | 0 | 0 |
| 0 | NA | NA | NA | NA | NA | Ighv1-19-1 | 0 | 0 | 0 | 0 | 0 | 0 | 0 | 0 | 0 | 0 | 0 |
| 0 | NA | NA | NA | NA | NA | Ighv1-21-1 | 0 | 0 | 0 | 0 | 0 | 0 | 0 | 0 | 0 | 0 | 0 |
| 0 | NA | NA | NA | NA | NA | Ighv1-23 | 0 | 0 | 0 | 0 | 0 | 0 | 0 | 0 | 0 | 0 | 0 |
| 0 | NA | NA | NA | NA | NA | Ighv1-24 | 0 | 0 | 0 | 0 | 0 | 0 | 0 | 0 | 0 | 0 | 0 |
| 0 | NA | NA | NA | NA | NA | Ighv1-27 | 0 | 0 | 0 | 0 | 0 | 0 | 0 | 0 | 0 | 0 | 0 |
| 0 | NA | NA | NA | NA | NA | Ighv1-28 | 0 | 0 | 0 | 0 | 0 | 0 | 0 | 0 | 0 | 0 | 0 |
| 0 | NA | NA | NA | NA | NA | Ighv1-29 | 0 | 0 | 0 | 0 | 0 | 0 | 0 | 0 | 0 | 0 | 0 |
| 0 | NA | NA | NA | NA | NA | Ighv1-30 | 0 | 0 | 0 | 0 | 0 | 0 | 0 | 0 | 0 | 0 | 0 |
| 0 | NA | NA | NA | NA | NA | Ighv1-32 | 0 | 0 | 0 | 0 | 0 | 0 | 0 | 0 | 0 | 0 | 0 |
| 0 | NA | NA | NA | NA | NA | Ighv1-35 | 0 | 0 | 0 | 0 | 0 | 0 | 0 | 0 | 0 | 0 | 0 |
| 0 | NA | NA | NA | NA | NA | Ighv1-36 | 0 | 0 | 0 | 0 | 0 | 0 | 0 | 0 | 0 | 0 | 0 |
| 0 | NA | NA | NA | NA | NA | Ighv1-37 | 0 | 0 | 0 | 0 | 0 | 0 | 0 | 0 | 0 | 0 | 0 |
| 0 | NA | NA | NA | NA | NA | Ighv1-38 | 0 | 0 | 0 | 0 | 0 | 0 | 0 | 0 | 0 | 0 | 0 |
| 0 | NA | NA | NA | NA | NA | Ighv1-40 | 0 | 0 | 0 | 0 | 0 | 0 | 0 | 0 | 0 | 0 | 0 |

| | | | | | | | | | | | | | | | | | |
|---|---|---|---|---|---|---|---|---|---|---|---|---|---|---|---|---|---|
| 0 | NA | NA | NA | NA | NA | Ighv1-41 | 0 | 0 | 0 | 0 | 0 | 0 | 0 | 0 | 0 | 0 | 0 |
| 0 | NA | NA | NA | NA | NA | Ighv1-44 | 0 | 0 | 0 | 0 | 0 | 0 | 0 | 0 | 0 | 0 | 0 |
| 0 | NA | NA | NA | NA | NA | Ighv1-45 | 0 | 0 | 0 | 0 | 0 | 0 | 0 | 0 | 0 | 0 | 0 |
| 0 | NA | NA | NA | NA | NA | Ighv1-46 | 0 | 0 | 0 | 0 | 0 | 0 | 0 | 0 | 0 | 0 | 0 |
| 0 | NA | NA | NA | NA | NA | Ighv8-3 | 0 | 0 | 0 | 0 | 0 | 0 | 0 | 0 | 0 | 0 | 0 |
| 0 | NA | NA | NA | NA | NA | Ighv8-4 | 0 | 0 | 0 | 0 | 0 | 0 | 0 | 0 | 0 | 0 | 0 |
| 0 | NA | NA | NA | NA | NA | Ighv1-48 | 0 | 0 | 0 | 0 | 0 | 0 | 0 | 0 | 0 | 0 | 0 |
| 0 | NA | NA | NA | NA | NA | Ighv1-49 | 0 | 0 | 0 | 0 | 0 | 0 | 0 | 0 | 0 | 0 | 0 |
| 0 | NA | NA | NA | NA | NA | Ighv1-51 | 0 | 0 | 0 | 0 | 0 | 0 | 0 | 0 | 0 | 0 | 0 |
| 0 | NA | NA | NA | NA | NA | Ighv8-6 | 0 | 0 | 0 | 0 | 0 | 0 | 0 | 0 | 0 | 0 | 0 |
| 0 | NA | NA | NA | NA | NA | Gm9227 | 0 | 0 | 0 | 0 | 0 | 0 | 0 | 0 | 0 | 0 | 0 |
| 0 | NA | NA | NA | NA | NA | Gm7142 | 0 | 0 | 0 | 0 | 0 | 0 | 0 | 0 | 0 | 0 | 0 |
| 0 | NA | NA | NA | NA | NA | Ighv8-7 | 0 | 0 | 0 | 0 | 0 | 0 | 0 | 0 | 0 | 0 | 0 |
| 0 | NA | NA | NA | NA | NA | Gm30948 | 0 | 0 | 0 | 0 | 0 | 0 | 0 | 0 | 0 | 0 | 0 |
| 0 | NA | NA | NA | NA | NA | Ighv1-57 | 0 | 0 | 0 | 0 | 0 | 0 | 0 | 0 | 0 | 0 | 0 |
| 0 | NA | NA | NA | NA | NA | Ighv1-58 | 0 | 0 | 0 | 0 | 0 | 0 | 0 | 0 | 0 | 0 | 0 |
| 0 | NA | NA | NA | NA | NA | Ighv1-60 | 0 | 0 | 0 | 0 | 0 | 0 | 0 | 0 | 0 | 0 | 0 |
| 0 | NA | NA | NA | NA | NA | Ighv1-62-1 | 0 | 0 | 0 | 0 | 0 | 0 | 0 | 0 | 0 | 0 | 0 |
| 0 | NA | NA | NA | NA | NA | Gm19331 | 0 | 0 | 0 | 0 | 0 | 0 | 0 | 0 | 0 | 0 | 0 |
| 0 | NA | NA | NA | NA | NA | Gm37511 | 0 | 0 | 0 | 0 | 0 | 0 | 0 | 0 | 0 | 0 | 0 |
| 0 | NA | NA | NA | NA | NA | Gm38184 | 0 | 0 | 0 | 0 | 0 | 0 | 0 | 0 | 0 | 0 | 0 |
| 0 | NA | NA | NA | NA | NA | Ighv8-9 | 0 | 0 | 0 | 0 | 0 | 0 | 0 | 0 | 0 | 0 | 0 |
| 0 | NA | NA | NA | NA | NA | Gm9236 | 0 | 0 | 0 | 0 | 0 | 0 | 0 | 0 | 0 | 0 | 0 |
| 0 | NA | NA | NA | NA | NA | Ighv1-63 | 0 | 0 | 0 | 0 | 0 | 0 | 0 | 0 | 0 | 0 | 0 |
| 0 | NA | NA | NA | NA | NA | Gm27722 | 0 | 0 | 0 | 0 | 0 | 0 | 0 | 0 | 0 | 0 | 0 |
| 0 | NA | NA | NA | NA | NA | Gm18340 | 0 | 0 | 0 | 0 | 0 | 0 | 0 | 0 | 0 | 0 | 0 |
| 0 | NA | NA | NA | NA | NA | Ighv1-65 | 0 | 0 | 0 | 0 | 0 | 0 | 0 | 0 | 0 | 0 | 0 |
| 0 | NA | NA | NA | NA | NA | Ighv1-66 | 0 | 0 | 0 | 0 | 0 | 0 | 0 | 0 | 0 | 0 | 0 |
| 0 | NA | NA | NA | NA | NA | Ighv1-67 | 0 | 0 | 0 | 0 | 0 | 0 | 0 | 0 | 0 | 0 | 0 |
| 0 | NA | NA | NA | NA | NA | Ighv1-68 | 0 | 0 | 0 | 0 | 0 | 0 | 0 | 0 | 0 | 0 | 0 |
| 0 | NA | NA | NA | NA | NA | Gm5660 | 0 | 0 | 0 | 0 | 0 | 0 | 0 | 0 | 0 | 0 | 0 |
| 0 | NA | NA | NA | NA | NA | Gm9245 | 0 | 0 | 0 | 0 | 0 | 0 | 0 | 0 | 0 | 0 | 0 |
| 0 | NA | NA | NA | NA | NA | Gm44782 | 0 | 0 | 0 | 0 | 0 | 0 | 0 | 0 | 0 | 0 | 0 |
| 0 | NA | NA | NA | NA | NA | Ighv1-70 | 0 | 0 | 0 | 0 | 0 | 0 | 0 | 0 | 0 | 0 | 0 |
| 0 | NA | NA | NA | NA | NA | Mir6388 | 0 | 0 | 0 | 0 | 0 | 0 | 0 | 0 | 0 | 0 | 0 |
| 0 | NA | NA | NA | NA | NA | Ighv8-13 | 0 | 0 | 0 | 0 | 0 | 0 | 0 | 0 | 0 | 0 | 0 |
| 0 | NA | NA | NA | NA | NA | Gm9251 | 0 | 0 | 0 | 0 | 0 | 0 | 0 | 0 | 0 | 0 | 0 |
| 0 | NA | NA | NA | NA | NA | Ighv1-73 | 0 | 0 | 0 | 0 | 0 | 0 | 0 | 0 | 0 | 0 | 0 |
| 0 | NA | NA | NA | NA | NA | Ighv8-14 | 0 | 0 | 0 | 0 | 0 | 0 | 0 | 0 | 0 | 0 | 0 |
| 0 | NA | NA | NA | NA | NA | Gm18341 | 0 | 0 | 0 | 0 | 0 | 0 | 0 | 0 | 0 | 0 | 0 |
| 0 | NA | NA | NA | NA | NA | Ighv1-75 | 0 | 0 | 0 | 0 | 0 | 0 | 0 | 0 | 0 | 0 | 0 |
| 0 | NA | NA | NA | NA | NA | Ighv8-15 | 0 | 0 | 0 | 0 | 0 | 0 | 0 | 0 | 0 | 0 | 0 |
| 0 | NA | NA | NA | NA | NA | Ighv1-76 | 0 | 0 | 0 | 0 | 0 | 0 | 0 | 0 | 0 | 0 | 0 |
| 0 | NA | NA | NA | NA | NA | Ighv8-16 | 0 | 0 | 0 | 0 | 0 | 0 | 0 | 0 | 0 | 0 | 0 |
| 0 | NA | NA | NA | NA | NA | Ighv1-77 | 0 | 0 | 0 | 0 | 0 | 0 | 0 | 0 | 0 | 0 | 0 |
| 0 | NA | NA | NA | NA | NA | Ighv1-78 | 0 | 0 | 0 | 0 | 0 | 0 | 0 | 0 | 0 | 0 | 0 |
| 0 | NA | NA | NA | NA | NA | Gm42643 | 0 | 0 | 0 | 0 | 0 | 0 | 0 | 0 | 0 | 0 | 0 |
| 0 | NA | NA | NA | NA | NA | Gm5034 | 0 | 0 | 0 | 0 | 0 | 0 | 0 | 0 | 0 | 0 | 0 |
| 0 | NA | NA | NA | NA | NA | Ighv1-79 | 0 | 0 | 0 | 0 | 0 | 0 | 0 | 0 | 0 | 0 | 0 |
| 0 | NA | NA | NA | NA | NA | Gm42990 | 0 | 0 | 0 | 0 | 0 | 0 | 0 | 0 | 0 | 0 | 0 |
| 0 | NA | NA | NA | NA | NA | Olfr241-ps1 | 0 | 0 | 0 | 0 | 0 | 0 | 0 | 0 | 0 | 0 | 0 |
| 0 | NA | NA | NA | NA | NA | Ighv1-84 | 0 | 0 | 0 | 0 | 0 | 0 | 0 | 0 | 0 | 0 | 0 |
| 0 | NA | NA | NA | NA | NA | Ighv1-86 | 0 | 0 | 0 | 0 | 0 | 0 | 0 | 0 | 0 | 0 | 0 |
| 0 | NA | NA | NA | NA | NA | Gm20658 | 0 | 0 | 0 | 0 | 0 | 0 | 0 | 0 | 0 | 0 | 0 |
| 0 | NA | NA | NA | NA | NA | Gm48618 | 0 | 0 | 0 | 0 | 0 | 0 | 0 | 0 | 0 | 0 | 0 |
| 0 | NA | NA | NA | NA | NA | Gm44372 | 0 | 0 | 0 | 0 | 0 | 0 | 0 | 0 | 0 | 0 | 0 |
| 0 | NA | NA | NA | NA | NA | Gm23732 | 0 | 0 | 0 | 0 | 0 | 0 | 0 | 0 | 0 | 0 | 0 |
| 0 | NA | NA | NA | NA | NA | Gm23190 | 0 | 0 | 0 | 0 | 0 | 0 | 0 | 0 | 0 | 0 | 0 |
| 0 | NA | NA | NA | NA | NA | Gm11027 | 0 | 0 | 0 | 0 | 0 | 0 | 0 | 0 | 0 | 0 | 0 |
| 0 | NA | NA | NA | NA | NA | Gm48119 | 0 | 0 | 0 | 0 | 0 | 0 | 0 | 0 | 0 | 0 | 0 |

| | | | | | | | | | | | | | | | | | |
|---|---|---|---|---|---|---|---|---|---|---|---|---|---|---|---|---|---|
| 0 | NA | NA | NA | NA | NA | Gm10421 | | 0 | 0 | 0 | 0 | 0 | 0 | 0 | 0 | 0 | 0 | 0 |
| 0 | NA | NA | NA | NA | NA | Gm24741 | | 0 | 0 | 0 | 0 | 0 | 0 | 0 | 0 | 0 | 0 | 0 |
| 0 | NA | NA | NA | NA | NA | A930023M06Rik | | 0 | 0 | 0 | 0 | 0 | 0 | 0 | 0 | 0 | 0 | 0 |
| 0 | NA | NA | NA | NA | NA | Gm18955 | | 0 | 0 | 0 | 0 | 0 | 0 | 0 | 0 | 0 | 0 | 0 |
| 0 | NA | NA | NA | NA | NA | Gm20613 | | 0 | 0 | 0 | 0 | 0 | 0 | 0 | 0 | 0 | 0 | 0 |
| 0 | NA | NA | NA | NA | NA | Gm25577 | | 0 | 0 | 0 | 0 | 0 | 0 | 0 | 0 | 0 | 0 | 0 |
| 0 | NA | NA | NA | NA | NA | D230030E09Rik | | 0 | 0 | 0 | 0 | 0 | 0 | 0 | 0 | 0 | 0 | 0 |
| 0 | NA | NA | NA | NA | NA | Gm22537 | | 0 | 0 | 0 | 0 | 0 | 0 | 0 | 0 | 0 | 0 | 0 |
| 0 | NA | NA | NA | NA | NA | Gm6171 | | 0 | 0 | 0 | 0 | 0 | 0 | 0 | 0 | 0 | 0 | 0 |
| 0 | NA | NA | NA | NA | NA | Abcb5 | ATP-binding cassette, sub-family B (MDR | 0 | 0 | 0 | 0 | 0 | 0 | 0 | 0 | 0 | 0 | 0 |
| 0 | NA | NA | NA | NA | NA | Gm18921 | | 0 | 0 | 0 | 0 | 0 | 0 | 0 | 0 | 0 | 0 | 0 |
| 0 | NA | NA | NA | NA | NA | Gm48681 | | 0 | 0 | 0 | 0 | 0 | 0 | 0 | 0 | 0 | 0 | 0 |
| 0 | NA | NA | NA | NA | NA | Gm9279 | | 0 | 0 | 0 | 0 | 0 | 0 | 0 | 0 | 0 | 0 | 0 |
| 0 | NA | NA | NA | NA | NA | Gm48685 | | 0 | 0 | 0 | 0 | 0 | 0 | 0 | 0 | 0 | 0 | 0 |
| 0 | NA | NA | NA | NA | NA | Gm48686 | | 0 | 0 | 0 | 0 | 0 | 0 | 0 | 0 | 0 | 0 | 0 |
| 0 | NA | NA | NA | NA | NA | Gm25675 | | 0 | 0 | 0 | 0 | 0 | 0 | 0 | 0 | 0 | 0 | 0 |
| 0 | NA | NA | NA | NA | NA | Gm27236 | | 0 | 0 | 0 | 0 | 0 | 0 | 0 | 0 | 0 | 0 | 0 |
| 0 | NA | NA | NA | NA | NA | Gm49287 | | 0 | 0 | 0 | 0 | 0 | 0 | 0 | 0 | 0 | 0 | 0 |
| 0 | NA | NA | NA | NA | NA | Gm17177 | | 0 | 0 | 0 | 0 | 0 | 0 | 0 | 0 | 0 | 0 | 0 |
| 0 | NA | NA | NA | NA | NA | Gm46400 | | 0 | 0 | 0 | 0 | 0 | 0 | 0 | 0 | 0 | 0 | 0 |
| 0 | NA | NA | NA | NA | NA | Gm18783 | | 0 | 0 | 0 | 0 | 0 | 0 | 0 | 0 | 0 | 0 | 0 |
| 0 | NA | NA | NA | NA | NA | Gm48519 | | 0 | 0 | 0 | 0 | 0 | 0 | 0 | 0 | 0 | 0 | 0 |
| 0 | NA | NA | NA | NA | NA | Gm48520 | | 0 | 0 | 0 | 0 | 0 | 0 | 0 | 0 | 0 | 0 | 0 |
| 0 | NA | NA | NA | NA | NA | Speer6-ps1 | | 0 | 0 | 0 | 0 | 0 | 0 | 0 | 0 | 0 | 0 | 0 |
| 0 | NA | NA | NA | NA | NA | Gm40653 | | 0 | 0 | 0 | 0 | 0 | 0 | 0 | 0 | 0 | 0 | 0 |
| 0 | NA | NA | NA | NA | NA | Gm19163 | | 0 | 0 | 0 | 0 | 0 | 0 | 0 | 0 | 0 | 0 | 0 |
| 0 | NA | NA | NA | NA | NA | Gm47695 | | 0 | 0 | 0 | 0 | 0 | 0 | 0 | 0 | 0 | 0 | 0 |
| 0 | NA | NA | NA | NA | NA | Gm35190 | | 0 | 0 | 0 | 0 | 0 | 0 | 0 | 0 | 0 | 0 | 0 |
| 0 | NA | NA | NA | NA | NA | Calml3 | calmodulin-like 3 | 0 | 0 | 0 | 0 | 0 | 0 | 0 | 0 | 0 | 0 | 0 |
| 0 | NA | NA | NA | NA | NA | Gm40655 | | 0 | 0 | 0 | 0 | 0 | 0 | 0 | 0 | 0 | 0 | 0 |
| 0 | NA | NA | NA | NA | NA | Calm5 | calmodulin 5 | 0 | 0 | 0 | 0 | 0 | 0 | 0 | 0 | 0 | 0 | 0 |
| 0 | NA | NA | NA | NA | NA | Gm22460 | | 0 | 0 | 0 | 0 | 0 | 0 | 0 | 0 | 0 | 0 | 0 |
| 0 | NA | NA | NA | NA | NA | Gm47853 | | 0 | 0 | 0 | 0 | 0 | 0 | 0 | 0 | 0 | 0 | 0 |
| 0 | NA | NA | NA | NA | NA | Gm47925 | | 0 | 0 | 0 | 0 | 0 | 0 | 0 | 0 | 0 | 0 | 0 |
| 0 | NA | NA | NA | NA | NA | Gm2123 | | 0 | 0 | 0 | 0 | 0 | 0 | 0 | 0 | 0 | 0 | 0 |
| 0 | NA | NA | NA | NA | NA | Marcksl1-ps4 | | 0 | 0 | 0 | 0 | 0 | 0 | 0 | 0 | 0 | 0 | 0 |
| 0 | NA | NA | NA | NA | NA | Gm9973 | | 0 | 0 | 0 | 0 | 0 | 0 | 0 | 0 | 0 | 0 | 0 |
| 0 | NA | NA | NA | NA | NA | Gm47951 | | 0 | 0 | 0 | 0 | 0 | 0 | 0 | 0 | 0 | 0 | 0 |
| 0 | NA | NA | NA | NA | NA | Akr1c6 | aldo-keto reductase family 1, member C6 | 0 | 0 | 0 | 0 | 0 | 0 | 0 | 0 | 0 | 0 | 0 |
| 0 | NA | NA | NA | NA | NA | Gm48010 | | 0 | 0 | 0 | 0 | 0 | 0 | 0 | 0 | 0 | 0 | 0 |
| 0 | NA | NA | NA | NA | NA | Gm40658 | | 0 | 0 | 0 | 0 | 0 | 0 | 0 | 0 | 0 | 0 | 0 |
| 0 | NA | NA | NA | NA | NA | Gm5444 | | 0 | 0 | 0 | 0 | 0 | 0 | 0 | 0 | 0 | 0 | 0 |
| 0 | NA | NA | NA | NA | NA | Gm47450 | | 0 | 0 | 0 | 0 | 0 | 0 | 0 | 0 | 0 | 0 | 0 |
| 0 | NA | NA | NA | NA | NA | Gm24790 | | 0 | 0 | 0 | 0 | 0 | 0 | 0 | 0 | 0 | 0 | 0 |
| 0 | NA | NA | NA | NA | NA | Gm47451 | | 0 | 0 | 0 | 0 | 0 | 0 | 0 | 0 | 0 | 0 | 0 |
| 0 | NA | NA | NA | NA | NA | Gm40660 | | 0 | 0 | 0 | 0 | 0 | 0 | 0 | 0 | 0 | 0 | 0 |
| 0 | NA | NA | NA | NA | NA | Gm35330 | | 0 | 0 | 0 | 0 | 0 | 0 | 0 | 0 | 0 | 0 | 0 |
| 0 | NA | NA | NA | NA | NA | Gm26043 | | 0 | 0 | 0 | 0 | 0 | 0 | 0 | 0 | 0 | 0 | 0 |
| 0 | NA | NA | NA | NA | NA | 1700016G22Rik | | 0 | 0 | 0 | 0 | 0 | 0 | 0 | 0 | 0 | 0 | 0 |
| 0 | NA | NA | NA | NA | NA | Gm46401 | | 0 | 0 | 0 | 0 | 0 | 0 | 0 | 0 | 0 | 0 | 0 |
| 0 | NA | NA | NA | NA | NA | Gm26259 | | 0 | 0 | 0 | 0 | 0 | 0 | 0 | 0 | 0 | 0 | 0 |
| 0 | NA | NA | NA | NA | NA | Gm47514 | | 0 | 0 | 0 | 0 | 0 | 0 | 0 | 0 | 0 | 0 | 0 |
| 0 | NA | NA | NA | NA | NA | Gm47548 | | 0 | 0 | 0 | 0 | 0 | 0 | 0 | 0 | 0 | 0 | 0 |
| 0 | NA | NA | NA | NA | NA | Gm47549 | | 0 | 0 | 0 | 0 | 0 | 0 | 0 | 0 | 0 | 0 | 0 |
| 0 | NA | NA | NA | NA | NA | Gm35615 | | 0 | 0 | 0 | 0 | 0 | 0 | 0 | 0 | 0 | 0 | 0 |
| 0 | NA | NA | NA | NA | NA | Gm28465 | | 0 | 0 | 0 | 0 | 0 | 0 | 0 | 0 | 0 | 0 | 0 |
| 0 | NA | NA | NA | NA | NA | 2900042G08Rik | | 0 | 0 | 0 | 0 | 0 | 0 | 0 | 0 | 0 | 0 | 0 |
| 0 | NA | NA | NA | NA | NA | Gm48238 | | 0 | 0 | 0 | 0 | 0 | 0 | 0 | 0 | 0 | 0 | 0 |
| 0 | NA | NA | NA | NA | NA | 2900024D18Rik | | 0 | 0 | 0 | 0 | 0 | 0 | 0 | 0 | 0 | 0 | 0 |
| 0 | NA | NA | NA | NA | NA | Gm40662 | | 0 | 0 | 0 | 0 | 0 | 0 | 0 | 0 | 0 | 0 | 0 |

| | | | | | | | | | | | | | | | | | |
|---|---|---|---|---|---|---|---|---|---|---|---|---|---|---|---|---|---|
| 0 | NA | NA | NA | NA | NA | Gm22689 | | 0 | 0 | 0 | 0 | 0 | 0 | 0 | 0 | 0 | 0 | 0 |
| 0 | NA | NA | NA | NA | NA | Gm40663 | | 0 | 0 | 0 | 0 | 0 | 0 | 0 | 0 | 0 | 0 | 0 |
| 0 | NA | NA | NA | NA | NA | Gm8725 | | 0 | 0 | 0 | 0 | 0 | 0 | 0 | 0 | 0 | 0 | 0 |
| 0 | NA | NA | NA | NA | NA | Gm36074 | | 0 | 0 | 0 | 0 | 0 | 0 | 0 | 0 | 0 | 0 | 0 |
| 0 | NA | NA | NA | NA | NA | Gm48251 | | 0 | 0 | 0 | 0 | 0 | 0 | 0 | 0 | 0 | 0 | 0 |
| 0 | NA | NA | NA | NA | NA | Gm48252 | | 0 | 0 | 0 | 0 | 0 | 0 | 0 | 0 | 0 | 0 | 0 |
| 0 | NA | NA | NA | NA | NA | Gm48253 | | 0 | 0 | 0 | 0 | 0 | 0 | 0 | 0 | 0 | 0 | 0 |
| 0 | NA | NA | NA | NA | NA | Gm9742 | | 0 | 0 | 0 | 0 | 0 | 0 | 0 | 0 | 0 | 0 | 0 |
| 0 | NA | NA | NA | NA | NA | Gm48257 | | 0 | 0 | 0 | 0 | 0 | 0 | 0 | 0 | 0 | 0 | 0 |
| 0 | NA | NA | NA | NA | NA | Gm48259 | | 0 | 0 | 0 | 0 | 0 | 0 | 0 | 0 | 0 | 0 | 0 |
| 0 | NA | NA | NA | NA | NA | Gm48261 | | 0 | 0 | 0 | 0 | 0 | 0 | 0 | 0 | 0 | 0 | 0 |
| 0 | NA | NA | NA | NA | NA | Gm48260 | | 0 | 0 | 0 | 0 | 0 | 0 | 0 | 0 | 0 | 0 | 0 |
| 0 | NA | NA | NA | NA | NA | Gm48262 | | 0 | 0 | 0 | 0 | 0 | 0 | 0 | 0 | 0 | 0 | 0 |
| 0 | NA | NA | NA | NA | NA | Gm48327 | | 0 | 0 | 0 | 0 | 0 | 0 | 0 | 0 | 0 | 0 | 0 |
| 0 | NA | NA | NA | NA | NA | Gm22750 | | 0 | 0 | 0 | 0 | 0 | 0 | 0 | 0 | 0 | 0 | 0 |
| 0 | NA | NA | NA | NA | NA | Rpl10a-ps2 | | 0 | 0 | 0 | 0 | 0 | 0 | 0 | 0 | 0 | 0 | 0 |
| 0 | NA | NA | NA | NA | NA | Gm9745 | | 0 | 0 | 0 | 0 | 0 | 0 | 0 | 0 | 0 | 0 | 0 |
| 0 | NA | NA | NA | NA | NA | Gm25274 | | 0 | 0 | 0 | 0 | 0 | 0 | 0 | 0 | 0 | 0 | 0 |
| 0 | NA | NA | NA | NA | NA | Gm22115 | | 0 | 0 | 0 | 0 | 0 | 0 | 0 | 0 | 0 | 0 | 0 |
| 0 | NA | NA | NA | NA | NA | Idi2 | isopentenyl-diphosphate delta isomerase | 0 | 0 | 0 | 0 | 0 | 0 | 0 | 0 | 0 | 0 | 0 |
| 0 | NA | NA | NA | NA | NA | Gm23923 | | 0 | 0 | 0 | 0 | 0 | 0 | 0 | 0 | 0 | 0 | 0 |
| 0 | NA | NA | NA | NA | NA | Gm48649 | | 0 | 0 | 0 | 0 | 0 | 0 | 0 | 0 | 0 | 0 | 0 |
| 0 | NA | NA | NA | NA | NA | Gm48650 | | 0 | 0 | 0 | 0 | 0 | 0 | 0 | 0 | 0 | 0 | 0 |
| 0 | NA | NA | NA | NA | NA | Gm36264 | | 0 | 0 | 0 | 0 | 0 | 0 | 0 | 0 | 0 | 0 | 0 |
| 0 | NA | NA | NA | NA | NA | Gm28155 | | 0 | 0 | 0 | 0 | 0 | 0 | 0 | 0 | 0 | 0 | 0 |
| 0 | NA | NA | NA | NA | NA | Gm48825 | | 0 | 0 | 0 | 0 | 0 | 0 | 0 | 0 | 0 | 0 | 0 |
| 0 | NA | NA | NA | NA | NA | Gm23496 | | 0 | 0 | 0 | 0 | 0 | 0 | 0 | 0 | 0 | 0 | 0 |
| 0 | NA | NA | NA | NA | NA | Gm48888 | | 0 | 0 | 0 | 0 | 0 | 0 | 0 | 0 | 0 | 0 | 0 |
| 0 | NA | NA | NA | NA | NA | Gm47802 | | 0 | 0 | 0 | 0 | 0 | 0 | 0 | 0 | 0 | 0 | 0 |
| 0 | NA | NA | NA | NA | NA | Gm47401 | | 0 | 0 | 0 | 0 | 0 | 0 | 0 | 0 | 0 | 0 | 0 |
| 0 | NA | NA | NA | NA | NA | Gm47406 | | 0 | 0 | 0 | 0 | 0 | 0 | 0 | 0 | 0 | 0 | 0 |
| 0 | NA | NA | NA | NA | NA | Gm47404 | | 0 | 0 | 0 | 0 | 0 | 0 | 0 | 0 | 0 | 0 | 0 |
| 0 | NA | NA | NA | NA | NA | Gm47407 | | 0 | 0 | 0 | 0 | 0 | 0 | 0 | 0 | 0 | 0 | 0 |
| 0 | NA | NA | NA | NA | NA | Gm36525 | | 0 | 0 | 0 | 0 | 0 | 0 | 0 | 0 | 0 | 0 | 0 |
| 0 | NA | NA | NA | NA | NA | Gm23415 | | 0 | 0 | 0 | 0 | 0 | 0 | 0 | 0 | 0 | 0 | 0 |
| 0 | NA | NA | NA | NA | NA | Gm47408 | | 0 | 0 | 0 | 0 | 0 | 0 | 0 | 0 | 0 | 0 | 0 |
| 0 | NA | NA | NA | NA | NA | Gm47486 | | 0 | 0 | 0 | 0 | 0 | 0 | 0 | 0 | 0 | 0 | 0 |
| 0 | NA | NA | NA | NA | NA | Gm25496 | | 0 | 0 | 0 | 0 | 0 | 0 | 0 | 0 | 0 | 0 | 0 |
| 0 | NA | NA | NA | NA | NA | Zp4-ps | | 0 | 0 | 0 | 0 | 0 | 0 | 0 | 0 | 0 | 0 | 0 |
| 0 | NA | NA | NA | NA | NA | Gm47487 | | 0 | 0 | 0 | 0 | 0 | 0 | 0 | 0 | 0 | 0 | 0 |
| 0 | NA | NA | NA | NA | NA | Gm47494 | | 0 | 0 | 0 | 0 | 0 | 0 | 0 | 0 | 0 | 0 | 0 |
| 0 | NA | NA | NA | NA | NA | Gm26956 | | 0 | 0 | 0 | 0 | 0 | 0 | 0 | 0 | 0 | 0 | 0 |
| 0 | NA | NA | NA | NA | NA | Gm47493 | | 0 | 0 | 0 | 0 | 0 | 0 | 0 | 0 | 0 | 0 | 0 |
| 0 | NA | NA | NA | NA | NA | Gm47323 | | 0 | 0 | 0 | 0 | 0 | 0 | 0 | 0 | 0 | 0 | 0 |
| 0 | NA | NA | NA | NA | NA | Gm16419 | | 0 | 0 | 0 | 0 | 0 | 0 | 0 | 0 | 0 | 0 | 0 |
| 0 | NA | NA | NA | NA | NA | Gm24065 | | 0 | 0 | 0 | 0 | 0 | 0 | 0 | 0 | 0 | 0 | 0 |
| 0 | NA | NA | NA | NA | NA | Gm48440 | | 0 | 0 | 0 | 0 | 0 | 0 | 0 | 0 | 0 | 0 | 0 |
| 0 | NA | NA | NA | NA | NA | Gm48445 | | 0 | 0 | 0 | 0 | 0 | 0 | 0 | 0 | 0 | 0 | 0 |
| 0 | NA | NA | NA | NA | NA | Gm48456 | | 0 | 0 | 0 | 0 | 0 | 0 | 0 | 0 | 0 | 0 | 0 |
| 0 | NA | NA | NA | NA | NA | Prl2c3 | prolactin family 2, subfamily c, member 3 | 0 | 0 | 0 | 0 | 0 | 0 | 0 | 0 | 0 | 0 | 0 |
| 0 | NA | NA | NA | NA | NA | Gm18262 | | 0 | 0 | 0 | 0 | 0 | 0 | 0 | 0 | 0 | 0 | 0 |
| 0 | NA | NA | NA | NA | NA | Gm2544 | | 0 | 0 | 0 | 0 | 0 | 0 | 0 | 0 | 0 | 0 | 0 |
| 0 | NA | NA | NA | NA | NA | Gm18878 | | 0 | 0 | 0 | 0 | 0 | 0 | 0 | 0 | 0 | 0 | 0 |
| 0 | NA | NA | NA | NA | NA | Gm48527 | | 0 | 0 | 0 | 0 | 0 | 0 | 0 | 0 | 0 | 0 | 0 |
| 0 | NA | NA | NA | NA | NA | Prl2c2 | prolactin family 2, subfamily c, member 2 | 0 | 0 | 0 | 0 | 0 | 0 | 0 | 0 | 0 | 0 | 0 |
| 0 | NA | NA | NA | NA | NA | Gm18263 | | 0 | 0 | 0 | 0 | 0 | 0 | 0 | 0 | 0 | 0 | 0 |
| 0 | NA | NA | NA | NA | NA | Gm32300 | | 0 | 0 | 0 | 0 | 0 | 0 | 0 | 0 | 0 | 0 | 0 |
| 0 | NA | NA | NA | NA | NA | Gm18879 | | 0 | 0 | 0 | 0 | 0 | 0 | 0 | 0 | 0 | 0 | 0 |
| 0 | NA | NA | NA | NA | NA | Gm9628 | | 0 | 0 | 0 | 0 | 0 | 0 | 0 | 0 | 0 | 0 | 0 |
| 0 | NA | NA | NA | NA | NA | Gm18259 | | 0 | 0 | 0 | 0 | 0 | 0 | 0 | 0 | 0 | 0 | 0 |

| | | | | | | | | | | | | | | | | | | |
|---|---|---|---|---|---|---|---|---|---|---|---|---|---|---|---|---|---|---|
| 0 | NA | NA | NA | NA | NA | Gm10811 | | 0 | 0 | 0 | 0 | 0 | 0 | 0 | 0 | 0 | 0 | 0 |
| 0 | NA | NA | NA | NA | NA | Prl2c5 | prolactin family 2, subfamily c, member 5 | 0 | 0 | 0 | 0 | 0 | 0 | 0 | 0 | 0 | 0 | 0 |
| 0 | NA | NA | NA | NA | NA | Gm18880 | | 0 | 0 | 0 | 0 | 0 | 0 | 0 | 0 | 0 | 0 | 0 |
| 0 | NA | NA | NA | NA | NA | Gm48588 | | 0 | 0 | 0 | 0 | 0 | 0 | 0 | 0 | 0 | 0 | 0 |
| 0 | NA | NA | NA | NA | NA | Gm48589 | | 0 | 0 | 0 | 0 | 0 | 0 | 0 | 0 | 0 | 0 | 0 |
| 0 | NA | NA | NA | NA | NA | Gm48590 | | 0 | 0 | 0 | 0 | 0 | 0 | 0 | 0 | 0 | 0 | 0 |
| 0 | NA | NA | NA | NA | NA | Gm26133 | | 0 | 0 | 0 | 0 | 0 | 0 | 0 | 0 | 0 | 0 | 0 |
| 0 | NA | NA | NA | NA | NA | Gm7040 | | 0 | 0 | 0 | 0 | 0 | 0 | 0 | 0 | 0 | 0 | 0 |
| 0 | NA | NA | NA | NA | NA | Gm25804 | | 0 | 0 | 0 | 0 | 0 | 0 | 0 | 0 | 0 | 0 | 0 |
| 0 | NA | NA | NA | NA | NA | Gm23081 | | 0 | 0 | 0 | 0 | 0 | 0 | 0 | 0 | 0 | 0 | 0 |
| 0 | NA | NA | NA | NA | NA | Gm48661 | | 0 | 0 | 0 | 0 | 0 | 0 | 0 | 0 | 0 | 0 | 0 |
| 0 | NA | NA | NA | NA | NA | Gm48672 | | 0 | 0 | 0 | 0 | 0 | 0 | 0 | 0 | 0 | 0 | 0 |
| 0 | NA | NA | NA | NA | NA | Gm7426 | | 0 | 0 | 0 | 0 | 0 | 0 | 0 | 0 | 0 | 0 | 0 |
| 0 | NA | NA | NA | NA | NA | Gm48682 | | 0 | 0 | 0 | 0 | 0 | 0 | 0 | 0 | 0 | 0 | 0 |
| 0 | NA | NA | NA | NA | NA | Gm7046 | | 0 | 0 | 0 | 0 | 0 | 0 | 0 | 0 | 0 | 0 | 0 |
| 0 | NA | NA | NA | NA | NA | Gm47884 | | 0 | 0 | 0 | 0 | 0 | 0 | 0 | 0 | 0 | 0 | 0 |
| 0 | NA | NA | NA | NA | NA | Gm18856 | | 0 | 0 | 0 | 0 | 0 | 0 | 0 | 0 | 0 | 0 | 0 |
| 0 | NA | NA | NA | NA | NA | Gm26129 | | 0 | 0 | 0 | 0 | 0 | 0 | 0 | 0 | 0 | 0 | 0 |
| 0 | NA | NA | NA | NA | NA | n-R5s52 | | 0 | 0 | 0 | 0 | 0 | 0 | 0 | 0 | 0 | 0 | 0 |
| 0 | NA | NA | NA | NA | NA | Gm2492 | | 0 | 0 | 0 | 0 | 0 | 0 | 0 | 0 | 0 | 0 | 0 |
| 0 | NA | NA | NA | NA | NA | Gm47011 | | 0 | 0 | 0 | 0 | 0 | 0 | 0 | 0 | 0 | 0 | 0 |
| 0 | NA | NA | NA | NA | NA | Gm30893 | | 0 | 0 | 0 | 0 | 0 | 0 | 0 | 0 | 0 | 0 | 0 |
| 0 | NA | NA | NA | NA | NA | Gm48096 | | 0 | 0 | 0 | 0 | 0 | 0 | 0 | 0 | 0 | 0 | 0 |
| 0 | NA | NA | NA | NA | NA | Gm32526 | | 0 | 0 | 0 | 0 | 0 | 0 | 0 | 0 | 0 | 0 | 0 |
| 0 | NA | NA | NA | NA | NA | Gm25444 | | 0 | 0 | 0 | 0 | 0 | 0 | 0 | 0 | 0 | 0 | 0 |
| 0 | NA | NA | NA | NA | NA | Gm19189 | | 0 | 0 | 0 | 0 | 0 | 0 | 0 | 0 | 0 | 0 | 0 |
| 0 | NA | NA | NA | NA | NA | Gm48329 | | 0 | 0 | 0 | 0 | 0 | 0 | 0 | 0 | 0 | 0 | 0 |
| 0 | NA | NA | NA | NA | NA | Gm48343 | | 0 | 0 | 0 | 0 | 0 | 0 | 0 | 0 | 0 | 0 | 0 |
| 0 | NA | NA | NA | NA | NA | A530046M15Rik | | 0 | 0 | 0 | 0 | 0 | 0 | 0 | 0 | 0 | 0 | 0 |
| 0 | NA | NA | NA | NA | NA | A530058O07Rik | | 0 | 0 | 0 | 0 | 0 | 0 | 0 | 0 | 0 | 0 | 0 |
| 0 | NA | NA | NA | NA | NA | Gm31450 | | 0 | 0 | 0 | 0 | 0 | 0 | 0 | 0 | 0 | 0 | 0 |
| 0 | NA | NA | NA | NA | NA | Gm48444 | | 0 | 0 | 0 | 0 | 0 | 0 | 0 | 0 | 0 | 0 | 0 |
| 0 | NA | NA | NA | NA | NA | Gm22628 | | 0 | 0 | 0 | 0 | 0 | 0 | 0 | 0 | 0 | 0 | 0 |
| 0 | NA | NA | NA | NA | NA | Gm6562 | | 0 | 0 | 0 | 0 | 0 | 0 | 0 | 0 | 0 | 0 | 0 |
| 0 | NA | NA | NA | NA | NA | Gm31711 | | 0 | 0 | 0 | 0 | 0 | 0 | 0 | 0 | 0 | 0 | 0 |
| 0 | NA | NA | NA | NA | NA | Gm48491 | | 0 | 0 | 0 | 0 | 0 | 0 | 0 | 0 | 0 | 0 | 0 |
| 0 | NA | NA | NA | NA | NA | Gm48492 | | 0 | 0 | 0 | 0 | 0 | 0 | 0 | 0 | 0 | 0 | 0 |
| 0 | NA | NA | NA | NA | NA | Gm48493 | | 0 | 0 | 0 | 0 | 0 | 0 | 0 | 0 | 0 | 0 | 0 |
| 0 | NA | NA | NA | NA | NA | Gm48494 | | 0 | 0 | 0 | 0 | 0 | 0 | 0 | 0 | 0 | 0 | 0 |
| 0 | NA | NA | NA | NA | NA | Gm48497 | | 0 | 0 | 0 | 0 | 0 | 0 | 0 | 0 | 0 | 0 | 0 |
| 0 | NA | NA | NA | NA | NA | Gm7537 | | 0 | 0 | 0 | 0 | 0 | 0 | 0 | 0 | 0 | 0 | 0 |
| 0 | NA | NA | NA | NA | NA | Gm26098 | | 0 | 0 | 0 | 0 | 0 | 0 | 0 | 0 | 0 | 0 | 0 |
| 0 | NA | NA | NA | NA | NA | Gm48503 | | 0 | 0 | 0 | 0 | 0 | 0 | 0 | 0 | 0 | 0 | 0 |
| 0 | NA | NA | NA | NA | NA | Gm48499 | | 0 | 0 | 0 | 0 | 0 | 0 | 0 | 0 | 0 | 0 | 0 |
| 0 | NA | NA | NA | NA | NA | Gm27043 | | 0 | 0 | 0 | 0 | 0 | 0 | 0 | 0 | 0 | 0 | 0 |
| 0 | NA | NA | NA | NA | NA | Gm25020 | | 0 | 0 | 0 | 0 | 0 | 0 | 0 | 0 | 0 | 0 | 0 |
| 0 | NA | NA | NA | NA | NA | Gm48502 | | 0 | 0 | 0 | 0 | 0 | 0 | 0 | 0 | 0 | 0 | 0 |
| 0 | NA | NA | NA | NA | NA | Gm48501 | | 0 | 0 | 0 | 0 | 0 | 0 | 0 | 0 | 0 | 0 | 0 |
| 0 | NA | NA | NA | NA | NA | Mir466i | | 0 | 0 | 0 | 0 | 0 | 0 | 0 | 0 | 0 | 0 | 0 |
| 0 | NA | NA | NA | NA | NA | Gm24018 | | 0 | 0 | 0 | 0 | 0 | 0 | 0 | 0 | 0 | 0 | 0 |
| 0 | NA | NA | NA | NA | NA | Gm18362 | | 0 | 0 | 0 | 0 | 0 | 0 | 0 | 0 | 0 | 0 | 0 |
| 0 | NA | NA | NA | NA | NA | Gm48754 | | 0 | 0 | 0 | 0 | 0 | 0 | 0 | 0 | 0 | 0 | 0 |
| 0 | NA | NA | NA | NA | NA | Gm48776 | | 0 | 0 | 0 | 0 | 0 | 0 | 0 | 0 | 0 | 0 | 0 |
| 0 | NA | NA | NA | NA | NA | Gm48778 | | 0 | 0 | 0 | 0 | 0 | 0 | 0 | 0 | 0 | 0 | 0 |
| 0 | NA | NA | NA | NA | NA | Gm48799 | | 0 | 0 | 0 | 0 | 0 | 0 | 0 | 0 | 0 | 0 | 0 |
| 0 | NA | NA | NA | NA | NA | Gm6575 | | 0 | 0 | 0 | 0 | 0 | 0 | 0 | 0 | 0 | 0 | 0 |
| 0 | NA | NA | NA | NA | NA | 4930448F12Rik | | 0 | 0 | 0 | 0 | 0 | 0 | 0 | 0 | 0 | 0 | 0 |
| 0 | NA | NA | NA | NA | NA | Gm48800 | | 0 | 0 | 0 | 0 | 0 | 0 | 0 | 0 | 0 | 0 | 0 |
| 0 | NA | NA | NA | NA | NA | Gm48829 | | 0 | 0 | 0 | 0 | 0 | 0 | 0 | 0 | 0 | 0 | 0 |
| 0 | NA | NA | NA | NA | NA | Gm26172 | | 0 | 0 | 0 | 0 | 0 | 0 | 0 | 0 | 0 | 0 | 0 |

| | | | | | | | | | | | | | | | | | | |
|---|---|---|---|---|---|---|---|---|---|---|---|---|---|---|---|---|---|---|
| 0 | NA | NA | NA | NA | NA | Gm18065 | | 0 | 0 | 0 | 0 | 0 | 0 | 0 | 0 | 0 | 0 | 0 |
| 0 | NA | NA | NA | NA | NA | Gm47661 | | 0 | 0 | 0 | 0 | 0 | 0 | 0 | 0 | 0 | 0 | 0 |
| 0 | NA | NA | NA | NA | NA | Gm47660 | | 0 | 0 | 0 | 0 | 0 | 0 | 0 | 0 | 0 | 0 | 0 |
| 0 | NA | NA | NA | NA | NA | Gm31887 | | 0 | 0 | 0 | 0 | 0 | 0 | 0 | 0 | 0 | 0 | 0 |
| 0 | NA | NA | NA | NA | NA | Trgv7 | | 0 | 0 | 0 | 0 | 0 | 0 | 0 | 0 | 0 | 0 | 0 |
| 0 | NA | NA | NA | NA | NA | Trgv4 | | 0 | 0 | 0 | 0 | 0 | 0 | 0 | 0 | 0 | 0 | 0 |
| 0 | NA | NA | NA | NA | NA | Trgv6 | | 0 | 0 | 0 | 0 | 0 | 0 | 0 | 0 | 0 | 0 | 0 |
| 0 | NA | NA | NA | NA | NA | Trgv5 | | 0 | 0 | 0 | 0 | 0 | 0 | 0 | 0 | 0 | 0 | 0 |
| 0 | NA | NA | NA | NA | NA | Trgj1 | | 0 | 0 | 0 | 0 | 0 | 0 | 0 | 0 | 0 | 0 | 0 |
| 0 | NA | NA | NA | NA | NA | Trgc1 | | 0 | 0 | 0 | 0 | 0 | 0 | 0 | 0 | 0 | 0 | 0 |
| 0 | NA | NA | NA | NA | NA | Trgv3 | | 0 | 0 | 0 | 0 | 0 | 0 | 0 | 0 | 0 | 0 | 0 |
| 0 | NA | NA | NA | NA | NA | Trgj3 | | 0 | 0 | 0 | 0 | 0 | 0 | 0 | 0 | 0 | 0 | 0 |
| 0 | NA | NA | NA | NA | NA | Trgc3 | | 0 | 0 | 0 | 0 | 0 | 0 | 0 | 0 | 0 | 0 | 0 |
| 0 | NA | NA | NA | NA | NA | Trgc2 | | 0 | 0 | 0 | 0 | 0 | 0 | 0 | 0 | 0 | 0 | 0 |
| 0 | NA | NA | NA | NA | NA | Trgj2 | | 0 | 0 | 0 | 0 | 0 | 0 | 0 | 0 | 0 | 0 | 0 |
| 0 | NA | NA | NA | NA | NA | Trgv2 | | 0 | 0 | 0 | 0 | 0 | 0 | 0 | 0 | 0 | 0 | 0 |
| 0 | NA | NA | NA | NA | NA | Trgv1 | | 0 | 0 | 0 | 0 | 0 | 0 | 0 | 0 | 0 | 0 | 0 |
| 0 | NA | NA | NA | NA | NA | Trgj4 | | 0 | 0 | 0 | 0 | 0 | 0 | 0 | 0 | 0 | 0 | 0 |
| 0 | NA | NA | NA | NA | NA | Trgc4 | | 0 | 0 | 0 | 0 | 0 | 0 | 0 | 0 | 0 | 0 | 0 |
| 0 | NA | NA | NA | NA | NA | Gm18131 | | 0 | 0 | 0 | 0 | 0 | 0 | 0 | 0 | 0 | 0 | 0 |
| 0 | NA | NA | NA | NA | NA | 9330199G10Rik | | 0 | 0 | 0 | 0 | 0 | 0 | 0 | 0 | 0 | 0 | 0 |
| 0 | NA | NA | NA | NA | NA | Gm18363 | | 0 | 0 | 0 | 0 | 0 | 0 | 0 | 0 | 0 | 0 | 0 |
| 0 | NA | NA | NA | NA | NA | Nme8 | NME/NM23 family member 8 | 0 | 0 | 0 | 0 | 0 | 0 | 0 | 0 | 0 | 0 | 0 |
| 0 | NA | NA | NA | NA | NA | Gm47606 | | 0 | 0 | 0 | 0 | 0 | 0 | 0 | 0 | 0 | 0 | 0 |
| 0 | NA | NA | NA | NA | NA | Gm19154 | | 0 | 0 | 0 | 0 | 0 | 0 | 0 | 0 | 0 | 0 | 0 |
| 0 | NA | NA | NA | NA | NA | Gm32036 | | 0 | 0 | 0 | 0 | 0 | 0 | 0 | 0 | 0 | 0 | 0 |
| 0 | NA | NA | NA | NA | NA | Gm47727 | | 0 | 0 | 0 | 0 | 0 | 0 | 0 | 0 | 0 | 0 | 0 |
| 0 | NA | NA | NA | NA | NA | Gm25605 | | 0 | 0 | 0 | 0 | 0 | 0 | 0 | 0 | 0 | 0 | 0 |
| 0 | NA | NA | NA | NA | NA | Olfr1370 | olfactory receptor 1370 | 0 | 0 | 0 | 0 | 0 | 0 | 0 | 0 | 0 | 0 | 0 |
| 0 | NA | NA | NA | NA | NA | Olfr1369-ps1 | | 0 | 0 | 0 | 0 | 0 | 0 | 0 | 0 | 0 | 0 | 0 |
| 0 | NA | NA | NA | NA | NA | Olfr263 | olfactory receptor 263 | 0 | 0 | 0 | 0 | 0 | 0 | 0 | 0 | 0 | 0 | 0 |
| 0 | NA | NA | NA | NA | NA | Olfr1368 | olfactory receptor 1368 | 0 | 0 | 0 | 0 | 0 | 0 | 0 | 0 | 0 | 0 | 0 |
| 0 | NA | NA | NA | NA | NA | Gm47806 | | 0 | 0 | 0 | 0 | 0 | 0 | 0 | 0 | 0 | 0 | 0 |
| 0 | NA | NA | NA | NA | NA | Gpx5 | glutathione peroxidase 5 | 0 | 0 | 0 | 0 | 0 | 0 | 0 | 0 | 0 | 0 | 0 |
| 0 | NA | NA | NA | NA | NA | Gm11270 | | 0 | 0 | 0 | 0 | 0 | 0 | 0 | 0 | 0 | 0 | 0 |
| 0 | NA | NA | NA | NA | NA | Olfr1367 | olfactory receptor 1367 | 0 | 0 | 0 | 0 | 0 | 0 | 0 | 0 | 0 | 0 | 0 |
| 0 | NA | NA | NA | NA | NA | Gm11271 | | 0 | 0 | 0 | 0 | 0 | 0 | 0 | 0 | 0 | 0 | 0 |
| 0 | NA | NA | NA | NA | NA | Mir1896 | | 0 | 0 | 0 | 0 | 0 | 0 | 0 | 0 | 0 | 0 | 0 |
| 0 | NA | NA | NA | NA | NA | Gm23779 | | 0 | 0 | 0 | 0 | 0 | 0 | 0 | 0 | 0 | 0 | 0 |
| 0 | NA | NA | NA | NA | NA | Olfr1366 | olfactory receptor 1366 | 0 | 0 | 0 | 0 | 0 | 0 | 0 | 0 | 0 | 0 | 0 |
| 0 | NA | NA | NA | NA | NA | Olfr1535 | olfactory receptor 1535 | 0 | 0 | 0 | 0 | 0 | 0 | 0 | 0 | 0 | 0 | 0 |
| 0 | NA | NA | NA | NA | NA | Olfr1364 | olfactory receptor 1364 | 0 | 0 | 0 | 0 | 0 | 0 | 0 | 0 | 0 | 0 | 0 |
| 0 | NA | NA | NA | NA | NA | Olfr1362 | olfactory receptor 1362 | 0 | 0 | 0 | 0 | 0 | 0 | 0 | 0 | 0 | 0 | 0 |
| 0 | NA | NA | NA | NA | NA | Olfr11 | olfactory receptor 11 | 0 | 0 | 0 | 0 | 0 | 0 | 0 | 0 | 0 | 0 | 0 |
| 0 | NA | NA | NA | NA | NA | Olfr1361 | olfactory receptor 1361 | 0 | 0 | 0 | 0 | 0 | 0 | 0 | 0 | 0 | 0 | 0 |
| 0 | NA | NA | NA | NA | NA | Olfr1360 | olfactory receptor 1360 | 0 | 0 | 0 | 0 | 0 | 0 | 0 | 0 | 0 | 0 | 0 |
| 0 | NA | NA | NA | NA | NA | Olfr1359 | olfactory receptor 1359 | 0 | 0 | 0 | 0 | 0 | 0 | 0 | 0 | 0 | 0 | 0 |
| 0 | NA | NA | NA | NA | NA | Gm44367 | | 0 | 0 | 0 | 0 | 0 | 0 | 0 | 0 | 0 | 0 | 0 |
| 0 | NA | NA | NA | NA | NA | Gm44357 | | 0 | 0 | 0 | 0 | 0 | 0 | 0 | 0 | 0 | 0 | 0 |
| 0 | NA | NA | NA | NA | NA | H2ac14 | | 0 | 0 | 0 | 0 | 0 | 0 | 0 | 0 | 0 | 0 | 0 |
| 0 | NA | NA | NA | NA | NA | Gm44454 | | 0 | 0 | 0 | 0 | 0 | 0 | 0 | 0 | 0 | 0 | 0 |
| 0 | NA | NA | NA | NA | NA | Hist1h2an | histone cluster 1, H2an | 0 | 0 | 0 | 0 | 0 | 0 | 0 | 0 | 0 | 0 | 0 |
| 0 | NA | NA | NA | NA | NA | Hist1h2ao | histone cluster 1, H2ao | 0 | 0 | 0 | 0 | 0 | 0 | 0 | 0 | 0 | 0 | 0 |
| 0 | NA | NA | NA | NA | NA | Gm19658 | | 0 | 0 | 0 | 0 | 0 | 0 | 0 | 0 | 0 | 0 | 0 |
| 0 | NA | NA | NA | NA | NA | Hist1h2ap | histone cluster 1, H2ap | 0 | 0 | 0 | 0 | 0 | 0 | 0 | 0 | 0 | 0 | 0 |
| 0 | NA | NA | NA | NA | NA | Gm44466 | | 0 | 0 | 0 | 0 | 0 | 0 | 0 | 0 | 0 | 0 | 0 |
| 0 | NA | NA | NA | NA | NA | Hist1h2br | histone cluster 1 H2br | 0 | 0 | 0 | 0 | 0 | 0 | 0 | 0 | 0 | 0 | 0 |
| 0 | NA | NA | NA | NA | NA | Gm11278 | | 0 | 0 | 0 | 0 | 0 | 0 | 0 | 0 | 0 | 0 | 0 |
| 0 | NA | NA | NA | NA | NA | Mir1983 | | 0 | 0 | 0 | 0 | 0 | 0 | 0 | 0 | 0 | 0 | 0 |
| 0 | NA | NA | NA | NA | NA | Gm11279 | | 0 | 0 | 0 | 0 | 0 | 0 | 0 | 0 | 0 | 0 | 0 |

| | | | | | | | | | | | | | | | | | | |
|---|---|---|---|---|---|---|---|---|---|---|---|---|---|---|---|---|---|---|
| 0 | NA | NA | NA | NA | NA | Gm44456 | | 0 | 0 | 0 | 0 | 0 | 0 | 0 | 0 | 0 | 0 | 0 |
| 0 | NA | NA | NA | NA | NA | Gm11280 | | 0 | 0 | 0 | 0 | 0 | 0 | 0 | 0 | 0 | 0 | 0 |
| 0 | NA | NA | NA | NA | NA | Gm11282 | | 0 | 0 | 0 | 0 | 0 | 0 | 0 | 0 | 0 | 0 | 0 |
| 0 | NA | NA | NA | NA | NA | Pom121l2 | POM121 membrane glycoprotein-like 2 (r | 0 | 0 | 0 | 0 | 0 | 0 | 0 | 0 | 0 | 0 | 0 |
| 0 | NA | NA | NA | NA | NA | Gm11292 | | 0 | 0 | 0 | 0 | 0 | 0 | 0 | 0 | 0 | 0 | 0 |
| 0 | NA | NA | NA | NA | NA | Gm44390 | | 0 | 0 | 0 | 0 | 0 | 0 | 0 | 0 | 0 | 0 | 0 |
| 0 | NA | NA | NA | NA | NA | H2ac12 | H2A clustered histone 12 | 0 | 0 | 0 | 0 | 0 | 0 | 0 | 0 | 0 | 0 | 0 |
| 0 | NA | NA | NA | NA | NA | H2ac11 | H2A clustered histone 11 | 0 | 0 | 0 | 0 | 0 | 0 | 0 | 0 | 0 | 0 | 0 |
| 0 | NA | NA | NA | NA | NA | Gm44486 | | 0 | 0 | 0 | 0 | 0 | 0 | 0 | 0 | 0 | 0 | 0 |
| 0 | NA | NA | NA | NA | NA | Vmn1r-ps92 | | 0 | 0 | 0 | 0 | 0 | 0 | 0 | 0 | 0 | 0 | 0 |
| 0 | NA | NA | NA | NA | NA | Vmn1r-ps93 | | 0 | 0 | 0 | 0 | 0 | 0 | 0 | 0 | 0 | 0 | 0 |
| 0 | NA | NA | NA | NA | NA | Gm26300 | | 0 | 0 | 0 | 0 | 0 | 0 | 0 | 0 | 0 | 0 | 0 |
| 0 | NA | NA | NA | NA | NA | Vmn1r188 | vomeronasal 1 receptor 188 | 0 | 0 | 0 | 0 | 0 | 0 | 0 | 0 | 0 | 0 | 0 |
| 0 | NA | NA | NA | NA | NA | Vmn1r-ps94 | | 0 | 0 | 0 | 0 | 0 | 0 | 0 | 0 | 0 | 0 | 0 |
| 0 | NA | NA | NA | NA | NA | Vmn1r189 | vomeronasal 1 receptor 189 | 0 | 0 | 0 | 0 | 0 | 0 | 0 | 0 | 0 | 0 | 0 |
| 0 | NA | NA | NA | NA | NA | Vmn1r-ps95 | | 0 | 0 | 0 | 0 | 0 | 0 | 0 | 0 | 0 | 0 | 0 |
| 0 | NA | NA | NA | NA | NA | Vmn1r-ps96 | | 0 | 0 | 0 | 0 | 0 | 0 | 0 | 0 | 0 | 0 | 0 |
| 0 | NA | NA | NA | NA | NA | Vmn1r-ps97 | | 0 | 0 | 0 | 0 | 0 | 0 | 0 | 0 | 0 | 0 | 0 |
| 0 | NA | NA | NA | NA | NA | Vmn1r190-ps | | 0 | 0 | 0 | 0 | 0 | 0 | 0 | 0 | 0 | 0 | 0 |
| 0 | NA | NA | NA | NA | NA | Vmn1r191 | vomeronasal 1 receptor 191 | 0 | 0 | 0 | 0 | 0 | 0 | 0 | 0 | 0 | 0 | 0 |
| 0 | NA | NA | NA | NA | NA | Vmn1r-ps99 | | 0 | 0 | 0 | 0 | 0 | 0 | 0 | 0 | 0 | 0 | 0 |
| 0 | NA | NA | NA | NA | NA | Vmn1r192 | vomeronasal 1 receptor 192 | 0 | 0 | 0 | 0 | 0 | 0 | 0 | 0 | 0 | 0 | 0 |
| 0 | NA | NA | NA | NA | NA | Gm23282 | | 0 | 0 | 0 | 0 | 0 | 0 | 0 | 0 | 0 | 0 | 0 |
| 0 | NA | NA | NA | NA | NA | Vmn1r193 | vomeronasal 1 receptor 193 | 0 | 0 | 0 | 0 | 0 | 0 | 0 | 0 | 0 | 0 | 0 |
| 0 | NA | NA | NA | NA | NA | Vmn1r194 | vomeronasal 1 receptor 194 | 0 | 0 | 0 | 0 | 0 | 0 | 0 | 0 | 0 | 0 | 0 |
| 0 | NA | NA | NA | NA | NA | Vmn1r195 | vomeronasal 1 receptor 195 | 0 | 0 | 0 | 0 | 0 | 0 | 0 | 0 | 0 | 0 | 0 |
| 0 | NA | NA | NA | NA | NA | Vmn1r196 | vomeronasal 1 receptor 196 | 0 | 0 | 0 | 0 | 0 | 0 | 0 | 0 | 0 | 0 | 0 |
| 0 | NA | NA | NA | NA | NA | Vmn1r-ps100 | | 0 | 0 | 0 | 0 | 0 | 0 | 0 | 0 | 0 | 0 | 0 |
| 0 | NA | NA | NA | NA | NA | Vmn1r197 | vomeronasal 1 receptor 197 | 0 | 0 | 0 | 0 | 0 | 0 | 0 | 0 | 0 | 0 | 0 |
| 0 | NA | NA | NA | NA | NA | Gm48489 | | 0 | 0 | 0 | 0 | 0 | 0 | 0 | 0 | 0 | 0 | 0 |
| 0 | NA | NA | NA | NA | NA | Vmn1r198 | vomeronasal 1 receptor 198 | 0 | 0 | 0 | 0 | 0 | 0 | 0 | 0 | 0 | 0 | 0 |
| 0 | NA | NA | NA | NA | NA | Vmn1r-ps101 | | 0 | 0 | 0 | 0 | 0 | 0 | 0 | 0 | 0 | 0 | 0 |
| 0 | NA | NA | NA | NA | NA | Vmn1r199 | vomeronasal 1 receptor 199 | 0 | 0 | 0 | 0 | 0 | 0 | 0 | 0 | 0 | 0 | 0 |
| 0 | NA | NA | NA | NA | NA | Vmn1r-ps102 | | 0 | 0 | 0 | 0 | 0 | 0 | 0 | 0 | 0 | 0 | 0 |
| 0 | NA | NA | NA | NA | NA | Vmn1r200 | vomeronasal 1 receptor 200 | 0 | 0 | 0 | 0 | 0 | 0 | 0 | 0 | 0 | 0 | 0 |
| 0 | NA | NA | NA | NA | NA | Vmn1r-ps103 | vomeronasal 1 receptor, pseudogene 10: | 0 | 0 | 0 | 0 | 0 | 0 | 0 | 0 | 0 | 0 | 0 |
| 0 | NA | NA | NA | NA | NA | Vmn1r-ps104 | | 0 | 0 | 0 | 0 | 0 | 0 | 0 | 0 | 0 | 0 | 0 |
| 0 | NA | NA | NA | NA | NA | Gm11298 | | 0 | 0 | 0 | 0 | 0 | 0 | 0 | 0 | 0 | 0 | 0 |
| 0 | NA | NA | NA | NA | NA | Vmn1r201 | vomeronasal 1 receptor 201 | 0 | 0 | 0 | 0 | 0 | 0 | 0 | 0 | 0 | 0 | 0 |
| 0 | NA | NA | NA | NA | NA | Vmn1r-ps106 | | 0 | 0 | 0 | 0 | 0 | 0 | 0 | 0 | 0 | 0 | 0 |
| 0 | NA | NA | NA | NA | NA | Vmn1r-ps107 | | 0 | 0 | 0 | 0 | 0 | 0 | 0 | 0 | 0 | 0 | 0 |
| 0 | NA | NA | NA | NA | NA | Vmn1r-ps108 | | 0 | 0 | 0 | 0 | 0 | 0 | 0 | 0 | 0 | 0 | 0 |
| 0 | NA | NA | NA | NA | NA | Gm11300 | | 0 | 0 | 0 | 0 | 0 | 0 | 0 | 0 | 0 | 0 | 0 |
| 0 | NA | NA | NA | NA | NA | Vmn1r204 | vomeronasal 1 receptor 204 | 0 | 0 | 0 | 0 | 0 | 0 | 0 | 0 | 0 | 0 | 0 |
| 0 | NA | NA | NA | NA | NA | Vmn1r205 | vomeronasal 1 receptor 205 | 0 | 0 | 0 | 0 | 0 | 0 | 0 | 0 | 0 | 0 | 0 |
| 0 | NA | NA | NA | NA | NA | Vmn1r206 | vomeronasal 1 receptor 206 | 0 | 0 | 0 | 0 | 0 | 0 | 0 | 0 | 0 | 0 | 0 |
| 0 | NA | NA | NA | NA | NA | Vmn1r-ps109 | | 0 | 0 | 0 | 0 | 0 | 0 | 0 | 0 | 0 | 0 | 0 |
| 0 | NA | NA | NA | NA | NA | Vmn1r-ps110 | | 0 | 0 | 0 | 0 | 0 | 0 | 0 | 0 | 0 | 0 | 0 |
| 0 | NA | NA | NA | NA | NA | Vmn1r-ps111 | | 0 | 0 | 0 | 0 | 0 | 0 | 0 | 0 | 0 | 0 | 0 |
| 0 | NA | NA | NA | NA | NA | Gm11309 | | 0 | 0 | 0 | 0 | 0 | 0 | 0 | 0 | 0 | 0 | 0 |
| 0 | NA | NA | NA | NA | NA | Vmn1r207 | vomeronasal 1 receptor 207 | 0 | 0 | 0 | 0 | 0 | 0 | 0 | 0 | 0 | 0 | 0 |
| 0 | NA | NA | NA | NA | NA | Vmn1r-ps112 | | 0 | 0 | 0 | 0 | 0 | 0 | 0 | 0 | 0 | 0 | 0 |
| 0 | NA | NA | NA | NA | NA | Vmn1r-ps113 | | 0 | 0 | 0 | 0 | 0 | 0 | 0 | 0 | 0 | 0 | 0 |
| 0 | NA | NA | NA | NA | NA | Gm11313 | | 0 | 0 | 0 | 0 | 0 | 0 | 0 | 0 | 0 | 0 | 0 |
| 0 | NA | NA | NA | NA | NA | Vmn1r208 | vomeronasal 1 receptor 208 | 0 | 0 | 0 | 0 | 0 | 0 | 0 | 0 | 0 | 0 | 0 |
| 0 | NA | NA | NA | NA | NA | Gm11314 | | 0 | 0 | 0 | 0 | 0 | 0 | 0 | 0 | 0 | 0 | 0 |
| 0 | NA | NA | NA | NA | NA | Vmn1r209 | vomeronasal 1 receptor 209 | 0 | 0 | 0 | 0 | 0 | 0 | 0 | 0 | 0 | 0 | 0 |
| 0 | NA | NA | NA | NA | NA | Vmn1r210 | vomeronasal 1 receptor 210 | 0 | 0 | 0 | 0 | 0 | 0 | 0 | 0 | 0 | 0 | 0 |
| 0 | NA | NA | NA | NA | NA | Vmn1r-ps114 | | 0 | 0 | 0 | 0 | 0 | 0 | 0 | 0 | 0 | 0 | 0 |
| 0 | NA | NA | NA | NA | NA | Vmn1r211 | vomeronasal 1 receptor 211 | 0 | 0 | 0 | 0 | 0 | 0 | 0 | 0 | 0 | 0 | 0 |

| | | | | | | | | | | | | | | | | | | |
|---|---|---|---|---|---|---|---|---|---|---|---|---|---|---|---|---|---|---|
| 0 | NA | NA | NA | NA | NA | Vmn1r-ps116 | | 0 | 0 | 0 | 0 | 0 | 0 | 0 | 0 | 0 | 0 | 0 |
| 0 | NA | NA | NA | NA | NA | Gm23774 | | 0 | 0 | 0 | 0 | 0 | 0 | 0 | 0 | 0 | 0 | 0 |
| 0 | NA | NA | NA | NA | NA | Vmn1r212 | vomeronasal 1 receptor 212 | 0 | 0 | 0 | 0 | 0 | 0 | 0 | 0 | 0 | 0 | 0 |
| 0 | NA | NA | NA | NA | NA | Vmn1r-ps117 | | 0 | 0 | 0 | 0 | 0 | 0 | 0 | 0 | 0 | 0 | 0 |
| 0 | NA | NA | NA | NA | NA | Gm11320 | | 0 | 0 | 0 | 0 | 0 | 0 | 0 | 0 | 0 | 0 | 0 |
| 0 | NA | NA | NA | NA | NA | Gm11321 | | 0 | 0 | 0 | 0 | 0 | 0 | 0 | 0 | 0 | 0 | 0 |
| 0 | NA | NA | NA | NA | NA | Vmn1r-ps120 | | 0 | 0 | 0 | 0 | 0 | 0 | 0 | 0 | 0 | 0 | 0 |
| 0 | NA | NA | NA | NA | NA | Gm11322 | | 0 | 0 | 0 | 0 | 0 | 0 | 0 | 0 | 0 | 0 | 0 |
| 0 | NA | NA | NA | NA | NA | Gm11318 | | 0 | 0 | 0 | 0 | 0 | 0 | 0 | 0 | 0 | 0 | 0 |
| 0 | NA | NA | NA | NA | NA | Vmn1r-ps123 | | 0 | 0 | 0 | 0 | 0 | 0 | 0 | 0 | 0 | 0 | 0 |
| 0 | NA | NA | NA | NA | NA | Vmn1r213 | vomeronasal 1 receptor 213 | 0 | 0 | 0 | 0 | 0 | 0 | 0 | 0 | 0 | 0 | 0 |
| 0 | NA | NA | NA | NA | NA | Vmn1r-ps125 | | 0 | 0 | 0 | 0 | 0 | 0 | 0 | 0 | 0 | 0 | 0 |
| 0 | NA | NA | NA | NA | NA | Vmn1r214 | vomeronasal 1 receptor 214 | 0 | 0 | 0 | 0 | 0 | 0 | 0 | 0 | 0 | 0 | 0 |
| 0 | NA | NA | NA | NA | NA | Gm19153 | | 0 | 0 | 0 | 0 | 0 | 0 | 0 | 0 | 0 | 0 | 0 |
| 0 | NA | NA | NA | NA | NA | Vmn1r-ps126 | | 0 | 0 | 0 | 0 | 0 | 0 | 0 | 0 | 0 | 0 | 0 |
| 0 | NA | NA | NA | NA | NA | Vmn1r215 | vomeronasal 1 receptor 215 | 0 | 0 | 0 | 0 | 0 | 0 | 0 | 0 | 0 | 0 | 0 |
| 0 | NA | NA | NA | NA | NA | Vmn1r216 | | 0 | 0 | 0 | 0 | 0 | 0 | 0 | 0 | 0 | 0 | 0 |
| 0 | NA | NA | NA | NA | NA | Vmn1r216 | vomeronasal 1 receptor 216 | 0 | 0 | 0 | 0 | 0 | 0 | 0 | 0 | 0 | 0 | 0 |
| 0 | NA | NA | NA | NA | NA | Gm11325 | | 0 | 0 | 0 | 0 | 0 | 0 | 0 | 0 | 0 | 0 | 0 |
| 0 | NA | NA | NA | NA | NA | Vmn1r-ps127 | | 0 | 0 | 0 | 0 | 0 | 0 | 0 | 0 | 0 | 0 | 0 |
| 0 | NA | NA | NA | NA | NA | Vmn1r217 | vomeronasal 1 receptor 217 | 0 | 0 | 0 | 0 | 0 | 0 | 0 | 0 | 0 | 0 | 0 |
| 0 | NA | NA | NA | NA | NA | Vmn1r-ps128 | | 0 | 0 | 0 | 0 | 0 | 0 | 0 | 0 | 0 | 0 | 0 |
| 0 | NA | NA | NA | NA | NA | Vmn1r218 | vomeronasal 1 receptor 218 | 0 | 0 | 0 | 0 | 0 | 0 | 0 | 0 | 0 | 0 | 0 |
| 0 | NA | NA | NA | NA | NA | Vmn1r219 | vomeronasal 1 receptor 219 | 0 | 0 | 0 | 0 | 0 | 0 | 0 | 0 | 0 | 0 | 0 |
| 0 | NA | NA | NA | NA | NA | Vmn1r220 | vomeronasal 1 receptor 220 | 0 | 0 | 0 | 0 | 0 | 0 | 0 | 0 | 0 | 0 | 0 |
| 0 | NA | NA | NA | NA | NA | Vmn1r-ps131 | | 0 | 0 | 0 | 0 | 0 | 0 | 0 | 0 | 0 | 0 | 0 |
| 0 | NA | NA | NA | NA | NA | Vmn1r-ps132 | | 0 | 0 | 0 | 0 | 0 | 0 | 0 | 0 | 0 | 0 | 0 |
| 0 | NA | NA | NA | NA | NA | Vmn1r221 | vomeronasal 1 receptor 221 | 0 | 0 | 0 | 0 | 0 | 0 | 0 | 0 | 0 | 0 | 0 |
| 0 | NA | NA | NA | NA | NA | Vmn1r-ps133 | | 0 | 0 | 0 | 0 | 0 | 0 | 0 | 0 | 0 | 0 | 0 |
| 0 | NA | NA | NA | NA | NA | Vmn1r222 | vomeronasal 1 receptor 222 | 0 | 0 | 0 | 0 | 0 | 0 | 0 | 0 | 0 | 0 | 0 |
| 0 | NA | NA | NA | NA | NA | Vmn1r223 | vomeronasal 1 receptor 223 | 0 | 0 | 0 | 0 | 0 | 0 | 0 | 0 | 0 | 0 | 0 |
| 0 | NA | NA | NA | NA | NA | Gm11331 | | 0 | 0 | 0 | 0 | 0 | 0 | 0 | 0 | 0 | 0 | 0 |
| 0 | NA | NA | NA | NA | NA | Vmn1r-ps135 | | 0 | 0 | 0 | 0 | 0 | 0 | 0 | 0 | 0 | 0 | 0 |
| 0 | NA | NA | NA | NA | NA | 4930557F10Rik | | 0 | 0 | 0 | 0 | 0 | 0 | 0 | 0 | 0 | 0 | 0 |
| 0 | NA | NA | NA | NA | NA | Gm11333 | | 0 | 0 | 0 | 0 | 0 | 0 | 0 | 0 | 0 | 0 | 0 |
| 0 | NA | NA | NA | NA | NA | Gm11334 | | 0 | 0 | 0 | 0 | 0 | 0 | 0 | 0 | 0 | 0 | 0 |
| 0 | NA | NA | NA | NA | NA | n-TStga1 | | 0 | 0 | 0 | 0 | 0 | 0 | 0 | 0 | 0 | 0 | 0 |
| 0 | NA | NA | NA | NA | NA | Gm44297 | | 0 | 0 | 0 | 0 | 0 | 0 | 0 | 0 | 0 | 0 | 0 |
| 0 | NA | NA | NA | NA | NA | Gm44359 | | 0 | 0 | 0 | 0 | 0 | 0 | 0 | 0 | 0 | 0 | 0 |
| 0 | NA | NA | NA | NA | NA | Gm11398 | | 0 | 0 | 0 | 0 | 0 | 0 | 0 | 0 | 0 | 0 | 0 |
| 0 | NA | NA | NA | NA | NA | Gm22452 | | 0 | 0 | 0 | 0 | 0 | 0 | 0 | 0 | 0 | 0 | 0 |
| 0 | NA | NA | NA | NA | NA | Gm24991 | | 0 | 0 | 0 | 0 | 0 | 0 | 0 | 0 | 0 | 0 | 0 |
| 0 | NA | NA | NA | NA | NA | 4930558J22Rik | | 0 | 0 | 0 | 0 | 0 | 0 | 0 | 0 | 0 | 0 | 0 |
| 0 | NA | NA | NA | NA | NA | Gm11337 | | 0 | 0 | 0 | 0 | 0 | 0 | 0 | 0 | 0 | 0 | 0 |
| 0 | NA | NA | NA | NA | NA | Slc17a3 | solute carrier family 17 (sodium phosphat | 0 | 0 | 0 | 0 | 0 | 0 | 0 | 0 | 0 | 0 | 0 |
| 0 | NA | NA | NA | NA | NA | Slc17a1 | solute carrier family 17 (sodium phosphat | 0 | 0 | 0 | 0 | 0 | 0 | 0 | 0 | 0 | 0 | 0 |
| 0 | NA | NA | NA | NA | NA | Snrp1c-ps2 | | 0 | 0 | 0 | 0 | 0 | 0 | 0 | 0 | 0 | 0 | 0 |
| 0 | NA | NA | NA | NA | NA | H2ac1 | H2A clustered histone 1 | 0 | 0 | 0 | 0 | 0 | 0 | 0 | 0 | 0 | 0 | 0 |
| 0 | NA | NA | NA | NA | NA | Gm44344 | | 0 | 0 | 0 | 0 | 0 | 0 | 0 | 0 | 0 | 0 | 0 |
| 0 | NA | NA | NA | NA | NA | Gm11339 | | 0 | 0 | 0 | 0 | 0 | 0 | 0 | 0 | 0 | 0 | 0 |
| 0 | NA | NA | NA | NA | NA | Gm11340 | | 0 | 0 | 0 | 0 | 0 | 0 | 0 | 0 | 0 | 0 | 0 |
| 0 | NA | NA | NA | NA | NA | Gm23340 | | 0 | 0 | 0 | 0 | 0 | 0 | 0 | 0 | 0 | 0 | 0 |
| 0 | NA | NA | NA | NA | NA | Gm11341 | | 0 | 0 | 0 | 0 | 0 | 0 | 0 | 0 | 0 | 0 | 0 |
| 0 | NA | NA | NA | NA | NA | | | 0 | 0 | 0 | 0 | 0 | 0 | 0 | 0 | 0 | 0 | 0 |
| 0 | NA | NA | NA | NA | NA | Gm11342 | | 0 | 0 | 0 | 0 | 0 | 0 | 0 | 0 | 0 | 0 | 0 |
| 0 | NA | NA | NA | NA | NA | Gm11344 | | 0 | 0 | 0 | 0 | 0 | 0 | 0 | 0 | 0 | 0 | 0 |
| 0 | NA | NA | NA | NA | NA | Gm22013 | | 0 | 0 | 0 | 0 | 0 | 0 | 0 | 0 | 0 | 0 | 0 |
| 0 | NA | NA | NA | NA | NA | Armh2 | armadillo-like helical domain containing 2 | 0 | 0 | 0 | 0 | 0 | 0 | 0 | 0 | 0 | 0 | 0 |
| 0 | NA | NA | NA | NA | NA | Gm22358 | | 0 | 0 | 0 | 0 | 0 | 0 | 0 | 0 | 0 | 0 | 0 |
| 0 | NA | NA | NA | NA | NA | Gm47605 | | 0 | 0 | 0 | 0 | 0 | 0 | 0 | 0 | 0 | 0 | 0 |

| | | | | | | | | | | | | | | | | | | |
|---|---|---|---|---|---|---|---|---|---|---|---|---|---|---|---|---|---|---|
| 0 | NA | NA | NA | NA | NA | Gm11348 | | 0 | 0 | 0 | 0 | 0 | 0 | 0 | 0 | 0 | 0 | 0 |
| 0 | NA | NA | NA | NA | NA | F830002E08Rik | | 0 | 0 | 0 | 0 | 0 | 0 | 0 | 0 | 0 | 0 | 0 |
| 0 | NA | NA | NA | NA | NA | Tpi-rs8 | | 0 | 0 | 0 | 0 | 0 | 0 | 0 | 0 | 0 | 0 | 0 |
| 0 | NA | NA | NA | NA | NA | Gm11350 | | 0 | 0 | 0 | 0 | 0 | 0 | 0 | 0 | 0 | 0 | 0 |
| 0 | NA | NA | NA | NA | NA | Gm11351 | | 0 | 0 | 0 | 0 | 0 | 0 | 0 | 0 | 0 | 0 | 0 |
| 0 | NA | NA | NA | NA | NA | Gm11352 | | 0 | 0 | 0 | 0 | 0 | 0 | 0 | 0 | 0 | 0 | 0 |
| 0 | NA | NA | NA | NA | NA | 1700092E19Rik | | 0 | 0 | 0 | 0 | 0 | 0 | 0 | 0 | 0 | 0 | 0 |
| 0 | NA | NA | NA | NA | NA | Gm47881 | | 0 | 0 | 0 | 0 | 0 | 0 | 0 | 0 | 0 | 0 | 0 |
| 0 | NA | NA | NA | NA | NA | Gm47883 | | 0 | 0 | 0 | 0 | 0 | 0 | 0 | 0 | 0 | 0 | 0 |
| 0 | NA | NA | NA | NA | NA | Hdgfl1 | HDGF like 1 | 0 | 0 | 0 | 0 | 0 | 0 | 0 | 0 | 0 | 0 | 0 |
| 0 | NA | NA | NA | NA | NA | 4921520N01Rik | | 0 | 0 | 0 | 0 | 0 | 0 | 0 | 0 | 0 | 0 | 0 |
| 0 | NA | NA | NA | NA | NA | Gm11354 | | 0 | 0 | 0 | 0 | 0 | 0 | 0 | 0 | 0 | 0 | 0 |
| 0 | NA | NA | NA | NA | NA | Gm11355 | | 0 | 0 | 0 | 0 | 0 | 0 | 0 | 0 | 0 | 0 | 0 |
| 0 | NA | NA | NA | NA | NA | Gm47888 | | 0 | 0 | 0 | 0 | 0 | 0 | 0 | 0 | 0 | 0 | 0 |
| 0 | NA | NA | NA | NA | NA | Prl | prolactin | 0 | 0 | 0 | 0 | 0 | 0 | 0 | 0 | 0 | 0 | 0 |
| 0 | NA | NA | NA | NA | NA | Prl3d1 | prolactin family 3, subfamily d, member 1 | 0 | 0 | 0 | 0 | 0 | 0 | 0 | 0 | 0 | 0 | 0 |
| 0 | NA | NA | NA | NA | NA | Prl3d2 | prolactin family 3, subfamily d, member 1 | 0 | 0 | 0 | 0 | 0 | 0 | 0 | 0 | 0 | 0 | 0 |
| 0 | NA | NA | NA | NA | NA | Prl3d3 | prolactin family 3, subfamily d, member 3 | 0 | 0 | 0 | 0 | 0 | 0 | 0 | 0 | 0 | 0 | 0 |
| 0 | NA | NA | NA | NA | NA | Prl3c1 | prolactin family 3, subfamily c, member 1 | 0 | 0 | 0 | 0 | 0 | 0 | 0 | 0 | 0 | 0 | 0 |
| 0 | NA | NA | NA | NA | NA | Prl3b1 | prolactin family 3, subfamily b, member 1 | 0 | 0 | 0 | 0 | 0 | 0 | 0 | 0 | 0 | 0 | 0 |
| 0 | NA | NA | NA | NA | NA | Prl3a1 | prolactin family 3, subfamily a, member 1 | 0 | 0 | 0 | 0 | 0 | 0 | 0 | 0 | 0 | 0 | 0 |
| 0 | NA | NA | NA | NA | NA | 4930511O05Rik | | 0 | 0 | 0 | 0 | 0 | 0 | 0 | 0 | 0 | 0 | 0 |
| 0 | NA | NA | NA | NA | NA | | | 0 | 0 | 0 | 0 | 0 | 0 | 0 | 0 | 0 | 0 | 0 |
| 0 | NA | NA | NA | NA | NA | Gm11356 | | 0 | 0 | 0 | 0 | 0 | 0 | 0 | 0 | 0 | 0 | 0 |
| 0 | NA | NA | NA | NA | NA | Prl6a1 | prolactin family 6, subfamily a, member 1 | 0 | 0 | 0 | 0 | 0 | 0 | 0 | 0 | 0 | 0 | 0 |
| 0 | NA | NA | NA | NA | NA | Gm11357 | | 0 | 0 | 0 | 0 | 0 | 0 | 0 | 0 | 0 | 0 | 0 |
| 0 | NA | NA | NA | NA | NA | Gm11358 | | 0 | 0 | 0 | 0 | 0 | 0 | 0 | 0 | 0 | 0 | 0 |
| 0 | NA | NA | NA | NA | NA | Prl8a2 | prolactin family 8, subfamily a, member 2 | 0 | 0 | 0 | 0 | 0 | 0 | 0 | 0 | 0 | 0 | 0 |
| 0 | NA | NA | NA | NA | NA | Prl2b1 | prolactin family 2, subfamily b, member 1 | 0 | 0 | 0 | 0 | 0 | 0 | 0 | 0 | 0 | 0 | 0 |
| 0 | NA | NA | NA | NA | NA | Prl8a6 | prolactin family 8, subfamily a, member 6 | 0 | 0 | 0 | 0 | 0 | 0 | 0 | 0 | 0 | 0 | 0 |
| 0 | NA | NA | NA | NA | NA | Prl8a8 | prolactin family 8, subfamily a, member 8 | 0 | 0 | 0 | 0 | 0 | 0 | 0 | 0 | 0 | 0 | 0 |
| 0 | NA | NA | NA | NA | NA | Prl8a9 | prolactin family8, subfamily a, member 9 | 0 | 0 | 0 | 0 | 0 | 0 | 0 | 0 | 0 | 0 | 0 |
| 0 | NA | NA | NA | NA | NA | Prl8a1 | prolactin family 8, subfamily a, member 1 | 0 | 0 | 0 | 0 | 0 | 0 | 0 | 0 | 0 | 0 | 0 |
| 0 | NA | NA | NA | NA | NA | Prl7b1 | prolactin family 7, subfamily b, member 1 | 0 | 0 | 0 | 0 | 0 | 0 | 0 | 0 | 0 | 0 | 0 |
| 0 | NA | NA | NA | NA | NA | Prl7a1 | prolactin family 7, subfamily a, member 1 | 0 | 0 | 0 | 0 | 0 | 0 | 0 | 0 | 0 | 0 | 0 |
| 0 | NA | NA | NA | NA | NA | Prl7a2 | prolactin family 7, subfamily a, member 2 | 0 | 0 | 0 | 0 | 0 | 0 | 0 | 0 | 0 | 0 | 0 |
| 0 | NA | NA | NA | NA | NA | Gm11359 | | 0 | 0 | 0 | 0 | 0 | 0 | 0 | 0 | 0 | 0 | 0 |
| 0 | NA | NA | NA | NA | NA | Prl7d1 | prolactin family 7, subfamily d, member 1 | 0 | 0 | 0 | 0 | 0 | 0 | 0 | 0 | 0 | 0 | 0 |
| 0 | NA | NA | NA | NA | NA | Prl7c1 | prolactin family 7, subfamily c, member 1 | 0 | 0 | 0 | 0 | 0 | 0 | 0 | 0 | 0 | 0 | 0 |
| 0 | NA | NA | NA | NA | NA | Prl2a1 | prolactin family 2, subfamily a, member 1 | 0 | 0 | 0 | 0 | 0 | 0 | 0 | 0 | 0 | 0 | 0 |
| 0 | NA | NA | NA | NA | NA | Prl2c1 | Prolactin family 2, subfamily c, member 1 | 0 | 0 | 0 | 0 | 0 | 0 | 0 | 0 | 0 | 0 | 0 |
| 0 | NA | NA | NA | NA | NA | Gm22476 | | 0 | 0 | 0 | 0 | 0 | 0 | 0 | 0 | 0 | 0 | 0 |
| 0 | NA | NA | NA | NA | NA | Gm34165 | | 0 | 0 | 0 | 0 | 0 | 0 | 0 | 0 | 0 | 0 | 0 |
| 0 | NA | NA | NA | NA | NA | Gm11360 | | 0 | 0 | 0 | 0 | 0 | 0 | 0 | 0 | 0 | 0 | 0 |
| 0 | NA | NA | NA | NA | NA | Gm18881 | | 0 | 0 | 0 | 0 | 0 | 0 | 0 | 0 | 0 | 0 | 0 |
| 0 | NA | NA | NA | NA | NA | Prl4a1 | prolactin family 4, subfamily a, member 1 | 0 | 0 | 0 | 0 | 0 | 0 | 0 | 0 | 0 | 0 | 0 |
| 0 | NA | NA | NA | NA | NA | Gm47019 | | 0 | 0 | 0 | 0 | 0 | 0 | 0 | 0 | 0 | 0 | 0 |
| 0 | NA | NA | NA | NA | NA | Prl5a1 | prolactin family 5, subfamily a, member 1 | 0 | 0 | 0 | 0 | 0 | 0 | 0 | 0 | 0 | 0 | 0 |
| 0 | NA | NA | NA | NA | NA | Gm47172 | | 0 | 0 | 0 | 0 | 0 | 0 | 0 | 0 | 0 | 0 | 0 |
| 0 | NA | NA | NA | NA | NA | Gm40841 | | 0 | 0 | 0 | 0 | 0 | 0 | 0 | 0 | 0 | 0 | 0 |
| 0 | NA | NA | NA | NA | NA | Gm6081 | | 0 | 0 | 0 | 0 | 0 | 0 | 0 | 0 | 0 | 0 | 0 |
| 0 | NA | NA | NA | NA | NA | Mir6368 | | 0 | 0 | 0 | 0 | 0 | 0 | 0 | 0 | 0 | 0 | 0 |
| 0 | NA | NA | NA | NA | NA | Gm17528 | | 0 | 0 | 0 | 0 | 0 | 0 | 0 | 0 | 0 | 0 | 0 |
| 0 | NA | NA | NA | NA | NA | Gm34639 | | 0 | 0 | 0 | 0 | 0 | 0 | 0 | 0 | 0 | 0 | 0 |
| 0 | NA | NA | NA | NA | NA | Gm11364 | | 0 | 0 | 0 | 0 | 0 | 0 | 0 | 0 | 0 | 0 | 0 |
| 0 | NA | NA | NA | NA | NA | Gm11365 | | 0 | 0 | 0 | 0 | 0 | 0 | 0 | 0 | 0 | 0 | 0 |
| 0 | NA | NA | NA | NA | NA | Gm11368 | | 0 | 0 | 0 | 0 | 0 | 0 | 0 | 0 | 0 | 0 | 0 |
| 0 | NA | NA | NA | NA | NA | Gm47259 | | 0 | 0 | 0 | 0 | 0 | 0 | 0 | 0 | 0 | 0 | 0 |
| 0 | NA | NA | NA | NA | NA | Gm23206 | | 0 | 0 | 0 | 0 | 0 | 0 | 0 | 0 | 0 | 0 | 0 |
| 0 | NA | NA | NA | NA | NA | Gm11366 | | 0 | 0 | 0 | 0 | 0 | 0 | 0 | 0 | 0 | 0 | 0 |

| | | | | | | | | | | | | | | | | | | |
|---|---|---|---|---|---|---|---|---|---|---|---|---|---|---|---|---|---|---|
| 0 | NA | NA | NA | NA | NA | Gm11369 | | 0 | 0 | 0 | 0 | 0 | 0 | 0 | 0 | 0 | 0 | 0 |
| 0 | NA | NA | NA | NA | NA | 4930519D14Rik | | 0 | 0 | 0 | 0 | 0 | 0 | 0 | 0 | 0 | 0 | 0 |
| 0 | NA | NA | NA | NA | NA | Gm11370 | | 0 | 0 | 0 | 0 | 0 | 0 | 0 | 0 | 0 | 0 | 0 |
| 0 | NA | NA | NA | NA | NA | Gm11371 | | 0 | 0 | 0 | 0 | 0 | 0 | 0 | 0 | 0 | 0 | 0 |
| 0 | NA | NA | NA | NA | NA | Gm48662 | | 0 | 0 | 0 | 0 | 0 | 0 | 0 | 0 | 0 | 0 | 0 |
| 0 | NA | NA | NA | NA | NA | Gm11372 | | 0 | 0 | 0 | 0 | 0 | 0 | 0 | 0 | 0 | 0 | 0 |
| 0 | NA | NA | NA | NA | NA | 4930401O12Rik | | 0 | 0 | 0 | 0 | 0 | 0 | 0 | 0 | 0 | 0 | 0 |
| 0 | NA | NA | NA | NA | NA | Gm11373 | | 0 | 0 | 0 | 0 | 0 | 0 | 0 | 0 | 0 | 0 | 0 |
| 0 | NA | NA | NA | NA | NA | Gm22330 | | 0 | 0 | 0 | 0 | 0 | 0 | 0 | 0 | 0 | 0 | 0 |
| 0 | NA | NA | NA | NA | NA | Gm11376 | | 0 | 0 | 0 | 0 | 0 | 0 | 0 | 0 | 0 | 0 | 0 |
| 0 | NA | NA | NA | NA | NA | Gm23351 | | 0 | 0 | 0 | 0 | 0 | 0 | 0 | 0 | 0 | 0 | 0 |
| 0 | NA | NA | NA | NA | NA | Gm11377 | | 0 | 0 | 0 | 0 | 0 | 0 | 0 | 0 | 0 | 0 | 0 |
| 0 | NA | NA | NA | NA | NA | 1700018A04Rik | | 0 | 0 | 0 | 0 | 0 | 0 | 0 | 0 | 0 | 0 | 0 |
| 0 | NA | NA | NA | NA | NA | Gm11378 | | 0 | 0 | 0 | 0 | 0 | 0 | 0 | 0 | 0 | 0 | 0 |
| 0 | NA | NA | NA | NA | NA | Gm27516 | | 0 | 0 | 0 | 0 | 0 | 0 | 0 | 0 | 0 | 0 | 0 |
| 0 | NA | NA | NA | NA | NA | Gm25186 | | 0 | 0 | 0 | 0 | 0 | 0 | 0 | 0 | 0 | 0 | 0 |
| 0 | NA | NA | NA | NA | NA | Gm11379 | | 0 | 0 | 0 | 0 | 0 | 0 | 0 | 0 | 0 | 0 | 0 |
| 0 | NA | NA | NA | NA | NA | Gm35732 | | 0 | 0 | 0 | 0 | 0 | 0 | 0 | 0 | 0 | 0 | 0 |
| 0 | NA | NA | NA | NA | NA | Gm11381 | | 0 | 0 | 0 | 0 | 0 | 0 | 0 | 0 | 0 | 0 | 0 |
| 0 | NA | NA | NA | NA | NA | | | 0 | 0 | 0 | 0 | 0 | 0 | 0 | 0 | 0 | 0 | 0 |
| 0 | NA | NA | NA | NA | NA | Gm27342 | | 0 | 0 | 0 | 0 | 0 | 0 | 0 | 0 | 0 | 0 | 0 |
| 0 | NA | NA | NA | NA | NA | Serpinb1-ps1 | | 0 | 0 | 0 | 0 | 0 | 0 | 0 | 0 | 0 | 0 | 0 |
| 0 | NA | NA | NA | NA | NA | Gm11382 | | 0 | 0 | 0 | 0 | 0 | 0 | 0 | 0 | 0 | 0 | 0 |
| 0 | NA | NA | NA | NA | NA | Gm36099 | | 0 | 0 | 0 | 0 | 0 | 0 | 0 | 0 | 0 | 0 | 0 |
| 0 | NA | NA | NA | NA | NA | Gm11384 | | 0 | 0 | 0 | 0 | 0 | 0 | 0 | 0 | 0 | 0 | 0 |
| 0 | NA | NA | NA | NA | NA | Gm11383 | | 0 | 0 | 0 | 0 | 0 | 0 | 0 | 0 | 0 | 0 | 0 |
| 0 | NA | NA | NA | NA | NA | Serpinb9c | serine (or cysteine) peptidase inhibitor, cl | 0 | 0 | 0 | 0 | 0 | 0 | 0 | 0 | 0 | 0 | 0 |
| 0 | NA | NA | NA | NA | NA | Serpinb9d | serine (or cysteine) peptidase inhibitor, cl | 0 | 0 | 0 | 0 | 0 | 0 | 0 | 0 | 0 | 0 | 0 |
| 0 | NA | NA | NA | NA | NA | Gm11385 | | 0 | 0 | 0 | 0 | 0 | 0 | 0 | 0 | 0 | 0 | 0 |
| 0 | NA | NA | NA | NA | NA | Gm47788 | | 0 | 0 | 0 | 0 | 0 | 0 | 0 | 0 | 0 | 0 | 0 |
| 0 | NA | NA | NA | NA | NA | Serpinb9e | serine (or cysteine) peptidase inhibitor, cl | 0 | 0 | 0 | 0 | 0 | 0 | 0 | 0 | 0 | 0 | 0 |
| 0 | NA | NA | NA | NA | NA | Serpinb9f | serine (or cysteine) peptidase inhibitor, cl | 0 | 0 | 0 | 0 | 0 | 0 | 0 | 0 | 0 | 0 | 0 |
| 0 | NA | NA | NA | NA | NA | Serpinb9h | serine (or cysteine) peptidase inhibitor, cl | 0 | 0 | 0 | 0 | 0 | 0 | 0 | 0 | 0 | 0 | 0 |
| 0 | NA | NA | NA | NA | NA | Serpinb9g | serine (or cysteine) peptidase inhibitor, cl | 0 | 0 | 0 | 0 | 0 | 0 | 0 | 0 | 0 | 0 | 0 |
| 0 | NA | NA | NA | NA | NA | Gm11387 | | 0 | 0 | 0 | 0 | 0 | 0 | 0 | 0 | 0 | 0 | 0 |
| 0 | NA | NA | NA | NA | NA | Gm11386 | | 0 | 0 | 0 | 0 | 0 | 0 | 0 | 0 | 0 | 0 | 0 |
| 0 | NA | NA | NA | NA | NA | Gm11393 | | 0 | 0 | 0 | 0 | 0 | 0 | 0 | 0 | 0 | 0 | 0 |
| 0 | NA | NA | NA | NA | NA | Gm6093 | | 0 | 0 | 0 | 0 | 0 | 0 | 0 | 0 | 0 | 0 | 0 |
| 0 | NA | NA | NA | NA | NA | Gm25043 | | 0 | 0 | 0 | 0 | 0 | 0 | 0 | 0 | 0 | 0 | 0 |
| 0 | NA | NA | NA | NA | NA | Gm47916 | | 0 | 0 | 0 | 0 | 0 | 0 | 0 | 0 | 0 | 0 | 0 |
| 0 | NA | NA | NA | NA | NA | Gm5194 | | 0 | 0 | 0 | 0 | 0 | 0 | 0 | 0 | 0 | 0 | 0 |
| 0 | NA | NA | NA | NA | NA | Gm11388 | | 0 | 0 | 0 | 0 | 0 | 0 | 0 | 0 | 0 | 0 | 0 |
| 0 | NA | NA | NA | NA | NA | Serpinb6d | serine (or cysteine) peptidase inhibitor, cl | 0 | 0 | 0 | 0 | 0 | 0 | 0 | 0 | 0 | 0 | 0 |
| 0 | NA | NA | NA | NA | NA | Gm11389 | | 0 | 0 | 0 | 0 | 0 | 0 | 0 | 0 | 0 | 0 | 0 |
| 0 | NA | NA | NA | NA | NA | Gm26701 | | 0 | 0 | 0 | 0 | 0 | 0 | 0 | 0 | 0 | 0 | 0 |
| 0 | NA | NA | NA | NA | NA | Gm11392 | | 0 | 0 | 0 | 0 | 0 | 0 | 0 | 0 | 0 | 0 | 0 |
| 0 | NA | NA | NA | NA | NA | Gm11394 | | 0 | 0 | 0 | 0 | 0 | 0 | 0 | 0 | 0 | 0 | 0 |
| 0 | NA | NA | NA | NA | NA | Gm11395 | | 0 | 0 | 0 | 0 | 0 | 0 | 0 | 0 | 0 | 0 | 0 |
| 0 | NA | NA | NA | NA | NA | Vmn2r-ps94 | | 0 | 0 | 0 | 0 | 0 | 0 | 0 | 0 | 0 | 0 | 0 |
| 0 | NA | NA | NA | NA | NA | Serpinb6e | serine (or cysteine) peptidase inhibitor, cl | 0 | 0 | 0 | 0 | 0 | 0 | 0 | 0 | 0 | 0 | 0 |
| 0 | NA | NA | NA | NA | NA | Gm17083 | | 0 | 0 | 0 | 0 | 0 | 0 | 0 | 0 | 0 | 0 | 0 |
| 0 | NA | NA | NA | NA | NA | Gm17084 | | 0 | 0 | 0 | 0 | 0 | 0 | 0 | 0 | 0 | 0 | 0 |
| 0 | NA | NA | NA | NA | NA | Gm47065 | | 0 | 0 | 0 | 0 | 0 | 0 | 0 | 0 | 0 | 0 | 0 |
| 0 | NA | NA | NA | NA | NA | Gm47076 | | 0 | 0 | 0 | 0 | 0 | 0 | 0 | 0 | 0 | 0 | 0 |
| 0 | NA | NA | NA | NA | NA | Gm24530 | | 0 | 0 | 0 | 0 | 0 | 0 | 0 | 0 | 0 | 0 | 0 |
| 0 | NA | NA | NA | NA | NA | Gm47086 | | 0 | 0 | 0 | 0 | 0 | 0 | 0 | 0 | 0 | 0 | 0 |
| 0 | NA | NA | NA | NA | NA | Gm47120 | | 0 | 0 | 0 | 0 | 0 | 0 | 0 | 0 | 0 | 0 | 0 |
| 0 | NA | NA | NA | NA | NA | Gm40909 | | 0 | 0 | 0 | 0 | 0 | 0 | 0 | 0 | 0 | 0 | 0 |
| 0 | NA | NA | NA | NA | NA | Gm47127 | | 0 | 0 | 0 | 0 | 0 | 0 | 0 | 0 | 0 | 0 | 0 |
| 0 | NA | NA | NA | NA | NA | Gm15908 | | 0 | 0 | 0 | 0 | 0 | 0 | 0 | 0 | 0 | 0 | 0 |

| | | | | | | | | | | | | | | | | | |
|---|---|---|---|---|---|---|---|---|---|---|---|---|---|---|---|---|---|
| 0 | NA | NA | NA | NA | NA | Gm47149 | | 0 | 0 | 0 | 0 | 0 | 0 | 0 | 0 | 0 | 0 | 0 |
| 0 | NA | NA | NA | NA | NA | Gm47150 | | 0 | 0 | 0 | 0 | 0 | 0 | 0 | 0 | 0 | 0 | 0 |
| 0 | NA | NA | NA | NA | NA | Gm47151 | | 0 | 0 | 0 | 0 | 0 | 0 | 0 | 0 | 0 | 0 | 0 |
| 0 | NA | NA | NA | NA | NA | Gm47152 | | 0 | 0 | 0 | 0 | 0 | 0 | 0 | 0 | 0 | 0 | 0 |
| 0 | NA | NA | NA | NA | NA | Fam50b | family with sequence similarity 50, membe | 0 | 0 | 0 | 0 | 0 | 0 | 0 | 0 | 0 | 0 | 0 |
| 0 | NA | NA | NA | NA | NA | Gm47157 | | 0 | 0 | 0 | 0 | 0 | 0 | 0 | 0 | 0 | 0 | 0 |
| 0 | NA | NA | NA | NA | NA | 4933417A18Ril | RIKEN cDNA 4933417A18 gene | 0 | 0 | 0 | 0 | 0 | 0 | 0 | 0 | 0 | 0 | 0 |
| 0 | NA | NA | NA | NA | NA | Gm8245 | | 0 | 0 | 0 | 0 | 0 | 0 | 0 | 0 | 0 | 0 | 0 |
| 0 | NA | NA | NA | NA | NA | Gm47217 | | 0 | 0 | 0 | 0 | 0 | 0 | 0 | 0 | 0 | 0 | 0 |
| 0 | NA | NA | NA | NA | NA | 4930529N20Rik | | 0 | 0 | 0 | 0 | 0 | 0 | 0 | 0 | 0 | 0 | 0 |
| 0 | NA | NA | NA | NA | NA | 1700019C18Rik | | 0 | 0 | 0 | 0 | 0 | 0 | 0 | 0 | 0 | 0 | 0 |
| 0 | NA | NA | NA | NA | NA | Gm48694 | | 0 | 0 | 0 | 0 | 0 | 0 | 0 | 0 | 0 | 0 | 0 |
| 0 | NA | NA | NA | NA | NA | Gm22674 | | 0 | 0 | 0 | 0 | 0 | 0 | 0 | 0 | 0 | 0 | 0 |
| 0 | NA | NA | NA | NA | NA | Gm48704 | | 0 | 0 | 0 | 0 | 0 | 0 | 0 | 0 | 0 | 0 | 0 |
| 0 | NA | NA | NA | NA | NA | Gm48707 | | 0 | 0 | 0 | 0 | 0 | 0 | 0 | 0 | 0 | 0 | 0 |
| 0 | NA | NA | NA | NA | NA | Gm22126 | | 0 | 0 | 0 | 0 | 0 | 0 | 0 | 0 | 0 | 0 | 0 |
| 0 | NA | NA | NA | NA | NA | Gm48708 | | 0 | 0 | 0 | 0 | 0 | 0 | 0 | 0 | 0 | 0 | 0 |
| 0 | NA | NA | NA | NA | NA | Gm48706 | | 0 | 0 | 0 | 0 | 0 | 0 | 0 | 0 | 0 | 0 | 0 |
| 0 | NA | NA | NA | NA | NA | Gm48756 | | 0 | 0 | 0 | 0 | 0 | 0 | 0 | 0 | 0 | 0 | 0 |
| 0 | NA | NA | NA | NA | NA | Gm30127 | | 0 | 0 | 0 | 0 | 0 | 0 | 0 | 0 | 0 | 0 | 0 |
| 0 | NA | NA | NA | NA | NA | Ppp1r3g | protein phosphatase 1, regulatory subuni | 0 | 0 | 0 | 0 | 0 | 0 | 0 | 0 | 0 | 0 | 0 |
| 0 | NA | NA | NA | NA | NA | Gm18132 | | 0 | 0 | 0 | 0 | 0 | 0 | 0 | 0 | 0 | 0 | 0 |
| 0 | NA | NA | NA | NA | NA | Gm48770 | | 0 | 0 | 0 | 0 | 0 | 0 | 0 | 0 | 0 | 0 | 0 |
| 0 | NA | NA | NA | NA | NA | Gm48765 | | 0 | 0 | 0 | 0 | 0 | 0 | 0 | 0 | 0 | 0 | 0 |
| 0 | NA | NA | NA | NA | NA | Gm48769 | | 0 | 0 | 0 | 0 | 0 | 0 | 0 | 0 | 0 | 0 | 0 |
| 0 | NA | NA | NA | NA | NA | Gm48763 | | 0 | 0 | 0 | 0 | 0 | 0 | 0 | 0 | 0 | 0 | 0 |
| 0 | NA | NA | NA | NA | NA | Gm48767 | | 0 | 0 | 0 | 0 | 0 | 0 | 0 | 0 | 0 | 0 | 0 |
| 0 | NA | NA | NA | NA | NA | 4933436N17Rik | | 0 | 0 | 0 | 0 | 0 | 0 | 0 | 0 | 0 | 0 | 0 |
| 0 | NA | NA | NA | NA | NA | Gm26395 | | 0 | 0 | 0 | 0 | 0 | 0 | 0 | 0 | 0 | 0 | 0 |
| 0 | NA | NA | NA | NA | NA | Gm49144 | | 0 | 0 | 0 | 0 | 0 | 0 | 0 | 0 | 0 | 0 | 0 |
| 0 | NA | NA | NA | NA | NA | Gm30489 | | 0 | 0 | 0 | 0 | 0 | 0 | 0 | 0 | 0 | 0 | 0 |
| 0 | NA | NA | NA | NA | NA | Gm18643 | | 0 | 0 | 0 | 0 | 0 | 0 | 0 | 0 | 0 | 0 | 0 |
| 0 | NA | NA | NA | NA | NA | Gm47707 | | 0 | 0 | 0 | 0 | 0 | 0 | 0 | 0 | 0 | 0 | 0 |
| 0 | NA | NA | NA | NA | NA | Gm30600 | | 0 | 0 | 0 | 0 | 0 | 0 | 0 | 0 | 0 | 0 | 0 |
| 0 | NA | NA | NA | NA | NA | Gm47711 | | 0 | 0 | 0 | 0 | 0 | 0 | 0 | 0 | 0 | 0 | 0 |
| 0 | NA | NA | NA | NA | NA | Gm4035 | | 0 | 0 | 0 | 0 | 0 | 0 | 0 | 0 | 0 | 0 | 0 |
| 0 | NA | NA | NA | NA | NA | Gm47730 | | 0 | 0 | 0 | 0 | 0 | 0 | 0 | 0 | 0 | 0 | 0 |
| 0 | NA | NA | NA | NA | NA | Gm47731 | | 0 | 0 | 0 | 0 | 0 | 0 | 0 | 0 | 0 | 0 | 0 |
| 0 | NA | NA | NA | NA | NA | Gm47732 | | 0 | 0 | 0 | 0 | 0 | 0 | 0 | 0 | 0 | 0 | 0 |
| 0 | NA | NA | NA | NA | NA | Gm47734 | | 0 | 0 | 0 | 0 | 0 | 0 | 0 | 0 | 0 | 0 | 0 |
| 0 | NA | NA | NA | NA | NA | Gm47751 | | 0 | 0 | 0 | 0 | 0 | 0 | 0 | 0 | 0 | 0 | 0 |
| 0 | NA | NA | NA | NA | NA | Gm47753 | | 0 | 0 | 0 | 0 | 0 | 0 | 0 | 0 | 0 | 0 | 0 |
| 0 | NA | NA | NA | NA | NA | Gm40915 | | 0 | 0 | 0 | 0 | 0 | 0 | 0 | 0 | 0 | 0 | 0 |
| 0 | NA | NA | NA | NA | NA | Gm40916 | | 0 | 0 | 0 | 0 | 0 | 0 | 0 | 0 | 0 | 0 | 0 |
| 0 | NA | NA | NA | NA | NA | Gm49350 | | 0 | 0 | 0 | 0 | 0 | 0 | 0 | 0 | 0 | 0 | 0 |
| 0 | NA | NA | NA | NA | NA | Gm49351 | | 0 | 0 | 0 | 0 | 0 | 0 | 0 | 0 | 0 | 0 | 0 |
| 0 | NA | NA | NA | NA | NA | Gm31600 | | 0 | 0 | 0 | 0 | 0 | 0 | 0 | 0 | 0 | 0 | 0 |
| 0 | NA | NA | NA | NA | NA | Gm46411 | | 0 | 0 | 0 | 0 | 0 | 0 | 0 | 0 | 0 | 0 | 0 |
| 0 | NA | NA | NA | NA | NA | Gm27387 | | 0 | 0 | 0 | 0 | 0 | 0 | 0 | 0 | 0 | 0 | 0 |
| 0 | NA | NA | NA | NA | NA | Gm47978 | predicted gene, 47978 | 0 | 0 | 0 | 0 | 0 | 0 | 0 | 0 | 0 | 0 | 0 |
| 0 | NA | NA | NA | NA | NA | Gm47991 | | 0 | 0 | 0 | 0 | 0 | 0 | 0 | 0 | 0 | 0 | 0 |
| 0 | NA | NA | NA | NA | NA | 4930579J19Rik | | 0 | 0 | 0 | 0 | 0 | 0 | 0 | 0 | 0 | 0 | 0 |
| 0 | NA | NA | NA | NA | NA | Gm47994 | | 0 | 0 | 0 | 0 | 0 | 0 | 0 | 0 | 0 | 0 | 0 |
| 0 | NA | NA | NA | NA | NA | Gm48056 | | 0 | 0 | 0 | 0 | 0 | 0 | 0 | 0 | 0 | 0 | 0 |
| 0 | NA | NA | NA | NA | NA | Gm40922 | | 0 | 0 | 0 | 0 | 0 | 0 | 0 | 0 | 0 | 0 | 0 |
| 0 | NA | NA | NA | NA | NA | Gm6200 | | 0 | 0 | 0 | 0 | 0 | 0 | 0 | 0 | 0 | 0 | 0 |
| 0 | NA | NA | NA | NA | NA | Gm46392 | | 0 | 0 | 0 | 0 | 0 | 0 | 0 | 0 | 0 | 0 | 0 |
| 0 | NA | NA | NA | NA | NA | Gm47510 | | 0 | 0 | 0 | 0 | 0 | 0 | 0 | 0 | 0 | 0 | 0 |
| 0 | NA | NA | NA | NA | NA | Gm22035 | | 0 | 0 | 0 | 0 | 0 | 0 | 0 | 0 | 0 | 0 | 0 |
| 0 | NA | NA | NA | NA | NA | Gm47530 | | 0 | 0 | 0 | 0 | 0 | 0 | 0 | 0 | 0 | 0 | 0 |

| | | | | | | | | | | | | | | | | | | |
|---|---|---|---|---|---|---|---|---|---|---|---|---|---|---|---|---|---|---|
| 0 | NA | NA | NA | NA | NA | Gm47348 | | 0 | 0 | 0 | 0 | 0 | 0 | 0 | 0 | 0 | 0 | 0 |
| 0 | NA | NA | NA | NA | NA | Gm3509 | | 0 | 0 | 0 | 0 | 0 | 0 | 0 | 0 | 0 | 0 | 0 |
| 0 | NA | NA | NA | NA | NA | Gm47349 | | 0 | 0 | 0 | 0 | 0 | 0 | 0 | 0 | 0 | 0 | 0 |
| 0 | NA | NA | NA | NA | NA | Gm47351 | | 0 | 0 | 0 | 0 | 0 | 0 | 0 | 0 | 0 | 0 | 0 |
| 0 | NA | NA | NA | NA | NA | Gm47352 | | 0 | 0 | 0 | 0 | 0 | 0 | 0 | 0 | 0 | 0 | 0 |
| 0 | NA | NA | NA | NA | NA | Gm32243 | | 0 | 0 | 0 | 0 | 0 | 0 | 0 | 0 | 0 | 0 | 0 |
| 0 | NA | NA | NA | NA | NA | Gm5195 | | 0 | 0 | 0 | 0 | 0 | 0 | 0 | 0 | 0 | 0 | 0 |
| 0 | NA | NA | NA | NA | NA | Gm40923 | | 0 | 0 | 0 | 0 | 0 | 0 | 0 | 0 | 0 | 0 | 0 |
| 0 | NA | NA | NA | NA | NA | A230103O09Rik | | 0 | 0 | 0 | 0 | 0 | 0 | 0 | 0 | 0 | 0 | 0 |
| 0 | NA | NA | NA | NA | NA | Gm47361 | | 0 | 0 | 0 | 0 | 0 | 0 | 0 | 0 | 0 | 0 | 0 |
| 0 | NA | NA | NA | NA | NA | Gm23972 | | 0 | 0 | 0 | 0 | 0 | 0 | 0 | 0 | 0 | 0 | 0 |
| 0 | NA | NA | NA | NA | NA | Gm47316 | | 0 | 0 | 0 | 0 | 0 | 0 | 0 | 0 | 0 | 0 | 0 |
| 0 | NA | NA | NA | NA | NA | Gm8352 | | 0 | 0 | 0 | 0 | 0 | 0 | 0 | 0 | 0 | 0 | 0 |
| 0 | NA | NA | NA | NA | NA | Gm47038 | | 0 | 0 | 0 | 0 | 0 | 0 | 0 | 0 | 0 | 0 | 0 |
| 0 | NA | NA | NA | NA | NA | Gm47039 | | 0 | 0 | 0 | 0 | 0 | 0 | 0 | 0 | 0 | 0 | 0 |
| 0 | NA | NA | NA | NA | NA | Gm47041 | | 0 | 0 | 0 | 0 | 0 | 0 | 0 | 0 | 0 | 0 | 0 |
| 0 | NA | NA | NA | NA | NA | Gm9979 | | 0 | 0 | 0 | 0 | 0 | 0 | 0 | 0 | 0 | 0 | 0 |
| 0 | NA | NA | NA | NA | NA | Gm26688 | | 0 | 0 | 0 | 0 | 0 | 0 | 0 | 0 | 0 | 0 | 0 |
| 0 | NA | NA | NA | NA | NA | Gm47067 | | 0 | 0 | 0 | 0 | 0 | 0 | 0 | 0 | 0 | 0 | 0 |
| 0 | NA | NA | NA | NA | NA | Gm48107 | | 0 | 0 | 0 | 0 | 0 | 0 | 0 | 0 | 0 | 0 | 0 |
| 0 | NA | NA | NA | NA | NA | Gm7201 | | 0 | 0 | 0 | 0 | 0 | 0 | 0 | 0 | 0 | 0 | 0 |
| 0 | NA | NA | NA | NA | NA | Gm35160 | | 0 | 0 | 0 | 0 | 0 | 0 | 0 | 0 | 0 | 0 | 0 |
| 0 | NA | NA | NA | NA | NA | Gm48192 | | 0 | 0 | 0 | 0 | 0 | 0 | 0 | 0 | 0 | 0 | 0 |
| 0 | NA | NA | NA | NA | NA | 5530402G07Rik | | 0 | 0 | 0 | 0 | 0 | 0 | 0 | 0 | 0 | 0 | 0 |
| 0 | NA | NA | NA | NA | NA | Gm48344 | | 0 | 0 | 0 | 0 | 0 | 0 | 0 | 0 | 0 | 0 | 0 |
| 0 | NA | NA | NA | NA | NA | Gcm2 | glial cells missing homolog 2 | 0 | 0 | 0 | 0 | 0 | 0 | 0 | 0 | 0 | 0 | 0 |
| 0 | NA | NA | NA | NA | NA | Gm48510 | | 0 | 0 | 0 | 0 | 0 | 0 | 0 | 0 | 0 | 0 | 0 |
| 0 | NA | NA | NA | NA | NA | Gm17364 | | 0 | 0 | 0 | 0 | 0 | 0 | 0 | 0 | 0 | 0 | 0 |
| 0 | NA | NA | NA | NA | NA | Gm48571 | | 0 | 0 | 0 | 0 | 0 | 0 | 0 | 0 | 0 | 0 | 0 |
| 0 | NA | NA | NA | NA | NA | Gm48570 | | 0 | 0 | 0 | 0 | 0 | 0 | 0 | 0 | 0 | 0 | 0 |
| 0 | NA | NA | NA | NA | NA | Gm48569 | | 0 | 0 | 0 | 0 | 0 | 0 | 0 | 0 | 0 | 0 | 0 |
| 0 | NA | NA | NA | NA | NA | Gm24819 | | 0 | 0 | 0 | 0 | 0 | 0 | 0 | 0 | 0 | 0 | 0 |
| 0 | NA | NA | NA | NA | NA | Gm5082 | | 0 | 0 | 0 | 0 | 0 | 0 | 0 | 0 | 0 | 0 | 0 |
| 0 | NA | NA | NA | NA | NA | 1700061E18Rik | | 0 | 0 | 0 | 0 | 0 | 0 | 0 | 0 | 0 | 0 | 0 |
| 0 | NA | NA | NA | NA | NA | Gm28564 | | 0 | 0 | 0 | 0 | 0 | 0 | 0 | 0 | 0 | 0 | 0 |
| 0 | NA | NA | NA | NA | NA | Gm37698 | | 0 | 0 | 0 | 0 | 0 | 0 | 0 | 0 | 0 | 0 | 0 |
| 0 | NA | NA | NA | NA | NA | Gm22157 | | 0 | 0 | 0 | 0 | 0 | 0 | 0 | 0 | 0 | 0 | 0 |
| 0 | NA | NA | NA | NA | NA | Gm47118 | | 0 | 0 | 0 | 0 | 0 | 0 | 0 | 0 | 0 | 0 | 0 |
| 0 | NA | NA | NA | NA | NA | Gm6049 | | 0 | 0 | 0 | 0 | 0 | 0 | 0 | 0 | 0 | 0 | 0 |
| 0 | NA | NA | NA | NA | NA | Gm47125 | | 0 | 0 | 0 | 0 | 0 | 0 | 0 | 0 | 0 | 0 | 0 |
| 0 | NA | NA | NA | NA | NA | Gm23086 | | 0 | 0 | 0 | 0 | 0 | 0 | 0 | 0 | 0 | 0 | 0 |
| 0 | NA | NA | NA | NA | NA | Gm15810 | | 0 | 0 | 0 | 0 | 0 | 0 | 0 | 0 | 0 | 0 | 0 |
| 0 | NA | NA | NA | NA | NA | Gm15812 | | 0 | 0 | 0 | 0 | 0 | 0 | 0 | 0 | 0 | 0 | 0 |
| 0 | NA | NA | NA | NA | NA | Gm26086 | | 0 | 0 | 0 | 0 | 0 | 0 | 0 | 0 | 0 | 0 | 0 |
| 0 | NA | NA | NA | NA | NA | Gm47597 | | 0 | 0 | 0 | 0 | 0 | 0 | 0 | 0 | 0 | 0 | 0 |
| 0 | NA | NA | NA | NA | NA | Gm47683 | | 0 | 0 | 0 | 0 | 0 | 0 | 0 | 0 | 0 | 0 | 0 |
| 0 | NA | NA | NA | NA | NA | Gm7214 | | 0 | 0 | 0 | 0 | 0 | 0 | 0 | 0 | 0 | 0 | 0 |
| 0 | NA | NA | NA | NA | NA | Gm32939 | | 0 | 0 | 0 | 0 | 0 | 0 | 0 | 0 | 0 | 0 | 0 |
| 0 | NA | NA | NA | NA | NA | Gm33115 | | 0 | 0 | 0 | 0 | 0 | 0 | 0 | 0 | 0 | 0 | 0 |
| 0 | NA | NA | NA | NA | NA | Gm23117 | | 0 | 0 | 0 | 0 | 0 | 0 | 0 | 0 | 0 | 0 | 0 |
| 0 | NA | NA | NA | NA | NA | Gm2233 | | 0 | 0 | 0 | 0 | 0 | 0 | 0 | 0 | 0 | 0 | 0 |
| 0 | NA | NA | NA | NA | NA | Gm33489 | | 0 | 0 | 0 | 0 | 0 | 0 | 0 | 0 | 0 | 0 | 0 |
| 0 | NA | NA | NA | NA | NA | Gm33630 | | 0 | 0 | 0 | 0 | 0 | 0 | 0 | 0 | 0 | 0 | 0 |
| 0 | NA | NA | NA | NA | NA | Gm33684 | | 0 | 0 | 0 | 0 | 0 | 0 | 0 | 0 | 0 | 0 | 0 |
| 0 | NA | NA | NA | NA | NA | A330076C08Rik | | 0 | 0 | 0 | 0 | 0 | 0 | 0 | 0 | 0 | 0 | 0 |
| 0 | NA | NA | NA | NA | NA | Gm47781 | | 0 | 0 | 0 | 0 | 0 | 0 | 0 | 0 | 0 | 0 | 0 |
| 0 | NA | NA | NA | NA | NA | Gm29676 | | 0 | 0 | 0 | 0 | 0 | 0 | 0 | 0 | 0 | 0 | 0 |
| 0 | NA | NA | NA | NA | NA | Gm33958 | | 0 | 0 | 0 | 0 | 0 | 0 | 0 | 0 | 0 | 0 | 0 |
| 0 | NA | NA | NA | NA | NA | Gm27007 | | 0 | 0 | 0 | 0 | 0 | 0 | 0 | 0 | 0 | 0 | 0 |
| 0 | NA | NA | NA | NA | NA | Gm27036 | | 0 | 0 | 0 | 0 | 0 | 0 | 0 | 0 | 0 | 0 | 0 |

| | | | | | | | | | | | | | | | | | | |
|---|---|---|---|---|---|---|---|---|---|---|---|---|---|---|---|---|---|---|
| 0 | NA | NA | NA | NA | NA | 1700029N11Rik | | 0 | 0 | 0 | 0 | 0 | 0 | 0 | 0 | 0 | 0 | 0 |
| 0 | NA | NA | NA | NA | NA | Gm47792 | | 0 | 0 | 0 | 0 | 0 | 0 | 0 | 0 | 0 | 0 | 0 |
| 0 | NA | NA | NA | NA | NA | Gm34276 | | 0 | 0 | 0 | 0 | 0 | 0 | 0 | 0 | 0 | 0 | 0 |
| 0 | NA | NA | NA | NA | NA | Gm47805 | | 0 | 0 | 0 | 0 | 0 | 0 | 0 | 0 | 0 | 0 | 0 |
| 0 | NA | NA | NA | NA | NA | Gm22213 | | 0 | 0 | 0 | 0 | 0 | 0 | 0 | 0 | 0 | 0 | 0 |
| 0 | NA | NA | NA | NA | NA | Gm23104 | | 0 | 0 | 0 | 0 | 0 | 0 | 0 | 0 | 0 | 0 | 0 |
| 0 | NA | NA | NA | NA | NA | Gm9817 | | 0 | 0 | 0 | 0 | 0 | 0 | 0 | 0 | 0 | 0 | 0 |
| 0 | NA | NA | NA | NA | NA | Gm40932 | | 0 | 0 | 0 | 0 | 0 | 0 | 0 | 0 | 0 | 0 | 0 |
| 0 | NA | NA | NA | NA | NA | Gm47448 | | 0 | 0 | 0 | 0 | 0 | 0 | 0 | 0 | 0 | 0 | 0 |
| 0 | NA | NA | NA | NA | NA | Gm34466 | | 0 | 0 | 0 | 0 | 0 | 0 | 0 | 0 | 0 | 0 | 0 |
| 0 | NA | NA | NA | NA | NA | Gm23755 | | 0 | 0 | 0 | 0 | 0 | 0 | 0 | 0 | 0 | 0 | 0 |
| 0 | NA | NA | NA | NA | NA | Gm47455 | | 0 | 0 | 0 | 0 | 0 | 0 | 0 | 0 | 0 | 0 | 0 |
| 0 | NA | NA | NA | NA | NA | Gm23387 | | 0 | 0 | 0 | 0 | 0 | 0 | 0 | 0 | 0 | 0 | 0 |
| 0 | NA | NA | NA | NA | NA | Gm47460 | | 0 | 0 | 0 | 0 | 0 | 0 | 0 | 0 | 0 | 0 | 0 |
| 0 | NA | NA | NA | NA | NA | 4930453C13Rik | | 0 | 0 | 0 | 0 | 0 | 0 | 0 | 0 | 0 | 0 | 0 |
| 0 | NA | NA | NA | NA | NA | Gm35110 | | 0 | 0 | 0 | 0 | 0 | 0 | 0 | 0 | 0 | 0 | 0 |
| 0 | NA | NA | NA | NA | NA | Gm24915 | | 0 | 0 | 0 | 0 | 0 | 0 | 0 | 0 | 0 | 0 | 0 |
| 0 | NA | NA | NA | NA | NA | Gm47523 | | 0 | 0 | 0 | 0 | 0 | 0 | 0 | 0 | 0 | 0 | 0 |
| 0 | NA | NA | NA | NA | NA | Gm48250 | | 0 | 0 | 0 | 0 | 0 | 0 | 0 | 0 | 0 | 0 | 0 |
| 0 | NA | NA | NA | NA | NA | Gm22960 | | 0 | 0 | 0 | 0 | 0 | 0 | 0 | 0 | 0 | 0 | 0 |
| 0 | NA | NA | NA | NA | NA | Gm48325 | | 0 | 0 | 0 | 0 | 0 | 0 | 0 | 0 | 0 | 0 | 0 |
| 0 | NA | NA | NA | NA | NA | Gm24769 | | 0 | 0 | 0 | 0 | 0 | 0 | 0 | 0 | 0 | 0 | 0 |
| 0 | NA | NA | NA | NA | NA | Gm18807 | | 0 | 0 | 0 | 0 | 0 | 0 | 0 | 0 | 0 | 0 | 0 |
| 0 | NA | NA | NA | NA | NA | Gm48581 | | 0 | 0 | 0 | 0 | 0 | 0 | 0 | 0 | 0 | 0 | 0 |
| 0 | NA | NA | NA | NA | NA | Gm48612 | | 0 | 0 | 0 | 0 | 0 | 0 | 0 | 0 | 0 | 0 | 0 |
| 0 | NA | NA | NA | NA | NA | Gm35733 | | 0 | 0 | 0 | 0 | 0 | 0 | 0 | 0 | 0 | 0 | 0 |
| 0 | NA | NA | NA | NA | NA | 4930471G24Rik | | 0 | 0 | 0 | 0 | 0 | 0 | 0 | 0 | 0 | 0 | 0 |
| 0 | NA | NA | NA | NA | NA | G630093K05Rik | | 0 | 0 | 0 | 0 | 0 | 0 | 0 | 0 | 0 | 0 | 0 |
| 0 | NA | NA | NA | NA | NA | A330033J07Rik | | 0 | 0 | 0 | 0 | 0 | 0 | 0 | 0 | 0 | 0 | 0 |
| 0 | NA | NA | NA | NA | NA | Gm36346 | | 0 | 0 | 0 | 0 | 0 | 0 | 0 | 0 | 0 | 0 | 0 |
| 0 | NA | NA | NA | NA | NA | Gm49291 | | 0 | 0 | 0 | 0 | 0 | 0 | 0 | 0 | 0 | 0 | 0 |
| 0 | NA | NA | NA | NA | NA | A330048O09Rik | | 0 | 0 | 0 | 0 | 0 | 0 | 0 | 0 | 0 | 0 | 0 |
| 0 | NA | NA | NA | NA | NA | Gm8588 | | 0 | 0 | 0 | 0 | 0 | 0 | 0 | 0 | 0 | 0 | 0 |
| 0 | NA | NA | NA | NA | NA | Gm48648 | | 0 | 0 | 0 | 0 | 0 | 0 | 0 | 0 | 0 | 0 | 0 |
| 0 | NA | NA | NA | NA | NA | Mirlet7d | | 0 | 0 | 0 | 0 | 0 | 0 | 0 | 0 | 0 | 0 | 0 |
| 0 | NA | NA | NA | NA | NA | Gm24111 | | 0 | 0 | 0 | 0 | 0 | 0 | 0 | 0 | 0 | 0 | 0 |
| 0 | NA | NA | NA | NA | NA | Mirlet7f-1 | | 0 | 0 | 0 | 0 | 0 | 0 | 0 | 0 | 0 | 0 | 0 |
| 0 | NA | NA | NA | NA | NA | Mirlet7a-1 | | 0 | 0 | 0 | 0 | 0 | 0 | 0 | 0 | 0 | 0 | 0 |
| 0 | NA | NA | NA | NA | NA | Gm25232 | | 0 | 0 | 0 | 0 | 0 | 0 | 0 | 0 | 0 | 0 | 0 |
| 0 | NA | NA | NA | NA | NA | Gm36550 | | 0 | 0 | 0 | 0 | 0 | 0 | 0 | 0 | 0 | 0 | 0 |
| 0 | NA | NA | NA | NA | NA | Rpl17-ps6 | | 0 | 0 | 0 | 0 | 0 | 0 | 0 | 0 | 0 | 0 | 0 |
| 0 | NA | NA | NA | NA | NA | Gm19334 | | 0 | 0 | 0 | 0 | 0 | 0 | 0 | 0 | 0 | 0 | 0 |
| 0 | NA | NA | NA | NA | NA | Gm270 | | 0 | 0 | 0 | 0 | 0 | 0 | 0 | 0 | 0 | 0 | 0 |
| 0 | NA | NA | NA | NA | NA | Gm30302 | predicted gene, 30302 | 0 | 0 | 0 | 0 | 0 | 0 | 0 | 0 | 0 | 0 | 0 |
| 0 | NA | NA | NA | NA | NA | Gm47977 | | 0 | 0 | 0 | 0 | 0 | 0 | 0 | 0 | 0 | 0 | 0 |
| 0 | NA | NA | NA | NA | NA | Gm272 | | 0 | 0 | 0 | 0 | 0 | 0 | 0 | 0 | 0 | 0 | 0 |
| 0 | NA | NA | NA | NA | NA | Gm8672 | | 0 | 0 | 0 | 0 | 0 | 0 | 0 | 0 | 0 | 0 | 0 |
| 0 | NA | NA | NA | NA | NA | Gm19011 | | 0 | 0 | 0 | 0 | 0 | 0 | 0 | 0 | 0 | 0 | 0 |
| 0 | NA | NA | NA | NA | NA | Gm48009 | | 0 | 0 | 0 | 0 | 0 | 0 | 0 | 0 | 0 | 0 | 0 |
| 0 | NA | NA | NA | NA | NA | Gm8674 | | 0 | 0 | 0 | 0 | 0 | 0 | 0 | 0 | 0 | 0 | 0 |
| 0 | NA | NA | NA | NA | NA | Gm48051 | | 0 | 0 | 0 | 0 | 0 | 0 | 0 | 0 | 0 | 0 | 0 |
| 0 | NA | NA | NA | NA | NA | Gm17611 | | 0 | 0 | 0 | 0 | 0 | 0 | 0 | 0 | 0 | 0 | 0 |
| 0 | NA | NA | NA | NA | NA | Gm6554 | | 0 | 0 | 0 | 0 | 0 | 0 | 0 | 0 | 0 | 0 | 0 |
| 0 | NA | NA | NA | NA | NA | Gm48052 | | 0 | 0 | 0 | 0 | 0 | 0 | 0 | 0 | 0 | 0 | 0 |
| 0 | NA | NA | NA | NA | NA | LOC667541 | | 0 | 0 | 0 | 0 | 0 | 0 | 0 | 0 | 0 | 0 | 0 |
| 0 | NA | NA | NA | NA | NA | Gm8700 | | 0 | 0 | 0 | 0 | 0 | 0 | 0 | 0 | 0 | 0 | 0 |
| 0 | NA | NA | NA | NA | NA | Gm24300 | | 0 | 0 | 0 | 0 | 0 | 0 | 0 | 0 | 0 | 0 | 0 |
| 0 | NA | NA | NA | NA | NA | Gm48054 | | 0 | 0 | 0 | 0 | 0 | 0 | 0 | 0 | 0 | 0 | 0 |
| 0 | NA | NA | NA | NA | NA | 4933433N18Rik | | 0 | 0 | 0 | 0 | 0 | 0 | 0 | 0 | 0 | 0 | 0 |
| 0 | NA | NA | NA | NA | NA | Gm47412 | | 0 | 0 | 0 | 0 | 0 | 0 | 0 | 0 | 0 | 0 | 0 |

| | | | | | | | | | | | | | | | | | |
|---|---|---|---|---|---|---|---|---|---|---|---|---|---|---|---|---|---|
| 0 | NA | NA | NA | NA | NA | Gm47413 | | 0 | 0 | 0 | 0 | 0 | 0 | 0 | 0 | 0 | 0 | 0 |
| 0 | NA | NA | NA | NA | NA | Gm906 | predicted gene 906 | 0 | 0 | 0 | 0 | 0 | 0 | 0 | 0 | 0 | 0 | 0 |
| 0 | NA | NA | NA | NA | NA | Gm24017 | | 0 | 0 | 0 | 0 | 0 | 0 | 0 | 0 | 0 | 0 | 0 |
| 0 | NA | NA | NA | NA | NA | Gm31126 | | 0 | 0 | 0 | 0 | 0 | 0 | 0 | 0 | 0 | 0 | 0 |
| 0 | NA | NA | NA | NA | NA | Gm8739 | | 0 | 0 | 0 | 0 | 0 | 0 | 0 | 0 | 0 | 0 | 0 |
| 0 | NA | NA | NA | NA | NA | Gm47429 | | 0 | 0 | 0 | 0 | 0 | 0 | 0 | 0 | 0 | 0 | 0 |
| 0 | NA | NA | NA | NA | NA | Nutm2 | NUT family member 2 | 0 | 0 | 0 | 0 | 0 | 0 | 0 | 0 | 0 | 0 | 0 |
| 0 | NA | NA | NA | NA | NA | Mir683-1 | | 0 | 0 | 0 | 0 | 0 | 0 | 0 | 0 | 0 | 0 | 0 |
| 0 | NA | NA | NA | NA | NA | Gm20789 | | 0 | 0 | 0 | 0 | 0 | 0 | 0 | 0 | 0 | 0 | 0 |
| 0 | NA | NA | NA | NA | NA | Mir683-2 | | 0 | 0 | 0 | 0 | 0 | 0 | 0 | 0 | 0 | 0 | 0 |
| 0 | NA | NA | NA | NA | NA | Gm904 | predicted gene 904 | 0 | 0 | 0 | 0 | 0 | 0 | 0 | 0 | 0 | 0 | 0 |
| 0 | NA | NA | NA | NA | NA | Gm23952 | | 0 | 0 | 0 | 0 | 0 | 0 | 0 | 0 | 0 | 0 | 0 |
| 0 | NA | NA | NA | NA | NA | Gm8765 | predicted gene 8765 | 0 | 0 | 0 | 0 | 0 | 0 | 0 | 0 | 0 | 0 | 0 |
| 0 | NA | NA | NA | NA | NA | 4930518P08Rik | | 0 | 0 | 0 | 0 | 0 | 0 | 0 | 0 | 0 | 0 | 0 |
| 0 | NA | NA | NA | NA | NA | Gm19009 | | 0 | 0 | 0 | 0 | 0 | 0 | 0 | 0 | 0 | 0 | 0 |
| 0 | NA | NA | NA | NA | NA | Gm22199 | | 0 | 0 | 0 | 0 | 0 | 0 | 0 | 0 | 0 | 0 | 0 |
| 0 | NA | NA | NA | NA | NA | Gm6056 | | 0 | 0 | 0 | 0 | 0 | 0 | 0 | 0 | 0 | 0 | 0 |
| 0 | NA | NA | NA | NA | NA | Gm47554 | | 0 | 0 | 0 | 0 | 0 | 0 | 0 | 0 | 0 | 0 | 0 |
| 0 | NA | NA | NA | NA | NA | Gm23566 | | 0 | 0 | 0 | 0 | 0 | 0 | 0 | 0 | 0 | 0 | 0 |
| 0 | NA | NA | NA | NA | NA | Gm26651 | | 0 | 0 | 0 | 0 | 0 | 0 | 0 | 0 | 0 | 0 | 0 |
| 0 | NA | NA | NA | NA | NA | Gm37872 | | 0 | 0 | 0 | 0 | 0 | 0 | 0 | 0 | 0 | 0 | 0 |
| 0 | NA | NA | NA | NA | NA | Gm48190 | | 0 | 0 | 0 | 0 | 0 | 0 | 0 | 0 | 0 | 0 | 0 |
| 0 | NA | NA | NA | NA | NA | Gm48199 | | 0 | 0 | 0 | 0 | 0 | 0 | 0 | 0 | 0 | 0 | 0 |
| 0 | NA | NA | NA | NA | NA | n-R5s54 | | 0 | 0 | 0 | 0 | 0 | 0 | 0 | 0 | 0 | 0 | 0 |
| 0 | NA | NA | NA | NA | NA | Gm33424 | | 0 | 0 | 0 | 0 | 0 | 0 | 0 | 0 | 0 | 0 | 0 |
| 0 | NA | NA | NA | NA | NA | Gm48336 | | 0 | 0 | 0 | 0 | 0 | 0 | 0 | 0 | 0 | 0 | 0 |
| 0 | NA | NA | NA | NA | NA | Gm43262 | | 0 | 0 | 0 | 0 | 0 | 0 | 0 | 0 | 0 | 0 | 0 |
| 0 | NA | NA | NA | NA | NA | Gm34022 | | 0 | 0 | 0 | 0 | 0 | 0 | 0 | 0 | 0 | 0 | 0 |
| 0 | NA | NA | NA | NA | NA | 9530014B07Rik | | 0 | 0 | 0 | 0 | 0 | 0 | 0 | 0 | 0 | 0 | 0 |
| 0 | NA | NA | NA | NA | NA | 4930555G21Rik | | 0 | 0 | 0 | 0 | 0 | 0 | 0 | 0 | 0 | 0 | 0 |
| 0 | NA | NA | NA | NA | NA | Gm34278 | | 0 | 0 | 0 | 0 | 0 | 0 | 0 | 0 | 0 | 0 | 0 |
| 0 | NA | NA | NA | NA | NA | Gm48548 | | 0 | 0 | 0 | 0 | 0 | 0 | 0 | 0 | 0 | 0 | 0 |
| 0 | NA | NA | NA | NA | NA | Gm18974 | | 0 | 0 | 0 | 0 | 0 | 0 | 0 | 0 | 0 | 0 | 0 |
| 0 | NA | NA | NA | NA | NA | E130119H09Rik | | 0 | 0 | 0 | 0 | 0 | 0 | 0 | 0 | 0 | 0 | 0 |
| 0 | NA | NA | NA | NA | NA | Gm34557 | | 0 | 0 | 0 | 0 | 0 | 0 | 0 | 0 | 0 | 0 | 0 |
| 0 | NA | NA | NA | NA | NA | Gm48550 | | 0 | 0 | 0 | 0 | 0 | 0 | 0 | 0 | 0 | 0 | 0 |
| 0 | NA | NA | NA | NA | NA | Gm18473 | | 0 | 0 | 0 | 0 | 0 | 0 | 0 | 0 | 0 | 0 | 0 |
| 0 | NA | NA | NA | NA | NA | Gm16578 | | 0 | 0 | 0 | 0 | 0 | 0 | 0 | 0 | 0 | 0 | 0 |
| 0 | NA | NA | NA | NA | NA | Gm48615 | | 0 | 0 | 0 | 0 | 0 | 0 | 0 | 0 | 0 | 0 | 0 |
| 0 | NA | NA | NA | NA | NA | Gm48622 | | 0 | 0 | 0 | 0 | 0 | 0 | 0 | 0 | 0 | 0 | 0 |
| 0 | NA | NA | NA | NA | NA | Gm22668 | | 0 | 0 | 0 | 0 | 0 | 0 | 0 | 0 | 0 | 0 | 0 |
| 0 | NA | NA | NA | NA | NA | Gm16249 | | 0 | 0 | 0 | 0 | 0 | 0 | 0 | 0 | 0 | 0 | 0 |
| 0 | NA | NA | NA | NA | NA | Gm16250 | | 0 | 0 | 0 | 0 | 0 | 0 | 0 | 0 | 0 | 0 | 0 |
| 0 | NA | NA | NA | NA | NA | Gm29431 | | 0 | 0 | 0 | 0 | 0 | 0 | 0 | 0 | 0 | 0 | 0 |
| 0 | NA | NA | NA | NA | NA | 5330429C05Rik | | 0 | 0 | 0 | 0 | 0 | 0 | 0 | 0 | 0 | 0 | 0 |
| 0 | NA | NA | NA | NA | NA | 4930526F13Rik | | 0 | 0 | 0 | 0 | 0 | 0 | 0 | 0 | 0 | 0 | 0 |
| 0 | NA | NA | NA | NA | NA | Gm17617 | | 0 | 0 | 0 | 0 | 0 | 0 | 0 | 0 | 0 | 0 | 0 |
| 0 | NA | NA | NA | NA | NA | Gm47816 | | 0 | 0 | 0 | 0 | 0 | 0 | 0 | 0 | 0 | 0 | 0 |
| 0 | NA | NA | NA | NA | NA | Mir6944 | | 0 | 0 | 0 | 0 | 0 | 0 | 0 | 0 | 0 | 0 | 0 |
| 0 | NA | NA | NA | NA | NA | Mir6945 | | 0 | 0 | 0 | 0 | 0 | 0 | 0 | 0 | 0 | 0 | 0 |
| 0 | NA | NA | NA | NA | NA | Gm15911 | | 0 | 0 | 0 | 0 | 0 | 0 | 0 | 0 | 0 | 0 | 0 |
| 0 | NA | NA | NA | NA | NA | Gm28760 | | 0 | 0 | 0 | 0 | 0 | 0 | 0 | 0 | 0 | 0 | 0 |
| 0 | NA | NA | NA | NA | NA | Gm47072 | | 0 | 0 | 0 | 0 | 0 | 0 | 0 | 0 | 0 | 0 | 0 |
| 0 | NA | NA | NA | NA | NA | Gm25148 | | 0 | 0 | 0 | 0 | 0 | 0 | 0 | 0 | 0 | 0 | 0 |
| 0 | NA | NA | NA | NA | NA | Gm47104 | | 0 | 0 | 0 | 0 | 0 | 0 | 0 | 0 | 0 | 0 | 0 |
| 0 | NA | NA | NA | NA | NA | 4930550C17Rik | | 0 | 0 | 0 | 0 | 0 | 0 | 0 | 0 | 0 | 0 | 0 |
| 0 | NA | NA | NA | NA | NA | Neurog1 | neurogenin 1 | 0 | 0 | 0 | 0 | 0 | 0 | 0 | 0 | 0 | 0 | 0 |
| 0 | NA | NA | NA | NA | NA | Gm17878 | | 0 | 0 | 0 | 0 | 0 | 0 | 0 | 0 | 0 | 0 | 0 |
| 0 | NA | NA | NA | NA | NA | Gm24716 | | 0 | 0 | 0 | 0 | 0 | 0 | 0 | 0 | 0 | 0 | 0 |
| 0 | NA | NA | NA | NA | NA | Gm10782 | | 0 | 0 | 0 | 0 | 0 | 0 | 0 | 0 | 0 | 0 | 0 |

| | | | | | | | | | | | | | | | | | |
|---|---|---|---|---|---|---|---|---|---|---|---|---|---|---|---|---|---|
| 0 | NA | NA | NA | NA | NA | Il9 | interleukin 9 | 0 | 0 | 0 | 0 | 0 | 0 | 0 | 0 | 0 | 0 | 0 |
| 0 | NA | NA | NA | NA | NA | Lect2 | leukocyte cell-derived chemotaxin 2 | 0 | 0 | 0 | 0 | 0 | 0 | 0 | 0 | 0 | 0 | 0 |
| 0 | NA | NA | NA | NA | NA | 2010203P06Rik | | 0 | 0 | 0 | 0 | 0 | 0 | 0 | 0 | 0 | 0 | 0 |
| 0 | NA | NA | NA | NA | NA | Gm27698 | | 0 | 0 | 0 | 0 | 0 | 0 | 0 | 0 | 0 | 0 | 0 |
| 0 | NA | NA | NA | NA | NA | Gm27301 | | 0 | 0 | 0 | 0 | 0 | 0 | 0 | 0 | 0 | 0 | 0 |
| 0 | NA | NA | NA | NA | NA | Gm27834 | | 0 | 0 | 0 | 0 | 0 | 0 | 0 | 0 | 0 | 0 | 0 |
| 0 | NA | NA | NA | NA | NA | Gm22777 | | 0 | 0 | 0 | 0 | 0 | 0 | 0 | 0 | 0 | 0 | 0 |
| 0 | NA | NA | NA | NA | NA | Gm48174 | | 0 | 0 | 0 | 0 | 0 | 0 | 0 | 0 | 0 | 0 | 0 |
| 0 | NA | NA | NA | NA | NA | Gm48176 | | 0 | 0 | 0 | 0 | 0 | 0 | 0 | 0 | 0 | 0 | 0 |
| 0 | NA | NA | NA | NA | NA | Mir874 | | 0 | 0 | 0 | 0 | 0 | 0 | 0 | 0 | 0 | 0 | 0 |
| 0 | NA | NA | NA | NA | NA | Gm26555 | | 0 | 0 | 0 | 0 | 0 | 0 | 0 | 0 | 0 | 0 | 0 |
| 0 | NA | NA | NA | NA | NA | Mir6369 | | 0 | 0 | 0 | 0 | 0 | 0 | 0 | 0 | 0 | 0 | 0 |
| 0 | NA | NA | NA | NA | NA | Gm24284 | | 0 | 0 | 0 | 0 | 0 | 0 | 0 | 0 | 0 | 0 | 0 |
| 0 | NA | NA | NA | NA | NA | Gm3131 | | 0 | 0 | 0 | 0 | 0 | 0 | 0 | 0 | 0 | 0 | 0 |
| 0 | NA | NA | NA | NA | NA | 4930455J16Rik | | 0 | 0 | 0 | 0 | 0 | 0 | 0 | 0 | 0 | 0 | 0 |
| 0 | NA | NA | NA | NA | NA | Gm47947 | | 0 | 0 | 0 | 0 | 0 | 0 | 0 | 0 | 0 | 0 | 0 |
| 0 | NA | NA | NA | NA | NA | Gm40968 | | 0 | 0 | 0 | 0 | 0 | 0 | 0 | 0 | 0 | 0 | 0 |
| 0 | NA | NA | NA | NA | NA | Gm47423 | | 0 | 0 | 0 | 0 | 0 | 0 | 0 | 0 | 0 | 0 | 0 |
| 0 | NA | NA | NA | NA | NA | 4930415C11Rik | | 0 | 0 | 0 | 0 | 0 | 0 | 0 | 0 | 0 | 0 | 0 |
| 0 | NA | NA | NA | NA | NA | Gm34245 | | 0 | 0 | 0 | 0 | 0 | 0 | 0 | 0 | 0 | 0 | 0 |
| 0 | NA | NA | NA | NA | NA | Gm34558 | | 0 | 0 | 0 | 0 | 0 | 0 | 0 | 0 | 0 | 0 | 0 |
| 0 | NA | NA | NA | NA | NA | Gm34672 | | 0 | 0 | 0 | 0 | 0 | 0 | 0 | 0 | 0 | 0 | 0 |
| 0 | NA | NA | NA | NA | NA | Gm47360 | | 0 | 0 | 0 | 0 | 0 | 0 | 0 | 0 | 0 | 0 | 0 |
| 0 | NA | NA | NA | NA | NA | Gm34721 | | 0 | 0 | 0 | 0 | 0 | 0 | 0 | 0 | 0 | 0 | 0 |
| 0 | NA | NA | NA | NA | NA | Gm34788 | | 0 | 0 | 0 | 0 | 0 | 0 | 0 | 0 | 0 | 0 | 0 |
| 0 | NA | NA | NA | NA | NA | 4930455M05Rik | | 0 | 0 | 0 | 0 | 0 | 0 | 0 | 0 | 0 | 0 | 0 |
| 0 | NA | NA | NA | NA | NA | 4921517D22Ril | RIKEN cDNA 4921517D22 gene | 0 | 0 | 0 | 0 | 0 | 0 | 0 | 0 | 0 | 0 | 0 |
| 0 | NA | NA | NA | NA | NA | Gm47269 | | 0 | 0 | 0 | 0 | 0 | 0 | 0 | 0 | 0 | 0 | 0 |
| 0 | NA | NA | NA | NA | NA | Spata31d1a | spermatogenesis associated 31 subfamily | 0 | 0 | 0 | 0 | 0 | 0 | 0 | 0 | 0 | 0 | 0 |
| 0 | NA | NA | NA | NA | NA | Spata31d1b | spermatogenesis associated 31 subfamily | 0 | 0 | 0 | 0 | 0 | 0 | 0 | 0 | 0 | 0 | 0 |
| 0 | NA | NA | NA | NA | NA | Spata31d1d | spermatogenesis associated 31 subfamily | 0 | 0 | 0 | 0 | 0 | 0 | 0 | 0 | 0 | 0 | 0 |
| 0 | NA | NA | NA | NA | NA | Gm46419 | | 0 | 0 | 0 | 0 | 0 | 0 | 0 | 0 | 0 | 0 | 0 |
| 0 | NA | NA | NA | NA | NA | 4930528D03Rik | | 0 | 0 | 0 | 0 | 0 | 0 | 0 | 0 | 0 | 0 | 0 |
| 0 | NA | NA | NA | NA | NA | Gm49354 | | 0 | 0 | 0 | 0 | 0 | 0 | 0 | 0 | 0 | 0 | 0 |
| 0 | NA | NA | NA | NA | NA | Gm48384 | | 0 | 0 | 0 | 0 | 0 | 0 | 0 | 0 | 0 | 0 | 0 |
| 0 | NA | NA | NA | NA | NA | Gm19866 | | 0 | 0 | 0 | 0 | 0 | 0 | 0 | 0 | 0 | 0 | 0 |
| 0 | NA | NA | NA | NA | NA | Gm48390 | | 0 | 0 | 0 | 0 | 0 | 0 | 0 | 0 | 0 | 0 | 0 |
| 0 | NA | NA | NA | NA | NA | A530065N20Rik | | 0 | 0 | 0 | 0 | 0 | 0 | 0 | 0 | 0 | 0 | 0 |
| 0 | NA | NA | NA | NA | NA | Gm48396 | | 0 | 0 | 0 | 0 | 0 | 0 | 0 | 0 | 0 | 0 | 0 |
| 0 | NA | NA | NA | NA | NA | Gm48397 | | 0 | 0 | 0 | 0 | 0 | 0 | 0 | 0 | 0 | 0 | 0 |
| 0 | NA | NA | NA | NA | NA | Gm48488 | | 0 | 0 | 0 | 0 | 0 | 0 | 0 | 0 | 0 | 0 | 0 |
| 0 | NA | NA | NA | NA | NA | Gm24999 | | 0 | 0 | 0 | 0 | 0 | 0 | 0 | 0 | 0 | 0 | 0 |
| 0 | NA | NA | NA | NA | NA | Gm10779 | | 0 | 0 | 0 | 0 | 0 | 0 | 0 | 0 | 0 | 0 | 0 |
| 0 | NA | NA | NA | NA | NA | Gm35333 | | 0 | 0 | 0 | 0 | 0 | 0 | 0 | 0 | 0 | 0 | 0 |
| 0 | NA | NA | NA | NA | NA | Gm25000 | | 0 | 0 | 0 | 0 | 0 | 0 | 0 | 0 | 0 | 0 | 0 |
| 0 | NA | NA | NA | NA | NA | Gm10312 | | 0 | 0 | 0 | 0 | 0 | 0 | 0 | 0 | 0 | 0 | 0 |
| 0 | NA | NA | NA | NA | NA | BC051665 | cDNA sequence BC051665 | 0 | 0 | 0 | 0 | 0 | 0 | 0 | 0 | 0 | 0 | 0 |
| 0 | NA | NA | NA | NA | NA | Ctsll3 | cathepsin L-like 3 | 0 | 0 | 0 | 0 | 0 | 0 | 0 | 0 | 0 | 0 | 0 |
| 0 | NA | NA | NA | NA | NA | 4930486L24Ril | RIKEN cDNA 4930486L24 gene | 0 | 0 | 0 | 0 | 0 | 0 | 0 | 0 | 0 | 0 | 0 |
| 0 | NA | NA | NA | NA | NA | Gm49392 | | 0 | 0 | 0 | 0 | 0 | 0 | 0 | 0 | 0 | 0 | 0 |
| 0 | NA | NA | NA | NA | NA | Gm49393 | | 0 | 0 | 0 | 0 | 0 | 0 | 0 | 0 | 0 | 0 | 0 |
| 0 | NA | NA | NA | NA | NA | Ctla2b | cytotoxic T lymphocyte-associated protein | 0 | 0 | 0 | 0 | 0 | 0 | 0 | 0 | 0 | 0 | 0 |
| 0 | NA | NA | NA | NA | NA | Gm49391 | | 0 | 0 | 0 | 0 | 0 | 0 | 0 | 0 | 0 | 0 | 0 |
| 0 | NA | NA | NA | NA | NA | Tpbpb | trophoblast specific protein beta | 0 | 0 | 0 | 0 | 0 | 0 | 0 | 0 | 0 | 0 | 0 |
| 0 | NA | NA | NA | NA | NA | Gm24594 | | 0 | 0 | 0 | 0 | 0 | 0 | 0 | 0 | 0 | 0 | 0 |
| 0 | NA | NA | NA | NA | NA | Tpbpa | trophoblast specific protein alpha | 0 | 0 | 0 | 0 | 0 | 0 | 0 | 0 | 0 | 0 | 0 |
| 0 | NA | NA | NA | NA | NA | Gm40975 | | 0 | 0 | 0 | 0 | 0 | 0 | 0 | 0 | 0 | 0 | 0 |
| 0 | NA | NA | NA | NA | NA | Gm40976 | | 0 | 0 | 0 | 0 | 0 | 0 | 0 | 0 | 0 | 0 | 0 |
| 0 | NA | NA | NA | NA | NA | Ctsj | cathepsin J | 0 | 0 | 0 | 0 | 0 | 0 | 0 | 0 | 0 | 0 | 0 |
| 0 | NA | NA | NA | NA | NA | Ctsq | cathepsin Q | 0 | 0 | 0 | 0 | 0 | 0 | 0 | 0 | 0 | 0 | 0 |

| | | | | | | | | | | | | | | | | | | |
|---|---|---|---|---|---|---|---|---|---|---|---|---|---|---|---|---|---|---|
| 0 | NA | NA | NA | NA | NA | Gm18064 | | 0 | 0 | 0 | 0 | 0 | 0 | 0 | 0 | 0 | 0 | 0 |
| 0 | NA | NA | NA | NA | NA | Ctsr | cathepsin R | 0 | 0 | 0 | 0 | 0 | 0 | 0 | 0 | 0 | 0 | 0 |
| 0 | NA | NA | NA | NA | NA | Cts6 | cathepsin 6 | 0 | 0 | 0 | 0 | 0 | 0 | 0 | 0 | 0 | 0 | 0 |
| 0 | NA | NA | NA | NA | NA | Cts8 | cathepsin 8 | 0 | 0 | 0 | 0 | 0 | 0 | 0 | 0 | 0 | 0 | 0 |
| 0 | NA | NA | NA | NA | NA | Gm49398 | | 0 | 0 | 0 | 0 | 0 | 0 | 0 | 0 | 0 | 0 | 0 |
| 0 | NA | NA | NA | NA | NA | Cts8-ps | | 0 | 0 | 0 | 0 | 0 | 0 | 0 | 0 | 0 | 0 | 0 |
| 0 | NA | NA | NA | NA | NA | Gm5664 | | 0 | 0 | 0 | 0 | 0 | 0 | 0 | 0 | 0 | 0 | 0 |
| 0 | NA | NA | NA | NA | NA | Gm27704 | | 0 | 0 | 0 | 0 | 0 | 0 | 0 | 0 | 0 | 0 | 0 |
| 0 | NA | NA | NA | NA | NA | Cts7 | cathepsin 7 | 0 | 0 | 0 | 0 | 0 | 0 | 0 | 0 | 0 | 0 | 0 |
| 0 | NA | NA | NA | NA | NA | Cts7-ps | | 0 | 0 | 0 | 0 | 0 | 0 | 0 | 0 | 0 | 0 | 0 |
| 0 | NA | NA | NA | NA | NA | Gm47825 | | 0 | 0 | 0 | 0 | 0 | 0 | 0 | 0 | 0 | 0 | 0 |
| 0 | NA | NA | NA | NA | NA | Ctsm-ps2 | | 0 | 0 | 0 | 0 | 0 | 0 | 0 | 0 | 0 | 0 | 0 |
| 0 | NA | NA | NA | NA | NA | Gm47848 | | 0 | 0 | 0 | 0 | 0 | 0 | 0 | 0 | 0 | 0 | 0 |
| 0 | NA | NA | NA | NA | NA | Ctsm | cathepsin M | 0 | 0 | 0 | 0 | 0 | 0 | 0 | 0 | 0 | 0 | 0 |
| 0 | NA | NA | NA | NA | NA | Gm49352 | | 0 | 0 | 0 | 0 | 0 | 0 | 0 | 0 | 0 | 0 | 0 |
| 0 | NA | NA | NA | NA | NA | Cts3 | cathepsin 3 | 0 | 0 | 0 | 0 | 0 | 0 | 0 | 0 | 0 | 0 | 0 |
| 0 | NA | NA | NA | NA | NA | Gm49357 | | 0 | 0 | 0 | 0 | 0 | 0 | 0 | 0 | 0 | 0 | 0 |
| 0 | NA | NA | NA | NA | NA | Gm48116 | | 0 | 0 | 0 | 0 | 0 | 0 | 0 | 0 | 0 | 0 | 0 |
| 0 | NA | NA | NA | NA | NA | Gm23739 | | 0 | 0 | 0 | 0 | 0 | 0 | 0 | 0 | 0 | 0 | 0 |
| 0 | NA | NA | NA | NA | NA | Gm24193 | | 0 | 0 | 0 | 0 | 0 | 0 | 0 | 0 | 0 | 0 | 0 |
| 0 | NA | NA | NA | NA | NA | Gm35619 | | 0 | 0 | 0 | 0 | 0 | 0 | 0 | 0 | 0 | 0 | 0 |
| 0 | NA | NA | NA | NA | NA | Gm48228 | | 0 | 0 | 0 | 0 | 0 | 0 | 0 | 0 | 0 | 0 | 0 |
| 0 | NA | NA | NA | NA | NA | Gm25649 | | 0 | 0 | 0 | 0 | 0 | 0 | 0 | 0 | 0 | 0 | 0 |
| 0 | NA | NA | NA | NA | NA | Gm40977 | | 0 | 0 | 0 | 0 | 0 | 0 | 0 | 0 | 0 | 0 | 0 |
| 0 | NA | NA | NA | NA | NA | Gm23713 | | 0 | 0 | 0 | 0 | 0 | 0 | 0 | 0 | 0 | 0 | 0 |
| 0 | NA | NA | NA | NA | NA | Gm24539 | | 0 | 0 | 0 | 0 | 0 | 0 | 0 | 0 | 0 | 0 | 0 |
| 0 | NA | NA | NA | NA | NA | Gm25933 | | 0 | 0 | 0 | 0 | 0 | 0 | 0 | 0 | 0 | 0 | 0 |
| 0 | NA | NA | NA | NA | NA | Gm23855 | | 0 | 0 | 0 | 0 | 0 | 0 | 0 | 0 | 0 | 0 | 0 |
| 0 | NA | NA | NA | NA | NA | Gm48699 | | 0 | 0 | 0 | 0 | 0 | 0 | 0 | 0 | 0 | 0 | 0 |
| 0 | NA | NA | NA | NA | NA | Zfp935 | | 0 | 0 | 0 | 0 | 0 | 0 | 0 | 0 | 0 | 0 | 0 |
| 0 | NA | NA | NA | NA | NA | Gm22256 | | 0 | 0 | 0 | 0 | 0 | 0 | 0 | 0 | 0 | 0 | 0 |
| 0 | NA | NA | NA | NA | NA | Gm24095 | | 0 | 0 | 0 | 0 | 0 | 0 | 0 | 0 | 0 | 0 | 0 |
| 0 | NA | NA | NA | NA | NA | Gm22615 | | 0 | 0 | 0 | 0 | 0 | 0 | 0 | 0 | 0 | 0 | 0 |
| 0 | NA | NA | NA | NA | NA | Gm24020 | | 0 | 0 | 0 | 0 | 0 | 0 | 0 | 0 | 0 | 0 | 0 |
| 0 | NA | NA | NA | NA | NA | Mir713 | | 0 | 0 | 0 | 0 | 0 | 0 | 0 | 0 | 0 | 0 | 0 |
| 0 | NA | NA | NA | NA | NA | Gm48795 | | 0 | 0 | 0 | 0 | 0 | 0 | 0 | 0 | 0 | 0 | 0 |
| 0 | NA | NA | NA | NA | NA | Gm48812 | | 0 | 0 | 0 | 0 | 0 | 0 | 0 | 0 | 0 | 0 | 0 |
| 0 | NA | NA | NA | NA | NA | Gm47584 | | 0 | 0 | 0 | 0 | 0 | 0 | 0 | 0 | 0 | 0 | 0 |
| 0 | NA | NA | NA | NA | NA | Gm47587 | | 0 | 0 | 0 | 0 | 0 | 0 | 0 | 0 | 0 | 0 | 0 |
| 0 | NA | NA | NA | NA | NA | Gm47603 | | 0 | 0 | 0 | 0 | 0 | 0 | 0 | 0 | 0 | 0 | 0 |
| 0 | NA | NA | NA | NA | NA | Gm47602 | | 0 | 0 | 0 | 0 | 0 | 0 | 0 | 0 | 0 | 0 | 0 |
| 0 | NA | NA | NA | NA | NA | Mir23b | | 0 | 0 | 0 | 0 | 0 | 0 | 0 | 0 | 0 | 0 | 0 |
| 0 | NA | NA | NA | NA | NA | Mir27b | | 0 | 0 | 0 | 0 | 0 | 0 | 0 | 0 | 0 | 0 | 0 |
| 0 | NA | NA | NA | NA | NA | Mir3074-1 | | 0 | 0 | 0 | 0 | 0 | 0 | 0 | 0 | 0 | 0 | 0 |
| 0 | NA | NA | NA | NA | NA | Mir24-1 | | 0 | 0 | 0 | 0 | 0 | 0 | 0 | 0 | 0 | 0 | 0 |
| 0 | NA | NA | NA | NA | NA | Gm16132 | | 0 | 0 | 0 | 0 | 0 | 0 | 0 | 0 | 0 | 0 | 0 |
| 0 | NA | NA | NA | NA | NA | Gm16134 | | 0 | 0 | 0 | 0 | 0 | 0 | 0 | 0 | 0 | 0 | 0 |
| 0 | NA | NA | NA | NA | NA | A930032L01Rik | | 0 | 0 | 0 | 0 | 0 | 0 | 0 | 0 | 0 | 0 | 0 |
| 0 | NA | NA | NA | NA | NA | Gm30655 | | 0 | 0 | 0 | 0 | 0 | 0 | 0 | 0 | 0 | 0 | 0 |
| 0 | NA | NA | NA | NA | NA | Gm30709 | | 0 | 0 | 0 | 0 | 0 | 0 | 0 | 0 | 0 | 0 | 0 |
| 0 | NA | NA | NA | NA | NA | Gm47387 | | 0 | 0 | 0 | 0 | 0 | 0 | 0 | 0 | 0 | 0 | 0 |
| 0 | NA | NA | NA | NA | NA | 1700024I08Rik | | 0 | 0 | 0 | 0 | 0 | 0 | 0 | 0 | 0 | 0 | 0 |
| 0 | NA | NA | NA | NA | NA | Gm47390 | | 0 | 0 | 0 | 0 | 0 | 0 | 0 | 0 | 0 | 0 | 0 |
| 0 | NA | NA | NA | NA | NA | Gm47417 | | 0 | 0 | 0 | 0 | 0 | 0 | 0 | 0 | 0 | 0 | 0 |
| 0 | NA | NA | NA | NA | NA | Gm47418 | | 0 | 0 | 0 | 0 | 0 | 0 | 0 | 0 | 0 | 0 | 0 |
| 0 | NA | NA | NA | NA | NA | Gm7695 | | 0 | 0 | 0 | 0 | 0 | 0 | 0 | 0 | 0 | 0 | 0 |
| 0 | NA | NA | NA | NA | NA | Gm46424 | | 0 | 0 | 0 | 0 | 0 | 0 | 0 | 0 | 0 | 0 | 0 |
| 0 | NA | NA | NA | NA | NA | | | 0 | 0 | 0 | 0 | 0 | 0 | 0 | 0 | 0 | 0 | 0 |
| 0 | NA | NA | NA | NA | NA | Gm47123 | | 0 | 0 | 0 | 0 | 0 | 0 | 0 | 0 | 0 | 0 | 0 |
| 0 | NA | NA | NA | NA | NA | Gm49230 | | 0 | 0 | 0 | 0 | 0 | 0 | 0 | 0 | 0 | 0 | 0 |

| | | | | | | | | | | | | | | | | | | |
|---|---|---|---|---|---|---|---|---|---|---|---|---|---|---|---|---|---|---|
| 0 | NA | NA | NA | NA | NA | Fam240b | family with sequence similarity 240 memb | 0 | 0 | 0 | 0 | 0 | 0 | 0 | 0 | 0 | 0 | 0 |
| 0 | NA | NA | NA | NA | NA | Gm47190 | | 0 | 0 | 0 | 0 | 0 | 0 | 0 | 0 | 0 | 0 | 0 |
| 0 | NA | NA | NA | NA | NA | Gm47191 | | 0 | 0 | 0 | 0 | 0 | 0 | 0 | 0 | 0 | 0 | 0 |
| 0 | NA | NA | NA | NA | NA | Gm47194 | | 0 | 0 | 0 | 0 | 0 | 0 | 0 | 0 | 0 | 0 | 0 |
| 0 | NA | NA | NA | NA | NA | Spata31 | spermatogenesis associated 31 | 0 | 0 | 0 | 0 | 0 | 0 | 0 | 0 | 0 | 0 | 0 |
| 0 | NA | NA | NA | NA | NA | Gm6888 | | 0 | 0 | 0 | 0 | 0 | 0 | 0 | 0 | 0 | 0 | 0 |
| 0 | NA | NA | NA | NA | NA | Spata31d1c | spermatogenesis associated 31 subfamily | 0 | 0 | 0 | 0 | 0 | 0 | 0 | 0 | 0 | 0 | 0 |
| 0 | NA | NA | NA | NA | NA | Prss47 | protease, serine 47 | 0 | 0 | 0 | 0 | 0 | 0 | 0 | 0 | 0 | 0 | 0 |
| 0 | NA | NA | NA | NA | NA | Olfr465-ps1 | | 0 | 0 | 0 | 0 | 0 | 0 | 0 | 0 | 0 | 0 | 0 |
| 0 | NA | NA | NA | NA | NA | Olfr466 | olfactory receptor 466 | 0 | 0 | 0 | 0 | 0 | 0 | 0 | 0 | 0 | 0 | 0 |
| 0 | NA | NA | NA | NA | NA | Gm24130 | | 0 | 0 | 0 | 0 | 0 | 0 | 0 | 0 | 0 | 0 | 0 |
| 0 | NA | NA | NA | NA | NA | Gm47249 | | 0 | 0 | 0 | 0 | 0 | 0 | 0 | 0 | 0 | 0 | 0 |
| 0 | NA | NA | NA | NA | NA | Gm47250 | | 0 | 0 | 0 | 0 | 0 | 0 | 0 | 0 | 0 | 0 | 0 |
| 0 | NA | NA | NA | NA | NA | Gm47251 | | 0 | 0 | 0 | 0 | 0 | 0 | 0 | 0 | 0 | 0 | 0 |
| 0 | NA | NA | NA | NA | NA | Gm47263 | | 0 | 0 | 0 | 0 | 0 | 0 | 0 | 0 | 0 | 0 | 0 |
| 0 | NA | NA | NA | NA | NA | Gm7762 | predicted gene 7762 | 0 | 0 | 0 | 0 | 0 | 0 | 0 | 0 | 0 | 0 | 0 |
| 0 | NA | NA | NA | NA | NA | Gm47264 | | 0 | 0 | 0 | 0 | 0 | 0 | 0 | 0 | 0 | 0 | 0 |
| 0 | NA | NA | NA | NA | NA | Vmn2r-ps95 | | 0 | 0 | 0 | 0 | 0 | 0 | 0 | 0 | 0 | 0 | 0 |
| 0 | NA | NA | NA | NA | NA | Gm7768 | | 0 | 0 | 0 | 0 | 0 | 0 | 0 | 0 | 0 | 0 | 0 |
| 0 | NA | NA | NA | NA | NA | Gm40983 | | 0 | 0 | 0 | 0 | 0 | 0 | 0 | 0 | 0 | 0 | 0 |
| 0 | NA | NA | NA | NA | NA | Gm48103 | | 0 | 0 | 0 | 0 | 0 | 0 | 0 | 0 | 0 | 0 | 0 |
| 0 | NA | NA | NA | NA | NA | Gm21930 | | 0 | 0 | 0 | 0 | 0 | 0 | 0 | 0 | 0 | 0 | 0 |
| 0 | NA | NA | NA | NA | NA | Gm48111 | | 0 | 0 | 0 | 0 | 0 | 0 | 0 | 0 | 0 | 0 | 0 |
| 0 | NA | NA | NA | NA | NA | Cbx3-ps5 | | 0 | 0 | 0 | 0 | 0 | 0 | 0 | 0 | 0 | 0 | 0 |
| 0 | NA | NA | NA | NA | NA | Vmn2r-ps96 | | 0 | 0 | 0 | 0 | 0 | 0 | 0 | 0 | 0 | 0 | 0 |
| 0 | NA | NA | NA | NA | NA | Hmgb1-ps11 | | 0 | 0 | 0 | 0 | 0 | 0 | 0 | 0 | 0 | 0 | 0 |
| 0 | NA | NA | NA | NA | NA | Gm48136 | | 0 | 0 | 0 | 0 | 0 | 0 | 0 | 0 | 0 | 0 | 0 |
| 0 | NA | NA | NA | NA | NA | Gm48165 | | 0 | 0 | 0 | 0 | 0 | 0 | 0 | 0 | 0 | 0 | 0 |
| 0 | NA | NA | NA | NA | NA | Gm48168 | | 0 | 0 | 0 | 0 | 0 | 0 | 0 | 0 | 0 | 0 | 0 |
| 0 | NA | NA | NA | NA | NA | Gm48166 | | 0 | 0 | 0 | 0 | 0 | 0 | 0 | 0 | 0 | 0 | 0 |
| 0 | NA | NA | NA | NA | NA | Gm10139 | | 0 | 0 | 0 | 0 | 0 | 0 | 0 | 0 | 0 | 0 | 0 |
| 0 | NA | NA | NA | NA | NA | Gm48171 | | 0 | 0 | 0 | 0 | 0 | 0 | 0 | 0 | 0 | 0 | 0 |
| 0 | NA | NA | NA | NA | NA | Gm48169 | | 0 | 0 | 0 | 0 | 0 | 0 | 0 | 0 | 0 | 0 | 0 |
| 0 | NA | NA | NA | NA | NA | Gm48167 | | 0 | 0 | 0 | 0 | 0 | 0 | 0 | 0 | 0 | 0 | 0 |
| 0 | NA | NA | NA | NA | NA | Cbx3-ps2 | | 0 | 0 | 0 | 0 | 0 | 0 | 0 | 0 | 0 | 0 | 0 |
| 0 | NA | NA | NA | NA | NA | Vmn2r-ps97 | | 0 | 0 | 0 | 0 | 0 | 0 | 0 | 0 | 0 | 0 | 0 |
| 0 | NA | NA | NA | NA | NA | Gm3829 | | 0 | 0 | 0 | 0 | 0 | 0 | 0 | 0 | 0 | 0 | 0 |
| 0 | NA | NA | NA | NA | NA | Gm26639 | | 0 | 0 | 0 | 0 | 0 | 0 | 0 | 0 | 0 | 0 | 0 |
| 0 | NA | NA | NA | NA | NA | Gm48170 | | 0 | 0 | 0 | 0 | 0 | 0 | 0 | 0 | 0 | 0 | 0 |
| 0 | NA | NA | NA | NA | NA | Gm48164 | | 0 | 0 | 0 | 0 | 0 | 0 | 0 | 0 | 0 | 0 | 0 |
| 0 | NA | NA | NA | NA | NA | Gm48185 | | 0 | 0 | 0 | 0 | 0 | 0 | 0 | 0 | 0 | 0 | 0 |
| 0 | NA | NA | NA | NA | NA | Gm48186 | | 0 | 0 | 0 | 0 | 0 | 0 | 0 | 0 | 0 | 0 | 0 |
| 0 | NA | NA | NA | NA | NA | Gm19428 | | 0 | 0 | 0 | 0 | 0 | 0 | 0 | 0 | 0 | 0 | 0 |
| 0 | NA | NA | NA | NA | NA | Gm19118 | | 0 | 0 | 0 | 0 | 0 | 0 | 0 | 0 | 0 | 0 | 0 |
| 0 | NA | NA | NA | NA | NA | Vmn2r-ps98 | | 0 | 0 | 0 | 0 | 0 | 0 | 0 | 0 | 0 | 0 | 0 |
| 0 | NA | NA | NA | NA | NA | Gm38428 | | 0 | 0 | 0 | 0 | 0 | 0 | 0 | 0 | 0 | 0 | 0 |
| 0 | NA | NA | NA | NA | NA | Gm7822 | | 0 | 0 | 0 | 0 | 0 | 0 | 0 | 0 | 0 | 0 | 0 |
| 0 | NA | NA | NA | NA | NA | Vmn2r-ps99 | | 0 | 0 | 0 | 0 | 0 | 0 | 0 | 0 | 0 | 0 | 0 |
| 0 | NA | NA | NA | NA | NA | Cbx3-ps1 | | 0 | 0 | 0 | 0 | 0 | 0 | 0 | 0 | 0 | 0 | 0 |
| 0 | NA | NA | NA | NA | NA | Gm48218 | | 0 | 0 | 0 | 0 | 0 | 0 | 0 | 0 | 0 | 0 | 0 |
| 0 | NA | NA | NA | NA | NA | Gm48219 | | 0 | 0 | 0 | 0 | 0 | 0 | 0 | 0 | 0 | 0 | 0 |
| 0 | NA | NA | NA | NA | NA | Vmn2r-ps100 | | 0 | 0 | 0 | 0 | 0 | 0 | 0 | 0 | 0 | 0 | 0 |
| 0 | NA | NA | NA | NA | NA | Gm40987 | | 0 | 0 | 0 | 0 | 0 | 0 | 0 | 0 | 0 | 0 | 0 |
| 0 | NA | NA | NA | NA | NA | Gm48220 | | 0 | 0 | 0 | 0 | 0 | 0 | 0 | 0 | 0 | 0 | 0 |
| 0 | NA | NA | NA | NA | NA | Gm48221 | | 0 | 0 | 0 | 0 | 0 | 0 | 0 | 0 | 0 | 0 | 0 |
| 0 | NA | NA | NA | NA | NA | Gm48222 | | 0 | 0 | 0 | 0 | 0 | 0 | 0 | 0 | 0 | 0 | 0 |
| 0 | NA | NA | NA | NA | NA | Gm10324 | predicted gene 10324 | 0 | 0 | 0 | 0 | 0 | 0 | 0 | 0 | 0 | 0 | 0 |
| 0 | NA | NA | NA | NA | NA | Gm50489 | | 0 | 0 | 0 | 0 | 0 | 0 | 0 | 0 | 0 | 0 | 0 |
| 0 | NA | NA | NA | NA | NA | Gm45965 | | 0 | 0 | 0 | 0 | 0 | 0 | 0 | 0 | 0 | 0 | 0 |
| 0 | NA | NA | NA | NA | NA | Gm48223 | | 0 | 0 | 0 | 0 | 0 | 0 | 0 | 0 | 0 | 0 | 0 |

| | | | | | | | | | | | | | | | | | | |
|---|---|---|---|---|---|---|---|---|---|---|---|---|---|---|---|---|---|---|
| 0 | NA | NA | NA | NA | NA | Vmn2r-ps102 | | 0 | 0 | 0 | 0 | 0 | 0 | 0 | 0 | 0 | 0 | 0 |
| 0 | NA | NA | NA | NA | NA | Gm48240 | | 0 | 0 | 0 | 0 | 0 | 0 | 0 | 0 | 0 | 0 | 0 |
| 0 | NA | NA | NA | NA | NA | Gm10772 | | 0 | 0 | 0 | 0 | 0 | 0 | 0 | 0 | 0 | 0 | 0 |
| 0 | NA | NA | NA | NA | NA | Gm48241 | | 0 | 0 | 0 | 0 | 0 | 0 | 0 | 0 | 0 | 0 | 0 |
| 0 | NA | NA | NA | NA | NA | Gm26754 | | 0 | 0 | 0 | 0 | 0 | 0 | 0 | 0 | 0 | 0 | 0 |
| 0 | NA | NA | NA | NA | NA | Vmn2r-ps103 | | 0 | 0 | 0 | 0 | 0 | 0 | 0 | 0 | 0 | 0 | 0 |
| 0 | NA | NA | NA | NA | NA | Gm48245 | | 0 | 0 | 0 | 0 | 0 | 0 | 0 | 0 | 0 | 0 | 0 |
| 0 | NA | NA | NA | NA | NA | Gm46426 | | 0 | 0 | 0 | 0 | 0 | 0 | 0 | 0 | 0 | 0 | 0 |
| 0 | NA | NA | NA | NA | NA | Gm18260 | | 0 | 0 | 0 | 0 | 0 | 0 | 0 | 0 | 0 | 0 | 0 |
| 0 | NA | NA | NA | NA | NA | Gm48246 | | 0 | 0 | 0 | 0 | 0 | 0 | 0 | 0 | 0 | 0 | 0 |
| 0 | NA | NA | NA | NA | NA | C330022B21Rik | | 0 | 0 | 0 | 0 | 0 | 0 | 0 | 0 | 0 | 0 | 0 |
| 0 | NA | NA | NA | NA | NA | Vmn2r-ps104 | | 0 | 0 | 0 | 0 | 0 | 0 | 0 | 0 | 0 | 0 | 0 |
| 0 | NA | NA | NA | NA | NA | Gm48247 | | 0 | 0 | 0 | 0 | 0 | 0 | 0 | 0 | 0 | 0 | 0 |
| 0 | NA | NA | NA | NA | NA | Gm48244 | | 0 | 0 | 0 | 0 | 0 | 0 | 0 | 0 | 0 | 0 | 0 |
| 0 | NA | NA | NA | NA | NA | Gm19119 | | 0 | 0 | 0 | 0 | 0 | 0 | 0 | 0 | 0 | 0 | 0 |
| 0 | NA | NA | NA | NA | NA | Gm48243 | | 0 | 0 | 0 | 0 | 0 | 0 | 0 | 0 | 0 | 0 | 0 |
| 0 | NA | NA | NA | NA | NA | 2410141K09Ril | RIKEN cDNA 2410141K09 gene | 0 | 0 | 0 | 0 | 0 | 0 | 0 | 0 | 0 | 0 | 0 |
| 0 | NA | NA | NA | NA | NA | Gm48322 | | 0 | 0 | 0 | 0 | 0 | 0 | 0 | 0 | 0 | 0 | 0 |
| 0 | NA | NA | NA | NA | NA | Gm48323 | | 0 | 0 | 0 | 0 | 0 | 0 | 0 | 0 | 0 | 0 | 0 |
| 0 | NA | NA | NA | NA | NA | Gm26806 | | 0 | 0 | 0 | 0 | 0 | 0 | 0 | 0 | 0 | 0 | 0 |
| 0 | NA | NA | NA | NA | NA | Vmn2r-ps105 | | 0 | 0 | 0 | 0 | 0 | 0 | 0 | 0 | 0 | 0 | 0 |
| 0 | NA | NA | NA | NA | NA | Gm46427 | | 0 | 0 | 0 | 0 | 0 | 0 | 0 | 0 | 0 | 0 | 0 |
| 0 | NA | NA | NA | NA | NA | Gm48351 | | 0 | 0 | 0 | 0 | 0 | 0 | 0 | 0 | 0 | 0 | 0 |
| 0 | NA | NA | NA | NA | NA | Gm48360 | | 0 | 0 | 0 | 0 | 0 | 0 | 0 | 0 | 0 | 0 | 0 |
| 0 | NA | NA | NA | NA | NA | Gm49531 | | 0 | 0 | 0 | 0 | 0 | 0 | 0 | 0 | 0 | 0 | 0 |
| 0 | NA | NA | NA | NA | NA | Gm10769 | | 0 | 0 | 0 | 0 | 0 | 0 | 0 | 0 | 0 | 0 | 0 |
| 0 | NA | NA | NA | NA | NA | Gm27501 | | 0 | 0 | 0 | 0 | 0 | 0 | 0 | 0 | 0 | 0 | 0 |
| 0 | NA | NA | NA | NA | NA | Gm49532 | | 0 | 0 | 0 | 0 | 0 | 0 | 0 | 0 | 0 | 0 | 0 |
| 0 | NA | NA | NA | NA | NA | Gm48361 | | 0 | 0 | 0 | 0 | 0 | 0 | 0 | 0 | 0 | 0 | 0 |
| 0 | NA | NA | NA | NA | NA | Zfp640 | | 0 | 0 | 0 | 0 | 0 | 0 | 0 | 0 | 0 | 0 | 0 |
| 0 | NA | NA | NA | NA | NA | Gm40988 | | 0 | 0 | 0 | 0 | 0 | 0 | 0 | 0 | 0 | 0 | 0 |
| 0 | NA | NA | NA | NA | NA | Vmn2r-ps106 | | 0 | 0 | 0 | 0 | 0 | 0 | 0 | 0 | 0 | 0 | 0 |
| 0 | NA | NA | NA | NA | NA | Gm40989 | | 0 | 0 | 0 | 0 | 0 | 0 | 0 | 0 | 0 | 0 | 0 |
| 0 | NA | NA | NA | NA | NA | Gm48404 | | 0 | 0 | 0 | 0 | 0 | 0 | 0 | 0 | 0 | 0 | 0 |
| 0 | NA | NA | NA | NA | NA | Gm48412 | | 0 | 0 | 0 | 0 | 0 | 0 | 0 | 0 | 0 | 0 | 0 |
| 0 | NA | NA | NA | NA | NA | Gm7896 | predicted gene 7869 | 0 | 0 | 0 | 0 | 0 | 0 | 0 | 0 | 0 | 0 | 0 |
| 0 | NA | NA | NA | NA | NA | Gm48413 | | 0 | 0 | 0 | 0 | 0 | 0 | 0 | 0 | 0 | 0 | 0 |
| 0 | NA | NA | NA | NA | NA | Vmn2r-ps108 | | 0 | 0 | 0 | 0 | 0 | 0 | 0 | 0 | 0 | 0 | 0 |
| 0 | NA | NA | NA | NA | NA | Gm48414 | | 0 | 0 | 0 | 0 | 0 | 0 | 0 | 0 | 0 | 0 | 0 |
| 0 | NA | NA | NA | NA | NA | Gm10323 | predicted gene 10323 | 0 | 0 | 0 | 0 | 0 | 0 | 0 | 0 | 0 | 0 | 0 |
| 0 | NA | NA | NA | NA | NA | Cbx3-ps4 | | 0 | 0 | 0 | 0 | 0 | 0 | 0 | 0 | 0 | 0 | 0 |
| 0 | NA | NA | NA | NA | NA | Vmn2r-ps109 | | 0 | 0 | 0 | 0 | 0 | 0 | 0 | 0 | 0 | 0 | 0 |
| 0 | NA | NA | NA | NA | NA | Gm48705 | | 0 | 0 | 0 | 0 | 0 | 0 | 0 | 0 | 0 | 0 | 0 |
| 0 | NA | NA | NA | NA | NA | Zfp708 | zinc finger protein 708 | 0 | 0 | 0 | 0 | 0 | 0 | 0 | 0 | 0 | 0 | 0 |
| 0 | NA | NA | NA | NA | NA | Gm48732 | | 0 | 0 | 0 | 0 | 0 | 0 | 0 | 0 | 0 | 0 | 0 |
| 0 | NA | NA | NA | NA | NA | Gm28044 | | 0 | 0 | 0 | 0 | 0 | 0 | 0 | 0 | 0 | 0 | 0 |
| 0 | NA | NA | NA | NA | NA | Gm28041 | | 0 | 0 | 0 | 0 | 0 | 0 | 0 | 0 | 0 | 0 | 0 |
| 0 | NA | NA | NA | NA | NA | Gm48900 | | 0 | 0 | 0 | 0 | 0 | 0 | 0 | 0 | 0 | 0 | 0 |
| 0 | NA | NA | NA | NA | NA | Gm7928 | | 0 | 0 | 0 | 0 | 0 | 0 | 0 | 0 | 0 | 0 | 0 |
| 0 | NA | NA | NA | NA | NA | Gm26965 | | 0 | 0 | 0 | 0 | 0 | 0 | 0 | 0 | 0 | 0 | 0 |
| 0 | NA | NA | NA | NA | NA | Gm49345 | | 0 | 0 | 0 | 0 | 0 | 0 | 0 | 0 | 0 | 0 | 0 |
| 0 | NA | NA | NA | NA | NA | Gm26587 | | 0 | 0 | 0 | 0 | 0 | 0 | 0 | 0 | 0 | 0 | 0 |
| 0 | NA | NA | NA | NA | NA | Gm26875 | | 0 | 0 | 0 | 0 | 0 | 0 | 0 | 0 | 0 | 0 | 0 |
| 0 | NA | NA | NA | NA | NA | Gm9894 | | 0 | 0 | 0 | 0 | 0 | 0 | 0 | 0 | 0 | 0 | 0 |
| 0 | NA | NA | NA | NA | NA | Gm7969 | | 0 | 0 | 0 | 0 | 0 | 0 | 0 | 0 | 0 | 0 | 0 |
| 0 | NA | NA | NA | NA | NA | Gm48419 | | 0 | 0 | 0 | 0 | 0 | 0 | 0 | 0 | 0 | 0 | 0 |
| 0 | NA | NA | NA | NA | NA | Gm48423 | | 0 | 0 | 0 | 0 | 0 | 0 | 0 | 0 | 0 | 0 | 0 |
| 0 | NA | NA | NA | NA | NA | Gm48424 | | 0 | 0 | 0 | 0 | 0 | 0 | 0 | 0 | 0 | 0 | 0 |
| 0 | NA | NA | NA | NA | NA | AA414992 | | 0 | 0 | 0 | 0 | 0 | 0 | 0 | 0 | 0 | 0 | 0 |
| 0 | NA | NA | NA | NA | NA | Gm8016 | | 0 | 0 | 0 | 0 | 0 | 0 | 0 | 0 | 0 | 0 | 0 |

| | | | | | | | | | | | | | | | | | | |
|---|---|---|---|---|---|---|---|---|---|---|---|---|---|---|---|---|---|---|
| 0 | NA | NA | NA | NA | NA | Gm48429 | | 0 | 0 | 0 | 0 | 0 | 0 | 0 | 0 | 0 | 0 | 0 |
| 0 | NA | NA | NA | NA | NA | Gm48436 | | 0 | 0 | 0 | 0 | 0 | 0 | 0 | 0 | 0 | 0 | 0 |
| 0 | NA | NA | NA | NA | NA | Gm48556 | | 0 | 0 | 0 | 0 | 0 | 0 | 0 | 0 | 0 | 0 | 0 |
| 0 | NA | NA | NA | NA | NA | Gm26929 | | 0 | 0 | 0 | 0 | 0 | 0 | 0 | 0 | 0 | 0 | 0 |
| 0 | NA | NA | NA | NA | NA | Gm48593 | | 0 | 0 | 0 | 0 | 0 | 0 | 0 | 0 | 0 | 0 | 0 |
| 0 | NA | NA | NA | NA | NA | Gm37122 | | 0 | 0 | 0 | 0 | 0 | 0 | 0 | 0 | 0 | 0 | 0 |
| 0 | NA | NA | NA | NA | NA | Gm26844 | | 0 | 0 | 0 | 0 | 0 | 0 | 0 | 0 | 0 | 0 | 0 |
| 0 | NA | NA | NA | NA | NA | Gm35514 | | 0 | 0 | 0 | 0 | 0 | 0 | 0 | 0 | 0 | 0 | 0 |
| 0 | NA | NA | NA | NA | NA | Gm44375 | | 0 | 0 | 0 | 0 | 0 | 0 | 0 | 0 | 0 | 0 | 0 |
| 0 | NA | NA | NA | NA | NA | Gm48676 | | 0 | 0 | 0 | 0 | 0 | 0 | 0 | 0 | 0 | 0 | 0 |
| 0 | NA | NA | NA | NA | NA | Gm17872 | | 0 | 0 | 0 | 0 | 0 | 0 | 0 | 0 | 0 | 0 | 0 |
| 0 | NA | NA | NA | NA | NA | Gm48816 | | 0 | 0 | 0 | 0 | 0 | 0 | 0 | 0 | 0 | 0 | 0 |
| 0 | NA | NA | NA | NA | NA | Gm48817 | | 0 | 0 | 0 | 0 | 0 | 0 | 0 | 0 | 0 | 0 | 0 |
| 0 | NA | NA | NA | NA | NA | Gm18313 | | 0 | 0 | 0 | 0 | 0 | 0 | 0 | 0 | 0 | 0 | 0 |
| 0 | NA | NA | NA | NA | NA | Gm35618 | | 0 | 0 | 0 | 0 | 0 | 0 | 0 | 0 | 0 | 0 | 0 |
| 0 | NA | NA | NA | NA | NA | Gm48819 | | 0 | 0 | 0 | 0 | 0 | 0 | 0 | 0 | 0 | 0 | 0 |
| 0 | NA | NA | NA | NA | NA | 4930547H16Rik | | 0 | 0 | 0 | 0 | 0 | 0 | 0 | 0 | 0 | 0 | 0 |
| 0 | NA | NA | NA | NA | NA | Gm20288 | | 0 | 0 | 0 | 0 | 0 | 0 | 0 | 0 | 0 | 0 | 0 |
| 0 | NA | NA | NA | NA | NA | 4933416O17Rik | | 0 | 0 | 0 | 0 | 0 | 0 | 0 | 0 | 0 | 0 | 0 |
| 0 | NA | NA | NA | NA | NA | 1700084F23Rik | | 0 | 0 | 0 | 0 | 0 | 0 | 0 | 0 | 0 | 0 | 0 |
| 0 | NA | NA | NA | NA | NA | Gm47656 | | 0 | 0 | 0 | 0 | 0 | 0 | 0 | 0 | 0 | 0 | 0 |
| 0 | NA | NA | NA | NA | NA | Gm26018 | | 0 | 0 | 0 | 0 | 0 | 0 | 0 | 0 | 0 | 0 | 0 |
| 0 | NA | NA | NA | NA | NA | 1700100L14Rik | | 0 | 0 | 0 | 0 | 0 | 0 | 0 | 0 | 0 | 0 | 0 |
| 0 | NA | NA | NA | NA | NA | Gm36426 | | 0 | 0 | 0 | 0 | 0 | 0 | 0 | 0 | 0 | 0 | 0 |
| 0 | NA | NA | NA | NA | NA | Gm36607 | | 0 | 0 | 0 | 0 | 0 | 0 | 0 | 0 | 0 | 0 | 0 |
| 0 | NA | NA | NA | NA | NA | Gm36529 | | 0 | 0 | 0 | 0 | 0 | 0 | 0 | 0 | 0 | 0 | 0 |
| 0 | NA | NA | NA | NA | NA | 8030423J24Rik | | 0 | 0 | 0 | 0 | 0 | 0 | 0 | 0 | 0 | 0 | 0 |
| 0 | NA | NA | NA | NA | NA | Gm38191 | | 0 | 0 | 0 | 0 | 0 | 0 | 0 | 0 | 0 | 0 | 0 |
| 0 | NA | NA | NA | NA | NA | Mir466f-4 | | 0 | 0 | 0 | 0 | 0 | 0 | 0 | 0 | 0 | 0 | 0 |
| 0 | NA | NA | NA | NA | NA | 1700112M02Rik | | 0 | 0 | 0 | 0 | 0 | 0 | 0 | 0 | 0 | 0 | 0 |
| 0 | NA | NA | NA | NA | NA | Gm47811 | | 0 | 0 | 0 | 0 | 0 | 0 | 0 | 0 | 0 | 0 | 0 |
| 0 | NA | NA | NA | NA | NA | Gm47812 | | 0 | 0 | 0 | 0 | 0 | 0 | 0 | 0 | 0 | 0 | 0 |
| 0 | NA | NA | NA | NA | NA | Gm47838 | | 0 | 0 | 0 | 0 | 0 | 0 | 0 | 0 | 0 | 0 | 0 |
| 0 | NA | NA | NA | NA | NA | Gm47890 | | 0 | 0 | 0 | 0 | 0 | 0 | 0 | 0 | 0 | 0 | 0 |
| 0 | NA | NA | NA | NA | NA | Gm47902 | | 0 | 0 | 0 | 0 | 0 | 0 | 0 | 0 | 0 | 0 | 0 |
| 0 | NA | NA | NA | NA | NA | D730050B12Rik | | 0 | 0 | 0 | 0 | 0 | 0 | 0 | 0 | 0 | 0 | 0 |
| 0 | NA | NA | NA | NA | NA | Rpl31-ps2 | | 0 | 0 | 0 | 0 | 0 | 0 | 0 | 0 | 0 | 0 | 0 |
| 0 | NA | NA | NA | NA | NA | Gm48000 | | 0 | 0 | 0 | 0 | 0 | 0 | 0 | 0 | 0 | 0 | 0 |
| 0 | NA | NA | NA | NA | NA | Slc6a3 | solute carrier family 6 (neurotransmitter tra | 0 | 0 | 0 | 0 | 0 | 0 | 0 | 0 | 0 | 0 | 0 |
| 0 | NA | NA | NA | NA | NA | Gm25197 | | 0 | 0 | 0 | 0 | 0 | 0 | 0 | 0 | 0 | 0 | 0 |
| 0 | NA | NA | NA | NA | NA | Gm41002 | | 0 | 0 | 0 | 0 | 0 | 0 | 0 | 0 | 0 | 0 | 0 |
| 0 | NA | NA | NA | NA | NA | Zdhhc11 | zinc finger, DHHC domain containing 11 | 0 | 0 | 0 | 0 | 0 | 0 | 0 | 0 | 0 | 0 | 0 |
| 0 | NA | NA | NA | NA | NA | Gm10126 | | 0 | 0 | 0 | 0 | 0 | 0 | 0 | 0 | 0 | 0 | 0 |
| 0 | NA | NA | NA | NA | NA | Slc9a3 | solute carrier family 9 (sodium/hydrogen e | 0 | 0 | 0 | 0 | 0 | 0 | 0 | 0 | 0 | 0 | 0 |
| 0 | NA | NA | NA | NA | NA | Gm8227 | | 0 | 0 | 0 | 0 | 0 | 0 | 0 | 0 | 0 | 0 | 0 |
| 0 | NA | NA | NA | NA | NA | Mir692-3 | | 0 | 0 | 0 | 0 | 0 | 0 | 0 | 0 | 0 | 0 | 0 |
| 0 | NA | NA | NA | NA | NA | Gm49762 | | 0 | 0 | 0 | 0 | 0 | 0 | 0 | 0 | 0 | 0 | 0 |
| 0 | NA | NA | NA | NA | NA | Gm49763 | | 0 | 0 | 0 | 0 | 0 | 0 | 0 | 0 | 0 | 0 | 0 |
| 0 | NA | NA | NA | NA | NA | Gm47470 | | 0 | 0 | 0 | 0 | 0 | 0 | 0 | 0 | 0 | 0 | 0 |
| 0 | NA | NA | NA | NA | NA | Gm19095 | | 0 | 0 | 0 | 0 | 0 | 0 | 0 | 0 | 0 | 0 | 0 |
| 0 | NA | NA | NA | NA | NA | Gm48235 | | 0 | 0 | 0 | 0 | 0 | 0 | 0 | 0 | 0 | 0 | 0 |
| 0 | NA | NA | NA | NA | NA | Gm3926 | | 0 | 0 | 0 | 0 | 0 | 0 | 0 | 0 | 0 | 0 | 0 |
| 0 | NA | NA | NA | NA | NA | Mir682 | | 0 | 0 | 0 | 0 | 0 | 0 | 0 | 0 | 0 | 0 | 0 |
| 0 | NA | NA | NA | NA | NA | Gm23012 | | 0 | 0 | 0 | 0 | 0 | 0 | 0 | 0 | 0 | 0 | 0 |
| 0 | NA | NA | NA | NA | NA | Gm23127 | | 0 | 0 | 0 | 0 | 0 | 0 | 0 | 0 | 0 | 0 | 0 |
| 0 | NA | NA | NA | NA | NA | Gm24163 | | 0 | 0 | 0 | 0 | 0 | 0 | 0 | 0 | 0 | 0 | 0 |
| 0 | NA | NA | NA | NA | NA | Gm23319 | | 0 | 0 | 0 | 0 | 0 | 0 | 0 | 0 | 0 | 0 | 0 |
| 0 | NA | NA | NA | NA | NA | Gm4165 | | 0 | 0 | 0 | 0 | 0 | 0 | 0 | 0 | 0 | 0 | 0 |
| 0 | NA | NA | NA | NA | NA | Fam81b | family with sequence similarity 81, membe | 0 | 0 | 0 | 0 | 0 | 0 | 0 | 0 | 0 | 0 | 0 |
| 0 | NA | NA | NA | NA | NA | Gm31219 | | 0 | 0 | 0 | 0 | 0 | 0 | 0 | 0 | 0 | 0 | 0 |

| | | | | | | | | | | | | | | | | | |
|---|---|---|---|---|---|---|---|---|---|---|---|---|---|---|---|---|---|
| 0 | NA | NA | NA | NA | NA | Gm24025 | 0 | 0 | 0 | 0 | 0 | 0 | 0 | 0 | 0 | 0 | 0 |
| 0 | NA | NA | NA | NA | NA | Gm24519 | 0 | 0 | 0 | 0 | 0 | 0 | 0 | 0 | 0 | 0 | 0 |
| 0 | NA | NA | NA | NA | NA | Gm29844 | 0 | 0 | 0 | 0 | 0 | 0 | 0 | 0 | 0 | 0 | 0 |
| 0 | NA | NA | NA | NA | NA | Gm9634 | 0 | 0 | 0 | 0 | 0 | 0 | 0 | 0 | 0 | 0 | 0 |
| 0 | NA | NA | NA | NA | NA | Gm26088 | 0 | 0 | 0 | 0 | 0 | 0 | 0 | 0 | 0 | 0 | 0 |
| 0 | NA | NA | NA | NA | NA | Gm38604 | 0 | 0 | 0 | 0 | 0 | 0 | 0 | 0 | 0 | 0 | 0 |
| 0 | NA | NA | NA | NA | NA | Gm46387 | 0 | 0 | 0 | 0 | 0 | 0 | 0 | 0 | 0 | 0 | 0 |
| 0 | NA | NA | NA | NA | NA | Gm3963 | 0 | 0 | 0 | 0 | 0 | 0 | 0 | 0 | 0 | 0 | 0 |
| 0 | NA | NA | NA | NA | NA | Gm48399 | 0 | 0 | 0 | 0 | 0 | 0 | 0 | 0 | 0 | 0 | 0 |
| 0 | NA | NA | NA | NA | NA | Gm48402 | 0 | 0 | 0 | 0 | 0 | 0 | 0 | 0 | 0 | 0 | 0 |
| 0 | NA | NA | NA | NA | NA | Gm25383 | 0 | 0 | 0 | 0 | 0 | 0 | 0 | 0 | 0 | 0 | 0 |
| 0 | NA | NA | NA | NA | NA | Gm29318 | 0 | 0 | 0 | 0 | 0 | 0 | 0 | 0 | 0 | 0 | 0 |
| 0 | NA | NA | NA | NA | NA | Gm38349 | 0 | 0 | 0 | 0 | 0 | 0 | 0 | 0 | 0 | 0 | 0 |
| 0 | NA | NA | NA | NA | NA | Gm48470 | 0 | 0 | 0 | 0 | 0 | 0 | 0 | 0 | 0 | 0 | 0 |
| 0 | NA | NA | NA | NA | NA | Gm24863 | 0 | 0 | 0 | 0 | 0 | 0 | 0 | 0 | 0 | 0 | 0 |
| 0 | NA | NA | NA | NA | NA | Gm48471 | 0 | 0 | 0 | 0 | 0 | 0 | 0 | 0 | 0 | 0 | 0 |
| 0 | NA | NA | NA | NA | NA | Gm28526 | 0 | 0 | 0 | 0 | 0 | 0 | 0 | 0 | 0 | 0 | 0 |
| 0 | NA | NA | NA | NA | NA | Gm19320 | 0 | 0 | 0 | 0 | 0 | 0 | 0 | 0 | 0 | 0 | 0 |
| 0 | NA | NA | NA | NA | NA | Gm5197 | 0 | 0 | 0 | 0 | 0 | 0 | 0 | 0 | 0 | 0 | 0 |
| 0 | NA | NA | NA | NA | NA | Gm24635 | 0 | 0 | 0 | 0 | 0 | 0 | 0 | 0 | 0 | 0 | 0 |
| 0 | NA | NA | NA | NA | NA | Gm46388 | 0 | 0 | 0 | 0 | 0 | 0 | 0 | 0 | 0 | 0 | 0 |
| 0 | NA | NA | NA | NA | NA | Gm49375 | 0 | 0 | 0 | 0 | 0 | 0 | 0 | 0 | 0 | 0 | 0 |
| 0 | NA | NA | NA | NA | NA | Gm48568 | 0 | 0 | 0 | 0 | 0 | 0 | 0 | 0 | 0 | 0 | 0 |
| 0 | NA | NA | NA | NA | NA | Gm48549 | 0 | 0 | 0 | 0 | 0 | 0 | 0 | 0 | 0 | 0 | 0 |
| 0 | NA | NA | NA | NA | NA | Gm25241 | 0 | 0 | 0 | 0 | 0 | 0 | 0 | 0 | 0 | 0 | 0 |
| 0 | NA | NA | NA | NA | NA | Gm18517 | 0 | 0 | 0 | 0 | 0 | 0 | 0 | 0 | 0 | 0 | 0 |
| 0 | NA | NA | NA | NA | NA | Gm48567 | 0 | 0 | 0 | 0 | 0 | 0 | 0 | 0 | 0 | 0 | 0 |
| 0 | NA | NA | NA | NA | NA | Gm48113 | 0 | 0 | 0 | 0 | 0 | 0 | 0 | 0 | 0 | 0 | 0 |
| 0 | NA | NA | NA | NA | NA | F830210D05Rik | 0 | 0 | 0 | 0 | 0 | 0 | 0 | 0 | 0 | 0 | 0 |
| 0 | NA | NA | NA | NA | NA | Gm24295 | 0 | 0 | 0 | 0 | 0 | 0 | 0 | 0 | 0 | 0 | 0 |
| 0 | NA | NA | NA | NA | NA | Gm48155 | 0 | 0 | 0 | 0 | 0 | 0 | 0 | 0 | 0 | 0 | 0 |
| 0 | NA | NA | NA | NA | NA | Mir3961 | 0 | 0 | 0 | 0 | 0 | 0 | 0 | 0 | 0 | 0 | 0 |
| 0 | NA | NA | NA | NA | NA | Gm48156 | 0 | 0 | 0 | 0 | 0 | 0 | 0 | 0 | 0 | 0 | 0 |
| 0 | NA | NA | NA | NA | NA | Gm26803 | 0 | 0 | 0 | 0 | 0 | 0 | 0 | 0 | 0 | 0 | 0 |
| 0 | NA | NA | NA | NA | NA | Gm33366 | 0 | 0 | 0 | 0 | 0 | 0 | 0 | 0 | 0 | 0 | 0 |
| 0 | NA | NA | NA | NA | NA | Mir9-2 | 0 | 0 | 0 | 0 | 0 | 0 | 0 | 0 | 0 | 0 | 0 |
| 0 | NA | NA | NA | NA | NA | Gm6411 | 0 | 0 | 0 | 0 | 0 | 0 | 0 | 0 | 0 | 0 | 0 |
| 0 | NA | NA | NA | NA | NA | 2810049E08Rik | 0 | 0 | 0 | 0 | 0 | 0 | 0 | 0 | 0 | 0 | 0 |
| 0 | NA | NA | NA | NA | NA | Gm4241 | 0 | 0 | 0 | 0 | 0 | 0 | 0 | 0 | 0 | 0 | 0 |
| 0 | NA | NA | NA | NA | NA | Gm8447 | 0 | 0 | 0 | 0 | 0 | 0 | 0 | 0 | 0 | 0 | 0 |
| 0 | NA | NA | NA | NA | NA | Gm17750 | 0 | 0 | 0 | 0 | 0 | 0 | 0 | 0 | 0 | 0 | 0 |
| 0 | NA | NA | NA | NA | NA | Gm8456 | 0 | 0 | 0 | 0 | 0 | 0 | 0 | 0 | 0 | 0 | 0 |
| 0 | NA | NA | NA | NA | NA | Gm24331 | 0 | 0 | 0 | 0 | 0 | 0 | 0 | 0 | 0 | 0 | 0 |
| 0 | NA | NA | NA | NA | NA | Gm26927 | 0 | 0 | 0 | 0 | 0 | 0 | 0 | 0 | 0 | 0 | 0 |
| 0 | NA | NA | NA | NA | NA | Gm26913 | 0 | 0 | 0 | 0 | 0 | 0 | 0 | 0 | 0 | 0 | 0 |
| 0 | NA | NA | NA | NA | NA | Gm4059 | 0 | 0 | 0 | 0 | 0 | 0 | 0 | 0 | 0 | 0 | 0 |
| 0 | NA | NA | NA | NA | NA | Gm47745 | 0 | 0 | 0 | 0 | 0 | 0 | 0 | 0 | 0 | 0 | 0 |
| 0 | NA | NA | NA | NA | NA | Gm47746 | 0 | 0 | 0 | 0 | 0 | 0 | 0 | 0 | 0 | 0 | 0 |
| 0 | NA | NA | NA | NA | NA | Gm37641 | 0 | 0 | 0 | 0 | 0 | 0 | 0 | 0 | 0 | 0 | 0 |
| 0 | NA | NA | NA | NA | NA | Gm47635 | 0 | 0 | 0 | 0 | 0 | 0 | 0 | 0 | 0 | 0 | 0 |
| 0 | NA | NA | NA | NA | NA | Gm5666 | 0 | 0 | 0 | 0 | 0 | 0 | 0 | 0 | 0 | 0 | 0 |
| 0 | NA | NA | NA | NA | NA | Gm27025 | 0 | 0 | 0 | 0 | 0 | 0 | 0 | 0 | 0 | 0 | 0 |
| 0 | NA | NA | NA | NA | NA | Gm25700 | 0 | 0 | 0 | 0 | 0 | 0 | 0 | 0 | 0 | 0 | 0 |
| 0 | NA | NA | NA | NA | NA | Gm47642 | 0 | 0 | 0 | 0 | 0 | 0 | 0 | 0 | 0 | 0 | 0 |
| 0 | NA | NA | NA | NA | NA | Gm18265 | 0 | 0 | 0 | 0 | 0 | 0 | 0 | 0 | 0 | 0 | 0 |
| 0 | NA | NA | NA | NA | NA | Gm24935 | 0 | 0 | 0 | 0 | 0 | 0 | 0 | 0 | 0 | 0 | 0 |
| 0 | NA | NA | NA | NA | NA | Gm19239 | 0 | 0 | 0 | 0 | 0 | 0 | 0 | 0 | 0 | 0 | 0 |
| 0 | NA | NA | NA | NA | NA | Gm47649 | 0 | 0 | 0 | 0 | 0 | 0 | 0 | 0 | 0 | 0 | 0 |
| 0 | NA | NA | NA | NA | NA | Gm27044 | 0 | 0 | 0 | 0 | 0 | 0 | 0 | 0 | 0 | 0 | 0 |
| 0 | NA | NA | NA | NA | NA | Gm47651 | 0 | 0 | 0 | 0 | 0 | 0 | 0 | 0 | 0 | 0 | 0 |

| | | | | | | | | | | | | | | | | | | |
|---|---|---|---|---|---|---|---|---|---|---|---|---|---|---|---|---|---|---|
| 0 | NA | NA | NA | NA | NA | Gm47653 | | 0 | 0 | 0 | 0 | 0 | 0 | 0 | 0 | 0 | 0 | 0 |
| 0 | NA | NA | NA | NA | NA | Gm19139 | | 0 | 0 | 0 | 0 | 0 | 0 | 0 | 0 | 0 | 0 | 0 |
| 0 | NA | NA | NA | NA | NA | Gm34585 | | 0 | 0 | 0 | 0 | 0 | 0 | 0 | 0 | 0 | 0 | 0 |
| 0 | NA | NA | NA | NA | NA | Gm5667 | | 0 | 0 | 0 | 0 | 0 | 0 | 0 | 0 | 0 | 0 | 0 |
| 0 | NA | NA | NA | NA | NA | Gm16318 | | 0 | 0 | 0 | 0 | 0 | 0 | 0 | 0 | 0 | 0 | 0 |
| 0 | NA | NA | NA | NA | NA | Gm24498 | | 0 | 0 | 0 | 0 | 0 | 0 | 0 | 0 | 0 | 0 | 0 |
| 0 | NA | NA | NA | NA | NA | Gm37727 | | 0 | 0 | 0 | 0 | 0 | 0 | 0 | 0 | 0 | 0 | 0 |
| 0 | NA | NA | NA | NA | NA | Gm29680 | | 0 | 0 | 0 | 0 | 0 | 0 | 0 | 0 | 0 | 0 | 0 |
| 0 | NA | NA | NA | NA | NA | Gm37054 | | 0 | 0 | 0 | 0 | 0 | 0 | 0 | 0 | 0 | 0 | 0 |
| 0 | NA | NA | NA | NA | NA | Gm47520 | | 0 | 0 | 0 | 0 | 0 | 0 | 0 | 0 | 0 | 0 | 0 |
| 0 | NA | NA | NA | NA | NA | Gm37708 | | 0 | 0 | 0 | 0 | 0 | 0 | 0 | 0 | 0 | 0 | 0 |
| 0 | NA | NA | NA | NA | NA | Gm36966 | | 0 | 0 | 0 | 0 | 0 | 0 | 0 | 0 | 0 | 0 | 0 |
| 0 | NA | NA | NA | NA | NA | Gm44338 | | 0 | 0 | 0 | 0 | 0 | 0 | 0 | 0 | 0 | 0 | 0 |
| 0 | NA | NA | NA | NA | NA | Gm47521 | | 0 | 0 | 0 | 0 | 0 | 0 | 0 | 0 | 0 | 0 | 0 |
| 0 | NA | NA | NA | NA | NA | Gm18518 | | 0 | 0 | 0 | 0 | 0 | 0 | 0 | 0 | 0 | 0 | 0 |
| 0 | NA | NA | NA | NA | NA | Atp6ap1l | ATPase, H+ transporting, lysosomal acce | 0 | 0 | 0 | 0 | 0 | 0 | 0 | 0 | 0 | 0 | 0 |
| 0 | NA | NA | NA | NA | NA | Gm47381 | | 0 | 0 | 0 | 0 | 0 | 0 | 0 | 0 | 0 | 0 | 0 |
| 0 | NA | NA | NA | NA | NA | Gm29540 | | 0 | 0 | 0 | 0 | 0 | 0 | 0 | 0 | 0 | 0 | 0 |
| 0 | NA | NA | NA | NA | NA | Gm22778 | | 0 | 0 | 0 | 0 | 0 | 0 | 0 | 0 | 0 | 0 | 0 |
| 0 | NA | NA | NA | NA | NA | Gm24597 | | 0 | 0 | 0 | 0 | 0 | 0 | 0 | 0 | 0 | 0 | 0 |
| 0 | NA | NA | NA | NA | NA | Gm27656 | | 0 | 0 | 0 | 0 | 0 | 0 | 0 | 0 | 0 | 0 | 0 |
| 0 | NA | NA | NA | NA | NA | Ckmt2 | creatine kinase, mitochondrial 2 | 0 | 0 | 0 | 0 | 0 | 0 | 0 | 0 | 0 | 0 | 0 |
| 0 | NA | NA | NA | NA | NA | 1700119I11Rik | | 0 | 0 | 0 | 0 | 0 | 0 | 0 | 0 | 0 | 0 | 0 |
| 0 | NA | NA | NA | NA | NA | Spz1 | spermatogenic leucine zipper 1 | 0 | 0 | 0 | 0 | 0 | 0 | 0 | 0 | 0 | 0 | 0 |
| 0 | NA | NA | NA | NA | NA | Gm5199 | | 0 | 0 | 0 | 0 | 0 | 0 | 0 | 0 | 0 | 0 | 0 |
| 0 | NA | NA | NA | NA | NA | Gm31630 | | 0 | 0 | 0 | 0 | 0 | 0 | 0 | 0 | 0 | 0 | 0 |
| 0 | NA | NA | NA | NA | NA | A630019I02Rik | | 0 | 0 | 0 | 0 | 0 | 0 | 0 | 0 | 0 | 0 | 0 |
| 0 | NA | NA | NA | NA | NA | Gm47088 | | 0 | 0 | 0 | 0 | 0 | 0 | 0 | 0 | 0 | 0 | 0 |
| 0 | NA | NA | NA | NA | NA | Gm18960 | | 0 | 0 | 0 | 0 | 0 | 0 | 0 | 0 | 0 | 0 | 0 |
| 0 | NA | NA | NA | NA | NA | Gm30155 | | 0 | 0 | 0 | 0 | 0 | 0 | 0 | 0 | 0 | 0 | 0 |
| 0 | NA | NA | NA | NA | NA | Gm15622 | | 0 | 0 | 0 | 0 | 0 | 0 | 0 | 0 | 0 | 0 | 0 |
| 0 | NA | NA | NA | NA | NA | Bhmt2 | betaine-homocysteine methyltransferase | 0 | 0 | 0 | 0 | 0 | 0 | 0 | 0 | 0 | 0 | 0 |
| 0 | NA | NA | NA | NA | NA | Dmgdh | dimethylglycine dehydrogenase precursc | 0 | 0 | 0 | 0 | 0 | 0 | 0 | 0 | 0 | 0 | 0 |
| 0 | NA | NA | NA | NA | NA | Gm15620 | | 0 | 0 | 0 | 0 | 0 | 0 | 0 | 0 | 0 | 0 | 0 |
| 0 | NA | NA | NA | NA | NA | Mir5624 | | 0 | 0 | 0 | 0 | 0 | 0 | 0 | 0 | 0 | 0 | 0 |
| 0 | NA | NA | NA | NA | NA | Gm25534 | | 0 | 0 | 0 | 0 | 0 | 0 | 0 | 0 | 0 | 0 | 0 |
| 0 | NA | NA | NA | NA | NA | Gm24737 | | 0 | 0 | 0 | 0 | 0 | 0 | 0 | 0 | 0 | 0 | 0 |
| 0 | NA | NA | NA | NA | NA | Gm47216 | | 0 | 0 | 0 | 0 | 0 | 0 | 0 | 0 | 0 | 0 | 0 |
| 0 | NA | NA | NA | NA | NA | Cycs-ps3 | | 0 | 0 | 0 | 0 | 0 | 0 | 0 | 0 | 0 | 0 | 0 |
| 0 | NA | NA | NA | NA | NA | Gm15907 | | 0 | 0 | 0 | 0 | 0 | 0 | 0 | 0 | 0 | 0 | 0 |
| 0 | NA | NA | NA | NA | NA | Gm48102 | | 0 | 0 | 0 | 0 | 0 | 0 | 0 | 0 | 0 | 0 | 0 |
| 0 | NA | NA | NA | NA | NA | Gm25109 | | 0 | 0 | 0 | 0 | 0 | 0 | 0 | 0 | 0 | 0 | 0 |
| 0 | NA | NA | NA | NA | NA | Gm36991 | | 0 | 0 | 0 | 0 | 0 | 0 | 0 | 0 | 0 | 0 | 0 |
| 0 | NA | NA | NA | NA | NA | Gm48287 | | 0 | 0 | 0 | 0 | 0 | 0 | 0 | 0 | 0 | 0 | 0 |
| 0 | NA | NA | NA | NA | NA | Gm32305 | | 0 | 0 | 0 | 0 | 0 | 0 | 0 | 0 | 0 | 0 | 0 |
| 0 | NA | NA | NA | NA | NA | Gm32351 | | 0 | 0 | 0 | 0 | 0 | 0 | 0 | 0 | 0 | 0 | 0 |
| 0 | NA | NA | NA | NA | NA | Gm48316 | | 0 | 0 | 0 | 0 | 0 | 0 | 0 | 0 | 0 | 0 | 0 |
| 0 | NA | NA | NA | NA | NA | Gm46389 | | 0 | 0 | 0 | 0 | 0 | 0 | 0 | 0 | 0 | 0 | 0 |
| 0 | NA | NA | NA | NA | NA | Otp | orthopedia homeobox | 0 | 0 | 0 | 0 | 0 | 0 | 0 | 0 | 0 | 0 | 0 |
| 0 | NA | NA | NA | NA | NA | Gm48350 | | 0 | 0 | 0 | 0 | 0 | 0 | 0 | 0 | 0 | 0 | 0 |
| 0 | NA | NA | NA | NA | NA | Gm16243 | | 0 | 0 | 0 | 0 | 0 | 0 | 0 | 0 | 0 | 0 | 0 |
| 0 | NA | NA | NA | NA | NA | Gm48712 | | 0 | 0 | 0 | 0 | 0 | 0 | 0 | 0 | 0 | 0 | 0 |
| 0 | NA | NA | NA | NA | NA | Gm48730 | | 0 | 0 | 0 | 0 | 0 | 0 | 0 | 0 | 0 | 0 | 0 |
| 0 | NA | NA | NA | NA | NA | Gm48731 | | 0 | 0 | 0 | 0 | 0 | 0 | 0 | 0 | 0 | 0 | 0 |
| 0 | NA | NA | NA | NA | NA | Gm48745 | | 0 | 0 | 0 | 0 | 0 | 0 | 0 | 0 | 0 | 0 | 0 |
| 0 | NA | NA | NA | NA | NA | Gm32880 | | 0 | 0 | 0 | 0 | 0 | 0 | 0 | 0 | 0 | 0 | 0 |
| 0 | NA | NA | NA | NA | NA | Gm24032 | | 0 | 0 | 0 | 0 | 0 | 0 | 0 | 0 | 0 | 0 | 0 |
| 0 | NA | NA | NA | NA | NA | Gm23955 | | 0 | 0 | 0 | 0 | 0 | 0 | 0 | 0 | 0 | 0 | 0 |
| 0 | NA | NA | NA | NA | NA | Gm29543 | | 0 | 0 | 0 | 0 | 0 | 0 | 0 | 0 | 0 | 0 | 0 |
| 0 | NA | NA | NA | NA | NA | Gm25213 | | 0 | 0 | 0 | 0 | 0 | 0 | 0 | 0 | 0 | 0 | 0 |

| | | | | | | | | | | | | | | | | | | |
|---|---|---|---|---|---|---|---|---|---|---|---|---|---|---|---|---|---|---|
| 0 | NA | NA | NA | NA | NA | Gm17622 | | 0 | 0 | 0 | 0 | 0 | 0 | 0 | 0 | 0 | 0 | 0 |
| 0 | NA | NA | NA | NA | NA | Gm48575 | | 0 | 0 | 0 | 0 | 0 | 0 | 0 | 0 | 0 | 0 | 0 |
| 0 | NA | NA | NA | NA | NA | Gm48580 | | 0 | 0 | 0 | 0 | 0 | 0 | 0 | 0 | 0 | 0 | 0 |
| 0 | NA | NA | NA | NA | NA | Gm48579 | | 0 | 0 | 0 | 0 | 0 | 0 | 0 | 0 | 0 | 0 | 0 |
| 0 | NA | NA | NA | NA | NA | Gm27792 | | 0 | 0 | 0 | 0 | 0 | 0 | 0 | 0 | 0 | 0 | 0 |
| 0 | NA | NA | NA | NA | NA | Gm48597 | | 0 | 0 | 0 | 0 | 0 | 0 | 0 | 0 | 0 | 0 | 0 |
| 0 | NA | NA | NA | NA | NA | Gm48609 | | 0 | 0 | 0 | 0 | 0 | 0 | 0 | 0 | 0 | 0 | 0 |
| 0 | NA | NA | NA | NA | NA | Gm48881 | | 0 | 0 | 0 | 0 | 0 | 0 | 0 | 0 | 0 | 0 | 0 |
| 0 | NA | NA | NA | NA | NA | C430039J16Rik | | 0 | 0 | 0 | 0 | 0 | 0 | 0 | 0 | 0 | 0 | 0 |
| 0 | NA | NA | NA | NA | NA | Gm48090 | | 0 | 0 | 0 | 0 | 0 | 0 | 0 | 0 | 0 | 0 | 0 |
| 0 | NA | NA | NA | NA | NA | Gm48070 | | 0 | 0 | 0 | 0 | 0 | 0 | 0 | 0 | 0 | 0 | 0 |
| 0 | NA | NA | NA | NA | NA | Gm47577 | | 0 | 0 | 0 | 0 | 0 | 0 | 0 | 0 | 0 | 0 | 0 |
| 0 | NA | NA | NA | NA | NA | Gm21976 | predicted gene 21976 | 0 | 0 | 0 | 0 | 0 | 0 | 0 | 0 | 0 | 0 | 0 |
| 0 | NA | NA | NA | NA | NA | 2310005E17Rik | | 0 | 0 | 0 | 0 | 0 | 0 | 0 | 0 | 0 | 0 | 0 |
| 0 | NA | NA | NA | NA | NA | Gm29501 | | 0 | 0 | 0 | 0 | 0 | 0 | 0 | 0 | 0 | 0 | 0 |
| 0 | NA | NA | NA | NA | NA | Gm37425 | | 0 | 0 | 0 | 0 | 0 | 0 | 0 | 0 | 0 | 0 | 0 |
| 0 | NA | NA | NA | NA | NA | Gm48049 | | 0 | 0 | 0 | 0 | 0 | 0 | 0 | 0 | 0 | 0 | 0 |
| 0 | NA | NA | NA | NA | NA | Gm22698 | | 0 | 0 | 0 | 0 | 0 | 0 | 0 | 0 | 0 | 0 | 0 |
| 0 | NA | NA | NA | NA | NA | Gm48050 | | 0 | 0 | 0 | 0 | 0 | 0 | 0 | 0 | 0 | 0 | 0 |
| 0 | NA | NA | NA | NA | NA | Tmem174 | transmembrane protein 174 | 0 | 0 | 0 | 0 | 0 | 0 | 0 | 0 | 0 | 0 | 0 |
| 0 | NA | NA | NA | NA | NA | Gm47550 | | 0 | 0 | 0 | 0 | 0 | 0 | 0 | 0 | 0 | 0 | 0 |
| 0 | NA | NA | NA | NA | NA | Gm47551 | | 0 | 0 | 0 | 0 | 0 | 0 | 0 | 0 | 0 | 0 | 0 |
| 0 | NA | NA | NA | NA | NA | H2bl1 | H2B.L histone variant 1 | 0 | 0 | 0 | 0 | 0 | 0 | 0 | 0 | 0 | 0 | 0 |
| 0 | NA | NA | NA | NA | NA | A930014D07Rik | | 0 | 0 | 0 | 0 | 0 | 0 | 0 | 0 | 0 | 0 | 0 |
| 0 | NA | NA | NA | NA | NA | Gm47057 | | 0 | 0 | 0 | 0 | 0 | 0 | 0 | 0 | 0 | 0 | 0 |
| 0 | NA | NA | NA | NA | NA | Gm24733 | | 0 | 0 | 0 | 0 | 0 | 0 | 0 | 0 | 0 | 0 | 0 |
| 0 | NA | NA | NA | NA | NA | Gm47158 | | 0 | 0 | 0 | 0 | 0 | 0 | 0 | 0 | 0 | 0 | 0 |
| 0 | NA | NA | NA | NA | NA | 1700012J22Rik | | 0 | 0 | 0 | 0 | 0 | 0 | 0 | 0 | 0 | 0 | 0 |
| 0 | NA | NA | NA | NA | NA | Gm24471 | | 0 | 0 | 0 | 0 | 0 | 0 | 0 | 0 | 0 | 0 | 0 |
| 0 | NA | NA | NA | NA | NA | Gm18601 | | 0 | 0 | 0 | 0 | 0 | 0 | 0 | 0 | 0 | 0 | 0 |
| 0 | NA | NA | NA | NA | NA | Gm48452 | | 0 | 0 | 0 | 0 | 0 | 0 | 0 | 0 | 0 | 0 | 0 |
| 0 | NA | NA | NA | NA | NA | Gm8847 | | 0 | 0 | 0 | 0 | 0 | 0 | 0 | 0 | 0 | 0 | 0 |
| 0 | NA | NA | NA | NA | NA | Gm37830 | | 0 | 0 | 0 | 0 | 0 | 0 | 0 | 0 | 0 | 0 | 0 |
| 0 | NA | NA | NA | NA | NA | Gm6114 | | 0 | 0 | 0 | 0 | 0 | 0 | 0 | 0 | 0 | 0 | 0 |
| 0 | NA | NA | NA | NA | NA | Gm36994 | | 0 | 0 | 0 | 0 | 0 | 0 | 0 | 0 | 0 | 0 | 0 |
| 0 | NA | NA | NA | NA | NA | Gm41043 | | 0 | 0 | 0 | 0 | 0 | 0 | 0 | 0 | 0 | 0 | 0 |
| 0 | NA | NA | NA | NA | NA | Gm19010 | | 0 | 0 | 0 | 0 | 0 | 0 | 0 | 0 | 0 | 0 | 0 |
| 0 | NA | NA | NA | NA | NA | Gm29341 | | 0 | 0 | 0 | 0 | 0 | 0 | 0 | 0 | 0 | 0 | 0 |
| 0 | NA | NA | NA | NA | NA | Gm19108 | | 0 | 0 | 0 | 0 | 0 | 0 | 0 | 0 | 0 | 0 | 0 |
| 0 | NA | NA | NA | NA | NA | Gm47007 | | 0 | 0 | 0 | 0 | 0 | 0 | 0 | 0 | 0 | 0 | 0 |
| 0 | NA | NA | NA | NA | NA | Gm17832 | | 0 | 0 | 0 | 0 | 0 | 0 | 0 | 0 | 0 | 0 | 0 |
| 0 | NA | NA | NA | NA | NA | Gm38133 | | 0 | 0 | 0 | 0 | 0 | 0 | 0 | 0 | 0 | 0 | 0 |
| 0 | NA | NA | NA | NA | NA | Gm25808 | | 0 | 0 | 0 | 0 | 0 | 0 | 0 | 0 | 0 | 0 | 0 |
| 0 | NA | NA | NA | NA | NA | Gm47008 | | 0 | 0 | 0 | 0 | 0 | 0 | 0 | 0 | 0 | 0 | 0 |
| 0 | NA | NA | NA | NA | NA | Gm29927 | | 0 | 0 | 0 | 0 | 0 | 0 | 0 | 0 | 0 | 0 | 0 |
| 0 | NA | NA | NA | NA | NA | Gm47014 | | 0 | 0 | 0 | 0 | 0 | 0 | 0 | 0 | 0 | 0 | 0 |
| 0 | NA | NA | NA | NA | NA | Gm17160 | | 0 | 0 | 0 | 0 | 0 | 0 | 0 | 0 | 0 | 0 | 0 |
| 0 | NA | NA | NA | NA | NA | n-R5s56 | | 0 | 0 | 0 | 0 | 0 | 0 | 0 | 0 | 0 | 0 | 0 |
| 0 | NA | NA | NA | NA | NA | Gm24441 | | 0 | 0 | 0 | 0 | 0 | 0 | 0 | 0 | 0 | 0 | 0 |
| 0 | NA | NA | NA | NA | NA | Gm24870 | | 0 | 0 | 0 | 0 | 0 | 0 | 0 | 0 | 0 | 0 | 0 |
| 0 | NA | NA | NA | NA | NA | Gm47851 | | 0 | 0 | 0 | 0 | 0 | 0 | 0 | 0 | 0 | 0 | 0 |
| 0 | NA | NA | NA | NA | NA | 2610204G07Rik | | 0 | 0 | 0 | 0 | 0 | 0 | 0 | 0 | 0 | 0 | 0 |
| 0 | NA | NA | NA | NA | NA | Nt5el | 5' nucleotidase, ecto-like | 0 | 0 | 0 | 0 | 0 | 0 | 0 | 0 | 0 | 0 | 0 |
| 0 | NA | NA | NA | NA | NA | Gm25631 | | 0 | 0 | 0 | 0 | 0 | 0 | 0 | 0 | 0 | 0 | 0 |
| 0 | NA | NA | NA | NA | NA | Htr1a | 5-hydroxytryptamine (serotonin) receptor | 0 | 0 | 0 | 0 | 0 | 0 | 0 | 0 | 0 | 0 | 0 |
| 0 | NA | NA | NA | NA | NA | Gm48208 | | 0 | 0 | 0 | 0 | 0 | 0 | 0 | 0 | 0 | 0 | 0 |
| 0 | NA | NA | NA | NA | NA | Gm48211 | | 0 | 0 | 0 | 0 | 0 | 0 | 0 | 0 | 0 | 0 | 0 |
| 0 | NA | NA | NA | NA | NA | Gm10739 | | 0 | 0 | 0 | 0 | 0 | 0 | 0 | 0 | 0 | 0 | 0 |
| 0 | NA | NA | NA | NA | NA | Gm48212 | | 0 | 0 | 0 | 0 | 0 | 0 | 0 | 0 | 0 | 0 | 0 |
| 0 | NA | NA | NA | NA | NA | Gm48214 | | 0 | 0 | 0 | 0 | 0 | 0 | 0 | 0 | 0 | 0 | 0 |

| | | | | | | | | | | | | | | | | | | |
|---|---|---|---|---|---|---|---|---|---|---|---|---|---|---|---|---|---|---|
| 0 | NA | NA | NA | NA | NA | Olfr717-ps1 | | 0 | 0 | 0 | 0 | 0 | 0 | 0 | 0 | 0 | 0 | 0 |
| 0 | NA | NA | NA | NA | NA | Gm48216 | | 0 | 0 | 0 | 0 | 0 | 0 | 0 | 0 | 0 | 0 | 0 |
| 0 | NA | NA | NA | NA | NA | Dph3b-ps | | 0 | 0 | 0 | 0 | 0 | 0 | 0 | 0 | 0 | 0 | 0 |
| 0 | NA | NA | NA | NA | NA | Gm30411 | | 0 | 0 | 0 | 0 | 0 | 0 | 0 | 0 | 0 | 0 | 0 |
| 0 | NA | NA | NA | NA | NA | Gm24527 | | 0 | 0 | 0 | 0 | 0 | 0 | 0 | 0 | 0 | 0 | 0 |
| 0 | NA | NA | NA | NA | NA | Gm28988 | | 0 | 0 | 0 | 0 | 0 | 0 | 0 | 0 | 0 | 0 | 0 |
| 0 | NA | NA | NA | NA | NA | Gm48457 | | 0 | 0 | 0 | 0 | 0 | 0 | 0 | 0 | 0 | 0 | 0 |
| 0 | NA | NA | NA | NA | NA | Gm24431 | | 0 | 0 | 0 | 0 | 0 | 0 | 0 | 0 | 0 | 0 | 0 |
| 0 | NA | NA | NA | NA | NA | Gm31452 | | 0 | 0 | 0 | 0 | 0 | 0 | 0 | 0 | 0 | 0 | 0 |
| 0 | NA | NA | NA | NA | NA | Gm2726 | | 0 | 0 | 0 | 0 | 0 | 0 | 0 | 0 | 0 | 0 | 0 |
| 0 | NA | NA | NA | NA | NA | Gm2736 | | 0 | 0 | 0 | 0 | 0 | 0 | 0 | 0 | 0 | 0 | 0 |
| 0 | NA | NA | NA | NA | NA | Gm32004 | | 0 | 0 | 0 | 0 | 0 | 0 | 0 | 0 | 0 | 0 | 0 |
| 0 | NA | NA | NA | NA | NA | Gm32090 | | 0 | 0 | 0 | 0 | 0 | 0 | 0 | 0 | 0 | 0 | 0 |
| 0 | NA | NA | NA | NA | NA | 1700006H21Rik | | 0 | 0 | 0 | 0 | 0 | 0 | 0 | 0 | 0 | 0 | 0 |
| 0 | NA | NA | NA | NA | NA | Gm22118 | | 0 | 0 | 0 | 0 | 0 | 0 | 0 | 0 | 0 | 0 | 0 |
| 0 | NA | NA | NA | NA | NA | Gm27572 | | 0 | 0 | 0 | 0 | 0 | 0 | 0 | 0 | 0 | 0 | 0 |
| 0 | NA | NA | NA | NA | NA | Gm28016 | | 0 | 0 | 0 | 0 | 0 | 0 | 0 | 0 | 0 | 0 | 0 |
| 0 | NA | NA | NA | NA | NA | Mir582 | | 0 | 0 | 0 | 0 | 0 | 0 | 0 | 0 | 0 | 0 | 0 |
| 0 | NA | NA | NA | NA | NA | Mir1904 | | 0 | 0 | 0 | 0 | 0 | 0 | 0 | 0 | 0 | 0 | 0 |
| 0 | NA | NA | NA | NA | NA | Gm33045 | | 0 | 0 | 0 | 0 | 0 | 0 | 0 | 0 | 0 | 0 | 0 |
| 0 | NA | NA | NA | NA | NA | Gm48533 | | 0 | 0 | 0 | 0 | 0 | 0 | 0 | 0 | 0 | 0 | 0 |
| 0 | NA | NA | NA | NA | NA | Gm48534 | | 0 | 0 | 0 | 0 | 0 | 0 | 0 | 0 | 0 | 0 | 0 |
| 0 | NA | NA | NA | NA | NA | Gm30839 | | 0 | 0 | 0 | 0 | 0 | 0 | 0 | 0 | 0 | 0 | 0 |
| 0 | NA | NA | NA | NA | NA | 4930526H09Rik | | 0 | 0 | 0 | 0 | 0 | 0 | 0 | 0 | 0 | 0 | 0 |
| 0 | NA | NA | NA | NA | NA | Gm38397 | | 0 | 0 | 0 | 0 | 0 | 0 | 0 | 0 | 0 | 0 | 0 |
| 0 | NA | NA | NA | NA | NA | Gm48547 | | 0 | 0 | 0 | 0 | 0 | 0 | 0 | 0 | 0 | 0 | 0 |
| 0 | NA | NA | NA | NA | NA | 3110015C05Rik | | 0 | 0 | 0 | 0 | 0 | 0 | 0 | 0 | 0 | 0 | 0 |
| 0 | NA | NA | NA | NA | NA | Actbl2 | actin, beta-like 2 | 0 | 0 | 0 | 0 | 0 | 0 | 0 | 0 | 0 | 0 | 0 |
| 0 | NA | NA | NA | NA | NA | Gm15289 | | 0 | 0 | 0 | 0 | 0 | 0 | 0 | 0 | 0 | 0 | 0 |
| 0 | NA | NA | NA | NA | NA | Gm6270 | | 0 | 0 | 0 | 0 | 0 | 0 | 0 | 0 | 0 | 0 | 0 |
| 0 | NA | NA | NA | NA | NA | Gm49496 | | 0 | 0 | 0 | 0 | 0 | 0 | 0 | 0 | 0 | 0 | 0 |
| 0 | NA | NA | NA | NA | NA | Gm15290 | | 0 | 0 | 0 | 0 | 0 | 0 | 0 | 0 | 0 | 0 | 0 |
| 0 | NA | NA | NA | NA | NA | Gm15288 | | 0 | 0 | 0 | 0 | 0 | 0 | 0 | 0 | 0 | 0 | 0 |
| 0 | NA | NA | NA | NA | NA | Gm15286 | | 0 | 0 | 0 | 0 | 0 | 0 | 0 | 0 | 0 | 0 | 0 |
| 0 | NA | NA | NA | NA | NA | Gm10198 | | 0 | 0 | 0 | 0 | 0 | 0 | 0 | 0 | 0 | 0 | 0 |
| 0 | NA | NA | NA | NA | NA | Gm15488 | | 0 | 0 | 0 | 0 | 0 | 0 | 0 | 0 | 0 | 0 | 0 |
| 0 | NA | NA | NA | NA | NA | Gm15287 | | 0 | 0 | 0 | 0 | 0 | 0 | 0 | 0 | 0 | 0 | 0 |
| 0 | NA | NA | NA | NA | NA | Gm15325 | | 0 | 0 | 0 | 0 | 0 | 0 | 0 | 0 | 0 | 0 | 0 |
| 0 | NA | NA | NA | NA | NA | Gm15324 | | 0 | 0 | 0 | 0 | 0 | 0 | 0 | 0 | 0 | 0 | 0 |
| 0 | NA | NA | NA | NA | NA | Gm9025 | | 0 | 0 | 0 | 0 | 0 | 0 | 0 | 0 | 0 | 0 | 0 |
| 0 | NA | NA | NA | NA | NA | Gm9025 | | 0 | 0 | 0 | 0 | 0 | 0 | 0 | 0 | 0 | 0 | 0 |
| 0 | NA | NA | NA | NA | NA | Gm15322 | | 0 | 0 | 0 | 0 | 0 | 0 | 0 | 0 | 0 | 0 | 0 |
| 0 | NA | NA | NA | NA | NA | Gm15323 | | 0 | 0 | 0 | 0 | 0 | 0 | 0 | 0 | 0 | 0 | 0 |
| 0 | NA | NA | NA | NA | NA | Gm31104 | | 0 | 0 | 0 | 0 | 0 | 0 | 0 | 0 | 0 | 0 | 0 |
| 0 | NA | NA | NA | NA | NA | Gm48802 | | 0 | 0 | 0 | 0 | 0 | 0 | 0 | 0 | 0 | 0 | 0 |
| 0 | NA | NA | NA | NA | NA | Gm37427 | | 0 | 0 | 0 | 0 | 0 | 0 | 0 | 0 | 0 | 0 | 0 |
| 0 | NA | NA | NA | NA | NA | Gm22768 | | 0 | 0 | 0 | 0 | 0 | 0 | 0 | 0 | 0 | 0 | 0 |
| 0 | NA | NA | NA | NA | NA | 2810403G07Rik | | 0 | 0 | 0 | 0 | 0 | 0 | 0 | 0 | 0 | 0 | 0 |
| 0 | NA | NA | NA | NA | NA | Gm48876 | | 0 | 0 | 0 | 0 | 0 | 0 | 0 | 0 | 0 | 0 | 0 |
| 0 | NA | NA | NA | NA | NA | Gm48879 | | 0 | 0 | 0 | 0 | 0 | 0 | 0 | 0 | 0 | 0 | 0 |
| 0 | NA | NA | NA | NA | NA | Gm47827 | | 0 | 0 | 0 | 0 | 0 | 0 | 0 | 0 | 0 | 0 | 0 |
| 0 | NA | NA | NA | NA | NA | Gm47850 | | 0 | 0 | 0 | 0 | 0 | 0 | 0 | 0 | 0 | 0 | 0 |
| 0 | NA | NA | NA | NA | NA | Gm26943 | | 0 | 0 | 0 | 0 | 0 | 0 | 0 | 0 | 0 | 0 | 0 |
| 0 | NA | NA | NA | NA | NA | Mcidas | multiciliate differentiation and DNA synthe | 0 | 0 | 0 | 0 | 0 | 0 | 0 | 0 | 0 | 0 | 0 |
| 0 | NA | NA | NA | NA | NA | Cdc20b | cell division cycle 20B | 0 | 0 | 0 | 0 | 0 | 0 | 0 | 0 | 0 | 0 | 0 |
| 0 | NA | NA | NA | NA | NA | Mir449c | | 0 | 0 | 0 | 0 | 0 | 0 | 0 | 0 | 0 | 0 | 0 |
| 0 | NA | NA | NA | NA | NA | Mir449b | | 0 | 0 | 0 | 0 | 0 | 0 | 0 | 0 | 0 | 0 | 0 |
| 0 | NA | NA | NA | NA | NA | Mir449a | | 0 | 0 | 0 | 0 | 0 | 0 | 0 | 0 | 0 | 0 | 0 |
| 0 | NA | NA | NA | NA | NA | 4930563N14Rik | | 0 | 0 | 0 | 0 | 0 | 0 | 0 | 0 | 0 | 0 | 0 |
| 0 | NA | NA | NA | NA | NA | Gm9848 | | 0 | 0 | 0 | 0 | 0 | 0 | 0 | 0 | 0 | 0 | 0 |

| | | | | | | | | | | | | | | | | | | |
|---|---|---|---|---|---|---|---|---|---|---|---|---|---|---|---|---|---|---|
| 0 | NA | NA | NA | NA | NA | 1700084D21Rik | | 0 | 0 | 0 | 0 | 0 | 0 | 0 | 0 | 0 | 0 | 0 |
| 0 | NA | NA | NA | NA | NA | Gm49564 | | 0 | 0 | 0 | 0 | 0 | 0 | 0 | 0 | 0 | 0 | 0 |
| 0 | NA | NA | NA | NA | NA | Gm22385 | | 0 | 0 | 0 | 0 | 0 | 0 | 0 | 0 | 0 | 0 | 0 |
| 0 | NA | NA | NA | NA | NA | Gm41071 | | 0 | 0 | 0 | 0 | 0 | 0 | 0 | 0 | 0 | 0 | 0 |
| 0 | NA | NA | NA | NA | NA | 4921509O07Rik | | 0 | 0 | 0 | 0 | 0 | 0 | 0 | 0 | 0 | 0 | 0 |
| 0 | NA | NA | NA | NA | NA | Gm47426 | | 0 | 0 | 0 | 0 | 0 | 0 | 0 | 0 | 0 | 0 | 0 |
| 0 | NA | NA | NA | NA | NA | Hspb3 | heat shock protein 3 | 0 | 0 | 0 | 0 | 0 | 0 | 0 | 0 | 0 | 0 | 0 |
| 0 | NA | NA | NA | NA | NA | Gm34586 | | 0 | 0 | 0 | 0 | 0 | 0 | 0 | 0 | 0 | 0 | 0 |
| 0 | NA | NA | NA | NA | NA | Gm47462 | | 0 | 0 | 0 | 0 | 0 | 0 | 0 | 0 | 0 | 0 | 0 |
| 0 | NA | NA | NA | NA | NA | Gm47464 | | 0 | 0 | 0 | 0 | 0 | 0 | 0 | 0 | 0 | 0 | 0 |
| 0 | NA | NA | NA | NA | NA | Gm47479 | | 0 | 0 | 0 | 0 | 0 | 0 | 0 | 0 | 0 | 0 | 0 |
| 0 | NA | NA | NA | NA | NA | A430090L17Rik | | 0 | 0 | 0 | 0 | 0 | 0 | 0 | 0 | 0 | 0 | 0 |
| 0 | NA | NA | NA | NA | NA | Gm37907 | | 0 | 0 | 0 | 0 | 0 | 0 | 0 | 0 | 0 | 0 | 0 |
| 0 | NA | NA | NA | NA | NA | 4631422I05Rik | | 0 | 0 | 0 | 0 | 0 | 0 | 0 | 0 | 0 | 0 | 0 |
| 0 | NA | NA | NA | NA | NA | Gm47569 | | 0 | 0 | 0 | 0 | 0 | 0 | 0 | 0 | 0 | 0 | 0 |
| 0 | NA | NA | NA | NA | NA | Gm25950 | | 0 | 0 | 0 | 0 | 0 | 0 | 0 | 0 | 0 | 0 | 0 |
| 0 | NA | NA | NA | NA | NA | 4930467J12Rik | | 0 | 0 | 0 | 0 | 0 | 0 | 0 | 0 | 0 | 0 | 0 |
| 0 | NA | NA | NA | NA | NA | Gm47776 | | 0 | 0 | 0 | 0 | 0 | 0 | 0 | 0 | 0 | 0 | 0 |
| 0 | NA | NA | NA | NA | NA | Gm49395 | | 0 | 0 | 0 | 0 | 0 | 0 | 0 | 0 | 0 | 0 | 0 |
| 0 | NA | NA | NA | NA | NA | Gm25985 | | 0 | 0 | 0 | 0 | 0 | 0 | 0 | 0 | 0 | 0 | 0 |
| 0 | NA | NA | NA | NA | NA | Gm18759 | | 0 | 0 | 0 | 0 | 0 | 0 | 0 | 0 | 0 | 0 | 0 |
| 0 | NA | NA | NA | NA | NA | Gm47891 | | 0 | 0 | 0 | 0 | 0 | 0 | 0 | 0 | 0 | 0 | 0 |
| 0 | NA | NA | NA | NA | NA | Gm47892 | | 0 | 0 | 0 | 0 | 0 | 0 | 0 | 0 | 0 | 0 | 0 |
| 0 | NA | NA | NA | NA | NA | Gm18135 | | 0 | 0 | 0 | 0 | 0 | 0 | 0 | 0 | 0 | 0 | 0 |
| 0 | NA | NA | NA | NA | NA | Gm47893 | | 0 | 0 | 0 | 0 | 0 | 0 | 0 | 0 | 0 | 0 | 0 |
| 0 | NA | NA | NA | NA | NA | Gm29764 | | 0 | 0 | 0 | 0 | 0 | 0 | 0 | 0 | 0 | 0 | 0 |
| 0 | NA | NA | NA | NA | NA | Gm47912 | | 0 | 0 | 0 | 0 | 0 | 0 | 0 | 0 | 0 | 0 | 0 |
| 0 | NA | NA | NA | NA | NA | Gm47913 | | 0 | 0 | 0 | 0 | 0 | 0 | 0 | 0 | 0 | 0 | 0 |
| 0 | NA | NA | NA | NA | NA | Gm23198 | | 0 | 0 | 0 | 0 | 0 | 0 | 0 | 0 | 0 | 0 | 0 |
| 0 | NA | NA | NA | NA | NA | 4930435F18Rik | | 0 | 0 | 0 | 0 | 0 | 0 | 0 | 0 | 0 | 0 | 0 |
| 0 | NA | NA | NA | NA | NA | Gm48026 | | 0 | 0 | 0 | 0 | 0 | 0 | 0 | 0 | 0 | 0 | 0 |
| 0 | NA | NA | NA | NA | NA | Gm6416 | | 0 | 0 | 0 | 0 | 0 | 0 | 0 | 0 | 0 | 0 | 0 |
| 0 | NA | NA | NA | NA | NA | Gm18945 | | 0 | 0 | 0 | 0 | 0 | 0 | 0 | 0 | 0 | 0 | 0 |
| 0 | NA | NA | NA | NA | NA | 4933413L06Rik | | 0 | 0 | 0 | 0 | 0 | 0 | 0 | 0 | 0 | 0 | 0 |
| 0 | NA | NA | NA | NA | NA | Gm26349 | | 0 | 0 | 0 | 0 | 0 | 0 | 0 | 0 | 0 | 0 | 0 |
| 0 | NA | NA | NA | NA | NA | Gm9633 | | 0 | 0 | 0 | 0 | 0 | 0 | 0 | 0 | 0 | 0 | 0 |
| 0 | NA | NA | NA | NA | NA | Gm47563 | | 0 | 0 | 0 | 0 | 0 | 0 | 0 | 0 | 0 | 0 | 0 |
| 0 | NA | NA | NA | NA | NA | 1700003P14Rik | | 0 | 0 | 0 | 0 | 0 | 0 | 0 | 0 | 0 | 0 | 0 |
| 0 | NA | NA | NA | NA | NA | Gm9098 | | 0 | 0 | 0 | 0 | 0 | 0 | 0 | 0 | 0 | 0 | 0 |
| 0 | NA | NA | NA | NA | NA | Gm47330 | | 0 | 0 | 0 | 0 | 0 | 0 | 0 | 0 | 0 | 0 | 0 |
| 0 | NA | NA | NA | NA | NA | Gm18361 | | 0 | 0 | 0 | 0 | 0 | 0 | 0 | 0 | 0 | 0 | 0 |
| 0 | NA | NA | NA | NA | NA | Gm10732 | | 0 | 0 | 0 | 0 | 0 | 0 | 0 | 0 | 0 | 0 | 0 |
| 0 | NA | NA | NA | NA | NA | Gm47335 | | 0 | 0 | 0 | 0 | 0 | 0 | 0 | 0 | 0 | 0 | 0 |
| 0 | NA | NA | NA | NA | NA | Gm44488 | | 0 | 0 | 0 | 0 | 0 | 0 | 0 | 0 | 0 | 0 | 0 |
| 0 | NA | NA | NA | NA | NA | Gm47336 | | 0 | 0 | 0 | 0 | 0 | 0 | 0 | 0 | 0 | 0 | 0 |
| 0 | NA | NA | NA | NA | NA | Nnt | | 0 | 0 | 0 | 0 | 0 | 0 | 0 | 0 | 0 | 0 | 0 |
| 0 | NA | NA | NA | NA | NA | 1700074H08Rik | | 0 | 0 | 0 | 0 | 0 | 0 | 0 | 0 | 0 | 0 | 0 |
| 0 | NA | NA | NA | NA | NA | Gm23308 | | 0 | 0 | 0 | 0 | 0 | 0 | 0 | 0 | 0 | 0 | 0 |
| 0 | NA | NA | NA | NA | NA | AF067063 | cDNA sequence AF067063 | 0 | 0 | 0 | 0 | 0 | 0 | 0 | 0 | 0 | 0 | 0 |
| 0 | NA | NA | NA | NA | NA | D13Ertd608e | | 0 | 0 | 0 | 0 | 0 | 0 | 0 | 0 | 0 | 0 | 0 |
| 0 | NA | NA | NA | NA | NA | Gm21358 | | 0 | 0 | 0 | 0 | 0 | 0 | 0 | 0 | 0 | 0 | 0 |
| 0 | NA | NA | NA | NA | NA | Gm37619 | | 0 | 0 | 0 | 0 | 0 | 0 | 0 | 0 | 0 | 0 | 0 |
| 0 | NA | NA | NA | NA | NA | Tcstv1 | 2-cell-stage, variable group, member 1 | 0 | 0 | 0 | 0 | 0 | 0 | 0 | 0 | 0 | 0 | 0 |
| 0 | NA | NA | NA | NA | NA | Gm21761 | predicted gene, 21761 | 0 | 0 | 0 | 0 | 0 | 0 | 0 | 0 | 0 | 0 | 0 |
| 0 | NA | NA | NA | NA | NA | Gm21181 | | 0 | 0 | 0 | 0 | 0 | 0 | 0 | 0 | 0 | 0 | 0 |
| 0 | NA | NA | NA | NA | NA | B020031M17Ri | RIKEN cDNA B020031M17 gene | 0 | 0 | 0 | 0 | 0 | 0 | 0 | 0 | 0 | 0 | 0 |
| 0 | NA | NA | NA | NA | NA | Gm22108 | | 0 | 0 | 0 | 0 | 0 | 0 | 0 | 0 | 0 | 0 | 0 |
| 0 | NA | NA | NA | NA | NA | Gm36079 | predicted gene, 36079 | 0 | 0 | 0 | 0 | 0 | 0 | 0 | 0 | 0 | 0 | 0 |
| 0 | NA | NA | NA | NA | NA | Gm24798 | | 0 | 0 | 0 | 0 | 0 | 0 | 0 | 0 | 0 | 0 | 0 |
| 0 | NA | NA | NA | NA | NA | Gm20767 | predicted gene, 20767 | 0 | 0 | 0 | 0 | 0 | 0 | 0 | 0 | 0 | 0 | 0 |

| | | | | | | | | | | | | | | | | | |
|---|---|---|---|---|---|---|---|---|---|---|---|---|---|---|---|---|---|
| 0 | NA | NA | NA | NA | NA | Gm20769 | | 0 | 0 | 0 | 0 | 0 | 0 | 0 | 0 | 0 | 0 | 0 |
| 0 | NA | NA | NA | NA | NA | Gm21818 | predicted gene, 21818 | 0 | 0 | 0 | 0 | 0 | 0 | 0 | 0 | 0 | 0 | 0 |
| 0 | NA | NA | NA | NA | NA | Gm20763 | | 0 | 0 | 0 | 0 | 0 | 0 | 0 | 0 | 0 | 0 | 0 |
| 0 | NA | NA | NA | NA | NA | Gm21731 | | 0 | 0 | 0 | 0 | 0 | 0 | 0 | 0 | 0 | 0 | 0 |
| 0 | NA | NA | NA | NA | NA | AF067061 | cDNA sequence AF067061 | 0 | 0 | 0 | 0 | 0 | 0 | 0 | 0 | 0 | 0 | 0 |
| 0 | NA | NA | NA | NA | NA | Tcstv3 | 2-cell-stage, variable group, member 3 | 0 | 0 | 0 | 0 | 0 | 0 | 0 | 0 | 0 | 0 | 0 |
| 0 | NA | NA | NA | NA | NA | Gm48109 | | 0 | 0 | 0 | 0 | 0 | 0 | 0 | 0 | 0 | 0 | 0 |
| 0 | NA | NA | NA | NA | NA | Gm2888 | predicted gene 2888 | 0 | 0 | 0 | 0 | 0 | 0 | 0 | 0 | 0 | 0 | 0 |
| 0 | NA | NA | NA | NA | NA | Gm17051 | | 0 | 0 | 0 | 0 | 0 | 0 | 0 | 0 | 0 | 0 | 0 |
| 0 | NA | NA | NA | NA | NA | Gm10340 | predicted gene 10340 | 0 | 0 | 0 | 0 | 0 | 0 | 0 | 0 | 0 | 0 | 0 |
| 0 | NA | NA | NA | NA | NA | Gm17221 | | 0 | 0 | 0 | 0 | 0 | 0 | 0 | 0 | 0 | 0 | 0 |
| 0 | NA | NA | NA | NA | NA | Gm17188 | | 0 | 0 | 0 | 0 | 0 | 0 | 0 | 0 | 0 | 0 | 0 |
| 0 | NA | NA | NA | NA | NA | Gm5795 | predicted gene 5795 | 0 | 0 | 0 | 0 | 0 | 0 | 0 | 0 | 0 | 0 | 0 |
| 0 | NA | NA | NA | NA | NA | D830030K20Ri | RIKEN cDNA D830030K20 gene | 0 | 0 | 0 | 0 | 0 | 0 | 0 | 0 | 0 | 0 | 0 |
| 0 | NA | NA | NA | NA | NA | Gm17222 | | 0 | 0 | 0 | 0 | 0 | 0 | 0 | 0 | 0 | 0 | 0 |
| 0 | NA | NA | NA | NA | NA | Gm4904 | | 0 | 0 | 0 | 0 | 0 | 0 | 0 | 0 | 0 | 0 | 0 |
| 0 | NA | NA | NA | NA | NA | 1700110I01Rik | | 0 | 0 | 0 | 0 | 0 | 0 | 0 | 0 | 0 | 0 | 0 |
| 0 | NA | NA | NA | NA | NA | Gm2956 | | 0 | 0 | 0 | 0 | 0 | 0 | 0 | 0 | 0 | 0 | 0 |
| 0 | NA | NA | NA | NA | NA | Gm17223 | | 0 | 0 | 0 | 0 | 0 | 0 | 0 | 0 | 0 | 0 | 0 |
| 0 | NA | NA | NA | NA | NA | Gm10413 | predicted gene 10413 | 0 | 0 | 0 | 0 | 0 | 0 | 0 | 0 | 0 | 0 | 0 |
| 0 | NA | NA | NA | NA | NA | Gm10409 | predicted gene 3667 | 0 | 0 | 0 | 0 | 0 | 0 | 0 | 0 | 0 | 0 | 0 |
| 0 | NA | NA | NA | NA | NA | Gm3005 | | 0 | 0 | 0 | 0 | 0 | 0 | 0 | 0 | 0 | 0 | 0 |
| 0 | NA | NA | NA | NA | NA | Gm26817 | | 0 | 0 | 0 | 0 | 0 | 0 | 0 | 0 | 0 | 0 | 0 |
| 0 | NA | NA | NA | NA | NA | Gm10408 | predicted gene 10408 | 0 | 0 | 0 | 0 | 0 | 0 | 0 | 0 | 0 | 0 | 0 |
| 0 | NA | NA | NA | NA | NA | Gm8323 | | 0 | 0 | 0 | 0 | 0 | 0 | 0 | 0 | 0 | 0 | 0 |
| 0 | NA | NA | NA | NA | NA | Gm8324 | | 0 | 0 | 0 | 0 | 0 | 0 | 0 | 0 | 0 | 0 | 0 |
| 0 | NA | NA | NA | NA | NA | Gm11109 | | 0 | 0 | 0 | 0 | 0 | 0 | 0 | 0 | 0 | 0 | 0 |
| 0 | NA | NA | NA | NA | NA | Gm17107 | | 0 | 0 | 0 | 0 | 0 | 0 | 0 | 0 | 0 | 0 | 0 |
| 0 | NA | NA | NA | NA | NA | Gm3012 | predicted gene 3012 | 0 | 0 | 0 | 0 | 0 | 0 | 0 | 0 | 0 | 0 | 0 |
| 0 | NA | NA | NA | NA | NA | Gm3020 | predicted gene 3667 | 0 | 0 | 0 | 0 | 0 | 0 | 0 | 0 | 0 | 0 | 0 |
| 0 | NA | NA | NA | NA | NA | Gm2866 | | 0 | 0 | 0 | 0 | 0 | 0 | 0 | 0 | 0 | 0 | 0 |
| 0 | NA | NA | NA | NA | NA | Gm3043 | predicted gene 3043 | 0 | 0 | 0 | 0 | 0 | 0 | 0 | 0 | 0 | 0 | 0 |
| 0 | NA | NA | NA | NA | NA | Gm2871 | | 0 | 0 | 0 | 0 | 0 | 0 | 0 | 0 | 0 | 0 | 0 |
| 0 | NA | NA | NA | NA | NA | Gm2877 | | 0 | 0 | 0 | 0 | 0 | 0 | 0 | 0 | 0 | 0 | 0 |
| 0 | NA | NA | NA | NA | NA | Gm21965 | | 0 | 0 | 0 | 0 | 0 | 0 | 0 | 0 | 0 | 0 | 0 |
| 0 | NA | NA | NA | NA | NA | Gm3002 | predicted gene 3667 | 0 | 0 | 0 | 0 | 0 | 0 | 0 | 0 | 0 | 0 | 0 |
| 0 | NA | NA | NA | NA | NA | Gm26886 | | 0 | 0 | 0 | 0 | 0 | 0 | 0 | 0 | 0 | 0 | 0 |
| 0 | NA | NA | NA | NA | NA | Gm26570 | | 0 | 0 | 0 | 0 | 0 | 0 | 0 | 0 | 0 | 0 | 0 |
| 0 | NA | NA | NA | NA | NA | Gm3033 | | 0 | 0 | 0 | 0 | 0 | 0 | 0 | 0 | 0 | 0 | 0 |
| 0 | NA | NA | NA | NA | NA | Gm2916 | | 0 | 0 | 0 | 0 | 0 | 0 | 0 | 0 | 0 | 0 | 0 |
| 0 | NA | NA | NA | NA | NA | Gm3029 | | 0 | 0 | 0 | 0 | 0 | 0 | 0 | 0 | 0 | 0 | 0 |
| 0 | NA | NA | NA | NA | NA | Gm3095 | predicted gene 3095 | 0 | 0 | 0 | 0 | 0 | 0 | 0 | 0 | 0 | 0 | 0 |
| 0 | NA | NA | NA | NA | NA | Gm17129 | | 0 | 0 | 0 | 0 | 0 | 0 | 0 | 0 | 0 | 0 | 0 |
| 0 | NA | NA | NA | NA | NA | Gm3099 | | 0 | 0 | 0 | 0 | 0 | 0 | 0 | 0 | 0 | 0 | 0 |
| 0 | NA | NA | NA | NA | NA | Gm11107 | | 0 | 0 | 0 | 0 | 0 | 0 | 0 | 0 | 0 | 0 | 0 |
| 0 | NA | NA | NA | NA | NA | Gm3115 | predicted gene 3115 | 0 | 0 | 0 | 0 | 0 | 0 | 0 | 0 | 0 | 0 | 0 |
| 0 | NA | NA | NA | NA | NA | Gm8108 | predicted gene 8108 | 0 | 0 | 0 | 0 | 0 | 0 | 0 | 0 | 0 | 0 | 0 |
| 0 | NA | NA | NA | NA | NA | Gm17152 | | 0 | 0 | 0 | 0 | 0 | 0 | 0 | 0 | 0 | 0 | 0 |
| 0 | NA | NA | NA | NA | NA | Gm3127 | predicted gene 3127 | 0 | 0 | 0 | 0 | 0 | 0 | 0 | 0 | 0 | 0 | 0 |
| 0 | NA | NA | NA | NA | NA | Gm2974 | predicted gene 2974 | 0 | 0 | 0 | 0 | 0 | 0 | 0 | 0 | 0 | 0 | 0 |
| 0 | NA | NA | NA | NA | NA | Gm11105 | | 0 | 0 | 0 | 0 | 0 | 0 | 0 | 0 | 0 | 0 | 0 |
| 0 | NA | NA | NA | NA | NA | Gm3138 | | 0 | 0 | 0 | 0 | 0 | 0 | 0 | 0 | 0 | 0 | 0 |
| 0 | NA | NA | NA | NA | NA | Gm17085 | | 0 | 0 | 0 | 0 | 0 | 0 | 0 | 0 | 0 | 0 | 0 |
| 0 | NA | NA | NA | NA | NA | Gm3149 | | 0 | 0 | 0 | 0 | 0 | 0 | 0 | 0 | 0 | 0 | 0 |
| 0 | NA | NA | NA | NA | NA | Gm17086 | | 0 | 0 | 0 | 0 | 0 | 0 | 0 | 0 | 0 | 0 | 0 |
| 0 | NA | NA | NA | NA | NA | Gm3159 | predicted gene 3159 | 0 | 0 | 0 | 0 | 0 | 0 | 0 | 0 | 0 | 0 | 0 |
| 0 | NA | NA | NA | NA | NA | Gm3164 | | 0 | 0 | 0 | 0 | 0 | 0 | 0 | 0 | 0 | 0 | 0 |
| 0 | NA | NA | NA | NA | NA | Gm3173 | predicted gene 3173 | 0 | 0 | 0 | 0 | 0 | 0 | 0 | 0 | 0 | 0 | 0 |
| 0 | NA | NA | NA | NA | NA | Gm17170 | | 0 | 0 | 0 | 0 | 0 | 0 | 0 | 0 | 0 | 0 | 0 |
| 0 | NA | NA | NA | NA | NA | Gm3182 | predicted gene 3182 | 0 | 0 | 0 | 0 | 0 | 0 | 0 | 0 | 0 | 0 | 0 |

| | | | | | | | | | | | | | | | | | | |
|---|---|---|---|---|---|---|---|---|---|---|---|---|---|---|---|---|---|---|
| 0 | NA | NA | NA | NA | NA | Gm17074 | | 0 | 0 | 0 | 0 | 0 | 0 | 0 | 0 | 0 | 0 | 0 |
| 0 | NA | NA | NA | NA | NA | Gm3030 | | 0 | 0 | 0 | 0 | 0 | 0 | 0 | 0 | 0 | 0 | 0 |
| 0 | NA | NA | NA | NA | NA | Gm26655 | | 0 | 0 | 0 | 0 | 0 | 0 | 0 | 0 | 0 | 0 | 0 |
| 0 | NA | NA | NA | NA | NA | Gm3047 | | 0 | 0 | 0 | 0 | 0 | 0 | 0 | 0 | 0 | 0 | 0 |
| 0 | NA | NA | NA | NA | NA | Gm8164 | | 0 | 0 | 0 | 0 | 0 | 0 | 0 | 0 | 0 | 0 | 0 |
| 0 | NA | NA | NA | NA | NA | Gm26898 | | 0 | 0 | 0 | 0 | 0 | 0 | 0 | 0 | 0 | 0 | 0 |
| 0 | NA | NA | NA | NA | NA | Gm8159 | | 0 | 0 | 0 | 0 | 0 | 0 | 0 | 0 | 0 | 0 | 0 |
| 0 | NA | NA | NA | NA | NA | Gm3239 | | 0 | 0 | 0 | 0 | 0 | 0 | 0 | 0 | 0 | 0 | 0 |
| 0 | NA | NA | NA | NA | NA | Gm17073 | | 0 | 0 | 0 | 0 | 0 | 0 | 0 | 0 | 0 | 0 | 0 |
| 0 | NA | NA | NA | NA | NA | Gm7876 | predicted gene 7876 | 0 | 0 | 0 | 0 | 0 | 0 | 0 | 0 | 0 | 0 | 0 |
| 0 | NA | NA | NA | NA | NA | Gm3252 | | 0 | 0 | 0 | 0 | 0 | 0 | 0 | 0 | 0 | 0 | 0 |
| 0 | NA | NA | NA | NA | NA | Gm3084 | | 0 | 0 | 0 | 0 | 0 | 0 | 0 | 0 | 0 | 0 | 0 |
| 0 | NA | NA | NA | NA | NA | Gm9602 | predicted gene 9602 | 0 | 0 | 0 | 0 | 0 | 0 | 0 | 0 | 0 | 0 | 0 |
| 0 | NA | NA | NA | NA | NA | Gm3091 | | 0 | 0 | 0 | 0 | 0 | 0 | 0 | 0 | 0 | 0 | 0 |
| 0 | NA | NA | NA | NA | NA | Gm3269 | | 0 | 0 | 0 | 0 | 0 | 0 | 0 | 0 | 0 | 0 | 0 |
| 0 | NA | NA | NA | NA | NA | Gm3264 | predicted gene 3264 | 0 | 0 | 0 | 0 | 0 | 0 | 0 | 0 | 0 | 0 | 0 |
| 0 | NA | NA | NA | NA | NA | Gm26650 | | 0 | 0 | 0 | 0 | 0 | 0 | 0 | 0 | 0 | 0 | 0 |
| 0 | NA | NA | NA | NA | NA | Gm3278 | predicted gene 3278 | 0 | 0 | 0 | 0 | 0 | 0 | 0 | 0 | 0 | 0 | 0 |
| 0 | NA | NA | NA | NA | NA | Gm8305 | | 0 | 0 | 0 | 0 | 0 | 0 | 0 | 0 | 0 | 0 | 0 |
| 0 | NA | NA | NA | NA | NA | Gm10410 | | 0 | 0 | 0 | 0 | 0 | 0 | 0 | 0 | 0 | 0 | 0 |
| 0 | NA | NA | NA | NA | NA | Gm8297 | | 0 | 0 | 0 | 0 | 0 | 0 | 0 | 0 | 0 | 0 | 0 |
| 0 | NA | NA | NA | NA | NA | 4930555G01Ril | RIKEN cDNA 4930555G01 gene | 0 | 0 | 0 | 0 | 0 | 0 | 0 | 0 | 0 | 0 | 0 |
| 0 | NA | NA | NA | NA | NA | Gm17069 | | 0 | 0 | 0 | 0 | 0 | 0 | 0 | 0 | 0 | 0 | 0 |
| 0 | NA | NA | NA | NA | NA | Gm8279 | predicted gene 8279 | 0 | 0 | 0 | 0 | 0 | 0 | 0 | 0 | 0 | 0 | 0 |
| 0 | NA | NA | NA | NA | NA | Gm26552 | | 0 | 0 | 0 | 0 | 0 | 0 | 0 | 0 | 0 | 0 | 0 |
| 0 | NA | NA | NA | NA | NA | Gm3317 | predicted gene 3317 | 0 | 0 | 0 | 0 | 0 | 0 | 0 | 0 | 0 | 0 | 0 |
| 0 | NA | NA | NA | NA | NA | Gm3140 | | 0 | 0 | 0 | 0 | 0 | 0 | 0 | 0 | 0 | 0 | 0 |
| 0 | NA | NA | NA | NA | NA | Gm8271 | predicted gene 8271 | 0 | 0 | 0 | 0 | 0 | 0 | 0 | 0 | 0 | 0 | 0 |
| 0 | NA | NA | NA | NA | NA | Gm3326 | | 0 | 0 | 0 | 0 | 0 | 0 | 0 | 0 | 0 | 0 | 0 |
| 0 | NA | NA | NA | NA | NA | Gm3526 | | 0 | 0 | 0 | 0 | 0 | 0 | 0 | 0 | 0 | 0 | 0 |
| 0 | NA | NA | NA | NA | NA | Gm3339 | | 0 | 0 | 0 | 0 | 0 | 0 | 0 | 0 | 0 | 0 | 0 |
| 0 | NA | NA | NA | NA | NA | Gm3542 | | 0 | 0 | 0 | 0 | 0 | 0 | 0 | 0 | 0 | 0 | 0 |
| 0 | NA | NA | NA | NA | NA | Gm17126 | | 0 | 0 | 0 | 0 | 0 | 0 | 0 | 0 | 0 | 0 | 0 |
| 0 | NA | NA | NA | NA | NA | Gm3348 | | 0 | 0 | 0 | 0 | 0 | 0 | 0 | 0 | 0 | 0 | 0 |
| 0 | NA | NA | NA | NA | NA | Gm3194 | | 0 | 0 | 0 | 0 | 0 | 0 | 0 | 0 | 0 | 0 | 0 |
| 0 | NA | NA | NA | NA | NA | Gm8246 | predicted gene 8246 | 0 | 0 | 0 | 0 | 0 | 0 | 0 | 0 | 0 | 0 | 0 |
| 0 | NA | NA | NA | NA | NA | Gm3187 | predicted gene 3187 | 0 | 0 | 0 | 0 | 0 | 0 | 0 | 0 | 0 | 0 | 0 |
| 0 | NA | NA | NA | NA | NA | Gm3488 | predicted gene, 3488 | 0 | 0 | 0 | 0 | 0 | 0 | 0 | 0 | 0 | 0 | 0 |
| 0 | NA | NA | NA | NA | NA | Gm26893 | | 0 | 0 | 0 | 0 | 0 | 0 | 0 | 0 | 0 | 0 | 0 |
| 0 | NA | NA | NA | NA | NA | Gm26746 | | 0 | 0 | 0 | 0 | 0 | 0 | 0 | 0 | 0 | 0 | 0 |
| 0 | NA | NA | NA | NA | NA | Gm3532 | | 0 | 0 | 0 | 0 | 0 | 0 | 0 | 0 | 0 | 0 | 0 |
| 0 | NA | NA | NA | NA | NA | Gm8266 | | 0 | 0 | 0 | 0 | 0 | 0 | 0 | 0 | 0 | 0 | 0 |
| 0 | NA | NA | NA | NA | NA | Gm26731 | | 0 | 0 | 0 | 0 | 0 | 0 | 0 | 0 | 0 | 0 | 0 |
| 0 | NA | NA | NA | NA | NA | Gm3383 | predicted gene 3383 | 0 | 0 | 0 | 0 | 0 | 0 | 0 | 0 | 0 | 0 | 0 |
| 0 | NA | NA | NA | NA | NA | Gm3344 | | 0 | 0 | 0 | 0 | 0 | 0 | 0 | 0 | 0 | 0 | 0 |
| 0 | NA | NA | NA | NA | NA | Gm8265 | | 0 | 0 | 0 | 0 | 0 | 0 | 0 | 0 | 0 | 0 | 0 |
| 0 | NA | NA | NA | NA | NA | Gm9603 | | 0 | 0 | 0 | 0 | 0 | 0 | 0 | 0 | 0 | 0 | 0 |
| 0 | NA | NA | NA | NA | NA | Gm3373 | predicted gene 3373 | 0 | 0 | 0 | 0 | 0 | 0 | 0 | 0 | 0 | 0 | 0 |
| 0 | NA | NA | NA | NA | NA | Gm3170 | | 0 | 0 | 0 | 0 | 0 | 0 | 0 | 0 | 0 | 0 | 0 |
| 0 | NA | NA | NA | NA | NA | Gm26557 | | 0 | 0 | 0 | 0 | 0 | 0 | 0 | 0 | 0 | 0 | 0 |
| 0 | NA | NA | NA | NA | NA | Gm3424 | predicted gene 3424 | 0 | 0 | 0 | 0 | 0 | 0 | 0 | 0 | 0 | 0 | 0 |
| 0 | NA | NA | NA | NA | NA | Gm17158 | | 0 | 0 | 0 | 0 | 0 | 0 | 0 | 0 | 0 | 0 | 0 |
| 0 | NA | NA | NA | NA | NA | Gm17046 | | 0 | 0 | 0 | 0 | 0 | 0 | 0 | 0 | 0 | 0 | 0 |
| 0 | NA | NA | NA | NA | NA | Gm3242 | | 0 | 0 | 0 | 0 | 0 | 0 | 0 | 0 | 0 | 0 | 0 |
| 0 | NA | NA | NA | NA | NA | Gm17047 | | 0 | 0 | 0 | 0 | 0 | 0 | 0 | 0 | 0 | 0 | 0 |
| 0 | NA | NA | NA | NA | NA | Gm17048 | | 0 | 0 | 0 | 0 | 0 | 0 | 0 | 0 | 0 | 0 | 0 |
| 0 | NA | NA | NA | NA | NA | Gm3453 | | 0 | 0 | 0 | 0 | 0 | 0 | 0 | 0 | 0 | 0 | 0 |
| 0 | NA | NA | NA | NA | NA | Gm3261 | | 0 | 0 | 0 | 0 | 0 | 0 | 0 | 0 | 0 | 0 | 0 |
| 0 | NA | NA | NA | NA | NA | Gm6337 | predicted gene 6337 | 0 | 0 | 0 | 0 | 0 | 0 | 0 | 0 | 0 | 0 | 0 |
| 0 | NA | NA | NA | NA | NA | Gm3275 | | 0 | 0 | 0 | 0 | 0 | 0 | 0 | 0 | 0 | 0 | 0 |

| | | | | | | | | | | | | | | | | | | |
|---|---|---|---|---|---|---|---|---|---|---|---|---|---|---|---|---|---|---|
| 0 | NA | NA | NA | NA | NA | Gm3476 | predicted gene 3476 | 0 | 0 | 0 | 0 | 0 | 0 | 0 | 0 | 0 | 0 | 0 |
| 0 | NA | NA | NA | NA | NA | Gm26991 | | 0 | 0 | 0 | 0 | 0 | 0 | 0 | 0 | 0 | 0 | 0 |
| 0 | NA | NA | NA | NA | NA | Gm3208 | | 0 | 0 | 0 | 0 | 0 | 0 | 0 | 0 | 0 | 0 | 0 |
| 0 | NA | NA | NA | NA | NA | Gm21560 | uncharacterized LOC105245675 | 0 | 0 | 0 | 0 | 0 | 0 | 0 | 0 | 0 | 0 | 0 |
| 0 | NA | NA | NA | NA | NA | Gm21772 | | 0 | 0 | 0 | 0 | 0 | 0 | 0 | 0 | 0 | 0 | 0 |
| 0 | NA | NA | NA | NA | NA | Gm3411 | predicted gene 3411 | 0 | 0 | 0 | 0 | 0 | 0 | 0 | 0 | 0 | 0 | 0 |
| 0 | NA | NA | NA | NA | NA | Gm21103 | | 0 | 0 | 0 | 0 | 0 | 0 | 0 | 0 | 0 | 0 | 0 |
| 0 | NA | NA | NA | NA | NA | Gm3618 | | 0 | 0 | 0 | 0 | 0 | 0 | 0 | 0 | 0 | 0 | 0 |
| 0 | NA | NA | NA | NA | NA | Gm26630 | | 0 | 0 | 0 | 0 | 0 | 0 | 0 | 0 | 0 | 0 | 0 |
| 0 | NA | NA | NA | NA | NA | Gm8340 | | 0 | 0 | 0 | 0 | 0 | 0 | 0 | 0 | 0 | 0 | 0 |
| 0 | NA | NA | NA | NA | NA | Gm16440 | predicted gene 16440 | 0 | 0 | 0 | 0 | 0 | 0 | 0 | 0 | 0 | 0 | 0 |
| 0 | NA | NA | NA | NA | NA | Lamtor3-ps | late endosomal/lysosomal adaptor, MAPK | 0 | 0 | 0 | 0 | 0 | 0 | 0 | 0 | 0 | 0 | 0 |
| 0 | NA | NA | NA | NA | NA | Gm3594 | predicted gene 3594 | 0 | 0 | 0 | 0 | 0 | 0 | 0 | 0 | 0 | 0 | 0 |
| 0 | NA | NA | NA | NA | NA | Gm8356 | uncharacterized LOC108168187 | 0 | 0 | 0 | 0 | 0 | 0 | 0 | 0 | 0 | 0 | 0 |
| 0 | NA | NA | NA | NA | NA | Gm3460 | | 0 | 0 | 0 | 0 | 0 | 0 | 0 | 0 | 0 | 0 | 0 |
| 0 | NA | NA | NA | NA | NA | Gm10251 | predicted gene 3629 | 0 | 0 | 0 | 0 | 0 | 0 | 0 | 0 | 0 | 0 | 0 |
| 0 | NA | NA | NA | NA | NA | Gm26680 | | 0 | 0 | 0 | 0 | 0 | 0 | 0 | 0 | 0 | 0 | 0 |
| 0 | NA | NA | NA | NA | NA | Gm3466 | | 0 | 0 | 0 | 0 | 0 | 0 | 0 | 0 | 0 | 0 | 0 |
| 0 | NA | NA | NA | NA | NA | Gm8050 | predicted gene 8050 | 0 | 0 | 0 | 0 | 0 | 0 | 0 | 0 | 0 | 0 | 0 |
| 0 | NA | NA | NA | NA | NA | Gm8362 | | 0 | 0 | 0 | 0 | 0 | 0 | 0 | 0 | 0 | 0 | 0 |
| 0 | NA | NA | NA | NA | NA | Gm3642 | predicted gene 3642 | 0 | 0 | 0 | 0 | 0 | 0 | 0 | 0 | 0 | 0 | 0 |
| 0 | NA | NA | NA | NA | NA | Gm3667 | predicted gene 3667 | 0 | 0 | 0 | 0 | 0 | 0 | 0 | 0 | 0 | 0 | 0 |
| 0 | NA | NA | NA | NA | NA | Gm47747 | | 0 | 0 | 0 | 0 | 0 | 0 | 0 | 0 | 0 | 0 | 0 |
| 0 | NA | NA | NA | NA | NA | Gm47749 | | 0 | 0 | 0 | 0 | 0 | 0 | 0 | 0 | 0 | 0 | 0 |
| 0 | NA | NA | NA | NA | NA | Gm3685 | predicted gene 3685 | 0 | 0 | 0 | 0 | 0 | 0 | 0 | 0 | 0 | 0 | 0 |
| 0 | NA | NA | NA | NA | NA | Gm47764 | | 0 | 0 | 0 | 0 | 0 | 0 | 0 | 0 | 0 | 0 | 0 |
| 0 | NA | NA | NA | NA | NA | Gm6676 | predicted gene 6676 | 0 | 0 | 0 | 0 | 0 | 0 | 0 | 0 | 0 | 0 | 0 |
| 0 | NA | NA | NA | NA | NA | Gm16434 | predicted gene 16434 | 0 | 0 | 0 | 0 | 0 | 0 | 0 | 0 | 0 | 0 | 0 |
| 0 | NA | NA | NA | NA | NA | Gm3727 | predicted gene 3727 | 0 | 0 | 0 | 0 | 0 | 0 | 0 | 0 | 0 | 0 | 0 |
| 0 | NA | NA | NA | NA | NA | Gm5797 | predicted gene 5797 | 0 | 0 | 0 | 0 | 0 | 0 | 0 | 0 | 0 | 0 | 0 |
| 0 | NA | NA | NA | NA | NA | Gm8374 | | 0 | 0 | 0 | 0 | 0 | 0 | 0 | 0 | 0 | 0 | 0 |
| 0 | NA | NA | NA | NA | NA | Gm10339 | | 0 | 0 | 0 | 0 | 0 | 0 | 0 | 0 | 0 | 0 | 0 |
| 0 | NA | NA | NA | NA | NA | 4933406F09Rik | | 0 | 0 | 0 | 0 | 0 | 0 | 0 | 0 | 0 | 0 | 0 |
| 0 | NA | NA | NA | NA | NA | Gm8379 | | 0 | 0 | 0 | 0 | 0 | 0 | 0 | 0 | 0 | 0 | 0 |
| 0 | NA | NA | NA | NA | NA | Gm3755 | | 0 | 0 | 0 | 0 | 0 | 0 | 0 | 0 | 0 | 0 | 0 |
| 0 | NA | NA | NA | NA | NA | Gm3752 | predicted gene 3005 | 0 | 0 | 0 | 0 | 0 | 0 | 0 | 0 | 0 | 0 | 0 |
| 0 | NA | NA | NA | NA | NA | Gm3760 | | 0 | 0 | 0 | 0 | 0 | 0 | 0 | 0 | 0 | 0 | 0 |
| 0 | NA | NA | NA | NA | NA | Gm17121 | | 0 | 0 | 0 | 0 | 0 | 0 | 0 | 0 | 0 | 0 | 0 |
| 0 | NA | NA | NA | NA | NA | Gm10338 | predicted gene 10338 | 0 | 0 | 0 | 0 | 0 | 0 | 0 | 0 | 0 | 0 | 0 |
| 0 | NA | NA | NA | NA | NA | Gm47876 | | 0 | 0 | 0 | 0 | 0 | 0 | 0 | 0 | 0 | 0 | 0 |
| 0 | NA | NA | NA | NA | NA | Gm10128 | | 0 | 0 | 0 | 0 | 0 | 0 | 0 | 0 | 0 | 0 | 0 |
| 0 | NA | NA | NA | NA | NA | Gm26661 | | 0 | 0 | 0 | 0 | 0 | 0 | 0 | 0 | 0 | 0 | 0 |
| 0 | NA | NA | NA | NA | NA | Gm8396 | | 0 | 0 | 0 | 0 | 0 | 0 | 0 | 0 | 0 | 0 | 0 |
| 0 | NA | NA | NA | NA | NA | Gm45521 | predicted gene 45521 | 0 | 0 | 0 | 0 | 0 | 0 | 0 | 0 | 0 | 0 | 0 |
| 0 | NA | NA | NA | NA | NA | Gm8416 | | 0 | 0 | 0 | 0 | 0 | 0 | 0 | 0 | 0 | 0 | 0 |
| 0 | NA | NA | NA | NA | NA | 4930455B14Rik | | 0 | 0 | 0 | 0 | 0 | 0 | 0 | 0 | 0 | 0 | 0 |
| 0 | NA | NA | NA | NA | NA | Gm20241 | | 0 | 0 | 0 | 0 | 0 | 0 | 0 | 0 | 0 | 0 | 0 |
| 0 | NA | NA | NA | NA | NA | Gm48364 | | 0 | 0 | 0 | 0 | 0 | 0 | 0 | 0 | 0 | 0 | 0 |
| 0 | NA | NA | NA | NA | NA | Gm18071 | | 0 | 0 | 0 | 0 | 0 | 0 | 0 | 0 | 0 | 0 | 0 |
| 0 | NA | NA | NA | NA | NA | Gm48370 | | 0 | 0 | 0 | 0 | 0 | 0 | 0 | 0 | 0 | 0 | 0 |
| 0 | NA | NA | NA | NA | NA | Hmgb3-ps | | 0 | 0 | 0 | 0 | 0 | 0 | 0 | 0 | 0 | 0 | 0 |
| 0 | NA | NA | NA | NA | NA | Gm48439 | | 0 | 0 | 0 | 0 | 0 | 0 | 0 | 0 | 0 | 0 | 0 |
| 0 | NA | NA | NA | NA | NA | Gm48442 | | 0 | 0 | 0 | 0 | 0 | 0 | 0 | 0 | 0 | 0 | 0 |
| 0 | NA | NA | NA | NA | NA | Gm22053 | | 0 | 0 | 0 | 0 | 0 | 0 | 0 | 0 | 0 | 0 | 0 |
| 0 | NA | NA | NA | NA | NA | Fhitos | | 0 | 0 | 0 | 0 | 0 | 0 | 0 | 0 | 0 | 0 | 0 |
| 0 | NA | NA | NA | NA | NA | Mir5124b | | 0 | 0 | 0 | 0 | 0 | 0 | 0 | 0 | 0 | 0 | 0 |
| 0 | NA | NA | NA | NA | NA | Gm25466 | | 0 | 0 | 0 | 0 | 0 | 0 | 0 | 0 | 0 | 0 | 0 |
| 0 | NA | NA | NA | NA | NA | Gm48592 | | 0 | 0 | 0 | 0 | 0 | 0 | 0 | 0 | 0 | 0 | 0 |
| 0 | NA | NA | NA | NA | NA | Gm48594 | | 0 | 0 | 0 | 0 | 0 | 0 | 0 | 0 | 0 | 0 | 0 |
| 0 | NA | NA | NA | NA | NA | Gm48595 | | 0 | 0 | 0 | 0 | 0 | 0 | 0 | 0 | 0 | 0 | 0 |

| | | | | | | | | | | | | | | | | | |
|---|---|---|---|---|---|---|---|---|---|---|---|---|---|---|---|---|---|
| 0 | NA | NA | NA | NA | NA | Gm48602 | | 0 | 0 | 0 | 0 | 0 | 0 | 0 | 0 | 0 | 0 | 0 |
| 0 | NA | NA | NA | NA | NA | Gm48600 | | 0 | 0 | 0 | 0 | 0 | 0 | 0 | 0 | 0 | 0 | 0 |
| 0 | NA | NA | NA | NA | NA | Gm3848 | | 0 | 0 | 0 | 0 | 0 | 0 | 0 | 0 | 0 | 0 | 0 |
| 0 | NA | NA | NA | NA | NA | Gm48603 | | 0 | 0 | 0 | 0 | 0 | 0 | 0 | 0 | 0 | 0 | 0 |
| 0 | NA | NA | NA | NA | NA | D030051J21Rik | | 0 | 0 | 0 | 0 | 0 | 0 | 0 | 0 | 0 | 0 | 0 |
| 0 | NA | NA | NA | NA | NA | Gm24578 | | 0 | 0 | 0 | 0 | 0 | 0 | 0 | 0 | 0 | 0 | 0 |
| 0 | NA | NA | NA | NA | NA | Gm18077 | | 0 | 0 | 0 | 0 | 0 | 0 | 0 | 0 | 0 | 0 | 0 |
| 0 | NA | NA | NA | NA | NA | Gm48266 | | 0 | 0 | 0 | 0 | 0 | 0 | 0 | 0 | 0 | 0 | 0 |
| 0 | NA | NA | NA | NA | NA | Gm48515 | | 0 | 0 | 0 | 0 | 0 | 0 | 0 | 0 | 0 | 0 | 0 |
| 0 | NA | NA | NA | NA | NA | Gm15913 | | 0 | 0 | 0 | 0 | 0 | 0 | 0 | 0 | 0 | 0 | 0 |
| 0 | NA | NA | NA | NA | NA | Sntn | sentan, cilia apical structure protein | 0 | 0 | 0 | 0 | 0 | 0 | 0 | 0 | 0 | 0 | 0 |
| 0 | NA | NA | NA | NA | NA | Gm27506 | | 0 | 0 | 0 | 0 | 0 | 0 | 0 | 0 | 0 | 0 | 0 |
| 0 | NA | NA | NA | NA | NA | Gm25964 | | 0 | 0 | 0 | 0 | 0 | 0 | 0 | 0 | 0 | 0 | 0 |
| 0 | NA | NA | NA | NA | NA | Gm48841 | | 0 | 0 | 0 | 0 | 0 | 0 | 0 | 0 | 0 | 0 | 0 |
| 0 | NA | NA | NA | NA | NA | Gm48852 | | 0 | 0 | 0 | 0 | 0 | 0 | 0 | 0 | 0 | 0 | 0 |
| 0 | NA | NA | NA | NA | NA | Gm3895 | | 0 | 0 | 0 | 0 | 0 | 0 | 0 | 0 | 0 | 0 | 0 |
| 0 | NA | NA | NA | NA | NA | Gm48859 | | 0 | 0 | 0 | 0 | 0 | 0 | 0 | 0 | 0 | 0 | 0 |
| 0 | NA | NA | NA | NA | NA | Olfr31 | olfactory receptor 31 | 0 | 0 | 0 | 0 | 0 | 0 | 0 | 0 | 0 | 0 | 0 |
| 0 | NA | NA | NA | NA | NA | Gm6734 | | 0 | 0 | 0 | 0 | 0 | 0 | 0 | 0 | 0 | 0 | 0 |
| 0 | NA | NA | NA | NA | NA | Olfr721-ps1 | | 0 | 0 | 0 | 0 | 0 | 0 | 0 | 0 | 0 | 0 | 0 |
| 0 | NA | NA | NA | NA | NA | Gm47654 | | 0 | 0 | 0 | 0 | 0 | 0 | 0 | 0 | 0 | 0 | 0 |
| 0 | NA | NA | NA | NA | NA | Gm21063 | | 0 | 0 | 0 | 0 | 0 | 0 | 0 | 0 | 0 | 0 | 0 |
| 0 | NA | NA | NA | NA | NA | Gm47678 | | 0 | 0 | 0 | 0 | 0 | 0 | 0 | 0 | 0 | 0 | 0 |
| 0 | NA | NA | NA | NA | NA | Gm47685 | | 0 | 0 | 0 | 0 | 0 | 0 | 0 | 0 | 0 | 0 | 0 |
| 0 | NA | NA | NA | NA | NA | Gm19018 | | 0 | 0 | 0 | 0 | 0 | 0 | 0 | 0 | 0 | 0 | 0 |
| 0 | NA | NA | NA | NA | NA | Gm18142 | | 0 | 0 | 0 | 0 | 0 | 0 | 0 | 0 | 0 | 0 | 0 |
| 0 | NA | NA | NA | NA | NA | 4921522A10Rik | | 0 | 0 | 0 | 0 | 0 | 0 | 0 | 0 | 0 | 0 | 0 |
| 0 | NA | NA | NA | NA | NA | Gm47756 | | 0 | 0 | 0 | 0 | 0 | 0 | 0 | 0 | 0 | 0 | 0 |
| 0 | NA | NA | NA | NA | NA | Gm47758 | | 0 | 0 | 0 | 0 | 0 | 0 | 0 | 0 | 0 | 0 | 0 |
| 0 | NA | NA | NA | NA | NA | Gm47759 | | 0 | 0 | 0 | 0 | 0 | 0 | 0 | 0 | 0 | 0 | 0 |
| 0 | NA | NA | NA | NA | NA | Gm47779 | | 0 | 0 | 0 | 0 | 0 | 0 | 0 | 0 | 0 | 0 | 0 |
| 0 | NA | NA | NA | NA | NA | Gm47782 | | 0 | 0 | 0 | 0 | 0 | 0 | 0 | 0 | 0 | 0 | 0 |
| 0 | NA | NA | NA | NA | NA | Gm31804 | | 0 | 0 | 0 | 0 | 0 | 0 | 0 | 0 | 0 | 0 | 0 |
| 0 | NA | NA | NA | NA | NA | Gm47780 | | 0 | 0 | 0 | 0 | 0 | 0 | 0 | 0 | 0 | 0 | 0 |
| 0 | NA | NA | NA | NA | NA | 1700062C10Rik | | 0 | 0 | 0 | 0 | 0 | 0 | 0 | 0 | 0 | 0 | 0 |
| 0 | NA | NA | NA | NA | NA | Gm48162 | | 0 | 0 | 0 | 0 | 0 | 0 | 0 | 0 | 0 | 0 | 0 |
| 0 | NA | NA | NA | NA | NA | Tpi-rs9 | | 0 | 0 | 0 | 0 | 0 | 0 | 0 | 0 | 0 | 0 | 0 |
| 0 | NA | NA | NA | NA | NA | Gm18927 | | 0 | 0 | 0 | 0 | 0 | 0 | 0 | 0 | 0 | 0 | 0 |
| 0 | NA | NA | NA | NA | NA | Gm48320 | | 0 | 0 | 0 | 0 | 0 | 0 | 0 | 0 | 0 | 0 | 0 |
| 0 | NA | NA | NA | NA | NA | Gm24140 | | 0 | 0 | 0 | 0 | 0 | 0 | 0 | 0 | 0 | 0 | 0 |
| 0 | NA | NA | NA | NA | NA | Gm48321 | | 0 | 0 | 0 | 0 | 0 | 0 | 0 | 0 | 0 | 0 | 0 |
| 0 | NA | NA | NA | NA | NA | Gm27851 | | 0 | 0 | 0 | 0 | 0 | 0 | 0 | 0 | 0 | 0 | 0 |
| 0 | NA | NA | NA | NA | NA | Gm19290 | | 0 | 0 | 0 | 0 | 0 | 0 | 0 | 0 | 0 | 0 | 0 |
| 0 | NA | NA | NA | NA | NA | Gm6773 | | 0 | 0 | 0 | 0 | 0 | 0 | 0 | 0 | 0 | 0 | 0 |
| 0 | NA | NA | NA | NA | NA | Gm48481 | | 0 | 0 | 0 | 0 | 0 | 0 | 0 | 0 | 0 | 0 | 0 |
| 0 | NA | NA | NA | NA | NA | Gm44469 | | 0 | 0 | 0 | 0 | 0 | 0 | 0 | 0 | 0 | 0 | 0 |
| 0 | NA | NA | NA | NA | NA | Gm7084 | | 0 | 0 | 0 | 0 | 0 | 0 | 0 | 0 | 0 | 0 | 0 |
| 0 | NA | NA | NA | NA | NA | Gm22350 | | 0 | 0 | 0 | 0 | 0 | 0 | 0 | 0 | 0 | 0 | 0 |
| 0 | NA | NA | NA | NA | NA | Gm22499 | | 0 | 0 | 0 | 0 | 0 | 0 | 0 | 0 | 0 | 0 | 0 |
| 0 | NA | NA | NA | NA | NA | Gm15916 | | 0 | 0 | 0 | 0 | 0 | 0 | 0 | 0 | 0 | 0 | 0 |
| 0 | NA | NA | NA | NA | NA | Gm6785 | | 0 | 0 | 0 | 0 | 0 | 0 | 0 | 0 | 0 | 0 | 0 |
| 0 | NA | NA | NA | NA | NA | Gm3778 | | 0 | 0 | 0 | 0 | 0 | 0 | 0 | 0 | 0 | 0 | 0 |
| 0 | NA | NA | NA | NA | NA | Gm34440 | | 0 | 0 | 0 | 0 | 0 | 0 | 0 | 0 | 0 | 0 | 0 |
| 0 | NA | NA | NA | NA | NA | Gm48782 | | 0 | 0 | 0 | 0 | 0 | 0 | 0 | 0 | 0 | 0 | 0 |
| 0 | NA | NA | NA | NA | NA | Gm18275 | | 0 | 0 | 0 | 0 | 0 | 0 | 0 | 0 | 0 | 0 | 0 |
| 0 | NA | NA | NA | NA | NA | Gm7089 | | 0 | 0 | 0 | 0 | 0 | 0 | 0 | 0 | 0 | 0 | 0 |
| 0 | NA | NA | NA | NA | NA | Gm48785 | | 0 | 0 | 0 | 0 | 0 | 0 | 0 | 0 | 0 | 0 | 0 |
| 0 | NA | NA | NA | NA | NA | Gm8582 | | 0 | 0 | 0 | 0 | 0 | 0 | 0 | 0 | 0 | 0 | 0 |
| 0 | NA | NA | NA | NA | NA | Gm8584 | | 0 | 0 | 0 | 0 | 0 | 0 | 0 | 0 | 0 | 0 | 0 |
| 0 | NA | NA | NA | NA | NA | Gm48098 | | 0 | 0 | 0 | 0 | 0 | 0 | 0 | 0 | 0 | 0 | 0 |

| | | | | | | | | | | | | | | | | | | |
|---|---|---|---|---|---|---|---|---|---|---|---|---|---|---|---|---|---|---|
| 0 | NA | NA | NA | NA | NA | Gm48099 | | 0 | 0 | 0 | 0 | 0 | 0 | 0 | 0 | 0 | 0 | 0 |
| 0 | NA | NA | NA | NA | NA | Gm6122 | | 0 | 0 | 0 | 0 | 0 | 0 | 0 | 0 | 0 | 0 | 0 |
| 0 | NA | NA | NA | NA | NA | Gm33022 | | 0 | 0 | 0 | 0 | 0 | 0 | 0 | 0 | 0 | 0 | 0 |
| 0 | NA | NA | NA | NA | NA | Gm5630 | | 0 | 0 | 0 | 0 | 0 | 0 | 0 | 0 | 0 | 0 | 0 |
| 0 | NA | NA | NA | NA | NA | Gm9637 | | 0 | 0 | 0 | 0 | 0 | 0 | 0 | 0 | 0 | 0 | 0 |
| 0 | NA | NA | NA | NA | NA | Gm36704 | | 0 | 0 | 0 | 0 | 0 | 0 | 0 | 0 | 0 | 0 | 0 |
| 0 | NA | NA | NA | NA | NA | Gm48105 | | 0 | 0 | 0 | 0 | 0 | 0 | 0 | 0 | 0 | 0 | 0 |
| 0 | NA | NA | NA | NA | NA | Gm18577 | | 0 | 0 | 0 | 0 | 0 | 0 | 0 | 0 | 0 | 0 | 0 |
| 0 | NA | NA | NA | NA | NA | Gm5458 | predicted gene 5458 | 0 | 0 | 0 | 0 | 0 | 0 | 0 | 0 | 0 | 0 | 0 |
| 0 | NA | NA | NA | NA | NA | Gm41102 | | 0 | 0 | 0 | 0 | 0 | 0 | 0 | 0 | 0 | 0 | 0 |
| 0 | NA | NA | NA | NA | NA | 4930469B13Rik | | 0 | 0 | 0 | 0 | 0 | 0 | 0 | 0 | 0 | 0 | 0 |
| 0 | NA | NA | NA | NA | NA | Gm49800 | | 0 | 0 | 0 | 0 | 0 | 0 | 0 | 0 | 0 | 0 | 0 |
| 0 | NA | NA | NA | NA | NA | Gm48157 | | 0 | 0 | 0 | 0 | 0 | 0 | 0 | 0 | 0 | 0 | 0 |
| 0 | NA | NA | NA | NA | NA | Gm17030 | | 0 | 0 | 0 | 0 | 0 | 0 | 0 | 0 | 0 | 0 | 0 |
| 0 | NA | NA | NA | NA | NA | Gm7371 | | 0 | 0 | 0 | 0 | 0 | 0 | 0 | 0 | 0 | 0 | 0 |
| 0 | NA | NA | NA | NA | NA | Gm49341 | | 0 | 0 | 0 | 0 | 0 | 0 | 0 | 0 | 0 | 0 | 0 |
| 0 | NA | NA | NA | NA | NA | D14Ertd670e | | 0 | 0 | 0 | 0 | 0 | 0 | 0 | 0 | 0 | 0 | 0 |
| 0 | NA | NA | NA | NA | NA | Gm48283 | | 0 | 0 | 0 | 0 | 0 | 0 | 0 | 0 | 0 | 0 | 0 |
| 0 | NA | NA | NA | NA | NA | Gm48339 | | 0 | 0 | 0 | 0 | 0 | 0 | 0 | 0 | 0 | 0 | 0 |
| 0 | NA | NA | NA | NA | NA | Gm48394 | | 0 | 0 | 0 | 0 | 0 | 0 | 0 | 0 | 0 | 0 | 0 |
| 0 | NA | NA | NA | NA | NA | Gm24912 | | 0 | 0 | 0 | 0 | 0 | 0 | 0 | 0 | 0 | 0 | 0 |
| 0 | NA | NA | NA | NA | NA | Gm48395 | | 0 | 0 | 0 | 0 | 0 | 0 | 0 | 0 | 0 | 0 | 0 |
| 0 | NA | NA | NA | NA | NA | Mir6946 | | 0 | 0 | 0 | 0 | 0 | 0 | 0 | 0 | 0 | 0 | 0 |
| 0 | NA | NA | NA | NA | NA | Gm25328 | | 0 | 0 | 0 | 0 | 0 | 0 | 0 | 0 | 0 | 0 | 0 |
| 0 | NA | NA | NA | NA | NA | Gm26372 | | 0 | 0 | 0 | 0 | 0 | 0 | 0 | 0 | 0 | 0 | 0 |
| 0 | NA | NA | NA | NA | NA | Gm25864 | | 0 | 0 | 0 | 0 | 0 | 0 | 0 | 0 | 0 | 0 | 0 |
| 0 | NA | NA | NA | NA | NA | Gm15935 | | 0 | 0 | 0 | 0 | 0 | 0 | 0 | 0 | 0 | 0 | 0 |
| 0 | NA | NA | NA | NA | NA | Gm23502 | | 0 | 0 | 0 | 0 | 0 | 0 | 0 | 0 | 0 | 0 | 0 |
| 0 | NA | NA | NA | NA | NA | 4931407E12Rik | | 0 | 0 | 0 | 0 | 0 | 0 | 0 | 0 | 0 | 0 | 0 |
| 0 | NA | NA | NA | NA | NA | Gm37590 | | 0 | 0 | 0 | 0 | 0 | 0 | 0 | 0 | 0 | 0 | 0 |
| 0 | NA | NA | NA | NA | NA | Gm27532 | | 0 | 0 | 0 | 0 | 0 | 0 | 0 | 0 | 0 | 0 | 0 |
| 0 | NA | NA | NA | NA | NA | Gm7480 | | 0 | 0 | 0 | 0 | 0 | 0 | 0 | 0 | 0 | 0 | 0 |
| 0 | NA | NA | NA | NA | NA | Gm5670 | | 0 | 0 | 0 | 0 | 0 | 0 | 0 | 0 | 0 | 0 | 0 |
| 0 | NA | NA | NA | NA | NA | Gm47601 | | 0 | 0 | 0 | 0 | 0 | 0 | 0 | 0 | 0 | 0 | 0 |
| 0 | NA | NA | NA | NA | NA | Gm10248 | | 0 | 0 | 0 | 0 | 0 | 0 | 0 | 0 | 0 | 0 | 0 |
| 0 | NA | NA | NA | NA | NA | Gm46443 | | 0 | 0 | 0 | 0 | 0 | 0 | 0 | 0 | 0 | 0 | 0 |
| 0 | NA | NA | NA | NA | NA | 5430425E15Rik | | 0 | 0 | 0 | 0 | 0 | 0 | 0 | 0 | 0 | 0 | 0 |
| 0 | NA | NA | NA | NA | NA | Gm29626 | | 0 | 0 | 0 | 0 | 0 | 0 | 0 | 0 | 0 | 0 | 0 |
| 0 | NA | NA | NA | NA | NA | Gm31392 | | 0 | 0 | 0 | 0 | 0 | 0 | 0 | 0 | 0 | 0 | 0 |
| 0 | NA | NA | NA | NA | NA | | | 0 | 0 | 0 | 0 | 0 | 0 | 0 | 0 | 0 | 0 | 0 |
| 0 | NA | NA | NA | NA | NA | Mir7210 | | 0 | 0 | 0 | 0 | 0 | 0 | 0 | 0 | 0 | 0 | 0 |
| 0 | NA | NA | NA | NA | NA | | | 0 | 0 | 0 | 0 | 0 | 0 | 0 | 0 | 0 | 0 | 0 |
| 0 | NA | NA | NA | NA | NA | Gm17105 | | 0 | 0 | 0 | 0 | 0 | 0 | 0 | 0 | 0 | 0 | 0 |
| 0 | NA | NA | NA | NA | NA | | | 0 | 0 | 0 | 0 | 0 | 0 | 0 | 0 | 0 | 0 | 0 |
| 0 | NA | NA | NA | NA | NA | Gm19034 | | 0 | 0 | 0 | 0 | 0 | 0 | 0 | 0 | 0 | 0 | 0 |
| 0 | NA | NA | NA | NA | NA | E330034G19Ri | RIKEN cDNA E330034G19 gene | 0 | 0 | 0 | 0 | 0 | 0 | 0 | 0 | 0 | 0 | 0 |
| 0 | NA | NA | NA | NA | NA | | | 0 | 0 | 0 | 0 | 0 | 0 | 0 | 0 | 0 | 0 | 0 |
| 0 | NA | NA | NA | NA | NA | | | 0 | 0 | 0 | 0 | 0 | 0 | 0 | 0 | 0 | 0 | 0 |
| 0 | NA | NA | NA | NA | NA | Gm18276 | | 0 | 0 | 0 | 0 | 0 | 0 | 0 | 0 | 0 | 0 | 0 |
| 0 | NA | NA | NA | NA | NA | | | 0 | 0 | 0 | 0 | 0 | 0 | 0 | 0 | 0 | 0 | 0 |
| 0 | NA | NA | NA | NA | NA | Gm47774 | | 0 | 0 | 0 | 0 | 0 | 0 | 0 | 0 | 0 | 0 | 0 |
| 0 | NA | NA | NA | NA | NA | | | 0 | 0 | 0 | 0 | 0 | 0 | 0 | 0 | 0 | 0 | 0 |
| 0 | NA | NA | NA | NA | NA | 4930542C16Rik | | 0 | 0 | 0 | 0 | 0 | 0 | 0 | 0 | 0 | 0 | 0 |
| 0 | NA | NA | NA | NA | NA | Gm47906 | | 0 | 0 | 0 | 0 | 0 | 0 | 0 | 0 | 0 | 0 | 0 |
| 0 | NA | NA | NA | NA | NA | Gm10398 | | 0 | 0 | 0 | 0 | 0 | 0 | 0 | 0 | 0 | 0 | 0 |
| 0 | NA | NA | NA | NA | NA | Gm47907 | | 0 | 0 | 0 | 0 | 0 | 0 | 0 | 0 | 0 | 0 | 0 |
| 0 | NA | NA | NA | NA | NA | Gm47909 | | 0 | 0 | 0 | 0 | 0 | 0 | 0 | 0 | 0 | 0 | 0 |
| 0 | NA | NA | NA | NA | NA | 4930572O13Rik | | 0 | 0 | 0 | 0 | 0 | 0 | 0 | 0 | 0 | 0 | 0 |
| 0 | NA | NA | NA | NA | NA | Gm17747 | | 0 | 0 | 0 | 0 | 0 | 0 | 0 | 0 | 0 | 0 | 0 |
| 0 | NA | NA | NA | NA | NA | Mir3075 | | 0 | 0 | 0 | 0 | 0 | 0 | 0 | 0 | 0 | 0 | 0 |

| | | | | | | | | | | | | | | | | | |
|---|---|---|---|---|---|---|---|---|---|---|---|---|---|---|---|---|---|
| 0 | NA | NA | NA | NA | NA | Ppifos | | 0 | 0 | 0 | 0 | 0 | 0 | 0 | 0 | 0 | 0 | 0 |
| 0 | NA | NA | NA | NA | NA | Gm27185 | | 0 | 0 | 0 | 0 | 0 | 0 | 0 | 0 | 0 | 0 | 0 |
| 0 | NA | NA | NA | NA | NA | Gm23701 | | 0 | 0 | 0 | 0 | 0 | 0 | 0 | 0 | 0 | 0 | 0 |
| 0 | NA | NA | NA | NA | NA | Anxa11os | | 0 | 0 | 0 | 0 | 0 | 0 | 0 | 0 | 0 | 0 | 0 |
| 0 | NA | NA | NA | NA | NA | Plac9a | placenta specific 9a | 0 | 0 | 0 | 0 | 0 | 0 | 0 | 0 | 0 | 0 | 0 |
| 0 | NA | NA | NA | NA | NA | Tmem254a | transmembrane protein 254a | 0 | 0 | 0 | 0 | 0 | 0 | 0 | 0 | 0 | 0 | 0 |
| 0 | NA | NA | NA | NA | NA | Cphx1 | cytoplasmic polyadenylated homeobox 1 | 0 | 0 | 0 | 0 | 0 | 0 | 0 | 0 | 0 | 0 | 0 |
| 0 | NA | NA | NA | NA | NA | Gm19013 | | 0 | 0 | 0 | 0 | 0 | 0 | 0 | 0 | 0 | 0 | 0 |
| 0 | NA | NA | NA | NA | NA | Duxbl1 | double homeobox B-like 1 | 0 | 0 | 0 | 0 | 0 | 0 | 0 | 0 | 0 | 0 | 0 |
| 0 | NA | NA | NA | NA | NA | Gm2260 | | 0 | 0 | 0 | 0 | 0 | 0 | 0 | 0 | 0 | 0 | 0 |
| 0 | NA | NA | NA | NA | NA | Gm9780 | predicted gene 9780 | 0 | 0 | 0 | 0 | 0 | 0 | 0 | 0 | 0 | 0 | 0 |
| 0 | NA | NA | NA | NA | NA | Tmem254c | transmembrane protein 254c | 0 | 0 | 0 | 0 | 0 | 0 | 0 | 0 | 0 | 0 | 0 |
| 0 | NA | NA | NA | NA | NA | Gm19014 | | 0 | 0 | 0 | 0 | 0 | 0 | 0 | 0 | 0 | 0 | 0 |
| 0 | NA | NA | NA | NA | NA | Cphx2 | cytoplasmic polyadenylated homeobox 2 | 0 | 0 | 0 | 0 | 0 | 0 | 0 | 0 | 0 | 0 | 0 |
| 0 | NA | NA | NA | NA | NA | Cphx3 | cytoplasmic polyadenylated homeobox 3 | 0 | 0 | 0 | 0 | 0 | 0 | 0 | 0 | 0 | 0 | 0 |
| 0 | NA | NA | NA | NA | NA | Duxbl2 | double homeobox B-like 2 | 0 | 0 | 0 | 0 | 0 | 0 | 0 | 0 | 0 | 0 | 0 |
| 0 | NA | NA | NA | NA | NA | Gm2274 | | 0 | 0 | 0 | 0 | 0 | 0 | 0 | 0 | 0 | 0 | 0 |
| 0 | NA | NA | NA | NA | NA | Plac9b | predicted gene 9780 | 0 | 0 | 0 | 0 | 0 | 0 | 0 | 0 | 0 | 0 | 0 |
| 0 | NA | NA | NA | NA | NA | Tmem254b | transmembrane protein 254b | 0 | 0 | 0 | 0 | 0 | 0 | 0 | 0 | 0 | 0 | 0 |
| 0 | NA | NA | NA | NA | NA | Gm19015 | | 0 | 0 | 0 | 0 | 0 | 0 | 0 | 0 | 0 | 0 | 0 |
| 0 | NA | NA | NA | NA | NA | Duxbl3 | double homeobox B-like 3 | 0 | 0 | 0 | 0 | 0 | 0 | 0 | 0 | 0 | 0 | 0 |
| 0 | NA | NA | NA | NA | NA | Gm18578 | | 0 | 0 | 0 | 0 | 0 | 0 | 0 | 0 | 0 | 0 | 0 |
| 0 | NA | NA | NA | NA | NA | Gm34059 | | 0 | 0 | 0 | 0 | 0 | 0 | 0 | 0 | 0 | 0 | 0 |
| 0 | NA | NA | NA | NA | NA | 4933413J09Rik | | 0 | 0 | 0 | 0 | 0 | 0 | 0 | 0 | 0 | 0 | 0 |
| 0 | NA | NA | NA | NA | NA | Crb1-ps | | 0 | 0 | 0 | 0 | 0 | 0 | 0 | 0 | 0 | 0 | 0 |
| 0 | NA | NA | NA | NA | NA | Slmapos2 | | 0 | 0 | 0 | 0 | 0 | 0 | 0 | 0 | 0 | 0 | 0 |
| 0 | NA | NA | NA | NA | NA | Gm24031 | | 0 | 0 | 0 | 0 | 0 | 0 | 0 | 0 | 0 | 0 | 0 |
| 0 | NA | NA | NA | NA | NA | Gm2178 | | 0 | 0 | 0 | 0 | 0 | 0 | 0 | 0 | 0 | 0 | 0 |
| 0 | NA | NA | NA | NA | NA | Gm18215 | | 0 | 0 | 0 | 0 | 0 | 0 | 0 | 0 | 0 | 0 | 0 |
| 0 | NA | NA | NA | NA | NA | Mir7672 | | 0 | 0 | 0 | 0 | 0 | 0 | 0 | 0 | 0 | 0 | 0 |
| 0 | NA | NA | NA | NA | NA | Asb14 | ankyrin repeat and SOCS box-containing | 0 | 0 | 0 | 0 | 0 | 0 | 0 | 0 | 0 | 0 | 0 |
| 0 | NA | NA | NA | NA | NA | Gm41115 | | 0 | 0 | 0 | 0 | 0 | 0 | 0 | 0 | 0 | 0 | 0 |
| 0 | NA | NA | NA | NA | NA | Gm23633 | | 0 | 0 | 0 | 0 | 0 | 0 | 0 | 0 | 0 | 0 | 0 |
| 0 | NA | NA | NA | NA | NA | Gm22485 | | 0 | 0 | 0 | 0 | 0 | 0 | 0 | 0 | 0 | 0 | 0 |
| 0 | NA | NA | NA | NA | NA | Gm25697 | | 0 | 0 | 0 | 0 | 0 | 0 | 0 | 0 | 0 | 0 | 0 |
| 0 | NA | NA | NA | NA | NA | Gm45645 | | 0 | 0 | 0 | 0 | 0 | 0 | 0 | 0 | 0 | 0 | 0 |
| 0 | NA | NA | NA | NA | NA | Gm35164 | | 0 | 0 | 0 | 0 | 0 | 0 | 0 | 0 | 0 | 0 | 0 |
| 0 | NA | NA | NA | NA | NA | Gm35217 | | 0 | 0 | 0 | 0 | 0 | 0 | 0 | 0 | 0 | 0 | 0 |
| 0 | NA | NA | NA | NA | NA | Gm23999 | | 0 | 0 | 0 | 0 | 0 | 0 | 0 | 0 | 0 | 0 | 0 |
| 0 | NA | NA | NA | NA | NA | Lrtm1 | leucine-rich repeats and transmembrane | 0 | 0 | 0 | 0 | 0 | 0 | 0 | 0 | 0 | 0 | 0 |
| 0 | NA | NA | NA | NA | NA | Gm48004 | | 0 | 0 | 0 | 0 | 0 | 0 | 0 | 0 | 0 | 0 | 0 |
| 0 | NA | NA | NA | NA | NA | Gm48006 | | 0 | 0 | 0 | 0 | 0 | 0 | 0 | 0 | 0 | 0 | 0 |
| 0 | NA | NA | NA | NA | NA | Gm48005 | | 0 | 0 | 0 | 0 | 0 | 0 | 0 | 0 | 0 | 0 | 0 |
| 0 | NA | NA | NA | NA | NA | Gm48003 | | 0 | 0 | 0 | 0 | 0 | 0 | 0 | 0 | 0 | 0 | 0 |
| 0 | NA | NA | NA | NA | NA | Gm48058 | | 0 | 0 | 0 | 0 | 0 | 0 | 0 | 0 | 0 | 0 | 0 |
| 0 | NA | NA | NA | NA | NA | Gm41118 | | 0 | 0 | 0 | 0 | 0 | 0 | 0 | 0 | 0 | 0 | 0 |
| 0 | NA | NA | NA | NA | NA | Gm35281 | | 0 | 0 | 0 | 0 | 0 | 0 | 0 | 0 | 0 | 0 | 0 |
| 0 | NA | NA | NA | NA | NA | Gm7633 | | 0 | 0 | 0 | 0 | 0 | 0 | 0 | 0 | 0 | 0 | 0 |
| 0 | NA | NA | NA | NA | NA | Gm18760 | | 0 | 0 | 0 | 0 | 0 | 0 | 0 | 0 | 0 | 0 | 0 |
| 0 | NA | NA | NA | NA | NA | Gm50467 | | 0 | 0 | 0 | 0 | 0 | 0 | 0 | 0 | 0 | 0 | 0 |
| 0 | NA | NA | NA | NA | NA | Gm47570 | | 0 | 0 | 0 | 0 | 0 | 0 | 0 | 0 | 0 | 0 | 0 |
| 0 | NA | NA | NA | NA | NA | Gm18072 | | 0 | 0 | 0 | 0 | 0 | 0 | 0 | 0 | 0 | 0 | 0 |
| 0 | NA | NA | NA | NA | NA | Gm47576 | | 0 | 0 | 0 | 0 | 0 | 0 | 0 | 0 | 0 | 0 | 0 |
| 0 | NA | NA | NA | NA | NA | Gm18681 | | 0 | 0 | 0 | 0 | 0 | 0 | 0 | 0 | 0 | 0 | 0 |
| 0 | NA | NA | NA | NA | NA | Mir3076 | | 0 | 0 | 0 | 0 | 0 | 0 | 0 | 0 | 0 | 0 | 0 |
| 0 | NA | NA | NA | NA | NA | Gm7644 | | 0 | 0 | 0 | 0 | 0 | 0 | 0 | 0 | 0 | 0 | 0 |
| 0 | NA | NA | NA | NA | NA | 1700087M22Rik | | 0 | 0 | 0 | 0 | 0 | 0 | 0 | 0 | 0 | 0 | 0 |
| 0 | NA | NA | NA | NA | NA | Snord69 | | 0 | 0 | 0 | 0 | 0 | 0 | 0 | 0 | 0 | 0 | 0 |
| 0 | NA | NA | NA | NA | NA | Gm24916 | | 0 | 0 | 0 | 0 | 0 | 0 | 0 | 0 | 0 | 0 | 0 |
| 0 | NA | NA | NA | NA | NA | Snord19 | | 0 | 0 | 0 | 0 | 0 | 0 | 0 | 0 | 0 | 0 | 0 |

| | | | | | | | | | | | | | | | | | | |
|---|---|---|---|---|---|---|---|---|---|---|---|---|---|---|---|---|---|---|
| 0 | NA | NA | NA | NA | NA | Gm49094 | | 0 | 0 | 0 | 0 | 0 | 0 | 0 | 0 | 0 | 0 | 0 |
| 0 | NA | NA | NA | NA | NA | Gm35823 | | 0 | 0 | 0 | 0 | 0 | 0 | 0 | 0 | 0 | 0 | 0 |
| 0 | NA | NA | NA | NA | NA | Gm49217 | | 0 | 0 | 0 | 0 | 0 | 0 | 0 | 0 | 0 | 0 | 0 |
| 0 | NA | NA | NA | NA | NA | Gm49791 | | 0 | 0 | 0 | 0 | 0 | 0 | 0 | 0 | 0 | 0 | 0 |
| 0 | NA | NA | NA | NA | NA | Gm24057 | | 0 | 0 | 0 | 0 | 0 | 0 | 0 | 0 | 0 | 0 | 0 |
| 0 | NA | NA | NA | NA | NA | Gm49387 | | 0 | 0 | 0 | 0 | 0 | 0 | 0 | 0 | 0 | 0 | 0 |
| 0 | NA | NA | NA | NA | NA | Gm49309 | | 0 | 0 | 0 | 0 | 0 | 0 | 0 | 0 | 0 | 0 | 0 |
| 0 | NA | NA | NA | NA | NA | Gm22876 | | 0 | 0 | 0 | 0 | 0 | 0 | 0 | 0 | 0 | 0 | 0 |
| 0 | NA | NA | NA | NA | NA | D830044D21Rik | | 0 | 0 | 0 | 0 | 0 | 0 | 0 | 0 | 0 | 0 | 0 |
| 0 | NA | NA | NA | NA | NA | 4930425P05Rik | | 0 | 0 | 0 | 0 | 0 | 0 | 0 | 0 | 0 | 0 | 0 |
| 0 | NA | NA | NA | NA | NA | Gm26045 | | 0 | 0 | 0 | 0 | 0 | 0 | 0 | 0 | 0 | 0 | 0 |
| 0 | NA | NA | NA | NA | NA | Gm49084 | | 0 | 0 | 0 | 0 | 0 | 0 | 0 | 0 | 0 | 0 | 0 |
| 0 | NA | NA | NA | NA | NA | Msmb | beta-microseminoprotein | 0 | 0 | 0 | 0 | 0 | 0 | 0 | 0 | 0 | 0 | 0 |
| 0 | NA | NA | NA | NA | NA | Gm18909 | | 0 | 0 | 0 | 0 | 0 | 0 | 0 | 0 | 0 | 0 | 0 |
| 0 | NA | NA | NA | NA | NA | Gm23946 | | 0 | 0 | 0 | 0 | 0 | 0 | 0 | 0 | 0 | 0 | 0 |
| 0 | NA | NA | NA | NA | NA | Gm17210 | | 0 | 0 | 0 | 0 | 0 | 0 | 0 | 0 | 0 | 0 | 0 |
| 0 | NA | NA | NA | NA | NA | 1700024G13Ril | RIKEN cDNA 1700024G13 gene | 0 | 0 | 0 | 0 | 0 | 0 | 0 | 0 | 0 | 0 | 0 |
| 0 | NA | NA | NA | NA | NA | Slc18a3 | solute carrier family 18 (vesicular monoar | 0 | 0 | 0 | 0 | 0 | 0 | 0 | 0 | 0 | 0 | 0 |
| 0 | NA | NA | NA | NA | NA | Fam170b | family with sequence similarity 170, meml | 0 | 0 | 0 | 0 | 0 | 0 | 0 | 0 | 0 | 0 | 0 |
| 0 | NA | NA | NA | NA | NA | Gm7734 | | 0 | 0 | 0 | 0 | 0 | 0 | 0 | 0 | 0 | 0 | 0 |
| 0 | NA | NA | NA | NA | NA | Gm49029 | | 0 | 0 | 0 | 0 | 0 | 0 | 0 | 0 | 0 | 0 | 0 |
| 0 | NA | NA | NA | NA | NA | Gm25498 | | 0 | 0 | 0 | 0 | 0 | 0 | 0 | 0 | 0 | 0 | 0 |
| 0 | NA | NA | NA | NA | NA | Gm6340 | | 0 | 0 | 0 | 0 | 0 | 0 | 0 | 0 | 0 | 0 | 0 |
| 0 | NA | NA | NA | NA | NA | Gm49030 | | 0 | 0 | 0 | 0 | 0 | 0 | 0 | 0 | 0 | 0 | 0 |
| 0 | NA | NA | NA | NA | NA | Gm49037 | | 0 | 0 | 0 | 0 | 0 | 0 | 0 | 0 | 0 | 0 | 0 |
| 0 | NA | NA | NA | NA | NA | Ptpn20 | protein tyrosine phosphatase, non-recept | 0 | 0 | 0 | 0 | 0 | 0 | 0 | 0 | 0 | 0 | 0 |
| 0 | NA | NA | NA | NA | NA | Gm26228 | | 0 | 0 | 0 | 0 | 0 | 0 | 0 | 0 | 0 | 0 | 0 |
| 0 | NA | NA | NA | NA | NA | Gm49108 | | 0 | 0 | 0 | 0 | 0 | 0 | 0 | 0 | 0 | 0 | 0 |
| 0 | NA | NA | NA | NA | NA | 4930503F20Rik | | 0 | 0 | 0 | 0 | 0 | 0 | 0 | 0 | 0 | 0 | 0 |
| 0 | NA | NA | NA | NA | NA | Zfp488 | zinc finger protein 488 | 0 | 0 | 0 | 0 | 0 | 0 | 0 | 0 | 0 | 0 | 0 |
| 0 | NA | NA | NA | NA | NA | Gm30083 | predicted gene, 30083 | 0 | 0 | 0 | 0 | 0 | 0 | 0 | 0 | 0 | 0 | 0 |
| 0 | NA | NA | NA | NA | NA | Gm5460 | predicted gene 5460 | 0 | 0 | 0 | 0 | 0 | 0 | 0 | 0 | 0 | 0 | 0 |
| 0 | NA | NA | NA | NA | NA | A630023A22Ril | RIKEN cDNA A630023A22 gene | 0 | 0 | 0 | 0 | 0 | 0 | 0 | 0 | 0 | 0 | 0 |
| 0 | NA | NA | NA | NA | NA | Antxrl | anthrax toxin receptor-like | 0 | 0 | 0 | 0 | 0 | 0 | 0 | 0 | 0 | 0 | 0 |
| 0 | NA | NA | NA | NA | NA | Gm49129 | | 0 | 0 | 0 | 0 | 0 | 0 | 0 | 0 | 0 | 0 | 0 |
| 0 | NA | NA | NA | NA | NA | Gm24998 | | 0 | 0 | 0 | 0 | 0 | 0 | 0 | 0 | 0 | 0 | 0 |
| 0 | NA | NA | NA | NA | NA | Gm30556 | | 0 | 0 | 0 | 0 | 0 | 0 | 0 | 0 | 0 | 0 | 0 |
| 0 | NA | NA | NA | NA | NA | Gm7778 | | 0 | 0 | 0 | 0 | 0 | 0 | 0 | 0 | 0 | 0 | 0 |
| 0 | NA | NA | NA | NA | NA | Gm18884 | | 0 | 0 | 0 | 0 | 0 | 0 | 0 | 0 | 0 | 0 | 0 |
| 0 | NA | NA | NA | NA | NA | Gm18813 | | 0 | 0 | 0 | 0 | 0 | 0 | 0 | 0 | 0 | 0 | 0 |
| 0 | NA | NA | NA | NA | NA | Gm49203 | | 0 | 0 | 0 | 0 | 0 | 0 | 0 | 0 | 0 | 0 | 0 |
| 0 | NA | NA | NA | NA | NA | Gm49202 | | 0 | 0 | 0 | 0 | 0 | 0 | 0 | 0 | 0 | 0 | 0 |
| 0 | NA | NA | NA | NA | NA | 9230112D13Ril | RIKEN cDNA 9230112D13 gene | 0 | 0 | 0 | 0 | 0 | 0 | 0 | 0 | 0 | 0 | 0 |
| 0 | NA | NA | NA | NA | NA | Gm48953 | | 0 | 0 | 0 | 0 | 0 | 0 | 0 | 0 | 0 | 0 | 0 |
| 0 | NA | NA | NA | NA | NA | Opn4 | opsin 4 (melanopsin) | 0 | 0 | 0 | 0 | 0 | 0 | 0 | 0 | 0 | 0 | 0 |
| 0 | NA | NA | NA | NA | NA | Gm49012 | | 0 | 0 | 0 | 0 | 0 | 0 | 0 | 0 | 0 | 0 | 0 |
| 0 | NA | NA | NA | NA | NA | Mir346 | | 0 | 0 | 0 | 0 | 0 | 0 | 0 | 0 | 0 | 0 | 0 |
| 0 | NA | NA | NA | NA | NA | Gm49034 | | 0 | 0 | 0 | 0 | 0 | 0 | 0 | 0 | 0 | 0 | 0 |
| 0 | NA | NA | NA | NA | NA | Gm19016 | | 0 | 0 | 0 | 0 | 0 | 0 | 0 | 0 | 0 | 0 | 0 |
| 0 | NA | NA | NA | NA | NA | Gm27617 | | 0 | 0 | 0 | 0 | 0 | 0 | 0 | 0 | 0 | 0 | 0 |
| 0 | NA | NA | NA | NA | NA | 4930596D02Ril | RIKEN cDNA 4930596D02 gene | 0 | 0 | 0 | 0 | 0 | 0 | 0 | 0 | 0 | 0 | 0 |
| 0 | NA | NA | NA | NA | NA | Gm49039 | | 0 | 0 | 0 | 0 | 0 | 0 | 0 | 0 | 0 | 0 | 0 |
| 0 | NA | NA | NA | NA | NA | Gm49040 | | 0 | 0 | 0 | 0 | 0 | 0 | 0 | 0 | 0 | 0 | 0 |
| 0 | NA | NA | NA | NA | NA | Gm46474 | | 0 | 0 | 0 | 0 | 0 | 0 | 0 | 0 | 0 | 0 | 0 |
| 0 | NA | NA | NA | NA | NA | Gm19129 | | 0 | 0 | 0 | 0 | 0 | 0 | 0 | 0 | 0 | 0 | 0 |
| 0 | NA | NA | NA | NA | NA | Gm7853 | | 0 | 0 | 0 | 0 | 0 | 0 | 0 | 0 | 0 | 0 | 0 |
| 0 | NA | NA | NA | NA | NA | 4930474N05Ril | RIKEN cDNA 4930474N05 gene | 0 | 0 | 0 | 0 | 0 | 0 | 0 | 0 | 0 | 0 | 0 |
| 0 | NA | NA | NA | NA | NA | Gm48663 | | 0 | 0 | 0 | 0 | 0 | 0 | 0 | 0 | 0 | 0 | 0 |
| 0 | NA | NA | NA | NA | NA | Gm24394 | | 0 | 0 | 0 | 0 | 0 | 0 | 0 | 0 | 0 | 0 | 0 |
| 0 | NA | NA | NA | NA | NA | Gm17940 | | 0 | 0 | 0 | 0 | 0 | 0 | 0 | 0 | 0 | 0 | 0 |

| | | | | | | | | | | | | | | | | | | |
|---|---|---|---|---|---|---|---|---|---|---|---|---|---|---|---|---|---|---|
| 0 | NA | NA | NA | NA | NA | Gm25204 | | 0 | 0 | 0 | 0 | 0 | 0 | 0 | 0 | 0 | 0 | 0 |
| 0 | NA | NA | NA | NA | NA | n-R5s46 | | 0 | 0 | 0 | 0 | 0 | 0 | 0 | 0 | 0 | 0 | 0 |
| 0 | NA | NA | NA | NA | NA | Rgr | retinal G protein coupled receptor | 0 | 0 | 0 | 0 | 0 | 0 | 0 | 0 | 0 | 0 | 0 |
| 0 | NA | NA | NA | NA | NA | Lrit1 | leucine-rich repeat, immunoglobulin-like a | 0 | 0 | 0 | 0 | 0 | 0 | 0 | 0 | 0 | 0 | 0 |
| 0 | NA | NA | NA | NA | NA | Lrit2 | leucine-rich repeat, immunoglobulin-like a | 0 | 0 | 0 | 0 | 0 | 0 | 0 | 0 | 0 | 0 | 0 |
| 0 | NA | NA | NA | NA | NA | Gm18834 | | 0 | 0 | 0 | 0 | 0 | 0 | 0 | 0 | 0 | 0 | 0 |
| 0 | NA | NA | NA | NA | NA | Gm22469 | | 0 | 0 | 0 | 0 | 0 | 0 | 0 | 0 | 0 | 0 | 0 |
| 0 | NA | NA | NA | NA | NA | Gm23909 | | 0 | 0 | 0 | 0 | 0 | 0 | 0 | 0 | 0 | 0 | 0 |
| 0 | NA | NA | NA | NA | NA | Gm47971 | | 0 | 0 | 0 | 0 | 0 | 0 | 0 | 0 | 0 | 0 | 0 |
| 0 | NA | NA | NA | NA | NA | Gm31393 | | 0 | 0 | 0 | 0 | 0 | 0 | 0 | 0 | 0 | 0 | 0 |
| 0 | NA | NA | NA | NA | NA | Gm5204 | | 0 | 0 | 0 | 0 | 0 | 0 | 0 | 0 | 0 | 0 | 0 |
| 0 | NA | NA | NA | NA | NA | Gm47974 | | 0 | 0 | 0 | 0 | 0 | 0 | 0 | 0 | 0 | 0 | 0 |
| 0 | NA | NA | NA | NA | NA | Gm18523 | | 0 | 0 | 0 | 0 | 0 | 0 | 0 | 0 | 0 | 0 | 0 |
| 0 | NA | NA | NA | NA | NA | Gm31456 | | 0 | 0 | 0 | 0 | 0 | 0 | 0 | 0 | 0 | 0 | 0 |
| 0 | NA | NA | NA | NA | NA | Gm20641 | | 0 | 0 | 0 | 0 | 0 | 0 | 0 | 0 | 0 | 0 | 0 |
| 0 | NA | NA | NA | NA | NA | Nrg3os | | 0 | 0 | 0 | 0 | 0 | 0 | 0 | 0 | 0 | 0 | 0 |
| 0 | NA | NA | NA | NA | NA | Gm27332 | | 0 | 0 | 0 | 0 | 0 | 0 | 0 | 0 | 0 | 0 | 0 |
| 0 | NA | NA | NA | NA | NA | Gm20642 | | 0 | 0 | 0 | 0 | 0 | 0 | 0 | 0 | 0 | 0 | 0 |
| 0 | NA | NA | NA | NA | NA | 1700109I08Rik | | 0 | 0 | 0 | 0 | 0 | 0 | 0 | 0 | 0 | 0 | 0 |
| 0 | NA | NA | NA | NA | NA | Gm35389 | | 0 | 0 | 0 | 0 | 0 | 0 | 0 | 0 | 0 | 0 | 0 |
| 0 | NA | NA | NA | NA | NA | Gm25012 | | 0 | 0 | 0 | 0 | 0 | 0 | 0 | 0 | 0 | 0 | 0 |
| 0 | NA | NA | NA | NA | NA | Gm22191 | | 0 | 0 | 0 | 0 | 0 | 0 | 0 | 0 | 0 | 0 | 0 |
| 0 | NA | NA | NA | NA | NA | Gm22726 | | 0 | 0 | 0 | 0 | 0 | 0 | 0 | 0 | 0 | 0 | 0 |
| 0 | NA | NA | NA | NA | NA | Gm48028 | | 0 | 0 | 0 | 0 | 0 | 0 | 0 | 0 | 0 | 0 | 0 |
| 0 | NA | NA | NA | NA | NA | Gm24388 | | 0 | 0 | 0 | 0 | 0 | 0 | 0 | 0 | 0 | 0 | 0 |
| 0 | NA | NA | NA | NA | NA | Gm48040 | | 0 | 0 | 0 | 0 | 0 | 0 | 0 | 0 | 0 | 0 | 0 |
| 0 | NA | NA | NA | NA | NA | Gm7920 | | 0 | 0 | 0 | 0 | 0 | 0 | 0 | 0 | 0 | 0 | 0 |
| 0 | NA | NA | NA | NA | NA | Gm47114 | | 0 | 0 | 0 | 0 | 0 | 0 | 0 | 0 | 0 | 0 | 0 |
| 0 | NA | NA | NA | NA | NA | Gm7925 | | 0 | 0 | 0 | 0 | 0 | 0 | 0 | 0 | 0 | 0 | 0 |
| 0 | NA | NA | NA | NA | NA | Gm22823 | | 0 | 0 | 0 | 0 | 0 | 0 | 0 | 0 | 0 | 0 | 0 |
| 0 | NA | NA | NA | NA | NA | Gm2832 | predicted gene 2832 | 0 | 0 | 0 | 0 | 0 | 0 | 0 | 0 | 0 | 0 | 0 |
| 0 | NA | NA | NA | NA | NA | Gm9671 | | 0 | 0 | 0 | 0 | 0 | 0 | 0 | 0 | 0 | 0 | 0 |
| 0 | NA | NA | NA | NA | NA | Gm5798 | predicted gene 5798 | 0 | 0 | 0 | 0 | 0 | 0 | 0 | 0 | 0 | 0 | 0 |
| 0 | NA | NA | NA | NA | NA | Gm2855 | | 0 | 0 | 0 | 0 | 0 | 0 | 0 | 0 | 0 | 0 | 0 |
| 0 | NA | NA | NA | NA | NA | Gm7945 | predicted gene 7945 | 0 | 0 | 0 | 0 | 0 | 0 | 0 | 0 | 0 | 0 | 0 |
| 0 | NA | NA | NA | NA | NA | Gm6482 | | 0 | 0 | 0 | 0 | 0 | 0 | 0 | 0 | 0 | 0 | 0 |
| 0 | NA | NA | NA | NA | NA | Gm7953 | | 0 | 0 | 0 | 0 | 0 | 0 | 0 | 0 | 0 | 0 | 0 |
| 0 | NA | NA | NA | NA | NA | Gm7954 | | 0 | 0 | 0 | 0 | 0 | 0 | 0 | 0 | 0 | 0 | 0 |
| 0 | NA | NA | NA | NA | NA | Gm2878 | | 0 | 0 | 0 | 0 | 0 | 0 | 0 | 0 | 0 | 0 | 0 |
| 0 | NA | NA | NA | NA | NA | Gm3486 | | 0 | 0 | 0 | 0 | 0 | 0 | 0 | 0 | 0 | 0 | 0 |
| 0 | NA | NA | NA | NA | NA | Gm6490 | | 0 | 0 | 0 | 0 | 0 | 0 | 0 | 0 | 0 | 0 | 0 |
| 0 | NA | NA | NA | NA | NA | Gm17061 | | 0 | 0 | 0 | 0 | 0 | 0 | 0 | 0 | 0 | 0 | 0 |
| 0 | NA | NA | NA | NA | NA | Gm7970 | predicted gene 7970 | 0 | 0 | 0 | 0 | 0 | 0 | 0 | 0 | 0 | 0 | 0 |
| 0 | NA | NA | NA | NA | NA | 1700049E17Ril | tandem duplication of RIKEN cDNA 1700 | 0 | 0 | 0 | 0 | 0 | 0 | 0 | 0 | 0 | 0 | 0 |
| 0 | NA | NA | NA | NA | NA | Gm17159 | | 0 | 0 | 0 | 0 | 0 | 0 | 0 | 0 | 0 | 0 | 0 |
| 0 | NA | NA | NA | NA | NA | Gm3072 | | 0 | 0 | 0 | 0 | 0 | 0 | 0 | 0 | 0 | 0 | 0 |
| 0 | NA | NA | NA | NA | NA | Gm3077 | | 0 | 0 | 0 | 0 | 0 | 0 | 0 | 0 | 0 | 0 | 0 |
| 0 | NA | NA | NA | NA | NA | Gm3676 | | 0 | 0 | 0 | 0 | 0 | 0 | 0 | 0 | 0 | 0 | 0 |
| 0 | NA | NA | NA | NA | NA | Gm8065 | | 0 | 0 | 0 | 0 | 0 | 0 | 0 | 0 | 0 | 0 | 0 |
| 0 | NA | NA | NA | NA | NA | Gm8068 | | 0 | 0 | 0 | 0 | 0 | 0 | 0 | 0 | 0 | 0 | 0 |
| 0 | NA | NA | NA | NA | NA | Gm7929 | | 0 | 0 | 0 | 0 | 0 | 0 | 0 | 0 | 0 | 0 | 0 |
| 0 | NA | NA | NA | NA | NA | Gm47189 | | 0 | 0 | 0 | 0 | 0 | 0 | 0 | 0 | 0 | 0 | 0 |
| 0 | NA | NA | NA | NA | NA | Gm17062 | | 0 | 0 | 0 | 0 | 0 | 0 | 0 | 0 | 0 | 0 | 0 |
| 0 | NA | NA | NA | NA | NA | Gm7980 | predicted gene 7980 | 0 | 0 | 0 | 0 | 0 | 0 | 0 | 0 | 0 | 0 | 0 |
| 0 | NA | NA | NA | NA | NA | 1700049E17Ril | RIKEN cDNA 1700049E17 gene, gene 1 | 0 | 0 | 0 | 0 | 0 | 0 | 0 | 0 | 0 | 0 | 0 |
| 0 | NA | NA | NA | NA | NA | 1700024B05Rik | | 0 | 0 | 0 | 0 | 0 | 0 | 0 | 0 | 0 | 0 | 0 |
| 0 | NA | NA | NA | NA | NA | Gm9890 | | 0 | 0 | 0 | 0 | 0 | 0 | 0 | 0 | 0 | 0 | 0 |
| 0 | NA | NA | NA | NA | NA | Gm6401 | disks large homolog 5-like | 0 | 0 | 0 | 0 | 0 | 0 | 0 | 0 | 0 | 0 | 0 |
| 0 | NA | NA | NA | NA | NA | Gm3543 | | 0 | 0 | 0 | 0 | 0 | 0 | 0 | 0 | 0 | 0 | 0 |
| 0 | NA | NA | NA | NA | NA | Gm17063 | | 0 | 0 | 0 | 0 | 0 | 0 | 0 | 0 | 0 | 0 | 0 |

| | | | | | | | | | | | | | | | | | | |
|---|---|---|---|---|---|---|---|---|---|---|---|---|---|---|---|---|---|---|
| 0 | NA | NA | NA | NA | NA | Gm2930 | | 0 | 0 | 0 | 0 | 0 | 0 | 0 | 0 | 0 | 0 | 0 |
| 0 | NA | NA | NA | NA | NA | Gm7991 | | 0 | 0 | 0 | 0 | 0 | 0 | 0 | 0 | 0 | 0 | 0 |
| 0 | NA | NA | NA | NA | NA | 1700091H14Rik | | 0 | 0 | 0 | 0 | 0 | 0 | 0 | 0 | 0 | 0 | 0 |
| 0 | NA | NA | NA | NA | NA | Gm7951 | | 0 | 0 | 0 | 0 | 0 | 0 | 0 | 0 | 0 | 0 | 0 |
| 0 | NA | NA | NA | NA | NA | Gm17064 | | 0 | 0 | 0 | 0 | 0 | 0 | 0 | 0 | 0 | 0 | 0 |
| 0 | NA | NA | NA | NA | NA | Gm17027 | | 0 | 0 | 0 | 0 | 0 | 0 | 0 | 0 | 0 | 0 | 0 |
| 0 | NA | NA | NA | NA | NA | Gm17166 | | 0 | 0 | 0 | 0 | 0 | 0 | 0 | 0 | 0 | 0 | 0 |
| 0 | NA | NA | NA | NA | NA | Gm3573 | | 0 | 0 | 0 | 0 | 0 | 0 | 0 | 0 | 0 | 0 | 0 |
| 0 | NA | NA | NA | NA | NA | Gm17165 | | 0 | 0 | 0 | 0 | 0 | 0 | 0 | 0 | 0 | 0 | 0 |
| 0 | NA | NA | NA | NA | NA | Gm17038 | | 0 | 0 | 0 | 0 | 0 | 0 | 0 | 0 | 0 | 0 | 0 |
| 0 | NA | NA | NA | NA | NA | Gm17026 | | 0 | 0 | 0 | 0 | 0 | 0 | 0 | 0 | 0 | 0 | 0 |
| 0 | NA | NA | NA | NA | NA | Gm25208 | | 0 | 0 | 0 | 0 | 0 | 0 | 0 | 0 | 0 | 0 | 0 |
| 0 | NA | NA | NA | NA | NA | Gm25892 | | 0 | 0 | 0 | 0 | 0 | 0 | 0 | 0 | 0 | 0 | 0 |
| 0 | NA | NA | NA | NA | NA | Gm9611 | predicted gene 9611 | 0 | 0 | 0 | 0 | 0 | 0 | 0 | 0 | 0 | 0 | 0 |
| 0 | NA | NA | NA | NA | NA | Gm7995 | predicted gene 7995 | 0 | 0 | 0 | 0 | 0 | 0 | 0 | 0 | 0 | 0 | 0 |
| 0 | NA | NA | NA | NA | NA | Gm17164 | | 0 | 0 | 0 | 0 | 0 | 0 | 0 | 0 | 0 | 0 | 0 |
| 0 | NA | NA | NA | NA | NA | Gm2951 | | 0 | 0 | 0 | 0 | 0 | 0 | 0 | 0 | 0 | 0 | 0 |
| 0 | NA | NA | NA | NA | NA | Gm2959 | | 0 | 0 | 0 | 0 | 0 | 0 | 0 | 0 | 0 | 0 | 0 |
| 0 | NA | NA | NA | NA | NA | Gm8005 | predicted gene 8005 | 0 | 0 | 0 | 0 | 0 | 0 | 0 | 0 | 0 | 0 | 0 |
| 0 | NA | NA | NA | NA | NA | Gm49360 | | 0 | 0 | 0 | 0 | 0 | 0 | 0 | 0 | 0 | 0 | 0 |
| 0 | NA | NA | NA | NA | NA | Gm8011 | predicted gene 2959 | 0 | 0 | 0 | 0 | 0 | 0 | 0 | 0 | 0 | 0 | 0 |
| 0 | NA | NA | NA | NA | NA | Gm2966 | | 0 | 0 | 0 | 0 | 0 | 0 | 0 | 0 | 0 | 0 | 0 |
| 0 | NA | NA | NA | NA | NA | Gm8020 | predicted gene 8020 | 0 | 0 | 0 | 0 | 0 | 0 | 0 | 0 | 0 | 0 | 0 |
| 0 | NA | NA | NA | NA | NA | Gm8024 | | 0 | 0 | 0 | 0 | 0 | 0 | 0 | 0 | 0 | 0 | 0 |
| 0 | NA | NA | NA | NA | NA | Gm48134 | | 0 | 0 | 0 | 0 | 0 | 0 | 0 | 0 | 0 | 0 | 0 |
| 0 | NA | NA | NA | NA | NA | Gm8032 | | 0 | 0 | 0 | 0 | 0 | 0 | 0 | 0 | 0 | 0 | 0 |
| 0 | NA | NA | NA | NA | NA | Gm17204 | | 0 | 0 | 0 | 0 | 0 | 0 | 0 | 0 | 0 | 0 | 0 |
| 0 | NA | NA | NA | NA | NA | Gm17124 | predicted gene 17124 | 0 | 0 | 0 | 0 | 0 | 0 | 0 | 0 | 0 | 0 | 0 |
| 0 | NA | NA | NA | NA | NA | Gm25228 | | 0 | 0 | 0 | 0 | 0 | 0 | 0 | 0 | 0 | 0 | 0 |
| 0 | NA | NA | NA | NA | NA | Gm23620 | | 0 | 0 | 0 | 0 | 0 | 0 | 0 | 0 | 0 | 0 | 0 |
| 0 | NA | NA | NA | NA | NA | Gm3633 | | 0 | 0 | 0 | 0 | 0 | 0 | 0 | 0 | 0 | 0 | 0 |
| 0 | NA | NA | NA | NA | NA | Gm10378 | | 0 | 0 | 0 | 0 | 0 | 0 | 0 | 0 | 0 | 0 | 0 |
| 0 | NA | NA | NA | NA | NA | Gm17162 | | 0 | 0 | 0 | 0 | 0 | 0 | 0 | 0 | 0 | 0 | 0 |
| 0 | NA | NA | NA | NA | NA | Gm5929 | | 0 | 0 | 0 | 0 | 0 | 0 | 0 | 0 | 0 | 0 | 0 |
| 0 | NA | NA | NA | NA | NA | Gm3008 | | 0 | 0 | 0 | 0 | 0 | 0 | 0 | 0 | 0 | 0 | 0 |
| 0 | NA | NA | NA | NA | NA | Gm17161 | | 0 | 0 | 0 | 0 | 0 | 0 | 0 | 0 | 0 | 0 | 0 |
| 0 | NA | NA | NA | NA | NA | Gm3015 | | 0 | 0 | 0 | 0 | 0 | 0 | 0 | 0 | 0 | 0 | 0 |
| 0 | NA | NA | NA | NA | NA | Gm3025 | | 0 | 0 | 0 | 0 | 0 | 0 | 0 | 0 | 0 | 0 | 0 |
| 0 | NA | NA | NA | NA | NA | Gm10377 | predicted gene 10377 | 0 | 0 | 0 | 0 | 0 | 0 | 0 | 0 | 0 | 0 | 0 |
| 0 | NA | NA | NA | NA | NA | Gm7152 | | 0 | 0 | 0 | 0 | 0 | 0 | 0 | 0 | 0 | 0 | 0 |
| 0 | NA | NA | NA | NA | NA | Gm21977 | | 0 | 0 | 0 | 0 | 0 | 0 | 0 | 0 | 0 | 0 | 0 |
| 0 | NA | NA | NA | NA | NA | Gm48217 | | 0 | 0 | 0 | 0 | 0 | 0 | 0 | 0 | 0 | 0 | 0 |
| 0 | NA | NA | NA | NA | NA | Gm3102 | | 0 | 0 | 0 | 0 | 0 | 0 | 0 | 0 | 0 | 0 | 0 |
| 0 | NA | NA | NA | NA | NA | Gm8082 | | 0 | 0 | 0 | 0 | 0 | 0 | 0 | 0 | 0 | 0 | 0 |
| 0 | NA | NA | NA | NA | NA | Gm3111 | | 0 | 0 | 0 | 0 | 0 | 0 | 0 | 0 | 0 | 0 | 0 |
| 0 | NA | NA | NA | NA | NA | Gm10376 | | 0 | 0 | 0 | 0 | 0 | 0 | 0 | 0 | 0 | 0 | 0 |
| 0 | NA | NA | NA | NA | NA | Gm6536 | | 0 | 0 | 0 | 0 | 0 | 0 | 0 | 0 | 0 | 0 | 0 |
| 0 | NA | NA | NA | NA | NA | Gm3123 | | 0 | 0 | 0 | 0 | 0 | 0 | 0 | 0 | 0 | 0 | 0 |
| 0 | NA | NA | NA | NA | NA | Gm8094 | predicted gene 8094 | 0 | 0 | 0 | 0 | 0 | 0 | 0 | 0 | 0 | 0 | 0 |
| 0 | NA | NA | NA | NA | NA | Gm49873 | | 0 | 0 | 0 | 0 | 0 | 0 | 0 | 0 | 0 | 0 | 0 |
| 0 | NA | NA | NA | NA | NA | Gm8104 | predicted gene 8104 | 0 | 0 | 0 | 0 | 0 | 0 | 0 | 0 | 0 | 0 | 0 |
| 0 | NA | NA | NA | NA | NA | Gm49135 | | 0 | 0 | 0 | 0 | 0 | 0 | 0 | 0 | 0 | 0 | 0 |
| 0 | NA | NA | NA | NA | NA | Gm8114 | | 0 | 0 | 0 | 0 | 0 | 0 | 0 | 0 | 0 | 0 | 0 |
| 0 | NA | NA | NA | NA | NA | Gm3141 | | 0 | 0 | 0 | 0 | 0 | 0 | 0 | 0 | 0 | 0 | 0 |
| 0 | NA | NA | NA | NA | NA | Gm49141 | | 0 | 0 | 0 | 0 | 0 | 0 | 0 | 0 | 0 | 0 | 0 |
| 0 | NA | NA | NA | NA | NA | Gm7233 | | 0 | 0 | 0 | 0 | 0 | 0 | 0 | 0 | 0 | 0 | 0 |
| 0 | NA | NA | NA | NA | NA | Gm7316 | | 0 | 0 | 0 | 0 | 0 | 0 | 0 | 0 | 0 | 0 | 0 |
| 0 | NA | NA | NA | NA | NA | Gm8122 | | 0 | 0 | 0 | 0 | 0 | 0 | 0 | 0 | 0 | 0 | 0 |
| 0 | NA | NA | NA | NA | NA | Gm49146 | | 0 | 0 | 0 | 0 | 0 | 0 | 0 | 0 | 0 | 0 | 0 |
| 0 | NA | NA | NA | NA | NA | Gm8126 | predicted gene 8126 | 0 | 0 | 0 | 0 | 0 | 0 | 0 | 0 | 0 | 0 | 0 |

| | | | | | | | | | | | | | | | | | |
|---|---|---|---|---|---|---|---|---|---|---|---|---|---|---|---|---|---|
| 0 | NA | NA | NA | NA | NA | Gm9732 | | 0 | 0 | 0 | 0 | 0 | 0 | 0 | 0 | 0 | 0 | 0 |
| 0 | NA | NA | NA | NA | NA | Gm17153 | | 0 | 0 | 0 | 0 | 0 | 0 | 0 | 0 | 0 | 0 | 0 |
| 0 | NA | NA | NA | NA | NA | Gm8127 | predicted gene 8127 | 0 | 0 | 0 | 0 | 0 | 0 | 0 | 0 | 0 | 0 | 0 |
| 0 | NA | NA | NA | NA | NA | Gm17155 | | 0 | 0 | 0 | 0 | 0 | 0 | 0 | 0 | 0 | 0 | 0 |
| 0 | NA | NA | NA | NA | NA | Gm17154 | | 0 | 0 | 0 | 0 | 0 | 0 | 0 | 0 | 0 | 0 | 0 |
| 0 | NA | NA | NA | NA | NA | 1700001F09Rik | RIKEN cDNA 1700001F09 gene | 0 | 0 | 0 | 0 | 0 | 0 | 0 | 0 | 0 | 0 | 0 |
| 0 | NA | NA | NA | NA | NA | Gm9597 | | 0 | 0 | 0 | 0 | 0 | 0 | 0 | 0 | 0 | 0 | 0 |
| 0 | NA | NA | NA | NA | NA | Gm49315 | | 0 | 0 | 0 | 0 | 0 | 0 | 0 | 0 | 0 | 0 | 0 |
| 0 | NA | NA | NA | NA | NA | Gm8138 | predicted gene 8138 | 0 | 0 | 0 | 0 | 0 | 0 | 0 | 0 | 0 | 0 | 0 |
| 0 | NA | NA | NA | NA | NA | Gm49072 | | 0 | 0 | 0 | 0 | 0 | 0 | 0 | 0 | 0 | 0 | 0 |
| 0 | NA | NA | NA | NA | NA | Gm21162 | | 0 | 0 | 0 | 0 | 0 | 0 | 0 | 0 | 0 | 0 | 0 |
| 0 | NA | NA | NA | NA | NA | Gm5799 | predicted gene 8126 | 0 | 0 | 0 | 0 | 0 | 0 | 0 | 0 | 0 | 0 | 0 |
| 0 | NA | NA | NA | NA | NA | Gm3216 | | 0 | 0 | 0 | 0 | 0 | 0 | 0 | 0 | 0 | 0 | 0 |
| 0 | NA | NA | NA | NA | NA | Gm17654 | | 0 | 0 | 0 | 0 | 0 | 0 | 0 | 0 | 0 | 0 | 0 |
| 0 | NA | NA | NA | NA | NA | Gm3221 | | 0 | 0 | 0 | 0 | 0 | 0 | 0 | 0 | 0 | 0 | 0 |
| 0 | NA | NA | NA | NA | NA | Gm10375 | predicted gene 10375 | 0 | 0 | 0 | 0 | 0 | 0 | 0 | 0 | 0 | 0 | 0 |
| 0 | NA | NA | NA | NA | NA | Gm8046 | | 0 | 0 | 0 | 0 | 0 | 0 | 0 | 0 | 0 | 0 | 0 |
| 0 | NA | NA | NA | NA | NA | Gm21154 | | 0 | 0 | 0 | 0 | 0 | 0 | 0 | 0 | 0 | 0 | 0 |
| 0 | NA | NA | NA | NA | NA | Gm8161 | | 0 | 0 | 0 | 0 | 0 | 0 | 0 | 0 | 0 | 0 | 0 |
| 0 | NA | NA | NA | NA | NA | Gm8165 | predicted gene 8165 | 0 | 0 | 0 | 0 | 0 | 0 | 0 | 0 | 0 | 0 | 0 |
| 0 | NA | NA | NA | NA | NA | Gm9672 | | 0 | 0 | 0 | 0 | 0 | 0 | 0 | 0 | 0 | 0 | 0 |
| 0 | NA | NA | NA | NA | NA | Gm17208 | | 0 | 0 | 0 | 0 | 0 | 0 | 0 | 0 | 0 | 0 | 0 |
| 0 | NA | NA | NA | NA | NA | Gm16506 | predicted gene 16506 | 0 | 0 | 0 | 0 | 0 | 0 | 0 | 0 | 0 | 0 | 0 |
| 0 | NA | NA | NA | NA | NA | Gm49078 | | 0 | 0 | 0 | 0 | 0 | 0 | 0 | 0 | 0 | 0 | 0 |
| 0 | NA | NA | NA | NA | NA | Gm49080 | | 0 | 0 | 0 | 0 | 0 | 0 | 0 | 0 | 0 | 0 | 0 |
| 0 | NA | NA | NA | NA | NA | Gm6526 | | 0 | 0 | 0 | 0 | 0 | 0 | 0 | 0 | 0 | 0 | 0 |
| 0 | NA | NA | NA | NA | NA | Gm49965 | | 0 | 0 | 0 | 0 | 0 | 0 | 0 | 0 | 0 | 0 | 0 |
| 0 | NA | NA | NA | NA | NA | Gm9587 | | 0 | 0 | 0 | 0 | 0 | 0 | 0 | 0 | 0 | 0 | 0 |
| 0 | NA | NA | NA | NA | NA | Gm8180 | | 0 | 0 | 0 | 0 | 0 | 0 | 0 | 0 | 0 | 0 | 0 |
| 0 | NA | NA | NA | NA | NA | Gm49082 | | 0 | 0 | 0 | 0 | 0 | 0 | 0 | 0 | 0 | 0 | 0 |
| 0 | NA | NA | NA | NA | NA | Gm49081 | | 0 | 0 | 0 | 0 | 0 | 0 | 0 | 0 | 0 | 0 | 0 |
| 0 | NA | NA | NA | NA | NA | Gm41138 | | 0 | 0 | 0 | 0 | 0 | 0 | 0 | 0 | 0 | 0 | 0 |
| 0 | NA | NA | NA | NA | NA | Ear-ps12 | | 0 | 0 | 0 | 0 | 0 | 0 | 0 | 0 | 0 | 0 | 0 |
| 0 | NA | NA | NA | NA | NA | Ang-ps2 | | 0 | 0 | 0 | 0 | 0 | 0 | 0 | 0 | 0 | 0 | 0 |
| 0 | NA | NA | NA | NA | NA | Ear-ps9 | | 0 | 0 | 0 | 0 | 0 | 0 | 0 | 0 | 0 | 0 | 0 |
| 0 | NA | NA | NA | NA | NA | Ear-ps8 | | 0 | 0 | 0 | 0 | 0 | 0 | 0 | 0 | 0 | 0 | 0 |
| 0 | NA | NA | NA | NA | NA | Ang-ps3 | | 0 | 0 | 0 | 0 | 0 | 0 | 0 | 0 | 0 | 0 | 0 |
| 0 | NA | NA | NA | NA | NA | Ear10 | eosinophil-associated, ribonuclease A fan | 0 | 0 | 0 | 0 | 0 | 0 | 0 | 0 | 0 | 0 | 0 |
| 0 | NA | NA | NA | NA | NA | Gm9550 | | 0 | 0 | 0 | 0 | 0 | 0 | 0 | 0 | 0 | 0 | 0 |
| 0 | NA | NA | NA | NA | NA | Ang5 | angiogenin, ribonuclease A family, memb | 0 | 0 | 0 | 0 | 0 | 0 | 0 | 0 | 0 | 0 | 0 |
| 0 | NA | NA | NA | NA | NA | Ear-ps7 | | 0 | 0 | 0 | 0 | 0 | 0 | 0 | 0 | 0 | 0 | 0 |
| 0 | NA | NA | NA | NA | NA | Gm49087 | | 0 | 0 | 0 | 0 | 0 | 0 | 0 | 0 | 0 | 0 | 0 |
| 0 | NA | NA | NA | NA | NA | Ear-ps10 | | 0 | 0 | 0 | 0 | 0 | 0 | 0 | 0 | 0 | 0 | 0 |
| 0 | NA | NA | NA | NA | NA | Ang6 | angiogenin, ribonuclease A family, memb | 0 | 0 | 0 | 0 | 0 | 0 | 0 | 0 | 0 | 0 | 0 |
| 0 | NA | NA | NA | NA | NA | Gm8194 | | 0 | 0 | 0 | 0 | 0 | 0 | 0 | 0 | 0 | 0 | 0 |
| 0 | NA | NA | NA | NA | NA | Gm3287 | | 0 | 0 | 0 | 0 | 0 | 0 | 0 | 0 | 0 | 0 | 0 |
| 0 | NA | NA | NA | NA | NA | Ang-ps1 | | 0 | 0 | 0 | 0 | 0 | 0 | 0 | 0 | 0 | 0 | 0 |
| 0 | NA | NA | NA | NA | NA | Gm49099 | | 0 | 0 | 0 | 0 | 0 | 0 | 0 | 0 | 0 | 0 | 0 |
| 0 | NA | NA | NA | NA | NA | Gm16545 | | 0 | 0 | 0 | 0 | 0 | 0 | 0 | 0 | 0 | 0 | 0 |
| 0 | NA | NA | NA | NA | NA | Gm49101 | | 0 | 0 | 0 | 0 | 0 | 0 | 0 | 0 | 0 | 0 | 0 |
| 0 | NA | NA | NA | NA | NA | Gm49102 | | 0 | 0 | 0 | 0 | 0 | 0 | 0 | 0 | 0 | 0 | 0 |
| 0 | NA | NA | NA | NA | NA | Gm49103 | | 0 | 0 | 0 | 0 | 0 | 0 | 0 | 0 | 0 | 0 | 0 |
| 0 | NA | NA | NA | NA | NA | Gm18216 | | 0 | 0 | 0 | 0 | 0 | 0 | 0 | 0 | 0 | 0 | 0 |
| 0 | NA | NA | NA | NA | NA | Gm3327 | | 0 | 0 | 0 | 0 | 0 | 0 | 0 | 0 | 0 | 0 | 0 |
| 0 | NA | NA | NA | NA | NA | Gm8207 | | 0 | 0 | 0 | 0 | 0 | 0 | 0 | 0 | 0 | 0 | 0 |
| 0 | NA | NA | NA | NA | NA | 4930503E14Ril | RIKEN cDNA 4930503E14 gene | 0 | 0 | 0 | 0 | 0 | 0 | 0 | 0 | 0 | 0 | 0 |
| 0 | NA | NA | NA | NA | NA | 1700047E10Rik | | 0 | 0 | 0 | 0 | 0 | 0 | 0 | 0 | 0 | 0 | 0 |
| 0 | NA | NA | NA | NA | NA | Gm8212 | | 0 | 0 | 0 | 0 | 0 | 0 | 0 | 0 | 0 | 0 | 0 |
| 0 | NA | NA | NA | NA | NA | Gm49104 | | 0 | 0 | 0 | 0 | 0 | 0 | 0 | 0 | 0 | 0 | 0 |
| 0 | NA | NA | NA | NA | NA | Gm32857 | | 0 | 0 | 0 | 0 | 0 | 0 | 0 | 0 | 0 | 0 | 0 |

| | | | | | | | | | | | | | | | | | | |
|---|---|---|---|---|---|---|---|---|---|---|---|---|---|---|---|---|---|---|
| 0 | NA | NA | NA | NA | NA | Gm49110 | | 0 | 0 | 0 | 0 | 0 | 0 | 0 | 0 | 0 | 0 | 0 |
| 0 | NA | NA | NA | NA | NA | Gm49111 | | 0 | 0 | 0 | 0 | 0 | 0 | 0 | 0 | 0 | 0 | 0 |
| 0 | NA | NA | NA | NA | NA | Gm8220 | predicted gene 8220 | 0 | 0 | 0 | 0 | 0 | 0 | 0 | 0 | 0 | 0 | 0 |
| 0 | NA | NA | NA | NA | NA | Gm5930 | | 0 | 0 | 0 | 0 | 0 | 0 | 0 | 0 | 0 | 0 | 0 |
| 0 | NA | NA | NA | NA | NA | Gm8229 | | 0 | 0 | 0 | 0 | 0 | 0 | 0 | 0 | 0 | 0 | 0 |
| 0 | NA | NA | NA | NA | NA | Gm3370 | | 0 | 0 | 0 | 0 | 0 | 0 | 0 | 0 | 0 | 0 | 0 |
| 0 | NA | NA | NA | NA | NA | Gm3371 | | 0 | 0 | 0 | 0 | 0 | 0 | 0 | 0 | 0 | 0 | 0 |
| 0 | NA | NA | NA | NA | NA | Gm8232 | predicted gene 8232 | 0 | 0 | 0 | 0 | 0 | 0 | 0 | 0 | 0 | 0 | 0 |
| 0 | NA | NA | NA | NA | NA | Gm49779 | | 0 | 0 | 0 | 0 | 0 | 0 | 0 | 0 | 0 | 0 | 0 |
| 0 | NA | NA | NA | NA | NA | Gm21991 | | 0 | 0 | 0 | 0 | 0 | 0 | 0 | 0 | 0 | 0 | 0 |
| 0 | NA | NA | NA | NA | NA | Gm17184 | | 0 | 0 | 0 | 0 | 0 | 0 | 0 | 0 | 0 | 0 | 0 |
| 0 | NA | NA | NA | NA | NA | BC061237 | cDNA sequence BC061237 | 0 | 0 | 0 | 0 | 0 | 0 | 0 | 0 | 0 | 0 | 0 |
| 0 | NA | NA | NA | NA | NA | Gm17093 | | 0 | 0 | 0 | 0 | 0 | 0 | 0 | 0 | 0 | 0 | 0 |
| 0 | NA | NA | NA | NA | NA | Gm17183 | | 0 | 0 | 0 | 0 | 0 | 0 | 0 | 0 | 0 | 0 | 0 |
| 0 | NA | NA | NA | NA | NA | Gm5624 | predicted gene 5930 | 0 | 0 | 0 | 0 | 0 | 0 | 0 | 0 | 0 | 0 | 0 |
| 0 | NA | NA | NA | NA | NA | Gm8247 | | 0 | 0 | 0 | 0 | 0 | 0 | 0 | 0 | 0 | 0 | 0 |
| 0 | NA | NA | NA | NA | NA | Gm8256 | | 0 | 0 | 0 | 0 | 0 | 0 | 0 | 0 | 0 | 0 | 0 |
| 0 | NA | NA | NA | NA | NA | Gm9593 | | 0 | 0 | 0 | 0 | 0 | 0 | 0 | 0 | 0 | 0 | 0 |
| 0 | NA | NA | NA | NA | NA | Gm8257 | | 0 | 0 | 0 | 0 | 0 | 0 | 0 | 0 | 0 | 0 | 0 |
| 0 | NA | NA | NA | NA | NA | Gm21754 | | 0 | 0 | 0 | 0 | 0 | 0 | 0 | 0 | 0 | 0 | 0 |
| 0 | NA | NA | NA | NA | NA | Gm6541 | | 0 | 0 | 0 | 0 | 0 | 0 | 0 | 0 | 0 | 0 | 0 |
| 0 | NA | NA | NA | NA | NA | Gm8267 | predicted gene 8267 | 0 | 0 | 0 | 0 | 0 | 0 | 0 | 0 | 0 | 0 | 0 |
| 0 | NA | NA | NA | NA | NA | Gm34060 | | 0 | 0 | 0 | 0 | 0 | 0 | 0 | 0 | 0 | 0 | 0 |
| 0 | NA | NA | NA | NA | NA | Gm49142 | | 0 | 0 | 0 | 0 | 0 | 0 | 0 | 0 | 0 | 0 | 0 |
| 0 | NA | NA | NA | NA | NA | Gm17187 | | 0 | 0 | 0 | 0 | 0 | 0 | 0 | 0 | 0 | 0 | 0 |
| 0 | NA | NA | NA | NA | NA | Gm49143 | | 0 | 0 | 0 | 0 | 0 | 0 | 0 | 0 | 0 | 0 | 0 |
| 0 | NA | NA | NA | NA | NA | Gm17070 | | 0 | 0 | 0 | 0 | 0 | 0 | 0 | 0 | 0 | 0 | 0 |
| 0 | NA | NA | NA | NA | NA | Gm49145 | | 0 | 0 | 0 | 0 | 0 | 0 | 0 | 0 | 0 | 0 | 0 |
| 0 | NA | NA | NA | NA | NA | Gm34198 | | 0 | 0 | 0 | 0 | 0 | 0 | 0 | 0 | 0 | 0 | 0 |
| 0 | NA | NA | NA | NA | NA | Ptgdr | prostaglandin D receptor | 0 | 0 | 0 | 0 | 0 | 0 | 0 | 0 | 0 | 0 | 0 |
| 0 | NA | NA | NA | NA | NA | Gm16976 | | 0 | 0 | 0 | 0 | 0 | 0 | 0 | 0 | 0 | 0 | 0 |
| 0 | NA | NA | NA | NA | NA | Gm18524 | | 0 | 0 | 0 | 0 | 0 | 0 | 0 | 0 | 0 | 0 | 0 |
| 0 | NA | NA | NA | NA | NA | Gm49148 | | 0 | 0 | 0 | 0 | 0 | 0 | 0 | 0 | 0 | 0 | 0 |
| 0 | NA | NA | NA | NA | NA | Gm49147 | | 0 | 0 | 0 | 0 | 0 | 0 | 0 | 0 | 0 | 0 | 0 |
| 0 | NA | NA | NA | NA | NA | Mir327 | | 0 | 0 | 0 | 0 | 0 | 0 | 0 | 0 | 0 | 0 | 0 |
| 0 | NA | NA | NA | NA | NA | 4930431P03Rik | | 0 | 0 | 0 | 0 | 0 | 0 | 0 | 0 | 0 | 0 | 0 |
| 0 | NA | NA | NA | NA | NA | Gm49364 | | 0 | 0 | 0 | 0 | 0 | 0 | 0 | 0 | 0 | 0 | 0 |
| 0 | NA | NA | NA | NA | NA | Gm49180 | | 0 | 0 | 0 | 0 | 0 | 0 | 0 | 0 | 0 | 0 | 0 |
| 0 | NA | NA | NA | NA | NA | Gm21010 | | 0 | 0 | 0 | 0 | 0 | 0 | 0 | 0 | 0 | 0 | 0 |
| 0 | NA | NA | NA | NA | NA | Gm49181 | | 0 | 0 | 0 | 0 | 0 | 0 | 0 | 0 | 0 | 0 | 0 |
| 0 | NA | NA | NA | NA | NA | Gm49183 | | 0 | 0 | 0 | 0 | 0 | 0 | 0 | 0 | 0 | 0 | 0 |
| 0 | NA | NA | NA | NA | NA | Gm49184 | | 0 | 0 | 0 | 0 | 0 | 0 | 0 | 0 | 0 | 0 | 0 |
| 0 | NA | NA | NA | NA | NA | Gm15765 | | 0 | 0 | 0 | 0 | 0 | 0 | 0 | 0 | 0 | 0 | 0 |
| 0 | NA | NA | NA | NA | NA | Gm34250 | | 0 | 0 | 0 | 0 | 0 | 0 | 0 | 0 | 0 | 0 | 0 |
| 0 | NA | NA | NA | NA | NA | Gm41144 | | 0 | 0 | 0 | 0 | 0 | 0 | 0 | 0 | 0 | 0 | 0 |
| 0 | NA | NA | NA | NA | NA | Gm15601 | | 0 | 0 | 0 | 0 | 0 | 0 | 0 | 0 | 0 | 0 | 0 |
| 0 | NA | NA | NA | NA | NA | Mir5131 | | 0 | 0 | 0 | 0 | 0 | 0 | 0 | 0 | 0 | 0 | 0 |
| 0 | NA | NA | NA | NA | NA | Gm15218 | | 0 | 0 | 0 | 0 | 0 | 0 | 0 | 0 | 0 | 0 | 0 |
| 0 | NA | NA | NA | NA | NA | Gm15220 | | 0 | 0 | 0 | 0 | 0 | 0 | 0 | 0 | 0 | 0 | 0 |
| 0 | NA | NA | NA | NA | NA | Gm46475 | | 0 | 0 | 0 | 0 | 0 | 0 | 0 | 0 | 0 | 0 | 0 |
| 0 | NA | NA | NA | NA | NA | Gm49192 | | 0 | 0 | 0 | 0 | 0 | 0 | 0 | 0 | 0 | 0 | 0 |
| 0 | NA | NA | NA | NA | NA | Gm15217 | | 0 | 0 | 0 | 0 | 0 | 0 | 0 | 0 | 0 | 0 | 0 |
| 0 | NA | NA | NA | NA | NA | Gm15221 | | 0 | 0 | 0 | 0 | 0 | 0 | 0 | 0 | 0 | 0 | 0 |
| 0 | NA | NA | NA | NA | NA | Gm48910 | | 0 | 0 | 0 | 0 | 0 | 0 | 0 | 0 | 0 | 0 | 0 |
| 0 | NA | NA | NA | NA | NA | Gm48911 | | 0 | 0 | 0 | 0 | 0 | 0 | 0 | 0 | 0 | 0 | 0 |
| 0 | NA | NA | NA | NA | NA | Gm34756 | | 0 | 0 | 0 | 0 | 0 | 0 | 0 | 0 | 0 | 0 | 0 |
| 0 | NA | NA | NA | NA | NA | Gm48912 | | 0 | 0 | 0 | 0 | 0 | 0 | 0 | 0 | 0 | 0 | 0 |
| 0 | NA | NA | NA | NA | NA | Gm49319 | | 0 | 0 | 0 | 0 | 0 | 0 | 0 | 0 | 0 | 0 | 0 |
| 0 | NA | NA | NA | NA | NA | E130120K24Rik | | 0 | 0 | 0 | 0 | 0 | 0 | 0 | 0 | 0 | 0 | 0 |
| 0 | NA | NA | NA | NA | NA | Rubie | | 0 | 0 | 0 | 0 | 0 | 0 | 0 | 0 | 0 | 0 | 0 |

| | | | | | | | | | | | | | | | | | | |
|---|---|---|---|---|---|---|---|---|---|---|---|---|---|---|---|---|---|---|
| 0 | NA | NA | NA | NA | NA | 4933425B07Rik | | 0 | 0 | 0 | 0 | 0 | 0 | 0 | 0 | 0 | 0 | 0 |
| 0 | NA | NA | NA | NA | NA | Gm18761 | | 0 | 0 | 0 | 0 | 0 | 0 | 0 | 0 | 0 | 0 | 0 |
| 0 | NA | NA | NA | NA | NA | D330046F09Rik | | 0 | 0 | 0 | 0 | 0 | 0 | 0 | 0 | 0 | 0 | 0 |
| 0 | NA | NA | NA | NA | NA | Gm48917 | | 0 | 0 | 0 | 0 | 0 | 0 | 0 | 0 | 0 | 0 | 0 |
| 0 | NA | NA | NA | NA | NA | Gm48924 | | 0 | 0 | 0 | 0 | 0 | 0 | 0 | 0 | 0 | 0 | 0 |
| 0 | NA | NA | NA | NA | NA | Gm48958 | | 0 | 0 | 0 | 0 | 0 | 0 | 0 | 0 | 0 | 0 | 0 |
| 0 | NA | NA | NA | NA | NA | Mir378c | | 0 | 0 | 0 | 0 | 0 | 0 | 0 | 0 | 0 | 0 | 0 |
| 0 | NA | NA | NA | NA | NA | Gm15562 | | 0 | 0 | 0 | 0 | 0 | 0 | 0 | 0 | 0 | 0 | 0 |
| 0 | NA | NA | NA | NA | NA | Gm49136 | | 0 | 0 | 0 | 0 | 0 | 0 | 0 | 0 | 0 | 0 | 0 |
| 0 | NA | NA | NA | NA | NA | Gm49137 | | 0 | 0 | 0 | 0 | 0 | 0 | 0 | 0 | 0 | 0 | 0 |
| 0 | NA | NA | NA | NA | NA | Gm49149 | | 0 | 0 | 0 | 0 | 0 | 0 | 0 | 0 | 0 | 0 | 0 |
| 0 | NA | NA | NA | NA | NA | Gm24378 | | 0 | 0 | 0 | 0 | 0 | 0 | 0 | 0 | 0 | 0 | 0 |
| 0 | NA | NA | NA | NA | NA | Gm49150 | | 0 | 0 | 0 | 0 | 0 | 0 | 0 | 0 | 0 | 0 | 0 |
| 0 | NA | NA | NA | NA | NA | Gm49190 | | 0 | 0 | 0 | 0 | 0 | 0 | 0 | 0 | 0 | 0 | 0 |
| 0 | NA | NA | NA | NA | NA | Gm8353 | | 0 | 0 | 0 | 0 | 0 | 0 | 0 | 0 | 0 | 0 | 0 |
| 0 | NA | NA | NA | NA | NA | Gm35166 | | 0 | 0 | 0 | 0 | 0 | 0 | 0 | 0 | 0 | 0 | 0 |
| 0 | NA | NA | NA | NA | NA | Gm49004 | | 0 | 0 | 0 | 0 | 0 | 0 | 0 | 0 | 0 | 0 | 0 |
| 0 | NA | NA | NA | NA | NA | Gm6616 | | 0 | 0 | 0 | 0 | 0 | 0 | 0 | 0 | 0 | 0 | 0 |
| 0 | NA | NA | NA | NA | NA | 4930538L07Rik | | 0 | 0 | 0 | 0 | 0 | 0 | 0 | 0 | 0 | 0 | 0 |
| 0 | NA | NA | NA | NA | NA | Gm41148 | | 0 | 0 | 0 | 0 | 0 | 0 | 0 | 0 | 0 | 0 | 0 |
| 0 | NA | NA | NA | NA | NA | Gm49302 | | 0 | 0 | 0 | 0 | 0 | 0 | 0 | 0 | 0 | 0 | 0 |
| 0 | NA | NA | NA | NA | NA | Gm35360 | | 0 | 0 | 0 | 0 | 0 | 0 | 0 | 0 | 0 | 0 | 0 |
| 0 | NA | NA | NA | NA | NA | 4930447J18Rik | | 0 | 0 | 0 | 0 | 0 | 0 | 0 | 0 | 0 | 0 | 0 |
| 0 | NA | NA | NA | NA | NA | Gm49303 | | 0 | 0 | 0 | 0 | 0 | 0 | 0 | 0 | 0 | 0 | 0 |
| 0 | NA | NA | NA | NA | NA | Gm49304 | | 0 | 0 | 0 | 0 | 0 | 0 | 0 | 0 | 0 | 0 | 0 |
| 0 | NA | NA | NA | NA | NA | Gm49305 | | 0 | 0 | 0 | 0 | 0 | 0 | 0 | 0 | 0 | 0 | 0 |
| 0 | NA | NA | NA | NA | NA | Gm49306 | | 0 | 0 | 0 | 0 | 0 | 0 | 0 | 0 | 0 | 0 | 0 |
| 0 | NA | NA | NA | NA | NA | 9530086P17Rik | | 0 | 0 | 0 | 0 | 0 | 0 | 0 | 0 | 0 | 0 | 0 |
| 0 | NA | NA | NA | NA | NA | Gm49310 | | 0 | 0 | 0 | 0 | 0 | 0 | 0 | 0 | 0 | 0 | 0 |
| 0 | NA | NA | NA | NA | NA | Gm49307 | | 0 | 0 | 0 | 0 | 0 | 0 | 0 | 0 | 0 | 0 | 0 |
| 0 | NA | NA | NA | NA | NA | Gm6498 | | 0 | 0 | 0 | 0 | 0 | 0 | 0 | 0 | 0 | 0 | 0 |
| 0 | NA | NA | NA | NA | NA | 1700128I11Rik | | 0 | 0 | 0 | 0 | 0 | 0 | 0 | 0 | 0 | 0 | 0 |
| 0 | NA | NA | NA | NA | NA | Gm23881 | | 0 | 0 | 0 | 0 | 0 | 0 | 0 | 0 | 0 | 0 | 0 |
| 0 | NA | NA | NA | NA | NA | Gm18962 | | 0 | 0 | 0 | 0 | 0 | 0 | 0 | 0 | 0 | 0 | 0 |
| 0 | NA | NA | NA | NA | NA | Gm49120 | | 0 | 0 | 0 | 0 | 0 | 0 | 0 | 0 | 0 | 0 | 0 |
| 0 | NA | NA | NA | NA | NA | Gm35766 | | 0 | 0 | 0 | 0 | 0 | 0 | 0 | 0 | 0 | 0 | 0 |
| 0 | NA | NA | NA | NA | NA | Gm49151 | | 0 | 0 | 0 | 0 | 0 | 0 | 0 | 0 | 0 | 0 | 0 |
| 0 | NA | NA | NA | NA | NA | Gm23469 | | 0 | 0 | 0 | 0 | 0 | 0 | 0 | 0 | 0 | 0 | 0 |
| 0 | NA | NA | NA | NA | NA | Otx2os1 | | 0 | 0 | 0 | 0 | 0 | 0 | 0 | 0 | 0 | 0 | 0 |
| 0 | NA | NA | NA | NA | NA | Gm49154 | | 0 | 0 | 0 | 0 | 0 | 0 | 0 | 0 | 0 | 0 | 0 |
| 0 | NA | NA | NA | NA | NA | Gm25380 | | 0 | 0 | 0 | 0 | 0 | 0 | 0 | 0 | 0 | 0 | 0 |
| 0 | NA | NA | NA | NA | NA | Gm49153 | | 0 | 0 | 0 | 0 | 0 | 0 | 0 | 0 | 0 | 0 | 0 |
| 0 | NA | NA | NA | NA | NA | Gm49156 | | 0 | 0 | 0 | 0 | 0 | 0 | 0 | 0 | 0 | 0 | 0 |
| 0 | NA | NA | NA | NA | NA | Gm22699 | | 0 | 0 | 0 | 0 | 0 | 0 | 0 | 0 | 0 | 0 | 0 |
| 0 | NA | NA | NA | NA | NA | Gm49152 | | 0 | 0 | 0 | 0 | 0 | 0 | 0 | 0 | 0 | 0 | 0 |
| 0 | NA | NA | NA | NA | NA | Gm49155 | | 0 | 0 | 0 | 0 | 0 | 0 | 0 | 0 | 0 | 0 | 0 |
| 0 | NA | NA | NA | NA | NA | Gm534 | | 0 | 0 | 0 | 0 | 0 | 0 | 0 | 0 | 0 | 0 | 0 |
| 0 | NA | NA | NA | NA | NA | 4933429O19Rik | | 0 | 0 | 0 | 0 | 0 | 0 | 0 | 0 | 0 | 0 | 0 |
| 0 | NA | NA | NA | NA | NA | Gm23470 | | 0 | 0 | 0 | 0 | 0 | 0 | 0 | 0 | 0 | 0 | 0 |
| 0 | NA | NA | NA | NA | NA | Gm48935 | | 0 | 0 | 0 | 0 | 0 | 0 | 0 | 0 | 0 | 0 | 0 |
| 0 | NA | NA | NA | NA | NA | Gm22116 | | 0 | 0 | 0 | 0 | 0 | 0 | 0 | 0 | 0 | 0 | 0 |
| 0 | NA | NA | NA | NA | NA | Gm22949 | | 0 | 0 | 0 | 0 | 0 | 0 | 0 | 0 | 0 | 0 | 0 |
| 0 | NA | NA | NA | NA | NA | Gm36146 | | 0 | 0 | 0 | 0 | 0 | 0 | 0 | 0 | 0 | 0 | 0 |
| 0 | NA | NA | NA | NA | NA | Gm48950 | | 0 | 0 | 0 | 0 | 0 | 0 | 0 | 0 | 0 | 0 | 0 |
| 0 | NA | NA | NA | NA | NA | Gm48948 | | 0 | 0 | 0 | 0 | 0 | 0 | 0 | 0 | 0 | 0 | 0 |
| 0 | NA | NA | NA | NA | NA | Gm22989 | | 0 | 0 | 0 | 0 | 0 | 0 | 0 | 0 | 0 | 0 | 0 |
| 0 | NA | NA | NA | NA | NA | Olfr722 | olfactory receptor 722 | 0 | 0 | 0 | 0 | 0 | 0 | 0 | 0 | 0 | 0 | 0 |
| 0 | NA | NA | NA | NA | NA | Olfr723 | olfactory receptor 723 | 0 | 0 | 0 | 0 | 0 | 0 | 0 | 0 | 0 | 0 | 0 |
| 0 | NA | NA | NA | NA | NA | Olfr724 | olfactory receptor 724 | 0 | 0 | 0 | 0 | 0 | 0 | 0 | 0 | 0 | 0 | 0 |
| 0 | NA | NA | NA | NA | NA | Gm49133 | | 0 | 0 | 0 | 0 | 0 | 0 | 0 | 0 | 0 | 0 | 0 |

| | | | | | | | | | | | | | | | | | |
|---|---|---|---|---|---|---|---|---|---|---|---|---|---|---|---|---|---|
| 0 | NA | NA | NA | NA | NA | Olfr725 | olfactory receptor 725 | 0 | 0 | 0 | 0 | 0 | 0 | 0 | 0 | 0 | 0 | 0 |
| 0 | NA | NA | NA | NA | NA | Olfr726 | olfactory receptor 726 | 0 | 0 | 0 | 0 | 0 | 0 | 0 | 0 | 0 | 0 | 0 |
| 0 | NA | NA | NA | NA | NA | Olfr727 | olfactory receptor 727 | 0 | 0 | 0 | 0 | 0 | 0 | 0 | 0 | 0 | 0 | 0 |
| 0 | NA | NA | NA | NA | NA | Olfr728 | olfactory receptor 728 | 0 | 0 | 0 | 0 | 0 | 0 | 0 | 0 | 0 | 0 | 0 |
| 0 | NA | NA | NA | NA | NA | Olfr729 | olfactory receptor 729 | 0 | 0 | 0 | 0 | 0 | 0 | 0 | 0 | 0 | 0 | 0 |
| 0 | NA | NA | NA | NA | NA | Olfr730 | olfactory receptor 730 | 0 | 0 | 0 | 0 | 0 | 0 | 0 | 0 | 0 | 0 | 0 |
| 0 | NA | NA | NA | NA | NA | Olfr731 | olfactory receptor 731 | 0 | 0 | 0 | 0 | 0 | 0 | 0 | 0 | 0 | 0 | 0 |
| 0 | NA | NA | NA | NA | NA | Olfr732 | olfactory receptor 732 | 0 | 0 | 0 | 0 | 0 | 0 | 0 | 0 | 0 | 0 | 0 |
| 0 | NA | NA | NA | NA | NA | Olfr733 | olfactory receptor 733 | 0 | 0 | 0 | 0 | 0 | 0 | 0 | 0 | 0 | 0 | 0 |
| 0 | NA | NA | NA | NA | NA | Olfr734 | olfactory receptor 734 | 0 | 0 | 0 | 0 | 0 | 0 | 0 | 0 | 0 | 0 | 0 |
| 0 | NA | NA | NA | NA | NA | Olfr735 | olfactory receptor 735 | 0 | 0 | 0 | 0 | 0 | 0 | 0 | 0 | 0 | 0 | 0 |
| 0 | NA | NA | NA | NA | NA | Olfr736 | olfactory receptor 736 | 0 | 0 | 0 | 0 | 0 | 0 | 0 | 0 | 0 | 0 | 0 |
| 0 | NA | NA | NA | NA | NA | Olfr738 | olfactory receptor 738 | 0 | 0 | 0 | 0 | 0 | 0 | 0 | 0 | 0 | 0 | 0 |
| 0 | NA | NA | NA | NA | NA | Olfr737-ps1 | | 0 | 0 | 0 | 0 | 0 | 0 | 0 | 0 | 0 | 0 | 0 |
| 0 | NA | NA | NA | NA | NA | Olfr739 | olfactory receptor 739 | 0 | 0 | 0 | 0 | 0 | 0 | 0 | 0 | 0 | 0 | 0 |
| 0 | NA | NA | NA | NA | NA | Olfr740 | olfactory receptor 740 | 0 | 0 | 0 | 0 | 0 | 0 | 0 | 0 | 0 | 0 | 0 |
| 0 | NA | NA | NA | NA | NA | Olfr741 | olfactory receptor 741 | 0 | 0 | 0 | 0 | 0 | 0 | 0 | 0 | 0 | 0 | 0 |
| 0 | NA | NA | NA | NA | NA | Olfr742 | olfactory receptor 742 | 0 | 0 | 0 | 0 | 0 | 0 | 0 | 0 | 0 | 0 | 0 |
| 0 | NA | NA | NA | NA | NA | Olfr743 | olfactory receptor 743 | 0 | 0 | 0 | 0 | 0 | 0 | 0 | 0 | 0 | 0 | 0 |
| 0 | NA | NA | NA | NA | NA | Gm19109 | | 0 | 0 | 0 | 0 | 0 | 0 | 0 | 0 | 0 | 0 | 0 |
| 0 | NA | NA | NA | NA | NA | Olfr744 | olfactory receptor 744 | 0 | 0 | 0 | 0 | 0 | 0 | 0 | 0 | 0 | 0 | 0 |
| 0 | NA | NA | NA | NA | NA | Olfr745 | olfactory receptor 745 | 0 | 0 | 0 | 0 | 0 | 0 | 0 | 0 | 0 | 0 | 0 |
| 0 | NA | NA | NA | NA | NA | Gm45192 | | 0 | 0 | 0 | 0 | 0 | 0 | 0 | 0 | 0 | 0 | 0 |
| 0 | NA | NA | NA | NA | NA | Olfr746 | olfactory receptor 746 | 0 | 0 | 0 | 0 | 0 | 0 | 0 | 0 | 0 | 0 | 0 |
| 0 | NA | NA | NA | NA | NA | Gm18762 | | 0 | 0 | 0 | 0 | 0 | 0 | 0 | 0 | 0 | 0 | 0 |
| 0 | NA | NA | NA | NA | NA | Olfr747 | olfactory receptor 747 | 0 | 0 | 0 | 0 | 0 | 0 | 0 | 0 | 0 | 0 | 0 |
| 0 | NA | NA | NA | NA | NA | Olfr748 | olfactory receptor 748 | 0 | 0 | 0 | 0 | 0 | 0 | 0 | 0 | 0 | 0 | 0 |
| 0 | NA | NA | NA | NA | NA | Gm24316 | | 0 | 0 | 0 | 0 | 0 | 0 | 0 | 0 | 0 | 0 | 0 |
| 0 | NA | NA | NA | NA | NA | Olfr749 | olfactory receptor 749 | 0 | 0 | 0 | 0 | 0 | 0 | 0 | 0 | 0 | 0 | 0 |
| 0 | NA | NA | NA | NA | NA | Gm48938 | | 0 | 0 | 0 | 0 | 0 | 0 | 0 | 0 | 0 | 0 | 0 |
| 0 | NA | NA | NA | NA | NA | Rpph1 | | 0 | 0 | 0 | 0 | 0 | 0 | 0 | 0 | 0 | 0 | 0 |
| 0 | NA | NA | NA | NA | NA | Gm24689 | | 0 | 0 | 0 | 0 | 0 | 0 | 0 | 0 | 0 | 0 | 0 |
| 0 | NA | NA | NA | NA | NA | Gm49342 | | 0 | 0 | 0 | 0 | 0 | 0 | 0 | 0 | 0 | 0 | 0 |
| 0 | NA | NA | NA | NA | NA | Gm38316 | | 0 | 0 | 0 | 0 | 0 | 0 | 0 | 0 | 0 | 0 | 0 |
| 0 | NA | NA | NA | NA | NA | Gm49038 | | 0 | 0 | 0 | 0 | 0 | 0 | 0 | 0 | 0 | 0 | 0 |
| 0 | NA | NA | NA | NA | NA | Rnase10 | ribonuclease, RNase A family, 10 (non-ac | 0 | 0 | 0 | 0 | 0 | 0 | 0 | 0 | 0 | 0 | 0 |
| 0 | NA | NA | NA | NA | NA | Rnase9 | ribonuclease, RNase A family, 9 (non-acti | 0 | 0 | 0 | 0 | 0 | 0 | 0 | 0 | 0 | 0 | 0 |
| 0 | NA | NA | NA | NA | NA | Rnase11 | ribonuclease, RNase A family, 11 (non-ac | 0 | 0 | 0 | 0 | 0 | 0 | 0 | 0 | 0 | 0 | 0 |
| 0 | NA | NA | NA | NA | NA | Rnase12 | ribonuclease, RNase A family, 12 (non-ac | 0 | 0 | 0 | 0 | 0 | 0 | 0 | 0 | 0 | 0 | 0 |
| 0 | NA | NA | NA | NA | NA | Olfr750 | olfactory receptor 750 | 0 | 0 | 0 | 0 | 0 | 0 | 0 | 0 | 0 | 0 | 0 |
| 0 | NA | NA | NA | NA | NA | Rnase2b | ribonuclease, RNase A family, 2B (liver, e | 0 | 0 | 0 | 0 | 0 | 0 | 0 | 0 | 0 | 0 | 0 |
| 0 | NA | NA | NA | NA | NA | Ear-ps1 | | 0 | 0 | 0 | 0 | 0 | 0 | 0 | 0 | 0 | 0 | 0 |
| 0 | NA | NA | NA | NA | NA | Ang2 | angiogenin, ribonuclease A family, memb | 0 | 0 | 0 | 0 | 0 | 0 | 0 | 0 | 0 | 0 | 0 |
| 0 | NA | NA | NA | NA | NA | Ear14 | eosinophil-associated, ribonuclease A fa | 0 | 0 | 0 | 0 | 0 | 0 | 0 | 0 | 0 | 0 | 0 |
| 0 | NA | NA | NA | NA | NA | Gm49245 | | 0 | 0 | 0 | 0 | 0 | 0 | 0 | 0 | 0 | 0 | 0 |
| 0 | NA | NA | NA | NA | NA | Gm21718 | | 0 | 0 | 0 | 0 | 0 | 0 | 0 | 0 | 0 | 0 | 0 |
| 0 | NA | NA | NA | NA | NA | 4930597G03Rik | | 0 | 0 | 0 | 0 | 0 | 0 | 0 | 0 | 0 | 0 | 0 |
| 0 | NA | NA | NA | NA | NA | Gm16289 | | 0 | 0 | 0 | 0 | 0 | 0 | 0 | 0 | 0 | 0 | 0 |
| 0 | NA | NA | NA | NA | NA | Gm7247 | predicted gene 7247 | 0 | 0 | 0 | 0 | 0 | 0 | 0 | 0 | 0 | 0 | 0 |
| 0 | NA | NA | NA | NA | NA | Vmn2r-ps110 | | 0 | 0 | 0 | 0 | 0 | 0 | 0 | 0 | 0 | 0 | 0 |
| 0 | NA | NA | NA | NA | NA | Vmn2r-ps111 | | 0 | 0 | 0 | 0 | 0 | 0 | 0 | 0 | 0 | 0 | 0 |
| 0 | NA | NA | NA | NA | NA | Vmn2r88 | vomeronasal 2, receptor 88 | 0 | 0 | 0 | 0 | 0 | 0 | 0 | 0 | 0 | 0 | 0 |
| 0 | NA | NA | NA | NA | NA | Vmn2r-ps111 | | 0 | 0 | 0 | 0 | 0 | 0 | 0 | 0 | 0 | 0 | 0 |
| 0 | NA | NA | NA | NA | NA | Vmn2r89 | vomeronasal 2, receptor 89 | 0 | 0 | 0 | 0 | 0 | 0 | 0 | 0 | 0 | 0 | 0 |
| 0 | NA | NA | NA | NA | NA | Gm5622 | predicted gene 5622 | 0 | 0 | 0 | 0 | 0 | 0 | 0 | 0 | 0 | 0 | 0 |
| 0 | NA | NA | NA | NA | NA | Gm16290 | | 0 | 0 | 0 | 0 | 0 | 0 | 0 | 0 | 0 | 0 | 0 |
| 0 | NA | NA | NA | NA | NA | Gm17175 | predicted gene 17175 | 0 | 0 | 0 | 0 | 0 | 0 | 0 | 0 | 0 | 0 | 0 |
| 0 | NA | NA | NA | NA | NA | Gm19156 | | 0 | 0 | 0 | 0 | 0 | 0 | 0 | 0 | 0 | 0 | 0 |
| 0 | NA | NA | NA | NA | NA | Gm17174 | | 0 | 0 | 0 | 0 | 0 | 0 | 0 | 0 | 0 | 0 | 0 |
| 0 | NA | NA | NA | NA | NA | Gm19157 | | 0 | 0 | 0 | 0 | 0 | 0 | 0 | 0 | 0 | 0 | 0 |

| | | | | | | | | | | | | | | | | | | |
|---|---|---|---|---|---|---|---|---|---|---|---|---|---|---|---|---|---|---|
| 0 | NA | NA | NA | NA | NA | Gm17078 | | 0 | 0 | 0 | 0 | 0 | 0 | 0 | 0 | 0 | 0 | 0 |
| 0 | NA | NA | NA | NA | NA | Gm19158 | | 0 | 0 | 0 | 0 | 0 | 0 | 0 | 0 | 0 | 0 | 0 |
| 0 | NA | NA | NA | NA | NA | Gm4181 | | 0 | 0 | 0 | 0 | 0 | 0 | 0 | 0 | 0 | 0 | 0 |
| 0 | NA | NA | NA | NA | NA | Gm17186 | | 0 | 0 | 0 | 0 | 0 | 0 | 0 | 0 | 0 | 0 | 0 |
| 0 | NA | NA | NA | NA | NA | Gm17079 | | 0 | 0 | 0 | 0 | 0 | 0 | 0 | 0 | 0 | 0 | 0 |
| 0 | NA | NA | NA | NA | NA | Gm5800 | predicted gene 5800 | 0 | 0 | 0 | 0 | 0 | 0 | 0 | 0 | 0 | 0 | 0 |
| 0 | NA | NA | NA | NA | NA | Gm49261 | | 0 | 0 | 0 | 0 | 0 | 0 | 0 | 0 | 0 | 0 | 0 |
| 0 | NA | NA | NA | NA | NA | Ear-ps3 | | 0 | 0 | 0 | 0 | 0 | 0 | 0 | 0 | 0 | 0 | 0 |
| 0 | NA | NA | NA | NA | NA | Ang4 | angiogenin, ribonuclease A family, memb | 0 | 0 | 0 | 0 | 0 | 0 | 0 | 0 | 0 | 0 | 0 |
| 0 | NA | NA | NA | NA | NA | Ear-ps4 | | 0 | 0 | 0 | 0 | 0 | 0 | 0 | 0 | 0 | 0 | 0 |
| 0 | NA | NA | NA | NA | NA | AY358078 | cDNA sequence AY358078 | 0 | 0 | 0 | 0 | 0 | 0 | 0 | 0 | 0 | 0 | 0 |
| 0 | NA | NA | NA | NA | NA | Gm27375 | | 0 | 0 | 0 | 0 | 0 | 0 | 0 | 0 | 0 | 0 | 0 |
| 0 | NA | NA | NA | NA | NA | Ear-ps5 | | 0 | 0 | 0 | 0 | 0 | 0 | 0 | 0 | 0 | 0 | 0 |
| 0 | NA | NA | NA | NA | NA | Ear6 | eosinophil-associated, ribonuclease A far | 0 | 0 | 0 | 0 | 0 | 0 | 0 | 0 | 0 | 0 | 0 |
| 0 | NA | NA | NA | NA | NA | Tppp2 | tubulin polymerization-promoting protein | 0 | 0 | 0 | 0 | 0 | 0 | 0 | 0 | 0 | 0 | 0 |
| 0 | NA | NA | NA | NA | NA | Rnase13 | ribonuclease, RNase A family, 13 (non-ac | 0 | 0 | 0 | 0 | 0 | 0 | 0 | 0 | 0 | 0 | 0 |
| 0 | NA | NA | NA | NA | NA | Gm49076 | | 0 | 0 | 0 | 0 | 0 | 0 | 0 | 0 | 0 | 0 | 0 |
| 0 | NA | NA | NA | NA | NA | Gm16617 | | 0 | 0 | 0 | 0 | 0 | 0 | 0 | 0 | 0 | 0 | 0 |
| 0 | NA | NA | NA | NA | NA | Olfr221 | olfactory receptor 221 | 0 | 0 | 0 | 0 | 0 | 0 | 0 | 0 | 0 | 0 | 0 |
| 0 | NA | NA | NA | NA | NA | Gm22354 | | 0 | 0 | 0 | 0 | 0 | 0 | 0 | 0 | 0 | 0 | 0 |
| 0 | NA | NA | NA | NA | NA | Gm49258 | | 0 | 0 | 0 | 0 | 0 | 0 | 0 | 0 | 0 | 0 | 0 |
| 0 | NA | NA | NA | NA | NA | Gm23758 | | 0 | 0 | 0 | 0 | 0 | 0 | 0 | 0 | 0 | 0 | 0 |
| 0 | NA | NA | NA | NA | NA | Gm26328 | | 0 | 0 | 0 | 0 | 0 | 0 | 0 | 0 | 0 | 0 | 0 |
| 0 | NA | NA | NA | NA | NA | Olfr1513 | olfactory receptor 1513 | 0 | 0 | 0 | 0 | 0 | 0 | 0 | 0 | 0 | 0 | 0 |
| 0 | NA | NA | NA | NA | NA | Olfr1512 | olfactory receptor 1512 | 0 | 0 | 0 | 0 | 0 | 0 | 0 | 0 | 0 | 0 | 0 |
| 0 | NA | NA | NA | NA | NA | Gm37504 | | 0 | 0 | 0 | 0 | 0 | 0 | 0 | 0 | 0 | 0 | 0 |
| 0 | NA | NA | NA | NA | NA | Olfr1511 | olfactory receptor 1511 | 0 | 0 | 0 | 0 | 0 | 0 | 0 | 0 | 0 | 0 | 0 |
| 0 | NA | NA | NA | NA | NA | Olfr1510 | olfactory receptor 1510 | 0 | 0 | 0 | 0 | 0 | 0 | 0 | 0 | 0 | 0 | 0 |
| 0 | NA | NA | NA | NA | NA | Gm43250 | | 0 | 0 | 0 | 0 | 0 | 0 | 0 | 0 | 0 | 0 | 0 |
| 0 | NA | NA | NA | NA | NA | Olfr1509 | olfactory receptor 1509 | 0 | 0 | 0 | 0 | 0 | 0 | 0 | 0 | 0 | 0 | 0 |
| 0 | NA | NA | NA | NA | NA | Olfr1508 | olfactory receptor 1508 | 0 | 0 | 0 | 0 | 0 | 0 | 0 | 0 | 0 | 0 | 0 |
| 0 | NA | NA | NA | NA | NA | Gm43512 | | 0 | 0 | 0 | 0 | 0 | 0 | 0 | 0 | 0 | 0 | 0 |
| 0 | NA | NA | NA | NA | NA | Gm36382 | | 0 | 0 | 0 | 0 | 0 | 0 | 0 | 0 | 0 | 0 | 0 |
| 0 | NA | NA | NA | NA | NA | Trav2 | | 0 | 0 | 0 | 0 | 0 | 0 | 0 | 0 | 0 | 0 | 0 |
| 0 | NA | NA | NA | NA | NA | Trav4-1 | | 0 | 0 | 0 | 0 | 0 | 0 | 0 | 0 | 0 | 0 | 0 |
| 0 | NA | NA | NA | NA | NA | Trav5-1 | | 0 | 0 | 0 | 0 | 0 | 0 | 0 | 0 | 0 | 0 | 0 |
| 0 | NA | NA | NA | NA | NA | Trav6-1 | | 0 | 0 | 0 | 0 | 0 | 0 | 0 | 0 | 0 | 0 | 0 |
| 0 | NA | NA | NA | NA | NA | Gm42988 | | 0 | 0 | 0 | 0 | 0 | 0 | 0 | 0 | 0 | 0 | 0 |
| 0 | NA | NA | NA | NA | NA | Trav7-1 | | 0 | 0 | 0 | 0 | 0 | 0 | 0 | 0 | 0 | 0 | 0 |
| 0 | NA | NA | NA | NA | NA | Trav6-2 | | 0 | 0 | 0 | 0 | 0 | 0 | 0 | 0 | 0 | 0 | 0 |
| 0 | NA | NA | NA | NA | NA | Trav7d-2 | | 0 | 0 | 0 | 0 | 0 | 0 | 0 | 0 | 0 | 0 | 0 |
| 0 | NA | NA | NA | NA | NA | Trav4d-2 | | 0 | 0 | 0 | 0 | 0 | 0 | 0 | 0 | 0 | 0 | 0 |
| 0 | NA | NA | NA | NA | NA | Trav6d-3 | | 0 | 0 | 0 | 0 | 0 | 0 | 0 | 0 | 0 | 0 | 0 |
| 0 | NA | NA | NA | NA | NA | Trav7d-3 | | 0 | 0 | 0 | 0 | 0 | 0 | 0 | 0 | 0 | 0 | 0 |
| 0 | NA | NA | NA | NA | NA | Trav6d-4 | | 0 | 0 | 0 | 0 | 0 | 0 | 0 | 0 | 0 | 0 | 0 |
| 0 | NA | NA | NA | NA | NA | Trav7d-4 | | 0 | 0 | 0 | 0 | 0 | 0 | 0 | 0 | 0 | 0 | 0 |
| 0 | NA | NA | NA | NA | NA | Trav9d-1 | | 0 | 0 | 0 | 0 | 0 | 0 | 0 | 0 | 0 | 0 | 0 |
| 0 | NA | NA | NA | NA | NA | Trav6d-5 | | 0 | 0 | 0 | 0 | 0 | 0 | 0 | 0 | 0 | 0 | 0 |
| 0 | NA | NA | NA | NA | NA | Trav10d | | 0 | 0 | 0 | 0 | 0 | 0 | 0 | 0 | 0 | 0 | 0 |
| 0 | NA | NA | NA | NA | NA | Trav6d-6 | | 0 | 0 | 0 | 0 | 0 | 0 | 0 | 0 | 0 | 0 | 0 |
| 0 | NA | NA | NA | NA | NA | Trav11d | | 0 | 0 | 0 | 0 | 0 | 0 | 0 | 0 | 0 | 0 | 0 |
| 0 | NA | NA | NA | NA | NA | Trav7d-5 | | 0 | 0 | 0 | 0 | 0 | 0 | 0 | 0 | 0 | 0 | 0 |
| 0 | NA | NA | NA | NA | NA | Trav12d-1 | | 0 | 0 | 0 | 0 | 0 | 0 | 0 | 0 | 0 | 0 | 0 |
| 0 | NA | NA | NA | NA | NA | Trav13d-1 | | 0 | 0 | 0 | 0 | 0 | 0 | 0 | 0 | 0 | 0 | 0 |
| 0 | NA | NA | NA | NA | NA | Trav14d-1 | | 0 | 0 | 0 | 0 | 0 | 0 | 0 | 0 | 0 | 0 | 0 |
| 0 | NA | NA | NA | NA | NA | Gm6740 | | 0 | 0 | 0 | 0 | 0 | 0 | 0 | 0 | 0 | 0 | 0 |
| 0 | NA | NA | NA | NA | NA | Trav15d-1-dv6d-1 | | 0 | 0 | 0 | 0 | 0 | 0 | 0 | 0 | 0 | 0 | 0 |
| 0 | NA | NA | NA | NA | NA | Trav3d-2 | | 0 | 0 | 0 | 0 | 0 | 0 | 0 | 0 | 0 | 0 | 0 |
| 0 | NA | NA | NA | NA | NA | Trav9d-2 | | 0 | 0 | 0 | 0 | 0 | 0 | 0 | 0 | 0 | 0 | 0 |
| 0 | NA | NA | NA | NA | NA | Trav4d-3 | | 0 | 0 | 0 | 0 | 0 | 0 | 0 | 0 | 0 | 0 | 0 |

| | | | | | | | | | | | | | | | | | |
|---|---|---|---|---|---|---|---|---|---|---|---|---|---|---|---|---|---|
| 0 | NA | NA | NA | NA | NA | Rps19-ps2 | 0 | 0 | 0 | 0 | 0 | 0 | 0 | 0 | 0 | 0 | 0 |
| 0 | NA | NA | NA | NA | NA | Trav5d-2 | 0 | 0 | 0 | 0 | 0 | 0 | 0 | 0 | 0 | 0 | 0 |
| 0 | NA | NA | NA | NA | NA | Trav12d-2 | 0 | 0 | 0 | 0 | 0 | 0 | 0 | 0 | 0 | 0 | 0 |
| 0 | NA | NA | NA | NA | NA | Trav9d-3 | 0 | 0 | 0 | 0 | 0 | 0 | 0 | 0 | 0 | 0 | 0 |
| 0 | NA | NA | NA | NA | NA | Trav5d-3 | 0 | 0 | 0 | 0 | 0 | 0 | 0 | 0 | 0 | 0 | 0 |
| 0 | NA | NA | NA | NA | NA | Trav12d-3 | 0 | 0 | 0 | 0 | 0 | 0 | 0 | 0 | 0 | 0 | 0 |
| 0 | NA | NA | NA | NA | NA | Trav13d-2 | 0 | 0 | 0 | 0 | 0 | 0 | 0 | 0 | 0 | 0 | 0 |
| 0 | NA | NA | NA | NA | NA | Trav14d-2 | 0 | 0 | 0 | 0 | 0 | 0 | 0 | 0 | 0 | 0 | 0 |
| 0 | NA | NA | NA | NA | NA | Trav15d-2-dv6d-2 | 0 | 0 | 0 | 0 | 0 | 0 | 0 | 0 | 0 | 0 | 0 |
| 0 | NA | NA | NA | NA | NA | Gm8704 | 0 | 0 | 0 | 0 | 0 | 0 | 0 | 0 | 0 | 0 | 0 |
| 0 | NA | NA | NA | NA | NA | Trav3d-3 | 0 | 0 | 0 | 0 | 0 | 0 | 0 | 0 | 0 | 0 | 0 |
| 0 | NA | NA | NA | NA | NA | Trav9d-4 | 0 | 0 | 0 | 0 | 0 | 0 | 0 | 0 | 0 | 0 | 0 |
| 0 | NA | NA | NA | NA | NA | Trav4d-4 | 0 | 0 | 0 | 0 | 0 | 0 | 0 | 0 | 0 | 0 | 0 |
| 0 | NA | NA | NA | NA | NA | Gm43104 | 0 | 0 | 0 | 0 | 0 | 0 | 0 | 0 | 0 | 0 | 0 |
| 0 | NA | NA | NA | NA | NA | Trav5d-4 | 0 | 0 | 0 | 0 | 0 | 0 | 0 | 0 | 0 | 0 | 0 |
| 0 | NA | NA | NA | NA | NA | Trav6d-7 | 0 | 0 | 0 | 0 | 0 | 0 | 0 | 0 | 0 | 0 | 0 |
| 0 | NA | NA | NA | NA | NA | Trav7d-6 | 0 | 0 | 0 | 0 | 0 | 0 | 0 | 0 | 0 | 0 | 0 |
| 0 | NA | NA | NA | NA | NA | Gm30214 | 0 | 0 | 0 | 0 | 0 | 0 | 0 | 0 | 0 | 0 | 0 |
| 0 | NA | NA | NA | NA | NA | Trav13d-3 | 0 | 0 | 0 | 0 | 0 | 0 | 0 | 0 | 0 | 0 | 0 |
| 0 | NA | NA | NA | NA | NA | Trav8d-2 | 0 | 0 | 0 | 0 | 0 | 0 | 0 | 0 | 0 | 0 | 0 |
| 0 | NA | NA | NA | NA | NA | Trav16d-dv11 | 0 | 0 | 0 | 0 | 0 | 0 | 0 | 0 | 0 | 0 | 0 |
| 0 | NA | NA | NA | NA | NA | Trav13d-4 | 0 | 0 | 0 | 0 | 0 | 0 | 0 | 0 | 0 | 0 | 0 |
| 0 | NA | NA | NA | NA | NA | Trav15d-3 | 0 | 0 | 0 | 0 | 0 | 0 | 0 | 0 | 0 | 0 | 0 |
| 0 | NA | NA | NA | NA | NA | Trav7n-4 | 0 | 0 | 0 | 0 | 0 | 0 | 0 | 0 | 0 | 0 | 0 |
| 0 | NA | NA | NA | NA | NA | Trav9n-1 | 0 | 0 | 0 | 0 | 0 | 0 | 0 | 0 | 0 | 0 | 0 |
| 0 | NA | NA | NA | NA | NA | Trav6n-5 | 0 | 0 | 0 | 0 | 0 | 0 | 0 | 0 | 0 | 0 | 0 |
| 0 | NA | NA | NA | NA | NA | Trav10n | 0 | 0 | 0 | 0 | 0 | 0 | 0 | 0 | 0 | 0 | 0 |
| 0 | NA | NA | NA | NA | NA | Trav6n-6 | 0 | 0 | 0 | 0 | 0 | 0 | 0 | 0 | 0 | 0 | 0 |
| 0 | NA | NA | NA | NA | NA | Trav11n | 0 | 0 | 0 | 0 | 0 | 0 | 0 | 0 | 0 | 0 | 0 |
| 0 | NA | NA | NA | NA | NA | Trav7n-5 | 0 | 0 | 0 | 0 | 0 | 0 | 0 | 0 | 0 | 0 | 0 |
| 0 | NA | NA | NA | NA | NA | Trav12n-1 | 0 | 0 | 0 | 0 | 0 | 0 | 0 | 0 | 0 | 0 | 0 |
| 0 | NA | NA | NA | NA | NA | Trav13n-1 | 0 | 0 | 0 | 0 | 0 | 0 | 0 | 0 | 0 | 0 | 0 |
| 0 | NA | NA | NA | NA | NA | Trav14n-1 | 0 | 0 | 0 | 0 | 0 | 0 | 0 | 0 | 0 | 0 | 0 |
| 0 | NA | NA | NA | NA | NA | Gm6755 | 0 | 0 | 0 | 0 | 0 | 0 | 0 | 0 | 0 | 0 | 0 |
| 0 | NA | NA | NA | NA | NA | Trav15n-1 | 0 | 0 | 0 | 0 | 0 | 0 | 0 | 0 | 0 | 0 | 0 |
| 0 | NA | NA | NA | NA | NA | Trav3n-2 | 0 | 0 | 0 | 0 | 0 | 0 | 0 | 0 | 0 | 0 | 0 |
| 0 | NA | NA | NA | NA | NA | Trav9n-2 | 0 | 0 | 0 | 0 | 0 | 0 | 0 | 0 | 0 | 0 | 0 |
| 0 | NA | NA | NA | NA | NA | Trav4n-3 | 0 | 0 | 0 | 0 | 0 | 0 | 0 | 0 | 0 | 0 | 0 |
| 0 | NA | NA | NA | NA | NA | Rps19-ps1 | 0 | 0 | 0 | 0 | 0 | 0 | 0 | 0 | 0 | 0 | 0 |
| 0 | NA | NA | NA | NA | NA | Trav5n-2 | 0 | 0 | 0 | 0 | 0 | 0 | 0 | 0 | 0 | 0 | 0 |
| 0 | NA | NA | NA | NA | NA | Trav12n-2 | 0 | 0 | 0 | 0 | 0 | 0 | 0 | 0 | 0 | 0 | 0 |
| 0 | NA | NA | NA | NA | NA | Trav9n-3 | 0 | 0 | 0 | 0 | 0 | 0 | 0 | 0 | 0 | 0 | 0 |
| 0 | NA | NA | NA | NA | NA | Trav5n-3 | 0 | 0 | 0 | 0 | 0 | 0 | 0 | 0 | 0 | 0 | 0 |
| 0 | NA | NA | NA | NA | NA | Trav12n-3 | 0 | 0 | 0 | 0 | 0 | 0 | 0 | 0 | 0 | 0 | 0 |
| 0 | NA | NA | NA | NA | NA | Trav13n-2 | 0 | 0 | 0 | 0 | 0 | 0 | 0 | 0 | 0 | 0 | 0 |
| 0 | NA | NA | NA | NA | NA | Trav14n-2 | 0 | 0 | 0 | 0 | 0 | 0 | 0 | 0 | 0 | 0 | 0 |
| 0 | NA | NA | NA | NA | NA | Trav15n-2 | 0 | 0 | 0 | 0 | 0 | 0 | 0 | 0 | 0 | 0 | 0 |
| 0 | NA | NA | NA | NA | NA | Gm3922 | 0 | 0 | 0 | 0 | 0 | 0 | 0 | 0 | 0 | 0 | 0 |
| 0 | NA | NA | NA | NA | NA | Trav3n-3 | 0 | 0 | 0 | 0 | 0 | 0 | 0 | 0 | 0 | 0 | 0 |
| 0 | NA | NA | NA | NA | NA | Trav9n-4 | 0 | 0 | 0 | 0 | 0 | 0 | 0 | 0 | 0 | 0 | 0 |
| 0 | NA | NA | NA | NA | NA | Trav4n-4 | 0 | 0 | 0 | 0 | 0 | 0 | 0 | 0 | 0 | 0 | 0 |
| 0 | NA | NA | NA | NA | NA | Trav5n-4 | 0 | 0 | 0 | 0 | 0 | 0 | 0 | 0 | 0 | 0 | 0 |
| 0 | NA | NA | NA | NA | NA | Trav6n-7 | 0 | 0 | 0 | 0 | 0 | 0 | 0 | 0 | 0 | 0 | 0 |
| 0 | NA | NA | NA | NA | NA | Trav7n-6 | 0 | 0 | 0 | 0 | 0 | 0 | 0 | 0 | 0 | 0 | 0 |
| 0 | NA | NA | NA | NA | NA | Gm43650 | 0 | 0 | 0 | 0 | 0 | 0 | 0 | 0 | 0 | 0 | 0 |
| 0 | NA | NA | NA | NA | NA | Trav13n-3 | 0 | 0 | 0 | 0 | 0 | 0 | 0 | 0 | 0 | 0 | 0 |
| 0 | NA | NA | NA | NA | NA | Trav16n | 0 | 0 | 0 | 0 | 0 | 0 | 0 | 0 | 0 | 0 | 0 |
| 0 | NA | NA | NA | NA | NA | Trav13n-4 | 0 | 0 | 0 | 0 | 0 | 0 | 0 | 0 | 0 | 0 | 0 |
| 0 | NA | NA | NA | NA | NA | Trav14n-3 | 0 | 0 | 0 | 0 | 0 | 0 | 0 | 0 | 0 | 0 | 0 |
| 0 | NA | NA | NA | NA | NA | Trav15n-3 | 0 | 0 | 0 | 0 | 0 | 0 | 0 | 0 | 0 | 0 | 0 |

| | | | | | | | | | | | | | | | | | |
|---|---|---|---|---|---|---|---|---|---|---|---|---|---|---|---|---|---|
| 0 | NA | NA | NA | NA | NA | Trav7-2 | 0 | 0 | 0 | 0 | 0 | 0 | 0 | 0 | 0 | 0 | 0 |
| 0 | NA | NA | NA | NA | NA | Trav6-3 | 0 | 0 | 0 | 0 | 0 | 0 | 0 | 0 | 0 | 0 | 0 |
| 0 | NA | NA | NA | NA | NA | Trav7-3 | 0 | 0 | 0 | 0 | 0 | 0 | 0 | 0 | 0 | 0 | 0 |
| 0 | NA | NA | NA | NA | NA | Trav6-4 | 0 | 0 | 0 | 0 | 0 | 0 | 0 | 0 | 0 | 0 | 0 |
| 0 | NA | NA | NA | NA | NA | Trav7-4 | 0 | 0 | 0 | 0 | 0 | 0 | 0 | 0 | 0 | 0 | 0 |
| 0 | NA | NA | NA | NA | NA | Gm18711 | 0 | 0 | 0 | 0 | 0 | 0 | 0 | 0 | 0 | 0 | 0 |
| 0 | NA | NA | NA | NA | NA | Trav8-1 | 0 | 0 | 0 | 0 | 0 | 0 | 0 | 0 | 0 | 0 | 0 |
| 0 | NA | NA | NA | NA | NA | Trav9-1 | 0 | 0 | 0 | 0 | 0 | 0 | 0 | 0 | 0 | 0 | 0 |
| 0 | NA | NA | NA | NA | NA | Trav10 | 0 | 0 | 0 | 0 | 0 | 0 | 0 | 0 | 0 | 0 | 0 |
| 0 | NA | NA | NA | NA | NA | Trav6-6 | 0 | 0 | 0 | 0 | 0 | 0 | 0 | 0 | 0 | 0 | 0 |
| 0 | NA | NA | NA | NA | NA | Trav11 | 0 | 0 | 0 | 0 | 0 | 0 | 0 | 0 | 0 | 0 | 0 |
| 0 | NA | NA | NA | NA | NA | Trav7-5 | 0 | 0 | 0 | 0 | 0 | 0 | 0 | 0 | 0 | 0 | 0 |
| 0 | NA | NA | NA | NA | NA | Trav15-1-dv6-1 | 0 | 0 | 0 | 0 | 0 | 0 | 0 | 0 | 0 | 0 | 0 |
| 0 | NA | NA | NA | NA | NA | Trav3-2 | 0 | 0 | 0 | 0 | 0 | 0 | 0 | 0 | 0 | 0 | 0 |
| 0 | NA | NA | NA | NA | NA | Trav9-2 | 0 | 0 | 0 | 0 | 0 | 0 | 0 | 0 | 0 | 0 | 0 |
| 0 | NA | NA | NA | NA | NA | Trav5-2 | 0 | 0 | 0 | 0 | 0 | 0 | 0 | 0 | 0 | 0 | 0 |
| 0 | NA | NA | NA | NA | NA | Trav12-2 | 0 | 0 | 0 | 0 | 0 | 0 | 0 | 0 | 0 | 0 | 0 |
| 0 | NA | NA | NA | NA | NA | Trav12-3 | 0 | 0 | 0 | 0 | 0 | 0 | 0 | 0 | 0 | 0 | 0 |
| 0 | NA | NA | NA | NA | NA | Trav13-2 | 0 | 0 | 0 | 0 | 0 | 0 | 0 | 0 | 0 | 0 | 0 |
| 0 | NA | NA | NA | NA | NA | Trav14-2 | 0 | 0 | 0 | 0 | 0 | 0 | 0 | 0 | 0 | 0 | 0 |
| 0 | NA | NA | NA | NA | NA | Trav15-2-dv6-2 | 0 | 0 | 0 | 0 | 0 | 0 | 0 | 0 | 0 | 0 | 0 |
| 0 | NA | NA | NA | NA | NA | Gm43551 | 0 | 0 | 0 | 0 | 0 | 0 | 0 | 0 | 0 | 0 | 0 |
| 0 | NA | NA | NA | NA | NA | Trav9-4 | 0 | 0 | 0 | 0 | 0 | 0 | 0 | 0 | 0 | 0 | 0 |
| 0 | NA | NA | NA | NA | NA | Trav4-4-dv10 | 0 | 0 | 0 | 0 | 0 | 0 | 0 | 0 | 0 | 0 | 0 |
| 0 | NA | NA | NA | NA | NA | Gm43550 | 0 | 0 | 0 | 0 | 0 | 0 | 0 | 0 | 0 | 0 | 0 |
| 0 | NA | NA | NA | NA | NA | Trav5-4 | 0 | 0 | 0 | 0 | 0 | 0 | 0 | 0 | 0 | 0 | 0 |
| 0 | NA | NA | NA | NA | NA | Trav6-7-dv9 | 0 | 0 | 0 | 0 | 0 | 0 | 0 | 0 | 0 | 0 | 0 |
| 0 | NA | NA | NA | NA | NA | Trav13-3 | 0 | 0 | 0 | 0 | 0 | 0 | 0 | 0 | 0 | 0 | 0 |
| 0 | NA | NA | NA | NA | NA | Trav8-2 | 0 | 0 | 0 | 0 | 0 | 0 | 0 | 0 | 0 | 0 | 0 |
| 0 | NA | NA | NA | NA | NA | Trav16 | 0 | 0 | 0 | 0 | 0 | 0 | 0 | 0 | 0 | 0 | 0 |
| 0 | NA | NA | NA | NA | NA | Trav13-4-dv7 | 0 | 0 | 0 | 0 | 0 | 0 | 0 | 0 | 0 | 0 | 0 |
| 0 | NA | NA | NA | NA | NA | Trav14-3 | 0 | 0 | 0 | 0 | 0 | 0 | 0 | 0 | 0 | 0 | 0 |
| 0 | NA | NA | NA | NA | NA | Trav15-3 | 0 | 0 | 0 | 0 | 0 | 0 | 0 | 0 | 0 | 0 | 0 |
| 0 | NA | NA | NA | NA | NA | Trav3-4 | 0 | 0 | 0 | 0 | 0 | 0 | 0 | 0 | 0 | 0 | 0 |
| 0 | NA | NA | NA | NA | NA | Trav12-4 | 0 | 0 | 0 | 0 | 0 | 0 | 0 | 0 | 0 | 0 | 0 |
| 0 | NA | NA | NA | NA | NA | Trav17 | 0 | 0 | 0 | 0 | 0 | 0 | 0 | 0 | 0 | 0 | 0 |
| 0 | NA | NA | NA | NA | NA | Trav18 | 0 | 0 | 0 | 0 | 0 | 0 | 0 | 0 | 0 | 0 | 0 |
| 0 | NA | NA | NA | NA | NA | Trav19 | 0 | 0 | 0 | 0 | 0 | 0 | 0 | 0 | 0 | 0 | 0 |
| 0 | NA | NA | NA | NA | NA | Trav20 | 0 | 0 | 0 | 0 | 0 | 0 | 0 | 0 | 0 | 0 | 0 |
| 0 | NA | NA | NA | NA | NA | Trav21-dv12 | 0 | 0 | 0 | 0 | 0 | 0 | 0 | 0 | 0 | 0 | 0 |
| 0 | NA | NA | NA | NA | NA | Trdv1 | 0 | 0 | 0 | 0 | 0 | 0 | 0 | 0 | 0 | 0 | 0 |
| 0 | NA | NA | NA | NA | NA | Gm43647 | 0 | 0 | 0 | 0 | 0 | 0 | 0 | 0 | 0 | 0 | 0 |
| 0 | NA | NA | NA | NA | NA | Trav22 | 0 | 0 | 0 | 0 | 0 | 0 | 0 | 0 | 0 | 0 | 0 |
| 0 | NA | NA | NA | NA | NA | Trdv2-1 | 0 | 0 | 0 | 0 | 0 | 0 | 0 | 0 | 0 | 0 | 0 |
| 0 | NA | NA | NA | NA | NA | Trdv2-2 | 0 | 0 | 0 | 0 | 0 | 0 | 0 | 0 | 0 | 0 | 0 |
| 0 | NA | NA | NA | NA | NA | Trav23 | 0 | 0 | 0 | 0 | 0 | 0 | 0 | 0 | 0 | 0 | 0 |
| 0 | NA | NA | NA | NA | NA | Trdv3 | 0 | 0 | 0 | 0 | 0 | 0 | 0 | 0 | 0 | 0 | 0 |
| 0 | NA | NA | NA | NA | NA | Gm30275 | 0 | 0 | 0 | 0 | 0 | 0 | 0 | 0 | 0 | 0 | 0 |
| 0 | NA | NA | NA | NA | NA | Trdd1 | 0 | 0 | 0 | 0 | 0 | 0 | 0 | 0 | 0 | 0 | 0 |
| 0 | NA | NA | NA | NA | NA | Gm23704 | 0 | 0 | 0 | 0 | 0 | 0 | 0 | 0 | 0 | 0 | 0 |
| 0 | NA | NA | NA | NA | NA | Trdd2 | 0 | 0 | 0 | 0 | 0 | 0 | 0 | 0 | 0 | 0 | 0 |
| 0 | NA | NA | NA | NA | NA | Trdj1 | 0 | 0 | 0 | 0 | 0 | 0 | 0 | 0 | 0 | 0 | 0 |
| 0 | NA | NA | NA | NA | NA | Gm43434 | 0 | 0 | 0 | 0 | 0 | 0 | 0 | 0 | 0 | 0 | 0 |
| 0 | NA | NA | NA | NA | NA | Trdj2 | 0 | 0 | 0 | 0 | 0 | 0 | 0 | 0 | 0 | 0 | 0 |
| 0 | NA | NA | NA | NA | NA | Trdv5 | 0 | 0 | 0 | 0 | 0 | 0 | 0 | 0 | 0 | 0 | 0 |
| 0 | NA | NA | NA | NA | NA | Traj61 | 0 | 0 | 0 | 0 | 0 | 0 | 0 | 0 | 0 | 0 | 0 |
| 0 | NA | NA | NA | NA | NA | Traj60 | 0 | 0 | 0 | 0 | 0 | 0 | 0 | 0 | 0 | 0 | 0 |
| 0 | NA | NA | NA | NA | NA | Traj59 | 0 | 0 | 0 | 0 | 0 | 0 | 0 | 0 | 0 | 0 | 0 |
| 0 | NA | NA | NA | NA | NA | Traj58 | 0 | 0 | 0 | 0 | 0 | 0 | 0 | 0 | 0 | 0 | 0 |
| 0 | NA | NA | NA | NA | NA | Traj57 | 0 | 0 | 0 | 0 | 0 | 0 | 0 | 0 | 0 | 0 | 0 |

| | | | | | | | | | | | | | | | | | | |
|---|---|---|---|---|---|---|---|---|---|---|---|---|---|---|---|---|---|---|
| 0 | NA | NA | NA | NA | NA | Traj56 | | 0 | 0 | 0 | 0 | 0 | 0 | 0 | 0 | 0 | 0 | 0 |
| 0 | NA | NA | NA | NA | NA | Traj55 | | 0 | 0 | 0 | 0 | 0 | 0 | 0 | 0 | 0 | 0 | 0 |
| 0 | NA | NA | NA | NA | NA | Traj54 | | 0 | 0 | 0 | 0 | 0 | 0 | 0 | 0 | 0 | 0 | 0 |
| 0 | NA | NA | NA | NA | NA | Traj53 | | 0 | 0 | 0 | 0 | 0 | 0 | 0 | 0 | 0 | 0 | 0 |
| 0 | NA | NA | NA | NA | NA | Traj52 | | 0 | 0 | 0 | 0 | 0 | 0 | 0 | 0 | 0 | 0 | 0 |
| 0 | NA | NA | NA | NA | NA | Traj51 | | 0 | 0 | 0 | 0 | 0 | 0 | 0 | 0 | 0 | 0 | 0 |
| 0 | NA | NA | NA | NA | NA | Traj50 | | 0 | 0 | 0 | 0 | 0 | 0 | 0 | 0 | 0 | 0 | 0 |
| 0 | NA | NA | NA | NA | NA | Traj49 | | 0 | 0 | 0 | 0 | 0 | 0 | 0 | 0 | 0 | 0 | 0 |
| 0 | NA | NA | NA | NA | NA | Traj48 | | 0 | 0 | 0 | 0 | 0 | 0 | 0 | 0 | 0 | 0 | 0 |
| 0 | NA | NA | NA | NA | NA | Traj47 | | 0 | 0 | 0 | 0 | 0 | 0 | 0 | 0 | 0 | 0 | 0 |
| 0 | NA | NA | NA | NA | NA | Traj46 | | 0 | 0 | 0 | 0 | 0 | 0 | 0 | 0 | 0 | 0 | 0 |
| 0 | NA | NA | NA | NA | NA | Traj45 | | 0 | 0 | 0 | 0 | 0 | 0 | 0 | 0 | 0 | 0 | 0 |
| 0 | NA | NA | NA | NA | NA | Traj43 | | 0 | 0 | 0 | 0 | 0 | 0 | 0 | 0 | 0 | 0 | 0 |
| 0 | NA | NA | NA | NA | NA | Traj42 | | 0 | 0 | 0 | 0 | 0 | 0 | 0 | 0 | 0 | 0 | 0 |
| 0 | NA | NA | NA | NA | NA | Traj41 | | 0 | 0 | 0 | 0 | 0 | 0 | 0 | 0 | 0 | 0 | 0 |
| 0 | NA | NA | NA | NA | NA | Traj40 | | 0 | 0 | 0 | 0 | 0 | 0 | 0 | 0 | 0 | 0 | 0 |
| 0 | NA | NA | NA | NA | NA | Traj39 | | 0 | 0 | 0 | 0 | 0 | 0 | 0 | 0 | 0 | 0 | 0 |
| 0 | NA | NA | NA | NA | NA | Traj38 | | 0 | 0 | 0 | 0 | 0 | 0 | 0 | 0 | 0 | 0 | 0 |
| 0 | NA | NA | NA | NA | NA | Traj37 | | 0 | 0 | 0 | 0 | 0 | 0 | 0 | 0 | 0 | 0 | 0 |
| 0 | NA | NA | NA | NA | NA | Traj36 | | 0 | 0 | 0 | 0 | 0 | 0 | 0 | 0 | 0 | 0 | 0 |
| 0 | NA | NA | NA | NA | NA | Traj35 | | 0 | 0 | 0 | 0 | 0 | 0 | 0 | 0 | 0 | 0 | 0 |
| 0 | NA | NA | NA | NA | NA | Traj34 | | 0 | 0 | 0 | 0 | 0 | 0 | 0 | 0 | 0 | 0 | 0 |
| 0 | NA | NA | NA | NA | NA | Traj33 | | 0 | 0 | 0 | 0 | 0 | 0 | 0 | 0 | 0 | 0 | 0 |
| 0 | NA | NA | NA | NA | NA | Traj32 | | 0 | 0 | 0 | 0 | 0 | 0 | 0 | 0 | 0 | 0 | 0 |
| 0 | NA | NA | NA | NA | NA | Traj31 | | 0 | 0 | 0 | 0 | 0 | 0 | 0 | 0 | 0 | 0 | 0 |
| 0 | NA | NA | NA | NA | NA | Traj30 | | 0 | 0 | 0 | 0 | 0 | 0 | 0 | 0 | 0 | 0 | 0 |
| 0 | NA | NA | NA | NA | NA | Traj29 | | 0 | 0 | 0 | 0 | 0 | 0 | 0 | 0 | 0 | 0 | 0 |
| 0 | NA | NA | NA | NA | NA | Traj28 | | 0 | 0 | 0 | 0 | 0 | 0 | 0 | 0 | 0 | 0 | 0 |
| 0 | NA | NA | NA | NA | NA | Traj27 | | 0 | 0 | 0 | 0 | 0 | 0 | 0 | 0 | 0 | 0 | 0 |
| 0 | NA | NA | NA | NA | NA | Traj26 | | 0 | 0 | 0 | 0 | 0 | 0 | 0 | 0 | 0 | 0 | 0 |
| 0 | NA | NA | NA | NA | NA | Traj25 | | 0 | 0 | 0 | 0 | 0 | 0 | 0 | 0 | 0 | 0 | 0 |
| 0 | NA | NA | NA | NA | NA | Traj24 | | 0 | 0 | 0 | 0 | 0 | 0 | 0 | 0 | 0 | 0 | 0 |
| 0 | NA | NA | NA | NA | NA | Traj23 | | 0 | 0 | 0 | 0 | 0 | 0 | 0 | 0 | 0 | 0 | 0 |
| 0 | NA | NA | NA | NA | NA | Traj22 | | 0 | 0 | 0 | 0 | 0 | 0 | 0 | 0 | 0 | 0 | 0 |
| 0 | NA | NA | NA | NA | NA | Traj21 | | 0 | 0 | 0 | 0 | 0 | 0 | 0 | 0 | 0 | 0 | 0 |
| 0 | NA | NA | NA | NA | NA | Traj20 | | 0 | 0 | 0 | 0 | 0 | 0 | 0 | 0 | 0 | 0 | 0 |
| 0 | NA | NA | NA | NA | NA | Traj19 | | 0 | 0 | 0 | 0 | 0 | 0 | 0 | 0 | 0 | 0 | 0 |
| 0 | NA | NA | NA | NA | NA | Traj18 | | 0 | 0 | 0 | 0 | 0 | 0 | 0 | 0 | 0 | 0 | 0 |
| 0 | NA | NA | NA | NA | NA | Traj17 | | 0 | 0 | 0 | 0 | 0 | 0 | 0 | 0 | 0 | 0 | 0 |
| 0 | NA | NA | NA | NA | NA | Traj16 | | 0 | 0 | 0 | 0 | 0 | 0 | 0 | 0 | 0 | 0 | 0 |
| 0 | NA | NA | NA | NA | NA | Traj15 | | 0 | 0 | 0 | 0 | 0 | 0 | 0 | 0 | 0 | 0 | 0 |
| 0 | NA | NA | NA | NA | NA | Traj14 | | 0 | 0 | 0 | 0 | 0 | 0 | 0 | 0 | 0 | 0 | 0 |
| 0 | NA | NA | NA | NA | NA | Traj13 | | 0 | 0 | 0 | 0 | 0 | 0 | 0 | 0 | 0 | 0 | 0 |
| 0 | NA | NA | NA | NA | NA | Traj12 | | 0 | 0 | 0 | 0 | 0 | 0 | 0 | 0 | 0 | 0 | 0 |
| 0 | NA | NA | NA | NA | NA | Traj11 | | 0 | 0 | 0 | 0 | 0 | 0 | 0 | 0 | 0 | 0 | 0 |
| 0 | NA | NA | NA | NA | NA | Traj9 | | 0 | 0 | 0 | 0 | 0 | 0 | 0 | 0 | 0 | 0 | 0 |
| 0 | NA | NA | NA | NA | NA | Traj8 | | 0 | 0 | 0 | 0 | 0 | 0 | 0 | 0 | 0 | 0 | 0 |
| 0 | NA | NA | NA | NA | NA | Traj6 | | 0 | 0 | 0 | 0 | 0 | 0 | 0 | 0 | 0 | 0 | 0 |
| 0 | NA | NA | NA | NA | NA | Traj5 | | 0 | 0 | 0 | 0 | 0 | 0 | 0 | 0 | 0 | 0 | 0 |
| 0 | NA | NA | NA | NA | NA | Traj4 | | 0 | 0 | 0 | 0 | 0 | 0 | 0 | 0 | 0 | 0 | 0 |
| 0 | NA | NA | NA | NA | NA | Traj3 | | 0 | 0 | 0 | 0 | 0 | 0 | 0 | 0 | 0 | 0 | 0 |
| 0 | NA | NA | NA | NA | NA | Traj2 | | 0 | 0 | 0 | 0 | 0 | 0 | 0 | 0 | 0 | 0 | 0 |
| 0 | NA | NA | NA | NA | NA | Traj1 | | 0 | 0 | 0 | 0 | 0 | 0 | 0 | 0 | 0 | 0 | 0 |
| 0 | NA | NA | NA | NA | NA | Olfr49 | olfactory receptor 49 | 0 | 0 | 0 | 0 | 0 | 0 | 0 | 0 | 0 | 0 | 0 |
| 0 | NA | NA | NA | NA | NA | Gm43304 | | 0 | 0 | 0 | 0 | 0 | 0 | 0 | 0 | 0 | 0 | 0 |
| 0 | NA | NA | NA | NA | NA | Gm3945 | | 0 | 0 | 0 | 0 | 0 | 0 | 0 | 0 | 0 | 0 | 0 |
| 0 | NA | NA | NA | NA | NA | 5330426L24Rik | | 0 | 0 | 0 | 0 | 0 | 0 | 0 | 0 | 0 | 0 | 0 |
| 0 | NA | NA | NA | NA | NA | 5430430K15Rik | | 0 | 0 | 0 | 0 | 0 | 0 | 0 | 0 | 0 | 0 | 0 |
| 0 | NA | NA | NA | NA | NA | Gm24424 | | 0 | 0 | 0 | 0 | 0 | 0 | 0 | 0 | 0 | 0 | 0 |
| 0 | NA | NA | NA | NA | NA | Gm24705 | | 0 | 0 | 0 | 0 | 0 | 0 | 0 | 0 | 0 | 0 | 0 |

| | | | | | | | | | | | | | | | | | | |
|---|---|---|---|---|---|---|---|---|---|---|---|---|---|---|---|---|---|---|
| 0 | NA | NA | NA | NA | NA | Mir686 | | 0 | 0 | 0 | 0 | 0 | 0 | 0 | 0 | 0 | 0 | 0 |
| 0 | NA | NA | NA | NA | NA | Psmb11 | proteasome (prosome, macropain) subun | 0 | 0 | 0 | 0 | 0 | 0 | 0 | 0 | 0 | 0 | 0 |
| 0 | NA | NA | NA | NA | NA | Gm17606 | | 0 | 0 | 0 | 0 | 0 | 0 | 0 | 0 | 0 | 0 | 0 |
| 0 | NA | NA | NA | NA | NA | Mir6948 | | 0 | 0 | 0 | 0 | 0 | 0 | 0 | 0 | 0 | 0 | 0 |
| 0 | NA | NA | NA | NA | NA | Gm29776 | predicted gene, 29776 | 0 | 0 | 0 | 0 | 0 | 0 | 0 | 0 | 0 | 0 | 0 |
| 0 | NA | NA | NA | NA | NA | Gm8865 | | 0 | 0 | 0 | 0 | 0 | 0 | 0 | 0 | 0 | 0 | 0 |
| 0 | NA | NA | NA | NA | NA | Gm20521 | | 0 | 0 | 0 | 0 | 0 | 0 | 0 | 0 | 0 | 0 | 0 |
| 0 | NA | NA | NA | NA | NA | Il25 | interleukin 25 | 0 | 0 | 0 | 0 | 0 | 0 | 0 | 0 | 0 | 0 | 0 |
| 0 | NA | NA | NA | NA | NA | Gm49130 | | 0 | 0 | 0 | 0 | 0 | 0 | 0 | 0 | 0 | 0 | 0 |
| 0 | NA | NA | NA | NA | NA | Mir208a | | 0 | 0 | 0 | 0 | 0 | 0 | 0 | 0 | 0 | 0 | 0 |
| 0 | NA | NA | NA | NA | NA | Mhrt | | 0 | 0 | 0 | 0 | 0 | 0 | 0 | 0 | 0 | 0 | 0 |
| 0 | NA | NA | NA | NA | NA | Gm29015 | | 0 | 0 | 0 | 0 | 0 | 0 | 0 | 0 | 0 | 0 | 0 |
| 0 | NA | NA | NA | NA | NA | Mir208b | | 0 | 0 | 0 | 0 | 0 | 0 | 0 | 0 | 0 | 0 | 0 |
| 0 | NA | NA | NA | NA | NA | Gm17428 | | 0 | 0 | 0 | 0 | 0 | 0 | 0 | 0 | 0 | 0 | 0 |
| 0 | NA | NA | NA | NA | NA | Gm31251 | | 0 | 0 | 0 | 0 | 0 | 0 | 0 | 0 | 0 | 0 | 0 |
| 0 | NA | NA | NA | NA | NA | Gm20687 | | 0 | 0 | 0 | 0 | 0 | 0 | 0 | 0 | 0 | 0 | 0 |
| 0 | NA | NA | NA | NA | NA | Gm49020 | | 0 | 0 | 0 | 0 | 0 | 0 | 0 | 0 | 0 | 0 | 0 |
| 0 | NA | NA | NA | NA | NA | Gm23596 | | 0 | 0 | 0 | 0 | 0 | 0 | 0 | 0 | 0 | 0 | 0 |
| 0 | NA | NA | NA | NA | NA | Dhrs2 | dehydrogenase/reductase member 2 | 0 | 0 | 0 | 0 | 0 | 0 | 0 | 0 | 0 | 0 | 0 |
| 0 | NA | NA | NA | NA | NA | Gm49021 | | 0 | 0 | 0 | 0 | 0 | 0 | 0 | 0 | 0 | 0 | 0 |
| 0 | NA | NA | NA | NA | NA | Gm49022 | | 0 | 0 | 0 | 0 | 0 | 0 | 0 | 0 | 0 | 0 | 0 |
| 0 | NA | NA | NA | NA | NA | Gm49023 | | 0 | 0 | 0 | 0 | 0 | 0 | 0 | 0 | 0 | 0 | 0 |
| 0 | NA | NA | NA | NA | NA | Gm10876 | | 0 | 0 | 0 | 0 | 0 | 0 | 0 | 0 | 0 | 0 | 0 |
| 0 | NA | NA | NA | NA | NA | Gm49117 | | 0 | 0 | 0 | 0 | 0 | 0 | 0 | 0 | 0 | 0 | 0 |
| 0 | NA | NA | NA | NA | NA | Gm49378 | | 0 | 0 | 0 | 0 | 0 | 0 | 0 | 0 | 0 | 0 | 0 |
| 0 | NA | NA | NA | NA | NA | Gm49299 | | 0 | 0 | 0 | 0 | 0 | 0 | 0 | 0 | 0 | 0 | 0 |
| 0 | NA | NA | NA | NA | NA | Gm25887 | | 0 | 0 | 0 | 0 | 0 | 0 | 0 | 0 | 0 | 0 | 0 |
| 0 | NA | NA | NA | NA | NA | Gm48925 | | 0 | 0 | 0 | 0 | 0 | 0 | 0 | 0 | 0 | 0 | 0 |
| 0 | NA | NA | NA | NA | NA | Mcpt-ps1 | | 0 | 0 | 0 | 0 | 0 | 0 | 0 | 0 | 0 | 0 | 0 |
| 0 | NA | NA | NA | NA | NA | Gm48927 | | 0 | 0 | 0 | 0 | 0 | 0 | 0 | 0 | 0 | 0 | 0 |
| 0 | NA | NA | NA | NA | NA | Mcpt1 | mast cell protease 1 | 0 | 0 | 0 | 0 | 0 | 0 | 0 | 0 | 0 | 0 | 0 |
| 0 | NA | NA | NA | NA | NA | Mcpt9 | mast cell protease 9 | 0 | 0 | 0 | 0 | 0 | 0 | 0 | 0 | 0 | 0 | 0 |
| 0 | NA | NA | NA | NA | NA | Mcpt2 | mast cell protease 2 | 0 | 0 | 0 | 0 | 0 | 0 | 0 | 0 | 0 | 0 | 0 |
| 0 | NA | NA | NA | NA | NA | Mcpt8 | mast cell protease 8 | 0 | 0 | 0 | 0 | 0 | 0 | 0 | 0 | 0 | 0 | 0 |
| 0 | NA | NA | NA | NA | NA | Mir6949 | | 0 | 0 | 0 | 0 | 0 | 0 | 0 | 0 | 0 | 0 | 0 |
| 0 | NA | NA | NA | NA | NA | Gzme | granzyme E | 0 | 0 | 0 | 0 | 0 | 0 | 0 | 0 | 0 | 0 | 0 |
| 0 | NA | NA | NA | NA | NA | Gzmd | granzyme D | 0 | 0 | 0 | 0 | 0 | 0 | 0 | 0 | 0 | 0 | 0 |
| 0 | NA | NA | NA | NA | NA | Gzmg | granzyme G | 0 | 0 | 0 | 0 | 0 | 0 | 0 | 0 | 0 | 0 | 0 |
| 0 | NA | NA | NA | NA | NA | Gm9547 | | 0 | 0 | 0 | 0 | 0 | 0 | 0 | 0 | 0 | 0 | 0 |
| 0 | NA | NA | NA | NA | NA | Gzmn | granzyme N | 0 | 0 | 0 | 0 | 0 | 0 | 0 | 0 | 0 | 0 | 0 |
| 0 | NA | NA | NA | NA | NA | Gm4022 | | 0 | 0 | 0 | 0 | 0 | 0 | 0 | 0 | 0 | 0 | 0 |
| 0 | NA | NA | NA | NA | NA | Gm8961 | | 0 | 0 | 0 | 0 | 0 | 0 | 0 | 0 | 0 | 0 | 0 |
| 0 | NA | NA | NA | NA | NA | Gm8963 | | 0 | 0 | 0 | 0 | 0 | 0 | 0 | 0 | 0 | 0 | 0 |
| 0 | NA | NA | NA | NA | NA | Gzmf | granzyme F | 0 | 0 | 0 | 0 | 0 | 0 | 0 | 0 | 0 | 0 | 0 |
| 0 | NA | NA | NA | NA | NA | Gzmc | granzyme C | 0 | 0 | 0 | 0 | 0 | 0 | 0 | 0 | 0 | 0 | 0 |
| 0 | NA | NA | NA | NA | NA | Gm24118 | | 0 | 0 | 0 | 0 | 0 | 0 | 0 | 0 | 0 | 0 | 0 |
| 0 | NA | NA | NA | NA | NA | Gm4819 | | 0 | 0 | 0 | 0 | 0 | 0 | 0 | 0 | 0 | 0 | 0 |
| 0 | NA | NA | NA | NA | NA | Atp12a | ATPase, H+/K+ transporting, nongastric, a | 0 | 0 | 0 | 0 | 0 | 0 | 0 | 0 | 0 | 0 | 0 |
| 0 | NA | NA | NA | NA | NA | Gm10873 | | 0 | 0 | 0 | 0 | 0 | 0 | 0 | 0 | 0 | 0 | 0 |
| 0 | NA | NA | NA | NA | NA | Gm24392 | | 0 | 0 | 0 | 0 | 0 | 0 | 0 | 0 | 0 | 0 | 0 |
| 0 | NA | NA | NA | NA | NA | Gm22431 | | 0 | 0 | 0 | 0 | 0 | 0 | 0 | 0 | 0 | 0 | 0 |
| 0 | NA | NA | NA | NA | NA | Gm4479 | | 0 | 0 | 0 | 0 | 0 | 0 | 0 | 0 | 0 | 0 | 0 |
| 0 | NA | NA | NA | NA | NA | Gm48906 | | 0 | 0 | 0 | 0 | 0 | 0 | 0 | 0 | 0 | 0 | 0 |
| 0 | NA | NA | NA | NA | NA | Gm22498 | | 0 | 0 | 0 | 0 | 0 | 0 | 0 | 0 | 0 | 0 | 0 |
| 0 | NA | NA | NA | NA | NA | Gm27635 | | 0 | 0 | 0 | 0 | 0 | 0 | 0 | 0 | 0 | 0 | 0 |
| 0 | NA | NA | NA | NA | NA | Gm22218 | | 0 | 0 | 0 | 0 | 0 | 0 | 0 | 0 | 0 | 0 | 0 |
| 0 | NA | NA | NA | NA | NA | B020004C17Ri | RIKEN cDNA B020004C17 gene | 0 | 0 | 0 | 0 | 0 | 0 | 0 | 0 | 0 | 0 | 0 |
| 0 | NA | NA | NA | NA | NA | Gja3 | gap junction protein, alpha 3 | 0 | 0 | 0 | 0 | 0 | 0 | 0 | 0 | 0 | 0 | 0 |
| 0 | NA | NA | NA | NA | NA | Gm4491 | | 0 | 0 | 0 | 0 | 0 | 0 | 0 | 0 | 0 | 0 | 0 |
| 0 | NA | NA | NA | NA | NA | Gjb6 | gap junction protein, beta 6 | 0 | 0 | 0 | 0 | 0 | 0 | 0 | 0 | 0 | 0 | 0 |

| | | | | | | | | | | | | | | | | | | |
|---|---|---|---|---|---|---|---|---|---|---|---|---|---|---|---|---|---|---|
| 0 | NA | NA | NA | NA | NA | Gm48725 | | 0 | 0 | 0 | 0 | 0 | 0 | 0 | 0 | 0 | 0 | 0 |
| 0 | NA | NA | NA | NA | NA | Gm8996 | | 0 | 0 | 0 | 0 | 0 | 0 | 0 | 0 | 0 | 0 | 0 |
| 0 | NA | NA | NA | NA | NA | Gm15836 | | 0 | 0 | 0 | 0 | 0 | 0 | 0 | 0 | 0 | 0 | 0 |
| 0 | NA | NA | NA | NA | NA | 1700039M10Rik | | 0 | 0 | 0 | 0 | 0 | 0 | 0 | 0 | 0 | 0 | 0 |
| 0 | NA | NA | NA | NA | NA | Gm49361 | | 0 | 0 | 0 | 0 | 0 | 0 | 0 | 0 | 0 | 0 | 0 |
| 0 | NA | NA | NA | NA | NA | Gm26440 | | 0 | 0 | 0 | 0 | 0 | 0 | 0 | 0 | 0 | 0 | 0 |
| 0 | NA | NA | NA | NA | NA | Mir3077 | | 0 | 0 | 0 | 0 | 0 | 0 | 0 | 0 | 0 | 0 | 0 |
| 0 | NA | NA | NA | NA | NA | Gm22738 | | 0 | 0 | 0 | 0 | 0 | 0 | 0 | 0 | 0 | 0 | 0 |
| 0 | NA | NA | NA | NA | NA | Gm17109 | | 0 | 0 | 0 | 0 | 0 | 0 | 0 | 0 | 0 | 0 | 0 |
| 0 | NA | NA | NA | NA | NA | Gm18080 | | 0 | 0 | 0 | 0 | 0 | 0 | 0 | 0 | 0 | 0 | 0 |
| 0 | NA | NA | NA | NA | NA | Gm33321 | | 0 | 0 | 0 | 0 | 0 | 0 | 0 | 0 | 0 | 0 | 0 |
| 0 | NA | NA | NA | NA | NA | Gm25614 | | 0 | 0 | 0 | 0 | 0 | 0 | 0 | 0 | 0 | 0 | 0 |
| 0 | NA | NA | NA | NA | NA | Gm48447 | | 0 | 0 | 0 | 0 | 0 | 0 | 0 | 0 | 0 | 0 | 0 |
| 0 | NA | NA | NA | NA | NA | Gm33472 | | 0 | 0 | 0 | 0 | 0 | 0 | 0 | 0 | 0 | 0 | 0 |
| 0 | NA | NA | NA | NA | NA | Gm41162 | | 0 | 0 | 0 | 0 | 0 | 0 | 0 | 0 | 0 | 0 | 0 |
| 0 | NA | NA | NA | NA | NA | Gm48453 | | 0 | 0 | 0 | 0 | 0 | 0 | 0 | 0 | 0 | 0 | 0 |
| 0 | NA | NA | NA | NA | NA | 1700129C05Ril | RIKEN cDNA 1700129C05 gene | 0 | 0 | 0 | 0 | 0 | 0 | 0 | 0 | 0 | 0 | 0 |
| 0 | NA | NA | NA | NA | NA | Gm5142 | predicted gene 5142 | 0 | 0 | 0 | 0 | 0 | 0 | 0 | 0 | 0 | 0 | 0 |
| 0 | NA | NA | NA | NA | NA | Gm19716 | | 0 | 0 | 0 | 0 | 0 | 0 | 0 | 0 | 0 | 0 | 0 |
| 0 | NA | NA | NA | NA | NA | Mir719 | | 0 | 0 | 0 | 0 | 0 | 0 | 0 | 0 | 0 | 0 | 0 |
| 0 | NA | NA | NA | NA | NA | Gm6066 | | 0 | 0 | 0 | 0 | 0 | 0 | 0 | 0 | 0 | 0 | 0 |
| 0 | NA | NA | NA | NA | NA | Gm47810 | | 0 | 0 | 0 | 0 | 0 | 0 | 0 | 0 | 0 | 0 | 0 |
| 0 | NA | NA | NA | NA | NA | Mipepos | | 0 | 0 | 0 | 0 | 0 | 0 | 0 | 0 | 0 | 0 | 0 |
| 0 | NA | NA | NA | NA | NA | Gm41168 | | 0 | 0 | 0 | 0 | 0 | 0 | 0 | 0 | 0 | 0 | 0 |
| 0 | NA | NA | NA | NA | NA | Sgcg | sarcoglycan, gamma (dystrophin-associa | 0 | 0 | 0 | 0 | 0 | 0 | 0 | 0 | 0 | 0 | 0 |
| 0 | NA | NA | NA | NA | NA | Gm19097 | | 0 | 0 | 0 | 0 | 0 | 0 | 0 | 0 | 0 | 0 | 0 |
| 0 | NA | NA | NA | NA | NA | Gm25716 | | 0 | 0 | 0 | 0 | 0 | 0 | 0 | 0 | 0 | 0 | 0 |
| 0 | NA | NA | NA | NA | NA | Vmn1r-ps136 | | 0 | 0 | 0 | 0 | 0 | 0 | 0 | 0 | 0 | 0 | 0 |
| 0 | NA | NA | NA | NA | NA | Gm27845 | | 0 | 0 | 0 | 0 | 0 | 0 | 0 | 0 | 0 | 0 | 0 |
| 0 | NA | NA | NA | NA | NA | Gm26916 | | 0 | 0 | 0 | 0 | 0 | 0 | 0 | 0 | 0 | 0 | 0 |
| 0 | NA | NA | NA | NA | NA | Mir16-1 | | 0 | 0 | 0 | 0 | 0 | 0 | 0 | 0 | 0 | 0 | 0 |
| 0 | NA | NA | NA | NA | NA | Mir15a | | 0 | 0 | 0 | 0 | 0 | 0 | 0 | 0 | 0 | 0 | 0 |
| 0 | NA | NA | NA | NA | NA | Gm27747 | | 0 | 0 | 0 | 0 | 0 | 0 | 0 | 0 | 0 | 0 | 0 |
| 0 | NA | NA | NA | NA | NA | Gm27928 | | 0 | 0 | 0 | 0 | 0 | 0 | 0 | 0 | 0 | 0 | 0 |
| 0 | NA | NA | NA | NA | NA | Gm47952 | | 0 | 0 | 0 | 0 | 0 | 0 | 0 | 0 | 0 | 0 | 0 |
| 0 | NA | NA | NA | NA | NA | Gm47953 | | 0 | 0 | 0 | 0 | 0 | 0 | 0 | 0 | 0 | 0 | 0 |
| 0 | NA | NA | NA | NA | NA | Gm35419 | | 0 | 0 | 0 | 0 | 0 | 0 | 0 | 0 | 0 | 0 | 0 |
| 0 | NA | NA | NA | NA | NA | Gm26969 | | 0 | 0 | 0 | 0 | 0 | 0 | 0 | 0 | 0 | 0 | 0 |
| 0 | NA | NA | NA | NA | NA | Gm4131 | predicted gene 4131 | 0 | 0 | 0 | 0 | 0 | 0 | 0 | 0 | 0 | 0 | 0 |
| 0 | NA | NA | NA | NA | NA | Gm6076 | | 0 | 0 | 0 | 0 | 0 | 0 | 0 | 0 | 0 | 0 | 0 |
| 0 | NA | NA | NA | NA | NA | Gm24625 | | 0 | 0 | 0 | 0 | 0 | 0 | 0 | 0 | 0 | 0 | 0 |
| 0 | NA | NA | NA | NA | NA | Gm23816 | | 0 | 0 | 0 | 0 | 0 | 0 | 0 | 0 | 0 | 0 | 0 |
| 0 | NA | NA | NA | NA | NA | Gm25162 | | 0 | 0 | 0 | 0 | 0 | 0 | 0 | 0 | 0 | 0 | 0 |
| 0 | NA | NA | NA | NA | NA | Gm18146 | | 0 | 0 | 0 | 0 | 0 | 0 | 0 | 0 | 0 | 0 | 0 |
| 0 | NA | NA | NA | NA | NA | Mir8098 | | 0 | 0 | 0 | 0 | 0 | 0 | 0 | 0 | 0 | 0 | 0 |
| 0 | NA | NA | NA | NA | NA | Mir6541 | | 0 | 0 | 0 | 0 | 0 | 0 | 0 | 0 | 0 | 0 | 0 |
| 0 | NA | NA | NA | NA | NA | n-R5s47 | | 0 | 0 | 0 | 0 | 0 | 0 | 0 | 0 | 0 | 0 | 0 |
| 0 | NA | NA | NA | NA | NA | Gm21021 | | 0 | 0 | 0 | 0 | 0 | 0 | 0 | 0 | 0 | 0 | 0 |
| 0 | NA | NA | NA | NA | NA | Defb48 | defensin beta 48 | 0 | 0 | 0 | 0 | 0 | 0 | 0 | 0 | 0 | 0 | 0 |
| 0 | NA | NA | NA | NA | NA | Defb47 | defensin beta 47 | 0 | 0 | 0 | 0 | 0 | 0 | 0 | 0 | 0 | 0 | 0 |
| 0 | NA | NA | NA | NA | NA | Defb43 | defensin beta 43 | 0 | 0 | 0 | 0 | 0 | 0 | 0 | 0 | 0 | 0 | 0 |
| 0 | NA | NA | NA | NA | NA | Gm19198 | | 0 | 0 | 0 | 0 | 0 | 0 | 0 | 0 | 0 | 0 | 0 |
| 0 | NA | NA | NA | NA | NA | Defb30 | defensin beta 30 | 0 | 0 | 0 | 0 | 0 | 0 | 0 | 0 | 0 | 0 | 0 |
| 0 | NA | NA | NA | NA | NA | Defb42 | defensin beta 42 | 0 | 0 | 0 | 0 | 0 | 0 | 0 | 0 | 0 | 0 | 0 |
| 0 | NA | NA | NA | NA | NA | Gm17941 | | 0 | 0 | 0 | 0 | 0 | 0 | 0 | 0 | 0 | 0 | 0 |
| 0 | NA | NA | NA | NA | NA | Gm23629 | | 0 | 0 | 0 | 0 | 0 | 0 | 0 | 0 | 0 | 0 | 0 |
| 0 | NA | NA | NA | NA | NA | Gm49390 | | 0 | 0 | 0 | 0 | 0 | 0 | 0 | 0 | 0 | 0 | 0 |
| 0 | NA | NA | NA | NA | NA | Gm5463 | | 0 | 0 | 0 | 0 | 0 | 0 | 0 | 0 | 0 | 0 | 0 |
| 0 | NA | NA | NA | NA | NA | Gm47074 | | 0 | 0 | 0 | 0 | 0 | 0 | 0 | 0 | 0 | 0 | 0 |
| 0 | NA | NA | NA | NA | NA | Mir598 | | 0 | 0 | 0 | 0 | 0 | 0 | 0 | 0 | 0 | 0 | 0 |

| | | | | | | | | | | | | | | | | | | |
|---|---|---|---|---|---|---|---|---|---|---|---|---|---|---|---|---|---|---|
| 0 | NA | NA | NA | NA | NA | 4930578I06Rik | RIKEN cDNA 4930578I06 gene | 0 | 0 | 0 | 0 | 0 | 0 | 0 | 0 | 0 | 0 | 0 |
| 0 | NA | NA | NA | NA | NA | Rp1l1 | retinitis pigmentosa 1 homolog like 1 | 0 | 0 | 0 | 0 | 0 | 0 | 0 | 0 | 0 | 0 | 0 |
| 0 | NA | NA | NA | NA | NA | Prss55 | protease, serine 55 | 0 | 0 | 0 | 0 | 0 | 0 | 0 | 0 | 0 | 0 | 0 |
| 0 | NA | NA | NA | NA | NA | Prss51 | protease, serine 51 | 0 | 0 | 0 | 0 | 0 | 0 | 0 | 0 | 0 | 0 | 0 |
| 0 | NA | NA | NA | NA | NA | 4930471C04Rik | | 0 | 0 | 0 | 0 | 0 | 0 | 0 | 0 | 0 | 0 | 0 |
| 0 | NA | NA | NA | NA | NA | Prss52 | protease, serine 52 | 0 | 0 | 0 | 0 | 0 | 0 | 0 | 0 | 0 | 0 | 0 |
| 0 | NA | NA | NA | NA | NA | Gm17116 | | 0 | 0 | 0 | 0 | 0 | 0 | 0 | 0 | 0 | 0 | 0 |
| 0 | NA | NA | NA | NA | NA | 9630015K15Rik | | 0 | 0 | 0 | 0 | 0 | 0 | 0 | 0 | 0 | 0 | 0 |
| 0 | NA | NA | NA | NA | NA | Mir124a-1hg | | 0 | 0 | 0 | 0 | 0 | 0 | 0 | 0 | 0 | 0 | 0 |
| 0 | NA | NA | NA | NA | NA | Mir124a-1 | | 0 | 0 | 0 | 0 | 0 | 0 | 0 | 0 | 0 | 0 | 0 |
| 0 | NA | NA | NA | NA | NA | Mir3078 | | 0 | 0 | 0 | 0 | 0 | 0 | 0 | 0 | 0 | 0 | 0 |
| 0 | NA | NA | NA | NA | NA | Gm26225 | | 0 | 0 | 0 | 0 | 0 | 0 | 0 | 0 | 0 | 0 | 0 |
| 0 | NA | NA | NA | NA | NA | Gm20111 | | 0 | 0 | 0 | 0 | 0 | 0 | 0 | 0 | 0 | 0 | 0 |
| 0 | NA | NA | NA | NA | NA | Gm4573 | | 0 | 0 | 0 | 0 | 0 | 0 | 0 | 0 | 0 | 0 | 0 |
| 0 | NA | NA | NA | NA | NA | Gm48299 | | 0 | 0 | 0 | 0 | 0 | 0 | 0 | 0 | 0 | 0 | 0 |
| 0 | NA | NA | NA | NA | NA | Gm30008 | | 0 | 0 | 0 | 0 | 0 | 0 | 0 | 0 | 0 | 0 | 0 |
| 0 | NA | NA | NA | NA | NA | Gm27263 | | 0 | 0 | 0 | 0 | 0 | 0 | 0 | 0 | 0 | 0 | 0 |
| 0 | NA | NA | NA | NA | NA | Gm9086 | | 0 | 0 | 0 | 0 | 0 | 0 | 0 | 0 | 0 | 0 | 0 |
| 0 | NA | NA | NA | NA | NA | Adam2 | a disintegrin and metallopeptidase domai | 0 | 0 | 0 | 0 | 0 | 0 | 0 | 0 | 0 | 0 | 0 |
| 0 | NA | NA | NA | NA | NA | Chrna2 | cholinergic receptor, nicotinic, alpha poly | 0 | 0 | 0 | 0 | 0 | 0 | 0 | 0 | 0 | 0 | 0 |
| 0 | NA | NA | NA | NA | NA | Gm41183 | | 0 | 0 | 0 | 0 | 0 | 0 | 0 | 0 | 0 | 0 | 0 |
| 0 | NA | NA | NA | NA | NA | Gm23899 | | 0 | 0 | 0 | 0 | 0 | 0 | 0 | 0 | 0 | 0 | 0 |
| 0 | NA | NA | NA | NA | NA | Gm10032 | | 0 | 0 | 0 | 0 | 0 | 0 | 0 | 0 | 0 | 0 | 0 |
| 0 | NA | NA | NA | NA | NA | 4930578I07Rik | | 0 | 0 | 0 | 0 | 0 | 0 | 0 | 0 | 0 | 0 | 0 |
| 0 | NA | NA | NA | NA | NA | Gm24258 | | 0 | 0 | 0 | 0 | 0 | 0 | 0 | 0 | 0 | 0 | 0 |
| 0 | NA | NA | NA | NA | NA | Gm27647 | | 0 | 0 | 0 | 0 | 0 | 0 | 0 | 0 | 0 | 0 | 0 |
| 0 | NA | NA | NA | NA | NA | Gm30806 | | 0 | 0 | 0 | 0 | 0 | 0 | 0 | 0 | 0 | 0 | 0 |
| 0 | NA | NA | NA | NA | NA | Gm20675 | | 0 | 0 | 0 | 0 | 0 | 0 | 0 | 0 | 0 | 0 | 0 |
| 0 | NA | NA | NA | NA | NA | Gm23178 | | 0 | 0 | 0 | 0 | 0 | 0 | 0 | 0 | 0 | 0 | 0 |
| 0 | NA | NA | NA | NA | NA | Gm6878 | predicted gene 6878 | 0 | 0 | 0 | 0 | 0 | 0 | 0 | 0 | 0 | 0 | 0 |
| 0 | NA | NA | NA | NA | NA | 1700120O09Rik | | 0 | 0 | 0 | 0 | 0 | 0 | 0 | 0 | 0 | 0 | 0 |
| 0 | NA | NA | NA | NA | NA | 4930438E09Rik | | 0 | 0 | 0 | 0 | 0 | 0 | 0 | 0 | 0 | 0 | 0 |
| 0 | NA | NA | NA | NA | NA | Gm10860 | | 0 | 0 | 0 | 0 | 0 | 0 | 0 | 0 | 0 | 0 | 0 |
| 0 | NA | NA | NA | NA | NA | Gm38409 | | 0 | 0 | 0 | 0 | 0 | 0 | 0 | 0 | 0 | 0 | 0 |
| 0 | NA | NA | NA | NA | NA | Gm24981 | | 0 | 0 | 0 | 0 | 0 | 0 | 0 | 0 | 0 | 0 | 0 |
| 0 | NA | NA | NA | NA | NA | Gm47010 | | 0 | 0 | 0 | 0 | 0 | 0 | 0 | 0 | 0 | 0 | 0 |
| 0 | NA | NA | NA | NA | NA | Mir6539 | | 0 | 0 | 0 | 0 | 0 | 0 | 0 | 0 | 0 | 0 | 0 |
| 0 | NA | NA | NA | NA | NA | Gm19216 | | 0 | 0 | 0 | 0 | 0 | 0 | 0 | 0 | 0 | 0 | 0 |
| 0 | NA | NA | NA | NA | NA | Gm22319 | | 0 | 0 | 0 | 0 | 0 | 0 | 0 | 0 | 0 | 0 | 0 |
| 0 | NA | NA | NA | NA | NA | Adam7 | a disintegrin and metallopeptidase domai | 0 | 0 | 0 | 0 | 0 | 0 | 0 | 0 | 0 | 0 | 0 |
| 0 | NA | NA | NA | NA | NA | Adam28 | a disintegrin and metallopeptidase domai | 0 | 0 | 0 | 0 | 0 | 0 | 0 | 0 | 0 | 0 | 0 |
| 0 | NA | NA | NA | NA | NA | Gm46476 | | 0 | 0 | 0 | 0 | 0 | 0 | 0 | 0 | 0 | 0 | 0 |
| 0 | NA | NA | NA | NA | NA | Gm47256 | | 0 | 0 | 0 | 0 | 0 | 0 | 0 | 0 | 0 | 0 | 0 |
| 0 | NA | NA | NA | NA | NA | Gm41192 | | 0 | 0 | 0 | 0 | 0 | 0 | 0 | 0 | 0 | 0 | 0 |
| 0 | NA | NA | NA | NA | NA | 1700092C10Rik | | 0 | 0 | 0 | 0 | 0 | 0 | 0 | 0 | 0 | 0 | 0 |
| 0 | NA | NA | NA | NA | NA | Nkx2-6 | NK2 homeobox 6 | 0 | 0 | 0 | 0 | 0 | 0 | 0 | 0 | 0 | 0 | 0 |
| 0 | NA | NA | NA | NA | NA | Nkx3-1 | NK3 homeobox 1 | 0 | 0 | 0 | 0 | 0 | 0 | 0 | 0 | 0 | 0 | 0 |
| 0 | NA | NA | NA | NA | NA | Gm37762 | | 0 | 0 | 0 | 0 | 0 | 0 | 0 | 0 | 0 | 0 | 0 |
| 0 | NA | NA | NA | NA | NA | Gm27222 | | 0 | 0 | 0 | 0 | 0 | 0 | 0 | 0 | 0 | 0 | 0 |
| 0 | NA | NA | NA | NA | NA | Gm20236 | | 0 | 0 | 0 | 0 | 0 | 0 | 0 | 0 | 0 | 0 | 0 |
| 0 | NA | NA | NA | NA | NA | Gm31748 | | 0 | 0 | 0 | 0 | 0 | 0 | 0 | 0 | 0 | 0 | 0 |
| 0 | NA | NA | NA | NA | NA | Gm27223 | | 0 | 0 | 0 | 0 | 0 | 0 | 0 | 0 | 0 | 0 | 0 |
| 0 | NA | NA | NA | NA | NA | Synb | syncytin b | 0 | 0 | 0 | 0 | 0 | 0 | 0 | 0 | 0 | 0 | 0 |
| 0 | NA | NA | NA | NA | NA | Entpd4 | ectonucleoside triphosphate diphosphohy | 0 | 0 | 0 | 0 | 0 | 0 | 0 | 0 | 0 | 0 | 0 |
| 0 | NA | NA | NA | NA | NA | Gm27221 | | 0 | 0 | 0 | 0 | 0 | 0 | 0 | 0 | 0 | 0 | 0 |
| 0 | NA | NA | NA | NA | NA | Gm38378 | | 0 | 0 | 0 | 0 | 0 | 0 | 0 | 0 | 0 | 0 | 0 |
| 0 | NA | NA | NA | NA | NA | Gm37094 | | 0 | 0 | 0 | 0 | 0 | 0 | 0 | 0 | 0 | 0 | 0 |
| 0 | NA | NA | NA | NA | NA | Gm37847 | | 0 | 0 | 0 | 0 | 0 | 0 | 0 | 0 | 0 | 0 | 0 |
| 0 | NA | NA | NA | NA | NA | Gm27175 | | 0 | 0 | 0 | 0 | 0 | 0 | 0 | 0 | 0 | 0 | 0 |
| 0 | NA | NA | NA | NA | NA | Gm27179 | syncytin-B | 0 | 0 | 0 | 0 | 0 | 0 | 0 | 0 | 0 | 0 | 0 |

| | | | | | | | | | | | | | | | | | |
|---|---|---|---|---|---|---|---|---|---|---|---|---|---|---|---|---|---|
| 0 | NA | NA | NA | NA | NA | Gm27174 | | 0 | 0 | 0 | 0 | 0 | 0 | 0 | 0 | 0 | 0 | 0 |
| 0 | NA | NA | NA | NA | NA | Entpd4b | ectonucleoside triphosphate diphosphohy | 0 | 0 | 0 | 0 | 0 | 0 | 0 | 0 | 0 | 0 | 0 |
| 0 | NA | NA | NA | NA | NA | Gm27176 | | 0 | 0 | 0 | 0 | 0 | 0 | 0 | 0 | 0 | 0 | 0 |
| 0 | NA | NA | NA | NA | NA | Gm27177 | | 0 | 0 | 0 | 0 | 0 | 0 | 0 | 0 | 0 | 0 | 0 |
| 0 | NA | NA | NA | NA | NA | Gm37513 | | 0 | 0 | 0 | 0 | 0 | 0 | 0 | 0 | 0 | 0 | 0 |
| 0 | NA | NA | NA | NA | NA | Mir6950 | | 0 | 0 | 0 | 0 | 0 | 0 | 0 | 0 | 0 | 0 | 0 |
| 0 | NA | NA | NA | NA | NA | Gm17740 | | 0 | 0 | 0 | 0 | 0 | 0 | 0 | 0 | 0 | 0 | 0 |
| 0 | NA | NA | NA | NA | NA | Pebp4 | phosphatidylethanolamine binding protei | 0 | 0 | 0 | 0 | 0 | 0 | 0 | 0 | 0 | 0 | 0 |
| 0 | NA | NA | NA | NA | NA | Gm33524 | | 0 | 0 | 0 | 0 | 0 | 0 | 0 | 0 | 0 | 0 | 0 |
| 0 | NA | NA | NA | NA | NA | Gm44340 | | 0 | 0 | 0 | 0 | 0 | 0 | 0 | 0 | 0 | 0 | 0 |
| 0 | NA | NA | NA | NA | NA | Gm4240 | | 0 | 0 | 0 | 0 | 0 | 0 | 0 | 0 | 0 | 0 | 0 |
| 0 | NA | NA | NA | NA | NA | Gm22725 | | 0 | 0 | 0 | 0 | 0 | 0 | 0 | 0 | 0 | 0 | 0 |
| 0 | NA | NA | NA | NA | NA | Gm24890 | | 0 | 0 | 0 | 0 | 0 | 0 | 0 | 0 | 0 | 0 | 0 |
| 0 | NA | NA | NA | NA | NA | Sftpc | surfactant associated protein C | 0 | 0 | 0 | 0 | 0 | 0 | 0 | 0 | 0 | 0 | 0 |
| 0 | NA | NA | NA | NA | NA | 2410012E07Rik | | 0 | 0 | 0 | 0 | 0 | 0 | 0 | 0 | 0 | 0 | 0 |
| 0 | NA | NA | NA | NA | NA | Gm49289 | | 0 | 0 | 0 | 0 | 0 | 0 | 0 | 0 | 0 | 0 | 0 |
| 0 | NA | NA | NA | NA | NA | Gm4251 | | 0 | 0 | 0 | 0 | 0 | 0 | 0 | 0 | 0 | 0 | 0 |
| 0 | NA | NA | NA | NA | NA | Gm37455 | | 0 | 0 | 0 | 0 | 0 | 0 | 0 | 0 | 0 | 0 | 0 |
| 0 | NA | NA | NA | NA | NA | 4930429C20Rik | | 0 | 0 | 0 | 0 | 0 | 0 | 0 | 0 | 0 | 0 | 0 |
| 0 | NA | NA | NA | NA | NA | Gm9192 | | 0 | 0 | 0 | 0 | 0 | 0 | 0 | 0 | 0 | 0 | 0 |
| 0 | NA | NA | NA | NA | NA | 4930434J06Rik | | 0 | 0 | 0 | 0 | 0 | 0 | 0 | 0 | 0 | 0 | 0 |
| 0 | NA | NA | NA | NA | NA | Gm23735 | | 0 | 0 | 0 | 0 | 0 | 0 | 0 | 0 | 0 | 0 | 0 |
| 0 | NA | NA | NA | NA | NA | Gm49290 | | 0 | 0 | 0 | 0 | 0 | 0 | 0 | 0 | 0 | 0 | 0 |
| 0 | NA | NA | NA | NA | NA | Mlnr-ps | | 0 | 0 | 0 | 0 | 0 | 0 | 0 | 0 | 0 | 0 | 0 |
| 0 | NA | NA | NA | NA | NA | Gm49293 | | 0 | 0 | 0 | 0 | 0 | 0 | 0 | 0 | 0 | 0 | 0 |
| 0 | NA | NA | NA | NA | NA | Gm41206 | | 0 | 0 | 0 | 0 | 0 | 0 | 0 | 0 | 0 | 0 | 0 |
| 0 | NA | NA | NA | NA | NA | Gm49295 | | 0 | 0 | 0 | 0 | 0 | 0 | 0 | 0 | 0 | 0 | 0 |
| 0 | NA | NA | NA | NA | NA | Gm9198 | | 0 | 0 | 0 | 0 | 0 | 0 | 0 | 0 | 0 | 0 | 0 |
| 0 | NA | NA | NA | NA | NA | Gm17233 | | 0 | 0 | 0 | 0 | 0 | 0 | 0 | 0 | 0 | 0 | 0 |
| 0 | NA | NA | NA | NA | NA | Mir687 | | 0 | 0 | 0 | 0 | 0 | 0 | 0 | 0 | 0 | 0 | 0 |
| 0 | NA | NA | NA | NA | NA | Gm49131 | | 0 | 0 | 0 | 0 | 0 | 0 | 0 | 0 | 0 | 0 | 0 |
| 0 | NA | NA | NA | NA | NA | Gm4266 | | 0 | 0 | 0 | 0 | 0 | 0 | 0 | 0 | 0 | 0 | 0 |
| 0 | NA | NA | NA | NA | NA | Gm21750 | | 0 | 0 | 0 | 0 | 0 | 0 | 0 | 0 | 0 | 0 | 0 |
| 0 | NA | NA | NA | NA | NA | Gm49165 | | 0 | 0 | 0 | 0 | 0 | 0 | 0 | 0 | 0 | 0 | 0 |
| 0 | NA | NA | NA | NA | NA | Gm49164 | | 0 | 0 | 0 | 0 | 0 | 0 | 0 | 0 | 0 | 0 | 0 |
| 0 | NA | NA | NA | NA | NA | Gm18683 | | 0 | 0 | 0 | 0 | 0 | 0 | 0 | 0 | 0 | 0 | 0 |
| 0 | NA | NA | NA | NA | NA | Gm22164 | | 0 | 0 | 0 | 0 | 0 | 0 | 0 | 0 | 0 | 0 | 0 |
| 0 | NA | NA | NA | NA | NA | Gm49170 | | 0 | 0 | 0 | 0 | 0 | 0 | 0 | 0 | 0 | 0 | 0 |
| 0 | NA | NA | NA | NA | NA | Gm49171 | | 0 | 0 | 0 | 0 | 0 | 0 | 0 | 0 | 0 | 0 | 0 |
| 0 | NA | NA | NA | NA | NA | 4933402J15Rik | | 0 | 0 | 0 | 0 | 0 | 0 | 0 | 0 | 0 | 0 | 0 |
| 0 | NA | NA | NA | NA | NA | Gm49174 | | 0 | 0 | 0 | 0 | 0 | 0 | 0 | 0 | 0 | 0 | 0 |
| 0 | NA | NA | NA | NA | NA | Gm49175 | | 0 | 0 | 0 | 0 | 0 | 0 | 0 | 0 | 0 | 0 | 0 |
| 0 | NA | NA | NA | NA | NA | Gm9212 | | 0 | 0 | 0 | 0 | 0 | 0 | 0 | 0 | 0 | 0 | 0 |
| 0 | NA | NA | NA | NA | NA | Gm10847 | | 0 | 0 | 0 | 0 | 0 | 0 | 0 | 0 | 0 | 0 | 0 |
| 0 | NA | NA | NA | NA | NA | Gm4278 | | 0 | 0 | 0 | 0 | 0 | 0 | 0 | 0 | 0 | 0 | 0 |
| 0 | NA | NA | NA | NA | NA | Gm15628 | | 0 | 0 | 0 | 0 | 0 | 0 | 0 | 0 | 0 | 0 | 0 |
| 0 | NA | NA | NA | NA | NA | Gm15629 | | 0 | 0 | 0 | 0 | 0 | 0 | 0 | 0 | 0 | 0 | 0 |
| 0 | NA | NA | NA | NA | NA | Gm44482 | | 0 | 0 | 0 | 0 | 0 | 0 | 0 | 0 | 0 | 0 | 0 |
| 0 | NA | NA | NA | NA | NA | Gm49200 | | 0 | 0 | 0 | 0 | 0 | 0 | 0 | 0 | 0 | 0 | 0 |
| 0 | NA | NA | NA | NA | NA | Cby2 | chibby family member 2 | 0 | 0 | 0 | 0 | 0 | 0 | 0 | 0 | 0 | 0 | 0 |
| 0 | NA | NA | NA | NA | NA | Erich6b | glutamate rich 6B | 0 | 0 | 0 | 0 | 0 | 0 | 0 | 0 | 0 | 0 | 0 |
| 0 | NA | NA | NA | NA | NA | Gm23622 | | 0 | 0 | 0 | 0 | 0 | 0 | 0 | 0 | 0 | 0 | 0 |
| 0 | NA | NA | NA | NA | NA | Gm24876 | | 0 | 0 | 0 | 0 | 0 | 0 | 0 | 0 | 0 | 0 | 0 |
| 0 | NA | NA | NA | NA | NA | Kctd4 | potassium channel tetramerisation domai | 0 | 0 | 0 | 0 | 0 | 0 | 0 | 0 | 0 | 0 | 0 |
| 0 | NA | NA | NA | NA | NA | Gm25517 | | 0 | 0 | 0 | 0 | 0 | 0 | 0 | 0 | 0 | 0 | 0 |
| 0 | NA | NA | NA | NA | NA | E130202H07Rik | | 0 | 0 | 0 | 0 | 0 | 0 | 0 | 0 | 0 | 0 | 0 |
| 0 | NA | NA | NA | NA | NA | Gm30246 | | 0 | 0 | 0 | 0 | 0 | 0 | 0 | 0 | 0 | 0 | 0 |
| 0 | NA | NA | NA | NA | NA | Gm48968 | | 0 | 0 | 0 | 0 | 0 | 0 | 0 | 0 | 0 | 0 | 0 |
| 0 | NA | NA | NA | NA | NA | Gm48969 | | 0 | 0 | 0 | 0 | 0 | 0 | 0 | 0 | 0 | 0 | 0 |
| 0 | NA | NA | NA | NA | NA | Gm49001 | | 0 | 0 | 0 | 0 | 0 | 0 | 0 | 0 | 0 | 0 | 0 |

| | | | | | | | | | | | | | | | | | | |
|---|---|---|---|---|---|---|---|---|---|---|---|---|---|---|---|---|---|---|
| 0 | NA | NA | NA | NA | NA | Gm49002 | | 0 | 0 | 0 | 0 | 0 | 0 | 0 | 0 | 0 | 0 | 0 |
| 0 | NA | NA | NA | NA | NA | Gm6994 | | 0 | 0 | 0 | 0 | 0 | 0 | 0 | 0 | 0 | 0 | 0 |
| 0 | NA | NA | NA | NA | NA | n-R5s48 | | 0 | 0 | 0 | 0 | 0 | 0 | 0 | 0 | 0 | 0 | 0 |
| 0 | NA | NA | NA | NA | NA | Gm23823 | | 0 | 0 | 0 | 0 | 0 | 0 | 0 | 0 | 0 | 0 | 0 |
| 0 | NA | NA | NA | NA | NA | Gm49003 | | 0 | 0 | 0 | 0 | 0 | 0 | 0 | 0 | 0 | 0 | 0 |
| 0 | NA | NA | NA | NA | NA | Gm30467 | | 0 | 0 | 0 | 0 | 0 | 0 | 0 | 0 | 0 | 0 | 0 |
| 0 | NA | NA | NA | NA | NA | Gm1587 | | 0 | 0 | 0 | 0 | 0 | 0 | 0 | 0 | 0 | 0 | 0 |
| 0 | NA | NA | NA | NA | NA | Gm48954 | | 0 | 0 | 0 | 0 | 0 | 0 | 0 | 0 | 0 | 0 | 0 |
| 0 | NA | NA | NA | NA | NA | Gm19301 | | 0 | 0 | 0 | 0 | 0 | 0 | 0 | 0 | 0 | 0 | 0 |
| 0 | NA | NA | NA | NA | NA | Gm48955 | | 0 | 0 | 0 | 0 | 0 | 0 | 0 | 0 | 0 | 0 | 0 |
| 0 | NA | NA | NA | NA | NA | Mir1971 | | 0 | 0 | 0 | 0 | 0 | 0 | 0 | 0 | 0 | 0 | 0 |
| 0 | NA | NA | NA | NA | NA | Gm48959 | | 0 | 0 | 0 | 0 | 0 | 0 | 0 | 0 | 0 | 0 | 0 |
| 0 | NA | NA | NA | NA | NA | Gm23652 | | 0 | 0 | 0 | 0 | 0 | 0 | 0 | 0 | 0 | 0 | 0 |
| 0 | NA | NA | NA | NA | NA | Gm26197 | | 0 | 0 | 0 | 0 | 0 | 0 | 0 | 0 | 0 | 0 | 0 |
| 0 | NA | NA | NA | NA | NA | Gm26251 | | 0 | 0 | 0 | 0 | 0 | 0 | 0 | 0 | 0 | 0 | 0 |
| 0 | NA | NA | NA | NA | NA | Gm18368 | | 0 | 0 | 0 | 0 | 0 | 0 | 0 | 0 | 0 | 0 | 0 |
| 0 | NA | NA | NA | NA | NA | Gm19098 | | 0 | 0 | 0 | 0 | 0 | 0 | 0 | 0 | 0 | 0 | 0 |
| 0 | NA | NA | NA | NA | NA | Gm49016 | | 0 | 0 | 0 | 0 | 0 | 0 | 0 | 0 | 0 | 0 | 0 |
| 0 | NA | NA | NA | NA | NA | Gm49011 | | 0 | 0 | 0 | 0 | 0 | 0 | 0 | 0 | 0 | 0 | 0 |
| 0 | NA | NA | NA | NA | NA | Zfp957 | zinc finger protein 957 | 0 | 0 | 0 | 0 | 0 | 0 | 0 | 0 | 0 | 0 | 0 |
| 0 | NA | NA | NA | NA | NA | Gm49028 | | 0 | 0 | 0 | 0 | 0 | 0 | 0 | 0 | 0 | 0 | 0 |
| 0 | NA | NA | NA | NA | NA | Gm27857 | | 0 | 0 | 0 | 0 | 0 | 0 | 0 | 0 | 0 | 0 | 0 |
| 0 | NA | NA | NA | NA | NA | Gm49042 | | 0 | 0 | 0 | 0 | 0 | 0 | 0 | 0 | 0 | 0 | 0 |
| 0 | NA | NA | NA | NA | NA | 4930452G13Rik | | 0 | 0 | 0 | 0 | 0 | 0 | 0 | 0 | 0 | 0 | 0 |
| 0 | NA | NA | NA | NA | NA | Mir759 | | 0 | 0 | 0 | 0 | 0 | 0 | 0 | 0 | 0 | 0 | 0 |
| 0 | NA | NA | NA | NA | NA | Gm17923 | | 0 | 0 | 0 | 0 | 0 | 0 | 0 | 0 | 0 | 0 | 0 |
| 0 | NA | NA | NA | NA | NA | Gm9578 | | 0 | 0 | 0 | 0 | 0 | 0 | 0 | 0 | 0 | 0 | 0 |
| 0 | NA | NA | NA | NA | NA | Gm49043 | | 0 | 0 | 0 | 0 | 0 | 0 | 0 | 0 | 0 | 0 | 0 |
| 0 | NA | NA | NA | NA | NA | Gm19120 | | 0 | 0 | 0 | 0 | 0 | 0 | 0 | 0 | 0 | 0 | 0 |
| 0 | NA | NA | NA | NA | NA | Gm49044 | | 0 | 0 | 0 | 0 | 0 | 0 | 0 | 0 | 0 | 0 | 0 |
| 0 | NA | NA | NA | NA | NA | Gm49045 | | 0 | 0 | 0 | 0 | 0 | 0 | 0 | 0 | 0 | 0 | 0 |
| 0 | NA | NA | NA | NA | NA | Gm49046 | | 0 | 0 | 0 | 0 | 0 | 0 | 0 | 0 | 0 | 0 | 0 |
| 0 | NA | NA | NA | NA | NA | Cyp2c73-ps | | 0 | 0 | 0 | 0 | 0 | 0 | 0 | 0 | 0 | 0 | 0 |
| 0 | NA | NA | NA | NA | NA | Gm41219 | | 0 | 0 | 0 | 0 | 0 | 0 | 0 | 0 | 0 | 0 | 0 |
| 0 | NA | NA | NA | NA | NA | Gm19311 | | 0 | 0 | 0 | 0 | 0 | 0 | 0 | 0 | 0 | 0 | 0 |
| 0 | NA | NA | NA | NA | NA | Gm18525 | | 0 | 0 | 0 | 0 | 0 | 0 | 0 | 0 | 0 | 0 | 0 |
| 0 | NA | NA | NA | NA | NA | Gm19050 | | 0 | 0 | 0 | 0 | 0 | 0 | 0 | 0 | 0 | 0 | 0 |
| 0 | NA | NA | NA | NA | NA | Gm49047 | | 0 | 0 | 0 | 0 | 0 | 0 | 0 | 0 | 0 | 0 | 0 |
| 0 | NA | NA | NA | NA | NA | Gm49048 | | 0 | 0 | 0 | 0 | 0 | 0 | 0 | 0 | 0 | 0 | 0 |
| 0 | NA | NA | NA | NA | NA | Gm22891 | | 0 | 0 | 0 | 0 | 0 | 0 | 0 | 0 | 0 | 0 | 0 |
| 0 | NA | NA | NA | NA | NA | Gm49049 | | 0 | 0 | 0 | 0 | 0 | 0 | 0 | 0 | 0 | 0 | 0 |
| 0 | NA | NA | NA | NA | NA | Gm49050 | | 0 | 0 | 0 | 0 | 0 | 0 | 0 | 0 | 0 | 0 | 0 |
| 0 | NA | NA | NA | NA | NA | Gm49051 | | 0 | 0 | 0 | 0 | 0 | 0 | 0 | 0 | 0 | 0 | 0 |
| 0 | NA | NA | NA | NA | NA | Gm24774 | | 0 | 0 | 0 | 0 | 0 | 0 | 0 | 0 | 0 | 0 | 0 |
| 0 | NA | NA | NA | NA | NA | Gm49052 | | 0 | 0 | 0 | 0 | 0 | 0 | 0 | 0 | 0 | 0 | 0 |
| 0 | NA | NA | NA | NA | NA | Gm49053 | | 0 | 0 | 0 | 0 | 0 | 0 | 0 | 0 | 0 | 0 | 0 |
| 0 | NA | NA | NA | NA | NA | Gm49054 | | 0 | 0 | 0 | 0 | 0 | 0 | 0 | 0 | 0 | 0 | 0 |
| 0 | NA | NA | NA | NA | NA | Gm49055 | | 0 | 0 | 0 | 0 | 0 | 0 | 0 | 0 | 0 | 0 | 0 |
| 0 | NA | NA | NA | NA | NA | Gm32455 | | 0 | 0 | 0 | 0 | 0 | 0 | 0 | 0 | 0 | 0 | 0 |
| 0 | NA | NA | NA | NA | NA | Gm49056 | | 0 | 0 | 0 | 0 | 0 | 0 | 0 | 0 | 0 | 0 | 0 |
| 0 | NA | NA | NA | NA | NA | Gm29800 | | 0 | 0 | 0 | 0 | 0 | 0 | 0 | 0 | 0 | 0 | 0 |
| 0 | NA | NA | NA | NA | NA | Gm49057 | | 0 | 0 | 0 | 0 | 0 | 0 | 0 | 0 | 0 | 0 | 0 |
| 0 | NA | NA | NA | NA | NA | 9630013A20Rik | | 0 | 0 | 0 | 0 | 0 | 0 | 0 | 0 | 0 | 0 | 0 |
| 0 | NA | NA | NA | NA | NA | Gm9264 | | 0 | 0 | 0 | 0 | 0 | 0 | 0 | 0 | 0 | 0 | 0 |
| 0 | NA | NA | NA | NA | NA | Gm49059 | | 0 | 0 | 0 | 0 | 0 | 0 | 0 | 0 | 0 | 0 | 0 |
| 0 | NA | NA | NA | NA | NA | Gm32729 | | 0 | 0 | 0 | 0 | 0 | 0 | 0 | 0 | 0 | 0 | 0 |
| 0 | NA | NA | NA | NA | NA | Gm49060 | | 0 | 0 | 0 | 0 | 0 | 0 | 0 | 0 | 0 | 0 | 0 |
| 0 | NA | NA | NA | NA | NA | Gm49061 | | 0 | 0 | 0 | 0 | 0 | 0 | 0 | 0 | 0 | 0 | 0 |
| 0 | NA | NA | NA | NA | NA | Gm49062 | | 0 | 0 | 0 | 0 | 0 | 0 | 0 | 0 | 0 | 0 | 0 |
| 0 | NA | NA | NA | NA | NA | 4930529K09Rik | | 0 | 0 | 0 | 0 | 0 | 0 | 0 | 0 | 0 | 0 | 0 |

| | | | | | | | | | | | | | | | | | | |
|---|---|---|---|---|---|---|---|---|---|---|---|---|---|---|---|---|---|---|
| 0 | NA | NA | NA | NA | NA | Gm49063 | | 0 | 0 | 0 | 0 | 0 | 0 | 0 | 0 | 0 | 0 | 0 |
| 0 | NA | NA | NA | NA | NA | Gm23926 | | 0 | 0 | 0 | 0 | 0 | 0 | 0 | 0 | 0 | 0 | 0 |
| 0 | NA | NA | NA | NA | NA | Gm4350 | | 0 | 0 | 0 | 0 | 0 | 0 | 0 | 0 | 0 | 0 | 0 |
| 0 | NA | NA | NA | NA | NA | Gm32913 | | 0 | 0 | 0 | 0 | 0 | 0 | 0 | 0 | 0 | 0 | 0 |
| 0 | NA | NA | NA | NA | NA | Gm25115 | | 0 | 0 | 0 | 0 | 0 | 0 | 0 | 0 | 0 | 0 | 0 |
| 0 | NA | NA | NA | NA | NA | Gm49157 | | 0 | 0 | 0 | 0 | 0 | 0 | 0 | 0 | 0 | 0 | 0 |
| 0 | NA | NA | NA | NA | NA | Gm49158 | | 0 | 0 | 0 | 0 | 0 | 0 | 0 | 0 | 0 | 0 | 0 |
| 0 | NA | NA | NA | NA | NA | Gm49159 | | 0 | 0 | 0 | 0 | 0 | 0 | 0 | 0 | 0 | 0 | 0 |
| 0 | NA | NA | NA | NA | NA | Gm23438 | | 0 | 0 | 0 | 0 | 0 | 0 | 0 | 0 | 0 | 0 | 0 |
| 0 | NA | NA | NA | NA | NA | Gm49193 | | 0 | 0 | 0 | 0 | 0 | 0 | 0 | 0 | 0 | 0 | 0 |
| 0 | NA | NA | NA | NA | NA | Gm37002 | | 0 | 0 | 0 | 0 | 0 | 0 | 0 | 0 | 0 | 0 | 0 |
| 0 | NA | NA | NA | NA | NA | Gm23278 | | 0 | 0 | 0 | 0 | 0 | 0 | 0 | 0 | 0 | 0 | 0 |
| 0 | NA | NA | NA | NA | NA | Gm48922 | | 0 | 0 | 0 | 0 | 0 | 0 | 0 | 0 | 0 | 0 | 0 |
| 0 | NA | NA | NA | NA | NA | Gm18812 | | 0 | 0 | 0 | 0 | 0 | 0 | 0 | 0 | 0 | 0 | 0 |
| 0 | NA | NA | NA | NA | NA | Gm4355 | | 0 | 0 | 0 | 0 | 0 | 0 | 0 | 0 | 0 | 0 | 0 |
| 0 | NA | NA | NA | NA | NA | Gm26405 | | 0 | 0 | 0 | 0 | 0 | 0 | 0 | 0 | 0 | 0 | 0 |
| 0 | NA | NA | NA | NA | NA | Gm33203 | | 0 | 0 | 0 | 0 | 0 | 0 | 0 | 0 | 0 | 0 | 0 |
| 0 | NA | NA | NA | NA | NA | Gm9274 | | 0 | 0 | 0 | 0 | 0 | 0 | 0 | 0 | 0 | 0 | 0 |
| 0 | NA | NA | NA | NA | NA | Gm48929 | | 0 | 0 | 0 | 0 | 0 | 0 | 0 | 0 | 0 | 0 | 0 |
| 0 | NA | NA | NA | NA | NA | Gm9275 | | 0 | 0 | 0 | 0 | 0 | 0 | 0 | 0 | 0 | 0 | 0 |
| 0 | NA | NA | NA | NA | NA | Gm48930 | | 0 | 0 | 0 | 0 | 0 | 0 | 0 | 0 | 0 | 0 | 0 |
| 0 | NA | NA | NA | NA | NA | Gm41225 | | 0 | 0 | 0 | 0 | 0 | 0 | 0 | 0 | 0 | 0 | 0 |
| 0 | NA | NA | NA | NA | NA | Gm36946 | | 0 | 0 | 0 | 0 | 0 | 0 | 0 | 0 | 0 | 0 | 0 |
| 0 | NA | NA | NA | NA | NA | Gm17882 | | 0 | 0 | 0 | 0 | 0 | 0 | 0 | 0 | 0 | 0 | 0 |
| 0 | NA | NA | NA | NA | NA | Gm25886 | | 0 | 0 | 0 | 0 | 0 | 0 | 0 | 0 | 0 | 0 | 0 |
| 0 | NA | NA | NA | NA | NA | Gm48941 | | 0 | 0 | 0 | 0 | 0 | 0 | 0 | 0 | 0 | 0 | 0 |
| 0 | NA | NA | NA | NA | NA | 4930474H20Rik | | 0 | 0 | 0 | 0 | 0 | 0 | 0 | 0 | 0 | 0 | 0 |
| 0 | NA | NA | NA | NA | NA | Gm25495 | | 0 | 0 | 0 | 0 | 0 | 0 | 0 | 0 | 0 | 0 | 0 |
| 0 | NA | NA | NA | NA | NA | Gm33246 | | 0 | 0 | 0 | 0 | 0 | 0 | 0 | 0 | 0 | 0 | 0 |
| 0 | NA | NA | NA | NA | NA | Gm18082 | | 0 | 0 | 0 | 0 | 0 | 0 | 0 | 0 | 0 | 0 | 0 |
| 0 | NA | NA | NA | NA | NA | Gm23225 | | 0 | 0 | 0 | 0 | 0 | 0 | 0 | 0 | 0 | 0 | 0 |
| 0 | NA | NA | NA | NA | NA | Gm7272 | | 0 | 0 | 0 | 0 | 0 | 0 | 0 | 0 | 0 | 0 | 0 |
| 0 | NA | NA | NA | NA | NA | Gm48942 | | 0 | 0 | 0 | 0 | 0 | 0 | 0 | 0 | 0 | 0 | 0 |
| 0 | NA | NA | NA | NA | NA | Gm48943 | | 0 | 0 | 0 | 0 | 0 | 0 | 0 | 0 | 0 | 0 | 0 |
| 0 | NA | NA | NA | NA | NA | Gm48944 | | 0 | 0 | 0 | 0 | 0 | 0 | 0 | 0 | 0 | 0 | 0 |
| 0 | NA | NA | NA | NA | NA | Gm24127 | | 0 | 0 | 0 | 0 | 0 | 0 | 0 | 0 | 0 | 0 | 0 |
| 0 | NA | NA | NA | NA | NA | Gm37341 | | 0 | 0 | 0 | 0 | 0 | 0 | 0 | 0 | 0 | 0 | 0 |
| 0 | NA | NA | NA | NA | NA | Gm48962 | | 0 | 0 | 0 | 0 | 0 | 0 | 0 | 0 | 0 | 0 | 0 |
| 0 | NA | NA | NA | NA | NA | 4930433E05Rik | | 0 | 0 | 0 | 0 | 0 | 0 | 0 | 0 | 0 | 0 | 0 |
| 0 | NA | NA | NA | NA | NA | Gm48963 | | 0 | 0 | 0 | 0 | 0 | 0 | 0 | 0 | 0 | 0 | 0 |
| 0 | NA | NA | NA | NA | NA | Gm23509 | | 0 | 0 | 0 | 0 | 0 | 0 | 0 | 0 | 0 | 0 | 0 |
| 0 | NA | NA | NA | NA | NA | Gm17922 | | 0 | 0 | 0 | 0 | 0 | 0 | 0 | 0 | 0 | 0 | 0 |
| 0 | NA | NA | NA | NA | NA | Gm48964 | | 0 | 0 | 0 | 0 | 0 | 0 | 0 | 0 | 0 | 0 | 0 |
| 0 | NA | NA | NA | NA | NA | Gm48967 | | 0 | 0 | 0 | 0 | 0 | 0 | 0 | 0 | 0 | 0 | 0 |
| 0 | NA | NA | NA | NA | NA | C230086J09Rik | | 0 | 0 | 0 | 0 | 0 | 0 | 0 | 0 | 0 | 0 | 0 |
| 0 | NA | NA | NA | NA | NA | Gm48981 | | 0 | 0 | 0 | 0 | 0 | 0 | 0 | 0 | 0 | 0 | 0 |
| 0 | NA | NA | NA | NA | NA | Gm48982 | | 0 | 0 | 0 | 0 | 0 | 0 | 0 | 0 | 0 | 0 | 0 |
| 0 | NA | NA | NA | NA | NA | Gm23324 | | 0 | 0 | 0 | 0 | 0 | 0 | 0 | 0 | 0 | 0 | 0 |
| 0 | NA | NA | NA | NA | NA | Gm48983 | | 0 | 0 | 0 | 0 | 0 | 0 | 0 | 0 | 0 | 0 | 0 |
| 0 | NA | NA | NA | NA | NA | Gm18074 | | 0 | 0 | 0 | 0 | 0 | 0 | 0 | 0 | 0 | 0 | 0 |
| 0 | NA | NA | NA | NA | NA | Gm33450 | | 0 | 0 | 0 | 0 | 0 | 0 | 0 | 0 | 0 | 0 | 0 |
| 0 | NA | NA | NA | NA | NA | Gm24680 | | 0 | 0 | 0 | 0 | 0 | 0 | 0 | 0 | 0 | 0 | 0 |
| 0 | NA | NA | NA | NA | NA | Gm9721 | | 0 | 0 | 0 | 0 | 0 | 0 | 0 | 0 | 0 | 0 | 0 |
| 0 | NA | NA | NA | NA | NA | Gm48989 | | 0 | 0 | 0 | 0 | 0 | 0 | 0 | 0 | 0 | 0 | 0 |
| 0 | NA | NA | NA | NA | NA | Gm48991 | | 0 | 0 | 0 | 0 | 0 | 0 | 0 | 0 | 0 | 0 | 0 |
| 0 | NA | NA | NA | NA | NA | Gm48994 | | 0 | 0 | 0 | 0 | 0 | 0 | 0 | 0 | 0 | 0 | 0 |
| 0 | NA | NA | NA | NA | NA | Gm48995 | | 0 | 0 | 0 | 0 | 0 | 0 | 0 | 0 | 0 | 0 | 0 |
| 0 | NA | NA | NA | NA | NA | Spertl | spermatid associated like | 0 | 0 | 0 | 0 | 0 | 0 | 0 | 0 | 0 | 0 | 0 |
| 0 | NA | NA | NA | NA | NA | Gm18885 | | 0 | 0 | 0 | 0 | 0 | 0 | 0 | 0 | 0 | 0 | 0 |
| 0 | NA | NA | NA | NA | NA | Gm19099 | | 0 | 0 | 0 | 0 | 0 | 0 | 0 | 0 | 0 | 0 | 0 |

| | | | | | | | | | | | | | | | | | | |
|---|---|---|---|---|---|---|---|---|---|---|---|---|---|---|---|---|---|---|
| 0 | NA | NA | NA | NA | NA | Gm15515 | | 0 | 0 | 0 | 0 | 0 | 0 | 0 | 0 | 0 | 0 | 0 |
| 0 | NA | NA | NA | NA | NA | Gm24043 | | 0 | 0 | 0 | 0 | 0 | 0 | 0 | 0 | 0 | 0 | 0 |
| 0 | NA | NA | NA | NA | NA | Gm25133 | | 0 | 0 | 0 | 0 | 0 | 0 | 0 | 0 | 0 | 0 | 0 |
| 0 | NA | NA | NA | NA | NA | Gm49006 | | 0 | 0 | 0 | 0 | 0 | 0 | 0 | 0 | 0 | 0 | 0 |
| 0 | NA | NA | NA | NA | NA | Gm18926 | | 0 | 0 | 0 | 0 | 0 | 0 | 0 | 0 | 0 | 0 | 0 |
| 0 | NA | NA | NA | NA | NA | Gm49007 | | 0 | 0 | 0 | 0 | 0 | 0 | 0 | 0 | 0 | 0 | 0 |
| 0 | NA | NA | NA | NA | NA | Gm24770 | | 0 | 0 | 0 | 0 | 0 | 0 | 0 | 0 | 0 | 0 | 0 |
| 0 | NA | NA | NA | NA | NA | Gm9287 | | 0 | 0 | 0 | 0 | 0 | 0 | 0 | 0 | 0 | 0 | 0 |
| 0 | NA | NA | NA | NA | NA | Gm33525 | | 0 | 0 | 0 | 0 | 0 | 0 | 0 | 0 | 0 | 0 | 0 |
| 0 | NA | NA | NA | NA | NA | Gm27441 | | 0 | 0 | 0 | 0 | 0 | 0 | 0 | 0 | 0 | 0 | 0 |
| 0 | NA | NA | NA | NA | NA | Gm23276 | | 0 | 0 | 0 | 0 | 0 | 0 | 0 | 0 | 0 | 0 | 0 |
| 0 | NA | NA | NA | NA | NA | Gm25831 | | 0 | 0 | 0 | 0 | 0 | 0 | 0 | 0 | 0 | 0 | 0 |
| 0 | NA | NA | NA | NA | NA | Gm16331 | | 0 | 0 | 0 | 0 | 0 | 0 | 0 | 0 | 0 | 0 | 0 |
| 0 | NA | NA | NA | NA | NA | Gm26006 | | 0 | 0 | 0 | 0 | 0 | 0 | 0 | 0 | 0 | 0 | 0 |
| 0 | NA | NA | NA | NA | NA | Gm46477 | | 0 | 0 | 0 | 0 | 0 | 0 | 0 | 0 | 0 | 0 | 0 |
| 0 | NA | NA | NA | NA | NA | Gm26139 | | 0 | 0 | 0 | 0 | 0 | 0 | 0 | 0 | 0 | 0 | 0 |
| 0 | NA | NA | NA | NA | NA | Gm27034 | | 0 | 0 | 0 | 0 | 0 | 0 | 0 | 0 | 0 | 0 | 0 |
| 0 | NA | NA | NA | NA | NA | Gm49018 | | 0 | 0 | 0 | 0 | 0 | 0 | 0 | 0 | 0 | 0 | 0 |
| 0 | NA | NA | NA | NA | NA | Gm18148 | | 0 | 0 | 0 | 0 | 0 | 0 | 0 | 0 | 0 | 0 | 0 |
| 0 | NA | NA | NA | NA | NA | Gm4412 | | 0 | 0 | 0 | 0 | 0 | 0 | 0 | 0 | 0 | 0 | 0 |
| 0 | NA | NA | NA | NA | NA | Gm22984 | | 0 | 0 | 0 | 0 | 0 | 0 | 0 | 0 | 0 | 0 | 0 |
| 0 | NA | NA | NA | NA | NA | Gm25130 | | 0 | 0 | 0 | 0 | 0 | 0 | 0 | 0 | 0 | 0 | 0 |
| 0 | NA | NA | NA | NA | NA | Gm49219 | | 0 | 0 | 0 | 0 | 0 | 0 | 0 | 0 | 0 | 0 | 0 |
| 0 | NA | NA | NA | NA | NA | Gm49220 | | 0 | 0 | 0 | 0 | 0 | 0 | 0 | 0 | 0 | 0 | 0 |
| 0 | NA | NA | NA | NA | NA | Gm41230 | | 0 | 0 | 0 | 0 | 0 | 0 | 0 | 0 | 0 | 0 | 0 |
| 0 | NA | NA | NA | NA | NA | Gm49225 | | 0 | 0 | 0 | 0 | 0 | 0 | 0 | 0 | 0 | 0 | 0 |
| 0 | NA | NA | NA | NA | NA | Gm9298 | | 0 | 0 | 0 | 0 | 0 | 0 | 0 | 0 | 0 | 0 | 0 |
| 0 | NA | NA | NA | NA | NA | Gm37093 | | 0 | 0 | 0 | 0 | 0 | 0 | 0 | 0 | 0 | 0 | 0 |
| 0 | NA | NA | NA | NA | NA | Gm22970 | | 0 | 0 | 0 | 0 | 0 | 0 | 0 | 0 | 0 | 0 | 0 |
| 0 | NA | NA | NA | NA | NA | Gm5958 | | 0 | 0 | 0 | 0 | 0 | 0 | 0 | 0 | 0 | 0 | 0 |
| 0 | NA | NA | NA | NA | NA | 4930517O19Rik | | 0 | 0 | 0 | 0 | 0 | 0 | 0 | 0 | 0 | 0 | 0 |
| 0 | NA | NA | NA | NA | NA | Gm16260 | | 0 | 0 | 0 | 0 | 0 | 0 | 0 | 0 | 0 | 0 | 0 |
| 0 | NA | NA | NA | NA | NA | Gm26367 | | 0 | 0 | 0 | 0 | 0 | 0 | 0 | 0 | 0 | 0 | 0 |
| 0 | NA | NA | NA | NA | NA | Gm9302 | | 0 | 0 | 0 | 0 | 0 | 0 | 0 | 0 | 0 | 0 | 0 |
| 0 | NA | NA | NA | NA | NA | Gm24345 | | 0 | 0 | 0 | 0 | 0 | 0 | 0 | 0 | 0 | 0 | 0 |
| 0 | NA | NA | NA | NA | NA | Gm49237 | | 0 | 0 | 0 | 0 | 0 | 0 | 0 | 0 | 0 | 0 | 0 |
| 0 | NA | NA | NA | NA | NA | Gm49238 | | 0 | 0 | 0 | 0 | 0 | 0 | 0 | 0 | 0 | 0 | 0 |
| 0 | NA | NA | NA | NA | NA | Gm22401 | | 0 | 0 | 0 | 0 | 0 | 0 | 0 | 0 | 0 | 0 | 0 |
| 0 | NA | NA | NA | NA | NA | Gm23538 | | 0 | 0 | 0 | 0 | 0 | 0 | 0 | 0 | 0 | 0 | 0 |
| 0 | NA | NA | NA | NA | NA | Gm49239 | | 0 | 0 | 0 | 0 | 0 | 0 | 0 | 0 | 0 | 0 | 0 |
| 0 | NA | NA | NA | NA | NA | Prr30 | proline rich 30 | 0 | 0 | 0 | 0 | 0 | 0 | 0 | 0 | 0 | 0 | 0 |
| 0 | NA | NA | NA | NA | NA | Gm26173 | | 0 | 0 | 0 | 0 | 0 | 0 | 0 | 0 | 0 | 0 | 0 |
| 0 | NA | NA | NA | NA | NA | Gm34417 | | 0 | 0 | 0 | 0 | 0 | 0 | 0 | 0 | 0 | 0 | 0 |
| 0 | NA | NA | NA | NA | NA | Gm18075 | | 0 | 0 | 0 | 0 | 0 | 0 | 0 | 0 | 0 | 0 | 0 |
| 0 | NA | NA | NA | NA | NA | Gm49269 | | 0 | 0 | 0 | 0 | 0 | 0 | 0 | 0 | 0 | 0 | 0 |
| 0 | NA | NA | NA | NA | NA | Gm22347 | | 0 | 0 | 0 | 0 | 0 | 0 | 0 | 0 | 0 | 0 | 0 |
| 0 | NA | NA | NA | NA | NA | Gm49292 | | 0 | 0 | 0 | 0 | 0 | 0 | 0 | 0 | 0 | 0 | 0 |
| 0 | NA | NA | NA | NA | NA | Gm23764 | | 0 | 0 | 0 | 0 | 0 | 0 | 0 | 0 | 0 | 0 | 0 |
| 0 | NA | NA | NA | NA | NA | Mir5130 | | 0 | 0 | 0 | 0 | 0 | 0 | 0 | 0 | 0 | 0 | 0 |
| 0 | NA | NA | NA | NA | NA | 4933432I03Rik | | 0 | 0 | 0 | 0 | 0 | 0 | 0 | 0 | 0 | 0 | 0 |
| 0 | NA | NA | NA | NA | NA | Gm37910 | | 0 | 0 | 0 | 0 | 0 | 0 | 0 | 0 | 0 | 0 | 0 |
| 0 | NA | NA | NA | NA | NA | 4930432J09Rik | | 0 | 0 | 0 | 0 | 0 | 0 | 0 | 0 | 0 | 0 | 0 |
| 0 | NA | NA | NA | NA | NA | D130079A08Rik | | 0 | 0 | 0 | 0 | 0 | 0 | 0 | 0 | 0 | 0 | 0 |
| 0 | NA | NA | NA | NA | NA | Gm22021 | | 0 | 0 | 0 | 0 | 0 | 0 | 0 | 0 | 0 | 0 | 0 |
| 0 | NA | NA | NA | NA | NA | D130009I18Rik | | 0 | 0 | 0 | 0 | 0 | 0 | 0 | 0 | 0 | 0 | 0 |
| 0 | NA | NA | NA | NA | NA | Gm25724 | | 0 | 0 | 0 | 0 | 0 | 0 | 0 | 0 | 0 | 0 | 0 |
| 0 | NA | NA | NA | NA | NA | Gm24280 | | 0 | 0 | 0 | 0 | 0 | 0 | 0 | 0 | 0 | 0 | 0 |
| 0 | NA | NA | NA | NA | NA | Gm5671 | | 0 | 0 | 0 | 0 | 0 | 0 | 0 | 0 | 0 | 0 | 0 |
| 0 | NA | NA | NA | NA | NA | Gm48937 | | 0 | 0 | 0 | 0 | 0 | 0 | 0 | 0 | 0 | 0 | 0 |
| 0 | NA | NA | NA | NA | NA | Gm22290 | | 0 | 0 | 0 | 0 | 0 | 0 | 0 | 0 | 0 | 0 | 0 |

| | | | | | | | | | | | | | | | | | | |
|---|---|---|---|---|---|---|---|---|---|---|---|---|---|---|---|---|---|---|
| 0 | NA | NA | NA | NA | NA | Gm24207 | | 0 | 0 | 0 | 0 | 0 | 0 | 0 | 0 | 0 | 0 | 0 |
| 0 | NA | NA | NA | NA | NA | Gm16259 | | 0 | 0 | 0 | 0 | 0 | 0 | 0 | 0 | 0 | 0 | 0 |
| 0 | NA | NA | NA | NA | NA | 4930449E01Rik | | 0 | 0 | 0 | 0 | 0 | 0 | 0 | 0 | 0 | 0 | 0 |
| 0 | NA | NA | NA | NA | NA | 9330188P03Rik | | 0 | 0 | 0 | 0 | 0 | 0 | 0 | 0 | 0 | 0 | 0 |
| 0 | NA | NA | NA | NA | NA | Gm6145 | | 0 | 0 | 0 | 0 | 0 | 0 | 0 | 0 | 0 | 0 | 0 |
| 0 | NA | NA | NA | NA | NA | Mir6390 | | 0 | 0 | 0 | 0 | 0 | 0 | 0 | 0 | 0 | 0 | 0 |
| 0 | NA | NA | NA | NA | NA | Gm22406 | | 0 | 0 | 0 | 0 | 0 | 0 | 0 | 0 | 0 | 0 | 0 |
| 0 | NA | NA | NA | NA | NA | Gm48970 | | 0 | 0 | 0 | 0 | 0 | 0 | 0 | 0 | 0 | 0 | 0 |
| 0 | NA | NA | NA | NA | NA | Frg2f4 | | 0 | 0 | 0 | 0 | 0 | 0 | 0 | 0 | 0 | 0 | 0 |
| 0 | NA | NA | NA | NA | NA | Trim52 | tripartite motif-containing 52 | 0 | 0 | 0 | 0 | 0 | 0 | 0 | 0 | 0 | 0 | 0 |
| 0 | NA | NA | NA | NA | NA | Gm48971 | | 0 | 0 | 0 | 0 | 0 | 0 | 0 | 0 | 0 | 0 | 0 |
| 0 | NA | NA | NA | NA | NA | Gm48972 | | 0 | 0 | 0 | 0 | 0 | 0 | 0 | 0 | 0 | 0 | 0 |
| 0 | NA | NA | NA | NA | NA | Gm23459 | | 0 | 0 | 0 | 0 | 0 | 0 | 0 | 0 | 0 | 0 | 0 |
| 0 | NA | NA | NA | NA | NA | Gm36107 | | 0 | 0 | 0 | 0 | 0 | 0 | 0 | 0 | 0 | 0 | 0 |
| 0 | NA | NA | NA | NA | NA | 1700128A07Rik | | 0 | 0 | 0 | 0 | 0 | 0 | 0 | 0 | 0 | 0 | 0 |
| 0 | NA | NA | NA | NA | NA | Gm24634 | | 0 | 0 | 0 | 0 | 0 | 0 | 0 | 0 | 0 | 0 | 0 |
| 0 | NA | NA | NA | NA | NA | Gm48973 | | 0 | 0 | 0 | 0 | 0 | 0 | 0 | 0 | 0 | 0 | 0 |
| 0 | NA | NA | NA | NA | NA | Gm17366 | | 0 | 0 | 0 | 0 | 0 | 0 | 0 | 0 | 0 | 0 | 0 |
| 0 | NA | NA | NA | NA | NA | Gm23437 | | 0 | 0 | 0 | 0 | 0 | 0 | 0 | 0 | 0 | 0 | 0 |
| 0 | NA | NA | NA | NA | NA | Gm9334 | | 0 | 0 | 0 | 0 | 0 | 0 | 0 | 0 | 0 | 0 | 0 |
| 0 | NA | NA | NA | NA | NA | Gm19146 | | 0 | 0 | 0 | 0 | 0 | 0 | 0 | 0 | 0 | 0 | 0 |
| 0 | NA | NA | NA | NA | NA | Gm9335 | | 0 | 0 | 0 | 0 | 0 | 0 | 0 | 0 | 0 | 0 | 0 |
| 0 | NA | NA | NA | NA | NA | Gm48976 | | 0 | 0 | 0 | 0 | 0 | 0 | 0 | 0 | 0 | 0 | 0 |
| 0 | NA | NA | NA | NA | NA | Gm48977 | | 0 | 0 | 0 | 0 | 0 | 0 | 0 | 0 | 0 | 0 | 0 |
| 0 | NA | NA | NA | NA | NA | Gm24818 | | 0 | 0 | 0 | 0 | 0 | 0 | 0 | 0 | 0 | 0 | 0 |
| 0 | NA | NA | NA | NA | NA | Gm24229 | | 0 | 0 | 0 | 0 | 0 | 0 | 0 | 0 | 0 | 0 | 0 |
| 0 | NA | NA | NA | NA | NA | Gm18977 | | 0 | 0 | 0 | 0 | 0 | 0 | 0 | 0 | 0 | 0 | 0 |
| 0 | NA | NA | NA | NA | NA | Gm7313 | | 0 | 0 | 0 | 0 | 0 | 0 | 0 | 0 | 0 | 0 | 0 |
| 0 | NA | NA | NA | NA | NA | Gm25901 | | 0 | 0 | 0 | 0 | 0 | 0 | 0 | 0 | 0 | 0 | 0 |
| 0 | NA | NA | NA | NA | NA | Gm24369 | | 0 | 0 | 0 | 0 | 0 | 0 | 0 | 0 | 0 | 0 | 0 |
| 0 | NA | NA | NA | NA | NA | n-R5s50 | | 0 | 0 | 0 | 0 | 0 | 0 | 0 | 0 | 0 | 0 | 0 |
| 0 | NA | NA | NA | NA | NA | Gm23242 | | 0 | 0 | 0 | 0 | 0 | 0 | 0 | 0 | 0 | 0 | 0 |
| 0 | NA | NA | NA | NA | NA | Gm48980 | | 0 | 0 | 0 | 0 | 0 | 0 | 0 | 0 | 0 | 0 | 0 |
| 0 | NA | NA | NA | NA | NA | Gm17932 | | 0 | 0 | 0 | 0 | 0 | 0 | 0 | 0 | 0 | 0 | 0 |
| 0 | NA | NA | NA | NA | NA | Gm25670 | | 0 | 0 | 0 | 0 | 0 | 0 | 0 | 0 | 0 | 0 | 0 |
| 0 | NA | NA | NA | NA | NA | Gm26255 | | 0 | 0 | 0 | 0 | 0 | 0 | 0 | 0 | 0 | 0 | 0 |
| 0 | NA | NA | NA | NA | NA | Gm48987 | | 0 | 0 | 0 | 0 | 0 | 0 | 0 | 0 | 0 | 0 | 0 |
| 0 | NA | NA | NA | NA | NA | Gm9348 | | 0 | 0 | 0 | 0 | 0 | 0 | 0 | 0 | 0 | 0 | 0 |
| 0 | NA | NA | NA | NA | NA | Gm22802 | | 0 | 0 | 0 | 0 | 0 | 0 | 0 | 0 | 0 | 0 | 0 |
| 0 | NA | NA | NA | NA | NA | Gm48988 | | 0 | 0 | 0 | 0 | 0 | 0 | 0 | 0 | 0 | 0 | 0 |
| 0 | NA | NA | NA | NA | NA | Gm48992 | | 0 | 0 | 0 | 0 | 0 | 0 | 0 | 0 | 0 | 0 | 0 |
| 0 | NA | NA | NA | NA | NA | Gm48993 | | 0 | 0 | 0 | 0 | 0 | 0 | 0 | 0 | 0 | 0 | 0 |
| 0 | NA | NA | NA | NA | NA | Gm18580 | | 0 | 0 | 0 | 0 | 0 | 0 | 0 | 0 | 0 | 0 | 0 |
| 0 | NA | NA | NA | NA | NA | | | 0 | 0 | 0 | 0 | 0 | 0 | 0 | 0 | 0 | 0 | 0 |
| 0 | NA | NA | NA | NA | NA | | | 0 | 0 | 0 | 0 | 0 | 0 | 0 | 0 | 0 | 0 | 0 |
| 0 | NA | NA | NA | NA | NA | Gm48999 | | 0 | 0 | 0 | 0 | 0 | 0 | 0 | 0 | 0 | 0 | 0 |
| 0 | NA | NA | NA | NA | NA | Gm22764 | | 0 | 0 | 0 | 0 | 0 | 0 | 0 | 0 | 0 | 0 | 0 |
| 0 | NA | NA | NA | NA | NA | Gm49000 | | 0 | 0 | 0 | 0 | 0 | 0 | 0 | 0 | 0 | 0 | 0 |
| 0 | NA | NA | NA | NA | NA | Gm20127 | | 0 | 0 | 0 | 0 | 0 | 0 | 0 | 0 | 0 | 0 | 0 |
| 0 | NA | NA | NA | NA | NA | Gm4822 | | 0 | 0 | 0 | 0 | 0 | 0 | 0 | 0 | 0 | 0 | 0 |
| 0 | NA | NA | NA | NA | NA | Gm19223 | | 0 | 0 | 0 | 0 | 0 | 0 | 0 | 0 | 0 | 0 | 0 |
| 0 | NA | NA | NA | NA | NA | Gm4486 | | 0 | 0 | 0 | 0 | 0 | 0 | 0 | 0 | 0 | 0 | 0 |
| 0 | NA | NA | NA | NA | NA | Gm49005 | | 0 | 0 | 0 | 0 | 0 | 0 | 0 | 0 | 0 | 0 | 0 |
| 0 | NA | NA | NA | NA | NA | Gm4487 | | 0 | 0 | 0 | 0 | 0 | 0 | 0 | 0 | 0 | 0 | 0 |
| 0 | NA | NA | NA | NA | NA | Gm18369 | | 0 | 0 | 0 | 0 | 0 | 0 | 0 | 0 | 0 | 0 | 0 |
| 0 | NA | NA | NA | NA | NA | Gm19829 | | 0 | 0 | 0 | 0 | 0 | 0 | 0 | 0 | 0 | 0 | 0 |
| 0 | NA | NA | NA | NA | NA | Gm37495 | | 0 | 0 | 0 | 0 | 0 | 0 | 0 | 0 | 0 | 0 | 0 |
| 0 | NA | NA | NA | NA | NA | 4930524C18Rik | | 0 | 0 | 0 | 0 | 0 | 0 | 0 | 0 | 0 | 0 | 0 |
| 0 | NA | NA | NA | NA | NA | Gm49010 | | 0 | 0 | 0 | 0 | 0 | 0 | 0 | 0 | 0 | 0 | 0 |
| 0 | NA | NA | NA | NA | NA | 1700100I10Rik | | 0 | 0 | 0 | 0 | 0 | 0 | 0 | 0 | 0 | 0 | 0 |

| | | | | | | | | | | | | | | | | | | |
|---|---|---|---|---|---|---|---|---|---|---|---|---|---|---|---|---|---|---|
| 0 | NA | NA | NA | NA | NA | Gm31072 | | 0 | 0 | 0 | 0 | 0 | 0 | 0 | 0 | 0 | 0 | 0 |
| 0 | NA | NA | NA | NA | NA | Gm24073 | | 0 | 0 | 0 | 0 | 0 | 0 | 0 | 0 | 0 | 0 | 0 |
| 0 | NA | NA | NA | NA | NA | Mir17 | | 0 | 0 | 0 | 0 | 0 | 0 | 0 | 0 | 0 | 0 | 0 |
| 0 | NA | NA | NA | NA | NA | Mir18 | | 0 | 0 | 0 | 0 | 0 | 0 | 0 | 0 | 0 | 0 | 0 |
| 0 | NA | NA | NA | NA | NA | Mir19a | | 0 | 0 | 0 | 0 | 0 | 0 | 0 | 0 | 0 | 0 | 0 |
| 0 | NA | NA | NA | NA | NA | Mir20a | | 0 | 0 | 0 | 0 | 0 | 0 | 0 | 0 | 0 | 0 | 0 |
| 0 | NA | NA | NA | NA | NA | Mir19b-1 | | 0 | 0 | 0 | 0 | 0 | 0 | 0 | 0 | 0 | 0 | 0 |
| 0 | NA | NA | NA | NA | NA | Mir92-1 | | 0 | 0 | 0 | 0 | 0 | 0 | 0 | 0 | 0 | 0 | 0 |
| 0 | NA | NA | NA | NA | NA | 4930505G20Rik | | 0 | 0 | 0 | 0 | 0 | 0 | 0 | 0 | 0 | 0 | 0 |
| 0 | NA | NA | NA | NA | NA | Gm20713 | | 0 | 0 | 0 | 0 | 0 | 0 | 0 | 0 | 0 | 0 | 0 |
| 0 | NA | NA | NA | NA | NA | Gm38045 | | 0 | 0 | 0 | 0 | 0 | 0 | 0 | 0 | 0 | 0 | 0 |
| 0 | NA | NA | NA | NA | NA | Mir6239 | | 0 | 0 | 0 | 0 | 0 | 0 | 0 | 0 | 0 | 0 | 0 |
| 0 | NA | NA | NA | NA | NA | 4930435M08Rik | | 0 | 0 | 0 | 0 | 0 | 0 | 0 | 0 | 0 | 0 | 0 |
| 0 | NA | NA | NA | NA | NA | 4933431J24Rik | | 0 | 0 | 0 | 0 | 0 | 0 | 0 | 0 | 0 | 0 | 0 |
| 0 | NA | NA | NA | NA | NA | Sox21 | SRY (sex determining region Y)-box 21 | 0 | 0 | 0 | 0 | 0 | 0 | 0 | 0 | 0 | 0 | 0 |
| 0 | NA | NA | NA | NA | NA | Gm4675 | | 0 | 0 | 0 | 0 | 0 | 0 | 0 | 0 | 0 | 0 | 0 |
| 0 | NA | NA | NA | NA | NA | Mir6241 | | 0 | 0 | 0 | 0 | 0 | 0 | 0 | 0 | 0 | 0 | 0 |
| 0 | NA | NA | NA | NA | NA | Gm9376 | predicted gene 9376 | 0 | 0 | 0 | 0 | 0 | 0 | 0 | 0 | 0 | 0 | 0 |
| 0 | NA | NA | NA | NA | NA | Gm23669 | | 0 | 0 | 0 | 0 | 0 | 0 | 0 | 0 | 0 | 0 | 0 |
| 0 | NA | NA | NA | NA | NA | Gm32093 | | 0 | 0 | 0 | 0 | 0 | 0 | 0 | 0 | 0 | 0 | 0 |
| 0 | NA | NA | NA | NA | NA | Gm26791 | | 0 | 0 | 0 | 0 | 0 | 0 | 0 | 0 | 0 | 0 | 0 |
| 0 | NA | NA | NA | NA | NA | 1700044C05Rik | | 0 | 0 | 0 | 0 | 0 | 0 | 0 | 0 | 0 | 0 | 0 |
| 0 | NA | NA | NA | NA | NA | Gm5672 | | 0 | 0 | 0 | 0 | 0 | 0 | 0 | 0 | 0 | 0 | 0 |
| 0 | NA | NA | NA | NA | NA | Gm49033 | | 0 | 0 | 0 | 0 | 0 | 0 | 0 | 0 | 0 | 0 | 0 |
| 0 | NA | NA | NA | NA | NA | Gm23302 | | 0 | 0 | 0 | 0 | 0 | 0 | 0 | 0 | 0 | 0 | 0 |
| 0 | NA | NA | NA | NA | NA | Mir6391 | | 0 | 0 | 0 | 0 | 0 | 0 | 0 | 0 | 0 | 0 | 0 |
| 0 | NA | NA | NA | NA | NA | Gm27198 | | 0 | 0 | 0 | 0 | 0 | 0 | 0 | 0 | 0 | 0 | 0 |
| 0 | NA | NA | NA | NA | NA | Gm22379 | | 0 | 0 | 0 | 0 | 0 | 0 | 0 | 0 | 0 | 0 | 0 |
| 0 | NA | NA | NA | NA | NA | n-R5s51 | | 0 | 0 | 0 | 0 | 0 | 0 | 0 | 0 | 0 | 0 | 0 |
| 0 | NA | NA | NA | NA | NA | Gm49179 | | 0 | 0 | 0 | 0 | 0 | 0 | 0 | 0 | 0 | 0 | 0 |
| 0 | NA | NA | NA | NA | NA | 4930404K13Rik | | 0 | 0 | 0 | 0 | 0 | 0 | 0 | 0 | 0 | 0 | 0 |
| 0 | NA | NA | NA | NA | NA | Oxgr1 | oxoglutarate (alpha-ketoglutarate) recept | 0 | 0 | 0 | 0 | 0 | 0 | 0 | 0 | 0 | 0 | 0 |
| 0 | NA | NA | NA | NA | NA | Gm20167 | | 0 | 0 | 0 | 0 | 0 | 0 | 0 | 0 | 0 | 0 | 0 |
| 0 | NA | NA | NA | NA | NA | Gm48997 | | 0 | 0 | 0 | 0 | 0 | 0 | 0 | 0 | 0 | 0 | 0 |
| 0 | NA | NA | NA | NA | NA | Gm49008 | | 0 | 0 | 0 | 0 | 0 | 0 | 0 | 0 | 0 | 0 | 0 |
| 0 | NA | NA | NA | NA | NA | Gm25781 | | 0 | 0 | 0 | 0 | 0 | 0 | 0 | 0 | 0 | 0 | 0 |
| 0 | NA | NA | NA | NA | NA | Gm49031 | | 0 | 0 | 0 | 0 | 0 | 0 | 0 | 0 | 0 | 0 | 0 |
| 0 | NA | NA | NA | NA | NA | Gm33299 | | 0 | 0 | 0 | 0 | 0 | 0 | 0 | 0 | 0 | 0 | 0 |
| 0 | NA | NA | NA | NA | NA | Gm49035 | | 0 | 0 | 0 | 0 | 0 | 0 | 0 | 0 | 0 | 0 | 0 |
| 0 | NA | NA | NA | NA | NA | Gm4529 | | 0 | 0 | 0 | 0 | 0 | 0 | 0 | 0 | 0 | 0 | 0 |
| 0 | NA | NA | NA | NA | NA | Gm9395 | | 0 | 0 | 0 | 0 | 0 | 0 | 0 | 0 | 0 | 0 | 0 |
| 0 | NA | NA | NA | NA | NA | 1700108J01Rik | | 0 | 0 | 0 | 0 | 0 | 0 | 0 | 0 | 0 | 0 | 0 |
| 0 | NA | NA | NA | NA | NA | Gm5089 | | 0 | 0 | 0 | 0 | 0 | 0 | 0 | 0 | 0 | 0 | 0 |
| 0 | NA | NA | NA | NA | NA | Gm25464 | | 0 | 0 | 0 | 0 | 0 | 0 | 0 | 0 | 0 | 0 | 0 |
| 0 | NA | NA | NA | NA | NA | Gm5089 | | 0 | 0 | 0 | 0 | 0 | 0 | 0 | 0 | 0 | 0 | 0 |
| 0 | NA | NA | NA | NA | NA | 2610035F20Rik | | 0 | 0 | 0 | 0 | 0 | 0 | 0 | 0 | 0 | 0 | 0 |
| 0 | NA | NA | NA | NA | NA | Zic2 | zinc finger protein of the cerebellum 2 | 0 | 0 | 0 | 0 | 0 | 0 | 0 | 0 | 0 | 0 | 0 |
| 0 | NA | NA | NA | NA | NA | Gm10837 | | 0 | 0 | 0 | 0 | 0 | 0 | 0 | 0 | 0 | 0 | 0 |
| 0 | NA | NA | NA | NA | NA | Gm9399 | | 0 | 0 | 0 | 0 | 0 | 0 | 0 | 0 | 0 | 0 | 0 |
| 0 | NA | NA | NA | NA | NA | Gm18143 | | 0 | 0 | 0 | 0 | 0 | 0 | 0 | 0 | 0 | 0 | 0 |
| 0 | NA | NA | NA | NA | NA | Gm49259 | | 0 | 0 | 0 | 0 | 0 | 0 | 0 | 0 | 0 | 0 | 0 |
| 0 | NA | NA | NA | NA | NA | Gm49294 | | 0 | 0 | 0 | 0 | 0 | 0 | 0 | 0 | 0 | 0 | 0 |
| 0 | NA | NA | NA | NA | NA | Gm25272 | | 0 | 0 | 0 | 0 | 0 | 0 | 0 | 0 | 0 | 0 | 0 |
| 0 | NA | NA | NA | NA | NA | 1700024B18Rik | | 0 | 0 | 0 | 0 | 0 | 0 | 0 | 0 | 0 | 0 | 0 |
| 0 | NA | NA | NA | NA | NA | Gm49070 | | 0 | 0 | 0 | 0 | 0 | 0 | 0 | 0 | 0 | 0 | 0 |
| 0 | NA | NA | NA | NA | NA | Gm49071 | | 0 | 0 | 0 | 0 | 0 | 0 | 0 | 0 | 0 | 0 | 0 |
| 0 | NA | NA | NA | NA | NA | Olfr277-ps1 | | 0 | 0 | 0 | 0 | 0 | 0 | 0 | 0 | 0 | 0 | 0 |
| 0 | NA | NA | NA | NA | NA | Gm49069 | | 0 | 0 | 0 | 0 | 0 | 0 | 0 | 0 | 0 | 0 | 0 |
| 0 | NA | NA | NA | NA | NA | 4930404O17Rik | | 0 | 0 | 0 | 0 | 0 | 0 | 0 | 0 | 0 | 0 | 0 |
| 0 | NA | NA | NA | NA | NA | Gm9614 | | 0 | 0 | 0 | 0 | 0 | 0 | 0 | 0 | 0 | 0 | 0 |

| | | | | | | | | | | | | | | | | | | |
|---|---|---|---|---|---|---|---|---|---|---|---|---|---|---|---|---|---|---|
| 0 | NA | NA | NA | NA | NA | Gm49222 | | 0 | 0 | 0 | 0 | 0 | 0 | 0 | 0 | 0 | 0 | 0 |
| 0 | NA | NA | NA | NA | NA | Gm49231 | | 0 | 0 | 0 | 0 | 0 | 0 | 0 | 0 | 0 | 0 | 0 |
| 0 | NA | NA | NA | NA | NA | Gm49232 | | 0 | 0 | 0 | 0 | 0 | 0 | 0 | 0 | 0 | 0 | 0 |
| 0 | NA | NA | NA | NA | NA | Gm22031 | | 0 | 0 | 0 | 0 | 0 | 0 | 0 | 0 | 0 | 0 | 0 |
| 0 | NA | NA | NA | NA | NA | Gm23228 | | 0 | 0 | 0 | 0 | 0 | 0 | 0 | 0 | 0 | 0 | 0 |
| 0 | NA | NA | NA | NA | NA | Gm15632 | | 0 | 0 | 0 | 0 | 0 | 0 | 0 | 0 | 0 | 0 | 0 |
| 0 | NA | NA | NA | NA | NA | Gm49073 | | 0 | 0 | 0 | 0 | 0 | 0 | 0 | 0 | 0 | 0 | 0 |
| 0 | NA | NA | NA | NA | NA | Gm49074 | | 0 | 0 | 0 | 0 | 0 | 0 | 0 | 0 | 0 | 0 | 0 |
| 0 | NA | NA | NA | NA | NA | Mroh2b | maestro heat-like repeat family member 2 | 0 | 0 | 0 | 0 | 0 | 0 | 0 | 0 | 0 | 0 | 0 |
| 0 | NA | NA | NA | NA | NA | Gm25652 | | 0 | 0 | 0 | 0 | 0 | 0 | 0 | 0 | 0 | 0 | 0 |
| 0 | NA | NA | NA | NA | NA | Gm16077 | | 0 | 0 | 0 | 0 | 0 | 0 | 0 | 0 | 0 | 0 | 0 |
| 0 | NA | NA | NA | NA | NA | Gm49093 | | 0 | 0 | 0 | 0 | 0 | 0 | 0 | 0 | 0 | 0 | 0 |
| 0 | NA | NA | NA | NA | NA | Gm2150 | | 0 | 0 | 0 | 0 | 0 | 0 | 0 | 0 | 0 | 0 | 0 |
| 0 | NA | NA | NA | NA | NA | Gm18715 | | 0 | 0 | 0 | 0 | 0 | 0 | 0 | 0 | 0 | 0 | 0 |
| 0 | NA | NA | NA | NA | NA | Gm8043 | | 0 | 0 | 0 | 0 | 0 | 0 | 0 | 0 | 0 | 0 | 0 |
| 0 | NA | NA | NA | NA | NA | Gm49095 | | 0 | 0 | 0 | 0 | 0 | 0 | 0 | 0 | 0 | 0 | 0 |
| 0 | NA | NA | NA | NA | NA | Gm8047 | | 0 | 0 | 0 | 0 | 0 | 0 | 0 | 0 | 0 | 0 | 0 |
| 0 | NA | NA | NA | NA | NA | Gm23139 | | 0 | 0 | 0 | 0 | 0 | 0 | 0 | 0 | 0 | 0 | 0 |
| 0 | NA | NA | NA | NA | NA | Gm16311 | | 0 | 0 | 0 | 0 | 0 | 0 | 0 | 0 | 0 | 0 | 0 |
| 0 | NA | NA | NA | NA | NA | Gm18888 | | 0 | 0 | 0 | 0 | 0 | 0 | 0 | 0 | 0 | 0 | 0 |
| 0 | NA | NA | NA | NA | NA | C9 | complement component 9 | 0 | 0 | 0 | 0 | 0 | 0 | 0 | 0 | 0 | 0 | 0 |
| 0 | NA | NA | NA | NA | NA | Gm23218 | | 0 | 0 | 0 | 0 | 0 | 0 | 0 | 0 | 0 | 0 | 0 |
| 0 | NA | NA | NA | NA | NA | Gm38282 | | 0 | 0 | 0 | 0 | 0 | 0 | 0 | 0 | 0 | 0 | 0 |
| 0 | NA | NA | NA | NA | NA | Gm16030 | | 0 | 0 | 0 | 0 | 0 | 0 | 0 | 0 | 0 | 0 | 0 |
| 0 | NA | NA | NA | NA | NA | Gm49206 | | 0 | 0 | 0 | 0 | 0 | 0 | 0 | 0 | 0 | 0 | 0 |
| 0 | NA | NA | NA | NA | NA | Gm37743 | | 0 | 0 | 0 | 0 | 0 | 0 | 0 | 0 | 0 | 0 | 0 |
| 0 | NA | NA | NA | NA | NA | Gm19227 | | 0 | 0 | 0 | 0 | 0 | 0 | 0 | 0 | 0 | 0 | 0 |
| 0 | NA | NA | NA | NA | NA | Gm27529 | | 0 | 0 | 0 | 0 | 0 | 0 | 0 | 0 | 0 | 0 | 0 |
| 0 | NA | NA | NA | NA | NA | Gm49247 | | 0 | 0 | 0 | 0 | 0 | 0 | 0 | 0 | 0 | 0 | 0 |
| 0 | NA | NA | NA | NA | NA | Gm49249 | | 0 | 0 | 0 | 0 | 0 | 0 | 0 | 0 | 0 | 0 | 0 |
| 0 | NA | NA | NA | NA | NA | Gm2310 | | 0 | 0 | 0 | 0 | 0 | 0 | 0 | 0 | 0 | 0 | 0 |
| 0 | NA | NA | NA | NA | NA | Gm2348 | | 0 | 0 | 0 | 0 | 0 | 0 | 0 | 0 | 0 | 0 | 0 |
| 0 | NA | NA | NA | NA | NA | Gm49250 | | 0 | 0 | 0 | 0 | 0 | 0 | 0 | 0 | 0 | 0 | 0 |
| 0 | NA | NA | NA | NA | NA | Gm37310 | | 0 | 0 | 0 | 0 | 0 | 0 | 0 | 0 | 0 | 0 | 0 |
| 0 | NA | NA | NA | NA | NA | Gm8139 | | 0 | 0 | 0 | 0 | 0 | 0 | 0 | 0 | 0 | 0 | 0 |
| 0 | NA | NA | NA | NA | NA | Ugt3a1 | UDP glycosyltransferases 3 family, polype | 0 | 0 | 0 | 0 | 0 | 0 | 0 | 0 | 0 | 0 | 0 |
| 0 | NA | NA | NA | NA | NA | Gm5219 | | 0 | 0 | 0 | 0 | 0 | 0 | 0 | 0 | 0 | 0 | 0 |
| 0 | NA | NA | NA | NA | NA | Gm26350 | | 0 | 0 | 0 | 0 | 0 | 0 | 0 | 0 | 0 | 0 | 0 |
| 0 | NA | NA | NA | NA | NA | Gm21973 | | 0 | 0 | 0 | 0 | 0 | 0 | 0 | 0 | 0 | 0 | 0 |
| 0 | NA | NA | NA | NA | NA | Gm17044 | | 0 | 0 | 0 | 0 | 0 | 0 | 0 | 0 | 0 | 0 | 0 |
| 0 | NA | NA | NA | NA | NA | Gm10389 | | 0 | 0 | 0 | 0 | 0 | 0 | 0 | 0 | 0 | 0 | 0 |
| 0 | NA | NA | NA | NA | NA | Gm19276 | | 0 | 0 | 0 | 0 | 0 | 0 | 0 | 0 | 0 | 0 | 0 |
| 0 | NA | NA | NA | NA | NA | Gm29742 | | 0 | 0 | 0 | 0 | 0 | 0 | 0 | 0 | 0 | 0 | 0 |
| 0 | NA | NA | NA | NA | NA | Gm49068 | | 0 | 0 | 0 | 0 | 0 | 0 | 0 | 0 | 0 | 0 | 0 |
| 0 | NA | NA | NA | NA | NA | Slc45a2 | solute carrier family 45, member 2 | 0 | 0 | 0 | 0 | 0 | 0 | 0 | 0 | 0 | 0 | 0 |
| 0 | NA | NA | NA | NA | NA | Gm24844 | | 0 | 0 | 0 | 0 | 0 | 0 | 0 | 0 | 0 | 0 | 0 |
| 0 | NA | NA | NA | NA | NA | Gm49106 | | 0 | 0 | 0 | 0 | 0 | 0 | 0 | 0 | 0 | 0 | 0 |
| 0 | NA | NA | NA | NA | NA | Gm49107 | | 0 | 0 | 0 | 0 | 0 | 0 | 0 | 0 | 0 | 0 | 0 |
| 0 | NA | NA | NA | NA | NA | Gm41271 | | 0 | 0 | 0 | 0 | 0 | 0 | 0 | 0 | 0 | 0 | 0 |
| 0 | NA | NA | NA | NA | NA | 1700047G03Rik | | 0 | 0 | 0 | 0 | 0 | 0 | 0 | 0 | 0 | 0 | 0 |
| 0 | NA | NA | NA | NA | NA | Gm24074 | | 0 | 0 | 0 | 0 | 0 | 0 | 0 | 0 | 0 | 0 | 0 |
| 0 | NA | NA | NA | NA | NA | Gm2559 | | 0 | 0 | 0 | 0 | 0 | 0 | 0 | 0 | 0 | 0 | 0 |
| 0 | NA | NA | NA | NA | NA | Gm25713 | | 0 | 0 | 0 | 0 | 0 | 0 | 0 | 0 | 0 | 0 | 0 |
| 0 | NA | NA | NA | NA | NA | Gm49240 | | 0 | 0 | 0 | 0 | 0 | 0 | 0 | 0 | 0 | 0 | 0 |
| 0 | NA | NA | NA | NA | NA | Gm2581 | | 0 | 0 | 0 | 0 | 0 | 0 | 0 | 0 | 0 | 0 | 0 |
| 0 | NA | NA | NA | NA | NA | Gm46497 | | 0 | 0 | 0 | 0 | 0 | 0 | 0 | 0 | 0 | 0 | 0 |
| 0 | NA | NA | NA | NA | NA | Gm24135 | | 0 | 0 | 0 | 0 | 0 | 0 | 0 | 0 | 0 | 0 | 0 |
| 0 | NA | NA | NA | NA | NA | F830212C03Rik | | 0 | 0 | 0 | 0 | 0 | 0 | 0 | 0 | 0 | 0 | 0 |
| 0 | NA | NA | NA | NA | NA | Gm24302 | | 0 | 0 | 0 | 0 | 0 | 0 | 0 | 0 | 0 | 0 | 0 |
| 0 | NA | NA | NA | NA | NA | Gm49276 | | 0 | 0 | 0 | 0 | 0 | 0 | 0 | 0 | 0 | 0 | 0 |

| | | | | | | | | | | | | | | | | | |
|---|---|---|---|---|---|---|---|---|---|---|---|---|---|---|---|---|---|
| 0 | NA | NA | NA | NA | NA | Gm17756 | 0 | 0 | 0 | 0 | 0 | 0 | 0 | 0 | 0 | 0 | 0 |
| 0 | NA | NA | NA | NA | NA | Gm49277 | 0 | 0 | 0 | 0 | 0 | 0 | 0 | 0 | 0 | 0 | 0 |
| 0 | NA | NA | NA | NA | NA | Gm47288 | 0 | 0 | 0 | 0 | 0 | 0 | 0 | 0 | 0 | 0 | 0 |
| 0 | NA | NA | NA | NA | NA | Gm49100 | 0 | 0 | 0 | 0 | 0 | 0 | 0 | 0 | 0 | 0 | 0 |
| 0 | NA | NA | NA | NA | NA | Gm24664 | 0 | 0 | 0 | 0 | 0 | 0 | 0 | 0 | 0 | 0 | 0 |
| 0 | NA | NA | NA | NA | NA | Gm49112 | 0 | 0 | 0 | 0 | 0 | 0 | 0 | 0 | 0 | 0 | 0 |
| 0 | NA | NA | NA | NA | NA | Gm18714 | 0 | 0 | 0 | 0 | 0 | 0 | 0 | 0 | 0 | 0 | 0 |
| 0 | NA | NA | NA | NA | NA | Gm49113 | 0 | 0 | 0 | 0 | 0 | 0 | 0 | 0 | 0 | 0 | 0 |
| 0 | NA | NA | NA | NA | NA | Gm49114 | 0 | 0 | 0 | 0 | 0 | 0 | 0 | 0 | 0 | 0 | 0 |
| 0 | NA | NA | NA | NA | NA | Gm49115 | 0 | 0 | 0 | 0 | 0 | 0 | 0 | 0 | 0 | 0 | 0 |
| 0 | NA | NA | NA | NA | NA | Gm49116 | 0 | 0 | 0 | 0 | 0 | 0 | 0 | 0 | 0 | 0 | 0 |
| 0 | NA | NA | NA | NA | NA | Gm9630 | 0 | 0 | 0 | 0 | 0 | 0 | 0 | 0 | 0 | 0 | 0 |
| 0 | NA | NA | NA | NA | NA | Gm24080 | 0 | 0 | 0 | 0 | 0 | 0 | 0 | 0 | 0 | 0 | 0 |
| 0 | NA | NA | NA | NA | NA | Gm49118 | 0 | 0 | 0 | 0 | 0 | 0 | 0 | 0 | 0 | 0 | 0 |
| 0 | NA | NA | NA | NA | NA | 4930557F08Rik | 0 | 0 | 0 | 0 | 0 | 0 | 0 | 0 | 0 | 0 | 0 |
| 0 | NA | NA | NA | NA | NA | Gm25550 | 0 | 0 | 0 | 0 | 0 | 0 | 0 | 0 | 0 | 0 | 0 |
| 0 | NA | NA | NA | NA | NA | Gm49122 | 0 | 0 | 0 | 0 | 0 | 0 | 0 | 0 | 0 | 0 | 0 |
| 0 | NA | NA | NA | NA | NA | Gm6479 | 0 | 0 | 0 | 0 | 0 | 0 | 0 | 0 | 0 | 0 | 0 |
| 0 | NA | NA | NA | NA | NA | Gm41276 | 0 | 0 | 0 | 0 | 0 | 0 | 0 | 0 | 0 | 0 | 0 |
| 0 | NA | NA | NA | NA | NA | Gm49126 | 0 | 0 | 0 | 0 | 0 | 0 | 0 | 0 | 0 | 0 | 0 |
| 0 | NA | NA | NA | NA | NA | Gm2611 | 0 | 0 | 0 | 0 | 0 | 0 | 0 | 0 | 0 | 0 | 0 |
| 0 | NA | NA | NA | NA | NA | Gm49127 | 0 | 0 | 0 | 0 | 0 | 0 | 0 | 0 | 0 | 0 | 0 |
| 0 | NA | NA | NA | NA | NA | Gm49128 | 0 | 0 | 0 | 0 | 0 | 0 | 0 | 0 | 0 | 0 | 0 |
| 0 | NA | NA | NA | NA | NA | Gm22032 | 0 | 0 | 0 | 0 | 0 | 0 | 0 | 0 | 0 | 0 | 0 |
| 0 | NA | NA | NA | NA | NA | Gm2638 | 0 | 0 | 0 | 0 | 0 | 0 | 0 | 0 | 0 | 0 | 0 |
| 0 | NA | NA | NA | NA | NA | Gm22895 | 0 | 0 | 0 | 0 | 0 | 0 | 0 | 0 | 0 | 0 | 0 |
| 0 | NA | NA | NA | NA | NA | Gm18532 | 0 | 0 | 0 | 0 | 0 | 0 | 0 | 0 | 0 | 0 | 0 |
| 0 | NA | NA | NA | NA | NA | Gm24713 | 0 | 0 | 0 | 0 | 0 | 0 | 0 | 0 | 0 | 0 | 0 |
| 0 | NA | NA | NA | NA | NA | Gm41277 | 0 | 0 | 0 | 0 | 0 | 0 | 0 | 0 | 0 | 0 | 0 |
| 0 | NA | NA | NA | NA | NA | 4921515E04Rik | 0 | 0 | 0 | 0 | 0 | 0 | 0 | 0 | 0 | 0 | 0 |
| 0 | NA | NA | NA | NA | NA | Gm49138 | 0 | 0 | 0 | 0 | 0 | 0 | 0 | 0 | 0 | 0 | 0 |
| 0 | NA | NA | NA | NA | NA | Gm49139 | 0 | 0 | 0 | 0 | 0 | 0 | 0 | 0 | 0 | 0 | 0 |
| 0 | NA | NA | NA | NA | NA | Gm49140 | 0 | 0 | 0 | 0 | 0 | 0 | 0 | 0 | 0 | 0 | 0 |
| 0 | NA | NA | NA | NA | NA | C030047K22Rik | 0 | 0 | 0 | 0 | 0 | 0 | 0 | 0 | 0 | 0 | 0 |
| 0 | NA | NA | NA | NA | NA | Gm8328 | 0 | 0 | 0 | 0 | 0 | 0 | 0 | 0 | 0 | 0 | 0 |
| 0 | NA | NA | NA | NA | NA | Gm35496 | 0 | 0 | 0 | 0 | 0 | 0 | 0 | 0 | 0 | 0 | 0 |
| 0 | NA | NA | NA | NA | NA | Gm18533 | 0 | 0 | 0 | 0 | 0 | 0 | 0 | 0 | 0 | 0 | 0 |
| 0 | NA | NA | NA | NA | NA | Gm49160 | 0 | 0 | 0 | 0 | 0 | 0 | 0 | 0 | 0 | 0 | 0 |
| 0 | NA | NA | NA | NA | NA | Gm49161 | 0 | 0 | 0 | 0 | 0 | 0 | 0 | 0 | 0 | 0 | 0 |
| 0 | NA | NA | NA | NA | NA | Gm49166 | 0 | 0 | 0 | 0 | 0 | 0 | 0 | 0 | 0 | 0 | 0 |
| 0 | NA | NA | NA | NA | NA | Gm18741 | 0 | 0 | 0 | 0 | 0 | 0 | 0 | 0 | 0 | 0 | 0 |
| 0 | NA | NA | NA | NA | NA | Gm46498 | 0 | 0 | 0 | 0 | 0 | 0 | 0 | 0 | 0 | 0 | 0 |
| 0 | NA | NA | NA | NA | NA | Gm49176 | 0 | 0 | 0 | 0 | 0 | 0 | 0 | 0 | 0 | 0 | 0 |
| 0 | NA | NA | NA | NA | NA | Gm25976 | 0 | 0 | 0 | 0 | 0 | 0 | 0 | 0 | 0 | 0 | 0 |
| 0 | NA | NA | NA | NA | NA | Gm37815 | 0 | 0 | 0 | 0 | 0 | 0 | 0 | 0 | 0 | 0 | 0 |
| 0 | NA | NA | NA | NA | NA | Gm19207 | 0 | 0 | 0 | 0 | 0 | 0 | 0 | 0 | 0 | 0 | 0 |
| 0 | NA | NA | NA | NA | NA | Gm49177 | 0 | 0 | 0 | 0 | 0 | 0 | 0 | 0 | 0 | 0 | 0 |
| 0 | NA | NA | NA | NA | NA | Gm49178 | 0 | 0 | 0 | 0 | 0 | 0 | 0 | 0 | 0 | 0 | 0 |
| 0 | NA | NA | NA | NA | NA | Gm49685 | 0 | 0 | 0 | 0 | 0 | 0 | 0 | 0 | 0 | 0 | 0 |
| 0 | NA | NA | NA | NA | NA | Gm49182 | 0 | 0 | 0 | 0 | 0 | 0 | 0 | 0 | 0 | 0 | 0 |
| 0 | NA | NA | NA | NA | NA | Gm25711 | 0 | 0 | 0 | 0 | 0 | 0 | 0 | 0 | 0 | 0 | 0 |
| 0 | NA | NA | NA | NA | NA | Gm27397 | 0 | 0 | 0 | 0 | 0 | 0 | 0 | 0 | 0 | 0 | 0 |
| 0 | NA | NA | NA | NA | NA | Gm19371 | 0 | 0 | 0 | 0 | 0 | 0 | 0 | 0 | 0 | 0 | 0 |
| 0 | NA | NA | NA | NA | NA | Gm25101 | 0 | 0 | 0 | 0 | 0 | 0 | 0 | 0 | 0 | 0 | 0 |
| 0 | NA | NA | NA | NA | NA | Gm38234 | 0 | 0 | 0 | 0 | 0 | 0 | 0 | 0 | 0 | 0 | 0 |
| 0 | NA | NA | NA | NA | NA | Gm44373 | 0 | 0 | 0 | 0 | 0 | 0 | 0 | 0 | 0 | 0 | 0 |
| 0 | NA | NA | NA | NA | NA | Gm49187 | 0 | 0 | 0 | 0 | 0 | 0 | 0 | 0 | 0 | 0 | 0 |
| 0 | NA | NA | NA | NA | NA | Gm27525 | 0 | 0 | 0 | 0 | 0 | 0 | 0 | 0 | 0 | 0 | 0 |
| 0 | NA | NA | NA | NA | NA | Gm24656 | 0 | 0 | 0 | 0 | 0 | 0 | 0 | 0 | 0 | 0 | 0 |
| 0 | NA | NA | NA | NA | NA | Gm49191 | 0 | 0 | 0 | 0 | 0 | 0 | 0 | 0 | 0 | 0 | 0 |

| | | | | | | | | | | | | | | | | | |
|---|---|---|---|---|---|---|---|---|---|---|---|---|---|---|---|---|---|
| 0 | NA | NA | NA | NA | NA | Gm48915 | 0 | 0 | 0 | 0 | 0 | 0 | 0 | 0 | 0 | 0 | 0 |
| 0 | NA | NA | NA | NA | NA | Gm23980 | 0 | 0 | 0 | 0 | 0 | 0 | 0 | 0 | 0 | 0 | 0 |
| 0 | NA | NA | NA | NA | NA | Gm48916 | 0 | 0 | 0 | 0 | 0 | 0 | 0 | 0 | 0 | 0 | 0 |
| 0 | NA | NA | NA | NA | NA | Gm18582 | 0 | 0 | 0 | 0 | 0 | 0 | 0 | 0 | 0 | 0 | 0 |
| 0 | NA | NA | NA | NA | NA | Gm48918 | 0 | 0 | 0 | 0 | 0 | 0 | 0 | 0 | 0 | 0 | 0 |
| 0 | NA | NA | NA | NA | NA | Gm48919 | 0 | 0 | 0 | 0 | 0 | 0 | 0 | 0 | 0 | 0 | 0 |
| 0 | NA | NA | NA | NA | NA | Gm8408 | 0 | 0 | 0 | 0 | 0 | 0 | 0 | 0 | 0 | 0 | 0 |
| 0 | NA | NA | NA | NA | NA | Gm49729 | 0 | 0 | 0 | 0 | 0 | 0 | 0 | 0 | 0 | 0 | 0 |
| 0 | NA | NA | NA | NA | NA | Gm36147 | 0 | 0 | 0 | 0 | 0 | 0 | 0 | 0 | 0 | 0 | 0 |
| 0 | NA | NA | NA | NA | NA | 4930540I17Rik | 0 | 0 | 0 | 0 | 0 | 0 | 0 | 0 | 0 | 0 | 0 |
| 0 | NA | NA | NA | NA | NA | Gm48920 | 0 | 0 | 0 | 0 | 0 | 0 | 0 | 0 | 0 | 0 | 0 |
| 0 | NA | NA | NA | NA | NA | Gm48921 | 0 | 0 | 0 | 0 | 0 | 0 | 0 | 0 | 0 | 0 | 0 |
| 0 | NA | NA | NA | NA | NA | Gm2803 | 0 | 0 | 0 | 0 | 0 | 0 | 0 | 0 | 0 | 0 | 0 |
| 0 | NA | NA | NA | NA | NA | Gm22810 | 0 | 0 | 0 | 0 | 0 | 0 | 0 | 0 | 0 | 0 | 0 |
| 0 | NA | NA | NA | NA | NA | Gm35769 | 0 | 0 | 0 | 0 | 0 | 0 | 0 | 0 | 0 | 0 | 0 |
| 0 | NA | NA | NA | NA | NA | Gm35996 | 0 | 0 | 0 | 0 | 0 | 0 | 0 | 0 | 0 | 0 | 0 |
| 0 | NA | NA | NA | NA | NA | 5830426I08Rik | 0 | 0 | 0 | 0 | 0 | 0 | 0 | 0 | 0 | 0 | 0 |
| 0 | NA | NA | NA | NA | NA | Gm38341 | 0 | 0 | 0 | 0 | 0 | 0 | 0 | 0 | 0 | 0 | 0 |
| 0 | NA | NA | NA | NA | NA | 4930445E18Rik | 0 | 0 | 0 | 0 | 0 | 0 | 0 | 0 | 0 | 0 | 0 |
| 0 | NA | NA | NA | NA | NA | Gm5468 | 0 | 0 | 0 | 0 | 0 | 0 | 0 | 0 | 0 | 0 | 0 |
| 0 | NA | NA | NA | NA | NA | Gm48956 | 0 | 0 | 0 | 0 | 0 | 0 | 0 | 0 | 0 | 0 | 0 |
| 0 | NA | NA | NA | NA | NA | Gm48957 | 0 | 0 | 0 | 0 | 0 | 0 | 0 | 0 | 0 | 0 | 0 |
| 0 | NA | NA | NA | NA | NA | Gm36642 | 0 | 0 | 0 | 0 | 0 | 0 | 0 | 0 | 0 | 0 | 0 |
| 0 | NA | NA | NA | NA | NA | Mir7212 | 0 | 0 | 0 | 0 | 0 | 0 | 0 | 0 | 0 | 0 | 0 |
| 0 | NA | NA | NA | NA | NA | Gm41279 | 0 | 0 | 0 | 0 | 0 | 0 | 0 | 0 | 0 | 0 | 0 |
| 0 | NA | NA | NA | NA | NA | Gm49267 | 0 | 0 | 0 | 0 | 0 | 0 | 0 | 0 | 0 | 0 | 0 |
| 0 | NA | NA | NA | NA | NA | 9430068D22Rik | 0 | 0 | 0 | 0 | 0 | 0 | 0 | 0 | 0 | 0 | 0 |
| 0 | NA | NA | NA | NA | NA | Gm49265 | 0 | 0 | 0 | 0 | 0 | 0 | 0 | 0 | 0 | 0 | 0 |
| 0 | NA | NA | NA | NA | NA | Gm49268 | 0 | 0 | 0 | 0 | 0 | 0 | 0 | 0 | 0 | 0 | 0 |
| 0 | NA | NA | NA | NA | NA | Gm6330 | 0 | 0 | 0 | 0 | 0 | 0 | 0 | 0 | 0 | 0 | 0 |
| 0 | NA | NA | NA | NA | NA | Gm49266 | 0 | 0 | 0 | 0 | 0 | 0 | 0 | 0 | 0 | 0 | 0 |
| 0 | NA | NA | NA | NA | NA | Gm23903 | 0 | 0 | 0 | 0 | 0 | 0 | 0 | 0 | 0 | 0 | 0 |
| 0 | NA | NA | NA | NA | NA | Gm19111 | 0 | 0 | 0 | 0 | 0 | 0 | 0 | 0 | 0 | 0 | 0 |
| 0 | NA | NA | NA | NA | NA | Mir7117 | 0 | 0 | 0 | 0 | 0 | 0 | 0 | 0 | 0 | 0 | 0 |
| 0 | NA | NA | NA | NA | NA | Gm46506 | 0 | 0 | 0 | 0 | 0 | 0 | 0 | 0 | 0 | 0 | 0 |
| 0 | NA | NA | NA | NA | NA | Gm9891 | 0 | 0 | 0 | 0 | 0 | 0 | 0 | 0 | 0 | 0 | 0 |
| 0 | NA | NA | NA | NA | NA | Gm19883 | 0 | 0 | 0 | 0 | 0 | 0 | 0 | 0 | 0 | 0 | 0 |
| 0 | NA | NA | NA | NA | NA | Gm31458 | 0 | 0 | 0 | 0 | 0 | 0 | 0 | 0 | 0 | 0 | 0 |
| 0 | NA | NA | NA | NA | NA | Gm8472 | 0 | 0 | 0 | 0 | 0 | 0 | 0 | 0 | 0 | 0 | 0 |
| 0 | NA | NA | NA | NA | NA | Gm49233 | 0 | 0 | 0 | 0 | 0 | 0 | 0 | 0 | 0 | 0 | 0 |
| 0 | NA | NA | NA | NA | NA | Gm49234 | 0 | 0 | 0 | 0 | 0 | 0 | 0 | 0 | 0 | 0 | 0 |
| 0 | NA | NA | NA | NA | NA | Gm49278 | 0 | 0 | 0 | 0 | 0 | 0 | 0 | 0 | 0 | 0 | 0 |
| 0 | NA | NA | NA | NA | NA | Gm49279 | 0 | 0 | 0 | 0 | 0 | 0 | 0 | 0 | 0 | 0 | 0 |
| 0 | NA | NA | NA | NA | NA | Gm8484 | 0 | 0 | 0 | 0 | 0 | 0 | 0 | 0 | 0 | 0 | 0 |
| 0 | NA | NA | NA | NA | NA | Gm49280 | 0 | 0 | 0 | 0 | 0 | 0 | 0 | 0 | 0 | 0 | 0 |
| 0 | NA | NA | NA | NA | NA | Mir3964 | 0 | 0 | 0 | 0 | 0 | 0 | 0 | 0 | 0 | 0 | 0 |
| 0 | NA | NA | NA | NA | NA | Gm23033 | 0 | 0 | 0 | 0 | 0 | 0 | 0 | 0 | 0 | 0 | 0 |
| 0 | NA | NA | NA | NA | NA | Gm49281 | 0 | 0 | 0 | 0 | 0 | 0 | 0 | 0 | 0 | 0 | 0 |
| 0 | NA | NA | NA | NA | NA | Gm24554 | 0 | 0 | 0 | 0 | 0 | 0 | 0 | 0 | 0 | 0 | 0 |
| 0 | NA | NA | NA | NA | NA | Gm23327 | 0 | 0 | 0 | 0 | 0 | 0 | 0 | 0 | 0 | 0 | 0 |
| 0 | NA | NA | NA | NA | NA | Gm5212 | 0 | 0 | 0 | 0 | 0 | 0 | 0 | 0 | 0 | 0 | 0 |
| 0 | NA | NA | NA | NA | NA | Gm26163 | 0 | 0 | 0 | 0 | 0 | 0 | 0 | 0 | 0 | 0 | 0 |
| 0 | NA | NA | NA | NA | NA | Gm49283 | 0 | 0 | 0 | 0 | 0 | 0 | 0 | 0 | 0 | 0 | 0 |
| 0 | NA | NA | NA | NA | NA | Gm23956 | 0 | 0 | 0 | 0 | 0 | 0 | 0 | 0 | 0 | 0 | 0 |
| 0 | NA | NA | NA | NA | NA | 4930570B17Rik | 0 | 0 | 0 | 0 | 0 | 0 | 0 | 0 | 0 | 0 | 0 |
| 0 | NA | NA | NA | NA | NA | Gm24487 | 0 | 0 | 0 | 0 | 0 | 0 | 0 | 0 | 0 | 0 | 0 |
| 0 | NA | NA | NA | NA | NA | 9630009A06Rik | 0 | 0 | 0 | 0 | 0 | 0 | 0 | 0 | 0 | 0 | 0 |
| 0 | NA | NA | NA | NA | NA | 4930430F21Rik | 0 | 0 | 0 | 0 | 0 | 0 | 0 | 0 | 0 | 0 | 0 |
| 0 | NA | NA | NA | NA | NA | Gm26416 | 0 | 0 | 0 | 0 | 0 | 0 | 0 | 0 | 0 | 0 | 0 |
| 0 | NA | NA | NA | NA | NA | Gm49296 | 0 | 0 | 0 | 0 | 0 | 0 | 0 | 0 | 0 | 0 | 0 |

| | | | | | | | | | | | | | | | | | | |
|---|---|---|---|---|---|---|---|---|---|---|---|---|---|---|---|---|---|---|
| 0 | NA | NA | NA | NA | NA | Gm49208 | | 0 | 0 | 0 | 0 | 0 | 0 | 0 | 0 | 0 | 0 | 0 |
| 0 | NA | NA | NA | NA | NA | Gm49214 | | 0 | 0 | 0 | 0 | 0 | 0 | 0 | 0 | 0 | 0 | 0 |
| 0 | NA | NA | NA | NA | NA | Gm49215 | | 0 | 0 | 0 | 0 | 0 | 0 | 0 | 0 | 0 | 0 | 0 |
| 0 | NA | NA | NA | NA | NA | Gm18003 | | 0 | 0 | 0 | 0 | 0 | 0 | 0 | 0 | 0 | 0 | 0 |
| 0 | NA | NA | NA | NA | NA | Gm49285 | | 0 | 0 | 0 | 0 | 0 | 0 | 0 | 0 | 0 | 0 | 0 |
| 0 | NA | NA | NA | NA | NA | Tas2r119 | taste receptor, type 2, member 119 | 0 | 0 | 0 | 0 | 0 | 0 | 0 | 0 | 0 | 0 | 0 |
| 0 | NA | NA | NA | NA | NA | 4930518I17Rik | | 0 | 0 | 0 | 0 | 0 | 0 | 0 | 0 | 0 | 0 | 0 |
| 0 | NA | NA | NA | NA | NA | 4930465K09Rik | | 0 | 0 | 0 | 0 | 0 | 0 | 0 | 0 | 0 | 0 | 0 |
| 0 | NA | NA | NA | NA | NA | Gm41289 | | 0 | 0 | 0 | 0 | 0 | 0 | 0 | 0 | 0 | 0 | 0 |
| 0 | NA | NA | NA | NA | NA | Gm32764 | | 0 | 0 | 0 | 0 | 0 | 0 | 0 | 0 | 0 | 0 | 0 |
| 0 | NA | NA | NA | NA | NA | 1700084J12Rik | | 0 | 0 | 0 | 0 | 0 | 0 | 0 | 0 | 0 | 0 | 0 |
| 0 | NA | NA | NA | NA | NA | Gm24476 | | 0 | 0 | 0 | 0 | 0 | 0 | 0 | 0 | 0 | 0 | 0 |
| 0 | NA | NA | NA | NA | NA | Gm49088 | | 0 | 0 | 0 | 0 | 0 | 0 | 0 | 0 | 0 | 0 | 0 |
| 0 | NA | NA | NA | NA | NA | Gm9643 | | 0 | 0 | 0 | 0 | 0 | 0 | 0 | 0 | 0 | 0 | 0 |
| 0 | NA | NA | NA | NA | NA | Gm41290 | | 0 | 0 | 0 | 0 | 0 | 0 | 0 | 0 | 0 | 0 | 0 |
| 0 | NA | NA | NA | NA | NA | Gm49091 | | 0 | 0 | 0 | 0 | 0 | 0 | 0 | 0 | 0 | 0 | 0 |
| 0 | NA | NA | NA | NA | NA | Gm24500 | | 0 | 0 | 0 | 0 | 0 | 0 | 0 | 0 | 0 | 0 | 0 |
| 0 | NA | NA | NA | NA | NA | Gm25809 | | 0 | 0 | 0 | 0 | 0 | 0 | 0 | 0 | 0 | 0 | 0 |
| 0 | NA | NA | NA | NA | NA | Gm24523 | | 0 | 0 | 0 | 0 | 0 | 0 | 0 | 0 | 0 | 0 | 0 |
| 0 | NA | NA | NA | NA | NA | Gm49188 | | 0 | 0 | 0 | 0 | 0 | 0 | 0 | 0 | 0 | 0 | 0 |
| 0 | NA | NA | NA | NA | NA | Gm48932 | | 0 | 0 | 0 | 0 | 0 | 0 | 0 | 0 | 0 | 0 | 0 |
| 0 | NA | NA | NA | NA | NA | BC048602 | | 0 | 0 | 0 | 0 | 0 | 0 | 0 | 0 | 0 | 0 | 0 |
| 0 | NA | NA | NA | NA | NA | Gm24771 | | 0 | 0 | 0 | 0 | 0 | 0 | 0 | 0 | 0 | 0 | 0 |
| 0 | NA | NA | NA | NA | NA | Mir599 | | 0 | 0 | 0 | 0 | 0 | 0 | 0 | 0 | 0 | 0 | 0 |
| 0 | NA | NA | NA | NA | NA | Mir875 | | 0 | 0 | 0 | 0 | 0 | 0 | 0 | 0 | 0 | 0 | 0 |
| 0 | NA | NA | NA | NA | NA | Gm25372 | | 0 | 0 | 0 | 0 | 0 | 0 | 0 | 0 | 0 | 0 | 0 |
| 0 | NA | NA | NA | NA | NA | Gm22979 | | 0 | 0 | 0 | 0 | 0 | 0 | 0 | 0 | 0 | 0 | 0 |
| 0 | NA | NA | NA | NA | NA | Mir8097 | | 0 | 0 | 0 | 0 | 0 | 0 | 0 | 0 | 0 | 0 | 0 |
| 0 | NA | NA | NA | NA | NA | Gm9501 | | 0 | 0 | 0 | 0 | 0 | 0 | 0 | 0 | 0 | 0 | 0 |
| 0 | NA | NA | NA | NA | NA | Gm49224 | | 0 | 0 | 0 | 0 | 0 | 0 | 0 | 0 | 0 | 0 | 0 |
| 0 | NA | NA | NA | NA | NA | Gm49246 | | 0 | 0 | 0 | 0 | 0 | 0 | 0 | 0 | 0 | 0 | 0 |
| 0 | NA | NA | NA | NA | NA | Gm44310 | | 0 | 0 | 0 | 0 | 0 | 0 | 0 | 0 | 0 | 0 | 0 |
| 0 | NA | NA | NA | NA | NA | Gm49262 | | 0 | 0 | 0 | 0 | 0 | 0 | 0 | 0 | 0 | 0 | 0 |
| 0 | NA | NA | NA | NA | NA | Gm49263 | | 0 | 0 | 0 | 0 | 0 | 0 | 0 | 0 | 0 | 0 | 0 |
| 0 | NA | NA | NA | NA | NA | Gm25303 | | 0 | 0 | 0 | 0 | 0 | 0 | 0 | 0 | 0 | 0 | 0 |
| 0 | NA | NA | NA | NA | NA | Gm41300 | | 0 | 0 | 0 | 0 | 0 | 0 | 0 | 0 | 0 | 0 | 0 |
| 0 | NA | NA | NA | NA | NA | Gm8664 | | 0 | 0 | 0 | 0 | 0 | 0 | 0 | 0 | 0 | 0 | 0 |
| 0 | NA | NA | NA | NA | NA | Gm16137 | | 0 | 0 | 0 | 0 | 0 | 0 | 0 | 0 | 0 | 0 | 0 |
| 0 | NA | NA | NA | NA | NA | Gm16138 | | 0 | 0 | 0 | 0 | 0 | 0 | 0 | 0 | 0 | 0 | 0 |
| 0 | NA | NA | NA | NA | NA | Gm8668 | | 0 | 0 | 0 | 0 | 0 | 0 | 0 | 0 | 0 | 0 | 0 |
| 0 | NA | NA | NA | NA | NA | 4930447A16Ril | RIKEN cDNA 4930447A16 gene | 0 | 0 | 0 | 0 | 0 | 0 | 0 | 0 | 0 | 0 | 0 |
| 0 | NA | NA | NA | NA | NA | Gm3267 | | 0 | 0 | 0 | 0 | 0 | 0 | 0 | 0 | 0 | 0 | 0 |
| 0 | NA | NA | NA | NA | NA | Gm49311 | | 0 | 0 | 0 | 0 | 0 | 0 | 0 | 0 | 0 | 0 | 0 |
| 0 | NA | NA | NA | NA | NA | Gm24833 | | 0 | 0 | 0 | 0 | 0 | 0 | 0 | 0 | 0 | 0 | 0 |
| 0 | NA | NA | NA | NA | NA | Gm15942 | | 0 | 0 | 0 | 0 | 0 | 0 | 0 | 0 | 0 | 0 | 0 |
| 0 | NA | NA | NA | NA | NA | Gm15940 | | 0 | 0 | 0 | 0 | 0 | 0 | 0 | 0 | 0 | 0 | 0 |
| 0 | NA | NA | NA | NA | NA | Gm22353 | | 0 | 0 | 0 | 0 | 0 | 0 | 0 | 0 | 0 | 0 | 0 |
| 0 | NA | NA | NA | NA | NA | Odf1 | outer dense fiber of sperm tails 1 | 0 | 0 | 0 | 0 | 0 | 0 | 0 | 0 | 0 | 0 | 0 |
| 0 | NA | NA | NA | NA | NA | Gm24098 | | 0 | 0 | 0 | 0 | 0 | 0 | 0 | 0 | 0 | 0 | 0 |
| 0 | NA | NA | NA | NA | NA | Gm49314 | | 0 | 0 | 0 | 0 | 0 | 0 | 0 | 0 | 0 | 0 | 0 |
| 0 | NA | NA | NA | NA | NA | 2310043O21Rik | | 0 | 0 | 0 | 0 | 0 | 0 | 0 | 0 | 0 | 0 | 0 |
| 0 | NA | NA | NA | NA | NA | Gm49090 | | 0 | 0 | 0 | 0 | 0 | 0 | 0 | 0 | 0 | 0 | 0 |
| 0 | NA | NA | NA | NA | NA | Siah1-ps2 | | 0 | 0 | 0 | 0 | 0 | 0 | 0 | 0 | 0 | 0 | 0 |
| 0 | NA | NA | NA | NA | NA | Gm45924 | | 0 | 0 | 0 | 0 | 0 | 0 | 0 | 0 | 0 | 0 | 0 |
| 0 | NA | NA | NA | NA | NA | Gm24789 | | 0 | 0 | 0 | 0 | 0 | 0 | 0 | 0 | 0 | 0 | 0 |
| 0 | NA | NA | NA | NA | NA | n-R5s39 | | 0 | 0 | 0 | 0 | 0 | 0 | 0 | 0 | 0 | 0 | 0 |
| 0 | NA | NA | NA | NA | NA | Gm25381 | | 0 | 0 | 0 | 0 | 0 | 0 | 0 | 0 | 0 | 0 | 0 |
| 0 | NA | NA | NA | NA | NA | Gm16291 | | 0 | 0 | 0 | 0 | 0 | 0 | 0 | 0 | 0 | 0 | 0 |
| 0 | NA | NA | NA | NA | NA | Gm49253 | | 0 | 0 | 0 | 0 | 0 | 0 | 0 | 0 | 0 | 0 | 0 |
| 0 | NA | NA | NA | NA | NA | Gm49254 | | 0 | 0 | 0 | 0 | 0 | 0 | 0 | 0 | 0 | 0 | 0 |

| | | | | | | | | | | | | | | | | | | |
|---|---|---|---|---|---|---|---|---|---|---|---|---|---|---|---|---|---|---|
| 0 | NA | NA | NA | NA | NA | Gm33251 | | 0 | 0 | 0 | 0 | 0 | 0 | 0 | 0 | 0 | 0 | 0 |
| 0 | NA | NA | NA | NA | NA | Gm25958 | | 0 | 0 | 0 | 0 | 0 | 0 | 0 | 0 | 0 | 0 | 0 |
| 0 | NA | NA | NA | NA | NA | Gm41311 | | 0 | 0 | 0 | 0 | 0 | 0 | 0 | 0 | 0 | 0 | 0 |
| 0 | NA | NA | NA | NA | NA | Gm16294 | | 0 | 0 | 0 | 0 | 0 | 0 | 0 | 0 | 0 | 0 | 0 |
| 0 | NA | NA | NA | NA | NA | Gm7382 | | 0 | 0 | 0 | 0 | 0 | 0 | 0 | 0 | 0 | 0 | 0 |
| 0 | NA | NA | NA | NA | NA | Gm49525 | | 0 | 0 | 0 | 0 | 0 | 0 | 0 | 0 | 0 | 0 | 0 |
| 0 | NA | NA | NA | NA | NA | Gm49526 | | 0 | 0 | 0 | 0 | 0 | 0 | 0 | 0 | 0 | 0 | 0 |
| 0 | NA | NA | NA | NA | NA | Gm49524 | | 0 | 0 | 0 | 0 | 0 | 0 | 0 | 0 | 0 | 0 | 0 |
| 0 | NA | NA | NA | NA | NA | 4930555K19Rik | | 0 | 0 | 0 | 0 | 0 | 0 | 0 | 0 | 0 | 0 | 0 |
| 0 | NA | NA | NA | NA | NA | Gm49529 | | 0 | 0 | 0 | 0 | 0 | 0 | 0 | 0 | 0 | 0 | 0 |
| 0 | NA | NA | NA | NA | NA | Gm26854 | | 0 | 0 | 0 | 0 | 0 | 0 | 0 | 0 | 0 | 0 | 0 |
| 0 | NA | NA | NA | NA | NA | Gm24751 | | 0 | 0 | 0 | 0 | 0 | 0 | 0 | 0 | 0 | 0 | 0 |
| 0 | NA | NA | NA | NA | NA | Gm49439 | | 0 | 0 | 0 | 0 | 0 | 0 | 0 | 0 | 0 | 0 | 0 |
| 0 | NA | NA | NA | NA | NA | Gm49438 | | 0 | 0 | 0 | 0 | 0 | 0 | 0 | 0 | 0 | 0 | 0 |
| 0 | NA | NA | NA | NA | NA | Gm49440 | | 0 | 0 | 0 | 0 | 0 | 0 | 0 | 0 | 0 | 0 | 0 |
| 0 | NA | NA | NA | NA | NA | Gm49441 | | 0 | 0 | 0 | 0 | 0 | 0 | 0 | 0 | 0 | 0 | 0 |
| 0 | NA | NA | NA | NA | NA | Gm18531 | | 0 | 0 | 0 | 0 | 0 | 0 | 0 | 0 | 0 | 0 | 0 |
| 0 | NA | NA | NA | NA | NA | Gm49452 | | 0 | 0 | 0 | 0 | 0 | 0 | 0 | 0 | 0 | 0 | 0 |
| 0 | NA | NA | NA | NA | NA | Gm18092 | | 0 | 0 | 0 | 0 | 0 | 0 | 0 | 0 | 0 | 0 | 0 |
| 0 | NA | NA | NA | NA | NA | Gm49453 | | 0 | 0 | 0 | 0 | 0 | 0 | 0 | 0 | 0 | 0 | 0 |
| 0 | NA | NA | NA | NA | NA | Gm22037 | | 0 | 0 | 0 | 0 | 0 | 0 | 0 | 0 | 0 | 0 | 0 |
| 0 | NA | NA | NA | NA | NA | Gm2140 | | 0 | 0 | 0 | 0 | 0 | 0 | 0 | 0 | 0 | 0 | 0 |
| 0 | NA | NA | NA | NA | NA | Trhr | thyrotropin releasing hormone receptor | 0 | 0 | 0 | 0 | 0 | 0 | 0 | 0 | 0 | 0 | 0 |
| 0 | NA | NA | NA | NA | NA | Gm19235 | | 0 | 0 | 0 | 0 | 0 | 0 | 0 | 0 | 0 | 0 | 0 |
| 0 | NA | NA | NA | NA | NA | Gm24903 | | 0 | 0 | 0 | 0 | 0 | 0 | 0 | 0 | 0 | 0 | 0 |
| 0 | NA | NA | NA | NA | NA | Gm25093 | | 0 | 0 | 0 | 0 | 0 | 0 | 0 | 0 | 0 | 0 | 0 |
| 0 | NA | NA | NA | NA | NA | Gm22392 | | 0 | 0 | 0 | 0 | 0 | 0 | 0 | 0 | 0 | 0 | 0 |
| 0 | NA | NA | NA | NA | NA | Gm24768 | | 0 | 0 | 0 | 0 | 0 | 0 | 0 | 0 | 0 | 0 | 0 |
| 0 | NA | NA | NA | NA | NA | Gm23343 | | 0 | 0 | 0 | 0 | 0 | 0 | 0 | 0 | 0 | 0 | 0 |
| 0 | NA | NA | NA | NA | NA | Gm18153 | | 0 | 0 | 0 | 0 | 0 | 0 | 0 | 0 | 0 | 0 | 0 |
| 0 | NA | NA | NA | NA | NA | | | 0 | 0 | 0 | 0 | 0 | 0 | 0 | 0 | 0 | 0 | 0 |
| 0 | NA | NA | NA | NA | NA | Gm2203 | | 0 | 0 | 0 | 0 | 0 | 0 | 0 | 0 | 0 | 0 | 0 |
| 0 | NA | NA | NA | NA | NA | Gm26063 | | 0 | 0 | 0 | 0 | 0 | 0 | 0 | 0 | 0 | 0 | 0 |
| 0 | NA | NA | NA | NA | NA | Gm49075 | | 0 | 0 | 0 | 0 | 0 | 0 | 0 | 0 | 0 | 0 | 0 |
| 0 | NA | NA | NA | NA | NA | 1700022A22Rik | | 0 | 0 | 0 | 0 | 0 | 0 | 0 | 0 | 0 | 0 | 0 |
| 0 | NA | NA | NA | NA | NA | Gm23562 | | 0 | 0 | 0 | 0 | 0 | 0 | 0 | 0 | 0 | 0 | 0 |
| 0 | NA | NA | NA | NA | NA | 4930523O13Rik | | 0 | 0 | 0 | 0 | 0 | 0 | 0 | 0 | 0 | 0 | 0 |
| 0 | NA | NA | NA | NA | NA | Gm49079 | | 0 | 0 | 0 | 0 | 0 | 0 | 0 | 0 | 0 | 0 | 0 |
| 0 | NA | NA | NA | NA | NA | 4930548G14Rik | | 0 | 0 | 0 | 0 | 0 | 0 | 0 | 0 | 0 | 0 | 0 |
| 0 | NA | NA | NA | NA | NA | Gm38541 | | 0 | 0 | 0 | 0 | 0 | 0 | 0 | 0 | 0 | 0 | 0 |
| 0 | NA | NA | NA | NA | NA | Gm23665 | | 0 | 0 | 0 | 0 | 0 | 0 | 0 | 0 | 0 | 0 | 0 |
| 0 | NA | NA | NA | NA | NA | Gm7508 | | 0 | 0 | 0 | 0 | 0 | 0 | 0 | 0 | 0 | 0 | 0 |
| 0 | NA | NA | NA | NA | NA | Gm2267 | | 0 | 0 | 0 | 0 | 0 | 0 | 0 | 0 | 0 | 0 | 0 |
| 0 | NA | NA | NA | NA | NA | Gm16300 | | 0 | 0 | 0 | 0 | 0 | 0 | 0 | 0 | 0 | 0 | 0 |
| 0 | NA | NA | NA | NA | NA | Gm23530 | | 0 | 0 | 0 | 0 | 0 | 0 | 0 | 0 | 0 | 0 | 0 |
| 0 | NA | NA | NA | NA | NA | Gm25007 | | 0 | 0 | 0 | 0 | 0 | 0 | 0 | 0 | 0 | 0 | 0 |
| 0 | NA | NA | NA | NA | NA | Gm49185 | | 0 | 0 | 0 | 0 | 0 | 0 | 0 | 0 | 0 | 0 | 0 |
| 0 | NA | NA | NA | NA | NA | Gm23185 | | 0 | 0 | 0 | 0 | 0 | 0 | 0 | 0 | 0 | 0 | 0 |
| 0 | NA | NA | NA | NA | NA | Gm2333 | | 0 | 0 | 0 | 0 | 0 | 0 | 0 | 0 | 0 | 0 | 0 |
| 0 | NA | NA | NA | NA | NA | Gm41318 | | 0 | 0 | 0 | 0 | 0 | 0 | 0 | 0 | 0 | 0 | 0 |
| 0 | NA | NA | NA | NA | NA | Gm23848 | | 0 | 0 | 0 | 0 | 0 | 0 | 0 | 0 | 0 | 0 | 0 |
| 0 | NA | NA | NA | NA | NA | Gm49186 | | 0 | 0 | 0 | 0 | 0 | 0 | 0 | 0 | 0 | 0 | 0 |
| 0 | NA | NA | NA | NA | NA | Gm49198 | | 0 | 0 | 0 | 0 | 0 | 0 | 0 | 0 | 0 | 0 | 0 |
| 0 | NA | NA | NA | NA | NA | Gm19303 | | 0 | 0 | 0 | 0 | 0 | 0 | 0 | 0 | 0 | 0 | 0 |
| 0 | NA | NA | NA | NA | NA | Gm2361 | | 0 | 0 | 0 | 0 | 0 | 0 | 0 | 0 | 0 | 0 | 0 |
| 0 | NA | NA | NA | NA | NA | Gm34562 | | 0 | 0 | 0 | 0 | 0 | 0 | 0 | 0 | 0 | 0 | 0 |
| 0 | NA | NA | NA | NA | NA | Gm34678 | | 0 | 0 | 0 | 0 | 0 | 0 | 0 | 0 | 0 | 0 | 0 |
| 0 | NA | NA | NA | NA | NA | Gm48923 | | 0 | 0 | 0 | 0 | 0 | 0 | 0 | 0 | 0 | 0 | 0 |
| 0 | NA | NA | NA | NA | NA | Gm48926 | | 0 | 0 | 0 | 0 | 0 | 0 | 0 | 0 | 0 | 0 | 0 |
| 0 | NA | NA | NA | NA | NA | Gm41322 | | 0 | 0 | 0 | 0 | 0 | 0 | 0 | 0 | 0 | 0 | 0 |

| | | | | | | | | | | | | | | | | | | |
|---|---|---|---|---|---|---|---|---|---|---|---|---|---|---|---|---|---|---|
| 0 | NA | NA | NA | NA | NA | Gm23545 | | 0 | 0 | 0 | 0 | 0 | 0 | 0 | 0 | 0 | 0 | 0 |
| 0 | NA | NA | NA | NA | NA | Gm23267 | | 0 | 0 | 0 | 0 | 0 | 0 | 0 | 0 | 0 | 0 | 0 |
| 0 | NA | NA | NA | NA | NA | Gm34794 | | 0 | 0 | 0 | 0 | 0 | 0 | 0 | 0 | 0 | 0 | 0 |
| 0 | NA | NA | NA | NA | NA | Gm27926 | | 0 | 0 | 0 | 0 | 0 | 0 | 0 | 0 | 0 | 0 | 0 |
| 0 | NA | NA | NA | NA | NA | Gm23200 | | 0 | 0 | 0 | 0 | 0 | 0 | 0 | 0 | 0 | 0 | 0 |
| 0 | NA | NA | NA | NA | NA | Mir28c | | 0 | 0 | 0 | 0 | 0 | 0 | 0 | 0 | 0 | 0 | 0 |
| 0 | NA | NA | NA | NA | NA | 1700015H07Rik | | 0 | 0 | 0 | 0 | 0 | 0 | 0 | 0 | 0 | 0 | 0 |
| 0 | NA | NA | NA | NA | NA | Gm7489 | | 0 | 0 | 0 | 0 | 0 | 0 | 0 | 0 | 0 | 0 | 0 |
| 0 | NA | NA | NA | NA | NA | Gm26933 | | 0 | 0 | 0 | 0 | 0 | 0 | 0 | 0 | 0 | 0 | 0 |
| 0 | NA | NA | NA | NA | NA | Gm7566 | | 0 | 0 | 0 | 0 | 0 | 0 | 0 | 0 | 0 | 0 | 0 |
| 0 | NA | NA | NA | NA | NA | Gm20070 | | 0 | 0 | 0 | 0 | 0 | 0 | 0 | 0 | 0 | 0 | 0 |
| 0 | NA | NA | NA | NA | NA | Gm7577 | | 0 | 0 | 0 | 0 | 0 | 0 | 0 | 0 | 0 | 0 | 0 |
| 0 | NA | NA | NA | NA | NA | 4930478E11Rik | | 0 | 0 | 0 | 0 | 0 | 0 | 0 | 0 | 0 | 0 | 0 |
| 0 | NA | NA | NA | NA | NA | Gm49036 | | 0 | 0 | 0 | 0 | 0 | 0 | 0 | 0 | 0 | 0 | 0 |
| 0 | NA | NA | NA | NA | NA | Gm41325 | | 0 | 0 | 0 | 0 | 0 | 0 | 0 | 0 | 0 | 0 | 0 |
| 0 | NA | NA | NA | NA | NA | Gm26296 | | 0 | 0 | 0 | 0 | 0 | 0 | 0 | 0 | 0 | 0 | 0 |
| 0 | NA | NA | NA | NA | NA | 1700040F17Rik | | 0 | 0 | 0 | 0 | 0 | 0 | 0 | 0 | 0 | 0 | 0 |
| 0 | NA | NA | NA | NA | NA | Gm18787 | | 0 | 0 | 0 | 0 | 0 | 0 | 0 | 0 | 0 | 0 | 0 |
| 0 | NA | NA | NA | NA | NA | Gm49257 | | 0 | 0 | 0 | 0 | 0 | 0 | 0 | 0 | 0 | 0 | 0 |
| 0 | NA | NA | NA | NA | NA | Gm46516 | | 0 | 0 | 0 | 0 | 0 | 0 | 0 | 0 | 0 | 0 | 0 |
| 0 | NA | NA | NA | NA | NA | Gm10370 | | 0 | 0 | 0 | 0 | 0 | 0 | 0 | 0 | 0 | 0 | 0 |
| 0 | NA | NA | NA | NA | NA | Gm7595 | | 0 | 0 | 0 | 0 | 0 | 0 | 0 | 0 | 0 | 0 | 0 |
| 0 | NA | NA | NA | NA | NA | Gm49210 | | 0 | 0 | 0 | 0 | 0 | 0 | 0 | 0 | 0 | 0 | 0 |
| 0 | NA | NA | NA | NA | NA | Gm49211 | | 0 | 0 | 0 | 0 | 0 | 0 | 0 | 0 | 0 | 0 | 0 |
| 0 | NA | NA | NA | NA | NA | Hba-ps3 | | 0 | 0 | 0 | 0 | 0 | 0 | 0 | 0 | 0 | 0 | 0 |
| 0 | NA | NA | NA | NA | NA | Gm49213 | | 0 | 0 | 0 | 0 | 0 | 0 | 0 | 0 | 0 | 0 | 0 |
| 0 | NA | NA | NA | NA | NA | Gm2582 | | 0 | 0 | 0 | 0 | 0 | 0 | 0 | 0 | 0 | 0 | 0 |
| 0 | NA | NA | NA | NA | NA | Gm26178 | | 0 | 0 | 0 | 0 | 0 | 0 | 0 | 0 | 0 | 0 | 0 |
| 0 | NA | NA | NA | NA | NA | Slc22a22 | solute carrier family 22 (organic cation tra | 0 | 0 | 0 | 0 | 0 | 0 | 0 | 0 | 0 | 0 | 0 |
| 0 | NA | NA | NA | NA | NA | Gm49229 | | 0 | 0 | 0 | 0 | 0 | 0 | 0 | 0 | 0 | 0 | 0 |
| 0 | NA | NA | NA | NA | NA | Gm18094 | | 0 | 0 | 0 | 0 | 0 | 0 | 0 | 0 | 0 | 0 | 0 |
| 0 | NA | NA | NA | NA | NA | Gm49235 | | 0 | 0 | 0 | 0 | 0 | 0 | 0 | 0 | 0 | 0 | 0 |
| 0 | NA | NA | NA | NA | NA | 9330154K18Rik | | 0 | 0 | 0 | 0 | 0 | 0 | 0 | 0 | 0 | 0 | 0 |
| 0 | NA | NA | NA | NA | NA | Gm22299 | | 0 | 0 | 0 | 0 | 0 | 0 | 0 | 0 | 0 | 0 | 0 |
| 0 | NA | NA | NA | NA | NA | Gm23475 | | 0 | 0 | 0 | 0 | 0 | 0 | 0 | 0 | 0 | 0 | 0 |
| 0 | NA | NA | NA | NA | NA | Gm23789 | | 0 | 0 | 0 | 0 | 0 | 0 | 0 | 0 | 0 | 0 | 0 |
| 0 | NA | NA | NA | NA | NA | Ube2d4 | | 0 | 0 | 0 | 0 | 0 | 0 | 0 | 0 | 0 | 0 | 0 |
| 0 | NA | NA | NA | NA | NA | Gm49356 | | 0 | 0 | 0 | 0 | 0 | 0 | 0 | 0 | 0 | 0 | 0 |
| 0 | NA | NA | NA | NA | NA | Zfp572 | | 0 | 0 | 0 | 0 | 0 | 0 | 0 | 0 | 0 | 0 | 0 |
| 0 | NA | NA | NA | NA | NA | Gm7708 | | 0 | 0 | 0 | 0 | 0 | 0 | 0 | 0 | 0 | 0 | 0 |
| 0 | NA | NA | NA | NA | NA | Gm7713 | | 0 | 0 | 0 | 0 | 0 | 0 | 0 | 0 | 0 | 0 | 0 |
| 0 | NA | NA | NA | NA | NA | 4933412E24Ril | RIKEN cDNA 4933412E24 gene | 0 | 0 | 0 | 0 | 0 | 0 | 0 | 0 | 0 | 0 | 0 |
| 0 | NA | NA | NA | NA | NA | Gm25875 | | 0 | 0 | 0 | 0 | 0 | 0 | 0 | 0 | 0 | 0 | 0 |
| 0 | NA | NA | NA | NA | NA | Gm2682 | | 0 | 0 | 0 | 0 | 0 | 0 | 0 | 0 | 0 | 0 | 0 |
| 0 | NA | NA | NA | NA | NA | Gm46507 | | 0 | 0 | 0 | 0 | 0 | 0 | 0 | 0 | 0 | 0 | 0 |
| 0 | NA | NA | NA | NA | NA | Gm4774 | | 0 | 0 | 0 | 0 | 0 | 0 | 0 | 0 | 0 | 0 | 0 |
| 0 | NA | NA | NA | NA | NA | Gm17892 | | 0 | 0 | 0 | 0 | 0 | 0 | 0 | 0 | 0 | 0 | 0 |
| 0 | NA | NA | NA | NA | NA | Gm48946 | | 0 | 0 | 0 | 0 | 0 | 0 | 0 | 0 | 0 | 0 | 0 |
| 0 | NA | NA | NA | NA | NA | Gm23835 | | 0 | 0 | 0 | 0 | 0 | 0 | 0 | 0 | 0 | 0 | 0 |
| 0 | NA | NA | NA | NA | NA | Gm7086 | | 0 | 0 | 0 | 0 | 0 | 0 | 0 | 0 | 0 | 0 | 0 |
| 0 | NA | NA | NA | NA | NA | Gm48947 | | 0 | 0 | 0 | 0 | 0 | 0 | 0 | 0 | 0 | 0 | 0 |
| 0 | NA | NA | NA | NA | NA | A1bg | alpha-1-B glycoprotein | 0 | 0 | 0 | 0 | 0 | 0 | 0 | 0 | 0 | 0 | 0 |
| 0 | NA | NA | NA | NA | NA | 4930402D18Rik | | 0 | 0 | 0 | 0 | 0 | 0 | 0 | 0 | 0 | 0 | 0 |
| 0 | NA | NA | NA | NA | NA | Gm48951 | | 0 | 0 | 0 | 0 | 0 | 0 | 0 | 0 | 0 | 0 | 0 |
| 0 | NA | NA | NA | NA | NA | Gm49497 | | 0 | 0 | 0 | 0 | 0 | 0 | 0 | 0 | 0 | 0 | 0 |
| 0 | NA | NA | NA | NA | NA | Gm29904 | | 0 | 0 | 0 | 0 | 0 | 0 | 0 | 0 | 0 | 0 | 0 |
| 0 | NA | NA | NA | NA | NA | Gm24696 | | 0 | 0 | 0 | 0 | 0 | 0 | 0 | 0 | 0 | 0 | 0 |
| 0 | NA | NA | NA | NA | NA | Gm49498 | | 0 | 0 | 0 | 0 | 0 | 0 | 0 | 0 | 0 | 0 | 0 |
| 0 | NA | NA | NA | NA | NA | Gm18154 | | 0 | 0 | 0 | 0 | 0 | 0 | 0 | 0 | 0 | 0 | 0 |
| 0 | NA | NA | NA | NA | NA | Gm37947 | | 0 | 0 | 0 | 0 | 0 | 0 | 0 | 0 | 0 | 0 | 0 |

| | | | | | | | | | | | | | | | | | | |
|---|---|---|---|---|---|---|---|---|---|---|---|---|---|---|---|---|---|---|
| 0 | NA | NA | NA | NA | NA | Gm27913 | | 0 | 0 | 0 | 0 | 0 | 0 | 0 | 0 | 0 | 0 | 0 |
| 0 | NA | NA | NA | NA | NA | Gm27782 | | 0 | 0 | 0 | 0 | 0 | 0 | 0 | 0 | 0 | 0 | 0 |
| 0 | NA | NA | NA | NA | NA | Gm27718 | | 0 | 0 | 0 | 0 | 0 | 0 | 0 | 0 | 0 | 0 | 0 |
| 0 | NA | NA | NA | NA | NA | Gm27957 | | 0 | 0 | 0 | 0 | 0 | 0 | 0 | 0 | 0 | 0 | 0 |
| 0 | NA | NA | NA | NA | NA | Gm41333 | | 0 | 0 | 0 | 0 | 0 | 0 | 0 | 0 | 0 | 0 | 0 |
| 0 | NA | NA | NA | NA | NA | Gm24810 | | 0 | 0 | 0 | 0 | 0 | 0 | 0 | 0 | 0 | 0 | 0 |
| 0 | NA | NA | NA | NA | NA | Gm22521 | | 0 | 0 | 0 | 0 | 0 | 0 | 0 | 0 | 0 | 0 | 0 |
| 0 | NA | NA | NA | NA | NA | Gm49013 | | 0 | 0 | 0 | 0 | 0 | 0 | 0 | 0 | 0 | 0 | 0 |
| 0 | NA | NA | NA | NA | NA | Gm41335 | | 0 | 0 | 0 | 0 | 0 | 0 | 0 | 0 | 0 | 0 | 0 |
| 0 | NA | NA | NA | NA | NA | Gm49014 | | 0 | 0 | 0 | 0 | 0 | 0 | 0 | 0 | 0 | 0 | 0 |
| 0 | NA | NA | NA | NA | NA | Gm25702 | | 0 | 0 | 0 | 0 | 0 | 0 | 0 | 0 | 0 | 0 | 0 |
| 0 | NA | NA | NA | NA | NA | Gm4942 | | 0 | 0 | 0 | 0 | 0 | 0 | 0 | 0 | 0 | 0 | 0 |
| 0 | NA | NA | NA | NA | NA | 4930449C09Rik | | 0 | 0 | 0 | 0 | 0 | 0 | 0 | 0 | 0 | 0 | 0 |
| 0 | NA | NA | NA | NA | NA | Gm41336 | | 0 | 0 | 0 | 0 | 0 | 0 | 0 | 0 | 0 | 0 | 0 |
| 0 | NA | NA | NA | NA | NA | Gm49019 | | 0 | 0 | 0 | 0 | 0 | 0 | 0 | 0 | 0 | 0 | 0 |
| 0 | NA | NA | NA | NA | NA | Gm46499 | | 0 | 0 | 0 | 0 | 0 | 0 | 0 | 0 | 0 | 0 | 0 |
| 0 | NA | NA | NA | NA | NA | Gsdmc | gasdermin C | 0 | 0 | 0 | 0 | 0 | 0 | 0 | 0 | 0 | 0 | 0 |
| 0 | NA | NA | NA | NA | NA | 1700065I16Rik | | 0 | 0 | 0 | 0 | 0 | 0 | 0 | 0 | 0 | 0 | 0 |
| 0 | NA | NA | NA | NA | NA | Gsdmcl1 | | 0 | 0 | 0 | 0 | 0 | 0 | 0 | 0 | 0 | 0 | 0 |
| 0 | NA | NA | NA | NA | NA | Gsdmcl2 | | 0 | 0 | 0 | 0 | 0 | 0 | 0 | 0 | 0 | 0 | 0 |
| 0 | NA | NA | NA | NA | NA | Gsdmcl-ps | | 0 | 0 | 0 | 0 | 0 | 0 | 0 | 0 | 0 | 0 | 0 |
| 0 | NA | NA | NA | NA | NA | Gm25628 | | 0 | 0 | 0 | 0 | 0 | 0 | 0 | 0 | 0 | 0 | 0 |
| 0 | NA | NA | NA | NA | NA | Gm20717 | | 0 | 0 | 0 | 0 | 0 | 0 | 0 | 0 | 0 | 0 | 0 |
| 0 | NA | NA | NA | NA | NA | Gm10926 | | 0 | 0 | 0 | 0 | 0 | 0 | 0 | 0 | 0 | 0 | 0 |
| 0 | NA | NA | NA | NA | NA | Gm30563 | | 0 | 0 | 0 | 0 | 0 | 0 | 0 | 0 | 0 | 0 | 0 |
| 0 | NA | NA | NA | NA | NA | Gm30691 | | 0 | 0 | 0 | 0 | 0 | 0 | 0 | 0 | 0 | 0 | 0 |
| 0 | NA | NA | NA | NA | NA | Gm21798 | | 0 | 0 | 0 | 0 | 0 | 0 | 0 | 0 | 0 | 0 | 0 |
| 0 | NA | NA | NA | NA | NA | Gm21961 | | 0 | 0 | 0 | 0 | 0 | 0 | 0 | 0 | 0 | 0 | 0 |
| 0 | NA | NA | NA | NA | NA | Gm5046 | | 0 | 0 | 0 | 0 | 0 | 0 | 0 | 0 | 0 | 0 | 0 |
| 0 | NA | NA | NA | NA | NA | Gm49241 | | 0 | 0 | 0 | 0 | 0 | 0 | 0 | 0 | 0 | 0 | 0 |
| 0 | NA | NA | NA | NA | NA | Gm7814 | | 0 | 0 | 0 | 0 | 0 | 0 | 0 | 0 | 0 | 0 | 0 |
| 0 | NA | NA | NA | NA | NA | Gm49242 | | 0 | 0 | 0 | 0 | 0 | 0 | 0 | 0 | 0 | 0 | 0 |
| 0 | NA | NA | NA | NA | NA | Gm49243 | | 0 | 0 | 0 | 0 | 0 | 0 | 0 | 0 | 0 | 0 | 0 |
| 0 | NA | NA | NA | NA | NA | Gm23907 | | 0 | 0 | 0 | 0 | 0 | 0 | 0 | 0 | 0 | 0 | 0 |
| 0 | NA | NA | NA | NA | NA | Oc90 | otoconin 90 | 0 | 0 | 0 | 0 | 0 | 0 | 0 | 0 | 0 | 0 | 0 |
| 0 | NA | NA | NA | NA | NA | Hhla1 | HERV-H LTR-associating 1 | 0 | 0 | 0 | 0 | 0 | 0 | 0 | 0 | 0 | 0 | 0 |
| 0 | NA | NA | NA | NA | NA | Gm27153 | | 0 | 0 | 0 | 0 | 0 | 0 | 0 | 0 | 0 | 0 | 0 |
| 0 | NA | NA | NA | NA | NA | Gm27242 | | 0 | 0 | 0 | 0 | 0 | 0 | 0 | 0 | 0 | 0 | 0 |
| 0 | NA | NA | NA | NA | NA | Gm17140 | | 0 | 0 | 0 | 0 | 0 | 0 | 0 | 0 | 0 | 0 | 0 |
| 0 | NA | NA | NA | NA | NA | 1700012I11Rik | | 0 | 0 | 0 | 0 | 0 | 0 | 0 | 0 | 0 | 0 | 0 |
| 0 | NA | NA | NA | NA | NA | Gm49408 | | 0 | 0 | 0 | 0 | 0 | 0 | 0 | 0 | 0 | 0 | 0 |
| 0 | NA | NA | NA | NA | NA | Gm49409 | | 0 | 0 | 0 | 0 | 0 | 0 | 0 | 0 | 0 | 0 | 0 |
| 0 | NA | NA | NA | NA | NA | Mir30b | | 0 | 0 | 0 | 0 | 0 | 0 | 0 | 0 | 0 | 0 | 0 |
| 0 | NA | NA | NA | NA | NA | Mir30d | | 0 | 0 | 0 | 0 | 0 | 0 | 0 | 0 | 0 | 0 | 0 |
| 0 | NA | NA | NA | NA | NA | Gm49414 | | 0 | 0 | 0 | 0 | 0 | 0 | 0 | 0 | 0 | 0 | 0 |
| 0 | NA | NA | NA | NA | NA | Gm25987 | | 0 | 0 | 0 | 0 | 0 | 0 | 0 | 0 | 0 | 0 | 0 |
| 0 | NA | NA | NA | NA | NA | Gm49415 | | 0 | 0 | 0 | 0 | 0 | 0 | 0 | 0 | 0 | 0 | 0 |
| 0 | NA | NA | NA | NA | NA | Gm18930 | | 0 | 0 | 0 | 0 | 0 | 0 | 0 | 0 | 0 | 0 | 0 |
| 0 | NA | NA | NA | NA | NA | Gm49422 | | 0 | 0 | 0 | 0 | 0 | 0 | 0 | 0 | 0 | 0 | 0 |
| 0 | NA | NA | NA | NA | NA | 4930573C08Rik | | 0 | 0 | 0 | 0 | 0 | 0 | 0 | 0 | 0 | 0 | 0 |
| 0 | NA | NA | NA | NA | NA | 4930504C09Rik | | 0 | 0 | 0 | 0 | 0 | 0 | 0 | 0 | 0 | 0 | 0 |
| 0 | NA | NA | NA | NA | NA | Gm24232 | | 0 | 0 | 0 | 0 | 0 | 0 | 0 | 0 | 0 | 0 | 0 |
| 0 | NA | NA | NA | NA | NA | Gm23987 | | 0 | 0 | 0 | 0 | 0 | 0 | 0 | 0 | 0 | 0 | 0 |
| 0 | NA | NA | NA | NA | NA | Gm19782 | | 0 | 0 | 0 | 0 | 0 | 0 | 0 | 0 | 0 | 0 | 0 |
| 0 | NA | NA | NA | NA | NA | Gm18155 | | 0 | 0 | 0 | 0 | 0 | 0 | 0 | 0 | 0 | 0 | 0 |
| 0 | NA | NA | NA | NA | NA | Gm49430 | | 0 | 0 | 0 | 0 | 0 | 0 | 0 | 0 | 0 | 0 | 0 |
| 0 | NA | NA | NA | NA | NA | Gm19199 | | 0 | 0 | 0 | 0 | 0 | 0 | 0 | 0 | 0 | 0 | 0 |
| 0 | NA | NA | NA | NA | NA | Gm47309 | | 0 | 0 | 0 | 0 | 0 | 0 | 0 | 0 | 0 | 0 | 0 |
| 0 | NA | NA | NA | NA | NA | Gm22519 | | 0 | 0 | 0 | 0 | 0 | 0 | 0 | 0 | 0 | 0 | 0 |
| 0 | NA | NA | NA | NA | NA | Gm49457 | | 0 | 0 | 0 | 0 | 0 | 0 | 0 | 0 | 0 | 0 | 0 |

| | | | | | | | | | | | | | | | | | | |
|---|---|---|---|---|---|---|---|---|---|---|---|---|---|---|---|---|---|---|
| 0 | NA | NA | NA | NA | NA | Gm23217 | | 0 | 0 | 0 | 0 | 0 | 0 | 0 | 0 | 0 | 0 | 0 |
| 0 | NA | NA | NA | NA | NA | Gm24390 | | 0 | 0 | 0 | 0 | 0 | 0 | 0 | 0 | 0 | 0 | 0 |
| 0 | NA | NA | NA | NA | NA | Gm49460 | | 0 | 0 | 0 | 0 | 0 | 0 | 0 | 0 | 0 | 0 | 0 |
| 0 | NA | NA | NA | NA | NA | Gm28020 | | 0 | 0 | 0 | 0 | 0 | 0 | 0 | 0 | 0 | 0 | 0 |
| 0 | NA | NA | NA | NA | NA | Gm49459 | | 0 | 0 | 0 | 0 | 0 | 0 | 0 | 0 | 0 | 0 | 0 |
| 0 | NA | NA | NA | NA | NA | 1700085D07Rik | | 0 | 0 | 0 | 0 | 0 | 0 | 0 | 0 | 0 | 0 | 0 |
| 0 | NA | NA | NA | NA | NA | Mir151 | | 0 | 0 | 0 | 0 | 0 | 0 | 0 | 0 | 0 | 0 | 0 |
| 0 | NA | NA | NA | NA | NA | Gm24787 | | 0 | 0 | 0 | 0 | 0 | 0 | 0 | 0 | 0 | 0 | 0 |
| 0 | NA | NA | NA | NA | NA | Gm19811 | | 0 | 0 | 0 | 0 | 0 | 0 | 0 | 0 | 0 | 0 | 0 |
| 0 | NA | NA | NA | NA | NA | Gm32405 | | 0 | 0 | 0 | 0 | 0 | 0 | 0 | 0 | 0 | 0 | 0 |
| 0 | NA | NA | NA | NA | NA | Gm26117 | | 0 | 0 | 0 | 0 | 0 | 0 | 0 | 0 | 0 | 0 | 0 |
| 0 | NA | NA | NA | NA | NA | Mroh4 | maestro heat-like repeat family member 4 | 0 | 0 | 0 | 0 | 0 | 0 | 0 | 0 | 0 | 0 | 0 |
| 0 | NA | NA | NA | NA | NA | Gm28976 | | 0 | 0 | 0 | 0 | 0 | 0 | 0 | 0 | 0 | 0 | 0 |
| 0 | NA | NA | NA | NA | NA | Slurp1 | secreted Ly6/Plaur domain containing 1 | 0 | 0 | 0 | 0 | 0 | 0 | 0 | 0 | 0 | 0 | 0 |
| 0 | NA | NA | NA | NA | NA | Lypd2 | Ly6/Plaur domain containing 2 | 0 | 0 | 0 | 0 | 0 | 0 | 0 | 0 | 0 | 0 | 0 |
| 0 | NA | NA | NA | NA | NA | Slurp2 | secreted Ly6/Plaur domain containing 2 | 0 | 0 | 0 | 0 | 0 | 0 | 0 | 0 | 0 | 0 | 0 |
| 0 | NA | NA | NA | NA | NA | Ly6g6g | | 0 | 0 | 0 | 0 | 0 | 0 | 0 | 0 | 0 | 0 | 0 |
| 0 | NA | NA | NA | NA | NA | Gm17189 | | 0 | 0 | 0 | 0 | 0 | 0 | 0 | 0 | 0 | 0 | 0 |
| 0 | NA | NA | NA | NA | NA | Gml | glycosylphosphatidylinositol anchored mo | 0 | 0 | 0 | 0 | 0 | 0 | 0 | 0 | 0 | 0 | 0 |
| 0 | NA | NA | NA | NA | NA | Gml2 | glycosylphosphatidylinositol anchored mo | 0 | 0 | 0 | 0 | 0 | 0 | 0 | 0 | 0 | 0 | 0 |
| 0 | NA | NA | NA | NA | NA | Cyp11b1 | cytochrome P450, family 11, subfamily b, | 0 | 0 | 0 | 0 | 0 | 0 | 0 | 0 | 0 | 0 | 0 |
| 0 | NA | NA | NA | NA | NA | Cyp11b2 | cytochrome P450, family 11, subfamily b, | 0 | 0 | 0 | 0 | 0 | 0 | 0 | 0 | 0 | 0 | 0 |
| 0 | NA | NA | NA | NA | NA | Ly6m | lymphocyte antigen 6 complex, locus M | 0 | 0 | 0 | 0 | 0 | 0 | 0 | 0 | 0 | 0 | 0 |
| 0 | NA | NA | NA | NA | NA | Gm6610 | | 0 | 0 | 0 | 0 | 0 | 0 | 0 | 0 | 0 | 0 | 0 |
| 0 | NA | NA | NA | NA | NA | Gm29126 | | 0 | 0 | 0 | 0 | 0 | 0 | 0 | 0 | 0 | 0 | 0 |
| 0 | NA | NA | NA | NA | NA | Gm37042 | | 0 | 0 | 0 | 0 | 0 | 0 | 0 | 0 | 0 | 0 | 0 |
| 0 | NA | NA | NA | NA | NA | Gm28068 | | 0 | 0 | 0 | 0 | 0 | 0 | 0 | 0 | 0 | 0 | 0 |
| 0 | NA | NA | NA | NA | NA | Gm28069 | | 0 | 0 | 0 | 0 | 0 | 0 | 0 | 0 | 0 | 0 | 0 |
| 0 | NA | NA | NA | NA | NA | Gm5960 | | 0 | 0 | 0 | 0 | 0 | 0 | 0 | 0 | 0 | 0 | 0 |
| 0 | NA | NA | NA | NA | NA | Gm28502 | | 0 | 0 | 0 | 0 | 0 | 0 | 0 | 0 | 0 | 0 | 0 |
| 0 | NA | NA | NA | NA | NA | Gm28117 | | 0 | 0 | 0 | 0 | 0 | 0 | 0 | 0 | 0 | 0 | 0 |
| 0 | NA | NA | NA | NA | NA | Gm10238 | | 0 | 0 | 0 | 0 | 0 | 0 | 0 | 0 | 0 | 0 | 0 |
| 0 | NA | NA | NA | NA | NA | Gm28115 | | 0 | 0 | 0 | 0 | 0 | 0 | 0 | 0 | 0 | 0 | 0 |
| 0 | NA | NA | NA | NA | NA | Ly6f | lymphocyte antigen 6 complex, locus F | 0 | 0 | 0 | 0 | 0 | 0 | 0 | 0 | 0 | 0 | 0 |
| 0 | NA | NA | NA | NA | NA | Gm28116 | | 0 | 0 | 0 | 0 | 0 | 0 | 0 | 0 | 0 | 0 | 0 |
| 0 | NA | NA | NA | NA | NA | Gm3454 | | 0 | 0 | 0 | 0 | 0 | 0 | 0 | 0 | 0 | 0 | 0 |
| 0 | NA | NA | NA | NA | NA | Gm28113 | | 0 | 0 | 0 | 0 | 0 | 0 | 0 | 0 | 0 | 0 | 0 |
| 0 | NA | NA | NA | NA | NA | Gm29517 | | 0 | 0 | 0 | 0 | 0 | 0 | 0 | 0 | 0 | 0 | 0 |
| 0 | NA | NA | NA | NA | NA | Gm28114 | | 0 | 0 | 0 | 0 | 0 | 0 | 0 | 0 | 0 | 0 | 0 |
| 0 | NA | NA | NA | NA | NA | Gm3142 | | 0 | 0 | 0 | 0 | 0 | 0 | 0 | 0 | 0 | 0 | 0 |
| 0 | NA | NA | NA | NA | NA | 9030619P08Rik | | 0 | 0 | 0 | 0 | 0 | 0 | 0 | 0 | 0 | 0 | 0 |
| 0 | NA | NA | NA | NA | NA | Ly6l | lymphocyte antigen 6 complex, locus L | 0 | 0 | 0 | 0 | 0 | 0 | 0 | 0 | 0 | 0 | 0 |
| 0 | NA | NA | NA | NA | NA | Gm34531 | | 0 | 0 | 0 | 0 | 0 | 0 | 0 | 0 | 0 | 0 | 0 |
| 0 | NA | NA | NA | NA | NA | Gm34593 | | 0 | 0 | 0 | 0 | 0 | 0 | 0 | 0 | 0 | 0 | 0 |
| 0 | NA | NA | NA | NA | NA | Gm34646 | predicted gene, 34646 | 0 | 0 | 0 | 0 | 0 | 0 | 0 | 0 | 0 | 0 | 0 |
| 0 | NA | NA | NA | NA | NA | Gm46519 | | 0 | 0 | 0 | 0 | 0 | 0 | 0 | 0 | 0 | 0 | 0 |
| 0 | NA | NA | NA | NA | NA | Gm23747 | | 0 | 0 | 0 | 0 | 0 | 0 | 0 | 0 | 0 | 0 | 0 |
| 0 | NA | NA | NA | NA | NA | Mir6952 | | 0 | 0 | 0 | 0 | 0 | 0 | 0 | 0 | 0 | 0 | 0 |
| 0 | NA | NA | NA | NA | NA | Gm19945 | | 0 | 0 | 0 | 0 | 0 | 0 | 0 | 0 | 0 | 0 | 0 |
| 0 | NA | NA | NA | NA | NA | Gm25720 | | 0 | 0 | 0 | 0 | 0 | 0 | 0 | 0 | 0 | 0 | 0 |
| 0 | NA | NA | NA | NA | NA | Mir1942 | | 0 | 0 | 0 | 0 | 0 | 0 | 0 | 0 | 0 | 0 | 0 |
| 0 | NA | NA | NA | NA | NA | Mir6953 | | 0 | 0 | 0 | 0 | 0 | 0 | 0 | 0 | 0 | 0 | 0 |
| 0 | NA | NA | NA | NA | NA | Mirt2 | | 0 | 0 | 0 | 0 | 0 | 0 | 0 | 0 | 0 | 0 | 0 |
| 0 | NA | NA | NA | NA | NA | Mirt2 | | 0 | 0 | 0 | 0 | 0 | 0 | 0 | 0 | 0 | 0 | 0 |
| 0 | NA | NA | NA | NA | NA | Spatc1 | spermatogenesis and centriole associate | 0 | 0 | 0 | 0 | 0 | 0 | 0 | 0 | 0 | 0 | 0 |
| 0 | NA | NA | NA | NA | NA | Mir3079 | | 0 | 0 | 0 | 0 | 0 | 0 | 0 | 0 | 0 | 0 | 0 |
| 0 | NA | NA | NA | NA | NA | Tssk5 | testis-specific serine kinase 5 | 0 | 0 | 0 | 0 | 0 | 0 | 0 | 0 | 0 | 0 | 0 |
| 0 | NA | NA | NA | NA | NA | Mir6954 | | 0 | 0 | 0 | 0 | 0 | 0 | 0 | 0 | 0 | 0 | 0 |
| 0 | NA | NA | NA | NA | NA | Tmem249 | transmembrane protein 249 | 0 | 0 | 0 | 0 | 0 | 0 | 0 | 0 | 0 | 0 | 0 |
| 0 | NA | NA | NA | NA | NA | Gm17271 | | 0 | 0 | 0 | 0 | 0 | 0 | 0 | 0 | 0 | 0 | 0 |

| | | | | | | | | | | | | | | | | | |
|---|---|---|---|---|---|---|---|---|---|---|---|---|---|---|---|---|---|
| 0 | NA | NA | NA | NA | NA | Gm49527 | | 0 | 0 | 0 | 0 | 0 | 0 | 0 | 0 | 0 | 0 | 0 |
| 0 | NA | NA | NA | NA | NA | Gm35933 | | 0 | 0 | 0 | 0 | 0 | 0 | 0 | 0 | 0 | 0 | 0 |
| 0 | NA | NA | NA | NA | NA | Mb | myoglobin | 0 | 0 | 0 | 0 | 0 | 0 | 0 | 0 | 0 | 0 | 0 |
| 0 | NA | NA | NA | NA | NA | Gm24753 | | 0 | 0 | 0 | 0 | 0 | 0 | 0 | 0 | 0 | 0 | 0 |
| 0 | NA | NA | NA | NA | NA | 1700109K24Rik | | 0 | 0 | 0 | 0 | 0 | 0 | 0 | 0 | 0 | 0 | 0 |
| 0 | NA | NA | NA | NA | NA | Gm49411 | | 0 | 0 | 0 | 0 | 0 | 0 | 0 | 0 | 0 | 0 | 0 |
| 0 | NA | NA | NA | NA | NA | Gm24056 | | 0 | 0 | 0 | 0 | 0 | 0 | 0 | 0 | 0 | 0 | 0 |
| 0 | NA | NA | NA | NA | NA | Gm20688 | | 0 | 0 | 0 | 0 | 0 | 0 | 0 | 0 | 0 | 0 | 0 |
| 0 | NA | NA | NA | NA | NA | Apol7b | apolipoprotein L 7b | 0 | 0 | 0 | 0 | 0 | 0 | 0 | 0 | 0 | 0 | 0 |
| 0 | NA | NA | NA | NA | NA | 4930571N24Rik | | 0 | 0 | 0 | 0 | 0 | 0 | 0 | 0 | 0 | 0 | 0 |
| 0 | NA | NA | NA | NA | NA | Gm49551 | | 0 | 0 | 0 | 0 | 0 | 0 | 0 | 0 | 0 | 0 | 0 |
| 0 | NA | NA | NA | NA | NA | Apol10a | apolipoprotein L 10A | 0 | 0 | 0 | 0 | 0 | 0 | 0 | 0 | 0 | 0 | 0 |
| 0 | NA | NA | NA | NA | NA | 1700025B11Rik | | 0 | 0 | 0 | 0 | 0 | 0 | 0 | 0 | 0 | 0 | 0 |
| 0 | NA | NA | NA | NA | NA | Gm36245 | | 0 | 0 | 0 | 0 | 0 | 0 | 0 | 0 | 0 | 0 | 0 |
| 0 | NA | NA | NA | NA | NA | Gm49407 | | 0 | 0 | 0 | 0 | 0 | 0 | 0 | 0 | 0 | 0 | 0 |
| 0 | NA | NA | NA | NA | NA | Gm8233 | | 0 | 0 | 0 | 0 | 0 | 0 | 0 | 0 | 0 | 0 | 0 |
| 0 | NA | NA | NA | NA | NA | Gm36329 | | 0 | 0 | 0 | 0 | 0 | 0 | 0 | 0 | 0 | 0 | 0 |
| 0 | NA | NA | NA | NA | NA | Mir7676-1 | | 0 | 0 | 0 | 0 | 0 | 0 | 0 | 0 | 0 | 0 | 0 |
| 0 | NA | NA | NA | NA | NA | Mir7676-2 | | 0 | 0 | 0 | 0 | 0 | 0 | 0 | 0 | 0 | 0 | 0 |
| 0 | NA | NA | NA | NA | NA | Gm26329 | | 0 | 0 | 0 | 0 | 0 | 0 | 0 | 0 | 0 | 0 | 0 |
| 0 | NA | NA | NA | NA | NA | Tex33 | testis expressed 33 | 0 | 0 | 0 | 0 | 0 | 0 | 0 | 0 | 0 | 0 | 0 |
| 0 | NA | NA | NA | NA | NA | Gm49481 | | 0 | 0 | 0 | 0 | 0 | 0 | 0 | 0 | 0 | 0 | 0 |
| 0 | NA | NA | NA | NA | NA | Gm36644 | | 0 | 0 | 0 | 0 | 0 | 0 | 0 | 0 | 0 | 0 | 0 |
| 0 | NA | NA | NA | NA | NA | 1700041B01Rik | | 0 | 0 | 0 | 0 | 0 | 0 | 0 | 0 | 0 | 0 | 0 |
| 0 | NA | NA | NA | NA | NA | Gm49510 | | 0 | 0 | 0 | 0 | 0 | 0 | 0 | 0 | 0 | 0 | 0 |
| 0 | NA | NA | NA | NA | NA | 1700027A07Rik | | 0 | 0 | 0 | 0 | 0 | 0 | 0 | 0 | 0 | 0 | 0 |
| 0 | NA | NA | NA | NA | NA | Mir6956 | | 0 | 0 | 0 | 0 | 0 | 0 | 0 | 0 | 0 | 0 | 0 |
| 0 | NA | NA | NA | NA | NA | Gm49471 | | 0 | 0 | 0 | 0 | 0 | 0 | 0 | 0 | 0 | 0 | 0 |
| 0 | NA | NA | NA | NA | NA | Gm25072 | | 0 | 0 | 0 | 0 | 0 | 0 | 0 | 0 | 0 | 0 | 0 |
| 0 | NA | NA | NA | NA | NA | Lgals1-ps2 | | 0 | 0 | 0 | 0 | 0 | 0 | 0 | 0 | 0 | 0 | 0 |
| 0 | NA | NA | NA | NA | NA | Gm20420 | | 0 | 0 | 0 | 0 | 0 | 0 | 0 | 0 | 0 | 0 | 0 |
| 0 | NA | NA | NA | NA | NA | Gm44302 | | 0 | 0 | 0 | 0 | 0 | 0 | 0 | 0 | 0 | 0 | 0 |
| 0 | NA | NA | NA | NA | NA | Gm27903 | | 0 | 0 | 0 | 0 | 0 | 0 | 0 | 0 | 0 | 0 | 0 |
| 0 | NA | NA | NA | NA | NA | Npcd | neuronal pentraxin chromo domain | 0 | 0 | 0 | 0 | 0 | 0 | 0 | 0 | 0 | 0 | 0 |
| 0 | NA | NA | NA | NA | NA | Gm10856 | | 0 | 0 | 0 | 0 | 0 | 0 | 0 | 0 | 0 | 0 | 0 |
| 0 | NA | NA | NA | NA | NA | Gm49412 | | 0 | 0 | 0 | 0 | 0 | 0 | 0 | 0 | 0 | 0 | 0 |
| 0 | NA | NA | NA | NA | NA | Mir7213 | | 0 | 0 | 0 | 0 | 0 | 0 | 0 | 0 | 0 | 0 | 0 |
| 0 | NA | NA | NA | NA | NA | Gm22077 | | 0 | 0 | 0 | 0 | 0 | 0 | 0 | 0 | 0 | 0 | 0 |
| 0 | NA | NA | NA | NA | NA | Gm24204 | | 0 | 0 | 0 | 0 | 0 | 0 | 0 | 0 | 0 | 0 | 0 |
| 0 | NA | NA | NA | NA | NA | 4930588D02Rik | | 0 | 0 | 0 | 0 | 0 | 0 | 0 | 0 | 0 | 0 | 0 |
| 0 | NA | NA | NA | NA | NA | | | 0 | 0 | 0 | 0 | 0 | 0 | 0 | 0 | 0 | 0 | 0 |
| 0 | NA | NA | NA | NA | NA | Gm31369 | | 0 | 0 | 0 | 0 | 0 | 0 | 0 | 0 | 0 | 0 | 0 |
| 0 | NA | NA | NA | NA | NA | Gm23220 | | 0 | 0 | 0 | 0 | 0 | 0 | 0 | 0 | 0 | 0 | 0 |
| 0 | NA | NA | NA | NA | NA | Gm19141 | | 0 | 0 | 0 | 0 | 0 | 0 | 0 | 0 | 0 | 0 | 0 |
| 0 | NA | NA | NA | NA | NA | Mir6957 | | 0 | 0 | 0 | 0 | 0 | 0 | 0 | 0 | 0 | 0 | 0 |
| 0 | NA | NA | NA | NA | NA | A430088P11Rik | | 0 | 0 | 0 | 0 | 0 | 0 | 0 | 0 | 0 | 0 | 0 |
| 0 | NA | NA | NA | NA | NA | Mir5113 | | 0 | 0 | 0 | 0 | 0 | 0 | 0 | 0 | 0 | 0 | 0 |
| 0 | NA | NA | NA | NA | NA | Gm25131 | | 0 | 0 | 0 | 0 | 0 | 0 | 0 | 0 | 0 | 0 | 0 |
| 0 | NA | NA | NA | NA | NA | Gm49522 | | 0 | 0 | 0 | 0 | 0 | 0 | 0 | 0 | 0 | 0 | 0 |
| 0 | NA | NA | NA | NA | NA | Gm49537 | | 0 | 0 | 0 | 0 | 0 | 0 | 0 | 0 | 0 | 0 | 0 |
| 0 | NA | NA | NA | NA | NA | Dnajb7 | DnaJ heat shock protein family (Hsp40) m | 0 | 0 | 0 | 0 | 0 | 0 | 0 | 0 | 0 | 0 | 0 |
| 0 | NA | NA | NA | NA | NA | Gm49406 | | 0 | 0 | 0 | 0 | 0 | 0 | 0 | 0 | 0 | 0 | 0 |
| 0 | NA | NA | NA | NA | NA | Gm23880 | | 0 | 0 | 0 | 0 | 0 | 0 | 0 | 0 | 0 | 0 | 0 |
| 0 | NA | NA | NA | NA | NA | Gm22067 | | 0 | 0 | 0 | 0 | 0 | 0 | 0 | 0 | 0 | 0 | 0 |
| 0 | NA | NA | NA | NA | NA | Gm49444 | | 0 | 0 | 0 | 0 | 0 | 0 | 0 | 0 | 0 | 0 | 0 |
| 0 | NA | NA | NA | NA | NA | Cyp2d67-ps | | 0 | 0 | 0 | 0 | 0 | 0 | 0 | 0 | 0 | 0 | 0 |
| 0 | NA | NA | NA | NA | NA | 1700029P11Ril | RIKEN cDNA 1700029P11 gene | 0 | 0 | 0 | 0 | 0 | 0 | 0 | 0 | 0 | 0 | 0 |
| 0 | NA | NA | NA | NA | NA | Gm49487 | | 0 | 0 | 0 | 0 | 0 | 0 | 0 | 0 | 0 | 0 | 0 |
| 0 | NA | NA | NA | NA | NA | Gm49488 | | 0 | 0 | 0 | 0 | 0 | 0 | 0 | 0 | 0 | 0 | 0 |
| 0 | NA | NA | NA | NA | NA | Gm49489 | | 0 | 0 | 0 | 0 | 0 | 0 | 0 | 0 | 0 | 0 | 0 |

| | | | | | | | | | | | | | | | | | | |
|---|---|---|---|---|---|---|---|---|---|---|---|---|---|---|---|---|---|---|
| 0 | NA | NA | NA | NA | NA | 4930407I10Rik | RIKEN cDNA 4930407I10 gene | 0 | 0 | 0 | 0 | 0 | 0 | 0 | 0 | 0 | 0 | 0 |
| 0 | NA | NA | NA | NA | NA | Mei1 | meiotic double-stranded break formation | 0 | 0 | 0 | 0 | 0 | 0 | 0 | 0 | 0 | 0 | 0 |
| 0 | NA | NA | NA | NA | NA | Mir33 | | 0 | 0 | 0 | 0 | 0 | 0 | 0 | 0 | 0 | 0 | 0 |
| 0 | NA | NA | NA | NA | NA | 1500009C09Rik | | 0 | 0 | 0 | 0 | 0 | 0 | 0 | 0 | 0 | 0 | 0 |
| 0 | NA | NA | NA | NA | NA | Cyp2d11 | cytochrome P450, family 2, subfamily d, p | 0 | 0 | 0 | 0 | 0 | 0 | 0 | 0 | 0 | 0 | 0 |
| 0 | NA | NA | NA | NA | NA | Mir7684 | | 0 | 0 | 0 | 0 | 0 | 0 | 0 | 0 | 0 | 0 | 0 |
| 0 | NA | NA | NA | NA | NA | Cyp2d10 | cytochrome P450, family 2, subfamily d, p | 0 | 0 | 0 | 0 | 0 | 0 | 0 | 0 | 0 | 0 | 0 |
| 0 | NA | NA | NA | NA | NA | Gm27331 | | 0 | 0 | 0 | 0 | 0 | 0 | 0 | 0 | 0 | 0 | 0 |
| 0 | NA | NA | NA | NA | NA | Gm25990 | | 0 | 0 | 0 | 0 | 0 | 0 | 0 | 0 | 0 | 0 | 0 |
| 0 | NA | NA | NA | NA | NA | Gm19267 | | 0 | 0 | 0 | 0 | 0 | 0 | 0 | 0 | 0 | 0 | 0 |
| 0 | NA | NA | NA | NA | NA | Gm46501 | | 0 | 0 | 0 | 0 | 0 | 0 | 0 | 0 | 0 | 0 | 0 |
| 0 | NA | NA | NA | NA | NA | Gm49421 | | 0 | 0 | 0 | 0 | 0 | 0 | 0 | 0 | 0 | 0 | 0 |
| 0 | NA | NA | NA | NA | NA | Gm49419 | | 0 | 0 | 0 | 0 | 0 | 0 | 0 | 0 | 0 | 0 | 0 |
| 0 | NA | NA | NA | NA | NA | Gm27622 | | 0 | 0 | 0 | 0 | 0 | 0 | 0 | 0 | 0 | 0 | 0 |
| 0 | NA | NA | NA | NA | NA | Cyp2d32-ps | | 0 | 0 | 0 | 0 | 0 | 0 | 0 | 0 | 0 | 0 | 0 |
| 0 | NA | NA | NA | NA | NA | Gm27825 | | 0 | 0 | 0 | 0 | 0 | 0 | 0 | 0 | 0 | 0 | 0 |
| 0 | NA | NA | NA | NA | NA | Gm27825 | | 0 | 0 | 0 | 0 | 0 | 0 | 0 | 0 | 0 | 0 | 0 |
| 0 | NA | NA | NA | NA | NA | Gm46525 | | 0 | 0 | 0 | 0 | 0 | 0 | 0 | 0 | 0 | 0 | 0 |
| 0 | NA | NA | NA | NA | NA | Gm46524 | | 0 | 0 | 0 | 0 | 0 | 0 | 0 | 0 | 0 | 0 | 0 |
| 0 | NA | NA | NA | NA | NA | Gm28023 | | 0 | 0 | 0 | 0 | 0 | 0 | 0 | 0 | 0 | 0 | 0 |
| 0 | NA | NA | NA | NA | NA | Gm28023 | | 0 | 0 | 0 | 0 | 0 | 0 | 0 | 0 | 0 | 0 | 0 |
| 0 | NA | NA | NA | NA | NA | Cyp2d12 | cytochrome P450, family 2, subfamily d, p | 0 | 0 | 0 | 0 | 0 | 0 | 0 | 0 | 0 | 0 | 0 |
| 0 | NA | NA | NA | NA | NA | Gm27924 | | 0 | 0 | 0 | 0 | 0 | 0 | 0 | 0 | 0 | 0 | 0 |
| 0 | NA | NA | NA | NA | NA | Cyp2d33-ps | | 0 | 0 | 0 | 0 | 0 | 0 | 0 | 0 | 0 | 0 | 0 |
| 0 | NA | NA | NA | NA | NA | Gm49429 | | 0 | 0 | 0 | 0 | 0 | 0 | 0 | 0 | 0 | 0 | 0 |
| 0 | NA | NA | NA | NA | NA | Cyp2d34-ps | | 0 | 0 | 0 | 0 | 0 | 0 | 0 | 0 | 0 | 0 | 0 |
| 0 | NA | NA | NA | NA | NA | Gm27742 | | 0 | 0 | 0 | 0 | 0 | 0 | 0 | 0 | 0 | 0 | 0 |
| 0 | NA | NA | NA | NA | NA | Gm49431 | | 0 | 0 | 0 | 0 | 0 | 0 | 0 | 0 | 0 | 0 | 0 |
| 0 | NA | NA | NA | NA | NA | Gm27463 | | 0 | 0 | 0 | 0 | 0 | 0 | 0 | 0 | 0 | 0 | 0 |
| 0 | NA | NA | NA | NA | NA | Cyp2d35-ps | | 0 | 0 | 0 | 0 | 0 | 0 | 0 | 0 | 0 | 0 | 0 |
| 0 | NA | NA | NA | NA | NA | Cyp2d36-ps | | 0 | 0 | 0 | 0 | 0 | 0 | 0 | 0 | 0 | 0 | 0 |
| 0 | NA | NA | NA | NA | NA | Cyp2d38-ps | | 0 | 0 | 0 | 0 | 0 | 0 | 0 | 0 | 0 | 0 | 0 |
| 0 | NA | NA | NA | NA | NA | Cyp2d39-ps | | 0 | 0 | 0 | 0 | 0 | 0 | 0 | 0 | 0 | 0 | 0 |
| 0 | NA | NA | NA | NA | NA | Gm49433 | | 0 | 0 | 0 | 0 | 0 | 0 | 0 | 0 | 0 | 0 | 0 |
| 0 | NA | NA | NA | NA | NA | Cyp2d40 | cytochrome P450, family 2, subfamily d, p | 0 | 0 | 0 | 0 | 0 | 0 | 0 | 0 | 0 | 0 | 0 |
| 0 | NA | NA | NA | NA | NA | Cyp2d41-ps | | 0 | 0 | 0 | 0 | 0 | 0 | 0 | 0 | 0 | 0 | 0 |
| 0 | NA | NA | NA | NA | NA | Gm27618 | | 0 | 0 | 0 | 0 | 0 | 0 | 0 | 0 | 0 | 0 | 0 |
| 0 | NA | NA | NA | NA | NA | Mir3080 | | 0 | 0 | 0 | 0 | 0 | 0 | 0 | 0 | 0 | 0 | 0 |
| 0 | NA | NA | NA | NA | NA | Gm17206 | | 0 | 0 | 0 | 0 | 0 | 0 | 0 | 0 | 0 | 0 | 0 |
| 0 | NA | NA | NA | NA | NA | Gm24575 | | 0 | 0 | 0 | 0 | 0 | 0 | 0 | 0 | 0 | 0 | 0 |
| 0 | NA | NA | NA | NA | NA | Gm46495 | | 0 | 0 | 0 | 0 | 0 | 0 | 0 | 0 | 0 | 0 | 0 |
| 0 | NA | NA | NA | NA | NA | Mpped1 | metallophosphoesterase domain containi | 0 | 0 | 0 | 0 | 0 | 0 | 0 | 0 | 0 | 0 | 0 |
| 0 | NA | NA | NA | NA | NA | Rpl31-ps22 | | 0 | 0 | 0 | 0 | 0 | 0 | 0 | 0 | 0 | 0 | 0 |
| 0 | NA | NA | NA | NA | NA | Gm33432 | | 0 | 0 | 0 | 0 | 0 | 0 | 0 | 0 | 0 | 0 | 0 |
| 0 | NA | NA | NA | NA | NA | Gm49513 | | 0 | 0 | 0 | 0 | 0 | 0 | 0 | 0 | 0 | 0 | 0 |
| 0 | NA | NA | NA | NA | NA | Gm22890 | | 0 | 0 | 0 | 0 | 0 | 0 | 0 | 0 | 0 | 0 | 0 |
| 0 | NA | NA | NA | NA | NA | Gm5214 | | 0 | 0 | 0 | 0 | 0 | 0 | 0 | 0 | 0 | 0 | 0 |
| 0 | NA | NA | NA | NA | NA | Gm34095 | | 0 | 0 | 0 | 0 | 0 | 0 | 0 | 0 | 0 | 0 | 0 |
| 0 | NA | NA | NA | NA | NA | 4930513L16Rik | | 0 | 0 | 0 | 0 | 0 | 0 | 0 | 0 | 0 | 0 | 0 |
| 0 | NA | NA | NA | NA | NA | Mir1249 | | 0 | 0 | 0 | 0 | 0 | 0 | 0 | 0 | 0 | 0 | 0 |
| 0 | NA | NA | NA | NA | NA | Gm49523 | | 0 | 0 | 0 | 0 | 0 | 0 | 0 | 0 | 0 | 0 | 0 |
| 0 | NA | NA | NA | NA | NA | Gm23517 | | 0 | 0 | 0 | 0 | 0 | 0 | 0 | 0 | 0 | 0 | 0 |
| 0 | NA | NA | NA | NA | NA | 7530416G11Ril | RIKEN cDNA 7530416G11 gene | 0 | 0 | 0 | 0 | 0 | 0 | 0 | 0 | 0 | 0 | 0 |
| 0 | NA | NA | NA | NA | NA | Mirlet7c-2 | | 0 | 0 | 0 | 0 | 0 | 0 | 0 | 0 | 0 | 0 | 0 |
| 0 | NA | NA | NA | NA | NA | Mirlet7b | | 0 | 0 | 0 | 0 | 0 | 0 | 0 | 0 | 0 | 0 | 0 |
| 0 | NA | NA | NA | NA | NA | Gm24011 | | 0 | 0 | 0 | 0 | 0 | 0 | 0 | 0 | 0 | 0 | 0 |
| 0 | NA | NA | NA | NA | NA | A930001M01Rik | | 0 | 0 | 0 | 0 | 0 | 0 | 0 | 0 | 0 | 0 | 0 |
| 0 | NA | NA | NA | NA | NA | Gm49464 | | 0 | 0 | 0 | 0 | 0 | 0 | 0 | 0 | 0 | 0 | 0 |
| 0 | NA | NA | NA | NA | NA | Gm49465 | | 0 | 0 | 0 | 0 | 0 | 0 | 0 | 0 | 0 | 0 | 0 |
| 0 | NA | NA | NA | NA | NA | Gm49466 | | 0 | 0 | 0 | 0 | 0 | 0 | 0 | 0 | 0 | 0 | 0 |

| | | | | | | | | | | | | | | | | | | |
|---|---|---|---|---|---|---|---|---|---|---|---|---|---|---|---|---|---|---|
| 0 | NA | NA | NA | NA | NA | Gm49467 | | 0 | 0 | 0 | 0 | 0 | 0 | 0 | 0 | 0 | 0 | 0 |
| 0 | NA | NA | NA | NA | NA | Gm49468 | | 0 | 0 | 0 | 0 | 0 | 0 | 0 | 0 | 0 | 0 | 0 |
| 0 | NA | NA | NA | NA | NA | Gm23416 | | 0 | 0 | 0 | 0 | 0 | 0 | 0 | 0 | 0 | 0 | 0 |
| 0 | NA | NA | NA | NA | NA | Gm49469 | | 0 | 0 | 0 | 0 | 0 | 0 | 0 | 0 | 0 | 0 | 0 |
| 0 | NA | NA | NA | NA | NA | Gm49470 | | 0 | 0 | 0 | 0 | 0 | 0 | 0 | 0 | 0 | 0 | 0 |
| 0 | NA | NA | NA | NA | NA | n-R5s42 | | 0 | 0 | 0 | 0 | 0 | 0 | 0 | 0 | 0 | 0 | 0 |
| 0 | NA | NA | NA | NA | NA | A930027H12Rik | | 0 | 0 | 0 | 0 | 0 | 0 | 0 | 0 | 0 | 0 | 0 |
| 0 | NA | NA | NA | NA | NA | Mir6393 | | 0 | 0 | 0 | 0 | 0 | 0 | 0 | 0 | 0 | 0 | 0 |
| 0 | NA | NA | NA | NA | NA | Gm49472 | | 0 | 0 | 0 | 0 | 0 | 0 | 0 | 0 | 0 | 0 | 0 |
| 0 | NA | NA | NA | NA | NA | Gm35772 | | 0 | 0 | 0 | 0 | 0 | 0 | 0 | 0 | 0 | 0 | 0 |
| 0 | NA | NA | NA | NA | NA | 4930445N06Rik | | 0 | 0 | 0 | 0 | 0 | 0 | 0 | 0 | 0 | 0 | 0 |
| 0 | NA | NA | NA | NA | NA | Zdhhc25 | zinc finger, DHHC domain containing 25 | 0 | 0 | 0 | 0 | 0 | 0 | 0 | 0 | 0 | 0 | 0 |
| 0 | NA | NA | NA | NA | NA | Gm23144 | | 0 | 0 | 0 | 0 | 0 | 0 | 0 | 0 | 0 | 0 | 0 |
| 0 | NA | NA | NA | NA | NA | Ttll8 | tubulin tyrosine ligase-like family, membe | 0 | 0 | 0 | 0 | 0 | 0 | 0 | 0 | 0 | 0 | 0 |
| 0 | NA | NA | NA | NA | NA | Mov10l1 | Mov10 like RISC complex RNA helicase | 0 | 0 | 0 | 0 | 0 | 0 | 0 | 0 | 0 | 0 | 0 |
| 0 | NA | NA | NA | NA | NA | Gm23737 | | 0 | 0 | 0 | 0 | 0 | 0 | 0 | 0 | 0 | 0 | 0 |
| 0 | NA | NA | NA | NA | NA | Mir7118 | | 0 | 0 | 0 | 0 | 0 | 0 | 0 | 0 | 0 | 0 | 0 |
| 0 | NA | NA | NA | NA | NA | Mir6958 | | 0 | 0 | 0 | 0 | 0 | 0 | 0 | 0 | 0 | 0 | 0 |
| 0 | NA | NA | NA | NA | NA | Klhdc7b | kelch domain containing 7B | 0 | 0 | 0 | 0 | 0 | 0 | 0 | 0 | 0 | 0 | 0 |
| 0 | NA | NA | NA | NA | NA | Syce3 | synaptonemal complex central element p | 0 | 0 | 0 | 0 | 0 | 0 | 0 | 0 | 0 | 0 | 0 |
| 0 | NA | NA | NA | NA | NA | Gm44502 | | 0 | 0 | 0 | 0 | 0 | 0 | 0 | 0 | 0 | 0 | 0 |
| 0 | NA | NA | NA | NA | NA | Gm15609 | | 0 | 0 | 0 | 0 | 0 | 0 | 0 | 0 | 0 | 0 | 0 |
| 0 | NA | NA | NA | NA | NA | Gm18476 | | 0 | 0 | 0 | 0 | 0 | 0 | 0 | 0 | 0 | 0 | 0 |
| 0 | NA | NA | NA | NA | NA | Gm24643 | | 0 | 0 | 0 | 0 | 0 | 0 | 0 | 0 | 0 | 0 | 0 |
| 0 | NA | NA | NA | NA | NA | Gm24415 | | 0 | 0 | 0 | 0 | 0 | 0 | 0 | 0 | 0 | 0 | 0 |
| 0 | NA | NA | NA | NA | NA | Gm36330 | | 0 | 0 | 0 | 0 | 0 | 0 | 0 | 0 | 0 | 0 | 0 |
| 0 | NA | NA | NA | NA | NA | Mir7648 | | 0 | 0 | 0 | 0 | 0 | 0 | 0 | 0 | 0 | 0 | 0 |
| 0 | NA | NA | NA | NA | NA | Gm36480 | | 0 | 0 | 0 | 0 | 0 | 0 | 0 | 0 | 0 | 0 | 0 |
| 0 | NA | NA | NA | NA | NA | Gm6746 | | 0 | 0 | 0 | 0 | 0 | 0 | 0 | 0 | 0 | 0 | 0 |
| 0 | NA | NA | NA | NA | NA | Gm49541 | | 0 | 0 | 0 | 0 | 0 | 0 | 0 | 0 | 0 | 0 | 0 |
| 0 | NA | NA | NA | NA | NA | Gm3861 | | 0 | 0 | 0 | 0 | 0 | 0 | 0 | 0 | 0 | 0 | 0 |
| 0 | NA | NA | NA | NA | NA | Spn-ps | | 0 | 0 | 0 | 0 | 0 | 0 | 0 | 0 | 0 | 0 | 0 |
| 0 | NA | NA | NA | NA | NA | Gm49545 | | 0 | 0 | 0 | 0 | 0 | 0 | 0 | 0 | 0 | 0 | 0 |
| 0 | NA | NA | NA | NA | NA | Gm18814 | | 0 | 0 | 0 | 0 | 0 | 0 | 0 | 0 | 0 | 0 | 0 |
| 0 | NA | NA | NA | NA | NA | Gm7167 | | 0 | 0 | 0 | 0 | 0 | 0 | 0 | 0 | 0 | 0 | 0 |
| 0 | NA | NA | NA | NA | NA | CN725425 | cDNA sequence CN725425 | 0 | 0 | 0 | 0 | 0 | 0 | 0 | 0 | 0 | 0 | 0 |
| 0 | NA | NA | NA | NA | NA | 4933438A12Rik | | 0 | 0 | 0 | 0 | 0 | 0 | 0 | 0 | 0 | 0 | 0 |
| 0 | NA | NA | NA | NA | NA | Gm25389 | | 0 | 0 | 0 | 0 | 0 | 0 | 0 | 0 | 0 | 0 | 0 |
| 0 | NA | NA | NA | NA | NA | Smgc | submandibular gland protein C | 0 | 0 | 0 | 0 | 0 | 0 | 0 | 0 | 0 | 0 | 0 |
| 0 | NA | NA | NA | NA | NA | Muc19 | | 0 | 0 | 0 | 0 | 0 | 0 | 0 | 0 | 0 | 0 | 0 |
| 0 | NA | NA | NA | NA | NA | Gm18979 | | 0 | 0 | 0 | 0 | 0 | 0 | 0 | 0 | 0 | 0 | 0 |
| 0 | NA | NA | NA | NA | NA | 4930588J15Rik | | 0 | 0 | 0 | 0 | 0 | 0 | 0 | 0 | 0 | 0 | 0 |
| 0 | NA | NA | NA | NA | NA | Gm49418 | | 0 | 0 | 0 | 0 | 0 | 0 | 0 | 0 | 0 | 0 | 0 |
| 0 | NA | NA | NA | NA | NA | Gm18815 | | 0 | 0 | 0 | 0 | 0 | 0 | 0 | 0 | 0 | 0 | 0 |
| 0 | NA | NA | NA | NA | NA | Gm30085 | | 0 | 0 | 0 | 0 | 0 | 0 | 0 | 0 | 0 | 0 | 0 |
| 0 | NA | NA | NA | NA | NA | Gm18685 | | 0 | 0 | 0 | 0 | 0 | 0 | 0 | 0 | 0 | 0 | 0 |
| 0 | NA | NA | NA | NA | NA | Gm49420 | | 0 | 0 | 0 | 0 | 0 | 0 | 0 | 0 | 0 | 0 | 0 |
| 0 | NA | NA | NA | NA | NA | Gm24447 | | 0 | 0 | 0 | 0 | 0 | 0 | 0 | 0 | 0 | 0 | 0 |
| 0 | NA | NA | NA | NA | NA | Gm26371 | | 0 | 0 | 0 | 0 | 0 | 0 | 0 | 0 | 0 | 0 | 0 |
| 0 | NA | NA | NA | NA | NA | Zfp42-ps1 | | 0 | 0 | 0 | 0 | 0 | 0 | 0 | 0 | 0 | 0 | 0 |
| 0 | NA | NA | NA | NA | NA | Gm41386 | | 0 | 0 | 0 | 0 | 0 | 0 | 0 | 0 | 0 | 0 | 0 |
| 0 | NA | NA | NA | NA | NA | Gm30339 | | 0 | 0 | 0 | 0 | 0 | 0 | 0 | 0 | 0 | 0 | 0 |
| 0 | NA | NA | NA | NA | NA | Gm49445 | | 0 | 0 | 0 | 0 | 0 | 0 | 0 | 0 | 0 | 0 | 0 |
| 0 | NA | NA | NA | NA | NA | 9430014N10Rik | | 0 | 0 | 0 | 0 | 0 | 0 | 0 | 0 | 0 | 0 | 0 |
| 0 | NA | NA | NA | NA | NA | Gm30564 | | 0 | 0 | 0 | 0 | 0 | 0 | 0 | 0 | 0 | 0 | 0 |
| 0 | NA | NA | NA | NA | NA | Gm23129 | | 0 | 0 | 0 | 0 | 0 | 0 | 0 | 0 | 0 | 0 | 0 |
| 0 | NA | NA | NA | NA | NA | 1700129L04Rik | | 0 | 0 | 0 | 0 | 0 | 0 | 0 | 0 | 0 | 0 | 0 |
| 0 | NA | NA | NA | NA | NA | Gm24668 | | 0 | 0 | 0 | 0 | 0 | 0 | 0 | 0 | 0 | 0 | 0 |
| 0 | NA | NA | NA | NA | NA | Gm8843 | | 0 | 0 | 0 | 0 | 0 | 0 | 0 | 0 | 0 | 0 | 0 |
| 0 | NA | NA | NA | NA | NA | Gm25070 | | 0 | 0 | 0 | 0 | 0 | 0 | 0 | 0 | 0 | 0 | 0 |

| | | | | | | | | | | | | | | | | | |
|---|---|---|---|---|---|---|---|---|---|---|---|---|---|---|---|---|---|
| 0 | NA | NA | NA | NA | NA | Gm25397 | | 0 | 0 | 0 | 0 | 0 | 0 | 0 | 0 | 0 | 0 | 0 |
| 0 | NA | NA | NA | NA | NA | Gm41392 | | 0 | 0 | 0 | 0 | 0 | 0 | 0 | 0 | 0 | 0 | 0 |
| 0 | NA | NA | NA | NA | NA | Gm22045 | | 0 | 0 | 0 | 0 | 0 | 0 | 0 | 0 | 0 | 0 | 0 |
| 0 | NA | NA | NA | NA | NA | Gm38144 | | 0 | 0 | 0 | 0 | 0 | 0 | 0 | 0 | 0 | 0 | 0 |
| 0 | NA | NA | NA | NA | NA | Gm8888 | | 0 | 0 | 0 | 0 | 0 | 0 | 0 | 0 | 0 | 0 | 0 |
| 0 | NA | NA | NA | NA | NA | Rpl10a-ps3 | | 0 | 0 | 0 | 0 | 0 | 0 | 0 | 0 | 0 | 0 | 0 |
| 0 | NA | NA | NA | NA | NA | Rapgef3os1 | | 0 | 0 | 0 | 0 | 0 | 0 | 0 | 0 | 0 | 0 | 0 |
| 0 | NA | NA | NA | NA | NA | Gm22202 | | 0 | 0 | 0 | 0 | 0 | 0 | 0 | 0 | 0 | 0 | 0 |
| 0 | NA | NA | NA | NA | NA | Gm25668 | | 0 | 0 | 0 | 0 | 0 | 0 | 0 | 0 | 0 | 0 | 0 |
| 0 | NA | NA | NA | NA | NA | Olfr288 | olfactory receptor 288 | 0 | 0 | 0 | 0 | 0 | 0 | 0 | 0 | 0 | 0 | 0 |
| 0 | NA | NA | NA | NA | NA | Gm44579 | | 0 | 0 | 0 | 0 | 0 | 0 | 0 | 0 | 0 | 0 | 0 |
| 0 | NA | NA | NA | NA | NA | Olfr287 | olfactory receptor 287 | 0 | 0 | 0 | 0 | 0 | 0 | 0 | 0 | 0 | 0 | 0 |
| 0 | NA | NA | NA | NA | NA | Olfr286 | olfactory receptor 286 | 0 | 0 | 0 | 0 | 0 | 0 | 0 | 0 | 0 | 0 | 0 |
| 0 | NA | NA | NA | NA | NA | H1f7 | H1.7 linker histone | 0 | 0 | 0 | 0 | 0 | 0 | 0 | 0 | 0 | 0 | 0 |
| 0 | NA | NA | NA | NA | NA | Olfr285 | olfactory receptor 285 | 0 | 0 | 0 | 0 | 0 | 0 | 0 | 0 | 0 | 0 | 0 |
| 0 | NA | NA | NA | NA | NA | Olfr284 | olfactory receptor 284 | 0 | 0 | 0 | 0 | 0 | 0 | 0 | 0 | 0 | 0 | 0 |
| 0 | NA | NA | NA | NA | NA | Olfr283 | olfactory receptor 283 | 0 | 0 | 0 | 0 | 0 | 0 | 0 | 0 | 0 | 0 | 0 |
| 0 | NA | NA | NA | NA | NA | Gm49443 | | 0 | 0 | 0 | 0 | 0 | 0 | 0 | 0 | 0 | 0 | 0 |
| 0 | NA | NA | NA | NA | NA | 1700031M16Rik | | 0 | 0 | 0 | 0 | 0 | 0 | 0 | 0 | 0 | 0 | 0 |
| 0 | NA | NA | NA | NA | NA | Olfr282 | olfactory receptor 282 | 0 | 0 | 0 | 0 | 0 | 0 | 0 | 0 | 0 | 0 | 0 |
| 0 | NA | NA | NA | NA | NA | Gm6979 | | 0 | 0 | 0 | 0 | 0 | 0 | 0 | 0 | 0 | 0 | 0 |
| 0 | NA | NA | NA | NA | NA | Olfr281 | olfactory receptor 281 | 0 | 0 | 0 | 0 | 0 | 0 | 0 | 0 | 0 | 0 | 0 |
| 0 | NA | NA | NA | NA | NA | | | 0 | 0 | 0 | 0 | 0 | 0 | 0 | 0 | 0 | 0 | 0 |
| 0 | NA | NA | NA | NA | NA | Lalba | lactalbumin, alpha | 0 | 0 | 0 | 0 | 0 | 0 | 0 | 0 | 0 | 0 | 0 |
| 0 | NA | NA | NA | NA | NA | Olfr279 | olfactory receptor 279 | 0 | 0 | 0 | 0 | 0 | 0 | 0 | 0 | 0 | 0 | 0 |
| 0 | NA | NA | NA | NA | NA | Olfr278-ps1 | | 0 | 0 | 0 | 0 | 0 | 0 | 0 | 0 | 0 | 0 | 0 |
| 0 | NA | NA | NA | NA | NA | Snora2b | | 0 | 0 | 0 | 0 | 0 | 0 | 0 | 0 | 0 | 0 | 0 |
| 0 | NA | NA | NA | NA | NA | Gm34152 | | 0 | 0 | 0 | 0 | 0 | 0 | 0 | 0 | 0 | 0 | 0 |
| 0 | NA | NA | NA | NA | NA | Gm49437 | | 0 | 0 | 0 | 0 | 0 | 0 | 0 | 0 | 0 | 0 | 0 |
| 0 | NA | NA | NA | NA | NA | Gm49450 | | 0 | 0 | 0 | 0 | 0 | 0 | 0 | 0 | 0 | 0 | 0 |
| 0 | NA | NA | NA | NA | NA | Gm8973 | | 0 | 0 | 0 | 0 | 0 | 0 | 0 | 0 | 0 | 0 | 0 |
| 0 | NA | NA | NA | NA | NA | Gm34284 | | 0 | 0 | 0 | 0 | 0 | 0 | 0 | 0 | 0 | 0 | 0 |
| 0 | NA | NA | NA | NA | NA | Mir6960 | | 0 | 0 | 0 | 0 | 0 | 0 | 0 | 0 | 0 | 0 | 0 |
| 0 | NA | NA | NA | NA | NA | Gm25183 | | 0 | 0 | 0 | 0 | 0 | 0 | 0 | 0 | 0 | 0 | 0 |
| 0 | NA | NA | NA | NA | NA | Gm25057 | | 0 | 0 | 0 | 0 | 0 | 0 | 0 | 0 | 0 | 0 | 0 |
| 0 | NA | NA | NA | NA | NA | Gm49458 | | 0 | 0 | 0 | 0 | 0 | 0 | 0 | 0 | 0 | 0 | 0 |
| 0 | NA | NA | NA | NA | NA | Gm25999 | | 0 | 0 | 0 | 0 | 0 | 0 | 0 | 0 | 0 | 0 | 0 |
| 0 | NA | NA | NA | NA | NA | Gm49478 | | 0 | 0 | 0 | 0 | 0 | 0 | 0 | 0 | 0 | 0 | 0 |
| 0 | NA | NA | NA | NA | NA | Gm22951 | | 0 | 0 | 0 | 0 | 0 | 0 | 0 | 0 | 0 | 0 | 0 |
| 0 | NA | NA | NA | NA | NA | Gm49483 | | 0 | 0 | 0 | 0 | 0 | 0 | 0 | 0 | 0 | 0 | 0 |
| 0 | NA | NA | NA | NA | NA | Nckap5los | | 0 | 0 | 0 | 0 | 0 | 0 | 0 | 0 | 0 | 0 | 0 |
| 0 | NA | NA | NA | NA | NA | Gm49484 | | 0 | 0 | 0 | 0 | 0 | 0 | 0 | 0 | 0 | 0 | 0 |
| 0 | NA | NA | NA | NA | NA | Gm49485 | | 0 | 0 | 0 | 0 | 0 | 0 | 0 | 0 | 0 | 0 | 0 |
| 0 | NA | NA | NA | NA | NA | Aqp2 | aquaporin 2 | 0 | 0 | 0 | 0 | 0 | 0 | 0 | 0 | 0 | 0 | 0 |
| 0 | NA | NA | NA | NA | NA | Aqp6 | aquaporin 6 | 0 | 0 | 0 | 0 | 0 | 0 | 0 | 0 | 0 | 0 | 0 |
| 0 | NA | NA | NA | NA | NA | Gm17058 | | 0 | 0 | 0 | 0 | 0 | 0 | 0 | 0 | 0 | 0 | 0 |
| 0 | NA | NA | NA | NA | NA | Gm25897 | | 0 | 0 | 0 | 0 | 0 | 0 | 0 | 0 | 0 | 0 | 0 |
| 0 | NA | NA | NA | NA | NA | Gm17057 | | 0 | 0 | 0 | 0 | 0 | 0 | 0 | 0 | 0 | 0 | 0 |
| 0 | NA | NA | NA | NA | NA | Gm20564 | | 0 | 0 | 0 | 0 | 0 | 0 | 0 | 0 | 0 | 0 | 0 |
| 0 | NA | NA | NA | NA | NA | Gm23133 | | 0 | 0 | 0 | 0 | 0 | 0 | 0 | 0 | 0 | 0 | 0 |
| 0 | NA | NA | NA | NA | NA | Gm49474 | | 0 | 0 | 0 | 0 | 0 | 0 | 0 | 0 | 0 | 0 | 0 |
| 0 | NA | NA | NA | NA | NA | 4930478M13Rik | | 0 | 0 | 0 | 0 | 0 | 0 | 0 | 0 | 0 | 0 | 0 |
| 0 | NA | NA | NA | NA | NA | Gm49475 | | 0 | 0 | 0 | 0 | 0 | 0 | 0 | 0 | 0 | 0 | 0 |
| 0 | NA | NA | NA | NA | NA | Tmprss12 | transmembrane (C-terminal) protease, se | 0 | 0 | 0 | 0 | 0 | 0 | 0 | 0 | 0 | 0 | 0 |
| 0 | NA | NA | NA | NA | NA | Methig1 | methyltransferase hypoxia inducible dom | 0 | 0 | 0 | 0 | 0 | 0 | 0 | 0 | 0 | 0 | 0 |
| 0 | NA | NA | NA | NA | NA | Higd1c | HIG1 domain family, member 1C | 0 | 0 | 0 | 0 | 0 | 0 | 0 | 0 | 0 | 0 | 0 |
| 0 | NA | NA | NA | NA | NA | n-R5s43 | | 0 | 0 | 0 | 0 | 0 | 0 | 0 | 0 | 0 | 0 | 0 |
| 0 | NA | NA | NA | NA | NA | Gm27209 | | 0 | 0 | 0 | 0 | 0 | 0 | 0 | 0 | 0 | 0 | 0 |
| 0 | NA | NA | NA | NA | NA | Mir6961 | | 0 | 0 | 0 | 0 | 0 | 0 | 0 | 0 | 0 | 0 | 0 |
| 0 | NA | NA | NA | NA | NA | Gm35569 | | 0 | 0 | 0 | 0 | 0 | 0 | 0 | 0 | 0 | 0 | 0 |

| | | | | | | | | | | | | | | | | | | |
|---|---|---|---|---|---|---|---|---|---|---|---|---|---|---|---|---|---|---|
| 0 | NA | NA | NA | NA | NA | Gm16031 | | 0 | 0 | 0 | 0 | 0 | 0 | 0 | 0 | 0 | 0 | 0 |
| 0 | NA | NA | NA | NA | NA | Mir6962 | | 0 | 0 | 0 | 0 | 0 | 0 | 0 | 0 | 0 | 0 | 0 |
| 0 | NA | NA | NA | NA | NA | AU021063 | | 0 | 0 | 0 | 0 | 0 | 0 | 0 | 0 | 0 | 0 | 0 |
| 0 | NA | NA | NA | NA | NA | Gm49499 | | 0 | 0 | 0 | 0 | 0 | 0 | 0 | 0 | 0 | 0 | 0 |
| 0 | NA | NA | NA | NA | NA | Mir1941 | | 0 | 0 | 0 | 0 | 0 | 0 | 0 | 0 | 0 | 0 | 0 |
| 0 | NA | NA | NA | NA | NA | Gm49501 | | 0 | 0 | 0 | 0 | 0 | 0 | 0 | 0 | 0 | 0 | 0 |
| 0 | NA | NA | NA | NA | NA | Krt83 | keratin 83 | 0 | 0 | 0 | 0 | 0 | 0 | 0 | 0 | 0 | 0 | 0 |
| 0 | NA | NA | NA | NA | NA | Gm49423 | | 0 | 0 | 0 | 0 | 0 | 0 | 0 | 0 | 0 | 0 | 0 |
| 0 | NA | NA | NA | NA | NA | Gm49424 | | 0 | 0 | 0 | 0 | 0 | 0 | 0 | 0 | 0 | 0 | 0 |
| 0 | NA | NA | NA | NA | NA | Gm49425 | | 0 | 0 | 0 | 0 | 0 | 0 | 0 | 0 | 0 | 0 | 0 |
| 0 | NA | NA | NA | NA | NA | Krt84 | keratin 84 | 0 | 0 | 0 | 0 | 0 | 0 | 0 | 0 | 0 | 0 | 0 |
| 0 | NA | NA | NA | NA | NA | Krt82 | keratin 82 | 0 | 0 | 0 | 0 | 0 | 0 | 0 | 0 | 0 | 0 | 0 |
| 0 | NA | NA | NA | NA | NA | Gm44393 | | 0 | 0 | 0 | 0 | 0 | 0 | 0 | 0 | 0 | 0 | 0 |
| 0 | NA | NA | NA | NA | NA | Krt75 | keratin 75 | 0 | 0 | 0 | 0 | 0 | 0 | 0 | 0 | 0 | 0 | 0 |
| 0 | NA | NA | NA | NA | NA | Gm44460 | | 0 | 0 | 0 | 0 | 0 | 0 | 0 | 0 | 0 | 0 | 0 |
| 0 | NA | NA | NA | NA | NA | Gm5476 | | 0 | 0 | 0 | 0 | 0 | 0 | 0 | 0 | 0 | 0 | 0 |
| 0 | NA | NA | NA | NA | NA | Gm5477 | | 0 | 0 | 0 | 0 | 0 | 0 | 0 | 0 | 0 | 0 | 0 |
| 0 | NA | NA | NA | NA | NA | Gm49426 | | 0 | 0 | 0 | 0 | 0 | 0 | 0 | 0 | 0 | 0 | 0 |
| 0 | NA | NA | NA | NA | NA | Gm5414 | predicted gene 5414 | 0 | 0 | 0 | 0 | 0 | 0 | 0 | 0 | 0 | 0 | 0 |
| 0 | NA | NA | NA | NA | NA | Gm17851 | | 0 | 0 | 0 | 0 | 0 | 0 | 0 | 0 | 0 | 0 | 0 |
| 0 | NA | NA | NA | NA | NA | Gm5478 | | 0 | 0 | 0 | 0 | 0 | 0 | 0 | 0 | 0 | 0 | 0 |
| 0 | NA | NA | NA | NA | NA | Gm44479 | | 0 | 0 | 0 | 0 | 0 | 0 | 0 | 0 | 0 | 0 | 0 |
| 0 | NA | NA | NA | NA | NA | | | 0 | 0 | 0 | 0 | 0 | 0 | 0 | 0 | 0 | 0 | 0 |
| 0 | NA | NA | NA | NA | NA | Krt6b | keratin 6B | 0 | 0 | 0 | 0 | 0 | 0 | 0 | 0 | 0 | 0 | 0 |
| 0 | NA | NA | NA | NA | NA | Gm44383 | | 0 | 0 | 0 | 0 | 0 | 0 | 0 | 0 | 0 | 0 | 0 |
| 0 | NA | NA | NA | NA | NA | Krt6a | keratin 6A | 0 | 0 | 0 | 0 | 0 | 0 | 0 | 0 | 0 | 0 | 0 |
| 0 | NA | NA | NA | NA | NA | Gm49428 | | 0 | 0 | 0 | 0 | 0 | 0 | 0 | 0 | 0 | 0 | 0 |
| 0 | NA | NA | NA | NA | NA | Gm44342 | | 0 | 0 | 0 | 0 | 0 | 0 | 0 | 0 | 0 | 0 | 0 |
| 0 | NA | NA | NA | NA | NA | Gm44323 | | 0 | 0 | 0 | 0 | 0 | 0 | 0 | 0 | 0 | 0 | 0 |
| 0 | NA | NA | NA | NA | NA | Krt73 | keratin 73 | 0 | 0 | 0 | 0 | 0 | 0 | 0 | 0 | 0 | 0 | 0 |
| 0 | NA | NA | NA | NA | NA | Krt2 | keratin 2 | 0 | 0 | 0 | 0 | 0 | 0 | 0 | 0 | 0 | 0 | 0 |
| 0 | NA | NA | NA | NA | NA | Gm44356 | | 0 | 0 | 0 | 0 | 0 | 0 | 0 | 0 | 0 | 0 | 0 |
| 0 | NA | NA | NA | NA | NA | Gm49434 | | 0 | 0 | 0 | 0 | 0 | 0 | 0 | 0 | 0 | 0 | 0 |
| 0 | NA | NA | NA | NA | NA | Krt4 | keratin 4 | 0 | 0 | 0 | 0 | 0 | 0 | 0 | 0 | 0 | 0 | 0 |
| 0 | NA | NA | NA | NA | NA | Krt79 | keratin 79 | 0 | 0 | 0 | 0 | 0 | 0 | 0 | 0 | 0 | 0 | 0 |
| 0 | NA | NA | NA | NA | NA | Gm49479 | | 0 | 0 | 0 | 0 | 0 | 0 | 0 | 0 | 0 | 0 | 0 |
| 0 | NA | NA | NA | NA | NA | Gm49480 | | 0 | 0 | 0 | 0 | 0 | 0 | 0 | 0 | 0 | 0 | 0 |
| 0 | NA | NA | NA | NA | NA | Gm23590 | | 0 | 0 | 0 | 0 | 0 | 0 | 0 | 0 | 0 | 0 | 0 |
| 0 | NA | NA | NA | NA | NA | Gm24128 | | 0 | 0 | 0 | 0 | 0 | 0 | 0 | 0 | 0 | 0 | 0 |
| 0 | NA | NA | NA | NA | NA | Gm27406 | | 0 | 0 | 0 | 0 | 0 | 0 | 0 | 0 | 0 | 0 | 0 |
| 0 | NA | NA | NA | NA | NA | Gm10337 | | 0 | 0 | 0 | 0 | 0 | 0 | 0 | 0 | 0 | 0 | 0 |
| 0 | NA | NA | NA | NA | NA | Gm28047 | | 0 | 0 | 0 | 0 | 0 | 0 | 0 | 0 | 0 | 0 | 0 |
| 0 | NA | NA | NA | NA | NA | Gm25141 | | 0 | 0 | 0 | 0 | 0 | 0 | 0 | 0 | 0 | 0 | 0 |
| 0 | NA | NA | NA | NA | NA | Gm4544 | | 0 | 0 | 0 | 0 | 0 | 0 | 0 | 0 | 0 | 0 | 0 |
| 0 | NA | NA | NA | NA | NA | Mir688 | | 0 | 0 | 0 | 0 | 0 | 0 | 0 | 0 | 0 | 0 | 0 |
| 0 | NA | NA | NA | NA | NA | Gm49473 | | 0 | 0 | 0 | 0 | 0 | 0 | 0 | 0 | 0 | 0 | 0 |
| 0 | NA | NA | NA | NA | NA | Gm36560 | | 0 | 0 | 0 | 0 | 0 | 0 | 0 | 0 | 0 | 0 | 0 |
| 0 | NA | NA | NA | NA | NA | Hoxc13 | homeobox C13 | 0 | 0 | 0 | 0 | 0 | 0 | 0 | 0 | 0 | 0 | 0 |
| 0 | NA | NA | NA | NA | NA | Hoxc12 | homeobox C12 | 0 | 0 | 0 | 0 | 0 | 0 | 0 | 0 | 0 | 0 | 0 |
| 0 | NA | NA | NA | NA | NA | Hotair | | 0 | 0 | 0 | 0 | 0 | 0 | 0 | 0 | 0 | 0 | 0 |
| 0 | NA | NA | NA | NA | NA | Gm27337 | | 0 | 0 | 0 | 0 | 0 | 0 | 0 | 0 | 0 | 0 | 0 |
| 0 | NA | NA | NA | NA | NA | Gm27443 | | 0 | 0 | 0 | 0 | 0 | 0 | 0 | 0 | 0 | 0 | 0 |
| 0 | NA | NA | NA | NA | NA | Gm28011 | | 0 | 0 | 0 | 0 | 0 | 0 | 0 | 0 | 0 | 0 | 0 |
| 0 | NA | NA | NA | NA | NA | Gm27535 | | 0 | 0 | 0 | 0 | 0 | 0 | 0 | 0 | 0 | 0 | 0 |
| 0 | NA | NA | NA | NA | NA | Hoxc11 | homeobox C11 | 0 | 0 | 0 | 0 | 0 | 0 | 0 | 0 | 0 | 0 | 0 |
| 0 | NA | NA | NA | NA | NA | Hoxc10 | homeobox C10 | 0 | 0 | 0 | 0 | 0 | 0 | 0 | 0 | 0 | 0 | 0 |
| 0 | NA | NA | NA | NA | NA | Mir196a-2 | | 0 | 0 | 0 | 0 | 0 | 0 | 0 | 0 | 0 | 0 | 0 |
| 0 | NA | NA | NA | NA | NA | Hoxc9 | homeobox C9 | 0 | 0 | 0 | 0 | 0 | 0 | 0 | 0 | 0 | 0 | 0 |
| 0 | NA | NA | NA | NA | NA | Hoxc8 | homeobox C8 | 0 | 0 | 0 | 0 | 0 | 0 | 0 | 0 | 0 | 0 | 0 |
| 0 | NA | NA | NA | NA | NA | Mir615 | | 0 | 0 | 0 | 0 | 0 | 0 | 0 | 0 | 0 | 0 | 0 |

| | | | | | | | | | | | | | | | | | | |
|---|---|---|---|---|---|---|---|---|---|---|---|---|---|---|---|---|---|---|
| 0 | NA | NA | NA | NA | NA | 5730585A16Rik | | 0 | 0 | 0 | 0 | 0 | 0 | 0 | 0 | 0 | 0 | 0 |
| 0 | NA | NA | NA | NA | NA | Gm10830 | | 0 | 0 | 0 | 0 | 0 | 0 | 0 | 0 | 0 | 0 | 0 |
| 0 | NA | NA | NA | NA | NA | Gm28265 | | 0 | 0 | 0 | 0 | 0 | 0 | 0 | 0 | 0 | 0 | 0 |
| 0 | NA | NA | NA | NA | NA | Mir148b | | 0 | 0 | 0 | 0 | 0 | 0 | 0 | 0 | 0 | 0 | 0 |
| 0 | NA | NA | NA | NA | NA | Mir6963 | | 0 | 0 | 0 | 0 | 0 | 0 | 0 | 0 | 0 | 0 | 0 |
| 0 | NA | NA | NA | NA | NA | Gm37322 | | 0 | 0 | 0 | 0 | 0 | 0 | 0 | 0 | 0 | 0 | 0 |
| 0 | NA | NA | NA | NA | NA | Gtsf1 | gametocyte specific factor 1 | 0 | 0 | 0 | 0 | 0 | 0 | 0 | 0 | 0 | 0 | 0 |
| 0 | NA | NA | NA | NA | NA | Gtsf2 | gametocyte specific factor 2 | 0 | 0 | 0 | 0 | 0 | 0 | 0 | 0 | 0 | 0 | 0 |
| 0 | NA | NA | NA | NA | NA | Glycam1 | glycosylation dependent cell adhesion m | 0 | 0 | 0 | 0 | 0 | 0 | 0 | 0 | 0 | 0 | 0 |
| 0 | NA | NA | NA | NA | NA | Gm49274 | | 0 | 0 | 0 | 0 | 0 | 0 | 0 | 0 | 0 | 0 | 0 |
| 0 | NA | NA | NA | NA | NA | Gm49275 | | 0 | 0 | 0 | 0 | 0 | 0 | 0 | 0 | 0 | 0 | 0 |
| 0 | NA | NA | NA | NA | NA | Gm49491 | | 0 | 0 | 0 | 0 | 0 | 0 | 0 | 0 | 0 | 0 | 0 |
| 0 | NA | NA | NA | NA | NA | Gm22852 | | 0 | 0 | 0 | 0 | 0 | 0 | 0 | 0 | 0 | 0 | 0 |
| 0 | NA | NA | NA | NA | NA | Mucl2 | mucin-like 2 | 0 | 0 | 0 | 0 | 0 | 0 | 0 | 0 | 0 | 0 | 0 |
| 0 | NA | NA | NA | NA | NA | Gm35912 | | 0 | 0 | 0 | 0 | 0 | 0 | 0 | 0 | 0 | 0 | 0 |
| 0 | NA | NA | NA | NA | NA | Vmn1r-ps137 | | 0 | 0 | 0 | 0 | 0 | 0 | 0 | 0 | 0 | 0 | 0 |
| 0 | NA | NA | NA | NA | NA | Gm49554 | | 0 | 0 | 0 | 0 | 0 | 0 | 0 | 0 | 0 | 0 | 0 |
| 0 | NA | NA | NA | NA | NA | Gm4106 | | 0 | 0 | 0 | 0 | 0 | 0 | 0 | 0 | 0 | 0 | 0 |
| 0 | NA | NA | NA | NA | NA | Gm3711 | | 0 | 0 | 0 | 0 | 0 | 0 | 0 | 0 | 0 | 0 | 0 |
| 0 | NA | NA | NA | NA | NA | Gm23215 | | 0 | 0 | 0 | 0 | 0 | 0 | 0 | 0 | 0 | 0 | 0 |
| 0 | NA | NA | NA | NA | NA | Gm3749 | | 0 | 0 | 0 | 0 | 0 | 0 | 0 | 0 | 0 | 0 | 0 |
| 0 | NA | NA | NA | NA | NA | Gm4111 | | 0 | 0 | 0 | 0 | 0 | 0 | 0 | 0 | 0 | 0 | 0 |
| 0 | NA | NA | NA | NA | NA | Gm49504 | | 0 | 0 | 0 | 0 | 0 | 0 | 0 | 0 | 0 | 0 | 0 |
| 0 | NA | NA | NA | NA | NA | Gm22862 | | 0 | 0 | 0 | 0 | 0 | 0 | 0 | 0 | 0 | 0 | 0 |
| 0 | NA | NA | NA | NA | NA | Gm49505 | | 0 | 0 | 0 | 0 | 0 | 0 | 0 | 0 | 0 | 0 | 0 |
| 0 | NA | NA | NA | NA | NA | Gm49507 | | 0 | 0 | 0 | 0 | 0 | 0 | 0 | 0 | 0 | 0 | 0 |
| 0 | NA | NA | NA | NA | NA | Gm35974 | | 0 | 0 | 0 | 0 | 0 | 0 | 0 | 0 | 0 | 0 | 0 |
| 0 | NA | NA | NA | NA | NA | Gm49508 | | 0 | 0 | 0 | 0 | 0 | 0 | 0 | 0 | 0 | 0 | 0 |
| 0 | NA | NA | NA | NA | NA | Gm49509 | | 0 | 0 | 0 | 0 | 0 | 0 | 0 | 0 | 0 | 0 | 0 |
| 0 | NA | NA | NA | NA | NA | Gm4116 | | 0 | 0 | 0 | 0 | 0 | 0 | 0 | 0 | 0 | 0 | 0 |
| 0 | NA | NA | NA | NA | NA | Gm22598 | | 0 | 0 | 0 | 0 | 0 | 0 | 0 | 0 | 0 | 0 | 0 |
| 0 | NA | NA | NA | NA | NA | Olfr161 | olfactory receptor 161 | 0 | 0 | 0 | 0 | 0 | 0 | 0 | 0 | 0 | 0 | 0 |
| 0 | NA | NA | NA | NA | NA | Olfr162-ps1 | | 0 | 0 | 0 | 0 | 0 | 0 | 0 | 0 | 0 | 0 | 0 |
| 0 | NA | NA | NA | NA | NA | Olfr163-ps1 | | 0 | 0 | 0 | 0 | 0 | 0 | 0 | 0 | 0 | 0 | 0 |
| 0 | NA | NA | NA | NA | NA | | | 0 | 0 | 0 | 0 | 0 | 0 | 0 | 0 | 0 | 0 | 0 |
| 0 | NA | NA | NA | NA | NA | Tdgf1-ps1 | | 0 | 0 | 0 | 0 | 0 | 0 | 0 | 0 | 0 | 0 | 0 |
| 0 | NA | NA | NA | NA | NA | Olfr15 | olfactory receptor 15 | 0 | 0 | 0 | 0 | 0 | 0 | 0 | 0 | 0 | 0 | 0 |
| 0 | NA | NA | NA | NA | NA | 1700016D08Rik | | 0 | 0 | 0 | 0 | 0 | 0 | 0 | 0 | 0 | 0 | 0 |
| 0 | NA | NA | NA | NA | NA | Gm23413 | | 0 | 0 | 0 | 0 | 0 | 0 | 0 | 0 | 0 | 0 | 0 |
| 0 | NA | NA | NA | NA | NA | Gm49547 | | 0 | 0 | 0 | 0 | 0 | 0 | 0 | 0 | 0 | 0 | 0 |
| 0 | NA | NA | NA | NA | NA | Gm15885 | | 0 | 0 | 0 | 0 | 0 | 0 | 0 | 0 | 0 | 0 | 0 |
| 0 | NA | NA | NA | NA | NA | Gm15835 | | 0 | 0 | 0 | 0 | 0 | 0 | 0 | 0 | 0 | 0 | 0 |
| 0 | NA | NA | NA | NA | NA | 4930562C15Ril | RIKEN cDNA 4930562C15 gene | 0 | 0 | 0 | 0 | 0 | 0 | 0 | 0 | 0 | 0 | 0 |
| 0 | NA | NA | NA | NA | NA | Septin12 | septin 12 | 0 | 0 | 0 | 0 | 0 | 0 | 0 | 0 | 0 | 0 | 0 |
| 0 | NA | NA | NA | NA | NA | Gm23483 | | 0 | 0 | 0 | 0 | 0 | 0 | 0 | 0 | 0 | 0 | 0 |
| 0 | NA | NA | NA | NA | NA | Mir6364 | | 0 | 0 | 0 | 0 | 0 | 0 | 0 | 0 | 0 | 0 | 0 |
| 0 | NA | NA | NA | NA | NA | Gm23689 | | 0 | 0 | 0 | 0 | 0 | 0 | 0 | 0 | 0 | 0 | 0 |
| 0 | NA | NA | NA | NA | NA | Gm26095 | | 0 | 0 | 0 | 0 | 0 | 0 | 0 | 0 | 0 | 0 | 0 |
| 0 | NA | NA | NA | NA | NA | n-R5s30 | | 0 | 0 | 0 | 0 | 0 | 0 | 0 | 0 | 0 | 0 | 0 |
| 0 | NA | NA | NA | NA | NA | 1700123O21Rik | | 0 | 0 | 0 | 0 | 0 | 0 | 0 | 0 | 0 | 0 | 0 |
| 0 | NA | NA | NA | NA | NA | Gm49533 | | 0 | 0 | 0 | 0 | 0 | 0 | 0 | 0 | 0 | 0 | 0 |
| 0 | NA | NA | NA | NA | NA | Gm49534 | | 0 | 0 | 0 | 0 | 0 | 0 | 0 | 0 | 0 | 0 | 0 |
| 0 | NA | NA | NA | NA | NA | Gm49535 | | 0 | 0 | 0 | 0 | 0 | 0 | 0 | 0 | 0 | 0 | 0 |
| 0 | NA | NA | NA | NA | NA | Gm23476 | | 0 | 0 | 0 | 0 | 0 | 0 | 0 | 0 | 0 | 0 | 0 |
| 0 | NA | NA | NA | NA | NA | Gm26159 | | 0 | 0 | 0 | 0 | 0 | 0 | 0 | 0 | 0 | 0 | 0 |
| 0 | NA | NA | NA | NA | NA | Gm24579 | | 0 | 0 | 0 | 0 | 0 | 0 | 0 | 0 | 0 | 0 | 0 |
| 0 | NA | NA | NA | NA | NA | Gm25782 | | 0 | 0 | 0 | 0 | 0 | 0 | 0 | 0 | 0 | 0 | 0 |
| 0 | NA | NA | NA | NA | NA | Gm23706 | | 0 | 0 | 0 | 0 | 0 | 0 | 0 | 0 | 0 | 0 | 0 |
| 0 | NA | NA | NA | NA | NA | Gm22137 | | 0 | 0 | 0 | 0 | 0 | 0 | 0 | 0 | 0 | 0 | 0 |
| 0 | NA | NA | NA | NA | NA | Gm49549 | | 0 | 0 | 0 | 0 | 0 | 0 | 0 | 0 | 0 | 0 | 0 |

| | | | | | | | | | | | | | | | | | | |
|---|---|---|---|---|---|---|---|---|---|---|---|---|---|---|---|---|---|---|
| 0 | NA | NA | NA | NA | NA | Gm25805 | | 0 | 0 | 0 | 0 | 0 | 0 | 0 | 0 | 0 | 0 | 0 |
| 0 | NA | NA | NA | NA | NA | Gm22224 | | 0 | 0 | 0 | 0 | 0 | 0 | 0 | 0 | 0 | 0 | 0 |
| 0 | NA | NA | NA | NA | NA | Gm30371 | | 0 | 0 | 0 | 0 | 0 | 0 | 0 | 0 | 0 | 0 | 0 |
| 0 | NA | NA | NA | NA | NA | Gm49447 | | 0 | 0 | 0 | 0 | 0 | 0 | 0 | 0 | 0 | 0 | 0 |
| 0 | NA | NA | NA | NA | NA | Gm30567 | | 0 | 0 | 0 | 0 | 0 | 0 | 0 | 0 | 0 | 0 | 0 |
| 0 | NA | NA | NA | NA | NA | Gm49448 | | 0 | 0 | 0 | 0 | 0 | 0 | 0 | 0 | 0 | 0 | 0 |
| 0 | NA | NA | NA | NA | NA | Gm49451 | | 0 | 0 | 0 | 0 | 0 | 0 | 0 | 0 | 0 | 0 | 0 |
| 0 | NA | NA | NA | NA | NA | Grin2a | glutamate receptor, ionotropic, NMDA2A ( | 0 | 0 | 0 | 0 | 0 | 0 | 0 | 0 | 0 | 0 | 0 |
| 0 | NA | NA | NA | NA | NA | Gm3890 | | 0 | 0 | 0 | 0 | 0 | 0 | 0 | 0 | 0 | 0 | 0 |
| 0 | NA | NA | NA | NA | NA | Gm49455 | | 0 | 0 | 0 | 0 | 0 | 0 | 0 | 0 | 0 | 0 | 0 |
| 0 | NA | NA | NA | NA | NA | Gm24667 | | 0 | 0 | 0 | 0 | 0 | 0 | 0 | 0 | 0 | 0 | 0 |
| 0 | NA | NA | NA | NA | NA | Gm49456 | | 0 | 0 | 0 | 0 | 0 | 0 | 0 | 0 | 0 | 0 | 0 |
| 0 | NA | NA | NA | NA | NA | Gm22932 | | 0 | 0 | 0 | 0 | 0 | 0 | 0 | 0 | 0 | 0 | 0 |
| 0 | NA | NA | NA | NA | NA | Gm1600 | | 0 | 0 | 0 | 0 | 0 | 0 | 0 | 0 | 0 | 0 | 0 |
| 0 | NA | NA | NA | NA | NA | Gm15558 | | 0 | 0 | 0 | 0 | 0 | 0 | 0 | 0 | 0 | 0 | 0 |
| 0 | NA | NA | NA | NA | NA | Tnp2 | transition protein 2 | 0 | 0 | 0 | 0 | 0 | 0 | 0 | 0 | 0 | 0 | 0 |
| 0 | NA | NA | NA | NA | NA | Prm3 | protamine 3 | 0 | 0 | 0 | 0 | 0 | 0 | 0 | 0 | 0 | 0 | 0 |
| 0 | NA | NA | NA | NA | NA | Prm2 | protamine 2 | 0 | 0 | 0 | 0 | 0 | 0 | 0 | 0 | 0 | 0 | 0 |
| 0 | NA | NA | NA | NA | NA | Prm1 | protamine 1 | 0 | 0 | 0 | 0 | 0 | 0 | 0 | 0 | 0 | 0 | 0 |
| 0 | NA | NA | NA | NA | NA | Gm11172 | | 0 | 0 | 0 | 0 | 0 | 0 | 0 | 0 | 0 | 0 | 0 |
| 0 | NA | NA | NA | NA | NA | Gm26822 | | 0 | 0 | 0 | 0 | 0 | 0 | 0 | 0 | 0 | 0 | 0 |
| 0 | NA | NA | NA | NA | NA | Gm49960 | | 0 | 0 | 0 | 0 | 0 | 0 | 0 | 0 | 0 | 0 | 0 |
| 0 | NA | NA | NA | NA | NA | Gm26268 | | 0 | 0 | 0 | 0 | 0 | 0 | 0 | 0 | 0 | 0 | 0 |
| 0 | NA | NA | NA | NA | NA | Gm24961 | | 0 | 0 | 0 | 0 | 0 | 0 | 0 | 0 | 0 | 0 | 0 |
| 0 | NA | NA | NA | NA | NA | Gm10832 | | 0 | 0 | 0 | 0 | 0 | 0 | 0 | 0 | 0 | 0 | 0 |
| 0 | NA | NA | NA | NA | NA | Gm3919 | | 0 | 0 | 0 | 0 | 0 | 0 | 0 | 0 | 0 | 0 | 0 |
| 0 | NA | NA | NA | NA | NA | Gm18724 | | 0 | 0 | 0 | 0 | 0 | 0 | 0 | 0 | 0 | 0 | 0 |
| 0 | NA | NA | NA | NA | NA | Mir1945 | | 0 | 0 | 0 | 0 | 0 | 0 | 0 | 0 | 0 | 0 | 0 |
| 0 | NA | NA | NA | NA | NA | Gm4279 | | 0 | 0 | 0 | 0 | 0 | 0 | 0 | 0 | 0 | 0 | 0 |
| 0 | NA | NA | NA | NA | NA | Gm9961 | | 0 | 0 | 0 | 0 | 0 | 0 | 0 | 0 | 0 | 0 | 0 |
| 0 | NA | NA | NA | NA | NA | C87114 | | 0 | 0 | 0 | 0 | 0 | 0 | 0 | 0 | 0 | 0 | 0 |
| 0 | NA | NA | NA | NA | NA | 4930414F18Rik | | 0 | 0 | 0 | 0 | 0 | 0 | 0 | 0 | 0 | 0 | 0 |
| 0 | NA | NA | NA | NA | NA | Gm38619 | | 0 | 0 | 0 | 0 | 0 | 0 | 0 | 0 | 0 | 0 | 0 |
| 0 | NA | NA | NA | NA | NA | Gm38619 | | 0 | 0 | 0 | 0 | 0 | 0 | 0 | 0 | 0 | 0 | 0 |
| 0 | NA | NA | NA | NA | NA | Gm6327 | | 0 | 0 | 0 | 0 | 0 | 0 | 0 | 0 | 0 | 0 | 0 |
| 0 | NA | NA | NA | NA | NA | 1700003L19Rik | | 0 | 0 | 0 | 0 | 0 | 0 | 0 | 0 | 0 | 0 | 0 |
| 0 | NA | NA | NA | NA | NA | Gm6332 | | 0 | 0 | 0 | 0 | 0 | 0 | 0 | 0 | 0 | 0 | 0 |
| 0 | NA | NA | NA | NA | NA | Mir193b | | 0 | 0 | 0 | 0 | 0 | 0 | 0 | 0 | 0 | 0 | 0 |
| 0 | NA | NA | NA | NA | NA | Mir365-1 | | 0 | 0 | 0 | 0 | 0 | 0 | 0 | 0 | 0 | 0 | 0 |
| 0 | NA | NA | NA | NA | NA | Gm32817 | | 0 | 0 | 0 | 0 | 0 | 0 | 0 | 0 | 0 | 0 | 0 |
| 0 | NA | NA | NA | NA | NA | Mir6365 | | 0 | 0 | 0 | 0 | 0 | 0 | 0 | 0 | 0 | 0 | 0 |
| 0 | NA | NA | NA | NA | NA | Gm25276 | | 0 | 0 | 0 | 0 | 0 | 0 | 0 | 0 | 0 | 0 | 0 |
| 0 | NA | NA | NA | NA | NA | Gm49432 | | 0 | 0 | 0 | 0 | 0 | 0 | 0 | 0 | 0 | 0 | 0 |
| 0 | NA | NA | NA | NA | NA | Gm27734 | | 0 | 0 | 0 | 0 | 0 | 0 | 0 | 0 | 0 | 0 | 0 |
| 0 | NA | NA | NA | NA | NA | Gm26213 | | 0 | 0 | 0 | 0 | 0 | 0 | 0 | 0 | 0 | 0 | 0 |
| 0 | NA | NA | NA | NA | NA | Gm27671 | | 0 | 0 | 0 | 0 | 0 | 0 | 0 | 0 | 0 | 0 | 0 |
| 0 | NA | NA | NA | NA | NA | Gm49517 | | 0 | 0 | 0 | 0 | 0 | 0 | 0 | 0 | 0 | 0 | 0 |
| 0 | NA | NA | NA | NA | NA | Gm23116 | | 0 | 0 | 0 | 0 | 0 | 0 | 0 | 0 | 0 | 0 | 0 |
| 0 | NA | NA | NA | NA | NA | Gm49528 | | 0 | 0 | 0 | 0 | 0 | 0 | 0 | 0 | 0 | 0 | 0 |
| 0 | NA | NA | NA | NA | NA | Gm1758 | | 0 | 0 | 0 | 0 | 0 | 0 | 0 | 0 | 0 | 0 | 0 |
| 0 | NA | NA | NA | NA | NA | A630010A05Rik | | 0 | 0 | 0 | 0 | 0 | 0 | 0 | 0 | 0 | 0 | 0 |
| 0 | NA | NA | NA | NA | NA | Gm49530 | | 0 | 0 | 0 | 0 | 0 | 0 | 0 | 0 | 0 | 0 | 0 |
| 0 | NA | NA | NA | NA | NA | Gm33696 | | 0 | 0 | 0 | 0 | 0 | 0 | 0 | 0 | 0 | 0 | 0 |
| 0 | NA | NA | NA | NA | NA | Gm49536 | | 0 | 0 | 0 | 0 | 0 | 0 | 0 | 0 | 0 | 0 | 0 |
| 0 | NA | NA | NA | NA | NA | Gm7731 | | 0 | 0 | 0 | 0 | 0 | 0 | 0 | 0 | 0 | 0 | 0 |
| 0 | NA | NA | NA | NA | NA | Gm5404 | | 0 | 0 | 0 | 0 | 0 | 0 | 0 | 0 | 0 | 0 | 0 |
| 0 | NA | NA | NA | NA | NA | Gm21897 | | 0 | 0 | 0 | 0 | 0 | 0 | 0 | 0 | 0 | 0 | 0 |
| 0 | NA | NA | NA | NA | NA | Gm49538 | | 0 | 0 | 0 | 0 | 0 | 0 | 0 | 0 | 0 | 0 | 0 |
| 0 | NA | NA | NA | NA | NA | Gm49511 | | 0 | 0 | 0 | 0 | 0 | 0 | 0 | 0 | 0 | 0 | 0 |
| 0 | NA | NA | NA | NA | NA | n-R5s31 | | 0 | 0 | 0 | 0 | 0 | 0 | 0 | 0 | 0 | 0 | 0 |

| | | | | | | | | | | | | | | | | | | |
|---|---|---|---|---|---|---|---|---|---|---|---|---|---|---|---|---|---|---|
| 0 | NA | NA | NA | NA | NA | Gm7765 | | 0 | 0 | 0 | 0 | 0 | 0 | 0 | 0 | 0 | 0 | 0 |
| 0 | NA | NA | NA | NA | NA | Gm18869 | | 0 | 0 | 0 | 0 | 0 | 0 | 0 | 0 | 0 | 0 | 0 |
| 0 | NA | NA | NA | NA | NA | | | 0 | 0 | 0 | 0 | 0 | 0 | 0 | 0 | 0 | 0 | 0 |
| 0 | NA | NA | NA | NA | NA | Gm16139 | | 0 | 0 | 0 | 0 | 0 | 0 | 0 | 0 | 0 | 0 | 0 |
| 0 | NA | NA | NA | NA | NA | Olfr19 | olfactory receptor 19 | 0 | 0 | 0 | 0 | 0 | 0 | 0 | 0 | 0 | 0 | 0 |
| 0 | NA | NA | NA | NA | NA | Gm44932 | | 0 | 0 | 0 | 0 | 0 | 0 | 0 | 0 | 0 | 0 | 0 |
| 0 | NA | NA | NA | NA | NA | 4933404G15Rik | | 0 | 0 | 0 | 0 | 0 | 0 | 0 | 0 | 0 | 0 | 0 |
| 0 | NA | NA | NA | NA | NA | Gm49644 | | 0 | 0 | 0 | 0 | 0 | 0 | 0 | 0 | 0 | 0 | 0 |
| 0 | NA | NA | NA | NA | NA | Igll1 | immunoglobulin lambda-like polypeptide | 0 | 0 | 0 | 0 | 0 | 0 | 0 | 0 | 0 | 0 | 0 |
| 0 | NA | NA | NA | NA | NA | Gm6438 | | 0 | 0 | 0 | 0 | 0 | 0 | 0 | 0 | 0 | 0 | 0 |
| 0 | NA | NA | NA | NA | NA | Gm22558 | | 0 | 0 | 0 | 0 | 0 | 0 | 0 | 0 | 0 | 0 | 0 |
| 0 | NA | NA | NA | NA | NA | Mir301b | | 0 | 0 | 0 | 0 | 0 | 0 | 0 | 0 | 0 | 0 | 0 |
| 0 | NA | NA | NA | NA | NA | Gm22129 | | 0 | 0 | 0 | 0 | 0 | 0 | 0 | 0 | 0 | 0 | 0 |
| 0 | NA | NA | NA | NA | NA | Gm49573 | | 0 | 0 | 0 | 0 | 0 | 0 | 0 | 0 | 0 | 0 | 0 |
| 0 | NA | NA | NA | NA | NA | Gm22770 | | 0 | 0 | 0 | 0 | 0 | 0 | 0 | 0 | 0 | 0 | 0 |
| 0 | NA | NA | NA | NA | NA | Gm49684 | | 0 | 0 | 0 | 0 | 0 | 0 | 0 | 0 | 0 | 0 | 0 |
| 0 | NA | NA | NA | NA | NA | Gm49683 | | 0 | 0 | 0 | 0 | 0 | 0 | 0 | 0 | 0 | 0 | 0 |
| 0 | NA | NA | NA | NA | NA | Gm6048 | | 0 | 0 | 0 | 0 | 0 | 0 | 0 | 0 | 0 | 0 | 0 |
| 0 | NA | NA | NA | NA | NA | Tssk2 | testis-specific serine kinase 2 | 0 | 0 | 0 | 0 | 0 | 0 | 0 | 0 | 0 | 0 | 0 |
| 0 | NA | NA | NA | NA | NA | Gsc2 | goosecoid homebox 2 | 0 | 0 | 0 | 0 | 0 | 0 | 0 | 0 | 0 | 0 | 0 |
| 0 | NA | NA | NA | NA | NA | Vpreb2 | pre-B lymphocyte gene 2 | 0 | 0 | 0 | 0 | 0 | 0 | 0 | 0 | 0 | 0 | 0 |
| 0 | NA | NA | NA | NA | NA | Gm49580 | | 0 | 0 | 0 | 0 | 0 | 0 | 0 | 0 | 0 | 0 | 0 |
| 0 | NA | NA | NA | NA | NA | Mir6366 | | 0 | 0 | 0 | 0 | 0 | 0 | 0 | 0 | 0 | 0 | 0 |
| 0 | NA | NA | NA | NA | NA | 4930483P17Rik | | 0 | 0 | 0 | 0 | 0 | 0 | 0 | 0 | 0 | 0 | 0 |
| 0 | NA | NA | NA | NA | NA | Ccdc188 | coiled-coil domain containing 188 | 0 | 0 | 0 | 0 | 0 | 0 | 0 | 0 | 0 | 0 | 0 |
| 0 | NA | NA | NA | NA | NA | Gm49776 | | 0 | 0 | 0 | 0 | 0 | 0 | 0 | 0 | 0 | 0 | 0 |
| 0 | NA | NA | NA | NA | NA | Mir1306 | | 0 | 0 | 0 | 0 | 0 | 0 | 0 | 0 | 0 | 0 | 0 |
| 0 | NA | NA | NA | NA | NA | Gm24572 | | 0 | 0 | 0 | 0 | 0 | 0 | 0 | 0 | 0 | 0 | 0 |
| 0 | NA | NA | NA | NA | NA | Mir185 | | 0 | 0 | 0 | 0 | 0 | 0 | 0 | 0 | 0 | 0 | 0 |
| 0 | NA | NA | NA | NA | NA | Gm26219 | | 0 | 0 | 0 | 0 | 0 | 0 | 0 | 0 | 0 | 0 | 0 |
| 0 | NA | NA | NA | NA | NA | 4930588K23Rik | | 0 | 0 | 0 | 0 | 0 | 0 | 0 | 0 | 0 | 0 | 0 |
| 0 | NA | NA | NA | NA | NA | Gp1bb | glycoprotein Ib, beta polypeptide | 0 | 0 | 0 | 0 | 0 | 0 | 0 | 0 | 0 | 0 | 0 |
| 0 | NA | NA | NA | NA | NA | Gm28539 | | 0 | 0 | 0 | 0 | 0 | 0 | 0 | 0 | 0 | 0 | 0 |
| 0 | NA | NA | NA | NA | NA | Gm18747 | | 0 | 0 | 0 | 0 | 0 | 0 | 0 | 0 | 0 | 0 | 0 |
| 0 | NA | NA | NA | NA | NA | Gm10088 | | 0 | 0 | 0 | 0 | 0 | 0 | 0 | 0 | 0 | 0 | 0 |
| 0 | NA | NA | NA | NA | NA | Iglj3p | | 0 | 0 | 0 | 0 | 0 | 0 | 0 | 0 | 0 | 0 | 0 |
| 0 | NA | NA | NA | NA | NA | Iglj3 | | 0 | 0 | 0 | 0 | 0 | 0 | 0 | 0 | 0 | 0 | 0 |
| 0 | NA | NA | NA | NA | NA | Gm18748 | | 0 | 0 | 0 | 0 | 0 | 0 | 0 | 0 | 0 | 0 | 0 |
| 0 | NA | NA | NA | NA | NA | Iglc4 | | 0 | 0 | 0 | 0 | 0 | 0 | 0 | 0 | 0 | 0 | 0 |
| 0 | NA | NA | NA | NA | NA | Iglj4 | | 0 | 0 | 0 | 0 | 0 | 0 | 0 | 0 | 0 | 0 | 0 |
| 0 | NA | NA | NA | NA | NA | Iglj2 | | 0 | 0 | 0 | 0 | 0 | 0 | 0 | 0 | 0 | 0 | 0 |
| 0 | NA | NA | NA | NA | NA | Olfr164 | olfactory receptor 164 | 0 | 0 | 0 | 0 | 0 | 0 | 0 | 0 | 0 | 0 | 0 |
| 0 | NA | NA | NA | NA | NA | Gm22158 | | 0 | 0 | 0 | 0 | 0 | 0 | 0 | 0 | 0 | 0 | 0 |
| 0 | NA | NA | NA | NA | NA | Gm23612 | | 0 | 0 | 0 | 0 | 0 | 0 | 0 | 0 | 0 | 0 | 0 |
| 0 | NA | NA | NA | NA | NA | Olfr165 | olfactory receptor 165 | 0 | 0 | 0 | 0 | 0 | 0 | 0 | 0 | 0 | 0 | 0 |
| 0 | NA | NA | NA | NA | NA | Gm25762 | | 0 | 0 | 0 | 0 | 0 | 0 | 0 | 0 | 0 | 0 | 0 |
| 0 | NA | NA | NA | NA | NA | Olfr167 | olfactory receptor 167 | 0 | 0 | 0 | 0 | 0 | 0 | 0 | 0 | 0 | 0 | 0 |
| 0 | NA | NA | NA | NA | NA | Olfr168 | olfactory receptor 168 | 0 | 0 | 0 | 0 | 0 | 0 | 0 | 0 | 0 | 0 | 0 |
| 0 | NA | NA | NA | NA | NA | Olfr169 | olfactory receptor 169 | 0 | 0 | 0 | 0 | 0 | 0 | 0 | 0 | 0 | 0 | 0 |
| 0 | NA | NA | NA | NA | NA | Olfr170 | olfactory receptor 170 | 0 | 0 | 0 | 0 | 0 | 0 | 0 | 0 | 0 | 0 | 0 |
| 0 | NA | NA | NA | NA | NA | Olfr171 | olfactory receptor 171 | 0 | 0 | 0 | 0 | 0 | 0 | 0 | 0 | 0 | 0 | 0 |
| 0 | NA | NA | NA | NA | NA | Gm49596 | | 0 | 0 | 0 | 0 | 0 | 0 | 0 | 0 | 0 | 0 | 0 |
| 0 | NA | NA | NA | NA | NA | Gm49597 | | 0 | 0 | 0 | 0 | 0 | 0 | 0 | 0 | 0 | 0 | 0 |
| 0 | NA | NA | NA | NA | NA | A930003A15Rik | | 0 | 0 | 0 | 0 | 0 | 0 | 0 | 0 | 0 | 0 | 0 |
| 0 | NA | NA | NA | NA | NA | Gm49598 | | 0 | 0 | 0 | 0 | 0 | 0 | 0 | 0 | 0 | 0 | 0 |
| 0 | NA | NA | NA | NA | NA | Gm27883 | | 0 | 0 | 0 | 0 | 0 | 0 | 0 | 0 | 0 | 0 | 0 |
| 0 | NA | NA | NA | NA | NA | Gm49566 | | 0 | 0 | 0 | 0 | 0 | 0 | 0 | 0 | 0 | 0 | 0 |
| 0 | NA | NA | NA | NA | NA | Gm20614 | | 0 | 0 | 0 | 0 | 0 | 0 | 0 | 0 | 0 | 0 | 0 |
| 0 | NA | NA | NA | NA | NA | Gm23332 | | 0 | 0 | 0 | 0 | 0 | 0 | 0 | 0 | 0 | 0 | 0 |
| 0 | NA | NA | NA | NA | NA | Gm20615 | | 0 | 0 | 0 | 0 | 0 | 0 | 0 | 0 | 0 | 0 | 0 |

| | | | | | | | | | | | | | | | | | | |
|---|---|---|---|---|---|---|---|---|---|---|---|---|---|---|---|---|---|---|
| 0 | NA | NA | NA | NA | NA | Gm49744 | | 0 | 0 | 0 | 0 | 0 | 0 | 0 | 0 | 0 | 0 | 0 |
| 0 | NA | NA | NA | NA | NA | Gm49746 | | 0 | 0 | 0 | 0 | 0 | 0 | 0 | 0 | 0 | 0 | 0 |
| 0 | NA | NA | NA | NA | NA | Gm20164 | | 0 | 0 | 0 | 0 | 0 | 0 | 0 | 0 | 0 | 0 | 0 |
| 0 | NA | NA | NA | NA | NA | Mir1224 | | 0 | 0 | 0 | 0 | 0 | 0 | 0 | 0 | 0 | 0 | 0 |
| 0 | NA | NA | NA | NA | NA | Gm49333 | Eef1akmt4-Ece2 readthrough | 0 | 0 | 0 | 0 | 0 | 0 | 0 | 0 | 0 | 0 | 0 |
| 0 | NA | NA | NA | NA | NA | Snord66 | | 0 | 0 | 0 | 0 | 0 | 0 | 0 | 0 | 0 | 0 | 0 |
| 0 | NA | NA | NA | NA | NA | Gm22741 | | 0 | 0 | 0 | 0 | 0 | 0 | 0 | 0 | 0 | 0 | 0 |
| 0 | NA | NA | NA | NA | NA | Gm49742 | | 0 | 0 | 0 | 0 | 0 | 0 | 0 | 0 | 0 | 0 | 0 |
| 0 | NA | NA | NA | NA | NA | Gm6551 | | 0 | 0 | 0 | 0 | 0 | 0 | 0 | 0 | 0 | 0 | 0 |
| 0 | NA | NA | NA | NA | NA | Gm18769 | | 0 | 0 | 0 | 0 | 0 | 0 | 0 | 0 | 0 | 0 | 0 |
| 0 | NA | NA | NA | NA | NA | Gm18770 | | 0 | 0 | 0 | 0 | 0 | 0 | 0 | 0 | 0 | 0 | 0 |
| 0 | NA | NA | NA | NA | NA | Gm6557 | | 0 | 0 | 0 | 0 | 0 | 0 | 0 | 0 | 0 | 0 | 0 |
| 0 | NA | NA | NA | NA | NA | Gm9697 | | 0 | 0 | 0 | 0 | 0 | 0 | 0 | 0 | 0 | 0 | 0 |
| 0 | NA | NA | NA | NA | NA | Gm18771 | | 0 | 0 | 0 | 0 | 0 | 0 | 0 | 0 | 0 | 0 | 0 |
| 0 | NA | NA | NA | NA | NA | Teddm3 | transmembrane epididymal family membe | 0 | 0 | 0 | 0 | 0 | 0 | 0 | 0 | 0 | 0 | 0 |
| 0 | NA | NA | NA | NA | NA | Gm49750 | | 0 | 0 | 0 | 0 | 0 | 0 | 0 | 0 | 0 | 0 | 0 |
| 0 | NA | NA | NA | NA | NA | Gm16863 | | 0 | 0 | 0 | 0 | 0 | 0 | 0 | 0 | 0 | 0 | 0 |
| 0 | NA | NA | NA | NA | NA | Mir7680 | | 0 | 0 | 0 | 0 | 0 | 0 | 0 | 0 | 0 | 0 | 0 |
| 0 | NA | NA | NA | NA | NA | Gm24898 | | 0 | 0 | 0 | 0 | 0 | 0 | 0 | 0 | 0 | 0 | 0 |
| 0 | NA | NA | NA | NA | NA | Gm49607 | | 0 | 0 | 0 | 0 | 0 | 0 | 0 | 0 | 0 | 0 | 0 |
| 0 | NA | NA | NA | NA | NA | Gm5962 | | 0 | 0 | 0 | 0 | 0 | 0 | 0 | 0 | 0 | 0 | 0 |
| 0 | NA | NA | NA | NA | NA | Ndufa11b | | 0 | 0 | 0 | 0 | 0 | 0 | 0 | 0 | 0 | 0 | 0 |
| 0 | NA | NA | NA | NA | NA | Gm25810 | | 0 | 0 | 0 | 0 | 0 | 0 | 0 | 0 | 0 | 0 | 0 |
| 0 | NA | NA | NA | NA | NA | Gm36796 | | 0 | 0 | 0 | 0 | 0 | 0 | 0 | 0 | 0 | 0 | 0 |
| 0 | NA | NA | NA | NA | NA | Gm49582 | | 0 | 0 | 0 | 0 | 0 | 0 | 0 | 0 | 0 | 0 | 0 |
| 0 | NA | NA | NA | NA | NA | Mir8095 | | 0 | 0 | 0 | 0 | 0 | 0 | 0 | 0 | 0 | 0 | 0 |
| 0 | NA | NA | NA | NA | NA | Gm49565 | | 0 | 0 | 0 | 0 | 0 | 0 | 0 | 0 | 0 | 0 | 0 |
| 0 | NA | NA | NA | NA | NA | Gm22481 | | 0 | 0 | 0 | 0 | 0 | 0 | 0 | 0 | 0 | 0 | 0 |
| 0 | NA | NA | NA | NA | NA | Mir6362 | | 0 | 0 | 0 | 0 | 0 | 0 | 0 | 0 | 0 | 0 | 0 |
| 0 | NA | NA | NA | NA | NA | Gm30505 | | 0 | 0 | 0 | 0 | 0 | 0 | 0 | 0 | 0 | 0 | 0 |
| 0 | NA | NA | NA | NA | NA | Gm8128 | | 0 | 0 | 0 | 0 | 0 | 0 | 0 | 0 | 0 | 0 | 0 |
| 0 | NA | NA | NA | NA | NA | Gm23715 | | 0 | 0 | 0 | 0 | 0 | 0 | 0 | 0 | 0 | 0 | 0 |
| 0 | NA | NA | NA | NA | NA | Snord2 | | 0 | 0 | 0 | 0 | 0 | 0 | 0 | 0 | 0 | 0 | 0 |
| 0 | NA | NA | NA | NA | NA | Gm24201 | | 0 | 0 | 0 | 0 | 0 | 0 | 0 | 0 | 0 | 0 | 0 |
| 0 | NA | NA | NA | NA | NA | Gm49346 | | 0 | 0 | 0 | 0 | 0 | 0 | 0 | 0 | 0 | 0 | 0 |
| 0 | NA | NA | NA | NA | NA | Snora81 | | 0 | 0 | 0 | 0 | 0 | 0 | 0 | 0 | 0 | 0 | 0 |
| 0 | NA | NA | NA | NA | NA | Gm26447 | | 0 | 0 | 0 | 0 | 0 | 0 | 0 | 0 | 0 | 0 | 0 |
| 0 | NA | NA | NA | NA | NA | Gm49624 | | 0 | 0 | 0 | 0 | 0 | 0 | 0 | 0 | 0 | 0 | 0 |
| 0 | NA | NA | NA | NA | NA | Gm15746 | | 0 | 0 | 0 | 0 | 0 | 0 | 0 | 0 | 0 | 0 | 0 |
| 0 | NA | NA | NA | NA | NA | Gm23430 | | 0 | 0 | 0 | 0 | 0 | 0 | 0 | 0 | 0 | 0 | 0 |
| 0 | NA | NA | NA | NA | NA | Gm23066 | | 0 | 0 | 0 | 0 | 0 | 0 | 0 | 0 | 0 | 0 | 0 |
| 0 | NA | NA | NA | NA | NA | Rtp1 | receptor transporter protein 1 | 0 | 0 | 0 | 0 | 0 | 0 | 0 | 0 | 0 | 0 | 0 |
| 0 | NA | NA | NA | NA | NA | Gm49515 | | 0 | 0 | 0 | 0 | 0 | 0 | 0 | 0 | 0 | 0 | 0 |
| 0 | NA | NA | NA | NA | NA | Gm49516 | | 0 | 0 | 0 | 0 | 0 | 0 | 0 | 0 | 0 | 0 | 0 |
| 0 | NA | NA | NA | NA | NA | Rtp2 | receptor transporter protein 2 | 0 | 0 | 0 | 0 | 0 | 0 | 0 | 0 | 0 | 0 | 0 |
| 0 | NA | NA | NA | NA | NA | Gm46545 | | 0 | 0 | 0 | 0 | 0 | 0 | 0 | 0 | 0 | 0 | 0 |
| 0 | NA | NA | NA | NA | NA | Gm31583 | | 0 | 0 | 0 | 0 | 0 | 0 | 0 | 0 | 0 | 0 | 0 |
| 0 | NA | NA | NA | NA | NA | Gm32126 | | 0 | 0 | 0 | 0 | 0 | 0 | 0 | 0 | 0 | 0 | 0 |
| 0 | NA | NA | NA | NA | NA | Gm31814 | | 0 | 0 | 0 | 0 | 0 | 0 | 0 | 0 | 0 | 0 | 0 |
| 0 | NA | NA | NA | NA | NA | Gm41434 | | 0 | 0 | 0 | 0 | 0 | 0 | 0 | 0 | 0 | 0 | 0 |
| 0 | NA | NA | NA | NA | NA | Gm24440 | | 0 | 0 | 0 | 0 | 0 | 0 | 0 | 0 | 0 | 0 | 0 |
| 0 | NA | NA | NA | NA | NA | Mir28a | | 0 | 0 | 0 | 0 | 0 | 0 | 0 | 0 | 0 | 0 | 0 |
| 0 | NA | NA | NA | NA | NA | Gm22672 | | 0 | 0 | 0 | 0 | 0 | 0 | 0 | 0 | 0 | 0 | 0 |
| 0 | NA | NA | NA | NA | NA | Gm18896 | | 0 | 0 | 0 | 0 | 0 | 0 | 0 | 0 | 0 | 0 | 0 |
| 0 | NA | NA | NA | NA | NA | Gm24928 | | 0 | 0 | 0 | 0 | 0 | 0 | 0 | 0 | 0 | 0 | 0 |
| 0 | NA | NA | NA | NA | NA | Gm27005 | | 0 | 0 | 0 | 0 | 0 | 0 | 0 | 0 | 0 | 0 | 0 |
| 0 | NA | NA | NA | NA | NA | Gm4524 | | 0 | 0 | 0 | 0 | 0 | 0 | 0 | 0 | 0 | 0 | 0 |
| 0 | NA | NA | NA | NA | NA | AU015336 | | 0 | 0 | 0 | 0 | 0 | 0 | 0 | 0 | 0 | 0 | 0 |
| 0 | NA | NA | NA | NA | NA | Cldn16 | claudin 16 | 0 | 0 | 0 | 0 | 0 | 0 | 0 | 0 | 0 | 0 | 0 |
| 0 | NA | NA | NA | NA | NA | Tmem207 | transmembrane protein 207 | 0 | 0 | 0 | 0 | 0 | 0 | 0 | 0 | 0 | 0 | 0 |

| | | | | | | | | | | | | | | | | | | |
|---|---|---|---|---|---|---|---|---|---|---|---|---|---|---|---|---|---|---|
| 0 | NA | NA | NA | NA | NA | Gm20319 | | 0 | 0 | 0 | 0 | 0 | 0 | 0 | 0 | 0 | 0 | 0 |
| 0 | NA | NA | NA | NA | NA | Gmnc | geminin coiled-coil domain containing | 0 | 0 | 0 | 0 | 0 | 0 | 0 | 0 | 0 | 0 | 0 |
| 0 | NA | NA | NA | NA | NA | Gm32312 | | 0 | 0 | 0 | 0 | 0 | 0 | 0 | 0 | 0 | 0 | 0 |
| 0 | NA | NA | NA | NA | NA | Gm15970 | | 0 | 0 | 0 | 0 | 0 | 0 | 0 | 0 | 0 | 0 | 0 |
| 0 | NA | NA | NA | NA | NA | Uts2b | urotensin 2B | 0 | 0 | 0 | 0 | 0 | 0 | 0 | 0 | 0 | 0 | 0 |
| 0 | NA | NA | NA | NA | NA | Gm10823 | | 0 | 0 | 0 | 0 | 0 | 0 | 0 | 0 | 0 | 0 | 0 |
| 0 | NA | NA | NA | NA | NA | Gm49765 | | 0 | 0 | 0 | 0 | 0 | 0 | 0 | 0 | 0 | 0 | 0 |
| 0 | NA | NA | NA | NA | NA | Mir690 | | 0 | 0 | 0 | 0 | 0 | 0 | 0 | 0 | 0 | 0 | 0 |
| 0 | NA | NA | NA | NA | NA | Gm15752 | | 0 | 0 | 0 | 0 | 0 | 0 | 0 | 0 | 0 | 0 | 0 |
| 0 | NA | NA | NA | NA | NA | | | 0 | 0 | 0 | 0 | 0 | 0 | 0 | 0 | 0 | 0 | 0 |
| 0 | NA | NA | NA | NA | NA | Gm32679 | | 0 | 0 | 0 | 0 | 0 | 0 | 0 | 0 | 0 | 0 | 0 |
| 0 | NA | NA | NA | NA | NA | Gm20040 | | 0 | 0 | 0 | 0 | 0 | 0 | 0 | 0 | 0 | 0 | 0 |
| 0 | NA | NA | NA | NA | NA | n-R5s32 | | 0 | 0 | 0 | 0 | 0 | 0 | 0 | 0 | 0 | 0 | 0 |
| 0 | NA | NA | NA | NA | NA | 1700025H01Rik | | 0 | 0 | 0 | 0 | 0 | 0 | 0 | 0 | 0 | 0 | 0 |
| 0 | NA | NA | NA | NA | NA | Gm49645 | | 0 | 0 | 0 | 0 | 0 | 0 | 0 | 0 | 0 | 0 | 0 |
| 0 | NA | NA | NA | NA | NA | Cpn2 | carboxypeptidase N, polypeptide 2 | 0 | 0 | 0 | 0 | 0 | 0 | 0 | 0 | 0 | 0 | 0 |
| 0 | NA | NA | NA | NA | NA | Gm49679 | | 0 | 0 | 0 | 0 | 0 | 0 | 0 | 0 | 0 | 0 | 0 |
| 0 | NA | NA | NA | NA | NA | Gm26295 | | 0 | 0 | 0 | 0 | 0 | 0 | 0 | 0 | 0 | 0 | 0 |
| 0 | NA | NA | NA | NA | NA | Gm49755 | | 0 | 0 | 0 | 0 | 0 | 0 | 0 | 0 | 0 | 0 | 0 |
| 0 | NA | NA | NA | NA | NA | Gm49756 | | 0 | 0 | 0 | 0 | 0 | 0 | 0 | 0 | 0 | 0 | 0 |
| 0 | NA | NA | NA | NA | NA | Gm49757 | | 0 | 0 | 0 | 0 | 0 | 0 | 0 | 0 | 0 | 0 | 0 |
| 0 | NA | NA | NA | NA | NA | Gm25375 | | 0 | 0 | 0 | 0 | 0 | 0 | 0 | 0 | 0 | 0 | 0 |
| 0 | NA | NA | NA | NA | NA | Gm18237 | | 0 | 0 | 0 | 0 | 0 | 0 | 0 | 0 | 0 | 0 | 0 |
| 0 | NA | NA | NA | NA | NA | Gm27415 | | 0 | 0 | 0 | 0 | 0 | 0 | 0 | 0 | 0 | 0 | 0 |
| 0 | NA | NA | NA | NA | NA | Gm49546 | | 0 | 0 | 0 | 0 | 0 | 0 | 0 | 0 | 0 | 0 | 0 |
| 0 | NA | NA | NA | NA | NA | AV205837 | | 0 | 0 | 0 | 0 | 0 | 0 | 0 | 0 | 0 | 0 | 0 |
| 0 | NA | NA | NA | NA | NA | Gm49404 | | 0 | 0 | 0 | 0 | 0 | 0 | 0 | 0 | 0 | 0 | 0 |
| 0 | NA | NA | NA | NA | NA | 1700007E05Rik | | 0 | 0 | 0 | 0 | 0 | 0 | 0 | 0 | 0 | 0 | 0 |
| 0 | NA | NA | NA | NA | NA | Gm49736 | | 0 | 0 | 0 | 0 | 0 | 0 | 0 | 0 | 0 | 0 | 0 |
| 0 | NA | NA | NA | NA | NA | Gm49695 | | 0 | 0 | 0 | 0 | 0 | 0 | 0 | 0 | 0 | 0 | 0 |
| 0 | NA | NA | NA | NA | NA | Gm49712 | | 0 | 0 | 0 | 0 | 0 | 0 | 0 | 0 | 0 | 0 | 0 |
| 0 | NA | NA | NA | NA | NA | Gm24879 | | 0 | 0 | 0 | 0 | 0 | 0 | 0 | 0 | 0 | 0 | 0 |
| 0 | NA | NA | NA | NA | NA | Gm49731 | | 0 | 0 | 0 | 0 | 0 | 0 | 0 | 0 | 0 | 0 | 0 |
| 0 | NA | NA | NA | NA | NA | Gm26269 | | 0 | 0 | 0 | 0 | 0 | 0 | 0 | 0 | 0 | 0 | 0 |
| 0 | NA | NA | NA | NA | NA | Gm26419 | | 0 | 0 | 0 | 0 | 0 | 0 | 0 | 0 | 0 | 0 | 0 |
| 0 | NA | NA | NA | NA | NA | Mir1946a | | 0 | 0 | 0 | 0 | 0 | 0 | 0 | 0 | 0 | 0 | 0 |
| 0 | NA | NA | NA | NA | NA | Smco1 | single-pass membrane protein with coilec | 0 | 0 | 0 | 0 | 0 | 0 | 0 | 0 | 0 | 0 | 0 |
| 0 | NA | NA | NA | NA | NA | 2210020O09Rik | | 0 | 0 | 0 | 0 | 0 | 0 | 0 | 0 | 0 | 0 | 0 |
| 0 | NA | NA | NA | NA | NA | Zdhhc19 | zinc finger, DHHC domain containing 19 | 0 | 0 | 0 | 0 | 0 | 0 | 0 | 0 | 0 | 0 | 0 |
| 0 | NA | NA | NA | NA | NA | Gm5405 | | 0 | 0 | 0 | 0 | 0 | 0 | 0 | 0 | 0 | 0 | 0 |
| 0 | NA | NA | NA | NA | NA | Tnk2os | | 0 | 0 | 0 | 0 | 0 | 0 | 0 | 0 | 0 | 0 | 0 |
| 0 | NA | NA | NA | NA | NA | Gm9556 | | 0 | 0 | 0 | 0 | 0 | 0 | 0 | 0 | 0 | 0 | 0 |
| 0 | NA | NA | NA | NA | NA | Gm26769 | | 0 | 0 | 0 | 0 | 0 | 0 | 0 | 0 | 0 | 0 | 0 |
| 0 | NA | NA | NA | NA | NA | Gm17106 | | 0 | 0 | 0 | 0 | 0 | 0 | 0 | 0 | 0 | 0 | 0 |
| 0 | NA | NA | NA | NA | NA | Gm22231 | | 0 | 0 | 0 | 0 | 0 | 0 | 0 | 0 | 0 | 0 | 0 |
| 0 | NA | NA | NA | NA | NA | Gm23916 | | 0 | 0 | 0 | 0 | 0 | 0 | 0 | 0 | 0 | 0 | 0 |
| 0 | NA | NA | NA | NA | NA | Gm15657 | | 0 | 0 | 0 | 0 | 0 | 0 | 0 | 0 | 0 | 0 | 0 |
| 0 | NA | NA | NA | NA | NA | Ropn1 | ropporin, rhophilin associated protein 1 | 0 | 0 | 0 | 0 | 0 | 0 | 0 | 0 | 0 | 0 | 0 |
| 0 | NA | NA | NA | NA | NA | 1700119H24Rik | | 0 | 0 | 0 | 0 | 0 | 0 | 0 | 0 | 0 | 0 | 0 |
| 0 | NA | NA | NA | NA | NA | E130310I04Rik | | 0 | 0 | 0 | 0 | 0 | 0 | 0 | 0 | 0 | 0 | 0 |
| 0 | NA | NA | NA | NA | NA | Gm25179 | | 0 | 0 | 0 | 0 | 0 | 0 | 0 | 0 | 0 | 0 | 0 |
| 0 | NA | NA | NA | NA | NA | Gm25079 | | 0 | 0 | 0 | 0 | 0 | 0 | 0 | 0 | 0 | 0 | 0 |
| 0 | NA | NA | NA | NA | NA | Gm23518 | | 0 | 0 | 0 | 0 | 0 | 0 | 0 | 0 | 0 | 0 | 0 |
| 0 | NA | NA | NA | NA | NA | Gm25548 | | 0 | 0 | 0 | 0 | 0 | 0 | 0 | 0 | 0 | 0 | 0 |
| 0 | NA | NA | NA | NA | NA | 1600019K03Rik | | 0 | 0 | 0 | 0 | 0 | 0 | 0 | 0 | 0 | 0 | 0 |
| 0 | NA | NA | NA | NA | NA | Gm25967 | | 0 | 0 | 0 | 0 | 0 | 0 | 0 | 0 | 0 | 0 | 0 |
| 0 | NA | NA | NA | NA | NA | Gm26838 | | 0 | 0 | 0 | 0 | 0 | 0 | 0 | 0 | 0 | 0 | 0 |
| 0 | NA | NA | NA | NA | NA | Gm49728 | | 0 | 0 | 0 | 0 | 0 | 0 | 0 | 0 | 0 | 0 | 0 |
| 0 | NA | NA | NA | NA | NA | Gm41467 | | 0 | 0 | 0 | 0 | 0 | 0 | 0 | 0 | 0 | 0 | 0 |
| 0 | NA | NA | NA | NA | NA | Csta1 | cystatin A1 | 0 | 0 | 0 | 0 | 0 | 0 | 0 | 0 | 0 | 0 | 0 |

| | | | | | | | | | | | | | | | | | | |
|---|---|---|---|---|---|---|---|---|---|---|---|---|---|---|---|---|---|---|
| 0 | NA | NA | NA | NA | NA | Gm10092 | | 0 | 0 | 0 | 0 | 0 | 0 | 0 | 0 | 0 | 0 | 0 |
| 0 | NA | NA | NA | NA | NA | Gm23080 | | 0 | 0 | 0 | 0 | 0 | 0 | 0 | 0 | 0 | 0 | 0 |
| 0 | NA | NA | NA | NA | NA | Stfa2l1 | stefin A2 like 1 | 0 | 0 | 0 | 0 | 0 | 0 | 0 | 0 | 0 | 0 | 0 |
| 0 | NA | NA | NA | NA | NA | Gm22885 | | 0 | 0 | 0 | 0 | 0 | 0 | 0 | 0 | 0 | 0 | 0 |
| 0 | NA | NA | NA | NA | NA | Cstdc4 | cystatin domain containing 4 | 0 | 0 | 0 | 0 | 0 | 0 | 0 | 0 | 0 | 0 | 0 |
| 0 | NA | NA | NA | NA | NA | Gm25024 | | 0 | 0 | 0 | 0 | 0 | 0 | 0 | 0 | 0 | 0 | 0 |
| 0 | NA | NA | NA | NA | NA | Gm6815 | | 0 | 0 | 0 | 0 | 0 | 0 | 0 | 0 | 0 | 0 | 0 |
| 0 | NA | NA | NA | NA | NA | Csta3 | cystatin A family member 3 | 0 | 0 | 0 | 0 | 0 | 0 | 0 | 0 | 0 | 0 | 0 |
| 0 | NA | NA | NA | NA | NA | Gm24801 | | 0 | 0 | 0 | 0 | 0 | 0 | 0 | 0 | 0 | 0 | 0 |
| 0 | NA | NA | NA | NA | NA | Gm10813 | | 0 | 0 | 0 | 0 | 0 | 0 | 0 | 0 | 0 | 0 | 0 |
| 0 | NA | NA | NA | NA | NA | Gm24569 | | 0 | 0 | 0 | 0 | 0 | 0 | 0 | 0 | 0 | 0 | 0 |
| 0 | NA | NA | NA | NA | NA | Stfa1 | stefin A1 | 0 | 0 | 0 | 0 | 0 | 0 | 0 | 0 | 0 | 0 | 0 |
| 0 | NA | NA | NA | NA | NA | Gm23306 | | 0 | 0 | 0 | 0 | 0 | 0 | 0 | 0 | 0 | 0 | 0 |
| 0 | NA | NA | NA | NA | NA | Cstdc3 | | 0 | 0 | 0 | 0 | 0 | 0 | 0 | 0 | 0 | 0 | 0 |
| 0 | NA | NA | NA | NA | NA | Gm25254 | | 0 | 0 | 0 | 0 | 0 | 0 | 0 | 0 | 0 | 0 | 0 |
| 0 | NA | NA | NA | NA | NA | Gm49647 | | 0 | 0 | 0 | 0 | 0 | 0 | 0 | 0 | 0 | 0 | 0 |
| 0 | NA | NA | NA | NA | NA | Gm24437 | | 0 | 0 | 0 | 0 | 0 | 0 | 0 | 0 | 0 | 0 | 0 |
| 0 | NA | NA | NA | NA | NA | Cstdc6 | cystatin domain containing 6 | 0 | 0 | 0 | 0 | 0 | 0 | 0 | 0 | 0 | 0 | 0 |
| 0 | NA | NA | NA | NA | NA | Gm25465 | | 0 | 0 | 0 | 0 | 0 | 0 | 0 | 0 | 0 | 0 | 0 |
| 0 | NA | NA | NA | NA | NA | Cstdc5 | cystatin domain containing 5 | 0 | 0 | 0 | 0 | 0 | 0 | 0 | 0 | 0 | 0 | 0 |
| 0 | NA | NA | NA | NA | NA | Gm15845 | | 0 | 0 | 0 | 0 | 0 | 0 | 0 | 0 | 0 | 0 | 0 |
| 0 | NA | NA | NA | NA | NA | Gm24218 | | 0 | 0 | 0 | 0 | 0 | 0 | 0 | 0 | 0 | 0 | 0 |
| 0 | NA | NA | NA | NA | NA | Gm7260 | | 0 | 0 | 0 | 0 | 0 | 0 | 0 | 0 | 0 | 0 | 0 |
| 0 | NA | NA | NA | NA | NA | Gm22349 | | 0 | 0 | 0 | 0 | 0 | 0 | 0 | 0 | 0 | 0 | 0 |
| 0 | NA | NA | NA | NA | NA | Stfa2 | stefin A2 | 0 | 0 | 0 | 0 | 0 | 0 | 0 | 0 | 0 | 0 | 0 |
| 0 | NA | NA | NA | NA | NA | Gm22263 | | 0 | 0 | 0 | 0 | 0 | 0 | 0 | 0 | 0 | 0 | 0 |
| 0 | NA | NA | NA | NA | NA | Gm49652 | | 0 | 0 | 0 | 0 | 0 | 0 | 0 | 0 | 0 | 0 | 0 |
| 0 | NA | NA | NA | NA | NA | Gm25903 | | 0 | 0 | 0 | 0 | 0 | 0 | 0 | 0 | 0 | 0 | 0 |
| 0 | NA | NA | NA | NA | NA | Stfa3 | stefin A3 | 0 | 0 | 0 | 0 | 0 | 0 | 0 | 0 | 0 | 0 | 0 |
| 0 | NA | NA | NA | NA | NA | Gm49653 | | 0 | 0 | 0 | 0 | 0 | 0 | 0 | 0 | 0 | 0 | 0 |
| 0 | NA | NA | NA | NA | NA | Gm49662 | | 0 | 0 | 0 | 0 | 0 | 0 | 0 | 0 | 0 | 0 | 0 |
| 0 | NA | NA | NA | NA | NA | Gm49664 | | 0 | 0 | 0 | 0 | 0 | 0 | 0 | 0 | 0 | 0 | 0 |
| 0 | NA | NA | NA | NA | NA | Gm22617 | | 0 | 0 | 0 | 0 | 0 | 0 | 0 | 0 | 0 | 0 | 0 |
| 0 | NA | NA | NA | NA | NA | 4930565N06Rik | | 0 | 0 | 0 | 0 | 0 | 0 | 0 | 0 | 0 | 0 | 0 |
| 0 | NA | NA | NA | NA | NA | Gm49605 | | 0 | 0 | 0 | 0 | 0 | 0 | 0 | 0 | 0 | 0 | 0 |
| 0 | NA | NA | NA | NA | NA | Gm49567 | | 0 | 0 | 0 | 0 | 0 | 0 | 0 | 0 | 0 | 0 | 0 |
| 0 | NA | NA | NA | NA | NA | Gm49569 | | 0 | 0 | 0 | 0 | 0 | 0 | 0 | 0 | 0 | 0 | 0 |
| 0 | NA | NA | NA | NA | NA | Gm46559 | | 0 | 0 | 0 | 0 | 0 | 0 | 0 | 0 | 0 | 0 | 0 |
| 0 | NA | NA | NA | NA | NA | Gm36028 | predicted gene, 36028 | 0 | 0 | 0 | 0 | 0 | 0 | 0 | 0 | 0 | 0 | 0 |
| 0 | NA | NA | NA | NA | NA | n-R5s34 | | 0 | 0 | 0 | 0 | 0 | 0 | 0 | 0 | 0 | 0 | 0 |
| 0 | NA | NA | NA | NA | NA | Gm15725 | | 0 | 0 | 0 | 0 | 0 | 0 | 0 | 0 | 0 | 0 | 0 |
| 0 | NA | NA | NA | NA | NA | Gm25140 | | 0 | 0 | 0 | 0 | 0 | 0 | 0 | 0 | 0 | 0 | 0 |
| 0 | NA | NA | NA | NA | NA | Gm25617 | | 0 | 0 | 0 | 0 | 0 | 0 | 0 | 0 | 0 | 0 | 0 |
| 0 | NA | NA | NA | NA | NA | Gm25617 | | 0 | 0 | 0 | 0 | 0 | 0 | 0 | 0 | 0 | 0 | 0 |
| 0 | NA | NA | NA | NA | NA | Gm27931 | | 0 | 0 | 0 | 0 | 0 | 0 | 0 | 0 | 0 | 0 | 0 |
| 0 | NA | NA | NA | NA | NA | Maats1os | | 0 | 0 | 0 | 0 | 0 | 0 | 0 | 0 | 0 | 0 | 0 |
| 0 | NA | NA | NA | NA | NA | Gm21987 | | 0 | 0 | 0 | 0 | 0 | 0 | 0 | 0 | 0 | 0 | 0 |
| 0 | NA | NA | NA | NA | NA | Gm15953 | | 0 | 0 | 0 | 0 | 0 | 0 | 0 | 0 | 0 | 0 | 0 |
| 0 | NA | NA | NA | NA | NA | Tex55 | testis expressed 55 | 0 | 0 | 0 | 0 | 0 | 0 | 0 | 0 | 0 | 0 | 0 |
| 0 | NA | NA | NA | NA | NA | Gm15803 | | 0 | 0 | 0 | 0 | 0 | 0 | 0 | 0 | 0 | 0 | 0 |
| 0 | NA | NA | NA | NA | NA | Gm22500 | | 0 | 0 | 0 | 0 | 0 | 0 | 0 | 0 | 0 | 0 | 0 |
| 0 | NA | NA | NA | NA | NA | Gm36903 | | 0 | 0 | 0 | 0 | 0 | 0 | 0 | 0 | 0 | 0 | 0 |
| 0 | NA | NA | NA | NA | NA | Gm24795 | | 0 | 0 | 0 | 0 | 0 | 0 | 0 | 0 | 0 | 0 | 0 |
| 0 | NA | NA | NA | NA | NA | Gm8609 | | 0 | 0 | 0 | 0 | 0 | 0 | 0 | 0 | 0 | 0 | 0 |
| 0 | NA | NA | NA | NA | NA | Gm37508 | | 0 | 0 | 0 | 0 | 0 | 0 | 0 | 0 | 0 | 0 | 0 |
| 0 | NA | NA | NA | NA | NA | Gm22543 | | 0 | 0 | 0 | 0 | 0 | 0 | 0 | 0 | 0 | 0 | 0 |
| 0 | NA | NA | NA | NA | NA | Gm36939 | | 0 | 0 | 0 | 0 | 0 | 0 | 0 | 0 | 0 | 0 | 0 |
| 0 | NA | NA | NA | NA | NA | Gm27887 | | 0 | 0 | 0 | 0 | 0 | 0 | 0 | 0 | 0 | 0 | 0 |
| 0 | NA | NA | NA | NA | NA | Gm27717 | | 0 | 0 | 0 | 0 | 0 | 0 | 0 | 0 | 0 | 0 | 0 |
| 0 | NA | NA | NA | NA | NA | Gm26381 | | 0 | 0 | 0 | 0 | 0 | 0 | 0 | 0 | 0 | 0 | 0 |

| | | | | | | | | | | | | | | | | | |
|---|---|---|---|---|---|---|---|---|---|---|---|---|---|---|---|---|---|
| 0 | NA | NA | NA | NA | NA | Gm47276 | | 0 | 0 | 0 | 0 | 0 | 0 | 0 | 0 | 0 | 0 | 0 |
| 0 | NA | NA | NA | NA | NA | Gm27587 | | 0 | 0 | 0 | 0 | 0 | 0 | 0 | 0 | 0 | 0 | 0 |
| 0 | NA | NA | NA | NA | NA | Gm28750 | | 0 | 0 | 0 | 0 | 0 | 0 | 0 | 0 | 0 | 0 | 0 |
| 0 | NA | NA | NA | NA | NA | Gm49737 | | 0 | 0 | 0 | 0 | 0 | 0 | 0 | 0 | 0 | 0 | 0 |
| 0 | NA | NA | NA | NA | NA | Gm49738 | | 0 | 0 | 0 | 0 | 0 | 0 | 0 | 0 | 0 | 0 | 0 |
| 0 | NA | NA | NA | NA | NA | Mir6540 | | 0 | 0 | 0 | 0 | 0 | 0 | 0 | 0 | 0 | 0 | 0 |
| 0 | NA | NA | NA | NA | NA | Gm25873 | | 0 | 0 | 0 | 0 | 0 | 0 | 0 | 0 | 0 | 0 | 0 |
| 0 | NA | NA | NA | NA | NA | Gm37946 | | 0 | 0 | 0 | 0 | 0 | 0 | 0 | 0 | 0 | 0 | 0 |
| 0 | NA | NA | NA | NA | NA | Gm15713 | | 0 | 0 | 0 | 0 | 0 | 0 | 0 | 0 | 0 | 0 | 0 |
| 0 | NA | NA | NA | NA | NA | Mir568 | | 0 | 0 | 0 | 0 | 0 | 0 | 0 | 0 | 0 | 0 | 0 |
| 0 | NA | NA | NA | NA | NA | Gm49734 | | 0 | 0 | 0 | 0 | 0 | 0 | 0 | 0 | 0 | 0 | 0 |
| 0 | NA | NA | NA | NA | NA | Gm49735 | | 0 | 0 | 0 | 0 | 0 | 0 | 0 | 0 | 0 | 0 | 0 |
| 0 | NA | NA | NA | NA | NA | Gm25996 | | 0 | 0 | 0 | 0 | 0 | 0 | 0 | 0 | 0 | 0 | 0 |
| 0 | NA | NA | NA | NA | NA | Gm26074 | | 0 | 0 | 0 | 0 | 0 | 0 | 0 | 0 | 0 | 0 | 0 |
| 0 | NA | NA | NA | NA | NA | Gm25989 | | 0 | 0 | 0 | 0 | 0 | 0 | 0 | 0 | 0 | 0 | 0 |
| 0 | NA | NA | NA | NA | NA | Mir3470b | | 0 | 0 | 0 | 0 | 0 | 0 | 0 | 0 | 0 | 0 | 0 |
| 0 | NA | NA | NA | NA | NA | Mir3081 | | 0 | 0 | 0 | 0 | 0 | 0 | 0 | 0 | 0 | 0 | 0 |
| 0 | NA | NA | NA | NA | NA | Gm5063 | | 0 | 0 | 0 | 0 | 0 | 0 | 0 | 0 | 0 | 0 | 0 |
| 0 | NA | NA | NA | NA | NA | Gm19142 | | 0 | 0 | 0 | 0 | 0 | 0 | 0 | 0 | 0 | 0 | 0 |
| 0 | NA | NA | NA | NA | NA | Gm5964 | | 0 | 0 | 0 | 0 | 0 | 0 | 0 | 0 | 0 | 0 | 0 |
| 0 | NA | NA | NA | NA | NA | Cd200r2 | Cd200 receptor 2 | 0 | 0 | 0 | 0 | 0 | 0 | 0 | 0 | 0 | 0 | 0 |
| 0 | NA | NA | NA | NA | NA | Gm17928 | | 0 | 0 | 0 | 0 | 0 | 0 | 0 | 0 | 0 | 0 | 0 |
| 0 | NA | NA | NA | NA | NA | Cd200r3 | CD200 receptor 3 | 0 | 0 | 0 | 0 | 0 | 0 | 0 | 0 | 0 | 0 | 0 |
| 0 | NA | NA | NA | NA | NA | Gm6903 | | 0 | 0 | 0 | 0 | 0 | 0 | 0 | 0 | 0 | 0 | 0 |
| 0 | NA | NA | NA | NA | NA | Gm49585 | | 0 | 0 | 0 | 0 | 0 | 0 | 0 | 0 | 0 | 0 | 0 |
| 0 | NA | NA | NA | NA | NA | Gm15640 | | 0 | 0 | 0 | 0 | 0 | 0 | 0 | 0 | 0 | 0 | 0 |
| 0 | NA | NA | NA | NA | NA | Gm24754 | | 0 | 0 | 0 | 0 | 0 | 0 | 0 | 0 | 0 | 0 | 0 |
| 0 | NA | NA | NA | NA | NA | Gm50487 | | 0 | 0 | 0 | 0 | 0 | 0 | 0 | 0 | 0 | 0 | 0 |
| 0 | NA | NA | NA | NA | NA | Gm26297 | | 0 | 0 | 0 | 0 | 0 | 0 | 0 | 0 | 0 | 0 | 0 |
| 0 | NA | NA | NA | NA | NA | Gm17900 | | 0 | 0 | 0 | 0 | 0 | 0 | 0 | 0 | 0 | 0 | 0 |
| 0 | NA | NA | NA | NA | NA | Gm6912 | | 0 | 0 | 0 | 0 | 0 | 0 | 0 | 0 | 0 | 0 | 0 |
| 0 | NA | NA | NA | NA | NA | Gm24651 | | 0 | 0 | 0 | 0 | 0 | 0 | 0 | 0 | 0 | 0 | 0 |
| 0 | NA | NA | NA | NA | NA | Gm25318 | | 0 | 0 | 0 | 0 | 0 | 0 | 0 | 0 | 0 | 0 | 0 |
| 0 | NA | NA | NA | NA | NA | Gm6914 | | 0 | 0 | 0 | 0 | 0 | 0 | 0 | 0 | 0 | 0 | 0 |
| 0 | NA | NA | NA | NA | NA | Gm25339 | | 0 | 0 | 0 | 0 | 0 | 0 | 0 | 0 | 0 | 0 | 0 |
| 0 | NA | NA | NA | NA | NA | Gm7275 | | 0 | 0 | 0 | 0 | 0 | 0 | 0 | 0 | 0 | 0 | 0 |
| 0 | NA | NA | NA | NA | NA | Gm17804 | | 0 | 0 | 0 | 0 | 0 | 0 | 0 | 0 | 0 | 0 | 0 |
| 0 | NA | NA | NA | NA | NA | 6430553K19Rik | | 0 | 0 | 0 | 0 | 0 | 0 | 0 | 0 | 0 | 0 | 0 |
| 0 | NA | NA | NA | NA | NA | Gm49770 | | 0 | 0 | 0 | 0 | 0 | 0 | 0 | 0 | 0 | 0 | 0 |
| 0 | NA | NA | NA | NA | NA | Dppa4 | developmental pluripotency associated 4 | 0 | 0 | 0 | 0 | 0 | 0 | 0 | 0 | 0 | 0 | 0 |
| 0 | NA | NA | NA | NA | NA | Gm49771 | | 0 | 0 | 0 | 0 | 0 | 0 | 0 | 0 | 0 | 0 | 0 |
| 0 | NA | NA | NA | NA | NA | n-R5s36 | | 0 | 0 | 0 | 0 | 0 | 0 | 0 | 0 | 0 | 0 | 0 |
| 0 | NA | NA | NA | NA | NA | Retnlb | resistin like beta | 0 | 0 | 0 | 0 | 0 | 0 | 0 | 0 | 0 | 0 | 0 |
| 0 | NA | NA | NA | NA | NA | Myh15 | myosin, heavy chain 15 | 0 | 0 | 0 | 0 | 0 | 0 | 0 | 0 | 0 | 0 | 0 |
| 0 | NA | NA | NA | NA | NA | 1700026J12Rik | | 0 | 0 | 0 | 0 | 0 | 0 | 0 | 0 | 0 | 0 | 0 |
| 0 | NA | NA | NA | NA | NA | Gm5407 | | 0 | 0 | 0 | 0 | 0 | 0 | 0 | 0 | 0 | 0 | 0 |
| 0 | NA | NA | NA | NA | NA | Gm18694 | | 0 | 0 | 0 | 0 | 0 | 0 | 0 | 0 | 0 | 0 | 0 |
| 0 | NA | NA | NA | NA | NA | Gm4802 | predicted gene 4802 | 0 | 0 | 0 | 0 | 0 | 0 | 0 | 0 | 0 | 0 | 0 |
| 0 | NA | NA | NA | NA | NA | G730013B05Rik | | 0 | 0 | 0 | 0 | 0 | 0 | 0 | 0 | 0 | 0 | 0 |
| 0 | NA | NA | NA | NA | NA | Ccdc54 | coiled-coil domain containing 54 | 0 | 0 | 0 | 0 | 0 | 0 | 0 | 0 | 0 | 0 | 0 |
| 0 | NA | NA | NA | NA | NA | Mir6363 | | 0 | 0 | 0 | 0 | 0 | 0 | 0 | 0 | 0 | 0 | 0 |
| 0 | NA | NA | NA | NA | NA | Gm24925 | | 0 | 0 | 0 | 0 | 0 | 0 | 0 | 0 | 0 | 0 | 0 |
| 0 | NA | NA | NA | NA | NA | Gm32431 | | 0 | 0 | 0 | 0 | 0 | 0 | 0 | 0 | 0 | 0 | 0 |
| 0 | NA | NA | NA | NA | NA | Gm49608 | | 0 | 0 | 0 | 0 | 0 | 0 | 0 | 0 | 0 | 0 | 0 |
| 0 | NA | NA | NA | NA | NA | Gm23513 | | 0 | 0 | 0 | 0 | 0 | 0 | 0 | 0 | 0 | 0 | 0 |
| 0 | NA | NA | NA | NA | NA | Gm49634 | | 0 | 0 | 0 | 0 | 0 | 0 | 0 | 0 | 0 | 0 | 0 |
| 0 | NA | NA | NA | NA | NA | 4930404A05Rik | | 0 | 0 | 0 | 0 | 0 | 0 | 0 | 0 | 0 | 0 | 0 |
| 0 | NA | NA | NA | NA | NA | 4930405D01Rik | | 0 | 0 | 0 | 0 | 0 | 0 | 0 | 0 | 0 | 0 | 0 |
| 0 | NA | NA | NA | NA | NA | Gm25723 | | 0 | 0 | 0 | 0 | 0 | 0 | 0 | 0 | 0 | 0 | 0 |
| 0 | NA | NA | NA | NA | NA | Gm22459 | | 0 | 0 | 0 | 0 | 0 | 0 | 0 | 0 | 0 | 0 | 0 |

| | | | | | | | | | | | | | | | | | |
|---|---|---|---|---|---|---|---|---|---|---|---|---|---|---|---|---|---|
| 0 | NA | NA | NA | NA | NA | Gm49637 | | 0 | 0 | 0 | 0 | 0 | 0 | 0 | 0 | 0 | 0 | 0 |
| 0 | NA | NA | NA | NA | NA | Gm49638 | | 0 | 0 | 0 | 0 | 0 | 0 | 0 | 0 | 0 | 0 | 0 |
| 0 | NA | NA | NA | NA | NA | Gm25041 | | 0 | 0 | 0 | 0 | 0 | 0 | 0 | 0 | 0 | 0 | 0 |
| 0 | NA | NA | NA | NA | NA | Gm49639 | | 0 | 0 | 0 | 0 | 0 | 0 | 0 | 0 | 0 | 0 | 0 |
| 0 | NA | NA | NA | NA | NA | Gm25853 | | 0 | 0 | 0 | 0 | 0 | 0 | 0 | 0 | 0 | 0 | 0 |
| 0 | NA | NA | NA | NA | NA | Gm46541 | | 0 | 0 | 0 | 0 | 0 | 0 | 0 | 0 | 0 | 0 | 0 |
| 0 | NA | NA | NA | NA | NA | Gm23180 | | 0 | 0 | 0 | 0 | 0 | 0 | 0 | 0 | 0 | 0 | 0 |
| 0 | NA | NA | NA | NA | NA | Gm8881 | | 0 | 0 | 0 | 0 | 0 | 0 | 0 | 0 | 0 | 0 | 0 |
| 0 | NA | NA | NA | NA | NA | Gm22977 | | 0 | 0 | 0 | 0 | 0 | 0 | 0 | 0 | 0 | 0 | 0 |
| 0 | NA | NA | NA | NA | NA | Mir5118 | | 0 | 0 | 0 | 0 | 0 | 0 | 0 | 0 | 0 | 0 | 0 |
| 0 | NA | NA | NA | NA | NA | Gm23003 | | 0 | 0 | 0 | 0 | 0 | 0 | 0 | 0 | 0 | 0 | 0 |
| 0 | NA | NA | NA | NA | NA | Gm19771 | | 0 | 0 | 0 | 0 | 0 | 0 | 0 | 0 | 0 | 0 | 0 |
| 0 | NA | NA | NA | NA | NA | Gm26369 | | 0 | 0 | 0 | 0 | 0 | 0 | 0 | 0 | 0 | 0 | 0 |
| 0 | NA | NA | NA | NA | NA | Gm8900 | | 0 | 0 | 0 | 0 | 0 | 0 | 0 | 0 | 0 | 0 | 0 |
| 0 | NA | NA | NA | NA | NA | Gm28037 | | 0 | 0 | 0 | 0 | 0 | 0 | 0 | 0 | 0 | 0 | 0 |
| 0 | NA | NA | NA | NA | NA | Gm24047 | | 0 | 0 | 0 | 0 | 0 | 0 | 0 | 0 | 0 | 0 | 0 |
| 0 | NA | NA | NA | NA | NA | Gm18723 | | 0 | 0 | 0 | 0 | 0 | 0 | 0 | 0 | 0 | 0 | 0 |
| 0 | NA | NA | NA | NA | NA | Gm49682 | | 0 | 0 | 0 | 0 | 0 | 0 | 0 | 0 | 0 | 0 | 0 |
| 0 | NA | NA | NA | NA | NA | Gm21050 | | 0 | 0 | 0 | 0 | 0 | 0 | 0 | 0 | 0 | 0 | 0 |
| 0 | NA | NA | NA | NA | NA | Gm15693 | | 0 | 0 | 0 | 0 | 0 | 0 | 0 | 0 | 0 | 0 | 0 |
| 0 | NA | NA | NA | NA | NA | Gm15839 | | 0 | 0 | 0 | 0 | 0 | 0 | 0 | 0 | 0 | 0 | 0 |
| 0 | NA | NA | NA | NA | NA | Gm15840 | | 0 | 0 | 0 | 0 | 0 | 0 | 0 | 0 | 0 | 0 | 0 |
| 0 | NA | NA | NA | NA | NA | Lnp1 | leukemia NUP98 fusion partner 1 | 0 | 0 | 0 | 0 | 0 | 0 | 0 | 0 | 0 | 0 | 0 |
| 0 | NA | NA | NA | NA | NA | Gm8938 | | 0 | 0 | 0 | 0 | 0 | 0 | 0 | 0 | 0 | 0 | 0 |
| 0 | NA | NA | NA | NA | NA | Gm49581 | | 0 | 0 | 0 | 0 | 0 | 0 | 0 | 0 | 0 | 0 | 0 |
| 0 | NA | NA | NA | NA | NA | 4921517D16Rik | | 0 | 0 | 0 | 0 | 0 | 0 | 0 | 0 | 0 | 0 | 0 |
| 0 | NA | NA | NA | NA | NA | Gm49579 | | 0 | 0 | 0 | 0 | 0 | 0 | 0 | 0 | 0 | 0 | 0 |
| 0 | NA | NA | NA | NA | NA | Gm5674 | | 0 | 0 | 0 | 0 | 0 | 0 | 0 | 0 | 0 | 0 | 0 |
| 0 | NA | NA | NA | NA | NA | Gm49575 | | 0 | 0 | 0 | 0 | 0 | 0 | 0 | 0 | 0 | 0 | 0 |
| 0 | NA | NA | NA | NA | NA | Gm24039 | | 0 | 0 | 0 | 0 | 0 | 0 | 0 | 0 | 0 | 0 | 0 |
| 0 | NA | NA | NA | NA | NA | Rpl7a-ps4 | | 0 | 0 | 0 | 0 | 0 | 0 | 0 | 0 | 0 | 0 | 0 |
| 0 | NA | NA | NA | NA | NA | Gm37458 | | 0 | 0 | 0 | 0 | 0 | 0 | 0 | 0 | 0 | 0 | 0 |
| 0 | NA | NA | NA | NA | NA | Gm49576 | | 0 | 0 | 0 | 0 | 0 | 0 | 0 | 0 | 0 | 0 | 0 |
| 0 | NA | NA | NA | NA | NA | Gm49574 | | 0 | 0 | 0 | 0 | 0 | 0 | 0 | 0 | 0 | 0 | 0 |
| 0 | NA | NA | NA | NA | NA | Gm37825 | | 0 | 0 | 0 | 0 | 0 | 0 | 0 | 0 | 0 | 0 | 0 |
| 0 | NA | NA | NA | NA | NA | Gm20942 | | 0 | 0 | 0 | 0 | 0 | 0 | 0 | 0 | 0 | 0 | 0 |
| 0 | NA | NA | NA | NA | NA | Ftdc1 | ferritin domain containing 1 | 0 | 0 | 0 | 0 | 0 | 0 | 0 | 0 | 0 | 0 | 0 |
| 0 | NA | NA | NA | NA | NA | Ftdc2 | ferritin domain containing 2 | 0 | 0 | 0 | 0 | 0 | 0 | 0 | 0 | 0 | 0 | 0 |
| 0 | NA | NA | NA | NA | NA | Gm27564 | | 0 | 0 | 0 | 0 | 0 | 0 | 0 | 0 | 0 | 0 | 0 |
| 0 | NA | NA | NA | NA | NA | Gm26019 | | 0 | 0 | 0 | 0 | 0 | 0 | 0 | 0 | 0 | 0 | 0 |
| 0 | NA | NA | NA | NA | NA | Olfr172 | olfactory receptor 172 | 0 | 0 | 0 | 0 | 0 | 0 | 0 | 0 | 0 | 0 | 0 |
| 0 | NA | NA | NA | NA | NA | Olfr173 | olfactory receptor 173 | 0 | 0 | 0 | 0 | 0 | 0 | 0 | 0 | 0 | 0 | 0 |
| 0 | NA | NA | NA | NA | NA | Olfr175 | olfactory receptor 175 | 0 | 0 | 0 | 0 | 0 | 0 | 0 | 0 | 0 | 0 | 0 |
| 0 | NA | NA | NA | NA | NA | Olfr177 | olfactory receptor 176 | 0 | 0 | 0 | 0 | 0 | 0 | 0 | 0 | 0 | 0 | 0 |
| 0 | NA | NA | NA | NA | NA | Olfr178 | olfactory receptor 178 | 0 | 0 | 0 | 0 | 0 | 0 | 0 | 0 | 0 | 0 | 0 |
| 0 | NA | NA | NA | NA | NA | Olfr180 | olfactory receptor 180 | 0 | 0 | 0 | 0 | 0 | 0 | 0 | 0 | 0 | 0 | 0 |
| 0 | NA | NA | NA | NA | NA | Olfr181 | olfactory receptor 181 | 0 | 0 | 0 | 0 | 0 | 0 | 0 | 0 | 0 | 0 | 0 |
| 0 | NA | NA | NA | NA | NA | Gm45071 | | 0 | 0 | 0 | 0 | 0 | 0 | 0 | 0 | 0 | 0 | 0 |
| 0 | NA | NA | NA | NA | NA | Olfr183 | olfactory receptor 183 | 0 | 0 | 0 | 0 | 0 | 0 | 0 | 0 | 0 | 0 | 0 |
| 0 | NA | NA | NA | NA | NA | Olfr184-ps1 | | 0 | 0 | 0 | 0 | 0 | 0 | 0 | 0 | 0 | 0 | 0 |
| 0 | NA | NA | NA | NA | NA | Olfr185-ps1 | | 0 | 0 | 0 | 0 | 0 | 0 | 0 | 0 | 0 | 0 | 0 |
| 0 | NA | NA | NA | NA | NA | Olfr186 | olfactory receptor 186 | 0 | 0 | 0 | 0 | 0 | 0 | 0 | 0 | 0 | 0 | 0 |
| 0 | NA | NA | NA | NA | NA | Olfr188-ps1 | | 0 | 0 | 0 | 0 | 0 | 0 | 0 | 0 | 0 | 0 | 0 |
| 0 | NA | NA | NA | NA | NA | Olfr189-ps1 | | 0 | 0 | 0 | 0 | 0 | 0 | 0 | 0 | 0 | 0 | 0 |
| 0 | NA | NA | NA | NA | NA | Olfr190 | olfactory receptor 190 | 0 | 0 | 0 | 0 | 0 | 0 | 0 | 0 | 0 | 0 | 0 |
| 0 | NA | NA | NA | NA | NA | Olfr191 | olfactory receptor 191 | 0 | 0 | 0 | 0 | 0 | 0 | 0 | 0 | 0 | 0 | 0 |
| 0 | NA | NA | NA | NA | NA | Olfr192 | | 0 | 0 | 0 | 0 | 0 | 0 | 0 | 0 | 0 | 0 | 0 |
| 0 | NA | NA | NA | NA | NA | Olfr193 | olfactory receptor 193 | 0 | 0 | 0 | 0 | 0 | 0 | 0 | 0 | 0 | 0 | 0 |
| 0 | NA | NA | NA | NA | NA | Olfr194 | olfactory receptor 194 | 0 | 0 | 0 | 0 | 0 | 0 | 0 | 0 | 0 | 0 | 0 |
| 0 | NA | NA | NA | NA | NA | Olfr195 | olfactory receptor 195 | 0 | 0 | 0 | 0 | 0 | 0 | 0 | 0 | 0 | 0 | 0 |

| | | | | | | | | | | | | | | | | | | |
|---|---|---|---|---|---|---|---|---|---|---|---|---|---|---|---|---|---|---|
| 0 | NA | NA | NA | NA | NA | Olfr196 | olfactory receptor 196 | 0 | 0 | 0 | 0 | 0 | 0 | 0 | 0 | 0 | 0 | 0 |
| 0 | NA | NA | NA | NA | NA | Olfr197 | | 0 | 0 | 0 | 0 | 0 | 0 | 0 | 0 | 0 | 0 | 0 |
| 0 | NA | NA | NA | NA | NA | Olfr198 | olfactory receptor 198 | 0 | 0 | 0 | 0 | 0 | 0 | 0 | 0 | 0 | 0 | 0 |
| 0 | NA | NA | NA | NA | NA | Olfr199 | olfactory receptor 199 | 0 | 0 | 0 | 0 | 0 | 0 | 0 | 0 | 0 | 0 | 0 |
| 0 | NA | NA | NA | NA | NA | Olfr200-ps1 | | 0 | 0 | 0 | 0 | 0 | 0 | 0 | 0 | 0 | 0 | 0 |
| 0 | NA | NA | NA | NA | NA | Olfr201 | olfactory receptor 201 | 0 | 0 | 0 | 0 | 0 | 0 | 0 | 0 | 0 | 0 | 0 |
| 0 | NA | NA | NA | NA | NA | Olfr202 | olfactory receptor 202 | 0 | 0 | 0 | 0 | 0 | 0 | 0 | 0 | 0 | 0 | 0 |
| 0 | NA | NA | NA | NA | NA | Olfr203 | olfactory receptor 203 | 0 | 0 | 0 | 0 | 0 | 0 | 0 | 0 | 0 | 0 | 0 |
| 0 | NA | NA | NA | NA | NA | Olfr205 | olfactory receptor 205 | 0 | 0 | 0 | 0 | 0 | 0 | 0 | 0 | 0 | 0 | 0 |
| 0 | NA | NA | NA | NA | NA | Olfr206 | olfactory receptor 206 | 0 | 0 | 0 | 0 | 0 | 0 | 0 | 0 | 0 | 0 | 0 |
| 0 | NA | NA | NA | NA | NA | Olfr209 | olfactory receptor 207 | 0 | 0 | 0 | 0 | 0 | 0 | 0 | 0 | 0 | 0 | 0 |
| 0 | NA | NA | NA | NA | NA | Gabrr3 | gamma-aminobutyric acid (GABA) recept | 0 | 0 | 0 | 0 | 0 | 0 | 0 | 0 | 0 | 0 | 0 |
| 0 | NA | NA | NA | NA | NA | 1700022E09Rik | | 0 | 0 | 0 | 0 | 0 | 0 | 0 | 0 | 0 | 0 | 0 |
| 0 | NA | NA | NA | NA | NA | Epha6 | Eph receptor A6 | 0 | 0 | 0 | 0 | 0 | 0 | 0 | 0 | 0 | 0 | 0 |
| 0 | NA | NA | NA | NA | NA | Gm24755 | | 0 | 0 | 0 | 0 | 0 | 0 | 0 | 0 | 0 | 0 | 0 |
| 0 | NA | NA | NA | NA | NA | Gm4660 | | 0 | 0 | 0 | 0 | 0 | 0 | 0 | 0 | 0 | 0 | 0 |
| 0 | NA | NA | NA | NA | NA | Gm9017 | | 0 | 0 | 0 | 0 | 0 | 0 | 0 | 0 | 0 | 0 | 0 |
| 0 | NA | NA | NA | NA | NA | Gm27001 | | 0 | 0 | 0 | 0 | 0 | 0 | 0 | 0 | 0 | 0 | 0 |
| 0 | NA | NA | NA | NA | NA | Gm49609 | | 0 | 0 | 0 | 0 | 0 | 0 | 0 | 0 | 0 | 0 | 0 |
| 0 | NA | NA | NA | NA | NA | Gm27784 | | 0 | 0 | 0 | 0 | 0 | 0 | 0 | 0 | 0 | 0 | 0 |
| 0 | NA | NA | NA | NA | NA | Gm17065 | | 0 | 0 | 0 | 0 | 0 | 0 | 0 | 0 | 0 | 0 | 0 |
| 0 | NA | NA | NA | NA | NA | Gm23852 | | 0 | 0 | 0 | 0 | 0 | 0 | 0 | 0 | 0 | 0 | 0 |
| 0 | NA | NA | NA | NA | NA | Gm17054 | | 0 | 0 | 0 | 0 | 0 | 0 | 0 | 0 | 0 | 0 | 0 |
| 0 | NA | NA | NA | NA | NA | Gm49610 | | 0 | 0 | 0 | 0 | 0 | 0 | 0 | 0 | 0 | 0 | 0 |
| 0 | NA | NA | NA | NA | NA | n-R5s37 | | 0 | 0 | 0 | 0 | 0 | 0 | 0 | 0 | 0 | 0 | 0 |
| 0 | NA | NA | NA | NA | NA | Gm17192 | | 0 | 0 | 0 | 0 | 0 | 0 | 0 | 0 | 0 | 0 | 0 |
| 0 | NA | NA | NA | NA | NA | Gm49615 | | 0 | 0 | 0 | 0 | 0 | 0 | 0 | 0 | 0 | 0 | 0 |
| 0 | NA | NA | NA | NA | NA | | | 0 | 0 | 0 | 0 | 0 | 0 | 0 | 0 | 0 | 0 | 0 |
| 0 | NA | NA | NA | NA | NA | Gm24164 | | 0 | 0 | 0 | 0 | 0 | 0 | 0 | 0 | 0 | 0 | 0 |
| 0 | NA | NA | NA | NA | NA | Gm19000 | | 0 | 0 | 0 | 0 | 0 | 0 | 0 | 0 | 0 | 0 | 0 |
| 0 | NA | NA | NA | NA | NA | Gm25986 | | 0 | 0 | 0 | 0 | 0 | 0 | 0 | 0 | 0 | 0 | 0 |
| 0 | NA | NA | NA | NA | NA | Gm18870 | | 0 | 0 | 0 | 0 | 0 | 0 | 0 | 0 | 0 | 0 | 0 |
| 0 | NA | NA | NA | NA | NA | Gm49621 | | 0 | 0 | 0 | 0 | 0 | 0 | 0 | 0 | 0 | 0 | 0 |
| 0 | NA | NA | NA | NA | NA | Gm33912 | | 0 | 0 | 0 | 0 | 0 | 0 | 0 | 0 | 0 | 0 | 0 |
| 0 | NA | NA | NA | NA | NA | Gm49622 | | 0 | 0 | 0 | 0 | 0 | 0 | 0 | 0 | 0 | 0 | 0 |
| 0 | NA | NA | NA | NA | NA | Gm22769 | | 0 | 0 | 0 | 0 | 0 | 0 | 0 | 0 | 0 | 0 | 0 |
| 0 | NA | NA | NA | NA | NA | Gm49627 | | 0 | 0 | 0 | 0 | 0 | 0 | 0 | 0 | 0 | 0 | 0 |
| 0 | NA | NA | NA | NA | NA | Gm23374 | | 0 | 0 | 0 | 0 | 0 | 0 | 0 | 0 | 0 | 0 | 0 |
| 0 | NA | NA | NA | NA | NA | Htr1f | 5-hydroxytryptamine (serotonin) receptor | 0 | 0 | 0 | 0 | 0 | 0 | 0 | 0 | 0 | 0 | 0 |
| 0 | NA | NA | NA | NA | NA | Gm22429 | | 0 | 0 | 0 | 0 | 0 | 0 | 0 | 0 | 0 | 0 | 0 |
| 0 | NA | NA | NA | NA | NA | Gm9584 | | 0 | 0 | 0 | 0 | 0 | 0 | 0 | 0 | 0 | 0 | 0 |
| 0 | NA | NA | NA | NA | NA | Gm49629 | | 0 | 0 | 0 | 0 | 0 | 0 | 0 | 0 | 0 | 0 | 0 |
| 0 | NA | NA | NA | NA | NA | Gm23659 | | 0 | 0 | 0 | 0 | 0 | 0 | 0 | 0 | 0 | 0 | 0 |
| 0 | NA | NA | NA | NA | NA | Gm49631 | | 0 | 0 | 0 | 0 | 0 | 0 | 0 | 0 | 0 | 0 | 0 |
| 0 | NA | NA | NA | NA | NA | Pou1f1 | POU domain, class 1, transcription factor | 0 | 0 | 0 | 0 | 0 | 0 | 0 | 0 | 0 | 0 | 0 |
| 0 | NA | NA | NA | NA | NA | Gm49633 | | 0 | 0 | 0 | 0 | 0 | 0 | 0 | 0 | 0 | 0 | 0 |
| 0 | NA | NA | NA | NA | NA | Gm5221 | | 0 | 0 | 0 | 0 | 0 | 0 | 0 | 0 | 0 | 0 | 0 |
| 0 | NA | NA | NA | NA | NA | Gm7366 | | 0 | 0 | 0 | 0 | 0 | 0 | 0 | 0 | 0 | 0 | 0 |
| 0 | NA | NA | NA | NA | NA | Gm49635 | | 0 | 0 | 0 | 0 | 0 | 0 | 0 | 0 | 0 | 0 | 0 |
| 0 | NA | NA | NA | NA | NA | Gm22217 | | 0 | 0 | 0 | 0 | 0 | 0 | 0 | 0 | 0 | 0 | 0 |
| 0 | NA | NA | NA | NA | NA | Gm22448 | | 0 | 0 | 0 | 0 | 0 | 0 | 0 | 0 | 0 | 0 | 0 |
| 0 | NA | NA | NA | NA | NA | 1700010K23Rik | | 0 | 0 | 0 | 0 | 0 | 0 | 0 | 0 | 0 | 0 | 0 |
| 0 | NA | NA | NA | NA | NA | 4933411O13Rik | | 0 | 0 | 0 | 0 | 0 | 0 | 0 | 0 | 0 | 0 | 0 |
| 0 | NA | NA | NA | NA | NA | Gm49640 | | 0 | 0 | 0 | 0 | 0 | 0 | 0 | 0 | 0 | 0 | 0 |
| 0 | NA | NA | NA | NA | NA | Gm15828 | | 0 | 0 | 0 | 0 | 0 | 0 | 0 | 0 | 0 | 0 | 0 |
| 0 | NA | NA | NA | NA | NA | Gm49648 | | 0 | 0 | 0 | 0 | 0 | 0 | 0 | 0 | 0 | 0 | 0 |
| 0 | NA | NA | NA | NA | NA | Gm49649 | | 0 | 0 | 0 | 0 | 0 | 0 | 0 | 0 | 0 | 0 | 0 |
| 0 | NA | NA | NA | NA | NA | Gm18495 | | 0 | 0 | 0 | 0 | 0 | 0 | 0 | 0 | 0 | 0 | 0 |
| 0 | NA | NA | NA | NA | NA | Gm49650 | | 0 | 0 | 0 | 0 | 0 | 0 | 0 | 0 | 0 | 0 | 0 |
| 0 | NA | NA | NA | NA | NA | Gm24681 | | 0 | 0 | 0 | 0 | 0 | 0 | 0 | 0 | 0 | 0 | 0 |

| | | | | | | | | | | | | | | | | | | |
|---|---|---|---|---|---|---|---|---|---|---|---|---|---|---|---|---|---|---|
| 0 | NA | NA | NA | NA | NA | Gm23591 | | 0 | 0 | 0 | 0 | 0 | 0 | 0 | 0 | 0 | 0 | 0 |
| 0 | NA | NA | NA | NA | NA | Gm18496 | | 0 | 0 | 0 | 0 | 0 | 0 | 0 | 0 | 0 | 0 | 0 |
| 0 | NA | NA | NA | NA | NA | 4930428D20Rik | | 0 | 0 | 0 | 0 | 0 | 0 | 0 | 0 | 0 | 0 | 0 |
| 0 | NA | NA | NA | NA | NA | Gm49655 | | 0 | 0 | 0 | 0 | 0 | 0 | 0 | 0 | 0 | 0 | 0 |
| 0 | NA | NA | NA | NA | NA | Gm19082 | | 0 | 0 | 0 | 0 | 0 | 0 | 0 | 0 | 0 | 0 | 0 |
| 0 | NA | NA | NA | NA | NA | Gm49656 | | 0 | 0 | 0 | 0 | 0 | 0 | 0 | 0 | 0 | 0 | 0 |
| 0 | NA | NA | NA | NA | NA | D16Ertd519e | | 0 | 0 | 0 | 0 | 0 | 0 | 0 | 0 | 0 | 0 | 0 |
| 0 | NA | NA | NA | NA | NA | Gm19291 | | 0 | 0 | 0 | 0 | 0 | 0 | 0 | 0 | 0 | 0 | 0 |
| 0 | NA | NA | NA | NA | NA | Gm49665 | | 0 | 0 | 0 | 0 | 0 | 0 | 0 | 0 | 0 | 0 | 0 |
| 0 | NA | NA | NA | NA | NA | Gm29823 | | 0 | 0 | 0 | 0 | 0 | 0 | 0 | 0 | 0 | 0 | 0 |
| 0 | NA | NA | NA | NA | NA | 4931420L22Rik | | 0 | 0 | 0 | 0 | 0 | 0 | 0 | 0 | 0 | 0 | 0 |
| 0 | NA | NA | NA | NA | NA | Gm24968 | | 0 | 0 | 0 | 0 | 0 | 0 | 0 | 0 | 0 | 0 | 0 |
| 0 | NA | NA | NA | NA | NA | Gm18918 | | 0 | 0 | 0 | 0 | 0 | 0 | 0 | 0 | 0 | 0 | 0 |
| 0 | NA | NA | NA | NA | NA | Gm49666 | | 0 | 0 | 0 | 0 | 0 | 0 | 0 | 0 | 0 | 0 | 0 |
| 0 | NA | NA | NA | NA | NA | 4930567J20Rik | | 0 | 0 | 0 | 0 | 0 | 0 | 0 | 0 | 0 | 0 | 0 |
| 0 | NA | NA | NA | NA | NA | Gm22797 | | 0 | 0 | 0 | 0 | 0 | 0 | 0 | 0 | 0 | 0 | 0 |
| 0 | NA | NA | NA | NA | NA | Gm18549 | | 0 | 0 | 0 | 0 | 0 | 0 | 0 | 0 | 0 | 0 | 0 |
| 0 | NA | NA | NA | NA | NA | Gm18550 | | 0 | 0 | 0 | 0 | 0 | 0 | 0 | 0 | 0 | 0 | 0 |
| 0 | NA | NA | NA | NA | NA | Gm49667 | | 0 | 0 | 0 | 0 | 0 | 0 | 0 | 0 | 0 | 0 | 0 |
| 0 | NA | NA | NA | NA | NA | 8030451O07Rik | | 0 | 0 | 0 | 0 | 0 | 0 | 0 | 0 | 0 | 0 | 0 |
| 0 | NA | NA | NA | NA | NA | Gm49670 | | 0 | 0 | 0 | 0 | 0 | 0 | 0 | 0 | 0 | 0 | 0 |
| 0 | NA | NA | NA | NA | NA | 4930500H12Rik | | 0 | 0 | 0 | 0 | 0 | 0 | 0 | 0 | 0 | 0 | 0 |
| 0 | NA | NA | NA | NA | NA | Gm25699 | | 0 | 0 | 0 | 0 | 0 | 0 | 0 | 0 | 0 | 0 | 0 |
| 0 | NA | NA | NA | NA | NA | Gm18825 | | 0 | 0 | 0 | 0 | 0 | 0 | 0 | 0 | 0 | 0 | 0 |
| 0 | NA | NA | NA | NA | NA | Gm49680 | | 0 | 0 | 0 | 0 | 0 | 0 | 0 | 0 | 0 | 0 | 0 |
| 0 | NA | NA | NA | NA | NA | Gm18168 | | 0 | 0 | 0 | 0 | 0 | 0 | 0 | 0 | 0 | 0 | 0 |
| 0 | NA | NA | NA | NA | NA | Gm24272 | | 0 | 0 | 0 | 0 | 0 | 0 | 0 | 0 | 0 | 0 | 0 |
| 0 | NA | NA | NA | NA | NA | Gm22163 | | 0 | 0 | 0 | 0 | 0 | 0 | 0 | 0 | 0 | 0 | 0 |
| 0 | NA | NA | NA | NA | NA | Gm18021 | | 0 | 0 | 0 | 0 | 0 | 0 | 0 | 0 | 0 | 0 | 0 |
| 0 | NA | NA | NA | NA | NA | Gm49671 | | 0 | 0 | 0 | 0 | 0 | 0 | 0 | 0 | 0 | 0 | 0 |
| 0 | NA | NA | NA | NA | NA | Gm26307 | | 0 | 0 | 0 | 0 | 0 | 0 | 0 | 0 | 0 | 0 | 0 |
| 0 | NA | NA | NA | NA | NA | Gm49674 | | 0 | 0 | 0 | 0 | 0 | 0 | 0 | 0 | 0 | 0 | 0 |
| 0 | NA | NA | NA | NA | NA | Gm26937 | | 0 | 0 | 0 | 0 | 0 | 0 | 0 | 0 | 0 | 0 | 0 |
| 0 | NA | NA | NA | NA | NA | A630036G19Rik | | 0 | 0 | 0 | 0 | 0 | 0 | 0 | 0 | 0 | 0 | 0 |
| 0 | NA | NA | NA | NA | NA | Gm49676 | | 0 | 0 | 0 | 0 | 0 | 0 | 0 | 0 | 0 | 0 | 0 |
| 0 | NA | NA | NA | NA | NA | Gm49678 | | 0 | 0 | 0 | 0 | 0 | 0 | 0 | 0 | 0 | 0 | 0 |
| 0 | NA | NA | NA | NA | NA | Gm49677 | | 0 | 0 | 0 | 0 | 0 | 0 | 0 | 0 | 0 | 0 | 0 |
| 0 | NA | NA | NA | NA | NA | Lipi | lipase, member I | 0 | 0 | 0 | 0 | 0 | 0 | 0 | 0 | 0 | 0 | 0 |
| 0 | NA | NA | NA | NA | NA | Gm30162 | | 0 | 0 | 0 | 0 | 0 | 0 | 0 | 0 | 0 | 0 | 0 |
| 0 | NA | NA | NA | NA | NA | Gm15553 | | 0 | 0 | 0 | 0 | 0 | 0 | 0 | 0 | 0 | 0 | 0 |
| 0 | NA | NA | NA | NA | NA | Gm22923 | | 0 | 0 | 0 | 0 | 0 | 0 | 0 | 0 | 0 | 0 | 0 |
| 0 | NA | NA | NA | NA | NA | Gm15554 | | 0 | 0 | 0 | 0 | 0 | 0 | 0 | 0 | 0 | 0 | 0 |
| 0 | NA | NA | NA | NA | NA | Gm15555 | | 0 | 0 | 0 | 0 | 0 | 0 | 0 | 0 | 0 | 0 | 0 |
| 0 | NA | NA | NA | NA | NA | 4930578N18Rik | | 0 | 0 | 0 | 0 | 0 | 0 | 0 | 0 | 0 | 0 | 0 |
| 0 | NA | NA | NA | NA | NA | Gm22268 | | 0 | 0 | 0 | 0 | 0 | 0 | 0 | 0 | 0 | 0 | 0 |
| 0 | NA | NA | NA | NA | NA | Gm26915 | | 0 | 0 | 0 | 0 | 0 | 0 | 0 | 0 | 0 | 0 | 0 |
| 0 | NA | NA | NA | NA | NA | Gm45030 | | 0 | 0 | 0 | 0 | 0 | 0 | 0 | 0 | 0 | 0 | 0 |
| 0 | NA | NA | NA | NA | NA | Gm30726 | | 0 | 0 | 0 | 0 | 0 | 0 | 0 | 0 | 0 | 0 | 0 |
| 0 | NA | NA | NA | NA | NA | Gm45029 | | 0 | 0 | 0 | 0 | 0 | 0 | 0 | 0 | 0 | 0 | 0 |
| 0 | NA | NA | NA | NA | NA | Gm30790 | | 0 | 0 | 0 | 0 | 0 | 0 | 0 | 0 | 0 | 0 | 0 |
| 0 | NA | NA | NA | NA | NA | 1700041M19Rik | | 0 | 0 | 0 | 0 | 0 | 0 | 0 | 0 | 0 | 0 | 0 |
| 0 | NA | NA | NA | NA | NA | 1700063K16Rik | | 0 | 0 | 0 | 0 | 0 | 0 | 0 | 0 | 0 | 0 | 0 |
| 0 | NA | NA | NA | NA | NA | Gm7501 | | 0 | 0 | 0 | 0 | 0 | 0 | 0 | 0 | 0 | 0 | 0 |
| 0 | NA | NA | NA | NA | NA | Gm31258 | | 0 | 0 | 0 | 0 | 0 | 0 | 0 | 0 | 0 | 0 | 0 |
| 0 | NA | NA | NA | NA | NA | Gm37466 | | 0 | 0 | 0 | 0 | 0 | 0 | 0 | 0 | 0 | 0 | 0 |
| 0 | NA | NA | NA | NA | NA | 9430092D12Rik | | 0 | 0 | 0 | 0 | 0 | 0 | 0 | 0 | 0 | 0 | 0 |
| 0 | NA | NA | NA | NA | NA | Gm38071 | | 0 | 0 | 0 | 0 | 0 | 0 | 0 | 0 | 0 | 0 | 0 |
| 0 | NA | NA | NA | NA | NA | 9430053O09Rik | | 0 | 0 | 0 | 0 | 0 | 0 | 0 | 0 | 0 | 0 | 0 |
| 0 | NA | NA | NA | NA | NA | Gm37063 | | 0 | 0 | 0 | 0 | 0 | 0 | 0 | 0 | 0 | 0 | 0 |
| 0 | NA | NA | NA | NA | NA | Gm37241 | | 0 | 0 | 0 | 0 | 0 | 0 | 0 | 0 | 0 | 0 | 0 |

| | | | | | | | | | | | | | | | | | |
|---|---|---|---|---|---|---|---|---|---|---|---|---|---|---|---|---|---|
| 0 | NA | NA | NA | NA | NA | Gm36963 | | 0 | 0 | 0 | 0 | 0 | 0 | 0 | 0 | 0 | 0 | 0 |
| 0 | NA | NA | NA | NA | NA | Gm11146 | | 0 | 0 | 0 | 0 | 0 | 0 | 0 | 0 | 0 | 0 | 0 |
| 0 | NA | NA | NA | NA | NA | Mir99a | | 0 | 0 | 0 | 0 | 0 | 0 | 0 | 0 | 0 | 0 | 0 |
| 0 | NA | NA | NA | NA | NA | Mirlet7c-1 | | 0 | 0 | 0 | 0 | 0 | 0 | 0 | 0 | 0 | 0 | 0 |
| 0 | NA | NA | NA | NA | NA | Gm37311 | | 0 | 0 | 0 | 0 | 0 | 0 | 0 | 0 | 0 | 0 | 0 |
| 0 | NA | NA | NA | NA | NA | Mir125b-2 | | 0 | 0 | 0 | 0 | 0 | 0 | 0 | 0 | 0 | 0 | 0 |
| 0 | NA | NA | NA | NA | NA | 9530003O04Rik | | 0 | 0 | 0 | 0 | 0 | 0 | 0 | 0 | 0 | 0 | 0 |
| 0 | NA | NA | NA | NA | NA | Gm17333 | | 0 | 0 | 0 | 0 | 0 | 0 | 0 | 0 | 0 | 0 | 0 |
| 0 | NA | NA | NA | NA | NA | Gm49589 | | 0 | 0 | 0 | 0 | 0 | 0 | 0 | 0 | 0 | 0 | 0 |
| 0 | NA | NA | NA | NA | NA | Gm25025 | | 0 | 0 | 0 | 0 | 0 | 0 | 0 | 0 | 0 | 0 | 0 |
| 0 | NA | NA | NA | NA | NA | Gm5675 | | 0 | 0 | 0 | 0 | 0 | 0 | 0 | 0 | 0 | 0 | 0 |
| 0 | NA | NA | NA | NA | NA | Gm25916 | | 0 | 0 | 0 | 0 | 0 | 0 | 0 | 0 | 0 | 0 | 0 |
| 0 | NA | NA | NA | NA | NA | Gm49591 | | 0 | 0 | 0 | 0 | 0 | 0 | 0 | 0 | 0 | 0 | 0 |
| 0 | NA | NA | NA | NA | NA | Gm25038 | | 0 | 0 | 0 | 0 | 0 | 0 | 0 | 0 | 0 | 0 | 0 |
| 0 | NA | NA | NA | NA | NA | Gm49603 | | 0 | 0 | 0 | 0 | 0 | 0 | 0 | 0 | 0 | 0 | 0 |
| 0 | NA | NA | NA | NA | NA | Gm49604 | | 0 | 0 | 0 | 0 | 0 | 0 | 0 | 0 | 0 | 0 | 0 |
| 0 | NA | NA | NA | NA | NA | 4930478L05Rik | | 0 | 0 | 0 | 0 | 0 | 0 | 0 | 0 | 0 | 0 | 0 |
| 0 | NA | NA | NA | NA | NA | Gm24861 | | 0 | 0 | 0 | 0 | 0 | 0 | 0 | 0 | 0 | 0 | 0 |
| 0 | NA | NA | NA | NA | NA | 4930570E03Rik | | 0 | 0 | 0 | 0 | 0 | 0 | 0 | 0 | 0 | 0 | 0 |
| 0 | NA | NA | NA | NA | NA | 4930403O18Rik | | 0 | 0 | 0 | 0 | 0 | 0 | 0 | 0 | 0 | 0 | 0 |
| 0 | NA | NA | NA | NA | NA | Tmprss15 | transmembrane protease, serine 15 | 0 | 0 | 0 | 0 | 0 | 0 | 0 | 0 | 0 | 0 | 0 |
| 0 | NA | NA | NA | NA | NA | Gm37709 | | 0 | 0 | 0 | 0 | 0 | 0 | 0 | 0 | 0 | 0 | 0 |
| 0 | NA | NA | NA | NA | NA | 1700066C05Ril | RIKEN cDNA 1700066C05 gene | 0 | 0 | 0 | 0 | 0 | 0 | 0 | 0 | 0 | 0 | 0 |
| 0 | NA | NA | NA | NA | NA | Gm23083 | | 0 | 0 | 0 | 0 | 0 | 0 | 0 | 0 | 0 | 0 | 0 |
| 0 | NA | NA | NA | NA | NA | Gm24773 | | 0 | 0 | 0 | 0 | 0 | 0 | 0 | 0 | 0 | 0 | 0 |
| 0 | NA | NA | NA | NA | NA | Gm49578 | | 0 | 0 | 0 | 0 | 0 | 0 | 0 | 0 | 0 | 0 | 0 |
| 0 | NA | NA | NA | NA | NA | Gm19867 | | 0 | 0 | 0 | 0 | 0 | 0 | 0 | 0 | 0 | 0 | 0 |
| 0 | NA | NA | NA | NA | NA | Fkbp1a-ps3 | | 0 | 0 | 0 | 0 | 0 | 0 | 0 | 0 | 0 | 0 | 0 |
| 0 | NA | NA | NA | NA | NA | Gm26023 | | 0 | 0 | 0 | 0 | 0 | 0 | 0 | 0 | 0 | 0 | 0 |
| 0 | NA | NA | NA | NA | NA | 7120432I05Rik | | 0 | 0 | 0 | 0 | 0 | 0 | 0 | 0 | 0 | 0 | 0 |
| 0 | NA | NA | NA | NA | NA | Gm49555 | | 0 | 0 | 0 | 0 | 0 | 0 | 0 | 0 | 0 | 0 | 0 |
| 0 | NA | NA | NA | NA | NA | Gm49558 | | 0 | 0 | 0 | 0 | 0 | 0 | 0 | 0 | 0 | 0 | 0 |
| 0 | NA | NA | NA | NA | NA | Gm5676 | | 0 | 0 | 0 | 0 | 0 | 0 | 0 | 0 | 0 | 0 | 0 |
| 0 | NA | NA | NA | NA | NA | Gm49559 | | 0 | 0 | 0 | 0 | 0 | 0 | 0 | 0 | 0 | 0 | 0 |
| 0 | NA | NA | NA | NA | NA | Gm49560 | | 0 | 0 | 0 | 0 | 0 | 0 | 0 | 0 | 0 | 0 | 0 |
| 0 | NA | NA | NA | NA | NA | Pgk1-ps1 | | 0 | 0 | 0 | 0 | 0 | 0 | 0 | 0 | 0 | 0 | 0 |
| 0 | NA | NA | NA | NA | NA | Gm41480 | | 0 | 0 | 0 | 0 | 0 | 0 | 0 | 0 | 0 | 0 | 0 |
| 0 | NA | NA | NA | NA | NA | Gm23355 | | 0 | 0 | 0 | 0 | 0 | 0 | 0 | 0 | 0 | 0 | 0 |
| 0 | NA | NA | NA | NA | NA | Gm7559 | | 0 | 0 | 0 | 0 | 0 | 0 | 0 | 0 | 0 | 0 | 0 |
| 0 | NA | NA | NA | NA | NA | Gm49561 | | 0 | 0 | 0 | 0 | 0 | 0 | 0 | 0 | 0 | 0 | 0 |
| 0 | NA | NA | NA | NA | NA | Gm41481 | | 0 | 0 | 0 | 0 | 0 | 0 | 0 | 0 | 0 | 0 | 0 |
| 0 | NA | NA | NA | NA | NA | Gm49562 | | 0 | 0 | 0 | 0 | 0 | 0 | 0 | 0 | 0 | 0 | 0 |
| 0 | NA | NA | NA | NA | NA | Gm7295 | | 0 | 0 | 0 | 0 | 0 | 0 | 0 | 0 | 0 | 0 | 0 |
| 0 | NA | NA | NA | NA | NA | Gm24508 | | 0 | 0 | 0 | 0 | 0 | 0 | 0 | 0 | 0 | 0 | 0 |
| 0 | NA | NA | NA | NA | NA | 4930553E22Rik | | 0 | 0 | 0 | 0 | 0 | 0 | 0 | 0 | 0 | 0 | 0 |
| 0 | NA | NA | NA | NA | NA | A730009L09Rik | | 0 | 0 | 0 | 0 | 0 | 0 | 0 | 0 | 0 | 0 | 0 |
| 0 | NA | NA | NA | NA | NA | Gm2477 | | 0 | 0 | 0 | 0 | 0 | 0 | 0 | 0 | 0 | 0 | 0 |
| 0 | NA | NA | NA | NA | NA | Gm25292 | | 0 | 0 | 0 | 0 | 0 | 0 | 0 | 0 | 0 | 0 | 0 |
| 0 | NA | NA | NA | NA | NA | Gm2349 | | 0 | 0 | 0 | 0 | 0 | 0 | 0 | 0 | 0 | 0 | 0 |
| 0 | NA | NA | NA | NA | NA | 4930551I23Rik | | 0 | 0 | 0 | 0 | 0 | 0 | 0 | 0 | 0 | 0 | 0 |
| 0 | NA | NA | NA | NA | NA | Mir155 | | 0 | 0 | 0 | 0 | 0 | 0 | 0 | 0 | 0 | 0 | 0 |
| 0 | NA | NA | NA | NA | NA | Gm49568 | | 0 | 0 | 0 | 0 | 0 | 0 | 0 | 0 | 0 | 0 | 0 |
| 0 | NA | NA | NA | NA | NA | Gm49228 | | 0 | 0 | 0 | 0 | 0 | 0 | 0 | 0 | 0 | 0 | 0 |
| 0 | NA | NA | NA | NA | NA | Gm27295 | | 0 | 0 | 0 | 0 | 0 | 0 | 0 | 0 | 0 | 0 | 0 |
| 0 | NA | NA | NA | NA | NA | 4930556C24Rik | | 0 | 0 | 0 | 0 | 0 | 0 | 0 | 0 | 0 | 0 | 0 |
| 0 | NA | NA | NA | NA | NA | Gm49570 | | 0 | 0 | 0 | 0 | 0 | 0 | 0 | 0 | 0 | 0 | 0 |
| 0 | NA | NA | NA | NA | NA | 1700007H22Rik | | 0 | 0 | 0 | 0 | 0 | 0 | 0 | 0 | 0 | 0 | 0 |
| 0 | NA | NA | NA | NA | NA | Gm32357 | | 0 | 0 | 0 | 0 | 0 | 0 | 0 | 0 | 0 | 0 | 0 |
| 0 | NA | NA | NA | NA | NA | Gm49572 | | 0 | 0 | 0 | 0 | 0 | 0 | 0 | 0 | 0 | 0 | 0 |
| 0 | NA | NA | NA | NA | NA | Gm25715 | | 0 | 0 | 0 | 0 | 0 | 0 | 0 | 0 | 0 | 0 | 0 |

| | | | | | | | | | | | | | | | | | | |
|---|---|---|---|---|---|---|---|---|---|---|---|---|---|---|---|---|---|---|
| 0 | NA | NA | NA | NA | NA | Gm38477 | | 0 | 0 | 0 | 0 | 0 | 0 | 0 | 0 | 0 | 0 | 0 |
| 0 | NA | NA | NA | NA | NA | Gm32865 | | 0 | 0 | 0 | 0 | 0 | 0 | 0 | 0 | 0 | 0 | 0 |
| 0 | NA | NA | NA | NA | NA | Gm25896 | | 0 | 0 | 0 | 0 | 0 | 0 | 0 | 0 | 0 | 0 | 0 |
| 0 | NA | NA | NA | NA | NA | Gm24891 | | 0 | 0 | 0 | 0 | 0 | 0 | 0 | 0 | 0 | 0 | 0 |
| 0 | NA | NA | NA | NA | NA | Gm22808 | | 0 | 0 | 0 | 0 | 0 | 0 | 0 | 0 | 0 | 0 | 0 |
| 0 | NA | NA | NA | NA | NA | Gm23994 | | 0 | 0 | 0 | 0 | 0 | 0 | 0 | 0 | 0 | 0 | 0 |
| 0 | NA | NA | NA | NA | NA | Grik1 | glutamate receptor, ionotropic, kainate 1 | 0 | 0 | 0 | 0 | 0 | 0 | 0 | 0 | 0 | 0 | 0 |
| 0 | NA | NA | NA | NA | NA | Gm33255 | | 0 | 0 | 0 | 0 | 0 | 0 | 0 | 0 | 0 | 0 | 0 |
| 0 | NA | NA | NA | NA | NA | 4930420G21Rik | | 0 | 0 | 0 | 0 | 0 | 0 | 0 | 0 | 0 | 0 | 0 |
| 0 | NA | NA | NA | NA | NA | Gm49688 | | 0 | 0 | 0 | 0 | 0 | 0 | 0 | 0 | 0 | 0 | 0 |
| 0 | NA | NA | NA | NA | NA | Krtap24-1 | keratin associated protein 24-1 | 0 | 0 | 0 | 0 | 0 | 0 | 0 | 0 | 0 | 0 | 0 |
| 0 | NA | NA | NA | NA | NA | Gm23587 | | 0 | 0 | 0 | 0 | 0 | 0 | 0 | 0 | 0 | 0 | 0 |
| 0 | NA | NA | NA | NA | NA | 2310079G19Ril | RIKEN cDNA 2310079G19 gene | 0 | 0 | 0 | 0 | 0 | 0 | 0 | 0 | 0 | 0 | 0 |
| 0 | NA | NA | NA | NA | NA | Krtap26-1 | keratin associated protein 26-1 | 0 | 0 | 0 | 0 | 0 | 0 | 0 | 0 | 0 | 0 | 0 |
| 0 | NA | NA | NA | NA | NA | Krtap27-1 | keratin associated protein 27-1 | 0 | 0 | 0 | 0 | 0 | 0 | 0 | 0 | 0 | 0 | 0 |
| 0 | NA | NA | NA | NA | NA | 2310061N02Ril | RIKEN cDNA 2310061N02 gene | 0 | 0 | 0 | 0 | 0 | 0 | 0 | 0 | 0 | 0 | 0 |
| 0 | NA | NA | NA | NA | NA | Krtap13-1 | keratin associated protein 13-1 | 0 | 0 | 0 | 0 | 0 | 0 | 0 | 0 | 0 | 0 | 0 |
| 0 | NA | NA | NA | NA | NA | Krtap13 | keratin associated protein 13 | 0 | 0 | 0 | 0 | 0 | 0 | 0 | 0 | 0 | 0 | 0 |
| 0 | NA | NA | NA | NA | NA | 2310034C09Ril | RIKEN cDNA 2310034C09 gene | 0 | 0 | 0 | 0 | 0 | 0 | 0 | 0 | 0 | 0 | 0 |
| 0 | NA | NA | NA | NA | NA | 2310057N15Ril | RIKEN cDNA 2310057N15 gene | 0 | 0 | 0 | 0 | 0 | 0 | 0 | 0 | 0 | 0 | 0 |
| 0 | NA | NA | NA | NA | NA | Gm5965 | predicted gene 5965 | 0 | 0 | 0 | 0 | 0 | 0 | 0 | 0 | 0 | 0 | 0 |
| 0 | NA | NA | NA | NA | NA | 4930553J12Rik | | 0 | 0 | 0 | 0 | 0 | 0 | 0 | 0 | 0 | 0 | 0 |
| 0 | NA | NA | NA | NA | NA | Gm20741 | | 0 | 0 | 0 | 0 | 0 | 0 | 0 | 0 | 0 | 0 | 0 |
| 0 | NA | NA | NA | NA | NA | Krtap14 | keratin associated protein 14 | 0 | 0 | 0 | 0 | 0 | 0 | 0 | 0 | 0 | 0 | 0 |
| 0 | NA | NA | NA | NA | NA | Krtap15 | keratin associated protein 15 | 0 | 0 | 0 | 0 | 0 | 0 | 0 | 0 | 0 | 0 | 0 |
| 0 | NA | NA | NA | NA | NA | Krtap19-1 | keratin associated protein 19-1 | 0 | 0 | 0 | 0 | 0 | 0 | 0 | 0 | 0 | 0 | 0 |
| 0 | NA | NA | NA | NA | NA | Krtap19-2 | keratin associated protein 19-2 | 0 | 0 | 0 | 0 | 0 | 0 | 0 | 0 | 0 | 0 | 0 |
| 0 | NA | NA | NA | NA | NA | Krtap19-3 | keratin associated protein 19-3 | 0 | 0 | 0 | 0 | 0 | 0 | 0 | 0 | 0 | 0 | 0 |
| 0 | NA | NA | NA | NA | NA | Krtap19-4 | keratin associated protein 19-4 | 0 | 0 | 0 | 0 | 0 | 0 | 0 | 0 | 0 | 0 | 0 |
| 0 | NA | NA | NA | NA | NA | Krtap19-5 | keratin associated protein 19-5 | 0 | 0 | 0 | 0 | 0 | 0 | 0 | 0 | 0 | 0 | 0 |
| 0 | NA | NA | NA | NA | NA | Krtap19-9a | | 0 | 0 | 0 | 0 | 0 | 0 | 0 | 0 | 0 | 0 | 0 |
| 0 | NA | NA | NA | NA | NA | Krtap19-9b | keratin associated protein 19-9B | 0 | 0 | 0 | 0 | 0 | 0 | 0 | 0 | 0 | 0 | 0 |
| 0 | NA | NA | NA | NA | NA | Krtap16-3 | keratin associated protein 16-3 | 0 | 0 | 0 | 0 | 0 | 0 | 0 | 0 | 0 | 0 | 0 |
| 0 | NA | NA | NA | NA | NA | Krtap22-2 | keratin associated protein 22-2 | 0 | 0 | 0 | 0 | 0 | 0 | 0 | 0 | 0 | 0 | 0 |
| 0 | NA | NA | NA | NA | NA | Gm10229 | predicted gene 10229 | 0 | 0 | 0 | 0 | 0 | 0 | 0 | 0 | 0 | 0 | 0 |
| 0 | NA | NA | NA | NA | NA | Krtap6-1 | keratin associated protein 6-1 | 0 | 0 | 0 | 0 | 0 | 0 | 0 | 0 | 0 | 0 | 0 |
| 0 | NA | NA | NA | NA | NA | Gm10228 | predicted gene 10228 | 0 | 0 | 0 | 0 | 0 | 0 | 0 | 0 | 0 | 0 | 0 |
| 0 | NA | NA | NA | NA | NA | Krtap6-5 | keratin associated protein 6-5 | 0 | 0 | 0 | 0 | 0 | 0 | 0 | 0 | 0 | 0 | 0 |
| 0 | NA | NA | NA | NA | NA | 1110025L11Rik | RIKEN cDNA 1110025L11 gene | 0 | 0 | 0 | 0 | 0 | 0 | 0 | 0 | 0 | 0 | 0 |
| 0 | NA | NA | NA | NA | NA | Krtap6-3 | | 0 | 0 | 0 | 0 | 0 | 0 | 0 | 0 | 0 | 0 | 0 |
| 0 | NA | NA | NA | NA | NA | Gm6358 | predicted gene, 33798 | 0 | 0 | 0 | 0 | 0 | 0 | 0 | 0 | 0 | 0 | 0 |
| 0 | NA | NA | NA | NA | NA | Gm10061 | predicted gene 10061 | 0 | 0 | 0 | 0 | 0 | 0 | 0 | 0 | 0 | 0 | 0 |
| 0 | NA | NA | NA | NA | NA | Gm9789 | predicted gene 9789 | 0 | 0 | 0 | 0 | 0 | 0 | 0 | 0 | 0 | 0 | 0 |
| 0 | NA | NA | NA | NA | NA | Gm7735 | predicted gene 7735 | 0 | 0 | 0 | 0 | 0 | 0 | 0 | 0 | 0 | 0 | 0 |
| 0 | NA | NA | NA | NA | NA | 1110057P08Ril | keratin-associated protein 20-2-like | 0 | 0 | 0 | 0 | 0 | 0 | 0 | 0 | 0 | 0 | 0 |
| 0 | NA | NA | NA | NA | NA | Gm34396 | predicted gene, 34396 | 0 | 0 | 0 | 0 | 0 | 0 | 0 | 0 | 0 | 0 | 0 |
| 0 | NA | NA | NA | NA | NA | Krtap20-2 | keratin associated protein 20-2 | 0 | 0 | 0 | 0 | 0 | 0 | 0 | 0 | 0 | 0 | 0 |
| 0 | NA | NA | NA | NA | NA | Krtap21-1 | keratin associated protein 21-1 | 0 | 0 | 0 | 0 | 0 | 0 | 0 | 0 | 0 | 0 | 0 |
| 0 | NA | NA | NA | NA | NA | Krtap6-2 | keratin associated protein 6-2 | 0 | 0 | 0 | 0 | 0 | 0 | 0 | 0 | 0 | 0 | 0 |
| 0 | NA | NA | NA | NA | NA | Gm23655 | | 0 | 0 | 0 | 0 | 0 | 0 | 0 | 0 | 0 | 0 | 0 |
| 0 | NA | NA | NA | NA | NA | Krtap8-1 | keratin associated protein 8-1 | 0 | 0 | 0 | 0 | 0 | 0 | 0 | 0 | 0 | 0 | 0 |
| 0 | NA | NA | NA | NA | NA | Krtap7-1 | keratin associated protein 7-1 | 0 | 0 | 0 | 0 | 0 | 0 | 0 | 0 | 0 | 0 | 0 |
| 0 | NA | NA | NA | NA | NA | Krtap11-1 | keratin associated protein 11-1 | 0 | 0 | 0 | 0 | 0 | 0 | 0 | 0 | 0 | 0 | 0 |
| 0 | NA | NA | NA | NA | NA | Gm49696 | | 0 | 0 | 0 | 0 | 0 | 0 | 0 | 0 | 0 | 0 | 0 |
| 0 | NA | NA | NA | NA | NA | Gm17844 | | 0 | 0 | 0 | 0 | 0 | 0 | 0 | 0 | 0 | 0 | 0 |
| 0 | NA | NA | NA | NA | NA | Gm7307 | | 0 | 0 | 0 | 0 | 0 | 0 | 0 | 0 | 0 | 0 | 0 |
| 0 | NA | NA | NA | NA | NA | Gm49707 | | 0 | 0 | 0 | 0 | 0 | 0 | 0 | 0 | 0 | 0 | 0 |
| 0 | NA | NA | NA | NA | NA | Gm49709 | | 0 | 0 | 0 | 0 | 0 | 0 | 0 | 0 | 0 | 0 | 0 |
| 0 | NA | NA | NA | NA | NA | Gm49710 | | 0 | 0 | 0 | 0 | 0 | 0 | 0 | 0 | 0 | 0 | 0 |
| 0 | NA | NA | NA | NA | NA | Gm36001 | | 0 | 0 | 0 | 0 | 0 | 0 | 0 | 0 | 0 | 0 | 0 |

| | | | | | | | | | | | | | | | | | | |
|---|---|---|---|---|---|---|---|---|---|---|---|---|---|---|---|---|---|---|
| 0 | NA | NA | NA | NA | NA | 1110008E08Rik | | 0 | 0 | 0 | 0 | 0 | 0 | 0 | 0 | 0 | 0 | 0 |
| 0 | NA | NA | NA | NA | NA | Gm49714 | | 0 | 0 | 0 | 0 | 0 | 0 | 0 | 0 | 0 | 0 | 0 |
| 0 | NA | NA | NA | NA | NA | Gm23406 | | 0 | 0 | 0 | 0 | 0 | 0 | 0 | 0 | 0 | 0 | 0 |
| 0 | NA | NA | NA | NA | NA | Gm49715 | | 0 | 0 | 0 | 0 | 0 | 0 | 0 | 0 | 0 | 0 | 0 |
| 0 | NA | NA | NA | NA | NA | Gm36363 | | 0 | 0 | 0 | 0 | 0 | 0 | 0 | 0 | 0 | 0 | 0 |
| 0 | NA | NA | NA | NA | NA | Gm22933 | | 0 | 0 | 0 | 0 | 0 | 0 | 0 | 0 | 0 | 0 | 0 |
| 0 | NA | NA | NA | NA | NA | Gm15965 | | 0 | 0 | 0 | 0 | 0 | 0 | 0 | 0 | 0 | 0 | 0 |
| 0 | NA | NA | NA | NA | NA | Gm49611 | | 0 | 0 | 0 | 0 | 0 | 0 | 0 | 0 | 0 | 0 | 0 |
| 0 | NA | NA | NA | NA | NA | Gm49612 | | 0 | 0 | 0 | 0 | 0 | 0 | 0 | 0 | 0 | 0 | 0 |
| 0 | NA | NA | NA | NA | NA | Gm49613 | | 0 | 0 | 0 | 0 | 0 | 0 | 0 | 0 | 0 | 0 | 0 |
| 0 | NA | NA | NA | NA | NA | Gm49614 | | 0 | 0 | 0 | 0 | 0 | 0 | 0 | 0 | 0 | 0 | 0 |
| 0 | NA | NA | NA | NA | NA | Olig2 | oligodendrocyte transcription factor 2 | 0 | 0 | 0 | 0 | 0 | 0 | 0 | 0 | 0 | 0 | 0 |
| 0 | NA | NA | NA | NA | NA | Gm37016 | | 0 | 0 | 0 | 0 | 0 | 0 | 0 | 0 | 0 | 0 | 0 |
| 0 | NA | NA | NA | NA | NA | Gm15966 | | 0 | 0 | 0 | 0 | 0 | 0 | 0 | 0 | 0 | 0 | 0 |
| 0 | NA | NA | NA | NA | NA | Gm24695 | | 0 | 0 | 0 | 0 | 0 | 0 | 0 | 0 | 0 | 0 | 0 |
| 0 | NA | NA | NA | NA | NA | Gm21970 | | 0 | 0 | 0 | 0 | 0 | 0 | 0 | 0 | 0 | 0 | 0 |
| 0 | NA | NA | NA | NA | NA | Gm46562 | | 0 | 0 | 0 | 0 | 0 | 0 | 0 | 0 | 0 | 0 | 0 |
| 0 | NA | NA | NA | NA | NA | Gm49626 | | 0 | 0 | 0 | 0 | 0 | 0 | 0 | 0 | 0 | 0 | 0 |
| 0 | NA | NA | NA | NA | NA | Mir6367 | | 0 | 0 | 0 | 0 | 0 | 0 | 0 | 0 | 0 | 0 | 0 |
| 0 | NA | NA | NA | NA | NA | Atp5o | | 0 | 0 | 0 | 0 | 0 | 0 | 0 | 0 | 0 | 0 | 0 |
| 0 | NA | NA | NA | NA | NA | Gm25245 | | 0 | 0 | 0 | 0 | 0 | 0 | 0 | 0 | 0 | 0 | 0 |
| 0 | NA | NA | NA | NA | NA | Gm15976 | | 0 | 0 | 0 | 0 | 0 | 0 | 0 | 0 | 0 | 0 | 0 |
| 0 | NA | NA | NA | NA | NA | Gm27773 | | 0 | 0 | 0 | 0 | 0 | 0 | 0 | 0 | 0 | 0 | 0 |
| 0 | NA | NA | NA | NA | NA | Gm18020 | | 0 | 0 | 0 | 0 | 0 | 0 | 0 | 0 | 0 | 0 | 0 |
| 0 | NA | NA | NA | NA | NA | Gm23210 | | 0 | 0 | 0 | 0 | 0 | 0 | 0 | 0 | 0 | 0 | 0 |
| 0 | NA | NA | NA | NA | NA | Gm49711 | | 0 | 0 | 0 | 0 | 0 | 0 | 0 | 0 | 0 | 0 | 0 |
| 0 | NA | NA | NA | NA | NA | Gm41496 | | 0 | 0 | 0 | 0 | 0 | 0 | 0 | 0 | 0 | 0 | 0 |
| 0 | NA | NA | NA | NA | NA | Gm41495 | | 0 | 0 | 0 | 0 | 0 | 0 | 0 | 0 | 0 | 0 | 0 |
| 0 | NA | NA | NA | NA | NA | Gm49716 | | 0 | 0 | 0 | 0 | 0 | 0 | 0 | 0 | 0 | 0 | 0 |
| 0 | NA | NA | NA | NA | NA | Gm29880 | | 0 | 0 | 0 | 0 | 0 | 0 | 0 | 0 | 0 | 0 | 0 |
| 0 | NA | NA | NA | NA | NA | Kcne2 | potassium voltage-gated channel, Isk-rela | 0 | 0 | 0 | 0 | 0 | 0 | 0 | 0 | 0 | 0 | 0 |
| 0 | NA | NA | NA | NA | NA | 1700048M11Rik | | 0 | 0 | 0 | 0 | 0 | 0 | 0 | 0 | 0 | 0 | 0 |
| 0 | NA | NA | NA | NA | NA | Smim34 | | 0 | 0 | 0 | 0 | 0 | 0 | 0 | 0 | 0 | 0 | 0 |
| 0 | NA | NA | NA | NA | NA | Kcne1 | potassium voltage-gated channel, Isk-rela | 0 | 0 | 0 | 0 | 0 | 0 | 0 | 0 | 0 | 0 | 0 |
| 0 | NA | NA | NA | NA | NA | Gm29805 | | 0 | 0 | 0 | 0 | 0 | 0 | 0 | 0 | 0 | 0 | 0 |
| 0 | NA | NA | NA | NA | NA | 2410124H12Ri | RIKEN cDNA 2410124H12 gene | 0 | 0 | 0 | 0 | 0 | 0 | 0 | 0 | 0 | 0 | 0 |
| 0 | NA | NA | NA | NA | NA | Gm49722 | | 0 | 0 | 0 | 0 | 0 | 0 | 0 | 0 | 0 | 0 | 0 |
| 0 | NA | NA | NA | NA | NA | Gm26626 | | 0 | 0 | 0 | 0 | 0 | 0 | 0 | 0 | 0 | 0 | 0 |
| 0 | NA | NA | NA | NA | NA | Gm49723 | | 0 | 0 | 0 | 0 | 0 | 0 | 0 | 0 | 0 | 0 | 0 |
| 0 | NA | NA | NA | NA | NA | Gm28003 | | 0 | 0 | 0 | 0 | 0 | 0 | 0 | 0 | 0 | 0 | 0 |
| 0 | NA | NA | NA | NA | NA | 1700029J03Rik | | 0 | 0 | 0 | 0 | 0 | 0 | 0 | 0 | 0 | 0 | 0 |
| 0 | NA | NA | NA | NA | NA | Gm30695 | | 0 | 0 | 0 | 0 | 0 | 0 | 0 | 0 | 0 | 0 | 0 |
| 0 | NA | NA | NA | NA | NA | Gm49725 | | 0 | 0 | 0 | 0 | 0 | 0 | 0 | 0 | 0 | 0 | 0 |
| 0 | NA | NA | NA | NA | NA | Gm49961 | | 0 | 0 | 0 | 0 | 0 | 0 | 0 | 0 | 0 | 0 | 0 |
| 0 | NA | NA | NA | NA | NA | Gm45384 | | 0 | 0 | 0 | 0 | 0 | 0 | 0 | 0 | 0 | 0 | 0 |
| 0 | NA | NA | NA | NA | NA | Gm22344 | | 0 | 0 | 0 | 0 | 0 | 0 | 0 | 0 | 0 | 0 | 0 |
| 0 | NA | NA | NA | NA | NA | | | 0 | 0 | 0 | 0 | 0 | 0 | 0 | 0 | 0 | 0 | 0 |
| 0 | NA | NA | NA | NA | NA | Gm49617 | | 0 | 0 | 0 | 0 | 0 | 0 | 0 | 0 | 0 | 0 | 0 |
| 0 | NA | NA | NA | NA | NA | Gm49618 | | 0 | 0 | 0 | 0 | 0 | 0 | 0 | 0 | 0 | 0 | 0 |
| 0 | NA | NA | NA | NA | NA | 2410003I16Rik | | 0 | 0 | 0 | 0 | 0 | 0 | 0 | 0 | 0 | 0 | 0 |
| 0 | NA | NA | NA | NA | NA | Gm49619 | | 0 | 0 | 0 | 0 | 0 | 0 | 0 | 0 | 0 | 0 | 0 |
| 0 | NA | NA | NA | NA | NA | Gm49620 | | 0 | 0 | 0 | 0 | 0 | 0 | 0 | 0 | 0 | 0 | 0 |
| 0 | NA | NA | NA | NA | NA | Gm23062 | | 0 | 0 | 0 | 0 | 0 | 0 | 0 | 0 | 0 | 0 | 0 |
| 0 | NA | NA | NA | NA | NA | Gm48979 | | 0 | 0 | 0 | 0 | 0 | 0 | 0 | 0 | 0 | 0 | 0 |
| 0 | NA | NA | NA | NA | NA | Gm48984 | | 0 | 0 | 0 | 0 | 0 | 0 | 0 | 0 | 0 | 0 | 0 |
| 0 | NA | NA | NA | NA | NA | Gm15310 | | 0 | 0 | 0 | 0 | 0 | 0 | 0 | 0 | 0 | 0 | 0 |
| 0 | NA | NA | NA | NA | NA | Gm23692 | | 0 | 0 | 0 | 0 | 0 | 0 | 0 | 0 | 0 | 0 | 0 |
| 0 | NA | NA | NA | NA | NA | Gm15971 | | 0 | 0 | 0 | 0 | 0 | 0 | 0 | 0 | 0 | 0 | 0 |
| 0 | NA | NA | NA | NA | NA | Gm22745 | | 0 | 0 | 0 | 0 | 0 | 0 | 0 | 0 | 0 | 0 | 0 |
| 0 | NA | NA | NA | NA | NA | Gm49641 | | 0 | 0 | 0 | 0 | 0 | 0 | 0 | 0 | 0 | 0 | 0 |

| | | | | | | | | | | | | | | | | | | |
|---|---|---|---|---|---|---|---|---|---|---|---|---|---|---|---|---|---|---|
| 0 | NA | NA | NA | NA | NA | Gm49642 | | 0 | 0 | 0 | 0 | 0 | 0 | 0 | 0 | 0 | 0 | 0 |
| 0 | NA | NA | NA | NA | NA | Gm31641 | | 0 | 0 | 0 | 0 | 0 | 0 | 0 | 0 | 0 | 0 | 0 |
| 0 | NA | NA | NA | NA | NA | Gm49643 | | 0 | 0 | 0 | 0 | 0 | 0 | 0 | 0 | 0 | 0 | 0 |
| 0 | NA | NA | NA | NA | NA | 1600002D24Rik | | 0 | 0 | 0 | 0 | 0 | 0 | 0 | 0 | 0 | 0 | 0 |
| 0 | NA | NA | NA | NA | NA | Gm37259 | | 0 | 0 | 0 | 0 | 0 | 0 | 0 | 0 | 0 | 0 | 0 |
| 0 | NA | NA | NA | NA | NA | 2810404F17Rik | | 0 | 0 | 0 | 0 | 0 | 0 | 0 | 0 | 0 | 0 | 0 |
| 0 | NA | NA | NA | NA | NA | Gm15340 | | 0 | 0 | 0 | 0 | 0 | 0 | 0 | 0 | 0 | 0 | 0 |
| 0 | NA | NA | NA | NA | NA | Gm27833 | | 0 | 0 | 0 | 0 | 0 | 0 | 0 | 0 | 0 | 0 | 0 |
| 0 | NA | NA | NA | NA | NA | Gm49953 | | 0 | 0 | 0 | 0 | 0 | 0 | 0 | 0 | 0 | 0 | 0 |
| 0 | NA | NA | NA | NA | NA | Gm49928 | | 0 | 0 | 0 | 0 | 0 | 0 | 0 | 0 | 0 | 0 | 0 |
| 0 | NA | NA | NA | NA | NA | Gm49948 | | 0 | 0 | 0 | 0 | 0 | 0 | 0 | 0 | 0 | 0 | 0 |
| 0 | NA | NA | NA | NA | NA | Itgb2l | integrin beta 2-like | 0 | 0 | 0 | 0 | 0 | 0 | 0 | 0 | 0 | 0 | 0 |
| 0 | NA | NA | NA | NA | NA | Gm49907 | | 0 | 0 | 0 | 0 | 0 | 0 | 0 | 0 | 0 | 0 | 0 |
| 0 | NA | NA | NA | NA | NA | Gm32509 | | 0 | 0 | 0 | 0 | 0 | 0 | 0 | 0 | 0 | 0 | 0 |
| 0 | NA | NA | NA | NA | NA | Gm32432 | | 0 | 0 | 0 | 0 | 0 | 0 | 0 | 0 | 0 | 0 | 0 |
| 0 | NA | NA | NA | NA | NA | 2210009P08Rik | | 0 | 0 | 0 | 0 | 0 | 0 | 0 | 0 | 0 | 0 | 0 |
| 0 | NA | NA | NA | NA | NA | Gm24777 | | 0 | 0 | 0 | 0 | 0 | 0 | 0 | 0 | 0 | 0 | 0 |
| 0 | NA | NA | NA | NA | NA | Gm49592 | | 0 | 0 | 0 | 0 | 0 | 0 | 0 | 0 | 0 | 0 | 0 |
| 0 | NA | NA | NA | NA | NA | Mir6964 | | 0 | 0 | 0 | 0 | 0 | 0 | 0 | 0 | 0 | 0 | 0 |
| 0 | NA | NA | NA | NA | NA | Gm49792 | | 0 | 0 | 0 | 0 | 0 | 0 | 0 | 0 | 0 | 0 | 0 |
| 0 | NA | NA | NA | NA | NA | Gm16569 | | 0 | 0 | 0 | 0 | 0 | 0 | 0 | 0 | 0 | 0 | 0 |
| 0 | NA | NA | NA | NA | NA | Gm25909 | | 0 | 0 | 0 | 0 | 0 | 0 | 0 | 0 | 0 | 0 | 0 |
| 0 | NA | NA | NA | NA | NA | Gm5679 | | 0 | 0 | 0 | 0 | 0 | 0 | 0 | 0 | 0 | 0 | 0 |
| 0 | NA | NA | NA | NA | NA | Gm49798 | | 0 | 0 | 0 | 0 | 0 | 0 | 0 | 0 | 0 | 0 | 0 |
| 0 | NA | NA | NA | NA | NA | Gm34035 | | 0 | 0 | 0 | 0 | 0 | 0 | 0 | 0 | 0 | 0 | 0 |
| 0 | NA | NA | NA | NA | NA | Gm49221 | | 0 | 0 | 0 | 0 | 0 | 0 | 0 | 0 | 0 | 0 | 0 |
| 0 | NA | NA | NA | NA | NA | Cldn20 | claudin 20 | 0 | 0 | 0 | 0 | 0 | 0 | 0 | 0 | 0 | 0 | 0 |
| 0 | NA | NA | NA | NA | NA | Nox3 | NADPH oxidase 3 | 0 | 0 | 0 | 0 | 0 | 0 | 0 | 0 | 0 | 0 | 0 |
| 0 | NA | NA | NA | NA | NA | 4930517M08Rik | | 0 | 0 | 0 | 0 | 0 | 0 | 0 | 0 | 0 | 0 | 0 |
| 0 | NA | NA | NA | NA | NA | Gm23186 | | 0 | 0 | 0 | 0 | 0 | 0 | 0 | 0 | 0 | 0 | 0 |
| 0 | NA | NA | NA | NA | NA | Gm49780 | | 0 | 0 | 0 | 0 | 0 | 0 | 0 | 0 | 0 | 0 | 0 |
| 0 | NA | NA | NA | NA | NA | 4930579D07Rik | | 0 | 0 | 0 | 0 | 0 | 0 | 0 | 0 | 0 | 0 | 0 |
| 0 | NA | NA | NA | NA | NA | Gm29050 | | 0 | 0 | 0 | 0 | 0 | 0 | 0 | 0 | 0 | 0 | 0 |
| 0 | NA | NA | NA | NA | NA | Gm22475 | | 0 | 0 | 0 | 0 | 0 | 0 | 0 | 0 | 0 | 0 | 0 |
| 0 | NA | NA | NA | NA | NA | Gm8363 | | 0 | 0 | 0 | 0 | 0 | 0 | 0 | 0 | 0 | 0 | 0 |
| 0 | NA | NA | NA | NA | NA | Gm26595 | | 0 | 0 | 0 | 0 | 0 | 0 | 0 | 0 | 0 | 0 | 0 |
| 0 | NA | NA | NA | NA | NA | Gm26622 | | 0 | 0 | 0 | 0 | 0 | 0 | 0 | 0 | 0 | 0 | 0 |
| 0 | NA | NA | NA | NA | NA | Gm8376 | | 0 | 0 | 0 | 0 | 0 | 0 | 0 | 0 | 0 | 0 | 0 |
| 0 | NA | NA | NA | NA | NA | Gm22774 | | 0 | 0 | 0 | 0 | 0 | 0 | 0 | 0 | 0 | 0 | 0 |
| 0 | NA | NA | NA | NA | NA | n-R5s26 | | 0 | 0 | 0 | 0 | 0 | 0 | 0 | 0 | 0 | 0 | 0 |
| 0 | NA | NA | NA | NA | NA | Ppp1r2-ps6 | | 0 | 0 | 0 | 0 | 0 | 0 | 0 | 0 | 0 | 0 | 0 |
| 0 | NA | NA | NA | NA | NA | | | 0 | 0 | 0 | 0 | 0 | 0 | 0 | 0 | 0 | 0 | 0 |
| 0 | NA | NA | NA | NA | NA | Gm49693 | | 0 | 0 | 0 | 0 | 0 | 0 | 0 | 0 | 0 | 0 | 0 |
| 0 | NA | NA | NA | NA | NA | Gm29719 | | 0 | 0 | 0 | 0 | 0 | 0 | 0 | 0 | 0 | 0 | 0 |
| 0 | NA | NA | NA | NA | NA | Gm29719 | | 0 | 0 | 0 | 0 | 0 | 0 | 0 | 0 | 0 | 0 | 0 |
| 0 | NA | NA | NA | NA | NA | Gm29721 | | 0 | 0 | 0 | 0 | 0 | 0 | 0 | 0 | 0 | 0 | 0 |
| 0 | NA | NA | NA | NA | NA | Gm49717 | | 0 | 0 | 0 | 0 | 0 | 0 | 0 | 0 | 0 | 0 | 0 |
| 0 | NA | NA | NA | NA | NA | 4933426B08Rik | | 0 | 0 | 0 | 0 | 0 | 0 | 0 | 0 | 0 | 0 | 0 |
| 0 | NA | NA | NA | NA | NA | 1700122H20Rik | | 0 | 0 | 0 | 0 | 0 | 0 | 0 | 0 | 0 | 0 | 0 |
| 0 | NA | NA | NA | NA | NA | Gm49721 | | 0 | 0 | 0 | 0 | 0 | 0 | 0 | 0 | 0 | 0 | 0 |
| 0 | NA | NA | NA | NA | NA | Gm1604b | predicted gene 1604b | 0 | 0 | 0 | 0 | 0 | 0 | 0 | 0 | 0 | 0 | 0 |
| 0 | NA | NA | NA | NA | NA | Gm25119 | | 0 | 0 | 0 | 0 | 0 | 0 | 0 | 0 | 0 | 0 | 0 |
| 0 | NA | NA | NA | NA | NA | Ttll2 | tubulin tyrosine ligase-like family, membe | 0 | 0 | 0 | 0 | 0 | 0 | 0 | 0 | 0 | 0 | 0 |
| 0 | NA | NA | NA | NA | NA | Gm49630 | | 0 | 0 | 0 | 0 | 0 | 0 | 0 | 0 | 0 | 0 | 0 |
| 0 | NA | NA | NA | NA | NA | Gm26057 | | 0 | 0 | 0 | 0 | 0 | 0 | 0 | 0 | 0 | 0 | 0 |
| 0 | NA | NA | NA | NA | NA | Gm33086 | | 0 | 0 | 0 | 0 | 0 | 0 | 0 | 0 | 0 | 0 | 0 |
| 0 | NA | NA | NA | NA | NA | Gm49632 | | 0 | 0 | 0 | 0 | 0 | 0 | 0 | 0 | 0 | 0 | 0 |
| 0 | NA | NA | NA | NA | NA | Gm25872 | | 0 | 0 | 0 | 0 | 0 | 0 | 0 | 0 | 0 | 0 | 0 |
| 0 | NA | NA | NA | NA | NA | Gm22416 | | 0 | 0 | 0 | 0 | 0 | 0 | 0 | 0 | 0 | 0 | 0 |
| 0 | NA | NA | NA | NA | NA | Gm49987 | | 0 | 0 | 0 | 0 | 0 | 0 | 0 | 0 | 0 | 0 | 0 |

| | | | | | | | | | | | | | | | | | |
|---|---|---|---|---|---|---|---|---|---|---|---|---|---|---|---|---|---|
| 0 | NA | NA | NA | NA | NA | Gm15426 | | 0 | 0 | 0 | 0 | 0 | 0 | 0 | 0 | 0 | 0 | 0 |
| 0 | NA | NA | NA | NA | NA | Gm29711 | | 0 | 0 | 0 | 0 | 0 | 0 | 0 | 0 | 0 | 0 | 0 |
| 0 | NA | NA | NA | NA | NA | 6530411M01Rik | | 0 | 0 | 0 | 0 | 0 | 0 | 0 | 0 | 0 | 0 | 0 |
| 0 | NA | NA | NA | NA | NA | C87487 | | 0 | 0 | 0 | 0 | 0 | 0 | 0 | 0 | 0 | 0 | 0 |
| 0 | NA | NA | NA | NA | NA | Gm49806 | | 0 | 0 | 0 | 0 | 0 | 0 | 0 | 0 | 0 | 0 | 0 |
| 0 | NA | NA | NA | NA | NA | Gm7947 | | 0 | 0 | 0 | 0 | 0 | 0 | 0 | 0 | 0 | 0 | 0 |
| 0 | NA | NA | NA | NA | NA | Gm17728 | | 0 | 0 | 0 | 0 | 0 | 0 | 0 | 0 | 0 | 0 | 0 |
| 0 | NA | NA | NA | NA | NA | Gm49807 | | 0 | 0 | 0 | 0 | 0 | 0 | 0 | 0 | 0 | 0 | 0 |
| 0 | NA | NA | NA | NA | NA | Pabpc6 | poly(A) binding protein, cytoplasmic 6 | 0 | 0 | 0 | 0 | 0 | 0 | 0 | 0 | 0 | 0 | 0 |
| 0 | NA | NA | NA | NA | NA | Gm34684 | | 0 | 0 | 0 | 0 | 0 | 0 | 0 | 0 | 0 | 0 | 0 |
| 0 | NA | NA | NA | NA | NA | 4930452A19Rik | | 0 | 0 | 0 | 0 | 0 | 0 | 0 | 0 | 0 | 0 | 0 |
| 0 | NA | NA | NA | NA | NA | Gm49808 | | 0 | 0 | 0 | 0 | 0 | 0 | 0 | 0 | 0 | 0 | 0 |
| 0 | NA | NA | NA | NA | NA | Gm49809 | | 0 | 0 | 0 | 0 | 0 | 0 | 0 | 0 | 0 | 0 | 0 |
| 0 | NA | NA | NA | NA | NA | Gm16168 | | 0 | 0 | 0 | 0 | 0 | 0 | 0 | 0 | 0 | 0 | 0 |
| 0 | NA | NA | NA | NA | NA | Gm16169 | | 0 | 0 | 0 | 0 | 0 | 0 | 0 | 0 | 0 | 0 | 0 |
| 0 | NA | NA | NA | NA | NA | Gm37332 | | 0 | 0 | 0 | 0 | 0 | 0 | 0 | 0 | 0 | 0 | 0 |
| 0 | NA | NA | NA | NA | NA | Gm28118 | | 0 | 0 | 0 | 0 | 0 | 0 | 0 | 0 | 0 | 0 | 0 |
| 0 | NA | NA | NA | NA | NA | Gm10513 | | 0 | 0 | 0 | 0 | 0 | 0 | 0 | 0 | 0 | 0 | 0 |
| 0 | NA | NA | NA | NA | NA | Tdgf1-ps2 | | 0 | 0 | 0 | 0 | 0 | 0 | 0 | 0 | 0 | 0 | 0 |
| 0 | NA | NA | NA | NA | NA | Gm49962 | | 0 | 0 | 0 | 0 | 0 | 0 | 0 | 0 | 0 | 0 | 0 |
| 0 | NA | NA | NA | NA | NA | Gm24794 | | 0 | 0 | 0 | 0 | 0 | 0 | 0 | 0 | 0 | 0 | 0 |
| 0 | NA | NA | NA | NA | NA | Gm49964 | | 0 | 0 | 0 | 0 | 0 | 0 | 0 | 0 | 0 | 0 | 0 |
| 0 | NA | NA | NA | NA | NA | C030013G03Rik | | 0 | 0 | 0 | 0 | 0 | 0 | 0 | 0 | 0 | 0 | 0 |
| 0 | NA | NA | NA | NA | NA | Gm49958 | | 0 | 0 | 0 | 0 | 0 | 0 | 0 | 0 | 0 | 0 | 0 |
| 0 | NA | NA | NA | NA | NA | Gm24085 | | 0 | 0 | 0 | 0 | 0 | 0 | 0 | 0 | 0 | 0 | 0 |
| 0 | NA | NA | NA | NA | NA | Gm49926 | | 0 | 0 | 0 | 0 | 0 | 0 | 0 | 0 | 0 | 0 | 0 |
| 0 | NA | NA | NA | NA | NA | Gm26130 | | 0 | 0 | 0 | 0 | 0 | 0 | 0 | 0 | 0 | 0 | 0 |
| 0 | NA | NA | NA | NA | NA | Gm11166 | | 0 | 0 | 0 | 0 | 0 | 0 | 0 | 0 | 0 | 0 | 0 |
| 0 | NA | NA | NA | NA | NA | Unc93a | unc-93 homolog A | 0 | 0 | 0 | 0 | 0 | 0 | 0 | 0 | 0 | 0 | 0 |
| 0 | NA | NA | NA | NA | NA | Gm23050 | | 0 | 0 | 0 | 0 | 0 | 0 | 0 | 0 | 0 | 0 | 0 |
| 0 | NA | NA | NA | NA | NA | Gm49898 | | 0 | 0 | 0 | 0 | 0 | 0 | 0 | 0 | 0 | 0 | 0 |
| 0 | NA | NA | NA | NA | NA | Gm10512 | | 0 | 0 | 0 | 0 | 0 | 0 | 0 | 0 | 0 | 0 | 0 |
| 0 | NA | NA | NA | NA | NA | Smok2a | sperm motility kinase 2A | 0 | 0 | 0 | 0 | 0 | 0 | 0 | 0 | 0 | 0 | 0 |
| 0 | NA | NA | NA | NA | NA | Smok2b | sperm motility kinase 2B | 0 | 0 | 0 | 0 | 0 | 0 | 0 | 0 | 0 | 0 | 0 |
| 0 | NA | NA | NA | NA | NA | Gm26500 | | 0 | 0 | 0 | 0 | 0 | 0 | 0 | 0 | 0 | 0 | 0 |
| 0 | NA | NA | NA | NA | NA | Tcp10c | t-complex protein 10c | 0 | 0 | 0 | 0 | 0 | 0 | 0 | 0 | 0 | 0 | 0 |
| 0 | NA | NA | NA | NA | NA | Gm49934 | | 0 | 0 | 0 | 0 | 0 | 0 | 0 | 0 | 0 | 0 | 0 |
| 0 | NA | NA | NA | NA | NA | Gm25730 | | 0 | 0 | 0 | 0 | 0 | 0 | 0 | 0 | 0 | 0 | 0 |
| 0 | NA | NA | NA | NA | NA | Gm49936 | | 0 | 0 | 0 | 0 | 0 | 0 | 0 | 0 | 0 | 0 | 0 |
| 0 | NA | NA | NA | NA | NA | Gm16050 | | 0 | 0 | 0 | 0 | 0 | 0 | 0 | 0 | 0 | 0 | 0 |
| 0 | NA | NA | NA | NA | NA | Gm8603 | | 0 | 0 | 0 | 0 | 0 | 0 | 0 | 0 | 0 | 0 | 0 |
| 0 | NA | NA | NA | NA | NA | Smok4a | | 0 | 0 | 0 | 0 | 0 | 0 | 0 | 0 | 0 | 0 | 0 |
| 0 | NA | NA | NA | NA | NA | Gm26375 | | 0 | 0 | 0 | 0 | 0 | 0 | 0 | 0 | 0 | 0 | 0 |
| 0 | NA | NA | NA | NA | NA | Gm24867 | | 0 | 0 | 0 | 0 | 0 | 0 | 0 | 0 | 0 | 0 | 0 |
| 0 | NA | NA | NA | NA | NA | Gm49121 | | 0 | 0 | 0 | 0 | 0 | 0 | 0 | 0 | 0 | 0 | 0 |
| 0 | NA | NA | NA | NA | NA | Gm7356 | predicted gene 7356 | 0 | 0 | 0 | 0 | 0 | 0 | 0 | 0 | 0 | 0 | 0 |
| 0 | NA | NA | NA | NA | NA | Gm7358 | | 0 | 0 | 0 | 0 | 0 | 0 | 0 | 0 | 0 | 0 | 0 |
| 0 | NA | NA | NA | NA | NA | Gm49903 | | 0 | 0 | 0 | 0 | 0 | 0 | 0 | 0 | 0 | 0 | 0 |
| 0 | NA | NA | NA | NA | NA | 4930474M22Rik | | 0 | 0 | 0 | 0 | 0 | 0 | 0 | 0 | 0 | 0 | 0 |
| 0 | NA | NA | NA | NA | NA | Gm38551 | | 0 | 0 | 0 | 0 | 0 | 0 | 0 | 0 | 0 | 0 | 0 |
| 0 | NA | NA | NA | NA | NA | Gm23352 | | 0 | 0 | 0 | 0 | 0 | 0 | 0 | 0 | 0 | 0 | 0 |
| 0 | NA | NA | NA | NA | NA | Gm3417 | predicted gene 3417 | 0 | 0 | 0 | 0 | 0 | 0 | 0 | 0 | 0 | 0 | 0 |
| 0 | NA | NA | NA | NA | NA | Gm3448 | predicted gene 3448 | 0 | 0 | 0 | 0 | 0 | 0 | 0 | 0 | 0 | 0 | 0 |
| 0 | NA | NA | NA | NA | NA | Tcte3 | t-complex-associated testis expressed 3 | 0 | 0 | 0 | 0 | 0 | 0 | 0 | 0 | 0 | 0 | 0 |
| 0 | NA | NA | NA | NA | NA | Gm5091 | | 0 | 0 | 0 | 0 | 0 | 0 | 0 | 0 | 0 | 0 | 0 |
| 0 | NA | NA | NA | NA | NA | Gm35576 | | 0 | 0 | 0 | 0 | 0 | 0 | 0 | 0 | 0 | 0 | 0 |
| 0 | NA | NA | NA | NA | NA | Gm7423 | | 0 | 0 | 0 | 0 | 0 | 0 | 0 | 0 | 0 | 0 | 0 |
| 0 | NA | NA | NA | NA | NA | Gm49669 | | 0 | 0 | 0 | 0 | 0 | 0 | 0 | 0 | 0 | 0 | 0 |
| 0 | NA | NA | NA | NA | NA | Gm7177 | | 0 | 0 | 0 | 0 | 0 | 0 | 0 | 0 | 0 | 0 | 0 |
| 0 | NA | NA | NA | NA | NA | Gm6686 | | 0 | 0 | 0 | 0 | 0 | 0 | 0 | 0 | 0 | 0 | 0 |

| | | | | | | | | | | | | | | | | | | |
|---|---|---|---|---|---|---|---|---|---|---|---|---|---|---|---|---|---|---|
| 0 | NA | NA | NA | NA | NA | 4933401D09Rik | | 0 | 0 | 0 | 0 | 0 | 0 | 0 | 0 | 0 | 0 | 0 |
| 0 | NA | NA | NA | NA | NA | Gm3292 | | 0 | 0 | 0 | 0 | 0 | 0 | 0 | 0 | 0 | 0 | 0 |
| 0 | NA | NA | NA | NA | NA | Gm25598 | | 0 | 0 | 0 | 0 | 0 | 0 | 0 | 0 | 0 | 0 | 0 |
| 0 | NA | NA | NA | NA | NA | Gm49777 | | 0 | 0 | 0 | 0 | 0 | 0 | 0 | 0 | 0 | 0 | 0 |
| 0 | NA | NA | NA | NA | NA | Gm49778 | | 0 | 0 | 0 | 0 | 0 | 0 | 0 | 0 | 0 | 0 | 0 |
| 0 | NA | NA | NA | NA | NA | Gm46610 | | 0 | 0 | 0 | 0 | 0 | 0 | 0 | 0 | 0 | 0 | 0 |
| 0 | NA | NA | NA | NA | NA | Gm7477 | | 0 | 0 | 0 | 0 | 0 | 0 | 0 | 0 | 0 | 0 | 0 |
| 0 | NA | NA | NA | NA | NA | Gm5223 | | 0 | 0 | 0 | 0 | 0 | 0 | 0 | 0 | 0 | 0 | 0 |
| 0 | NA | NA | NA | NA | NA | Gm49775 | | 0 | 0 | 0 | 0 | 0 | 0 | 0 | 0 | 0 | 0 | 0 |
| 0 | NA | NA | NA | NA | NA | Gm7483 | | 0 | 0 | 0 | 0 | 0 | 0 | 0 | 0 | 0 | 0 | 0 |
| 0 | NA | NA | NA | NA | NA | Gm5232 | | 0 | 0 | 0 | 0 | 0 | 0 | 0 | 0 | 0 | 0 | 0 |
| 0 | NA | NA | NA | NA | NA | Gm49784 | | 0 | 0 | 0 | 0 | 0 | 0 | 0 | 0 | 0 | 0 | 0 |
| 0 | NA | NA | NA | NA | NA | Gm7513 | | 0 | 0 | 0 | 0 | 0 | 0 | 0 | 0 | 0 | 0 | 0 |
| 0 | NA | NA | NA | NA | NA | Gm49787 | | 0 | 0 | 0 | 0 | 0 | 0 | 0 | 0 | 0 | 0 | 0 |
| 0 | NA | NA | NA | NA | NA | Vmn2r90 | vomeronasal 2, receptor 90 | 0 | 0 | 0 | 0 | 0 | 0 | 0 | 0 | 0 | 0 | 0 |
| 0 | NA | NA | NA | NA | NA | Gm23837 | | 0 | 0 | 0 | 0 | 0 | 0 | 0 | 0 | 0 | 0 | 0 |
| 0 | NA | NA | NA | NA | NA | Mir99b | | 0 | 0 | 0 | 0 | 0 | 0 | 0 | 0 | 0 | 0 | 0 |
| 0 | NA | NA | NA | NA | NA | Gm25927 | | 0 | 0 | 0 | 0 | 0 | 0 | 0 | 0 | 0 | 0 | 0 |
| 0 | NA | NA | NA | NA | NA | Mirlet7e | | 0 | 0 | 0 | 0 | 0 | 0 | 0 | 0 | 0 | 0 | 0 |
| 0 | NA | NA | NA | NA | NA | Mir125a | | 0 | 0 | 0 | 0 | 0 | 0 | 0 | 0 | 0 | 0 | 0 |
| 0 | NA | NA | NA | NA | NA | Fpr3 | formyl peptide receptor 3 | 0 | 0 | 0 | 0 | 0 | 0 | 0 | 0 | 0 | 0 | 0 |
| 0 | NA | NA | NA | NA | NA | Gm7531 | | 0 | 0 | 0 | 0 | 0 | 0 | 0 | 0 | 0 | 0 | 0 |
| 0 | NA | NA | NA | NA | NA | Gm5966 | | 0 | 0 | 0 | 0 | 0 | 0 | 0 | 0 | 0 | 0 | 0 |
| 0 | NA | NA | NA | NA | NA | Gm5966 | | 0 | 0 | 0 | 0 | 0 | 0 | 0 | 0 | 0 | 0 | 0 |
| 0 | NA | NA | NA | NA | NA | Fpr-rs5 | | 0 | 0 | 0 | 0 | 0 | 0 | 0 | 0 | 0 | 0 | 0 |
| 0 | NA | NA | NA | NA | NA | Fpr-rs4 | formyl peptide receptor, related sequence | 0 | 0 | 0 | 0 | 0 | 0 | 0 | 0 | 0 | 0 | 0 |
| 0 | NA | NA | NA | NA | NA | Vmn2r124 | vomeronasal 2, receptor 124 | 0 | 0 | 0 | 0 | 0 | 0 | 0 | 0 | 0 | 0 | 0 |
| 0 | NA | NA | NA | NA | NA | Vmn2r91 | vomeronasal 2, receptor 91 | 0 | 0 | 0 | 0 | 0 | 0 | 0 | 0 | 0 | 0 | 0 |
| 0 | NA | NA | NA | NA | NA | Vmn2r92 | vomeronasal 2, receptor 92 | 0 | 0 | 0 | 0 | 0 | 0 | 0 | 0 | 0 | 0 | 0 |
| 0 | NA | NA | NA | NA | NA | Vmn2r94 | vomeronasal 2, receptor 94 | 0 | 0 | 0 | 0 | 0 | 0 | 0 | 0 | 0 | 0 | 0 |
| 0 | NA | NA | NA | NA | NA | Vmn2r93 | vomeronasal 2, receptor 93 | 0 | 0 | 0 | 0 | 0 | 0 | 0 | 0 | 0 | 0 | 0 |
| 0 | NA | NA | NA | NA | NA | Vmn2r95 | vomeronasal 2, receptor 95 | 0 | 0 | 0 | 0 | 0 | 0 | 0 | 0 | 0 | 0 | 0 |
| 0 | NA | NA | NA | NA | NA | Gm49963 | | 0 | 0 | 0 | 0 | 0 | 0 | 0 | 0 | 0 | 0 | 0 |
| 0 | NA | NA | NA | NA | NA | Vmn2r-ps114 | | 0 | 0 | 0 | 0 | 0 | 0 | 0 | 0 | 0 | 0 | 0 |
| 0 | NA | NA | NA | NA | NA | Vmn2r-ps115 | | 0 | 0 | 0 | 0 | 0 | 0 | 0 | 0 | 0 | 0 | 0 |
| 0 | NA | NA | NA | NA | NA | Vmn2r-ps116 | | 0 | 0 | 0 | 0 | 0 | 0 | 0 | 0 | 0 | 0 | 0 |
| 0 | NA | NA | NA | NA | NA | Vmn2r-ps117 | | 0 | 0 | 0 | 0 | 0 | 0 | 0 | 0 | 0 | 0 | 0 |
| 0 | NA | NA | NA | NA | NA | Vmn2r97 | vomeronasal 2, receptor 97 | 0 | 0 | 0 | 0 | 0 | 0 | 0 | 0 | 0 | 0 | 0 |
| 0 | NA | NA | NA | NA | NA | Vmn2r-ps118 | | 0 | 0 | 0 | 0 | 0 | 0 | 0 | 0 | 0 | 0 | 0 |
| 0 | NA | NA | NA | NA | NA | Vmn2r98 | vomeronasal 2, receptor 98 | 0 | 0 | 0 | 0 | 0 | 0 | 0 | 0 | 0 | 0 | 0 |
| 0 | NA | NA | NA | NA | NA | Vmn2r127 | | 0 | 0 | 0 | 0 | 0 | 0 | 0 | 0 | 0 | 0 | 0 |
| 0 | NA | NA | NA | NA | NA | Gm49753 | | 0 | 0 | 0 | 0 | 0 | 0 | 0 | 0 | 0 | 0 | 0 |
| 0 | NA | NA | NA | NA | NA | Gm49752 | | 0 | 0 | 0 | 0 | 0 | 0 | 0 | 0 | 0 | 0 | 0 |
| 0 | NA | NA | NA | NA | NA | Vmn2r-ps120 | | 0 | 0 | 0 | 0 | 0 | 0 | 0 | 0 | 0 | 0 | 0 |
| 0 | NA | NA | NA | NA | NA | Vmn2r99 | vomeronasal 2, receptor 99 | 0 | 0 | 0 | 0 | 0 | 0 | 0 | 0 | 0 | 0 | 0 |
| 0 | NA | NA | NA | NA | NA | 4930549P19Rik | | 0 | 0 | 0 | 0 | 0 | 0 | 0 | 0 | 0 | 0 | 0 |
| 0 | NA | NA | NA | NA | NA | Vmn2r-ps121 | | 0 | 0 | 0 | 0 | 0 | 0 | 0 | 0 | 0 | 0 | 0 |
| 0 | NA | NA | NA | NA | NA | Vmn2r101 | vomeronasal 2, receptor 101 | 0 | 0 | 0 | 0 | 0 | 0 | 0 | 0 | 0 | 0 | 0 |
| 0 | NA | NA | NA | NA | NA | Vmn2r102 | vomeronasal 2, receptor 102 | 0 | 0 | 0 | 0 | 0 | 0 | 0 | 0 | 0 | 0 | 0 |
| 0 | NA | NA | NA | NA | NA | Vmn2r-ps122 | | 0 | 0 | 0 | 0 | 0 | 0 | 0 | 0 | 0 | 0 | 0 |
| 0 | NA | NA | NA | NA | NA | Vmn2r103 | vomeronasal 2, receptor 103 | 0 | 0 | 0 | 0 | 0 | 0 | 0 | 0 | 0 | 0 | 0 |
| 0 | NA | NA | NA | NA | NA | Vmn2r-ps123 | | 0 | 0 | 0 | 0 | 0 | 0 | 0 | 0 | 0 | 0 | 0 |
| 0 | NA | NA | NA | NA | NA | Vmn2r-ps124 | | 0 | 0 | 0 | 0 | 0 | 0 | 0 | 0 | 0 | 0 | 0 |
| 0 | NA | NA | NA | NA | NA | Vmn2r-ps125 | | 0 | 0 | 0 | 0 | 0 | 0 | 0 | 0 | 0 | 0 | 0 |
| 0 | NA | NA | NA | NA | NA | Vmn2r104 | vomeronasal 2, receptor 104 | 0 | 0 | 0 | 0 | 0 | 0 | 0 | 0 | 0 | 0 | 0 |
| 0 | NA | NA | NA | NA | NA | Gm3547 | | 0 | 0 | 0 | 0 | 0 | 0 | 0 | 0 | 0 | 0 | 0 |
| 0 | NA | NA | NA | NA | NA | Vmn2r-ps126 | | 0 | 0 | 0 | 0 | 0 | 0 | 0 | 0 | 0 | 0 | 0 |
| 0 | NA | NA | NA | NA | NA | Fpr-rs7 | formyl peptide receptor, related sequence | 0 | 0 | 0 | 0 | 0 | 0 | 0 | 0 | 0 | 0 | 0 |
| 0 | NA | NA | NA | NA | NA | Vmn1r-ps139 | | 0 | 0 | 0 | 0 | 0 | 0 | 0 | 0 | 0 | 0 | 0 |
| 0 | NA | NA | NA | NA | NA | Fpr-rs6 | formyl peptide receptor, related sequence | 0 | 0 | 0 | 0 | 0 | 0 | 0 | 0 | 0 | 0 | 0 |

| | | | | | | | | | | | | | | | | | | |
|---|---|---|---|---|---|---|---|---|---|---|---|---|---|---|---|---|---|---|
| 0 | NA | NA | NA | NA | NA | Vmn1r-ps140 | | 0 | 0 | 0 | 0 | 0 | 0 | 0 | 0 | 0 | 0 | 0 |
| 0 | NA | NA | NA | NA | NA | Vmn2r105 | vomeronasal 2, receptor 105 | 0 | 0 | 0 | 0 | 0 | 0 | 0 | 0 | 0 | 0 | 0 |
| 0 | NA | NA | NA | NA | NA | Vmn2r106 | vomeronasal 2, receptor 106 | 0 | 0 | 0 | 0 | 0 | 0 | 0 | 0 | 0 | 0 | 0 |
| 0 | NA | NA | NA | NA | NA | Vmn2r-ps127 | | 0 | 0 | 0 | 0 | 0 | 0 | 0 | 0 | 0 | 0 | 0 |
| 0 | NA | NA | NA | NA | NA | Vmn2r107 | vomeronasal 2, receptor 107 | 0 | 0 | 0 | 0 | 0 | 0 | 0 | 0 | 0 | 0 | 0 |
| 0 | NA | NA | NA | NA | NA | Gm7217 | | 0 | 0 | 0 | 0 | 0 | 0 | 0 | 0 | 0 | 0 | 0 |
| 0 | NA | NA | NA | NA | NA | Vmn1r-ps141 | | 0 | 0 | 0 | 0 | 0 | 0 | 0 | 0 | 0 | 0 | 0 |
| 0 | NA | NA | NA | NA | NA | Vmn1r224 | vomeronasal 1 receptor 224 | 0 | 0 | 0 | 0 | 0 | 0 | 0 | 0 | 0 | 0 | 0 |
| 0 | NA | NA | NA | NA | NA | Vmn1r-ps142 | | 0 | 0 | 0 | 0 | 0 | 0 | 0 | 0 | 0 | 0 | 0 |
| 0 | NA | NA | NA | NA | NA | Vmn1r-ps143 | | 0 | 0 | 0 | 0 | 0 | 0 | 0 | 0 | 0 | 0 | 0 |
| 0 | NA | NA | NA | NA | NA | Vmn2r108 | vomeronasal 2, receptor 108 | 0 | 0 | 0 | 0 | 0 | 0 | 0 | 0 | 0 | 0 | 0 |
| 0 | NA | NA | NA | NA | NA | Vmn1r225 | vomeronasal 1 receptor 225 | 0 | 0 | 0 | 0 | 0 | 0 | 0 | 0 | 0 | 0 | 0 |
| 0 | NA | NA | NA | NA | NA | Vmn2r109 | vomeronasal 2, receptor 109 | 0 | 0 | 0 | 0 | 0 | 0 | 0 | 0 | 0 | 0 | 0 |
| 0 | NA | NA | NA | NA | NA | Vmn2r110 | vomeronasal 2, receptor 110 | 0 | 0 | 0 | 0 | 0 | 0 | 0 | 0 | 0 | 0 | 0 |
| 0 | NA | NA | NA | NA | NA | Fpr-rs3 | formyl peptide receptor, related sequence | 0 | 0 | 0 | 0 | 0 | 0 | 0 | 0 | 0 | 0 | 0 |
| 0 | NA | NA | NA | NA | NA | Vmn1r226 | vomeronasal 1 receptor 226 | 0 | 0 | 0 | 0 | 0 | 0 | 0 | 0 | 0 | 0 | 0 |
| 0 | NA | NA | NA | NA | NA | Vmn1r227 | vomeronasal 1 receptor 227 | 0 | 0 | 0 | 0 | 0 | 0 | 0 | 0 | 0 | 0 | 0 |
| 0 | NA | NA | NA | NA | NA | Vmn1r228 | vomeronasal 1 receptor 228 | 0 | 0 | 0 | 0 | 0 | 0 | 0 | 0 | 0 | 0 | 0 |
| 0 | NA | NA | NA | NA | NA | Vmn1r229 | vomeronasal 1 receptor 229 | 0 | 0 | 0 | 0 | 0 | 0 | 0 | 0 | 0 | 0 | 0 |
| 0 | NA | NA | NA | NA | NA | Vmn1r230 | vomeronasal 1 receptor 230 | 0 | 0 | 0 | 0 | 0 | 0 | 0 | 0 | 0 | 0 | 0 |
| 0 | NA | NA | NA | NA | NA | Vmn1r231 | vomeronasal 1 receptor 231 | 0 | 0 | 0 | 0 | 0 | 0 | 0 | 0 | 0 | 0 | 0 |
| 0 | NA | NA | NA | NA | NA | Vmn1r232 | vomeronasal 1 receptor 232 | 0 | 0 | 0 | 0 | 0 | 0 | 0 | 0 | 0 | 0 | 0 |
| 0 | NA | NA | NA | NA | NA | Vmn1r-ps144 | | 0 | 0 | 0 | 0 | 0 | 0 | 0 | 0 | 0 | 0 | 0 |
| 0 | NA | NA | NA | NA | NA | Vmn1r-ps145 | | 0 | 0 | 0 | 0 | 0 | 0 | 0 | 0 | 0 | 0 | 0 |
| 0 | NA | NA | NA | NA | NA | Vmn1r233 | vomeronasal 1 receptor 233 | 0 | 0 | 0 | 0 | 0 | 0 | 0 | 0 | 0 | 0 | 0 |
| 0 | NA | NA | NA | NA | NA | Gm5681 | | 0 | 0 | 0 | 0 | 0 | 0 | 0 | 0 | 0 | 0 | 0 |
| 0 | NA | NA | NA | NA | NA | Vmn1r-ps146 | | 0 | 0 | 0 | 0 | 0 | 0 | 0 | 0 | 0 | 0 | 0 |
| 0 | NA | NA | NA | NA | NA | Gm7732 | | 0 | 0 | 0 | 0 | 0 | 0 | 0 | 0 | 0 | 0 | 0 |
| 0 | NA | NA | NA | NA | NA | Gm7736 | | 0 | 0 | 0 | 0 | 0 | 0 | 0 | 0 | 0 | 0 | 0 |
| 0 | NA | NA | NA | NA | NA | Gm7740 | | 0 | 0 | 0 | 0 | 0 | 0 | 0 | 0 | 0 | 0 | 0 |
| 0 | NA | NA | NA | NA | NA | Gm7742 | | 0 | 0 | 0 | 0 | 0 | 0 | 0 | 0 | 0 | 0 | 0 |
| 0 | NA | NA | NA | NA | NA | Vmn1r-ps147 | | 0 | 0 | 0 | 0 | 0 | 0 | 0 | 0 | 0 | 0 | 0 |
| 0 | NA | NA | NA | NA | NA | Vmn1r234 | vomeronasal 1 receptor 234 | 0 | 0 | 0 | 0 | 0 | 0 | 0 | 0 | 0 | 0 | 0 |
| 0 | NA | NA | NA | NA | NA | Vmn1r-ps148 | | 0 | 0 | 0 | 0 | 0 | 0 | 0 | 0 | 0 | 0 | 0 |
| 0 | NA | NA | NA | NA | NA | Vmn1r235 | vomeronasal 1 receptor 235 | 0 | 0 | 0 | 0 | 0 | 0 | 0 | 0 | 0 | 0 | 0 |
| 0 | NA | NA | NA | NA | NA | Vmn1r-ps149 | | 0 | 0 | 0 | 0 | 0 | 0 | 0 | 0 | 0 | 0 | 0 |
| 0 | NA | NA | NA | NA | NA | Vmn1r236 | vomeronasal 1 receptor 236 | 0 | 0 | 0 | 0 | 0 | 0 | 0 | 0 | 0 | 0 | 0 |
| 0 | NA | NA | NA | NA | NA | Vmn1r237 | vomeronasal 1 receptor 237 | 0 | 0 | 0 | 0 | 0 | 0 | 0 | 0 | 0 | 0 | 0 |
| 0 | NA | NA | NA | NA | NA | Vmn1r-ps150 | | 0 | 0 | 0 | 0 | 0 | 0 | 0 | 0 | 0 | 0 | 0 |
| 0 | NA | NA | NA | NA | NA | Gm18269 | | 0 | 0 | 0 | 0 | 0 | 0 | 0 | 0 | 0 | 0 | 0 |
| 0 | NA | NA | NA | NA | NA | Gm18519 | | 0 | 0 | 0 | 0 | 0 | 0 | 0 | 0 | 0 | 0 | 0 |
| 0 | NA | NA | NA | NA | NA | 4930515G13Rik | | 0 | 0 | 0 | 0 | 0 | 0 | 0 | 0 | 0 | 0 | 0 |
| 0 | NA | NA | NA | NA | NA | Gm9818 | | 0 | 0 | 0 | 0 | 0 | 0 | 0 | 0 | 0 | 0 | 0 |
| 0 | NA | NA | NA | NA | NA | Gm7805 | | 0 | 0 | 0 | 0 | 0 | 0 | 0 | 0 | 0 | 0 | 0 |
| 0 | NA | NA | NA | NA | NA | Gm49950 | | 0 | 0 | 0 | 0 | 0 | 0 | 0 | 0 | 0 | 0 | 0 |
| 0 | NA | NA | NA | NA | NA | Gm9772 | | 0 | 0 | 0 | 0 | 0 | 0 | 0 | 0 | 0 | 0 | 0 |
| 0 | NA | NA | NA | NA | NA | Gm7052 | | 0 | 0 | 0 | 0 | 0 | 0 | 0 | 0 | 0 | 0 | 0 |
| 0 | NA | NA | NA | NA | NA | Gm49951 | | 0 | 0 | 0 | 0 | 0 | 0 | 0 | 0 | 0 | 0 | 0 |
| 0 | NA | NA | NA | NA | NA | Gm49955 | | 0 | 0 | 0 | 0 | 0 | 0 | 0 | 0 | 0 | 0 | 0 |
| 0 | NA | NA | NA | NA | NA | Gm49956 | | 0 | 0 | 0 | 0 | 0 | 0 | 0 | 0 | 0 | 0 | 0 |
| 0 | NA | NA | NA | NA | NA | Gm19012 | | 0 | 0 | 0 | 0 | 0 | 0 | 0 | 0 | 0 | 0 | 0 |
| 0 | NA | NA | NA | NA | NA | Vmn2r111 | vomeronasal 2, receptor 111 | 0 | 0 | 0 | 0 | 0 | 0 | 0 | 0 | 0 | 0 | 0 |
| 0 | NA | NA | NA | NA | NA | Vmn2r112 | vomeronasal 2, receptor 112 | 0 | 0 | 0 | 0 | 0 | 0 | 0 | 0 | 0 | 0 | 0 |
| 0 | NA | NA | NA | NA | NA | Gm5224 | | 0 | 0 | 0 | 0 | 0 | 0 | 0 | 0 | 0 | 0 | 0 |
| 0 | NA | NA | NA | NA | NA | Gm32892 | | 0 | 0 | 0 | 0 | 0 | 0 | 0 | 0 | 0 | 0 | 0 |
| 0 | NA | NA | NA | NA | NA | Gm18214 | | 0 | 0 | 0 | 0 | 0 | 0 | 0 | 0 | 0 | 0 | 0 |
| 0 | NA | NA | NA | NA | NA | Vmn2r-ps128 | | 0 | 0 | 0 | 0 | 0 | 0 | 0 | 0 | 0 | 0 | 0 |
| 0 | NA | NA | NA | NA | NA | Vmn2r113 | vomeronasal 2, receptor 113 | 0 | 0 | 0 | 0 | 0 | 0 | 0 | 0 | 0 | 0 | 0 |
| 0 | NA | NA | NA | NA | NA | Vmn2r-ps129 | | 0 | 0 | 0 | 0 | 0 | 0 | 0 | 0 | 0 | 0 | 0 |
| 0 | NA | NA | NA | NA | NA | Gm18415 | | 0 | 0 | 0 | 0 | 0 | 0 | 0 | 0 | 0 | 0 | 0 |

| | | | | | | | | | | | | | | | | | |
|---|---|---|---|---|---|---|---|---|---|---|---|---|---|---|---|---|---|
| 0 | NA | NA | NA | NA | NA | Gm18396 | | 0 | 0 | 0 | 0 | 0 | 0 | 0 | 0 | 0 | 0 | 0 |
| 0 | NA | NA | NA | NA | NA | Vmn2r-ps130 | | 0 | 0 | 0 | 0 | 0 | 0 | 0 | 0 | 0 | 0 | 0 |
| 0 | NA | NA | NA | NA | NA | Gm18397 | | 0 | 0 | 0 | 0 | 0 | 0 | 0 | 0 | 0 | 0 | 0 |
| 0 | NA | NA | NA | NA | NA | Gm18416 | | 0 | 0 | 0 | 0 | 0 | 0 | 0 | 0 | 0 | 0 | 0 |
| 0 | NA | NA | NA | NA | NA | Vmn2r-ps131 | | 0 | 0 | 0 | 0 | 0 | 0 | 0 | 0 | 0 | 0 | 0 |
| 0 | NA | NA | NA | NA | NA | Vmn2r-ps132 | | 0 | 0 | 0 | 0 | 0 | 0 | 0 | 0 | 0 | 0 | 0 |
| 0 | NA | NA | NA | NA | NA | Vmn2r-ps133 | | 0 | 0 | 0 | 0 | 0 | 0 | 0 | 0 | 0 | 0 | 0 |
| 0 | NA | NA | NA | NA | NA | Vmn2r114 | vomeronasal 2, receptor 114 | 0 | 0 | 0 | 0 | 0 | 0 | 0 | 0 | 0 | 0 | 0 |
| 0 | NA | NA | NA | NA | NA | Vmn2r115 | vomeronasal 2, receptor 115 | 0 | 0 | 0 | 0 | 0 | 0 | 0 | 0 | 0 | 0 | 0 |
| 0 | NA | NA | NA | NA | NA | Vmn2r116 | vomeronasal 2, receptor 116 | 0 | 0 | 0 | 0 | 0 | 0 | 0 | 0 | 0 | 0 | 0 |
| 0 | NA | NA | NA | NA | NA | Gm49952 | | 0 | 0 | 0 | 0 | 0 | 0 | 0 | 0 | 0 | 0 | 0 |
| 0 | NA | NA | NA | NA | NA | Vmn2r-ps134 | | 0 | 0 | 0 | 0 | 0 | 0 | 0 | 0 | 0 | 0 | 0 |
| 0 | NA | NA | NA | NA | NA | Vmn2r117 | vomeronasal 2, receptor 117 | 0 | 0 | 0 | 0 | 0 | 0 | 0 | 0 | 0 | 0 | 0 |
| 0 | NA | NA | NA | NA | NA | Olfr752-ps1 | | 0 | 0 | 0 | 0 | 0 | 0 | 0 | 0 | 0 | 0 | 0 |
| 0 | NA | NA | NA | NA | NA | Gm7898 | | 0 | 0 | 0 | 0 | 0 | 0 | 0 | 0 | 0 | 0 | 0 |
| 0 | NA | NA | NA | NA | NA | Gm7773 | | 0 | 0 | 0 | 0 | 0 | 0 | 0 | 0 | 0 | 0 | 0 |
| 0 | NA | NA | NA | NA | NA | Gm49163 | | 0 | 0 | 0 | 0 | 0 | 0 | 0 | 0 | 0 | 0 | 0 |
| 0 | NA | NA | NA | NA | NA | Gm49911 | | 0 | 0 | 0 | 0 | 0 | 0 | 0 | 0 | 0 | 0 | 0 |
| 0 | NA | NA | NA | NA | NA | Prss33 | protease, serine 33 | 0 | 0 | 0 | 0 | 0 | 0 | 0 | 0 | 0 | 0 | 0 |
| 0 | NA | NA | NA | NA | NA | Gm15947 | | 0 | 0 | 0 | 0 | 0 | 0 | 0 | 0 | 0 | 0 | 0 |
| 0 | NA | NA | NA | NA | NA | Prss21 | protease, serine 21 | 0 | 0 | 0 | 0 | 0 | 0 | 0 | 0 | 0 | 0 | 0 |
| 0 | NA | NA | NA | NA | NA | Dcpp1 | demilune cell and parotid protein 1 | 0 | 0 | 0 | 0 | 0 | 0 | 0 | 0 | 0 | 0 | 0 |
| 0 | NA | NA | NA | NA | NA | Dcpp2 | demilune cell and parotid protein 2 | 0 | 0 | 0 | 0 | 0 | 0 | 0 | 0 | 0 | 0 | 0 |
| 0 | NA | NA | NA | NA | NA | Gm50057 | | 0 | 0 | 0 | 0 | 0 | 0 | 0 | 0 | 0 | 0 | 0 |
| 0 | NA | NA | NA | NA | NA | Dcpp3 | demilune cell and parotid protein 3 | 0 | 0 | 0 | 0 | 0 | 0 | 0 | 0 | 0 | 0 | 0 |
| 0 | NA | NA | NA | NA | NA | Sbp | spermine binding protein | 0 | 0 | 0 | 0 | 0 | 0 | 0 | 0 | 0 | 0 | 0 |
| 0 | NA | NA | NA | NA | NA | Prss22 | protease, serine 22 | 0 | 0 | 0 | 0 | 0 | 0 | 0 | 0 | 0 | 0 | 0 |
| 0 | NA | NA | NA | NA | NA | Gm50066 | | 0 | 0 | 0 | 0 | 0 | 0 | 0 | 0 | 0 | 0 | 0 |
| 0 | NA | NA | NA | NA | NA | Gm27700 | | 0 | 0 | 0 | 0 | 0 | 0 | 0 | 0 | 0 | 0 | 0 |
| 0 | NA | NA | NA | NA | NA | Gm25092 | | 0 | 0 | 0 | 0 | 0 | 0 | 0 | 0 | 0 | 0 | 0 |
| 0 | NA | NA | NA | NA | NA | Gm5225 | | 0 | 0 | 0 | 0 | 0 | 0 | 0 | 0 | 0 | 0 | 0 |
| 0 | NA | NA | NA | NA | NA | Prss27 | protease, serine 27 | 0 | 0 | 0 | 0 | 0 | 0 | 0 | 0 | 0 | 0 | 0 |
| 0 | NA | NA | NA | NA | NA | Gm50071 | | 0 | 0 | 0 | 0 | 0 | 0 | 0 | 0 | 0 | 0 | 0 |
| 0 | NA | NA | NA | NA | NA | Mir5134 | | 0 | 0 | 0 | 0 | 0 | 0 | 0 | 0 | 0 | 0 | 0 |
| 0 | NA | NA | NA | NA | NA | Mir6965 | | 0 | 0 | 0 | 0 | 0 | 0 | 0 | 0 | 0 | 0 | 0 |
| 0 | NA | NA | NA | NA | NA | Gm24427 | | 0 | 0 | 0 | 0 | 0 | 0 | 0 | 0 | 0 | 0 | 0 |
| 0 | NA | NA | NA | NA | NA | Gm25618 | | 0 | 0 | 0 | 0 | 0 | 0 | 0 | 0 | 0 | 0 | 0 |
| 0 | NA | NA | NA | NA | NA | Gm50062 | | 0 | 0 | 0 | 0 | 0 | 0 | 0 | 0 | 0 | 0 | 0 |
| 0 | NA | NA | NA | NA | NA | Snord60 | | 0 | 0 | 0 | 0 | 0 | 0 | 0 | 0 | 0 | 0 | 0 |
| 0 | NA | NA | NA | NA | NA | Gm33508 | | 0 | 0 | 0 | 0 | 0 | 0 | 0 | 0 | 0 | 0 | 0 |
| 0 | NA | NA | NA | NA | NA | Gm24589 | | 0 | 0 | 0 | 0 | 0 | 0 | 0 | 0 | 0 | 0 | 0 |
| 0 | NA | NA | NA | NA | NA | Snora64 | | 0 | 0 | 0 | 0 | 0 | 0 | 0 | 0 | 0 | 0 | 0 |
| 0 | NA | NA | NA | NA | NA | Rpl3l | ribosomal protein L3-like | 0 | 0 | 0 | 0 | 0 | 0 | 0 | 0 | 0 | 0 | 0 |
| 0 | NA | NA | NA | NA | NA | Ptx4 | pentraxin 4 | 0 | 0 | 0 | 0 | 0 | 0 | 0 | 0 | 0 | 0 | 0 |
| 0 | NA | NA | NA | NA | NA | Ccdc154 | coiled-coil domain containing 154 | 0 | 0 | 0 | 0 | 0 | 0 | 0 | 0 | 0 | 0 | 0 |
| 0 | NA | NA | NA | NA | NA | Mir3547 | | 0 | 0 | 0 | 0 | 0 | 0 | 0 | 0 | 0 | 0 | 0 |
| 0 | NA | NA | NA | NA | NA | Gm25921 | | 0 | 0 | 0 | 0 | 0 | 0 | 0 | 0 | 0 | 0 | 0 |
| 0 | NA | NA | NA | NA | NA | Prss28 | protease, serine 28 | 0 | 0 | 0 | 0 | 0 | 0 | 0 | 0 | 0 | 0 | 0 |
| 0 | NA | NA | NA | NA | NA | Prss29 | protease, serine 29 | 0 | 0 | 0 | 0 | 0 | 0 | 0 | 0 | 0 | 0 | 0 |
| 0 | NA | NA | NA | NA | NA | Gm50085 | | 0 | 0 | 0 | 0 | 0 | 0 | 0 | 0 | 0 | 0 | 0 |
| 0 | NA | NA | NA | NA | NA | Tpsg1 | tryptase gamma 1 | 0 | 0 | 0 | 0 | 0 | 0 | 0 | 0 | 0 | 0 | 0 |
| 0 | NA | NA | NA | NA | NA | Gm33727 | | 0 | 0 | 0 | 0 | 0 | 0 | 0 | 0 | 0 | 0 | 0 |
| 0 | NA | NA | NA | NA | NA | Gm50025 | | 0 | 0 | 0 | 0 | 0 | 0 | 0 | 0 | 0 | 0 | 0 |
| 0 | NA | NA | NA | NA | NA | Rpsa-ps6 | | 0 | 0 | 0 | 0 | 0 | 0 | 0 | 0 | 0 | 0 | 0 |
| 0 | NA | NA | NA | NA | NA | Gm50026 | | 0 | 0 | 0 | 0 | 0 | 0 | 0 | 0 | 0 | 0 | 0 |
| 0 | NA | NA | NA | NA | NA | Gm22645 | | 0 | 0 | 0 | 0 | 0 | 0 | 0 | 0 | 0 | 0 | 0 |
| 0 | NA | NA | NA | NA | NA | 4930541O19Rik | | 0 | 0 | 0 | 0 | 0 | 0 | 0 | 0 | 0 | 0 | 0 |
| 0 | NA | NA | NA | NA | NA | Gm50312 | | 0 | 0 | 0 | 0 | 0 | 0 | 0 | 0 | 0 | 0 | 0 |
| 0 | NA | NA | NA | NA | NA | Gm5836 | | 0 | 0 | 0 | 0 | 0 | 0 | 0 | 0 | 0 | 0 | 0 |
| 0 | NA | NA | NA | NA | NA | Gm25083 | | 0 | 0 | 0 | 0 | 0 | 0 | 0 | 0 | 0 | 0 | 0 |

| | | | | | | | | | | | | | | | | | |
|---|---|---|---|---|---|---|---|---|---|---|---|---|---|---|---|---|---|
| 0 | NA | NA | NA | NA | NA | 1700080C20Rik | | 0 | 0 | 0 | 0 | 0 | 0 | 0 | 0 | 0 | 0 | 0 |
| 0 | NA | NA | NA | NA | NA | Mir6966 | | 0 | 0 | 0 | 0 | 0 | 0 | 0 | 0 | 0 | 0 | 0 |
| 0 | NA | NA | NA | NA | NA | Gm50367 | | 0 | 0 | 0 | 0 | 0 | 0 | 0 | 0 | 0 | 0 | 0 |
| 0 | NA | NA | NA | NA | NA | Mir3082 | | 0 | 0 | 0 | 0 | 0 | 0 | 0 | 0 | 0 | 0 | 0 |
| 0 | NA | NA | NA | NA | NA | Nhlrc4 | NHL repeat containing 4 | 0 | 0 | 0 | 0 | 0 | 0 | 0 | 0 | 0 | 0 | 0 |
| 0 | NA | NA | NA | NA | NA | A930017K11Ril | RIKEN cDNA A930017K11 gene | 0 | 0 | 0 | 0 | 0 | 0 | 0 | 0 | 0 | 0 | 0 |
| 0 | NA | NA | NA | NA | NA | D630044L22Rik | | 0 | 0 | 0 | 0 | 0 | 0 | 0 | 0 | 0 | 0 | 0 |
| 0 | NA | NA | NA | NA | NA | Gm17814 | | 0 | 0 | 0 | 0 | 0 | 0 | 0 | 0 | 0 | 0 | 0 |
| 0 | NA | NA | NA | NA | NA | Gm49983 | | 0 | 0 | 0 | 0 | 0 | 0 | 0 | 0 | 0 | 0 | 0 |
| 0 | NA | NA | NA | NA | NA | Gm23123 | | 0 | 0 | 0 | 0 | 0 | 0 | 0 | 0 | 0 | 0 | 0 |
| 0 | NA | NA | NA | NA | NA | Gm50007 | | 0 | 0 | 0 | 0 | 0 | 0 | 0 | 0 | 0 | 0 | 0 |
| 0 | NA | NA | NA | NA | NA | Gm50013 | | 0 | 0 | 0 | 0 | 0 | 0 | 0 | 0 | 0 | 0 | 0 |
| 0 | NA | NA | NA | NA | NA | Hba-ps4 | | 0 | 0 | 0 | 0 | 0 | 0 | 0 | 0 | 0 | 0 | 0 |
| 0 | NA | NA | NA | NA | NA | Gm50274 | | 0 | 0 | 0 | 0 | 0 | 0 | 0 | 0 | 0 | 0 | 0 |
| 0 | NA | NA | NA | NA | NA | Gm22420 | | 0 | 0 | 0 | 0 | 0 | 0 | 0 | 0 | 0 | 0 | 0 |
| 0 | NA | NA | NA | NA | NA | Gm17218 | | 0 | 0 | 0 | 0 | 0 | 0 | 0 | 0 | 0 | 0 | 0 |
| 0 | NA | NA | NA | NA | NA | Gm41555 | | 0 | 0 | 0 | 0 | 0 | 0 | 0 | 0 | 0 | 0 | 0 |
| 0 | NA | NA | NA | NA | NA | Gm27775 | | 0 | 0 | 0 | 0 | 0 | 0 | 0 | 0 | 0 | 0 | 0 |
| 0 | NA | NA | NA | NA | NA | Mir3083 | | 0 | 0 | 0 | 0 | 0 | 0 | 0 | 0 | 0 | 0 | 0 |
| 0 | NA | NA | NA | NA | NA | Mir7677 | | 0 | 0 | 0 | 0 | 0 | 0 | 0 | 0 | 0 | 0 | 0 |
| 0 | NA | NA | NA | NA | NA | | | 0 | 0 | 0 | 0 | 0 | 0 | 0 | 0 | 0 | 0 | 0 |
| 0 | NA | NA | NA | NA | NA | Gm50328 | | 0 | 0 | 0 | 0 | 0 | 0 | 0 | 0 | 0 | 0 | 0 |
| 0 | NA | NA | NA | NA | NA | Gm49793 | | 0 | 0 | 0 | 0 | 0 | 0 | 0 | 0 | 0 | 0 | 0 |
| 0 | NA | NA | NA | NA | NA | Mir7214 | | 0 | 0 | 0 | 0 | 0 | 0 | 0 | 0 | 0 | 0 | 0 |
| 0 | NA | NA | NA | NA | NA | Gm10505 | | 0 | 0 | 0 | 0 | 0 | 0 | 0 | 0 | 0 | 0 | 0 |
| 0 | NA | NA | NA | NA | NA | Mir7216 | | 0 | 0 | 0 | 0 | 0 | 0 | 0 | 0 | 0 | 0 | 0 |
| 0 | NA | NA | NA | NA | NA | Gm49794 | | 0 | 0 | 0 | 0 | 0 | 0 | 0 | 0 | 0 | 0 | 0 |
| 0 | NA | NA | NA | NA | NA | Mir7217 | | 0 | 0 | 0 | 0 | 0 | 0 | 0 | 0 | 0 | 0 | 0 |
| 0 | NA | NA | NA | NA | NA | Mir7218 | | 0 | 0 | 0 | 0 | 0 | 0 | 0 | 0 | 0 | 0 | 0 |
| 0 | NA | NA | NA | NA | NA | Gm49796 | | 0 | 0 | 0 | 0 | 0 | 0 | 0 | 0 | 0 | 0 | 0 |
| 0 | NA | NA | NA | NA | NA | Gm49789 | | 0 | 0 | 0 | 0 | 0 | 0 | 0 | 0 | 0 | 0 | 0 |
| 0 | NA | NA | NA | NA | NA | Gm49790 | | 0 | 0 | 0 | 0 | 0 | 0 | 0 | 0 | 0 | 0 | 0 |
| 0 | NA | NA | NA | NA | NA | Gm49773 | | 0 | 0 | 0 | 0 | 0 | 0 | 0 | 0 | 0 | 0 | 0 |
| 0 | NA | NA | NA | NA | NA | Gm49804 | | 0 | 0 | 0 | 0 | 0 | 0 | 0 | 0 | 0 | 0 | 0 |
| 0 | NA | NA | NA | NA | NA | Gm15458 | | 0 | 0 | 0 | 0 | 0 | 0 | 0 | 0 | 0 | 0 | 0 |
| 0 | NA | NA | NA | NA | NA | Gm49803 | | 0 | 0 | 0 | 0 | 0 | 0 | 0 | 0 | 0 | 0 | 0 |
| 0 | NA | NA | NA | NA | NA | Gm35290 | | 0 | 0 | 0 | 0 | 0 | 0 | 0 | 0 | 0 | 0 | 0 |
| 0 | NA | NA | NA | NA | NA | Gm50238 | | 0 | 0 | 0 | 0 | 0 | 0 | 0 | 0 | 0 | 0 | 0 |
| 0 | NA | NA | NA | NA | NA | Gm49897 | | 0 | 0 | 0 | 0 | 0 | 0 | 0 | 0 | 0 | 0 | 0 |
| 0 | NA | NA | NA | NA | NA | Gm49886 | | 0 | 0 | 0 | 0 | 0 | 0 | 0 | 0 | 0 | 0 | 0 |
| 0 | NA | NA | NA | NA | NA | Gm49799 | | 0 | 0 | 0 | 0 | 0 | 0 | 0 | 0 | 0 | 0 | 0 |
| 0 | NA | NA | NA | NA | NA | Gm49861 | | 0 | 0 | 0 | 0 | 0 | 0 | 0 | 0 | 0 | 0 | 0 |
| 0 | NA | NA | NA | NA | NA | Gm25195 | | 0 | 0 | 0 | 0 | 0 | 0 | 0 | 0 | 0 | 0 | 0 |
| 0 | NA | NA | NA | NA | NA | Gm41506 | | 0 | 0 | 0 | 0 | 0 | 0 | 0 | 0 | 0 | 0 | 0 |
| 0 | NA | NA | NA | NA | NA | Clpsl2 | colipase-like 2 | 0 | 0 | 0 | 0 | 0 | 0 | 0 | 0 | 0 | 0 | 0 |
| 0 | NA | NA | NA | NA | NA | Mir6969 | | 0 | 0 | 0 | 0 | 0 | 0 | 0 | 0 | 0 | 0 | 0 |
| 0 | NA | NA | NA | NA | NA | Gm35883 | | 0 | 0 | 0 | 0 | 0 | 0 | 0 | 0 | 0 | 0 | 0 |
| 0 | NA | NA | NA | NA | NA | 1700030A11Rik | | 0 | 0 | 0 | 0 | 0 | 0 | 0 | 0 | 0 | 0 | 0 |
| 0 | NA | NA | NA | NA | NA | Gm49868 | | 0 | 0 | 0 | 0 | 0 | 0 | 0 | 0 | 0 | 0 | 0 |
| 0 | NA | NA | NA | NA | NA | Gm16195 | | 0 | 0 | 0 | 0 | 0 | 0 | 0 | 0 | 0 | 0 | 0 |
| 0 | NA | NA | NA | NA | NA | Gm27599 | | 0 | 0 | 0 | 0 | 0 | 0 | 0 | 0 | 0 | 0 | 0 |
| 0 | NA | NA | NA | NA | NA | Rab44 | RAB44, member RAS oncogene family | 0 | 0 | 0 | 0 | 0 | 0 | 0 | 0 | 0 | 0 | 0 |
| 0 | NA | NA | NA | NA | NA | Gm18271 | | 0 | 0 | 0 | 0 | 0 | 0 | 0 | 0 | 0 | 0 | 0 |
| 0 | NA | NA | NA | NA | NA | Gm50054 | | 0 | 0 | 0 | 0 | 0 | 0 | 0 | 0 | 0 | 0 | 0 |
| 0 | NA | NA | NA | NA | NA | Gm10045 | | 0 | 0 | 0 | 0 | 0 | 0 | 0 | 0 | 0 | 0 | 0 |
| 0 | NA | NA | NA | NA | NA | Gm36199 | | 0 | 0 | 0 | 0 | 0 | 0 | 0 | 0 | 0 | 0 | 0 |
| 0 | NA | NA | NA | NA | NA | Gm17657 | transmembrane protein 217 | 0 | 0 | 0 | 0 | 0 | 0 | 0 | 0 | 0 | 0 | 0 |
| 0 | NA | NA | NA | NA | NA | Gm28052 | | 0 | 0 | 0 | 0 | 0 | 0 | 0 | 0 | 0 | 0 | 0 |
| 0 | NA | NA | NA | NA | NA | Tbc1d22bos | | 0 | 0 | 0 | 0 | 0 | 0 | 0 | 0 | 0 | 0 | 0 |
| 0 | NA | NA | NA | NA | NA | Mir7667 | | 0 | 0 | 0 | 0 | 0 | 0 | 0 | 0 | 0 | 0 | 0 |

| | | | | | | | | | | | | | | | | | |
|---|---|---|---|---|---|---|---|---|---|---|---|---|---|---|---|---|---|
| 0 | NA | NA | NA | NA | NA | Gm20161 | | 0 | 0 | 0 | 0 | 0 | 0 | 0 | 0 | 0 | 0 | 0 |
| 0 | NA | NA | NA | NA | NA | Gm50016 | | 0 | 0 | 0 | 0 | 0 | 0 | 0 | 0 | 0 | 0 | 0 |
| 0 | NA | NA | NA | NA | NA | Gm50017 | | 0 | 0 | 0 | 0 | 0 | 0 | 0 | 0 | 0 | 0 | 0 |
| 0 | NA | NA | NA | NA | NA | Gm50018 | | 0 | 0 | 0 | 0 | 0 | 0 | 0 | 0 | 0 | 0 | 0 |
| 0 | NA | NA | NA | NA | NA | Gm22327 | | 0 | 0 | 0 | 0 | 0 | 0 | 0 | 0 | 0 | 0 | 0 |
| 0 | NA | NA | NA | NA | NA | Gm25049 | | 0 | 0 | 0 | 0 | 0 | 0 | 0 | 0 | 0 | 0 | 0 |
| 0 | NA | NA | NA | NA | NA | Gm50250 | | 0 | 0 | 0 | 0 | 0 | 0 | 0 | 0 | 0 | 0 | 0 |
| 0 | NA | NA | NA | NA | NA | Gm50252 | | 0 | 0 | 0 | 0 | 0 | 0 | 0 | 0 | 0 | 0 | 0 |
| 0 | NA | NA | NA | NA | NA | Gm9874 | | 0 | 0 | 0 | 0 | 0 | 0 | 0 | 0 | 0 | 0 | 0 |
| 0 | NA | NA | NA | NA | NA | Gm50245 | | 0 | 0 | 0 | 0 | 0 | 0 | 0 | 0 | 0 | 0 | 0 |
| 0 | NA | NA | NA | NA | NA | Gm24661 | | 0 | 0 | 0 | 0 | 0 | 0 | 0 | 0 | 0 | 0 | 0 |
| 0 | NA | NA | NA | NA | NA | Gm41561 | | 0 | 0 | 0 | 0 | 0 | 0 | 0 | 0 | 0 | 0 | 0 |
| 0 | NA | NA | NA | NA | NA | Gm10503 | | 0 | 0 | 0 | 0 | 0 | 0 | 0 | 0 | 0 | 0 | 0 |
| 0 | NA | NA | NA | NA | NA | Gm24026 | | 0 | 0 | 0 | 0 | 0 | 0 | 0 | 0 | 0 | 0 | 0 |
| 0 | NA | NA | NA | NA | NA | Gm50249 | | 0 | 0 | 0 | 0 | 0 | 0 | 0 | 0 | 0 | 0 | 0 |
| 0 | NA | NA | NA | NA | NA | Gm25447 | | 0 | 0 | 0 | 0 | 0 | 0 | 0 | 0 | 0 | 0 | 0 |
| 0 | NA | NA | NA | NA | NA | Tff1 | trefoil factor 1 | 0 | 0 | 0 | 0 | 0 | 0 | 0 | 0 | 0 | 0 | 0 |
| 0 | NA | NA | NA | NA | NA | Gm50138 | | 0 | 0 | 0 | 0 | 0 | 0 | 0 | 0 | 0 | 0 | 0 |
| 0 | NA | NA | NA | NA | NA | Gm50102 | | 0 | 0 | 0 | 0 | 0 | 0 | 0 | 0 | 0 | 0 | 0 |
| 0 | NA | NA | NA | NA | NA | Gm50108 | | 0 | 0 | 0 | 0 | 0 | 0 | 0 | 0 | 0 | 0 | 0 |
| 0 | NA | NA | NA | NA | NA | Gm24970 | | 0 | 0 | 0 | 0 | 0 | 0 | 0 | 0 | 0 | 0 | 0 |
| 0 | NA | NA | NA | NA | NA | Gm50225 | | 0 | 0 | 0 | 0 | 0 | 0 | 0 | 0 | 0 | 0 | 0 |
| 0 | NA | NA | NA | NA | NA | Gm49999 | | 0 | 0 | 0 | 0 | 0 | 0 | 0 | 0 | 0 | 0 | 0 |
| 0 | NA | NA | NA | NA | NA | Gm50000 | | 0 | 0 | 0 | 0 | 0 | 0 | 0 | 0 | 0 | 0 | 0 |
| 0 | NA | NA | NA | NA | NA | Gm17572 | | 0 | 0 | 0 | 0 | 0 | 0 | 0 | 0 | 0 | 0 | 0 |
| 0 | NA | NA | NA | NA | NA | Gm25883 | | 0 | 0 | 0 | 0 | 0 | 0 | 0 | 0 | 0 | 0 | 0 |
| 0 | NA | NA | NA | NA | NA | Gm4432 | | 0 | 0 | 0 | 0 | 0 | 0 | 0 | 0 | 0 | 0 | 0 |
| 0 | NA | NA | NA | NA | NA | Gm22201 | | 0 | 0 | 0 | 0 | 0 | 0 | 0 | 0 | 0 | 0 | 0 |
| 0 | NA | NA | NA | NA | NA | Gm50036 | | 0 | 0 | 0 | 0 | 0 | 0 | 0 | 0 | 0 | 0 | 0 |
| 0 | NA | NA | NA | NA | NA | Gm50038 | | 0 | 0 | 0 | 0 | 0 | 0 | 0 | 0 | 0 | 0 | 0 |
| 0 | NA | NA | NA | NA | NA | Cyp4f38-ps | | 0 | 0 | 0 | 0 | 0 | 0 | 0 | 0 | 0 | 0 | 0 |
| 0 | NA | NA | NA | NA | NA | Gm50046 | | 0 | 0 | 0 | 0 | 0 | 0 | 0 | 0 | 0 | 0 | 0 |
| 0 | NA | NA | NA | NA | NA | Gm6452 | | 0 | 0 | 0 | 0 | 0 | 0 | 0 | 0 | 0 | 0 | 0 |
| 0 | NA | NA | NA | NA | NA | Gm17948 | | 0 | 0 | 0 | 0 | 0 | 0 | 0 | 0 | 0 | 0 | 0 |
| 0 | NA | NA | NA | NA | NA | Gm25555 | | 0 | 0 | 0 | 0 | 0 | 0 | 0 | 0 | 0 | 0 | 0 |
| 0 | NA | NA | NA | NA | NA | Gm26693 | | 0 | 0 | 0 | 0 | 0 | 0 | 0 | 0 | 0 | 0 | 0 |
| 0 | NA | NA | NA | NA | NA | Gm18272 | | 0 | 0 | 0 | 0 | 0 | 0 | 0 | 0 | 0 | 0 | 0 |
| 0 | NA | NA | NA | NA | NA | Gm18273 | | 0 | 0 | 0 | 0 | 0 | 0 | 0 | 0 | 0 | 0 | 0 |
| 0 | NA | NA | NA | NA | NA | Gm50051 | | 0 | 0 | 0 | 0 | 0 | 0 | 0 | 0 | 0 | 0 | 0 |
| 0 | NA | NA | NA | NA | NA | Phf8-ps | | 0 | 0 | 0 | 0 | 0 | 0 | 0 | 0 | 0 | 0 | 0 |
| 0 | NA | NA | NA | NA | NA | Morc2b | microrchidia 2B | 0 | 0 | 0 | 0 | 0 | 0 | 0 | 0 | 0 | 0 | 0 |
| 0 | NA | NA | NA | NA | NA | Olfr55 | olfactory receptor 55 | 0 | 0 | 0 | 0 | 0 | 0 | 0 | 0 | 0 | 0 | 0 |
| 0 | NA | NA | NA | NA | NA | Olfr239 | olfactory receptor 239 | 0 | 0 | 0 | 0 | 0 | 0 | 0 | 0 | 0 | 0 | 0 |
| 0 | NA | NA | NA | NA | NA | Olfr1564 | olfactory receptor 1564 | 0 | 0 | 0 | 0 | 0 | 0 | 0 | 0 | 0 | 0 | 0 |
| 0 | NA | NA | NA | NA | NA | Olfr63 | olfactory receptor 63 | 0 | 0 | 0 | 0 | 0 | 0 | 0 | 0 | 0 | 0 | 0 |
| 0 | NA | NA | NA | NA | NA | Actl9 | actin-like 9 | 0 | 0 | 0 | 0 | 0 | 0 | 0 | 0 | 0 | 0 | 0 |
| 0 | NA | NA | NA | NA | NA | Gm4125 | | 0 | 0 | 0 | 0 | 0 | 0 | 0 | 0 | 0 | 0 | 0 |
| 0 | NA | NA | NA | NA | NA | Gm20507 | | 0 | 0 | 0 | 0 | 0 | 0 | 0 | 0 | 0 | 0 | 0 |
| 0 | NA | NA | NA | NA | NA | Gm22943 | | 0 | 0 | 0 | 0 | 0 | 0 | 0 | 0 | 0 | 0 | 0 |
| 0 | NA | NA | NA | NA | NA | 4931413I07Rik | | 0 | 0 | 0 | 0 | 0 | 0 | 0 | 0 | 0 | 0 | 0 |
| 0 | NA | NA | NA | NA | NA | Smim40-ps | | 0 | 0 | 0 | 0 | 0 | 0 | 0 | 0 | 0 | 0 | 0 |
| 0 | NA | NA | NA | NA | NA | Gm27740 | | 0 | 0 | 0 | 0 | 0 | 0 | 0 | 0 | 0 | 0 | 0 |
| 0 | NA | NA | NA | NA | NA | Gm26940 | | 0 | 0 | 0 | 0 | 0 | 0 | 0 | 0 | 0 | 0 | 0 |
| 0 | NA | NA | NA | NA | NA | Mir219a-1 | | 0 | 0 | 0 | 0 | 0 | 0 | 0 | 0 | 0 | 0 | 0 |
| 0 | NA | NA | NA | NA | NA | Mir219c | | 0 | 0 | 0 | 0 | 0 | 0 | 0 | 0 | 0 | 0 | 0 |
| 0 | NA | NA | NA | NA | NA | H2-Pb | | 0 | 0 | 0 | 0 | 0 | 0 | 0 | 0 | 0 | 0 | 0 |
| 0 | NA | NA | NA | NA | NA | BC051537 | | 0 | 0 | 0 | 0 | 0 | 0 | 0 | 0 | 0 | 0 | 0 |
| 0 | NA | NA | NA | NA | NA | H2-Pa | | 0 | 0 | 0 | 0 | 0 | 0 | 0 | 0 | 0 | 0 | 0 |
| 0 | NA | NA | NA | NA | NA | Gm50333 | | 0 | 0 | 0 | 0 | 0 | 0 | 0 | 0 | 0 | 0 | 0 |
| 0 | NA | NA | NA | NA | NA | Gm50335 | | 0 | 0 | 0 | 0 | 0 | 0 | 0 | 0 | 0 | 0 | 0 |

| | | | | | | | | | | | | | | | | | | |
|---|---|---|---|---|---|---|---|---|---|---|---|---|---|---|---|---|---|---|
| 0 | NA | NA | NA | NA | NA | Gm20493 | | 0 | 0 | 0 | 0 | 0 | 0 | 0 | 0 | 0 | 0 | 0 |
| 0 | NA | NA | NA | NA | NA | Gm15332 | | 0 | 0 | 0 | 0 | 0 | 0 | 0 | 0 | 0 | 0 | 0 |
| 0 | NA | NA | NA | NA | NA | Gm15331 | | 0 | 0 | 0 | 0 | 0 | 0 | 0 | 0 | 0 | 0 | 0 |
| 0 | NA | NA | NA | NA | NA | Btnl6 | butyrophilin-like 6 | 0 | 0 | 0 | 0 | 0 | 0 | 0 | 0 | 0 | 0 | 0 |
| 0 | NA | NA | NA | NA | NA | Gm15329 | | 0 | 0 | 0 | 0 | 0 | 0 | 0 | 0 | 0 | 0 | 0 |
| 0 | NA | NA | NA | NA | NA | Gm20462 | | 0 | 0 | 0 | 0 | 0 | 0 | 0 | 0 | 0 | 0 | 0 |
| 0 | NA | NA | NA | NA | NA | Gm20460 | | 0 | 0 | 0 | 0 | 0 | 0 | 0 | 0 | 0 | 0 | 0 |
| 0 | NA | NA | NA | NA | NA | Gm20461 | | 0 | 0 | 0 | 0 | 0 | 0 | 0 | 0 | 0 | 0 | 0 |
| 0 | NA | NA | NA | NA | NA | Gm24389 | | 0 | 0 | 0 | 0 | 0 | 0 | 0 | 0 | 0 | 0 | 0 |
| 0 | NA | NA | NA | NA | NA | Cyp21a2-ps | | 0 | 0 | 0 | 0 | 0 | 0 | 0 | 0 | 0 | 0 | 0 |
| 0 | NA | NA | NA | NA | NA | Stk19-ps1 | | 0 | 0 | 0 | 0 | 0 | 0 | 0 | 0 | 0 | 0 | 0 |
| 0 | NA | NA | NA | NA | NA | Gm27471 | | 0 | 0 | 0 | 0 | 0 | 0 | 0 | 0 | 0 | 0 | 0 |
| 0 | NA | NA | NA | NA | NA | Tnxa | | 0 | 0 | 0 | 0 | 0 | 0 | 0 | 0 | 0 | 0 | 0 |
| 0 | NA | NA | NA | NA | NA | Cyp21a1 | cytochrome P450, family 21, subfamily a, | 0 | 0 | 0 | 0 | 0 | 0 | 0 | 0 | 0 | 0 | 0 |
| 0 | NA | NA | NA | NA | NA | Mir6971 | | 0 | 0 | 0 | 0 | 0 | 0 | 0 | 0 | 0 | 0 | 0 |
| 0 | NA | NA | NA | NA | NA | Mir6970 | | 0 | 0 | 0 | 0 | 0 | 0 | 0 | 0 | 0 | 0 | 0 |
| 0 | NA | NA | NA | NA | NA | Gm20547 | | 0 | 0 | 0 | 0 | 0 | 0 | 0 | 0 | 0 | 0 | 0 |
| 0 | NA | NA | NA | NA | NA | Mir6972 | | 0 | 0 | 0 | 0 | 0 | 0 | 0 | 0 | 0 | 0 | 0 |
| 0 | NA | NA | NA | NA | NA | Snord52 | | 0 | 0 | 0 | 0 | 0 | 0 | 0 | 0 | 0 | 0 | 0 |
| 0 | NA | NA | NA | NA | NA | Gm25744 | | 0 | 0 | 0 | 0 | 0 | 0 | 0 | 0 | 0 | 0 | 0 |
| 0 | NA | NA | NA | NA | NA | Ly6g6c | lymphocyte antigen 6 complex, locus G6( | 0 | 0 | 0 | 0 | 0 | 0 | 0 | 0 | 0 | 0 | 0 |
| 0 | NA | NA | NA | NA | NA | Ly6g5c | lymphocyte antigen 6 complex, locus G5( | 0 | 0 | 0 | 0 | 0 | 0 | 0 | 0 | 0 | 0 | 0 |
| 0 | NA | NA | NA | NA | NA | Ncr3-ps | | 0 | 0 | 0 | 0 | 0 | 0 | 0 | 0 | 0 | 0 | 0 |
| 0 | NA | NA | NA | NA | NA | Mir6973a | | 0 | 0 | 0 | 0 | 0 | 0 | 0 | 0 | 0 | 0 | 0 |
| 0 | NA | NA | NA | NA | NA | Mir6974 | | 0 | 0 | 0 | 0 | 0 | 0 | 0 | 0 | 0 | 0 | 0 |
| 0 | NA | NA | NA | NA | NA | Gm16181 | | 0 | 0 | 0 | 0 | 0 | 0 | 0 | 0 | 0 | 0 | 0 |
| 0 | NA | NA | NA | NA | NA | Gm22589 | | 0 | 0 | 0 | 0 | 0 | 0 | 0 | 0 | 0 | 0 | 0 |
| 0 | NA | NA | NA | NA | NA | Gm8750 | | 0 | 0 | 0 | 0 | 0 | 0 | 0 | 0 | 0 | 0 | 0 |
| 0 | NA | NA | NA | NA | NA | Pou5f1 | POU domain, class 5, transcription factor | 0 | 0 | 0 | 0 | 0 | 0 | 0 | 0 | 0 | 0 | 0 |
| 0 | NA | NA | NA | NA | NA | Gm19553 | | 0 | 0 | 0 | 0 | 0 | 0 | 0 | 0 | 0 | 0 | 0 |
| 0 | NA | NA | NA | NA | NA | | | 0 | 0 | 0 | 0 | 0 | 0 | 0 | 0 | 0 | 0 | 0 |
| 0 | NA | NA | NA | NA | NA | Psors1c2 | psoriasis susceptibility 1 candidate 2 (hur | 0 | 0 | 0 | 0 | 0 | 0 | 0 | 0 | 0 | 0 | 0 |
| 0 | NA | NA | NA | NA | NA | Sfta2 | surfactant associated 2 | 0 | 0 | 0 | 0 | 0 | 0 | 0 | 0 | 0 | 0 | 0 |
| 0 | NA | NA | NA | NA | NA | Gm9573 | predicted gene 9573 | 0 | 0 | 0 | 0 | 0 | 0 | 0 | 0 | 0 | 0 | 0 |
| 0 | NA | NA | NA | NA | NA | Gm20442 | | 0 | 0 | 0 | 0 | 0 | 0 | 0 | 0 | 0 | 0 | 0 |
| 0 | NA | NA | NA | NA | NA | 4833427F10Rik | | 0 | 0 | 0 | 0 | 0 | 0 | 0 | 0 | 0 | 0 | 0 |
| 0 | NA | NA | NA | NA | NA | Mir1894 | | 0 | 0 | 0 | 0 | 0 | 0 | 0 | 0 | 0 | 0 | 0 |
| 0 | NA | NA | NA | NA | NA | Mir877 | | 0 | 0 | 0 | 0 | 0 | 0 | 0 | 0 | 0 | 0 | 0 |
| 0 | NA | NA | NA | NA | NA | Gm8810 | | 0 | 0 | 0 | 0 | 0 | 0 | 0 | 0 | 0 | 0 | 0 |
| 0 | NA | NA | NA | NA | NA | Gm20530 | | 0 | 0 | 0 | 0 | 0 | 0 | 0 | 0 | 0 | 0 | 0 |
| 0 | NA | NA | NA | NA | NA | Gm8815 | | 0 | 0 | 0 | 0 | 0 | 0 | 0 | 0 | 0 | 0 | 0 |
| 0 | NA | NA | NA | NA | NA | Gm9574 | | 0 | 0 | 0 | 0 | 0 | 0 | 0 | 0 | 0 | 0 | 0 |
| 0 | NA | NA | NA | NA | NA | Gm17782 | | 0 | 0 | 0 | 0 | 0 | 0 | 0 | 0 | 0 | 0 | 0 |
| 0 | NA | NA | NA | NA | NA | Gm20495 | | 0 | 0 | 0 | 0 | 0 | 0 | 0 | 0 | 0 | 0 | 0 |
| 0 | NA | NA | NA | NA | NA | Gm20447 | | 0 | 0 | 0 | 0 | 0 | 0 | 0 | 0 | 0 | 0 | 0 |
| 0 | NA | NA | NA | NA | NA | Gm8835 | | 0 | 0 | 0 | 0 | 0 | 0 | 0 | 0 | 0 | 0 | 0 |
| 0 | NA | NA | NA | NA | NA | Gm5682 | | 0 | 0 | 0 | 0 | 0 | 0 | 0 | 0 | 0 | 0 | 0 |
| 0 | NA | NA | NA | NA | NA | Gm20392 | | 0 | 0 | 0 | 0 | 0 | 0 | 0 | 0 | 0 | 0 | 0 |
| 0 | NA | NA | NA | NA | NA | Gm29402 | | 0 | 0 | 0 | 0 | 0 | 0 | 0 | 0 | 0 | 0 | 0 |
| 0 | NA | NA | NA | NA | NA | Gm18604 | | 0 | 0 | 0 | 0 | 0 | 0 | 0 | 0 | 0 | 0 | 0 |
| 0 | NA | NA | NA | NA | NA | Gm20391 | | 0 | 0 | 0 | 0 | 0 | 0 | 0 | 0 | 0 | 0 | 0 |
| 0 | NA | NA | NA | NA | NA | Gm22453 | | 0 | 0 | 0 | 0 | 0 | 0 | 0 | 0 | 0 | 0 | 0 |
| 0 | NA | NA | NA | NA | NA | Gm4252 | | 0 | 0 | 0 | 0 | 0 | 0 | 0 | 0 | 0 | 0 | 0 |
| 0 | NA | NA | NA | NA | NA | Gm20546 | | 0 | 0 | 0 | 0 | 0 | 0 | 0 | 0 | 0 | 0 | 0 |
| 0 | NA | NA | NA | NA | NA | Gm6633 | | 0 | 0 | 0 | 0 | 0 | 0 | 0 | 0 | 0 | 0 | 0 |
| 0 | NA | NA | NA | NA | NA | H2-M10.2 | histocompatibility 2, M region locus 10.2 | 0 | 0 | 0 | 0 | 0 | 0 | 0 | 0 | 0 | 0 | 0 |
| 0 | NA | NA | NA | NA | NA | H2-M10.1 | histocompatibility 2, M region locus 10.1 | 0 | 0 | 0 | 0 | 0 | 0 | 0 | 0 | 0 | 0 | 0 |
| 0 | NA | NA | NA | NA | NA | H2-M10.3 | histocompatibility 2, M region locus 10.3 | 0 | 0 | 0 | 0 | 0 | 0 | 0 | 0 | 0 | 0 | 0 |
| 0 | NA | NA | NA | NA | NA | Stk-ps1 | | 0 | 0 | 0 | 0 | 0 | 0 | 0 | 0 | 0 | 0 | 0 |
| 0 | NA | NA | NA | NA | NA | H2-M10.5-ps1 | | 0 | 0 | 0 | 0 | 0 | 0 | 0 | 0 | 0 | 0 | 0 |

| | | | | | | | | | | | | | | | | | | |
|---|---|---|---|---|---|---|---|---|---|---|---|---|---|---|---|---|---|---|
| 0 | NA | NA | NA | NA | NA | Rpl9-ps9 | | 0 | 0 | 0 | 0 | 0 | 0 | 0 | 0 | 0 | 0 | 0 |
| 0 | NA | NA | NA | NA | NA | Obox3-ps8 | | 0 | 0 | 0 | 0 | 0 | 0 | 0 | 0 | 0 | 0 | 0 |
| 0 | NA | NA | NA | NA | NA | H2-M10.4 | histocompatibility 2, M region locus 10.4 | 0 | 0 | 0 | 0 | 0 | 0 | 0 | 0 | 0 | 0 | 0 |
| 0 | NA | NA | NA | NA | NA | Gm20452 | | 0 | 0 | 0 | 0 | 0 | 0 | 0 | 0 | 0 | 0 | 0 |
| 0 | NA | NA | NA | NA | NA | Gm20451 | | 0 | 0 | 0 | 0 | 0 | 0 | 0 | 0 | 0 | 0 | 0 |
| 0 | NA | NA | NA | NA | NA | Gm20454 | | 0 | 0 | 0 | 0 | 0 | 0 | 0 | 0 | 0 | 0 | 0 |
| 0 | NA | NA | NA | NA | NA | Gm20453 | | 0 | 0 | 0 | 0 | 0 | 0 | 0 | 0 | 0 | 0 | 0 |
| 0 | NA | NA | NA | NA | NA | Gm8868 | | 0 | 0 | 0 | 0 | 0 | 0 | 0 | 0 | 0 | 0 | 0 |
| 0 | NA | NA | NA | NA | NA | Gm20450 | | 0 | 0 | 0 | 0 | 0 | 0 | 0 | 0 | 0 | 0 | 0 |
| 0 | NA | NA | NA | NA | NA | H2-M1 | histocompatibility 2, M region locus 1 | 0 | 0 | 0 | 0 | 0 | 0 | 0 | 0 | 0 | 0 | 0 |
| 0 | NA | NA | NA | NA | NA | Gm8877 | | 0 | 0 | 0 | 0 | 0 | 0 | 0 | 0 | 0 | 0 | 0 |
| 0 | NA | NA | NA | NA | NA | Gm20549 | | 0 | 0 | 0 | 0 | 0 | 0 | 0 | 0 | 0 | 0 | 0 |
| 0 | NA | NA | NA | NA | NA | Gm20523 | | 0 | 0 | 0 | 0 | 0 | 0 | 0 | 0 | 0 | 0 | 0 |
| 0 | NA | NA | NA | NA | NA | Gm20525 | | 0 | 0 | 0 | 0 | 0 | 0 | 0 | 0 | 0 | 0 | 0 |
| 0 | NA | NA | NA | NA | NA | Gm20526 | | 0 | 0 | 0 | 0 | 0 | 0 | 0 | 0 | 0 | 0 | 0 |
| 0 | NA | NA | NA | NA | NA | Gm4271 | | 0 | 0 | 0 | 0 | 0 | 0 | 0 | 0 | 0 | 0 | 0 |
| 0 | NA | NA | NA | NA | NA | Gm20401 | | 0 | 0 | 0 | 0 | 0 | 0 | 0 | 0 | 0 | 0 | 0 |
| 0 | NA | NA | NA | NA | NA | H2-M10.5 | histocompatibility 2, M region locus 10.5 | 0 | 0 | 0 | 0 | 0 | 0 | 0 | 0 | 0 | 0 | 0 |
| 0 | NA | NA | NA | NA | NA | H2-M10.6 | histocompatibility 2, M region locus 10.6 | 0 | 0 | 0 | 0 | 0 | 0 | 0 | 0 | 0 | 0 | 0 |
| 0 | NA | NA | NA | NA | NA | Gm25973 | | 0 | 0 | 0 | 0 | 0 | 0 | 0 | 0 | 0 | 0 | 0 |
| 0 | NA | NA | NA | NA | NA | Trim15 | tripartite motif-containing 15 | 0 | 0 | 0 | 0 | 0 | 0 | 0 | 0 | 0 | 0 | 0 |
| 0 | NA | NA | NA | NA | NA | Trim10 | tripartite motif-containing 10 | 0 | 0 | 0 | 0 | 0 | 0 | 0 | 0 | 0 | 0 | 0 |
| 0 | NA | NA | NA | NA | NA | Trim40 | tripartite motif-containing 40 | 0 | 0 | 0 | 0 | 0 | 0 | 0 | 0 | 0 | 0 | 0 |
| 0 | NA | NA | NA | NA | NA | 1700031A10Rik | | 0 | 0 | 0 | 0 | 0 | 0 | 0 | 0 | 0 | 0 | 0 |
| 0 | NA | NA | NA | NA | NA | Gm27735 | | 0 | 0 | 0 | 0 | 0 | 0 | 0 | 0 | 0 | 0 | 0 |
| 0 | NA | NA | NA | NA | NA | H2-M6-ps | | 0 | 0 | 0 | 0 | 0 | 0 | 0 | 0 | 0 | 0 | 0 |
| 0 | NA | NA | NA | NA | NA | 2410137M14Ri | RIKEN cDNA 2410137M14 gene | 0 | 0 | 0 | 0 | 0 | 0 | 0 | 0 | 0 | 0 | 0 |
| 0 | NA | NA | NA | NA | NA | Mog | myelin oligodendrocyte glycoprotein | 0 | 0 | 0 | 0 | 0 | 0 | 0 | 0 | 0 | 0 | 0 |
| 0 | NA | NA | NA | NA | NA | Gm22388 | | 0 | 0 | 0 | 0 | 0 | 0 | 0 | 0 | 0 | 0 | 0 |
| 0 | NA | NA | NA | NA | NA | Olfr90 | olfactory receptor 90 | 0 | 0 | 0 | 0 | 0 | 0 | 0 | 0 | 0 | 0 | 0 |
| 0 | NA | NA | NA | NA | NA | Olfr91 | olfactory receptor 91 | 0 | 0 | 0 | 0 | 0 | 0 | 0 | 0 | 0 | 0 | 0 |
| 0 | NA | NA | NA | NA | NA | Olfr92 | olfactory receptor 92 | 0 | 0 | 0 | 0 | 0 | 0 | 0 | 0 | 0 | 0 | 0 |
| 0 | NA | NA | NA | NA | NA | Gm4831 | | 0 | 0 | 0 | 0 | 0 | 0 | 0 | 0 | 0 | 0 | 0 |
| 0 | NA | NA | NA | NA | NA | Olfr93 | olfactory receptor 93 | 0 | 0 | 0 | 0 | 0 | 0 | 0 | 0 | 0 | 0 | 0 |
| 0 | NA | NA | NA | NA | NA | Vhl-ps1 | | 0 | 0 | 0 | 0 | 0 | 0 | 0 | 0 | 0 | 0 | 0 |
| 0 | NA | NA | NA | NA | NA | Olfr753-ps1 | | 0 | 0 | 0 | 0 | 0 | 0 | 0 | 0 | 0 | 0 | 0 |
| 0 | NA | NA | NA | NA | NA | Gm7031 | | 0 | 0 | 0 | 0 | 0 | 0 | 0 | 0 | 0 | 0 | 0 |
| 0 | NA | NA | NA | NA | NA | Olfr94 | olfactory receptor 94 | 0 | 0 | 0 | 0 | 0 | 0 | 0 | 0 | 0 | 0 | 0 |
| 0 | NA | NA | NA | NA | NA | Olfr95 | olfactory receptor 95 | 0 | 0 | 0 | 0 | 0 | 0 | 0 | 0 | 0 | 0 | 0 |
| 0 | NA | NA | NA | NA | NA | Olfr96 | olfactory receptor 96 | 0 | 0 | 0 | 0 | 0 | 0 | 0 | 0 | 0 | 0 | 0 |
| 0 | NA | NA | NA | NA | NA | Olfr97 | olfactory receptor 97 | 0 | 0 | 0 | 0 | 0 | 0 | 0 | 0 | 0 | 0 | 0 |
| 0 | NA | NA | NA | NA | NA | Gm20429 | | 0 | 0 | 0 | 0 | 0 | 0 | 0 | 0 | 0 | 0 | 0 |
| 0 | NA | NA | NA | NA | NA | Olfr754-ps1 | | 0 | 0 | 0 | 0 | 0 | 0 | 0 | 0 | 0 | 0 | 0 |
| 0 | NA | NA | NA | NA | NA | Olfr98 | olfactory receptor 98 | 0 | 0 | 0 | 0 | 0 | 0 | 0 | 0 | 0 | 0 | 0 |
| 0 | NA | NA | NA | NA | NA | Gm25104 | | 0 | 0 | 0 | 0 | 0 | 0 | 0 | 0 | 0 | 0 | 0 |
| 0 | NA | NA | NA | NA | NA | Olfr99 | olfactory receptor 99 | 0 | 0 | 0 | 0 | 0 | 0 | 0 | 0 | 0 | 0 | 0 |
| 0 | NA | NA | NA | NA | NA | Olfr756-ps1 | | 0 | 0 | 0 | 0 | 0 | 0 | 0 | 0 | 0 | 0 | 0 |
| 0 | NA | NA | NA | NA | NA | Olfr101 | olfactory receptor 101 | 0 | 0 | 0 | 0 | 0 | 0 | 0 | 0 | 0 | 0 | 0 |
| 0 | NA | NA | NA | NA | NA | Olfr102 | olfactory receptor 102 | 0 | 0 | 0 | 0 | 0 | 0 | 0 | 0 | 0 | 0 | 0 |
| 0 | NA | NA | NA | NA | NA | Olfr103 | olfactory receptor 103 | 0 | 0 | 0 | 0 | 0 | 0 | 0 | 0 | 0 | 0 | 0 |
| 0 | NA | NA | NA | NA | NA | Olfr104-ps | | 0 | 0 | 0 | 0 | 0 | 0 | 0 | 0 | 0 | 0 | 0 |
| 0 | NA | NA | NA | NA | NA | Olfr757-ps1 | | 0 | 0 | 0 | 0 | 0 | 0 | 0 | 0 | 0 | 0 | 0 |
| 0 | NA | NA | NA | NA | NA | Olfr105-ps | | 0 | 0 | 0 | 0 | 0 | 0 | 0 | 0 | 0 | 0 | 0 |
| 0 | NA | NA | NA | NA | NA | Olfr758-ps1 | | 0 | 0 | 0 | 0 | 0 | 0 | 0 | 0 | 0 | 0 | 0 |
| 0 | NA | NA | NA | NA | NA | Olfr106-ps | | 0 | 0 | 0 | 0 | 0 | 0 | 0 | 0 | 0 | 0 | 0 |
| 0 | NA | NA | NA | NA | NA | Olfr107 | olfactory receptor 107 | 0 | 0 | 0 | 0 | 0 | 0 | 0 | 0 | 0 | 0 | 0 |
| 0 | NA | NA | NA | NA | NA | Olfr759-ps1 | | 0 | 0 | 0 | 0 | 0 | 0 | 0 | 0 | 0 | 0 | 0 |
| 0 | NA | NA | NA | NA | NA | Gm18808 | | 0 | 0 | 0 | 0 | 0 | 0 | 0 | 0 | 0 | 0 | 0 |
| 0 | NA | NA | NA | NA | NA | Olfr108 | olfactory receptor 108 | 0 | 0 | 0 | 0 | 0 | 0 | 0 | 0 | 0 | 0 | 0 |
| 0 | NA | NA | NA | NA | NA | Olfr760-ps1 | | 0 | 0 | 0 | 0 | 0 | 0 | 0 | 0 | 0 | 0 | 0 |

| | | | | | | | | | | | | | | | | | | |
|---|---|---|---|---|---|---|---|---|---|---|---|---|---|---|---|---|---|---|
| 0 | NA | NA | NA | NA | NA | Olfr109 | olfactory receptor 109 | 0 | 0 | 0 | 0 | 0 | 0 | 0 | 0 | 0 | 0 | 0 |
| 0 | NA | NA | NA | NA | NA | Olfr110 | olfactory receptor 110 | 0 | 0 | 0 | 0 | 0 | 0 | 0 | 0 | 0 | 0 | 0 |
| 0 | NA | NA | NA | NA | NA | Olfr112 | olfactory receptor 112 | 0 | 0 | 0 | 0 | 0 | 0 | 0 | 0 | 0 | 0 | 0 |
| 0 | NA | NA | NA | NA | NA | Olfr113 | olfactory receptor 113 | 0 | 0 | 0 | 0 | 0 | 0 | 0 | 0 | 0 | 0 | 0 |
| 0 | NA | NA | NA | NA | NA | Olfr114 | olfactory receptor 114 | 0 | 0 | 0 | 0 | 0 | 0 | 0 | 0 | 0 | 0 | 0 |
| 0 | NA | NA | NA | NA | NA | Gm18213 | | 0 | 0 | 0 | 0 | 0 | 0 | 0 | 0 | 0 | 0 | 0 |
| 0 | NA | NA | NA | NA | NA | Olfr115 | olfactory receptor 115 | 0 | 0 | 0 | 0 | 0 | 0 | 0 | 0 | 0 | 0 | 0 |
| 0 | NA | NA | NA | NA | NA | Olfr116 | olfactory receptor 116 | 0 | 0 | 0 | 0 | 0 | 0 | 0 | 0 | 0 | 0 | 0 |
| 0 | NA | NA | NA | NA | NA | Olfr117 | olfactory receptor 117 | 0 | 0 | 0 | 0 | 0 | 0 | 0 | 0 | 0 | 0 | 0 |
| 0 | NA | NA | NA | NA | NA | Olfr118 | olfactory receptor 118 | 0 | 0 | 0 | 0 | 0 | 0 | 0 | 0 | 0 | 0 | 0 |
| 0 | NA | NA | NA | NA | NA | Olfr119 | olfactory receptor 119 | 0 | 0 | 0 | 0 | 0 | 0 | 0 | 0 | 0 | 0 | 0 |
| 0 | NA | NA | NA | NA | NA | Olfr120 | olfactory receptor 120 | 0 | 0 | 0 | 0 | 0 | 0 | 0 | 0 | 0 | 0 | 0 |
| 0 | NA | NA | NA | NA | NA | Olfr121 | olfactory receptor 121 | 0 | 0 | 0 | 0 | 0 | 0 | 0 | 0 | 0 | 0 | 0 |
| 0 | NA | NA | NA | NA | NA | Olfr122 | olfactory receptor 122 | 0 | 0 | 0 | 0 | 0 | 0 | 0 | 0 | 0 | 0 | 0 |
| 0 | NA | NA | NA | NA | NA | Gm18268 | | 0 | 0 | 0 | 0 | 0 | 0 | 0 | 0 | 0 | 0 | 0 |
| 0 | NA | NA | NA | NA | NA | Olfr123 | olfactory receptor 123 | 0 | 0 | 0 | 0 | 0 | 0 | 0 | 0 | 0 | 0 | 0 |
| 0 | NA | NA | NA | NA | NA | Olfr124 | olfactory receptor 124 | 0 | 0 | 0 | 0 | 0 | 0 | 0 | 0 | 0 | 0 | 0 |
| 0 | NA | NA | NA | NA | NA | Olfr125 | olfactory receptor 125 | 0 | 0 | 0 | 0 | 0 | 0 | 0 | 0 | 0 | 0 | 0 |
| 0 | NA | NA | NA | NA | NA | Gm9577 | | 0 | 0 | 0 | 0 | 0 | 0 | 0 | 0 | 0 | 0 | 0 |
| 0 | NA | NA | NA | NA | NA | Olfr126 | olfactory receptor 126 | 0 | 0 | 0 | 0 | 0 | 0 | 0 | 0 | 0 | 0 | 0 |
| 0 | NA | NA | NA | NA | NA | Gm18520 | | 0 | 0 | 0 | 0 | 0 | 0 | 0 | 0 | 0 | 0 | 0 |
| 0 | NA | NA | NA | NA | NA | Olfr127 | olfactory receptor 127 | 0 | 0 | 0 | 0 | 0 | 0 | 0 | 0 | 0 | 0 | 0 |
| 0 | NA | NA | NA | NA | NA | Olfr128 | olfactory receptor 128 | 0 | 0 | 0 | 0 | 0 | 0 | 0 | 0 | 0 | 0 | 0 |
| 0 | NA | NA | NA | NA | NA | Gm20543 | | 0 | 0 | 0 | 0 | 0 | 0 | 0 | 0 | 0 | 0 | 0 |
| 0 | NA | NA | NA | NA | NA | Gm20542 | | 0 | 0 | 0 | 0 | 0 | 0 | 0 | 0 | 0 | 0 | 0 |
| 0 | NA | NA | NA | NA | NA | Olfr761 | olfactory receptor 761 | 0 | 0 | 0 | 0 | 0 | 0 | 0 | 0 | 0 | 0 | 0 |
| 0 | NA | NA | NA | NA | NA | Gm18522 | | 0 | 0 | 0 | 0 | 0 | 0 | 0 | 0 | 0 | 0 | 0 |
| 0 | NA | NA | NA | NA | NA | Olfr129 | olfactory receptor 129 | 0 | 0 | 0 | 0 | 0 | 0 | 0 | 0 | 0 | 0 | 0 |
| 0 | NA | NA | NA | NA | NA | Olfr130 | olfactory receptor 130 | 0 | 0 | 0 | 0 | 0 | 0 | 0 | 0 | 0 | 0 | 0 |
| 0 | NA | NA | NA | NA | NA | Gm18833 | | 0 | 0 | 0 | 0 | 0 | 0 | 0 | 0 | 0 | 0 | 0 |
| 0 | NA | NA | NA | NA | NA | Olfr131 | olfactory receptor 131 | 0 | 0 | 0 | 0 | 0 | 0 | 0 | 0 | 0 | 0 | 0 |
| 0 | NA | NA | NA | NA | NA | Olfr133 | olfactory receptor 133 | 0 | 0 | 0 | 0 | 0 | 0 | 0 | 0 | 0 | 0 | 0 |
| 0 | NA | NA | NA | NA | NA | Olfr134 | olfactory receptor 134 | 0 | 0 | 0 | 0 | 0 | 0 | 0 | 0 | 0 | 0 | 0 |
| 0 | NA | NA | NA | NA | NA | Olfr135 | olfactory receptor 135 | 0 | 0 | 0 | 0 | 0 | 0 | 0 | 0 | 0 | 0 | 0 |
| 0 | NA | NA | NA | NA | NA | Olfr138 | olfactory receptor 138 | 0 | 0 | 0 | 0 | 0 | 0 | 0 | 0 | 0 | 0 | 0 |
| 0 | NA | NA | NA | NA | NA | Olfr137 | olfactory receptor 137 | 0 | 0 | 0 | 0 | 0 | 0 | 0 | 0 | 0 | 0 | 0 |
| 0 | NA | NA | NA | NA | NA | Olfr762-ps1 | | 0 | 0 | 0 | 0 | 0 | 0 | 0 | 0 | 0 | 0 | 0 |
| 0 | NA | NA | NA | NA | NA | Olfr136 | olfactory receptor 136 | 0 | 0 | 0 | 0 | 0 | 0 | 0 | 0 | 0 | 0 | 0 |
| 0 | NA | NA | NA | NA | NA | Gm49810 | | 0 | 0 | 0 | 0 | 0 | 0 | 0 | 0 | 0 | 0 | 0 |
| 0 | NA | NA | NA | NA | NA | Gm49811 | | 0 | 0 | 0 | 0 | 0 | 0 | 0 | 0 | 0 | 0 | 0 |
| 0 | NA | NA | NA | NA | NA | Esp36 | exocrine gland secreted peptide 36 | 0 | 0 | 0 | 0 | 0 | 0 | 0 | 0 | 0 | 0 | 0 |
| 0 | NA | NA | NA | NA | NA | Gm20409 | | 0 | 0 | 0 | 0 | 0 | 0 | 0 | 0 | 0 | 0 | 0 |
| 0 | NA | NA | NA | NA | NA | Gm6726 | | 0 | 0 | 0 | 0 | 0 | 0 | 0 | 0 | 0 | 0 | 0 |
| 0 | NA | NA | NA | NA | NA | Gm49812 | | 0 | 0 | 0 | 0 | 0 | 0 | 0 | 0 | 0 | 0 | 0 |
| 0 | NA | NA | NA | NA | NA | Esp34 | exocrine gland secreted peptide 34 | 0 | 0 | 0 | 0 | 0 | 0 | 0 | 0 | 0 | 0 | 0 |
| 0 | NA | NA | NA | NA | NA | Gm49813 | | 0 | 0 | 0 | 0 | 0 | 0 | 0 | 0 | 0 | 0 | 0 |
| 0 | NA | NA | NA | NA | NA | Gm49814 | | 0 | 0 | 0 | 0 | 0 | 0 | 0 | 0 | 0 | 0 | 0 |
| 0 | NA | NA | NA | NA | NA | Esp31 | exocrine gland secreted peptide 31 | 0 | 0 | 0 | 0 | 0 | 0 | 0 | 0 | 0 | 0 | 0 |
| 0 | NA | NA | NA | NA | NA | Gm49815 | | 0 | 0 | 0 | 0 | 0 | 0 | 0 | 0 | 0 | 0 | 0 |
| 0 | NA | NA | NA | NA | NA | Gm49816 | | 0 | 0 | 0 | 0 | 0 | 0 | 0 | 0 | 0 | 0 | 0 |
| 0 | NA | NA | NA | NA | NA | Gm49818 | | 0 | 0 | 0 | 0 | 0 | 0 | 0 | 0 | 0 | 0 | 0 |
| 0 | NA | NA | NA | NA | NA | Gm49819 | | 0 | 0 | 0 | 0 | 0 | 0 | 0 | 0 | 0 | 0 | 0 |
| 0 | NA | NA | NA | NA | NA | Gm49820 | | 0 | 0 | 0 | 0 | 0 | 0 | 0 | 0 | 0 | 0 | 0 |
| 0 | NA | NA | NA | NA | NA | Gm49821 | | 0 | 0 | 0 | 0 | 0 | 0 | 0 | 0 | 0 | 0 | 0 |
| 0 | NA | NA | NA | NA | NA | Gm49822 | | 0 | 0 | 0 | 0 | 0 | 0 | 0 | 0 | 0 | 0 | 0 |
| 0 | NA | NA | NA | NA | NA | Gm49823 | | 0 | 0 | 0 | 0 | 0 | 0 | 0 | 0 | 0 | 0 | 0 |
| 0 | NA | NA | NA | NA | NA | Gm49824 | | 0 | 0 | 0 | 0 | 0 | 0 | 0 | 0 | 0 | 0 | 0 |
| 0 | NA | NA | NA | NA | NA | Gm49825 | | 0 | 0 | 0 | 0 | 0 | 0 | 0 | 0 | 0 | 0 | 0 |
| 0 | NA | NA | NA | NA | NA | Gm49826 | | 0 | 0 | 0 | 0 | 0 | 0 | 0 | 0 | 0 | 0 | 0 |
| 0 | NA | NA | NA | NA | NA | Esp24 | exocrine gland secreted peptide 24 | 0 | 0 | 0 | 0 | 0 | 0 | 0 | 0 | 0 | 0 | 0 |

| | | | | | | | | | | | | | | | | | |
|---|---|---|---|---|---|---|---|---|---|---|---|---|---|---|---|---|---|
| 0 | NA | NA | NA | NA | NA | Gm21903 | | 0 | 0 | 0 | 0 | 0 | 0 | 0 | 0 | 0 | 0 | 0 |
| 0 | NA | NA | NA | NA | NA | Esp23 | exocrine gland secreted peptide 23 | 0 | 0 | 0 | 0 | 0 | 0 | 0 | 0 | 0 | 0 | 0 |
| 0 | NA | NA | NA | NA | NA | Esp22 | | 0 | 0 | 0 | 0 | 0 | 0 | 0 | 0 | 0 | 0 | 0 |
| 0 | NA | NA | NA | NA | NA | Gm18784 | | 0 | 0 | 0 | 0 | 0 | 0 | 0 | 0 | 0 | 0 | 0 |
| 0 | NA | NA | NA | NA | NA | Gm49827 | | 0 | 0 | 0 | 0 | 0 | 0 | 0 | 0 | 0 | 0 | 0 |
| 0 | NA | NA | NA | NA | NA | Gm49828 | | 0 | 0 | 0 | 0 | 0 | 0 | 0 | 0 | 0 | 0 | 0 |
| 0 | NA | NA | NA | NA | NA | Gm49829 | | 0 | 0 | 0 | 0 | 0 | 0 | 0 | 0 | 0 | 0 | 0 |
| 0 | NA | NA | NA | NA | NA | Gm49830 | | 0 | 0 | 0 | 0 | 0 | 0 | 0 | 0 | 0 | 0 | 0 |
| 0 | NA | NA | NA | NA | NA | Esp18 | exocrine gland secreted peptide 18 | 0 | 0 | 0 | 0 | 0 | 0 | 0 | 0 | 0 | 0 | 0 |
| 0 | NA | NA | NA | NA | NA | Gm49831 | | 0 | 0 | 0 | 0 | 0 | 0 | 0 | 0 | 0 | 0 | 0 |
| 0 | NA | NA | NA | NA | NA | Gm26429 | | 0 | 0 | 0 | 0 | 0 | 0 | 0 | 0 | 0 | 0 | 0 |
| 0 | NA | NA | NA | NA | NA | Esp16 | exocrine gland secreted peptide 16 | 0 | 0 | 0 | 0 | 0 | 0 | 0 | 0 | 0 | 0 | 0 |
| 0 | NA | NA | NA | NA | NA | Gm49832 | | 0 | 0 | 0 | 0 | 0 | 0 | 0 | 0 | 0 | 0 | 0 |
| 0 | NA | NA | NA | NA | NA | Esp15 | exocrine gland secreted peptide 15 | 0 | 0 | 0 | 0 | 0 | 0 | 0 | 0 | 0 | 0 | 0 |
| 0 | NA | NA | NA | NA | NA | Gm49833 | | 0 | 0 | 0 | 0 | 0 | 0 | 0 | 0 | 0 | 0 | 0 |
| 0 | NA | NA | NA | NA | NA | Esp38 | exocrine gland secreted peptide 38 | 0 | 0 | 0 | 0 | 0 | 0 | 0 | 0 | 0 | 0 | 0 |
| 0 | NA | NA | NA | NA | NA | Gm26376 | | 0 | 0 | 0 | 0 | 0 | 0 | 0 | 0 | 0 | 0 | 0 |
| 0 | NA | NA | NA | NA | NA | Gm49835 | | 0 | 0 | 0 | 0 | 0 | 0 | 0 | 0 | 0 | 0 | 0 |
| 0 | NA | NA | NA | NA | NA | Tgif2-ps2 | | 0 | 0 | 0 | 0 | 0 | 0 | 0 | 0 | 0 | 0 | 0 |
| 0 | NA | NA | NA | NA | NA | Gm49836 | | 0 | 0 | 0 | 0 | 0 | 0 | 0 | 0 | 0 | 0 | 0 |
| 0 | NA | NA | NA | NA | NA | Gm49837 | | 0 | 0 | 0 | 0 | 0 | 0 | 0 | 0 | 0 | 0 | 0 |
| 0 | NA | NA | NA | NA | NA | Pgk2 | phosphoglycerate kinase 2 | 0 | 0 | 0 | 0 | 0 | 0 | 0 | 0 | 0 | 0 | 0 |
| 0 | NA | NA | NA | NA | NA | Crisp3 | cysteine-rich secretory protein 3 | 0 | 0 | 0 | 0 | 0 | 0 | 0 | 0 | 0 | 0 | 0 |
| 0 | NA | NA | NA | NA | NA | Crisp1 | cysteine-rich secretory protein 1 | 0 | 0 | 0 | 0 | 0 | 0 | 0 | 0 | 0 | 0 | 0 |
| 0 | NA | NA | NA | NA | NA | Gm49840 | | 0 | 0 | 0 | 0 | 0 | 0 | 0 | 0 | 0 | 0 | 0 |
| 0 | NA | NA | NA | NA | NA | Gm49841 | | 0 | 0 | 0 | 0 | 0 | 0 | 0 | 0 | 0 | 0 | 0 |
| 0 | NA | NA | NA | NA | NA | Gm49843 | | 0 | 0 | 0 | 0 | 0 | 0 | 0 | 0 | 0 | 0 | 0 |
| 0 | NA | NA | NA | NA | NA | Gm45926 | | 0 | 0 | 0 | 0 | 0 | 0 | 0 | 0 | 0 | 0 | 0 |
| 0 | NA | NA | NA | NA | NA | Gm34266 | | 0 | 0 | 0 | 0 | 0 | 0 | 0 | 0 | 0 | 0 | 0 |
| 0 | NA | NA | NA | NA | NA | Gm49844 | | 0 | 0 | 0 | 0 | 0 | 0 | 0 | 0 | 0 | 0 | 0 |
| 0 | NA | NA | NA | NA | NA | Gm49845 | | 0 | 0 | 0 | 0 | 0 | 0 | 0 | 0 | 0 | 0 | 0 |
| 0 | NA | NA | NA | NA | NA | Gm49846 | | 0 | 0 | 0 | 0 | 0 | 0 | 0 | 0 | 0 | 0 | 0 |
| 0 | NA | NA | NA | NA | NA | Esp8 | exocrine gland secreted peptide 8 | 0 | 0 | 0 | 0 | 0 | 0 | 0 | 0 | 0 | 0 | 0 |
| 0 | NA | NA | NA | NA | NA | Gm49847 | | 0 | 0 | 0 | 0 | 0 | 0 | 0 | 0 | 0 | 0 | 0 |
| 0 | NA | NA | NA | NA | NA | Gm29748 | | 0 | 0 | 0 | 0 | 0 | 0 | 0 | 0 | 0 | 0 | 0 |
| 0 | NA | NA | NA | NA | NA | Gm46573 | | 0 | 0 | 0 | 0 | 0 | 0 | 0 | 0 | 0 | 0 | 0 |
| 0 | NA | NA | NA | NA | NA | Esp6 | exocrine gland secreted peptide 6 | 0 | 0 | 0 | 0 | 0 | 0 | 0 | 0 | 0 | 0 | 0 |
| 0 | NA | NA | NA | NA | NA | Gm44501 | predicted readthrough transcript, 44501 | 0 | 0 | 0 | 0 | 0 | 0 | 0 | 0 | 0 | 0 | 0 |
| 0 | NA | NA | NA | NA | NA | Esp5 | exocrine gland secreted peptide 5 | 0 | 0 | 0 | 0 | 0 | 0 | 0 | 0 | 0 | 0 | 0 |
| 0 | NA | NA | NA | NA | NA | Esp4 | exocrine gland secreted peptide 4 | 0 | 0 | 0 | 0 | 0 | 0 | 0 | 0 | 0 | 0 | 0 |
| 0 | NA | NA | NA | NA | NA | Esp3 | exocrine gland secreted peptide 3 | 0 | 0 | 0 | 0 | 0 | 0 | 0 | 0 | 0 | 0 | 0 |
| 0 | NA | NA | NA | NA | NA | Gm49850 | | 0 | 0 | 0 | 0 | 0 | 0 | 0 | 0 | 0 | 0 | 0 |
| 0 | NA | NA | NA | NA | NA | Esp1 | exocrine gland secreted peptide 1 | 0 | 0 | 0 | 0 | 0 | 0 | 0 | 0 | 0 | 0 | 0 |
| 0 | NA | NA | NA | NA | NA | Rhag | Rhesus blood group-associated A glycop | 0 | 0 | 0 | 0 | 0 | 0 | 0 | 0 | 0 | 0 | 0 |
| 0 | NA | NA | NA | NA | NA | Gm20574 | | 0 | 0 | 0 | 0 | 0 | 0 | 0 | 0 | 0 | 0 | 0 |
| 0 | NA | NA | NA | NA | NA | Glyatl3 | glycine-N-acyltransferase-like 3 | 0 | 0 | 0 | 0 | 0 | 0 | 0 | 0 | 0 | 0 | 0 |
| 0 | NA | NA | NA | NA | NA | Gm23078 | | 0 | 0 | 0 | 0 | 0 | 0 | 0 | 0 | 0 | 0 | 0 |
| 0 | NA | NA | NA | NA | NA | Gm49854 | | 0 | 0 | 0 | 0 | 0 | 0 | 0 | 0 | 0 | 0 | 0 |
| 0 | NA | NA | NA | NA | NA | Gm49855 | | 0 | 0 | 0 | 0 | 0 | 0 | 0 | 0 | 0 | 0 | 0 |
| 0 | NA | NA | NA | NA | NA | Gm49857 | | 0 | 0 | 0 | 0 | 0 | 0 | 0 | 0 | 0 | 0 | 0 |
| 0 | NA | NA | NA | NA | NA | Gm49858 | | 0 | 0 | 0 | 0 | 0 | 0 | 0 | 0 | 0 | 0 | 0 |
| 0 | NA | NA | NA | NA | NA | Opn5 | opsin 5 | 0 | 0 | 0 | 0 | 0 | 0 | 0 | 0 | 0 | 0 | 0 |
| 0 | NA | NA | NA | NA | NA | Gm25135 | | 0 | 0 | 0 | 0 | 0 | 0 | 0 | 0 | 0 | 0 | 0 |
| 0 | NA | NA | NA | NA | NA | 1700071M16Rik | | 0 | 0 | 0 | 0 | 0 | 0 | 0 | 0 | 0 | 0 | 0 |
| 0 | NA | NA | NA | NA | NA | Gm25783 | | 0 | 0 | 0 | 0 | 0 | 0 | 0 | 0 | 0 | 0 | 0 |
| 0 | NA | NA | NA | NA | NA | Gm24532 | | 0 | 0 | 0 | 0 | 0 | 0 | 0 | 0 | 0 | 0 | 0 |
| 0 | NA | NA | NA | NA | NA | Gm49872 | | 0 | 0 | 0 | 0 | 0 | 0 | 0 | 0 | 0 | 0 | 0 |
| 0 | NA | NA | NA | NA | NA | Runx2os3 | | 0 | 0 | 0 | 0 | 0 | 0 | 0 | 0 | 0 | 0 | 0 |
| 0 | NA | NA | NA | NA | NA | n-R5s27 | | 0 | 0 | 0 | 0 | 0 | 0 | 0 | 0 | 0 | 0 | 0 |
| 0 | NA | NA | NA | NA | NA | Runx2os2 | | 0 | 0 | 0 | 0 | 0 | 0 | 0 | 0 | 0 | 0 | 0 |

| | | | | | | | | | | | | | | | | | |
|---|---|---|---|---|---|---|---|---|---|---|---|---|---|---|---|---|---|
| 0 | NA | NA | NA | NA | NA | 4930564C03Rik | | 0 | 0 | 0 | 0 | 0 | 0 | 0 | 0 | 0 | 0 | 0 |
| 0 | NA | NA | NA | NA | NA | Gm4766 | | 0 | 0 | 0 | 0 | 0 | 0 | 0 | 0 | 0 | 0 | 0 |
| 0 | NA | NA | NA | NA | NA | Gm24979 | | 0 | 0 | 0 | 0 | 0 | 0 | 0 | 0 | 0 | 0 | 0 |
| 0 | NA | NA | NA | NA | NA | Mymx | myomixer, myoblast fusion factor | 0 | 0 | 0 | 0 | 0 | 0 | 0 | 0 | 0 | 0 | 0 |
| 0 | NA | NA | NA | NA | NA | Gm25008 | | 0 | 0 | 0 | 0 | 0 | 0 | 0 | 0 | 0 | 0 | 0 |
| 0 | NA | NA | NA | NA | NA | Gm35692 | | 0 | 0 | 0 | 0 | 0 | 0 | 0 | 0 | 0 | 0 | 0 |
| 0 | NA | NA | NA | NA | NA | Gm19969 | | 0 | 0 | 0 | 0 | 0 | 0 | 0 | 0 | 0 | 0 | 0 |
| 0 | NA | NA | NA | NA | NA | Mir693 | | 0 | 0 | 0 | 0 | 0 | 0 | 0 | 0 | 0 | 0 | 0 |
| 0 | NA | NA | NA | NA | NA | Gm22474 | | 0 | 0 | 0 | 0 | 0 | 0 | 0 | 0 | 0 | 0 | 0 |
| 0 | NA | NA | NA | NA | NA | Gm9706 | | 0 | 0 | 0 | 0 | 0 | 0 | 0 | 0 | 0 | 0 | 0 |
| 0 | NA | NA | NA | NA | NA | Gm47119 | | 0 | 0 | 0 | 0 | 0 | 0 | 0 | 0 | 0 | 0 | 0 |
| 0 | NA | NA | NA | NA | NA | Ptcra | pre T cell antigen receptor alpha | 0 | 0 | 0 | 0 | 0 | 0 | 0 | 0 | 0 | 0 | 0 |
| 0 | NA | NA | NA | NA | NA | Prph2 | peripherin 2 | 0 | 0 | 0 | 0 | 0 | 0 | 0 | 0 | 0 | 0 | 0 |
| 0 | NA | NA | NA | NA | NA | Gm23657 | | 0 | 0 | 0 | 0 | 0 | 0 | 0 | 0 | 0 | 0 | 0 |
| 0 | NA | NA | NA | NA | NA | Gm26216 | | 0 | 0 | 0 | 0 | 0 | 0 | 0 | 0 | 0 | 0 | 0 |
| 0 | NA | NA | NA | NA | NA | Guca1a | guanylate cyclase activator 1a (retina) | 0 | 0 | 0 | 0 | 0 | 0 | 0 | 0 | 0 | 0 | 0 |
| 0 | NA | NA | NA | NA | NA | Gm50461 | | 0 | 0 | 0 | 0 | 0 | 0 | 0 | 0 | 0 | 0 | 0 |
| 0 | NA | NA | NA | NA | NA | Gm21981 | | 0 | 0 | 0 | 0 | 0 | 0 | 0 | 0 | 0 | 0 | 0 |
| 0 | NA | NA | NA | NA | NA | Frs3os | | 0 | 0 | 0 | 0 | 0 | 0 | 0 | 0 | 0 | 0 | 0 |
| 0 | NA | NA | NA | NA | NA | Pgc | progastricsin (pepsinogen C) | 0 | 0 | 0 | 0 | 0 | 0 | 0 | 0 | 0 | 0 | 0 |
| 0 | NA | NA | NA | NA | NA | Gm25201 | | 0 | 0 | 0 | 0 | 0 | 0 | 0 | 0 | 0 | 0 | 0 |
| 0 | NA | NA | NA | NA | NA | n-R5s28 | | 0 | 0 | 0 | 0 | 0 | 0 | 0 | 0 | 0 | 0 | 0 |
| 0 | NA | NA | NA | NA | NA | Gm5228 | | 0 | 0 | 0 | 0 | 0 | 0 | 0 | 0 | 0 | 0 | 0 |
| 0 | NA | NA | NA | NA | NA | Gm14871 | | 0 | 0 | 0 | 0 | 0 | 0 | 0 | 0 | 0 | 0 | 0 |
| 0 | NA | NA | NA | NA | NA | Gm20596 | | 0 | 0 | 0 | 0 | 0 | 0 | 0 | 0 | 0 | 0 | 0 |
| 0 | NA | NA | NA | NA | NA | A730006G06Rik | | 0 | 0 | 0 | 0 | 0 | 0 | 0 | 0 | 0 | 0 | 0 |
| 0 | NA | NA | NA | NA | NA | Gm49946 | | 0 | 0 | 0 | 0 | 0 | 0 | 0 | 0 | 0 | 0 | 0 |
| 0 | NA | NA | NA | NA | NA | Gm49947 | | 0 | 0 | 0 | 0 | 0 | 0 | 0 | 0 | 0 | 0 | 0 |
| 0 | NA | NA | NA | NA | NA | Gm18679 | | 0 | 0 | 0 | 0 | 0 | 0 | 0 | 0 | 0 | 0 | 0 |
| 0 | NA | NA | NA | NA | NA | Tspo2 | translocator protein 2 | 0 | 0 | 0 | 0 | 0 | 0 | 0 | 0 | 0 | 0 | 0 |
| 0 | NA | NA | NA | NA | NA | Gm24071 | | 0 | 0 | 0 | 0 | 0 | 0 | 0 | 0 | 0 | 0 | 0 |
| 0 | NA | NA | NA | NA | NA | Gm23642 | | 0 | 0 | 0 | 0 | 0 | 0 | 0 | 0 | 0 | 0 | 0 |
| 0 | NA | NA | NA | NA | NA | Gm44304 | | 0 | 0 | 0 | 0 | 0 | 0 | 0 | 0 | 0 | 0 | 0 |
| 0 | NA | NA | NA | NA | NA | 1700008K24Rik | | 0 | 0 | 0 | 0 | 0 | 0 | 0 | 0 | 0 | 0 | 0 |
| 0 | NA | NA | NA | NA | NA | Gm9586 | | 0 | 0 | 0 | 0 | 0 | 0 | 0 | 0 | 0 | 0 | 0 |
| 0 | NA | NA | NA | NA | NA | Gm22978 | | 0 | 0 | 0 | 0 | 0 | 0 | 0 | 0 | 0 | 0 | 0 |
| 0 | NA | NA | NA | NA | NA | Gm17830 | | 0 | 0 | 0 | 0 | 0 | 0 | 0 | 0 | 0 | 0 | 0 |
| 0 | NA | NA | NA | NA | NA | Gm18067 | | 0 | 0 | 0 | 0 | 0 | 0 | 0 | 0 | 0 | 0 | 0 |
| 0 | NA | NA | NA | NA | NA | Gm16555 | | 0 | 0 | 0 | 0 | 0 | 0 | 0 | 0 | 0 | 0 | 0 |
| 0 | NA | NA | NA | NA | NA | Gm16554 | | 0 | 0 | 0 | 0 | 0 | 0 | 0 | 0 | 0 | 0 | 0 |
| 0 | NA | NA | NA | NA | NA | | | 0 | 0 | 0 | 0 | 0 | 0 | 0 | 0 | 0 | 0 | 0 |
| 0 | NA | NA | NA | NA | NA | Gm49901 | | 0 | 0 | 0 | 0 | 0 | 0 | 0 | 0 | 0 | 0 | 0 |
| 0 | NA | NA | NA | NA | NA | Gm37173 | | 0 | 0 | 0 | 0 | 0 | 0 | 0 | 0 | 0 | 0 | 0 |
| 0 | NA | NA | NA | NA | NA | Gm9191 | | 0 | 0 | 0 | 0 | 0 | 0 | 0 | 0 | 0 | 0 | 0 |
| 0 | NA | NA | NA | NA | NA | Gm18735 | | 0 | 0 | 0 | 0 | 0 | 0 | 0 | 0 | 0 | 0 | 0 |
| 0 | NA | NA | NA | NA | NA | Gm41597 | | 0 | 0 | 0 | 0 | 0 | 0 | 0 | 0 | 0 | 0 | 0 |
| 0 | NA | NA | NA | NA | NA | Gm49906 | | 0 | 0 | 0 | 0 | 0 | 0 | 0 | 0 | 0 | 0 | 0 |
| 0 | NA | NA | NA | NA | NA | Gm25177 | | 0 | 0 | 0 | 0 | 0 | 0 | 0 | 0 | 0 | 0 | 0 |
| 0 | NA | NA | NA | NA | NA | Gm49908 | | 0 | 0 | 0 | 0 | 0 | 0 | 0 | 0 | 0 | 0 | 0 |
| 0 | NA | NA | NA | NA | NA | C330011F03Rik | | 0 | 0 | 0 | 0 | 0 | 0 | 0 | 0 | 0 | 0 | 0 |
| 0 | NA | NA | NA | NA | NA | Gm46574 | | 0 | 0 | 0 | 0 | 0 | 0 | 0 | 0 | 0 | 0 | 0 |
| 0 | NA | NA | NA | NA | NA | Gm31143 | | 0 | 0 | 0 | 0 | 0 | 0 | 0 | 0 | 0 | 0 | 0 |
| 0 | NA | NA | NA | NA | NA | 5830444F18Rik | | 0 | 0 | 0 | 0 | 0 | 0 | 0 | 0 | 0 | 0 | 0 |
| 0 | NA | NA | NA | NA | NA | Gm20098 | | 0 | 0 | 0 | 0 | 0 | 0 | 0 | 0 | 0 | 0 | 0 |
| 0 | NA | NA | NA | NA | NA | Gm19197 | | 0 | 0 | 0 | 0 | 0 | 0 | 0 | 0 | 0 | 0 | 0 |
| 0 | NA | NA | NA | NA | NA | Gm41600 | | 0 | 0 | 0 | 0 | 0 | 0 | 0 | 0 | 0 | 0 | 0 |
| 0 | NA | NA | NA | NA | NA | Gm49912 | | 0 | 0 | 0 | 0 | 0 | 0 | 0 | 0 | 0 | 0 | 0 |
| 0 | NA | NA | NA | NA | NA | Gm5094 | | 0 | 0 | 0 | 0 | 0 | 0 | 0 | 0 | 0 | 0 | 0 |
| 0 | NA | NA | NA | NA | NA | Gm49913 | | 0 | 0 | 0 | 0 | 0 | 0 | 0 | 0 | 0 | 0 | 0 |
| 0 | NA | NA | NA | NA | NA | Gm49915 | | 0 | 0 | 0 | 0 | 0 | 0 | 0 | 0 | 0 | 0 | 0 |

| | | | | | | | | | | | | | | | | | | |
|---|---|---|---|---|---|---|---|---|---|---|---|---|---|---|---|---|---|---|
| 0 | NA | NA | NA | NA | NA | Pp2d1 | protein phosphatase 2C-like domain cont | 0 | 0 | 0 | 0 | 0 | 0 | 0 | 0 | 0 | 0 | 0 |
| 0 | NA | NA | NA | NA | NA | 4932415M13Rik | | 0 | 0 | 0 | 0 | 0 | 0 | 0 | 0 | 0 | 0 | 0 |
| 0 | NA | NA | NA | NA | NA | Gm49917 | | 0 | 0 | 0 | 0 | 0 | 0 | 0 | 0 | 0 | 0 | 0 |
| 0 | NA | NA | NA | NA | NA | Mark1-ps2 | | 0 | 0 | 0 | 0 | 0 | 0 | 0 | 0 | 0 | 0 | 0 |
| 0 | NA | NA | NA | NA | NA | Hcfc1r1-ps2 | | 0 | 0 | 0 | 0 | 0 | 0 | 0 | 0 | 0 | 0 | 0 |
| 0 | NA | NA | NA | NA | NA | Sult1c2-ps2 | | 0 | 0 | 0 | 0 | 0 | 0 | 0 | 0 | 0 | 0 | 0 |
| 0 | NA | NA | NA | NA | NA | Sult1c2-ps1 | | 0 | 0 | 0 | 0 | 0 | 0 | 0 | 0 | 0 | 0 | 0 |
| 0 | NA | NA | NA | NA | NA | Hcfc1r1-ps1 | | 0 | 0 | 0 | 0 | 0 | 0 | 0 | 0 | 0 | 0 | 0 |
| 0 | NA | NA | NA | NA | NA | Mark4-ps | | 0 | 0 | 0 | 0 | 0 | 0 | 0 | 0 | 0 | 0 | 0 |
| 0 | NA | NA | NA | NA | NA | Mark1-ps1 | | 0 | 0 | 0 | 0 | 0 | 0 | 0 | 0 | 0 | 0 | 0 |
| 0 | NA | NA | NA | NA | NA | Gm37966 | | 0 | 0 | 0 | 0 | 0 | 0 | 0 | 0 | 0 | 0 | 0 |
| 0 | NA | NA | NA | NA | NA | 1700025K24Rik | | 0 | 0 | 0 | 0 | 0 | 0 | 0 | 0 | 0 | 0 | 0 |
| 0 | NA | NA | NA | NA | NA | Gm49919 | | 0 | 0 | 0 | 0 | 0 | 0 | 0 | 0 | 0 | 0 | 0 |
| 0 | NA | NA | NA | NA | NA | Gm4946 | | 0 | 0 | 0 | 0 | 0 | 0 | 0 | 0 | 0 | 0 | 0 |
| 0 | NA | NA | NA | NA | NA | Gm26291 | | 0 | 0 | 0 | 0 | 0 | 0 | 0 | 0 | 0 | 0 | 0 |
| 0 | NA | NA | NA | NA | NA | Gm32055 | | 0 | 0 | 0 | 0 | 0 | 0 | 0 | 0 | 0 | 0 | 0 |
| 0 | NA | NA | NA | NA | NA | Gm49920 | | 0 | 0 | 0 | 0 | 0 | 0 | 0 | 0 | 0 | 0 | 0 |
| 0 | NA | NA | NA | NA | NA | Gm49921 | | 0 | 0 | 0 | 0 | 0 | 0 | 0 | 0 | 0 | 0 | 0 |
| 0 | NA | NA | NA | NA | NA | Vmn2r118 | vomeronasal 2, receptor 118 | 0 | 0 | 0 | 0 | 0 | 0 | 0 | 0 | 0 | 0 | 0 |
| 0 | NA | NA | NA | NA | NA | Gm22654 | | 0 | 0 | 0 | 0 | 0 | 0 | 0 | 0 | 0 | 0 | 0 |
| 0 | NA | NA | NA | NA | NA | Gm18648 | | 0 | 0 | 0 | 0 | 0 | 0 | 0 | 0 | 0 | 0 | 0 |
| 0 | NA | NA | NA | NA | NA | Gm18365 | | 0 | 0 | 0 | 0 | 0 | 0 | 0 | 0 | 0 | 0 | 0 |
| 0 | NA | NA | NA | NA | NA | Gm44397 | | 0 | 0 | 0 | 0 | 0 | 0 | 0 | 0 | 0 | 0 | 0 |
| 0 | NA | NA | NA | NA | NA | Arrdc5 | arrestin domain containing 5 | 0 | 0 | 0 | 0 | 0 | 0 | 0 | 0 | 0 | 0 | 0 |
| 0 | NA | NA | NA | NA | NA | Mir6977 | | 0 | 0 | 0 | 0 | 0 | 0 | 0 | 0 | 0 | 0 | 0 |
| 0 | NA | NA | NA | NA | NA | Acsbg2 | acyl-CoA synthetase bubblegum family m | 0 | 0 | 0 | 0 | 0 | 0 | 0 | 0 | 0 | 0 | 0 |
| 0 | NA | NA | NA | NA | NA | Gm49887 | | 0 | 0 | 0 | 0 | 0 | 0 | 0 | 0 | 0 | 0 | 0 |
| 0 | NA | NA | NA | NA | NA | Pspn | persephin | 0 | 0 | 0 | 0 | 0 | 0 | 0 | 0 | 0 | 0 | 0 |
| 0 | NA | NA | NA | NA | NA | Slc25a41 | solute carrier family 25, member 41 | 0 | 0 | 0 | 0 | 0 | 0 | 0 | 0 | 0 | 0 | 0 |
| 0 | NA | NA | NA | NA | NA | Gm49892 | | 0 | 0 | 0 | 0 | 0 | 0 | 0 | 0 | 0 | 0 | 0 |
| 0 | NA | NA | NA | NA | NA | Mir6978 | | 0 | 0 | 0 | 0 | 0 | 0 | 0 | 0 | 0 | 0 | 0 |
| 0 | NA | NA | NA | NA | NA | Gm46575 | | 0 | 0 | 0 | 0 | 0 | 0 | 0 | 0 | 0 | 0 | 0 |
| 0 | NA | NA | NA | NA | NA | Vmn2r120 | vomeronasal 2, receptor 120 | 0 | 0 | 0 | 0 | 0 | 0 | 0 | 0 | 0 | 0 | 0 |
| 0 | NA | NA | NA | NA | NA | Gm49842 | | 0 | 0 | 0 | 0 | 0 | 0 | 0 | 0 | 0 | 0 | 0 |
| 0 | NA | NA | NA | NA | NA | Gm21834 | | 0 | 0 | 0 | 0 | 0 | 0 | 0 | 0 | 0 | 0 | 0 |
| 0 | NA | NA | NA | NA | NA | Gm18139 | | 0 | 0 | 0 | 0 | 0 | 0 | 0 | 0 | 0 | 0 | 0 |
| 0 | NA | NA | NA | NA | NA | Gm25102 | | 0 | 0 | 0 | 0 | 0 | 0 | 0 | 0 | 0 | 0 | 0 |
| 0 | NA | NA | NA | NA | NA | Gm17920 | | 0 | 0 | 0 | 0 | 0 | 0 | 0 | 0 | 0 | 0 | 0 |
| 0 | NA | NA | NA | NA | NA | Cntnap5c | contactin associated protein-like 5C | 0 | 0 | 0 | 0 | 0 | 0 | 0 | 0 | 0 | 0 | 0 |
| 0 | NA | NA | NA | NA | NA | Gm6149 | | 0 | 0 | 0 | 0 | 0 | 0 | 0 | 0 | 0 | 0 | 0 |
| 0 | NA | NA | NA | NA | NA | Gm49849 | | 0 | 0 | 0 | 0 | 0 | 0 | 0 | 0 | 0 | 0 | 0 |
| 0 | NA | NA | NA | NA | NA | A330072L02Rik | | 0 | 0 | 0 | 0 | 0 | 0 | 0 | 0 | 0 | 0 | 0 |
| 0 | NA | NA | NA | NA | NA | Pdzph1 | PDZ and pleckstrin homology domains 1 | 0 | 0 | 0 | 0 | 0 | 0 | 0 | 0 | 0 | 0 | 0 |
| 0 | NA | NA | NA | NA | NA | Gm23769 | | 0 | 0 | 0 | 0 | 0 | 0 | 0 | 0 | 0 | 0 | 0 |
| 0 | NA | NA | NA | NA | NA | Gm23203 | | 0 | 0 | 0 | 0 | 0 | 0 | 0 | 0 | 0 | 0 | 0 |
| 0 | NA | NA | NA | NA | NA | Gm49851 | | 0 | 0 | 0 | 0 | 0 | 0 | 0 | 0 | 0 | 0 | 0 |
| 0 | NA | NA | NA | NA | NA | Gm49852 | | 0 | 0 | 0 | 0 | 0 | 0 | 0 | 0 | 0 | 0 | 0 |
| 0 | NA | NA | NA | NA | NA | Gm49853 | | 0 | 0 | 0 | 0 | 0 | 0 | 0 | 0 | 0 | 0 | 0 |
| 0 | NA | NA | NA | NA | NA | Gm29051 | | 0 | 0 | 0 | 0 | 0 | 0 | 0 | 0 | 0 | 0 | 0 |
| 0 | NA | NA | NA | NA | NA | Gm23447 | | 0 | 0 | 0 | 0 | 0 | 0 | 0 | 0 | 0 | 0 | 0 |
| 0 | NA | NA | NA | NA | NA | Gm4512 | | 0 | 0 | 0 | 0 | 0 | 0 | 0 | 0 | 0 | 0 | 0 |
| 0 | NA | NA | NA | NA | NA | Gm18521 | | 0 | 0 | 0 | 0 | 0 | 0 | 0 | 0 | 0 | 0 | 0 |
| 0 | NA | NA | NA | NA | NA | Gm49856 | | 0 | 0 | 0 | 0 | 0 | 0 | 0 | 0 | 0 | 0 | 0 |
| 0 | NA | NA | NA | NA | NA | Gm33890 | | 0 | 0 | 0 | 0 | 0 | 0 | 0 | 0 | 0 | 0 | 0 |
| 0 | NA | NA | NA | NA | NA | Gm18070 | | 0 | 0 | 0 | 0 | 0 | 0 | 0 | 0 | 0 | 0 | 0 |
| 0 | NA | NA | NA | NA | NA | Gm23773 | | 0 | 0 | 0 | 0 | 0 | 0 | 0 | 0 | 0 | 0 | 0 |
| 0 | NA | NA | NA | NA | NA | Obox6-ps1 | | 0 | 0 | 0 | 0 | 0 | 0 | 0 | 0 | 0 | 0 | 0 |
| 0 | NA | NA | NA | NA | NA | Gm6174 | | 0 | 0 | 0 | 0 | 0 | 0 | 0 | 0 | 0 | 0 | 0 |
| 0 | NA | NA | NA | NA | NA | Gm49859 | | 0 | 0 | 0 | 0 | 0 | 0 | 0 | 0 | 0 | 0 | 0 |
| 0 | NA | NA | NA | NA | NA | Gm49860 | | 0 | 0 | 0 | 0 | 0 | 0 | 0 | 0 | 0 | 0 | 0 |

| | | | | | | | | | | | | | | | | | |
|---|---|---|---|---|---|---|---|---|---|---|---|---|---|---|---|---|---|
| 0 | NA | NA | NA | NA | NA | Gm25800 | | 0 | 0 | 0 | 0 | 0 | 0 | 0 | 0 | 0 | 0 | 0 |
| 0 | NA | NA | NA | NA | NA | Gm24730 | | 0 | 0 | 0 | 0 | 0 | 0 | 0 | 0 | 0 | 0 | 0 |
| 0 | NA | NA | NA | NA | NA | Gm49862 | | 0 | 0 | 0 | 0 | 0 | 0 | 0 | 0 | 0 | 0 | 0 |
| 0 | NA | NA | NA | NA | NA | Gm9300 | | 0 | 0 | 0 | 0 | 0 | 0 | 0 | 0 | 0 | 0 | 0 |
| 0 | NA | NA | NA | NA | NA | Gm24194 | | 0 | 0 | 0 | 0 | 0 | 0 | 0 | 0 | 0 | 0 | 0 |
| 0 | NA | NA | NA | NA | NA | Mir6420 | | 0 | 0 | 0 | 0 | 0 | 0 | 0 | 0 | 0 | 0 | 0 |
| 0 | NA | NA | NA | NA | NA | AU016765 | | 0 | 0 | 0 | 0 | 0 | 0 | 0 | 0 | 0 | 0 | 0 |
| 0 | NA | NA | NA | NA | NA | Gm17133 | | 0 | 0 | 0 | 0 | 0 | 0 | 0 | 0 | 0 | 0 | 0 |
| 0 | NA | NA | NA | NA | NA | Dreh | | 0 | 0 | 0 | 0 | 0 | 0 | 0 | 0 | 0 | 0 | 0 |
| 0 | NA | NA | NA | NA | NA | 4930583I09Rik | RIKEN cDNA 4930583I09 gene | 0 | 0 | 0 | 0 | 0 | 0 | 0 | 0 | 0 | 0 | 0 |
| 0 | NA | NA | NA | NA | NA | Gm4701 | | 0 | 0 | 0 | 0 | 0 | 0 | 0 | 0 | 0 | 0 | 0 |
| 0 | NA | NA | NA | NA | NA | Gm25344 | | 0 | 0 | 0 | 0 | 0 | 0 | 0 | 0 | 0 | 0 | 0 |
| 0 | NA | NA | NA | NA | NA | Gm49865 | | 0 | 0 | 0 | 0 | 0 | 0 | 0 | 0 | 0 | 0 | 0 |
| 0 | NA | NA | NA | NA | NA | Gm18992 | | 0 | 0 | 0 | 0 | 0 | 0 | 0 | 0 | 0 | 0 | 0 |
| 0 | NA | NA | NA | NA | NA | Gm9288 | | 0 | 0 | 0 | 0 | 0 | 0 | 0 | 0 | 0 | 0 | 0 |
| 0 | NA | NA | NA | NA | NA | Gm25731 | | 0 | 0 | 0 | 0 | 0 | 0 | 0 | 0 | 0 | 0 | 0 |
| 0 | NA | NA | NA | NA | NA | Gm18138 | | 0 | 0 | 0 | 0 | 0 | 0 | 0 | 0 | 0 | 0 | 0 |
| 0 | NA | NA | NA | NA | NA | Gm49866 | | 0 | 0 | 0 | 0 | 0 | 0 | 0 | 0 | 0 | 0 | 0 |
| 0 | NA | NA | NA | NA | NA | Txndc2 | thioredoxin domain containing 2 (sperma | 0 | 0 | 0 | 0 | 0 | 0 | 0 | 0 | 0 | 0 | 0 |
| 0 | NA | NA | NA | NA | NA | Gm35550 | | 0 | 0 | 0 | 0 | 0 | 0 | 0 | 0 | 0 | 0 | 0 |
| 0 | NA | NA | NA | NA | NA | Gm23264 | | 0 | 0 | 0 | 0 | 0 | 0 | 0 | 0 | 0 | 0 | 0 |
| 0 | NA | NA | NA | NA | NA | Gm49927 | | 0 | 0 | 0 | 0 | 0 | 0 | 0 | 0 | 0 | 0 | 0 |
| 0 | NA | NA | NA | NA | NA | Gm49930 | | 0 | 0 | 0 | 0 | 0 | 0 | 0 | 0 | 0 | 0 | 0 |
| 0 | NA | NA | NA | NA | NA | Gm49931 | | 0 | 0 | 0 | 0 | 0 | 0 | 0 | 0 | 0 | 0 | 0 |
| 0 | NA | NA | NA | NA | NA | Gm20742 | | 0 | 0 | 0 | 0 | 0 | 0 | 0 | 0 | 0 | 0 | 0 |
| 0 | NA | NA | NA | NA | NA | Gm49938 | | 0 | 0 | 0 | 0 | 0 | 0 | 0 | 0 | 0 | 0 | 0 |
| 0 | NA | NA | NA | NA | NA | Gm46576 | | 0 | 0 | 0 | 0 | 0 | 0 | 0 | 0 | 0 | 0 | 0 |
| 0 | NA | NA | NA | NA | NA | Gm49941 | | 0 | 0 | 0 | 0 | 0 | 0 | 0 | 0 | 0 | 0 | 0 |
| 0 | NA | NA | NA | NA | NA | 1700016K05Rik | | 0 | 0 | 0 | 0 | 0 | 0 | 0 | 0 | 0 | 0 | 0 |
| 0 | NA | NA | NA | NA | NA | Gm18961 | | 0 | 0 | 0 | 0 | 0 | 0 | 0 | 0 | 0 | 0 | 0 |
| 0 | NA | NA | NA | NA | NA | Gm49943 | | 0 | 0 | 0 | 0 | 0 | 0 | 0 | 0 | 0 | 0 | 0 |
| 0 | NA | NA | NA | NA | NA | Gm49940 | | 0 | 0 | 0 | 0 | 0 | 0 | 0 | 0 | 0 | 0 | 0 |
| 0 | NA | NA | NA | NA | NA | Gm49942 | | 0 | 0 | 0 | 0 | 0 | 0 | 0 | 0 | 0 | 0 | 0 |
| 0 | NA | NA | NA | NA | NA | Gm9984 | | 0 | 0 | 0 | 0 | 0 | 0 | 0 | 0 | 0 | 0 | 0 |
| 0 | NA | NA | NA | NA | NA | Mir5709 | | 0 | 0 | 0 | 0 | 0 | 0 | 0 | 0 | 0 | 0 | 0 |
| 0 | NA | NA | NA | NA | NA | Gm22277 | | 0 | 0 | 0 | 0 | 0 | 0 | 0 | 0 | 0 | 0 | 0 |
| 0 | NA | NA | NA | NA | NA | Lrrc30 | leucine rich repeat containing 30 | 0 | 0 | 0 | 0 | 0 | 0 | 0 | 0 | 0 | 0 | 0 |
| 0 | NA | NA | NA | NA | NA | Gm49944 | | 0 | 0 | 0 | 0 | 0 | 0 | 0 | 0 | 0 | 0 | 0 |
| 0 | NA | NA | NA | NA | NA | Gm49945 | | 0 | 0 | 0 | 0 | 0 | 0 | 0 | 0 | 0 | 0 | 0 |
| 0 | NA | NA | NA | NA | NA | Gm36252 | | 0 | 0 | 0 | 0 | 0 | 0 | 0 | 0 | 0 | 0 | 0 |
| 0 | NA | NA | NA | NA | NA | Gm15974 | | 0 | 0 | 0 | 0 | 0 | 0 | 0 | 0 | 0 | 0 | 0 |
| 0 | NA | NA | NA | NA | NA | Gm17921 | | 0 | 0 | 0 | 0 | 0 | 0 | 0 | 0 | 0 | 0 | 0 |
| 0 | NA | NA | NA | NA | NA | Gm36487 | | 0 | 0 | 0 | 0 | 0 | 0 | 0 | 0 | 0 | 0 | 0 |
| 0 | NA | NA | NA | NA | NA | A730049N16Rik | | 0 | 0 | 0 | 0 | 0 | 0 | 0 | 0 | 0 | 0 | 0 |
| 0 | NA | NA | NA | NA | NA | Gm9319 | | 0 | 0 | 0 | 0 | 0 | 0 | 0 | 0 | 0 | 0 | 0 |
| 0 | NA | NA | NA | NA | NA | Gm22967 | | 0 | 0 | 0 | 0 | 0 | 0 | 0 | 0 | 0 | 0 | 0 |
| 0 | NA | NA | NA | NA | NA | Gm49895 | | 0 | 0 | 0 | 0 | 0 | 0 | 0 | 0 | 0 | 0 | 0 |
| 0 | NA | NA | NA | NA | NA | Gm4561 | | 0 | 0 | 0 | 0 | 0 | 0 | 0 | 0 | 0 | 0 | 0 |
| 0 | NA | NA | NA | NA | NA | 5031415H12Rik | | 0 | 0 | 0 | 0 | 0 | 0 | 0 | 0 | 0 | 0 | 0 |
| 0 | NA | NA | NA | NA | NA | Gm41609 | | 0 | 0 | 0 | 0 | 0 | 0 | 0 | 0 | 0 | 0 | 0 |
| 0 | NA | NA | NA | NA | NA | Mir1195 | | 0 | 0 | 0 | 0 | 0 | 0 | 0 | 0 | 0 | 0 | 0 |
| 0 | NA | NA | NA | NA | NA | Gm49902 | | 0 | 0 | 0 | 0 | 0 | 0 | 0 | 0 | 0 | 0 | 0 |
| 0 | NA | NA | NA | NA | NA | Gm49904 | | 0 | 0 | 0 | 0 | 0 | 0 | 0 | 0 | 0 | 0 | 0 |
| 0 | NA | NA | NA | NA | NA | Gm49905 | | 0 | 0 | 0 | 0 | 0 | 0 | 0 | 0 | 0 | 0 | 0 |
| 0 | NA | NA | NA | NA | NA | Gm49909 | | 0 | 0 | 0 | 0 | 0 | 0 | 0 | 0 | 0 | 0 | 0 |
| 0 | NA | NA | NA | NA | NA | 1600022D10Rik | | 0 | 0 | 0 | 0 | 0 | 0 | 0 | 0 | 0 | 0 | 0 |
| 0 | NA | NA | NA | NA | NA | Gm49910 | | 0 | 0 | 0 | 0 | 0 | 0 | 0 | 0 | 0 | 0 | 0 |
| 0 | NA | NA | NA | NA | NA | 4930471L23Rik | | 0 | 0 | 0 | 0 | 0 | 0 | 0 | 0 | 0 | 0 | 0 |
| 0 | NA | NA | NA | NA | NA | Gm49916 | | 0 | 0 | 0 | 0 | 0 | 0 | 0 | 0 | 0 | 0 | 0 |
| 0 | NA | NA | NA | NA | NA | Gm18738 | | 0 | 0 | 0 | 0 | 0 | 0 | 0 | 0 | 0 | 0 | 0 |

| | | | | | | | | | | | | | | | | | | |
|---|---|---|---|---|---|---|---|---|---|---|---|---|---|---|---|---|---|---|
| 0 | NA | NA | NA | NA | NA | Gm23115 | | 0 | 0 | 0 | 0 | 0 | 0 | 0 | 0 | 0 | 0 | 0 |
| 0 | NA | NA | NA | NA | NA | Snord92 | | 0 | 0 | 0 | 0 | 0 | 0 | 0 | 0 | 0 | 0 | 0 |
| 0 | NA | NA | NA | NA | NA | Gm15641 | | 0 | 0 | 0 | 0 | 0 | 0 | 0 | 0 | 0 | 0 | 0 |
| 0 | NA | NA | NA | NA | NA | Pcare | photoreceptor cilium actin regulator | 0 | 0 | 0 | 0 | 0 | 0 | 0 | 0 | 0 | 0 | 0 |
| 0 | NA | NA | NA | NA | NA | Gm49922 | | 0 | 0 | 0 | 0 | 0 | 0 | 0 | 0 | 0 | 0 | 0 |
| 0 | NA | NA | NA | NA | NA | Gm24736 | | 0 | 0 | 0 | 0 | 0 | 0 | 0 | 0 | 0 | 0 | 0 |
| 0 | NA | NA | NA | NA | NA | Gm49923 | | 0 | 0 | 0 | 0 | 0 | 0 | 0 | 0 | 0 | 0 | 0 |
| 0 | NA | NA | NA | NA | NA | Gm26963 | | 0 | 0 | 0 | 0 | 0 | 0 | 0 | 0 | 0 | 0 | 0 |
| 0 | NA | NA | NA | NA | NA | Gm23649 | | 0 | 0 | 0 | 0 | 0 | 0 | 0 | 0 | 0 | 0 | 0 |
| 0 | NA | NA | NA | NA | NA | Gm30420 | | 0 | 0 | 0 | 0 | 0 | 0 | 0 | 0 | 0 | 0 | 0 |
| 0 | NA | NA | NA | NA | NA | Gm49875 | | 0 | 0 | 0 | 0 | 0 | 0 | 0 | 0 | 0 | 0 | 0 |
| 0 | NA | NA | NA | NA | NA | Gm49876 | | 0 | 0 | 0 | 0 | 0 | 0 | 0 | 0 | 0 | 0 | 0 |
| 0 | NA | NA | NA | NA | NA | Gm49878 | | 0 | 0 | 0 | 0 | 0 | 0 | 0 | 0 | 0 | 0 | 0 |
| 0 | NA | NA | NA | NA | NA | Gm18068 | | 0 | 0 | 0 | 0 | 0 | 0 | 0 | 0 | 0 | 0 | 0 |
| 0 | NA | NA | NA | NA | NA | Gm24475 | | 0 | 0 | 0 | 0 | 0 | 0 | 0 | 0 | 0 | 0 | 0 |
| 0 | NA | NA | NA | NA | NA | Srd5a2 | steroid 5 alpha-reductase 2 | 0 | 0 | 0 | 0 | 0 | 0 | 0 | 0 | 0 | 0 | 0 |
| 0 | NA | NA | NA | NA | NA | Gm49880 | | 0 | 0 | 0 | 0 | 0 | 0 | 0 | 0 | 0 | 0 | 0 |
| 0 | NA | NA | NA | NA | NA | Gm31084 | | 0 | 0 | 0 | 0 | 0 | 0 | 0 | 0 | 0 | 0 | 0 |
| 0 | NA | NA | NA | NA | NA | Gm49881 | | 0 | 0 | 0 | 0 | 0 | 0 | 0 | 0 | 0 | 0 | 0 |
| 0 | NA | NA | NA | NA | NA | Gm49882 | | 0 | 0 | 0 | 0 | 0 | 0 | 0 | 0 | 0 | 0 | 0 |
| 0 | NA | NA | NA | NA | NA | Gm31328 | | 0 | 0 | 0 | 0 | 0 | 0 | 0 | 0 | 0 | 0 | 0 |
| 0 | NA | NA | NA | NA | NA | Gm9351 | | 0 | 0 | 0 | 0 | 0 | 0 | 0 | 0 | 0 | 0 | 0 |
| 0 | NA | NA | NA | NA | NA | Gm50020 | | 0 | 0 | 0 | 0 | 0 | 0 | 0 | 0 | 0 | 0 | 0 |
| 0 | NA | NA | NA | NA | NA | Gm50021 | | 0 | 0 | 0 | 0 | 0 | 0 | 0 | 0 | 0 | 0 | 0 |
| 0 | NA | NA | NA | NA | NA | Gm31759 | | 0 | 0 | 0 | 0 | 0 | 0 | 0 | 0 | 0 | 0 | 0 |
| 0 | NA | NA | NA | NA | NA | Gm31818 | | 0 | 0 | 0 | 0 | 0 | 0 | 0 | 0 | 0 | 0 | 0 |
| 0 | NA | NA | NA | NA | NA | Gm9360 | | 0 | 0 | 0 | 0 | 0 | 0 | 0 | 0 | 0 | 0 | 0 |
| 0 | NA | NA | NA | NA | NA | Gm50099 | | 0 | 0 | 0 | 0 | 0 | 0 | 0 | 0 | 0 | 0 | 0 |
| 0 | NA | NA | NA | NA | NA | Gm24126 | | 0 | 0 | 0 | 0 | 0 | 0 | 0 | 0 | 0 | 0 | 0 |
| 0 | NA | NA | NA | NA | NA | Gm26124 | | 0 | 0 | 0 | 0 | 0 | 0 | 0 | 0 | 0 | 0 | 0 |
| 0 | NA | NA | NA | NA | NA | Gm16391 | | 0 | 0 | 0 | 0 | 0 | 0 | 0 | 0 | 0 | 0 | 0 |
| 0 | NA | NA | NA | NA | NA | Gm50028 | | 0 | 0 | 0 | 0 | 0 | 0 | 0 | 0 | 0 | 0 | 0 |
| 0 | NA | NA | NA | NA | NA | Gm18366 | | 0 | 0 | 0 | 0 | 0 | 0 | 0 | 0 | 0 | 0 | 0 |
| 0 | NA | NA | NA | NA | NA | Gm6496 | | 0 | 0 | 0 | 0 | 0 | 0 | 0 | 0 | 0 | 0 | 0 |
| 0 | NA | NA | NA | NA | NA | Gm4711 | | 0 | 0 | 0 | 0 | 0 | 0 | 0 | 0 | 0 | 0 | 0 |
| 0 | NA | NA | NA | NA | NA | Gm50030 | | 0 | 0 | 0 | 0 | 0 | 0 | 0 | 0 | 0 | 0 | 0 |
| 0 | NA | NA | NA | NA | NA | Gm5229 | | 0 | 0 | 0 | 0 | 0 | 0 | 0 | 0 | 0 | 0 | 0 |
| 0 | NA | NA | NA | NA | NA | Gm50031 | | 0 | 0 | 0 | 0 | 0 | 0 | 0 | 0 | 0 | 0 | 0 |
| 0 | NA | NA | NA | NA | NA | Gm25069 | | 0 | 0 | 0 | 0 | 0 | 0 | 0 | 0 | 0 | 0 | 0 |
| 0 | NA | NA | NA | NA | NA | Gm50034 | | 0 | 0 | 0 | 0 | 0 | 0 | 0 | 0 | 0 | 0 | 0 |
| 0 | NA | NA | NA | NA | NA | Gm22562 | | 0 | 0 | 0 | 0 | 0 | 0 | 0 | 0 | 0 | 0 | 0 |
| 0 | NA | NA | NA | NA | NA | Gm49863 | | 0 | 0 | 0 | 0 | 0 | 0 | 0 | 0 | 0 | 0 | 0 |
| 0 | NA | NA | NA | NA | NA | Gm18649 | | 0 | 0 | 0 | 0 | 0 | 0 | 0 | 0 | 0 | 0 | 0 |
| 0 | NA | NA | NA | NA | NA | Gm26957 | | 0 | 0 | 0 | 0 | 0 | 0 | 0 | 0 | 0 | 0 | 0 |
| 0 | NA | NA | NA | NA | NA | Gm49869 | | 0 | 0 | 0 | 0 | 0 | 0 | 0 | 0 | 0 | 0 | 0 |
| 0 | NA | NA | NA | NA | NA | Gm49984 | | 0 | 0 | 0 | 0 | 0 | 0 | 0 | 0 | 0 | 0 | 0 |
| 0 | NA | NA | NA | NA | NA | Gm37555 | | 0 | 0 | 0 | 0 | 0 | 0 | 0 | 0 | 0 | 0 | 0 |
| 0 | NA | NA | NA | NA | NA | 1700106O07Rik | | 0 | 0 | 0 | 0 | 0 | 0 | 0 | 0 | 0 | 0 | 0 |
| 0 | NA | NA | NA | NA | NA | Gm49985 | | 0 | 0 | 0 | 0 | 0 | 0 | 0 | 0 | 0 | 0 | 0 |
| 0 | NA | NA | NA | NA | NA | Gm41625 | | 0 | 0 | 0 | 0 | 0 | 0 | 0 | 0 | 0 | 0 | 0 |
| 0 | NA | NA | NA | NA | NA | Gm22215 | | 0 | 0 | 0 | 0 | 0 | 0 | 0 | 0 | 0 | 0 | 0 |
| 0 | NA | NA | NA | NA | NA | Gm23125 | | 0 | 0 | 0 | 0 | 0 | 0 | 0 | 0 | 0 | 0 | 0 |
| 0 | NA | NA | NA | NA | NA | Gm25706 | | 0 | 0 | 0 | 0 | 0 | 0 | 0 | 0 | 0 | 0 | 0 |
| 0 | NA | NA | NA | NA | NA | C230072F16Rik | | 0 | 0 | 0 | 0 | 0 | 0 | 0 | 0 | 0 | 0 | 0 |
| 0 | NA | NA | NA | NA | NA | Gm50042 | | 0 | 0 | 0 | 0 | 0 | 0 | 0 | 0 | 0 | 0 | 0 |
| 0 | NA | NA | NA | NA | NA | Gm11096 | | 0 | 0 | 0 | 0 | 0 | 0 | 0 | 0 | 0 | 0 | 0 |
| 0 | NA | NA | NA | NA | NA | Rpl31-ps25 | | 0 | 0 | 0 | 0 | 0 | 0 | 0 | 0 | 0 | 0 | 0 |
| 0 | NA | NA | NA | NA | NA | Gm4599 | | 0 | 0 | 0 | 0 | 0 | 0 | 0 | 0 | 0 | 0 | 0 |
| 0 | NA | NA | NA | NA | NA | Gm27359 | | 0 | 0 | 0 | 0 | 0 | 0 | 0 | 0 | 0 | 0 | 0 |
| 0 | NA | NA | NA | NA | NA | Gm22386 | | 0 | 0 | 0 | 0 | 0 | 0 | 0 | 0 | 0 | 0 | 0 |

| | | | | | | | | | | | | | | | | | |
|---|---|---|---|---|---|---|---|---|---|---|---|---|---|---|---|---|---|
| 0 | NA | NA | NA | NA | NA | Gm23400 | | 0 | 0 | 0 | 0 | 0 | 0 | 0 | 0 | 0 | 0 | 0 |
| 0 | NA | NA | NA | NA | NA | Gm9390 | | 0 | 0 | 0 | 0 | 0 | 0 | 0 | 0 | 0 | 0 | 0 |
| 0 | NA | NA | NA | NA | NA | Gm4601 | | 0 | 0 | 0 | 0 | 0 | 0 | 0 | 0 | 0 | 0 | 0 |
| 0 | NA | NA | NA | NA | NA | Gm19689 | | 0 | 0 | 0 | 0 | 0 | 0 | 0 | 0 | 0 | 0 | 0 |
| 0 | NA | NA | NA | NA | NA | Gm29052 | | 0 | 0 | 0 | 0 | 0 | 0 | 0 | 0 | 0 | 0 | 0 |
| 0 | NA | NA | NA | NA | NA | Gm24240 | | 0 | 0 | 0 | 0 | 0 | 0 | 0 | 0 | 0 | 0 | 0 |
| 0 | NA | NA | NA | NA | NA | Gm23905 | | 0 | 0 | 0 | 0 | 0 | 0 | 0 | 0 | 0 | 0 | 0 |
| 0 | NA | NA | NA | NA | NA | Gm41638 | | 0 | 0 | 0 | 0 | 0 | 0 | 0 | 0 | 0 | 0 | 0 |
| 0 | NA | NA | NA | NA | NA | Gm37549 | | 0 | 0 | 0 | 0 | 0 | 0 | 0 | 0 | 0 | 0 | 0 |
| 0 | NA | NA | NA | NA | NA | Gm49967 | | 0 | 0 | 0 | 0 | 0 | 0 | 0 | 0 | 0 | 0 | 0 |
| 0 | NA | NA | NA | NA | NA | 4933433H22Rik | | 0 | 0 | 0 | 0 | 0 | 0 | 0 | 0 | 0 | 0 | 0 |
| 0 | NA | NA | NA | NA | NA | Gm24492 | | 0 | 0 | 0 | 0 | 0 | 0 | 0 | 0 | 0 | 0 | 0 |
| 0 | NA | NA | NA | NA | NA | Gm24722 | | 0 | 0 | 0 | 0 | 0 | 0 | 0 | 0 | 0 | 0 | 0 |
| 0 | NA | NA | NA | NA | NA | 4930500A05Rik | | 0 | 0 | 0 | 0 | 0 | 0 | 0 | 0 | 0 | 0 | 0 |
| 0 | NA | NA | NA | NA | NA | Gm49969 | | 0 | 0 | 0 | 0 | 0 | 0 | 0 | 0 | 0 | 0 | 0 |
| 0 | NA | NA | NA | NA | NA | Gm26066 | | 0 | 0 | 0 | 0 | 0 | 0 | 0 | 0 | 0 | 0 | 0 |
| 0 | NA | NA | NA | NA | NA | Gm49979 | | 0 | 0 | 0 | 0 | 0 | 0 | 0 | 0 | 0 | 0 | 0 |
| 0 | NA | NA | NA | NA | NA | Gm49981 | | 0 | 0 | 0 | 0 | 0 | 0 | 0 | 0 | 0 | 0 | 0 |
| 0 | NA | NA | NA | NA | NA | Gm9400 | | 0 | 0 | 0 | 0 | 0 | 0 | 0 | 0 | 0 | 0 | 0 |
| 0 | NA | NA | NA | NA | NA | Gm49982 | | 0 | 0 | 0 | 0 | 0 | 0 | 0 | 0 | 0 | 0 | 0 |
| 0 | NA | NA | NA | NA | NA | Gm5231 | | 0 | 0 | 0 | 0 | 0 | 0 | 0 | 0 | 0 | 0 | 0 |
| 0 | NA | NA | NA | NA | NA | Gm27178 | | 0 | 0 | 0 | 0 | 0 | 0 | 0 | 0 | 0 | 0 | 0 |
| 0 | NA | NA | NA | NA | NA | Six3os1 | | 0 | 0 | 0 | 0 | 0 | 0 | 0 | 0 | 0 | 0 | 0 |
| 0 | NA | NA | NA | NA | NA | Gm27435 | | 0 | 0 | 0 | 0 | 0 | 0 | 0 | 0 | 0 | 0 | 0 |
| 0 | NA | NA | NA | NA | NA | Gm27801 | | 0 | 0 | 0 | 0 | 0 | 0 | 0 | 0 | 0 | 0 | 0 |
| 0 | NA | NA | NA | NA | NA | Gm27610 | | 0 | 0 | 0 | 0 | 0 | 0 | 0 | 0 | 0 | 0 | 0 |
| 0 | NA | NA | NA | NA | NA | Gm27283 | | 0 | 0 | 0 | 0 | 0 | 0 | 0 | 0 | 0 | 0 | 0 |
| 0 | NA | NA | NA | NA | NA | Gm27275 | | 0 | 0 | 0 | 0 | 0 | 0 | 0 | 0 | 0 | 0 | 0 |
| 0 | NA | NA | NA | NA | NA | Gm27502 | | 0 | 0 | 0 | 0 | 0 | 0 | 0 | 0 | 0 | 0 | 0 |
| 0 | NA | NA | NA | NA | NA | Gm27427 | | 0 | 0 | 0 | 0 | 0 | 0 | 0 | 0 | 0 | 0 | 0 |
| 0 | NA | NA | NA | NA | NA | Gm28528 | | 0 | 0 | 0 | 0 | 0 | 0 | 0 | 0 | 0 | 0 | 0 |
| 0 | NA | NA | NA | NA | NA | Gm29418 | | 0 | 0 | 0 | 0 | 0 | 0 | 0 | 0 | 0 | 0 | 0 |
| 0 | NA | NA | NA | NA | NA | Gm30117 | | 0 | 0 | 0 | 0 | 0 | 0 | 0 | 0 | 0 | 0 | 0 |
| 0 | NA | NA | NA | NA | NA | 2010106C02Rik | | 0 | 0 | 0 | 0 | 0 | 0 | 0 | 0 | 0 | 0 | 0 |
| 0 | NA | NA | NA | NA | NA | Gm18832 | | 0 | 0 | 0 | 0 | 0 | 0 | 0 | 0 | 0 | 0 | 0 |
| 0 | NA | NA | NA | NA | NA | Gm22235 | | 0 | 0 | 0 | 0 | 0 | 0 | 0 | 0 | 0 | 0 | 0 |
| 0 | NA | NA | NA | NA | NA | Tmem247 | transmembrane protein 247 | 0 | 0 | 0 | 0 | 0 | 0 | 0 | 0 | 0 | 0 | 0 |
| 0 | NA | NA | NA | NA | NA | Gm50011 | | 0 | 0 | 0 | 0 | 0 | 0 | 0 | 0 | 0 | 0 | 0 |
| 0 | NA | NA | NA | NA | NA | Atp6v1e2 | ATPase, H+ transporting, lysosomal V1 su | 0 | 0 | 0 | 0 | 0 | 0 | 0 | 0 | 0 | 0 | 0 |
| 0 | NA | NA | NA | NA | NA | Gm24648 | | 0 | 0 | 0 | 0 | 0 | 0 | 0 | 0 | 0 | 0 | 0 |
| 0 | NA | NA | NA | NA | NA | Gm22346 | | 0 | 0 | 0 | 0 | 0 | 0 | 0 | 0 | 0 | 0 | 0 |
| 0 | NA | NA | NA | NA | NA | C330024C12Rik | | 0 | 0 | 0 | 0 | 0 | 0 | 0 | 0 | 0 | 0 | 0 |
| 0 | NA | NA | NA | NA | NA | 0610012D04Rik | | 0 | 0 | 0 | 0 | 0 | 0 | 0 | 0 | 0 | 0 | 0 |
| 0 | NA | NA | NA | NA | NA | Stpg4 | sperm tail PG rich repeat containing 4 | 0 | 0 | 0 | 0 | 0 | 0 | 0 | 0 | 0 | 0 | 0 |
| 0 | NA | NA | NA | NA | NA | Gm26005 | | 0 | 0 | 0 | 0 | 0 | 0 | 0 | 0 | 0 | 0 | 0 |
| 0 | NA | NA | NA | NA | NA | Gm28676 | | 0 | 0 | 0 | 0 | 0 | 0 | 0 | 0 | 0 | 0 | 0 |
| 0 | NA | NA | NA | NA | NA | Gm46587 | | 0 | 0 | 0 | 0 | 0 | 0 | 0 | 0 | 0 | 0 | 0 |
| 0 | NA | NA | NA | NA | NA | Gm50040 | | 0 | 0 | 0 | 0 | 0 | 0 | 0 | 0 | 0 | 0 | 0 |
| 0 | NA | NA | NA | NA | NA | Gm31615 | | 0 | 0 | 0 | 0 | 0 | 0 | 0 | 0 | 0 | 0 | 0 |
| 0 | NA | NA | NA | NA | NA | 1700116H05Rik | | 0 | 0 | 0 | 0 | 0 | 0 | 0 | 0 | 0 | 0 | 0 |
| 0 | NA | NA | NA | NA | NA | Gm22542 | | 0 | 0 | 0 | 0 | 0 | 0 | 0 | 0 | 0 | 0 | 0 |
| 0 | NA | NA | NA | NA | NA | Gm23947 | | 0 | 0 | 0 | 0 | 0 | 0 | 0 | 0 | 0 | 0 | 0 |
| 0 | NA | NA | NA | NA | NA | Gm9406 | | 0 | 0 | 0 | 0 | 0 | 0 | 0 | 0 | 0 | 0 | 0 |
| 0 | NA | NA | NA | NA | NA | Gm50056 | | 0 | 0 | 0 | 0 | 0 | 0 | 0 | 0 | 0 | 0 | 0 |
| 0 | NA | NA | NA | NA | NA | Fshr | follicle stimulating hormone receptor | 0 | 0 | 0 | 0 | 0 | 0 | 0 | 0 | 0 | 0 | 0 |
| 0 | NA | NA | NA | NA | NA | 4933400B14Rik | | 0 | 0 | 0 | 0 | 0 | 0 | 0 | 0 | 0 | 0 | 0 |
| 0 | NA | NA | NA | NA | NA | Gm4719 | | 0 | 0 | 0 | 0 | 0 | 0 | 0 | 0 | 0 | 0 | 0 |
| 0 | NA | NA | NA | NA | NA | Gm50061 | | 0 | 0 | 0 | 0 | 0 | 0 | 0 | 0 | 0 | 0 | 0 |
| 0 | NA | NA | NA | NA | NA | Gm41653 | | 0 | 0 | 0 | 0 | 0 | 0 | 0 | 0 | 0 | 0 | 0 |
| 0 | NA | NA | NA | NA | NA | Gm24510 | | 0 | 0 | 0 | 0 | 0 | 0 | 0 | 0 | 0 | 0 | 0 |

| | | | | | | | | | | | | | | | | | |
|---|---|---|---|---|---|---|---|---|---|---|---|---|---|---|---|---|---|
| 0 | NA | NA | NA | NA | NA | Gm10493 | | 0 | 0 | 0 | 0 | 0 | 0 | 0 | 0 | 0 | 0 | 0 |
| 0 | NA | NA | NA | NA | NA | Gm20491 | | 0 | 0 | 0 | 0 | 0 | 0 | 0 | 0 | 0 | 0 | 0 |
| 0 | NA | NA | NA | NA | NA | 4930480K15Rik | | 0 | 0 | 0 | 0 | 0 | 0 | 0 | 0 | 0 | 0 | 0 |
| 0 | NA | NA | NA | NA | NA | Gm47307 | | 0 | 0 | 0 | 0 | 0 | 0 | 0 | 0 | 0 | 0 | 0 |
| 0 | NA | NA | NA | NA | NA | Gm27475 | | 0 | 0 | 0 | 0 | 0 | 0 | 0 | 0 | 0 | 0 | 0 |
| 0 | NA | NA | NA | NA | NA | Gm9410 | | 0 | 0 | 0 | 0 | 0 | 0 | 0 | 0 | 0 | 0 | 0 |
| 0 | NA | NA | NA | NA | NA | Gm9411 | | 0 | 0 | 0 | 0 | 0 | 0 | 0 | 0 | 0 | 0 | 0 |
| 0 | NA | NA | NA | NA | NA | Gm6741 | | 0 | 0 | 0 | 0 | 0 | 0 | 0 | 0 | 0 | 0 | 0 |
| 0 | NA | NA | NA | NA | NA | Gm49988 | | 0 | 0 | 0 | 0 | 0 | 0 | 0 | 0 | 0 | 0 | 0 |
| 0 | NA | NA | NA | NA | NA | Gm49989 | | 0 | 0 | 0 | 0 | 0 | 0 | 0 | 0 | 0 | 0 | 0 |
| 0 | NA | NA | NA | NA | NA | Gm15404 | | 0 | 0 | 0 | 0 | 0 | 0 | 0 | 0 | 0 | 0 | 0 |
| 0 | NA | NA | NA | NA | NA | Gm49991 | | 0 | 0 | 0 | 0 | 0 | 0 | 0 | 0 | 0 | 0 | 0 |
| 0 | NA | NA | NA | NA | NA | Gm15403 | | 0 | 0 | 0 | 0 | 0 | 0 | 0 | 0 | 0 | 0 | 0 |
| 0 | NA | NA | NA | NA | NA | Gm49992 | | 0 | 0 | 0 | 0 | 0 | 0 | 0 | 0 | 0 | 0 | 0 |
| 0 | NA | NA | NA | NA | NA | Gm49993 | | 0 | 0 | 0 | 0 | 0 | 0 | 0 | 0 | 0 | 0 | 0 |
| 0 | NA | NA | NA | NA | NA | Gm41654 | | 0 | 0 | 0 | 0 | 0 | 0 | 0 | 0 | 0 | 0 | 0 |
| 0 | NA | NA | NA | NA | NA | Gm24244 | | 0 | 0 | 0 | 0 | 0 | 0 | 0 | 0 | 0 | 0 | 0 |
| 0 | NA | NA | NA | NA | NA | Gm44345 | | 0 | 0 | 0 | 0 | 0 | 0 | 0 | 0 | 0 | 0 | 0 |
| 0 | NA | NA | NA | NA | NA | Gm49994 | | 0 | 0 | 0 | 0 | 0 | 0 | 0 | 0 | 0 | 0 | 0 |
| 0 | NA | NA | NA | NA | NA | Gm19145 | | 0 | 0 | 0 | 0 | 0 | 0 | 0 | 0 | 0 | 0 | 0 |
| 0 | NA | NA | NA | NA | NA | Gm6751 | | 0 | 0 | 0 | 0 | 0 | 0 | 0 | 0 | 0 | 0 | 0 |
| 0 | NA | NA | NA | NA | NA | Gm27039 | | 0 | 0 | 0 | 0 | 0 | 0 | 0 | 0 | 0 | 0 | 0 |
| 0 | NA | NA | NA | NA | NA | Gm49995 | | 0 | 0 | 0 | 0 | 0 | 0 | 0 | 0 | 0 | 0 | 0 |
| 0 | NA | NA | NA | NA | NA | Gm21018 | | 0 | 0 | 0 | 0 | 0 | 0 | 0 | 0 | 0 | 0 | 0 |
| 0 | NA | NA | NA | NA | NA | Gm49997 | | 0 | 0 | 0 | 0 | 0 | 0 | 0 | 0 | 0 | 0 | 0 |
| 0 | NA | NA | NA | NA | NA | Gm49998 | | 0 | 0 | 0 | 0 | 0 | 0 | 0 | 0 | 0 | 0 | 0 |
| 0 | NA | NA | NA | NA | NA | Adcyap1 | adenylate cyclase activating polypeptide | 0 | 0 | 0 | 0 | 0 | 0 | 0 | 0 | 0 | 0 | 0 |
| 0 | NA | NA | NA | NA | NA | Gm25935 | | 0 | 0 | 0 | 0 | 0 | 0 | 0 | 0 | 0 | 0 | 0 |
| 0 | NA | NA | NA | NA | NA | Gm9665 | | 0 | 0 | 0 | 0 | 0 | 0 | 0 | 0 | 0 | 0 | 0 |
| 0 | NA | NA | NA | NA | NA | Gm6764 | | 0 | 0 | 0 | 0 | 0 | 0 | 0 | 0 | 0 | 0 | 0 |
| 0 | NA | NA | NA | NA | NA | Gm50001 | | 0 | 0 | 0 | 0 | 0 | 0 | 0 | 0 | 0 | 0 | 0 |
| 0 | NA | NA | NA | NA | NA | Gm50002 | | 0 | 0 | 0 | 0 | 0 | 0 | 0 | 0 | 0 | 0 | 0 |
| 0 | NA | NA | NA | NA | NA | Gm50003 | | 0 | 0 | 0 | 0 | 0 | 0 | 0 | 0 | 0 | 0 | 0 |
| 0 | NA | NA | NA | NA | NA | Gm50004 | | 0 | 0 | 0 | 0 | 0 | 0 | 0 | 0 | 0 | 0 | 0 |
| 0 | NA | NA | NA | NA | NA | Gm22593 | | 0 | 0 | 0 | 0 | 0 | 0 | 0 | 0 | 0 | 0 | 0 |
| 0 | NA | NA | NA | NA | NA | Gm50005 | | 0 | 0 | 0 | 0 | 0 | 0 | 0 | 0 | 0 | 0 | 0 |
| 0 | NA | NA | NA | NA | NA | Gm23964 | | 0 | 0 | 0 | 0 | 0 | 0 | 0 | 0 | 0 | 0 | 0 |
| 0 | NA | NA | NA | NA | NA | Gm23102 | | 0 | 0 | 0 | 0 | 0 | 0 | 0 | 0 | 0 | 0 | 0 |
| 0 | NA | NA | NA | NA | NA | Gm50072 | | 0 | 0 | 0 | 0 | 0 | 0 | 0 | 0 | 0 | 0 | 0 |
| 0 | NA | NA | NA | NA | NA | Vmn1r-ps151 | | 0 | 0 | 0 | 0 | 0 | 0 | 0 | 0 | 0 | 0 | 0 |
| 0 | NA | NA | NA | NA | NA | Vmn1r-ps152 | | 0 | 0 | 0 | 0 | 0 | 0 | 0 | 0 | 0 | 0 | 0 |
| 0 | NA | NA | NA | NA | NA | Gm50073 | | 0 | 0 | 0 | 0 | 0 | 0 | 0 | 0 | 0 | 0 | 0 |
| 0 | NA | NA | NA | NA | NA | Vmn1r238 | vomeronasal 1 receptor, 238 | 0 | 0 | 0 | 0 | 0 | 0 | 0 | 0 | 0 | 0 | 0 |
| 0 | NA | NA | NA | NA | NA | Gm50074 | | 0 | 0 | 0 | 0 | 0 | 0 | 0 | 0 | 0 | 0 | 0 |
| 0 | NA | NA | NA | NA | NA | Gm50075 | | 0 | 0 | 0 | 0 | 0 | 0 | 0 | 0 | 0 | 0 | 0 |
| 0 | NA | NA | NA | NA | NA | Gm50088 | | 0 | 0 | 0 | 0 | 0 | 0 | 0 | 0 | 0 | 0 | 0 |
| 0 | NA | NA | NA | NA | NA | G430049J08Rik | | 0 | 0 | 0 | 0 | 0 | 0 | 0 | 0 | 0 | 0 | 0 |
| 0 | NA | NA | NA | NA | NA | Gm50089 | | 0 | 0 | 0 | 0 | 0 | 0 | 0 | 0 | 0 | 0 | 0 |
| 0 | NA | NA | NA | NA | NA | Gm50090 | | 0 | 0 | 0 | 0 | 0 | 0 | 0 | 0 | 0 | 0 | 0 |
| 0 | NA | NA | NA | NA | NA | Gm50091 | | 0 | 0 | 0 | 0 | 0 | 0 | 0 | 0 | 0 | 0 | 0 |
| 0 | NA | NA | NA | NA | NA | Gm23643 | | 0 | 0 | 0 | 0 | 0 | 0 | 0 | 0 | 0 | 0 | 0 |
| 0 | NA | NA | NA | NA | NA | Lyzl1 | lysozyme-like 1 | 0 | 0 | 0 | 0 | 0 | 0 | 0 | 0 | 0 | 0 | 0 |
| 0 | NA | NA | NA | NA | NA | Gm7400 | | 0 | 0 | 0 | 0 | 0 | 0 | 0 | 0 | 0 | 0 | 0 |
| 0 | NA | NA | NA | NA | NA | Gm50096 | | 0 | 0 | 0 | 0 | 0 | 0 | 0 | 0 | 0 | 0 | 0 |
| 0 | NA | NA | NA | NA | NA | Gm26865 | | 0 | 0 | 0 | 0 | 0 | 0 | 0 | 0 | 0 | 0 | 0 |
| 0 | NA | NA | NA | NA | NA | Gm50024 | | 0 | 0 | 0 | 0 | 0 | 0 | 0 | 0 | 0 | 0 | 0 |
| 0 | NA | NA | NA | NA | NA | Gm10556 | | 0 | 0 | 0 | 0 | 0 | 0 | 0 | 0 | 0 | 0 | 0 |
| 0 | NA | NA | NA | NA | NA | Gm50064 | | 0 | 0 | 0 | 0 | 0 | 0 | 0 | 0 | 0 | 0 | 0 |
| 0 | NA | NA | NA | NA | NA | Gm50065 | | 0 | 0 | 0 | 0 | 0 | 0 | 0 | 0 | 0 | 0 | 0 |
| 0 | NA | NA | NA | NA | NA | Gm16147 | | 0 | 0 | 0 | 0 | 0 | 0 | 0 | 0 | 0 | 0 | 0 |

| | | | | | | | | | | | | | | | | | |
|---|---|---|---|---|---|---|---|---|---|---|---|---|---|---|---|---|---|
| 0 | NA | NA | NA | NA | NA | Gm22709 | | 0 | 0 | 0 | 0 | 0 | 0 | 0 | 0 | 0 | 0 | 0 |
| 0 | NA | NA | NA | NA | NA | Gm34804 | | 0 | 0 | 0 | 0 | 0 | 0 | 0 | 0 | 0 | 0 | 0 |
| 0 | NA | NA | NA | NA | NA | Gm49986 | | 0 | 0 | 0 | 0 | 0 | 0 | 0 | 0 | 0 | 0 | 0 |
| 0 | NA | NA | NA | NA | NA | Mir1893 | | 0 | 0 | 0 | 0 | 0 | 0 | 0 | 0 | 0 | 0 | 0 |
| 0 | NA | NA | NA | NA | NA | 4921524L21Rik | RIKEN cDNA 4921524L21 gene | 0 | 0 | 0 | 0 | 0 | 0 | 0 | 0 | 0 | 0 | 0 |
| 0 | NA | NA | NA | NA | NA | Gm50077 | | 0 | 0 | 0 | 0 | 0 | 0 | 0 | 0 | 0 | 0 | 0 |
| 0 | NA | NA | NA | NA | NA | Gm50081 | | 0 | 0 | 0 | 0 | 0 | 0 | 0 | 0 | 0 | 0 | 0 |
| 0 | NA | NA | NA | NA | NA | Gm2350 | | 0 | 0 | 0 | 0 | 0 | 0 | 0 | 0 | 0 | 0 | 0 |
| 0 | NA | NA | NA | NA | NA | Fabp5l2 | | 0 | 0 | 0 | 0 | 0 | 0 | 0 | 0 | 0 | 0 | 0 |
| 0 | NA | NA | NA | NA | NA | Gm24292 | | 0 | 0 | 0 | 0 | 0 | 0 | 0 | 0 | 0 | 0 | 0 |
| 0 | NA | NA | NA | NA | NA | Gm50027 | | 0 | 0 | 0 | 0 | 0 | 0 | 0 | 0 | 0 | 0 | 0 |
| 0 | NA | NA | NA | NA | NA | Gm35031 | | 0 | 0 | 0 | 0 | 0 | 0 | 0 | 0 | 0 | 0 | 0 |
| 0 | NA | NA | NA | NA | NA | Gm35232 | | 0 | 0 | 0 | 0 | 0 | 0 | 0 | 0 | 0 | 0 | 0 |
| 0 | NA | NA | NA | NA | NA | Gm26119 | | 0 | 0 | 0 | 0 | 0 | 0 | 0 | 0 | 0 | 0 | 0 |
| 0 | NA | NA | NA | NA | NA | Gm37148 | | 0 | 0 | 0 | 0 | 0 | 0 | 0 | 0 | 0 | 0 | 0 |
| 0 | NA | NA | NA | NA | NA | Gm7523 | | 0 | 0 | 0 | 0 | 0 | 0 | 0 | 0 | 0 | 0 | 0 |
| 0 | NA | NA | NA | NA | NA | Gm18526 | | 0 | 0 | 0 | 0 | 0 | 0 | 0 | 0 | 0 | 0 | 0 |
| 0 | NA | NA | NA | NA | NA | Gjd4 | gap junction protein, delta 4 | 0 | 0 | 0 | 0 | 0 | 0 | 0 | 0 | 0 | 0 | 0 |
| 0 | NA | NA | NA | NA | NA | Gm50053 | | 0 | 0 | 0 | 0 | 0 | 0 | 0 | 0 | 0 | 0 | 0 |
| 0 | NA | NA | NA | NA | NA | Gm7274 | | 0 | 0 | 0 | 0 | 0 | 0 | 0 | 0 | 0 | 0 | 0 |
| 0 | NA | NA | NA | NA | NA | Gm46641 | | 0 | 0 | 0 | 0 | 0 | 0 | 0 | 0 | 0 | 0 | 0 |
| 0 | NA | NA | NA | NA | NA | Gm22765 | | 0 | 0 | 0 | 0 | 0 | 0 | 0 | 0 | 0 | 0 | 0 |
| 0 | NA | NA | NA | NA | NA | Cetn1 | centrin 1 | 0 | 0 | 0 | 0 | 0 | 0 | 0 | 0 | 0 | 0 | 0 |
| 0 | NA | NA | NA | NA | NA | Gm50080 | | 0 | 0 | 0 | 0 | 0 | 0 | 0 | 0 | 0 | 0 | 0 |
| 0 | NA | NA | NA | NA | NA | Gm50078 | | 0 | 0 | 0 | 0 | 0 | 0 | 0 | 0 | 0 | 0 | 0 |
| 0 | NA | NA | NA | NA | NA | Gm23350 | | 0 | 0 | 0 | 0 | 0 | 0 | 0 | 0 | 0 | 0 | 0 |
| 0 | NA | NA | NA | NA | NA | Gm50079 | | 0 | 0 | 0 | 0 | 0 | 0 | 0 | 0 | 0 | 0 | 0 |
| 0 | NA | NA | NA | NA | NA | Gm23637 | | 0 | 0 | 0 | 0 | 0 | 0 | 0 | 0 | 0 | 0 | 0 |
| 0 | NA | NA | NA | NA | NA | Gm2424 | | 0 | 0 | 0 | 0 | 0 | 0 | 0 | 0 | 0 | 0 | 0 |
| 0 | NA | NA | NA | NA | NA | Gm5501 | | 0 | 0 | 0 | 0 | 0 | 0 | 0 | 0 | 0 | 0 | 0 |
| 0 | NA | NA | NA | NA | NA | Gm26244 | | 0 | 0 | 0 | 0 | 0 | 0 | 0 | 0 | 0 | 0 | 0 |
| 0 | NA | NA | NA | NA | NA | Gm50032 | | 0 | 0 | 0 | 0 | 0 | 0 | 0 | 0 | 0 | 0 | 0 |
| 0 | NA | NA | NA | NA | NA | Gm7156 | | 0 | 0 | 0 | 0 | 0 | 0 | 0 | 0 | 0 | 0 | 0 |
| 0 | NA | NA | NA | NA | NA | Gm19052 | | 0 | 0 | 0 | 0 | 0 | 0 | 0 | 0 | 0 | 0 | 0 |
| 0 | NA | NA | NA | NA | NA | Gm50033 | | 0 | 0 | 0 | 0 | 0 | 0 | 0 | 0 | 0 | 0 | 0 |
| 0 | NA | NA | NA | NA | NA | Gm44409 | | 0 | 0 | 0 | 0 | 0 | 0 | 0 | 0 | 0 | 0 | 0 |
| 0 | NA | NA | NA | NA | NA | Nutf2-ps2 | | 0 | 0 | 0 | 0 | 0 | 0 | 0 | 0 | 0 | 0 | 0 |
| 0 | NA | NA | NA | NA | NA | Gm24894 | | 0 | 0 | 0 | 0 | 0 | 0 | 0 | 0 | 0 | 0 | 0 |
| 0 | NA | NA | NA | NA | NA | Gm25116 | | 0 | 0 | 0 | 0 | 0 | 0 | 0 | 0 | 0 | 0 | 0 |
| 0 | NA | NA | NA | NA | NA | Mir133a-1hg | | 0 | 0 | 0 | 0 | 0 | 0 | 0 | 0 | 0 | 0 | 0 |
| 0 | NA | NA | NA | NA | NA | Mir133a-1 | | 0 | 0 | 0 | 0 | 0 | 0 | 0 | 0 | 0 | 0 | 0 |
| 0 | NA | NA | NA | NA | NA | Mir1b | | 0 | 0 | 0 | 0 | 0 | 0 | 0 | 0 | 0 | 0 | 0 |
| 0 | NA | NA | NA | NA | NA | Mir1a-2 | | 0 | 0 | 0 | 0 | 0 | 0 | 0 | 0 | 0 | 0 | 0 |
| 0 | NA | NA | NA | NA | NA | Gm18764 | | 0 | 0 | 0 | 0 | 0 | 0 | 0 | 0 | 0 | 0 | 0 |
| 0 | NA | NA | NA | NA | NA | Gm18713 | | 0 | 0 | 0 | 0 | 0 | 0 | 0 | 0 | 0 | 0 | 0 |
| 0 | NA | NA | NA | NA | NA | Mir1901 | | 0 | 0 | 0 | 0 | 0 | 0 | 0 | 0 | 0 | 0 | 0 |
| 0 | NA | NA | NA | NA | NA | Gm25289 | | 0 | 0 | 0 | 0 | 0 | 0 | 0 | 0 | 0 | 0 | 0 |
| 0 | NA | NA | NA | NA | NA | Gm29201 | | 0 | 0 | 0 | 0 | 0 | 0 | 0 | 0 | 0 | 0 | 0 |
| 0 | NA | NA | NA | NA | NA | Gm29199 | | 0 | 0 | 0 | 0 | 0 | 0 | 0 | 0 | 0 | 0 | 0 |
| 0 | NA | NA | NA | NA | NA | Gm29200 | | 0 | 0 | 0 | 0 | 0 | 0 | 0 | 0 | 0 | 0 | 0 |
| 0 | NA | NA | NA | NA | NA | Gm22340 | | 0 | 0 | 0 | 0 | 0 | 0 | 0 | 0 | 0 | 0 | 0 |
| 0 | NA | NA | NA | NA | NA | Gm50063 | | 0 | 0 | 0 | 0 | 0 | 0 | 0 | 0 | 0 | 0 | 0 |
| 0 | NA | NA | NA | NA | NA | Gm41668 | | 0 | 0 | 0 | 0 | 0 | 0 | 0 | 0 | 0 | 0 | 0 |
| 0 | NA | NA | NA | NA | NA | Mir1948 | | 0 | 0 | 0 | 0 | 0 | 0 | 0 | 0 | 0 | 0 | 0 |
| 0 | NA | NA | NA | NA | NA | Gm7599 | | 0 | 0 | 0 | 0 | 0 | 0 | 0 | 0 | 0 | 0 | 0 |
| 0 | NA | NA | NA | NA | NA | Gm18083 | | 0 | 0 | 0 | 0 | 0 | 0 | 0 | 0 | 0 | 0 | 0 |
| 0 | NA | NA | NA | NA | NA | Gm22251 | | 0 | 0 | 0 | 0 | 0 | 0 | 0 | 0 | 0 | 0 | 0 |
| 0 | NA | NA | NA | NA | NA | Gm41669 | | 0 | 0 | 0 | 0 | 0 | 0 | 0 | 0 | 0 | 0 | 0 |
| 0 | NA | NA | NA | NA | NA | Gm50094 | | 0 | 0 | 0 | 0 | 0 | 0 | 0 | 0 | 0 | 0 | 0 |
| 0 | NA | NA | NA | NA | NA | 8430422H06Rik | | 0 | 0 | 0 | 0 | 0 | 0 | 0 | 0 | 0 | 0 | 0 |

| | | | | | | | | | | | | | | | | | | |
|---|---|---|---|---|---|---|---|---|---|---|---|---|---|---|---|---|---|---|
| 0 | NA | NA | NA | NA | NA | Gm50095 | | 0 | 0 | 0 | 0 | 0 | 0 | 0 | 0 | 0 | 0 | 0 |
| 0 | NA | NA | NA | NA | NA | Gm50097 | | 0 | 0 | 0 | 0 | 0 | 0 | 0 | 0 | 0 | 0 | 0 |
| 0 | NA | NA | NA | NA | NA | Gm24924 | | 0 | 0 | 0 | 0 | 0 | 0 | 0 | 0 | 0 | 0 | 0 |
| 0 | NA | NA | NA | NA | NA | Gm21055 | | 0 | 0 | 0 | 0 | 0 | 0 | 0 | 0 | 0 | 0 | 0 |
| 0 | NA | NA | NA | NA | NA | Gm9474 | | 0 | 0 | 0 | 0 | 0 | 0 | 0 | 0 | 0 | 0 | 0 |
| 0 | NA | NA | NA | NA | NA | 4933424G05Rik | | 0 | 0 | 0 | 0 | 0 | 0 | 0 | 0 | 0 | 0 | 0 |
| 0 | NA | NA | NA | NA | NA | Gm50035 | | 0 | 0 | 0 | 0 | 0 | 0 | 0 | 0 | 0 | 0 | 0 |
| 0 | NA | NA | NA | NA | NA | E430002N23Rik | | 0 | 0 | 0 | 0 | 0 | 0 | 0 | 0 | 0 | 0 | 0 |
| 0 | NA | NA | NA | NA | NA | A830021F12Rik | | 0 | 0 | 0 | 0 | 0 | 0 | 0 | 0 | 0 | 0 | 0 |
| 0 | NA | NA | NA | NA | NA | Gm24228 | | 0 | 0 | 0 | 0 | 0 | 0 | 0 | 0 | 0 | 0 | 0 |
| 0 | NA | NA | NA | NA | NA | Gm22389 | | 0 | 0 | 0 | 0 | 0 | 0 | 0 | 0 | 0 | 0 | 0 |
| 0 | NA | NA | NA | NA | NA | Gm50076 | | 0 | 0 | 0 | 0 | 0 | 0 | 0 | 0 | 0 | 0 | 0 |
| 0 | NA | NA | NA | NA | NA | Gm24385 | | 0 | 0 | 0 | 0 | 0 | 0 | 0 | 0 | 0 | 0 | 0 |
| 0 | NA | NA | NA | NA | NA | Gm15485 | | 0 | 0 | 0 | 0 | 0 | 0 | 0 | 0 | 0 | 0 | 0 |
| 0 | NA | NA | NA | NA | NA | 1700001G01Rik | | 0 | 0 | 0 | 0 | 0 | 0 | 0 | 0 | 0 | 0 | 0 |
| 0 | NA | NA | NA | NA | NA | Gm22470 | | 0 | 0 | 0 | 0 | 0 | 0 | 0 | 0 | 0 | 0 | 0 |
| 0 | NA | NA | NA | NA | NA | Gm24015 | | 0 | 0 | 0 | 0 | 0 | 0 | 0 | 0 | 0 | 0 | 0 |
| 0 | NA | NA | NA | NA | NA | 4921533I20Rik | | 0 | 0 | 0 | 0 | 0 | 0 | 0 | 0 | 0 | 0 | 0 |
| 0 | NA | NA | NA | NA | NA | 4930545E07Rik | | 0 | 0 | 0 | 0 | 0 | 0 | 0 | 0 | 0 | 0 | 0 |
| 0 | NA | NA | NA | NA | NA | Gm50082 | | 0 | 0 | 0 | 0 | 0 | 0 | 0 | 0 | 0 | 0 | 0 |
| 0 | NA | NA | NA | NA | NA | Gm50083 | | 0 | 0 | 0 | 0 | 0 | 0 | 0 | 0 | 0 | 0 | 0 |
| 0 | NA | NA | NA | NA | NA | Gm26235 | | 0 | 0 | 0 | 0 | 0 | 0 | 0 | 0 | 0 | 0 | 0 |
| 0 | NA | NA | NA | NA | NA | Gm50084 | | 0 | 0 | 0 | 0 | 0 | 0 | 0 | 0 | 0 | 0 | 0 |
| 0 | NA | NA | NA | NA | NA | Gm31086 | | 0 | 0 | 0 | 0 | 0 | 0 | 0 | 0 | 0 | 0 | 0 |
| 0 | NA | NA | NA | NA | NA | Gm38432 | | 0 | 0 | 0 | 0 | 0 | 0 | 0 | 0 | 0 | 0 | 0 |
| 0 | NA | NA | NA | NA | NA | Gm7720 | | 0 | 0 | 0 | 0 | 0 | 0 | 0 | 0 | 0 | 0 | 0 |
| 0 | NA | NA | NA | NA | NA | Gm25392 | | 0 | 0 | 0 | 0 | 0 | 0 | 0 | 0 | 0 | 0 | 0 |
| 0 | NA | NA | NA | NA | NA | Gm24238 | | 0 | 0 | 0 | 0 | 0 | 0 | 0 | 0 | 0 | 0 | 0 |
| 0 | NA | NA | NA | NA | NA | Dsg1a | desmoglein 1 alpha | 0 | 0 | 0 | 0 | 0 | 0 | 0 | 0 | 0 | 0 | 0 |
| 0 | NA | NA | NA | NA | NA | Dsg4 | desmoglein 4 | 0 | 0 | 0 | 0 | 0 | 0 | 0 | 0 | 0 | 0 | 0 |
| 0 | NA | NA | NA | NA | NA | Mep1b | meprin 1 beta | 0 | 0 | 0 | 0 | 0 | 0 | 0 | 0 | 0 | 0 | 0 |
| 0 | NA | NA | NA | NA | NA | Gm24372 | | 0 | 0 | 0 | 0 | 0 | 0 | 0 | 0 | 0 | 0 | 0 |
| 0 | NA | NA | NA | NA | NA | Gm50049 | | 0 | 0 | 0 | 0 | 0 | 0 | 0 | 0 | 0 | 0 | 0 |
| 0 | NA | NA | NA | NA | NA | Gm50052 | | 0 | 0 | 0 | 0 | 0 | 0 | 0 | 0 | 0 | 0 | 0 |
| 0 | NA | NA | NA | NA | NA | Gm22886 | | 0 | 0 | 0 | 0 | 0 | 0 | 0 | 0 | 0 | 0 | 0 |
| 0 | NA | NA | NA | NA | NA | Gm22495 | | 0 | 0 | 0 | 0 | 0 | 0 | 0 | 0 | 0 | 0 | 0 |
| 0 | NA | NA | NA | NA | NA | Mir6360 | | 0 | 0 | 0 | 0 | 0 | 0 | 0 | 0 | 0 | 0 | 0 |
| 0 | NA | NA | NA | NA | NA | Gm23207 | | 0 | 0 | 0 | 0 | 0 | 0 | 0 | 0 | 0 | 0 | 0 |
| 0 | NA | NA | NA | NA | NA | Gm18085 | | 0 | 0 | 0 | 0 | 0 | 0 | 0 | 0 | 0 | 0 | 0 |
| 0 | NA | NA | NA | NA | NA | Mir187 | | 0 | 0 | 0 | 0 | 0 | 0 | 0 | 0 | 0 | 0 | 0 |
| 0 | NA | NA | NA | NA | NA | Gm23606 | | 0 | 0 | 0 | 0 | 0 | 0 | 0 | 0 | 0 | 0 | 0 |
| 0 | NA | NA | NA | NA | NA | n-R5s23 | | 0 | 0 | 0 | 0 | 0 | 0 | 0 | 0 | 0 | 0 | 0 |
| 0 | NA | NA | NA | NA | NA | Gm24294 | | 0 | 0 | 0 | 0 | 0 | 0 | 0 | 0 | 0 | 0 | 0 |
| 0 | NA | NA | NA | NA | NA | Gm23674 | | 0 | 0 | 0 | 0 | 0 | 0 | 0 | 0 | 0 | 0 | 0 |
| 0 | NA | NA | NA | NA | NA | Gm49976 | | 0 | 0 | 0 | 0 | 0 | 0 | 0 | 0 | 0 | 0 | 0 |
| 0 | NA | NA | NA | NA | NA | Gm26106 | | 0 | 0 | 0 | 0 | 0 | 0 | 0 | 0 | 0 | 0 | 0 |
| 0 | NA | NA | NA | NA | NA | Gm18089 | | 0 | 0 | 0 | 0 | 0 | 0 | 0 | 0 | 0 | 0 | 0 |
| 0 | NA | NA | NA | NA | NA | Gm7874 | | 0 | 0 | 0 | 0 | 0 | 0 | 0 | 0 | 0 | 0 | 0 |
| 0 | NA | NA | NA | NA | NA | Gm49977 | | 0 | 0 | 0 | 0 | 0 | 0 | 0 | 0 | 0 | 0 | 0 |
| 0 | NA | NA | NA | NA | NA | Gm49978 | | 0 | 0 | 0 | 0 | 0 | 0 | 0 | 0 | 0 | 0 | 0 |
| 0 | NA | NA | NA | NA | NA | Gm33674 | | 0 | 0 | 0 | 0 | 0 | 0 | 0 | 0 | 0 | 0 | 0 |
| 0 | NA | NA | NA | NA | NA | Gm24246 | | 0 | 0 | 0 | 0 | 0 | 0 | 0 | 0 | 0 | 0 | 0 |
| 0 | NA | NA | NA | NA | NA | Gm34658 | | 0 | 0 | 0 | 0 | 0 | 0 | 0 | 0 | 0 | 0 | 0 |
| 0 | NA | NA | NA | NA | NA | Gm20096 | | 0 | 0 | 0 | 0 | 0 | 0 | 0 | 0 | 0 | 0 | 0 |
| 0 | NA | NA | NA | NA | NA | Gm33948 | | 0 | 0 | 0 | 0 | 0 | 0 | 0 | 0 | 0 | 0 | 0 |
| 0 | NA | NA | NA | NA | NA | 4930578E11Rik | | 0 | 0 | 0 | 0 | 0 | 0 | 0 | 0 | 0 | 0 | 0 |
| 0 | NA | NA | NA | NA | NA | 4930527G23Rik | | 0 | 0 | 0 | 0 | 0 | 0 | 0 | 0 | 0 | 0 | 0 |
| 0 | NA | NA | NA | NA | NA | Gm7917 | | 0 | 0 | 0 | 0 | 0 | 0 | 0 | 0 | 0 | 0 | 0 |
| 0 | NA | NA | NA | NA | NA | Gm7930 | | 0 | 0 | 0 | 0 | 0 | 0 | 0 | 0 | 0 | 0 | 0 |
| 0 | NA | NA | NA | NA | NA | Gm22870 | | 0 | 0 | 0 | 0 | 0 | 0 | 0 | 0 | 0 | 0 | 0 |

| | | | | | | | | | | | | | | | | | |
|---|---|---|---|---|---|---|---|---|---|---|---|---|---|---|---|---|---|
| 0 | NA | NA | NA | NA | NA | Gm7941 | | 0 | 0 | 0 | 0 | 0 | 0 | 0 | 0 | 0 | 0 | 0 |
| 0 | NA | NA | NA | NA | NA | Gm23283 | | 0 | 0 | 0 | 0 | 0 | 0 | 0 | 0 | 0 | 0 | 0 |
| 0 | NA | NA | NA | NA | NA | Gm25396 | | 0 | 0 | 0 | 0 | 0 | 0 | 0 | 0 | 0 | 0 | 0 |
| 0 | NA | NA | NA | NA | NA | Gm5241 | | 0 | 0 | 0 | 0 | 0 | 0 | 0 | 0 | 0 | 0 | 0 |
| 0 | NA | NA | NA | NA | NA | Gm16345 | | 0 | 0 | 0 | 0 | 0 | 0 | 0 | 0 | 0 | 0 | 0 |
| 0 | NA | NA | NA | NA | NA | 4933435E02Rik | | 0 | 0 | 0 | 0 | 0 | 0 | 0 | 0 | 0 | 0 | 0 |
| 0 | NA | NA | NA | NA | NA | Gm22325 | | 0 | 0 | 0 | 0 | 0 | 0 | 0 | 0 | 0 | 0 | 0 |
| 0 | NA | NA | NA | NA | NA | Gm35060 | predicted gene, 35060 | 0 | 0 | 0 | 0 | 0 | 0 | 0 | 0 | 0 | 0 | 0 |
| 0 | NA | NA | NA | NA | NA | Gm20136 | | 0 | 0 | 0 | 0 | 0 | 0 | 0 | 0 | 0 | 0 | 0 |
| 0 | NA | NA | NA | NA | NA | Gm25826 | | 0 | 0 | 0 | 0 | 0 | 0 | 0 | 0 | 0 | 0 | 0 |
| 0 | NA | NA | NA | NA | NA | Gm26378 | | 0 | 0 | 0 | 0 | 0 | 0 | 0 | 0 | 0 | 0 | 0 |
| 0 | NA | NA | NA | NA | NA | Mir6361 | | 0 | 0 | 0 | 0 | 0 | 0 | 0 | 0 | 0 | 0 | 0 |
| 0 | NA | NA | NA | NA | NA | Gm22814 | | 0 | 0 | 0 | 0 | 0 | 0 | 0 | 0 | 0 | 0 | 0 |
| 0 | NA | NA | NA | NA | NA | Gm50392 | | 0 | 0 | 0 | 0 | 0 | 0 | 0 | 0 | 0 | 0 | 0 |
| 0 | NA | NA | NA | NA | NA | Gm22788 | | 0 | 0 | 0 | 0 | 0 | 0 | 0 | 0 | 0 | 0 | 0 |
| 0 | NA | NA | NA | NA | NA | Gm50265 | | 0 | 0 | 0 | 0 | 0 | 0 | 0 | 0 | 0 | 0 | 0 |
| 0 | NA | NA | NA | NA | NA | Gm24432 | | 0 | 0 | 0 | 0 | 0 | 0 | 0 | 0 | 0 | 0 | 0 |
| 0 | NA | NA | NA | NA | NA | Gm5238 | | 0 | 0 | 0 | 0 | 0 | 0 | 0 | 0 | 0 | 0 | 0 |
| 0 | NA | NA | NA | NA | NA | Gm23411 | | 0 | 0 | 0 | 0 | 0 | 0 | 0 | 0 | 0 | 0 | 0 |
| 0 | NA | NA | NA | NA | NA | n-R5s24 | | 0 | 0 | 0 | 0 | 0 | 0 | 0 | 0 | 0 | 0 | 0 |
| 0 | NA | NA | NA | NA | NA | Wnt8a | wingless-type MMTV integration site famil | 0 | 0 | 0 | 0 | 0 | 0 | 0 | 0 | 0 | 0 | 0 |
| 0 | NA | NA | NA | NA | NA | n-R5s25 | | 0 | 0 | 0 | 0 | 0 | 0 | 0 | 0 | 0 | 0 | 0 |
| 0 | NA | NA | NA | NA | NA | Gm36037 | | 0 | 0 | 0 | 0 | 0 | 0 | 0 | 0 | 0 | 0 | 0 |
| 0 | NA | NA | NA | NA | NA | Gm3587 | | 0 | 0 | 0 | 0 | 0 | 0 | 0 | 0 | 0 | 0 | 0 |
| 0 | NA | NA | NA | NA | NA | Gm50146 | | 0 | 0 | 0 | 0 | 0 | 0 | 0 | 0 | 0 | 0 | 0 |
| 0 | NA | NA | NA | NA | NA | Snhg4 | | 0 | 0 | 0 | 0 | 0 | 0 | 0 | 0 | 0 | 0 | 0 |
| 0 | NA | NA | NA | NA | NA | Mir1949 | | 0 | 0 | 0 | 0 | 0 | 0 | 0 | 0 | 0 | 0 | 0 |
| 0 | NA | NA | NA | NA | NA | Mir1949 | | 0 | 0 | 0 | 0 | 0 | 0 | 0 | 0 | 0 | 0 | 0 |
| 0 | NA | NA | NA | NA | NA | Snora74a | | 0 | 0 | 0 | 0 | 0 | 0 | 0 | 0 | 0 | 0 | 0 |
| 0 | NA | NA | NA | NA | NA | Gm49347 | | 0 | 0 | 0 | 0 | 0 | 0 | 0 | 0 | 0 | 0 | 0 |
| 0 | NA | NA | NA | NA | NA | Slc23a1 | solute carrier family 23 (nucleobase trans | 0 | 0 | 0 | 0 | 0 | 0 | 0 | 0 | 0 | 0 | 0 |
| 0 | NA | NA | NA | NA | NA | Gm50318 | | 0 | 0 | 0 | 0 | 0 | 0 | 0 | 0 | 0 | 0 | 0 |
| 0 | NA | NA | NA | NA | NA | Gm50371 | | 0 | 0 | 0 | 0 | 0 | 0 | 0 | 0 | 0 | 0 | 0 |
| 0 | NA | NA | NA | NA | NA | C330008A17Rik | | 0 | 0 | 0 | 0 | 0 | 0 | 0 | 0 | 0 | 0 | 0 |
| 0 | NA | NA | NA | NA | NA | Gm23882 | | 0 | 0 | 0 | 0 | 0 | 0 | 0 | 0 | 0 | 0 | 0 |
| 0 | NA | NA | NA | NA | NA | 4930471G03Rik | | 0 | 0 | 0 | 0 | 0 | 0 | 0 | 0 | 0 | 0 | 0 |
| 0 | NA | NA | NA | NA | NA | Gm50199 | | 0 | 0 | 0 | 0 | 0 | 0 | 0 | 0 | 0 | 0 | 0 |
| 0 | NA | NA | NA | NA | NA | Slc4a9 | solute carrier family 4, sodium bicarbonat | 0 | 0 | 0 | 0 | 0 | 0 | 0 | 0 | 0 | 0 | 0 |
| 0 | NA | NA | NA | NA | NA | Gm23424 | | 0 | 0 | 0 | 0 | 0 | 0 | 0 | 0 | 0 | 0 | 0 |
| 0 | NA | NA | NA | NA | NA | Gm37751 | | 0 | 0 | 0 | 0 | 0 | 0 | 0 | 0 | 0 | 0 | 0 |
| 0 | NA | NA | NA | NA | NA | Pcdha4 | protocadherin alpha 4 | 0 | 0 | 0 | 0 | 0 | 0 | 0 | 0 | 0 | 0 | 0 |
| 0 | NA | NA | NA | NA | NA | Gm38225 | | 0 | 0 | 0 | 0 | 0 | 0 | 0 | 0 | 0 | 0 | 0 |
| 0 | NA | NA | NA | NA | NA | Gm42416 | predicted gene, 37013 | 0 | 0 | 0 | 0 | 0 | 0 | 0 | 0 | 0 | 0 | 0 |
| 0 | NA | NA | NA | NA | NA | Gm37013 | | 0 | 0 | 0 | 0 | 0 | 0 | 0 | 0 | 0 | 0 | 0 |
| 0 | NA | NA | NA | NA | NA | Gm37388 | | 0 | 0 | 0 | 0 | 0 | 0 | 0 | 0 | 0 | 0 | 0 |
| 0 | NA | NA | NA | NA | NA | Gm18150 | | 0 | 0 | 0 | 0 | 0 | 0 | 0 | 0 | 0 | 0 | 0 |
| 0 | NA | NA | NA | NA | NA | Gm38097 | | 0 | 0 | 0 | 0 | 0 | 0 | 0 | 0 | 0 | 0 | 0 |
| 0 | NA | NA | NA | NA | NA | Gm18529 | | 0 | 0 | 0 | 0 | 0 | 0 | 0 | 0 | 0 | 0 | 0 |
| 0 | NA | NA | NA | NA | NA | 4930517L18Rik | | 0 | 0 | 0 | 0 | 0 | 0 | 0 | 0 | 0 | 0 | 0 |
| 0 | NA | NA | NA | NA | NA | Gm37118 | | 0 | 0 | 0 | 0 | 0 | 0 | 0 | 0 | 0 | 0 | 0 |
| 0 | NA | NA | NA | NA | NA | Gm24401 | | 0 | 0 | 0 | 0 | 0 | 0 | 0 | 0 | 0 | 0 | 0 |
| 0 | NA | NA | NA | NA | NA | Gm38182 | | 0 | 0 | 0 | 0 | 0 | 0 | 0 | 0 | 0 | 0 | 0 |
| 0 | NA | NA | NA | NA | NA | Mir6979 | | 0 | 0 | 0 | 0 | 0 | 0 | 0 | 0 | 0 | 0 | 0 |
| 0 | NA | NA | NA | NA | NA | Gm50122 | | 0 | 0 | 0 | 0 | 0 | 0 | 0 | 0 | 0 | 0 | 0 |
| 0 | NA | NA | NA | NA | NA | Mir6981 | | 0 | 0 | 0 | 0 | 0 | 0 | 0 | 0 | 0 | 0 | 0 |
| 0 | NA | NA | NA | NA | NA | Mir6980 | | 0 | 0 | 0 | 0 | 0 | 0 | 0 | 0 | 0 | 0 | 0 |
| 0 | NA | NA | NA | NA | NA | Gm24690 | | 0 | 0 | 0 | 0 | 0 | 0 | 0 | 0 | 0 | 0 | 0 |
| 0 | NA | NA | NA | NA | NA | Gm50344 | | 0 | 0 | 0 | 0 | 0 | 0 | 0 | 0 | 0 | 0 | 0 |
| 0 | NA | NA | NA | NA | NA | Gm50349 | | 0 | 0 | 0 | 0 | 0 | 0 | 0 | 0 | 0 | 0 | 0 |
| 0 | NA | NA | NA | NA | NA | Gm19100 | | 0 | 0 | 0 | 0 | 0 | 0 | 0 | 0 | 0 | 0 | 0 |

| | | | | | | | | | | | | | | | | | |
|---|---|---|---|---|---|---|---|---|---|---|---|---|---|---|---|---|---|
| 0 | NA | NA | NA | NA | NA | Gm27369 | | 0 | 0 | 0 | 0 | 0 | 0 | 0 | 0 | 0 | 0 | 0 |
| 0 | NA | NA | NA | NA | NA | Gm27804 | | 0 | 0 | 0 | 0 | 0 | 0 | 0 | 0 | 0 | 0 | 0 |
| 0 | NA | NA | NA | NA | NA | Arhgap26 | | 0 | 0 | 0 | 0 | 0 | 0 | 0 | 0 | 0 | 0 | 0 |
| 0 | NA | NA | NA | NA | NA | Gm15335 | | 0 | 0 | 0 | 0 | 0 | 0 | 0 | 0 | 0 | 0 | 0 |
| 0 | NA | NA | NA | NA | NA | Gm15333 | | 0 | 0 | 0 | 0 | 0 | 0 | 0 | 0 | 0 | 0 | 0 |
| 0 | NA | NA | NA | NA | NA | Gm50397 | | 0 | 0 | 0 | 0 | 0 | 0 | 0 | 0 | 0 | 0 | 0 |
| 0 | NA | NA | NA | NA | NA | Gm50398 | | 0 | 0 | 0 | 0 | 0 | 0 | 0 | 0 | 0 | 0 | 0 |
| 0 | NA | NA | NA | NA | NA | Gm41708 | | 0 | 0 | 0 | 0 | 0 | 0 | 0 | 0 | 0 | 0 | 0 |
| 0 | NA | NA | NA | NA | NA | Gm50395 | | 0 | 0 | 0 | 0 | 0 | 0 | 0 | 0 | 0 | 0 | 0 |
| 0 | NA | NA | NA | NA | NA | Gm50396 | | 0 | 0 | 0 | 0 | 0 | 0 | 0 | 0 | 0 | 0 | 0 |
| 0 | NA | NA | NA | NA | NA | Gm50403 | | 0 | 0 | 0 | 0 | 0 | 0 | 0 | 0 | 0 | 0 | 0 |
| 0 | NA | NA | NA | NA | NA | Gm31205 | | 0 | 0 | 0 | 0 | 0 | 0 | 0 | 0 | 0 | 0 | 0 |
| 0 | NA | NA | NA | NA | NA | Gm22515 | | 0 | 0 | 0 | 0 | 0 | 0 | 0 | 0 | 0 | 0 | 0 |
| 0 | NA | NA | NA | NA | NA | Gm50407 | | 0 | 0 | 0 | 0 | 0 | 0 | 0 | 0 | 0 | 0 | 0 |
| 0 | NA | NA | NA | NA | NA | Gm22483 | | 0 | 0 | 0 | 0 | 0 | 0 | 0 | 0 | 0 | 0 | 0 |
| 0 | NA | NA | NA | NA | NA | Gm44476 | | 0 | 0 | 0 | 0 | 0 | 0 | 0 | 0 | 0 | 0 | 0 |
| 0 | NA | NA | NA | NA | NA | Gm50408 | | 0 | 0 | 0 | 0 | 0 | 0 | 0 | 0 | 0 | 0 | 0 |
| 0 | NA | NA | NA | NA | NA | Gm50409 | | 0 | 0 | 0 | 0 | 0 | 0 | 0 | 0 | 0 | 0 | 0 |
| 0 | NA | NA | NA | NA | NA | Gm18530 | | 0 | 0 | 0 | 0 | 0 | 0 | 0 | 0 | 0 | 0 | 0 |
| 0 | NA | NA | NA | NA | NA | Gm50410 | | 0 | 0 | 0 | 0 | 0 | 0 | 0 | 0 | 0 | 0 | 0 |
| 0 | NA | NA | NA | NA | NA | Gm18371 | | 0 | 0 | 0 | 0 | 0 | 0 | 0 | 0 | 0 | 0 | 0 |
| 0 | NA | NA | NA | NA | NA | Grxcr2 | glutaredoxin, cysteine rich 2 | 0 | 0 | 0 | 0 | 0 | 0 | 0 | 0 | 0 | 0 | 0 |
| 0 | NA | NA | NA | NA | NA | Cstdc7 | | 0 | 0 | 0 | 0 | 0 | 0 | 0 | 0 | 0 | 0 | 0 |
| 0 | NA | NA | NA | NA | NA | Plac8l1 | PLAC8-like 1 | 0 | 0 | 0 | 0 | 0 | 0 | 0 | 0 | 0 | 0 | 0 |
| 0 | NA | NA | NA | NA | NA | Gm22697 | | 0 | 0 | 0 | 0 | 0 | 0 | 0 | 0 | 0 | 0 | 0 |
| 0 | NA | NA | NA | NA | NA | Pou4f3 | POU domain, class 4, transcription factor | 0 | 0 | 0 | 0 | 0 | 0 | 0 | 0 | 0 | 0 | 0 |
| 0 | NA | NA | NA | NA | NA | Gm3631 | | 0 | 0 | 0 | 0 | 0 | 0 | 0 | 0 | 0 | 0 | 0 |
| 0 | NA | NA | NA | NA | NA | Gm50214 | | 0 | 0 | 0 | 0 | 0 | 0 | 0 | 0 | 0 | 0 | 0 |
| 0 | NA | NA | NA | NA | NA | Gm50213 | | 0 | 0 | 0 | 0 | 0 | 0 | 0 | 0 | 0 | 0 | 0 |
| 0 | NA | NA | NA | NA | NA | C030004G16Rik | | 0 | 0 | 0 | 0 | 0 | 0 | 0 | 0 | 0 | 0 | 0 |
| 0 | NA | NA | NA | NA | NA | 4933407I08Rik | | 0 | 0 | 0 | 0 | 0 | 0 | 0 | 0 | 0 | 0 | 0 |
| 0 | NA | NA | NA | NA | NA | 4930588A03Rik | | 0 | 0 | 0 | 0 | 0 | 0 | 0 | 0 | 0 | 0 | 0 |
| 0 | NA | NA | NA | NA | NA | Gm22264 | | 0 | 0 | 0 | 0 | 0 | 0 | 0 | 0 | 0 | 0 | 0 |
| 0 | NA | NA | NA | NA | NA | Gm27325 | | 0 | 0 | 0 | 0 | 0 | 0 | 0 | 0 | 0 | 0 | 0 |
| 0 | NA | NA | NA | NA | NA | Gm23894 | | 0 | 0 | 0 | 0 | 0 | 0 | 0 | 0 | 0 | 0 | 0 |
| 0 | NA | NA | NA | NA | NA | Scgb3a2 | secretoglobin, family 3A, member 2 | 0 | 0 | 0 | 0 | 0 | 0 | 0 | 0 | 0 | 0 | 0 |
| 0 | NA | NA | NA | NA | NA | Gm94 | predicted gene 94 | 0 | 0 | 0 | 0 | 0 | 0 | 0 | 0 | 0 | 0 | 0 |
| 0 | NA | NA | NA | NA | NA | Gm41715 | | 0 | 0 | 0 | 0 | 0 | 0 | 0 | 0 | 0 | 0 | 0 |
| 0 | NA | NA | NA | NA | NA | Gm50206 | | 0 | 0 | 0 | 0 | 0 | 0 | 0 | 0 | 0 | 0 | 0 |
| 0 | NA | NA | NA | NA | NA | Gm50207 | | 0 | 0 | 0 | 0 | 0 | 0 | 0 | 0 | 0 | 0 | 0 |
| 0 | NA | NA | NA | NA | NA | Spink5 | serine peptidase inhibitor, Kazal type 5 | 0 | 0 | 0 | 0 | 0 | 0 | 0 | 0 | 0 | 0 | 0 |
| 0 | NA | NA | NA | NA | NA | Spink14 | serine peptidase inhibitor, Kazal type 14 | 0 | 0 | 0 | 0 | 0 | 0 | 0 | 0 | 0 | 0 | 0 |
| 0 | NA | NA | NA | NA | NA | Spink6 | serine peptidase inhibitor, Kazal type 6 | 0 | 0 | 0 | 0 | 0 | 0 | 0 | 0 | 0 | 0 | 0 |
| 0 | NA | NA | NA | NA | NA | Spink12 | serine peptidase inhibitor, Kazal type 12 | 0 | 0 | 0 | 0 | 0 | 0 | 0 | 0 | 0 | 0 | 0 |
| 0 | NA | NA | NA | NA | NA | Gm37797 | | 0 | 0 | 0 | 0 | 0 | 0 | 0 | 0 | 0 | 0 | 0 |
| 0 | NA | NA | NA | NA | NA | Gm10267 | predicted gene 10267 | 0 | 0 | 0 | 0 | 0 | 0 | 0 | 0 | 0 | 0 | 0 |
| 0 | NA | NA | NA | NA | NA | Spinkl | serine protease inhibitor, Kazal type-like | 0 | 0 | 0 | 0 | 0 | 0 | 0 | 0 | 0 | 0 | 0 |
| 0 | NA | NA | NA | NA | NA | Spink11 | serine peptidase inhibitor, Kazal type 11 | 0 | 0 | 0 | 0 | 0 | 0 | 0 | 0 | 0 | 0 | 0 |
| 0 | NA | NA | NA | NA | NA | Gm10542 | | 0 | 0 | 0 | 0 | 0 | 0 | 0 | 0 | 0 | 0 | 0 |
| 0 | NA | NA | NA | NA | NA | Npy6r | neuropeptide Y receptor Y6 | 0 | 0 | 0 | 0 | 0 | 0 | 0 | 0 | 0 | 0 | 0 |
| 0 | NA | NA | NA | NA | NA | Gm41717 | | 0 | 0 | 0 | 0 | 0 | 0 | 0 | 0 | 0 | 0 | 0 |
| 0 | NA | NA | NA | NA | NA | Myot | myotilin | 0 | 0 | 0 | 0 | 0 | 0 | 0 | 0 | 0 | 0 | 0 |
| 0 | NA | NA | NA | NA | NA | Gm26928 | | 0 | 0 | 0 | 0 | 0 | 0 | 0 | 0 | 0 | 0 | 0 |
| 0 | NA | NA | NA | NA | NA | Gm4839 | | 0 | 0 | 0 | 0 | 0 | 0 | 0 | 0 | 0 | 0 | 0 |
| 0 | NA | NA | NA | NA | NA | Gm18652 | | 0 | 0 | 0 | 0 | 0 | 0 | 0 | 0 | 0 | 0 | 0 |
| 0 | NA | NA | NA | NA | NA | A330093E20Rik | | 0 | 0 | 0 | 0 | 0 | 0 | 0 | 0 | 0 | 0 | 0 |
| 0 | NA | NA | NA | NA | NA | Gm50125 | | 0 | 0 | 0 | 0 | 0 | 0 | 0 | 0 | 0 | 0 | 0 |
| 0 | NA | NA | NA | NA | NA | Gm26262 | | 0 | 0 | 0 | 0 | 0 | 0 | 0 | 0 | 0 | 0 | 0 |
| 0 | NA | NA | NA | NA | NA | Gm38337 | | 0 | 0 | 0 | 0 | 0 | 0 | 0 | 0 | 0 | 0 | 0 |
| 0 | NA | NA | NA | NA | NA | 1700018A14Rik | | 0 | 0 | 0 | 0 | 0 | 0 | 0 | 0 | 0 | 0 | 0 |

| | | | | | | | | | | | | | | | | | | |
|---|---|---|---|---|---|---|---|---|---|---|---|---|---|---|---|---|---|---|
| 0 | NA | NA | NA | NA | NA | 4930415P13Rik | | 0 | 0 | 0 | 0 | 0 | 0 | 0 | 0 | 0 | 0 | 0 |
| 0 | NA | NA | NA | NA | NA | Gm4107 | | 0 | 0 | 0 | 0 | 0 | 0 | 0 | 0 | 0 | 0 | 0 |
| 0 | NA | NA | NA | NA | NA | Gm24617 | | 0 | 0 | 0 | 0 | 0 | 0 | 0 | 0 | 0 | 0 | 0 |
| 0 | NA | NA | NA | NA | NA | Gm24076 | | 0 | 0 | 0 | 0 | 0 | 0 | 0 | 0 | 0 | 0 | 0 |
| 0 | NA | NA | NA | NA | NA | Gm3720 | | 0 | 0 | 0 | 0 | 0 | 0 | 0 | 0 | 0 | 0 | 0 |
| 0 | NA | NA | NA | NA | NA | Gm23610 | | 0 | 0 | 0 | 0 | 0 | 0 | 0 | 0 | 0 | 0 | 0 |
| 0 | NA | NA | NA | NA | NA | Gm49973 | | 0 | 0 | 0 | 0 | 0 | 0 | 0 | 0 | 0 | 0 | 0 |
| 0 | NA | NA | NA | NA | NA | Gm3744 | | 0 | 0 | 0 | 0 | 0 | 0 | 0 | 0 | 0 | 0 | 0 |
| 0 | NA | NA | NA | NA | NA | Gm22791 | | 0 | 0 | 0 | 0 | 0 | 0 | 0 | 0 | 0 | 0 | 0 |
| 0 | NA | NA | NA | NA | NA | Gm8468 | | 0 | 0 | 0 | 0 | 0 | 0 | 0 | 0 | 0 | 0 | 0 |
| 0 | NA | NA | NA | NA | NA | Gm25036 | | 0 | 0 | 0 | 0 | 0 | 0 | 0 | 0 | 0 | 0 | 0 |
| 0 | NA | NA | NA | NA | NA | Gm50086 | | 0 | 0 | 0 | 0 | 0 | 0 | 0 | 0 | 0 | 0 | 0 |
| 0 | NA | NA | NA | NA | NA | 9130209A04Rik | | 0 | 0 | 0 | 0 | 0 | 0 | 0 | 0 | 0 | 0 | 0 |
| 0 | NA | NA | NA | NA | NA | Gm5095 | | 0 | 0 | 0 | 0 | 0 | 0 | 0 | 0 | 0 | 0 | 0 |
| 0 | NA | NA | NA | NA | NA | Gm5236 | | 0 | 0 | 0 | 0 | 0 | 0 | 0 | 0 | 0 | 0 | 0 |
| 0 | NA | NA | NA | NA | NA | Gm4146 | | 0 | 0 | 0 | 0 | 0 | 0 | 0 | 0 | 0 | 0 | 0 |
| 0 | NA | NA | NA | NA | NA | Gm50224 | | 0 | 0 | 0 | 0 | 0 | 0 | 0 | 0 | 0 | 0 | 0 |
| 0 | NA | NA | NA | NA | NA | G630055G22Rik | | 0 | 0 | 0 | 0 | 0 | 0 | 0 | 0 | 0 | 0 | 0 |
| 0 | NA | NA | NA | NA | NA | Gm5237 | | 0 | 0 | 0 | 0 | 0 | 0 | 0 | 0 | 0 | 0 | 0 |
| 0 | NA | NA | NA | NA | NA | Gm5839 | | 0 | 0 | 0 | 0 | 0 | 0 | 0 | 0 | 0 | 0 | 0 |
| 0 | NA | NA | NA | NA | NA | Gm50229 | | 0 | 0 | 0 | 0 | 0 | 0 | 0 | 0 | 0 | 0 | 0 |
| 0 | NA | NA | NA | NA | NA | Gm18993 | | 0 | 0 | 0 | 0 | 0 | 0 | 0 | 0 | 0 | 0 | 0 |
| 0 | NA | NA | NA | NA | NA | Gm50231 | | 0 | 0 | 0 | 0 | 0 | 0 | 0 | 0 | 0 | 0 | 0 |
| 0 | NA | NA | NA | NA | NA | 1700044K03Rik | | 0 | 0 | 0 | 0 | 0 | 0 | 0 | 0 | 0 | 0 | 0 |
| 0 | NA | NA | NA | NA | NA | Gm25440 | | 0 | 0 | 0 | 0 | 0 | 0 | 0 | 0 | 0 | 0 | 0 |
| 0 | NA | NA | NA | NA | NA | | | 0 | 0 | 0 | 0 | 0 | 0 | 0 | 0 | 0 | 0 | 0 |
| 0 | NA | NA | NA | NA | NA | Fam170a | family with sequence similarity 170, meml | 0 | 0 | 0 | 0 | 0 | 0 | 0 | 0 | 0 | 0 | 0 |
| 0 | NA | NA | NA | NA | NA | Pudp | pseudouridine 5'-phosphatase | 0 | 0 | 0 | 0 | 0 | 0 | 0 | 0 | 0 | 0 | 0 |
| 0 | NA | NA | NA | NA | NA | Gm25739 | | 0 | 0 | 0 | 0 | 0 | 0 | 0 | 0 | 0 | 0 | 0 |
| 0 | NA | NA | NA | NA | NA | Gm25785 | | 0 | 0 | 0 | 0 | 0 | 0 | 0 | 0 | 0 | 0 | 0 |
| 0 | NA | NA | NA | NA | NA | Gm50446 | | 0 | 0 | 0 | 0 | 0 | 0 | 0 | 0 | 0 | 0 | 0 |
| 0 | NA | NA | NA | NA | NA | Gm50447 | | 0 | 0 | 0 | 0 | 0 | 0 | 0 | 0 | 0 | 0 | 0 |
| 0 | NA | NA | NA | NA | NA | Gm50448 | | 0 | 0 | 0 | 0 | 0 | 0 | 0 | 0 | 0 | 0 | 0 |
| 0 | NA | NA | NA | NA | NA | Gm22259 | | 0 | 0 | 0 | 0 | 0 | 0 | 0 | 0 | 0 | 0 | 0 |
| 0 | NA | NA | NA | NA | NA | Ftmt | ferritin mitochondrial | 0 | 0 | 0 | 0 | 0 | 0 | 0 | 0 | 0 | 0 | 0 |
| 0 | NA | NA | NA | NA | NA | Gm27026 | | 0 | 0 | 0 | 0 | 0 | 0 | 0 | 0 | 0 | 0 | 0 |
| 0 | NA | NA | NA | NA | NA | Gm41724 | | 0 | 0 | 0 | 0 | 0 | 0 | 0 | 0 | 0 | 0 | 0 |
| 0 | NA | NA | NA | NA | NA | 1700034E13Ril | RIKEN cDNA 1700034E13 gene | 0 | 0 | 0 | 0 | 0 | 0 | 0 | 0 | 0 | 0 | 0 |
| 0 | NA | NA | NA | NA | NA | Gm8572 | | 0 | 0 | 0 | 0 | 0 | 0 | 0 | 0 | 0 | 0 | 0 |
| 0 | NA | NA | NA | NA | NA | Gm1859 | | 0 | 0 | 0 | 0 | 0 | 0 | 0 | 0 | 0 | 0 | 0 |
| 0 | NA | NA | NA | NA | NA | Gm19466 | | 0 | 0 | 0 | 0 | 0 | 0 | 0 | 0 | 0 | 0 | 0 |
| 0 | NA | NA | NA | NA | NA | Gm8594 | | 0 | 0 | 0 | 0 | 0 | 0 | 0 | 0 | 0 | 0 | 0 |
| 0 | NA | NA | NA | NA | NA | Redrum | | 0 | 0 | 0 | 0 | 0 | 0 | 0 | 0 | 0 | 0 | 0 |
| 0 | NA | NA | NA | NA | NA | Gm50361 | | 0 | 0 | 0 | 0 | 0 | 0 | 0 | 0 | 0 | 0 | 0 |
| 0 | NA | NA | NA | NA | NA | 4933434P08Rik | | 0 | 0 | 0 | 0 | 0 | 0 | 0 | 0 | 0 | 0 | 0 |
| 0 | NA | NA | NA | NA | NA | Gm50365 | | 0 | 0 | 0 | 0 | 0 | 0 | 0 | 0 | 0 | 0 | 0 |
| 0 | NA | NA | NA | NA | NA | Gm50366 | | 0 | 0 | 0 | 0 | 0 | 0 | 0 | 0 | 0 | 0 | 0 |
| 0 | NA | NA | NA | NA | NA | Gm24183 | | 0 | 0 | 0 | 0 | 0 | 0 | 0 | 0 | 0 | 0 | 0 |
| 0 | NA | NA | NA | NA | NA | | | 0 | 0 | 0 | 0 | 0 | 0 | 0 | 0 | 0 | 0 | 0 |
| 0 | NA | NA | NA | NA | NA | Gm34073 | | 0 | 0 | 0 | 0 | 0 | 0 | 0 | 0 | 0 | 0 | 0 |
| 0 | NA | NA | NA | NA | NA | Gm22597 | | 0 | 0 | 0 | 0 | 0 | 0 | 0 | 0 | 0 | 0 | 0 |
| 0 | NA | NA | NA | NA | NA | Gm37828 | | 0 | 0 | 0 | 0 | 0 | 0 | 0 | 0 | 0 | 0 | 0 |
| 0 | NA | NA | NA | NA | NA | Gm37337 | | 0 | 0 | 0 | 0 | 0 | 0 | 0 | 0 | 0 | 0 | 0 |
| 0 | NA | NA | NA | NA | NA | Gm24514 | | 0 | 0 | 0 | 0 | 0 | 0 | 0 | 0 | 0 | 0 | 0 |
| 0 | NA | NA | NA | NA | NA | Gm26959 | | 0 | 0 | 0 | 0 | 0 | 0 | 0 | 0 | 0 | 0 | 0 |
| 0 | NA | NA | NA | NA | NA | Gm8614 | | 0 | 0 | 0 | 0 | 0 | 0 | 0 | 0 | 0 | 0 | 0 |
| 0 | NA | NA | NA | NA | NA | Gm6942 | | 0 | 0 | 0 | 0 | 0 | 0 | 0 | 0 | 0 | 0 | 0 |
| 0 | NA | NA | NA | NA | NA | Gm4230 | | 0 | 0 | 0 | 0 | 0 | 0 | 0 | 0 | 0 | 0 | 0 |
| 0 | NA | NA | NA | NA | NA | Gm23335 | | 0 | 0 | 0 | 0 | 0 | 0 | 0 | 0 | 0 | 0 | 0 |
| 0 | NA | NA | NA | NA | NA | Gm22484 | | 0 | 0 | 0 | 0 | 0 | 0 | 0 | 0 | 0 | 0 | 0 |

| | | | | | | | | | | | | | | | | | | |
|---|---|---|---|---|---|---|---|---|---|---|---|---|---|---|---|---|---|---|
| 0 | NA | NA | NA | NA | NA | Gm23555 | | 0 | 0 | 0 | 0 | 0 | 0 | 0 | 0 | 0 | 0 | 0 |
| 0 | NA | NA | NA | NA | NA | Gm25476 | | 0 | 0 | 0 | 0 | 0 | 0 | 0 | 0 | 0 | 0 | 0 |
| 0 | NA | NA | NA | NA | NA | Gm50385 | | 0 | 0 | 0 | 0 | 0 | 0 | 0 | 0 | 0 | 0 | 0 |
| 0 | NA | NA | NA | NA | NA | Mir1258 | | 0 | 0 | 0 | 0 | 0 | 0 | 0 | 0 | 0 | 0 | 0 |
| 0 | NA | NA | NA | NA | NA | Tex43 | testis expressed 43 | 0 | 0 | 0 | 0 | 0 | 0 | 0 | 0 | 0 | 0 | 0 |
| 0 | NA | NA | NA | NA | NA | Gm50283 | | 0 | 0 | 0 | 0 | 0 | 0 | 0 | 0 | 0 | 0 | 0 |
| 0 | NA | NA | NA | NA | NA | Gm50289 | | 0 | 0 | 0 | 0 | 0 | 0 | 0 | 0 | 0 | 0 | 0 |
| 0 | NA | NA | NA | NA | NA | Gm18739 | | 0 | 0 | 0 | 0 | 0 | 0 | 0 | 0 | 0 | 0 | 0 |
| 0 | NA | NA | NA | NA | NA | Gm19486 | | 0 | 0 | 0 | 0 | 0 | 0 | 0 | 0 | 0 | 0 | 0 |
| 0 | NA | NA | NA | NA | NA | Gm50200 | | 0 | 0 | 0 | 0 | 0 | 0 | 0 | 0 | 0 | 0 | 0 |
| 0 | NA | NA | NA | NA | NA | Gm50201 | | 0 | 0 | 0 | 0 | 0 | 0 | 0 | 0 | 0 | 0 | 0 |
| 0 | NA | NA | NA | NA | NA | Gm18929 | | 0 | 0 | 0 | 0 | 0 | 0 | 0 | 0 | 0 | 0 | 0 |
| 0 | NA | NA | NA | NA | NA | 1700066O22Rik | | 0 | 0 | 0 | 0 | 0 | 0 | 0 | 0 | 0 | 0 | 0 |
| 0 | NA | NA | NA | NA | NA | 2210409D07Rik | | 0 | 0 | 0 | 0 | 0 | 0 | 0 | 0 | 0 | 0 | 0 |
| 0 | NA | NA | NA | NA | NA | Gm26038 | | 0 | 0 | 0 | 0 | 0 | 0 | 0 | 0 | 0 | 0 | 0 |
| 0 | NA | NA | NA | NA | NA | Gm44491 | | 0 | 0 | 0 | 0 | 0 | 0 | 0 | 0 | 0 | 0 | 0 |
| 0 | NA | NA | NA | NA | NA | Gm19519 | | 0 | 0 | 0 | 0 | 0 | 0 | 0 | 0 | 0 | 0 | 0 |
| 0 | NA | NA | NA | NA | NA | Gm25660 | | 0 | 0 | 0 | 0 | 0 | 0 | 0 | 0 | 0 | 0 | 0 |
| 0 | NA | NA | NA | NA | NA | Gm23270 | | 0 | 0 | 0 | 0 | 0 | 0 | 0 | 0 | 0 | 0 | 0 |
| 0 | NA | NA | NA | NA | NA | Gm50227 | | 0 | 0 | 0 | 0 | 0 | 0 | 0 | 0 | 0 | 0 | 0 |
| 0 | NA | NA | NA | NA | NA | Mir6355 | | 0 | 0 | 0 | 0 | 0 | 0 | 0 | 0 | 0 | 0 | 0 |
| 0 | NA | NA | NA | NA | NA | Gm26028 | | 0 | 0 | 0 | 0 | 0 | 0 | 0 | 0 | 0 | 0 | 0 |
| 0 | NA | NA | NA | NA | NA | Gm38695 | | 0 | 0 | 0 | 0 | 0 | 0 | 0 | 0 | 0 | 0 | 0 |
| 0 | NA | NA | NA | NA | NA | Gm50235 | | 0 | 0 | 0 | 0 | 0 | 0 | 0 | 0 | 0 | 0 | 0 |
| 0 | NA | NA | NA | NA | NA | Gm18217 | | 0 | 0 | 0 | 0 | 0 | 0 | 0 | 0 | 0 | 0 | 0 |
| 0 | NA | NA | NA | NA | NA | Gm23576 | | 0 | 0 | 0 | 0 | 0 | 0 | 0 | 0 | 0 | 0 | 0 |
| 0 | NA | NA | NA | NA | NA | | | 0 | 0 | 0 | 0 | 0 | 0 | 0 | 0 | 0 | 0 | 0 |
| 0 | NA | NA | NA | NA | NA | Gm50237 | | 0 | 0 | 0 | 0 | 0 | 0 | 0 | 0 | 0 | 0 | 0 |
| 0 | NA | NA | NA | NA | NA | Gm8714 | | 0 | 0 | 0 | 0 | 0 | 0 | 0 | 0 | 0 | 0 | 0 |
| 0 | NA | NA | NA | NA | NA | Gm6960 | | 0 | 0 | 0 | 0 | 0 | 0 | 0 | 0 | 0 | 0 | 0 |
| 0 | NA | NA | NA | NA | NA | Myoz3 | myozenin 3 | 0 | 0 | 0 | 0 | 0 | 0 | 0 | 0 | 0 | 0 | 0 |
| 0 | NA | NA | NA | NA | NA | Mir7220 | | 0 | 0 | 0 | 0 | 0 | 0 | 0 | 0 | 0 | 0 | 0 |
| 0 | NA | NA | NA | NA | NA | Mir6982 | | 0 | 0 | 0 | 0 | 0 | 0 | 0 | 0 | 0 | 0 | 0 |
| 0 | NA | NA | NA | NA | NA | Mir6983 | | 0 | 0 | 0 | 0 | 0 | 0 | 0 | 0 | 0 | 0 | 0 |
| 0 | NA | NA | NA | NA | NA | Gm25301 | | 0 | 0 | 0 | 0 | 0 | 0 | 0 | 0 | 0 | 0 | 0 |
| 0 | NA | NA | NA | NA | NA | Mir378a | | 0 | 0 | 0 | 0 | 0 | 0 | 0 | 0 | 0 | 0 | 0 |
| 0 | NA | NA | NA | NA | NA | Gm8755 | | 0 | 0 | 0 | 0 | 0 | 0 | 0 | 0 | 0 | 0 | 0 |
| 0 | NA | NA | NA | NA | NA | Gm25834 | | 0 | 0 | 0 | 0 | 0 | 0 | 0 | 0 | 0 | 0 | 0 |
| 0 | NA | NA | NA | NA | NA | Lncbate10 | | 0 | 0 | 0 | 0 | 0 | 0 | 0 | 0 | 0 | 0 | 0 |
| 0 | NA | NA | NA | NA | NA | Gm46637 | | 0 | 0 | 0 | 0 | 0 | 0 | 0 | 0 | 0 | 0 | 0 |
| 0 | NA | NA | NA | NA | NA | Mir145a | | 0 | 0 | 0 | 0 | 0 | 0 | 0 | 0 | 0 | 0 | 0 |
| 0 | NA | NA | NA | NA | NA | Mir143 | | 0 | 0 | 0 | 0 | 0 | 0 | 0 | 0 | 0 | 0 | 0 |
| 0 | NA | NA | NA | NA | NA | Il17b | interleukin 17B | 0 | 0 | 0 | 0 | 0 | 0 | 0 | 0 | 0 | 0 | 0 |
| 0 | NA | NA | NA | NA | NA | Gm38165 | | 0 | 0 | 0 | 0 | 0 | 0 | 0 | 0 | 0 | 0 | 0 |
| 0 | NA | NA | NA | NA | NA | Gm37774 | | 0 | 0 | 0 | 0 | 0 | 0 | 0 | 0 | 0 | 0 | 0 |
| 0 | NA | NA | NA | NA | NA | Gm36368 | | 0 | 0 | 0 | 0 | 0 | 0 | 0 | 0 | 0 | 0 | 0 |
| 0 | NA | NA | NA | NA | NA | Spink7 | serine peptidase inhibitor, Kazal type 7 (p | 0 | 0 | 0 | 0 | 0 | 0 | 0 | 0 | 0 | 0 | 0 |
| 0 | NA | NA | NA | NA | NA | Gm19159 | | 0 | 0 | 0 | 0 | 0 | 0 | 0 | 0 | 0 | 0 | 0 |
| 0 | NA | NA | NA | NA | NA | Gm50171 | | 0 | 0 | 0 | 0 | 0 | 0 | 0 | 0 | 0 | 0 | 0 |
| 0 | NA | NA | NA | NA | NA | Gm50172 | | 0 | 0 | 0 | 0 | 0 | 0 | 0 | 0 | 0 | 0 | 0 |
| 0 | NA | NA | NA | NA | NA | Gm26972 | | 0 | 0 | 0 | 0 | 0 | 0 | 0 | 0 | 0 | 0 | 0 |
| 0 | NA | NA | NA | NA | NA | Gm23581 | | 0 | 0 | 0 | 0 | 0 | 0 | 0 | 0 | 0 | 0 | 0 |
| 0 | NA | NA | NA | NA | NA | Gm22560 | | 0 | 0 | 0 | 0 | 0 | 0 | 0 | 0 | 0 | 0 | 0 |
| 0 | NA | NA | NA | NA | NA | Gm22210 | | 0 | 0 | 0 | 0 | 0 | 0 | 0 | 0 | 0 | 0 | 0 |
| 0 | NA | NA | NA | NA | NA | Gm24848 | | 0 | 0 | 0 | 0 | 0 | 0 | 0 | 0 | 0 | 0 | 0 |
| 0 | NA | NA | NA | NA | NA | Gm50104 | | 0 | 0 | 0 | 0 | 0 | 0 | 0 | 0 | 0 | 0 | 0 |
| 0 | NA | NA | NA | NA | NA | Gm50106 | | 0 | 0 | 0 | 0 | 0 | 0 | 0 | 0 | 0 | 0 | 0 |
| 0 | NA | NA | NA | NA | NA | Gm19076 | | 0 | 0 | 0 | 0 | 0 | 0 | 0 | 0 | 0 | 0 | 0 |
| 0 | NA | NA | NA | NA | NA | Gm24384 | | 0 | 0 | 0 | 0 | 0 | 0 | 0 | 0 | 0 | 0 | 0 |
| 0 | NA | NA | NA | NA | NA | Gm50109 | | 0 | 0 | 0 | 0 | 0 | 0 | 0 | 0 | 0 | 0 | 0 |

| | | | | | | | | | | | | | | | | | | |
|---|---|---|---|---|---|---|---|---|---|---|---|---|---|---|---|---|---|---|
| 0 | NA | NA | NA | NA | NA | Gm45934 | | 0 | 0 | 0 | 0 | 0 | 0 | 0 | 0 | 0 | 0 | 0 |
| 0 | NA | NA | NA | NA | NA | Prdx1-ps | | 0 | 0 | 0 | 0 | 0 | 0 | 0 | 0 | 0 | 0 | 0 |
| 0 | NA | NA | NA | NA | NA | Gm23016 | | 0 | 0 | 0 | 0 | 0 | 0 | 0 | 0 | 0 | 0 | 0 |
| 0 | NA | NA | NA | NA | NA | Gm24504 | | 0 | 0 | 0 | 0 | 0 | 0 | 0 | 0 | 0 | 0 | 0 |
| 0 | NA | NA | NA | NA | NA | Gm29860 | | 0 | 0 | 0 | 0 | 0 | 0 | 0 | 0 | 0 | 0 | 0 |
| 0 | NA | NA | NA | NA | NA | Gm50368 | | 0 | 0 | 0 | 0 | 0 | 0 | 0 | 0 | 0 | 0 | 0 |
| 0 | NA | NA | NA | NA | NA | Gm29966 | | 0 | 0 | 0 | 0 | 0 | 0 | 0 | 0 | 0 | 0 | 0 |
| 0 | NA | NA | NA | NA | NA | Mir122 | | 0 | 0 | 0 | 0 | 0 | 0 | 0 | 0 | 0 | 0 | 0 |
| 0 | NA | NA | NA | NA | NA | F730048M01Rik | | 0 | 0 | 0 | 0 | 0 | 0 | 0 | 0 | 0 | 0 | 0 |
| 0 | NA | NA | NA | NA | NA | Gm50220 | | 0 | 0 | 0 | 0 | 0 | 0 | 0 | 0 | 0 | 0 | 0 |
| 0 | NA | NA | NA | NA | NA | Gm41757 | | 0 | 0 | 0 | 0 | 0 | 0 | 0 | 0 | 0 | 0 | 0 |
| 0 | NA | NA | NA | NA | NA | Gm50221 | | 0 | 0 | 0 | 0 | 0 | 0 | 0 | 0 | 0 | 0 | 0 |
| 0 | NA | NA | NA | NA | NA | Gm50228 | | 0 | 0 | 0 | 0 | 0 | 0 | 0 | 0 | 0 | 0 | 0 |
| 0 | NA | NA | NA | NA | NA | Gm46618 | | 0 | 0 | 0 | 0 | 0 | 0 | 0 | 0 | 0 | 0 | 0 |
| 0 | NA | NA | NA | NA | NA | Gm50246 | | 0 | 0 | 0 | 0 | 0 | 0 | 0 | 0 | 0 | 0 | 0 |
| 0 | NA | NA | NA | NA | NA | Gm30018 | | 0 | 0 | 0 | 0 | 0 | 0 | 0 | 0 | 0 | 0 | 0 |
| 0 | NA | NA | NA | NA | NA | Gm50247 | | 0 | 0 | 0 | 0 | 0 | 0 | 0 | 0 | 0 | 0 | 0 |
| 0 | NA | NA | NA | NA | NA | Gm50115 | | 0 | 0 | 0 | 0 | 0 | 0 | 0 | 0 | 0 | 0 | 0 |
| 0 | NA | NA | NA | NA | NA | Gm50234 | | 0 | 0 | 0 | 0 | 0 | 0 | 0 | 0 | 0 | 0 | 0 |
| 0 | NA | NA | NA | NA | NA | Gm50256 | | 0 | 0 | 0 | 0 | 0 | 0 | 0 | 0 | 0 | 0 | 0 |
| 0 | NA | NA | NA | NA | NA | Gm50175 | | 0 | 0 | 0 | 0 | 0 | 0 | 0 | 0 | 0 | 0 | 0 |
| 0 | NA | NA | NA | NA | NA | Gm30094 | | 0 | 0 | 0 | 0 | 0 | 0 | 0 | 0 | 0 | 0 | 0 |
| 0 | NA | NA | NA | NA | NA | Gm25150 | | 0 | 0 | 0 | 0 | 0 | 0 | 0 | 0 | 0 | 0 | 0 |
| 0 | NA | NA | NA | NA | NA | Gm50117 | | 0 | 0 | 0 | 0 | 0 | 0 | 0 | 0 | 0 | 0 | 0 |
| 0 | NA | NA | NA | NA | NA | Gm50118 | | 0 | 0 | 0 | 0 | 0 | 0 | 0 | 0 | 0 | 0 | 0 |
| 0 | NA | NA | NA | NA | NA | Gm50119 | | 0 | 0 | 0 | 0 | 0 | 0 | 0 | 0 | 0 | 0 | 0 |
| 0 | NA | NA | NA | NA | NA | Rax | retina and anterior neural fold homeobox | 0 | 0 | 0 | 0 | 0 | 0 | 0 | 0 | 0 | 0 | 0 |
| 0 | NA | NA | NA | NA | NA | Raxos1 | | 0 | 0 | 0 | 0 | 0 | 0 | 0 | 0 | 0 | 0 | 0 |
| 0 | NA | NA | NA | NA | NA | Gm50142 | | 0 | 0 | 0 | 0 | 0 | 0 | 0 | 0 | 0 | 0 | 0 |
| 0 | NA | NA | NA | NA | NA | Gm50143 | | 0 | 0 | 0 | 0 | 0 | 0 | 0 | 0 | 0 | 0 | 0 |
| 0 | NA | NA | NA | NA | NA | Mir694 | | 0 | 0 | 0 | 0 | 0 | 0 | 0 | 0 | 0 | 0 | 0 |
| 0 | NA | NA | NA | NA | NA | Gm50148 | | 0 | 0 | 0 | 0 | 0 | 0 | 0 | 0 | 0 | 0 | 0 |
| 0 | NA | NA | NA | NA | NA | Gm50153 | | 0 | 0 | 0 | 0 | 0 | 0 | 0 | 0 | 0 | 0 | 0 |
| 0 | NA | NA | NA | NA | NA | Gm20294 | | 0 | 0 | 0 | 0 | 0 | 0 | 0 | 0 | 0 | 0 | 0 |
| 0 | NA | NA | NA | NA | NA | Gm50154 | | 0 | 0 | 0 | 0 | 0 | 0 | 0 | 0 | 0 | 0 | 0 |
| 0 | NA | NA | NA | NA | NA | Gm50157 | | 0 | 0 | 0 | 0 | 0 | 0 | 0 | 0 | 0 | 0 | 0 |
| 0 | NA | NA | NA | NA | NA | Gm50158 | | 0 | 0 | 0 | 0 | 0 | 0 | 0 | 0 | 0 | 0 | 0 |
| 0 | NA | NA | NA | NA | NA | Gm24679 | | 0 | 0 | 0 | 0 | 0 | 0 | 0 | 0 | 0 | 0 | 0 |
| 0 | NA | NA | NA | NA | NA | Gm36092 | | 0 | 0 | 0 | 0 | 0 | 0 | 0 | 0 | 0 | 0 | 0 |
| 0 | NA | NA | NA | NA | NA | Gm50159 | | 0 | 0 | 0 | 0 | 0 | 0 | 0 | 0 | 0 | 0 | 0 |
| 0 | NA | NA | NA | NA | NA | Gm50160 | | 0 | 0 | 0 | 0 | 0 | 0 | 0 | 0 | 0 | 0 | 0 |
| 0 | NA | NA | NA | NA | NA | Gm50161 | | 0 | 0 | 0 | 0 | 0 | 0 | 0 | 0 | 0 | 0 | 0 |
| 0 | NA | NA | NA | NA | NA | Gm30190 | | 0 | 0 | 0 | 0 | 0 | 0 | 0 | 0 | 0 | 0 | 0 |
| 0 | NA | NA | NA | NA | NA | Gm22219 | | 0 | 0 | 0 | 0 | 0 | 0 | 0 | 0 | 0 | 0 | 0 |
| 0 | NA | NA | NA | NA | NA | Gm50110 | | 0 | 0 | 0 | 0 | 0 | 0 | 0 | 0 | 0 | 0 | 0 |
| 0 | NA | NA | NA | NA | NA | Gm50162 | | 0 | 0 | 0 | 0 | 0 | 0 | 0 | 0 | 0 | 0 | 0 |
| 0 | NA | NA | NA | NA | NA | Gm50164 | | 0 | 0 | 0 | 0 | 0 | 0 | 0 | 0 | 0 | 0 | 0 |
| 0 | NA | NA | NA | NA | NA | Gm50165 | | 0 | 0 | 0 | 0 | 0 | 0 | 0 | 0 | 0 | 0 | 0 |
| 0 | NA | NA | NA | NA | NA | Gm50166 | | 0 | 0 | 0 | 0 | 0 | 0 | 0 | 0 | 0 | 0 | 0 |
| 0 | NA | NA | NA | NA | NA | Gm50170 | | 0 | 0 | 0 | 0 | 0 | 0 | 0 | 0 | 0 | 0 | 0 |
| 0 | NA | NA | NA | NA | NA | Gm50167 | | 0 | 0 | 0 | 0 | 0 | 0 | 0 | 0 | 0 | 0 | 0 |
| 0 | NA | NA | NA | NA | NA | Gm50169 | | 0 | 0 | 0 | 0 | 0 | 0 | 0 | 0 | 0 | 0 | 0 |
| 0 | NA | NA | NA | NA | NA | Gm24900 | | 0 | 0 | 0 | 0 | 0 | 0 | 0 | 0 | 0 | 0 | 0 |
| 0 | NA | NA | NA | NA | NA | Gm8887 | | 0 | 0 | 0 | 0 | 0 | 0 | 0 | 0 | 0 | 0 | 0 |
| 0 | NA | NA | NA | NA | NA | Gm41764 | | 0 | 0 | 0 | 0 | 0 | 0 | 0 | 0 | 0 | 0 | 0 |
| 0 | NA | NA | NA | NA | NA | Gm18149 | | 0 | 0 | 0 | 0 | 0 | 0 | 0 | 0 | 0 | 0 | 0 |
| 0 | NA | NA | NA | NA | NA | Mir7219 | | 0 | 0 | 0 | 0 | 0 | 0 | 0 | 0 | 0 | 0 | 0 |
| 0 | NA | NA | NA | NA | NA | Gm50258 | | 0 | 0 | 0 | 0 | 0 | 0 | 0 | 0 | 0 | 0 | 0 |
| 0 | NA | NA | NA | NA | NA | Gm29677 | | 0 | 0 | 0 | 0 | 0 | 0 | 0 | 0 | 0 | 0 | 0 |
| 0 | NA | NA | NA | NA | NA | Gm50259 | | 0 | 0 | 0 | 0 | 0 | 0 | 0 | 0 | 0 | 0 | 0 |

| | | | | | | | | | | | | | | | | | |
|---|---|---|---|---|---|---|---|---|---|---|---|---|---|---|---|---|---|
| 0 | NA | NA | NA | NA | NA | Gm30641 | | 0 | 0 | 0 | 0 | 0 | 0 | 0 | 0 | 0 | 0 | 0 |
| 0 | NA | NA | NA | NA | NA | Mir6356 | | 0 | 0 | 0 | 0 | 0 | 0 | 0 | 0 | 0 | 0 | 0 |
| 0 | NA | NA | NA | NA | NA | Gm50260 | | 0 | 0 | 0 | 0 | 0 | 0 | 0 | 0 | 0 | 0 | 0 |
| 0 | NA | NA | NA | NA | NA | 4930546C10Rik | | 0 | 0 | 0 | 0 | 0 | 0 | 0 | 0 | 0 | 0 | 0 |
| 0 | NA | NA | NA | NA | NA | Mir145b | | 0 | 0 | 0 | 0 | 0 | 0 | 0 | 0 | 0 | 0 | 0 |
| 0 | NA | NA | NA | NA | NA | Gm23536 | | 0 | 0 | 0 | 0 | 0 | 0 | 0 | 0 | 0 | 0 | 0 |
| 0 | NA | NA | NA | NA | NA | Gm31294 | | 0 | 0 | 0 | 0 | 0 | 0 | 0 | 0 | 0 | 0 | 0 |
| 0 | NA | NA | NA | NA | NA | Gm4842 | | 0 | 0 | 0 | 0 | 0 | 0 | 0 | 0 | 0 | 0 | 0 |
| 0 | NA | NA | NA | NA | NA | Gm24845 | | 0 | 0 | 0 | 0 | 0 | 0 | 0 | 0 | 0 | 0 | 0 |
| 0 | NA | NA | NA | NA | NA | Gm22508 | | 0 | 0 | 0 | 0 | 0 | 0 | 0 | 0 | 0 | 0 | 0 |
| 0 | NA | NA | NA | NA | NA | 1700061H18Rik | | 0 | 0 | 0 | 0 | 0 | 0 | 0 | 0 | 0 | 0 | 0 |
| 0 | NA | NA | NA | NA | NA | Gm50219 | | 0 | 0 | 0 | 0 | 0 | 0 | 0 | 0 | 0 | 0 | 0 |
| 0 | NA | NA | NA | NA | NA | Dynap | dynactin associated protein | 0 | 0 | 0 | 0 | 0 | 0 | 0 | 0 | 0 | 0 | 0 |
| 0 | NA | NA | NA | NA | NA | Mir6357 | | 0 | 0 | 0 | 0 | 0 | 0 | 0 | 0 | 0 | 0 | 0 |
| 0 | NA | NA | NA | NA | NA | Gm50261 | | 0 | 0 | 0 | 0 | 0 | 0 | 0 | 0 | 0 | 0 | 0 |
| 0 | NA | NA | NA | NA | NA | Gm18963 | | 0 | 0 | 0 | 0 | 0 | 0 | 0 | 0 | 0 | 0 | 0 |
| 0 | NA | NA | NA | NA | NA | Gm21046 | | 0 | 0 | 0 | 0 | 0 | 0 | 0 | 0 | 0 | 0 | 0 |
| 0 | NA | NA | NA | NA | NA | Gm50262 | | 0 | 0 | 0 | 0 | 0 | 0 | 0 | 0 | 0 | 0 | 0 |
| 0 | NA | NA | NA | NA | NA | Gm25509 | | 0 | 0 | 0 | 0 | 0 | 0 | 0 | 0 | 0 | 0 | 0 |
| 0 | NA | NA | NA | NA | NA | Gm31908 | | 0 | 0 | 0 | 0 | 0 | 0 | 0 | 0 | 0 | 0 | 0 |
| 0 | NA | NA | NA | NA | NA | Gm26438 | | 0 | 0 | 0 | 0 | 0 | 0 | 0 | 0 | 0 | 0 | 0 |
| 0 | NA | NA | NA | NA | NA | Gm22065 | | 0 | 0 | 0 | 0 | 0 | 0 | 0 | 0 | 0 | 0 | 0 |
| 0 | NA | NA | NA | NA | NA | Gm50126 | | 0 | 0 | 0 | 0 | 0 | 0 | 0 | 0 | 0 | 0 | 0 |
| 0 | NA | NA | NA | NA | NA | Gm32548 | | 0 | 0 | 0 | 0 | 0 | 0 | 0 | 0 | 0 | 0 | 0 |
| 0 | NA | NA | NA | NA | NA | Gm17943 | | 0 | 0 | 0 | 0 | 0 | 0 | 0 | 0 | 0 | 0 | 0 |
| 0 | NA | NA | NA | NA | NA | Gm5690 | | 0 | 0 | 0 | 0 | 0 | 0 | 0 | 0 | 0 | 0 | 0 |
| 0 | NA | NA | NA | NA | NA | Gm50168 | | 0 | 0 | 0 | 0 | 0 | 0 | 0 | 0 | 0 | 0 | 0 |
| 0 | NA | NA | NA | NA | NA | 1700120E14Rik | | 0 | 0 | 0 | 0 | 0 | 0 | 0 | 0 | 0 | 0 | 0 |
| 0 | NA | NA | NA | NA | NA | Gm24559 | | 0 | 0 | 0 | 0 | 0 | 0 | 0 | 0 | 0 | 0 | 0 |
| 0 | NA | NA | NA | NA | NA | Rpl37a-ps2 | | 0 | 0 | 0 | 0 | 0 | 0 | 0 | 0 | 0 | 0 | 0 |
| 0 | NA | NA | NA | NA | NA | Gm25594 | | 0 | 0 | 0 | 0 | 0 | 0 | 0 | 0 | 0 | 0 | 0 |
| 0 | NA | NA | NA | NA | NA | Gm18786 | | 0 | 0 | 0 | 0 | 0 | 0 | 0 | 0 | 0 | 0 | 0 |
| 0 | NA | NA | NA | NA | NA | 9030625G05Rik | | 0 | 0 | 0 | 0 | 0 | 0 | 0 | 0 | 0 | 0 | 0 |
| 0 | NA | NA | NA | NA | NA | Gm27781 | | 0 | 0 | 0 | 0 | 0 | 0 | 0 | 0 | 0 | 0 | 0 |
| 0 | NA | NA | NA | NA | NA | 2010010A06Rik | | 0 | 0 | 0 | 0 | 0 | 0 | 0 | 0 | 0 | 0 | 0 |
| 0 | NA | NA | NA | NA | NA | Gm10532 | | 0 | 0 | 0 | 0 | 0 | 0 | 0 | 0 | 0 | 0 | 0 |
| 0 | NA | NA | NA | NA | NA | 1700034B16Rik | | 0 | 0 | 0 | 0 | 0 | 0 | 0 | 0 | 0 | 0 | 0 |
| 0 | NA | NA | NA | NA | NA | 2900057B20Rik | | 0 | 0 | 0 | 0 | 0 | 0 | 0 | 0 | 0 | 0 | 0 |
| 0 | NA | NA | NA | NA | NA | Mir6358 | | 0 | 0 | 0 | 0 | 0 | 0 | 0 | 0 | 0 | 0 | 0 |
| 0 | NA | NA | NA | NA | NA | Gm31933 | | 0 | 0 | 0 | 0 | 0 | 0 | 0 | 0 | 0 | 0 | 0 |
| 0 | NA | NA | NA | NA | NA | Skor2 | SKI family transcriptional corepressor 2 | 0 | 0 | 0 | 0 | 0 | 0 | 0 | 0 | 0 | 0 | 0 |
| 0 | NA | NA | NA | NA | NA | Gm15957 | | 0 | 0 | 0 | 0 | 0 | 0 | 0 | 0 | 0 | 0 | 0 |
| 0 | NA | NA | NA | NA | NA | Gm50284 | | 0 | 0 | 0 | 0 | 0 | 0 | 0 | 0 | 0 | 0 | 0 |
| 0 | NA | NA | NA | NA | NA | Loxhd1 | lipoxygenase homology domains 1 | 0 | 0 | 0 | 0 | 0 | 0 | 0 | 0 | 0 | 0 | 0 |
| 0 | NA | NA | NA | NA | NA | Gm50197 | | 0 | 0 | 0 | 0 | 0 | 0 | 0 | 0 | 0 | 0 | 0 |
| 0 | NA | NA | NA | NA | NA | Gm26014 | | 0 | 0 | 0 | 0 | 0 | 0 | 0 | 0 | 0 | 0 | 0 |
| 0 | NA | NA | NA | NA | NA | Gm23954 | | 0 | 0 | 0 | 0 | 0 | 0 | 0 | 0 | 0 | 0 | 0 |
| 0 | NA | NA | NA | NA | NA | Siglec15 | sialic acid binding Ig-like lectin 15 | 0 | 0 | 0 | 0 | 0 | 0 | 0 | 0 | 0 | 0 | 0 |
| 0 | NA | NA | NA | NA | NA | 4931439C15Rik | | 0 | 0 | 0 | 0 | 0 | 0 | 0 | 0 | 0 | 0 | 0 |
| 0 | NA | NA | NA | NA | NA | Gm26430 | | 0 | 0 | 0 | 0 | 0 | 0 | 0 | 0 | 0 | 0 | 0 |
| 0 | NA | NA | NA | NA | NA | Gm23895 | | 0 | 0 | 0 | 0 | 0 | 0 | 0 | 0 | 0 | 0 | 0 |
| 0 | NA | NA | NA | NA | NA | Gm18978 | | 0 | 0 | 0 | 0 | 0 | 0 | 0 | 0 | 0 | 0 | 0 |
| 0 | NA | NA | NA | NA | NA | Gm2116 | | 0 | 0 | 0 | 0 | 0 | 0 | 0 | 0 | 0 | 0 | 0 |
| 0 | NA | NA | NA | NA | NA | Gm20593 | | 0 | 0 | 0 | 0 | 0 | 0 | 0 | 0 | 0 | 0 | 0 |
| 0 | NA | NA | NA | NA | NA | Gm25824 | | 0 | 0 | 0 | 0 | 0 | 0 | 0 | 0 | 0 | 0 | 0 |
| 0 | NA | NA | NA | NA | NA | Gm50176 | | 0 | 0 | 0 | 0 | 0 | 0 | 0 | 0 | 0 | 0 | 0 |
| 0 | NA | NA | NA | NA | NA | Gm50177 | | 0 | 0 | 0 | 0 | 0 | 0 | 0 | 0 | 0 | 0 | 0 |
| 0 | NA | NA | NA | NA | NA | Gm50178 | | 0 | 0 | 0 | 0 | 0 | 0 | 0 | 0 | 0 | 0 | 0 |
| 0 | NA | NA | NA | NA | NA | Gm19062 | | 0 | 0 | 0 | 0 | 0 | 0 | 0 | 0 | 0 | 0 | 0 |
| 0 | NA | NA | NA | NA | NA | Gm18910 | | 0 | 0 | 0 | 0 | 0 | 0 | 0 | 0 | 0 | 0 | 0 |

| | | | | | | | | | | | | | | | | | | |
|---|---|---|---|---|---|---|---|---|---|---|---|---|---|---|---|---|---|---|
| 0 | NA | NA | NA | NA | NA | AA684185 | | 0 | 0 | 0 | 0 | 0 | 0 | 0 | 0 | 0 | 0 | 0 |
| 0 | NA | NA | NA | NA | NA | Gm50179 | | 0 | 0 | 0 | 0 | 0 | 0 | 0 | 0 | 0 | 0 | 0 |
| 0 | NA | NA | NA | NA | NA | Gm36627 | | 0 | 0 | 0 | 0 | 0 | 0 | 0 | 0 | 0 | 0 | 0 |
| 0 | NA | NA | NA | NA | NA | Gm36718 | | 0 | 0 | 0 | 0 | 0 | 0 | 0 | 0 | 0 | 0 | 0 |
| 0 | NA | NA | NA | NA | NA | Gm50180 | | 0 | 0 | 0 | 0 | 0 | 0 | 0 | 0 | 0 | 0 | 0 |
| 0 | NA | NA | NA | NA | NA | Gm50190 | | 0 | 0 | 0 | 0 | 0 | 0 | 0 | 0 | 0 | 0 | 0 |
| 0 | NA | NA | NA | NA | NA | Gm50192 | | 0 | 0 | 0 | 0 | 0 | 0 | 0 | 0 | 0 | 0 | 0 |
| 0 | NA | NA | NA | NA | NA | Gm50193 | | 0 | 0 | 0 | 0 | 0 | 0 | 0 | 0 | 0 | 0 | 0 |
| 0 | NA | NA | NA | NA | NA | Gm50195 | | 0 | 0 | 0 | 0 | 0 | 0 | 0 | 0 | 0 | 0 | 0 |
| 0 | NA | NA | NA | NA | NA | Gm21886 | | 0 | 0 | 0 | 0 | 0 | 0 | 0 | 0 | 0 | 0 | 0 |
| 0 | NA | NA | NA | NA | NA | Gm41787 | | 0 | 0 | 0 | 0 | 0 | 0 | 0 | 0 | 0 | 0 | 0 |
| 0 | NA | NA | NA | NA | NA | Gm7447 | | 0 | 0 | 0 | 0 | 0 | 0 | 0 | 0 | 0 | 0 | 0 |
| 0 | NA | NA | NA | NA | NA | Gm46629 | | 0 | 0 | 0 | 0 | 0 | 0 | 0 | 0 | 0 | 0 | 0 |
| 0 | NA | NA | NA | NA | NA | Gm37015 | | 0 | 0 | 0 | 0 | 0 | 0 | 0 | 0 | 0 | 0 | 0 |
| 0 | NA | NA | NA | NA | NA | Gm26676 | | 0 | 0 | 0 | 0 | 0 | 0 | 0 | 0 | 0 | 0 | 0 |
| 0 | NA | NA | NA | NA | NA | D330025C20Rik | | 0 | 0 | 0 | 0 | 0 | 0 | 0 | 0 | 0 | 0 | 0 |
| 0 | NA | NA | NA | NA | NA | Gm50211 | | 0 | 0 | 0 | 0 | 0 | 0 | 0 | 0 | 0 | 0 | 0 |
| 0 | NA | NA | NA | NA | NA | Gm47272 | | 0 | 0 | 0 | 0 | 0 | 0 | 0 | 0 | 0 | 0 | 0 |
| 0 | NA | NA | NA | NA | NA | Gm25071 | | 0 | 0 | 0 | 0 | 0 | 0 | 0 | 0 | 0 | 0 | 0 |
| 0 | NA | NA | NA | NA | NA | Sall3 | spalt like transcription factor 3 | 0 | 0 | 0 | 0 | 0 | 0 | 0 | 0 | 0 | 0 | 0 |
| 0 | NA | NA | NA | NA | NA | Gm27239 | | 0 | 0 | 0 | 0 | 0 | 0 | 0 | 0 | 0 | 0 | 0 |
| 0 | NA | NA | NA | NA | NA | Gm7503 | | 0 | 0 | 0 | 0 | 0 | 0 | 0 | 0 | 0 | 0 | 0 |
| 0 | NA | NA | NA | NA | NA | Gm25718 | | 0 | 0 | 0 | 0 | 0 | 0 | 0 | 0 | 0 | 0 | 0 |
| 0 | NA | NA | NA | NA | NA | Gm50424 | | 0 | 0 | 0 | 0 | 0 | 0 | 0 | 0 | 0 | 0 | 0 |
| 0 | NA | NA | NA | NA | NA | Gm50423 | | 0 | 0 | 0 | 0 | 0 | 0 | 0 | 0 | 0 | 0 | 0 |
| 0 | NA | NA | NA | NA | NA | 4930594M17Rik | | 0 | 0 | 0 | 0 | 0 | 0 | 0 | 0 | 0 | 0 | 0 |
| 0 | NA | NA | NA | NA | NA | Gm30192 | | 0 | 0 | 0 | 0 | 0 | 0 | 0 | 0 | 0 | 0 | 0 |
| 0 | NA | NA | NA | NA | NA | Gm30288 | | 0 | 0 | 0 | 0 | 0 | 0 | 0 | 0 | 0 | 0 | 0 |
| 0 | NA | NA | NA | NA | NA | Gm50412 | | 0 | 0 | 0 | 0 | 0 | 0 | 0 | 0 | 0 | 0 | 0 |
| 0 | NA | NA | NA | NA | NA | Gm41792 | | 0 | 0 | 0 | 0 | 0 | 0 | 0 | 0 | 0 | 0 | 0 |
| 0 | NA | NA | NA | NA | NA | Gm50414 | | 0 | 0 | 0 | 0 | 0 | 0 | 0 | 0 | 0 | 0 | 0 |
| 0 | NA | NA | NA | NA | NA | Gm50290 | | 0 | 0 | 0 | 0 | 0 | 0 | 0 | 0 | 0 | 0 | 0 |
| 0 | NA | NA | NA | NA | NA | Gm50291 | | 0 | 0 | 0 | 0 | 0 | 0 | 0 | 0 | 0 | 0 | 0 |
| 0 | NA | NA | NA | NA | NA | Gm46614 | | 0 | 0 | 0 | 0 | 0 | 0 | 0 | 0 | 0 | 0 | 0 |
| 0 | NA | NA | NA | NA | NA | Gm30454 | | 0 | 0 | 0 | 0 | 0 | 0 | 0 | 0 | 0 | 0 | 0 |
| 0 | NA | NA | NA | NA | NA | Gm50292 | | 0 | 0 | 0 | 0 | 0 | 0 | 0 | 0 | 0 | 0 | 0 |
| 0 | NA | NA | NA | NA | NA | Gm30593 | | 0 | 0 | 0 | 0 | 0 | 0 | 0 | 0 | 0 | 0 | 0 |
| 0 | NA | NA | NA | NA | NA | Gm26012 | | 0 | 0 | 0 | 0 | 0 | 0 | 0 | 0 | 0 | 0 | 0 |
| 0 | NA | NA | NA | NA | NA | Mir5127 | | 0 | 0 | 0 | 0 | 0 | 0 | 0 | 0 | 0 | 0 | 0 |
| 0 | NA | NA | NA | NA | NA | Gm50293 | | 0 | 0 | 0 | 0 | 0 | 0 | 0 | 0 | 0 | 0 | 0 |
| 0 | NA | NA | NA | NA | NA | Gm50294 | | 0 | 0 | 0 | 0 | 0 | 0 | 0 | 0 | 0 | 0 | 0 |
| 0 | NA | NA | NA | NA | NA | Gm50297 | | 0 | 0 | 0 | 0 | 0 | 0 | 0 | 0 | 0 | 0 | 0 |
| 0 | NA | NA | NA | NA | NA | Gm50298 | | 0 | 0 | 0 | 0 | 0 | 0 | 0 | 0 | 0 | 0 | 0 |
| 0 | NA | NA | NA | NA | NA | B020010K11Rik | | 0 | 0 | 0 | 0 | 0 | 0 | 0 | 0 | 0 | 0 | 0 |
| 0 | NA | NA | NA | NA | NA | Gm25060 | | 0 | 0 | 0 | 0 | 0 | 0 | 0 | 0 | 0 | 0 | 0 |
| 0 | NA | NA | NA | NA | NA | Gm50347 | | 0 | 0 | 0 | 0 | 0 | 0 | 0 | 0 | 0 | 0 | 0 |
| 0 | NA | NA | NA | NA | NA | Gm50350 | | 0 | 0 | 0 | 0 | 0 | 0 | 0 | 0 | 0 | 0 | 0 |
| 0 | NA | NA | NA | NA | NA | Gm41793 | | 0 | 0 | 0 | 0 | 0 | 0 | 0 | 0 | 0 | 0 | 0 |
| 0 | NA | NA | NA | NA | NA | Mir5112 | | 0 | 0 | 0 | 0 | 0 | 0 | 0 | 0 | 0 | 0 | 0 |
| 0 | NA | NA | NA | NA | NA | Gm50355 | | 0 | 0 | 0 | 0 | 0 | 0 | 0 | 0 | 0 | 0 | 0 |
| 0 | NA | NA | NA | NA | NA | 2210420H20Rik | | 0 | 0 | 0 | 0 | 0 | 0 | 0 | 0 | 0 | 0 | 0 |
| 0 | NA | NA | NA | NA | NA | Gm50356 | | 0 | 0 | 0 | 0 | 0 | 0 | 0 | 0 | 0 | 0 | 0 |
| 0 | NA | NA | NA | NA | NA | Gm50437 | | 0 | 0 | 0 | 0 | 0 | 0 | 0 | 0 | 0 | 0 | 0 |
| 0 | NA | NA | NA | NA | NA | 1700095A13Rik | | 0 | 0 | 0 | 0 | 0 | 0 | 0 | 0 | 0 | 0 | 0 |
| 0 | NA | NA | NA | NA | NA | 4921531P14Rik | | 0 | 0 | 0 | 0 | 0 | 0 | 0 | 0 | 0 | 0 | 0 |
| 0 | NA | NA | NA | NA | NA | Gm24720 | | 0 | 0 | 0 | 0 | 0 | 0 | 0 | 0 | 0 | 0 | 0 |
| 0 | NA | NA | NA | NA | NA | Gm31265 | | 0 | 0 | 0 | 0 | 0 | 0 | 0 | 0 | 0 | 0 | 0 |
| 0 | NA | NA | NA | NA | NA | Gm31503 | | 0 | 0 | 0 | 0 | 0 | 0 | 0 | 0 | 0 | 0 | 0 |
| 0 | NA | NA | NA | NA | NA | Gm50413 | | 0 | 0 | 0 | 0 | 0 | 0 | 0 | 0 | 0 | 0 | 0 |
| 0 | NA | NA | NA | NA | NA | 4933401L05Rik | | 0 | 0 | 0 | 0 | 0 | 0 | 0 | 0 | 0 | 0 | 0 |

| | | | | | | | | | | | | | | | | | | |
|---|---|---|---|---|---|---|---|---|---|---|---|---|---|---|---|---|---|---|
| 0 | NA | NA | NA | NA | NA | Gm50415 | | 0 | 0 | 0 | 0 | 0 | 0 | 0 | 0 | 0 | 0 | 0 |
| 0 | NA | NA | NA | NA | NA | Gm50416 | | 0 | 0 | 0 | 0 | 0 | 0 | 0 | 0 | 0 | 0 | 0 |
| 0 | NA | NA | NA | NA | NA | Gm50417 | | 0 | 0 | 0 | 0 | 0 | 0 | 0 | 0 | 0 | 0 | 0 |
| 0 | NA | NA | NA | NA | NA | Gm50418 | | 0 | 0 | 0 | 0 | 0 | 0 | 0 | 0 | 0 | 0 | 0 |
| 0 | NA | NA | NA | NA | NA | Gm50419 | | 0 | 0 | 0 | 0 | 0 | 0 | 0 | 0 | 0 | 0 | 0 |
| 0 | NA | NA | NA | NA | NA | Gm50420 | | 0 | 0 | 0 | 0 | 0 | 0 | 0 | 0 | 0 | 0 | 0 |
| 0 | NA | NA | NA | NA | NA | Gm50421 | | 0 | 0 | 0 | 0 | 0 | 0 | 0 | 0 | 0 | 0 | 0 |
| 0 | NA | NA | NA | NA | NA | Gm31854 | | 0 | 0 | 0 | 0 | 0 | 0 | 0 | 0 | 0 | 0 | 0 |
| 0 | NA | NA | NA | NA | NA | Gm50309 | | 0 | 0 | 0 | 0 | 0 | 0 | 0 | 0 | 0 | 0 | 0 |
| 0 | NA | NA | NA | NA | NA | Gm37216 | | 0 | 0 | 0 | 0 | 0 | 0 | 0 | 0 | 0 | 0 | 0 |
| 0 | NA | NA | NA | NA | NA | Cndp1 | carnosine dipeptidase 1 (metallopeptidas | 0 | 0 | 0 | 0 | 0 | 0 | 0 | 0 | 0 | 0 | 0 |
| 0 | NA | NA | NA | NA | NA | Gm50314 | | 0 | 0 | 0 | 0 | 0 | 0 | 0 | 0 | 0 | 0 | 0 |
| 0 | NA | NA | NA | NA | NA | D030046N08Rik | | 0 | 0 | 0 | 0 | 0 | 0 | 0 | 0 | 0 | 0 | 0 |
| 0 | NA | NA | NA | NA | NA | Gm22014 | | 0 | 0 | 0 | 0 | 0 | 0 | 0 | 0 | 0 | 0 | 0 |
| 0 | NA | NA | NA | NA | NA | Gm25005 | | 0 | 0 | 0 | 0 | 0 | 0 | 0 | 0 | 0 | 0 | 0 |
| 0 | NA | NA | NA | NA | NA | Gm38576 | | 0 | 0 | 0 | 0 | 0 | 0 | 0 | 0 | 0 | 0 | 0 |
| 0 | NA | NA | NA | NA | NA | Gm17266 | predicted gene, 17266 | 0 | 0 | 0 | 0 | 0 | 0 | 0 | 0 | 0 | 0 | 0 |
| 0 | NA | NA | NA | NA | NA | Gm22819 | | 0 | 0 | 0 | 0 | 0 | 0 | 0 | 0 | 0 | 0 | 0 |
| 0 | NA | NA | NA | NA | NA | Gm16146 | | 0 | 0 | 0 | 0 | 0 | 0 | 0 | 0 | 0 | 0 | 0 |
| 0 | NA | NA | NA | NA | NA | Gm50369 | | 0 | 0 | 0 | 0 | 0 | 0 | 0 | 0 | 0 | 0 | 0 |
| 0 | NA | NA | NA | NA | NA | Gm50370 | | 0 | 0 | 0 | 0 | 0 | 0 | 0 | 0 | 0 | 0 | 0 |
| 0 | NA | NA | NA | NA | NA | Gm7612 | | 0 | 0 | 0 | 0 | 0 | 0 | 0 | 0 | 0 | 0 | 0 |
| 0 | NA | NA | NA | NA | NA | Gm50381 | | 0 | 0 | 0 | 0 | 0 | 0 | 0 | 0 | 0 | 0 | 0 |
| 0 | NA | NA | NA | NA | NA | Gm25453 | | 0 | 0 | 0 | 0 | 0 | 0 | 0 | 0 | 0 | 0 | 0 |
| 0 | NA | NA | NA | NA | NA | Neto1 | neuropilin (NRP) and tolloid (TLL)-like 1 | 0 | 0 | 0 | 0 | 0 | 0 | 0 | 0 | 0 | 0 | 0 |
| 0 | NA | NA | NA | NA | NA | Gm50384 | | 0 | 0 | 0 | 0 | 0 | 0 | 0 | 0 | 0 | 0 | 0 |
| 0 | NA | NA | NA | NA | NA | Gm50386 | | 0 | 0 | 0 | 0 | 0 | 0 | 0 | 0 | 0 | 0 | 0 |
| 0 | NA | NA | NA | NA | NA | Gm46615 | | 0 | 0 | 0 | 0 | 0 | 0 | 0 | 0 | 0 | 0 | 0 |
| 0 | NA | NA | NA | NA | NA | Gm5691 | | 0 | 0 | 0 | 0 | 0 | 0 | 0 | 0 | 0 | 0 | 0 |
| 0 | NA | NA | NA | NA | NA | Gm5971 | | 0 | 0 | 0 | 0 | 0 | 0 | 0 | 0 | 0 | 0 | 0 |
| 0 | NA | NA | NA | NA | NA | Gm50389 | | 0 | 0 | 0 | 0 | 0 | 0 | 0 | 0 | 0 | 0 | 0 |
| 0 | NA | NA | NA | NA | NA | Gm5096 | | 0 | 0 | 0 | 0 | 0 | 0 | 0 | 0 | 0 | 0 | 0 |
| 0 | NA | NA | NA | NA | NA | Gm24987 | | 0 | 0 | 0 | 0 | 0 | 0 | 0 | 0 | 0 | 0 | 0 |
| 0 | NA | NA | NA | NA | NA | Gm50391 | | 0 | 0 | 0 | 0 | 0 | 0 | 0 | 0 | 0 | 0 | 0 |
| 0 | NA | NA | NA | NA | NA | Gm50400 | | 0 | 0 | 0 | 0 | 0 | 0 | 0 | 0 | 0 | 0 | 0 |
| 0 | NA | NA | NA | NA | NA | Gm50401 | | 0 | 0 | 0 | 0 | 0 | 0 | 0 | 0 | 0 | 0 | 0 |
| 0 | NA | NA | NA | NA | NA | Gm7662 | | 0 | 0 | 0 | 0 | 0 | 0 | 0 | 0 | 0 | 0 | 0 |
| 0 | NA | NA | NA | NA | NA | Gm50404 | | 0 | 0 | 0 | 0 | 0 | 0 | 0 | 0 | 0 | 0 | 0 |
| 0 | NA | NA | NA | NA | NA | Gm50406 | | 0 | 0 | 0 | 0 | 0 | 0 | 0 | 0 | 0 | 0 | 0 |
| 0 | NA | NA | NA | NA | NA | Gm50267 | | 0 | 0 | 0 | 0 | 0 | 0 | 0 | 0 | 0 | 0 | 0 |
| 0 | NA | NA | NA | NA | NA | Mir6359 | | 0 | 0 | 0 | 0 | 0 | 0 | 0 | 0 | 0 | 0 | 0 |
| 0 | NA | NA | NA | NA | NA | Gm16296 | | 0 | 0 | 0 | 0 | 0 | 0 | 0 | 0 | 0 | 0 | 0 |
| 0 | NA | NA | NA | NA | NA | Gm22694 | | 0 | 0 | 0 | 0 | 0 | 0 | 0 | 0 | 0 | 0 | 0 |
| 0 | NA | NA | NA | NA | NA | Gm6173 | | 0 | 0 | 0 | 0 | 0 | 0 | 0 | 0 | 0 | 0 | 0 |
| 0 | NA | NA | NA | NA | NA | Gm23859 | | 0 | 0 | 0 | 0 | 0 | 0 | 0 | 0 | 0 | 0 | 0 |
| 0 | NA | NA | NA | NA | NA | Gm7674 | | 0 | 0 | 0 | 0 | 0 | 0 | 0 | 0 | 0 | 0 | 0 |
| 0 | NA | NA | NA | NA | NA | Gm25418 | | 0 | 0 | 0 | 0 | 0 | 0 | 0 | 0 | 0 | 0 | 0 |
| 0 | NA | NA | NA | NA | NA | Gm20039 | | 0 | 0 | 0 | 0 | 0 | 0 | 0 | 0 | 0 | 0 | 0 |
| 0 | NA | NA | NA | NA | NA | 1700030N03Rik | | 0 | 0 | 0 | 0 | 0 | 0 | 0 | 0 | 0 | 0 | 0 |
| 0 | NA | NA | NA | NA | NA | Gm25431 | | 0 | 0 | 0 | 0 | 0 | 0 | 0 | 0 | 0 | 0 | 0 |
| 0 | NA | NA | NA | NA | NA | Gm50427 | | 0 | 0 | 0 | 0 | 0 | 0 | 0 | 0 | 0 | 0 | 0 |
| 0 | NA | NA | NA | NA | NA | Gm50426 | | 0 | 0 | 0 | 0 | 0 | 0 | 0 | 0 | 0 | 0 | 0 |
| 0 | NA | NA | NA | NA | NA | Gm7798 | | 0 | 0 | 0 | 0 | 0 | 0 | 0 | 0 | 0 | 0 | 0 |
| 0 | NA | NA | NA | NA | NA | Mir6984 | | 0 | 0 | 0 | 0 | 0 | 0 | 0 | 0 | 0 | 0 | 0 |
| 0 | NA | NA | NA | NA | NA | Gm48696 | | 0 | 0 | 0 | 0 | 0 | 0 | 0 | 0 | 0 | 0 | 0 |
| 0 | NA | NA | NA | NA | NA | Gm23940 | | 0 | 0 | 0 | 0 | 0 | 0 | 0 | 0 | 0 | 0 | 0 |
| 0 | NA | NA | NA | NA | NA | Gm48683 | | 0 | 0 | 0 | 0 | 0 | 0 | 0 | 0 | 0 | 0 | 0 |
| 0 | NA | NA | NA | NA | NA | Gm2141 | | 0 | 0 | 0 | 0 | 0 | 0 | 0 | 0 | 0 | 0 | 0 |
| 0 | NA | NA | NA | NA | NA | Gm50374 | | 0 | 0 | 0 | 0 | 0 | 0 | 0 | 0 | 0 | 0 | 0 |
| 0 | NA | NA | NA | NA | NA | Tbx10 | T-box 10 | 0 | 0 | 0 | 0 | 0 | 0 | 0 | 0 | 0 | 0 | 0 |

| | | | | | | | | | | | | | | | | | | |
|---|---|---|---|---|---|---|---|---|---|---|---|---|---|---|---|---|---|---|
| 0 | NA | NA | NA | NA | NA | Gm23927 | | 0 | 0 | 0 | 0 | 0 | 0 | 0 | 0 | 0 | 0 | 0 |
| 0 | NA | NA | NA | NA | NA | Gm23927 | | 0 | 0 | 0 | 0 | 0 | 0 | 0 | 0 | 0 | 0 | 0 |
| 0 | NA | NA | NA | NA | NA | Gpr152 | G protein-coupled receptor 152 | 0 | 0 | 0 | 0 | 0 | 0 | 0 | 0 | 0 | 0 | 0 |
| 0 | NA | NA | NA | NA | NA | Gm45928 | | 0 | 0 | 0 | 0 | 0 | 0 | 0 | 0 | 0 | 0 | 0 |
| 0 | NA | NA | NA | NA | NA | Mir6985 | | 0 | 0 | 0 | 0 | 0 | 0 | 0 | 0 | 0 | 0 | 0 |
| 0 | NA | NA | NA | NA | NA | Gm50152 | | 0 | 0 | 0 | 0 | 0 | 0 | 0 | 0 | 0 | 0 | 0 |
| 0 | NA | NA | NA | NA | NA | 2010003K11Ril | RIKEN cDNA 2010003K11 gene | 0 | 0 | 0 | 0 | 0 | 0 | 0 | 0 | 0 | 0 | 0 |
| 0 | NA | NA | NA | NA | NA | Gm50372 | | 0 | 0 | 0 | 0 | 0 | 0 | 0 | 0 | 0 | 0 | 0 |
| 0 | NA | NA | NA | NA | NA | Gm21844 | | 0 | 0 | 0 | 0 | 0 | 0 | 0 | 0 | 0 | 0 | 0 |
| 0 | NA | NA | NA | NA | NA | Gm25334 | | 0 | 0 | 0 | 0 | 0 | 0 | 0 | 0 | 0 | 0 | 0 |
| 0 | NA | NA | NA | NA | NA | Gm50388 | | 0 | 0 | 0 | 0 | 0 | 0 | 0 | 0 | 0 | 0 | 0 |
| 0 | NA | NA | NA | NA | NA | Gm27468 | | 0 | 0 | 0 | 0 | 0 | 0 | 0 | 0 | 0 | 0 | 0 |
| 0 | NA | NA | NA | NA | NA | Gm50189 | | 0 | 0 | 0 | 0 | 0 | 0 | 0 | 0 | 0 | 0 | 0 |
| 0 | NA | NA | NA | NA | NA | Gm50208 | | 0 | 0 | 0 | 0 | 0 | 0 | 0 | 0 | 0 | 0 | 0 |
| 0 | NA | NA | NA | NA | NA | Catsper1 | cation channel, sperm associated 1 | 0 | 0 | 0 | 0 | 0 | 0 | 0 | 0 | 0 | 0 | 0 |
| 0 | NA | NA | NA | NA | NA | | | 0 | 0 | 0 | 0 | 0 | 0 | 0 | 0 | 0 | 0 | 0 |
| 0 | NA | NA | NA | NA | NA | Gm50111 | | 0 | 0 | 0 | 0 | 0 | 0 | 0 | 0 | 0 | 0 | 0 |
| 0 | NA | NA | NA | NA | NA | Gm31070 | | 0 | 0 | 0 | 0 | 0 | 0 | 0 | 0 | 0 | 0 | 0 |
| 0 | NA | NA | NA | NA | NA | Mir6987 | | 0 | 0 | 0 | 0 | 0 | 0 | 0 | 0 | 0 | 0 | 0 |
| 0 | NA | NA | NA | NA | NA | Kcnk7 | potassium channel, subfamily K, member | 0 | 0 | 0 | 0 | 0 | 0 | 0 | 0 | 0 | 0 | 0 |
| 0 | NA | NA | NA | NA | NA | Gm16538 | | 0 | 0 | 0 | 0 | 0 | 0 | 0 | 0 | 0 | 0 | 0 |
| 0 | NA | NA | NA | NA | NA | Fam89b | family with sequence similarity 89, memb | 0 | 0 | 0 | 0 | 0 | 0 | 0 | 0 | 0 | 0 | 0 |
| 0 | NA | NA | NA | NA | NA | Gm27694 | | 0 | 0 | 0 | 0 | 0 | 0 | 0 | 0 | 0 | 0 | 0 |
| 0 | NA | NA | NA | NA | NA | Gm27702 | | 0 | 0 | 0 | 0 | 0 | 0 | 0 | 0 | 0 | 0 | 0 |
| 0 | NA | NA | NA | NA | NA | Gm27533 | | 0 | 0 | 0 | 0 | 0 | 0 | 0 | 0 | 0 | 0 | 0 |
| 0 | NA | NA | NA | NA | NA | Gm27505 | | 0 | 0 | 0 | 0 | 0 | 0 | 0 | 0 | 0 | 0 | 0 |
| 0 | NA | NA | NA | NA | NA | Gm28006 | | 0 | 0 | 0 | 0 | 0 | 0 | 0 | 0 | 0 | 0 | 0 |
| 0 | NA | NA | NA | NA | NA | Slc22a20 | solute carrier family 22 (organic anion tra | 0 | 0 | 0 | 0 | 0 | 0 | 0 | 0 | 0 | 0 | 0 |
| 0 | NA | NA | NA | NA | NA | Gm19505 | | 0 | 0 | 0 | 0 | 0 | 0 | 0 | 0 | 0 | 0 | 0 |
| 0 | NA | NA | NA | NA | NA | Mir6988 | | 0 | 0 | 0 | 0 | 0 | 0 | 0 | 0 | 0 | 0 | 0 |
| 0 | NA | NA | NA | NA | NA | | | 0 | 0 | 0 | 0 | 0 | 0 | 0 | 0 | 0 | 0 | 0 |
| 0 | NA | NA | NA | NA | NA | n-R5s19 | | 0 | 0 | 0 | 0 | 0 | 0 | 0 | 0 | 0 | 0 | 0 |
| 0 | NA | NA | NA | NA | NA | Gpha2 | glycoprotein hormone alpha 2 | 0 | 0 | 0 | 0 | 0 | 0 | 0 | 0 | 0 | 0 | 0 |
| 0 | NA | NA | NA | NA | NA | Mir194-2 | | 0 | 0 | 0 | 0 | 0 | 0 | 0 | 0 | 0 | 0 | 0 |
| 0 | NA | NA | NA | NA | NA | Mir6989 | | 0 | 0 | 0 | 0 | 0 | 0 | 0 | 0 | 0 | 0 | 0 |
| 0 | NA | NA | NA | NA | NA | Gm50151 | | 0 | 0 | 0 | 0 | 0 | 0 | 0 | 0 | 0 | 0 | 0 |
| 0 | NA | NA | NA | NA | NA | Slc22a12 | solute carrier family 22 (organic anion/cat | 0 | 0 | 0 | 0 | 0 | 0 | 0 | 0 | 0 | 0 | 0 |
| 0 | NA | NA | NA | NA | NA | Gm26460 | | 0 | 0 | 0 | 0 | 0 | 0 | 0 | 0 | 0 | 0 | 0 |
| 0 | NA | NA | NA | NA | NA | Gm50435 | | 0 | 0 | 0 | 0 | 0 | 0 | 0 | 0 | 0 | 0 | 0 |
| 0 | NA | NA | NA | NA | NA | 2410152P15Rik | | 0 | 0 | 0 | 0 | 0 | 0 | 0 | 0 | 0 | 0 | 0 |
| 0 | NA | NA | NA | NA | NA | Gm14968 | | 0 | 0 | 0 | 0 | 0 | 0 | 0 | 0 | 0 | 0 | 0 |
| 0 | NA | NA | NA | NA | NA | | | 0 | 0 | 0 | 0 | 0 | 0 | 0 | 0 | 0 | 0 | 0 |
| 0 | NA | NA | NA | NA | NA | Mir5046 | | 0 | 0 | 0 | 0 | 0 | 0 | 0 | 0 | 0 | 0 | 0 |
| 0 | NA | NA | NA | NA | NA | Mir6990 | | 0 | 0 | 0 | 0 | 0 | 0 | 0 | 0 | 0 | 0 | 0 |
| 0 | NA | NA | NA | NA | NA | Catsperz | cation channel sperm associated auxiliar | 0 | 0 | 0 | 0 | 0 | 0 | 0 | 0 | 0 | 0 | 0 |
| 0 | NA | NA | NA | NA | NA | Gm22113 | | 0 | 0 | 0 | 0 | 0 | 0 | 0 | 0 | 0 | 0 | 0 |
| 0 | NA | NA | NA | NA | NA | Gm23997 | | 0 | 0 | 0 | 0 | 0 | 0 | 0 | 0 | 0 | 0 | 0 |
| 0 | NA | NA | NA | NA | NA | Gm50232 | | 0 | 0 | 0 | 0 | 0 | 0 | 0 | 0 | 0 | 0 | 0 |
| 0 | NA | NA | NA | NA | NA | Gm18842 | | 0 | 0 | 0 | 0 | 0 | 0 | 0 | 0 | 0 | 0 | 0 |
| 0 | NA | NA | NA | NA | NA | Plaat5 | phospholipase A and acyltransferase 5 | 0 | 0 | 0 | 0 | 0 | 0 | 0 | 0 | 0 | 0 | 0 |
| 0 | NA | NA | NA | NA | NA | Slc22a19 | solute carrier family 22 (organic anion tra | 0 | 0 | 0 | 0 | 0 | 0 | 0 | 0 | 0 | 0 | 0 |
| 0 | NA | NA | NA | NA | NA | Gm15967 | | 0 | 0 | 0 | 0 | 0 | 0 | 0 | 0 | 0 | 0 | 0 |
| 0 | NA | NA | NA | NA | NA | Gm15968 | | 0 | 0 | 0 | 0 | 0 | 0 | 0 | 0 | 0 | 0 | 0 |
| 0 | NA | NA | NA | NA | NA | Gm7096 | | 0 | 0 | 0 | 0 | 0 | 0 | 0 | 0 | 0 | 0 | 0 |
| 0 | NA | NA | NA | NA | NA | Slc22a30 | solute carrier family 22, member 30 | 0 | 0 | 0 | 0 | 0 | 0 | 0 | 0 | 0 | 0 | 0 |
| 0 | NA | NA | NA | NA | NA | Slc22a6 | solute carrier family 22 (organic anion tra | 0 | 0 | 0 | 0 | 0 | 0 | 0 | 0 | 0 | 0 | 0 |
| 0 | NA | NA | NA | NA | NA | 9830166K06Rik | | 0 | 0 | 0 | 0 | 0 | 0 | 0 | 0 | 0 | 0 | 0 |
| 0 | NA | NA | NA | NA | NA | Gm22820 | | 0 | 0 | 0 | 0 | 0 | 0 | 0 | 0 | 0 | 0 | 0 |
| 0 | NA | NA | NA | NA | NA | Gm25855 | | 0 | 0 | 0 | 0 | 0 | 0 | 0 | 0 | 0 | 0 | 0 |
| 0 | NA | NA | NA | NA | NA | Gm25894 | | 0 | 0 | 0 | 0 | 0 | 0 | 0 | 0 | 0 | 0 | 0 |

| | | | | | | | | | | | | | | | | | | |
|---|---|---|---|---|---|---|---|---|---|---|---|---|---|---|---|---|---|---|
| 0 | NA | NA | NA | NA | NA | Gm24452 | | 0 | 0 | 0 | 0 | 0 | 0 | 0 | 0 | 0 | 0 | 0 |
| 0 | NA | NA | NA | NA | NA | Gm24453 | | 0 | 0 | 0 | 0 | 0 | 0 | 0 | 0 | 0 | 0 | 0 |
| 0 | NA | NA | NA | NA | NA | Gm22744 | | 0 | 0 | 0 | 0 | 0 | 0 | 0 | 0 | 0 | 0 | 0 |
| 0 | NA | NA | NA | NA | NA | Gm22680 | | 0 | 0 | 0 | 0 | 0 | 0 | 0 | 0 | 0 | 0 | 0 |
| 0 | NA | NA | NA | NA | NA | Gm23246 | | 0 | 0 | 0 | 0 | 0 | 0 | 0 | 0 | 0 | 0 | 0 |
| 0 | NA | NA | NA | NA | NA | Snord22 | | 0 | 0 | 0 | 0 | 0 | 0 | 0 | 0 | 0 | 0 | 0 |
| 0 | NA | NA | NA | NA | NA | Mir6992 | | 0 | 0 | 0 | 0 | 0 | 0 | 0 | 0 | 0 | 0 | 0 |
| 0 | NA | NA | NA | NA | NA | Gm50131 | | 0 | 0 | 0 | 0 | 0 | 0 | 0 | 0 | 0 | 0 | 0 |
| 0 | NA | NA | NA | NA | NA | Gm50139 | | 0 | 0 | 0 | 0 | 0 | 0 | 0 | 0 | 0 | 0 | 0 |
| 0 | NA | NA | NA | NA | NA | Gm49403 | | 0 | 0 | 0 | 0 | 0 | 0 | 0 | 0 | 0 | 0 | 0 |
| 0 | NA | NA | NA | NA | NA | Gm49416 | | 0 | 0 | 0 | 0 | 0 | 0 | 0 | 0 | 0 | 0 | 0 |
| 0 | NA | NA | NA | NA | NA | Scgb1a1 | secretoglobin, family 1A, member 1 (uterc | 0 | 0 | 0 | 0 | 0 | 0 | 0 | 0 | 0 | 0 | 0 |
| 0 | NA | NA | NA | NA | NA | Gm24543 | | 0 | 0 | 0 | 0 | 0 | 0 | 0 | 0 | 0 | 0 | 0 |
| 0 | NA | NA | NA | NA | NA | Gm2617 | | 0 | 0 | 0 | 0 | 0 | 0 | 0 | 0 | 0 | 0 | 0 |
| 0 | NA | NA | NA | NA | NA | Gm29231 | | 0 | 0 | 0 | 0 | 0 | 0 | 0 | 0 | 0 | 0 | 0 |
| 0 | NA | NA | NA | NA | NA | Gm50320 | | 0 | 0 | 0 | 0 | 0 | 0 | 0 | 0 | 0 | 0 | 0 |
| 0 | NA | NA | NA | NA | NA | Scgb2a2 | secretoglobin, family 2A, member 2 | 0 | 0 | 0 | 0 | 0 | 0 | 0 | 0 | 0 | 0 | 0 |
| 0 | NA | NA | NA | NA | NA | Gm22369 | | 0 | 0 | 0 | 0 | 0 | 0 | 0 | 0 | 0 | 0 | 0 |
| 0 | NA | NA | NA | NA | NA | Gm50324 | | 0 | 0 | 0 | 0 | 0 | 0 | 0 | 0 | 0 | 0 | 0 |
| 0 | NA | NA | NA | NA | NA | Gm24484 | | 0 | 0 | 0 | 0 | 0 | 0 | 0 | 0 | 0 | 0 | 0 |
| 0 | NA | NA | NA | NA | NA | Gm36913 | | 0 | 0 | 0 | 0 | 0 | 0 | 0 | 0 | 0 | 0 | 0 |
| 0 | NA | NA | NA | NA | NA | Gm50343 | | 0 | 0 | 0 | 0 | 0 | 0 | 0 | 0 | 0 | 0 | 0 |
| 0 | NA | NA | NA | NA | NA | Gm50345 | | 0 | 0 | 0 | 0 | 0 | 0 | 0 | 0 | 0 | 0 | 0 |
| 0 | NA | NA | NA | NA | NA | Gm50346 | | 0 | 0 | 0 | 0 | 0 | 0 | 0 | 0 | 0 | 0 | 0 |
| 0 | NA | NA | NA | NA | NA | Gm50359 | | 0 | 0 | 0 | 0 | 0 | 0 | 0 | 0 | 0 | 0 | 0 |
| 0 | NA | NA | NA | NA | NA | Gm50205 | | 0 | 0 | 0 | 0 | 0 | 0 | 0 | 0 | 0 | 0 | 0 |
| 0 | NA | NA | NA | NA | NA | 4930524O05Rik | | 0 | 0 | 0 | 0 | 0 | 0 | 0 | 0 | 0 | 0 | 0 |
| 0 | NA | NA | NA | NA | NA | 2210404E10Rik | | 0 | 0 | 0 | 0 | 0 | 0 | 0 | 0 | 0 | 0 | 0 |
| 0 | NA | NA | NA | NA | NA | Gm10802 | | 0 | 0 | 0 | 0 | 0 | 0 | 0 | 0 | 0 | 0 | 0 |
| 0 | NA | NA | NA | NA | NA | Zp1 | zona pellucida glycoprotein 1 | 0 | 0 | 0 | 0 | 0 | 0 | 0 | 0 | 0 | 0 | 0 |
| 0 | NA | NA | NA | NA | NA | Ms4a15 | membrane-spanning 4-domains, subfami | 0 | 0 | 0 | 0 | 0 | 0 | 0 | 0 | 0 | 0 | 0 |
| 0 | NA | NA | NA | NA | NA | Ms4a18 | membrane-spanning 4-domains, subfami | 0 | 0 | 0 | 0 | 0 | 0 | 0 | 0 | 0 | 0 | 0 |
| 0 | NA | NA | NA | NA | NA | Ms4a20 | membrane-spanning 4-domains, subfami | 0 | 0 | 0 | 0 | 0 | 0 | 0 | 0 | 0 | 0 | 0 |
| 0 | NA | NA | NA | NA | NA | Ms4a19 | membrane-spanning 4-domains, subfami | 0 | 0 | 0 | 0 | 0 | 0 | 0 | 0 | 0 | 0 | 0 |
| 0 | NA | NA | NA | NA | NA | Ms4a13 | membrane-spanning 4-domains, subfami | 0 | 0 | 0 | 0 | 0 | 0 | 0 | 0 | 0 | 0 | 0 |
| 0 | NA | NA | NA | NA | NA | 4930526L06Rik | | 0 | 0 | 0 | 0 | 0 | 0 | 0 | 0 | 0 | 0 | 0 |
| 0 | NA | NA | NA | NA | NA | Ms4a12 | membrane-spanning 4-domains, subfami | 0 | 0 | 0 | 0 | 0 | 0 | 0 | 0 | 0 | 0 | 0 |
| 0 | NA | NA | NA | NA | NA | Ms4a5 | membrane-spanning 4-domains, subfami | 0 | 0 | 0 | 0 | 0 | 0 | 0 | 0 | 0 | 0 | 0 |
| 0 | NA | NA | NA | NA | NA | Gm28935 | | 0 | 0 | 0 | 0 | 0 | 0 | 0 | 0 | 0 | 0 | 0 |
| 0 | NA | NA | NA | NA | NA | Gm5692 | | 0 | 0 | 0 | 0 | 0 | 0 | 0 | 0 | 0 | 0 | 0 |
| 0 | NA | NA | NA | NA | NA | Gm19261 | | 0 | 0 | 0 | 0 | 0 | 0 | 0 | 0 | 0 | 0 | 0 |
| 0 | NA | NA | NA | NA | NA | Gm10212 | | 0 | 0 | 0 | 0 | 0 | 0 | 0 | 0 | 0 | 0 | 0 |
| 0 | NA | NA | NA | NA | NA | Gm38275 | | 0 | 0 | 0 | 0 | 0 | 0 | 0 | 0 | 0 | 0 | 0 |
| 0 | NA | NA | NA | NA | NA | Gm24599 | | 0 | 0 | 0 | 0 | 0 | 0 | 0 | 0 | 0 | 0 | 0 |
| 0 | NA | NA | NA | NA | NA | Ms4a3 | membrane-spanning 4-domains, subfami | 0 | 0 | 0 | 0 | 0 | 0 | 0 | 0 | 0 | 0 | 0 |
| 0 | NA | NA | NA | NA | NA | Gm26183 | | 0 | 0 | 0 | 0 | 0 | 0 | 0 | 0 | 0 | 0 | 0 |
| 0 | NA | NA | NA | NA | NA | Gm25443 | | 0 | 0 | 0 | 0 | 0 | 0 | 0 | 0 | 0 | 0 | 0 |
| 0 | NA | NA | NA | NA | NA | Oosp3 | oocyte secreted protein 3 | 0 | 0 | 0 | 0 | 0 | 0 | 0 | 0 | 0 | 0 | 0 |
| 0 | NA | NA | NA | NA | NA | Gm19224 | | 0 | 0 | 0 | 0 | 0 | 0 | 0 | 0 | 0 | 0 | 0 |
| 0 | NA | NA | NA | NA | NA | Cblif | cobalamin binding intrinsic factor | 0 | 0 | 0 | 0 | 0 | 0 | 0 | 0 | 0 | 0 | 0 |
| 0 | NA | NA | NA | NA | NA | Olfr1417 | olfactory receptor 1417 | 0 | 0 | 0 | 0 | 0 | 0 | 0 | 0 | 0 | 0 | 0 |
| 0 | NA | NA | NA | NA | NA | Olfr1418 | olfactory receptor 1418 | 0 | 0 | 0 | 0 | 0 | 0 | 0 | 0 | 0 | 0 | 0 |
| 0 | NA | NA | NA | NA | NA | Gm45119 | | 0 | 0 | 0 | 0 | 0 | 0 | 0 | 0 | 0 | 0 | 0 |
| 0 | NA | NA | NA | NA | NA | Olfr1419 | olfactory receptor 1419 | 0 | 0 | 0 | 0 | 0 | 0 | 0 | 0 | 0 | 0 | 0 |
| 0 | NA | NA | NA | NA | NA | Olfr1420 | olfactory receptor 1420 | 0 | 0 | 0 | 0 | 0 | 0 | 0 | 0 | 0 | 0 | 0 |
| 0 | NA | NA | NA | NA | NA | Mir6994 | | 0 | 0 | 0 | 0 | 0 | 0 | 0 | 0 | 0 | 0 | 0 |
| 0 | NA | NA | NA | NA | NA | Gm22272 | | 0 | 0 | 0 | 0 | 0 | 0 | 0 | 0 | 0 | 0 | 0 |
| 0 | NA | NA | NA | NA | NA | Olfr1421-ps1 | | 0 | 0 | 0 | 0 | 0 | 0 | 0 | 0 | 0 | 0 | 0 |
| 0 | NA | NA | NA | NA | NA | Gm45115 | | 0 | 0 | 0 | 0 | 0 | 0 | 0 | 0 | 0 | 0 | 0 |
| 0 | NA | NA | NA | NA | NA | Olfr1422-ps1 | | 0 | 0 | 0 | 0 | 0 | 0 | 0 | 0 | 0 | 0 | 0 |

| | | | | | | | | | | | | | | | | | | |
|---|---|---|---|---|---|---|---|---|---|---|---|---|---|---|---|---|---|---|
| 0 | NA | NA | NA | NA | NA | Gm45116 | | 0 | 0 | 0 | 0 | 0 | 0 | 0 | 0 | 0 | 0 | 0 |
| 0 | NA | NA | NA | NA | NA | Olfr1423 | olfactory receptor 1423 | 0 | 0 | 0 | 0 | 0 | 0 | 0 | 0 | 0 | 0 | 0 |
| 0 | NA | NA | NA | NA | NA | Gm44859 | | 0 | 0 | 0 | 0 | 0 | 0 | 0 | 0 | 0 | 0 | 0 |
| 0 | NA | NA | NA | NA | NA | Olfr1424 | olfactory receptor 1424 | 0 | 0 | 0 | 0 | 0 | 0 | 0 | 0 | 0 | 0 | 0 |
| 0 | NA | NA | NA | NA | NA | Olfr1425 | olfactory receptor 1425 | 0 | 0 | 0 | 0 | 0 | 0 | 0 | 0 | 0 | 0 | 0 |
| 0 | NA | NA | NA | NA | NA | Olfr1426 | olfactory receptor 1426 | 0 | 0 | 0 | 0 | 0 | 0 | 0 | 0 | 0 | 0 | 0 |
| 0 | NA | NA | NA | NA | NA | Olfr1427 | olfactory receptor 1427 | 0 | 0 | 0 | 0 | 0 | 0 | 0 | 0 | 0 | 0 | 0 |
| 0 | NA | NA | NA | NA | NA | Olfr1428 | olfactory receptor 1428 | 0 | 0 | 0 | 0 | 0 | 0 | 0 | 0 | 0 | 0 | 0 |
| 0 | NA | NA | NA | NA | NA | Olfr76 | olfactory receptor 76 | 0 | 0 | 0 | 0 | 0 | 0 | 0 | 0 | 0 | 0 | 0 |
| 0 | NA | NA | NA | NA | NA | Olfr1555-ps1 | | 0 | 0 | 0 | 0 | 0 | 0 | 0 | 0 | 0 | 0 | 0 |
| 0 | NA | NA | NA | NA | NA | Olfr1430-ps1 | | 0 | 0 | 0 | 0 | 0 | 0 | 0 | 0 | 0 | 0 | 0 |
| 0 | NA | NA | NA | NA | NA | Olfr1431 | olfactory receptor 1431 | 0 | 0 | 0 | 0 | 0 | 0 | 0 | 0 | 0 | 0 | 0 |
| 0 | NA | NA | NA | NA | NA | Olfr1432 | olfactory receptor 1432 | 0 | 0 | 0 | 0 | 0 | 0 | 0 | 0 | 0 | 0 | 0 |
| 0 | NA | NA | NA | NA | NA | Olfr262 | olfactory receptor 262 | 0 | 0 | 0 | 0 | 0 | 0 | 0 | 0 | 0 | 0 | 0 |
| 0 | NA | NA | NA | NA | NA | Gm26864 | | 0 | 0 | 0 | 0 | 0 | 0 | 0 | 0 | 0 | 0 | 0 |
| 0 | NA | NA | NA | NA | NA | Olfr235 | olfactory receptor 235 | 0 | 0 | 0 | 0 | 0 | 0 | 0 | 0 | 0 | 0 | 0 |
| 0 | NA | NA | NA | NA | NA | Olfr1434 | olfactory receptor 1434 | 0 | 0 | 0 | 0 | 0 | 0 | 0 | 0 | 0 | 0 | 0 |
| 0 | NA | NA | NA | NA | NA | Olfr1435-ps1 | | 0 | 0 | 0 | 0 | 0 | 0 | 0 | 0 | 0 | 0 | 0 |
| 0 | NA | NA | NA | NA | NA | Olfr1436 | olfactory receptor 1436 | 0 | 0 | 0 | 0 | 0 | 0 | 0 | 0 | 0 | 0 | 0 |
| 0 | NA | NA | NA | NA | NA | Gm44858 | | 0 | 0 | 0 | 0 | 0 | 0 | 0 | 0 | 0 | 0 | 0 |
| 0 | NA | NA | NA | NA | NA | Olfr1437 | olfactory receptor 1437 | 0 | 0 | 0 | 0 | 0 | 0 | 0 | 0 | 0 | 0 | 0 |
| 0 | NA | NA | NA | NA | NA | Olfr1438-ps1 | | 0 | 0 | 0 | 0 | 0 | 0 | 0 | 0 | 0 | 0 | 0 |
| 0 | NA | NA | NA | NA | NA | Olfr1439-ps1 | | 0 | 0 | 0 | 0 | 0 | 0 | 0 | 0 | 0 | 0 | 0 |
| 0 | NA | NA | NA | NA | NA | Olfr1440 | olfactory receptor 1440 | 0 | 0 | 0 | 0 | 0 | 0 | 0 | 0 | 0 | 0 | 0 |
| 0 | NA | NA | NA | NA | NA | Olfr1441 | olfactory receptor 1441 | 0 | 0 | 0 | 0 | 0 | 0 | 0 | 0 | 0 | 0 | 0 |
| 0 | NA | NA | NA | NA | NA | Pfpl | pore forming protein-like | 0 | 0 | 0 | 0 | 0 | 0 | 0 | 0 | 0 | 0 | 0 |
| 0 | NA | NA | NA | NA | NA | Gm47243 | | 0 | 0 | 0 | 0 | 0 | 0 | 0 | 0 | 0 | 0 | 0 |
| 0 | NA | NA | NA | NA | NA | Gm4952 | predicted gene 4952 | 0 | 0 | 0 | 0 | 0 | 0 | 0 | 0 | 0 | 0 | 0 |
| 0 | NA | NA | NA | NA | NA | Gm24521 | | 0 | 0 | 0 | 0 | 0 | 0 | 0 | 0 | 0 | 0 | 0 |
| 0 | NA | NA | NA | NA | NA | Olfr1442 | olfactory receptor 1442 | 0 | 0 | 0 | 0 | 0 | 0 | 0 | 0 | 0 | 0 | 0 |
| 0 | NA | NA | NA | NA | NA | Olfr1443 | olfactory receptor 1443 | 0 | 0 | 0 | 0 | 0 | 0 | 0 | 0 | 0 | 0 | 0 |
| 0 | NA | NA | NA | NA | NA | Gm49772 | | 0 | 0 | 0 | 0 | 0 | 0 | 0 | 0 | 0 | 0 | 0 |
| 0 | NA | NA | NA | NA | NA | Gm15962 | | 0 | 0 | 0 | 0 | 0 | 0 | 0 | 0 | 0 | 0 | 0 |
| 0 | NA | NA | NA | NA | NA | Zfp91 | | 0 | 0 | 0 | 0 | 0 | 0 | 0 | 0 | 0 | 0 | 0 |
| 0 | NA | NA | NA | NA | NA | Gm5244 | | 0 | 0 | 0 | 0 | 0 | 0 | 0 | 0 | 0 | 0 | 0 |
| 0 | NA | NA | NA | NA | NA | Olfr1444 | olfactory receptor 1444 | 0 | 0 | 0 | 0 | 0 | 0 | 0 | 0 | 0 | 0 | 0 |
| 0 | NA | NA | NA | NA | NA | Olfr1445 | olfactory receptor 1445 | 0 | 0 | 0 | 0 | 0 | 0 | 0 | 0 | 0 | 0 | 0 |
| 0 | NA | NA | NA | NA | NA | Olfr1446 | olfactory receptor 1446 | 0 | 0 | 0 | 0 | 0 | 0 | 0 | 0 | 0 | 0 | 0 |
| 0 | NA | NA | NA | NA | NA | Olfr1447 | olfactory receptor 1447 | 0 | 0 | 0 | 0 | 0 | 0 | 0 | 0 | 0 | 0 | 0 |
| 0 | NA | NA | NA | NA | NA | Olfr1448 | olfactory receptor 1448 | 0 | 0 | 0 | 0 | 0 | 0 | 0 | 0 | 0 | 0 | 0 |
| 0 | NA | NA | NA | NA | NA | Olfr1449 | olfactory receptor 1449 | 0 | 0 | 0 | 0 | 0 | 0 | 0 | 0 | 0 | 0 | 0 |
| 0 | NA | NA | NA | NA | NA | Olfr1450 | olfactory receptor 1450 | 0 | 0 | 0 | 0 | 0 | 0 | 0 | 0 | 0 | 0 | 0 |
| 0 | NA | NA | NA | NA | NA | Olfr1451 | olfactory receptor 1451 | 0 | 0 | 0 | 0 | 0 | 0 | 0 | 0 | 0 | 0 | 0 |
| 0 | NA | NA | NA | NA | NA | Olfr1452-ps1 | | 0 | 0 | 0 | 0 | 0 | 0 | 0 | 0 | 0 | 0 | 0 |
| 0 | NA | NA | NA | NA | NA | Olfr1453 | olfactory receptor 1453 | 0 | 0 | 0 | 0 | 0 | 0 | 0 | 0 | 0 | 0 | 0 |
| 0 | NA | NA | NA | NA | NA | Olfr1454 | olfactory receptor 1454 | 0 | 0 | 0 | 0 | 0 | 0 | 0 | 0 | 0 | 0 | 0 |
| 0 | NA | NA | NA | NA | NA | Olfr1455-ps1 | | 0 | 0 | 0 | 0 | 0 | 0 | 0 | 0 | 0 | 0 | 0 |
| 0 | NA | NA | NA | NA | NA | Olfr1456-ps1 | | 0 | 0 | 0 | 0 | 0 | 0 | 0 | 0 | 0 | 0 | 0 |
| 0 | NA | NA | NA | NA | NA | Olfr1457 | olfactory receptor 1457 | 0 | 0 | 0 | 0 | 0 | 0 | 0 | 0 | 0 | 0 | 0 |
| 0 | NA | NA | NA | NA | NA | Olfr1458 | olfactory receptor 1458 | 0 | 0 | 0 | 0 | 0 | 0 | 0 | 0 | 0 | 0 | 0 |
| 0 | NA | NA | NA | NA | NA | Olfr1459 | olfactory receptor 1459 | 0 | 0 | 0 | 0 | 0 | 0 | 0 | 0 | 0 | 0 | 0 |
| 0 | NA | NA | NA | NA | NA | Olfr1460-ps1 | | 0 | 0 | 0 | 0 | 0 | 0 | 0 | 0 | 0 | 0 | 0 |
| 0 | NA | NA | NA | NA | NA | Olfr1461 | olfactory receptor 1461 | 0 | 0 | 0 | 0 | 0 | 0 | 0 | 0 | 0 | 0 | 0 |
| 0 | NA | NA | NA | NA | NA | Olfr1462 | olfactory receptor 1462 | 0 | 0 | 0 | 0 | 0 | 0 | 0 | 0 | 0 | 0 | 0 |
| 0 | NA | NA | NA | NA | NA | Olfr1463 | olfactory receptor 1463 | 0 | 0 | 0 | 0 | 0 | 0 | 0 | 0 | 0 | 0 | 0 |
| 0 | NA | NA | NA | NA | NA | Olfr1464-ps1 | | 0 | 0 | 0 | 0 | 0 | 0 | 0 | 0 | 0 | 0 | 0 |
| 0 | NA | NA | NA | NA | NA | Gm22443 | | 0 | 0 | 0 | 0 | 0 | 0 | 0 | 0 | 0 | 0 | 0 |
| 0 | NA | NA | NA | NA | NA | Olfr1465 | olfactory receptor 1465 | 0 | 0 | 0 | 0 | 0 | 0 | 0 | 0 | 0 | 0 | 0 |
| 0 | NA | NA | NA | NA | NA | Olfr1466 | olfactory receptor 1466 | 0 | 0 | 0 | 0 | 0 | 0 | 0 | 0 | 0 | 0 | 0 |
| 0 | NA | NA | NA | NA | NA | Olfr1467 | olfactory receptor 1467 | 0 | 0 | 0 | 0 | 0 | 0 | 0 | 0 | 0 | 0 | 0 |

| | | | | | | | | | | | | | | | | | | |
|---|---|---|---|---|---|---|---|---|---|---|---|---|---|---|---|---|---|---|
| 0 | NA | NA | NA | NA | NA | Olfr1468-ps1 | | 0 | 0 | 0 | 0 | 0 | 0 | 0 | 0 | 0 | 0 | 0 |
| 0 | NA | NA | NA | NA | NA | Gm44849 | | 0 | 0 | 0 | 0 | 0 | 0 | 0 | 0 | 0 | 0 | 0 |
| 0 | NA | NA | NA | NA | NA | Olfr1469 | olfactory receptor 1469 | 0 | 0 | 0 | 0 | 0 | 0 | 0 | 0 | 0 | 0 | 0 |
| 0 | NA | NA | NA | NA | NA | Olfr1470-ps1 | | 0 | 0 | 0 | 0 | 0 | 0 | 0 | 0 | 0 | 0 | 0 |
| 0 | NA | NA | NA | NA | NA | Olfr1471 | olfactory receptor 1471 | 0 | 0 | 0 | 0 | 0 | 0 | 0 | 0 | 0 | 0 | 0 |
| 0 | NA | NA | NA | NA | NA | Olfr1472 | olfactory receptor 1472 | 0 | 0 | 0 | 0 | 0 | 0 | 0 | 0 | 0 | 0 | 0 |
| 0 | NA | NA | NA | NA | NA | Olfr1473-ps1 | | 0 | 0 | 0 | 0 | 0 | 0 | 0 | 0 | 0 | 0 | 0 |
| 0 | NA | NA | NA | NA | NA | Olfr1474 | olfactory receptor 1474 | 0 | 0 | 0 | 0 | 0 | 0 | 0 | 0 | 0 | 0 | 0 |
| 0 | NA | NA | NA | NA | NA | Olfr1475 | olfactory receptor 1475 | 0 | 0 | 0 | 0 | 0 | 0 | 0 | 0 | 0 | 0 | 0 |
| 0 | NA | NA | NA | NA | NA | Olfr1476-ps1 | | 0 | 0 | 0 | 0 | 0 | 0 | 0 | 0 | 0 | 0 | 0 |
| 0 | NA | NA | NA | NA | NA | Olfr1477 | olfactory receptor 1477 | 0 | 0 | 0 | 0 | 0 | 0 | 0 | 0 | 0 | 0 | 0 |
| 0 | NA | NA | NA | NA | NA | Olfr1478-ps1 | | 0 | 0 | 0 | 0 | 0 | 0 | 0 | 0 | 0 | 0 | 0 |
| 0 | NA | NA | NA | NA | NA | Olfr1479-ps1 | | 0 | 0 | 0 | 0 | 0 | 0 | 0 | 0 | 0 | 0 | 0 |
| 0 | NA | NA | NA | NA | NA | Gm3188 | | 0 | 0 | 0 | 0 | 0 | 0 | 0 | 0 | 0 | 0 | 0 |
| 0 | NA | NA | NA | NA | NA | Olfr1480 | olfactory receptor 1480 | 0 | 0 | 0 | 0 | 0 | 0 | 0 | 0 | 0 | 0 | 0 |
| 0 | NA | NA | NA | NA | NA | Olfr1481-ps1 | | 0 | 0 | 0 | 0 | 0 | 0 | 0 | 0 | 0 | 0 | 0 |
| 0 | NA | NA | NA | NA | NA | Olfr1482-ps1 | | 0 | 0 | 0 | 0 | 0 | 0 | 0 | 0 | 0 | 0 | 0 |
| 0 | NA | NA | NA | NA | NA | Olfr1483-ps1 | | 0 | 0 | 0 | 0 | 0 | 0 | 0 | 0 | 0 | 0 | 0 |
| 0 | NA | NA | NA | NA | NA | Olfr1484 | olfactory receptor 1484 | 0 | 0 | 0 | 0 | 0 | 0 | 0 | 0 | 0 | 0 | 0 |
| 0 | NA | NA | NA | NA | NA | Olfr1485-ps1 | | 0 | 0 | 0 | 0 | 0 | 0 | 0 | 0 | 0 | 0 | 0 |
| 0 | NA | NA | NA | NA | NA | Olfr1486-ps1 | | 0 | 0 | 0 | 0 | 0 | 0 | 0 | 0 | 0 | 0 | 0 |
| 0 | NA | NA | NA | NA | NA | Gm22323 | | 0 | 0 | 0 | 0 | 0 | 0 | 0 | 0 | 0 | 0 | 0 |
| 0 | NA | NA | NA | NA | NA | Gm23931 | | 0 | 0 | 0 | 0 | 0 | 0 | 0 | 0 | 0 | 0 | 0 |
| 0 | NA | NA | NA | NA | NA | Olfr1487 | olfactory receptor 1487 | 0 | 0 | 0 | 0 | 0 | 0 | 0 | 0 | 0 | 0 | 0 |
| 0 | NA | NA | NA | NA | NA | Olfr1488-ps1 | | 0 | 0 | 0 | 0 | 0 | 0 | 0 | 0 | 0 | 0 | 0 |
| 0 | NA | NA | NA | NA | NA | Olfr1489 | olfactory receptor 1489 | 0 | 0 | 0 | 0 | 0 | 0 | 0 | 0 | 0 | 0 | 0 |
| 0 | NA | NA | NA | NA | NA | Gm24124 | | 0 | 0 | 0 | 0 | 0 | 0 | 0 | 0 | 0 | 0 | 0 |
| 0 | NA | NA | NA | NA | NA | Olfr1490 | olfactory receptor 1490 | 0 | 0 | 0 | 0 | 0 | 0 | 0 | 0 | 0 | 0 | 0 |
| 0 | NA | NA | NA | NA | NA | Olfr1491 | olfactory receptor 1491 | 0 | 0 | 0 | 0 | 0 | 0 | 0 | 0 | 0 | 0 | 0 |
| 0 | NA | NA | NA | NA | NA | Olfr1492-ps1 | | 0 | 0 | 0 | 0 | 0 | 0 | 0 | 0 | 0 | 0 | 0 |
| 0 | NA | NA | NA | NA | NA | Olfr1493-ps1 | | 0 | 0 | 0 | 0 | 0 | 0 | 0 | 0 | 0 | 0 | 0 |
| 0 | NA | NA | NA | NA | NA | Olfr1494 | olfactory receptor 1494 | 0 | 0 | 0 | 0 | 0 | 0 | 0 | 0 | 0 | 0 | 0 |
| 0 | NA | NA | NA | NA | NA | Olfr1495 | olfactory receptor 1495 | 0 | 0 | 0 | 0 | 0 | 0 | 0 | 0 | 0 | 0 | 0 |
| 0 | NA | NA | NA | NA | NA | Olfr1496 | olfactory receptor 1496 | 0 | 0 | 0 | 0 | 0 | 0 | 0 | 0 | 0 | 0 | 0 |
| 0 | NA | NA | NA | NA | NA | Olfr1497 | olfactory receptor 1497 | 0 | 0 | 0 | 0 | 0 | 0 | 0 | 0 | 0 | 0 | 0 |
| 0 | NA | NA | NA | NA | NA | Olfr1498-ps1 | | 0 | 0 | 0 | 0 | 0 | 0 | 0 | 0 | 0 | 0 | 0 |
| 0 | NA | NA | NA | NA | NA | Olfr1499 | olfactory receptor 1499 | 0 | 0 | 0 | 0 | 0 | 0 | 0 | 0 | 0 | 0 | 0 |
| 0 | NA | NA | NA | NA | NA | Olfr1500 | olfactory receptor 1500 | 0 | 0 | 0 | 0 | 0 | 0 | 0 | 0 | 0 | 0 | 0 |
| 0 | NA | NA | NA | NA | NA | Olfr1501 | olfactory receptor 1501 | 0 | 0 | 0 | 0 | 0 | 0 | 0 | 0 | 0 | 0 | 0 |
| 0 | NA | NA | NA | NA | NA | Olfr1502 | olfactory receptor 1502 | 0 | 0 | 0 | 0 | 0 | 0 | 0 | 0 | 0 | 0 | 0 |
| 0 | NA | NA | NA | NA | NA | Olfr1503-ps1 | | 0 | 0 | 0 | 0 | 0 | 0 | 0 | 0 | 0 | 0 | 0 |
| 0 | NA | NA | NA | NA | NA | Olfr1504 | olfactory receptor 1504 | 0 | 0 | 0 | 0 | 0 | 0 | 0 | 0 | 0 | 0 | 0 |
| 0 | NA | NA | NA | NA | NA | Olfr1505 | olfactory receptor 1505 | 0 | 0 | 0 | 0 | 0 | 0 | 0 | 0 | 0 | 0 | 0 |
| 0 | NA | NA | NA | NA | NA | Gm26993 | | 0 | 0 | 0 | 0 | 0 | 0 | 0 | 0 | 0 | 0 | 0 |
| 0 | NA | NA | NA | NA | NA | Gm37997 | | 0 | 0 | 0 | 0 | 0 | 0 | 0 | 0 | 0 | 0 | 0 |
| 0 | NA | NA | NA | NA | NA | Gm26026 | | 0 | 0 | 0 | 0 | 0 | 0 | 0 | 0 | 0 | 0 | 0 |
| 0 | NA | NA | NA | NA | NA | Gm24319 | | 0 | 0 | 0 | 0 | 0 | 0 | 0 | 0 | 0 | 0 | 0 |
| 0 | NA | NA | NA | NA | NA | Gm3329 | | 0 | 0 | 0 | 0 | 0 | 0 | 0 | 0 | 0 | 0 | 0 |
| 0 | NA | NA | NA | NA | NA | Rpl37-ps1 | | 0 | 0 | 0 | 0 | 0 | 0 | 0 | 0 | 0 | 0 | 0 |
| 0 | NA | NA | NA | NA | NA | Mir496b | | 0 | 0 | 0 | 0 | 0 | 0 | 0 | 0 | 0 | 0 | 0 |
| 0 | NA | NA | NA | NA | NA | Gm19312 | | 0 | 0 | 0 | 0 | 0 | 0 | 0 | 0 | 0 | 0 | 0 |
| 0 | NA | NA | NA | NA | NA | Gm50362 | | 0 | 0 | 0 | 0 | 0 | 0 | 0 | 0 | 0 | 0 | 0 |
| 0 | NA | NA | NA | NA | NA | Gm8222 | | 0 | 0 | 0 | 0 | 0 | 0 | 0 | 0 | 0 | 0 | 0 |
| 0 | NA | NA | NA | NA | NA | Gm50363 | | 0 | 0 | 0 | 0 | 0 | 0 | 0 | 0 | 0 | 0 | 0 |
| 0 | NA | NA | NA | NA | NA | Gm9806 | | 0 | 0 | 0 | 0 | 0 | 0 | 0 | 0 | 0 | 0 | 0 |
| 0 | NA | NA | NA | NA | NA | Gm32341 | | 0 | 0 | 0 | 0 | 0 | 0 | 0 | 0 | 0 | 0 | 0 |
| 0 | NA | NA | NA | NA | NA | Foxb2 | forkhead box B2 | 0 | 0 | 0 | 0 | 0 | 0 | 0 | 0 | 0 | 0 | 0 |
| 0 | NA | NA | NA | NA | NA | Gm27970 | | 0 | 0 | 0 | 0 | 0 | 0 | 0 | 0 | 0 | 0 | 0 |
| 0 | NA | NA | NA | NA | NA | Gm27796 | | 0 | 0 | 0 | 0 | 0 | 0 | 0 | 0 | 0 | 0 | 0 |
| 0 | NA | NA | NA | NA | NA | Gm50276 | | 0 | 0 | 0 | 0 | 0 | 0 | 0 | 0 | 0 | 0 | 0 |

| | | | | | | | | | | | | | | | | | | |
|---|---|---|---|---|---|---|---|---|---|---|---|---|---|---|---|---|---|---|
| 0 | NA | NA | NA | NA | NA | Gm50277 | | 0 | 0 | 0 | 0 | 0 | 0 | 0 | 0 | 0 | 0 | 0 |
| 0 | NA | NA | NA | NA | NA | Gm50280 | | 0 | 0 | 0 | 0 | 0 | 0 | 0 | 0 | 0 | 0 | 0 |
| 0 | NA | NA | NA | NA | NA | Gm22880 | | 0 | 0 | 0 | 0 | 0 | 0 | 0 | 0 | 0 | 0 | 0 |
| 0 | NA | NA | NA | NA | NA | Gm50181 | | 0 | 0 | 0 | 0 | 0 | 0 | 0 | 0 | 0 | 0 | 0 |
| 0 | NA | NA | NA | NA | NA | Gm23063 | | 0 | 0 | 0 | 0 | 0 | 0 | 0 | 0 | 0 | 0 | 0 |
| 0 | NA | NA | NA | NA | NA | Gm50182 | | 0 | 0 | 0 | 0 | 0 | 0 | 0 | 0 | 0 | 0 | 0 |
| 0 | NA | NA | NA | NA | NA | Gm18610 | | 0 | 0 | 0 | 0 | 0 | 0 | 0 | 0 | 0 | 0 | 0 |
| 0 | NA | NA | NA | NA | NA | Gm50183 | | 0 | 0 | 0 | 0 | 0 | 0 | 0 | 0 | 0 | 0 | 0 |
| 0 | NA | NA | NA | NA | NA | Gm50202 | | 0 | 0 | 0 | 0 | 0 | 0 | 0 | 0 | 0 | 0 | 0 |
| 0 | NA | NA | NA | NA | NA | Gm23238 | | 0 | 0 | 0 | 0 | 0 | 0 | 0 | 0 | 0 | 0 | 0 |
| 0 | NA | NA | NA | NA | NA | Gm24190 | | 0 | 0 | 0 | 0 | 0 | 0 | 0 | 0 | 0 | 0 | 0 |
| 0 | NA | NA | NA | NA | NA | Gm50212 | | 0 | 0 | 0 | 0 | 0 | 0 | 0 | 0 | 0 | 0 | 0 |
| 0 | NA | NA | NA | NA | NA | Gm25437 | | 0 | 0 | 0 | 0 | 0 | 0 | 0 | 0 | 0 | 0 | 0 |
| 0 | NA | NA | NA | NA | NA | Gm41815 | | 0 | 0 | 0 | 0 | 0 | 0 | 0 | 0 | 0 | 0 | 0 |
| 0 | NA | NA | NA | NA | NA | Gm50216 | | 0 | 0 | 0 | 0 | 0 | 0 | 0 | 0 | 0 | 0 | 0 |
| 0 | NA | NA | NA | NA | NA | Gm3728 | | 0 | 0 | 0 | 0 | 0 | 0 | 0 | 0 | 0 | 0 | 0 |
| 0 | NA | NA | NA | NA | NA | Gm22684 | | 0 | 0 | 0 | 0 | 0 | 0 | 0 | 0 | 0 | 0 | 0 |
| 0 | NA | NA | NA | NA | NA | Gm32629 | | 0 | 0 | 0 | 0 | 0 | 0 | 0 | 0 | 0 | 0 | 0 |
| 0 | NA | NA | NA | NA | NA | 1500015L24Rik | | 0 | 0 | 0 | 0 | 0 | 0 | 0 | 0 | 0 | 0 | 0 |
| 0 | NA | NA | NA | NA | NA | C730002L08Rik | | 0 | 0 | 0 | 0 | 0 | 0 | 0 | 0 | 0 | 0 | 0 |
| 0 | NA | NA | NA | NA | NA | Tmc1 | transmembrane channel-like gene family | 0 | 0 | 0 | 0 | 0 | 0 | 0 | 0 | 0 | 0 | 0 |
| 0 | NA | NA | NA | NA | NA | Gm32750 | | 0 | 0 | 0 | 0 | 0 | 0 | 0 | 0 | 0 | 0 | 0 |
| 0 | NA | NA | NA | NA | NA | 4930554I06Rik | | 0 | 0 | 0 | 0 | 0 | 0 | 0 | 0 | 0 | 0 | 0 |
| 0 | NA | NA | NA | NA | NA | | | 0 | 0 | 0 | 0 | 0 | 0 | 0 | 0 | 0 | 0 | 0 |
| 0 | NA | NA | NA | NA | NA | Gm50236 | | 0 | 0 | 0 | 0 | 0 | 0 | 0 | 0 | 0 | 0 | 0 |
| 0 | NA | NA | NA | NA | NA | Gm50243 | | 0 | 0 | 0 | 0 | 0 | 0 | 0 | 0 | 0 | 0 | 0 |
| 0 | NA | NA | NA | NA | NA | Gm50255 | | 0 | 0 | 0 | 0 | 0 | 0 | 0 | 0 | 0 | 0 | 0 |
| 0 | NA | NA | NA | NA | NA | Gm3443 | | 0 | 0 | 0 | 0 | 0 | 0 | 0 | 0 | 0 | 0 | 0 |
| 0 | NA | NA | NA | NA | NA | Gm50129 | | 0 | 0 | 0 | 0 | 0 | 0 | 0 | 0 | 0 | 0 | 0 |
| 0 | NA | NA | NA | NA | NA | Gm50133 | | 0 | 0 | 0 | 0 | 0 | 0 | 0 | 0 | 0 | 0 | 0 |
| 0 | NA | NA | NA | NA | NA | Gm41819 | | 0 | 0 | 0 | 0 | 0 | 0 | 0 | 0 | 0 | 0 | 0 |
| 0 | NA | NA | NA | NA | NA | Gm50134 | | 0 | 0 | 0 | 0 | 0 | 0 | 0 | 0 | 0 | 0 | 0 |
| 0 | NA | NA | NA | NA | NA | Gm22506 | | 0 | 0 | 0 | 0 | 0 | 0 | 0 | 0 | 0 | 0 | 0 |
| 0 | NA | NA | NA | NA | NA | Gm27151 | | 0 | 0 | 0 | 0 | 0 | 0 | 0 | 0 | 0 | 0 | 0 |
| 0 | NA | NA | NA | NA | NA | Mir204 | | 0 | 0 | 0 | 0 | 0 | 0 | 0 | 0 | 0 | 0 | 0 |
| 0 | NA | NA | NA | NA | NA | Gm41821 | | 0 | 0 | 0 | 0 | 0 | 0 | 0 | 0 | 0 | 0 | 0 |
| 0 | NA | NA | NA | NA | NA | Gm24263 | | 0 | 0 | 0 | 0 | 0 | 0 | 0 | 0 | 0 | 0 | 0 |
| 0 | NA | NA | NA | NA | NA | Gm50445 | | 0 | 0 | 0 | 0 | 0 | 0 | 0 | 0 | 0 | 0 | 0 |
| 0 | NA | NA | NA | NA | NA | Gm50443 | | 0 | 0 | 0 | 0 | 0 | 0 | 0 | 0 | 0 | 0 | 0 |
| 0 | NA | NA | NA | NA | NA | 1700028P14Ril | RIKEN cDNA 1700028P14 gene | 0 | 0 | 0 | 0 | 0 | 0 | 0 | 0 | 0 | 0 | 0 |
| 0 | NA | NA | NA | NA | NA | Gm50422 | | 0 | 0 | 0 | 0 | 0 | 0 | 0 | 0 | 0 | 0 | 0 |
| 0 | NA | NA | NA | NA | NA | Gm22144 | | 0 | 0 | 0 | 0 | 0 | 0 | 0 | 0 | 0 | 0 | 0 |
| 0 | NA | NA | NA | NA | NA | Gm21542 | | 0 | 0 | 0 | 0 | 0 | 0 | 0 | 0 | 0 | 0 | 0 |
| 0 | NA | NA | NA | NA | NA | Gm50299 | | 0 | 0 | 0 | 0 | 0 | 0 | 0 | 0 | 0 | 0 | 0 |
| 0 | NA | NA | NA | NA | NA | Gm50300 | | 0 | 0 | 0 | 0 | 0 | 0 | 0 | 0 | 0 | 0 | 0 |
| 0 | NA | NA | NA | NA | NA | Gm50305 | | 0 | 0 | 0 | 0 | 0 | 0 | 0 | 0 | 0 | 0 | 0 |
| 0 | NA | NA | NA | NA | NA | 1700021P04Rik | | 0 | 0 | 0 | 0 | 0 | 0 | 0 | 0 | 0 | 0 | 0 |
| 0 | NA | NA | NA | NA | NA | Gm50310 | | 0 | 0 | 0 | 0 | 0 | 0 | 0 | 0 | 0 | 0 | 0 |
| 0 | NA | NA | NA | NA | NA | Gm50308 | | 0 | 0 | 0 | 0 | 0 | 0 | 0 | 0 | 0 | 0 | 0 |
| 0 | NA | NA | NA | NA | NA | Gm16216 | | 0 | 0 | 0 | 0 | 0 | 0 | 0 | 0 | 0 | 0 | 0 |
| 0 | NA | NA | NA | NA | NA | Gm17800 | | 0 | 0 | 0 | 0 | 0 | 0 | 0 | 0 | 0 | 0 | 0 |
| 0 | NA | NA | NA | NA | NA | Gm50342 | | 0 | 0 | 0 | 0 | 0 | 0 | 0 | 0 | 0 | 0 | 0 |
| 0 | NA | NA | NA | NA | NA | Mir3084-1 | | 0 | 0 | 0 | 0 | 0 | 0 | 0 | 0 | 0 | 0 | 0 |
| 0 | NA | NA | NA | NA | NA | Gm24252 | | 0 | 0 | 0 | 0 | 0 | 0 | 0 | 0 | 0 | 0 | 0 |
| 0 | NA | NA | NA | NA | NA | Gm26207 | | 0 | 0 | 0 | 0 | 0 | 0 | 0 | 0 | 0 | 0 | 0 |
| 0 | NA | NA | NA | NA | NA | Dmrt1 | doublesex and mab-3 related transcriptio | 0 | 0 | 0 | 0 | 0 | 0 | 0 | 0 | 0 | 0 | 0 |
| 0 | NA | NA | NA | NA | NA | 2610016A17Rik | | 0 | 0 | 0 | 0 | 0 | 0 | 0 | 0 | 0 | 0 | 0 |
| 0 | NA | NA | NA | NA | NA | 1700048O14Rik | | 0 | 0 | 0 | 0 | 0 | 0 | 0 | 0 | 0 | 0 | 0 |
| 0 | NA | NA | NA | NA | NA | Gm50376 | | 0 | 0 | 0 | 0 | 0 | 0 | 0 | 0 | 0 | 0 | 0 |
| 0 | NA | NA | NA | NA | NA | Gm25176 | | 0 | 0 | 0 | 0 | 0 | 0 | 0 | 0 | 0 | 0 | 0 |

| | | | | | | | | | | | | | | | | | |
|---|---|---|---|---|---|---|---|---|---|---|---|---|---|---|---|---|---|
| 0 | NA | NA | NA | NA | NA | Gm50377 | | 0 | 0 | 0 | 0 | 0 | 0 | 0 | 0 | 0 | 0 | 0 |
| 0 | NA | NA | NA | NA | NA | Gm50378 | | 0 | 0 | 0 | 0 | 0 | 0 | 0 | 0 | 0 | 0 | 0 |
| 0 | NA | NA | NA | NA | NA | Gm50379 | | 0 | 0 | 0 | 0 | 0 | 0 | 0 | 0 | 0 | 0 | 0 |
| 0 | NA | NA | NA | NA | NA | Gm48775 | | 0 | 0 | 0 | 0 | 0 | 0 | 0 | 0 | 0 | 0 | 0 |
| 0 | NA | NA | NA | NA | NA | Gm18380 | | 0 | 0 | 0 | 0 | 0 | 0 | 0 | 0 | 0 | 0 | 0 |
| 0 | NA | NA | NA | NA | NA | Gm815 | predicted gene 815 | 0 | 0 | 0 | 0 | 0 | 0 | 0 | 0 | 0 | 0 | 0 |
| 0 | NA | NA | NA | NA | NA | Gm50121 | | 0 | 0 | 0 | 0 | 0 | 0 | 0 | 0 | 0 | 0 | 0 |
| 0 | NA | NA | NA | NA | NA | Gm35438 | | 0 | 0 | 0 | 0 | 0 | 0 | 0 | 0 | 0 | 0 | 0 |
| 0 | NA | NA | NA | NA | NA | Gm35438 | | 0 | 0 | 0 | 0 | 0 | 0 | 0 | 0 | 0 | 0 | 0 |
| 0 | NA | NA | NA | NA | NA | Gm50114 | | 0 | 0 | 0 | 0 | 0 | 0 | 0 | 0 | 0 | 0 | 0 |
| 0 | NA | NA | NA | NA | NA | Kcnv2 | potassium channel, subfamily V, member | 0 | 0 | 0 | 0 | 0 | 0 | 0 | 0 | 0 | 0 | 0 |
| 0 | NA | NA | NA | NA | NA | Gm35554 | | 0 | 0 | 0 | 0 | 0 | 0 | 0 | 0 | 0 | 0 | 0 |
| 0 | NA | NA | NA | NA | NA | Gm50113 | | 0 | 0 | 0 | 0 | 0 | 0 | 0 | 0 | 0 | 0 | 0 |
| 0 | NA | NA | NA | NA | NA | Gm50101 | | 0 | 0 | 0 | 0 | 0 | 0 | 0 | 0 | 0 | 0 | 0 |
| 0 | NA | NA | NA | NA | NA | Gm24782 | | 0 | 0 | 0 | 0 | 0 | 0 | 0 | 0 | 0 | 0 | 0 |
| 0 | NA | NA | NA | NA | NA | Gm20407 | | 0 | 0 | 0 | 0 | 0 | 0 | 0 | 0 | 0 | 0 | 0 |
| 0 | NA | NA | NA | NA | NA | Gm28228 | | 0 | 0 | 0 | 0 | 0 | 0 | 0 | 0 | 0 | 0 | 0 |
| 0 | NA | NA | NA | NA | NA | Gm26942 | | 0 | 0 | 0 | 0 | 0 | 0 | 0 | 0 | 0 | 0 | 0 |
| 0 | NA | NA | NA | NA | NA | Gm50425 | | 0 | 0 | 0 | 0 | 0 | 0 | 0 | 0 | 0 | 0 | 0 |
| 0 | NA | NA | NA | NA | NA | Rln1 | relaxin 1 | 0 | 0 | 0 | 0 | 0 | 0 | 0 | 0 | 0 | 0 | 0 |
| 0 | NA | NA | NA | NA | NA | Gm50380 | | 0 | 0 | 0 | 0 | 0 | 0 | 0 | 0 | 0 | 0 | 0 |
| 0 | NA | NA | NA | NA | NA | Gm26046 | | 0 | 0 | 0 | 0 | 0 | 0 | 0 | 0 | 0 | 0 | 0 |
| 0 | NA | NA | NA | NA | NA | Trpd52l3 | tumor protein D52-like 3 | 0 | 0 | 0 | 0 | 0 | 0 | 0 | 0 | 0 | 0 | 0 |
| 0 | NA | NA | NA | NA | NA | Gm24108 | | 0 | 0 | 0 | 0 | 0 | 0 | 0 | 0 | 0 | 0 | 0 |
| 0 | NA | NA | NA | NA | NA | Gm24883 | | 0 | 0 | 0 | 0 | 0 | 0 | 0 | 0 | 0 | 0 | 0 |
| 0 | NA | NA | NA | NA | NA | Gm17885 | | 0 | 0 | 0 | 0 | 0 | 0 | 0 | 0 | 0 | 0 | 0 |
| 0 | NA | NA | NA | NA | NA | Dkk1 | dickkopf WNT signaling pathway inhibitor | 0 | 0 | 0 | 0 | 0 | 0 | 0 | 0 | 0 | 0 | 0 |
| 0 | NA | NA | NA | NA | NA | Gm23426 | | 0 | 0 | 0 | 0 | 0 | 0 | 0 | 0 | 0 | 0 | 0 |
| 0 | NA | NA | NA | NA | NA | Gm50266 | | 0 | 0 | 0 | 0 | 0 | 0 | 0 | 0 | 0 | 0 | 0 |
| 0 | NA | NA | NA | NA | NA | Gm50210 | | 0 | 0 | 0 | 0 | 0 | 0 | 0 | 0 | 0 | 0 | 0 |
| 0 | NA | NA | NA | NA | NA | Gm38616 | | 0 | 0 | 0 | 0 | 0 | 0 | 0 | 0 | 0 | 0 | 0 |
| 0 | NA | NA | NA | NA | NA | Gm9563 | | 0 | 0 | 0 | 0 | 0 | 0 | 0 | 0 | 0 | 0 | 0 |
| 0 | NA | NA | NA | NA | NA | Gm22482 | | 0 | 0 | 0 | 0 | 0 | 0 | 0 | 0 | 0 | 0 | 0 |
| 0 | NA | NA | NA | NA | NA | Gm36860 | | 0 | 0 | 0 | 0 | 0 | 0 | 0 | 0 | 0 | 0 | 0 |
| 0 | NA | NA | NA | NA | NA | Gm29863 | | 0 | 0 | 0 | 0 | 0 | 0 | 0 | 0 | 0 | 0 | 0 |
| 0 | NA | NA | NA | NA | NA | Gm29946 | | 0 | 0 | 0 | 0 | 0 | 0 | 0 | 0 | 0 | 0 | 0 |
| 0 | NA | NA | NA | NA | NA | Mir3970 | | 0 | 0 | 0 | 0 | 0 | 0 | 0 | 0 | 0 | 0 | 0 |
| 0 | NA | NA | NA | NA | NA | Gm7237 | | 0 | 0 | 0 | 0 | 0 | 0 | 0 | 0 | 0 | 0 | 0 |
| 0 | NA | NA | NA | NA | NA | Gm19724 | | 0 | 0 | 0 | 0 | 0 | 0 | 0 | 0 | 0 | 0 | 0 |
| 0 | NA | NA | NA | NA | NA | Gm50233 | | 0 | 0 | 0 | 0 | 0 | 0 | 0 | 0 | 0 | 0 | 0 |
| 0 | NA | NA | NA | NA | NA | | | 0 | 0 | 0 | 0 | 0 | 0 | 0 | 0 | 0 | 0 | 0 |
| 0 | NA | NA | NA | NA | NA | Gm27440 | | 0 | 0 | 0 | 0 | 0 | 0 | 0 | 0 | 0 | 0 | 0 |
| 0 | NA | NA | NA | NA | NA | Gm8978 | predicted gene 8978 | 0 | 0 | 0 | 0 | 0 | 0 | 0 | 0 | 0 | 0 | 0 |
| 0 | NA | NA | NA | NA | NA | Gm19755 | | 0 | 0 | 0 | 0 | 0 | 0 | 0 | 0 | 0 | 0 | 0 |
| 0 | NA | NA | NA | NA | NA | Gm18293 | | 0 | 0 | 0 | 0 | 0 | 0 | 0 | 0 | 0 | 0 | 0 |
| 0 | NA | NA | NA | NA | NA | Lipo1 | lipase, member O1 | 0 | 0 | 0 | 0 | 0 | 0 | 0 | 0 | 0 | 0 | 0 |
| 0 | NA | NA | NA | NA | NA | Gm5519 | | 0 | 0 | 0 | 0 | 0 | 0 | 0 | 0 | 0 | 0 | 0 |
| 0 | NA | NA | NA | NA | NA | Gm8988 | | 0 | 0 | 0 | 0 | 0 | 0 | 0 | 0 | 0 | 0 | 0 |
| 0 | NA | NA | NA | NA | NA | Gm50120 | | 0 | 0 | 0 | 0 | 0 | 0 | 0 | 0 | 0 | 0 | 0 |
| 0 | NA | NA | NA | NA | NA | Lipf | lipase, gastric | 0 | 0 | 0 | 0 | 0 | 0 | 0 | 0 | 0 | 0 | 0 |
| 0 | NA | NA | NA | NA | NA | Lipk | lipase, family member K | 0 | 0 | 0 | 0 | 0 | 0 | 0 | 0 | 0 | 0 | 0 |
| 0 | NA | NA | NA | NA | NA | Lipn | lipase, family member N | 0 | 0 | 0 | 0 | 0 | 0 | 0 | 0 | 0 | 0 | 0 |
| 0 | NA | NA | NA | NA | NA | Gm23334 | | 0 | 0 | 0 | 0 | 0 | 0 | 0 | 0 | 0 | 0 | 0 |
| 0 | NA | NA | NA | NA | NA | Lipm | lipase, family member M | 0 | 0 | 0 | 0 | 0 | 0 | 0 | 0 | 0 | 0 | 0 |
| 0 | NA | NA | NA | NA | NA | Gm50239 | | 0 | 0 | 0 | 0 | 0 | 0 | 0 | 0 | 0 | 0 | 0 |
| 0 | NA | NA | NA | NA | NA | Gm27818 | | 0 | 0 | 0 | 0 | 0 | 0 | 0 | 0 | 0 | 0 | 0 |
| 0 | NA | NA | NA | NA | NA | Gm23060 | | 0 | 0 | 0 | 0 | 0 | 0 | 0 | 0 | 0 | 0 | 0 |
| 0 | NA | NA | NA | NA | NA | Gm50257 | | 0 | 0 | 0 | 0 | 0 | 0 | 0 | 0 | 0 | 0 | 0 |
| 0 | NA | NA | NA | NA | NA | Cnn2-ps | | 0 | 0 | 0 | 0 | 0 | 0 | 0 | 0 | 0 | 0 | 0 |
| 0 | NA | NA | NA | NA | NA | Gm27476 | | 0 | 0 | 0 | 0 | 0 | 0 | 0 | 0 | 0 | 0 | 0 |

| | | | | | | | | | | | | | | | | | | |
|---|---|---|---|---|---|---|---|---|---|---|---|---|---|---|---|---|---|---|
| 0 | NA | NA | NA | NA | NA | Gm50173 | | 0 | 0 | 0 | 0 | 0 | 0 | 0 | 0 | 0 | 0 | 0 |
| 0 | NA | NA | NA | NA | NA | Gm22082 | | 0 | 0 | 0 | 0 | 0 | 0 | 0 | 0 | 0 | 0 | 0 |
| 0 | NA | NA | NA | NA | NA | Gm50116 | | 0 | 0 | 0 | 0 | 0 | 0 | 0 | 0 | 0 | 0 | 0 |
| 0 | NA | NA | NA | NA | NA | Mir107 | | 0 | 0 | 0 | 0 | 0 | 0 | 0 | 0 | 0 | 0 | 0 |
| 0 | NA | NA | NA | NA | NA | Mir1950 | | 0 | 0 | 0 | 0 | 0 | 0 | 0 | 0 | 0 | 0 | 0 |
| 0 | NA | NA | NA | NA | NA | Gm5248 | | 0 | 0 | 0 | 0 | 0 | 0 | 0 | 0 | 0 | 0 | 0 |
| 0 | NA | NA | NA | NA | NA | Gm50137 | | 0 | 0 | 0 | 0 | 0 | 0 | 0 | 0 | 0 | 0 | 0 |
| 0 | NA | NA | NA | NA | NA | Gm50140 | | 0 | 0 | 0 | 0 | 0 | 0 | 0 | 0 | 0 | 0 | 0 |
| 0 | NA | NA | NA | NA | NA | Gm24006 | | 0 | 0 | 0 | 0 | 0 | 0 | 0 | 0 | 0 | 0 | 0 |
| 0 | NA | NA | NA | NA | NA | Gm23075 | | 0 | 0 | 0 | 0 | 0 | 0 | 0 | 0 | 0 | 0 | 0 |
| 0 | NA | NA | NA | NA | NA | Gm22853 | | 0 | 0 | 0 | 0 | 0 | 0 | 0 | 0 | 0 | 0 | 0 |
| 0 | NA | NA | NA | NA | NA | Gm25611 | | 0 | 0 | 0 | 0 | 0 | 0 | 0 | 0 | 0 | 0 | 0 |
| 0 | NA | NA | NA | NA | NA | Gm32081 | | 0 | 0 | 0 | 0 | 0 | 0 | 0 | 0 | 0 | 0 | 0 |
| 0 | NA | NA | NA | NA | NA | Gm50147 | | 0 | 0 | 0 | 0 | 0 | 0 | 0 | 0 | 0 | 0 | 0 |
| 0 | NA | NA | NA | NA | NA | Gm47773 | | 0 | 0 | 0 | 0 | 0 | 0 | 0 | 0 | 0 | 0 | 0 |
| 0 | NA | NA | NA | NA | NA | Gm23170 | | 0 | 0 | 0 | 0 | 0 | 0 | 0 | 0 | 0 | 0 | 0 |
| 0 | NA | NA | NA | NA | NA | F530104D19Rik | | 0 | 0 | 0 | 0 | 0 | 0 | 0 | 0 | 0 | 0 | 0 |
| 0 | NA | NA | NA | NA | NA | Gm50112 | | 0 | 0 | 0 | 0 | 0 | 0 | 0 | 0 | 0 | 0 | 0 |
| 0 | NA | NA | NA | NA | NA | Gm22714 | | 0 | 0 | 0 | 0 | 0 | 0 | 0 | 0 | 0 | 0 | 0 |
| 0 | NA | NA | NA | NA | NA | Gm25268 | | 0 | 0 | 0 | 0 | 0 | 0 | 0 | 0 | 0 | 0 | 0 |
| 0 | NA | NA | NA | NA | NA | Gm23026 | | 0 | 0 | 0 | 0 | 0 | 0 | 0 | 0 | 0 | 0 | 0 |
| 0 | NA | NA | NA | NA | NA | 1700122C19Rik | | 0 | 0 | 0 | 0 | 0 | 0 | 0 | 0 | 0 | 0 | 0 |
| 0 | NA | NA | NA | NA | NA | Cyp26c1 | cytochrome P450, family 26, subfamily c, | 0 | 0 | 0 | 0 | 0 | 0 | 0 | 0 | 0 | 0 | 0 |
| 0 | NA | NA | NA | NA | NA | Gm9066 | | 0 | 0 | 0 | 0 | 0 | 0 | 0 | 0 | 0 | 0 | 0 |
| 0 | NA | NA | NA | NA | NA | Gm4757 | | 0 | 0 | 0 | 0 | 0 | 0 | 0 | 0 | 0 | 0 | 0 |
| 0 | NA | NA | NA | NA | NA | Gm23300 | | 0 | 0 | 0 | 0 | 0 | 0 | 0 | 0 | 0 | 0 | 0 |
| 0 | NA | NA | NA | NA | NA | Gm50150 | | 0 | 0 | 0 | 0 | 0 | 0 | 0 | 0 | 0 | 0 | 0 |
| 0 | NA | NA | NA | NA | NA | Gm50155 | | 0 | 0 | 0 | 0 | 0 | 0 | 0 | 0 | 0 | 0 | 0 |
| 0 | NA | NA | NA | NA | NA | Gm18936 | | 0 | 0 | 0 | 0 | 0 | 0 | 0 | 0 | 0 | 0 | 0 |
| 0 | NA | NA | NA | NA | NA | Gm24930 | | 0 | 0 | 0 | 0 | 0 | 0 | 0 | 0 | 0 | 0 | 0 |
| 0 | NA | NA | NA | NA | NA | Gm28991 | | 0 | 0 | 0 | 0 | 0 | 0 | 0 | 0 | 0 | 0 | 0 |
| 0 | NA | NA | NA | NA | NA | Gm25838 | | 0 | 0 | 0 | 0 | 0 | 0 | 0 | 0 | 0 | 0 | 0 |
| 0 | NA | NA | NA | NA | NA | Gm23800 | | 0 | 0 | 0 | 0 | 0 | 0 | 0 | 0 | 0 | 0 | 0 |
| 0 | NA | NA | NA | NA | NA | Cyp2c65 | cytochrome P450, family 2, subfamily c, p | 0 | 0 | 0 | 0 | 0 | 0 | 0 | 0 | 0 | 0 | 0 |
| 0 | NA | NA | NA | NA | NA | Cyp2c66 | cytochrome P450, family 2, subfamily c, p | 0 | 0 | 0 | 0 | 0 | 0 | 0 | 0 | 0 | 0 | 0 |
| 0 | NA | NA | NA | NA | NA | Cyp2c53-ps | | 0 | 0 | 0 | 0 | 0 | 0 | 0 | 0 | 0 | 0 | 0 |
| 0 | NA | NA | NA | NA | NA | Cyp2c29 | cytochrome P450, family 2, subfamily c, p | 0 | 0 | 0 | 0 | 0 | 0 | 0 | 0 | 0 | 0 | 0 |
| 0 | NA | NA | NA | NA | NA | Cyp2c72-ps | | 0 | 0 | 0 | 0 | 0 | 0 | 0 | 0 | 0 | 0 | 0 |
| 0 | NA | NA | NA | NA | NA | Cyp2c38 | cytochrome P450, family 2, subfamily c, p | 0 | 0 | 0 | 0 | 0 | 0 | 0 | 0 | 0 | 0 | 0 |
| 0 | NA | NA | NA | NA | NA | Cyp2c39 | cytochrome P450, family 2, subfamily c, p | 0 | 0 | 0 | 0 | 0 | 0 | 0 | 0 | 0 | 0 | 0 |
| 0 | NA | NA | NA | NA | NA | Cyp2c67 | cytochrome P450, family 2, subfamily c, p | 0 | 0 | 0 | 0 | 0 | 0 | 0 | 0 | 0 | 0 | 0 |
| 0 | NA | NA | NA | NA | NA | Cyp2c68 | cytochrome P450, family 2, subfamily c, p | 0 | 0 | 0 | 0 | 0 | 0 | 0 | 0 | 0 | 0 | 0 |
| 0 | NA | NA | NA | NA | NA | Cyp2c40 | cytochrome P450, family 2, subfamily c, p | 0 | 0 | 0 | 0 | 0 | 0 | 0 | 0 | 0 | 0 | 0 |
| 0 | NA | NA | NA | NA | NA | Cyp2c71-ps | | 0 | 0 | 0 | 0 | 0 | 0 | 0 | 0 | 0 | 0 | 0 |
| 0 | NA | NA | NA | NA | NA | Cyp2c37 | cytochrome P450, family 2. subfamily c, p | 0 | 0 | 0 | 0 | 0 | 0 | 0 | 0 | 0 | 0 | 0 |
| 0 | NA | NA | NA | NA | NA | Cyp2c54 | cytochrome P450, family 2, subfamily c, p | 0 | 0 | 0 | 0 | 0 | 0 | 0 | 0 | 0 | 0 | 0 |
| 0 | NA | NA | NA | NA | NA | Gm5693 | | 0 | 0 | 0 | 0 | 0 | 0 | 0 | 0 | 0 | 0 | 0 |
| 0 | NA | NA | NA | NA | NA | A930028N01Rik | | 0 | 0 | 0 | 0 | 0 | 0 | 0 | 0 | 0 | 0 | 0 |
| 0 | NA | NA | NA | NA | NA | Gm8783 | | 0 | 0 | 0 | 0 | 0 | 0 | 0 | 0 | 0 | 0 | 0 |
| 0 | NA | NA | NA | NA | NA | Gm50132 | | 0 | 0 | 0 | 0 | 0 | 0 | 0 | 0 | 0 | 0 | 0 |
| 0 | NA | NA | NA | NA | NA | Mir8092 | | 0 | 0 | 0 | 0 | 0 | 0 | 0 | 0 | 0 | 0 | 0 |
| 0 | NA | NA | NA | NA | NA | Opalin | oligodendrocytic myelin paranodal and in | 0 | 0 | 0 | 0 | 0 | 0 | 0 | 0 | 0 | 0 | 0 |
| 0 | NA | NA | NA | NA | NA | Gm9788 | | 0 | 0 | 0 | 0 | 0 | 0 | 0 | 0 | 0 | 0 | 0 |
| 0 | NA | NA | NA | NA | NA | Gm22601 | | 0 | 0 | 0 | 0 | 0 | 0 | 0 | 0 | 0 | 0 | 0 |
| 0 | NA | NA | NA | NA | NA | Gm46644 | | 0 | 0 | 0 | 0 | 0 | 0 | 0 | 0 | 0 | 0 | 0 |
| 0 | NA | NA | NA | NA | NA | Gm23113 | | 0 | 0 | 0 | 0 | 0 | 0 | 0 | 0 | 0 | 0 | 0 |
| 0 | NA | NA | NA | NA | NA | Gm26339 | | 0 | 0 | 0 | 0 | 0 | 0 | 0 | 0 | 0 | 0 | 0 |
| 0 | NA | NA | NA | NA | NA | Mir3085 | | 0 | 0 | 0 | 0 | 0 | 0 | 0 | 0 | 0 | 0 | 0 |
| 0 | NA | NA | NA | NA | NA | Gm50184 | | 0 | 0 | 0 | 0 | 0 | 0 | 0 | 0 | 0 | 0 | 0 |
| 0 | NA | NA | NA | NA | NA | Gm16541 | | 0 | 0 | 0 | 0 | 0 | 0 | 0 | 0 | 0 | 0 | 0 |

| | | | | | | | | | | | | | | | | | |
|---|---|---|---|---|---|---|---|---|---|---|---|---|---|---|---|---|---|
| 0 | NA | NA | NA | NA | NA | Gm25216 | | 0 | 0 | 0 | 0 | 0 | 0 | 0 | 0 | 0 | 0 | 0 |
| 0 | NA | NA | NA | NA | NA | Gm22135 | | 0 | 0 | 0 | 0 | 0 | 0 | 0 | 0 | 0 | 0 | 0 |
| 0 | NA | NA | NA | NA | NA | Gm34517 | | 0 | 0 | 0 | 0 | 0 | 0 | 0 | 0 | 0 | 0 | 0 |
| 0 | NA | NA | NA | NA | NA | Gm50194 | | 0 | 0 | 0 | 0 | 0 | 0 | 0 | 0 | 0 | 0 | 0 |
| 0 | NA | NA | NA | NA | NA | Gm19179 | | 0 | 0 | 0 | 0 | 0 | 0 | 0 | 0 | 0 | 0 | 0 |
| 0 | NA | NA | NA | NA | NA | Gm34636 | | 0 | 0 | 0 | 0 | 0 | 0 | 0 | 0 | 0 | 0 | 0 |
| 0 | NA | NA | NA | NA | NA | Gm22646 | | 0 | 0 | 0 | 0 | 0 | 0 | 0 | 0 | 0 | 0 | 0 |
| 0 | NA | NA | NA | NA | NA | Gm34922 | | 0 | 0 | 0 | 0 | 0 | 0 | 0 | 0 | 0 | 0 | 0 |
| 0 | NA | NA | NA | NA | NA | n-R5s21 | | 0 | 0 | 0 | 0 | 0 | 0 | 0 | 0 | 0 | 0 | 0 |
| 0 | NA | NA | NA | NA | NA | Gm24336 | | 0 | 0 | 0 | 0 | 0 | 0 | 0 | 0 | 0 | 0 | 0 |
| 0 | NA | NA | NA | NA | NA | Gm6951 | | 0 | 0 | 0 | 0 | 0 | 0 | 0 | 0 | 0 | 0 | 0 |
| 0 | NA | NA | NA | NA | NA | Gm35183 | | 0 | 0 | 0 | 0 | 0 | 0 | 0 | 0 | 0 | 0 | 0 |
| 0 | NA | NA | NA | NA | NA | Mir5114 | | 0 | 0 | 0 | 0 | 0 | 0 | 0 | 0 | 0 | 0 | 0 |
| 0 | NA | NA | NA | NA | NA | Gm50337 | | 0 | 0 | 0 | 0 | 0 | 0 | 0 | 0 | 0 | 0 | 0 |
| 0 | NA | NA | NA | NA | NA | Gm35406 | | 0 | 0 | 0 | 0 | 0 | 0 | 0 | 0 | 0 | 0 | 0 |
| 0 | NA | NA | NA | NA | NA | Gm20538 | | 0 | 0 | 0 | 0 | 0 | 0 | 0 | 0 | 0 | 0 | 0 |
| 0 | NA | NA | NA | NA | NA | Gm35460 | | 0 | 0 | 0 | 0 | 0 | 0 | 0 | 0 | 0 | 0 | 0 |
| 0 | NA | NA | NA | NA | NA | Gm26644 | | 0 | 0 | 0 | 0 | 0 | 0 | 0 | 0 | 0 | 0 | 0 |
| 0 | NA | NA | NA | NA | NA | Gm35610 | | 0 | 0 | 0 | 0 | 0 | 0 | 0 | 0 | 0 | 0 | 0 |
| 0 | NA | NA | NA | NA | NA | Mir6405 | | 0 | 0 | 0 | 0 | 0 | 0 | 0 | 0 | 0 | 0 | 0 |
| 0 | NA | NA | NA | NA | NA | 1700039E22Rik | | 0 | 0 | 0 | 0 | 0 | 0 | 0 | 0 | 0 | 0 | 0 |
| 0 | NA | NA | NA | NA | NA | Gm5246 | | 0 | 0 | 0 | 0 | 0 | 0 | 0 | 0 | 0 | 0 | 0 |
| 0 | NA | NA | NA | NA | NA | Gm25482 | | 0 | 0 | 0 | 0 | 0 | 0 | 0 | 0 | 0 | 0 | 0 |
| 0 | NA | NA | NA | NA | NA | Tlx1os | | 0 | 0 | 0 | 0 | 0 | 0 | 0 | 0 | 0 | 0 | 0 |
| 0 | NA | NA | NA | NA | NA | Gm29594 | | 0 | 0 | 0 | 0 | 0 | 0 | 0 | 0 | 0 | 0 | 0 |
| 0 | NA | NA | NA | NA | NA | 1700016H03Rik | | 0 | 0 | 0 | 0 | 0 | 0 | 0 | 0 | 0 | 0 | 0 |
| 0 | NA | NA | NA | NA | NA | Fgf8 | fibroblast growth factor 8 | 0 | 0 | 0 | 0 | 0 | 0 | 0 | 0 | 0 | 0 | 0 |
| 0 | NA | NA | NA | NA | NA | Gm50196 | | 0 | 0 | 0 | 0 | 0 | 0 | 0 | 0 | 0 | 0 | 0 |
| 0 | NA | NA | NA | NA | NA | Gm50306 | | 0 | 0 | 0 | 0 | 0 | 0 | 0 | 0 | 0 | 0 | 0 |
| 0 | NA | NA | NA | NA | NA | Elovl3 | elongation of very long chain fatty acids (I | 0 | 0 | 0 | 0 | 0 | 0 | 0 | 0 | 0 | 0 | 0 |
| 0 | NA | NA | NA | NA | NA | Mir146b | | 0 | 0 | 0 | 0 | 0 | 0 | 0 | 0 | 0 | 0 | 0 |
| 0 | NA | NA | NA | NA | NA | Gm36602 | | 0 | 0 | 0 | 0 | 0 | 0 | 0 | 0 | 0 | 0 | 0 |
| 0 | NA | NA | NA | NA | NA | Gm50319 | | 0 | 0 | 0 | 0 | 0 | 0 | 0 | 0 | 0 | 0 | 0 |
| 0 | NA | NA | NA | NA | NA | Calhm1 | calcium homeostasis modulator 1 | 0 | 0 | 0 | 0 | 0 | 0 | 0 | 0 | 0 | 0 | 0 |
| 0 | NA | NA | NA | NA | NA | Gm36693 | | 0 | 0 | 0 | 0 | 0 | 0 | 0 | 0 | 0 | 0 | 0 |
| 0 | NA | NA | NA | NA | NA | Gm50339 | | 0 | 0 | 0 | 0 | 0 | 0 | 0 | 0 | 0 | 0 | 0 |
| 0 | NA | NA | NA | NA | NA | Mir6995 | | 0 | 0 | 0 | 0 | 0 | 0 | 0 | 0 | 0 | 0 | 0 |
| 0 | NA | NA | NA | NA | NA | Gm50358 | | 0 | 0 | 0 | 0 | 0 | 0 | 0 | 0 | 0 | 0 | 0 |
| 0 | NA | NA | NA | NA | NA | Gm16068 | | 0 | 0 | 0 | 0 | 0 | 0 | 0 | 0 | 0 | 0 | 0 |
| 0 | NA | NA | NA | NA | NA | Gm50281 | | 0 | 0 | 0 | 0 | 0 | 0 | 0 | 0 | 0 | 0 | 0 |
| 0 | NA | NA | NA | NA | NA | Gm26085 | | 0 | 0 | 0 | 0 | 0 | 0 | 0 | 0 | 0 | 0 | 0 |
| 0 | NA | NA | NA | NA | NA | Gm24805 | | 0 | 0 | 0 | 0 | 0 | 0 | 0 | 0 | 0 | 0 | 0 |
| 0 | NA | NA | NA | NA | NA | Gm50282 | | 0 | 0 | 0 | 0 | 0 | 0 | 0 | 0 | 0 | 0 | 0 |
| 0 | NA | NA | NA | NA | NA | Mir8090 | | 0 | 0 | 0 | 0 | 0 | 0 | 0 | 0 | 0 | 0 | 0 |
| 0 | NA | NA | NA | NA | NA | Gm22240 | | 0 | 0 | 0 | 0 | 0 | 0 | 0 | 0 | 0 | 0 | 0 |
| 0 | NA | NA | NA | NA | NA | Gm50430 | | 0 | 0 | 0 | 0 | 0 | 0 | 0 | 0 | 0 | 0 | 0 |
| 0 | NA | NA | NA | NA | NA | Gm23857 | | 0 | 0 | 0 | 0 | 0 | 0 | 0 | 0 | 0 | 0 | 0 |
| 0 | NA | NA | NA | NA | NA | Gm50436 | | 0 | 0 | 0 | 0 | 0 | 0 | 0 | 0 | 0 | 0 | 0 |
| 0 | NA | NA | NA | NA | NA | 4930535F04Rik | | 0 | 0 | 0 | 0 | 0 | 0 | 0 | 0 | 0 | 0 | 0 |
| 0 | NA | NA | NA | NA | NA | Gm50441 | | 0 | 0 | 0 | 0 | 0 | 0 | 0 | 0 | 0 | 0 | 0 |
| 0 | NA | NA | NA | NA | NA | Gm50442 | | 0 | 0 | 0 | 0 | 0 | 0 | 0 | 0 | 0 | 0 | 0 |
| 0 | NA | NA | NA | NA | NA | Gm50444 | | 0 | 0 | 0 | 0 | 0 | 0 | 0 | 0 | 0 | 0 | 0 |
| 0 | NA | NA | NA | NA | NA | Gm27935 | | 0 | 0 | 0 | 0 | 0 | 0 | 0 | 0 | 0 | 0 | 0 |
| 0 | NA | NA | NA | NA | NA | Siah1-ps1 | | 0 | 0 | 0 | 0 | 0 | 0 | 0 | 0 | 0 | 0 | 0 |
| 0 | NA | NA | NA | NA | NA | Gm27749 | | 0 | 0 | 0 | 0 | 0 | 0 | 0 | 0 | 0 | 0 | 0 |
| 0 | NA | NA | NA | NA | NA | Gm26629 | | 0 | 0 | 0 | 0 | 0 | 0 | 0 | 0 | 0 | 0 | 0 |
| 0 | NA | NA | NA | NA | NA | Gm50302 | | 0 | 0 | 0 | 0 | 0 | 0 | 0 | 0 | 0 | 0 | 0 |
| 0 | NA | NA | NA | NA | NA | Gm25659 | | 0 | 0 | 0 | 0 | 0 | 0 | 0 | 0 | 0 | 0 | 0 |
| 0 | NA | NA | NA | NA | NA | Gm50317 | | 0 | 0 | 0 | 0 | 0 | 0 | 0 | 0 | 0 | 0 | 0 |
| 0 | NA | NA | NA | NA | NA | Gm25644 | | 0 | 0 | 0 | 0 | 0 | 0 | 0 | 0 | 0 | 0 | 0 |

| | | | | | | | | | | | | | | | | | | |
|---|---|---|---|---|---|---|---|---|---|---|---|---|---|---|---|---|---|---|
| 0 | NA | NA | NA | NA | NA | Gm50325 | | 0 | 0 | 0 | 0 | 0 | 0 | 0 | 0 | 0 | 0 | 0 |
| 0 | NA | NA | NA | NA | NA | Gm50326 | | 0 | 0 | 0 | 0 | 0 | 0 | 0 | 0 | 0 | 0 | 0 |
| 0 | NA | NA | NA | NA | NA | Gm50330 | | 0 | 0 | 0 | 0 | 0 | 0 | 0 | 0 | 0 | 0 | 0 |
| 0 | NA | NA | NA | NA | NA | Gm50329 | | 0 | 0 | 0 | 0 | 0 | 0 | 0 | 0 | 0 | 0 | 0 |
| 0 | NA | NA | NA | NA | NA | Gm50331 | | 0 | 0 | 0 | 0 | 0 | 0 | 0 | 0 | 0 | 0 | 0 |
| 0 | NA | NA | NA | NA | NA | Gm50332 | | 0 | 0 | 0 | 0 | 0 | 0 | 0 | 0 | 0 | 0 | 0 |
| 0 | NA | NA | NA | NA | NA | Mir6407 | | 0 | 0 | 0 | 0 | 0 | 0 | 0 | 0 | 0 | 0 | 0 |
| 0 | NA | NA | NA | NA | NA | Gm30381 | | 0 | 0 | 0 | 0 | 0 | 0 | 0 | 0 | 0 | 0 | 0 |
| 0 | NA | NA | NA | NA | NA | Gm9618 | | 0 | 0 | 0 | 0 | 0 | 0 | 0 | 0 | 0 | 0 | 0 |
| 0 | NA | NA | NA | NA | NA | Gm24724 | | 0 | 0 | 0 | 0 | 0 | 0 | 0 | 0 | 0 | 0 | 0 |
| 0 | NA | NA | NA | NA | NA | 1700001K23Rik | | 0 | 0 | 0 | 0 | 0 | 0 | 0 | 0 | 0 | 0 | 0 |
| 0 | NA | NA | NA | NA | NA | Gm36339 | | 0 | 0 | 0 | 0 | 0 | 0 | 0 | 0 | 0 | 0 | 0 |
| 0 | NA | NA | NA | NA | NA | Gm10197 | | 0 | 0 | 0 | 0 | 0 | 0 | 0 | 0 | 0 | 0 | 0 |
| 0 | NA | NA | NA | NA | NA | Gm50393 | | 0 | 0 | 0 | 0 | 0 | 0 | 0 | 0 | 0 | 0 | 0 |
| 0 | NA | NA | NA | NA | NA | Gm50272 | | 0 | 0 | 0 | 0 | 0 | 0 | 0 | 0 | 0 | 0 | 0 |
| 0 | NA | NA | NA | NA | NA | Gm50273 | | 0 | 0 | 0 | 0 | 0 | 0 | 0 | 0 | 0 | 0 | 0 |
| 0 | NA | NA | NA | NA | NA | 4930484I04Rik | | 0 | 0 | 0 | 0 | 0 | 0 | 0 | 0 | 0 | 0 | 0 |
| 0 | NA | NA | NA | NA | NA | Gm50187 | | 0 | 0 | 0 | 0 | 0 | 0 | 0 | 0 | 0 | 0 | 0 |
| 0 | NA | NA | NA | NA | NA | Gm50188 | | 0 | 0 | 0 | 0 | 0 | 0 | 0 | 0 | 0 | 0 | 0 |
| 0 | NA | NA | NA | NA | NA | Tectb | tectorin beta | 0 | 0 | 0 | 0 | 0 | 0 | 0 | 0 | 0 | 0 | 0 |
| 0 | NA | NA | NA | NA | NA | Mir6715 | | 0 | 0 | 0 | 0 | 0 | 0 | 0 | 0 | 0 | 0 | 0 |
| 0 | NA | NA | NA | NA | NA | Gm50135 | | 0 | 0 | 0 | 0 | 0 | 0 | 0 | 0 | 0 | 0 | 0 |
| 0 | NA | NA | NA | NA | NA | D730002M21Rik | | 0 | 0 | 0 | 0 | 0 | 0 | 0 | 0 | 0 | 0 | 0 |
| 0 | NA | NA | NA | NA | NA | Gm31912 | | 0 | 0 | 0 | 0 | 0 | 0 | 0 | 0 | 0 | 0 | 0 |
| 0 | NA | NA | NA | NA | NA | Gm22271 | | 0 | 0 | 0 | 0 | 0 | 0 | 0 | 0 | 0 | 0 | 0 |
| 0 | NA | NA | NA | NA | NA | Gm17197 | | 0 | 0 | 0 | 0 | 0 | 0 | 0 | 0 | 0 | 0 | 0 |
| 0 | NA | NA | NA | NA | NA | Gm32441 | | 0 | 0 | 0 | 0 | 0 | 0 | 0 | 0 | 0 | 0 | 0 |
| 0 | NA | NA | NA | NA | NA | Gm32492 | | 0 | 0 | 0 | 0 | 0 | 0 | 0 | 0 | 0 | 0 | 0 |
| 0 | NA | NA | NA | NA | NA | Tdrd1 | tudor domain containing 1 | 0 | 0 | 0 | 0 | 0 | 0 | 0 | 0 | 0 | 0 | 0 |
| 0 | NA | NA | NA | NA | NA | Gm50270 | | 0 | 0 | 0 | 0 | 0 | 0 | 0 | 0 | 0 | 0 | 0 |
| 0 | NA | NA | NA | NA | NA | Gm32932 | | 0 | 0 | 0 | 0 | 0 | 0 | 0 | 0 | 0 | 0 | 0 |
| 0 | NA | NA | NA | NA | NA | Gm50285 | | 0 | 0 | 0 | 0 | 0 | 0 | 0 | 0 | 0 | 0 | 0 |
| 0 | NA | NA | NA | NA | NA | Gm50286 | | 0 | 0 | 0 | 0 | 0 | 0 | 0 | 0 | 0 | 0 | 0 |
| 0 | NA | NA | NA | NA | NA | Mir5623 | | 0 | 0 | 0 | 0 | 0 | 0 | 0 | 0 | 0 | 0 | 0 |
| 0 | NA | NA | NA | NA | NA | Gm50287 | | 0 | 0 | 0 | 0 | 0 | 0 | 0 | 0 | 0 | 0 | 0 |
| 0 | NA | NA | NA | NA | NA | Gm16277 | | 0 | 0 | 0 | 0 | 0 | 0 | 0 | 0 | 0 | 0 | 0 |
| 0 | NA | NA | NA | NA | NA | Ccdc172 | coiled-coil domain containing 172 | 0 | 0 | 0 | 0 | 0 | 0 | 0 | 0 | 0 | 0 | 0 |
| 0 | NA | NA | NA | NA | NA | 1700019N19Ril | RIKEN cDNA 1700019N19 gene | 0 | 0 | 0 | 0 | 0 | 0 | 0 | 0 | 0 | 0 | 0 |
| 0 | NA | NA | NA | NA | NA | Gm22561 | | 0 | 0 | 0 | 0 | 0 | 0 | 0 | 0 | 0 | 0 | 0 |
| 0 | NA | NA | NA | NA | NA | Mir3086 | | 0 | 0 | 0 | 0 | 0 | 0 | 0 | 0 | 0 | 0 | 0 |
| 0 | NA | NA | NA | NA | NA | Vax1 | ventral anterior homeobox 1 | 0 | 0 | 0 | 0 | 0 | 0 | 0 | 0 | 0 | 0 | 0 |
| 0 | NA | NA | NA | NA | NA | Kcnk18 | potassium channel, subfamily K, member | 0 | 0 | 0 | 0 | 0 | 0 | 0 | 0 | 0 | 0 | 0 |
| 0 | NA | NA | NA | NA | NA | Gm29261 | | 0 | 0 | 0 | 0 | 0 | 0 | 0 | 0 | 0 | 0 | 0 |
| 0 | NA | NA | NA | NA | NA | Gm50440 | | 0 | 0 | 0 | 0 | 0 | 0 | 0 | 0 | 0 | 0 | 0 |
| 0 | NA | NA | NA | NA | NA | Gm33756 | | 0 | 0 | 0 | 0 | 0 | 0 | 0 | 0 | 0 | 0 | 0 |
| 0 | NA | NA | NA | NA | NA | 2700089I24Rik | | 0 | 0 | 0 | 0 | 0 | 0 | 0 | 0 | 0 | 0 | 0 |
| 0 | NA | NA | NA | NA | NA | Gm18161 | | 0 | 0 | 0 | 0 | 0 | 0 | 0 | 0 | 0 | 0 | 0 |
| 0 | NA | NA | NA | NA | NA | Gm19956 | | 0 | 0 | 0 | 0 | 0 | 0 | 0 | 0 | 0 | 0 | 0 |
| 0 | NA | NA | NA | NA | NA | Gm50411 | | 0 | 0 | 0 | 0 | 0 | 0 | 0 | 0 | 0 | 0 | 0 |
| 0 | NA | NA | NA | NA | NA | Csf1r-ps | | 0 | 0 | 0 | 0 | 0 | 0 | 0 | 0 | 0 | 0 | 0 |
| 0 | NA | NA | NA | NA | NA | Gm50296 | | 0 | 0 | 0 | 0 | 0 | 0 | 0 | 0 | 0 | 0 | 0 |
| 0 | NA | NA | NA | NA | NA | Gm28351 | | 0 | 0 | 0 | 0 | 0 | 0 | 0 | 0 | 0 | 0 | 0 |
| 0 | NA | NA | NA | NA | NA | Gm46652 | | 0 | 0 | 0 | 0 | 0 | 0 | 0 | 0 | 0 | 0 | 0 |
| 0 | NA | NA | NA | NA | NA | Mir3084-2 | | 0 | 0 | 0 | 0 | 0 | 0 | 0 | 0 | 0 | 0 | 0 |
| 0 | NA | NA | NA | NA | NA | Gm15959 | | 0 | 0 | 0 | 0 | 0 | 0 | 0 | 0 | 0 | 0 | 0 |
| 0 | NA | NA | NA | NA | NA | Gm9529 | | 0 | 0 | 0 | 0 | 0 | 0 | 0 | 0 | 0 | 0 | 0 |
| 0 | NA | NA | NA | NA | NA | Gm22520 | | 0 | 0 | 0 | 0 | 0 | 0 | 0 | 0 | 0 | 0 | 0 |
| 0 | NA | NA | NA | NA | NA | Gm22365 | | 0 | 0 | 0 | 0 | 0 | 0 | 0 | 0 | 0 | 0 | 0 |
| 0 | NA | NA | NA | NA | NA | Gm4242 | | 0 | 0 | 0 | 0 | 0 | 0 | 0 | 0 | 0 | 0 | 0 |
| 0 | NA | NA | NA | NA | NA | Gm18999 | | 0 | 0 | 0 | 0 | 0 | 0 | 0 | 0 | 0 | 0 | 0 |

| | | | | | | | | | | | | | | | | | | |
|---|---|---|---|---|---|---|---|---|---|---|---|---|---|---|---|---|---|---|
| 0 | NA | NA | NA | NA | NA | Gm21060 | | 0 | 0 | 0 | 0 | 0 | 0 | 0 | 0 | 0 | 0 | 0 |
| 0 | NA | NA | NA | NA | NA | Btbd35f23 | BTB domain containing 35, family membe | 0 | 0 | 0 | 0 | 0 | 0 | 0 | 0 | 0 | 0 | 0 |
| 0 | NA | NA | NA | NA | NA | Btbd35f24 | | 0 | 0 | 0 | 0 | 0 | 0 | 0 | 0 | 0 | 0 | 0 |
| 0 | NA | NA | NA | NA | NA | Gm14331 | | 0 | 0 | 0 | 0 | 0 | 0 | 0 | 0 | 0 | 0 | 0 |
| 0 | NA | NA | NA | NA | NA | Gm14333 | | 0 | 0 | 0 | 0 | 0 | 0 | 0 | 0 | 0 | 0 | 0 |
| 0 | NA | NA | NA | NA | NA | Gm14332 | | 0 | 0 | 0 | 0 | 0 | 0 | 0 | 0 | 0 | 0 | 0 |
| 0 | NA | NA | NA | NA | NA | Gm14334 | | 0 | 0 | 0 | 0 | 0 | 0 | 0 | 0 | 0 | 0 | 0 |
| 0 | NA | NA | NA | NA | NA | Gm14352 | | 0 | 0 | 0 | 0 | 0 | 0 | 0 | 0 | 0 | 0 | 0 |
| 0 | NA | NA | NA | NA | NA | Btbd35f11 | BTB domain containing 35, family membe | 0 | 0 | 0 | 0 | 0 | 0 | 0 | 0 | 0 | 0 | 0 |
| 0 | NA | NA | NA | NA | NA | Gm14353 | | 0 | 0 | 0 | 0 | 0 | 0 | 0 | 0 | 0 | 0 | 0 |
| 0 | NA | NA | NA | NA | NA | Gm14360 | | 0 | 0 | 0 | 0 | 0 | 0 | 0 | 0 | 0 | 0 | 0 |
| 0 | NA | NA | NA | NA | NA | Gm14361 | | 0 | 0 | 0 | 0 | 0 | 0 | 0 | 0 | 0 | 0 | 0 |
| 0 | NA | NA | NA | NA | NA | Btbd35f18 | BTB domain containing 35, family membe | 0 | 0 | 0 | 0 | 0 | 0 | 0 | 0 | 0 | 0 | 0 |
| 0 | NA | NA | NA | NA | NA | Btbd35f10 | BTB domain containing 35, family membe | 0 | 0 | 0 | 0 | 0 | 0 | 0 | 0 | 0 | 0 | 0 |
| 0 | NA | NA | NA | NA | NA | Gm14357 | | 0 | 0 | 0 | 0 | 0 | 0 | 0 | 0 | 0 | 0 | 0 |
| 0 | NA | NA | NA | NA | NA | Gm14359 | | 0 | 0 | 0 | 0 | 0 | 0 | 0 | 0 | 0 | 0 | 0 |
| 0 | NA | NA | NA | NA | NA | Gm14355 | | 0 | 0 | 0 | 0 | 0 | 0 | 0 | 0 | 0 | 0 | 0 |
| 0 | NA | NA | NA | NA | NA | Spin2-ps1 | | 0 | 0 | 0 | 0 | 0 | 0 | 0 | 0 | 0 | 0 | 0 |
| 0 | NA | NA | NA | NA | NA | Btbd35f16 | BTB domain containing 35, family membe | 0 | 0 | 0 | 0 | 0 | 0 | 0 | 0 | 0 | 0 | 0 |
| 0 | NA | NA | NA | NA | NA | Gm15251 | | 0 | 0 | 0 | 0 | 0 | 0 | 0 | 0 | 0 | 0 | 0 |
| 0 | NA | NA | NA | NA | NA | Gm15257 | | 0 | 0 | 0 | 0 | 0 | 0 | 0 | 0 | 0 | 0 | 0 |
| 0 | NA | NA | NA | NA | NA | Btbd35f3 | BTB domain containing 35, family membe | 0 | 0 | 0 | 0 | 0 | 0 | 0 | 0 | 0 | 0 | 0 |
| 0 | NA | NA | NA | NA | NA | Gm15258 | | 0 | 0 | 0 | 0 | 0 | 0 | 0 | 0 | 0 | 0 | 0 |
| 0 | NA | NA | NA | NA | NA | Gm14350 | | 0 | 0 | 0 | 0 | 0 | 0 | 0 | 0 | 0 | 0 | 0 |
| 0 | NA | NA | NA | NA | NA | Gm14349 | | 0 | 0 | 0 | 0 | 0 | 0 | 0 | 0 | 0 | 0 | 0 |
| 0 | NA | NA | NA | NA | NA | Gm14358 | | 0 | 0 | 0 | 0 | 0 | 0 | 0 | 0 | 0 | 0 | 0 |
| 0 | NA | NA | NA | NA | NA | Gm14362 | | 0 | 0 | 0 | 0 | 0 | 0 | 0 | 0 | 0 | 0 | 0 |
| 0 | NA | NA | NA | NA | NA | Gm14356 | | 0 | 0 | 0 | 0 | 0 | 0 | 0 | 0 | 0 | 0 | 0 |
| 0 | NA | NA | NA | NA | NA | Btbd35f28 | BTB domain containing 35, family membe | 0 | 0 | 0 | 0 | 0 | 0 | 0 | 0 | 0 | 0 | 0 |
| 0 | NA | NA | NA | NA | NA | Btbd35f20 | BTB domain containing 35, family membe | 0 | 0 | 0 | 0 | 0 | 0 | 0 | 0 | 0 | 0 | 0 |
| 0 | NA | NA | NA | NA | NA | Gm14348 | | 0 | 0 | 0 | 0 | 0 | 0 | 0 | 0 | 0 | 0 | 0 |
| 0 | NA | NA | NA | NA | NA | Gm14364 | | 0 | 0 | 0 | 0 | 0 | 0 | 0 | 0 | 0 | 0 | 0 |
| 0 | NA | NA | NA | NA | NA | Gm14368 | | 0 | 0 | 0 | 0 | 0 | 0 | 0 | 0 | 0 | 0 | 0 |
| 0 | NA | NA | NA | NA | NA | Gm15260 | | 0 | 0 | 0 | 0 | 0 | 0 | 0 | 0 | 0 | 0 | 0 |
| 0 | NA | NA | NA | NA | NA | Btbd35f17 | BTB domain containing 35, family membe | 0 | 0 | 0 | 0 | 0 | 0 | 0 | 0 | 0 | 0 | 0 |
| 0 | NA | NA | NA | NA | NA | Gm6019 | | 0 | 0 | 0 | 0 | 0 | 0 | 0 | 0 | 0 | 0 | 0 |
| 0 | NA | NA | NA | NA | NA | Gm9427 | | 0 | 0 | 0 | 0 | 0 | 0 | 0 | 0 | 0 | 0 | 0 |
| 0 | NA | NA | NA | NA | NA | Gm15256 | | 0 | 0 | 0 | 0 | 0 | 0 | 0 | 0 | 0 | 0 | 0 |
| 0 | NA | NA | NA | NA | NA | Gm5925 | | 0 | 0 | 0 | 0 | 0 | 0 | 0 | 0 | 0 | 0 | 0 |
| 0 | NA | NA | NA | NA | NA | Gm4725 | | 0 | 0 | 0 | 0 | 0 | 0 | 0 | 0 | 0 | 0 | 0 |
| 0 | NA | NA | NA | NA | NA | Gm9498 | | 0 | 0 | 0 | 0 | 0 | 0 | 0 | 0 | 0 | 0 | 0 |
| 0 | NA | NA | NA | NA | NA | Gm15252 | | 0 | 0 | 0 | 0 | 0 | 0 | 0 | 0 | 0 | 0 | 0 |
| 0 | NA | NA | NA | NA | NA | Btbd35f4 | BTB domain containing 35, family membe | 0 | 0 | 0 | 0 | 0 | 0 | 0 | 0 | 0 | 0 | 0 |
| 0 | NA | NA | NA | NA | NA | Gm4726 | | 0 | 0 | 0 | 0 | 0 | 0 | 0 | 0 | 0 | 0 | 0 |
| 0 | NA | NA | NA | NA | NA | Gm3757 | | 0 | 0 | 0 | 0 | 0 | 0 | 0 | 0 | 0 | 0 | 0 |
| 0 | NA | NA | NA | NA | NA | Gm15259 | | 0 | 0 | 0 | 0 | 0 | 0 | 0 | 0 | 0 | 0 | 0 |
| 0 | NA | NA | NA | NA | NA | Gm15254 | | 0 | 0 | 0 | 0 | 0 | 0 | 0 | 0 | 0 | 0 | 0 |
| 0 | NA | NA | NA | NA | NA | Btbd35f27 | BTB domain containing 35, family membe | 0 | 0 | 0 | 0 | 0 | 0 | 0 | 0 | 0 | 0 | 0 |
| 0 | NA | NA | NA | NA | NA | Gm14370 | | 0 | 0 | 0 | 0 | 0 | 0 | 0 | 0 | 0 | 0 | 0 |
| 0 | NA | NA | NA | NA | NA | Gm14367 | | 0 | 0 | 0 | 0 | 0 | 0 | 0 | 0 | 0 | 0 | 0 |
| 0 | NA | NA | NA | NA | NA | Gm14365 | | 0 | 0 | 0 | 0 | 0 | 0 | 0 | 0 | 0 | 0 | 0 |
| 0 | NA | NA | NA | NA | NA | Gm9428 | | 0 | 0 | 0 | 0 | 0 | 0 | 0 | 0 | 0 | 0 | 0 |
| 0 | NA | NA | NA | NA | NA | Gm5924 | | 0 | 0 | 0 | 0 | 0 | 0 | 0 | 0 | 0 | 0 | 0 |
| 0 | NA | NA | NA | NA | NA | Gm14369 | | 0 | 0 | 0 | 0 | 0 | 0 | 0 | 0 | 0 | 0 | 0 |
| 0 | NA | NA | NA | NA | NA | Gm14363 | | 0 | 0 | 0 | 0 | 0 | 0 | 0 | 0 | 0 | 0 | 0 |
| 0 | NA | NA | NA | NA | NA | Gm14366 | | 0 | 0 | 0 | 0 | 0 | 0 | 0 | 0 | 0 | 0 | 0 |
| 0 | NA | NA | NA | NA | NA | Gm14375 | | 0 | 0 | 0 | 0 | 0 | 0 | 0 | 0 | 0 | 0 | 0 |
| 0 | NA | NA | NA | NA | NA | Btbd35f7 | BTB domain containing 35, family membe | 0 | 0 | 0 | 0 | 0 | 0 | 0 | 0 | 0 | 0 | 0 |
| 0 | NA | NA | NA | NA | NA | Gm14452 | | 0 | 0 | 0 | 0 | 0 | 0 | 0 | 0 | 0 | 0 | 0 |
| 0 | NA | NA | NA | NA | NA | Gm14451 | | 0 | 0 | 0 | 0 | 0 | 0 | 0 | 0 | 0 | 0 | 0 |

| | | | | | | | | | | | | | | | | | |
|---|---|---|---|---|---|---|---|---|---|---|---|---|---|---|---|---|---|
| 0 | NA | NA | NA | NA | NA | Gm27360 | | 0 | 0 | 0 | 0 | 0 | 0 | 0 | 0 | 0 | 0 | 0 |
| 0 | NA | NA | NA | NA | NA | n-R5s2 | | 0 | 0 | 0 | 0 | 0 | 0 | 0 | 0 | 0 | 0 | 0 |
| 0 | NA | NA | NA | NA | NA | Ccnb3 | cyclin B3 | 0 | 0 | 0 | 0 | 0 | 0 | 0 | 0 | 0 | 0 | 0 |
| 0 | NA | NA | NA | NA | NA | Akap4 | A kinase (PRKA) anchor protein 4 | 0 | 0 | 0 | 0 | 0 | 0 | 0 | 0 | 0 | 0 | 0 |
| 0 | NA | NA | NA | NA | NA | Gm14380 | | 0 | 0 | 0 | 0 | 0 | 0 | 0 | 0 | 0 | 0 | 0 |
| 0 | NA | NA | NA | NA | NA | Gm26401 | | 0 | 0 | 0 | 0 | 0 | 0 | 0 | 0 | 0 | 0 | 0 |
| 0 | NA | NA | NA | NA | NA | Mir500 | | 0 | 0 | 0 | 0 | 0 | 0 | 0 | 0 | 0 | 0 | 0 |
| 0 | NA | NA | NA | NA | NA | Mir501 | | 0 | 0 | 0 | 0 | 0 | 0 | 0 | 0 | 0 | 0 | 0 |
| 0 | NA | NA | NA | NA | NA | Mir362 | | 0 | 0 | 0 | 0 | 0 | 0 | 0 | 0 | 0 | 0 | 0 |
| 0 | NA | NA | NA | NA | NA | Mir188 | | 0 | 0 | 0 | 0 | 0 | 0 | 0 | 0 | 0 | 0 | 0 |
| 0 | NA | NA | NA | NA | NA | Mir532 | | 0 | 0 | 0 | 0 | 0 | 0 | 0 | 0 | 0 | 0 | 0 |
| 0 | NA | NA | NA | NA | NA | Gm14381 | | 0 | 0 | 0 | 0 | 0 | 0 | 0 | 0 | 0 | 0 | 0 |
| 0 | NA | NA | NA | NA | NA | Gm14395 | | 0 | 0 | 0 | 0 | 0 | 0 | 0 | 0 | 0 | 0 | 0 |
| 0 | NA | NA | NA | NA | NA | Gm14394 | | 0 | 0 | 0 | 0 | 0 | 0 | 0 | 0 | 0 | 0 | 0 |
| 0 | NA | NA | NA | NA | NA | Gm23082 | | 0 | 0 | 0 | 0 | 0 | 0 | 0 | 0 | 0 | 0 | 0 |
| 0 | NA | NA | NA | NA | NA | Gm6079 | | 0 | 0 | 0 | 0 | 0 | 0 | 0 | 0 | 0 | 0 | 0 |
| 0 | NA | NA | NA | NA | NA | Gm45208 | | 0 | 0 | 0 | 0 | 0 | 0 | 0 | 0 | 0 | 0 | 0 |
| 0 | NA | NA | NA | NA | NA | Gm10491 | | 0 | 0 | 0 | 0 | 0 | 0 | 0 | 0 | 0 | 0 | 0 |
| 0 | NA | NA | NA | NA | NA | Eras | ES cell-expressed Ras | 0 | 0 | 0 | 0 | 0 | 0 | 0 | 0 | 0 | 0 | 0 |
| 0 | NA | NA | NA | NA | NA | Psmb6-ps | | 0 | 0 | 0 | 0 | 0 | 0 | 0 | 0 | 0 | 0 | 0 |
| 0 | NA | NA | NA | NA | NA | Gm14799 | | 0 | 0 | 0 | 0 | 0 | 0 | 0 | 0 | 0 | 0 | 0 |
| 0 | NA | NA | NA | NA | NA | Gm22083 | | 0 | 0 | 0 | 0 | 0 | 0 | 0 | 0 | 0 | 0 | 0 |
| 0 | NA | NA | NA | NA | NA | Ssxb10 | synovial sarcoma, X member B10 | 0 | 0 | 0 | 0 | 0 | 0 | 0 | 0 | 0 | 0 | 0 |
| 0 | NA | NA | NA | NA | NA | Ssxb9 | synovial sarcoma, X member B9 | 0 | 0 | 0 | 0 | 0 | 0 | 0 | 0 | 0 | 0 | 0 |
| 0 | NA | NA | NA | NA | NA | Ssxb1 | synovial sarcoma, X member B1 | 0 | 0 | 0 | 0 | 0 | 0 | 0 | 0 | 0 | 0 | 0 |
| 0 | NA | NA | NA | NA | NA | Ssxb2 | synovial sarcoma, X member B2 | 0 | 0 | 0 | 0 | 0 | 0 | 0 | 0 | 0 | 0 | 0 |
| 0 | NA | NA | NA | NA | NA | Gm14459 | predicted gene 14459 | 0 | 0 | 0 | 0 | 0 | 0 | 0 | 0 | 0 | 0 | 0 |
| 0 | NA | NA | NA | NA | NA | Ssxb6 | synovial sarcoma, X member B6 | 0 | 0 | 0 | 0 | 0 | 0 | 0 | 0 | 0 | 0 | 0 |
| 0 | NA | NA | NA | NA | NA | Ssxb3 | synovial sarcoma, X member B3 | 0 | 0 | 0 | 0 | 0 | 0 | 0 | 0 | 0 | 0 | 0 |
| 0 | NA | NA | NA | NA | NA | Gm14500 | | 0 | 0 | 0 | 0 | 0 | 0 | 0 | 0 | 0 | 0 | 0 |
| 0 | NA | NA | NA | NA | NA | Gm6797 | synovial sarcoma, X member B7 | 0 | 0 | 0 | 0 | 0 | 0 | 0 | 0 | 0 | 0 | 0 |
| 0 | NA | NA | NA | NA | NA | Gm6798 | | 0 | 0 | 0 | 0 | 0 | 0 | 0 | 0 | 0 | 0 | 0 |
| 0 | NA | NA | NA | NA | NA | Gm9430 | | 0 | 0 | 0 | 0 | 0 | 0 | 0 | 0 | 0 | 0 | 0 |
| 0 | NA | NA | NA | NA | NA | Ssxb8 | synovial sarcoma, X member B8 | 0 | 0 | 0 | 0 | 0 | 0 | 0 | 0 | 0 | 0 | 0 |
| 0 | NA | NA | NA | NA | NA | Ssx9 | synovial sarcoma, X 9 | 0 | 0 | 0 | 0 | 0 | 0 | 0 | 0 | 0 | 0 | 0 |
| 0 | NA | NA | NA | NA | NA | Ssxb5 | synovial sarcoma, X member B5 | 0 | 0 | 0 | 0 | 0 | 0 | 0 | 0 | 0 | 0 | 0 |
| 0 | NA | NA | NA | NA | NA | Gm6592 | predicted gene 6592 | 0 | 0 | 0 | 0 | 0 | 0 | 0 | 0 | 0 | 0 | 0 |
| 0 | NA | NA | NA | NA | NA | Gm5751 | | 0 | 0 | 0 | 0 | 0 | 0 | 0 | 0 | 0 | 0 | 0 |
| 0 | NA | NA | NA | NA | NA | Gm5752 | | 0 | 0 | 0 | 0 | 0 | 0 | 0 | 0 | 0 | 0 | 0 |
| 0 | NA | NA | NA | NA | NA | Gm16479 | | 0 | 0 | 0 | 0 | 0 | 0 | 0 | 0 | 0 | 0 | 0 |
| 0 | NA | NA | NA | NA | NA | Gm14510 | | 0 | 0 | 0 | 0 | 0 | 0 | 0 | 0 | 0 | 0 | 0 |
| 0 | NA | NA | NA | NA | NA | Fthl17b | ferritin, heavy polypeptide-like 17, membe | 0 | 0 | 0 | 0 | 0 | 0 | 0 | 0 | 0 | 0 | 0 |
| 0 | NA | NA | NA | NA | NA | Gm14509 | | 0 | 0 | 0 | 0 | 0 | 0 | 0 | 0 | 0 | 0 | 0 |
| 0 | NA | NA | NA | NA | NA | Fthl17c | ferritin, heavy polypeptide-like 17, membe | 0 | 0 | 0 | 0 | 0 | 0 | 0 | 0 | 0 | 0 | 0 |
| 0 | NA | NA | NA | NA | NA | Gm14457 | | 0 | 0 | 0 | 0 | 0 | 0 | 0 | 0 | 0 | 0 | 0 |
| 0 | NA | NA | NA | NA | NA | Fthl17d | ferritin, heavy polypeptide-like 17, membe | 0 | 0 | 0 | 0 | 0 | 0 | 0 | 0 | 0 | 0 | 0 |
| 0 | NA | NA | NA | NA | NA | Gm14865 | | 0 | 0 | 0 | 0 | 0 | 0 | 0 | 0 | 0 | 0 | 0 |
| 0 | NA | NA | NA | NA | NA | Fthl17e | ferritin, heavy polypeptide-like 17, membe | 0 | 0 | 0 | 0 | 0 | 0 | 0 | 0 | 0 | 0 | 0 |
| 0 | NA | NA | NA | NA | NA | Gm14503 | | 0 | 0 | 0 | 0 | 0 | 0 | 0 | 0 | 0 | 0 | 0 |
| 0 | NA | NA | NA | NA | NA | Fthl17-ps1 | | 0 | 0 | 0 | 0 | 0 | 0 | 0 | 0 | 0 | 0 | 0 |
| 0 | NA | NA | NA | NA | NA | Gm14864 | | 0 | 0 | 0 | 0 | 0 | 0 | 0 | 0 | 0 | 0 | 0 |
| 0 | NA | NA | NA | NA | NA | Gm5379 | | 0 | 0 | 0 | 0 | 0 | 0 | 0 | 0 | 0 | 0 | 0 |
| 0 | NA | NA | NA | NA | NA | Fthl17f | ferritin, heavy polypeptide-like 17, membe | 0 | 0 | 0 | 0 | 0 | 0 | 0 | 0 | 0 | 0 | 0 |
| 0 | NA | NA | NA | NA | NA | Gm16480 | | 0 | 0 | 0 | 0 | 0 | 0 | 0 | 0 | 0 | 0 | 0 |
| 0 | NA | NA | NA | NA | NA | Fthl17-ps2 | | 0 | 0 | 0 | 0 | 0 | 0 | 0 | 0 | 0 | 0 | 0 |
| 0 | NA | NA | NA | NA | NA | 4930402K13Ril | RIKEN cDNA 4930402K13 gene | 0 | 0 | 0 | 0 | 0 | 0 | 0 | 0 | 0 | 0 | 0 |
| 0 | NA | NA | NA | NA | NA | Fthl17-ps3 | | 0 | 0 | 0 | 0 | 0 | 0 | 0 | 0 | 0 | 0 | 0 |
| 0 | NA | NA | NA | NA | NA | Gm14862 | | 0 | 0 | 0 | 0 | 0 | 0 | 0 | 0 | 0 | 0 | 0 |
| 0 | NA | NA | NA | NA | NA | Gm25481 | | 0 | 0 | 0 | 0 | 0 | 0 | 0 | 0 | 0 | 0 | 0 |
| 0 | NA | NA | NA | NA | NA | H2al1k | H2A histone family member L1K | 0 | 0 | 0 | 0 | 0 | 0 | 0 | 0 | 0 | 0 | 0 |

| | | | | | | | | | | | | | | | | | | |
|---|---|---|---|---|---|---|---|---|---|---|---|---|---|---|---|---|---|---|
| 0 | NA | NA | NA | NA | NA | Gm25728 | | 0 | 0 | 0 | 0 | 0 | 0 | 0 | 0 | 0 | 0 | 0 |
| 0 | NA | NA | NA | NA | NA | H2al1o | H2A histone family member L1O | 0 | 0 | 0 | 0 | 0 | 0 | 0 | 0 | 0 | 0 | 0 |
| 0 | NA | NA | NA | NA | NA | | | 0 | 0 | 0 | 0 | 0 | 0 | 0 | 0 | 0 | 0 | 0 |
| 0 | NA | NA | NA | NA | NA | H2al3 | H2A histone family member L3 | 0 | 0 | 0 | 0 | 0 | 0 | 0 | 0 | 0 | 0 | 0 |
| 0 | NA | NA | NA | NA | NA | Otc | ornithine transcarbamylase | 0 | 0 | 0 | 0 | 0 | 0 | 0 | 0 | 0 | 0 | 0 |
| 0 | NA | NA | NA | NA | NA | 4933416E14Rik | | 0 | 0 | 0 | 0 | 0 | 0 | 0 | 0 | 0 | 0 | 0 |
| 0 | NA | NA | NA | NA | NA | Gm25885 | | 0 | 0 | 0 | 0 | 0 | 0 | 0 | 0 | 0 | 0 | 0 |
| 0 | NA | NA | NA | NA | NA | Gm14470 | | 0 | 0 | 0 | 0 | 0 | 0 | 0 | 0 | 0 | 0 | 0 |
| 0 | NA | NA | NA | NA | NA | Gm14485 | | 0 | 0 | 0 | 0 | 0 | 0 | 0 | 0 | 0 | 0 | 0 |
| 0 | NA | NA | NA | NA | NA | Gm14481 | | 0 | 0 | 0 | 0 | 0 | 0 | 0 | 0 | 0 | 0 | 0 |
| 0 | NA | NA | NA | NA | NA | H2al1a | H2A histone family member L1A | 0 | 0 | 0 | 0 | 0 | 0 | 0 | 0 | 0 | 0 | 0 |
| 0 | NA | NA | NA | NA | NA | H2al1b | H2A histone family member L1B | 0 | 0 | 0 | 0 | 0 | 0 | 0 | 0 | 0 | 0 | 0 |
| 0 | NA | NA | NA | NA | NA | H2al1c | H2A histone family member L1C | 0 | 0 | 0 | 0 | 0 | 0 | 0 | 0 | 0 | 0 | 0 |
| 0 | NA | NA | NA | NA | NA | H2al1d | H2A histone family member L1D | 0 | 0 | 0 | 0 | 0 | 0 | 0 | 0 | 0 | 0 | 0 |
| 0 | NA | NA | NA | NA | NA | H2al1e | H2A histone family member L1E | 0 | 0 | 0 | 0 | 0 | 0 | 0 | 0 | 0 | 0 | 0 |
| 0 | NA | NA | NA | NA | NA | H2al1f | H2A histone family member L1F | 0 | 0 | 0 | 0 | 0 | 0 | 0 | 0 | 0 | 0 | 0 |
| 0 | NA | NA | NA | NA | NA | H2al1g | H2A histone family member L1G | 0 | 0 | 0 | 0 | 0 | 0 | 0 | 0 | 0 | 0 | 0 |
| 0 | NA | NA | NA | NA | NA | H2al1h | H2A histone family member L1H | 0 | 0 | 0 | 0 | 0 | 0 | 0 | 0 | 0 | 0 | 0 |
| 0 | NA | NA | NA | NA | NA | H2al1i | H2A histone family member L1I | 0 | 0 | 0 | 0 | 0 | 0 | 0 | 0 | 0 | 0 | 0 |
| 0 | NA | NA | NA | NA | NA | H2al1j | H2A histone family member L1J | 0 | 0 | 0 | 0 | 0 | 0 | 0 | 0 | 0 | 0 | 0 |
| 0 | NA | NA | NA | NA | NA | Gm14515 | | 0 | 0 | 0 | 0 | 0 | 0 | 0 | 0 | 0 | 0 | 0 |
| 0 | NA | NA | NA | NA | NA | Gm26314 | | 0 | 0 | 0 | 0 | 0 | 0 | 0 | 0 | 0 | 0 | 0 |
| 0 | NA | NA | NA | NA | NA | Rpl23a-ps13 | | 0 | 0 | 0 | 0 | 0 | 0 | 0 | 0 | 0 | 0 | 0 |
| 0 | NA | NA | NA | NA | NA | Gm14491 | | 0 | 0 | 0 | 0 | 0 | 0 | 0 | 0 | 0 | 0 | 0 |
| 0 | NA | NA | NA | NA | NA | Gm14512 | | 0 | 0 | 0 | 0 | 0 | 0 | 0 | 0 | 0 | 0 | 0 |
| 0 | NA | NA | NA | NA | NA | Gm14521 | | 0 | 0 | 0 | 0 | 0 | 0 | 0 | 0 | 0 | 0 | 0 |
| 0 | NA | NA | NA | NA | NA | Gm14635 | | 0 | 0 | 0 | 0 | 0 | 0 | 0 | 0 | 0 | 0 | 0 |
| 0 | NA | NA | NA | NA | NA | Gm4984 | | 0 | 0 | 0 | 0 | 0 | 0 | 0 | 0 | 0 | 0 | 0 |
| 0 | NA | NA | NA | NA | NA | Gm25063 | | 0 | 0 | 0 | 0 | 0 | 0 | 0 | 0 | 0 | 0 | 0 |
| 0 | NA | NA | NA | NA | NA | Gm14524 | | 0 | 0 | 0 | 0 | 0 | 0 | 0 | 0 | 0 | 0 | 0 |
| 0 | NA | NA | NA | NA | NA | Gm14534 | | 0 | 0 | 0 | 0 | 0 | 0 | 0 | 0 | 0 | 0 | 0 |
| 0 | NA | NA | NA | NA | NA | Gm15027 | | 0 | 0 | 0 | 0 | 0 | 0 | 0 | 0 | 0 | 0 | 0 |
| 0 | NA | NA | NA | NA | NA | Gm14536 | | 0 | 0 | 0 | 0 | 0 | 0 | 0 | 0 | 0 | 0 | 0 |
| 0 | NA | NA | NA | NA | NA | n-R5s3 | | 0 | 0 | 0 | 0 | 0 | 0 | 0 | 0 | 0 | 0 | 0 |
| 0 | NA | NA | NA | NA | NA | Gm16481 | | 0 | 0 | 0 | 0 | 0 | 0 | 0 | 0 | 0 | 0 | 0 |
| 0 | NA | NA | NA | NA | NA | Gm25246 | | 0 | 0 | 0 | 0 | 0 | 0 | 0 | 0 | 0 | 0 | 0 |
| 0 | NA | NA | NA | NA | NA | Gm5381 | | 0 | 0 | 0 | 0 | 0 | 0 | 0 | 0 | 0 | 0 | 0 |
| 0 | NA | NA | NA | NA | NA | Gm25552 | | 0 | 0 | 0 | 0 | 0 | 0 | 0 | 0 | 0 | 0 | 0 |
| 0 | NA | NA | NA | NA | NA | H2al1n | H2A histone family member L1N | 0 | 0 | 0 | 0 | 0 | 0 | 0 | 0 | 0 | 0 | 0 |
| 0 | NA | NA | NA | NA | NA | Gm14867 | | 0 | 0 | 0 | 0 | 0 | 0 | 0 | 0 | 0 | 0 | 0 |
| 0 | NA | NA | NA | NA | NA | Gm25202 | | 0 | 0 | 0 | 0 | 0 | 0 | 0 | 0 | 0 | 0 | 0 |
| 0 | NA | NA | NA | NA | NA | Gm14505 | | 0 | 0 | 0 | 0 | 0 | 0 | 0 | 0 | 0 | 0 | 0 |
| 0 | NA | NA | NA | NA | NA | Ppp1r2-ps9 | | 0 | 0 | 0 | 0 | 0 | 0 | 0 | 0 | 0 | 0 | 0 |
| 0 | NA | NA | NA | NA | NA | Gm14518 | | 0 | 0 | 0 | 0 | 0 | 0 | 0 | 0 | 0 | 0 | 0 |
| 0 | NA | NA | NA | NA | NA | Gm14517 | | 0 | 0 | 0 | 0 | 0 | 0 | 0 | 0 | 0 | 0 | 0 |
| 0 | NA | NA | NA | NA | NA | Gm25207 | | 0 | 0 | 0 | 0 | 0 | 0 | 0 | 0 | 0 | 0 | 0 |
| 0 | NA | NA | NA | NA | NA | Gm22464 | | 0 | 0 | 0 | 0 | 0 | 0 | 0 | 0 | 0 | 0 | 0 |
| 0 | NA | NA | NA | NA | NA | Gm23516 | | 0 | 0 | 0 | 0 | 0 | 0 | 0 | 0 | 0 | 0 | 0 |
| 0 | NA | NA | NA | NA | NA | Drr1 | | 0 | 0 | 0 | 0 | 0 | 0 | 0 | 0 | 0 | 0 | 0 |
| 0 | NA | NA | NA | NA | NA | Gm14520 | | 0 | 0 | 0 | 0 | 0 | 0 | 0 | 0 | 0 | 0 | 0 |
| 0 | NA | NA | NA | NA | NA | Gm14516 | | 0 | 0 | 0 | 0 | 0 | 0 | 0 | 0 | 0 | 0 | 0 |
| 0 | NA | NA | NA | NA | NA | Cypt1 | cysteine-rich perinuclear theca 1 | 0 | 0 | 0 | 0 | 0 | 0 | 0 | 0 | 0 | 0 | 0 |
| 0 | NA | NA | NA | NA | NA | Gm14514 | | 0 | 0 | 0 | 0 | 0 | 0 | 0 | 0 | 0 | 0 | 0 |
| 0 | NA | NA | NA | NA | NA | Gm22294 | | 0 | 0 | 0 | 0 | 0 | 0 | 0 | 0 | 0 | 0 | 0 |
| 0 | NA | NA | NA | NA | NA | Gm5073 | | 0 | 0 | 0 | 0 | 0 | 0 | 0 | 0 | 0 | 0 | 0 |
| 0 | NA | NA | NA | NA | NA | Gm5755 | | 0 | 0 | 0 | 0 | 0 | 0 | 0 | 0 | 0 | 0 | 0 |
| 0 | NA | NA | NA | NA | NA | Gm5123 | | 0 | 0 | 0 | 0 | 0 | 0 | 0 | 0 | 0 | 0 | 0 |
| 0 | NA | NA | NA | NA | NA | Dusp21 | dual specificity phosphatase 21 | 0 | 0 | 0 | 0 | 0 | 0 | 0 | 0 | 0 | 0 | 0 |
| 0 | NA | NA | NA | NA | NA | n-R5s5 | | 0 | 0 | 0 | 0 | 0 | 0 | 0 | 0 | 0 | 0 | 0 |
| 0 | NA | NA | NA | NA | NA | Gm14883 | | 0 | 0 | 0 | 0 | 0 | 0 | 0 | 0 | 0 | 0 | 0 |

| | | | | | | | | | | | | | | | | | |
|---|---|---|---|---|---|---|---|---|---|---|---|---|---|---|---|---|---|
| 0 | NA | NA | NA | NA | NA | Gm6923 | | 0 | 0 | 0 | 0 | 0 | 0 | 0 | 0 | 0 | 0 | 0 |
| 0 | NA | NA | NA | NA | NA | Gm5383 | | 0 | 0 | 0 | 0 | 0 | 0 | 0 | 0 | 0 | 0 | 0 |
| 0 | NA | NA | NA | NA | NA | Mir222 | | 0 | 0 | 0 | 0 | 0 | 0 | 0 | 0 | 0 | 0 | 0 |
| 0 | NA | NA | NA | NA | NA | Gm22783 | | 0 | 0 | 0 | 0 | 0 | 0 | 0 | 0 | 0 | 0 | 0 |
| 0 | NA | NA | NA | NA | NA | Gm14885 | | 0 | 0 | 0 | 0 | 0 | 0 | 0 | 0 | 0 | 0 | 0 |
| 0 | NA | NA | NA | NA | NA | Tex13c3 | TEX13 family member C3 | 0 | 0 | 0 | 0 | 0 | 0 | 0 | 0 | 0 | 0 | 0 |
| 0 | NA | NA | NA | NA | NA | Gm14531 | | 0 | 0 | 0 | 0 | 0 | 0 | 0 | 0 | 0 | 0 | 0 |
| 0 | NA | NA | NA | NA | NA | Gm23678 | | 0 | 0 | 0 | 0 | 0 | 0 | 0 | 0 | 0 | 0 | 0 |
| 0 | NA | NA | NA | NA | NA | Gm9085 | | 0 | 0 | 0 | 0 | 0 | 0 | 0 | 0 | 0 | 0 | 0 |
| 0 | NA | NA | NA | NA | NA | Gm14884 | | 0 | 0 | 0 | 0 | 0 | 0 | 0 | 0 | 0 | 0 | 0 |
| 0 | NA | NA | NA | NA | NA | | | 0 | 0 | 0 | 0 | 0 | 0 | 0 | 0 | 0 | 0 | 0 |
| 0 | NA | NA | NA | NA | NA | Gm26065 | | 0 | 0 | 0 | 0 | 0 | 0 | 0 | 0 | 0 | 0 | 0 |
| 0 | NA | NA | NA | NA | NA | Gm24824 | | 0 | 0 | 0 | 0 | 0 | 0 | 0 | 0 | 0 | 0 | 0 |
| 0 | NA | NA | NA | NA | NA | Gm15028 | | 0 | 0 | 0 | 0 | 0 | 0 | 0 | 0 | 0 | 0 | 0 |
| 0 | NA | NA | NA | NA | NA | Mir5617 | | 0 | 0 | 0 | 0 | 0 | 0 | 0 | 0 | 0 | 0 | 0 |
| 0 | NA | NA | NA | NA | NA | Gm15029 | | 0 | 0 | 0 | 0 | 0 | 0 | 0 | 0 | 0 | 0 | 0 |
| 0 | NA | NA | NA | NA | NA | Spaca5 | sperm acrosome associated 5 | 0 | 0 | 0 | 0 | 0 | 0 | 0 | 0 | 0 | 0 | 0 |
| 0 | NA | NA | NA | NA | NA | Ssxa1 | | 0 | 0 | 0 | 0 | 0 | 0 | 0 | 0 | 0 | 0 | 0 |
| 0 | NA | NA | NA | NA | NA | Gm21876 | | 0 | 0 | 0 | 0 | 0 | 0 | 0 | 0 | 0 | 0 | 0 |
| 0 | NA | NA | NA | NA | NA | 4930453H23Ril | RIKEN cDNA 4930453H23 gene | 0 | 0 | 0 | 0 | 0 | 0 | 0 | 0 | 0 | 0 | 0 |
| 0 | NA | NA | NA | NA | NA | Agtr2 | angiotensin II receptor, type 2 | 0 | 0 | 0 | 0 | 0 | 0 | 0 | 0 | 0 | 0 | 0 |
| 0 | NA | NA | NA | NA | NA | Gm22592 | | 0 | 0 | 0 | 0 | 0 | 0 | 0 | 0 | 0 | 0 | 0 |
| 0 | NA | NA | NA | NA | NA | Gm14558 | | 0 | 0 | 0 | 0 | 0 | 0 | 0 | 0 | 0 | 0 | 0 |
| 0 | NA | NA | NA | NA | NA | Gm25553 | | 0 | 0 | 0 | 0 | 0 | 0 | 0 | 0 | 0 | 0 | 0 |
| 0 | NA | NA | NA | NA | NA | Gm28269 | | 0 | 0 | 0 | 0 | 0 | 0 | 0 | 0 | 0 | 0 | 0 |
| 0 | NA | NA | NA | NA | NA | Gm28268 | | 0 | 0 | 0 | 0 | 0 | 0 | 0 | 0 | 0 | 0 | 0 |
| 0 | NA | NA | NA | NA | NA | Gm24535 | | 0 | 0 | 0 | 0 | 0 | 0 | 0 | 0 | 0 | 0 | 0 |
| 0 | NA | NA | NA | NA | NA | Gm14563 | | 0 | 0 | 0 | 0 | 0 | 0 | 0 | 0 | 0 | 0 | 0 |
| 0 | NA | NA | NA | NA | NA | Gm14561 | | 0 | 0 | 0 | 0 | 0 | 0 | 0 | 0 | 0 | 0 | 0 |
| 0 | NA | NA | NA | NA | NA | | | 0 | 0 | 0 | 0 | 0 | 0 | 0 | 0 | 0 | 0 | 0 |
| 0 | NA | NA | NA | NA | NA | Gm6071 | | 0 | 0 | 0 | 0 | 0 | 0 | 0 | 0 | 0 | 0 | 0 |
| 0 | NA | NA | NA | NA | NA | Gm25812 | | 0 | 0 | 0 | 0 | 0 | 0 | 0 | 0 | 0 | 0 | 0 |
| 0 | NA | NA | NA | NA | NA | Gm14562 | | 0 | 0 | 0 | 0 | 0 | 0 | 0 | 0 | 0 | 0 | 0 |
| 0 | NA | NA | NA | NA | NA | Gm15030 | | 0 | 0 | 0 | 0 | 0 | 0 | 0 | 0 | 0 | 0 | 0 |
| 0 | NA | NA | NA | NA | NA | Ppp1r2-ps7 | | 0 | 0 | 0 | 0 | 0 | 0 | 0 | 0 | 0 | 0 | 0 |
| 0 | NA | NA | NA | NA | NA | Gm26131 | | 0 | 0 | 0 | 0 | 0 | 0 | 0 | 0 | 0 | 0 | 0 |
| 0 | NA | NA | NA | NA | NA | Gm7164 | | 0 | 0 | 0 | 0 | 0 | 0 | 0 | 0 | 0 | 0 | 0 |
| 0 | NA | NA | NA | NA | NA | Gm4789 | | 0 | 0 | 0 | 0 | 0 | 0 | 0 | 0 | 0 | 0 | 0 |
| 0 | NA | NA | NA | NA | NA | Gm4907 | predicted gene 4907 | 0 | 0 | 0 | 0 | 0 | 0 | 0 | 0 | 0 | 0 | 0 |
| 0 | NA | NA | NA | NA | NA | Gm4985 | | 0 | 0 | 0 | 0 | 0 | 0 | 0 | 0 | 0 | 0 | 0 |
| 0 | NA | NA | NA | NA | NA | Gm25299 | | 0 | 0 | 0 | 0 | 0 | 0 | 0 | 0 | 0 | 0 | 0 |
| 0 | NA | NA | NA | NA | NA | Gm4732 | | 0 | 0 | 0 | 0 | 0 | 0 | 0 | 0 | 0 | 0 | 0 |
| 0 | NA | NA | NA | NA | NA | Gm9440 | | 0 | 0 | 0 | 0 | 0 | 0 | 0 | 0 | 0 | 0 | 0 |
| 0 | NA | NA | NA | NA | NA | Gm6956 | | 0 | 0 | 0 | 0 | 0 | 0 | 0 | 0 | 0 | 0 | 0 |
| 0 | NA | NA | NA | NA | NA | Gm27192 | | 0 | 0 | 0 | 0 | 0 | 0 | 0 | 0 | 0 | 0 | 0 |
| 0 | NA | NA | NA | NA | NA | Gm5934 | predicted gene 5934 | 0 | 0 | 0 | 0 | 0 | 0 | 0 | 0 | 0 | 0 | 0 |
| 0 | NA | NA | NA | NA | NA | Gm4297 | predicted gene 4297 | 0 | 0 | 0 | 0 | 0 | 0 | 0 | 0 | 0 | 0 | 0 |
| 0 | NA | NA | NA | NA | NA | Gm27910 | | 0 | 0 | 0 | 0 | 0 | 0 | 0 | 0 | 0 | 0 | 0 |
| 0 | NA | NA | NA | NA | NA | Gm27839 | | 0 | 0 | 0 | 0 | 0 | 0 | 0 | 0 | 0 | 0 | 0 |
| 0 | NA | NA | NA | NA | NA | Gm5935 | predicted gene 5935 | 0 | 0 | 0 | 0 | 0 | 0 | 0 | 0 | 0 | 0 | 0 |
| 0 | NA | NA | NA | NA | NA | Gm2759 | | 0 | 0 | 0 | 0 | 0 | 0 | 0 | 0 | 0 | 0 | 0 |
| 0 | NA | NA | NA | NA | NA | Gm44480 | | 0 | 0 | 0 | 0 | 0 | 0 | 0 | 0 | 0 | 0 | 0 |
| 0 | NA | NA | NA | NA | NA | Gm2768 | | 0 | 0 | 0 | 0 | 0 | 0 | 0 | 0 | 0 | 0 | 0 |
| 0 | NA | NA | NA | NA | NA | Gm5169 | predicted gene 5169 | 0 | 0 | 0 | 0 | 0 | 0 | 0 | 0 | 0 | 0 | 0 |
| 0 | NA | NA | NA | NA | NA | Gm1989 | | 0 | 0 | 0 | 0 | 0 | 0 | 0 | 0 | 0 | 0 | 0 |
| 0 | NA | NA | NA | NA | NA | Gm1993 | predicted gene 1993 | 0 | 0 | 0 | 0 | 0 | 0 | 0 | 0 | 0 | 0 | 0 |
| 0 | NA | NA | NA | NA | NA | Gm2003 | predicted pseudogene 2003 | 0 | 0 | 0 | 0 | 0 | 0 | 0 | 0 | 0 | 0 | 0 |
| 0 | NA | NA | NA | NA | NA | Gm27892 | | 0 | 0 | 0 | 0 | 0 | 0 | 0 | 0 | 0 | 0 | 0 |
| 0 | NA | NA | NA | NA | NA | Gm7350 | | 0 | 0 | 0 | 0 | 0 | 0 | 0 | 0 | 0 | 0 | 0 |
| 0 | NA | NA | NA | NA | NA | E330010L02Rik | | 0 | 0 | 0 | 0 | 0 | 0 | 0 | 0 | 0 | 0 | 0 |

| | | | | | | | | | | | | | | | | | | |
|---|---|---|---|---|---|---|---|---|---|---|---|---|---|---|---|---|---|---|
| 0 | NA | NA | NA | NA | NA | Gm14533 | | 0 | 0 | 0 | 0 | 0 | 0 | 0 | 0 | 0 | 0 | 0 |
| 0 | NA | NA | NA | NA | NA | Gm5168 | predicted gene 5168 | 0 | 0 | 0 | 0 | 0 | 0 | 0 | 0 | 0 | 0 | 0 |
| 0 | NA | NA | NA | NA | NA | Gm2012 | predicted gene 2012 | 0 | 0 | 0 | 0 | 0 | 0 | 0 | 0 | 0 | 0 | 0 |
| 0 | NA | NA | NA | NA | NA | Gm26302 | | 0 | 0 | 0 | 0 | 0 | 0 | 0 | 0 | 0 | 0 | 0 |
| 0 | NA | NA | NA | NA | NA | Gm2030 | predicted gene 2030 | 0 | 0 | 0 | 0 | 0 | 0 | 0 | 0 | 0 | 0 | 0 |
| 0 | NA | NA | NA | NA | NA | Gm2005 | | 0 | 0 | 0 | 0 | 0 | 0 | 0 | 0 | 0 | 0 | 0 |
| 0 | NA | NA | NA | NA | NA | Slx | Sycp3 like X-linked | 0 | 0 | 0 | 0 | 0 | 0 | 0 | 0 | 0 | 0 | 0 |
| 0 | NA | NA | NA | NA | NA | Gm27341 | | 0 | 0 | 0 | 0 | 0 | 0 | 0 | 0 | 0 | 0 | 0 |
| 0 | NA | NA | NA | NA | NA | Gm14525 | predicted gene 14525 | 0 | 0 | 0 | 0 | 0 | 0 | 0 | 0 | 0 | 0 | 0 |
| 0 | NA | NA | NA | NA | NA | Gm6113 | | 0 | 0 | 0 | 0 | 0 | 0 | 0 | 0 | 0 | 0 | 0 |
| 0 | NA | NA | NA | NA | NA | Gm6121 | predicted gene 6121 | 0 | 0 | 0 | 0 | 0 | 0 | 0 | 0 | 0 | 0 | 0 |
| 0 | NA | NA | NA | NA | NA | Gm7365 | | 0 | 0 | 0 | 0 | 0 | 0 | 0 | 0 | 0 | 0 | 0 |
| 0 | NA | NA | NA | NA | NA | Gm7375 | | 0 | 0 | 0 | 0 | 0 | 0 | 0 | 0 | 0 | 0 | 0 |
| 0 | NA | NA | NA | NA | NA | Gm10230 | predicted gene 10230 | 0 | 0 | 0 | 0 | 0 | 0 | 0 | 0 | 0 | 0 | 0 |
| 0 | NA | NA | NA | NA | NA | Gm44452 | | 0 | 0 | 0 | 0 | 0 | 0 | 0 | 0 | 0 | 0 | 0 |
| 0 | NA | NA | NA | NA | NA | Gm2092 | | 0 | 0 | 0 | 0 | 0 | 0 | 0 | 0 | 0 | 0 | 0 |
| 0 | NA | NA | NA | NA | NA | Gm44407 | | 0 | 0 | 0 | 0 | 0 | 0 | 0 | 0 | 0 | 0 | 0 |
| 0 | NA | NA | NA | NA | NA | Gm2101 | | 0 | 0 | 0 | 0 | 0 | 0 | 0 | 0 | 0 | 0 | 0 |
| 0 | NA | NA | NA | NA | NA | Gm7386 | | 0 | 0 | 0 | 0 | 0 | 0 | 0 | 0 | 0 | 0 | 0 |
| 0 | NA | NA | NA | NA | NA | Gm10058 | predicted gene 10058 | 0 | 0 | 0 | 0 | 0 | 0 | 0 | 0 | 0 | 0 | 0 |
| 0 | NA | NA | NA | NA | NA | Gm2117 | | 0 | 0 | 0 | 0 | 0 | 0 | 0 | 0 | 0 | 0 | 0 |
| 0 | NA | NA | NA | NA | NA | Gm7391 | | 0 | 0 | 0 | 0 | 0 | 0 | 0 | 0 | 0 | 0 | 0 |
| 0 | NA | NA | NA | NA | NA | Gm4836 | predicted gene 4836 | 0 | 0 | 0 | 0 | 0 | 0 | 0 | 0 | 0 | 0 | 0 |
| 0 | NA | NA | NA | NA | NA | Gm7398 | | 0 | 0 | 0 | 0 | 0 | 0 | 0 | 0 | 0 | 0 | 0 |
| 0 | NA | NA | NA | NA | NA | Gm10147 | predicted gene 10147 | 0 | 0 | 0 | 0 | 0 | 0 | 0 | 0 | 0 | 0 | 0 |
| 0 | NA | NA | NA | NA | NA | Gm44499 | | 0 | 0 | 0 | 0 | 0 | 0 | 0 | 0 | 0 | 0 | 0 |
| 0 | NA | NA | NA | NA | NA | Gm2165 | | 0 | 0 | 0 | 0 | 0 | 0 | 0 | 0 | 0 | 0 | 0 |
| 0 | NA | NA | NA | NA | NA | Gm7408 | | 0 | 0 | 0 | 0 | 0 | 0 | 0 | 0 | 0 | 0 | 0 |
| 0 | NA | NA | NA | NA | NA | Gm10096 | predicted gene 10096 | 0 | 0 | 0 | 0 | 0 | 0 | 0 | 0 | 0 | 0 | 0 |
| 0 | NA | NA | NA | NA | NA | Gm2200 | | 0 | 0 | 0 | 0 | 0 | 0 | 0 | 0 | 0 | 0 | 0 |
| 0 | NA | NA | NA | NA | NA | Gm7415 | | 0 | 0 | 0 | 0 | 0 | 0 | 0 | 0 | 0 | 0 | 0 |
| 0 | NA | NA | NA | NA | NA | Gm2066 | | 0 | 0 | 0 | 0 | 0 | 0 | 0 | 0 | 0 | 0 | 0 |
| 0 | NA | NA | NA | NA | NA | Gm7421 | | 0 | 0 | 0 | 0 | 0 | 0 | 0 | 0 | 0 | 0 | 0 |
| 0 | NA | NA | NA | NA | NA | Gm14527 | | 0 | 0 | 0 | 0 | 0 | 0 | 0 | 0 | 0 | 0 | 0 |
| 0 | NA | NA | NA | NA | NA | Gm44496 | | 0 | 0 | 0 | 0 | 0 | 0 | 0 | 0 | 0 | 0 | 0 |
| 0 | NA | NA | NA | NA | NA | Gm3657 | | 0 | 0 | 0 | 0 | 0 | 0 | 0 | 0 | 0 | 0 | 0 |
| 0 | NA | NA | NA | NA | NA | Gm3669 | | 0 | 0 | 0 | 0 | 0 | 0 | 0 | 0 | 0 | 0 | 0 |
| 0 | NA | NA | NA | NA | NA | Gm10488 | predicted gene 10488 | 0 | 0 | 0 | 0 | 0 | 0 | 0 | 0 | 0 | 0 | 0 |
| 0 | NA | NA | NA | NA | NA | Gm44298 | | 0 | 0 | 0 | 0 | 0 | 0 | 0 | 0 | 0 | 0 | 0 |
| 0 | NA | NA | NA | NA | NA | E330016L19Rik | | 0 | 0 | 0 | 0 | 0 | 0 | 0 | 0 | 0 | 0 | 0 |
| 0 | NA | NA | NA | NA | NA | Gm3681 | | 0 | 0 | 0 | 0 | 0 | 0 | 0 | 0 | 0 | 0 | 0 |
| 0 | NA | NA | NA | NA | NA | Gm14632 | predicted gene 14632 | 0 | 0 | 0 | 0 | 0 | 0 | 0 | 0 | 0 | 0 | 0 |
| 0 | NA | NA | NA | NA | NA | Gm7437 | | 0 | 0 | 0 | 0 | 0 | 0 | 0 | 0 | 0 | 0 | 0 |
| 0 | NA | NA | NA | NA | NA | Gm14974 | | 0 | 0 | 0 | 0 | 0 | 0 | 0 | 0 | 0 | 0 | 0 |
| 0 | NA | NA | NA | NA | NA | Gm7438 | | 0 | 0 | 0 | 0 | 0 | 0 | 0 | 0 | 0 | 0 | 0 |
| 0 | NA | NA | NA | NA | NA | Gm10487 | predicted gene 10487 | 0 | 0 | 0 | 0 | 0 | 0 | 0 | 0 | 0 | 0 | 0 |
| 0 | NA | NA | NA | NA | NA | Btbd35f29 | BTB domain containing 35, family membe | 0 | 0 | 0 | 0 | 0 | 0 | 0 | 0 | 0 | 0 | 0 |
| 0 | NA | NA | NA | NA | NA | Spin2f | spindlin-2A-like | 0 | 0 | 0 | 0 | 0 | 0 | 0 | 0 | 0 | 0 | 0 |
| 0 | NA | NA | NA | NA | NA | Btbd35f26 | BTB domain containing 35, family membe | 0 | 0 | 0 | 0 | 0 | 0 | 0 | 0 | 0 | 0 | 0 |
| 0 | NA | NA | NA | NA | NA | Gm15280 | | 0 | 0 | 0 | 0 | 0 | 0 | 0 | 0 | 0 | 0 | 0 |
| 0 | NA | NA | NA | NA | NA | Gm15276 | | 0 | 0 | 0 | 0 | 0 | 0 | 0 | 0 | 0 | 0 | 0 |
| 0 | NA | NA | NA | NA | NA | Btbd35f19 | BTB domain containing 35, family membe | 0 | 0 | 0 | 0 | 0 | 0 | 0 | 0 | 0 | 0 | 0 |
| 0 | NA | NA | NA | NA | NA | Gm21883 | | 0 | 0 | 0 | 0 | 0 | 0 | 0 | 0 | 0 | 0 | 0 |
| 0 | NA | NA | NA | NA | NA | Spin2e | predicted gene 5923 | 0 | 0 | 0 | 0 | 0 | 0 | 0 | 0 | 0 | 0 | 0 |
| 0 | NA | NA | NA | NA | NA | Btbd35f25 | BTB domain containing 35, family membe | 0 | 0 | 0 | 0 | 0 | 0 | 0 | 0 | 0 | 0 | 0 |
| 0 | NA | NA | NA | NA | NA | Gm21637 | spindlin family, member 2, pseudogene 1 | 0 | 0 | 0 | 0 | 0 | 0 | 0 | 0 | 0 | 0 | 0 |
| 0 | NA | NA | NA | NA | NA | Btbd35f22 | BTB domain containing 35, family membe | 0 | 0 | 0 | 0 | 0 | 0 | 0 | 0 | 0 | 0 | 0 |
| 0 | NA | NA | NA | NA | NA | Gm5923 | | 0 | 0 | 0 | 0 | 0 | 0 | 0 | 0 | 0 | 0 | 0 |
| 0 | NA | NA | NA | NA | NA | Btbd35f2 | BTB domain containing 35, family membe | 0 | 0 | 0 | 0 | 0 | 0 | 0 | 0 | 0 | 0 | 0 |
| 0 | NA | NA | NA | NA | NA | Btbd35f1 | BTB domain containing 35, family membe | 0 | 0 | 0 | 0 | 0 | 0 | 0 | 0 | 0 | 0 | 0 |

| | | | | | | | | | | | | | | | | | | |
|---|---|---|---|---|---|---|---|---|---|---|---|---|---|---|---|---|---|---|
| 0 | NA | NA | NA | NA | NA | Gm5926 | predicted gene 5926 | 0 | 0 | 0 | 0 | 0 | 0 | 0 | 0 | 0 | 0 | 0 |
| 0 | NA | NA | NA | NA | NA | Btbd35f21 | BTB domain containing 35, family membe | 0 | 0 | 0 | 0 | 0 | 0 | 0 | 0 | 0 | 0 | 0 |
| 0 | NA | NA | NA | NA | NA | Btbd35f30 | BTB domain containing 35, family membe | 0 | 0 | 0 | 0 | 0 | 0 | 0 | 0 | 0 | 0 | 0 |
| 0 | NA | NA | NA | NA | NA | Gm21789 | | 0 | 0 | 0 | 0 | 0 | 0 | 0 | 0 | 0 | 0 | 0 |
| 0 | NA | NA | NA | NA | NA | Btbd35f5 | BTB domain containing 35, family membe | 0 | 0 | 0 | 0 | 0 | 0 | 0 | 0 | 0 | 0 | 0 |
| 0 | NA | NA | NA | NA | NA | Spin2-ps6 | | 0 | 0 | 0 | 0 | 0 | 0 | 0 | 0 | 0 | 0 | 0 |
| 0 | NA | NA | NA | NA | NA | Gm15275 | | 0 | 0 | 0 | 0 | 0 | 0 | 0 | 0 | 0 | 0 | 0 |
| 0 | NA | NA | NA | NA | NA | Gm15274 | | 0 | 0 | 0 | 0 | 0 | 0 | 0 | 0 | 0 | 0 | 0 |
| 0 | NA | NA | NA | NA | NA | Btbd35f12 | BTB domain containing 35, family membe | 0 | 0 | 0 | 0 | 0 | 0 | 0 | 0 | 0 | 0 | 0 |
| 0 | NA | NA | NA | NA | NA | Gm2182 | | 0 | 0 | 0 | 0 | 0 | 0 | 0 | 0 | 0 | 0 | 0 |
| 0 | NA | NA | NA | NA | NA | Gm2854 | | 0 | 0 | 0 | 0 | 0 | 0 | 0 | 0 | 0 | 0 | 0 |
| 0 | NA | NA | NA | NA | NA | Gm21979 | | 0 | 0 | 0 | 0 | 0 | 0 | 0 | 0 | 0 | 0 | 0 |
| 0 | NA | NA | NA | NA | NA | Gm21990 | | 0 | 0 | 0 | 0 | 0 | 0 | 0 | 0 | 0 | 0 | 0 |
| 0 | NA | NA | NA | NA | NA | Gm2892 | | 0 | 0 | 0 | 0 | 0 | 0 | 0 | 0 | 0 | 0 | 0 |
| 0 | NA | NA | NA | NA | NA | Gm2231 | | 0 | 0 | 0 | 0 | 0 | 0 | 0 | 0 | 0 | 0 | 0 |
| 0 | NA | NA | NA | NA | NA | Btbd35f15 | BTB domain containing 35, family membe | 0 | 0 | 0 | 0 | 0 | 0 | 0 | 0 | 0 | 0 | 0 |
| 0 | NA | NA | NA | NA | NA | Gm15273 | | 0 | 0 | 0 | 0 | 0 | 0 | 0 | 0 | 0 | 0 | 0 |
| 0 | NA | NA | NA | NA | NA | Gm15281 | | 0 | 0 | 0 | 0 | 0 | 0 | 0 | 0 | 0 | 0 | 0 |
| 0 | NA | NA | NA | NA | NA | Gm15279 | | 0 | 0 | 0 | 0 | 0 | 0 | 0 | 0 | 0 | 0 | 0 |
| 0 | NA | NA | NA | NA | NA | Gm7023 | | 0 | 0 | 0 | 0 | 0 | 0 | 0 | 0 | 0 | 0 | 0 |
| 0 | NA | NA | NA | NA | NA | Btbd35f13 | BTB domain containing 35, family membe | 0 | 0 | 0 | 0 | 0 | 0 | 0 | 0 | 0 | 0 | 0 |
| 0 | NA | NA | NA | NA | NA | Btbd35f14 | BTB domain containing 35, family membe | 0 | 0 | 0 | 0 | 0 | 0 | 0 | 0 | 0 | 0 | 0 |
| 0 | NA | NA | NA | NA | NA | Btbd35f6 | BTB domain containing 35, family membe | 0 | 0 | 0 | 0 | 0 | 0 | 0 | 0 | 0 | 0 | 0 |
| 0 | NA | NA | NA | NA | NA | Gm21870 | | 0 | 0 | 0 | 0 | 0 | 0 | 0 | 0 | 0 | 0 | 0 |
| 0 | NA | NA | NA | NA | NA | Btbd35f9 | BTB domain containing 35, family membe | 0 | 0 | 0 | 0 | 0 | 0 | 0 | 0 | 0 | 0 | 0 |
| 0 | NA | NA | NA | NA | NA | Spin2g | spindlin family, member 2G | 0 | 0 | 0 | 0 | 0 | 0 | 0 | 0 | 0 | 0 | 0 |
| 0 | NA | NA | NA | NA | NA | Btbd35f8 | BTB domain containing 35, family membe | 0 | 0 | 0 | 0 | 0 | 0 | 0 | 0 | 0 | 0 | 0 |
| 0 | NA | NA | NA | NA | NA | Gm2288 | | 0 | 0 | 0 | 0 | 0 | 0 | 0 | 0 | 0 | 0 | 0 |
| 0 | NA | NA | NA | NA | NA | Gm14552 | | 0 | 0 | 0 | 0 | 0 | 0 | 0 | 0 | 0 | 0 | 0 |
| 0 | NA | NA | NA | NA | NA | Gm10486 | predicted gene 10486 | 0 | 0 | 0 | 0 | 0 | 0 | 0 | 0 | 0 | 0 | 0 |
| 0 | NA | NA | NA | NA | NA | Gm2309 | | 0 | 0 | 0 | 0 | 0 | 0 | 0 | 0 | 0 | 0 | 0 |
| 0 | NA | NA | NA | NA | NA | Gm2318 | | 0 | 0 | 0 | 0 | 0 | 0 | 0 | 0 | 0 | 0 | 0 |
| 0 | NA | NA | NA | NA | NA | Gm14553 | | 0 | 0 | 0 | 0 | 0 | 0 | 0 | 0 | 0 | 0 | 0 |
| 0 | NA | NA | NA | NA | NA | Gm14819 | predicted gene 14819 | 0 | 0 | 0 | 0 | 0 | 0 | 0 | 0 | 0 | 0 | 0 |
| 0 | NA | NA | NA | NA | NA | Gm14554 | | 0 | 0 | 0 | 0 | 0 | 0 | 0 | 0 | 0 | 0 | 0 |
| 0 | NA | NA | NA | NA | NA | Gm25624 | | 0 | 0 | 0 | 0 | 0 | 0 | 0 | 0 | 0 | 0 | 0 |
| 0 | NA | NA | NA | NA | NA | Gm14560 | | 0 | 0 | 0 | 0 | 0 | 0 | 0 | 0 | 0 | 0 | 0 |
| 0 | NA | NA | NA | NA | NA | Gm14559 | | 0 | 0 | 0 | 0 | 0 | 0 | 0 | 0 | 0 | 0 | 0 |
| 0 | NA | NA | NA | NA | NA | Gm14568 | | 0 | 0 | 0 | 0 | 0 | 0 | 0 | 0 | 0 | 0 | 0 |
| 0 | NA | NA | NA | NA | NA | Gm44485 | | 0 | 0 | 0 | 0 | 0 | 0 | 0 | 0 | 0 | 0 | 0 |
| 0 | NA | NA | NA | NA | NA | Gm6268 | | 0 | 0 | 0 | 0 | 0 | 0 | 0 | 0 | 0 | 0 | 0 |
| 0 | NA | NA | NA | NA | NA | Gm22364 | | 0 | 0 | 0 | 0 | 0 | 0 | 0 | 0 | 0 | 0 | 0 |
| 0 | NA | NA | NA | NA | NA | Gm14569 | predicted gene 14569 | 0 | 0 | 0 | 0 | 0 | 0 | 0 | 0 | 0 | 0 | 0 |
| 0 | NA | NA | NA | NA | NA | Gm23613 | | 0 | 0 | 0 | 0 | 0 | 0 | 0 | 0 | 0 | 0 | 0 |
| 0 | NA | NA | NA | NA | NA | Gm14549 | | 0 | 0 | 0 | 0 | 0 | 0 | 0 | 0 | 0 | 0 | 0 |
| 0 | NA | NA | NA | NA | NA | Gm14542 | | 0 | 0 | 0 | 0 | 0 | 0 | 0 | 0 | 0 | 0 | 0 |
| 0 | NA | NA | NA | NA | NA | Sowahd | sosondowah ankyrin repeat domain famil | 0 | 0 | 0 | 0 | 0 | 0 | 0 | 0 | 0 | 0 | 0 |
| 0 | NA | NA | NA | NA | NA | Gm26099 | | 0 | 0 | 0 | 0 | 0 | 0 | 0 | 0 | 0 | 0 | 0 |
| 0 | NA | NA | NA | NA | NA | Gm24809 | | 0 | 0 | 0 | 0 | 0 | 0 | 0 | 0 | 0 | 0 | 0 |
| 0 | NA | NA | NA | NA | NA | Gm9 | predicted gene 9 | 0 | 0 | 0 | 0 | 0 | 0 | 0 | 0 | 0 | 0 | 0 |
| 0 | NA | NA | NA | NA | NA | Rhox1 | reproductive homeobox 1 | 0 | 0 | 0 | 0 | 0 | 0 | 0 | 0 | 0 | 0 | 0 |
| 0 | NA | NA | NA | NA | NA | Rhox2a | reproductive homeobox 2A | 0 | 0 | 0 | 0 | 0 | 0 | 0 | 0 | 0 | 0 | 0 |
| 0 | NA | NA | NA | NA | NA | Rhox3a | reproductive homeobox 3A | 0 | 0 | 0 | 0 | 0 | 0 | 0 | 0 | 0 | 0 | 0 |
| 0 | NA | NA | NA | NA | NA | Rhox3a2 | reproductive homeobox 3A2 | 0 | 0 | 0 | 0 | 0 | 0 | 0 | 0 | 0 | 0 | 0 |
| 0 | NA | NA | NA | NA | NA | Rhox4a2 | reproductive homeobox 4A2 | 0 | 0 | 0 | 0 | 0 | 0 | 0 | 0 | 0 | 0 | 0 |
| 0 | NA | NA | NA | NA | NA | Rhox2b | reproductive homeobox 2B | 0 | 0 | 0 | 0 | 0 | 0 | 0 | 0 | 0 | 0 | 0 |
| 0 | NA | NA | NA | NA | NA | Rhox2c | reproductive homeobox 2C | 0 | 0 | 0 | 0 | 0 | 0 | 0 | 0 | 0 | 0 | 0 |
| 0 | NA | NA | NA | NA | NA | Rhox3c | reproductive homeobox 3C | 0 | 0 | 0 | 0 | 0 | 0 | 0 | 0 | 0 | 0 | 0 |
| 0 | NA | NA | NA | NA | NA | Rhox4c | reproductive homeobox 4C | 0 | 0 | 0 | 0 | 0 | 0 | 0 | 0 | 0 | 0 | 0 |
| 0 | NA | NA | NA | NA | NA | Rhox2d | reproductive homeobox 2D | 0 | 0 | 0 | 0 | 0 | 0 | 0 | 0 | 0 | 0 | 0 |

| | | | | | | | | | | | | | | | | | | |
|---|---|---|---|---|---|---|---|---|---|---|---|---|---|---|---|---|---|---|
| 0 | NA | NA | NA | NA | NA | Gm26494 | | 0 | 0 | 0 | 0 | 0 | 0 | 0 | 0 | 0 | 0 | 0 |
| 0 | NA | NA | NA | NA | NA | Rhox3d-ps | | 0 | 0 | 0 | 0 | 0 | 0 | 0 | 0 | 0 | 0 | 0 |
| 0 | NA | NA | NA | NA | NA | Rhox4d | reproductive homeobox 4D | 0 | 0 | 0 | 0 | 0 | 0 | 0 | 0 | 0 | 0 | 0 |
| 0 | NA | NA | NA | NA | NA | Rhox2e | reproductive homeobox 2E | 0 | 0 | 0 | 0 | 0 | 0 | 0 | 0 | 0 | 0 | 0 |
| 0 | NA | NA | NA | NA | NA | Rhox3e | reproductive homeobox 3E | 0 | 0 | 0 | 0 | 0 | 0 | 0 | 0 | 0 | 0 | 0 |
| 0 | NA | NA | NA | NA | NA | Rhox4e | reproductive homeobox 4E | 0 | 0 | 0 | 0 | 0 | 0 | 0 | 0 | 0 | 0 | 0 |
| 0 | NA | NA | NA | NA | NA | Rhox2f | reproductive homeobox 2F | 0 | 0 | 0 | 0 | 0 | 0 | 0 | 0 | 0 | 0 | 0 |
| 0 | NA | NA | NA | NA | NA | Rhox4f | reproductive homeobox 4F | 0 | 0 | 0 | 0 | 0 | 0 | 0 | 0 | 0 | 0 | 0 |
| 0 | NA | NA | NA | NA | NA | Gm24493 | | 0 | 0 | 0 | 0 | 0 | 0 | 0 | 0 | 0 | 0 | 0 |
| 0 | NA | NA | NA | NA | NA | Rhox2g | reproductive homeobox 2G | 0 | 0 | 0 | 0 | 0 | 0 | 0 | 0 | 0 | 0 | 0 |
| 0 | NA | NA | NA | NA | NA | Rhox4g | reproductive homeobox 4G | 0 | 0 | 0 | 0 | 0 | 0 | 0 | 0 | 0 | 0 | 0 |
| 0 | NA | NA | NA | NA | NA | Gm24147 | | 0 | 0 | 0 | 0 | 0 | 0 | 0 | 0 | 0 | 0 | 0 |
| 0 | NA | NA | NA | NA | NA | Rhox2h | reproductive homeobox 2H | 0 | 0 | 0 | 0 | 0 | 0 | 0 | 0 | 0 | 0 | 0 |
| 0 | NA | NA | NA | NA | NA | Rhox7-ps2 | | 0 | 0 | 0 | 0 | 0 | 0 | 0 | 0 | 0 | 0 | 0 |
| 0 | NA | NA | NA | NA | NA | Rhox2-ps | | 0 | 0 | 0 | 0 | 0 | 0 | 0 | 0 | 0 | 0 | 0 |
| 0 | NA | NA | NA | NA | NA | Rhox6 | reproductive homeobox 6 | 0 | 0 | 0 | 0 | 0 | 0 | 0 | 0 | 0 | 0 | 0 |
| 0 | NA | NA | NA | NA | NA | Rhox7a | reproductive homeobox 7A | 0 | 0 | 0 | 0 | 0 | 0 | 0 | 0 | 0 | 0 | 0 |
| 0 | NA | NA | NA | NA | NA | Rhox8 | reproductive homeobox 8 | 0 | 0 | 0 | 0 | 0 | 0 | 0 | 0 | 0 | 0 | 0 |
| 0 | NA | NA | NA | NA | NA | Rhox7b | | 0 | 0 | 0 | 0 | 0 | 0 | 0 | 0 | 0 | 0 | 0 |
| 0 | NA | NA | NA | NA | NA | Rhox9 | reproductive homeobox 9 | 0 | 0 | 0 | 0 | 0 | 0 | 0 | 0 | 0 | 0 | 0 |
| 0 | NA | NA | NA | NA | NA | Rhox11-ps1 | | 0 | 0 | 0 | 0 | 0 | 0 | 0 | 0 | 0 | 0 | 0 |
| 0 | NA | NA | NA | NA | NA | Rhox11-ps2 | | 0 | 0 | 0 | 0 | 0 | 0 | 0 | 0 | 0 | 0 | 0 |
| 0 | NA | NA | NA | NA | NA | Btg1b | BTG anti-proliferation factor 1B | 0 | 0 | 0 | 0 | 0 | 0 | 0 | 0 | 0 | 0 | 0 |
| 0 | NA | NA | NA | NA | NA | Btg1c | BTG anti-proliferation factor 1C | 0 | 0 | 0 | 0 | 0 | 0 | 0 | 0 | 0 | 0 | 0 |
| 0 | NA | NA | NA | NA | NA | Rhox10 | reproductive homeobox 10 | 0 | 0 | 0 | 0 | 0 | 0 | 0 | 0 | 0 | 0 | 0 |
| 0 | NA | NA | NA | NA | NA | Rhox11 | reproductive homeobox 11 | 0 | 0 | 0 | 0 | 0 | 0 | 0 | 0 | 0 | 0 | 0 |
| 0 | NA | NA | NA | NA | NA | Rhox11-ps3 | | 0 | 0 | 0 | 0 | 0 | 0 | 0 | 0 | 0 | 0 | 0 |
| 0 | NA | NA | NA | NA | NA | Rhox12 | reproductive homeobox 12 | 0 | 0 | 0 | 0 | 0 | 0 | 0 | 0 | 0 | 0 | 0 |
| 0 | NA | NA | NA | NA | NA | Rhox13 | reproductive homeobox 13 | 0 | 0 | 0 | 0 | 0 | 0 | 0 | 0 | 0 | 0 | 0 |
| 0 | NA | NA | NA | NA | NA | Mir3110 | | 0 | 0 | 0 | 0 | 0 | 0 | 0 | 0 | 0 | 0 | 0 |
| 0 | NA | NA | NA | NA | NA | Rhox7-ps1 | | 0 | 0 | 0 | 0 | 0 | 0 | 0 | 0 | 0 | 0 | 0 |
| 0 | NA | NA | NA | NA | NA | Gm40 | | 0 | 0 | 0 | 0 | 0 | 0 | 0 | 0 | 0 | 0 | 0 |
| 0 | NA | NA | NA | NA | NA | Gm25642 | | 0 | 0 | 0 | 0 | 0 | 0 | 0 | 0 | 0 | 0 | 0 |
| 0 | NA | NA | NA | NA | NA | Gm14564 | | 0 | 0 | 0 | 0 | 0 | 0 | 0 | 0 | 0 | 0 | 0 |
| 0 | NA | NA | NA | NA | NA | Gm16423 | | 0 | 0 | 0 | 0 | 0 | 0 | 0 | 0 | 0 | 0 | 0 |
| 0 | NA | NA | NA | NA | NA | Gm14551 | | 0 | 0 | 0 | 0 | 0 | 0 | 0 | 0 | 0 | 0 | 0 |
| 0 | NA | NA | NA | NA | NA | Gm14566 | | 0 | 0 | 0 | 0 | 0 | 0 | 0 | 0 | 0 | 0 | 0 |
| 0 | NA | NA | NA | NA | NA | Gm26437 | | 0 | 0 | 0 | 0 | 0 | 0 | 0 | 0 | 0 | 0 | 0 |
| 0 | NA | NA | NA | NA | NA | Cypt15 | cysteine-rich perinuclear theca 15 | 0 | 0 | 0 | 0 | 0 | 0 | 0 | 0 | 0 | 0 | 0 |
| 0 | NA | NA | NA | NA | NA | Gm14550 | | 0 | 0 | 0 | 0 | 0 | 0 | 0 | 0 | 0 | 0 | 0 |
| 0 | NA | NA | NA | NA | NA | Gm14570 | | 0 | 0 | 0 | 0 | 0 | 0 | 0 | 0 | 0 | 0 | 0 |
| 0 | NA | NA | NA | NA | NA | Gm14571 | | 0 | 0 | 0 | 0 | 0 | 0 | 0 | 0 | 0 | 0 | 0 |
| 0 | NA | NA | NA | NA | NA | Cypt14 | cysteine-rich perinuclear theca 14 | 0 | 0 | 0 | 0 | 0 | 0 | 0 | 0 | 0 | 0 | 0 |
| 0 | NA | NA | NA | NA | NA | Gm26194 | | 0 | 0 | 0 | 0 | 0 | 0 | 0 | 0 | 0 | 0 | 0 |
| 0 | NA | NA | NA | NA | NA | Gm16383 | | 0 | 0 | 0 | 0 | 0 | 0 | 0 | 0 | 0 | 0 | 0 |
| 0 | NA | NA | NA | NA | NA | Gm14575 | | 0 | 0 | 0 | 0 | 0 | 0 | 0 | 0 | 0 | 0 | 0 |
| 0 | NA | NA | NA | NA | NA | Gm14574 | | 0 | 0 | 0 | 0 | 0 | 0 | 0 | 0 | 0 | 0 | 0 |
| 0 | NA | NA | NA | NA | NA | Gm14577 | | 0 | 0 | 0 | 0 | 0 | 0 | 0 | 0 | 0 | 0 | 0 |
| 0 | NA | NA | NA | NA | NA | Gm22413 | | 0 | 0 | 0 | 0 | 0 | 0 | 0 | 0 | 0 | 0 | 0 |
| 0 | NA | NA | NA | NA | NA | Gm14614 | | 0 | 0 | 0 | 0 | 0 | 0 | 0 | 0 | 0 | 0 | 0 |
| 0 | NA | NA | NA | NA | NA | Gm14613 | | 0 | 0 | 0 | 0 | 0 | 0 | 0 | 0 | 0 | 0 | 0 |
| 0 | NA | NA | NA | NA | NA | Gm7657 | | 0 | 0 | 0 | 0 | 0 | 0 | 0 | 0 | 0 | 0 | 0 |
| 0 | NA | NA | NA | NA | NA | Gm5135 | | 0 | 0 | 0 | 0 | 0 | 0 | 0 | 0 | 0 | 0 | 0 |
| 0 | NA | NA | NA | NA | NA | Gm44490 | | 0 | 0 | 0 | 0 | 0 | 0 | 0 | 0 | 0 | 0 | 0 |
| 0 | NA | NA | NA | NA | NA | Gm14615 | | 0 | 0 | 0 | 0 | 0 | 0 | 0 | 0 | 0 | 0 | 0 |
| 0 | NA | NA | NA | NA | NA | Gm37564 | | 0 | 0 | 0 | 0 | 0 | 0 | 0 | 0 | 0 | 0 | 0 |
| 0 | NA | NA | NA | NA | NA | Gm14619 | | 0 | 0 | 0 | 0 | 0 | 0 | 0 | 0 | 0 | 0 | 0 |
| 0 | NA | NA | NA | NA | NA | Tex13c2 | TEX13 family member D | 0 | 0 | 0 | 0 | 0 | 0 | 0 | 0 | 0 | 0 | 0 |
| 0 | NA | NA | NA | NA | NA | Sh2d1a | SH2 domain containing 1A | 0 | 0 | 0 | 0 | 0 | 0 | 0 | 0 | 0 | 0 | 0 |
| 0 | NA | NA | NA | NA | NA | Gm23705 | | 0 | 0 | 0 | 0 | 0 | 0 | 0 | 0 | 0 | 0 | 0 |

| | | | | | | | | | | | | | | | | | | |
|---|---|---|---|---|---|---|---|---|---|---|---|---|---|---|---|---|---|---|
| 0 | NA | NA | NA | NA | NA | Gm14578 | | 0 | 0 | 0 | 0 | 0 | 0 | 0 | 0 | 0 | 0 | 0 |
| 0 | NA | NA | NA | NA | NA | Gm24763 | | 0 | 0 | 0 | 0 | 0 | 0 | 0 | 0 | 0 | 0 | 0 |
| 0 | NA | NA | NA | NA | NA | Gm14587 | | 0 | 0 | 0 | 0 | 0 | 0 | 0 | 0 | 0 | 0 | 0 |
| 0 | NA | NA | NA | NA | NA | Gm14588 | | 0 | 0 | 0 | 0 | 0 | 0 | 0 | 0 | 0 | 0 | 0 |
| 0 | NA | NA | NA | NA | NA | Gm14579 | | 0 | 0 | 0 | 0 | 0 | 0 | 0 | 0 | 0 | 0 | 0 |
| 0 | NA | NA | NA | NA | NA | Tex13c1 | TEX13 family member C1 | 0 | 0 | 0 | 0 | 0 | 0 | 0 | 0 | 0 | 0 | 0 |
| 0 | NA | NA | NA | NA | NA | Gm14580 | | 0 | 0 | 0 | 0 | 0 | 0 | 0 | 0 | 0 | 0 | 0 |
| 0 | NA | NA | NA | NA | NA | Gm14628 | | 0 | 0 | 0 | 0 | 0 | 0 | 0 | 0 | 0 | 0 | 0 |
| 0 | NA | NA | NA | NA | NA | Gm14603 | | 0 | 0 | 0 | 0 | 0 | 0 | 0 | 0 | 0 | 0 | 0 |
| 0 | NA | NA | NA | NA | NA | Itpa-ps3 | | 0 | 0 | 0 | 0 | 0 | 0 | 0 | 0 | 0 | 0 | 0 |
| 0 | NA | NA | NA | NA | NA | Gm14602 | | 0 | 0 | 0 | 0 | 0 | 0 | 0 | 0 | 0 | 0 | 0 |
| 0 | NA | NA | NA | NA | NA | Gm7700 | | 0 | 0 | 0 | 0 | 0 | 0 | 0 | 0 | 0 | 0 | 0 |
| 0 | NA | NA | NA | NA | NA | Prr32 | proline rich 32 | 0 | 0 | 0 | 0 | 0 | 0 | 0 | 0 | 0 | 0 | 0 |
| 0 | NA | NA | NA | NA | NA | Gm14606 | | 0 | 0 | 0 | 0 | 0 | 0 | 0 | 0 | 0 | 0 | 0 |
| 0 | NA | NA | NA | NA | NA | Gm14605 | | 0 | 0 | 0 | 0 | 0 | 0 | 0 | 0 | 0 | 0 | 0 |
| 0 | NA | NA | NA | NA | NA | Gm5386 | | 0 | 0 | 0 | 0 | 0 | 0 | 0 | 0 | 0 | 0 | 0 |
| 0 | NA | NA | NA | NA | NA | Gm25078 | | 0 | 0 | 0 | 0 | 0 | 0 | 0 | 0 | 0 | 0 | 0 |
| 0 | NA | NA | NA | NA | NA | Gm14653 | | 0 | 0 | 0 | 0 | 0 | 0 | 0 | 0 | 0 | 0 | 0 |
| 0 | NA | NA | NA | NA | NA | 4930515L19Rik | | 0 | 0 | 0 | 0 | 0 | 0 | 0 | 0 | 0 | 0 | 0 |
| 0 | NA | NA | NA | NA | NA | Actrt1 | actin-related protein T1 | 0 | 0 | 0 | 0 | 0 | 0 | 0 | 0 | 0 | 0 | 0 |
| 0 | NA | NA | NA | NA | NA | Gm14654 | | 0 | 0 | 0 | 0 | 0 | 0 | 0 | 0 | 0 | 0 | 0 |
| 0 | NA | NA | NA | NA | NA | Gm29242 | | 0 | 0 | 0 | 0 | 0 | 0 | 0 | 0 | 0 | 0 | 0 |
| 0 | NA | NA | NA | NA | NA | Gm14609 | | 0 | 0 | 0 | 0 | 0 | 0 | 0 | 0 | 0 | 0 | 0 |
| 0 | NA | NA | NA | NA | NA | Gm14610 | | 0 | 0 | 0 | 0 | 0 | 0 | 0 | 0 | 0 | 0 | 0 |
| 0 | NA | NA | NA | NA | NA | Gm4908 | | 0 | 0 | 0 | 0 | 0 | 0 | 0 | 0 | 0 | 0 | 0 |
| 0 | NA | NA | NA | NA | NA | Gm14608 | | 0 | 0 | 0 | 0 | 0 | 0 | 0 | 0 | 0 | 0 | 0 |
| 0 | NA | NA | NA | NA | NA | Gm6480 | | 0 | 0 | 0 | 0 | 0 | 0 | 0 | 0 | 0 | 0 | 0 |
| 0 | NA | NA | NA | NA | NA | Gm24038 | | 0 | 0 | 0 | 0 | 0 | 0 | 0 | 0 | 0 | 0 | 0 |
| 0 | NA | NA | NA | NA | NA | Gm22528 | | 0 | 0 | 0 | 0 | 0 | 0 | 0 | 0 | 0 | 0 | 0 |
| 0 | NA | NA | NA | NA | NA | Gm7212 | | 0 | 0 | 0 | 0 | 0 | 0 | 0 | 0 | 0 | 0 | 0 |
| 0 | NA | NA | NA | NA | NA | Gm22612 | | 0 | 0 | 0 | 0 | 0 | 0 | 0 | 0 | 0 | 0 | 0 |
| 0 | NA | NA | NA | NA | NA | Gm27828 | | 0 | 0 | 0 | 0 | 0 | 0 | 0 | 0 | 0 | 0 | 0 |
| 0 | NA | NA | NA | NA | NA | Gm7219 | | 0 | 0 | 0 | 0 | 0 | 0 | 0 | 0 | 0 | 0 | 0 |
| 0 | NA | NA | NA | NA | NA | Gm15034 | | 0 | 0 | 0 | 0 | 0 | 0 | 0 | 0 | 0 | 0 | 0 |
| 0 | NA | NA | NA | NA | NA | Gm15036 | | 0 | 0 | 0 | 0 | 0 | 0 | 0 | 0 | 0 | 0 | 0 |
| 0 | NA | NA | NA | NA | NA | Gm14696 | | 0 | 0 | 0 | 0 | 0 | 0 | 0 | 0 | 0 | 0 | 0 |
| 0 | NA | NA | NA | NA | NA | Gm15037 | | 0 | 0 | 0 | 0 | 0 | 0 | 0 | 0 | 0 | 0 | 0 |
| 0 | NA | NA | NA | NA | NA | Olfr1320 | olfactory receptor 1320 | 0 | 0 | 0 | 0 | 0 | 0 | 0 | 0 | 0 | 0 | 0 |
| 0 | NA | NA | NA | NA | NA | Olfr1321 | olfactory receptor 1321 | 0 | 0 | 0 | 0 | 0 | 0 | 0 | 0 | 0 | 0 | 0 |
| 0 | NA | NA | NA | NA | NA | n-R5s6 | | 0 | 0 | 0 | 0 | 0 | 0 | 0 | 0 | 0 | 0 | 0 |
| 0 | NA | NA | NA | NA | NA | Olfr1323 | olfactory receptor 1323 | 0 | 0 | 0 | 0 | 0 | 0 | 0 | 0 | 0 | 0 | 0 |
| 0 | NA | NA | NA | NA | NA | Gm23385 | | 0 | 0 | 0 | 0 | 0 | 0 | 0 | 0 | 0 | 0 | 0 |
| 0 | NA | NA | NA | NA | NA | | | 0 | 0 | 0 | 0 | 0 | 0 | 0 | 0 | 0 | 0 | 0 |
| 0 | NA | NA | NA | NA | NA | Gm22288 | | 0 | 0 | 0 | 0 | 0 | 0 | 0 | 0 | 0 | 0 | 0 |
| 0 | NA | NA | NA | NA | NA | Olfr1324 | olfactory receptor 1324 | 0 | 0 | 0 | 0 | 0 | 0 | 0 | 0 | 0 | 0 | 0 |
| 0 | NA | NA | NA | NA | NA | Gm5387 | | 0 | 0 | 0 | 0 | 0 | 0 | 0 | 0 | 0 | 0 | 0 |
| 0 | NA | NA | NA | NA | NA | Gm7834 | | 0 | 0 | 0 | 0 | 0 | 0 | 0 | 0 | 0 | 0 | 0 |
| 0 | NA | NA | NA | NA | NA | Gm23417 | | 0 | 0 | 0 | 0 | 0 | 0 | 0 | 0 | 0 | 0 | 0 |
| 0 | NA | NA | NA | NA | NA | Gm14621 | | 0 | 0 | 0 | 0 | 0 | 0 | 0 | 0 | 0 | 0 | 0 |
| 0 | NA | NA | NA | NA | NA | Gm14581 | | 0 | 0 | 0 | 0 | 0 | 0 | 0 | 0 | 0 | 0 | 0 |
| 0 | NA | NA | NA | NA | NA | Usp26 | ubiquitin specific peptidase 26 | 0 | 0 | 0 | 0 | 0 | 0 | 0 | 0 | 0 | 0 | 0 |
| 0 | NA | NA | NA | NA | NA | Gm7840 | | 0 | 0 | 0 | 0 | 0 | 0 | 0 | 0 | 0 | 0 | 0 |
| 0 | NA | NA | NA | NA | NA | Gm22539 | | 0 | 0 | 0 | 0 | 0 | 0 | 0 | 0 | 0 | 0 | 0 |
| 0 | NA | NA | NA | NA | NA | 1700080O16Ril | RIKEN cDNA 1700080O16 gene | 0 | 0 | 0 | 0 | 0 | 0 | 0 | 0 | 0 | 0 | 0 |
| 0 | NA | NA | NA | NA | NA | Gm14582 | | 0 | 0 | 0 | 0 | 0 | 0 | 0 | 0 | 0 | 0 | 0 |
| 0 | NA | NA | NA | NA | NA | Mir6384 | | 0 | 0 | 0 | 0 | 0 | 0 | 0 | 0 | 0 | 0 | 0 |
| 0 | NA | NA | NA | NA | NA | Mir717 | | 0 | 0 | 0 | 0 | 0 | 0 | 0 | 0 | 0 | 0 | 0 |
| 0 | NA | NA | NA | NA | NA | Mir363 | | 0 | 0 | 0 | 0 | 0 | 0 | 0 | 0 | 0 | 0 | 0 |
| 0 | NA | NA | NA | NA | NA | Mir92-2 | | 0 | 0 | 0 | 0 | 0 | 0 | 0 | 0 | 0 | 0 | 0 |
| 0 | NA | NA | NA | NA | NA | Mir19b-2 | | 0 | 0 | 0 | 0 | 0 | 0 | 0 | 0 | 0 | 0 | 0 |

| | | | | | | | | | | | | | | | | | | |
|---|---|---|---|---|---|---|---|---|---|---|---|---|---|---|---|---|---|---|
| 0 | NA | NA | NA | NA | NA | Mir20b | | 0 | 0 | 0 | 0 | 0 | 0 | 0 | 0 | 0 | 0 | 0 |
| 0 | NA | NA | NA | NA | NA | Mir18b | | 0 | 0 | 0 | 0 | 0 | 0 | 0 | 0 | 0 | 0 | 0 |
| 0 | NA | NA | NA | NA | NA | Mir106a | | 0 | 0 | 0 | 0 | 0 | 0 | 0 | 0 | 0 | 0 | 0 |
| 0 | NA | NA | NA | NA | NA | Kis2 | | 0 | 0 | 0 | 0 | 0 | 0 | 0 | 0 | 0 | 0 | 0 |
| 0 | NA | NA | NA | NA | NA | Gm7851 | | 0 | 0 | 0 | 0 | 0 | 0 | 0 | 0 | 0 | 0 | 0 |
| 0 | NA | NA | NA | NA | NA | Gm14607 | | 0 | 0 | 0 | 0 | 0 | 0 | 0 | 0 | 0 | 0 | 0 |
| 0 | NA | NA | NA | NA | NA | Gm6539 | | 0 | 0 | 0 | 0 | 0 | 0 | 0 | 0 | 0 | 0 | 0 |
| 0 | NA | NA | NA | NA | NA | Mir450b | | 0 | 0 | 0 | 0 | 0 | 0 | 0 | 0 | 0 | 0 | 0 |
| 0 | NA | NA | NA | NA | NA | Mir450-1 | | 0 | 0 | 0 | 0 | 0 | 0 | 0 | 0 | 0 | 0 | 0 |
| 0 | NA | NA | NA | NA | NA | Mir450-2 | | 0 | 0 | 0 | 0 | 0 | 0 | 0 | 0 | 0 | 0 | 0 |
| 0 | NA | NA | NA | NA | NA | Mir542 | | 0 | 0 | 0 | 0 | 0 | 0 | 0 | 0 | 0 | 0 | 0 |
| 0 | NA | NA | NA | NA | NA | Gm28730 | | 0 | 0 | 0 | 0 | 0 | 0 | 0 | 0 | 0 | 0 | 0 |
| 0 | NA | NA | NA | NA | NA | Mir351 | | 0 | 0 | 0 | 0 | 0 | 0 | 0 | 0 | 0 | 0 | 0 |
| 0 | NA | NA | NA | NA | NA | Mir503 | | 0 | 0 | 0 | 0 | 0 | 0 | 0 | 0 | 0 | 0 | 0 |
| 0 | NA | NA | NA | NA | NA | Mir322 | | 0 | 0 | 0 | 0 | 0 | 0 | 0 | 0 | 0 | 0 | 0 |
| 0 | NA | NA | NA | NA | NA | Plac1 | placental specific protein 1 | 0 | 0 | 0 | 0 | 0 | 0 | 0 | 0 | 0 | 0 | 0 |
| 0 | NA | NA | NA | NA | NA | Fam122c | family with sequence similarity 122, meml | 0 | 0 | 0 | 0 | 0 | 0 | 0 | 0 | 0 | 0 | 0 |
| 0 | NA | NA | NA | NA | NA | Gm23718 | | 0 | 0 | 0 | 0 | 0 | 0 | 0 | 0 | 0 | 0 | 0 |
| 0 | NA | NA | NA | NA | NA | Gm14585 | | 0 | 0 | 0 | 0 | 0 | 0 | 0 | 0 | 0 | 0 | 0 |
| 0 | NA | NA | NA | NA | NA | Etd | embryonic testis differentiation | 0 | 0 | 0 | 0 | 0 | 0 | 0 | 0 | 0 | 0 | 0 |
| 0 | NA | NA | NA | NA | NA | Gm14597 | RIKEN cDNA 4933416I08 gene | 0 | 0 | 0 | 0 | 0 | 0 | 0 | 0 | 0 | 0 | 0 |
| 0 | NA | NA | NA | NA | NA | Gm14592 | | 0 | 0 | 0 | 0 | 0 | 0 | 0 | 0 | 0 | 0 | 0 |
| 0 | NA | NA | NA | NA | NA | Gm14600 | | 0 | 0 | 0 | 0 | 0 | 0 | 0 | 0 | 0 | 0 | 0 |
| 0 | NA | NA | NA | NA | NA | 4933416I08Rik | | 0 | 0 | 0 | 0 | 0 | 0 | 0 | 0 | 0 | 0 | 0 |
| 0 | NA | NA | NA | NA | NA | 1700013H16Ril | RIKEN cDNA 1700013H16 gene | 0 | 0 | 0 | 0 | 0 | 0 | 0 | 0 | 0 | 0 | 0 |
| 0 | NA | NA | NA | NA | NA | Gm6025 | | 0 | 0 | 0 | 0 | 0 | 0 | 0 | 0 | 0 | 0 | 0 |
| 0 | NA | NA | NA | NA | NA | Gm14593 | | 0 | 0 | 0 | 0 | 0 | 0 | 0 | 0 | 0 | 0 | 0 |
| 0 | NA | NA | NA | NA | NA | Zfp36l3 | zinc finger protein 36, C3H type-like 3 | 0 | 0 | 0 | 0 | 0 | 0 | 0 | 0 | 0 | 0 | 0 |
| 0 | NA | NA | NA | NA | NA | Gm14599 | | 0 | 0 | 0 | 0 | 0 | 0 | 0 | 0 | 0 | 0 | 0 |
| 0 | NA | NA | NA | NA | NA | Gm14589 | | 0 | 0 | 0 | 0 | 0 | 0 | 0 | 0 | 0 | 0 | 0 |
| 0 | NA | NA | NA | NA | NA | Gm15482 | | 0 | 0 | 0 | 0 | 0 | 0 | 0 | 0 | 0 | 0 | 0 |
| 0 | NA | NA | NA | NA | NA | Gm14601 | | 0 | 0 | 0 | 0 | 0 | 0 | 0 | 0 | 0 | 0 | 0 |
| 0 | NA | NA | NA | NA | NA | Gm14644 | | 0 | 0 | 0 | 0 | 0 | 0 | 0 | 0 | 0 | 0 | 0 |
| 0 | NA | NA | NA | NA | NA | Gm22550 | | 0 | 0 | 0 | 0 | 0 | 0 | 0 | 0 | 0 | 0 | 0 |
| 0 | NA | NA | NA | NA | NA | Gm16428 | | 0 | 0 | 0 | 0 | 0 | 0 | 0 | 0 | 0 | 0 | 0 |
| 0 | NA | NA | NA | NA | NA | Gm7916 | | 0 | 0 | 0 | 0 | 0 | 0 | 0 | 0 | 0 | 0 | 0 |
| 0 | NA | NA | NA | NA | NA | Gm14598 | | 0 | 0 | 0 | 0 | 0 | 0 | 0 | 0 | 0 | 0 | 0 |
| 0 | NA | NA | NA | NA | NA | Gm6591 | | 0 | 0 | 0 | 0 | 0 | 0 | 0 | 0 | 0 | 0 | 0 |
| 0 | NA | NA | NA | NA | NA | Gm14596 | predicted gene 14596 | 0 | 0 | 0 | 0 | 0 | 0 | 0 | 0 | 0 | 0 | 0 |
| 0 | NA | NA | NA | NA | NA | Gm14594 | predicted gene 14594 | 0 | 0 | 0 | 0 | 0 | 0 | 0 | 0 | 0 | 0 | 0 |
| 0 | NA | NA | NA | NA | NA | Gm16405 | predicted gene 14625 | 0 | 0 | 0 | 0 | 0 | 0 | 0 | 0 | 0 | 0 | 0 |
| 0 | NA | NA | NA | NA | NA | Gm16430 | predicted gene 14625 | 0 | 0 | 0 | 0 | 0 | 0 | 0 | 0 | 0 | 0 | 0 |
| 0 | NA | NA | NA | NA | NA | Gm14595 | predicted gene 14595 | 0 | 0 | 0 | 0 | 0 | 0 | 0 | 0 | 0 | 0 | 0 |
| 0 | NA | NA | NA | NA | NA | Gm14590 | predicted gene 14590 | 0 | 0 | 0 | 0 | 0 | 0 | 0 | 0 | 0 | 0 | 0 |
| 0 | NA | NA | NA | NA | NA | Gm16431 | | 0 | 0 | 0 | 0 | 0 | 0 | 0 | 0 | 0 | 0 | 0 |
| 0 | NA | NA | NA | NA | NA | Gm7950 | | 0 | 0 | 0 | 0 | 0 | 0 | 0 | 0 | 0 | 0 | 0 |
| 0 | NA | NA | NA | NA | NA | Gm7958 | | 0 | 0 | 0 | 0 | 0 | 0 | 0 | 0 | 0 | 0 | 0 |
| 0 | NA | NA | NA | NA | NA | Gm7974 | | 0 | 0 | 0 | 0 | 0 | 0 | 0 | 0 | 0 | 0 | 0 |
| 0 | NA | NA | NA | NA | NA | Slxl1 | Slx-like 1 | 0 | 0 | 0 | 0 | 0 | 0 | 0 | 0 | 0 | 0 | 0 |
| 0 | NA | NA | NA | NA | NA | Gm6660 | | 0 | 0 | 0 | 0 | 0 | 0 | 0 | 0 | 0 | 0 | 0 |
| 0 | NA | NA | NA | NA | NA | Gm14625 | predicted gene 14625 | 0 | 0 | 0 | 0 | 0 | 0 | 0 | 0 | 0 | 0 | 0 |
| 0 | NA | NA | NA | NA | NA | Gm16404 | predicted gene 16404 | 0 | 0 | 0 | 0 | 0 | 0 | 0 | 0 | 0 | 0 | 0 |
| 0 | NA | NA | NA | NA | NA | Gm14622 | | 0 | 0 | 0 | 0 | 0 | 0 | 0 | 0 | 0 | 0 | 0 |
| 0 | NA | NA | NA | NA | NA | Gm6664 | predicted gene 6664 | 0 | 0 | 0 | 0 | 0 | 0 | 0 | 0 | 0 | 0 | 0 |
| 0 | NA | NA | NA | NA | NA | Gm14626 | | 0 | 0 | 0 | 0 | 0 | 0 | 0 | 0 | 0 | 0 | 0 |
| 0 | NA | NA | NA | NA | NA | Gm14623 | | 0 | 0 | 0 | 0 | 0 | 0 | 0 | 0 | 0 | 0 | 0 |
| 0 | NA | NA | NA | NA | NA | Gm14624 | | 0 | 0 | 0 | 0 | 0 | 0 | 0 | 0 | 0 | 0 | 0 |
| 0 | NA | NA | NA | NA | NA | | | 0 | 0 | 0 | 0 | 0 | 0 | 0 | 0 | 0 | 0 | 0 |
| 0 | NA | NA | NA | NA | NA | Gm22103 | | 0 | 0 | 0 | 0 | 0 | 0 | 0 | 0 | 0 | 0 | 0 |
| 0 | NA | NA | NA | NA | NA | Gm14638 | | 0 | 0 | 0 | 0 | 0 | 0 | 0 | 0 | 0 | 0 | 0 |

| | | | | | | | | | | | | | | | | | | |
|---|---|---|---|---|---|---|---|---|---|---|---|---|---|---|---|---|---|---|
| 0 | NA | NA | NA | NA | NA | Gm773 | predicted gene 773 | 0 | 0 | 0 | 0 | 0 | 0 | 0 | 0 | 0 | 0 | 0 |
| 0 | NA | NA | NA | NA | NA | Gm14627 | | 0 | 0 | 0 | 0 | 0 | 0 | 0 | 0 | 0 | 0 | 0 |
| 0 | NA | NA | NA | NA | NA | 1600025M17Rik | | 0 | 0 | 0 | 0 | 0 | 0 | 0 | 0 | 0 | 0 | 0 |
| 0 | NA | NA | NA | NA | NA | Gm10477 | | 0 | 0 | 0 | 0 | 0 | 0 | 0 | 0 | 0 | 0 | 0 |
| 0 | NA | NA | NA | NA | NA | Mir5116 | | 0 | 0 | 0 | 0 | 0 | 0 | 0 | 0 | 0 | 0 | 0 |
| 0 | NA | NA | NA | NA | NA | Gm26312 | | 0 | 0 | 0 | 0 | 0 | 0 | 0 | 0 | 0 | 0 | 0 |
| 0 | NA | NA | NA | NA | NA | Gm23258 | | 0 | 0 | 0 | 0 | 0 | 0 | 0 | 0 | 0 | 0 | 0 |
| 0 | NA | NA | NA | NA | NA | Vgll1 | vestigial like family member 1 | 0 | 0 | 0 | 0 | 0 | 0 | 0 | 0 | 0 | 0 | 0 |
| 0 | NA | NA | NA | NA | NA | Gm14718 | | 0 | 0 | 0 | 0 | 0 | 0 | 0 | 0 | 0 | 0 | 0 |
| 0 | NA | NA | NA | NA | NA | Snord61 | | 0 | 0 | 0 | 0 | 0 | 0 | 0 | 0 | 0 | 0 | 0 |
| 0 | NA | NA | NA | NA | NA | Gm364 | predicted gene 364 | 0 | 0 | 0 | 0 | 0 | 0 | 0 | 0 | 0 | 0 | 0 |
| 0 | NA | NA | NA | NA | NA | Gpr101 | G protein-coupled receptor 101 | 0 | 0 | 0 | 0 | 0 | 0 | 0 | 0 | 0 | 0 | 0 |
| 0 | NA | NA | NA | NA | NA | Gm14629 | | 0 | 0 | 0 | 0 | 0 | 0 | 0 | 0 | 0 | 0 | 0 |
| 0 | NA | NA | NA | NA | NA | Gm8107 | | 0 | 0 | 0 | 0 | 0 | 0 | 0 | 0 | 0 | 0 | 0 |
| 0 | NA | NA | NA | NA | NA | Gm8111 | | 0 | 0 | 0 | 0 | 0 | 0 | 0 | 0 | 0 | 0 | 0 |
| 0 | NA | NA | NA | NA | NA | Gm14631 | | 0 | 0 | 0 | 0 | 0 | 0 | 0 | 0 | 0 | 0 | 0 |
| 0 | NA | NA | NA | NA | NA | Gm25023 | | 0 | 0 | 0 | 0 | 0 | 0 | 0 | 0 | 0 | 0 | 0 |
| 0 | NA | NA | NA | NA | NA | Zic3 | zinc finger protein of the cerebellum 3 | 0 | 0 | 0 | 0 | 0 | 0 | 0 | 0 | 0 | 0 | 0 |
| 0 | NA | NA | NA | NA | NA | Gm14649 | | 0 | 0 | 0 | 0 | 0 | 0 | 0 | 0 | 0 | 0 | 0 |
| 0 | NA | NA | NA | NA | NA | Gm14647 | | 0 | 0 | 0 | 0 | 0 | 0 | 0 | 0 | 0 | 0 | 0 |
| 0 | NA | NA | NA | NA | NA | Gm25851 | | 0 | 0 | 0 | 0 | 0 | 0 | 0 | 0 | 0 | 0 | 0 |
| 0 | NA | NA | NA | NA | NA | Gm14645 | | 0 | 0 | 0 | 0 | 0 | 0 | 0 | 0 | 0 | 0 | 0 |
| 0 | NA | NA | NA | NA | NA | Gm7071 | | 0 | 0 | 0 | 0 | 0 | 0 | 0 | 0 | 0 | 0 | 0 |
| 0 | NA | NA | NA | NA | NA | Gm44333 | | 0 | 0 | 0 | 0 | 0 | 0 | 0 | 0 | 0 | 0 | 0 |
| 0 | NA | NA | NA | NA | NA | 4930550L24Rik | RIKEN cDNA 4930550L24 gene | 0 | 0 | 0 | 0 | 0 | 0 | 0 | 0 | 0 | 0 | 0 |
| 0 | NA | NA | NA | NA | NA | Gm26351 | | 0 | 0 | 0 | 0 | 0 | 0 | 0 | 0 | 0 | 0 | 0 |
| 0 | NA | NA | NA | NA | NA | Mir504 | | 0 | 0 | 0 | 0 | 0 | 0 | 0 | 0 | 0 | 0 | 0 |
| 0 | NA | NA | NA | NA | NA | Gm26487 | | 0 | 0 | 0 | 0 | 0 | 0 | 0 | 0 | 0 | 0 | 0 |
| 0 | NA | NA | NA | NA | NA | Gm44500 | | 0 | 0 | 0 | 0 | 0 | 0 | 0 | 0 | 0 | 0 | 0 |
| 0 | NA | NA | NA | NA | NA | Gm44318 | | 0 | 0 | 0 | 0 | 0 | 0 | 0 | 0 | 0 | 0 | 0 |
| 0 | NA | NA | NA | NA | NA | Gm14887 | | 0 | 0 | 0 | 0 | 0 | 0 | 0 | 0 | 0 | 0 | 0 |
| 0 | NA | NA | NA | NA | NA | Gm14891 | | 0 | 0 | 0 | 0 | 0 | 0 | 0 | 0 | 0 | 0 | 0 |
| 0 | NA | NA | NA | NA | NA | Gm22287 | | 0 | 0 | 0 | 0 | 0 | 0 | 0 | 0 | 0 | 0 | 0 |
| 0 | NA | NA | NA | NA | NA | Mir505 | | 0 | 0 | 0 | 0 | 0 | 0 | 0 | 0 | 0 | 0 | 0 |
| 0 | NA | NA | NA | NA | NA | Gm7073 | predicted gene 7073 | 0 | 0 | 0 | 0 | 0 | 0 | 0 | 0 | 0 | 0 | 0 |
| 0 | NA | NA | NA | NA | NA | Gm14661 | | 0 | 0 | 0 | 0 | 0 | 0 | 0 | 0 | 0 | 0 | 0 |
| 0 | NA | NA | NA | NA | NA | Gm14651 | | 0 | 0 | 0 | 0 | 0 | 0 | 0 | 0 | 0 | 0 | 0 |
| 0 | NA | NA | NA | NA | NA | Gm4909 | | 0 | 0 | 0 | 0 | 0 | 0 | 0 | 0 | 0 | 0 | 0 |
| 0 | NA | NA | NA | NA | NA | Gm14660 | | 0 | 0 | 0 | 0 | 0 | 0 | 0 | 0 | 0 | 0 | 0 |
| 0 | NA | NA | NA | NA | NA | Sox3 | SRY (sex determining region Y)-box 3 | 0 | 0 | 0 | 0 | 0 | 0 | 0 | 0 | 0 | 0 | 0 |
| 0 | NA | NA | NA | NA | NA | Gm14662 | | 0 | 0 | 0 | 0 | 0 | 0 | 0 | 0 | 0 | 0 | 0 |
| 0 | NA | NA | NA | NA | NA | Rpl7l1-ps1 | | 0 | 0 | 0 | 0 | 0 | 0 | 0 | 0 | 0 | 0 | 0 |
| 0 | NA | NA | NA | NA | NA | Gm14664 | | 0 | 0 | 0 | 0 | 0 | 0 | 0 | 0 | 0 | 0 | 0 |
| 0 | NA | NA | NA | NA | NA | Gm14663 | | 0 | 0 | 0 | 0 | 0 | 0 | 0 | 0 | 0 | 0 | 0 |
| 0 | NA | NA | NA | NA | NA | | | 0 | 0 | 0 | 0 | 0 | 0 | 0 | 0 | 0 | 0 | 0 |
| 0 | NA | NA | NA | NA | NA | Gm24396 | | 0 | 0 | 0 | 0 | 0 | 0 | 0 | 0 | 0 | 0 | 0 |
| 0 | NA | NA | NA | NA | NA | Cdr1 | cerebellar degeneration related antigen 1 | 0 | 0 | 0 | 0 | 0 | 0 | 0 | 0 | 0 | 0 | 0 |
| 0 | NA | NA | NA | NA | NA | | predicted gene 4988 | 0 | 0 | 0 | 0 | 0 | 0 | 0 | 0 | 0 | 0 | 0 |
| 0 | NA | NA | NA | NA | NA | Gm14665 | | 0 | 0 | 0 | 0 | 0 | 0 | 0 | 0 | 0 | 0 | 0 |
| 0 | NA | NA | NA | NA | NA | Gm14666 | | 0 | 0 | 0 | 0 | 0 | 0 | 0 | 0 | 0 | 0 | 0 |
| 0 | NA | NA | NA | NA | NA | Gm14821 | | 0 | 0 | 0 | 0 | 0 | 0 | 0 | 0 | 0 | 0 | 0 |
| 0 | NA | NA | NA | NA | NA | Gm14655 | | 0 | 0 | 0 | 0 | 0 | 0 | 0 | 0 | 0 | 0 | 0 |
| 0 | NA | NA | NA | NA | NA | Ldoc1 | regulator of NFKB signaling | 0 | 0 | 0 | 0 | 0 | 0 | 0 | 0 | 0 | 0 | 0 |
| 0 | NA | NA | NA | NA | NA | Gm25412 | | 0 | 0 | 0 | 0 | 0 | 0 | 0 | 0 | 0 | 0 | 0 |
| 0 | NA | NA | NA | NA | NA | Gm8215 | | 0 | 0 | 0 | 0 | 0 | 0 | 0 | 0 | 0 | 0 | 0 |
| 0 | NA | NA | NA | NA | NA | Gm5390 | | 0 | 0 | 0 | 0 | 0 | 0 | 0 | 0 | 0 | 0 | 0 |
| 0 | NA | NA | NA | NA | NA | Gm14656 | | 0 | 0 | 0 | 0 | 0 | 0 | 0 | 0 | 0 | 0 | 0 |
| 0 | NA | NA | NA | NA | NA | Gm16411 | | 0 | 0 | 0 | 0 | 0 | 0 | 0 | 0 | 0 | 0 | 0 |
| 0 | NA | NA | NA | NA | NA | 4933402E13Ril | RIKEN cDNA 4933402E13 gene | 0 | 0 | 0 | 0 | 0 | 0 | 0 | 0 | 0 | 0 | 0 |
| 0 | NA | NA | NA | NA | NA | Gm14679 | | 0 | 0 | 0 | 0 | 0 | 0 | 0 | 0 | 0 | 0 | 0 |

| | | | | | | | | | | | | | | | | | | |
|---|---|---|---|---|---|---|---|---|---|---|---|---|---|---|---|---|---|---|
| 0 | NA | NA | NA | NA | NA | 4931400O07Rik | | 0 | 0 | 0 | 0 | 0 | 0 | 0 | 0 | 0 | 0 | 0 |
| 0 | NA | NA | NA | NA | NA | Gm14678 | | 0 | 0 | 0 | 0 | 0 | 0 | 0 | 0 | 0 | 0 | 0 |
| 0 | NA | NA | NA | NA | NA | Gm14677 | | 0 | 0 | 0 | 0 | 0 | 0 | 0 | 0 | 0 | 0 | 0 |
| 0 | NA | NA | NA | NA | NA | Gm8244 | | 0 | 0 | 0 | 0 | 0 | 0 | 0 | 0 | 0 | 0 | 0 |
| 0 | NA | NA | NA | NA | NA | Gm6754 | | 0 | 0 | 0 | 0 | 0 | 0 | 0 | 0 | 0 | 0 | 0 |
| 0 | NA | NA | NA | NA | NA | Gm14659 | | 0 | 0 | 0 | 0 | 0 | 0 | 0 | 0 | 0 | 0 | 0 |
| 0 | NA | NA | NA | NA | NA | Gm14657 | | 0 | 0 | 0 | 0 | 0 | 0 | 0 | 0 | 0 | 0 | 0 |
| 0 | NA | NA | NA | NA | NA | Gm8260 | | 0 | 0 | 0 | 0 | 0 | 0 | 0 | 0 | 0 | 0 | 0 |
| 0 | NA | NA | NA | NA | NA | Gm6760 | predicted gene 6760 | 0 | 0 | 0 | 0 | 0 | 0 | 0 | 0 | 0 | 0 | 0 |
| 0 | NA | NA | NA | NA | NA | 3830417A13Ril | RIKEN cDNA 3830417A13 gene | 0 | 0 | 0 | 0 | 0 | 0 | 0 | 0 | 0 | 0 | 0 |
| 0 | NA | NA | NA | NA | NA | Gm14676 | | 0 | 0 | 0 | 0 | 0 | 0 | 0 | 0 | 0 | 0 | 0 |
| 0 | NA | NA | NA | NA | NA | Gm26111 | | 0 | 0 | 0 | 0 | 0 | 0 | 0 | 0 | 0 | 0 | 0 |
| 0 | NA | NA | NA | NA | NA | Gm14667 | | 0 | 0 | 0 | 0 | 0 | 0 | 0 | 0 | 0 | 0 | 0 |
| 0 | NA | NA | NA | NA | NA | Gm14673 | | 0 | 0 | 0 | 0 | 0 | 0 | 0 | 0 | 0 | 0 | 0 |
| 0 | NA | NA | NA | NA | NA | Ctag2 | cancer/testis antigen 2 | 0 | 0 | 0 | 0 | 0 | 0 | 0 | 0 | 0 | 0 | 0 |
| 0 | NA | NA | NA | NA | NA | Gm14669 | | 0 | 0 | 0 | 0 | 0 | 0 | 0 | 0 | 0 | 0 | 0 |
| 0 | NA | NA | NA | NA | NA | Gm14670 | | 0 | 0 | 0 | 0 | 0 | 0 | 0 | 0 | 0 | 0 | 0 |
| 0 | NA | NA | NA | NA | NA | Gm14668 | | 0 | 0 | 0 | 0 | 0 | 0 | 0 | 0 | 0 | 0 | 0 |
| 0 | NA | NA | NA | NA | NA | Gm14671 | | 0 | 0 | 0 | 0 | 0 | 0 | 0 | 0 | 0 | 0 | 0 |
| 0 | NA | NA | NA | NA | NA | Gm14674 | | 0 | 0 | 0 | 0 | 0 | 0 | 0 | 0 | 0 | 0 | 0 |
| 0 | NA | NA | NA | NA | NA | 4930447F04Rik | | 0 | 0 | 0 | 0 | 0 | 0 | 0 | 0 | 0 | 0 | 0 |
| 0 | NA | NA | NA | NA | NA | Gm23028 | | 0 | 0 | 0 | 0 | 0 | 0 | 0 | 0 | 0 | 0 | 0 |
| 0 | NA | NA | NA | NA | NA | Gm23000 | | 0 | 0 | 0 | 0 | 0 | 0 | 0 | 0 | 0 | 0 | 0 |
| 0 | NA | NA | NA | NA | NA | Gm8310 | | 0 | 0 | 0 | 0 | 0 | 0 | 0 | 0 | 0 | 0 | 0 |
| 0 | NA | NA | NA | NA | NA | Mir743 | | 0 | 0 | 0 | 0 | 0 | 0 | 0 | 0 | 0 | 0 | 0 |
| 0 | NA | NA | NA | NA | NA | Mir743b | | 0 | 0 | 0 | 0 | 0 | 0 | 0 | 0 | 0 | 0 | 0 |
| 0 | NA | NA | NA | NA | NA | Mir742 | | 0 | 0 | 0 | 0 | 0 | 0 | 0 | 0 | 0 | 0 | 0 |
| 0 | NA | NA | NA | NA | NA | Mir883a | | 0 | 0 | 0 | 0 | 0 | 0 | 0 | 0 | 0 | 0 | 0 |
| 0 | NA | NA | NA | NA | NA | Mir883b | | 0 | 0 | 0 | 0 | 0 | 0 | 0 | 0 | 0 | 0 | 0 |
| 0 | NA | NA | NA | NA | NA | Mir471 | | 0 | 0 | 0 | 0 | 0 | 0 | 0 | 0 | 0 | 0 | 0 |
| 0 | NA | NA | NA | NA | NA | Mir741 | | 0 | 0 | 0 | 0 | 0 | 0 | 0 | 0 | 0 | 0 | 0 |
| 0 | NA | NA | NA | NA | NA | Mir463 | | 0 | 0 | 0 | 0 | 0 | 0 | 0 | 0 | 0 | 0 | 0 |
| 0 | NA | NA | NA | NA | NA | Mir880 | | 0 | 0 | 0 | 0 | 0 | 0 | 0 | 0 | 0 | 0 | 0 |
| 0 | NA | NA | NA | NA | NA | Mir878 | | 0 | 0 | 0 | 0 | 0 | 0 | 0 | 0 | 0 | 0 | 0 |
| 0 | NA | NA | NA | NA | NA | Mir881 | | 0 | 0 | 0 | 0 | 0 | 0 | 0 | 0 | 0 | 0 | 0 |
| 0 | NA | NA | NA | NA | NA | Mir871 | | 0 | 0 | 0 | 0 | 0 | 0 | 0 | 0 | 0 | 0 | 0 |
| 0 | NA | NA | NA | NA | NA | Mir470 | | 0 | 0 | 0 | 0 | 0 | 0 | 0 | 0 | 0 | 0 | 0 |
| 0 | NA | NA | NA | NA | NA | Mir465d | | 0 | 0 | 0 | 0 | 0 | 0 | 0 | 0 | 0 | 0 | 0 |
| 0 | NA | NA | NA | NA | NA | Mir465c-1 | | 0 | 0 | 0 | 0 | 0 | 0 | 0 | 0 | 0 | 0 | 0 |
| 0 | NA | NA | NA | NA | NA | Mir465b-1 | | 0 | 0 | 0 | 0 | 0 | 0 | 0 | 0 | 0 | 0 | 0 |
| 0 | NA | NA | NA | NA | NA | Mir465c-2 | | 0 | 0 | 0 | 0 | 0 | 0 | 0 | 0 | 0 | 0 | 0 |
| 0 | NA | NA | NA | NA | NA | Mir465b-2 | | 0 | 0 | 0 | 0 | 0 | 0 | 0 | 0 | 0 | 0 | 0 |
| 0 | NA | NA | NA | NA | NA | Mir465 | | 0 | 0 | 0 | 0 | 0 | 0 | 0 | 0 | 0 | 0 | 0 |
| 0 | NA | NA | NA | NA | NA | Gm14682 | | 0 | 0 | 0 | 0 | 0 | 0 | 0 | 0 | 0 | 0 | 0 |
| 0 | NA | NA | NA | NA | NA | Gm8312 | | 0 | 0 | 0 | 0 | 0 | 0 | 0 | 0 | 0 | 0 | 0 |
| 0 | NA | NA | NA | NA | NA | 1700036O09Rik | | 0 | 0 | 0 | 0 | 0 | 0 | 0 | 0 | 0 | 0 | 0 |
| 0 | NA | NA | NA | NA | NA | Gm26276 | | 0 | 0 | 0 | 0 | 0 | 0 | 0 | 0 | 0 | 0 | 0 |
| 0 | NA | NA | NA | NA | NA | Gm14687 | | 0 | 0 | 0 | 0 | 0 | 0 | 0 | 0 | 0 | 0 | 0 |
| 0 | NA | NA | NA | NA | NA | Gm1140 | predicted gene 1140 | 0 | 0 | 0 | 0 | 0 | 0 | 0 | 0 | 0 | 0 | 0 |
| 0 | NA | NA | NA | NA | NA | Gm14692 | predicted gene 14692 | 0 | 0 | 0 | 0 | 0 | 0 | 0 | 0 | 0 | 0 | 0 |
| 0 | NA | NA | NA | NA | NA | 4933436I01Rik | RIKEN cDNA 4933436I01 gene | 0 | 0 | 0 | 0 | 0 | 0 | 0 | 0 | 0 | 0 | 0 |
| 0 | NA | NA | NA | NA | NA | Mir201 | | 0 | 0 | 0 | 0 | 0 | 0 | 0 | 0 | 0 | 0 | 0 |
| 0 | NA | NA | NA | NA | NA | Mir547 | | 0 | 0 | 0 | 0 | 0 | 0 | 0 | 0 | 0 | 0 | 0 |
| 0 | NA | NA | NA | NA | NA | Gm14688 | | 0 | 0 | 0 | 0 | 0 | 0 | 0 | 0 | 0 | 0 | 0 |
| 0 | NA | NA | NA | NA | NA | Mir509 | | 0 | 0 | 0 | 0 | 0 | 0 | 0 | 0 | 0 | 0 | 0 |
| 0 | NA | NA | NA | NA | NA | Gm6806 | | 0 | 0 | 0 | 0 | 0 | 0 | 0 | 0 | 0 | 0 | 0 |
| 0 | NA | NA | NA | NA | NA | Gm22332 | | 0 | 0 | 0 | 0 | 0 | 0 | 0 | 0 | 0 | 0 | 0 |
| 0 | NA | NA | NA | NA | NA | Gm14701 | | 0 | 0 | 0 | 0 | 0 | 0 | 0 | 0 | 0 | 0 | 0 |
| 0 | NA | NA | NA | NA | NA | Gm27979 | | 0 | 0 | 0 | 0 | 0 | 0 | 0 | 0 | 0 | 0 | 0 |
| 0 | NA | NA | NA | NA | NA | Gm27838 | | 0 | 0 | 0 | 0 | 0 | 0 | 0 | 0 | 0 | 0 | 0 |

| | | | | | | | | | | | | | | | | | |
|---|---|---|---|---|---|---|---|---|---|---|---|---|---|---|---|---|---|
| 0 | NA | NA | NA | NA | NA | Gm22435 | | 0 | 0 | 0 | 0 | 0 | 0 | 0 | 0 | 0 | 0 | 0 |
| 0 | NA | NA | NA | NA | NA | Gm6812 | predicted gene 6812 | 0 | 0 | 0 | 0 | 0 | 0 | 0 | 0 | 0 | 0 | 0 |
| 0 | NA | NA | NA | NA | NA | Gm8344 | | 0 | 0 | 0 | 0 | 0 | 0 | 0 | 0 | 0 | 0 | 0 |
| 0 | NA | NA | NA | NA | NA | n-R5s7 | | 0 | 0 | 0 | 0 | 0 | 0 | 0 | 0 | 0 | 0 | 0 |
| 0 | NA | NA | NA | NA | NA | 1700020N15Ril | RIKEN cDNA 1700020N15 gene | 0 | 0 | 0 | 0 | 0 | 0 | 0 | 0 | 0 | 0 | 0 |
| 0 | NA | NA | NA | NA | NA | 4930567H17Rik | | 0 | 0 | 0 | 0 | 0 | 0 | 0 | 0 | 0 | 0 | 0 |
| 0 | NA | NA | NA | NA | NA | Gm16407 | | 0 | 0 | 0 | 0 | 0 | 0 | 0 | 0 | 0 | 0 | 0 |
| 0 | NA | NA | NA | NA | NA | Gm14704 | | 0 | 0 | 0 | 0 | 0 | 0 | 0 | 0 | 0 | 0 | 0 |
| 0 | NA | NA | NA | NA | NA | Gm6823 | | 0 | 0 | 0 | 0 | 0 | 0 | 0 | 0 | 0 | 0 | 0 |
| 0 | NA | NA | NA | NA | NA | Gm16441 | | 0 | 0 | 0 | 0 | 0 | 0 | 0 | 0 | 0 | 0 | 0 |
| 0 | NA | NA | NA | NA | NA | Gm14722 | | 0 | 0 | 0 | 0 | 0 | 0 | 0 | 0 | 0 | 0 | 0 |
| 0 | NA | NA | NA | NA | NA | Gm14721 | | 0 | 0 | 0 | 0 | 0 | 0 | 0 | 0 | 0 | 0 | 0 |
| 0 | NA | NA | NA | NA | NA | Gm14727 | | 0 | 0 | 0 | 0 | 0 | 0 | 0 | 0 | 0 | 0 | 0 |
| 0 | NA | NA | NA | NA | NA | Gm14729 | | 0 | 0 | 0 | 0 | 0 | 0 | 0 | 0 | 0 | 0 | 0 |
| 0 | NA | NA | NA | NA | NA | Gm6038 | | 0 | 0 | 0 | 0 | 0 | 0 | 0 | 0 | 0 | 0 | 0 |
| 0 | NA | NA | NA | NA | NA | Gm14726 | | 0 | 0 | 0 | 0 | 0 | 0 | 0 | 0 | 0 | 0 | 0 |
| 0 | NA | NA | NA | NA | NA | Gm14724 | | 0 | 0 | 0 | 0 | 0 | 0 | 0 | 0 | 0 | 0 | 0 |
| 0 | NA | NA | NA | NA | NA | Gm14725 | predicted gene 14725 | 0 | 0 | 0 | 0 | 0 | 0 | 0 | 0 | 0 | 0 | 0 |
| 0 | NA | NA | NA | NA | NA | Gm14728 | | 0 | 0 | 0 | 0 | 0 | 0 | 0 | 0 | 0 | 0 | 0 |
| 0 | NA | NA | NA | NA | NA | Gm14732 | | 0 | 0 | 0 | 0 | 0 | 0 | 0 | 0 | 0 | 0 | 0 |
| 0 | NA | NA | NA | NA | NA | Gm14731 | | 0 | 0 | 0 | 0 | 0 | 0 | 0 | 0 | 0 | 0 | 0 |
| 0 | NA | NA | NA | NA | NA | Gm22785 | | 0 | 0 | 0 | 0 | 0 | 0 | 0 | 0 | 0 | 0 | 0 |
| 0 | NA | NA | NA | NA | NA | Gm8410 | | 0 | 0 | 0 | 0 | 0 | 0 | 0 | 0 | 0 | 0 | 0 |
| 0 | NA | NA | NA | NA | NA | Gm6062 | | 0 | 0 | 0 | 0 | 0 | 0 | 0 | 0 | 0 | 0 | 0 |
| 0 | NA | NA | NA | NA | NA | Gm16189 | | 0 | 0 | 0 | 0 | 0 | 0 | 0 | 0 | 0 | 0 | 0 |
| 0 | NA | NA | NA | NA | NA | Gm8421 | | 0 | 0 | 0 | 0 | 0 | 0 | 0 | 0 | 0 | 0 | 0 |
| 0 | NA | NA | NA | NA | NA | Gm23945 | | 0 | 0 | 0 | 0 | 0 | 0 | 0 | 0 | 0 | 0 | 0 |
| 0 | NA | NA | NA | NA | NA | Gpr50 | G-protein-coupled receptor 50 | 0 | 0 | 0 | 0 | 0 | 0 | 0 | 0 | 0 | 0 | 0 |
| 0 | NA | NA | NA | NA | NA | Gm14723 | | 0 | 0 | 0 | 0 | 0 | 0 | 0 | 0 | 0 | 0 | 0 |
| 0 | NA | NA | NA | NA | NA | Fate1 | | 0 | 0 | 0 | 0 | 0 | 0 | 0 | 0 | 0 | 0 | 0 |
| 0 | NA | NA | NA | NA | NA | Cnga2 | cyclic nucleotide gated channel alpha 2 | 0 | 0 | 0 | 0 | 0 | 0 | 0 | 0 | 0 | 0 | 0 |
| 0 | NA | NA | NA | NA | NA | Magea4 | melanoma antigen, family A, 4 | 0 | 0 | 0 | 0 | 0 | 0 | 0 | 0 | 0 | 0 | 0 |
| 0 | NA | NA | NA | NA | NA | Gm47265 | | 0 | 0 | 0 | 0 | 0 | 0 | 0 | 0 | 0 | 0 | 0 |
| 0 | NA | NA | NA | NA | NA | Mir224 | | 0 | 0 | 0 | 0 | 0 | 0 | 0 | 0 | 0 | 0 | 0 |
| 0 | NA | NA | NA | NA | NA | Mir452 | | 0 | 0 | 0 | 0 | 0 | 0 | 0 | 0 | 0 | 0 | 0 |
| 0 | NA | NA | NA | NA | NA | Magea10 | melanoma antigen family A, 10 | 0 | 0 | 0 | 0 | 0 | 0 | 0 | 0 | 0 | 0 | 0 |
| 0 | NA | NA | NA | NA | NA | Gm14735 | | 0 | 0 | 0 | 0 | 0 | 0 | 0 | 0 | 0 | 0 | 0 |
| 0 | NA | NA | NA | NA | NA | Mir767 | | 0 | 0 | 0 | 0 | 0 | 0 | 0 | 0 | 0 | 0 | 0 |
| 0 | NA | NA | NA | NA | NA | Mir105 | | 0 | 0 | 0 | 0 | 0 | 0 | 0 | 0 | 0 | 0 | 0 |
| 0 | NA | NA | NA | NA | NA | Gm23461 | | 0 | 0 | 0 | 0 | 0 | 0 | 0 | 0 | 0 | 0 | 0 |
| 0 | NA | NA | NA | NA | NA | Gm14736 | | 0 | 0 | 0 | 0 | 0 | 0 | 0 | 0 | 0 | 0 | 0 |
| 0 | NA | NA | NA | NA | NA | Magea9-ps | | 0 | 0 | 0 | 0 | 0 | 0 | 0 | 0 | 0 | 0 | 0 |
| 0 | NA | NA | NA | NA | NA | Gm14684 | | 0 | 0 | 0 | 0 | 0 | 0 | 0 | 0 | 0 | 0 | 0 |
| 0 | NA | NA | NA | NA | NA | Mir2137 | | 0 | 0 | 0 | 0 | 0 | 0 | 0 | 0 | 0 | 0 | 0 |
| 0 | NA | NA | NA | NA | NA | Gm6858 | | 0 | 0 | 0 | 0 | 0 | 0 | 0 | 0 | 0 | 0 | 0 |
| 0 | NA | NA | NA | NA | NA | Gm20455 | | 0 | 0 | 0 | 0 | 0 | 0 | 0 | 0 | 0 | 0 | 0 |
| 0 | NA | NA | NA | NA | NA | Xlr5a | X-linked lymphocyte-regulated 5A | 0 | 0 | 0 | 0 | 0 | 0 | 0 | 0 | 0 | 0 | 0 |
| 0 | NA | NA | NA | NA | NA | Spin2d | spindlin family, member 2D | 0 | 0 | 0 | 0 | 0 | 0 | 0 | 0 | 0 | 0 | 0 |
| 0 | NA | NA | NA | NA | NA | Xlr5c | X-linked lymphocyte-regulated 5C | 0 | 0 | 0 | 0 | 0 | 0 | 0 | 0 | 0 | 0 | 0 |
| 0 | NA | NA | NA | NA | NA | Gm8501 | | 0 | 0 | 0 | 0 | 0 | 0 | 0 | 0 | 0 | 0 | 0 |
| 0 | NA | NA | NA | NA | NA | Gm45015 | | 0 | 0 | 0 | 0 | 0 | 0 | 0 | 0 | 0 | 0 | 0 |
| 0 | NA | NA | NA | NA | NA | Gm37741 | | 0 | 0 | 0 | 0 | 0 | 0 | 0 | 0 | 0 | 0 | 0 |
| 0 | NA | NA | NA | NA | NA | Gm9730 | | 0 | 0 | 0 | 0 | 0 | 0 | 0 | 0 | 0 | 0 | 0 |
| 0 | NA | NA | NA | NA | NA | Gm25226 | | 0 | 0 | 0 | 0 | 0 | 0 | 0 | 0 | 0 | 0 | 0 |
| 0 | NA | NA | NA | NA | NA | Gm23615 | | 0 | 0 | 0 | 0 | 0 | 0 | 0 | 0 | 0 | 0 | 0 |
| 0 | NA | NA | NA | NA | NA | Avpr2 | arginine vasopressin receptor 2 | 0 | 0 | 0 | 0 | 0 | 0 | 0 | 0 | 0 | 0 | 0 |
| 0 | NA | NA | NA | NA | NA | Mir5132 | | 0 | 0 | 0 | 0 | 0 | 0 | 0 | 0 | 0 | 0 | 0 |
| 0 | NA | NA | NA | NA | NA | Mir718 | | 0 | 0 | 0 | 0 | 0 | 0 | 0 | 0 | 0 | 0 | 0 |
| 0 | NA | NA | NA | NA | NA | Tex28 | testis expressed 28 | 0 | 0 | 0 | 0 | 0 | 0 | 0 | 0 | 0 | 0 | 0 |
| 0 | NA | NA | NA | NA | NA | Tktl1 | transketolase-like 1 | 0 | 0 | 0 | 0 | 0 | 0 | 0 | 0 | 0 | 0 | 0 |

| | | | | | | | | | | | | | | | | | | |
|---|---|---|---|---|---|---|---|---|---|---|---|---|---|---|---|---|---|---|
| 0 | NA | NA | NA | NA | NA | Gm14686 | | 0 | 0 | 0 | 0 | 0 | 0 | 0 | 0 | 0 | 0 | 0 |
| 0 | NA | NA | NA | NA | NA | Snora70 | | 0 | 0 | 0 | 0 | 0 | 0 | 0 | 0 | 0 | 0 | 0 |
| 0 | NA | NA | NA | NA | NA | Mir7091 | | 0 | 0 | 0 | 0 | 0 | 0 | 0 | 0 | 0 | 0 | 0 |
| 0 | NA | NA | NA | NA | NA | Gm14846 | | 0 | 0 | 0 | 0 | 0 | 0 | 0 | 0 | 0 | 0 | 0 |
| 0 | NA | NA | NA | NA | NA | Mir7092 | | 0 | 0 | 0 | 0 | 0 | 0 | 0 | 0 | 0 | 0 | 0 |
| 0 | NA | NA | NA | NA | NA | Ctag2l1 | CTAG2 like 1 | 0 | 0 | 0 | 0 | 0 | 0 | 0 | 0 | 0 | 0 | 0 |
| 0 | NA | NA | NA | NA | NA | Gm15362 | | 0 | 0 | 0 | 0 | 0 | 0 | 0 | 0 | 0 | 0 | 0 |
| 0 | NA | NA | NA | NA | NA | Gm15364 | | 0 | 0 | 0 | 0 | 0 | 0 | 0 | 0 | 0 | 0 | 0 |
| 0 | NA | NA | NA | NA | NA | Gm26125 | | 0 | 0 | 0 | 0 | 0 | 0 | 0 | 0 | 0 | 0 | 0 |
| 0 | NA | NA | NA | NA | NA | Olfr1326-ps1 | | 0 | 0 | 0 | 0 | 0 | 0 | 0 | 0 | 0 | 0 | 0 |
| 0 | NA | NA | NA | NA | NA | Olfr1325 | olfactory receptor 1325 | 0 | 0 | 0 | 0 | 0 | 0 | 0 | 0 | 0 | 0 | 0 |
| 0 | NA | NA | NA | NA | NA | Gm26417 | | 0 | 0 | 0 | 0 | 0 | 0 | 0 | 0 | 0 | 0 | 0 |
| 0 | NA | NA | NA | NA | NA | Xlr5d-ps | | 0 | 0 | 0 | 0 | 0 | 0 | 0 | 0 | 0 | 0 | 0 |
| 0 | NA | NA | NA | NA | NA | Spin2-ps2 | | 0 | 0 | 0 | 0 | 0 | 0 | 0 | 0 | 0 | 0 | 0 |
| 0 | NA | NA | NA | NA | NA | Xlr3d-ps | | 0 | 0 | 0 | 0 | 0 | 0 | 0 | 0 | 0 | 0 | 0 |
| 0 | NA | NA | NA | NA | NA | Gm22351 | | 0 | 0 | 0 | 0 | 0 | 0 | 0 | 0 | 0 | 0 | 0 |
| 0 | NA | NA | NA | NA | NA | Olfr1327-ps1 | | 0 | 0 | 0 | 0 | 0 | 0 | 0 | 0 | 0 | 0 | 0 |
| 0 | NA | NA | NA | NA | NA | Gm14758 | | 0 | 0 | 0 | 0 | 0 | 0 | 0 | 0 | 0 | 0 | 0 |
| 0 | NA | NA | NA | NA | NA | Gm14695 | | 0 | 0 | 0 | 0 | 0 | 0 | 0 | 0 | 0 | 0 | 0 |
| 0 | NA | NA | NA | NA | NA | Gm8617 | | 0 | 0 | 0 | 0 | 0 | 0 | 0 | 0 | 0 | 0 | 0 |
| 0 | NA | NA | NA | NA | NA | Spin2-ps3 | | 0 | 0 | 0 | 0 | 0 | 0 | 0 | 0 | 0 | 0 | 0 |
| 0 | NA | NA | NA | NA | NA | Xlr5e-ps | | 0 | 0 | 0 | 0 | 0 | 0 | 0 | 0 | 0 | 0 | 0 |
| 0 | NA | NA | NA | NA | NA | Gm14694 | | 0 | 0 | 0 | 0 | 0 | 0 | 0 | 0 | 0 | 0 | 0 |
| 0 | NA | NA | NA | NA | NA | Gm6897 | | 0 | 0 | 0 | 0 | 0 | 0 | 0 | 0 | 0 | 0 | 0 |
| 0 | NA | NA | NA | NA | NA | Gm14756 | | 0 | 0 | 0 | 0 | 0 | 0 | 0 | 0 | 0 | 0 | 0 |
| 0 | NA | NA | NA | NA | NA | Spin2-ps4 | | 0 | 0 | 0 | 0 | 0 | 0 | 0 | 0 | 0 | 0 | 0 |
| 0 | NA | NA | NA | NA | NA | Xlr3e-ps | | 0 | 0 | 0 | 0 | 0 | 0 | 0 | 0 | 0 | 0 | 0 |
| 0 | NA | NA | NA | NA | NA | Xlr4e-ps | | 0 | 0 | 0 | 0 | 0 | 0 | 0 | 0 | 0 | 0 | 0 |
| 0 | NA | NA | NA | NA | NA | Gm15363 | | 0 | 0 | 0 | 0 | 0 | 0 | 0 | 0 | 0 | 0 | 0 |
| 0 | NA | NA | NA | NA | NA | Gm25629 | | 0 | 0 | 0 | 0 | 0 | 0 | 0 | 0 | 0 | 0 | 0 |
| 0 | NA | NA | NA | NA | NA | Gm15367 | | 0 | 0 | 0 | 0 | 0 | 0 | 0 | 0 | 0 | 0 | 0 |
| 0 | NA | NA | NA | NA | NA | Gm15378 | | 0 | 0 | 0 | 0 | 0 | 0 | 0 | 0 | 0 | 0 | 0 |
| 0 | NA | NA | NA | NA | NA | Fam220-ps | | 0 | 0 | 0 | 0 | 0 | 0 | 0 | 0 | 0 | 0 | 0 |
| 0 | NA | NA | NA | NA | NA | Gm25421 | | 0 | 0 | 0 | 0 | 0 | 0 | 0 | 0 | 0 | 0 | 0 |
| 0 | NA | NA | NA | NA | NA | Gm15065 | | 0 | 0 | 0 | 0 | 0 | 0 | 0 | 0 | 0 | 0 | 0 |
| 0 | NA | NA | NA | NA | NA | Gm23320 | | 0 | 0 | 0 | 0 | 0 | 0 | 0 | 0 | 0 | 0 | 0 |
| 0 | NA | NA | NA | NA | NA | Oat-rs1 | | 0 | 0 | 0 | 0 | 0 | 0 | 0 | 0 | 0 | 0 | 0 |
| 0 | NA | NA | NA | NA | NA | Gm14715 | | 0 | 0 | 0 | 0 | 0 | 0 | 0 | 0 | 0 | 0 | 0 |
| 0 | NA | NA | NA | NA | NA | Gm7155 | | 0 | 0 | 0 | 0 | 0 | 0 | 0 | 0 | 0 | 0 | 0 |
| 0 | NA | NA | NA | NA | NA | Gm14710 | | 0 | 0 | 0 | 0 | 0 | 0 | 0 | 0 | 0 | 0 | 0 |
| 0 | NA | NA | NA | NA | NA | Gm5397 | | 0 | 0 | 0 | 0 | 0 | 0 | 0 | 0 | 0 | 0 | 0 |
| 0 | NA | NA | NA | NA | NA | Gm14711 | | 0 | 0 | 0 | 0 | 0 | 0 | 0 | 0 | 0 | 0 | 0 |
| 0 | NA | NA | NA | NA | NA | Gm14707 | | 0 | 0 | 0 | 0 | 0 | 0 | 0 | 0 | 0 | 0 | 0 |
| 0 | NA | NA | NA | NA | NA | Gm14717 | | 0 | 0 | 0 | 0 | 0 | 0 | 0 | 0 | 0 | 0 | 0 |
| 0 | NA | NA | NA | NA | NA | Cldn34b3 | claudin 34B3 | 0 | 0 | 0 | 0 | 0 | 0 | 0 | 0 | 0 | 0 | 0 |
| 0 | NA | NA | NA | NA | NA | Gm14712 | | 0 | 0 | 0 | 0 | 0 | 0 | 0 | 0 | 0 | 0 | 0 |
| 0 | NA | NA | NA | NA | NA | Magea7-ps | | 0 | 0 | 0 | 0 | 0 | 0 | 0 | 0 | 0 | 0 | 0 |
| 0 | NA | NA | NA | NA | NA | Gm14709 | | 0 | 0 | 0 | 0 | 0 | 0 | 0 | 0 | 0 | 0 | 0 |
| 0 | NA | NA | NA | NA | NA | Cldn34b4 | claudin 34B4 | 0 | 0 | 0 | 0 | 0 | 0 | 0 | 0 | 0 | 0 | 0 |
| 0 | NA | NA | NA | NA | NA | Gm5793 | | 0 | 0 | 0 | 0 | 0 | 0 | 0 | 0 | 0 | 0 | 0 |
| 0 | NA | NA | NA | NA | NA | Cldn34d | claudin 34D | 0 | 0 | 0 | 0 | 0 | 0 | 0 | 0 | 0 | 0 | 0 |
| 0 | NA | NA | NA | NA | NA | Gm14716 | | 0 | 0 | 0 | 0 | 0 | 0 | 0 | 0 | 0 | 0 | 0 |
| 0 | NA | NA | NA | NA | NA | Gm14713 | | 0 | 0 | 0 | 0 | 0 | 0 | 0 | 0 | 0 | 0 | 0 |
| 0 | NA | NA | NA | NA | NA | Gm26041 | | 0 | 0 | 0 | 0 | 0 | 0 | 0 | 0 | 0 | 0 | 0 |
| 0 | NA | NA | NA | NA | NA | Gm23280 | | 0 | 0 | 0 | 0 | 0 | 0 | 0 | 0 | 0 | 0 | 0 |
| 0 | NA | NA | NA | NA | NA | Gm14714 | | 0 | 0 | 0 | 0 | 0 | 0 | 0 | 0 | 0 | 0 | 0 |
| 0 | NA | NA | NA | NA | NA | Gm5398 | | 0 | 0 | 0 | 0 | 0 | 0 | 0 | 0 | 0 | 0 | 0 |
| 0 | NA | NA | NA | NA | NA | Gm5757 | | 0 | 0 | 0 | 0 | 0 | 0 | 0 | 0 | 0 | 0 | 0 |
| 0 | NA | NA | NA | NA | NA | Gm23121 | | 0 | 0 | 0 | 0 | 0 | 0 | 0 | 0 | 0 | 0 | 0 |
| 0 | NA | NA | NA | NA | NA | Cldn34-ps | claudin 34, pseudogene | 0 | 0 | 0 | 0 | 0 | 0 | 0 | 0 | 0 | 0 | 0 |

| | | | | | | | | | | | | | | | | | |
|---|---|---|---|---|---|---|---|---|---|---|---|---|---|---|---|---|---|
| 0 | NA | NA | NA | NA | NA | Gm14746 | | 0 | 0 | 0 | 0 | 0 | 0 | 0 | 0 | 0 | 0 | 0 |
| 0 | NA | NA | NA | NA | NA | Gm14748 | | 0 | 0 | 0 | 0 | 0 | 0 | 0 | 0 | 0 | 0 | 0 |
| 0 | NA | NA | NA | NA | NA | Gm14742 | | 0 | 0 | 0 | 0 | 0 | 0 | 0 | 0 | 0 | 0 | 0 |
| 0 | NA | NA | NA | NA | NA | Pbsn | probasin | 0 | 0 | 0 | 0 | 0 | 0 | 0 | 0 | 0 | 0 | 0 |
| 0 | NA | NA | NA | NA | NA | Gm14744 | predicted gene 14744 | 0 | 0 | 0 | 0 | 0 | 0 | 0 | 0 | 0 | 0 | 0 |
| 0 | NA | NA | NA | NA | NA | 5430402E10Ril | RIKEN cDNA 5430402E10 gene | 0 | 0 | 0 | 0 | 0 | 0 | 0 | 0 | 0 | 0 | 0 |
| 0 | NA | NA | NA | NA | NA | Gm14750 | | 0 | 0 | 0 | 0 | 0 | 0 | 0 | 0 | 0 | 0 | 0 |
| 0 | NA | NA | NA | NA | NA | Gm16458 | | 0 | 0 | 0 | 0 | 0 | 0 | 0 | 0 | 0 | 0 | 0 |
| 0 | NA | NA | NA | NA | NA | Obp1a | odorant binding protein IA | 0 | 0 | 0 | 0 | 0 | 0 | 0 | 0 | 0 | 0 | 0 |
| 0 | NA | NA | NA | NA | NA | Gm5938 | predicted gene 5938 | 0 | 0 | 0 | 0 | 0 | 0 | 0 | 0 | 0 | 0 | 0 |
| 0 | NA | NA | NA | NA | NA | Obp1b | odorant binding protein IB | 0 | 0 | 0 | 0 | 0 | 0 | 0 | 0 | 0 | 0 | 0 |
| 0 | NA | NA | NA | NA | NA | Gm14743 | predicted gene 14743 | 0 | 0 | 0 | 0 | 0 | 0 | 0 | 0 | 0 | 0 | 0 |
| 0 | NA | NA | NA | NA | NA | Gm8749 | | 0 | 0 | 0 | 0 | 0 | 0 | 0 | 0 | 0 | 0 | 0 |
| 0 | NA | NA | NA | NA | NA | 4930480E11Ril | RIKEN cDNA 4930480E11 gene | 0 | 0 | 0 | 0 | 0 | 0 | 0 | 0 | 0 | 0 | 0 |
| 0 | NA | NA | NA | NA | NA | Gm14749 | | 0 | 0 | 0 | 0 | 0 | 0 | 0 | 0 | 0 | 0 | 0 |
| 0 | NA | NA | NA | NA | NA | Gm8754 | | 0 | 0 | 0 | 0 | 0 | 0 | 0 | 0 | 0 | 0 | 0 |
| 0 | NA | NA | NA | NA | NA | Gm14752 | | 0 | 0 | 0 | 0 | 0 | 0 | 0 | 0 | 0 | 0 | 0 |
| 0 | NA | NA | NA | NA | NA | Gm22843 | | 0 | 0 | 0 | 0 | 0 | 0 | 0 | 0 | 0 | 0 | 0 |
| 0 | NA | NA | NA | NA | NA | Gm8761 | | 0 | 0 | 0 | 0 | 0 | 0 | 0 | 0 | 0 | 0 | 0 |
| 0 | NA | NA | NA | NA | NA | Gm14747 | | 0 | 0 | 0 | 0 | 0 | 0 | 0 | 0 | 0 | 0 | 0 |
| 0 | NA | NA | NA | NA | NA | Fam47c | family with sequence similarity 47, membe | 0 | 0 | 0 | 0 | 0 | 0 | 0 | 0 | 0 | 0 | 0 |
| 0 | NA | NA | NA | NA | NA | Gm8770 | | 0 | 0 | 0 | 0 | 0 | 0 | 0 | 0 | 0 | 0 | 0 |
| 0 | NA | NA | NA | NA | NA | Gm23556 | | 0 | 0 | 0 | 0 | 0 | 0 | 0 | 0 | 0 | 0 | 0 |
| 0 | NA | NA | NA | NA | NA | Gm14751 | | 0 | 0 | 0 | 0 | 0 | 0 | 0 | 0 | 0 | 0 | 0 |
| 0 | NA | NA | NA | NA | NA | Mageb16 | melanoma antigen family B, 16 | 0 | 0 | 0 | 0 | 0 | 0 | 0 | 0 | 0 | 0 | 0 |
| 0 | NA | NA | NA | NA | NA | Gm4772 | | 0 | 0 | 0 | 0 | 0 | 0 | 0 | 0 | 0 | 0 | 0 |
| 0 | NA | NA | NA | NA | NA | Gm4773 | | 0 | 0 | 0 | 0 | 0 | 0 | 0 | 0 | 0 | 0 | 0 |
| 0 | NA | NA | NA | NA | NA | Gm6952 | | 0 | 0 | 0 | 0 | 0 | 0 | 0 | 0 | 0 | 0 | 0 |
| 0 | NA | NA | NA | NA | NA | Gm14768 | | 0 | 0 | 0 | 0 | 0 | 0 | 0 | 0 | 0 | 0 | 0 |
| 0 | NA | NA | NA | NA | NA | Gm5940 | | 0 | 0 | 0 | 0 | 0 | 0 | 0 | 0 | 0 | 0 | 0 |
| 0 | NA | NA | NA | NA | NA | Gm6954 | | 0 | 0 | 0 | 0 | 0 | 0 | 0 | 0 | 0 | 0 | 0 |
| 0 | NA | NA | NA | NA | NA | Gm14733 | | 0 | 0 | 0 | 0 | 0 | 0 | 0 | 0 | 0 | 0 | 0 |
| 0 | NA | NA | NA | NA | NA | Gm14754 | | 0 | 0 | 0 | 0 | 0 | 0 | 0 | 0 | 0 | 0 | 0 |
| 0 | NA | NA | NA | NA | NA | 4930595M18Ri | RIKEN cDNA 4930595M18 gene | 0 | 0 | 0 | 0 | 0 | 0 | 0 | 0 | 0 | 0 | 0 |
| 0 | NA | NA | NA | NA | NA | Gm26174 | | 0 | 0 | 0 | 0 | 0 | 0 | 0 | 0 | 0 | 0 | 0 |
| 0 | NA | NA | NA | NA | NA | Gm22686 | | 0 | 0 | 0 | 0 | 0 | 0 | 0 | 0 | 0 | 0 | 0 |
| 0 | NA | NA | NA | NA | NA | Rpl10-ps4 | | 0 | 0 | 0 | 0 | 0 | 0 | 0 | 0 | 0 | 0 | 0 |
| 0 | NA | NA | NA | NA | NA | Gm24706 | | 0 | 0 | 0 | 0 | 0 | 0 | 0 | 0 | 0 | 0 | 0 |
| 0 | NA | NA | NA | NA | NA | Gm14740 | | 0 | 0 | 0 | 0 | 0 | 0 | 0 | 0 | 0 | 0 | 0 |
| 0 | NA | NA | NA | NA | NA | Gm14739 | | 0 | 0 | 0 | 0 | 0 | 0 | 0 | 0 | 0 | 0 | 0 |
| 0 | NA | NA | NA | NA | NA | Gm24470 | | 0 | 0 | 0 | 0 | 0 | 0 | 0 | 0 | 0 | 0 | 0 |
| 0 | NA | NA | NA | NA | NA | Gm14741 | | 0 | 0 | 0 | 0 | 0 | 0 | 0 | 0 | 0 | 0 | 0 |
| 0 | NA | NA | NA | NA | NA | Gm23671 | | 0 | 0 | 0 | 0 | 0 | 0 | 0 | 0 | 0 | 0 | 0 |
| 0 | NA | NA | NA | NA | NA | n-R5s9 | | 0 | 0 | 0 | 0 | 0 | 0 | 0 | 0 | 0 | 0 | 0 |
| 0 | NA | NA | NA | NA | NA | Tsga8 | testis specific gene A8 | 0 | 0 | 0 | 0 | 0 | 0 | 0 | 0 | 0 | 0 | 0 |
| 0 | NA | NA | NA | NA | NA | Gm22580 | | 0 | 0 | 0 | 0 | 0 | 0 | 0 | 0 | 0 | 0 | 0 |
| 0 | NA | NA | NA | NA | NA | Gm14770 | | 0 | 0 | 0 | 0 | 0 | 0 | 0 | 0 | 0 | 0 | 0 |
| 0 | NA | NA | NA | NA | NA | Gm14771 | | 0 | 0 | 0 | 0 | 0 | 0 | 0 | 0 | 0 | 0 | 0 |
| 0 | NA | NA | NA | NA | NA | Fthl17a | ferritin, heavy polypeptide-like 17, membe | 0 | 0 | 0 | 0 | 0 | 0 | 0 | 0 | 0 | 0 | 0 |
| 0 | NA | NA | NA | NA | NA | Gm14765 | | 0 | 0 | 0 | 0 | 0 | 0 | 0 | 0 | 0 | 0 | 0 |
| 0 | NA | NA | NA | NA | NA | Gm44378 | | 0 | 0 | 0 | 0 | 0 | 0 | 0 | 0 | 0 | 0 | 0 |
| 0 | NA | NA | NA | NA | NA | Gm14767 | | 0 | 0 | 0 | 0 | 0 | 0 | 0 | 0 | 0 | 0 | 0 |
| 0 | NA | NA | NA | NA | NA | Gm14764 | | 0 | 0 | 0 | 0 | 0 | 0 | 0 | 0 | 0 | 0 | 0 |
| 0 | NA | NA | NA | NA | NA | Gm6964 | | 0 | 0 | 0 | 0 | 0 | 0 | 0 | 0 | 0 | 0 | 0 |
| 0 | NA | NA | NA | NA | NA | Gm14761 | | 0 | 0 | 0 | 0 | 0 | 0 | 0 | 0 | 0 | 0 | 0 |
| 0 | NA | NA | NA | NA | NA | Gm14762 | | 0 | 0 | 0 | 0 | 0 | 0 | 0 | 0 | 0 | 0 | 0 |
| 0 | NA | NA | NA | NA | NA | Gm14772 | | 0 | 0 | 0 | 0 | 0 | 0 | 0 | 0 | 0 | 0 | 0 |
| 0 | NA | NA | NA | NA | NA | Samt3 | spermatogenesis associated multipass tra | 0 | 0 | 0 | 0 | 0 | 0 | 0 | 0 | 0 | 0 | 0 |
| 0 | NA | NA | NA | NA | NA | Mageb4 | melanoma antigen, family B, 4 | 0 | 0 | 0 | 0 | 0 | 0 | 0 | 0 | 0 | 0 | 0 |
| 0 | NA | NA | NA | NA | NA | Mageb10-ps | | 0 | 0 | 0 | 0 | 0 | 0 | 0 | 0 | 0 | 0 | 0 |

| | | | | | | | | | | | | | | | | | | |
|---|---|---|---|---|---|---|---|---|---|---|---|---|---|---|---|---|---|---|
| 0 | NA | NA | NA | NA | NA | Mageb8-ps | | 0 | 0 | 0 | 0 | 0 | 0 | 0 | 0 | 0 | 0 | 0 |
| 0 | NA | NA | NA | NA | NA | Gm41 | | 0 | 0 | 0 | 0 | 0 | 0 | 0 | 0 | 0 | 0 | 0 |
| 0 | NA | NA | NA | NA | NA | Mageb6-ps | | 0 | 0 | 0 | 0 | 0 | 0 | 0 | 0 | 0 | 0 | 0 |
| 0 | NA | NA | NA | NA | NA | | | 0 | 0 | 0 | 0 | 0 | 0 | 0 | 0 | 0 | 0 | 0 |
| 0 | NA | NA | NA | NA | NA | Mageb7-ps | | 0 | 0 | 0 | 0 | 0 | 0 | 0 | 0 | 0 | 0 | 0 |
| 0 | NA | NA | NA | NA | NA | n-R5s10 | | 0 | 0 | 0 | 0 | 0 | 0 | 0 | 0 | 0 | 0 | 0 |
| 0 | NA | NA | NA | NA | NA | Gm14775 | | 0 | 0 | 0 | 0 | 0 | 0 | 0 | 0 | 0 | 0 | 0 |
| 0 | NA | NA | NA | NA | NA | Gm8855 | | 0 | 0 | 0 | 0 | 0 | 0 | 0 | 0 | 0 | 0 | 0 |
| 0 | NA | NA | NA | NA | NA | Gm24812 | | 0 | 0 | 0 | 0 | 0 | 0 | 0 | 0 | 0 | 0 | 0 |
| 0 | NA | NA | NA | NA | NA | Gm14774 | | 0 | 0 | 0 | 0 | 0 | 0 | 0 | 0 | 0 | 0 | 0 |
| 0 | NA | NA | NA | NA | NA | Gm25006 | | 0 | 0 | 0 | 0 | 0 | 0 | 0 | 0 | 0 | 0 | 0 |
| 0 | NA | NA | NA | NA | NA | Gm14793 | | 0 | 0 | 0 | 0 | 0 | 0 | 0 | 0 | 0 | 0 | 0 |
| 0 | NA | NA | NA | NA | NA | Gm14795 | | 0 | 0 | 0 | 0 | 0 | 0 | 0 | 0 | 0 | 0 | 0 |
| 0 | NA | NA | NA | NA | NA | Gm14792 | | 0 | 0 | 0 | 0 | 0 | 0 | 0 | 0 | 0 | 0 | 0 |
| 0 | NA | NA | NA | NA | NA | Ppp4r3c-ps | | 0 | 0 | 0 | 0 | 0 | 0 | 0 | 0 | 0 | 0 | 0 |
| 0 | NA | NA | NA | NA | NA | Mir1906-2 | | 0 | 0 | 0 | 0 | 0 | 0 | 0 | 0 | 0 | 0 | 0 |
| 0 | NA | NA | NA | NA | NA | Gm27335 | | 0 | 0 | 0 | 0 | 0 | 0 | 0 | 0 | 0 | 0 | 0 |
| 0 | NA | NA | NA | NA | NA | Gm5759 | | 0 | 0 | 0 | 0 | 0 | 0 | 0 | 0 | 0 | 0 | 0 |
| 0 | NA | NA | NA | NA | NA | Gm14791 | | 0 | 0 | 0 | 0 | 0 | 0 | 0 | 0 | 0 | 0 | 0 |
| 0 | NA | NA | NA | NA | NA | Gm14796 | | 0 | 0 | 0 | 0 | 0 | 0 | 0 | 0 | 0 | 0 | 0 |
| 0 | NA | NA | NA | NA | NA | Gm14797 | | 0 | 0 | 0 | 0 | 0 | 0 | 0 | 0 | 0 | 0 | 0 |
| 0 | NA | NA | NA | NA | NA | Gm649 | | 0 | 0 | 0 | 0 | 0 | 0 | 0 | 0 | 0 | 0 | 0 |
| 0 | NA | NA | NA | NA | NA | | | 0 | 0 | 0 | 0 | 0 | 0 | 0 | 0 | 0 | 0 | 0 |
| 0 | NA | NA | NA | NA | NA | Gm14805 | | 0 | 0 | 0 | 0 | 0 | 0 | 0 | 0 | 0 | 0 | 0 |
| 0 | NA | NA | NA | NA | NA | Dcaf8l | DDB1 and CUL4 associated factor 8 like | 0 | 0 | 0 | 0 | 0 | 0 | 0 | 0 | 0 | 0 | 0 |
| 0 | NA | NA | NA | NA | NA | Gm14804 | | 0 | 0 | 0 | 0 | 0 | 0 | 0 | 0 | 0 | 0 | 0 |
| 0 | NA | NA | NA | NA | NA | Gm6973 | | 0 | 0 | 0 | 0 | 0 | 0 | 0 | 0 | 0 | 0 | 0 |
| 0 | NA | NA | NA | NA | NA | 4932429P05Ril | RIKEN cDNA 4932429P05 gene | 0 | 0 | 0 | 0 | 0 | 0 | 0 | 0 | 0 | 0 | 0 |
| 0 | NA | NA | NA | NA | NA | Gm24954 | | 0 | 0 | 0 | 0 | 0 | 0 | 0 | 0 | 0 | 0 | 0 |
| 0 | NA | NA | NA | NA | NA | Nanog-ps1 | | 0 | 0 | 0 | 0 | 0 | 0 | 0 | 0 | 0 | 0 | 0 |
| 0 | NA | NA | NA | NA | NA | 4930415L06Ril | RIKEN cDNA 4930415L06 gene | 0 | 0 | 0 | 0 | 0 | 0 | 0 | 0 | 0 | 0 | 0 |
| 0 | NA | NA | NA | NA | NA | Gm6027 | | 0 | 0 | 0 | 0 | 0 | 0 | 0 | 0 | 0 | 0 | 0 |
| 0 | NA | NA | NA | NA | NA | Gm4746 | | 0 | 0 | 0 | 0 | 0 | 0 | 0 | 0 | 0 | 0 | 0 |
| 0 | NA | NA | NA | NA | NA | Gm44 | predicted gene 44 | 0 | 0 | 0 | 0 | 0 | 0 | 0 | 0 | 0 | 0 | 0 |
| 0 | NA | NA | NA | NA | NA | Gm14773 | | 0 | 0 | 0 | 0 | 0 | 0 | 0 | 0 | 0 | 0 | 0 |
| 0 | NA | NA | NA | NA | NA | Gm24835 | | 0 | 0 | 0 | 0 | 0 | 0 | 0 | 0 | 0 | 0 | 0 |
| 0 | NA | NA | NA | NA | NA | Gm14786 | | 0 | 0 | 0 | 0 | 0 | 0 | 0 | 0 | 0 | 0 | 0 |
| 0 | NA | NA | NA | NA | NA | Gm8908 | | 0 | 0 | 0 | 0 | 0 | 0 | 0 | 0 | 0 | 0 | 0 |
| 0 | NA | NA | NA | NA | NA | Mageb2 | melanoma antigen, family B, 2 | 0 | 0 | 0 | 0 | 0 | 0 | 0 | 0 | 0 | 0 | 0 |
| 0 | NA | NA | NA | NA | NA | Gm14783 | | 0 | 0 | 0 | 0 | 0 | 0 | 0 | 0 | 0 | 0 | 0 |
| 0 | NA | NA | NA | NA | NA | Gm5072 | predicted gene 5072 | 0 | 0 | 0 | 0 | 0 | 0 | 0 | 0 | 0 | 0 | 0 |
| 0 | NA | NA | NA | NA | NA | Gm8914 | predicted gene 8914 | 0 | 0 | 0 | 0 | 0 | 0 | 0 | 0 | 0 | 0 | 0 |
| 0 | NA | NA | NA | NA | NA | Gm14787 | | 0 | 0 | 0 | 0 | 0 | 0 | 0 | 0 | 0 | 0 | 0 |
| 0 | NA | NA | NA | NA | NA | Gm8916 | | 0 | 0 | 0 | 0 | 0 | 0 | 0 | 0 | 0 | 0 | 0 |
| 0 | NA | NA | NA | NA | NA | Gm14785 | | 0 | 0 | 0 | 0 | 0 | 0 | 0 | 0 | 0 | 0 | 0 |
| 0 | NA | NA | NA | NA | NA | 1700084M14Rik | | 0 | 0 | 0 | 0 | 0 | 0 | 0 | 0 | 0 | 0 | 0 |
| 0 | NA | NA | NA | NA | NA | Mageb17-ps | melanoma antigen family B, 17, pseudog | 0 | 0 | 0 | 0 | 0 | 0 | 0 | 0 | 0 | 0 | 0 |
| 0 | NA | NA | NA | NA | NA | Gm14781 | predicted gene 14781 | 0 | 0 | 0 | 0 | 0 | 0 | 0 | 0 | 0 | 0 | 0 |
| 0 | NA | NA | NA | NA | NA | Mageb5 | melanoma antigen, family B, 5 | 0 | 0 | 0 | 0 | 0 | 0 | 0 | 0 | 0 | 0 | 0 |
| 0 | NA | NA | NA | NA | NA | Gm14782 | | 0 | 0 | 0 | 0 | 0 | 0 | 0 | 0 | 0 | 0 | 0 |
| 0 | NA | NA | NA | NA | NA | Gm7113 | | 0 | 0 | 0 | 0 | 0 | 0 | 0 | 0 | 0 | 0 | 0 |
| 0 | NA | NA | NA | NA | NA | Gm5071 | predicted pseudogene 5071 | 0 | 0 | 0 | 0 | 0 | 0 | 0 | 0 | 0 | 0 | 0 |
| 0 | NA | NA | NA | NA | NA | Gm14784 | | 0 | 0 | 0 | 0 | 0 | 0 | 0 | 0 | 0 | 0 | 0 |
| 0 | NA | NA | NA | NA | NA | Mageb1 | melanoma antigen, family B, 1 | 0 | 0 | 0 | 0 | 0 | 0 | 0 | 0 | 0 | 0 | 0 |
| 0 | NA | NA | NA | NA | NA | Gm4911 | | 0 | 0 | 0 | 0 | 0 | 0 | 0 | 0 | 0 | 0 | 0 |
| 0 | NA | NA | NA | NA | NA | Gm4912 | | 0 | 0 | 0 | 0 | 0 | 0 | 0 | 0 | 0 | 0 | 0 |
| 0 | NA | NA | NA | NA | NA | Mageb18 | melanoma antigen family B, 18 | 0 | 0 | 0 | 0 | 0 | 0 | 0 | 0 | 0 | 0 | 0 |
| 0 | NA | NA | NA | NA | NA | Gm14777 | | 0 | 0 | 0 | 0 | 0 | 0 | 0 | 0 | 0 | 0 | 0 |
| 0 | NA | NA | NA | NA | NA | Gm5941 | predicted gene 5941 | 0 | 0 | 0 | 0 | 0 | 0 | 0 | 0 | 0 | 0 | 0 |
| 0 | NA | NA | NA | NA | NA | Gm6982 | | 0 | 0 | 0 | 0 | 0 | 0 | 0 | 0 | 0 | 0 | 0 |

| | | | | | | | | | | | | | | | | | | |
|---|---|---|---|---|---|---|---|---|---|---|---|---|---|---|---|---|---|---|
| 0 | NA | NA | NA | NA | NA | Gm25332 | | 0 | 0 | 0 | 0 | 0 | 0 | 0 | 0 | 0 | 0 | 0 |
| 0 | NA | NA | NA | NA | NA | Gm24973 | | 0 | 0 | 0 | 0 | 0 | 0 | 0 | 0 | 0 | 0 | 0 |
| 0 | NA | NA | NA | NA | NA | 1700003E24Rik | | 0 | 0 | 0 | 0 | 0 | 0 | 0 | 0 | 0 | 0 | 0 |
| 0 | NA | NA | NA | NA | NA | Gm24123 | | 0 | 0 | 0 | 0 | 0 | 0 | 0 | 0 | 0 | 0 | 0 |
| 0 | NA | NA | NA | NA | NA | BC061195 | | 0 | 0 | 0 | 0 | 0 | 0 | 0 | 0 | 0 | 0 | 0 |
| 0 | NA | NA | NA | NA | NA | Gm23454 | | 0 | 0 | 0 | 0 | 0 | 0 | 0 | 0 | 0 | 0 | 0 |
| 0 | NA | NA | NA | NA | NA | Gm23557 | | 0 | 0 | 0 | 0 | 0 | 0 | 0 | 0 | 0 | 0 | 0 |
| 0 | NA | NA | NA | NA | NA | Gm6985 | | 0 | 0 | 0 | 0 | 0 | 0 | 0 | 0 | 0 | 0 | 0 |
| 0 | NA | NA | NA | NA | NA | Supt20-ps | | 0 | 0 | 0 | 0 | 0 | 0 | 0 | 0 | 0 | 0 | 0 |
| 0 | NA | NA | NA | NA | NA | Gm15164 | | 0 | 0 | 0 | 0 | 0 | 0 | 0 | 0 | 0 | 0 | 0 |
| 0 | NA | NA | NA | NA | NA | Gm14790 | | 0 | 0 | 0 | 0 | 0 | 0 | 0 | 0 | 0 | 0 | 0 |
| 0 | NA | NA | NA | NA | NA | Gm6987 | | 0 | 0 | 0 | 0 | 0 | 0 | 0 | 0 | 0 | 0 | 0 |
| 0 | NA | NA | NA | NA | NA | Gm14798 | | 0 | 0 | 0 | 0 | 0 | 0 | 0 | 0 | 0 | 0 | 0 |
| 0 | NA | NA | NA | NA | NA | Mir7673 | | 0 | 0 | 0 | 0 | 0 | 0 | 0 | 0 | 0 | 0 | 0 |
| 0 | NA | NA | NA | NA | NA | Gm5400 | | 0 | 0 | 0 | 0 | 0 | 0 | 0 | 0 | 0 | 0 | 0 |
| 0 | NA | NA | NA | NA | NA | Gm14826 | | 0 | 0 | 0 | 0 | 0 | 0 | 0 | 0 | 0 | 0 | 0 |
| 0 | NA | NA | NA | NA | NA | Gm14825 | | 0 | 0 | 0 | 0 | 0 | 0 | 0 | 0 | 0 | 0 | 0 |
| 0 | NA | NA | NA | NA | NA | | family with sequence similarity 90, memb | 0 | 0 | 0 | 0 | 0 | 0 | 0 | 0 | 0 | 0 | 0 |
| 0 | NA | NA | NA | NA | NA | Gm44477 | | 0 | 0 | 0 | 0 | 0 | 0 | 0 | 0 | 0 | 0 | 0 |
| 0 | NA | NA | NA | NA | NA | Gm9003 | | 0 | 0 | 0 | 0 | 0 | 0 | 0 | 0 | 0 | 0 | 0 |
| 0 | NA | NA | NA | NA | NA | | | 0 | 0 | 0 | 0 | 0 | 0 | 0 | 0 | 0 | 0 | 0 |
| 0 | NA | NA | NA | NA | NA | Gm24595 | | 0 | 0 | 0 | 0 | 0 | 0 | 0 | 0 | 0 | 0 | 0 |
| 0 | NA | NA | NA | NA | NA | Gm9005 | | 0 | 0 | 0 | 0 | 0 | 0 | 0 | 0 | 0 | 0 | 0 |
| 0 | NA | NA | NA | NA | NA | Gm14831 | | 0 | 0 | 0 | 0 | 0 | 0 | 0 | 0 | 0 | 0 | 0 |
| 0 | NA | NA | NA | NA | NA | Gm5401 | | 0 | 0 | 0 | 0 | 0 | 0 | 0 | 0 | 0 | 0 | 0 |
| 0 | NA | NA | NA | NA | NA | Gm5402 | | 0 | 0 | 0 | 0 | 0 | 0 | 0 | 0 | 0 | 0 | 0 |
| 0 | NA | NA | NA | NA | NA | Rpl13-ps5 | | 0 | 0 | 0 | 0 | 0 | 0 | 0 | 0 | 0 | 0 | 0 |
| 0 | NA | NA | NA | NA | NA | Gm371 | | 0 | 0 | 0 | 0 | 0 | 0 | 0 | 0 | 0 | 0 | 0 |
| 0 | NA | NA | NA | NA | NA | Gm6092 | | 0 | 0 | 0 | 0 | 0 | 0 | 0 | 0 | 0 | 0 | 0 |
| 0 | NA | NA | NA | NA | NA | Gm3712 | | 0 | 0 | 0 | 0 | 0 | 0 | 0 | 0 | 0 | 0 | 0 |
| 0 | NA | NA | NA | NA | NA | Gm14806 | | 0 | 0 | 0 | 0 | 0 | 0 | 0 | 0 | 0 | 0 | 0 |
| 0 | NA | NA | NA | NA | NA | Asb12 | ankyrin repeat and SOCS box-containing | 0 | 0 | 0 | 0 | 0 | 0 | 0 | 0 | 0 | 0 | 0 |
| 0 | NA | NA | NA | NA | NA | Pfn5 | | 0 | 0 | 0 | 0 | 0 | 0 | 0 | 0 | 0 | 0 | 0 |
| 0 | NA | NA | NA | NA | NA | Gm9043 | | 0 | 0 | 0 | 0 | 0 | 0 | 0 | 0 | 0 | 0 | 0 |
| 0 | NA | NA | NA | NA | NA | 1700010D01Ri | RIKEN cDNA 1700010D01 gene | 0 | 0 | 0 | 0 | 0 | 0 | 0 | 0 | 0 | 0 | 0 |
| 0 | NA | NA | NA | NA | NA | Gm24718 | | 0 | 0 | 0 | 0 | 0 | 0 | 0 | 0 | 0 | 0 | 0 |
| 0 | NA | NA | NA | NA | NA | Mir223 | | 0 | 0 | 0 | 0 | 0 | 0 | 0 | 0 | 0 | 0 | 0 |
| 0 | NA | NA | NA | NA | NA | Hsf3 | heat shock transcription factor 3 | 0 | 0 | 0 | 0 | 0 | 0 | 0 | 0 | 0 | 0 | 0 |
| 0 | NA | NA | NA | NA | NA | Gm14933 | | 0 | 0 | 0 | 0 | 0 | 0 | 0 | 0 | 0 | 0 | 0 |
| 0 | NA | NA | NA | NA | NA | Gm14935 | | 0 | 0 | 0 | 0 | 0 | 0 | 0 | 0 | 0 | 0 | 0 |
| 0 | NA | NA | NA | NA | NA | Gm16394 | | 0 | 0 | 0 | 0 | 0 | 0 | 0 | 0 | 0 | 0 | 0 |
| 0 | NA | NA | NA | NA | NA | Gm14803 | | 0 | 0 | 0 | 0 | 0 | 0 | 0 | 0 | 0 | 0 | 0 |
| 0 | NA | NA | NA | NA | NA | Gm9059 | | 0 | 0 | 0 | 0 | 0 | 0 | 0 | 0 | 0 | 0 | 0 |
| 0 | NA | NA | NA | NA | NA | Gm9060 | | 0 | 0 | 0 | 0 | 0 | 0 | 0 | 0 | 0 | 0 | 0 |
| 0 | NA | NA | NA | NA | NA | Pgr15l | G protein-coupled receptor 15-like | 0 | 0 | 0 | 0 | 0 | 0 | 0 | 0 | 0 | 0 | 0 |
| 0 | NA | NA | NA | NA | NA | Gm14800 | | 0 | 0 | 0 | 0 | 0 | 0 | 0 | 0 | 0 | 0 | 0 |
| 0 | NA | NA | NA | NA | NA | Gm9062 | | 0 | 0 | 0 | 0 | 0 | 0 | 0 | 0 | 0 | 0 | 0 |
| 0 | NA | NA | NA | NA | NA | Gm14802 | | 0 | 0 | 0 | 0 | 0 | 0 | 0 | 0 | 0 | 0 | 0 |
| 0 | NA | NA | NA | NA | NA | Gm14811 | | 0 | 0 | 0 | 0 | 0 | 0 | 0 | 0 | 0 | 0 | 0 |
| 0 | NA | NA | NA | NA | NA | Gm14815 | | 0 | 0 | 0 | 0 | 0 | 0 | 0 | 0 | 0 | 0 | 0 |
| 0 | NA | NA | NA | NA | NA | Gm27554 | | 0 | 0 | 0 | 0 | 0 | 0 | 0 | 0 | 0 | 0 | 0 |
| 0 | NA | NA | NA | NA | NA | Gm26000 | | 0 | 0 | 0 | 0 | 0 | 0 | 0 | 0 | 0 | 0 | 0 |
| 0 | NA | NA | NA | NA | NA | Gm14809 | | 0 | 0 | 0 | 0 | 0 | 0 | 0 | 0 | 0 | 0 | 0 |
| 0 | NA | NA | NA | NA | NA | Gm14808 | | 0 | 0 | 0 | 0 | 0 | 0 | 0 | 0 | 0 | 0 | 0 |
| 0 | NA | NA | NA | NA | NA | Gm14816 | | 0 | 0 | 0 | 0 | 0 | 0 | 0 | 0 | 0 | 0 | 0 |
| 0 | NA | NA | NA | NA | NA | Pcna-ps1 | | 0 | 0 | 0 | 0 | 0 | 0 | 0 | 0 | 0 | 0 | 0 |
| 0 | NA | NA | NA | NA | NA | Gm14832 | | 0 | 0 | 0 | 0 | 0 | 0 | 0 | 0 | 0 | 0 | 0 |
| 0 | NA | NA | NA | NA | NA | Mir676 | | 0 | 0 | 0 | 0 | 0 | 0 | 0 | 0 | 0 | 0 | 0 |
| 0 | NA | NA | NA | NA | NA | Otud6a | OTU domain containing 6A | 0 | 0 | 0 | 0 | 0 | 0 | 0 | 0 | 0 | 0 | 0 |
| 0 | NA | NA | NA | NA | NA | Dgat2l6 | diacylglycerol O-acyltransferase 2-like 6 | 0 | 0 | 0 | 0 | 0 | 0 | 0 | 0 | 0 | 0 | 0 |

| | | | | | | | | | | | | | | | | | | |
|---|---|---|---|---|---|---|---|---|---|---|---|---|---|---|---|---|---|---|
| 0 | NA | NA | NA | NA | NA | Awat1 | acyl-CoA wax alcohol acyltransferase 1 | 0 | 0 | 0 | 0 | 0 | 0 | 0 | 0 | 0 | 0 | 0 |
| 0 | NA | NA | NA | NA | NA | Gm6175 | | 0 | 0 | 0 | 0 | 0 | 0 | 0 | 0 | 0 | 0 | 0 |
| 0 | NA | NA | NA | NA | NA | P2ry4 | pyrimidinergic receptor P2Y, G-protein co | 0 | 0 | 0 | 0 | 0 | 0 | 0 | 0 | 0 | 0 | 0 |
| 0 | NA | NA | NA | NA | NA | Arr3 | arrestin 3, retinal | 0 | 0 | 0 | 0 | 0 | 0 | 0 | 0 | 0 | 0 | 0 |
| 0 | NA | NA | NA | NA | NA | Gm14866 | | 0 | 0 | 0 | 0 | 0 | 0 | 0 | 0 | 0 | 0 | 0 |
| 0 | NA | NA | NA | NA | NA | Gm14902 | | 0 | 0 | 0 | 0 | 0 | 0 | 0 | 0 | 0 | 0 | 0 |
| 0 | NA | NA | NA | NA | NA | Gm22160 | | 0 | 0 | 0 | 0 | 0 | 0 | 0 | 0 | 0 | 0 | 0 |
| 0 | NA | NA | NA | NA | NA | Gm24577 | | 0 | 0 | 0 | 0 | 0 | 0 | 0 | 0 | 0 | 0 | 0 |
| 0 | NA | NA | NA | NA | NA | Gm27510 | | 0 | 0 | 0 | 0 | 0 | 0 | 0 | 0 | 0 | 0 | 0 |
| 0 | NA | NA | NA | NA | NA | Gm14901 | | 0 | 0 | 0 | 0 | 0 | 0 | 0 | 0 | 0 | 0 | 0 |
| 0 | NA | NA | NA | NA | NA | Gm24061 | | 0 | 0 | 0 | 0 | 0 | 0 | 0 | 0 | 0 | 0 | 0 |
| 0 | NA | NA | NA | NA | NA | Gm14842 | | 0 | 0 | 0 | 0 | 0 | 0 | 0 | 0 | 0 | 0 | 0 |
| 0 | NA | NA | NA | NA | NA | Gm22472 | | 0 | 0 | 0 | 0 | 0 | 0 | 0 | 0 | 0 | 0 | 0 |
| 0 | NA | NA | NA | NA | NA | Gm614 | predicted gene 614 | 0 | 0 | 0 | 0 | 0 | 0 | 0 | 0 | 0 | 0 | 0 |
| 0 | NA | NA | NA | NA | NA | Gm20489 | | 0 | 0 | 0 | 0 | 0 | 0 | 0 | 0 | 0 | 0 | 0 |
| 0 | NA | NA | NA | NA | NA | Gm50476 | | 0 | 0 | 0 | 0 | 0 | 0 | 0 | 0 | 0 | 0 | 0 |
| 0 | NA | NA | NA | NA | NA | Gm9089 | | 0 | 0 | 0 | 0 | 0 | 0 | 0 | 0 | 0 | 0 | 0 |
| 0 | NA | NA | NA | NA | NA | Gm9097 | | 0 | 0 | 0 | 0 | 0 | 0 | 0 | 0 | 0 | 0 | 0 |
| 0 | NA | NA | NA | NA | NA | | | 0 | 0 | 0 | 0 | 0 | 0 | 0 | 0 | 0 | 0 | 0 |
| 0 | NA | NA | NA | NA | NA | Gm25828 | | 0 | 0 | 0 | 0 | 0 | 0 | 0 | 0 | 0 | 0 | 0 |
| 0 | NA | NA | NA | NA | NA | Gm27843 | | 0 | 0 | 0 | 0 | 0 | 0 | 0 | 0 | 0 | 0 | 0 |
| 0 | NA | NA | NA | NA | NA | Gm5125 | | 0 | 0 | 0 | 0 | 0 | 0 | 0 | 0 | 0 | 0 | 0 |
| 0 | NA | NA | NA | NA | NA | Gm26318 | | 0 | 0 | 0 | 0 | 0 | 0 | 0 | 0 | 0 | 0 | 0 |
| 0 | NA | NA | NA | NA | NA | Gm14840 | | 0 | 0 | 0 | 0 | 0 | 0 | 0 | 0 | 0 | 0 | 0 |
| 0 | NA | NA | NA | NA | NA | Gm3880 | | 0 | 0 | 0 | 0 | 0 | 0 | 0 | 0 | 0 | 0 | 0 |
| 0 | NA | NA | NA | NA | NA | Gm9112 | predicted gene 9112 | 0 | 0 | 0 | 0 | 0 | 0 | 0 | 0 | 0 | 0 | 0 |
| 0 | NA | NA | NA | NA | NA | Dmrtc1b | DMRT-like family C1b | 0 | 0 | 0 | 0 | 0 | 0 | 0 | 0 | 0 | 0 | 0 |
| 0 | NA | NA | NA | NA | NA | Gm14839 | | 0 | 0 | 0 | 0 | 0 | 0 | 0 | 0 | 0 | 0 | 0 |
| 0 | NA | NA | NA | NA | NA | Gm4234 | | 0 | 0 | 0 | 0 | 0 | 0 | 0 | 0 | 0 | 0 | 0 |
| 0 | NA | NA | NA | NA | NA | Gm9119 | | 0 | 0 | 0 | 0 | 0 | 0 | 0 | 0 | 0 | 0 | 0 |
| 0 | NA | NA | NA | NA | NA | Gm5166 | | 0 | 0 | 0 | 0 | 0 | 0 | 0 | 0 | 0 | 0 | 0 |
| 0 | NA | NA | NA | NA | NA | Gm25358 | | 0 | 0 | 0 | 0 | 0 | 0 | 0 | 0 | 0 | 0 | 0 |
| 0 | NA | NA | NA | NA | NA | Dmrtc1c1 | DMRT-like family C1c1 | 0 | 0 | 0 | 0 | 0 | 0 | 0 | 0 | 0 | 0 | 0 |
| 0 | NA | NA | NA | NA | NA | Gm9126 | | 0 | 0 | 0 | 0 | 0 | 0 | 0 | 0 | 0 | 0 | 0 |
| 0 | NA | NA | NA | NA | NA | Gm16511 | | 0 | 0 | 0 | 0 | 0 | 0 | 0 | 0 | 0 | 0 | 0 |
| 0 | NA | NA | NA | NA | NA | Gm24509 | | 0 | 0 | 0 | 0 | 0 | 0 | 0 | 0 | 0 | 0 | 0 |
| 0 | NA | NA | NA | NA | NA | Dmrtc1c2 | DMRT-like family C1c2 | 0 | 0 | 0 | 0 | 0 | 0 | 0 | 0 | 0 | 0 | 0 |
| 0 | NA | NA | NA | NA | NA | Gm26152 | | 0 | 0 | 0 | 0 | 0 | 0 | 0 | 0 | 0 | 0 | 0 |
| 0 | NA | NA | NA | NA | NA | Gm9133 | | 0 | 0 | 0 | 0 | 0 | 0 | 0 | 0 | 0 | 0 | 0 |
| 0 | NA | NA | NA | NA | NA | 1700011M02Rik | | 0 | 0 | 0 | 0 | 0 | 0 | 0 | 0 | 0 | 0 | 0 |
| 0 | NA | NA | NA | NA | NA | 1700018G05Rik | | 0 | 0 | 0 | 0 | 0 | 0 | 0 | 0 | 0 | 0 | 0 |
| 0 | NA | NA | NA | NA | NA | Gm14841 | | 0 | 0 | 0 | 0 | 0 | 0 | 0 | 0 | 0 | 0 | 0 |
| 0 | NA | NA | NA | NA | NA | Gm14837 | | 0 | 0 | 0 | 0 | 0 | 0 | 0 | 0 | 0 | 0 | 0 |
| 0 | NA | NA | NA | NA | NA | Pabpc1l2b-ps | | 0 | 0 | 0 | 0 | 0 | 0 | 0 | 0 | 0 | 0 | 0 |
| 0 | NA | NA | NA | NA | NA | Pabpc1l2a-ps | | 0 | 0 | 0 | 0 | 0 | 0 | 0 | 0 | 0 | 0 | 0 |
| 0 | NA | NA | NA | NA | NA | Gm14838 | | 0 | 0 | 0 | 0 | 0 | 0 | 0 | 0 | 0 | 0 | 0 |
| 0 | NA | NA | NA | NA | NA | Gm9050 | | 0 | 0 | 0 | 0 | 0 | 0 | 0 | 0 | 0 | 0 | 0 |
| 0 | NA | NA | NA | NA | NA | Gm14847 | | 0 | 0 | 0 | 0 | 0 | 0 | 0 | 0 | 0 | 0 | 0 |
| 0 | NA | NA | NA | NA | NA | Gm14848 | | 0 | 0 | 0 | 0 | 0 | 0 | 0 | 0 | 0 | 0 | 0 |
| 0 | NA | NA | NA | NA | NA | Gm3931 | | 0 | 0 | 0 | 0 | 0 | 0 | 0 | 0 | 0 | 0 | 0 |
| 0 | NA | NA | NA | NA | NA | Gm9516 | | 0 | 0 | 0 | 0 | 0 | 0 | 0 | 0 | 0 | 0 | 0 |
| 0 | NA | NA | NA | NA | NA | Gm26952 | | 0 | 0 | 0 | 0 | 0 | 0 | 0 | 0 | 0 | 0 | 0 |
| 0 | NA | NA | NA | NA | NA | Gm26992 | | 0 | 0 | 0 | 0 | 0 | 0 | 0 | 0 | 0 | 0 | 0 |
| 0 | NA | NA | NA | NA | NA | Gm27733 | | 0 | 0 | 0 | 0 | 0 | 0 | 0 | 0 | 0 | 0 | 0 |
| 0 | NA | NA | NA | NA | NA | Gm27927 | | 0 | 0 | 0 | 0 | 0 | 0 | 0 | 0 | 0 | 0 | 0 |
| 0 | NA | NA | NA | NA | NA | Gm27520 | | 0 | 0 | 0 | 0 | 0 | 0 | 0 | 0 | 0 | 0 | 0 |
| 0 | NA | NA | NA | NA | NA | Gm27875 | | 0 | 0 | 0 | 0 | 0 | 0 | 0 | 0 | 0 | 0 | 0 |
| 0 | NA | NA | NA | NA | NA | Gm14860 | | 0 | 0 | 0 | 0 | 0 | 0 | 0 | 0 | 0 | 0 | 0 |
| 0 | NA | NA | NA | NA | NA | Gm9785 | | 0 | 0 | 0 | 0 | 0 | 0 | 0 | 0 | 0 | 0 | 0 |
| 0 | NA | NA | NA | NA | NA | Mir374c | | 0 | 0 | 0 | 0 | 0 | 0 | 0 | 0 | 0 | 0 | 0 |

| | | | | | | | | | | | | | | | | | | |
|---|---|---|---|---|---|---|---|---|---|---|---|---|---|---|---|---|---|---|
| 0 | NA | NA | NA | NA | NA | Gm27798 | | 0 | 0 | 0 | 0 | 0 | 0 | 0 | 0 | 0 | 0 | 0 |
| 0 | NA | NA | NA | NA | NA | Gm9159 | | 0 | 0 | 0 | 0 | 0 | 0 | 0 | 0 | 0 | 0 | 0 |
| 0 | NA | NA | NA | NA | NA | Gm27987 | | 0 | 0 | 0 | 0 | 0 | 0 | 0 | 0 | 0 | 0 | 0 |
| 0 | NA | NA | NA | NA | NA | Gm27285 | | 0 | 0 | 0 | 0 | 0 | 0 | 0 | 0 | 0 | 0 | 0 |
| 0 | NA | NA | NA | NA | NA | Gm14874 | | 0 | 0 | 0 | 0 | 0 | 0 | 0 | 0 | 0 | 0 | 0 |
| 0 | NA | NA | NA | NA | NA | Gm14875 | | 0 | 0 | 0 | 0 | 0 | 0 | 0 | 0 | 0 | 0 | 0 |
| 0 | NA | NA | NA | NA | NA | Gm3943 | | 0 | 0 | 0 | 0 | 0 | 0 | 0 | 0 | 0 | 0 | 0 |
| 0 | NA | NA | NA | NA | NA | Gm37956 | | 0 | 0 | 0 | 0 | 0 | 0 | 0 | 0 | 0 | 0 | 0 |
| 0 | NA | NA | NA | NA | NA | Gm14893 | | 0 | 0 | 0 | 0 | 0 | 0 | 0 | 0 | 0 | 0 | 0 |
| 0 | NA | NA | NA | NA | NA | Gm9166 | | 0 | 0 | 0 | 0 | 0 | 0 | 0 | 0 | 0 | 0 | 0 |
| 0 | NA | NA | NA | NA | NA | Gm14892 | | 0 | 0 | 0 | 0 | 0 | 0 | 0 | 0 | 0 | 0 | 0 |
| 0 | NA | NA | NA | NA | NA | Gm23378 | | 0 | 0 | 0 | 0 | 0 | 0 | 0 | 0 | 0 | 0 | 0 |
| 0 | NA | NA | NA | NA | NA | Gm22266 | | 0 | 0 | 0 | 0 | 0 | 0 | 0 | 0 | 0 | 0 | 0 |
| 0 | NA | NA | NA | NA | NA | Gm24993 | | 0 | 0 | 0 | 0 | 0 | 0 | 0 | 0 | 0 | 0 | 0 |
| 0 | NA | NA | NA | NA | NA | Mir672 | | 0 | 0 | 0 | 0 | 0 | 0 | 0 | 0 | 0 | 0 | 0 |
| 0 | NA | NA | NA | NA | NA | Gm22430 | | 0 | 0 | 0 | 0 | 0 | 0 | 0 | 0 | 0 | 0 | 0 |
| 0 | NA | NA | NA | NA | NA | 1700121L16Rik | | 0 | 0 | 0 | 0 | 0 | 0 | 0 | 0 | 0 | 0 | 0 |
| 0 | NA | NA | NA | NA | NA | Gm22090 | | 0 | 0 | 0 | 0 | 0 | 0 | 0 | 0 | 0 | 0 | 0 |
| 0 | NA | NA | NA | NA | NA | Gm9673 | | 0 | 0 | 0 | 0 | 0 | 0 | 0 | 0 | 0 | 0 | 0 |
| 0 | NA | NA | NA | NA | NA | Mir325 | | 0 | 0 | 0 | 0 | 0 | 0 | 0 | 0 | 0 | 0 | 0 |
| 0 | NA | NA | NA | NA | NA | Rab9b-ps1 | | 0 | 0 | 0 | 0 | 0 | 0 | 0 | 0 | 0 | 0 | 0 |
| 0 | NA | NA | NA | NA | NA | Cypt2 | cysteine-rich perinuclear theca 2 | 0 | 0 | 0 | 0 | 0 | 0 | 0 | 0 | 0 | 0 | 0 |
| 0 | NA | NA | NA | NA | NA | Gm14823 | | 0 | 0 | 0 | 0 | 0 | 0 | 0 | 0 | 0 | 0 | 0 |
| 0 | NA | NA | NA | NA | NA | Gm14855 | | 0 | 0 | 0 | 0 | 0 | 0 | 0 | 0 | 0 | 0 | 0 |
| 0 | NA | NA | NA | NA | NA | Gm14854 | | 0 | 0 | 0 | 0 | 0 | 0 | 0 | 0 | 0 | 0 | 0 |
| 0 | NA | NA | NA | NA | NA | Gm26020 | | 0 | 0 | 0 | 0 | 0 | 0 | 0 | 0 | 0 | 0 | 0 |
| 0 | NA | NA | NA | NA | NA | Gm23656 | | 0 | 0 | 0 | 0 | 0 | 0 | 0 | 0 | 0 | 0 | 0 |
| 0 | NA | NA | NA | NA | NA | Gm14856 | | 0 | 0 | 0 | 0 | 0 | 0 | 0 | 0 | 0 | 0 | 0 |
| 0 | NA | NA | NA | NA | NA | Fnd3c2 | fibronectin type III domain containing 3C2 | 0 | 0 | 0 | 0 | 0 | 0 | 0 | 0 | 0 | 0 | 0 |
| 0 | NA | NA | NA | NA | NA | Gm14852 | | 0 | 0 | 0 | 0 | 0 | 0 | 0 | 0 | 0 | 0 | 0 |
| 0 | NA | NA | NA | NA | NA | Gm5133 | | 0 | 0 | 0 | 0 | 0 | 0 | 0 | 0 | 0 | 0 | 0 |
| 0 | NA | NA | NA | NA | NA | Mir6382 | | 0 | 0 | 0 | 0 | 0 | 0 | 0 | 0 | 0 | 0 | 0 |
| 0 | NA | NA | NA | NA | NA | Gm14876 | | 0 | 0 | 0 | 0 | 0 | 0 | 0 | 0 | 0 | 0 | 0 |
| 0 | NA | NA | NA | NA | NA | Gm6325 | | 0 | 0 | 0 | 0 | 0 | 0 | 0 | 0 | 0 | 0 | 0 |
| 0 | NA | NA | NA | NA | NA | Fndc3c1 | fibronectin type III domain containing 3C1 | 0 | 0 | 0 | 0 | 0 | 0 | 0 | 0 | 0 | 0 | 0 |
| 0 | NA | NA | NA | NA | NA | Gm27428 | | 0 | 0 | 0 | 0 | 0 | 0 | 0 | 0 | 0 | 0 | 0 |
| 0 | NA | NA | NA | NA | NA | Gm24622 | | 0 | 0 | 0 | 0 | 0 | 0 | 0 | 0 | 0 | 0 | 0 |
| 0 | NA | NA | NA | NA | NA | Gm14877 | | 0 | 0 | 0 | 0 | 0 | 0 | 0 | 0 | 0 | 0 | 0 |
| 0 | NA | NA | NA | NA | NA | Gm14878 | | 0 | 0 | 0 | 0 | 0 | 0 | 0 | 0 | 0 | 0 | 0 |
| 0 | NA | NA | NA | NA | NA | Gm14879 | | 0 | 0 | 0 | 0 | 0 | 0 | 0 | 0 | 0 | 0 | 0 |
| 0 | NA | NA | NA | NA | NA | Gm24491 | | 0 | 0 | 0 | 0 | 0 | 0 | 0 | 0 | 0 | 0 | 0 |
| 0 | NA | NA | NA | NA | NA | Gm22102 | | 0 | 0 | 0 | 0 | 0 | 0 | 0 | 0 | 0 | 0 | 0 |
| 0 | NA | NA | NA | NA | NA | Gm14882 | | 0 | 0 | 0 | 0 | 0 | 0 | 0 | 0 | 0 | 0 | 0 |
| 0 | NA | NA | NA | NA | NA | Gm14881 | | 0 | 0 | 0 | 0 | 0 | 0 | 0 | 0 | 0 | 0 | 0 |
| 0 | NA | NA | NA | NA | NA | Gm14880 | | 0 | 0 | 0 | 0 | 0 | 0 | 0 | 0 | 0 | 0 | 0 |
| 0 | NA | NA | NA | NA | NA | Gm14834 | | 0 | 0 | 0 | 0 | 0 | 0 | 0 | 0 | 0 | 0 | 0 |
| 0 | NA | NA | NA | NA | NA | Gm14836 | | 0 | 0 | 0 | 0 | 0 | 0 | 0 | 0 | 0 | 0 | 0 |
| 0 | NA | NA | NA | NA | NA | Gm9208 | | 0 | 0 | 0 | 0 | 0 | 0 | 0 | 0 | 0 | 0 | 0 |
| 0 | NA | NA | NA | NA | NA | Gm14833 | | 0 | 0 | 0 | 0 | 0 | 0 | 0 | 0 | 0 | 0 | 0 |
| 0 | NA | NA | NA | NA | NA | Gm14835 | | 0 | 0 | 0 | 0 | 0 | 0 | 0 | 0 | 0 | 0 | 0 |
| 0 | NA | NA | NA | NA | NA | 4933401B06Rik | | 0 | 0 | 0 | 0 | 0 | 0 | 0 | 0 | 0 | 0 | 0 |
| 0 | NA | NA | NA | NA | NA | Tbx22 | T-box 22 | 0 | 0 | 0 | 0 | 0 | 0 | 0 | 0 | 0 | 0 | 0 |
| 0 | NA | NA | NA | NA | NA | Gm732 | predicted gene 732 | 0 | 0 | 0 | 0 | 0 | 0 | 0 | 0 | 0 | 0 | 0 |
| 0 | NA | NA | NA | NA | NA | Gm14849 | | 0 | 0 | 0 | 0 | 0 | 0 | 0 | 0 | 0 | 0 | 0 |
| 0 | NA | NA | NA | NA | NA | Gm378 | | 0 | 0 | 0 | 0 | 0 | 0 | 0 | 0 | 0 | 0 | 0 |
| 0 | NA | NA | NA | NA | NA | Gm9218 | | 0 | 0 | 0 | 0 | 0 | 0 | 0 | 0 | 0 | 0 | 0 |
| 0 | NA | NA | NA | NA | NA | Gm25797 | | 0 | 0 | 0 | 0 | 0 | 0 | 0 | 0 | 0 | 0 | 0 |
| 0 | NA | NA | NA | NA | NA | Gm26415 | | 0 | 0 | 0 | 0 | 0 | 0 | 0 | 0 | 0 | 0 | 0 |
| 0 | NA | NA | NA | NA | NA | Gm14869 | | 0 | 0 | 0 | 0 | 0 | 0 | 0 | 0 | 0 | 0 | 0 |
| 0 | NA | NA | NA | NA | NA | Gm14868 | | 0 | 0 | 0 | 0 | 0 | 0 | 0 | 0 | 0 | 0 | 0 |

| | | | | | | | | | | | | | | | | | | |
|---|---|---|---|---|---|---|---|---|---|---|---|---|---|---|---|---|---|---|
| 0 | NA | NA | NA | NA | NA | Gm4991 | | 0 | 0 | 0 | 0 | 0 | 0 | 0 | 0 | 0 | 0 | 0 |
| 0 | NA | NA | NA | NA | NA | Gm7134 | | 0 | 0 | 0 | 0 | 0 | 0 | 0 | 0 | 0 | 0 | 0 |
| 0 | NA | NA | NA | NA | NA | Gm9225 | | 0 | 0 | 0 | 0 | 0 | 0 | 0 | 0 | 0 | 0 | 0 |
| 0 | NA | NA | NA | NA | NA | Gm14886 | | 0 | 0 | 0 | 0 | 0 | 0 | 0 | 0 | 0 | 0 | 0 |
| 0 | NA | NA | NA | NA | NA | Gm26108 | | 0 | 0 | 0 | 0 | 0 | 0 | 0 | 0 | 0 | 0 | 0 |
| 0 | NA | NA | NA | NA | NA | Gm44593 | | 0 | 0 | 0 | 0 | 0 | 0 | 0 | 0 | 0 | 0 | 0 |
| 0 | NA | NA | NA | NA | NA | Cylc1 | cylicin, basic protein of sperm head cytosl | 0 | 0 | 0 | 0 | 0 | 0 | 0 | 0 | 0 | 0 | 0 |
| 0 | NA | NA | NA | NA | NA | Gm10112 | | 0 | 0 | 0 | 0 | 0 | 0 | 0 | 0 | 0 | 0 | 0 |
| 0 | NA | NA | NA | NA | NA | Gm7340 | | 0 | 0 | 0 | 0 | 0 | 0 | 0 | 0 | 0 | 0 | 0 |
| 0 | NA | NA | NA | NA | NA | Gm14898 | | 0 | 0 | 0 | 0 | 0 | 0 | 0 | 0 | 0 | 0 | 0 |
| 0 | NA | NA | NA | NA | NA | Gm14899 | | 0 | 0 | 0 | 0 | 0 | 0 | 0 | 0 | 0 | 0 | 0 |
| 0 | NA | NA | NA | NA | NA | Gm45194 | | 0 | 0 | 0 | 0 | 0 | 0 | 0 | 0 | 0 | 0 | 0 |
| 0 | NA | NA | NA | NA | NA | Gm14900 | | 0 | 0 | 0 | 0 | 0 | 0 | 0 | 0 | 0 | 0 | 0 |
| 0 | NA | NA | NA | NA | NA | 4933403O08Ril | RIKEN cDNA 4933403O08 gene | 0 | 0 | 0 | 0 | 0 | 0 | 0 | 0 | 0 | 0 | 0 |
| 0 | NA | NA | NA | NA | NA | Gm5393 | | 0 | 0 | 0 | 0 | 0 | 0 | 0 | 0 | 0 | 0 | 0 |
| 0 | NA | NA | NA | NA | NA | Satl1 | spermidine/spermine N1-acetyl transferas | 0 | 0 | 0 | 0 | 0 | 0 | 0 | 0 | 0 | 0 | 0 |
| 0 | NA | NA | NA | NA | NA | Gm6418 | | 0 | 0 | 0 | 0 | 0 | 0 | 0 | 0 | 0 | 0 | 0 |
| 0 | NA | NA | NA | NA | NA | Gm14908 | | 0 | 0 | 0 | 0 | 0 | 0 | 0 | 0 | 0 | 0 | 0 |
| 0 | NA | NA | NA | NA | NA | 2010106E10Ril | RIKEN cDNA 2010106E10 gene | 0 | 0 | 0 | 0 | 0 | 0 | 0 | 0 | 0 | 0 | 0 |
| 0 | NA | NA | NA | NA | NA | Gm14921 | | 0 | 0 | 0 | 0 | 0 | 0 | 0 | 0 | 0 | 0 | 0 |
| 0 | NA | NA | NA | NA | NA | Gm14922 | | 0 | 0 | 0 | 0 | 0 | 0 | 0 | 0 | 0 | 0 | 0 |
| 0 | NA | NA | NA | NA | NA | Gm14923 | | 0 | 0 | 0 | 0 | 0 | 0 | 0 | 0 | 0 | 0 | 0 |
| 0 | NA | NA | NA | NA | NA | Gm6427 | | 0 | 0 | 0 | 0 | 0 | 0 | 0 | 0 | 0 | 0 | 0 |
| 0 | NA | NA | NA | NA | NA | Gm6429 | | 0 | 0 | 0 | 0 | 0 | 0 | 0 | 0 | 0 | 0 | 0 |
| 0 | NA | NA | NA | NA | NA | Gm14937 | | 0 | 0 | 0 | 0 | 0 | 0 | 0 | 0 | 0 | 0 | 0 |
| 0 | NA | NA | NA | NA | NA | Gm14938 | | 0 | 0 | 0 | 0 | 0 | 0 | 0 | 0 | 0 | 0 | 0 |
| 0 | NA | NA | NA | NA | NA | Gm14936 | | 0 | 0 | 0 | 0 | 0 | 0 | 0 | 0 | 0 | 0 | 0 |
| 0 | NA | NA | NA | NA | NA | Gm14939 | | 0 | 0 | 0 | 0 | 0 | 0 | 0 | 0 | 0 | 0 | 0 |
| 0 | NA | NA | NA | NA | NA | Mir361 | | 0 | 0 | 0 | 0 | 0 | 0 | 0 | 0 | 0 | 0 | 0 |
| 0 | NA | NA | NA | NA | NA | Gm7405 | | 0 | 0 | 0 | 0 | 0 | 0 | 0 | 0 | 0 | 0 | 0 |
| 0 | NA | NA | NA | NA | NA | Gm24831 | | 0 | 0 | 0 | 0 | 0 | 0 | 0 | 0 | 0 | 0 | 0 |
| 0 | NA | NA | NA | NA | NA | Gm23985 | | 0 | 0 | 0 | 0 | 0 | 0 | 0 | 0 | 0 | 0 | 0 |
| 0 | NA | NA | NA | NA | NA | Gm4915 | | 0 | 0 | 0 | 0 | 0 | 0 | 0 | 0 | 0 | 0 | 0 |
| 0 | NA | NA | NA | NA | NA | Gm14896 | | 0 | 0 | 0 | 0 | 0 | 0 | 0 | 0 | 0 | 0 | 0 |
| 0 | NA | NA | NA | NA | NA | Gm7146 | | 0 | 0 | 0 | 0 | 0 | 0 | 0 | 0 | 0 | 0 | 0 |
| 0 | NA | NA | NA | NA | NA | Gm14895 | | 0 | 0 | 0 | 0 | 0 | 0 | 0 | 0 | 0 | 0 | 0 |
| 0 | NA | NA | NA | NA | NA | Gm14894 | | 0 | 0 | 0 | 0 | 0 | 0 | 0 | 0 | 0 | 0 | 0 |
| 0 | NA | NA | NA | NA | NA | Gm5942 | | 0 | 0 | 0 | 0 | 0 | 0 | 0 | 0 | 0 | 0 | 0 |
| 0 | NA | NA | NA | NA | NA | Gm14897 | | 0 | 0 | 0 | 0 | 0 | 0 | 0 | 0 | 0 | 0 | 0 |
| 0 | NA | NA | NA | NA | NA | Ube2dnl1 | ubiquitin-conjugating enzyme E2D N-tern | 0 | 0 | 0 | 0 | 0 | 0 | 0 | 0 | 0 | 0 | 0 |
| 0 | NA | NA | NA | NA | NA | Ube2dnl2 | ubiquitin-conjugating enzyme E2D N-tern | 0 | 0 | 0 | 0 | 0 | 0 | 0 | 0 | 0 | 0 | 0 |
| 0 | NA | NA | NA | NA | NA | 4930555B12Rik | | 0 | 0 | 0 | 0 | 0 | 0 | 0 | 0 | 0 | 0 | 0 |
| 0 | NA | NA | NA | NA | NA | Gm4914 | | 0 | 0 | 0 | 0 | 0 | 0 | 0 | 0 | 0 | 0 | 0 |
| 0 | NA | NA | NA | NA | NA | Gm14905 | | 0 | 0 | 0 | 0 | 0 | 0 | 0 | 0 | 0 | 0 | 0 |
| 0 | NA | NA | NA | NA | NA | Gm25366 | | 0 | 0 | 0 | 0 | 0 | 0 | 0 | 0 | 0 | 0 | 0 |
| 0 | NA | NA | NA | NA | NA | Gm27414 | | 0 | 0 | 0 | 0 | 0 | 0 | 0 | 0 | 0 | 0 | 0 |
| 0 | NA | NA | NA | NA | NA | Gm14903 | | 0 | 0 | 0 | 0 | 0 | 0 | 0 | 0 | 0 | 0 | 0 |
| 0 | NA | NA | NA | NA | NA | Gm14906 | | 0 | 0 | 0 | 0 | 0 | 0 | 0 | 0 | 0 | 0 | 0 |
| 0 | NA | NA | NA | NA | NA | Gm14907 | | 0 | 0 | 0 | 0 | 0 | 0 | 0 | 0 | 0 | 0 | 0 |
| 0 | NA | NA | NA | NA | NA | Gm24683 | | 0 | 0 | 0 | 0 | 0 | 0 | 0 | 0 | 0 | 0 | 0 |
| 0 | NA | NA | NA | NA | NA | Gm14904 | | 0 | 0 | 0 | 0 | 0 | 0 | 0 | 0 | 0 | 0 | 0 |
| 0 | NA | NA | NA | NA | NA | Gm24624 | | 0 | 0 | 0 | 0 | 0 | 0 | 0 | 0 | 0 | 0 | 0 |
| 0 | NA | NA | NA | NA | NA | Cpxcr1 | CPX chromosome region, candidate 1 | 0 | 0 | 0 | 0 | 0 | 0 | 0 | 0 | 0 | 0 | 0 |
| 0 | NA | NA | NA | NA | NA | H2ab2 | H2A.B variant histone 2 | 0 | 0 | 0 | 0 | 0 | 0 | 0 | 0 | 0 | 0 | 0 |
| 0 | NA | NA | NA | NA | NA | Gm24438 | | 0 | 0 | 0 | 0 | 0 | 0 | 0 | 0 | 0 | 0 | 0 |
| 0 | NA | NA | NA | NA | NA | Gm14919 | | 0 | 0 | 0 | 0 | 0 | 0 | 0 | 0 | 0 | 0 | 0 |
| 0 | NA | NA | NA | NA | NA | H2ab1 | H2A.B variant histone 1 | 0 | 0 | 0 | 0 | 0 | 0 | 0 | 0 | 0 | 0 | 0 |
| 0 | NA | NA | NA | NA | NA | Gm14918 | | 0 | 0 | 0 | 0 | 0 | 0 | 0 | 0 | 0 | 0 | 0 |
| 0 | NA | NA | NA | NA | NA | Gm14913 | | 0 | 0 | 0 | 0 | 0 | 0 | 0 | 0 | 0 | 0 | 0 |
| 0 | NA | NA | NA | NA | NA | Gm28579 | | 0 | 0 | 0 | 0 | 0 | 0 | 0 | 0 | 0 | 0 | 0 |

| | | | | | | | | | | | | | | | | | | |
|---|---|---|---|---|---|---|---|---|---|---|---|---|---|---|---|---|---|---|
| 0 | NA | NA | NA | NA | NA | Gm14911 | | 0 | 0 | 0 | 0 | 0 | 0 | 0 | 0 | 0 | 0 | 0 |
| 0 | NA | NA | NA | NA | NA | Gm14914 | | 0 | 0 | 0 | 0 | 0 | 0 | 0 | 0 | 0 | 0 | 0 |
| 0 | NA | NA | NA | NA | NA | Gm14917 | | 0 | 0 | 0 | 0 | 0 | 0 | 0 | 0 | 0 | 0 | 0 |
| 0 | NA | NA | NA | NA | NA | Gm14916 | | 0 | 0 | 0 | 0 | 0 | 0 | 0 | 0 | 0 | 0 | 0 |
| 0 | NA | NA | NA | NA | NA | Gm14912 | | 0 | 0 | 0 | 0 | 0 | 0 | 0 | 0 | 0 | 0 | 0 |
| 0 | NA | NA | NA | NA | NA | Gm14915 | | 0 | 0 | 0 | 0 | 0 | 0 | 0 | 0 | 0 | 0 | 0 |
| 0 | NA | NA | NA | NA | NA | Gm14909 | | 0 | 0 | 0 | 0 | 0 | 0 | 0 | 0 | 0 | 0 | 0 |
| 0 | NA | NA | NA | NA | NA | Gm14910 | | 0 | 0 | 0 | 0 | 0 | 0 | 0 | 0 | 0 | 0 | 0 |
| 0 | NA | NA | NA | NA | NA | Tgif2lx2 | TGFB-induced factor homeobox 2-like, X- | 0 | 0 | 0 | 0 | 0 | 0 | 0 | 0 | 0 | 0 | 0 |
| 0 | NA | NA | NA | NA | NA | Tgif2lx1 | TGFB-induced factor homeobox 2-like, X- | 0 | 0 | 0 | 0 | 0 | 0 | 0 | 0 | 0 | 0 | 0 |
| 0 | NA | NA | NA | NA | NA | Gm7485 | | 0 | 0 | 0 | 0 | 0 | 0 | 0 | 0 | 0 | 0 | 0 |
| 0 | NA | NA | NA | NA | NA | Gm14926 | | 0 | 0 | 0 | 0 | 0 | 0 | 0 | 0 | 0 | 0 | 0 |
| 0 | NA | NA | NA | NA | NA | Gm26182 | | 0 | 0 | 0 | 0 | 0 | 0 | 0 | 0 | 0 | 0 | 0 |
| 0 | NA | NA | NA | NA | NA | Gm14929 | | 0 | 0 | 0 | 0 | 0 | 0 | 0 | 0 | 0 | 0 | 0 |
| 0 | NA | NA | NA | NA | NA | Gm16420 | | 0 | 0 | 0 | 0 | 0 | 0 | 0 | 0 | 0 | 0 | 0 |
| 0 | NA | NA | NA | NA | NA | H2ab3 | H2A.B variant histone 3 | 0 | 0 | 0 | 0 | 0 | 0 | 0 | 0 | 0 | 0 | 0 |
| 0 | NA | NA | NA | NA | NA | Gm14931 | | 0 | 0 | 0 | 0 | 0 | 0 | 0 | 0 | 0 | 0 | 0 |
| 0 | NA | NA | NA | NA | NA | Gm14930 | | 0 | 0 | 0 | 0 | 0 | 0 | 0 | 0 | 0 | 0 | 0 |
| 0 | NA | NA | NA | NA | NA | Gm4993 | | 0 | 0 | 0 | 0 | 0 | 0 | 0 | 0 | 0 | 0 | 0 |
| 0 | NA | NA | NA | NA | NA | Gm22590 | | 0 | 0 | 0 | 0 | 0 | 0 | 0 | 0 | 0 | 0 | 0 |
| 0 | NA | NA | NA | NA | NA | Gm5394 | | 0 | 0 | 0 | 0 | 0 | 0 | 0 | 0 | 0 | 0 | 0 |
| 0 | NA | NA | NA | NA | NA | Gm25825 | | 0 | 0 | 0 | 0 | 0 | 0 | 0 | 0 | 0 | 0 | 0 |
| 0 | NA | NA | NA | NA | NA | Gm14932 | | 0 | 0 | 0 | 0 | 0 | 0 | 0 | 0 | 0 | 0 | 0 |
| 0 | NA | NA | NA | NA | NA | Gm23277 | | 0 | 0 | 0 | 0 | 0 | 0 | 0 | 0 | 0 | 0 | 0 |
| 0 | NA | NA | NA | NA | NA | Gm14943 | | 0 | 0 | 0 | 0 | 0 | 0 | 0 | 0 | 0 | 0 | 0 |
| 0 | NA | NA | NA | NA | NA | Gm14945 | | 0 | 0 | 0 | 0 | 0 | 0 | 0 | 0 | 0 | 0 | 0 |
| 0 | NA | NA | NA | NA | NA | Gm4992 | | 0 | 0 | 0 | 0 | 0 | 0 | 0 | 0 | 0 | 0 | 0 |
| 0 | NA | NA | NA | NA | NA | Gm24783 | | 0 | 0 | 0 | 0 | 0 | 0 | 0 | 0 | 0 | 0 | 0 |
| 0 | NA | NA | NA | NA | NA | Gm16421 | | 0 | 0 | 0 | 0 | 0 | 0 | 0 | 0 | 0 | 0 | 0 |
| 0 | NA | NA | NA | NA | NA | Gm14946 | | 0 | 0 | 0 | 0 | 0 | 0 | 0 | 0 | 0 | 0 | 0 |
| 0 | NA | NA | NA | NA | NA | Gm14947 | | 0 | 0 | 0 | 0 | 0 | 0 | 0 | 0 | 0 | 0 | 0 |
| 0 | NA | NA | NA | NA | NA | Gm14940 | | 0 | 0 | 0 | 0 | 0 | 0 | 0 | 0 | 0 | 0 | 0 |
| 0 | NA | NA | NA | NA | NA | Gm17521 | | 0 | 0 | 0 | 0 | 0 | 0 | 0 | 0 | 0 | 0 | 0 |
| 0 | NA | NA | NA | NA | NA | Astx6 | | 0 | 0 | 0 | 0 | 0 | 0 | 0 | 0 | 0 | 0 | 0 |
| 0 | NA | NA | NA | NA | NA | Srsx | serine-rich, secreted, X-linked | 0 | 0 | 0 | 0 | 0 | 0 | 0 | 0 | 0 | 0 | 0 |
| 0 | NA | NA | NA | NA | NA | Gm17577 | | 0 | 0 | 0 | 0 | 0 | 0 | 0 | 0 | 0 | 0 | 0 |
| 0 | NA | NA | NA | NA | NA | Gm14951 | | 0 | 0 | 0 | 0 | 0 | 0 | 0 | 0 | 0 | 0 | 0 |
| 0 | NA | NA | NA | NA | NA | Rps12-ps21 | | 0 | 0 | 0 | 0 | 0 | 0 | 0 | 0 | 0 | 0 | 0 |
| 0 | NA | NA | NA | NA | NA | Gm2411 | predicted gene 2411 | 0 | 0 | 0 | 0 | 0 | 0 | 0 | 0 | 0 | 0 | 0 |
| 0 | NA | NA | NA | NA | NA | Astx2 | | 0 | 0 | 0 | 0 | 0 | 0 | 0 | 0 | 0 | 0 | 0 |
| 0 | NA | NA | NA | NA | NA | Gm17412 | | 0 | 0 | 0 | 0 | 0 | 0 | 0 | 0 | 0 | 0 | 0 |
| 0 | NA | NA | NA | NA | NA | Rps12-ps22 | | 0 | 0 | 0 | 0 | 0 | 0 | 0 | 0 | 0 | 0 | 0 |
| 0 | NA | NA | NA | NA | NA | Cldn34c2 | claudin 34C2 | 0 | 0 | 0 | 0 | 0 | 0 | 0 | 0 | 0 | 0 | 0 |
| 0 | NA | NA | NA | NA | NA | Gm14950 | | 0 | 0 | 0 | 0 | 0 | 0 | 0 | 0 | 0 | 0 | 0 |
| 0 | NA | NA | NA | NA | NA | Gm17467 | | 0 | 0 | 0 | 0 | 0 | 0 | 0 | 0 | 0 | 0 | 0 |
| 0 | NA | NA | NA | NA | NA | Rps12-ps23 | | 0 | 0 | 0 | 0 | 0 | 0 | 0 | 0 | 0 | 0 | 0 |
| 0 | NA | NA | NA | NA | NA | Cldn34c3 | | 0 | 0 | 0 | 0 | 0 | 0 | 0 | 0 | 0 | 0 | 0 |
| 0 | NA | NA | NA | NA | NA | Astx5 | | 0 | 0 | 0 | 0 | 0 | 0 | 0 | 0 | 0 | 0 | 0 |
| 0 | NA | NA | NA | NA | NA | Rps12-ps20 | | 0 | 0 | 0 | 0 | 0 | 0 | 0 | 0 | 0 | 0 | 0 |
| 0 | NA | NA | NA | NA | NA | Gm5945 | claudin 34C3 | 0 | 0 | 0 | 0 | 0 | 0 | 0 | 0 | 0 | 0 | 0 |
| 0 | NA | NA | NA | NA | NA | Vmn2r121 | vomeronasal 2, receptor 121 | 0 | 0 | 0 | 0 | 0 | 0 | 0 | 0 | 0 | 0 | 0 |
| 0 | NA | NA | NA | NA | NA | Rps12-ps6 | | 0 | 0 | 0 | 0 | 0 | 0 | 0 | 0 | 0 | 0 | 0 |
| 0 | NA | NA | NA | NA | NA | Astx1a | | 0 | 0 | 0 | 0 | 0 | 0 | 0 | 0 | 0 | 0 | 0 |
| 0 | NA | NA | NA | NA | NA | Rps12-ps11 | | 0 | 0 | 0 | 0 | 0 | 0 | 0 | 0 | 0 | 0 | 0 |
| 0 | NA | NA | NA | NA | NA | Gm17584 | | 0 | 0 | 0 | 0 | 0 | 0 | 0 | 0 | 0 | 0 | 0 |
| 0 | NA | NA | NA | NA | NA | Astx4a | | 0 | 0 | 0 | 0 | 0 | 0 | 0 | 0 | 0 | 0 | 0 |
| 0 | NA | NA | NA | NA | NA | Rps12-ps12 | | 0 | 0 | 0 | 0 | 0 | 0 | 0 | 0 | 0 | 0 | 0 |
| 0 | NA | NA | NA | NA | NA | Gm17469 | | 0 | 0 | 0 | 0 | 0 | 0 | 0 | 0 | 0 | 0 | 0 |
| 0 | NA | NA | NA | NA | NA | Astx4b | | 0 | 0 | 0 | 0 | 0 | 0 | 0 | 0 | 0 | 0 | 0 |
| 0 | NA | NA | NA | NA | NA | Rps12-ps18 | | 0 | 0 | 0 | 0 | 0 | 0 | 0 | 0 | 0 | 0 | 0 |

| | | | | | | | | | | | | | | | | | | |
|---|---|---|---|---|---|---|---|---|---|---|---|---|---|---|---|---|---|---|
| 0 | NA | NA | NA | NA | NA | Gm50465 | | 0 | 0 | 0 | 0 | 0 | 0 | 0 | 0 | 0 | 0 | 0 |
| 0 | NA | NA | NA | NA | NA | Astx1b | | 0 | 0 | 0 | 0 | 0 | 0 | 0 | 0 | 0 | 0 | 0 |
| 0 | NA | NA | NA | NA | NA | Rps12-ps15 | | 0 | 0 | 0 | 0 | 0 | 0 | 0 | 0 | 0 | 0 | 0 |
| 0 | NA | NA | NA | NA | NA | Gm17361 | | 0 | 0 | 0 | 0 | 0 | 0 | 0 | 0 | 0 | 0 | 0 |
| 0 | NA | NA | NA | NA | NA | Gm50485 | | 0 | 0 | 0 | 0 | 0 | 0 | 0 | 0 | 0 | 0 | 0 |
| 0 | NA | NA | NA | NA | NA | Gm21616 | | 0 | 0 | 0 | 0 | 0 | 0 | 0 | 0 | 0 | 0 | 0 |
| 0 | NA | NA | NA | NA | NA | Rps12-ps17 | | 0 | 0 | 0 | 0 | 0 | 0 | 0 | 0 | 0 | 0 | 0 |
| 0 | NA | NA | NA | NA | NA | Gm50486 | | 0 | 0 | 0 | 0 | 0 | 0 | 0 | 0 | 0 | 0 | 0 |
| 0 | NA | NA | NA | NA | NA | Astx4c | | 0 | 0 | 0 | 0 | 0 | 0 | 0 | 0 | 0 | 0 | 0 |
| 0 | NA | NA | NA | NA | NA | Rps12-ps13 | | 0 | 0 | 0 | 0 | 0 | 0 | 0 | 0 | 0 | 0 | 0 |
| 0 | NA | NA | NA | NA | NA | Gm17693 | | 0 | 0 | 0 | 0 | 0 | 0 | 0 | 0 | 0 | 0 | 0 |
| 0 | NA | NA | NA | NA | NA | Astx1c | | 0 | 0 | 0 | 0 | 0 | 0 | 0 | 0 | 0 | 0 | 0 |
| 0 | NA | NA | NA | NA | NA | Rps12-ps14 | | 0 | 0 | 0 | 0 | 0 | 0 | 0 | 0 | 0 | 0 | 0 |
| 0 | NA | NA | NA | NA | NA | Gm17522 | | 0 | 0 | 0 | 0 | 0 | 0 | 0 | 0 | 0 | 0 | 0 |
| 0 | NA | NA | NA | NA | NA | Gm17522 | | 0 | 0 | 0 | 0 | 0 | 0 | 0 | 0 | 0 | 0 | 0 |
| 0 | NA | NA | NA | NA | NA | Astx4d | | 0 | 0 | 0 | 0 | 0 | 0 | 0 | 0 | 0 | 0 | 0 |
| 0 | NA | NA | NA | NA | NA | Rps12-ps16 | | 0 | 0 | 0 | 0 | 0 | 0 | 0 | 0 | 0 | 0 | 0 |
| 0 | NA | NA | NA | NA | NA | Gm17267 | | 0 | 0 | 0 | 0 | 0 | 0 | 0 | 0 | 0 | 0 | 0 |
| 0 | NA | NA | NA | NA | NA | Astx3 | | 0 | 0 | 0 | 0 | 0 | 0 | 0 | 0 | 0 | 0 | 0 |
| 0 | NA | NA | NA | NA | NA | Gm14957 | | 0 | 0 | 0 | 0 | 0 | 0 | 0 | 0 | 0 | 0 | 0 |
| 0 | NA | NA | NA | NA | NA | Gm14971 | | 0 | 0 | 0 | 0 | 0 | 0 | 0 | 0 | 0 | 0 | 0 |
| 0 | NA | NA | NA | NA | NA | Gm7746 | | 0 | 0 | 0 | 0 | 0 | 0 | 0 | 0 | 0 | 0 | 0 |
| 0 | NA | NA | NA | NA | NA | 4932411N23Ri | RIKEN cDNA 4932411N23 gene | 0 | 0 | 0 | 0 | 0 | 0 | 0 | 0 | 0 | 0 | 0 |
| 0 | NA | NA | NA | NA | NA | Gm23586 | | 0 | 0 | 0 | 0 | 0 | 0 | 0 | 0 | 0 | 0 | 0 |
| 0 | NA | NA | NA | NA | NA | Gm381 | | 0 | 0 | 0 | 0 | 0 | 0 | 0 | 0 | 0 | 0 | 0 |
| 0 | NA | NA | NA | NA | NA | Gm14973 | | 0 | 0 | 0 | 0 | 0 | 0 | 0 | 0 | 0 | 0 | 0 |
| 0 | NA | NA | NA | NA | NA | Gm14972 | | 0 | 0 | 0 | 0 | 0 | 0 | 0 | 0 | 0 | 0 | 0 |
| 0 | NA | NA | NA | NA | NA | Gm382 | predicted gene 382 | 0 | 0 | 0 | 0 | 0 | 0 | 0 | 0 | 0 | 0 | 0 |
| 0 | NA | NA | NA | NA | NA | Gm22139 | | 0 | 0 | 0 | 0 | 0 | 0 | 0 | 0 | 0 | 0 | 0 |
| 0 | NA | NA | NA | NA | NA | 4921511C20Rik | | 0 | 0 | 0 | 0 | 0 | 0 | 0 | 0 | 0 | 0 | 0 |
| 0 | NA | NA | NA | NA | NA | Gm22636 | | 0 | 0 | 0 | 0 | 0 | 0 | 0 | 0 | 0 | 0 | 0 |
| 0 | NA | NA | NA | NA | NA | Cldn34c4 | claudin 34C4 | 0 | 0 | 0 | 0 | 0 | 0 | 0 | 0 | 0 | 0 | 0 |
| 0 | NA | NA | NA | NA | NA | Gm25061 | | 0 | 0 | 0 | 0 | 0 | 0 | 0 | 0 | 0 | 0 | 0 |
| 0 | NA | NA | NA | NA | NA | Gm14975 | | 0 | 0 | 0 | 0 | 0 | 0 | 0 | 0 | 0 | 0 | 0 |
| 0 | NA | NA | NA | NA | NA | Gm4916 | | 0 | 0 | 0 | 0 | 0 | 0 | 0 | 0 | 0 | 0 | 0 |
| 0 | NA | NA | NA | NA | NA | Gm38390 | | 0 | 0 | 0 | 0 | 0 | 0 | 0 | 0 | 0 | 0 | 0 |
| 0 | NA | NA | NA | NA | NA | Gm26475 | | 0 | 0 | 0 | 0 | 0 | 0 | 0 | 0 | 0 | 0 | 0 |
| 0 | NA | NA | NA | NA | NA | Gm24522 | | 0 | 0 | 0 | 0 | 0 | 0 | 0 | 0 | 0 | 0 | 0 |
| 0 | NA | NA | NA | NA | NA | 4930558G05Rik | | 0 | 0 | 0 | 0 | 0 | 0 | 0 | 0 | 0 | 0 | 0 |
| 0 | NA | NA | NA | NA | NA | Gm26029 | | 0 | 0 | 0 | 0 | 0 | 0 | 0 | 0 | 0 | 0 | 0 |
| 0 | NA | NA | NA | NA | NA | Gm22225 | | 0 | 0 | 0 | 0 | 0 | 0 | 0 | 0 | 0 | 0 | 0 |
| 0 | NA | NA | NA | NA | NA | Gm14986 | | 0 | 0 | 0 | 0 | 0 | 0 | 0 | 0 | 0 | 0 | 0 |
| 0 | NA | NA | NA | NA | NA | Gm14985 | | 0 | 0 | 0 | 0 | 0 | 0 | 0 | 0 | 0 | 0 | 0 |
| 0 | NA | NA | NA | NA | NA | Gm14983 | | 0 | 0 | 0 | 0 | 0 | 0 | 0 | 0 | 0 | 0 | 0 |
| 0 | NA | NA | NA | NA | NA | Gm14984 | | 0 | 0 | 0 | 0 | 0 | 0 | 0 | 0 | 0 | 0 | 0 |
| 0 | NA | NA | NA | NA | NA | Gm14987 | | 0 | 0 | 0 | 0 | 0 | 0 | 0 | 0 | 0 | 0 | 0 |
| 0 | NA | NA | NA | NA | NA | Gm14982 | | 0 | 0 | 0 | 0 | 0 | 0 | 0 | 0 | 0 | 0 | 0 |
| 0 | NA | NA | NA | NA | NA | Gm26209 | | 0 | 0 | 0 | 0 | 0 | 0 | 0 | 0 | 0 | 0 | 0 |
| 0 | NA | NA | NA | NA | NA | Gm25170 | | 0 | 0 | 0 | 0 | 0 | 0 | 0 | 0 | 0 | 0 | 0 |
| 0 | NA | NA | NA | NA | NA | Gm14976 | | 0 | 0 | 0 | 0 | 0 | 0 | 0 | 0 | 0 | 0 | 0 |
| 0 | NA | NA | NA | NA | NA | Gm7790 | | 0 | 0 | 0 | 0 | 0 | 0 | 0 | 0 | 0 | 0 | 0 |
| 0 | NA | NA | NA | NA | NA | Gm14977 | | 0 | 0 | 0 | 0 | 0 | 0 | 0 | 0 | 0 | 0 | 0 |
| 0 | NA | NA | NA | NA | NA | Gm4994 | | 0 | 0 | 0 | 0 | 0 | 0 | 0 | 0 | 0 | 0 | 0 |
| 0 | NA | NA | NA | NA | NA | Gm7058 | | 0 | 0 | 0 | 0 | 0 | 0 | 0 | 0 | 0 | 0 | 0 |
| 0 | NA | NA | NA | NA | NA | Gm14981 | | 0 | 0 | 0 | 0 | 0 | 0 | 0 | 0 | 0 | 0 | 0 |
| 0 | NA | NA | NA | NA | NA | Gm26406 | | 0 | 0 | 0 | 0 | 0 | 0 | 0 | 0 | 0 | 0 | 0 |
| 0 | NA | NA | NA | NA | NA | Gm23768 | | 0 | 0 | 0 | 0 | 0 | 0 | 0 | 0 | 0 | 0 | 0 |
| 0 | NA | NA | NA | NA | NA | Gm22143 | | 0 | 0 | 0 | 0 | 0 | 0 | 0 | 0 | 0 | 0 | 0 |
| 0 | NA | NA | NA | NA | NA | Gm7821 | | 0 | 0 | 0 | 0 | 0 | 0 | 0 | 0 | 0 | 0 | 0 |
| 0 | NA | NA | NA | NA | NA | Gm14979 | | 0 | 0 | 0 | 0 | 0 | 0 | 0 | 0 | 0 | 0 | 0 |

| | | | | | | | | | | | | | | | | | | |
|---|---|---|---|---|---|---|---|---|---|---|---|---|---|---|---|---|---|---|
| 0 | NA | NA | NA | NA | NA | Gm22552 | | 0 | 0 | 0 | 0 | 0 | 0 | 0 | 0 | 0 | 0 | 0 |
| 0 | NA | NA | NA | NA | NA | Gm14980 | | 0 | 0 | 0 | 0 | 0 | 0 | 0 | 0 | 0 | 0 | 0 |
| 0 | NA | NA | NA | NA | NA | Tnmd | tenomodulin | 0 | 0 | 0 | 0 | 0 | 0 | 0 | 0 | 0 | 0 | 0 |
| 0 | NA | NA | NA | NA | NA | 4930570D08Rik | | 0 | 0 | 0 | 0 | 0 | 0 | 0 | 0 | 0 | 0 | 0 |
| 0 | NA | NA | NA | NA | NA | Gm7841 | | 0 | 0 | 0 | 0 | 0 | 0 | 0 | 0 | 0 | 0 | 0 |
| 0 | NA | NA | NA | NA | NA | Mir3112 | | 0 | 0 | 0 | 0 | 0 | 0 | 0 | 0 | 0 | 0 | 0 |
| 0 | NA | NA | NA | NA | NA | Arl13a | ADP-ribosylation factor-like 13A | 0 | 0 | 0 | 0 | 0 | 0 | 0 | 0 | 0 | 0 | 0 |
| 0 | NA | NA | NA | NA | NA | Taf7l | TATA-box binding protein associated fact | 0 | 0 | 0 | 0 | 0 | 0 | 0 | 0 | 0 | 0 | 0 |
| 0 | NA | NA | NA | NA | NA | Gm23124 | | 0 | 0 | 0 | 0 | 0 | 0 | 0 | 0 | 0 | 0 | 0 |
| 0 | NA | NA | NA | NA | NA | Gm15154 | | 0 | 0 | 0 | 0 | 0 | 0 | 0 | 0 | 0 | 0 | 0 |
| 0 | NA | NA | NA | NA | NA | Gm7855 | | 0 | 0 | 0 | 0 | 0 | 0 | 0 | 0 | 0 | 0 | 0 |
| 0 | NA | NA | NA | NA | NA | Gm7857 | | 0 | 0 | 0 | 0 | 0 | 0 | 0 | 0 | 0 | 0 | 0 |
| 0 | NA | NA | NA | NA | NA | Gm16397 | | 0 | 0 | 0 | 0 | 0 | 0 | 0 | 0 | 0 | 0 | 0 |
| 0 | NA | NA | NA | NA | NA | Gm15018 | | 0 | 0 | 0 | 0 | 0 | 0 | 0 | 0 | 0 | 0 | 0 |
| 0 | NA | NA | NA | NA | NA | Gm6207 | | 0 | 0 | 0 | 0 | 0 | 0 | 0 | 0 | 0 | 0 | 0 |
| 0 | NA | NA | NA | NA | NA | Gm15020 | | 0 | 0 | 0 | 0 | 0 | 0 | 0 | 0 | 0 | 0 | 0 |
| 0 | NA | NA | NA | NA | NA | Gm25868 | | 0 | 0 | 0 | 0 | 0 | 0 | 0 | 0 | 0 | 0 | 0 |
| 0 | NA | NA | NA | NA | NA | Gm15023 | predicted gene 15023 | 0 | 0 | 0 | 0 | 0 | 0 | 0 | 0 | 0 | 0 | 0 |
| 0 | NA | NA | NA | NA | NA | Gm15022 | | 0 | 0 | 0 | 0 | 0 | 0 | 0 | 0 | 0 | 0 | 0 |
| 0 | NA | NA | NA | NA | NA | Pramel | preferentially expressed antigen in melan | 0 | 0 | 0 | 0 | 0 | 0 | 0 | 0 | 0 | 0 | 0 |
| 0 | NA | NA | NA | NA | NA | Pramel3 | preferentially expressed antigen in melan | 0 | 0 | 0 | 0 | 0 | 0 | 0 | 0 | 0 | 0 | 0 |
| 0 | NA | NA | NA | NA | NA | Gm25936 | | 0 | 0 | 0 | 0 | 0 | 0 | 0 | 0 | 0 | 0 | 0 |
| 0 | NA | NA | NA | NA | NA | Gm15017 | | 0 | 0 | 0 | 0 | 0 | 0 | 0 | 0 | 0 | 0 | 0 |
| 0 | NA | NA | NA | NA | NA | Gm6215 | | 0 | 0 | 0 | 0 | 0 | 0 | 0 | 0 | 0 | 0 | 0 |
| 0 | NA | NA | NA | NA | NA | Gm5128 | predicted gene 5128 | 0 | 0 | 0 | 0 | 0 | 0 | 0 | 0 | 0 | 0 | 0 |
| 0 | NA | NA | NA | NA | NA | Gm22006 | | 0 | 0 | 0 | 0 | 0 | 0 | 0 | 0 | 0 | 0 | 0 |
| 0 | NA | NA | NA | NA | NA | Gm6221 | | 0 | 0 | 0 | 0 | 0 | 0 | 0 | 0 | 0 | 0 | 0 |
| 0 | NA | NA | NA | NA | NA | Gm7903 | predicted gene 7903 | 0 | 0 | 0 | 0 | 0 | 0 | 0 | 0 | 0 | 0 | 0 |
| 0 | NA | NA | NA | NA | NA | Gm24416 | | 0 | 0 | 0 | 0 | 0 | 0 | 0 | 0 | 0 | 0 | 0 |
| 0 | NA | NA | NA | NA | NA | Gm7905 | | 0 | 0 | 0 | 0 | 0 | 0 | 0 | 0 | 0 | 0 | 0 |
| 0 | NA | NA | NA | NA | NA | AV320801 | expressed sequence AV320801 | 0 | 0 | 0 | 0 | 0 | 0 | 0 | 0 | 0 | 0 | 0 |
| 0 | NA | NA | NA | NA | NA | Gm22233 | | 0 | 0 | 0 | 0 | 0 | 0 | 0 | 0 | 0 | 0 | 0 |
| 0 | NA | NA | NA | NA | NA | Gm15015 | | 0 | 0 | 0 | 0 | 0 | 0 | 0 | 0 | 0 | 0 | 0 |
| 0 | NA | NA | NA | NA | NA | Gm6228 | | 0 | 0 | 0 | 0 | 0 | 0 | 0 | 0 | 0 | 0 | 0 |
| 0 | NA | NA | NA | NA | NA | Prame | preferentially expressed antigen in melan | 0 | 0 | 0 | 0 | 0 | 0 | 0 | 0 | 0 | 0 | 0 |
| 0 | NA | NA | NA | NA | NA | Gm15021 | | 0 | 0 | 0 | 0 | 0 | 0 | 0 | 0 | 0 | 0 | 0 |
| 0 | NA | NA | NA | NA | NA | Tcp11x2 | t-complex 11 family, X-linked 2 | 0 | 0 | 0 | 0 | 0 | 0 | 0 | 0 | 0 | 0 | 0 |
| 0 | NA | NA | NA | NA | NA | Tmsb15a | thymosin beta 15a | 0 | 0 | 0 | 0 | 0 | 0 | 0 | 0 | 0 | 0 | 0 |
| 0 | NA | NA | NA | NA | NA | Gm15019 | | 0 | 0 | 0 | 0 | 0 | 0 | 0 | 0 | 0 | 0 | 0 |
| 0 | NA | NA | NA | NA | NA | Mir7646 | | 0 | 0 | 0 | 0 | 0 | 0 | 0 | 0 | 0 | 0 | 0 |
| 0 | NA | NA | NA | NA | NA | Mir1970 | | 0 | 0 | 0 | 0 | 0 | 0 | 0 | 0 | 0 | 0 | 0 |
| 0 | NA | NA | NA | NA | NA | 5730412P04Rik | | 0 | 0 | 0 | 0 | 0 | 0 | 0 | 0 | 0 | 0 | 0 |
| 0 | NA | NA | NA | NA | NA | Mir6383 | | 0 | 0 | 0 | 0 | 0 | 0 | 0 | 0 | 0 | 0 | 0 |
| 0 | NA | NA | NA | NA | NA | n-R5s12 | | 0 | 0 | 0 | 0 | 0 | 0 | 0 | 0 | 0 | 0 | 0 |
| 0 | NA | NA | NA | NA | NA | Gm14997 | | 0 | 0 | 0 | 0 | 0 | 0 | 0 | 0 | 0 | 0 | 0 |
| 0 | NA | NA | NA | NA | NA | Gm4917 | | 0 | 0 | 0 | 0 | 0 | 0 | 0 | 0 | 0 | 0 | 0 |
| 0 | NA | NA | NA | NA | NA | Gm14994 | | 0 | 0 | 0 | 0 | 0 | 0 | 0 | 0 | 0 | 0 | 0 |
| 0 | NA | NA | NA | NA | NA | Gm14999 | | 0 | 0 | 0 | 0 | 0 | 0 | 0 | 0 | 0 | 0 | 0 |
| 0 | NA | NA | NA | NA | NA | Gm14998 | | 0 | 0 | 0 | 0 | 0 | 0 | 0 | 0 | 0 | 0 | 0 |
| 0 | NA | NA | NA | NA | NA | Gm14993 | | 0 | 0 | 0 | 0 | 0 | 0 | 0 | 0 | 0 | 0 | 0 |
| 0 | NA | NA | NA | NA | NA | Gm14992 | | 0 | 0 | 0 | 0 | 0 | 0 | 0 | 0 | 0 | 0 | 0 |
| 0 | NA | NA | NA | NA | NA | Gm14996 | | 0 | 0 | 0 | 0 | 0 | 0 | 0 | 0 | 0 | 0 | 0 |
| 0 | NA | NA | NA | NA | NA | Gm14990 | | 0 | 0 | 0 | 0 | 0 | 0 | 0 | 0 | 0 | 0 | 0 |
| 0 | NA | NA | NA | NA | NA | Gm14995 | | 0 | 0 | 0 | 0 | 0 | 0 | 0 | 0 | 0 | 0 | 0 |
| 0 | NA | NA | NA | NA | NA | Kir3dl2 | killer cell immunoglobulin-like receptor, th | 0 | 0 | 0 | 0 | 0 | 0 | 0 | 0 | 0 | 0 | 0 |
| 0 | NA | NA | NA | NA | NA | Gm14991 | | 0 | 0 | 0 | 0 | 0 | 0 | 0 | 0 | 0 | 0 | 0 |
| 0 | NA | NA | NA | NA | NA | Kir3dl1 | killer cell immunoglobulin-like receptor, th | 0 | 0 | 0 | 0 | 0 | 0 | 0 | 0 | 0 | 0 | 0 |
| 0 | NA | NA | NA | NA | NA | Gm15006 | | 0 | 0 | 0 | 0 | 0 | 0 | 0 | 0 | 0 | 0 | 0 |
| 0 | NA | NA | NA | NA | NA | Gm15003 | | 0 | 0 | 0 | 0 | 0 | 0 | 0 | 0 | 0 | 0 | 0 |
| 0 | NA | NA | NA | NA | NA | Gm15005 | | 0 | 0 | 0 | 0 | 0 | 0 | 0 | 0 | 0 | 0 | 0 |

| | | | | | | | | | | | | | | | | | | |
|---|---|---|---|---|---|---|---|---|---|---|---|---|---|---|---|---|---|---|
| 0 | NA | NA | NA | NA | NA | Gm15004 | | 0 | 0 | 0 | 0 | 0 | 0 | 0 | 0 | 0 | 0 | 0 |
| 0 | NA | NA | NA | NA | NA | Gm15010 | | 0 | 0 | 0 | 0 | 0 | 0 | 0 | 0 | 0 | 0 | 0 |
| 0 | NA | NA | NA | NA | NA | Gm23068 | | 0 | 0 | 0 | 0 | 0 | 0 | 0 | 0 | 0 | 0 | 0 |
| 0 | NA | NA | NA | NA | NA | Gm15009 | | 0 | 0 | 0 | 0 | 0 | 0 | 0 | 0 | 0 | 0 | 0 |
| 0 | NA | NA | NA | NA | NA | Gm15026 | | 0 | 0 | 0 | 0 | 0 | 0 | 0 | 0 | 0 | 0 | 0 |
| 0 | NA | NA | NA | NA | NA | H2bfm | H2B histone family, member M | 0 | 0 | 0 | 0 | 0 | 0 | 0 | 0 | 0 | 0 | 0 |
| 0 | NA | NA | NA | NA | NA | Gm15038 | | 0 | 0 | 0 | 0 | 0 | 0 | 0 | 0 | 0 | 0 | 0 |
| 0 | NA | NA | NA | NA | NA | Gm25770 | | 0 | 0 | 0 | 0 | 0 | 0 | 0 | 0 | 0 | 0 | 0 |
| 0 | NA | NA | NA | NA | NA | Gm15039 | | 0 | 0 | 0 | 0 | 0 | 0 | 0 | 0 | 0 | 0 | 0 |
| 0 | NA | NA | NA | NA | NA | Gm15042 | | 0 | 0 | 0 | 0 | 0 | 0 | 0 | 0 | 0 | 0 | 0 |
| 0 | NA | NA | NA | NA | NA | Gm15045 | | 0 | 0 | 0 | 0 | 0 | 0 | 0 | 0 | 0 | 0 | 0 |
| 0 | NA | NA | NA | NA | NA | Gm15043 | | 0 | 0 | 0 | 0 | 0 | 0 | 0 | 0 | 0 | 0 | 0 |
| 0 | NA | NA | NA | NA | NA | Gm15044 | | 0 | 0 | 0 | 0 | 0 | 0 | 0 | 0 | 0 | 0 | 0 |
| 0 | NA | NA | NA | NA | NA | Gm15040 | | 0 | 0 | 0 | 0 | 0 | 0 | 0 | 0 | 0 | 0 | 0 |
| 0 | NA | NA | NA | NA | NA | Gm15041 | | 0 | 0 | 0 | 0 | 0 | 0 | 0 | 0 | 0 | 0 | 0 |
| 0 | NA | NA | NA | NA | NA | Gm23679 | | 0 | 0 | 0 | 0 | 0 | 0 | 0 | 0 | 0 | 0 | 0 |
| 0 | NA | NA | NA | NA | NA | Il1rapl2 | interleukin 1 receptor accessory protein-li | 0 | 0 | 0 | 0 | 0 | 0 | 0 | 0 | 0 | 0 | 0 |
| 0 | NA | NA | NA | NA | NA | Gm14988 | | 0 | 0 | 0 | 0 | 0 | 0 | 0 | 0 | 0 | 0 | 0 |
| 0 | NA | NA | NA | NA | NA | Gm25393 | | 0 | 0 | 0 | 0 | 0 | 0 | 0 | 0 | 0 | 0 | 0 |
| 0 | NA | NA | NA | NA | NA | Gm14989 | | 0 | 0 | 0 | 0 | 0 | 0 | 0 | 0 | 0 | 0 | 0 |
| 0 | NA | NA | NA | NA | NA | Gm24675 | | 0 | 0 | 0 | 0 | 0 | 0 | 0 | 0 | 0 | 0 | 0 |
| 0 | NA | NA | NA | NA | NA | Tex13a | testis expressed 13A | 0 | 0 | 0 | 0 | 0 | 0 | 0 | 0 | 0 | 0 | 0 |
| 0 | NA | NA | NA | NA | NA | Gm22057 | | 0 | 0 | 0 | 0 | 0 | 0 | 0 | 0 | 0 | 0 | 0 |
| 0 | NA | NA | NA | NA | NA | Gm23958 | | 0 | 0 | 0 | 0 | 0 | 0 | 0 | 0 | 0 | 0 | 0 |
| 0 | NA | NA | NA | NA | NA | Gm15000 | | 0 | 0 | 0 | 0 | 0 | 0 | 0 | 0 | 0 | 0 | 0 |
| 0 | NA | NA | NA | NA | NA | Gm15001 | | 0 | 0 | 0 | 0 | 0 | 0 | 0 | 0 | 0 | 0 | 0 |
| 0 | NA | NA | NA | NA | NA | 4933428M09Ri | RIKEN cDNA 4933428M09 gene | 0 | 0 | 0 | 0 | 0 | 0 | 0 | 0 | 0 | 0 | 0 |
| 0 | NA | NA | NA | NA | NA | Platr21 | | 0 | 0 | 0 | 0 | 0 | 0 | 0 | 0 | 0 | 0 | 0 |
| 0 | NA | NA | NA | NA | NA | Gm15002 | | 0 | 0 | 0 | 0 | 0 | 0 | 0 | 0 | 0 | 0 | 0 |
| 0 | NA | NA | NA | NA | NA | Trap1a | tumor rejection antigen P1A | 0 | 0 | 0 | 0 | 0 | 0 | 0 | 0 | 0 | 0 | 0 |
| 0 | NA | NA | NA | NA | NA | Gm8061 | | 0 | 0 | 0 | 0 | 0 | 0 | 0 | 0 | 0 | 0 | 0 |
| 0 | NA | NA | NA | NA | NA | Gm24380 | | 0 | 0 | 0 | 0 | 0 | 0 | 0 | 0 | 0 | 0 | 0 |
| 0 | NA | NA | NA | NA | NA | Gm6322 | | 0 | 0 | 0 | 0 | 0 | 0 | 0 | 0 | 0 | 0 | 0 |
| 0 | NA | NA | NA | NA | NA | Ripply1 | ripply transcriptional repressor 1 | 0 | 0 | 0 | 0 | 0 | 0 | 0 | 0 | 0 | 0 | 0 |
| 0 | NA | NA | NA | NA | NA | Gm15012 | | 0 | 0 | 0 | 0 | 0 | 0 | 0 | 0 | 0 | 0 | 0 |
| 0 | NA | NA | NA | NA | NA | Gm15011 | | 0 | 0 | 0 | 0 | 0 | 0 | 0 | 0 | 0 | 0 | 0 |
| 0 | NA | NA | NA | NA | NA | Gm15046 | | 0 | 0 | 0 | 0 | 0 | 0 | 0 | 0 | 0 | 0 | 0 |
| 0 | NA | NA | NA | NA | NA | Mir3475 | | 0 | 0 | 0 | 0 | 0 | 0 | 0 | 0 | 0 | 0 | 0 |
| 0 | NA | NA | NA | NA | NA | | | 0 | 0 | 0 | 0 | 0 | 0 | 0 | 0 | 0 | 0 | 0 |
| 0 | NA | NA | NA | NA | NA | Gm15047 | | 0 | 0 | 0 | 0 | 0 | 0 | 0 | 0 | 0 | 0 | 0 |
| 0 | NA | NA | NA | NA | NA | Gm22025 | | 0 | 0 | 0 | 0 | 0 | 0 | 0 | 0 | 0 | 0 | 0 |
| 0 | NA | NA | NA | NA | NA | Gm23199 | | 0 | 0 | 0 | 0 | 0 | 0 | 0 | 0 | 0 | 0 | 0 |
| 0 | NA | NA | NA | NA | NA | Eif2c5 | | 0 | 0 | 0 | 0 | 0 | 0 | 0 | 0 | 0 | 0 | 0 |
| 0 | NA | NA | NA | NA | NA | Tex13b | testis expressed 13B | 0 | 0 | 0 | 0 | 0 | 0 | 0 | 0 | 0 | 0 | 0 |
| 0 | NA | NA | NA | NA | NA | Gm15077 | | 0 | 0 | 0 | 0 | 0 | 0 | 0 | 0 | 0 | 0 | 0 |
| 0 | NA | NA | NA | NA | NA | Gm15078 | | 0 | 0 | 0 | 0 | 0 | 0 | 0 | 0 | 0 | 0 | 0 |
| 0 | NA | NA | NA | NA | NA | Gm15079 | | 0 | 0 | 0 | 0 | 0 | 0 | 0 | 0 | 0 | 0 | 0 |
| 0 | NA | NA | NA | NA | NA | Gm15268 | | 0 | 0 | 0 | 0 | 0 | 0 | 0 | 0 | 0 | 0 | 0 |
| 0 | NA | NA | NA | NA | NA | Gm15282 | | 0 | 0 | 0 | 0 | 0 | 0 | 0 | 0 | 0 | 0 | 0 |
| 0 | NA | NA | NA | NA | NA | Gm15295 | | 0 | 0 | 0 | 0 | 0 | 0 | 0 | 0 | 0 | 0 | 0 |
| 0 | NA | NA | NA | NA | NA | Gm15297 | | 0 | 0 | 0 | 0 | 0 | 0 | 0 | 0 | 0 | 0 | 0 |
| 0 | NA | NA | NA | NA | NA | Hmgn2l6 | | 0 | 0 | 0 | 0 | 0 | 0 | 0 | 0 | 0 | 0 | 0 |
| 0 | NA | NA | NA | NA | NA | Gm15298 | | 0 | 0 | 0 | 0 | 0 | 0 | 0 | 0 | 0 | 0 | 0 |
| 0 | NA | NA | NA | NA | NA | Gm5762 | | 0 | 0 | 0 | 0 | 0 | 0 | 0 | 0 | 0 | 0 | 0 |
| 0 | NA | NA | NA | NA | NA | Gm6373 | | 0 | 0 | 0 | 0 | 0 | 0 | 0 | 0 | 0 | 0 | 0 |
| 0 | NA | NA | NA | NA | NA | Gm15014 | | 0 | 0 | 0 | 0 | 0 | 0 | 0 | 0 | 0 | 0 | 0 |
| 0 | NA | NA | NA | NA | NA | Gm6067 | | 0 | 0 | 0 | 0 | 0 | 0 | 0 | 0 | 0 | 0 | 0 |
| 0 | NA | NA | NA | NA | NA | Gm15031 | | 0 | 0 | 0 | 0 | 0 | 0 | 0 | 0 | 0 | 0 | 0 |
| 0 | NA | NA | NA | NA | NA | Gm25915 | | 0 | 0 | 0 | 0 | 0 | 0 | 0 | 0 | 0 | 0 | 0 |
| 0 | NA | NA | NA | NA | NA | Gm15049 | | 0 | 0 | 0 | 0 | 0 | 0 | 0 | 0 | 0 | 0 | 0 |

| | | | | | | | | | | | | | | | | | | |
|---|---|---|---|---|---|---|---|---|---|---|---|---|---|---|---|---|---|---|
| 0 | NA | NA | NA | NA | NA | Gm44351 | | 0 | 0 | 0 | 0 | 0 | 0 | 0 | 0 | 0 | 0 | 0 |
| 0 | NA | NA | NA | NA | NA | Gm23325 | | 0 | 0 | 0 | 0 | 0 | 0 | 0 | 0 | 0 | 0 | 0 |
| 0 | NA | NA | NA | NA | NA | Gm47298 | | 0 | 0 | 0 | 0 | 0 | 0 | 0 | 0 | 0 | 0 | 0 |
| 0 | NA | NA | NA | NA | NA | Gm15048 | | 0 | 0 | 0 | 0 | 0 | 0 | 0 | 0 | 0 | 0 | 0 |
| 0 | NA | NA | NA | NA | NA | Gm8202 | | 0 | 0 | 0 | 0 | 0 | 0 | 0 | 0 | 0 | 0 | 0 |
| 0 | NA | NA | NA | NA | NA | Gm15067 | | 0 | 0 | 0 | 0 | 0 | 0 | 0 | 0 | 0 | 0 | 0 |
| 0 | NA | NA | NA | NA | NA | Mir680-2 | | 0 | 0 | 0 | 0 | 0 | 0 | 0 | 0 | 0 | 0 | 0 |
| 0 | NA | NA | NA | NA | NA | Trpc5os | transient receptor potential cation channe | 0 | 0 | 0 | 0 | 0 | 0 | 0 | 0 | 0 | 0 | 0 |
| 0 | NA | NA | NA | NA | NA | Gm15071 | | 0 | 0 | 0 | 0 | 0 | 0 | 0 | 0 | 0 | 0 | 0 |
| 0 | NA | NA | NA | NA | NA | Mageb16-ps1 | | 0 | 0 | 0 | 0 | 0 | 0 | 0 | 0 | 0 | 0 | 0 |
| 0 | NA | NA | NA | NA | NA | Gm26024 | | 0 | 0 | 0 | 0 | 0 | 0 | 0 | 0 | 0 | 0 | 0 |
| 0 | NA | NA | NA | NA | NA | Mir3472 | | 0 | 0 | 0 | 0 | 0 | 0 | 0 | 0 | 0 | 0 | 0 |
| 0 | NA | NA | NA | NA | NA | Gm25097 | | 0 | 0 | 0 | 0 | 0 | 0 | 0 | 0 | 0 | 0 | 0 |
| 0 | NA | NA | NA | NA | NA | Gm15073 | | 0 | 0 | 0 | 0 | 0 | 0 | 0 | 0 | 0 | 0 | 0 |
| 0 | NA | NA | NA | NA | NA | Gm6441 | | 0 | 0 | 0 | 0 | 0 | 0 | 0 | 0 | 0 | 0 | 0 |
| 0 | NA | NA | NA | NA | NA | Gm15060 | | 0 | 0 | 0 | 0 | 0 | 0 | 0 | 0 | 0 | 0 | 0 |
| 0 | NA | NA | NA | NA | NA | Gm27659 | | 0 | 0 | 0 | 0 | 0 | 0 | 0 | 0 | 0 | 0 | 0 |
| 0 | NA | NA | NA | NA | NA | Gm15058 | | 0 | 0 | 0 | 0 | 0 | 0 | 0 | 0 | 0 | 0 | 0 |
| 0 | NA | NA | NA | NA | NA | Gm6446 | | 0 | 0 | 0 | 0 | 0 | 0 | 0 | 0 | 0 | 0 | 0 |
| 0 | NA | NA | NA | NA | NA | Gm6447 | | 0 | 0 | 0 | 0 | 0 | 0 | 0 | 0 | 0 | 0 | 0 |
| 0 | NA | NA | NA | NA | NA | Gm15062 | | 0 | 0 | 0 | 0 | 0 | 0 | 0 | 0 | 0 | 0 | 0 |
| 0 | NA | NA | NA | NA | NA | Gm4918 | | 0 | 0 | 0 | 0 | 0 | 0 | 0 | 0 | 0 | 0 | 0 |
| 0 | NA | NA | NA | NA | NA | Gm15061 | | 0 | 0 | 0 | 0 | 0 | 0 | 0 | 0 | 0 | 0 | 0 |
| 0 | NA | NA | NA | NA | NA | Gm26151 | | 0 | 0 | 0 | 0 | 0 | 0 | 0 | 0 | 0 | 0 | 0 |
| 0 | NA | NA | NA | NA | NA | Gm15076 | | 0 | 0 | 0 | 0 | 0 | 0 | 0 | 0 | 0 | 0 | 0 |
| 0 | NA | NA | NA | NA | NA | Gm15075 | | 0 | 0 | 0 | 0 | 0 | 0 | 0 | 0 | 0 | 0 | 0 |
| 0 | NA | NA | NA | NA | NA | Gm15074 | | 0 | 0 | 0 | 0 | 0 | 0 | 0 | 0 | 0 | 0 | 0 |
| 0 | NA | NA | NA | NA | NA | Gm4996 | | 0 | 0 | 0 | 0 | 0 | 0 | 0 | 0 | 0 | 0 | 0 |
| 0 | NA | NA | NA | NA | NA | Htr2c | 5-hydroxytryptamine (serotonin) receptor | 0 | 0 | 0 | 0 | 0 | 0 | 0 | 0 | 0 | 0 | 0 |
| 0 | NA | NA | NA | NA | NA | Snora35 | | 0 | 0 | 0 | 0 | 0 | 0 | 0 | 0 | 0 | 0 | 0 |
| 0 | NA | NA | NA | NA | NA | Gm22650 | | 0 | 0 | 0 | 0 | 0 | 0 | 0 | 0 | 0 | 0 | 0 |
| 0 | NA | NA | NA | NA | NA | Mir764 | | 0 | 0 | 0 | 0 | 0 | 0 | 0 | 0 | 0 | 0 | 0 |
| 0 | NA | NA | NA | NA | NA | Mir1912 | | 0 | 0 | 0 | 0 | 0 | 0 | 0 | 0 | 0 | 0 | 0 |
| 0 | NA | NA | NA | NA | NA | Mir1264 | | 0 | 0 | 0 | 0 | 0 | 0 | 0 | 0 | 0 | 0 | 0 |
| 0 | NA | NA | NA | NA | NA | Mir1298 | | 0 | 0 | 0 | 0 | 0 | 0 | 0 | 0 | 0 | 0 | 0 |
| 0 | NA | NA | NA | NA | NA | Gm24627 | | 0 | 0 | 0 | 0 | 0 | 0 | 0 | 0 | 0 | 0 | 0 |
| 0 | NA | NA | NA | NA | NA | Mir448 | | 0 | 0 | 0 | 0 | 0 | 0 | 0 | 0 | 0 | 0 | 0 |
| 0 | NA | NA | NA | NA | NA | Vmn1r239-ps | | 0 | 0 | 0 | 0 | 0 | 0 | 0 | 0 | 0 | 0 | 0 |
| 0 | NA | NA | NA | NA | NA | Gm16400 | | 0 | 0 | 0 | 0 | 0 | 0 | 0 | 0 | 0 | 0 | 0 |
| 0 | NA | NA | NA | NA | NA | Gm15167 | | 0 | 0 | 0 | 0 | 0 | 0 | 0 | 0 | 0 | 0 | 0 |
| 0 | NA | NA | NA | NA | NA | Gm8289 | | 0 | 0 | 0 | 0 | 0 | 0 | 0 | 0 | 0 | 0 | 0 |
| 0 | NA | NA | NA | NA | NA | Gm15128 | predicted gene 15128 | 0 | 0 | 0 | 0 | 0 | 0 | 0 | 0 | 0 | 0 | 0 |
| 0 | NA | NA | NA | NA | NA | Gm15081 | | 0 | 0 | 0 | 0 | 0 | 0 | 0 | 0 | 0 | 0 | 0 |
| 0 | NA | NA | NA | NA | NA | Gm15080 | predicted gene 15080 | 0 | 0 | 0 | 0 | 0 | 0 | 0 | 0 | 0 | 0 | 0 |
| 0 | NA | NA | NA | NA | NA | Gm15108 | | 0 | 0 | 0 | 0 | 0 | 0 | 0 | 0 | 0 | 0 | 0 |
| 0 | NA | NA | NA | NA | NA | Gm15110 | | 0 | 0 | 0 | 0 | 0 | 0 | 0 | 0 | 0 | 0 | 0 |
| 0 | NA | NA | NA | NA | NA | Gm8303 | | 0 | 0 | 0 | 0 | 0 | 0 | 0 | 0 | 0 | 0 | 0 |
| 0 | NA | NA | NA | NA | NA | Gm15107 | predicted gene 15107 | 0 | 0 | 0 | 0 | 0 | 0 | 0 | 0 | 0 | 0 | 0 |
| 0 | NA | NA | NA | NA | NA | Gm15109 | | 0 | 0 | 0 | 0 | 0 | 0 | 0 | 0 | 0 | 0 | 0 |
| 0 | NA | NA | NA | NA | NA | Gm15111 | | 0 | 0 | 0 | 0 | 0 | 0 | 0 | 0 | 0 | 0 | 0 |
| 0 | NA | NA | NA | NA | NA | Gm15113 | | 0 | 0 | 0 | 0 | 0 | 0 | 0 | 0 | 0 | 0 | 0 |
| 0 | NA | NA | NA | NA | NA | Gm8320 | | 0 | 0 | 0 | 0 | 0 | 0 | 0 | 0 | 0 | 0 | 0 |
| 0 | NA | NA | NA | NA | NA | Gm15114 | predicted gene 15114 | 0 | 0 | 0 | 0 | 0 | 0 | 0 | 0 | 0 | 0 | 0 |
| 0 | NA | NA | NA | NA | NA | Gm15106 | | 0 | 0 | 0 | 0 | 0 | 0 | 0 | 0 | 0 | 0 | 0 |
| 0 | NA | NA | NA | NA | NA | Gm15112 | | 0 | 0 | 0 | 0 | 0 | 0 | 0 | 0 | 0 | 0 | 0 |
| 0 | NA | NA | NA | NA | NA | Gm8331 | | 0 | 0 | 0 | 0 | 0 | 0 | 0 | 0 | 0 | 0 | 0 |
| 0 | NA | NA | NA | NA | NA | Gm8334 | | 0 | 0 | 0 | 0 | 0 | 0 | 0 | 0 | 0 | 0 | 0 |
| 0 | NA | NA | NA | NA | NA | Gm15127 | predicted gene 15127 | 0 | 0 | 0 | 0 | 0 | 0 | 0 | 0 | 0 | 0 | 0 |
| 0 | NA | NA | NA | NA | NA | Gm15125 | | 0 | 0 | 0 | 0 | 0 | 0 | 0 | 0 | 0 | 0 | 0 |
| 0 | NA | NA | NA | NA | NA | Gm15126 | | 0 | 0 | 0 | 0 | 0 | 0 | 0 | 0 | 0 | 0 | 0 |

| | | | | | | | | | | | | | | | | | | |
|---|---|---|---|---|---|---|---|---|---|---|---|---|---|---|---|---|---|---|
| 0 | NA | NA | NA | NA | NA | Kars-ps1 | | 0 | 0 | 0 | 0 | 0 | 0 | 0 | 0 | 0 | 0 | 0 |
| 0 | NA | NA | NA | NA | NA | Luzp4 | leucine zipper protein 4 | 0 | 0 | 0 | 0 | 0 | 0 | 0 | 0 | 0 | 0 | 0 |
| 0 | NA | NA | NA | NA | NA | Gm15099 | | 0 | 0 | 0 | 0 | 0 | 0 | 0 | 0 | 0 | 0 | 0 |
| 0 | NA | NA | NA | NA | NA | Gm15084 | | 0 | 0 | 0 | 0 | 0 | 0 | 0 | 0 | 0 | 0 | 0 |
| 0 | NA | NA | NA | NA | NA | Gm15094 | | 0 | 0 | 0 | 0 | 0 | 0 | 0 | 0 | 0 | 0 | 0 |
| 0 | NA | NA | NA | NA | NA | Gm15096 | | 0 | 0 | 0 | 0 | 0 | 0 | 0 | 0 | 0 | 0 | 0 |
| 0 | NA | NA | NA | NA | NA | Ott | ovary testis transcribed | 0 | 0 | 0 | 0 | 0 | 0 | 0 | 0 | 0 | 0 | 0 |
| 0 | NA | NA | NA | NA | NA | Gm15092 | | 0 | 0 | 0 | 0 | 0 | 0 | 0 | 0 | 0 | 0 | 0 |
| 0 | NA | NA | NA | NA | NA | Gm15088 | | 0 | 0 | 0 | 0 | 0 | 0 | 0 | 0 | 0 | 0 | 0 |
| 0 | NA | NA | NA | NA | NA | Gm15090 | | 0 | 0 | 0 | 0 | 0 | 0 | 0 | 0 | 0 | 0 | 0 |
| 0 | NA | NA | NA | NA | NA | Gm8346 | | 0 | 0 | 0 | 0 | 0 | 0 | 0 | 0 | 0 | 0 | 0 |
| 0 | NA | NA | NA | NA | NA | Gm15093 | predicted gene 15093 | 0 | 0 | 0 | 0 | 0 | 0 | 0 | 0 | 0 | 0 | 0 |
| 0 | NA | NA | NA | NA | NA | Gm15098 | | 0 | 0 | 0 | 0 | 0 | 0 | 0 | 0 | 0 | 0 | 0 |
| 0 | NA | NA | NA | NA | NA | Gm15089 | | 0 | 0 | 0 | 0 | 0 | 0 | 0 | 0 | 0 | 0 | 0 |
| 0 | NA | NA | NA | NA | NA | Gm8366 | | 0 | 0 | 0 | 0 | 0 | 0 | 0 | 0 | 0 | 0 | 0 |
| 0 | NA | NA | NA | NA | NA | Gm15100 | | 0 | 0 | 0 | 0 | 0 | 0 | 0 | 0 | 0 | 0 | 0 |
| 0 | NA | NA | NA | NA | NA | Gm15085 | predicted gene 15085 | 0 | 0 | 0 | 0 | 0 | 0 | 0 | 0 | 0 | 0 | 0 |
| 0 | NA | NA | NA | NA | NA | Gm15086 | | 0 | 0 | 0 | 0 | 0 | 0 | 0 | 0 | 0 | 0 | 0 |
| 0 | NA | NA | NA | NA | NA | Gm10439 | predicted gene 10439 | 0 | 0 | 0 | 0 | 0 | 0 | 0 | 0 | 0 | 0 | 0 |
| 0 | NA | NA | NA | NA | NA | Gm15101 | | 0 | 0 | 0 | 0 | 0 | 0 | 0 | 0 | 0 | 0 | 0 |
| 0 | NA | NA | NA | NA | NA | Gm8383 | | 0 | 0 | 0 | 0 | 0 | 0 | 0 | 0 | 0 | 0 | 0 |
| 0 | NA | NA | NA | NA | NA | Gm15097 | predicted gene 15097 | 0 | 0 | 0 | 0 | 0 | 0 | 0 | 0 | 0 | 0 | 0 |
| 0 | NA | NA | NA | NA | NA | Gm15095 | | 0 | 0 | 0 | 0 | 0 | 0 | 0 | 0 | 0 | 0 | 0 |
| 0 | NA | NA | NA | NA | NA | Gm15087 | | 0 | 0 | 0 | 0 | 0 | 0 | 0 | 0 | 0 | 0 | 0 |
| 0 | NA | NA | NA | NA | NA | Gm15091 | predicted gene 15091 | 0 | 0 | 0 | 0 | 0 | 0 | 0 | 0 | 0 | 0 | 0 |
| 0 | NA | NA | NA | NA | NA | Gm15102 | | 0 | 0 | 0 | 0 | 0 | 0 | 0 | 0 | 0 | 0 | 0 |
| 0 | NA | NA | NA | NA | NA | Gm15103 | | 0 | 0 | 0 | 0 | 0 | 0 | 0 | 0 | 0 | 0 | 0 |
| 0 | NA | NA | NA | NA | NA | Gm8402 | | 0 | 0 | 0 | 0 | 0 | 0 | 0 | 0 | 0 | 0 | 0 |
| 0 | NA | NA | NA | NA | NA | Gm15104 | predicted gene 15104 | 0 | 0 | 0 | 0 | 0 | 0 | 0 | 0 | 0 | 0 | 0 |
| 0 | NA | NA | NA | NA | NA | Gm15105 | | 0 | 0 | 0 | 0 | 0 | 0 | 0 | 0 | 0 | 0 | 0 |
| 0 | NA | NA | NA | NA | NA | Gm15129 | | 0 | 0 | 0 | 0 | 0 | 0 | 0 | 0 | 0 | 0 | 0 |
| 0 | NA | NA | NA | NA | NA | Gm15132 | | 0 | 0 | 0 | 0 | 0 | 0 | 0 | 0 | 0 | 0 | 0 |
| 0 | NA | NA | NA | NA | NA | Gm27191 | | 0 | 0 | 0 | 0 | 0 | 0 | 0 | 0 | 0 | 0 | 0 |
| 0 | NA | NA | NA | NA | NA | Gm8424 | | 0 | 0 | 0 | 0 | 0 | 0 | 0 | 0 | 0 | 0 | 0 |
| 0 | NA | NA | NA | NA | NA | Gm15152 | | 0 | 0 | 0 | 0 | 0 | 0 | 0 | 0 | 0 | 0 | 0 |
| 0 | NA | NA | NA | NA | NA | Rpl21-ps15 | | 0 | 0 | 0 | 0 | 0 | 0 | 0 | 0 | 0 | 0 | 0 |
| 0 | NA | NA | NA | NA | NA | Gm26441 | | 0 | 0 | 0 | 0 | 0 | 0 | 0 | 0 | 0 | 0 | 0 |
| 0 | NA | NA | NA | NA | NA | Gm22248 | | 0 | 0 | 0 | 0 | 0 | 0 | 0 | 0 | 0 | 0 | 0 |
| 0 | NA | NA | NA | NA | NA | Gm24460 | | 0 | 0 | 0 | 0 | 0 | 0 | 0 | 0 | 0 | 0 | 0 |
| 0 | NA | NA | NA | NA | NA | n-R5s13 | | 0 | 0 | 0 | 0 | 0 | 0 | 0 | 0 | 0 | 0 | 0 |
| 0 | NA | NA | NA | NA | NA | Gm15161 | | 0 | 0 | 0 | 0 | 0 | 0 | 0 | 0 | 0 | 0 | 0 |
| 0 | NA | NA | NA | NA | NA | Mir3113 | | 0 | 0 | 0 | 0 | 0 | 0 | 0 | 0 | 0 | 0 | 0 |
| 0 | NA | NA | NA | NA | NA | Mirlet7f-2 | | 0 | 0 | 0 | 0 | 0 | 0 | 0 | 0 | 0 | 0 | 0 |
| 0 | NA | NA | NA | NA | NA | Mir98 | | 0 | 0 | 0 | 0 | 0 | 0 | 0 | 0 | 0 | 0 | 0 |
| 0 | NA | NA | NA | NA | NA | n-R5s14 | | 0 | 0 | 0 | 0 | 0 | 0 | 0 | 0 | 0 | 0 | 0 |
| 0 | NA | NA | NA | NA | NA | Gm15266 | predicted gene 15266 | 0 | 0 | 0 | 0 | 0 | 0 | 0 | 0 | 0 | 0 | 0 |
| 0 | NA | NA | NA | NA | NA | n-R5s15 | | 0 | 0 | 0 | 0 | 0 | 0 | 0 | 0 | 0 | 0 | 0 |
| 0 | NA | NA | NA | NA | NA | Gm4750 | | 0 | 0 | 0 | 0 | 0 | 0 | 0 | 0 | 0 | 0 | 0 |
| 0 | NA | NA | NA | NA | NA | Cldn34a | claudin 34A | 0 | 0 | 0 | 0 | 0 | 0 | 0 | 0 | 0 | 0 | 0 |
| 0 | NA | NA | NA | NA | NA | Gm7158 | | 0 | 0 | 0 | 0 | 0 | 0 | 0 | 0 | 0 | 0 | 0 |
| 0 | NA | NA | NA | NA | NA | Gm8475 | | 0 | 0 | 0 | 0 | 0 | 0 | 0 | 0 | 0 | 0 | 0 |
| 0 | NA | NA | NA | NA | NA | Gpr143 | G protein-coupled receptor 143 | 0 | 0 | 0 | 0 | 0 | 0 | 0 | 0 | 0 | 0 | 0 |
| 0 | NA | NA | NA | NA | NA | Gm24687 | | 0 | 0 | 0 | 0 | 0 | 0 | 0 | 0 | 0 | 0 | 0 |
| 0 | NA | NA | NA | NA | NA | Gm26035 | | 0 | 0 | 0 | 0 | 0 | 0 | 0 | 0 | 0 | 0 | 0 |
| 0 | NA | NA | NA | NA | NA | Gm7150 | | 0 | 0 | 0 | 0 | 0 | 0 | 0 | 0 | 0 | 0 | 0 |
| 0 | NA | NA | NA | NA | NA | Gm15166 | | 0 | 0 | 0 | 0 | 0 | 0 | 0 | 0 | 0 | 0 | 0 |
| 0 | NA | NA | NA | NA | NA | Cypt3 | cysteine-rich perinuclear theca 3 | 0 | 0 | 0 | 0 | 0 | 0 | 0 | 0 | 0 | 0 | 0 |
| 0 | NA | NA | NA | NA | NA | Gm25895 | | 0 | 0 | 0 | 0 | 0 | 0 | 0 | 0 | 0 | 0 | 0 |
| 0 | NA | NA | NA | NA | NA | Samt1 | spermatogenesis associated multipass tra | 0 | 0 | 0 | 0 | 0 | 0 | 0 | 0 | 0 | 0 | 0 |
| 0 | NA | NA | NA | NA | NA | Samt1b | predicted pseudogene 5645 | 0 | 0 | 0 | 0 | 0 | 0 | 0 | 0 | 0 | 0 | 0 |

| | | | | | | | | | | | | | | | | | |
|---|---|---|---|---|---|---|---|---|---|---|---|---|---|---|---|---|---|
| 0 | NA | NA | NA | NA | NA | Samt1c | predicted pseudogene 5646 | 0 | 0 | 0 | 0 | 0 | 0 | 0 | 0 | 0 | 0 | 0 |
| 0 | NA | NA | NA | NA | NA | Gm15143 | | 0 | 0 | 0 | 0 | 0 | 0 | 0 | 0 | 0 | 0 | 0 |
| 0 | NA | NA | NA | NA | NA | Gm5947 | | 0 | 0 | 0 | 0 | 0 | 0 | 0 | 0 | 0 | 0 | 0 |
| 0 | NA | NA | NA | NA | NA | Samt1d | spermatogenesis associated multipass tra | 0 | 0 | 0 | 0 | 0 | 0 | 0 | 0 | 0 | 0 | 0 |
| 0 | NA | NA | NA | NA | NA | Gm5645 | | 0 | 0 | 0 | 0 | 0 | 0 | 0 | 0 | 0 | 0 | 0 |
| 0 | NA | NA | NA | NA | NA | Gm16445 | | 0 | 0 | 0 | 0 | 0 | 0 | 0 | 0 | 0 | 0 | 0 |
| 0 | NA | NA | NA | NA | NA | Gm6620 | | 0 | 0 | 0 | 0 | 0 | 0 | 0 | 0 | 0 | 0 | 0 |
| 0 | NA | NA | NA | NA | NA | Gm15147 | | 0 | 0 | 0 | 0 | 0 | 0 | 0 | 0 | 0 | 0 | 0 |
| 0 | NA | NA | NA | NA | NA | Gm5060 | | 0 | 0 | 0 | 0 | 0 | 0 | 0 | 0 | 0 | 0 | 0 |
| 0 | NA | NA | NA | NA | NA | Gm15141 | | 0 | 0 | 0 | 0 | 0 | 0 | 0 | 0 | 0 | 0 | 0 |
| 0 | NA | NA | NA | NA | NA | Samt2b | spermatogenesis associated multipass tra | 0 | 0 | 0 | 0 | 0 | 0 | 0 | 0 | 0 | 0 | 0 |
| 0 | NA | NA | NA | NA | NA | Gm5646 | | 0 | 0 | 0 | 0 | 0 | 0 | 0 | 0 | 0 | 0 | 0 |
| 0 | NA | NA | NA | NA | NA | Gm8569 | | 0 | 0 | 0 | 0 | 0 | 0 | 0 | 0 | 0 | 0 | 0 |
| 0 | NA | NA | NA | NA | NA | Samt4 | spermatogenesis associated multipass tra | 0 | 0 | 0 | 0 | 0 | 0 | 0 | 0 | 0 | 0 | 0 |
| 0 | NA | NA | NA | NA | NA | Samt2 | spermatogenesis associated multipass tra | 0 | 0 | 0 | 0 | 0 | 0 | 0 | 0 | 0 | 0 | 0 |
| 0 | NA | NA | NA | NA | NA | Gm15142 | predicted gene 15142 | 0 | 0 | 0 | 0 | 0 | 0 | 0 | 0 | 0 | 0 | 0 |
| 0 | NA | NA | NA | NA | NA | Gm22359 | | 0 | 0 | 0 | 0 | 0 | 0 | 0 | 0 | 0 | 0 | 0 |
| 0 | NA | NA | NA | NA | NA | Cldn34b1 | claudin 34B1 | 0 | 0 | 0 | 0 | 0 | 0 | 0 | 0 | 0 | 0 | 0 |
| 0 | NA | NA | NA | NA | NA | Magea6 | melanoma antigen, family A, 6 | 0 | 0 | 0 | 0 | 0 | 0 | 0 | 0 | 0 | 0 | 0 |
| 0 | NA | NA | NA | NA | NA | Magea3 | melanoma antigen, family A, 3 | 0 | 0 | 0 | 0 | 0 | 0 | 0 | 0 | 0 | 0 | 0 |
| 0 | NA | NA | NA | NA | NA | Gm25257 | | 0 | 0 | 0 | 0 | 0 | 0 | 0 | 0 | 0 | 0 | 0 |
| 0 | NA | NA | NA | NA | NA | Magea8 | melanoma antigen, family A, 8 | 0 | 0 | 0 | 0 | 0 | 0 | 0 | 0 | 0 | 0 | 0 |
| 0 | NA | NA | NA | NA | NA | Magea2 | melanoma antigen, family A, 2 | 0 | 0 | 0 | 0 | 0 | 0 | 0 | 0 | 0 | 0 | 0 |
| 0 | NA | NA | NA | NA | NA | Gm24268 | | 0 | 0 | 0 | 0 | 0 | 0 | 0 | 0 | 0 | 0 | 0 |
| 0 | NA | NA | NA | NA | NA | Magea5 | melanoma antigen, family A, 5 | 0 | 0 | 0 | 0 | 0 | 0 | 0 | 0 | 0 | 0 | 0 |
| 0 | NA | NA | NA | NA | NA | Magea1 | melanoma antigen, family A, 1 | 0 | 0 | 0 | 0 | 0 | 0 | 0 | 0 | 0 | 0 | 0 |
| 0 | NA | NA | NA | NA | NA | Cldn34b2 | claudin 34B2 | 0 | 0 | 0 | 0 | 0 | 0 | 0 | 0 | 0 | 0 | 0 |
| 0 | NA | NA | NA | NA | NA | Gm25429 | | 0 | 0 | 0 | 0 | 0 | 0 | 0 | 0 | 0 | 0 | 0 |
| 0 | NA | NA | NA | NA | NA | Gm8595 | | 0 | 0 | 0 | 0 | 0 | 0 | 0 | 0 | 0 | 0 | 0 |
| 0 | NA | NA | NA | NA | NA | Gm15159 | | 0 | 0 | 0 | 0 | 0 | 0 | 0 | 0 | 0 | 0 | 0 |
| 0 | NA | NA | NA | NA | NA | Gm650 | | 0 | 0 | 0 | 0 | 0 | 0 | 0 | 0 | 0 | 0 | 0 |
| 0 | NA | NA | NA | NA | NA | Gm15158 | | 0 | 0 | 0 | 0 | 0 | 0 | 0 | 0 | 0 | 0 | 0 |
| 0 | NA | NA | NA | NA | NA | n-R5s16 | | 0 | 0 | 0 | 0 | 0 | 0 | 0 | 0 | 0 | 0 | 0 |
| 0 | NA | NA | NA | NA | NA | Gm15155 | | 0 | 0 | 0 | 0 | 0 | 0 | 0 | 0 | 0 | 0 | 0 |
| 0 | NA | NA | NA | NA | NA | Gm15160 | | 0 | 0 | 0 | 0 | 0 | 0 | 0 | 0 | 0 | 0 | 0 |
| 0 | NA | NA | NA | NA | NA | Gm23404 | | 0 | 0 | 0 | 0 | 0 | 0 | 0 | 0 | 0 | 0 | 0 |
| 0 | NA | NA | NA | NA | NA | 4930503H13Rik | | 0 | 0 | 0 | 0 | 0 | 0 | 0 | 0 | 0 | 0 | 0 |
| 0 | NA | NA | NA | NA | NA | Gm7312 | | 0 | 0 | 0 | 0 | 0 | 0 | 0 | 0 | 0 | 0 | 0 |
| 0 | NA | NA | NA | NA | NA | Gm15162 | | 0 | 0 | 0 | 0 | 0 | 0 | 0 | 0 | 0 | 0 | 0 |
| 0 | NA | NA | NA | NA | NA | Gm22666 | | 0 | 0 | 0 | 0 | 0 | 0 | 0 | 0 | 0 | 0 | 0 |
| 0 | NA | NA | NA | NA | NA | Gm23798 | | 0 | 0 | 0 | 0 | 0 | 0 | 0 | 0 | 0 | 0 | 0 |
| 0 | NA | NA | NA | NA | NA | Gm25795 | | 0 | 0 | 0 | 0 | 0 | 0 | 0 | 0 | 0 | 0 | 0 |
| 0 | NA | NA | NA | NA | NA | Gm15172 | | 0 | 0 | 0 | 0 | 0 | 0 | 0 | 0 | 0 | 0 | 0 |
| 0 | NA | NA | NA | NA | NA | Gm1947 | | 0 | 0 | 0 | 0 | 0 | 0 | 0 | 0 | 0 | 0 | 0 |
| 0 | NA | NA | NA | NA | NA | Gm24281 | | 0 | 0 | 0 | 0 | 0 | 0 | 0 | 0 | 0 | 0 | 0 |
| 0 | NA | NA | NA | NA | NA | Yy2 | Yy2 transcription factor | 0 | 0 | 0 | 0 | 0 | 0 | 0 | 0 | 0 | 0 | 0 |
| 0 | NA | NA | NA | NA | NA | Gm15176 | | 0 | 0 | 0 | 0 | 0 | 0 | 0 | 0 | 0 | 0 | 0 |
| 0 | NA | NA | NA | NA | NA | Gm15175 | | 0 | 0 | 0 | 0 | 0 | 0 | 0 | 0 | 0 | 0 | 0 |
| 0 | NA | NA | NA | NA | NA | Gm15173 | | 0 | 0 | 0 | 0 | 0 | 0 | 0 | 0 | 0 | 0 | 0 |
| 0 | NA | NA | NA | NA | NA | Gm23381 | | 0 | 0 | 0 | 0 | 0 | 0 | 0 | 0 | 0 | 0 | 0 |
| 0 | NA | NA | NA | NA | NA | Gm8644 | | 0 | 0 | 0 | 0 | 0 | 0 | 0 | 0 | 0 | 0 | 0 |
| 0 | NA | NA | NA | NA | NA | Gm15171 | | 0 | 0 | 0 | 0 | 0 | 0 | 0 | 0 | 0 | 0 | 0 |
| 0 | NA | NA | NA | NA | NA | Smpx | small muscle protein, X-linked | 0 | 0 | 0 | 0 | 0 | 0 | 0 | 0 | 0 | 0 | 0 |
| 0 | NA | NA | NA | NA | NA | Klhl34 | kelch-like 34 | 0 | 0 | 0 | 0 | 0 | 0 | 0 | 0 | 0 | 0 | 0 |
| 0 | NA | NA | NA | NA | NA | Gm15170 | | 0 | 0 | 0 | 0 | 0 | 0 | 0 | 0 | 0 | 0 | 0 |
| 0 | NA | NA | NA | NA | NA | Gm15177 | | 0 | 0 | 0 | 0 | 0 | 0 | 0 | 0 | 0 | 0 | 0 |
| 0 | NA | NA | NA | NA | NA | Gm22522 | | 0 | 0 | 0 | 0 | 0 | 0 | 0 | 0 | 0 | 0 | 0 |
| 0 | NA | NA | NA | NA | NA | Gm15168 | | 0 | 0 | 0 | 0 | 0 | 0 | 0 | 0 | 0 | 0 | 0 |
| 0 | NA | NA | NA | NA | NA | Gm15185 | | 0 | 0 | 0 | 0 | 0 | 0 | 0 | 0 | 0 | 0 | 0 |
| 0 | NA | NA | NA | NA | NA | Gm15188 | | 0 | 0 | 0 | 0 | 0 | 0 | 0 | 0 | 0 | 0 | 0 |

| | | | | | | | | | | | | | | | | | | |
|---|---|---|---|---|---|---|---|---|---|---|---|---|---|---|---|---|---|---|
| 0 | NA | NA | NA | NA | NA | Gm15189 | | 0 | 0 | 0 | 0 | 0 | 0 | 0 | 0 | 0 | 0 | 0 |
| 0 | NA | NA | NA | NA | NA | Gm15187 | | 0 | 0 | 0 | 0 | 0 | 0 | 0 | 0 | 0 | 0 | 0 |
| 0 | NA | NA | NA | NA | NA | Gm15194 | | 0 | 0 | 0 | 0 | 0 | 0 | 0 | 0 | 0 | 0 | 0 |
| 0 | NA | NA | NA | NA | NA | Gm15192 | | 0 | 0 | 0 | 0 | 0 | 0 | 0 | 0 | 0 | 0 | 0 |
| 0 | NA | NA | NA | NA | NA | Gm15196 | | 0 | 0 | 0 | 0 | 0 | 0 | 0 | 0 | 0 | 0 | 0 |
| 0 | NA | NA | NA | NA | NA | Gm15193 | | 0 | 0 | 0 | 0 | 0 | 0 | 0 | 0 | 0 | 0 | 0 |
| 0 | NA | NA | NA | NA | NA | Gm15241 | | 0 | 0 | 0 | 0 | 0 | 0 | 0 | 0 | 0 | 0 | 0 |
| 0 | NA | NA | NA | NA | NA | Gm15242 | | 0 | 0 | 0 | 0 | 0 | 0 | 0 | 0 | 0 | 0 | 0 |
| 0 | NA | NA | NA | NA | NA | Gm15244 | | 0 | 0 | 0 | 0 | 0 | 0 | 0 | 0 | 0 | 0 | 0 |
| 0 | NA | NA | NA | NA | NA | Rs1 | retinoschisis (X-linked, juvenile) 1 (humar | 0 | 0 | 0 | 0 | 0 | 0 | 0 | 0 | 0 | 0 | 0 |
| 0 | NA | NA | NA | NA | NA | Gm5419 | | 0 | 0 | 0 | 0 | 0 | 0 | 0 | 0 | 0 | 0 | 0 |
| 0 | NA | NA | NA | NA | NA | Gm15262 | predicted gene 15262 | 0 | 0 | 0 | 0 | 0 | 0 | 0 | 0 | 0 | 0 | 0 |
| 0 | NA | NA | NA | NA | NA | Scml1 | | 0 | 0 | 0 | 0 | 0 | 0 | 0 | 0 | 0 | 0 | 0 |
| 0 | NA | NA | NA | NA | NA | Gm15202 | | 0 | 0 | 0 | 0 | 0 | 0 | 0 | 0 | 0 | 0 | 0 |
| 0 | NA | NA | NA | NA | NA | Gm26317 | | 0 | 0 | 0 | 0 | 0 | 0 | 0 | 0 | 0 | 0 | 0 |
| 0 | NA | NA | NA | NA | NA | Gm23794 | | 0 | 0 | 0 | 0 | 0 | 0 | 0 | 0 | 0 | 0 | 0 |
| 0 | NA | NA | NA | NA | NA | Gm27490 | | 0 | 0 | 0 | 0 | 0 | 0 | 0 | 0 | 0 | 0 | 0 |
| 0 | NA | NA | NA | NA | NA | Gm23867 | | 0 | 0 | 0 | 0 | 0 | 0 | 0 | 0 | 0 | 0 | 0 |
| 0 | NA | NA | NA | NA | NA | Gm15206 | | 0 | 0 | 0 | 0 | 0 | 0 | 0 | 0 | 0 | 0 | 0 |
| 0 | NA | NA | NA | NA | NA | Gm15203 | | 0 | 0 | 0 | 0 | 0 | 0 | 0 | 0 | 0 | 0 | 0 |
| 0 | NA | NA | NA | NA | NA | Gm22257 | | 0 | 0 | 0 | 0 | 0 | 0 | 0 | 0 | 0 | 0 | 0 |
| 0 | NA | NA | NA | NA | NA | Gm25557 | | 0 | 0 | 0 | 0 | 0 | 0 | 0 | 0 | 0 | 0 | 0 |
| 0 | NA | NA | NA | NA | NA | Gm23786 | | 0 | 0 | 0 | 0 | 0 | 0 | 0 | 0 | 0 | 0 | 0 |
| 0 | NA | NA | NA | NA | NA | Gm23786 | | 0 | 0 | 0 | 0 | 0 | 0 | 0 | 0 | 0 | 0 | 0 |
| 0 | NA | NA | NA | NA | NA | Gm15207 | | 0 | 0 | 0 | 0 | 0 | 0 | 0 | 0 | 0 | 0 | 0 |
| 0 | NA | NA | NA | NA | NA | Gm23901 | | 0 | 0 | 0 | 0 | 0 | 0 | 0 | 0 | 0 | 0 | 0 |
| 0 | NA | NA | NA | NA | NA | Gm15209 | | 0 | 0 | 0 | 0 | 0 | 0 | 0 | 0 | 0 | 0 | 0 |
| 0 | NA | NA | NA | NA | NA | S100g | S100 calcium binding protein G | 0 | 0 | 0 | 0 | 0 | 0 | 0 | 0 | 0 | 0 | 0 |
| 0 | NA | NA | NA | NA | NA | Gm23501 | | 0 | 0 | 0 | 0 | 0 | 0 | 0 | 0 | 0 | 0 | 0 |
| 0 | NA | NA | NA | NA | NA | Gm22746 | | 0 | 0 | 0 | 0 | 0 | 0 | 0 | 0 | 0 | 0 | 0 |
| 0 | NA | NA | NA | NA | NA | Gm15215 | | 0 | 0 | 0 | 0 | 0 | 0 | 0 | 0 | 0 | 0 | 0 |
| 0 | NA | NA | NA | NA | NA | Gm15214 | | 0 | 0 | 0 | 0 | 0 | 0 | 0 | 0 | 0 | 0 | 0 |
| 0 | NA | NA | NA | NA | NA | Gm26007 | | 0 | 0 | 0 | 0 | 0 | 0 | 0 | 0 | 0 | 0 | 0 |
| 0 | NA | NA | NA | NA | NA | Gm7199 | | 0 | 0 | 0 | 0 | 0 | 0 | 0 | 0 | 0 | 0 | 0 |
| 0 | NA | NA | NA | NA | NA | Gm15213 | | 0 | 0 | 0 | 0 | 0 | 0 | 0 | 0 | 0 | 0 | 0 |
| 0 | NA | NA | NA | NA | NA | n-R5s18 | | 0 | 0 | 0 | 0 | 0 | 0 | 0 | 0 | 0 | 0 | 0 |
| 0 | NA | NA | NA | NA | NA | Gm25367 | | 0 | 0 | 0 | 0 | 0 | 0 | 0 | 0 | 0 | 0 | 0 |
| 0 | NA | NA | NA | NA | NA | Gm22023 | | 0 | 0 | 0 | 0 | 0 | 0 | 0 | 0 | 0 | 0 | 0 |
| 0 | NA | NA | NA | NA | NA | Gm15216 | | 0 | 0 | 0 | 0 | 0 | 0 | 0 | 0 | 0 | 0 | 0 |
| 0 | NA | NA | NA | NA | NA | Gm17604 | | 0 | 0 | 0 | 0 | 0 | 0 | 0 | 0 | 0 | 0 | 0 |
| 0 | NA | NA | NA | NA | NA | Gm15236 | | 0 | 0 | 0 | 0 | 0 | 0 | 0 | 0 | 0 | 0 | 0 |
| 0 | NA | NA | NA | NA | NA | Gm22062 | | 0 | 0 | 0 | 0 | 0 | 0 | 0 | 0 | 0 | 0 | 0 |
| 0 | NA | NA | NA | NA | NA | Gm15224 | | 0 | 0 | 0 | 0 | 0 | 0 | 0 | 0 | 0 | 0 | 0 |
| 0 | NA | NA | NA | NA | NA | Gm15225 | | 0 | 0 | 0 | 0 | 0 | 0 | 0 | 0 | 0 | 0 | 0 |
| 0 | NA | NA | NA | NA | NA | Gm15227 | | 0 | 0 | 0 | 0 | 0 | 0 | 0 | 0 | 0 | 0 | 0 |
| 0 | NA | NA | NA | NA | NA | Gm15229 | | 0 | 0 | 0 | 0 | 0 | 0 | 0 | 0 | 0 | 0 | 0 |
| 0 | NA | NA | NA | NA | NA | Gm15226 | | 0 | 0 | 0 | 0 | 0 | 0 | 0 | 0 | 0 | 0 | 0 |
| 0 | NA | NA | NA | NA | NA | Gm1720 | | 0 | 0 | 0 | 0 | 0 | 0 | 0 | 0 | 0 | 0 | 0 |
| 0 | NA | NA | NA | NA | NA | Rpl7a-ps11 | | 0 | 0 | 0 | 0 | 0 | 0 | 0 | 0 | 0 | 0 | 0 |
| 0 | NA | NA | NA | NA | NA | Gm15230 | | 0 | 0 | 0 | 0 | 0 | 0 | 0 | 0 | 0 | 0 | 0 |
| 0 | NA | NA | NA | NA | NA | Gm15231 | | 0 | 0 | 0 | 0 | 0 | 0 | 0 | 0 | 0 | 0 | 0 |
| 0 | NA | NA | NA | NA | NA | Gm44376 | | 0 | 0 | 0 | 0 | 0 | 0 | 0 | 0 | 0 | 0 | 0 |
| 0 | NA | NA | NA | NA | NA | Gm22368 | | 0 | 0 | 0 | 0 | 0 | 0 | 0 | 0 | 0 | 0 | 0 |
| 0 | NA | NA | NA | NA | NA | Gm8817 | predicted gene 8817 | 0 | 0 | 0 | 0 | 0 | 0 | 0 | 0 | 0 | 0 | 0 |
| 0 | NA | NA | NA | NA | NA | Gm15228 | | 0 | 0 | 0 | 0 | 0 | 0 | 0 | 0 | 0 | 0 | 0 |
| 0 | NA | NA | NA | NA | NA | Gm15239 | | 0 | 0 | 0 | 0 | 0 | 0 | 0 | 0 | 0 | 0 | 0 |
| 0 | NA | NA | NA | NA | NA | Gm26368 | | 0 | 0 | 0 | 0 | 0 | 0 | 0 | 0 | 0 | 0 | 0 |
| 0 | NA | NA | NA | NA | NA | Gm6744 | | 0 | 0 | 0 | 0 | 0 | 0 | 0 | 0 | 0 | 0 | 0 |
| 0 | NA | NA | NA | NA | NA | Gm15240 | | 0 | 0 | 0 | 0 | 0 | 0 | 0 | 0 | 0 | 0 | 0 |
| 0 | NA | NA | NA | NA | NA | Gm15237 | | 0 | 0 | 0 | 0 | 0 | 0 | 0 | 0 | 0 | 0 | 0 |

| | | | | | | | | | | | | | | | | | |
|---|---|---|---|---|---|---|---|---|---|---|---|---|---|---|---|---|---|
| 0 | NA | NA | NA | NA | NA | Gm7209 | | 0 | 0 | 0 | 0 | 0 | 0 | 0 | 0 | 0 | 0 | 0 |
| 0 | NA | NA | NA | NA | NA | Gm23197 | | 0 | 0 | 0 | 0 | 0 | 0 | 0 | 0 | 0 | 0 | 0 |
| 0 | NA | NA | NA | NA | NA | Gm15233 | | 0 | 0 | 0 | 0 | 0 | 0 | 0 | 0 | 0 | 0 | 0 |
| 0 | NA | NA | NA | NA | NA | Gm15245 | | 0 | 0 | 0 | 0 | 0 | 0 | 0 | 0 | 0 | 0 | 0 |
| 0 | NA | NA | NA | NA | NA | Gm25651 | | 0 | 0 | 0 | 0 | 0 | 0 | 0 | 0 | 0 | 0 | 0 |
| 0 | NA | NA | NA | NA | NA | 4933400A11Rik | | 0 | 0 | 0 | 0 | 0 | 0 | 0 | 0 | 0 | 0 | 0 |
| 0 | NA | NA | NA | NA | NA | Gm15069 | | 0 | 0 | 0 | 0 | 0 | 0 | 0 | 0 | 0 | 0 | 0 |
| 0 | NA | NA | NA | NA | NA | Gm15068 | | 0 | 0 | 0 | 0 | 0 | 0 | 0 | 0 | 0 | 0 | 0 |
| 0 | NA | NA | NA | NA | NA | Gm29277 | | 0 | 0 | 0 | 0 | 0 | 0 | 0 | 0 | 0 | 0 | 0 |
| 0 | NA | NA | NA | NA | NA | Gm29089 | | 0 | 0 | 0 | 0 | 0 | 0 | 0 | 0 | 0 | 0 | 0 |
| 0 | NA | NA | NA | NA | NA | Gm2098 | | 0 | 0 | 0 | 0 | 0 | 0 | 0 | 0 | 0 | 0 | 0 |
| 0 | NA | NA | NA | NA | NA | Zfy1 | zinc finger protein 1, Y-linked | 0 | 0 | 0 | 0 | 0 | 0 | 0 | 0 | 0 | 0 | 0 |
| 0 | NA | NA | NA | NA | NA | Uba1y | ubiquitin-activating enzyme, Chr Y | 0 | 0 | 0 | 0 | 0 | 0 | 0 | 0 | 0 | 0 | 0 |
| 0 | NA | NA | NA | NA | NA | Gm28588 | | 0 | 0 | 0 | 0 | 0 | 0 | 0 | 0 | 0 | 0 | 0 |
| 0 | NA | NA | NA | NA | NA | Gm28587 | | 0 | 0 | 0 | 0 | 0 | 0 | 0 | 0 | 0 | 0 | 0 |
| 0 | NA | NA | NA | NA | NA | n-R5s1 | | 0 | 0 | 0 | 0 | 0 | 0 | 0 | 0 | 0 | 0 | 0 |
| 0 | NA | NA | NA | NA | NA | Gm47281 | | 0 | 0 | 0 | 0 | 0 | 0 | 0 | 0 | 0 | 0 | 0 |
| 0 | NA | NA | NA | NA | NA | Usp9y | ubiquitin specific peptidase 9, Y chromos | 0 | 0 | 0 | 0 | 0 | 0 | 0 | 0 | 0 | 0 | 0 |
| 0 | NA | NA | NA | NA | NA | Gm25565 | | 0 | 0 | 0 | 0 | 0 | 0 | 0 | 0 | 0 | 0 | 0 |
| 0 | NA | NA | NA | NA | NA | Gm17790 | | 0 | 0 | 0 | 0 | 0 | 0 | 0 | 0 | 0 | 0 | 0 |
| 0 | NA | NA | NA | NA | NA | Gm29167 | | 0 | 0 | 0 | 0 | 0 | 0 | 0 | 0 | 0 | 0 | 0 |
| 0 | NA | NA | NA | NA | NA | Gm8446 | | 0 | 0 | 0 | 0 | 0 | 0 | 0 | 0 | 0 | 0 | 0 |
| 0 | NA | NA | NA | NA | NA | Gm28534 | | 0 | 0 | 0 | 0 | 0 | 0 | 0 | 0 | 0 | 0 | 0 |
| 0 | NA | NA | NA | NA | NA | Gm28975 | | 0 | 0 | 0 | 0 | 0 | 0 | 0 | 0 | 0 | 0 | 0 |
| 0 | NA | NA | NA | NA | NA | Gm28974 | | 0 | 0 | 0 | 0 | 0 | 0 | 0 | 0 | 0 | 0 | 0 |
| 0 | NA | NA | NA | NA | NA | Gm18797 | | 0 | 0 | 0 | 0 | 0 | 0 | 0 | 0 | 0 | 0 | 0 |
| 0 | NA | NA | NA | NA | NA | Rhoay-ps1 | | 0 | 0 | 0 | 0 | 0 | 0 | 0 | 0 | 0 | 0 | 0 |
| 0 | NA | NA | NA | NA | NA | Zfy2 | zinc finger protein 2, Y-linked | 0 | 0 | 0 | 0 | 0 | 0 | 0 | 0 | 0 | 0 | 0 |
| 0 | NA | NA | NA | NA | NA | Uba1y-ps2 | | 0 | 0 | 0 | 0 | 0 | 0 | 0 | 0 | 0 | 0 | 0 |
| 0 | NA | NA | NA | NA | NA | Gm18796 | | 0 | 0 | 0 | 0 | 0 | 0 | 0 | 0 | 0 | 0 | 0 |
| 0 | NA | NA | NA | NA | NA | Gm29552 | | 0 | 0 | 0 | 0 | 0 | 0 | 0 | 0 | 0 | 0 | 0 |
| 0 | NA | NA | NA | NA | NA | Gm29550 | | 0 | 0 | 0 | 0 | 0 | 0 | 0 | 0 | 0 | 0 | 0 |
| 0 | NA | NA | NA | NA | NA | Gm29551 | | 0 | 0 | 0 | 0 | 0 | 0 | 0 | 0 | 0 | 0 | 0 |
| 0 | NA | NA | NA | NA | NA | Gm8480 | | 0 | 0 | 0 | 0 | 0 | 0 | 0 | 0 | 0 | 0 | 0 |
| 0 | NA | NA | NA | NA | NA | Gm29553 | | 0 | 0 | 0 | 0 | 0 | 0 | 0 | 0 | 0 | 0 | 0 |
| 0 | NA | NA | NA | NA | NA | Gm22119 | | 0 | 0 | 0 | 0 | 0 | 0 | 0 | 0 | 0 | 0 | 0 |
| 0 | NA | NA | NA | NA | NA | Gm28630 | | 0 | 0 | 0 | 0 | 0 | 0 | 0 | 0 | 0 | 0 | 0 |
| 0 | NA | NA | NA | NA | NA | Gm28629 | | 0 | 0 | 0 | 0 | 0 | 0 | 0 | 0 | 0 | 0 | 0 |
| 0 | NA | NA | NA | NA | NA | H2al2c | H2A histone family member L2C | 0 | 0 | 0 | 0 | 0 | 0 | 0 | 0 | 0 | 0 | 0 |
| 0 | NA | NA | NA | NA | NA | Uba1y-ps1 | | 0 | 0 | 0 | 0 | 0 | 0 | 0 | 0 | 0 | 0 | 0 |
| 0 | NA | NA | NA | NA | NA | Sry | sex determining region of Chr Y | 0 | 0 | 0 | 0 | 0 | 0 | 0 | 0 | 0 | 0 | 0 |
| 0 | NA | NA | NA | NA | NA | Gm18798 | | 0 | 0 | 0 | 0 | 0 | 0 | 0 | 0 | 0 | 0 | 0 |
| 0 | NA | NA | NA | NA | NA | H2al2b | H2A histone family member L2B | 0 | 0 | 0 | 0 | 0 | 0 | 0 | 0 | 0 | 0 | 0 |
| 0 | NA | NA | NA | NA | NA | Gm28628 | | 0 | 0 | 0 | 0 | 0 | 0 | 0 | 0 | 0 | 0 | 0 |
| 0 | NA | NA | NA | NA | NA | Gm4064 | predicted gene 4064 | 0 | 0 | 0 | 0 | 0 | 0 | 0 | 0 | 0 | 0 | 0 |
| 0 | NA | NA | NA | NA | NA | Gm6137 | | 0 | 0 | 0 | 0 | 0 | 0 | 0 | 0 | 0 | 0 | 0 |
| 0 | NA | NA | NA | NA | NA | Gm8498 | | 0 | 0 | 0 | 0 | 0 | 0 | 0 | 0 | 0 | 0 | 0 |
| 0 | NA | NA | NA | NA | NA | Rbmy | RNA binding motif protein, Y chromosom | 0 | 0 | 0 | 0 | 0 | 0 | 0 | 0 | 0 | 0 | 0 |
| 0 | NA | NA | NA | NA | NA | Gm10256 | predicted gene 10256 | 0 | 0 | 0 | 0 | 0 | 0 | 0 | 0 | 0 | 0 | 0 |
| 0 | NA | NA | NA | NA | NA | Gm8502 | | 0 | 0 | 0 | 0 | 0 | 0 | 0 | 0 | 0 | 0 | 0 |
| 0 | NA | NA | NA | NA | NA | Gm8506 | | 0 | 0 | 0 | 0 | 0 | 0 | 0 | 0 | 0 | 0 | 0 |
| 0 | NA | NA | NA | NA | NA | Gm10352 | predicted gene 10352 | 0 | 0 | 0 | 0 | 0 | 0 | 0 | 0 | 0 | 0 | 0 |
| 0 | NA | NA | NA | NA | NA | Gm8510 | | 0 | 0 | 0 | 0 | 0 | 0 | 0 | 0 | 0 | 0 | 0 |
| 0 | NA | NA | NA | NA | NA | Gm29289 | | 0 | 0 | 0 | 0 | 0 | 0 | 0 | 0 | 0 | 0 | 0 |
| 0 | NA | NA | NA | NA | NA | Gm21677 | predicted gene, 21677 | 0 | 0 | 0 | 0 | 0 | 0 | 0 | 0 | 0 | 0 | 0 |
| 0 | NA | NA | NA | NA | NA | Gm29288 | | 0 | 0 | 0 | 0 | 0 | 0 | 0 | 0 | 0 | 0 | 0 |
| 0 | NA | NA | NA | NA | NA | Gm21693 | predicted gene, 21693 | 0 | 0 | 0 | 0 | 0 | 0 | 0 | 0 | 0 | 0 | 0 |
| 0 | NA | NA | NA | NA | NA | Gm21139 | | 0 | 0 | 0 | 0 | 0 | 0 | 0 | 0 | 0 | 0 | 0 |
| 0 | NA | NA | NA | NA | NA | Gm21704 | predicted gene, 21704 | 0 | 0 | 0 | 0 | 0 | 0 | 0 | 0 | 0 | 0 | 0 |
| 0 | NA | NA | NA | NA | NA | Gm21147 | | 0 | 0 | 0 | 0 | 0 | 0 | 0 | 0 | 0 | 0 | 0 |

| | | | | | | | | | | | | | | | | | | |
|---|---|---|---|---|---|---|---|---|---|---|---|---|---|---|---|---|---|---|
| 0 | NA | NA | NA | NA | NA | Gm21708 | predicted gene, 21708 | 0 | 0 | 0 | 0 | 0 | 0 | 0 | 0 | 0 | 0 | 0 |
| 0 | NA | NA | NA | NA | NA | Gm21156 | | 0 | 0 | 0 | 0 | 0 | 0 | 0 | 0 | 0 | 0 | 0 |
| 0 | NA | NA | NA | NA | NA | Rhoay-ps2 | | 0 | 0 | 0 | 0 | 0 | 0 | 0 | 0 | 0 | 0 | 0 |
| 0 | NA | NA | NA | NA | NA | Gm2316 | | 0 | 0 | 0 | 0 | 0 | 0 | 0 | 0 | 0 | 0 | 0 |
| 0 | NA | NA | NA | NA | NA | Gm3376 | predicted gene 3376 | 0 | 0 | 0 | 0 | 0 | 0 | 0 | 0 | 0 | 0 | 0 |
| 0 | NA | NA | NA | NA | NA | Gm2325 | | 0 | 0 | 0 | 0 | 0 | 0 | 0 | 0 | 0 | 0 | 0 |
| 0 | NA | NA | NA | NA | NA | Gm28649 | | 0 | 0 | 0 | 0 | 0 | 0 | 0 | 0 | 0 | 0 | 0 |
| 0 | NA | NA | NA | NA | NA | Gm8521 | | 0 | 0 | 0 | 0 | 0 | 0 | 0 | 0 | 0 | 0 | 0 |
| 0 | NA | NA | NA | NA | NA | Gm18177 | | 0 | 0 | 0 | 0 | 0 | 0 | 0 | 0 | 0 | 0 | 0 |
| 0 | NA | NA | NA | NA | NA | Gm28429 | | 0 | 0 | 0 | 0 | 0 | 0 | 0 | 0 | 0 | 0 | 0 |
| 0 | NA | NA | NA | NA | NA | Gm28413 | | 0 | 0 | 0 | 0 | 0 | 0 | 0 | 0 | 0 | 0 | 0 |
| 0 | NA | NA | NA | NA | NA | Gm28412 | | 0 | 0 | 0 | 0 | 0 | 0 | 0 | 0 | 0 | 0 | 0 |
| 0 | NA | NA | NA | NA | NA | Gm8525 | | 0 | 0 | 0 | 0 | 0 | 0 | 0 | 0 | 0 | 0 | 0 |
| 0 | NA | NA | NA | NA | NA | Gm28414 | | 0 | 0 | 0 | 0 | 0 | 0 | 0 | 0 | 0 | 0 | 0 |
| 0 | NA | NA | NA | NA | NA | Gm28648 | | 0 | 0 | 0 | 0 | 0 | 0 | 0 | 0 | 0 | 0 | 0 |
| 0 | NA | NA | NA | NA | NA | Gm28647 | | 0 | 0 | 0 | 0 | 0 | 0 | 0 | 0 | 0 | 0 | 0 |
| 0 | NA | NA | NA | NA | NA | Gm29435 | | 0 | 0 | 0 | 0 | 0 | 0 | 0 | 0 | 0 | 0 | 0 |
| 0 | NA | NA | NA | NA | NA | Vmn2r-ps139 | | 0 | 0 | 0 | 0 | 0 | 0 | 0 | 0 | 0 | 0 | 0 |
| 0 | NA | NA | NA | NA | NA | Gm29038 | | 0 | 0 | 0 | 0 | 0 | 0 | 0 | 0 | 0 | 0 | 0 |
| 0 | NA | NA | NA | NA | NA | Gm28191 | | 0 | 0 | 0 | 0 | 0 | 0 | 0 | 0 | 0 | 0 | 0 |
| 0 | NA | NA | NA | NA | NA | Gm28186 | | 0 | 0 | 0 | 0 | 0 | 0 | 0 | 0 | 0 | 0 | 0 |
| 0 | NA | NA | NA | NA | NA | Gm21064 | | 0 | 0 | 0 | 0 | 0 | 0 | 0 | 0 | 0 | 0 | 0 |
| 0 | NA | NA | NA | NA | NA | Gm28241 | | 0 | 0 | 0 | 0 | 0 | 0 | 0 | 0 | 0 | 0 | 0 |
| 0 | NA | NA | NA | NA | NA | Gm28242 | | 0 | 0 | 0 | 0 | 0 | 0 | 0 | 0 | 0 | 0 | 0 |
| 0 | NA | NA | NA | NA | NA | Gm37222 | | 0 | 0 | 0 | 0 | 0 | 0 | 0 | 0 | 0 | 0 | 0 |
| 0 | NA | NA | NA | NA | NA | Gm28356 | | 0 | 0 | 0 | 0 | 0 | 0 | 0 | 0 | 0 | 0 | 0 |
| 0 | NA | NA | NA | NA | NA | Gm28357 | | 0 | 0 | 0 | 0 | 0 | 0 | 0 | 0 | 0 | 0 | 0 |
| 0 | NA | NA | NA | NA | NA | Gm29352 | | 0 | 0 | 0 | 0 | 0 | 0 | 0 | 0 | 0 | 0 | 0 |
| 0 | NA | NA | NA | NA | NA | Gm29351 | | 0 | 0 | 0 | 0 | 0 | 0 | 0 | 0 | 0 | 0 | 0 |
| 0 | NA | NA | NA | NA | NA | Gm29350 | | 0 | 0 | 0 | 0 | 0 | 0 | 0 | 0 | 0 | 0 | 0 |
| 0 | NA | NA | NA | NA | NA | Gm29349 | | 0 | 0 | 0 | 0 | 0 | 0 | 0 | 0 | 0 | 0 | 0 |
| 0 | NA | NA | NA | NA | NA | Gm29355 | | 0 | 0 | 0 | 0 | 0 | 0 | 0 | 0 | 0 | 0 | 0 |
| 0 | NA | NA | NA | NA | NA | Gm20918 | predicted gene, 20918 | 0 | 0 | 0 | 0 | 0 | 0 | 0 | 0 | 0 | 0 | 0 |
| 0 | NA | NA | NA | NA | NA | Gm29354 | | 0 | 0 | 0 | 0 | 0 | 0 | 0 | 0 | 0 | 0 | 0 |
| 0 | NA | NA | NA | NA | NA | Gm29353 | | 0 | 0 | 0 | 0 | 0 | 0 | 0 | 0 | 0 | 0 | 0 |
| 0 | NA | NA | NA | NA | NA | Gm50460 | | 0 | 0 | 0 | 0 | 0 | 0 | 0 | 0 | 0 | 0 | 0 |
| 0 | NA | NA | NA | NA | NA | Gm21820 | | 0 | 0 | 0 | 0 | 0 | 0 | 0 | 0 | 0 | 0 | 0 |
| 0 | NA | NA | NA | NA | NA | Gm29357 | | 0 | 0 | 0 | 0 | 0 | 0 | 0 | 0 | 0 | 0 | 0 |
| 0 | NA | NA | NA | NA | NA | Gm29356 | | 0 | 0 | 0 | 0 | 0 | 0 | 0 | 0 | 0 | 0 | 0 |
| 0 | NA | NA | NA | NA | NA | Gm29161 | | 0 | 0 | 0 | 0 | 0 | 0 | 0 | 0 | 0 | 0 | 0 |
| 0 | NA | NA | NA | NA | NA | Gm28767 | | 0 | 0 | 0 | 0 | 0 | 0 | 0 | 0 | 0 | 0 | 0 |
| 0 | NA | NA | NA | NA | NA | Gm29160 | | 0 | 0 | 0 | 0 | 0 | 0 | 0 | 0 | 0 | 0 | 0 |
| 0 | NA | NA | NA | NA | NA | Gm20915 | | 0 | 0 | 0 | 0 | 0 | 0 | 0 | 0 | 0 | 0 | 0 |
| 0 | NA | NA | NA | NA | NA | Gm28766 | | 0 | 0 | 0 | 0 | 0 | 0 | 0 | 0 | 0 | 0 | 0 |
| 0 | NA | NA | NA | NA | NA | Gm29159 | | 0 | 0 | 0 | 0 | 0 | 0 | 0 | 0 | 0 | 0 | 0 |
| 0 | NA | NA | NA | NA | NA | Gm27663 | | 0 | 0 | 0 | 0 | 0 | 0 | 0 | 0 | 0 | 0 | 0 |
| 0 | NA | NA | NA | NA | NA | Gm29158 | | 0 | 0 | 0 | 0 | 0 | 0 | 0 | 0 | 0 | 0 | 0 |
| 0 | NA | NA | NA | NA | NA | Gm21854 | | 0 | 0 | 0 | 0 | 0 | 0 | 0 | 0 | 0 | 0 | 0 |
| 0 | NA | NA | NA | NA | NA | Gm28768 | | 0 | 0 | 0 | 0 | 0 | 0 | 0 | 0 | 0 | 0 | 0 |
| 0 | NA | NA | NA | NA | NA | Gm28769 | | 0 | 0 | 0 | 0 | 0 | 0 | 0 | 0 | 0 | 0 | 0 |
| 0 | NA | NA | NA | NA | NA | Gm20914 | predicted gene, 20914 | 0 | 0 | 0 | 0 | 0 | 0 | 0 | 0 | 0 | 0 | 0 |
| 0 | NA | NA | NA | NA | NA | Gm28171 | | 0 | 0 | 0 | 0 | 0 | 0 | 0 | 0 | 0 | 0 | 0 |
| 0 | NA | NA | NA | NA | NA | Gm28173 | | 0 | 0 | 0 | 0 | 0 | 0 | 0 | 0 | 0 | 0 | 0 |
| 0 | NA | NA | NA | NA | NA | Gm37236 | | 0 | 0 | 0 | 0 | 0 | 0 | 0 | 0 | 0 | 0 | 0 |
| 0 | NA | NA | NA | NA | NA | Gm29194 | | 0 | 0 | 0 | 0 | 0 | 0 | 0 | 0 | 0 | 0 | 0 |
| 0 | NA | NA | NA | NA | NA | Gm21778 | | 0 | 0 | 0 | 0 | 0 | 0 | 0 | 0 | 0 | 0 | 0 |
| 0 | NA | NA | NA | NA | NA | Gm29195 | | 0 | 0 | 0 | 0 | 0 | 0 | 0 | 0 | 0 | 0 | 0 |
| 0 | NA | NA | NA | NA | NA | Gm29196 | | 0 | 0 | 0 | 0 | 0 | 0 | 0 | 0 | 0 | 0 | 0 |
| 0 | NA | NA | NA | NA | NA | Gm28435 | | 0 | 0 | 0 | 0 | 0 | 0 | 0 | 0 | 0 | 0 | 0 |
| 0 | NA | NA | NA | NA | NA | Gm28436 | | 0 | 0 | 0 | 0 | 0 | 0 | 0 | 0 | 0 | 0 | 0 |

| | | | | | | | | | | | | | | | | | |
|---|---|---|---|---|---|---|---|---|---|---|---|---|---|---|---|---|---|
| 0 | NA | NA | NA | NA | NA | Gm29197 | | 0 | 0 | 0 | 0 | 0 | 0 | 0 | 0 | 0 | 0 | 0 |
| 0 | NA | NA | NA | NA | NA | Gm29198 | | 0 | 0 | 0 | 0 | 0 | 0 | 0 | 0 | 0 | 0 | 0 |
| 0 | NA | NA | NA | NA | NA | Gm29193 | | 0 | 0 | 0 | 0 | 0 | 0 | 0 | 0 | 0 | 0 | 0 |
| 0 | NA | NA | NA | NA | NA | Gm28430 | | 0 | 0 | 0 | 0 | 0 | 0 | 0 | 0 | 0 | 0 | 0 |
| 0 | NA | NA | NA | NA | NA | Gm21746 | | 0 | 0 | 0 | 0 | 0 | 0 | 0 | 0 | 0 | 0 | 0 |
| 0 | NA | NA | NA | NA | NA | Gm28593 | | 0 | 0 | 0 | 0 | 0 | 0 | 0 | 0 | 0 | 0 | 0 |
| 0 | NA | NA | NA | NA | NA | Gm20873 | predicted gene, 20873 | 0 | 0 | 0 | 0 | 0 | 0 | 0 | 0 | 0 | 0 | 0 |
| 0 | NA | NA | NA | NA | NA | Gm28916 | | 0 | 0 | 0 | 0 | 0 | 0 | 0 | 0 | 0 | 0 | 0 |
| 0 | NA | NA | NA | NA | NA | Gm28915 | | 0 | 0 | 0 | 0 | 0 | 0 | 0 | 0 | 0 | 0 | 0 |
| 0 | NA | NA | NA | NA | NA | Gm28918 | | 0 | 0 | 0 | 0 | 0 | 0 | 0 | 0 | 0 | 0 | 0 |
| 0 | NA | NA | NA | NA | NA | Gm28917 | | 0 | 0 | 0 | 0 | 0 | 0 | 0 | 0 | 0 | 0 | 0 |
| 0 | NA | NA | NA | NA | NA | Gm28919 | | 0 | 0 | 0 | 0 | 0 | 0 | 0 | 0 | 0 | 0 | 0 |
| 0 | NA | NA | NA | NA | NA | Gm21719 | | 0 | 0 | 0 | 0 | 0 | 0 | 0 | 0 | 0 | 0 | 0 |
| 0 | NA | NA | NA | NA | NA | Gm28921 | | 0 | 0 | 0 | 0 | 0 | 0 | 0 | 0 | 0 | 0 | 0 |
| 0 | NA | NA | NA | NA | NA | Gm28920 | | 0 | 0 | 0 | 0 | 0 | 0 | 0 | 0 | 0 | 0 | 0 |
| 0 | NA | NA | NA | NA | NA | Gm28444 | | 0 | 0 | 0 | 0 | 0 | 0 | 0 | 0 | 0 | 0 | 0 |
| 0 | NA | NA | NA | NA | NA | Gm28443 | | 0 | 0 | 0 | 0 | 0 | 0 | 0 | 0 | 0 | 0 | 0 |
| 0 | NA | NA | NA | NA | NA | Gm20830 | predicted gene, 20830 | 0 | 0 | 0 | 0 | 0 | 0 | 0 | 0 | 0 | 0 | 0 |
| 0 | NA | NA | NA | NA | NA | Gm28442 | | 0 | 0 | 0 | 0 | 0 | 0 | 0 | 0 | 0 | 0 | 0 |
| 0 | NA | NA | NA | NA | NA | Gm28445 | | 0 | 0 | 0 | 0 | 0 | 0 | 0 | 0 | 0 | 0 | 0 |
| 0 | NA | NA | NA | NA | NA | Gm20788 | | 0 | 0 | 0 | 0 | 0 | 0 | 0 | 0 | 0 | 0 | 0 |
| 0 | NA | NA | NA | NA | NA | Gm28999 | | 0 | 0 | 0 | 0 | 0 | 0 | 0 | 0 | 0 | 0 | 0 |
| 0 | NA | NA | NA | NA | NA | Gm21244 | predicted gene, 21244 | 0 | 0 | 0 | 0 | 0 | 0 | 0 | 0 | 0 | 0 | 0 |
| 0 | NA | NA | NA | NA | NA | Gm28997 | | 0 | 0 | 0 | 0 | 0 | 0 | 0 | 0 | 0 | 0 | 0 |
| 0 | NA | NA | NA | NA | NA | Gm28998 | predicted gene 28998 | 0 | 0 | 0 | 0 | 0 | 0 | 0 | 0 | 0 | 0 | 0 |
| 0 | NA | NA | NA | NA | NA | Gm28398 | | 0 | 0 | 0 | 0 | 0 | 0 | 0 | 0 | 0 | 0 | 0 |
| 0 | NA | NA | NA | NA | NA | Gm28397 | | 0 | 0 | 0 | 0 | 0 | 0 | 0 | 0 | 0 | 0 | 0 |
| 0 | NA | NA | NA | NA | NA | Gm28400 | | 0 | 0 | 0 | 0 | 0 | 0 | 0 | 0 | 0 | 0 | 0 |
| 0 | NA | NA | NA | NA | NA | Gm28399 | | 0 | 0 | 0 | 0 | 0 | 0 | 0 | 0 | 0 | 0 | 0 |
| 0 | NA | NA | NA | NA | NA | Gm28395 | | 0 | 0 | 0 | 0 | 0 | 0 | 0 | 0 | 0 | 0 | 0 |
| 0 | NA | NA | NA | NA | NA | Gm21788 | | 0 | 0 | 0 | 0 | 0 | 0 | 0 | 0 | 0 | 0 | 0 |
| 0 | NA | NA | NA | NA | NA | Gm28394 | | 0 | 0 | 0 | 0 | 0 | 0 | 0 | 0 | 0 | 0 | 0 |
| 0 | NA | NA | NA | NA | NA | Gm28396 | | 0 | 0 | 0 | 0 | 0 | 0 | 0 | 0 | 0 | 0 | 0 |
| 0 | NA | NA | NA | NA | NA | Gm28393 | | 0 | 0 | 0 | 0 | 0 | 0 | 0 | 0 | 0 | 0 | 0 |
| 0 | NA | NA | NA | NA | NA | Gm20826 | predicted gene, 20826 | 0 | 0 | 0 | 0 | 0 | 0 | 0 | 0 | 0 | 0 | 0 |
| 0 | NA | NA | NA | NA | NA | Vmn2r-ps149 | | 0 | 0 | 0 | 0 | 0 | 0 | 0 | 0 | 0 | 0 | 0 |
| 0 | NA | NA | NA | NA | NA | Gm28576 | predicted gene 28576 | 0 | 0 | 0 | 0 | 0 | 0 | 0 | 0 | 0 | 0 | 0 |
| 0 | NA | NA | NA | NA | NA | Gm28575 | | 0 | 0 | 0 | 0 | 0 | 0 | 0 | 0 | 0 | 0 | 0 |
| 0 | NA | NA | NA | NA | NA | Gm28574 | | 0 | 0 | 0 | 0 | 0 | 0 | 0 | 0 | 0 | 0 | 0 |
| 0 | NA | NA | NA | NA | NA | Gm28573 | | 0 | 0 | 0 | 0 | 0 | 0 | 0 | 0 | 0 | 0 | 0 |
| 0 | NA | NA | NA | NA | NA | Gm28572 | | 0 | 0 | 0 | 0 | 0 | 0 | 0 | 0 | 0 | 0 | 0 |
| 0 | NA | NA | NA | NA | NA | Gm28571 | | 0 | 0 | 0 | 0 | 0 | 0 | 0 | 0 | 0 | 0 | 0 |
| 0 | NA | NA | NA | NA | NA | Gm28570 | | 0 | 0 | 0 | 0 | 0 | 0 | 0 | 0 | 0 | 0 | 0 |
| 0 | NA | NA | NA | NA | NA | Gm29049 | | 0 | 0 | 0 | 0 | 0 | 0 | 0 | 0 | 0 | 0 | 0 |
| 0 | NA | NA | NA | NA | NA | Gm20825 | predicted gene, 20825 | 0 | 0 | 0 | 0 | 0 | 0 | 0 | 0 | 0 | 0 | 0 |
| 0 | NA | NA | NA | NA | NA | Gm29047 | | 0 | 0 | 0 | 0 | 0 | 0 | 0 | 0 | 0 | 0 | 0 |
| 0 | NA | NA | NA | NA | NA | Gm29048 | | 0 | 0 | 0 | 0 | 0 | 0 | 0 | 0 | 0 | 0 | 0 |
| 0 | NA | NA | NA | NA | NA | Gm29045 | | 0 | 0 | 0 | 0 | 0 | 0 | 0 | 0 | 0 | 0 | 0 |
| 0 | NA | NA | NA | NA | NA | Gm29046 | | 0 | 0 | 0 | 0 | 0 | 0 | 0 | 0 | 0 | 0 | 0 |
| 0 | NA | NA | NA | NA | NA | Gm29043 | | 0 | 0 | 0 | 0 | 0 | 0 | 0 | 0 | 0 | 0 | 0 |
| 0 | NA | NA | NA | NA | NA | Gm29044 | | 0 | 0 | 0 | 0 | 0 | 0 | 0 | 0 | 0 | 0 | 0 |
| 0 | NA | NA | NA | NA | NA | Gm37951 | | 0 | 0 | 0 | 0 | 0 | 0 | 0 | 0 | 0 | 0 | 0 |
| 0 | NA | NA | NA | NA | NA | Gm20824 | | 0 | 0 | 0 | 0 | 0 | 0 | 0 | 0 | 0 | 0 | 0 |
| 0 | NA | NA | NA | NA | NA | Gm28146 | | 0 | 0 | 0 | 0 | 0 | 0 | 0 | 0 | 0 | 0 | 0 |
| 0 | NA | NA | NA | NA | NA | Gm28145 | | 0 | 0 | 0 | 0 | 0 | 0 | 0 | 0 | 0 | 0 | 0 |
| 0 | NA | NA | NA | NA | NA | Gm28148 | | 0 | 0 | 0 | 0 | 0 | 0 | 0 | 0 | 0 | 0 | 0 |
| 0 | NA | NA | NA | NA | NA | Gm28147 | | 0 | 0 | 0 | 0 | 0 | 0 | 0 | 0 | 0 | 0 | 0 |
| 0 | NA | NA | NA | NA | NA | Gm28142 | | 0 | 0 | 0 | 0 | 0 | 0 | 0 | 0 | 0 | 0 | 0 |
| 0 | NA | NA | NA | NA | NA | Gm28141 | | 0 | 0 | 0 | 0 | 0 | 0 | 0 | 0 | 0 | 0 | 0 |
| 0 | NA | NA | NA | NA | NA | Gm28144 | | 0 | 0 | 0 | 0 | 0 | 0 | 0 | 0 | 0 | 0 | 0 |

| | | | | | | | | | | | | | | | | | | |
|---|---|---|---|---|---|---|---|---|---|---|---|---|---|---|---|---|---|---|
| 0 | NA | NA | NA | NA | NA | Gm28143 | | 0 | 0 | 0 | 0 | 0 | 0 | 0 | 0 | 0 | 0 | 0 |
| 0 | NA | NA | NA | NA | NA | Gm28150 | | 0 | 0 | 0 | 0 | 0 | 0 | 0 | 0 | 0 | 0 | 0 |
| 0 | NA | NA | NA | NA | NA | Gm28149 | | 0 | 0 | 0 | 0 | 0 | 0 | 0 | 0 | 0 | 0 | 0 |
| 0 | NA | NA | NA | NA | NA | Gm28098 | | 0 | 0 | 0 | 0 | 0 | 0 | 0 | 0 | 0 | 0 | 0 |
| 0 | NA | NA | NA | NA | NA | Gm28330 | | 0 | 0 | 0 | 0 | 0 | 0 | 0 | 0 | 0 | 0 | 0 |
| 0 | NA | NA | NA | NA | NA | Gm28331 | | 0 | 0 | 0 | 0 | 0 | 0 | 0 | 0 | 0 | 0 | 0 |
| 0 | NA | NA | NA | NA | NA | Gm28099 | | 0 | 0 | 0 | 0 | 0 | 0 | 0 | 0 | 0 | 0 | 0 |
| 0 | NA | NA | NA | NA | NA | Gm20815 | predicted gene, 20815 | 0 | 0 | 0 | 0 | 0 | 0 | 0 | 0 | 0 | 0 | 0 |
| 0 | NA | NA | NA | NA | NA | Gm28328 | | 0 | 0 | 0 | 0 | 0 | 0 | 0 | 0 | 0 | 0 | 0 |
| 0 | NA | NA | NA | NA | NA | Gm28329 | | 0 | 0 | 0 | 0 | 0 | 0 | 0 | 0 | 0 | 0 | 0 |
| 0 | NA | NA | NA | NA | NA | Gm28270 | | 0 | 0 | 0 | 0 | 0 | 0 | 0 | 0 | 0 | 0 | 0 |
| 0 | NA | NA | NA | NA | NA | Gm28327 | | 0 | 0 | 0 | 0 | 0 | 0 | 0 | 0 | 0 | 0 | 0 |
| 0 | NA | NA | NA | NA | NA | Gm28097 | | 0 | 0 | 0 | 0 | 0 | 0 | 0 | 0 | 0 | 0 | 0 |
| 0 | NA | NA | NA | NA | NA | Gm29365 | | 0 | 0 | 0 | 0 | 0 | 0 | 0 | 0 | 0 | 0 | 0 |
| 0 | NA | NA | NA | NA | NA | Gm29364 | | 0 | 0 | 0 | 0 | 0 | 0 | 0 | 0 | 0 | 0 | 0 |
| 0 | NA | NA | NA | NA | NA | Gm29363 | | 0 | 0 | 0 | 0 | 0 | 0 | 0 | 0 | 0 | 0 | 0 |
| 0 | NA | NA | NA | NA | NA | Gm21292 | predicted gene, 21292 | 0 | 0 | 0 | 0 | 0 | 0 | 0 | 0 | 0 | 0 | 0 |
| 0 | NA | NA | NA | NA | NA | Gm21721 | | 0 | 0 | 0 | 0 | 0 | 0 | 0 | 0 | 0 | 0 | 0 |
| 0 | NA | NA | NA | NA | NA | Gm29366 | | 0 | 0 | 0 | 0 | 0 | 0 | 0 | 0 | 0 | 0 | 0 |
| 0 | NA | NA | NA | NA | NA | Gm29281 | | 0 | 0 | 0 | 0 | 0 | 0 | 0 | 0 | 0 | 0 | 0 |
| 0 | NA | NA | NA | NA | NA | Gm29330 | | 0 | 0 | 0 | 0 | 0 | 0 | 0 | 0 | 0 | 0 | 0 |
| 0 | NA | NA | NA | NA | NA | Gm29329 | | 0 | 0 | 0 | 0 | 0 | 0 | 0 | 0 | 0 | 0 | 0 |
| 0 | NA | NA | NA | NA | NA | Gm21812 | | 0 | 0 | 0 | 0 | 0 | 0 | 0 | 0 | 0 | 0 | 0 |
| 0 | NA | NA | NA | NA | NA | Gm29535 | | 0 | 0 | 0 | 0 | 0 | 0 | 0 | 0 | 0 | 0 | 0 |
| 0 | NA | NA | NA | NA | NA | Gm29533 | | 0 | 0 | 0 | 0 | 0 | 0 | 0 | 0 | 0 | 0 | 0 |
| 0 | NA | NA | NA | NA | NA | Gm29534 | | 0 | 0 | 0 | 0 | 0 | 0 | 0 | 0 | 0 | 0 | 0 |
| 0 | NA | NA | NA | NA | NA | Gm21874 | | 0 | 0 | 0 | 0 | 0 | 0 | 0 | 0 | 0 | 0 | 0 |
| 0 | NA | NA | NA | NA | NA | Gm28849 | | 0 | 0 | 0 | 0 | 0 | 0 | 0 | 0 | 0 | 0 | 0 |
| 0 | NA | NA | NA | NA | NA | Gm28847 | | 0 | 0 | 0 | 0 | 0 | 0 | 0 | 0 | 0 | 0 | 0 |
| 0 | NA | NA | NA | NA | NA | Gm28848 | | 0 | 0 | 0 | 0 | 0 | 0 | 0 | 0 | 0 | 0 | 0 |
| 0 | NA | NA | NA | NA | NA | Gm28846 | | 0 | 0 | 0 | 0 | 0 | 0 | 0 | 0 | 0 | 0 | 0 |
| 0 | NA | NA | NA | NA | NA | Gm29471 | | 0 | 0 | 0 | 0 | 0 | 0 | 0 | 0 | 0 | 0 | 0 |
| 0 | NA | NA | NA | NA | NA | Gm29474 | | 0 | 0 | 0 | 0 | 0 | 0 | 0 | 0 | 0 | 0 | 0 |
| 0 | NA | NA | NA | NA | NA | Gm29475 | | 0 | 0 | 0 | 0 | 0 | 0 | 0 | 0 | 0 | 0 | 0 |
| 0 | NA | NA | NA | NA | NA | Gm20821 | | 0 | 0 | 0 | 0 | 0 | 0 | 0 | 0 | 0 | 0 | 0 |
| 0 | NA | NA | NA | NA | NA | Gm28506 | | 0 | 0 | 0 | 0 | 0 | 0 | 0 | 0 | 0 | 0 | 0 |
| 0 | NA | NA | NA | NA | NA | Gm28507 | | 0 | 0 | 0 | 0 | 0 | 0 | 0 | 0 | 0 | 0 | 0 |
| 0 | NA | NA | NA | NA | NA | Gm21301 | predicted gene, 21301 | 0 | 0 | 0 | 0 | 0 | 0 | 0 | 0 | 0 | 0 | 0 |
| 0 | NA | NA | NA | NA | NA | Gm29142 | | 0 | 0 | 0 | 0 | 0 | 0 | 0 | 0 | 0 | 0 | 0 |
| 0 | NA | NA | NA | NA | NA | Gm21310 | predicted gene, 21310 | 0 | 0 | 0 | 0 | 0 | 0 | 0 | 0 | 0 | 0 | 0 |
| 0 | NA | NA | NA | NA | NA | Gm28509 | | 0 | 0 | 0 | 0 | 0 | 0 | 0 | 0 | 0 | 0 | 0 |
| 0 | NA | NA | NA | NA | NA | Gm28508 | | 0 | 0 | 0 | 0 | 0 | 0 | 0 | 0 | 0 | 0 | 0 |
| 0 | NA | NA | NA | NA | NA | Gm20834 | predicted gene, 20834 | 0 | 0 | 0 | 0 | 0 | 0 | 0 | 0 | 0 | 0 | 0 |
| 0 | NA | NA | NA | NA | NA | Gm28510 | | 0 | 0 | 0 | 0 | 0 | 0 | 0 | 0 | 0 | 0 | 0 |
| 0 | NA | NA | NA | NA | NA | Gm29525 | | 0 | 0 | 0 | 0 | 0 | 0 | 0 | 0 | 0 | 0 | 0 |
| 0 | NA | NA | NA | NA | NA | Gm29526 | | 0 | 0 | 0 | 0 | 0 | 0 | 0 | 0 | 0 | 0 | 0 |
| 0 | NA | NA | NA | NA | NA | Gm20737 | predicted gene, 20737 | 0 | 0 | 0 | 0 | 0 | 0 | 0 | 0 | 0 | 0 | 0 |
| 0 | NA | NA | NA | NA | NA | Gm21775 | | 0 | 0 | 0 | 0 | 0 | 0 | 0 | 0 | 0 | 0 | 0 |
| 0 | NA | NA | NA | NA | NA | Gm21900 | | 0 | 0 | 0 | 0 | 0 | 0 | 0 | 0 | 0 | 0 | 0 |
| 0 | NA | NA | NA | NA | NA | Gm29527 | | 0 | 0 | 0 | 0 | 0 | 0 | 0 | 0 | 0 | 0 | 0 |
| 0 | NA | NA | NA | NA | NA | Gm29522 | | 0 | 0 | 0 | 0 | 0 | 0 | 0 | 0 | 0 | 0 | 0 |
| 0 | NA | NA | NA | NA | NA | Gm29523 | | 0 | 0 | 0 | 0 | 0 | 0 | 0 | 0 | 0 | 0 | 0 |
| 0 | NA | NA | NA | NA | NA | Gm29524 | | 0 | 0 | 0 | 0 | 0 | 0 | 0 | 0 | 0 | 0 | 0 |
| 0 | NA | NA | NA | NA | NA | Gm29528 | | 0 | 0 | 0 | 0 | 0 | 0 | 0 | 0 | 0 | 0 | 0 |
| 0 | NA | NA | NA | NA | NA | Gm20777 | predicted gene, 20777 | 0 | 0 | 0 | 0 | 0 | 0 | 0 | 0 | 0 | 0 | 0 |
| 0 | NA | NA | NA | NA | NA | Gm28955 | | 0 | 0 | 0 | 0 | 0 | 0 | 0 | 0 | 0 | 0 | 0 |
| 0 | NA | NA | NA | NA | NA | Gm28954 | | 0 | 0 | 0 | 0 | 0 | 0 | 0 | 0 | 0 | 0 | 0 |
| 0 | NA | NA | NA | NA | NA | Gm28953 | | 0 | 0 | 0 | 0 | 0 | 0 | 0 | 0 | 0 | 0 | 0 |
| 0 | NA | NA | NA | NA | NA | Gm20776 | | 0 | 0 | 0 | 0 | 0 | 0 | 0 | 0 | 0 | 0 | 0 |
| 0 | NA | NA | NA | NA | NA | Gm28958 | | 0 | 0 | 0 | 0 | 0 | 0 | 0 | 0 | 0 | 0 | 0 |

| | | | | | | | | | | | | | | | | | | |
|---|---|---|---|---|---|---|---|---|---|---|---|---|---|---|---|---|---|---|
| 0 | NA | NA | NA | NA | NA | Gm20828 | predicted gene, 20828 | 0 | 0 | 0 | 0 | 0 | 0 | 0 | 0 | 0 | 0 | 0 |
| 0 | NA | NA | NA | NA | NA | Gm28957 | | 0 | 0 | 0 | 0 | 0 | 0 | 0 | 0 | 0 | 0 | 0 |
| 0 | NA | NA | NA | NA | NA | Gm28956 | | 0 | 0 | 0 | 0 | 0 | 0 | 0 | 0 | 0 | 0 | 0 |
| 0 | NA | NA | NA | NA | NA | Gm28952 | | 0 | 0 | 0 | 0 | 0 | 0 | 0 | 0 | 0 | 0 | 0 |
| 0 | NA | NA | NA | NA | NA | Gm28951 | | 0 | 0 | 0 | 0 | 0 | 0 | 0 | 0 | 0 | 0 | 0 |
| 0 | NA | NA | NA | NA | NA | Gm28390 | | 0 | 0 | 0 | 0 | 0 | 0 | 0 | 0 | 0 | 0 | 0 |
| 0 | NA | NA | NA | NA | NA | Gm28391 | | 0 | 0 | 0 | 0 | 0 | 0 | 0 | 0 | 0 | 0 | 0 |
| 0 | NA | NA | NA | NA | NA | Gm28389 | | 0 | 0 | 0 | 0 | 0 | 0 | 0 | 0 | 0 | 0 | 0 |
| 0 | NA | NA | NA | NA | NA | Gm28655 | | 0 | 0 | 0 | 0 | 0 | 0 | 0 | 0 | 0 | 0 | 0 |
| 0 | NA | NA | NA | NA | NA | Gm28654 | | 0 | 0 | 0 | 0 | 0 | 0 | 0 | 0 | 0 | 0 | 0 |
| 0 | NA | NA | NA | NA | NA | Gm28657 | | 0 | 0 | 0 | 0 | 0 | 0 | 0 | 0 | 0 | 0 | 0 |
| 0 | NA | NA | NA | NA | NA | Gm28656 | | 0 | 0 | 0 | 0 | 0 | 0 | 0 | 0 | 0 | 0 | 0 |
| 0 | NA | NA | NA | NA | NA | Gm29554 | | 0 | 0 | 0 | 0 | 0 | 0 | 0 | 0 | 0 | 0 | 0 |
| 0 | NA | NA | NA | NA | NA | Gm29556 | | 0 | 0 | 0 | 0 | 0 | 0 | 0 | 0 | 0 | 0 | 0 |
| 0 | NA | NA | NA | NA | NA | Gm20812 | predicted gene, 20812 | 0 | 0 | 0 | 0 | 0 | 0 | 0 | 0 | 0 | 0 | 0 |
| 0 | NA | NA | NA | NA | NA | Gm35108 | | 0 | 0 | 0 | 0 | 0 | 0 | 0 | 0 | 0 | 0 | 0 |
| 0 | NA | NA | NA | NA | NA | Gm37739 | | 0 | 0 | 0 | 0 | 0 | 0 | 0 | 0 | 0 | 0 | 0 |
| 0 | NA | NA | NA | NA | NA | Gm21736 | | 0 | 0 | 0 | 0 | 0 | 0 | 0 | 0 | 0 | 0 | 0 |
| 0 | NA | NA | NA | NA | NA | Gm21379 | predicted gene, 21379 | 0 | 0 | 0 | 0 | 0 | 0 | 0 | 0 | 0 | 0 | 0 |
| 0 | NA | NA | NA | NA | NA | Gm36928 | | 0 | 0 | 0 | 0 | 0 | 0 | 0 | 0 | 0 | 0 | 0 |
| 0 | NA | NA | NA | NA | NA | Gm20807 | predicted gene, 20807 | 0 | 0 | 0 | 0 | 0 | 0 | 0 | 0 | 0 | 0 | 0 |
| 0 | NA | NA | NA | NA | NA | Gm38084 | | 0 | 0 | 0 | 0 | 0 | 0 | 0 | 0 | 0 | 0 | 0 |
| 0 | NA | NA | NA | NA | NA | Gm37231 | | 0 | 0 | 0 | 0 | 0 | 0 | 0 | 0 | 0 | 0 | 0 |
| 0 | NA | NA | NA | NA | NA | Gm21891 | | 0 | 0 | 0 | 0 | 0 | 0 | 0 | 0 | 0 | 0 | 0 |
| 0 | NA | NA | NA | NA | NA | Gm21425 | predicted gene, 20793 | 0 | 0 | 0 | 0 | 0 | 0 | 0 | 0 | 0 | 0 | 0 |
| 0 | NA | NA | NA | NA | NA | Gm37252 | | 0 | 0 | 0 | 0 | 0 | 0 | 0 | 0 | 0 | 0 | 0 |
| 0 | NA | NA | NA | NA | NA | Gm21828 | | 0 | 0 | 0 | 0 | 0 | 0 | 0 | 0 | 0 | 0 | 0 |
| 0 | NA | NA | NA | NA | NA | Gm21440 | predicted gene, 20793 | 0 | 0 | 0 | 0 | 0 | 0 | 0 | 0 | 0 | 0 | 0 |
| 0 | NA | NA | NA | NA | NA | Gm21454 | predicted gene, 20910 | 0 | 0 | 0 | 0 | 0 | 0 | 0 | 0 | 0 | 0 | 0 |
| 0 | NA | NA | NA | NA | NA | Gm37147 | | 0 | 0 | 0 | 0 | 0 | 0 | 0 | 0 | 0 | 0 | 0 |
| 0 | NA | NA | NA | NA | NA | Gm38159 | | 0 | 0 | 0 | 0 | 0 | 0 | 0 | 0 | 0 | 0 | 0 |
| 0 | NA | NA | NA | NA | NA | Gm21725 | | 0 | 0 | 0 | 0 | 0 | 0 | 0 | 0 | 0 | 0 | 0 |
| 0 | NA | NA | NA | NA | NA | Gm37870 | | 0 | 0 | 0 | 0 | 0 | 0 | 0 | 0 | 0 | 0 | 0 |
| 0 | NA | NA | NA | NA | NA | Gm37378 | | 0 | 0 | 0 | 0 | 0 | 0 | 0 | 0 | 0 | 0 | 0 |
| 0 | NA | NA | NA | NA | NA | Gm21866 | | 0 | 0 | 0 | 0 | 0 | 0 | 0 | 0 | 0 | 0 | 0 |
| 0 | NA | NA | NA | NA | NA | Gm20773 | predicted gene, 20773 | 0 | 0 | 0 | 0 | 0 | 0 | 0 | 0 | 0 | 0 | 0 |
| 0 | NA | NA | NA | NA | NA | Gm36398 | | 0 | 0 | 0 | 0 | 0 | 0 | 0 | 0 | 0 | 0 | 0 |
| 0 | NA | NA | NA | NA | NA | Gm37441 | | 0 | 0 | 0 | 0 | 0 | 0 | 0 | 0 | 0 | 0 | 0 |
| 0 | NA | NA | NA | NA | NA | Gm37561 | | 0 | 0 | 0 | 0 | 0 | 0 | 0 | 0 | 0 | 0 | 0 |
| 0 | NA | NA | NA | NA | NA | Gm36728 | | 0 | 0 | 0 | 0 | 0 | 0 | 0 | 0 | 0 | 0 | 0 |
| 0 | NA | NA | NA | NA | NA | Gm32295 | | 0 | 0 | 0 | 0 | 0 | 0 | 0 | 0 | 0 | 0 | 0 |
| 0 | NA | NA | NA | NA | NA | Gm20836 | predicted gene, 20836 | 0 | 0 | 0 | 0 | 0 | 0 | 0 | 0 | 0 | 0 | 0 |
| 0 | NA | NA | NA | NA | NA | Gm37263 | | 0 | 0 | 0 | 0 | 0 | 0 | 0 | 0 | 0 | 0 | 0 |
| 0 | NA | NA | NA | NA | NA | Gm21767 | | 0 | 0 | 0 | 0 | 0 | 0 | 0 | 0 | 0 | 0 | 0 |
| 0 | NA | NA | NA | NA | NA | Gm37130 | | 0 | 0 | 0 | 0 | 0 | 0 | 0 | 0 | 0 | 0 | 0 |
| 0 | NA | NA | NA | NA | NA | Gm37467 | | 0 | 0 | 0 | 0 | 0 | 0 | 0 | 0 | 0 | 0 | 0 |
| 0 | NA | NA | NA | NA | NA | Gm20928 | predicted gene, 20928 | 0 | 0 | 0 | 0 | 0 | 0 | 0 | 0 | 0 | 0 | 0 |
| 0 | NA | NA | NA | NA | NA | Gm38363 | | 0 | 0 | 0 | 0 | 0 | 0 | 0 | 0 | 0 | 0 | 0 |
| 0 | NA | NA | NA | NA | NA | Gm30174 | | 0 | 0 | 0 | 0 | 0 | 0 | 0 | 0 | 0 | 0 | 0 |
| 0 | NA | NA | NA | NA | NA | Gm30353 | | 0 | 0 | 0 | 0 | 0 | 0 | 0 | 0 | 0 | 0 | 0 |
| 0 | NA | NA | NA | NA | NA | Gm30686 | | 0 | 0 | 0 | 0 | 0 | 0 | 0 | 0 | 0 | 0 | 0 |
| 0 | NA | NA | NA | NA | NA | Gm30737 | predicted gene, 30737 | 0 | 0 | 0 | 0 | 0 | 0 | 0 | 0 | 0 | 0 | 0 |
| 0 | NA | NA | NA | NA | NA | Gm36987 | | 0 | 0 | 0 | 0 | 0 | 0 | 0 | 0 | 0 | 0 | 0 |
| 0 | NA | NA | NA | NA | NA | Gm31186 | | 0 | 0 | 0 | 0 | 0 | 0 | 0 | 0 | 0 | 0 | 0 |
| 0 | NA | NA | NA | NA | NA | Gm37654 | | 0 | 0 | 0 | 0 | 0 | 0 | 0 | 0 | 0 | 0 | 0 |
| 0 | NA | NA | NA | NA | NA | Gm37044 | | 0 | 0 | 0 | 0 | 0 | 0 | 0 | 0 | 0 | 0 | 0 |
| 0 | NA | NA | NA | NA | NA | Gm31511 | | 0 | 0 | 0 | 0 | 0 | 0 | 0 | 0 | 0 | 0 | 0 |
| 0 | NA | NA | NA | NA | NA | Gm37572 | | 0 | 0 | 0 | 0 | 0 | 0 | 0 | 0 | 0 | 0 | 0 |
| 0 | NA | NA | NA | NA | NA | Gm37344 | | 0 | 0 | 0 | 0 | 0 | 0 | 0 | 0 | 0 | 0 | 0 |
| 0 | NA | NA | NA | NA | NA | Gm37059 | | 0 | 0 | 0 | 0 | 0 | 0 | 0 | 0 | 0 | 0 | 0 |

| | | | | | | | | | | | | | | | | | | |
|---|---|---|---|---|---|---|---|---|---|---|---|---|---|---|---|---|---|---|
| 0 | NA | NA | NA | NA | NA | Gm31942 | | 0 | 0 | 0 | 0 | 0 | 0 | 0 | 0 | 0 | 0 | 0 |
| 0 | NA | NA | NA | NA | NA | Gm37440 | | 0 | 0 | 0 | 0 | 0 | 0 | 0 | 0 | 0 | 0 | 0 |
| 0 | NA | NA | NA | NA | NA | Gm38185 | | 0 | 0 | 0 | 0 | 0 | 0 | 0 | 0 | 0 | 0 | 0 |
| 0 | NA | NA | NA | NA | NA | Gm37395 | | 0 | 0 | 0 | 0 | 0 | 0 | 0 | 0 | 0 | 0 | 0 |
| 0 | NA | NA | NA | NA | NA | Gm37577 | | 0 | 0 | 0 | 0 | 0 | 0 | 0 | 0 | 0 | 0 | 0 |
| 0 | NA | NA | NA | NA | NA | Gm20822 | predicted gene, 20822 | 0 | 0 | 0 | 0 | 0 | 0 | 0 | 0 | 0 | 0 | 0 |
| 0 | NA | NA | NA | NA | NA | Gm38183 | | 0 | 0 | 0 | 0 | 0 | 0 | 0 | 0 | 0 | 0 | 0 |
| 0 | NA | NA | NA | NA | NA | Gm38149 | | 0 | 0 | 0 | 0 | 0 | 0 | 0 | 0 | 0 | 0 | 0 |
| 0 | NA | NA | NA | NA | NA | Gm38145 | | 0 | 0 | 0 | 0 | 0 | 0 | 0 | 0 | 0 | 0 | 0 |
| 0 | NA | NA | NA | NA | NA | Gm32033 | | 0 | 0 | 0 | 0 | 0 | 0 | 0 | 0 | 0 | 0 | 0 |
| 0 | NA | NA | NA | NA | NA | Gm32447 | | 0 | 0 | 0 | 0 | 0 | 0 | 0 | 0 | 0 | 0 | 0 |
| 0 | NA | NA | NA | NA | NA | Gm20770 | | 0 | 0 | 0 | 0 | 0 | 0 | 0 | 0 | 0 | 0 | 0 |
| 0 | NA | NA | NA | NA | NA | Gm38016 | | 0 | 0 | 0 | 0 | 0 | 0 | 0 | 0 | 0 | 0 | 0 |
| 0 | NA | NA | NA | NA | NA | Gm37158 | | 0 | 0 | 0 | 0 | 0 | 0 | 0 | 0 | 0 | 0 | 0 |
| 0 | NA | NA | NA | NA | NA | Gm36998 | | 0 | 0 | 0 | 0 | 0 | 0 | 0 | 0 | 0 | 0 | 0 |
| 0 | NA | NA | NA | NA | NA | Gm33112 | | 0 | 0 | 0 | 0 | 0 | 0 | 0 | 0 | 0 | 0 | 0 |
| 0 | NA | NA | NA | NA | NA | Gm21809 | | 0 | 0 | 0 | 0 | 0 | 0 | 0 | 0 | 0 | 0 | 0 |
| 0 | NA | NA | NA | NA | NA | Gm34015 | | 0 | 0 | 0 | 0 | 0 | 0 | 0 | 0 | 0 | 0 | 0 |
| 0 | NA | NA | NA | NA | NA | Gm20877 | predicted gene, 20877 | 0 | 0 | 0 | 0 | 0 | 0 | 0 | 0 | 0 | 0 | 0 |
| 0 | NA | NA | NA | NA | NA | Gm33191 | | 0 | 0 | 0 | 0 | 0 | 0 | 0 | 0 | 0 | 0 | 0 |
| 0 | NA | NA | NA | NA | NA | Gm38003 | | 0 | 0 | 0 | 0 | 0 | 0 | 0 | 0 | 0 | 0 | 0 |
| 0 | NA | NA | NA | NA | NA | Gm20781 | | 0 | 0 | 0 | 0 | 0 | 0 | 0 | 0 | 0 | 0 | 0 |
| 0 | NA | NA | NA | NA | NA | Rbm31y | RNA binding motif 31, Y-linked | 0 | 0 | 0 | 0 | 0 | 0 | 0 | 0 | 0 | 0 | 0 |
| 0 | NA | NA | NA | NA | NA | Gm37325 | | 0 | 0 | 0 | 0 | 0 | 0 | 0 | 0 | 0 | 0 | 0 |
| 0 | NA | NA | NA | NA | NA | Gm20833 | predicted gene, 20833 | 0 | 0 | 0 | 0 | 0 | 0 | 0 | 0 | 0 | 0 | 0 |
| 0 | NA | NA | NA | NA | NA | Gm37865 | | 0 | 0 | 0 | 0 | 0 | 0 | 0 | 0 | 0 | 0 | 0 |
| 0 | NA | NA | NA | NA | NA | Gm36941 | | 0 | 0 | 0 | 0 | 0 | 0 | 0 | 0 | 0 | 0 | 0 |
| 0 | NA | NA | NA | NA | NA | Gm21904 | | 0 | 0 | 0 | 0 | 0 | 0 | 0 | 0 | 0 | 0 | 0 |
| 0 | NA | NA | NA | NA | NA | Gm37937 | | 0 | 0 | 0 | 0 | 0 | 0 | 0 | 0 | 0 | 0 | 0 |
| 0 | NA | NA | NA | NA | NA | Gm20772 | predicted gene, 20772 | 0 | 0 | 0 | 0 | 0 | 0 | 0 | 0 | 0 | 0 | 0 |
| 0 | NA | NA | NA | NA | NA | Gm20831 | predicted gene, 20793 | 0 | 0 | 0 | 0 | 0 | 0 | 0 | 0 | 0 | 0 | 0 |
| 0 | NA | NA | NA | NA | NA | Gm37286 | | 0 | 0 | 0 | 0 | 0 | 0 | 0 | 0 | 0 | 0 | 0 |
| 0 | NA | NA | NA | NA | NA | Srsy | serine-rich, secreted, Y-linked | 0 | 0 | 0 | 0 | 0 | 0 | 0 | 0 | 0 | 0 | 0 |
| 0 | NA | NA | NA | NA | NA | Gm38028 | | 0 | 0 | 0 | 0 | 0 | 0 | 0 | 0 | 0 | 0 | 0 |
| 0 | NA | NA | NA | NA | NA | Gm21764 | | 0 | 0 | 0 | 0 | 0 | 0 | 0 | 0 | 0 | 0 | 0 |
| 0 | NA | NA | NA | NA | NA | Ssty1 | spermiogenesis specific transcript on the | 0 | 0 | 0 | 0 | 0 | 0 | 0 | 0 | 0 | 0 | 0 |
| 0 | NA | NA | NA | NA | NA | Gm38072 | | 0 | 0 | 0 | 0 | 0 | 0 | 0 | 0 | 0 | 0 | 0 |
| 0 | NA | NA | NA | NA | NA | Gm20851 | predicted gene, 20851 | 0 | 0 | 0 | 0 | 0 | 0 | 0 | 0 | 0 | 0 | 0 |
| 0 | NA | NA | NA | NA | NA | Gm30312 | | 0 | 0 | 0 | 0 | 0 | 0 | 0 | 0 | 0 | 0 | 0 |
| 0 | NA | NA | NA | NA | NA | Gm37433 | | 0 | 0 | 0 | 0 | 0 | 0 | 0 | 0 | 0 | 0 | 0 |
| 0 | NA | NA | NA | NA | NA | Gm37473 | | 0 | 0 | 0 | 0 | 0 | 0 | 0 | 0 | 0 | 0 | 0 |
| 0 | NA | NA | NA | NA | NA | Gm37656 | | 0 | 0 | 0 | 0 | 0 | 0 | 0 | 0 | 0 | 0 | 0 |
| 0 | NA | NA | NA | NA | NA | Gm38127 | | 0 | 0 | 0 | 0 | 0 | 0 | 0 | 0 | 0 | 0 | 0 |
| 0 | NA | NA | NA | NA | NA | Gm30638 | | 0 | 0 | 0 | 0 | 0 | 0 | 0 | 0 | 0 | 0 | 0 |
| 0 | NA | NA | NA | NA | NA | Gm38361 | | 0 | 0 | 0 | 0 | 0 | 0 | 0 | 0 | 0 | 0 | 0 |
| 0 | NA | NA | NA | NA | NA | Gm31260 | | 0 | 0 | 0 | 0 | 0 | 0 | 0 | 0 | 0 | 0 | 0 |
| 0 | NA | NA | NA | NA | NA | Gm21660 | predicted gene, 21660 | 0 | 0 | 0 | 0 | 0 | 0 | 0 | 0 | 0 | 0 | 0 |
| 0 | NA | NA | NA | NA | NA | Gm38209 | | 0 | 0 | 0 | 0 | 0 | 0 | 0 | 0 | 0 | 0 | 0 |
| 0 | NA | NA | NA | NA | NA | Gm21822 | | 0 | 0 | 0 | 0 | 0 | 0 | 0 | 0 | 0 | 0 | 0 |
| 0 | NA | NA | NA | NA | NA | Gm37734 | | 0 | 0 | 0 | 0 | 0 | 0 | 0 | 0 | 0 | 0 | 0 |
| 0 | NA | NA | NA | NA | NA | Gm38013 | | 0 | 0 | 0 | 0 | 0 | 0 | 0 | 0 | 0 | 0 | 0 |
| 0 | NA | NA | NA | NA | NA | Gm21678 | predicted gene, 21678 | 0 | 0 | 0 | 0 | 0 | 0 | 0 | 0 | 0 | 0 | 0 |
| 0 | NA | NA | NA | NA | NA | Gm37948 | | 0 | 0 | 0 | 0 | 0 | 0 | 0 | 0 | 0 | 0 | 0 |
| 0 | NA | NA | NA | NA | NA | Gm37442 | | 0 | 0 | 0 | 0 | 0 | 0 | 0 | 0 | 0 | 0 | 0 |
| 0 | NA | NA | NA | NA | NA | Gm34550 | | 0 | 0 | 0 | 0 | 0 | 0 | 0 | 0 | 0 | 0 | 0 |
| 0 | NA | NA | NA | NA | NA | Gm37544 | | 0 | 0 | 0 | 0 | 0 | 0 | 0 | 0 | 0 | 0 | 0 |
| 0 | NA | NA | NA | NA | NA | Gm34716 | | 0 | 0 | 0 | 0 | 0 | 0 | 0 | 0 | 0 | 0 | 0 |
| 0 | NA | NA | NA | NA | NA | Gm37635 | | 0 | 0 | 0 | 0 | 0 | 0 | 0 | 0 | 0 | 0 | 0 |
| 0 | NA | NA | NA | NA | NA | Gm35070 | | 0 | 0 | 0 | 0 | 0 | 0 | 0 | 0 | 0 | 0 | 0 |
| 0 | NA | NA | NA | NA | NA | Gm35134 | predicted gene, 35134 | 0 | 0 | 0 | 0 | 0 | 0 | 0 | 0 | 0 | 0 | 0 |

| | | | | | | | | | | | | | | | | | | |
|---|---|---|---|---|---|---|---|---|---|---|---|---|---|---|---|---|---|---|
| 0 | NA | NA | NA | NA | NA | Gm37998 | | 0 | 0 | 0 | 0 | 0 | 0 | 0 | 0 | 0 | 0 | 0 |
| 0 | NA | NA | NA | NA | NA | Gm38094 | | 0 | 0 | 0 | 0 | 0 | 0 | 0 | 0 | 0 | 0 | 0 |
| 0 | NA | NA | NA | NA | NA | Gm21763 | | 0 | 0 | 0 | 0 | 0 | 0 | 0 | 0 | 0 | 0 | 0 |
| 0 | NA | NA | NA | NA | NA | Gm38369 | | 0 | 0 | 0 | 0 | 0 | 0 | 0 | 0 | 0 | 0 | 0 |
| 0 | NA | NA | NA | NA | NA | Gm35843 | | 0 | 0 | 0 | 0 | 0 | 0 | 0 | 0 | 0 | 0 | 0 |
| 0 | NA | NA | NA | NA | NA | Gm38168 | | 0 | 0 | 0 | 0 | 0 | 0 | 0 | 0 | 0 | 0 | 0 |
| 0 | NA | NA | NA | NA | NA | Gm37740 | | 0 | 0 | 0 | 0 | 0 | 0 | 0 | 0 | 0 | 0 | 0 |
| 0 | NA | NA | NA | NA | NA | Gm27701 | | 0 | 0 | 0 | 0 | 0 | 0 | 0 | 0 | 0 | 0 | 0 |
| 0 | NA | NA | NA | NA | NA | Gm37723 | | 0 | 0 | 0 | 0 | 0 | 0 | 0 | 0 | 0 | 0 | 0 |
| 0 | NA | NA | NA | NA | NA | Gm36261 | | 0 | 0 | 0 | 0 | 0 | 0 | 0 | 0 | 0 | 0 | 0 |
| 0 | NA | NA | NA | NA | NA | Gm37898 | | 0 | 0 | 0 | 0 | 0 | 0 | 0 | 0 | 0 | 0 | 0 |
| 0 | NA | NA | NA | NA | NA | Gm36467 | | 0 | 0 | 0 | 0 | 0 | 0 | 0 | 0 | 0 | 0 | 0 |
| 0 | NA | NA | NA | NA | NA | Gm37927 | | 0 | 0 | 0 | 0 | 0 | 0 | 0 | 0 | 0 | 0 | 0 |
| 0 | NA | NA | NA | NA | NA | Gm36891 | | 0 | 0 | 0 | 0 | 0 | 0 | 0 | 0 | 0 | 0 | 0 |
| 0 | NA | NA | NA | NA | NA | Gm36929 | | 0 | 0 | 0 | 0 | 0 | 0 | 0 | 0 | 0 | 0 | 0 |
| 0 | NA | NA | NA | NA | NA | Gm38174 | | 0 | 0 | 0 | 0 | 0 | 0 | 0 | 0 | 0 | 0 | 0 |
| 0 | NA | NA | NA | NA | NA | Gm29866 | | 0 | 0 | 0 | 0 | 0 | 0 | 0 | 0 | 0 | 0 | 0 |
| 0 | NA | NA | NA | NA | NA | Gm36950 | | 0 | 0 | 0 | 0 | 0 | 0 | 0 | 0 | 0 | 0 | 0 |
| 0 | NA | NA | NA | NA | NA | Gm20910 | predicted gene, 20910 | 0 | 0 | 0 | 0 | 0 | 0 | 0 | 0 | 0 | 0 | 0 |
| 0 | NA | NA | NA | NA | NA | Gm30598 | | 0 | 0 | 0 | 0 | 0 | 0 | 0 | 0 | 0 | 0 | 0 |
| 0 | NA | NA | NA | NA | NA | Gm20909 | predicted gene, 20868 | 0 | 0 | 0 | 0 | 0 | 0 | 0 | 0 | 0 | 0 | 0 |
| 0 | NA | NA | NA | NA | NA | Gm20865 | predicted gene, 20865 | 0 | 0 | 0 | 0 | 0 | 0 | 0 | 0 | 0 | 0 | 0 |
| 0 | NA | NA | NA | NA | NA | Gm37462 | | 0 | 0 | 0 | 0 | 0 | 0 | 0 | 0 | 0 | 0 | 0 |
| 0 | NA | NA | NA | NA | NA | Gm37658 | | 0 | 0 | 0 | 0 | 0 | 0 | 0 | 0 | 0 | 0 | 0 |
| 0 | NA | NA | NA | NA | NA | Gm32347 | | 0 | 0 | 0 | 0 | 0 | 0 | 0 | 0 | 0 | 0 | 0 |
| 0 | NA | NA | NA | NA | NA | Gm31123 | | 0 | 0 | 0 | 0 | 0 | 0 | 0 | 0 | 0 | 0 | 0 |
| 0 | NA | NA | NA | NA | NA | Gm38370 | | 0 | 0 | 0 | 0 | 0 | 0 | 0 | 0 | 0 | 0 | 0 |
| 0 | NA | NA | NA | NA | NA | Gm38054 | | 0 | 0 | 0 | 0 | 0 | 0 | 0 | 0 | 0 | 0 | 0 |
| 0 | NA | NA | NA | NA | NA | Gm37476 | | 0 | 0 | 0 | 0 | 0 | 0 | 0 | 0 | 0 | 0 | 0 |
| 0 | NA | NA | NA | NA | NA | Gm37369 | | 0 | 0 | 0 | 0 | 0 | 0 | 0 | 0 | 0 | 0 | 0 |
| 0 | NA | NA | NA | NA | NA | Gm38181 | | 0 | 0 | 0 | 0 | 0 | 0 | 0 | 0 | 0 | 0 | 0 |
| 0 | NA | NA | NA | NA | NA | Gm31335 | | 0 | 0 | 0 | 0 | 0 | 0 | 0 | 0 | 0 | 0 | 0 |
| 0 | NA | NA | NA | NA | NA | Gm37690 | | 0 | 0 | 0 | 0 | 0 | 0 | 0 | 0 | 0 | 0 | 0 |
| 0 | NA | NA | NA | NA | NA | Gm38107 | | 0 | 0 | 0 | 0 | 0 | 0 | 0 | 0 | 0 | 0 | 0 |
| 0 | NA | NA | NA | NA | NA | Gm31571 | | 0 | 0 | 0 | 0 | 0 | 0 | 0 | 0 | 0 | 0 | 0 |
| 0 | NA | NA | NA | NA | NA | Gm32114 | | 0 | 0 | 0 | 0 | 0 | 0 | 0 | 0 | 0 | 0 | 0 |
| 0 | NA | NA | NA | NA | NA | Gm37075 | | 0 | 0 | 0 | 0 | 0 | 0 | 0 | 0 | 0 | 0 | 0 |
| 0 | NA | NA | NA | NA | NA | Gm37921 | | 0 | 0 | 0 | 0 | 0 | 0 | 0 | 0 | 0 | 0 | 0 |
| 0 | NA | NA | NA | NA | NA | Gm32180 | | 0 | 0 | 0 | 0 | 0 | 0 | 0 | 0 | 0 | 0 | 0 |
| 0 | NA | NA | NA | NA | NA | Gm38371 | | 0 | 0 | 0 | 0 | 0 | 0 | 0 | 0 | 0 | 0 | 0 |
| 0 | NA | NA | NA | NA | NA | Gm37657 | | 0 | 0 | 0 | 0 | 0 | 0 | 0 | 0 | 0 | 0 | 0 |
| 0 | NA | NA | NA | NA | NA | Gm37448 | | 0 | 0 | 0 | 0 | 0 | 0 | 0 | 0 | 0 | 0 | 0 |
| 0 | NA | NA | NA | NA | NA | Gm33163 | | 0 | 0 | 0 | 0 | 0 | 0 | 0 | 0 | 0 | 0 | 0 |
| 0 | NA | NA | NA | NA | NA | Gm21065 | predicted gene, 21065 | 0 | 0 | 0 | 0 | 0 | 0 | 0 | 0 | 0 | 0 | 0 |
| 0 | NA | NA | NA | NA | NA | Gm36978 | | 0 | 0 | 0 | 0 | 0 | 0 | 0 | 0 | 0 | 0 | 0 |
| 0 | NA | NA | NA | NA | NA | Gm37721 | | 0 | 0 | 0 | 0 | 0 | 0 | 0 | 0 | 0 | 0 | 0 |
| 0 | NA | NA | NA | NA | NA | Gm37129 | | 0 | 0 | 0 | 0 | 0 | 0 | 0 | 0 | 0 | 0 | 0 |
| 0 | NA | NA | NA | NA | NA | Gm37990 | | 0 | 0 | 0 | 0 | 0 | 0 | 0 | 0 | 0 | 0 | 0 |
| 0 | NA | NA | NA | NA | NA | Gm21366 | X-linked lymphocyte-regulated protein PM | 0 | 0 | 0 | 0 | 0 | 0 | 0 | 0 | 0 | 0 | 0 |
| 0 | NA | NA | NA | NA | NA | Gm33954 | | 0 | 0 | 0 | 0 | 0 | 0 | 0 | 0 | 0 | 0 | 0 |
| 0 | NA | NA | NA | NA | NA | Gm37538 | | 0 | 0 | 0 | 0 | 0 | 0 | 0 | 0 | 0 | 0 | 0 |
| 0 | NA | NA | NA | NA | NA | Gm21871 | | 0 | 0 | 0 | 0 | 0 | 0 | 0 | 0 | 0 | 0 | 0 |
| 0 | NA | NA | NA | NA | NA | Gm34274 | | 0 | 0 | 0 | 0 | 0 | 0 | 0 | 0 | 0 | 0 | 0 |
| 0 | NA | NA | NA | NA | NA | Gm34217 | | 0 | 0 | 0 | 0 | 0 | 0 | 0 | 0 | 0 | 0 | 0 |
| 0 | NA | NA | NA | NA | NA | Gm21469 | | 0 | 0 | 0 | 0 | 0 | 0 | 0 | 0 | 0 | 0 | 0 |
| 0 | NA | NA | NA | NA | NA | Gm21661 | Y-linked testis-specific protein 1-like | 0 | 0 | 0 | 0 | 0 | 0 | 0 | 0 | 0 | 0 | 0 |
| 0 | NA | NA | NA | NA | NA | Gm37952 | | 0 | 0 | 0 | 0 | 0 | 0 | 0 | 0 | 0 | 0 | 0 |
| 0 | NA | NA | NA | NA | NA | Gm21907 | | 0 | 0 | 0 | 0 | 0 | 0 | 0 | 0 | 0 | 0 | 0 |
| 0 | NA | NA | NA | NA | NA | Gm37627 | | 0 | 0 | 0 | 0 | 0 | 0 | 0 | 0 | 0 | 0 | 0 |
| 0 | NA | NA | NA | NA | NA | Gm20925 | predicted gene, 20925 | 0 | 0 | 0 | 0 | 0 | 0 | 0 | 0 | 0 | 0 | 0 |

| | | | | | | | | | | | | | | | | | | |
|---|---|---|---|---|---|---|---|---|---|---|---|---|---|---|---|---|---|---|
| 0 | NA | NA | NA | NA | NA | Gm20926 | predicted gene, 20926 | 0 | 0 | 0 | 0 | 0 | 0 | 0 | 0 | 0 | 0 | 0 |
| 0 | NA | NA | NA | NA | NA | Gm37454 | | 0 | 0 | 0 | 0 | 0 | 0 | 0 | 0 | 0 | 0 | 0 |
| 0 | NA | NA | NA | NA | NA | Gm38138 | | 0 | 0 | 0 | 0 | 0 | 0 | 0 | 0 | 0 | 0 | 0 |
| 0 | NA | NA | NA | NA | NA | Gm29740 | predicted gene, 29740 | 0 | 0 | 0 | 0 | 0 | 0 | 0 | 0 | 0 | 0 | 0 |
| 0 | NA | NA | NA | NA | NA | Gm37840 | | 0 | 0 | 0 | 0 | 0 | 0 | 0 | 0 | 0 | 0 | 0 |
| 0 | NA | NA | NA | NA | NA | Gm37574 | | 0 | 0 | 0 | 0 | 0 | 0 | 0 | 0 | 0 | 0 | 0 |
| 0 | NA | NA | NA | NA | NA | Gm21768 | | 0 | 0 | 0 | 0 | 0 | 0 | 0 | 0 | 0 | 0 | 0 |
| 0 | NA | NA | NA | NA | NA | Gm37172 | | 0 | 0 | 0 | 0 | 0 | 0 | 0 | 0 | 0 | 0 | 0 |
| 0 | NA | NA | NA | NA | NA | Gm35670 | | 0 | 0 | 0 | 0 | 0 | 0 | 0 | 0 | 0 | 0 | 0 |
| 0 | NA | NA | NA | NA | NA | Gm20840 | predicted gene, 20840 | 0 | 0 | 0 | 0 | 0 | 0 | 0 | 0 | 0 | 0 | 0 |
| 0 | NA | NA | NA | NA | NA | Gm35955 | | 0 | 0 | 0 | 0 | 0 | 0 | 0 | 0 | 0 | 0 | 0 |
| 0 | NA | NA | NA | NA | NA | Gm37687 | | 0 | 0 | 0 | 0 | 0 | 0 | 0 | 0 | 0 | 0 | 0 |
| 0 | NA | NA | NA | NA | NA | Gm38296 | | 0 | 0 | 0 | 0 | 0 | 0 | 0 | 0 | 0 | 0 | 0 |
| 0 | NA | NA | NA | NA | NA | Gm38290 | | 0 | 0 | 0 | 0 | 0 | 0 | 0 | 0 | 0 | 0 | 0 |
| 0 | NA | NA | NA | NA | NA | Gm38121 | | 0 | 0 | 0 | 0 | 0 | 0 | 0 | 0 | 0 | 0 | 0 |
| 0 | NA | NA | NA | NA | NA | Gm36047 | | 0 | 0 | 0 | 0 | 0 | 0 | 0 | 0 | 0 | 0 | 0 |
| 0 | NA | NA | NA | NA | NA | Gm36345 | | 0 | 0 | 0 | 0 | 0 | 0 | 0 | 0 | 0 | 0 | 0 |
| 0 | NA | NA | NA | NA | NA | Gm37157 | | 0 | 0 | 0 | 0 | 0 | 0 | 0 | 0 | 0 | 0 | 0 |
| 0 | NA | NA | NA | NA | NA | Gm38017 | | 0 | 0 | 0 | 0 | 0 | 0 | 0 | 0 | 0 | 0 | 0 |
| 0 | NA | NA | NA | NA | NA | Gm36468 | | 0 | 0 | 0 | 0 | 0 | 0 | 0 | 0 | 0 | 0 | 0 |
| 0 | NA | NA | NA | NA | NA | Gm21845 | | 0 | 0 | 0 | 0 | 0 | 0 | 0 | 0 | 0 | 0 | 0 |
| 0 | NA | NA | NA | NA | NA | Gm20809 | predicted gene, 20809 | 0 | 0 | 0 | 0 | 0 | 0 | 0 | 0 | 0 | 0 | 0 |
| 0 | NA | NA | NA | NA | NA | Gm37434 | | 0 | 0 | 0 | 0 | 0 | 0 | 0 | 0 | 0 | 0 | 0 |
| 0 | NA | NA | NA | NA | NA | Gm37765 | | 0 | 0 | 0 | 0 | 0 | 0 | 0 | 0 | 0 | 0 | 0 |
| 0 | NA | NA | NA | NA | NA | Gm38229 | | 0 | 0 | 0 | 0 | 0 | 0 | 0 | 0 | 0 | 0 | 0 |
| 0 | NA | NA | NA | NA | NA | Gm36866 | | 0 | 0 | 0 | 0 | 0 | 0 | 0 | 0 | 0 | 0 | 0 |
| 0 | NA | NA | NA | NA | NA | Gm30045 | | 0 | 0 | 0 | 0 | 0 | 0 | 0 | 0 | 0 | 0 | 0 |
| 0 | NA | NA | NA | NA | NA | Gm21921 | | 0 | 0 | 0 | 0 | 0 | 0 | 0 | 0 | 0 | 0 | 0 |
| 0 | NA | NA | NA | NA | NA | Gm37112 | | 0 | 0 | 0 | 0 | 0 | 0 | 0 | 0 | 0 | 0 | 0 |
| 0 | NA | NA | NA | NA | NA | Gm37287 | | 0 | 0 | 0 | 0 | 0 | 0 | 0 | 0 | 0 | 0 | 0 |
| 0 | NA | NA | NA | NA | NA | Gm30175 | | 0 | 0 | 0 | 0 | 0 | 0 | 0 | 0 | 0 | 0 | 0 |
| 0 | NA | NA | NA | NA | NA | Gm21350 | | 0 | 0 | 0 | 0 | 0 | 0 | 0 | 0 | 0 | 0 | 0 |
| 0 | NA | NA | NA | NA | NA | Gm21412 | Y-linked testis-specific protein 1-like | 0 | 0 | 0 | 0 | 0 | 0 | 0 | 0 | 0 | 0 | 0 |
| 0 | NA | NA | NA | NA | NA | Gm37071 | | 0 | 0 | 0 | 0 | 0 | 0 | 0 | 0 | 0 | 0 | 0 |
| 0 | NA | NA | NA | NA | NA | Gm37547 | | 0 | 0 | 0 | 0 | 0 | 0 | 0 | 0 | 0 | 0 | 0 |
| 0 | NA | NA | NA | NA | NA | Gm36968 | | 0 | 0 | 0 | 0 | 0 | 0 | 0 | 0 | 0 | 0 | 0 |
| 0 | NA | NA | NA | NA | NA | Gm38035 | | 0 | 0 | 0 | 0 | 0 | 0 | 0 | 0 | 0 | 0 | 0 |
| 0 | NA | NA | NA | NA | NA | Gm20857 | | 0 | 0 | 0 | 0 | 0 | 0 | 0 | 0 | 0 | 0 | 0 |
| 0 | NA | NA | NA | NA | NA | Gm30705 | | 0 | 0 | 0 | 0 | 0 | 0 | 0 | 0 | 0 | 0 | 0 |
| 0 | NA | NA | NA | NA | NA | Gm21773 | | 0 | 0 | 0 | 0 | 0 | 0 | 0 | 0 | 0 | 0 | 0 |
| 0 | NA | NA | NA | NA | NA | Gm37451 | | 0 | 0 | 0 | 0 | 0 | 0 | 0 | 0 | 0 | 0 | 0 |
| 0 | NA | NA | NA | NA | NA | Gm37793 | | 0 | 0 | 0 | 0 | 0 | 0 | 0 | 0 | 0 | 0 | 0 |
| 0 | NA | NA | NA | NA | NA | Gm30858 | | 0 | 0 | 0 | 0 | 0 | 0 | 0 | 0 | 0 | 0 | 0 |
| 0 | NA | NA | NA | NA | NA | Gm21791 | | 0 | 0 | 0 | 0 | 0 | 0 | 0 | 0 | 0 | 0 | 0 |
| 0 | NA | NA | NA | NA | NA | Gm21427 | Y-linked testis-specific protein 1-like | 0 | 0 | 0 | 0 | 0 | 0 | 0 | 0 | 0 | 0 | 0 |
| 0 | NA | NA | NA | NA | NA | Gm37875 | | 0 | 0 | 0 | 0 | 0 | 0 | 0 | 0 | 0 | 0 | 0 |
| 0 | NA | NA | NA | NA | NA | Gm37237 | | 0 | 0 | 0 | 0 | 0 | 0 | 0 | 0 | 0 | 0 | 0 |
| 0 | NA | NA | NA | NA | NA | Gm38350 | | 0 | 0 | 0 | 0 | 0 | 0 | 0 | 0 | 0 | 0 | 0 |
| 0 | NA | NA | NA | NA | NA | Gm20858 | | 0 | 0 | 0 | 0 | 0 | 0 | 0 | 0 | 0 | 0 | 0 |
| 0 | NA | NA | NA | NA | NA | Gm31422 | | 0 | 0 | 0 | 0 | 0 | 0 | 0 | 0 | 0 | 0 | 0 |
| 0 | NA | NA | NA | NA | NA | Gm37798 | | 0 | 0 | 0 | 0 | 0 | 0 | 0 | 0 | 0 | 0 | 0 |
| 0 | NA | NA | NA | NA | NA | Gm37318 | | 0 | 0 | 0 | 0 | 0 | 0 | 0 | 0 | 0 | 0 | 0 |
| 0 | NA | NA | NA | NA | NA | Gm31572 | | 0 | 0 | 0 | 0 | 0 | 0 | 0 | 0 | 0 | 0 | 0 |
| 0 | NA | NA | NA | NA | NA | Gm21890 | | 0 | 0 | 0 | 0 | 0 | 0 | 0 | 0 | 0 | 0 | 0 |
| 0 | NA | NA | NA | NA | NA | Gm21443 | Y-linked testis-specific protein 1-like | 0 | 0 | 0 | 0 | 0 | 0 | 0 | 0 | 0 | 0 | 0 |
| 0 | NA | NA | NA | NA | NA | Gm37346 | | 0 | 0 | 0 | 0 | 0 | 0 | 0 | 0 | 0 | 0 | 0 |
| 0 | NA | NA | NA | NA | NA | Gm38136 | | 0 | 0 | 0 | 0 | 0 | 0 | 0 | 0 | 0 | 0 | 0 |
| 0 | NA | NA | NA | NA | NA | Gm37991 | | 0 | 0 | 0 | 0 | 0 | 0 | 0 | 0 | 0 | 0 | 0 |
| 0 | NA | NA | NA | NA | NA | Gm37808 | | 0 | 0 | 0 | 0 | 0 | 0 | 0 | 0 | 0 | 0 | 0 |
| 0 | NA | NA | NA | NA | NA | Gm33571 | | 0 | 0 | 0 | 0 | 0 | 0 | 0 | 0 | 0 | 0 | 0 |

| | | | | | | | | | | | | | | | | | |
|---|---|---|---|---|---|---|---|---|---|---|---|---|---|---|---|---|---|
| 0 | NA | NA | NA | NA | NA | Gm32181 | | 0 | 0 | 0 | 0 | 0 | 0 | 0 | 0 | 0 | 0 | 0 |
| 0 | NA | NA | NA | NA | NA | Gm28662 | | 0 | 0 | 0 | 0 | 0 | 0 | 0 | 0 | 0 | 0 | 0 |
| 0 | NA | NA | NA | NA | NA | Gm28780 | | 0 | 0 | 0 | 0 | 0 | 0 | 0 | 0 | 0 | 0 | 0 |
| 0 | NA | NA | NA | NA | NA | Gm21755 | | 0 | 0 | 0 | 0 | 0 | 0 | 0 | 0 | 0 | 0 | 0 |
| 0 | NA | NA | NA | NA | NA | Gm20895 | predicted gene, 20895 | 0 | 0 | 0 | 0 | 0 | 0 | 0 | 0 | 0 | 0 | 0 |
| 0 | NA | NA | NA | NA | NA | Gm28853 | | 0 | 0 | 0 | 0 | 0 | 0 | 0 | 0 | 0 | 0 | 0 |
| 0 | NA | NA | NA | NA | NA | Gm28852 | | 0 | 0 | 0 | 0 | 0 | 0 | 0 | 0 | 0 | 0 | 0 |
| 0 | NA | NA | NA | NA | NA | Gm28855 | | 0 | 0 | 0 | 0 | 0 | 0 | 0 | 0 | 0 | 0 | 0 |
| 0 | NA | NA | NA | NA | NA | Gm28854 | | 0 | 0 | 0 | 0 | 0 | 0 | 0 | 0 | 0 | 0 | 0 |
| 0 | NA | NA | NA | NA | NA | Gm28858 | | 0 | 0 | 0 | 0 | 0 | 0 | 0 | 0 | 0 | 0 | 0 |
| 0 | NA | NA | NA | NA | NA | Gm20890 | predicted gene, 20890 | 0 | 0 | 0 | 0 | 0 | 0 | 0 | 0 | 0 | 0 | 0 |
| 0 | NA | NA | NA | NA | NA | Gm28245 | | 0 | 0 | 0 | 0 | 0 | 0 | 0 | 0 | 0 | 0 | 0 |
| 0 | NA | NA | NA | NA | NA | Gm28246 | | 0 | 0 | 0 | 0 | 0 | 0 | 0 | 0 | 0 | 0 | 0 |
| 0 | NA | NA | NA | NA | NA | Gm28247 | | 0 | 0 | 0 | 0 | 0 | 0 | 0 | 0 | 0 | 0 | 0 |
| 0 | NA | NA | NA | NA | NA | Gm28248 | | 0 | 0 | 0 | 0 | 0 | 0 | 0 | 0 | 0 | 0 | 0 |
| 0 | NA | NA | NA | NA | NA | Gm20892 | predicted gene, 20892 | 0 | 0 | 0 | 0 | 0 | 0 | 0 | 0 | 0 | 0 | 0 |
| 0 | NA | NA | NA | NA | NA | Gm28249 | | 0 | 0 | 0 | 0 | 0 | 0 | 0 | 0 | 0 | 0 | 0 |
| 0 | NA | NA | NA | NA | NA | Gm28250 | | 0 | 0 | 0 | 0 | 0 | 0 | 0 | 0 | 0 | 0 | 0 |
| 0 | NA | NA | NA | NA | NA | Gm28251 | | 0 | 0 | 0 | 0 | 0 | 0 | 0 | 0 | 0 | 0 | 0 |
| 0 | NA | NA | NA | NA | NA | Gm28243 | | 0 | 0 | 0 | 0 | 0 | 0 | 0 | 0 | 0 | 0 | 0 |
| 0 | NA | NA | NA | NA | NA | Gm28244 | | 0 | 0 | 0 | 0 | 0 | 0 | 0 | 0 | 0 | 0 | 0 |
| 0 | NA | NA | NA | NA | NA | Gm21771 | | 0 | 0 | 0 | 0 | 0 | 0 | 0 | 0 | 0 | 0 | 0 |
| 0 | NA | NA | NA | NA | NA | Gm21476 | predicted gene, 21476 | 0 | 0 | 0 | 0 | 0 | 0 | 0 | 0 | 0 | 0 | 0 |
| 0 | NA | NA | NA | NA | NA | Gm29021 | | 0 | 0 | 0 | 0 | 0 | 0 | 0 | 0 | 0 | 0 | 0 |
| 0 | NA | NA | NA | NA | NA | Gm28880 | | 0 | 0 | 0 | 0 | 0 | 0 | 0 | 0 | 0 | 0 | 0 |
| 0 | NA | NA | NA | NA | NA | Gm28452 | | 0 | 0 | 0 | 0 | 0 | 0 | 0 | 0 | 0 | 0 | 0 |
| 0 | NA | NA | NA | NA | NA | Gm21488 | predicted gene, 21488 | 0 | 0 | 0 | 0 | 0 | 0 | 0 | 0 | 0 | 0 | 0 |
| 0 | NA | NA | NA | NA | NA | Gm28454 | | 0 | 0 | 0 | 0 | 0 | 0 | 0 | 0 | 0 | 0 | 0 |
| 0 | NA | NA | NA | NA | NA | Gm28878 | | 0 | 0 | 0 | 0 | 0 | 0 | 0 | 0 | 0 | 0 | 0 |
| 0 | NA | NA | NA | NA | NA | Gm28879 | | 0 | 0 | 0 | 0 | 0 | 0 | 0 | 0 | 0 | 0 | 0 |
| 0 | NA | NA | NA | NA | NA | Gm29369 | | 0 | 0 | 0 | 0 | 0 | 0 | 0 | 0 | 0 | 0 | 0 |
| 0 | NA | NA | NA | NA | NA | Gm21506 | predicted gene, 21506 | 0 | 0 | 0 | 0 | 0 | 0 | 0 | 0 | 0 | 0 | 0 |
| 0 | NA | NA | NA | NA | NA | Gm29399 | | 0 | 0 | 0 | 0 | 0 | 0 | 0 | 0 | 0 | 0 | 0 |
| 0 | NA | NA | NA | NA | NA | Gm29400 | | 0 | 0 | 0 | 0 | 0 | 0 | 0 | 0 | 0 | 0 | 0 |
| 0 | NA | NA | NA | NA | NA | Gm29401 | | 0 | 0 | 0 | 0 | 0 | 0 | 0 | 0 | 0 | 0 | 0 |
| 0 | NA | NA | NA | NA | NA | Gm20891 | predicted gene, 20891 | 0 | 0 | 0 | 0 | 0 | 0 | 0 | 0 | 0 | 0 | 0 |
| 0 | NA | NA | NA | NA | NA | Gm20893 | predicted gene, 20893 | 0 | 0 | 0 | 0 | 0 | 0 | 0 | 0 | 0 | 0 | 0 |
| 0 | NA | NA | NA | NA | NA | Gm28799 | | 0 | 0 | 0 | 0 | 0 | 0 | 0 | 0 | 0 | 0 | 0 |
| 0 | NA | NA | NA | NA | NA | Gm28798 | | 0 | 0 | 0 | 0 | 0 | 0 | 0 | 0 | 0 | 0 | 0 |
| 0 | NA | NA | NA | NA | NA | Gm28797 | | 0 | 0 | 0 | 0 | 0 | 0 | 0 | 0 | 0 | 0 | 0 |
| 0 | NA | NA | NA | NA | NA | Gm28796 | | 0 | 0 | 0 | 0 | 0 | 0 | 0 | 0 | 0 | 0 | 0 |
| 0 | NA | NA | NA | NA | NA | Gm20894 | X-linked lymphocyte-regulated protein PM | 0 | 0 | 0 | 0 | 0 | 0 | 0 | 0 | 0 | 0 | 0 |
| 0 | NA | NA | NA | NA | NA | Gm28938 | | 0 | 0 | 0 | 0 | 0 | 0 | 0 | 0 | 0 | 0 | 0 |
| 0 | NA | NA | NA | NA | NA | Gm28937 | | 0 | 0 | 0 | 0 | 0 | 0 | 0 | 0 | 0 | 0 | 0 |
| 0 | NA | NA | NA | NA | NA | Gm28936 | | 0 | 0 | 0 | 0 | 0 | 0 | 0 | 0 | 0 | 0 | 0 |
| 0 | NA | NA | NA | NA | NA | Gm21642 | predicted gene, 21642 | 0 | 0 | 0 | 0 | 0 | 0 | 0 | 0 | 0 | 0 | 0 |
| 0 | NA | NA | NA | NA | NA | Gm21804 | | 0 | 0 | 0 | 0 | 0 | 0 | 0 | 0 | 0 | 0 | 0 |
| 0 | NA | NA | NA | NA | NA | Gm29122 | | 0 | 0 | 0 | 0 | 0 | 0 | 0 | 0 | 0 | 0 | 0 |
| 0 | NA | NA | NA | NA | NA | Gm28939 | | 0 | 0 | 0 | 0 | 0 | 0 | 0 | 0 | 0 | 0 | 0 |
| 0 | NA | NA | NA | NA | NA | Gm28983 | | 0 | 0 | 0 | 0 | 0 | 0 | 0 | 0 | 0 | 0 | 0 |
| 0 | NA | NA | NA | NA | NA | Gm28985 | | 0 | 0 | 0 | 0 | 0 | 0 | 0 | 0 | 0 | 0 | 0 |
| 0 | NA | NA | NA | NA | NA | Gm20810 | predicted gene, 20810 | 0 | 0 | 0 | 0 | 0 | 0 | 0 | 0 | 0 | 0 | 0 |
| 0 | NA | NA | NA | NA | NA | Gm20811 | | 0 | 0 | 0 | 0 | 0 | 0 | 0 | 0 | 0 | 0 | 0 |
| 0 | NA | NA | NA | NA | NA | Gm28725 | | 0 | 0 | 0 | 0 | 0 | 0 | 0 | 0 | 0 | 0 | 0 |
| 0 | NA | NA | NA | NA | NA | Gm28984 | | 0 | 0 | 0 | 0 | 0 | 0 | 0 | 0 | 0 | 0 | 0 |
| 0 | NA | NA | NA | NA | NA | Gm28987 | | 0 | 0 | 0 | 0 | 0 | 0 | 0 | 0 | 0 | 0 | 0 |
| 0 | NA | NA | NA | NA | NA | Gm28986 | | 0 | 0 | 0 | 0 | 0 | 0 | 0 | 0 | 0 | 0 | 0 |
| 0 | NA | NA | NA | NA | NA | Gm29171 | | 0 | 0 | 0 | 0 | 0 | 0 | 0 | 0 | 0 | 0 | 0 |
| 0 | NA | NA | NA | NA | NA | Gm28129 | | 0 | 0 | 0 | 0 | 0 | 0 | 0 | 0 | 0 | 0 | 0 |
| 0 | NA | NA | NA | NA | NA | Gm28127 | | 0 | 0 | 0 | 0 | 0 | 0 | 0 | 0 | 0 | 0 | 0 |

| | | | | | | | | | | | | | | | | | |
|---|---|---|---|---|---|---|---|---|---|---|---|---|---|---|---|---|---|
| 0 | NA | NA | NA | NA | NA | Gm28128 | 0 | 0 | 0 | 0 | 0 | 0 | 0 | 0 | 0 | 0 | 0 |
| 0 | NA | NA | NA | NA | NA | Gm21853 | 0 | 0 | 0 | 0 | 0 | 0 | 0 | 0 | 0 | 0 | 0 |
| 0 | NA | NA | NA | NA | NA | Gm28132 | 0 | 0 | 0 | 0 | 0 | 0 | 0 | 0 | 0 | 0 | 0 |
| 0 | NA | NA | NA | NA | NA | Gm28130 | 0 | 0 | 0 | 0 | 0 | 0 | 0 | 0 | 0 | 0 | 0 |
| 0 | NA | NA | NA | NA | NA | Gm28131 | 0 | 0 | 0 | 0 | 0 | 0 | 0 | 0 | 0 | 0 | 0 |
| 0 | NA | NA | NA | NA | NA | Gm21679 | 0 | 0 | 0 | 0 | 0 | 0 | 0 | 0 | 0 | 0 | 0 |
| 0 | NA | NA | NA | NA | NA | Gm29317 | 0 | 0 | 0 | 0 | 0 | 0 | 0 | 0 | 0 | 0 | 0 |
| 0 | NA | NA | NA | NA | NA | Gm29316 | 0 | 0 | 0 | 0 | 0 | 0 | 0 | 0 | 0 | 0 | 0 |
| 0 | NA | NA | NA | NA | NA | Gm29315 | 0 | 0 | 0 | 0 | 0 | 0 | 0 | 0 | 0 | 0 | 0 |
| 0 | NA | NA | NA | NA | NA | Gm28216 | 0 | 0 | 0 | 0 | 0 | 0 | 0 | 0 | 0 | 0 | 0 |
| 0 | NA | NA | NA | NA | NA | Gm28217 | 0 | 0 | 0 | 0 | 0 | 0 | 0 | 0 | 0 | 0 | 0 |
| 0 | NA | NA | NA | NA | NA | Gm28093 | 0 | 0 | 0 | 0 | 0 | 0 | 0 | 0 | 0 | 0 | 0 |
| 0 | NA | NA | NA | NA | NA | Gm28092 | 0 | 0 | 0 | 0 | 0 | 0 | 0 | 0 | 0 | 0 | 0 |
| 0 | NA | NA | NA | NA | NA | Gm28091 | 0 | 0 | 0 | 0 | 0 | 0 | 0 | 0 | 0 | 0 | 0 |
| 0 | NA | NA | NA | NA | NA | Gm28090 | 0 | 0 | 0 | 0 | 0 | 0 | 0 | 0 | 0 | 0 | 0 |
| 0 | NA | NA | NA | NA | NA | Gm29580 | 0 | 0 | 0 | 0 | 0 | 0 | 0 | 0 | 0 | 0 | 0 |
| 0 | NA | NA | NA | NA | NA | Gm28089 | 0 | 0 | 0 | 0 | 0 | 0 | 0 | 0 | 0 | 0 | 0 |
| 0 | NA | NA | NA | NA | NA | Gm28088 | 0 | 0 | 0 | 0 | 0 | 0 | 0 | 0 | 0 | 0 | 0 |
| 0 | NA | NA | NA | NA | NA | Gm29579 | 0 | 0 | 0 | 0 | 0 | 0 | 0 | 0 | 0 | 0 | 0 |
| 0 | NA | NA | NA | NA | NA | Gm29582 | 0 | 0 | 0 | 0 | 0 | 0 | 0 | 0 | 0 | 0 | 0 |
| 0 | NA | NA | NA | NA | NA | Gm21529 | 0 | 0 | 0 | 0 | 0 | 0 | 0 | 0 | 0 | 0 | 0 |
| 0 | NA | NA | NA | NA | NA | Gm29581 | 0 | 0 | 0 | 0 | 0 | 0 | 0 | 0 | 0 | 0 | 0 |
| 0 | NA | NA | NA | NA | NA | Gm21539 | 0 | 0 | 0 | 0 | 0 | 0 | 0 | 0 | 0 | 0 | 0 |
| 0 | NA | NA | NA | NA | NA | Gm29578 | 0 | 0 | 0 | 0 | 0 | 0 | 0 | 0 | 0 | 0 | 0 |
| 0 | NA | NA | NA | NA | NA | Gm21908 | 0 | 0 | 0 | 0 | 0 | 0 | 0 | 0 | 0 | 0 | 0 |
| 0 | NA | NA | NA | NA | NA | Gm21780 | 0 | 0 | 0 | 0 | 0 | 0 | 0 | 0 | 0 | 0 | 0 |
| 0 | NA | NA | NA | NA | NA | Gm28486 | 0 | 0 | 0 | 0 | 0 | 0 | 0 | 0 | 0 | 0 | 0 |
| 0 | NA | NA | NA | NA | NA | Gm21737 | 0 | 0 | 0 | 0 | 0 | 0 | 0 | 0 | 0 | 0 | 0 |
| 0 | NA | NA | NA | NA | NA | Gm28487 | 0 | 0 | 0 | 0 | 0 | 0 | 0 | 0 | 0 | 0 | 0 |
| 0 | NA | NA | NA | NA | NA | Gm28489 | 0 | 0 | 0 | 0 | 0 | 0 | 0 | 0 | 0 | 0 | 0 |
| 0 | NA | NA | NA | NA | NA | Gm28488 | 0 | 0 | 0 | 0 | 0 | 0 | 0 | 0 | 0 | 0 | 0 |
| 0 | NA | NA | NA | NA | NA | Gm28491 | 0 | 0 | 0 | 0 | 0 | 0 | 0 | 0 | 0 | 0 | 0 |
| 0 | NA | NA | NA | NA | NA | Gm28490 | 0 | 0 | 0 | 0 | 0 | 0 | 0 | 0 | 0 | 0 | 0 |
| 0 | NA | NA | NA | NA | NA | Gm21562 | 0 | 0 | 0 | 0 | 0 | 0 | 0 | 0 | 0 | 0 | 0 |
| 0 | NA | NA | NA | NA | NA | Gm28298 | 0 | 0 | 0 | 0 | 0 | 0 | 0 | 0 | 0 | 0 | 0 |
| 0 | NA | NA | NA | NA | NA | Gm21572 | 0 | 0 | 0 | 0 | 0 | 0 | 0 | 0 | 0 | 0 | 0 |
| 0 | NA | NA | NA | NA | NA | Gm28297 | 0 | 0 | 0 | 0 | 0 | 0 | 0 | 0 | 0 | 0 | 0 |
| 0 | NA | NA | NA | NA | NA | Gm21753 | 0 | 0 | 0 | 0 | 0 | 0 | 0 | 0 | 0 | 0 | 0 |
| 0 | NA | NA | NA | NA | NA | Gm21723 | 0 | 0 | 0 | 0 | 0 | 0 | 0 | 0 | 0 | 0 | 0 |
| 0 | NA | NA | NA | NA | NA | Gm28296 | 0 | 0 | 0 | 0 | 0 | 0 | 0 | 0 | 0 | 0 | 0 |
| 0 | NA | NA | NA | NA | NA | Gm21821 | 0 | 0 | 0 | 0 | 0 | 0 | 0 | 0 | 0 | 0 | 0 |
| 0 | NA | NA | NA | NA | NA | Gm28295 | 0 | 0 | 0 | 0 | 0 | 0 | 0 | 0 | 0 | 0 | 0 |
| 0 | NA | NA | NA | NA | NA | Gm21842 | 0 | 0 | 0 | 0 | 0 | 0 | 0 | 0 | 0 | 0 | 0 |
| 0 | NA | NA | NA | NA | NA | Gm28170 | 0 | 0 | 0 | 0 | 0 | 0 | 0 | 0 | 0 | 0 | 0 |
| 0 | NA | NA | NA | NA | NA | Gm21744 | 0 | 0 | 0 | 0 | 0 | 0 | 0 | 0 | 0 | 0 | 0 |
| 0 | NA | NA | NA | NA | NA | Gm28764 | 0 | 0 | 0 | 0 | 0 | 0 | 0 | 0 | 0 | 0 | 0 |
| 0 | NA | NA | NA | NA | NA | Gm28762 | 0 | 0 | 0 | 0 | 0 | 0 | 0 | 0 | 0 | 0 | 0 |
| 0 | NA | NA | NA | NA | NA | Gm28763 | 0 | 0 | 0 | 0 | 0 | 0 | 0 | 0 | 0 | 0 | 0 |
| 0 | NA | NA | NA | NA | NA | Gm21094 | 0 | 0 | 0 | 0 | 0 | 0 | 0 | 0 | 0 | 0 | 0 |
| 0 | NA | NA | NA | NA | NA | Gm21588 | 0 | 0 | 0 | 0 | 0 | 0 | 0 | 0 | 0 | 0 | 0 |
| 0 | NA | NA | NA | NA | NA | Gm28761 | 0 | 0 | 0 | 0 | 0 | 0 | 0 | 0 | 0 | 0 | 0 |
| 0 | NA | NA | NA | NA | NA | Gm21599 | 0 | 0 | 0 | 0 | 0 | 0 | 0 | 0 | 0 | 0 | 0 |
| 0 | NA | NA | NA | NA | NA | Gm28765 | 0 | 0 | 0 | 0 | 0 | 0 | 0 | 0 | 0 | 0 | 0 |
| 0 | NA | NA | NA | NA | NA | Gm21810 | 0 | 0 | 0 | 0 | 0 | 0 | 0 | 0 | 0 | 0 | 0 |
| 0 | NA | NA | NA | NA | NA | Gm21916 | 0 | 0 | 0 | 0 | 0 | 0 | 0 | 0 | 0 | 0 | 0 |
| 0 | NA | NA | NA | NA | NA | Gm29191 | 0 | 0 | 0 | 0 | 0 | 0 | 0 | 0 | 0 | 0 | 0 |
| 0 | NA | NA | NA | NA | NA | Gm21756 | 0 | 0 | 0 | 0 | 0 | 0 | 0 | 0 | 0 | 0 | 0 |
| 0 | NA | NA | NA | NA | NA | Gm28210 | 0 | 0 | 0 | 0 | 0 | 0 | 0 | 0 | 0 | 0 | 0 |
| 0 | NA | NA | NA | NA | NA | Gm29074 | 0 | 0 | 0 | 0 | 0 | 0 | 0 | 0 | 0 | 0 | 0 |
| 0 | NA | NA | NA | NA | NA | Gm28211 | 0 | 0 | 0 | 0 | 0 | 0 | 0 | 0 | 0 | 0 | 0 |

| | | | | | | | | | | | | | | | | | |
|---|---|---|---|---|---|---|---|---|---|---|---|---|---|---|---|---|---|
| 0 | NA | NA | NA | NA | NA | Gm28518 | | 0 | 0 | 0 | 0 | 0 | 0 | 0 | 0 | 0 | 0 | 0 |
| 0 | NA | NA | NA | NA | NA | Gm28519 | | 0 | 0 | 0 | 0 | 0 | 0 | 0 | 0 | 0 | 0 | 0 |
| 0 | NA | NA | NA | NA | NA | Gm28520 | | 0 | 0 | 0 | 0 | 0 | 0 | 0 | 0 | 0 | 0 | 0 |
| 0 | NA | NA | NA | NA | NA | Gm20838 | | 0 | 0 | 0 | 0 | 0 | 0 | 0 | 0 | 0 | 0 | 0 |
| 0 | NA | NA | NA | NA | NA | Gm21626 | | 0 | 0 | 0 | 0 | 0 | 0 | 0 | 0 | 0 | 0 | 0 |
| 0 | NA | NA | NA | NA | NA | Gm28521 | | 0 | 0 | 0 | 0 | 0 | 0 | 0 | 0 | 0 | 0 | 0 |
| 0 | NA | NA | NA | NA | NA | Gm21633 | | 0 | 0 | 0 | 0 | 0 | 0 | 0 | 0 | 0 | 0 | 0 |
| 0 | NA | NA | NA | NA | NA | Gm28522 | | 0 | 0 | 0 | 0 | 0 | 0 | 0 | 0 | 0 | 0 | 0 |
| 0 | NA | NA | NA | NA | NA | Gm21745 | | 0 | 0 | 0 | 0 | 0 | 0 | 0 | 0 | 0 | 0 | 0 |
| 0 | NA | NA | NA | NA | NA | Gm28517 | | 0 | 0 | 0 | 0 | 0 | 0 | 0 | 0 | 0 | 0 | 0 |
| 0 | NA | NA | NA | NA | NA | Gm29532 | | 0 | 0 | 0 | 0 | 0 | 0 | 0 | 0 | 0 | 0 | 0 |
| 0 | NA | NA | NA | NA | NA | Gm29531 | | 0 | 0 | 0 | 0 | 0 | 0 | 0 | 0 | 0 | 0 | 0 |
| 0 | NA | NA | NA | NA | NA | Gm28464 | | 0 | 0 | 0 | 0 | 0 | 0 | 0 | 0 | 0 | 0 | 0 |
| 0 | NA | NA | NA | NA | NA | Gm29656 | | 0 | 0 | 0 | 0 | 0 | 0 | 0 | 0 | 0 | 0 | 0 |
| 0 | NA | NA | NA | NA | NA | Gm29655 | | 0 | 0 | 0 | 0 | 0 | 0 | 0 | 0 | 0 | 0 | 0 |
| 0 | NA | NA | NA | NA | NA | Gm21647 | | 0 | 0 | 0 | 0 | 0 | 0 | 0 | 0 | 0 | 0 | 0 |
| 0 | NA | NA | NA | NA | NA | Gm29653 | | 0 | 0 | 0 | 0 | 0 | 0 | 0 | 0 | 0 | 0 | 0 |
| 0 | NA | NA | NA | NA | NA | Gm29654 | | 0 | 0 | 0 | 0 | 0 | 0 | 0 | 0 | 0 | 0 | 0 |
| 0 | NA | NA | NA | NA | NA | Gm29651 | | 0 | 0 | 0 | 0 | 0 | 0 | 0 | 0 | 0 | 0 | 0 |
| 0 | NA | NA | NA | NA | NA | Gm29652 | | 0 | 0 | 0 | 0 | 0 | 0 | 0 | 0 | 0 | 0 | 0 |
| 0 | NA | NA | NA | NA | NA | Gm29080 | | 0 | 0 | 0 | 0 | 0 | 0 | 0 | 0 | 0 | 0 | 0 |
| 0 | NA | NA | NA | NA | NA | Gm21184 | Y-linked testis-specific protein 1-like | 0 | 0 | 0 | 0 | 0 | 0 | 0 | 0 | 0 | 0 | 0 |
| 0 | NA | NA | NA | NA | NA | Gm21201 | Y-linked testis-specific protein 1-like | 0 | 0 | 0 | 0 | 0 | 0 | 0 | 0 | 0 | 0 | 0 |
| 0 | NA | NA | NA | NA | NA | Gm29081 | | 0 | 0 | 0 | 0 | 0 | 0 | 0 | 0 | 0 | 0 | 0 |
| 0 | NA | NA | NA | NA | NA | Gm29077 | | 0 | 0 | 0 | 0 | 0 | 0 | 0 | 0 | 0 | 0 | 0 |
| 0 | NA | NA | NA | NA | NA | Gm29076 | | 0 | 0 | 0 | 0 | 0 | 0 | 0 | 0 | 0 | 0 | 0 |
| 0 | NA | NA | NA | NA | NA | Gm29079 | | 0 | 0 | 0 | 0 | 0 | 0 | 0 | 0 | 0 | 0 | 0 |
| 0 | NA | NA | NA | NA | NA | Gm29078 | | 0 | 0 | 0 | 0 | 0 | 0 | 0 | 0 | 0 | 0 | 0 |
| 0 | NA | NA | NA | NA | NA | Gm20853 | predicted gene, 20853 | 0 | 0 | 0 | 0 | 0 | 0 | 0 | 0 | 0 | 0 | 0 |
| 0 | NA | NA | NA | NA | NA | Gm29075 | | 0 | 0 | 0 | 0 | 0 | 0 | 0 | 0 | 0 | 0 | 0 |
| 0 | NA | NA | NA | NA | NA | Gm29225 | | 0 | 0 | 0 | 0 | 0 | 0 | 0 | 0 | 0 | 0 | 0 |
| 0 | NA | NA | NA | NA | NA | Gm29226 | | 0 | 0 | 0 | 0 | 0 | 0 | 0 | 0 | 0 | 0 | 0 |
| 0 | NA | NA | NA | NA | NA | Gm29227 | | 0 | 0 | 0 | 0 | 0 | 0 | 0 | 0 | 0 | 0 | 0 |
| 0 | NA | NA | NA | NA | NA | Gm28339 | | 0 | 0 | 0 | 0 | 0 | 0 | 0 | 0 | 0 | 0 | 0 |
| 0 | NA | NA | NA | NA | NA | Gm20855 | X-linked lymphocyte-regulated protein PM | 0 | 0 | 0 | 0 | 0 | 0 | 0 | 0 | 0 | 0 | 0 |
| 0 | NA | NA | NA | NA | NA | Gm29386 | | 0 | 0 | 0 | 0 | 0 | 0 | 0 | 0 | 0 | 0 | 0 |
| 0 | NA | NA | NA | NA | NA | Gm29384 | | 0 | 0 | 0 | 0 | 0 | 0 | 0 | 0 | 0 | 0 | 0 |
| 0 | NA | NA | NA | NA | NA | Gm21896 | | 0 | 0 | 0 | 0 | 0 | 0 | 0 | 0 | 0 | 0 | 0 |
| 0 | NA | NA | NA | NA | NA | Gm29280 | | 0 | 0 | 0 | 0 | 0 | 0 | 0 | 0 | 0 | 0 | 0 |
| 0 | NA | NA | NA | NA | NA | Gm20799 | | 0 | 0 | 0 | 0 | 0 | 0 | 0 | 0 | 0 | 0 | 0 |
| 0 | NA | NA | NA | NA | NA | Gm29279 | | 0 | 0 | 0 | 0 | 0 | 0 | 0 | 0 | 0 | 0 | 0 |
| 0 | NA | NA | NA | NA | NA | Gm28260 | | 0 | 0 | 0 | 0 | 0 | 0 | 0 | 0 | 0 | 0 | 0 |
| 0 | NA | NA | NA | NA | NA | Gm21795 | | 0 | 0 | 0 | 0 | 0 | 0 | 0 | 0 | 0 | 0 | 0 |
| 0 | NA | NA | NA | NA | NA | Gm28261 | | 0 | 0 | 0 | 0 | 0 | 0 | 0 | 0 | 0 | 0 | 0 |
| 0 | NA | NA | NA | NA | NA | Gm28264 | | 0 | 0 | 0 | 0 | 0 | 0 | 0 | 0 | 0 | 0 | 0 |
| 0 | NA | NA | NA | NA | NA | Gm21872 | | 0 | 0 | 0 | 0 | 0 | 0 | 0 | 0 | 0 | 0 | 0 |
| 0 | NA | NA | NA | NA | NA | Gm20798 | predicted gene, 20798 | 0 | 0 | 0 | 0 | 0 | 0 | 0 | 0 | 0 | 0 | 0 |
| 0 | NA | NA | NA | NA | NA | Gm28463 | | 0 | 0 | 0 | 0 | 0 | 0 | 0 | 0 | 0 | 0 | 0 |
| 0 | NA | NA | NA | NA | NA | Gm28584 | | 0 | 0 | 0 | 0 | 0 | 0 | 0 | 0 | 0 | 0 | 0 |
| 0 | NA | NA | NA | NA | NA | Gm28585 | | 0 | 0 | 0 | 0 | 0 | 0 | 0 | 0 | 0 | 0 | 0 |
| 0 | NA | NA | NA | NA | NA | Gm20896 | X-linked lymphocyte-regulated protein PM | 0 | 0 | 0 | 0 | 0 | 0 | 0 | 0 | 0 | 0 | 0 |
| 0 | NA | NA | NA | NA | NA | Gm28470 | | 0 | 0 | 0 | 0 | 0 | 0 | 0 | 0 | 0 | 0 | 0 |
| 0 | NA | NA | NA | NA | NA | Gm28469 | | 0 | 0 | 0 | 0 | 0 | 0 | 0 | 0 | 0 | 0 | 0 |
| 0 | NA | NA | NA | NA | NA | Gm21898 | | 0 | 0 | 0 | 0 | 0 | 0 | 0 | 0 | 0 | 0 | 0 |
| 0 | NA | NA | NA | NA | NA | Gm28468 | | 0 | 0 | 0 | 0 | 0 | 0 | 0 | 0 | 0 | 0 | 0 |
| 0 | NA | NA | NA | NA | NA | Gm20898 | predicted gene, 20898 | 0 | 0 | 0 | 0 | 0 | 0 | 0 | 0 | 0 | 0 | 0 |
| 0 | NA | NA | NA | NA | NA | Gm29333 | | 0 | 0 | 0 | 0 | 0 | 0 | 0 | 0 | 0 | 0 | 0 |
| 0 | NA | NA | NA | NA | NA | Gm28332 | | 0 | 0 | 0 | 0 | 0 | 0 | 0 | 0 | 0 | 0 | 0 |
| 0 | NA | NA | NA | NA | NA | Gm28333 | | 0 | 0 | 0 | 0 | 0 | 0 | 0 | 0 | 0 | 0 | 0 |
| 0 | NA | NA | NA | NA | NA | Gm28334 | | 0 | 0 | 0 | 0 | 0 | 0 | 0 | 0 | 0 | 0 | 0 |

| | | | | | | | | | | | | | | | | | | |
|---|---|---|---|---|---|---|---|---|---|---|---|---|---|---|---|---|---|---|
| 0 | NA | NA | NA | NA | NA | Gm28335 | | 0 | 0 | 0 | 0 | 0 | 0 | 0 | 0 | 0 | 0 | 0 |
| 0 | NA | NA | NA | NA | NA | Gm28336 | | 0 | 0 | 0 | 0 | 0 | 0 | 0 | 0 | 0 | 0 | 0 |
| 0 | NA | NA | NA | NA | NA | Gm28337 | | 0 | 0 | 0 | 0 | 0 | 0 | 0 | 0 | 0 | 0 | 0 |
| 0 | NA | NA | NA | NA | NA | Gm28338 | | 0 | 0 | 0 | 0 | 0 | 0 | 0 | 0 | 0 | 0 | 0 |
| 0 | NA | NA | NA | NA | NA | Gm28812 | | 0 | 0 | 0 | 0 | 0 | 0 | 0 | 0 | 0 | 0 | 0 |
| 0 | NA | NA | NA | NA | NA | Gm28811 | | 0 | 0 | 0 | 0 | 0 | 0 | 0 | 0 | 0 | 0 | 0 |
| 0 | NA | NA | NA | NA | NA | Gm28810 | | 0 | 0 | 0 | 0 | 0 | 0 | 0 | 0 | 0 | 0 | 0 |
| 0 | NA | NA | NA | NA | NA | Gm20835 | X-linked lymphocyte-regulated protein PM | 0 | 0 | 0 | 0 | 0 | 0 | 0 | 0 | 0 | 0 | 0 |
| 0 | NA | NA | NA | NA | NA | Gm29027 | | 0 | 0 | 0 | 0 | 0 | 0 | 0 | 0 | 0 | 0 | 0 |
| 0 | NA | NA | NA | NA | NA | Gm29028 | | 0 | 0 | 0 | 0 | 0 | 0 | 0 | 0 | 0 | 0 | 0 |
| 0 | NA | NA | NA | NA | NA | Gm29029 | | 0 | 0 | 0 | 0 | 0 | 0 | 0 | 0 | 0 | 0 | 0 |
| 0 | NA | NA | NA | NA | NA | Gm29030 | | 0 | 0 | 0 | 0 | 0 | 0 | 0 | 0 | 0 | 0 | 0 |
| 0 | NA | NA | NA | NA | NA | Gm29023 | | 0 | 0 | 0 | 0 | 0 | 0 | 0 | 0 | 0 | 0 | 0 |
| 0 | NA | NA | NA | NA | NA | Gm29024 | | 0 | 0 | 0 | 0 | 0 | 0 | 0 | 0 | 0 | 0 | 0 |
| 0 | NA | NA | NA | NA | NA | Gm29025 | | 0 | 0 | 0 | 0 | 0 | 0 | 0 | 0 | 0 | 0 | 0 |
| 0 | NA | NA | NA | NA | NA | Gm29026 | | 0 | 0 | 0 | 0 | 0 | 0 | 0 | 0 | 0 | 0 | 0 |
| 0 | NA | NA | NA | NA | NA | Gm28213 | | 0 | 0 | 0 | 0 | 0 | 0 | 0 | 0 | 0 | 0 | 0 |
| 0 | NA | NA | NA | NA | NA | Gm28212 | | 0 | 0 | 0 | 0 | 0 | 0 | 0 | 0 | 0 | 0 | 0 |
| 0 | NA | NA | NA | NA | NA | Gm21799 | | 0 | 0 | 0 | 0 | 0 | 0 | 0 | 0 | 0 | 0 | 0 |
| 0 | NA | NA | NA | NA | NA | Gm28227 | | 0 | 0 | 0 | 0 | 0 | 0 | 0 | 0 | 0 | 0 | 0 |
| 0 | NA | NA | NA | NA | NA | Gm20905 | X-linked lymphocyte-regulated protein PM | 0 | 0 | 0 | 0 | 0 | 0 | 0 | 0 | 0 | 0 | 0 |
| 0 | NA | NA | NA | NA | NA | Gm29411 | | 0 | 0 | 0 | 0 | 0 | 0 | 0 | 0 | 0 | 0 | 0 |
| 0 | NA | NA | NA | NA | NA | Gm28222 | | 0 | 0 | 0 | 0 | 0 | 0 | 0 | 0 | 0 | 0 | 0 |
| 0 | NA | NA | NA | NA | NA | Gm21835 | | 0 | 0 | 0 | 0 | 0 | 0 | 0 | 0 | 0 | 0 | 0 |
| 0 | NA | NA | NA | NA | NA | Gm28820 | | 0 | 0 | 0 | 0 | 0 | 0 | 0 | 0 | 0 | 0 | 0 |
| 0 | NA | NA | NA | NA | NA | Gm28223 | | 0 | 0 | 0 | 0 | 0 | 0 | 0 | 0 | 0 | 0 | 0 |
| 0 | NA | NA | NA | NA | NA | Gm29412 | | 0 | 0 | 0 | 0 | 0 | 0 | 0 | 0 | 0 | 0 | 0 |
| 0 | NA | NA | NA | NA | NA | Gm29413 | | 0 | 0 | 0 | 0 | 0 | 0 | 0 | 0 | 0 | 0 | 0 |
| 0 | NA | NA | NA | NA | NA | Gm29671 | | 0 | 0 | 0 | 0 | 0 | 0 | 0 | 0 | 0 | 0 | 0 |
| 0 | NA | NA | NA | NA | NA | Gm21878 | | 0 | 0 | 0 | 0 | 0 | 0 | 0 | 0 | 0 | 0 | 0 |
| 0 | NA | NA | NA | NA | NA | Gm28162 | | 0 | 0 | 0 | 0 | 0 | 0 | 0 | 0 | 0 | 0 | 0 |
| 0 | NA | NA | NA | NA | NA | Gm20922 | | 0 | 0 | 0 | 0 | 0 | 0 | 0 | 0 | 0 | 0 | 0 |
| 0 | NA | NA | NA | NA | NA | Gm20738 | predicted gene, 20738 | 0 | 0 | 0 | 0 | 0 | 0 | 0 | 0 | 0 | 0 | 0 |
| 0 | NA | NA | NA | NA | NA | Gm28545 | | 0 | 0 | 0 | 0 | 0 | 0 | 0 | 0 | 0 | 0 | 0 |
| 0 | NA | NA | NA | NA | NA | Gm28547 | | 0 | 0 | 0 | 0 | 0 | 0 | 0 | 0 | 0 | 0 | 0 |
| 0 | NA | NA | NA | NA | NA | Gm28546 | | 0 | 0 | 0 | 0 | 0 | 0 | 0 | 0 | 0 | 0 | 0 |
| 0 | NA | NA | NA | NA | NA | Gm21924 | | 0 | 0 | 0 | 0 | 0 | 0 | 0 | 0 | 0 | 0 | 0 |
| 0 | NA | NA | NA | NA | NA | Gm21735 | | 0 | 0 | 0 | 0 | 0 | 0 | 0 | 0 | 0 | 0 | 0 |
| 0 | NA | NA | NA | NA | NA | Gm28771 | | 0 | 0 | 0 | 0 | 0 | 0 | 0 | 0 | 0 | 0 | 0 |
| 0 | NA | NA | NA | NA | NA | Gm28772 | | 0 | 0 | 0 | 0 | 0 | 0 | 0 | 0 | 0 | 0 | 0 |
| 0 | NA | NA | NA | NA | NA | Gm28773 | | 0 | 0 | 0 | 0 | 0 | 0 | 0 | 0 | 0 | 0 | 0 |
| 0 | NA | NA | NA | NA | NA | Gm20897 | X-linked lymphocyte-regulated protein PM | 0 | 0 | 0 | 0 | 0 | 0 | 0 | 0 | 0 | 0 | 0 |
| 0 | NA | NA | NA | NA | NA | Gm28774 | | 0 | 0 | 0 | 0 | 0 | 0 | 0 | 0 | 0 | 0 | 0 |
| 0 | NA | NA | NA | NA | NA | Gm28775 | | 0 | 0 | 0 | 0 | 0 | 0 | 0 | 0 | 0 | 0 | 0 |
| 0 | NA | NA | NA | NA | NA | Gm21765 | | 0 | 0 | 0 | 0 | 0 | 0 | 0 | 0 | 0 | 0 | 0 |
| 0 | NA | NA | NA | NA | NA | Gm28776 | | 0 | 0 | 0 | 0 | 0 | 0 | 0 | 0 | 0 | 0 | 0 |
| 0 | NA | NA | NA | NA | NA | Gm21787 | | 0 | 0 | 0 | 0 | 0 | 0 | 0 | 0 | 0 | 0 | 0 |
| 0 | NA | NA | NA | NA | NA | Gm28770 | | 0 | 0 | 0 | 0 | 0 | 0 | 0 | 0 | 0 | 0 | 0 |
| 0 | NA | NA | NA | NA | NA | Gm20901 | | 0 | 0 | 0 | 0 | 0 | 0 | 0 | 0 | 0 | 0 | 0 |
| 0 | NA | NA | NA | NA | NA | Gm28176 | | 0 | 0 | 0 | 0 | 0 | 0 | 0 | 0 | 0 | 0 | 0 |
| 0 | NA | NA | NA | NA | NA | Gm28786 | | 0 | 0 | 0 | 0 | 0 | 0 | 0 | 0 | 0 | 0 | 0 |
| 0 | NA | NA | NA | NA | NA | Gm29360 | | 0 | 0 | 0 | 0 | 0 | 0 | 0 | 0 | 0 | 0 | 0 |
| 0 | NA | NA | NA | NA | NA | Gm29496 | | 0 | 0 | 0 | 0 | 0 | 0 | 0 | 0 | 0 | 0 | 0 |
| 0 | NA | NA | NA | NA | NA | Gm29497 | | 0 | 0 | 0 | 0 | 0 | 0 | 0 | 0 | 0 | 0 | 0 |
| 0 | NA | NA | NA | NA | NA | Gm29499 | | 0 | 0 | 0 | 0 | 0 | 0 | 0 | 0 | 0 | 0 | 0 |
| 0 | NA | NA | NA | NA | NA | Gm29498 | | 0 | 0 | 0 | 0 | 0 | 0 | 0 | 0 | 0 | 0 | 0 |
| 0 | NA | NA | NA | NA | NA | Gm36782 | | 0 | 0 | 0 | 0 | 0 | 0 | 0 | 0 | 0 | 0 | 0 |
| 0 | NA | NA | NA | NA | NA | Gm29839 | | 0 | 0 | 0 | 0 | 0 | 0 | 0 | 0 | 0 | 0 | 0 |
| 0 | NA | NA | NA | NA | NA | Gm29495 | | 0 | 0 | 0 | 0 | 0 | 0 | 0 | 0 | 0 | 0 | 0 |
| 0 | NA | NA | NA | NA | NA | Gm28371 | | 0 | 0 | 0 | 0 | 0 | 0 | 0 | 0 | 0 | 0 | 0 |

| | | | | | | | | | | | | | | | | | | |
|---|---|---|---|---|---|---|---|---|---|---|---|---|---|---|---|---|---|---|
| 0 | NA | NA | NA | NA | NA | Gm28595 | | 0 | 0 | 0 | 0 | 0 | 0 | 0 | 0 | 0 | 0 | 0 |
| 0 | NA | NA | NA | NA | NA | Gm37032 | | 0 | 0 | 0 | 0 | 0 | 0 | 0 | 0 | 0 | 0 | 0 |
| 0 | NA | NA | NA | NA | NA | Gm21865 | predicted gene, 21865 | 0 | 0 | 0 | 0 | 0 | 0 | 0 | 0 | 0 | 0 | 0 |
| 0 | NA | NA | NA | NA | NA | Gm29433 | | 0 | 0 | 0 | 0 | 0 | 0 | 0 | 0 | 0 | 0 | 0 |
| 0 | NA | NA | NA | NA | NA | Gm29215 | | 0 | 0 | 0 | 0 | 0 | 0 | 0 | 0 | 0 | 0 | 0 |
| 0 | NA | NA | NA | NA | NA | Gm21882 | | 0 | 0 | 0 | 0 | 0 | 0 | 0 | 0 | 0 | 0 | 0 |
| 0 | NA | NA | NA | NA | NA | Gm21801 | | 0 | 0 | 0 | 0 | 0 | 0 | 0 | 0 | 0 | 0 | 0 |
| 0 | NA | NA | NA | NA | NA | Gm28850 | | 0 | 0 | 0 | 0 | 0 | 0 | 0 | 0 | 0 | 0 | 0 |
| 0 | NA | NA | NA | NA | NA | Gm21914 | | 0 | 0 | 0 | 0 | 0 | 0 | 0 | 0 | 0 | 0 | 0 |
| 0 | NA | NA | NA | NA | NA | Gm28617 | | 0 | 0 | 0 | 0 | 0 | 0 | 0 | 0 | 0 | 0 | 0 |
| 0 | NA | NA | NA | NA | NA | Gm28618 | | 0 | 0 | 0 | 0 | 0 | 0 | 0 | 0 | 0 | 0 | 0 |
| 0 | NA | NA | NA | NA | NA | Gm28619 | | 0 | 0 | 0 | 0 | 0 | 0 | 0 | 0 | 0 | 0 | 0 |
| 0 | NA | NA | NA | NA | NA | Gm20793 | predicted gene, 20793 | 0 | 0 | 0 | 0 | 0 | 0 | 0 | 0 | 0 | 0 | 0 |
| 0 | NA | NA | NA | NA | NA | Gm27466 | | 0 | 0 | 0 | 0 | 0 | 0 | 0 | 0 | 0 | 0 | 0 |
| 0 | NA | NA | NA | NA | NA | Gm28018 | | 0 | 0 | 0 | 0 | 0 | 0 | 0 | 0 | 0 | 0 | 0 |
| 0 | NA | NA | NA | NA | NA | Gm28993 | | 0 | 0 | 0 | 0 | 0 | 0 | 0 | 0 | 0 | 0 | 0 |
| 0 | NA | NA | NA | NA | NA | Gm28994 | | 0 | 0 | 0 | 0 | 0 | 0 | 0 | 0 | 0 | 0 | 0 |
| 0 | NA | NA | NA | NA | NA | Gm21797 | | 0 | 0 | 0 | 0 | 0 | 0 | 0 | 0 | 0 | 0 | 0 |
| 0 | NA | NA | NA | NA | NA | Gm20795 | predicted gene, 20795 | 0 | 0 | 0 | 0 | 0 | 0 | 0 | 0 | 0 | 0 | 0 |
| 0 | NA | NA | NA | NA | NA | Gm28596 | | 0 | 0 | 0 | 0 | 0 | 0 | 0 | 0 | 0 | 0 | 0 |
| 0 | NA | NA | NA | NA | NA | Gm20832 | | 0 | 0 | 0 | 0 | 0 | 0 | 0 | 0 | 0 | 0 | 0 |
| 0 | NA | NA | NA | NA | NA | Gm28597 | | 0 | 0 | 0 | 0 | 0 | 0 | 0 | 0 | 0 | 0 | 0 |
| 0 | NA | NA | NA | NA | NA | Gm28428 | | 0 | 0 | 0 | 0 | 0 | 0 | 0 | 0 | 0 | 0 | 0 |
| 0 | NA | NA | NA | NA | NA | Gm20827 | predicted gene, 20827 | 0 | 0 | 0 | 0 | 0 | 0 | 0 | 0 | 0 | 0 | 0 |
| 0 | NA | NA | NA | NA | NA | Gm28427 | | 0 | 0 | 0 | 0 | 0 | 0 | 0 | 0 | 0 | 0 | 0 |
| 0 | NA | NA | NA | NA | NA | Gm21800 | | 0 | 0 | 0 | 0 | 0 | 0 | 0 | 0 | 0 | 0 | 0 |
| 0 | NA | NA | NA | NA | NA | Gm28426 | | 0 | 0 | 0 | 0 | 0 | 0 | 0 | 0 | 0 | 0 | 0 |
| 0 | NA | NA | NA | NA | NA | Gm28425 | | 0 | 0 | 0 | 0 | 0 | 0 | 0 | 0 | 0 | 0 | 0 |
| 0 | NA | NA | NA | NA | NA | Gm21729 | | 0 | 0 | 0 | 0 | 0 | 0 | 0 | 0 | 0 | 0 | 0 |
| 0 | NA | NA | NA | NA | NA | Gm28834 | | 0 | 0 | 0 | 0 | 0 | 0 | 0 | 0 | 0 | 0 | 0 |
| 0 | NA | NA | NA | NA | NA | Gm28833 | | 0 | 0 | 0 | 0 | 0 | 0 | 0 | 0 | 0 | 0 | 0 |
| 0 | NA | NA | NA | NA | NA | Gm21774 | | 0 | 0 | 0 | 0 | 0 | 0 | 0 | 0 | 0 | 0 | 0 |
| 0 | NA | NA | NA | NA | NA | Gm29537 | | 0 | 0 | 0 | 0 | 0 | 0 | 0 | 0 | 0 | 0 | 0 |
| 0 | NA | NA | NA | NA | NA | Gm29274 | | 0 | 0 | 0 | 0 | 0 | 0 | 0 | 0 | 0 | 0 | 0 |
| 0 | NA | NA | NA | NA | NA | Gm29275 | | 0 | 0 | 0 | 0 | 0 | 0 | 0 | 0 | 0 | 0 | 0 |
| 0 | NA | NA | NA | NA | NA | Gm29276 | | 0 | 0 | 0 | 0 | 0 | 0 | 0 | 0 | 0 | 0 | 0 |
| 0 | NA | NA | NA | NA | NA | Gm21805 | | 0 | 0 | 0 | 0 | 0 | 0 | 0 | 0 | 0 | 0 | 0 |
| 0 | NA | NA | NA | NA | NA | Gm29645 | | 0 | 0 | 0 | 0 | 0 | 0 | 0 | 0 | 0 | 0 | 0 |
| 0 | NA | NA | NA | NA | NA | Gm28312 | | 0 | 0 | 0 | 0 | 0 | 0 | 0 | 0 | 0 | 0 | 0 |
| 0 | NA | NA | NA | NA | NA | Gm29165 | | 0 | 0 | 0 | 0 | 0 | 0 | 0 | 0 | 0 | 0 | 0 |
| 0 | NA | NA | NA | NA | NA | Gm29042 | | 0 | 0 | 0 | 0 | 0 | 0 | 0 | 0 | 0 | 0 | 0 |
| 0 | NA | NA | NA | NA | NA | Gm28311 | | 0 | 0 | 0 | 0 | 0 | 0 | 0 | 0 | 0 | 0 | 0 |
| 0 | NA | NA | NA | NA | NA | Gm28310 | | 0 | 0 | 0 | 0 | 0 | 0 | 0 | 0 | 0 | 0 | 0 |
| 0 | NA | NA | NA | NA | NA | Gm28504 | | 0 | 0 | 0 | 0 | 0 | 0 | 0 | 0 | 0 | 0 | 0 |
| 0 | NA | NA | NA | NA | NA | Gm28567 | | 0 | 0 | 0 | 0 | 0 | 0 | 0 | 0 | 0 | 0 | 0 |
| 0 | NA | NA | NA | NA | NA | Gm28568 | | 0 | 0 | 0 | 0 | 0 | 0 | 0 | 0 | 0 | 0 | 0 |
| 0 | NA | NA | NA | NA | NA | Gm29286 | | 0 | 0 | 0 | 0 | 0 | 0 | 0 | 0 | 0 | 0 | 0 |
| 0 | NA | NA | NA | NA | NA | Gm29285 | | 0 | 0 | 0 | 0 | 0 | 0 | 0 | 0 | 0 | 0 | 0 |
| 0 | NA | NA | NA | NA | NA | Gm28280 | | 0 | 0 | 0 | 0 | 0 | 0 | 0 | 0 | 0 | 0 | 0 |
| 0 | NA | NA | NA | NA | NA | Gm21241 | | 0 | 0 | 0 | 0 | 0 | 0 | 0 | 0 | 0 | 0 | 0 |
| 0 | NA | NA | NA | NA | NA | Gm28278 | | 0 | 0 | 0 | 0 | 0 | 0 | 0 | 0 | 0 | 0 | 0 |
| 0 | NA | NA | NA | NA | NA | Gm28279 | | 0 | 0 | 0 | 0 | 0 | 0 | 0 | 0 | 0 | 0 | 0 |
| 0 | NA | NA | NA | NA | NA | Gm21739 | | 0 | 0 | 0 | 0 | 0 | 0 | 0 | 0 | 0 | 0 | 0 |
| 0 | NA | NA | NA | NA | NA | Gm28886 | | 0 | 0 | 0 | 0 | 0 | 0 | 0 | 0 | 0 | 0 | 0 |
| 0 | NA | NA | NA | NA | NA | Gm29426 | | 0 | 0 | 0 | 0 | 0 | 0 | 0 | 0 | 0 | 0 | 0 |
| 0 | NA | NA | NA | NA | NA | Gm28885 | | 0 | 0 | 0 | 0 | 0 | 0 | 0 | 0 | 0 | 0 | 0 |
| 0 | NA | NA | NA | NA | NA | Gm21856 | | 0 | 0 | 0 | 0 | 0 | 0 | 0 | 0 | 0 | 0 | 0 |
| 0 | NA | NA | NA | NA | NA | Gm28887 | | 0 | 0 | 0 | 0 | 0 | 0 | 0 | 0 | 0 | 0 | 0 |
| 0 | NA | NA | NA | NA | NA | Gm29444 | | 0 | 0 | 0 | 0 | 0 | 0 | 0 | 0 | 0 | 0 | 0 |
| 0 | NA | NA | NA | NA | NA | Gm21783 | | 0 | 0 | 0 | 0 | 0 | 0 | 0 | 0 | 0 | 0 | 0 |

| | | | | | | | | | | | | | | | | | | |
|---|---|---|---|---|---|---|---|---|---|---|---|---|---|---|---|---|---|---|
| 0 | NA | NA | NA | NA | NA | Gm29445 | | 0 | 0 | 0 | 0 | 0 | 0 | 0 | 0 | 0 | 0 | 0 |
| 0 | NA | NA | NA | NA | NA | Gm28457 | | 0 | 0 | 0 | 0 | 0 | 0 | 0 | 0 | 0 | 0 | 0 |
| 0 | NA | NA | NA | NA | NA | Gm21779 | | 0 | 0 | 0 | 0 | 0 | 0 | 0 | 0 | 0 | 0 | 0 |
| 0 | NA | NA | NA | NA | NA | Gm28458 | | 0 | 0 | 0 | 0 | 0 | 0 | 0 | 0 | 0 | 0 | 0 |
| 0 | NA | NA | NA | NA | NA | Gm28459 | | 0 | 0 | 0 | 0 | 0 | 0 | 0 | 0 | 0 | 0 | 0 |
| 0 | NA | NA | NA | NA | NA | Gm28461 | | 0 | 0 | 0 | 0 | 0 | 0 | 0 | 0 | 0 | 0 | 0 |
| 0 | NA | NA | NA | NA | NA | Gm28460 | | 0 | 0 | 0 | 0 | 0 | 0 | 0 | 0 | 0 | 0 | 0 |
| 0 | NA | NA | NA | NA | NA | Gm28851 | | 0 | 0 | 0 | 0 | 0 | 0 | 0 | 0 | 0 | 0 | 0 |
| 0 | NA | NA | NA | NA | NA | Gm28456 | | 0 | 0 | 0 | 0 | 0 | 0 | 0 | 0 | 0 | 0 | 0 |
| 0 | NA | NA | NA | NA | NA | Gm29073 | | 0 | 0 | 0 | 0 | 0 | 0 | 0 | 0 | 0 | 0 | 0 |
| 0 | NA | NA | NA | NA | NA | Gm29648 | | 0 | 0 | 0 | 0 | 0 | 0 | 0 | 0 | 0 | 0 | 0 |
| 0 | NA | NA | NA | NA | NA | Gm29647 | | 0 | 0 | 0 | 0 | 0 | 0 | 0 | 0 | 0 | 0 | 0 |
| 0 | NA | NA | NA | NA | NA | Gm29646 | | 0 | 0 | 0 | 0 | 0 | 0 | 0 | 0 | 0 | 0 | 0 |
| 0 | NA | NA | NA | NA | NA | Gm28549 | | 0 | 0 | 0 | 0 | 0 | 0 | 0 | 0 | 0 | 0 | 0 |
| 0 | NA | NA | NA | NA | NA | Gm29211 | | 0 | 0 | 0 | 0 | 0 | 0 | 0 | 0 | 0 | 0 | 0 |
| 0 | NA | NA | NA | NA | NA | Gm29212 | | 0 | 0 | 0 | 0 | 0 | 0 | 0 | 0 | 0 | 0 | 0 |
| 0 | NA | NA | NA | NA | NA | Gm21247 | Y-linked testis-specific protein 1-like | 0 | 0 | 0 | 0 | 0 | 0 | 0 | 0 | 0 | 0 | 0 |
| 0 | NA | NA | NA | NA | NA | Gm29210 | | 0 | 0 | 0 | 0 | 0 | 0 | 0 | 0 | 0 | 0 | 0 |
| 0 | NA | NA | NA | NA | NA | Gm29213 | | 0 | 0 | 0 | 0 | 0 | 0 | 0 | 0 | 0 | 0 | 0 |
| 0 | NA | NA | NA | NA | NA | Gm28550 | | 0 | 0 | 0 | 0 | 0 | 0 | 0 | 0 | 0 | 0 | 0 |
| 0 | NA | NA | NA | NA | NA | Gm28604 | | 0 | 0 | 0 | 0 | 0 | 0 | 0 | 0 | 0 | 0 | 0 |
| 0 | NA | NA | NA | NA | NA | Gm28603 | | 0 | 0 | 0 | 0 | 0 | 0 | 0 | 0 | 0 | 0 | 0 |
| 0 | NA | NA | NA | NA | NA | Gm28565 | | 0 | 0 | 0 | 0 | 0 | 0 | 0 | 0 | 0 | 0 | 0 |
| 0 | NA | NA | NA | NA | NA | Gm28566 | | 0 | 0 | 0 | 0 | 0 | 0 | 0 | 0 | 0 | 0 | 0 |
| 0 | NA | NA | NA | NA | NA | Gm28697 | | 0 | 0 | 0 | 0 | 0 | 0 | 0 | 0 | 0 | 0 | 0 |
| 0 | NA | NA | NA | NA | NA | Gm28554 | | 0 | 0 | 0 | 0 | 0 | 0 | 0 | 0 | 0 | 0 | 0 |
| 0 | NA | NA | NA | NA | NA | Gm21256 | | 0 | 0 | 0 | 0 | 0 | 0 | 0 | 0 | 0 | 0 | 0 |
| 0 | NA | NA | NA | NA | NA | Gm29056 | | 0 | 0 | 0 | 0 | 0 | 0 | 0 | 0 | 0 | 0 | 0 |
| 0 | NA | NA | NA | NA | NA | Gm29098 | | 0 | 0 | 0 | 0 | 0 | 0 | 0 | 0 | 0 | 0 | 0 |
| 0 | NA | NA | NA | NA | NA | Gm21170 | | 0 | 0 | 0 | 0 | 0 | 0 | 0 | 0 | 0 | 0 | 0 |
| 0 | NA | NA | NA | NA | NA | Gm29373 | | 0 | 0 | 0 | 0 | 0 | 0 | 0 | 0 | 0 | 0 | 0 |
| 0 | NA | NA | NA | NA | NA | Gm29370 | | 0 | 0 | 0 | 0 | 0 | 0 | 0 | 0 | 0 | 0 | 0 |
| 0 | NA | NA | NA | NA | NA | Gm21751 | | 0 | 0 | 0 | 0 | 0 | 0 | 0 | 0 | 0 | 0 | 0 |
| 0 | NA | NA | NA | NA | NA | Gm29166 | | 0 | 0 | 0 | 0 | 0 | 0 | 0 | 0 | 0 | 0 | 0 |
| 0 | NA | NA | NA | NA | NA | Gm28194 | | 0 | 0 | 0 | 0 | 0 | 0 | 0 | 0 | 0 | 0 | 0 |
| 0 | NA | NA | NA | NA | NA | Gm20792 | | 0 | 0 | 0 | 0 | 0 | 0 | 0 | 0 | 0 | 0 | 0 |
| 0 | NA | NA | NA | NA | NA | Gm28754 | | 0 | 0 | 0 | 0 | 0 | 0 | 0 | 0 | 0 | 0 | 0 |
| 0 | NA | NA | NA | NA | NA | Gm28753 | | 0 | 0 | 0 | 0 | 0 | 0 | 0 | 0 | 0 | 0 | 0 |
| 0 | NA | NA | NA | NA | NA | Gm21191 | | 0 | 0 | 0 | 0 | 0 | 0 | 0 | 0 | 0 | 0 | 0 |
| 0 | NA | NA | NA | NA | NA | Gm21271 | | 0 | 0 | 0 | 0 | 0 | 0 | 0 | 0 | 0 | 0 | 0 |
| 0 | NA | NA | NA | NA | NA | Gm21281 | Y-linked testis-specific protein 1-like | 0 | 0 | 0 | 0 | 0 | 0 | 0 | 0 | 0 | 0 | 0 |
| 0 | NA | NA | NA | NA | NA | Gm28193 | | 0 | 0 | 0 | 0 | 0 | 0 | 0 | 0 | 0 | 0 | 0 |
| 0 | NA | NA | NA | NA | NA | Gm20802 | | 0 | 0 | 0 | 0 | 0 | 0 | 0 | 0 | 0 | 0 | 0 |
| 0 | NA | NA | NA | NA | NA | Gm29070 | | 0 | 0 | 0 | 0 | 0 | 0 | 0 | 0 | 0 | 0 | 0 |
| 0 | NA | NA | NA | NA | NA | Gm28553 | predicted gene 28553 | 0 | 0 | 0 | 0 | 0 | 0 | 0 | 0 | 0 | 0 | 0 |
| 0 | NA | NA | NA | NA | NA | Gm28751 | | 0 | 0 | 0 | 0 | 0 | 0 | 0 | 0 | 0 | 0 | 0 |
| 0 | NA | NA | NA | NA | NA | Gm28752 | | 0 | 0 | 0 | 0 | 0 | 0 | 0 | 0 | 0 | 0 | 0 |
| 0 | NA | NA | NA | NA | NA | Gm28726 | | 0 | 0 | 0 | 0 | 0 | 0 | 0 | 0 | 0 | 0 | 0 |
| 0 | NA | NA | NA | NA | NA | Gm21308 | | 0 | 0 | 0 | 0 | 0 | 0 | 0 | 0 | 0 | 0 | 0 |
| 0 | NA | NA | NA | NA | NA | Gm29267 | | 0 | 0 | 0 | 0 | 0 | 0 | 0 | 0 | 0 | 0 | 0 |
| 0 | NA | NA | NA | NA | NA | Gm29268 | | 0 | 0 | 0 | 0 | 0 | 0 | 0 | 0 | 0 | 0 | 0 |
| 0 | NA | NA | NA | NA | NA | Gm29269 | | 0 | 0 | 0 | 0 | 0 | 0 | 0 | 0 | 0 | 0 | 0 |
| 0 | NA | NA | NA | NA | NA | Gm29270 | | 0 | 0 | 0 | 0 | 0 | 0 | 0 | 0 | 0 | 0 | 0 |
| 0 | NA | NA | NA | NA | NA | Gm29271 | | 0 | 0 | 0 | 0 | 0 | 0 | 0 | 0 | 0 | 0 | 0 |
| 0 | NA | NA | NA | NA | NA | Gm21209 | predicted gene, 21209 | 0 | 0 | 0 | 0 | 0 | 0 | 0 | 0 | 0 | 0 | 0 |
| 0 | NA | NA | NA | NA | NA | Gm28682 | | 0 | 0 | 0 | 0 | 0 | 0 | 0 | 0 | 0 | 0 | 0 |
| 0 | NA | NA | NA | NA | NA | Gm28569 | | 0 | 0 | 0 | 0 | 0 | 0 | 0 | 0 | 0 | 0 | 0 |
| 0 | NA | NA | NA | NA | NA | Gm28681 | | 0 | 0 | 0 | 0 | 0 | 0 | 0 | 0 | 0 | 0 | 0 |
| 0 | NA | NA | NA | NA | NA | Gm28684 | | 0 | 0 | 0 | 0 | 0 | 0 | 0 | 0 | 0 | 0 | 0 |
| 0 | NA | NA | NA | NA | NA | Gm21316 | | 0 | 0 | 0 | 0 | 0 | 0 | 0 | 0 | 0 | 0 | 0 |

| | | | | | | | | | | | | | | | | | | |
|---|---|---|---|---|---|---|---|---|---|---|---|---|---|---|---|---|---|---|
| 0 | NA | NA | NA | NA | NA | Gm21923 | | 0 | 0 | 0 | 0 | 0 | 0 | 0 | 0 | 0 | 0 | 0 |
| 0 | NA | NA | NA | NA | NA | Gm28683 | | 0 | 0 | 0 | 0 | 0 | 0 | 0 | 0 | 0 | 0 | 0 |
| 0 | NA | NA | NA | NA | NA | Gm28680 | | 0 | 0 | 0 | 0 | 0 | 0 | 0 | 0 | 0 | 0 | 0 |
| 0 | NA | NA | NA | NA | NA | Gm28679 | | 0 | 0 | 0 | 0 | 0 | 0 | 0 | 0 | 0 | 0 | 0 |
| 0 | NA | NA | NA | NA | NA | Gm28870 | | 0 | 0 | 0 | 0 | 0 | 0 | 0 | 0 | 0 | 0 | 0 |
| 0 | NA | NA | NA | NA | NA | Gm28871 | | 0 | 0 | 0 | 0 | 0 | 0 | 0 | 0 | 0 | 0 | 0 |
| 0 | NA | NA | NA | NA | NA | Gm29512 | | 0 | 0 | 0 | 0 | 0 | 0 | 0 | 0 | 0 | 0 | 0 |
| 0 | NA | NA | NA | NA | NA | Gm29511 | | 0 | 0 | 0 | 0 | 0 | 0 | 0 | 0 | 0 | 0 | 0 |
| 0 | NA | NA | NA | NA | NA | Gm20880 | predicted gene, 20880 | 0 | 0 | 0 | 0 | 0 | 0 | 0 | 0 | 0 | 0 | 0 |
| 0 | NA | NA | NA | NA | NA | Gm20881 | predicted gene, 20881 | 0 | 0 | 0 | 0 | 0 | 0 | 0 | 0 | 0 | 0 | 0 |
| 0 | NA | NA | NA | NA | NA | Gm29121 | | 0 | 0 | 0 | 0 | 0 | 0 | 0 | 0 | 0 | 0 | 0 |
| 0 | NA | NA | NA | NA | NA | Gm29120 | | 0 | 0 | 0 | 0 | 0 | 0 | 0 | 0 | 0 | 0 | 0 |
| 0 | NA | NA | NA | NA | NA | Gm29117 | | 0 | 0 | 0 | 0 | 0 | 0 | 0 | 0 | 0 | 0 | 0 |
| 0 | NA | NA | NA | NA | NA | Gm20883 | X-linked lymphocyte-regulated protein PM | 0 | 0 | 0 | 0 | 0 | 0 | 0 | 0 | 0 | 0 | 0 |
| 0 | NA | NA | NA | NA | NA | Gm29119 | | 0 | 0 | 0 | 0 | 0 | 0 | 0 | 0 | 0 | 0 | 0 |
| 0 | NA | NA | NA | NA | NA | Gm29118 | | 0 | 0 | 0 | 0 | 0 | 0 | 0 | 0 | 0 | 0 | 0 |
| 0 | NA | NA | NA | NA | NA | Gm29116 | | 0 | 0 | 0 | 0 | 0 | 0 | 0 | 0 | 0 | 0 | 0 |
| 0 | NA | NA | NA | NA | NA | Gm20889 | predicted gene, 20889 | 0 | 0 | 0 | 0 | 0 | 0 | 0 | 0 | 0 | 0 | 0 |
| 0 | NA | NA | NA | NA | NA | Gm20887 | | 0 | 0 | 0 | 0 | 0 | 0 | 0 | 0 | 0 | 0 | 0 |
| 0 | NA | NA | NA | NA | NA | Gm29320 | | 0 | 0 | 0 | 0 | 0 | 0 | 0 | 0 | 0 | 0 | 0 |
| 0 | NA | NA | NA | NA | NA | Gm21849 | | 0 | 0 | 0 | 0 | 0 | 0 | 0 | 0 | 0 | 0 | 0 |
| 0 | NA | NA | NA | NA | NA | Gm29321 | | 0 | 0 | 0 | 0 | 0 | 0 | 0 | 0 | 0 | 0 | 0 |
| 0 | NA | NA | NA | NA | NA | Gm21117 | predicted gene, 21117 | 0 | 0 | 0 | 0 | 0 | 0 | 0 | 0 | 0 | 0 | 0 |
| 0 | NA | NA | NA | NA | NA | Gm29319 | | 0 | 0 | 0 | 0 | 0 | 0 | 0 | 0 | 0 | 0 | 0 |
| 0 | NA | NA | NA | NA | NA | Gm28317 | | 0 | 0 | 0 | 0 | 0 | 0 | 0 | 0 | 0 | 0 | 0 |
| 0 | NA | NA | NA | NA | NA | Gm28318 | | 0 | 0 | 0 | 0 | 0 | 0 | 0 | 0 | 0 | 0 | 0 |
| 0 | NA | NA | NA | NA | NA | Gm20884 | predicted gene, 20884 | 0 | 0 | 0 | 0 | 0 | 0 | 0 | 0 | 0 | 0 | 0 |
| 0 | NA | NA | NA | NA | NA | Gm28313 | | 0 | 0 | 0 | 0 | 0 | 0 | 0 | 0 | 0 | 0 | 0 |
| 0 | NA | NA | NA | NA | NA | Gm28314 | | 0 | 0 | 0 | 0 | 0 | 0 | 0 | 0 | 0 | 0 | 0 |
| 0 | NA | NA | NA | NA | NA | Gm28315 | | 0 | 0 | 0 | 0 | 0 | 0 | 0 | 0 | 0 | 0 | 0 |
| 0 | NA | NA | NA | NA | NA | Gm28316 | | 0 | 0 | 0 | 0 | 0 | 0 | 0 | 0 | 0 | 0 | 0 |
| 0 | NA | NA | NA | NA | NA | Gm21387 | predicted gene, 21387 | 0 | 0 | 0 | 0 | 0 | 0 | 0 | 0 | 0 | 0 | 0 |
| 0 | NA | NA | NA | NA | NA | Gm21396 | predicted gene, 21396 | 0 | 0 | 0 | 0 | 0 | 0 | 0 | 0 | 0 | 0 | 0 |
| 0 | NA | NA | NA | NA | NA | Gm28944 | | 0 | 0 | 0 | 0 | 0 | 0 | 0 | 0 | 0 | 0 | 0 |
| 0 | NA | NA | NA | NA | NA | Gm28943 | | 0 | 0 | 0 | 0 | 0 | 0 | 0 | 0 | 0 | 0 | 0 |
| 0 | NA | NA | NA | NA | NA | Gm28946 | | 0 | 0 | 0 | 0 | 0 | 0 | 0 | 0 | 0 | 0 | 0 |
| 0 | NA | NA | NA | NA | NA | Gm28945 | | 0 | 0 | 0 | 0 | 0 | 0 | 0 | 0 | 0 | 0 | 0 |
| 0 | NA | NA | NA | NA | NA | Gm28947 | | 0 | 0 | 0 | 0 | 0 | 0 | 0 | 0 | 0 | 0 | 0 |
| 0 | NA | NA | NA | NA | NA | Gm20920 | X-linked lymphocyte-regulated protein PM | 0 | 0 | 0 | 0 | 0 | 0 | 0 | 0 | 0 | 0 | 0 |
| 0 | NA | NA | NA | NA | NA | Gm28949 | | 0 | 0 | 0 | 0 | 0 | 0 | 0 | 0 | 0 | 0 | 0 |
| 0 | NA | NA | NA | NA | NA | Gm28948 | | 0 | 0 | 0 | 0 | 0 | 0 | 0 | 0 | 0 | 0 | 0 |
| 0 | NA | NA | NA | NA | NA | Gm28950 | | 0 | 0 | 0 | 0 | 0 | 0 | 0 | 0 | 0 | 0 | 0 |
| 0 | NA | NA | NA | NA | NA | Gm21245 | predicted gene, 21245 | 0 | 0 | 0 | 0 | 0 | 0 | 0 | 0 | 0 | 0 | 0 |
| 0 | NA | NA | NA | NA | NA | Gm29169 | | 0 | 0 | 0 | 0 | 0 | 0 | 0 | 0 | 0 | 0 | 0 |
| 0 | NA | NA | NA | NA | NA | Gm21899 | | 0 | 0 | 0 | 0 | 0 | 0 | 0 | 0 | 0 | 0 | 0 |
| 0 | NA | NA | NA | NA | NA | Gm29063 | | 0 | 0 | 0 | 0 | 0 | 0 | 0 | 0 | 0 | 0 | 0 |
| 0 | NA | NA | NA | NA | NA | Gm28737 | | 0 | 0 | 0 | 0 | 0 | 0 | 0 | 0 | 0 | 0 | 0 |
| 0 | NA | NA | NA | NA | NA | Gm28738 | | 0 | 0 | 0 | 0 | 0 | 0 | 0 | 0 | 0 | 0 | 0 |
| 0 | NA | NA | NA | NA | NA | Gm20923 | | 0 | 0 | 0 | 0 | 0 | 0 | 0 | 0 | 0 | 0 | 0 |
| 0 | NA | NA | NA | NA | NA | Gm28197 | | 0 | 0 | 0 | 0 | 0 | 0 | 0 | 0 | 0 | 0 | 0 |
| 0 | NA | NA | NA | NA | NA | Gm21249 | Y-linked testis-specific protein 1-like | 0 | 0 | 0 | 0 | 0 | 0 | 0 | 0 | 0 | 0 | 0 |
| 0 | NA | NA | NA | NA | NA | Gm28736 | | 0 | 0 | 0 | 0 | 0 | 0 | 0 | 0 | 0 | 0 | 0 |
| 0 | NA | NA | NA | NA | NA | Gm21840 | | 0 | 0 | 0 | 0 | 0 | 0 | 0 | 0 | 0 | 0 | 0 |
| 0 | NA | NA | NA | NA | NA | Gm28735 | | 0 | 0 | 0 | 0 | 0 | 0 | 0 | 0 | 0 | 0 | 0 |
| 0 | NA | NA | NA | NA | NA | Gm21258 | | 0 | 0 | 0 | 0 | 0 | 0 | 0 | 0 | 0 | 0 | 0 |
| 0 | NA | NA | NA | NA | NA | Gm28734 | | 0 | 0 | 0 | 0 | 0 | 0 | 0 | 0 | 0 | 0 | 0 |
| 0 | NA | NA | NA | NA | NA | Gm28232 | | 0 | 0 | 0 | 0 | 0 | 0 | 0 | 0 | 0 | 0 | 0 |
| 0 | NA | NA | NA | NA | NA | Gm28732 | | 0 | 0 | 0 | 0 | 0 | 0 | 0 | 0 | 0 | 0 | 0 |
| 0 | NA | NA | NA | NA | NA | Gm28733 | | 0 | 0 | 0 | 0 | 0 | 0 | 0 | 0 | 0 | 0 | 0 |
| 0 | NA | NA | NA | NA | NA | Gm21268 | predicted gene, 21268 | 0 | 0 | 0 | 0 | 0 | 0 | 0 | 0 | 0 | 0 | 0 |

| | | | | | | | | | | | | | | | | | | |
|---|---|---|---|---|---|---|---|---|---|---|---|---|---|---|---|---|---|---|
| 0 | NA | NA | NA | NA | NA | Gm21275 | predicted gene, 21275 | 0 | 0 | 0 | 0 | 0 | 0 | 0 | 0 | 0 | 0 | 0 |
| 0 | NA | NA | NA | NA | NA | Gm28233 | | 0 | 0 | 0 | 0 | 0 | 0 | 0 | 0 | 0 | 0 | 0 |
| 0 | NA | NA | NA | NA | NA | Gm28235 | | 0 | 0 | 0 | 0 | 0 | 0 | 0 | 0 | 0 | 0 | 0 |
| 0 | NA | NA | NA | NA | NA | Gm28234 | | 0 | 0 | 0 | 0 | 0 | 0 | 0 | 0 | 0 | 0 | 0 |
| 0 | NA | NA | NA | NA | NA | Gm21902 | | 0 | 0 | 0 | 0 | 0 | 0 | 0 | 0 | 0 | 0 | 0 |
| 0 | NA | NA | NA | NA | NA | Gm28236 | | 0 | 0 | 0 | 0 | 0 | 0 | 0 | 0 | 0 | 0 | 0 |
| 0 | NA | NA | NA | NA | NA | Gm21285 | predicted gene, 21285 | 0 | 0 | 0 | 0 | 0 | 0 | 0 | 0 | 0 | 0 | 0 |
| 0 | NA | NA | NA | NA | NA | Gm20747 | predicted gene, 20747 | 0 | 0 | 0 | 0 | 0 | 0 | 0 | 0 | 0 | 0 | 0 |
| 0 | NA | NA | NA | NA | NA | Gm29250 | | 0 | 0 | 0 | 0 | 0 | 0 | 0 | 0 | 0 | 0 | 0 |
| 0 | NA | NA | NA | NA | NA | Gm29251 | | 0 | 0 | 0 | 0 | 0 | 0 | 0 | 0 | 0 | 0 | 0 |
| 0 | NA | NA | NA | NA | NA | Gm21906 | | 0 | 0 | 0 | 0 | 0 | 0 | 0 | 0 | 0 | 0 | 0 |
| 0 | NA | NA | NA | NA | NA | Gm29252 | | 0 | 0 | 0 | 0 | 0 | 0 | 0 | 0 | 0 | 0 | 0 |
| 0 | NA | NA | NA | NA | NA | Gm29248 | | 0 | 0 | 0 | 0 | 0 | 0 | 0 | 0 | 0 | 0 | 0 |
| 0 | NA | NA | NA | NA | NA | Gm29249 | | 0 | 0 | 0 | 0 | 0 | 0 | 0 | 0 | 0 | 0 | 0 |
| 0 | NA | NA | NA | NA | NA | Gm29484 | | 0 | 0 | 0 | 0 | 0 | 0 | 0 | 0 | 0 | 0 | 0 |
| 0 | NA | NA | NA | NA | NA | Gm21920 | | 0 | 0 | 0 | 0 | 0 | 0 | 0 | 0 | 0 | 0 | 0 |
| 0 | NA | NA | NA | NA | NA | Gm28908 | | 0 | 0 | 0 | 0 | 0 | 0 | 0 | 0 | 0 | 0 | 0 |
| 0 | NA | NA | NA | NA | NA | Gm28907 | | 0 | 0 | 0 | 0 | 0 | 0 | 0 | 0 | 0 | 0 | 0 |
| 0 | NA | NA | NA | NA | NA | Gm21749 | | 0 | 0 | 0 | 0 | 0 | 0 | 0 | 0 | 0 | 0 | 0 |
| 0 | NA | NA | NA | NA | NA | Gm20929 | predicted gene, 20929 | 0 | 0 | 0 | 0 | 0 | 0 | 0 | 0 | 0 | 0 | 0 |
| 0 | NA | NA | NA | NA | NA | Gm28910 | | 0 | 0 | 0 | 0 | 0 | 0 | 0 | 0 | 0 | 0 | 0 |
| 0 | NA | NA | NA | NA | NA | Gm28909 | | 0 | 0 | 0 | 0 | 0 | 0 | 0 | 0 | 0 | 0 | 0 |
| 0 | NA | NA | NA | NA | NA | Gm28276 | | 0 | 0 | 0 | 0 | 0 | 0 | 0 | 0 | 0 | 0 | 0 |
| 0 | NA | NA | NA | NA | NA | Gm28274 | | 0 | 0 | 0 | 0 | 0 | 0 | 0 | 0 | 0 | 0 | 0 |
| 0 | NA | NA | NA | NA | NA | Gm28275 | | 0 | 0 | 0 | 0 | 0 | 0 | 0 | 0 | 0 | 0 | 0 |
| 0 | NA | NA | NA | NA | NA | Gm21302 | | 0 | 0 | 0 | 0 | 0 | 0 | 0 | 0 | 0 | 0 | 0 |
| 0 | NA | NA | NA | NA | NA | Gm28290 | | 0 | 0 | 0 | 0 | 0 | 0 | 0 | 0 | 0 | 0 | 0 |
| 0 | NA | NA | NA | NA | NA | Gm28292 | | 0 | 0 | 0 | 0 | 0 | 0 | 0 | 0 | 0 | 0 | 0 |
| 0 | NA | NA | NA | NA | NA | Gm28291 | | 0 | 0 | 0 | 0 | 0 | 0 | 0 | 0 | 0 | 0 | 0 |
| 0 | NA | NA | NA | NA | NA | Gm28293 | | 0 | 0 | 0 | 0 | 0 | 0 | 0 | 0 | 0 | 0 | 0 |
| 0 | NA | NA | NA | NA | NA | Gm21740 | | 0 | 0 | 0 | 0 | 0 | 0 | 0 | 0 | 0 | 0 | 0 |
| 0 | NA | NA | NA | NA | NA | Gm21912 | | 0 | 0 | 0 | 0 | 0 | 0 | 0 | 0 | 0 | 0 | 0 |
| 0 | NA | NA | NA | NA | NA | Gm28698 | | 0 | 0 | 0 | 0 | 0 | 0 | 0 | 0 | 0 | 0 | 0 |
| 0 | NA | NA | NA | NA | NA | Gm29304 | | 0 | 0 | 0 | 0 | 0 | 0 | 0 | 0 | 0 | 0 | 0 |
| 0 | NA | NA | NA | NA | NA | Gm29305 | | 0 | 0 | 0 | 0 | 0 | 0 | 0 | 0 | 0 | 0 | 0 |
| 0 | NA | NA | NA | NA | NA | Gm29303 | | 0 | 0 | 0 | 0 | 0 | 0 | 0 | 0 | 0 | 0 | 0 |
| 0 | NA | NA | NA | NA | NA | Gm21894 | | 0 | 0 | 0 | 0 | 0 | 0 | 0 | 0 | 0 | 0 | 0 |
| 0 | NA | NA | NA | NA | NA | Gm29307 | | 0 | 0 | 0 | 0 | 0 | 0 | 0 | 0 | 0 | 0 | 0 |
| 0 | NA | NA | NA | NA | NA | Gm29557 | | 0 | 0 | 0 | 0 | 0 | 0 | 0 | 0 | 0 | 0 | 0 |
| 0 | NA | NA | NA | NA | NA | Gm23052 | | 0 | 0 | 0 | 0 | 0 | 0 | 0 | 0 | 0 | 0 | 0 |
| 0 | NA | NA | NA | NA | NA | Gm20927 | Y-linked testis-specific protein 1-like | 0 | 0 | 0 | 0 | 0 | 0 | 0 | 0 | 0 | 0 | 0 |
| 0 | NA | NA | NA | NA | NA | Gm20930 | predicted gene, 20930 | 0 | 0 | 0 | 0 | 0 | 0 | 0 | 0 | 0 | 0 | 0 |
| 0 | NA | NA | NA | NA | NA | Gm29306 | | 0 | 0 | 0 | 0 | 0 | 0 | 0 | 0 | 0 | 0 | 0 |
| 0 | NA | NA | NA | NA | NA | Gm29300 | | 0 | 0 | 0 | 0 | 0 | 0 | 0 | 0 | 0 | 0 | 0 |
| 0 | NA | NA | NA | NA | NA | Gm29301 | | 0 | 0 | 0 | 0 | 0 | 0 | 0 | 0 | 0 | 0 | 0 |
| 0 | NA | NA | NA | NA | NA | Gm20931 | predicted gene, 20931 | 0 | 0 | 0 | 0 | 0 | 0 | 0 | 0 | 0 | 0 | 0 |
| 0 | NA | NA | NA | NA | NA | Gm29559 | | 0 | 0 | 0 | 0 | 0 | 0 | 0 | 0 | 0 | 0 | 0 |
| 0 | NA | NA | NA | NA | NA | Gm29558 | | 0 | 0 | 0 | 0 | 0 | 0 | 0 | 0 | 0 | 0 | 0 |
| 0 | NA | NA | NA | NA | NA | Gm29555 | | 0 | 0 | 0 | 0 | 0 | 0 | 0 | 0 | 0 | 0 | 0 |
| 0 | NA | NA | NA | NA | NA | Gm21733 | | 0 | 0 | 0 | 0 | 0 | 0 | 0 | 0 | 0 | 0 | 0 |
| 0 | NA | NA | NA | NA | NA | Gm21330 | | 0 | 0 | 0 | 0 | 0 | 0 | 0 | 0 | 0 | 0 | 0 |
| 0 | NA | NA | NA | NA | NA | Gm28355 | | 0 | 0 | 0 | 0 | 0 | 0 | 0 | 0 | 0 | 0 | 0 |
| 0 | NA | NA | NA | NA | NA | Gm28354 | | 0 | 0 | 0 | 0 | 0 | 0 | 0 | 0 | 0 | 0 | 0 |
| 0 | NA | NA | NA | NA | NA | Gm28353 | | 0 | 0 | 0 | 0 | 0 | 0 | 0 | 0 | 0 | 0 | 0 |
| 0 | NA | NA | NA | NA | NA | Gm28352 | | 0 | 0 | 0 | 0 | 0 | 0 | 0 | 0 | 0 | 0 | 0 |
| 0 | NA | NA | NA | NA | NA | Gm21858 | | 0 | 0 | 0 | 0 | 0 | 0 | 0 | 0 | 0 | 0 | 0 |
| 0 | NA | NA | NA | NA | NA | Gm29244 | | 0 | 0 | 0 | 0 | 0 | 0 | 0 | 0 | 0 | 0 | 0 |
| 0 | NA | NA | NA | NA | NA | Gm28200 | | 0 | 0 | 0 | 0 | 0 | 0 | 0 | 0 | 0 | 0 | 0 |
| 0 | NA | NA | NA | NA | NA | Gm28201 | | 0 | 0 | 0 | 0 | 0 | 0 | 0 | 0 | 0 | 0 | 0 |
| 0 | NA | NA | NA | NA | NA | Gm28202 | | 0 | 0 | 0 | 0 | 0 | 0 | 0 | 0 | 0 | 0 | 0 |

| | | | | | | | | | | | | | | | | | | |
|---|---|---|---|---|---|---|---|---|---|---|---|---|---|---|---|---|---|---|
| 0 | NA | NA | NA | NA | NA | Gm28203 | | 0 | 0 | 0 | 0 | 0 | 0 | 0 | 0 | 0 | 0 | 0 |
| 0 | NA | NA | NA | NA | NA | Gm21340 | predicted gene, 21340 | 0 | 0 | 0 | 0 | 0 | 0 | 0 | 0 | 0 | 0 | 0 |
| 0 | NA | NA | NA | NA | NA | Gm28204 | | 0 | 0 | 0 | 0 | 0 | 0 | 0 | 0 | 0 | 0 | 0 |
| 0 | NA | NA | NA | NA | NA | Gm28205 | | 0 | 0 | 0 | 0 | 0 | 0 | 0 | 0 | 0 | 0 | 0 |
| 0 | NA | NA | NA | NA | NA | Gm28206 | | 0 | 0 | 0 | 0 | 0 | 0 | 0 | 0 | 0 | 0 | 0 |
| 0 | NA | NA | NA | NA | NA | Gm28207 | | 0 | 0 | 0 | 0 | 0 | 0 | 0 | 0 | 0 | 0 | 0 |
| 0 | NA | NA | NA | NA | NA | Gm28208 | | 0 | 0 | 0 | 0 | 0 | 0 | 0 | 0 | 0 | 0 | 0 |
| 0 | NA | NA | NA | NA | NA | Gm21851 | | 0 | 0 | 0 | 0 | 0 | 0 | 0 | 0 | 0 | 0 | 0 |
| 0 | NA | NA | NA | NA | NA | Gm21819 | | 0 | 0 | 0 | 0 | 0 | 0 | 0 | 0 | 0 | 0 | 0 |
| 0 | NA | NA | NA | NA | NA | Gm29224 | | 0 | 0 | 0 | 0 | 0 | 0 | 0 | 0 | 0 | 0 | 0 |
| 0 | NA | NA | NA | NA | NA | Gm29223 | | 0 | 0 | 0 | 0 | 0 | 0 | 0 | 0 | 0 | 0 | 0 |
| 0 | NA | NA | NA | NA | NA | Gm29222 | | 0 | 0 | 0 | 0 | 0 | 0 | 0 | 0 | 0 | 0 | 0 |
| 0 | NA | NA | NA | NA | NA | Gm29221 | | 0 | 0 | 0 | 0 | 0 | 0 | 0 | 0 | 0 | 0 | 0 |
| 0 | NA | NA | NA | NA | NA | Gm29220 | | 0 | 0 | 0 | 0 | 0 | 0 | 0 | 0 | 0 | 0 | 0 |
| 0 | NA | NA | NA | NA | NA | Gm21815 | | 0 | 0 | 0 | 0 | 0 | 0 | 0 | 0 | 0 | 0 | 0 |
| 0 | NA | NA | NA | NA | NA | Gm29407 | | 0 | 0 | 0 | 0 | 0 | 0 | 0 | 0 | 0 | 0 | 0 |
| 0 | NA | NA | NA | NA | NA | Gm29405 | | 0 | 0 | 0 | 0 | 0 | 0 | 0 | 0 | 0 | 0 | 0 |
| 0 | NA | NA | NA | NA | NA | Gm29406 | | 0 | 0 | 0 | 0 | 0 | 0 | 0 | 0 | 0 | 0 | 0 |
| 0 | NA | NA | NA | NA | NA | Gm20913 | predicted gene, 20913 | 0 | 0 | 0 | 0 | 0 | 0 | 0 | 0 | 0 | 0 | 0 |
| 0 | NA | NA | NA | NA | NA | Gm29404 | | 0 | 0 | 0 | 0 | 0 | 0 | 0 | 0 | 0 | 0 | 0 |
| 0 | NA | NA | NA | NA | NA | Gm29403 | | 0 | 0 | 0 | 0 | 0 | 0 | 0 | 0 | 0 | 0 | 0 |
| 0 | NA | NA | NA | NA | NA | Gm28350 | | 0 | 0 | 0 | 0 | 0 | 0 | 0 | 0 | 0 | 0 | 0 |
| 0 | NA | NA | NA | NA | NA | Gm28964 | | 0 | 0 | 0 | 0 | 0 | 0 | 0 | 0 | 0 | 0 | 0 |
| 0 | NA | NA | NA | NA | NA | Sly | Sycp3 like Y-linked | 0 | 0 | 0 | 0 | 0 | 0 | 0 | 0 | 0 | 0 | 0 |
| 0 | NA | NA | NA | NA | NA | Gm28165 | | 0 | 0 | 0 | 0 | 0 | 0 | 0 | 0 | 0 | 0 | 0 |
| 0 | NA | NA | NA | NA | NA | Gm28709 | | 0 | 0 | 0 | 0 | 0 | 0 | 0 | 0 | 0 | 0 | 0 |
| 0 | NA | NA | NA | NA | NA | Gm28965 | | 0 | 0 | 0 | 0 | 0 | 0 | 0 | 0 | 0 | 0 | 0 |
| 0 | NA | NA | NA | NA | NA | Gm21868 | | 0 | 0 | 0 | 0 | 0 | 0 | 0 | 0 | 0 | 0 | 0 |
| 0 | NA | NA | NA | NA | NA | Gm21943 | predicted gene, 21943 | 0 | 0 | 0 | 0 | 0 | 0 | 0 | 0 | 0 | 0 | 0 |
| 0 | NA | NA | NA | NA | NA | Gm28532 | | 0 | 0 | 0 | 0 | 0 | 0 | 0 | 0 | 0 | 0 | 0 |
| 0 | NA | NA | NA | NA | NA | Gm28533 | | 0 | 0 | 0 | 0 | 0 | 0 | 0 | 0 | 0 | 0 | 0 |
| 0 | NA | NA | NA | NA | NA | Gm28966 | | 0 | 0 | 0 | 0 | 0 | 0 | 0 | 0 | 0 | 0 | 0 |
| 0 | NA | NA | NA | NA | NA | Gm28961 | | 0 | 0 | 0 | 0 | 0 | 0 | 0 | 0 | 0 | 0 | 0 |
| 0 | NA | NA | NA | NA | NA | Gm28962 | | 0 | 0 | 0 | 0 | 0 | 0 | 0 | 0 | 0 | 0 | 0 |
| 0 | NA | NA | NA | NA | NA | Gm28963 | | 0 | 0 | 0 | 0 | 0 | 0 | 0 | 0 | 0 | 0 | 0 |
| 0 | NA | NA | NA | NA | NA | Gm29466 | | 0 | 0 | 0 | 0 | 0 | 0 | 0 | 0 | 0 | 0 | 0 |
| 0 | NA | NA | NA | NA | NA | Gm29467 | | 0 | 0 | 0 | 0 | 0 | 0 | 0 | 0 | 0 | 0 | 0 |
| 0 | NA | NA | NA | NA | NA | Gm21724 | | 0 | 0 | 0 | 0 | 0 | 0 | 0 | 0 | 0 | 0 | 0 |
| 0 | NA | NA | NA | NA | NA | Gm20963 | | 0 | 0 | 0 | 0 | 0 | 0 | 0 | 0 | 0 | 0 | 0 |
| 0 | NA | NA | NA | NA | NA | Gm28326 | | 0 | 0 | 0 | 0 | 0 | 0 | 0 | 0 | 0 | 0 | 0 |
| 0 | NA | NA | NA | NA | NA | Gm28325 | | 0 | 0 | 0 | 0 | 0 | 0 | 0 | 0 | 0 | 0 | 0 |
| 0 | NA | NA | NA | NA | NA | Gm21405 | | 0 | 0 | 0 | 0 | 0 | 0 | 0 | 0 | 0 | 0 | 0 |
| 0 | NA | NA | NA | NA | NA | Gm28284 | | 0 | 0 | 0 | 0 | 0 | 0 | 0 | 0 | 0 | 0 | 0 |
| 0 | NA | NA | NA | NA | NA | Gm29368 | | 0 | 0 | 0 | 0 | 0 | 0 | 0 | 0 | 0 | 0 | 0 |
| 0 | NA | NA | NA | NA | NA | Gm21861 | | 0 | 0 | 0 | 0 | 0 | 0 | 0 | 0 | 0 | 0 | 0 |
| 0 | NA | NA | NA | NA | NA | Gm21413 | | 0 | 0 | 0 | 0 | 0 | 0 | 0 | 0 | 0 | 0 | 0 |
| 0 | NA | NA | NA | NA | NA | Gm28073 | | 0 | 0 | 0 | 0 | 0 | 0 | 0 | 0 | 0 | 0 | 0 |
| 0 | NA | NA | NA | NA | NA | Gm21419 | | 0 | 0 | 0 | 0 | 0 | 0 | 0 | 0 | 0 | 0 | 0 |
| 0 | NA | NA | NA | NA | NA | Gm28074 | | 0 | 0 | 0 | 0 | 0 | 0 | 0 | 0 | 0 | 0 | 0 |
| 0 | NA | NA | NA | NA | NA | Gm21428 | | 0 | 0 | 0 | 0 | 0 | 0 | 0 | 0 | 0 | 0 | 0 |
| 0 | NA | NA | NA | NA | NA | Gm28072 | | 0 | 0 | 0 | 0 | 0 | 0 | 0 | 0 | 0 | 0 | 0 |
| 0 | NA | NA | NA | NA | NA | Gm21758 | | 0 | 0 | 0 | 0 | 0 | 0 | 0 | 0 | 0 | 0 | 0 |
| 0 | NA | NA | NA | NA | NA | Gm29672 | | 0 | 0 | 0 | 0 | 0 | 0 | 0 | 0 | 0 | 0 | 0 |
| 0 | NA | NA | NA | NA | NA | Gm29131 | | 0 | 0 | 0 | 0 | 0 | 0 | 0 | 0 | 0 | 0 | 0 |
| 0 | NA | NA | NA | NA | NA | Gm20871 | | 0 | 0 | 0 | 0 | 0 | 0 | 0 | 0 | 0 | 0 | 0 |
| 0 | NA | NA | NA | NA | NA | Gm29130 | | 0 | 0 | 0 | 0 | 0 | 0 | 0 | 0 | 0 | 0 | 0 |
| 0 | NA | NA | NA | NA | NA | Gm20978 | | 0 | 0 | 0 | 0 | 0 | 0 | 0 | 0 | 0 | 0 | 0 |
| 0 | NA | NA | NA | NA | NA | Gm29132 | | 0 | 0 | 0 | 0 | 0 | 0 | 0 | 0 | 0 | 0 | 0 |
| 0 | NA | NA | NA | NA | NA | Gm21450 | | 0 | 0 | 0 | 0 | 0 | 0 | 0 | 0 | 0 | 0 | 0 |
| 0 | NA | NA | NA | NA | NA | Gm21843 | | 0 | 0 | 0 | 0 | 0 | 0 | 0 | 0 | 0 | 0 | 0 |

| | | | | | | | | | | | | | | | | | | |
|---|---|---|---|---|---|---|---|---|---|---|---|---|---|---|---|---|---|---|
| 0 | NA | NA | NA | NA | NA | Gm28889 | | 0 | 0 | 0 | 0 | 0 | 0 | 0 | 0 | 0 | 0 | 0 |
| 0 | NA | NA | NA | NA | NA | Gm28890 | | 0 | 0 | 0 | 0 | 0 | 0 | 0 | 0 | 0 | 0 | 0 |
| 0 | NA | NA | NA | NA | NA | Gm28891 | | 0 | 0 | 0 | 0 | 0 | 0 | 0 | 0 | 0 | 0 | 0 |
| 0 | NA | NA | NA | NA | NA | Gm28691 | | 0 | 0 | 0 | 0 | 0 | 0 | 0 | 0 | 0 | 0 | 0 |
| 0 | NA | NA | NA | NA | NA | Gm28692 | | 0 | 0 | 0 | 0 | 0 | 0 | 0 | 0 | 0 | 0 | 0 |
| 0 | NA | NA | NA | NA | NA | Gm20794 | | 0 | 0 | 0 | 0 | 0 | 0 | 0 | 0 | 0 | 0 | 0 |
| 0 | NA | NA | NA | NA | NA | Gm28690 | | 0 | 0 | 0 | 0 | 0 | 0 | 0 | 0 | 0 | 0 | 0 |
| 0 | NA | NA | NA | NA | NA | Gm28689 | | 0 | 0 | 0 | 0 | 0 | 0 | 0 | 0 | 0 | 0 | 0 |
| 0 | NA | NA | NA | NA | NA | Gm28687 | | 0 | 0 | 0 | 0 | 0 | 0 | 0 | 0 | 0 | 0 | 0 |
| 0 | NA | NA | NA | NA | NA | Gm20987 | | 0 | 0 | 0 | 0 | 0 | 0 | 0 | 0 | 0 | 0 | 0 |
| 0 | NA | NA | NA | NA | NA | Gm21918 | | 0 | 0 | 0 | 0 | 0 | 0 | 0 | 0 | 0 | 0 | 0 |
| 0 | NA | NA | NA | NA | NA | 1700040F15Rik | | 0 | 0 | 0 | 0 | 0 | 0 | 0 | 0 | 0 | 0 | 0 |
| 0 | NA | NA | NA | NA | NA | Gm29207 | | 0 | 0 | 0 | 0 | 0 | 0 | 0 | 0 | 0 | 0 | 0 |
| 0 | NA | NA | NA | NA | NA | Gm29204 | | 0 | 0 | 0 | 0 | 0 | 0 | 0 | 0 | 0 | 0 | 0 |
| 0 | NA | NA | NA | NA | NA | Gm21489 | | 0 | 0 | 0 | 0 | 0 | 0 | 0 | 0 | 0 | 0 | 0 |
| 0 | NA | NA | NA | NA | NA | Gm21832 | | 0 | 0 | 0 | 0 | 0 | 0 | 0 | 0 | 0 | 0 | 0 |
| 0 | NA | NA | NA | NA | NA | Gm21573 | Y-linked testis-specific protein 1-like | 0 | 0 | 0 | 0 | 0 | 0 | 0 | 0 | 0 | 0 | 0 |
| 0 | NA | NA | NA | NA | NA | Gm28179 | | 0 | 0 | 0 | 0 | 0 | 0 | 0 | 0 | 0 | 0 | 0 |
| 0 | NA | NA | NA | NA | NA | Gm21577 | | 0 | 0 | 0 | 0 | 0 | 0 | 0 | 0 | 0 | 0 | 0 |
| 0 | NA | NA | NA | NA | NA | Gm29206 | | 0 | 0 | 0 | 0 | 0 | 0 | 0 | 0 | 0 | 0 | 0 |
| 0 | NA | NA | NA | NA | NA | Gm29203 | | 0 | 0 | 0 | 0 | 0 | 0 | 0 | 0 | 0 | 0 | 0 |
| 0 | NA | NA | NA | NA | NA | Gm21497 | | 0 | 0 | 0 | 0 | 0 | 0 | 0 | 0 | 0 | 0 | 0 |
| 0 | NA | NA | NA | NA | NA | Gm29614 | | 0 | 0 | 0 | 0 | 0 | 0 | 0 | 0 | 0 | 0 | 0 |
| 0 | NA | NA | NA | NA | NA | Gm29613 | | 0 | 0 | 0 | 0 | 0 | 0 | 0 | 0 | 0 | 0 | 0 |
| 0 | NA | NA | NA | NA | NA | Gm29612 | | 0 | 0 | 0 | 0 | 0 | 0 | 0 | 0 | 0 | 0 | 0 |
| 0 | NA | NA | NA | NA | NA | Gm29343 | | 0 | 0 | 0 | 0 | 0 | 0 | 0 | 0 | 0 | 0 | 0 |
| 0 | NA | NA | NA | NA | NA | Gm29342 | | 0 | 0 | 0 | 0 | 0 | 0 | 0 | 0 | 0 | 0 | 0 |
| 0 | NA | NA | NA | NA | NA | Gm20803 | predicted gene, 20803 | 0 | 0 | 0 | 0 | 0 | 0 | 0 | 0 | 0 | 0 | 0 |
| 0 | NA | NA | NA | NA | NA | Gm20804 | predicted gene, 20804 | 0 | 0 | 0 | 0 | 0 | 0 | 0 | 0 | 0 | 0 | 0 |
| 0 | NA | NA | NA | NA | NA | Gm28598 | | 0 | 0 | 0 | 0 | 0 | 0 | 0 | 0 | 0 | 0 | 0 |
| 0 | NA | NA | NA | NA | NA | Gm28599 | | 0 | 0 | 0 | 0 | 0 | 0 | 0 | 0 | 0 | 0 | 0 |
| 0 | NA | NA | NA | NA | NA | Gm28600 | | 0 | 0 | 0 | 0 | 0 | 0 | 0 | 0 | 0 | 0 | 0 |
| 0 | NA | NA | NA | NA | NA | Gm21518 | predicted gene, 21518 | 0 | 0 | 0 | 0 | 0 | 0 | 0 | 0 | 0 | 0 | 0 |
| 0 | NA | NA | NA | NA | NA | Gm28185 | | 0 | 0 | 0 | 0 | 0 | 0 | 0 | 0 | 0 | 0 | 0 |
| 0 | NA | NA | NA | NA | NA | Gm29037 | | 0 | 0 | 0 | 0 | 0 | 0 | 0 | 0 | 0 | 0 | 0 |
| 0 | NA | NA | NA | NA | NA | Gm20823 | predicted gene, 20823 | 0 | 0 | 0 | 0 | 0 | 0 | 0 | 0 | 0 | 0 | 0 |
| 0 | NA | NA | NA | NA | NA | Gm21877 | | 0 | 0 | 0 | 0 | 0 | 0 | 0 | 0 | 0 | 0 | 0 |
| 0 | NA | NA | NA | NA | NA | Gm28835 | | 0 | 0 | 0 | 0 | 0 | 0 | 0 | 0 | 0 | 0 | 0 |
| 0 | NA | NA | NA | NA | NA | Gm29548 | | 0 | 0 | 0 | 0 | 0 | 0 | 0 | 0 | 0 | 0 | 0 |
| 0 | NA | NA | NA | NA | NA | Gm29547 | | 0 | 0 | 0 | 0 | 0 | 0 | 0 | 0 | 0 | 0 | 0 |
| 0 | NA | NA | NA | NA | NA | Gm29546 | | 0 | 0 | 0 | 0 | 0 | 0 | 0 | 0 | 0 | 0 | 0 |
| 0 | NA | NA | NA | NA | NA | Gm29545 | | 0 | 0 | 0 | 0 | 0 | 0 | 0 | 0 | 0 | 0 | 0 |
| 0 | NA | NA | NA | NA | NA | Gm29544 | | 0 | 0 | 0 | 0 | 0 | 0 | 0 | 0 | 0 | 0 | 0 |
| 0 | NA | NA | NA | NA | NA | Gm29209 | | 0 | 0 | 0 | 0 | 0 | 0 | 0 | 0 | 0 | 0 | 0 |
| 0 | NA | NA | NA | NA | NA | Gm29549 | | 0 | 0 | 0 | 0 | 0 | 0 | 0 | 0 | 0 | 0 | 0 |
| 0 | NA | NA | NA | NA | NA | Gm21530 | predicted gene, 21530 | 0 | 0 | 0 | 0 | 0 | 0 | 0 | 0 | 0 | 0 | 0 |
| 0 | NA | NA | NA | NA | NA | Gm21617 | predicted gene, 21617 | 0 | 0 | 0 | 0 | 0 | 0 | 0 | 0 | 0 | 0 | 0 |
| 0 | NA | NA | NA | NA | NA | Gm28589 | | 0 | 0 | 0 | 0 | 0 | 0 | 0 | 0 | 0 | 0 | 0 |
| 0 | NA | NA | NA | NA | NA | Gm29420 | | 0 | 0 | 0 | 0 | 0 | 0 | 0 | 0 | 0 | 0 | 0 |
| 0 | NA | NA | NA | NA | NA | Gm29421 | | 0 | 0 | 0 | 0 | 0 | 0 | 0 | 0 | 0 | 0 | 0 |
| 0 | NA | NA | NA | NA | NA | Gm21627 | predicted gene, 21627 | 0 | 0 | 0 | 0 | 0 | 0 | 0 | 0 | 0 | 0 | 0 |
| 0 | NA | NA | NA | NA | NA | Gm29422 | | 0 | 0 | 0 | 0 | 0 | 0 | 0 | 0 | 0 | 0 | 0 |
| 0 | NA | NA | NA | NA | NA | Gm28808 | | 0 | 0 | 0 | 0 | 0 | 0 | 0 | 0 | 0 | 0 | 0 |
| 0 | NA | NA | NA | NA | NA | Gm21634 | predicted gene, 21634 | 0 | 0 | 0 | 0 | 0 | 0 | 0 | 0 | 0 | 0 | 0 |
| 0 | NA | NA | NA | NA | NA | Gm28446 | | 0 | 0 | 0 | 0 | 0 | 0 | 0 | 0 | 0 | 0 | 0 |
| 0 | NA | NA | NA | NA | NA | Gm29060 | | 0 | 0 | 0 | 0 | 0 | 0 | 0 | 0 | 0 | 0 | 0 |
| 0 | NA | NA | NA | NA | NA | Gm29061 | | 0 | 0 | 0 | 0 | 0 | 0 | 0 | 0 | 0 | 0 | 0 |
| 0 | NA | NA | NA | NA | NA | Gm20885 | predicted gene, 20885 | 0 | 0 | 0 | 0 | 0 | 0 | 0 | 0 | 0 | 0 | 0 |
| 0 | NA | NA | NA | NA | NA | Gm29062 | | 0 | 0 | 0 | 0 | 0 | 0 | 0 | 0 | 0 | 0 | 0 |
| 0 | NA | NA | NA | NA | NA | Gm29057 | | 0 | 0 | 0 | 0 | 0 | 0 | 0 | 0 | 0 | 0 | 0 |

| | | | | | | | | | | | | | | | | | | |
|---|---|---|---|---|---|---|---|---|---|---|---|---|---|---|---|---|---|---|
| 0 | NA | NA | NA | NA | NA | Gm29625 | | 0 | 0 | 0 | 0 | 0 | 0 | 0 | 0 | 0 | 0 | 0 |
| 0 | NA | NA | NA | NA | NA | Gm21654 | predicted gene, 21654 | 0 | 0 | 0 | 0 | 0 | 0 | 0 | 0 | 0 | 0 | 0 |
| 0 | NA | NA | NA | NA | NA | Gm29058 | | 0 | 0 | 0 | 0 | 0 | 0 | 0 | 0 | 0 | 0 | 0 |
| 0 | NA | NA | NA | NA | NA | Gm29059 | | 0 | 0 | 0 | 0 | 0 | 0 | 0 | 0 | 0 | 0 | 0 |
| 0 | NA | NA | NA | NA | NA | Gm29628 | | 0 | 0 | 0 | 0 | 0 | 0 | 0 | 0 | 0 | 0 | 0 |
| 0 | NA | NA | NA | NA | NA | Gm20882 | predicted gene, 20882 | 0 | 0 | 0 | 0 | 0 | 0 | 0 | 0 | 0 | 0 | 0 |
| 0 | NA | NA | NA | NA | NA | Gm20907 | predicted gene, 20907 | 0 | 0 | 0 | 0 | 0 | 0 | 0 | 0 | 0 | 0 | 0 |
| 0 | NA | NA | NA | NA | NA | Gm29624 | | 0 | 0 | 0 | 0 | 0 | 0 | 0 | 0 | 0 | 0 | 0 |
| 0 | NA | NA | NA | NA | NA | Gm29623 | | 0 | 0 | 0 | 0 | 0 | 0 | 0 | 0 | 0 | 0 | 0 |
| 0 | NA | NA | NA | NA | NA | Gm29622 | | 0 | 0 | 0 | 0 | 0 | 0 | 0 | 0 | 0 | 0 | 0 |
| 0 | NA | NA | NA | NA | NA | Gm29265 | | 0 | 0 | 0 | 0 | 0 | 0 | 0 | 0 | 0 | 0 | 0 |
| 0 | NA | NA | NA | NA | NA | Gm20908 | predicted gene, 20908 | 0 | 0 | 0 | 0 | 0 | 0 | 0 | 0 | 0 | 0 | 0 |
| 0 | NA | NA | NA | NA | NA | Gm29259 | | 0 | 0 | 0 | 0 | 0 | 0 | 0 | 0 | 0 | 0 | 0 |
| 0 | NA | NA | NA | NA | NA | Gm28860 | | 0 | 0 | 0 | 0 | 0 | 0 | 0 | 0 | 0 | 0 | 0 |
| 0 | NA | NA | NA | NA | NA | Gm28252 | | 0 | 0 | 0 | 0 | 0 | 0 | 0 | 0 | 0 | 0 | 0 |
| 0 | NA | NA | NA | NA | NA | Gm28859 | | 0 | 0 | 0 | 0 | 0 | 0 | 0 | 0 | 0 | 0 | 0 |
| 0 | NA | NA | NA | NA | NA | Gm20921 | predicted gene, 20921 | 0 | 0 | 0 | 0 | 0 | 0 | 0 | 0 | 0 | 0 | 0 |
| 0 | NA | NA | NA | NA | NA | Gm28253 | | 0 | 0 | 0 | 0 | 0 | 0 | 0 | 0 | 0 | 0 | 0 |
| 0 | NA | NA | NA | NA | NA | Gm21826 | | 0 | 0 | 0 | 0 | 0 | 0 | 0 | 0 | 0 | 0 | 0 |
| 0 | NA | NA | NA | NA | NA | Gm28254 | | 0 | 0 | 0 | 0 | 0 | 0 | 0 | 0 | 0 | 0 | 0 |
| 0 | NA | NA | NA | NA | NA | Gm28861 | | 0 | 0 | 0 | 0 | 0 | 0 | 0 | 0 | 0 | 0 | 0 |
| 0 | NA | NA | NA | NA | NA | Gm28255 | | 0 | 0 | 0 | 0 | 0 | 0 | 0 | 0 | 0 | 0 | 0 |
| 0 | NA | NA | NA | NA | NA | Gm21171 | predicted gene, 21171 | 0 | 0 | 0 | 0 | 0 | 0 | 0 | 0 | 0 | 0 | 0 |
| 0 | NA | NA | NA | NA | NA | Gm20924 | Y-linked testis-specific protein 1-like | 0 | 0 | 0 | 0 | 0 | 0 | 0 | 0 | 0 | 0 | 0 |
| 0 | NA | NA | NA | NA | NA | Gm28990 | | 0 | 0 | 0 | 0 | 0 | 0 | 0 | 0 | 0 | 0 | 0 |
| 0 | NA | NA | NA | NA | NA | Gm21848 | | 0 | 0 | 0 | 0 | 0 | 0 | 0 | 0 | 0 | 0 | 0 |
| 0 | NA | NA | NA | NA | NA | Gm29457 | | 0 | 0 | 0 | 0 | 0 | 0 | 0 | 0 | 0 | 0 | 0 |
| 0 | NA | NA | NA | NA | NA | Gm20736 | predicted gene, 20736 | 0 | 0 | 0 | 0 | 0 | 0 | 0 | 0 | 0 | 0 | 0 |
| 0 | NA | NA | NA | NA | NA | Gm28558 | | 0 | 0 | 0 | 0 | 0 | 0 | 0 | 0 | 0 | 0 | 0 |
| 0 | NA | NA | NA | NA | NA | Gm28560 | | 0 | 0 | 0 | 0 | 0 | 0 | 0 | 0 | 0 | 0 | 0 |
| 0 | NA | NA | NA | NA | NA | Gm28561 | | 0 | 0 | 0 | 0 | 0 | 0 | 0 | 0 | 0 | 0 | 0 |
| 0 | NA | NA | NA | NA | NA | Gm29584 | | 0 | 0 | 0 | 0 | 0 | 0 | 0 | 0 | 0 | 0 | 0 |
| 0 | NA | NA | NA | NA | NA | Gm20936 | predicted gene, 20936 | 0 | 0 | 0 | 0 | 0 | 0 | 0 | 0 | 0 | 0 | 0 |
| 0 | NA | NA | NA | NA | NA | Gm20934 | predicted gene, 20934 | 0 | 0 | 0 | 0 | 0 | 0 | 0 | 0 | 0 | 0 | 0 |
| 0 | NA | NA | NA | NA | NA | Gm29586 | | 0 | 0 | 0 | 0 | 0 | 0 | 0 | 0 | 0 | 0 | 0 |
| 0 | NA | NA | NA | NA | NA | Gm21895 | | 0 | 0 | 0 | 0 | 0 | 0 | 0 | 0 | 0 | 0 | 0 |
| 0 | NA | NA | NA | NA | NA | Gm28134 | | 0 | 0 | 0 | 0 | 0 | 0 | 0 | 0 | 0 | 0 | 0 |
| 0 | NA | NA | NA | NA | NA | Gm28135 | | 0 | 0 | 0 | 0 | 0 | 0 | 0 | 0 | 0 | 0 | 0 |
| 0 | NA | NA | NA | NA | NA | Gm20933 | predicted gene, 20933 | 0 | 0 | 0 | 0 | 0 | 0 | 0 | 0 | 0 | 0 | 0 |
| 0 | NA | NA | NA | NA | NA | Gm28133 | | 0 | 0 | 0 | 0 | 0 | 0 | 0 | 0 | 0 | 0 | 0 |
| 0 | NA | NA | NA | NA | NA | Gm28137 | | 0 | 0 | 0 | 0 | 0 | 0 | 0 | 0 | 0 | 0 | 0 |
| 0 | NA | NA | NA | NA | NA | Gm28138 | | 0 | 0 | 0 | 0 | 0 | 0 | 0 | 0 | 0 | 0 | 0 |
| 0 | NA | NA | NA | NA | NA | Gm21837 | | 0 | 0 | 0 | 0 | 0 | 0 | 0 | 0 | 0 | 0 | 0 |
| 0 | NA | NA | NA | NA | NA | Gm28136 | | 0 | 0 | 0 | 0 | 0 | 0 | 0 | 0 | 0 | 0 | 0 |
| 0 | NA | NA | NA | NA | NA | Gm21910 | | 0 | 0 | 0 | 0 | 0 | 0 | 0 | 0 | 0 | 0 | 0 |
| 0 | NA | NA | NA | NA | NA | Gm29377 | | 0 | 0 | 0 | 0 | 0 | 0 | 0 | 0 | 0 | 0 | 0 |
| 0 | NA | NA | NA | NA | NA | Gm21889 | | 0 | 0 | 0 | 0 | 0 | 0 | 0 | 0 | 0 | 0 | 0 |
| 0 | NA | NA | NA | NA | NA | Gm29378 | | 0 | 0 | 0 | 0 | 0 | 0 | 0 | 0 | 0 | 0 | 0 |
| 0 | NA | NA | NA | NA | NA | Gm29379 | | 0 | 0 | 0 | 0 | 0 | 0 | 0 | 0 | 0 | 0 | 0 |
| 0 | NA | NA | NA | NA | NA | Gm29380 | | 0 | 0 | 0 | 0 | 0 | 0 | 0 | 0 | 0 | 0 | 0 |
| 0 | NA | NA | NA | NA | NA | Gm29381 | | 0 | 0 | 0 | 0 | 0 | 0 | 0 | 0 | 0 | 0 | 0 |
| 0 | NA | NA | NA | NA | NA | Gm28795 | | 0 | 0 | 0 | 0 | 0 | 0 | 0 | 0 | 0 | 0 | 0 |
| 0 | NA | NA | NA | NA | NA | Gm20852 | predicted gene, 20852 | 0 | 0 | 0 | 0 | 0 | 0 | 0 | 0 | 0 | 0 | 0 |
| 0 | NA | NA | NA | NA | NA | Gm29003 | | 0 | 0 | 0 | 0 | 0 | 0 | 0 | 0 | 0 | 0 | 0 |
| 0 | NA | NA | NA | NA | NA | Gm28749 | | 0 | 0 | 0 | 0 | 0 | 0 | 0 | 0 | 0 | 0 | 0 |
| 0 | NA | NA | NA | NA | NA | Gm29001 | | 0 | 0 | 0 | 0 | 0 | 0 | 0 | 0 | 0 | 0 | 0 |
| 0 | NA | NA | NA | NA | NA | Gm29002 | | 0 | 0 | 0 | 0 | 0 | 0 | 0 | 0 | 0 | 0 | 0 |
| 0 | NA | NA | NA | NA | NA | Ssty2 | spermiogenesis specific transcript on the | 0 | 0 | 0 | 0 | 0 | 0 | 0 | 0 | 0 | 0 | 0 |
| 0 | NA | NA | NA | NA | NA | Gm29000 | | 0 | 0 | 0 | 0 | 0 | 0 | 0 | 0 | 0 | 0 | 0 |
| 0 | NA | NA | NA | NA | NA | Gm28646 | | 0 | 0 | 0 | 0 | 0 | 0 | 0 | 0 | 0 | 0 | 0 |

| | | | | | | | | | | | | | | | | | | |
|---|---|---|---|---|---|---|---|---|---|---|---|---|---|---|---|---|---|---|
| 0 | NA | NA | NA | NA | NA | Gm28612 | | 0 | 0 | 0 | 0 | 0 | 0 | 0 | 0 | 0 | 0 | 0 |
| 0 | NA | NA | NA | NA | NA | Gm28613 | | 0 | 0 | 0 | 0 | 0 | 0 | 0 | 0 | 0 | 0 | 0 |
| 0 | NA | NA | NA | NA | NA | Gm28611 | | 0 | 0 | 0 | 0 | 0 | 0 | 0 | 0 | 0 | 0 | 0 |
| 0 | NA | NA | NA | NA | NA | Gm21114 | predicted gene, 21114 | 0 | 0 | 0 | 0 | 0 | 0 | 0 | 0 | 0 | 0 | 0 |
| 0 | NA | NA | NA | NA | NA | Gm29633 | | 0 | 0 | 0 | 0 | 0 | 0 | 0 | 0 | 0 | 0 | 0 |
| 0 | NA | NA | NA | NA | NA | Gm21794 | predicted gene, 21794 | 0 | 0 | 0 | 0 | 0 | 0 | 0 | 0 | 0 | 0 | 0 |
| 0 | NA | NA | NA | NA | NA | Gm29219 | | 0 | 0 | 0 | 0 | 0 | 0 | 0 | 0 | 0 | 0 | 0 |
| 0 | NA | NA | NA | NA | NA | Gm29632 | | 0 | 0 | 0 | 0 | 0 | 0 | 0 | 0 | 0 | 0 | 0 |
| 0 | NA | NA | NA | NA | NA | Gm29631 | | 0 | 0 | 0 | 0 | 0 | 0 | 0 | 0 | 0 | 0 | 0 |
| 0 | NA | NA | NA | NA | NA | Gm29637 | | 0 | 0 | 0 | 0 | 0 | 0 | 0 | 0 | 0 | 0 | 0 |
| 0 | NA | NA | NA | NA | NA | Gm20874 | predicted gene, 20874 | 0 | 0 | 0 | 0 | 0 | 0 | 0 | 0 | 0 | 0 | 0 |
| 0 | NA | NA | NA | NA | NA | Gm29082 | | 0 | 0 | 0 | 0 | 0 | 0 | 0 | 0 | 0 | 0 | 0 |
| 0 | NA | NA | NA | NA | NA | Gm29217 | | 0 | 0 | 0 | 0 | 0 | 0 | 0 | 0 | 0 | 0 | 0 |
| 0 | NA | NA | NA | NA | NA | Gm29636 | | 0 | 0 | 0 | 0 | 0 | 0 | 0 | 0 | 0 | 0 | 0 |
| 0 | NA | NA | NA | NA | NA | Gm29635 | | 0 | 0 | 0 | 0 | 0 | 0 | 0 | 0 | 0 | 0 | 0 |
| 0 | NA | NA | NA | NA | NA | Gm29634 | | 0 | 0 | 0 | 0 | 0 | 0 | 0 | 0 | 0 | 0 | 0 |
| 0 | NA | NA | NA | NA | NA | Gm29639 | | 0 | 0 | 0 | 0 | 0 | 0 | 0 | 0 | 0 | 0 | 0 |
| 0 | NA | NA | NA | NA | NA | Gm29218 | | 0 | 0 | 0 | 0 | 0 | 0 | 0 | 0 | 0 | 0 | 0 |
| 0 | NA | NA | NA | NA | NA | Gm21806 | predicted gene, 21806 | 0 | 0 | 0 | 0 | 0 | 0 | 0 | 0 | 0 | 0 | 0 |
| 0 | NA | NA | NA | NA | NA | Gm20935 | predicted gene, 20935 | 0 | 0 | 0 | 0 | 0 | 0 | 0 | 0 | 0 | 0 | 0 |
| 0 | NA | NA | NA | NA | NA | Gm29662 | | 0 | 0 | 0 | 0 | 0 | 0 | 0 | 0 | 0 | 0 | 0 |
| 0 | NA | NA | NA | NA | NA | Gm29660 | | 0 | 0 | 0 | 0 | 0 | 0 | 0 | 0 | 0 | 0 | 0 |
| 0 | NA | NA | NA | NA | NA | Gm29661 | | 0 | 0 | 0 | 0 | 0 | 0 | 0 | 0 | 0 | 0 | 0 |
| 0 | NA | NA | NA | NA | NA | Gm29659 | | 0 | 0 | 0 | 0 | 0 | 0 | 0 | 0 | 0 | 0 | 0 |
| 0 | NA | NA | NA | NA | NA | Gm20937 | predicted gene, 20874 | 0 | 0 | 0 | 0 | 0 | 0 | 0 | 0 | 0 | 0 | 0 |
| 0 | NA | NA | NA | NA | NA | Gm29450 | | 0 | 0 | 0 | 0 | 0 | 0 | 0 | 0 | 0 | 0 | 0 |
| 0 | NA | NA | NA | NA | NA | Gm29449 | | 0 | 0 | 0 | 0 | 0 | 0 | 0 | 0 | 0 | 0 | 0 |
| 0 | NA | NA | NA | NA | NA | Gm21867 | | 0 | 0 | 0 | 0 | 0 | 0 | 0 | 0 | 0 | 0 | 0 |
| 0 | NA | NA | NA | NA | NA | Gm29448 | | 0 | 0 | 0 | 0 | 0 | 0 | 0 | 0 | 0 | 0 | 0 |
| 0 | NA | NA | NA | NA | NA | Gm21127 | predicted gene, 21127 | 0 | 0 | 0 | 0 | 0 | 0 | 0 | 0 | 0 | 0 | 0 |
| 0 | NA | NA | NA | NA | NA | Gm20816 | predicted gene, 20816 | 0 | 0 | 0 | 0 | 0 | 0 | 0 | 0 | 0 | 0 | 0 |
| 0 | NA | NA | NA | NA | NA | Gm29447 | | 0 | 0 | 0 | 0 | 0 | 0 | 0 | 0 | 0 | 0 | 0 |
| 0 | NA | NA | NA | NA | NA | Gm29446 | | 0 | 0 | 0 | 0 | 0 | 0 | 0 | 0 | 0 | 0 | 0 |
| 0 | NA | NA | NA | NA | NA | Gm29452 | | 0 | 0 | 0 | 0 | 0 | 0 | 0 | 0 | 0 | 0 | 0 |
| 0 | NA | NA | NA | NA | NA | Gm29451 | | 0 | 0 | 0 | 0 | 0 | 0 | 0 | 0 | 0 | 0 | 0 |
| 0 | NA | NA | NA | NA | NA | Gm20817 | | 0 | 0 | 0 | 0 | 0 | 0 | 0 | 0 | 0 | 0 | 0 |
| 0 | NA | NA | NA | NA | NA | Gm28633 | | 0 | 0 | 0 | 0 | 0 | 0 | 0 | 0 | 0 | 0 | 0 |
| 0 | NA | NA | NA | NA | NA | Gm29410 | | 0 | 0 | 0 | 0 | 0 | 0 | 0 | 0 | 0 | 0 | 0 |
| 0 | NA | NA | NA | NA | NA | Gm28819 | | 0 | 0 | 0 | 0 | 0 | 0 | 0 | 0 | 0 | 0 | 0 |
| 0 | NA | NA | NA | NA | NA | Gm28462 | | 0 | 0 | 0 | 0 | 0 | 0 | 0 | 0 | 0 | 0 | 0 |
| 0 | NA | NA | NA | NA | NA | Gm20818 | Y-linked testis-specific protein 1-like | 0 | 0 | 0 | 0 | 0 | 0 | 0 | 0 | 0 | 0 | 0 |
| 0 | NA | NA | NA | NA | NA | Gm21151 | Y-linked testis-specific protein 1-like | 0 | 0 | 0 | 0 | 0 | 0 | 0 | 0 | 0 | 0 | 0 |
| 0 | NA | NA | NA | NA | NA | Gm29409 | | 0 | 0 | 0 | 0 | 0 | 0 | 0 | 0 | 0 | 0 | 0 |
| 0 | NA | NA | NA | NA | NA | Gm28632 | | 0 | 0 | 0 | 0 | 0 | 0 | 0 | 0 | 0 | 0 | 0 |
| 0 | NA | NA | NA | NA | NA | Gm28101 | | 0 | 0 | 0 | 0 | 0 | 0 | 0 | 0 | 0 | 0 | 0 |
| 0 | NA | NA | NA | NA | NA | Gm33815 | | 0 | 0 | 0 | 0 | 0 | 0 | 0 | 0 | 0 | 0 | 0 |
| 0 | NA | NA | NA | NA | NA | Gm21160 | predicted gene, 21160 | 0 | 0 | 0 | 0 | 0 | 0 | 0 | 0 | 0 | 0 | 0 |
| 0 | NA | NA | NA | NA | NA | Gm28637 | | 0 | 0 | 0 | 0 | 0 | 0 | 0 | 0 | 0 | 0 | 0 |
| 0 | NA | NA | NA | NA | NA | Gm28636 | | 0 | 0 | 0 | 0 | 0 | 0 | 0 | 0 | 0 | 0 | 0 |
| 0 | NA | NA | NA | NA | NA | Gm20932 | Y-linked testis-specific protein 1-like | 0 | 0 | 0 | 0 | 0 | 0 | 0 | 0 | 0 | 0 | 0 |
| 0 | NA | NA | NA | NA | NA | Gm29134 | | 0 | 0 | 0 | 0 | 0 | 0 | 0 | 0 | 0 | 0 | 0 |
| 0 | NA | NA | NA | NA | NA | Gm28303 | | 0 | 0 | 0 | 0 | 0 | 0 | 0 | 0 | 0 | 0 | 0 |
| 0 | NA | NA | NA | NA | NA | Gm28304 | | 0 | 0 | 0 | 0 | 0 | 0 | 0 | 0 | 0 | 0 | 0 |
| 0 | NA | NA | NA | NA | NA | Gm28824 | | 0 | 0 | 0 | 0 | 0 | 0 | 0 | 0 | 0 | 0 | 0 |
| 0 | NA | NA | NA | NA | NA | Gm28081 | | 0 | 0 | 0 | 0 | 0 | 0 | 0 | 0 | 0 | 0 | 0 |
| 0 | NA | NA | NA | NA | NA | Gm28080 | | 0 | 0 | 0 | 0 | 0 | 0 | 0 | 0 | 0 | 0 | 0 |
| 0 | NA | NA | NA | NA | NA | Gm21524 | predicted gene, 21524 | 0 | 0 | 0 | 0 | 0 | 0 | 0 | 0 | 0 | 0 | 0 |
| 0 | NA | NA | NA | NA | NA | Gm28079 | | 0 | 0 | 0 | 0 | 0 | 0 | 0 | 0 | 0 | 0 | 0 |
| 0 | NA | NA | NA | NA | NA | Gm28302 | | 0 | 0 | 0 | 0 | 0 | 0 | 0 | 0 | 0 | 0 | 0 |
| 0 | NA | NA | NA | NA | NA | Gm28082 | | 0 | 0 | 0 | 0 | 0 | 0 | 0 | 0 | 0 | 0 | 0 |

| | | | | | | | | | | | | | | | | | | |
|---|---|---|---|---|---|---|---|---|---|---|---|---|---|---|---|---|---|---|
| 0 | NA | NA | NA | NA | NA | Gm21672 | predicted gene, 21672 | 0 | 0 | 0 | 0 | 0 | 0 | 0 | 0 | 0 | 0 | 0 |
| 0 | NA | NA | NA | NA | NA | Gm21118 | predicted gene, 21118 | 0 | 0 | 0 | 0 | 0 | 0 | 0 | 0 | 0 | 0 | 0 |
| 0 | NA | NA | NA | NA | NA | Gm29644 | | 0 | 0 | 0 | 0 | 0 | 0 | 0 | 0 | 0 | 0 | 0 |
| 0 | NA | NA | NA | NA | NA | Gm29423 | | 0 | 0 | 0 | 0 | 0 | 0 | 0 | 0 | 0 | 0 | 0 |
| 0 | NA | NA | NA | NA | NA | Gm29424 | | 0 | 0 | 0 | 0 | 0 | 0 | 0 | 0 | 0 | 0 | 0 |
| 0 | NA | NA | NA | NA | NA | 4932431L22Rik | | 0 | 0 | 0 | 0 | 0 | 0 | 0 | 0 | 0 | 0 | 0 |
| 0 | NA | NA | NA | NA | NA | Gm29425 | | 0 | 0 | 0 | 0 | 0 | 0 | 0 | 0 | 0 | 0 | 0 |
| 0 | NA | NA | NA | NA | NA | Gm20888 | predicted gene, 20874 | 0 | 0 | 0 | 0 | 0 | 0 | 0 | 0 | 0 | 0 | 0 |
| 0 | NA | NA | NA | NA | NA | Gm28238 | | 0 | 0 | 0 | 0 | 0 | 0 | 0 | 0 | 0 | 0 | 0 |
| 0 | NA | NA | NA | NA | NA | Gm28433 | | 0 | 0 | 0 | 0 | 0 | 0 | 0 | 0 | 0 | 0 | 0 |
| 0 | NA | NA | NA | NA | NA | Gm28431 | | 0 | 0 | 0 | 0 | 0 | 0 | 0 | 0 | 0 | 0 | 0 |
| 0 | NA | NA | NA | NA | NA | Gm28432 | | 0 | 0 | 0 | 0 | 0 | 0 | 0 | 0 | 0 | 0 | 0 |
| 0 | NA | NA | NA | NA | NA | Gm20886 | | 0 | 0 | 0 | 0 | 0 | 0 | 0 | 0 | 0 | 0 | 0 |
| 0 | NA | NA | NA | NA | NA | Gm21808 | | 0 | 0 | 0 | 0 | 0 | 0 | 0 | 0 | 0 | 0 | 0 |
| 0 | NA | NA | NA | NA | NA | Gm28840 | | 0 | 0 | 0 | 0 | 0 | 0 | 0 | 0 | 0 | 0 | 0 |
| 0 | NA | NA | NA | NA | NA | Gm28841 | | 0 | 0 | 0 | 0 | 0 | 0 | 0 | 0 | 0 | 0 | 0 |
| 0 | NA | NA | NA | NA | NA | Gm28842 | | 0 | 0 | 0 | 0 | 0 | 0 | 0 | 0 | 0 | 0 | 0 |
| 0 | NA | NA | NA | NA | NA | Gm28843 | | 0 | 0 | 0 | 0 | 0 | 0 | 0 | 0 | 0 | 0 | 0 |
| 0 | NA | NA | NA | NA | NA | Gm28837 | | 0 | 0 | 0 | 0 | 0 | 0 | 0 | 0 | 0 | 0 | 0 |
| 0 | NA | NA | NA | NA | NA | Gm28838 | | 0 | 0 | 0 | 0 | 0 | 0 | 0 | 0 | 0 | 0 | 0 |
| 0 | NA | NA | NA | NA | NA | Gm28839 | | 0 | 0 | 0 | 0 | 0 | 0 | 0 | 0 | 0 | 0 | 0 |
| 0 | NA | NA | NA | NA | NA | Gm20866 | predicted gene, 20866 | 0 | 0 | 0 | 0 | 0 | 0 | 0 | 0 | 0 | 0 | 0 |
| 0 | NA | NA | NA | NA | NA | Gm20868 | | 0 | 0 | 0 | 0 | 0 | 0 | 0 | 0 | 0 | 0 | 0 |
| 0 | NA | NA | NA | NA | NA | Gm28844 | | 0 | 0 | 0 | 0 | 0 | 0 | 0 | 0 | 0 | 0 | 0 |
| 0 | NA | NA | NA | NA | NA | Gm21841 | | 0 | 0 | 0 | 0 | 0 | 0 | 0 | 0 | 0 | 0 | 0 |
| 0 | NA | NA | NA | NA | NA | Gm28668 | | 0 | 0 | 0 | 0 | 0 | 0 | 0 | 0 | 0 | 0 | 0 |
| 0 | NA | NA | NA | NA | NA | Gm20869 | X-linked lymphocyte-regulated protein PM | 0 | 0 | 0 | 0 | 0 | 0 | 0 | 0 | 0 | 0 | 0 |
| 0 | NA | NA | NA | NA | NA | Gm28481 | | 0 | 0 | 0 | 0 | 0 | 0 | 0 | 0 | 0 | 0 | 0 |
| 0 | NA | NA | NA | NA | NA | Gm28482 | | 0 | 0 | 0 | 0 | 0 | 0 | 0 | 0 | 0 | 0 | 0 |
| 0 | NA | NA | NA | NA | NA | Gm28485 | | 0 | 0 | 0 | 0 | 0 | 0 | 0 | 0 | 0 | 0 | 0 |
| 0 | NA | NA | NA | NA | NA | Gm20870 | RIKEN cDNA 1700040F15 gene | 0 | 0 | 0 | 0 | 0 | 0 | 0 | 0 | 0 | 0 | 0 |
| 0 | NA | NA | NA | NA | NA | Gm28483 | | 0 | 0 | 0 | 0 | 0 | 0 | 0 | 0 | 0 | 0 | 0 |
| 0 | NA | NA | NA | NA | NA | Gm28484 | | 0 | 0 | 0 | 0 | 0 | 0 | 0 | 0 | 0 | 0 | 0 |
| 0 | NA | NA | NA | NA | NA | Gm28480 | | 0 | 0 | 0 | 0 | 0 | 0 | 0 | 0 | 0 | 0 | 0 |
| 0 | NA | NA | NA | NA | NA | Gm20842 | predicted gene, 20842 | 0 | 0 | 0 | 0 | 0 | 0 | 0 | 0 | 0 | 0 | 0 |
| 0 | NA | NA | NA | NA | NA | Gm29179 | | 0 | 0 | 0 | 0 | 0 | 0 | 0 | 0 | 0 | 0 | 0 |
| 0 | NA | NA | NA | NA | NA | Gm21880 | | 0 | 0 | 0 | 0 | 0 | 0 | 0 | 0 | 0 | 0 | 0 |
| 0 | NA | NA | NA | NA | NA | Gm29180 | | 0 | 0 | 0 | 0 | 0 | 0 | 0 | 0 | 0 | 0 | 0 |
| 0 | NA | NA | NA | NA | NA | Gm29181 | | 0 | 0 | 0 | 0 | 0 | 0 | 0 | 0 | 0 | 0 | 0 |
| 0 | NA | NA | NA | NA | NA | Gm20843 | predicted gene, 20874 | 0 | 0 | 0 | 0 | 0 | 0 | 0 | 0 | 0 | 0 | 0 |
| 0 | NA | NA | NA | NA | NA | Gm29182 | | 0 | 0 | 0 | 0 | 0 | 0 | 0 | 0 | 0 | 0 | 0 |
| 0 | NA | NA | NA | NA | NA | Gm20844 | predicted gene, 20844 | 0 | 0 | 0 | 0 | 0 | 0 | 0 | 0 | 0 | 0 | 0 |
| 0 | NA | NA | NA | NA | NA | Gm29178 | | 0 | 0 | 0 | 0 | 0 | 0 | 0 | 0 | 0 | 0 | 0 |
| 0 | NA | NA | NA | NA | NA | Gm21757 | | 0 | 0 | 0 | 0 | 0 | 0 | 0 | 0 | 0 | 0 | 0 |
| 0 | NA | NA | NA | NA | NA | Gm28968 | | 0 | 0 | 0 | 0 | 0 | 0 | 0 | 0 | 0 | 0 | 0 |
| 0 | NA | NA | NA | NA | NA | Gm28495 | | 0 | 0 | 0 | 0 | 0 | 0 | 0 | 0 | 0 | 0 | 0 |
| 0 | NA | NA | NA | NA | NA | Gm20846 | predicted gene, 20846 | 0 | 0 | 0 | 0 | 0 | 0 | 0 | 0 | 0 | 0 | 0 |
| 0 | NA | NA | NA | NA | NA | Gm20847 | predicted gene, 20847 | 0 | 0 | 0 | 0 | 0 | 0 | 0 | 0 | 0 | 0 | 0 |
| 0 | NA | NA | NA | NA | NA | Gm20848 | predicted gene, 20848 | 0 | 0 | 0 | 0 | 0 | 0 | 0 | 0 | 0 | 0 | 0 |
| 0 | NA | NA | NA | NA | NA | Gm28969 | | 0 | 0 | 0 | 0 | 0 | 0 | 0 | 0 | 0 | 0 | 0 |
| 0 | NA | NA | NA | NA | NA | Gm21766 | | 0 | 0 | 0 | 0 | 0 | 0 | 0 | 0 | 0 | 0 | 0 |
| 0 | NA | NA | NA | NA | NA | Gm20800 | predicted gene, 20800 | 0 | 0 | 0 | 0 | 0 | 0 | 0 | 0 | 0 | 0 | 0 |
| 0 | NA | NA | NA | NA | NA | Gm21683 | predicted gene, 21683 | 0 | 0 | 0 | 0 | 0 | 0 | 0 | 0 | 0 | 0 | 0 |
| 0 | NA | NA | NA | NA | NA | Gm28677 | | 0 | 0 | 0 | 0 | 0 | 0 | 0 | 0 | 0 | 0 | 0 |
| 0 | NA | NA | NA | NA | NA | Gm21855 | | 0 | 0 | 0 | 0 | 0 | 0 | 0 | 0 | 0 | 0 | 0 |
| 0 | NA | NA | NA | NA | NA | Gm29255 | | 0 | 0 | 0 | 0 | 0 | 0 | 0 | 0 | 0 | 0 | 0 |
| 0 | NA | NA | NA | NA | NA | Gm20902 | predicted gene, 20902 | 0 | 0 | 0 | 0 | 0 | 0 | 0 | 0 | 0 | 0 | 0 |
| 0 | NA | NA | NA | NA | NA | Gm28678 | | 0 | 0 | 0 | 0 | 0 | 0 | 0 | 0 | 0 | 0 | 0 |
| 0 | NA | NA | NA | NA | NA | Gm20903 | predicted gene, 20903 | 0 | 0 | 0 | 0 | 0 | 0 | 0 | 0 | 0 | 0 | 0 |
| 0 | NA | NA | NA | NA | NA | Gm28272 | | 0 | 0 | 0 | 0 | 0 | 0 | 0 | 0 | 0 | 0 | 0 |

| | | | | | | | | | | | | | | | | | |
|---|---|---|---|---|---|---|---|---|---|---|---|---|---|---|---|---|---|
| 0 | NA | NA | NA | NA | NA | Gm29256 | | 0 | 0 | 0 | 0 | 0 | 0 | 0 | 0 | 0 | 0 | 0 |
| 0 | NA | NA | NA | NA | NA | Gm28259 | | 0 | 0 | 0 | 0 | 0 | 0 | 0 | 0 | 0 | 0 | 0 |
| 0 | NA | NA | NA | NA | NA | Gm28273 | | 0 | 0 | 0 | 0 | 0 | 0 | 0 | 0 | 0 | 0 | 0 |
| 0 | NA | NA | NA | NA | NA | Gm21715 | | 0 | 0 | 0 | 0 | 0 | 0 | 0 | 0 | 0 | 0 | 0 |
| 0 | NA | NA | NA | NA | NA | Gm29617 | | 0 | 0 | 0 | 0 | 0 | 0 | 0 | 0 | 0 | 0 | 0 |
| 0 | NA | NA | NA | NA | NA | Gm29615 | | 0 | 0 | 0 | 0 | 0 | 0 | 0 | 0 | 0 | 0 | 0 |
| 0 | NA | NA | NA | NA | NA | Gm29616 | | 0 | 0 | 0 | 0 | 0 | 0 | 0 | 0 | 0 | 0 | 0 |
| 0 | NA | NA | NA | NA | NA | Gm21784 | | 0 | 0 | 0 | 0 | 0 | 0 | 0 | 0 | 0 | 0 | 0 |
| 0 | NA | NA | NA | NA | NA | Gm20805 | predicted gene, 20805 | 0 | 0 | 0 | 0 | 0 | 0 | 0 | 0 | 0 | 0 | 0 |
| 0 | NA | NA | NA | NA | NA | Gm29618 | | 0 | 0 | 0 | 0 | 0 | 0 | 0 | 0 | 0 | 0 | 0 |
| 0 | NA | NA | NA | NA | NA | Gm29109 | | 0 | 0 | 0 | 0 | 0 | 0 | 0 | 0 | 0 | 0 | 0 |
| 0 | NA | NA | NA | NA | NA | Gm29108 | | 0 | 0 | 0 | 0 | 0 | 0 | 0 | 0 | 0 | 0 | 0 |
| 0 | NA | NA | NA | NA | NA | Gm29110 | | 0 | 0 | 0 | 0 | 0 | 0 | 0 | 0 | 0 | 0 | 0 |
| 0 | NA | NA | NA | NA | NA | Gm28781 | | 0 | 0 | 0 | 0 | 0 | 0 | 0 | 0 | 0 | 0 | 0 |
| 0 | NA | NA | NA | NA | NA | Gm21913 | | 0 | 0 | 0 | 0 | 0 | 0 | 0 | 0 | 0 | 0 | 0 |
| 0 | NA | NA | NA | NA | NA | Gm28174 | | 0 | 0 | 0 | 0 | 0 | 0 | 0 | 0 | 0 | 0 | 0 |
| 0 | NA | NA | NA | NA | NA | Gm22832 | | 0 | 0 | 0 | 0 | 0 | 0 | 0 | 0 | 0 | 0 | 0 |
| 0 | NA | NA | NA | NA | NA | Gm21076 | Y-linked testis-specific protein 1-like | 0 | 0 | 0 | 0 | 0 | 0 | 0 | 0 | 0 | 0 | 0 |
| 0 | NA | NA | NA | NA | NA | Gm29416 | | 0 | 0 | 0 | 0 | 0 | 0 | 0 | 0 | 0 | 0 | 0 |
| 0 | NA | NA | NA | NA | NA | Gm25359 | | 0 | 0 | 0 | 0 | 0 | 0 | 0 | 0 | 0 | 0 | 0 |
| 0 | NA | NA | NA | NA | NA | Gm21869 | | 0 | 0 | 0 | 0 | 0 | 0 | 0 | 0 | 0 | 0 | 0 |
| 0 | NA | NA | NA | NA | NA | Gm20813 | predicted gene, 20813 | 0 | 0 | 0 | 0 | 0 | 0 | 0 | 0 | 0 | 0 | 0 |
| 0 | NA | NA | NA | NA | NA | Gm28790 | | 0 | 0 | 0 | 0 | 0 | 0 | 0 | 0 | 0 | 0 | 0 |
| 0 | NA | NA | NA | NA | NA | Gm29272 | | 0 | 0 | 0 | 0 | 0 | 0 | 0 | 0 | 0 | 0 | 0 |
| 0 | NA | NA | NA | NA | NA | Gm28789 | | 0 | 0 | 0 | 0 | 0 | 0 | 0 | 0 | 0 | 0 | 0 |
| 0 | NA | NA | NA | NA | NA | Gm20814 | X-linked lymphocyte-regulated protein PM | 0 | 0 | 0 | 0 | 0 | 0 | 0 | 0 | 0 | 0 | 0 |
| 0 | NA | NA | NA | NA | NA | Gm28787 | | 0 | 0 | 0 | 0 | 0 | 0 | 0 | 0 | 0 | 0 | 0 |
| 0 | NA | NA | NA | NA | NA | Gm28788 | | 0 | 0 | 0 | 0 | 0 | 0 | 0 | 0 | 0 | 0 | 0 |
| 0 | NA | NA | NA | NA | NA | Gm29278 | | 0 | 0 | 0 | 0 | 0 | 0 | 0 | 0 | 0 | 0 | 0 |
| 0 | NA | NA | NA | NA | NA | Gm21922 | | 0 | 0 | 0 | 0 | 0 | 0 | 0 | 0 | 0 | 0 | 0 |
| 0 | NA | NA | NA | NA | NA | Gm21111 | predicted gene, 21111 | 0 | 0 | 0 | 0 | 0 | 0 | 0 | 0 | 0 | 0 | 0 |
| 0 | NA | NA | NA | NA | NA | Gm28608 | | 0 | 0 | 0 | 0 | 0 | 0 | 0 | 0 | 0 | 0 | 0 |
| 0 | NA | NA | NA | NA | NA | Gm28607 | | 0 | 0 | 0 | 0 | 0 | 0 | 0 | 0 | 0 | 0 | 0 |
| 0 | NA | NA | NA | NA | NA | Gm28606 | | 0 | 0 | 0 | 0 | 0 | 0 | 0 | 0 | 0 | 0 | 0 |
| 0 | NA | NA | NA | NA | NA | Gm20850 | predicted gene, 20850 | 0 | 0 | 0 | 0 | 0 | 0 | 0 | 0 | 0 | 0 | 0 |
| 0 | NA | NA | NA | NA | NA | Gm29574 | | 0 | 0 | 0 | 0 | 0 | 0 | 0 | 0 | 0 | 0 | 0 |
| 0 | NA | NA | NA | NA | NA | Gm29575 | | 0 | 0 | 0 | 0 | 0 | 0 | 0 | 0 | 0 | 0 | 0 |
| 0 | NA | NA | NA | NA | NA | Gm21875 | | 0 | 0 | 0 | 0 | 0 | 0 | 0 | 0 | 0 | 0 | 0 |
| 0 | NA | NA | NA | NA | NA | Gm20904 | | 0 | 0 | 0 | 0 | 0 | 0 | 0 | 0 | 0 | 0 | 0 |
| 0 | NA | NA | NA | NA | NA | Gm29576 | | 0 | 0 | 0 | 0 | 0 | 0 | 0 | 0 | 0 | 0 | 0 |
| 0 | NA | NA | NA | NA | NA | Gm29577 | | 0 | 0 | 0 | 0 | 0 | 0 | 0 | 0 | 0 | 0 | 0 |
| 0 | NA | NA | NA | NA | NA | Gm29573 | | 0 | 0 | 0 | 0 | 0 | 0 | 0 | 0 | 0 | 0 | 0 |
| 0 | NA | NA | NA | NA | NA | Gm28377 | | 0 | 0 | 0 | 0 | 0 | 0 | 0 | 0 | 0 | 0 | 0 |
| 0 | NA | NA | NA | NA | NA | Gm21776 | | 0 | 0 | 0 | 0 | 0 | 0 | 0 | 0 | 0 | 0 | 0 |
| 0 | NA | NA | NA | NA | NA | Gm21155 | Y-linked testis-specific protein 1-like | 0 | 0 | 0 | 0 | 0 | 0 | 0 | 0 | 0 | 0 | 0 |
| 0 | NA | NA | NA | NA | NA | Gm21163 | Y-linked testis-specific protein 1-like | 0 | 0 | 0 | 0 | 0 | 0 | 0 | 0 | 0 | 0 | 0 |
| 0 | NA | NA | NA | NA | NA | Gm28122 | | 0 | 0 | 0 | 0 | 0 | 0 | 0 | 0 | 0 | 0 | 0 |
| 0 | NA | NA | NA | NA | NA | Gm28121 | | 0 | 0 | 0 | 0 | 0 | 0 | 0 | 0 | 0 | 0 | 0 |
| 0 | NA | NA | NA | NA | NA | Gm21173 | | 0 | 0 | 0 | 0 | 0 | 0 | 0 | 0 | 0 | 0 | 0 |
| 0 | NA | NA | NA | NA | NA | Gm28123 | | 0 | 0 | 0 | 0 | 0 | 0 | 0 | 0 | 0 | 0 | 0 |
| 0 | NA | NA | NA | NA | NA | Gm28120 | | 0 | 0 | 0 | 0 | 0 | 0 | 0 | 0 | 0 | 0 | 0 |
| 0 | NA | NA | NA | NA | NA | Gm29313 | | 0 | 0 | 0 | 0 | 0 | 0 | 0 | 0 | 0 | 0 | 0 |
| 0 | NA | NA | NA | NA | NA | Gm21180 | predicted gene, 21180 | 0 | 0 | 0 | 0 | 0 | 0 | 0 | 0 | 0 | 0 | 0 |
| 0 | NA | NA | NA | NA | NA | Gm20859 | predicted gene, 20859 | 0 | 0 | 0 | 0 | 0 | 0 | 0 | 0 | 0 | 0 | 0 |
| 0 | NA | NA | NA | NA | NA | Gm29314 | | 0 | 0 | 0 | 0 | 0 | 0 | 0 | 0 | 0 | 0 | 0 |
| 0 | NA | NA | NA | NA | NA | Gm29310 | | 0 | 0 | 0 | 0 | 0 | 0 | 0 | 0 | 0 | 0 | 0 |
| 0 | NA | NA | NA | NA | NA | Gm29311 | | 0 | 0 | 0 | 0 | 0 | 0 | 0 | 0 | 0 | 0 | 0 |
| 0 | NA | NA | NA | NA | NA | Gm21638 | predicted gene, 21638 | 0 | 0 | 0 | 0 | 0 | 0 | 0 | 0 | 0 | 0 | 0 |
| 0 | NA | NA | NA | NA | NA | Gm29312 | | 0 | 0 | 0 | 0 | 0 | 0 | 0 | 0 | 0 | 0 | 0 |
| 0 | NA | NA | NA | NA | NA | Gm29308 | | 0 | 0 | 0 | 0 | 0 | 0 | 0 | 0 | 0 | 0 | 0 |

| | | | | | | | | | | | | | | | | | |
|---|---|---|---|---|---|---|---|---|---|---|---|---|---|---|---|---|---|
| 0 | NA | NA | NA | NA | NA | Gm29309 | | 0 | 0 | 0 | 0 | 0 | 0 | 0 | 0 | 0 | 0 | 0 |
| 0 | NA | NA | NA | NA | NA | Gm28702 | | 0 | 0 | 0 | 0 | 0 | 0 | 0 | 0 | 0 | 0 | 0 |
| 0 | NA | NA | NA | NA | NA | Gm28701 | | 0 | 0 | 0 | 0 | 0 | 0 | 0 | 0 | 0 | 0 | 0 |
| 0 | NA | NA | NA | NA | NA | Gm28700 | | 0 | 0 | 0 | 0 | 0 | 0 | 0 | 0 | 0 | 0 | 0 |
| 0 | NA | NA | NA | NA | NA | Gm28699 | | 0 | 0 | 0 | 0 | 0 | 0 | 0 | 0 | 0 | 0 | 0 |
| 0 | NA | NA | NA | NA | NA | Gm28705 | | 0 | 0 | 0 | 0 | 0 | 0 | 0 | 0 | 0 | 0 | 0 |
| 0 | NA | NA | NA | NA | NA | Gm28704 | | 0 | 0 | 0 | 0 | 0 | 0 | 0 | 0 | 0 | 0 | 0 |
| 0 | NA | NA | NA | NA | NA | Gm28540 | | 0 | 0 | 0 | 0 | 0 | 0 | 0 | 0 | 0 | 0 | 0 |
| 0 | NA | NA | NA | NA | NA | Gm21650 | predicted gene, 20874 | 0 | 0 | 0 | 0 | 0 | 0 | 0 | 0 | 0 | 0 | 0 |
| 0 | NA | NA | NA | NA | NA | Gm28541 | | 0 | 0 | 0 | 0 | 0 | 0 | 0 | 0 | 0 | 0 | 0 |
| 0 | NA | NA | NA | NA | NA | Gm28542 | | 0 | 0 | 0 | 0 | 0 | 0 | 0 | 0 | 0 | 0 | 0 |
| 0 | NA | NA | NA | NA | NA | Gm28537 | | 0 | 0 | 0 | 0 | 0 | 0 | 0 | 0 | 0 | 0 | 0 |
| 0 | NA | NA | NA | NA | NA | Gm28538 | | 0 | 0 | 0 | 0 | 0 | 0 | 0 | 0 | 0 | 0 | 0 |
| 0 | NA | NA | NA | NA | NA | Gm29569 | | 0 | 0 | 0 | 0 | 0 | 0 | 0 | 0 | 0 | 0 | 0 |
| 0 | NA | NA | NA | NA | NA | Gm20867 | predicted gene, 20867 | 0 | 0 | 0 | 0 | 0 | 0 | 0 | 0 | 0 | 0 | 0 |
| 0 | NA | NA | NA | NA | NA | Gm21658 | | 0 | 0 | 0 | 0 | 0 | 0 | 0 | 0 | 0 | 0 | 0 |
| 0 | NA | NA | NA | NA | NA | Gm29568 | | 0 | 0 | 0 | 0 | 0 | 0 | 0 | 0 | 0 | 0 | 0 |
| 0 | NA | NA | NA | NA | NA | Gm29567 | | 0 | 0 | 0 | 0 | 0 | 0 | 0 | 0 | 0 | 0 | 0 |
| 0 | NA | NA | NA | NA | NA | Gm21198 | predicted gene, 21198 | 0 | 0 | 0 | 0 | 0 | 0 | 0 | 0 | 0 | 0 | 0 |
| 0 | NA | NA | NA | NA | NA | Gm29566 | | 0 | 0 | 0 | 0 | 0 | 0 | 0 | 0 | 0 | 0 | 0 |
| 0 | NA | NA | NA | NA | NA | Gm29565 | | 0 | 0 | 0 | 0 | 0 | 0 | 0 | 0 | 0 | 0 | 0 |
| 0 | NA | NA | NA | NA | NA | Gm29564 | | 0 | 0 | 0 | 0 | 0 | 0 | 0 | 0 | 0 | 0 | 0 |
| 0 | NA | NA | NA | NA | NA | Gm29147 | | 0 | 0 | 0 | 0 | 0 | 0 | 0 | 0 | 0 | 0 | 0 |
| 0 | NA | NA | NA | NA | NA | Gm29146 | | 0 | 0 | 0 | 0 | 0 | 0 | 0 | 0 | 0 | 0 | 0 |
| 0 | NA | NA | NA | NA | NA | Gm28664 | | 0 | 0 | 0 | 0 | 0 | 0 | 0 | 0 | 0 | 0 | 0 |
| 0 | NA | NA | NA | NA | NA | Gm28663 | | 0 | 0 | 0 | 0 | 0 | 0 | 0 | 0 | 0 | 0 | 0 |
| 0 | NA | NA | NA | NA | NA | Gm29149 | | 0 | 0 | 0 | 0 | 0 | 0 | 0 | 0 | 0 | 0 | 0 |
| 0 | NA | NA | NA | NA | NA | Gm29148 | | 0 | 0 | 0 | 0 | 0 | 0 | 0 | 0 | 0 | 0 | 0 |
| 0 | NA | NA | NA | NA | NA | Gm29145 | | 0 | 0 | 0 | 0 | 0 | 0 | 0 | 0 | 0 | 0 | 0 |
| 0 | NA | NA | NA | NA | NA | Gm29144 | | 0 | 0 | 0 | 0 | 0 | 0 | 0 | 0 | 0 | 0 | 0 |
| 0 | NA | NA | NA | NA | NA | Gm28817 | | 0 | 0 | 0 | 0 | 0 | 0 | 0 | 0 | 0 | 0 | 0 |
| 0 | NA | NA | NA | NA | NA | Gm28816 | | 0 | 0 | 0 | 0 | 0 | 0 | 0 | 0 | 0 | 0 | 0 |
| 0 | NA | NA | NA | NA | NA | Gm20808 | Y-linked testis-specific protein 1-like | 0 | 0 | 0 | 0 | 0 | 0 | 0 | 0 | 0 | 0 | 0 |
| 0 | NA | NA | NA | NA | NA | Gm20806 | spermiogenesis specific transcript on the | 0 | 0 | 0 | 0 | 0 | 0 | 0 | 0 | 0 | 0 | 0 |
| 0 | NA | NA | NA | NA | NA | Gm28815 | | 0 | 0 | 0 | 0 | 0 | 0 | 0 | 0 | 0 | 0 | 0 |
| 0 | NA | NA | NA | NA | NA | Gm28814 | | 0 | 0 | 0 | 0 | 0 | 0 | 0 | 0 | 0 | 0 | 0 |
| 0 | NA | NA | NA | NA | NA | Gm28813 | | 0 | 0 | 0 | 0 | 0 | 0 | 0 | 0 | 0 | 0 | 0 |
| 0 | NA | NA | NA | NA | NA | Gm20919 | Y-linked testis-specific protein 1-like | 0 | 0 | 0 | 0 | 0 | 0 | 0 | 0 | 0 | 0 | 0 |
| 0 | NA | NA | NA | NA | NA | Gm20917 | predicted gene, 20917 | 0 | 0 | 0 | 0 | 0 | 0 | 0 | 0 | 0 | 0 | 0 |
| 0 | NA | NA | NA | NA | NA | Gm28421 | | 0 | 0 | 0 | 0 | 0 | 0 | 0 | 0 | 0 | 0 | 0 |
| 0 | NA | NA | NA | NA | NA | Gm28420 | | 0 | 0 | 0 | 0 | 0 | 0 | 0 | 0 | 0 | 0 | 0 |
| 0 | NA | NA | NA | NA | NA | Gm28423 | | 0 | 0 | 0 | 0 | 0 | 0 | 0 | 0 | 0 | 0 | 0 |
| 0 | NA | NA | NA | NA | NA | Gm28422 | | 0 | 0 | 0 | 0 | 0 | 0 | 0 | 0 | 0 | 0 | 0 |
| 0 | NA | NA | NA | NA | NA | Gm20916 | predicted gene, 20916 | 0 | 0 | 0 | 0 | 0 | 0 | 0 | 0 | 0 | 0 | 0 |
| 0 | NA | NA | NA | NA | NA | Gm29033 | | 0 | 0 | 0 | 0 | 0 | 0 | 0 | 0 | 0 | 0 | 0 |
| 0 | NA | NA | NA | NA | NA | Gm29034 | | 0 | 0 | 0 | 0 | 0 | 0 | 0 | 0 | 0 | 0 | 0 |
| 0 | NA | NA | NA | NA | NA | Gm21823 | | 0 | 0 | 0 | 0 | 0 | 0 | 0 | 0 | 0 | 0 | 0 |
| 0 | NA | NA | NA | NA | NA | Gm21242 | | 0 | 0 | 0 | 0 | 0 | 0 | 0 | 0 | 0 | 0 | 0 |
| 0 | NA | NA | NA | NA | NA | Gm29031 | | 0 | 0 | 0 | 0 | 0 | 0 | 0 | 0 | 0 | 0 | 0 |
| 0 | NA | NA | NA | NA | NA | Gm29032 | | 0 | 0 | 0 | 0 | 0 | 0 | 0 | 0 | 0 | 0 | 0 |
| 0 | NA | NA | NA | NA | NA | Gm29606 | | 0 | 0 | 0 | 0 | 0 | 0 | 0 | 0 | 0 | 0 | 0 |
| 0 | NA | NA | NA | NA | NA | Gm29607 | | 0 | 0 | 0 | 0 | 0 | 0 | 0 | 0 | 0 | 0 | 0 |
| 0 | NA | NA | NA | NA | NA | Gm21248 | | 0 | 0 | 0 | 0 | 0 | 0 | 0 | 0 | 0 | 0 | 0 |
| 0 | NA | NA | NA | NA | NA | Gm29189 | | 0 | 0 | 0 | 0 | 0 | 0 | 0 | 0 | 0 | 0 | 0 |
| 0 | NA | NA | NA | NA | NA | Gm21873 | | 0 | 0 | 0 | 0 | 0 | 0 | 0 | 0 | 0 | 0 | 0 |
| 0 | NA | NA | NA | NA | NA | Gm29190 | | 0 | 0 | 0 | 0 | 0 | 0 | 0 | 0 | 0 | 0 | 0 |
| 0 | NA | NA | NA | NA | NA | Gm21760 | | 0 | 0 | 0 | 0 | 0 | 0 | 0 | 0 | 0 | 0 | 0 |
| 0 | NA | NA | NA | NA | NA | Gm28159 | | 0 | 0 | 0 | 0 | 0 | 0 | 0 | 0 | 0 | 0 | 0 |
| 0 | NA | NA | NA | NA | NA | Gm28160 | | 0 | 0 | 0 | 0 | 0 | 0 | 0 | 0 | 0 | 0 | 0 |
| 0 | NA | NA | NA | NA | NA | Gm28157 | | 0 | 0 | 0 | 0 | 0 | 0 | 0 | 0 | 0 | 0 | 0 |

| | | | | | | | | | | | | | | | | | | |
|---|---|---|---|---|---|---|---|---|---|---|---|---|---|---|---|---|---|---|
| 0 | NA | NA | NA | NA | NA | Gm28158 | | 0 | 0 | 0 | 0 | 0 | 0 | 0 | 0 | 0 | 0 | 0 |
| 0 | NA | NA | NA | NA | NA | Gm21257 | predicted gene, 21257 | 0 | 0 | 0 | 0 | 0 | 0 | 0 | 0 | 0 | 0 | 0 |
| 0 | NA | NA | NA | NA | NA | Gm28161 | | 0 | 0 | 0 | 0 | 0 | 0 | 0 | 0 | 0 | 0 | 0 |
| 0 | NA | NA | NA | NA | NA | Gm28695 | | 0 | 0 | 0 | 0 | 0 | 0 | 0 | 0 | 0 | 0 | 0 |
| 0 | NA | NA | NA | NA | NA | Gm28696 | | 0 | 0 | 0 | 0 | 0 | 0 | 0 | 0 | 0 | 0 | 0 |
| 0 | NA | NA | NA | NA | NA | Gm20791 | | 0 | 0 | 0 | 0 | 0 | 0 | 0 | 0 | 0 | 0 | 0 |
| 0 | NA | NA | NA | NA | NA | Gm20841 | predicted gene, 20841 | 0 | 0 | 0 | 0 | 0 | 0 | 0 | 0 | 0 | 0 | 0 |
| 0 | NA | NA | NA | NA | NA | Gm28899 | | 0 | 0 | 0 | 0 | 0 | 0 | 0 | 0 | 0 | 0 | 0 |
| 0 | NA | NA | NA | NA | NA | Gm21722 | | 0 | 0 | 0 | 0 | 0 | 0 | 0 | 0 | 0 | 0 | 0 |
| 0 | NA | NA | NA | NA | NA | Gm28898 | | 0 | 0 | 0 | 0 | 0 | 0 | 0 | 0 | 0 | 0 | 0 |
| 0 | NA | NA | NA | NA | NA | Gm28897 | | 0 | 0 | 0 | 0 | 0 | 0 | 0 | 0 | 0 | 0 | 0 |
| 0 | NA | NA | NA | NA | NA | Gm28896 | | 0 | 0 | 0 | 0 | 0 | 0 | 0 | 0 | 0 | 0 | 0 |
| 0 | NA | NA | NA | NA | NA | Gm21282 | predicted gene, 21282 | 0 | 0 | 0 | 0 | 0 | 0 | 0 | 0 | 0 | 0 | 0 |
| 0 | NA | NA | NA | NA | NA | Gm28895 | | 0 | 0 | 0 | 0 | 0 | 0 | 0 | 0 | 0 | 0 | 0 |
| 0 | NA | NA | NA | NA | NA | Gm28894 | | 0 | 0 | 0 | 0 | 0 | 0 | 0 | 0 | 0 | 0 | 0 |
| 0 | NA | NA | NA | NA | NA | Gm21829 | | 0 | 0 | 0 | 0 | 0 | 0 | 0 | 0 | 0 | 0 | 0 |
| 0 | NA | NA | NA | NA | NA | Gm29324 | | 0 | 0 | 0 | 0 | 0 | 0 | 0 | 0 | 0 | 0 | 0 |
| 0 | NA | NA | NA | NA | NA | Gm20911 | predicted gene, 20911 | 0 | 0 | 0 | 0 | 0 | 0 | 0 | 0 | 0 | 0 | 0 |
| 0 | NA | NA | NA | NA | NA | Gm28739 | | 0 | 0 | 0 | 0 | 0 | 0 | 0 | 0 | 0 | 0 | 0 |
| 0 | NA | NA | NA | NA | NA | Gm28741 | | 0 | 0 | 0 | 0 | 0 | 0 | 0 | 0 | 0 | 0 | 0 |
| 0 | NA | NA | NA | NA | NA | Gm20912 | predicted gene, 20912 | 0 | 0 | 0 | 0 | 0 | 0 | 0 | 0 | 0 | 0 | 0 |
| 0 | NA | NA | NA | NA | NA | Gm20801 | Y-linked testis-specific protein 1-like | 0 | 0 | 0 | 0 | 0 | 0 | 0 | 0 | 0 | 0 | 0 |
| 0 | NA | NA | NA | NA | NA | Gm28742 | | 0 | 0 | 0 | 0 | 0 | 0 | 0 | 0 | 0 | 0 | 0 |
| 0 | NA | NA | NA | NA | NA | Gm21919 | | 0 | 0 | 0 | 0 | 0 | 0 | 0 | 0 | 0 | 0 | 0 |
| 0 | NA | NA | NA | NA | NA | Gm28743 | | 0 | 0 | 0 | 0 | 0 | 0 | 0 | 0 | 0 | 0 | 0 |
| 0 | NA | NA | NA | NA | NA | Gm28124 | | 0 | 0 | 0 | 0 | 0 | 0 | 0 | 0 | 0 | 0 | 0 |
| 0 | NA | NA | NA | NA | NA | Gm21790 | | 0 | 0 | 0 | 0 | 0 | 0 | 0 | 0 | 0 | 0 | 0 |
| 0 | NA | NA | NA | NA | NA | Gm21728 | | 0 | 0 | 0 | 0 | 0 | 0 | 0 | 0 | 0 | 0 | 0 |
| 0 | NA | NA | NA | NA | NA | Gm28126 | | 0 | 0 | 0 | 0 | 0 | 0 | 0 | 0 | 0 | 0 | 0 |
| 0 | NA | NA | NA | NA | NA | Gm28125 | | 0 | 0 | 0 | 0 | 0 | 0 | 0 | 0 | 0 | 0 | 0 |
| 0 | NA | NA | NA | NA | NA | Gm21825 | | 0 | 0 | 0 | 0 | 0 | 0 | 0 | 0 | 0 | 0 | 0 |
| 0 | NA | NA | NA | NA | NA | Gm21317 | X-linked lymphocyte-regulated protein PM | 0 | 0 | 0 | 0 | 0 | 0 | 0 | 0 | 0 | 0 | 0 |
| 0 | NA | NA | NA | NA | NA | Gm29468 | | 0 | 0 | 0 | 0 | 0 | 0 | 0 | 0 | 0 | 0 | 0 |
| 0 | NA | NA | NA | NA | NA | Gm29469 | | 0 | 0 | 0 | 0 | 0 | 0 | 0 | 0 | 0 | 0 | 0 |
| 0 | NA | NA | NA | NA | NA | Gm29472 | | 0 | 0 | 0 | 0 | 0 | 0 | 0 | 0 | 0 | 0 | 0 |
| 0 | NA | NA | NA | NA | NA | Gm29473 | | 0 | 0 | 0 | 0 | 0 | 0 | 0 | 0 | 0 | 0 | 0 |
| 0 | NA | NA | NA | NA | NA | Gm29470 | | 0 | 0 | 0 | 0 | 0 | 0 | 0 | 0 | 0 | 0 | 0 |
| 0 | NA | NA | NA | NA | NA | Gm21333 | predicted gene, 21333 | 0 | 0 | 0 | 0 | 0 | 0 | 0 | 0 | 0 | 0 | 0 |
| 0 | NA | NA | NA | NA | NA | Gm29382 | | 0 | 0 | 0 | 0 | 0 | 0 | 0 | 0 | 0 | 0 | 0 |
| 0 | NA | NA | NA | NA | NA | Gm28224 | | 0 | 0 | 0 | 0 | 0 | 0 | 0 | 0 | 0 | 0 | 0 |
| 0 | NA | NA | NA | NA | NA | Gm28226 | | 0 | 0 | 0 | 0 | 0 | 0 | 0 | 0 | 0 | 0 | 0 |
| 0 | NA | NA | NA | NA | NA | Gm28225 | | 0 | 0 | 0 | 0 | 0 | 0 | 0 | 0 | 0 | 0 | 0 |
| 0 | NA | NA | NA | NA | NA | Gm21344 | Y-linked testis-specific protein 1-like | 0 | 0 | 0 | 0 | 0 | 0 | 0 | 0 | 0 | 0 | 0 |
| 0 | NA | NA | NA | NA | NA | Gm20849 | | 0 | 0 | 0 | 0 | 0 | 0 | 0 | 0 | 0 | 0 | 0 |
| 0 | NA | NA | NA | NA | NA | Gm28829 | | 0 | 0 | 0 | 0 | 0 | 0 | 0 | 0 | 0 | 0 | 0 |
| 0 | NA | NA | NA | NA | NA | Gm21901 | | 0 | 0 | 0 | 0 | 0 | 0 | 0 | 0 | 0 | 0 | 0 |
| 0 | NA | NA | NA | NA | NA | Gm28827 | | 0 | 0 | 0 | 0 | 0 | 0 | 0 | 0 | 0 | 0 | 0 |
| 0 | NA | NA | NA | NA | NA | Gm28828 | | 0 | 0 | 0 | 0 | 0 | 0 | 0 | 0 | 0 | 0 | 0 |
| 0 | NA | NA | NA | NA | NA | Gm28831 | | 0 | 0 | 0 | 0 | 0 | 0 | 0 | 0 | 0 | 0 | 0 |
| 0 | NA | NA | NA | NA | NA | Gm28832 | | 0 | 0 | 0 | 0 | 0 | 0 | 0 | 0 | 0 | 0 | 0 |
| 0 | NA | NA | NA | NA | NA | Gm21864 | | 0 | 0 | 0 | 0 | 0 | 0 | 0 | 0 | 0 | 0 | 0 |
| 0 | NA | NA | NA | NA | NA | Gm21802 | | 0 | 0 | 0 | 0 | 0 | 0 | 0 | 0 | 0 | 0 | 0 |
| 0 | NA | NA | NA | NA | NA | Gm28658 | | 0 | 0 | 0 | 0 | 0 | 0 | 0 | 0 | 0 | 0 | 0 |
| 0 | NA | NA | NA | NA | NA | Gm29238 | | 0 | 0 | 0 | 0 | 0 | 0 | 0 | 0 | 0 | 0 | 0 |
| 0 | NA | NA | NA | NA | NA | Gm21088 | | 0 | 0 | 0 | 0 | 0 | 0 | 0 | 0 | 0 | 0 | 0 |
| 0 | NA | NA | NA | NA | NA | Gm29239 | | 0 | 0 | 0 | 0 | 0 | 0 | 0 | 0 | 0 | 0 | 0 |
| 0 | NA | NA | NA | NA | NA | Gm25729 | | 0 | 0 | 0 | 0 | 0 | 0 | 0 | 0 | 0 | 0 | 0 |
| 0 | NA | NA | NA | NA | NA | Gm21741 | | 0 | 0 | 0 | 0 | 0 | 0 | 0 | 0 | 0 | 0 | 0 |
| 0 | NA | NA | NA | NA | NA | Gm20862 | predicted gene, 20862 | 0 | 0 | 0 | 0 | 0 | 0 | 0 | 0 | 0 | 0 | 0 |
| 0 | NA | NA | NA | NA | NA | Gm22624 | | 0 | 0 | 0 | 0 | 0 | 0 | 0 | 0 | 0 | 0 | 0 |

| | | | | | | | | | | | | | | | | | |
|---|---|---|---|---|---|---|---|---|---|---|---|---|---|---|---|---|---|
| 0 | NA | NA | NA | NA | NA | Gm20863 | Y-linked testis-specific protein 1-like | 0 | 0 | 0 | 0 | 0 | 0 | 0 | 0 | 0 | 0 | 0 |
| 0 | NA | NA | NA | NA | NA | Gm20864 | | 0 | 0 | 0 | 0 | 0 | 0 | 0 | 0 | 0 | 0 | 0 |
| 0 | NA | NA | NA | NA | NA | Gm29476 | | 0 | 0 | 0 | 0 | 0 | 0 | 0 | 0 | 0 | 0 | 0 |
| 0 | NA | NA | NA | NA | NA | Gm21717 | | 0 | 0 | 0 | 0 | 0 | 0 | 0 | 0 | 0 | 0 | 0 |
| 0 | NA | NA | NA | NA | NA | Gm28472 | | 0 | 0 | 0 | 0 | 0 | 0 | 0 | 0 | 0 | 0 | 0 |
| 0 | NA | NA | NA | NA | NA | Gm21095 | | 0 | 0 | 0 | 0 | 0 | 0 | 0 | 0 | 0 | 0 | 0 |
| 0 | NA | NA | NA | NA | NA | Gm28473 | | 0 | 0 | 0 | 0 | 0 | 0 | 0 | 0 | 0 | 0 | 0 |
| 0 | NA | NA | NA | NA | NA | Gm28474 | | 0 | 0 | 0 | 0 | 0 | 0 | 0 | 0 | 0 | 0 | 0 |
| 0 | NA | NA | NA | NA | NA | Gm28475 | | 0 | 0 | 0 | 0 | 0 | 0 | 0 | 0 | 0 | 0 | 0 |
| 0 | NA | NA | NA | NA | NA | Gm28476 | | 0 | 0 | 0 | 0 | 0 | 0 | 0 | 0 | 0 | 0 | 0 |
| 0 | NA | NA | NA | NA | NA | Gm21380 | predicted gene, 21380 | 0 | 0 | 0 | 0 | 0 | 0 | 0 | 0 | 0 | 0 | 0 |
| 0 | NA | NA | NA | NA | NA | Gm21782 | | 0 | 0 | 0 | 0 | 0 | 0 | 0 | 0 | 0 | 0 | 0 |
| 0 | NA | NA | NA | NA | NA | Gm21394 | Y-linked testis-specific protein 1-like | 0 | 0 | 0 | 0 | 0 | 0 | 0 | 0 | 0 | 0 | 0 |
| 0 | NA | NA | NA | NA | NA | Gm28471 | | 0 | 0 | 0 | 0 | 0 | 0 | 0 | 0 | 0 | 0 | 0 |
| 0 | NA | NA | NA | NA | NA | Gm21824 | | 0 | 0 | 0 | 0 | 0 | 0 | 0 | 0 | 0 | 0 | 0 |
| 0 | NA | NA | NA | NA | NA | Gm29338 | | 0 | 0 | 0 | 0 | 0 | 0 | 0 | 0 | 0 | 0 | 0 |
| 0 | NA | NA | NA | NA | NA | Gm21409 | X-linked lymphocyte-regulated protein PM | 0 | 0 | 0 | 0 | 0 | 0 | 0 | 0 | 0 | 0 | 0 |
| 0 | NA | NA | NA | NA | NA | Gm29339 | | 0 | 0 | 0 | 0 | 0 | 0 | 0 | 0 | 0 | 0 | 0 |
| 0 | NA | NA | NA | NA | NA | Gm28759 | | 0 | 0 | 0 | 0 | 0 | 0 | 0 | 0 | 0 | 0 | 0 |
| 0 | NA | NA | NA | NA | NA | Gm28758 | | 0 | 0 | 0 | 0 | 0 | 0 | 0 | 0 | 0 | 0 | 0 |
| 0 | NA | NA | NA | NA | NA | Gm29071 | | 0 | 0 | 0 | 0 | 0 | 0 | 0 | 0 | 0 | 0 | 0 |
| 0 | NA | NA | NA | NA | NA | Gm21879 | | 0 | 0 | 0 | 0 | 0 | 0 | 0 | 0 | 0 | 0 | 0 |
| 0 | NA | NA | NA | NA | NA | Gm28757 | | 0 | 0 | 0 | 0 | 0 | 0 | 0 | 0 | 0 | 0 | 0 |
| 0 | NA | NA | NA | NA | NA | Gm20856 | | 0 | 0 | 0 | 0 | 0 | 0 | 0 | 0 | 0 | 0 | 0 |
| 0 | NA | NA | NA | NA | NA | Gm28348 | | 0 | 0 | 0 | 0 | 0 | 0 | 0 | 0 | 0 | 0 | 0 |
| 0 | NA | NA | NA | NA | NA | Gm28973 | | 0 | 0 | 0 | 0 | 0 | 0 | 0 | 0 | 0 | 0 | 0 |
| 0 | NA | NA | NA | NA | NA | Gm21734 | | 0 | 0 | 0 | 0 | 0 | 0 | 0 | 0 | 0 | 0 | 0 |
| 0 | NA | NA | NA | NA | NA | Gm28972 | | 0 | 0 | 0 | 0 | 0 | 0 | 0 | 0 | 0 | 0 | 0 |
| 0 | NA | NA | NA | NA | NA | Gm28970 | | 0 | 0 | 0 | 0 | 0 | 0 | 0 | 0 | 0 | 0 | 0 |
| 0 | NA | NA | NA | NA | NA | Gm28971 | | 0 | 0 | 0 | 0 | 0 | 0 | 0 | 0 | 0 | 0 | 0 |
| 0 | NA | NA | NA | NA | NA | Gm24828 | | 0 | 0 | 0 | 0 | 0 | 0 | 0 | 0 | 0 | 0 | 0 |
| 0 | NA | NA | NA | NA | NA | Gm21785 | | 0 | 0 | 0 | 0 | 0 | 0 | 0 | 0 | 0 | 0 | 0 |
| 0 | NA | NA | NA | NA | NA | Gm20854 | predicted gene, 20854 | 0 | 0 | 0 | 0 | 0 | 0 | 0 | 0 | 0 | 0 | 0 |
| 0 | NA | NA | NA | NA | NA | Gm28349 | | 0 | 0 | 0 | 0 | 0 | 0 | 0 | 0 | 0 | 0 | 0 |
| 0 | NA | NA | NA | NA | NA | Gm21892 | | 0 | 0 | 0 | 0 | 0 | 0 | 0 | 0 | 0 | 0 | 0 |
| 0 | NA | NA | NA | NA | NA | Gm28346 | | 0 | 0 | 0 | 0 | 0 | 0 | 0 | 0 | 0 | 0 | 0 |
| 0 | NA | NA | NA | NA | NA | Gm28345 | | 0 | 0 | 0 | 0 | 0 | 0 | 0 | 0 | 0 | 0 | 0 |
| 0 | NA | NA | NA | NA | NA | Gm29090 | | 0 | 0 | 0 | 0 | 0 | 0 | 0 | 0 | 0 | 0 | 0 |
| 0 | NA | NA | NA | NA | NA | Gm28152 | | 0 | 0 | 0 | 0 | 0 | 0 | 0 | 0 | 0 | 0 | 0 |
| 0 | NA | NA | NA | NA | NA | Gm23770 | | 0 | 0 | 0 | 0 | 0 | 0 | 0 | 0 | 0 | 0 | 0 |
| 0 | NA | NA | NA | NA | NA | Gm21881 | | 0 | 0 | 0 | 0 | 0 | 0 | 0 | 0 | 0 | 0 | 0 |
| 0 | NA | NA | NA | NA | NA | Gm21435 | predicted gene, 21435 | 0 | 0 | 0 | 0 | 0 | 0 | 0 | 0 | 0 | 0 | 0 |
| 0 | NA | NA | NA | NA | NA | Gm29505 | | 0 | 0 | 0 | 0 | 0 | 0 | 0 | 0 | 0 | 0 | 0 |
| 0 | NA | NA | NA | NA | NA | Gm21862 | | 0 | 0 | 0 | 0 | 0 | 0 | 0 | 0 | 0 | 0 | 0 |
| 0 | NA | NA | NA | NA | NA | Gm28823 | | 0 | 0 | 0 | 0 | 0 | 0 | 0 | 0 | 0 | 0 | 0 |
| 0 | NA | NA | NA | NA | NA | Gm28220 | | 0 | 0 | 0 | 0 | 0 | 0 | 0 | 0 | 0 | 0 | 0 |
| 0 | NA | NA | NA | NA | NA | Gm20820 | | 0 | 0 | 0 | 0 | 0 | 0 | 0 | 0 | 0 | 0 | 0 |
| 0 | NA | NA | NA | NA | NA | Gm28219 | | 0 | 0 | 0 | 0 | 0 | 0 | 0 | 0 | 0 | 0 | 0 |
| 0 | NA | NA | NA | NA | NA | Gm28218 | | 0 | 0 | 0 | 0 | 0 | 0 | 0 | 0 | 0 | 0 | 0 |
| 0 | NA | NA | NA | NA | NA | Gm23863 | | 0 | 0 | 0 | 0 | 0 | 0 | 0 | 0 | 0 | 0 | 0 |
| 0 | NA | NA | NA | NA | NA | Gm20861 | Y-linked testis-specific protein 1-like | 0 | 0 | 0 | 0 | 0 | 0 | 0 | 0 | 0 | 0 | 0 |
| 0 | NA | NA | NA | NA | NA | Gm20860 | predicted gene, 20860 | 0 | 0 | 0 | 0 | 0 | 0 | 0 | 0 | 0 | 0 | 0 |
| 0 | NA | NA | NA | NA | NA | Gm28057 | | 0 | 0 | 0 | 0 | 0 | 0 | 0 | 0 | 0 | 0 | 0 |
| 0 | NA | NA | NA | NA | NA | Gm21796 | | 0 | 0 | 0 | 0 | 0 | 0 | 0 | 0 | 0 | 0 | 0 |
| 0 | NA | NA | NA | NA | NA | Gm29302 | | 0 | 0 | 0 | 0 | 0 | 0 | 0 | 0 | 0 | 0 | 0 |
| 0 | NA | NA | NA | NA | NA | Gm28440 | | 0 | 0 | 0 | 0 | 0 | 0 | 0 | 0 | 0 | 0 | 0 |
| 0 | NA | NA | NA | NA | NA | Gm29299 | | 0 | 0 | 0 | 0 | 0 | 0 | 0 | 0 | 0 | 0 | 0 |
| 0 | NA | NA | NA | NA | NA | Gm29298 | | 0 | 0 | 0 | 0 | 0 | 0 | 0 | 0 | 0 | 0 | 0 |
| 0 | NA | NA | NA | NA | NA | Gm29092 | | 0 | 0 | 0 | 0 | 0 | 0 | 0 | 0 | 0 | 0 | 0 |
| 0 | NA | NA | NA | NA | NA | Gm29091 | | 0 | 0 | 0 | 0 | 0 | 0 | 0 | 0 | 0 | 0 | 0 |

| | | | | | | | | | | | | | | | | | | |
|---|---|---|---|---|---|---|---|---|---|---|---|---|---|---|---|---|---|---|
| 0 | NA | NA | NA | NA | NA | Gm29297 | | 0 | 0 | 0 | 0 | 0 | 0 | 0 | 0 | 0 | 0 | 0 |
| 0 | NA | NA | NA | NA | NA | Gm21462 | Y-linked testis-specific protein 1-like | 0 | 0 | 0 | 0 | 0 | 0 | 0 | 0 | 0 | 0 | 0 |
| 0 | NA | NA | NA | NA | NA | Gm21470 | predicted gene, 21470 | 0 | 0 | 0 | 0 | 0 | 0 | 0 | 0 | 0 | 0 | 0 |
| 0 | NA | NA | NA | NA | NA | Gm28366 | | 0 | 0 | 0 | 0 | 0 | 0 | 0 | 0 | 0 | 0 | 0 |
| 0 | NA | NA | NA | NA | NA | Gm21752 | | 0 | 0 | 0 | 0 | 0 | 0 | 0 | 0 | 0 | 0 | 0 |
| 0 | NA | NA | NA | NA | NA | Gm28365 | | 0 | 0 | 0 | 0 | 0 | 0 | 0 | 0 | 0 | 0 | 0 |
| 0 | NA | NA | NA | NA | NA | Gm21477 | X-linked lymphocyte-regulated protein PM | 0 | 0 | 0 | 0 | 0 | 0 | 0 | 0 | 0 | 0 | 0 |
| 0 | NA | NA | NA | NA | NA | Gm28368 | | 0 | 0 | 0 | 0 | 0 | 0 | 0 | 0 | 0 | 0 | 0 |
| 0 | NA | NA | NA | NA | NA | Gm28369 | | 0 | 0 | 0 | 0 | 0 | 0 | 0 | 0 | 0 | 0 | 0 |
| 0 | NA | NA | NA | NA | NA | Gm28367 | | 0 | 0 | 0 | 0 | 0 | 0 | 0 | 0 | 0 | 0 | 0 |
| 0 | NA | NA | NA | NA | NA | Gm29115 | | 0 | 0 | 0 | 0 | 0 | 0 | 0 | 0 | 0 | 0 | 0 |
| 0 | NA | NA | NA | NA | NA | Gm21813 | | 0 | 0 | 0 | 0 | 0 | 0 | 0 | 0 | 0 | 0 | 0 |
| 0 | NA | NA | NA | NA | NA | Gm20906 | predicted gene, 20906 | 0 | 0 | 0 | 0 | 0 | 0 | 0 | 0 | 0 | 0 | 0 |
| 0 | NA | NA | NA | NA | NA | Gm29111 | | 0 | 0 | 0 | 0 | 0 | 0 | 0 | 0 | 0 | 0 | 0 |
| 0 | NA | NA | NA | NA | NA | Gm28110 | | 0 | 0 | 0 | 0 | 0 | 0 | 0 | 0 | 0 | 0 | 0 |
| 0 | NA | NA | NA | NA | NA | Gm28109 | | 0 | 0 | 0 | 0 | 0 | 0 | 0 | 0 | 0 | 0 | 0 |
| 0 | NA | NA | NA | NA | NA | Gm28108 | | 0 | 0 | 0 | 0 | 0 | 0 | 0 | 0 | 0 | 0 | 0 |
| 0 | NA | NA | NA | NA | NA | Gm28107 | | 0 | 0 | 0 | 0 | 0 | 0 | 0 | 0 | 0 | 0 | 0 |
| 0 | NA | NA | NA | NA | NA | Gm28106 | | 0 | 0 | 0 | 0 | 0 | 0 | 0 | 0 | 0 | 0 | 0 |
| 0 | NA | NA | NA | NA | NA | Gm28105 | | 0 | 0 | 0 | 0 | 0 | 0 | 0 | 0 | 0 | 0 | 0 |
| 0 | NA | NA | NA | NA | NA | Gm28104 | | 0 | 0 | 0 | 0 | 0 | 0 | 0 | 0 | 0 | 0 | 0 |
| 0 | NA | NA | NA | NA | NA | Gm28103 | | 0 | 0 | 0 | 0 | 0 | 0 | 0 | 0 | 0 | 0 | 0 |
| 0 | NA | NA | NA | NA | NA | Gm28102 | | 0 | 0 | 0 | 0 | 0 | 0 | 0 | 0 | 0 | 0 | 0 |
| 0 | NA | NA | NA | NA | NA | Gm28716 | | 0 | 0 | 0 | 0 | 0 | 0 | 0 | 0 | 0 | 0 | 0 |
| 0 | NA | NA | NA | NA | NA | Gm28717 | | 0 | 0 | 0 | 0 | 0 | 0 | 0 | 0 | 0 | 0 | 0 |
| 0 | NA | NA | NA | NA | NA | Gm28718 | | 0 | 0 | 0 | 0 | 0 | 0 | 0 | 0 | 0 | 0 | 0 |
| 0 | NA | NA | NA | NA | NA | Gm29162 | | 0 | 0 | 0 | 0 | 0 | 0 | 0 | 0 | 0 | 0 | 0 |
| 0 | NA | NA | NA | NA | NA | Gm21777 | | 0 | 0 | 0 | 0 | 0 | 0 | 0 | 0 | 0 | 0 | 0 |
| 0 | NA | NA | NA | NA | NA | Gm28866 | | 0 | 0 | 0 | 0 | 0 | 0 | 0 | 0 | 0 | 0 | 0 |
| 0 | NA | NA | NA | NA | NA | Gm28405 | | 0 | 0 | 0 | 0 | 0 | 0 | 0 | 0 | 0 | 0 | 0 |
| 0 | NA | NA | NA | NA | NA | Gm21792 | | 0 | 0 | 0 | 0 | 0 | 0 | 0 | 0 | 0 | 0 | 0 |
| 0 | NA | NA | NA | NA | NA | Gm28406 | | 0 | 0 | 0 | 0 | 0 | 0 | 0 | 0 | 0 | 0 | 0 |
| 0 | NA | NA | NA | NA | NA | Gm28978 | | 0 | 0 | 0 | 0 | 0 | 0 | 0 | 0 | 0 | 0 | 0 |
| 0 | NA | NA | NA | NA | NA | Gm29436 | | 0 | 0 | 0 | 0 | 0 | 0 | 0 | 0 | 0 | 0 | 0 |
| 0 | NA | NA | NA | NA | NA | Gm20819 | | 0 | 0 | 0 | 0 | 0 | 0 | 0 | 0 | 0 | 0 | 0 |
| 0 | NA | NA | NA | NA | NA | Gm29437 | | 0 | 0 | 0 | 0 | 0 | 0 | 0 | 0 | 0 | 0 | 0 |
| 0 | NA | NA | NA | NA | NA | Gm28977 | | 0 | 0 | 0 | 0 | 0 | 0 | 0 | 0 | 0 | 0 | 0 |
| 0 | NA | NA | NA | NA | NA | Gm28407 | | 0 | 0 | 0 | 0 | 0 | 0 | 0 | 0 | 0 | 0 | 0 |
| 0 | NA | NA | NA | NA | NA | Gm21287 | predicted gene, 21287 | 0 | 0 | 0 | 0 | 0 | 0 | 0 | 0 | 0 | 0 | 0 |
| 0 | NA | NA | NA | NA | NA | Gm29390 | | 0 | 0 | 0 | 0 | 0 | 0 | 0 | 0 | 0 | 0 | 0 |
| 0 | NA | NA | NA | NA | NA | Gm29391 | | 0 | 0 | 0 | 0 | 0 | 0 | 0 | 0 | 0 | 0 | 0 |
| 0 | NA | NA | NA | NA | NA | Gm21720 | | 0 | 0 | 0 | 0 | 0 | 0 | 0 | 0 | 0 | 0 | 0 |
| 0 | NA | NA | NA | NA | NA | Gm29392 | | 0 | 0 | 0 | 0 | 0 | 0 | 0 | 0 | 0 | 0 | 0 |
| 0 | NA | NA | NA | NA | NA | Gm29393 | | 0 | 0 | 0 | 0 | 0 | 0 | 0 | 0 | 0 | 0 | 0 |
| 0 | NA | NA | NA | NA | NA | Gm21294 | | 0 | 0 | 0 | 0 | 0 | 0 | 0 | 0 | 0 | 0 | 0 |
| 0 | NA | NA | NA | NA | NA | Gm28672 | | 0 | 0 | 0 | 0 | 0 | 0 | 0 | 0 | 0 | 0 | 0 |
| 0 | NA | NA | NA | NA | NA | Gm21838 | | 0 | 0 | 0 | 0 | 0 | 0 | 0 | 0 | 0 | 0 | 0 |
| 0 | NA | NA | NA | NA | NA | Gm28670 | | 0 | 0 | 0 | 0 | 0 | 0 | 0 | 0 | 0 | 0 | 0 |
| 0 | NA | NA | NA | NA | NA | Gm28671 | | 0 | 0 | 0 | 0 | 0 | 0 | 0 | 0 | 0 | 0 | 0 |
| 0 | NA | NA | NA | NA | NA | Gm28673 | | 0 | 0 | 0 | 0 | 0 | 0 | 0 | 0 | 0 | 0 | 0 |
| 0 | NA | NA | NA | NA | NA | Gm28674 | | 0 | 0 | 0 | 0 | 0 | 0 | 0 | 0 | 0 | 0 | 0 |
| 0 | NA | NA | NA | NA | NA | Gm28531 | | 0 | 0 | 0 | 0 | 0 | 0 | 0 | 0 | 0 | 0 | 0 |
| 0 | NA | NA | NA | NA | NA | Gm21996 | | 0 | 0 | 0 | 0 | 0 | 0 | 0 | 0 | 0 | 0 | 0 |
| 0 | NA | NA | NA | NA | NA | Gm21803 | | 0 | 0 | 0 | 0 | 0 | 0 | 0 | 0 | 0 | 0 | 0 |
| 0 | NA | NA | NA | NA | NA | Gm28930 | | 0 | 0 | 0 | 0 | 0 | 0 | 0 | 0 | 0 | 0 | 0 |
| 0 | NA | NA | NA | NA | NA | Gm20879 | | 0 | 0 | 0 | 0 | 0 | 0 | 0 | 0 | 0 | 0 | 0 |
| 0 | NA | NA | NA | NA | NA | Gm29504 | | 0 | 0 | 0 | 0 | 0 | 0 | 0 | 0 | 0 | 0 | 0 |
| 0 | NA | NA | NA | NA | NA | Gm20837 | | 0 | 0 | 0 | 0 | 0 | 0 | 0 | 0 | 0 | 0 | 0 |
| 0 | NA | NA | NA | NA | NA | Gm29503 | | 0 | 0 | 0 | 0 | 0 | 0 | 0 | 0 | 0 | 0 | 0 |
| 0 | NA | NA | NA | NA | NA | Gm28300 | | 0 | 0 | 0 | 0 | 0 | 0 | 0 | 0 | 0 | 0 | 0 |

| | | | | | | | | | | | | | | | | | | |
|---|---|---|---|---|---|---|---|---|---|---|---|---|---|---|---|---|---|---|
| 0 | NA | NA | NA | NA | NA | Gm28301 | | 0 | 0 | 0 | 0 | 0 | 0 | 0 | 0 | 0 | 0 | 0 |
| 0 | NA | NA | NA | NA | NA | Gm21860 | | 0 | 0 | 0 | 0 | 0 | 0 | 0 | 0 | 0 | 0 | 0 |
| 0 | NA | NA | NA | NA | NA | Gm21742 | | 0 | 0 | 0 | 0 | 0 | 0 | 0 | 0 | 0 | 0 | 0 |
| 0 | NA | NA | NA | NA | NA | Gm21748 | | 0 | 0 | 0 | 0 | 0 | 0 | 0 | 0 | 0 | 0 | 0 |
| 0 | NA | NA | NA | NA | NA | mt-Tf | | 0 | 0 | 0 | 0 | 0 | 0 | 0 | 0 | 0 | 0 | 0 |
| 0 | NA | NA | NA | NA | NA | mt-Tv | | 0 | 0 | 0 | 0 | 0 | 0 | 0 | 0 | 0 | 0 | 0 |
| 0 | NA | NA | NA | NA | NA | mt-Ta | | 0 | 0 | 0 | 0 | 0 | 0 | 0 | 0 | 0 | 0 | 0 |
| 0 | NA | NA | NA | NA | NA | mt-Tn | | 0 | 0 | 0 | 0 | 0 | 0 | 0 | 0 | 0 | 0 | 0 |
| 0 | NA | NA | NA | NA | NA | mt-Tc | | 0 | 0 | 0 | 0 | 0 | 0 | 0 | 0 | 0 | 0 | 0 |
| 0 | NA | NA | NA | NA | NA | mt-Ty | | 0 | 0 | 0 | 0 | 0 | 0 | 0 | 0 | 0 | 0 | 0 |
| 0 | NA | NA | NA | NA | NA | mt-Ts1 | | 0 | 0 | 0 | 0 | 0 | 0 | 0 | 0 | 0 | 0 | 0 |
| 0 | NA | NA | NA | NA | NA | mt-Td | | 0 | 0 | 0 | 0 | 0 | 0 | 0 | 0 | 0 | 0 | 0 |
| 0 | NA | NA | NA | NA | NA | mt-Co2 | cytochrome c oxidase subunit II | 0 | 0 | 0 | 0 | 0 | 0 | 0 | 0 | 0 | 0 | 0 |
| 0 | NA | NA | NA | NA | NA | mt-Tk | | 0 | 0 | 0 | 0 | 0 | 0 | 0 | 0 | 0 | 0 | 0 |
| 0 | NA | NA | NA | NA | NA | mt-Atp8 | ATP synthase F0 subunit 8 | 0 | 0 | 0 | 0 | 0 | 0 | 0 | 0 | 0 | 0 | 0 |
| 0 | NA | NA | NA | NA | NA | mt-Tg | | 0 | 0 | 0 | 0 | 0 | 0 | 0 | 0 | 0 | 0 | 0 |
| 0 | NA | NA | NA | NA | NA | mt-Tr | | 0 | 0 | 0 | 0 | 0 | 0 | 0 | 0 | 0 | 0 | 0 |
| 0 | NA | NA | NA | NA | NA | mt-Nd4l | NADH dehydrogenase subunit 4L | 0 | 0 | 0 | 0 | 0 | 0 | 0 | 0 | 0 | 0 | 0 |
| 0 | NA | NA | NA | NA | NA | mt-Th | | 0 | 0 | 0 | 0 | 0 | 0 | 0 | 0 | 0 | 0 | 0 |
| 0 | NA | NA | NA | NA | NA | mt-Ts2 | | 0 | 0 | 0 | 0 | 0 | 0 | 0 | 0 | 0 | 0 | 0 |
| 0 | NA | NA | NA | NA | NA | mt-Tl2 | | 0 | 0 | 0 | 0 | 0 | 0 | 0 | 0 | 0 | 0 | 0 |
| 0 | NA | NA | NA | NA | NA | mt-Te | | 0 | 0 | 0 | 0 | 0 | 0 | 0 | 0 | 0 | 0 | 0 |
